## Supplementary material for "Completing the BASEL phage collection to unlock hidden diversity for systematic exploration of phage-host interactions": Table S3

### S3 Table. List of all bacterial strains used in this study

| **Strain** | **Genotype** | **relevant plasmid(s)** | **Selection** | **Source/Description** |
| --- | --- | --- | --- | --- |
| AH-E02-148 | *Escherichia coli* K-12 MG1655  F^–^ λ^–^ *ilvG*^–^ *rfb-50 rph-1* | none | none | our laboratory collection; *E. coli* K-12 laboratory wildtype strain |
| AH-E03-169 | *Escherichia coli* K-12 EMG2  F^+^ λ^+^ |  |  | old variant of the *E. coli* K-12 laboratory strain; obtained from the Coli Genetic Stock Center (CGSC #4401) |
| AH-E03-200 | *E. coli* K-12 MG1655 ΔRM  MG1655 Δ*mrr-hsdRMS-mcrBC* Δ*mcrA* | none | none | our laboratory collection [1]; strain lacking all known restriction systems of *E. coli* K-12 |
| AH-E03-217 | *E. coli* K-12 MG1655 ΔRM | pBR322_ΔP*tet*  F(*pifA::zeoR*) | Amp50  Zeo50 | our laboratory collection [1]; control for phenotyping experiments (without O-antigen, with K-12 cryptic prophages) |
| AH-E03-243 | *E. coli* K-12 MG1655 ΔRM *wbbL(+)* | none | none | our laboratory collection [1] |
| AH-E04-280 | *E. coli* K-12 MG1655 ΔRM *wbbL(+)* | pBR322_ΔP*tet*  F(*pifA::zeoR*) | Amp50  Zeo50 | our laboratory collection [1]; host for phenotypic experiments (with O-antigen, with K-12 cryptic prophages); for characterization of phage sensitivity or resistance to bacterial immunity, the pBR322_ΔP*tet* empty vector was replaced with one of diverse plasmids encoding antiviral defenses (see list in Table S5) |
| AH-E08-621 | *E. coli* K-12 MG1655 ΔRM *wbbL(+)* | pBR322_ΔP*tet*  pAH200e | Amp50  Kan25 | this study; pAH200e is an F-plasmid variant (encoding *pifA*) tagged with kanamycin resistance) [1] |
| AH-E01-047 | *Escherichia coli* K-12 BW25113  F^-^ Δ(*araD-araB*)567, Δ*lacZ4787(::rrnB-3*), λ^-^, *rph-1*, Δ(*rhaD-rhaB*)568, *hsdR514* | none | none | our laboratory collection [2] |
| AH-E10-740 | *E. coli* K-12 BW25113 *wbbL(+)* | none | none | this study |
| DH-E01-039 | *E. coli* K-12 BW25113 *wbbL(+)* | pBR322_ΔP*tet*  F(*pifA::zeoR*) | Amp50  Zeo50 | this study; control for phenotyping experiments (with O-antigen, with K-12 cryptic prophages) |
| AH-E01-029 | *Escherichia coli* K-12 BW25113 Δ9CP  BW25113 Δ*rac* Δ*CP4-57* Δ*CPS-53* Δ*DLP12* Δ*Qin* Δ*e14* Δ*CP4-6* Δ*CPZ-55* Δ*CP4-44* | none | none | our laboratory collection (originally created by Wang et al. [3]) |
| AH-E09-711 | *E. coli* K-12 BW25113 Δ9CP | pBR322_ΔP*tet*  F(*pifA::zeoR*) | Amp50  Zeo50 | this study; control for phenotyping experiments (without O-antigen, without K-12 cryptic prophages) |
| AH-E10-744 | *E. coli* K-12 BW25113 Δ9CP *wbbL(+)* | none | none | this study |
| DH-E01-025 | *E. coli* K-12 BW25113 Δ9CP *wbbL(+)* | pBR322_ΔP*tet*  F(*pifA::zeoR*) | Amp50  Zeo50 | this study; host for phenotypic experiments (with O-antigen, without K-12 cryptic prophages);for characterization of phage sensitivity or resistance to bacterial immunity, the pBR322_ΔP*tet* empty vector was replaced with one of diverse plasmids encoding antiviral defenses (see list in Table S5) |
| DH-E01-032 | *E. coli* K-12 BW25113 Δ9CP *wbbL(+)* | pBR322_ΔP*tet*  pAH200e | Amp50  Kan25 | this study; pAH200e is an F-plasmid variant (encoding *pifA*) tagged with kanamycin resistance) [1] |
| AH-E08-634 | *Escherichia coli* K-12 BW25113 *nfrA::kanR* | pAS001 | Kan25, Cam25 | our laboratory collection (Keio collection mutant [4]); strain lacking the NGR glycan transformed with pAS001 to restore O16-type O-antigen expression |
| DP-E03-152 | *Escherichia coli* K-12 BW25113 Δ*gtrS* | none | none | this study; clean deletion of the *gtrS* gene (encoding the glucosyltransferase of the GtrABS O-antigen glycosylation system) |
| DP-E03-170 | *Escherichia coli* K-12 BW25113 Δ*gtrS wbbL(+)* | none | none | this study; *gtrS* mutant with restored O16-type O-antigen expression |
| AH-E01-044 | *E. coli* B REL606 | none | none | our laboratory collection |
| AH-E03-168 | *E. coli* UTI89 | none | none | our laboratory collection |
| AH-E04-284 | *E. coli* CFT073 *rpoS(+)* | none | none | our laboratory collection |
| AH-E06-481 | *E. coli* 55989 | none | none | our laboratory collection |
| AH-E04-297 | *Salmonella enterica* subsp. *enterica* serovar Typhimurium 12023s (also known as ATCC 14028) | none | none | our laboratory collection |
| AH-E06-438 | *S.* Typhimurium SL1344 | none | none | our laboratory collection |
| AH-E08-577 | *E. coli* isolate 720834-18 | none | none | collection of Prof. Adrian Egli (phylogroup B2, ST131, O16-type O-antigen) [5] |
| AH-E08-578 | *E. coli* isolate 714478-19 | none | none | collection of Prof. Adrian Egli (phylogroup B2, ST144, O16-type O-antigen) [5] |
| AH-E08-576 | *E. coli* isolate 711043-19 | none | none | collection of Prof. Adrian Egli (phylogroup B2, ST8281, O16-type O-antigen) [5] |

### References (Table S3)
