## Supplementary material for "Completing the BASEL phage collection to unlock hidden diversity for systematic exploration of phage-host interactions": Table S4

### S4 Table. List of all oligonucleotide primers used in this study

| ***Primer name*** | ***Sequence (5'-3')*** |
| --- | --- |
| prAH2151 | CATTCATCCGCTTATTATCACTTA |
| prAH2152 | GTAATGACCTCAGAACTCCATCT |
| prAS0001 | ATAAGTGATAATAAGCGGATGAATGTAAGGAGGAACAATATGGTATATATAATAATCGTTTCCCAC |
| prAS0002 | CCAGATGGAGTTCTGAGGTCATTACGCTTTATATTACGGGTGAAAAACT |
| prDP0079 | TTGAAACCAAAAAACGCCCGAAATACATCATCAAGAGAGTCAAAAAATGACGTAATTTTTTTAAGGCAGTTATTG |
| prDP0080 | TAGGTTGGTATTATAGCTTGTGCGCGCCATGATTGGCGCGCAATTTAAACCTTACTGTCCCTAGTGCTTGG |
| prDP0081 | AAAACGCCCGAAATACATCATCAAGAGAGTCAAAAAATGAGTTTAAATTGCGCGCCAATCATGGCGCGCACAAGCTATAA |
| prDP0082 | TTATAGCTTGTGCGCGCCATGATTGGCGCGCAATTTAAACTCATTTTTTGACTCTCTTGATGATGTATTTCGGGCGTTTT |
| prDP0083 | TTAGTGGTAGCCAGTGTAGC |
| prDP0084 | CATTTGATAGTCAATACCGC |
