## Supplementary material for "Completing the BASEL phage collection to unlock hidden diversity for systematic exploration of phage-host interactions": Table S5

### S5 Table. List of all plasmids used in this study

| **name** | **Selection** | **Description** | **Source** |
| --- | --- | --- | --- |
| pWRG99 | Amp100 | lambda red recombineering plasmid with inducible I-SceI | our laboratory collection [1] |
| pWRG100 | Cam25 | template plasmid for the recombineering double-selectable cassette encoding chloramphenicol resistance and an I-SceI recognition site | our laboratory collection [1] |
| pUA139_cat-sacB_v3 | Cam25, Kan25 | template plasmid for recombineering double-selectable cassettes encoding *sacB* and either chloramphenicol or kanamycin resistance | our laboratory collection [2] |
| pAS001 | Cam25 | constitutive expression of *wbbL* to restore the O16-type O-antigen of K-12 strains | this study |
| pBR322_ΔP*tet* | Amp50 | variant of pBR322 in which the tetracycline resistance cassette and its promoter have been deleted; empty-vector control for immunity experiments | our laboratory collection [3] |
| pAH213_EcoKI | Amp50 | pBR322 derivative expressing type I RM system EcoKI of *E. coli* K‑12 | our laboratory collection [2] |
| pAH213_EcoCFT_I | Amp50 | pAH186_SC101e derivative expressing type I RM system EcoCFT_I of *E. coli* CFT073 | our laboratory collection [2] |
| pEcoRI | Amp50 | pBR322 derivative expressing type II RM system EcoRI | our laboratory collection [3] |
| pEcoRV | Amp50 | pBR322 derivative expressing type II RM system EcoRII | our laboratory collection [3] |
| pAH213_EcoCFT_II | Amp50 | pBR322 derivative expressing type III RM system EcoCFT_II of *E. coli* CFT073 | our laboratory collection [2] |
| pAH213_EcoP1_I | Amp50 | pAH186_SC101e derivative expressing type I RM system EcoP1_I of *E. coli* phage P1 | our laboratory collection [2] |
| pAH213_RexAB | Amp50 | pBR322 derivative expressing the RexAB Abi system of *E. coli* phage lambda | our laboratory collection [2] |
| pAH200e | Kan25 | F-plasmid in which the *tn1000* locus was replaced with a kanamycin resistance cassette by recombineering | our laboratory collection [2] |
| pAH213_Fun/Z | Amp50 | pBR322 derivative expressing the Fun/Z Abi system of *E. coli* phage P2 | our laboratory collection [2] |
| pAH213_Old | Amp50 | pBR322 derivative expressing the Old Abi system of *E. coli* phage P2 | our laboratory collection [2] |
| pAH213_Tin | Amp50 | pBR322 derivative expressing the Tin Abi system of *E. coli* phage P2 | our laboratory collection [2] |

### References (S5 Table)

1.	Blank K, Hensel M, Gerlach RG. Rapid and highly efficient method for scarless mutagenesis within the *Salmonella enterica* chromosome. PLoS One. 2011;6(1):e15763. doi: 10.1371/journal.pone.0015763. PubMed PMID: 21264289; PubMed Central PMCID: PMCPMC3021506.

2.	Maffei E, Shaidullina A, Burkolter M, Heyer Y, Estermann F, Druelle V, et al. Systematic exploration of *Escherichia coli* phage-host interactions with the BASEL phage collection. PLoS Biol. 2021;19(11):e3001424. Epub 2021/11/17. doi: 10.1371/journal.pbio.3001424. PubMed PMID: 34784345; PubMed Central PMCID: PMCPMC8594841.

3.	Pleska M, Qian L, Okura R, Bergmiller T, Wakamoto Y, Kussell E, et al. Bacterial Autoimmunity Due to a Restriction-Modification System. Curr Biol. 2016;26(3):404-9. Epub 2016/01/26. doi: 10.1016/j.cub.2015.12.041. PubMed PMID: 26804559.
