## Supplementary material for "Completing the BASEL phage collection to unlock hidden diversity for systematic exploration of phage-host interactions": Data S2: 1.html

FANPEZAQ\_CDS\_0001


Return to summary | Go to next

|  |  |
| --- | --- |
| FANPEZAQ\_CDS\_0001  Page creation date: 02 Sep 2024, 12:00  Project folder: n/a  Input sequences file: Escherichia\_virus\_HeidiAbel.gb | terminase small packaging dna nu1 phage hypothetical domain\_containing fragment helix\_turn\_helix duf1441 hth merr\_type dna\_packaging elements external origin putative dna\_binding transcriptional regulator |

### Sequence information

|  |  |
| --- | --- |
| Name | FANPEZAQ\_CDS\_0001  01\_FANPEZAQ\_CDS\_0001 (pipeline id) |
| Imported annotations | Escherichia\_virus\_HeidiAbel Bas97 |
| Protein sequence | MWRKDGCPVVTMGGRGKEWEFDTAEVANWLRDRDVRAATGDKVTDQEELKRRKLAAETEK AELELAKAKGEVAPLDQVERAVARAFAEVRAGMRNLPQRTVSMLIGETDERRFKSVMMGE IDEVLKSLVTASLLDGYDEGGDGDSEE |
| Number of residues | 147 |
| Molecular weight (Da) | 16389.26 |
| Output files | ../../query\_sequences/01\_FANPEZAQ\_CDS\_0001.fasta |

### Putative domain architecture and protein family

#### Search results (HHblits)^1^

|  |  |
| --- | --- |
| Domain family databases searched | Pfam, Ncbi-cd, Cath, Phrogs |
| Results, scheme(s)  (Top layers only; threshold 1.00e-03 (evalue)) | xml version="1.0" encoding="utf-8" standalone="no"?       2024-09-02T21:08:11.598965 image/svg+xml   Matplotlib v3.7.2, https://matplotlib.org/ |
| Results, table  (E-value ≤ 1.00e-03 (evalue)) | db \| id \| prob \| evalue \| pvalue \| score \| cols \| query \| query\_len \| template \| template\_len \| name \| description; pfam \| PF07278 \| 99.2 \| 5.9e-16 \| 1e-19 \| 94.3 \| 113 \| (16, 133) \| 147 \| (33, 145) \| 147 \| DUF1441 \| Protein of unknown function (DUF1441); pfam \| PF07471 \| 99.0 \| 1.8e-14 \| 3e-18 \| 88.3 \| 124 \| (2, 127) \| 147 \| (22, 161) \| 162 \| Phage\_Nu1 \| Phage DNA packaging protein Nu1; phrogs \| 141 \| 99.8 \| 2.1e-24 \| 2.5e-28 \| 150.6 \| 121 \| (17, 145) \| 147 \| (41, 161) \| 163 \| terminase small subunit \| terminase small subunit; Category: head and packaging; p69558 VI\_02299; phrogs \| 57 \| 99.7 \| 9.1e-23 \| 1.2e-26 \| 143.9 \| 128 \| (2, 136) \| 147 \| (24, 154) \| 173 \| terminase small subunit \| terminase small subunit; Category: head and packaging; p277333 VI\_02593; phrogs \| 21782 \| 99.6 \| 1.2e-20 \| 1.3e-24 \| 121.2 \| 126 \| (17, 145) \| 147 \| (7, 132) \| 136 \| NA \| NA; Category: unknown function; p302384 VI\_08027; phrogs \| 1790 \| 98.6 \| 7.5e-12 \| 8.4e-16 \| 77.4 \| 86 \| (44, 131) \| 147 \| (12, 97) \| 114 \| terminase small subunit \| terminase small subunit; Category: head and packaging; p410932 VI\_07810; phrogs \| 5710 \| 97.2 \| 4.1e-07 \| 4.7e-11 \| 55.2 \| 62 \| (16, 79) \| 147 \| (41, 102) \| 104 \| NA \| NA; Category: unknown function; p435585 VI\_08997; phrogs \| 6039 \| 97.1 \| 5.1e-07 \| 5.7e-11 \| 48.2 \| 54 \| (91, 147) \| 147 \| (2, 55) \| 55 \| NA \| NA; Category: unknown function; p386363 VI\_02030 |
| Top keywords  (threshold 1.00e-03 (evalue)) | **packaging, terminase, small, head, and, DUF1441, Phage, DNA, Nu1, p69558** |
| Output files | ../../domain\_architecture/01\_FANPEZAQ\_CDS\_0001\_cath.hhr ../../domain\_architecture/01\_FANPEZAQ\_CDS\_0001\_merged.svg ../../domain\_architecture/01\_FANPEZAQ\_CDS\_0001\_ncbi-cd.hhr ../../domain\_architecture/01\_FANPEZAQ\_CDS\_0001\_pfam.hhr ../../domain\_architecture/01\_FANPEZAQ\_CDS\_0001\_phrogs.hhr |

### Identical protein sequences/structures

#### Search results

|  |  |
| --- | --- |
| Protein sequence databases searched | Pdb, Swissprot, Refseq |
| Identical proteins found | -- |
| Top keywords | -- |
| Output files | -- |

### Similar protein sequences/structures

#### Sequence similarity search results (HHblits)^1^

|  |  |
| --- | --- |
| Sequence databases searched | Uniclust, Pdb70 |
| Results, scheme(s)  (Top layers only, threshold 1.00e-03 (evalue)) | xml version="1.0" encoding="utf-8" standalone="no"?       2024-09-02T21:08:27.846833 image/svg+xml   Matplotlib v3.7.2, https://matplotlib.org/ |
| Results, table(s)  (threshold 1.00e-03 (evalue)) | db \| id \| prob \| evalue \| pvalue \| score \| cols \| query \| query\_len \| template \| template\_len \| name \| description; uniclust \| UniRef100\_A0A017HCB4 \| 99.9 \| 1.7e-28 \| 3.6e-34 \| 182.6 \| 142 \| (1, 142) \| 147 \| (57, 198) \| 231 \| Phage DNA packaging Nu1 \| Phage DNA packaging Nu1; uniclust \| UniRef100\_A0A073IUP9 \| 99.9 \| 4.4e-25 \| 9.4e-31 \| 164.7 \| 130 \| (3, 137) \| 147 \| (59, 189) \| 229 \| Terminase small subunit \| Terminase small subunit; uniclust \| UniRef100\_A0A103EHP4 \| 99.8 \| 2.6e-22 \| 5e-28 \| 138.7 \| 142 \| (1, 142) \| 147 \| (25, 166) \| 175 \| Terminase small subunit \| Terminase small subunit; uniclust \| UniRef100\_A0A3B8QTC1 \| 99.8 \| 3.2e-22 \| 6.2e-28 \| 140.3 \| 136 \| (1, 139) \| 147 \| (27, 164) \| 180 \| DNA packaging protein \| DNA packaging protein; uniclust \| UniRef100\_A0A192IJE2 \| 99.8 \| 5.8e-22 \| 1.2e-27 \| 144.2 \| 125 \| (2, 130) \| 147 \| (34, 161) \| 189 \| Terminase small subunit \| Terminase small subunit; uniclust \| UniRef100\_A0A137SPY3 \| 99.8 \| 7.7e-22 \| 1.5e-27 \| 143.2 \| 134 \| (3, 141) \| 147 \| (35, 169) \| 216 \| Phage DNA packaging protein Nu1 \| Phage DNA packaging protein Nu1; uniclust \| UniRef100\_A0A0C9PQA6 \| 99.8 \| 1.2e-21 \| 2.3e-27 \| 139.5 \| 130 \| (1, 134) \| 147 \| (49, 178) \| 188 \| Uncharacterized protein \| Uncharacterized protein; uniclust \| UniRef100\_A0A031G4Z9 \| 99.7 \| 3.1e-21 \| 6.3e-27 \| 139.8 \| 127 \| (1, 133) \| 147 \| (44, 172) \| 208 \| Phage DNA packaging protein Nu1 \| Phage DNA packaging protein Nu1; uniclust \| UniRef100\_A0A081RFK1 \| 99.7 \| 6.2e-21 \| 1.3e-26 \| 140.8 \| 133 \| (2, 141) \| 147 \| (61, 197) \| 220 \| Phage DNA packaging protein Nu1 \| Phage DNA packaging protein Nu1; uniclust \| UniRef100\_A0A0B1Q7B2 \| 99.7 \| 6.3e-21 \| 1.3e-26 \| 139.1 \| 123 \| (2, 130) \| 147 \| (37, 164) \| 196 \| Uncharacterized protein \| Uncharacterized protein; uniclust \| UniRef100\_A0A061NPR6 \| 99.7 \| 1.2e-20 \| 2.4e-26 \| 140.5 \| 134 \| (2, 140) \| 147 \| (58, 197) \| 229 \| Uncharacterized protein \| Uncharacterized protein; uniclust \| UniRef100\_A0A0R3MP30 \| 99.7 \| 1.2e-20 \| 2.5e-26 \| 138.4 \| 129 \| (1, 129) \| 147 \| (50, 180) \| 212 \| DNA packaging protein \| DNA packaging protein; uniclust \| UniRef100\_A0A062V3Q6 \| 99.7 \| 2.3e-20 \| 4.7e-26 \| 131.5 \| 124 \| (2, 132) \| 147 \| (44, 169) \| 172 \| Phage DNA packaging protein Nu1 \| Phage DNA packaging protein Nu1; uniclust \| UniRef100\_A0A0A8K3F1 \| 99.7 \| 4.5e-20 \| 9.4e-26 \| 135.6 \| 124 \| (2, 132) \| 147 \| (54, 181) \| 205 \| Phage DNA packaging protein, Nu1 subunit of terminase \| Phage DNA packaging protein, Nu1 subunit of terminase; uniclust \| UniRef100\_A0A1X7L0X5 \| 99.7 \| 5.4e-20 \| 1e-25 \| 127.1 \| 131 \| (1, 137) \| 147 \| (32, 165) \| 168 \| Phage DNA packaging protein, Nu1 subunit of terminase \| Phage DNA packaging protein, Nu1 subunit of terminase; uniclust \| UniRef100\_A0A2D8RC70 \| 99.7 \| 6.6e-20 \| 1.3e-25 \| 129.5 \| 141 \| (2, 142) \| 147 \| (23, 165) \| 174 \| Terminase small subunit \| Terminase small subunit; uniclust \| UniRef100\_A0A1M6SLK9 \| 99.7 \| 7.8e-20 \| 1.6e-25 \| 133.0 \| 135 \| (2, 142) \| 147 \| (34, 175) \| 205 \| Phage DNA packaging protein, Nu1 subunit of terminase \| Phage DNA packaging protein, Nu1 subunit of terminase; uniclust \| UniRef100\_A0A069I596 \| 99.7 \| 2e-19 \| 4e-25 \| 128.0 \| 130 \| (1, 136) \| 147 \| (25, 155) \| 178 \| DNA packaging protein \| DNA packaging protein; uniclust \| UniRef100\_A0A126NZA6 \| 99.7 \| 2.2e-19 \| 4.3e-25 \| 123.6 \| 132 \| (1, 133) \| 147 \| (28, 159) \| 162 \| Terminase small subunit \| Terminase small subunit; uniclust \| UniRef100\_A0A1Q3WRG7 \| 99.6 \| 3.3e-19 \| 6.6e-25 \| 126.8 \| 132 \| (1, 139) \| 147 \| (35, 166) \| 184 \| Terminase small subunit \| Terminase small subunit; uniclust \| UniRef100\_A0A1C3WPE9 \| 99.6 \| 4e-19 \| 8e-25 \| 129.9 \| 126 \| (3, 134) \| 147 \| (33, 160) \| 215 \| Phage DNA packaging protein, Nu1 subunit of terminase \| Phage DNA packaging protein, Nu1 subunit of terminase; uniclust \| UniRef100\_A0A078MJZ8 \| 99.6 \| 4.7e-19 \| 9.4e-25 \| 130.7 \| 130 \| (3, 137) \| 147 \| (38, 169) \| 224 \| Phage DNA packaging protein Nu1 \| Phage DNA packaging protein Nu1; uniclust \| UniRef100\_A0A1H8QIJ8 \| 99.6 \| 5.5e-19 \| 1.1e-24 \| 125.0 \| 130 \| (2, 138) \| 147 \| (32, 163) \| 187 \| Uncharacterized protein \| Uncharacterized protein; uniclust \| UniRef100\_A0A8J7MTI5 \| 99.6 \| 8.8e-19 \| 1.7e-24 \| 119.4 \| 132 \| (1, 133) \| 147 \| (22, 153) \| 159 \| Terminase small subunit \| Terminase small subunit; uniclust \| UniRef100\_A0A2D5VNR8 \| 99.6 \| 2.6e-18 \| 5e-24 \| 119.6 \| 133 \| (2, 134) \| 147 \| (22, 160) \| 169 \| Terminase small subunit \| Terminase small subunit; uniclust \| UniRef100\_A0A088C3V1 \| 99.6 \| 3.1e-18 \| 6e-24 \| 122.2 \| 136 \| (1, 139) \| 147 \| (44, 182) \| 202 \| Terminase small subunit \| Terminase small subunit; uniclust \| UniRef100\_A0A0D0QHJ8 \| 99.6 \| 6.7e-18 \| 1.4e-23 \| 123.0 \| 126 \| (2, 135) \| 147 \| (49, 176) \| 196 \| Phage DNA packaging protein, Nu1 subunit of terminase \| Phage DNA packaging protein, Nu1 subunit of terminase; uniclust \| UniRef100\_A0A1H5XMK5 \| 99.6 \| 9.8e-18 \| 2e-23 \| 122.1 \| 128 \| (2, 131) \| 147 \| (40, 170) \| 201 \| Phage DNA packaging protein Nu1 \| Phage DNA packaging protein Nu1; uniclust \| UniRef100\_UPI00174BBF1A \| 99.5 \| 1.4e-17 \| 2.6e-23 \| 117.6 \| 135 \| (1, 135) \| 147 \| (28, 165) \| 200 \| terminase small subunit \| terminase small subunit; uniclust \| UniRef100\_A0A240TSP7 \| 99.5 \| 1.4e-17 \| 2.8e-23 \| 118.8 \| 127 \| (3, 135) \| 147 \| (51, 178) \| 202 \| Terminase small subunit, Nu1 \| Terminase small subunit, Nu1; uniclust \| UniRef100\_A0A1Q6PWQ4 \| 99.5 \| 1.6e-17 \| 3.2e-23 \| 117.1 \| 130 \| (7, 142) \| 147 \| (35, 164) \| 176 \| Terminase small subunit \| Terminase small subunit; uniclust \| UniRef100\_A0A081NI16 \| 99.5 \| 1.5e-17 \| 3.2e-23 \| 124.8 \| 131 \| (3, 138) \| 147 \| (42, 195) \| 216 \| Terminase small subunit \| Terminase small subunit; uniclust \| UniRef100\_A0A0Q4E400 \| 99.5 \| 6.7e-17 \| 1.3e-22 \| 115.4 \| 128 \| (2, 135) \| 147 \| (37, 168) \| 175 \| HTH merR-type domain-containing protein \| HTH merR-type domain-containing protein; uniclust \| UniRef100\_A0A1G1HG69 \| 99.5 \| 9.5e-17 \| 1.7e-22 \| 110.4 \| 132 \| (1, 134) \| 147 \| (19, 150) \| 185 \| Uncharacterized protein \| Uncharacterized protein; uniclust \| UniRef100\_A0A087M4C5 \| 99.5 \| 9.1e-17 \| 1.9e-22 \| 118.0 \| 126 \| (1, 133) \| 147 \| (42, 171) \| 204 \| DNA packaging protein \| DNA packaging protein; uniclust \| UniRef100\_A0A165IZT2 \| 99.5 \| 9.8e-17 \| 2e-22 \| 119.7 \| 131 \| (1, 137) \| 147 \| (37, 192) \| 219 \| Terminase small subunit \| Terminase small subunit; uniclust \| UniRef100\_A0A248UJZ5 \| 99.5 \| 1.4e-16 \| 2.6e-22 \| 119.8 \| 129 \| (2, 137) \| 147 \| (148, 279) \| 309 \| Homeo-like domain protein \| Homeo-like domain protein; uniclust \| UniRef100\_A0A1U7MFY0 \| 99.5 \| 1.4e-16 \| 2.9e-22 \| 115.6 \| 133 \| (1, 141) \| 147 \| (42, 176) \| 188 \| Phage DNA packaging protein Nu1 \| Phage DNA packaging protein Nu1; uniclust \| UniRef100\_A0A7C6RL71 \| 99.5 \| 2.1e-16 \| 4.1e-22 \| 112.2 \| 138 \| (1, 138) \| 147 \| (23, 172) \| 181 \| Terminase small subunit \| Terminase small subunit; uniclust \| UniRef100\_A0A7V5XCS4 \| 99.5 \| 2.2e-16 \| 4.2e-22 \| 112.6 \| 124 \| (1, 128) \| 147 \| (48, 171) \| 202 \| Terminase small subunit \| Terminase small subunit; uniclust \| UniRef100\_A0A068SKV9 \| 99.5 \| 2.1e-16 \| 4.2e-22 \| 118.3 \| 131 \| (1, 132) \| 147 \| (94, 231) \| 251 \| Phage DNA packaging Nu1 \| Phage DNA packaging Nu1; uniclust \| UniRef100\_A0A0K1NDZ1 \| 99.5 \| 2.2e-16 \| 4.5e-22 \| 116.9 \| 125 \| (2, 131) \| 147 \| (38, 184) \| 214 \| Uncharacterized protein \| Uncharacterized protein; uniclust \| UniRef100\_A0A1G1JJM4 \| 99.4 \| 2.9e-16 \| 5.8e-22 \| 116.2 \| 125 \| (1, 134) \| 147 \| (77, 201) \| 230 \| HTH merR-type domain-containing protein \| HTH merR-type domain-containing protein; uniclust \| UniRef100\_A0A6L4BEY6 \| 99.4 \| 3.4e-16 \| 6.3e-22 \| 108.5 \| 134 \| (1, 136) \| 147 \| (44, 177) \| 193 \| Terminase small subunit \| Terminase small subunit; uniclust \| UniRef100\_A0A127CGJ9 \| 99.4 \| 3.1e-16 \| 6.4e-22 \| 114.3 \| 135 \| (2, 142) \| 147 \| (45, 190) \| 196 \| DNA packaging protein \| DNA packaging protein; uniclust \| UniRef100\_A0A086Y826 \| 99.4 \| 3.2e-16 \| 6.5e-22 \| 118.6 \| 126 \| (1, 129) \| 147 \| (74, 219) \| 252 \| DNA packaging protein \| DNA packaging protein; uniclust \| UniRef100\_A0A0Q7XWC0 \| 99.4 \| 3.8e-16 \| 7.4e-22 \| 114.8 \| 133 \| (1, 134) \| 147 \| (73, 214) \| 228 \| Terminase small subunit \| Terminase small subunit; uniclust \| UniRef100\_UPI0021CCE98E \| 99.4 \| 5.2e-16 \| 9.6e-22 \| 110.2 \| 135 \| (1, 135) \| 147 \| (29, 163) \| 231 \| terminase small subunit \| terminase small subunit; uniclust \| UniRef100\_A0A073J8E6 \| 99.4 \| 6.3e-16 \| 1.3e-21 \| 118.0 \| 106 \| (1, 106) \| 147 \| (75, 201) \| 261 \| Phage DNA packaging protein Nu1 \| Phage DNA packaging protein Nu1; uniclust \| UniRef100\_A0A2R3Q9L1 \| 99.4 \| 7.5e-16 \| 1.5e-21 \| 111.0 \| 123 \| (2, 135) \| 147 \| (31, 183) \| 196 \| Terminase small subunit \| Terminase small subunit; uniclust \| UniRef100\_A0A0H3ZUV8 \| 99.4 \| 1.1e-15 \| 2.3e-21 \| 111.6 \| 128 \| (1, 128) \| 147 \| (39, 172) \| 211 \| Phage DNA packaging \| Phage DNA packaging; uniclust \| UniRef100\_A0A510E7W2 \| 99.4 \| 1.4e-15 \| 2.7e-21 \| 106.8 \| 127 \| (1, 134) \| 147 \| (24, 150) \| 174 \| Uncharacterized protein \| Uncharacterized protein; uniclust \| UniRef100\_A0A2T5K719 \| 99.4 \| 1.6e-15 \| 3e-21 \| 109.9 \| 134 \| (1, 134) \| 147 \| (44, 180) \| 215 \| Phage terminase Nu1 subunit (DNA packaging protein) \| Phage terminase Nu1 subunit (DNA packaging protein); uniclust \| UniRef100\_A0A136M3Y9 \| 99.4 \| 1.7e-15 \| 3.3e-21 \| 98.9 \| 91 \| (43, 133) \| 147 \| (11, 101) \| 109 \| Tetrapyrrole biosynthesis glutamyl-tRNA reductase dimerisation domain-containing protein (Fragment) \| Tetrapyrrole biosynthesis glutamyl-tRNA reductase dimerisation domain-containing protein (Fragment); uniclust \| UniRef100\_A0A235H584 \| 99.4 \| 1.9e-15 \| 3.8e-21 \| 113.4 \| 135 \| (1, 136) \| 147 \| (82, 224) \| 240 \| Terminase small subunit (DNA packaging protein Nu1) \| Terminase small subunit (DNA packaging protein Nu1); uniclust \| UniRef100\_A0A2N2QIM9 \| 99.4 \| 2.9e-15 \| 5.8e-21 \| 109.5 \| 89 \| (44, 132) \| 147 \| (89, 177) \| 209 \| Terminase \| Terminase; uniclust \| UniRef100\_A0A0E1N6N6 \| 99.3 \| 4.1e-15 \| 7.9e-21 \| 105.5 \| 129 \| (2, 135) \| 147 \| (27, 159) \| 174 \| Phage DNA packaging Nu1 family protein \| Phage DNA packaging Nu1 family protein; uniclust \| UniRef100\_A0A0F9D3W0 \| 99.3 \| 3.9e-15 \| 8e-21 \| 108.3 \| 136 \| (1, 138) \| 147 \| (23, 173) \| 190 \| Terminase small subunit (Fragment) \| Terminase small subunit (Fragment); uniclust \| UniRef100\_A0A071LTT6 \| 99.3 \| 5.3e-15 \| 1.1e-20 \| 112.1 \| 128 \| (1, 133) \| 147 \| (63, 191) \| 240 \| Uncharacterized protein \| Uncharacterized protein; uniclust \| UniRef100\_A0A318U841 \| 99.3 \| 6.4e-15 \| 1.2e-20 \| 107.4 \| 121 \| (4, 131) \| 147 \| (103, 228) \| 230 \| Phage terminase Nu1 subunit (DNA packaging protein) \| Phage terminase Nu1 subunit (DNA packaging protein); uniclust \| UniRef100\_A0A1M7CWV3 \| 99.3 \| 6.9e-15 \| 1.4e-20 \| 107.6 \| 130 \| (2, 137) \| 147 \| (31, 169) \| 197 \| Phage DNA packaging protein, Nu1 subunit of terminase \| Phage DNA packaging protein, Nu1 subunit of terminase; uniclust \| UniRef100\_A0A1F9UST3 \| 99.3 \| 8.2e-15 \| 1.5e-20 \| 100.6 \| 126 \| (1, 131) \| 147 \| (47, 172) \| 176 \| Helix-turn-helix domain-containing protein \| Helix-turn-helix domain-containing protein; uniclust \| UniRef100\_A0A1F9I5E8 \| 99.3 \| 8e-15 \| 1.6e-20 \| 106.7 \| 94 \| (44, 137) \| 147 \| (101, 194) \| 200 \| Terminase small subunit \| Terminase small subunit; uniclust \| UniRef100\_A0A315EM69 \| 99.3 \| 1e-14 \| 2e-20 \| 102.3 \| 135 \| (1, 135) \| 147 \| (32, 166) \| 182 \| DNA-packaging protein \| DNA-packaging protein; uniclust \| UniRef100\_A0A024HPJ0 \| 99.3 \| 1.1e-14 \| 2.1e-20 \| 107.8 \| 127 \| (2, 135) \| 147 \| (42, 182) \| 213 \| p21 prophage-derived terminase small subunit \| p21 prophage-derived terminase small subunit; uniclust \| UniRef100\_A0A3N2E0T8 \| 99.3 \| 1.7e-14 \| 3e-20 \| 98.7 \| 138 \| (1, 138) \| 147 \| (27, 165) \| 170 \| Phage terminase Nu1 subunit (DNA packaging protein) \| Phage terminase Nu1 subunit (DNA packaging protein); uniclust \| UniRef100\_A0A0C2YAI7 \| 99.3 \| 1.8e-14 \| 3.6e-20 \| 103.8 \| 133 \| (2, 140) \| 147 \| (34, 179) \| 186 \| Uncharacterized protein \| Uncharacterized protein; uniclust \| UniRef100\_A0A031IUW6 \| 99.3 \| 1.8e-14 \| 3.9e-20 \| 109.0 \| 95 \| (44, 138) \| 147 \| (99, 193) \| 230 \| Terminase small subunit \| Terminase small subunit; uniclust \| UniRef100\_A0A4Y3TG37 \| 99.3 \| 2.2e-14 \| 4e-20 \| 100.7 \| 132 \| (1, 133) \| 147 \| (27, 160) \| 203 \| Terminase small subunit \| Terminase small subunit; uniclust \| UniRef100\_A0A098F576 \| 99.3 \| 2.3e-14 \| 4.7e-20 \| 105.0 \| 129 \| (1, 135) \| 147 \| (50, 187) \| 200 \| Phage DNA packaging protein, Nu1 subunit of terminase \| Phage DNA packaging protein, Nu1 subunit of terminase; uniclust \| UniRef100\_A0A7G8BY94 \| 99.2 \| 4.3e-14 \| 8e-20 \| 99.1 \| 127 \| (2, 131) \| 147 \| (28, 165) \| 183 \| Terminase small subunit \| Terminase small subunit; uniclust \| UniRef100\_A0A3B1BSM9 \| 99.2 \| 4.7e-14 \| 8.6e-20 \| 92.3 \| 109 \| (21, 135) \| 147 \| (9, 117) \| 126 \| Uncharacterized protein \| Uncharacterized protein; uniclust \| UniRef100\_A0A0E4BPR8 \| 99.2 \| 4.8e-14 \| 1e-19 \| 106.7 \| 115 \| (2, 131) \| 147 \| (71, 185) \| 229 \| Terminase small subunit \| Terminase small subunit; uniclust \| UniRef100\_A0A0F8WMT9 \| 99.2 \| 5.7e-14 \| 1.1e-19 \| 99.0 \| 128 \| (3, 135) \| 147 \| (28, 158) \| 196 \| Uncharacterized protein \| Uncharacterized protein; uniclust \| UniRef100\_A0A021X9M2 \| 99.2 \| 8.2e-14 \| 1.7e-19 \| 106.9 \| 94 \| (45, 138) \| 147 \| (119, 212) \| 242 \| Terminase small subunit \| Terminase small subunit; uniclust \| UniRef100\_A0A957S2F1 \| 99.2 \| 1.5e-13 \| 2.9e-19 \| 98.6 \| 131 \| (3, 135) \| 147 \| (25, 186) \| 192 \| Uncharacterized protein \| Uncharacterized protein; uniclust \| UniRef100\_A0A075KE40 \| 99.2 \| 1.6e-13 \| 3.3e-19 \| 100.0 \| 127 \| (1, 136) \| 147 \| (44, 173) \| 183 \| Terminase small subunit \| Terminase small subunit; uniclust \| UniRef100\_A0A024E8L0 \| 99.1 \| 2.7e-13 \| 5.6e-19 \| 106.2 \| 93 \| (44, 136) \| 147 \| (136, 228) \| 291 \| Terminase small subunit \| Terminase small subunit; uniclust \| UniRef100\_A0A5K8AH95 \| 99.1 \| 3.8e-13 \| 7.2e-19 \| 93.6 \| 132 \| (1, 132) \| 147 \| (19, 155) \| 157 \| Terminase small subunit \| Terminase small subunit; uniclust \| UniRef100\_A0A1A8TAI5 \| 99.1 \| 4.1e-13 \| 7.7e-19 \| 93.8 \| 128 \| (1, 131) \| 147 \| (23, 156) \| 166 \| Phage DNA packaging protein Nu1 \| Phage DNA packaging protein Nu1; uniclust \| UniRef100\_E8X2N4 \| 99.1 \| 5.3e-13 \| 9.7e-19 \| 93.1 \| 132 \| (1, 139) \| 147 \| (31, 169) \| 189 \| Putative terminase small subunit, Nu1 \| Putative terminase small subunit, Nu1; uniclust \| UniRef100\_A0A0G3XF80 \| 99.1 \| 4.9e-13 \| 1e-18 \| 100.9 \| 125 \| (2, 129) \| 147 \| (46, 194) \| 225 \| Putative transcriptional regulator \| Putative transcriptional regulator; uniclust \| UniRef100\_A0A0S6UUS0 \| 99.1 \| 7e-13 \| 1.3e-18 \| 86.3 \| 87 \| (44, 132) \| 147 \| (13, 99) \| 102 \| Uncharacterized protein \| Uncharacterized protein; uniclust \| UniRef100\_A0A011QU19 \| 99.1 \| 7.5e-13 \| 1.5e-18 \| 100.3 \| 95 \| (43, 137) \| 147 \| (111, 207) \| 233 \| Elements of external origin \| Elements of external origin; uniclust \| UniRef100\_A0A1M3NUS7 \| 99.1 \| 8.4e-13 \| 1.7e-18 \| 99.1 \| 97 \| (2, 106) \| 147 \| (53, 149) \| 212 \| Terminase small subunit \| Terminase small subunit; uniclust \| UniRef100\_UPI0013FDE30D \| 99.1 \| 9.5e-13 \| 1.7e-18 \| 92.4 \| 128 \| (1, 129) \| 147 \| (37, 169) \| 196 \| terminase small subunit \| terminase small subunit; uniclust \| UniRef100\_A0A066T0V2 \| 99.1 \| 1e-12 \| 2e-18 \| 101.9 \| 131 \| (1, 133) \| 147 \| (52, 198) \| 313 \| Terminase small subunit \| Terminase small subunit; uniclust \| UniRef100\_UPI00177B5103 \| 99.0 \| 1.3e-12 \| 2.4e-18 \| 92.9 \| 122 \| (3, 131) \| 147 \| (89, 211) \| 214 \| terminase small subunit \| terminase small subunit; uniclust \| UniRef100\_A0A3M2C0J2 \| 99.0 \| 1.6e-12 \| 3e-18 \| 87.0 \| 124 \| (3, 131) \| 147 \| (13, 139) \| 141 \| Terminase \| Terminase; uniclust \| UniRef100\_A0A0A0F9C3 \| 99.0 \| 1.6e-12 \| 3.1e-18 \| 94.1 \| 132 \| (1, 142) \| 147 \| (32, 164) \| 205 \| Terminase small subunit \| Terminase small subunit; uniclust \| UniRef100\_A0A1F2RY64 \| 99.0 \| 1.6e-12 \| 3.2e-18 \| 92.4 \| 89 \| (43, 131) \| 147 \| (80, 168) \| 174 \| Uncharacterized protein \| Uncharacterized protein; uniclust \| UniRef100\_A0A0F8W5B6 \| 99.0 \| 1.7e-12 \| 3.3e-18 \| 93.9 \| 132 \| (1, 135) \| 147 \| (34, 166) \| 198 \| Terminase small subunit (Fragment) \| Terminase small subunit (Fragment); uniclust \| UniRef100\_A0A0E4FVD0 \| 99.0 \| 1.8e-12 \| 3.5e-18 \| 91.8 \| 127 \| (3, 135) \| 147 \| (26, 156) \| 177 \| Uncharacterized protein \| Uncharacterized protein; uniclust \| UniRef100\_A0A1C3XV01 \| 99.0 \| 1.9e-12 \| 3.5e-18 \| 93.4 \| 126 \| (2, 133) \| 147 \| (43, 171) \| 203 \| Phage DNA packaging protein, Nu1 subunit of terminase \| Phage DNA packaging protein, Nu1 subunit of terminase; uniclust \| UniRef100\_A0A081MYL8 \| 99.0 \| 2.4e-12 \| 4.6e-18 \| 93.2 \| 136 \| (1, 136) \| 147 \| (24, 171) \| 188 \| Terminase small subunit \| Terminase small subunit; uniclust \| UniRef100\_A0A1S6TPB0 \| 99.0 \| 2.5e-12 \| 5e-18 \| 93.2 \| 123 \| (3, 135) \| 147 \| (25, 147) \| 184 \| Phage DNA packaging protein, Nu1 subunit of terminase \| Phage DNA packaging protein, Nu1 subunit of terminase; uniclust \| UniRef100\_A0A9D5VVB4 \| 99.0 \| 3e-12 \| 5.5e-18 \| 87.2 \| 125 \| (1, 131) \| 147 \| (26, 151) \| 157 \| Uncharacterized protein \| Uncharacterized protein; uniclust \| UniRef100\_A0A3A8WV56 \| 99.0 \| 3.4e-12 \| 6.4e-18 \| 88.2 \| 132 \| (3, 139) \| 147 \| (24, 156) \| 164 \| Terminase small subunit \| Terminase small subunit; uniclust \| UniRef100\_A0A023XKN2 \| 99.0 \| 3.3e-12 \| 6.5e-18 \| 94.1 \| 111 \| (3, 129) \| 147 \| (54, 164) \| 199 \| Uncharacterized protein \| Uncharacterized protein; uniclust \| UniRef100\_A0A1V1UM65 \| 99.0 \| 3.4e-12 \| 6.7e-18 \| 94.9 \| 127 \| (2, 133) \| 147 \| (44, 174) \| 213 \| Phage DNA packaging protein Nu1 \| Phage DNA packaging protein Nu1; uniclust \| UniRef100\_A0A133ZYV3 \| 99.0 \| 3.6e-12 \| 7.1e-18 \| 96.3 \| 133 \| (2, 134) \| 147 \| (84, 227) \| 248 \| Phage DNA packaging protein Nu1 \| Phage DNA packaging protein Nu1; uniclust \| UniRef100\_A0A1H5S2V5 \| 99.0 \| 4.1e-12 \| 8.3e-18 \| 93.5 \| 119 \| (23, 141) \| 147 \| (45, 166) \| 184 \| Phage DNA packaging protein, Nu1 subunit of terminase \| Phage DNA packaging protein, Nu1 subunit of terminase; uniclust \| UniRef100\_A0A2E5KQ39 \| 99.0 \| 4.7e-12 \| 8.9e-18 \| 89.8 \| 121 \| (3, 135) \| 147 \| (27, 147) \| 175 \| Terminase small subunit, Nu1 \| Terminase small subunit, Nu1; uniclust \| UniRef100\_A0A1G1KXY1 \| 99.0 \| 5.1e-12 \| 9.4e-18 \| 87.6 \| 120 \| (1, 130) \| 147 \| (34, 153) \| 176 \| Terminase small subunit \| Terminase small subunit; uniclust \| UniRef100\_K2J734 \| 99.0 \| 5.1e-12 \| 9.4e-18 \| 89.3 \| 131 \| (1, 134) \| 147 \| (32, 187) \| 202 \| Terminase small subunit \| Terminase small subunit; uniclust \| UniRef100\_A0A962SUK3 \| 99.0 \| 5.2e-12 \| 9.6e-18 \| 87.8 \| 126 \| (1, 129) \| 147 \| (25, 155) \| 166 \| DUF1441 family protein \| DUF1441 family protein; uniclust \| UniRef100\_A0A540V7P3 \| 98.9 \| 5.8e-12 \| 1.1e-17 \| 86.1 \| 121 \| (1, 130) \| 147 \| (26, 146) \| 160 \| Terminase small subunit \| Terminase small subunit; uniclust \| UniRef100\_A0A0A1YW04 \| 98.9 \| 6e-12 \| 1.2e-17 \| 94.8 \| 122 \| (21, 142) \| 147 \| (61, 185) \| 234 \| Uncharacterized protein \| Uncharacterized protein; uniclust \| UniRef100\_A0A176Z207 \| 98.9 \| 7.1e-12 \| 1.4e-17 \| 90.0 \| 93 \| (43, 135) \| 147 \| (74, 166) \| 175 \| Uncharacterized protein \| Uncharacterized protein; uniclust \| UniRef100\_A0A175RTU2 \| 98.9 \| 7.2e-12 \| 1.4e-17 \| 95.1 \| 128 \| (1, 129) \| 147 \| (60, 198) \| 248 \| Terminase small subunit \| Terminase small subunit; uniclust \| UniRef100\_A0A379CXL1 \| 98.9 \| 9.5e-12 \| 1.8e-17 \| 90.4 \| 132 \| (1, 140) \| 147 \| (40, 172) \| 225 \| Phage DNA packaging protein Nu1 \| Phage DNA packaging protein Nu1; uniclust \| UniRef100\_A0A088FV50 \| 98.9 \| 9.7e-12 \| 1.8e-17 \| 80.9 \| 99 \| (48, 146) \| 147 \| (6, 104) \| 108 \| Small terminase subunit \| Small terminase subunit; uniclust \| UniRef100\_A0A928NFM1 \| 98.9 \| 1.1e-11 \| 1.9e-17 \| 79.7 \| 90 \| (43, 132) \| 147 \| (14, 103) \| 107 \| Terminase small subunit \| Terminase small subunit; uniclust \| UniRef100\_A0A024L3J4 \| 98.9 \| 1.1e-11 \| 2e-17 \| 92.6 \| 129 \| (1, 131) \| 147 \| (36, 180) \| 278 \| DNA-packaging protein \| DNA-packaging protein; uniclust \| UniRef100\_A0A1H7YHA6 \| 98.9 \| 1.1e-11 \| 2.2e-17 \| 90.6 \| 130 \| (1, 136) \| 147 \| (43, 184) \| 192 \| Phage DNA packaging protein, Nu1 subunit of terminase \| Phage DNA packaging protein, Nu1 subunit of terminase; uniclust \| UniRef100\_A0A1B7L4T6 \| 98.9 \| 1.2e-11 \| 2.4e-17 \| 89.1 \| 116 \| (21, 136) \| 147 \| (41, 158) \| 185 \| Phage DNA packaging protein Nu1 \| Phage DNA packaging protein Nu1; uniclust \| UniRef100\_A0A946DB58 \| 98.9 \| 1.3e-11 \| 2.5e-17 \| 87.9 \| 125 \| (2, 131) \| 147 \| (32, 157) \| 192 \| Uncharacterized protein \| Uncharacterized protein; uniclust \| UniRef100\_UPI0018E13DD1 \| 98.9 \| 1.5e-11 \| 2.7e-17 \| 89.6 \| 124 \| (5, 134) \| 147 \| (101, 225) \| 251 \| type IV toxin-antitoxin system AbiEi family antitoxin domain-containing protein \| type IV toxin-antitoxin system AbiEi family antitoxin domain-containing protein; uniclust \| UniRef100\_A0A316SBE2 \| 98.9 \| 1.6e-11 \| 2.9e-17 \| 88.2 \| 130 \| (1, 137) \| 147 \| (74, 206) \| 223 \| Terminase small subunit \| Terminase small subunit; uniclust \| UniRef100\_A0A933XTS0 \| 98.9 \| 1.7e-11 \| 3e-17 \| 81.8 \| 92 \| (42, 133) \| 147 \| (30, 121) \| 134 \| Uncharacterized protein \| Uncharacterized protein; uniclust \| UniRef100\_A0A935WCC7 \| 98.9 \| 1.7e-11 \| 3e-17 \| 84.6 \| 129 \| (2, 136) \| 147 \| (23, 159) \| 168 \| Uncharacterized protein \| Uncharacterized protein; uniclust \| UniRef100\_A0A2A5EK29 \| 98.9 \| 1.7e-11 \| 3.1e-17 \| 87.6 \| 131 \| (2, 132) \| 147 \| (44, 185) \| 216 \| Terminase small subunit \| Terminase small subunit; uniclust \| UniRef100\_A0A0A8IL49 \| 98.9 \| 1.6e-11 \| 3.3e-17 \| 91.6 \| 95 \| (43, 137) \| 147 \| (91, 187) \| 199 \| Terminase small subunit \| Terminase small subunit; uniclust \| UniRef100\_A0A0Q6MN60 \| 98.9 \| 1.7e-11 \| 3.5e-17 \| 93.0 \| 96 \| (45, 140) \| 147 \| (108, 204) \| 231 \| Uncharacterized protein \| Uncharacterized protein; uniclust \| UniRef100\_UPI00041AA28B \| 98.8 \| 2.2e-11 \| 4e-17 \| 87.4 \| 126 \| (3, 134) \| 147 \| (51, 177) \| 213 \| terminase small subunit \| terminase small subunit; uniclust \| UniRef100\_A0A158L615 \| 98.8 \| 2.2e-11 \| 4.5e-17 \| 91.8 \| 88 \| (44, 131) \| 147 \| (96, 186) \| 213 \| Putative bacteriophage--like protein \| Putative bacteriophage--like protein; uniclust \| UniRef100\_A0A7L5ZY90 \| 98.8 \| 2.5e-11 \| 4.7e-17 \| 79.7 \| 87 \| (55, 141) \| 147 \| (4, 90) \| 121 \| Uncharacterized protein \| Uncharacterized protein; uniclust \| UniRef100\_UPI00223107AD \| 98.8 \| 2.8e-11 \| 5.2e-17 \| 82.5 \| 109 \| (29, 137) \| 147 \| (3, 113) \| 154 \| terminase small subunit, Nu1 \| terminase small subunit, Nu1; uniclust \| UniRef100\_A0A0S4V076 \| 98.8 \| 2.8e-11 \| 5.5e-17 \| 80.9 \| 94 \| (43, 136) \| 147 \| (7, 102) \| 107 \| Bacteriophage-related protein \| Bacteriophage-related protein; uniclust \| UniRef100\_A0A0J6WV29 \| 98.8 \| 3e-11 \| 5.6e-17 \| 85.1 \| 133 \| (2, 140) \| 147 \| (31, 163) \| 173 \| DNA-binding protein \| DNA-binding protein; uniclust \| UniRef100\_A0A1L3ZRR3 \| 98.8 \| 2.9e-11 \| 5.7e-17 \| 92.0 \| 126 \| (1, 129) \| 147 \| (44, 193) \| 244 \| Terminase small subunit \| Terminase small subunit; uniclust \| UniRef100\_A0A2X2UEZ9 \| 98.8 \| 3.5e-11 \| 6.4e-17 \| 87.0 \| 120 \| (18, 137) \| 147 \| (97, 216) \| 233 \| Phage DNA packaging protein, Nu1 subunit of terminase \| Phage DNA packaging protein, Nu1 subunit of terminase; uniclust \| UniRef100\_H8YXV1 \| 98.8 \| 3.7e-11 \| 6.8e-17 \| 81.8 \| 117 \| (15, 134) \| 147 \| (32, 148) \| 152 \| Phage DNA packaging protein, Nu1 subunit of terminase \| Phage DNA packaging protein, Nu1 subunit of terminase; uniclust \| UniRef100\_UPI0022B810E0 \| 98.8 \| 3.9e-11 \| 7.1e-17 \| 85.2 \| 130 \| (2, 137) \| 147 \| (55, 196) \| 203 \| hypothetical protein \| hypothetical protein; uniclust \| UniRef100\_A0A7V9ZEA9 \| 98.8 \| 3.7e-11 \| 7.2e-17 \| 88.1 \| 96 \| (44, 141) \| 147 \| (100, 195) \| 201 \| Terminase small subunit \| Terminase small subunit; uniclust \| UniRef100\_A0A0Q4ZGW7 \| 98.8 \| 3.8e-11 \| 7.4e-17 \| 89.3 \| 109 \| (1, 109) \| 147 \| (57, 173) \| 218 \| Uncharacterized protein \| Uncharacterized protein; uniclust \| UniRef100\_A0A9D1S5U4 \| 98.8 \| 4.2e-11 \| 7.7e-17 \| 85.0 \| 126 \| (1, 132) \| 147 \| (32, 172) \| 202 \| Uncharacterized protein \| Uncharacterized protein; uniclust \| UniRef100\_A0A0X8CI04 \| 98.8 \| 4.1e-11 \| 7.9e-17 \| 87.8 \| 130 \| (3, 132) \| 147 \| (64, 198) \| 207 \| Terminase small subunit \| Terminase small subunit; uniclust \| UniRef100\_D1Y312 \| 98.8 \| 4.6e-11 \| 8.5e-17 \| 82.3 \| 133 \| (3, 140) \| 147 \| (10, 147) \| 164 \| Phage DNA packaging protein Nu1 \| Phage DNA packaging protein Nu1; uniclust \| UniRef100\_A0A014MCD6 \| 98.8 \| 5.3e-11 \| 1.1e-16 \| 88.8 \| 114 \| (2, 126) \| 147 \| (39, 156) \| 197 \| Terminase small subunit \| Terminase small subunit; uniclust \| UniRef100\_A0A023E0R0 \| 98.8 \| 5.9e-11 \| 1.1e-16 \| 86.1 \| 87 \| (48, 134) \| 147 \| (103, 189) \| 196 \| Terminase small subunit \| Terminase small subunit; uniclust \| UniRef100\_A0A6I6DNA2 \| 98.7 \| 7.9e-11 \| 1.5e-16 \| 85.4 \| 128 \| (2, 140) \| 147 \| (70, 198) \| 231 \| Terminase small subunit \| Terminase small subunit; uniclust \| UniRef100\_A0A060BBC7 \| 98.7 \| 8.1e-11 \| 1.7e-16 \| 90.6 \| 91 \| (13, 105) \| 147 \| (76, 166) \| 234 \| Terminase small subunit \| Terminase small subunit; uniclust \| UniRef100\_UPI0009DC06AE \| 98.7 \| 1e-10 \| 2e-16 \| 84.9 \| 127 \| (1, 134) \| 147 \| (68, 198) \| 212 \| terminase small subunit \| terminase small subunit; uniclust \| UniRef100\_A0A7Z8R5N7 \| 98.7 \| 1e-10 \| 2e-16 \| 83.8 \| 126 \| (1, 126) \| 147 \| (22, 154) \| 184 \| Terminase small subunit \| Terminase small subunit; uniclust \| UniRef100\_B0U9E3 \| 98.7 \| 1.1e-10 \| 2.1e-16 \| 76.1 \| 89 \| (44, 132) \| 147 \| (7, 95) \| 115 \| Uncharacterized protein \| Uncharacterized protein; uniclust \| UniRef100\_A0A0H3ZKI4 \| 98.7 \| 1e-10 \| 2.1e-16 \| 86.4 \| 131 \| (2, 136) \| 147 \| (30, 166) \| 190 \| Terminase small subunit \| Terminase small subunit; uniclust \| UniRef100\_UPI0006B47411 \| 98.7 \| 1.2e-10 \| 2.2e-16 \| 81.3 \| 132 \| (3, 139) \| 147 \| (37, 172) \| 177 \| hypothetical protein \| hypothetical protein; uniclust \| UniRef100\_A0A017HTQ6 \| 98.7 \| 1.2e-10 \| 2.2e-16 \| 80.1 \| 137 \| (1, 137) \| 147 \| (10, 148) \| 160 \| Phage DNA packaging protein Nu1 \| Phage DNA packaging protein Nu1; uniclust \| UniRef100\_A0A1H8NQA9 \| 98.7 \| 1.2e-10 \| 2.3e-16 \| 84.8 \| 87 \| (46, 132) \| 147 \| (98, 184) \| 190 \| Phage DNA packaging protein, Nu1 subunit of terminase \| Phage DNA packaging protein, Nu1 subunit of terminase; uniclust \| UniRef100\_A0A071M1J9 \| 98.7 \| 1.2e-10 \| 2.4e-16 \| 91.8 \| 98 \| (45, 142) \| 147 \| (154, 253) \| 291 \| Uncharacterized protein \| Uncharacterized protein; uniclust \| UniRef100\_A0A8J6NMN9 \| 98.7 \| 1.4e-10 \| 2.6e-16 \| 79.0 \| 123 \| (2, 131) \| 147 \| (26, 148) \| 151 \| Uncharacterized protein \| Uncharacterized protein; uniclust \| UniRef100\_A0A6N6MDM0 \| 98.7 \| 1.3e-10 \| 2.7e-16 \| 84.3 \| 97 \| (2, 106) \| 147 \| (36, 132) \| 175 \| Terminase small subunit, Nu1 \| Terminase small subunit, Nu1; uniclust \| UniRef100\_UPI0012E9DC02 \| 98.7 \| 1.5e-10 \| 2.7e-16 \| 82.8 \| 122 \| (2, 130) \| 147 \| (86, 207) \| 209 \| hypothetical protein \| hypothetical protein; uniclust \| UniRef100\_A0A0Q7F4I9 \| 98.7 \| 1.4e-10 \| 2.8e-16 \| 84.4 \| 119 \| (6, 131) \| 147 \| (30, 170) \| 185 \| Terminase small subunit \| Terminase small subunit; uniclust \| UniRef100\_A0A1V5Z7S9 \| 98.7 \| 1.6e-10 \| 3e-16 \| 80.4 \| 91 \| (43, 133) \| 147 \| (82, 172) \| 174 \| Phage DNA packaging protein Nu1 \| Phage DNA packaging protein Nu1; uniclust \| UniRef100\_A0A7T1HQ48 \| 98.7 \| 1.8e-10 \| 3.3e-16 \| 82.7 \| 87 \| (48, 134) \| 147 \| (87, 179) \| 190 \| Terminase small subunit \| Terminase small subunit; uniclust \| UniRef100\_A0A0K6HIX8 \| 98.7 \| 1.7e-10 \| 3.4e-16 \| 86.8 \| 96 \| (2, 105) \| 147 \| (64, 159) \| 221 \| Phage DNA packaging protein, Nu1 subunit of terminase \| Phage DNA packaging protein, Nu1 subunit of terminase; uniclust \| UniRef100\_A0A0F2PHE5 \| 98.7 \| 2.3e-10 \| 4.4e-16 \| 81.0 \| 127 \| (3, 136) \| 147 \| (32, 159) \| 174 \| HTH merR-type domain-containing protein \| HTH merR-type domain-containing protein; uniclust \| UniRef100\_A0A0C5XPQ2 \| 98.7 \| 2.3e-10 \| 4.5e-16 \| 83.9 \| 112 \| (16, 134) \| 147 \| (62, 173) \| 193 \| Terminase \| Terminase; uniclust \| UniRef100\_A0A6L9M419 \| 98.7 \| 2.5e-10 \| 4.7e-16 \| 79.3 \| 91 \| (44, 134) \| 147 \| (69, 159) \| 170 \| Terminase small subunit \| Terminase small subunit; uniclust \| UniRef100\_A0A1G2ZKR4 \| 98.6 \| 2.8e-10 \| 5.1e-16 \| 84.4 \| 130 \| (2, 138) \| 147 \| (117, 246) \| 276 \| Helix-turn-helix domain-containing protein \| Helix-turn-helix domain-containing protein; uniclust \| UniRef100\_A0A517Q5A6 \| 98.6 \| 3e-10 \| 5.8e-16 \| 82.4 \| 127 \| (1, 132) \| 147 \| (60, 189) \| 195 \| Phage DNA packaging protein Nu1 \| Phage DNA packaging protein Nu1; uniclust \| UniRef100\_A0A956GQK7 \| 98.6 \| 3.2e-10 \| 5.9e-16 \| 81.7 \| 122 \| (1, 129) \| 147 \| (37, 159) \| 220 \| Uncharacterized protein \| Uncharacterized protein; uniclust \| UniRef100\_A0A059KUP0 \| 98.6 \| 3.2e-10 \| 6.3e-16 \| 86.5 \| 129 \| (4, 136) \| 147 \| (51, 185) \| 237 \| DNA packaging protein \| DNA packaging protein; uniclust \| UniRef100\_A0A064A069 \| 98.6 \| 3.4e-10 \| 6.4e-16 \| 82.1 \| 106 \| (21, 136) \| 147 \| (65, 170) \| 186 \| Terminase \| Terminase; uniclust \| UniRef100\_A0A099EVD2 \| 98.6 \| 3.2e-10 \| 6.4e-16 \| 86.1 \| 96 \| (43, 138) \| 147 \| (112, 209) \| 225 \| Elements of external origin \| Elements of external origin; uniclust \| UniRef100\_A0L5S8 \| 98.6 \| 3.6e-10 \| 6.6e-16 \| 84.4 \| 114 \| (21, 134) \| 147 \| (167, 283) \| 292 \| Transcriptional regulator, Fis family \| Transcriptional regulator, Fis family; uniclust \| UniRef100\_A0A0U2S671 \| 98.6 \| 4.6e-10 \| 8.5e-16 \| 79.8 \| 126 \| (1, 128) \| 147 \| (21, 162) \| 181 \| Terminase small subunit \| Terminase small subunit; uniclust \| UniRef100\_A0A0M9GBF1 \| 98.6 \| 5.6e-10 \| 1.1e-15 \| 75.6 \| 91 \| (43, 133) \| 147 \| (22, 112) \| 122 \| Uncharacterized protein (Fragment) \| Uncharacterized protein (Fragment); uniclust \| UniRef100\_A0A064AK97 \| 98.6 \| 5.4e-10 \| 1.1e-15 \| 83.6 \| 122 \| (2, 142) \| 147 \| (78, 199) \| 210 \| Helix-turn-helix domain-containing protein \| Helix-turn-helix domain-containing protein; uniclust \| UniRef100\_A0A076GEC0 \| 98.6 \| 6.1e-10 \| 1.2e-15 \| 80.6 \| 133 \| (1, 134) \| 147 \| (54, 195) \| 200 \| Small terminase subunit \| Small terminase subunit; uniclust \| UniRef100\_UPI0020034B75 \| 98.6 \| 6.3e-10 \| 1.2e-15 \| 78.7 \| 126 \| (2, 131) \| 147 \| (63, 189) \| 190 \| hypothetical protein \| hypothetical protein; uniclust \| UniRef100\_A0A8S7YJX2 \| 98.6 \| 6.5e-10 \| 1.2e-15 \| 87.7 \| 127 \| (1, 129) \| 147 \| (21, 163) \| 473 \| Phage tail collar domain-containing protein \| Phage tail collar domain-containing protein; uniclust \| UniRef100\_E7C6A2 \| 98.6 \| 7.3e-10 \| 1.3e-15 \| 72.4 \| 90 \| (45, 134) \| 147 \| (3, 92) \| 113 \| Uncharacterized protein \| Uncharacterized protein; uniclust \| UniRef100\_A0A1T4PVA8 \| 98.6 \| 7.4e-10 \| 1.4e-15 \| 80.8 \| 126 \| (3, 135) \| 147 \| (83, 208) \| 213 \| Phage DNA packaging protein, Nu1 subunit of terminase \| Phage DNA packaging protein, Nu1 subunit of terminase; uniclust \| UniRef100\_A0A4R6UIY8 \| 98.5 \| 7.8e-10 \| 1.5e-15 \| 69.3 \| 72 \| (63, 134) \| 147 \| (2, 73) \| 81 \| Uncharacterized protein \| Uncharacterized protein; uniclust \| UniRef100\_A0A945GZD5 \| 98.5 \| 8.3e-10 \| 1.5e-15 \| 71.4 \| 89 \| (44, 132) \| 147 \| (13, 101) \| 106 \| Uncharacterized protein \| Uncharacterized protein; uniclust \| UniRef100\_UPI0020CD6DAF \| 98.5 \| 8.4e-10 \| 1.6e-15 \| 81.1 \| 90 \| (43, 132) \| 147 \| (128, 217) \| 229 \| hypothetical protein \| hypothetical protein; uniclust \| UniRef100\_A0A1E7PYA9 \| 98.5 \| 8.9e-10 \| 1.8e-15 \| 79.5 \| 91 \| (43, 136) \| 147 \| (62, 152) \| 162 \| Terminase small subunit \| Terminase small subunit; uniclust \| UniRef100\_UPI001EE2AD5A \| 98.5 \| 1e-09 \| 1.8e-15 \| 82.8 \| 127 \| (2, 134) \| 147 \| (23, 175) \| 310 \| hypothetical protein \| hypothetical protein; uniclust \| UniRef100\_A0A151FHD5 \| 98.5 \| 1e-09 \| 1.9e-15 \| 78.7 \| 92 \| (43, 134) \| 147 \| (71, 162) \| 172 \| Terminase \| Terminase; uniclust \| UniRef100\_A0A2T5IIL6 \| 98.5 \| 1e-09 \| 2e-15 \| 69.2 \| 75 \| (60, 134) \| 147 \| (2, 76) \| 84 \| Uncharacterized protein \| Uncharacterized protein; uniclust \| UniRef100\_A0A1G7GZW8 \| 98.5 \| 1.3e-09 \| 2.5e-15 \| 79.4 \| 115 \| (2, 131) \| 147 \| (30, 144) \| 168 \| Uncharacterized protein \| Uncharacterized protein; uniclust \| UniRef100\_E2CDD4 \| 98.5 \| 1.4e-09 \| 2.7e-15 \| 79.3 \| 87 \| (45, 131) \| 147 \| (118, 204) \| 217 \| Uncharacterized protein \| Uncharacterized protein; uniclust \| UniRef100\_A0A6J5EMI8 \| 98.5 \| 1.5e-09 \| 2.8e-15 \| 72.1 \| 93 \| (44, 136) \| 147 \| (8, 102) \| 106 \| Uncharacterized protein \| Uncharacterized protein; uniclust \| UniRef100\_UPI00223516E7 \| 98.5 \| 1.6e-09 \| 2.9e-15 \| 85.3 \| 127 \| (1, 129) \| 147 \| (21, 163) \| 452 \| terminase small subunit \| terminase small subunit; uniclust \| UniRef100\_UPI001FB86660 \| 98.5 \| 1.7e-09 \| 3.1e-15 \| 73.4 \| 122 \| (6, 132) \| 147 \| (4, 126) \| 141 \| hypothetical protein \| hypothetical protein; uniclust \| UniRef100\_D8F202 \| 98.5 \| 2e-09 \| 3.6e-15 \| 74.4 \| 127 \| (1, 132) \| 147 \| (22, 149) \| 159 \| Uncharacterized protein \| Uncharacterized protein; uniclust \| UniRef100\_UPI001C52BE35 \| 98.5 \| 2e-09 \| 3.7e-15 \| 75.9 \| 115 \| (20, 134) \| 147 \| (46, 163) \| 172 \| hypothetical protein \| hypothetical protein; uniclust \| UniRef100\_A0A6I1JC22 \| 98.5 \| 2e-09 \| 3.8e-15 \| 76.5 \| 97 \| (2, 106) \| 147 \| (29, 125) \| 165 \| Helix-turn-helix domain-containing protein \| Helix-turn-helix domain-containing protein; uniclust \| UniRef100\_A0A3S0BZF6 \| 98.4 \| 2.2e-09 \| 4.1e-15 \| 75.6 \| 129 \| (3, 134) \| 147 \| (21, 152) \| 164 \| Terminase small subunit \| Terminase small subunit; uniclust \| UniRef100\_UPI0012ED759B \| 98.4 \| 2.4e-09 \| 4.4e-15 \| 71.4 \| 91 \| (44, 134) \| 147 \| (32, 122) \| 126 \| hypothetical protein \| hypothetical protein; uniclust \| UniRef100\_A0A094YII6 \| 98.4 \| 2.3e-09 \| 4.5e-15 \| 80.5 \| 88 \| (45, 132) \| 147 \| (102, 189) \| 212 \| Helix-turn-helix domain-containing protein \| Helix-turn-helix domain-containing protein; uniclust \| UniRef100\_A0A167H541 \| 98.4 \| 2.5e-09 \| 5e-15 \| 79.4 \| 127 \| (1, 134) \| 147 \| (20, 161) \| 189 \| Uncharacterized protein \| Uncharacterized protein; uniclust \| UniRef100\_A0A8I0KDZ5 \| 98.4 \| 3.1e-09 \| 5.7e-15 \| 64.0 \| 64 \| (66, 129) \| 147 \| (6, 69) \| 70 \| Uncharacterized protein \| Uncharacterized protein; uniclust \| UniRef100\_A0A077NSP9 \| 98.4 \| 3.2e-09 \| 6.2e-15 \| 78.2 \| 127 \| (1, 132) \| 147 \| (22, 187) \| 196 \| Uncharacterized protein \| Uncharacterized protein; uniclust \| UniRef100\_A0A9E8X8G8 \| 98.4 \| 3.4e-09 \| 6.3e-15 \| 76.0 \| 116 \| (8, 130) \| 147 \| (84, 202) \| 205 \| Uncharacterized protein \| Uncharacterized protein; uniclust \| UniRef100\_A0A060HBH3 \| 98.4 \| 3.3e-09 \| 6.5e-15 \| 79.5 \| 92 \| (44, 135) \| 147 \| (100, 191) \| 219 \| Phage-related protein \| Phage-related protein; uniclust \| UniRef100\_UPI0012F7E1C9 \| 98.4 \| 3.5e-09 \| 6.5e-15 \| 72.8 \| 110 \| (17, 130) \| 147 \| (40, 149) \| 153 \| DUF1441 family protein \| DUF1441 family protein; uniclust \| UniRef100\_A0A929F7T2 \| 98.4 \| 3.6e-09 \| 6.7e-15 \| 75.3 \| 90 \| (43, 132) \| 147 \| (102, 191) \| 193 \| Uncharacterized protein \| Uncharacterized protein; uniclust \| UniRef100\_UPI001688EBE8 \| 98.4 \| 3.9e-09 \| 7.1e-15 \| 73.7 \| 125 \| (2, 131) \| 147 \| (32, 163) \| 169 \| hypothetical protein \| hypothetical protein; uniclust \| UniRef100\_H3RLL8 \| 98.4 \| 3.8e-09 \| 7.1e-15 \| 69.4 \| 94 \| (44, 137) \| 147 \| (3, 99) \| 103 \| DNA-directed RNA polymerase sigma subunit \| DNA-directed RNA polymerase sigma subunit; uniclust \| UniRef100\_A0A7C3DJT8 \| 98.4 \| 3.8e-09 \| 7.1e-15 \| 68.2 \| 84 \| (45, 129) \| 147 \| (12, 95) \| 98 \| Tetrapyrrole biosynthesis glutamyl-tRNA reductase dimerisation domain-containing protein \| Tetrapyrrole biosynthesis glutamyl-tRNA reductase dimerisation domain-containing protein; uniclust \| UniRef100\_A0A0J6K589 \| 98.4 \| 3.7e-09 \| 7.1e-15 \| 78.9 \| 92 \| (44, 135) \| 147 \| (114, 207) \| 213 \| Elements of external origin \| Elements of external origin; uniclust \| UniRef100\_A0A8G2CJ81 \| 98.4 \| 4.4e-09 \| 8.1e-15 \| 76.2 \| 126 \| (2, 132) \| 147 \| (21, 152) \| 219 \| Phage DNA packaging protein, Nu1 subunit of terminase \| Phage DNA packaging protein, Nu1 subunit of terminase; uniclust \| UniRef100\_A0A9E5R9F2 \| 98.4 \| 4.6e-09 \| 8.4e-15 \| 72.0 \| 121 \| (1, 131) \| 147 \| (19, 147) \| 149 \| Terminase small subunit \| Terminase small subunit; uniclust \| UniRef100\_A0A0A1AKA0 \| 98.4 \| 4.5e-09 \| 8.5e-15 \| 76.0 \| 123 \| (4, 132) \| 147 \| (26, 178) \| 187 \| Terminase small subunit \| Terminase small subunit; uniclust \| UniRef100\_A0A109J6S7 \| 98.4 \| 4.6e-09 \| 8.6e-15 \| 77.9 \| 127 \| (2, 134) \| 147 \| (84, 212) \| 233 \| Phage terminase Nu1 subunit (DNA packaging protein) \| Phage terminase Nu1 subunit (DNA packaging protein); uniclust \| UniRef100\_A0A6I6MLK3 \| 98.4 \| 4.7e-09 \| 8.8e-15 \| 71.1 \| 85 \| (46, 132) \| 147 \| (28, 112) \| 125 \| Uncharacterized protein \| Uncharacterized protein; uniclust \| UniRef100\_UPI00135677F5 \| 98.4 \| 5.1e-09 \| 9.4e-15 \| 68.8 \| 96 \| (4, 104) \| 147 \| (17, 112) \| 114 \| hypothetical protein \| hypothetical protein; uniclust \| UniRef100\_A0A562RXT0 \| 98.4 \| 4.9e-09 \| 9.4e-15 \| 78.9 \| 125 \| (3, 132) \| 147 \| (35, 204) \| 228 \| Phage terminase small subunit \| Phage terminase small subunit; uniclust \| UniRef100\_A0A7Y3AH85 \| 98.4 \| 5.6e-09 \| 1e-14 \| 72.6 \| 110 \| (17, 134) \| 147 \| (39, 148) \| 164 \| DUF1441 family protein \| DUF1441 family protein; uniclust \| UniRef100\_A0A7C4DTE2 \| 98.4 \| 5.6e-09 \| 1e-14 \| 73.0 \| 123 \| (3, 132) \| 147 \| (28, 160) \| 169 \| DNA packaging protein \| DNA packaging protein; uniclust \| UniRef100\_A0A376VNK9 \| 98.3 \| 6e-09 \| 1.1e-14 \| 73.5 \| 107 \| (1, 107) \| 147 \| (25, 147) \| 166 \| DNA packaging protein \| DNA packaging protein; uniclust \| UniRef100\_A0A1Q6UMH2 \| 98.3 \| 5.7e-09 \| 1.1e-14 \| 80.0 \| 128 \| (1, 138) \| 147 \| (82, 214) \| 232 \| Terminase small subunit \| Terminase small subunit; uniclust \| UniRef100\_A0A974SWB6 \| 98.3 \| 6.1e-09 \| 1.2e-14 \| 75.7 \| 117 \| (14, 133) \| 147 \| (41, 157) \| 179 \| DUF1441 family protein \| DUF1441 family protein; uniclust \| UniRef100\_UPI0020003892 \| 98.3 \| 7e-09 \| 1.3e-14 \| 69.3 \| 95 \| (1, 95) \| 147 \| (23, 125) \| 126 \| terminase small subunit \| terminase small subunit; uniclust \| UniRef100\_UPI001FF8B96B \| 98.3 \| 6.8e-09 \| 1.3e-14 \| 64.3 \| 70 \| (61, 132) \| 147 \| (2, 71) \| 73 \| hypothetical protein \| hypothetical protein; uniclust \| UniRef100\_A0A3D1NTE5 \| 98.3 \| 7.2e-09 \| 1.3e-14 \| 75.1 \| 90 \| (44, 133) \| 147 \| (106, 195) \| 199 \| Terminase small subunit \| Terminase small subunit; uniclust \| UniRef100\_A0A7L6A869 \| 98.3 \| 8.7e-09 \| 1.6e-14 \| 71.6 \| 103 \| (28, 130) \| 147 \| (41, 143) \| 162 \| MarR family transcriptional regulator \| MarR family transcriptional regulator; uniclust \| UniRef100\_A0A0P6VRG9 \| 98.3 \| 8.2e-09 \| 1.6e-14 \| 75.7 \| 128 \| (3, 135) \| 147 \| (25, 168) \| 178 \| Elements of external origin \| Elements of external origin; uniclust \| UniRef100\_A0A2D2C217 \| 98.3 \| 8.7e-09 \| 1.6e-14 \| 76.6 \| 121 \| (12, 132) \| 147 \| (61, 203) \| 230 \| Terminase small subunit \| Terminase small subunit; uniclust \| UniRef100\_A0A1J5SR61 \| 98.3 \| 8.4e-09 \| 1.6e-14 \| 76.8 \| 124 \| (16, 139) \| 147 \| (28, 155) \| 207 \| Uncharacterized protein \| Uncharacterized protein; uniclust \| UniRef100\_A0A1G0DQX9 \| 98.3 \| 7.7e-09 \| 1.6e-14 \| 79.5 \| 89 \| (45, 133) \| 147 \| (109, 197) \| 212 \| Terminase small subunit \| Terminase small subunit; uniclust \| UniRef100\_A0A165RXZ6 \| 98.3 \| 9.1e-09 \| 1.7e-14 \| 75.1 \| 95 \| (43, 137) \| 147 \| (96, 192) \| 195 \| Elements of external origin \| Elements of external origin; uniclust \| UniRef100\_A0A3T0L2W5 \| 98.3 \| 9.2e-09 \| 1.7e-14 \| 76.0 \| 88 \| (44, 131) \| 147 \| (126, 213) \| 217 \| Terminase \| Terminase; uniclust \| UniRef100\_A0A1I3TC35 \| 98.3 \| 1e-08 \| 1.9e-14 \| 73.6 \| 123 \| (2, 134) \| 147 \| (21, 147) \| 177 \| Uncharacterized protein \| Uncharacterized protein; uniclust \| UniRef100\_UPI00137A5DD6 \| 98.3 \| 1.2e-08 \| 2.1e-14 \| 75.2 \| 124 \| (14, 137) \| 147 \| (86, 210) \| 243 \| terminase small subunit \| terminase small subunit; uniclust \| UniRef100\_A0A7W6H5E1 \| 98.3 \| 1.1e-08 \| 2.1e-14 \| 68.5 \| 83 \| (44, 130) \| 147 \| (30, 112) \| 113 \| Phage terminase Nu1 subunit (DNA packaging protein) \| Phage terminase Nu1 subunit (DNA packaging protein); uniclust \| UniRef100\_UPI0020049ABE \| 98.3 \| 1.2e-08 \| 2.3e-14 \| 70.2 \| 110 \| (1, 118) \| 147 \| (24, 134) \| 147 \| terminase small subunit \| terminase small subunit; uniclust \| UniRef100\_A0A2M8EDC0 \| 98.3 \| 1.3e-08 \| 2.4e-14 \| 71.5 \| 94 \| (45, 138) \| 147 \| (64, 157) \| 172 \| Terminase small subunit (Fragment) \| Terminase small subunit (Fragment); uniclust \| UniRef100\_A0A0A0FF05 \| 98.3 \| 1.3e-08 \| 2.6e-14 \| 71.3 \| 89 \| (1, 89) \| 147 \| (24, 128) \| 136 \| DNA-packaging protein \| DNA-packaging protein; uniclust \| UniRef100\_UPI00036DAC41 \| 98.2 \| 1.5e-08 \| 2.8e-14 \| 73.0 \| 129 \| (1, 129) \| 147 \| (32, 184) \| 205 \| DUF1441 family protein \| DUF1441 family protein; uniclust \| UniRef100\_UPI000A1E6997 \| 98.2 \| 1.5e-08 \| 2.8e-14 \| 67.5 \| 96 \| (1, 102) \| 147 \| (23, 120) \| 122 \| terminase small subunit \| terminase small subunit; uniclust \| UniRef100\_A0A973GUH3 \| 98.2 \| 1.5e-08 \| 2.9e-14 \| 70.1 \| 90 \| (44, 134) \| 147 \| (32, 121) \| 144 \| Uncharacterized protein \| Uncharacterized protein; uniclust \| UniRef100\_A0A1M7R7I9 \| 98.2 \| 1.4e-08 \| 2.9e-14 \| 75.5 \| 93 \| (44, 136) \| 147 \| (94, 186) \| 192 \| Phage terminase small subunit \| Phage terminase small subunit; uniclust \| UniRef100\_UPI00045BE5E8 \| 98.2 \| 1.6e-08 \| 3e-14 \| 72.4 \| 135 \| (4, 140) \| 147 \| (33, 181) \| 198 \| hypothetical protein \| hypothetical protein; uniclust \| UniRef100\_UPI001FAD9689 \| 98.2 \| 1.7e-08 \| 3.1e-14 \| 69.2 \| 114 \| (21, 134) \| 147 \| (5, 122) \| 146 \| hypothetical protein \| hypothetical protein; uniclust \| UniRef100\_A0A418VWY4 \| 98.2 \| 1.7e-08 \| 3.2e-14 \| 69.7 \| 92 \| (43, 134) \| 147 \| (36, 127) \| 133 \| Uncharacterized protein \| Uncharacterized protein; uniclust \| UniRef100\_A0A6M0K1R5 \| 98.2 \| 1.8e-08 \| 3.3e-14 \| 69.3 \| 128 \| (7, 134) \| 147 \| (1, 132) \| 150 \| Terminase small subunit \| Terminase small subunit; uniclust \| UniRef100\_A0A661I0F5 \| 98.2 \| 1.9e-08 \| 3.4e-14 \| 71.6 \| 90 \| (44, 133) \| 147 \| (97, 186) \| 188 \| Uncharacterized protein \| Uncharacterized protein; uniclust \| UniRef100\_A0A8S5NY96 \| 98.2 \| 2e-08 \| 3.6e-14 \| 73.1 \| 100 \| (42, 142) \| 147 \| (112, 211) \| 220 \| Uncharacterized protein \| Uncharacterized protein; uniclust \| UniRef100\_A0A067ZIZ9 \| 98.2 \| 1.9e-08 \| 3.7e-14 \| 73.5 \| 123 \| (4, 133) \| 147 \| (26, 176) \| 190 \| Terminase small subunit \| Terminase small subunit; uniclust \| UniRef100\_A0A952DRL5 \| 98.2 \| 2.1e-08 \| 3.9e-14 \| 73.0 \| 87 \| (46, 132) \| 147 \| (97, 183) \| 187 \| Uncharacterized protein \| Uncharacterized protein; uniclust \| UniRef100\_UPI0015E45FFC \| 98.2 \| 2.2e-08 \| 4e-14 \| 74.1 \| 105 \| (1, 105) \| 147 \| (58, 183) \| 249 \| terminase small subunit \| terminase small subunit; uniclust \| UniRef100\_A0A6H1ZN68 \| 98.2 \| 2.8e-08 \| 5.1e-14 \| 61.9 \| 77 \| (54, 130) \| 147 \| (2, 78) \| 81 \| Uncharacterized protein (Fragment) \| Uncharacterized protein (Fragment); uniclust \| UniRef100\_A0A0F7L3Q2 \| 98.2 \| 2.8e-08 \| 5.1e-14 \| 71.9 \| 129 \| (1, 132) \| 147 \| (36, 202) \| 209 \| MerR family transcriptional regulator \| MerR family transcriptional regulator; uniclust \| UniRef100\_A0A560D0R5 \| 98.2 \| 2.6e-08 \| 5.1e-14 \| 75.7 \| 125 \| (1, 132) \| 147 \| (26, 208) \| 237 \| Terminase small subunit \| Terminase small subunit; uniclust \| UniRef100\_A0A1F9DZH8 \| 98.2 \| 2.9e-08 \| 5.3e-14 \| 72.0 \| 105 \| (1, 107) \| 147 \| (23, 127) \| 213 \| HTH merR-type domain-containing protein \| HTH merR-type domain-containing protein; uniclust \| UniRef100\_A0A2E1R925 \| 98.2 \| 2.8e-08 \| 5.4e-14 \| 73.6 \| 132 \| (4, 135) \| 147 \| (33, 190) \| 202 \| Terminase small subunit \| Terminase small subunit; uniclust \| UniRef100\_A0A5E8GTT9 \| 98.2 \| 3.1e-08 \| 5.7e-14 \| 70.1 \| 128 \| (3, 134) \| 147 \| (37, 167) \| 180 \| Phage DNA packaging protein, Nu1 subunit of terminase \| Phage DNA packaging protein, Nu1 subunit of terminase; uniclust \| UniRef100\_A0A0F9RRF6 \| 98.2 \| 3.1e-08 \| 6e-14 \| 74.7 \| 127 \| (4, 136) \| 147 \| (89, 218) \| 236 \| Helix-turn-helix domain-containing protein \| Helix-turn-helix domain-containing protein; uniclust \| UniRef100\_F8KQH1 \| 98.2 \| 3.1e-08 \| 6e-14 \| 80.6 \| 122 \| (4, 137) \| 147 \| (234, 355) \| 392 \| Uncharacterized protein \| Uncharacterized protein; uniclust \| UniRef100\_A0A0Q5ZNV5 \| 98.2 \| 3.1e-08 \| 6e-14 \| 69.6 \| 92 \| (43, 134) \| 147 \| (43, 136) \| 140 \| Elements of external origin \| Elements of external origin; uniclust \| UniRef100\_X1GDY6 \| 98.1 \| 3.4e-08 \| 6.3e-14 \| 68.0 \| 125 \| (1, 130) \| 147 \| (24, 148) \| 149 \| Helix-turn-helix domain-containing protein (Fragment) \| Helix-turn-helix domain-containing protein (Fragment); uniclust \| UniRef100\_A0A2J0PYP4 \| 98.1 \| 4e-08 \| 8e-14 \| 70.1 \| 63 \| (44, 106) \| 147 \| (41, 103) \| 141 \| Terminase \| Terminase; uniclust \| UniRef100\_A0A2D9DW54 \| 98.1 \| 4e-08 \| 8.3e-14 \| 73.8 \| 119 \| (24, 142) \| 147 \| (43, 165) \| 181 \| Terminase small subunit \| Terminase small subunit; uniclust \| UniRef100\_UPI001032D246 \| 98.1 \| 4.6e-08 \| 8.5e-14 \| 71.3 \| 126 \| (1, 132) \| 147 \| (87, 217) \| 220 \| hypothetical protein \| hypothetical protein; uniclust \| UniRef100\_A0A4P7WC83 \| 98.1 \| 4.5e-08 \| 9e-14 \| 74.8 \| 105 \| (1, 105) \| 147 \| (39, 165) \| 216 \| Terminase small subunit \| Terminase small subunit; uniclust \| UniRef100\_UPI00186BB4F4 \| 98.1 \| 5e-08 \| 9.2e-14 \| 62.4 \| 83 \| (47, 131) \| 147 \| (10, 92) \| 94 \| hypothetical protein \| hypothetical protein; uniclust \| UniRef100\_UPI0021AED1F4 \| 98.1 \| 5.4e-08 \| 9.9e-14 \| 63.9 \| 102 \| (28, 131) \| 147 \| (2, 106) \| 109 \| hypothetical protein \| hypothetical protein; uniclust \| UniRef100\_A0A2I5AR94 \| 98.1 \| 5.6e-08 \| 1.1e-13 \| 71.2 \| 87 \| (44, 132) \| 147 \| (102, 188) \| 191 \| Terminase small subunit \| Terminase small subunit; uniclust \| UniRef100\_UPI0021A78783 \| 98.1 \| 5.9e-08 \| 1.1e-13 \| 64.3 \| 89 \| (44, 132) \| 147 \| (18, 106) \| 116 \| hypothetical protein \| hypothetical protein; uniclust \| UniRef100\_A0A7C7KJQ7 \| 98.1 \| 6.2e-08 \| 1.1e-13 \| 69.6 \| 128 \| (1, 131) \| 147 \| (32, 183) \| 197 \| Terminase small subunit \| Terminase small subunit; uniclust \| UniRef100\_A0A0F8X8C5 \| 98.1 \| 6.2e-08 \| 1.1e-13 \| 66.1 \| 97 \| (3, 106) \| 147 \| (39, 135) \| 138 \| Uncharacterized protein \| Uncharacterized protein; uniclust \| UniRef100\_A0A8J7C8B9 \| 98.1 \| 6.5e-08 \| 1.2e-13 \| 67.9 \| 111 \| (21, 135) \| 147 \| (54, 164) \| 167 \| Uncharacterized protein \| Uncharacterized protein; uniclust \| UniRef100\_UPI0018C227CA \| 98.1 \| 6.5e-08 \| 1.2e-13 \| 71.1 \| 90 \| (46, 135) \| 147 \| (114, 203) \| 208 \| hypothetical protein \| hypothetical protein; uniclust \| UniRef100\_UPI0012DF7B95 \| 98.1 \| 7e-08 \| 1.3e-13 \| 66.9 \| 122 \| (2, 128) \| 147 \| (23, 144) \| 153 \| hypothetical protein \| hypothetical protein; uniclust \| UniRef100\_A0A2A4V1Q2 \| 98.1 \| 7e-08 \| 1.3e-13 \| 69.1 \| 87 \| (44, 130) \| 147 \| (98, 184) \| 191 \| Uncharacterized protein \| Uncharacterized protein; uniclust \| UniRef100\_A0A166JZ29 \| 98.1 \| 6.7e-08 \| 1.3e-13 \| 72.4 \| 123 \| (4, 134) \| 147 \| (23, 163) \| 196 \| DNA packaging protein \| DNA packaging protein; uniclust \| UniRef100\_UPI001C90BF93 \| 98.1 \| 7.5e-08 \| 1.4e-13 \| 70.6 \| 131 \| (1, 132) \| 147 \| (59, 198) \| 209 \| terminase small subunit \| terminase small subunit; uniclust \| UniRef100\_F5R896 \| 98.1 \| 7.7e-08 \| 1.4e-13 \| 67.5 \| 124 \| (1, 132) \| 147 \| (25, 148) \| 166 \| Uncharacterized protein \| Uncharacterized protein; uniclust \| UniRef100\_A0A6G5Y155 \| 98.1 \| 7.4e-08 \| 1.4e-13 \| 70.7 \| 85 \| (45, 129) \| 147 \| (101, 185) \| 194 \| Terminase small subunit \| Terminase small subunit; uniclust \| UniRef100\_A0A0G9K8N3 \| 98.0 \| 7.4e-08 \| 1.4e-13 \| 73.2 \| 89 \| (45, 133) \| 147 \| (137, 225) \| 230 \| Uncharacterized protein \| Uncharacterized protein; uniclust \| UniRef100\_A0A085EXP4 \| 98.0 \| 7.1e-08 \| 1.5e-13 \| 75.0 \| 132 \| (1, 133) \| 147 \| (43, 198) \| 237 \| MarR family protein \| MarR family protein; uniclust \| UniRef100\_A0A3G8M604 \| 98.0 \| 7.7e-08 \| 1.5e-13 \| 70.9 \| 89 \| (46, 134) \| 147 \| (111, 199) \| 207 \| Uncharacterized protein \| Uncharacterized protein; uniclust \| UniRef100\_A0A934VIF6 \| 98.0 \| 8.1e-08 \| 1.5e-13 \| 69.7 \| 88 \| (42, 129) \| 147 \| (118, 205) \| 211 \| Uncharacterized protein \| Uncharacterized protein; uniclust \| UniRef100\_A0A934FD39 \| 98.0 \| 8.3e-08 \| 1.5e-13 \| 62.5 \| 77 \| (1, 83) \| 147 \| (22, 98) \| 103 \| Terminase small subunit (Fragment) \| Terminase small subunit (Fragment); uniclust \| UniRef100\_A0A258L5M1 \| 98.0 \| 8.6e-08 \| 1.6e-13 \| 68.7 \| 114 \| (4, 130) \| 147 \| (55, 168) \| 193 \| Terminase small subunit \| Terminase small subunit; uniclust \| UniRef100\_A0A7V0QL40 \| 98.0 \| 8.6e-08 \| 1.6e-13 \| 69.6 \| 124 \| (1, 132) \| 147 \| (79, 203) \| 211 \| DNA-binding protein \| DNA-binding protein; uniclust \| UniRef100\_A0A0F9MYQ4 \| 98.0 \| 9.7e-08 \| 1.8e-13 \| 67.8 \| 126 \| (1, 132) \| 147 \| (30, 160) \| 180 \| HTH merR-type domain-containing protein (Fragment) \| HTH merR-type domain-containing protein (Fragment); uniclust \| UniRef100\_UPI001E5479A9 \| 98.0 \| 9.8e-08 \| 1.8e-13 \| 67.9 \| 81 \| (47, 129) \| 147 \| (99, 179) \| 181 \| hypothetical protein \| hypothetical protein; uniclust \| UniRef100\_A0A963KDJ3 \| 98.0 \| 1.1e-07 \| 2e-13 \| 66.8 \| 94 \| (44, 137) \| 147 \| (70, 163) \| 167 \| Uncharacterized protein \| Uncharacterized protein; uniclust \| UniRef100\_D8F5N7 \| 98.0 \| 1.1e-07 \| 2.1e-13 \| 63.3 \| 105 \| (27, 132) \| 147 \| (3, 108) \| 118 \| Uncharacterized protein \| Uncharacterized protein; uniclust \| UniRef100\_A0A071MGC1 \| 98.0 \| 1.3e-07 \| 2.5e-13 \| 73.1 \| 102 \| (5, 106) \| 147 \| (69, 203) \| 250 \| Uncharacterized protein \| Uncharacterized protein; uniclust \| UniRef100\_UPI001F38D2C2 \| 98.0 \| 1.3e-07 \| 2.5e-13 \| 68.6 \| 126 \| (4, 129) \| 147 \| (72, 206) \| 210 \| terminase small subunit \| terminase small subunit; uniclust \| UniRef100\_UPI000AA827A0 \| 98.0 \| 1.4e-07 \| 2.5e-13 \| 66.6 \| 111 \| (21, 131) \| 147 \| (45, 163) \| 171 \| hypothetical protein \| hypothetical protein; uniclust \| UniRef100\_A0A951ZIQ2 \| 98.0 \| 1.5e-07 \| 2.7e-13 \| 67.1 \| 121 \| (2, 129) \| 147 \| (42, 162) \| 182 \| Uncharacterized protein \| Uncharacterized protein; uniclust \| UniRef100\_A0A2V7CIN0 \| 98.0 \| 1.5e-07 \| 2.7e-13 \| 66.2 \| 122 \| (6, 133) \| 147 \| (34, 156) \| 167 \| Terminase small subunit \| Terminase small subunit; uniclust \| UniRef100\_A0A1Q8Q2C1 \| 98.0 \| 1.5e-07 \| 2.8e-13 \| 65.8 \| 87 \| (48, 134) \| 147 \| (29, 118) \| 128 \| Uncharacterized protein \| Uncharacterized protein; uniclust \| UniRef100\_A0A1L8CPS1 \| 98.0 \| 1.6e-07 \| 3e-13 \| 67.0 \| 118 \| (13, 136) \| 147 \| (33, 150) \| 161 \| Phage DNA packaging protein Nu1 \| Phage DNA packaging protein Nu1; uniclust \| UniRef100\_A0A501WV30 \| 98.0 \| 1.7e-07 \| 3e-13 \| 74.1 \| 105 \| (1, 105) \| 147 \| (224, 349) \| 415 \| Terminase small subunit \| Terminase small subunit; uniclust \| UniRef100\_A0A1I3KS91 \| 97.9 \| 1.7e-07 \| 3.1e-13 \| 66.2 \| 122 \| (4, 130) \| 147 \| (9, 149) \| 155 \| Uncharacterized protein \| Uncharacterized protein; uniclust \| UniRef100\_A0A973E0Z4 \| 97.9 \| 1.7e-07 \| 3.2e-13 \| 67.4 \| 88 \| (44, 131) \| 147 \| (74, 162) \| 174 \| Uncharacterized protein \| Uncharacterized protein; uniclust \| UniRef100\_UPI001359000C \| 97.9 \| 1.9e-07 \| 3.5e-13 \| 59.9 \| 75 \| (58, 133) \| 147 \| (15, 89) \| 94 \| hypothetical protein \| hypothetical protein; uniclust \| UniRef100\_UPI000BFF95C5 \| 97.9 \| 2e-07 \| 3.6e-13 \| 67.7 \| 86 \| (45, 130) \| 147 \| (117, 202) \| 206 \| hypothetical protein \| hypothetical protein; uniclust \| UniRef100\_A0A835YX25 \| 97.9 \| 2.4e-07 \| 4.4e-13 \| 79.1 \| 123 \| (3, 133) \| 147 \| (258, 381) \| 962 \| Phage terminase large subunit-domain-containing protein \| Phage terminase large subunit-domain-containing protein; uniclust \| UniRef100\_A0A254T9S7 \| 97.9 \| 2.4e-07 \| 4.5e-13 \| 66.0 \| 105 \| (24, 130) \| 147 \| (25, 132) \| 159 \| DNA packaging Nu1 \| DNA packaging Nu1; uniclust \| UniRef100\_UPI000F3E537D \| 97.9 \| 2.5e-07 \| 4.6e-13 \| 59.8 \| 69 \| (1, 75) \| 147 \| (23, 92) \| 97 \| terminase small subunit \| terminase small subunit; uniclust \| UniRef100\_A0A5E4NWJ3 \| 97.9 \| 2.5e-07 \| 4.6e-13 \| 67.3 \| 122 \| (1, 132) \| 147 \| (55, 186) \| 209 \| Uncharacterized protein \| Uncharacterized protein; uniclust \| UniRef100\_UPI00209DF2FA \| 97.9 \| 2.5e-07 \| 4.6e-13 \| 68.6 \| 125 \| (6, 135) \| 147 \| (82, 216) \| 240 \| hypothetical protein \| hypothetical protein; uniclust \| UniRef100\_A0A1H5Z527 \| 97.9 \| 2.4e-07 \| 4.7e-13 \| 71.6 \| 107 \| (20, 128) \| 147 \| (99, 209) \| 258 \| DNA packaging protein \| DNA packaging protein; uniclust \| UniRef100\_M5PQI4 \| 97.9 \| 2.9e-07 \| 5.3e-13 \| 65.7 \| 91 \| (44, 134) \| 147 \| (73, 163) \| 181 \| Terminase small subunit \| Terminase small subunit; uniclust \| UniRef100\_UPI001AE9382A \| 97.9 \| 3e-07 \| 5.5e-13 \| 58.7 \| 80 \| (55, 134) \| 147 \| (3, 84) \| 90 \| hypothetical protein \| hypothetical protein; uniclust \| UniRef100\_UPI001E317423 \| 97.9 \| 3.1e-07 \| 5.6e-13 \| 62.7 \| 83 \| (46, 131) \| 147 \| (51, 133) \| 135 \| terminase small subunit \| terminase small subunit; uniclust \| UniRef100\_A0A0F9PXT5 \| 97.9 \| 2.9e-07 \| 5.7e-13 \| 69.9 \| 91 \| (45, 135) \| 147 \| (113, 203) \| 222 \| HTH merR-type domain-containing protein \| HTH merR-type domain-containing protein; uniclust \| UniRef100\_A0A6H1ZMV7 \| 97.9 \| 3.2e-07 \| 5.9e-13 \| 64.0 \| 119 \| (5, 129) \| 147 \| (33, 152) \| 155 \| Putative terminase \| Putative terminase; uniclust \| UniRef100\_A0A2X1LFL7 \| 97.9 \| 3.2e-07 \| 6e-13 \| 62.0 \| 86 \| (45, 131) \| 147 \| (32, 117) \| 124 \| Uncharacterized protein \| Uncharacterized protein; uniclust \| UniRef100\_A0A327KXV4 \| 97.9 \| 3.4e-07 \| 6.2e-13 \| 62.9 \| 91 \| (45, 136) \| 147 \| (41, 131) \| 140 \| Tetrapyrrole biosynthesis glutamyl-tRNA reductase dimerisation domain-containing protein \| Tetrapyrrole biosynthesis glutamyl-tRNA reductase dimerisation domain-containing protein; uniclust \| UniRef100\_UPI001FDEB81B \| 97.9 \| 3.5e-07 \| 6.4e-13 \| 64.6 \| 102 \| (4, 105) \| 147 \| (2, 123) \| 168 \| DUF1441 family protein \| DUF1441 family protein; uniclust \| UniRef100\_A0A2X1K069 \| 97.8 \| 3.3e-07 \| 6.5e-13 \| 68.2 \| 96 \| (6, 103) \| 147 \| (49, 145) \| 187 \| Prophage protein \| Prophage protein; uniclust \| UniRef100\_A0A2J7TM92 \| 97.8 \| 3.6e-07 \| 6.7e-13 \| 59.4 \| 87 \| (46, 133) \| 147 \| (3, 89) \| 100 \| Uncharacterized protein \| Uncharacterized protein; uniclust \| UniRef100\_A0A1W1CCQ2 \| 97.8 \| 3.9e-07 \| 7.1e-13 \| 63.6 \| 125 \| (2, 131) \| 147 \| (22, 152) \| 155 \| HTH merR-type domain-containing protein \| HTH merR-type domain-containing protein; uniclust \| UniRef100\_UPI0012EB806B \| 97.8 \| 3.9e-07 \| 7.2e-13 \| 62.3 \| 90 \| (45, 134) \| 147 \| (28, 119) \| 136 \| hypothetical protein \| hypothetical protein; uniclust \| UniRef100\_A0A7C7JGL7 \| 97.8 \| 4.1e-07 \| 7.5e-13 \| 60.8 \| 87 \| (44, 131) \| 147 \| (28, 114) \| 118 \| Host attachment protein \| Host attachment protein; uniclust \| UniRef100\_A0A256F2V5 \| 97.8 \| 4.1e-07 \| 7.9e-13 \| 66.4 \| 113 \| (3, 129) \| 147 \| (40, 154) \| 173 \| Terminase small subunit \| Terminase small subunit; uniclust \| UniRef100\_A0A5E7Q645 \| 97.8 \| 4.3e-07 \| 7.9e-13 \| 65.5 \| 87 \| (45, 131) \| 147 \| (101, 187) \| 195 \| Terminase small subunit \| Terminase small subunit; uniclust \| UniRef100\_UPI00235EB4F2 \| 97.8 \| 4.5e-07 \| 8.3e-13 \| 65.6 \| 92 \| (44, 135) \| 147 \| (104, 195) \| 198 \| hypothetical protein \| hypothetical protein; uniclust \| UniRef100\_UPI001CD487CF \| 97.8 \| 4.4e-07 \| 8.4e-13 \| 68.1 \| 116 \| (2, 132) \| 147 \| (63, 178) \| 220 \| hypothetical protein \| hypothetical protein; uniclust \| UniRef100\_A0A369R942 \| 97.8 \| 4.5e-07 \| 8.4e-13 \| 65.4 \| 121 \| (3, 132) \| 147 \| (36, 156) \| 180 \| DNA-binding protein \| DNA-binding protein; uniclust \| UniRef100\_A0A0H4A1N2 \| 97.8 \| 4.3e-07 \| 8.5e-13 \| 68.5 \| 130 \| (1, 136) \| 147 \| (30, 172) \| 202 \| Phage protein \| Phage protein; uniclust \| UniRef100\_UPI0011982A0B \| 97.8 \| 4.7e-07 \| 8.7e-13 \| 62.7 \| 89 \| (44, 132) \| 147 \| (37, 127) \| 135 \| hypothetical protein \| hypothetical protein; uniclust \| UniRef100\_M5T875 \| 97.8 \| 4.8e-07 \| 9e-13 \| 64.8 \| 122 \| (1, 129) \| 147 \| (23, 153) \| 165 \| Host attachment protein \| Host attachment protein; uniclust \| UniRef100\_UPI001FA8243C \| 97.8 \| 4.8e-07 \| 9e-13 \| 60.6 \| 77 \| (3, 84) \| 147 \| (26, 104) \| 108 \| hypothetical protein \| hypothetical protein; uniclust \| UniRef100\_UPI001E589372 \| 97.8 \| 5e-07 \| 9.2e-13 \| 58.4 \| 71 \| (1, 74) \| 147 \| (26, 96) \| 96 \| terminase small subunit \| terminase small subunit; uniclust \| UniRef100\_E3H7D0 \| 97.8 \| 5.5e-07 \| 1e-12 \| 63.1 \| 118 \| (20, 142) \| 147 \| (35, 152) \| 159 \| Uncharacterized protein \| Uncharacterized protein; uniclust \| UniRef100\_A0A358JR36 \| 97.8 \| 5.5e-07 \| 1e-12 \| 61.9 \| 88 \| (44, 131) \| 147 \| (43, 130) \| 139 \| Uncharacterized protein \| Uncharacterized protein; uniclust \| UniRef100\_A0A3D4L0I6 \| 97.8 \| 5.5e-07 \| 1e-12 \| 69.7 \| 131 \| (2, 134) \| 147 \| (171, 309) \| 336 \| Uncharacterized protein \| Uncharacterized protein; uniclust \| UniRef100\_A0A1S1TKW3 \| 97.8 \| 5.7e-07 \| 1.1e-12 \| 66.9 \| 88 \| (50, 137) \| 147 \| (90, 179) \| 187 \| Uncharacterized protein \| Uncharacterized protein; uniclust \| UniRef100\_A0A345DE68 \| 97.8 \| 6.6e-07 \| 1.2e-12 \| 64.6 \| 124 \| (2, 131) \| 147 \| (32, 162) \| 193 \| Terminase small subunit \| Terminase small subunit; uniclust \| UniRef100\_UPI00098D0DFA \| 97.8 \| 6.7e-07 \| 1.2e-12 \| 61.3 \| 124 \| (7, 132) \| 147 \| (1, 126) \| 137 \| hypothetical protein \| hypothetical protein; uniclust \| UniRef100\_A0A651HJ89 \| 97.8 \| 6.7e-07 \| 1.2e-12 \| 65.6 \| 105 \| (4, 111) \| 147 \| (53, 158) \| 217 \| Helix-turn-helix domain-containing protein \| Helix-turn-helix domain-containing protein; uniclust \| UniRef100\_A0A369QU30 \| 97.8 \| 6.8e-07 \| 1.2e-12 \| 63.4 \| 87 \| (44, 130) \| 147 \| (77, 163) \| 170 \| Terminase small subunit \| Terminase small subunit; uniclust \| UniRef100\_A0A5C7Q6H9 \| 97.8 \| 7.1e-07 \| 1.3e-12 \| 64.9 \| 132 \| (2, 133) \| 147 \| (25, 196) \| 203 \| Terminase small subunit \| Terminase small subunit; uniclust \| UniRef100\_UPI000B983DC3 \| 97.7 \| 7.1e-07 \| 1.3e-12 \| 60.4 \| 84 \| (48, 131) \| 147 \| (40, 123) \| 126 \| hypothetical protein \| hypothetical protein; uniclust \| UniRef100\_A0A0S9PYS1 \| 97.7 \| 7e-07 \| 1.3e-12 \| 66.7 \| 125 \| (2, 133) \| 147 \| (24, 192) \| 195 \| Uncharacterized protein \| Uncharacterized protein; uniclust \| UniRef100\_A0A978BM29 \| 97.7 \| 7.7e-07 \| 1.4e-12 \| 61.8 \| 121 \| (1, 131) \| 147 \| (21, 141) \| 148 \| DNA packaging protein \| DNA packaging protein; uniclust \| UniRef100\_A0A149VFT2 \| 97.7 \| 7.5e-07 \| 1.4e-12 \| 63.0 \| 89 \| (45, 133) \| 147 \| (48, 136) \| 144 \| Uncharacterized protein \| Uncharacterized protein; uniclust \| UniRef100\_A0A7V8JF95 \| 97.7 \| 7.8e-07 \| 1.5e-12 \| 62.5 \| 93 \| (43, 135) \| 147 \| (36, 130) \| 136 \| Uncharacterized protein \| Uncharacterized protein; uniclust \| UniRef100\_A0A812JEV6 \| 97.7 \| 8.1e-07 \| 1.5e-12 \| 82.6 \| 90 \| (43, 132) \| 147 \| (273, 362) \| 4451 \| Phage portal protein \| Phage portal protein; uniclust \| UniRef100\_A0A2U3QE23 \| 97.7 \| 8.4e-07 \| 1.5e-12 \| 63.0 \| 92 \| (43, 134) \| 147 \| (62, 153) \| 171 \| Terminase small subunit \| Terminase small subunit; uniclust \| UniRef100\_A0A272EML8 \| 97.7 \| 9.1e-07 \| 1.7e-12 \| 55.9 \| 70 \| (62, 131) \| 147 \| (6, 75) \| 84 \| Uncharacterized protein \| Uncharacterized protein; uniclust \| UniRef100\_A0A3A0EF07 \| 97.7 \| 9.2e-07 \| 1.7e-12 \| 64.8 \| 89 \| (44, 132) \| 147 \| (83, 171) \| 194 \| Terminase small subunit \| Terminase small subunit; uniclust \| UniRef100\_A0A965H8A5 \| 97.7 \| 9.5e-07 \| 1.7e-12 \| 56.3 \| 84 \| (46, 131) \| 147 \| (2, 85) \| 88 \| Uncharacterized protein \| Uncharacterized protein; uniclust \| UniRef100\_A0A937TUA9 \| 97.7 \| 9.5e-07 \| 1.7e-12 \| 62.3 \| 124 \| (1, 133) \| 147 \| (29, 153) \| 163 \| Uncharacterized protein \| Uncharacterized protein; uniclust \| UniRef100\_UPI0021F8C375 \| 97.7 \| 1e-06 \| 1.9e-12 \| 59.4 \| 83 \| (50, 132) \| 147 \| (37, 119) \| 122 \| hypothetical protein \| hypothetical protein; uniclust \| UniRef100\_A0A3D9Z369 \| 97.7 \| 1.1e-06 \| 1.9e-12 \| 61.2 \| 107 \| (22, 128) \| 147 \| (2, 113) \| 149 \| Uncharacterized protein \| Uncharacterized protein; uniclust \| UniRef100\_A0A176FBY6 \| 97.7 \| 1.1e-06 \| 2.1e-12 \| 64.4 \| 113 \| (3, 130) \| 147 \| (34, 146) \| 170 \| Uncharacterized protein \| Uncharacterized protein; uniclust \| UniRef100\_A0A0F9KLK2 \| 97.7 \| 1.2e-06 \| 2.2e-12 \| 59.4 \| 84 \| (47, 130) \| 147 \| (34, 119) \| 126 \| Uncharacterized protein (Fragment) \| Uncharacterized protein (Fragment); uniclust \| UniRef100\_A0A8S0GWK0 \| 97.7 \| 1.2e-06 \| 2.3e-12 \| 56.8 \| 71 \| (63, 133) \| 147 \| (1, 71) \| 91 \| Uncharacterized protein \| Uncharacterized protein; uniclust \| UniRef100\_A0A0F9I3X3 \| 97.7 \| 1.3e-06 \| 2.3e-12 \| 59.4 \| 88 \| (2, 96) \| 147 \| (34, 124) \| 127 \| DNA-binding protein (Fragment) \| DNA-binding protein (Fragment); uniclust \| UniRef100\_UPI0011E5ABD1 \| 97.7 \| 1.3e-06 \| 2.4e-12 \| 59.2 \| 74 \| (61, 134) \| 147 \| (20, 93) \| 126 \| hypothetical protein \| hypothetical protein; uniclust \| UniRef100\_A0A0Q1ADF8 \| 97.7 \| 1.2e-06 \| 2.4e-12 \| 67.0 \| 87 \| (46, 132) \| 147 \| (124, 210) \| 218 \| Terminase \| Terminase; uniclust \| UniRef100\_UPI0006948996 \| 97.7 \| 1.3e-06 \| 2.5e-12 \| 62.6 \| 86 \| (1, 86) \| 147 \| (48, 155) \| 156 \| hypothetical protein \| hypothetical protein; uniclust \| UniRef100\_UPI002306CCEF \| 97.7 \| 1.3e-06 \| 2.5e-12 \| 64.9 \| 96 \| (1, 102) \| 147 \| (44, 139) \| 238 \| hypothetical protein \| hypothetical protein; uniclust \| UniRef100\_UPI0010B5C855 \| 97.7 \| 1.4e-06 \| 2.5e-12 \| 65.1 \| 115 \| (3, 132) \| 147 \| (69, 183) \| 225 \| hypothetical protein \| hypothetical protein; uniclust \| UniRef100\_A0A964PCS7 \| 97.6 \| 1.4e-06 \| 2.6e-12 \| 62.4 \| 132 \| (5, 139) \| 147 \| (45, 177) \| 180 \| Uncharacterized protein \| Uncharacterized protein; uniclust \| UniRef100\_A0A179D3Q0 \| 97.6 \| 1.4e-06 \| 2.7e-12 \| 64.7 \| 129 \| (1, 135) \| 147 \| (27, 156) \| 207 \| HTH merR-type domain-containing protein \| HTH merR-type domain-containing protein; uniclust \| UniRef100\_UPI0009E87574 \| 97.6 \| 1.5e-06 \| 2.8e-12 \| 56.5 \| 88 \| (48, 135) \| 147 \| (4, 91) \| 98 \| hypothetical protein \| hypothetical protein; uniclust \| UniRef100\_R7UYV5 \| 97.6 \| 1.6e-06 \| 2.8e-12 \| 66.8 \| 80 \| (54, 133) \| 147 \| (3, 82) \| 310 \| Phage terminase large subunit GpA ATPase domain-containing protein (Fragment) \| Phage terminase large subunit GpA ATPase domain-containing protein (Fragment); uniclust \| UniRef100\_UPI0004D72F9B \| 97.6 \| 1.7e-06 \| 3.1e-12 \| 72.5 \| 107 \| (1, 107) \| 147 \| (278, 400) \| 697 \| terminase small subunit \| terminase small subunit; uniclust \| UniRef100\_A0A124V088 \| 97.6 \| 1.7e-06 \| 3.1e-12 \| 55.0 \| 70 \| (63, 132) \| 147 \| (9, 78) \| 86 \| Uncharacterized protein \| Uncharacterized protein; uniclust \| UniRef100\_A0A090R9J5 \| 97.6 \| 1.7e-06 \| 3.3e-12 \| 59.8 \| 62 \| (45, 106) \| 147 \| (15, 76) \| 115 \| Phage protein \| Phage protein; uniclust \| UniRef100\_A0A935IH67 \| 97.6 \| 1.9e-06 \| 3.5e-12 \| 57.5 \| 85 \| (45, 129) \| 147 \| (28, 112) \| 113 \| Uncharacterized protein \| Uncharacterized protein; uniclust \| UniRef100\_UPI0019149DFE \| 97.6 \| 1.9e-06 \| 3.5e-12 \| 60.3 \| 84 \| (46, 129) \| 147 \| (63, 146) \| 152 \| hypothetical protein \| hypothetical protein; uniclust \| UniRef100\_A0A847LN12 \| 97.6 \| 2e-06 \| 3.6e-12 \| 61.6 \| 125 \| (1, 131) \| 147 \| (42, 169) \| 178 \| Terminase small subunit \| Terminase small subunit; uniclust \| UniRef100\_A0A432RHB9 \| 97.6 \| 2e-06 \| 3.6e-12 \| 59.7 \| 87 \| (45, 132) \| 147 \| (50, 136) \| 144 \| Host attachment protein \| Host attachment protein; uniclust \| UniRef100\_A0A351U3V4 \| 97.6 \| 2.1e-06 \| 3.9e-12 \| 60.4 \| 115 \| (1, 129) \| 147 \| (25, 139) \| 149 \| Helix-turn-helix domain-containing protein (Fragment) \| Helix-turn-helix domain-containing protein (Fragment); uniclust \| UniRef100\_UPI000C997E74 \| 97.6 \| 2.3e-06 \| 4.1e-12 \| 62.2 \| 110 \| (21, 131) \| 147 \| (59, 169) \| 197 \| hypothetical protein \| hypothetical protein; uniclust \| UniRef100\_A0A3G2R721 \| 97.6 \| 2.4e-06 \| 4.4e-12 \| 53.6 \| 59 \| (74, 132) \| 147 \| (3, 61) \| 79 \| Uncharacterized protein \| Uncharacterized protein; uniclust \| UniRef100\_A0A2E9M7Y8 \| 97.6 \| 2.3e-06 \| 4.4e-12 \| 61.1 \| 107 \| (14, 130) \| 147 \| (36, 143) \| 144 \| Uncharacterized protein \| Uncharacterized protein; uniclust \| UniRef100\_A0A7J6YMH7 \| 97.6 \| 2.4e-06 \| 4.5e-12 \| 70.6 \| 83 \| (49, 131) \| 147 \| (447, 529) \| 531 \| ParB/Sulfiredoxin domain-containing protein \| ParB/Sulfiredoxin domain-containing protein; uniclust \| UniRef100\_A0A090DYF1 \| 97.6 \| 2.3e-06 \| 4.6e-12 \| 65.6 \| 93 \| (3, 103) \| 147 \| (62, 154) \| 199 \| Uncharacterized protein \| Uncharacterized protein; uniclust \| UniRef100\_UPI00125E4FA9 \| 97.6 \| 2.5e-06 \| 4.7e-12 \| 55.4 \| 81 \| (50, 130) \| 147 \| (9, 89) \| 96 \| hypothetical protein \| hypothetical protein; uniclust \| UniRef100\_A0A849W6W6 \| 97.6 \| 2.6e-06 \| 4.7e-12 \| 58.7 \| 128 \| (4, 132) \| 147 \| (4, 135) \| 137 \| Uncharacterized protein \| Uncharacterized protein; uniclust \| UniRef100\_UPI000F0FA4D5 \| 97.6 \| 2.5e-06 \| 4.7e-12 \| 59.8 \| 80 \| (21, 100) \| 147 \| (41, 121) \| 131 \| terminase small subunit \| terminase small subunit; uniclust \| UniRef100\_A0A1F9BH28 \| 97.6 \| 2.6e-06 \| 4.8e-12 \| 62.1 \| 130 \| (1, 135) \| 147 \| (39, 182) \| 201 \| Helix-turn-helix domain-containing protein \| Helix-turn-helix domain-containing protein; uniclust \| UniRef100\_A0A8X6HP51 \| 97.5 \| 2.7e-06 \| 5e-12 \| 66.6 \| 80 \| (50, 129) \| 147 \| (269, 348) \| 350 \| DNA methylase-like protein \| DNA methylase-like protein; uniclust \| UniRef100\_A0A9D5VQR4 \| 97.5 \| 2.7e-06 \| 5e-12 \| 61.3 \| 83 \| (46, 129) \| 147 \| (103, 185) \| 186 \| Uncharacterized protein \| Uncharacterized protein; uniclust \| UniRef100\_A0A850RI89 \| 97.5 \| 2.9e-06 \| 5.3e-12 \| 61.1 \| 127 \| (6, 133) \| 147 \| (43, 170) \| 184 \| Terminase small subunit \| Terminase small subunit; uniclust \| UniRef100\_UPI001FBB6124 \| 97.5 \| 2.9e-06 \| 5.3e-12 \| 56.3 \| 88 \| (45, 132) \| 147 \| (5, 92) \| 108 \| hypothetical protein \| hypothetical protein; uniclust \| UniRef100\_A0A0B0VTU2 \| 97.5 \| 2.9e-06 \| 5.4e-12 \| 61.8 \| 128 \| (1, 130) \| 147 \| (24, 167) \| 185 \| Terminase \| Terminase; uniclust \| UniRef100\_A0A1G3M3Z5 \| 97.5 \| 3e-06 \| 5.6e-12 \| 59.9 \| 83 \| (44, 129) \| 147 \| (75, 157) \| 162 \| Uncharacterized protein \| Uncharacterized protein; uniclust \| UniRef100\_UPI0018800AA8 \| 97.5 \| 3.2e-06 \| 5.8e-12 \| 60.3 \| 88 \| (44, 131) \| 147 \| (79, 166) \| 172 \| hypothetical protein \| hypothetical protein; uniclust \| UniRef100\_A0A352L2D0 \| 97.5 \| 3.2e-06 \| 5.9e-12 \| 62.6 \| 106 \| (21, 130) \| 147 \| (71, 176) \| 194 \| Terminase small subunit \| Terminase small subunit; uniclust \| UniRef100\_A0A965BIW1 \| 97.5 \| 3.3e-06 \| 6.1e-12 \| 55.6 \| 87 \| (46, 132) \| 147 \| (11, 97) \| 103 \| Uncharacterized protein \| Uncharacterized protein; uniclust \| UniRef100\_A0A066Q1C8 \| 97.5 \| 3.2e-06 \| 6.1e-12 \| 64.2 \| 124 \| (4, 131) \| 147 \| (58, 186) \| 209 \| Prophage Qin DNA packaging protein NU1-like protein \| Prophage Qin DNA packaging protein NU1-like protein; uniclust \| UniRef100\_A0A0F9B6S6 \| 97.5 \| 3.5e-06 \| 6.3e-12 \| 64.0 \| 110 \| (21, 131) \| 147 \| (130, 239) \| 270 \| Uncharacterized protein \| Uncharacterized protein; uniclust \| UniRef100\_A0A953PU68 \| 97.5 \| 3.6e-06 \| 6.6e-12 \| 61.3 \| 92 \| (44, 137) \| 147 \| (99, 190) \| 198 \| Uncharacterized protein \| Uncharacterized protein; uniclust \| UniRef100\_UPI0013EA2CF4 \| 97.5 \| 3.6e-06 \| 6.7e-12 \| 62.3 \| 87 \| (45, 131) \| 147 \| (126, 212) \| 222 \| hypothetical protein \| hypothetical protein; uniclust \| UniRef100\_UPI00039DC22E \| 97.5 \| 3.7e-06 \| 6.8e-12 \| 59.8 \| 88 \| (44, 131) \| 147 \| (72, 159) \| 168 \| hypothetical protein \| hypothetical protein; uniclust \| UniRef100\_A0A348YJM0 \| 97.5 \| 4.3e-06 \| 8e-12 \| 61.5 \| 86 \| (44, 130) \| 147 \| (103, 188) \| 191 \| Terminase small subunit \| Terminase small subunit; uniclust \| UniRef100\_A0A286GNB3 \| 97.5 \| 4.4e-06 \| 8.1e-12 \| 58.5 \| 118 \| (6, 129) \| 147 \| (17, 144) \| 150 \| Uncharacterized protein \| Uncharacterized protein; uniclust \| UniRef100\_A0A2T2PZJ4 \| 97.5 \| 4.6e-06 \| 8.4e-12 \| 62.6 \| 94 \| (2, 103) \| 147 \| (85, 178) \| 233 \| Uncharacterized protein \| Uncharacterized protein; uniclust \| UniRef100\_A0A8I2ABK2 \| 97.5 \| 4.7e-06 \| 8.8e-12 \| 59.6 \| 121 \| (4, 132) \| 147 \| (27, 147) \| 153 \| Uncharacterized protein \| Uncharacterized protein; uniclust \| UniRef100\_A0A5C8B3W1 \| 97.5 \| 4.8e-06 \| 8.9e-12 \| 59.9 \| 87 \| (45, 132) \| 147 \| (84, 170) \| 181 \| DNA packaging protein \| DNA packaging protein; uniclust \| UniRef100\_UPI001421EE26 \| 97.4 \| 5.1e-06 \| 9.4e-12 \| 58.7 \| 123 \| (2, 130) \| 147 \| (24, 156) \| 159 \| terminase small subunit \| terminase small subunit; uniclust \| UniRef100\_UPI001008D493 \| 97.4 \| 5.2e-06 \| 9.6e-12 \| 59.8 \| 113 \| (21, 133) \| 147 \| (27, 147) \| 181 \| hypothetical protein \| hypothetical protein; uniclust \| UniRef100\_A0A177QEF8 \| 97.4 \| 5.3e-06 \| 9.8e-12 \| 52.4 \| 67 \| (66, 132) \| 147 \| (2, 68) \| 81 \| Uncharacterized protein \| Uncharacterized protein; uniclust \| UniRef100\_D8JWB5 \| 97.4 \| 5.2e-06 \| 9.8e-12 \| 61.8 \| 85 \| (44, 133) \| 147 \| (111, 195) \| 203 \| Terminase small subunit \| Terminase small subunit; uniclust \| UniRef100\_A0A4V2SN76 \| 97.4 \| 5.5e-06 \| 1e-11 \| 63.1 \| 83 \| (47, 129) \| 147 \| (187, 269) \| 275 \| Phage terminase Nu1 subunit (DNA packaging protein) (Fragment) \| Phage terminase Nu1 subunit (DNA packaging protein) (Fragment); uniclust \| UniRef100\_A0A2V9B9X4 \| 97.4 \| 5.5e-06 \| 1e-11 \| 54.6 \| 85 \| (25, 109) \| 147 \| (5, 96) \| 103 \| Uncharacterized protein \| Uncharacterized protein; uniclust \| UniRef100\_A0A0J1LWF8 \| 97.4 \| 5.6e-06 \| 1.1e-11 \| 60.7 \| 93 \| (12, 106) \| 147 \| (41, 133) \| 176 \| Terminase \| Terminase; uniclust \| UniRef100\_A0A969QZA3 \| 97.4 \| 5.8e-06 \| 1.1e-11 \| 59.1 \| 84 \| (47, 130) \| 147 \| (87, 170) \| 171 \| Uncharacterized protein \| Uncharacterized protein; uniclust \| UniRef100\_A0A7C1MJZ5 \| 97.4 \| 5.8e-06 \| 1.1e-11 \| 60.6 \| 113 \| (17, 134) \| 147 \| (68, 180) \| 204 \| DUF1441 family protein \| DUF1441 family protein; uniclust \| UniRef100\_A0A536XED3 \| 97.4 \| 5.8e-06 \| 1.1e-11 \| 57.0 \| 86 \| (44, 130) \| 147 \| (48, 133) \| 135 \| Uncharacterized protein \| Uncharacterized protein; uniclust \| UniRef100\_A0A0D6PUD1 \| 97.4 \| 5.8e-06 \| 1.1e-11 \| 65.2 \| 104 \| (1, 104) \| 147 \| (77, 207) \| 265 \| Phage DNA packaging protein \| Phage DNA packaging protein; uniclust \| UniRef100\_A0A437MJQ9 \| 97.4 \| 5.9e-06 \| 1.1e-11 \| 62.1 \| 60 \| (42, 101) \| 147 \| (85, 144) \| 197 \| DNA packaging protein \| DNA packaging protein; uniclust \| UniRef100\_A0A5E4N3A7 \| 97.4 \| 6.6e-06 \| 1.2e-11 \| 64.8 \| 84 \| (50, 133) \| 147 \| (39, 122) \| 359 \| Ankyrin repeat-containing domain,Ankyrin repeat \| Ankyrin repeat-containing domain,Ankyrin repeat; uniclust \| UniRef100\_A0A518LAV0 \| 97.4 \| 6.7e-06 \| 1.3e-11 \| 60.6 \| 125 \| (1, 130) \| 147 \| (27, 154) \| 197 \| Phage DNA packaging protein Nu1 \| Phage DNA packaging protein Nu1; uniclust \| UniRef100\_E6X1P9 \| 97.4 \| 7e-06 \| 1.3e-11 \| 59.6 \| 84 \| (48, 131) \| 147 \| (106, 189) \| 191 \| Terminase small subunit \| Terminase small subunit; uniclust \| UniRef100\_A0A106BVR3 \| 97.4 \| 7.1e-06 \| 1.3e-11 \| 60.2 \| 79 \| (62, 140) \| 147 \| (98, 176) \| 196 \| Terminase small subunit \| Terminase small subunit; uniclust \| UniRef100\_A0A6G5R4R9 \| 97.4 \| 7.5e-06 \| 1.4e-11 \| 55.0 \| 81 \| (46, 129) \| 147 \| (21, 101) \| 115 \| Uncharacterized protein \| Uncharacterized protein; uniclust \| UniRef100\_UPI001EE44BAD \| 97.4 \| 7.5e-06 \| 1.4e-11 \| 52.9 \| 79 \| (47, 125) \| 147 \| (10, 88) \| 91 \| terminase small subunit \| terminase small subunit; uniclust \| UniRef100\_A0A1V5JDJ1 \| 97.4 \| 7.6e-06 \| 1.4e-11 \| 59.0 \| 79 \| (1, 91) \| 147 \| (40, 118) \| 181 \| Phage DNA packaging protein Nu1 \| Phage DNA packaging protein Nu1; uniclust \| UniRef100\_A0A517V825 \| 97.4 \| 7.2e-06 \| 1.4e-11 \| 60.9 \| 117 \| (1, 133) \| 147 \| (33, 150) \| 169 \| Uncharacterized protein \| Uncharacterized protein; uniclust \| UniRef100\_A0A354SKF5 \| 97.4 \| 7.8e-06 \| 1.4e-11 \| 61.1 \| 81 \| (51, 131) \| 147 \| (147, 227) \| 234 \| Uncharacterized protein \| Uncharacterized protein; uniclust \| UniRef100\_A0A7W2H2Q2 \| 97.4 \| 8.4e-06 \| 1.5e-11 \| 53.5 \| 74 \| (62, 136) \| 147 \| (2, 75) \| 99 \| Uncharacterized protein \| Uncharacterized protein; uniclust \| UniRef100\_A0A316FGX7 \| 97.3 \| 8.5e-06 \| 1.7e-11 \| 54.7 \| 34 \| (1, 34) \| 147 \| (50, 83) \| 90 \| Excisionase DUF1233 \| Excisionase DUF1233; uniclust \| UniRef100\_UPI0015923F61 \| 97.3 \| 9.3e-06 \| 1.7e-11 \| 56.4 \| 90 \| (45, 134) \| 147 \| (45, 136) \| 140 \| hypothetical protein \| hypothetical protein; uniclust \| UniRef100\_A0A9E1EL47 \| 97.3 \| 9.4e-06 \| 1.8e-11 \| 57.9 \| 97 \| (3, 104) \| 147 \| (52, 149) \| 153 \| Uncharacterized protein (Fragment) \| Uncharacterized protein (Fragment); uniclust \| UniRef100\_A0A6J7X1I5 \| 97.3 \| 9.6e-06 \| 1.8e-11 \| 59.4 \| 122 \| (1, 129) \| 147 \| (41, 165) \| 200 \| Terminase small subunit \| Terminase small subunit; uniclust \| UniRef100\_A0A1I3HJ39 \| 97.3 \| 9.1e-06 \| 1.8e-11 \| 61.3 \| 90 \| (44, 133) \| 147 \| (89, 178) \| 191 \| Homeodomain-like domain-containing protein \| Homeodomain-like domain-containing protein; uniclust \| UniRef100\_A0A353ZI83 \| 97.3 \| 1e-05 \| 1.9e-11 \| 59.0 \| 118 \| (5, 134) \| 147 \| (59, 178) \| 194 \| Uncharacterized protein \| Uncharacterized protein; uniclust \| UniRef100\_UPI000D30D27B \| 97.3 \| 1.1e-05 \| 2e-11 \| 54.4 \| 86 \| (45, 130) \| 147 \| (25, 110) \| 116 \| hypothetical protein \| hypothetical protein; uniclust \| UniRef100\_A0A0S9S7I7 \| 97.3 \| 1.1e-05 \| 2.1e-11 \| 61.8 \| 105 \| (1, 106) \| 147 \| (52, 171) \| 283 \| HTH merR-type domain-containing protein \| HTH merR-type domain-containing protein; uniclust \| UniRef100\_A0A177RGU4 \| 97.3 \| 1e-05 \| 2.1e-11 \| 63.9 \| 91 \| (6, 103) \| 147 \| (93, 191) \| 242 \| Terminase small subunit \| Terminase small subunit; uniclust \| UniRef100\_A0A7X8UB87 \| 97.3 \| 1.2e-05 \| 2.1e-11 \| 57.3 \| 122 \| (1, 134) \| 147 \| (26, 147) \| 164 \| Terminase small subunit \| Terminase small subunit; uniclust \| UniRef100\_UPI000D0BA657 \| 97.3 \| 1.2e-05 \| 2.1e-11 \| 55.5 \| 107 \| (25, 131) \| 147 \| (10, 118) \| 133 \| hypothetical protein \| hypothetical protein; uniclust \| UniRef100\_A0A2Z6DY60 \| 97.3 \| 1.2e-05 \| 2.1e-11 \| 57.8 \| 85 \| (46, 131) \| 147 \| (87, 171) \| 174 \| Uncharacterized protein \| Uncharacterized protein; uniclust \| UniRef100\_A0A5C7PCC4 \| 97.3 \| 1.2e-05 \| 2.1e-11 \| 59.5 \| 101 \| (1, 106) \| 147 \| (87, 193) \| 213 \| Terminase small subunit \| Terminase small subunit; uniclust \| UniRef100\_UPI001FBB0D88 \| 97.3 \| 1.2e-05 \| 2.2e-11 \| 55.2 \| 78 \| (1, 78) \| 147 \| (29, 111) \| 129 \| terminase small subunit \| terminase small subunit; uniclust \| UniRef100\_UPI00223AE246 \| 97.3 \| 1.2e-05 \| 2.3e-11 \| 59.8 \| 82 \| (50, 131) \| 147 \| (141, 222) \| 225 \| hypothetical protein \| hypothetical protein; uniclust \| UniRef100\_A0A1X7JJP1 \| 97.3 \| 1.3e-05 \| 2.4e-11 \| 54.7 \| 93 \| (43, 135) \| 147 \| (19, 111) \| 124 \| Uncharacterized protein \| Uncharacterized protein; uniclust \| UniRef100\_UPI001FB94040 \| 97.3 \| 1.3e-05 \| 2.4e-11 \| 59.4 \| 89 \| (45, 134) \| 147 \| (62, 150) \| 217 \| hypothetical protein \| hypothetical protein; uniclust \| UniRef100\_A0A0Q5HEW5 \| 97.3 \| 1.4e-05 \| 2.5e-11 \| 57.2 \| 104 \| (1, 107) \| 147 \| (53, 162) \| 169 \| Terminase small subunit \| Terminase small subunit; uniclust \| UniRef100\_A0A7V3AH88 \| 97.3 \| 1.4e-05 \| 2.6e-11 \| 57.9 \| 95 \| (6, 105) \| 147 \| (27, 127) \| 183 \| Elements of external origin \| Elements of external origin; uniclust \| UniRef100\_UPI00211ECC39 \| 97.3 \| 1.4e-05 \| 2.6e-11 \| 56.0 \| 108 \| (21, 136) \| 147 \| (36, 143) \| 147 \| hypothetical protein \| hypothetical protein; uniclust \| UniRef100\_A0A4Q3DN64 \| 97.3 \| 1.3e-05 \| 2.6e-11 \| 64.2 \| 91 \| (44, 135) \| 147 \| (74, 164) \| 284 \| Uncharacterized protein \| Uncharacterized protein; uniclust \| UniRef100\_A0A1Y6C3X7 \| 97.3 \| 1.4e-05 \| 2.6e-11 \| 56.3 \| 88 \| (43, 130) \| 147 \| (60, 147) \| 152 \| Phage DNA packaging protein, Nu1 subunit of terminase \| Phage DNA packaging protein, Nu1 subunit of terminase; uniclust \| UniRef100\_A0A356R6T8 \| 97.3 \| 1.5e-05 \| 2.7e-11 \| 56.2 \| 111 \| (16, 131) \| 147 \| (39, 149) \| 152 \| Terminase small subunit \| Terminase small subunit; uniclust \| UniRef100\_A0A1Q8YAQ8 \| 97.3 \| 1.5e-05 \| 2.7e-11 \| 55.7 \| 89 \| (42, 130) \| 147 \| (37, 125) \| 143 \| Uncharacterized protein \| Uncharacterized protein; uniclust \| UniRef100\_A0A966W431 \| 97.3 \| 1.5e-05 \| 2.7e-11 \| 53.4 \| 107 \| (21, 129) \| 147 \| (2, 108) \| 111 \| Uncharacterized protein \| Uncharacterized protein; uniclust \| UniRef100\_A0A0P1F0B1 \| 97.3 \| 1.5e-05 \| 2.7e-11 \| 53.8 \| 84 \| (2, 90) \| 147 \| (21, 107) \| 111 \| Uncharacterized protein \| Uncharacterized protein; uniclust \| UniRef100\_A0A9D2GWA4 \| 97.2 \| 1.4e-05 \| 2.8e-11 \| 61.8 \| 123 \| (3, 132) \| 147 \| (97, 221) \| 235 \| Uncharacterized protein \| Uncharacterized protein; uniclust \| UniRef100\_UPI001F0AC2C8 \| 97.2 \| 1.6e-05 \| 2.8e-11 \| 65.2 \| 96 \| (46, 141) \| 147 \| (64, 159) \| 506 \| phage terminase large subunit family protein \| phage terminase large subunit family protein; uniclust \| UniRef100\_A0A1I4VF06 \| 97.2 \| 1.6e-05 \| 3e-11 \| 58.9 \| 111 \| (21, 132) \| 147 \| (80, 193) \| 217 \| Phage DNA packaging protein, Nu1 subunit of terminase \| Phage DNA packaging protein, Nu1 subunit of terminase; uniclust \| UniRef100\_A0A068YVF4 \| 97.2 \| 1.7e-05 \| 3.1e-11 \| 59.5 \| 90 \| (44, 133) \| 147 \| (101, 193) \| 205 \| RNA polymerase subunit sigma-70 \| RNA polymerase subunit sigma-70; uniclust \| UniRef100\_I6APC4 \| 97.2 \| 1.8e-05 \| 3.2e-11 \| 59.8 \| 123 \| (4, 131) \| 147 \| (86, 210) \| 237 \| Uncharacterized protein \| Uncharacterized protein; uniclust \| UniRef100\_A0A2H0LPX9 \| 97.2 \| 1.8e-05 \| 3.3e-11 \| 56.7 \| 122 \| (2, 131) \| 147 \| (39, 160) \| 170 \| HTH merR-type domain-containing protein \| HTH merR-type domain-containing protein; uniclust \| UniRef100\_UPI001AB0391E \| 97.2 \| 1.8e-05 \| 3.3e-11 \| 58.3 \| 88 \| (45, 132) \| 147 \| (103, 190) \| 206 \| hypothetical protein \| hypothetical protein; uniclust \| UniRef100\_A0A3G6WFJ2 \| 97.2 \| 1.8e-05 \| 3.3e-11 \| 62.2 \| 103 \| (1, 103) \| 147 \| (47, 171) \| 342 \| Uncharacterized protein \| Uncharacterized protein; uniclust \| UniRef100\_UPI002004F906 \| 97.2 \| 1.8e-05 \| 3.4e-11 \| 58.7 \| 113 \| (3, 116) \| 147 \| (53, 174) \| 217 \| hypothetical protein \| hypothetical protein; uniclust \| UniRef100\_N9NDI8 \| 97.2 \| 1.9e-05 \| 3.4e-11 \| 54.9 \| 81 \| (50, 130) \| 147 \| (52, 132) \| 139 \| Uncharacterized protein \| Uncharacterized protein; uniclust \| UniRef100\_A0A8S5U955 \| 97.2 \| 1.9e-05 \| 3.5e-11 \| 59.5 \| 125 \| (1, 131) \| 147 \| (99, 235) \| 242 \| DNA packaging protein \| DNA packaging protein; uniclust \| UniRef100\_UPI00137449B1 \| 97.2 \| 2e-05 \| 3.6e-11 \| 59.5 \| 115 \| (17, 131) \| 147 \| (40, 181) \| 244 \| hypothetical protein \| hypothetical protein; uniclust \| UniRef100\_A0A978BJL5 \| 97.2 \| 2e-05 \| 3.7e-11 \| 55.7 \| 88 \| (43, 130) \| 147 \| (65, 152) \| 154 \| Uncharacterized protein \| Uncharacterized protein; uniclust \| UniRef100\_A0A2V5U841 \| 97.2 \| 2.2e-05 \| 4e-11 \| 56.2 \| 122 \| (5, 132) \| 147 \| (38, 159) \| 167 \| DUF4332 domain-containing protein \| DUF4332 domain-containing protein; uniclust \| UniRef100\_UPI0018655FAC \| 97.2 \| 2.2e-05 \| 4.1e-11 \| 54.2 \| 89 \| (45, 133) \| 147 \| (23, 114) \| 121 \| hypothetical protein \| hypothetical protein; uniclust \| UniRef100\_A0A4Q2XV87 \| 97.2 \| 2.3e-05 \| 4.3e-11 \| 52.5 \| 58 \| (43, 100) \| 147 \| (53, 110) \| 110 \| Uncharacterized protein \| Uncharacterized protein; uniclust \| UniRef100\_UPI00142DB13B \| 97.2 \| 2.5e-05 \| 4.6e-11 \| 55.2 \| 88 \| (43, 132) \| 147 \| (25, 112) \| 153 \| terminase small subunit \| terminase small subunit; uniclust \| UniRef100\_UPI00130115E7 \| 97.1 \| 2.6e-05 \| 4.8e-11 \| 54.2 \| 85 \| (44, 128) \| 147 \| (14, 98) \| 137 \| hypothetical protein \| hypothetical protein; uniclust \| UniRef100\_A0A248JSB1 \| 97.1 \| 2.6e-05 \| 5e-11 \| 60.0 \| 87 \| (45, 131) \| 147 \| (105, 191) \| 216 \| Terminase small subunit \| Terminase small subunit; uniclust \| UniRef100\_UPI00069BD42E \| 97.1 \| 2.8e-05 \| 5.2e-11 \| 56.4 \| 118 \| (5, 126) \| 147 \| (25, 146) \| 182 \| terminase small subunit \| terminase small subunit; uniclust \| UniRef100\_UPI002240E942 \| 97.1 \| 2.8e-05 \| 5.2e-11 \| 56.7 \| 121 \| (2, 129) \| 147 \| (36, 157) \| 189 \| hypothetical protein \| hypothetical protein; uniclust \| UniRef100\_A0A847HLK2 \| 97.1 \| 2.8e-05 \| 5.2e-11 \| 56.8 \| 122 \| (1, 132) \| 147 \| (34, 155) \| 193 \| Terminase small subunit \| Terminase small subunit; uniclust \| UniRef100\_A0A930QBP3 \| 97.1 \| 2.8e-05 \| 5.2e-11 \| 52.4 \| 86 \| (48, 134) \| 147 \| (22, 110) \| 114 \| Uncharacterized protein \| Uncharacterized protein; uniclust \| UniRef100\_A0A0Q7W4M6 \| 97.1 \| 2.8e-05 \| 5.5e-11 \| 60.9 \| 85 \| (50, 134) \| 147 \| (115, 199) \| 234 \| Terminase small subunit \| Terminase small subunit; uniclust \| UniRef100\_A0A2T5K4X9 \| 97.1 \| 3e-05 \| 5.5e-11 \| 57.3 \| 91 \| (44, 134) \| 147 \| (103, 193) \| 207 \| Phage terminase Nu1 subunit (DNA packaging protein) \| Phage terminase Nu1 subunit (DNA packaging protein); uniclust \| UniRef100\_A0A379V2M5 \| 97.1 \| 3e-05 \| 5.5e-11 \| 54.1 \| 74 \| (1, 74) \| 147 \| (21, 110) \| 140 \| Terminase \| Terminase; uniclust \| UniRef100\_A0A0F9I7J9 \| 97.1 \| 3.1e-05 \| 5.7e-11 \| 55.7 \| 125 \| (2, 132) \| 147 \| (38, 163) \| 172 \| Helix-turn-helix domain-containing protein \| Helix-turn-helix domain-containing protein; uniclust \| UniRef100\_A0A376MST9 \| 97.1 \| 3.1e-05 \| 5.7e-11 \| 52.4 \| 88 \| (20, 107) \| 147 \| (2, 105) \| 112 \| Terminase small subunit \| Terminase small subunit; uniclust \| UniRef100\_A0A929Y0V3 \| 97.1 \| 3.2e-05 \| 5.8e-11 \| 53.5 \| 100 \| (6, 110) \| 147 \| (27, 127) \| 132 \| DNA-packaging protein (Fragment) \| DNA-packaging protein (Fragment); uniclust \| UniRef100\_A0A936H996 \| 97.1 \| 3.1e-05 \| 5.9e-11 \| 53.3 \| 83 \| (43, 128) \| 147 \| (13, 95) \| 116 \| DUF1441 family protein \| DUF1441 family protein; uniclust \| UniRef100\_A0A1D8IV69 \| 97.1 \| 3.3e-05 \| 6e-11 \| 52.6 \| 81 \| (48, 131) \| 147 \| (31, 111) \| 120 \| DUF1441 family protein \| DUF1441 family protein; uniclust \| UniRef100\_UPI000BE6EFA5 \| 97.1 \| 3.3e-05 \| 6e-11 \| 49.4 \| 35 \| (1, 35) \| 147 \| (19, 53) \| 83 \| terminase small subunit \| terminase small subunit; uniclust \| UniRef100\_UPI0002E51B9A \| 97.1 \| 3.4e-05 \| 6.2e-11 \| 56.5 \| 92 \| (45, 137) \| 147 \| (72, 166) \| 193 \| hypothetical protein \| hypothetical protein; uniclust \| UniRef100\_A0A4Q3LZ14 \| 97.1 \| 3.5e-05 \| 6.5e-11 \| 55.2 \| 106 \| (29, 134) \| 147 \| (47, 159) \| 166 \| DUF892 family protein \| DUF892 family protein; uniclust \| UniRef100\_A0A1V6E2G2 \| 97.1 \| 3.6e-05 \| 6.6e-11 \| 54.6 \| 96 \| (1, 106) \| 147 \| (40, 135) \| 156 \| Uncharacterized protein \| Uncharacterized protein; uniclust \| UniRef100\_A0A0F9FRE9 \| 97.1 \| 3.8e-05 \| 6.9e-11 \| 55.3 \| 116 \| (1, 123) \| 147 \| (23, 140) \| 171 \| Terminase small subunit (Fragment) \| Terminase small subunit (Fragment); uniclust \| UniRef100\_A0A382JWV4 \| 97.1 \| 3.8e-05 \| 7e-11 \| 53.2 \| 65 \| (43, 107) \| 147 \| (57, 121) \| 134 \| Helix-turn-helix domain-containing protein \| Helix-turn-helix domain-containing protein; uniclust \| UniRef100\_A0A934R2B7 \| 97.1 \| 3.9e-05 \| 7.1e-11 \| 54.7 \| 110 \| (21, 131) \| 147 \| (37, 147) \| 160 \| Uncharacterized protein \| Uncharacterized protein; uniclust \| UniRef100\_UPI001CF1D285 \| 97.1 \| 4.1e-05 \| 7.5e-11 \| 49.3 \| 36 \| (1, 36) \| 147 \| (21, 56) \| 86 \| terminase small subunit \| terminase small subunit; uniclust \| UniRef100\_UPI0013598FD9 \| 97.1 \| 4.1e-05 \| 7.7e-11 \| 57.8 \| 112 \| (16, 129) \| 147 \| (65, 182) \| 203 \| hypothetical protein \| hypothetical protein; uniclust \| UniRef100\_A0A376PB86 \| 97.1 \| 4.1e-05 \| 7.8e-11 \| 51.9 \| 35 \| (1, 35) \| 147 \| (21, 55) \| 98 \| DNA packaging protein \| DNA packaging protein; uniclust \| UniRef100\_A0A2M7YS23 \| 97.0 \| 4.7e-05 \| 9.1e-11 \| 55.5 \| 57 \| (43, 99) \| 147 \| (90, 146) \| 146 \| Uncharacterized protein (Fragment) \| Uncharacterized protein (Fragment); uniclust \| UniRef100\_UPI0005F41AB5 \| 97.0 \| 5e-05 \| 9.2e-11 \| 67.2 \| 83 \| (49, 131) \| 147 \| (289, 371) \| 1163 \| uncharacterized protein LOC105557561 \| uncharacterized protein LOC105557561; uniclust \| UniRef100\_A0A389MXY8 \| 97.0 \| 5e-05 \| 9.2e-11 \| 57.6 \| 129 \| (1, 130) \| 147 \| (63, 214) \| 246 \| Terminase small subunit \| Terminase small subunit; uniclust \| UniRef100\_X1DAN0 \| 97.0 \| 5.1e-05 \| 9.3e-11 \| 50.1 \| 86 \| (45, 130) \| 147 \| (5, 90) \| 99 \| Uncharacterized protein (Fragment) \| Uncharacterized protein (Fragment); uniclust \| UniRef100\_A0A7X6FQV0 \| 97.0 \| 5.1e-05 \| 9.4e-11 \| 49.6 \| 74 \| (21, 98) \| 147 \| (19, 92) \| 93 \| Uncharacterized protein \| Uncharacterized protein; uniclust \| UniRef100\_A0A376RGB0 \| 97.0 \| 5.2e-05 \| 9.6e-11 \| 50.8 \| 35 \| (1, 35) \| 147 \| (21, 55) \| 106 \| DNA packaging protein \| DNA packaging protein; uniclust \| UniRef100\_A0A3D0I2E3 \| 97.0 \| 5.3e-05 \| 9.7e-11 \| 54.1 \| 115 \| (15, 134) \| 147 \| (34, 152) \| 160 \| Terminase small subunit \| Terminase small subunit; uniclust \| UniRef100\_UPI002023B37C \| 97.0 \| 5.5e-05 \| 1e-10 \| 57.4 \| 87 \| (45, 131) \| 147 \| (148, 237) \| 246 \| hypothetical protein \| hypothetical protein; uniclust \| UniRef100\_UPI0002DEF5FE \| 97.0 \| 5.8e-05 \| 1.1e-10 \| 52.0 \| 93 \| (44, 136) \| 147 \| (14, 106) \| 127 \| terminase small subunit \| terminase small subunit; uniclust \| UniRef100\_A0A7X7H556 \| 97.0 \| 5.9e-05 \| 1.1e-10 \| 52.7 \| 118 \| (1, 127) \| 147 \| (8, 130) \| 138 \| Terminase small subunit \| Terminase small subunit; uniclust \| UniRef100\_UPI00209FEA06 \| 97.0 \| 6.8e-05 \| 1.2e-10 \| 50.9 \| 85 \| (48, 132) \| 147 \| (24, 108) \| 116 \| hypothetical protein \| hypothetical protein; uniclust \| UniRef100\_A0A517X1A7 \| 97.0 \| 6.8e-05 \| 1.2e-10 \| 53.3 \| 125 \| (1, 130) \| 147 \| (21, 148) \| 155 \| Phage DNA packaging protein Nu1 \| Phage DNA packaging protein Nu1; uniclust \| UniRef100\_UPI001BCF985F \| 97.0 \| 6.8e-05 \| 1.3e-10 \| 54.9 \| 91 \| (43, 133) \| 147 \| (70, 160) \| 189 \| hypothetical protein \| hypothetical protein; uniclust \| UniRef100\_A0A426QG20 \| 96.9 \| 6.9e-05 \| 1.3e-10 \| 52.6 \| 112 \| (23, 134) \| 147 \| (12, 126) \| 142 \| Uncharacterized protein \| Uncharacterized protein |
| Top keywords  (threshold 1.00e-03 (evalue)) | **Terminase, small, DNA, packaging, hypothetical, Phage, Nu1, domain\_containing, Fragment, Helix\_turn\_helix** |
| Output files | ../../similar\_sequences/01\_FANPEZAQ\_CDS\_0001\_merged.svg ../../similar\_sequences/01\_FANPEZAQ\_CDS\_0001\_pdb70.a3m ../../similar\_sequences/01\_FANPEZAQ\_CDS\_0001\_pdb70.hhr ../../similar\_sequences/01\_FANPEZAQ\_CDS\_0001\_uniclust.a3m ../../similar\_sequences/01\_FANPEZAQ\_CDS\_0001\_uniclust.hhr |

#### Structure prediction (AlphaFold)^2^

|  |  |
| --- | --- |
| Stats | xml version="1.0" encoding="utf-8" standalone="no"?       2024-09-02T21:08:57.193908 image/svg+xml   Matplotlib v3.7.2, https://matplotlib.org/ |
| Predicted structure | **NGL Viewer Controls:**  - Center: *Left-Click* - Rotate: *Left-Click + Drag* - Translate: *Right-Click + Drag* - Zoom: *Shift + Left-Click + Drag* |
| Output files | ../../predicted\_structures/01\_FANPEZAQ\_CDS\_0001/features.pkl ../../predicted\_structures/01\_FANPEZAQ\_CDS\_0001/ranked\_0.pdb ../../predicted\_structures/01\_FANPEZAQ\_CDS\_0001/ranked\_0\_plots.svg ../../predicted\_structures/01\_FANPEZAQ\_CDS\_0001/result\_model\_1\_ptm\_pred\_0.pkl |

#### Structure similarity search results (Foldseek)^3^

|  |  |
| --- | --- |
| Structure databases searched | Pdb, Afdb-proteome, Afdb-uniprot50 |
| Results, scheme(s)  (Top layers only, threshold 1.00e-02 (evalue)) | xml version="1.0" encoding="utf-8" standalone="no"?       2024-09-02T21:10:02.435958 image/svg+xml   Matplotlib v3.7.2, https://matplotlib.org/ |
| Results, table  (threshold 1.00e-02 (evalue)) | db \| id \| prob \| evalue \| bits \| fident \| alnlen \| mismatch \| gapopen \| qstart \| qend \| tstart \| tend \| name \| description; afdb-proteome \| AF-P31062-F1-MODEL\_V4 \| 0.981 \| 0.005043 \| 79 \| 0.194 \| 159 \| 92 \| 5 \| 2 \| 130 \| 22 \| 174 \| DNA-packaging protein NU1 homolog \| DNA-packaging protein NU1 homolog; afdb-uniprot50 \| AF-A0A5Y3B0P0-F1-MODEL\_V4 \| 1.0 \| 6.8e-12 \| 421 \| 0.605 \| 147 \| 57 \| 1 \| 2 \| 147 \| 29 \| 175 \| Terminase small subunit \| Terminase small subunit; afdb-uniprot50 \| AF-A0A2Z3I612-F1-MODEL\_V4 \| 1.0 \| 4.42e-12 \| 419 \| 0.567 \| 141 \| 61 \| 0 \| 2 \| 142 \| 26 \| 166 \| DNA-packaging protein \| DNA-packaging protein; afdb-uniprot50 \| AF-A0A410UF08-F1-MODEL\_V4 \| 1.0 \| 7.975e-11 \| 387 \| 0.527 \| 146 \| 69 \| 0 \| 2 \| 147 \| 28 \| 173 \| Terminase small subunit \| Terminase small subunit; afdb-uniprot50 \| AF-A0A0Q2LYU8-F1-MODEL\_V4 \| 1.0 \| 2.477e-11 \| 381 \| 0.554 \| 146 \| 62 \| 1 \| 2 \| 147 \| 30 \| 172 \| DNA-packaging protein \| DNA-packaging protein; afdb-uniprot50 \| AF-A0A3G6WMJ0-F1-MODEL\_V4 \| 1.0 \| 4.752e-10 \| 356 \| 0.475 \| 145 \| 76 \| 0 \| 2 \| 146 \| 58 \| 202 \| Helix-turn-helix domain-containing protein \| Helix-turn-helix domain-containing protein; afdb-uniprot50 \| AF-A0A149SVI6-F1-MODEL\_V4 \| 1.0 \| 5.572e-09 \| 336 \| 0.443 \| 133 \| 73 \| 1 \| 2 \| 134 \| 30 \| 161 \| Uncharacterized protein \| Uncharacterized protein; afdb-uniprot50 \| AF-A0A6A4RCC3-F1-MODEL\_V4 \| 1.0 \| 3.951e-10 \| 336 \| 0.516 \| 151 \| 67 \| 5 \| 2 \| 147 \| 32 \| 181 \| Terminase small subunit \| Terminase small subunit; afdb-uniprot50 \| AF-A0A1R7Q9M5-F1-MODEL\_V4 \| 1.0 \| 5.926e-09 \| 332 \| 0.496 \| 131 \| 66 \| 0 \| 2 \| 132 \| 29 \| 159 \| Phage DNA packaging protein Nu1 \| Phage DNA packaging protein Nu1; afdb-uniprot50 \| AF-A0A291LYW6-F1-MODEL\_V4 \| 1.0 \| 4.752e-10 \| 327 \| 0.493 \| 148 \| 72 \| 3 \| 2 \| 147 \| 30 \| 176 \| DNA-packaging protein \| DNA-packaging protein; afdb-uniprot50 \| AF-A0A7J0BX75-F1-MODEL\_V4 \| 1.0 \| 1.166e-08 \| 319 \| 0.407 \| 140 \| 71 \| 2 \| 2 \| 134 \| 43 \| 177 \| Uncharacterized protein \| Uncharacterized protein; afdb-uniprot50 \| AF-A0A8A6KGW4-F1-MODEL\_V4 \| 1.0 \| 6.302e-09 \| 314 \| 0.446 \| 150 \| 74 \| 5 \| 2 \| 147 \| 28 \| 172 \| Terminase small subunit \| Terminase small subunit; afdb-uniprot50 \| AF-A0A1I1F6U7-F1-MODEL\_V4 \| 1.0 \| 2.354e-09 \| 307 \| 0.387 \| 147 \| 88 \| 2 \| 2 \| 147 \| 20 \| 165 \| Phage DNA packaging protein, Nu1 subunit of terminase \| Phage DNA packaging protein, Nu1 subunit of terminase; afdb-uniprot50 \| AF-A0A844HQQ0-F1-MODEL\_V4 \| 1.0 \| 7.86e-08 \| 297 \| 0.41 \| 146 \| 85 \| 1 \| 2 \| 147 \| 27 \| 171 \| Terminase small subunit \| Terminase small subunit; afdb-uniprot50 \| AF-A0A6I6JFF5-F1-MODEL\_V4 \| 1.0 \| 1.794e-08 \| 292 \| 0.386 \| 150 \| 82 \| 3 \| 2 \| 147 \| 29 \| 172 \| Terminase small subunit \| Terminase small subunit; afdb-uniprot50 \| AF-A0A103EHP4-F1-MODEL\_V4 \| 1.0 \| 3.755e-08 \| 290 \| 0.449 \| 138 \| 75 \| 1 \| 11 \| 147 \| 3 \| 140 \| Uncharacterized protein \| Uncharacterized protein; afdb-uniprot50 \| AF-A0A0A6SUS5-F1-MODEL\_V4 \| 1.0 \| 1.86e-07 \| 274 \| 0.321 \| 146 \| 97 \| 2 \| 2 \| 147 \| 33 \| 176 \| Uncharacterized protein \| Uncharacterized protein; afdb-uniprot50 \| AF-A0A017HTQ6-F1-MODEL\_V4 \| 1.0 \| 2.104e-07 \| 270 \| 0.31 \| 148 \| 99 \| 3 \| 2 \| 147 \| 11 \| 157 \| Uncharacterized protein \| Uncharacterized protein; afdb-uniprot50 \| AF-A0A2D7GUJ0-F1-MODEL\_V4 \| 1.0 \| 7.86e-08 \| 265 \| 0.392 \| 130 \| 78 \| 1 \| 2 \| 130 \| 22 \| 151 \| Uncharacterized protein \| Uncharacterized protein; afdb-uniprot50 \| AF-A0A7C8HWC0-F1-MODEL\_V4 \| 1.0 \| 6.776e-07 \| 261 \| 0.359 \| 142 \| 84 \| 3 \| 2 \| 138 \| 40 \| 179 \| Uncharacterized protein \| Uncharacterized protein; afdb-uniprot50 \| AF-A0A3N2E0T8-F1-MODEL\_V4 \| 1.0 \| 3.443e-07 \| 259 \| 0.356 \| 146 \| 88 \| 3 \| 2 \| 144 \| 28 \| 170 \| Phage terminase Nu1 subunit (DNA packaging protein) \| Phage terminase Nu1 subunit (DNA packaging protein); afdb-uniprot50 \| AF-A0A126NZA6-F1-MODEL\_V4 \| 1.0 \| 5.991e-07 \| 252 \| 0.333 \| 132 \| 85 \| 3 \| 2 \| 132 \| 9 \| 138 \| Uncharacterized protein \| Uncharacterized protein; afdb-uniprot50 \| AF-A0A5P9IWW9-F1-MODEL\_V4 \| 1.0 \| 4.141e-07 \| 252 \| 0.342 \| 140 \| 81 \| 1 \| 2 \| 130 \| 22 \| 161 \| Phage DNA packaging protein Nu1 \| Phage DNA packaging protein Nu1; afdb-uniprot50 \| AF-A0A7W6S2H6-F1-MODEL\_V4 \| 1.0 \| 3.443e-07 \| 239 \| 0.328 \| 149 \| 95 \| 3 \| 2 \| 146 \| 25 \| 172 \| Phage terminase Nu1 subunit (DNA packaging protein) \| Phage terminase Nu1 subunit (DNA packaging protein); afdb-uniprot50 \| AF-A0A315EM69-F1-MODEL\_V4 \| 1.0 \| 3.994e-08 \| 238 \| 0.4 \| 155 \| 84 \| 3 \| 2 \| 147 \| 23 \| 177 \| Uncharacterized protein \| Uncharacterized protein; afdb-uniprot50 \| AF-A0A1W1Z3G8-F1-MODEL\_V4 \| 1.0 \| 8.667e-07 \| 237 \| 0.324 \| 148 \| 96 \| 4 \| 2 \| 147 \| 24 \| 169 \| Phage DNA packaging protein Nu1 \| Phage DNA packaging protein Nu1; afdb-uniprot50 \| AF-A0A7V5XCS4-F1-MODEL\_V4 \| 1.0 \| 6.606e-06 \| 223 \| 0.273 \| 146 \| 98 \| 3 \| 2 \| 143 \| 49 \| 190 \| Uncharacterized protein \| Uncharacterized protein; afdb-uniprot50 \| AF-A0A178LGQ7-F1-MODEL\_V4 \| 1.0 \| 1.47e-05 \| 216 \| 0.282 \| 145 \| 92 \| 5 \| 3 \| 147 \| 10 \| 142 \| Uncharacterized protein \| Uncharacterized protein; afdb-uniprot50 \| AF-A0A2E0ENP5-F1-MODEL\_V4 \| 1.0 \| 1.222e-05 \| 216 \| 0.251 \| 139 \| 93 \| 2 \| 2 \| 140 \| 24 \| 151 \| Uncharacterized protein \| Uncharacterized protein; afdb-uniprot50 \| AF-A0A6A8A3Z2-F1-MODEL\_V4 \| 1.0 \| 2.968e-06 \| 216 \| 0.298 \| 151 \| 101 \| 2 \| 2 \| 147 \| 32 \| 182 \| Uncharacterized protein \| Uncharacterized protein; afdb-uniprot50 \| AF-A0A1G1HG69-F1-MODEL\_V4 \| 1.0 \| 1.47e-05 \| 211 \| 0.237 \| 135 \| 101 \| 1 \| 2 \| 136 \| 20 \| 152 \| Uncharacterized protein \| Uncharacterized protein; afdb-uniprot50 \| AF-A0A386UIG2-F1-MODEL\_V4 \| 1.0 \| 5.492e-06 \| 210 \| 0.283 \| 148 \| 102 \| 3 \| 2 \| 147 \| 29 \| 174 \| Uncharacterized protein \| Uncharacterized protein; afdb-uniprot50 \| AF-A0A165U8L6-F1-MODEL\_V4 \| 1.0 \| 2.721e-05 \| 202 \| 0.29 \| 131 \| 92 \| 1 \| 2 \| 132 \| 24 \| 153 \| Phage DNA packaging protein Nu1 \| Phage DNA packaging protein Nu1; afdb-uniprot50 \| AF-A0A7X2NZP6-F1-MODEL\_V4 \| 1.0 \| 8.761e-05 \| 201 \| 0.412 \| 126 \| 69 \| 2 \| 22 \| 147 \| 1 \| 121 \| Uncharacterized protein \| Uncharacterized protein; afdb-uniprot50 \| AF-A0A7Y7QTY3-F1-MODEL\_V4 \| 1.0 \| 2.721e-05 \| 200 \| 0.291 \| 144 \| 95 \| 2 \| 4 \| 147 \| 33 \| 169 \| Terminase small subunit \| Terminase small subunit; afdb-uniprot50 \| AF-A0A1Q5SRW0-F1-MODEL\_V4 \| 1.0 \| 2.558e-05 \| 197 \| 0.344 \| 145 \| 82 \| 4 \| 3 \| 147 \| 29 \| 160 \| Uncharacterized protein \| Uncharacterized protein; afdb-uniprot50 \| AF-A0A2A5EK29-F1-MODEL\_V4 \| 1.0 \| 2e-05 \| 197 \| 0.212 \| 165 \| 111 \| 3 \| 2 \| 147 \| 44 \| 208 \| Uncharacterized protein \| Uncharacterized protein; afdb-uniprot50 \| AF-A0A4R2YRS0-F1-MODEL\_V4 \| 1.0 \| 4.735e-05 \| 195 \| 0.289 \| 145 \| 91 \| 6 \| 3 \| 147 \| 34 \| 166 \| Uncharacterized protein \| Uncharacterized protein; afdb-uniprot50 \| AF-A0A235H584-F1-MODEL\_V4 \| 1.0 \| 2.558e-05 \| 194 \| 0.258 \| 155 \| 106 \| 4 \| 2 \| 147 \| 25 \| 179 \| Uncharacterized protein \| Uncharacterized protein; afdb-uniprot50 \| AF-A0A2U2N0D2-F1-MODEL\_V4 \| 1.0 \| 0.0001621 \| 193 \| 0.244 \| 139 \| 99 \| 3 \| 2 \| 136 \| 22 \| 158 \| Uncharacterized protein \| Uncharacterized protein; afdb-uniprot50 \| AF-A0A433J1A9-F1-MODEL\_V4 \| 1.0 \| 4.452e-05 \| 192 \| 0.282 \| 138 \| 93 \| 2 \| 2 \| 134 \| 27 \| 163 \| Terminase small subunit (DNA packaging protein Nu1) \| Terminase small subunit (DNA packaging protein Nu1); afdb-uniprot50 \| AF-A0A1X7JJP1-F1-MODEL\_V4 \| 1.0 \| 0.0002074 \| 191 \| 0.401 \| 107 \| 61 \| 2 \| 22 \| 128 \| 1 \| 104 \| Uncharacterized protein \| Uncharacterized protein; afdb-uniprot50 \| AF-A0A3M6EKX5-F1-MODEL\_V4 \| 1.0 \| 0.0003393 \| 191 \| 0.305 \| 118 \| 80 \| 1 \| 16 \| 133 \| 36 \| 151 \| Uncharacterized protein \| Uncharacterized protein; afdb-uniprot50 \| AF-A0A2V5D2Z0-F1-MODEL\_V4 \| 1.0 \| 0.0001192 \| 188 \| 0.315 \| 133 \| 84 \| 5 \| 16 \| 147 \| 38 \| 164 \| Uncharacterized protein \| Uncharacterized protein; afdb-uniprot50 \| AF-G4KQ86-F1-MODEL\_V4 \| 1.0 \| 0.0001348 \| 187 \| 0.21 \| 147 \| 109 \| 5 \| 3 \| 147 \| 23 \| 164 \| Putative terminase small subunit \| Putative terminase small subunit; afdb-uniprot50 \| AF-A0A1G3BHK7-F1-MODEL\_V4 \| 1.0 \| 7.284e-05 \| 186 \| 0.224 \| 129 \| 94 \| 2 \| 2 \| 130 \| 23 \| 145 \| Uncharacterized protein \| Uncharacterized protein; afdb-uniprot50 \| AF-A0A4S2H4R7-F1-MODEL\_V4 \| 1.0 \| 0.0001724 \| 185 \| 0.232 \| 146 \| 106 \| 3 \| 3 \| 145 \| 27 \| 169 \| Terminase small subunit, Nu1 \| Terminase small subunit, Nu1; afdb-uniprot50 \| AF-A0A8B2NVC2-F1-MODEL\_V4 \| 1.0 \| 3.48e-05 \| 183 \| 0.31 \| 148 \| 99 \| 3 \| 2 \| 147 \| 41 \| 187 \| Uncharacterized protein \| Uncharacterized protein; afdb-uniprot50 \| AF-A9IRX2-F1-MODEL\_V4 \| 1.0 \| 2.721e-05 \| 182 \| 0.31 \| 145 \| 83 \| 6 \| 3 \| 147 \| 18 \| 145 \| Uncharacterized protein \| Uncharacterized protein; afdb-uniprot50 \| AF-A0A2E1HAF3-F1-MODEL\_V4 \| 1.0 \| 0.0001434 \| 181 \| 0.234 \| 141 \| 97 \| 3 \| 10 \| 147 \| 27 \| 159 \| Uncharacterized protein \| Uncharacterized protein; afdb-uniprot50 \| AF-A0A6L4BEY6-F1-MODEL\_V4 \| 1.0 \| 0.0001192 \| 179 \| 0.288 \| 142 \| 97 \| 2 \| 2 \| 143 \| 45 \| 182 \| Terminase small subunit \| Terminase small subunit; afdb-uniprot50 \| AF-A0A6L3T3B7-F1-MODEL\_V4 \| 1.0 \| 0.0002821 \| 177 \| 0.281 \| 135 \| 90 \| 2 \| 2 \| 130 \| 30 \| 163 \| Terminase small subunit \| Terminase small subunit; afdb-uniprot50 \| AF-A0A2D2C217-F1-MODEL\_V4 \| 1.0 \| 0.0001267 \| 177 \| 0.231 \| 160 \| 106 \| 4 \| 3 \| 147 \| 46 \| 203 \| Uncharacterized protein \| Uncharacterized protein; afdb-uniprot50 \| AF-A0A2A5A3W5-F1-MODEL\_V4 \| 1.0 \| 7.284e-05 \| 176 \| 0.333 \| 135 \| 82 \| 4 \| 2 \| 132 \| 26 \| 156 \| Uncharacterized protein \| Uncharacterized protein; afdb-uniprot50 \| AF-A0A1C3WPE9-F1-MODEL\_V4 \| 1.0 \| 0.0001054 \| 176 \| 0.285 \| 147 \| 91 \| 4 \| 10 \| 147 \| 27 \| 168 \| Phage DNA packaging protein, Nu1 subunit of terminase \| Phage DNA packaging protein, Nu1 subunit of terminase; afdb-uniprot50 \| AF-G2IX52-F1-MODEL\_V4 \| 1.0 \| 0.0001054 \| 173 \| 0.205 \| 146 \| 104 \| 2 \| 2 \| 147 \| 24 \| 157 \| Phage terminase small subunit gpNu1 \| Phage terminase small subunit gpNu1; afdb-uniprot50 \| AF-A0A4Y3TG37-F1-MODEL\_V4 \| 1.0 \| 0.0004909 \| 173 \| 0.26 \| 150 \| 106 \| 4 \| 2 \| 147 \| 28 \| 176 \| Uncharacterized protein \| Uncharacterized protein; afdb-uniprot50 \| AF-A0A2T7TXU6-F1-MODEL\_V4 \| 1.0 \| 0.0002653 \| 170 \| 0.22 \| 136 \| 99 \| 2 \| 2 \| 130 \| 27 \| 162 \| Terminase small subunit (DNA packaging protein Nu1) \| Terminase small subunit (DNA packaging protein Nu1); afdb-uniprot50 \| AF-X1HCJ8-F1-MODEL\_V4 \| 1.0 \| 0.0001724 \| 170 \| 0.246 \| 154 \| 102 \| 4 \| 2 \| 147 \| 23 \| 170 \| Uncharacterized protein \| Uncharacterized protein; afdb-uniprot50 \| AF-A0A3U4W908-F1-MODEL\_V4 \| 1.0 \| 0.0003191 \| 167 \| 0.251 \| 135 \| 100 \| 1 \| 14 \| 147 \| 33 \| 167 \| Uncharacterized protein \| Uncharacterized protein; afdb-uniprot50 \| AF-A0A1S7LLS5-F1-MODEL\_V4 \| 1.0 \| 0.002022 \| 166 \| 0.237 \| 118 \| 87 \| 2 \| 20 \| 134 \| 31 \| 148 \| Uncharacterized protein \| Uncharacterized protein; afdb-uniprot50 \| AF-A0A192IJE2-F1-MODEL\_V4 \| 1.0 \| 0.0003393 \| 166 \| 0.255 \| 133 \| 97 \| 2 \| 3 \| 135 \| 22 \| 152 \| Uncharacterized protein \| Uncharacterized protein; afdb-uniprot50 \| AF-A0A3B8QTC1-F1-MODEL\_V4 \| 1.0 \| 0.0001434 \| 166 \| 0.25 \| 128 \| 85 \| 4 \| 2 \| 128 \| 22 \| 139 \| Uncharacterized protein \| Uncharacterized protein; afdb-uniprot50 \| AF-A0A7G8BY94-F1-MODEL\_V4 \| 1.0 \| 0.0002494 \| 166 \| 0.272 \| 158 \| 96 \| 5 \| 2 \| 146 \| 28 \| 179 \| Terminase small subunit \| Terminase small subunit; afdb-uniprot50 \| AF-A0A7Z8R5N7-F1-MODEL\_V4 \| 1.0 \| 0.0003191 \| 165 \| 0.205 \| 136 \| 101 \| 3 \| 2 \| 130 \| 17 \| 152 \| Terminase small subunit \| Terminase small subunit; afdb-uniprot50 \| AF-A0A1V5Z7S9-F1-MODEL\_V4 \| 1.0 \| 0.001314 \| 165 \| 0.259 \| 108 \| 78 \| 1 \| 22 \| 129 \| 63 \| 168 \| Phage DNA packaging protein Nu1 \| Phage DNA packaging protein Nu1; afdb-uniprot50 \| AF-A0A0C9PQA6-F1-MODEL\_V4 \| 1.0 \| 0.000195 \| 164 \| 0.234 \| 149 \| 101 \| 5 \| 2 \| 144 \| 45 \| 186 \| Uncharacterized protein \| Uncharacterized protein; afdb-uniprot50 \| AF-A0A2D8RC70-F1-MODEL\_V4 \| 1.0 \| 0.0003838 \| 162 \| 0.222 \| 148 \| 109 \| 4 \| 5 \| 147 \| 18 \| 164 \| Uncharacterized protein \| Uncharacterized protein; afdb-uniprot50 \| AF-A0A7X1W7E9-F1-MODEL\_V4 \| 1.0 \| 0.001398 \| 161 \| 0.287 \| 132 \| 88 \| 4 \| 4 \| 135 \| 25 \| 150 \| Terminase small subunit, Nu1 \| Terminase small subunit, Nu1; afdb-uniprot50 \| AF-A0A7C6RL71-F1-MODEL\_V4 \| 1.0 \| 8.238e-05 \| 161 \| 0.261 \| 157 \| 87 \| 4 \| 2 \| 134 \| 20 \| 171 \| Terminase small subunit \| Terminase small subunit; afdb-uniprot50 \| AF-A0A7Z0KXQ8-F1-MODEL\_V4 \| 1.0 \| 0.000434 \| 159 \| 0.279 \| 143 \| 93 \| 5 \| 3 \| 143 \| 27 \| 161 \| Terminase small subunit, Nu1 \| Terminase small subunit, Nu1; afdb-uniprot50 \| AF-A0A7U9IVZ5-F1-MODEL\_V4 \| 1.0 \| 0.0003393 \| 159 \| 0.238 \| 147 \| 99 \| 4 \| 2 \| 147 \| 25 \| 159 \| Uncharacterized protein \| Uncharacterized protein; afdb-uniprot50 \| AF-A0A6A4R6A4-F1-MODEL\_V4 \| 1.0 \| 0.0005552 \| 158 \| 0.27 \| 137 \| 92 \| 1 \| 2 \| 130 \| 24 \| 160 \| Uncharacterized protein \| Uncharacterized protein; afdb-uniprot50 \| AF-A0A8A5HVC6-F1-MODEL\_V4 \| 1.0 \| 0.0004081 \| 156 \| 0.238 \| 147 \| 99 \| 4 \| 2 \| 147 \| 12 \| 146 \| Uncharacterized protein \| Uncharacterized protein; afdb-uniprot50 \| AF-A0A7L6A869-F1-MODEL\_V4 \| 1.0 \| 0.001681 \| 156 \| 0.304 \| 125 \| 84 \| 2 \| 24 \| 147 \| 38 \| 160 \| Uncharacterized protein \| Uncharacterized protein; afdb-uniprot50 \| AF-A0A7C5CB12-F1-MODEL\_V4 \| 1.0 \| 0.002586 \| 156 \| 0.214 \| 126 \| 93 \| 2 \| 24 \| 147 \| 46 \| 167 \| Terminase small subunit, Nu1 \| Terminase small subunit, Nu1; afdb-uniprot50 \| AF-A0A3B1BSM9-F1-MODEL\_V4 \| 1.0 \| 0.002022 \| 155 \| 0.248 \| 137 \| 89 \| 3 \| 11 \| 147 \| 3 \| 125 \| Uncharacterized protein \| Uncharacterized protein; afdb-uniprot50 \| AF-A0A6N6NL26-F1-MODEL\_V4 \| 1.0 \| 0.0005221 \| 155 \| 0.211 \| 151 \| 104 \| 5 \| 2 \| 147 \| 54 \| 194 \| Terminase small subunit \| Terminase small subunit; afdb-uniprot50 \| AF-A0A1I5MM69-F1-MODEL\_V4 \| 1.0 \| 0.004786 \| 154 \| 0.22 \| 127 \| 99 \| 0 \| 21 \| 147 \| 46 \| 172 \| Phage DNA packaging protein, Nu1 subunit of terminase \| Phage DNA packaging protein, Nu1 subunit of terminase; afdb-uniprot50 \| AF-E8V6R1-F1-MODEL\_V4 \| 1.0 \| 0.0006678 \| 153 \| 0.266 \| 150 \| 87 \| 4 \| 2 \| 134 \| 13 \| 156 \| Uncharacterized protein \| Uncharacterized protein; afdb-uniprot50 \| AF-A0A0S9LPH7-F1-MODEL\_V4 \| 1.0 \| 0.003111 \| 151 \| 0.347 \| 118 \| 73 \| 2 \| 24 \| 140 \| 44 \| 158 \| Uncharacterized protein \| Uncharacterized protein; afdb-uniprot50 \| AF-A0A3M2C0J2-F1-MODEL\_V4 \| 1.0 \| 0.002586 \| 150 \| 0.233 \| 133 \| 94 \| 3 \| 3 \| 132 \| 13 \| 140 \| Uncharacterized protein \| Uncharacterized protein; afdb-uniprot50 \| AF-A0A1H9NYA0-F1-MODEL\_V4 \| 1.0 \| 0.0005904 \| 150 \| 0.258 \| 139 \| 89 \| 2 \| 2 \| 130 \| 24 \| 158 \| Phage DNA packaging protein, Nu1 subunit of terminase \| Phage DNA packaging protein, Nu1 subunit of terminase; afdb-uniprot50 \| AF-U7T5L9-F1-MODEL\_V4 \| 1.0 \| 0.001398 \| 150 \| 0.183 \| 147 \| 114 \| 4 \| 3 \| 147 \| 28 \| 170 \| Uncharacterized protein \| Uncharacterized protein; afdb-uniprot50 \| AF-A0A2T5K719-F1-MODEL\_V4 \| 1.0 \| 0.0003838 \| 150 \| 0.186 \| 166 \| 115 \| 4 \| 2 \| 147 \| 44 \| 209 \| Phage terminase Nu1 subunit (DNA packaging protein) \| Phage terminase Nu1 subunit (DNA packaging protein); afdb-uniprot50 \| AF-A0A075KAS7-F1-MODEL\_V4 \| 1.0 \| 0.005413 \| 149 \| 0.223 \| 121 \| 86 \| 2 \| 21 \| 139 \| 42 \| 156 \| Uncharacterized protein \| Uncharacterized protein; afdb-uniprot50 \| AF-A0A661I0F5-F1-MODEL\_V4 \| 1.0 \| 0.001027 \| 148 \| 0.218 \| 133 \| 102 \| 1 \| 2 \| 132 \| 53 \| 185 \| Uncharacterized protein \| Uncharacterized protein; afdb-uniprot50 \| AF-A0A4Q2B780-F1-MODEL\_V4 \| 1.0 \| 0.002751 \| 147 \| 0.174 \| 143 \| 108 \| 4 \| 3 \| 139 \| 38 \| 176 \| Protoporphyrinogen oxidase \| Protoporphyrinogen oxidase; afdb-uniprot50 \| AF-A0A8A5CXW5-F1-MODEL\_V4 \| 1.0 \| 0.003308 \| 146 \| 0.207 \| 135 \| 103 \| 3 \| 14 \| 147 \| 38 \| 169 \| Uncharacterized protein \| Uncharacterized protein; afdb-uniprot50 \| AF-A0A2V8RWY8-F1-MODEL\_V4 \| 1.0 \| 0.003979 \| 146 \| 0.211 \| 123 \| 86 \| 2 \| 25 \| 147 \| 46 \| 157 \| Uncharacterized protein \| Uncharacterized protein; afdb-uniprot50 \| AF-A0A2U1XZ16-F1-MODEL\_V4 \| 1.0 \| 0.0001192 \| 146 \| 0.3 \| 150 \| 99 \| 4 \| 2 \| 147 \| 24 \| 171 \| Uncharacterized protein \| Uncharacterized protein; afdb-uniprot50 \| AF-A0A7G9WG98-F1-MODEL\_V4 \| 1.0 \| 0.002432 \| 146 \| 0.211 \| 142 \| 100 \| 4 \| 3 \| 139 \| 24 \| 158 \| Excisionase family DNA-binding protein \| Excisionase family DNA-binding protein; afdb-uniprot50 \| AF-A0A1Q9PIU8-F1-MODEL\_V4 \| 1.0 \| 0.003518 \| 146 \| 0.216 \| 143 \| 95 \| 2 \| 16 \| 147 \| 41 \| 177 \| Uncharacterized protein \| Uncharacterized protein; afdb-uniprot50 \| AF-A0A853K663-F1-MODEL\_V4 \| 1.0 \| 0.003742 \| 145 \| 0.321 \| 112 \| 74 \| 1 \| 36 \| 147 \| 2 \| 111 \| Terminase small subunit \| Terminase small subunit; afdb-uniprot50 \| AF-A0A2I8QTZ5-F1-MODEL\_V4 \| 1.0 \| 0.002287 \| 145 \| 0.231 \| 138 \| 101 \| 2 \| 11 \| 147 \| 1 \| 134 \| Uncharacterized protein \| Uncharacterized protein; afdb-uniprot50 \| AF-A0A1M6SLK9-F1-MODEL\_V4 \| 1.0 \| 0.004232 \| 145 \| 0.248 \| 129 \| 91 \| 4 \| 22 \| 147 \| 30 \| 155 \| Uncharacterized protein \| Uncharacterized protein; afdb-uniprot50 \| AF-A0A752JIU9-F1-MODEL\_V4 \| 1.0 \| 0.001162 \| 145 \| 0.193 \| 150 \| 107 \| 3 \| 2 \| 147 \| 8 \| 147 \| Terminase small subunit \| Terminase small subunit; afdb-uniprot50 \| AF-A0A5B9Y7V3-F1-MODEL\_V4 \| 1.0 \| 0.001901 \| 145 \| 0.234 \| 141 \| 98 \| 5 \| 3 \| 138 \| 36 \| 171 \| Uncharacterized protein \| Uncharacterized protein; afdb-uniprot50 \| AF-A0A846KET6-F1-MODEL\_V4 \| 1.0 \| 0.00509 \| 144 \| 0.205 \| 146 \| 110 \| 4 \| 3 \| 147 \| 23 \| 163 \| DNA-packaging protein \| DNA-packaging protein; afdb-uniprot50 \| AF-A0A098AXF9-F1-MODEL\_V4 \| 1.0 \| 0.002287 \| 144 \| 0.215 \| 158 \| 108 \| 4 \| 3 \| 147 \| 34 \| 188 \| Uncharacterized protein \| Uncharacterized protein; afdb-uniprot50 \| AF-A0A854GZ82-F1-MODEL\_V4 \| 1.0 \| 0.002287 \| 143 \| 0.215 \| 153 \| 108 \| 3 \| 3 \| 147 \| 29 \| 177 \| Uncharacterized protein \| Uncharacterized protein; afdb-uniprot50 \| AF-A0A7V2AKR6-F1-MODEL\_V4 \| 1.0 \| 0.003979 \| 142 \| 0.208 \| 149 \| 104 \| 5 \| 2 \| 146 \| 26 \| 164 \| Terminase small subunit, Nu1 \| Terminase small subunit, Nu1; afdb-uniprot50 \| AF-A0A3B9ZIR9-F1-MODEL\_V4 \| 1.0 \| 0.001901 \| 142 \| 0.231 \| 147 \| 100 \| 6 \| 2 \| 147 \| 73 \| 207 \| Uncharacterized protein \| Uncharacterized protein; afdb-uniprot50 \| AF-A0A7C9M626-F1-MODEL\_V4 \| 1.0 \| 0.009419 \| 141 \| 0.204 \| 127 \| 97 \| 1 \| 16 \| 138 \| 44 \| 170 \| Uncharacterized protein \| Uncharacterized protein; afdb-uniprot50 \| AF-A0A4R2YZM0-F1-MODEL\_V4 \| 1.0 \| 0.00215 \| 140 \| 0.177 \| 152 \| 114 \| 5 \| 2 \| 147 \| 29 \| 175 \| Uncharacterized protein \| Uncharacterized protein; afdb-uniprot50 \| AF-A0A2S0MHN3-F1-MODEL\_V4 \| 1.0 \| 0.00509 \| 139 \| 0.284 \| 109 \| 75 \| 2 \| 23 \| 130 \| 39 \| 145 \| Uncharacterized protein \| Uncharacterized protein; afdb-uniprot50 \| AF-A0A139GP13-F1-MODEL\_V4 \| 1.0 \| 0.0009084 \| 139 \| 0.212 \| 146 \| 104 \| 4 \| 2 \| 147 \| 21 \| 155 \| Uncharacterized protein \| Uncharacterized protein; afdb-uniprot50 \| AF-A0A1V5QJC7-F1-MODEL\_V4 \| 1.0 \| 0.009419 \| 139 \| 0.215 \| 116 \| 84 \| 2 \| 21 \| 130 \| 50 \| 164 \| Uncharacterized protein \| Uncharacterized protein; afdb-uniprot50 \| AF-A0A442XVU9-F1-MODEL\_V4 \| 1.0 \| 0.0045 \| 139 \| 0.244 \| 147 \| 103 \| 4 \| 4 \| 147 \| 27 \| 168 \| Terminase small subunit, Nu1 \| Terminase small subunit, Nu1; afdb-uniprot50 \| AF-A0A510KMJ0-F1-MODEL\_V4 \| 1.0 \| 0.005757 \| 139 \| 0.147 \| 136 \| 112 \| 2 \| 16 \| 147 \| 38 \| 173 \| Uncharacterized protein \| Uncharacterized protein; afdb-uniprot50 \| AF-A0A2I8QQ64-F1-MODEL\_V4 \| 1.0 \| 0.002586 \| 138 \| 0.238 \| 151 \| 99 \| 3 \| 2 \| 147 \| 25 \| 164 \| Uncharacterized protein \| Uncharacterized protein; afdb-uniprot50 \| AF-A0A516SJA0-F1-MODEL\_V4 \| 1.0 \| 0.007831 \| 138 \| 0.205 \| 146 \| 110 \| 4 \| 3 \| 147 \| 24 \| 164 \| Uncharacterized protein \| Uncharacterized protein; afdb-uniprot50 \| AF-A0A2U3QE23-F1-MODEL\_V4 \| 1.0 \| 0.002432 \| 137 \| 0.261 \| 134 \| 87 \| 4 \| 4 \| 133 \| 27 \| 152 \| Uncharacterized protein \| Uncharacterized protein; afdb-uniprot50 \| AF-A0A0B6D5F6-F1-MODEL\_V4 \| 1.0 \| 0.00215 \| 137 \| 0.21 \| 114 \| 86 \| 3 \| 19 \| 130 \| 55 \| 166 \| Uncharacterized protein \| Uncharacterized protein; afdb-uniprot50 \| AF-A0A7R8B6S8-F1-MODEL\_V4 \| 1.0 \| 0.0005904 \| 137 \| 0.312 \| 147 \| 78 \| 5 \| 2 \| 130 \| 30 \| 171 \| Uncharacterized protein \| Uncharacterized protein; afdb-uniprot50 \| AF-A0A7V8TMM0-F1-MODEL\_V4 \| 1.0 \| 0.001681 \| 136 \| 0.231 \| 147 \| 100 \| 3 \| 2 \| 147 \| 23 \| 157 \| Uncharacterized protein \| Uncharacterized protein; afdb-uniprot50 \| AF-A0A2X2Q8M2-F1-MODEL\_V4 \| 1.0 \| 0.007831 \| 136 \| 0.246 \| 146 \| 98 \| 6 \| 4 \| 147 \| 30 \| 165 \| Uncharacterized protein \| Uncharacterized protein; afdb-uniprot50 \| AF-A0A317PTI1-F1-MODEL\_V4 \| 1.0 \| 0.001901 \| 136 \| 0.24 \| 129 \| 89 \| 3 \| 22 \| 147 \| 32 \| 154 \| Uncharacterized protein \| Uncharacterized protein; afdb-uniprot50 \| AF-A0A0H3ZUV8-F1-MODEL\_V4 \| 1.0 \| 0.0045 \| 136 \| 0.266 \| 139 \| 90 \| 2 \| 2 \| 130 \| 23 \| 159 \| Phage DNA packaging \| Phage DNA packaging; afdb-uniprot50 \| AF-A0A843B8J5-F1-MODEL\_V4 \| 1.0 \| 0.005757 \| 136 \| 0.224 \| 125 \| 82 \| 1 \| 6 \| 130 \| 69 \| 178 \| Uncharacterized protein \| Uncharacterized protein; afdb-uniprot50 \| AF-A0A1F9DZH8-F1-MODEL\_V4 \| 1.0 \| 0.0005221 \| 136 \| 0.264 \| 136 \| 87 \| 4 \| 2 \| 130 \| 24 \| 153 \| Uncharacterized protein \| Uncharacterized protein; afdb-uniprot50 \| AF-A0A090UYZ9-F1-MODEL\_V4 \| 1.0 \| 0.001581 \| 135 \| 0.234 \| 132 \| 85 \| 3 \| 2 \| 130 \| 22 \| 140 \| Uncharacterized protein \| Uncharacterized protein; afdb-uniprot50 \| AF-A0A371JBP9-F1-MODEL\_V4 \| 1.0 \| 0.004786 \| 134 \| 0.19 \| 142 \| 103 \| 5 \| 2 \| 139 \| 28 \| 161 \| Protoporphyrinogen oxidase \| Protoporphyrinogen oxidase; afdb-uniprot50 \| AF-A0A2E9M7Y8-F1-MODEL\_V4 \| 1.0 \| 0.005413 \| 133 \| 0.2 \| 120 \| 89 \| 3 \| 11 \| 130 \| 30 \| 142 \| Uncharacterized protein \| Uncharacterized protein; afdb-uniprot50 \| AF-A0A2E1P375-F1-MODEL\_V4 \| 1.0 \| 0.006924 \| 133 \| 0.198 \| 131 \| 96 \| 3 \| 3 \| 129 \| 22 \| 147 \| Uncharacterized protein \| Uncharacterized protein; afdb-uniprot50 \| AF-A0A3P6JQE4-F1-MODEL\_V4 \| 1.0 \| 0.005757 \| 133 \| 0.22 \| 154 \| 107 \| 5 \| 3 \| 146 \| 37 \| 187 \| Phage DNA packaging protein, Nu1 subunit of terminase \| Phage DNA packaging protein, Nu1 subunit of terminase; afdb-uniprot50 \| AF-A0A096AL02-F1-MODEL\_V4 \| 1.0 \| 0.0045 \| 132 \| 0.178 \| 140 \| 101 \| 4 \| 3 \| 139 \| 31 \| 159 \| Uncharacterized protein \| Uncharacterized protein; afdb-uniprot50 \| AF-A0A0E4G066-F1-MODEL\_V4 \| 1.0 \| 0.001681 \| 132 \| 0.209 \| 162 \| 106 \| 5 \| 2 \| 147 \| 21 \| 176 \| Uncharacterized protein \| Uncharacterized protein; afdb-uniprot50 \| AF-A0A554W5S1-F1-MODEL\_V4 \| 1.0 \| 0.002022 \| 131 \| 0.232 \| 146 \| 101 \| 2 \| 3 \| 138 \| 26 \| 170 \| Uncharacterized protein \| Uncharacterized protein; afdb-uniprot50 \| AF-A0A1B1C3B3-F1-MODEL\_V4 \| 1.0 \| 0.0009084 \| 131 \| 0.253 \| 150 \| 91 \| 5 \| 2 \| 144 \| 45 \| 180 \| Uncharacterized protein \| Uncharacterized protein; afdb-uniprot50 \| AF-A0A2A5ADY0-F1-MODEL\_V4 \| 1.0 \| 0.008857 \| 129 \| 0.198 \| 131 \| 95 \| 4 \| 3 \| 129 \| 22 \| 146 \| Uncharacterized protein \| Uncharacterized protein; afdb-uniprot50 \| AF-A0A841I4T7-F1-MODEL\_V4 \| 1.0 \| 0.003518 \| 129 \| 0.228 \| 118 \| 86 \| 2 \| 15 \| 129 \| 36 \| 151 \| Uncharacterized protein \| Uncharacterized protein; afdb-uniprot50 \| AF-A0A6I4RSJ3-F1-MODEL\_V4 \| 1.0 \| 0.003979 \| 129 \| 0.233 \| 124 \| 88 \| 2 \| 12 \| 130 \| 33 \| 154 \| Uncharacterized protein \| Uncharacterized protein; afdb-uniprot50 \| AF-A0A1I0D922-F1-MODEL\_V4 \| 1.0 \| 0.006924 \| 129 \| 0.231 \| 138 \| 90 \| 4 \| 4 \| 139 \| 26 \| 149 \| Phage DNA packaging protein, Nu1 subunit of terminase \| Phage DNA packaging protein, Nu1 subunit of terminase; afdb-uniprot50 \| AF-A0A160U3F2-F1-MODEL\_V4 \| 1.0 \| 0.003308 \| 129 \| 0.18 \| 133 \| 104 \| 1 \| 3 \| 130 \| 53 \| 185 \| Uncharacterized protein \| Uncharacterized protein; afdb-uniprot50 \| AF-X1GDY6-F1-MODEL\_V4 \| 1.0 \| 0.006511 \| 128 \| 0.165 \| 133 \| 100 \| 3 \| 2 \| 131 \| 25 \| 149 \| Uncharacterized protein \| Uncharacterized protein; afdb-uniprot50 \| AF-A0A316DDI3-F1-MODEL\_V4 \| 1.0 \| 0.00509 \| 128 \| 0.2 \| 155 \| 105 \| 7 \| 3 \| 147 \| 29 \| 174 \| Phage terminase Nu1 subunit (DNA packaging protein) \| Phage terminase Nu1 subunit (DNA packaging protein); afdb-uniprot50 \| AF-A0A7S7J7F6-F1-MODEL\_V4 \| 1.0 \| 0.003308 \| 128 \| 0.189 \| 148 \| 110 \| 5 \| 2 \| 147 \| 26 \| 165 \| Uncharacterized protein \| Uncharacterized protein; afdb-uniprot50 \| AF-A4J3T3-F1-MODEL\_V4 \| 1.0 \| 0.004786 \| 128 \| 0.209 \| 148 \| 100 \| 4 \| 4 \| 147 \| 30 \| 164 \| Uncharacterized protein \| Uncharacterized protein; afdb-uniprot50 \| AF-A0A358PNV0-F1-MODEL\_V4 \| 1.0 \| 0.005413 \| 127 \| 0.215 \| 139 \| 98 \| 4 \| 3 \| 131 \| 32 \| 169 \| Protoporphyrinogen oxidase \| Protoporphyrinogen oxidase; afdb-uniprot50 \| AF-E7C7Y8-F1-MODEL\_V4 \| 1.0 \| 0.006122 \| 126 \| 0.289 \| 152 \| 91 \| 5 \| 3 \| 147 \| 25 \| 166 \| Uncharacterized protein \| Uncharacterized protein; afdb-uniprot50 \| AF-A0A0U9HMP1-F1-MODEL\_V4 \| 1.0 \| 0.0045 \| 126 \| 0.262 \| 141 \| 90 \| 5 \| 3 \| 139 \| 33 \| 163 \| Phage DNA packaging protein, Nu1 subunit of terminase \| Phage DNA packaging protein, Nu1 subunit of terminase; afdb-uniprot50 \| AF-A0A553SNI5-F1-MODEL\_V4 \| 1.0 \| 0.003979 \| 126 \| 0.232 \| 159 \| 104 \| 6 \| 2 \| 147 \| 51 \| 204 \| Uncharacterized protein \| Uncharacterized protein; afdb-uniprot50 \| AF-A0A2S4QSD8-F1-MODEL\_V4 \| 1.0 \| 0.006122 \| 125 \| 0.185 \| 135 \| 105 \| 2 \| 14 \| 147 \| 31 \| 161 \| Uncharacterized protein \| Uncharacterized protein; afdb-uniprot50 \| AF-A0A382N6L8-F1-MODEL\_V4 \| 1.0 \| 0.007831 \| 124 \| 0.191 \| 136 \| 94 \| 3 \| 20 \| 147 \| 34 \| 161 \| Uncharacterized protein \| Uncharacterized protein; afdb-uniprot50 \| AF-A0A8B0KWW7-F1-MODEL\_V4 \| 1.0 \| 0.007364 \| 124 \| 0.221 \| 149 \| 100 \| 5 \| 2 \| 147 \| 21 \| 156 \| Uncharacterized protein \| Uncharacterized protein; afdb-uniprot50 \| AF-A0A2N8HQG2-F1-MODEL\_V4 \| 1.0 \| 0.008328 \| 124 \| 0.207 \| 140 \| 101 \| 4 \| 2 \| 139 \| 42 \| 173 \| Uncharacterized protein \| Uncharacterized protein; afdb-uniprot50 \| AF-G1WGD7-F1-MODEL\_V4 \| 1.0 \| 0.008857 \| 123 \| 0.209 \| 143 \| 99 \| 5 \| 3 \| 139 \| 25 \| 159 \| Uncharacterized protein \| Uncharacterized protein; afdb-uniprot50 \| AF-A0A7I8DI47-F1-MODEL\_V4 \| 1.0 \| 0.009419 \| 123 \| 0.212 \| 155 \| 109 \| 7 \| 3 \| 147 \| 35 \| 186 \| Uncharacterized protein \| Uncharacterized protein; afdb-uniprot50 \| AF-Q89JP3-F1-MODEL\_V4 \| 1.0 \| 0.003979 \| 123 \| 0.208 \| 144 \| 97 \| 3 \| 20 \| 147 \| 56 \| 198 \| Bll5240 protein \| Bll5240 protein; afdb-uniprot50 \| AF-A0A7T5R3N3-F1-MODEL\_V4 \| 1.0 \| 0.008328 \| 120 \| 0.173 \| 121 \| 81 \| 2 \| 10 \| 130 \| 70 \| 171 \| Uncharacterized protein \| Uncharacterized protein; afdb-uniprot50 \| AF-A0A839W5R0-F1-MODEL\_V4 \| 1.0 \| 0.003742 \| 120 \| 0.202 \| 143 \| 101 \| 6 \| 2 \| 138 \| 25 \| 160 \| Phage terminase Nu1 subunit (DNA packaging protein) \| Phage terminase Nu1 subunit (DNA packaging protein); afdb-uniprot50 \| AF-A0A1E7WZ77-F1-MODEL\_V4 \| 1.0 \| 0.006924 \| 118 \| 0.221 \| 122 \| 79 \| 2 \| 11 \| 130 \| 77 \| 184 \| Uncharacterized protein \| Uncharacterized protein; afdb-uniprot50 \| AF-A0A1C6BMG4-F1-MODEL\_V4 \| 1.0 \| 0.00509 \| 117 \| 0.164 \| 158 \| 110 \| 6 \| 2 \| 147 \| 22 \| 169 \| Uncharacterized protein \| Uncharacterized protein; afdb-uniprot50 \| AF-D7XY77-F1-MODEL\_V4 \| 1.0 \| 0.007364 \| 117 \| 0.181 \| 154 \| 112 \| 6 \| 2 \| 147 \| 26 \| 173 \| Uncharacterized protein \| Uncharacterized protein; afdb-uniprot50 \| AF-A0A4D7QKX8-F1-MODEL\_V4 \| 1.0 \| 0.006924 \| 117 \| 0.207 \| 130 \| 89 \| 3 \| 3 \| 132 \| 82 \| 197 \| Uncharacterized protein \| Uncharacterized protein; afdb-uniprot50 \| AF-A0A554WU06-F1-MODEL\_V4 \| 1.0 \| 0.006924 \| 115 \| 0.226 \| 137 \| 84 \| 1 \| 2 \| 138 \| 16 \| 130 \| Uncharacterized protein \| Uncharacterized protein; afdb-uniprot50 \| AF-A0A7L6ENJ2-F1-MODEL\_V4 \| 1.0 \| 0.004232 \| 115 \| 0.207 \| 159 \| 100 \| 5 \| 2 \| 147 \| 22 \| 167 \| Uncharacterized protein \| Uncharacterized protein; afdb-uniprot50 \| AF-A0A413EDA4-F1-MODEL\_V4 \| 1.0 \| 0.008328 \| 115 \| 0.191 \| 157 \| 100 \| 6 \| 3 \| 147 \| 31 \| 172 \| Uncharacterized protein \| Uncharacterized protein; afdb-uniprot50 \| AF-A0A4Q8WZ92-F1-MODEL\_V4 \| 1.0 \| 0.007364 \| 115 \| 0.216 \| 143 \| 94 \| 5 \| 21 \| 147 \| 40 \| 180 \| Uncharacterized protein \| Uncharacterized protein; afdb-uniprot50 \| AF-A0A496KCG1-F1-MODEL\_V4 \| 1.0 \| 0.006924 \| 114 \| 0.225 \| 151 \| 102 \| 4 \| 2 \| 147 \| 22 \| 162 \| DNA packaging protein \| DNA packaging protein; afdb-uniprot50 \| AF-A0A2D5TTA5-F1-MODEL\_V4 \| 1.0 \| 0.001788 \| 113 \| 0.189 \| 164 \| 105 \| 4 \| 2 \| 147 \| 25 \| 178 \| Uncharacterized protein \| Uncharacterized protein; afdb-uniprot50 \| AF-A0A081MYL8-F1-MODEL\_V4 \| 1.0 \| 0.004786 \| 113 \| 0.171 \| 163 \| 118 \| 4 \| 2 \| 147 \| 24 \| 186 \| Uncharacterized protein \| Uncharacterized protein; afdb-uniprot50 \| AF-A0A844Q904-F1-MODEL\_V4 \| 1.0 \| 0.008328 \| 113 \| 0.205 \| 151 \| 91 \| 5 \| 2 \| 139 \| 33 \| 167 \| Uncharacterized protein \| Uncharacterized protein; afdb-uniprot50 \| AF-A0A3D4L0I6-F1-MODEL\_V4 \| 1.0 \| 0.007364 \| 113 \| 0.216 \| 148 \| 98 \| 6 \| 2 \| 139 \| 171 \| 310 \| Uncharacterized protein \| Uncharacterized protein; afdb-uniprot50 \| AF-A0A7U0PNM6-F1-MODEL\_V4 \| 1.0 \| 0.0045 \| 111 \| 0.198 \| 151 \| 106 \| 4 \| 2 \| 147 \| 25 \| 165 \| Uncharacterized protein \| Uncharacterized protein; afdb-uniprot50 \| AF-A0A4Y8EEM3-F1-MODEL\_V4 \| 1.0 \| 0.008328 \| 111 \| 0.189 \| 148 \| 111 \| 5 \| 2 \| 147 \| 12 \| 152 \| Uncharacterized protein \| Uncharacterized protein |
| Top keywords  (threshold 1.00e-02 (evalue)) | **Terminase, Nu1, small, DNA, packaging, Phage, DNA\_packaging, Protoporphyrinogen, oxidase, homolog** |
| Output files | ../../similar\_structures/01\_FANPEZAQ\_CDS\_0001\_afdb-proteome\_foldseek.tsv ../../similar\_structures/01\_FANPEZAQ\_CDS\_0001\_afdb-uniprot50\_foldseek.tsv ../../similar\_structures/01\_FANPEZAQ\_CDS\_0001\_merged.svg ../../similar\_structures/01\_FANPEZAQ\_CDS\_0001\_pdb\_foldseek.tsv |

  
  
  

Return to summary | Go to next

  


---

**Sequence/structure alignments coloring**  
Each object in the alignment figures is colored according to its E-value following this color coding:

1e-100
10
