## Supplementary material for "Completing the BASEL phage collection to unlock hidden diversity for systematic exploration of phage-host interactions": Data S2: 2.html

FANPEZAQ\_CDS\_0002


Return to summary | Go to previous | Go to next

|  |  |
| --- | --- |
| FANPEZAQ\_CDS\_0002  Page creation date: 02 Sep 2024, 12:00  Project folder: n/a  Input sequences file: Escherichia\_virus\_HeidiAbel.gb | terminase large phage helicase gpa rna dna fragment a atp\_dependent domain\_containing atpase in c complex e and yes adp tail i hydrolase engineered the dead assembly fold escherichia coli beta box bacteriophage alpha containing gpa\_like organism\_taxid nucleotide dead\_box expressed arch 3\_layer aba sandwich rossmann gene expression\_system\_taxid triphosphate p\_loop hydrolases factor |

### Sequence information

|  |  |
| --- | --- |
| Name | FANPEZAQ\_CDS\_0002  02\_FANPEZAQ\_CDS\_0002 (pipeline id) |
| Imported annotations | Escherichia\_virus\_HeidiAbel Bas97 |
| Protein sequence | MAMTKGATVTAKNNPFANIEGIDQAIRRAQQQLLPPPDMKPSEWAQANIRIPSGNAVPGP LRLDNAPYQREPMDMLVDPDCYRVTLMWGAQVGKTMLALCVQAYSVAAEPRSQMMMQPSQ DDLRTWLETKFNPLAEECAPVNNAIAKPRGRDGVNNQKMKSYPGGFLMFAWSGSPKTMRG RSAPLIVCDEVDGYERTSEGHPVGLLWQRSATFGDQRFLLEISTPTLKGESYIEKAYEMG DQRHFFVCCPHCDDHVTLEWENIKWDGQRETDAETLAAIEEQKPETAHLYCPSCGAQWND GERIAAIRQAEGKGAGWKASRPFEGHASYHLNELYSTFRKVPAIVRDYLDKLKTDDLQTF TNVSLARTWEEEGDKVDPDSLMARREEYAAPVPAGGLYITAGIDMQVDRLECEVVAWGVG EESWSLGYHVLWGDPLTPDPWDALDDLLAETYQHETGAIMPIMAACLDTGGTSGYTQAAY EYARGRTGRRLFAIKGVGGWGRPIVEKPQRKQSGKDARKVDLFLVGTDEAKLITARRLNK TAPGPGYCHFPADRDEEWFKQITAEKLTMRYVKGQPIREWRKPDKARNEALDCRNYALAA LKIMQPSFRRLAERFGVANDNAPPERRKRLPNVKPIVKPVEVPQPANDKPQEEEKQQVTE TKPIKRSARASKRRGGGWVSRW |
| Number of residues | 682 |
| Molecular weight (Da) | 76687.12 |
| Output files | ../../query\_sequences/02\_FANPEZAQ\_CDS\_0002.fasta |

### Putative domain architecture and protein family

#### Search results (HHblits)^1^

|  |  |
| --- | --- |
| Domain family databases searched | Pfam, Ncbi-cd, Cath, Phrogs |
| Results, scheme(s)  (Top layers only; threshold 1.00e-03 (evalue)) | xml version="1.0" encoding="utf-8" standalone="no"?       2024-09-02T21:08:11.971900 image/svg+xml   Matplotlib v3.7.2, https://matplotlib.org/ |
| Results, table  (E-value ≤ 1.00e-03 (evalue)) | db \| id \| prob \| evalue \| pvalue \| score \| cols \| query \| query\_len \| template \| template\_len \| name \| description; pfam \| PF20454 \| 100.0 \| 1.4e-41 \| 2.6e-45 \| 349.3 \| 274 \| (326, 605) \| 682 \| (1, 297) \| 306 \| GpA\_nuclease \| Terminase large subunit gpA, endonuclease domain; pfam \| PF05876 \| 99.7 \| 1.8e-23 \| 3.4e-27 \| 203.7 \| 240 \| (56, 309) \| 682 \| (2, 244) \| 247 \| GpA\_ATPase \| Phage terminase large subunit gpA, ATPase domain; pfam \| PF03354 \| 97.8 \| 3.5e-09 \| 6.7e-13 \| 92.7 \| 162 \| (67, 240) \| 682 \| (1, 173) \| 175 \| TerL\_ATPase \| Terminase large subunit, ATPase domain; pfam \| PF04466 \| 97.0 \| 4.1e-07 \| 7.9e-11 \| 81.0 \| 146 \| (82, 240) \| 682 \| (2, 154) \| 202 \| Terminase\_3 \| Phage terminase large subunit; pfam \| PF02562 \| 96.9 \| 7.9e-07 \| 1.5e-10 \| 77.5 \| 153 \| (64, 228) \| 682 \| (4, 158) \| 205 \| PhoH \| PhoH-like protein; pfam \| PF04851 \| 96.5 \| 4.8e-06 \| 9.4e-10 \| 67.3 \| 62 \| (63, 125) \| 682 \| (2, 66) \| 151 \| ResIII \| Type III restriction enzyme, res subunit; pfam \| PF03237 \| 96.0 \| 2.3e-05 \| 4.3e-09 \| 70.5 \| 142 \| (85, 240) \| 682 \| (1, 157) \| 234 \| Terminase\_6N \| Terminase large subunit, T4likevirus-type, N-terminal; pfam \| PF05127 \| 95.3 \| 0.00012 \| 2.3e-08 \| 60.1 \| 117 \| (85, 228) \| 682 \| (1, 118) \| 168 \| Helicase\_RecD \| Helicase; pfam \| PF13245 \| 93.7 \| 0.00094 \| 1.8e-07 \| 51.6 \| 36 \| (69, 106) \| 682 \| (1, 36) \| 141 \| AAA\_19 \| AAA domain; ncbi-cd \| cd18021 \| 96.7 \| 1.1e-06 \| 3.7e-10 \| 63.0 \| 175 \| (65, 248) \| 682 \| (4, 191) \| 191 \| DEXHc\_Brr2\_2 \| cd18021 DEXHc\_Brr2\_2; C-terminal D[D/E]X[H/Q]-box helicase domain of spliceosomal Brr2 RNA helicase. Brr2 is a type II DEAD box helicase that mediates spliceosome catalytic activation.; ncbi-cd \| cd17974 \| 95.9 \| 1.9e-05 \| 6.5e-09 \| 52.4 \| 60 \| (64, 125) \| 682 \| (2, 62) \| 174 \| DEXHc\_DHX16 \| cd17974 DEXHc\_DHX16; DEXH-box helicase domain of DEAH-box helicase 16. DEAH-box helicase 16 (DHX16) is probably involved in pre-mRNA splicing.; ncbi-cd \| cd18026 \| 95.8 \| 2.1e-05 \| 6.7e-09 \| 55.3 \| 65 \| (64, 128) \| 682 \| (16, 80) \| 202 \| DEXHc\_POLQ-like \| cd18026 DEXHc\_POLQ-like; DEXH-box helicase domain of DNA polymerase theta. DNA polymerase theta (POLQ) is important in the repair of genomic double-strand breaks (DSBs).; ncbi-cd \| cd18028 \| 95.8 \| 2.1e-05 \| 6.7e-09 \| 54.2 \| 62 \| (65, 128) \| 682 \| (2, 63) \| 177 \| DEXHc\_archSki2 \| cd18028 DEXHc\_archSki2; DEXH-box helicase domain of archaeal Ski2-type helicase. Archaeal Ski2-type RNA helicases play an important role in RNA degradation, processing and splicing pathways.; ncbi-cd \| cd17940 \| 95.7 \| 2.5e-05 \| 8.5e-09 \| 53.2 \| 60 \| (64, 125) \| 682 \| (21, 82) \| 201 \| DEADc\_DDX6 \| cd17940 DEADc\_DDX6; DEAD-box helicase domain of DEAD box protein 6. DEAD box protein 6 (DDX6, also known as Rck or p54) participates in mRNA regulation mediated by miRNA-mediated silencing.; ncbi-cd \| cd17927 \| 95.7 \| 2.8e-05 \| 9.1e-09 \| 55.5 \| 60 \| (64, 125) \| 682 \| (2, 65) \| 201 \| DEXHc\_RIG-I \| cd17927 DEXHc\_RIG-I; DEXH-box helicase domain of DEAD-like helicase RIG-I family proteins. Members of the RIG-I family include FANCM, dicer, Hef, and the RIG-I-like receptors.; ncbi-cd \| cd18016 \| 95.6 \| 3.5e-05 \| 1.1e-08 \| 55.3 \| 60 \| (64, 129) \| 682 \| (17, 76) \| 208 \| DEXHc\_RecQ2\_BLM \| cd18016 DEXHc\_RecQ2\_BLM; DEAH-box helicase domain of RecQ2. ATP-dependent DNA helicase Q2 (RecQ2, also called Bloom syndrome protein homolog or BLM) is part of the RecQ family of highly conserved DNA repair helicases that is part of the type II DEAD box helicase superfamily, a diverse family of proteins involved in ATP-dependent RNA or DNA unwinding.; ncbi-cd \| cd18011 \| 95.6 \| 3.6e-05 \| 1.2e-08 \| 55.4 \| 63 \| (66, 129) \| 682 \| (2, 65) \| 207 \| DEXDc\_RapA \| cd18011 DEXDc\_RapA; DEXH-box helicase domain of RapA. In bacteria, RapA is an RNA polymerase (RNAP)-associated SWI2/SNF2 (switch/sucrose non-fermentable) protein that mediates RNAP recycling during transcription.; ncbi-cd \| cd18007 \| 95.4 \| 5.5e-05 \| 1.7e-08 \| 57.8 \| 64 \| (65, 129) \| 682 \| (1, 75) \| 239 \| DEXHc\_ATRX-like \| cd18007 DEXHc\_ATRX-like; DEXH-box helicase domain of ATRX-like proteins. This family includes ATRX-like members such as transcriptional regulator ATRX (also called alpha thalassemia/mental retardation syndrome X-linked and X-linked nuclear protein or XNP) which is involved in transcriptional regulation and chromatin remodeling, and ARIP4 (also called androgen receptor-interacting protein 4, RAD54 like 2 or RAD54L2) which modulates androgen receptor (AR)-dependent transactivation in a promoter-dependent manner.; ncbi-cd \| cd18034 \| 95.4 \| 5.4e-05 \| 1.7e-08 \| 54.9 \| 62 \| (64, 128) \| 682 \| (2, 69) \| 200 \| DEXHc\_dicer \| cd18034 DEXHc\_dicer; DEXH-box helicase domain of endoribonuclease Dicer. Dicer ribonucleases cleave double-stranded RNA (dsRNA) precursors to generate microRNAs (miRNAs) and small interfering RNAs (siRNAs).; ncbi-cd \| cd17921 \| 95.3 \| 6.3e-05 \| 2.1e-08 \| 50.9 \| 62 \| (65, 128) \| 682 \| (2, 64) \| 181 \| DEXHc\_Ski2 \| cd17921 DEXHc\_Ski2; DEXH-box helicase domain of DEAD-like helicase Ski2 family proteins. Ski2-like RNA helicases play an important role in RNA degradation, processing, and splicing pathways.; ncbi-cd \| cd17981 \| 95.3 \| 6.6e-05 \| 2.2e-08 \| 51.2 \| 64 \| (64, 129) \| 682 \| (2, 69) \| 180 \| DEXHc\_DHX36 \| cd17981 DEXHc\_DHX36; DEXH-box helicase domain of DEAH-box helicase 36. DEAH-box helicase 36 (DHX36, also known as G4-resolvase 1 or G4R1, MLE-like protein 1 and RNA helicase associated with AU-rich element or RHAU) unwinds a G4-quadruplex in human telomerase RNA.; ncbi-cd \| cd17998 \| 95.3 \| 7.1e-05 \| 2.3e-08 \| 52.9 \| 64 \| (65, 129) \| 682 \| (1, 67) \| 187 \| DEXHc\_SMARCAD1 \| cd17998 DEXHc\_SMARCAD1; DEXH-box helicase domain of SMARCAD1. SWI/SNF-related matrix-associated actin-dependent regulator of chromatin subfamily A containing DEAD/H box 1 (SMARCAD1, also known as ATP-dependent helicase 1 or Hel1) possesses intrinsic ATP-dependent nucleosome-remodeling activity and is required for both DNA repair and heterochromatin organization.; ncbi-cd \| cd17984 \| 95.3 \| 6.9e-05 \| 2.3e-08 \| 50.8 \| 57 \| (64, 122) \| 682 \| (2, 58) \| 178 \| DEXHc\_DHX40 \| cd17984 DEXHc\_DHX40; DEXH-box helicase domain of DEAH-box helicase 40. DEAH-box helicase 40 (DHX40) belongs to the DEAD-like helicase superfamily, a diverse family of proteins involved in ATP-dependent RNA or DNA unwinding.; ncbi-cd \| cd18022 \| 95.3 \| 6.9e-05 \| 2.3e-08 \| 51.1 \| 64 \| (65, 129) \| 682 \| (2, 66) \| 189 \| DEXHc\_ASCC3\_2 \| cd18022 DEXHc\_ASCC3\_2; C-terminal DEXH-box helicase domain of Activating signal cointegrator 1 complex subunit 3.; ncbi-cd \| cd09710 \| 95.3 \| 7.4e-05 \| 2.4e-08 \| 56.5 \| 58 \| (68, 129) \| 682 \| (1, 58) \| 353 \| Cas3\_I-D \| cd09710 Cas3\_I-D; CRISPR/Cas system-associated protein Cas3; Distinct diverged subfamily of Cas3 helicase domain.; ncbi-cd \| cd17988 \| 95.2 \| 7.7e-05 \| 2.5e-08 \| 50.7 \| 59 \| (65, 125) \| 682 \| (3, 63) \| 180 \| DEXHc\_TDRD9 \| cd17988 DEXHc\_TDRD9; DEXH-box helicase domain of tudor domain containing 9. Tudor domain containing 9 (TDRD9, also known as HIG-1or NET54 or C14orf75) is a part of the nuclear PIWI-interacting RNA (piRNA) pathway essential for transposon silencing and male fertility TDRD9 belongs to the DEAD-like helicase superfamily, a diverse family of proteins involved in ATP-dependent RNA or DNA unwinding.; ncbi-cd \| cd17989 \| 95.2 \| 8.1e-05 \| 2.7e-08 \| 49.7 \| 55 \| (65, 121) \| 682 \| (3, 57) \| 173 \| DEXHc\_HrpA \| cd17989 DEXHc\_HrpA; DEXH-box helicase domain of ATP-dependent RNA helicase HrpA. HrpA is part of the HrpB-HrpA two-partner secretion (TPS) system, a secretion pathway important to the secretion of large virulence-associated proteins.; ncbi-cd \| cd17987 \| 95.1 \| 9.9e-05 \| 3.2e-08 \| 50.4 \| 60 \| (64, 125) \| 682 \| (2, 63) \| 176 \| DEXHc\_YTHDC2 \| cd17987 DEXHc\_YTHDC2; DEXH-box helicase domain of YTH domain containing 2. YTH domain containing 2 (YTHDC2) regulates mRNA translation and stability via binding to N6-methyladenosine, a modified RNA nucleotide enriched in the stop codons and 3' UTRs of eukaryotic messenger RNAs.; ncbi-cd \| cd18015 \| 95.1 \| 0.0001 \| 3.4e-08 \| 52.2 \| 57 \| (64, 126) \| 682 \| (18, 74) \| 209 \| DEXHc\_RecQ1 \| cd18015 DEXHc\_RecQ1; DEXH-box helicase domain of RecQ1. ATP-dependent DNA helicase Q1 (RecQ1) is part of the RecQ family of highly conserved DNA repair helicases that is part of the type II DEAD box helicase superfamily, a diverse family of proteins involved in ATP-dependent RNA or DNA unwinding.; ncbi-cd \| cd17923 \| 95.1 \| 0.00011 \| 3.5e-08 \| 50.5 \| 60 \| (66, 127) \| 682 \| (2, 62) \| 182 \| DEXHc\_Hrq1-like \| cd17923 DEXHc\_Hrq1-like; DEAH-box helicase domain of Hrq1 and similar proteins. Yeast Hrq1, similar to RecQ4, plays a role in DNA inter-strand crosslink (ICL) repair and in telomere maintenance.; ncbi-cd \| cd18025 \| 94.8 \| 0.00015 \| 5.1e-08 \| 48.9 \| 61 \| (65, 127) \| 682 \| (2, 63) \| 192 \| DEXHc\_DDX60 \| cd18025 DEXHc\_DDX60; DEXH-box helicase domain of DEAD box protein 60. DEAD box protein 60 (DDX60) is an IFN-inducible cytoplasmic helicase that plays a role in RIG-I-mediated type I interferon (IFN) nuclease-mediated viral RNA degradation.; ncbi-cd \| cd18075 \| 94.8 \| 0.00016 \| 5.1e-08 \| 52.1 \| 65 \| (64, 130) \| 682 \| (2, 67) \| 200 \| DEXHc\_RLR-3 \| cd18075 DEXHc\_RLR-3; DEXH-box helicase domain of RLR-3. RIG-I-like receptor 3 (RLR-3, also known as laboratory of genetics and physiology 2 or LGP2 and DHX58) appears to positively and negatively regulate MDA5 and RIG-I signaling, respectively.; ncbi-cd \| cd17995 \| 94.7 \| 0.00018 \| 5.6e-08 \| 52.8 \| 64 \| (65, 129) \| 682 \| (1, 68) \| 223 \| DEXHc\_CHD6\_7\_8\_9 \| cd17995 DEXHc\_CHD6\_7\_8\_9; DEXH-box helicase domain of the chromodomain helicase DNA binding protein 6, 7, 8 and 9.; ncbi-cd \| cd18032 \| 94.7 \| 0.00018 \| 6e-08 \| 48.3 \| 61 \| (66, 126) \| 682 \| (2, 66) \| 163 \| DEXHc\_RE\_I\_III\_res \| cd18032 DEXHc\_RE\_I\_III\_res; DEXH-box helicase domain of type III restriction enzyme res subunit. Members of this cd includes both type I and type III restriction enzymes.; ncbi-cd \| cd17991 \| 94.7 \| 0.00019 \| 6.1e-08 \| 49.8 \| 66 \| (62, 128) \| 682 \| (13, 82) \| 193 \| DEXHc\_TRCF \| cd17991 DEXHc\_TRCF; DEXH/Q-box helicase domain of the transcription-repair coupling factor. Transcription-repair coupling factor (TrcF) dissociates transcription elongation complexes blocked at nonpairing lesions and mediates recruitment of DNA repair proteins.; ncbi-cd \| cd18006 \| 94.7 \| 0.00019 \| 6.1e-08 \| 51.4 \| 63 \| (66, 129) \| 682 \| (2, 68) \| 216 \| DEXHc\_CHD1L \| cd18006 DEXHc\_CHD1L; DEAH/Q-box helicase domain of CHD1L. Chromodomain helicase DNA binding protein 1 like (CHD1L, also known as ALC1) is involved in DNA repair by regulating chromatin relaxation following DNA damage.; ncbi-cd \| cd18033 \| 94.7 \| 0.00019 \| 6.2e-08 \| 48.2 \| 60 \| (64, 126) \| 682 \| (2, 62) \| 182 \| DEXDc\_FANCM \| cd18033 DEXDc\_FANCM; DEAH-box helicase domain of FANCM. Fanconi anemia group M (FANCM) protein is a DNA-dependent ATPase component of the Fanconi anemia (FA) core complex.; ncbi-cd \| cd18014 \| 94.7 \| 0.0002 \| 6.4e-08 \| 51.1 \| 58 \| (66, 128) \| 682 \| (15, 72) \| 205 \| DEXHc\_RecQ5 \| cd18014 DEXHc\_RecQ5; DEAH-box helicase domain of RecQ5. ATP-dependent DNA helicase Q5 (RecQ5) is part of the RecQ family of highly conserved DNA repair helicases that is part of the type II DEAD box helicase superfamily, a diverse family of proteins involved in ATP-dependent RNA or DNA unwinding.; ncbi-cd \| cd17924 \| 94.6 \| 0.0002 \| 6.4e-08 \| 49.7 \| 62 \| (63, 127) \| 682 \| (16, 77) \| 189 \| DDXDc\_reverse\_gyrase \| cd17924 DDXDc\_reverse\_gyrase; DDXD-box helicase domain of reverse gyrase. Reverse gyrase modifies the topological state of DNA by introducing positive supercoils in an ATP-dependent process.; ncbi-cd \| cd18035 \| 94.6 \| 0.0002 \| 6.5e-08 \| 49.6 \| 59 \| (67, 127) \| 682 \| (4, 62) \| 181 \| DEXHc\_Hef \| cd18035 DEXHc\_Hef; DEXH-box helicase domain of Hef. Hef (helicase-associated endonuclease fork-structure) belongs to the XPF/MUS81/FANCM family of endonucleases and is involved in stalled replication fork repair.; ncbi-cd \| cd18031 \| 94.6 \| 0.0002 \| 6.6e-08 \| 47.5 \| 61 \| (66, 128) \| 682 \| (2, 62) \| 161 \| DEXHc\_UvsW \| cd18031 DEXHc\_UvsW; DEXH-box helicase domain of bacteriophage UvsW. Bacteriophage UvsW is part of the WXY system that repairs DNA damage by a process that involves homologous recombination.; ncbi-cd \| cd18076 \| 94.6 \| 0.0002 \| 6.7e-08 \| 50.4 \| 68 \| (65, 132) \| 682 \| (2, 74) \| 230 \| DEXXQc\_HELZ2-N \| cd18076 DEXXQc\_HELZ2-N; N-terminal DEXXQ-box helicase domain of HELZ2. Helicase with zinc finger 2 (HELZ2, also known as PPAR-alpha-interacting complex protein 285 or PRIC285 and PPAR-gamma DBD-interacting protein 1 or PDIP1) acts as a transcriptional coactivator for a number of nuclear receptors including PPARA, PPARG, THRA, THRB, and RXRA.; ncbi-cd \| cd17978 \| 94.5 \| 0.00023 \| 7.8e-08 \| 46.1 \| 53 \| (65, 119) \| 682 \| (3, 56) \| 179 \| DEXHc\_DHX33 \| cd17978 DEXHc\_DHX33; DEXH-box helicase domain of DEAH-box helicase 33. DEAH-box helicase 33 (DHX33) stimulates RNA polymerase I transcription of the 47S precursor rRNA.; ncbi-cd \| cd18066 \| 94.5 \| 0.00027 \| 8e-08 \| 54.0 \| 64 \| (65, 129) \| 682 \| (1, 78) \| 235 \| DEXHc\_RAD54B \| cd18066 DEXHc\_RAD54B; DEXH-box helicase domain of RAD54B. DNA repair and recombination protein RAD54B, also known as RDH54, binds to double-stranded DNA, displays ATPase activity in the presence of DNA, and may have a role in meiotic and mitotic recombination.; ncbi-cd \| cd18073 \| 94.4 \| 0.00029 \| 9.3e-08 \| 50.0 \| 62 \| (64, 127) \| 682 \| (2, 67) \| 202 \| DEXHc\_RIG-I\_DDX58 \| cd18073 DEXHc\_RIG-I\_DDX58; DEXH-box helicase domain of RIG-I. RIG-I (Retinoic acid-inducible gene I protein), also called DEAD box protein 58 (DDX58), is a pathogen-recognition receptor that recognizes viral 5'-triphosphates carrying double-stranded RNA.; ncbi-cd \| cd17941 \| 94.3 \| 0.00029 \| 9.8e-08 \| 47.5 \| 62 \| (64, 127) \| 682 \| (12, 79) \| 198 \| DEADc\_DDX10 \| cd17941 DEADc\_DDX10; DEAD-box helicase domain of DEAD box protein 10. Fusion of the DDX10 gene and the nucleoporin gene, NUP98, by inversion 11 (p15q22) chromosome translocation is found in the patients with de novo or therapy-related myeloid malignancies.; ncbi-cd \| cd17950 \| 94.3 \| 0.0003 \| 9.9e-08 \| 48.6 \| 62 \| (64, 127) \| 682 \| (24, 87) \| 208 \| DEADc\_DDX39 \| cd17950 DEADc\_DDX39; DEAD-box helicase domain of DEAD box protein 39. DDX39A is involved in pre-mRNA splicing and is required for the export of mRNA out of the nucleus.; ncbi-cd \| cd18036 \| 94.2 \| 0.00034 \| 1.1e-07 \| 48.8 \| 60 \| (64, 125) \| 682 \| (2, 66) \| 204 \| DEXHc\_RLR \| cd18036 DEXHc\_RLR; DEXH-box helicase domain of RIG-I-like receptors. RIG-I-like receptors (RLRs) sense cytoplasmic viral RNA and comprise RIG-I, RLR-2/MDA5 (melanoma differentiation-associated protein 5) and RLR-3/LGP2 (laboratory of genetics and physiology 2).; ncbi-cd \| cd18013 \| 94.2 \| 0.00039 \| 1.1e-07 \| 53.8 \| 61 \| (65, 127) \| 682 \| (1, 62) \| 218 \| DEXQc\_bact\_SNF2 \| cd18013 DEXQc\_bact\_SNF2; DEXQ-box helicase domain of bacterial SNF2 family proteins. Proteins belonging to the SNF2 family of DNA dependent ATPases are important members of the chromatin remodeling complexes that are implicated in epigenetic control of gene expression.; ncbi-cd \| cd17983 \| 94.2 \| 0.00035 \| 1.2e-07 \| 45.8 \| 53 \| (66, 120) \| 682 \| (4, 56) \| 173 \| DEXHc\_DHX38 \| cd17983 DEXHc\_DHX38; DEXH-box helicase domain of DEAH-box helicase 38. DEAH-box helicase 38 (DHX38, also known as PRP16) is involved in pre-mRNA splicing.; ncbi-cd \| cd18027 \| 94.2 \| 0.00036 \| 1.2e-07 \| 47.2 \| 60 \| (63, 125) \| 682 \| (7, 66) \| 179 \| DEXHc\_SKIV2L \| cd18027 DEXHc\_SKIV2L; DEXH-box helicase domain of SKIV2L. Superkiller viralicidic activity 2-like (SKIV2L, also called SKI2 or DHX13) plays a role in a number of cellular processes involving alteration of RNA secondary structure such as translation initiation, nuclear and mitochondrial splicing, and ribosome and spliceosome assembly.; ncbi-cd \| cd18038 \| 94.2 \| 0.00037 \| 1.2e-07 \| 50.4 \| 65 \| (65, 129) \| 682 \| (2, 69) \| 229 \| DEXXQc\_Helz-like \| cd18038 DEXXQc\_Helz-like; DEXXQ/H-box helicase domain of Helz-like helicase. This subfamily contains HELZ, Mov10L1, and similar proteins.; ncbi-cd \| cd17975 \| 94.1 \| 0.00039 \| 1.3e-07 \| 46.5 \| 61 \| (64, 126) \| 682 \| (2, 67) \| 183 \| DEXHc\_DHX29 \| cd17975 DEXHc\_DHX29; DEXH-box helicase domain of DEAH-box helicase 29. DEAH-box helicase 29 (DHX29) is a part of the 43S pre-initiation complex involved in translation initiation of mRNAs with structured 5'-UTRs.; ncbi-cd \| cd17955 \| 94.0 \| 0.00043 \| 1.4e-07 \| 46.9 \| 62 \| (64, 127) \| 682 \| (21, 84) \| 204 \| DEADc\_DDX49 \| cd17955 DEADc\_DDX49; DEAD-box helicase domain of DEAD box protein 49. DDX49 (also called Dbp8) is a member of the DEAD-box helicases, a diverse family of proteins involved in ATP-dependent RNA unwinding, needed in a variety of cellular processes including splicing, ribosome biogenesis and RNA degradation.; ncbi-cd \| cd17999 \| 94.0 \| 0.00047 \| 1.5e-07 \| 50.0 \| 64 \| (65, 129) \| 682 \| (1, 73) \| 232 \| DEXHc\_Mot1 \| cd17999 DEXHc\_Mot1; DEXH-box helicase domain of Mot1. Modifier of transcription 1 (Mot1, also known as TAF172 in eukaryotes) regulates transcription in association with TATA binding protein (TBP).; ncbi-cd \| cd17938 \| 93.9 \| 0.00045 \| 1.5e-07 \| 47.4 \| 55 \| (64, 125) \| 682 \| (21, 75) \| 204 \| DEADc\_DDX1 \| cd17938 DEADc\_DDX1; DEAD-box helicase domain of DEAD box protein 1. DEAD box protein 1 (DDX1) acts as an ATP-dependent RNA helicase, able to unwind both RNA-RNA and RNA-DNA duplexes.; ncbi-cd \| cd17977 \| 93.9 \| 0.00045 \| 1.5e-07 \| 45.6 \| 60 \| (64, 125) \| 682 \| (2, 64) \| 176 \| DEXHc\_DHX32 \| cd17977 DEXHc\_DHX32; DEXH-box helicase domain of DEAH-box helicase 32. DEAH-box helicase 32 (DHX32) belongs to the DEAD-like helicase superfamily, a diverse family of proteins involved in ATP-dependent RNA or DNA unwinding.; ncbi-cd \| cd17992 \| 93.9 \| 0.00051 \| 1.6e-07 \| 50.4 \| 71 \| (56, 127) \| 682 \| (37, 111) \| 225 \| DEXHc\_RecG \| cd17992 DEXHc\_RecG; DEXH/Q-box helicase domain of RecG. ATP-dependent DNA helicase RecG plays a critical role in recombination and DNA repair.; ncbi-cd \| cd18074 \| 93.8 \| 0.00052 \| 1.7e-07 \| 49.3 \| 67 \| (64, 132) \| 682 \| (2, 73) \| 216 \| DEXHc\_RLR-2 \| cd18074 DEXHc\_RLR-2; DEXH-box helicase domain of RLR-2. RIG-I-like receptor 2 (RLR-2, also known as melanoma differentiation-associated protein 5 or Mda5 and IFIH1) is a viral double-stranded RNA (dsRNA) receptor that shares sequence similarity and signaling pathways with RIG-I, yet plays essential functions in antiviral immunity through distinct specificity for viral RNA.; ncbi-cd \| cd17943 \| 93.8 \| 0.00049 \| 1.7e-07 \| 45.8 \| 61 \| (64, 126) \| 682 \| (12, 74) \| 192 \| DEADc\_DDX20 \| cd17943 DEADc\_DDX20; DEAD-box helicase domain of DEAD box protein 20. DDX20 (also called DEAD Box Protein DP 103, Component Of Gems 3, Gemin-3, and SMN-Interacting Protein) interacts directly with SMN (survival of motor neurons), the spinal muscular atrophy gene product, and may play a catalytic role in the function of the SMN complex on ribonucleoproteins.; ncbi-cd \| cd18024 \| 93.8 \| 0.00052 \| 1.7e-07 \| 47.8 \| 66 \| (60, 128) \| 682 \| (28, 93) \| 205 \| DEXHc\_Mtr4-like \| cd18024 DEXHc\_Mtr4-like; DEXH-box helicase domain of ATP-dependent RNA helicase Mtr4. Mtr4 (also known as DOB1 or SKIV2L2) is a type II DEAD box helicase that plays a role in the processing of structured RNAs, including the maturation of 5.; ncbi-cd \| cd18029 \| 93.8 \| 0.00053 \| 1.7e-07 \| 45.8 \| 62 \| (63, 128) \| 682 \| (7, 69) \| 169 \| DEXHc\_XPB \| cd18029 DEXHc\_XPB; DEXH-box helicase domain of TFIIH XPB subunit and similar proteins. TFIIH basal transcription factor complex helicase XPB subunit (also known as DNA excision repair protein ERCC-3 or TFIIH 89 kDa subunit) is the ATP-dependent 3'-5' DNA helicase component of the core-TFIIH basal transcription factor, involved in nucleotide excision repair (NER) of DNA and, when complexed to CAK, in RNA transcription by RNA polymerase II.; ncbi-cd \| cd18077 \| 93.7 \| 0.00058 \| 1.9e-07 \| 48.3 \| 65 \| (65, 129) \| 682 \| (2, 69) \| 226 \| DEXXQc\_HELZ \| cd18077 DEXXQc\_HELZ; DEXXQ-box helicase domain of HELZ. Helicase with zinc finger (HELZ) acts as a helicase that plays a role in RNA metabolism during development.; ncbi-cd \| cd17961 \| 93.7 \| 0.00057 \| 1.9e-07 \| 46.3 \| 62 \| (64, 127) \| 682 \| (16, 85) \| 206 \| DEADc\_DDX56 \| cd17961 DEADc\_DDX56; DEAD-box helicase domain of DEAD box protein 56. DDX56 is a helicase required for assembly of infectious West Nile virus particles.; ncbi-cd \| cd17942 \| 93.7 \| 0.00058 \| 1.9e-07 \| 46.4 \| 60 \| (65, 126) \| 682 \| (13, 78) \| 198 \| DEADc\_DDX18 \| cd17942 DEADc\_DDX18; DEAD-box helicase domain of DEAD box protein 18. This DDX18 gene encodes a DEAD box protein and is activated by Myc protein.; ncbi-cd \| cd17963 \| 93.7 \| 0.00059 \| 2e-07 \| 46.5 \| 65 \| (64, 128) \| 682 \| (16, 82) \| 196 \| DEADc\_DDX19\_DDX25 \| cd17963 DEADc\_DDX19\_DDX25; DEAD-box helicase domain of ATP-dependent RNA helicases DDX19 and DDX25. DDX19 (also called DEAD box RNA helicase DEAD5) and DDX25 (also called gonadotropin-regulated testicular RNA helicase (GRTH)) are members of the DEAD-box helicases, a diverse family of proteins involved in ATP-dependent RNA unwinding, needed in a variety of cellular processes including splicing, ribosome biogenesis and RNA degradation.; ncbi-cd \| cd18018 \| 93.6 \| 0.00063 \| 2e-07 \| 48.3 \| 61 \| (64, 126) \| 682 \| (12, 72) \| 201 \| DEXHc\_RecQ4-like \| cd18018 DEXHc\_RecQ4-like; DEAH-box helicase domain of RecQ4 and similar proteins. ATP-dependent DNA helicase Q4 (RecQ4) is part of the RecQ family of highly conserved DNA repair helicases that is part of the type II DEAD box helicase superfamily, a diverse family of proteins involved in ATP-dependent RNA or DNA unwinding.; ncbi-cd \| cd18019 \| 93.6 \| 0.00061 \| 2e-07 \| 47.3 \| 65 \| (63, 128) \| 682 \| (16, 90) \| 214 \| DEXHc\_Brr2\_1 \| cd18019 DEXHc\_Brr2\_1; N-terminal DEXH-box helicase domain of spliceosomal Brr2 RNA helicase. Brr2 is a type II DEAD box helicase that mediates spliceosome catalytic activation.; ncbi-cd \| cd17954 \| 93.6 \| 0.0006 \| 2e-07 \| 45.6 \| 60 \| (64, 125) \| 682 \| (22, 83) \| 203 \| DEADc\_DDX47 \| cd17954 DEADc\_DDX47; DEAD-box helicase domain of DEAD box protein 47. DDX47 (also called E4-DEAD box protein) can shuttle between the nucleus and the cytoplasm, and has an RNA-independent ATPase activity.; ncbi-cd \| cd18000 \| 93.6 \| 0.00062 \| 2e-07 \| 46.9 \| 63 \| (65, 128) \| 682 \| (1, 67) \| 193 \| DEXHc\_ERCC6 \| cd18000 DEXHc\_ERCC6; DEXH-box helicase domain of ERCC6. ERCC excision repair 6, chromatin remodeling factor (ERCC6, also known Cockayne syndrome group B (CSB), Rad26 in Saccharomyces cerevisiae, and Rhp26 in Schizosaccharomyces pombe) is a DNA-binding protein that is important in transcription-coupled excision repair.; ncbi-cd \| cd18045 \| 93.6 \| 0.00062 \| 2.1e-07 \| 46.6 \| 62 \| (64, 127) \| 682 \| (21, 84) \| 201 \| DEADc\_EIF4AIII\_DDX48 \| cd18045 DEADc\_EIF4AIII\_DDX48; DEAD-box helicase domain of eukaryotic initiation factor 4A-III. Eukaryotic initiation factor 4A-III (EIF4AIII, also known as DDX48) is part of the exon junction complex (EJC) that plays a major role in posttranscriptional regulation of mRNA.; ncbi-cd \| cd17976 \| 93.6 \| 0.00065 \| 2.1e-07 \| 46.0 \| 44 \| (82, 125) \| 682 \| (18, 65) \| 178 \| DEXHc\_DHX30 \| cd17976 DEXHc\_DHX30; DEXH-box helicase domain of DEAH-box helicase 30. DEAH-box helicase 30 (DHX30) plays an important role in the assembly of the mitochondrial large ribosomal subunit.; ncbi-cd \| cd17964 \| 93.5 \| 0.00067 \| 2.2e-07 \| 46.3 \| 63 \| (64, 127) \| 682 \| (16, 85) \| 211 \| DEADc\_MSS116 \| cd17964 DEADc\_MSS116; DEAD-box helicase domain of DEAD-box helicase Mss116. Mss116 is an RNA chaperone important for mitochondrial group I and II intron splicing, translational activation, and RNA end processing.; ncbi-cd \| cd17939 \| 93.5 \| 0.00067 \| 2.3e-07 \| 44.9 \| 62 \| (64, 127) \| 682 \| (19, 82) \| 199 \| DEADc\_EIF4A \| cd17939 DEADc\_EIF4A; DEAD-box helicase domain of eukaryotic initiation factor 4A. The eukaryotic initiation factor-4A (eIF4A) family consists of 3 proteins EIF4A1, EIF4A2, and EIF4A3.; ncbi-cd \| cd18001 \| 93.5 \| 0.00073 \| 2.3e-07 \| 48.6 \| 63 \| (66, 129) \| 682 \| (2, 67) \| 232 \| DEXHc\_ERCC6L \| cd18001 DEXHc\_ERCC6L; DEXH-box helicase domain of ERCC6L. ERCC excision repair 6 like, spindle assembly checkpoint helicase (ERCC6L, also known as RAD26L) is an essential component of the mitotic spindle assembly checkpoint, by acting as a tension sensor that associates with catenated DNA which is stretched under tension until it is resolved during anaphase.; ncbi-cd \| cd17926 \| 93.4 \| 0.00072 \| 2.4e-07 \| 43.7 \| 60 \| (66, 128) \| 682 \| (2, 62) \| 146 \| DEXHc\_RE \| cd17926 DEXHc\_RE; DEXH-box helicase domain of DEAD-like helicase restriction enzyme family proteins.; ncbi-cd \| cd18058 \| 93.3 \| 0.0008 \| 2.6e-07 \| 46.7 \| 63 \| (66, 129) \| 682 \| (2, 67) \| 222 \| DEXHc\_CHD6 \| cd18058 DEXHc\_CHD6; DEAH-box helicase domain of the chromodomain helicase DNA binding protein 6. Chromodomain-helicase-DNA-binding protein 6 (CHD6) is a DNA-dependent ATPase that plays a role in chromatin remodeling.; ncbi-cd \| cd17957 \| 93.2 \| 0.00086 \| 2.9e-07 \| 44.8 \| 62 \| (64, 127) \| 682 \| (12, 77) \| 198 \| DEADc\_DDX52 \| cd17957 DEADc\_DDX52; DEAD-box helicase domain of DEAD box protein 52. DDX52 (also called ROK1 and HUSSY19) is ubiquitously expressed in testis, endometrium, and other tissues in humans.; ncbi-cd \| cd18017 \| 93.2 \| 0.00091 \| 2.9e-07 \| 46.8 \| 60 \| (64, 128) \| 682 \| (12, 71) \| 193 \| DEXHc\_RecQ3 \| cd18017 DEXHc\_RecQ3; DEAH-box helicase domain of RecQ3. DEAD-like helicase RecQ3 (also called Werner syndrome ATP-dependent helicase or WRN) is part of the RecQ family of highly conserved DNA repair helicases that is part of the type II DEAD box helicase superfamily, a diverse family of proteins involved in ATP-dependent RNA or DNA unwinding.; ncbi-cd \| cd17918 \| 93.1 \| 0.00096 \| 3.2e-07 \| 44.9 \| 63 \| (64, 127) \| 682 \| (15, 81) \| 180 \| DEXHc\_RecG \| cd17918 DEXHc\_RecG; DEXH/Q-box helicase domain of DEAD-like helicase RecG family proteins. The DEAD-like helicase RecG family is part of the DEAD-like helicases superfamily, a diverse family of proteins involved in ATP-dependent RNA or DNA unwinding.; ncbi-cd \| cd17967 \| 93.1 \| 0.00099 \| 3.2e-07 \| 47.0 \| 63 \| (64, 128) \| 682 \| (22, 96) \| 221 \| DEADc\_DDX3\_DDX4 \| cd17967 DEADc\_DDX3\_DDX4; DEAD-box helicase domain of ATP-dependent RNA helicases DDX3 and DDX4. This subfamily includes Drosophila melanogaster Vasa, which is essential for development.; ncbi-cd \| cd18004 \| 93.1 \| 0.001 \| 3.2e-07 \| 48.3 \| 64 \| (66, 130) \| 682 \| (2, 78) \| 240 \| DEXHc\_RAD54 \| cd18004 DEXHc\_RAD54; DEXH-box helicase domain of RAD54. RAD54 proteins play a role in recombination. They are members of the DEAD-like helicase superfamily, a diverse family of proteins involved in ATP-dependent RNA or DNA unwinding.; ncbi-cd \| cd17951 \| 93.0 \| 0.001 \| 3.4e-07 \| 46.0 \| 62 \| (64, 127) \| 682 \| (12, 83) \| 206 \| DEADc\_DDX41 \| cd17951 DEADc\_DDX41; DEAD-box helicase domain of DEAD box protein 41. DDX41 (also called ABS and MPLPF) interacts with several spliceosomal proteins and may recognize the bacterial second messengers cyclic di-GMP and cyclic di-AMP, resulting in the induction of genes involved in the innate immune response.; ncbi-cd \| cd17947 \| 93.0 \| 0.001 \| 3.4e-07 \| 44.3 \| 63 \| (64, 128) \| 682 \| (12, 79) \| 196 \| DEADc\_DDX27 \| cd17947 DEADc\_DDX27; DEAD-box helicase domain of DEAD box protein 27. DDX27 (also called RHLP, deficiency of ribosomal subunits protein 1 homolog, and probable ATP-dependent RNA helicase DDX27) is involved in the processing of 5.; cath \| 2o0jA02 \| 98.1 \| 4.2e-10 \| 8.3e-14 \| 102.0 \| 196 \| (29, 240) \| 682 \| (46, 242) \| 254 \| Dna packaging protein gp17 \| CATHCODE: 3.40.50.300 NAME: Dna packaging protein gp17. Chain: a. Fragment: n-terminal atpase domain. Synonym: terminase. Engineered: yes. Mutation: yes SOURCE: Enterobacteria phage t4. Organism\_taxid: 10665. Gene: 17. Expressed in: escherichia coli. Expression\_system\_taxid: 562. CLASS: Alpha Beta, ARCH: 3-Layer(aba) Sandwich, TOPOL: Rossmann fold, HOMOL: P-loop containing nucleotide triphosphate hydrolases; cath \| 4idhA01 \| 97.4 \| 5e-08 \| 1e-11 \| 83.4 \| 154 \| (65, 240) \| 682 \| (2, 160) \| 208 \| Gene 2 protein \| CATHCODE: 3.40.50.300 NAME: Gene 2 protein. Chain: a. Engineered: yes SOURCE: Shigella phage sf6. Shigella flexneri bacteriophage vi. Organism\_taxid: 10761. Expressed in: escherichia coli. Expression\_system\_taxid: 469008. CLASS: Alpha Beta, ARCH: 3-Layer(aba) Sandwich, TOPOL: Rossmann fold, HOMOL: P-loop containing nucleotide triphosphate hydrolases; cath \| 1wp9A01 \| 97.4 \| 7.5e-08 \| 1.5e-11 \| 80.0 \| 177 \| (63, 252) \| 682 \| (8, 192) \| 199 \| Atp-dependent RNA helicase, putative \| CATHCODE: 3.40.50.300 NAME: Atp-dependent RNA helicase, putative. Chain: a, b, c, d, e, f. Fragment: residues 2-495. Synonym: hef helicase, nuclease. Engineered: yes SOURCE: Pyrococcus furiosus. Organism\_taxid: 186497. Strain: dsm 3638. Expressed in: escherichia coli bl21(de3). Expression\_system\_taxid: 469008. CLASS: Alpha Beta, ARCH: 3-Layer(aba) Sandwich, TOPOL: Rossmann fold, HOMOL: P-loop containing nucleotide triphosphate hydrolases; cath \| 3b85A00 \| 97.3 \| 1.1e-07 \| 2.3e-11 \| 80.0 \| 156 \| (63, 229) \| 682 \| (6, 162) \| 208 \| Phosphate starvation-inducible protein \| CATHCODE: 3.40.50.300 NAME: Phosphate starvation-inducible protein. Chain: a, b. Fragment: residues 116-320. Engineered: yes SOURCE: Corynebacterium glutamicum atcc 13032. Organism\_taxid: 196627. Strain:dsm 20300 / jcm 1318 / lmg 3730 / ncimb 10025. Atcc: 13032. Gene: phoh2, cg2513. Expressed in: escherichia coli bl21(de3). Expression\_system\_taxid: 469008. CLASS: Alpha Beta, ARCH: 3-Layer(aba) Sandwich, TOPOL: Rossmann fold, HOMOL: P-loop containing nucleotide triphosphate hydrolases; cath \| 1rifA02 \| 97.0 \| 3.9e-07 \| 8.1e-11 \| 73.9 \| 150 \| (63, 229) \| 682 \| (22, 176) \| 192 \| Dna helicase uvsw \| CATHCODE: 3.40.50.300 NAME: Dna helicase uvsw. Chain: a, b. Synonym: dar protein. Engineered: yes.Mutation: yes SOURCE: Enterobacteria phage t4. Organism\_taxid: 10665. Gene: uvsw, dar. Expressed in: escherichia coli bl21(de3). Expression\_system\_taxid: 469008. CLASS: Alpha Beta, ARCH: 3-Layer(aba) Sandwich, TOPOL: Rossmann fold, HOMOL: P-loop containing nucleotide triphosphate hydrolases; cath \| 1hv8A01 \| 97.0 \| 4.1e-07 \| 8.8e-11 \| 74.5 \| 157 \| (63, 229) \| 682 \| (27, 192) \| 210 \| Putative atp-dependent RNA helicase mj0669 \| CATHCODE: 3.40.50.300 NAME: Putative atp-dependent RNA helicase mj0669. Chain: a, b. Synonym: deadbox helicase. Engineered: yes SOURCE: Methanocaldococcus jannaschii. Organism\_taxid: 2190. Expressed in: escherichia coli. Expression\_system\_taxid: 562 CLASS: Alpha Beta, ARCH: 3-Layer(aba) Sandwich, TOPOL: Rossmann fold, HOMOL: P-loop containing nucleotide triphosphate hydrolases; cath \| 2xgjB01 \| 97.0 \| 5.3e-07 \| 1.1e-10 \| 76.7 \| 156 \| (60, 239) \| 682 \| (29, 192) \| 206 \| Atp-dependent RNA helicase dob1 \| CATHCODE: 3.40.50.300 NAME: Atp-dependent RNA helicase dob1. Chain: a, b. Fragment: residues 81-1073. Synonym: mtr4p, mRNA transport regulator mtr4., 3.6.1.-. Rna (5'-(\*ap\*ap\*ap\*ap\*a)-3'). Chain: c, d SOURCE: Saccharomyces cerevisiae. Baker's yeast. Organism\_taxid: 4932. CLASS: Alpha Beta, ARCH: 3-Layer(aba) Sandwich, TOPOL: Rossmann fold, HOMOL: P-loop containing nucleotide triphosphate hydrolases; cath \| 3berA00 \| 97.0 \| 5.7e-07 \| 1.2e-10 \| 75.2 \| 157 \| (64, 229) \| 682 \| (36, 202) \| 220 \| Probable atp-dependent RNA helicase ddx47 \| CATHCODE: 3.40.50.300 NAME: Probable atp-dependent RNA helicase ddx47. Chain: a. Fragment: conserved domain i (dead): residues 5-230. Synonym: dead box protein 47. Engineered: yes SOURCE: Homo sapiens. Human. Organism\_taxid: 9606. Gene: ddx47. Expressed in:escherichia coli. Expression\_system\_taxid: 562. CLASS: Alpha Beta, ARCH: 3-Layer(aba) Sandwich, TOPOL: Rossmann fold, HOMOL: P-loop containing nucleotide triphosphate hydrolases; cath \| 1fuuA00 \| 96.9 \| 6.7e-07 \| 1.4e-10 \| 74.3 \| 67 \| (64, 133) \| 682 \| (43, 111) \| 225 \| Yeast initiation factor 4a \| CATHCODE: 3.40.50.300 NAME: Yeast initiation factor 4a. Chain: a, b. Fragment: mRNA helicase. Synonym: eukaryotic initiation factor 4a. Engineered: yes SOURCE: Saccharomyces cerevisiae. Baker's yeast. Organism\_taxid: 4932. Expressed in: escherichia coli. Expression\_system\_taxid: 562 CLASS: Alpha Beta, ARCH: 3-Layer(aba) Sandwich, TOPOL: Rossmann fold, HOMOL: P-loop containing nucleotide triphosphate hydrolases; cath \| 1vecA00 \| 96.8 \| 1.2e-06 \| 2.5e-10 \| 70.9 \| 67 \| (63, 132) \| 682 \| (24, 92) \| 206 \| Atp-dependent RNA helicase p54 \| CATHCODE: 3.40.50.300 NAME: Atp-dependent RNA helicase p54. Chain: a, b. Fragment: n-terminal domain. Synonym: rck, dead-box protein 6. Engineered: yes SOURCE: Homo sapiens. Human. Organism\_taxid: 9606. Gene: humrck. Expressed in: escherichia coli. Expression\_system\_taxid: 562. CLASS: Alpha Beta, ARCH: 3-Layer(aba) Sandwich, TOPOL: Rossmann fold, HOMOL: P-loop containing nucleotide triphosphate hydrolases; cath \| 1q0uA00 \| 96.8 \| 1.2e-06 \| 2.6e-10 \| 73.2 \| 69 \| (63, 134) \| 682 \| (25, 95) \| 219 \| Bstdead \| CATHCODE: 3.40.50.300 NAME: Bstdead. Chain: a, b. Fragment: n-terminal domain. Engineered: yes SOURCE: Geobacillus stearothermophilus. Organism\_taxid: 1422. Expressed in: escherichia coli. Expression\_system\_taxid: 562. CLASS: Alpha Beta, ARCH: 3-Layer(aba) Sandwich, TOPOL: Rossmann fold, HOMOL: P-loop containing nucleotide triphosphate hydrolases; cath \| 2p6rA01 \| 96.8 \| 1.4e-06 \| 2.8e-10 \| 71.9 \| 60 \| (64, 127) \| 682 \| (19, 79) \| 197 \| 25-mer \| CATHCODE: 3.40.50.300 NAME: 25-mer. Chain: x. Engineered: yes. 5'- d(\*cp\*tp\*ap\*gp\*ap\*gp\*ap\*cp\*tp\*ap\*tp\*cp\*gp\*ap\*t)-3'. Chain: y. Engineered: yes. Afuhel308 helicase. Chain: a. Engineered: yes SOURCE: Yes. Yes. CLASS: Alpha Beta, ARCH: 3-Layer(aba) Sandwich, TOPOL: Rossmann fold, HOMOL: P-loop containing nucleotide triphosphate hydrolases; cath \| 5lstA01 \| 96.8 \| 1.4e-06 \| 3e-10 \| 72.6 \| 68 \| (63, 134) \| 682 \| (31, 99) \| 215 \| Atp-dependent DNA helicase q4 \| CATHCODE: 3.40.50.300 NAME: Atp-dependent DNA helicase q4. Chain: a. Synonym: DNA helicase,recq-like type 4,recq4,rts,recq protein-like 4. Engineered: yes SOURCE: Homo sapiens. Human. Organism\_taxid: 9606. Gene: recql4, recq4. Expressed in: escherichia coli. Expression\_system\_taxid: 469008. CLASS: Alpha Beta, ARCH: 3-Layer(aba) Sandwich, TOPOL: Rossmann fold, HOMOL: P-loop containing nucleotide triphosphate hydrolases; cath \| 2oxcA00 \| 96.7 \| 2.1e-06 \| 4.6e-10 \| 69.5 \| 68 \| (63, 133) \| 682 \| (22, 91) \| 207 \| Probable atp-dependent RNA helicase ddx20 \| CATHCODE: 3.40.50.300 NAME: Probable atp-dependent RNA helicase ddx20. Chain: a, b. Fragment: deaddomain. Synonym: dead box protein 20, dead box protein dp 103, component of gems 3, gemin-3. Engineered: yes SOURCE: Homo sapiens. Human. Organism\_taxid: 9606. Gene: ddx20, dp103, gemin3.Expressed in: escherichia coli. Expression\_system\_taxid: 562. CLASS: Alpha Beta, ARCH: 3-Layer(aba) Sandwich, TOPOL: Rossmann fold, HOMOL: P-loop containing nucleotide triphosphate hydrolases; cath \| 5gvrA00 \| 96.6 \| 2.4e-06 \| 5e-10 \| 71.9 \| 67 \| (63, 132) \| 682 \| (34, 110) \| 234 \| Probable atp-dependent RNA helicase ddx41 \| CATHCODE: 3.40.50.300 NAME: Probable atp-dependent RNA helicase ddx41. Chain: a. Fragment: unp residues 169-402. Synonym: dead box protein 41,dead box protein abstrakt homolog. Engineered: yes SOURCE: Homo sapiens. Human. Organism\_taxid: 9606. Gene: ddx41, abs. Expressed in: escherichia coli. Expression\_system\_taxid: 562. Expression\_system\_vector\_type: plasmid CLASS: Alpha Beta, ARCH: 3-Layer(aba) Sandwich, TOPOL: Rossmann fold, HOMOL: P-loop containing nucleotide triphosphate hydrolases; cath \| 2w00A03 \| 96.6 \| 2.7e-06 \| 5.7e-10 \| 68.2 \| 51 \| (81, 132) \| 682 \| (30, 81) \| 191 \| Hsdr \| CATHCODE: 3.40.50.300 NAME: Hsdr. Chain: a, b. Synonym: r.ecor124i. Engineered: yes SOURCE: Escherichia coli. Organism\_taxid: 562. Strain: b834(de3). Expressed in: escherichia coli. Expression\_system\_taxid: 562. CLASS: Alpha Beta, ARCH: 3-Layer(aba) Sandwich, TOPOL: Rossmann fold, HOMOL: P-loop containing nucleotide triphosphate hydrolases; cath \| 1gm5A04 \| 96.6 \| 2.9e-06 \| 5.9e-10 \| 71.0 \| 152 \| (62, 229) \| 682 \| (18, 182) \| 204 \| Recg \| CATHCODE: 3.40.50.300 NAME: Recg. Chain: a. Engineered: yes. Dna (5'-(\*cp\*ap\*gp\*cp\*tp\*cp\*cp\*ap\*tp\*gp\*ap\*tp\* cp\*ap\*tp\*tp\*gp\*gp\*cp\*a)-3'). Chain: x. Dna (5'-(\*gp\*cp\*ap\*gp\*tp\*gp\*cp\*tp\*cp\*gp\*cp\*ap\* tp\*gp\*gp\*ap\*gp\*cp\*tp\*g)-3'). Chain: y. Dna(5'-(\*gp\*ap\*gp\*cp\*ap\*cp\*tp\*gp\*c)-3'). Chain: z SOURCE: Thermotoga maritima. Organism\_taxid: 2336. Expressed in: escherichia coli. Expression\_system\_taxid: 562. CLASS: Alpha Beta, ARCH: 3-Layer(aba) Sandwich, TOPOL: Rossmann fold, HOMOL: P-loop containing nucleotide triphosphate hydrolases; cath \| 4a15A01 \| 96.6 \| 2.9e-06 \| 6.1e-10 \| 68.5 \| 69 \| (63, 133) \| 682 \| (2, 73) \| 188 \| Atp-dependent DNA helicase ta0057 \| CATHCODE: 3.40.50.300 NAME: Atp-dependent DNA helicase ta0057. Chain: a. Synonym: xpd helicase. Engineered: yes. 5'-d(\*dtp\*ap\*cp\*gp)-3'. Chain: e SOURCE: Thermoplasma acidophilum. Organism\_taxid: 2303. Expressed in: escherichia coli. Expression\_system\_taxid: 469008. CLASS: Alpha Beta, ARCH: 3-Layer(aba) Sandwich, TOPOL: Rossmann fold, HOMOL: P-loop containing nucleotide triphosphate hydrolases; cath \| 2pl3A00 \| 96.5 \| 4.3e-06 \| 9e-10 \| 69.3 \| 67 \| (63, 132) \| 682 \| (46, 118) \| 236 \| Probable atp-dependent RNA helicase ddx10 \| CATHCODE: 3.40.50.300 NAME: Probable atp-dependent RNA helicase ddx10. Chain: a. Fragment: dead domain. Synonym: dead box protein 10. Engineered: yes SOURCE: Homo sapiens. Human. Organism\_taxid: 9606. Gene: ddx10. Expressed in:escherichia coli. Expression\_system\_taxid: 562. CLASS: Alpha Beta, ARCH: 3-Layer(aba) Sandwich, TOPOL: Rossmann fold, HOMOL: P-loop containing nucleotide triphosphate hydrolases; cath \| 3h1tA02 \| 96.5 \| 4.6e-06 \| 9.5e-10 \| 68.1 \| 64 \| (62, 127) \| 682 \| (19, 95) \| 202 \| Type i site-specific restriction-modification system, r (restriction) subunit \| CATHCODE: 3.40.50.300 NAME: Type i site-specific restriction-modification system, r (restriction) subunit. Chain: a. Fragment: unp residues 1-590. Engineered: yes SOURCE: Vibrio vulnificus. Organism\_taxid: 196600. Strain: yj016. Gene: vv0265. Expressed in: escherichia coli. Expression\_system\_taxid: 562. CLASS: Alpha Beta, ARCH: 3-Layer(aba) Sandwich, TOPOL: Rossmann fold, HOMOL: P-loop containing nucleotide triphosphate hydrolases; cath \| 1w36D02 \| 96.5 \| 5.1e-06 \| 9.9e-10 \| 73.2 \| 98 \| (16, 127) \| 682 \| (3, 103) \| 245 \| Exodeoxyribonuclease v beta chain \| CATHCODE: 3.40.50.300 NAME: Exodeoxyribonuclease v beta chain. Chain: b, e. Synonym: recb. Engineered: yes. Exodeoxyribonuclease v gamma chain. Chain: c, f. Synonym: recc. Engineered: yes. Exodeoxyribonuclease v alpha chain. Chain: d, g. Synonym: recd. Engineered: yes. Dna hairpin. Chain: y, z SOURCE: Escherichia coli. Organism\_taxid: 562. Expressed in: escherichia coli. Expression\_system\_taxid: 562. Escherichia coli. Organism\_taxid: 562.Expressed in: escherichia coli. Expression\_system\_taxid: 562. Escherichia coli. Organism\_taxid: 562. Expressed in: escherichia coli. Expression\_system\_taxid: 562. CLASS: Alpha Beta, ARCH: 3-Layer(aba) Sandwich, TOPOL: Rossmann fold, HOMOL: P-loop containing nucleotide triphosphate hydrolases; cath \| 4f92B02 \| 96.4 \| 5.4e-06 \| 1.1e-09 \| 69.3 \| 68 \| (63, 133) \| 682 \| (26, 104) \| 228 \| U5 small nuclear ribonucleoprotein 200 kda helicase \| CATHCODE: 3.40.50.300 NAME: U5 small nuclear ribonucleoprotein 200 kda helicase. Chain: b. Fragment: brr2 helicase region. Synonym: activating signal cointegrator 1 complex subunit 3-like 1, brr2 homolog, u5 snrnp-specific 200 kda protein, u5-200kd. Engineered: yes. Mutation: yes SOURCE: Homo sapiens. Human. Organism\_taxid: 9606. Gene: snrnp200, ascc3l1, helic2, kiaa0788. Expressed in: spodoptera frugiperda. Expression\_system\_taxid: 7108 CLASS: Alpha Beta, ARCH: 3-Layer(aba) Sandwich, TOPOL: Rossmann fold, HOMOL: P-loop containing nucleotide triphosphate hydrolases; cath \| 2db3A01 \| 96.3 \| 9.5e-06 \| 1.9e-09 \| 70.3 \| 68 \| (63, 133) \| 682 \| (77, 151) \| 267 \| 5'-r(\*up\*up\*up\*up\*up\*up\*up\*up\*up\*u)-3' \| CATHCODE: 3.40.50.300 NAME: 5'-r(\*up\*up\*up\*up\*up\*up\*up\*up\*up\*u)-3'. Chain: e, f, g, h. Engineered:yes. Atp-dependent RNA helicase vasa. Chain: a, b, c, d. Fragment: residues 200-623. Synonym: vasa protein, antigen mab46f11. Engineered: yes SOURCE: Yes. CLASS: Alpha Beta, ARCH: 3-Layer(aba) Sandwich, TOPOL: Rossmann fold, HOMOL: P-loop containing nucleotide triphosphate hydrolases; cath \| 5supC01 \| 96.3 \| 9.3e-06 \| 1.9e-09 \| 66.8 \| 68 \| (63, 133) \| 682 \| (27, 96) \| 220 \| Atp-dependent RNA helicase sub2 \| CATHCODE: 3.40.50.300 NAME: Atp-dependent RNA helicase sub2. Chain: a, b, c. Fragment: residues 61-446. Synonym: suppressor of brr1 protein 2. Engineered: yes. Rna annealing protein yra1. Chain: g, h, i. Fragment: residues 200-226. Engineered: yes. Rna (5'-r(p\*up\*up\*up\*up\*up\*u)-3'). Chain: d, e, f. Engineered: yes SOURCE: Saccharomyces cerevisiae. Baker's yeast. Organism\_taxid: 559292. Strain: atcc 204508 / s288c. Gene: sub2, ydl084w. Expressed in: escherichia coli. Expression\_system\_taxid: 562. Saccharomyces cerevisiae. Baker's yeast. Organism\_taxid: 559292. Strain: atcc 204508 / s288c. Gene: yra1, ydr381w, d9481.2, d9509.1. Expressed in: escherichia coli. Expression\_system\_taxid: 562. CLASS: Alpha Beta, ARCH: 3-Layer(aba) Sandwich, TOPOL: Rossmann fold, HOMOL: P-loop containing nucleotide triphosphate hydrolases; cath \| 4ljyA01 \| 96.3 \| 9.6e-06 \| 2e-09 \| 70.4 \| 67 \| (64, 133) \| 682 \| (73, 147) \| 265 \| Pre-mRNA-processing atp-dependent RNA helicase prp5 \| CATHCODE: 3.40.50.300 NAME: Pre-mRNA-processing atp-dependent RNA helicase prp5. Chain: a. Fragment: unp residues 206-698. Engineered: yes SOURCE: Saccharomyces cerevisiae. Yeast. Organism\_taxid: 559292. Strain: atcc 204508 / s288c. Gene: prp5, RNA5, ybr237w, ybr1603. Expressed in: escherichia coli. Expression\_system\_taxid: 562. Expression\_system\_vector\_type: plasmid CLASS: Alpha Beta, ARCH: 3-Layer(aba) Sandwich, TOPOL: Rossmann fold, HOMOL: P-loop containing nucleotide triphosphate hydrolases; cath \| 4w7sA01 \| 96.2 \| 1.1e-05 \| 2.1e-09 \| 71.5 \| 69 \| (63, 134) \| 682 \| (70, 152) \| 277 \| Pre-mRNA-splicing atp-dependent RNA helicase prp28 \| CATHCODE: 3.40.50.300 NAME: Pre-mRNA-splicing atp-dependent RNA helicase prp28. Chain: a, b. Synonym: helicase ca8. Engineered: yes SOURCE: Saccharomyces cerevisiae. Baker's yeast. Organism\_taxid: 559292. Strain: atcc 204508 / s288c. Gene: prp28, ydr243c, yd8419.10c. Expressed in: escherichia coli. Expression\_system\_taxid: 562. CLASS: Alpha Beta, ARCH: 3-Layer(aba) Sandwich, TOPOL: Rossmann fold, HOMOL: P-loop containing nucleotide triphosphate hydrolases; cath \| 4f92B07 \| 96.2 \| 1.1e-05 \| 2.4e-09 \| 67.0 \| 63 \| (63, 127) \| 682 \| (37, 101) \| 230 \| U5 small nuclear ribonucleoprotein 200 kda helicase \| CATHCODE: 3.40.50.300 NAME: U5 small nuclear ribonucleoprotein 200 kda helicase. Chain: b. Fragment: brr2 helicase region. Synonym: activating signal cointegrator 1 complex subunit 3-like 1, brr2 homolog, u5 snrnp-specific 200 kda protein, u5-200kd. Engineered: yes. Mutation: yes SOURCE: Homo sapiens. Human. Organism\_taxid: 9606. Gene: snrnp200, ascc3l1, helic2, kiaa0788. Expressed in: spodoptera frugiperda. Expression\_system\_taxid: 7108 CLASS: Alpha Beta, ARCH: 3-Layer(aba) Sandwich, TOPOL: Rossmann fold, HOMOL: P-loop containing nucleotide triphosphate hydrolases; cath \| 4q48A01 \| 96.2 \| 1.2e-05 \| 2.4e-09 \| 66.3 \| 63 \| (63, 133) \| 682 \| (23, 86) \| 207 \| Dna helicase recq \| CATHCODE: 3.40.50.300 NAME: Dna helicase recq. Chain: a, b. Fragment: unp residues 1-517. Engineered: yes SOURCE: Deinococcus radiodurans. Organism\_taxid: 243230. Strain: atcc 13939 / dsm 20539 / jcm 16871 / lmg 4051 / nbrc 15346 / ncimb 9279 / r1 / vkm b-1422. Gene: dr\_1289. Expressed in: escherichia coli. Expression\_system\_taxid: 562 CLASS: Alpha Beta, ARCH: 3-Layer(aba) Sandwich, TOPOL: Rossmann fold, HOMOL: P-loop containing nucleotide triphosphate hydrolases; cath \| 2ykgA01 \| 96.1 \| 1.4e-05 \| 2.9e-09 \| 65.5 \| 68 \| (62, 132) \| 682 \| (11, 82) \| 224 \| Probable atp-dependent RNA helicase ddx58 \| CATHCODE: 3.40.50.300 NAME: Probable atp-dependent RNA helicase ddx58. Chain: a. Fragment: residues 230-925. Synonym: dead box protein 58, retinoic acid-inducible gene 1 protein, rig-1, retinoic acid-inducible gene i protein, rig-i. 5'-r(\*gp\*cp\*gp\*cp\*gp\*cp\*gp\*cp\*gp\*cp)-3'. Chain: c, d SOURCE: Homo sapiens. Organism\_taxid: 9606. CLASS: Alpha Beta, ARCH: 3-Layer(aba) Sandwich, TOPOL: Rossmann fold, HOMOL: P-loop containing nucleotide triphosphate hydrolases; cath \| 3dmqA04 \| 96.1 \| 1.9e-05 \| 3.7e-09 \| 71.9 \| 62 \| (63, 127) \| 682 \| (3, 67) \| 302 \| Rna polymerase-associated protein rapa \| CATHCODE: 3.40.50.10810 NAME: Rna polymerase-associated protein rapa. Chain: a, b. Synonym: atp-dependent helicase hepa. Engineered: yes. Mutation: yes SOURCE: Escherichia coli k12. Organism\_taxid: 83333. Strain: k12 / mg1655. Atcc: 47076. Gene: rapa, hepa, yaba, b0059, jw0058. Expressed in: escherichia coli. Expression\_system\_taxid: 562 CLASS: Alpha Beta, ARCH: 3-Layer(aba) Sandwich, TOPOL: Rossmann fold, HOMOL: Tandem AAA-ATPase domain; cath \| 2fwrA02 \| 96.0 \| 2e-05 \| 4.1e-09 \| 63.8 \| 136 \| (62, 228) \| 682 \| (12, 152) \| 178 \| Dna repair protein rad25 \| CATHCODE: 3.40.50.300 NAME: Dna repair protein rad25. Chain: a, b, c, d. Engineered: yes SOURCE: Archaeoglobus fulgidus. Organism\_taxid: 2234. Expressed in: escherichia coli bl21(de3). Expression\_system\_taxid: 469008. CLASS: Alpha Beta, ARCH: 3-Layer(aba) Sandwich, TOPOL: Rossmann fold, HOMOL: P-loop containing nucleotide triphosphate hydrolases; cath \| 5jajA01 \| 95.9 \| 2.6e-05 \| 5.4e-09 \| 66.5 \| 63 \| (63, 127) \| 682 \| (8, 71) \| 269 \| Lgp2 \| CATHCODE: 3.40.50.300 NAME: Lgp2. Chain: a. Engineered: yes. Mutation: yes. Rna (5'-r(p\*gp\*gp\*up\*ap\*cp\*gp\*up\*ap\*cp\*cp\*c)-3'). Chain: x. Engineered: yes. Rna (5'-r(p\*gp\*gp\*up\*ap\*cp\*gp\*up\*ap\*cp\*c)-3'). Chain: y. Engineered: yes SOURCE: Gallus gallus. Chicken. Organism\_taxid: 9031. Expressed in: escherichia coli bl21(de3). Expression\_system\_taxid: 469008. Expression\_system\_variant: rosetta 2. CLASS: Alpha Beta, ARCH: 3-Layer(aba) Sandwich, TOPOL: Rossmann fold, HOMOL: P-loop containing nucleotide triphosphate hydrolases; cath \| 3dkpA00 \| 95.8 \| 3.5e-05 \| 7.3e-09 \| 64.9 \| 69 \| (63, 134) \| 682 \| (50, 121) \| 245 \| Probable atp-dependent RNA helicase ddx52 \| CATHCODE: 3.40.50.300 NAME: Probable atp-dependent RNA helicase ddx52. Chain: a. Fragment: conserved domain i: residues 139-381. Synonym: dead box protein 52, atp-dependent RNA helicase rok1-like. Engineered: yes SOURCE: Homo sapiens. Organism\_taxid: 9606. Gene: ddx52, rok1. Expressed in: escherichia coli. CLASS: Alpha Beta, ARCH: 3-Layer(aba) Sandwich, TOPOL: Rossmann fold, HOMOL: P-loop containing nucleotide triphosphate hydrolases; cath \| 3llmA01 \| 95.7 \| 5e-05 \| 9.8e-09 \| 65.3 \| 63 \| (63, 127) \| 682 \| (50, 116) \| 225 \| Atp-dependent RNA helicase a \| CATHCODE: 3.40.50.300 NAME: Atp-dependent RNA helicase a. Chain: a, b. Fragment: nucleotide binding domain (unp residues 329-563). Synonym: nuclear DNA helicase ii, ndhii, deah box protein 9. Engineered: yes SOURCE: Homo sapiens. Human. Organism\_taxid: 9606. Gene: ddx9, dhx9, lkp, ndh2. Expressed in: escherichia coli. Expression\_system\_taxid: 562. CLASS: Alpha Beta, ARCH: 3-Layer(aba) Sandwich, TOPOL: Rossmann fold, HOMOL: P-loop containing nucleotide triphosphate hydrolases; cath \| 1gkuB02 \| 95.6 \| 5.6e-05 \| 1.1e-08 \| 63.2 \| 65 \| (64, 133) \| 682 \| (24, 89) \| 213 \| Reverse gyrase \| CATHCODE: 3.40.50.300 NAME: Reverse gyrase. Chain: b. Synonym: top-rg. Engineered: yes. Mutation: yes SOURCE: Archaeoglobus fulgidus. Organism\_taxid: 224325. Strain: vc-16. Atcc: 49558. Expressed in: escherichia coli bl21(de3). Expression\_system\_taxid: 469008. Expression\_system\_variant: c41. CLASS: Alpha Beta, ARCH: 3-Layer(aba) Sandwich, TOPOL: Rossmann fold, HOMOL: P-loop containing nucleotide triphosphate hydrolases; cath \| 1w36B01 \| 95.6 \| 5.6e-05 \| 1.2e-08 \| 63.4 \| 61 \| (64, 127) \| 682 \| (2, 73) \| 248 \| Exodeoxyribonuclease v beta chain \| CATHCODE: 3.40.50.300 NAME: Exodeoxyribonuclease v beta chain. Chain: b, e. Synonym: recb. Engineered: yes. Exodeoxyribonuclease v gamma chain. Chain: c, f. Synonym: recc. Engineered: yes. Exodeoxyribonuclease v alpha chain. Chain: d, g. Synonym: recd. Engineered: yes. Dna hairpin. Chain: y, z SOURCE: Escherichia coli. Organism\_taxid: 562. Expressed in: escherichia coli. Expression\_system\_taxid: 562. Escherichia coli. Organism\_taxid: 562.Expressed in: escherichia coli. Expression\_system\_taxid: 562. Escherichia coli. Organism\_taxid: 562. Expressed in: escherichia coli. Expression\_system\_taxid: 562. CLASS: Alpha Beta, ARCH: 3-Layer(aba) Sandwich, TOPOL: Rossmann fold, HOMOL: P-loop containing nucleotide triphosphate hydrolases; cath \| 3crvA01 \| 95.6 \| 6.3e-05 \| 1.3e-08 \| 64.1 \| 66 \| (63, 133) \| 682 \| (2, 69) \| 246 \| Xpd/rad3 related DNA helicase \| CATHCODE: 3.40.50.300 NAME: Xpd/rad3 related DNA helicase. Chain: a. Ec: 3.-.-.-. Engineered: yes SOURCE: Sulfolobus acidocaldarius. Organism\_taxid: 2285. Gene: saci\_0192. Expressed in: escherichia coli. Expression\_system\_taxid: 562. Expression\_system\_vector\_type: plasmid. CLASS: Alpha Beta, ARCH: 3-Layer(aba) Sandwich, TOPOL: Rossmann fold, HOMOL: P-loop containing nucleotide triphosphate hydrolases; cath \| 4nl4H03 \| 95.5 \| 6.7e-05 \| 1.4e-08 \| 60.1 \| 62 \| (62, 127) \| 682 \| (13, 79) \| 196 \| Primosome assembly protein pria \| CATHCODE: 3.40.50.300 NAME: Primosome assembly protein pria. Chain: h. Engineered: yes SOURCE: Klebsiella pneumoniae subsp. Pneumoniae. Organism\_taxid: 272620. Strain: mgh 78578. Gene: kpn78578\_41850, kpn\_04230, pria. Expressed in: escherichia coli. Expression\_system\_taxid: 562. Expression\_system\_vector\_type: plasmid. CLASS: Alpha Beta, ARCH: 3-Layer(aba) Sandwich, TOPOL: Rossmann fold, HOMOL: P-loop containing nucleotide triphosphate hydrolases; cath \| 1m6nA01 \| 95.2 \| 0.00013 \| 2.5e-08 \| 66.4 \| 66 \| (62, 134) \| 682 \| (78, 144) \| 286 \| Preprotein translocase seca \| CATHCODE: 3.40.50.300 NAME: Preprotein translocase seca. Chain: a. Synonym: seca. Engineered: yes SOURCE: Bacillus subtilis. Organism\_taxid: 1423. Gene: div. Expressed in: escherichia coli bl21(de3). Expression\_system\_taxid: 469008. CLASS: Alpha Beta, ARCH: 3-Layer(aba) Sandwich, TOPOL: Rossmann fold, HOMOL: P-loop containing nucleotide triphosphate hydrolases; cath \| 2vl7A01 \| 95.2 \| 0.00013 \| 2.6e-08 \| 58.7 \| 60 \| (63, 127) \| 682 \| (6, 68) \| 179 \| Xpd \| CATHCODE: 3.40.50.300 NAME: Xpd. Chain: a. Synonym: uncharacterized protein st1307. Engineered: yes SOURCE: Sulfolobus tokodaii. Organism\_taxid: 111955. Strain: 7. Expressed in:escherichia coli. Expression\_system\_taxid: 562. Expression\_system\_vector\_type: plasmid. CLASS: Alpha Beta, ARCH: 3-Layer(aba) Sandwich, TOPOL: Rossmann fold, HOMOL: P-loop containing nucleotide triphosphate hydrolases; cath \| 3fhtB01 \| 95.1 \| 0.00016 \| 3.3e-08 \| 60.5 \| 63 \| (63, 127) \| 682 \| (46, 112) \| 235 \| Atp-dependent RNA helicase ddx19b \| CATHCODE: 3.40.50.300 NAME: Atp-dependent RNA helicase ddx19b. Chain: a, b. Fragment: helicase atp-binding domain, c-terminal domain, residues 68-479. Synonym: dead-boxhelicase 5, dbp5, dead box protein 19b, dead box RNA helicase dead5. Engineered: yes. Rna (5'-r(\*up\*up\*up\*up\*up\*up\*up\*up\*up\*u)-3'). Chain: c, d. Engineered: yes SOURCE: Homo sapiens. Human. Organism\_taxid: 9606. Gene: ddx19b (dbp5). Expressed in: escherichia coli. Expression\_system\_taxid: 511693. CLASS: Alpha Beta, ARCH: 3-Layer(aba) Sandwich, TOPOL: Rossmann fold, HOMOL: P-loop containing nucleotide triphosphate hydrolases; cath \| 1pjrA01 \| 95.0 \| 0.00017 \| 3.6e-08 \| 56.9 \| 61 \| (63, 127) \| 682 \| (8, 71) \| 202 \| Pcra \| CATHCODE: 3.40.50.300 NAME: Pcra. Chain: a. Engineered: yes SOURCE: Geobacillus stearothermophilus. Organism\_taxid: 1422. Strain: nca1503.Gene: pcra. Expressed in: escherichia coli. Expression\_system\_taxid:562. Expression\_system\_vector\_type: t7 promoter. CLASS: Alpha Beta, ARCH: 3-Layer(aba) Sandwich, TOPOL: Rossmann fold, HOMOL: P-loop containing nucleotide triphosphate hydrolases; cath \| 3i5xA01 \| 94.9 \| 0.0002 \| 4.2e-08 \| 60.3 \| 70 \| (64, 134) \| 682 \| (40, 115) \| 247 \| Atp-dependent RNA helicase mss116 \| CATHCODE: 3.40.50.300 NAME: Atp-dependent RNA helicase mss116. Chain: a. Fragment: unp residues 37to 597. Engineered: yes. 5'-r(\*up\*up\*up\*up\*up\*up\*up\*up\*up\*u)-3'. Chain: b. Engineered: yes SOURCE: Saccharomyces cerevisiae. Yeast. Organism\_taxid: 4932. Gene: mss116, yd9346.05c, ydr194c. Expressed in: escherichia coli. Expression\_system\_taxid: 562. CLASS: Alpha Beta, ARCH: 3-Layer(aba) Sandwich, TOPOL: Rossmann fold, HOMOL: P-loop containing nucleotide triphosphate hydrolases; cath \| 2v1xA01 \| 94.9 \| 0.0002 \| 4.2e-08 \| 59.2 \| 57 \| (64, 127) \| 682 \| (44, 101) \| 235 \| Atp-dependent DNA helicase q1 \| CATHCODE: 3.40.50.300 NAME: Atp-dependent DNA helicase q1. Chain: a, b. Fragment: residues 49-616.Synonym: DNA-dependent atpase q1, recq DNA helicase. Engineered: yes SOURCE: Homo sapiens. Human. Organism\_taxid: 9606. Expressed in: escherichia coli. Expression\_system\_taxid: 469008. CLASS: Alpha Beta, ARCH: 3-Layer(aba) Sandwich, TOPOL: Rossmann fold, HOMOL: P-loop containing nucleotide triphosphate hydrolases; cath \| 1a1vA01 \| 94.8 \| 0.00025 \| 4.9e-08 \| 56.0 \| 42 \| (81, 127) \| 682 \| (8, 50) \| 135 \| Protein (ns3 protein) \| CATHCODE: 3.40.50.300 NAME: Protein (ns3 protein). Chain: a. Fragment: helicase domain. Engineered: yes. Mutation: yes. Dna (5'-d(\*up\*up\*up\*up\*up\*up\*up\*u)-3'). Chain: b. Engineered: yes. Other\_details: single stranded DNA SOURCE: Hepatitis c virus (isolate h). Organism\_taxid: 11108. Strain: h. Gene:ns3. Expressed in: escherichia coli. Expression\_system\_taxid: 562. CLASS: Alpha Beta, ARCH: 3-Layer(aba) Sandwich, TOPOL: Rossmann fold, HOMOL: P-loop containing nucleotide triphosphate hydrolases; cath \| 3upuA01 \| 94.8 \| 0.00025 \| 5.2e-08 \| 56.0 \| 40 \| (182, 228) \| 682 \| (106, 145) \| 176 \| Atp-dependent DNA helicase dda \| CATHCODE: 3.40.50.300 NAME: Atp-dependent DNA helicase dda. Chain: a, b, c. Synonym: sf1b helicasedda. Engineered: yes. Mutation: yes. 5'-d(\*tp\*tp\*tp\*tp\*tp\*tp\*tp\*t)-3'. Chain: d, e, f. Engineered: yes SOURCE: Enterobacteria phage t4. Organism\_taxid: 10665. Gene: dda, sud. Expressed in: escherichia coli. Expression\_system\_taxid: 562. CLASS: Alpha Beta, ARCH: 3-Layer(aba) Sandwich, TOPOL: Rossmann fold, HOMOL: P-loop containing nucleotide triphosphate hydrolases; cath \| 3mwyW03 \| 94.3 \| 0.00049 \| 1e-07 \| 58.8 \| 65 \| (62, 127) \| 682 \| (32, 100) \| 282 \| Chromo domain-containing protein 1 \| CATHCODE: 3.40.50.10810 NAME: Chromo domain-containing protein 1. Chain: w. Fragment: double chromodomains and atpase motor (unp residues 142- 939). Synonym: atp-dependent helicase chd1. Engineered: yes SOURCE: Saccharomyces cerevisiae. Brewer's yeast,lager beer yeast,yeast. Organism\_taxid: 4932. Gene: chd1, sygp-orf4, yer164w. Expressed in: escherichia coli. Expression\_system\_taxid: 469008. CLASS: Alpha Beta, ARCH: 3-Layer(aba) Sandwich, TOPOL: Rossmann fold, HOMOL: Tandem AAA-ATPase domain; cath \| 2gk6A01 \| 94.0 \| 0.00072 \| 1.4e-07 \| 59.2 \| 62 \| (63, 127) \| 682 \| (89, 151) \| 267 \| Regulator of nonsense transcripts 1 \| CATHCODE: 3.40.50.300 NAME: Regulator of nonsense transcripts 1. Chain: a, b. Fragment: helicase core domain(residues 295-914). Synonym: atp-dependent helicase rent1, nonsense mRNA reducing factor 1, norf1, up-frameshift suppressor 1 homolog, hupf1. Engineered: yes SOURCE: Homo sapiens. Human. Organism\_taxid: 9606. Expressed in: escherichia coli. Expression\_system\_taxid: 562. CLASS: Alpha Beta, ARCH: 3-Layer(aba) Sandwich, TOPOL: Rossmann fold, HOMOL: P-loop containing nucleotide triphosphate hydrolases; cath \| 2xauA02 \| 93.8 \| 0.00082 \| 1.7e-07 \| 55.4 \| 60 \| (65, 126) \| 682 \| (57, 118) \| 232 \| Pre-mRNA-splicing factor atp-dependent RNA helicase prp43 \| CATHCODE: 3.40.50.300 NAME: Pre-mRNA-splicing factor atp-dependent RNA helicase prp43. Chain: a, b. Synonym: helicase ja1, deah-box RNA helicase prp43p. Engineered: yes SOURCE: Saccharomyces cerevisiae. Baker's yeast. Organism\_taxid: 4932. Expressed in: escherichia coli. Expression\_system\_taxid: 562. CLASS: Alpha Beta, ARCH: 3-Layer(aba) Sandwich, TOPOL: Rossmann fold, HOMOL: P-loop containing nucleotide triphosphate hydrolases; cath \| 1z63A01 \| 93.7 \| 0.0009 \| 1.9e-07 \| 51.7 \| 45 \| (82, 127) \| 682 \| (19, 64) \| 206 \| 5'-d(\*ap\*ap\*ap\*ap\*ap\*a\*ap\*tp\*tp\*gp\*cp\*cp\*gp\*ap\*ap\*gp\*ap\*cp\* gp\*ap\*ap\*ap\*ap\*ap\*a)-3' \| CATHCODE: 3.40.50.10810 NAME: 5'-d(\*ap\*ap\*ap\*ap\*ap\*a\*ap\*tp\*tp\*gp\*cp\*cp\*gp\*ap\*ap\*gp\*ap\*cp\* gp\*ap\*ap\*ap\*ap\*ap\*a)-3'. Chain: c, e. Engineered: yes. 5'-d(\*tp\*tp\*tp\*tp\*tp\*tp\*tp\*cp\*gp\*tp\*cp\*tp\*tp\*cp\*gp\*gp\*cp\*ap \*ap\*tp\*tp\*tp\*tp\*tp\*t)-3'. Chain: d,f. Engineered: yes. Helicase of the snf2/rad54 family. Chain: a, b. Fragment: residues 407-902. Engineered: yes SOURCE: Yes. Yes. CLASS: Alpha Beta, ARCH: 3-Layer(aba) Sandwich, TOPOL: Rossmann fold, HOMOL: Tandem AAA-ATPase domain; phrogs \| 15 \| 100.0 \| 1e-128 \| 1e-132 \| 1098.8 \| 574 \| (13, 619) \| 682 \| (2, 616) \| 641 \| terminase large subunit \| terminase large subunit; Category: head and packaging; p173497 VI\_06341; phrogs \| 675 \| 100.0 \| 1.3e-89 \| 1.7e-93 \| 765.8 \| 510 \| (31, 607) \| 682 \| (26, 568) \| 598 \| terminase large subunit \| terminase large subunit; Category: head and packaging; MF288921\_p63; phrogs \| 12876 \| 100.0 \| 3.7e-47 \| 4.2e-51 \| 359.0 \| 252 \| (125, 407) \| 682 \| (1, 262) \| 262 \| terminase large subunit \| terminase large subunit; Category: head and packaging; NC\_027374\_p36; phrogs \| 3028 \| 100.0 \| 3.9e-42 \| 4.5e-46 \| 326.0 \| 215 \| (396, 617) \| 682 \| (1, 222) \| 249 \| NA \| NA; Category: unknown function; p399924 VI\_07252; phrogs \| 15657 \| 100.0 \| 9.9e-42 \| 1.1e-45 \| 319.0 \| 210 \| (322, 537) \| 682 \| (23, 234) \| 234 \| NA \| NA; Category: unknown function; p353452 VI\_08078; phrogs \| 21311 \| 100.0 \| 4.8e-39 \| 5.5e-43 \| 293.7 \| 183 \| (127, 331) \| 682 \| (1, 194) \| 199 \| terminase large subunit \| terminase large subunit; Category: head and packaging; p353453 VI\_08078; phrogs \| 4145 \| 100.0 \| 5.3e-37 \| 6e-41 \| 282.2 \| 220 \| (18, 240) \| 682 \| (7, 230) \| 230 \| terminase large subunit \| terminase large subunit; Category: head and packaging; p354851 VI\_02330; phrogs \| 3925 \| 99.8 \| 1.2e-23 \| 1.4e-27 \| 177.3 \| 104 \| (21, 126) \| 682 \| (10, 115) \| 115 \| terminase large subunit \| terminase large subunit; Category: head and packaging; p10469 VI\_06522; phrogs \| 16406 \| 99.6 \| 5.2e-20 \| 5.8e-24 \| 156.0 \| 125 \| (382, 508) \| 682 \| (2, 126) \| 136 \| NA \| NA; Category: unknown function; p397021 VI\_02904; phrogs \| 23163 \| 99.5 \| 9e-19 \| 1e-22 \| 147.7 \| 125 \| (18, 142) \| 682 \| (3, 128) \| 130 \| NA \| NA; Category: unknown function; p282317 VI\_07250; phrogs \| 33134 \| 99.2 \| 1.2e-15 \| 1.3e-19 \| 127.1 \| 90 \| (278, 372) \| 682 \| (13, 122) \| 132 \| terminase large subunit \| terminase large subunit; Category: head and packaging; KR296689\_p12; phrogs \| 12089 \| 98.6 \| 5.3e-12 \| 5.9e-16 \| 110.0 \| 104 \| (506, 616) \| 682 \| (12, 122) \| 173 \| NA \| NA; Category: unknown function; p27927 VI\_03821; phrogs \| 7579 \| 97.7 \| 1.7e-08 \| 2e-12 \| 94.5 \| 147 \| (80, 243) \| 682 \| (33, 182) \| 246 \| terminase large subunit \| terminase large subunit; Category: head and packaging; JF937090\_p2; phrogs \| 9315 \| 97.4 \| 9e-08 \| 1e-11 \| 78.9 \| 69 \| (32, 109) \| 682 \| (27, 106) \| 111 \| terminase large subunit \| terminase large subunit; Category: head and packaging; NC\_027352\_p8; phrogs \| 5989 \| 97.2 \| 3.5e-07 \| 4.3e-11 \| 86.6 \| 144 \| (40, 194) \| 682 \| (67, 211) \| 238 \| terminase large subunit \| terminase large subunit; Category: head and packaging; NC\_018861\_p56; phrogs \| 20035 \| 96.7 \| 3.1e-06 \| 3.5e-10 \| 64.3 \| 71 \| (19, 89) \| 682 \| (2, 72) \| 76 \| NA \| NA; Category: unknown function; p329195 VI\_06416; phrogs \| 37509 \| 96.5 \| 6.9e-06 \| 7.7e-10 \| 70.7 \| 80 \| (528, 607) \| 682 \| (40, 121) \| 189 \| NA \| NA; Category: unknown function; p186239 VI\_04222; phrogs \| 2022 \| 95.3 \| 0.00018 \| 2e-08 \| 53.6 \| 38 \| (578, 616) \| 682 \| (3, 40) \| 68 \| NA \| NA; Category: unknown function; p413704 VI\_07200; phrogs \| 34048 \| 95.3 \| 0.0002 \| 2.2e-08 \| 50.1 \| 53 \| (123, 181) \| 682 \| (1, 53) \| 55 \| terminase large subunit \| terminase large subunit; Category: head and packaging; KX532239\_p29 |
| Top keywords  (threshold 1.00e-03 (evalue)) | **helicase, a, in, yes, RNA, and, Engineered, the, DNA, i** |
| Output files | ../../domain\_architecture/02\_FANPEZAQ\_CDS\_0002\_cath.hhr ../../domain\_architecture/02\_FANPEZAQ\_CDS\_0002\_merged.svg ../../domain\_architecture/02\_FANPEZAQ\_CDS\_0002\_ncbi-cd.hhr ../../domain\_architecture/02\_FANPEZAQ\_CDS\_0002\_pfam.hhr ../../domain\_architecture/02\_FANPEZAQ\_CDS\_0002\_phrogs.hhr |

### Identical protein sequences/structures

#### Search results

|  |  |
| --- | --- |
| Protein sequence databases searched | Pdb, Swissprot, Refseq |
| Identical proteins found | -- |
| Top keywords | -- |
| Output files | -- |

### Similar protein sequences/structures

#### Sequence similarity search results (HHblits)^1^

|  |  |
| --- | --- |
| Sequence databases searched | Uniclust, Pdb70 |
| Results, scheme(s)  (Top layers only, threshold 1.00e-03 (evalue)) | xml version="1.0" encoding="utf-8" standalone="no"?       2024-09-02T21:08:28.584782 image/svg+xml   Matplotlib v3.7.2, https://matplotlib.org/ |
| Results, table(s)  (threshold 1.00e-03 (evalue)) | db \| id \| prob \| evalue \| pvalue \| score \| cols \| query \| query\_len \| template \| template\_len \| name \| description; uniclust \| UniRef100\_A0A060H2N6 \| 100.0 \| 9e-146 \| 2e-151 \| 1132.9 \| 655 \| (5, 682) \| 682 \| (21, 676) \| 678 \| Terminase \| Terminase; uniclust \| UniRef100\_A0A021X9J1 \| 100.0 \| 1e-133 \| 3e-139 \| 1077.1 \| 561 \| (27, 608) \| 682 \| (28, 607) \| 707 \| Bacteriophage tail assembly protein \| Bacteriophage tail assembly protein; uniclust \| UniRef100\_A0A059ZLR8 \| 100.0 \| 5e-129 \| 1e-134 \| 1041.3 \| 558 \| (26, 608) \| 682 \| (37, 611) \| 764 \| Phage terminase, large subunit \| Phage terminase, large subunit; uniclust \| UniRef100\_A0A0A1VHF5 \| 100.0 \| 8e-128 \| 2e-133 \| 1013.0 \| 575 \| (19, 616) \| 682 \| (23, 618) \| 685 \| Bacteriophage tail assembly protein \| Bacteriophage tail assembly protein; uniclust \| UniRef100\_A0A024E947 \| 100.0 \| 1e-127 \| 3e-133 \| 1068.0 \| 566 \| (19, 607) \| 682 \| (90, 669) \| 828 \| Terminase, large subunit \| Terminase, large subunit; uniclust \| UniRef100\_A0A016XHY7 \| 100.0 \| 3e-127 \| 7e-133 \| 1076.8 \| 570 \| (22, 615) \| 682 \| (138, 714) \| 841 \| Terminase \| Terminase; uniclust \| UniRef100\_A0A011TXQ2 \| 100.0 \| 2e-126 \| 4e-132 \| 1006.6 \| 571 \| (22, 616) \| 682 \| (196, 796) \| 966 \| Phage tail protein \| Phage tail protein; uniclust \| UniRef100\_A0A061JGL3 \| 100.0 \| 4e-124 \| 7e-130 \| 976.6 \| 557 \| (23, 606) \| 682 \| (57, 624) \| 684 \| Large terminase protein \| Large terminase protein; uniclust \| UniRef100\_A0A011PCR3 \| 100.0 \| 3e-120 \| 6e-126 \| 938.9 \| 566 \| (17, 608) \| 682 \| (150, 744) \| 826 \| Bacteriophage tail assembly protein \| Bacteriophage tail assembly protein; uniclust \| UniRef100\_A0A096GRB5 \| 100.0 \| 6e-117 \| 1e-122 \| 945.8 \| 556 \| (25, 609) \| 682 \| (108, 681) \| 813 \| Terminase \| Terminase; uniclust \| UniRef100\_A0A081RFK0 \| 100.0 \| 1e-116 \| 3e-122 \| 939.4 \| 564 \| (20, 616) \| 682 \| (46, 616) \| 686 \| Phage terminase, large subunit \| Phage terminase, large subunit; uniclust \| UniRef100\_A0A0P1FAJ6 \| 100.0 \| 2e-112 \| 3e-118 \| 928.7 \| 491 \| (93, 608) \| 682 \| (492, 1001) \| 1064 \| Bacteriophage tail assembly protein \| Bacteriophage tail assembly protein; uniclust \| UniRef100\_A0A011UNI9 \| 100.0 \| 5e-110 \| 1e-115 \| 927.9 \| 559 \| (24, 608) \| 682 \| (132, 735) \| 823 \| Terminase GpA \| Terminase GpA; uniclust \| UniRef100\_A0A014Q8E3 \| 100.0 \| 2e-107 \| 3e-113 \| 883.3 \| 561 \| (19, 608) \| 682 \| (79, 678) \| 834 \| Terminase \| Terminase; uniclust \| UniRef100\_A0A074TCV6 \| 100.0 \| 1e-104 \| 3e-110 \| 841.3 \| 568 \| (26, 616) \| 682 \| (21, 606) \| 689 \| Terminase \| Terminase; uniclust \| UniRef100\_A0A1T4W516 \| 100.0 \| 4e-104 \| 8e-110 \| 796.1 \| 553 \| (30, 608) \| 682 \| (12, 584) \| 645 \| Phage terminase, large subunit GpA \| Phage terminase, large subunit GpA; uniclust \| UniRef100\_A0A031G5Z5 \| 100.0 \| 3e-102 \| 6e-108 \| 832.5 \| 586 \| (22, 618) \| 682 \| (61, 678) \| 801 \| Terminase GpA \| Terminase GpA; uniclust \| UniRef100\_A0A1F7S212 \| 100.0 \| 5.8e-99 \| 1e-104 \| 785.7 \| 572 \| (27, 616) \| 682 \| (31, 621) \| 657 \| Terminase \| Terminase; uniclust \| UniRef100\_A0A0H3ZME8 \| 100.0 \| 4.6e-98 \| 9e-104 \| 804.9 \| 556 \| (21, 608) \| 682 \| (125, 732) \| 799 \| Phage terminase large subunit GpA \| Phage terminase large subunit GpA; uniclust \| UniRef100\_A0A1I6AD23 \| 100.0 \| 7.9e-98 \| 2e-103 \| 742.2 \| 552 \| (28, 604) \| 682 \| (12, 570) \| 623 \| Phage terminase, large subunit GpA \| Phage terminase, large subunit GpA; uniclust \| UniRef100\_A0A071M5H9 \| 100.0 \| 3.1e-97 \| 6e-103 \| 784.0 \| 572 \| (5, 604) \| 682 \| (37, 621) \| 717 \| Terminase \| Terminase; uniclust \| UniRef100\_A0A0F2N5L2 \| 100.0 \| 3e-96 \| 7e-102 \| 813.1 \| 553 \| (25, 608) \| 682 \| (39, 624) \| 692 \| Terminase \| Terminase; uniclust \| UniRef100\_A0A2G6EYH6 \| 100.0 \| 4.9e-96 \| 1e-101 \| 747.5 \| 458 \| (23, 505) \| 682 \| (31, 496) \| 507 \| Terminase (Fragment) \| Terminase (Fragment); uniclust \| UniRef100\_A0A0F9R794 \| 100.0 \| 5.2e-96 \| 1e-101 \| 767.0 \| 544 \| (28, 608) \| 682 \| (67, 644) \| 686 \| Terminase large subunit gp17-like C-terminal domain-containing protein \| Terminase large subunit gp17-like C-terminal domain-containing protein; uniclust \| UniRef100\_A0A7J6YKX9 \| 100.0 \| 1.7e-95 \| 3e-101 \| 739.1 \| 555 \| (28, 608) \| 682 \| (6, 584) \| 991 \| Uncharacterized protein \| Uncharacterized protein; uniclust \| UniRef100\_UPI0005F41AB5 \| 100.0 \| 1.8e-94 \| 3e-100 \| 734.4 \| 555 \| (28, 608) \| 682 \| (383, 953) \| 1163 \| uncharacterized protein LOC105557561 \| uncharacterized protein LOC105557561; uniclust \| UniRef100\_A0A0F8ZCZ7 \| 100.0 \| 8.2e-92 \| 1.6e-97 \| 698.3 \| 463 \| (134, 616) \| 682 \| (1, 487) \| 534 \| Phage terminase large subunit N-terminal domain-containing protein (Fragment) \| Phage terminase large subunit N-terminal domain-containing protein (Fragment); uniclust \| UniRef100\_A0A0E2EB01 \| 100.0 \| 9.4e-92 \| 1.9e-97 \| 744.3 \| 562 \| (21, 607) \| 682 \| (22, 640) \| 689 \| Phage terminase large subunit \| Phage terminase large subunit; uniclust \| UniRef100\_A0A660NKM7 \| 100.0 \| 1.4e-91 \| 2.6e-97 \| 675.4 \| 493 \| (11, 515) \| 682 \| (5, 497) \| 501 \| Phage terminase large subunit family protein (Fragment) \| Phage terminase large subunit family protein (Fragment); uniclust \| UniRef100\_A0A2V6CXN5 \| 100.0 \| 2.2e-91 \| 4.4e-97 \| 712.8 \| 520 \| (28, 560) \| 682 \| (29, 571) \| 571 \| Terminase \| Terminase; uniclust \| UniRef100\_A0A257ERR4 \| 100.0 \| 4.3e-91 \| 8.3e-97 \| 701.8 \| 456 \| (28, 504) \| 682 \| (12, 479) \| 569 \| Phage tail protein (Fragment) \| Phage tail protein (Fragment); uniclust \| UniRef100\_A0A0S9PWZ7 \| 100.0 \| 7.3e-90 \| 1.4e-95 \| 713.2 \| 574 \| (12, 607) \| 682 \| (14, 612) \| 756 \| Terminase \| Terminase; uniclust \| UniRef100\_UPI00147320C4 \| 100.0 \| 2.3e-89 \| 4.3e-95 \| 684.4 \| 556 \| (28, 607) \| 682 \| (15, 594) \| 886 \| phage terminase large subunit family protein \| phage terminase large subunit family protein; uniclust \| UniRef100\_A0A060BGK3 \| 100.0 \| 3.5e-89 \| 7.5e-95 \| 758.2 \| 571 \| (13, 607) \| 682 \| (60, 702) \| 793 \| DNA packaging protein \| DNA packaging protein; uniclust \| UniRef100\_A0A0H2UY08 \| 100.0 \| 1.3e-88 \| 2.6e-94 \| 688.3 \| 545 \| (33, 608) \| 682 \| (22, 606) \| 649 \| Putative DNA packaging protein of prophage CP-933R terminase large subunit \| Putative DNA packaging protein of prophage CP-933R terminase large subunit; uniclust \| UniRef100\_A0A3G8M2W2 \| 100.0 \| 8.7e-88 \| 1.7e-93 \| 682.3 \| 553 \| (23, 607) \| 682 \| (18, 581) \| 616 \| Terminase \| Terminase; uniclust \| UniRef100\_A0A1A9VKE5 \| 100.0 \| 1.3e-87 \| 2.4e-93 \| 676.6 \| 517 \| (28, 569) \| 682 \| (434, 973) \| 974 \| ParB/Sulfiredoxin domain-containing protein \| ParB/Sulfiredoxin domain-containing protein; uniclust \| UniRef100\_A0A7J6YL28 \| 100.0 \| 1.6e-87 \| 2.9e-93 \| 692.1 \| 551 \| (32, 608) \| 682 \| (490, 1058) \| 1419 \| DUF1016 family protein \| DUF1016 family protein; uniclust \| UniRef100\_A0A1Y6CNV9 \| 100.0 \| 8.8e-87 \| 1.7e-92 \| 671.2 \| 564 \| (24, 616) \| 682 \| (15, 613) \| 621 \| Phage terminase, large subunit GpA \| Phage terminase, large subunit GpA; uniclust \| UniRef100\_A0A1X0SVE4 \| 100.0 \| 1.2e-86 \| 2.3e-92 \| 653.2 \| 420 \| (167, 608) \| 682 \| (10, 446) \| 492 \| Phage terminase GpA \| Phage terminase GpA; uniclust \| UniRef100\_A0A0A0YNL4 \| 100.0 \| 2.1e-84 \| 4.4e-90 \| 694.8 \| 546 \| (37, 604) \| 682 \| (32, 654) \| 684 \| Terminase, large subunit \| Terminase, large subunit; uniclust \| UniRef100\_A0A227JQ12 \| 100.0 \| 5.5e-84 \| 1e-89 \| 628.0 \| 453 \| (134, 609) \| 682 \| (1, 466) \| 629 \| Uncharacterized protein \| Uncharacterized protein; uniclust \| UniRef100\_A0A916FSY5 \| 100.0 \| 8.1e-83 \| 1.5e-88 \| 631.5 \| 554 \| (25, 613) \| 682 \| (6, 570) \| 779 \| Uncharacterized protein \| Uncharacterized protein; uniclust \| UniRef100\_A0A073IUG7 \| 100.0 \| 7.4e-83 \| 1.5e-88 \| 690.7 \| 565 \| (16, 607) \| 682 \| (38, 674) \| 734 \| Terminase \| Terminase; uniclust \| UniRef100\_A0A0A2V4M4 \| 100.0 \| 1.2e-81 \| 2.2e-87 \| 664.6 \| 572 \| (28, 616) \| 682 \| (434, 1068) \| 1696 \| Portal protein B \| Portal protein B; uniclust \| UniRef100\_A0A812JEV6 \| 100.0 \| 1.5e-81 \| 2.8e-87 \| 674.5 \| 525 \| (65, 615) \| 682 \| (376, 919) \| 4451 \| Phage portal protein \| Phage portal protein; uniclust \| UniRef100\_A0A061NN40 \| 100.0 \| 2e-81 \| 4e-87 \| 634.8 \| 379 \| (25, 429) \| 682 \| (21, 405) \| 456 \| Phage terminase, large subunit \| Phage terminase, large subunit; uniclust \| UniRef100\_A0A9D2GTG3 \| 100.0 \| 4.3e-81 \| 8.2e-87 \| 631.0 \| 556 \| (26, 616) \| 682 \| (31, 607) \| 648 \| Phage terminase large subunit family protein \| Phage terminase large subunit family protein; uniclust \| UniRef100\_A0A1M3KG66 \| 100.0 \| 2.1e-80 \| 3.8e-86 \| 623.9 \| 563 \| (29, 603) \| 682 \| (286, 901) \| 948 \| Terminase \| Terminase; uniclust \| UniRef100\_A0A061QCF2 \| 100.0 \| 3.6e-80 \| 7.2e-86 \| 652.2 \| 561 \| (20, 603) \| 682 \| (39, 647) \| 723 \| Large terminase protein \| Large terminase protein; uniclust \| UniRef100\_A0A1E3GXY7 \| 100.0 \| 6.4e-80 \| 1.3e-85 \| 609.4 \| 365 \| (24, 407) \| 682 \| (72, 445) \| 445 \| Phage terminase large subunit (GpA) \| Phage terminase large subunit (GpA); uniclust \| UniRef100\_A0A0D6PTK1 \| 100.0 \| 1.4e-79 \| 2.8e-85 \| 643.6 \| 563 \| (19, 607) \| 682 \| (25, 657) \| 745 \| Bacteriophage terminase large subunit \| Bacteriophage terminase large subunit; uniclust \| UniRef100\_A0A068SM63 \| 100.0 \| 1.7e-79 \| 3.5e-85 \| 647.7 \| 547 \| (32, 608) \| 682 \| (44, 625) \| 730 \| Terminase large subunit (Gp2) \| Terminase large subunit (Gp2); uniclust \| UniRef100\_A0A0A0FB79 \| 100.0 \| 1.7e-78 \| 3.2e-84 \| 611.2 \| 555 \| (17, 603) \| 682 \| (3, 573) \| 635 \| Terminase large subunit \| Terminase large subunit; uniclust \| UniRef100\_A0A953ZK54 \| 100.0 \| 2.2e-78 \| 4.1e-84 \| 605.0 \| 552 \| (25, 605) \| 682 \| (16, 605) \| 649 \| Phage terminase large subunit family protein \| Phage terminase large subunit family protein; uniclust \| UniRef100\_A0A1V1UMI0 \| 100.0 \| 2.8e-78 \| 5.5e-84 \| 618.9 \| 544 \| (28, 604) \| 682 \| (25, 615) \| 687 \| Phage terminase large subunit \| Phage terminase large subunit; uniclust \| UniRef100\_A0A1V0BGV6 \| 100.0 \| 4.5e-78 \| 8.5e-84 \| 629.1 \| 543 \| (25, 595) \| 682 \| (44, 631) \| 1084 \| Terminase \| Terminase; uniclust \| UniRef100\_UPI00034C077C \| 100.0 \| 1.6e-77 \| 3e-83 \| 596.7 \| 544 \| (32, 605) \| 682 \| (4, 588) \| 661 \| phage terminase large subunit family protein \| phage terminase large subunit family protein; uniclust \| UniRef100\_A0A2N3B688 \| 100.0 \| 1.7e-77 \| 3.2e-83 \| 612.4 \| 544 \| (24, 590) \| 682 \| (12, 626) \| 626 \| Terminase (Fragment) \| Terminase (Fragment); uniclust \| UniRef100\_A0A0S8GHT9 \| 100.0 \| 2e-77 \| 3.9e-83 \| 598.0 \| 547 \| (28, 606) \| 682 \| (18, 610) \| 656 \| Terminase \| Terminase; uniclust \| UniRef100\_A0A063ZY87 \| 100.0 \| 3.1e-76 \| 5.7e-82 \| 599.3 \| 547 \| (28, 604) \| 682 \| (14, 574) \| 1075 \| Terminase \| Terminase; uniclust \| UniRef100\_UPI001D18FAC7 \| 100.0 \| 6.8e-76 \| 1.3e-81 \| 572.9 \| 408 \| (80, 505) \| 682 \| (4, 472) \| 493 \| phage terminase large subunit family protein \| phage terminase large subunit family protein; uniclust \| UniRef100\_A0A1Y6CQE5 \| 100.0 \| 1.5e-75 \| 3e-81 \| 589.6 \| 402 \| (186, 616) \| 682 \| (2, 415) \| 494 \| Phage terminase large subunit (GpA) \| Phage terminase large subunit (GpA); uniclust \| UniRef100\_A0A0F9GJQ5 \| 100.0 \| 2.8e-75 \| 5.2e-81 \| 564.6 \| 415 \| (27, 462) \| 682 \| (14, 447) \| 449 \| Uncharacterized protein (Fragment) \| Uncharacterized protein (Fragment); uniclust \| UniRef100\_A0A257VFA9 \| 100.0 \| 3.1e-75 \| 5.8e-81 \| 556.1 \| 373 \| (20, 410) \| 682 \| (14, 394) \| 394 \| Phage tail protein (Fragment) \| Phage tail protein (Fragment); uniclust \| UniRef100\_A0A1V6JK58 \| 100.0 \| 5.9e-75 \| 1.1e-80 \| 590.7 \| 537 \| (26, 605) \| 682 \| (15, 605) \| 623 \| Phage terminase large subunit (GpA) \| Phage terminase large subunit (GpA); uniclust \| UniRef100\_H1D5G9 \| 100.0 \| 2.4e-74 \| 4.5e-80 \| 553.9 \| 414 \| (178, 613) \| 682 \| (2, 426) \| 455 \| Terminase \| Terminase; uniclust \| UniRef100\_A0A091AQ85 \| 100.0 \| 5.3e-74 \| 1e-79 \| 581.0 \| 539 \| (27, 603) \| 682 \| (22, 625) \| 697 \| Terminase \| Terminase; uniclust \| UniRef100\_A0A0Q4Z1A9 \| 100.0 \| 1.6e-73 \| 3.2e-79 \| 615.1 \| 550 \| (26, 604) \| 682 \| (42, 651) \| 797 \| Terminase \| Terminase; uniclust \| UniRef100\_UPI001F05D527 \| 100.0 \| 2.4e-73 \| 4.4e-79 \| 538.9 \| 428 \| (28, 470) \| 682 \| (9, 449) \| 470 \| phage terminase large subunit family protein \| phage terminase large subunit family protein; uniclust \| UniRef100\_A0A1V6HQW6 \| 100.0 \| 3.7e-73 \| 7.6e-79 \| 599.7 \| 542 \| (26, 607) \| 682 \| (22, 605) \| 640 \| Phage terminase large subunit (GpA) \| Phage terminase large subunit (GpA); uniclust \| UniRef100\_A0A3S3RLY6 \| 100.0 \| 9.8e-73 \| 1.9e-78 \| 562.2 \| 459 \| (28, 509) \| 682 \| (68, 540) \| 560 \| Terminase (Fragment) \| Terminase (Fragment); uniclust \| UniRef100\_A0A166JZ20 \| 100.0 \| 2.1e-72 \| 4e-78 \| 567.7 \| 538 \| (27, 608) \| 682 \| (12, 599) \| 639 \| Terminase \| Terminase; uniclust \| UniRef100\_UPI00186BB8CC \| 100.0 \| 4e-72 \| 7.4e-78 \| 563.3 \| 488 \| (39, 553) \| 682 \| (6, 519) \| 917 \| phage terminase large subunit family protein \| phage terminase large subunit family protein; uniclust \| UniRef100\_A0A1Q6UMF3 \| 100.0 \| 8.3e-72 \| 1.7e-77 \| 592.6 \| 534 \| (24, 606) \| 682 \| (32, 606) \| 635 \| Terminase \| Terminase; uniclust \| UniRef100\_A0A0F9BAB4 \| 100.0 \| 1.3e-71 \| 2.5e-77 \| 539.5 \| 338 \| (246, 608) \| 682 \| (1, 351) \| 399 \| Terminase large subunit GpA endonuclease domain-containing protein (Fragment) \| Terminase large subunit GpA endonuclease domain-containing protein (Fragment); uniclust \| UniRef100\_A0A0F8X804 \| 100.0 \| 7.3e-71 \| 1.4e-76 \| 530.1 \| 348 \| (105, 472) \| 682 \| (11, 367) \| 373 \| Phage terminase large subunit (GpA) (Fragment) \| Phage terminase large subunit (GpA) (Fragment); uniclust \| UniRef100\_A0A059KU81 \| 100.0 \| 2.8e-70 \| 5.2e-76 \| 545.3 \| 569 \| (22, 615) \| 682 \| (11, 623) \| 681 \| Terminase \| Terminase; uniclust \| UniRef100\_A0A2H0PF84 \| 100.0 \| 5.2e-70 \| 9.8e-76 \| 504.8 \| 333 \| (152, 506) \| 682 \| (2, 337) \| 343 \| Terminase (Fragment) \| Terminase (Fragment); uniclust \| UniRef100\_A0A835YX25 \| 100.0 \| 1.3e-69 \| 2.4e-75 \| 547.2 \| 525 \| (57, 616) \| 682 \| (393, 937) \| 962 \| Phage terminase large subunit-domain-containing protein \| Phage terminase large subunit-domain-containing protein; uniclust \| UniRef100\_Q1QI78 \| 100.0 \| 5.9e-69 \| 1.1e-74 \| 550.1 \| 535 \| (34, 602) \| 682 \| (2, 552) \| 1179 \| Phage terminase GpA \| Phage terminase GpA; uniclust \| UniRef100\_A0A285D3Y8 \| 100.0 \| 7.2e-69 \| 1.3e-74 \| 519.9 \| 426 \| (157, 604) \| 682 \| (59, 505) \| 543 \| Transposase IS166 family protein \| Transposase IS166 family protein; uniclust \| UniRef100\_A0A1G1LJQ7 \| 100.0 \| 1.1e-68 \| 2.2e-74 \| 533.8 \| 413 \| (167, 602) \| 682 \| (3, 438) \| 492 \| Terminase (Fragment) \| Terminase (Fragment); uniclust \| UniRef100\_A0A3C1F771 \| 100.0 \| 2.5e-68 \| 5e-74 \| 547.7 \| 453 \| (26, 498) \| 682 \| (54, 536) \| 554 \| Terminase \| Terminase; uniclust \| UniRef100\_A0A090VRU1 \| 100.0 \| 6e-68 \| 1.1e-73 \| 530.6 \| 496 \| (79, 604) \| 682 \| (7, 521) \| 580 \| Putative terminase large subunit \| Putative terminase large subunit; uniclust \| UniRef100\_A0A0P6WGK7 \| 100.0 \| 6.5e-68 \| 1.3e-73 \| 527.3 \| 422 \| (165, 608) \| 682 \| (5, 450) \| 526 \| Terminase large subunit GpA endonuclease domain-containing protein \| Terminase large subunit GpA endonuclease domain-containing protein; uniclust \| UniRef100\_A0A0F9JWS4 \| 100.0 \| 8.1e-68 \| 1.7e-73 \| 572.9 \| 572 \| (22, 608) \| 682 \| (59, 680) \| 726 \| Phage terminase large subunit (GpA) \| Phage terminase large subunit (GpA); uniclust \| UniRef100\_A0A376PVS1 \| 100.0 \| 1.9e-67 \| 3.6e-73 \| 518.9 \| 449 \| (24, 498) \| 682 \| (13, 493) \| 535 \| Terminase large subunit (Gp2) \| Terminase large subunit (Gp2); uniclust \| UniRef100\_A0A2K3TKA3 \| 100.0 \| 2e-67 \| 3.8e-73 \| 515.0 \| 366 \| (216, 609) \| 682 \| (11, 411) \| 453 \| DNA packaging protein \| DNA packaging protein; uniclust \| UniRef100\_A0A2N9AI26 \| 100.0 \| 2.3e-67 \| 4.3e-73 \| 540.9 \| 548 \| (42, 607) \| 682 \| (2, 598) \| 1265 \| Terminase GpA (Modular protein) \| Terminase GpA (Modular protein); uniclust \| UniRef100\_A0A2N1TNL2 \| 100.0 \| 5.5e-67 \| 1e-72 \| 524.5 \| 564 \| (35, 608) \| 682 \| (34, 635) \| 670 \| Terminase \| Terminase; uniclust \| UniRef100\_UPI001184B44D \| 100.0 \| 1.2e-66 \| 2.1e-72 \| 491.8 \| 416 \| (179, 616) \| 682 \| (2, 419) \| 451 \| phage terminase large subunit family protein \| phage terminase large subunit family protein; uniclust \| UniRef100\_A0A1G8VFE7 \| 100.0 \| 1.2e-66 \| 2.3e-72 \| 512.9 \| 454 \| (27, 502) \| 682 \| (13, 510) \| 514 \| Phage terminase large subunit (GpA) (Fragment) \| Phage terminase large subunit (GpA) (Fragment); uniclust \| UniRef100\_A0A1H8P223 \| 100.0 \| 1.3e-66 \| 2.4e-72 \| 505.2 \| 427 \| (155, 616) \| 682 \| (56, 488) \| 508 \| Phage terminase, large subunit GpA \| Phage terminase, large subunit GpA; uniclust \| UniRef100\_A0A0D0Q7Y7 \| 100.0 \| 2.6e-66 \| 4.9e-72 \| 507.4 \| 549 \| (24, 609) \| 682 \| (9, 571) \| 597 \| Bacteriophage tail assembly protein \| Bacteriophage tail assembly protein; uniclust \| UniRef100\_UPI000753E5A0 \| 100.0 \| 2.9e-66 \| 5.5e-72 \| 509.2 \| 378 \| (218, 607) \| 682 \| (1, 399) \| 483 \| phage terminase large subunit family protein \| phage terminase large subunit family protein; uniclust \| UniRef100\_UPI000F7E4571 \| 100.0 \| 3.9e-66 \| 7.1e-72 \| 535.2 \| 537 \| (36, 603) \| 682 \| (25, 601) \| 1407 \| phage terminase large subunit family protein \| phage terminase large subunit family protein; uniclust \| UniRef100\_A0A1L8CRQ1 \| 100.0 \| 5.2e-66 \| 1e-71 \| 497.4 \| 294 \| (308, 613) \| 682 \| (6, 309) \| 346 \| Terminase \| Terminase; uniclust \| UniRef100\_UPI002041E7F5 \| 100.0 \| 8.8e-66 \| 1.6e-71 \| 523.2 \| 558 \| (27, 608) \| 682 \| (6, 595) \| 1052 \| phage portal protein \| phage portal protein; uniclust \| UniRef100\_A0A0F9MFA2 \| 100.0 \| 9.2e-66 \| 1.9e-71 \| 532.2 \| 507 \| (28, 604) \| 682 \| (14, 535) \| 552 \| Terminase \| Terminase; uniclust \| UniRef100\_A0A0F7LA42 \| 100.0 \| 1.6e-65 \| 3.2e-71 \| 528.3 \| 552 \| (27, 614) \| 682 \| (29, 634) \| 670 \| Phage terminase GpA \| Phage terminase GpA; uniclust \| UniRef100\_A0A4Q8X289 \| 100.0 \| 3.7e-65 \| 7.1e-71 \| 505.6 \| 457 \| (131, 613) \| 682 \| (35, 502) \| 554 \| Terminase (Fragment) \| Terminase (Fragment); uniclust \| UniRef100\_A0A949KD76 \| 100.0 \| 4.3e-65 \| 8e-71 \| 511.1 \| 576 \| (25, 620) \| 682 \| (24, 635) \| 659 \| Phage terminase large subunit family protein \| Phage terminase large subunit family protein; uniclust \| UniRef100\_A0A0Q4WDP7 \| 100.0 \| 5.2e-65 \| 9.6e-71 \| 514.5 \| 569 \| (16, 607) \| 682 \| (7, 605) \| 965 \| Uncharacterized protein \| Uncharacterized protein; uniclust \| UniRef100\_A0A2A2W7E6 \| 100.0 \| 9.2e-65 \| 1.8e-70 \| 523.1 \| 554 \| (27, 613) \| 682 \| (43, 642) \| 694 \| Terminase \| Terminase; uniclust \| UniRef100\_A0A6J4G269 \| 100.0 \| 1.4e-64 \| 2.7e-70 \| 482.5 \| 320 \| (198, 540) \| 682 \| (10, 344) \| 357 \| Bacteriophage tail assembly protein \| Bacteriophage tail assembly protein; uniclust \| UniRef100\_A0A518B2T0 \| 100.0 \| 1.6e-64 \| 3e-70 \| 490.8 \| 521 \| (55, 603) \| 682 \| (8, 565) \| 593 \| Phage terminase large subunit (GpA) \| Phage terminase large subunit (GpA); uniclust \| UniRef100\_A0A7C3C8G1 \| 100.0 \| 6.5e-64 \| 1.2e-69 \| 467.6 \| 308 \| (25, 352) \| 682 \| (28, 342) \| 343 \| Phage terminase large subunit family protein (Fragment) \| Phage terminase large subunit family protein (Fragment); uniclust \| UniRef100\_A0A0F9QMA2 \| 100.0 \| 1.2e-63 \| 2.6e-69 \| 531.0 \| 498 \| (33, 603) \| 682 \| (34, 548) \| 579 \| Phage terminase large subunit GpA ATPase domain-containing protein \| Phage terminase large subunit GpA ATPase domain-containing protein; uniclust \| UniRef100\_A0A1G8I030 \| 100.0 \| 1.5e-63 \| 2.8e-69 \| 497.1 \| 474 \| (114, 608) \| 682 \| (1, 516) \| 560 \| Phage terminase, large subunit GpA (Fragment) \| Phage terminase, large subunit GpA (Fragment); uniclust \| UniRef100\_A0A1X3LWX1 \| 100.0 \| 2.7e-63 \| 4.9e-69 \| 484.3 \| 498 \| (26, 553) \| 682 \| (26, 560) \| 570 \| Phage terminase large subunit (GpA) \| Phage terminase large subunit (GpA); uniclust \| UniRef100\_A0A1B7HMK9 \| 100.0 \| 4.1e-63 \| 8e-69 \| 504.4 \| 425 \| (160, 606) \| 682 \| (23, 519) \| 581 \| Phage terminase large subunit \| Phage terminase large subunit; uniclust \| UniRef100\_A0A1Y1R385 \| 100.0 \| 4.1e-63 \| 8.1e-69 \| 485.0 \| 293 \| (317, 616) \| 682 \| (17, 317) \| 356 \| Terminase large subunit GpA endonuclease domain-containing protein (Fragment) \| Terminase large subunit GpA endonuclease domain-containing protein (Fragment); uniclust \| UniRef100\_A0A371J7D6 \| 100.0 \| 6.4e-63 \| 1.2e-68 \| 472.0 \| 443 \| (155, 616) \| 682 \| (3, 471) \| 502 \| Phage terminase large subunit family protein (Fragment) \| Phage terminase large subunit family protein (Fragment); uniclust \| UniRef100\_A0A0P6X8Q6 \| 100.0 \| 1.2e-62 \| 2.4e-68 \| 522.4 \| 564 \| (26, 608) \| 682 \| (45, 677) \| 737 \| Terminase \| Terminase; uniclust \| UniRef100\_A0A1H5ZFL4 \| 100.0 \| 2.6e-62 \| 5e-68 \| 505.6 \| 332 \| (94, 444) \| 682 \| (452, 791) \| 792 \| Intein C-terminal splicing region (Fragment) \| Intein C-terminal splicing region (Fragment); uniclust \| UniRef100\_A0A0K1TMK8 \| 100.0 \| 3e-62 \| 5.8e-68 \| 459.6 \| 300 \| (84, 403) \| 682 \| (1, 307) \| 312 \| Large terminase subunit (Fragment) \| Large terminase subunit (Fragment); uniclust \| UniRef100\_A0A0B1U2X3 \| 100.0 \| 2.9e-62 \| 6.2e-68 \| 526.0 \| 478 \| (41, 607) \| 682 \| (40, 528) \| 573 \| Phage terminase, large subunit \| Phage terminase, large subunit; uniclust \| UniRef100\_A0A062VMA9 \| 100.0 \| 7.6e-62 \| 1.5e-67 \| 473.4 \| 257 \| (342, 604) \| 682 \| (2, 262) \| 310 \| Phage terminase GpA \| Phage terminase GpA; uniclust \| UniRef100\_C1A9W4 \| 100.0 \| 1.3e-61 \| 2.5e-67 \| 476.8 \| 569 \| (31, 608) \| 682 \| (25, 618) \| 683 \| Phage terminase GpA family protein \| Phage terminase GpA family protein; uniclust \| UniRef100\_A0A2W8GFJ1 \| 100.0 \| 2e-61 \| 3.8e-67 \| 471.0 \| 429 \| (26, 480) \| 682 \| (29, 489) \| 490 \| Phage terminase large subunit family protein (Fragment) \| Phage terminase large subunit family protein (Fragment); uniclust \| UniRef100\_UPI000D1F08C5 \| 100.0 \| 5.3e-61 \| 1e-66 \| 485.4 \| 546 \| (31, 608) \| 682 \| (37, 637) \| 740 \| phage terminase large subunit family protein \| phage terminase large subunit family protein; uniclust \| UniRef100\_A0A0F9GKY5 \| 100.0 \| 5.6e-61 \| 1.1e-66 \| 472.5 \| 435 \| (38, 487) \| 682 \| (51, 514) \| 528 \| Terminase large subunit gp17-like C-terminal domain-containing protein (Fragment) \| Terminase large subunit gp17-like C-terminal domain-containing protein (Fragment); uniclust \| UniRef100\_A0A431I9D4 \| 100.0 \| 1.5e-59 \| 2.9e-65 \| 476.9 \| 553 \| (27, 604) \| 682 \| (45, 644) \| 696 \| Terminase \| Terminase; uniclust \| UniRef100\_A0A0F2NJJ8 \| 100.0 \| 1.7e-59 \| 3.3e-65 \| 472.0 \| 542 \| (27, 608) \| 682 \| (52, 624) \| 647 \| Terminase \| Terminase; uniclust \| UniRef100\_A0A968KUZ0 \| 100.0 \| 3e-59 \| 5.6e-65 \| 465.8 \| 541 \| (36, 604) \| 682 \| (34, 635) \| 652 \| Uncharacterized protein \| Uncharacterized protein; uniclust \| UniRef100\_A0A4Q8X3C7 \| 100.0 \| 3.7e-59 \| 7.3e-65 \| 448.8 \| 336 \| (16, 365) \| 682 \| (4, 341) \| 345 \| Terminase (Fragment) \| Terminase (Fragment); uniclust \| UniRef100\_A0A7X7H546 \| 100.0 \| 1e-58 \| 1.9e-64 \| 454.7 \| 561 \| (26, 608) \| 682 \| (22, 619) \| 649 \| Phage terminase large subunit family protein \| Phage terminase large subunit family protein; uniclust \| UniRef100\_A0A5C7M1A7 \| 100.0 \| 1.9e-58 \| 3.6e-64 \| 488.9 \| 502 \| (85, 607) \| 682 \| (431, 1005) \| 1063 \| DOD-type homing endonuclease domain-containing protein \| DOD-type homing endonuclease domain-containing protein; uniclust \| UniRef100\_A0A376J619 \| 100.0 \| 1e-57 \| 1.9e-63 \| 443.1 \| 409 \| (159, 596) \| 682 \| (3, 449) \| 452 \| Terminase large subunit (Gp2) \| Terminase large subunit (Gp2); uniclust \| UniRef100\_A0A0Q0G0Q0 \| 100.0 \| 1.7e-57 \| 3.3e-63 \| 438.5 \| 356 \| (26, 408) \| 682 \| (13, 387) \| 388 \| Terminase, large subunit \| Terminase, large subunit; uniclust \| UniRef100\_UPI0022710B4D \| 100.0 \| 2.2e-57 \| 4.1e-63 \| 466.9 \| 559 \| (27, 608) \| 682 \| (13, 607) \| 1208 \| phage terminase large subunit family protein \| phage terminase large subunit family protein; uniclust \| UniRef100\_A0A2N2T364 \| 100.0 \| 3.7e-57 \| 6.9e-63 \| 426.7 \| 411 \| (139, 562) \| 682 \| (2, 437) \| 437 \| Terminase (Fragment) \| Terminase (Fragment); uniclust \| UniRef100\_A0A062V8V4 \| 100.0 \| 6.4e-57 \| 1.2e-62 \| 431.4 \| 323 \| (25, 372) \| 682 \| (8, 335) \| 361 \| Terminase GpA (Fragment) \| Terminase GpA (Fragment); uniclust \| UniRef100\_A0A060H3Y1 \| 100.0 \| 7.6e-57 \| 1.4e-62 \| 448.1 \| 536 \| (28, 601) \| 682 \| (21, 621) \| 699 \| Terminase \| Terminase; uniclust \| UniRef100\_A0A1G0RDU7 \| 100.0 \| 2.3e-56 \| 4.3e-62 \| 439.1 \| 573 \| (22, 616) \| 682 \| (22, 631) \| 662 \| Terminase \| Terminase; uniclust \| UniRef100\_A0A3D0SUB6 \| 100.0 \| 2.2e-56 \| 4.3e-62 \| 455.6 \| 408 \| (178, 605) \| 682 \| (2, 480) \| 539 \| Terminase large subunit GpA endonuclease domain-containing protein \| Terminase large subunit GpA endonuclease domain-containing protein; uniclust \| UniRef100\_A0A7R7A6L2 \| 100.0 \| 5.1e-56 \| 9.7e-62 \| 450.2 \| 554 \| (34, 608) \| 682 \| (15, 615) \| 642 \| Phage terminase large subunit \| Phage terminase large subunit; uniclust \| UniRef100\_A0A0E1M1W5 \| 100.0 \| 8.4e-56 \| 1.6e-61 \| 441.3 \| 559 \| (21, 603) \| 682 \| (10, 600) \| 602 \| Phage terminase large subunit (GpA) \| Phage terminase large subunit (GpA); uniclust \| UniRef100\_A0A8J7H8R6 \| 100.0 \| 9.4e-56 \| 1.8e-61 \| 414.9 \| 301 \| (24, 344) \| 682 \| (14, 320) \| 320 \| Phage terminase large subunit family protein (Fragment) \| Phage terminase large subunit family protein (Fragment); uniclust \| UniRef100\_A0A9E6BMG2 \| 100.0 \| 1.1e-55 \| 2e-61 \| 407.7 \| 351 \| (160, 540) \| 682 \| (3, 358) \| 359 \| Uncharacterized protein \| Uncharacterized protein; uniclust \| UniRef100\_A0A353ZI82 \| 100.0 \| 2.2e-55 \| 4.1e-61 \| 441.4 \| 548 \| (27, 604) \| 682 \| (14, 619) \| 653 \| Terminase \| Terminase; uniclust \| UniRef100\_UPI001CC6C82B \| 100.0 \| 7e-55 \| 1.3e-60 \| 428.4 \| 442 \| (26, 485) \| 682 \| (18, 495) \| 528 \| phage terminase large subunit family protein \| phage terminase large subunit family protein; uniclust \| UniRef100\_A0A894KFS4 \| 100.0 \| 1.1e-54 \| 2.1e-60 \| 422.9 \| 336 \| (25, 373) \| 682 \| (32, 377) \| 418 \| Terminase large subunit (Fragment) \| Terminase large subunit (Fragment); uniclust \| UniRef100\_UPI000A7FB1A1 \| 100.0 \| 1.2e-54 \| 2.1e-60 \| 410.1 \| 300 \| (62, 380) \| 682 \| (83, 390) \| 437 \| phage terminase large subunit family protein \| phage terminase large subunit family protein; uniclust \| UniRef100\_A0A1V6IVJ1 \| 100.0 \| 1.5e-54 \| 2.8e-60 \| 405.5 \| 298 \| (290, 609) \| 682 \| (1, 307) \| 350 \| Phage terminase large subunit (GpA) \| Phage terminase large subunit (GpA); uniclust \| UniRef100\_A0A2N3C6B8 \| 100.0 \| 1.7e-54 \| 3.3e-60 \| 418.3 \| 278 \| (326, 608) \| 682 \| (10, 309) \| 368 \| Terminase (Fragment) \| Terminase (Fragment); uniclust \| UniRef100\_A0A0Q6UXK2 \| 100.0 \| 2.4e-54 \| 4.3e-60 \| 425.0 \| 547 \| (25, 604) \| 682 \| (31, 610) \| 659 \| Terminase \| Terminase; uniclust \| UniRef100\_UPI00037AC9E0 \| 100.0 \| 8.4e-54 \| 1.5e-59 \| 423.3 \| 570 \| (26, 606) \| 682 \| (32, 648) \| 698 \| phage terminase large subunit family protein \| phage terminase large subunit family protein; uniclust \| UniRef100\_A0A0F8WXC1 \| 100.0 \| 7.9e-54 \| 1.6e-59 \| 408.5 \| 241 \| (26, 266) \| 682 \| (17, 261) \| 266 \| Phage terminase large subunit GpA ATPase domain-containing protein (Fragment) \| Phage terminase large subunit GpA ATPase domain-containing protein (Fragment); uniclust \| UniRef100\_A0A1B7L553 \| 100.0 \| 9.7e-54 \| 1.9e-59 \| 416.0 \| 401 \| (15, 431) \| 682 \| (4, 410) \| 411 \| Phage terminase large subunit GpA ATPase domain-containing protein \| Phage terminase large subunit GpA ATPase domain-containing protein; uniclust \| UniRef100\_A0A2I6PHS9 \| 100.0 \| 1.2e-53 \| 2.6e-59 \| 455.5 \| 523 \| (25, 606) \| 682 \| (22, 556) \| 599 \| Terminase large subunit \| Terminase large subunit; uniclust \| UniRef100\_A0A1A6FHD2 \| 100.0 \| 1.6e-53 \| 2.9e-59 \| 391.5 \| 274 \| (204, 498) \| 682 \| (2, 278) \| 343 \| Terminase \| Terminase; uniclust \| UniRef100\_A0A813BE77 \| 100.0 \| 2.4e-53 \| 4.4e-59 \| 457.2 \| 578 \| (18, 614) \| 682 \| (2650, 3275) \| 3302 \| PKD domain-containing protein \| PKD domain-containing protein; uniclust \| UniRef100\_A0A0E3VUL0 \| 100.0 \| 2.8e-53 \| 5.5e-59 \| 416.1 \| 375 \| (25, 414) \| 682 \| (8, 424) \| 424 \| Phage terminase large subunit GpA ATPase domain-containing protein \| Phage terminase large subunit GpA ATPase domain-containing protein; uniclust \| UniRef100\_A0A0F9S3N8 \| 100.0 \| 2.7e-53 \| 5.6e-59 \| 450.5 \| 475 \| (39, 603) \| 682 \| (45, 525) \| 583 \| Phage terminase large subunit GpA ATPase domain-containing protein \| Phage terminase large subunit GpA ATPase domain-containing protein; uniclust \| UniRef100\_A0A0F9FSR4 \| 100.0 \| 6.1e-53 \| 1.1e-58 \| 398.8 \| 318 \| (100, 437) \| 682 \| (101, 434) \| 438 \| Uncharacterized protein (Fragment) \| Uncharacterized protein (Fragment); uniclust \| UniRef100\_UPI001F53B0E2 \| 100.0 \| 6.4e-53 \| 1.2e-58 \| 426.7 \| 373 \| (29, 421) \| 682 \| (12, 394) \| 924 \| phage terminase large subunit family protein \| phage terminase large subunit family protein; uniclust \| UniRef100\_A0A7V3JJ92 \| 100.0 \| 6.7e-53 \| 1.2e-58 \| 380.9 \| 260 \| (73, 352) \| 682 \| (1, 262) \| 267 \| Phage terminase large subunit family protein (Fragment) \| Phage terminase large subunit family protein (Fragment); uniclust \| UniRef100\_A0A349EMJ1 \| 100.0 \| 1e-52 \| 2e-58 \| 417.4 \| 420 \| (162, 608) \| 682 \| (11, 449) \| 479 \| Terminase (Fragment) \| Terminase (Fragment); uniclust \| UniRef100\_A0A7X7KP84 \| 100.0 \| 1.3e-52 \| 2.4e-58 \| 410.8 \| 555 \| (23, 607) \| 682 \| (14, 598) \| 622 \| Terminase \| Terminase; uniclust \| UniRef100\_A0A890UZR5 \| 100.0 \| 1.5e-52 \| 2.9e-58 \| 386.4 \| 240 \| (73, 332) \| 682 \| (1, 244) \| 252 \| Terminase large subunit (Fragment) \| Terminase large subunit (Fragment); uniclust \| UniRef100\_A0A5C7P718 \| 100.0 \| 5.9e-52 \| 1.1e-57 \| 408.9 \| 551 \| (31, 604) \| 682 \| (14, 629) \| 668 \| Terminase \| Terminase; uniclust \| UniRef100\_A0A376U8Q0 \| 100.0 \| 1e-51 \| 1.9e-57 \| 393.7 \| 383 \| (32, 438) \| 682 \| (21, 435) \| 442 \| Terminase large subunit of prophage CP-933O \| Terminase large subunit of prophage CP-933O; uniclust \| UniRef100\_A0A4U9DA01 \| 100.0 \| 1.4e-51 \| 2.7e-57 \| 400.4 \| 386 \| (24, 419) \| 682 \| (21, 460) \| 478 \| Bacteriophage tail assembly protein \| Bacteriophage tail assembly protein; uniclust \| UniRef100\_UPI002074EEA0 \| 100.0 \| 3e-51 \| 5.5e-57 \| 391.6 \| 421 \| (42, 485) \| 682 \| (2, 453) \| 483 \| phage terminase large subunit family protein \| phage terminase large subunit family protein; uniclust \| UniRef100\_A0A7V2AL58 \| 100.0 \| 3.2e-51 \| 6e-57 \| 377.3 \| 273 \| (128, 412) \| 682 \| (7, 284) \| 349 \| Phage portal protein (Fragment) \| Phage portal protein (Fragment); uniclust \| UniRef100\_A0A0F9TJX8 \| 100.0 \| 4.8e-51 \| 1e-56 \| 441.3 \| 489 \| (35, 609) \| 682 \| (63, 576) \| 614 \| Phage terminase large subunit GpA ATPase domain-containing protein \| Phage terminase large subunit GpA ATPase domain-containing protein; uniclust \| UniRef100\_A0A1V5QJ79 \| 100.0 \| 6.6e-51 \| 1.4e-56 \| 393.9 \| 217 \| (390, 613) \| 682 \| (9, 226) \| 261 \| Phage terminase large subunit (GpA) \| Phage terminase large subunit (GpA); uniclust \| UniRef100\_A0A9E7APL5 \| 100.0 \| 1.5e-50 \| 2.7e-56 \| 387.6 \| 422 \| (26, 472) \| 682 \| (18, 482) \| 492 \| Terminase (Fragment) \| Terminase (Fragment); uniclust \| UniRef100\_UPI000FE04CD5 \| 100.0 \| 1.8e-50 \| 3.4e-56 \| 380.6 \| 388 \| (91, 497) \| 682 \| (1, 418) \| 421 \| phage terminase large subunit family protein \| phage terminase large subunit family protein; uniclust \| UniRef100\_A0A6I1JTB6 \| 100.0 \| 1.9e-50 \| 3.7e-56 \| 388.8 \| 309 \| (283, 616) \| 682 \| (7, 319) \| 343 \| Terminase large subunit GpA endonuclease domain-containing protein \| Terminase large subunit GpA endonuclease domain-containing protein; uniclust \| UniRef100\_UPI001D178A05 \| 100.0 \| 2.2e-50 \| 4e-56 \| 386.5 \| 424 \| (27, 479) \| 682 \| (26, 466) \| 493 \| phage terminase large subunit family protein \| phage terminase large subunit family protein; uniclust \| UniRef100\_A0A2W5ATV1 \| 100.0 \| 3.7e-50 \| 7.3e-56 \| 398.8 \| 384 \| (19, 418) \| 682 \| (13, 440) \| 440 \| Terminase (Fragment) \| Terminase (Fragment); uniclust \| UniRef100\_A0A966TY96 \| 100.0 \| 9.2e-50 \| 1.7e-55 \| 365.5 \| 276 \| (330, 609) \| 682 \| (1, 293) \| 331 \| Phage terminase large subunit family protein \| Phage terminase large subunit family protein; uniclust \| UniRef100\_A0A376TF77 \| 100.0 \| 3.2e-49 \| 6e-55 \| 389.8 \| 397 \| (85, 498) \| 682 \| (2, 464) \| 485 \| Terminase large subunit \| Terminase large subunit; uniclust \| UniRef100\_A0A8J3DC88 \| 100.0 \| 3.4e-49 \| 6.3e-55 \| 387.3 \| 552 \| (29, 604) \| 682 \| (22, 605) \| 625 \| Terminase \| Terminase; uniclust \| UniRef100\_A0A024B0A0 \| 100.0 \| 1e-48 \| 2.1e-54 \| 410.6 \| 500 \| (38, 604) \| 682 \| (42, 574) \| 637 \| Terminase, large subunit \| Terminase, large subunit; uniclust \| UniRef100\_A0A930EFS3 \| 100.0 \| 1.3e-48 \| 2.4e-54 \| 370.1 \| 309 \| (282, 608) \| 682 \| (9, 323) \| 358 \| Phage terminase large subunit family protein \| Phage terminase large subunit family protein; uniclust \| UniRef100\_A0A098MRD8 \| 100.0 \| 2.3e-48 \| 4.8e-54 \| 409.6 \| 485 \| (39, 603) \| 682 \| (62, 560) \| 589 \| Terminase \| Terminase; uniclust \| UniRef100\_UPI001E457DA2 \| 100.0 \| 3.3e-48 \| 6.1e-54 \| 352.6 \| 307 \| (78, 389) \| 682 \| (2, 308) \| 311 \| phage terminase large subunit family protein \| phage terminase large subunit family protein; uniclust \| UniRef100\_A0A329VRL4 \| 100.0 \| 3.8e-48 \| 7.3e-54 \| 353.8 \| 219 \| (346, 570) \| 682 \| (1, 223) \| 241 \| Terminase (Fragment) \| Terminase (Fragment); uniclust \| UniRef100\_UPI001F424B1F \| 100.0 \| 4.9e-48 \| 8.9e-54 \| 394.6 \| 429 \| (28, 486) \| 682 \| (21, 462) \| 1019 \| phage portal protein \| phage portal protein; uniclust \| UniRef100\_UPI001651FEFD \| 100.0 \| 1.2e-47 \| 2.3e-53 \| 391.6 \| 493 \| (93, 608) \| 682 \| (412, 955) \| 1015 \| phage terminase large subunit family protein \| phage terminase large subunit family protein; uniclust \| UniRef100\_A0A4R3NP09 \| 100.0 \| 2.1e-47 \| 4e-53 \| 359.6 \| 310 \| (27, 354) \| 682 \| (14, 331) \| 338 \| Phage terminase large subunit GpA (Fragment) \| Phage terminase large subunit GpA (Fragment); uniclust \| UniRef100\_A0A1V3W4P1 \| 100.0 \| 2.6e-47 \| 5e-53 \| 398.1 \| 404 \| (17, 452) \| 682 \| (6, 456) \| 793 \| Phage terminase large subunit family protein \| Phage terminase large subunit family protein; uniclust \| UniRef100\_A0A4Q6BLM2 \| 100.0 \| 5.3e-47 \| 9.8e-53 \| 355.5 \| 338 \| (153, 498) \| 682 \| (9, 373) \| 400 \| Terminase (Fragment) \| Terminase (Fragment); uniclust \| UniRef100\_A0A955CZT1 \| 100.0 \| 7.4e-47 \| 1.4e-52 \| 360.6 \| 393 \| (31, 432) \| 682 \| (15, 456) \| 466 \| Phage terminase large subunit family protein (Fragment) \| Phage terminase large subunit family protein (Fragment); uniclust \| UniRef100\_UPI0021BA7DFC \| 100.0 \| 7.9e-47 \| 1.5e-52 \| 378.9 \| 558 \| (28, 604) \| 682 \| (33, 700) \| 797 \| phage terminase large subunit family protein \| phage terminase large subunit family protein; uniclust \| UniRef100\_A0A2X0QVC0 \| 100.0 \| 1.2e-46 \| 2.2e-52 \| 350.3 \| 278 \| (28, 320) \| 682 \| (25, 306) \| 372 \| Phage terminase large subunit GpA ATPase domain-containing protein \| Phage terminase large subunit GpA ATPase domain-containing protein; uniclust \| UniRef100\_A0A0A0DDD5 \| 100.0 \| 1.4e-46 \| 2.9e-52 \| 395.7 \| 506 \| (36, 604) \| 682 \| (56, 572) \| 632 \| Phage terminase large subunit GpA ATPase domain-containing protein \| Phage terminase large subunit GpA ATPase domain-containing protein; uniclust \| UniRef100\_A0A0F9ENP6 \| 100.0 \| 2.3e-46 \| 4.4e-52 \| 356.0 \| 250 \| (353, 608) \| 682 \| (2, 266) \| 336 \| Terminase large subunit GpA endonuclease domain-containing protein (Fragment) \| Terminase large subunit GpA endonuclease domain-containing protein (Fragment); uniclust \| UniRef100\_A0A894KGS5 \| 100.0 \| 2.4e-46 \| 4.5e-52 \| 338.1 \| 237 \| (102, 353) \| 682 \| (2, 250) \| 253 \| Terminase large subunit (Fragment) \| Terminase large subunit (Fragment); uniclust \| UniRef100\_A0A0F8ZZL1 \| 100.0 \| 2.9e-46 \| 5.5e-52 \| 348.8 \| 261 \| (22, 302) \| 682 \| (24, 287) \| 297 \| Phage terminase large subunit GpA ATPase domain-containing protein (Fragment) \| Phage terminase large subunit GpA ATPase domain-containing protein (Fragment); uniclust \| UniRef100\_A0A367M3E6 \| 100.0 \| 3.2e-46 \| 6e-52 \| 350.1 \| 289 \| (40, 338) \| 682 \| (2, 297) \| 323 \| Phage terminase large subunit family protein (Fragment) \| Phage terminase large subunit family protein (Fragment); uniclust \| UniRef100\_A0A0F8WPC3 \| 100.0 \| 3.1e-46 \| 6.2e-52 \| 380.0 \| 393 \| (39, 498) \| 682 \| (43, 448) \| 467 \| Phage terminase large subunit GpA ATPase domain-containing protein (Fragment) \| Phage terminase large subunit GpA ATPase domain-containing protein (Fragment); uniclust \| UniRef100\_A0A1W6RGK6 \| 100.0 \| 4.7e-46 \| 8.8e-52 \| 349.0 \| 284 \| (25, 328) \| 682 \| (17, 304) \| 342 \| Phage terminase large subunit GpA ATPase domain-containing protein \| Phage terminase large subunit GpA ATPase domain-containing protein; uniclust \| UniRef100\_Q2W750 \| 100.0 \| 5.7e-46 \| 1.1e-51 \| 362.1 \| 392 \| (200, 608) \| 682 \| (28, 492) \| 573 \| Bacteriophage tail assembly protein \| Bacteriophage tail assembly protein; uniclust \| UniRef100\_UPI002279BDA3 \| 100.0 \| 6.7e-46 \| 1.2e-51 \| 364.6 \| 402 \| (170, 602) \| 682 \| (2, 466) \| 543 \| phage terminase large subunit family protein \| phage terminase large subunit family protein; uniclust \| UniRef100\_A0A193QJ57 \| 100.0 \| 8.3e-46 \| 1.6e-51 \| 365.1 \| 440 \| (27, 484) \| 682 \| (17, 523) \| 524 \| Phage terminase large subunit (GpA) \| Phage terminase large subunit (GpA); uniclust \| UniRef100\_A0A661I5B0 \| 100.0 \| 9.7e-46 \| 1.8e-51 \| 348.5 \| 311 \| (286, 613) \| 682 \| (7, 327) \| 359 \| Terminase large subunit GpA endonuclease domain-containing protein (Fragment) \| Terminase large subunit GpA endonuclease domain-containing protein (Fragment); uniclust \| UniRef100\_A0A2T6APL5 \| 100.0 \| 1.3e-45 \| 2.4e-51 \| 346.6 \| 278 \| (140, 436) \| 682 \| (3, 285) \| 403 \| Phage terminase large subunit GpA (Fragment) \| Phage terminase large subunit GpA (Fragment); uniclust \| UniRef100\_A0A1E3G618 \| 100.0 \| 2.3e-45 \| 4.3e-51 \| 353.1 \| 272 \| (324, 608) \| 682 \| (27, 312) \| 366 \| Terminase large subunit GpA endonuclease domain-containing protein \| Terminase large subunit GpA endonuclease domain-containing protein; uniclust \| UniRef100\_A0A0F9EXV6 \| 100.0 \| 2.3e-45 \| 4.4e-51 \| 357.0 \| 399 \| (198, 608) \| 682 \| (9, 431) \| 472 \| Terminase large subunit gp17-like C-terminal domain-containing protein (Fragment) \| Terminase large subunit gp17-like C-terminal domain-containing protein (Fragment); uniclust \| UniRef100\_UPI001F14FD3F \| 100.0 \| 2.5e-45 \| 4.7e-51 \| 356.2 \| 389 \| (27, 433) \| 682 \| (132, 525) \| 549 \| phage terminase large subunit family protein \| phage terminase large subunit family protein; uniclust \| UniRef100\_A0A0B0HWI4 \| 100.0 \| 5.2e-45 \| 9.6e-51 \| 342.5 \| 290 \| (114, 417) \| 682 \| (1, 295) \| 401 \| Phage terminase large subunit (GpA) \| Phage terminase large subunit (GpA); uniclust \| UniRef100\_A0A965PRW8 \| 100.0 \| 6.4e-45 \| 1.2e-50 \| 352.4 \| 376 \| (187, 605) \| 682 \| (1, 397) \| 416 \| Uncharacterized protein \| Uncharacterized protein; uniclust \| UniRef100\_A0A894KGQ0 \| 100.0 \| 7.2e-45 \| 1.3e-50 \| 330.3 \| 246 \| (284, 540) \| 682 \| (5, 258) \| 262 \| Terminase large subunit (Fragment) \| Terminase large subunit (Fragment); uniclust \| UniRef100\_A0A0F8XMS1 \| 100.0 \| 8.3e-45 \| 1.6e-50 \| 360.0 \| 475 \| (41, 602) \| 682 \| (3, 480) \| 500 \| Terminase large subunit gp17-like C-terminal domain-containing protein (Fragment) \| Terminase large subunit gp17-like C-terminal domain-containing protein (Fragment); uniclust \| UniRef100\_X0S680 \| 100.0 \| 8.8e-45 \| 1.6e-50 \| 348.2 \| 429 \| (162, 607) \| 682 \| (17, 477) \| 486 \| Phage terminase large subunit (GpA) (Fragment) \| Phage terminase large subunit (GpA) (Fragment); uniclust \| UniRef100\_A0A1H7AEM5 \| 100.0 \| 9.4e-45 \| 1.9e-50 \| 351.4 \| 225 \| (380, 608) \| 682 \| (9, 247) \| 327 \| Phage terminase large subunit (GpA) (Fragment) \| Phage terminase large subunit (GpA) (Fragment); uniclust \| UniRef100\_A0A7X1E4I2 \| 100.0 \| 1.1e-44 \| 2e-50 \| 354.3 \| 519 \| (57, 608) \| 682 \| (10, 549) \| 590 \| Phage terminase large subunit family protein \| Phage terminase large subunit family protein; uniclust \| UniRef100\_A0A2X1LDM4 \| 100.0 \| 1.3e-44 \| 2.4e-50 \| 328.8 \| 278 \| (255, 552) \| 682 \| (1, 285) \| 291 \| Phage terminase large subunit (GpA) \| Phage terminase large subunit (GpA); uniclust \| UniRef100\_A0A0F9FJK3 \| 100.0 \| 2.2e-44 \| 4.4e-50 \| 374.7 \| 475 \| (59, 604) \| 682 \| (50, 542) \| 605 \| Terminase large subunit gp17-like C-terminal domain-containing protein (Fragment) \| Terminase large subunit gp17-like C-terminal domain-containing protein (Fragment); uniclust \| UniRef100\_A0A2D6M8M9 \| 100.0 \| 4.6e-44 \| 8.5e-50 \| 355.6 \| 548 \| (31, 608) \| 682 \| (14, 650) \| 702 \| Terminase \| Terminase; uniclust \| UniRef100\_A0A158E7G5 \| 100.0 \| 4.7e-44 \| 9.6e-50 \| 385.7 \| 514 \| (34, 602) \| 682 \| (43, 600) \| 693 \| Phage terminase large subunit (GpA) \| Phage terminase large subunit (GpA); uniclust \| UniRef100\_A0A3R8ZAZ8 \| 100.0 \| 1.5e-43 \| 2.7e-49 \| 310.2 \| 217 \| (201, 437) \| 682 \| (6, 228) \| 231 \| Phage terminase large subunit family protein (Fragment) \| Phage terminase large subunit family protein (Fragment); uniclust \| UniRef100\_A0A1M6GTL4 \| 100.0 \| 1.5e-43 \| 2.9e-49 \| 343.1 \| 247 \| (365, 616) \| 682 \| (1, 267) \| 319 \| Phage terminase large subunit (GpA) \| Phage terminase large subunit (GpA); uniclust \| UniRef100\_R9TRM6 \| 100.0 \| 2e-43 \| 3.6e-49 \| 335.7 \| 356 \| (228, 608) \| 682 \| (12, 385) \| 443 \| Terminase large subunit (Fragment) \| Terminase large subunit (Fragment); uniclust \| UniRef100\_A0A2E9IKG8 \| 100.0 \| 2.1e-43 \| 4e-49 \| 358.2 \| 492 \| (60, 605) \| 682 \| (42, 547) \| 581 \| Terminase \| Terminase; uniclust \| UniRef100\_A0A3D3NRT9 \| 100.0 \| 3.5e-43 \| 6.6e-49 \| 357.7 \| 535 \| (32, 602) \| 682 \| (68, 666) \| 680 \| Terminase \| Terminase; uniclust \| UniRef100\_A0A072TET7 \| 100.0 \| 3.9e-43 \| 7.1e-49 \| 369.4 \| 515 \| (24, 552) \| 682 \| (16, 595) \| 1593 \| Phage portal protein, lambda family protein (Fragment) \| Phage portal protein, lambda family protein (Fragment); uniclust \| UniRef100\_A0A193QJW3 \| 100.0 \| 5.3e-43 \| 1e-48 \| 347.0 \| 311 \| (283, 605) \| 682 \| (16, 379) \| 441 \| Phage terminase large subunit (GpA) \| Phage terminase large subunit (GpA); uniclust \| UniRef100\_A0A061JHL7 \| 100.0 \| 6.6e-43 \| 1.3e-48 \| 328.8 \| 213 \| (390, 608) \| 682 \| (1, 224) \| 280 \| Large terminase protein \| Large terminase protein; uniclust \| UniRef100\_A0A963WE82 \| 100.0 \| 9.4e-43 \| 1.8e-48 \| 331.1 \| 304 \| (288, 605) \| 682 \| (1, 320) \| 380 \| Phage terminase large subunit family protein \| Phage terminase large subunit family protein; uniclust \| UniRef100\_A0A060AGH6 \| 100.0 \| 1.4e-42 \| 2.6e-48 \| 356.4 \| 480 \| (65, 604) \| 682 \| (63, 572) \| 616 \| Terminase large subunit \| Terminase large subunit; uniclust \| UniRef100\_UPI0004D72D9B \| 100.0 \| 2.3e-42 \| 4.3e-48 \| 331.7 \| 345 \| (38, 406) \| 682 \| (27, 403) \| 409 \| phage terminase large subunit family protein \| phage terminase large subunit family protein; uniclust \| UniRef100\_A0A3D0WC11 \| 100.0 \| 2.9e-42 \| 5.3e-48 \| 326.8 \| 354 \| (27, 408) \| 682 \| (28, 395) \| 398 \| Terminase (Fragment) \| Terminase (Fragment); uniclust \| UniRef100\_A0A2W6VA45 \| 100.0 \| 3.4e-42 \| 6.4e-48 \| 315.7 \| 254 \| (26, 296) \| 682 \| (8, 270) \| 272 \| Phage terminase large subunit GpA ATPase domain-containing protein \| Phage terminase large subunit GpA ATPase domain-containing protein; uniclust \| UniRef100\_A0A0D6AYL4 \| 100.0 \| 3.5e-42 \| 6.5e-48 \| 349.1 \| 562 \| (19, 604) \| 682 \| (21, 652) \| 722 \| Putative phage terminase large subunit \| Putative phage terminase large subunit; uniclust \| UniRef100\_A0A7X3QD08 \| 100.0 \| 4.4e-42 \| 8.1e-48 \| 336.9 \| 512 \| (34, 604) \| 682 \| (56, 582) \| 598 \| Terminase \| Terminase; uniclust \| UniRef100\_A0A893A8J2 \| 100.0 \| 5.2e-42 \| 1e-47 \| 299.9 \| 183 \| (183, 385) \| 682 \| (1, 186) \| 187 \| Terminase large subunit (Fragment) \| Terminase large subunit (Fragment); uniclust \| UniRef100\_UPI0022580570 \| 100.0 \| 8e-42 \| 1.5e-47 \| 316.6 \| 332 \| (187, 541) \| 682 \| (1, 337) \| 352 \| phage terminase large subunit family protein \| phage terminase large subunit family protein; uniclust \| UniRef100\_A0A9E4LFM8 \| 100.0 \| 9.7e-42 \| 1.8e-47 \| 335.2 \| 531 \| (41, 608) \| 682 \| (4, 594) \| 612 \| Phage terminase large subunit family protein \| Phage terminase large subunit family protein; uniclust \| UniRef100\_A0A0F8VYW0 \| 100.0 \| 1e-41 \| 1.9e-47 \| 322.0 \| 291 \| (306, 608) \| 682 \| (1, 310) \| 356 \| Terminase large subunit GpA endonuclease domain-containing protein (Fragment) \| Terminase large subunit GpA endonuclease domain-containing protein (Fragment); uniclust \| UniRef100\_A0A1S6J105 \| 100.0 \| 1.4e-41 \| 2.7e-47 \| 317.6 \| 246 \| (356, 608) \| 682 \| (6, 260) \| 300 \| Terminase large subunit \| Terminase large subunit; uniclust \| UniRef100\_A0A896ZAW1 \| 100.0 \| 3e-41 \| 5.7e-47 \| 343.6 \| 525 \| (35, 607) \| 682 \| (15, 587) \| 634 \| Terminase large subunit \| Terminase large subunit; uniclust \| UniRef100\_A0A1V5XPC1 \| 100.0 \| 4.4e-41 \| 8.7e-47 \| 347.1 \| 480 \| (39, 607) \| 682 \| (25, 533) \| 539 \| Phage terminase large subunit (GpA) \| Phage terminase large subunit (GpA); uniclust \| UniRef100\_A0A965PDD7 \| 100.0 \| 5.8e-41 \| 1.1e-46 \| 308.6 \| 314 \| (134, 485) \| 682 \| (1, 329) \| 331 \| Uncharacterized protein (Fragment) \| Uncharacterized protein (Fragment); uniclust \| UniRef100\_A0A7V0MM37 \| 100.0 \| 9.9e-41 \| 1.8e-46 \| 309.7 \| 305 \| (26, 350) \| 682 \| (14, 353) \| 354 \| Phage terminase large subunit GpA ATPase domain-containing protein (Fragment) \| Phage terminase large subunit GpA ATPase domain-containing protein (Fragment); uniclust \| UniRef100\_UPI001F06295E \| 100.0 \| 1.4e-40 \| 2.5e-46 \| 306.6 \| 242 \| (367, 615) \| 682 \| (5, 256) \| 305 \| phage terminase large subunit family protein \| phage terminase large subunit family protein; uniclust \| UniRef100\_A0A0D8BXD1 \| 100.0 \| 1.6e-40 \| 3.1e-46 \| 336.4 \| 492 \| (40, 608) \| 682 \| (19, 533) \| 565 \| Terminase-like family protein \| Terminase-like family protein; uniclust \| UniRef100\_A0A3N5QG09 \| 100.0 \| 2.2e-40 \| 4.2e-46 \| 325.7 \| 415 \| (24, 478) \| 682 \| (17, 445) \| 446 \| Terminase (Fragment) \| Terminase (Fragment); uniclust \| UniRef100\_A0A522X2A6 \| 100.0 \| 2.8e-40 \| 5.1e-46 \| 313.5 \| 215 \| (29, 243) \| 682 \| (5, 223) \| 427 \| Phage terminase large subunit GpA ATPase domain-containing protein \| Phage terminase large subunit GpA ATPase domain-containing protein; uniclust \| UniRef100\_A0A318EPN7 \| 100.0 \| 2.9e-40 \| 5.3e-46 \| 300.9 \| 277 \| (29, 321) \| 682 \| (20, 299) \| 304 \| Phage terminase large subunit GpA \| Phage terminase large subunit GpA; uniclust \| UniRef100\_A0A1Z9JFF5 \| 100.0 \| 4.3e-40 \| 7.9e-46 \| 322.0 \| 530 \| (31, 604) \| 682 \| (11, 569) \| 575 \| Terminase \| Terminase; uniclust \| UniRef100\_A0A3P5DNH7 \| 100.0 \| 4.7e-40 \| 9e-46 \| 322.3 \| 348 \| (25, 389) \| 682 \| (26, 414) \| 416 \| Phage terminase large subunit GpA ATPase domain-containing protein \| Phage terminase large subunit GpA ATPase domain-containing protein; uniclust \| UniRef100\_A0A068CB79 \| 100.0 \| 8.1e-40 \| 1.6e-45 \| 328.7 \| 366 \| (65, 470) \| 682 \| (73, 450) \| 461 \| Terminase \| Terminase; uniclust \| UniRef100\_UPI000CF61C34 \| 100.0 \| 1.3e-39 \| 2.4e-45 \| 310.0 \| 359 \| (249, 616) \| 682 \| (1, 401) \| 439 \| phage terminase large subunit family protein \| phage terminase large subunit family protein; uniclust \| UniRef100\_UPI001F3996E7 \| 100.0 \| 2.9e-39 \| 5.4e-45 \| 301.7 \| 305 \| (182, 504) \| 682 \| (2, 345) \| 370 \| phage terminase large subunit family protein \| phage terminase large subunit family protein; uniclust \| UniRef100\_A0A7C0ZXV4 \| 100.0 \| 3.1e-39 \| 5.8e-45 \| 326.6 \| 496 \| (41, 605) \| 682 \| (16, 542) \| 658 \| Phage terminase large subunit GpA ATPase domain-containing protein \| Phage terminase large subunit GpA ATPase domain-containing protein; uniclust \| UniRef100\_A0A0T7A6I5 \| 100.0 \| 4.3e-39 \| 7.9e-45 \| 329.4 \| 483 \| (82, 603) \| 682 \| (394, 909) \| 977 \| DOD-type homing endonuclease domain-containing protein \| DOD-type homing endonuclease domain-containing protein; uniclust \| UniRef100\_A0A497BF11 \| 100.0 \| 7.4e-39 \| 1.4e-44 \| 310.8 \| 482 \| (36, 603) \| 682 \| (21, 505) \| 526 \| Phage terminase large subunit GpA ATPase domain-containing protein \| Phage terminase large subunit GpA ATPase domain-containing protein; uniclust \| UniRef100\_A0A2A4RGG3 \| 100.0 \| 1e-38 \| 1.9e-44 \| 320.9 \| 552 \| (31, 607) \| 682 \| (16, 714) \| 774 \| Terminase \| Terminase; uniclust \| UniRef100\_UPI0021122F9B \| 100.0 \| 1.1e-38 \| 2e-44 \| 314.1 \| 355 \| (233, 607) \| 682 \| (13, 421) \| 610 \| phage terminase large subunit family protein \| phage terminase large subunit family protein; uniclust \| UniRef100\_A0A136Q9K5 \| 100.0 \| 1.2e-38 \| 2.4e-44 \| 310.6 \| 235 \| (177, 433) \| 682 \| (2, 266) \| 292 \| Terminase (Fragment) \| Terminase (Fragment); uniclust \| UniRef100\_A0A376ZEG5 \| 100.0 \| 1.4e-38 \| 2.6e-44 \| 297.8 \| 266 \| (250, 542) \| 682 \| (11, 303) \| 332 \| Phage terminase large subunit (GpA) \| Phage terminase large subunit (GpA); uniclust \| UniRef100\_A0A753AN12 \| 100.0 \| 1.4e-38 \| 2.6e-44 \| 337.9 \| 297 \| (33, 354) \| 682 \| (22, 349) \| 991 \| DUF1983 domain-containing protein \| DUF1983 domain-containing protein; uniclust \| UniRef100\_A0A0F9P2E4 \| 100.0 \| 1.4e-38 \| 2.6e-44 \| 321.1 \| 469 \| (36, 602) \| 682 \| (10, 485) \| 535 \| Phage terminase large subunit GpA ATPase domain-containing protein \| Phage terminase large subunit GpA ATPase domain-containing protein; uniclust \| UniRef100\_A0A0J5K4W6 \| 100.0 \| 1.4e-38 \| 2.7e-44 \| 298.8 \| 233 \| (369, 608) \| 682 \| (2, 242) \| 299 \| Terminase large subunit GpA endonuclease domain-containing protein \| Terminase large subunit GpA endonuclease domain-containing protein; uniclust \| UniRef100\_A0A0F8Y7S9 \| 100.0 \| 1.5e-38 \| 2.9e-44 \| 300.8 \| 191 \| (414, 608) \| 682 \| (2, 204) \| 272 \| Terminase large subunit GpA endonuclease domain-containing protein (Fragment) \| Terminase large subunit GpA endonuclease domain-containing protein (Fragment); uniclust \| UniRef100\_A0A2E3DCF8 \| 100.0 \| 1.8e-38 \| 3.4e-44 \| 320.6 \| 557 \| (31, 603) \| 682 \| (105, 747) \| 818 \| Terminase \| Terminase; uniclust \| UniRef100\_UPI001E522592 \| 100.0 \| 2.7e-38 \| 5e-44 \| 281.9 \| 224 \| (94, 322) \| 682 \| (2, 225) \| 226 \| phage terminase large subunit family protein \| phage terminase large subunit family protein; uniclust \| UniRef100\_A0A8T4YN14 \| 100.0 \| 2.8e-38 \| 5.2e-44 \| 284.4 \| 259 \| (257, 540) \| 682 \| (2, 268) \| 275 \| Phage terminase large subunit family protein \| Phage terminase large subunit family protein; uniclust \| UniRef100\_A0A7W8WPL2 \| 100.0 \| 3.4e-38 \| 6.2e-44 \| 295.9 \| 224 \| (326, 553) \| 682 \| (95, 325) \| 382 \| Phage terminase large subunit GpA-like protein \| Phage terminase large subunit GpA-like protein; uniclust \| UniRef100\_A0A2M7U537 \| 100.0 \| 3.4e-38 \| 6.7e-44 \| 324.3 \| 486 \| (38, 603) \| 682 \| (26, 532) \| 583 \| Phage terminase large subunit GpA ATPase domain-containing protein \| Phage terminase large subunit GpA ATPase domain-containing protein; uniclust \| UniRef100\_A0A954FR98 \| 100.0 \| 6.4e-38 \| 1.2e-43 \| 298.5 \| 374 \| (27, 422) \| 682 \| (14, 435) \| 437 \| Phage terminase large subunit family protein (Fragment) \| Phage terminase large subunit family protein (Fragment); uniclust \| UniRef100\_UPI0022C511CD \| 100.0 \| 6.8e-38 \| 1.2e-43 \| 314.6 \| 343 \| (25, 385) \| 682 \| (20, 397) \| 759 \| phage terminase large subunit family protein \| phage terminase large subunit family protein; uniclust \| UniRef100\_A0A0F9ARK8 \| 100.0 \| 8.4e-38 \| 1.5e-43 \| 297.6 \| 364 \| (220, 607) \| 682 \| (6, 389) \| 435 \| Terminase (Fragment) \| Terminase (Fragment); uniclust \| UniRef100\_A0A7W8DQ18 \| 100.0 \| 9.3e-38 \| 1.7e-43 \| 310.3 \| 555 \| (31, 604) \| 682 \| (14, 636) \| 670 \| Phage terminase large subunit GpA-like protein \| Phage terminase large subunit GpA-like protein; uniclust \| UniRef100\_UPI0014957A8C \| 100.0 \| 9.7e-38 \| 1.8e-43 \| 291.3 \| 315 \| (177, 511) \| 682 \| (5, 351) \| 365 \| phage terminase large subunit family protein \| phage terminase large subunit family protein; uniclust \| UniRef100\_A0A965JVY7 \| 100.0 \| 1.3e-37 \| 2.5e-43 \| 289.6 \| 296 \| (28, 335) \| 682 \| (11, 311) \| 311 \| Phage terminase large subunit GpA ATPase domain-containing protein (Fragment) \| Phage terminase large subunit GpA ATPase domain-containing protein (Fragment); uniclust \| UniRef100\_A0A827HT50 \| 100.0 \| 1.8e-37 \| 3.4e-43 \| 286.1 \| 251 \| (326, 582) \| 682 \| (26, 285) \| 287 \| Phage terminase large subunit family protein (Fragment) \| Phage terminase large subunit family protein (Fragment); uniclust \| UniRef100\_UPI000AE0F248 \| 100.0 \| 2.2e-37 \| 4e-43 \| 290.8 \| 326 \| (55, 395) \| 682 \| (6, 344) \| 345 \| phage terminase large subunit family protein \| phage terminase large subunit family protein; uniclust \| UniRef100\_UPI00203DBA6B \| 100.0 \| 2.2e-37 \| 4.1e-43 \| 272.5 \| 222 \| (159, 396) \| 682 \| (3, 231) \| 233 \| phage terminase large subunit family protein \| phage terminase large subunit family protein; uniclust \| UniRef100\_A0A8S4QPM0 \| 100.0 \| 2.6e-37 \| 4.7e-43 \| 320.3 \| 235 \| (31, 266) \| 682 \| (906, 1144) \| 1151 \| Jg24627 protein (Fragment) \| Jg24627 protein (Fragment); uniclust \| UniRef100\_A0A494X9H5 \| 100.0 \| 2.6e-37 \| 5.1e-43 \| 332.1 \| 479 \| (37, 571) \| 682 \| (229, 752) \| 868 \| Phage terminase large subunit GpA ATPase domain-containing protein \| Phage terminase large subunit GpA ATPase domain-containing protein; uniclust \| UniRef100\_D8F666 \| 100.0 \| 2.7e-37 \| 5.1e-43 \| 297.6 \| 302 \| (284, 602) \| 682 \| (21, 343) \| 390 \| Terminase large subunit GpA endonuclease domain-containing protein \| Terminase large subunit GpA endonuclease domain-containing protein; uniclust \| UniRef100\_B7WRM7 \| 100.0 \| 3.8e-37 \| 7.1e-43 \| 295.8 \| 304 \| (284, 608) \| 682 \| (89, 408) \| 472 \| Bacteriophage tail assembly protein-like protein \| Bacteriophage tail assembly protein-like protein; uniclust \| UniRef100\_A0A0F8VNF6 \| 100.0 \| 4.3e-37 \| 8.2e-43 \| 300.0 \| 356 \| (39, 470) \| 682 \| (17, 373) \| 388 \| Phage terminase large subunit GpA ATPase domain-containing protein (Fragment) \| Phage terminase large subunit GpA ATPase domain-containing protein (Fragment); uniclust \| UniRef100\_A0A5C5YUV7 \| 100.0 \| 4.8e-37 \| 8.7e-43 \| 299.4 \| 421 \| (167, 613) \| 682 \| (13, 479) \| 543 \| Phage terminase large subunit (GpA) \| Phage terminase large subunit (GpA); uniclust \| UniRef100\_A0A5A7MYI8 \| 100.0 \| 5.6e-37 \| 1e-42 \| 300.1 \| 337 \| (249, 603) \| 682 \| (3, 386) \| 473 \| Terminase \| Terminase; uniclust \| UniRef100\_A0A0M7BBY2 \| 100.0 \| 5.9e-37 \| 1.1e-42 \| 284.0 \| 213 \| (20, 235) \| 682 \| (27, 244) \| 252 \| Bacteriophage tail assembly protein \| Bacteriophage tail assembly protein; uniclust \| UniRef100\_A0A8D8A671 \| 100.0 \| 6.7e-37 \| 1.3e-42 \| 265.1 \| 183 \| (251, 452) \| 682 \| (1, 188) \| 188 \| Terminase, large subunit (Fragment) \| Terminase, large subunit (Fragment); uniclust \| UniRef100\_A0A0B0SDH7 \| 100.0 \| 1.1e-36 \| 2.3e-42 \| 322.8 \| 513 \| (39, 603) \| 682 \| (35, 597) \| 694 \| Phage terminase large subunit GpA ATPase domain-containing protein \| Phage terminase large subunit GpA ATPase domain-containing protein; uniclust \| UniRef100\_A0A894KQY8 \| 100.0 \| 1.3e-36 \| 2.4e-42 \| 263.1 \| 170 \| (326, 497) \| 682 \| (10, 184) \| 184 \| Terminase large subunit (Fragment) \| Terminase large subunit (Fragment); uniclust \| UniRef100\_UPI001F0AC2C8 \| 100.0 \| 1.4e-36 \| 2.6e-42 \| 294.1 \| 333 \| (23, 370) \| 682 \| (169, 505) \| 506 \| phage terminase large subunit family protein \| phage terminase large subunit family protein; uniclust \| UniRef100\_A0A0F9HNG5 \| 100.0 \| 2.2e-36 \| 4.4e-42 \| 312.2 \| 505 \| (38, 603) \| 682 \| (23, 577) \| 618 \| Phage terminase large subunit GpA ATPase domain-containing protein \| Phage terminase large subunit GpA ATPase domain-containing protein; uniclust \| UniRef100\_A0A0S9PYT1 \| 100.0 \| 3.1e-36 \| 5.7e-42 \| 285.1 \| 282 \| (23, 311) \| 682 \| (15, 297) \| 363 \| Phage terminase large subunit GpA ATPase domain-containing protein \| Phage terminase large subunit GpA ATPase domain-containing protein; uniclust \| UniRef100\_A0A1N7N106 \| 100.0 \| 3.3e-36 \| 6.4e-42 \| 274.3 \| 197 \| (18, 214) \| 682 \| (24, 224) \| 227 \| Phage terminase large subunit (GpA) \| Phage terminase large subunit (GpA); uniclust \| UniRef100\_A0A2D4TR23 \| 100.0 \| 3.6e-36 \| 6.8e-42 \| 272.6 \| 214 \| (395, 616) \| 682 \| (2, 215) \| 244 \| Terminase large subunit GpA endonuclease domain-containing protein \| Terminase large subunit GpA endonuclease domain-containing protein; uniclust \| UniRef100\_R7J609 \| 100.0 \| 6.4e-36 \| 1.2e-41 \| 280.4 \| 305 \| (284, 604) \| 682 \| (32, 350) \| 376 \| Phage terminase large subunit \| Phage terminase large subunit; uniclust \| UniRef100\_A0A3C0YP94 \| 100.0 \| 1.1e-35 \| 2.2e-41 \| 281.5 \| 221 \| (27, 250) \| 682 \| (36, 260) \| 260 \| Phage terminase large subunit GpA ATPase domain-containing protein (Fragment) \| Phage terminase large subunit GpA ATPase domain-containing protein (Fragment); uniclust \| UniRef100\_A0A8T6UVM0 \| 100.0 \| 1.8e-35 \| 3.4e-41 \| 280.1 \| 377 \| (28, 422) \| 682 \| (12, 406) \| 410 \| Terminase (Fragment) \| Terminase (Fragment); uniclust \| UniRef100\_A0A523S777 \| 100.0 \| 1.9e-35 \| 3.5e-41 \| 288.2 \| 400 \| (155, 603) \| 682 \| (9, 432) \| 446 \| Terminase (Fragment) \| Terminase (Fragment); uniclust \| UniRef100\_A0A0E3UJM4 \| 100.0 \| 2.2e-35 \| 4e-41 \| 296.1 \| 544 \| (37, 607) \| 682 \| (68, 710) \| 731 \| Terminase \| Terminase; uniclust \| UniRef100\_A0A2P9HPM1 \| 100.0 \| 2.6e-35 \| 4.8e-41 \| 283.1 \| 319 \| (31, 370) \| 682 \| (39, 375) \| 384 \| Phage terminase, large subunit \| Phage terminase, large subunit; uniclust \| UniRef100\_A0A376K1X9 \| 100.0 \| 2.5e-35 \| 5e-41 \| 289.2 \| 264 \| (327, 604) \| 682 \| (1, 277) \| 344 \| Phage terminase large subunit (GpA) \| Phage terminase large subunit (GpA); uniclust \| UniRef100\_UPI001FEFB116 \| 100.0 \| 3e-35 \| 5.5e-41 \| 273.5 \| 309 \| (32, 358) \| 682 \| (4, 347) \| 348 \| phage terminase large subunit family protein \| phage terminase large subunit family protein; uniclust \| UniRef100\_A0A5C5XIN6 \| 100.0 \| 5e-35 \| 9.4e-41 \| 287.9 \| 394 \| (28, 432) \| 682 \| (29, 446) \| 453 \| Phage terminase large subunit (GpA) \| Phage terminase large subunit (GpA); uniclust \| UniRef100\_A0A0F9SHF5 \| 100.0 \| 7.2e-35 \| 1.4e-40 \| 307.1 \| 491 \| (56, 612) \| 682 \| (56, 584) \| 615 \| Phage terminase large subunit GpA ATPase domain-containing protein \| Phage terminase large subunit GpA ATPase domain-containing protein; uniclust \| UniRef100\_A0A098N0R4 \| 100.0 \| 9.7e-35 \| 1.9e-40 \| 298.1 \| 509 \| (24, 605) \| 682 \| (20, 549) \| 561 \| Terminase \| Terminase; uniclust \| UniRef100\_A0A0F8W5R2 \| 100.0 \| 1.4e-34 \| 2.7e-40 \| 258.7 \| 186 \| (29, 214) \| 682 \| (7, 194) \| 196 \| Phage terminase large subunit GpA ATPase domain-containing protein (Fragment) \| Phage terminase large subunit GpA ATPase domain-containing protein (Fragment); uniclust \| UniRef100\_A0A6J5NTB7 \| 100.0 \| 1.9e-34 \| 3.5e-40 \| 284.1 \| 522 \| (39, 608) \| 682 \| (18, 576) \| 592 \| Bacteriophage lambda, GpA \| Bacteriophage lambda, GpA; uniclust \| UniRef100\_UPI0015B461E7 \| 100.0 \| 2.2e-34 \| 4e-40 \| 271.3 \| 271 \| (325, 607) \| 682 \| (47, 326) \| 388 \| phage terminase large subunit family protein \| phage terminase large subunit family protein; uniclust \| UniRef100\_UPI0022E897D2 \| 100.0 \| 2.2e-34 \| 4.1e-40 \| 252.1 \| 204 \| (154, 366) \| 682 \| (8, 221) \| 221 \| phage terminase large subunit family protein \| phage terminase large subunit family protein; uniclust \| UniRef100\_A0A0F9JW81 \| 100.0 \| 2.2e-34 \| 4.2e-40 \| 288.5 \| 480 \| (39, 604) \| 682 \| (7, 505) \| 525 \| Phage terminase large subunit GpA ATPase domain-containing protein (Fragment) \| Phage terminase large subunit GpA ATPase domain-containing protein (Fragment); uniclust \| UniRef100\_A0A956UQF2 \| 100.0 \| 2.5e-34 \| 4.6e-40 \| 280.6 \| 481 \| (31, 603) \| 682 \| (28, 530) \| 539 \| Phage terminase large subunit family protein \| Phage terminase large subunit family protein; uniclust \| UniRef100\_A0A077PUK0 \| 100.0 \| 2.5e-34 \| 5.1e-40 \| 262.8 \| 175 \| (20, 194) \| 682 \| (13, 188) \| 189 \| Putative phage tail assembly protein \| Putative phage tail assembly protein; uniclust \| UniRef100\_A0A2G4JNV2 \| 100.0 \| 7.6e-34 \| 1.4e-39 \| 274.9 \| 364 \| (114, 498) \| 682 \| (1, 431) \| 493 \| Terminase \| Terminase; uniclust \| UniRef100\_UPI001834D0C0 \| 100.0 \| 1.2e-33 \| 2.2e-39 \| 276.5 \| 240 \| (55, 309) \| 682 \| (8, 250) \| 549 \| phage tail tape measure protein \| phage tail tape measure protein; uniclust \| UniRef100\_A0A965LJP3 \| 100.0 \| 1.7e-33 \| 3.1e-39 \| 280.5 \| 544 \| (30, 605) \| 682 \| (13, 647) \| 665 \| Uncharacterized protein \| Uncharacterized protein; uniclust \| UniRef100\_UPI0021C412FA \| 100.0 \| 1.9e-33 \| 3.5e-39 \| 248.6 \| 201 \| (281, 497) \| 682 \| (27, 232) \| 235 \| phage terminase large subunit family protein \| phage terminase large subunit family protein; uniclust \| UniRef100\_A0A894KR06 \| 100.0 \| 2.2e-33 \| 4.2e-39 \| 268.2 \| 309 \| (25, 352) \| 682 \| (10, 322) \| 337 \| Terminase large subunit (Fragment) \| Terminase large subunit (Fragment); uniclust \| UniRef100\_S9S4Q4 \| 100.0 \| 2.9e-33 \| 5.4e-39 \| 260.8 \| 317 \| (28, 358) \| 682 \| (19, 342) \| 351 \| Terminase GpA (Fragment) \| Terminase GpA (Fragment); uniclust \| UniRef100\_A0A1B7HIA8 \| 100.0 \| 3.3e-33 \| 6.2e-39 \| 268.0 \| 284 \| (22, 322) \| 682 \| (17, 319) \| 353 \| Phage terminase large subunit (Fragment) \| Phage terminase large subunit (Fragment); uniclust \| UniRef100\_A0A4Q2YME8 \| 100.0 \| 1.1e-32 \| 2.1e-38 \| 276.1 \| 559 \| (25, 605) \| 682 \| (28, 691) \| 702 \| Phage terminase large subunit GpA ATPase domain-containing protein \| Phage terminase large subunit GpA ATPase domain-containing protein; uniclust \| UniRef100\_A0A1F7TIH1 \| 100.0 \| 1.2e-32 \| 2.3e-38 \| 275.3 \| 336 \| (40, 416) \| 682 \| (83, 432) \| 459 \| Phage terminase large subunit GpA ATPase domain-containing protein \| Phage terminase large subunit GpA ATPase domain-containing protein; uniclust \| UniRef100\_A0A0F9DJC4 \| 100.0 \| 1.6e-32 \| 3.3e-38 \| 309.5 \| 252 \| (356, 609) \| 682 \| (561, 859) \| 911 \| Terminase large subunit GpA endonuclease domain-containing protein \| Terminase large subunit GpA endonuclease domain-containing protein; uniclust \| UniRef100\_A0A3B9NUU6 \| 100.0 \| 1.8e-32 \| 3.5e-38 \| 246.9 \| 205 \| (408, 616) \| 682 \| (2, 213) \| 242 \| Terminase large subunit GpA endonuclease domain-containing protein \| Terminase large subunit GpA endonuclease domain-containing protein; uniclust \| UniRef100\_A0A4Q3NM87 \| 100.0 \| 1.9e-32 \| 3.5e-38 \| 256.7 \| 251 \| (353, 607) \| 682 \| (7, 271) \| 366 \| Terminase large subunit GpA endonuclease domain-containing protein (Fragment) \| Terminase large subunit GpA endonuclease domain-containing protein (Fragment); uniclust \| UniRef100\_UPI001F411F2F \| 100.0 \| 2.4e-32 \| 4.5e-38 \| 243.3 \| 164 \| (453, 616) \| 682 \| (11, 174) \| 218 \| phage terminase large subunit family protein \| phage terminase large subunit family protein; uniclust \| UniRef100\_A0A661TC80 \| 99.9 \| 2.7e-32 \| 5.1e-38 \| 264.2 \| 316 \| (39, 384) \| 682 \| (28, 352) \| 354 \| Phage terminase large subunit GpA ATPase domain-containing protein (Fragment) \| Phage terminase large subunit GpA ATPase domain-containing protein (Fragment); uniclust \| UniRef100\_UPI002101A6F7 \| 99.9 \| 3.2e-32 \| 6e-38 \| 251.3 \| 284 \| (198, 499) \| 682 \| (8, 301) \| 309 \| phage terminase large subunit family protein \| phage terminase large subunit family protein; uniclust \| UniRef100\_A0A7W6SXB3 \| 99.9 \| 3.2e-32 \| 6e-38 \| 247.4 \| 198 \| (415, 616) \| 682 \| (2, 207) \| 236 \| Phage terminase large subunit GpA-like protein \| Phage terminase large subunit GpA-like protein; uniclust \| UniRef100\_A0A2X3K131 \| 99.9 \| 3.2e-32 \| 6.3e-38 \| 255.9 \| 207 \| (395, 608) \| 682 \| (19, 232) \| 238 \| Phage terminase large subunit (GpA) \| Phage terminase large subunit (GpA); uniclust \| UniRef100\_UPI0015945207 \| 99.9 \| 3.3e-32 \| 6.5e-38 \| 259.2 \| 228 \| (24, 257) \| 682 \| (35, 286) \| 287 \| phage terminase large subunit family protein \| phage terminase large subunit family protein; uniclust \| UniRef100\_A0A0F9BBK9 \| 99.9 \| 4.9e-32 \| 9.4e-38 \| 254.5 \| 277 \| (30, 338) \| 682 \| (14, 293) \| 296 \| Phage terminase large subunit GpA ATPase domain-containing protein (Fragment) \| Phage terminase large subunit GpA ATPase domain-containing protein (Fragment); uniclust \| UniRef100\_A0A659YSG8 \| 99.9 \| 6.6e-32 \| 1.2e-37 \| 265.8 \| 352 \| (37, 419) \| 682 \| (25, 390) \| 455 \| Phage terminase large subunit GpA ATPase domain-containing protein \| Phage terminase large subunit GpA ATPase domain-containing protein; uniclust \| UniRef100\_A0A8D8WC28 \| 99.9 \| 7.5e-32 \| 1.4e-37 \| 233.2 \| 176 \| (326, 504) \| 682 \| (3, 183) \| 200 \| Terminase, large subunit (Fragment) \| Terminase, large subunit (Fragment); uniclust \| UniRef100\_UPI001C8BCF95 \| 99.9 \| 9.8e-32 \| 1.8e-37 \| 257.2 \| 353 \| (32, 386) \| 682 \| (7, 379) \| 436 \| phage terminase large subunit family protein \| phage terminase large subunit family protein; uniclust \| UniRef100\_A0A7W5C6Z2 \| 99.9 \| 1.1e-31 \| 2e-37 \| 240.0 \| 183 \| (217, 421) \| 682 \| (17, 202) \| 216 \| Phage terminase large subunit GpA-like protein \| Phage terminase large subunit GpA-like protein; uniclust \| UniRef100\_A0A381KV82 \| 99.9 \| 1.3e-31 \| 2.5e-37 \| 249.9 \| 239 \| (28, 266) \| 682 \| (28, 268) \| 288 \| Phage terminase large subunit \| Phage terminase large subunit; uniclust \| UniRef100\_A0A948S8L3 \| 99.9 \| 1.3e-31 \| 2.5e-37 \| 258.9 \| 406 \| (159, 604) \| 682 \| (10, 458) \| 479 \| Uncharacterized protein \| Uncharacterized protein; uniclust \| UniRef100\_UPI0008FD63AC \| 99.9 \| 1.6e-31 \| 3e-37 \| 231.0 \| 163 \| (218, 389) \| 682 \| (14, 177) \| 178 \| phage terminase large subunit family protein \| phage terminase large subunit family protein; uniclust \| UniRef100\_UPI001F50DAB2 \| 99.9 \| 2e-31 \| 3.6e-37 \| 242.9 \| 272 \| (74, 372) \| 682 \| (2, 287) \| 289 \| phage terminase large subunit family protein \| phage terminase large subunit family protein; uniclust \| UniRef100\_UPI001561DD3F \| 99.9 \| 3.1e-31 \| 5.8e-37 \| 239.4 \| 212 \| (355, 570) \| 682 \| (5, 234) \| 243 \| phage terminase large subunit family protein \| phage terminase large subunit family protein; uniclust \| UniRef100\_A0A1X0YW67 \| 99.9 \| 4.1e-31 \| 8.1e-37 \| 254.3 \| 240 \| (50, 301) \| 682 \| (3, 261) \| 281 \| Phage terminase large subunit GpA ATPase domain-containing protein (Fragment) \| Phage terminase large subunit GpA ATPase domain-containing protein (Fragment); uniclust \| UniRef100\_UPI000B18891F \| 99.9 \| 6.5e-31 \| 1.2e-36 \| 238.0 \| 244 \| (217, 479) \| 682 \| (3, 274) \| 275 \| phage terminase large subunit family protein \| phage terminase large subunit family protein; uniclust \| UniRef100\_R7UYV5 \| 99.9 \| 1.1e-30 \| 2.1e-36 \| 240.3 \| 193 \| (28, 220) \| 682 \| (113, 309) \| 310 \| Phage terminase large subunit GpA ATPase domain-containing protein (Fragment) \| Phage terminase large subunit GpA ATPase domain-containing protein (Fragment); uniclust \| UniRef100\_A0A945W5E5 \| 99.9 \| 1.2e-30 \| 2.3e-36 \| 257.6 \| 486 \| (60, 605) \| 682 \| (51, 555) \| 586 \| Uncharacterized protein \| Uncharacterized protein; uniclust \| UniRef100\_A0A5C7Q184 \| 99.9 \| 1.3e-30 \| 2.4e-36 \| 257.8 \| 533 \| (28, 605) \| 682 \| (6, 592) \| 594 \| Phage terminase large subunit GpA ATPase domain-containing protein \| Phage terminase large subunit GpA ATPase domain-containing protein; uniclust \| UniRef100\_UPI00095BDF8E \| 99.9 \| 1.8e-30 \| 3.4e-36 \| 247.6 \| 312 \| (22, 352) \| 682 \| (9, 362) \| 370 \| phage terminase large subunit family protein \| phage terminase large subunit family protein; uniclust \| UniRef100\_UPI00077C3110 \| 99.9 \| 2e-30 \| 3.7e-36 \| 245.5 \| 227 \| (386, 616) \| 682 \| (95, 331) \| 391 \| phage terminase large subunit family protein \| phage terminase large subunit family protein; uniclust \| UniRef100\_A0A2D5CRA2 \| 99.9 \| 2.5e-30 \| 4.6e-36 \| 252.7 \| 468 \| (106, 603) \| 682 \| (7, 516) \| 524 \| Terminase \| Terminase; uniclust \| UniRef100\_UPI001CC6F246 \| 99.9 \| 2.6e-30 \| 4.7e-36 \| 251.3 \| 312 \| (24, 351) \| 682 \| (10, 328) \| 499 \| phage terminase large subunit family protein \| phage terminase large subunit family protein; uniclust \| UniRef100\_A0A6M8BKE7 \| 99.9 \| 2.9e-30 \| 5.7e-36 \| 270.9 \| 506 \| (37, 603) \| 682 \| (57, 662) \| 686 \| Phage terminase large subunit GpA ATPase domain-containing protein \| Phage terminase large subunit GpA ATPase domain-containing protein; uniclust \| UniRef100\_A0A430BRK0 \| 99.9 \| 3.9e-30 \| 7.2e-36 \| 230.6 \| 232 \| (66, 319) \| 682 \| (16, 254) \| 254 \| Terminase (Fragment) \| Terminase (Fragment); uniclust \| UniRef100\_A0A1Y5U3Z7 \| 99.9 \| 3.9e-30 \| 7.5e-36 \| 231.7 \| 163 \| (447, 615) \| 682 \| (2, 166) \| 199 \| Phage terminase large subunit (GpA) \| Phage terminase large subunit (GpA); uniclust \| UniRef100\_A0A5T3NHC4 \| 99.9 \| 4e-30 \| 7.6e-36 \| 265.4 \| 337 \| (22, 375) \| 682 \| (11, 408) \| 753 \| Uncharacterized protein \| Uncharacterized protein; uniclust \| UniRef100\_UPI000BDE7CC4 \| 99.9 \| 4.6e-30 \| 8.8e-36 \| 236.7 \| 181 \| (249, 450) \| 682 \| (1, 209) \| 258 \| phage terminase large subunit family protein \| phage terminase large subunit family protein; uniclust \| UniRef100\_UPI00155FC26A \| 99.9 \| 5.2e-30 \| 9.6e-36 \| 252.8 \| 362 \| (27, 406) \| 682 \| (21, 427) \| 575 \| phage terminase large subunit family protein \| phage terminase large subunit family protein; uniclust \| UniRef100\_A0A894KG93 \| 99.9 \| 8e-30 \| 1.5e-35 \| 232.1 \| 181 \| (426, 609) \| 682 \| (1, 186) \| 249 \| Terminase large subunit \| Terminase large subunit; uniclust \| UniRef100\_A0A2N8K826 \| 99.9 \| 8.4e-30 \| 1.5e-35 \| 206.0 \| 127 \| (169, 310) \| 682 \| (2, 130) \| 130 \| Terminase (Fragment) \| Terminase (Fragment); uniclust \| UniRef100\_UPI000E1FC18F \| 99.9 \| 9e-30 \| 1.7e-35 \| 252.3 \| 316 \| (22, 354) \| 682 \| (11, 349) \| 601 \| phage terminase large subunit family protein \| phage terminase large subunit family protein; uniclust \| UniRef100\_A0A1G0XTS0 \| 99.9 \| 9.1e-30 \| 1.7e-35 \| 253.8 \| 453 \| (102, 604) \| 682 \| (3, 471) \| 518 \| Terminase \| Terminase; uniclust \| UniRef100\_A0A4Q8RL69 \| 99.9 \| 1.1e-29 \| 2.1e-35 \| 236.6 \| 285 \| (67, 366) \| 682 \| (2, 309) \| 309 \| Phage terminase large subunit family protein (Fragment) \| Phage terminase large subunit family protein (Fragment); uniclust \| UniRef100\_A0A062VB30 \| 99.9 \| 1.3e-29 \| 2.3e-35 \| 225.4 \| 213 \| (390, 616) \| 682 \| (6, 219) \| 238 \| Terminase GpA (Fragment) \| Terminase GpA (Fragment); uniclust \| UniRef100\_A0A928NBY3 \| 99.9 \| 1.3e-29 \| 2.5e-35 \| 223.2 \| 212 \| (26, 237) \| 682 \| (11, 222) \| 223 \| Phage terminase large subunit family protein (Fragment) \| Phage terminase large subunit family protein (Fragment); uniclust \| UniRef100\_UPI00187D135A \| 99.9 \| 1.4e-29 \| 2.6e-35 \| 236.2 \| 241 \| (245, 499) \| 682 \| (1, 285) \| 298 \| phage terminase large subunit family protein \| phage terminase large subunit family protein; uniclust \| UniRef100\_A0A3S1RU85 \| 99.9 \| 1.9e-29 \| 3.5e-35 \| 228.1 \| 253 \| (229, 503) \| 682 \| (9, 268) \| 269 \| Terminase (Fragment) \| Terminase (Fragment); uniclust \| UniRef100\_A0A343US01 \| 99.9 \| 2.1e-29 \| 3.9e-35 \| 251.5 \| 468 \| (68, 604) \| 682 \| (35, 512) \| 527 \| Phage terminase large subunit GpA-like protein \| Phage terminase large subunit GpA-like protein; uniclust \| UniRef100\_A0A021WYP0 \| 99.9 \| 2.1e-29 \| 4e-35 \| 246.1 \| 279 \| (21, 323) \| 682 \| (24, 313) \| 395 \| Bacteriophage tail assembly protein \| Bacteriophage tail assembly protein; uniclust \| UniRef100\_A0A5C7PZL4 \| 99.9 \| 3.3e-29 \| 6e-35 \| 248.9 \| 524 \| (34, 604) \| 682 \| (20, 601) \| 614 \| Phage terminase large subunit GpA ATPase domain-containing protein \| Phage terminase large subunit GpA ATPase domain-containing protein; uniclust \| UniRef100\_A0A1V6IIE5 \| 99.9 \| 3.6e-29 \| 6.5e-35 \| 245.9 \| 475 \| (63, 616) \| 682 \| (26, 514) \| 547 \| Phage terminase large subunit (GpA) \| Phage terminase large subunit (GpA); uniclust \| UniRef100\_A0A529QAD6 \| 99.9 \| 4.8e-29 \| 8.9e-35 \| 218.6 \| 173 \| (200, 387) \| 682 \| (2, 185) \| 193 \| Terminase (Fragment) \| Terminase (Fragment); uniclust \| UniRef100\_UPI001E56E5A7 \| 99.9 \| 5e-29 \| 9.2e-35 \| 209.0 \| 155 \| (286, 447) \| 682 \| (5, 161) \| 161 \| phage terminase large subunit family protein \| phage terminase large subunit family protein; uniclust \| UniRef100\_A0A4U9DCF0 \| 99.9 \| 5.2e-29 \| 9.8e-35 \| 227.7 \| 184 \| (428, 616) \| 682 \| (5, 200) \| 251 \| Bacteriophage tail assembly protein \| Bacteriophage tail assembly protein; uniclust \| UniRef100\_A0A3D1USL3 \| 99.9 \| 5e-29 \| 9.9e-35 \| 249.5 \| 233 \| (374, 608) \| 682 \| (46, 322) \| 368 \| Terminase large subunit GpA endonuclease domain-containing protein \| Terminase large subunit GpA endonuclease domain-containing protein; uniclust \| UniRef100\_A0A5C5VTC1 \| 99.9 \| 5.8e-29 \| 1.1e-34 \| 244.0 \| 395 \| (198, 608) \| 682 \| (49, 493) \| 536 \| Phage terminase large subunit (GpA) \| Phage terminase large subunit (GpA); uniclust \| UniRef100\_UPI0022E19F4F \| 99.9 \| 6.7e-29 \| 1.2e-34 \| 228.8 \| 297 \| (59, 400) \| 682 \| (1, 309) \| 309 \| phage terminase large subunit family protein \| phage terminase large subunit family protein; uniclust \| UniRef100\_A0A974WV16 \| 99.9 \| 7.1e-29 \| 1.3e-34 \| 248.3 \| 526 \| (30, 600) \| 682 \| (25, 613) \| 662 \| Phage terminase large subunit family protein \| Phage terminase large subunit family protein; uniclust \| UniRef100\_UPI0022E14ED5 \| 99.9 \| 7.6e-29 \| 1.4e-34 \| 231.5 \| 232 \| (31, 267) \| 682 \| (59, 293) \| 344 \| phage terminase large subunit family protein \| phage terminase large subunit family protein; uniclust \| UniRef100\_A0A440JQM9 \| 99.9 \| 7.8e-29 \| 1.4e-34 \| 233.4 \| 305 \| (28, 351) \| 682 \| (19, 340) \| 369 \| Phage terminase large subunit GpA ATPase domain-containing protein \| Phage terminase large subunit GpA ATPase domain-containing protein; uniclust \| UniRef100\_A0A0F9GI62 \| 99.9 \| 8.1e-29 \| 1.5e-34 \| 244.9 \| 495 \| (40, 607) \| 682 \| (23, 531) \| 582 \| Phage terminase large subunit GpA ATPase domain-containing protein (Fragment) \| Phage terminase large subunit GpA ATPase domain-containing protein (Fragment); uniclust \| UniRef100\_UPI001586C996 \| 99.9 \| 9.1e-29 \| 1.7e-34 \| 210.6 \| 175 \| (411, 592) \| 682 \| (1, 177) \| 177 \| phage terminase large subunit family protein \| phage terminase large subunit family protein; uniclust \| UniRef100\_A0A1V5IZJ9 \| 99.9 \| 9.6e-29 \| 1.9e-34 \| 249.8 \| 251 \| (356, 608) \| 682 \| (142, 439) \| 471 \| Phage terminase large subunit (GpA) \| Phage terminase large subunit (GpA); uniclust \| UniRef100\_A0A2V5PXK1 \| 99.9 \| 1.5e-28 \| 2.7e-34 \| 245.2 \| 538 \| (28, 601) \| 682 \| (12, 619) \| 636 \| Phage terminase large subunit GpA ATPase domain-containing protein \| Phage terminase large subunit GpA ATPase domain-containing protein; uniclust \| UniRef100\_UPI0021C46AE6 \| 99.9 \| 1.5e-28 \| 2.7e-34 \| 227.6 \| 237 \| (326, 568) \| 682 \| (24, 277) \| 320 \| phage terminase large subunit family protein \| phage terminase large subunit family protein; uniclust \| UniRef100\_UPI00210E3936 \| 99.9 \| 2.2e-28 \| 4e-34 \| 220.6 \| 233 \| (363, 603) \| 682 \| (13, 251) \| 262 \| phage terminase large subunit family protein \| phage terminase large subunit family protein; uniclust \| UniRef100\_UPI001CBE42DF \| 99.9 \| 2.3e-28 \| 4.2e-34 \| 225.8 \| 238 \| (357, 606) \| 682 \| (2, 250) \| 314 \| phage terminase large subunit family protein \| phage terminase large subunit family protein; uniclust \| UniRef100\_A0A059UUJ4 \| 99.9 \| 2.5e-28 \| 4.6e-34 \| 212.6 \| 160 \| (159, 338) \| 682 \| (3, 167) \| 204 \| Terminase, large subunit \| Terminase, large subunit; uniclust \| UniRef100\_UPI000F0B10AB \| 99.9 \| 2.7e-28 \| 5e-34 \| 234.9 \| 369 \| (37, 431) \| 682 \| (22, 405) \| 447 \| phage terminase large subunit family protein \| phage terminase large subunit family protein; uniclust \| UniRef100\_UPI00207C9BF4 \| 99.9 \| 3.2e-28 \| 5.9e-34 \| 226.3 \| 246 \| (353, 607) \| 682 \| (13, 280) \| 330 \| phage terminase large subunit family protein \| phage terminase large subunit family protein; uniclust \| UniRef100\_A0A661TT30 \| 99.9 \| 3.3e-28 \| 6e-34 \| 231.8 \| 311 \| (282, 608) \| 682 \| (30, 370) \| 405 \| Terminase large subunit GpA endonuclease domain-containing protein \| Terminase large subunit GpA endonuclease domain-containing protein; uniclust \| UniRef100\_A0A958D0X9 \| 99.9 \| 3.8e-28 \| 6.9e-34 \| 219.4 \| 206 \| (397, 608) \| 682 \| (2, 225) \| 264 \| Phage terminase large subunit family protein \| Phage terminase large subunit family protein; uniclust \| UniRef100\_UPI0013B8BA29 \| 99.9 \| 5e-28 \| 9.1e-34 \| 228.6 \| 247 \| (356, 605) \| 682 \| (54, 313) \| 376 \| phage terminase large subunit family protein \| phage terminase large subunit family protein; uniclust \| UniRef100\_UPI001C905274 \| 99.9 \| 5.6e-28 \| 1.1e-33 \| 229.7 \| 259 \| (34, 304) \| 682 \| (55, 332) \| 335 \| phage terminase large subunit family protein \| phage terminase large subunit family protein; uniclust \| UniRef100\_A0A349YR87 \| 99.9 \| 6.3e-28 \| 1.2e-33 \| 210.6 \| 202 \| (75, 291) \| 682 \| (3, 207) \| 207 \| Terminase (Fragment) \| Terminase (Fragment); uniclust \| UniRef100\_UPI0006F7D793 \| 99.9 \| 6.5e-28 \| 1.2e-33 \| 233.1 \| 299 \| (300, 607) \| 682 \| (3, 321) \| 459 \| phage terminase large subunit family protein \| phage terminase large subunit family protein; uniclust \| UniRef100\_UPI0018EFD575 \| 99.9 \| 7.2e-28 \| 1.3e-33 \| 214.2 \| 223 \| (222, 464) \| 682 \| (3, 234) \| 235 \| phage terminase large subunit family protein \| phage terminase large subunit family protein; uniclust \| UniRef100\_A0A2N8GT90 \| 99.9 \| 8e-28 \| 1.5e-33 \| 199.8 \| 134 \| (361, 496) \| 682 \| (1, 135) \| 136 \| Terminase (Fragment) \| Terminase (Fragment); uniclust \| UniRef100\_UPI0021565FA8 \| 99.9 \| 1.1e-27 \| 2e-33 \| 232.4 \| 237 \| (78, 324) \| 682 \| (5, 242) \| 474 \| phage terminase large subunit family protein \| phage terminase large subunit family protein; uniclust \| UniRef100\_UPI000B13FF44 \| 99.9 \| 1.5e-27 \| 2.7e-33 \| 237.4 \| 265 \| (33, 321) \| 682 \| (22, 297) \| 611 \| phage portal protein \| phage portal protein; uniclust \| UniRef100\_A0A2E1H9V8 \| 99.9 \| 2.8e-27 \| 5.2e-33 \| 208.8 \| 190 \| (308, 505) \| 682 \| (10, 204) \| 222 \| Terminase large subunit GpA endonuclease domain-containing protein \| Terminase large subunit GpA endonuclease domain-containing protein; uniclust \| UniRef100\_A0A0F9H7D9 \| 99.9 \| 3.2e-27 \| 6e-33 \| 225.2 \| 351 \| (182, 603) \| 682 \| (2, 356) \| 405 \| Phage terminase large subunit GpA ATPase domain-containing protein (Fragment) \| Phage terminase large subunit GpA ATPase domain-containing protein (Fragment); uniclust \| UniRef100\_A0A101IP78 \| 99.9 \| 3.5e-27 \| 6.5e-33 \| 229.1 \| 299 \| (58, 389) \| 682 \| (30, 334) \| 477 \| Gp3 \| Gp3; uniclust \| UniRef100\_A0A7W1RR82 \| 99.9 \| 4.2e-27 \| 7.8e-33 \| 236.0 \| 535 \| (32, 604) \| 682 \| (21, 632) \| 662 \| Phage terminase large subunit family protein \| Phage terminase large subunit family protein; uniclust \| UniRef100\_UPI0018EB5B5E \| 99.9 \| 5.3e-27 \| 9.8e-33 \| 224.6 \| 274 \| (326, 605) \| 682 \| (17, 324) \| 386 \| phage terminase large subunit family protein \| phage terminase large subunit family protein; uniclust \| UniRef100\_A0A2E6W638 \| 99.9 \| 5.4e-27 \| 1e-32 \| 235.8 \| 430 \| (60, 603) \| 682 \| (71, 529) \| 543 \| Terminase \| Terminase; uniclust \| UniRef100\_A0A1Y1R6T6 \| 99.9 \| 5.2e-27 \| 1.1e-32 \| 226.3 \| 159 \| (443, 608) \| 682 \| (25, 191) \| 229 \| Terminase large subunit GpA endonuclease domain-containing protein (Fragment) \| Terminase large subunit GpA endonuclease domain-containing protein (Fragment); uniclust \| UniRef100\_A0A971AUN1 \| 99.9 \| 6.6e-27 \| 1.2e-32 \| 231.7 \| 466 \| (68, 612) \| 682 \| (30, 521) \| 580 \| Phage terminase large subunit GpA ATPase domain-containing protein \| Phage terminase large subunit GpA ATPase domain-containing protein; uniclust \| UniRef100\_A0A497I8D6 \| 99.9 \| 8.6e-27 \| 1.6e-32 \| 229.8 \| 496 \| (37, 604) \| 682 \| (16, 528) \| 552 \| Phage terminase large subunit GpA ATPase domain-containing protein \| Phage terminase large subunit GpA ATPase domain-containing protein; uniclust \| UniRef100\_UPI001C92A03A \| 99.9 \| 8.7e-27 \| 1.6e-32 \| 222.8 \| 281 \| (307, 604) \| 682 \| (4, 308) \| 411 \| phage terminase large subunit family protein \| phage terminase large subunit family protein; uniclust \| UniRef100\_A0A376MST0 \| 99.9 \| 8.7e-27 \| 1.6e-32 \| 204.9 \| 164 \| (326, 491) \| 682 \| (31, 196) \| 198 \| Terminase GpA \| Terminase GpA; uniclust \| UniRef100\_UPI0022F0AE72 \| 99.9 \| 9.2e-27 \| 1.7e-32 \| 192.0 \| 140 \| (183, 337) \| 682 \| (1, 143) \| 143 \| phage terminase large subunit family protein \| phage terminase large subunit family protein; uniclust \| UniRef100\_A0A9C9TIT2 \| 99.9 \| 9.3e-27 \| 1.7e-32 \| 224.1 \| 315 \| (20, 352) \| 682 \| (5, 355) \| 437 \| Uncharacterized protein \| Uncharacterized protein; uniclust \| UniRef100\_UPI0018EACED0 \| 99.9 \| 1e-26 \| 1.9e-32 \| 197.7 \| 147 \| (139, 299) \| 682 \| (1, 149) \| 153 \| phage terminase large subunit family protein \| phage terminase large subunit family protein; uniclust \| UniRef100\_UPI00214843D1 \| 99.9 \| 1.7e-26 \| 3.2e-32 \| 193.4 \| 152 \| (178, 348) \| 682 \| (1, 157) \| 157 \| phage terminase large subunit family protein \| phage terminase large subunit family protein; uniclust \| UniRef100\_A0A3C0YPZ3 \| 99.9 \| 1.8e-26 \| 3.3e-32 \| 217.9 \| 275 \| (326, 608) \| 682 \| (51, 342) \| 368 \| Terminase large subunit GpA endonuclease domain-containing protein \| Terminase large subunit GpA endonuclease domain-containing protein; uniclust \| UniRef100\_A0A7M3MA58 \| 99.9 \| 2.3e-26 \| 4.2e-32 \| 206.8 \| 228 \| (109, 351) \| 682 \| (1, 249) \| 251 \| Phage terminase large subunit family protein (Fragment) \| Phage terminase large subunit family protein (Fragment); uniclust \| UniRef100\_UPI000ACF156D \| 99.9 \| 2.6e-26 \| 4.9e-32 \| 201.4 \| 159 \| (152, 324) \| 682 \| (12, 173) \| 190 \| phage terminase large subunit family protein \| phage terminase large subunit family protein; uniclust \| UniRef100\_UPI001BAE199D \| 99.9 \| 3.2e-26 \| 5.8e-32 \| 223.5 \| 326 \| (22, 367) \| 682 \| (15, 375) \| 495 \| phage terminase large subunit family protein \| phage terminase large subunit family protein; uniclust \| UniRef100\_A0A6M0HXK1 \| 99.9 \| 3.3e-26 \| 6e-32 \| 205.6 \| 207 \| (150, 370) \| 682 \| (6, 249) \| 249 \| Terminase (Fragment) \| Terminase (Fragment); uniclust \| UniRef100\_A0A2M9P6D8 \| 99.9 \| 6.5e-26 \| 1.2e-31 \| 193.0 \| 126 \| (306, 436) \| 682 \| (1, 137) \| 152 \| Phage tail protein (Fragment) \| Phage tail protein (Fragment); uniclust \| UniRef100\_UPI00106A2ED5 \| 99.9 \| 7.5e-26 \| 1.4e-31 \| 206.0 \| 182 \| (420, 608) \| 682 \| (2, 190) \| 237 \| phage terminase large subunit family protein \| phage terminase large subunit family protein; uniclust \| UniRef100\_UPI0020775569 \| 99.9 \| 8e-26 \| 1.5e-31 \| 214.4 \| 198 \| (326, 529) \| 682 \| (6, 205) \| 380 \| phage terminase large subunit family protein \| phage terminase large subunit family protein; uniclust \| UniRef100\_A0A6M3IK98 \| 99.9 \| 7.7e-26 \| 1.5e-31 \| 219.5 \| 305 \| (237, 602) \| 682 \| (2, 313) \| 332 \| Putative tail assembly (Fragment) \| Putative tail assembly (Fragment); uniclust \| UniRef100\_A0A0F9JEA4 \| 99.9 \| 9.5e-26 \| 1.7e-31 \| 192.8 \| 175 \| (73, 248) \| 682 \| (1, 177) \| 177 \| Phage terminase large subunit GpA ATPase domain-containing protein \| Phage terminase large subunit GpA ATPase domain-containing protein; uniclust \| UniRef100\_A0A0E3GM94 \| 99.9 \| 9.7e-26 \| 1.8e-31 \| 224.3 \| 523 \| (40, 604) \| 682 \| (12, 584) \| 595 \| Uncharacterized protein \| Uncharacterized protein; uniclust \| UniRef100\_UPI00207B9352 \| 99.9 \| 1e-25 \| 1.8e-31 \| 196.1 \| 176 \| (211, 397) \| 682 \| (9, 188) \| 199 \| phage terminase large subunit family protein \| phage terminase large subunit family protein; uniclust \| UniRef100\_UPI0015907DDA \| 99.9 \| 1.2e-25 \| 2.1e-31 \| 210.7 \| 240 \| (365, 608) \| 682 \| (1, 264) \| 342 \| phage terminase large subunit family protein \| phage terminase large subunit family protein; uniclust \| UniRef100\_A0A965PS09 \| 99.9 \| 1.3e-25 \| 2.4e-31 \| 217.3 \| 374 \| (36, 431) \| 682 \| (12, 413) \| 452 \| Phage terminase large subunit GpA ATPase domain-containing protein (Fragment) \| Phage terminase large subunit GpA ATPase domain-containing protein (Fragment); uniclust \| UniRef100\_A0A7C6PAD2 \| 99.9 \| 1.4e-25 \| 2.5e-31 \| 216.4 \| 293 \| (133, 496) \| 682 \| (1, 294) \| 438 \| Phage terminase large subunit GpA ATPase domain-containing protein \| Phage terminase large subunit GpA ATPase domain-containing protein; uniclust \| UniRef100\_D8A639 \| 99.9 \| 1.8e-25 \| 3.4e-31 \| 198.9 \| 178 \| (326, 505) \| 682 \| (23, 202) \| 232 \| Terminase large subunit GpA endonuclease domain-containing protein (Fragment) \| Terminase large subunit GpA endonuclease domain-containing protein (Fragment); uniclust \| UniRef100\_A0A7W4PGD7 \| 99.9 \| 1.8e-25 \| 3.4e-31 \| 214.0 \| 259 \| (55, 338) \| 682 \| (5, 293) \| 342 \| Phage terminase large subunit GpA ATPase domain-containing protein \| Phage terminase large subunit GpA ATPase domain-containing protein; uniclust \| UniRef100\_A0A1F9IYL2 \| 99.9 \| 2.1e-25 \| 4e-31 \| 204.4 \| 201 \| (387, 608) \| 682 \| (4, 222) \| 248 \| Terminase large subunit GpA endonuclease domain-containing protein \| Terminase large subunit GpA endonuclease domain-containing protein; uniclust \| UniRef100\_A0A496P6X6 \| 99.9 \| 2.7e-25 \| 5e-31 \| 206.7 \| 206 \| (382, 608) \| 682 \| (10, 240) \| 270 \| Terminase (Fragment) \| Terminase (Fragment); uniclust \| UniRef100\_A0A1V5N0R1 \| 99.9 \| 2.9e-25 \| 5.6e-31 \| 242.6 \| 252 \| (355, 608) \| 682 \| (802, 1099) \| 1127 \| Phage terminase large subunit (GpA) \| Phage terminase large subunit (GpA); uniclust \| UniRef100\_A0A0E4BQP1 \| 99.9 \| 3.2e-25 \| 5.9e-31 \| 203.4 \| 218 \| (377, 605) \| 682 \| (11, 240) \| 288 \| Terminase large subunit GpA endonuclease domain-containing protein \| Terminase large subunit GpA endonuclease domain-containing protein; uniclust \| UniRef100\_UPI0004D72F9B \| 99.9 \| 4.3e-25 \| 7.9e-31 \| 223.0 \| 230 \| (372, 608) \| 682 \| (424, 661) \| 697 \| terminase small subunit \| terminase small subunit; uniclust \| UniRef100\_A0A369QXE7 \| 99.9 \| 5.1e-25 \| 9.3e-31 \| 204.5 \| 215 \| (377, 604) \| 682 \| (20, 251) \| 315 \| Terminase large subunit GpA endonuclease domain-containing protein \| Terminase large subunit GpA endonuclease domain-containing protein; uniclust \| UniRef100\_A0A941P6F5 \| 99.9 \| 5.4e-25 \| 9.9e-31 \| 220.2 \| 538 \| (36, 606) \| 682 \| (15, 613) \| 625 \| Phage terminase large subunit family protein \| Phage terminase large subunit family protein; uniclust \| UniRef100\_A0A965UQ11 \| 99.8 \| 6e-25 \| 1.1e-30 \| 193.4 \| 162 \| (100, 266) \| 682 \| (35, 199) \| 213 \| Phage terminase large subunit GpA ATPase domain-containing protein (Fragment) \| Phage terminase large subunit GpA ATPase domain-containing protein (Fragment); uniclust \| UniRef100\_A0A894KLY6 \| 99.8 \| 6.4e-25 \| 1.2e-30 \| 191.1 \| 148 \| (217, 373) \| 682 \| (3, 155) \| 198 \| Terminase large subunit (Fragment) \| Terminase large subunit (Fragment); uniclust \| UniRef100\_A0A2D5TJ79 \| 99.8 \| 6.6e-25 \| 1.2e-30 \| 233.5 \| 446 \| (102, 603) \| 682 \| (489, 943) \| 988 \| DOD-type homing endonuclease domain-containing protein \| DOD-type homing endonuclease domain-containing protein; uniclust \| UniRef100\_A0A3M2D4J1 \| 99.8 \| 7.6e-25 \| 1.4e-30 \| 219.8 \| 496 \| (40, 602) \| 682 \| (98, 619) \| 646 \| Phage terminase large subunit GpA ATPase domain-containing protein \| Phage terminase large subunit GpA ATPase domain-containing protein; uniclust \| UniRef100\_A0A061NTX6 \| 99.8 \| 9.4e-25 \| 1.8e-30 \| 192.0 \| 139 \| (467, 613) \| 682 \| (1, 144) \| 172 \| Phage terminase, large subunit \| Phage terminase, large subunit; uniclust \| UniRef100\_A0A0P6WAH7 \| 99.8 \| 1.1e-24 \| 2.1e-30 \| 199.5 \| 201 \| (27, 229) \| 682 \| (14, 218) \| 226 \| Phage terminase large subunit GpA ATPase domain-containing protein \| Phage terminase large subunit GpA ATPase domain-containing protein; uniclust \| UniRef100\_A0A6M3XH57 \| 99.8 \| 1.2e-24 \| 2.2e-30 \| 224.3 \| 424 \| (109, 609) \| 682 \| (426, 871) \| 883 \| Putative terminase \| Putative terminase; uniclust \| UniRef100\_A0A969XTT3 \| 99.8 \| 1.3e-24 \| 2.3e-30 \| 201.3 \| 235 \| (27, 266) \| 682 \| (17, 255) \| 308 \| Phage terminase large subunit GpA ATPase domain-containing protein (Fragment) \| Phage terminase large subunit GpA ATPase domain-containing protein (Fragment); uniclust \| UniRef100\_A0A539CTX1 \| 99.8 \| 1.5e-24 \| 2.8e-30 \| 200.1 \| 225 \| (111, 354) \| 682 \| (46, 274) \| 299 \| Terminase \| Terminase; uniclust \| UniRef100\_A0A529NU64 \| 99.8 \| 1.9e-24 \| 3.5e-30 \| 189.2 \| 200 \| (61, 262) \| 682 \| (2, 204) \| 204 \| Terminase (Fragment) \| Terminase (Fragment); uniclust \| UniRef100\_UPI002283A5C9 \| 99.8 \| 2.8e-24 \| 5.1e-30 \| 175.0 \| 123 \| (306, 433) \| 682 \| (1, 127) \| 130 \| phage terminase large subunit family protein \| phage terminase large subunit family protein; uniclust \| UniRef100\_A0A0B8PLS3 \| 99.8 \| 2.7e-24 \| 5.2e-30 \| 197.5 \| 226 \| (58, 300) \| 682 \| (4, 239) \| 241 \| Phage terminase \| Phage terminase; uniclust \| UniRef100\_X0STT9 \| 99.8 \| 3.2e-24 \| 5.8e-30 \| 207.8 \| 388 \| (155, 608) \| 682 \| (11, 419) \| 449 \| Terminase (Fragment) \| Terminase (Fragment); uniclust \| UniRef100\_A0A956ER55 \| 99.8 \| 3.3e-24 \| 6e-30 \| 203.4 \| 275 \| (216, 498) \| 682 \| (6, 325) \| 373 \| Phage terminase large subunit family protein (Fragment) \| Phage terminase large subunit family protein (Fragment); uniclust \| UniRef100\_A0A178IH68 \| 99.8 \| 3.4e-24 \| 6.7e-30 \| 230.5 \| 527 \| (31, 604) \| 682 \| (112, 735) \| 770 \| Terminase \| Terminase; uniclust \| UniRef100\_A0A0F9MVA9 \| 99.8 \| 3.9e-24 \| 7.7e-30 \| 230.9 \| 228 \| (375, 608) \| 682 \| (444, 701) \| 743 \| Terminase large subunit GpA endonuclease domain-containing protein \| Terminase large subunit GpA endonuclease domain-containing protein; uniclust \| UniRef100\_A0A844G3X9 \| 99.8 \| 4.3e-24 \| 7.9e-30 \| 213.3 \| 430 \| (30, 499) \| 682 \| (26, 484) \| 605 \| Phage terminase large subunit GpA ATPase domain-containing protein \| Phage terminase large subunit GpA ATPase domain-containing protein; uniclust \| UniRef100\_A0A0M0C1A5 \| 99.8 \| 3.9e-24 \| 8.2e-30 \| 233.0 \| 164 \| (62, 241) \| 682 \| (101, 266) \| 573 \| Uncharacterized protein \| Uncharacterized protein; uniclust \| UniRef100\_A0A645HKJ2 \| 99.8 \| 5e-24 \| 9.7e-30 \| 193.0 \| 170 \| (422, 608) \| 682 \| (2, 173) \| 204 \| Terminase large subunit GpA endonuclease domain-containing protein \| Terminase large subunit GpA endonuclease domain-containing protein; uniclust \| UniRef100\_A0A936HC74 \| 99.8 \| 6e-24 \| 1.1e-29 \| 192.8 \| 183 \| (232, 428) \| 682 \| (9, 218) \| 219 \| Phage terminase large subunit family protein \| Phage terminase large subunit family protein; uniclust \| UniRef100\_A0A950WLK3 \| 99.8 \| 6.9e-24 \| 1.3e-29 \| 206.1 \| 413 \| (24, 470) \| 682 \| (21, 451) \| 461 \| Phage terminase large subunit family protein (Fragment) \| Phage terminase large subunit family protein (Fragment); uniclust \| UniRef100\_A0A2X3JNN5 \| 99.8 \| 6.8e-24 \| 1.3e-29 \| 185.2 \| 171 \| (351, 527) \| 682 \| (3, 175) \| 176 \| Phage terminase large subunit \| Phage terminase large subunit; uniclust \| UniRef100\_B4CXQ4 \| 99.8 \| 8.4e-24 \| 1.5e-29 \| 203.0 \| 319 \| (246, 604) \| 682 \| (21, 386) \| 411 \| Terminase large subunit GpA endonuclease domain-containing protein \| Terminase large subunit GpA endonuclease domain-containing protein; uniclust \| UniRef100\_A0A9D7JDZ3 \| 99.8 \| 8.7e-24 \| 1.6e-29 \| 193.8 \| 219 \| (327, 552) \| 682 \| (2, 234) \| 282 \| Phage terminase large subunit family protein \| Phage terminase large subunit family protein; uniclust \| UniRef100\_X0S7D8 \| 99.8 \| 9.1e-24 \| 1.7e-29 \| 193.4 \| 251 \| (45, 307) \| 682 \| (2, 271) \| 280 \| Phage terminase large subunit GpA ATPase domain-containing protein (Fragment) \| Phage terminase large subunit GpA ATPase domain-containing protein (Fragment); uniclust \| UniRef100\_A0A9E7YBD2 \| 99.8 \| 9.4e-24 \| 1.7e-29 \| 198.0 \| 240 \| (364, 608) \| 682 \| (1, 264) \| 337 \| Phage terminase large subunit family protein \| Phage terminase large subunit family protein; uniclust \| UniRef100\_A0A0S9QR24 \| 99.8 \| 9.5e-24 \| 1.7e-29 \| 200.3 \| 299 \| (166, 472) \| 682 \| (12, 351) \| 372 \| Terminase \| Terminase; uniclust \| UniRef100\_A0A660UHB7 \| 99.8 \| 1.3e-23 \| 2.5e-29 \| 203.4 \| 353 \| (234, 604) \| 682 \| (4, 407) \| 445 \| Terminase (Fragment) \| Terminase (Fragment); uniclust \| UniRef100\_A0A0E2L6E2 \| 99.8 \| 1.3e-23 \| 2.5e-29 \| 191.9 \| 202 \| (400, 608) \| 682 \| (1, 210) \| 246 \| Large terminase \| Large terminase; uniclust \| UniRef100\_A0A084XW67 \| 99.8 \| 1.5e-23 \| 2.7e-29 \| 216.0 \| 338 \| (39, 416) \| 682 \| (27, 373) \| 854 \| Bacteriophage tail assembly protein \| Bacteriophage tail assembly protein; uniclust \| UniRef100\_A0A0G0PPN9 \| 99.8 \| 1.5e-23 \| 2.7e-29 \| 215.3 \| 316 \| (38, 408) \| 682 \| (20, 335) \| 824 \| Bacteriophage tail assembly protein \| Bacteriophage tail assembly protein; uniclust \| UniRef100\_A0A1Y2KA79 \| 99.8 \| 1.6e-23 \| 3e-29 \| 181.8 \| 129 \| (19, 147) \| 682 \| (11, 141) \| 149 \| Phage terminase large subunit GpA ATPase domain-containing protein \| Phage terminase large subunit GpA ATPase domain-containing protein; uniclust \| UniRef100\_UPI0009DF5824 \| 99.8 \| 1.7e-23 \| 3.2e-29 \| 207.5 \| 178 \| (19, 196) \| 682 \| (29, 208) \| 557 \| phage terminase large subunit family protein \| phage terminase large subunit family protein; uniclust \| UniRef100\_A0A949VC24 \| 99.8 \| 1.8e-23 \| 3.4e-29 \| 198.1 \| 327 \| (81, 420) \| 682 \| (9, 364) \| 366 \| Phage terminase large subunit family protein (Fragment) \| Phage terminase large subunit family protein (Fragment); uniclust \| UniRef100\_A0A3S0BG77 \| 99.8 \| 1.9e-23 \| 3.6e-29 \| 211.0 \| 489 \| (42, 604) \| 682 \| (40, 553) \| 577 \| Phage terminase large subunit GpA ATPase domain-containing protein \| Phage terminase large subunit GpA ATPase domain-containing protein; uniclust \| UniRef100\_A0A7V4S3L3 \| 99.8 \| 2e-23 \| 3.7e-29 \| 208.6 \| 484 \| (37, 604) \| 682 \| (81, 581) \| 603 \| Phage terminase large subunit GpA ATPase domain-containing protein \| Phage terminase large subunit GpA ATPase domain-containing protein; uniclust \| UniRef100\_A0A7C7KMY0 \| 99.8 \| 2.3e-23 \| 4.1e-29 \| 190.4 \| 239 \| (24, 266) \| 682 \| (17, 259) \| 274 \| Phage terminase large subunit GpA ATPase domain-containing protein (Fragment) \| Phage terminase large subunit GpA ATPase domain-containing protein (Fragment); uniclust \| UniRef100\_A0A965UR17 \| 99.8 \| 2.4e-23 \| 4.4e-29 \| 200.0 \| 219 \| (27, 250) \| 682 \| (39, 260) \| 410 \| Phage terminase large subunit GpA ATPase domain-containing protein \| Phage terminase large subunit GpA ATPase domain-containing protein; uniclust \| UniRef100\_UPI001BAF8797 \| 99.8 \| 2.5e-23 \| 4.6e-29 \| 193.0 \| 247 \| (286, 553) \| 682 \| (48, 302) \| 308 \| phage terminase large subunit family protein \| phage terminase large subunit family protein; uniclust \| UniRef100\_A0A822IGS8 \| 99.8 \| 3.2e-23 \| 6.1e-29 \| 211.1 \| 444 \| (42, 600) \| 682 \| (41, 501) \| 523 \| Terminase \| Terminase; uniclust \| UniRef100\_A0A963I8E7 \| 99.8 \| 3.9e-23 \| 7.2e-29 \| 189.4 \| 243 \| (327, 570) \| 682 \| (1, 269) \| 280 \| Phage terminase large subunit family protein (Fragment) \| Phage terminase large subunit family protein (Fragment); uniclust \| UniRef100\_A0A376JFP1 \| 99.8 \| 3.9e-23 \| 7.4e-29 \| 180.9 \| 112 \| (326, 437) \| 682 \| (50, 163) \| 170 \| Terminase large subunit (Gp2) \| Terminase large subunit (Gp2); uniclust \| UniRef100\_A0A1H4FKT7 \| 99.8 \| 4.1e-23 \| 7.6e-29 \| 197.2 \| 182 \| (16, 197) \| 682 \| (22, 205) \| 390 \| Phage terminase large subunit (GpA) \| Phage terminase large subunit (GpA); uniclust \| UniRef100\_UPI0006F2C31D \| 99.8 \| 4.7e-23 \| 8.6e-29 \| 186.5 \| 169 \| (444, 616) \| 682 \| (8, 184) \| 254 \| phage terminase large subunit family protein \| phage terminase large subunit family protein; uniclust \| UniRef100\_A8TUC3 \| 99.8 \| 5.2e-23 \| 9.6e-29 \| 177.5 \| 139 \| (158, 311) \| 682 \| (2, 142) \| 183 \| Phage terminase GpA \| Phage terminase GpA; uniclust \| UniRef100\_A0A2E1H9V6 \| 99.8 \| 7.1e-23 \| 1.3e-28 \| 176.4 \| 164 \| (132, 311) \| 682 \| (2, 165) \| 181 \| Phage terminase large subunit GpA ATPase domain-containing protein \| Phage terminase large subunit GpA ATPase domain-containing protein; uniclust \| UniRef100\_A0A420Z3P3 \| 99.8 \| 7.2e-23 \| 1.3e-28 \| 219.3 \| 441 \| (93, 601) \| 682 \| (537, 1001) \| 1025 \| DOD-type homing endonuclease domain-containing protein \| DOD-type homing endonuclease domain-containing protein; uniclust \| UniRef100\_S9S9L7 \| 99.8 \| 1.1e-22 \| 2e-28 \| 188.2 \| 214 \| (390, 607) \| 682 \| (8, 230) \| 299 \| Terminase GpA (Fragment) \| Terminase GpA (Fragment); uniclust \| UniRef100\_A0A3B0M177 \| 99.8 \| 1.1e-22 \| 2e-28 \| 172.9 \| 144 \| (262, 419) \| 682 \| (9, 156) \| 166 \| Uncharacterized protein \| Uncharacterized protein; uniclust \| UniRef100\_A0A447UU55 \| 99.8 \| 1.1e-22 \| 2.1e-28 \| 193.8 \| 197 \| (392, 604) \| 682 \| (21, 227) \| 284 \| Phage terminase large subunit \| Phage terminase large subunit; uniclust \| UniRef100\_UPI00069CACCF \| 99.8 \| 1.3e-22 \| 2.4e-28 \| 183.1 \| 210 \| (102, 323) \| 682 \| (4, 217) \| 248 \| phage terminase large subunit family protein \| phage terminase large subunit family protein; uniclust \| UniRef100\_A0A087RM50 \| 99.8 \| 1.7e-22 \| 3.5e-28 \| 219.7 \| 491 \| (34, 603) \| 682 \| (47, 602) \| 621 \| Terminase-like domain containing protein \| Terminase-like domain containing protein; uniclust \| UniRef100\_UPI000810B029 \| 99.8 \| 2.8e-22 \| 5.1e-28 \| 180.3 \| 194 \| (106, 319) \| 682 \| (6, 224) \| 241 \| phage terminase large subunit family protein \| phage terminase large subunit family protein; uniclust \| UniRef100\_A0A022LQP3 \| 99.8 \| 2.6e-22 \| 5.3e-28 \| 215.7 \| 258 \| (40, 408) \| 682 \| (25, 285) \| 596 \| Terminase large subunit gp17-like C-terminal domain-containing protein \| Terminase large subunit gp17-like C-terminal domain-containing protein; uniclust \| UniRef100\_H0T4D4 \| 99.8 \| 3.2e-22 \| 5.9e-28 \| 181.1 \| 202 \| (394, 604) \| 682 \| (19, 222) \| 252 \| Terminase large subunit GpA endonuclease domain-containing protein \| Terminase large subunit GpA endonuclease domain-containing protein; uniclust \| UniRef100\_A0A7V4LST6 \| 99.8 \| 4.5e-22 \| 8.2e-28 \| 190.5 \| 165 \| (102, 267) \| 682 \| (2, 166) \| 392 \| Glycosyl hydrolase family 32 N-terminal domain-containing protein \| Glycosyl hydrolase family 32 N-terminal domain-containing protein; uniclust \| UniRef100\_A0A2X2UA18 \| 99.8 \| 4.5e-22 \| 8.8e-28 \| 188.0 \| 163 \| (437, 606) \| 682 \| (23, 198) \| 259 \| Bacteriophage tail assembly protein \| Bacteriophage tail assembly protein; uniclust \| UniRef100\_UPI0011BAB5E6 \| 99.8 \| 7.2e-22 \| 1.3e-27 \| 180.3 \| 216 \| (353, 568) \| 682 \| (13, 262) \| 267 \| phage terminase large subunit family protein \| phage terminase large subunit family protein; uniclust \| UniRef100\_A0A379V3E7 \| 99.8 \| 7.8e-22 \| 1.6e-27 \| 190.5 \| 189 \| (99, 302) \| 682 \| (26, 224) \| 248 \| Terminase \| Terminase; uniclust \| UniRef100\_A0A2E9Y9W4 \| 99.8 \| 8.2e-22 \| 1.6e-27 \| 175.4 \| 110 \| (490, 600) \| 682 \| (7, 120) \| 163 \| Terminase large subunit GpA endonuclease domain-containing protein \| Terminase large subunit GpA endonuclease domain-containing protein; uniclust \| UniRef100\_A0A0A0RP77 \| 99.8 \| 1.1e-21 \| 2e-27 \| 188.1 \| 258 \| (123, 406) \| 682 \| (12, 285) \| 295 \| Large terminase subunit \| Large terminase subunit; uniclust \| UniRef100\_A0A0F9CRH0 \| 99.8 \| 1.2e-21 \| 2.2e-27 \| 173.8 \| 211 \| (164, 407) \| 682 \| (3, 216) \| 217 \| Phage terminase large subunit GpA ATPase domain-containing protein (Fragment) \| Phage terminase large subunit GpA ATPase domain-containing protein (Fragment); uniclust \| UniRef100\_UPI0009657AEE \| 99.8 \| 1.2e-21 \| 2.3e-27 \| 163.6 \| 144 \| (123, 266) \| 682 \| (1, 145) \| 148 \| phage terminase large subunit family protein \| phage terminase large subunit family protein; uniclust \| UniRef100\_A0A0F7KJN6 \| 99.8 \| 1.3e-21 \| 2.4e-27 \| 180.6 \| 215 \| (383, 606) \| 682 \| (3, 230) \| 289 \| Phage terminase large subunit GpA \| Phage terminase large subunit GpA; uniclust \| UniRef100\_A0A1E3GZ05 \| 99.8 \| 1.4e-21 \| 2.5e-27 \| 168.0 \| 133 \| (256, 407) \| 682 \| (2, 140) \| 176 \| Phage terminase large subunit (GpA) \| Phage terminase large subunit (GpA); uniclust \| UniRef100\_UPI000D368E8B \| 99.7 \| 1.7e-21 \| 3.2e-27 \| 172.6 \| 200 \| (391, 606) \| 682 \| (9, 208) \| 215 \| phage terminase large subunit family protein \| phage terminase large subunit family protein; uniclust \| UniRef100\_UPI00190FBED9 \| 99.7 \| 1.8e-21 \| 3.4e-27 \| 159.3 \| 113 \| (390, 504) \| 682 \| (4, 116) \| 118 \| phage terminase large subunit family protein \| phage terminase large subunit family protein; pdb70 \| 2O0J\_A \| 98.7 \| 2.4e-12 \| 2.5e-16 \| 134.2 \| 195 \| (31, 241) \| 682 \| (132, 327) \| 385 \| DNA packaging protein Gp17 \| 2O0J\_A DNA packaging protein Gp17 nucleotide-binding fold, HYDROLASE HET: ADP; pdb70 \| 3EZK\_E \| 98.4 \| 4e-11 \| 4.6e-15 \| 128.0 \| 195 \| (30, 240) \| 682 \| (106, 301) \| 577 \| DNA packaging protein Gp17 \| 3EZK\_E DNA packaging protein Gp17 pentameric motor, DNA packaging, Alternative; pdb70 \| 3CPE\_A \| 98.4 \| 5e-11 \| 5.7e-15 \| 127.8 \| 196 \| (30, 241) \| 682 \| (131, 327) \| 592 \| DNA packaging protein Gp17 \| 3CPE\_A DNA packaging protein Gp17 large terminase, Alternative initiation, ATP-binding HET: PO4; pdb70 \| 4ZNK\_A \| 98.3 \| 9.5e-11 \| 1.2e-14 \| 110.2 \| 175 \| (38, 242) \| 682 \| (8, 200) \| 274 \| Phage terminase large subunit \| 4ZNK\_A Phage terminase large subunit DNA Translocation, VIRAL PROTEIN HET: SO4; pdb70 \| 4BIJ\_A \| 98.0 \| 2.1e-09 \| 2.6e-13 \| 109.4 \| 191 \| (30, 238) \| 682 \| (8, 214) \| 476 \| DNA MATURASE B \| 4BIJ\_A DNA MATURASE B HYDROLASE, ATPASE, DNA TRANSLOCATION, SINGLE-PARTICLE; pdb70 \| 4BIL\_C \| 98.0 \| 2.1e-09 \| 2.6e-13 \| 109.4 \| 191 \| (30, 238) \| 682 \| (8, 214) \| 476 \| DNA MATURASE B \| 4BIL\_C DNA MATURASE B HYDROLASE, PACKAGING MOTOR, CONNECTOR, DNA; pdb70 \| 5OEA\_B \| 97.8 \| 7.7e-09 \| 9.8e-13 \| 101.6 \| 165 \| (61, 240) \| 682 \| (12, 189) \| 420 \| Large subunit terminase \| 5OEA\_B Large subunit terminase large terminase, VIRAL PROTEIN HET: AGS; pdb70 \| 5OE8\_B \| 97.7 \| 1.1e-08 \| 1.5e-12 \| 100.9 \| 164 \| (62, 240) \| 682 \| (13, 189) \| 430 \| Large subunit terminase \| 5OE8\_B Large subunit terminase large terminase, VIRAL PROTEIN; pdb70 \| 5OE9\_A \| 97.7 \| 1.1e-08 \| 1.5e-12 \| 100.9 \| 164 \| (62, 240) \| 682 \| (13, 189) \| 430 \| Large subunit terminase \| 5OE9\_A Large subunit terminase large terminase, VIRAL PROTEIN HET: SO4; pdb70 \| 1RIF\_A \| 97.7 \| 2.1e-08 \| 2.4e-12 \| 96.6 \| 149 \| (62, 227) \| 682 \| (111, 264) \| 282 \| DNA helicase uvsW \| 1RIF\_A DNA helicase uvsW Bacteriophage, T4, Helicase, UvsW, RecG; pdb70 \| 2OCA\_A \| 97.4 \| 7.3e-08 \| 9e-12 \| 98.7 \| 149 \| (62, 227) \| 682 \| (111, 264) \| 510 \| ATP-dependent DNA helicase uvsW (E.C.3.6.1.8) \| 2OCA\_A ATP-dependent DNA helicase uvsW (E.C.3.6.1.8) ATP-dependant helicase, T4-bacteriophage, Recombination, HYDROLASE; pdb70 \| 3MWL\_A \| 97.4 \| 8.2e-08 \| 1.1e-11 \| 82.4 \| 70 \| (63, 135) \| 682 \| (22, 96) \| 207 \| Heat resistant RNA dependent ATPase \| 3MWL\_A Heat resistant RNA dependent ATPase RNA HELICASE, RIBOSOME BIOGENESIS, THERMOPHILIC HET: SO4, 8OX; pdb70 \| 4BUJ\_A \| 97.2 \| 2.8e-07 \| 3.1e-11 \| 104.7 \| 148 \| (61, 228) \| 682 \| (330, 485) \| 1044 \| ANTIVIRAL HELICASE SKI2 (E.C.3.6.4.13), SUPERKILLER \| 4BUJ\_A ANTIVIRAL HELICASE SKI2 (E.C.3.6.4.13), SUPERKILLER HYDROLASE, DEXH BOX HELICASE, RNA HET: SO4; pdb70 \| 2GXQ\_A \| 97.2 \| 2.4e-07 \| 3.3e-11 \| 79.0 \| 70 \| (63, 135) \| 682 \| (22, 96) \| 207 \| heat resistant RNA dependent ATPase \| 2GXQ\_A heat resistant RNA dependent ATPase RNA helicase, atomic resolution, AMP HET: AMP; pdb70 \| 3BER\_A \| 97.2 \| 2.7e-07 \| 3.3e-11 \| 85.1 \| 69 \| (63, 134) \| 682 \| (64, 134) \| 249 \| Probable ATP-dependent RNA helicase DDX47 \| 3BER\_A Probable ATP-dependent RNA helicase DDX47 RNA HELICASE, DEAD, AMP, Structural HET: PGE, AMP; pdb70 \| 4KBF\_A \| 97.2 \| 2.4e-07 \| 3.3e-11 \| 85.1 \| 70 \| (63, 135) \| 682 \| (22, 96) \| 365 \| Heat resistant RNA dependent ATPase \| 4KBF\_A Heat resistant RNA dependent ATPase DEAD BOX RNA HELICASE, DIMER HET: AMP; pdb70 \| 4BUJ\_E \| 97.2 \| 3.2e-07 \| 3.6e-11 \| 104.2 \| 149 \| (60, 228) \| 682 \| (329, 485) \| 1044 \| ANTIVIRAL HELICASE SKI2 (E.C.3.6.4.13), SUPERKILLER \| 4BUJ\_E ANTIVIRAL HELICASE SKI2 (E.C.3.6.4.13), SUPERKILLER HYDROLASE, DEXH BOX HELICASE, RNA HET: SO4; pdb70 \| 5MC6\_h \| 97.2 \| 3.7e-07 \| 4e-11 \| 106.7 \| 149 \| (60, 228) \| 682 \| (325, 481) \| 1287 \| 40S ribosomal protein S3, 40S \| 5MC6\_h 40S ribosomal protein S3, 40S cryo-EM, ribosome, RNA, helicase HET: 5CT; pdb70 \| 5F9F\_I \| 97.1 \| 4e-07 \| 4.9e-11 \| 96.6 \| 127 \| (61, 196) \| 682 \| (9, 149) \| 695 \| Probable ATP-dependent RNA helicase DDX58/RNA \| 5F9F\_I Probable ATP-dependent RNA helicase DDX58/RNA Complex, RIG-I, capped RNA, self HET: BU3, ETF; pdb70 \| 6FT6\_MM \| 97.1 \| 4.4e-07 \| 5.1e-11 \| 102.1 \| 145 \| (61, 227) \| 682 \| (84, 236) \| 1011 \| Ribosomal 60S subunit protein L2B \| 6FT6\_MM Ribosomal 60S subunit protein L2B RNA exosome, Ribosome, pre-ribosome, Mtr4 HET: GTP; pdb70 \| 5Z3G\_Y \| 97.1 \| 4.2e-07 \| 5.3e-11 \| 91.6 \| 69 \| (63, 134) \| 682 \| (63, 137) \| 505 \| Ribosome biogenesis protein RLP7, Ribosome \| 5Z3G\_Y Ribosome biogenesis protein RLP7, Ribosome ribosome, pre-60S, pre-ribosome, protein-RNA complex; pdb70 \| 6EM3\_D \| 97.1 \| 4.2e-07 \| 5.3e-11 \| 91.6 \| 69 \| (63, 134) \| 682 \| (63, 137) \| 505 \| Ribosome production factor 1, 60S \| 6EM3\_D Ribosome production factor 1, 60S Large Subunit Biogenesis Nucleolus, RIBOSOME; pdb70 \| 1Q0U\_B \| 97.1 \| 4.2e-07 \| 5.7e-11 \| 77.6 \| 69 \| (63, 134) \| 682 \| (25, 95) \| 219 \| BstDEAD \| 1Q0U\_B BstDEAD DEAD PROTEIN, RNA BINDING PROTEIN; pdb70 \| 6UV1\_A \| 97.1 \| 4.6e-07 \| 6.1e-11 \| 87.7 \| 69 \| (63, 134) \| 682 \| (84, 159) \| 448 \| Probable ATP-dependent RNA helicase DDX17/RNA \| 6UV1\_A Probable ATP-dependent RNA helicase DDX17/RNA DEAD-box ATPase, RNA helicase, RNA HET: MG, ADP; pdb70 \| 6UV2\_A \| 97.1 \| 4.6e-07 \| 6.1e-11 \| 87.7 \| 69 \| (63, 134) \| 682 \| (84, 159) \| 448 \| Probable ATP-dependent RNA helicase DDX17/RNA \| 6UV2\_A Probable ATP-dependent RNA helicase DDX17/RNA DEAD-box ATPase, RNA helicase, RNA HET: ADP; pdb70 \| 5E02\_A \| 97.1 \| 5.3e-07 \| 6.3e-11 \| 102.0 \| 145 \| (62, 228) \| 682 \| (178, 330) \| 1106 \| FRQ-interacting RNA helicase/RNA Complex \| 5E02\_A FRQ-interacting RNA helicase/RNA Complex circadian clock, RNA BINDING PROTEIN-RNA HET: ADP; pdb70 \| 6BB8\_A \| 97.1 \| 5.3e-07 \| 6.3e-11 \| 102.0 \| 145 \| (62, 228) \| 682 \| (178, 330) \| 1106 \| FRQ-interacting RNA helicase \| 6BB8\_A FRQ-interacting RNA helicase RNA helicase, RNA BINDING PROTEIN; pdb70 \| 5OF4\_A \| 97.1 \| 5.6e-07 \| 6.5e-11 \| 94.7 \| 145 \| (62, 229) \| 682 \| (84, 245) \| 553 \| TFIIH basal transcription factor complex \| 5OF4\_A TFIIH basal transcription factor complex transcription initiation, DNA repair, multiprotein; pdb70 \| 6UV3\_A \| 97.1 \| 5.4e-07 \| 7.1e-11 \| 87.3 \| 69 \| (63, 134) \| 682 \| (84, 159) \| 448 \| Probable ATP-dependent RNA helicase DDX17/RNA \| 6UV3\_A Probable ATP-dependent RNA helicase DDX17/RNA DEAD-box ATPase, RNA helicase, RNA HET: ADP; pdb70 \| 1Q0U\_A \| 97.1 \| 5.7e-07 \| 7.7e-11 \| 77.0 \| 69 \| (63, 134) \| 682 \| (25, 95) \| 219 \| BstDEAD \| 1Q0U\_A BstDEAD DEAD PROTEIN, RNA BINDING PROTEIN HET: MSE; pdb70 \| 1WP9\_A \| 97.0 \| 5.7e-07 \| 8.2e-11 \| 84.6 \| 125 \| (62, 196) \| 682 \| (7, 139) \| 494 \| Hef helicase/nuclease \| 1WP9\_A Hef helicase/nuclease Helicase, ATPase, DNA replication, DNA HET: PO4; pdb70 \| 1WP9\_C \| 97.0 \| 5.9e-07 \| 8.4e-11 \| 84.5 \| 124 \| (62, 196) \| 682 \| (7, 139) \| 494 \| Hef helicase/nuclease \| 1WP9\_C Hef helicase/nuclease Helicase, ATPase, DNA replication, DNA HET: PO4; pdb70 \| 1FUU\_B \| 97.0 \| 6.3e-07 \| 8.5e-11 \| 84.4 \| 70 \| (63, 135) \| 682 \| (42, 113) \| 394 \| INITIATION FACTOR 4A \| 1FUU\_B INITIATION FACTOR 4A IF4A, HELICASE, DEAD-BOX PROTEIN, TRANSLATION; pdb70 \| 4A4Z\_A \| 97.0 \| 7.7e-07 \| 8.6e-11 \| 100.6 \| 148 \| (61, 228) \| 682 \| (36, 191) \| 997 \| ANTIVIRAL HELICASE SKI2 (E.C.3.6.4.13) \| 4A4Z\_A ANTIVIRAL HELICASE SKI2 (E.C.3.6.4.13) HYDROLASE, ATPASE, MRNA DEGRADATION, EXOSOME HET: ANP, EDO; pdb70 \| 2VSO\_B \| 97.0 \| 6.7e-07 \| 8.9e-11 \| 84.3 \| 70 \| (63, 135) \| 682 \| (43, 114) \| 395 \| ATP-DEPENDENT RNA HELICASE EIF4A (E.C.3.6.1.-) \| 2VSO\_B ATP-DEPENDENT RNA HELICASE EIF4A (E.C.3.6.1.-) ACETYLATION, ATP-BINDING, PHOSPHOPROTEIN, PROTEIN BIOSYNTHESIS HET: AMP; pdb70 \| 1HV8\_A \| 97.0 \| 6.5e-07 \| 9e-11 \| 82.1 \| 71 \| (63, 135) \| 682 \| (27, 98) \| 367 \| DEAD BOX HELICASE \| 1HV8\_A DEAD BOX HELICASE Helicase, RNA-binding Protein, ATPase, RNA HET: MSE, SO4; pdb70 \| 4U4C\_A \| 97.0 \| 8.1e-07 \| 9.6e-11 \| 99.3 \| 145 \| (61, 227) \| 682 \| (71, 223) \| 998 \| Mtr4, Dob1 (E.C.3.6.4.13), Air2-Trf4 Fusion \| 4U4C\_A Mtr4, Dob1 (E.C.3.6.4.13), Air2-Trf4 Fusion helicase, ATPase, poly(A)polymerase, RNA degradation HET: PG4, SO4; pdb70 \| 1FUU\_A \| 97.0 \| 7.9e-07 \| 1.1e-10 \| 83.7 \| 70 \| (63, 135) \| 682 \| (42, 113) \| 394 \| INITIATION FACTOR 4A \| 1FUU\_A INITIATION FACTOR 4A IF4A, HELICASE, DEAD-BOX PROTEIN, TRANSLATION HET: MSE; pdb70 \| 5FMF\_1 \| 97.0 \| 8.6e-07 \| 1.1e-10 \| 89.4 \| 142 \| (63, 228) \| 682 \| (67, 227) \| 492 \| DNA REPAIR HELICASE RAD25, SSL2 \| 5FMF\_1 DNA REPAIR HELICASE RAD25, SSL2 TRANSCRIPTION, PRE-INITIATION COMPLEX, RNA POLYMERASE; pdb70 \| 6NE3\_W \| 97.0 \| 8e-07 \| 1.1e-10 \| 84.4 \| 152 \| (63, 228) \| 682 \| (13, 177) \| 467 \| Histone H3.2, Histone H4, Histone \| 6NE3\_W Histone H3.2, Histone H4, Histone ISWI, Chromatin, Nucleosome, DNA, SNF2h HET: ADP; pdb70 \| 4XGT\_A \| 97.0 \| 9.6e-07 \| 1.1e-10 \| 98.8 \| 146 \| (61, 228) \| 682 \| (64, 217) \| 993 \| FRQ-interacting RNA helicase \| 4XGT\_A FRQ-interacting RNA helicase ATPase, circadian clock, HYDROLASE; pdb70 \| 4BRW\_A \| 97.0 \| 8.8e-07 \| 1.2e-10 \| 82.0 \| 69 \| (63, 134) \| 682 \| (22, 92) \| 377 \| ATP-DEPENDENT RNA HELICASE DHH1 (E.C.3.6.4.13) \| 4BRW\_A ATP-DEPENDENT RNA HELICASE DHH1 (E.C.3.6.4.13) HYDROLASE, TRANSLATIONAL REPRESSION, MRNP REMODELING HET: 1PE; pdb70 \| 5IVL\_B \| 97.0 \| 9.2e-07 \| 1.2e-10 \| 85.0 \| 69 \| (64, 135) \| 682 \| (24, 94) \| 429 \| DEAD-box ATP-dependent RNA helicase CshA \| 5IVL\_B DEAD-box ATP-dependent RNA helicase CshA Rec-A like domain DEAD-box helicase HET: SO4; pdb70 \| 1Z6A\_A \| 97.0 \| 9.7e-07 \| 1.2e-10 \| 87.9 \| 148 \| (63, 228) \| 682 \| (36, 190) \| 500 \| Helicase of the snf2/rad54 family \| 1Z6A\_A Helicase of the snf2/rad54 family HYDROLASE, RECOMBINATION, HYDROLASE-RECOMBINATION COMPLEX HET: PO4; pdb70 \| 6AIB\_A \| 97.0 \| 9.6e-07 \| 1.3e-10 \| 76.5 \| 69 \| (64, 135) \| 682 \| (24, 93) \| 214 \| DEAD-box ATP-dependent RNA helicase CshA \| 6AIB\_A DEAD-box ATP-dependent RNA helicase CshA STRUCTURAL PROTEIN; pdb70 \| 5ELX\_A \| 97.0 \| 9.3e-07 \| 1.3e-10 \| 82.3 \| 71 \| (63, 134) \| 682 \| (23, 95) \| 391 \| ATP-dependent RNA helicase DBP5/RNA Complex \| 5ELX\_A ATP-dependent RNA helicase DBP5/RNA Complex Fluorescent, Nucleotide, Mant, ADP, RNA HET: M2A; pdb70 \| 2Z0M\_A \| 96.9 \| 9.3e-07 \| 1.3e-10 \| 79.5 \| 64 \| (63, 134) \| 682 \| (15, 79) \| 337 \| 337aa long hypothetical ATP-dependent RNA \| 2Z0M\_A 337aa long hypothetical ATP-dependent RNA helicase, ATP-binding, Hydrolase, Nucleotide-binding, RNA; pdb70 \| 5JAJ\_A \| 96.9 \| 1.1e-06 \| 1.3e-10 \| 92.5 \| 69 \| (63, 134) \| 682 \| (8, 77) \| 681 \| LGP2/RNA Complex \| 5JAJ\_A LGP2/RNA Complex Innate immune pattern recognition receptor HET: EDO, ADP; pdb70 \| 6QDV\_7 \| 96.9 \| 1e-06 \| 1.4e-10 \| 82.0 \| 69 \| (64, 135) \| 682 \| (39, 109) \| 390 \| Eukaryotic initiation factor 4A-III, RNA-binding \| 6QDV\_7 Eukaryotic initiation factor 4A-III, RNA-binding spliceosome, RNA, complex, SPLICING HET: GTP, I6P, ATP, SEP; pdb70 \| 5JB2\_A \| 96.9 \| 1.1e-06 \| 1.4e-10 \| 92.2 \| 69 \| (63, 134) \| 682 \| (7, 76) \| 680 \| LGP2/RNA Complex \| 5JB2\_A LGP2/RNA Complex Innate immune pattern recognition receptor HET: GTP, ADP; pdb70 \| 6G19\_A \| 96.9 \| 1.1e-06 \| 1.4e-10 \| 92.4 \| 62 \| (64, 127) \| 682 \| (2, 68) \| 696 \| Interferon-induced helicase C domain-containing protein \| 6G19\_A Interferon-induced helicase C domain-containing protein Protein-RNA complex, helical filament, ATPase HET: ANP; pdb70 \| 3FE2\_B \| 96.9 \| 1.1e-06 \| 1.4e-10 \| 77.4 \| 69 \| (63, 134) \| 682 \| (50, 125) \| 242 \| Probable ATP-dependent RNA helicase DDX5 \| 3FE2\_B Probable ATP-dependent RNA helicase DDX5 RNA HELICASE, DEAD, ADP, ATP-binding HET: SO4, ADP; pdb70 \| 1XTI\_A \| 96.9 \| 1.1e-06 \| 1.4e-10 \| 82.4 \| 70 \| (63, 135) \| 682 \| (29, 100) \| 391 \| Probable ATP-dependent RNA helicase p47 \| 1XTI\_A Probable ATP-dependent RNA helicase p47 alpha-beta fold, GENE REGULATION HET: IPA; pdb70 \| 4CT4\_B \| 96.9 \| 1.2e-06 \| 1.6e-10 \| 81.2 \| 69 \| (63, 134) \| 682 \| (26, 96) \| 378 \| CCR4-NOT TRANSCRIPTION COMPLEX SUBUNIT 1 \| 4CT4\_B CCR4-NOT TRANSCRIPTION COMPLEX SUBUNIT 1 RNA BINDING PROTEIN, DEADENYLATION, TRANSCRIPTION; pdb70 \| 1XTK\_A \| 96.9 \| 1.2e-06 \| 1.6e-10 \| 82.0 \| 70 \| (63, 135) \| 682 \| (28, 99) \| 390 \| Probable ATP-dependent RNA helicase p47 \| 1XTK\_A Probable ATP-dependent RNA helicase p47 alpha-beta fold, GENE REGULATION; pdb70 \| 1Z3I\_X \| 96.9 \| 1.3e-06 \| 1.7e-10 \| 90.6 \| 155 \| (62, 228) \| 682 \| (53, 233) \| 644 \| similar to RAD54-like \| 1Z3I\_X similar to RAD54-like Recombination ATPase helicase, Recombination-DNA binding HET: SO4; pdb70 \| 1S2M\_A \| 96.9 \| 1.2e-06 \| 1.7e-10 \| 82.5 \| 69 \| (63, 134) \| 682 \| (42, 112) \| 400 \| Putative ATP-dependent RNA helicase DHH1 \| 1S2M\_A Putative ATP-dependent RNA helicase DHH1 ATP-binding, RNA-binding, Helicase, RNA BINDING; pdb70 \| 3LY5\_B \| 96.9 \| 1.3e-06 \| 1.7e-10 \| 79.4 \| 69 \| (64, 135) \| 682 \| (76, 150) \| 262 \| ATP-dependent RNA helicase DDX18 (E.C.3.6.1.-) \| 3LY5\_B ATP-dependent RNA helicase DDX18 (E.C.3.6.1.-) alpha-beta, Structural Genomics, Structural Genomics; pdb70 \| 5TNU\_A \| 96.9 \| 1.4e-06 \| 1.7e-10 \| 86.2 \| 135 \| (63, 228) \| 682 \| (87, 226) \| 459 \| DNA-dependent ATPase XPBII \| 5TNU\_A DNA-dependent ATPase XPBII Helicase, NER, transcription HET: GOL, SO4; pdb70 \| 4LJY\_A \| 96.9 \| 1.4e-06 \| 1.8e-10 \| 86.0 \| 70 \| (63, 135) \| 682 \| (72, 149) \| 493 \| Pre-mRNA-processing ATP-dependent RNA helicase PRP5 \| 4LJY\_A Pre-mRNA-processing ATP-dependent RNA helicase PRP5 Prp5, DEAD box, RNA splicing HET: MRD, ADP; pdb70 \| 5GJU\_A \| 96.9 \| 1.4e-06 \| 1.8e-10 \| 73.5 \| 68 \| (63, 133) \| 682 \| (22, 91) \| 205 \| ATP-dependent RNA helicase DeaD (E.C.3.6.4.13) \| 5GJU\_A ATP-dependent RNA helicase DeaD (E.C.3.6.4.13) RecA-like, RNA helicase, HYDROLASE HET: AMP; pdb70 \| 2YKG\_A \| 96.9 \| 1.5e-06 \| 1.9e-10 \| 92.0 \| 71 \| (62, 135) \| 682 \| (11, 85) \| 696 \| PROBABLE ATP-DEPENDENT RNA HELICASE DDX58 \| 2YKG\_A PROBABLE ATP-DEPENDENT RNA HELICASE DDX58 HYDROLASE, INNATE IMMUNITY; pdb70 \| 5ANR\_B \| 96.9 \| 1.4e-06 \| 1.9e-10 \| 80.7 \| 69 \| (63, 134) \| 682 \| (26, 96) \| 378 \| CCR4-NOT TRANSCRIPTION COMPLEX SUBUNIT 1 \| 5ANR\_B CCR4-NOT TRANSCRIPTION COMPLEX SUBUNIT 1 RNA BINDING PROTEIN; pdb70 \| 1QDE\_A \| 96.9 \| 1.5e-06 \| 1.9e-10 \| 76.8 \| 68 \| (64, 134) \| 682 \| (36, 105) \| 224 \| TRANSLATION INITIATION FACTOR 4A \| 1QDE\_A TRANSLATION INITIATION FACTOR 4A TRANSLATION INITIATION, SACCHAROMYCES CEREVISIAE, DEAD; pdb70 \| 6JDE\_B \| 96.9 \| 1.7e-06 \| 1.9e-10 \| 91.6 \| 144 \| (63, 228) \| 682 \| (3, 156) \| 586 \| Putative DNA repair helicase RadD \| 6JDE\_B Putative DNA repair helicase RadD DNA repair protein, HYDROLASE; pdb70 \| 5SUQ\_C \| 96.9 \| 1.5e-06 \| 1.9e-10 \| 84.7 \| 70 \| (63, 135) \| 682 \| (83, 154) \| 446 \| ATP-dependent RNA helicase SUB2 (E.C.3.6.4.13) \| 5SUQ\_C ATP-dependent RNA helicase SUB2 (E.C.3.6.4.13) mRNA export, HYDROLASE HET: KEG; pdb70 \| 4IDH\_A \| 96.8 \| 1.7e-06 \| 2e-10 \| 88.8 \| 144 \| (80, 240) \| 682 \| (34, 182) \| 490 \| Gene 2 protein \| 4IDH\_A Gene 2 protein DNA packaging, terminase, ATPase, nuclease; pdb70 \| 4IEE\_A \| 96.8 \| 1.7e-06 \| 2e-10 \| 88.8 \| 144 \| (80, 240) \| 682 \| (34, 182) \| 490 \| Gene 2 protein \| 4IEE\_A Gene 2 protein DNA packaging, terminase, ATPase, nuclease HET: AGS; pdb70 \| 4BPB\_A \| 96.8 \| 1.7e-06 \| 2e-10 \| 92.0 \| 72 \| (61, 135) \| 682 \| (10, 85) \| 698 \| PROBABLE ATP-DEPENDENT RNA HELICASE DDX58 \| 4BPB\_A PROBABLE ATP-DEPENDENT RNA HELICASE DDX58 HYDROLASE-RNA COMPLEX, ADENOSINE TRIPHOSPHATE, DEAD-BOX; pdb70 \| 5SUP\_B \| 96.8 \| 1.6e-06 \| 2.1e-10 \| 81.5 \| 70 \| (63, 135) \| 682 \| (27, 98) \| 390 \| ATP-dependent RNA helicase SUB2, YRA1/RNA \| 5SUP\_B ATP-dependent RNA helicase SUB2, YRA1/RNA mRNA export, HYDROLASE-RNA complex HET: ADP, BEF; pdb70 \| 2FZ4\_A \| 96.8 \| 1.8e-06 \| 2.1e-10 \| 79.7 \| 135 \| (63, 228) \| 682 \| (92, 231) \| 237 \| DNA repair protein RAD25 \| 2FZ4\_A DNA repair protein RAD25 RecA-like domain, DNA damage recognition; pdb70 \| 3LY5\_A \| 96.8 \| 1.7e-06 \| 2.1e-10 \| 78.5 \| 69 \| (64, 135) \| 682 \| (76, 150) \| 262 \| ATP-dependent RNA helicase DDX18 (E.C.3.6.1.-) \| 3LY5\_A ATP-dependent RNA helicase DDX18 (E.C.3.6.1.-) alpha-beta, Structural Genomics, Structural Genomics; pdb70 \| 3B85\_A \| 96.8 \| 1.6e-06 \| 2.2e-10 \| 73.0 \| 60 \| (64, 125) \| 682 \| (7, 67) \| 208 \| Phosphate starvation-inducible protein \| 3B85\_A Phosphate starvation-inducible protein Corynebacterium glutamicum, PhoH2, ATPase, PFAM: HET: MSE, SO4; pdb70 \| 3PEY\_A \| 96.8 \| 1.6e-06 \| 2.2e-10 \| 81.0 \| 71 \| (63, 134) \| 682 \| (26, 98) \| 395 \| ATP-dependent RNA helicase DBP5 (E.C.3.6.4.13) \| 3PEY\_A ATP-dependent RNA helicase DBP5 (E.C.3.6.4.13) RecA, DEAD-box, ATPase, Helicase, mRNA-export HET: ADP, NO3; pdb70 \| 2FWR\_A \| 96.8 \| 1.8e-06 \| 2.2e-10 \| 86.4 \| 135 \| (63, 228) \| 682 \| (92, 231) \| 472 \| DNA repair protein RAD25 \| 2FWR\_A DNA repair protein RAD25 DNA Unwinding, DNA Repair, XPB HET: IPA, PO4; pdb70 \| 5JC3\_A \| 96.8 \| 1.9e-06 \| 2.3e-10 \| 91.0 \| 64 \| (62, 127) \| 682 \| (5, 73) \| 701 \| LGP2/RNA Complex \| 5JC3\_A LGP2/RNA Complex Innate immune pattern recognition receptor HET: ADP; pdb70 \| 5JCH\_A \| 96.8 \| 1.9e-06 \| 2.3e-10 \| 91.0 \| 64 \| (62, 127) \| 682 \| (5, 73) \| 701 \| LGP2/RNA Complex \| 5JCH\_A LGP2/RNA Complex Innate immune pattern recognition receptor HET: ADP; pdb70 \| 6AIC\_A \| 96.8 \| 1.8e-06 \| 2.3e-10 \| 74.7 \| 69 \| (64, 135) \| 682 \| (24, 93) \| 214 \| DEAD-box ATP-dependent RNA helicase CshA \| 6AIC\_A DEAD-box ATP-dependent RNA helicase CshA STRUCTURAL PROTEIN HET: AMP; pdb70 \| 3FMO\_B \| 96.8 \| 2.1e-06 \| 2.4e-10 \| 82.4 \| 69 \| (63, 134) \| 682 \| (113, 185) \| 300 \| Nuclear pore complex protein Nup214 \| 3FMO\_B Nuclear pore complex protein Nup214 nuclear porin, nuclear pore complex HET: ADP; pdb70 \| 5GVR\_A \| 96.8 \| 1.8e-06 \| 2.5e-10 \| 75.2 \| 68 \| (64, 134) \| 682 \| (35, 112) \| 234 \| Probable ATP-dependent RNA helicase DDX41 \| 5GVR\_A Probable ATP-dependent RNA helicase DDX41 ATPase, DEAD box protein, HYDROLASE; pdb70 \| 1Z63\_A \| 96.8 \| 2.1e-06 \| 2.7e-10 \| 85.3 \| 148 \| (63, 228) \| 682 \| (36, 190) \| 500 \| Helicase of the snf2/rad54 family/DNA \| 1Z63\_A Helicase of the snf2/rad54 family/DNA protein-DNA complex, HYDROLASE-DNA complex COMPLEX HET: MSE; pdb70 \| 3TBK\_A \| 96.8 \| 2.1e-06 \| 2.8e-10 \| 85.8 \| 70 \| (63, 135) \| 682 \| (3, 76) \| 555 \| RIG-I Helicase Domain (E.C.3.6.4.13) \| 3TBK\_A RIG-I Helicase Domain (E.C.3.6.4.13) DECH Helicase, Helicase, ATP binding HET: ANP; pdb70 \| 6C90\_A \| 96.8 \| 2.3e-06 \| 2.8e-10 \| 91.9 \| 145 \| (62, 228) \| 682 \| (70, 222) \| 734 \| Superkiller viralicidic activity 2-like 2 \| 6C90\_A Superkiller viralicidic activity 2-like 2 HYDROLASE, NEXT, EXOSOME, NUCLEOTIDE-BINDING, ATPASE HET: ADP, TLA, MLI; pdb70 \| 5DZR\_A \| 96.8 \| 2.5e-06 \| 2.9e-10 \| 95.4 \| 145 \| (62, 228) \| 682 \| (65, 217) \| 993 \| FRQ-interacting RNA helicase \| 5DZR\_A FRQ-interacting RNA helicase helicase, circadian clock, RNA BINDING; pdb70 \| 1QVA\_A \| 96.8 \| 2.3e-06 \| 3e-10 \| 75.6 \| 68 \| (64, 134) \| 682 \| (43, 112) \| 223 \| INITIATION FACTOR 4A \| 1QVA\_A INITIATION FACTOR 4A RNA HELICASE, DEAD BOX, EIF4A; pdb70 \| 4A36\_A \| 96.7 \| 2.4e-06 \| 3.1e-10 \| 85.4 \| 70 \| (63, 135) \| 682 \| (6, 79) \| 556 \| RETINOIC ACID INDUCIBLE PROTEIN I \| 4A36\_A RETINOIC ACID INDUCIBLE PROTEIN I RNA BINDING PROTEIN-RNA COMPLEX, SUPERFAMILY HET: ADP; pdb70 \| 6O9M\_7 \| 96.7 \| 2.9e-06 \| 3.1e-10 \| 94.1 \| 145 \| (62, 229) \| 682 \| (313, 474) \| 782 \| TFIIH basal transcription factor complex \| 6O9M\_7 TFIIH basal transcription factor complex Transcription initiation, Molecular dynamics, Gene; pdb70 \| 6RO4\_A \| 96.7 \| 2.9e-06 \| 3.1e-10 \| 94.1 \| 145 \| (62, 229) \| 682 \| (313, 474) \| 782 \| General transcription and DNA repair \| 6RO4\_A General transcription and DNA repair Complex, Helicase, Translocase, DNA repair; pdb70 \| 4A4D\_A \| 96.7 \| 2.4e-06 \| 3.2e-10 \| 75.8 \| 69 \| (63, 134) \| 682 \| (64, 139) \| 253 \| PROBABLE ATP-DEPENDENT RNA HELICASE DDX5 \| 4A4D\_A PROBABLE ATP-DEPENDENT RNA HELICASE DDX5 ATP-BINDING, HYDROLASE, RNA-BINDING; pdb70 \| 2J0S\_A \| 96.7 \| 2.5e-06 \| 3.3e-10 \| 80.9 \| 68 \| (64, 134) \| 682 \| (59, 128) \| 410 \| ATP-DEPENDENT RNA HELICASE DDX48, PROTEIN \| 2J0S\_A ATP-DEPENDENT RNA HELICASE DDX48, PROTEIN MRNA PROCESSING, PHOSPHORYLATION, RRNA PROCESSING HET: ANP; pdb70 \| 2PL3\_A \| 96.7 \| 2.6e-06 \| 3.3e-10 \| 75.8 \| 69 \| (63, 134) \| 682 \| (46, 120) \| 236 \| Probable ATP-dependent RNA helicase DDX10 \| 2PL3\_A Probable ATP-dependent RNA helicase DDX10 RNA, HELICASE, DEAD, STRUCTURAL GENOMICS HET: EOH, ADP; pdb70 \| 4W7S\_B \| 96.7 \| 2.6e-06 \| 3.4e-10 \| 82.8 \| 72 \| (63, 135) \| 682 \| (70, 153) \| 463 \| S.cerevisiae Prp28 (127-588 aa) \| 4W7S\_B S.cerevisiae Prp28 (127-588 aa) splicing factor, DEAD-box protein, ATPase HET: ANP, MSE, P6G; pdb70 \| 6GEJ\_M \| 96.7 \| 3.4e-06 \| 3.5e-10 \| 100.2 \| 154 \| (62, 229) \| 682 \| (693, 859) \| 1514 \| Vacuolar protein sorting-associated protein 72 \| 6GEJ\_M Vacuolar protein sorting-associated protein 72 Chromatin, Remodeller, ATPase, Histone, NUCLEAR HET: ADP; pdb70 \| 2I4I\_A \| 96.7 \| 2.6e-06 \| 3.5e-10 \| 80.2 \| 70 \| (63, 135) \| 682 \| (36, 125) \| 417 \| ATP-dependent RNA helicase DDX3X (E.C.3.6.1.-) \| 2I4I\_A ATP-dependent RNA helicase DDX3X (E.C.3.6.1.-) RNA, HELICASE, DEAD, STRUCTURAL GENOMICS HET: AMP; pdb70 \| 5OQJ\_7 \| 96.7 \| 3.5e-06 \| 3.6e-10 \| 94.6 \| 143 \| (62, 228) \| 682 \| (359, 520) \| 843 \| DNA-directed RNA polymerase II subunit \| 5OQJ\_7 DNA-directed RNA polymerase II subunit transcription initiation, TRANSCRIPTION, macromolecular complex HET: ZN; pdb70 \| 5HZR\_A \| 96.7 \| 3.1e-06 \| 3.7e-10 \| 90.8 \| 153 \| (63, 228) \| 682 \| (108, 272) \| 732 \| SNF2-family ATP dependent chromatin remodeling \| 5HZR\_A SNF2-family ATP dependent chromatin remodeling Swi2/Snf2, chromatin remodeling, TRANSCRIPTION HET: KH2, SO4; pdb70 \| 6PWF\_K \| 96.7 \| 3.6e-06 \| 4.1e-10 \| 89.7 \| 154 \| (62, 229) \| 682 \| (100, 266) \| 640 \| Histone H3, Histone H4, Histone \| 6PWF\_K Histone H3, Histone H4, Histone nucleosome, DNA-binding protein, ATP-dependent chromatin; pdb70 \| 5ZBZ\_A \| 96.7 \| 3.1e-06 \| 4.1e-10 \| 73.4 \| 68 \| (64, 134) \| 682 \| (36, 105) \| 220 \| Eukaryotic initiation factor 4A-I (E.C.3.6.4.13) \| 5ZBZ\_A Eukaryotic initiation factor 4A-I (E.C.3.6.4.13) eukaryotic translation initiation factor 4A HET: MLY, SAU; pdb70 \| 1VEC\_B \| 96.7 \| 3.3e-06 \| 4.2e-10 \| 73.9 \| 67 \| (63, 132) \| 682 \| (24, 92) \| 206 \| ATP-dependent RNA helicase p54 \| 1VEC\_B ATP-dependent RNA helicase p54 RNA HELICASE, DEAD-BOX PROTEIN, RNA HET: TLA; pdb70 \| 3IUY\_A \| 96.7 \| 3.2e-06 \| 4.2e-10 \| 74.2 \| 69 \| (63, 134) \| 682 \| (41, 117) \| 228 \| Probable ATP-dependent RNA helicase DDX53 \| 3IUY\_A Probable ATP-dependent RNA helicase DDX53 REC-A-like, DEAD-box, Structural Genomics, Structural HET: AMP; pdb70 \| 4A2P\_A \| 96.7 \| 3.3e-06 \| 4.2e-10 \| 84.6 \| 70 \| (63, 135) \| 682 \| (6, 79) \| 556 \| RETINOIC ACID INDUCIBLE PROTEIN I \| 4A2P\_A RETINOIC ACID INDUCIBLE PROTEIN I HYDROLASE, SUPERFAMILY 2 RNA HELICASE; pdb70 \| 2DB3\_D \| 96.7 \| 3.3e-06 \| 4.4e-10 \| 81.1 \| 69 \| (63, 134) \| 682 \| (77, 152) \| 434 \| ATP-dependent RNA helicase vasa(E.C.3.6.1.3)/RNA complex \| 2DB3\_D ATP-dependent RNA helicase vasa(E.C.3.6.1.3)/RNA complex DEAD-BOX, HELICASE, PROTEIN-RNA COMPLEX, ATPase HET: ANP; pdb70 \| 3FMP\_B \| 96.7 \| 3.4e-06 \| 4.4e-10 \| 83.7 \| 70 \| (64, 134) \| 682 \| (114, 185) \| 479 \| Nuclear pore complex protein Nup214 \| 3FMP\_B Nuclear pore complex protein Nup214 nuclear porin, nuclear pore complex HET: ADP; pdb70 \| 6NMI\_A \| 96.7 \| 3.8e-06 \| 4.4e-10 \| 89.6 \| 145 \| (62, 229) \| 682 \| (236, 397) \| 653 \| General transcription and DNA repair \| 6NMI\_A General transcription and DNA repair transcription, DNA repair, helicase, multiprotein; pdb70 \| 5ZC9\_A \| 96.7 \| 3.3e-06 \| 4.4e-10 \| 79.0 \| 69 \| (63, 134) \| 682 \| (41, 111) \| 394 \| Eukaryotic initiation factor 4A-I/RNA Complex \| 5ZC9\_A Eukaryotic initiation factor 4A-I/RNA Complex INITIATION FACTOR, DEAD-BOX, HELICASE, PROTEIN-RNA HET: ANP, RCG; pdb70 \| 5JXR\_A \| 96.7 \| 4.2e-06 \| 4.6e-10 \| 91.6 \| 153 \| (63, 229) \| 682 \| (104, 269) \| 723 \| Chromatin-remodeling complex ATPase-like protein \| 5JXR\_A Chromatin-remodeling complex ATPase-like protein chromatin remodeler, ISWI, TRANSCRIPTION; pdb70 \| 3P4X\_B \| 96.6 \| 3.8e-06 \| 4.8e-10 \| 82.3 \| 68 \| (63, 135) \| 682 \| (19, 87) \| 413 \| reverse gyrase helicase-like domain \| 3P4X\_B reverse gyrase helicase-like domain TOPOISOMERASE, DNA SUPERCOILING, ARCHAEA, HELICASE HET: ADP; pdb70 \| 5SVA\_Z \| 96.6 \| 4.7e-06 \| 4.9e-10 \| 93.6 \| 145 \| (63, 229) \| 682 \| (360, 521) \| 843 \| DNA-directed RNA polymerase II subunit \| 5SVA\_Z DNA-directed RNA polymerase II subunit Transcriptional Initiation, Mediator, Pre-Initiation Complex; pdb70 \| 4W7S\_A \| 96.6 \| 3.8e-06 \| 5e-10 \| 81.6 \| 72 \| (63, 135) \| 682 \| (70, 153) \| 463 \| S.cerevisiae Prp28 (127-588 aa) \| 4W7S\_A S.cerevisiae Prp28 (127-588 aa) splicing factor, DEAD-box protein, ATPase HET: P6G, ANP, MSE; pdb70 \| 3OIY\_A \| 96.6 \| 4e-06 \| 5e-10 \| 82.2 \| 68 \| (63, 135) \| 682 \| (20, 88) \| 414 \| reverse gyrase helicase domain \| 3OIY\_A reverse gyrase helicase domain TOPOISOMERASE, DNA SUPERCOILING, ARCHAEA, HELICASE; pdb70 \| 5H1Y\_B \| 96.6 \| 4.1e-06 \| 5.1e-10 \| 77.9 \| 69 \| (63, 134) \| 682 \| (73, 151) \| 281 \| Probable ATP-dependent RNA helicase DDX41 \| 5H1Y\_B Probable ATP-dependent RNA helicase DDX41 DNA sensor CDNs receptor ATPase HET: SO4; pdb70 \| 3FHT\_A \| 96.6 \| 4e-06 \| 5.3e-10 \| 79.6 \| 69 \| (64, 133) \| 682 \| (47, 117) \| 412 \| ATP-dependent RNA helicase DDX19B (E.C.3.6.1.-)/RNA \| 3FHT\_A ATP-dependent RNA helicase DDX19B (E.C.3.6.1.-)/RNA Dbp5, DEAD-box helicase, RNA dependent HET: GOL, ANP; pdb70 \| 6BOG\_A \| 96.6 \| 5e-06 \| 5.6e-10 \| 93.8 \| 157 \| (63, 228) \| 682 \| (152, 319) \| 968 \| RNA polymerase-associated protein RapA (E.C.3.6.4.-) \| 6BOG\_A RNA polymerase-associated protein RapA (E.C.3.6.4.-) hydrolase, TRANSCRIPTION HET: SO4, MSE; pdb70 \| 3FHC\_B \| 96.6 \| 4.4e-06 \| 5.9e-10 \| 72.9 \| 69 \| (63, 134) \| 682 \| (46, 118) \| 235 \| Nuclear pore complex protein Nup214 \| 3FHC\_B Nuclear pore complex protein Nup214 DEAD-box helicase, mRNA export, Nucleoporin; pdb70 \| 2OXC\_A \| 96.6 \| 4.4e-06 \| 5.9e-10 \| 72.0 \| 69 \| (63, 134) \| 682 \| (45, 115) \| 230 \| Probable ATP-dependent RNA helicase DDX20 \| 2OXC\_A Probable ATP-dependent RNA helicase DDX20 RNA, HELICASE, DEAD, STRUCTURAL GENOMICS HET: ADP; pdb70 \| 3B7G\_A \| 96.6 \| 4.4e-06 \| 5.9e-10 \| 72.0 \| 69 \| (63, 134) \| 682 \| (45, 115) \| 230 \| Probable ATP-dependent RNA helicase DDX20 \| 3B7G\_A Probable ATP-dependent RNA helicase DDX20 RNA, HELICASE, DEAD, CONSERVED DOMAIN HET: ANP; pdb70 \| 5E7I\_B \| 96.6 \| 4.5e-06 \| 5.9e-10 \| 80.8 \| 70 \| (63, 135) \| 682 \| (69, 158) \| 452 \| DEAD (Asp-Glu-Ala-Asp) box helicase 3 \| 5E7I\_B DEAD (Asp-Glu-Ala-Asp) box helicase 3 DEAD-box protein, RNA helicase, RecA; pdb70 \| 6IY2\_O \| 96.6 \| 5.8e-06 \| 6.6e-10 \| 89.1 \| 153 \| (63, 229) \| 682 \| (96, 261) \| 679 \| Transcription regulatory protein SNF2, Histone \| 6IY2\_O Transcription regulatory protein SNF2, Histone complex, nucleosome, chromatin remodeling, gene HET: ADP; pdb70 \| 3IUY\_B \| 96.6 \| 5.2e-06 \| 6.7e-10 \| 72.8 \| 69 \| (63, 134) \| 682 \| (41, 117) \| 228 \| Probable ATP-dependent RNA helicase DDX53 \| 3IUY\_B Probable ATP-dependent RNA helicase DDX53 REC-A-like, DEAD-box, Structural Genomics, Structural HET: AMP; pdb70 \| 4A2Q\_B \| 96.6 \| 6.1e-06 \| 7e-10 \| 90.3 \| 126 \| (63, 196) \| 682 \| (247, 385) \| 797 \| RETINOIC ACID INDUCIBLE PROTEIN I \| 4A2Q\_B RETINOIC ACID INDUCIBLE PROTEIN I HYDROLASE, SUPERFAMILY 2 RNA HELICASE; pdb70 \| 5ZWN\_y \| 96.5 \| 6.3e-06 \| 7.2e-10 \| 87.0 \| 71 \| (64, 135) \| 682 \| (196, 278) \| 588 \| U1 small nuclear ribonucleoprotein 70 \| 5ZWN\_y U1 small nuclear ribonucleoprotein 70 spliceosme, assemply, pre-B complex, U1; pdb70 \| 6IEH\_B \| 96.5 \| 6.3e-06 \| 7.3e-10 \| 92.0 \| 144 \| (62, 227) \| 682 \| (74, 225) \| 979 \| Exosome RNA helicase MTR4 (E.C.3.6.4.13) \| 6IEH\_B Exosome RNA helicase MTR4 (E.C.3.6.4.13) RNA helicase, MTR4, NRDE2, Complex HET: ATP; pdb70 \| 3MWY\_W \| 96.5 \| 6.2e-06 \| 7.5e-10 \| 88.7 \| 152 \| (63, 228) \| 682 \| (235, 408) \| 800 \| --REMARK 3 \| 3MWY\_W --REMARK 3 SWI2/SNF2 ATPase, double chromodomains, HYDROLASE HET: AGS; pdb70 \| 5DTU\_A \| 96.5 \| 5.8e-06 \| 7.8e-10 \| 79.8 \| 125 \| (63, 196) \| 682 \| (63, 206) \| 475 \| Putative uncharacterized protein \| 5DTU\_A Putative uncharacterized protein DEAD-box Protein, ATPase, RNA-helicase, DDX23 HET: ADP; pdb70 \| 6B4J\_F \| 96.5 \| 6.1e-06 \| 8e-10 \| 79.3 \| 69 \| (64, 133) \| 682 \| (65, 135) \| 430 \| Nucleoporin GLE1, Nucleoporin like 2 \| 6B4J\_F Nucleoporin GLE1, Nucleoporin like 2 Complex, Nuclear Pore Complex, mRNA HET: PO4, ANP; pdb70 \| 3H1T\_A \| 96.5 \| 7.1e-06 \| 8.6e-10 \| 85.0 \| 147 \| (63, 229) \| 682 \| (177, 345) \| 590 \| Type I site-specific restriction-modification system \| 3H1T\_A Type I site-specific restriction-modification system hydrolase, restriction enzyme hsdR, ATP-binding; pdb70 \| 6JYL\_K \| 96.5 \| 8.5e-06 \| 8.7e-10 \| 93.7 \| 153 \| (63, 229) \| 682 \| (126, 291) \| 1061 \| Histone H3, Histone H4, Histone \| 6JYL\_K Histone H3, Histone H4, Histone chromatin remodelling, single particle Cryo-EM HET: ADP; pdb70 \| 3FHO\_B \| 96.5 \| 8e-06 \| 9.2e-10 \| 84.5 \| 71 \| (64, 135) \| 682 \| (141, 213) \| 508 \| ATP-dependent RNA helicase dbp5 (E.C.3.6.1.-) \| 3FHO\_B ATP-dependent RNA helicase dbp5 (E.C.3.6.1.-) RNA helicase, mRNA export, ATPase; pdb70 \| 4A2W\_B \| 96.5 \| 8.7e-06 \| 9.7e-10 \| 91.3 \| 126 \| (63, 196) \| 682 \| (247, 385) \| 936 \| RETINOIC ACID INDUCIBLE PROTEIN I \| 4A2W\_B RETINOIC ACID INDUCIBLE PROTEIN I HYDROLASE, SUPERFAMILY 2 RNA HELICASE; pdb70 \| 6HTS\_G \| 96.4 \| 1e-05 \| 1e-09 \| 94.8 \| 153 \| (63, 229) \| 682 \| (250, 421) \| 1290 \| RuvB-like 1, RuvB-like 2, DNA \| 6HTS\_G RuvB-like 1, RuvB-like 2, DNA Chromatin, Remodeller, Nucleosome, DNA Binding HET: ADP; pdb70 \| 2KBE\_A \| 96.4 \| 7.6e-06 \| 1e-09 \| 70.4 \| 68 \| (63, 133) \| 682 \| (43, 114) \| 226 \| ATP-dependent RNA helicase DBP5 (E.C.3.6.1.-) \| 2KBE\_A ATP-dependent RNA helicase DBP5 (E.C.3.6.1.-) dbp5p, ATP-binding, Helicase, Hydrolase, Membrane; pdb70 \| 6QW6\_5X \| 96.4 \| 8.8e-06 \| 1e-09 \| 88.8 \| 124 \| (64, 196) \| 682 \| (413, 556) \| 820 \| Small nuclear ribonucleoprotein Sm D1 \| 6QW6\_5X Small nuclear ribonucleoprotein Sm D1 RNP complex, splicing, RNA, protein HET: IHP, M7M, GTP; pdb70 \| 6FML\_G \| 96.4 \| 1.1e-05 \| 1e-09 \| 96.8 \| 154 \| (62, 229) \| 682 \| (969, 1142) \| 1856 \| RuvB-like helicase (E.C.3.6.4.12), Ino80, Uncharacterized \| 6FML\_G RuvB-like helicase (E.C.3.6.4.12), Ino80, Uncharacterized INO80, Nucleosome, ATP dependent Chromatin HET: ATP, ADP; pdb70 \| 4A2W\_A \| 96.4 \| 9.9e-06 \| 1.1e-09 \| 90.8 \| 126 \| (63, 196) \| 682 \| (247, 385) \| 936 \| RETINOIC ACID INDUCIBLE PROTEIN I \| 4A2W\_A RETINOIC ACID INDUCIBLE PROTEIN I HYDROLASE, SUPERFAMILY 2 RNA HELICASE; pdb70 \| 4D25\_A \| 96.4 \| 8.6e-06 \| 1.2e-09 \| 77.6 \| 69 \| (63, 134) \| 682 \| (71, 150) \| 434 \| BMVLG PROTEIN \| 4D25\_A BMVLG PROTEIN HYDROLASE, PIRNA, AMPLIFIER COMPLEX, TRANSPOSON HET: ANP; pdb70 \| 4D26\_A \| 96.4 \| 9.2e-06 \| 1.2e-09 \| 77.4 \| 69 \| (63, 134) \| 682 \| (71, 150) \| 434 \| BMVLG PROTEIN \| 4D26\_A BMVLG PROTEIN HYDROLASE, PIRNA, AMPLIFIER COMPLEX, TRANSPOSON HET: ADP; pdb70 \| 2P6R\_A \| 96.4 \| 1.1e-05 \| 1.3e-09 \| 85.2 \| 67 \| (63, 134) \| 682 \| (24, 91) \| 702 \| afUHEL308 HELICASE/DNA Complex \| 2P6R\_A afUHEL308 HELICASE/DNA Complex PROTEIN-DNA COMPLEX, SF2 HELICASE, ARCHAEAL; pdb70 \| 6FTX\_W \| 96.4 \| 1.2e-05 \| 1.4e-09 \| 88.9 \| 152 \| (63, 228) \| 682 \| (192, 365) \| 878 \| Histone H3.3C, Histone H4, Histone \| 6FTX\_W Histone H3.3C, Histone H4, Histone Chromatin remodellers, MOTOR PROTEIN HET: ADP; pdb70 \| 6GKM\_A \| 96.4 \| 1.3e-05 \| 1.4e-09 \| 90.4 \| 64 \| (62, 127) \| 682 \| (306, 374) \| 1007 \| Interferon-induced helicase C domain-containing protein \| 6GKM\_A Interferon-induced helicase C domain-containing protein Protein-RNA complex, helical filament, ATPase HET: ATP; pdb70 \| 4CBG\_B \| 96.4 \| 1.2e-05 \| 1.4e-09 \| 81.6 \| 144 \| (64, 228) \| 682 \| (29, 181) \| 516 \| SERINE PROTEASE NS3 (E.C.3.4.21.113, 3.6.1.15 \| 4CBG\_B SERINE PROTEASE NS3 (E.C.3.4.21.113, 3.6.1.15 HYDROLASE, FLAVIVIRIDAE NS3, SAXS HET: MSE; pdb70 \| 4CBH\_A \| 96.4 \| 1.2e-05 \| 1.4e-09 \| 81.6 \| 144 \| (64, 228) \| 682 \| (29, 181) \| 516 \| SERINE PROTEASE NS3 (E.C.3.4.21.113, 3.6.1.15 \| 4CBH\_A SERINE PROTEASE NS3 (E.C.3.4.21.113, 3.6.1.15 HYDROLASE, SF2 HELICASES, FLAVIVIRIDAE NS3; pdb70 \| 5VVR\_M \| 96.4 \| 1.4e-05 \| 1.4e-09 \| 91.8 \| 72 \| (63, 136) \| 682 \| (295, 370) \| 1085 \| DNA-directed RNA polymerase II subunit \| 5VVR\_M DNA-directed RNA polymerase II subunit complex, RNA polymerase, CSB, transcription; pdb70 \| 6FHS\_G \| 96.3 \| 1.4e-05 \| 1.5e-09 \| 91.8 \| 154 \| (62, 229) \| 682 \| (220, 393) \| 1107 \| RuvB-like helicase (E.C.3.6.4.12), Ino80, Uncharacterized \| 6FHS\_G RuvB-like helicase (E.C.3.6.4.12), Ino80, Uncharacterized DNA BINDING PROTEIN HET: ADP, ATP; pdb70 \| 4NHO\_A \| 96.3 \| 1.1e-05 \| 1.5e-09 \| 78.7 \| 69 \| (63, 134) \| 682 \| (80, 159) \| 488 \| Probable ATP-dependent RNA helicase DDX23 \| 4NHO\_A Probable ATP-dependent RNA helicase DDX23 DEAD-BOX, HELICASE, HYDROLASE HET: GOL, CXS, SO4; pdb70 \| 3LLM\_B \| 96.3 \| 1.1e-05 \| 1.5e-09 \| 70.4 \| 69 \| (63, 134) \| 682 \| (60, 132) \| 235 \| ATP-dependent RNA helicase A (E.C.3.6.1.-) \| 3LLM\_B ATP-dependent RNA helicase A (E.C.3.6.1.-) alpha-beta-alpha, Structural Genomics, Structural Genomics HET: ADP, MSE; pdb70 \| 4PX9\_A \| 96.3 \| 1.3e-05 \| 1.5e-09 \| 75.8 \| 68 \| (64, 134) \| 682 \| (87, 174) \| 292 \| ATP-dependent RNA helicase DDX3X (E.C.3.6.4.13) \| 4PX9\_A ATP-dependent RNA helicase DDX3X (E.C.3.6.4.13) DEAD-box helicase, HYDROLASE, TRANSLATION, RNA HET: ADP; pdb70 \| 5V9X\_A \| 96.3 \| 1.3e-05 \| 1.5e-09 \| 88.6 \| 67 \| (64, 133) \| 682 \| (47, 119) \| 877 \| ATP-dependent DNA helicase/DNA Complex \| 5V9X\_A ATP-dependent DNA helicase/DNA Complex Helicase, HYDROLASE-DNA complex HET: GOL, ANP; pdb70 \| 6G0L\_W \| 96.3 \| 1.5e-05 \| 1.6e-09 \| 94.0 \| 152 \| (63, 228) \| 682 \| (374, 547) \| 1468 \| Histone H3.3C, Histone H4, Histone \| 6G0L\_W Histone H3.3C, Histone H4, Histone Chromatin remodellers, MOTOR PROTEIN HET: ADP; pdb70 \| 2ZJ8\_A \| 96.3 \| 1.3e-05 \| 1.6e-09 \| 84.4 \| 153 \| (63, 228) \| 682 \| (22, 182) \| 720 \| Putative ski2-type helicase (E.C.3.6.1.-) \| 2ZJ8\_A Putative ski2-type helicase (E.C.3.6.1.-) RecA fold, ATP-binding, Helicase, Hydrolase; pdb70 \| 2ZJA\_A \| 96.3 \| 1.3e-05 \| 1.6e-09 \| 84.4 \| 153 \| (63, 228) \| 682 \| (22, 182) \| 720 \| Putative ski2-type helicase (E.C.3.6.1.-) \| 2ZJA\_A Putative ski2-type helicase (E.C.3.6.1.-) RecA fold, ATP-binding, Helicase, Hydrolase HET: ACP; pdb70 \| 6K15\_J \| 96.3 \| 1.7e-05 \| 1.6e-09 \| 93.4 \| 153 \| (63, 229) \| 682 \| (468, 633) \| 1359 \| RSC7, Chromatin structure-remodeling complex protein \| 6K15\_J RSC7, Chromatin structure-remodeling complex protein chromatin remodeler, SWI/SNF family, DNA; pdb70 \| 6TDA\_S \| 96.3 \| 1.7e-05 \| 1.6e-09 \| 93.4 \| 153 \| (63, 229) \| 682 \| (468, 633) \| 1359 \| Histone H3.2, Histone H4, Histone \| 6TDA\_S Histone H3.2, Histone H4, Histone Chromatin remodeler DNA binding Nucleosome HET: MSE; pdb70 \| 6RFL\_Y \| 96.3 \| 1.4e-05 \| 1.7e-09 \| 83.6 \| 118 \| (63, 196) \| 682 \| (29, 148) \| 631 \| 15 kDa core protein \| 6RFL\_Y 15 kDa core protein Vaccinia, RNA polymerase, RNA Polymerase HET: SEP; pdb70 \| 6G7E\_B \| 96.3 \| 1.4e-05 \| 1.7e-09 \| 93.8 \| 154 \| (62, 229) \| 682 \| (1294, 1468) \| 1852 \| Helicase-like protein \| 6G7E\_B Helicase-like protein ATPase, Hydrolase, TRANSCRIPTION HET: MSE; pdb70 \| 6LTJ\_I \| 96.3 \| 1.6e-05 \| 1.7e-09 \| 94.7 \| 155 \| (62, 229) \| 682 \| (751, 917) \| 1647 \| Histone H3.2, Histone H4, Histone \| 6LTJ\_I Histone H3.2, Histone H4, Histone Chromatin remodeler, Complex, GENE REGULATION; pdb70 \| 5FFJ\_B \| 96.3 \| 1.5e-05 \| 1.8e-09 \| 92.5 \| 72 \| (62, 134) \| 682 \| (11, 86) \| 1406 \| LlaGI/DNA Complex \| 5FFJ\_B LlaGI/DNA Complex Helicase-like ATPase, methyltransferase, DNA-binding protein; pdb70 \| 3I62\_A \| 96.3 \| 1.4e-05 \| 1.8e-09 \| 81.5 \| 71 \| (63, 134) \| 682 \| (93, 169) \| 563 \| ATP-dependent RNA helicase MSS116 (E.C.3.6.1.-)/RNA \| 3I62\_A ATP-dependent RNA helicase MSS116 (E.C.3.6.1.-)/RNA Protein-RNA complex, RNA helicase, DEAD-box HET: ADP; pdb70 \| 5AGA\_A \| 96.3 \| 1.6e-05 \| 1.9e-09 \| 86.2 \| 69 \| (63, 135) \| 682 \| (26, 97) \| 830 \| DNA POLYMERASE THETA (E.C.2.7.7.7) \| 5AGA\_A DNA POLYMERASE THETA (E.C.2.7.7.7) TRANSFERASE, POLQ, DNA REPAIR HET: ANP, FLC; pdb70 \| 4PX9\_C \| 96.3 \| 1.6e-05 \| 1.9e-09 \| 74.9 \| 68 \| (64, 134) \| 682 \| (87, 174) \| 292 \| ATP-dependent RNA helicase DDX3X (E.C.3.6.4.13) \| 4PX9\_C ATP-dependent RNA helicase DDX3X (E.C.3.6.4.13) DEAD-box helicase, HYDROLASE, TRANSLATION, RNA HET: ADP; pdb70 \| 5O9G\_W \| 96.3 \| 1.8e-05 \| 1.9e-09 \| 93.2 \| 152 \| (63, 228) \| 682 \| (374, 547) \| 1468 \| Histone H3.2, Histone H4, Histone \| 5O9G\_W Histone H3.2, Histone H4, Histone ATPase, Complex, Nucleosome, DNA, DNA HET: ADP; pdb70 \| 2VA8\_B \| 96.3 \| 1.5e-05 \| 2e-09 \| 83.0 \| 152 \| (63, 228) \| 682 \| (29, 188) \| 715 \| SKI2-TYPE HELICASE \| 2VA8\_B SKI2-TYPE HELICASE HEL308, SSO2462, HELICASE, HYDROLASE, DNA HET: SO4; pdb70 \| 1OYW\_A \| 96.3 \| 1.6e-05 \| 2e-09 \| 80.2 \| 62 \| (64, 133) \| 682 \| (25, 87) \| 523 \| ATP-dependent DNA helicase (E.C.3.6.1.-) \| 1OYW\_A ATP-dependent DNA helicase (E.C.3.6.1.-) RecQ, helicase, Winged helix, Helix-turn-helix; pdb70 \| 6O5F\_A \| 96.2 \| 1.6e-05 \| 2.1e-09 \| 77.5 \| 70 \| (63, 135) \| 682 \| (70, 159) \| 476 \| Heterogeneous nuclear ribonucleoprotein A1/RNA Complex \| 6O5F\_A Heterogeneous nuclear ribonucleoprotein A1/RNA Complex DDX3X, DEAD-box, RNA helicase, HYDROLASE; pdb70 \| 4TYN\_A \| 96.2 \| 1.6e-05 \| 2.1e-09 \| 79.2 \| 71 \| (63, 134) \| 682 \| (40, 116) \| 509 \| Mss116 catalytic core/DNA Complex \| 4TYN\_A Mss116 catalytic core/DNA Complex DEAD-box, RNA helicase, hydrolase, RNA HET: ADP; pdb70 \| 3I5X\_A \| 96.2 \| 1.7e-05 \| 2.1e-09 \| 80.9 \| 71 \| (63, 134) \| 682 \| (93, 169) \| 563 \| ATP-dependent RNA helicase MSS116 (E.C.3.6.1.-)/RNA \| 3I5X\_A ATP-dependent RNA helicase MSS116 (E.C.3.6.1.-)/RNA Protein-RNA complex, RNA helicase, DEAD-box HET: ANP; pdb70 \| 5E7J\_A \| 96.2 \| 1.7e-05 \| 2.3e-09 \| 76.5 \| 70 \| (63, 135) \| 682 \| (69, 158) \| 452 \| DEAD (Asp-Glu-Ala-Asp) box helicase 3 \| 5E7J\_A DEAD (Asp-Glu-Ala-Asp) box helicase 3 DEAD-box protein, RNA helicase, RecA HET: AMP; pdb70 \| 2Z83\_A \| 96.2 \| 1.9e-05 \| 2.3e-09 \| 78.7 \| 66 \| (64, 133) \| 682 \| (6, 72) \| 459 \| Helicase/Nucleoside Triphosphatase \| 2Z83\_A Helicase/Nucleoside Triphosphatase HELICASE, Hydrolase, Membrane, Nucleotide-binding, RNA; pdb70 \| 2EYQ\_A \| 96.2 \| 2.1e-05 \| 2.3e-09 \| 90.1 \| 153 \| (61, 228) \| 682 \| (600, 764) \| 1151 \| Transcription-repair coupling factor \| 2EYQ\_A Transcription-repair coupling factor Mfd, SF2 ATPase, HYDROLASE HET: EPE, SO4; pdb70 \| 1GM5\_A \| 96.2 \| 2.2e-05 \| 2.5e-09 \| 86.4 \| 153 \| (61, 228) \| 682 \| (365, 529) \| 780 \| RECG \| 1GM5\_A RECG HELICASE, REPLICATION RESTART HET: ADP; pdb70 \| 6FF7\_q \| 96.2 \| 2.4e-05 \| 2.6e-09 \| 89.0 \| 69 \| (63, 134) \| 682 \| (399, 468) \| 1041 \| RNA-binding motif protein, X-linked 2 \| 6FF7\_q RNA-binding motif protein, X-linked 2 spliceosome, human, HELA, BACT, dynamics HET: GTP, IHP, ADP, ZN; pdb70 \| 6IGM\_H \| 96.2 \| 3.1e-05 \| 2.8e-09 \| 95.7 \| 152 \| (63, 229) \| 682 \| (616, 781) \| 3230 \| RuvB-like 1 (E.C.3.6.4.12), RuvB-like 2 \| 6IGM\_H RuvB-like 1 (E.C.3.6.4.12), RuvB-like 2 SRCAP complex, TRANSCRIPTION; pdb70 \| 2EYQ\_B \| 96.1 \| 2.6e-05 \| 2.8e-09 \| 89.4 \| 153 \| (61, 228) \| 682 \| (600, 764) \| 1151 \| Transcription-repair coupling factor \| 2EYQ\_B Transcription-repair coupling factor Mfd, SF2 ATPase, HYDROLASE HET: EPE, SO4; pdb70 \| 4U7D\_C \| 96.1 \| 2.5e-05 \| 2.9e-09 \| 82.0 \| 62 \| (64, 133) \| 682 \| (44, 106) \| 591 \| ATP-DEPENDENT DNA HELICASE Q1/DNA Complex \| 4U7D\_C ATP-DEPENDENT DNA HELICASE Q1/DNA Complex HYDROLASE-DNA COMPLEX, NUCLEAR PROTEIN, HYDROLASE; pdb70 \| 1W36\_D \| 96.1 \| 2.7e-05 \| 3.2e-09 \| 81.4 \| 145 \| (65, 228) \| 682 \| (150, 301) \| 608 \| EXODEOXYRIBONUCLEASE V BETA CHAIN (E.C.3.1.11.5) \| 1W36\_D EXODEOXYRIBONUCLEASE V BETA CHAIN (E.C.3.1.11.5) RECOMBINATION, HELICASE, NUCLEASE, HYDROLASE, DNA; pdb70 \| 3K70\_D \| 96.1 \| 2.7e-05 \| 3.2e-09 \| 81.4 \| 145 \| (65, 228) \| 682 \| (150, 301) \| 608 \| Exodeoxyribonuclease V beta chain (E.C.3.1.11.5) \| 3K70\_D Exodeoxyribonuclease V beta chain (E.C.3.1.11.5) RECOMBINATION, HELICASE, NUCLEASE, HYDROLASE, DNA HET: 5IU; pdb70 \| 6UXW\_A \| 96.1 \| 3.5e-05 \| 3.3e-09 \| 92.1 \| 154 \| (63, 229) \| 682 \| (765, 930) \| 1703 \| Histone H3.2, Histone H4, Histone \| 6UXW\_A Histone H3.2, Histone H4, Histone SWI/SNF, chromatin remodeler, TRANSCRIPTION, TRANSCRIPTION-DNA HET: ADP, PO4; pdb70 \| 5XC7\_A \| 96.1 \| 2.7e-05 \| 3.3e-09 \| 76.7 \| 66 \| (64, 133) \| 682 \| (4, 70) \| 451 \| NS3 Helicase \| 5XC7\_A NS3 Helicase Helicase, Dengue NS3, Hydrolase HET: GOL; pdb70 \| 4Q48\_A \| 96.1 \| 2.9e-05 \| 3.6e-09 \| 78.8 \| 62 \| (64, 133) \| 682 \| (24, 86) \| 525 \| DNA helicase RecQ \| 4Q48\_A DNA helicase RecQ DNA unwinding, helicase, DNA BINDING; pdb70 \| 6BU9\_A \| 96.1 \| 3.2e-05 \| 3.6e-09 \| 91.8 \| 154 \| (62, 228) \| 682 \| (7, 180) \| 1724 \| Dicer-2, isoform A/RNA Complex \| 6BU9\_A Dicer-2, isoform A/RNA Complex Dicer, Dcr2, Dcr-2, dmDcr-2, Dicer-2; pdb70 \| 6BUA\_A \| 96.1 \| 3.2e-05 \| 3.6e-09 \| 91.8 \| 154 \| (62, 228) \| 682 \| (7, 180) \| 1724 \| Drosophila Dicer-2 \| 6BUA\_A Drosophila Dicer-2 Dicer, dmDcr2, Dicer-2, helicase, platform; pdb70 \| 4Q47\_A \| 96.1 \| 3e-05 \| 3.6e-09 \| 78.7 \| 63 \| (64, 134) \| 682 \| (24, 87) \| 525 \| DNA helicase RecQ \| 4Q47\_A DNA helicase RecQ DNA unwinding, Topoisomerase, helicase, DNA HET: ADP; pdb70 \| 6FAC\_A \| 96.1 \| 2.9e-05 \| 3.7e-09 \| 80.1 \| 69 \| (63, 134) \| 682 \| (31, 100) \| 655 \| Putative mRNA splicing factor \| 6FAC\_A Putative mRNA splicing factor SPLICING, ATPASE, HELICASE, G-PATCH, HYDROLASE HET: ADP; pdb70 \| 3B6E\_A \| 96.0 \| 2.8e-05 \| 3.8e-09 \| 66.6 \| 64 \| (62, 127) \| 682 \| (31, 99) \| 216 \| Interferon-induced helicase C domain-containing protein \| 3B6E\_A Interferon-induced helicase C domain-containing protein DECH, DExD/H RNA-binding helicase, innate; pdb70 \| 2JLQ\_A \| 96.0 \| 3.1e-05 \| 3.9e-09 \| 76.1 \| 65 \| (65, 133) \| 682 \| (5, 70) \| 451 \| SERINE PROTEASE SUBUNIT NS3 (E.C.3.4.21.91) \| 2JLQ\_A SERINE PROTEASE SUBUNIT NS3 (E.C.3.4.21.91) RIBONUCLEOPROTEIN, NUCLEOTIDE-BINDING, VIRAL NUCLEOPROTEIN, ENDOPLASMIC; pdb70 \| 2V1X\_B \| 96.0 \| 3.4e-05 \| 3.9e-09 \| 81.2 \| 62 \| (64, 133) \| 682 \| (44, 106) \| 591 \| ATP-DEPENDENT DNA HELICASE Q1 (E.C.3.6.1.-) \| 2V1X\_B ATP-DEPENDENT DNA HELICASE Q1 (E.C.3.6.1.-) DNA STRAND ANNEALING, MISMATCH REPAIR HET: ADP; pdb70 \| 2WWY\_B \| 96.0 \| 3.4e-05 \| 3.9e-09 \| 81.2 \| 62 \| (64, 133) \| 682 \| (44, 106) \| 591 \| ATP-DEPENDENT DNA HELICASE Q1 (E.C.3.6.1.-) \| 2WWY\_B ATP-DEPENDENT DNA HELICASE Q1 (E.C.3.6.1.-) HYDROLASE-DNA COMPLEX, NUCLEAR PROTEIN, HYDROLASE HET: SO4; pdb70 \| 5WWP\_A \| 96.0 \| 4e-05 \| 4.3e-09 \| 82.2 \| 147 \| (63, 228) \| 682 \| (258, 407) \| 600 \| ORF1ab \| 5WWP\_A ORF1ab Middle East respiratory syndrome coronavirus HET: SO4; pdb70 \| 5XDR\_A \| 96.0 \| 3.5e-05 \| 4.3e-09 \| 80.9 \| 70 \| (63, 135) \| 682 \| (32, 104) \| 690 \| Pre-mRNA-splicing factor ATP-dependent RNA helicase \| 5XDR\_A Pre-mRNA-splicing factor ATP-dependent RNA helicase RNA helicase, DEAH-box, DHX15, Prp43 HET: ADP, SO4; pdb70 \| 4BGD\_A \| 96.0 \| 3.7e-05 \| 4.4e-09 \| 90.0 \| 71 \| (63, 135) \| 682 \| (56, 137) \| 1722 \| PRE-MRNA-SPLICING HELICASE BRR2 (E.C.3.6.4.13), PRE-MRNA-SPLICING \| 4BGD\_A PRE-MRNA-SPLICING HELICASE BRR2 (E.C.3.6.4.13), PRE-MRNA-SPLICING TRANSCRIPTION, SPLICEOSOME, RNA HELICASE, U5 HET: ADP, PE5; pdb70 \| 4XQK\_A \| 96.0 \| 4.3e-05 \| 4.7e-09 \| 90.3 \| 72 \| (62, 134) \| 682 \| (175, 250) \| 1578 \| LlaBIII/DNA Complex \| 4XQK\_A LlaBIII/DNA Complex ATP-dependent restriction-modification enzyme, Type ISP; pdb70 \| 6H2J\_B \| 96.0 \| 4.3e-05 \| 4.8e-09 \| 86.3 \| 150 \| (63, 228) \| 682 \| (270, 441) \| 1038 \| Type I restriction enzyme R \| 6H2J\_B Type I restriction enzyme R EcoR124, HsdR, C-terminal domain, restriction-modification HET: MSE, ATP; pdb70 \| 1T6N\_A \| 95.9 \| 4.1e-05 \| 5.2e-09 \| 67.2 \| 67 \| (64, 133) \| 682 \| (36, 104) \| 220 \| Probable ATP-dependent RNA helicase p47 \| 1T6N\_A Probable ATP-dependent RNA helicase p47 RecA-like fold, pre-mRNA processing protein HET: FLC; pdb70 \| 5LD2\_D \| 95.9 \| 5e-05 \| 5.8e-09 \| 80.2 \| 144 \| (66, 228) \| 682 \| (152, 302) \| 609 \| RecBCD enzyme subunit RecB, RecBCD \| 5LD2\_D RecBCD enzyme subunit RecB, RecBCD Helicase, Nuclease, SH3, Homologous Recombination HET: ANP; pdb70 \| 4BGD\_A \| 95.9 \| 4.9e-05 \| 5.8e-09 \| 88.9 \| 68 \| (64, 134) \| 682 \| (906, 974) \| 1722 \| PRE-MRNA-SPLICING HELICASE BRR2 (E.C.3.6.4.13), PRE-MRNA-SPLICING \| 4BGD\_A PRE-MRNA-SPLICING HELICASE BRR2 (E.C.3.6.4.13), PRE-MRNA-SPLICING TRANSCRIPTION, SPLICEOSOME, RNA HELICASE, U5 HET: ADP, PE5; pdb70 \| 4XJX\_A \| 95.9 \| 5.4e-05 \| 6e-09 \| 85.5 \| 150 \| (63, 228) \| 682 \| (270, 441) \| 1038 \| HSDR subunit of type I \| 4XJX\_A HSDR subunit of type I Restriction enzyme, ATP, hydrolase HET: ATP; pdb70 \| 4TMU\_A \| 95.9 \| 5.2e-05 \| 6.3e-09 \| 77.8 \| 62 \| (64, 133) \| 682 \| (45, 107) \| 541 \| Protein/DNA Complex \| 4TMU\_A Protein/DNA Complex RecQ, helicase, Winged helix, ATP; pdb70 \| 6EUD\_A \| 95.9 \| 5.4e-05 \| 6.3e-09 \| 82.6 \| 68 \| (63, 135) \| 682 \| (7, 75) \| 812 \| ATP-dependent RNA helicase HrpB (E.C.3.6.4.13) \| 6EUD\_A ATP-dependent RNA helicase HrpB (E.C.3.6.4.13) HrpB, DEAH/RHA helicase, bacterial helicase; pdb70 \| 5VHA\_A \| 95.9 \| 5.3e-05 \| 6.4e-09 \| 82.2 \| 69 \| (63, 134) \| 682 \| (60, 132) \| 870 \| DEAH (Asp-Glu-Ala-His) box polypeptide 36 \| 5VHA\_A DEAH (Asp-Glu-Ala-His) box polypeptide 36 HYDROLASE; pdb70 \| 5DCA\_A \| 95.8 \| 6.4e-05 \| 7.2e-09 \| 90.0 \| 71 \| (63, 135) \| 682 \| (294, 375) \| 1948 \| Pre-mRNA-splicing helicase BRR2 (E.C.3.6.4.13), Pre-mRNA-splicing \| 5DCA\_A Pre-mRNA-splicing helicase BRR2 (E.C.3.6.4.13), Pre-mRNA-splicing protein complex, helicase, RNP remodeling; pdb70 \| 3DKP\_A \| 95.8 \| 6.1e-05 \| 7.8e-09 \| 67.4 \| 69 \| (63, 134) \| 682 \| (50, 121) \| 245 \| Probable ATP-dependent RNA helicase DDX52 \| 3DKP\_A Probable ATP-dependent RNA helicase DDX52 RNA HELICASE, DEAD, ADP, Structural HET: ADP; pdb70 \| 3SQW\_A \| 95.7 \| 7.1e-05 \| 8.4e-09 \| 77.7 \| 72 \| (63, 135) \| 682 \| (42, 119) \| 579 \| ATP-dependent RNA helicase MSS116, mitochondrial \| 3SQW\_A ATP-dependent RNA helicase MSS116, mitochondrial RecA fold, RNA dependent ATPase HET: ANP; pdb70 \| 5N9D\_B \| 95.7 \| 7.4e-05 \| 8.8e-09 \| 82.3 \| 67 \| (63, 132) \| 682 \| (153, 223) \| 944 \| CG9323, isoform A/DNA Complex \| 5N9D\_B CG9323, isoform A/DNA Complex Helicase DExH ssDNA, hydrolase; pdb70 \| 5N8R\_B \| 95.7 \| 7.5e-05 \| 8.9e-09 \| 82.3 \| 67 \| (63, 132) \| 682 \| (153, 223) \| 944 \| CG9323, isoform A/DNA Complex \| 5N8R\_B CG9323, isoform A/DNA Complex Helicase, DExH, ssDNA, hydrolase; pdb70 \| 5VHD\_D \| 95.7 \| 7.5e-05 \| 9e-09 \| 81.0 \| 69 \| (63, 134) \| 682 \| (60, 132) \| 870 \| DEAH (Asp-Glu-Ala-His) box polypeptide 36 \| 5VHD\_D DEAH (Asp-Glu-Ala-His) box polypeptide 36 HYDROLASE HET: ADP; pdb70 \| 6ICZ\_Y \| 95.7 \| 8.1e-05 \| 9.1e-09 \| 85.3 \| 69 \| (63, 135) \| 682 \| (565, 634) \| 1220 \| Protein mago nashi homolog 2 \| 6ICZ\_Y Protein mago nashi homolog 2 Human Post-catalytic Spliceosome, SPLICING HET: GTP, ATP, SEP, I6P; pdb70 \| 6ID1\_V \| 95.7 \| 7.7e-05 \| 9.1e-09 \| 80.3 \| 69 \| (63, 134) \| 682 \| (137, 208) \| 795 \| Pre-mRNA-processing-splicing factor 8, 116 kDa \| 6ID1\_V Pre-mRNA-processing-splicing factor 8, 116 kDa Human Intron Lariat Spliceosome, SPLICING HET: GTP, SEP, I6P; pdb70 \| 6AC8\_A \| 95.7 \| 8.7e-05 \| 9.2e-09 \| 86.0 \| 153 \| (61, 228) \| 682 \| (659, 823) \| 1235 \| Mycobacterium smegmatis Mfd (E.C.3.6.4.-) \| 6AC8\_A Mycobacterium smegmatis Mfd (E.C.3.6.4.-) Transcription repair coupling factor, Mfd HET: SO4; pdb70 \| 6ACX\_A \| 95.7 \| 8.7e-05 \| 9.2e-09 \| 86.0 \| 153 \| (61, 228) \| 682 \| (659, 823) \| 1235 \| Mycobacterium smegmatis Mfd (E.C.3.6.4.-) \| 6ACX\_A Mycobacterium smegmatis Mfd (E.C.3.6.4.-) Transcription repair coupling factor, Mfd HET: ADP, SO4; pdb70 \| 5VHE\_A \| 95.7 \| 8.1e-05 \| 9.4e-09 \| 82.2 \| 69 \| (63, 134) \| 682 \| (123, 195) \| 933 \| DEAH (Asp-Glu-Ala-His) box polypeptide 36/DNA \| 5VHE\_A DEAH (Asp-Glu-Ala-His) box polypeptide 36/DNA HYDROLASE; pdb70 \| 5ZAM\_A \| 95.7 \| 9.4e-05 \| 1e-08 \| 89.0 \| 151 \| (63, 227) \| 682 \| (42, 211) \| 1922 \| Endoribonuclease Dicer, RISC-loading complex subunit \| 5ZAM\_A Endoribonuclease Dicer, RISC-loading complex subunit Dicer, TRBP, Cryo-EM, RNA interference; pdb70 \| 6RM9\_A \| 95.7 \| 8.1e-05 \| 1e-08 \| 76.7 \| 69 \| (63, 134) \| 682 \| (36, 105) \| 660 \| Putative mRNA splicing factor, Putative \| 6RM9\_A Putative mRNA splicing factor, Putative Prp2, DEAH-box ATPase, G-patch, spliceosome HET: GOL, ADP; pdb70 \| 5DCA\_A \| 95.7 \| 9e-05 \| 1e-08 \| 88.7 \| 69 \| (64, 135) \| 682 \| (1144, 1213) \| 1948 \| Pre-mRNA-splicing helicase BRR2 (E.C.3.6.4.13), Pre-mRNA-splicing \| 5DCA\_A Pre-mRNA-splicing helicase BRR2 (E.C.3.6.4.13), Pre-mRNA-splicing protein complex, helicase, RNP remodeling; pdb70 \| 5HD9\_A \| 95.6 \| 9.9e-05 \| 1.1e-08 \| 67.8 \| 146 \| (80, 240) \| 682 \| (13, 176) \| 194 \| Encapsidation protein \| 5HD9\_A Encapsidation protein ASCE fold, VIRAL PROTEIN HET: MSE; pdb70 \| 5MZ4\_B \| 95.6 \| 9.8e-05 \| 1.1e-08 \| 80.2 \| 130 \| (80, 228) \| 682 \| (279, 417) \| 752 \| NS3/NS4A (E.C.3.4.22.-,3.4.21.113,3.6.1.15,3.6.4.13,2.7.7.48) \| 5MZ4\_B NS3/NS4A (E.C.3.4.22.-,3.4.21.113,3.6.1.15,3.6.4.13,2.7.7.48) Flaviviridae, NS3/NS4A, Protease/Helicase, viral protein; pdb70 \| 5K9T\_A \| 95.6 \| 9.2e-05 \| 1.1e-08 \| 75.8 \| 65 \| (63, 134) \| 682 \| (69, 134) \| 578 \| Protein translocase subunit SecA \| 5K9T\_A Protein translocase subunit SecA preprotein translocase, SecA-N68, ATPase, C-terminal HET: ADP; pdb70 \| 1WRB\_B \| 95.6 \| 9.4e-05 \| 1.2e-08 \| 67.3 \| 67 \| (64, 133) \| 682 \| (45, 122) \| 253 \| DjVLGB \| 1WRB\_B DjVLGB RNA HELICASE, DEAD BOX, VASA HET: SO4; pdb70 \| 5GM6\_Y \| 95.6 \| 0.0001 \| 1.2e-08 \| 81.5 \| 68 \| (64, 134) \| 682 \| (224, 293) \| 876 \| Prp8, Brr2, Snu114, Rse1, Cus1 \| 5GM6\_Y Prp8, Brr2, Snu114, Rse1, Cus1 spliceosome, RNA splicing, Bact, Catalytically HET: GTP, ADP, ZN; pdb70 \| 5LQW\_O \| 95.6 \| 0.0001 \| 1.2e-08 \| 81.5 \| 68 \| (64, 134) \| 682 \| (224, 293) \| 876 \| Pre-mRNA-splicing factor 8, Pre-mRNA-splicing factor \| 5LQW\_O Pre-mRNA-splicing factor 8, Pre-mRNA-splicing factor activated spliceosome, spliceosome, pre-mRNA splicing; pdb70 \| 5YZG\_Z \| 95.6 \| 0.00011 \| 1.2e-08 \| 85.0 \| 68 \| (63, 134) \| 682 \| (532, 600) \| 1227 \| Pre-mRNA-processing-splicing factor 8, 116 kDa \| 5YZG\_Z Pre-mRNA-processing-splicing factor 8, 116 kDa Structure of a Human Catalytic HET: GTP, I6P, ATP, ADP, SEP; pdb70 \| 2V8O\_A \| 95.6 \| 0.00011 \| 1.2e-08 \| 75.2 \| 51 \| (80, 132) \| 682 \| (11, 62) \| 444 \| FLAVIVIRIN PROTEASE NS3 (E.C.3.4.21.91) \| 2V8O\_A FLAVIVIRIN PROTEASE NS3 (E.C.3.4.21.91) MURRAY VALLEY ENCEPHALITIS VIRUS, GLYCOPROTEIN; pdb70 \| 5AOR\_A \| 95.6 \| 0.00011 \| 1.2e-08 \| 84.2 \| 68 \| (63, 133) \| 682 \| (384, 455) \| 1293 \| DOSAGE COMPENSATION REGULATOR (E.C.3.6.4.13) \| 5AOR\_A DOSAGE COMPENSATION REGULATOR (E.C.3.6.4.13) HYDROLASE-RNA COMPLEX, HELICASE, DOSAGE COMPENSATION HET: ADP, ALF, GOL; pdb70 \| 6HYT\_B \| 95.6 \| 0.0001 \| 1.3e-08 \| 77.0 \| 69 \| (63, 135) \| 682 \| (18, 87) \| 673 \| ATP-dependent RNA helicase DHX8 (E.C.3.6.4.13) \| 6HYT\_B ATP-dependent RNA helicase DHX8 (E.C.3.6.4.13) helicase, splicing, RNA, RNA BINDING HET: ADP, EDO, DMS; pdb70 \| 6HYS\_A \| 95.6 \| 0.0001 \| 1.3e-08 \| 76.9 \| 69 \| (63, 135) \| 682 \| (18, 87) \| 673 \| ATP-dependent RNA helicase DHX8 (E.C.3.6.4.13) \| 6HYS\_A ATP-dependent RNA helicase DHX8 (E.C.3.6.4.13) Helicase, Splicing, RNA, RNA BINDING HET: ADP, DMS, EDO, ACT; pdb70 \| 6HYT\_A \| 95.6 \| 0.0001 \| 1.3e-08 \| 76.9 \| 69 \| (63, 135) \| 682 \| (18, 87) \| 673 \| ATP-dependent RNA helicase DHX8 (E.C.3.6.4.13) \| 6HYT\_A ATP-dependent RNA helicase DHX8 (E.C.3.6.4.13) helicase, splicing, RNA, RNA BINDING HET: ADP, DMS, EDO; pdb70 \| 5AOR\_B \| 95.6 \| 0.00011 \| 1.3e-08 \| 84.0 \| 68 \| (63, 133) \| 682 \| (384, 455) \| 1293 \| DOSAGE COMPENSATION REGULATOR (E.C.3.6.4.13) \| 5AOR\_B DOSAGE COMPENSATION REGULATOR (E.C.3.6.4.13) HYDROLASE-RNA COMPLEX, HELICASE, DOSAGE COMPENSATION HET: ADP, ALF, GOL; pdb70 \| 5LJ5\_Q \| 95.5 \| 0.00013 \| 1.4e-08 \| 83.1 \| 69 \| (63, 134) \| 682 \| (350, 420) \| 1071 \| Pre-mRNA-splicing factor 8, Pre-mRNA-splicing helicase \| 5LJ5\_Q Pre-mRNA-splicing factor 8, Pre-mRNA-splicing helicase spliceosome, snRNP, pre-mRNA splicing, trans-esterification HET: GTP; pdb70 \| 5WSG\_e \| 95.5 \| 0.00013 \| 1.4e-08 \| 83.1 \| 69 \| (63, 134) \| 682 \| (350, 420) \| 1071 \| Pre-mRNA-splicing factor 8, Pre-mRNA-splicing factor \| 5WSG\_e Pre-mRNA-splicing factor 8, Pre-mRNA-splicing factor Catalytic Step II spliceosome, C\* HET: GTP; pdb70 \| 3BXZ\_B \| 95.5 \| 0.00011 \| 1.4e-08 \| 72.0 \| 67 \| (63, 134) \| 682 \| (81, 148) \| 471 \| Preprotein translocase subunit secA \| 3BXZ\_B Preprotein translocase subunit secA Protein transport, Translocation, DEAD motor HET: SPD, ADP; pdb70 \| 2WV9\_A \| 95.5 \| 0.00013 \| 1.5e-08 \| 78.0 \| 67 \| (64, 133) \| 682 \| (225, 292) \| 673 \| FLAVIVIRIN PROTEASE NS2B REGULATORY SUBUNIT \| 2WV9\_A FLAVIVIRIN PROTEASE NS2B REGULATORY SUBUNIT NUCLEOTIDE-BINDING, CAPSID PROTEIN, RNA REPLICATION; pdb70 \| 2IBM\_A \| 95.5 \| 0.00013 \| 1.6e-08 \| 78.0 \| 67 \| (62, 135) \| 682 \| (78, 145) \| 780 \| Preprotein translocase secA subunit \| 2IBM\_A Preprotein translocase secA subunit protein translocation, SecA, signal peptide HET: ADP; pdb70 \| 6QV4\_A \| 95.4 \| 0.00013 \| 1.7e-08 \| 84.2 \| 70 \| (64, 135) \| 682 \| (910, 980) \| 1725 \| Pre-mRNA splicing helicase-like protein \| 6QV4\_A Pre-mRNA splicing helicase-like protein Helicase ATP-gamma-S Brr2 Ski2 Chaetomium HET: AGS; pdb70 \| 5LST\_A \| 95.4 \| 0.00015 \| 1.7e-08 \| 77.3 \| 67 \| (63, 133) \| 682 \| (56, 123) \| 693 \| ATP-dependent DNA helicase Q4 (E.C.3.6.4.12) \| 5LST\_A ATP-dependent DNA helicase Q4 (E.C.3.6.4.12) RecQ4, helicase, Rothmund-Thomson-Syndrome, RAPADILINO-Syndrome, hydrolase; pdb70 \| 5B7I\_A \| 95.4 \| 0.00018 \| 1.8e-08 \| 82.5 \| 72 \| (63, 135) \| 682 \| (395, 473) \| 1082 \| CRISPR-associated nuclease/helicase Cas3 subtype I-F/YPEST \| 5B7I\_A CRISPR-associated nuclease/helicase Cas3 subtype I-F/YPEST DNA nuclease, phagy protein, Anti-CRISPR HET: ADP, MSE; pdb70 \| 1YKS\_A \| 95.4 \| 0.00016 \| 2e-08 \| 70.5 \| 53 \| (80, 134) \| 682 \| (7, 60) \| 440 \| Genome polyprotein [contains: Flavivirin protease \| 1YKS\_A Genome polyprotein [contains: Flavivirin protease helicase, flavivirus, DEAD-box, ATPase, RTPase; pdb70 \| 3E1S\_A \| 95.4 \| 0.00017 \| 2e-08 \| 74.9 \| 130 \| (63, 228) \| 682 \| (188, 318) \| 574 \| Exodeoxyribonuclease V, subunit RecD \| 3E1S\_A Exodeoxyribonuclease V, subunit RecD alpha and beta protein, ATP-binding; pdb70 \| 3GP8\_A \| 95.4 \| 0.00017 \| 2e-08 \| 74.9 \| 130 \| (63, 228) \| 682 \| (188, 318) \| 574 \| Exodeoxyribonuclease V, subunit RecD, putative \| 3GP8\_A Exodeoxyribonuclease V, subunit RecD, putative ALPHA AND BETA PROTEIN, ATP-BINDING; pdb70 \| 5M59\_E \| 95.3 \| 0.00016 \| 2e-08 \| 83.8 \| 71 \| (63, 135) \| 682 \| (106, 186) \| 1772 \| Putative pre-mRNA splicing factor, Pre-mRNA \| 5M59\_E Putative pre-mRNA splicing factor, Pre-mRNA Brr2, pre-mRNA splicing, RNA-helicase, Prp8 HET: ACT; pdb70 \| 4Q2C\_A \| 95.3 \| 0.00018 \| 2.1e-08 \| 80.0 \| 72 \| (63, 135) \| 682 \| (306, 379) \| 949 \| CRISPR-associated helicase Cas3 \| 4Q2C\_A CRISPR-associated helicase Cas3 RecA, HD nuclease, Hydrolase; pdb70 \| 4Q2D\_A \| 95.3 \| 0.00018 \| 2.1e-08 \| 80.0 \| 72 \| (63, 135) \| 682 \| (306, 379) \| 949 \| CRISPR-associated helicase Cas3 \| 4Q2D\_A CRISPR-associated helicase Cas3 RecA, HD nuclease, Hydrolase HET: DTP; pdb70 \| 6QDV\_B \| 95.3 \| 0.00017 \| 2.1e-08 \| 83.3 \| 68 \| (64, 133) \| 682 \| (924, 992) \| 1722 \| Eukaryotic initiation factor 4A-III, RNA-binding \| 6QDV\_B Eukaryotic initiation factor 4A-III, RNA-binding spliceosome, RNA, complex, SPLICING HET: GTP, I6P, ATP, SEP; pdb70 \| 2VDA\_A \| 95.3 \| 0.00019 \| 2.2e-08 \| 78.4 \| 66 \| (62, 134) \| 682 \| (72, 138) \| 828 \| TRANSLOCASE SUBUNIT SECA, MALTOPORIN \| 2VDA\_A TRANSLOCASE SUBUNIT SECA, MALTOPORIN SUGAR TRANSPORT, PROTEIN TRANSPORT, PROTEIN; pdb70 \| 2FSF\_A \| 95.3 \| 0.00019 \| 2.2e-08 \| 78.4 \| 66 \| (62, 134) \| 682 \| (72, 138) \| 853 \| Preprotein translocase secA subunit \| 2FSF\_A Preprotein translocase secA subunit ATPase, DNA-RNA helicase, Protein translocation; pdb70 \| 2FSH\_A \| 95.3 \| 0.00019 \| 2.2e-08 \| 78.4 \| 66 \| (62, 134) \| 682 \| (72, 138) \| 853 \| Preprotein translocase secA subunit \| 2FSH\_A Preprotein translocase secA subunit ATPase, DNA-RNA helicase, Protein translocation HET: ANP; pdb70 \| 4KIT\_B \| 95.3 \| 0.00018 \| 2.2e-08 \| 83.2 \| 71 \| (63, 135) \| 682 \| (89, 169) \| 1739 \| U5 small nuclear ribonucleoprotein 200 \| 4KIT\_B U5 small nuclear ribonucleoprotein 200 RecA domain, winged helix domain HET: ADP; pdb70 \| 4KIT\_B \| 95.3 \| 0.00018 \| 2.2e-08 \| 83.2 \| 68 \| (64, 133) \| 682 \| (937, 1005) \| 1739 \| U5 small nuclear ribonucleoprotein 200 \| 4KIT\_B U5 small nuclear ribonucleoprotein 200 RecA domain, winged helix domain HET: ADP; pdb70 \| 4CGZ\_A \| 95.3 \| 0.00018 \| 2.2e-08 \| 75.3 \| 62 \| (64, 133) \| 682 \| (34, 96) \| 665 \| BLOOM'S SYNDROME HELICASE (E.C.3.6.4.12) \| 4CGZ\_A BLOOM'S SYNDROME HELICASE (E.C.3.6.4.12) HYDROLASE-DNA COMPLEX HET: ADP; pdb70 \| 3KQL\_A \| 95.3 \| 0.00018 \| 2.2e-08 \| 70.7 \| 97 \| (80, 196) \| 682 \| (8, 109) \| 437 \| Serine protease/NTPase/helicase NS3 (E.C.3.4.21.98, 3.6.1.15 \| 3KQL\_A Serine protease/NTPase/helicase NS3 (E.C.3.4.21.98, 3.6.1.15 helicase-substrate transition-state complex, HCV, NS3 HET: ADP, ALF; pdb70 \| 3KQU\_D \| 95.3 \| 0.00018 \| 2.2e-08 \| 70.7 \| 97 \| (80, 196) \| 682 \| (8, 109) \| 437 \| Serine protease/NTPase/helicase NS3 (E.C.3.4.21.98, 3.6.1.15 \| 3KQU\_D Serine protease/NTPase/helicase NS3 (E.C.3.4.21.98, 3.6.1.15 helicase-substrate ground-state complex, HCV, NS3 HET: ADP, BEF; pdb70 \| 4QQW\_A \| 95.3 \| 0.0002 \| 2.2e-08 \| 79.7 \| 72 \| (63, 135) \| 682 \| (298, 372) \| 964 \| CRISPR-associated helicase, Cas3 family/dna \| 4QQW\_A CRISPR-associated helicase, Cas3 family/dna CRISPR, Cas3, helicase, Hydrolase-DNA complex; pdb70 \| 4DDT\_A \| 95.3 \| 0.0002 \| 2.3e-08 \| 80.4 \| 69 \| (62, 135) \| 682 \| (76, 145) \| 1104 \| Reverse gyrase (E.C.3.6.4.12, 5.99.1.3) \| 4DDT\_A Reverse gyrase (E.C.3.6.4.12, 5.99.1.3) TOPOISOMERASE, DNA SUPERCOILING, ARCHAEA, HELICASE; pdb70 \| 4DDW\_A \| 95.3 \| 0.0002 \| 2.3e-08 \| 80.4 \| 69 \| (62, 135) \| 682 \| (76, 145) \| 1104 \| Reverse gyrase (E.C.3.6.4.12, 5.99.1.3) \| 4DDW\_A Reverse gyrase (E.C.3.6.4.12, 5.99.1.3) TOPOISOMERASE, DNA SUPERCOILING, ARCHAEA, HELICASE; pdb70 \| 6QV4\_A \| 95.3 \| 0.00019 \| 2.3e-08 \| 83.0 \| 71 \| (63, 135) \| 682 \| (59, 139) \| 1725 \| Pre-mRNA splicing helicase-like protein \| 6QV4\_A Pre-mRNA splicing helicase-like protein Helicase ATP-gamma-S Brr2 Ski2 Chaetomium HET: AGS; pdb70 \| 3KQN\_A \| 95.3 \| 0.00019 \| 2.3e-08 \| 70.4 \| 97 \| (80, 196) \| 682 \| (8, 109) \| 437 \| Serine protease/NTPase/helicase NS3 (E.C.3.4.21.98, 3.6.1.15 \| 3KQN\_A Serine protease/NTPase/helicase NS3 (E.C.3.4.21.98, 3.6.1.15 helicase-substrate transition-state complex, HCV, NS3 HET: ADP; pdb70 \| 6O16\_A \| 95.3 \| 0.00021 \| 2.4e-08 \| 79.5 \| 65 \| (67, 134) \| 682 \| (76, 142) \| 975 \| DEAH (Asp-Glu-Ala-His) box polypeptide 37/RNA \| 6O16\_A DEAH (Asp-Glu-Ala-His) box polypeptide 37/RNA RNA helicase, ribosome biogenesis, RNA-dependent; pdb70 \| 6S8Q\_B \| 95.3 \| 0.00019 \| 2.4e-08 \| 83.0 \| 71 \| (63, 135) \| 682 \| (90, 170) \| 1747 \| U5 small nuclear ribonucleoprotein 200 \| 6S8Q\_B U5 small nuclear ribonucleoprotein 200 RNP REMODELING, PRE-MRNA SPLICING, SPLICEOSOME HET: SO4; pdb70 \| 5WX1\_A \| 95.3 \| 0.00021 \| 2.4e-08 \| 77.2 \| 129 \| (81, 228) \| 682 \| (269, 406) \| 733 \| Serine protease NS3 \| 5WX1\_A Serine protease NS3 protease, RNA helicase, HYDROLASE; pdb70 \| 4DDU\_A \| 95.3 \| 0.00021 \| 2.4e-08 \| 80.3 \| 69 \| (62, 135) \| 682 \| (76, 145) \| 1104 \| Reverse gyrase (E.C.3.6.4.12, 5.99.1.3) \| 4DDU\_A Reverse gyrase (E.C.3.6.4.12, 5.99.1.3) TOPOISOMERASE, DNA SUPERCOILING, ARCHAEA, HELICASE; pdb70 \| 6S8Q\_B \| 95.3 \| 0.0002 \| 2.4e-08 \| 82.9 \| 68 \| (64, 133) \| 682 \| (938, 1006) \| 1747 \| U5 small nuclear ribonucleoprotein 200 \| 6S8Q\_B U5 small nuclear ribonucleoprotein 200 RNP REMODELING, PRE-MRNA SPLICING, SPLICEOSOME HET: SO4; pdb70 \| 4O3M\_A \| 95.3 \| 0.0002 \| 2.4e-08 \| 74.8 \| 63 \| (63, 133) \| 682 \| (27, 90) \| 659 \| Bloom syndrome protein (E.C.3.6.4.12) \| 4O3M\_A Bloom syndrome protein (E.C.3.6.4.12) Winged Helix, Helicase, Hydrolase-DNA complex HET: EDO, ADP; pdb70 \| 5M59\_E \| 95.2 \| 0.0002 \| 2.4e-08 \| 83.1 \| 70 \| (64, 135) \| 682 \| (957, 1027) \| 1772 \| Putative pre-mRNA splicing factor, Pre-mRNA \| 5M59\_E Putative pre-mRNA splicing factor, Pre-mRNA Brr2, pre-mRNA splicing, RNA-helicase, Prp8 HET: ACT; pdb70 \| 6QW6\_5B \| 95.2 \| 0.00021 \| 2.5e-08 \| 85.7 \| 70 \| (64, 135) \| 682 \| (480, 559) \| 2136 \| Small nuclear ribonucleoprotein Sm D1 \| 6QW6\_5B Small nuclear ribonucleoprotein Sm D1 RNP complex, splicing, RNA, protein HET: IHP, M7M, GTP; pdb70 \| 6QDV\_B \| 95.2 \| 0.0002 \| 2.5e-08 \| 82.7 \| 71 \| (63, 135) \| 682 \| (76, 156) \| 1722 \| Eukaryotic initiation factor 4A-III, RNA-binding \| 6QDV\_B Eukaryotic initiation factor 4A-III, RNA-binding spliceosome, RNA, complex, SPLICING HET: GTP, I6P, ATP, SEP; pdb70 \| 6QX9\_5B \| 95.2 \| 0.00022 \| 2.5e-08 \| 85.6 \| 71 \| (64, 136) \| 682 \| (480, 560) \| 2136 \| Splicing factor 3B subunit 4 \| 6QX9\_5B Splicing factor 3B subunit 4 RNP complex, splicing, RNA, protein HET: M7M, IHP, GTP; pdb70 \| 2V6I\_A \| 95.2 \| 0.0002 \| 2.6e-08 \| 69.4 \| 51 \| (81, 133) \| 682 \| (2, 53) \| 431 \| RNA HELICASE \| 2V6I\_A RNA HELICASE MEMBRANE, HELICASE, HYDROLASE, RNA HELICASE; pdb70 \| 5WSO\_A \| 95.2 \| 0.00024 \| 2.6e-08 \| 73.6 \| 60 \| (69, 133) \| 682 \| (2, 62) \| 476 \| NS3 helicase \| 5WSO\_A NS3 helicase BVDV, Bovine viral diarrhea virus; pdb70 \| 6RXU\_CS \| 95.2 \| 0.00026 \| 2.6e-08 \| 81.2 \| 70 \| (62, 133) \| 682 \| (254, 329) \| 1073 \| Periodic tryptophan protein 2-like protein \| 6RXU\_CS Periodic tryptophan protein 2-like protein ribosome, ribosome biogenesis, rRNA HET: GTP; pdb70 \| 3JV2\_A \| 95.2 \| 0.00023 \| 2.8e-08 \| 76.0 \| 66 \| (63, 135) \| 682 \| (82, 148) \| 783 \| Protein translocase subunit secA \| 3JV2\_A Protein translocase subunit secA protein translocation, ATPase, conformational change HET: ADP; pdb70 \| 6QW6\_5B \| 95.2 \| 0.00025 \| 2.9e-08 \| 85.1 \| 69 \| (64, 134) \| 682 \| (1327, 1396) \| 2136 \| Small nuclear ribonucleoprotein Sm D1 \| 6QW6\_5B Small nuclear ribonucleoprotein Sm D1 RNP complex, splicing, RNA, protein HET: IHP, M7M, GTP; pdb70 \| 6QX9\_5B \| 95.2 \| 0.00025 \| 2.9e-08 \| 85.1 \| 70 \| (64, 135) \| 682 \| (1327, 1397) \| 2136 \| Splicing factor 3B subunit 4 \| 6QX9\_5B Splicing factor 3B subunit 4 RNP complex, splicing, RNA, protein HET: M7M, IHP, GTP; pdb70 \| 6DGD\_A \| 95.1 \| 0.00025 \| 3.1e-08 \| 75.1 \| 125 \| (63, 195) \| 682 \| (216, 345) \| 751 \| Primosomal protein N'/DNA Complex \| 6DGD\_A Primosomal protein N'/DNA Complex DNA replication restart, PriA helicase HET: SO4; pdb70 \| 1CU1\_B \| 95.1 \| 0.00029 \| 3.3e-08 \| 75.1 \| 109 \| (66, 196) \| 682 \| (198, 311) \| 645 \| PROTEIN (HEPATITIS C VIRUS) \| 1CU1\_B PROTEIN (HEPATITIS C VIRUS) hepatitis C Virus, bifunctional, protease-helicase HET: PO4; pdb70 \| 6EG2\_A \| 95.1 \| 0.00025 \| 3.3e-08 \| 70.7 \| 157 \| (61, 229) \| 682 \| (386, 553) \| 621 \| Maltose/maltodextrin-binding periplasmic protein,Probable global transcription \| 6EG2\_A Maltose/maltodextrin-binding periplasmic protein,Probable global transcription Helicase, ATPase, Chromatin remodeling, inhibitor HET: J7J; pdb70 \| 6EG3\_A \| 95.1 \| 0.00025 \| 3.3e-08 \| 70.7 \| 157 \| (61, 229) \| 682 \| (386, 553) \| 621 \| Maltose/maltodextrin-binding periplasmic protein,Probable global transcription \| 6EG3\_A Maltose/maltodextrin-binding periplasmic protein,Probable global transcription Helicase, ATPase, Chromatin remodeling, inhibitor HET: EOH, J7G; pdb70 \| 5LB3\_B \| 95.1 \| 0.00026 \| 3.4e-08 \| 68.0 \| 60 \| (67, 133) \| 682 \| (24, 84) \| 445 \| ATP-dependent DNA helicase Q5 (E.C.3.6.4.12) \| 5LB3\_B ATP-dependent DNA helicase Q5 (E.C.3.6.4.12) Helicase, RecQ, Transcription, DNA repair HET: ADP; pdb70 \| 2V6J\_A \| 95.0 \| 0.00028 \| 3.6e-08 \| 68.2 \| 51 \| (81, 133) \| 682 \| (2, 53) \| 431 \| RNA HELICASE \| 2V6J\_A RNA HELICASE MEMBRANE, HELICASE, HYDROLASE, RNA HELICASE; pdb70 \| 5YVW\_B \| 95.0 \| 0.00031 \| 3.6e-08 \| 73.3 \| 52 \| (80, 133) \| 682 \| (166, 218) \| 599 \| Genome polyprotein \| 5YVW\_B Genome polyprotein SERINE PROTEASE, NON-STRUCTURAL PROTEIN 3 HET: EPE; pdb70 \| 4UAQ\_A \| 95.0 \| 0.00032 \| 3.7e-08 \| 75.7 \| 67 \| (64, 135) \| 682 \| (87, 154) \| 778 \| SecA2 \| 4UAQ\_A SecA2 Protein Transport, DEAD/DEAH box helicase HET: MSE; pdb70 \| 5EAN\_A \| 94.9 \| 0.00039 \| 4.4e-08 \| 78.2 \| 66 \| (63, 132) \| 682 \| (628, 694) \| 1059 \| DNA replication ATP-dependent helicase/nuclease DNA2/DNA \| 5EAN\_A DNA replication ATP-dependent helicase/nuclease DNA2/DNA DNA binding protein, Hydrolase-DNA complex HET: ADP; pdb70 \| 3O8B\_B \| 94.9 \| 0.0004 \| 4.5e-08 \| 74.5 \| 109 \| (66, 196) \| 682 \| (219, 332) \| 666 \| HCV NS3 protease/helicase (E.C.3.4.21.98, 3.6.1.15 \| 3O8B\_B HCV NS3 protease/helicase (E.C.3.4.21.98, 3.6.1.15 helicase, NTPase, HCV, RNA, translocation HET: SO4; pdb70 \| 5YW1\_B \| 94.9 \| 0.0004 \| 4.7e-08 \| 72.9 \| 52 \| (80, 133) \| 682 \| (185, 237) \| 618 \| Genome polyprotein, PTI protein \| 5YW1\_B Genome polyprotein, PTI protein SERINE PROTEASE, NON-STRUCTURAL PROTEIN 3; pdb70 \| 5MFX\_A \| 94.9 \| 0.00039 \| 4.8e-08 \| 68.3 \| 52 \| (80, 133) \| 682 \| (14, 66) \| 451 \| NS3 helicase/RNA Complex \| 5MFX\_A NS3 helicase/RNA Complex Helicase, RNA, hydrolase HET: FLC; pdb70 \| 6H57\_A \| 94.8 \| 0.00051 \| 5.3e-08 \| 79.7 \| 117 \| (66, 195) \| 682 \| (394, 522) \| 1267 \| Probable ATP-dependent RNA helicase DHR1 \| 6H57\_A Probable ATP-dependent RNA helicase DHR1 protein, HYDROLASE HET: EDO; pdb70 \| 1GKU\_B \| 94.8 \| 0.00048 \| 5.4e-08 \| 77.5 \| 68 \| (63, 135) \| 682 \| (55, 123) \| 1054 \| REVERSE GYRASE \| 1GKU\_B REVERSE GYRASE TOPOISOMERASE, DNA SUPERCOILING, ARCHAEA, HELICASE; pdb70 \| 2L8B\_A \| 94.8 \| 0.00045 \| 5.5e-08 \| 60.2 \| 57 \| (62, 120) \| 682 \| (32, 90) \| 189 \| Protein traI (E.C.3.6.4.12) \| 2L8B\_A Protein traI (E.C.3.6.4.12) RecD, HYDROLASE; pdb70 \| 5LB8\_A \| 94.7 \| 0.00052 \| 6.3e-08 \| 69.3 \| 60 \| (67, 133) \| 682 \| (24, 84) \| 518 \| ATP-dependent DNA helicase Q5 (E.C.3.6.4.12) \| 5LB8\_A ATP-dependent DNA helicase Q5 (E.C.3.6.4.12) Helicase, RecQ, Transcription, DNA repair; pdb70 \| 6DCR\_B \| 94.7 \| 0.00052 \| 6.3e-08 \| 71.6 \| 124 \| (64, 195) \| 682 \| (160, 288) \| 694 \| Primosomal protein N' (E.C.3.6.4.-) \| 6DCR\_B Primosomal protein N' (E.C.3.6.4.-) PriA, Helicase, DNA replication restart HET: SO4; pdb70 \| 2WHX\_A \| 94.7 \| 0.00055 \| 6.4e-08 \| 71.7 \| 52 \| (80, 133) \| 682 \| (185, 237) \| 618 \| SERINE PROTEASE/NTPASE/HELICASE NS3 (E.C.3.4.21.91, 3.6.1.15 \| 2WHX\_A SERINE PROTEASE/NTPASE/HELICASE NS3 (E.C.3.4.21.91, 3.6.1.15 TRANSCRIPTION, HYDROLASE, ATP-BINDING, RETICULUM, NUCLEOTIDYLTRANSFERASE HET: ADP; pdb70 \| 5Y4Z\_A \| 94.6 \| 0.00054 \| 6.9e-08 \| 66.1 \| 52 \| (81, 134) \| 682 \| (10, 62) \| 440 \| NS3 helicase (E.C.3.4.21.91,3.6.1.15,3.6.4.13) \| 5Y4Z\_A NS3 helicase (E.C.3.4.21.91,3.6.1.15,3.6.4.13) ZIKA virus, NS3 helicase, AMPPNP HET: ANP; pdb70 \| 1TF5\_A \| 94.6 \| 0.00058 \| 7.1e-08 \| 72.8 \| 66 \| (64, 134) \| 682 \| (81, 147) \| 844 \| Preprotein translocase secA subunit \| 1TF5\_A Preprotein translocase secA subunit ATPase, helicase, translocation, secretion, PROTEIN; pdb70 \| 6HPH\_A \| 94.5 \| 0.00058 \| 7.4e-08 \| 65.6 \| 55 \| (63, 123) \| 682 \| (3, 58) \| 418 \| ATP-dependent DNA helicase PIF1 (E.C.3.6.4.12) \| 6HPH\_A ATP-dependent DNA helicase PIF1 (E.C.3.6.4.12) Pif1, 5'-3' DNA helicase, Duplex HET: ANP; pdb70 \| 1GL9\_B \| 94.5 \| 0.00067 \| 7.6e-08 \| 76.1 \| 67 \| (64, 135) \| 682 \| (56, 123) \| 1054 \| REVERSE GYRASE \| 1GL9\_B REVERSE GYRASE TOPOISOMERASE, DNA SUPERCOILING, ARCHAEA, HELICASE HET: ANP; pdb70 \| 6JYT\_B \| 94.5 \| 0.00071 \| 7.8e-08 \| 71.8 \| 129 \| (81, 228) \| 682 \| (277, 407) \| 603 \| Helicase (E.C.3.6.4.12,3.6.4.13) \| 6JYT\_B Helicase (E.C.3.6.4.12,3.6.4.13) SARS-Nsp13, HYDROLASE HET: MSE; pdb70 \| 1NKT\_A \| 94.3 \| 0.00088 \| 9.6e-08 \| 74.8 \| 66 \| (64, 134) \| 682 \| (109, 175) \| 922 \| PREPROTEIN TRANSLOCASE SECA 1 SUBUNIT \| 1NKT\_A PREPROTEIN TRANSLOCASE SECA 1 SUBUNIT PREPROTEIN TRANSLOCATION, ATPASE, TRANSMEMBRANE TRANSPORT HET: ADP; pdb70 \| 1NL3\_A \| 94.3 \| 0.00088 \| 9.6e-08 \| 74.8 \| 66 \| (64, 134) \| 682 \| (109, 175) \| 922 \| PREPROTEIN TRANSLOCASE SECA 1 SUBUNIT \| 1NL3\_A PREPROTEIN TRANSLOCASE SECA 1 SUBUNIT PREPROTEIN TRANSLOCATION, ATPASE, TRANSMEMBRANE TRANSPORT; pdb70 \| 1NL3\_B \| 94.3 \| 0.00088 \| 9.6e-08 \| 74.8 \| 66 \| (64, 134) \| 682 \| (109, 175) \| 922 \| PREPROTEIN TRANSLOCASE SECA 1 SUBUNIT \| 1NL3\_B PREPROTEIN TRANSLOCASE SECA 1 SUBUNIT PREPROTEIN TRANSLOCATION, ATPASE, TRANSMEMBRANE TRANSPORT; pdb70 \| 6SXA\_F \| 94.2 \| 0.00091 \| 1.1e-07 \| 72.7 \| 62 \| (63, 131) \| 682 \| (13, 75) \| 916 \| DNA repair endonuclease XPF (E.C.3.1.-.-) \| 6SXA\_F DNA repair endonuclease XPF (E.C.3.1.-.-) DNA Repair enzyme. Nucleotide excision; pdb70 \| 6HPU\_A \| 94.2 \| 0.0009 \| 1.1e-07 \| 64.4 \| 54 \| (63, 122) \| 682 \| (3, 57) \| 418 \| ATP-dependent DNA helicase PIF1 (E.C.3.6.4.12) \| 6HPU\_A ATP-dependent DNA helicase PIF1 (E.C.3.6.4.12) Pif1, 5'-3' DNA helicase, Duplex HET: ADP, ALF |
| Top keywords  (threshold 1.00e-03 (evalue)) | **terminase, large, Phage, helicase, RNA, GpA, Fragment, ATPase, domain\_containing, DNA** |
| Output files | ../../similar\_sequences/02\_FANPEZAQ\_CDS\_0002\_merged.svg ../../similar\_sequences/02\_FANPEZAQ\_CDS\_0002\_pdb70.a3m ../../similar\_sequences/02\_FANPEZAQ\_CDS\_0002\_pdb70.hhr ../../similar\_sequences/02\_FANPEZAQ\_CDS\_0002\_uniclust.a3m ../../similar\_sequences/02\_FANPEZAQ\_CDS\_0002\_uniclust.hhr |

#### Structure prediction (AlphaFold)^2^

|  |  |
| --- | --- |
| Stats | xml version="1.0" encoding="utf-8" standalone="no"?       2024-09-02T21:08:59.285358 image/svg+xml   Matplotlib v3.7.2, https://matplotlib.org/ |
| Predicted structure | **NGL Viewer Controls:**  - Center: *Left-Click* - Rotate: *Left-Click + Drag* - Translate: *Right-Click + Drag* - Zoom: *Shift + Left-Click + Drag* |
| Output files | ../../predicted\_structures/02\_FANPEZAQ\_CDS\_0002/features.pkl ../../predicted\_structures/02\_FANPEZAQ\_CDS\_0002/ranked\_0.pdb ../../predicted\_structures/02\_FANPEZAQ\_CDS\_0002/ranked\_0\_plots.svg ../../predicted\_structures/02\_FANPEZAQ\_CDS\_0002/result\_model\_1\_ptm\_pred\_0.pkl |

#### Structure similarity search results (Foldseek)^3^

|  |  |
| --- | --- |
| Structure databases searched | Pdb, Afdb-proteome, Afdb-uniprot50 |
| Results, scheme(s)  (Top layers only, threshold 1.00e-02 (evalue)) | xml version="1.0" encoding="utf-8" standalone="no"?       2024-09-02T21:10:06.729080 image/svg+xml   Matplotlib v3.7.2, https://matplotlib.org/ |
| Results, table  (threshold 1.00e-02 (evalue)) | db \| id \| prob \| evalue \| bits \| fident \| alnlen \| mismatch \| gapopen \| qstart \| qend \| tstart \| tend \| name \| description; pdb \| 4IFE\_A \| 1.0 \| 1.299e-10 \| 197 \| 0.111 \| 572 \| 328 \| 39 \| 72 \| 604 \| 4 \| 433 \| Gene 2 protein \| Gene 2 protein; pdb \| 3EZK\_A \| 1.0 \| 1.154e-09 \| 193 \| 0.103 \| 600 \| 338 \| 36 \| 41 \| 602 \| 108 \| 545 \| DNA packaging protein Gp17 \| DNA packaging protein Gp17; pdb \| 3CPE\_A \| 1.0 \| 8.247e-10 \| 190 \| 0.106 \| 594 \| 339 \| 32 \| 44 \| 603 \| 111 \| 546 \| DNA packaging protein Gp17 \| DNA packaging protein Gp17; pdb \| 2O0K\_A \| 1.0 \| 8.989e-06 \| 188 \| 0.15 \| 200 \| 149 \| 11 \| 44 \| 240 \| 120 \| 301 \| DNA packaging protein Gp17 \| DNA packaging protein Gp17; pdb \| 4IDH\_A \| 1.0 \| 1.162e-10 \| 188 \| 0.115 \| 579 \| 317 \| 42 \| 72 \| 604 \| 5 \| 434 \| Gene 2 protein \| Gene 2 protein; pdb \| 4ZNJ\_A \| 1.0 \| 5.168e-07 \| 182 \| 0.146 \| 307 \| 157 \| 16 \| 83 \| 369 \| 29 \| 250 \| Phage terminase large subunit \| Phage terminase large subunit; pdb \| 4IEE\_A \| 1.0 \| 3.765e-10 \| 180 \| 0.113 \| 584 \| 314 \| 40 \| 72 \| 604 \| 4 \| 434 \| Gene 2 protein \| Gene 2 protein; pdb \| 6Z6D\_A \| 1.0 \| 1.084e-08 \| 171 \| 0.119 \| 634 \| 331 \| 46 \| 39 \| 600 \| 1 \| 478 \| Terminase large subunit \| Terminase large subunit; pdb \| 4ZNL\_C \| 1.0 \| 2.217e-06 \| 168 \| 0.155 \| 302 \| 160 \| 15 \| 83 \| 369 \| 30 \| 251 \| Phage terminase large subunit \| Phage terminase large subunit; pdb \| 4BIL\_A \| 1.0 \| 1.356e-08 \| 154 \| 0.102 \| 625 \| 371 \| 39 \| 23 \| 610 \| 3 \| 474 \| DNA MATURASE B \| DNA MATURASE B; pdb \| 4BIJ\_A \| 1.0 \| 6.153e-08 \| 139 \| 0.094 \| 624 \| 375 \| 41 \| 23 \| 610 \| 3 \| 472 \| DNA MATURASE B \| DNA MATURASE B; pdb \| 5C15\_A \| 1.0 \| 0.0005996 \| 103 \| 0.129 \| 262 \| 145 \| 21 \| 376 \| 598 \| 4 \| 221 \| Gene 2 protein \| Gene 2 protein; pdb \| 6M5S\_A \| 0.998 \| 1.197e-06 \| 94 \| 0.125 \| 589 \| 319 \| 36 \| 73 \| 607 \| 195 \| 641 \| Tripartite terminase subunit 3 \| Tripartite terminase subunit 3; pdb \| 4IOX\_C \| 0.996 \| 0.001944 \| 90 \| 0.111 \| 251 \| 154 \| 16 \| 378 \| 607 \| 8 \| 210 \| Tripartite terminase subunit UL15 \| Tripartite terminase subunit UL15; pdb \| 2P6R\_A \| 0.993 \| 0.005957 \| 86 \| 0.111 \| 179 \| 117 \| 11 \| 60 \| 223 \| 25 \| 176 \| afUHEL308 HELICASE \| afUHEL308 HELICASE; pdb \| 4LD0\_A \| 0.992 \| 0.008336 \| 85 \| 0.148 \| 108 \| 76 \| 5 \| 399 \| 495 \| 2 \| 104 \| Crossover junction endodeoxyribonuclease RuvC \| Crossover junction endodeoxyribonuclease RuvC; pdb \| 1RIF\_B \| 0.99 \| 0.005036 \| 83 \| 0.122 \| 245 \| 157 \| 19 \| 15 \| 235 \| 61 \| 271 \| DNA helicase uvsW \| DNA helicase uvsW; pdb \| 4EP5\_A \| 0.986 \| 0.007882 \| 81 \| 0.14 \| 213 \| 113 \| 14 \| 399 \| 604 \| 2 \| 151 \| Crossover junction endodeoxyribonuclease RuvC \| Crossover junction endodeoxyribonuclease RuvC; pdb \| 6S16\_B \| 0.984 \| 0.008336 \| 80 \| 0.181 \| 110 \| 72 \| 7 \| 398 \| 495 \| 2 \| 105 \| Crossover junction endodeoxyribonuclease RuvC \| Crossover junction endodeoxyribonuclease RuvC; pdb \| 3N4P\_C \| 0.984 \| 0.0063 \| 80 \| 0.114 \| 254 \| 144 \| 16 \| 378 \| 603 \| 7 \| 207 \| Terminase subunit UL89 protein \| Terminase subunit UL89 protein; pdb \| 4EP4\_B \| 0.981 \| 0.008816 \| 79 \| 0.127 \| 227 \| 111 \| 17 \| 399 \| 605 \| 2 \| 161 \| Crossover junction endodeoxyribonuclease RuvC \| Crossover junction endodeoxyribonuclease RuvC; pdb \| 4NL8\_B \| 0.961 \| 0.0001322 \| 74 \| 0.16 \| 387 \| 184 \| 34 \| 61 \| 349 \| 97 \| 440 \| Primosome assembly protein PriA \| Primosome assembly protein PriA; pdb \| 4NL8\_E \| 0.9 \| 0.0004051 \| 67 \| 0.15 \| 412 \| 193 \| 35 \| 61 \| 362 \| 90 \| 454 \| Primosome assembly protein PriA \| Primosome assembly protein PriA; pdb \| 6DCR\_A \| 0.9 \| 0.0008874 \| 67 \| 0.146 \| 383 \| 178 \| 33 \| 46 \| 331 \| 116 \| 446 \| Primosomal protein N' \| Primosomal protein N'; pdb \| 2XGJ\_A \| 0.855 \| 0.004257 \| 64 \| 0.145 \| 254 \| 152 \| 18 \| 34 \| 258 \| 45 \| 262 \| ATP-DEPENDENT RNA HELICASE DOB1 \| ATP-DEPENDENT RNA HELICASE DOB1; pdb \| 4NL8\_A \| 0.837 \| 0.0009385 \| 63 \| 0.152 \| 400 \| 175 \| 33 \| 61 \| 351 \| 97 \| 441 \| Primosome assembly protein PriA \| Primosome assembly protein PriA; pdb \| 4NL4\_H \| 0.772 \| 0.0004793 \| 60 \| 0.135 \| 444 \| 198 \| 36 \| 53 \| 370 \| 188 \| 571 \| Primosome assembly protein PriA \| Primosome assembly protein PriA; pdb \| 6DGD\_B \| 0.747 \| 0.0005996 \| 59 \| 0.142 \| 407 \| 192 \| 33 \| 61 \| 356 \| 195 \| 555 \| Primosomal protein N' \| Primosomal protein N'; pdb \| 8FAK\_H \| 0.569 \| 0.003403 \| 53 \| 0.125 \| 437 \| 207 \| 28 \| 64 \| 370 \| 113 \| 504 \| Primosomal protein N' \| Primosomal protein N'; pdb \| 7XSX\_M \| 0.505 \| 0.003042 \| 51 \| 0.133 \| 270 \| 151 \| 22 \| 300 \| 534 \| 382 \| 603 \| Transcription elongation factor Spt6 \| Transcription elongation factor Spt6; afdb-proteome \| AF-Q8ZQH1-F1-MODEL\_V4 \| 1.0 \| 3.644e-45 \| 1331 \| 0.213 \| 739 \| 469 \| 33 \| 14 \| 679 \| 4 \| 702 \| Fels-1 prophage protein \| Fels-1 prophage protein; afdb-proteome \| AF-Q8ZMZ3-F1-MODEL\_V4 \| 1.0 \| 5.534e-49 \| 1259 \| 0.253 \| 646 \| 414 \| 23 \| 19 \| 623 \| 8 \| 625 \| Gifsy-1 prophage protein \| Gifsy-1 prophage protein; afdb-proteome \| AF-Q8ZQ96-F1-MODEL\_V4 \| 1.0 \| 7.694e-48 \| 1112 \| 0.213 \| 753 \| 478 \| 33 \| 3 \| 682 \| 1 \| 711 \| Gifsy-2 prophage protein \| Gifsy-2 prophage protein; afdb-proteome \| AF-A0A077ZJY7-F1-MODEL\_V4 \| 0.998 \| 0.0005788 \| 93 \| 0.111 \| 288 \| 173 \| 20 \| 1 \| 223 \| 4 \| 273 \| Helicase POLQ-like \| Helicase POLQ-like; afdb-proteome \| AF-A0A0K0E265-F1-MODEL\_V4 \| 0.996 \| 0.001984 \| 90 \| 0.125 \| 270 \| 151 \| 18 \| 1 \| 237 \| 73 \| 290 \| Uncharacterized protein \| Uncharacterized protein; afdb-proteome \| AF-Q6ZR08-F4-MODEL\_V4 \| 0.986 \| 0.004597 \| 81 \| 0.128 \| 249 \| 169 \| 15 \| 5 \| 225 \| 1 \| 229 \| Dynein axonemal heavy chain 12 \| Dynein axonemal heavy chain 12; afdb-proteome \| AF-Q54CI4-F1-MODEL\_V4 \| 0.978 \| 0.004861 \| 78 \| 0.127 \| 267 \| 149 \| 19 \| 1 \| 236 \| 231 \| 444 \| Uncharacterized protein \| Uncharacterized protein; afdb-proteome \| AF-Q54X98-F1-MODEL\_V4 \| 0.975 \| 0.005438 \| 77 \| 0.123 \| 267 \| 151 \| 14 \| 1 \| 223 \| 386 \| 613 \| Uncharacterized protein \| Uncharacterized protein; afdb-proteome \| AF-H2KY86-F1-MODEL\_V4 \| 0.971 \| 0.002099 \| 76 \| 0.108 \| 287 \| 172 \| 18 \| 1 \| 223 \| 62 \| 328 \| Helicase C-terminal domain-containing protein \| Helicase C-terminal domain-containing protein; afdb-proteome \| AF-A0A3Q0KLA5-F1-MODEL\_V4 \| 0.971 \| 0.006082 \| 76 \| 0.129 \| 254 \| 153 \| 20 \| 19 \| 236 \| 446 \| 667 \| Putative chromodomain helicase DNA binding protein \| Putative chromodomain helicase DNA binding protein; afdb-proteome \| AF-F4KGU4-F1-MODEL\_V4 \| 0.473 \| 0.00952 \| 50 \| 0.093 \| 546 \| 252 \| 34 \| 63 \| 557 \| 306 \| 659 \| ATP-dependent RNA helicase DEAH12, chloroplastic \| ATP-dependent RNA helicase DEAH12, chloroplastic; afdb-uniprot50 \| AF-Q9P9Q0-F1-MODEL\_V4 \| 1.0 \| 5.809e-89 \| 3256 \| 0.618 \| 606 \| 205 \| 4 \| 77 \| 682 \| 1 \| 580 \| Phage-related terminase large subunit \| Phage-related terminase large subunit; afdb-uniprot50 \| AF-A0A1C3CVG6-F1-MODEL\_V4 \| 1.0 \| 2.147e-91 \| 3248 \| 0.542 \| 652 \| 278 \| 4 \| 33 \| 677 \| 4 \| 641 \| Terminase \| Terminase; afdb-uniprot50 \| AF-A0A7C8LYM0-F1-MODEL\_V4 \| 1.0 \| 7.036e-84 \| 3066 \| 0.497 \| 617 \| 285 \| 8 \| 4 \| 616 \| 1 \| 596 \| Uncharacterized protein \| Uncharacterized protein; afdb-uniprot50 \| AF-A0A1H2R4X1-F1-MODEL\_V4 \| 1.0 \| 7.939e-94 \| 3026 \| 0.523 \| 667 \| 286 \| 6 \| 15 \| 681 \| 4 \| 638 \| Phage terminase, large subunit GpA \| Phage terminase, large subunit GpA; afdb-uniprot50 \| AF-A0A238JBF0-F1-MODEL\_V4 \| 1.0 \| 3.199e-92 \| 2982 \| 0.524 \| 667 \| 282 \| 4 \| 15 \| 681 \| 4 \| 635 \| Phage terminase large subunit (GpA) \| Phage terminase large subunit (GpA); afdb-uniprot50 \| AF-A0A3N2E110-F1-MODEL\_V4 \| 1.0 \| 6.61e-83 \| 2942 \| 0.43 \| 700 \| 331 \| 17 \| 11 \| 681 \| 18 \| 678 \| Phage terminase large subunit GpA-like protein \| Phage terminase large subunit GpA-like protein; afdb-uniprot50 \| AF-A0A2S0MNT9-F1-MODEL\_V4 \| 1.0 \| 2.057e-93 \| 2910 \| 0.555 \| 680 \| 269 \| 8 \| 2 \| 678 \| 8 \| 656 \| Phage terminase large subunit family protein \| Phage terminase large subunit family protein; afdb-uniprot50 \| AF-A0A4Q3YY62-F1-MODEL\_V4 \| 1.0 \| 3.532e-90 \| 2839 \| 0.524 \| 664 \| 276 \| 6 \| 21 \| 678 \| 6 \| 635 \| Phage terminase large subunit family protein \| Phage terminase large subunit family protein; afdb-uniprot50 \| AF-A0A0C1G363-F1-MODEL\_V4 \| 1.0 \| 4.054e-94 \| 2764 \| 0.544 \| 689 \| 258 \| 8 \| 10 \| 681 \| 1 \| 650 \| Terminase \| Terminase; afdb-uniprot50 \| AF-A0A846VUS9-F1-MODEL\_V4 \| 1.0 \| 4.734e-92 \| 2736 \| 0.527 \| 667 \| 283 \| 7 \| 15 \| 681 \| 4 \| 638 \| Phage terminase large subunit family protein \| Phage terminase large subunit family protein; afdb-uniprot50 \| AF-A0A3B9NPU7-F1-MODEL\_V4 \| 1.0 \| 3.24e-68 \| 2442 \| 0.372 \| 639 \| 361 \| 16 \| 7 \| 634 \| 1 \| 610 \| Terminase \| Terminase; afdb-uniprot50 \| AF-A0A660NKM7-F1-MODEL\_V4 \| 1.0 \| 9.17e-73 \| 2402 \| 0.63 \| 484 \| 165 \| 3 \| 13 \| 495 \| 2 \| 472 \| Phage terminase large subunit family protein \| Phage terminase large subunit family protein; afdb-uniprot50 \| AF-M2Z026-F1-MODEL\_V4 \| 1.0 \| 3.339e-64 \| 2319 \| 0.387 \| 596 \| 325 \| 14 \| 17 \| 602 \| 5 \| 570 \| Phage terminase GpA \| Phage terminase GpA; afdb-uniprot50 \| AF-A0A442XW79-F1-MODEL\_V4 \| 1.0 \| 1.218e-64 \| 2298 \| 0.36 \| 654 \| 354 \| 16 \| 34 \| 681 \| 5 \| 599 \| Phage terminase large subunit family protein \| Phage terminase large subunit family protein; afdb-uniprot50 \| AF-A0A0B6D8Y0-F1-MODEL\_V4 \| 1.0 \| 7.937e-68 \| 2276 \| 0.357 \| 604 \| 359 \| 13 \| 20 \| 614 \| 3 \| 586 \| Phage terminase large subunit family protein \| Phage terminase large subunit family protein; afdb-uniprot50 \| AF-F4BFS2-F1-MODEL\_V4 \| 1.0 \| 4.514e-76 \| 2249 \| 0.453 \| 620 \| 309 \| 14 \| 19 \| 628 \| 3 \| 602 \| Terminase large subunit \| Terminase large subunit; afdb-uniprot50 \| AF-A0A7V2AL36-F1-MODEL\_V4 \| 1.0 \| 2.147e-65 \| 2246 \| 0.354 \| 643 \| 363 \| 18 \| 21 \| 633 \| 5 \| 625 \| Phage terminase large subunit family protein \| Phage terminase large subunit family protein; afdb-uniprot50 \| AF-A0A7X3ZN88-F1-MODEL\_V4 \| 1.0 \| 1.961e-77 \| 2238 \| 0.479 \| 607 \| 288 \| 9 \| 35 \| 633 \| 8 \| 594 \| Phage terminase large subunit family protein \| Phage terminase large subunit family protein; afdb-uniprot50 \| AF-R6ITZ2-F1-MODEL\_V4 \| 1.0 \| 2.966e-63 \| 2234 \| 0.354 \| 652 \| 357 \| 21 \| 15 \| 632 \| 2 \| 623 \| Phage terminase GpA \| Phage terminase GpA; afdb-uniprot50 \| AF-A0A0Q7VFI6-F1-MODEL\_V4 \| 1.0 \| 1.515e-63 \| 2222 \| 0.371 \| 624 \| 334 \| 16 \| 62 \| 681 \| 1 \| 570 \| Uncharacterized protein \| Uncharacterized protein; afdb-uniprot50 \| AF-A0A348G5S0-F1-MODEL\_V4 \| 1.0 \| 1.515e-63 \| 2201 \| 0.36 \| 630 \| 360 \| 17 \| 17 \| 631 \| 2 \| 603 \| Terminase \| Terminase; afdb-uniprot50 \| AF-A0A6G8F2D4-F1-MODEL\_V4 \| 1.0 \| 1.505e-62 \| 2194 \| 0.366 \| 638 \| 354 \| 21 \| 18 \| 632 \| 2 \| 612 \| Terminase \| Terminase; afdb-uniprot50 \| AF-A0A3R8IUB2-F1-MODEL\_V4 \| 1.0 \| 2.523e-64 \| 2185 \| 0.331 \| 621 \| 376 \| 15 \| 19 \| 614 \| 9 \| 615 \| Phage terminase large subunit family protein \| Phage terminase large subunit family protein; afdb-uniprot50 \| AF-A0A2A5DI28-F1-MODEL\_V4 \| 1.0 \| 7.219e-62 \| 2183 \| 0.309 \| 626 \| 395 \| 13 \| 19 \| 634 \| 2 \| 599 \| Uncharacterized protein \| Uncharacterized protein; afdb-uniprot50 \| AF-A0A1S6XRH5-F1-MODEL\_V4 \| 1.0 \| 1.017e-62 \| 2163 \| 0.34 \| 640 \| 369 \| 22 \| 19 \| 632 \| 3 \| 615 \| Phage terminase, large subunit GpA \| Phage terminase, large subunit GpA; afdb-uniprot50 \| AF-A0A4R3LGZ5-F1-MODEL\_V4 \| 1.0 \| 6.782e-61 \| 2161 \| 0.389 \| 595 \| 318 \| 18 \| 55 \| 632 \| 3 \| 568 \| Phage terminase large subunit GpA-like protein \| Phage terminase large subunit GpA-like protein; afdb-uniprot50 \| AF-A0A2E0ENC7-F1-MODEL\_V4 \| 1.0 \| 5.038e-67 \| 2147 \| 0.334 \| 676 \| 381 \| 16 \| 19 \| 682 \| 6 \| 624 \| Terminase \| Terminase; afdb-uniprot50 \| AF-A0A2W4R482-F1-MODEL\_V4 \| 1.0 \| 3.734e-64 \| 2125 \| 0.325 \| 697 \| 407 \| 19 \| 1 \| 679 \| 6 \| 657 \| Uncharacterized protein \| Uncharacterized protein; afdb-uniprot50 \| AF-A0A354SLA4-F1-MODEL\_V4 \| 1.0 \| 4.878e-62 \| 2112 \| 0.321 \| 638 \| 389 \| 19 \| 17 \| 632 \| 2 \| 617 \| Uncharacterized protein \| Uncharacterized protein; afdb-uniprot50 \| AF-A0A6M5G3M5-F1-MODEL\_V4 \| 1.0 \| 6.024e-60 \| 2082 \| 0.306 \| 679 \| 382 \| 18 \| 19 \| 679 \| 2 \| 609 \| Phage terminase large subunit family protein \| Phage terminase large subunit family protein; afdb-uniprot50 \| AF-A0A523CMU2-F1-MODEL\_V4 \| 1.0 \| 1.115e-59 \| 2047 \| 0.346 \| 641 \| 359 \| 21 \| 17 \| 632 \| 3 \| 608 \| Phage terminase large subunit family protein \| Phage terminase large subunit family protein; afdb-uniprot50 \| AF-G7ZAT3-F1-MODEL\_V4 \| 1.0 \| 4.612e-62 \| 2041 \| 0.325 \| 643 \| 384 \| 20 \| 19 \| 638 \| 4 \| 619 \| Putative phage terminase GpA \| Putative phage terminase GpA; afdb-uniprot50 \| AF-A0A1X7L0Y3-F1-MODEL\_V4 \| 1.0 \| 4.053e-68 \| 2023 \| 0.353 \| 677 \| 359 \| 18 \| 19 \| 682 \| 3 \| 613 \| Phage terminase, large subunit GpA \| Phage terminase, large subunit GpA; afdb-uniprot50 \| AF-A0A7Y0IMI9-F1-MODEL\_V4 \| 1.0 \| 3.639e-60 \| 2019 \| 0.296 \| 607 \| 382 \| 16 \| 22 \| 609 \| 3 \| 583 \| Phage terminase large subunit GpA-like protein \| Phage terminase large subunit GpA-like protein; afdb-uniprot50 \| AF-E8V6R2-F1-MODEL\_V4 \| 1.0 \| 1.395e-59 \| 2006 \| 0.285 \| 710 \| 418 \| 23 \| 4 \| 679 \| 1 \| 654 \| Terminase GpA \| Terminase GpA; afdb-uniprot50 \| AF-A0A3M9XNZ3-F1-MODEL\_V4 \| 1.0 \| 1.834e-58 \| 1997 \| 0.291 \| 627 \| 381 \| 21 \| 17 \| 614 \| 2 \| 593 \| Uncharacterized protein \| Uncharacterized protein; afdb-uniprot50 \| AF-A0A212RBV5-F1-MODEL\_V4 \| 1.0 \| 5.947e-58 \| 1991 \| 0.367 \| 590 \| 329 \| 15 \| 65 \| 637 \| 144 \| 706 \| Phage terminase, large subunit GpA \| Phage terminase, large subunit GpA; afdb-uniprot50 \| AF-A0A353UTN9-F1-MODEL\_V4 \| 1.0 \| 5.092e-60 \| 1990 \| 0.311 \| 710 \| 401 \| 26 \| 1 \| 674 \| 7 \| 664 \| Phage tail protein \| Phage tail protein; afdb-uniprot50 \| AF-A0A286IHK8-F1-MODEL\_V4 \| 1.0 \| 6.33e-59 \| 1979 \| 0.3 \| 629 \| 389 \| 22 \| 18 \| 632 \| 6 \| 597 \| Phage terminase large subunit GpA-like protein \| Phage terminase large subunit GpA-like protein; afdb-uniprot50 \| AF-A0A7V9YMV3-F1-MODEL\_V4 \| 1.0 \| 2.617e-61 \| 1970 \| 0.3 \| 686 \| 406 \| 21 \| 23 \| 679 \| 2 \| 642 \| Phage terminase large subunit GpA-like protein \| Phage terminase large subunit GpA-like protein; afdb-uniprot50 \| AF-A0A4Q0MAA4-F1-MODEL\_V4 \| 1.0 \| 1.087e-55 \| 1964 \| 0.319 \| 613 \| 365 \| 19 \| 31 \| 632 \| 2 \| 573 \| Phage terminase large subunit family protein \| Phage terminase large subunit family protein; afdb-uniprot50 \| AF-A0A564WH44-F1-MODEL\_V4 \| 1.0 \| 4.523e-59 \| 1963 \| 0.32 \| 621 \| 373 \| 17 \| 19 \| 619 \| 4 \| 595 \| Putative Bacteriophage tail assembly protein \| Putative Bacteriophage tail assembly protein; afdb-uniprot50 \| AF-M2ZRG0-F1-MODEL\_V4 \| 1.0 \| 7.919e-59 \| 1954 \| 0.328 \| 637 \| 362 \| 23 \| 37 \| 634 \| 5 \| 614 \| Bacteriophage tail assembly protein \| Bacteriophage tail assembly protein; afdb-uniprot50 \| AF-A0A6H1ZNN5-F1-MODEL\_V4 \| 1.0 \| 3.775e-57 \| 1922 \| 0.297 \| 646 \| 405 \| 23 \| 19 \| 632 \| 6 \| 634 \| Putative terminase \| Putative terminase; afdb-uniprot50 \| AF-A0A521H782-F1-MODEL\_V4 \| 1.0 \| 3.419e-59 \| 1918 \| 0.289 \| 714 \| 410 \| 26 \| 1 \| 682 \| 6 \| 653 \| Phage terminase large subunit family protein \| Phage terminase large subunit family protein; afdb-uniprot50 \| AF-D1ALM4-F1-MODEL\_V4 \| 1.0 \| 4.044e-59 \| 1911 \| 0.323 \| 633 \| 376 \| 18 \| 21 \| 641 \| 2 \| 593 \| Terminase GpA \| Terminase GpA; afdb-uniprot50 \| AF-A0A5N0TEJ8-F1-MODEL\_V4 \| 1.0 \| 8.763e-66 \| 1905 \| 0.369 \| 614 \| 352 \| 16 \| 14 \| 615 \| 2 \| 592 \| Phage terminase large subunit family protein \| Phage terminase large subunit family protein; afdb-uniprot50 \| AF-A0A7R7T6E2-F1-MODEL\_V4 \| 1.0 \| 1.302e-57 \| 1896 \| 0.278 \| 726 \| 410 \| 24 \| 3 \| 679 \| 1 \| 661 \| Terminase \| Terminase; afdb-uniprot50 \| AF-A0A839DEQ6-F1-MODEL\_V4 \| 1.0 \| 3.915e-54 \| 1894 \| 0.292 \| 606 \| 378 \| 18 \| 42 \| 640 \| 2 \| 563 \| Phage terminase large subunit GpA-like protein \| Phage terminase large subunit GpA-like protein; afdb-uniprot50 \| AF-V6F5I0-F1-MODEL\_V4 \| 1.0 \| 1.834e-58 \| 1894 \| 0.317 \| 658 \| 378 \| 22 \| 22 \| 633 \| 3 \| 635 \| Bacteriophage tail assembly protein \| Bacteriophage tail assembly protein; afdb-uniprot50 \| AF-A0A7C4G200-F1-MODEL\_V4 \| 1.0 \| 5.491e-63 \| 1893 \| 0.356 \| 623 \| 357 \| 16 \| 22 \| 631 \| 4 \| 595 \| Phage terminase large subunit family protein \| Phage terminase large subunit family protein; afdb-uniprot50 \| AF-A0A3D6C8I9-F1-MODEL\_V4 \| 1.0 \| 3.899e-62 \| 1892 \| 0.351 \| 645 \| 375 \| 17 \| 7 \| 634 \| 98 \| 715 \| Terminase \| Terminase; afdb-uniprot50 \| AF-A0A1Q3NUL5-F1-MODEL\_V4 \| 1.0 \| 6.944e-56 \| 1889 \| 0.298 \| 637 \| 391 \| 20 \| 17 \| 626 \| 3 \| 610 \| Uncharacterized protein \| Uncharacterized protein; afdb-uniprot50 \| AF-A0A0Q1F5T3-F1-MODEL\_V4 \| 1.0 \| 6.608e-57 \| 1889 \| 0.285 \| 665 \| 406 \| 25 \| 3 \| 632 \| 1 \| 631 \| Uncharacterized protein \| Uncharacterized protein; afdb-uniprot50 \| AF-A0A2V7CRE4-F1-MODEL\_V4 \| 1.0 \| 2.715e-58 \| 1862 \| 0.297 \| 636 \| 394 \| 20 \| 23 \| 632 \| 2 \| 610 \| Uncharacterized protein \| Uncharacterized protein; afdb-uniprot50 \| AF-A0A1S2FDW6-F1-MODEL\_V4 \| 1.0 \| 9.129e-55 \| 1857 \| 0.289 \| 642 \| 403 \| 19 \| 19 \| 632 \| 2 \| 618 \| Terminase \| Terminase; afdb-uniprot50 \| AF-A0A1Z9Q5D0-F1-MODEL\_V4 \| 1.0 \| 5.527e-64 \| 1842 \| 0.315 \| 622 \| 383 \| 15 \| 18 \| 632 \| 11 \| 596 \| Uncharacterized protein \| Uncharacterized protein; afdb-uniprot50 \| AF-A0A0J6IF00-F1-MODEL\_V4 \| 1.0 \| 1.551e-58 \| 1825 \| 0.285 \| 711 \| 433 \| 26 \| 5 \| 681 \| 1 \| 670 \| Terminase \| Terminase; afdb-uniprot50 \| AF-A0A3A0FKS0-F1-MODEL\_V4 \| 1.0 \| 1.272e-62 \| 1805 \| 0.308 \| 671 \| 382 \| 17 \| 11 \| 637 \| 10 \| 642 \| Uncharacterized protein \| Uncharacterized protein; afdb-uniprot50 \| AF-A0A849W7C2-F1-MODEL\_V4 \| 1.0 \| 1.115e-59 \| 1803 \| 0.279 \| 641 \| 411 \| 20 \| 20 \| 632 \| 2 \| 619 \| Phage terminase large subunit family protein \| Phage terminase large subunit family protein; afdb-uniprot50 \| AF-A0A2E7V2D5-F1-MODEL\_V4 \| 1.0 \| 8.005e-52 \| 1797 \| 0.399 \| 498 \| 258 \| 10 \| 147 \| 631 \| 1 \| 470 \| Terminase \| Terminase; afdb-uniprot50 \| AF-A0A5C8AYK7-F1-MODEL\_V4 \| 1.0 \| 4.553e-60 \| 1794 \| 0.29 \| 630 \| 400 \| 16 \| 23 \| 632 \| 3 \| 605 \| Phage terminase large subunit family protein \| Phage terminase large subunit family protein; afdb-uniprot50 \| AF-A0A0Q7A9C1-F1-MODEL\_V4 \| 1.0 \| 6.089e-53 \| 1793 \| 0.279 \| 618 \| 392 \| 21 \| 19 \| 615 \| 5 \| 589 \| Uncharacterized protein \| Uncharacterized protein; afdb-uniprot50 \| AF-A0A2E3Q9Q3-F1-MODEL\_V4 \| 1.0 \| 8.161e-55 \| 1780 \| 0.259 \| 718 \| 434 \| 26 \| 2 \| 679 \| 1 \| 660 \| Terminase \| Terminase; afdb-uniprot50 \| AF-A0A0F2PJS0-F1-MODEL\_V4 \| 1.0 \| 7.618e-53 \| 1777 \| 0.266 \| 631 \| 418 \| 21 \| 19 \| 626 \| 3 \| 611 \| Uncharacterized protein \| Uncharacterized protein; afdb-uniprot50 \| AF-A0A1F9USZ4-F1-MODEL\_V4 \| 1.0 \| 2.119e-63 \| 1774 \| 0.325 \| 672 \| 379 \| 20 \| 23 \| 682 \| 2 \| 610 \| Uncharacterized protein \| Uncharacterized protein; afdb-uniprot50 \| AF-A0A7K0GMS7-F1-MODEL\_V4 \| 1.0 \| 1.438e-55 \| 1764 \| 0.272 \| 631 \| 381 \| 18 \| 19 \| 608 \| 5 \| 598 \| Phage terminase large subunit family protein \| Phage terminase large subunit family protein; afdb-uniprot50 \| AF-D8JWB6-F1-MODEL\_V4 \| 1.0 \| 2.715e-58 \| 1763 \| 0.305 \| 635 \| 375 \| 22 \| 19 \| 616 \| 2 \| 607 \| Terminase GpA \| Terminase GpA; afdb-uniprot50 \| AF-A0A7V5JH40-F1-MODEL\_V4 \| 1.0 \| 2.427e-58 \| 1763 \| 0.339 \| 693 \| 352 \| 21 \| 34 \| 682 \| 13 \| 643 \| Phage terminase large subunit family protein \| Phage terminase large subunit family protein; afdb-uniprot50 \| AF-A0A3A6P726-F1-MODEL\_V4 \| 1.0 \| 3.975e-65 \| 1762 \| 0.342 \| 649 \| 383 \| 17 \| 22 \| 648 \| 3 \| 629 \| Phage terminase large subunit family protein \| Phage terminase large subunit family protein; afdb-uniprot50 \| AF-A0A2A4V2N5-F1-MODEL\_V4 \| 1.0 \| 2.959e-54 \| 1761 \| 0.273 \| 703 \| 424 \| 28 \| 15 \| 680 \| 2 \| 654 \| Terminase \| Terminase; afdb-uniprot50 \| AF-A0A0C2YP84-F1-MODEL\_V4 \| 1.0 \| 3.31e-54 \| 1756 \| 0.317 \| 623 \| 361 \| 19 \| 55 \| 637 \| 6 \| 603 \| Phage terminase large subunit \| Phage terminase large subunit; afdb-uniprot50 \| AF-A0A4V6IMC3-F1-MODEL\_V4 \| 1.0 \| 2.798e-54 \| 1755 \| 0.273 \| 692 \| 426 \| 26 \| 3 \| 653 \| 1 \| 656 \| Terminase GpA \| Terminase GpA; afdb-uniprot50 \| AF-A0A2E1HAG1-F1-MODEL\_V4 \| 1.0 \| 1.815e-65 \| 1753 \| 0.325 \| 658 \| 396 \| 17 \| 1 \| 637 \| 2 \| 632 \| Uncharacterized protein \| Uncharacterized protein; afdb-uniprot50 \| AF-A0A1G2ZN51-F1-MODEL\_V4 \| 1.0 \| 5.563e-65 \| 1750 \| 0.32 \| 687 \| 387 \| 21 \| 17 \| 682 \| 2 \| 629 \| Uncharacterized protein \| Uncharacterized protein; afdb-uniprot50 \| AF-A0A0Q6MRX7-F1-MODEL\_V4 \| 1.0 \| 4.114e-53 \| 1746 \| 0.25 \| 666 \| 428 \| 22 \| 7 \| 632 \| 1 \| 635 \| Terminase \| Terminase; afdb-uniprot50 \| AF-A0A3M6EXA9-F1-MODEL\_V4 \| 1.0 \| 1.048e-58 \| 1744 \| 0.289 \| 701 \| 424 \| 25 \| 15 \| 681 \| 2 \| 662 \| Terminase, large subunit \| Terminase, large subunit; afdb-uniprot50 \| AF-A0A840Q0P9-F1-MODEL\_V4 \| 1.0 \| 1.87e-61 \| 1736 \| 0.321 \| 631 \| 376 \| 15 \| 19 \| 633 \| 3 \| 597 \| Phage terminase large subunit GpA-like protein \| Phage terminase large subunit GpA-like protein; afdb-uniprot50 \| AF-A0A1V6DMU9-F1-MODEL\_V4 \| 1.0 \| 1.735e-58 \| 1729 \| 0.312 \| 689 \| 386 \| 23 \| 22 \| 681 \| 5 \| 634 \| Phage terminase large subunit (GpA) \| Phage terminase large subunit (GpA); afdb-uniprot50 \| AF-A0A3N6NUK1-F1-MODEL\_V4 \| 1.0 \| 1.145e-63 \| 1721 \| 0.319 \| 699 \| 398 \| 22 \| 1 \| 677 \| 1 \| 643 \| Phage terminase large subunit family protein \| Phage terminase large subunit family protein; afdb-uniprot50 \| AF-A8U464-F1-MODEL\_V4 \| 1.0 \| 6.222e-65 \| 1721 \| 0.361 \| 692 \| 365 \| 23 \| 1 \| 680 \| 8 \| 634 \| Phage terminase GpA \| Phage terminase GpA; afdb-uniprot50 \| AF-A0A7S7SL97-F1-MODEL\_V4 \| 1.0 \| 5.316e-58 \| 1719 \| 0.33 \| 593 \| 357 \| 14 \| 51 \| 632 \| 2 \| 565 \| Phage terminase large subunit family protein \| Phage terminase large subunit family protein; afdb-uniprot50 \| AF-A0A524RUS2-F1-MODEL\_V4 \| 1.0 \| 7.25e-54 \| 1718 \| 0.301 \| 641 \| 379 \| 26 \| 17 \| 616 \| 2 \| 614 \| Phage terminase large subunit family protein \| Phage terminase large subunit family protein; afdb-uniprot50 \| AF-A0A1M7PM43-F1-MODEL\_V4 \| 1.0 \| 1.014e-53 \| 1711 \| 0.238 \| 733 \| 451 \| 29 \| 7 \| 677 \| 1 \| 687 \| Phage terminase, large subunit GpA \| Phage terminase, large subunit GpA; afdb-uniprot50 \| AF-A0A4R3HZ06-F1-MODEL\_V4 \| 1.0 \| 2.54e-65 \| 1710 \| 0.33 \| 702 \| 410 \| 19 \| 3 \| 681 \| 1 \| 665 \| Phage terminase large subunit GpA-like protein \| Phage terminase large subunit GpA-like protein; afdb-uniprot50 \| AF-A0A1V5QS85-F1-MODEL\_V4 \| 1.0 \| 9.09e-63 \| 1709 \| 0.3 \| 686 \| 404 \| 15 \| 17 \| 679 \| 5 \| 637 \| Phage terminase large subunit (GpA) \| Phage terminase large subunit (GpA); afdb-uniprot50 \| AF-A0A075KAN7-F1-MODEL\_V4 \| 1.0 \| 6.454e-62 \| 1706 \| 0.341 \| 648 \| 378 \| 19 \| 10 \| 632 \| 1 \| 624 \| Terminase GpA \| Terminase GpA; afdb-uniprot50 \| AF-A0A6B3RPI9-F1-MODEL\_V4 \| 1.0 \| 2.256e-64 \| 1706 \| 0.34 \| 675 \| 381 \| 19 \| 1 \| 637 \| 1 \| 649 \| Phage terminase large subunit family protein \| Phage terminase large subunit family protein; afdb-uniprot50 \| AF-A0A6F8UND4-F1-MODEL\_V4 \| 1.0 \| 1.337e-61 \| 1701 \| 0.335 \| 640 \| 377 \| 17 \| 17 \| 632 \| 4 \| 618 \| Terminase \| Terminase; afdb-uniprot50 \| AF-A0A6H1ZMA4-F1-MODEL\_V4 \| 1.0 \| 6.011e-51 \| 1696 \| 0.285 \| 621 \| 392 \| 22 \| 42 \| 632 \| 2 \| 600 \| Putative terminase \| Putative terminase; afdb-uniprot50 \| AF-A0A5E9ZQA4-F1-MODEL\_V4 \| 1.0 \| 1.866e-52 \| 1691 \| 0.261 \| 691 \| 431 \| 28 \| 14 \| 682 \| 2 \| 634 \| Terminase \| Terminase; afdb-uniprot50 \| AF-A0A844IPM8-F1-MODEL\_V4 \| 1.0 \| 7.219e-62 \| 1687 \| 0.325 \| 632 \| 378 \| 19 \| 22 \| 634 \| 4 \| 605 \| Phage terminase large subunit family protein \| Phage terminase large subunit family protein; afdb-uniprot50 \| AF-A0A2G1CVR6-F1-MODEL\_V4 \| 1.0 \| 3.486e-62 \| 1682 \| 0.338 \| 642 \| 377 \| 20 \| 3 \| 632 \| 1 \| 606 \| Terminase \| Terminase; afdb-uniprot50 \| AF-A0A6L9LZ87-F1-MODEL\_V4 \| 1.0 \| 7.635e-62 \| 1680 \| 0.343 \| 666 \| 360 \| 16 \| 36 \| 682 \| 5 \| 611 \| Phage terminase large subunit family protein \| Phage terminase large subunit family protein; afdb-uniprot50 \| AF-A0A2K9JRQ1-F1-MODEL\_V4 \| 1.0 \| 1.075e-62 \| 1678 \| 0.362 \| 623 \| 346 \| 19 \| 39 \| 632 \| 1 \| 601 \| Phage tail protein \| Phage tail protein; afdb-uniprot50 \| AF-A0A735CWF0-F1-MODEL\_V4 \| 1.0 \| 2.35e-53 \| 1674 \| 0.254 \| 692 \| 438 \| 27 \| 12 \| 682 \| 2 \| 636 \| Terminase \| Terminase; afdb-uniprot50 \| AF-A0A533RCU9-F1-MODEL\_V4 \| 1.0 \| 1.135e-53 \| 1673 \| 0.277 \| 641 \| 411 \| 21 \| 1 \| 616 \| 2 \| 615 \| Uncharacterized protein \| Uncharacterized protein; afdb-uniprot50 \| AF-A0A4Y8GNU3-F1-MODEL\_V4 \| 1.0 \| 7.156e-52 \| 1667 \| 0.267 \| 688 \| 430 \| 28 \| 14 \| 682 \| 2 \| 634 \| Terminase \| Terminase; afdb-uniprot50 \| AF-A0A2D2C105-F1-MODEL\_V4 \| 1.0 \| 2.092e-61 \| 1667 \| 0.321 \| 685 \| 396 \| 23 \| 14 \| 680 \| 6 \| 639 \| Uncharacterized protein \| Uncharacterized protein; afdb-uniprot50 \| AF-A0A6M7VZA9-F1-MODEL\_V4 \| 1.0 \| 1.683e-62 \| 1662 \| 0.321 \| 690 \| 386 \| 22 \| 26 \| 682 \| 2 \| 641 \| Phage terminase large subunit family protein \| Phage terminase large subunit family protein; afdb-uniprot50 \| AF-A0A4R6RGI3-F1-MODEL\_V4 \| 1.0 \| 1.669e-52 \| 1657 \| 0.248 \| 716 \| 431 \| 26 \| 3 \| 680 \| 1 \| 647 \| Phage terminase large subunit GpA-like protein \| Phage terminase large subunit GpA-like protein; afdb-uniprot50 \| AF-G4KQ85-F1-MODEL\_V4 \| 1.0 \| 3.275e-61 \| 1654 \| 0.3 \| 658 \| 405 \| 21 \| 1 \| 633 \| 1 \| 627 \| Putative terminase large subunit \| Putative terminase large subunit; afdb-uniprot50 \| AF-A0A3S0BAZ2-F1-MODEL\_V4 \| 1.0 \| 6.87e-63 \| 1653 \| 0.29 \| 699 \| 425 \| 18 \| 1 \| 682 \| 112 \| 756 \| Phage terminase large subunit family protein \| Phage terminase large subunit family protein; afdb-uniprot50 \| AF-A0A081NI15-F1-MODEL\_V4 \| 1.0 \| 1.055e-59 \| 1651 \| 0.329 \| 668 \| 374 \| 20 \| 25 \| 675 \| 5 \| 615 \| Uncharacterized protein \| Uncharacterized protein; afdb-uniprot50 \| AF-A0A1B1C364-F1-MODEL\_V4 \| 1.0 \| 4.361e-62 \| 1645 \| 0.291 \| 714 \| 436 \| 22 \| 1 \| 681 \| 1 \| 677 \| Terminase \| Terminase; afdb-uniprot50 \| AF-A0A315EXH7-F1-MODEL\_V4 \| 1.0 \| 1.63e-57 \| 1644 \| 0.297 \| 696 \| 417 \| 25 \| 16 \| 679 \| 2 \| 657 \| Terminase \| Terminase; afdb-uniprot50 \| AF-A0A842J5L3-F1-MODEL\_V4 \| 1.0 \| 1.658e-51 \| 1642 \| 0.288 \| 590 \| 381 \| 17 \| 23 \| 605 \| 2 \| 559 \| Phage terminase large subunit family protein \| Phage terminase large subunit family protein; afdb-uniprot50 \| AF-A0A370UA85-F1-MODEL\_V4 \| 1.0 \| 2.34e-61 \| 1641 \| 0.297 \| 700 \| 436 \| 21 \| 1 \| 679 \| 1 \| 665 \| Uncharacterized protein \| Uncharacterized protein; afdb-uniprot50 \| AF-A0A5C1B6F6-F1-MODEL\_V4 \| 1.0 \| 2.744e-51 \| 1638 \| 0.248 \| 708 \| 438 \| 28 \| 18 \| 679 \| 6 \| 665 \| Terminase \| Terminase; afdb-uniprot50 \| AF-A0A7U3R0X7-F1-MODEL\_V4 \| 1.0 \| 2.355e-62 \| 1636 \| 0.341 \| 662 \| 365 \| 20 \| 39 \| 676 \| 1 \| 615 \| Terminase \| Terminase; afdb-uniprot50 \| AF-A0A6L4BC06-F1-MODEL\_V4 \| 1.0 \| 5.351e-59 \| 1632 \| 0.31 \| 625 \| 401 \| 13 \| 16 \| 632 \| 6 \| 608 \| Phage terminase large subunit family protein \| Phage terminase large subunit family protein; afdb-uniprot50 \| AF-W0J6N1-F1-MODEL\_V4 \| 1.0 \| 7.52e-51 \| 1630 \| 0.33 \| 571 \| 332 \| 16 \| 99 \| 632 \| 2 \| 558 \| Uncharacterized protein \| Uncharacterized protein; afdb-uniprot50 \| AF-A0A1Q5XTZ6-F1-MODEL\_V4 \| 1.0 \| 4.018e-58 \| 1629 \| 0.294 \| 612 \| 388 \| 15 \| 17 \| 608 \| 3 \| 590 \| Terminase \| Terminase; afdb-uniprot50 \| AF-A0A327Q9H2-F1-MODEL\_V4 \| 1.0 \| 1.788e-54 \| 1629 \| 0.275 \| 624 \| 400 \| 21 \| 19 \| 617 \| 2 \| 598 \| Phage terminase large subunit GpA-like protein \| Phage terminase large subunit GpA-like protein; afdb-uniprot50 \| AF-A0A5K8AHA3-F1-MODEL\_V4 \| 1.0 \| 8.801e-58 \| 1628 \| 0.288 \| 679 \| 396 \| 21 \| 20 \| 681 \| 4 \| 612 \| Terminase \| Terminase; afdb-uniprot50 \| AF-R5BHA3-F1-MODEL\_V4 \| 1.0 \| 1.757e-60 \| 1623 \| 0.312 \| 679 \| 388 \| 22 \| 20 \| 679 \| 2 \| 620 \| Phage terminase large subunit family protein \| Phage terminase large subunit family protein; afdb-uniprot50 \| AF-A0A7C5GLW2-F1-MODEL\_V4 \| 1.0 \| 3.941e-55 \| 1616 \| 0.363 \| 533 \| 309 \| 12 \| 23 \| 543 \| 2 \| 515 \| Phage terminase large subunit family protein \| Phage terminase large subunit family protein; afdb-uniprot50 \| AF-A0A1T4PVQ0-F1-MODEL\_V4 \| 1.0 \| 2.156e-57 \| 1613 \| 0.287 \| 609 \| 385 \| 16 \| 17 \| 613 \| 2 \| 573 \| Phage terminase, large subunit GpA \| Phage terminase, large subunit GpA; afdb-uniprot50 \| AF-A0A848NXM2-F1-MODEL\_V4 \| 1.0 \| 2.31e-59 \| 1613 \| 0.302 \| 645 \| 388 \| 25 \| 23 \| 632 \| 6 \| 623 \| Phage terminase large subunit family protein \| Phage terminase large subunit family protein; afdb-uniprot50 \| AF-A0A4R2P8A1-F1-MODEL\_V4 \| 1.0 \| 8.022e-61 \| 1608 \| 0.313 \| 714 \| 413 \| 29 \| 1 \| 679 \| 1 \| 671 \| Phage terminase large subunit GpA-like protein \| Phage terminase large subunit GpA-like protein; afdb-uniprot50 \| AF-A0A255YQE1-F1-MODEL\_V4 \| 1.0 \| 7.585e-61 \| 1608 \| 0.299 \| 664 \| 398 \| 16 \| 1 \| 614 \| 58 \| 704 \| Uncharacterized protein \| Uncharacterized protein; afdb-uniprot50 \| AF-A0A2P1VUZ1-F1-MODEL\_V4 \| 1.0 \| 2.034e-48 \| 1607 \| 0.265 \| 640 \| 399 \| 24 \| 19 \| 616 \| 8 \| 618 \| Terminase \| Terminase; afdb-uniprot50 \| AF-A0A1C0SI12-F1-MODEL\_V4 \| 1.0 \| 1.482e-51 \| 1607 \| 0.228 \| 761 \| 466 \| 32 \| 3 \| 681 \| 1 \| 722 \| Uncharacterized protein \| Uncharacterized protein; afdb-uniprot50 \| AF-A0A2E6KNG5-F1-MODEL\_V4 \| 1.0 \| 2.715e-58 \| 1604 \| 0.318 \| 635 \| 372 \| 24 \| 30 \| 632 \| 3 \| 608 \| Phage tail protein \| Phage tail protein; afdb-uniprot50 \| AF-A0A7W6RGT1-F1-MODEL\_V4 \| 1.0 \| 1.272e-62 \| 1604 \| 0.281 \| 746 \| 430 \| 21 \| 3 \| 679 \| 1 \| 709 \| Phage terminase large subunit GpA-like protein \| Phage terminase large subunit GpA-like protein; afdb-uniprot50 \| AF-A0A532DWA4-F1-MODEL\_V4 \| 1.0 \| 6.371e-60 \| 1603 \| 0.322 \| 691 \| 395 \| 23 \| 19 \| 682 \| 11 \| 655 \| Phage terminase large subunit family protein \| Phage terminase large subunit family protein; afdb-uniprot50 \| AF-A0A8B2NR07-F1-MODEL\_V4 \| 1.0 \| 4.333e-61 \| 1601 \| 0.291 \| 714 \| 420 \| 23 \| 14 \| 678 \| 2 \| 678 \| Terminase \| Terminase; afdb-uniprot50 \| AF-A0A2U1SE29-F1-MODEL\_V4 \| 1.0 \| 6.089e-53 \| 1596 \| 0.284 \| 623 \| 375 \| 22 \| 17 \| 628 \| 5 \| 567 \| Uncharacterized protein \| Uncharacterized protein; afdb-uniprot50 \| AF-A0A0F9L7P8-F1-MODEL\_V4 \| 1.0 \| 2.412e-57 \| 1595 \| 0.309 \| 626 \| 391 \| 17 \| 22 \| 632 \| 6 \| 604 \| Uncharacterized protein \| Uncharacterized protein; afdb-uniprot50 \| AF-R8AR60-F1-MODEL\_V4 \| 1.0 \| 3.766e-48 \| 1595 \| 0.265 \| 640 \| 399 \| 24 \| 19 \| 616 \| 8 \| 618 \| DNA packaging protein \| DNA packaging protein; afdb-uniprot50 \| AF-U1IPL3-F1-MODEL\_V4 \| 1.0 \| 5.515e-55 \| 1593 \| 0.277 \| 606 \| 385 \| 18 \| 23 \| 611 \| 4 \| 573 \| Uncharacterized protein \| Uncharacterized protein; afdb-uniprot50 \| AF-A0A1M6ZDE1-F1-MODEL\_V4 \| 1.0 \| 9.972e-60 \| 1593 \| 0.31 \| 648 \| 398 \| 24 \| 6 \| 637 \| 1 \| 615 \| Phage terminase, large subunit GpA \| Phage terminase, large subunit GpA; afdb-uniprot50 \| AF-A0A1G0RDU7-F1-MODEL\_V4 \| 1.0 \| 1.073e-53 \| 1592 \| 0.26 \| 652 \| 429 \| 19 \| 19 \| 634 \| 19 \| 653 \| Uncharacterized protein \| Uncharacterized protein; afdb-uniprot50 \| AF-A0A418VVC1-F1-MODEL\_V4 \| 1.0 \| 5.305e-49 \| 1591 \| 0.239 \| 701 \| 453 \| 24 \| 10 \| 682 \| 2 \| 650 \| Terminase \| Terminase; afdb-uniprot50 \| AF-A0A7W6S2Q9-F1-MODEL\_V4 \| 1.0 \| 1.846e-59 \| 1586 \| 0.292 \| 702 \| 405 \| 21 \| 23 \| 680 \| 2 \| 655 \| Phage terminase large subunit GpA-like protein \| Phage terminase large subunit GpA-like protein; afdb-uniprot50 \| AF-A0A1T4W516-F1-MODEL\_V4 \| 1.0 \| 2.909e-60 \| 1580 \| 0.298 \| 656 \| 397 \| 22 \| 16 \| 634 \| 2 \| 630 \| Phage terminase, large subunit GpA \| Phage terminase, large subunit GpA; afdb-uniprot50 \| AF-A0A2K9MBB9-F1-MODEL\_V4 \| 1.0 \| 2.534e-56 \| 1579 \| 0.315 \| 625 \| 381 \| 18 \| 18 \| 632 \| 2 \| 589 \| Terminase \| Terminase; afdb-uniprot50 \| AF-U5QDZ1-F1-MODEL\_V4 \| 1.0 \| 7.126e-60 \| 1579 \| 0.312 \| 662 \| 355 \| 19 \| 57 \| 682 \| 1 \| 598 \| Phage terminase GpA \| Phage terminase GpA; afdb-uniprot50 \| AF-A0A497V3E9-F1-MODEL\_V4 \| 1.0 \| 1.087e-55 \| 1579 \| 0.294 \| 636 \| 394 \| 23 \| 25 \| 633 \| 2 \| 609 \| Phage terminase large subunit GpA-like protein \| Phage terminase large subunit GpA-like protein; afdb-uniprot50 \| AF-A0A1Z8TD19-F1-MODEL\_V4 \| 1.0 \| 7.08e-59 \| 1577 \| 0.285 \| 645 \| 399 \| 22 \| 17 \| 632 \| 3 \| 614 \| Uncharacterized protein \| Uncharacterized protein; afdb-uniprot50 \| AF-A0A1H5XTY2-F1-MODEL\_V4 \| 1.0 \| 1.405e-60 \| 1565 \| 0.304 \| 690 \| 408 \| 25 \| 26 \| 681 \| 2 \| 653 \| Phage terminase, large subunit GpA \| Phage terminase, large subunit GpA; afdb-uniprot50 \| AF-A0A7W6KFB7-F1-MODEL\_V4 \| 1.0 \| 3.966e-56 \| 1563 \| 0.292 \| 619 \| 389 \| 19 \| 22 \| 632 \| 4 \| 581 \| Phage terminase large subunit GpA-like protein \| Phage terminase large subunit GpA-like protein; afdb-uniprot50 \| AF-A0A3B9VGB5-F1-MODEL\_V4 \| 1.0 \| 5.282e-57 \| 1559 \| 0.31 \| 644 \| 374 \| 24 \| 18 \| 615 \| 12 \| 631 \| Phage tail protein \| Phage tail protein; afdb-uniprot50 \| AF-A0A2E5F8C5-F1-MODEL\_V4 \| 1.0 \| 6.738e-60 \| 1559 \| 0.311 \| 715 \| 414 \| 23 \| 4 \| 681 \| 1 \| 673 \| Terminase \| Terminase; afdb-uniprot50 \| AF-A0A3M2YA84-F1-MODEL\_V4 \| 1.0 \| 8.858e-59 \| 1558 \| 0.285 \| 704 \| 423 \| 25 \| 15 \| 681 \| 2 \| 662 \| Uncharacterized protein \| Uncharacterized protein; afdb-uniprot50 \| AF-A0A2S6FJF2-F1-MODEL\_V4 \| 1.0 \| 2.065e-59 \| 1556 \| 0.294 \| 706 \| 426 \| 26 \| 5 \| 680 \| 1 \| 664 \| Terminase \| Terminase; afdb-uniprot50 \| AF-A0A177P6Y7-F1-MODEL\_V4 \| 1.0 \| 1.742e-50 \| 1554 \| 0.235 \| 671 \| 422 \| 28 \| 3 \| 621 \| 1 \| 632 \| Uncharacterized protein \| Uncharacterized protein; afdb-uniprot50 \| AF-A0A2D8RC69-F1-MODEL\_V4 \| 1.0 \| 4.465e-57 \| 1548 \| 0.298 \| 636 \| 396 \| 21 \| 13 \| 632 \| 2 \| 603 \| Terminase \| Terminase; afdb-uniprot50 \| AF-A0A4Q0MF59-F1-MODEL\_V4 \| 1.0 \| 2.422e-49 \| 1545 \| 0.241 \| 734 \| 446 \| 28 \| 19 \| 680 \| 9 \| 703 \| Uncharacterized protein \| Uncharacterized protein; afdb-uniprot50 \| AF-A0A3G8M2W2-F1-MODEL\_V4 \| 1.0 \| 2.124e-46 \| 1540 \| 0.252 \| 629 \| 395 \| 24 \| 16 \| 620 \| 3 \| 580 \| Uncharacterized protein \| Uncharacterized protein; afdb-uniprot50 \| AF-A0A7W6S4A2-F1-MODEL\_V4 \| 1.0 \| 1.055e-59 \| 1540 \| 0.32 \| 690 \| 399 \| 27 \| 19 \| 678 \| 3 \| 652 \| Phage terminase large subunit GpA-like protein \| Phage terminase large subunit GpA-like protein; afdb-uniprot50 \| AF-A0A2E3Q5K6-F1-MODEL\_V4 \| 1.0 \| 6.33e-59 \| 1540 \| 0.306 \| 698 \| 410 \| 25 \| 15 \| 681 \| 2 \| 656 \| Terminase \| Terminase; afdb-uniprot50 \| AF-A0A4Z0BA59-F1-MODEL\_V4 \| 1.0 \| 3.375e-57 \| 1540 \| 0.294 \| 703 \| 414 \| 28 \| 23 \| 681 \| 9 \| 673 \| Phage terminase large subunit family protein \| Phage terminase large subunit family protein; afdb-uniprot50 \| AF-A0A1Y2KA39-F1-MODEL\_V4 \| 1.0 \| 5.61e-49 \| 1539 \| 0.317 \| 607 \| 321 \| 18 \| 114 \| 681 \| 2 \| 553 \| Putative phage terminase large subunit \| Putative phage terminase large subunit; afdb-uniprot50 \| AF-R8VXB3-F1-MODEL\_V4 \| 1.0 \| 4.249e-58 \| 1539 \| 0.305 \| 658 \| 402 \| 20 \| 3 \| 632 \| 1 \| 631 \| Uncharacterized protein \| Uncharacterized protein; afdb-uniprot50 \| AF-A0A7J6YKX9-F1-MODEL\_V4 \| 1.0 \| 3.191e-57 \| 1535 \| 0.33 \| 626 \| 357 \| 24 \| 22 \| 615 \| 2 \| 597 \| Uncharacterized protein \| Uncharacterized protein; afdb-uniprot50 \| AF-A0A7U7FYJ0-F1-MODEL\_V4 \| 1.0 \| 8.321e-58 \| 1534 \| 0.322 \| 661 \| 392 \| 20 \| 1 \| 632 \| 1 \| 634 \| Putative phage terminase large subunit \| Putative phage terminase large subunit; afdb-uniprot50 \| AF-A0A099GM22-F1-MODEL\_V4 \| 1.0 \| 3.799e-58 \| 1534 \| 0.304 \| 677 \| 396 \| 22 \| 3 \| 632 \| 1 \| 649 \| Bacteriophage tail assembly protein \| Bacteriophage tail assembly protein; afdb-uniprot50 \| AF-A0A2W6SSS1-F1-MODEL\_V4 \| 1.0 \| 5.61e-49 \| 1533 \| 0.304 \| 610 \| 334 \| 21 \| 104 \| 680 \| 2 \| 553 \| Terminase \| Terminase; afdb-uniprot50 \| AF-L7U9X3-F1-MODEL\_V4 \| 1.0 \| 3.191e-57 \| 1533 \| 0.293 \| 655 \| 383 \| 24 \| 3 \| 628 \| 1 \| 604 \| Phage terminase, large subunit \| Phage terminase, large subunit; afdb-uniprot50 \| AF-A0A4P5VNP2-F1-MODEL\_V4 \| 1.0 \| 1.369e-56 \| 1532 \| 0.285 \| 649 \| 410 \| 23 \| 15 \| 632 \| 2 \| 627 \| Terminase \| Terminase; afdb-uniprot50 \| AF-A0A345DE67-F1-MODEL\_V4 \| 1.0 \| 8.375e-59 \| 1531 \| 0.282 \| 687 \| 416 \| 25 \| 19 \| 680 \| 9 \| 643 \| Uncharacterized protein \| Uncharacterized protein; afdb-uniprot50 \| AF-A0A1M7FP01-F1-MODEL\_V4 \| 1.0 \| 2.427e-58 \| 1528 \| 0.294 \| 675 \| 403 \| 26 \| 39 \| 682 \| 1 \| 632 \| Phage terminase, large subunit GpA \| Phage terminase, large subunit GpA; afdb-uniprot50 \| AF-H1PWD6-F1-MODEL\_V4 \| 1.0 \| 4.408e-55 \| 1527 \| 0.291 \| 603 \| 384 \| 17 \| 20 \| 608 \| 3 \| 575 \| Uncharacterized protein \| Uncharacterized protein; afdb-uniprot50 \| AF-A0A0U1HUR1-F1-MODEL\_V4 \| 1.0 \| 1.518e-46 \| 1526 \| 0.254 \| 675 \| 413 \| 25 \| 34 \| 680 \| 1 \| 612 \| Bacteriophage tail assembly protein \| Bacteriophage tail assembly protein; afdb-uniprot50 \| AF-A0A510JH28-F1-MODEL\_V4 \| 1.0 \| 3.15e-55 \| 1521 \| 0.271 \| 640 \| 401 \| 16 \| 12 \| 618 \| 2 \| 608 \| Phage terminase large subunit \| Phage terminase large subunit; afdb-uniprot50 \| AF-A0A358C8T4-F1-MODEL\_V4 \| 1.0 \| 2.534e-56 \| 1520 \| 0.312 \| 624 \| 370 \| 20 \| 1 \| 589 \| 10 \| 609 \| DOD-type homing endonuclease domain-containing protein \| DOD-type homing endonuclease domain-containing protein; afdb-uniprot50 \| AF-A0A3G6WID2-F1-MODEL\_V4 \| 1.0 \| 3.569e-57 \| 1519 \| 0.3 \| 679 \| 403 \| 21 \| 1 \| 632 \| 5 \| 658 \| Phage terminase large subunit family protein \| Phage terminase large subunit family protein; afdb-uniprot50 \| AF-A0A7G9GXH3-F1-MODEL\_V4 \| 1.0 \| 1.701e-55 \| 1518 \| 0.292 \| 626 \| 389 \| 22 \| 4 \| 613 \| 1 \| 588 \| Phage terminase large subunit family protein \| Phage terminase large subunit family protein; afdb-uniprot50 \| AF-A0A7C4ZUK4-F1-MODEL\_V4 \| 1.0 \| 5.757e-53 \| 1518 \| 0.297 \| 626 \| 389 \| 21 \| 16 \| 633 \| 18 \| 600 \| Phage terminase large subunit family protein \| Phage terminase large subunit family protein; afdb-uniprot50 \| AF-A0A1F2QZ43-F1-MODEL\_V4 \| 1.0 \| 3.017e-57 \| 1517 \| 0.312 \| 691 \| 391 \| 27 \| 21 \| 682 \| 4 \| 639 \| Uncharacterized protein \| Uncharacterized protein; afdb-uniprot50 \| AF-A0A2D2C208-F1-MODEL\_V4 \| 1.0 \| 2.443e-59 \| 1517 \| 0.283 \| 724 \| 424 \| 23 \| 15 \| 681 \| 4 \| 689 \| Uncharacterized protein \| Uncharacterized protein; afdb-uniprot50 \| AF-A8U0J2-F1-MODEL\_V4 \| 1.0 \| 1.974e-52 \| 1513 \| 0.365 \| 542 \| 302 \| 18 \| 1 \| 530 \| 7 \| 518 \| Phage terminase GpA \| Phage terminase GpA; afdb-uniprot50 \| AF-A0A7V1FWZ6-F1-MODEL\_V4 \| 1.0 \| 1.811e-56 \| 1513 \| 0.298 \| 654 \| 396 \| 21 \| 13 \| 632 \| 2 \| 626 \| Uncharacterized protein \| Uncharacterized protein; afdb-uniprot50 \| AF-A0A2D3WRN4-F1-MODEL\_V4 \| 1.0 \| 1.712e-56 \| 1511 \| 0.303 \| 663 \| 387 \| 21 \| 3 \| 632 \| 1 \| 621 \| Uncharacterized protein \| Uncharacterized protein; afdb-uniprot50 \| AF-A0A7G8X000-F1-MODEL\_V4 \| 1.0 \| 1.36e-55 \| 1511 \| 0.282 \| 623 \| 398 \| 19 \| 17 \| 619 \| 6 \| 599 \| Phage terminase large subunit family protein \| Phage terminase large subunit family protein; afdb-uniprot50 \| AF-N6USV6-F1-MODEL\_V4 \| 1.0 \| 9.844e-58 \| 1507 \| 0.332 \| 629 \| 340 \| 20 \| 74 \| 678 \| 2 \| 574 \| Phage terminase large subunit GpA \| Phage terminase large subunit GpA; afdb-uniprot50 \| AF-A0A1E3G716-F1-MODEL\_V4 \| 1.0 \| 2.142e-56 \| 1505 \| 0.307 \| 615 \| 379 \| 17 \| 39 \| 637 \| 1 \| 584 \| Uncharacterized protein \| Uncharacterized protein; afdb-uniprot50 \| AF-A0A7V8TMI0-F1-MODEL\_V4 \| 1.0 \| 8.668e-47 \| 1504 \| 0.25 \| 623 \| 407 \| 27 \| 19 \| 616 \| 16 \| 603 \| Phage terminase large subunit family protein \| Phage terminase large subunit family protein; afdb-uniprot50 \| AF-A0A1I5F2B5-F1-MODEL\_V4 \| 1.0 \| 2.518e-55 \| 1503 \| 0.289 \| 612 \| 381 \| 19 \| 22 \| 618 \| 2 \| 574 \| Phage terminase, large subunit GpA \| Phage terminase, large subunit GpA; afdb-uniprot50 \| AF-A0A7C1Y9M6-F1-MODEL\_V4 \| 1.0 \| 4.379e-54 \| 1503 \| 0.309 \| 644 \| 365 \| 23 \| 1 \| 614 \| 48 \| 641 \| Phage terminase large subunit family protein \| Phage terminase large subunit family protein; afdb-uniprot50 \| AF-A0A0K6HIC0-F1-MODEL\_V4 \| 1.0 \| 3.017e-57 \| 1502 \| 0.304 \| 670 \| 396 \| 21 \| 1 \| 632 \| 2 \| 639 \| Phage terminase, large subunit GpA \| Phage terminase, large subunit GpA; afdb-uniprot50 \| AF-A0A2J8GXT5-F1-MODEL\_V4 \| 1.0 \| 1.928e-57 \| 1500 \| 0.301 \| 664 \| 392 \| 22 \| 7 \| 632 \| 1 \| 630 \| Terminase \| Terminase; afdb-uniprot50 \| AF-A0A1I2E7I9-F1-MODEL\_V4 \| 1.0 \| 7.44e-58 \| 1500 \| 0.287 \| 724 \| 436 \| 26 \| 1 \| 680 \| 1 \| 688 \| Phage terminase, large subunit GpA \| Phage terminase, large subunit GpA; afdb-uniprot50 \| AF-A0A1S6XPU5-F1-MODEL\_V4 \| 1.0 \| 6.481e-54 \| 1499 \| 0.336 \| 567 \| 335 \| 18 \| 19 \| 572 \| 3 \| 541 \| Phage terminase, large subunit GpA \| Phage terminase, large subunit GpA; afdb-uniprot50 \| AF-Q1ILS3-F1-MODEL\_V4 \| 1.0 \| 1.378e-57 \| 1499 \| 0.275 \| 726 \| 445 \| 20 \| 4 \| 682 \| 1 \| 692 \| Phage terminase GpA \| Phage terminase GpA; afdb-uniprot50 \| AF-A0A011PCR3-F1-MODEL\_V4 \| 1.0 \| 1.578e-52 \| 1497 \| 0.257 \| 815 \| 378 \| 27 \| 10 \| 632 \| 2 \| 780 \| Bacteriophage tail assembly protein \| Bacteriophage tail assembly protein; afdb-uniprot50 \| AF-Q1QI78-F1-MODEL\_V4 \| 1.0 \| 1.334e-52 \| 1496 \| 0.315 \| 584 \| 346 \| 18 \| 34 \| 599 \| 2 \| 549 \| Phage terminase GpA \| Phage terminase GpA; afdb-uniprot50 \| AF-A0A1M7R7Y1-F1-MODEL\_V4 \| 1.0 \| 7.25e-54 \| 1495 \| 0.283 \| 648 \| 402 \| 24 \| 19 \| 636 \| 3 \| 617 \| Phage terminase, large subunit GpA \| Phage terminase, large subunit GpA; afdb-uniprot50 \| AF-A0A5B0KMA8-F1-MODEL\_V4 \| 1.0 \| 4.342e-44 \| 1492 \| 0.314 \| 518 \| 308 \| 15 \| 141 \| 646 \| 4 \| 486 \| Uncharacterized protein \| Uncharacterized protein; afdb-uniprot50 \| AF-A0A2J6HVG4-F1-MODEL\_V4 \| 1.0 \| 6.566e-56 \| 1492 \| 0.287 \| 598 \| 384 \| 18 \| 30 \| 608 \| 37 \| 611 \| Uncharacterized protein \| Uncharacterized protein; afdb-uniprot50 \| AF-A0A793I175-F1-MODEL\_V4 \| 1.0 \| 9.759e-48 \| 1491 \| 0.265 \| 633 \| 397 \| 25 \| 19 \| 611 \| 8 \| 612 \| Phage terminase large subunit family protein \| Phage terminase large subunit family protein; afdb-uniprot50 \| AF-A0A063ZY87-F1-MODEL\_V4 \| 1.0 \| 7.717e-55 \| 1490 \| 0.269 \| 624 \| 411 \| 18 \| 20 \| 632 \| 6 \| 595 \| Uncharacterized protein \| Uncharacterized protein; afdb-uniprot50 \| AF-A0A3N1MB28-F1-MODEL\_V4 \| 1.0 \| 1.034e-56 \| 1487 \| 0.273 \| 701 \| 425 \| 23 \| 17 \| 676 \| 4 \| 660 \| Phage terminase large subunit GpA-like protein \| Phage terminase large subunit GpA-like protein; afdb-uniprot50 \| AF-A0A1M7D8E3-F1-MODEL\_V4 \| 1.0 \| 5.502e-46 \| 1486 \| 0.259 \| 655 \| 428 \| 23 \| 3 \| 632 \| 1 \| 623 \| Phage terminase, large subunit GpA \| Phage terminase, large subunit GpA; afdb-uniprot50 \| AF-A0A2D8S0G5-F1-MODEL\_V4 \| 1.0 \| 1.701e-55 \| 1486 \| 0.27 \| 647 \| 398 \| 24 \| 21 \| 614 \| 17 \| 642 \| Terminase \| Terminase; afdb-uniprot50 \| AF-A0A2X2EBD0-F1-MODEL\_V4 \| 1.0 \| 3.17e-56 \| 1486 \| 0.319 \| 648 \| 382 \| 20 \| 18 \| 632 \| 196 \| 817 \| Bacteriophage tail assembly protein \| Bacteriophage tail assembly protein; afdb-uniprot50 \| AF-A0A4D8Q014-F1-MODEL\_V4 \| 1.0 \| 2.061e-50 \| 1484 \| 0.256 \| 716 \| 435 \| 30 \| 12 \| 677 \| 14 \| 681 \| Terminase \| Terminase; afdb-uniprot50 \| AF-A0A031IRQ6-F1-MODEL\_V4 \| 1.0 \| 9.248e-57 \| 1483 \| 0.278 \| 711 \| 423 \| 26 \| 6 \| 681 \| 1 \| 656 \| Phage terminase, large subunit \| Phage terminase, large subunit; afdb-uniprot50 \| AF-A3UIT1-F1-MODEL\_V4 \| 1.0 \| 2.021e-47 \| 1483 \| 0.226 \| 693 \| 431 \| 25 \| 3 \| 616 \| 1 \| 667 \| Prophage LambdaW1, terminase large subunit, putative \| Prophage LambdaW1, terminase large subunit, putative; afdb-uniprot50 \| AF-A0A7Y3JAE3-F1-MODEL\_V4 \| 1.0 \| 1.531e-56 \| 1479 \| 0.282 \| 702 \| 418 \| 26 \| 10 \| 682 \| 2 \| 646 \| Phage terminase large subunit family protein \| Phage terminase large subunit family protein; afdb-uniprot50 \| AF-F5R897-F1-MODEL\_V4 \| 1.0 \| 2.266e-56 \| 1477 \| 0.26 \| 722 \| 459 \| 27 \| 2 \| 677 \| 1 \| 693 \| Phage terminase GpA \| Phage terminase GpA; afdb-uniprot50 \| AF-A0A7X3UYU0-F1-MODEL\_V4 \| 1.0 \| 2.646e-54 \| 1476 \| 0.278 \| 618 \| 389 \| 20 \| 22 \| 630 \| 5 \| 574 \| Phage terminase large subunit family protein \| Phage terminase large subunit family protein; afdb-uniprot50 \| AF-A0A1M3KG66-F1-MODEL\_V4 \| 1.0 \| 1.834e-58 \| 1475 \| 0.299 \| 705 \| 403 \| 21 \| 3 \| 632 \| 258 \| 946 \| Uncharacterized protein \| Uncharacterized protein; afdb-uniprot50 \| AF-A0A1G2ZKT4-F1-MODEL\_V4 \| 1.0 \| 3.375e-57 \| 1474 \| 0.323 \| 624 \| 354 \| 23 \| 74 \| 682 \| 2 \| 572 \| Uncharacterized protein \| Uncharacterized protein; afdb-uniprot50 \| AF-A0A418VZ41-F1-MODEL\_V4 \| 1.0 \| 7.817e-57 \| 1474 \| 0.311 \| 671 \| 380 \| 22 \| 42 \| 682 \| 2 \| 620 \| Phage terminase large subunit family protein \| Phage terminase large subunit family protein; afdb-uniprot50 \| AF-A0A2N9AI26-F1-MODEL\_V4 \| 1.0 \| 7.44e-58 \| 1474 \| 0.263 \| 646 \| 401 \| 18 \| 44 \| 633 \| 4 \| 630 \| Terminase GpA (Modular protein) \| Terminase GpA (Modular protein); afdb-uniprot50 \| AF-A0A363UL95-F1-MODEL\_V4 \| 1.0 \| 5.87e-56 \| 1473 \| 0.252 \| 722 \| 454 \| 29 \| 3 \| 677 \| 1 \| 683 \| Terminase \| Terminase; afdb-uniprot50 \| AF-C1D856-F1-MODEL\_V4 \| 1.0 \| 1.619e-56 \| 1472 \| 0.278 \| 661 \| 415 \| 26 \| 21 \| 632 \| 4 \| 651 \| Phage terminase GpA \| Phage terminase GpA; afdb-uniprot50 \| AF-A0A1Y6CNV9-F1-MODEL\_V4 \| 1.0 \| 8.109e-54 \| 1470 \| 0.247 \| 655 \| 414 \| 21 \| 4 \| 616 \| 1 \| 618 \| Phage terminase, large subunit GpA \| Phage terminase, large subunit GpA; afdb-uniprot50 \| AF-A0A2D2M641-F1-MODEL\_V4 \| 1.0 \| 1.164e-57 \| 1470 \| 0.279 \| 697 \| 429 \| 22 \| 18 \| 681 \| 8 \| 663 \| Terminase \| Terminase; afdb-uniprot50 \| AF-A0A5E7Q6C7-F1-MODEL\_V4 \| 1.0 \| 1.208e-54 \| 1468 \| 0.264 \| 703 \| 422 \| 29 \| 22 \| 678 \| 6 \| 659 \| Uncharacterized protein \| Uncharacterized protein; afdb-uniprot50 \| AF-A0A2D5FQ80-F1-MODEL\_V4 \| 1.0 \| 3.742e-47 \| 1468 \| 0.215 \| 746 \| 472 \| 25 \| 3 \| 673 \| 1 \| 707 \| Uncharacterized protein \| Uncharacterized protein; afdb-uniprot50 \| AF-A0A0P7ZPK1-F1-MODEL\_V4 \| 1.0 \| 1.092e-47 \| 1467 \| 0.369 \| 487 \| 273 \| 11 \| 19 \| 497 \| 2 \| 462 \| Bacteriophage tail assembly protein \| Bacteriophage tail assembly protein; afdb-uniprot50 \| AF-A0A1I7L8M6-F1-MODEL\_V4 \| 1.0 \| 6.289e-58 \| 1467 \| 0.269 \| 654 \| 407 \| 20 \| 35 \| 633 \| 3 \| 640 \| Phage terminase, large subunit GpA \| Phage terminase, large subunit GpA; afdb-uniprot50 \| AF-A0A3S0TCK1-F1-MODEL\_V4 \| 1.0 \| 3.992e-57 \| 1467 \| 0.266 \| 746 \| 438 \| 17 \| 1 \| 677 \| 4 \| 708 \| Terminase \| Terminase; afdb-uniprot50 \| AF-A0A0F3IVH9-F1-MODEL\_V4 \| 1.0 \| 7.344e-56 \| 1465 \| 0.279 \| 694 \| 411 \| 24 \| 21 \| 679 \| 5 \| 644 \| Uncharacterized protein \| Uncharacterized protein; afdb-uniprot50 \| AF-A0A7Y4RQQ7-F1-MODEL\_V4 \| 1.0 \| 2.816e-55 \| 1461 \| 0.278 \| 656 \| 408 \| 27 \| 3 \| 633 \| 1 \| 615 \| Uncharacterized protein \| Uncharacterized protein; afdb-uniprot50 \| AF-A0A2S6H5F8-F1-MODEL\_V4 \| 1.0 \| 1.974e-52 \| 1461 \| 0.263 \| 641 \| 406 \| 22 \| 11 \| 624 \| 3 \| 604 \| Phage terminase large subunit GpA-like protein \| Phage terminase large subunit GpA-like protein; afdb-uniprot50 \| AF-A0A3N5Q187-F1-MODEL\_V4 \| 1.0 \| 1.374e-48 \| 1461 \| 0.222 \| 674 \| 458 \| 31 \| 3 \| 631 \| 1 \| 653 \| Uncharacterized protein \| Uncharacterized protein; afdb-uniprot50 \| AF-A0A6P2IYT3-F1-MODEL\_V4 \| 1.0 \| 1.63e-57 \| 1458 \| 0.288 \| 687 \| 428 \| 25 \| 33 \| 681 \| 2 \| 665 \| Terminase \| Terminase; afdb-uniprot50 \| AF-A0A377AKI7-F1-MODEL\_V4 \| 1.0 \| 1.094e-56 \| 1457 \| 0.31 \| 579 \| 359 \| 16 \| 74 \| 632 \| 2 \| 559 \| Phage terminase large subunit (GpA) \| Phage terminase large subunit (GpA); afdb-uniprot50 \| AF-A0A5N9IEX1-F1-MODEL\_V4 \| 1.0 \| 6.249e-57 \| 1457 \| 0.277 \| 718 \| 423 \| 26 \| 13 \| 680 \| 2 \| 673 \| Phage terminase large subunit family protein \| Phage terminase large subunit family protein; afdb-uniprot50 \| AF-A0A2T5K775-F1-MODEL\_V4 \| 1.0 \| 2.697e-57 \| 1457 \| 0.289 \| 721 \| 415 \| 23 \| 16 \| 680 \| 5 \| 683 \| Phage terminase large subunit GpA-like protein \| Phage terminase large subunit GpA-like protein; afdb-uniprot50 \| AF-A0A2A4WLK3-F1-MODEL\_V4 \| 1.0 \| 3.941e-55 \| 1456 \| 0.247 \| 687 \| 445 \| 25 \| 1 \| 640 \| 1 \| 662 \| Terminase \| Terminase; afdb-uniprot50 \| AF-A0A1C7BSH8-F1-MODEL\_V4 \| 1.0 \| 4.408e-55 \| 1455 \| 0.269 \| 650 \| 415 \| 23 \| 30 \| 633 \| 2 \| 637 \| Uncharacterized protein \| Uncharacterized protein; afdb-uniprot50 \| AF-A0A0N8REA1-F1-MODEL\_V4 \| 1.0 \| 5.55e-56 \| 1455 \| 0.274 \| 696 \| 427 \| 26 \| 19 \| 680 \| 2 \| 653 \| Phage terminase \| Phage terminase; afdb-uniprot50 \| AF-A0A2N3B951-F1-MODEL\_V4 \| 1.0 \| 5.114e-52 \| 1454 \| 0.308 \| 593 \| 356 \| 17 \| 1 \| 564 \| 7 \| 574 \| Phage tail protein \| Phage tail protein; afdb-uniprot50 \| AF-A0A557QLR6-F1-MODEL\_V4 \| 1.0 \| 1.028e-55 \| 1454 \| 0.276 \| 687 \| 411 \| 23 \| 22 \| 679 \| 5 \| 634 \| Phage terminase large subunit family protein \| Phage terminase large subunit family protein; afdb-uniprot50 \| AF-A0A5C8P8I7-F1-MODEL\_V4 \| 1.0 \| 1.302e-57 \| 1454 \| 0.272 \| 745 \| 446 \| 30 \| 1 \| 679 \| 1 \| 715 \| Terminase \| Terminase; afdb-uniprot50 \| AF-A0A1X0SVE4-F1-MODEL\_V4 \| 1.0 \| 8.196e-47 \| 1452 \| 0.332 \| 521 \| 271 \| 16 \| 172 \| 680 \| 6 \| 461 \| Phage terminase GpA \| Phage terminase GpA; afdb-uniprot50 \| AF-A0A1Y2K9P1-F1-MODEL\_V4 \| 1.0 \| 4.888e-45 \| 1451 \| 0.384 \| 502 \| 265 \| 14 \| 148 \| 632 \| 4 \| 478 \| Putative phage terminase GpA \| Putative phage terminase GpA; afdb-uniprot50 \| AF-A0A031G5Z5-F1-MODEL\_V4 \| 1.0 \| 1.788e-54 \| 1451 \| 0.269 \| 697 \| 425 \| 26 \| 19 \| 681 \| 2 \| 647 \| Terminase GpA \| Terminase GpA; afdb-uniprot50 \| AF-A0A437MJX3-F1-MODEL\_V4 \| 1.0 \| 1.334e-52 \| 1451 \| 0.269 \| 638 \| 404 \| 26 \| 19 \| 626 \| 5 \| 610 \| Uncharacterized protein \| Uncharacterized protein; afdb-uniprot50 \| AF-A0A4R1J3Z1-F1-MODEL\_V4 \| 1.0 \| 4.994e-57 \| 1450 \| 0.283 \| 698 \| 423 \| 24 \| 17 \| 679 \| 2 \| 657 \| Phage terminase large subunit GpA-like protein \| Phage terminase large subunit GpA-like protein; afdb-uniprot50 \| AF-W7DJG2-F1-MODEL\_V4 \| 1.0 \| 9.188e-56 \| 1449 \| 0.305 \| 651 \| 390 \| 23 \| 13 \| 632 \| 2 \| 621 \| Phage terminase large subunit \| Phage terminase large subunit; afdb-uniprot50 \| AF-A0A096GRB5-F1-MODEL\_V4 \| 1.0 \| 1.765e-52 \| 1449 \| 0.281 \| 667 \| 386 \| 27 \| 27 \| 642 \| 2 \| 625 \| Terminase \| Terminase; afdb-uniprot50 \| AF-A0A1W0CDL5-F1-MODEL\_V4 \| 1.0 \| 2.834e-56 \| 1449 \| 0.288 \| 690 \| 421 \| 23 \| 21 \| 681 \| 5 \| 653 \| Terminase \| Terminase; afdb-uniprot50 \| AF-A0A541CKZ6-F1-MODEL\_V4 \| 1.0 \| 5.515e-55 \| 1444 \| 0.313 \| 648 \| 369 \| 24 \| 33 \| 632 \| 5 \| 624 \| Phage terminase large subunit family protein \| Phage terminase large subunit family protein; afdb-uniprot50 \| AF-S1I5L9-F1-MODEL\_V4 \| 1.0 \| 1.447e-56 \| 1444 \| 0.268 \| 699 \| 425 \| 20 \| 3 \| 631 \| 1 \| 682 \| Uncharacterized protein \| Uncharacterized protein; afdb-uniprot50 \| AF-A0A1H9QET3-F1-MODEL\_V4 \| 1.0 \| 3.501e-54 \| 1443 \| 0.32 \| 612 \| 365 \| 23 \| 42 \| 632 \| 2 \| 583 \| Phage terminase, large subunit GpA \| Phage terminase, large subunit GpA; afdb-uniprot50 \| AF-A0A8A5CXP9-F1-MODEL\_V4 \| 1.0 \| 3.089e-52 \| 1443 \| 0.29 \| 629 \| 407 \| 16 \| 22 \| 633 \| 3 \| 608 \| Phage terminase large subunit family protein \| Phage terminase large subunit family protein; afdb-uniprot50 \| AF-A0A7Y5K1K5-F1-MODEL\_V4 \| 1.0 \| 3.726e-55 \| 1443 \| 0.286 \| 669 \| 393 \| 19 \| 51 \| 682 \| 2 \| 622 \| Phage terminase large subunit family protein \| Phage terminase large subunit family protein; afdb-uniprot50 \| AF-A0A1E3ZLB5-F1-MODEL\_V4 \| 1.0 \| 2.251e-55 \| 1442 \| 0.263 \| 698 \| 433 \| 25 \| 22 \| 681 \| 4 \| 658 \| Uncharacterized protein \| Uncharacterized protein; afdb-uniprot50 \| AF-A0A613PYZ4-F1-MODEL\_V4 \| 1.0 \| 3.13e-54 \| 1441 \| 0.332 \| 625 \| 359 \| 21 \| 39 \| 632 \| 1 \| 597 \| Phage terminase large subunit family protein \| Phage terminase large subunit family protein; afdb-uniprot50 \| AF-J2TG39-F1-MODEL\_V4 \| 1.0 \| 9.717e-56 \| 1441 \| 0.277 \| 656 \| 405 \| 23 \| 18 \| 632 \| 7 \| 634 \| Bacteriophage tail assembly protein \| Bacteriophage tail assembly protein; afdb-uniprot50 \| AF-A0A2X0QXE2-F1-MODEL\_V4 \| 1.0 \| 6.855e-54 \| 1440 \| 0.274 \| 704 \| 411 \| 23 \| 1 \| 681 \| 6 \| 632 \| Phage terminase GpA \| Phage terminase GpA; afdb-uniprot50 \| AF-A0A2G6T743-F1-MODEL\_V4 \| 1.0 \| 1.087e-55 \| 1439 \| 0.266 \| 693 \| 443 \| 27 \| 21 \| 680 \| 6 \| 665 \| Phage terminase large subunit GpA-like protein \| Phage terminase large subunit GpA-like protein; afdb-uniprot50 \| AF-A0A4D8PB68-F1-MODEL\_V4 \| 1.0 \| 1.69e-54 \| 1436 \| 0.301 \| 666 \| 387 \| 26 \| 3 \| 632 \| 1 \| 623 \| Phage terminase large subunit family protein \| Phage terminase large subunit family protein; afdb-uniprot50 \| AF-A0A495D1J5-F1-MODEL\_V4 \| 1.0 \| 3.718e-46 \| 1436 \| 0.221 \| 708 \| 430 \| 32 \| 3 \| 621 \| 1 \| 675 \| Phage terminase large subunit GpA-like protein \| Phage terminase large subunit GpA-like protein; afdb-uniprot50 \| AF-A0A521VMB5-F1-MODEL\_V4 \| 1.0 \| 7.344e-56 \| 1432 \| 0.268 \| 727 \| 458 \| 22 \| 3 \| 682 \| 8 \| 707 \| Phage terminase large subunit family protein \| Phage terminase large subunit family protein; afdb-uniprot50 \| AF-A0A3M5A852-F1-MODEL\_V4 \| 1.0 \| 4.408e-55 \| 1432 \| 0.291 \| 646 \| 380 \| 24 \| 1 \| 617 \| 1 \| 597 \| Portal protein \| Portal protein; afdb-uniprot50 \| AF-A0A6A1TPS3-F1-MODEL\_V4 \| 1.0 \| 1.261e-52 \| 1430 \| 0.267 \| 631 \| 408 \| 18 \| 22 \| 632 \| 4 \| 599 \| Phage terminase large subunit family protein \| Phage terminase large subunit family protein; afdb-uniprot50 \| AF-A0A842IZ28-F1-MODEL\_V4 \| 1.0 \| 1.999e-54 \| 1430 \| 0.264 \| 650 \| 422 \| 19 \| 1 \| 632 \| 2 \| 613 \| Phage terminase large subunit family protein \| Phage terminase large subunit family protein; afdb-uniprot50 \| AF-G2IX51-F1-MODEL\_V4 \| 1.0 \| 1.277e-54 \| 1429 \| 0.276 \| 688 \| 420 \| 25 \| 34 \| 672 \| 2 \| 660 \| Phage terminase large subunit GpA \| Phage terminase large subunit GpA; afdb-uniprot50 \| AF-A0A512JIP3-F1-MODEL\_V4 \| 1.0 \| 1.923e-48 \| 1429 \| 0.273 \| 719 \| 422 \| 34 \| 12 \| 679 \| 13 \| 681 \| Terminase \| Terminase; afdb-uniprot50 \| AF-A0A5C7NUV1-F1-MODEL\_V4 \| 1.0 \| 2.646e-54 \| 1426 \| 0.269 \| 687 \| 437 \| 26 \| 21 \| 675 \| 6 \| 659 \| Phage terminase large subunit family protein \| Phage terminase large subunit family protein; afdb-uniprot50 \| AF-A0A329VGS0-F1-MODEL\_V4 \| 1.0 \| 9.408e-51 \| 1424 \| 0.33 \| 581 \| 340 \| 19 \| 6 \| 561 \| 1 \| 557 \| Terminase \| Terminase; afdb-uniprot50 \| AF-A0A3B9NW07-F1-MODEL\_V4 \| 1.0 \| 7.618e-53 \| 1424 \| 0.273 \| 695 \| 418 \| 24 \| 13 \| 676 \| 2 \| 640 \| Terminase \| Terminase; afdb-uniprot50 \| AF-A0A0F9JB30-F1-MODEL\_V4 \| 1.0 \| 1.669e-52 \| 1422 \| 0.312 \| 566 \| 328 \| 15 \| 99 \| 632 \| 3 \| 539 \| Uncharacterized protein \| Uncharacterized protein; afdb-uniprot50 \| AF-N6V822-F1-MODEL\_V4 \| 1.0 \| 6.811e-53 \| 1421 \| 0.347 \| 549 \| 312 \| 19 \| 74 \| 607 \| 2 \| 519 \| Phage terminase large subunit GpA \| Phage terminase large subunit GpA; afdb-uniprot50 \| AF-A0A351RVU7-F1-MODEL\_V4 \| 1.0 \| 7.297e-55 \| 1420 \| 0.276 \| 655 \| 423 \| 23 \| 1 \| 633 \| 22 \| 647 \| Uncharacterized protein \| Uncharacterized protein; afdb-uniprot50 \| AF-A0A7C1RGY2-F1-MODEL\_V4 \| 1.0 \| 1.521e-55 \| 1420 \| 0.254 \| 742 \| 451 \| 25 \| 1 \| 682 \| 110 \| 808 \| Terminase \| Terminase; afdb-uniprot50 \| AF-A0A7Z0KZA8-F1-MODEL\_V4 \| 1.0 \| 2.335e-52 \| 1418 \| 0.317 \| 611 \| 361 \| 17 \| 2 \| 584 \| 6 \| 588 \| Phage terminase large subunit family protein \| Phage terminase large subunit family protein; afdb-uniprot50 \| AF-A0A3D4V590-F1-MODEL\_V4 \| 1.0 \| 1.501e-53 \| 1418 \| 0.244 \| 736 \| 454 \| 25 \| 1 \| 677 \| 98 \| 790 \| Uncharacterized protein \| Uncharacterized protein; afdb-uniprot50 \| AF-A0A386UIC0-F1-MODEL\_V4 \| 1.0 \| 2.012e-55 \| 1415 \| 0.309 \| 620 \| 363 \| 19 \| 77 \| 680 \| 2 \| 572 \| Phage terminase large subunit family protein \| Phage terminase large subunit family protein; afdb-uniprot50 \| AF-A0A7C2YSF8-F1-MODEL\_V4 \| 1.0 \| 2.485e-53 \| 1415 \| 0.291 \| 656 \| 398 \| 26 \| 1 \| 621 \| 1 \| 624 \| Phage terminase large subunit family protein \| Phage terminase large subunit family protein; afdb-uniprot50 \| AF-A0A4D7QHJ6-F1-MODEL\_V4 \| 1.0 \| 9.53e-53 \| 1414 \| 0.272 \| 656 \| 393 \| 24 \| 17 \| 632 \| 5 \| 615 \| Uncharacterized protein \| Uncharacterized protein; afdb-uniprot50 \| AF-A0A165RN54-F1-MODEL\_V4 \| 1.0 \| 6.84e-45 \| 1414 \| 0.234 \| 679 \| 441 \| 27 \| 34 \| 680 \| 1 \| 632 \| Phage terminase large subunit (GpA) \| Phage terminase large subunit (GpA); afdb-uniprot50 \| AF-A0A2N1TNL2-F1-MODEL\_V4 \| 1.0 \| 4.651e-46 \| 1412 \| 0.225 \| 684 \| 445 \| 29 \| 3 \| 619 \| 1 \| 666 \| Uncharacterized protein \| Uncharacterized protein; afdb-uniprot50 \| AF-A0A0Q0T412-F1-MODEL\_V4 \| 1.0 \| 6.011e-51 \| 1411 \| 0.298 \| 580 \| 349 \| 20 \| 10 \| 561 \| 1 \| 550 \| Terminase \| Terminase; afdb-uniprot50 \| AF-A0A0Q6UXK2-F1-MODEL\_V4 \| 1.0 \| 4.919e-46 \| 1411 \| 0.236 \| 676 \| 437 \| 29 \| 1 \| 632 \| 1 \| 641 \| Uncharacterized protein \| Uncharacterized protein; afdb-uniprot50 \| AF-A0A2K4MSU4-F1-MODEL\_V4 \| 1.0 \| 2.236e-54 \| 1411 \| 0.28 \| 702 \| 416 \| 26 \| 19 \| 679 \| 8 \| 661 \| Terminase \| Terminase; afdb-uniprot50 \| AF-A0A1L3ZRN2-F1-MODEL\_V4 \| 1.0 \| 3.932e-46 \| 1411 \| 0.233 \| 721 \| 456 \| 31 \| 2 \| 682 \| 6 \| 669 \| Uncharacterized protein \| Uncharacterized protein; afdb-uniprot50 \| AF-A0A0E1SI48-F1-MODEL\_V4 \| 1.0 \| 2.534e-56 \| 1411 \| 0.267 \| 722 \| 436 \| 26 \| 14 \| 681 \| 2 \| 684 \| Phage terminase large subunit (GpA) \| Phage terminase large subunit (GpA); afdb-uniprot50 \| AF-A0A6N7W1I2-F1-MODEL\_V4 \| 1.0 \| 1.2e-53 \| 1409 \| 0.263 \| 637 \| 414 \| 15 \| 24 \| 632 \| 7 \| 616 \| Terminase \| Terminase; afdb-uniprot50 \| AF-A0A225SLS1-F1-MODEL\_V4 \| 1.0 \| 1.712e-56 \| 1409 \| 0.27 \| 721 \| 457 \| 24 \| 1 \| 681 \| 1 \| 692 \| Terminase \| Terminase; afdb-uniprot50 \| AF-K2JJZ4-F1-MODEL\_V4 \| 1.0 \| 7.75e-47 \| 1409 \| 0.221 \| 748 \| 465 \| 33 \| 1 \| 679 \| 1 \| 699 \| Phage terminase large subunit \| Phage terminase large subunit; afdb-uniprot50 \| AF-A0A839Y4B2-F1-MODEL\_V4 \| 1.0 \| 4.379e-54 \| 1407 \| 0.306 \| 626 \| 369 \| 18 \| 74 \| 677 \| 2 \| 584 \| Phage terminase large subunit GpA-like protein \| Phage terminase large subunit GpA-like protein; afdb-uniprot50 \| AF-A0A2N8HQF8-F1-MODEL\_V4 \| 1.0 \| 1.878e-53 \| 1407 \| 0.255 \| 653 \| 425 \| 23 \| 19 \| 637 \| 6 \| 631 \| Terminase \| Terminase; afdb-uniprot50 \| AF-A0A369R412-F1-MODEL\_V4 \| 1.0 \| 1.776e-53 \| 1406 \| 0.279 \| 691 \| 407 \| 23 \| 19 \| 677 \| 7 \| 638 \| Uncharacterized protein \| Uncharacterized protein; afdb-uniprot50 \| AF-A0A0F9NFU4-F1-MODEL\_V4 \| 1.0 \| 9.592e-54 \| 1405 \| 0.229 \| 757 \| 459 \| 23 \| 3 \| 682 \| 1 \| 709 \| Uncharacterized protein \| Uncharacterized protein; afdb-uniprot50 \| AF-A0A1V0B9M4-F1-MODEL\_V4 \| 1.0 \| 2.78e-53 \| 1404 \| 0.263 \| 675 \| 416 \| 23 \| 39 \| 679 \| 1 \| 628 \| Terminase \| Terminase; afdb-uniprot50 \| AF-A0A2E5KQ34-F1-MODEL\_V4 \| 1.0 \| 3.302e-45 \| 1402 \| 0.358 \| 468 \| 262 \| 11 \| 42 \| 495 \| 2 \| 445 \| Terminase \| Terminase; afdb-uniprot50 \| AF-A0A2E0M071-F1-MODEL\_V4 \| 1.0 \| 2.115e-54 \| 1402 \| 0.269 \| 705 \| 420 \| 29 \| 16 \| 675 \| 2 \| 656 \| Uncharacterized protein \| Uncharacterized protein; afdb-uniprot50 \| AF-A0A661I0H7-F1-MODEL\_V4 \| 1.0 \| 3.84e-51 \| 1401 \| 0.273 \| 597 \| 393 \| 15 \| 24 \| 608 \| 5 \| 572 \| Uncharacterized protein \| Uncharacterized protein; afdb-uniprot50 \| AF-A0A370D963-F1-MODEL\_V4 \| 1.0 \| 1.334e-52 \| 1396 \| 0.266 \| 668 \| 413 \| 23 \| 16 \| 645 \| 9 \| 637 \| Uncharacterized protein \| Uncharacterized protein; afdb-uniprot50 \| AF-A0A1A9VKE5-F1-MODEL\_V4 \| 1.0 \| 2.335e-52 \| 1396 \| 0.314 \| 594 \| 345 \| 21 \| 3 \| 570 \| 417 \| 974 \| ParB domain-containing protein \| ParB domain-containing protein; afdb-uniprot50 \| AF-A0A1N6S847-F1-MODEL\_V4 \| 1.0 \| 8.576e-54 \| 1392 \| 0.28 \| 632 \| 400 \| 26 \| 39 \| 632 \| 1 \| 615 \| Phage terminase, large subunit GpA \| Phage terminase, large subunit GpA; afdb-uniprot50 \| AF-A0A3C0NSJ5-F1-MODEL\_V4 \| 1.0 \| 1.492e-52 \| 1392 \| 0.259 \| 709 \| 436 \| 26 \| 5 \| 682 \| 1 \| 651 \| Terminase \| Terminase; afdb-uniprot50 \| AF-A0A7X7H546-F1-MODEL\_V4 \| 1.0 \| 1.192e-52 \| 1389 \| 0.249 \| 657 \| 429 \| 24 \| 19 \| 632 \| 14 \| 649 \| Phage terminase large subunit family protein \| Phage terminase large subunit family protein; afdb-uniprot50 \| AF-A0A1V5WQS7-F1-MODEL\_V4 \| 1.0 \| 1.261e-52 \| 1389 \| 0.267 \| 681 \| 405 \| 21 \| 23 \| 677 \| 7 \| 619 \| Phage terminase large subunit (GpA) \| Phage terminase large subunit (GpA); afdb-uniprot50 \| AF-A0A0Q4WDP7-F1-MODEL\_V4 \| 1.0 \| 5.409e-52 \| 1389 \| 0.251 \| 648 \| 423 \| 24 \| 10 \| 621 \| 1 \| 622 \| Peptidase\_S49 domain-containing protein \| Peptidase\_S49 domain-containing protein; afdb-uniprot50 \| AF-A0A6S6LEK8-F1-MODEL\_V4 \| 1.0 \| 1.258e-43 \| 1385 \| 0.259 \| 597 \| 363 \| 22 \| 75 \| 623 \| 3 \| 568 \| Terminase \| Terminase; afdb-uniprot50 \| AF-A0A1E4Q6H8-F1-MODEL\_V4 \| 1.0 \| 4.351e-53 \| 1382 \| 0.268 \| 705 \| 437 \| 24 \| 7 \| 681 \| 1 \| 656 \| Uncharacterized protein \| Uncharacterized protein; afdb-uniprot50 \| AF-A0A3L7ALY9-F1-MODEL\_V4 \| 1.0 \| 2.865e-49 \| 1382 \| 0.245 \| 732 \| 446 \| 31 \| 1 \| 680 \| 1 \| 677 \| Terminase \| Terminase; afdb-uniprot50 \| AF-A0A286GNA0-F1-MODEL\_V4 \| 1.0 \| 1.567e-51 \| 1376 \| 0.253 \| 635 \| 418 \| 22 \| 3 \| 622 \| 1 \| 594 \| Phage terminase, large subunit GpA \| Phage terminase, large subunit GpA; afdb-uniprot50 \| AF-A0A2W7IT17-F1-MODEL\_V4 \| 1.0 \| 2.261e-47 \| 1374 \| 0.336 \| 511 \| 299 \| 14 \| 33 \| 526 \| 2 \| 489 \| Phage terminase large subunit GpA-like protein \| Phage terminase large subunit GpA-like protein; afdb-uniprot50 \| AF-A0A016XHY7-F1-MODEL\_V4 \| 1.0 \| 2.101e-53 \| 1371 \| 0.257 \| 698 \| 443 \| 25 \| 18 \| 679 \| 2 \| 659 \| Uncharacterized protein \| Uncharacterized protein; afdb-uniprot50 \| AF-C1A9W4-F1-MODEL\_V4 \| 1.0 \| 1.334e-52 \| 1370 \| 0.247 \| 708 \| 441 \| 24 \| 17 \| 677 \| 11 \| 673 \| Phage terminase GpA family protein \| Phage terminase GpA family protein; afdb-uniprot50 \| AF-A0A251ZT21-F1-MODEL\_V4 \| 1.0 \| 2.798e-54 \| 1369 \| 0.315 \| 612 \| 369 \| 19 \| 74 \| 659 \| 2 \| 589 \| Terminase \| Terminase; afdb-uniprot50 \| AF-A0A7V4IFC7-F1-MODEL\_V4 \| 1.0 \| 5.933e-49 \| 1369 \| 0.251 \| 685 \| 424 \| 33 \| 18 \| 682 \| 7 \| 622 \| Uncharacterized protein \| Uncharacterized protein; afdb-uniprot50 \| AF-A0A254T6Y7-F1-MODEL\_V4 \| 1.0 \| 4.168e-55 \| 1369 \| 0.272 \| 698 \| 423 \| 23 \| 36 \| 682 \| 7 \| 670 \| Uncharacterized protein \| Uncharacterized protein; afdb-uniprot50 \| AF-A0A6V8U717-F1-MODEL\_V4 \| 1.0 \| 1.237e-49 \| 1368 \| 0.261 \| 628 \| 404 \| 24 \| 19 \| 616 \| 10 \| 607 \| Terminase \| Terminase; afdb-uniprot50 \| AF-S9QNB7-F1-MODEL\_V4 \| 1.0 \| 2.991e-47 \| 1367 \| 0.296 \| 618 \| 342 \| 19 \| 93 \| 676 \| 284 \| 842 \| Bacteriophage tail assembly protein \| Bacteriophage tail assembly protein; afdb-uniprot50 \| AF-A0A522VF53-F1-MODEL\_V4 \| 1.0 \| 1.492e-52 \| 1365 \| 0.262 \| 662 \| 429 \| 21 \| 1 \| 634 \| 1 \| 630 \| Phage terminase large subunit family protein \| Phage terminase large subunit family protein; afdb-uniprot50 \| AF-A0A848JGX1-F1-MODEL\_V4 \| 1.0 \| 1.501e-53 \| 1364 \| 0.27 \| 735 \| 440 \| 27 \| 2 \| 682 \| 4 \| 695 \| Phage terminase large subunit GpA-like protein \| Phage terminase large subunit GpA-like protein; afdb-uniprot50 \| AF-A0A5M9I9K8-F1-MODEL\_V4 \| 1.0 \| 2.101e-53 \| 1363 \| 0.272 \| 694 \| 416 \| 25 \| 23 \| 632 \| 2 \| 690 \| Uncharacterized protein \| Uncharacterized protein; afdb-uniprot50 \| AF-A0A7C2H138-F1-MODEL\_V4 \| 1.0 \| 1.19e-43 \| 1361 \| 0.339 \| 466 \| 270 \| 15 \| 17 \| 470 \| 3 \| 442 \| Phage terminase large subunit family protein \| Phage terminase large subunit family protein; afdb-uniprot50 \| AF-A0A1Q6LLP8-F1-MODEL\_V4 \| 1.0 \| 1.669e-52 \| 1361 \| 0.272 \| 660 \| 409 \| 21 \| 10 \| 634 \| 4 \| 627 \| Uncharacterized protein \| Uncharacterized protein; afdb-uniprot50 \| AF-A0A7X5KSM0-F1-MODEL\_V4 \| 1.0 \| 9.408e-51 \| 1357 \| 0.242 \| 636 \| 420 \| 24 \| 13 \| 632 \| 3 \| 592 \| Phage terminase large subunit family protein \| Phage terminase large subunit family protein; afdb-uniprot50 \| AF-A0A1T2CHX3-F1-MODEL\_V4 \| 1.0 \| 4.898e-54 \| 1355 \| 0.311 \| 643 \| 347 \| 19 \| 76 \| 680 \| 6 \| 590 \| Phage tail protein \| Phage tail protein; afdb-uniprot50 \| AF-A0A285M891-F1-MODEL\_V4 \| 1.0 \| 3.184e-48 \| 1354 \| 0.241 \| 621 \| 422 \| 19 \| 21 \| 616 \| 7 \| 603 \| Phage terminase, large subunit GpA \| Phage terminase, large subunit GpA; afdb-uniprot50 \| AF-V0AGV8-F1-MODEL\_V4 \| 1.0 \| 1.891e-54 \| 1351 \| 0.282 \| 659 \| 389 \| 19 \| 42 \| 631 \| 2 \| 645 \| Phage terminase large subunit \| Phage terminase large subunit; afdb-uniprot50 \| AF-A0A4R4G7X9-F1-MODEL\_V4 \| 1.0 \| 1.765e-52 \| 1348 \| 0.307 \| 631 \| 355 \| 20 \| 67 \| 681 \| 2 \| 566 \| Phage terminase large subunit family protein \| Phage terminase large subunit family protein; afdb-uniprot50 \| AF-A0A6M4M8U3-F1-MODEL\_V4 \| 1.0 \| 3.03e-49 \| 1347 \| 0.267 \| 665 \| 407 \| 21 \| 19 \| 637 \| 8 \| 638 \| Phage terminase large subunit family protein \| Phage terminase large subunit family protein; afdb-uniprot50 \| AF-A0A370DPR9-F1-MODEL\_V4 \| 1.0 \| 2.595e-51 \| 1346 \| 0.253 \| 694 \| 402 \| 28 \| 20 \| 682 \| 97 \| 705 \| Uncharacterized protein \| Uncharacterized protein; afdb-uniprot50 \| AF-A0A1E3VZI0-F1-MODEL\_V4 \| 1.0 \| 3.218e-41 \| 1344 \| 0.243 \| 582 \| 384 \| 18 \| 87 \| 633 \| 2 \| 561 \| Uncharacterized protein \| Uncharacterized protein; afdb-uniprot50 \| AF-A0A1Y4JAM6-F1-MODEL\_V4 \| 1.0 \| 1.842e-50 \| 1343 \| 0.241 \| 621 \| 414 \| 22 \| 29 \| 635 \| 4 \| 581 \| Terminase \| Terminase; afdb-uniprot50 \| AF-A0A4Q3P8Y5-F1-MODEL\_V4 \| 1.0 \| 4.114e-53 \| 1343 \| 0.247 \| 727 \| 467 \| 25 \| 3 \| 682 \| 6 \| 699 \| Uncharacterized protein \| Uncharacterized protein; afdb-uniprot50 \| AF-A0A022G186-F1-MODEL\_V4 \| 1.0 \| 2.629e-53 \| 1342 \| 0.26 \| 718 \| 430 \| 27 \| 9 \| 678 \| 2 \| 666 \| Terminase \| Terminase; afdb-uniprot50 \| AF-A0A369T581-F1-MODEL\_V4 \| 1.0 \| 1.886e-45 \| 1342 \| 0.224 \| 744 \| 460 \| 34 \| 3 \| 679 \| 1 \| 694 \| Uncharacterized protein \| Uncharacterized protein; afdb-uniprot50 \| AF-A0A4V2KT04-F1-MODEL\_V4 \| 1.0 \| 6.05e-52 \| 1340 \| 0.291 \| 651 \| 392 \| 27 \| 2 \| 625 \| 1 \| 608 \| Uncharacterized protein \| Uncharacterized protein; afdb-uniprot50 \| AF-A0A447IRD4-F1-MODEL\_V4 \| 1.0 \| 1.588e-53 \| 1340 \| 0.251 \| 739 \| 457 \| 24 \| 3 \| 680 \| 1 \| 703 \| Phage terminase large subunit (GpA) \| Phage terminase large subunit (GpA); afdb-uniprot50 \| AF-A0A6G3I3U2-F1-MODEL\_V4 \| 1.0 \| 2.375e-46 \| 1339 \| 0.29 \| 523 \| 320 \| 15 \| 5 \| 511 \| 1 \| 488 \| Phage terminase large subunit family protein \| Phage terminase large subunit family protein; afdb-uniprot50 \| AF-A1VPI7-F1-MODEL\_V4 \| 1.0 \| 1.401e-51 \| 1339 \| 0.24 \| 720 \| 470 \| 23 \| 1 \| 682 \| 1 \| 681 \| Phage terminase GpA \| Phage terminase GpA; afdb-uniprot50 \| AF-A0A1E3LH66-F1-MODEL\_V4 \| 1.0 \| 5.181e-54 \| 1337 \| 0.282 \| 668 \| 416 \| 20 \| 55 \| 682 \| 7 \| 650 \| Uncharacterized protein \| Uncharacterized protein; afdb-uniprot50 \| AF-A0A1V5QV10-F1-MODEL\_V4 \| 1.0 \| 7.11e-51 \| 1336 \| 0.258 \| 662 \| 426 \| 26 \| 13 \| 633 \| 2 \| 639 \| Phage terminase large subunit (GpA) \| Phage terminase large subunit (GpA); afdb-uniprot50 \| AF-A0A248LMV6-F1-MODEL\_V4 \| 1.0 \| 9.011e-53 \| 1335 \| 0.302 \| 644 \| 356 \| 26 \| 76 \| 682 \| 6 \| 592 \| Phage terminase, large subunit \| Phage terminase, large subunit; afdb-uniprot50 \| AF-A0A1Y3TRI1-F1-MODEL\_V4 \| 1.0 \| 4.773e-50 \| 1335 \| 0.245 \| 618 \| 414 \| 22 \| 29 \| 632 \| 4 \| 582 \| Terminase \| Terminase; afdb-uniprot50 \| AF-A0A5U4F806-F1-MODEL\_V4 \| 1.0 \| 3.561e-48 \| 1335 \| 0.261 \| 627 \| 390 \| 20 \| 19 \| 604 \| 8 \| 602 \| Phage terminase large subunit family protein \| Phage terminase large subunit family protein; afdb-uniprot50 \| AF-A0A1I3KSI7-F1-MODEL\_V4 \| 1.0 \| 1.658e-51 \| 1335 \| 0.243 \| 707 \| 426 \| 22 \| 17 \| 677 \| 2 \| 645 \| Phage terminase, large subunit GpA \| Phage terminase, large subunit GpA; afdb-uniprot50 \| AF-A0A6N7AWA7-F1-MODEL\_V4 \| 1.0 \| 2.485e-53 \| 1334 \| 0.236 \| 668 \| 446 \| 27 \| 1 \| 632 \| 7 \| 646 \| Phage terminase GpA \| Phage terminase GpA; afdb-uniprot50 \| AF-A0A258LL44-F1-MODEL\_V4 \| 1.0 \| 1.83e-49 \| 1333 \| 0.283 \| 631 \| 394 \| 24 \| 24 \| 634 \| 100 \| 692 \| Uncharacterized protein \| Uncharacterized protein; afdb-uniprot50 \| AF-A0A839HET4-F1-MODEL\_V4 \| 1.0 \| 5.114e-52 \| 1328 \| 0.284 \| 611 \| 376 \| 20 \| 55 \| 632 \| 6 \| 588 \| Phage terminase large subunit family protein \| Phage terminase large subunit family protein; afdb-uniprot50 \| AF-A0A1J5T405-F1-MODEL\_V4 \| 1.0 \| 1.501e-53 \| 1327 \| 0.312 \| 634 \| 372 \| 23 \| 82 \| 682 \| 2 \| 604 \| Phage terminase large subunit (GpA) \| Phage terminase large subunit (GpA); afdb-uniprot50 \| AF-A0A829R1A8-F1-MODEL\_V4 \| 1.0 \| 3.815e-50 \| 1327 \| 0.256 \| 652 \| 402 \| 24 \| 9 \| 614 \| 2 \| 616 \| Phage terminase large subunit GpA \| Phage terminase large subunit GpA; afdb-uniprot50 \| AF-A0A5C7PVY5-F1-MODEL\_V4 \| 1.0 \| 2.464e-43 \| 1326 \| 0.261 \| 593 \| 368 \| 20 \| 114 \| 681 \| 2 \| 549 \| Phage terminase large subunit family protein \| Phage terminase large subunit family protein; afdb-uniprot50 \| AF-A0A2H4TLX0-F1-MODEL\_V4 \| 1.0 \| 3.865e-52 \| 1325 \| 0.322 \| 607 \| 357 \| 18 \| 74 \| 648 \| 2 \| 585 \| Terminase \| Terminase; afdb-uniprot50 \| AF-A0A4R9WUL6-F1-MODEL\_V4 \| 1.0 \| 1.753e-51 \| 1325 \| 0.254 \| 637 \| 411 \| 21 \| 28 \| 625 \| 2 \| 613 \| Terminase \| Terminase; afdb-uniprot50 \| AF-A0A7C6A5Y1-F1-MODEL\_V4 \| 1.0 \| 7.233e-45 \| 1324 \| 0.36 \| 466 \| 254 \| 13 \| 187 \| 637 \| 3 \| 439 \| Phage terminase large subunit family protein \| Phage terminase large subunit family protein; afdb-uniprot50 \| AF-A0A827C522-F1-MODEL\_V4 \| 1.0 \| 1.229e-48 \| 1324 \| 0.268 \| 648 \| 404 \| 28 \| 19 \| 623 \| 8 \| 628 \| Phage terminase large subunit family protein \| Phage terminase large subunit family protein; afdb-uniprot50 \| AF-A0A5C7Q4Y0-F1-MODEL\_V4 \| 1.0 \| 6.011e-51 \| 1324 \| 0.243 \| 674 \| 433 \| 22 \| 1 \| 632 \| 1 \| 639 \| Uncharacterized protein \| Uncharacterized protein; afdb-uniprot50 \| AF-W0E4D8-F1-MODEL\_V4 \| 1.0 \| 3.225e-50 \| 1323 \| 0.277 \| 699 \| 410 \| 27 \| 17 \| 682 \| 3 \| 639 \| Terminase \| Terminase; afdb-uniprot50 \| AF-A0A3D5MID3-F1-MODEL\_V4 \| 1.0 \| 1.482e-51 \| 1323 \| 0.246 \| 746 \| 431 \| 25 \| 7 \| 677 \| 1 \| 690 \| Terminase \| Terminase; afdb-uniprot50 \| AF-A0A248JSE1-F1-MODEL\_V4 \| 1.0 \| 3.267e-52 \| 1320 \| 0.245 \| 759 \| 460 \| 30 \| 4 \| 681 \| 1 \| 727 \| Uncharacterized protein \| Uncharacterized protein; afdb-uniprot50 \| AF-A0A4E0LAD2-F1-MODEL\_V4 \| 1.0 \| 5.574e-48 \| 1318 \| 0.259 \| 617 \| 384 \| 20 \| 19 \| 594 \| 8 \| 592 \| Phage terminase large subunit family protein \| Phage terminase large subunit family protein; afdb-uniprot50 \| AF-A0A5U4F844-F1-MODEL\_V4 \| 1.0 \| 6.974e-48 \| 1318 \| 0.251 \| 627 \| 397 \| 19 \| 19 \| 604 \| 8 \| 602 \| Phage terminase large subunit family protein \| Phage terminase large subunit family protein; afdb-uniprot50 \| AF-W9GZ58-F1-MODEL\_V4 \| 1.0 \| 2.744e-51 \| 1316 \| 0.226 \| 727 \| 463 \| 25 \| 4 \| 677 \| 1 \| 680 \| Uncharacterized protein \| Uncharacterized protein; afdb-uniprot50 \| AF-A0A838M0Z5-F1-MODEL\_V4 \| 1.0 \| 1.742e-50 \| 1315 \| 0.259 \| 705 \| 434 \| 24 \| 26 \| 680 \| 2 \| 668 \| Phage terminase large subunit family protein \| Phage terminase large subunit family protein; afdb-uniprot50 \| AF-A0A0K6IXN7-F1-MODEL\_V4 \| 1.0 \| 1.488e-43 \| 1311 \| 0.334 \| 499 \| 291 \| 14 \| 14 \| 497 \| 2 \| 474 \| Phage terminase, large subunit GpA \| Phage terminase, large subunit GpA; afdb-uniprot50 \| AF-A0A1M3KJ46-F1-MODEL\_V4 \| 1.0 \| 1.598e-54 \| 1308 \| 0.303 \| 626 \| 356 \| 16 \| 75 \| 632 \| 3 \| 616 \| Uncharacterized protein \| Uncharacterized protein; afdb-uniprot50 \| AF-A0A653RI20-F1-MODEL\_V4 \| 1.0 \| 4.484e-49 \| 1303 \| 0.239 \| 689 \| 418 \| 25 \| 23 \| 632 \| 12 \| 673 \| Terminase GpA \| Terminase GpA; afdb-uniprot50 \| AF-A0A0S9PWZ7-F1-MODEL\_V4 \| 1.0 \| 2.047e-49 \| 1300 \| 0.237 \| 653 \| 431 \| 25 \| 7 \| 620 \| 1 \| 625 \| Uncharacterized protein \| Uncharacterized protein; afdb-uniprot50 \| AF-E4LA55-F1-MODEL\_V4 \| 1.0 \| 2.261e-47 \| 1299 \| 0.211 \| 606 \| 421 \| 22 \| 19 \| 605 \| 24 \| 591 \| Phage terminase large subunit (GpA) \| Phage terminase large subunit (GpA); afdb-uniprot50 \| AF-F9Y4K8-F1-MODEL\_V4 \| 1.0 \| 8.039e-44 \| 1299 \| 0.195 \| 764 \| 479 \| 35 \| 1 \| 681 \| 1 \| 711 \| Phage terminase GpA \| Phage terminase GpA; afdb-uniprot50 \| AF-A0A1Q3WRM6-F1-MODEL\_V4 \| 1.0 \| 1.106e-49 \| 1299 \| 0.238 \| 726 \| 466 \| 22 \| 3 \| 679 \| 1 \| 688 \| Terminase \| Terminase; afdb-uniprot50 \| AF-D2U9G3-F1-MODEL\_V4 \| 1.0 \| 8.005e-52 \| 1295 \| 0.26 \| 663 \| 421 \| 25 \| 1 \| 632 \| 2 \| 625 \| Hypothetical phage terminase large subunit protein \| Hypothetical phage terminase large subunit protein; afdb-uniprot50 \| AF-A0A450WC01-F1-MODEL\_V4 \| 1.0 \| 5.374e-51 \| 1294 \| 0.275 \| 647 \| 415 \| 25 \| 39 \| 638 \| 1 \| 640 \| Phage terminase, large subunit GpA \| Phage terminase, large subunit GpA; afdb-uniprot50 \| AF-A0A831U048-F1-MODEL\_V4 \| 1.0 \| 1.059e-51 \| 1293 \| 0.253 \| 666 \| 426 \| 28 \| 39 \| 682 \| 19 \| 634 \| Uncharacterized protein \| Uncharacterized protein; afdb-uniprot50 \| AF-A0A371WUR5-F1-MODEL\_V4 \| 1.0 \| 1.245e-50 \| 1292 \| 0.259 \| 718 \| 439 \| 26 \| 1 \| 682 \| 1 \| 661 \| Uncharacterized protein \| Uncharacterized protein; afdb-uniprot50 \| AF-A5V8D6-F1-MODEL\_V4 \| 1.0 \| 2.33e-43 \| 1292 \| 0.218 \| 695 \| 446 \| 32 \| 26 \| 681 \| 2 \| 638 \| Phage terminase GpA \| Phage terminase GpA; afdb-uniprot50 \| AF-A0A1H5S0Q5-F1-MODEL\_V4 \| 1.0 \| 6.011e-51 \| 1290 \| 0.283 \| 638 \| 395 \| 27 \| 46 \| 632 \| 2 \| 628 \| Phage terminase, large subunit GpA \| Phage terminase, large subunit GpA; afdb-uniprot50 \| AF-A0A5N7WFR3-F1-MODEL\_V4 \| 1.0 \| 1.482e-51 \| 1290 \| 0.234 \| 734 \| 444 \| 30 \| 5 \| 679 \| 1 \| 675 \| Phage terminase large subunit family protein \| Phage terminase large subunit family protein; afdb-uniprot50 \| AF-S7HSN4-F1-MODEL\_V4 \| 1.0 \| 4.213e-48 \| 1289 \| 0.283 \| 589 \| 357 \| 18 \| 20 \| 572 \| 5 \| 564 \| Phage terminase large subunit (GpA) \| Phage terminase large subunit (GpA); afdb-uniprot50 \| AF-A0A5A7NAV3-F1-MODEL\_V4 \| 1.0 \| 9.571e-45 \| 1289 \| 0.232 \| 670 \| 415 \| 31 \| 75 \| 679 \| 3 \| 637 \| DNA packaging protein \| DNA packaging protein; afdb-uniprot50 \| AF-A0A1H2HHQ2-F1-MODEL\_V4 \| 1.0 \| 2.859e-40 \| 1286 \| 0.223 \| 644 \| 412 \| 29 \| 19 \| 616 \| 2 \| 603 \| Phage terminase, large subunit GpA \| Phage terminase, large subunit GpA; afdb-uniprot50 \| AF-E6X1P8-F1-MODEL\_V4 \| 1.0 \| 1.72e-48 \| 1286 \| 0.224 \| 687 \| 424 \| 27 \| 23 \| 677 \| 4 \| 613 \| Terminase GpA \| Terminase GpA; afdb-uniprot50 \| AF-A0A1I3H6J9-F1-MODEL\_V4 \| 1.0 \| 1.636e-49 \| 1286 \| 0.269 \| 652 \| 407 \| 27 \| 20 \| 632 \| 2 \| 622 \| Phage terminase, large subunit GpA \| Phage terminase, large subunit GpA; afdb-uniprot50 \| AF-A0A510IP07-F1-MODEL\_V4 \| 1.0 \| 3.433e-51 \| 1286 \| 0.258 \| 692 \| 426 \| 27 \| 39 \| 680 \| 1 \| 655 \| Terminase \| Terminase; afdb-uniprot50 \| AF-A0A480BWT3-F1-MODEL\_V4 \| 1.0 \| 1.72e-48 \| 1285 \| 0.285 \| 708 \| 390 \| 28 \| 5 \| 678 \| 1 \| 626 \| Uncharacterized protein \| Uncharacterized protein; afdb-uniprot50 \| AF-A0A2W4V675-F1-MODEL\_V4 \| 1.0 \| 5.574e-48 \| 1285 \| 0.255 \| 715 \| 410 \| 32 \| 10 \| 633 \| 1 \| 683 \| Uncharacterized protein \| Uncharacterized protein; afdb-uniprot50 \| AF-A0A420WD47-F1-MODEL\_V4 \| 1.0 \| 5.972e-50 \| 1284 \| 0.243 \| 669 \| 420 \| 23 \| 20 \| 630 \| 35 \| 675 \| Phage terminase large subunit GpA-like protein \| Phage terminase large subunit GpA-like protein; afdb-uniprot50 \| AF-A0A071M5H9-F1-MODEL\_V4 \| 1.0 \| 1.154e-47 \| 1283 \| 0.255 \| 622 \| 399 \| 24 \| 39 \| 632 \| 1 \| 586 \| Uncharacterized protein \| Uncharacterized protein; afdb-uniprot50 \| AF-A0A109K2Q7-F1-MODEL\_V4 \| 1.0 \| 1.765e-52 \| 1280 \| 0.291 \| 655 \| 370 \| 20 \| 75 \| 682 \| 3 \| 610 \| Uncharacterized protein \| Uncharacterized protein; afdb-uniprot50 \| AF-A0A1Z9VRR1-F1-MODEL\_V4 \| 1.0 \| 2.047e-49 \| 1280 \| 0.24 \| 683 \| 458 \| 25 \| 1 \| 630 \| 1 \| 675 \| Uncharacterized protein \| Uncharacterized protein; afdb-uniprot50 \| AF-A0A3S4FNV3-F1-MODEL\_V4 \| 1.0 \| 1.229e-48 \| 1279 \| 0.261 \| 670 \| 425 \| 26 \| 7 \| 637 \| 1 \| 639 \| Bacteriophage tail assembly protein \| Bacteriophage tail assembly protein; afdb-uniprot50 \| AF-A0A2P1VF79-F1-MODEL\_V4 \| 1.0 \| 2.744e-51 \| 1279 \| 0.246 \| 723 \| 462 \| 26 \| 3 \| 678 \| 1 \| 687 \| Terminase \| Terminase; afdb-uniprot50 \| AF-A0A2E3DCF8-F1-MODEL\_V4 \| 1.0 \| 2.083e-43 \| 1278 \| 0.211 \| 779 \| 469 \| 32 \| 2 \| 674 \| 79 \| 817 \| Uncharacterized protein \| Uncharacterized protein; afdb-uniprot50 \| AF-A0A3C1INY8-F1-MODEL\_V4 \| 1.0 \| 3.389e-49 \| 1277 \| 0.236 \| 661 \| 437 \| 26 \| 1 \| 632 \| 1 \| 622 \| Terminase \| Terminase; afdb-uniprot50 \| AF-A0A7C5KPE2-F1-MODEL\_V4 \| 1.0 \| 9.347e-50 \| 1277 \| 0.248 \| 713 \| 431 \| 32 \| 1 \| 679 \| 1 \| 642 \| Uncharacterized protein \| Uncharacterized protein; afdb-uniprot50 \| AF-A0A4Y1MW56-F1-MODEL\_V4 \| 1.0 \| 3.184e-48 \| 1275 \| 0.254 \| 645 \| 405 \| 25 \| 7 \| 616 \| 1 \| 604 \| Terminase large subunit \| Terminase large subunit; afdb-uniprot50 \| AF-A0A5E7QHL0-F1-MODEL\_V4 \| 1.0 \| 1.878e-53 \| 1274 \| 0.276 \| 670 \| 387 \| 20 \| 75 \| 681 \| 2 \| 636 \| Uncharacterized protein \| Uncharacterized protein; afdb-uniprot50 \| AF-G7ZCA7-F1-MODEL\_V4 \| 1.0 \| 3.03e-49 \| 1274 \| 0.226 \| 688 \| 459 \| 24 \| 14 \| 678 \| 2 \| 638 \| Terminase large subunit (DNA packaging protein A) (GpA) \| Terminase large subunit (DNA packaging protein A) (GpA); afdb-uniprot50 \| AF-A0A5E8H1L7-F1-MODEL\_V4 \| 1.0 \| 2.047e-49 \| 1272 \| 0.274 \| 638 \| 404 \| 24 \| 3 \| 615 \| 1 \| 604 \| Bacteriophage tail assembly protein \| Bacteriophage tail assembly protein; afdb-uniprot50 \| AF-F4V0R1-F1-MODEL\_V4 \| 1.0 \| 4.37e-45 \| 1272 \| 0.244 \| 663 \| 417 \| 30 \| 19 \| 633 \| 8 \| 634 \| Phage terminase large subunit (GpA) \| Phage terminase large subunit (GpA); afdb-uniprot50 \| AF-A0A5C8IZ84-F1-MODEL\_V4 \| 1.0 \| 4.268e-50 \| 1272 \| 0.24 \| 708 \| 440 \| 26 \| 14 \| 677 \| 2 \| 655 \| Phage terminase large subunit family protein \| Phage terminase large subunit family protein; afdb-uniprot50 \| AF-A0A5B8BZZ0-F1-MODEL\_V4 \| 1.0 \| 1.185e-51 \| 1272 \| 0.244 \| 741 \| 435 \| 35 \| 3 \| 677 \| 1 \| 682 \| Terminase \| Terminase; afdb-uniprot50 \| AF-A0A1A9R818-F1-MODEL\_V4 \| 1.0 \| 1.237e-49 \| 1270 \| 0.278 \| 643 \| 371 \| 25 \| 43 \| 632 \| 2 \| 604 \| Terminase \| Terminase; afdb-uniprot50 \| AF-A0A6V7DRP3-F1-MODEL\_V4 \| 1.0 \| 8.781e-49 \| 1270 \| 0.261 \| 680 \| 396 \| 25 \| 7 \| 615 \| 1 \| 644 \| Uncharacterized protein \| Uncharacterized protein; afdb-uniprot50 \| AF-A0A827AJ43-F1-MODEL\_V4 \| 1.0 \| 2.217e-44 \| 1269 \| 0.261 \| 592 \| 363 \| 18 \| 67 \| 614 \| 2 \| 563 \| Phage terminase large subunit family protein \| Phage terminase large subunit family protein; afdb-uniprot50 \| AF-A0A1V0BGV6-F1-MODEL\_V4 \| 1.0 \| 3.389e-49 \| 1268 \| 0.257 \| 622 \| 391 \| 21 \| 13 \| 589 \| 2 \| 597 \| Uncharacterized protein \| Uncharacterized protein; afdb-uniprot50 \| AF-S6SLZ1-F1-MODEL\_V4 \| 1.0 \| 2.275e-48 \| 1267 \| 0.294 \| 578 \| 346 \| 22 \| 88 \| 633 \| 2 \| 549 \| Terminase large subunit \| Terminase large subunit; afdb-uniprot50 \| AF-D1Y313-F1-MODEL\_V4 \| 1.0 \| 1.444e-47 \| 1266 \| 0.218 \| 635 \| 425 \| 24 \| 1 \| 605 \| 8 \| 600 \| Phage terminase large subunit (GpA) \| Phage terminase large subunit (GpA); afdb-uniprot50 \| AF-M3AEW4-F1-MODEL\_V4 \| 1.0 \| 6.384e-43 \| 1266 \| 0.222 \| 750 \| 443 \| 36 \| 3 \| 679 \| 1 \| 683 \| Terminase large subunit \| Terminase large subunit; afdb-uniprot50 \| AF-A0A840FD40-F1-MODEL\_V4 \| 1.0 \| 1.969e-43 \| 1266 \| 0.203 \| 738 \| 469 \| 37 \| 1 \| 677 \| 1 \| 680 \| Phage terminase large subunit GpA-like protein \| Phage terminase large subunit GpA-like protein; afdb-uniprot50 \| AF-A0A124UXC3-F1-MODEL\_V4 \| 1.0 \| 1.982e-44 \| 1265 \| 0.302 \| 529 \| 318 \| 17 \| 42 \| 543 \| 2 \| 506 \| Uncharacterized protein \| Uncharacterized protein; afdb-uniprot50 \| AF-A0A8B2PFX3-F1-MODEL\_V4 \| 1.0 \| 1.177e-50 \| 1261 \| 0.257 \| 766 \| 423 \| 28 \| 13 \| 677 \| 2 \| 722 \| Uncharacterized protein \| Uncharacterized protein; afdb-uniprot50 \| AF-W0BQE7-F1-MODEL\_V4 \| 1.0 \| 2.101e-53 \| 1260 \| 0.299 \| 638 \| 364 \| 17 \| 75 \| 644 \| 3 \| 625 \| Uncharacterized protein \| Uncharacterized protein; afdb-uniprot50 \| AF-A0A1Y4I4D2-F1-MODEL\_V4 \| 1.0 \| 1.357e-46 \| 1259 \| 0.223 \| 604 \| 415 \| 22 \| 25 \| 614 \| 4 \| 567 \| Uncharacterized protein \| Uncharacterized protein; afdb-uniprot50 \| AF-A0A4R3A929-F1-MODEL\_V4 \| 1.0 \| 4.323e-52 \| 1255 \| 0.268 \| 677 \| 387 \| 24 \| 75 \| 679 \| 3 \| 642 \| Phage terminase large subunit GpA-like protein \| Phage terminase large subunit GpA-like protein; afdb-uniprot50 \| AF-A0A531A524-F1-MODEL\_V4 \| 1.0 \| 7.471e-50 \| 1253 \| 0.243 \| 695 \| 421 \| 26 \| 7 \| 625 \| 1 \| 666 \| Terminase \| Terminase; afdb-uniprot50 \| AF-A0A1Y5TUH2-F1-MODEL\_V4 \| 1.0 \| 1.348e-45 \| 1252 \| 0.302 \| 556 \| 336 \| 17 \| 1 \| 540 \| 7 \| 526 \| Phage terminase large subunit (GpA) \| Phage terminase large subunit (GpA); afdb-uniprot50 \| AF-A0A3B9ZHZ9-F1-MODEL\_V4 \| 1.0 \| 7.851e-49 \| 1252 \| 0.217 \| 703 \| 445 \| 29 \| 5 \| 679 \| 1 \| 626 \| Terminase \| Terminase; afdb-uniprot50 \| AF-A0A377BJ29-F1-MODEL\_V4 \| 1.0 \| 1.174e-41 \| 1252 \| 0.264 \| 589 \| 346 \| 24 \| 87 \| 623 \| 2 \| 554 \| Phage terminase large subunit \| Phage terminase large subunit; afdb-uniprot50 \| AF-A0A1H3IRJ0-F1-MODEL\_V4 \| 1.0 \| 2.629e-53 \| 1250 \| 0.333 \| 612 \| 329 \| 17 \| 97 \| 682 \| 329 \| 887 \| Phage terminase, large subunit GpA \| Phage terminase, large subunit GpA; afdb-uniprot50 \| AF-G9PUH4-F1-MODEL\_V4 \| 1.0 \| 1.72e-48 \| 1248 \| 0.243 \| 645 \| 399 \| 31 \| 1 \| 604 \| 4 \| 600 \| Uncharacterized protein \| Uncharacterized protein; afdb-uniprot50 \| AF-A0A191WAA1-F1-MODEL\_V4 \| 1.0 \| 3.239e-42 \| 1248 \| 0.194 \| 657 \| 430 \| 25 \| 24 \| 605 \| 18 \| 649 \| DNA packaging protein \| DNA packaging protein; afdb-uniprot50 \| AF-A0A540VA48-F1-MODEL\_V4 \| 1.0 \| 2.246e-46 \| 1247 \| 0.292 \| 647 \| 360 \| 28 \| 2 \| 603 \| 1 \| 594 \| Uncharacterized protein \| Uncharacterized protein; afdb-uniprot50 \| AF-A0A076LPJ9-F1-MODEL\_V4 \| 1.0 \| 7.569e-52 \| 1244 \| 0.295 \| 622 \| 359 \| 17 \| 75 \| 632 \| 3 \| 608 \| Phage terminase large subunit (GpA) \| Phage terminase large subunit (GpA); afdb-uniprot50 \| AF-A0A2H4TZ33-F1-MODEL\_V4 \| 1.0 \| 2.612e-52 \| 1243 \| 0.295 \| 629 \| 358 \| 17 \| 75 \| 632 \| 3 \| 617 \| Uncharacterized protein \| Uncharacterized protein; afdb-uniprot50 \| AF-A0A1U7MFY3-F1-MODEL\_V4 \| 1.0 \| 1.615e-47 \| 1240 \| 0.226 \| 622 \| 406 \| 26 \| 39 \| 632 \| 1 \| 575 \| Phage terminase large subunit GpA \| Phage terminase large subunit GpA; afdb-uniprot50 \| AF-A0A7Y9NBJ2-F1-MODEL\_V4 \| 1.0 \| 2.865e-49 \| 1240 \| 0.246 \| 756 \| 428 \| 30 \| 1 \| 681 \| 1 \| 689 \| Phage terminase large subunit GpA-like protein \| Phage terminase large subunit GpA-like protein; afdb-uniprot50 \| AF-A0A7Z3TXV8-F1-MODEL\_V4 \| 1.0 \| 2.709e-49 \| 1239 \| 0.23 \| 681 \| 436 \| 27 \| 7 \| 633 \| 1 \| 647 \| Phage terminase large subunit family protein \| Phage terminase large subunit family protein; afdb-uniprot50 \| AF-A0A0S2F4Y2-F1-MODEL\_V4 \| 1.0 \| 4.743e-49 \| 1239 \| 0.311 \| 613 \| 335 \| 23 \| 98 \| 681 \| 446 \| 1000 \| Phage terminase large subunit family protein \| Phage terminase large subunit family protein; afdb-uniprot50 \| AF-B8IDR3-F1-MODEL\_V4 \| 1.0 \| 1.106e-49 \| 1238 \| 0.253 \| 741 \| 438 \| 30 \| 3 \| 676 \| 1 \| 692 \| Terminase GpA \| Terminase GpA; afdb-uniprot50 \| AF-A0A249MUF3-F1-MODEL\_V4 \| 1.0 \| 7.552e-43 \| 1237 \| 0.21 \| 693 \| 440 \| 36 \| 1 \| 634 \| 2 \| 646 \| Uncharacterized protein \| Uncharacterized protein; afdb-uniprot50 \| AF-A0A2P7RZY3-F1-MODEL\_V4 \| 1.0 \| 7.503e-42 \| 1236 \| 0.199 \| 753 \| 442 \| 34 \| 22 \| 679 \| 21 \| 707 \| Uncharacterized protein \| Uncharacterized protein; afdb-uniprot50 \| AF-A0A0S8GHT9-F1-MODEL\_V4 \| 1.0 \| 3.983e-48 \| 1235 \| 0.202 \| 722 \| 457 \| 25 \| 3 \| 680 \| 1 \| 647 \| Uncharacterized protein \| Uncharacterized protein; afdb-uniprot50 \| AF-A0A355TC58-F1-MODEL\_V4 \| 1.0 \| 7.733e-38 \| 1234 \| 0.332 \| 445 \| 262 \| 13 \| 18 \| 450 \| 2 \| 423 \| Phage tail protein \| Phage tail protein; afdb-uniprot50 \| AF-D8F5N8-F1-MODEL\_V4 \| 1.0 \| 6.194e-47 \| 1231 \| 0.236 \| 601 \| 414 \| 20 \| 24 \| 599 \| 5 \| 585 \| Phage terminase large subunit (GpA) \| Phage terminase large subunit (GpA); afdb-uniprot50 \| AF-A0A8B2T473-F1-MODEL\_V4 \| 1.0 \| 1.665e-43 \| 1229 \| 0.226 \| 636 \| 400 \| 27 \| 74 \| 681 \| 2 \| 573 \| Terminase \| Terminase; afdb-uniprot50 \| AF-A0A530QYK9-F1-MODEL\_V4 \| 1.0 \| 2.133e-38 \| 1226 \| 0.318 \| 449 \| 270 \| 12 \| 26 \| 457 \| 2 \| 431 \| Phage terminase large subunit family protein \| Phage terminase large subunit family protein; afdb-uniprot50 \| AF-A0A147HZ17-F1-MODEL\_V4 \| 1.0 \| 1.059e-51 \| 1226 \| 0.294 \| 670 \| 368 \| 17 \| 75 \| 680 \| 3 \| 631 \| Terminase \| Terminase; afdb-uniprot50 \| AF-A0A6M0JYU4-F1-MODEL\_V4 \| 1.0 \| 5.574e-48 \| 1224 \| 0.237 \| 706 \| 429 \| 28 \| 20 \| 677 \| 7 \| 650 \| Uncharacterized protein \| Uncharacterized protein; afdb-uniprot50 \| AF-A0A518LAV4-F1-MODEL\_V4 \| 1.0 \| 1.057e-42 \| 1224 \| 0.191 \| 712 \| 469 \| 30 \| 9 \| 681 \| 2 \| 645 \| Phage terminase large subunit (GpA) \| Phage terminase large subunit (GpA); afdb-uniprot50 \| AF-A0A2G6EYH6-F1-MODEL\_V4 \| 1.0 \| 4.258e-41 \| 1223 \| 0.3 \| 492 \| 313 \| 12 \| 20 \| 503 \| 10 \| 478 \| Terminase \| Terminase; afdb-uniprot50 \| AF-A0A520BGZ4-F1-MODEL\_V4 \| 1.0 \| 3.584e-49 \| 1223 \| 0.239 \| 705 \| 444 \| 29 \| 17 \| 675 \| 5 \| 662 \| Uncharacterized protein \| Uncharacterized protein; afdb-uniprot50 \| AF-A0A4Q3K4E2-F1-MODEL\_V4 \| 1.0 \| 9.509e-44 \| 1222 \| 0.263 \| 595 \| 357 \| 20 \| 7 \| 551 \| 1 \| 563 \| Phage terminase large subunit family protein \| Phage terminase large subunit family protein; afdb-uniprot50 \| AF-A0A2G2LIL6-F1-MODEL\_V4 \| 1.0 \| 1.159e-39 \| 1217 \| 0.211 \| 625 \| 394 \| 27 \| 50 \| 604 \| 2 \| 597 \| Terminase \| Terminase; afdb-uniprot50 \| AF-A0A7Z8LJQ9-F1-MODEL\_V4 \| 1.0 \| 4.009e-49 \| 1216 \| 0.229 \| 737 \| 454 \| 26 \| 18 \| 681 \| 28 \| 723 \| Phage terminase large subunit GpA-like protein \| Phage terminase large subunit GpA-like protein; afdb-uniprot50 \| AF-A0A0D0Q7Y7-F1-MODEL\_V4 \| 1.0 \| 3.382e-40 \| 1213 \| 0.294 \| 476 \| 293 \| 13 \| 7 \| 473 \| 1 \| 442 \| Bacteriophage tail assembly protein \| Bacteriophage tail assembly protein; afdb-uniprot50 \| AF-A0A1V6E218-F1-MODEL\_V4 \| 1.0 \| 3.766e-48 \| 1213 \| 0.253 \| 635 \| 398 \| 27 \| 37 \| 634 \| 2 \| 597 \| Phage terminase large subunit (GpA) \| Phage terminase large subunit (GpA); afdb-uniprot50 \| AF-A0A291AQS8-F1-MODEL\_V4 \| 1.0 \| 2.709e-49 \| 1212 \| 0.271 \| 675 \| 421 \| 23 \| 51 \| 679 \| 1 \| 650 \| Terminase \| Terminase; afdb-uniprot50 \| AF-D2Z2A1-F1-MODEL\_V4 \| 1.0 \| 1.911e-47 \| 1211 \| 0.234 \| 635 \| 428 \| 28 \| 30 \| 634 \| 10 \| 616 \| Terminase GpA \| Terminase GpA; afdb-uniprot50 \| AF-A0A1H3FGF1-F1-MODEL\_V4 \| 1.0 \| 2.261e-47 \| 1211 \| 0.229 \| 681 \| 453 \| 25 \| 24 \| 682 \| 2 \| 632 \| Phage terminase, large subunit GpA \| Phage terminase, large subunit GpA; afdb-uniprot50 \| AF-A0A2A2R1I2-F1-MODEL\_V4 \| 1.0 \| 5.037e-41 \| 1210 \| 0.207 \| 645 \| 435 \| 31 \| 1 \| 615 \| 6 \| 604 \| Uncharacterized protein \| Uncharacterized protein; afdb-uniprot50 \| AF-A0A1U9NPU3-F1-MODEL\_V4 \| 1.0 \| 1.213e-46 \| 1208 \| 0.224 \| 668 \| 450 \| 25 \| 1 \| 636 \| 3 \| 634 \| Bacteriophage tail assembly protein \| Bacteriophage tail assembly protein; afdb-uniprot50 \| AF-U2DIH0-F1-MODEL\_V4 \| 1.0 \| 6.384e-43 \| 1206 \| 0.228 \| 560 \| 383 \| 21 \| 10 \| 555 \| 2 \| 526 \| Phage terminase large subunit \| Phage terminase large subunit; afdb-uniprot50 \| AF-A0A6L9MJI6-F1-MODEL\_V4 \| 1.0 \| 4.825e-43 \| 1204 \| 0.255 \| 537 \| 354 \| 16 \| 14 \| 531 \| 2 \| 511 \| Terminase \| Terminase; afdb-uniprot50 \| AF-A0A835YX25-F1-MODEL\_V4 \| 1.0 \| 7.851e-49 \| 1204 \| 0.278 \| 640 \| 393 \| 18 \| 20 \| 632 \| 363 \| 960 \| Phage terminase large subunit-domain-containing protein \| Phage terminase large subunit-domain-containing protein; afdb-uniprot50 \| AF-A0A753AN12-F1-MODEL\_V4 \| 1.0 \| 6.884e-46 \| 1203 \| 0.261 \| 605 \| 374 \| 19 \| 44 \| 607 \| 3 \| 575 \| DUF1983 domain-containing protein \| DUF1983 domain-containing protein; afdb-uniprot50 \| AF-A0A7W6JZH5-F1-MODEL\_V4 \| 1.0 \| 1.819e-48 \| 1200 \| 0.22 \| 762 \| 457 \| 31 \| 3 \| 679 \| 1 \| 710 \| Phage terminase large subunit GpA-like protein \| Phage terminase large subunit GpA-like protein; afdb-uniprot50 \| AF-A0A521CPV3-F1-MODEL\_V4 \| 1.0 \| 2.496e-45 \| 1197 \| 0.216 \| 628 \| 429 \| 29 \| 27 \| 632 \| 6 \| 592 \| Phage terminase, large subunit GpA \| Phage terminase, large subunit GpA; afdb-uniprot50 \| AF-A0A7X7X896-F1-MODEL\_V4 \| 1.0 \| 9.696e-47 \| 1197 \| 0.234 \| 692 \| 416 \| 28 \| 39 \| 681 \| 45 \| 671 \| Terminase \| Terminase; afdb-uniprot50 \| AF-A0A518HJ51-F1-MODEL\_V4 \| 1.0 \| 4.258e-41 \| 1196 \| 0.182 \| 723 \| 477 \| 31 \| 1 \| 681 \| 1 \| 651 \| Phage terminase large subunit (GpA) \| Phage terminase large subunit (GpA); afdb-uniprot50 \| AF-A0A2S7YYZ0-F1-MODEL\_V4 \| 1.0 \| 1.357e-46 \| 1195 \| 0.216 \| 697 \| 445 \| 28 \| 1 \| 680 \| 15 \| 627 \| Terminase \| Terminase; afdb-uniprot50 \| AF-A0A853ND71-F1-MODEL\_V4 \| 1.0 \| 2.124e-46 \| 1195 \| 0.249 \| 662 \| 388 \| 28 \| 39 \| 638 \| 7 \| 621 \| Terminase \| Terminase; afdb-uniprot50 \| AF-A0A520YB68-F1-MODEL\_V4 \| 1.0 \| 1.911e-47 \| 1195 \| 0.218 \| 665 \| 437 \| 28 \| 2 \| 616 \| 8 \| 639 \| Uncharacterized protein \| Uncharacterized protein; afdb-uniprot50 \| AF-A0A1M7RIC4-F1-MODEL\_V4 \| 1.0 \| 7.359e-39 \| 1190 \| 0.278 \| 482 \| 303 \| 13 \| 2 \| 470 \| 1 \| 450 \| Phage terminase, large subunit GpA \| Phage terminase, large subunit GpA; afdb-uniprot50 \| AF-A0A6B3RJA5-F1-MODEL\_V4 \| 1.0 \| 6.275e-49 \| 1189 \| 0.245 \| 738 \| 431 \| 27 \| 5 \| 682 \| 1 \| 672 \| Terminase \| Terminase; afdb-uniprot50 \| AF-A0A1G3QLG8-F1-MODEL\_V4 \| 1.0 \| 1.092e-47 \| 1188 \| 0.239 \| 642 \| 395 \| 28 \| 17 \| 604 \| 3 \| 604 \| Uncharacterized protein \| Uncharacterized protein; afdb-uniprot50 \| AF-A0A1Y5U4W6-F1-MODEL\_V4 \| 1.0 \| 3.01e-48 \| 1187 \| 0.231 \| 740 \| 464 \| 36 \| 4 \| 682 \| 1 \| 696 \| Phage terminase large subunit (GpA) \| Phage terminase large subunit (GpA); afdb-uniprot50 \| AF-A0A833CCD2-F1-MODEL\_V4 \| 1.0 \| 5.502e-46 \| 1184 \| 0.231 \| 592 \| 408 \| 22 \| 54 \| 632 \| 5 \| 562 \| Phage terminase large subunit family protein \| Phage terminase large subunit family protein; afdb-uniprot50 \| AF-A0A0F8ZCZ7-F1-MODEL\_V4 \| 1.0 \| 8.392e-42 \| 1182 \| 0.338 \| 455 \| 259 \| 8 \| 225 \| 675 \| 1 \| 417 \| Uncharacterized protein \| Uncharacterized protein; afdb-uniprot50 \| AF-A0A560F1W1-F1-MODEL\_V4 \| 1.0 \| 3.561e-48 \| 1180 \| 0.228 \| 726 \| 459 \| 24 \| 1 \| 681 \| 1 \| 670 \| Phage terminase large subunit GpA-like protein \| Phage terminase large subunit GpA-like protein; afdb-uniprot50 \| AF-A0A1R1INW1-F1-MODEL\_V4 \| 1.0 \| 2.008e-46 \| 1180 \| 0.228 \| 708 \| 434 \| 33 \| 14 \| 632 \| 2 \| 685 \| Uncharacterized protein \| Uncharacterized protein; afdb-uniprot50 \| AF-A0A4P8IMH6-F1-MODEL\_V4 \| 1.0 \| 1.291e-47 \| 1179 \| 0.252 \| 685 \| 412 \| 28 \| 6 \| 620 \| 1 \| 655 \| Phage terminase large subunit family protein \| Phage terminase large subunit family protein; afdb-uniprot50 \| AF-A0A833GAI6-F1-MODEL\_V4 \| 1.0 \| 1.05e-41 \| 1176 \| 0.249 \| 546 \| 364 \| 17 \| 1 \| 525 \| 44 \| 564 \| Uncharacterized protein \| Uncharacterized protein; afdb-uniprot50 \| AF-A0A2W5LM25-F1-MODEL\_V4 \| 1.0 \| 5.857e-47 \| 1175 \| 0.251 \| 745 \| 437 \| 28 \| 6 \| 682 \| 1 \| 692 \| Terminase \| Terminase; afdb-uniprot50 \| AF-G0EQQ3-F1-MODEL\_V4 \| 1.0 \| 3.515e-46 \| 1174 \| 0.213 \| 651 \| 409 \| 32 \| 39 \| 616 \| 8 \| 628 \| Putative large terminase subunit \| Putative large terminase subunit; afdb-uniprot50 \| AF-A0A0F6MR09-F1-MODEL\_V4 \| 1.0 \| 1.899e-46 \| 1173 \| 0.216 \| 669 \| 428 \| 29 \| 18 \| 615 \| 6 \| 649 \| Uncharacterized protein \| Uncharacterized protein; afdb-uniprot50 \| AF-A0A7A6RPU2-F1-MODEL\_V4 \| 1.0 \| 1.697e-46 \| 1170 \| 0.262 \| 643 \| 381 \| 30 \| 39 \| 622 \| 1 \| 608 \| Phage terminase large subunit family protein \| Phage terminase large subunit family protein; afdb-uniprot50 \| AF-M5SXL0-F1-MODEL\_V4 \| 1.0 \| 3.218e-41 \| 1170 \| 0.192 \| 753 \| 465 \| 36 \| 3 \| 676 \| 1 \| 689 \| Phage terminase GpA \| Phage terminase GpA; afdb-uniprot50 \| AF-A0A7K3NN95-F1-MODEL\_V4 \| 1.0 \| 2.11e-45 \| 1168 \| 0.223 \| 617 \| 403 \| 30 \| 25 \| 605 \| 46 \| 622 \| Uncharacterized protein \| Uncharacterized protein; afdb-uniprot50 \| AF-A0A679GM58-F1-MODEL\_V4 \| 1.0 \| 9.108e-46 \| 1163 \| 0.241 \| 647 \| 409 \| 23 \| 39 \| 633 \| 1 \| 617 \| Terminase \| Terminase; afdb-uniprot50 \| AF-A0A537MC64-F1-MODEL\_V4 \| 1.0 \| 3.53e-38 \| 1161 \| 0.189 \| 670 \| 453 \| 33 \| 1 \| 615 \| 11 \| 645 \| Uncharacterized protein \| Uncharacterized protein; afdb-uniprot50 \| AF-A0A7V7NR71-F1-MODEL\_V4 \| 1.0 \| 2.36e-45 \| 1160 \| 0.268 \| 648 \| 387 \| 26 \| 39 \| 633 \| 7 \| 620 \| DNA packaging protein \| DNA packaging protein; afdb-uniprot50 \| AF-A4U1Y6-F1-MODEL\_V4 \| 1.0 \| 7.503e-42 \| 1157 \| 0.389 \| 429 \| 223 \| 14 \| 223 \| 640 \| 3 \| 403 \| Phage terminase GpA \| Phage terminase GpA; afdb-uniprot50 \| AF-A0A316NZJ1-F1-MODEL\_V4 \| 1.0 \| 9.05e-45 \| 1156 \| 0.243 \| 628 \| 411 \| 24 \| 19 \| 604 \| 7 \| 612 \| Uncharacterized protein \| Uncharacterized protein; afdb-uniprot50 \| AF-A0A7W7NR19-F1-MODEL\_V4 \| 1.0 \| 5.857e-47 \| 1156 \| 0.25 \| 726 \| 436 \| 35 \| 3 \| 681 \| 1 \| 665 \| Phage terminase large subunit GpA-like protein \| Phage terminase large subunit GpA-like protein; afdb-uniprot50 \| AF-A0A290MQW5-F1-MODEL\_V4 \| 1.0 \| 5.819e-46 \| 1156 \| 0.226 \| 754 \| 437 \| 30 \| 21 \| 677 \| 14 \| 717 \| Uncharacterized protein \| Uncharacterized protein; afdb-uniprot50 \| AF-A0A7W1G3V0-F1-MODEL\_V4 \| 1.0 \| 7.7e-46 \| 1154 \| 0.219 \| 651 \| 434 \| 31 \| 37 \| 634 \| 2 \| 631 \| Phage terminase large subunit family protein \| Phage terminase large subunit family protein; afdb-uniprot50 \| AF-A0A021X9J1-F1-MODEL\_V4 \| 1.0 \| 6.929e-47 \| 1153 \| 0.288 \| 611 \| 373 \| 20 \| 88 \| 678 \| 2 \| 570 \| Bacteriophage tail assembly protein \| Bacteriophage tail assembly protein; afdb-uniprot50 \| AF-A0A518B2T0-F1-MODEL\_V4 \| 1.0 \| 1.416e-44 \| 1153 \| 0.262 \| 621 \| 378 \| 24 \| 55 \| 632 \| 8 \| 591 \| Phage terminase large subunit (GpA) \| Phage terminase large subunit (GpA); afdb-uniprot50 \| AF-A0A1E5NH26-F1-MODEL\_V4 \| 1.0 \| 1.995e-45 \| 1153 \| 0.209 \| 679 \| 446 \| 31 \| 9 \| 619 \| 13 \| 668 \| Uncharacterized protein \| Uncharacterized protein; afdb-uniprot50 \| AF-A0A496X5L8-F1-MODEL\_V4 \| 1.0 \| 1.084e-46 \| 1153 \| 0.216 \| 659 \| 422 \| 28 \| 19 \| 604 \| 2 \| 638 \| Terminase \| Terminase; afdb-uniprot50 \| AF-A0A255XTH6-F1-MODEL\_V4 \| 1.0 \| 1.518e-46 \| 1153 \| 0.221 \| 735 \| 446 \| 35 \| 24 \| 682 \| 10 \| 693 \| Uncharacterized protein \| Uncharacterized protein; afdb-uniprot50 \| AF-A0A0Q4Z1A9-F1-MODEL\_V4 \| 1.0 \| 3.807e-41 \| 1153 \| 0.209 \| 779 \| 475 \| 40 \| 3 \| 682 \| 1 \| 737 \| Uncharacterized protein \| Uncharacterized protein; afdb-uniprot50 \| AF-A0A1J5JMJ2-F1-MODEL\_V4 \| 1.0 \| 4.417e-38 \| 1149 \| 0.306 \| 486 \| 283 \| 13 \| 4 \| 465 \| 1 \| 456 \| Phage tail protein \| Phage tail protein; afdb-uniprot50 \| AF-B8ICM2-F1-MODEL\_V4 \| 1.0 \| 1.795e-46 \| 1149 \| 0.228 \| 718 \| 452 \| 30 \| 10 \| 679 \| 2 \| 665 \| Terminase GpA \| Terminase GpA; afdb-uniprot50 \| AF-A0A1L8CRQ1-F1-MODEL\_V4 \| 1.0 \| 8.374e-33 \| 1148 \| 0.404 \| 319 \| 163 \| 7 \| 332 \| 637 \| 2 \| 306 \| Terminase \| Terminase; afdb-uniprot50 \| AF-A0A1Y5U0V6-F1-MODEL\_V4 \| 1.0 \| 1.147e-46 \| 1148 \| 0.237 \| 677 \| 416 \| 32 \| 1 \| 610 \| 2 \| 644 \| Phage terminase large subunit (GpA) \| Phage terminase large subunit (GpA); afdb-uniprot50 \| AF-A0A6M3J2A4-F1-MODEL\_V4 \| 1.0 \| 8.991e-44 \| 1146 \| 0.217 \| 662 \| 422 \| 26 \| 22 \| 634 \| 2 \| 616 \| Putative terminase \| Putative terminase; afdb-uniprot50 \| AF-A0A0H3A7D8-F1-MODEL\_V4 \| 1.0 \| 2.496e-45 \| 1145 \| 0.221 \| 618 \| 410 \| 27 \| 25 \| 605 \| 24 \| 607 \| Phage terminase GpA \| Phage terminase GpA; afdb-uniprot50 \| AF-A0A2G4KD28-F1-MODEL\_V4 \| 1.0 \| 2.375e-46 \| 1143 \| 0.23 \| 765 \| 444 \| 35 \| 3 \| 681 \| 1 \| 706 \| Uncharacterized protein \| Uncharacterized protein; afdb-uniprot50 \| AF-X6D541-F1-MODEL\_V4 \| 1.0 \| 1.705e-38 \| 1141 \| 0.254 \| 507 \| 330 \| 17 \| 7 \| 497 \| 1 \| 475 \| Uncharacterized protein \| Uncharacterized protein; afdb-uniprot50 \| AF-A0A0Q4XH76-F1-MODEL\_V4 \| 1.0 \| 1.899e-46 \| 1139 \| 0.218 \| 723 \| 459 \| 23 \| 10 \| 677 \| 2 \| 673 \| Terminase \| Terminase; afdb-uniprot50 \| AF-R7UZY0-F1-MODEL\_V4 \| 1.0 \| 2.03e-39 \| 1138 \| 0.317 \| 429 \| 263 \| 10 \| 76 \| 497 \| 6 \| 411 \| Uncharacterized protein \| Uncharacterized protein; afdb-uniprot50 \| AF-E1JR86-F1-MODEL\_V4 \| 1.0 \| 2.375e-46 \| 1138 \| 0.199 \| 693 \| 442 \| 32 \| 25 \| 681 \| 53 \| 668 \| Terminase GpA \| Terminase GpA; afdb-uniprot50 \| AF-A0A2G9X118-F1-MODEL\_V4 \| 1.0 \| 1.043e-40 \| 1138 \| 0.213 \| 720 \| 457 \| 35 \| 34 \| 682 \| 2 \| 682 \| Uncharacterized protein \| Uncharacterized protein; afdb-uniprot50 \| AF-A0A7W0C9U2-F1-MODEL\_V4 \| 1.0 \| 1.508e-45 \| 1137 \| 0.225 \| 638 \| 411 \| 31 \| 2 \| 597 \| 1 \| 597 \| Phage terminase large subunit GpA-like protein \| Phage terminase large subunit GpA-like protein; afdb-uniprot50 \| AF-A0A156IAQ8-F1-MODEL\_V4 \| 1.0 \| 3.907e-45 \| 1136 \| 0.235 \| 600 \| 391 \| 25 \| 68 \| 632 \| 10 \| 576 \| Phage terminase, large subunit \| Phage terminase, large subunit; afdb-uniprot50 \| AF-A0A212KXJ4-F1-MODEL\_V4 \| 1.0 \| 1.357e-46 \| 1136 \| 0.239 \| 711 \| 453 \| 31 \| 4 \| 678 \| 1 \| 659 \| Phage terminase large subunit \| Phage terminase large subunit; afdb-uniprot50 \| AF-A0A660W1Z2-F1-MODEL\_V4 \| 1.0 \| 1.14e-45 \| 1136 \| 0.195 \| 684 \| 447 \| 30 \| 22 \| 632 \| 5 \| 657 \| Uncharacterized protein \| Uncharacterized protein; afdb-uniprot50 \| AF-A0A2T5K4X7-F1-MODEL\_V4 \| 1.0 \| 6.723e-51 \| 1136 \| 0.26 \| 699 \| 394 \| 20 \| 80 \| 681 \| 9 \| 681 \| Phage terminase large subunit GpA-like protein \| Phage terminase large subunit GpA-like protein; afdb-uniprot50 \| AF-A0A4Q6FEN8-F1-MODEL\_V4 \| 1.0 \| 8.593e-37 \| 1133 \| 0.295 \| 464 \| 280 \| 16 \| 18 \| 460 \| 8 \| 445 \| Phage terminase large subunit family protein \| Phage terminase large subunit family protein; afdb-uniprot50 \| AF-F7XA70-F1-MODEL\_V4 \| 1.0 \| 4.621e-45 \| 1130 \| 0.22 \| 707 \| 447 \| 28 \| 21 \| 677 \| 7 \| 659 \| Phage terminase GpA \| Phage terminase GpA; afdb-uniprot50 \| AF-A0A1T4WWP7-F1-MODEL\_V4 \| 1.0 \| 4.203e-39 \| 1128 \| 0.217 \| 615 \| 412 \| 28 \| 31 \| 609 \| 13 \| 593 \| Phage terminase, large subunit GpA \| Phage terminase, large subunit GpA; afdb-uniprot50 \| AF-A0A2N9Y9A6-F1-MODEL\_V4 \| 1.0 \| 3.538e-47 \| 1125 \| 0.345 \| 561 \| 291 \| 19 \| 145 \| 678 \| 1 \| 512 \| Terminase \| Terminase; afdb-uniprot50 \| AF-A0A351WKF4-F1-MODEL\_V4 \| 1.0 \| 3.493e-45 \| 1125 \| 0.21 \| 698 \| 432 \| 29 \| 24 \| 679 \| 53 \| 673 \| Terminase \| Terminase; afdb-uniprot50 \| AF-A0A258VI65-F1-MODEL\_V4 \| 1.0 \| 3.907e-45 \| 1123 \| 0.213 \| 739 \| 470 \| 25 \| 1 \| 681 \| 2 \| 686 \| Uncharacterized protein \| Uncharacterized protein; afdb-uniprot50 \| AF-A0A5U8JIR8-F1-MODEL\_V4 \| 1.0 \| 2.137e-47 \| 1122 \| 0.202 \| 752 \| 479 \| 31 \| 1 \| 682 \| 2 \| 702 \| Phage terminase large subunit family protein \| Phage terminase large subunit family protein; afdb-uniprot50 \| AF-F7L006-F1-MODEL\_V4 \| 1.0 \| 7.633e-36 \| 1121 \| 0.273 \| 454 \| 300 \| 10 \| 6 \| 452 \| 1 \| 431 \| Phage terminase large subunit (GpA) \| Phage terminase large subunit (GpA); afdb-uniprot50 \| AF-A0A075KCW9-F1-MODEL\_V4 \| 1.0 \| 1.772e-44 \| 1119 \| 0.208 \| 624 \| 402 \| 27 \| 74 \| 681 \| 2 \| 549 \| Terminase GpA \| Terminase GpA; afdb-uniprot50 \| AF-A0A357ZWA1-F1-MODEL\_V4 \| 1.0 \| 3.923e-37 \| 1119 \| 0.195 \| 646 \| 414 \| 32 \| 2 \| 605 \| 1 \| 582 \| Uncharacterized protein \| Uncharacterized protein; afdb-uniprot50 \| AF-A0A1H5XMY4-F1-MODEL\_V4 \| 1.0 \| 8.781e-49 \| 1119 \| 0.26 \| 619 \| 373 \| 20 \| 98 \| 681 \| 469 \| 1037 \| Phage terminase, large subunit GpA \| Phage terminase, large subunit GpA; afdb-uniprot50 \| AF-A0A2D5TTG4-F1-MODEL\_V4 \| 1.0 \| 2.512e-46 \| 1118 \| 0.203 \| 706 \| 450 \| 27 \| 3 \| 632 \| 1 \| 669 \| Uncharacterized protein \| Uncharacterized protein; afdb-uniprot50 \| AF-A0A061NN40-F1-MODEL\_V4 \| 1.0 \| 8.021e-35 \| 1116 \| 0.289 \| 421 \| 261 \| 9 \| 22 \| 434 \| 4 \| 394 \| Phage terminase, large subunit \| Phage terminase, large subunit; afdb-uniprot50 \| AF-A0A2P2E5V7-F1-MODEL\_V4 \| 1.0 \| 4.132e-45 \| 1116 \| 0.212 \| 750 \| 461 \| 31 \| 4 \| 678 \| 1 \| 695 \| Uncharacterized protein \| Uncharacterized protein; afdb-uniprot50 \| AF-A0A258JF97-F1-MODEL\_V4 \| 1.0 \| 2.55e-31 \| 1113 \| 0.274 \| 386 \| 248 \| 11 \| 14 \| 394 \| 2 \| 360 \| Uncharacterized protein \| Uncharacterized protein; afdb-uniprot50 \| AF-A0A3S0WK41-F1-MODEL\_V4 \| 1.0 \| 1.043e-40 \| 1113 \| 0.252 \| 618 \| 383 \| 25 \| 95 \| 681 \| 6 \| 575 \| Terminase \| Terminase; afdb-uniprot50 \| AF-A0A1G0EIH4-F1-MODEL\_V4 \| 1.0 \| 2.512e-46 \| 1113 \| 0.278 \| 646 \| 382 \| 27 \| 82 \| 676 \| 2 \| 614 \| Uncharacterized protein \| Uncharacterized protein; afdb-uniprot50 \| AF-A0A5C7Q828-F1-MODEL\_V4 \| 1.0 \| 3.718e-46 \| 1111 \| 0.215 \| 701 \| 453 \| 33 \| 22 \| 675 \| 6 \| 656 \| Uncharacterized protein \| Uncharacterized protein; afdb-uniprot50 \| AF-A0A238KFY0-F1-MODEL\_V4 \| 1.0 \| 1.416e-44 \| 1107 \| 0.216 \| 765 \| 459 \| 37 \| 3 \| 678 \| 1 \| 713 \| Phage terminase large subunit (GpA) \| Phage terminase large subunit (GpA); afdb-uniprot50 \| AF-A0A1H2I406-F1-MODEL\_V4 \| 1.0 \| 1.676e-44 \| 1104 \| 0.222 \| 644 \| 419 \| 26 \| 4 \| 597 \| 6 \| 617 \| Phage terminase, large subunit GpA \| Phage terminase, large subunit GpA; afdb-uniprot50 \| AF-A0A7Y7I299-F1-MODEL\_V4 \| 1.0 \| 3.493e-45 \| 1104 \| 0.227 \| 695 \| 418 \| 30 \| 1 \| 621 \| 1 \| 650 \| Phage terminase large subunit GpA-like protein \| Phage terminase large subunit GpA-like protein; afdb-uniprot50 \| AF-A0A431JU41-F1-MODEL\_V4 \| 1.0 \| 3.281e-44 \| 1103 \| 0.217 \| 713 \| 442 \| 30 \| 10 \| 681 \| 1 \| 638 \| Uncharacterized protein \| Uncharacterized protein; afdb-uniprot50 \| AF-B6WS38-F1-MODEL\_V4 \| 1.0 \| 2.217e-44 \| 1102 \| 0.22 \| 648 \| 409 \| 34 \| 38 \| 632 \| 18 \| 622 \| Phage terminase large subunit (GpA) \| Phage terminase large subunit (GpA); afdb-uniprot50 \| AF-A0A5U4JN84-F1-MODEL\_V4 \| 1.0 \| 1.147e-46 \| 1102 \| 0.213 \| 751 \| 479 \| 31 \| 1 \| 679 \| 2 \| 712 \| Phage terminase large subunit family protein \| Phage terminase large subunit family protein; afdb-uniprot50 \| AF-A0A522MCR0-F1-MODEL\_V4 \| 1.0 \| 3.718e-46 \| 1101 \| 0.214 \| 746 \| 460 \| 31 \| 13 \| 682 \| 3 \| 698 \| Phage terminase large subunit family protein \| Phage terminase large subunit family protein; afdb-uniprot50 \| AF-A0A3D4DQL7-F1-MODEL\_V4 \| 1.0 \| 8.876e-42 \| 1099 \| 0.309 \| 542 \| 295 \| 19 \| 172 \| 681 \| 6 \| 499 \| Phage tail protein \| Phage tail protein; afdb-uniprot50 \| AF-A0A1V5D5G2-F1-MODEL\_V4 \| 1.0 \| 3.67e-44 \| 1099 \| 0.206 \| 711 \| 450 \| 32 \| 10 \| 679 \| 1 \| 637 \| Phage terminase large subunit (GpA) \| Phage terminase large subunit (GpA); afdb-uniprot50 \| AF-A0A3D6C8S5-F1-MODEL\_V4 \| 1.0 \| 2.953e-45 \| 1099 \| 0.2 \| 733 \| 466 \| 34 \| 1 \| 681 \| 12 \| 676 \| Terminase \| Terminase; afdb-uniprot50 \| AF-G5NAD3-F1-MODEL\_V4 \| 1.0 \| 9.509e-44 \| 1096 \| 0.338 \| 479 \| 276 \| 15 \| 172 \| 632 \| 9 \| 464 \| Phage terminase large subunit \| Phage terminase large subunit; afdb-uniprot50 \| AF-A0A7C1NNI0-F1-MODEL\_V4 \| 1.0 \| 1.749e-42 \| 1094 \| 0.212 \| 659 \| 421 \| 30 \| 10 \| 608 \| 1 \| 621 \| Uncharacterized protein \| Uncharacterized protein; afdb-uniprot50 \| AF-C7G5Z2-F1-MODEL\_V4 \| 1.0 \| 3.232e-33 \| 1092 \| 0.281 \| 416 \| 272 \| 11 \| 14 \| 418 \| 2 \| 401 \| Phage terminase large subunit (GpA) \| Phage terminase large subunit (GpA); afdb-uniprot50 \| AF-A0A7Y1BDW4-F1-MODEL\_V4 \| 1.0 \| 2.255e-38 \| 1092 \| 0.206 \| 684 \| 449 \| 22 \| 51 \| 680 \| 1 \| 644 \| Terminase \| Terminase; afdb-uniprot50 \| AF-A0A1H5ZMF5-F1-MODEL\_V4 \| 1.0 \| 1.294e-30 \| 1091 \| 0.297 \| 396 \| 246 \| 10 \| 1 \| 385 \| 4 \| 378 \| Phage terminase large subunit (GpA) \| Phage terminase large subunit (GpA); afdb-uniprot50 \| AF-A0A833NDZ4-F1-MODEL\_V4 \| 1.0 \| 6.84e-45 \| 1091 \| 0.215 \| 676 \| 431 \| 29 \| 3 \| 605 \| 1 \| 649 \| Phage terminase, large subunit \| Phage terminase, large subunit; afdb-uniprot50 \| AF-A0A660ULI2-F1-MODEL\_V4 \| 1.0 \| 1.25e-42 \| 1089 \| 0.227 \| 610 \| 409 \| 32 \| 35 \| 614 \| 19 \| 595 \| Uncharacterized protein \| Uncharacterized protein; afdb-uniprot50 \| AF-A0A6L9M202-F1-MODEL\_V4 \| 1.0 \| 2.774e-44 \| 1089 \| 0.216 \| 656 \| 417 \| 33 \| 29 \| 634 \| 30 \| 638 \| Phage terminase large subunit family protein \| Phage terminase large subunit family protein; afdb-uniprot50 \| AF-A0A496YPQ9-F1-MODEL\_V4 \| 1.0 \| 9.928e-42 \| 1088 \| 0.22 \| 607 \| 397 \| 26 \| 30 \| 596 \| 46 \| 616 \| Terminase \| Terminase; afdb-uniprot50 \| AF-A0A2M7FCS2-F1-MODEL\_V4 \| 1.0 \| 4.342e-44 \| 1088 \| 0.218 \| 692 \| 424 \| 35 \| 1 \| 634 \| 1 \| 633 \| Terminase \| Terminase; afdb-uniprot50 \| AF-A0A1B7JZW4-F1-MODEL\_V4 \| 1.0 \| 1.283e-46 \| 1086 \| 0.29 \| 612 \| 352 \| 17 \| 88 \| 631 \| 2 \| 598 \| Phage terminase large subunit \| Phage terminase large subunit; afdb-uniprot50 \| AF-G1V7I8-F1-MODEL\_V4 \| 1.0 \| 6.795e-44 \| 1086 \| 0.195 \| 720 \| 450 \| 31 \| 1 \| 677 \| 12 \| 644 \| Phage terminase large subunit \| Phage terminase large subunit; afdb-uniprot50 \| AF-A0A0F2NJJ8-F1-MODEL\_V4 \| 1.0 \| 5.959e-41 \| 1084 \| 0.205 \| 633 \| 422 \| 26 \| 35 \| 638 \| 3 \| 583 \| Uncharacterized protein \| Uncharacterized protein; afdb-uniprot50 \| AF-A0A1F9BJA2-F1-MODEL\_V4 \| 1.0 \| 7.14e-43 \| 1084 \| 0.226 \| 637 \| 407 \| 28 \| 3 \| 599 \| 1 \| 591 \| Uncharacterized protein \| Uncharacterized protein; afdb-uniprot50 \| AF-A0A165XGI1-F1-MODEL\_V4 \| 1.0 \| 6.795e-44 \| 1084 \| 0.216 \| 692 \| 432 \| 28 \| 39 \| 678 \| 1 \| 633 \| Phage terminase large subunit (GpA) \| Phage terminase large subunit (GpA); afdb-uniprot50 \| AF-A0A2T4JG50-F1-MODEL\_V4 \| 1.0 \| 9.633e-46 \| 1084 \| 0.227 \| 743 \| 442 \| 28 \| 3 \| 681 \| 1 \| 675 \| Terminase \| Terminase; afdb-uniprot50 \| AF-A0A0Q7XLX4-F1-MODEL\_V4 \| 1.0 \| 4e-40 \| 1084 \| 0.193 \| 723 \| 493 \| 31 \| 2 \| 677 \| 1 \| 680 \| Uncharacterized protein \| Uncharacterized protein; afdb-uniprot50 \| AF-A0A377BC22-F1-MODEL\_V4 \| 1.0 \| 7.438e-32 \| 1083 \| 0.275 \| 403 \| 239 \| 15 \| 5 \| 389 \| 1 \| 368 \| Phage terminase large subunit (GpA) \| Phage terminase large subunit (GpA); afdb-uniprot50 \| AF-A0A607GXG2-F1-MODEL\_V4 \| 1.0 \| 1.159e-39 \| 1082 \| 0.316 \| 467 \| 255 \| 14 \| 224 \| 678 \| 4 \| 418 \| Phage terminase large subunit family protein \| Phage terminase large subunit family protein; afdb-uniprot50 \| AF-A0A793N2K2-F1-MODEL\_V4 \| 1.0 \| 1.226e-39 \| 1082 \| 0.254 \| 566 \| 362 \| 21 \| 19 \| 551 \| 8 \| 546 \| Phage terminase large subunit family protein \| Phage terminase large subunit family protein; afdb-uniprot50 \| AF-A0A828NZJ9-F1-MODEL\_V4 \| 1.0 \| 5.502e-46 \| 1081 \| 0.203 \| 753 \| 467 \| 32 \| 10 \| 679 \| 2 \| 704 \| Phage terminase large subunit family protein \| Phage terminase large subunit family protein; afdb-uniprot50 \| AF-A0A2E9I639-F1-MODEL\_V4 \| 1.0 \| 2.33e-43 \| 1080 \| 0.208 \| 663 \| 429 \| 41 \| 19 \| 616 \| 9 \| 640 \| Uncharacterized protein \| Uncharacterized protein; afdb-uniprot50 \| AF-A0A7X3I7L0-F1-MODEL\_V4 \| 1.0 \| 2.915e-43 \| 1080 \| 0.23 \| 732 \| 447 \| 35 \| 19 \| 681 \| 19 \| 702 \| Uncharacterized protein \| Uncharacterized protein; afdb-uniprot50 \| AF-A0A5R1YK61-F1-MODEL\_V4 \| 1.0 \| 1.03e-38 \| 1077 \| 0.247 \| 569 \| 356 \| 20 \| 9 \| 542 \| 2 \| 533 \| Terminase \| Terminase; afdb-uniprot50 \| AF-A0A168N2Y6-F1-MODEL\_V4 \| 1.0 \| 8.557e-45 \| 1077 \| 0.257 \| 673 \| 393 \| 28 \| 62 \| 681 \| 1 \| 619 \| Uncharacterized protein \| Uncharacterized protein; afdb-uniprot50 \| AF-A0A2S1XLC7-F1-MODEL\_V4 \| 1.0 \| 4.105e-44 \| 1076 \| 0.213 \| 731 \| 475 \| 27 \| 1 \| 681 \| 2 \| 682 \| Uncharacterized protein \| Uncharacterized protein; afdb-uniprot50 \| AF-A0A7X8TUV2-F1-MODEL\_V4 \| 1.0 \| 5.49e-37 \| 1076 \| 0.167 \| 679 \| 437 \| 35 \| 24 \| 612 \| 7 \| 646 \| Uncharacterized protein \| Uncharacterized protein; afdb-uniprot50 \| AF-A0A134CAE4-F1-MODEL\_V4 \| 1.0 \| 6.194e-47 \| 1076 \| 0.256 \| 611 \| 365 \| 16 \| 97 \| 675 \| 426 \| 978 \| Phage terminase large subunit \| Phage terminase large subunit; afdb-uniprot50 \| AF-A0A158SPU5-F1-MODEL\_V4 \| 1.0 \| 5.744e-44 \| 1073 \| 0.211 \| 687 \| 434 \| 26 \| 5 \| 606 \| 1 \| 664 \| Uncharacterized protein \| Uncharacterized protein; afdb-uniprot50 \| AF-A0A2V4YP94-F1-MODEL\_V4 \| 1.0 \| 6.261e-40 \| 1072 \| 0.243 \| 587 \| 378 \| 23 \| 22 \| 565 \| 8 \| 571 \| Phage terminase large subunit GpA-like protein \| Phage terminase large subunit GpA-like protein; afdb-uniprot50 \| AF-A0A2D6FNL4-F1-MODEL\_V4 \| 1.0 \| 3.218e-41 \| 1071 \| 0.212 \| 644 \| 422 \| 29 \| 23 \| 633 \| 2 \| 593 \| Uncharacterized protein \| Uncharacterized protein; afdb-uniprot50 \| AF-A0A6B3LJH6-F1-MODEL\_V4 \| 1.0 \| 7.049e-41 \| 1071 \| 0.224 \| 656 \| 400 \| 25 \| 51 \| 632 \| 1 \| 621 \| Terminase \| Terminase; afdb-uniprot50 \| AF-A0A839H3C0-F1-MODEL\_V4 \| 1.0 \| 2.517e-29 \| 1069 \| 0.275 \| 363 \| 235 \| 11 \| 18 \| 372 \| 4 \| 346 \| Phage terminase large subunit GpA-like protein \| Phage terminase large subunit GpA-like protein; afdb-uniprot50 \| AF-A0A1V6JK58-F1-MODEL\_V4 \| 1.0 \| 2.448e-42 \| 1068 \| 0.226 \| 663 \| 400 \| 37 \| 3 \| 605 \| 1 \| 610 \| Phage terminase large subunit (GpA) \| Phage terminase large subunit (GpA); afdb-uniprot50 \| AF-A0A167H527-F1-MODEL\_V4 \| 1.0 \| 5.997e-42 \| 1068 \| 0.207 \| 645 \| 420 \| 28 \| 27 \| 614 \| 2 \| 612 \| Uncharacterized protein \| Uncharacterized protein; afdb-uniprot50 \| AF-A0A2E3H6Q3-F1-MODEL\_V4 \| 1.0 \| 2.64e-45 \| 1068 \| 0.228 \| 745 \| 441 \| 31 \| 1 \| 681 \| 1 \| 675 \| Terminase \| Terminase; afdb-uniprot50 \| AF-A0A161V854-F1-MODEL\_V4 \| 1.0 \| 3.143e-46 \| 1068 \| 0.216 \| 744 \| 468 \| 31 \| 23 \| 680 \| 22 \| 736 \| Phage terminase large subunit (GpA) \| Phage terminase large subunit (GpA); afdb-uniprot50 \| AF-A0A5F2KV72-F1-MODEL\_V4 \| 1.0 \| 1.371e-39 \| 1066 \| 0.26 \| 568 \| 354 \| 25 \| 19 \| 550 \| 8 \| 545 \| Phage terminase large subunit family protein \| Phage terminase large subunit family protein; afdb-uniprot50 \| AF-A0A5B9MCC3-F1-MODEL\_V4 \| 1.0 \| 1.944e-41 \| 1065 \| 0.187 \| 697 \| 457 \| 35 \| 1 \| 647 \| 1 \| 637 \| Phage terminase large subunit (GpA) \| Phage terminase large subunit (GpA); afdb-uniprot50 \| AF-A0A1M3E5M2-F1-MODEL\_V4 \| 1.0 \| 5.707e-43 \| 1064 \| 0.227 \| 693 \| 412 \| 38 \| 3 \| 616 \| 1 \| 648 \| Uncharacterized protein \| Uncharacterized protein; afdb-uniprot50 \| AF-E1JR22-F1-MODEL\_V4 \| 1.0 \| 1.957e-42 \| 1062 \| 0.223 \| 661 \| 437 \| 32 \| 1 \| 634 \| 1 \| 611 \| Terminase GpA \| Terminase GpA; afdb-uniprot50 \| AF-A0A227JQ12-F1-MODEL\_V4 \| 1.0 \| 6.509e-46 \| 1060 \| 0.351 \| 561 \| 295 \| 16 \| 135 \| 679 \| 2 \| 509 \| Uncharacterized protein \| Uncharacterized protein; afdb-uniprot50 \| AF-A0A4Q3P2E8-F1-MODEL\_V4 \| 1.0 \| 2.657e-46 \| 1059 \| 0.263 \| 603 \| 367 \| 25 \| 110 \| 681 \| 3 \| 559 \| Phage terminase large subunit family protein \| Phage terminase large subunit family protein; afdb-uniprot50 \| AF-A0A7W4IBA7-F1-MODEL\_V4 \| 1.0 \| 1.584e-44 \| 1059 \| 0.208 \| 762 \| 463 \| 33 \| 2 \| 680 \| 1 \| 704 \| Phage terminase large subunit family protein \| Phage terminase large subunit family protein; afdb-uniprot50 \| AF-A0A175RVL8-F1-MODEL\_V4 \| 1.0 \| 7.935e-42 \| 1059 \| 0.21 \| 655 \| 426 \| 31 \| 19 \| 615 \| 20 \| 641 \| Uncharacterized protein \| Uncharacterized protein; afdb-uniprot50 \| AF-A0A3M1QYL8-F1-MODEL\_V4 \| 1.0 \| 7.935e-42 \| 1058 \| 0.208 \| 666 \| 413 \| 29 \| 58 \| 682 \| 1 \| 593 \| Uncharacterized protein \| Uncharacterized protein; afdb-uniprot50 \| AF-A0A351VMH9-F1-MODEL\_V4 \| 1.0 \| 4.562e-43 \| 1058 \| 0.197 \| 669 \| 460 \| 29 \| 1 \| 633 \| 4 \| 631 \| Uncharacterized protein \| Uncharacterized protein; afdb-uniprot50 \| AF-A0A1W1Y9R2-F1-MODEL\_V4 \| 1.0 \| 5.397e-43 \| 1058 \| 0.196 \| 692 \| 447 \| 33 \| 19 \| 632 \| 22 \| 682 \| Phage terminase, large subunit GpA \| Phage terminase, large subunit GpA; afdb-uniprot50 \| AF-A0A662DNF2-F1-MODEL\_V4 \| 1.0 \| 2.651e-37 \| 1055 \| 0.249 \| 530 \| 329 \| 22 \| 1 \| 483 \| 13 \| 520 \| Large terminase subunit \| Large terminase subunit; afdb-uniprot50 \| AF-A0A2E6Y4R5-F1-MODEL\_V4 \| 1.0 \| 2.203e-43 \| 1054 \| 0.222 \| 653 \| 411 \| 25 \| 23 \| 605 \| 22 \| 647 \| Terminase \| Terminase; afdb-uniprot50 \| AF-A0A3Z9V4B3-F1-MODEL\_V4 \| 1.0 \| 2.36e-45 \| 1052 \| 0.204 \| 754 \| 453 \| 32 \| 16 \| 679 \| 3 \| 699 \| Phage terminase large subunit \| Phage terminase large subunit; afdb-uniprot50 \| AF-A0A4Z0F7F8-F1-MODEL\_V4 \| 1.0 \| 6.795e-44 \| 1049 \| 0.221 \| 735 \| 438 \| 29 \| 1 \| 630 \| 14 \| 718 \| Terminase \| Terminase; afdb-uniprot50 \| AF-A0A850I4C3-F1-MODEL\_V4 \| 1.0 \| 3.082e-43 \| 1048 \| 0.236 \| 694 \| 419 \| 27 \| 39 \| 677 \| 1 \| 638 \| Phage terminase large subunit family protein \| Phage terminase large subunit family protein; afdb-uniprot50 \| AF-A0A3E0WSL6-F1-MODEL\_V4 \| 1.0 \| 1.665e-43 \| 1048 \| 0.214 \| 668 \| 423 \| 31 \| 3 \| 605 \| 1 \| 631 \| Uncharacterized protein \| Uncharacterized protein; afdb-uniprot50 \| AF-A0A7G9NZU4-F1-MODEL\_V4 \| 1.0 \| 6.036e-43 \| 1047 \| 0.204 \| 698 \| 441 \| 31 \| 1 \| 632 \| 2 \| 650 \| Uncharacterized protein \| Uncharacterized protein; afdb-uniprot50 \| AF-A0A7V1K4K4-F1-MODEL\_V4 \| 1.0 \| 9.737e-39 \| 1044 \| 0.342 \| 441 \| 220 \| 15 \| 224 \| 615 \| 3 \| 422 \| Terminase \| Terminase; afdb-uniprot50 \| AF-A0A0M7BDF5-F1-MODEL\_V4 \| 1.0 \| 1.619e-30 \| 1044 \| 0.292 \| 404 \| 248 \| 13 \| 1 \| 390 \| 7 \| 386 \| Bacteriophage tail assembly protein \| Bacteriophage tail assembly protein; afdb-uniprot50 \| AF-W0DXT1-F1-MODEL\_V4 \| 1.0 \| 8.933e-43 \| 1044 \| 0.222 \| 656 \| 416 \| 30 \| 13 \| 605 \| 3 \| 627 \| DNA packaging protein \| DNA packaging protein; afdb-uniprot50 \| AF-A0A142BA63-F1-MODEL\_V4 \| 1.0 \| 1.125e-43 \| 1044 \| 0.23 \| 685 \| 413 \| 30 \| 1 \| 605 \| 1 \| 651 \| DNA packaging protein of prophage (Terminase large subunit) \| DNA packaging protein of prophage (Terminase large subunit); afdb-uniprot50 \| AF-A0A843EIY9-F1-MODEL\_V4 \| 1.0 \| 6.693e-33 \| 1043 \| 0.261 \| 439 \| 284 \| 14 \| 19 \| 435 \| 3 \| 423 \| Phage terminase large subunit family protein \| Phage terminase large subunit family protein; afdb-uniprot50 \| AF-A0A5S5MCE4-F1-MODEL\_V4 \| 1.0 \| 1.331e-43 \| 1042 \| 0.209 \| 741 \| 453 \| 37 \| 1 \| 682 \| 1 \| 667 \| Terminase \| Terminase; afdb-uniprot50 \| AF-F2K1G0-F1-MODEL\_V4 \| 1.0 \| 3.082e-43 \| 1041 \| 0.216 \| 655 \| 420 \| 31 \| 23 \| 605 \| 6 \| 638 \| Terminase GpA \| Terminase GpA; afdb-uniprot50 \| AF-A0A7Z6QWQ0-F1-MODEL\_V4 \| 1.0 \| 7.601e-44 \| 1039 \| 0.22 \| 718 \| 439 \| 30 \| 1 \| 631 \| 3 \| 686 \| Phage terminase large subunit family protein \| Phage terminase large subunit family protein; afdb-uniprot50 \| AF-A0A6B5HXD7-F1-MODEL\_V4 \| 1.0 \| 1.694e-37 \| 1038 \| 0.254 \| 555 \| 358 \| 20 \| 19 \| 544 \| 8 \| 535 \| Phage terminase large subunit family protein \| Phage terminase large subunit family protein; afdb-uniprot50 \| AF-A0A380N7T6-F1-MODEL\_V4 \| 1.0 \| 1.11e-41 \| 1038 \| 0.222 \| 576 \| 394 \| 26 \| 74 \| 632 \| 2 \| 540 \| Bacteriophage tail assembly protein \| Bacteriophage tail assembly protein; afdb-uniprot50 \| AF-A0A2V8S8T2-F1-MODEL\_V4 \| 1.0 \| 2.786e-36 \| 1038 \| 0.171 \| 663 \| 418 \| 31 \| 17 \| 607 \| 2 \| 604 \| Uncharacterized protein \| Uncharacterized protein; afdb-uniprot50 \| AF-A0A2W5TH91-F1-MODEL\_V4 \| 1.0 \| 8.991e-44 \| 1038 \| 0.236 \| 681 \| 413 \| 33 \| 3 \| 614 \| 1 \| 643 \| Terminase \| Terminase; afdb-uniprot50 \| AF-A0A5Z2FDT8-F1-MODEL\_V4 \| 1.0 \| 5.327e-41 \| 1037 \| 0.27 \| 569 \| 331 \| 24 \| 104 \| 623 \| 3 \| 536 \| Phage terminase large subunit family protein \| Phage terminase large subunit family protein; afdb-uniprot50 \| AF-A0A380T424-F1-MODEL\_V4 \| 1.0 \| 1.103e-40 \| 1036 \| 0.272 \| 506 \| 335 \| 14 \| 102 \| 585 \| 18 \| 512 \| Terminase, large subunit \| Terminase, large subunit; afdb-uniprot50 \| AF-A0A6G7TDA6-F1-MODEL\_V4 \| 1.0 \| 3.677e-27 \| 1034 \| 0.277 \| 353 \| 238 \| 7 \| 20 \| 369 \| 2 \| 340 \| Uncharacterized protein \| Uncharacterized protein; afdb-uniprot50 \| AF-A0A270BER1-F1-MODEL\_V4 \| 1.0 \| 2.432e-41 \| 1034 \| 0.216 \| 655 \| 409 \| 26 \| 22 \| 601 \| 6 \| 630 \| Uncharacterized protein \| Uncharacterized protein; afdb-uniprot50 \| AF-A0A0Q4QC50-F1-MODEL\_V4 \| 1.0 \| 1.676e-44 \| 1032 \| 0.246 \| 638 \| 373 \| 27 \| 74 \| 637 \| 2 \| 605 \| Terminase \| Terminase; afdb-uniprot50 \| AF-A0A3S3RLY6-F1-MODEL\_V4 \| 1.0 \| 8.971e-35 \| 1031 \| 0.254 \| 506 \| 320 \| 17 \| 3 \| 484 \| 1 \| 473 \| Terminase \| Terminase; afdb-uniprot50 \| AF-A0A7C1NH53-F1-MODEL\_V4 \| 1.0 \| 3.043e-41 \| 1031 \| 0.219 \| 630 \| 393 \| 30 \| 55 \| 633 \| 2 \| 583 \| Terminase \| Terminase; afdb-uniprot50 \| AF-A0A4U8YRI6-F1-MODEL\_V4 \| 1.0 \| 3.218e-41 \| 1029 \| 0.205 \| 653 \| 435 \| 35 \| 13 \| 632 \| 2 \| 603 \| Bacteriophage lambda gpa \| Bacteriophage lambda gpa; afdb-uniprot50 \| AF-A0A316N395-F1-MODEL\_V4 \| 1.0 \| 5.671e-42 \| 1029 \| 0.212 \| 672 \| 435 \| 35 \| 3 \| 632 \| 1 \| 619 \| Uncharacterized protein \| Uncharacterized protein; afdb-uniprot50 \| AF-A0A1G0ICW4-F1-MODEL\_V4 \| 1.0 \| 4.533e-42 \| 1029 \| 0.228 \| 664 \| 423 \| 36 \| 7 \| 626 \| 1 \| 618 \| Uncharacterized protein \| Uncharacterized protein; afdb-uniprot50 \| AF-A0A259DHP5-F1-MODEL\_V4 \| 1.0 \| 1.354e-37 \| 1028 \| 0.301 \| 488 \| 256 \| 17 \| 223 \| 680 \| 9 \| 441 \| Phage tail protein \| Phage tail protein; afdb-uniprot50 \| AF-I6B1C0-F1-MODEL\_V4 \| 1.0 \| 6.221e-39 \| 1028 \| 0.197 \| 633 \| 419 \| 32 \| 22 \| 605 \| 6 \| 598 \| Bacteriophage tail assembly protein \| Bacteriophage tail assembly protein; afdb-uniprot50 \| AF-A0A0F2N5L2-F1-MODEL\_V4 \| 1.0 \| 8.818e-41 \| 1027 \| 0.2 \| 604 \| 415 \| 23 \| 39 \| 608 \| 32 \| 601 \| Uncharacterized protein \| Uncharacterized protein; afdb-uniprot50 \| AF-A0A7V2AKH4-F1-MODEL\_V4 \| 1.0 \| 2.03e-39 \| 1026 \| 0.351 \| 444 \| 235 \| 18 \| 224 \| 632 \| 1 \| 426 \| Phage terminase large subunit family protein \| Phage terminase large subunit family protein; afdb-uniprot50 \| AF-A0A661RFV1-F1-MODEL\_V4 \| 1.0 \| 3.6e-41 \| 1025 \| 0.228 \| 660 \| 406 \| 34 \| 1 \| 598 \| 23 \| 640 \| Uncharacterized protein \| Uncharacterized protein; afdb-uniprot50 \| AF-A0A0F9R2K8-F1-MODEL\_V4 \| 1.0 \| 1.398e-42 \| 1025 \| 0.206 \| 712 \| 448 \| 31 \| 1 \| 629 \| 1 \| 678 \| Uncharacterized protein \| Uncharacterized protein; afdb-uniprot50 \| AF-A0A1X7MIT4-F1-MODEL\_V4 \| 1.0 \| 6.622e-40 \| 1024 \| 0.309 \| 511 \| 261 \| 16 \| 205 \| 679 \| 2 \| 456 \| Phage terminase large subunit (GpA) \| Phage terminase large subunit (GpA); afdb-uniprot50 \| AF-A0A5C7P718-F1-MODEL\_V4 \| 1.0 \| 1.314e-41 \| 1024 \| 0.19 \| 684 \| 453 \| 32 \| 19 \| 632 \| 2 \| 654 \| Uncharacterized protein \| Uncharacterized protein; afdb-uniprot50 \| AF-A0A5J6QMM2-F1-MODEL\_V4 \| 1.0 \| 2.448e-42 \| 1023 \| 0.216 \| 662 \| 418 \| 30 \| 23 \| 605 \| 6 \| 645 \| Phage terminase large subunit family protein \| Phage terminase large subunit family protein; afdb-uniprot50 \| AF-A0A540VMF1-F1-MODEL\_V4 \| 1.0 \| 4.661e-29 \| 1021 \| 0.408 \| 274 \| 149 \| 7 \| 365 \| 632 \| 6 \| 272 \| Phage terminase large subunit family protein \| Phage terminase large subunit family protein; afdb-uniprot50 \| AF-A0A2N8I4W4-F1-MODEL\_V4 \| 1.0 \| 2.03e-39 \| 1021 \| 0.225 \| 602 \| 394 \| 29 \| 30 \| 605 \| 2 \| 556 \| Uncharacterized protein \| Uncharacterized protein; afdb-uniprot50 \| AF-A0A4V2J7R8-F1-MODEL\_V4 \| 1.0 \| 3.102e-44 \| 1021 \| 0.206 \| 733 \| 460 \| 30 \| 24 \| 682 \| 8 \| 692 \| Phage terminase large subunit family protein \| Phage terminase large subunit family protein; afdb-uniprot50 \| AF-A0A1V3W4P1-F1-MODEL\_V4 \| 1.0 \| 8.338e-41 \| 1020 \| 0.244 \| 629 \| 382 \| 27 \| 19 \| 583 \| 8 \| 606 \| Uncharacterized protein \| Uncharacterized protein; afdb-uniprot50 \| AF-A0A0H3EJ51-F1-MODEL\_V4 \| 1.0 \| 1.498e-44 \| 1018 \| 0.207 \| 731 \| 436 \| 32 \| 39 \| 679 \| 1 \| 677 \| Putative terminase large subunit \| Putative terminase large subunit; afdb-uniprot50 \| AF-A0A4P7WC01-F1-MODEL\_V4 \| 1.0 \| 4.503e-41 \| 1018 \| 0.212 \| 683 \| 426 \| 33 \| 1 \| 605 \| 1 \| 649 \| Phage terminase large subunit family protein \| Phage terminase large subunit family protein; afdb-uniprot50 \| AF-A0A3A1YIE0-F1-MODEL\_V4 \| 1.0 \| 6.708e-42 \| 1018 \| 0.209 \| 662 \| 441 \| 29 \| 4 \| 605 \| 50 \| 688 \| Uncharacterized protein \| Uncharacterized protein; afdb-uniprot50 \| AF-A0A660LVS2-F1-MODEL\_V4 \| 1.0 \| 6.737e-34 \| 1017 \| 0.238 \| 473 \| 321 \| 16 \| 1 \| 462 \| 9 \| 453 \| Terminase \| Terminase; afdb-uniprot50 \| AF-A0A441I2M4-F1-MODEL\_V4 \| 1.0 \| 4.314e-43 \| 1015 \| 0.21 \| 693 \| 417 \| 35 \| 2 \| 599 \| 3 \| 660 \| Uncharacterized protein \| Uncharacterized protein; afdb-uniprot50 \| AF-A0A443J087-F1-MODEL\_V4 \| 1.0 \| 1.25e-42 \| 1014 \| 0.218 \| 676 \| 403 \| 31 \| 22 \| 602 \| 16 \| 660 \| Uncharacterized protein \| Uncharacterized protein; afdb-uniprot50 \| AF-A0A3D9BKG4-F1-MODEL\_V4 \| 1.0 \| 1.05e-41 \| 1010 \| 0.33 \| 541 \| 272 \| 21 \| 172 \| 677 \| 8 \| 493 \| Phage terminase large subunit family protein \| Phage terminase large subunit family protein; afdb-uniprot50 \| AF-A0A1H5Z1S1-F1-MODEL\_V4 \| 1.0 \| 1.197e-44 \| 1010 \| 0.267 \| 595 \| 361 \| 23 \| 99 \| 637 \| 451 \| 1026 \| Replicative DNA helicase (DnaB) \| Replicative DNA helicase (DnaB); afdb-uniprot50 \| AF-A0A1L3GDS2-F1-MODEL\_V4 \| 1.0 \| 1.46e-40 \| 1009 \| 0.218 \| 682 \| 439 \| 34 \| 3 \| 632 \| 1 \| 640 \| Uncharacterized protein \| Uncharacterized protein; afdb-uniprot50 \| AF-A0A2N5XX61-F1-MODEL\_V4 \| 1.0 \| 1.064e-43 \| 1009 \| 0.212 \| 762 \| 458 \| 39 \| 3 \| 679 \| 30 \| 734 \| Uncharacterized protein \| Uncharacterized protein; afdb-uniprot50 \| AF-A0A2D6XDS8-F1-MODEL\_V4 \| 1.0 \| 1.749e-42 \| 1008 \| 0.202 \| 734 \| 450 \| 32 \| 1 \| 677 \| 1 \| 655 \| Uncharacterized protein \| Uncharacterized protein; afdb-uniprot50 \| AF-F3Z2S1-F1-MODEL\_V4 \| 1.0 \| 1.554e-41 \| 1007 \| 0.216 \| 679 \| 448 \| 28 \| 5 \| 631 \| 1 \| 647 \| Uncharacterized protein \| Uncharacterized protein; afdb-uniprot50 \| AF-A0A350YH52-F1-MODEL\_V4 \| 1.0 \| 7.264e-37 \| 1004 \| 0.298 \| 509 \| 264 \| 18 \| 207 \| 680 \| 3 \| 453 \| Phage tail protein \| Phage tail protein; afdb-uniprot50 \| AF-A0A2D6M8M9-F1-MODEL\_V4 \| 1.0 \| 3.36e-39 \| 1004 \| 0.207 \| 684 \| 402 \| 33 \| 36 \| 616 \| 19 \| 665 \| Uncharacterized protein \| Uncharacterized protein; afdb-uniprot50 \| AF-A0A4Z1QY08-F1-MODEL\_V4 \| 1.0 \| 1.11e-41 \| 1003 \| 0.221 \| 673 \| 415 \| 32 \| 1 \| 605 \| 109 \| 740 \| Terminase \| Terminase; afdb-uniprot50 \| AF-A0A827VNM2-F1-MODEL\_V4 \| 1.0 \| 5.49e-37 \| 1002 \| 0.26 \| 538 \| 334 \| 25 \| 19 \| 523 \| 8 \| 514 \| Phage terminase large subunit family protein \| Phage terminase large subunit family protein; afdb-uniprot50 \| AF-A0A2E9SLK1-F1-MODEL\_V4 \| 1.0 \| 5.397e-43 \| 1002 \| 0.198 \| 709 \| 435 \| 32 \| 14 \| 632 \| 2 \| 666 \| DNA packaging protein \| DNA packaging protein; afdb-uniprot50 \| AF-A0A2S5JD74-F1-MODEL\_V4 \| 1.0 \| 1.218e-38 \| 1001 \| 0.307 \| 458 \| 276 \| 12 \| 98 \| 539 \| 470 \| 902 \| Intein \| Intein; afdb-uniprot50 \| AF-A0A6G8D7N8-F1-MODEL\_V4 \| 1.0 \| 3.082e-43 \| 998 \| 0.224 \| 707 \| 415 \| 38 \| 1 \| 624 \| 1 \| 656 \| Phage terminase large subunit family protein \| Phage terminase large subunit family protein; afdb-uniprot50 \| AF-A0A2W5JHQ6-F1-MODEL\_V4 \| 1.0 \| 1.089e-38 \| 997 \| 0.313 \| 527 \| 273 \| 24 \| 191 \| 682 \| 1 \| 473 \| Terminase \| Terminase; afdb-uniprot50 \| AF-A0A0F9GKY5-F1-MODEL\_V4 \| 1.0 \| 7.683e-37 \| 997 \| 0.234 \| 521 \| 351 \| 22 \| 7 \| 495 \| 23 \| 527 \| Uncharacterized protein \| Uncharacterized protein; afdb-uniprot50 \| AF-A0A1Z8QJ35-F1-MODEL\_V4 \| 1.0 \| 4.436e-30 \| 996 \| 0.398 \| 314 \| 148 \| 7 \| 369 \| 682 \| 31 \| 303 \| Uncharacterized protein \| Uncharacterized protein; afdb-uniprot50 \| AF-A0A6A8A390-F1-MODEL\_V4 \| 1.0 \| 5.478e-28 \| 996 \| 0.254 \| 385 \| 237 \| 9 \| 17 \| 364 \| 9 \| 380 \| Terminase \| Terminase; afdb-uniprot50 \| AF-A0A5C7QC12-F1-MODEL\_V4 \| 1.0 \| 7.094e-42 \| 996 \| 0.2 \| 724 \| 451 \| 32 \| 23 \| 649 \| 6 \| 698 \| Terminase \| Terminase; afdb-uniprot50 \| AF-A0A166JZ20-F1-MODEL\_V4 \| 1.0 \| 1.057e-42 \| 995 \| 0.242 \| 590 \| 348 \| 29 \| 77 \| 610 \| 2 \| 548 \| Uncharacterized protein \| Uncharacterized protein; afdb-uniprot50 \| AF-H1D5G9-F1-MODEL\_V4 \| 1.0 \| 7.584e-35 \| 993 \| 0.299 \| 394 \| 243 \| 11 \| 223 \| 608 \| 4 \| 372 \| Uncharacterized protein \| Uncharacterized protein; afdb-uniprot50 \| AF-A0A1V5E5H3-F1-MODEL\_V4 \| 1.0 \| 1.103e-40 \| 991 \| 0.227 \| 709 \| 404 \| 38 \| 39 \| 681 \| 1 \| 631 \| Phage terminase large subunit (GpA) \| Phage terminase large subunit (GpA); afdb-uniprot50 \| AF-A0A4Q2XVX2-F1-MODEL\_V4 \| 1.0 \| 4.35e-27 \| 990 \| 0.274 \| 364 \| 230 \| 13 \| 18 \| 372 \| 3 \| 341 \| Phage terminase large subunit family protein \| Phage terminase large subunit family protein; afdb-uniprot50 \| AF-A0A6C1RW32-F1-MODEL\_V4 \| 1.0 \| 2.859e-40 \| 988 \| 0.204 \| 660 \| 439 \| 30 \| 17 \| 610 \| 2 \| 641 \| Phage terminase large subunit family protein \| Phage terminase large subunit family protein; afdb-uniprot50 \| AF-A0A5P9EIV2-F1-MODEL\_V4 \| 1.0 \| 9.387e-42 \| 988 \| 0.184 \| 818 \| 425 \| 30 \| 1 \| 605 \| 1 \| 789 \| Phage terminase large subunit (GpA) \| Phage terminase large subunit (GpA); afdb-uniprot50 \| AF-A0A258G251-F1-MODEL\_V4 \| 1.0 \| 7.003e-40 \| 987 \| 0.326 \| 517 \| 284 \| 20 \| 161 \| 641 \| 1 \| 489 \| Phage tail protein \| Phage tail protein; afdb-uniprot50 \| AF-A0A2E3R5R9-F1-MODEL\_V4 \| 1.0 \| 3.623e-42 \| 987 \| 0.205 \| 691 \| 449 \| 29 \| 1 \| 632 \| 15 \| 664 \| Uncharacterized protein \| Uncharacterized protein; afdb-uniprot50 \| AF-A0A4Q5QYI3-F1-MODEL\_V4 \| 1.0 \| 2.908e-34 \| 986 \| 0.256 \| 515 \| 311 \| 20 \| 18 \| 487 \| 19 \| 506 \| Terminase \| Terminase; afdb-uniprot50 \| AF-Q6AMW9-F1-MODEL\_V4 \| 1.0 \| 9.387e-42 \| 986 \| 0.213 \| 669 \| 438 \| 31 \| 1 \| 614 \| 9 \| 643 \| Related to terminase large subunit of phage lambda \| Related to terminase large subunit of phage lambda; afdb-uniprot50 \| AF-A0A2T6LY95-F1-MODEL\_V4 \| 1.0 \| 6.708e-42 \| 985 \| 0.184 \| 753 \| 475 \| 31 \| 10 \| 681 \| 1 \| 695 \| Phage terminase large subunit GpA-like protein \| Phage terminase large subunit GpA-like protein; afdb-uniprot50 \| AF-A0A6S5X3W7-F1-MODEL\_V4 \| 1.0 \| 1.057e-42 \| 984 \| 0.232 \| 676 \| 400 \| 36 \| 33 \| 624 \| 18 \| 658 \| Phage terminase large subunit \| Phage terminase large subunit; afdb-uniprot50 \| AF-A0A7C5M9C6-F1-MODEL\_V4 \| 1.0 \| 5.293e-40 \| 983 \| 0.327 \| 483 \| 253 \| 21 \| 187 \| 632 \| 3 \| 450 \| Phage terminase large subunit family protein \| Phage terminase large subunit family protein; afdb-uniprot50 \| AF-A0A5C8SPM8-F1-MODEL\_V4 \| 1.0 \| 6.302e-41 \| 983 \| 0.209 \| 751 \| 469 \| 38 \| 1 \| 678 \| 1 \| 699 \| Uncharacterized protein \| Uncharacterized protein; afdb-uniprot50 \| AF-A0A4U9DA01-F1-MODEL\_V4 \| 1.0 \| 5.658e-33 \| 981 \| 0.235 \| 496 \| 296 \| 16 \| 3 \| 434 \| 1 \| 477 \| Bacteriophage tail assembly protein \| Bacteriophage tail assembly protein; afdb-uniprot50 \| AF-A0A7W6FW72-F1-MODEL\_V4 \| 1.0 \| 1.87e-35 \| 981 \| 0.235 \| 616 \| 337 \| 27 \| 145 \| 678 \| 3 \| 566 \| Phage terminase large subunit GpA-like protein \| Phage terminase large subunit GpA-like protein; afdb-uniprot50 \| AF-A0A6I2UCD1-F1-MODEL\_V4 \| 1.0 \| 6.795e-44 \| 981 \| 0.249 \| 633 \| 388 \| 17 \| 91 \| 681 \| 60 \| 647 \| Terminase \| Terminase; afdb-uniprot50 \| AF-A0A1G3M2E9-F1-MODEL\_V4 \| 1.0 \| 5.959e-41 \| 980 \| 0.216 \| 684 \| 434 \| 33 \| 1 \| 632 \| 6 \| 639 \| Uncharacterized protein \| Uncharacterized protein; afdb-uniprot50 \| AF-A0A2G0Q134-F1-MODEL\_V4 \| 1.0 \| 6.036e-43 \| 978 \| 0.345 \| 547 \| 293 \| 21 \| 140 \| 650 \| 6 \| 523 \| Terminase \| Terminase; afdb-uniprot50 \| AF-A0A379VTP8-F1-MODEL\_V4 \| 1.0 \| 2.056e-41 \| 978 \| 0.255 \| 607 \| 366 \| 26 \| 110 \| 677 \| 5 \| 564 \| Phage terminase large subunit \| Phage terminase large subunit; afdb-uniprot50 \| AF-A0A1V1UMI0-F1-MODEL\_V4 \| 1.0 \| 2.685e-39 \| 975 \| 0.217 \| 736 \| 438 \| 41 \| 17 \| 681 \| 2 \| 670 \| Phage terminase large subunit \| Phage terminase large subunit; afdb-uniprot50 \| AF-A0A454TI12-F1-MODEL\_V4 \| 1.0 \| 5.113e-26 \| 974 \| 0.314 \| 337 \| 200 \| 11 \| 17 \| 344 \| 3 \| 317 \| Phage terminase large subunit family protein \| Phage terminase large subunit family protein; afdb-uniprot50 \| AF-A0A2T5JQ45-F1-MODEL\_V4 \| 1.0 \| 5.562e-39 \| 974 \| 0.306 \| 489 \| 289 \| 18 \| 159 \| 640 \| 3 \| 448 \| Phage terminase large subunit GpA-like protein \| Phage terminase large subunit GpA-like protein; afdb-uniprot50 \| AF-A0A1V4WK69-F1-MODEL\_V4 \| 1.0 \| 3.063e-42 \| 974 \| 0.202 \| 712 \| 454 \| 37 \| 1 \| 636 \| 4 \| 677 \| Phage terminase large subunit (GpA) \| Phage terminase large subunit (GpA); afdb-uniprot50 \| AF-A0A0A0FB79-F1-MODEL\_V4 \| 1.0 \| 9.55e-36 \| 973 \| 0.238 \| 486 \| 326 \| 19 \| 68 \| 539 \| 18 \| 473 \| Terminase large subunit \| Terminase large subunit; afdb-uniprot50 \| AF-A0A285D3Y8-F1-MODEL\_V4 \| 1.0 \| 9.427e-34 \| 973 \| 0.372 \| 360 \| 207 \| 9 \| 282 \| 633 \| 189 \| 537 \| Transposase IS166 family protein \| Transposase IS166 family protein; afdb-uniprot50 \| AF-D7XY82-F1-MODEL\_V4 \| 1.0 \| 8.055e-27 \| 971 \| 0.301 \| 338 \| 217 \| 8 \| 18 \| 352 \| 3 \| 324 \| Uncharacterized protein \| Uncharacterized protein; afdb-uniprot50 \| AF-A0A1M5AMT0-F1-MODEL\_V4 \| 1.0 \| 2.611e-26 \| 971 \| 0.301 \| 348 \| 211 \| 11 \| 14 \| 352 \| 2 \| 326 \| Phage terminase large subunit (GpA) \| Phage terminase large subunit (GpA); afdb-uniprot50 \| AF-A0A3A1YAX6-F1-MODEL\_V4 \| 1.0 \| 2.133e-38 \| 970 \| 0.192 \| 728 \| 447 \| 39 \| 3 \| 621 \| 1 \| 696 \| Uncharacterized protein \| Uncharacterized protein; afdb-uniprot50 \| AF-F3G6T3-F1-MODEL\_V4 \| 1.0 \| 1.073e-27 \| 969 \| 0.26 \| 368 \| 243 \| 13 \| 39 \| 389 \| 1 \| 356 \| Phage terminase GpA \| Phage terminase GpA; afdb-uniprot50 \| AF-A0A6B9FZ59-F1-MODEL\_V4 \| 1.0 \| 3.592e-32 \| 965 \| 0.264 \| 477 \| 308 \| 19 \| 8 \| 462 \| 9 \| 464 \| Uncharacterized protein \| Uncharacterized protein; afdb-uniprot50 \| AF-A0A518ICL6-F1-MODEL\_V4 \| 1.0 \| 1.815e-39 \| 965 \| 0.203 \| 708 \| 460 \| 34 \| 15 \| 677 \| 4 \| 652 \| Phage terminase large subunit (GpA) \| Phage terminase large subunit (GpA); afdb-uniprot50 \| AF-A0A546X7W2-F1-MODEL\_V4 \| 1.0 \| 3.782e-40 \| 964 \| 0.202 \| 764 \| 484 \| 35 \| 1 \| 679 \| 1 \| 723 \| Uncharacterized protein \| Uncharacterized protein; afdb-uniprot50 \| AF-A0A1B9VJS7-F1-MODEL\_V4 \| 1.0 \| 1.85e-42 \| 963 \| 0.22 \| 697 \| 409 \| 35 \| 1 \| 605 \| 1 \| 654 \| Terminase GpA \| Terminase GpA; afdb-uniprot50 \| AF-A0A1C1YRY5-F1-MODEL\_V4 \| 1.0 \| 2.401e-39 \| 963 \| 0.197 \| 663 \| 462 \| 27 \| 12 \| 632 \| 17 \| 651 \| Uncharacterized protein \| Uncharacterized protein; afdb-uniprot50 \| AF-A0A1W2AC69-F1-MODEL\_V4 \| 1.0 \| 3.807e-41 \| 960 \| 0.183 \| 783 \| 488 \| 34 \| 4 \| 679 \| 11 \| 748 \| Phage terminase, large subunit GpA \| Phage terminase, large subunit GpA; afdb-uniprot50 \| AF-A0A7J0BHR8-F1-MODEL\_V4 \| 1.0 \| 2.84e-39 \| 959 \| 0.2 \| 629 \| 417 \| 28 \| 55 \| 632 \| 2 \| 595 \| Terminase \| Terminase; afdb-uniprot50 \| AF-A0A7R6V9X5-F1-MODEL\_V4 \| 1.0 \| 1.705e-38 \| 959 \| 0.168 \| 689 \| 453 \| 34 \| 27 \| 682 \| 8 \| 609 \| Phage terminase large subunit \| Phage terminase large subunit; afdb-uniprot50 \| AF-A0A431I9D4-F1-MODEL\_V4 \| 1.0 \| 2.255e-38 \| 958 \| 0.197 \| 699 \| 455 \| 40 \| 1 \| 633 \| 19 \| 677 \| Uncharacterized protein \| Uncharacterized protein; afdb-uniprot50 \| AF-A0A5Y1WM45-F1-MODEL\_V4 \| 1.0 \| 7.033e-32 \| 957 \| 0.236 \| 487 \| 309 \| 16 \| 7 \| 460 \| 1 \| 457 \| Phage terminase large subunit family protein \| Phage terminase large subunit family protein; afdb-uniprot50 \| AF-A0A2A3WZH3-F1-MODEL\_V4 \| 1.0 \| 6.494e-37 \| 957 \| 0.244 \| 553 \| 358 \| 24 \| 19 \| 537 \| 8 \| 534 \| Terminase \| Terminase; afdb-uniprot50 \| AF-A0A7T6AR28-F1-MODEL\_V4 \| 1.0 \| 7.783e-39 \| 956 \| 0.199 \| 638 \| 419 \| 30 \| 33 \| 633 \| 55 \| 637 \| Phage terminase large subunit family protein \| Phage terminase large subunit family protein; afdb-uniprot50 \| AF-A0A560D0T7-F1-MODEL\_V4 \| 1.0 \| 4.445e-39 \| 955 \| 0.222 \| 682 \| 413 \| 25 \| 51 \| 677 \| 2 \| 620 \| Phage terminase large subunit GpA-like protein \| Phage terminase large subunit GpA-like protein; afdb-uniprot50 \| AF-I6APC5-F1-MODEL\_V4 \| 1.0 \| 1.017e-36 \| 955 \| 0.186 \| 682 \| 434 \| 31 \| 23 \| 636 \| 2 \| 630 \| Bacteriophage tail assembly protein \| Bacteriophage tail assembly protein; afdb-uniprot50 \| AF-A0A7W8DQ18-F1-MODEL\_V4 \| 1.0 \| 1.023e-37 \| 953 \| 0.159 \| 688 \| 470 \| 32 \| 22 \| 635 \| 4 \| 656 \| Phage terminase large subunit GpA-like protein \| Phage terminase large subunit GpA-like protein; afdb-uniprot50 \| AF-A0A625VUZ6-F1-MODEL\_V4 \| 1.0 \| 1.144e-37 \| 952 \| 0.204 \| 598 \| 383 \| 23 \| 1 \| 531 \| 2 \| 573 \| Phage terminase large subunit family protein \| Phage terminase large subunit family protein; afdb-uniprot50 \| AF-A0A1G5T1A6-F1-MODEL\_V4 \| 1.0 \| 7.003e-40 \| 951 \| 0.192 \| 695 \| 463 \| 31 \| 23 \| 648 \| 24 \| 688 \| Phage terminase, large subunit GpA \| Phage terminase, large subunit GpA; afdb-uniprot50 \| AF-A0A4S1MLI7-F1-MODEL\_V4 \| 1.0 \| 1.043e-40 \| 948 \| 0.21 \| 694 \| 436 \| 30 \| 34 \| 640 \| 10 \| 678 \| Terminase \| Terminase; afdb-uniprot50 \| AF-A0A1D8A8W8-F1-MODEL\_V4 \| 1.0 \| 7.407e-40 \| 947 \| 0.191 \| 704 \| 439 \| 30 \| 11 \| 610 \| 3 \| 679 \| Uncharacterized protein \| Uncharacterized protein; afdb-uniprot50 \| AF-A0A353ZI82-F1-MODEL\_V4 \| 1.0 \| 7.733e-38 \| 946 \| 0.2 \| 644 \| 431 \| 27 \| 17 \| 604 \| 4 \| 619 \| Uncharacterized protein \| Uncharacterized protein; afdb-uniprot50 \| AF-A0A371JL63-F1-MODEL\_V4 \| 1.0 \| 1.45e-39 \| 945 \| 0.223 \| 690 \| 423 \| 36 \| 17 \| 618 \| 3 \| 667 \| Ig-like domain-containing protein \| Ig-like domain-containing protein; afdb-uniprot50 \| AF-A0A2V6CXN5-F1-MODEL\_V4 \| 1.0 \| 1.764e-26 \| 944 \| 0.223 \| 393 \| 263 \| 9 \| 7 \| 388 \| 9 \| 370 \| Uncharacterized protein \| Uncharacterized protein; afdb-uniprot50 \| AF-A0A1S7MA44-F1-MODEL\_V4 \| 1.0 \| 2.539e-39 \| 942 \| 0.191 \| 685 \| 448 \| 33 \| 3 \| 617 \| 1 \| 649 \| Putative Terminase large subunit GpA \| Putative Terminase large subunit GpA; afdb-uniprot50 \| AF-A0A6L9AAZ1-F1-MODEL\_V4 \| 1.0 \| 3.382e-40 \| 941 \| 0.263 \| 554 \| 340 \| 20 \| 111 \| 623 \| 4 \| 530 \| Phage terminase large subunit family protein \| Phage terminase large subunit family protein; afdb-uniprot50 \| AF-A0A0F2NFW3-F1-MODEL\_V4 \| 1.0 \| 3.004e-39 \| 940 \| 0.187 \| 650 \| 430 \| 28 \| 39 \| 633 \| 55 \| 661 \| Uncharacterized protein \| Uncharacterized protein; afdb-uniprot50 \| AF-A0A2W6X6N4-F1-MODEL\_V4 \| 1.0 \| 8.284e-40 \| 939 \| 0.195 \| 690 \| 447 \| 28 \| 7 \| 606 \| 1 \| 672 \| Uncharacterized protein \| Uncharacterized protein; afdb-uniprot50 \| AF-A0A1Z9JFF5-F1-MODEL\_V4 \| 1.0 \| 4.611e-36 \| 938 \| 0.187 \| 630 \| 408 \| 34 \| 19 \| 605 \| 2 \| 570 \| Uncharacterized protein \| Uncharacterized protein; afdb-uniprot50 \| AF-A0A5A9FAT1-F1-MODEL\_V4 \| 1.0 \| 3.197e-40 \| 935 \| 0.216 \| 629 \| 379 \| 30 \| 65 \| 632 \| 4 \| 579 \| Terminase \| Terminase; afdb-uniprot50 \| AF-A0A846QMH4-F1-MODEL\_V4 \| 1.0 \| 7.359e-39 \| 935 \| 0.212 \| 658 \| 434 \| 33 \| 4 \| 609 \| 1 \| 625 \| Phage terminase large subunit GpA-like protein \| Phage terminase large subunit GpA-like protein; afdb-uniprot50 \| AF-A0A7G6RHV7-F1-MODEL\_V4 \| 1.0 \| 2.49e-36 \| 931 \| 0.202 \| 618 \| 408 \| 28 \| 1 \| 563 \| 2 \| 589 \| Uncharacterized protein \| Uncharacterized protein; afdb-uniprot50 \| AF-A0A368TS88-F1-MODEL\_V4 \| 1.0 \| 9.8e-40 \| 930 \| 0.187 \| 775 \| 454 \| 36 \| 17 \| 675 \| 3 \| 717 \| Phage terminase large subunit family protein \| Phage terminase large subunit family protein; afdb-uniprot50 \| AF-A0A371J7D6-F1-MODEL\_V4 \| 1.0 \| 6.579e-39 \| 929 \| 0.284 \| 510 \| 310 \| 19 \| 155 \| 633 \| 3 \| 488 \| Phage terminase large subunit family protein \| Phage terminase large subunit family protein; afdb-uniprot50 \| AF-A0A6J7LP44-F1-MODEL\_V4 \| 1.0 \| 2.871e-32 \| 929 \| 0.147 \| 678 \| 454 \| 27 \| 17 \| 632 \| 2 \| 617 \| Unannotated protein \| Unannotated protein; afdb-uniprot50 \| AF-A0A2Y0RP32-F1-MODEL\_V4 \| 1.0 \| 1.447e-30 \| 924 \| 0.285 \| 421 \| 230 \| 15 \| 244 \| 623 \| 2 \| 392 \| DNA packaging protein of prophage terminase large subunit \| DNA packaging protein of prophage terminase large subunit; afdb-uniprot50 \| AF-A0A358B1W0-F1-MODEL\_V4 \| 1.0 \| 1.113e-24 \| 923 \| 0.297 \| 336 \| 208 \| 8 \| 22 \| 352 \| 4 \| 316 \| Terminase \| Terminase; afdb-uniprot50 \| AF-A0A3E0N1X8-F1-MODEL\_V4 \| 1.0 \| 3.096e-35 \| 922 \| 0.186 \| 626 \| 410 \| 36 \| 19 \| 621 \| 5 \| 553 \| Uncharacterized protein \| Uncharacterized protein; afdb-uniprot50 \| AF-A0A1Y1R385-F1-MODEL\_V4 \| 1.0 \| 3.522e-29 \| 919 \| 0.361 \| 340 \| 175 \| 10 \| 343 \| 675 \| 10 \| 314 \| Uncharacterized protein \| Uncharacterized protein; afdb-uniprot50 \| AF-A0A3D3NRT9-F1-MODEL\_V4 \| 1.0 \| 8.649e-38 \| 919 \| 0.166 \| 690 \| 465 \| 25 \| 1 \| 611 \| 24 \| 682 \| Uncharacterized protein \| Uncharacterized protein; afdb-uniprot50 \| AF-A0A1H0W316-F1-MODEL\_V4 \| 1.0 \| 6.181e-38 \| 916 \| 0.2 \| 578 \| 393 \| 25 \| 67 \| 608 \| 2 \| 546 \| Phage terminase, large subunit GpA \| Phage terminase, large subunit GpA; afdb-uniprot50 \| AF-A0A509DY50-F1-MODEL\_V4 \| 1.0 \| 2.685e-39 \| 913 \| 0.24 \| 606 \| 391 \| 26 \| 89 \| 632 \| 3 \| 600 \| Uncharacterized protein \| Uncharacterized protein; afdb-uniprot50 \| AF-A0A1I4C945-F1-MODEL\_V4 \| 1.0 \| 7.935e-42 \| 913 \| 0.228 \| 664 \| 389 \| 28 \| 81 \| 681 \| 13 \| 615 \| Phage terminase, large subunit GpA \| Phage terminase, large subunit GpA; afdb-uniprot50 \| AF-A0A257ERR4-F1-MODEL\_V4 \| 1.0 \| 2.078e-34 \| 912 \| 0.315 \| 419 \| 252 \| 11 \| 113 \| 520 \| 2 \| 396 \| Phage tail protein \| Phage tail protein; afdb-uniprot50 \| AF-A0A0P1EW26-F1-MODEL\_V4 \| 1.0 \| 7.049e-41 \| 911 \| 0.227 \| 667 \| 401 \| 30 \| 75 \| 681 \| 4 \| 615 \| Bacteriophage tail assembly protein \| Bacteriophage tail assembly protein; afdb-uniprot50 \| AF-A0A1K0IL49-F1-MODEL\_V4 \| 1.0 \| 3.576e-40 \| 910 \| 0.206 \| 721 \| 426 \| 34 \| 17 \| 632 \| 3 \| 682 \| Terminase GpA \| Terminase GpA; afdb-uniprot50 \| AF-A0A6N7ZQ45-F1-MODEL\_V4 \| 1.0 \| 3.116e-36 \| 909 \| 0.331 \| 435 \| 241 \| 13 \| 176 \| 583 \| 2 \| 413 \| Terminase \| Terminase; afdb-uniprot50 \| AF-K4W1X9-F1-MODEL\_V4 \| 1.0 \| 1.757e-34 \| 909 \| 0.25 \| 507 \| 329 \| 21 \| 48 \| 526 \| 1 \| 484 \| Terminase large subunit \| Terminase large subunit; afdb-uniprot50 \| AF-A0A1V5F3R7-F1-MODEL\_V4 \| 1.0 \| 5.682e-25 \| 905 \| 0.297 \| 339 \| 204 \| 10 \| 21 \| 347 \| 8 \| 324 \| Phage terminase large subunit (GpA) \| Phage terminase large subunit (GpA); afdb-uniprot50 \| AF-A0A766JRV8-F1-MODEL\_V4 \| 1.0 \| 2.128e-29 \| 905 \| 0.279 \| 429 \| 227 \| 19 \| 243 \| 623 \| 5 \| 399 \| Phage terminase large subunit family protein \| Phage terminase large subunit family protein; afdb-uniprot50 \| AF-A0A7V4LRV2-F1-MODEL\_V4 \| 1.0 \| 6.397e-26 \| 902 \| 0.46 \| 241 \| 114 \| 8 \| 392 \| 626 \| 2 \| 232 \| Phage terminase large subunit family protein \| Phage terminase large subunit family protein; afdb-uniprot50 \| AF-A0A0B0HFD7-F1-MODEL\_V4 \| 1.0 \| 1.438e-29 \| 901 \| 0.319 \| 357 \| 210 \| 9 \| 74 \| 425 \| 2 \| 330 \| Phage terminase large subunit (GpA) \| Phage terminase large subunit (GpA); afdb-uniprot50 \| AF-A0A838QRG0-F1-MODEL\_V4 \| 1.0 \| 3.734e-38 \| 901 \| 0.183 \| 685 \| 458 \| 27 \| 2 \| 608 \| 1 \| 661 \| Phage terminase large subunit family protein \| Phage terminase large subunit family protein; afdb-uniprot50 \| AF-A0A776QV88-F1-MODEL\_V4 \| 1.0 \| 1.87e-35 \| 900 \| 0.254 \| 511 \| 320 \| 17 \| 87 \| 561 \| 2 \| 487 \| Phage terminase large subunit family protein \| Phage terminase large subunit family protein; afdb-uniprot50 \| AF-A0A844G3X9-F1-MODEL\_V4 \| 1.0 \| 3.75e-30 \| 898 \| 0.153 \| 644 \| 444 \| 32 \| 6 \| 608 \| 1 \| 584 \| Uncharacterized protein \| Uncharacterized protein; afdb-uniprot50 \| AF-A0A0F9BAB4-F1-MODEL\_V4 \| 1.0 \| 5.443e-27 \| 897 \| 0.357 \| 274 \| 158 \| 10 \| 355 \| 619 \| 2 \| 266 \| Uncharacterized protein \| Uncharacterized protein; afdb-uniprot50 \| AF-A0A521CZS3-F1-MODEL\_V4 \| 1.0 \| 9.326e-41 \| 896 \| 0.237 \| 669 \| 391 \| 30 \| 75 \| 681 \| 4 \| 615 \| Phage terminase, large subunit GpA \| Phage terminase, large subunit GpA; afdb-uniprot50 \| AF-A0A378GMQ5-F1-MODEL\_V4 \| 1.0 \| 8.686e-30 \| 895 \| 0.238 \| 406 \| 283 \| 13 \| 68 \| 462 \| 18 \| 408 \| Bacteriophage tail assembly protein \| Bacteriophage tail assembly protein; afdb-uniprot50 \| AF-A0A7Z1KL01-F1-MODEL\_V4 \| 1.0 \| 3.274e-35 \| 895 \| 0.198 \| 578 \| 380 \| 25 \| 1 \| 516 \| 3 \| 558 \| DNA packaging protein \| DNA packaging protein; afdb-uniprot50 \| AF-A0A1Y6CQE5-F1-MODEL\_V4 \| 1.0 \| 1.187e-34 \| 894 \| 0.25 \| 456 \| 273 \| 18 \| 208 \| 633 \| 6 \| 422 \| Phage terminase large subunit (GpA) \| Phage terminase large subunit (GpA); afdb-uniprot50 \| AF-A0A2T6CPG6-F1-MODEL\_V4 \| 1.0 \| 1.061e-34 \| 894 \| 0.181 \| 661 \| 434 \| 28 \| 4 \| 605 \| 1 \| 613 \| Uncharacterized protein \| Uncharacterized protein; afdb-uniprot50 \| AF-A0A6I4TWI3-F1-MODEL\_V4 \| 1.0 \| 5.882e-39 \| 894 \| 0.2 \| 760 \| 473 \| 36 \| 4 \| 675 \| 1 \| 713 \| Uncharacterized protein \| Uncharacterized protein; afdb-uniprot50 \| AF-A0A1Y2XHW8-F1-MODEL\_V4 \| 1.0 \| 2.834e-30 \| 893 \| 0.245 \| 456 \| 294 \| 16 \| 36 \| 462 \| 1 \| 435 \| Terminase \| Terminase; afdb-uniprot50 \| AF-A0A0F9JWS4-F1-MODEL\_V4 \| 1.0 \| 2.017e-38 \| 893 \| 0.192 \| 659 \| 452 \| 28 \| 22 \| 612 \| 6 \| 652 \| Uncharacterized protein \| Uncharacterized protein; afdb-uniprot50 \| AF-A0A847BW98-F1-MODEL\_V4 \| 1.0 \| 3.232e-33 \| 891 \| 0.194 \| 556 \| 370 \| 23 \| 6 \| 518 \| 9 \| 529 \| Uncharacterized protein \| Uncharacterized protein; afdb-uniprot50 \| AF-A0A062VMA9-F1-MODEL\_V4 \| 1.0 \| 1.401e-25 \| 890 \| 0.402 \| 261 \| 138 \| 7 \| 392 \| 648 \| 2 \| 248 \| Phage terminase GpA \| Phage terminase GpA; afdb-uniprot50 \| AF-A0A842HKI1-F1-MODEL\_V4 \| 1.0 \| 6.062e-35 \| 890 \| 0.172 \| 648 \| 425 \| 31 \| 23 \| 614 \| 2 \| 593 \| Phage terminase large subunit family protein \| Phage terminase large subunit family protein; afdb-uniprot50 \| AF-A0A2A2RKY9-F1-MODEL\_V4 \| 1.0 \| 2.49e-36 \| 890 \| 0.172 \| 671 \| 432 \| 32 \| 18 \| 632 \| 3 \| 605 \| Uncharacterized protein \| Uncharacterized protein; afdb-uniprot50 \| AF-X0YXM7-F1-MODEL\_V4 \| 1.0 \| 8.686e-30 \| 888 \| 0.264 \| 443 \| 271 \| 24 \| 196 \| 604 \| 6 \| 427 \| Uncharacterized protein \| Uncharacterized protein; afdb-uniprot50 \| AF-A0A5C5WEI3-F1-MODEL\_V4 \| 1.0 \| 1.432e-37 \| 886 \| 0.195 \| 681 \| 431 \| 38 \| 28 \| 637 \| 9 \| 643 \| Phage terminase large subunit (GpA) \| Phage terminase large subunit (GpA); afdb-uniprot50 \| AF-A0A6N9PVQ4-F1-MODEL\_V4 \| 1.0 \| 7.616e-27 \| 885 \| 0.318 \| 352 \| 202 \| 11 \| 331 \| 672 \| 9 \| 332 \| Phage terminase large subunit family protein \| Phage terminase large subunit family protein; afdb-uniprot50 \| AF-A0A7V5UQF9-F1-MODEL\_V4 \| 1.0 \| 3.17e-30 \| 881 \| 0.363 \| 344 \| 198 \| 10 \| 298 \| 633 \| 2 \| 332 \| Phage terminase large subunit family protein \| Phage terminase large subunit family protein; afdb-uniprot50 \| AF-A0A2N3ATA9-F1-MODEL\_V4 \| 1.0 \| 3.774e-31 \| 881 \| 0.304 \| 411 \| 233 \| 15 \| 252 \| 632 \| 1 \| 388 \| Phage tail protein \| Phage tail protein; afdb-uniprot50 \| AF-A0A4Q3K415-F1-MODEL\_V4 \| 1.0 \| 1.087e-29 \| 878 \| 0.32 \| 353 \| 210 \| 12 \| 343 \| 677 \| 38 \| 378 \| Phage terminase large subunit family protein \| Phage terminase large subunit family protein; afdb-uniprot50 \| AF-A0A7X1E4I2-F1-MODEL\_V4 \| 1.0 \| 5.769e-36 \| 878 \| 0.187 \| 625 \| 414 \| 31 \| 48 \| 632 \| 2 \| 572 \| Phage terminase large subunit family protein \| Phage terminase large subunit family protein; afdb-uniprot50 \| AF-A0A1V6HQW6-F1-MODEL\_V4 \| 1.0 \| 6.411e-35 \| 878 \| 0.18 \| 743 \| 472 \| 41 \| 1 \| 678 \| 1 \| 671 \| Phage terminase large subunit (GpA) \| Phage terminase large subunit (GpA); afdb-uniprot50 \| AF-A0A212LQT7-F1-MODEL\_V4 \| 1.0 \| 7.783e-39 \| 877 \| 0.21 \| 670 \| 411 \| 31 \| 72 \| 682 \| 5 \| 615 \| Phage terminase GpA \| Phage terminase GpA; afdb-uniprot50 \| AF-A0A2D6EP56-F1-MODEL\_V4 \| 1.0 \| 3.815e-24 \| 875 \| 0.266 \| 360 \| 225 \| 13 \| 1 \| 350 \| 1 \| 331 \| Uncharacterized protein \| Uncharacterized protein; afdb-uniprot50 \| AF-A0A2E4W680-F1-MODEL\_V4 \| 1.0 \| 2.458e-34 \| 873 \| 0.171 \| 623 \| 419 \| 31 \| 19 \| 611 \| 3 \| 558 \| Uncharacterized protein \| Uncharacterized protein; afdb-uniprot50 \| AF-A0A5C7Q7J6-F1-MODEL\_V4 \| 1.0 \| 6.824e-36 \| 872 \| 0.204 \| 660 \| 435 \| 31 \| 3 \| 608 \| 1 \| 624 \| Uncharacterized protein \| Uncharacterized protein; afdb-uniprot50 \| AF-A0A527W6W1-F1-MODEL\_V4 \| 1.0 \| 3.823e-33 \| 870 \| 0.272 \| 418 \| 260 \| 14 \| 228 \| 626 \| 1 \| 393 \| Terminase \| Terminase; afdb-uniprot50 \| AF-A0A6N3UAX6-F1-MODEL\_V4 \| 1.0 \| 5.091e-34 \| 870 \| 0.311 \| 443 \| 253 \| 15 \| 223 \| 634 \| 16 \| 437 \| Terminase \| Terminase; afdb-uniprot50 \| AF-A0A0A1VHF5-F1-MODEL\_V4 \| 1.0 \| 1.413e-35 \| 868 \| 0.265 \| 584 \| 288 \| 20 \| 121 \| 680 \| 2 \| 468 \| Bacteriophage tail assembly protein \| Bacteriophage tail assembly protein; afdb-uniprot50 \| AF-A0A843PU22-F1-MODEL\_V4 \| 1.0 \| 9.406e-25 \| 863 \| 0.335 \| 295 \| 175 \| 8 \| 335 \| 615 \| 7 \| 294 \| Phage terminase large subunit family protein \| Phage terminase large subunit family protein; afdb-uniprot50 \| AF-A0A2N2T364-F1-MODEL\_V4 \| 1.0 \| 1.404e-34 \| 860 \| 0.293 \| 446 \| 277 \| 18 \| 141 \| 561 \| 4 \| 436 \| Terminase \| Terminase; afdb-uniprot50 \| AF-A0A4Q3JQ82-F1-MODEL\_V4 \| 1.0 \| 2.662e-29 \| 859 \| 0.31 \| 370 \| 211 \| 11 \| 333 \| 678 \| 2 \| 351 \| Phage terminase large subunit family protein \| Phage terminase large subunit family protein; afdb-uniprot50 \| AF-A0A1H7AEM5-F1-MODEL\_V4 \| 1.0 \| 5.719e-26 \| 858 \| 0.289 \| 314 \| 179 \| 9 \| 382 \| 678 \| 2 \| 288 \| Phage terminase large subunit (GpA) \| Phage terminase large subunit (GpA); afdb-uniprot50 \| AF-A0A2H0PF84-F1-MODEL\_V4 \| 1.0 \| 1.239e-32 \| 857 \| 0.364 \| 368 \| 205 \| 9 \| 151 \| 514 \| 1 \| 343 \| Terminase \| Terminase; afdb-uniprot50 \| AF-A0A826IDF5-F1-MODEL\_V4 \| 1.0 \| 1.69e-28 \| 855 \| 0.242 \| 445 \| 285 \| 17 \| 19 \| 433 \| 8 \| 430 \| Phage terminase large subunit family protein \| Phage terminase large subunit family protein; afdb-uniprot50 \| AF-A0A520NQB2-F1-MODEL\_V4 \| 1.0 \| 1.598e-28 \| 853 \| 0.354 \| 330 \| 190 \| 5 \| 111 \| 437 \| 17 \| 326 \| Uncharacterized protein \| Uncharacterized protein; afdb-uniprot50 \| AF-A0A4D8QI81-F1-MODEL\_V4 \| 1.0 \| 4.641e-37 \| 852 \| 0.213 \| 713 \| 393 \| 32 \| 1 \| 620 \| 1 \| 638 \| Phage portal protein \| Phage portal protein; afdb-uniprot50 \| AF-A0A1H0NRI3-F1-MODEL\_V4 \| 1.0 \| 1.413e-35 \| 849 \| 0.195 \| 630 \| 410 \| 34 \| 53 \| 632 \| 2 \| 584 \| Phage terminase, large subunit GpA \| Phage terminase, large subunit GpA; afdb-uniprot50 \| AF-A0A3N5Q4N6-F1-MODEL\_V4 \| 1.0 \| 3.823e-33 \| 847 \| 0.194 \| 576 \| 369 \| 28 \| 19 \| 554 \| 4 \| 524 \| Uncharacterized protein \| Uncharacterized protein; afdb-uniprot50 \| AF-A0A3B9NTQ1-F1-MODEL\_V4 \| 1.0 \| 3.288e-27 \| 846 \| 0.282 \| 368 \| 210 \| 18 \| 288 \| 615 \| 3 \| 356 \| Uncharacterized protein \| Uncharacterized protein; afdb-uniprot50 \| AF-A0A2U8G1G7-F1-MODEL\_V4 \| 1.0 \| 4.721e-31 \| 844 \| 0.215 \| 520 \| 319 \| 25 \| 31 \| 482 \| 5 \| 503 \| Uncharacterized protein \| Uncharacterized protein; afdb-uniprot50 \| AF-A0A5C7LMI1-F1-MODEL\_V4 \| 1.0 \| 3.508e-37 \| 844 \| 0.208 \| 724 \| 430 \| 45 \| 1 \| 637 \| 10 \| 677 \| Uncharacterized protein \| Uncharacterized protein; afdb-uniprot50 \| AF-A0A4R4G0L4-F1-MODEL\_V4 \| 1.0 \| 2.708e-23 \| 842 \| 0.294 \| 285 \| 186 \| 8 \| 51 \| 329 \| 1 \| 276 \| Phage terminase large subunit family protein \| Phage terminase large subunit family protein; afdb-uniprot50 \| AF-A0A3E0N1C3-F1-MODEL\_V4 \| 1.0 \| 5.124e-35 \| 842 \| 0.165 \| 684 \| 471 \| 26 \| 21 \| 637 \| 4 \| 654 \| Uncharacterized protein \| Uncharacterized protein; afdb-uniprot50 \| AF-A0A259JD23-F1-MODEL\_V4 \| 1.0 \| 2.902e-25 \| 841 \| 0.303 \| 363 \| 221 \| 10 \| 64 \| 411 \| 2 \| 347 \| Uncharacterized protein \| Uncharacterized protein; afdb-uniprot50 \| AF-A0A450TWC8-F1-MODEL\_V4 \| 1.0 \| 1.354e-37 \| 841 \| 0.28 \| 538 \| 338 \| 21 \| 141 \| 636 \| 10 \| 540 \| Phage terminase, large subunit GpA \| Phage terminase, large subunit GpA; afdb-uniprot50 \| AF-A0A327K9L2-F1-MODEL\_V4 \| 1.0 \| 1.636e-23 \| 840 \| 0.274 \| 350 \| 203 \| 9 \| 53 \| 365 \| 1 \| 336 \| Uncharacterized protein \| Uncharacterized protein; afdb-uniprot50 \| AF-A0A1M7U327-F1-MODEL\_V4 \| 1.0 \| 4.742e-23 \| 839 \| 0.203 \| 339 \| 223 \| 12 \| 39 \| 372 \| 4 \| 300 \| Phage terminase large subunit (GpA) \| Phage terminase large subunit (GpA); afdb-uniprot50 \| AF-A0A0F8VYW0-F1-MODEL\_V4 \| 1.0 \| 2.179e-24 \| 839 \| 0.275 \| 316 \| 200 \| 11 \| 328 \| 620 \| 2 \| 311 \| Uncharacterized protein \| Uncharacterized protein; afdb-uniprot50 \| AF-A0A7A6TWF3-F1-MODEL\_V4 \| 1.0 \| 1.101e-31 \| 838 \| 0.294 \| 435 \| 245 \| 17 \| 223 \| 616 \| 6 \| 419 \| Phage terminase large subunit family protein \| Phage terminase large subunit family protein; afdb-uniprot50 \| AF-A0A827QZE9-F1-MODEL\_V4 \| 1.0 \| 6.088e-27 \| 837 \| 0.296 \| 371 \| 220 \| 12 \| 283 \| 623 \| 1 \| 360 \| Phage terminase large subunit family protein \| Phage terminase large subunit family protein; afdb-uniprot50 \| AF-A0A853EE69-F1-MODEL\_V4 \| 1.0 \| 1.264e-35 \| 836 \| 0.285 \| 494 \| 289 \| 19 \| 149 \| 606 \| 20 \| 485 \| Phage terminase large subunit family protein \| Phage terminase large subunit family protein; afdb-uniprot50 \| AF-A0A7G6DUX0-F1-MODEL\_V4 \| 1.0 \| 2.065e-33 \| 836 \| 0.164 \| 615 \| 431 \| 28 \| 19 \| 605 \| 4 \| 563 \| Phage terminase large subunit family protein \| Phage terminase large subunit family protein; afdb-uniprot50 \| AF-A0A377M1M2-F1-MODEL\_V4 \| 1.0 \| 9.246e-31 \| 832 \| 0.205 \| 511 \| 330 \| 22 \| 24 \| 473 \| 8 \| 503 \| Phage terminase large subunit \| Phage terminase large subunit; afdb-uniprot50 \| AF-A0A645DZF6-F1-MODEL\_V4 \| 1.0 \| 8.894e-25 \| 831 \| 0.312 \| 307 \| 189 \| 6 \| 346 \| 632 \| 2 \| 306 \| Uncharacterized protein \| Uncharacterized protein; afdb-uniprot50 \| AF-A0A5Q3L868-F1-MODEL\_V4 \| 1.0 \| 4.332e-35 \| 828 \| 0.182 \| 668 \| 447 \| 34 \| 3 \| 608 \| 1 \| 631 \| Phage terminase large subunit (GpA) \| Phage terminase large subunit (GpA); afdb-uniprot50 \| AF-A0A2N3B688-F1-MODEL\_V4 \| 1.0 \| 3.75e-30 \| 827 \| 0.216 \| 500 \| 310 \| 21 \| 26 \| 465 \| 2 \| 479 \| Terminase \| Terminase; afdb-uniprot50 \| AF-Q24VG8-F1-MODEL\_V4 \| 1.0 \| 9.345e-24 \| 826 \| 0.227 \| 339 \| 215 \| 10 \| 39 \| 372 \| 4 \| 300 \| Uncharacterized protein \| Uncharacterized protein; afdb-uniprot50 \| AF-A0A853ECF6-F1-MODEL\_V4 \| 1.0 \| 3.584e-23 \| 825 \| 0.337 \| 270 \| 162 \| 7 \| 347 \| 606 \| 2 \| 264 \| Phage terminase large subunit family protein \| Phage terminase large subunit family protein; afdb-uniprot50 \| AF-B6XJF8-F1-MODEL\_V4 \| 1.0 \| 7.017e-23 \| 825 \| 0.319 \| 257 \| 156 \| 7 \| 375 \| 619 \| 2 \| 251 \| Uncharacterized protein \| Uncharacterized protein; afdb-uniprot50 \| AF-B3CPA2-F1-MODEL\_V4 \| 1.0 \| 4.294e-25 \| 824 \| 0.352 \| 284 \| 162 \| 12 \| 357 \| 626 \| 2 \| 277 \| Phage terminase large subunit GpA \| Phage terminase large subunit GpA; afdb-uniprot50 \| AF-A0A742P1A9-F1-MODEL\_V4 \| 1.0 \| 3.864e-26 \| 824 \| 0.247 \| 440 \| 273 \| 14 \| 19 \| 425 \| 8 \| 422 \| Phage terminase large subunit family protein \| Phage terminase large subunit family protein; afdb-uniprot50 \| AF-A0A3T1DR27-F1-MODEL\_V4 \| 1.0 \| 2.485e-27 \| 823 \| 0.371 \| 312 \| 166 \| 13 \| 336 \| 632 \| 3 \| 299 \| Uncharacterized protein \| Uncharacterized protein; afdb-uniprot50 \| AF-A0A369QUH9-F1-MODEL\_V4 \| 1.0 \| 9.884e-24 \| 823 \| 0.248 \| 382 \| 231 \| 22 \| 16 \| 371 \| 5 \| 356 \| Uncharacterized protein \| Uncharacterized protein; afdb-uniprot50 \| AF-A0A527ITJ7-F1-MODEL\_V4 \| 1.0 \| 9.406e-25 \| 823 \| 0.277 \| 396 \| 249 \| 15 \| 3 \| 384 \| 1 \| 373 \| Terminase \| Terminase; afdb-uniprot50 \| AF-A0A3U0PIZ9-F1-MODEL\_V4 \| 1.0 \| 7.518e-25 \| 822 \| 0.339 \| 306 \| 178 \| 10 \| 335 \| 623 \| 13 \| 311 \| Phage terminase large subunit family protein \| Phage terminase large subunit family protein; afdb-uniprot50 \| AF-A0A0F8X804-F1-MODEL\_V4 \| 1.0 \| 7.666e-28 \| 819 \| 0.307 \| 354 \| 209 \| 8 \| 99 \| 437 \| 2 \| 334 \| Uncharacterized protein \| Uncharacterized protein; afdb-uniprot50 \| AF-A0A6L3XYL3-F1-MODEL\_V4 \| 1.0 \| 3.19e-31 \| 818 \| 0.19 \| 529 \| 349 \| 22 \| 23 \| 489 \| 7 \| 518 \| Phage terminase large subunit family protein \| Phage terminase large subunit family protein; afdb-uniprot50 \| AF-A0A3M6FD43-F1-MODEL\_V4 \| 1.0 \| 9.285e-23 \| 817 \| 0.213 \| 351 \| 228 \| 15 \| 39 \| 371 \| 2 \| 322 \| Terminase, large subunit \| Terminase, large subunit; afdb-uniprot50 \| AF-A0A2U8VPY4-F1-MODEL\_V4 \| 1.0 \| 1.169e-23 \| 816 \| 0.279 \| 333 \| 217 \| 7 \| 1 \| 331 \| 1 \| 312 \| Uncharacterized protein \| Uncharacterized protein; afdb-uniprot50 \| AF-A0A3N5QG09-F1-MODEL\_V4 \| 1.0 \| 2.939e-27 \| 816 \| 0.216 \| 435 \| 285 \| 20 \| 19 \| 430 \| 12 \| 413 \| Uncharacterized protein \| Uncharacterized protein; afdb-uniprot50 \| AF-A0A454TI69-F1-MODEL\_V4 \| 1.0 \| 1.902e-29 \| 815 \| 0.316 \| 395 \| 208 \| 17 \| 317 \| 681 \| 9 \| 371 \| Phage terminase large subunit family protein \| Phage terminase large subunit family protein; afdb-uniprot50 \| AF-A0A3S1CDJ9-F1-MODEL\_V4 \| 1.0 \| 6.329e-33 \| 815 \| 0.176 \| 584 \| 377 \| 24 \| 13 \| 517 \| 2 \| 560 \| Uncharacterized protein \| Uncharacterized protein; afdb-uniprot50 \| AF-A0A2D4TR23-F1-MODEL\_V4 \| 1.0 \| 3.388e-23 \| 814 \| 0.348 \| 238 \| 144 \| 4 \| 395 \| 632 \| 2 \| 228 \| Uncharacterized protein \| Uncharacterized protein; afdb-uniprot50 \| AF-A0A4Q3AGK8-F1-MODEL\_V4 \| 1.0 \| 1.308e-23 \| 811 \| 0.23 \| 368 \| 250 \| 14 \| 17 \| 372 \| 6 \| 352 \| Uncharacterized protein \| Uncharacterized protein; afdb-uniprot50 \| AF-A0A1Q9KNH1-F1-MODEL\_V4 \| 1.0 \| 4.35e-27 \| 810 \| 0.25 \| 444 \| 275 \| 20 \| 19 \| 429 \| 8 \| 426 \| Terminase \| Terminase; afdb-uniprot50 \| AF-A0A839IWR0-F1-MODEL\_V4 \| 1.0 \| 1.137e-36 \| 810 \| 0.275 \| 570 \| 338 \| 24 \| 158 \| 682 \| 4 \| 543 \| Phage terminase large subunit family protein \| Phage terminase large subunit family protein; afdb-uniprot50 \| AF-A0A5C5VC85-F1-MODEL\_V4 \| 1.0 \| 9.488e-35 \| 810 \| 0.15 \| 755 \| 506 \| 32 \| 1 \| 646 \| 3 \| 730 \| Phage terminase large subunit (GpA) \| Phage terminase large subunit (GpA); afdb-uniprot50 \| AF-A0A6L5VNK7-F1-MODEL\_V4 \| 1.0 \| 1.054e-33 \| 806 \| 0.255 \| 543 \| 273 \| 21 \| 172 \| 614 \| 3 \| 513 \| Uncharacterized protein \| Uncharacterized protein; afdb-uniprot50 \| AF-S9RVT3-F1-MODEL\_V4 \| 1.0 \| 7.584e-35 \| 804 \| 0.235 \| 585 \| 349 \| 20 \| 141 \| 679 \| 8 \| 539 \| Uncharacterized protein \| Uncharacterized protein; afdb-uniprot50 \| AF-A0A1Y6C0B5-F1-MODEL\_V4 \| 1.0 \| 6.722e-25 \| 803 \| 0.31 \| 329 \| 196 \| 13 \| 317 \| 621 \| 9 \| 330 \| Phage terminase large subunit (GpA) \| Phage terminase large subunit (GpA); afdb-uniprot50 \| AF-A0A0S4VSP0-F1-MODEL\_V4 \| 1.0 \| 5.281e-31 \| 802 \| 0.297 \| 447 \| 262 \| 14 \| 262 \| 681 \| 11 \| 432 \| Terminase GpA \| Terminase GpA; afdb-uniprot50 \| AF-X0S680-F1-MODEL\_V4 \| 1.0 \| 3.991e-31 \| 802 \| 0.22 \| 481 \| 338 \| 18 \| 151 \| 605 \| 6 \| 475 \| Uncharacterized protein \| Uncharacterized protein; afdb-uniprot50 \| AF-A0A7C2B2N3-F1-MODEL\_V4 \| 1.0 \| 1.157e-30 \| 799 \| 0.352 \| 392 \| 214 \| 15 \| 283 \| 645 \| 15 \| 395 \| Phage terminase large subunit family protein \| Phage terminase large subunit family protein; afdb-uniprot50 \| AF-A0A4R3NP09-F1-MODEL\_V4 \| 1.0 \| 1.453e-22 \| 798 \| 0.292 \| 349 \| 219 \| 12 \| 14 \| 355 \| 3 \| 330 \| Phage terminase large subunit GpA \| Phage terminase large subunit GpA; afdb-uniprot50 \| AF-A0A522WV68-F1-MODEL\_V4 \| 1.0 \| 3.839e-25 \| 796 \| 0.333 \| 312 \| 174 \| 11 \| 355 \| 637 \| 11 \| 317 \| Phage terminase large subunit family protein \| Phage terminase large subunit family protein; afdb-uniprot50 \| AF-A0A0F9EXV6-F1-MODEL\_V4 \| 1.0 \| 1.577e-26 \| 796 \| 0.237 \| 442 \| 272 \| 22 \| 236 \| 628 \| 2 \| 427 \| Uncharacterized protein \| Uncharacterized protein; afdb-uniprot50 \| AF-A0A537X5I4-F1-MODEL\_V4 \| 1.0 \| 3.568e-31 \| 796 \| 0.195 \| 574 \| 378 \| 22 \| 38 \| 607 \| 10 \| 503 \| Uncharacterized protein \| Uncharacterized protein; afdb-uniprot50 \| AF-A0A5C5VSZ6-F1-MODEL\_V4 \| 1.0 \| 5.407e-26 \| 794 \| 0.225 \| 443 \| 295 \| 15 \| 3 \| 416 \| 1 \| 424 \| Phage terminase large subunit (GpA) \| Phage terminase large subunit (GpA); afdb-uniprot50 \| AF-A0A4Q3NM87-F1-MODEL\_V4 \| 1.0 \| 1.567e-25 \| 792 \| 0.306 \| 352 \| 199 \| 13 \| 350 \| 680 \| 2 \| 329 \| Uncharacterized protein \| Uncharacterized protein; afdb-uniprot50 \| AF-A0A3N9N400-F1-MODEL\_V4 \| 1.0 \| 1.466e-32 \| 791 \| 0.156 \| 702 \| 482 \| 30 \| 19 \| 647 \| 7 \| 671 \| Uncharacterized protein \| Uncharacterized protein; afdb-uniprot50 \| AF-E6BU52-F1-MODEL\_V4 \| 1.0 \| 2.179e-24 \| 789 \| 0.331 \| 323 \| 191 \| 12 \| 318 \| 623 \| 11 \| 325 \| Phage terminase large subunit (GpA) \| Phage terminase large subunit (GpA); afdb-uniprot50 \| AF-A0A381KUK4-F1-MODEL\_V4 \| 1.0 \| 3.477e-27 \| 789 \| 0.351 \| 341 \| 184 \| 9 \| 288 \| 618 \| 3 \| 316 \| Phage terminase large subunit \| Phage terminase large subunit; afdb-uniprot50 \| AF-A0A2X2RZY6-F1-MODEL\_V4 \| 1.0 \| 2.06e-24 \| 789 \| 0.197 \| 390 \| 254 \| 16 \| 24 \| 372 \| 8 \| 379 \| Phage terminase large subunit \| Phage terminase large subunit; afdb-uniprot50 \| AF-A0A3D0WC11-F1-MODEL\_V4 \| 1.0 \| 1.472e-24 \| 787 \| 0.251 \| 401 \| 253 \| 17 \| 3 \| 386 \| 1 \| 371 \| Terminase \| Terminase; afdb-uniprot50 \| AF-A0A0E2L746-F1-MODEL\_V4 \| 1.0 \| 2.708e-23 \| 786 \| 0.232 \| 408 \| 255 \| 14 \| 19 \| 393 \| 8 \| 390 \| Large terminase \| Large terminase; afdb-uniprot50 \| AF-A0A1T5CSF4-F1-MODEL\_V4 \| 1.0 \| 8.32e-32 \| 786 \| 0.151 \| 620 \| 435 \| 34 \| 38 \| 632 \| 19 \| 572 \| Phage terminase large subunit (GpA) \| Phage terminase large subunit (GpA); afdb-uniprot50 \| AF-A0A1G8I030-F1-MODEL\_V4 \| 1.0 \| 5.35e-33 \| 785 \| 0.279 \| 512 \| 283 \| 22 \| 178 \| 642 \| 2 \| 474 \| Phage terminase, large subunit GpA \| Phage terminase, large subunit GpA; afdb-uniprot50 \| AF-A0A8B3NLS1-F1-MODEL\_V4 \| 1.0 \| 1.169e-23 \| 784 \| 0.232 \| 409 \| 263 \| 15 \| 1 \| 389 \| 1 \| 378 \| Terminase \| Terminase; afdb-uniprot50 \| AF-A0A3R0XSB2-F1-MODEL\_V4 \| 1.0 \| 1.647e-24 \| 784 \| 0.235 \| 433 \| 273 \| 14 \| 19 \| 418 \| 8 \| 415 \| Phage terminase large subunit family protein \| Phage terminase large subunit family protein; afdb-uniprot50 \| AF-A0A2V5PXK1-F1-MODEL\_V4 \| 1.0 \| 9.366e-33 \| 783 \| 0.16 \| 680 \| 431 \| 41 \| 15 \| 607 \| 2 \| 628 \| Uncharacterized protein \| Uncharacterized protein; afdb-uniprot50 \| AF-A0A358B3M6-F1-MODEL\_V4 \| 1.0 \| 3.815e-24 \| 782 \| 0.321 \| 286 \| 171 \| 11 \| 355 \| 637 \| 9 \| 274 \| Terminase \| Terminase; afdb-uniprot50 \| AF-A0A6I1JTB6-F1-MODEL\_V4 \| 1.0 \| 6.678e-24 \| 780 \| 0.305 \| 291 \| 164 \| 10 \| 367 \| 637 \| 9 \| 281 \| Uncharacterized protein \| Uncharacterized protein; afdb-uniprot50 \| AF-A0A6I2KEC9-F1-MODEL\_V4 \| 1.0 \| 1.045e-23 \| 778 \| 0.52 \| 250 \| 95 \| 3 \| 435 \| 682 \| 1 \| 227 \| Phage terminase large subunit family protein \| Phage terminase large subunit family protein; afdb-uniprot50 \| AF-A0A5C7PZL4-F1-MODEL\_V4 \| 1.0 \| 1.723e-31 \| 778 \| 0.175 \| 666 \| 427 \| 40 \| 3 \| 607 \| 1 \| 605 \| Uncharacterized protein \| Uncharacterized protein; afdb-uniprot50 \| AF-A0A1X0Y5S6-F1-MODEL\_V4 \| 1.0 \| 8.779e-23 \| 776 \| 0.34 \| 264 \| 152 \| 8 \| 375 \| 623 \| 2 \| 258 \| Uncharacterized protein \| Uncharacterized protein; afdb-uniprot50 \| AF-A0A1V2N6H4-F1-MODEL\_V4 \| 1.0 \| 4.113e-27 \| 775 \| 0.337 \| 347 \| 194 \| 17 \| 317 \| 644 \| 7 \| 336 \| Phage terminase large subunit family protein \| Phage terminase large subunit family protein; afdb-uniprot50 \| AF-A0A3R0Z272-F1-MODEL\_V4 \| 1.0 \| 3.19e-31 \| 775 \| 0.255 \| 489 \| 286 \| 20 \| 104 \| 555 \| 3 \| 450 \| Phage terminase large subunit family protein \| Phage terminase large subunit family protein; afdb-uniprot50 \| AF-A0A376JNQ2-F1-MODEL\_V4 \| 1.0 \| 7.849e-23 \| 773 \| 0.329 \| 288 \| 170 \| 10 \| 344 \| 616 \| 16 \| 295 \| Phage terminase large subunit (GpA) \| Phage terminase large subunit (GpA); afdb-uniprot50 \| AF-A0A1M4LEQ2-F1-MODEL\_V4 \| 1.0 \| 4.35e-27 \| 773 \| 0.179 \| 458 \| 307 \| 17 \| 22 \| 430 \| 7 \| 444 \| Uncharacterized protein \| Uncharacterized protein; afdb-uniprot50 \| AF-A0A5C7Q184-F1-MODEL\_V4 \| 1.0 \| 5.281e-31 \| 769 \| 0.166 \| 650 \| 414 \| 39 \| 22 \| 604 \| 3 \| 591 \| Uncharacterized protein \| Uncharacterized protein; afdb-uniprot50 \| AF-A0A6H1ZZ96-F1-MODEL\_V4 \| 1.0 \| 3.352e-30 \| 767 \| 0.189 \| 618 \| 388 \| 32 \| 14 \| 610 \| 3 \| 528 \| Putative terminase \| Putative terminase; afdb-uniprot50 \| AF-A0A062VEY8-F1-MODEL\_V4 \| 1.0 \| 3.01e-22 \| 766 \| 0.295 \| 257 \| 157 \| 8 \| 385 \| 640 \| 1 \| 234 \| Terminase GpA \| Terminase GpA; afdb-uniprot50 \| AF-S9S4Q4-F1-MODEL\_V4 \| 1.0 \| 3.766e-22 \| 766 \| 0.23 \| 377 \| 255 \| 13 \| 1 \| 369 \| 1 \| 350 \| Terminase GpA \| Terminase GpA; afdb-uniprot50 \| AF-A0A6G3FD37-F1-MODEL\_V4 \| 1.0 \| 1.245e-24 \| 764 \| 0.313 \| 351 \| 204 \| 13 \| 280 \| 613 \| 3 \| 333 \| Phage terminase large subunit family protein \| Phage terminase large subunit family protein; afdb-uniprot50 \| AF-A0A847VIV5-F1-MODEL\_V4 \| 1.0 \| 1.605e-20 \| 763 \| 0.314 \| 283 \| 173 \| 6 \| 21 \| 298 \| 6 \| 272 \| Phage terminase large subunit family protein \| Phage terminase large subunit family protein; afdb-uniprot50 \| AF-A0A830WLI0-F1-MODEL\_V4 \| 1.0 \| 8.779e-23 \| 763 \| 0.353 \| 269 \| 158 \| 9 \| 335 \| 594 \| 12 \| 273 \| Phage terminase large subunit family protein \| Phage terminase large subunit family protein; afdb-uniprot50 \| AF-A0A4C3FSC6-F1-MODEL\_V4 \| 1.0 \| 1.277e-28 \| 763 \| 0.234 \| 457 \| 289 \| 15 \| 75 \| 497 \| 3 \| 432 \| Phage terminase large subunit \| Phage terminase large subunit; afdb-uniprot50 \| AF-A0A257VFA9-F1-MODEL\_V4 \| 1.0 \| 7.326e-21 \| 761 \| 0.332 \| 286 \| 171 \| 5 \| 16 \| 296 \| 10 \| 280 \| Phage tail protein \| Phage tail protein; afdb-uniprot50 \| AF-A0A7V0MM37-F1-MODEL\_V4 \| 1.0 \| 3.367e-22 \| 760 \| 0.251 \| 385 \| 219 \| 19 \| 3 \| 350 \| 1 \| 353 \| Uncharacterized protein \| Uncharacterized protein; afdb-uniprot50 \| AF-A0A1V6E2A3-F1-MODEL\_V4 \| 1.0 \| 5.658e-33 \| 760 \| 0.185 \| 695 \| 449 \| 33 \| 19 \| 632 \| 2 \| 660 \| Phage terminase large subunit (GpA) \| Phage terminase large subunit (GpA); afdb-uniprot50 \| AF-H0T4D4-F1-MODEL\_V4 \| 1.0 \| 6.972e-22 \| 756 \| 0.266 \| 255 \| 163 \| 5 \| 377 \| 616 \| 2 \| 247 \| Uncharacterized protein \| Uncharacterized protein; afdb-uniprot50 \| AF-A0A062VB30-F1-MODEL\_V4 \| 1.0 \| 1.356e-20 \| 753 \| 0.339 \| 215 \| 129 \| 5 \| 385 \| 599 \| 1 \| 202 \| Terminase GpA \| Terminase GpA; afdb-uniprot50 \| AF-A0A0F9ELH0-F1-MODEL\_V4 \| 1.0 \| 2.691e-22 \| 752 \| 0.342 \| 260 \| 142 \| 10 \| 381 \| 619 \| 2 \| 253 \| Uncharacterized protein \| Uncharacterized protein; afdb-uniprot50 \| AF-A0A4Q2YME8-F1-MODEL\_V4 \| 1.0 \| 2.427e-32 \| 751 \| 0.161 \| 729 \| 445 \| 41 \| 1 \| 603 \| 2 \| 689 \| Uncharacterized protein \| Uncharacterized protein; afdb-uniprot50 \| AF-A0A318EPN7-F1-MODEL\_V4 \| 1.0 \| 4.95e-21 \| 750 \| 0.237 \| 316 \| 215 \| 11 \| 13 \| 321 \| 3 \| 299 \| Phage terminase large subunit GpA \| Phage terminase large subunit GpA; afdb-uniprot50 \| AF-A0A0F9FSR4-F1-MODEL\_V4 \| 1.0 \| 5.113e-26 \| 750 \| 0.322 \| 350 \| 199 \| 10 \| 102 \| 434 \| 103 \| 431 \| Uncharacterized protein \| Uncharacterized protein; afdb-uniprot50 \| AF-A0A518G2X6-F1-MODEL\_V4 \| 1.0 \| 3.592e-32 \| 747 \| 0.142 \| 723 \| 501 \| 34 \| 1 \| 632 \| 37 \| 731 \| Phage terminase large subunit (GpA) \| Phage terminase large subunit (GpA); afdb-uniprot50 \| AF-A0A7C1N5P3-F1-MODEL\_V4 \| 1.0 \| 1.2e-27 \| 745 \| 0.204 \| 518 \| 325 \| 21 \| 2 \| 448 \| 22 \| 523 \| Uncharacterized protein \| Uncharacterized protein; afdb-uniprot50 \| AF-A0A430BX37-F1-MODEL\_V4 \| 1.0 \| 1.392e-24 \| 743 \| 0.251 \| 449 \| 273 \| 23 \| 259 \| 681 \| 1 \| 412 \| Terminase \| Terminase; afdb-uniprot50 \| AF-A0A7X5Z5Z9-F1-MODEL\_V4 \| 1.0 \| 2.797e-28 \| 742 \| 0.266 \| 450 \| 247 \| 18 \| 248 \| 679 \| 2 \| 386 \| Phage terminase large subunit family protein \| Phage terminase large subunit family protein; afdb-uniprot50 \| AF-S1IKM4-F1-MODEL\_V4 \| 1.0 \| 6.274e-23 \| 741 \| 0.218 \| 403 \| 265 \| 17 \| 19 \| 392 \| 8 \| 389 \| Large terminase \| Large terminase; afdb-uniprot50 \| AF-A0A827Z2U0-F1-MODEL\_V4 \| 1.0 \| 1.291e-21 \| 739 \| 0.211 \| 378 \| 257 \| 15 \| 19 \| 372 \| 8 \| 368 \| Phage terminase large subunit family protein \| Phage terminase large subunit family protein; afdb-uniprot50 \| AF-A0A1G8VFE7-F1-MODEL\_V4 \| 1.0 \| 5.047e-24 \| 737 \| 0.25 \| 439 \| 260 \| 14 \| 23 \| 416 \| 8 \| 422 \| Phage terminase large subunit (GpA) \| Phage terminase large subunit (GpA); afdb-uniprot50 \| AF-A0A6I1JCS5-F1-MODEL\_V4 \| 1.0 \| 9.82e-23 \| 736 \| 0.319 \| 310 \| 157 \| 12 \| 392 \| 678 \| 1 \| 279 \| Phage terminase large subunit (GpA) \| Phage terminase large subunit (GpA); afdb-uniprot50 \| AF-A0A2W6V870-F1-MODEL\_V4 \| 1.0 \| 1.134e-27 \| 736 \| 0.275 \| 378 \| 240 \| 13 \| 263 \| 631 \| 4 \| 356 \| Uncharacterized protein \| Uncharacterized protein; afdb-uniprot50 \| AF-A0A380TG39-F1-MODEL\_V4 \| 1.0 \| 3.388e-23 \| 734 \| 0.309 \| 304 \| 158 \| 9 \| 395 \| 680 \| 2 \| 271 \| Phage terminase large subunit (GpA) \| Phage terminase large subunit (GpA); afdb-uniprot50 \| AF-Q2W750-F1-MODEL\_V4 \| 1.0 \| 1.164e-31 \| 733 \| 0.229 \| 497 \| 288 \| 20 \| 198 \| 618 \| 26 \| 503 \| Bacteriophage tail assembly protein \| Bacteriophage tail assembly protein; afdb-uniprot50 \| AF-A0A329VGZ4-F1-MODEL\_V4 \| 1.0 \| 3.374e-31 \| 730 \| 0.372 \| 379 \| 197 \| 14 \| 203 \| 561 \| 5 \| 362 \| Terminase \| Terminase; afdb-uniprot50 \| AF-I6B2X7-F1-MODEL\_V4 \| 1.0 \| 1.54e-31 \| 729 \| 0.156 \| 685 \| 439 \| 35 \| 24 \| 633 \| 2 \| 622 \| Bacteriophage tail assembly protein \| Bacteriophage tail assembly protein; afdb-uniprot50 \| AF-A0A382VDL2-F1-MODEL\_V4 \| 1.0 \| 2.92e-26 \| 727 \| 0.366 \| 300 \| 170 \| 7 \| 176 \| 471 \| 2 \| 285 \| Uncharacterized protein \| Uncharacterized protein; afdb-uniprot50 \| AF-A0A661I5B0-F1-MODEL\_V4 \| 1.0 \| 1.557e-24 \| 726 \| 0.285 \| 336 \| 203 \| 14 \| 288 \| 608 \| 9 \| 322 \| Uncharacterized protein \| Uncharacterized protein; afdb-uniprot50 \| AF-A0A7A2LFN7-F1-MODEL\_V4 \| 1.0 \| 1.12e-25 \| 725 \| 0.294 \| 391 \| 215 \| 17 \| 263 \| 616 \| 17 \| 383 \| Phage terminase large subunit family protein \| Phage terminase large subunit family protein; afdb-uniprot50 \| AF-A0A2M9PAZ4-F1-MODEL\_V4 \| 1.0 \| 4.397e-20 \| 723 \| 0.33 \| 275 \| 173 \| 7 \| 1 \| 269 \| 8 \| 277 \| Phage tail protein \| Phage tail protein; afdb-uniprot50 \| AF-A0A6M3KIZ8-F1-MODEL\_V4 \| 1.0 \| 7.765e-30 \| 723 \| 0.168 \| 598 \| 386 \| 30 \| 39 \| 607 \| 11 \| 525 \| Putative terminase \| Putative terminase; afdb-uniprot50 \| AF-D7XLX5-F1-MODEL\_V4 \| 1.0 \| 6.972e-22 \| 722 \| 0.199 \| 371 \| 239 \| 15 \| 39 \| 369 \| 1 \| 353 \| Phage terminase large subunit (GpA) \| Phage terminase large subunit (GpA); afdb-uniprot50 \| AF-A0A4Y8EE10-F1-MODEL\_V4 \| 1.0 \| 2.577e-24 \| 719 \| 0.201 \| 436 \| 276 \| 19 \| 39 \| 424 \| 1 \| 414 \| DNA packaging protein \| DNA packaging protein; afdb-uniprot50 \| AF-A0A1H4SJ75-F1-MODEL\_V4 \| 1.0 \| 1.215e-29 \| 719 \| 0.157 \| 678 \| 460 \| 39 \| 14 \| 677 \| 2 \| 581 \| Phage terminase large subunit (GpA) \| Phage terminase large subunit (GpA); afdb-uniprot50 \| AF-A0A662JRG2-F1-MODEL\_V4 \| 1.0 \| 2.165e-23 \| 718 \| 0.307 \| 315 \| 179 \| 11 \| 344 \| 649 \| 23 \| 307 \| Uncharacterized protein \| Uncharacterized protein; afdb-uniprot50 \| AF-A0A4Q8X3C7-F1-MODEL\_V4 \| 1.0 \| 2.495e-19 \| 717 \| 0.3 \| 293 \| 188 \| 8 \| 18 \| 306 \| 3 \| 282 \| Terminase \| Terminase; afdb-uniprot50 \| AF-A0A440JQM9-F1-MODEL\_V4 \| 1.0 \| 3.163e-21 \| 717 \| 0.23 \| 364 \| 237 \| 13 \| 7 \| 351 \| 1 \| 340 \| Uncharacterized protein \| Uncharacterized protein; afdb-uniprot50 \| AF-A0A1H5ZFL4-F1-MODEL\_V4 \| 1.0 \| 1.708e-21 \| 717 \| 0.297 \| 296 \| 180 \| 7 \| 98 \| 385 \| 469 \| 744 \| Intein C-terminal splicing region \| Intein C-terminal splicing region; afdb-uniprot50 \| AF-A0A1I1MSX7-F1-MODEL\_V4 \| 1.0 \| 3.142e-20 \| 716 \| 0.301 \| 295 \| 185 \| 6 \| 1 \| 290 \| 54 \| 332 \| Phage terminase large subunit (GpA) \| Phage terminase large subunit (GpA); afdb-uniprot50 \| AF-A0A4Q6F2I6-F1-MODEL\_V4 \| 1.0 \| 9.59e-28 \| 716 \| 0.278 \| 399 \| 245 \| 15 \| 155 \| 541 \| 1 \| 368 \| Uncharacterized protein \| Uncharacterized protein; afdb-uniprot50 \| AF-A0A367M9T2-F1-MODEL\_V4 \| 1.0 \| 1.299e-22 \| 715 \| 0.304 \| 305 \| 179 \| 12 \| 359 \| 633 \| 9 \| 310 \| Phage terminase large subunit family protein \| Phage terminase large subunit family protein; afdb-uniprot50 \| AF-A0A1V5F4U4-F1-MODEL\_V4 \| 1.0 \| 1.162e-22 \| 713 \| 0.299 \| 307 \| 163 \| 13 \| 395 \| 680 \| 2 \| 277 \| Phage terminase large subunit (GpA) \| Phage terminase large subunit (GpA); afdb-uniprot50 \| AF-A0A6A8A1L0-F1-MODEL\_V4 \| 1.0 \| 7.017e-23 \| 713 \| 0.28 \| 325 \| 189 \| 15 \| 384 \| 679 \| 1 \| 309 \| Phage terminase large subunit family protein \| Phage terminase large subunit family protein; afdb-uniprot50 \| AF-A0A4S0VF86-F1-MODEL\_V4 \| 1.0 \| 2.25e-29 \| 712 \| 0.145 \| 632 \| 451 \| 34 \| 21 \| 632 \| 4 \| 566 \| Uncharacterized protein \| Uncharacterized protein; afdb-uniprot50 \| AF-A0A7W0A032-F1-MODEL\_V4 \| 1.0 \| 1.521e-29 \| 711 \| 0.17 \| 664 \| 450 \| 34 \| 1 \| 632 \| 1 \| 595 \| Phage terminase large subunit family protein \| Phage terminase large subunit family protein; afdb-uniprot50 \| AF-A0A6D0FA16-F1-MODEL\_V4 \| 1.0 \| 5.202e-20 \| 709 \| 0.366 \| 221 \| 126 \| 7 \| 386 \| 599 \| 1 \| 214 \| Phage terminase large subunit family protein \| Phage terminase large subunit family protein; afdb-uniprot50 \| AF-A0A2X0QVC0-F1-MODEL\_V4 \| 1.0 \| 3.957e-21 \| 709 \| 0.276 \| 329 \| 203 \| 11 \| 1 \| 319 \| 2 \| 305 \| Uncharacterized protein \| Uncharacterized protein; afdb-uniprot50 \| AF-A0A1H8WM18-F1-MODEL\_V4 \| 1.0 \| 1.866e-26 \| 706 \| 0.257 \| 407 \| 240 \| 15 \| 227 \| 596 \| 1 \| 382 \| Phage terminase large subunit (GpA) \| Phage terminase large subunit (GpA); afdb-uniprot50 \| AF-Q4EBR6-F1-MODEL\_V4 \| 1.0 \| 2.26e-21 \| 703 \| 0.352 \| 258 \| 138 \| 11 \| 398 \| 641 \| 2 \| 244 \| Uncharacterized protein \| Uncharacterized protein; afdb-uniprot50 \| AF-A0A827LNW8-F1-MODEL\_V4 \| 1.0 \| 1.025e-20 \| 702 \| 0.186 \| 364 \| 245 \| 12 \| 17 \| 347 \| 2 \| 347 \| Phage terminase large subunit family protein \| Phage terminase large subunit family protein; afdb-uniprot50 \| AF-A0A0F9SQU8-F1-MODEL\_V4 \| 1.0 \| 2.501e-28 \| 702 \| 0.166 \| 692 \| 435 \| 38 \| 3 \| 678 \| 1 \| 566 \| Uncharacterized protein \| Uncharacterized protein; afdb-uniprot50 \| AF-A0A327KGK2-F1-MODEL\_V4 \| 1.0 \| 6.635e-23 \| 700 \| 0.313 \| 344 \| 192 \| 15 \| 345 \| 679 \| 2 \| 310 \| Uncharacterized protein \| Uncharacterized protein; afdb-uniprot50 \| AF-A0A828GA55-F1-MODEL\_V4 \| 1.0 \| 1.025e-20 \| 697 \| 0.18 \| 372 \| 258 \| 11 \| 17 \| 357 \| 2 \| 357 \| Phage terminase large subunit family protein \| Phage terminase large subunit family protein; afdb-uniprot50 \| AF-A0A495FWY6-F1-MODEL\_V4 \| 1.0 \| 1.277e-28 \| 695 \| 0.155 \| 632 \| 445 \| 34 \| 13 \| 632 \| 2 \| 556 \| Phage terminase large subunit GpA \| Phage terminase large subunit GpA; afdb-uniprot50 \| AF-A0A1V6IVJ1-F1-MODEL\_V4 \| 1.0 \| 1.177e-24 \| 692 \| 0.322 \| 357 \| 188 \| 15 \| 291 \| 624 \| 2 \| 327 \| Phage terminase large subunit (GpA) \| Phage terminase large subunit (GpA); afdb-uniprot50 \| AF-A0A4Y7RJZ5-F1-MODEL\_V4 \| 1.0 \| 5.513e-29 \| 692 \| 0.155 \| 590 \| 399 \| 27 \| 38 \| 607 \| 14 \| 523 \| Uncharacterized protein \| Uncharacterized protein; afdb-uniprot50 \| AF-A0A521KKJ7-F1-MODEL\_V4 \| 1.0 \| 5.513e-29 \| 687 \| 0.18 \| 672 \| 378 \| 36 \| 4 \| 605 \| 1 \| 569 \| Uncharacterized protein \| Uncharacterized protein; afdb-uniprot50 \| AF-A0A2D6MBB3-F1-MODEL\_V4 \| 1.0 \| 4.691e-30 \| 686 \| 0.16 \| 662 \| 432 \| 35 \| 19 \| 634 \| 4 \| 587 \| Uncharacterized protein \| Uncharacterized protein; afdb-uniprot50 \| AF-A0A5T3NHC4-F1-MODEL\_V4 \| 1.0 \| 9.631e-20 \| 685 \| 0.211 \| 383 \| 243 \| 14 \| 19 \| 368 \| 8 \| 364 \| Collar domain-containing protein \| Collar domain-containing protein; afdb-uniprot50 \| AF-A0A376L7B0-F1-MODEL\_V4 \| 1.0 \| 1.567e-25 \| 683 \| 0.274 \| 372 \| 210 \| 14 \| 244 \| 583 \| 2 \| 345 \| Terminase large subunit (Gp2) \| Terminase large subunit (Gp2); afdb-uniprot50 \| AF-A0A5C5XIN6-F1-MODEL\_V4 \| 1.0 \| 2.809e-20 \| 683 \| 0.185 \| 367 \| 266 \| 15 \| 24 \| 369 \| 7 \| 361 \| Phage terminase large subunit (GpA) \| Phage terminase large subunit (GpA); afdb-uniprot50 \| AF-A0A349EMJ1-F1-MODEL\_V4 \| 1.0 \| 8.213e-30 \| 683 \| 0.22 \| 448 \| 296 \| 20 \| 187 \| 605 \| 4 \| 427 \| Uncharacterized protein \| Uncharacterized protein; afdb-uniprot50 \| AF-A0A2W5ATV1-F1-MODEL\_V4 \| 1.0 \| 6.927e-21 \| 682 \| 0.187 \| 394 \| 264 \| 17 \| 24 \| 372 \| 8 \| 390 \| Terminase \| Terminase; afdb-uniprot50 \| AF-A0A2P9HPM1-F1-MODEL\_V4 \| 1.0 \| 2.512e-20 \| 680 \| 0.216 \| 383 \| 255 \| 19 \| 12 \| 372 \| 8 \| 367 \| Phage terminase, large subunit \| Phage terminase, large subunit; afdb-uniprot50 \| AF-A0A7C1MEA8-F1-MODEL\_V4 \| 1.0 \| 4.681e-21 \| 679 \| 0.182 \| 379 \| 252 \| 14 \| 14 \| 357 \| 2 \| 357 \| Uncharacterized protein \| Uncharacterized protein; afdb-uniprot50 \| AF-A0A0F9N643-F1-MODEL\_V4 \| 1.0 \| 6.48e-28 \| 679 \| 0.153 \| 598 \| 384 \| 32 \| 39 \| 603 \| 11 \| 518 \| Uncharacterized protein \| Uncharacterized protein; afdb-uniprot50 \| AF-A0A4R8KN63-F1-MODEL\_V4 \| 1.0 \| 3.029e-23 \| 678 \| 0.244 \| 340 \| 226 \| 11 \| 107 \| 434 \| 2 \| 322 \| Phage terminase large subunit GpA \| Phage terminase large subunit GpA; afdb-uniprot50 \| AF-A0A0P6WGK7-F1-MODEL\_V4 \| 1.0 \| 4.294e-25 \| 677 \| 0.248 \| 410 \| 256 \| 15 \| 298 \| 680 \| 2 \| 386 \| Uncharacterized protein \| Uncharacterized protein; afdb-uniprot50 \| AF-A0A7C0ZXV4-F1-MODEL\_V4 \| 1.0 \| 2.939e-27 \| 677 \| 0.148 \| 618 \| 408 \| 35 \| 40 \| 632 \| 14 \| 537 \| Uncharacterized protein \| Uncharacterized protein; afdb-uniprot50 \| AF-A0A4C3GAR0-F1-MODEL\_V4 \| 1.0 \| 1.453e-22 \| 677 \| 0.204 \| 445 \| 270 \| 22 \| 19 \| 416 \| 8 \| 415 \| Phage terminase large subunit \| Phage terminase large subunit; afdb-uniprot50 \| AF-A0A1W6RGK6-F1-MODEL\_V4 \| 1.0 \| 6.838e-19 \| 672 \| 0.285 \| 305 \| 189 \| 10 \| 6 \| 301 \| 1 \| 285 \| Uncharacterized protein \| Uncharacterized protein; afdb-uniprot50 \| AF-A0A7W8UWC0-F1-MODEL\_V4 \| 1.0 \| 1.995e-19 \| 669 \| 0.426 \| 197 \| 104 \| 5 \| 371 \| 563 \| 2 \| 193 \| Phage terminase large subunit GpA-like protein \| Phage terminase large subunit GpA-like protein; afdb-uniprot50 \| AF-A0A1V6IIE5-F1-MODEL\_V4 \| 1.0 \| 2.101e-27 \| 669 \| 0.148 \| 594 \| 392 \| 31 \| 59 \| 631 \| 22 \| 522 \| Phage terminase large subunit (GpA) \| Phage terminase large subunit (GpA); afdb-uniprot50 \| AF-Z9JKT1-F1-MODEL\_V4 \| 1.0 \| 1.444e-21 \| 668 \| 0.539 \| 226 \| 83 \| 5 \| 451 \| 676 \| 9 \| 213 \| Uncharacterized protein \| Uncharacterized protein; afdb-uniprot50 \| AF-D7XM55-F1-MODEL\_V4 \| 1.0 \| 9.569e-19 \| 668 \| 0.347 \| 230 \| 132 \| 7 \| 398 \| 616 \| 1 \| 223 \| Uncharacterized protein \| Uncharacterized protein; afdb-uniprot50 \| AF-A0A530AH38-F1-MODEL\_V4 \| 1.0 \| 1.527e-21 \| 668 \| 0.241 \| 339 \| 215 \| 11 \| 354 \| 677 \| 8 \| 319 \| Terminase \| Terminase; afdb-uniprot50 \| AF-A0A529ZVC1-F1-MODEL\_V4 \| 1.0 \| 2.231e-19 \| 666 \| 0.293 \| 310 \| 191 \| 12 \| 35 \| 331 \| 3 \| 297 \| Terminase \| Terminase; afdb-uniprot50 \| AF-F3IRR5-F1-MODEL\_V4 \| 1.0 \| 6.508e-20 \| 666 \| 0.252 \| 313 \| 194 \| 11 \| 88 \| 384 \| 2 \| 290 \| Terminase, large subunit \| Terminase, large subunit; afdb-uniprot50 \| AF-A0A0E3UJM4-F1-MODEL\_V4 \| 1.0 \| 8.686e-30 \| 663 \| 0.148 \| 734 \| 475 \| 40 \| 1 \| 624 \| 35 \| 728 \| Uncharacterized protein \| Uncharacterized protein; afdb-uniprot50 \| AF-A0A1H8U4H6-F1-MODEL\_V4 \| 1.0 \| 5.478e-28 \| 659 \| 0.154 \| 620 \| 415 \| 32 \| 40 \| 634 \| 17 \| 551 \| Phage terminase large subunit (GpA) \| Phage terminase large subunit (GpA); afdb-uniprot50 \| AF-B7WRM7-F1-MODEL\_V4 \| 1.0 \| 4.087e-26 \| 657 \| 0.282 \| 417 \| 234 \| 16 \| 286 \| 680 \| 91 \| 464 \| Bacteriophage tail assembly protein-like protein \| Bacteriophage tail assembly protein-like protein; afdb-uniprot50 \| AF-A0A0S9PYT1-F1-MODEL\_V4 \| 1.0 \| 5.43e-18 \| 656 \| 0.305 \| 252 \| 169 \| 4 \| 19 \| 269 \| 2 \| 248 \| Uncharacterized protein \| Uncharacterized protein; afdb-uniprot50 \| AF-A0A5U5NLR7-F1-MODEL\_V4 \| 1.0 \| 1.995e-19 \| 654 \| 0.288 \| 225 \| 145 \| 6 \| 375 \| 591 \| 4 \| 221 \| Phage terminase large subunit family protein \| Phage terminase large subunit family protein; afdb-uniprot50 \| AF-A0A1S6J105-F1-MODEL\_V4 \| 1.0 \| 8.142e-20 \| 650 \| 0.289 \| 294 \| 156 \| 11 \| 397 \| 679 \| 2 \| 253 \| Terminase large subunit \| Terminase large subunit; afdb-uniprot50 \| AF-A0A0G1C921-F1-MODEL\_V4 \| 1.0 \| 2.087e-26 \| 648 \| 0.125 \| 599 \| 405 \| 31 \| 40 \| 628 \| 13 \| 502 \| Uncharacterized protein \| Uncharacterized protein; afdb-uniprot50 \| AF-A0A522VYJ8-F1-MODEL\_V4 \| 1.0 \| 5.501e-20 \| 643 \| 0.31 \| 296 \| 175 \| 8 \| 112 \| 398 \| 4 \| 279 \| Uncharacterized protein \| Uncharacterized protein; afdb-uniprot50 \| AF-A0A2W5RBW1-F1-MODEL\_V4 \| 1.0 \| 1.205e-19 \| 642 \| 0.211 \| 359 \| 223 \| 13 \| 39 \| 358 \| 3 \| 340 \| Terminase \| Terminase; afdb-uniprot50 \| AF-A0A7L6UGP5-F1-MODEL\_V4 \| 1.0 \| 2.577e-24 \| 641 \| 0.197 \| 612 \| 249 \| 14 \| 19 \| 616 \| 8 \| 391 \| Phage terminase large subunit family protein \| Phage terminase large subunit family protein; afdb-uniprot50 \| AF-A0A2T6APL5-F1-MODEL\_V4 \| 1.0 \| 3.79e-23 \| 640 \| 0.353 \| 300 \| 168 \| 7 \| 141 \| 434 \| 4 \| 283 \| Phage terminase large subunit GpA \| Phage terminase large subunit GpA; afdb-uniprot50 \| AF-A0A810UT14-F1-MODEL\_V4 \| 1.0 \| 9.106e-20 \| 639 \| 0.193 \| 357 \| 240 \| 14 \| 50 \| 371 \| 2 \| 345 \| Uncharacterized protein \| Uncharacterized protein; afdb-uniprot50 \| AF-A0A285D2I8-F1-MODEL\_V4 \| 1.0 \| 2.359e-19 \| 638 \| 0.425 \| 235 \| 113 \| 10 \| 405 \| 630 \| 1 \| 222 \| Phage terminase large subunit GpA \| Phage terminase large subunit GpA; afdb-uniprot50 \| AF-A0A765BM22-F1-MODEL\_V4 \| 1.0 \| 2.773e-18 \| 633 \| 0.214 \| 368 \| 224 \| 16 \| 7 \| 341 \| 1 \| 336 \| Phage terminase large subunit family protein \| Phage terminase large subunit family protein; afdb-uniprot50 \| AF-A0A2N3C6B8-F1-MODEL\_V4 \| 1.0 \| 1.229e-22 \| 632 \| 0.291 \| 391 \| 188 \| 16 \| 318 \| 682 \| 4 \| 331 \| Terminase \| Terminase; afdb-uniprot50 \| AF-A0A6J4G269-F1-MODEL\_V4 \| 1.0 \| 2.207e-26 \| 632 \| 0.336 \| 348 \| 194 \| 12 \| 204 \| 538 \| 2 \| 325 \| Bacteriophage tail assembly protein \| Bacteriophage tail assembly protein; afdb-uniprot50 \| AF-D8A4J1-F1-MODEL\_V4 \| 1.0 \| 8.037e-18 \| 631 \| 0.204 \| 352 \| 224 \| 12 \| 19 \| 339 \| 8 \| 334 \| Phage terminase large subunit (GpA) \| Phage terminase large subunit (GpA); afdb-uniprot50 \| AF-A0A776L0V5-F1-MODEL\_V4 \| 1.0 \| 1.266e-18 \| 628 \| 0.212 \| 362 \| 235 \| 17 \| 19 \| 352 \| 8 \| 347 \| Phage terminase large subunit family protein \| Phage terminase large subunit family protein; afdb-uniprot50 \| AF-A0A4Q6EZD1-F1-MODEL\_V4 \| 1.0 \| 1.383e-23 \| 624 \| 0.256 \| 362 \| 230 \| 12 \| 126 \| 472 \| 2 \| 339 \| Uncharacterized protein \| Uncharacterized protein; afdb-uniprot50 \| AF-A0A285CJV0-F1-MODEL\_V4 \| 1.0 \| 1.491e-26 \| 621 \| 0.15 \| 664 \| 404 \| 40 \| 1 \| 605 \| 1 \| 563 \| Phage terminase large subunit GpA \| Phage terminase large subunit GpA; afdb-uniprot50 \| AF-A8U1E7-F1-MODEL\_V4 \| 1.0 \| 3.082e-17 \| 619 \| 0.353 \| 249 \| 151 \| 6 \| 1 \| 241 \| 1 \| 247 \| Phage terminase GpA \| Phage terminase GpA; afdb-uniprot50 \| AF-A0A1V5XPC1-F1-MODEL\_V4 \| 1.0 \| 3.089e-26 \| 618 \| 0.153 \| 598 \| 393 \| 32 \| 39 \| 605 \| 1 \| 515 \| Phage terminase large subunit (GpA) \| Phage terminase large subunit (GpA); afdb-uniprot50 \| AF-A0A776QT91-F1-MODEL\_V4 \| 1.0 \| 1.419e-27 \| 617 \| 0.277 \| 422 \| 243 \| 17 \| 176 \| 561 \| 12 \| 407 \| Phage terminase large subunit family protein \| Phage terminase large subunit family protein; afdb-uniprot50 \| AF-A0A2W6VA45-F1-MODEL\_V4 \| 1.0 \| 7.985e-17 \| 612 \| 0.301 \| 282 \| 173 \| 6 \| 15 \| 293 \| 3 \| 263 \| Uncharacterized protein \| Uncharacterized protein; afdb-uniprot50 \| AF-A0A4T5F4C3-F1-MODEL\_V4 \| 1.0 \| 3.669e-18 \| 612 \| 0.213 \| 323 \| 207 \| 16 \| 39 \| 330 \| 1 \| 307 \| Phage terminase large subunit family protein \| Phage terminase large subunit family protein; afdb-uniprot50 \| AF-A0A2A6PYT6-F1-MODEL\_V4 \| 1.0 \| 1.425e-19 \| 607 \| 0.237 \| 374 \| 238 \| 14 \| 67 \| 412 \| 4 \| 358 \| Terminase \| Terminase; afdb-uniprot50 \| AF-D8F666-F1-MODEL\_V4 \| 1.0 \| 8.666e-21 \| 597 \| 0.232 \| 348 \| 220 \| 15 \| 277 \| 599 \| 2 \| 327 \| Uncharacterized protein \| Uncharacterized protein; afdb-uniprot50 \| AF-A0A6I1Z9L3-F1-MODEL\_V4 \| 1.0 \| 2.611e-26 \| 597 \| 0.164 \| 595 \| 389 \| 30 \| 38 \| 604 \| 68 \| 582 \| Uncharacterized protein \| Uncharacterized protein; afdb-uniprot50 \| AF-A0A3Z2C6Q0-F1-MODEL\_V4 \| 1.0 \| 3.122e-19 \| 597 \| 0.207 \| 410 \| 263 \| 18 \| 19 \| 395 \| 8 \| 388 \| Uncharacterized protein \| Uncharacterized protein; afdb-uniprot50 \| AF-A0A633N905-F1-MODEL\_V4 \| 1.0 \| 8.989e-18 \| 595 \| 0.223 \| 317 \| 207 \| 14 \| 24 \| 319 \| 10 \| 308 \| Phage terminase large subunit family protein \| Phage terminase large subunit family protein; afdb-uniprot50 \| AF-B6Y9I7-F1-MODEL\_V4 \| 1.0 \| 1.956e-16 \| 593 \| 0.302 \| 238 \| 141 \| 6 \| 103 \| 336 \| 2 \| 218 \| Phage terminase large subunit GpA \| Phage terminase large subunit GpA; afdb-uniprot50 \| AF-A0A7X7UKL1-F1-MODEL\_V4 \| 1.0 \| 1.886e-19 \| 593 \| 0.17 \| 399 \| 266 \| 15 \| 40 \| 437 \| 11 \| 345 \| Uncharacterized protein \| Uncharacterized protein; afdb-uniprot50 \| AF-A0A855W3M0-F1-MODEL\_V4 \| 1.0 \| 1.982e-18 \| 590 \| 0.227 \| 386 \| 248 \| 16 \| 50 \| 406 \| 2 \| 366 \| Terminase \| Terminase; afdb-uniprot50 \| AF-A0A786Q0T4-F1-MODEL\_V4 \| 1.0 \| 5.466e-19 \| 583 \| 0.328 \| 228 \| 137 \| 6 \| 344 \| 561 \| 6 \| 227 \| Phage terminase large subunit family protein \| Phage terminase large subunit family protein; afdb-uniprot50 \| AF-A0A827QIZ6-F1-MODEL\_V4 \| 1.0 \| 1.057e-16 \| 580 \| 0.3 \| 203 \| 128 \| 4 \| 373 \| 567 \| 3 \| 199 \| Phage terminase large subunit family protein \| Phage terminase large subunit family protein; afdb-uniprot50 \| AF-A0A2W5FWH4-F1-MODEL\_V4 \| 1.0 \| 2.329e-17 \| 579 \| 0.287 \| 261 \| 159 \| 9 \| 98 \| 353 \| 435 \| 673 \| DOD-type homing endonuclease domain-containing protein \| DOD-type homing endonuclease domain-containing protein; afdb-uniprot50 \| AF-A0A2V6CK06-F1-MODEL\_V4 \| 1.0 \| 2.344e-18 \| 577 \| 0.235 \| 310 \| 185 \| 11 \| 390 \| 682 \| 10 \| 284 \| Uncharacterized protein \| Uncharacterized protein; afdb-uniprot50 \| AF-A0A2N1GMP5-F1-MODEL\_V4 \| 1.0 \| 5.743e-18 \| 576 \| 0.208 \| 378 \| 239 \| 21 \| 3 \| 345 \| 1 \| 353 \| Terminase \| Terminase; afdb-uniprot50 \| AF-A0A0F9B9Q1-F1-MODEL\_V4 \| 1.0 \| 2.26e-21 \| 574 \| 0.162 \| 486 \| 305 \| 28 \| 40 \| 495 \| 10 \| 423 \| Uncharacterized protein \| Uncharacterized protein; afdb-uniprot50 \| AF-A0A524ITI1-F1-MODEL\_V4 \| 1.0 \| 8.5e-18 \| 570 \| 0.278 \| 291 \| 160 \| 13 \| 400 \| 670 \| 1 \| 261 \| Uncharacterized protein \| Uncharacterized protein; afdb-uniprot50 \| AF-A0A7B5N0Z9-F1-MODEL\_V4 \| 1.0 \| 1.818e-22 \| 569 \| 0.242 \| 412 \| 236 \| 20 \| 114 \| 484 \| 2 \| 378 \| Phage terminase large subunit family protein \| Phage terminase large subunit family protein; afdb-uniprot50 \| AF-S9S9L7-F1-MODEL\_V4 \| 1.0 \| 2.773e-18 \| 567 \| 0.245 \| 293 \| 188 \| 9 \| 386 \| 648 \| 4 \| 293 \| Terminase GpA \| Terminase GpA; afdb-uniprot50 \| AF-R9L5W8-F1-MODEL\_V4 \| 1.0 \| 6.592e-22 \| 564 \| 0.201 \| 412 \| 259 \| 20 \| 280 \| 677 \| 3 \| 358 \| Uncharacterized protein \| Uncharacterized protein; afdb-uniprot50 \| AF-A0A2N3CM29-F1-MODEL\_V4 \| 1.0 \| 1.931e-14 \| 563 \| 0.32 \| 215 \| 138 \| 5 \| 20 \| 228 \| 5 \| 217 \| Phage tail protein \| Phage tail protein; afdb-uniprot50 \| AF-A0A4Q3NU98-F1-MODEL\_V4 \| 1.0 \| 5.957e-15 \| 562 \| 0.251 \| 231 \| 168 \| 4 \| 27 \| 254 \| 2 \| 230 \| Phage terminase large subunit family protein \| Phage terminase large subunit family protein; afdb-uniprot50 \| AF-A0A7A6XDC4-F1-MODEL\_V4 \| 1.0 \| 6.838e-19 \| 561 \| 0.316 \| 237 \| 146 \| 6 \| 335 \| 561 \| 28 \| 258 \| Phage terminase large subunit family protein \| Phage terminase large subunit family protein; afdb-uniprot50 \| AF-A0A0F8VNF6-F1-MODEL\_V4 \| 1.0 \| 4.051e-16 \| 560 \| 0.146 \| 335 \| 229 \| 14 \| 40 \| 369 \| 12 \| 294 \| Uncharacterized protein \| Uncharacterized protein; afdb-uniprot50 \| AF-A0A1V6JWY6-F1-MODEL\_V4 \| 1.0 \| 2.611e-26 \| 560 \| 0.216 \| 532 \| 316 \| 33 \| 136 \| 608 \| 3 \| 492 \| Phage terminase large subunit (GpA) \| Phage terminase large subunit (GpA); afdb-uniprot50 \| AF-A0A522X2A6-F1-MODEL\_V4 \| 1.0 \| 1.162e-22 \| 559 \| 0.201 \| 641 \| 254 \| 17 \| 26 \| 632 \| 2 \| 418 \| Uncharacterized protein \| Uncharacterized protein; afdb-uniprot50 \| AF-A0A826Y2T5-F1-MODEL\_V4 \| 1.0 \| 3.622e-16 \| 556 \| 0.209 \| 305 \| 209 \| 10 \| 9 \| 302 \| 2 \| 285 \| Phage terminase large subunit family protein \| Phage terminase large subunit family protein; afdb-uniprot50 \| AF-A0A529QDC6-F1-MODEL\_V4 \| 1.0 \| 4.793e-16 \| 554 \| 0.253 \| 205 \| 142 \| 6 \| 385 \| 582 \| 1 \| 201 \| Terminase \| Terminase; afdb-uniprot50 \| AF-A0A520NQ90-F1-MODEL\_V4 \| 1.0 \| 6.074e-18 \| 554 \| 0.409 \| 242 \| 101 \| 6 \| 441 \| 682 \| 15 \| 214 \| Uncharacterized protein \| Uncharacterized protein; afdb-uniprot50 \| AF-A0A1A6FHD2-F1-MODEL\_V4 \| 1.0 \| 4.803e-25 \| 551 \| 0.376 \| 303 \| 159 \| 9 \| 223 \| 517 \| 21 \| 301 \| Uncharacterized protein \| Uncharacterized protein; afdb-uniprot50 \| AF-A0A826ZB13-F1-MODEL\_V4 \| 1.0 \| 6.274e-23 \| 551 \| 0.274 \| 371 \| 201 \| 17 \| 223 \| 555 \| 12 \| 352 \| Phage terminase large subunit family protein \| Phage terminase large subunit family protein; afdb-uniprot50 \| AF-A0A0J5K4V1-F1-MODEL\_V4 \| 1.0 \| 6.314e-24 \| 550 \| 0.254 \| 420 \| 240 \| 18 \| 148 \| 530 \| 1 \| 384 \| Terminase \| Terminase; afdb-uniprot50 \| AF-A0A539CTX1-F1-MODEL\_V4 \| 1.0 \| 1.182e-16 \| 546 \| 0.192 \| 306 \| 209 \| 10 \| 78 \| 369 \| 8 \| 289 \| Uncharacterized protein \| Uncharacterized protein; afdb-uniprot50 \| AF-A0A3S4UPH8-F1-MODEL\_V4 \| 1.0 \| 1.258e-17 \| 546 \| 0.22 \| 327 \| 206 \| 17 \| 17 \| 319 \| 3 \| 304 \| Bacteriophage terminase large subunit \| Bacteriophage terminase large subunit; afdb-uniprot50 \| AF-A0A2P9HNN7-F1-MODEL\_V4 \| 1.0 \| 1.982e-18 \| 544 \| 0.251 \| 354 \| 200 \| 16 \| 375 \| 682 \| 133 \| 467 \| Terminase large subunit \| Terminase large subunit; afdb-uniprot50 \| AF-A0A0F8WXC1-F1-MODEL\_V4 \| 1.0 \| 5.292e-14 \| 543 \| 0.277 \| 220 \| 150 \| 6 \| 4 \| 220 \| 1 \| 214 \| Uncharacterized protein \| Uncharacterized protein; afdb-uniprot50 \| AF-N6VN50-F1-MODEL\_V4 \| 1.0 \| 4.762e-15 \| 543 \| 0.267 \| 262 \| 172 \| 7 \| 2 \| 256 \| 1 \| 249 \| Phage terminase large subunit GpA2 \| Phage terminase large subunit GpA2; afdb-uniprot50 \| AF-A0A827LVK1-F1-MODEL\_V4 \| 1.0 \| 2.314e-16 \| 541 \| 0.223 \| 326 \| 206 \| 16 \| 16 \| 318 \| 3 \| 304 \| Phage terminase large subunit family protein \| Phage terminase large subunit family protein; afdb-uniprot50 \| AF-A0A827HT50-F1-MODEL\_V4 \| 1.0 \| 1.389e-15 \| 538 \| 0.336 \| 196 \| 117 \| 5 \| 377 \| 565 \| 3 \| 192 \| Phage terminase large subunit family protein \| Phage terminase large subunit family protein; afdb-uniprot50 \| AF-A0A329VSW5-F1-MODEL\_V4 \| 1.0 \| 2.702e-14 \| 536 \| 0.277 \| 238 \| 163 \| 6 \| 6 \| 238 \| 1 \| 234 \| Terminase \| Terminase; afdb-uniprot50 \| AF-A0A2X1IZC0-F1-MODEL\_V4 \| 1.0 \| 2.174e-15 \| 534 \| 0.221 \| 298 \| 198 \| 13 \| 19 \| 302 \| 8 \| 285 \| Terminase GpA \| Terminase GpA; afdb-uniprot50 \| AF-A0A777DWG3-F1-MODEL\_V4 \| 1.0 \| 4.257e-15 \| 533 \| 0.215 \| 293 \| 206 \| 9 \| 19 \| 302 \| 8 \| 285 \| Phage terminase large subunit family protein \| Phage terminase large subunit family protein; afdb-uniprot50 \| AF-A0A2D8LXN6-F1-MODEL\_V4 \| 1.0 \| 3.176e-13 \| 532 \| 0.303 \| 198 \| 132 \| 4 \| 41 \| 233 \| 2 \| 198 \| Phage tail protein \| Phage tail protein; afdb-uniprot50 \| AF-A0A381KV82-F1-MODEL\_V4 \| 1.0 \| 2.27e-13 \| 532 \| 0.253 \| 193 \| 143 \| 1 \| 34 \| 225 \| 13 \| 205 \| Phage terminase large subunit \| Phage terminase large subunit; afdb-uniprot50 \| AF-A0A2N9Y7Y3-F1-MODEL\_V4 \| 1.0 \| 3.023e-14 \| 532 \| 0.289 \| 211 \| 141 \| 6 \| 20 \| 224 \| 4 \| 211 \| Terminase \| Terminase; afdb-uniprot50 \| AF-B6XBA7-F1-MODEL\_V4 \| 1.0 \| 6.3e-15 \| 531 \| 0.223 \| 295 \| 197 \| 9 \| 9 \| 292 \| 2 \| 275 \| Uncharacterized protein \| Uncharacterized protein; afdb-uniprot50 \| AF-A0A356C2D8-F1-MODEL\_V4 \| 1.0 \| 2.447e-16 \| 529 \| 0.442 \| 199 \| 98 \| 4 \| 441 \| 632 \| 19 \| 211 \| Uncharacterized protein \| Uncharacterized protein; afdb-uniprot50 \| AF-A0A430BRK0-F1-MODEL\_V4 \| 1.0 \| 3.217e-15 \| 529 \| 0.244 \| 274 \| 172 \| 13 \| 55 \| 319 \| 7 \| 254 \| Terminase \| Terminase; afdb-uniprot50 \| AF-A0A540VIA7-F1-MODEL\_V4 \| 1.0 \| 9.264e-14 \| 526 \| 0.241 \| 232 \| 170 \| 4 \| 10 \| 238 \| 2 \| 230 \| Phage terminase large subunit family protein \| Phage terminase large subunit family protein; afdb-uniprot50 \| AF-A0A7C7KMY0-F1-MODEL\_V4 \| 1.0 \| 5.326e-15 \| 526 \| 0.219 \| 282 \| 200 \| 9 \| 15 \| 291 \| 8 \| 274 \| Uncharacterized protein \| Uncharacterized protein; afdb-uniprot50 \| AF-A0A4Q3I4G8-F1-MODEL\_V4 \| 1.0 \| 1.159e-13 \| 523 \| 0.243 \| 205 \| 150 \| 4 \| 35 \| 235 \| 9 \| 212 \| Phage terminase large subunit family protein \| Phage terminase large subunit family protein; afdb-uniprot50 \| AF-A0A0F8Y264-F1-MODEL\_V4 \| 1.0 \| 6.75e-17 \| 521 \| 0.224 \| 267 \| 160 \| 13 \| 386 \| 617 \| 7 \| 261 \| Uncharacterized protein \| Uncharacterized protein; afdb-uniprot50 \| AF-R7J609-F1-MODEL\_V4 \| 1.0 \| 1.274e-19 \| 520 \| 0.22 \| 359 \| 235 \| 15 \| 285 \| 617 \| 33 \| 372 \| Phage terminase large subunit \| Phage terminase large subunit; afdb-uniprot50 \| AF-A0A356HJD3-F1-MODEL\_V4 \| 1.0 \| 6.62e-14 \| 518 \| 0.273 \| 216 \| 147 \| 6 \| 24 \| 234 \| 2 \| 212 \| Terminase \| Terminase; afdb-uniprot50 \| AF-A0A8B5XJ81-F1-MODEL\_V4 \| 1.0 \| 5.919e-14 \| 518 \| 0.224 \| 245 \| 180 \| 7 \| 19 \| 257 \| 8 \| 248 \| Phage terminase large subunit family protein \| Phage terminase large subunit family protein; afdb-uniprot50 \| AF-A0A376JSG1-F1-MODEL\_V4 \| 1.0 \| 8.336e-15 \| 518 \| 0.227 \| 299 \| 195 \| 13 \| 39 \| 320 \| 7 \| 286 \| Phage terminase large subunit (GpA) \| Phage terminase large subunit (GpA); afdb-uniprot50 \| AF-A0A4Q6BLM2-F1-MODEL\_V4 \| 1.0 \| 4.711e-22 \| 516 \| 0.212 \| 410 \| 270 \| 12 \| 147 \| 520 \| 3 \| 395 \| Uncharacterized protein \| Uncharacterized protein; afdb-uniprot50 \| AF-A0A0E4BQP1-F1-MODEL\_V4 \| 1.0 \| 1.478e-16 \| 515 \| 0.233 \| 317 \| 190 \| 17 \| 377 \| 679 \| 3 \| 280 \| Uncharacterized protein \| Uncharacterized protein; afdb-uniprot50 \| AF-A0A829DWZ5-F1-MODEL\_V4 \| 1.0 \| 9.926e-16 \| 515 \| 0.216 \| 286 \| 187 \| 12 \| 33 \| 296 \| 18 \| 288 \| Phage terminase large subunit family protein \| Phage terminase large subunit family protein; afdb-uniprot50 \| AF-A0A601WLN8-F1-MODEL\_V4 \| 1.0 \| 3.023e-14 \| 514 \| 0.21 \| 299 \| 204 \| 9 \| 9 \| 296 \| 2 \| 279 \| Phage terminase large subunit family protein \| Phage terminase large subunit family protein; afdb-uniprot50 \| AF-A3VBF4-F1-MODEL\_V4 \| 1.0 \| 2.217e-18 \| 514 \| 0.201 \| 327 \| 213 \| 13 \| 318 \| 604 \| 28 \| 346 \| Putative phage terminase large subunit \| Putative phage terminase large subunit; afdb-uniprot50 \| AF-A0A377CJ19-F1-MODEL\_V4 \| 1.0 \| 8.817e-15 \| 513 \| 0.222 \| 265 \| 188 \| 9 \| 19 \| 271 \| 8 \| 266 \| Phage terminase, large subunit \| Phage terminase, large subunit; afdb-uniprot50 \| AF-A0A376JFJ4-F1-MODEL\_V4 \| 1.0 \| 4.051e-16 \| 513 \| 0.196 \| 346 \| 231 \| 13 \| 7 \| 331 \| 1 \| 320 \| Terminase large subunit (Gp2) \| Terminase large subunit (Gp2); afdb-uniprot50 \| AF-A0A496P6X6-F1-MODEL\_V4 \| 1.0 \| 4.502e-15 \| 508 \| 0.216 \| 226 \| 159 \| 10 \| 385 \| 604 \| 5 \| 218 \| Terminase \| Terminase; afdb-uniprot50 \| AF-A0A1F7TJM4-F1-MODEL\_V4 \| 1.0 \| 4.285e-16 \| 505 \| 0.188 \| 392 \| 233 \| 19 \| 3 \| 369 \| 1 \| 332 \| Uncharacterized protein \| Uncharacterized protein; afdb-uniprot50 \| AF-A0A0M7BBY2-F1-MODEL\_V4 \| 1.0 \| 2.84e-13 \| 504 \| 0.368 \| 179 \| 110 \| 2 \| 27 \| 202 \| 2 \| 180 \| Bacteriophage tail assembly protein \| Bacteriophage tail assembly protein; afdb-uniprot50 \| AF-A0A7W8WPL2-F1-MODEL\_V4 \| 1.0 \| 4.185e-21 \| 504 \| 0.28 \| 325 \| 171 \| 10 \| 288 \| 553 \| 5 \| 325 \| Phage terminase large subunit GpA-like protein \| Phage terminase large subunit GpA-like protein; afdb-uniprot50 \| AF-A0A416VMY3-F1-MODEL\_V4 \| 1.0 \| 2.146e-13 \| 500 \| 0.196 \| 229 \| 175 \| 5 \| 10 \| 235 \| 2 \| 224 \| Uncharacterized protein \| Uncharacterized protein; afdb-uniprot50 \| AF-A0A0P0YGY0-F1-MODEL\_V4 \| 1.0 \| 8.283e-14 \| 499 \| 0.202 \| 222 \| 166 \| 7 \| 4 \| 223 \| 1 \| 213 \| Bacteriophage terminase large subunit \| Bacteriophage terminase large subunit; afdb-uniprot50 \| AF-A0A0Q0XNC5-F1-MODEL\_V4 \| 1.0 \| 1.956e-16 \| 499 \| 0.314 \| 277 \| 171 \| 9 \| 415 \| 679 \| 3 \| 272 \| Uncharacterized protein \| Uncharacterized protein; afdb-uniprot50 \| AF-A0A6N9S2Q2-F1-MODEL\_V4 \| 1.0 \| 1.159e-13 \| 498 \| 0.24 \| 249 \| 177 \| 9 \| 19 \| 260 \| 8 \| 251 \| Phage terminase large subunit family protein \| Phage terminase large subunit family protein; afdb-uniprot50 \| AF-A0A355B5P3-F1-MODEL\_V4 \| 1.0 \| 4.051e-16 \| 498 \| 0.186 \| 397 \| 245 \| 20 \| 1 \| 369 \| 37 \| 383 \| Uncharacterized protein \| Uncharacterized protein; afdb-uniprot50 \| AF-A0A0F8YZ73-F1-MODEL\_V4 \| 1.0 \| 7.405e-14 \| 496 \| 0.228 \| 241 \| 169 \| 8 \| 6 \| 241 \| 1 \| 229 \| Uncharacterized protein \| Uncharacterized protein; afdb-uniprot50 \| AF-A0A377MYS1-F1-MODEL\_V4 \| 1.0 \| 1.534e-13 \| 495 \| 0.256 \| 203 \| 148 \| 3 \| 68 \| 269 \| 10 \| 210 \| Phage terminase large subunit (GpA) \| Phage terminase large subunit (GpA); afdb-uniprot50 \| AF-A0A1B7L553-F1-MODEL\_V4 \| 1.0 \| 1.167e-14 \| 494 \| 0.266 \| 240 \| 151 \| 8 \| 68 \| 298 \| 6 \| 229 \| Uncharacterized protein \| Uncharacterized protein; afdb-uniprot50 \| AF-A0A0F9CN62-F1-MODEL\_V4 \| 1.0 \| 2.029e-13 \| 488 \| 0.287 \| 195 \| 127 \| 7 \| 34 \| 225 \| 6 \| 191 \| Uncharacterized protein \| Uncharacterized protein; afdb-uniprot50 \| AF-A0A329VRL4-F1-MODEL\_V4 \| 1.0 \| 3.042e-15 \| 485 \| 0.409 \| 176 \| 92 \| 4 \| 398 \| 565 \| 2 \| 173 \| Terminase \| Terminase; afdb-uniprot50 \| AF-R7UYV5-F1-MODEL\_V4 \| 1.0 \| 4.416e-12 \| 485 \| 0.305 \| 193 \| 128 \| 4 \| 33 \| 220 \| 118 \| 309 \| Uncharacterized protein \| Uncharacterized protein; afdb-uniprot50 \| AF-A0A3R8ZB61-F1-MODEL\_V4 \| 1.0 \| 1.03e-12 \| 483 \| 0.272 \| 209 \| 143 \| 6 \| 17 \| 220 \| 2 \| 206 \| Phage terminase large subunit family protein \| Phage terminase large subunit family protein; afdb-uniprot50 \| AF-A0A796TC99-F1-MODEL\_V4 \| 1.0 \| 1.362e-12 \| 473 \| 0.23 \| 226 \| 165 \| 7 \| 19 \| 239 \| 8 \| 229 \| Phage terminase large subunit family protein \| Phage terminase large subunit family protein; afdb-uniprot50 \| AF-A0A3G2IBA0-F1-MODEL\_V4 \| 1.0 \| 7.002e-14 \| 473 \| 0.245 \| 293 \| 196 \| 12 \| 17 \| 298 \| 6 \| 284 \| Phage terminase large subunit family protein \| Phage terminase large subunit family protein; afdb-uniprot50 \| AF-A0A2N8K836-F1-MODEL\_V4 \| 1.0 \| 3.782e-14 \| 472 \| 0.352 \| 156 \| 90 \| 5 \| 377 \| 526 \| 3 \| 153 \| Terminase \| Terminase; afdb-uniprot50 \| AF-A0A0F9F3T8-F1-MODEL\_V4 \| 1.0 \| 4.473e-14 \| 471 \| 0.202 \| 326 \| 210 \| 14 \| 3 \| 296 \| 1 \| 308 \| Uncharacterized protein \| Uncharacterized protein; afdb-uniprot50 \| AF-A0A0F9JEA4-F1-MODEL\_V4 \| 1.0 \| 7.31e-12 \| 470 \| 0.276 \| 170 \| 120 \| 2 \| 74 \| 241 \| 2 \| 170 \| Uncharacterized protein \| Uncharacterized protein; afdb-uniprot50 \| AF-A0A5C5VTC1-F1-MODEL\_V4 \| 1.0 \| 1.537e-22 \| 468 \| 0.181 \| 486 \| 317 \| 23 \| 197 \| 629 \| 8 \| 465 \| Phage terminase large subunit (GpA) \| Phage terminase large subunit (GpA); afdb-uniprot50 \| AF-A0A7C2EB29-F1-MODEL\_V4 \| 1.0 \| 2.132e-12 \| 465 \| 0.317 \| 205 \| 129 \| 6 \| 17 \| 215 \| 3 \| 202 \| Phage terminase large subunit family protein \| Phage terminase large subunit family protein; afdb-uniprot50 \| AF-A0A853EDX0-F1-MODEL\_V4 \| 1.0 \| 3.381e-14 \| 465 \| 0.344 \| 212 \| 131 \| 5 \| 335 \| 544 \| 2 \| 207 \| Phage terminase large subunit family protein \| Phage terminase large subunit family protein; afdb-uniprot50 \| AF-A0A1H4FKT7-F1-MODEL\_V4 \| 1.0 \| 1.906e-12 \| 465 \| 0.246 \| 227 \| 164 \| 6 \| 1 \| 223 \| 7 \| 230 \| Phage terminase large subunit (GpA) \| Phage terminase large subunit (GpA); afdb-uniprot50 \| AF-A0A793I649-F1-MODEL\_V4 \| 1.0 \| 1.524e-12 \| 461 \| 0.206 \| 228 \| 162 \| 9 \| 3 \| 225 \| 1 \| 214 \| Phage terminase large subunit family protein \| Phage terminase large subunit family protein; afdb-uniprot50 \| AF-A0A810USD3-F1-MODEL\_V4 \| 1.0 \| 7.731e-12 \| 459 \| 0.262 \| 194 \| 136 \| 3 \| 19 \| 209 \| 6 \| 195 \| Uncharacterized protein \| Uncharacterized protein; afdb-uniprot50 \| AF-A0A455RRL4-F1-MODEL\_V4 \| 1.0 \| 4.67e-12 \| 455 \| 0.206 \| 237 \| 181 \| 5 \| 19 \| 251 \| 8 \| 241 \| Phage terminase, large subunit \| Phage terminase, large subunit; afdb-uniprot50 \| AF-I4SDW6-F1-MODEL\_V4 \| 1.0 \| 7.31e-12 \| 454 \| 0.222 \| 198 \| 144 \| 8 \| 33 \| 225 \| 18 \| 210 \| Phage terminase large subunit \| Phage terminase large subunit; afdb-uniprot50 \| AF-A0A828PQX7-F1-MODEL\_V4 \| 1.0 \| 5.224e-12 \| 452 \| 0.226 \| 243 \| 176 \| 7 \| 19 \| 254 \| 8 \| 245 \| Phage terminase large subunit family protein \| Phage terminase large subunit family protein; afdb-uniprot50 \| AF-A0A529SUP9-F1-MODEL\_V4 \| 1.0 \| 1.218e-12 \| 451 \| 0.276 \| 253 \| 162 \| 8 \| 60 \| 306 \| 1 \| 238 \| Terminase \| Terminase; afdb-uniprot50 \| AF-A0A0F9E166-F1-MODEL\_V4 \| 1.0 \| 6.957e-13 \| 449 \| 0.192 \| 250 \| 186 \| 9 \| 1 \| 241 \| 108 \| 350 \| Uncharacterized protein \| Uncharacterized protein; afdb-uniprot50 \| AF-A0A6D1D8D3-F1-MODEL\_V4 \| 1.0 \| 6.179e-12 \| 448 \| 0.215 \| 200 \| 143 \| 8 \| 33 \| 225 \| 18 \| 210 \| Phage terminase large subunit family protein \| Phage terminase large subunit family protein; afdb-uniprot50 \| AF-A0A742XX60-F1-MODEL\_V4 \| 1.0 \| 5.135e-18 \| 444 \| 0.316 \| 259 \| 154 \| 8 \| 258 \| 509 \| 1 \| 243 \| Phage terminase large subunit family protein \| Phage terminase large subunit family protein; afdb-uniprot50 \| AF-A0A5T8FU82-F1-MODEL\_V4 \| 1.0 \| 2.984e-12 \| 443 \| 0.198 \| 237 \| 175 \| 9 \| 1 \| 232 \| 2 \| 228 \| Phage terminase large subunit family protein \| Phage terminase large subunit family protein; afdb-uniprot50 \| AF-A3VBF5-F1-MODEL\_V4 \| 1.0 \| 9.735e-13 \| 442 \| 0.19 \| 236 \| 177 \| 9 \| 1 \| 228 \| 2 \| 231 \| Putative phage terminase large subunit \| Putative phage terminase large subunit; afdb-uniprot50 \| AF-A0A5Y3PSE3-F1-MODEL\_V4 \| 1.0 \| 1.144e-11 \| 441 \| 0.226 \| 243 \| 176 \| 7 \| 19 \| 254 \| 8 \| 245 \| Phage terminase large subunit family protein \| Phage terminase large subunit family protein; afdb-uniprot50 \| AF-A0A4Y5RQN5-F1-MODEL\_V4 \| 1.0 \| 7.405e-14 \| 439 \| 0.223 \| 269 \| 177 \| 6 \| 68 \| 330 \| 18 \| 260 \| Uncharacterized protein \| Uncharacterized protein; afdb-uniprot50 \| AF-A0A436S558-F1-MODEL\_V4 \| 1.0 \| 1.152e-12 \| 438 \| 0.252 \| 269 \| 179 \| 9 \| 10 \| 261 \| 1 \| 264 \| Terminase \| Terminase; afdb-uniprot50 \| AF-A0A826JJD2-F1-MODEL\_V4 \| 1.0 \| 2.755e-17 \| 436 \| 0.284 \| 309 \| 175 \| 12 \| 176 \| 457 \| 10 \| 299 \| Phage terminase large subunit family protein \| Phage terminase large subunit family protein; afdb-uniprot50 \| AF-A0A6N8NNT2-F1-MODEL\_V4 \| 1.0 \| 4.148e-11 \| 435 \| 0.215 \| 204 \| 159 \| 1 \| 4 \| 206 \| 1 \| 204 \| Terminase \| Terminase; afdb-uniprot50 \| AF-A0A3D5V0P4-F1-MODEL\_V4 \| 1.0 \| 7.263e-11 \| 430 \| 0.288 \| 177 \| 118 \| 5 \| 21 \| 191 \| 17 \| 191 \| Terminase \| Terminase; afdb-uniprot50 \| AF-V0YI02-F1-MODEL\_V4 \| 1.0 \| 8.445e-17 \| 425 \| 0.245 \| 346 \| 202 \| 13 \| 149 \| 460 \| 4 \| 324 \| Phage terminase large subunit \| Phage terminase large subunit; afdb-uniprot50 \| AF-A0A7V4LST6-F1-MODEL\_V4 \| 1.0 \| 7.731e-12 \| 425 \| 0.333 \| 165 \| 109 \| 1 \| 102 \| 266 \| 2 \| 165 \| Glyco\_hydro\_32N domain-containing protein \| Glyco\_hydro\_32N domain-containing protein; afdb-uniprot50 \| AF-A0A4U9DCF0-F1-MODEL\_V4 \| 1.0 \| 3.576e-14 \| 419 \| 0.328 \| 213 \| 126 \| 6 \| 431 \| 632 \| 8 \| 214 \| Bacteriophage tail assembly protein \| Bacteriophage tail assembly protein; afdb-uniprot50 \| AF-A0A7C7GM07-F1-MODEL\_V4 \| 1.0 \| 1.152e-12 \| 418 \| 0.333 \| 201 \| 116 \| 5 \| 445 \| 644 \| 2 \| 185 \| Uncharacterized protein \| Uncharacterized protein; afdb-uniprot50 \| AF-A7BN38-F1-MODEL\_V4 \| 1.0 \| 1.422e-10 \| 417 \| 0.2 \| 209 \| 160 \| 5 \| 19 \| 223 \| 8 \| 213 \| Phage terminase GpA \| Phage terminase GpA; afdb-uniprot50 \| AF-A0A777TUX5-F1-MODEL\_V4 \| 1.0 \| 4.416e-12 \| 413 \| 0.36 \| 161 \| 97 \| 4 \| 384 \| 544 \| 2 \| 156 \| Phage terminase large subunit family protein \| Phage terminase large subunit family protein; afdb-uniprot50 \| AF-A0A826S5Z3-F1-MODEL\_V4 \| 1.0 \| 8.123e-11 \| 410 \| 0.216 \| 212 \| 155 \| 9 \| 19 \| 224 \| 8 \| 214 \| Phage terminase large subunit family protein \| Phage terminase large subunit family protein; afdb-uniprot50 \| AF-A0A2W5G3V7-F1-MODEL\_V4 \| 1.0 \| 4.61e-10 \| 409 \| 0.314 \| 181 \| 120 \| 3 \| 20 \| 197 \| 7 \| 186 \| Terminase \| Terminase; afdb-uniprot50 \| AF-A0A1Z8QJE7-F1-MODEL\_V4 \| 1.0 \| 2.226e-10 \| 407 \| 0.266 \| 210 \| 150 \| 3 \| 1 \| 206 \| 2 \| 211 \| Uncharacterized protein \| Uncharacterized protein; afdb-uniprot50 \| AF-A0A367DAH4-F1-MODEL\_V4 \| 1.0 \| 1.682e-10 \| 406 \| 0.226 \| 212 \| 155 \| 7 \| 19 \| 225 \| 8 \| 215 \| Uncharacterized protein \| Uncharacterized protein; afdb-uniprot50 \| AF-A0A766PYH0-F1-MODEL\_V4 \| 1.0 \| 1.407e-17 \| 405 \| 0.25 \| 344 \| 192 \| 13 \| 190 \| 497 \| 1 \| 314 \| Phage terminase large subunit family protein \| Phage terminase large subunit family protein; afdb-uniprot50 \| AF-A0A5R2N7S2-F1-MODEL\_V4 \| 1.0 \| 1.144e-11 \| 403 \| 0.22 \| 227 \| 147 \| 12 \| 39 \| 242 \| 1 \| 220 \| Uncharacterized protein \| Uncharacterized protein; afdb-uniprot50 \| AF-A0A529NU64-F1-MODEL\_V4 \| 1.0 \| 1.075e-10 \| 402 \| 0.28 \| 207 \| 142 \| 6 \| 60 \| 262 \| 1 \| 204 \| Terminase \| Terminase; afdb-uniprot50 \| AF-A0A379V4Q9-F1-MODEL\_V4 \| 1.0 \| 3.259e-17 \| 401 \| 0.266 \| 327 \| 187 \| 14 \| 281 \| 584 \| 3 \| 299 \| Terminase \| Terminase; afdb-uniprot50 \| AF-A0A6P1B1H7-F1-MODEL\_V4 \| 1.0 \| 2.226e-10 \| 400 \| 0.666 \| 87 \| 29 \| 0 \| 392 \| 478 \| 1 \| 87 \| Phage terminase large subunit family protein \| Phage terminase large subunit family protein; afdb-uniprot50 \| AF-A0A1T1IZG5-F1-MODEL\_V4 \| 1.0 \| 1.218e-12 \| 399 \| 0.62 \| 137 \| 43 \| 1 \| 502 \| 638 \| 1 \| 128 \| Uncharacterized protein \| Uncharacterized protein; afdb-uniprot50 \| AF-Q9PBM0-F1-MODEL\_V4 \| 1.0 \| 4.025e-15 \| 398 \| 0.205 \| 350 \| 199 \| 15 \| 387 \| 680 \| 50 \| 376 \| Uncharacterized protein \| Uncharacterized protein; afdb-uniprot50 \| AF-A0A3S1PC79-F1-MODEL\_V4 \| 1.0 \| 8.123e-11 \| 396 \| 0.258 \| 224 \| 157 \| 6 \| 17 \| 232 \| 4 \| 226 \| Terminase \| Terminase; afdb-uniprot50 \| AF-A0A371JBP2-F1-MODEL\_V4 \| 1.0 \| 1.413e-09 \| 395 \| 0.256 \| 152 \| 112 \| 1 \| 19 \| 169 \| 11 \| 162 \| Phage terminase large subunit family protein \| Phage terminase large subunit family protein; afdb-uniprot50 \| AF-A0A6G4PKK6-F1-MODEL\_V4 \| 1.0 \| 1.226e-13 \| 393 \| 0.34 \| 191 \| 118 \| 5 \| 335 \| 523 \| 9 \| 193 \| Phage terminase large subunit family protein \| Phage terminase large subunit family protein; afdb-uniprot50 \| AF-A0A841I805-F1-MODEL\_V4 \| 1.0 \| 1.345e-10 \| 392 \| 0.193 \| 222 \| 165 \| 9 \| 17 \| 230 \| 3 \| 218 \| Phage terminase large subunit GpA-like protein \| Phage terminase large subunit GpA-like protein; afdb-uniprot50 \| AF-A0A855SXW0-F1-MODEL\_V4 \| 1.0 \| 1.043e-14 \| 390 \| 0.426 \| 190 \| 101 \| 5 \| 317 \| 501 \| 1 \| 187 \| Terminase \| Terminase; afdb-uniprot50 \| AF-A0A3D5V012-F1-MODEL\_V4 \| 1.0 \| 4.051e-16 \| 388 \| 0.262 \| 267 \| 160 \| 8 \| 223 \| 465 \| 27 \| 280 \| Uncharacterized protein \| Uncharacterized protein; afdb-uniprot50 \| AF-A0A377PIQ2-F1-MODEL\_V4 \| 1.0 \| 2.945e-10 \| 387 \| 0.193 \| 196 \| 141 \| 9 \| 33 \| 223 \| 18 \| 201 \| Bacteriophage tail assembly protein \| Bacteriophage tail assembly protein; afdb-uniprot50 \| AF-A0A447JGS3-F1-MODEL\_V4 \| 1.0 \| 1.85e-16 \| 387 \| 0.273 \| 274 \| 148 \| 9 \| 223 \| 467 \| 28 \| 279 \| Terminase \| Terminase; afdb-uniprot50 \| AF-A0A3C0YP94-F1-MODEL\_V4 \| 1.0 \| 5.19e-11 \| 384 \| 0.216 \| 236 \| 166 \| 7 \| 2 \| 230 \| 17 \| 240 \| Uncharacterized protein \| Uncharacterized protein; afdb-uniprot50 \| AF-A0A4Q6CYM5-F1-MODEL\_V4 \| 1.0 \| 1.01e-09 \| 383 \| 0.264 \| 178 \| 127 \| 2 \| 78 \| 251 \| 3 \| 180 \| Phage terminase large subunit family protein \| Phage terminase large subunit family protein; afdb-uniprot50 \| AF-A0A2A3WZ49-F1-MODEL\_V4 \| 1.0 \| 2.72e-15 \| 383 \| 0.275 \| 269 \| 144 \| 9 \| 223 \| 462 \| 6 \| 252 \| Terminase \| Terminase; afdb-uniprot50 \| AF-A0A826PUF0-F1-MODEL\_V4 \| 1.0 \| 7.501e-16 \| 383 \| 0.289 \| 266 \| 161 \| 10 \| 283 \| 530 \| 13 \| 268 \| Phage terminase large subunit family protein \| Phage terminase large subunit family protein; afdb-uniprot50 \| AF-A0A848YLQ0-F1-MODEL\_V4 \| 1.0 \| 8.123e-11 \| 382 \| 0.199 \| 246 \| 165 \| 8 \| 4 \| 232 \| 13 \| 243 \| Phage terminase large subunit family protein \| Phage terminase large subunit family protein; afdb-uniprot50 \| AF-A0A748BBD8-F1-MODEL\_V4 \| 1.0 \| 3.238e-16 \| 382 \| 0.278 \| 273 \| 161 \| 10 \| 283 \| 533 \| 2 \| 260 \| Phage terminase large subunit family protein \| Phage terminase large subunit family protein; afdb-uniprot50 \| AF-A0A376L8P3-F1-MODEL\_V4 \| 1.0 \| 7.631e-10 \| 380 \| 0.18 \| 221 \| 170 \| 5 \| 7 \| 223 \| 1 \| 214 \| Terminase large subunit of prophage CP-933O \| Terminase large subunit of prophage CP-933O; afdb-uniprot50 \| AF-A0A3A4P3Q9-F1-MODEL\_V4 \| 1.0 \| 6.867e-11 \| 379 \| 0.248 \| 201 \| 139 \| 2 \| 1 \| 189 \| 66 \| 266 \| Uncharacterized protein \| Uncharacterized protein; afdb-uniprot50 \| AF-A0A1V2N6H0-F1-MODEL\_V4 \| 1.0 \| 1.194e-09 \| 377 \| 0.274 \| 182 \| 124 \| 4 \| 24 \| 198 \| 2 \| 182 \| Uncharacterized protein \| Uncharacterized protein; afdb-uniprot50 \| AF-A0A2A5DAB2-F1-MODEL\_V4 \| 1.0 \| 1.194e-09 \| 374 \| 0.269 \| 193 \| 129 \| 5 \| 10 \| 200 \| 1 \| 183 \| Phage tail protein \| Phage tail protein; afdb-uniprot50 \| AF-A0A539CU23-F1-MODEL\_V4 \| 1.0 \| 7.263e-11 \| 373 \| 0.277 \| 144 \| 96 \| 3 \| 388 \| 526 \| 21 \| 161 \| Uncharacterized protein \| Uncharacterized protein; afdb-uniprot50 \| AF-A0A2N3B934-F1-MODEL\_V4 \| 1.0 \| 2.767e-09 \| 372 \| 0.302 \| 139 \| 95 \| 1 \| 99 \| 235 \| 398 \| 536 \| Terminase \| Terminase; afdb-uniprot50 \| AF-A0A5U5V548-F1-MODEL\_V4 \| 1.0 \| 1.99e-10 \| 371 \| 0.282 \| 202 \| 124 \| 5 \| 5 \| 200 \| 1 \| 187 \| Phage portal protein \| Phage portal protein; afdb-uniprot50 \| AF-A0A793H1F0-F1-MODEL\_V4 \| 1.0 \| 3.238e-16 \| 370 \| 0.26 \| 288 \| 159 \| 10 \| 243 \| 501 \| 5 \| 267 \| Phage terminase large subunit family protein \| Phage terminase large subunit family protein; afdb-uniprot50 \| AF-A0A1Y6C3G5-F1-MODEL\_V4 \| 1.0 \| 5.123e-09 \| 363 \| 0.23 \| 169 \| 124 \| 3 \| 22 \| 189 \| 3 \| 166 \| Phage terminase large subunit (GpA) \| Phage terminase large subunit (GpA); afdb-uniprot50 \| AF-A0A439EPL3-F1-MODEL\_V4 \| 1.0 \| 2.385e-12 \| 362 \| 0.21 \| 256 \| 145 \| 12 \| 374 \| 577 \| 2 \| 252 \| Terminase \| Terminase; afdb-uniprot50 \| AF-A0A8B2SLK2-F1-MODEL\_V4 \| 1.0 \| 2.49e-10 \| 362 \| 0.187 \| 262 \| 180 \| 9 \| 54 \| 302 \| 4 \| 245 \| Phage terminase large subunit family protein \| Phage terminase large subunit family protein; afdb-uniprot50 \| AF-A0A1V5NAS0-F1-MODEL\_V4 \| 1.0 \| 3.537e-21 \| 362 \| 0.138 \| 735 \| 443 \| 43 \| 17 \| 635 \| 2 \| 661 \| Phage terminase large subunit (GpA) \| Phage terminase large subunit (GpA); afdb-uniprot50 \| AF-A0A7U8Y278-F1-MODEL\_V4 \| 1.0 \| 4.148e-11 \| 359 \| 0.335 \| 146 \| 94 \| 2 \| 358 \| 502 \| 2 \| 145 \| Phage terminase large subunit family protein \| Phage terminase large subunit family protein; afdb-uniprot50 \| AF-A0A564UPM6-F1-MODEL\_V4 \| 1.0 \| 2.522e-12 \| 358 \| 0.377 \| 180 \| 93 \| 8 \| 463 \| 633 \| 53 \| 222 \| Phage terminase large subunit (GpA) \| Phage terminase large subunit (GpA); afdb-uniprot50 \| AF-E9CQP6-F1-MODEL\_V4 \| 1.0 \| 6.061e-09 \| 356 \| 0.241 \| 195 \| 143 \| 4 \| 5 \| 197 \| 1 \| 192 \| Putative phage tail assembly protein \| Putative phage tail assembly protein; afdb-uniprot50 \| AF-A0A7V8T050-F1-MODEL\_V4 \| 1.0 \| 5.768e-10 \| 356 \| 0.166 \| 265 \| 193 \| 9 \| 1 \| 257 \| 1 \| 245 \| Phage terminase large subunit family protein \| Phage terminase large subunit family protein; afdb-uniprot50 \| AF-A0A0F9G8T2-F1-MODEL\_V4 \| 1.0 \| 6.153e-20 \| 356 \| 0.158 \| 604 \| 352 \| 34 \| 112 \| 608 \| 13 \| 566 \| Uncharacterized protein \| Uncharacterized protein; afdb-uniprot50 \| AF-A0A271QD77-F1-MODEL\_V4 \| 1.0 \| 8.391e-16 \| 354 \| 0.268 \| 283 \| 146 \| 11 \| 223 \| 471 \| 14 \| 269 \| Terminase \| Terminase; afdb-uniprot50 \| AF-A0A786YKN4-F1-MODEL\_V4 \| 1.0 \| 6.341e-16 \| 351 \| 0.283 \| 268 \| 148 \| 10 \| 243 \| 486 \| 8 \| 255 \| Phage terminase large subunit family protein \| Phage terminase large subunit family protein; afdb-uniprot50 \| AF-A0A0P6WAH7-F1-MODEL\_V4 \| 1.0 \| 6.823e-10 \| 349 \| 0.247 \| 234 \| 156 \| 12 \| 6 \| 232 \| 1 \| 221 \| Uncharacterized protein \| Uncharacterized protein; afdb-uniprot50 \| AF-A0A6N1BJG2-F1-MODEL\_V4 \| 1.0 \| 2.212e-09 \| 349 \| 0.288 \| 177 \| 112 \| 6 \| 17 \| 181 \| 4 \| 178 \| Phage terminase large subunit family protein \| Phage terminase large subunit family protein; afdb-uniprot50 \| AF-A0A7X0Q037-F1-MODEL\_V4 \| 1.0 \| 2.324e-08 \| 346 \| 0.282 \| 163 \| 115 \| 2 \| 26 \| 186 \| 2 \| 164 \| Phage terminase large subunit family protein \| Phage terminase large subunit family protein; afdb-uniprot50 \| AF-A0A855SVD4-F1-MODEL\_V4 \| 1.0 \| 1.353e-11 \| 345 \| 0.325 \| 200 \| 118 \| 6 \| 463 \| 651 \| 1 \| 194 \| Terminase \| Terminase; afdb-uniprot50 \| AF-A0A660NIR9-F1-MODEL\_V4 \| 1.0 \| 1.814e-13 \| 345 \| 0.437 \| 231 \| 79 \| 4 \| 494 \| 682 \| 12 \| 233 \| Uncharacterized protein \| Uncharacterized protein; afdb-uniprot50 \| AF-A0A827LK07-F1-MODEL\_V4 \| 1.0 \| 1.858e-08 \| 344 \| 0.209 \| 196 \| 149 \| 4 \| 19 \| 210 \| 8 \| 201 \| Phage terminase large subunit family protein \| Phage terminase large subunit family protein; afdb-uniprot50 \| AF-A0A376ZGU3-F1-MODEL\_V4 \| 1.0 \| 3.661e-09 \| 344 \| 0.179 \| 229 \| 177 \| 8 \| 2 \| 222 \| 1 \| 226 \| Terminase large subunit (Gp2) \| Terminase large subunit (Gp2); afdb-uniprot50 \| AF-A0A173WP28-F1-MODEL\_V4 \| 1.0 \| 7.169e-09 \| 337 \| 0.209 \| 191 \| 129 \| 4 \| 1 \| 188 \| 1 \| 172 \| Bacteriophage tail assembly protein \| Bacteriophage tail assembly protein; afdb-uniprot50 \| AF-A0A518HU91-F1-MODEL\_V4 \| 1.0 \| 1.898e-20 \| 334 \| 0.123 \| 779 \| 429 \| 52 \| 19 \| 632 \| 2 \| 691 \| Phage terminase large subunit (GpA) \| Phage terminase large subunit (GpA); afdb-uniprot50 \| AF-A0A4R1YXE3-F1-MODEL\_V4 \| 1.0 \| 1.431e-11 \| 330 \| 0.169 \| 396 \| 251 \| 17 \| 4 \| 384 \| 1 \| 333 \| Mor transcription activator family protein \| Mor transcription activator family protein; afdb-uniprot50 \| AF-A0A2N8GT90-F1-MODEL\_V4 \| 1.0 \| 4.58e-09 \| 329 \| 0.447 \| 105 \| 58 \| 0 \| 369 \| 473 \| 1 \| 105 \| Terminase \| Terminase; afdb-uniprot50 \| AF-A0A0F8ZZL1-F1-MODEL\_V4 \| 1.0 \| 1.977e-09 \| 327 \| 0.236 \| 207 \| 133 \| 7 \| 1 \| 200 \| 1 \| 189 \| Uncharacterized protein \| Uncharacterized protein; afdb-uniprot50 \| AF-A0A811VMZ9-F1-MODEL\_V4 \| 1.0 \| 8.019e-09 \| 326 \| 0.317 \| 104 \| 67 \| 2 \| 388 \| 491 \| 7 \| 106 \| Phage terminase large subunit (GpA) \| Phage terminase large subunit (GpA); afdb-uniprot50 \| AF-A0A6G1XPZ7-F1-MODEL\_V4 \| 1.0 \| 1.661e-08 \| 320 \| 0.314 \| 124 \| 82 \| 2 \| 393 \| 515 \| 1 \| 122 \| Phage terminase large subunit family protein \| Phage terminase large subunit family protein; afdb-uniprot50 \| AF-A0A7Y6ERD3-F1-MODEL\_V4 \| 1.0 \| 4.069e-08 \| 317 \| 0.206 \| 194 \| 142 \| 6 \| 19 \| 205 \| 8 \| 196 \| Phage terminase large subunit family protein \| Phage terminase large subunit family protein; afdb-uniprot50 \| AF-A0A1V6CWB5-F1-MODEL\_V4 \| 1.0 \| 1.982e-18 \| 317 \| 0.131 \| 755 \| 449 \| 49 \| 17 \| 634 \| 47 \| 731 \| Phage terminase large subunit (GpA) \| Phage terminase large subunit (GpA); afdb-uniprot50 \| AF-A0A434AE22-F1-MODEL\_V4 \| 1.0 \| 4.844e-09 \| 316 \| 0.338 \| 127 \| 82 \| 1 \| 390 \| 516 \| 1 \| 125 \| Phage terminase large subunit family protein \| Phage terminase large subunit family protein; afdb-uniprot50 \| AF-A0A1Y5U6C2-F1-MODEL\_V4 \| 1.0 \| 4.61e-10 \| 316 \| 0.389 \| 159 \| 88 \| 4 \| 480 \| 633 \| 3 \| 157 \| Phage terminase large subunit (GpA) \| Phage terminase large subunit (GpA); afdb-uniprot50 \| AF-A0A6N7ZP18-F1-MODEL\_V4 \| 1.0 \| 1.952e-07 \| 314 \| 0.273 \| 161 \| 113 \| 3 \| 26 \| 183 \| 2 \| 161 \| Phage terminase large subunit family protein \| Phage terminase large subunit family protein; afdb-uniprot50 \| AF-A0A2V5RDT8-F1-MODEL\_V4 \| 1.0 \| 9.968e-08 \| 310 \| 0.213 \| 178 \| 137 \| 2 \| 7 \| 183 \| 7 \| 182 \| Uncharacterized protein \| Uncharacterized protein; afdb-uniprot50 \| AF-A0A1G0ZSU7-F1-MODEL\_V4 \| 1.0 \| 3.102e-18 \| 310 \| 0.135 \| 811 \| 458 \| 57 \| 1 \| 680 \| 1 \| 698 \| Uncharacterized protein \| Uncharacterized protein; afdb-uniprot50 \| AF-A0A356GP32-F1-MODEL\_V4 \| 1.0 \| 6.823e-10 \| 309 \| 0.369 \| 176 \| 93 \| 4 \| 468 \| 632 \| 2 \| 170 \| Uncharacterized protein \| Uncharacterized protein; afdb-uniprot50 \| AF-A0A7W8PL95-F1-MODEL\_V4 \| 1.0 \| 6.735e-08 \| 307 \| 0.274 \| 124 \| 88 \| 2 \| 104 \| 227 \| 3 \| 124 \| Phage terminase large subunit GpA-like protein \| Phage terminase large subunit GpA-like protein; afdb-uniprot50 \| AF-A0A627PGR2-F1-MODEL\_V4 \| 1.0 \| 1.846e-07 \| 306 \| 0.221 \| 185 \| 138 \| 4 \| 57 \| 238 \| 4 \| 185 \| Phage terminase large subunit family protein \| Phage terminase large subunit family protein; afdb-uniprot50 \| AF-A0A382RRM4-F1-MODEL\_V4 \| 1.0 \| 2.339e-09 \| 300 \| 0.366 \| 161 \| 95 \| 3 \| 480 \| 640 \| 1 \| 154 \| Uncharacterized protein \| Uncharacterized protein; afdb-uniprot50 \| AF-D8A639-F1-MODEL\_V4 \| 1.0 \| 2.945e-10 \| 300 \| 0.375 \| 149 \| 89 \| 3 \| 335 \| 481 \| 7 \| 153 \| Uncharacterized protein \| Uncharacterized protein; afdb-uniprot50 \| AF-A0A847I6D0-F1-MODEL\_V4 \| 1.0 \| 2.479e-18 \| 300 \| 0.135 \| 753 \| 428 \| 42 \| 19 \| 632 \| 5 \| 673 \| Uncharacterized protein \| Uncharacterized protein; afdb-uniprot50 \| AF-A0A377DG38-F1-MODEL\_V4 \| 1.0 \| 1.226e-13 \| 299 \| 0.238 \| 281 \| 161 \| 11 \| 243 \| 497 \| 7 \| 260 \| Terminase GpA \| Terminase GpA; afdb-uniprot50 \| AF-A0A0M9GD64-F1-MODEL\_V4 \| 1.0 \| 6.867e-11 \| 298 \| 0.284 \| 246 \| 138 \| 9 \| 460 \| 680 \| 3 \| 235 \| Bacteriophage tail assembly protein \| Bacteriophage tail assembly protein; afdb-uniprot50 \| AF-A0A376ZEG5-F1-MODEL\_V4 \| 1.0 \| 3.337e-12 \| 297 \| 0.298 \| 221 \| 134 \| 6 \| 281 \| 495 \| 6 \| 211 \| Phage terminase large subunit (GpA) \| Phage terminase large subunit (GpA); afdb-uniprot50 \| AF-A0A1A6FGJ8-F1-MODEL\_V4 \| 1.0 \| 3.822e-07 \| 296 \| 0.34 \| 147 \| 96 \| 1 \| 53 \| 198 \| 3 \| 149 \| Uncharacterized protein \| Uncharacterized protein; afdb-uniprot50 \| AF-A0A645HKJ2-F1-MODEL\_V4 \| 1.0 \| 7.169e-09 \| 293 \| 0.206 \| 174 \| 119 \| 9 \| 441 \| 605 \| 18 \| 181 \| Uncharacterized protein \| Uncharacterized protein; afdb-uniprot50 \| AF-A0A0J5K276-F1-MODEL\_V4 \| 1.0 \| 5.983e-07 \| 290 \| 0.2 \| 175 \| 136 \| 3 \| 17 \| 189 \| 6 \| 178 \| Terminase \| Terminase; afdb-uniprot50 \| AF-A0A090UZ83-F1-MODEL\_V4 \| 1.0 \| 3.614e-07 \| 288 \| 0.201 \| 174 \| 129 \| 3 \| 18 \| 186 \| 5 \| 173 \| Uncharacterized protein \| Uncharacterized protein; afdb-uniprot50 \| AF-A0A0K4ZPK0-F1-MODEL\_V4 \| 1.0 \| 7.967e-08 \| 286 \| 0.195 \| 200 \| 146 \| 6 \| 3 \| 200 \| 1 \| 187 \| Phage terminase large subunit \| Phage terminase large subunit; afdb-uniprot50 \| AF-A0A5C8XUH5-F1-MODEL\_V4 \| 1.0 \| 5.349e-07 \| 283 \| 0.185 \| 210 \| 154 \| 8 \| 7 \| 209 \| 1 \| 200 \| Phage terminase large subunit family protein \| Phage terminase large subunit family protein; afdb-uniprot50 \| AF-A0A830TE26-F1-MODEL\_V4 \| 1.0 \| 1.57e-08 \| 281 \| 0.362 \| 135 \| 83 \| 2 \| 354 \| 487 \| 7 \| 139 \| Phage terminase large subunit family protein \| Phage terminase large subunit family protein; afdb-uniprot50 \| AF-A0A0F8W5R2-F1-MODEL\_V4 \| 1.0 \| 1.55e-06 \| 279 \| 0.267 \| 146 \| 106 \| 1 \| 51 \| 195 \| 2 \| 147 \| Uncharacterized protein \| Uncharacterized protein; afdb-uniprot50 \| AF-A0A2W7VTJ7-F1-MODEL\_V4 \| 1.0 \| 3.439e-08 \| 278 \| 0.333 \| 132 \| 82 \| 3 \| 362 \| 492 \| 2 \| 128 \| Uncharacterized protein \| Uncharacterized protein; afdb-uniprot50 \| AF-A0A7U2BFP7-F1-MODEL\_V4 \| 1.0 \| 4.782e-07 \| 278 \| 0.228 \| 162 \| 122 \| 2 \| 19 \| 180 \| 7 \| 165 \| Phage terminase large subunit family protein \| Phage terminase large subunit family protein; afdb-uniprot50 \| AF-A0A827B2P6-F1-MODEL\_V4 \| 1.0 \| 2.616e-09 \| 276 \| 0.375 \| 141 \| 86 \| 2 \| 333 \| 471 \| 11 \| 151 \| Phage terminase large subunit family protein \| Phage terminase large subunit family protein; afdb-uniprot50 \| AF-X6D4R2-F1-MODEL\_V4 \| 1.0 \| 2.091e-09 \| 276 \| 0.247 \| 234 \| 132 \| 10 \| 466 \| 678 \| 3 \| 213 \| Uncharacterized protein \| Uncharacterized protein; afdb-uniprot50 \| AF-A0A607GXP9-F1-MODEL\_V4 \| 1.0 \| 9.364e-07 \| 271 \| 0.281 \| 167 \| 99 \| 5 \| 5 \| 165 \| 1 \| 152 \| Phage terminase large subunit family protein \| Phage terminase large subunit family protein; afdb-uniprot50 \| AF-A0A7U9B7N7-F1-MODEL\_V4 \| 1.0 \| 4.042e-07 \| 257 \| 0.355 \| 104 \| 65 \| 1 \| 384 \| 485 \| 4 \| 107 \| Terminase large subunit (DNA packaging protein A) (GpA) \| Terminase large subunit (DNA packaging protein A) (GpA); afdb-uniprot50 \| AF-A0A1E3G601-F1-MODEL\_V4 \| 1.0 \| 3.822e-07 \| 256 \| 0.182 \| 164 \| 115 \| 3 \| 94 \| 257 \| 3 \| 147 \| Uncharacterized protein \| Uncharacterized protein; afdb-uniprot50 \| AF-A0A781K8L4-F1-MODEL\_V4 \| 1.0 \| 1.041e-05 \| 248 \| 0.225 \| 160 \| 117 \| 4 \| 39 \| 195 \| 1 \| 156 \| Phage terminase large subunit family protein \| Phage terminase large subunit family protein; afdb-uniprot50 \| AF-A0A2X3JNN5-F1-MODEL\_V4 \| 1.0 \| 4.016e-06 \| 246 \| 0.358 \| 92 \| 57 \| 1 \| 396 \| 487 \| 2 \| 91 \| Phage terminase large subunit \| Phage terminase large subunit; afdb-uniprot50 \| AF-A0A2J0QJY4-F1-MODEL\_V4 \| 1.0 \| 4.844e-09 \| 243 \| 0.329 \| 158 \| 100 \| 4 \| 317 \| 468 \| 13 \| 170 \| Uncharacterized protein \| Uncharacterized protein; afdb-uniprot50 \| AF-A0A1X7MCA7-F1-MODEL\_V4 \| 1.0 \| 2.155e-05 \| 241 \| 0.285 \| 112 \| 80 \| 0 \| 89 \| 200 \| 3 \| 114 \| Phage terminase large subunit (GpA) \| Phage terminase large subunit (GpA); afdb-uniprot50 \| AF-A0A7Z0VHE6-F1-MODEL\_V4 \| 1.0 \| 4.751e-06 \| 237 \| 0.214 \| 168 \| 122 \| 6 \| 19 \| 181 \| 8 \| 170 \| Phage terminase large subunit (GpA) \| Phage terminase large subunit (GpA); afdb-uniprot50 \| AF-A0A447MUP3-F1-MODEL\_V4 \| 1.0 \| 1.822e-05 \| 233 \| 0.36 \| 75 \| 48 \| 0 \| 398 \| 472 \| 3 \| 77 \| Phage terminase large subunit \| Phage terminase large subunit; afdb-uniprot50 \| AF-A0A0F8YV96-F1-MODEL\_V4 \| 1.0 \| 3.373e-05 \| 231 \| 0.226 \| 115 \| 89 \| 0 \| 74 \| 188 \| 2 \| 116 \| Uncharacterized protein \| Uncharacterized protein; afdb-uniprot50 \| AF-A0A2E6PVU1-F1-MODEL\_V4 \| 1.0 \| 2.105e-10 \| 230 \| 0.117 \| 569 \| 307 \| 34 \| 64 \| 607 \| 25 \| 423 \| ATP-binding protein \| ATP-binding protein; afdb-uniprot50 \| AF-A0A418GMF3-F1-MODEL\_V4 \| 1.0 \| 5.906e-05 \| 222 \| 0.328 \| 73 \| 49 \| 0 \| 396 \| 468 \| 2 \| 74 \| Phage terminase large subunit family protein \| Phage terminase large subunit family protein; afdb-uniprot50 \| AF-A0A7C0WJF8-F1-MODEL\_V4 \| 1.0 \| 1.693e-11 \| 221 \| 0.107 \| 650 \| 401 \| 47 \| 40 \| 614 \| 16 \| 561 \| Uncharacterized protein \| Uncharacterized protein; afdb-uniprot50 \| AF-A0A7A6R3G1-F1-MODEL\_V4 \| 1.0 \| 8.74e-05 \| 219 \| 0.19 \| 152 \| 119 \| 3 \| 19 \| 168 \| 8 \| 157 \| Phage terminase large subunit family protein \| Phage terminase large subunit family protein; afdb-uniprot50 \| AF-A0A6H1ZTH9-F1-MODEL\_V4 \| 1.0 \| 5.157e-10 \| 219 \| 0.106 \| 566 \| 333 \| 30 \| 68 \| 608 \| 15 \| 432 \| Putative terminase \| Putative terminase; afdb-uniprot50 \| AF-A0A3A9EXD7-F1-MODEL\_V4 \| 1.0 \| 3.822e-07 \| 217 \| 0.228 \| 175 \| 117 \| 9 \| 463 \| 632 \| 1 \| 162 \| Uncharacterized protein \| Uncharacterized protein; afdb-uniprot50 \| AF-A0A7B8XUA3-F1-MODEL\_V4 \| 1.0 \| 2.852e-05 \| 215 \| 0.415 \| 77 \| 45 \| 0 \| 384 \| 460 \| 5 \| 81 \| Phage terminase large subunit family protein \| Phage terminase large subunit family protein; afdb-uniprot50 \| AF-A0A844W300-F1-MODEL\_V4 \| 1.0 \| 6.327e-07 \| 215 \| 0.289 \| 169 \| 107 \| 4 \| 492 \| 649 \| 2 \| 168 \| Terminase \| Terminase; afdb-uniprot50 \| AF-A0A0F9FUP5-F1-MODEL\_V4 \| 1.0 \| 3.798e-06 \| 202 \| 0.382 \| 123 \| 71 \| 2 \| 525 \| 644 \| 2 \| 122 \| Uncharacterized protein \| Uncharacterized protein; afdb-uniprot50 \| AF-A0A5B2VA92-F1-MODEL\_V4 \| 1.0 \| 8.426e-08 \| 202 \| 0.344 \| 151 \| 91 \| 4 \| 317 \| 460 \| 16 \| 165 \| Phage terminase large subunit family protein \| Phage terminase large subunit family protein; afdb-uniprot50 \| AF-A0A1M7PCZ5-F1-MODEL\_V4 \| 1.0 \| 1.734e-06 \| 199 \| 0.158 \| 164 \| 122 \| 5 \| 341 \| 492 \| 118 \| 277 \| Phage terminase large subunit (GpA) \| Phage terminase large subunit (GpA); afdb-uniprot50 \| AF-A0A258KXC3-F1-MODEL\_V4 \| 1.0 \| 0.0002025 \| 197 \| 0.288 \| 135 \| 90 \| 5 \| 51 \| 181 \| 2 \| 134 \| Uncharacterized protein \| Uncharacterized protein; afdb-uniprot50 \| AF-A0A1G8I1T2-F1-MODEL\_V4 \| 1.0 \| 0.000181 \| 196 \| 0.2 \| 145 \| 111 \| 4 \| 26 \| 168 \| 2 \| 143 \| Phage terminase large subunit (GpA) \| Phage terminase large subunit (GpA); afdb-uniprot50 \| AF-A0A2A3WXQ1-F1-MODEL\_V4 \| 1.0 \| 8.74e-05 \| 193 \| 0.333 \| 84 \| 56 \| 0 \| 379 \| 462 \| 18 \| 101 \| Terminase \| Terminase; afdb-uniprot50 \| AF-H6SLW0-F1-MODEL\_V4 \| 1.0 \| 3.035e-06 \| 191 \| 0.32 \| 178 \| 88 \| 6 \| 509 \| 678 \| 7 \| 159 \| Phage terminase GpA \| Phage terminase GpA; afdb-uniprot50 \| AF-A0A6P0ZEA2-F1-MODEL\_V4 \| 1.0 \| 1.65e-07 \| 191 \| 0.368 \| 190 \| 93 \| 6 \| 499 \| 682 \| 105 \| 273 \| Uncharacterized protein \| Uncharacterized protein; afdb-uniprot50 \| AF-A0A844YZ98-F1-MODEL\_V4 \| 1.0 \| 3.685e-10 \| 191 \| 0.092 \| 550 \| 321 \| 33 \| 82 \| 607 \| 28 \| 423 \| Uncharacterized protein \| Uncharacterized protein; afdb-uniprot50 \| AF-A0A524GG08-F1-MODEL\_V4 \| 1.0 \| 7.031e-06 \| 190 \| 0.134 \| 246 \| 184 \| 14 \| 2 \| 236 \| 1 \| 228 \| Uncharacterized protein \| Uncharacterized protein; afdb-uniprot50 \| AF-A0A317H0R0-F1-MODEL\_V4 \| 1.0 \| 8.797e-06 \| 190 \| 0.135 \| 228 \| 157 \| 15 \| 34 \| 238 \| 1 \| 211 \| Uncharacterized protein \| Uncharacterized protein; afdb-uniprot50 \| AF-A0A0F9MIU9-F1-MODEL\_V4 \| 1.0 \| 8.536e-10 \| 190 \| 0.124 \| 668 \| 371 \| 43 \| 41 \| 632 \| 19 \| 548 \| Uncharacterized protein \| Uncharacterized protein; afdb-uniprot50 \| AF-A0A2E0YGA1-F1-MODEL\_V4 \| 1.0 \| 1.768e-09 \| 181 \| 0.112 \| 700 \| 409 \| 44 \| 17 \| 631 \| 2 \| 573 \| Uncharacterized protein \| Uncharacterized protein; afdb-uniprot50 \| AF-A0A4P0YGW9-F1-MODEL\_V4 \| 1.0 \| 1.466e-06 \| 176 \| 0.326 \| 144 \| 94 \| 3 \| 318 \| 459 \| 11 \| 153 \| Terminase GpA \| Terminase GpA; afdb-uniprot50 \| AF-A0A4Z0P9X8-F1-MODEL\_V4 \| 1.0 \| 8.019e-09 \| 174 \| 0.115 \| 591 \| 320 \| 40 \| 68 \| 607 \| 6 \| 444 \| Terminase\_3 domain-containing protein \| Terminase\_3 domain-containing protein; afdb-uniprot50 \| AF-A0A2E7BN62-F1-MODEL\_V4 \| 1.0 \| 1.336e-09 \| 173 \| 0.105 \| 796 \| 441 \| 51 \| 11 \| 682 \| 61 \| 709 \| Uncharacterized protein \| Uncharacterized protein; afdb-uniprot50 \| AF-A0A844WAL0-F1-MODEL\_V4 \| 1.0 \| 0.002517 \| 168 \| 0.212 \| 127 \| 100 \| 0 \| 55 \| 181 \| 7 \| 133 \| Terminase \| Terminase; afdb-uniprot50 \| AF-A0A0F9G2T5-F1-MODEL\_V4 \| 1.0 \| 4.463e-05 \| 165 \| 0.125 \| 223 \| 155 \| 14 \| 39 \| 237 \| 2 \| 208 \| Uncharacterized protein \| Uncharacterized protein; afdb-uniprot50 \| AF-A0A1Y4DBY9-F1-MODEL\_V4 \| 1.0 \| 1.65e-07 \| 165 \| 0.091 \| 547 \| 315 \| 31 \| 76 \| 605 \| 16 \| 397 \| Terminase\_3 domain-containing protein \| Terminase\_3 domain-containing protein; afdb-uniprot50 \| AF-A0A2W5BGY9-F1-MODEL\_V4 \| 1.0 \| 5.584e-05 \| 159 \| 0.319 \| 141 \| 89 \| 3 \| 510 \| 644 \| 1 \| 140 \| Terminase \| Terminase; afdb-uniprot50 \| AF-A0A3D3HHB8-F1-MODEL\_V4 \| 1.0 \| 3.252e-08 \| 158 \| 0.082 \| 713 \| 370 \| 42 \| 20 \| 602 \| 86 \| 643 \| Uncharacterized protein \| Uncharacterized protein; afdb-uniprot50 \| AF-A0A3S0GVK1-F1-MODEL\_V4 \| 1.0 \| 9.776e-05 \| 149 \| 0.262 \| 156 \| 94 \| 7 \| 526 \| 678 \| 15 \| 152 \| Uncharacterized protein \| Uncharacterized protein; afdb-uniprot50 \| AF-A0A104MQ19-F1-MODEL\_V4 \| 1.0 \| 0.0001914 \| 140 \| 0.258 \| 170 \| 111 \| 4 \| 525 \| 680 \| 4 \| 172 \| Uncharacterized protein \| Uncharacterized protein; afdb-uniprot50 \| AF-A0A356HIS5-F1-MODEL\_V4 \| 1.0 \| 0.0007341 \| 130 \| 0.291 \| 144 \| 80 \| 3 \| 541 \| 681 \| 3 \| 127 \| Terminase \| Terminase; afdb-uniprot50 \| AF-A0A6N4T2G2-F1-MODEL\_V4 \| 1.0 \| 7.389e-05 \| 127 \| 0.273 \| 190 \| 105 \| 7 \| 507 \| 679 \| 31 \| 204 \| Uncharacterized protein \| Uncharacterized protein; afdb-uniprot50 \| AF-A0A0Q6MRT9-F1-MODEL\_V4 \| 1.0 \| 0.00238 \| 115 \| 0.258 \| 147 \| 82 \| 4 \| 534 \| 677 \| 2 \| 124 \| Uncharacterized protein \| Uncharacterized protein; afdb-uniprot50 \| AF-A0A381VRU7-F1-MODEL\_V4 \| 1.0 \| 0.0005868 \| 113 \| 0.111 \| 242 \| 147 \| 14 \| 23 \| 223 \| 11 \| 225 \| Terminase\_6 domain-containing protein \| Terminase\_6 domain-containing protein |
| Top keywords  (threshold 1.00e-02 (evalue)) | **terminase, Phage, large, GpA, tail, GpA\_like, assembly, Bacteriophage, DNA, Putative** |
| Output files | ../../similar\_structures/02\_FANPEZAQ\_CDS\_0002\_afdb-proteome\_foldseek.tsv ../../similar\_structures/02\_FANPEZAQ\_CDS\_0002\_afdb-uniprot50\_foldseek.tsv ../../similar\_structures/02\_FANPEZAQ\_CDS\_0002\_merged.svg ../../similar\_structures/02\_FANPEZAQ\_CDS\_0002\_pdb\_foldseek.tsv |

  
  
  

Return to summary | Go to previous | Go to next

  


---

**Sequence/structure alignments coloring**  
Each object in the alignment figures is colored according to its E-value following this color coding:

1e-100
10
