## Supplementary material for "Completing the BASEL phage collection to unlock hidden diversity for systematic exploration of phage-host interactions": Data S2: 3.html

FANPEZAQ\_CDS\_0003


Return to summary | Go to previous | Go to next

|  |  |
| --- | --- |
| FANPEZAQ\_CDS\_0003  Page creation date: 02 Sep 2024, 12:00  Project folder: n/a  Input sequences file: Escherichia\_virus\_HeidiAbel.gb | domain\_containing bppu\_n light phage receptor fab duf2479 hypothetical bppu dolichyl\_diphosphooligosaccharide\_\_protein glycosyltransferase n\_terminal immunoglobulin fibronectin motile sperm tail type repeat and ig\_like antibody heavy cell fragment copc sodium channel alpha leucine\_rich sequence putative region kda wall anchor type\_iii peptidylprolyl isomerase primosomal replication prib pric iii signal baseplate upper ig gamma transmembrane |

### Sequence information

|  |  |
| --- | --- |
| Name | FANPEZAQ\_CDS\_0003  03\_FANPEZAQ\_CDS\_0003 (pipeline id) |
| Imported annotations | Escherichia\_virus\_HeidiAbel Bas97 |
| Protein sequence | MKCSQLPQKITAGLTFESRLNLPDYPATEWAVRLLLRGPQSIDVPAVVDGKTSVIRAAAG VTGTWPAGEYWYSIRATRGSDVVEVGSGQITIMPDLAAAEAGYTAKTHAQRTLEAIEAVI EKRASMDQERYRINNRELYRTPIADLLKLRDLYRLEVAREQQAQRGCNPFGRKVRVSLR |
| Number of residues | 179 |
| Molecular weight (Da) | 19973.62 |
| Output files | ../../query\_sequences/03\_FANPEZAQ\_CDS\_0003.fasta |

### Putative domain architecture and protein family

#### Search results (HHblits)^1^

|  |  |
| --- | --- |
| Domain family databases searched | Pfam, Ncbi-cd, Cath, Phrogs |
| Results, scheme(s)  (Top layers only; threshold 1.00e-03 (evalue)) | xml version="1.0" encoding="utf-8" standalone="no"?       2024-09-02T21:08:12.337035 image/svg+xml   Matplotlib v3.7.2, https://matplotlib.org/ |
| Results, table  (E-value ≤ 1.00e-03 (evalue)) | db \| id \| prob \| evalue \| pvalue \| score \| cols \| query \| query\_len \| template \| template\_len \| name \| description; phrogs \| 5774 \| 100.0 \| 2.4e-42 \| 3e-46 \| 271.9 \| 171 \| (5, 178) \| 179 \| (2, 173) \| 176 \| head-tail joining \| head-tail joining; Category: connector; NC\_009016\_p5; phrogs \| 1627 \| 99.7 \| 5.8e-23 \| 7.5e-27 \| 149.9 \| 94 \| (3, 96) \| 179 \| (1, 107) \| 109 \| NA \| NA; Category: unknown function; NC\_024367\_p52; phrogs \| 118 \| 99.5 \| 4.9e-19 \| 6.4e-23 \| 119.2 \| 62 \| (106, 169) \| 179 \| (3, 64) \| 70 \| head-tail adaptor \| head-tail adaptor; Category: connector; p69560 VI\_02299; phrogs \| 35772 \| 97.3 \| 1.9e-07 \| 2.1e-11 \| 66.1 \| 56 \| (106, 163) \| 179 \| (66, 121) \| 136 \| NA \| NA; Category: unknown function; p130612 VI\_06329; phrogs \| 20307 \| 96.8 \| 2.2e-06 \| 2.5e-10 \| 54.6 \| 57 \| (107, 163) \| 179 \| (2, 58) \| 70 \| NA \| NA; Category: unknown function; p166236 VI\_12298 |
| Top keywords  (threshold 1.00e-03 (evalue)) | **head\_tail, connector, joining, NC\_009016\_p5, NC\_024367\_p52, adaptor, p69560, VI\_02299, p130612, VI\_06329** |
| Output files | ../../domain\_architecture/03\_FANPEZAQ\_CDS\_0003\_cath.hhr ../../domain\_architecture/03\_FANPEZAQ\_CDS\_0003\_merged.svg ../../domain\_architecture/03\_FANPEZAQ\_CDS\_0003\_ncbi-cd.hhr ../../domain\_architecture/03\_FANPEZAQ\_CDS\_0003\_pfam.hhr ../../domain\_architecture/03\_FANPEZAQ\_CDS\_0003\_phrogs.hhr |

### Identical protein sequences/structures

#### Search results

|  |  |
| --- | --- |
| Protein sequence databases searched | Pdb, Swissprot, Refseq |
| Identical proteins found | -- |
| Top keywords | -- |
| Output files | -- |

### Similar protein sequences/structures

#### Sequence similarity search results (HHblits)^1^

|  |  |
| --- | --- |
| Sequence databases searched | Uniclust, Pdb70 |
| Results, scheme(s)  (Top layers only, threshold 1.00e-03 (evalue)) | xml version="1.0" encoding="utf-8" standalone="no"?       2024-09-02T21:08:28.900931 image/svg+xml   Matplotlib v3.7.2, https://matplotlib.org/ |
| Results, table(s)  (threshold 1.00e-03 (evalue)) | db \| id \| prob \| evalue \| pvalue \| score \| cols \| query \| query\_len \| template \| template\_len \| name \| description; uniclust \| UniRef100\_A0A0A8TGY6 \| 100.0 \| 1.8e-51 \| 4.3e-57 \| 329.9 \| 176 \| (2, 179) \| 179 \| (24, 200) \| 218 \| Uncharacterized protein \| Uncharacterized protein; uniclust \| UniRef100\_A0A165R292 \| 100.0 \| 1.3e-45 \| 3e-51 \| 290.3 \| 175 \| (2, 179) \| 179 \| (10, 195) \| 206 \| Uncharacterized protein \| Uncharacterized protein; uniclust \| UniRef100\_A0A060H347 \| 100.0 \| 6.8e-45 \| 1.4e-50 \| 275.1 \| 179 \| (1, 179) \| 179 \| (5, 184) \| 185 \| Uncharacterized protein \| Uncharacterized protein; uniclust \| UniRef100\_A0A0S8DP01 \| 100.0 \| 7.6e-44 \| 1.6e-49 \| 279.5 \| 178 \| (2, 179) \| 179 \| (8, 198) \| 209 \| Uncharacterized protein \| Uncharacterized protein; uniclust \| UniRef100\_A0A2P1VF72 \| 100.0 \| 3e-38 \| 6.2e-44 \| 234.4 \| 144 \| (35, 179) \| 179 \| (4, 147) \| 150 \| Uncharacterized protein \| Uncharacterized protein; uniclust \| UniRef100\_A0A1W6TB99 \| 100.0 \| 3e-34 \| 5.6e-40 \| 214.5 \| 174 \| (4, 179) \| 179 \| (1, 175) \| 175 \| Uncharacterized protein \| Uncharacterized protein; uniclust \| UniRef100\_A0A0F9GVA5 \| 99.9 \| 2.8e-31 \| 5.4e-37 \| 201.5 \| 162 \| (1, 165) \| 179 \| (1, 164) \| 184 \| Uncharacterized protein (Fragment) \| Uncharacterized protein (Fragment); uniclust \| UniRef100\_A0A510E8K6 \| 99.9 \| 1.9e-30 \| 3.7e-36 \| 198.4 \| 174 \| (2, 179) \| 179 \| (1, 194) \| 195 \| Uncharacterized protein \| Uncharacterized protein; uniclust \| UniRef100\_A0A1F9VFN4 \| 99.9 \| 6.2e-29 \| 1.2e-34 \| 192.3 \| 170 \| (5, 179) \| 179 \| (20, 191) \| 197 \| Phage tail protein \| Phage tail protein; uniclust \| UniRef100\_A0A088FQK4 \| 99.9 \| 3.5e-28 \| 7.1e-34 \| 171.9 \| 101 \| (78, 179) \| 179 \| (2, 102) \| 103 \| Uncharacterized protein \| Uncharacterized protein; uniclust \| UniRef100\_A0A2D5ZTJ7 \| 99.9 \| 1.4e-27 \| 2.8e-33 \| 181.9 \| 170 \| (4, 179) \| 179 \| (1, 171) \| 171 \| Uncharacterized protein \| Uncharacterized protein; uniclust \| UniRef100\_A0A1J5SRA9 \| 99.9 \| 7.9e-27 \| 1.5e-32 \| 177.4 \| 175 \| (4, 179) \| 179 \| (1, 187) \| 187 \| Uncharacterized protein \| Uncharacterized protein; uniclust \| UniRef100\_Q1ILS4 \| 99.9 \| 3.2e-25 \| 5.8e-31 \| 167.9 \| 171 \| (8, 179) \| 179 \| (10, 182) \| 201 \| Uncharacterized protein \| Uncharacterized protein; uniclust \| UniRef100\_A0A510E7U2 \| 99.9 \| 3.9e-25 \| 7.2e-31 \| 163.9 \| 166 \| (4, 179) \| 179 \| (2, 168) \| 172 \| Uncharacterized protein \| Uncharacterized protein; uniclust \| UniRef100\_A0A0N8FSS6 \| 99.8 \| 1.1e-22 \| 2e-28 \| 154.6 \| 155 \| (6, 161) \| 179 \| (12, 171) \| 184 \| DUF1254 domain-containing protein (Fragment) \| DUF1254 domain-containing protein (Fragment); uniclust \| UniRef100\_A0A090V164 \| 99.8 \| 2.7e-22 \| 5.8e-28 \| 158.6 \| 161 \| (1, 175) \| 179 \| (4, 168) \| 185 \| Uncharacterized protein \| Uncharacterized protein; uniclust \| UniRef100\_A0A2E7ZMH9 \| 99.7 \| 3.5e-21 \| 6.7e-27 \| 138.7 \| 109 \| (69, 179) \| 179 \| (1, 114) \| 125 \| Uncharacterized protein (Fragment) \| Uncharacterized protein (Fragment); uniclust \| UniRef100\_A0A1V5CWH5 \| 99.7 \| 7.2e-21 \| 1.3e-26 \| 144.3 \| 174 \| (4, 179) \| 179 \| (1, 193) \| 194 \| Uncharacterized protein \| Uncharacterized protein; uniclust \| UniRef100\_A0A3B9Q8J6 \| 99.7 \| 8.9e-21 \| 1.8e-26 \| 147.3 \| 169 \| (7, 179) \| 179 \| (2, 181) \| 182 \| Uncharacterized protein \| Uncharacterized protein; uniclust \| UniRef100\_A0A958HLF9 \| 99.7 \| 2.7e-20 \| 4.9e-26 \| 144.8 \| 170 \| (5, 177) \| 179 \| (50, 230) \| 233 \| Uncharacterized protein \| Uncharacterized protein; uniclust \| UniRef100\_UPI0016849037 \| 99.7 \| 3.6e-20 \| 6.6e-26 \| 139.5 \| 170 \| (3, 179) \| 179 \| (10, 182) \| 182 \| hypothetical protein \| hypothetical protein; uniclust \| UniRef100\_A0A2D7GUN5 \| 99.7 \| 6.2e-20 \| 1.1e-25 \| 136.7 \| 165 \| (7, 178) \| 179 \| (3, 168) \| 168 \| Uncharacterized protein \| Uncharacterized protein; uniclust \| UniRef100\_A0A969NUQ2 \| 99.7 \| 1.8e-19 \| 3.3e-25 \| 135.3 \| 175 \| (1, 179) \| 179 \| (1, 176) \| 177 \| Uncharacterized protein \| Uncharacterized protein; uniclust \| UniRef100\_A0A286GPM6 \| 99.7 \| 2e-19 \| 3.7e-25 \| 133.3 \| 146 \| (9, 160) \| 179 \| (2, 147) \| 161 \| Uncharacterized protein \| Uncharacterized protein; uniclust \| UniRef100\_A0A812JEV6 \| 99.6 \| 4.7e-19 \| 8.6e-25 \| 169.0 \| 153 \| (8, 160) \| 179 \| (960, 1116) \| 4451 \| Phage portal protein \| Phage portal protein; uniclust \| UniRef100\_UPI00101302DA \| 99.6 \| 8.8e-19 \| 1.6e-24 \| 124.7 \| 120 \| (5, 126) \| 179 \| (1, 121) \| 122 \| hypothetical protein \| hypothetical protein; uniclust \| UniRef100\_A0A071LUC0 \| 99.6 \| 1.7e-18 \| 3.6e-24 \| 133.3 \| 139 \| (4, 164) \| 179 \| (1, 141) \| 161 \| Uncharacterized protein \| Uncharacterized protein; uniclust \| UniRef100\_UPI001982702E \| 99.6 \| 8.3e-18 \| 1.5e-23 \| 115.1 \| 95 \| (62, 157) \| 179 \| (1, 95) \| 95 \| hypothetical protein \| hypothetical protein; uniclust \| UniRef100\_A0A934RPA6 \| 99.6 \| 8.9e-18 \| 1.6e-23 \| 125.6 \| 145 \| (9, 156) \| 179 \| (2, 153) \| 168 \| Uncharacterized protein \| Uncharacterized protein; uniclust \| UniRef100\_A0A6H1ZNP5 \| 99.5 \| 1.5e-17 \| 2.8e-23 \| 126.3 \| 167 \| (6, 178) \| 179 \| (13, 182) \| 186 \| Uncharacterized protein \| Uncharacterized protein; uniclust \| UniRef100\_A0A0Q7SMP9 \| 99.5 \| 5.6e-17 \| 1.1e-22 \| 124.8 \| 176 \| (2, 178) \| 179 \| (1, 187) \| 187 \| Uncharacterized protein \| Uncharacterized protein; uniclust \| UniRef100\_A0A974WVA0 \| 99.5 \| 8.4e-17 \| 1.5e-22 \| 120.9 \| 156 \| (4, 161) \| 179 \| (2, 160) \| 170 \| Uncharacterized protein \| Uncharacterized protein; uniclust \| UniRef100\_A0A2U1B226 \| 99.5 \| 1.4e-16 \| 2.6e-22 \| 120.1 \| 134 \| (23, 165) \| 179 \| (12, 148) \| 159 \| Uncharacterized protein \| Uncharacterized protein; uniclust \| UniRef100\_A0A965HE76 \| 99.4 \| 1.6e-15 \| 2.9e-21 \| 108.3 \| 107 \| (71, 178) \| 179 \| (7, 119) \| 119 \| Uncharacterized protein \| Uncharacterized protein; uniclust \| UniRef100\_A0A6J5RTA8 \| 99.4 \| 2e-15 \| 3.7e-21 \| 115.0 \| 160 \| (4, 167) \| 179 \| (7, 170) \| 184 \| Uncharacterized protein \| Uncharacterized protein; uniclust \| UniRef100\_A0A968GWW4 \| 99.3 \| 4.5e-15 \| 8.2e-21 \| 111.7 \| 150 \| (6, 158) \| 179 \| (3, 155) \| 167 \| Uncharacterized protein \| Uncharacterized protein; uniclust \| UniRef100\_A0A2D9MYC7 \| 99.3 \| 5.8e-15 \| 1.1e-20 \| 107.3 \| 109 \| (5, 114) \| 179 \| (8, 118) \| 132 \| Uncharacterized protein \| Uncharacterized protein; uniclust \| UniRef100\_UPI0018F8B0B0 \| 99.3 \| 9.4e-15 \| 1.7e-20 \| 106.8 \| 135 \| (4, 139) \| 179 \| (1, 136) \| 136 \| hypothetical protein \| hypothetical protein; uniclust \| UniRef100\_A0A3S0DLF1 \| 99.3 \| 2.9e-14 \| 5.3e-20 \| 105.7 \| 128 \| (33, 161) \| 179 \| (2, 136) \| 148 \| Uncharacterized protein \| Uncharacterized protein; uniclust \| UniRef100\_A0A0F9JR02 \| 99.2 \| 2.7e-14 \| 5.9e-20 \| 109.7 \| 103 \| (1, 103) \| 179 \| (2, 118) \| 134 \| BppU N-terminal domain-containing protein \| BppU N-terminal domain-containing protein; uniclust \| UniRef100\_A0A8S5N661 \| 99.2 \| 3.7e-14 \| 7e-20 \| 108.4 \| 129 \| (31, 163) \| 179 \| (18, 148) \| 164 \| Uncharacterized protein \| Uncharacterized protein; uniclust \| UniRef100\_A0A0F9QC72 \| 99.2 \| 7.6e-14 \| 1.4e-19 \| 106.5 \| 156 \| (5, 161) \| 179 \| (10, 166) \| 180 \| Uncharacterized protein \| Uncharacterized protein; uniclust \| UniRef100\_A0A377P1N6 \| 99.1 \| 2e-13 \| 4.1e-19 \| 106.6 \| 155 \| (3, 179) \| 179 \| (6, 166) \| 170 \| Uncharacterized protein \| Uncharacterized protein; uniclust \| UniRef100\_UPI00035F2C0A \| 99.1 \| 3.8e-13 \| 7.1e-19 \| 101.8 \| 125 \| (35, 166) \| 179 \| (29, 154) \| 167 \| hypothetical protein \| hypothetical protein; uniclust \| UniRef100\_A0A1Z8QIZ8 \| 99.1 \| 4.5e-13 \| 8.6e-19 \| 100.5 \| 110 \| (3, 115) \| 179 \| (16, 131) \| 141 \| Uncharacterized protein \| Uncharacterized protein; uniclust \| UniRef100\_A0A2A4LXZ7 \| 99.1 \| 7e-13 \| 1.3e-18 \| 93.3 \| 67 \| (5, 71) \| 179 \| (8, 76) \| 99 \| Uncharacterized protein (Fragment) \| Uncharacterized protein (Fragment); uniclust \| UniRef100\_UPI0022F406F8 \| 99.0 \| 1.4e-12 \| 2.6e-18 \| 98.1 \| 141 \| (23, 163) \| 179 \| (2, 147) \| 158 \| hypothetical protein \| hypothetical protein; uniclust \| UniRef100\_A0A061NTW5 \| 99.0 \| 1.2e-12 \| 2.7e-18 \| 94.4 \| 57 \| (104, 162) \| 179 \| (13, 69) \| 88 \| Phage protein \| Phage protein; uniclust \| UniRef100\_A0A1B1IUI2 \| 99.0 \| 1.3e-12 \| 2.8e-18 \| 98.2 \| 99 \| (1, 99) \| 179 \| (2, 113) \| 117 \| Tail protein \| Tail protein; uniclust \| UniRef100\_A0A2E7ZNQ6 \| 99.0 \| 2.3e-12 \| 4.2e-18 \| 94.6 \| 118 \| (4, 123) \| 179 \| (1, 132) \| 133 \| Uncharacterized protein \| Uncharacterized protein; uniclust \| UniRef100\_A0A0F9ESU0 \| 99.0 \| 2.3e-12 \| 5e-18 \| 96.8 \| 96 \| (4, 99) \| 179 \| (6, 114) \| 117 \| Uncharacterized protein \| Uncharacterized protein; uniclust \| UniRef100\_A0A9E5RBB4 \| 99.0 \| 3.4e-12 \| 6.3e-18 \| 97.3 \| 153 \| (2, 163) \| 179 \| (1, 154) \| 172 \| Uncharacterized protein \| Uncharacterized protein; uniclust \| UniRef100\_A0A098F6N2 \| 99.0 \| 3.3e-12 \| 7.2e-18 \| 91.4 \| 56 \| (105, 161) \| 179 \| (13, 68) \| 87 \| Peptidylprolyl isomerase \| Peptidylprolyl isomerase; uniclust \| UniRef100\_A0A0B0HXU3 \| 99.0 \| 3.3e-12 \| 7.5e-18 \| 92.8 \| 57 \| (105, 163) \| 179 \| (15, 71) \| 92 \| Uncharacterized protein \| Uncharacterized protein; uniclust \| UniRef100\_A0A031G636 \| 99.0 \| 4.3e-12 \| 9e-18 \| 90.1 \| 60 \| (106, 166) \| 179 \| (9, 68) \| 86 \| Preprotein translocase subunit SecA \| Preprotein translocase subunit SecA; uniclust \| UniRef100\_A0A2D3WPF3 \| 98.9 \| 5.7e-12 \| 1.1e-17 \| 87.0 \| 76 \| (104, 179) \| 179 \| (5, 83) \| 84 \| Uncharacterized protein \| Uncharacterized protein; uniclust \| UniRef100\_A0A564WI89 \| 98.9 \| 6.7e-12 \| 1.2e-17 \| 94.8 \| 99 \| (63, 161) \| 179 \| (49, 147) \| 160 \| Uncharacterized protein \| Uncharacterized protein; uniclust \| UniRef100\_UPI001E2E0B13 \| 98.9 \| 8.6e-12 \| 1.6e-17 \| 85.7 \| 81 \| (41, 122) \| 179 \| (3, 83) \| 88 \| hypothetical protein \| hypothetical protein; uniclust \| UniRef100\_A0A8S0I2S7 \| 98.9 \| 9.8e-12 \| 1.8e-17 \| 90.1 \| 108 \| (4, 111) \| 179 \| (1, 120) \| 121 \| Uncharacterized protein \| Uncharacterized protein; uniclust \| UniRef100\_A0A5Q0TJP6 \| 98.9 \| 1.1e-11 \| 2.3e-17 \| 86.9 \| 73 \| (104, 178) \| 179 \| (9, 81) \| 81 \| Phage tail protein \| Phage tail protein; uniclust \| UniRef100\_A0A1C6BMD3 \| 98.8 \| 2.4e-11 \| 5.1e-17 \| 85.9 \| 57 \| (104, 161) \| 179 \| (5, 61) \| 80 \| Peptidylprolyl isomerase \| Peptidylprolyl isomerase; uniclust \| UniRef100\_A0A1G1JJJ1 \| 98.8 \| 2.9e-11 \| 5.6e-17 \| 81.3 \| 52 \| (108, 159) \| 179 \| (5, 57) \| 71 \| Phage tail protein \| Phage tail protein; uniclust \| UniRef100\_A0A073IS86 \| 98.8 \| 2.9e-11 \| 5.8e-17 \| 83.7 \| 54 \| (107, 161) \| 179 \| (5, 58) \| 77 \| Uncharacterized protein \| Uncharacterized protein; uniclust \| UniRef100\_A0A081NV45 \| 98.8 \| 2.8e-11 \| 6.2e-17 \| 88.3 \| 72 \| (104, 178) \| 179 \| (18, 89) \| 93 \| Uncharacterized protein \| Uncharacterized protein; uniclust \| UniRef100\_A0A3C0VX60 \| 98.8 \| 3.7e-11 \| 6.9e-17 \| 85.3 \| 89 \| (25, 114) \| 179 \| (1, 91) \| 105 \| Uncharacterized protein (Fragment) \| Uncharacterized protein (Fragment); uniclust \| UniRef100\_A0A0A8JM71 \| 98.8 \| 3.6e-11 \| 7.6e-17 \| 86.3 \| 72 \| (104, 178) \| 179 \| (16, 87) \| 90 \| Uncharacterized protein \| Uncharacterized protein; uniclust \| UniRef100\_A0A193LN67 \| 98.7 \| 9e-11 \| 1.7e-16 \| 92.6 \| 156 \| (1, 178) \| 179 \| (24, 185) \| 187 \| Uncharacterized protein \| Uncharacterized protein; uniclust \| UniRef100\_A0A0F9J4G4 \| 98.7 \| 1.1e-10 \| 2.5e-16 \| 91.4 \| 97 \| (3, 99) \| 179 \| (22, 137) \| 140 \| BppU N-terminal domain-containing protein \| BppU N-terminal domain-containing protein; uniclust \| UniRef100\_A0A1B7HN81 \| 98.6 \| 3e-10 \| 6.1e-16 \| 89.7 \| 160 \| (2, 179) \| 179 \| (1, 166) \| 168 \| Uncharacterized protein \| Uncharacterized protein; uniclust \| UniRef100\_A0A069I7I0 \| 98.6 \| 3.1e-10 \| 6.6e-16 \| 80.9 \| 59 \| (106, 166) \| 179 \| (14, 72) \| 84 \| Primosomal replication protein PriB/PriC domain protein \| Primosomal replication protein PriB/PriC domain protein; uniclust \| UniRef100\_UPI00197DA380 \| 98.6 \| 6.1e-10 \| 1.1e-15 \| 77.7 \| 89 \| (88, 179) \| 179 \| (1, 92) \| 92 \| hypothetical protein \| hypothetical protein; uniclust \| UniRef100\_A0A0K0KVG3 \| 98.5 \| 7.3e-10 \| 1.6e-15 \| 85.4 \| 99 \| (1, 99) \| 179 \| (13, 125) \| 128 \| Virion structural protein \| Virion structural protein; uniclust \| UniRef100\_A0A0T2MKH9 \| 98.4 \| 2e-09 \| 4.2e-15 \| 78.7 \| 58 \| (105, 164) \| 179 \| (9, 66) \| 95 \| Uncharacterized protein \| Uncharacterized protein; uniclust \| UniRef100\_A0A0Q4JQ41 \| 98.4 \| 2.3e-09 \| 4.9e-15 \| 77.4 \| 58 \| (104, 163) \| 179 \| (6, 63) \| 87 \| Uncharacterized protein \| Uncharacterized protein; uniclust \| UniRef100\_UPI000D3BEE15 \| 98.4 \| 3.2e-09 \| 5.8e-15 \| 71.4 \| 59 \| (104, 162) \| 179 \| (5, 63) \| 73 \| hypothetical protein \| hypothetical protein; uniclust \| UniRef100\_A0A3B9VER5 \| 98.4 \| 3.1e-09 \| 6.3e-15 \| 81.2 \| 97 \| (3, 99) \| 179 \| (5, 120) \| 127 \| BppU N-terminal domain-containing protein \| BppU N-terminal domain-containing protein; uniclust \| UniRef100\_A0A101WGN2 \| 98.4 \| 3.5e-09 \| 7.5e-15 \| 77.3 \| 63 \| (105, 167) \| 179 \| (15, 79) \| 92 \| Chemotaxis protein \| Chemotaxis protein; uniclust \| UniRef100\_A0A6J5QJF9 \| 98.3 \| 6.3e-09 \| 1.3e-14 \| 77.7 \| 89 \| (4, 93) \| 179 \| (10, 106) \| 107 \| Uncharacterized protein \| Uncharacterized protein; uniclust \| UniRef100\_A0A1G1LJR7 \| 98.3 \| 7.4e-09 \| 1.4e-14 \| 69.2 \| 53 \| (109, 161) \| 179 \| (4, 57) \| 70 \| Phage tail protein \| Phage tail protein; uniclust \| UniRef100\_A0A125WEA2 \| 98.3 \| 6.8e-09 \| 1.4e-14 \| 75.6 \| 74 \| (104, 179) \| 179 \| (17, 94) \| 95 \| Phage associated protein \| Phage associated protein; uniclust \| UniRef100\_A0A0A2SJL1 \| 98.2 \| 1.7e-08 \| 3.1e-14 \| 69.0 \| 54 \| (106, 161) \| 179 \| (6, 59) \| 76 \| Phage protein \| Phage protein; uniclust \| UniRef100\_A0A0A8TH18 \| 98.2 \| 1.4e-08 \| 3.2e-14 \| 81.7 \| 97 \| (3, 99) \| 179 \| (32, 146) \| 152 \| BppU N-terminal domain-containing protein \| BppU N-terminal domain-containing protein; uniclust \| UniRef100\_A0A072T1F6 \| 98.2 \| 1.4e-08 \| 3.4e-14 \| 82.6 \| 95 \| (5, 99) \| 179 \| (23, 137) \| 165 \| Auto-transporter adhesin head GIN domain-containing protein \| Auto-transporter adhesin head GIN domain-containing protein; uniclust \| UniRef100\_A0A142XVQ1 \| 98.2 \| 2.1e-08 \| 4.3e-14 \| 72.5 \| 49 \| (108, 157) \| 179 \| (5, 53) \| 87 \| Peptidylprolyl isomerase \| Peptidylprolyl isomerase; uniclust \| UniRef100\_A0A0F9I9A9 \| 98.2 \| 2.3e-08 \| 5e-14 \| 77.1 \| 98 \| (1, 99) \| 179 \| (5, 119) \| 123 \| Uncharacterized protein \| Uncharacterized protein; uniclust \| UniRef100\_A0A2D5YE29 \| 98.2 \| 2.9e-08 \| 6.1e-14 \| 75.7 \| 98 \| (2, 99) \| 179 \| (3, 113) \| 116 \| BppU N-terminal domain-containing protein \| BppU N-terminal domain-containing protein; uniclust \| UniRef100\_A0A2D6E192 \| 98.1 \| 4.5e-08 \| 9.4e-14 \| 76.1 \| 99 \| (1, 99) \| 179 \| (9, 124) \| 131 \| Uncharacterized protein \| Uncharacterized protein; uniclust \| UniRef100\_A0A432F9K8 \| 98.1 \| 5.4e-08 \| 9.9e-14 \| 65.4 \| 54 \| (5, 58) \| 179 \| (7, 62) \| 70 \| Uncharacterized protein \| Uncharacterized protein; uniclust \| UniRef100\_UPI000E2A48E6 \| 98.1 \| 5.5e-08 \| 1.1e-13 \| 74.2 \| 127 \| (2, 145) \| 179 \| (1, 132) \| 139 \| hypothetical protein \| hypothetical protein; uniclust \| UniRef100\_A0A1Q6KUC0 \| 98.1 \| 5.7e-08 \| 1.1e-13 \| 69.7 \| 62 \| (104, 166) \| 179 \| (14, 75) \| 87 \| Peptidylprolyl isomerase \| Peptidylprolyl isomerase; uniclust \| UniRef100\_A0A2E9Y0J9 \| 98.1 \| 6.9e-08 \| 1.3e-13 \| 66.5 \| 63 \| (115, 178) \| 179 \| (15, 80) \| 80 \| Uncharacterized protein (Fragment) \| Uncharacterized protein (Fragment); uniclust \| UniRef100\_A0A023D777 \| 98.0 \| 7.6e-08 \| 1.6e-13 \| 74.2 \| 99 \| (1, 100) \| 179 \| (3, 113) \| 120 \| Uncharacterized protein \| Uncharacterized protein; uniclust \| UniRef100\_A0A6G5Y298 \| 98.0 \| 8.6e-08 \| 1.6e-13 \| 67.2 \| 57 \| (104, 161) \| 179 \| (10, 66) \| 80 \| Phage protein \| Phage protein; uniclust \| UniRef100\_UPI002251F4A5 \| 98.0 \| 9.9e-08 \| 1.8e-13 \| 67.5 \| 87 \| (4, 90) \| 179 \| (5, 93) \| 93 \| hypothetical protein \| hypothetical protein; uniclust \| UniRef100\_A0A1J6Y2J5 \| 98.0 \| 9.4e-08 \| 1.9e-13 \| 73.4 \| 97 \| (3, 99) \| 179 \| (7, 116) \| 127 \| Uncharacterized protein \| Uncharacterized protein; uniclust \| UniRef100\_A0A3R8K1E5 \| 98.0 \| 1.4e-07 \| 2.6e-13 \| 68.1 \| 56 \| (108, 163) \| 179 \| (33, 88) \| 101 \| Uncharacterized protein \| Uncharacterized protein; uniclust \| UniRef100\_A0A075KCY2 \| 98.0 \| 1.4e-07 \| 3e-13 \| 66.2 \| 54 \| (107, 161) \| 179 \| (5, 58) \| 72 \| Uncharacterized protein \| Uncharacterized protein; uniclust \| UniRef100\_A0A7X3ZJZ6 \| 97.9 \| 2.1e-07 \| 3.9e-13 \| 62.9 \| 58 \| (104, 161) \| 179 \| (4, 61) \| 71 \| Uncharacterized protein \| Uncharacterized protein; uniclust \| UniRef100\_A0A3D2IPX1 \| 97.9 \| 2.4e-07 \| 4.4e-13 \| 64.9 \| 49 \| (109, 158) \| 179 \| (17, 65) \| 86 \| Peptidylprolyl isomerase \| Peptidylprolyl isomerase; uniclust \| UniRef100\_UPI001F055D47 \| 97.9 \| 2.4e-07 \| 4.5e-13 \| 73.3 \| 70 \| (106, 178) \| 179 \| (35, 105) \| 189 \| DUF6148 family protein \| DUF6148 family protein; uniclust \| UniRef100\_A0A0F9R4H3 \| 97.9 \| 2e-07 \| 4.6e-13 \| 72.4 \| 98 \| (1, 98) \| 179 \| (4, 119) \| 120 \| BppU N-terminal domain-containing protein \| BppU N-terminal domain-containing protein; uniclust \| UniRef100\_A0A060H7N8 \| 97.9 \| 2.9e-07 \| 5.8e-13 \| 71.9 \| 61 \| (104, 166) \| 179 \| (52, 112) \| 136 \| Phage-related protein \| Phage-related protein; uniclust \| UniRef100\_A0A974WXV3 \| 97.8 \| 3.9e-07 \| 7.2e-13 \| 66.0 \| 88 \| (71, 160) \| 179 \| (6, 93) \| 104 \| Uncharacterized protein \| Uncharacterized protein; uniclust \| UniRef100\_A0A661I002 \| 97.8 \| 3.5e-07 \| 7.2e-13 \| 65.5 \| 52 \| (109, 161) \| 179 \| (8, 59) \| 80 \| Uncharacterized protein \| Uncharacterized protein; uniclust \| UniRef100\_A0A1Q8QVH5 \| 97.8 \| 3.5e-07 \| 7.4e-13 \| 66.6 \| 57 \| (105, 163) \| 179 \| (8, 64) \| 85 \| PH domain-containing protein \| PH domain-containing protein; uniclust \| UniRef100\_A0A177QGG6 \| 97.8 \| 4.5e-07 \| 8.5e-13 \| 67.9 \| 96 \| (2, 97) \| 179 \| (12, 115) \| 116 \| CBM2 domain-containing protein \| CBM2 domain-containing protein; uniclust \| UniRef100\_A0A7V3MME9 \| 97.8 \| 5.1e-07 \| 1e-12 \| 68.6 \| 97 \| (1, 98) \| 179 \| (1, 109) \| 113 \| DUF2479 domain-containing protein \| DUF2479 domain-containing protein; uniclust \| UniRef100\_A0A1Q6PWT8 \| 97.8 \| 5.8e-07 \| 1.2e-12 \| 64.7 \| 62 \| (105, 166) \| 179 \| (10, 71) \| 82 \| Uncharacterized protein \| Uncharacterized protein; uniclust \| UniRef100\_A0A8J7TTR6 \| 97.8 \| 6.5e-07 \| 1.2e-12 \| 60.9 \| 56 \| (110, 166) \| 179 \| (5, 60) \| 72 \| Uncharacterized protein \| Uncharacterized protein; uniclust \| UniRef100\_A0A022G2W7 \| 97.8 \| 5.9e-07 \| 1.3e-12 \| 68.1 \| 60 \| (105, 166) \| 179 \| (18, 77) \| 106 \| Primosomal replication protein PriB/PriC domain protein \| Primosomal replication protein PriB/PriC domain protein; uniclust \| UniRef100\_A0A2A4ZZL1 \| 97.7 \| 7.6e-07 \| 1.5e-12 \| 61.8 \| 59 \| (107, 167) \| 179 \| (4, 62) \| 73 \| Primosomal replication protein PriB/PriC domain protein \| Primosomal replication protein PriB/PriC domain protein; uniclust \| UniRef100\_L8M8S2 \| 97.7 \| 8.5e-07 \| 1.6e-12 \| 61.2 \| 67 \| (108, 178) \| 179 \| (8, 75) \| 78 \| Uncharacterized protein \| Uncharacterized protein; uniclust \| UniRef100\_A0A090G6L9 \| 97.7 \| 7.2e-07 \| 1.7e-12 \| 69.6 \| 96 \| (1, 99) \| 179 \| (2, 116) \| 119 \| BppU N-terminal domain-containing protein \| BppU N-terminal domain-containing protein; uniclust \| UniRef100\_K2K034 \| 97.7 \| 1.2e-06 \| 2.2e-12 \| 59.5 \| 60 \| (105, 166) \| 179 \| (4, 63) \| 71 \| Uncharacterized protein \| Uncharacterized protein; uniclust \| UniRef100\_UPI001CCE3333 \| 97.7 \| 1.3e-06 \| 2.3e-12 \| 62.1 \| 57 \| (101, 158) \| 179 \| (19, 75) \| 91 \| hypothetical protein \| hypothetical protein; uniclust \| UniRef100\_A0A346JU94 \| 97.7 \| 1.2e-06 \| 2.3e-12 \| 62.1 \| 61 \| (101, 163) \| 179 \| (8, 68) \| 85 \| Uncharacterized protein \| Uncharacterized protein; uniclust \| UniRef100\_UPI001E29A3BE \| 97.6 \| 1.5e-06 \| 2.8e-12 \| 58.3 \| 60 \| (4, 63) \| 179 \| (6, 65) \| 66 \| hypothetical protein \| hypothetical protein; uniclust \| UniRef100\_A0A1M6NZM2 \| 97.6 \| 1.5e-06 \| 3.2e-12 \| 66.1 \| 92 \| (6, 99) \| 179 \| (7, 105) \| 107 \| BppU N-terminal domain-containing protein \| BppU N-terminal domain-containing protein; uniclust \| UniRef100\_A0A0K8PBJ4 \| 97.6 \| 1.6e-06 \| 3.3e-12 \| 66.7 \| 91 \| (7, 98) \| 179 \| (11, 116) \| 118 \| BppU N-terminal domain-containing protein \| BppU N-terminal domain-containing protein; uniclust \| UniRef100\_A0A1Y1R2U1 \| 97.6 \| 1.9e-06 \| 3.8e-12 \| 67.7 \| 91 \| (9, 99) \| 179 \| (13, 117) \| 134 \| Uncharacterized protein \| Uncharacterized protein; uniclust \| UniRef100\_A0A1Q8G4U9 \| 97.6 \| 2.1e-06 \| 3.9e-12 \| 58.8 \| 59 \| (107, 167) \| 179 \| (2, 60) \| 69 \| Primosomal replication protein PriB/PriC domain protein \| Primosomal replication protein PriB/PriC domain protein; uniclust \| UniRef100\_A0A315ENZ3 \| 97.6 \| 2.2e-06 \| 4.4e-12 \| 67.3 \| 103 \| (1, 103) \| 179 \| (8, 123) \| 135 \| Uncharacterized protein \| Uncharacterized protein; uniclust \| UniRef100\_UPI000A3F844A \| 97.6 \| 2.5e-06 \| 4.6e-12 \| 61.2 \| 75 \| (5, 79) \| 179 \| (12, 92) \| 95 \| hypothetical protein \| hypothetical protein; uniclust \| UniRef100\_A0A1V4WKC7 \| 97.5 \| 4e-06 \| 7.9e-12 \| 59.3 \| 51 \| (109, 161) \| 179 \| (6, 56) \| 76 \| Uncharacterized protein \| Uncharacterized protein; uniclust \| UniRef100\_A0A3S0P0G6 \| 97.5 \| 4.2e-06 \| 8.3e-12 \| 63.6 \| 90 \| (8, 97) \| 179 \| (15, 115) \| 116 \| Uncharacterized protein \| Uncharacterized protein; uniclust \| UniRef100\_UPI001C940115 \| 97.5 \| 4.8e-06 \| 8.8e-12 \| 58.3 \| 74 \| (102, 176) \| 179 \| (7, 80) \| 82 \| hypothetical protein \| hypothetical protein; uniclust \| UniRef100\_A0A023NI20 \| 97.4 \| 4.2e-06 \| 8.9e-12 \| 64.3 \| 98 \| (1, 98) \| 179 \| (1, 109) \| 111 \| BppU N-terminal domain-containing protein \| BppU N-terminal domain-containing protein; uniclust \| UniRef100\_UPI001880E38C \| 97.4 \| 5.7e-06 \| 1e-11 \| 59.4 \| 75 \| (105, 179) \| 179 \| (13, 92) \| 94 \| hypothetical protein \| hypothetical protein; uniclust \| UniRef100\_A0A268E9X3 \| 97.4 \| 5.2e-06 \| 1.1e-11 \| 58.8 \| 54 \| (106, 161) \| 179 \| (6, 59) \| 73 \| Uncharacterized protein \| Uncharacterized protein; uniclust \| UniRef100\_UPI0023578956 \| 97.4 \| 6.2e-06 \| 1.1e-11 \| 59.0 \| 59 \| (106, 166) \| 179 \| (22, 80) \| 92 \| DUF6148 family protein \| DUF6148 family protein; uniclust \| UniRef100\_UPI001FBB5195 \| 97.4 \| 6.6e-06 \| 1.2e-11 \| 53.2 \| 34 \| (128, 161) \| 179 \| (4, 37) \| 53 \| peptidylprolyl isomerase \| peptidylprolyl isomerase; uniclust \| UniRef100\_A0A1F7S430 \| 97.4 \| 6.4e-06 \| 1.2e-11 \| 57.3 \| 54 \| (106, 161) \| 179 \| (6, 59) \| 72 \| Uncharacterized protein \| Uncharacterized protein; uniclust \| UniRef100\_A0A2E5PQX4 \| 97.4 \| 6.6e-06 \| 1.2e-11 \| 59.8 \| 78 \| (3, 80) \| 179 \| (11, 99) \| 101 \| Uncharacterized protein \| Uncharacterized protein; uniclust \| UniRef100\_A0A7C9BLN2 \| 97.4 \| 7.1e-06 \| 1.4e-11 \| 61.8 \| 88 \| (9, 97) \| 179 \| (14, 108) \| 109 \| Uncharacterized protein \| Uncharacterized protein; uniclust \| UniRef100\_A0A2D6FIR8 \| 97.4 \| 6.9e-06 \| 1.5e-11 \| 65.5 \| 98 \| (2, 99) \| 179 \| (11, 126) \| 133 \| Cohesin domain-containing protein \| Cohesin domain-containing protein; uniclust \| UniRef100\_A0A496ZSP7 \| 97.3 \| 8.9e-06 \| 1.8e-11 \| 62.2 \| 94 \| (5, 99) \| 179 \| (2, 111) \| 114 \| BppU N-terminal domain-containing protein \| BppU N-terminal domain-containing protein; uniclust \| UniRef100\_A0A9D2L9Q1 \| 97.3 \| 1.1e-05 \| 2.1e-11 \| 58.4 \| 50 \| (110, 160) \| 179 \| (31, 80) \| 98 \| Peptidylprolyl isomerase \| Peptidylprolyl isomerase; uniclust \| UniRef100\_A0A1M5RH09 \| 97.3 \| 1.2e-05 \| 2.3e-11 \| 59.1 \| 60 \| (105, 166) \| 179 \| (29, 88) \| 98 \| Uncharacterized protein \| Uncharacterized protein; uniclust \| UniRef100\_A0A1V6E205 \| 97.3 \| 1.3e-05 \| 2.4e-11 \| 55.0 \| 52 \| (109, 160) \| 179 \| (4, 56) \| 71 \| Uncharacterized protein \| Uncharacterized protein; uniclust \| UniRef100\_A0A094YXQ6 \| 97.3 \| 1.1e-05 \| 2.4e-11 \| 62.7 \| 91 \| (8, 99) \| 179 \| (8, 109) \| 111 \| YtkA-like domain-containing protein \| YtkA-like domain-containing protein; uniclust \| UniRef100\_A0A948EQ09 \| 97.3 \| 1.4e-05 \| 2.7e-11 \| 55.4 \| 61 \| (105, 166) \| 179 \| (3, 63) \| 71 \| Uncharacterized protein \| Uncharacterized protein; uniclust \| UniRef100\_A0A6J5NE21 \| 97.2 \| 1.4e-05 \| 2.8e-11 \| 62.6 \| 95 \| (5, 99) \| 179 \| (6, 121) \| 125 \| Uncharacterized protein \| Uncharacterized protein; uniclust \| UniRef100\_A0A0B3S182 \| 97.2 \| 1.5e-05 \| 2.9e-11 \| 55.7 \| 53 \| (109, 161) \| 179 \| (6, 58) \| 70 \| Uncharacterized protein \| Uncharacterized protein; uniclust \| UniRef100\_A0A064AM50 \| 97.2 \| 1.6e-05 \| 3.2e-11 \| 57.5 \| 54 \| (105, 160) \| 179 \| (8, 61) \| 79 \| Uncharacterized protein \| Uncharacterized protein; uniclust \| UniRef100\_A0A3Q9BR07 \| 97.2 \| 1.9e-05 \| 3.7e-11 \| 55.7 \| 58 \| (107, 166) \| 179 \| (7, 64) \| 75 \| Uncharacterized protein \| Uncharacterized protein; uniclust \| UniRef100\_A0A959C7P8 \| 97.2 \| 2e-05 \| 3.8e-11 \| 60.5 \| 93 \| (4, 97) \| 179 \| (14, 125) \| 126 \| Uncharacterized protein \| Uncharacterized protein; uniclust \| UniRef100\_A0A2D7FMC3 \| 97.2 \| 2.1e-05 \| 3.9e-11 \| 57.2 \| 69 \| (3, 71) \| 179 \| (16, 90) \| 99 \| Uncharacterized protein \| Uncharacterized protein; uniclust \| UniRef100\_A0A0R3MN91 \| 97.2 \| 1.9e-05 \| 3.9e-11 \| 56.9 \| 52 \| (109, 161) \| 179 \| (5, 56) \| 79 \| Uncharacterized protein \| Uncharacterized protein; uniclust \| UniRef100\_A0A949G187 \| 97.2 \| 2e-05 \| 4.1e-11 \| 62.0 \| 90 \| (6, 95) \| 179 \| (13, 126) \| 126 \| Uncharacterized protein \| Uncharacterized protein; uniclust \| UniRef100\_UPI001431CBAE \| 97.2 \| 2.3e-05 \| 4.2e-11 \| 58.6 \| 107 \| (64, 178) \| 179 \| (3, 114) \| 116 \| hypothetical protein \| hypothetical protein; uniclust \| UniRef100\_A0A5B8C3L8 \| 97.2 \| 2.5e-05 \| 4.6e-11 \| 53.5 \| 53 \| (109, 161) \| 179 \| (6, 58) \| 70 \| Phage tail protein \| Phage tail protein; uniclust \| UniRef100\_A0A7V2T445 \| 97.1 \| 2.9e-05 \| 5.4e-11 \| 56.6 \| 55 \| (108, 163) \| 179 \| (3, 57) \| 93 \| Primosomal replication protein PriB/PriC domain protein \| Primosomal replication protein PriB/PriC domain protein; uniclust \| UniRef100\_A0A0F9BY64 \| 97.1 \| 3e-05 \| 5.6e-11 \| 53.8 \| 63 \| (3, 65) \| 179 \| (6, 75) \| 75 \| Uncharacterized protein (Fragment) \| Uncharacterized protein (Fragment); uniclust \| UniRef100\_A0A0F7L4W2 \| 97.1 \| 3.3e-05 \| 6e-11 \| 53.0 \| 47 \| (113, 161) \| 179 \| (9, 55) \| 70 \| Uncharacterized protein \| Uncharacterized protein; uniclust \| UniRef100\_A0A6M3KMZ3 \| 97.1 \| 3.1e-05 \| 6.2e-11 \| 60.7 \| 96 \| (4, 99) \| 179 \| (8, 120) \| 126 \| Putative tail protein \| Putative tail protein; uniclust \| UniRef100\_A0A850BA57 \| 97.1 \| 4e-05 \| 7.3e-11 \| 54.4 \| 57 \| (43, 99) \| 179 \| (22, 80) \| 84 \| MBG domain-containing protein \| MBG domain-containing protein; uniclust \| UniRef100\_UPI000CE90355 \| 97.0 \| 4.5e-05 \| 8.3e-11 \| 52.9 \| 59 \| (107, 167) \| 179 \| (7, 65) \| 74 \| DUF6148 family protein \| DUF6148 family protein; uniclust \| UniRef100\_E3HBM6 \| 97.0 \| 4.8e-05 \| 8.9e-11 \| 52.5 \| 55 \| (106, 161) \| 179 \| (5, 59) \| 72 \| Uncharacterized protein \| Uncharacterized protein; uniclust \| UniRef100\_A0A956K140 \| 97.0 \| 5e-05 \| 9.1e-11 \| 53.5 \| 62 \| (104, 167) \| 179 \| (3, 68) \| 80 \| Uncharacterized protein \| Uncharacterized protein; uniclust \| UniRef100\_A0A2W5LUH5 \| 97.0 \| 4.8e-05 \| 9.2e-11 \| 55.0 \| 58 \| (107, 166) \| 179 \| (13, 70) \| 84 \| Primosomal replication protein PriB/PriC domain protein \| Primosomal replication protein PriB/PriC domain protein; uniclust \| UniRef100\_A0A0S2ZD68 \| 97.0 \| 4.8e-05 \| 1e-10 \| 60.6 \| 52 \| (108, 161) \| 179 \| (67, 118) \| 135 \| Phage protein \| Phage protein; uniclust \| UniRef100\_A0A0F9B885 \| 97.0 \| 5.7e-05 \| 1.1e-10 \| 53.6 \| 54 \| (108, 161) \| 179 \| (7, 60) \| 77 \| Uncharacterized protein \| Uncharacterized protein; uniclust \| UniRef100\_A0A0E3F1B5 \| 97.0 \| 5.8e-05 \| 1.1e-10 \| 62.4 \| 94 \| (6, 99) \| 179 \| (7, 115) \| 191 \| WW domain-containing protein \| WW domain-containing protein; uniclust \| UniRef100\_A0A0A0YPM7 \| 96.9 \| 5.9e-05 \| 1.4e-10 \| 59.6 \| 98 \| (1, 99) \| 179 \| (1, 115) \| 118 \| Uncharacterized protein \| Uncharacterized protein; uniclust \| UniRef100\_A0A1T5JQ31 \| 96.9 \| 8.4e-05 \| 1.6e-10 \| 57.0 \| 89 \| (6, 97) \| 179 \| (12, 117) \| 118 \| Uncharacterized protein \| Uncharacterized protein; uniclust \| UniRef100\_UPI00063BFA67 \| 96.9 \| 9e-05 \| 1.6e-10 \| 59.3 \| 103 \| (53, 165) \| 179 \| (45, 148) \| 175 \| hypothetical protein \| hypothetical protein; uniclust \| UniRef100\_A0A4R5VYM7 \| 96.9 \| 8.2e-05 \| 1.7e-10 \| 58.0 \| 98 \| (1, 98) \| 179 \| (1, 117) \| 118 \| BppU N-terminal domain-containing protein \| BppU N-terminal domain-containing protein; uniclust \| UniRef100\_C6S6G3 \| 96.9 \| 9.7e-05 \| 1.8e-10 \| 56.2 \| 73 \| (104, 178) \| 179 \| (49, 125) \| 125 \| Putative phage associated protein \| Putative phage associated protein; uniclust \| UniRef100\_A0A955V1P7 \| 96.8 \| 0.00011 \| 2e-10 \| 49.6 \| 54 \| (5, 58) \| 179 \| (7, 61) \| 63 \| Uncharacterized protein (Fragment) \| Uncharacterized protein (Fragment); uniclust \| UniRef100\_UPI0009772D17 \| 96.8 \| 0.0001 \| 2e-10 \| 56.7 \| 91 \| (4, 94) \| 179 \| (5, 107) \| 111 \| hypothetical protein \| hypothetical protein; uniclust \| UniRef100\_A0A1G8EFZ3 \| 96.8 \| 0.00013 \| 2.7e-10 \| 57.4 \| 96 \| (1, 98) \| 179 \| (1, 115) \| 118 \| Uncharacterized protein \| Uncharacterized protein; uniclust \| UniRef100\_A0A965I963 \| 96.8 \| 0.00015 \| 2.9e-10 \| 53.2 \| 42 \| (118, 160) \| 179 \| (26, 67) \| 85 \| Uncharacterized protein \| Uncharacterized protein; uniclust \| UniRef100\_A0A6V8M2G4 \| 96.8 \| 0.00015 \| 3e-10 \| 56.0 \| 97 \| (1, 97) \| 179 \| (1, 110) \| 111 \| BppU N-terminal domain-containing protein \| BppU N-terminal domain-containing protein; uniclust \| UniRef100\_A0A075KJY6 \| 96.7 \| 0.00017 \| 3.4e-10 \| 56.9 \| 95 \| (3, 97) \| 179 \| (12, 121) \| 125 \| BppU N-terminal domain-containing protein \| BppU N-terminal domain-containing protein; uniclust \| UniRef100\_A0A6M3J343 \| 96.7 \| 0.0002 \| 3.9e-10 \| 50.8 \| 50 \| (110, 160) \| 179 \| (6, 55) \| 71 \| Uncharacterized protein \| Uncharacterized protein; uniclust \| UniRef100\_A0A173W983 \| 96.7 \| 0.00022 \| 4.6e-10 \| 57.4 \| 91 \| (6, 96) \| 179 \| (32, 133) \| 136 \| Uncharacterized protein \| Uncharacterized protein; uniclust \| UniRef100\_A0A9D8STH6 \| 96.6 \| 0.00033 \| 6e-10 \| 52.4 \| 63 \| (105, 172) \| 179 \| (37, 99) \| 109 \| Uncharacterized protein \| Uncharacterized protein; uniclust \| UniRef100\_A0A1U9NPV2 \| 96.6 \| 0.00032 \| 6.2e-10 \| 50.0 \| 52 \| (108, 160) \| 179 \| (4, 55) \| 72 \| Uncharacterized protein \| Uncharacterized protein; uniclust \| UniRef100\_A0A7J9RFL5 \| 96.6 \| 0.00034 \| 6.3e-10 \| 48.3 \| 57 \| (41, 97) \| 179 \| (9, 68) \| 69 \| Uncharacterized protein \| Uncharacterized protein; uniclust \| UniRef100\_UPI00187511A5 \| 96.6 \| 0.00037 \| 6.8e-10 \| 50.3 \| 72 \| (62, 143) \| 179 \| (15, 87) \| 88 \| hypothetical protein \| hypothetical protein; uniclust \| UniRef100\_A0A2D5MP89 \| 96.5 \| 0.00039 \| 7.2e-10 \| 52.7 \| 93 \| (4, 96) \| 179 \| (4, 110) \| 111 \| Uncharacterized protein \| Uncharacterized protein; uniclust \| UniRef100\_A0A0J6HKU7 \| 96.5 \| 0.00036 \| 7.4e-10 \| 54.3 \| 92 \| (4, 96) \| 179 \| (3, 106) \| 112 \| Uncharacterized protein \| Uncharacterized protein; uniclust \| UniRef100\_A0A0W1G8D0 \| 96.5 \| 0.00036 \| 7.6e-10 \| 55.2 \| 78 \| (2, 79) \| 179 \| (1, 97) \| 119 \| Uncharacterized protein \| Uncharacterized protein; uniclust \| UniRef100\_A0A081MYL7 \| 96.5 \| 0.00045 \| 8.7e-10 \| 49.1 \| 52 \| (106, 158) \| 179 \| (5, 60) \| 71 \| Phage tail protein \| Phage tail protein; uniclust \| UniRef100\_A0A7Y4VZ35 \| 96.5 \| 0.00046 \| 8.8e-10 \| 53.0 \| 65 \| (109, 178) \| 179 \| (46, 110) \| 115 \| Uncharacterized protein \| Uncharacterized protein; uniclust \| UniRef100\_A0A2E5T0B5 \| 96.5 \| 0.00044 \| 8.8e-10 \| 55.4 \| 96 \| (4, 99) \| 179 \| (5, 125) \| 136 \| Uncharacterized protein \| Uncharacterized protein; uniclust \| UniRef100\_A0A929FI38 \| 96.5 \| 0.00047 \| 9.1e-10 \| 53.4 \| 92 \| (7, 98) \| 179 \| (13, 117) \| 119 \| Uncharacterized protein \| Uncharacterized protein; uniclust \| UniRef100\_UPI00201A6BDC \| 96.5 \| 0.00054 \| 9.9e-10 \| 47.8 \| 63 \| (115, 178) \| 179 \| (5, 72) \| 72 \| hypothetical protein \| hypothetical protein; uniclust \| UniRef100\_A0A1V5IZW5 \| 96.4 \| 0.00055 \| 1e-09 \| 48.4 \| 59 \| (104, 163) \| 179 \| (2, 64) \| 77 \| GpW protein \| GpW protein; uniclust \| UniRef100\_A0A2V2FWH8 \| 96.4 \| 0.00058 \| 1.1e-09 \| 50.3 \| 64 \| (104, 167) \| 179 \| (13, 76) \| 89 \| Uncharacterized protein \| Uncharacterized protein; uniclust \| UniRef100\_A0A4U8YKJ0 \| 96.4 \| 0.00057 \| 1.1e-09 \| 49.2 \| 55 \| (105, 160) \| 179 \| (7, 61) \| 75 \| Uncharacterized protein \| Uncharacterized protein; uniclust \| UniRef100\_A0A2U0TA70 \| 96.4 \| 0.00053 \| 1.1e-09 \| 54.3 \| 94 \| (3, 99) \| 179 \| (5, 115) \| 118 \| Nuclear transport factor 2 family protein \| Nuclear transport factor 2 family protein; uniclust \| UniRef100\_A0A016XJH1 \| 96.4 \| 0.0005 \| 1.1e-09 \| 51.7 \| 50 \| (108, 161) \| 179 \| (16, 65) \| 85 \| Phage tail protein \| Phage tail protein; uniclust \| UniRef100\_A0A381C5X6 \| 96.4 \| 0.00059 \| 1.2e-09 \| 49.2 \| 52 \| (108, 161) \| 179 \| (5, 56) \| 72 \| Uncharacterized protein \| Uncharacterized protein; uniclust \| UniRef100\_A0A0F9H6E2 \| 96.4 \| 0.00062 \| 1.2e-09 \| 48.7 \| 53 \| (107, 161) \| 179 \| (5, 57) \| 74 \| Uncharacterized protein \| Uncharacterized protein; uniclust \| UniRef100\_A0A350IBU3 \| 96.4 \| 0.00064 \| 1.2e-09 \| 52.6 \| 91 \| (9, 99) \| 179 \| (10, 120) \| 122 \| Uncharacterized protein \| Uncharacterized protein; uniclust \| UniRef100\_A0A521CSU3 \| 96.4 \| 0.0007 \| 1.3e-09 \| 47.0 \| 64 \| (108, 179) \| 179 \| (5, 68) \| 69 \| GpW protein \| GpW protein; uniclust \| UniRef100\_A0A2W4YA14 \| 96.4 \| 0.00068 \| 1.3e-09 \| 49.3 \| 56 \| (109, 166) \| 179 \| (6, 61) \| 83 \| Primosomal replication protein PriB/PriC domain protein \| Primosomal replication protein PriB/PriC domain protein; uniclust \| UniRef100\_A0A009I3H9 \| 96.3 \| 0.00069 \| 1.5e-09 \| 53.9 \| 94 \| (2, 96) \| 179 \| (3, 117) \| 120 \| Uncharacterized protein \| Uncharacterized protein; uniclust \| UniRef100\_A0A061Y8V6 \| 96.3 \| 0.00078 \| 1.5e-09 \| 50.9 \| 59 \| (107, 167) \| 179 \| (16, 74) \| 100 \| Prophage protein \| Prophage protein; uniclust \| UniRef100\_A0A0F9D8Q2 \| 96.3 \| 0.00099 \| 1.8e-09 \| 50.0 \| 93 \| (5, 97) \| 179 \| (5, 107) \| 108 \| BppU N-terminal domain-containing protein \| BppU N-terminal domain-containing protein; uniclust \| UniRef100\_A0A846QP15 \| 96.3 \| 0.00098 \| 1.9e-09 \| 46.0 \| 48 \| (112, 160) \| 179 \| (4, 51) \| 63 \| GpW protein \| GpW protein; uniclust \| UniRef100\_A0A0J6WRI4 \| 96.3 \| 0.00084 \| 1.9e-09 \| 53.8 \| 92 \| (6, 98) \| 179 \| (11, 117) \| 121 \| BppU N-terminal domain-containing protein \| BppU N-terminal domain-containing protein |
| Top keywords  (threshold 1.00e-03 (evalue)) | **domain\_containing, hypothetical, BppU, N\_terminal, Phage, tail, Fragment, Peptidylprolyl, isomerase, Primosomal** |
| Output files | ../../similar\_sequences/03\_FANPEZAQ\_CDS\_0003\_merged.svg ../../similar\_sequences/03\_FANPEZAQ\_CDS\_0003\_pdb70.a3m ../../similar\_sequences/03\_FANPEZAQ\_CDS\_0003\_pdb70.hhr ../../similar\_sequences/03\_FANPEZAQ\_CDS\_0003\_uniclust.a3m ../../similar\_sequences/03\_FANPEZAQ\_CDS\_0003\_uniclust.hhr |

#### Structure prediction (AlphaFold)^2^

|  |  |
| --- | --- |
| Stats | xml version="1.0" encoding="utf-8" standalone="no"?       2024-09-02T21:09:00.613669 image/svg+xml   Matplotlib v3.7.2, https://matplotlib.org/ |
| Predicted structure | **NGL Viewer Controls:**  - Center: *Left-Click* - Rotate: *Left-Click + Drag* - Translate: *Right-Click + Drag* - Zoom: *Shift + Left-Click + Drag* |
| Output files | ../../predicted\_structures/03\_FANPEZAQ\_CDS\_0003/features.pkl ../../predicted\_structures/03\_FANPEZAQ\_CDS\_0003/ranked\_0.pdb ../../predicted\_structures/03\_FANPEZAQ\_CDS\_0003/ranked\_0\_plots.svg ../../predicted\_structures/03\_FANPEZAQ\_CDS\_0003/result\_model\_1\_ptm\_pred\_0.pkl |

#### Structure similarity search results (Foldseek)^3^

|  |  |
| --- | --- |
| Structure databases searched | Pdb, Afdb-proteome, Afdb-uniprot50 |
| Results, scheme(s)  (Top layers only, threshold 1.00e-02 (evalue)) | xml version="1.0" encoding="utf-8" standalone="no"?       2024-09-02T21:10:08.411433 image/svg+xml   Matplotlib v3.7.2, https://matplotlib.org/ |
| Results, table  (threshold 1.00e-02 (evalue)) | db \| id \| prob \| evalue \| bits \| fident \| alnlen \| mismatch \| gapopen \| qstart \| qend \| tstart \| tend \| name \| description; pdb \| 8GTD\_M \| 1.0 \| 5.143e-14 \| 410 \| 0.303 \| 181 \| 107 \| 8 \| 7 \| 178 \| 8 \| 178 \| Head-to-tail joining protein \| Head-to-tail joining protein; pdb \| 6SWS\_E \| 1.0 \| 0.0002162 \| 131 \| 0.159 \| 94 \| 67 \| 5 \| 7 \| 97 \| 6 \| 90 \| Phosphoinositide 3-kinase adapter protein 1 \| Phosphoinositide 3-kinase adapter protein 1; pdb \| 7KDD\_I \| 1.0 \| 0.0007933 \| 103 \| 0.12 \| 108 \| 69 \| 5 \| 7 \| 94 \| 7 \| 108 \| SM5-1 Fab antibody light chain \| SM5-1 Fab antibody light chain; pdb \| 4V96\_AN \| 1.0 \| 0.0001921 \| 103 \| 0.103 \| 126 \| 102 \| 5 \| 9 \| 124 \| 21 \| 145 \| ORF48 \| ORF48; pdb \| 1OAR\_O \| 0.999 \| 0.00152 \| 98 \| 0.112 \| 107 \| 70 \| 4 \| 7 \| 94 \| 5 \| 105 \| IMMUNOGLOBULING E \| IMMUNOGLOBULING E; pdb \| 7E5S\_Q \| 0.998 \| 0.003687 \| 93 \| 0.114 \| 105 \| 68 \| 5 \| 9 \| 94 \| 10 \| 108 \| FC05 light chain \| FC05 light chain; pdb \| 7XW6\_L \| 0.997 \| 0.002438 \| 92 \| 0.081 \| 111 \| 71 \| 6 \| 7 \| 94 \| 7 \| 109 \| M22 antibody light chain \| M22 antibody light chain; pdb \| 8DIM\_C \| 0.997 \| 0.0007933 \| 91 \| 0.099 \| 111 \| 70 \| 7 \| 7 \| 94 \| 5 \| 108 \| CR6261 Fab light chain \| CR6261 Fab light chain; pdb \| 7JG2\_E \| 0.997 \| 0.00415 \| 91 \| 0.164 \| 97 \| 63 \| 8 \| 6 \| 95 \| 449 \| 534 \| Polymeric immunoglobulin receptor \| Polymeric immunoglobulin receptor; pdb \| 1OAX\_O \| 0.996 \| 0.00415 \| 90 \| 0.112 \| 107 \| 70 \| 4 \| 7 \| 94 \| 6 \| 106 \| IMMUNOGLOBULIN E \| IMMUNOGLOBULIN E; pdb \| 7UR6\_K \| 0.996 \| 0.002298 \| 90 \| 0.083 \| 108 \| 72 \| 6 \| 7 \| 93 \| 6 \| 107 \| Light Chain \| Light Chain; pdb \| 6MID\_L \| 0.996 \| 0.003088 \| 90 \| 0.102 \| 107 \| 73 \| 3 \| 7 \| 94 \| 7 \| 109 \| monoclonal antibody ZIKV-195 light chain \| monoclonal antibody ZIKV-195 light chain; pdb \| 8JJ5\_B \| 0.996 \| 0.0008416 \| 90 \| 0.134 \| 134 \| 96 \| 8 \| 3 \| 124 \| 7 \| 132 \| Uroplakin 2 \| Uroplakin 2; pdb \| 7UAR\_L \| 0.996 \| 0.00135 \| 89 \| 0.104 \| 105 \| 69 \| 4 \| 9 \| 94 \| 11 \| 109 \| C1717 Fab Light Chain \| C1717 Fab Light Chain; pdb \| 7TGF\_B \| 0.996 \| 0.003476 \| 89 \| 0.075 \| 119 \| 71 \| 5 \| 9 \| 93 \| 10 \| 123 \| VHH camelid antibody \| VHH camelid antibody; pdb \| 5YAX\_A \| 0.996 \| 0.005256 \| 89 \| 0.103 \| 106 \| 68 \| 5 \| 9 \| 94 \| 129 \| 227 \| scFv1 antibody \| scFv1 antibody; pdb \| 1MH5\_H \| 0.995 \| 0.008433 \| 88 \| 0.108 \| 120 \| 70 \| 6 \| 9 \| 100 \| 10 \| 120 \| IMMUNOGLOBULIN MS6-164 \| IMMUNOGLOBULIN MS6-164; pdb \| 7T0W\_L \| 0.994 \| 0.006275 \| 87 \| 0.127 \| 110 \| 68 \| 5 \| 7 \| 94 \| 5 \| 108 \| Fab115 light chain, IgG1 \| Fab115 light chain, IgG1; pdb \| 4V96\_AK \| 0.994 \| 0.0006644 \| 87 \| 0.117 \| 128 \| 97 \| 7 \| 9 \| 124 \| 21 \| 144 \| ORF48 \| ORF48; pdb \| 7PA8\_KKK \| 0.993 \| 0.003687 \| 86 \| 0.117 \| 111 \| 67 \| 8 \| 7 \| 94 \| 6 \| 108 \| Fab 27C2 light chain \| Fab 27C2 light chain; pdb \| 6UYN\_L \| 0.993 \| 0.00152 \| 86 \| 0.107 \| 121 \| 77 \| 7 \| 7 \| 104 \| 5 \| 117 \| CR6261 Fab light chain \| CR6261 Fab light chain; pdb \| 5CUS\_O \| 0.993 \| 0.006658 \| 86 \| 0.116 \| 103 \| 70 \| 2 \| 9 \| 94 \| 10 \| 108 \| Fab LC region of KTN3379 \| Fab LC region of KTN3379; pdb \| 8TEA\_G \| 0.991 \| 0.003687 \| 84 \| 0.102 \| 107 \| 67 \| 6 \| 9 \| 94 \| 10 \| 108 \| CS2pt1p2\_A10L Fab light chain \| CS2pt1p2\_A10L Fab light chain; pdb \| 7XW7\_B \| 0.991 \| 0.002911 \| 84 \| 0.09 \| 110 \| 71 \| 5 \| 7 \| 94 \| 6 \| 108 \| K1-70 scFv light chain \| K1-70 scFv light chain; pdb \| 4GXU\_P \| 0.991 \| 0.001612 \| 84 \| 0.103 \| 106 \| 69 \| 6 \| 9 \| 94 \| 11 \| 110 \| Antibody 1F1, light chain \| Antibody 1F1, light chain; pdb \| 6WIT\_L \| 0.991 \| 0.00415 \| 84 \| 0.107 \| 112 \| 68 \| 7 \| 7 \| 94 \| 6 \| 109 \| NHP GN1-SD7 Fab Light Chain \| NHP GN1-SD7 Fab Light Chain; pdb \| 6WAS\_B \| 0.991 \| 0.006275 \| 84 \| 0.111 \| 117 \| 76 \| 6 \| 7 \| 101 \| 7 \| 117 \| GN1\_PA8 Fab Light chain \| GN1\_PA8 Fab Light chain; pdb \| 6UUP\_B \| 0.991 \| 0.005576 \| 84 \| 0.115 \| 104 \| 67 \| 5 \| 9 \| 94 \| 128 \| 224 \| Anti-CD33 conditional scFv \| Anti-CD33 conditional scFv; pdb \| 2A9N\_L \| 0.99 \| 0.005576 \| 83 \| 0.09 \| 110 \| 71 \| 7 \| 7 \| 94 \| 5 \| 107 \| fluorescein-scfv light chain \| fluorescein-scfv light chain; pdb \| 3T0W\_B \| 0.988 \| 0.009491 \| 82 \| 0.091 \| 109 \| 72 \| 5 \| 7 \| 94 \| 7 \| 109 \| immunoglobulin variable lambda domain \| immunoglobulin variable lambda domain; pdb \| 6OO0\_L \| 0.988 \| 0.007493 \| 82 \| 0.09 \| 110 \| 71 \| 5 \| 7 \| 94 \| 6 \| 108 \| NC-Cow1 light chain \| NC-Cow1 light chain; pdb \| 7U9G\_D \| 0.986 \| 0.005576 \| 81 \| 0.082 \| 109 \| 73 \| 5 \| 7 \| 94 \| 6 \| 108 \| RVA122 Fab Light Chain \| RVA122 Fab Light Chain; pdb \| 6U0L\_Q \| 0.986 \| 0.007493 \| 81 \| 0.116 \| 112 \| 66 \| 6 \| 7 \| 94 \| 7 \| 109 \| E51 Fab light chain \| E51 Fab light chain; pdb \| 6QN7\_L \| 0.986 \| 0.008946 \| 81 \| 0.121 \| 107 \| 71 \| 3 \| 7 \| 94 \| 5 \| 107 \| Light chain of bovine anti-RSV B13 \| Light chain of bovine anti-RSV B13; pdb \| 6EQC\_D \| 0.986 \| 0.008433 \| 81 \| 0.09 \| 111 \| 72 \| 5 \| 9 \| 95 \| 127 \| 232 \| scFv of 9C12 antibody \| scFv of 9C12 antibody; pdb \| 6CNV\_L \| 0.984 \| 0.007493 \| 80 \| 0.127 \| 110 \| 67 \| 7 \| 7 \| 94 \| 6 \| 108 \| CR9114 Light chain \| CR9114 Light chain; pdb \| 6WIT\_B \| 0.984 \| 0.002298 \| 80 \| 0.117 \| 136 \| 81 \| 9 \| 7 \| 115 \| 5 \| 128 \| NHP GN1-SD7 Fab Light Chain \| NHP GN1-SD7 Fab Light Chain; pdb \| 7FAU\_D \| 0.981 \| 0.008946 \| 79 \| 0.117 \| 119 \| 66 \| 6 \| 9 \| 92 \| 10 \| 124 \| NB\_1B11 \| NB\_1B11; pdb \| 7PA8\_III \| 0.981 \| 0.007493 \| 79 \| 0.117 \| 111 \| 67 \| 7 \| 7 \| 94 \| 6 \| 108 \| Fab 27C2 light chain \| Fab 27C2 light chain; pdb \| 7Q9F\_J \| 0.981 \| 0.007493 \| 79 \| 0.067 \| 118 \| 78 \| 5 \| 9 \| 100 \| 9 \| 120 \| Beta-50 heavy chain \| Beta-50 heavy chain; pdb \| 8DY5\_B \| 0.981 \| 0.009491 \| 79 \| 0.101 \| 118 \| 79 \| 5 \| 7 \| 103 \| 7 \| 118 \| spFv CAT2200 LH \| spFv CAT2200 LH; pdb \| 7T3M\_J \| 0.978 \| 0.009491 \| 78 \| 0.1 \| 109 \| 72 \| 6 \| 7 \| 94 \| 6 \| 109 \| Antibody 2-7 scFv, variable light chain VL \| Antibody 2-7 scFv, variable light chain VL; pdb \| 6IDI\_J \| 0.978 \| 0.008433 \| 78 \| 0.101 \| 108 \| 70 \| 5 \| 7 \| 93 \| 5 \| 106 \| Fab 1H10 light chain (V-region) \| Fab 1H10 light chain (V-region); pdb \| 8A67\_H \| 0.978 \| 0.00467 \| 78 \| 0.075 \| 120 \| 79 \| 5 \| 9 \| 100 \| 10 \| 125 \| Synthetic Nanobody NbSL3.3Q \| Synthetic Nanobody NbSL3.3Q; pdb \| 6V8I\_BJ \| 0.978 \| 0.001131 \| 78 \| 0.139 \| 158 \| 98 \| 10 \| 2 \| 128 \| 15 \| 165 \| Fiber Lower, gp62 \| Fiber Lower, gp62; pdb \| 3MLT\_A \| 0.978 \| 0.007949 \| 78 \| 0.126 \| 111 \| 68 \| 8 \| 7 \| 94 \| 7 \| 111 \| Human monoclonal anti-HIV-1 gp120 V3 antibody 2557 Fab light chain \| Human monoclonal anti-HIV-1 gp120 V3 antibody 2557 Fab light chain; pdb \| 7UP9\_H \| 0.975 \| 0.006658 \| 77 \| 0.081 \| 111 \| 74 \| 5 \| 7 \| 92 \| 9 \| 116 \| Fab 2D3 heavy chain \| Fab 2D3 heavy chain; pdb \| 4HKB\_B \| 0.975 \| 0.007063 \| 77 \| 0.158 \| 101 \| 63 \| 7 \| 7 \| 95 \| 6 \| 96 \| CH67 light chain \| CH67 light chain; pdb \| 5CUS\_L \| 0.975 \| 0.008433 \| 77 \| 0.114 \| 105 \| 68 \| 4 \| 9 \| 94 \| 10 \| 108 \| Fab LC region of KTN3379 \| Fab LC region of KTN3379; pdb \| 4Y5X\_B \| 0.971 \| 0.008433 \| 76 \| 0.117 \| 111 \| 67 \| 7 \| 7 \| 94 \| 7 \| 109 \| diabody 310 VH domain \| diabody 310 VH domain; pdb \| 7S2S\_A \| 0.971 \| 0.005915 \| 76 \| 0.09 \| 122 \| 69 \| 6 \| 9 \| 93 \| 10 \| 126 \| IL2Rb-binding nanobody \| IL2Rb-binding nanobody; pdb \| 7MLH\_E \| 0.971 \| 0.007063 \| 76 \| 0.076 \| 117 \| 79 \| 4 \| 8 \| 100 \| 10 \| 121 \| IgE Heavy chain \| IgE Heavy chain; pdb \| 5OCK\_L \| 0.971 \| 0.005915 \| 76 \| 0.145 \| 137 \| 76 \| 9 \| 7 \| 115 \| 7 \| 130 \| Human ACPA E4 Fab fragment - Light chain \| Human ACPA E4 Fab fragment - Light chain; pdb \| 7SJ0\_L \| 0.967 \| 0.004402 \| 75 \| 0.127 \| 110 \| 70 \| 6 \| 5 \| 94 \| 4 \| 107 \| A7V3 Fab light chain \| A7V3 Fab light chain; pdb \| 8DD3\_J \| 0.961 \| 0.005576 \| 74 \| 0.061 \| 113 \| 74 \| 6 \| 9 \| 95 \| 10 \| 116 \| IgG2b Fab Heavy Chain \| IgG2b Fab Heavy Chain; pdb \| 7AQG\_C \| 0.961 \| 0.007949 \| 74 \| 0.071 \| 112 \| 75 \| 5 \| 9 \| 93 \| 11 \| 120 \| VHH-2w-64 (Nb64) \| VHH-2w-64 (Nb64); pdb \| 1MH5\_B \| 0.961 \| 0.007493 \| 74 \| 0.093 \| 118 \| 74 \| 6 \| 9 \| 100 \| 10 \| 120 \| IMMUNOGLOBULIN MS6-164 \| IMMUNOGLOBULIN MS6-164; pdb \| 6U59\_H \| 0.956 \| 0.008433 \| 73 \| 0.059 \| 117 \| 74 \| 8 \| 9 \| 95 \| 8 \| 118 \| rabbit antibody 13B Fragment antigen binding heavy chain \| rabbit antibody 13B Fragment antigen binding heavy chain; pdb \| 6ZG3\_J \| 0.956 \| 0.003687 \| 73 \| 0.058 \| 120 \| 73 \| 5 \| 9 \| 93 \| 10 \| 124 \| CA14381 nanobody \| CA14381 nanobody; pdb \| 6LCS\_A \| 0.956 \| 0.007493 \| 73 \| 0.104 \| 144 \| 100 \| 6 \| 7 \| 124 \| 8 \| 148 \| VH-SARAH \| VH-SARAH; pdb \| 5HBV\_D \| 0.956 \| 0.003687 \| 73 \| 0.11 \| 136 \| 80 \| 6 \| 7 \| 107 \| 9 \| 138 \| Fab35, Heavy Chain \| Fab35, Heavy Chain; pdb \| 7RFC\_H \| 0.956 \| 0.009491 \| 73 \| 0.082 \| 121 \| 79 \| 6 \| 9 \| 100 \| 10 \| 127 \| mAb1382 Heavy Chain \| mAb1382 Heavy Chain; pdb \| 7FEJ\_L \| 0.949 \| 0.00415 \| 72 \| 0.119 \| 109 \| 67 \| 7 \| 7 \| 94 \| 6 \| 106 \| IG LAMDA CHAIN VARIABLE REGION \| IG LAMDA CHAIN VARIABLE REGION; pdb \| 7W9M\_B \| 0.949 \| 0.0004945 \| 72 \| 0.084 \| 189 \| 116 \| 8 \| 7 \| 162 \| 8 \| 172 \| Sodium channel subunit beta-1 \| Sodium channel subunit beta-1; pdb \| 6MEK\_E \| 0.949 \| 0.009491 \| 72 \| 0.117 \| 136 \| 83 \| 8 \| 7 \| 115 \| 6 \| 131 \| HEPC46 Light Chain \| HEPC46 Light Chain; pdb \| 7N4M\_L \| 0.949 \| 0.008946 \| 72 \| 0.12 \| 133 \| 85 \| 7 \| 7 \| 115 \| 7 \| 131 \| WRAIR-2151 antibody Fab light chain \| WRAIR-2151 antibody Fab light chain; pdb \| 7WQV\_N \| 0.949 \| 0.007063 \| 72 \| 0.126 \| 111 \| 67 \| 7 \| 7 \| 94 \| 129 \| 232 \| Ab08 \| Ab08; pdb \| 7M8J\_H \| 0.941 \| 0.004402 \| 71 \| 0.094 \| 117 \| 69 \| 7 \| 9 \| 95 \| 10 \| 119 \| CM25 Fab - Heavy Chain \| CM25 Fab - Heavy Chain; pdb \| 7TJ9\_B \| 0.941 \| 0.001432 \| 71 \| 0.107 \| 149 \| 94 \| 9 \| 7 \| 124 \| 8 \| 148 \| Sodium channel subunit beta-3 \| Sodium channel subunit beta-3; pdb \| 6II4\_L \| 0.941 \| 0.009491 \| 71 \| 0.111 \| 135 \| 84 \| 9 \| 7 \| 115 \| 7 \| 131 \| Light chain of L4A-14 Fab \| Light chain of L4A-14 Fab; pdb \| 7LFD\_H \| 0.941 \| 0.009491 \| 71 \| 0.106 \| 122 \| 76 \| 5 \| 7 \| 100 \| 9 \| 125 \| Fab 7D6 heavy chain \| Fab 7D6 heavy chain; pdb \| 7M1H\_B \| 0.933 \| 0.007949 \| 70 \| 0.102 \| 117 \| 68 \| 5 \| 9 \| 92 \| 10 \| 122 \| JPU-D12 \| JPU-D12; pdb \| 6MWR\_C \| 0.933 \| 0.005576 \| 70 \| 0.101 \| 138 \| 87 \| 8 \| 9 \| 115 \| 10 \| 141 \| G7 Gamma chain T cell receptor \| G7 Gamma chain T cell receptor; pdb \| 7JXD\_D \| 0.923 \| 0.009491 \| 69 \| 0.116 \| 137 \| 82 \| 11 \| 7 \| 115 \| 7 \| 132 \| S2A4 antigen-binding (Fab) fragment \| S2A4 antigen-binding (Fab) fragment; pdb \| 5XCV\_B \| 0.912 \| 0.007063 \| 68 \| 0.112 \| 142 \| 83 \| 9 \| 7 \| 124 \| 7 \| 129 \| VL-SARAH(S37C) chimera,VL-SARAH(S37C) chimera \| VL-SARAH(S37C) chimera,VL-SARAH(S37C) chimera; pdb \| 8F6O\_A \| 0.9 \| 0.008433 \| 67 \| 0.064 \| 124 \| 79 \| 7 \| 8 \| 100 \| 10 \| 127 \| h22B3 Fab heavy chain \| h22B3 Fab heavy chain; pdb \| 7T0R\_H \| 0.872 \| 0.008433 \| 65 \| 0.071 \| 126 \| 79 \| 6 \| 7 \| 101 \| 9 \| 127 \| Ibalizumab Heavy Chain \| Ibalizumab Heavy Chain; pdb \| 7K5Y\_N \| 0.855 \| 0.007063 \| 64 \| 0.073 \| 122 \| 69 \| 6 \| 7 \| 95 \| 117 \| 227 \| scFv20 \| scFv20; pdb \| 6DB6\_H \| 0.837 \| 0.008433 \| 63 \| 0.072 \| 125 \| 75 \| 6 \| 9 \| 100 \| 11 \| 127 \| Human monoclonal anti-HIV-1 gp120 V3 antibody 311-11D Fab heavy chain \| Human monoclonal anti-HIV-1 gp120 V3 antibody 311-11D Fab heavy chain; pdb \| 5L7U\_B \| 0.326 \| 0.009491 \| 45 \| 0.104 \| 143 \| 70 \| 12 \| 8 \| 96 \| 50 \| 188 \| Glycoside hydrolase \| Glycoside hydrolase; afdb-proteome \| AF-F1QIR0-F1-MODEL\_V4 \| 1.0 \| 0.0001196 \| 116 \| 0.112 \| 133 \| 84 \| 9 \| 7 \| 128 \| 201 \| 310 \| Phosphoinositide-3-kinase adaptor protein 1 \| Phosphoinositide-3-kinase adaptor protein 1; afdb-proteome \| AF-Q9EQ32-F1-MODEL\_V4 \| 1.0 \| 5.549e-05 \| 114 \| 0.137 \| 138 \| 80 \| 9 \| 7 \| 118 \| 184 \| 308 \| Phosphoinositide 3-kinase adapter protein 1 \| Phosphoinositide 3-kinase adapter protein 1; afdb-proteome \| AF-A0A044VIN6-F1-MODEL\_V4 \| 1.0 \| 0.00627 \| 111 \| 0.113 \| 97 \| 73 \| 5 \| 5 \| 96 \| 250 \| 338 \| Uncharacterized protein \| Uncharacterized protein; afdb-proteome \| AF-Q32J71-F1-MODEL\_V4 \| 1.0 \| 0.004666 \| 108 \| 0.157 \| 89 \| 66 \| 6 \| 8 \| 92 \| 684 \| 767 \| Uncharacterized protein \| Uncharacterized protein; afdb-proteome \| AF-Q6ZUJ8-F1-MODEL\_V4 \| 1.0 \| 7.029e-05 \| 103 \| 0.147 \| 156 \| 87 \| 10 \| 7 \| 129 \| 183 \| 325 \| Phosphoinositide 3-kinase adapter protein 1 \| Phosphoinositide 3-kinase adapter protein 1; afdb-proteome \| AF-Q32J72-F1-MODEL\_V4 \| 1.0 \| 0.004146 \| 101 \| 0.123 \| 97 \| 68 \| 7 \| 8 \| 96 \| 356 \| 443 \| Uncharacterized protein \| Uncharacterized protein; afdb-proteome \| AF-G4V6G9-F1-MODEL\_V4 \| 1.0 \| 0.0002903 \| 100 \| 0.17 \| 129 \| 93 \| 6 \| 9 \| 132 \| 441 \| 560 \| Putative titin \| Putative titin; afdb-proteome \| AF-Q2FX74-F1-MODEL\_V4 \| 0.999 \| 0.001004 \| 96 \| 0.125 \| 192 \| 121 \| 11 \| 9 \| 179 \| 24 \| 189 \| Phage tail fiber protein, putative \| Phage tail fiber protein, putative; afdb-proteome \| AF-Q641Y0-F1-MODEL\_V4 \| 0.999 \| 0.0009463 \| 96 \| 0.095 \| 147 \| 102 \| 9 \| 7 \| 134 \| 307 \| 441 \| Dolichyl-diphosphooligosaccharide--protein glycosyltransferase 48 kDa subunit \| Dolichyl-diphosphooligosaccharide--protein glycosyltransferase 48 kDa subunit; afdb-proteome \| AF-A0A1U7F3R9-F1-MODEL\_V4 \| 0.999 \| 0.008425 \| 96 \| 0.173 \| 92 \| 61 \| 5 \| 7 \| 95 \| 358 \| 437 \| Bm6990 \| Bm6990; afdb-proteome \| AF-O54734-F1-MODEL\_V4 \| 0.998 \| 0.002436 \| 94 \| 0.097 \| 133 \| 90 \| 8 \| 7 \| 124 \| 307 \| 424 \| Dolichyl-diphosphooligosaccharide--protein glycosyltransferase 48 kDa subunit \| Dolichyl-diphosphooligosaccharide--protein glycosyltransferase 48 kDa subunit; afdb-proteome \| AF-A0A044QQS6-F1-MODEL\_V4 \| 0.998 \| 0.002908 \| 93 \| 0.131 \| 145 \| 103 \| 12 \| 7 \| 136 \| 305 \| 441 \| Dolichyl-diphosphooligosaccharide--protein glycosyltransferase 48 kDa subunit \| Dolichyl-diphosphooligosaccharide--protein glycosyltransferase 48 kDa subunit; afdb-proteome \| AF-O95932-F1-MODEL\_V4 \| 0.998 \| 0.009482 \| 93 \| 0.139 \| 129 \| 84 \| 8 \| 9 \| 113 \| 31 \| 156 \| Protein-glutamine gamma-glutamyltransferase 6 \| Protein-glutamine gamma-glutamyltransferase 6; afdb-proteome \| AF-A0A077ZB36-F1-MODEL\_V4 \| 0.998 \| 0.005571 \| 93 \| 0.141 \| 92 \| 73 \| 5 \| 7 \| 93 \| 528 \| 618 \| Plexin A4 \| Plexin A4; afdb-proteome \| AF-A0A0H3GY50-F1-MODEL\_V4 \| 0.997 \| 0.0008409 \| 92 \| 0.117 \| 179 \| 112 \| 12 \| 6 \| 145 \| 3 \| 174 \| Uncharacterized protein \| Uncharacterized protein; afdb-proteome \| AF-A0A0G2K7L7-F1-MODEL\_V4 \| 0.997 \| 0.0004138 \| 91 \| 0.096 \| 155 \| 115 \| 9 \| 6 \| 142 \| 249 \| 396 \| ANTXR-like \| ANTXR-like; afdb-proteome \| AF-D3ZH36-F1-MODEL\_V4 \| 0.997 \| 0.000494 \| 91 \| 0.099 \| 171 \| 128 \| 8 \| 10 \| 178 \| 303 \| 449 \| Extracellular leucine-rich repeat and fibronectin type III domain-containing 2 \| Extracellular leucine-rich repeat and fibronectin type III domain-containing 2; afdb-proteome \| AF-A0A0G2JUH9-F1-MODEL\_V4 \| 0.996 \| 0.0006258 \| 90 \| 0.145 \| 186 \| 119 \| 10 \| 1 \| 166 \| 3 \| 168 \| SKICH domain-containing protein \| SKICH domain-containing protein; afdb-proteome \| AF-A0A0K0DYW0-F1-MODEL\_V4 \| 0.996 \| 0.002164 \| 90 \| 0.11 \| 154 \| 104 \| 9 \| 7 \| 140 \| 308 \| 448 \| Dolichyl-diphosphooligosaccharide--protein glycosyltransferase 48 kDa subunit \| Dolichyl-diphosphooligosaccharide--protein glycosyltransferase 48 kDa subunit; afdb-proteome \| AF-Q32J74-F1-MODEL\_V4 \| 0.996 \| 0.00591 \| 90 \| 0.177 \| 96 \| 62 \| 7 \| 8 \| 95 \| 1032 \| 1118 \| Uncharacterized protein \| Uncharacterized protein; afdb-proteome \| AF-P39656-F1-MODEL\_V4 \| 0.996 \| 0.002164 \| 89 \| 0.088 \| 147 \| 103 \| 9 \| 7 \| 134 \| 322 \| 456 \| Dolichyl-diphosphooligosaccharide--protein glycosyltransferase 48 kDa subunit \| Dolichyl-diphosphooligosaccharide--protein glycosyltransferase 48 kDa subunit; afdb-proteome \| AF-Q04864-F1-MODEL\_V4 \| 0.995 \| 0.002908 \| 88 \| 0.12 \| 141 \| 110 \| 8 \| 8 \| 142 \| 194 \| 326 \| Proto-oncogene c-Rel \| Proto-oncogene c-Rel; afdb-proteome \| AF-A0A5P3FZ45-F1-MODEL\_V4 \| 0.995 \| 0.002296 \| 88 \| 0.129 \| 177 \| 111 \| 12 \| 4 \| 164 \| 13 \| 162 \| Uncharacterized protein \| Uncharacterized protein; afdb-proteome \| AF-A0A3P7FSY3-F1-MODEL\_V4 \| 0.994 \| 0.003472 \| 87 \| 0.09 \| 154 \| 105 \| 9 \| 7 \| 141 \| 272 \| 409 \| Dolichyl-diphosphooligosaccharide--protein glycosyltransferase 48 kDa subunit \| Dolichyl-diphosphooligosaccharide--protein glycosyltransferase 48 kDa subunit; afdb-proteome \| AF-A0A2R8QBU7-F1-MODEL\_V4 \| 0.994 \| 0.0004657 \| 87 \| 0.113 \| 185 \| 124 \| 9 \| 10 \| 179 \| 297 \| 456 \| LRRCT domain-containing protein \| LRRCT domain-containing protein; afdb-proteome \| AF-Q9NZC2-F1-MODEL\_V4 \| 0.993 \| 0.0002903 \| 86 \| 0.14 \| 213 \| 116 \| 12 \| 9 \| 179 \| 25 \| 212 \| Triggering receptor expressed on myeloid cells 2 \| Triggering receptor expressed on myeloid cells 2; afdb-proteome \| AF-F1Q6X1-F1-MODEL\_V4 \| 0.993 \| 0.001272 \| 86 \| 0.096 \| 156 \| 103 \| 9 \| 7 \| 142 \| 225 \| 362 \| ANTXR cell adhesion molecule 1d \| ANTXR cell adhesion molecule 1d; afdb-proteome \| AF-G5EFS1-F1-MODEL\_V4 \| 0.993 \| 0.001431 \| 86 \| 0.112 \| 125 \| 89 \| 9 \| 7 \| 126 \| 49 \| 156 \| ANK\_REP\_REGION domain-containing protein \| ANK\_REP\_REGION domain-containing protein; afdb-proteome \| AF-A0A5K4F288-F1-MODEL\_V4 \| 0.993 \| 0.000892 \| 86 \| 0.164 \| 170 \| 102 \| 10 \| 9 \| 163 \| 708 \| 852 \| Kettin/titin-related protein \| Kettin/titin-related protein; afdb-proteome \| AF-A0A0N4UPP3-F1-MODEL\_V4 \| 0.993 \| 0.002584 \| 86 \| 0.114 \| 157 \| 103 \| 9 \| 4 \| 142 \| 779 \| 917 \| Uncharacterized protein \| Uncharacterized protein; afdb-proteome \| AF-P01740-F1-MODEL\_V4 \| 0.992 \| 0.008425 \| 85 \| 0.104 \| 115 \| 72 \| 7 \| 8 \| 95 \| 25 \| 135 \| T-cell receptor gamma chain V region V108A \| T-cell receptor gamma chain V region V108A; afdb-proteome \| AF-C6T7K0-F1-MODEL\_V4 \| 0.992 \| 0.001065 \| 85 \| 0.12 \| 150 \| 96 \| 12 \| 1 \| 126 \| 70 \| 207 \| Signal sequence receptor subunit alpha \| Signal sequence receptor subunit alpha; afdb-proteome \| AF-F1RDW2-F1-MODEL\_V4 \| 0.992 \| 0.00039 \| 85 \| 0.117 \| 162 \| 108 \| 9 \| 10 \| 146 \| 558 \| 709 \| Si:ch211-180f4.1 \| Si:ch211-180f4.1; afdb-proteome \| AF-Q68FM6-F1-MODEL\_V4 \| 0.992 \| 0.0005899 \| 85 \| 0.08 \| 175 \| 129 \| 8 \| 10 \| 178 \| 303 \| 451 \| Protein phosphatase 1 regulatory subunit 29 \| Protein phosphatase 1 regulatory subunit 29; afdb-proteome \| AF-Q9P7K7-F1-MODEL\_V4 \| 0.991 \| 0.001349 \| 84 \| 0.101 \| 168 \| 119 \| 7 \| 8 \| 165 \| 32 \| 177 \| Uncharacterized protein C21C3.17c \| Uncharacterized protein C21C3.17c; afdb-proteome \| AF-A0A175VWL7-F1-MODEL\_V4 \| 0.991 \| 0.002436 \| 84 \| 0.094 \| 201 \| 141 \| 10 \| 8 \| 179 \| 126 \| 314 \| Uncharacterized protein \| Uncharacterized protein; afdb-proteome \| AF-Q8BVM2-F1-MODEL\_V4 \| 0.991 \| 0.000892 \| 84 \| 0.084 \| 154 \| 112 \| 10 \| 7 \| 142 \| 260 \| 402 \| Anthrax toxin receptor-like \| Anthrax toxin receptor-like; afdb-proteome \| AF-Q61809-F1-MODEL\_V4 \| 0.991 \| 0.001349 \| 84 \| 0.104 \| 153 \| 108 \| 8 \| 10 \| 142 \| 533 \| 676 \| Leucine-rich repeat neuronal protein 1 \| Leucine-rich repeat neuronal protein 1; afdb-proteome \| AF-Q8BHK2-F1-MODEL\_V4 \| 0.99 \| 0.00039 \| 83 \| 0.115 \| 208 \| 124 \| 10 \| 7 \| 178 \| 32 \| 215 \| Sodium channel subunit beta-3 \| Sodium channel subunit beta-3; afdb-proteome \| AF-Q6QX36-F1-MODEL\_V4 \| 0.99 \| 0.000892 \| 83 \| 0.108 \| 212 \| 129 \| 13 \| 3 \| 179 \| 22 \| 208 \| CRKD-binding protein \| CRKD-binding protein; afdb-proteome \| AF-A6NF34-F1-MODEL\_V4 \| 0.99 \| 0.001349 \| 83 \| 0.14 \| 149 \| 103 \| 7 \| 7 \| 142 \| 259 \| 395 \| Anthrax toxin receptor-like \| Anthrax toxin receptor-like; afdb-proteome \| AF-A0A5K4F3S2-F1-MODEL\_V4 \| 0.99 \| 0.001431 \| 83 \| 0.161 \| 204 \| 109 \| 11 \| 9 \| 179 \| 708 \| 882 \| Kettin/titin-related protein \| Kettin/titin-related protein; afdb-proteome \| AF-A0A0N4UJG6-F1-MODEL\_V4 \| 0.988 \| 0.003085 \| 82 \| 0.083 \| 156 \| 105 \| 11 \| 7 \| 142 \| 305 \| 442 \| Dolichyl-diphosphooligosaccharide--protein glycosyltransferase 48 kDa subunit \| Dolichyl-diphosphooligosaccharide--protein glycosyltransferase 48 kDa subunit; afdb-proteome \| AF-Q6AWS7-F1-MODEL\_V4 \| 0.986 \| 0.001065 \| 81 \| 0.129 \| 154 \| 89 \| 11 \| 1 \| 126 \| 65 \| 201 \| Signal sequence receptor subunit alpha \| Signal sequence receptor subunit alpha; afdb-proteome \| AF-K0EVZ8-F1-MODEL\_V4 \| 0.986 \| 0.009482 \| 81 \| 0.125 \| 120 \| 87 \| 6 \| 7 \| 122 \| 543 \| 648 \| Non-specific serine/threonine protein kinase \| Non-specific serine/threonine protein kinase; afdb-proteome \| AF-P0C7U0-F1-MODEL\_V4 \| 0.986 \| 0.001199 \| 81 \| 0.135 \| 185 \| 121 \| 8 \| 5 \| 178 \| 323 \| 479 \| Protein ELFN1 \| Protein ELFN1; afdb-proteome \| AF-Q9JK00-F1-MODEL\_V4 \| 0.984 \| 0.0006639 \| 80 \| 0.115 \| 208 \| 124 \| 10 \| 7 \| 178 \| 32 \| 215 \| Sodium channel subunit beta-3 \| Sodium channel subunit beta-3; afdb-proteome \| AF-Q8BG84-F1-MODEL\_V4 \| 0.984 \| 0.001813 \| 80 \| 0.13 \| 176 \| 118 \| 8 \| 9 \| 179 \| 38 \| 183 \| Leukocyte-associated immunoglobulin-like receptor 1 \| Leukocyte-associated immunoglobulin-like receptor 1; afdb-proteome \| AF-Q7TQN4-F1-MODEL\_V4 \| 0.984 \| 0.00204 \| 80 \| 0.137 \| 145 \| 104 \| 9 \| 8 \| 142 \| 202 \| 335 \| Nuclear factor kappaB subunit p65 \| Nuclear factor kappaB subunit p65; afdb-proteome \| AF-Q09147-F1-MODEL\_V4 \| 0.984 \| 0.0004657 \| 80 \| 0.131 \| 175 \| 105 \| 9 \| 8 \| 164 \| 495 \| 640 \| Fibroblast growth factor receptor homolog 2 \| Fibroblast growth factor receptor homolog 2; afdb-proteome \| AF-Q9NY72-F1-MODEL\_V4 \| 0.981 \| 0.0007926 \| 79 \| 0.096 \| 217 \| 128 \| 11 \| 3 \| 179 \| 26 \| 214 \| Sodium channel subunit beta-3 \| Sodium channel subunit beta-3; afdb-proteome \| AF-B4FE49-F1-MODEL\_V4 \| 0.981 \| 0.0009463 \| 79 \| 0.128 \| 195 \| 115 \| 13 \| 8 \| 162 \| 78 \| 257 \| Signal sequence receptor subunit alpha \| Signal sequence receptor subunit alpha; afdb-proteome \| AF-Q6ZLK0-F1-MODEL\_V4 \| 0.981 \| 0.009482 \| 79 \| 0.111 \| 135 \| 87 \| 9 \| 7 \| 124 \| 310 \| 428 \| Dolichyl-diphosphooligosaccharide--protein glycosyltransferase 48 kDa subunit \| Dolichyl-diphosphooligosaccharide--protein glycosyltransferase 48 kDa subunit; afdb-proteome \| AF-Q9ESY6-F1-MODEL\_V4 \| 0.981 \| 0.001431 \| 79 \| 0.088 \| 147 \| 106 \| 8 \| 10 \| 142 \| 531 \| 663 \| Leucine-rich repeat neuronal protein 3 \| Leucine-rich repeat neuronal protein 3; afdb-proteome \| AF-Q8C8T7-F1-MODEL\_V4 \| 0.981 \| 0.001611 \| 79 \| 0.13 \| 184 \| 122 \| 8 \| 5 \| 178 \| 323 \| 478 \| Protein ELFN1 \| Protein ELFN1; afdb-proteome \| AF-A0A2R8RRG5-F1-MODEL\_V4 \| 0.978 \| 0.001709 \| 78 \| 0.092 \| 228 \| 128 \| 12 \| 3 \| 178 \| 24 \| 224 \| Si:rp71-81e14.2 \| Si:rp71-81e14.2; afdb-proteome \| AF-P15307-F1-MODEL\_V4 \| 0.978 \| 0.00627 \| 78 \| 0.112 \| 142 \| 107 \| 9 \| 8 \| 142 \| 194 \| 323 \| Proto-oncogene c-Rel \| Proto-oncogene c-Rel; afdb-proteome \| AF-Q93Z16-F1-MODEL\_V4 \| 0.978 \| 0.005571 \| 78 \| 0.115 \| 182 \| 122 \| 13 \| 9 \| 168 \| 455 \| 619 \| Dolichyl-diphosphooligosaccharide--protein glycosyltransferase subunit 2 \| Dolichyl-diphosphooligosaccharide--protein glycosyltransferase subunit 2; afdb-proteome \| AF-D3ZAV8-F1-MODEL\_V4 \| 0.978 \| 0.005571 \| 78 \| 0.112 \| 142 \| 109 \| 6 \| 10 \| 142 \| 535 \| 668 \| Leucine-rich repeat neuronal 2 \| Leucine-rich repeat neuronal 2; afdb-proteome \| AF-Q9P244-F1-MODEL\_V4 \| 0.978 \| 0.002741 \| 78 \| 0.14 \| 157 \| 99 \| 8 \| 12 \| 142 \| 434 \| 580 \| Leucine-rich repeat and fibronectin type III domain-containing protein 1 \| Leucine-rich repeat and fibronectin type III domain-containing protein 1; afdb-proteome \| AF-A9Q7H1-F1-MODEL\_V4 \| 0.975 \| 0.001272 \| 77 \| 0.114 \| 192 \| 111 \| 10 \| 3 \| 163 \| 22 \| 185 \| RIKEN cDNA 9830107B12 gene \| RIKEN cDNA 9830107B12 gene; afdb-proteome \| AF-F8W4B3-F1-MODEL\_V4 \| 0.975 \| 0.001709 \| 77 \| 0.105 \| 219 \| 127 \| 13 \| 3 \| 179 \| 47 \| 238 \| Sodium channel subunit beta-3 \| Sodium channel subunit beta-3; afdb-proteome \| AF-O59866-F1-MODEL\_V4 \| 0.975 \| 0.002584 \| 77 \| 0.089 \| 156 \| 103 \| 11 \| 7 \| 136 \| 290 \| 432 \| Dolichyl-diphosphooligosaccharide--protein glycosyltransferase subunit wbp1 \| Dolichyl-diphosphooligosaccharide--protein glycosyltransferase subunit wbp1; afdb-proteome \| AF-A0A0H2UKN7-F1-MODEL\_V4 \| 0.971 \| 0.0004138 \| 76 \| 0.091 \| 197 \| 123 \| 8 \| 3 \| 166 \| 8 \| 181 \| Sodium channel, voltage-gated, type I, beta b \| Sodium channel, voltage-gated, type I, beta b; afdb-proteome \| AF-Q07699-F1-MODEL\_V4 \| 0.971 \| 0.0006258 \| 76 \| 0.107 \| 195 \| 114 \| 9 \| 7 \| 166 \| 27 \| 196 \| Sodium channel subunit beta-1 \| Sodium channel subunit beta-1; afdb-proteome \| AF-F1QJZ8-F1-MODEL\_V4 \| 0.971 \| 0.002908 \| 76 \| 0.124 \| 177 \| 121 \| 8 \| 9 \| 179 \| 473 \| 621 \| Sema domain, immunoglobulin domain (Ig), transmembrane domain (TM) and short cytoplasmic domain, (semaphorin) 4Gb \| Sema domain, immunoglobulin domain (Ig), transmembrane domain (TM) and short cytoplasmic domain, (semaphorin) 4Gb; afdb-proteome \| AF-F1Q7Y5-F1-MODEL\_V4 \| 0.971 \| 0.002164 \| 76 \| 0.096 \| 177 \| 130 \| 7 \| 12 \| 179 \| 330 \| 485 \| Extracellular leucine-rich repeat and fibronectin type III domain-containing 1b \| Extracellular leucine-rich repeat and fibronectin type III domain-containing 1b; afdb-proteome \| AF-Q5R3F8-F1-MODEL\_V4 \| 0.971 \| 0.00204 \| 76 \| 0.093 \| 181 \| 127 \| 9 \| 10 \| 179 \| 303 \| 457 \| Protein phosphatase 1 regulatory subunit 29 \| Protein phosphatase 1 regulatory subunit 29; afdb-proteome \| AF-D3ZZ44-F1-MODEL\_V4 \| 0.971 \| 0.002164 \| 76 \| 0.129 \| 186 \| 120 \| 9 \| 5 \| 178 \| 323 \| 478 \| Extracellular leucine-rich repeat and fibronectin type III domain-containing 1 \| Extracellular leucine-rich repeat and fibronectin type III domain-containing 1; afdb-proteome \| AF-Q04207-F1-MODEL\_V4 \| 0.967 \| 0.003684 \| 75 \| 0.144 \| 145 \| 102 \| 10 \| 8 \| 142 \| 202 \| 334 \| Transcription factor p65 \| Transcription factor p65; afdb-proteome \| AF-A0A0D2DWU6-F1-MODEL\_V4 \| 0.961 \| 0.001518 \| 74 \| 0.151 \| 172 \| 99 \| 11 \| 9 \| 142 \| 57 \| 219 \| Signal sequence receptor subunit alpha \| Signal sequence receptor subunit alpha; afdb-proteome \| AF-A0A1C1C8U1-F1-MODEL\_V4 \| 0.961 \| 0.001199 \| 74 \| 0.146 \| 177 \| 99 \| 12 \| 9 \| 142 \| 57 \| 224 \| Signal sequence receptor subunit alpha \| Signal sequence receptor subunit alpha; afdb-proteome \| AF-Q8I3K4-F1-MODEL\_V4 \| 0.961 \| 0.005251 \| 74 \| 0.086 \| 184 \| 119 \| 6 \| 7 \| 141 \| 103 \| 286 \| Uncharacterized protein \| Uncharacterized protein; afdb-proteome \| AF-A0A2I3BPJ2-F1-MODEL\_V4 \| 0.961 \| 0.003908 \| 74 \| 0.115 \| 199 \| 128 \| 11 \| 8 \| 179 \| 258 \| 435 \| Predicted gene, 30083 \| Predicted gene, 30083; afdb-proteome \| AF-M9MM94-F1-MODEL\_V4 \| 0.961 \| 0.001923 \| 74 \| 0.093 \| 172 \| 110 \| 7 \| 8 \| 142 \| 577 \| 739 \| Sema domain, immunoglobulin domain (Ig), short basic domain, secreted, (semaphorin) 3B \| Sema domain, immunoglobulin domain (Ig), short basic domain, secreted, (semaphorin) 3B; afdb-proteome \| AF-F8W525-F1-MODEL\_V4 \| 0.956 \| 0.002296 \| 73 \| 0.115 \| 182 \| 115 \| 9 \| 9 \| 168 \| 33 \| 190 \| CD79a molecule, immunoglobulin-associated alpha \| CD79a molecule, immunoglobulin-associated alpha; afdb-proteome \| AF-Q00954-F1-MODEL\_V4 \| 0.956 \| 0.0008409 \| 73 \| 0.106 \| 198 \| 111 \| 11 \| 7 \| 166 \| 27 \| 196 \| Sodium channel subunit beta-1 \| Sodium channel subunit beta-1; afdb-proteome \| AF-P97952-F1-MODEL\_V4 \| 0.956 \| 0.000892 \| 73 \| 0.1 \| 199 \| 111 \| 11 \| 7 \| 166 \| 27 \| 196 \| Sodium channel subunit beta-1 \| Sodium channel subunit beta-1; afdb-proteome \| AF-Q6DFX2-F1-MODEL\_V4 \| 0.956 \| 0.005251 \| 73 \| 0.121 \| 156 \| 100 \| 8 \| 7 \| 142 \| 226 \| 364 \| Anthrax toxin receptor 2 \| Anthrax toxin receptor 2; afdb-proteome \| AF-A2AS37-F1-MODEL\_V4 \| 0.949 \| 0.002164 \| 72 \| 0.12 \| 191 \| 111 \| 14 \| 7 \| 153 \| 140 \| 317 \| Expressed sequence AI182371 \| Expressed sequence AI182371; afdb-proteome \| AF-B0S5A5-F1-MODEL\_V4 \| 0.949 \| 0.005571 \| 72 \| 0.123 \| 162 \| 105 \| 9 \| 6 \| 142 \| 250 \| 399 \| Si:ch211-150o23.3 \| Si:ch211-150o23.3; afdb-proteome \| AF-A0A2R8S050-F1-MODEL\_V4 \| 0.949 \| 0.004398 \| 72 \| 0.116 \| 154 \| 95 \| 10 \| 8 \| 142 \| 231 \| 362 \| RHD domain-containing protein \| RHD domain-containing protein; afdb-proteome \| AF-P48551-F1-MODEL\_V4 \| 0.949 \| 0.007942 \| 72 \| 0.101 \| 157 \| 108 \| 9 \| 10 \| 142 \| 142 \| 289 \| Interferon alpha/beta receptor 2 \| Interferon alpha/beta receptor 2; afdb-proteome \| AF-X1WEK5-F1-MODEL\_V4 \| 0.949 \| 0.003085 \| 72 \| 0.082 \| 145 \| 103 \| 10 \| 8 \| 142 \| 313 \| 437 \| V-rel avian reticuloendotheliosis viral oncogene homolog B \| V-rel avian reticuloendotheliosis viral oncogene homolog B; afdb-proteome \| AF-P0C7J6-F1-MODEL\_V4 \| 0.949 \| 0.00591 \| 72 \| 0.132 \| 158 \| 100 \| 8 \| 12 \| 142 \| 434 \| 581 \| Leucine-rich repeat and fibronectin type III domain-containing protein 1 \| Leucine-rich repeat and fibronectin type III domain-containing protein 1; afdb-proteome \| AF-Q4LFA9-F1-MODEL\_V4 \| 0.949 \| 0.004146 \| 72 \| 0.145 \| 192 \| 118 \| 11 \| 4 \| 179 \| 585 \| 746 \| Semaphorin-3G \| Semaphorin-3G; afdb-proteome \| AF-I1LIQ3-F1-MODEL\_V4 \| 0.941 \| 0.001611 \| 71 \| 0.145 \| 158 \| 85 \| 12 \| 8 \| 128 \| 80 \| 224 \| Signal sequence receptor subunit alpha \| Signal sequence receptor subunit alpha; afdb-proteome \| AF-Q7Z6A9-F1-MODEL\_V4 \| 0.941 \| 0.00113 \| 71 \| 0.137 \| 167 \| 98 \| 11 \| 9 \| 142 \| 47 \| 200 \| B- and T-lymphocyte attenuator \| B- and T-lymphocyte attenuator; afdb-proteome \| AF-Q9CZT5-F1-MODEL\_V4 \| 0.941 \| 0.003273 \| 71 \| 0.125 \| 160 \| 105 \| 7 \| 12 \| 142 \| 473 \| 626 \| Vasorin \| Vasorin; afdb-proteome \| AF-A0A0D2DMU8-F1-MODEL\_V4 \| 0.933 \| 0.002584 \| 70 \| 0.112 \| 240 \| 129 \| 12 \| 8 \| 179 \| 44 \| 267 \| Unplaced genomic scaffold supercont1.5, whole genome shotgun sequence \| Unplaced genomic scaffold supercont1.5, whole genome shotgun sequence; afdb-proteome \| AF-Q8NET5-F1-MODEL\_V4 \| 0.923 \| 0.003273 \| 69 \| 0.127 \| 172 \| 100 \| 12 \| 8 \| 142 \| 53 \| 211 \| NFAT activation molecule 1 \| NFAT activation molecule 1; afdb-proteome \| AF-Q63203-F1-MODEL\_V4 \| 0.923 \| 0.003472 \| 69 \| 0.12 \| 149 \| 102 \| 6 \| 7 \| 142 \| 127 \| 259 \| Low affinity immunoglobulin gamma Fc region receptor II \| Low affinity immunoglobulin gamma Fc region receptor II; afdb-proteome \| AF-F7FEU1-F1-MODEL\_V4 \| 0.923 \| 0.001709 \| 69 \| 0.105 \| 209 \| 116 \| 14 \| 7 \| 165 \| 26 \| 213 \| V-set and immunoglobulin domain-containing 4 \| V-set and immunoglobulin domain-containing 4; afdb-proteome \| AF-T1NXB5-F1-MODEL\_V4 \| 0.923 \| 0.003273 \| 69 \| 0.103 \| 213 \| 124 \| 13 \| 9 \| 178 \| 35 \| 223 \| V-set and transmembrane domain-containing protein 4 \| V-set and transmembrane domain-containing protein 4; afdb-proteome \| AF-Q96PJ5-F1-MODEL\_V4 \| 0.923 \| 0.007057 \| 69 \| 0.108 \| 148 \| 107 \| 9 \| 7 \| 142 \| 297 \| 431 \| Fc receptor-like protein 4 \| Fc receptor-like protein 4; afdb-proteome \| AF-P97484-F1-MODEL\_V4 \| 0.923 \| 0.001611 \| 69 \| 0.118 \| 203 \| 120 \| 11 \| 5 \| 179 \| 529 \| 700 \| Leukocyte immunoglobulin-like receptor subfamily B member 3 \| Leukocyte immunoglobulin-like receptor subfamily B member 3; afdb-proteome \| AF-O75325-F1-MODEL\_V4 \| 0.912 \| 0.003684 \| 68 \| 0.108 \| 147 \| 102 \| 7 \| 10 \| 142 \| 535 \| 666 \| Leucine-rich repeat neuronal protein 2 \| Leucine-rich repeat neuronal protein 2; afdb-proteome \| AF-Q14517-F9-MODEL\_V4 \| 0.912 \| 0.003908 \| 68 \| 0.155 \| 154 \| 96 \| 10 \| 6 \| 142 \| 629 \| 765 \| Protocadherin Fat 1 \| Protocadherin Fat 1; afdb-proteome \| AF-F1LU58-F1-MODEL\_V4 \| 0.9 \| 0.003684 \| 67 \| 0.095 \| 200 \| 137 \| 10 \| 3 \| 167 \| 9 \| 199 \| Triggering receptor-expressed on myeloid cells-like 4 \| Triggering receptor-expressed on myeloid cells-like 4; afdb-proteome \| AF-Q5Z9P6-F1-MODEL\_V4 \| 0.9 \| 0.002584 \| 67 \| 0.121 \| 181 \| 107 \| 14 \| 1 \| 145 \| 71 \| 235 \| Signal sequence receptor subunit alpha \| Signal sequence receptor subunit alpha; afdb-proteome \| AF-F6TUL9-F1-MODEL\_V4 \| 0.9 \| 0.001813 \| 67 \| 0.115 \| 216 \| 113 \| 15 \| 7 \| 170 \| 26 \| 215 \| V-set and immunoglobulin domain-containing 4 \| V-set and immunoglobulin domain-containing 4; afdb-proteome \| AF-Q8K558-F1-MODEL\_V4 \| 0.9 \| 0.00591 \| 67 \| 0.111 \| 225 \| 119 \| 13 \| 3 \| 179 \| 20 \| 211 \| Trem-like transcript 1 protein \| Trem-like transcript 1 protein; afdb-proteome \| AF-K7MP75-F1-MODEL\_V4 \| 0.9 \| 0.004146 \| 67 \| 0.125 \| 175 \| 118 \| 11 \| 7 \| 168 \| 392 \| 544 \| Uncharacterized protein \| Uncharacterized protein; afdb-proteome \| AF-Q9FYG2-F1-MODEL\_V4 \| 0.9 \| 0.00495 \| 67 \| 0.127 \| 173 \| 123 \| 11 \| 7 \| 168 \| 453 \| 608 \| Calmodulin-binding transcription activator 4 \| Calmodulin-binding transcription activator 4; afdb-proteome \| AF-A0A7I4KBK6-F1-MODEL\_V4 \| 0.9 \| 0.007486 \| 67 \| 0.107 \| 158 \| 106 \| 8 \| 8 \| 142 \| 703 \| 848 \| "BMA-FLN-2, isoform a \| "BMA-FLN-2, isoform a; afdb-proteome \| AF-P01732-F1-MODEL\_V4 \| 0.887 \| 0.003908 \| 66 \| 0.158 \| 221 \| 123 \| 13 \| 6 \| 179 \| 29 \| 233 \| T-cell surface glycoprotein CD8 alpha chain \| T-cell surface glycoprotein CD8 alpha chain; afdb-proteome \| AF-A0A0N4STN9-F1-MODEL\_V4 \| 0.887 \| 0.009482 \| 66 \| 0.082 \| 195 \| 121 \| 11 \| 6 \| 179 \| 144 \| 301 \| Novel immune-type receptor 1k \| Novel immune-type receptor 1k; afdb-proteome \| AF-B8A473-F1-MODEL\_V4 \| 0.887 \| 0.00591 \| 66 \| 0.086 \| 208 \| 130 \| 10 \| 9 \| 179 \| 34 \| 218 \| V-set and transmembrane domain-containing 4a \| V-set and transmembrane domain-containing 4a; afdb-proteome \| AF-P08101-F1-MODEL\_V4 \| 0.887 \| 0.00495 \| 66 \| 0.138 \| 144 \| 105 \| 7 \| 7 \| 142 \| 125 \| 257 \| Low affinity immunoglobulin gamma Fc region receptor II \| Low affinity immunoglobulin gamma Fc region receptor II; afdb-proteome \| AF-Q0JDP3-F1-MODEL\_V4 \| 0.887 \| 0.004398 \| 66 \| 0.149 \| 174 \| 119 \| 13 \| 7 \| 169 \| 456 \| 611 \| Os04g0388500 protein \| Os04g0388500 protein; afdb-proteome \| AF-Q8TBE3-F1-MODEL\_V4 \| 0.872 \| 0.005251 \| 65 \| 0.107 \| 167 \| 100 \| 10 \| 12 \| 142 \| 9 \| 162 \| Fibronectin type III domain-containing protein 9 \| Fibronectin type III domain-containing protein 9; afdb-proteome \| AF-Q57965-F1-MODEL\_V4 \| 0.872 \| 0.007057 \| 65 \| 0.13 \| 153 \| 108 \| 11 \| 7 \| 143 \| 34 \| 177 \| Uncharacterized protein MJ0545 \| Uncharacterized protein MJ0545; afdb-proteome \| AF-Q4QQR8-F1-MODEL\_V4 \| 0.872 \| 0.004398 \| 65 \| 0.09 \| 211 \| 129 \| 11 \| 9 \| 178 \| 35 \| 223 \| Uncharacterized protein RGD1310423 \| Uncharacterized protein RGD1310423; afdb-proteome \| AF-U7PRC0-F1-MODEL\_V4 \| 0.872 \| 0.009482 \| 65 \| 0.087 \| 137 \| 96 \| 6 \| 8 \| 124 \| 124 \| 251 \| Uncharacterized protein \| Uncharacterized protein; afdb-proteome \| AF-Q7TNJ4-F1-MODEL\_V4 \| 0.872 \| 0.00591 \| 65 \| 0.126 \| 150 \| 106 \| 9 \| 8 \| 142 \| 297 \| 436 \| Amphoterin-induced protein 2 \| Amphoterin-induced protein 2; afdb-proteome \| AF-Q1LVE1-F1-MODEL\_V4 \| 0.872 \| 0.00591 \| 65 \| 0.103 \| 193 \| 126 \| 11 \| 8 \| 179 \| 573 \| 739 \| Sema domain, immunoglobulin domain (Ig), short basic domain, secreted, (semaphorin) 3H \| Sema domain, immunoglobulin domain (Ig), short basic domain, secreted, (semaphorin) 3H; afdb-proteome \| AF-O75015-F1-MODEL\_V4 \| 0.855 \| 0.008938 \| 64 \| 0.12 \| 124 \| 89 \| 6 \| 7 \| 125 \| 115 \| 223 \| Low affinity immunoglobulin gamma Fc region receptor III-B \| Low affinity immunoglobulin gamma Fc region receptor III-B; afdb-proteome \| AF-A0A1D6ECQ6-F1-MODEL\_V4 \| 0.855 \| 0.007486 \| 64 \| 0.119 \| 176 \| 119 \| 12 \| 7 \| 168 \| 427 \| 580 \| Calmodulin-binding transcription activator 4 \| Calmodulin-binding transcription activator 4; afdb-proteome \| AF-Q4DYJ9-F1-MODEL\_V4 \| 0.855 \| 0.008938 \| 64 \| 0.102 \| 214 \| 126 \| 17 \| 9 \| 168 \| 1080 \| 1281 \| Uncharacterized protein \| Uncharacterized protein; afdb-proteome \| AF-Q9D659-F1-MODEL\_V4 \| 0.837 \| 0.00591 \| 63 \| 0.105 \| 228 \| 121 \| 14 \| 9 \| 179 \| 43 \| 244 \| V-type immunoglobulin domain-containing suppressor of T-cell activation \| V-type immunoglobulin domain-containing suppressor of T-cell activation; afdb-proteome \| AF-Q8IW00-F1-MODEL\_V4 \| 0.837 \| 0.007942 \| 63 \| 0.109 \| 220 \| 119 \| 14 \| 9 \| 179 \| 35 \| 226 \| V-set and transmembrane domain-containing protein 4 \| V-set and transmembrane domain-containing protein 4; afdb-proteome \| AF-O75022-F1-MODEL\_V4 \| 0.817 \| 0.003908 \| 62 \| 0.104 \| 192 \| 119 \| 11 \| 8 \| 179 \| 333 \| 491 \| Leukocyte immunoglobulin-like receptor subfamily B member 3 \| Leukocyte immunoglobulin-like receptor subfamily B member 3; afdb-proteome \| AF-Q18438-F1-MODEL\_V4 \| 0.795 \| 0.001813 \| 61 \| 0.075 \| 198 \| 127 \| 11 \| 7 \| 158 \| 15 \| 202 \| Motile sperm domain-containing protein 1 \| Motile sperm domain-containing protein 1; afdb-proteome \| AF-P08637-F1-MODEL\_V4 \| 0.795 \| 0.005571 \| 61 \| 0.096 \| 176 \| 118 \| 8 \| 7 \| 177 \| 115 \| 254 \| Low affinity immunoglobulin gamma Fc region receptor III-A \| Low affinity immunoglobulin gamma Fc region receptor III-A; afdb-proteome \| AF-O15041-F1-MODEL\_V4 \| 0.772 \| 0.007057 \| 60 \| 0.111 \| 198 \| 125 \| 12 \| 9 \| 179 \| 591 \| 764 \| Semaphorin-3E \| Semaphorin-3E; afdb-proteome \| AF-Q5RJS6-F1-MODEL\_V4 \| 0.747 \| 0.005251 \| 59 \| 0.086 \| 196 \| 128 \| 8 \| 8 \| 159 \| 24 \| 212 \| Motile sperm domain-containing protein 1 \| Motile sperm domain-containing protein 1; afdb-proteome \| AF-F1R9R3-F1-MODEL\_V4 \| 0.747 \| 0.003684 \| 59 \| 0.107 \| 205 \| 123 \| 11 \| 7 \| 159 \| 22 \| 218 \| Motile sperm domain-containing protein 1 \| Motile sperm domain-containing protein 1; afdb-proteome \| AF-Q9D5K1-F1-MODEL\_V4 \| 0.747 \| 0.004398 \| 59 \| 0.144 \| 201 \| 110 \| 17 \| 9 \| 178 \| 90 \| 259 \| Transmembrane protein 81 \| Transmembrane protein 81; afdb-proteome \| AF-G4LVE3-F1-MODEL\_V4 \| 0.747 \| 0.001349 \| 59 \| 0.137 \| 167 \| 102 \| 12 \| 7 \| 141 \| 13 \| 169 \| MSP domain-containing protein \| MSP domain-containing protein; afdb-proteome \| AF-Q9C0C4-F1-MODEL\_V4 \| 0.747 \| 0.007942 \| 59 \| 0.137 \| 203 \| 108 \| 13 \| 6 \| 179 \| 561 \| 725 \| Semaphorin-4C \| Semaphorin-4C; afdb-proteome \| AF-Q59YF4-F1-MODEL\_V4 \| 0.72 \| 0.00627 \| 58 \| 0.106 \| 187 \| 100 \| 12 \| 6 \| 142 \| 43 \| 212 \| Increased recombination centers protein 22-1 \| Increased recombination centers protein 22-1; afdb-proteome \| AF-A0A0G2KSM2-F1-MODEL\_V4 \| 0.72 \| 0.008938 \| 58 \| 0.097 \| 185 \| 111 \| 13 \| 8 \| 142 \| 794 \| 972 \| Uncharacterized protein \| Uncharacterized protein; afdb-proteome \| AF-Q59XW9-F1-MODEL\_V4 \| 0.692 \| 0.00627 \| 57 \| 0.112 \| 187 \| 99 \| 12 \| 6 \| 142 \| 43 \| 212 \| Increased recombination centers protein 22-2 \| Increased recombination centers protein 22-2; afdb-proteome \| AF-Q4CMW8-F1-MODEL\_V4 \| 0.692 \| 0.002164 \| 57 \| 0.114 \| 244 \| 131 \| 13 \| 9 \| 178 \| 679 \| 911 \| Uncharacterized protein \| Uncharacterized protein; afdb-proteome \| AF-D3ZAE6-F1-MODEL\_V4 \| 0.663 \| 0.007057 \| 56 \| 0.139 \| 186 \| 100 \| 8 \| 12 \| 178 \| 473 \| 617 \| RCG49849 \| RCG49849; afdb-proteome \| AF-F6PZL4-F1-MODEL\_V4 \| 0.663 \| 0.008938 \| 56 \| 0.089 \| 178 \| 107 \| 10 \| 7 \| 163 \| 533 \| 676 \| Predicted gene 15448 \| Predicted gene 15448; afdb-proteome \| AF-F8VQ94-F1-MODEL\_V4 \| 0.632 \| 0.009482 \| 55 \| 0.099 \| 181 \| 106 \| 11 \| 5 \| 163 \| 529 \| 674 \| Paired-Ig-like receptor A2 \| Paired-Ig-like receptor A2; afdb-proteome \| AF-Q9UJG1-F1-MODEL\_V4 \| 0.537 \| 0.007942 \| 52 \| 0.094 \| 201 \| 121 \| 11 \| 8 \| 159 \| 24 \| 212 \| Motile sperm domain-containing protein 1 \| Motile sperm domain-containing protein 1; afdb-proteome \| AF-F1QU86-F1-MODEL\_V4 \| 0.326 \| 0.008425 \| 45 \| 0.126 \| 197 \| 114 \| 10 \| 9 \| 179 \| 600 \| 764 \| Sema domain, immunoglobulin domain (Ig), transmembrane domain (TM) and short cytoplasmic domain, (semaphorin) 4Bb \| Sema domain, immunoglobulin domain (Ig), transmembrane domain (TM) and short cytoplasmic domain, (semaphorin) 4Bb; afdb-uniprot50 \| AF-E4PPT5-F1-MODEL\_V4 \| 1.0 \| 1.554e-22 \| 848 \| 0.514 \| 171 \| 83 \| 0 \| 4 \| 174 \| 2 \| 172 \| Uncharacterized protein \| Uncharacterized protein; afdb-uniprot50 \| AF-A0A515BE33-F1-MODEL\_V4 \| 1.0 \| 5.702e-22 \| 820 \| 0.502 \| 177 \| 86 \| 2 \| 5 \| 179 \| 3 \| 179 \| Uncharacterized protein \| Uncharacterized protein; afdb-uniprot50 \| AF-A0A291LZK4-F1-MODEL\_V4 \| 1.0 \| 3.35e-22 \| 816 \| 0.485 \| 173 \| 89 \| 0 \| 6 \| 178 \| 5 \| 177 \| Uncharacterized protein \| Uncharacterized protein; afdb-uniprot50 \| AF-A0A143DE12-F1-MODEL\_V4 \| 1.0 \| 1.752e-21 \| 799 \| 0.439 \| 173 \| 97 \| 0 \| 6 \| 178 \| 4 \| 176 \| Uncharacterized protein \| Uncharacterized protein; afdb-uniprot50 \| AF-A0A1H9YC26-F1-MODEL\_V4 \| 1.0 \| 2.228e-19 \| 752 \| 0.342 \| 175 \| 115 \| 0 \| 4 \| 178 \| 2 \| 176 \| Uncharacterized protein \| Uncharacterized protein; afdb-uniprot50 \| AF-A0A4V1V5L7-F1-MODEL\_V4 \| 1.0 \| 6.443e-20 \| 729 \| 0.404 \| 173 \| 103 \| 0 \| 6 \| 178 \| 24 \| 196 \| Uncharacterized protein \| Uncharacterized protein; afdb-uniprot50 \| AF-A0A846VV37-F1-MODEL\_V4 \| 1.0 \| 1.03e-21 \| 705 \| 0.477 \| 174 \| 91 \| 0 \| 5 \| 178 \| 4 \| 177 \| Uncharacterized protein \| Uncharacterized protein; afdb-uniprot50 \| AF-A0A3A0FKV7-F1-MODEL\_V4 \| 1.0 \| 5.736e-19 \| 698 \| 0.407 \| 179 \| 102 \| 2 \| 4 \| 178 \| 5 \| 183 \| Uncharacterized protein \| Uncharacterized protein; afdb-uniprot50 \| AF-A0A1L2B383-F1-MODEL\_V4 \| 1.0 \| 1.756e-20 \| 686 \| 0.408 \| 181 \| 95 \| 2 \| 3 \| 178 \| 7 \| 180 \| Uncharacterized protein \| Uncharacterized protein; afdb-uniprot50 \| AF-A0A6A9VNP5-F1-MODEL\_V4 \| 1.0 \| 1.984e-18 \| 681 \| 0.314 \| 175 \| 117 \| 3 \| 6 \| 179 \| 3 \| 175 \| Uncharacterized protein \| Uncharacterized protein; afdb-uniprot50 \| AF-A0A831XDR5-F1-MODEL\_V4 \| 1.0 \| 1.038e-17 \| 677 \| 0.307 \| 179 \| 118 \| 4 \| 5 \| 178 \| 3 \| 180 \| Uncharacterized protein \| Uncharacterized protein; afdb-uniprot50 \| AF-A0A2N1AP69-F1-MODEL\_V4 \| 1.0 \| 1.036e-18 \| 667 \| 0.518 \| 164 \| 78 \| 1 \| 16 \| 179 \| 1 \| 163 \| Uncharacterized protein \| Uncharacterized protein; afdb-uniprot50 \| AF-A0A0H3ZTY7-F1-MODEL\_V4 \| 1.0 \| 4.285e-17 \| 664 \| 0.345 \| 159 \| 103 \| 1 \| 5 \| 163 \| 2 \| 159 \| Uncharacterized protein \| Uncharacterized protein; afdb-uniprot50 \| AF-A0A4Y3TFT0-F1-MODEL\_V4 \| 1.0 \| 3.189e-17 \| 649 \| 0.35 \| 177 \| 110 \| 5 \| 7 \| 179 \| 4 \| 179 \| Uncharacterized protein \| Uncharacterized protein; afdb-uniprot50 \| AF-A0A359LUV7-F1-MODEL\_V4 \| 1.0 \| 3.189e-17 \| 647 \| 0.359 \| 181 \| 109 \| 4 \| 4 \| 178 \| 6 \| 185 \| Uncharacterized protein \| Uncharacterized protein; afdb-uniprot50 \| AF-A0A3C0NSL4-F1-MODEL\_V4 \| 1.0 \| 3.383e-17 \| 643 \| 0.297 \| 178 \| 120 \| 2 \| 6 \| 178 \| 1 \| 178 \| Uncharacterized protein \| Uncharacterized protein; afdb-uniprot50 \| AF-A0A5Y3B0U0-F1-MODEL\_V4 \| 1.0 \| 1.314e-17 \| 638 \| 0.503 \| 159 \| 79 \| 0 \| 21 \| 179 \| 4 \| 162 \| Uncharacterized protein \| Uncharacterized protein; afdb-uniprot50 \| AF-A0A1G2ZKV4-F1-MODEL\_V4 \| 1.0 \| 1.038e-17 \| 630 \| 0.311 \| 183 \| 116 \| 4 \| 4 \| 179 \| 5 \| 184 \| Uncharacterized protein \| Uncharacterized protein; afdb-uniprot50 \| AF-U5QDV1-F1-MODEL\_V4 \| 1.0 \| 7.294e-17 \| 624 \| 0.376 \| 170 \| 103 \| 3 \| 5 \| 171 \| 2 \| 171 \| Uncharacterized protein \| Uncharacterized protein; afdb-uniprot50 \| AF-A0A369R582-F1-MODEL\_V4 \| 1.0 \| 1.241e-16 \| 616 \| 0.308 \| 178 \| 119 \| 3 \| 4 \| 177 \| 6 \| 183 \| Uncharacterized protein \| Uncharacterized protein; afdb-uniprot50 \| AF-A0A0U5FTR8-F1-MODEL\_V4 \| 1.0 \| 8.209e-17 \| 601 \| 0.385 \| 184 \| 102 \| 7 \| 4 \| 179 \| 6 \| 186 \| Putative phage-related protein \| Putative phage-related protein; afdb-uniprot50 \| AF-A0A7J0BYR9-F1-MODEL\_V4 \| 1.0 \| 1.482e-16 \| 600 \| 0.327 \| 159 \| 101 \| 3 \| 6 \| 163 \| 4 \| 157 \| Uncharacterized protein \| Uncharacterized protein; afdb-uniprot50 \| AF-A0A7X2TJT6-F1-MODEL\_V4 \| 1.0 \| 1.763e-18 \| 599 \| 0.469 \| 164 \| 86 \| 1 \| 16 \| 178 \| 1 \| 164 \| Uncharacterized protein \| Uncharacterized protein; afdb-uniprot50 \| AF-A0A542RRX4-F1-MODEL\_V4 \| 1.0 \| 1.241e-16 \| 597 \| 0.312 \| 179 \| 116 \| 4 \| 4 \| 178 \| 6 \| 181 \| Uncharacterized protein \| Uncharacterized protein; afdb-uniprot50 \| AF-A0A7M1L5E3-F1-MODEL\_V4 \| 1.0 \| 3.012e-16 \| 595 \| 0.301 \| 176 \| 118 \| 3 \| 6 \| 179 \| 8 \| 180 \| Uncharacterized protein \| Uncharacterized protein; afdb-uniprot50 \| AF-A0A679JLL1-F1-MODEL\_V4 \| 1.0 \| 5.439e-16 \| 593 \| 0.312 \| 176 \| 113 \| 4 \| 6 \| 179 \| 9 \| 178 \| Uncharacterized protein \| Uncharacterized protein; afdb-uniprot50 \| AF-A0A6M3J2Y0-F1-MODEL\_V4 \| 1.0 \| 1.097e-19 \| 591 \| 0.386 \| 176 \| 104 \| 4 \| 6 \| 179 \| 2 \| 175 \| Uncharacterized protein \| Uncharacterized protein; afdb-uniprot50 \| AF-A0A2G6D355-F1-MODEL\_V4 \| 1.0 \| 2.383e-15 \| 591 \| 0.263 \| 167 \| 117 \| 4 \| 5 \| 166 \| 5 \| 170 \| Uncharacterized protein \| Uncharacterized protein; afdb-uniprot50 \| AF-A0A1F9B6L6-F1-MODEL\_V4 \| 1.0 \| 1.773e-15 \| 589 \| 0.282 \| 177 \| 122 \| 3 \| 6 \| 178 \| 8 \| 183 \| Uncharacterized protein \| Uncharacterized protein; afdb-uniprot50 \| AF-A0A2A5ADZ7-F1-MODEL\_V4 \| 1.0 \| 5.439e-16 \| 583 \| 0.3 \| 180 \| 116 \| 7 \| 7 \| 177 \| 10 \| 188 \| Uncharacterized protein \| Uncharacterized protein; afdb-uniprot50 \| AF-A0A3A4NQ75-F1-MODEL\_V4 \| 1.0 \| 1.32e-15 \| 580 \| 0.297 \| 178 \| 119 \| 4 \| 6 \| 179 \| 7 \| 182 \| Uncharacterized protein \| Uncharacterized protein; afdb-uniprot50 \| AF-A0A7C8LXQ6-F1-MODEL\_V4 \| 1.0 \| 5.77e-16 \| 578 \| 0.298 \| 174 \| 111 \| 4 \| 6 \| 178 \| 5 \| 168 \| Uncharacterized protein \| Uncharacterized protein; afdb-uniprot50 \| AF-A0A3B9NUU4-F1-MODEL\_V4 \| 1.0 \| 1.4e-15 \| 574 \| 0.306 \| 183 \| 113 \| 7 \| 6 \| 178 \| 8 \| 186 \| Uncharacterized protein \| Uncharacterized protein; afdb-uniprot50 \| AF-A0A5N0TG52-F1-MODEL\_V4 \| 1.0 \| 4.294e-16 \| 571 \| 0.287 \| 174 \| 118 \| 3 \| 7 \| 179 \| 5 \| 173 \| Uncharacterized protein \| Uncharacterized protein; afdb-uniprot50 \| AF-A0A1X7L1C8-F1-MODEL\_V4 \| 1.0 \| 1.77e-16 \| 569 \| 0.361 \| 177 \| 100 \| 4 \| 6 \| 178 \| 4 \| 171 \| Uncharacterized protein \| Uncharacterized protein; afdb-uniprot50 \| AF-A0A1V6GEE1-F1-MODEL\_V4 \| 1.0 \| 3.012e-16 \| 569 \| 0.27 \| 196 \| 121 \| 6 \| 5 \| 178 \| 4 \| 199 \| Uncharacterized protein \| Uncharacterized protein; afdb-uniprot50 \| AF-W0E3G9-F1-MODEL\_V4 \| 1.0 \| 4.048e-16 \| 568 \| 0.306 \| 186 \| 117 \| 6 \| 4 \| 179 \| 7 \| 190 \| Uncharacterized protein \| Uncharacterized protein; afdb-uniprot50 \| AF-A0A254T8E3-F1-MODEL\_V4 \| 1.0 \| 2.383e-15 \| 568 \| 0.234 \| 183 \| 130 \| 6 \| 4 \| 178 \| 15 \| 195 \| Uncharacterized protein \| Uncharacterized protein; afdb-uniprot50 \| AF-A0A6H2H8M8-F1-MODEL\_V4 \| 1.0 \| 1.576e-15 \| 566 \| 0.314 \| 178 \| 114 \| 5 \| 7 \| 178 \| 13 \| 188 \| Uncharacterized protein \| Uncharacterized protein; afdb-uniprot50 \| AF-A0A2W5J2E3-F1-MODEL\_V4 \| 1.0 \| 1.996e-15 \| 561 \| 0.284 \| 179 \| 119 \| 5 \| 8 \| 179 \| 4 \| 180 \| Uncharacterized protein \| Uncharacterized protein; afdb-uniprot50 \| AF-A0A3C1GA42-F1-MODEL\_V4 \| 1.0 \| 1.485e-15 \| 557 \| 0.275 \| 185 \| 122 \| 6 \| 6 \| 179 \| 7 \| 190 \| Uncharacterized protein \| Uncharacterized protein; afdb-uniprot50 \| AF-A0A2E1P353-F1-MODEL\_V4 \| 1.0 \| 1.485e-15 \| 552 \| 0.297 \| 188 \| 116 \| 7 \| 4 \| 178 \| 12 \| 196 \| Uncharacterized protein \| Uncharacterized protein; afdb-uniprot50 \| AF-A0A3S5CWM4-F1-MODEL\_V4 \| 1.0 \| 8.726e-16 \| 551 \| 0.267 \| 176 \| 121 \| 5 \| 5 \| 179 \| 2 \| 170 \| Uncharacterized protein \| Uncharacterized protein; afdb-uniprot50 \| AF-Z9JLC1-F1-MODEL\_V4 \| 1.0 \| 2.523e-16 \| 550 \| 0.386 \| 137 \| 84 \| 0 \| 6 \| 142 \| 10 \| 146 \| Uncharacterized protein \| Uncharacterized protein; afdb-uniprot50 \| AF-A0A2E2KNA8-F1-MODEL\_V4 \| 1.0 \| 3.808e-17 \| 549 \| 0.335 \| 176 \| 112 \| 4 \| 6 \| 179 \| 3 \| 175 \| Uncharacterized protein \| Uncharacterized protein; afdb-uniprot50 \| AF-A0A843YIL2-F1-MODEL\_V4 \| 1.0 \| 5.439e-16 \| 549 \| 0.302 \| 185 \| 117 \| 6 \| 1 \| 178 \| 2 \| 181 \| Uncharacterized protein \| Uncharacterized protein; afdb-uniprot50 \| AF-A0A1C2K2F5-F1-MODEL\_V4 \| 1.0 \| 1.101e-17 \| 544 \| 0.348 \| 175 \| 108 \| 4 \| 6 \| 179 \| 2 \| 171 \| Uncharacterized protein \| Uncharacterized protein; afdb-uniprot50 \| AF-A0A4T2A8A7-F1-MODEL\_V4 \| 1.0 \| 1.675e-14 \| 543 \| 0.321 \| 174 \| 108 \| 4 \| 6 \| 178 \| 5 \| 169 \| Uncharacterized protein \| Uncharacterized protein; afdb-uniprot50 \| AF-A0A1Y5TZC7-F1-MODEL\_V4 \| 1.0 \| 4.842e-15 \| 540 \| 0.316 \| 183 \| 116 \| 5 \| 4 \| 179 \| 6 \| 186 \| Uncharacterized protein \| Uncharacterized protein; afdb-uniprot50 \| AF-A0A0S9N2Z7-F1-MODEL\_V4 \| 1.0 \| 5.137e-15 \| 539 \| 0.284 \| 183 \| 119 \| 7 \| 6 \| 179 \| 6 \| 185 \| Uncharacterized protein \| Uncharacterized protein; afdb-uniprot50 \| AF-A0A423HS27-F1-MODEL\_V4 \| 1.0 \| 4.564e-15 \| 536 \| 0.287 \| 174 \| 118 \| 4 \| 6 \| 178 \| 2 \| 170 \| Uncharacterized protein \| Uncharacterized protein; afdb-uniprot50 \| AF-A0A2D5ZTJ7-F1-MODEL\_V4 \| 1.0 \| 1.885e-14 \| 531 \| 0.225 \| 173 \| 131 \| 3 \| 6 \| 177 \| 3 \| 173 \| Uncharacterized protein \| Uncharacterized protein; afdb-uniprot50 \| AF-A0A6L4BCL0-F1-MODEL\_V4 \| 1.0 \| 8.225e-16 \| 530 \| 0.302 \| 175 \| 112 \| 3 \| 5 \| 178 \| 2 \| 167 \| Uncharacterized protein \| Uncharacterized protein; afdb-uniprot50 \| AF-A0A368L7T0-F1-MODEL\_V4 \| 1.0 \| 1.246e-14 \| 524 \| 0.247 \| 186 \| 128 \| 6 \| 6 \| 179 \| 7 \| 192 \| Uncharacterized protein \| Uncharacterized protein; afdb-uniprot50 \| AF-Q1ILS4-F1-MODEL\_V4 \| 1.0 \| 6.481e-17 \| 524 \| 0.268 \| 175 \| 127 \| 1 \| 5 \| 178 \| 7 \| 181 \| Uncharacterized protein \| Uncharacterized protein; afdb-uniprot50 \| AF-A0A661FDS9-F1-MODEL\_V4 \| 1.0 \| 3.83e-14 \| 522 \| 0.259 \| 181 \| 125 \| 6 \| 6 \| 178 \| 3 \| 182 \| Uncharacterized protein \| Uncharacterized protein; afdb-uniprot50 \| AF-A0A1F9VFN4-F1-MODEL\_V4 \| 1.0 \| 4.833e-16 \| 520 \| 0.343 \| 160 \| 102 \| 3 \| 7 \| 164 \| 22 \| 180 \| Uncharacterized protein \| Uncharacterized protein; afdb-uniprot50 \| AF-E7C7Y6-F1-MODEL\_V4 \| 1.0 \| 3.611e-14 \| 518 \| 0.254 \| 185 \| 124 \| 6 \| 4 \| 176 \| 12 \| 194 \| Uncharacterized protein \| Uncharacterized protein; afdb-uniprot50 \| AF-A0A2V5PK36-F1-MODEL\_V4 \| 1.0 \| 1.877e-16 \| 518 \| 0.349 \| 163 \| 99 \| 5 \| 5 \| 161 \| 24 \| 185 \| Uncharacterized protein \| Uncharacterized protein; afdb-uniprot50 \| AF-A0A2Z6UL02-F1-MODEL\_V4 \| 1.0 \| 2.004e-13 \| 515 \| 0.258 \| 170 \| 121 \| 4 \| 9 \| 177 \| 5 \| 170 \| Phage-related protein \| Phage-related protein; afdb-uniprot50 \| AF-A0A0S8DP01-F1-MODEL\_V4 \| 1.0 \| 1.675e-14 \| 513 \| 0.225 \| 186 \| 132 \| 5 \| 6 \| 179 \| 3 \| 188 \| Uncharacterized protein \| Uncharacterized protein; afdb-uniprot50 \| AF-A0A2A5BDD7-F1-MODEL\_V4 \| 1.0 \| 9.859e-14 \| 511 \| 0.26 \| 165 \| 116 \| 4 \| 6 \| 166 \| 7 \| 169 \| Uncharacterized protein \| Uncharacterized protein; afdb-uniprot50 \| AF-A0A3B9NQ74-F1-MODEL\_V4 \| 1.0 \| 8.258e-14 \| 511 \| 0.26 \| 184 \| 121 \| 6 \| 6 \| 178 \| 14 \| 193 \| Uncharacterized protein \| Uncharacterized protein; afdb-uniprot50 \| AF-A0A838M0T7-F1-MODEL\_V4 \| 1.0 \| 1.044e-14 \| 509 \| 0.241 \| 182 \| 124 \| 6 \| 7 \| 179 \| 4 \| 180 \| Uncharacterized protein \| Uncharacterized protein; afdb-uniprot50 \| AF-A0A165R292-F1-MODEL\_V4 \| 1.0 \| 3.215e-13 \| 504 \| 0.26 \| 184 \| 126 \| 5 \| 4 \| 179 \| 11 \| 192 \| Uncharacterized protein \| Uncharacterized protein; afdb-uniprot50 \| AF-A0A6A4RDA2-F1-MODEL\_V4 \| 1.0 \| 3.823e-15 \| 503 \| 0.36 \| 147 \| 91 \| 3 \| 7 \| 150 \| 3 \| 149 \| Uncharacterized protein \| Uncharacterized protein; afdb-uniprot50 \| AF-A0A1T1IZJ5-F1-MODEL\_V4 \| 1.0 \| 7.769e-15 \| 498 \| 0.405 \| 143 \| 80 \| 2 \| 1 \| 141 \| 1 \| 140 \| Uncharacterized protein \| Uncharacterized protein; afdb-uniprot50 \| AF-A0A2E0ENC4-F1-MODEL\_V4 \| 1.0 \| 2.121e-14 \| 494 \| 0.248 \| 181 \| 127 \| 5 \| 4 \| 179 \| 6 \| 182 \| Uncharacterized protein \| Uncharacterized protein; afdb-uniprot50 \| AF-A0A7V8KQN1-F1-MODEL\_V4 \| 1.0 \| 3.403e-14 \| 492 \| 0.285 \| 182 \| 118 \| 6 \| 4 \| 178 \| 10 \| 186 \| Uncharacterized protein \| Uncharacterized protein; afdb-uniprot50 \| AF-A0A1Z9Q622-F1-MODEL\_V4 \| 1.0 \| 4.311e-14 \| 488 \| 0.226 \| 181 \| 128 \| 6 \| 7 \| 178 \| 15 \| 192 \| Uncharacterized protein \| Uncharacterized protein; afdb-uniprot50 \| AF-A0A2D7GUN5-F1-MODEL\_V4 \| 1.0 \| 1.678e-13 \| 487 \| 0.274 \| 175 \| 117 \| 6 \| 6 \| 178 \| 2 \| 168 \| Uncharacterized protein \| Uncharacterized protein; afdb-uniprot50 \| AF-A0A7G8BY96-F1-MODEL\_V4 \| 1.0 \| 1.406e-13 \| 487 \| 0.245 \| 179 \| 129 \| 4 \| 6 \| 179 \| 21 \| 198 \| Uncharacterized protein \| Uncharacterized protein; afdb-uniprot50 \| AF-A0A5P9IVF8-F1-MODEL\_V4 \| 1.0 \| 1.046e-13 \| 484 \| 0.264 \| 178 \| 125 \| 5 \| 1 \| 175 \| 1 \| 175 \| Uncharacterized protein \| Uncharacterized protein; afdb-uniprot50 \| AF-A0A7X8AH15-F1-MODEL\_V4 \| 1.0 \| 6.519e-14 \| 481 \| 0.308 \| 185 \| 107 \| 5 \| 14 \| 178 \| 1 \| 184 \| Uncharacterized protein \| Uncharacterized protein; afdb-uniprot50 \| AF-A0A653JDQ3-F1-MODEL\_V4 \| 1.0 \| 4.573e-14 \| 479 \| 0.274 \| 182 \| 119 \| 6 \| 4 \| 178 \| 6 \| 181 \| Uncharacterized protein \| Uncharacterized protein; afdb-uniprot50 \| AF-A0A1M7FPB6-F1-MODEL\_V4 \| 1.0 \| 6.507e-15 \| 477 \| 0.219 \| 182 \| 134 \| 3 \| 6 \| 179 \| 3 \| 184 \| Uncharacterized protein \| Uncharacterized protein; afdb-uniprot50 \| AF-A0A7T6ARA8-F1-MODEL\_V4 \| 1.0 \| 9.859e-14 \| 472 \| 0.237 \| 177 \| 125 \| 6 \| 6 \| 179 \| 8 \| 177 \| Uncharacterized protein \| Uncharacterized protein; afdb-uniprot50 \| AF-A0A1J5SRA9-F1-MODEL\_V4 \| 1.0 \| 3.403e-14 \| 471 \| 0.196 \| 183 \| 138 \| 4 \| 6 \| 179 \| 3 \| 185 \| Uncharacterized protein \| Uncharacterized protein; afdb-uniprot50 \| AF-A0A1B3WBU8-F1-MODEL\_V4 \| 1.0 \| 6.121e-16 \| 466 \| 0.341 \| 173 \| 106 \| 5 \| 7 \| 178 \| 6 \| 171 \| Uncharacterized protein \| Uncharacterized protein; afdb-uniprot50 \| AF-A0A2A5DGI2-F1-MODEL\_V4 \| 1.0 \| 4.582e-13 \| 466 \| 0.24 \| 166 \| 118 \| 5 \| 7 \| 166 \| 4 \| 167 \| Uncharacterized protein \| Uncharacterized protein; afdb-uniprot50 \| AF-A0A2E7WAD6-F1-MODEL\_V4 \| 1.0 \| 4.852e-14 \| 466 \| 0.225 \| 186 \| 131 \| 5 \| 7 \| 179 \| 4 \| 189 \| Uncharacterized protein \| Uncharacterized protein; afdb-uniprot50 \| AF-A0A1M7U224-F1-MODEL\_V4 \| 1.0 \| 3.215e-13 \| 465 \| 0.272 \| 180 \| 118 \| 6 \| 6 \| 179 \| 11 \| 183 \| Uncharacterized protein \| Uncharacterized protein; afdb-uniprot50 \| AF-A0A0Q7SMP9-F1-MODEL\_V4 \| 1.0 \| 1.78e-13 \| 461 \| 0.266 \| 180 \| 124 \| 5 \| 6 \| 177 \| 5 \| 184 \| Uncharacterized protein \| Uncharacterized protein; afdb-uniprot50 \| AF-A0A1L3ZRN8-F1-MODEL\_V4 \| 1.0 \| 4.072e-13 \| 461 \| 0.306 \| 189 \| 110 \| 8 \| 6 \| 178 \| 12 \| 195 \| Uncharacterized protein \| Uncharacterized protein; afdb-uniprot50 \| AF-A0A844IFC1-F1-MODEL\_V4 \| 1.0 \| 1.585e-12 \| 458 \| 0.214 \| 177 \| 120 \| 5 \| 6 \| 166 \| 5 \| 178 \| Uncharacterized protein \| Uncharacterized protein; afdb-uniprot50 \| AF-A0A2M7YUE3-F1-MODEL\_V4 \| 1.0 \| 1.322e-14 \| 456 \| 0.303 \| 178 \| 116 \| 5 \| 4 \| 178 \| 6 \| 178 \| Uncharacterized protein \| Uncharacterized protein; afdb-uniprot50 \| AF-A0A286GPM6-F1-MODEL\_V4 \| 1.0 \| 9.879e-13 \| 438 \| 0.263 \| 171 \| 115 \| 3 \| 9 \| 179 \| 2 \| 161 \| Uncharacterized protein \| Uncharacterized protein; afdb-uniprot50 \| AF-A0A1G0RDT9-F1-MODEL\_V4 \| 1.0 \| 2.004e-13 \| 436 \| 0.265 \| 177 \| 119 \| 7 \| 7 \| 179 \| 6 \| 175 \| Uncharacterized protein \| Uncharacterized protein; afdb-uniprot50 \| AF-A0A501WT91-F1-MODEL\_V4 \| 1.0 \| 3.625e-12 \| 423 \| 0.25 \| 180 \| 118 \| 6 \| 7 \| 179 \| 10 \| 179 \| Uncharacterized protein \| Uncharacterized protein; afdb-uniprot50 \| AF-A0A6H1ZNP5-F1-MODEL\_V4 \| 1.0 \| 7.815e-12 \| 407 \| 0.229 \| 174 \| 127 \| 4 \| 9 \| 179 \| 16 \| 185 \| Uncharacterized protein \| Uncharacterized protein; afdb-uniprot50 \| AF-F5R899-F1-MODEL\_V4 \| 1.0 \| 1.892e-12 \| 407 \| 0.272 \| 180 \| 122 \| 6 \| 6 \| 179 \| 8 \| 184 \| Phage-related protein \| Phage-related protein; afdb-uniprot50 \| AF-A0A6M4Y9K2-F1-MODEL\_V4 \| 1.0 \| 8.795e-12 \| 399 \| 0.558 \| 136 \| 58 \| 1 \| 46 \| 179 \| 3 \| 138 \| Uncharacterized protein \| Uncharacterized protein; afdb-uniprot50 \| AF-A0A7Y0NUN9-F1-MODEL\_V4 \| 1.0 \| 1.182e-11 \| 393 \| 0.144 \| 159 \| 130 \| 4 \| 6 \| 160 \| 8 \| 164 \| Uncharacterized protein \| Uncharacterized protein; afdb-uniprot50 \| AF-A0A0F9IDA6-F1-MODEL\_V4 \| 1.0 \| 7.352e-13 \| 389 \| 0.278 \| 187 \| 124 \| 5 \| 1 \| 179 \| 1 \| 184 \| Uncharacterized protein \| Uncharacterized protein; afdb-uniprot50 \| AF-A0A1V5CWH5-F1-MODEL\_V4 \| 1.0 \| 1.78e-13 \| 389 \| 0.211 \| 194 \| 130 \| 7 \| 5 \| 179 \| 2 \| 191 \| Uncharacterized protein \| Uncharacterized protein; afdb-uniprot50 \| AF-A0A0F9JRZ2-F1-MODEL\_V4 \| 1.0 \| 5.167e-12 \| 382 \| 0.234 \| 196 \| 124 \| 8 \| 6 \| 179 \| 9 \| 200 \| Uncharacterized protein \| Uncharacterized protein; afdb-uniprot50 \| AF-A0A358KKZ0-F1-MODEL\_V4 \| 1.0 \| 2.016e-10 \| 381 \| 0.232 \| 125 \| 93 \| 2 \| 6 \| 128 \| 9 \| 132 \| Uncharacterized protein \| Uncharacterized protein; afdb-uniprot50 \| AF-A0A6L6JB30-F1-MODEL\_V4 \| 1.0 \| 8.795e-12 \| 381 \| 0.485 \| 140 \| 71 \| 1 \| 40 \| 178 \| 1 \| 140 \| Uncharacterized protein \| Uncharacterized protein; afdb-uniprot50 \| AF-A0A0N8FSS6-F1-MODEL\_V4 \| 1.0 \| 2.402e-11 \| 381 \| 0.312 \| 144 \| 89 \| 4 \| 6 \| 142 \| 12 \| 152 \| Uncharacterized protein \| Uncharacterized protein; afdb-uniprot50 \| AF-A0A3B9Q8J6-F1-MODEL\_V4 \| 1.0 \| 6.558e-11 \| 364 \| 0.209 \| 181 \| 131 \| 6 \| 6 \| 178 \| 1 \| 177 \| Uncharacterized protein \| Uncharacterized protein; afdb-uniprot50 \| AF-A0A0F9QC72-F1-MODEL\_V4 \| 1.0 \| 4.591e-12 \| 362 \| 0.239 \| 171 \| 110 \| 5 \| 6 \| 166 \| 11 \| 171 \| Uncharacterized protein \| Uncharacterized protein; afdb-uniprot50 \| AF-A0A848V5M5-F1-MODEL\_V4 \| 1.0 \| 1.259e-09 \| 356 \| 0.251 \| 159 \| 107 \| 5 \| 29 \| 177 \| 3 \| 159 \| Uncharacterized protein \| Uncharacterized protein; afdb-uniprot50 \| AF-A0A2E7ZNQ6-F1-MODEL\_V4 \| 1.0 \| 1.691e-09 \| 340 \| 0.206 \| 131 \| 88 \| 5 \| 7 \| 123 \| 4 \| 132 \| Uncharacterized protein \| Uncharacterized protein; afdb-uniprot50 \| AF-A0A3B9NUI5-F1-MODEL\_V4 \| 1.0 \| 1.333e-10 \| 339 \| 0.185 \| 183 \| 130 \| 9 \| 6 \| 178 \| 9 \| 182 \| Uncharacterized protein \| Uncharacterized protein; afdb-uniprot50 \| AF-A0A1F9UT01-F1-MODEL\_V4 \| 1.0 \| 9.918e-11 \| 317 \| 0.275 \| 189 \| 119 \| 9 \| 6 \| 179 \| 8 \| 193 \| Uncharacterized protein \| Uncharacterized protein; afdb-uniprot50 \| AF-A0A7V8FKH2-F1-MODEL\_V4 \| 1.0 \| 6.985e-09 \| 313 \| 0.518 \| 108 \| 52 \| 0 \| 57 \| 164 \| 4 \| 111 \| Uncharacterized protein \| Uncharacterized protein; afdb-uniprot50 \| AF-A0A2P1VF72-F1-MODEL\_V4 \| 1.0 \| 3.444e-08 \| 302 \| 0.33 \| 142 \| 85 \| 3 \| 40 \| 179 \| 2 \| 135 \| Uncharacterized protein \| Uncharacterized protein; afdb-uniprot50 \| AF-A0A564WI89-F1-MODEL\_V4 \| 1.0 \| 3.246e-08 \| 299 \| 0.21 \| 171 \| 121 \| 5 \| 7 \| 176 \| 3 \| 160 \| Uncharacterized protein \| Uncharacterized protein; afdb-uniprot50 \| AF-A0A2V2GR66-F1-MODEL\_V4 \| 1.0 \| 1.338e-08 \| 284 \| 0.242 \| 165 \| 103 \| 8 \| 7 \| 165 \| 1 \| 149 \| Uncharacterized protein \| Uncharacterized protein; afdb-uniprot50 \| AF-A0A432FDA3-F1-MODEL\_V4 \| 1.0 \| 9.423e-07 \| 282 \| 0.311 \| 90 \| 58 \| 3 \| 6 \| 92 \| 9 \| 97 \| Uncharacterized protein \| Uncharacterized protein; afdb-uniprot50 \| AF-A0A0F9KXQ4-F1-MODEL\_V4 \| 1.0 \| 1.194e-06 \| 264 \| 0.091 \| 98 \| 84 \| 4 \| 9 \| 102 \| 5 \| 101 \| Uncharacterized protein \| Uncharacterized protein; afdb-uniprot50 \| AF-A0A3G9FQ52-F1-MODEL\_V4 \| 1.0 \| 3.072e-06 \| 260 \| 0.553 \| 94 \| 42 \| 0 \| 86 \| 179 \| 5 \| 98 \| Uncharacterized protein \| Uncharacterized protein; afdb-uniprot50 \| AF-A0A1F8QKN5-F1-MODEL\_V4 \| 1.0 \| 8.34e-09 \| 260 \| 0.287 \| 167 \| 109 \| 4 \| 22 \| 179 \| 1 \| 166 \| Uncharacterized protein \| Uncharacterized protein; afdb-uniprot50 \| AF-A0A382W2M1-F1-MODEL\_V4 \| 1.0 \| 1.702e-06 \| 247 \| 0.21 \| 100 \| 67 \| 4 \| 9 \| 96 \| 9 \| 108 \| Uncharacterized protein \| Uncharacterized protein; afdb-uniprot50 \| AF-A0A0A2Y143-F1-MODEL\_V4 \| 1.0 \| 1.343e-06 \| 246 \| 0.17 \| 100 \| 71 \| 3 \| 9 \| 96 \| 7 \| 106 \| Uncharacterized protein \| Uncharacterized protein; afdb-uniprot50 \| AF-A0A0E3UJT2-F1-MODEL\_V4 \| 1.0 \| 1.915e-06 \| 239 \| 0.2 \| 105 \| 69 \| 5 \| 9 \| 98 \| 18 \| 122 \| Uncharacterized protein \| Uncharacterized protein; afdb-uniprot50 \| AF-A0A2E3JRL0-F1-MODEL\_V4 \| 1.0 \| 3.668e-06 \| 237 \| 0.205 \| 102 \| 70 \| 4 \| 9 \| 99 \| 10 \| 111 \| Uncharacterized protein \| Uncharacterized protein; afdb-uniprot50 \| AF-A0A2D6TNC0-F1-MODEL\_V4 \| 1.0 \| 1.915e-06 \| 236 \| 0.173 \| 104 \| 74 \| 4 \| 9 \| 100 \| 8 \| 111 \| Uncharacterized protein \| Uncharacterized protein; afdb-uniprot50 \| AF-A0A2E5AWZ1-F1-MODEL\_V4 \| 1.0 \| 4.929e-06 \| 233 \| 0.198 \| 101 \| 68 \| 4 \| 9 \| 96 \| 9 \| 109 \| Uncharacterized protein \| Uncharacterized protein; afdb-uniprot50 \| AF-A6FS73-F1-MODEL\_V4 \| 1.0 \| 2.286e-06 \| 232 \| 0.156 \| 102 \| 75 \| 3 \| 9 \| 100 \| 5 \| 105 \| Uncharacterized protein \| Uncharacterized protein; afdb-uniprot50 \| AF-A0A0F9JR02-F1-MODEL\_V4 \| 1.0 \| 3.259e-06 \| 231 \| 0.247 \| 105 \| 65 \| 6 \| 6 \| 96 \| 5 \| 109 \| Uncharacterized protein \| Uncharacterized protein; afdb-uniprot50 \| AF-A0A0F9D8Q2-F1-MODEL\_V4 \| 1.0 \| 7.027e-06 \| 228 \| 0.176 \| 102 \| 74 \| 5 \| 5 \| 96 \| 5 \| 106 \| Uncharacterized protein \| Uncharacterized protein; afdb-uniprot50 \| AF-A0A2Z3H230-F1-MODEL\_V4 \| 1.0 \| 9.997e-07 \| 228 \| 0.174 \| 109 \| 72 \| 4 \| 10 \| 100 \| 1 \| 109 \| Uncharacterized protein \| Uncharacterized protein; afdb-uniprot50 \| AF-A0A3C1DS41-F1-MODEL\_V4 \| 1.0 \| 3.892e-06 \| 227 \| 0.198 \| 106 \| 70 \| 6 \| 6 \| 96 \| 7 \| 112 \| Uncharacterized protein \| Uncharacterized protein; afdb-uniprot50 \| AF-A0A3B0VIW3-F1-MODEL\_V4 \| 1.0 \| 8.883e-07 \| 225 \| 0.13 \| 107 \| 87 \| 5 \| 7 \| 108 \| 1 \| 106 \| Uncharacterized protein \| Uncharacterized protein; afdb-uniprot50 \| AF-A0A3N5X8Y3-F1-MODEL\_V4 \| 1.0 \| 5.548e-06 \| 224 \| 0.155 \| 103 \| 72 \| 6 \| 9 \| 96 \| 10 \| 112 \| Uncharacterized protein \| Uncharacterized protein; afdb-uniprot50 \| AF-A0A2E9KUT5-F1-MODEL\_V4 \| 1.0 \| 2.426e-06 \| 223 \| 0.203 \| 113 \| 75 \| 5 \| 9 \| 106 \| 8 \| 120 \| Uncharacterized protein \| Uncharacterized protein; afdb-uniprot50 \| AF-A0A381YBA8-F1-MODEL\_V4 \| 1.0 \| 9.442e-06 \| 220 \| 0.2 \| 100 \| 68 \| 5 \| 9 \| 96 \| 11 \| 110 \| Uncharacterized protein \| Uncharacterized protein; afdb-uniprot50 \| AF-A0A7X7UJS9-F1-MODEL\_V4 \| 1.0 \| 2.286e-06 \| 219 \| 0.152 \| 105 \| 80 \| 6 \| 9 \| 104 \| 8 \| 112 \| Uncharacterized protein \| Uncharacterized protein; afdb-uniprot50 \| AF-A0A5M8IDW4-F1-MODEL\_V4 \| 1.0 \| 3.892e-06 \| 214 \| 0.175 \| 108 \| 74 \| 6 \| 6 \| 98 \| 9 \| 116 \| Uncharacterized protein \| Uncharacterized protein; afdb-uniprot50 \| AF-A0A418M5Y9-F1-MODEL\_V4 \| 1.0 \| 3.465e-05 \| 213 \| 0.175 \| 91 \| 69 \| 4 \| 9 \| 96 \| 17 \| 104 \| Uncharacterized protein \| Uncharacterized protein; afdb-uniprot50 \| AF-A0A2D6E192-F1-MODEL\_V4 \| 1.0 \| 8.9e-06 \| 213 \| 0.194 \| 103 \| 68 \| 6 \| 9 \| 96 \| 8 \| 110 \| Uncharacterized protein \| Uncharacterized protein; afdb-uniprot50 \| AF-A0A382DI07-F1-MODEL\_V4 \| 1.0 \| 3.266e-05 \| 212 \| 0.13 \| 100 \| 77 \| 4 \| 9 \| 98 \| 21 \| 120 \| Uncharacterized protein \| Uncharacterized protein; afdb-uniprot50 \| AF-A0A2I1XBL1-F1-MODEL\_V4 \| 1.0 \| 3.465e-05 \| 209 \| 0.161 \| 99 \| 74 \| 3 \| 9 \| 98 \| 8 \| 106 \| Uncharacterized protein \| Uncharacterized protein; afdb-uniprot50 \| AF-A0A1H9GD31-F1-MODEL\_V4 \| 1.0 \| 6.243e-06 \| 206 \| 0.186 \| 118 \| 74 \| 7 \| 1 \| 96 \| 1 \| 118 \| Uncharacterized protein \| Uncharacterized protein; afdb-uniprot50 \| AF-A0A378PHR1-F1-MODEL\_V4 \| 1.0 \| 4.137e-05 \| 205 \| 0.126 \| 103 \| 78 \| 6 \| 9 \| 99 \| 6 \| 108 \| Uncharacterized protein \| Uncharacterized protein; afdb-uniprot50 \| AF-A0A1B7X5C5-F1-MODEL\_V4 \| 1.0 \| 2.43e-05 \| 203 \| 0.163 \| 104 \| 74 \| 5 \| 9 \| 99 \| 13 \| 116 \| Uncharacterized protein \| Uncharacterized protein; afdb-uniprot50 \| AF-A0A0F9R4H3-F1-MODEL\_V4 \| 1.0 \| 2.578e-05 \| 203 \| 0.157 \| 108 \| 73 \| 7 \| 9 \| 98 \| 12 \| 119 \| Uncharacterized protein \| Uncharacterized protein; afdb-uniprot50 \| AF-A0A849MUW1-F1-MODEL\_V4 \| 1.0 \| 6.243e-06 \| 203 \| 0.178 \| 123 \| 79 \| 5 \| 9 \| 109 \| 9 \| 131 \| Uncharacterized protein \| Uncharacterized protein; afdb-uniprot50 \| AF-A0A1T0ACG6-F1-MODEL\_V4 \| 1.0 \| 2.735e-05 \| 200 \| 0.166 \| 108 \| 76 \| 7 \| 9 \| 104 \| 5 \| 110 \| Uncharacterized protein \| Uncharacterized protein; afdb-uniprot50 \| AF-A0A3S0DMG9-F1-MODEL\_V4 \| 1.0 \| 7.469e-05 \| 200 \| 0.115 \| 104 \| 76 \| 5 \| 9 \| 96 \| 8 \| 111 \| Uncharacterized protein \| Uncharacterized protein; afdb-uniprot50 \| AF-A0A5C7PP54-F1-MODEL\_V4 \| 1.0 \| 1.607e-05 \| 199 \| 0.168 \| 101 \| 72 \| 3 \| 9 \| 97 \| 4 \| 104 \| Uncharacterized protein \| Uncharacterized protein; afdb-uniprot50 \| AF-A0A378UGU1-F1-MODEL\_V4 \| 1.0 \| 1.809e-05 \| 199 \| 0.183 \| 109 \| 80 \| 4 \| 9 \| 108 \| 7 \| 115 \| Uncharacterized protein \| Uncharacterized protein; afdb-uniprot50 \| AF-A0A2N0B444-F1-MODEL\_V4 \| 1.0 \| 1.512e-06 \| 198 \| 0.136 \| 139 \| 98 \| 7 \| 5 \| 130 \| 4 \| 133 \| Uncharacterized protein \| Uncharacterized protein; afdb-uniprot50 \| AF-A0A2N2D0B1-F1-MODEL\_V4 \| 1.0 \| 3.899e-05 \| 197 \| 0.149 \| 107 \| 76 \| 5 \| 9 \| 107 \| 2 \| 101 \| Uncharacterized protein \| Uncharacterized protein; afdb-uniprot50 \| AF-A0A257NKB8-F1-MODEL\_V4 \| 1.0 \| 1.269e-05 \| 197 \| 0.178 \| 112 \| 77 \| 8 \| 9 \| 107 \| 6 \| 115 \| Uncharacterized protein \| Uncharacterized protein; afdb-uniprot50 \| AF-A0A497T2U7-F1-MODEL\_V4 \| 1.0 \| 3.676e-05 \| 197 \| 0.138 \| 108 \| 73 \| 6 \| 9 \| 96 \| 9 \| 116 \| Uncharacterized protein \| Uncharacterized protein; afdb-uniprot50 \| AF-A0A3N4MWR2-F1-MODEL\_V4 \| 1.0 \| 1.127e-05 \| 196 \| 0.214 \| 112 \| 74 \| 6 \| 3 \| 100 \| 2 \| 113 \| Uncharacterized protein \| Uncharacterized protein; afdb-uniprot50 \| AF-A0A1Y3RG66-F1-MODEL\_V4 \| 1.0 \| 1.604e-06 \| 196 \| 0.177 \| 124 \| 87 \| 7 \| 7 \| 124 \| 13 \| 127 \| Uncharacterized protein \| Uncharacterized protein; afdb-uniprot50 \| AF-A0A1S1CDS1-F1-MODEL\_V4 \| 1.0 \| 5.537e-07 \| 196 \| 0.168 \| 166 \| 106 \| 3 \| 9 \| 142 \| 9 \| 174 \| Uncharacterized protein \| Uncharacterized protein; afdb-uniprot50 \| AF-A0A4Q4GW66-F1-MODEL\_V4 \| 1.0 \| 2.578e-05 \| 195 \| 0.13 \| 107 \| 78 \| 7 \| 6 \| 98 \| 1 \| 106 \| Uncharacterized protein \| Uncharacterized protein; afdb-uniprot50 \| AF-A0A351GJN3-F1-MODEL\_V4 \| 1.0 \| 5.548e-06 \| 194 \| 0.166 \| 114 \| 69 \| 7 \| 9 \| 96 \| 12 \| 125 \| Uncharacterized protein \| Uncharacterized protein; afdb-uniprot50 \| AF-A0A2D5QIA3-F1-MODEL\_V4 \| 1.0 \| 1.515e-05 \| 194 \| 0.151 \| 119 \| 72 \| 5 \| 9 \| 98 \| 9 \| 127 \| Uncharacterized protein \| Uncharacterized protein; afdb-uniprot50 \| AF-A0A090MR43-F1-MODEL\_V4 \| 1.0 \| 9.461e-05 \| 193 \| 0.118 \| 110 \| 82 \| 4 \| 5 \| 99 \| 4 \| 113 \| Uncharacterized protein \| Uncharacterized protein; afdb-uniprot50 \| AF-A0A838JK26-F1-MODEL\_V4 \| 1.0 \| 8.406e-05 \| 191 \| 0.095 \| 105 \| 78 \| 5 \| 9 \| 96 \| 4 \| 108 \| Uncharacterized protein \| Uncharacterized protein; afdb-uniprot50 \| AF-A0A3E0R3L4-F1-MODEL\_V4 \| 1.0 \| 7.469e-05 \| 189 \| 0.16 \| 106 \| 71 \| 4 \| 9 \| 96 \| 8 \| 113 \| Uncharacterized protein \| Uncharacterized protein; afdb-uniprot50 \| AF-A0A2D8B2N1-F1-MODEL\_V4 \| 1.0 \| 1.428e-05 \| 189 \| 0.136 \| 117 \| 81 \| 5 \| 4 \| 100 \| 12 \| 128 \| Uncharacterized protein \| Uncharacterized protein; afdb-uniprot50 \| AF-A0A1Q5PL77-F1-MODEL\_V4 \| 1.0 \| 1.705e-05 \| 188 \| 0.123 \| 105 \| 81 \| 6 \| 8 \| 107 \| 47 \| 145 \| Uncharacterized protein \| Uncharacterized protein; afdb-uniprot50 \| AF-A0A2L0W7Z5-F1-MODEL\_V4 \| 1.0 \| 0.0001518 \| 185 \| 0.123 \| 105 \| 75 \| 5 \| 9 \| 96 \| 8 \| 112 \| Uncharacterized protein \| Uncharacterized protein; afdb-uniprot50 \| AF-A0A524LAF8-F1-MODEL\_V4 \| 1.0 \| 1.915e-06 \| 182 \| 0.153 \| 143 \| 93 \| 6 \| 9 \| 142 \| 8 \| 131 \| Uncharacterized protein \| Uncharacterized protein; afdb-uniprot50 \| AF-A0A7C5R5E6-F1-MODEL\_V4 \| 1.0 \| 5.559e-05 \| 181 \| 0.154 \| 110 \| 75 \| 5 \| 9 \| 100 \| 10 \| 119 \| Uncharacterized protein \| Uncharacterized protein; afdb-uniprot50 \| AF-A0A1Y3SHK2-F1-MODEL\_V4 \| 1.0 \| 2.896e-06 \| 180 \| 0.157 \| 146 \| 104 \| 9 \| 6 \| 144 \| 12 \| 145 \| Uncharacterized protein \| Uncharacterized protein; afdb-uniprot50 \| AF-A0A397Q662-F1-MODEL\_V4 \| 1.0 \| 0.0001431 \| 179 \| 0.109 \| 110 \| 81 \| 7 \| 9 \| 101 \| 5 \| 114 \| Uncharacterized protein \| Uncharacterized protein; afdb-uniprot50 \| AF-A0A0F9S9C7-F1-MODEL\_V4 \| 1.0 \| 8.918e-05 \| 179 \| 0.165 \| 103 \| 68 \| 5 \| 11 \| 95 \| 14 \| 116 \| Uncharacterized protein \| Uncharacterized protein; afdb-uniprot50 \| AF-A0A495J3S7-F1-MODEL\_V4 \| 1.0 \| 2.159e-05 \| 179 \| 0.133 \| 120 \| 83 \| 6 \| 9 \| 107 \| 3 \| 122 \| Uncharacterized protein \| Uncharacterized protein; afdb-uniprot50 \| AF-A0A3C0VX60-F1-MODEL\_V4 \| 1.0 \| 7.924e-05 \| 178 \| 0.21 \| 119 \| 78 \| 3 \| 25 \| 141 \| 1 \| 105 \| Uncharacterized protein \| Uncharacterized protein; afdb-uniprot50 \| AF-A0A0F9HMU7-F1-MODEL\_V4 \| 1.0 \| 3.259e-06 \| 178 \| 0.136 \| 146 \| 101 \| 7 \| 3 \| 142 \| 2 \| 128 \| Uncharacterized protein \| Uncharacterized protein; afdb-uniprot50 \| AF-A0A447CZS3-F1-MODEL\_V4 \| 1.0 \| 0.0001065 \| 177 \| 0.14 \| 114 \| 75 \| 5 \| 9 \| 105 \| 8 \| 115 \| Uncharacterized protein \| Uncharacterized protein; afdb-uniprot50 \| AF-A0A1H4UM15-F1-MODEL\_V4 \| 1.0 \| 0.0001518 \| 177 \| 0.129 \| 116 \| 84 \| 6 \| 9 \| 107 \| 8 \| 123 \| Uncharacterized protein \| Uncharacterized protein; afdb-uniprot50 \| AF-A0A662BHI3-F1-MODEL\_V4 \| 1.0 \| 2.286e-06 \| 177 \| 0.125 \| 143 \| 107 \| 6 \| 9 \| 143 \| 6 \| 138 \| Uncharacterized protein \| Uncharacterized protein; afdb-uniprot50 \| AF-A0A1H6QBZ3-F1-MODEL\_V4 \| 1.0 \| 2.578e-05 \| 176 \| 0.169 \| 118 \| 71 \| 7 \| 6 \| 101 \| 12 \| 124 \| Uncharacterized protein \| Uncharacterized protein; afdb-uniprot50 \| AF-F9GSL4-F1-MODEL\_V4 \| 1.0 \| 3.676e-05 \| 176 \| 0.129 \| 116 \| 83 \| 7 \| 9 \| 108 \| 10 \| 123 \| Uncharacterized protein \| Uncharacterized protein; afdb-uniprot50 \| AF-A0A7C3YAM0-F1-MODEL\_V4 \| 1.0 \| 7.469e-05 \| 176 \| 0.168 \| 95 \| 65 \| 6 \| 8 \| 95 \| 66 \| 153 \| Uncharacterized protein \| Uncharacterized protein; afdb-uniprot50 \| AF-M5V2Z7-F1-MODEL\_V4 \| 1.0 \| 2.291e-05 \| 176 \| 0.124 \| 137 \| 99 \| 7 \| 5 \| 128 \| 4 \| 132 \| Uncharacterized protein \| Uncharacterized protein; afdb-uniprot50 \| AF-A0A432R0Z8-F1-MODEL\_V4 \| 1.0 \| 0.0001004 \| 175 \| 0.102 \| 107 \| 77 \| 5 \| 9 \| 96 \| 5 \| 111 \| Uncharacterized protein \| Uncharacterized protein; afdb-uniprot50 \| AF-A0A7W5UGR6-F1-MODEL\_V4 \| 1.0 \| 0.0001708 \| 175 \| 0.13 \| 115 \| 76 \| 7 \| 9 \| 106 \| 8 \| 115 \| Uncharacterized protein \| Uncharacterized protein; afdb-uniprot50 \| AF-A0A7V6LFQ4-F1-MODEL\_V4 \| 1.0 \| 3.458e-06 \| 175 \| 0.158 \| 164 \| 106 \| 8 \| 9 \| 142 \| 23 \| 184 \| Uncharacterized protein \| Uncharacterized protein; afdb-uniprot50 \| AF-R6F1T1-F1-MODEL\_V4 \| 1.0 \| 4.939e-05 \| 175 \| 0.158 \| 120 \| 94 \| 5 \| 9 \| 124 \| 20 \| 136 \| Uncharacterized protein \| Uncharacterized protein; afdb-uniprot50 \| AF-A0A418VEF3-F1-MODEL\_V4 \| 1.0 \| 4.939e-05 \| 171 \| 0.178 \| 112 \| 80 \| 5 \| 1 \| 100 \| 3 \| 114 \| Uncharacterized protein \| Uncharacterized protein; afdb-uniprot50 \| AF-A0A4R8HZ05-F1-MODEL\_V4 \| 1.0 \| 4.388e-05 \| 169 \| 0.14 \| 114 \| 81 \| 6 \| 9 \| 107 \| 5 \| 116 \| Uncharacterized protein \| Uncharacterized protein; afdb-uniprot50 \| AF-A0A417HAH9-F1-MODEL\_V4 \| 1.0 \| 3.266e-05 \| 169 \| 0.175 \| 120 \| 87 \| 5 \| 7 \| 120 \| 13 \| 126 \| Uncharacterized protein \| Uncharacterized protein; afdb-uniprot50 \| AF-A0A658JVP5-F1-MODEL\_V4 \| 1.0 \| 8.9e-06 \| 169 \| 0.103 \| 155 \| 118 \| 6 \| 8 \| 144 \| 13 \| 164 \| Uncharacterized protein \| Uncharacterized protein; afdb-uniprot50 \| AF-A0A1Z8QCH1-F1-MODEL\_V4 \| 1.0 \| 9.461e-05 \| 168 \| 0.14 \| 121 \| 75 \| 5 \| 9 \| 100 \| 21 \| 141 \| Uncharacterized protein \| Uncharacterized protein; afdb-uniprot50 \| AF-A0A1B1UD36-F1-MODEL\_V4 \| 1.0 \| 0.0002583 \| 167 \| 0.122 \| 106 \| 74 \| 5 \| 9 \| 95 \| 5 \| 110 \| Uncharacterized protein \| Uncharacterized protein; afdb-uniprot50 \| AF-A0A844AGF1-F1-MODEL\_V4 \| 1.0 \| 0.0001812 \| 167 \| 0.081 \| 111 \| 82 \| 6 \| 9 \| 99 \| 5 \| 115 \| Uncharacterized protein \| Uncharacterized protein; afdb-uniprot50 \| AF-A0A5C8SNL3-F1-MODEL\_V4 \| 1.0 \| 0.000113 \| 167 \| 0.159 \| 113 \| 77 \| 6 \| 5 \| 99 \| 9 \| 121 \| Uncharacterized protein \| Uncharacterized protein; afdb-uniprot50 \| AF-A0A382BVZ5-F1-MODEL\_V4 \| 1.0 \| 3.676e-05 \| 167 \| 0.152 \| 138 \| 72 \| 6 \| 9 \| 104 \| 13 \| 147 \| Uncharacterized protein \| Uncharacterized protein; afdb-uniprot50 \| AF-A0A1Q4CYS2-F1-MODEL\_V4 \| 1.0 \| 0.000525 \| 167 \| 0.191 \| 99 \| 70 \| 2 \| 8 \| 96 \| 110 \| 208 \| Uncharacterized protein \| Uncharacterized protein; afdb-uniprot50 \| AF-A0A1C5N9C8-F1-MODEL\_V4 \| 1.0 \| 4.646e-06 \| 167 \| 0.152 \| 144 \| 108 \| 5 \| 5 \| 144 \| 14 \| 147 \| Uncharacterized protein \| Uncharacterized protein; afdb-uniprot50 \| AF-A0A5P3MUE5-F1-MODEL\_V4 \| 1.0 \| 5.229e-06 \| 167 \| 0.127 \| 172 \| 117 \| 5 \| 9 \| 165 \| 9 \| 162 \| Uncharacterized protein \| Uncharacterized protein; afdb-uniprot50 \| AF-A0A1C5PNG8-F1-MODEL\_V4 \| 1.0 \| 0.0001431 \| 166 \| 0.11 \| 109 \| 86 \| 8 \| 7 \| 106 \| 1 \| 107 \| Uncharacterized protein \| Uncharacterized protein; afdb-uniprot50 \| AF-A0A1Y3XMP8-F1-MODEL\_V4 \| 1.0 \| 1.428e-05 \| 166 \| 0.146 \| 123 \| 90 \| 6 \| 9 \| 124 \| 14 \| 128 \| Uncharacterized protein \| Uncharacterized protein; afdb-uniprot50 \| AF-A0A651G663-F1-MODEL\_V4 \| 1.0 \| 0.0002295 \| 165 \| 0.145 \| 117 \| 76 \| 8 \| 9 \| 104 \| 5 \| 118 \| Uncharacterized protein \| Uncharacterized protein; afdb-uniprot50 \| AF-A0A1I3T8Y1-F1-MODEL\_V4 \| 1.0 \| 0.000204 \| 164 \| 0.095 \| 115 \| 77 \| 5 \| 9 \| 96 \| 15 \| 129 \| Uncharacterized protein \| Uncharacterized protein; afdb-uniprot50 \| AF-A0A5C7LQ67-F1-MODEL\_V4 \| 1.0 \| 5.874e-07 \| 163 \| 0.138 \| 188 \| 123 \| 7 \| 9 \| 158 \| 10 \| 196 \| Uncharacterized protein \| Uncharacterized protein; afdb-uniprot50 \| AF-A0A1U7GKV9-F1-MODEL\_V4 \| 1.0 \| 2.291e-05 \| 163 \| 0.182 \| 137 \| 94 \| 6 \| 9 \| 128 \| 7 \| 142 \| Uncharacterized protein \| Uncharacterized protein; afdb-uniprot50 \| AF-A0A7X5E0T7-F1-MODEL\_V4 \| 1.0 \| 3.892e-06 \| 162 \| 0.144 \| 166 \| 114 \| 5 \| 8 \| 166 \| 17 \| 161 \| Uncharacterized protein \| Uncharacterized protein; afdb-uniprot50 \| AF-A0A7J3I1Y6-F1-MODEL\_V4 \| 1.0 \| 2.43e-05 \| 160 \| 0.128 \| 132 \| 104 \| 7 \| 1 \| 124 \| 1 \| 129 \| Uncharacterized protein \| Uncharacterized protein; afdb-uniprot50 \| AF-A0A1Y3ZCV4-F1-MODEL\_V4 \| 1.0 \| 2.73e-06 \| 160 \| 0.152 \| 170 \| 114 \| 8 \| 6 \| 166 \| 10 \| 158 \| Uncharacterized protein \| Uncharacterized protein; afdb-uniprot50 \| AF-A0A5F2B0I3-F1-MODEL\_V4 \| 1.0 \| 1.127e-05 \| 160 \| 0.188 \| 143 \| 82 \| 9 \| 9 \| 129 \| 7 \| 137 \| Uncharacterized protein \| Uncharacterized protein; afdb-uniprot50 \| AF-A0A7M3JY29-F1-MODEL\_V4 \| 1.0 \| 4.929e-06 \| 160 \| 0.18 \| 144 \| 82 \| 8 \| 9 \| 129 \| 7 \| 137 \| Uncharacterized protein \| Uncharacterized protein; afdb-uniprot50 \| AF-A0A662SJ11-F1-MODEL\_V4 \| 1.0 \| 1.428e-05 \| 159 \| 0.138 \| 152 \| 97 \| 9 \| 7 \| 142 \| 736 \| 869 \| SBP\_bac\_5 domain-containing protein \| SBP\_bac\_5 domain-containing protein; afdb-uniprot50 \| AF-A0A443JEA3-F1-MODEL\_V4 \| 1.0 \| 0.0003683 \| 158 \| 0.188 \| 106 \| 74 \| 4 \| 3 \| 96 \| 108 \| 213 \| Uncharacterized protein \| Uncharacterized protein; afdb-uniprot50 \| AF-A0A353H0K0-F1-MODEL\_V4 \| 1.0 \| 5.559e-05 \| 158 \| 0.169 \| 124 \| 71 \| 6 \| 9 \| 101 \| 197 \| 319 \| Ig-like domain-containing protein \| Ig-like domain-containing protein; afdb-uniprot50 \| AF-J9UW90-F1-MODEL\_V4 \| 1.0 \| 0.0006268 \| 158 \| 0.128 \| 101 \| 71 \| 4 \| 7 \| 93 \| 718 \| 815 \| Solute-binding protein like protein \| Solute-binding protein like protein; afdb-uniprot50 \| AF-A0A518G2U2-F1-MODEL\_V4 \| 1.0 \| 0.0003272 \| 157 \| 0.111 \| 108 \| 78 \| 4 \| 8 \| 97 \| 24 \| 131 \| Uncharacterized protein \| Uncharacterized protein; afdb-uniprot50 \| AF-A0A1Y4LVL1-F1-MODEL\_V4 \| 1.0 \| 1.705e-05 \| 157 \| 0.164 \| 146 \| 103 \| 8 \| 7 \| 145 \| 16 \| 149 \| Uncharacterized protein \| Uncharacterized protein; afdb-uniprot50 \| AF-A0A1Y4HDA8-F1-MODEL\_V4 \| 1.0 \| 4.137e-05 \| 157 \| 0.14 \| 128 \| 95 \| 6 \| 9 \| 127 \| 2 \| 123 \| Uncharacterized protein \| Uncharacterized protein; afdb-uniprot50 \| AF-A0A243KKJ3-F1-MODEL\_V4 \| 1.0 \| 3.465e-05 \| 156 \| 0.106 \| 150 \| 117 \| 6 \| 6 \| 142 \| 1 \| 146 \| BppU\_N domain-containing protein \| BppU\_N domain-containing protein; afdb-uniprot50 \| AF-A0A562NRU4-F1-MODEL\_V4 \| 1.0 \| 0.000525 \| 155 \| 0.181 \| 99 \| 70 \| 4 \| 8 \| 96 \| 116 \| 213 \| Uncharacterized protein \| Uncharacterized protein; afdb-uniprot50 \| AF-A0A2R7Z2V5-F1-MODEL\_V4 \| 1.0 \| 6.256e-05 \| 155 \| 0.161 \| 105 \| 74 \| 6 \| 9 \| 107 \| 48 \| 144 \| ZnMc domain-containing protein \| ZnMc domain-containing protein; afdb-uniprot50 \| AF-A0A1Y3ZUZ1-F1-MODEL\_V4 \| 1.0 \| 4.38e-06 \| 154 \| 0.186 \| 161 \| 96 \| 9 \| 7 \| 142 \| 12 \| 162 \| Uncharacterized protein \| Uncharacterized protein; afdb-uniprot50 \| AF-A0A2N6C104-F1-MODEL\_V4 \| 1.0 \| 0.0006268 \| 153 \| 0.181 \| 99 \| 72 \| 6 \| 4 \| 96 \| 2 \| 97 \| Uncharacterized protein \| Uncharacterized protein; afdb-uniprot50 \| AF-A0A1V5X591-F1-MODEL\_V4 \| 1.0 \| 0.0008423 \| 153 \| 0.204 \| 83 \| 54 \| 4 \| 26 \| 96 \| 111 \| 193 \| Uncharacterized protein \| Uncharacterized protein; afdb-uniprot50 \| AF-A0A6M1VR84-F1-MODEL\_V4 \| 1.0 \| 5.24e-05 \| 152 \| 0.132 \| 143 \| 102 \| 9 \| 9 \| 134 \| 2 \| 139 \| BppU family phage baseplate upper protein \| BppU family phage baseplate upper protein; afdb-uniprot50 \| AF-A0A448VHT3-F1-MODEL\_V4 \| 1.0 \| 1.919e-05 \| 152 \| 0.157 \| 190 \| 116 \| 9 \| 9 \| 166 \| 8 \| 185 \| Uncharacterized protein \| Uncharacterized protein; afdb-uniprot50 \| AF-A0A6G7X1N8-F1-MODEL\_V4 \| 1.0 \| 7.454e-06 \| 151 \| 0.155 \| 167 \| 91 \| 7 \| 6 \| 129 \| 2 \| 161 \| Uncharacterized protein \| Uncharacterized protein; afdb-uniprot50 \| AF-A0A2B0J102-F1-MODEL\_V4 \| 1.0 \| 4.388e-05 \| 150 \| 0.128 \| 125 \| 102 \| 4 \| 8 \| 127 \| 5 \| 127 \| BppU\_N domain-containing protein \| BppU\_N domain-containing protein; afdb-uniprot50 \| AF-A0A1Y3ZD32-F1-MODEL\_V4 \| 1.0 \| 0.0001431 \| 150 \| 0.155 \| 109 \| 81 \| 5 \| 15 \| 122 \| 2 \| 100 \| Uncharacterized protein \| Uncharacterized protein; afdb-uniprot50 \| AF-A0A1Q6JK56-F1-MODEL\_V4 \| 1.0 \| 1.269e-05 \| 150 \| 0.191 \| 136 \| 92 \| 6 \| 5 \| 125 \| 18 \| 150 \| Uncharacterized protein \| Uncharacterized protein; afdb-uniprot50 \| AF-T3D9W5-F1-MODEL\_V4 \| 1.0 \| 9.442e-06 \| 149 \| 0.103 \| 164 \| 118 \| 7 \| 9 \| 166 \| 19 \| 159 \| Uncharacterized protein \| Uncharacterized protein; afdb-uniprot50 \| AF-A0A1Y4EZM3-F1-MODEL\_V4 \| 1.0 \| 4.939e-05 \| 149 \| 0.144 \| 152 \| 105 \| 7 \| 6 \| 136 \| 17 \| 164 \| Uncharacterized protein \| Uncharacterized protein; afdb-uniprot50 \| AF-A0A6C2CBK0-F1-MODEL\_V4 \| 1.0 \| 2.291e-05 \| 149 \| 0.146 \| 150 \| 107 \| 9 \| 8 \| 142 \| 22 \| 165 \| Collagen-like protein \| Collagen-like protein; afdb-uniprot50 \| AF-A0A487D5S2-F1-MODEL\_V4 \| 1.0 \| 1.346e-05 \| 149 \| 0.13 \| 168 \| 115 \| 8 \| 9 \| 166 \| 19 \| 165 \| Phage pre-neck appendage-like protein \| Phage pre-neck appendage-like protein; afdb-uniprot50 \| AF-G0J1Z2-F1-MODEL\_V4 \| 1.0 \| 0.0003085 \| 148 \| 0.118 \| 110 \| 79 \| 6 \| 9 \| 100 \| 12 \| 121 \| Uncharacterized protein \| Uncharacterized protein; afdb-uniprot50 \| AF-A0A7C5VHK6-F1-MODEL\_V4 \| 1.0 \| 0.0003085 \| 148 \| 0.12 \| 108 \| 80 \| 8 \| 9 \| 107 \| 700 \| 801 \| SBP\_bac\_5 domain-containing protein \| SBP\_bac\_5 domain-containing protein; afdb-uniprot50 \| AF-A0A3D6CSD6-F1-MODEL\_V4 \| 1.0 \| 0.0004145 \| 147 \| 0.204 \| 98 \| 65 \| 4 \| 6 \| 94 \| 1 \| 94 \| Uncharacterized protein \| Uncharacterized protein; afdb-uniprot50 \| AF-A0A089QFE0-F1-MODEL\_V4 \| 1.0 \| 3.266e-05 \| 147 \| 0.09 \| 154 \| 116 \| 9 \| 7 \| 142 \| 8 \| 155 \| Phage-associated protein, putative structural protein \| Phage-associated protein, putative structural protein; afdb-uniprot50 \| AF-A0A1Y3X1Y0-F1-MODEL\_V4 \| 1.0 \| 1.919e-05 \| 147 \| 0.142 \| 169 \| 117 \| 7 \| 6 \| 166 \| 10 \| 158 \| Uncharacterized protein \| Uncharacterized protein; afdb-uniprot50 \| AF-A0A2E6W1A1-F1-MODEL\_V4 \| 1.0 \| 0.0008423 \| 146 \| 0.193 \| 93 \| 61 \| 6 \| 9 \| 95 \| 7 \| 91 \| Uncharacterized protein \| Uncharacterized protein; afdb-uniprot50 \| AF-A0A4Q2YKC0-F1-MODEL\_V4 \| 1.0 \| 0.0001518 \| 146 \| 0.157 \| 121 \| 82 \| 7 \| 8 \| 108 \| 4 \| 124 \| Uncharacterized protein \| Uncharacterized protein; afdb-uniprot50 \| AF-A0A843HUJ8-F1-MODEL\_V4 \| 1.0 \| 3.676e-05 \| 146 \| 0.154 \| 123 \| 85 \| 6 \| 6 \| 109 \| 1 \| 123 \| Uncharacterized protein \| Uncharacterized protein; afdb-uniprot50 \| AF-A0A0Q8VPZ8-F1-MODEL\_V4 \| 1.0 \| 0.0001349 \| 146 \| 0.126 \| 119 \| 97 \| 5 \| 9 \| 124 \| 254 \| 368 \| Uncharacterized protein \| Uncharacterized protein; afdb-uniprot50 \| AF-A0A137PPG3-F1-MODEL\_V4 \| 1.0 \| 0.000794 \| 144 \| 0.096 \| 114 \| 93 \| 4 \| 3 \| 108 \| 15 \| 126 \| BppU\_N domain-containing protein \| BppU\_N domain-containing protein; afdb-uniprot50 \| AF-A0A4P9YNR0-F1-MODEL\_V4 \| 1.0 \| 2.291e-05 \| 144 \| 0.131 \| 182 \| 125 \| 9 \| 9 \| 178 \| 58 \| 218 \| Dolichyl-diphosphooligosaccharide--protein glycosyltransferase subunit 2 \| Dolichyl-diphosphooligosaccharide--protein glycosyltransferase subunit 2; afdb-uniprot50 \| AF-A0A3R6UUE4-F1-MODEL\_V4 \| 1.0 \| 7.454e-06 \| 144 \| 0.111 \| 171 \| 128 \| 5 \| 8 \| 164 \| 16 \| 176 \| Uncharacterized protein \| Uncharacterized protein; afdb-uniprot50 \| AF-A0A3C0GD26-F1-MODEL\_V4 \| 1.0 \| 6.256e-05 \| 143 \| 0.107 \| 140 \| 104 \| 6 \| 7 \| 125 \| 1 \| 140 \| Uncharacterized protein \| Uncharacterized protein; afdb-uniprot50 \| AF-Q6NTZ0-F1-MODEL\_V4 \| 1.0 \| 0.0002295 \| 143 \| 0.161 \| 130 \| 90 \| 7 \| 7 \| 126 \| 5 \| 125 \| MGC81410 protein \| MGC81410 protein; afdb-uniprot50 \| AF-A0A1K2HBG8-F1-MODEL\_V4 \| 1.0 \| 7.469e-05 \| 143 \| 0.106 \| 150 \| 114 \| 8 \| 8 \| 142 \| 10 \| 154 \| Collagen triple helix repeat-containing protein \| Collagen triple helix repeat-containing protein; afdb-uniprot50 \| AF-A0A3D2H3M5-F1-MODEL\_V4 \| 1.0 \| 0.000204 \| 143 \| 0.156 \| 128 \| 92 \| 7 \| 9 \| 124 \| 209 \| 332 \| Uncharacterized protein \| Uncharacterized protein; afdb-uniprot50 \| AF-A0A2D6EZF0-F1-MODEL\_V4 \| 1.0 \| 0.0001923 \| 142 \| 0.085 \| 140 \| 104 \| 6 \| 9 \| 142 \| 5 \| 126 \| Uncharacterized protein \| Uncharacterized protein; afdb-uniprot50 \| AF-A0A373N6D8-F1-MODEL\_V4 \| 1.0 \| 2.735e-05 \| 142 \| 0.115 \| 147 \| 106 \| 9 \| 8 \| 141 \| 11 \| 146 \| Uncharacterized protein \| Uncharacterized protein; afdb-uniprot50 \| AF-A0A3R6XI26-F1-MODEL\_V4 \| 1.0 \| 6.256e-05 \| 142 \| 0.13 \| 199 \| 130 \| 12 \| 8 \| 179 \| 16 \| 198 \| Collagen-like protein \| Collagen-like protein; afdb-uniprot50 \| AF-A0A7H9CMM2-F1-MODEL\_V4 \| 1.0 \| 9.461e-05 \| 142 \| 0.11 \| 136 \| 100 \| 7 \| 6 \| 129 \| 7 \| 133 \| Uncharacterized protein \| Uncharacterized protein; afdb-uniprot50 \| AF-U2L7Y7-F1-MODEL\_V4 \| 1.0 \| 7.908e-06 \| 142 \| 0.158 \| 158 \| 110 \| 8 \| 14 \| 163 \| 47 \| 189 \| Uncharacterized protein \| Uncharacterized protein; afdb-uniprot50 \| AF-D9SVZ5-F1-MODEL\_V4 \| 1.0 \| 0.0002435 \| 142 \| 0.1 \| 130 \| 98 \| 7 \| 8 \| 126 \| 18 \| 139 \| Uncharacterized protein \| Uncharacterized protein; afdb-uniprot50 \| AF-A0A6I4LXN3-F1-MODEL\_V4 \| 1.0 \| 0.0002295 \| 141 \| 0.159 \| 113 \| 74 \| 5 \| 6 \| 98 \| 2 \| 113 \| Uncharacterized protein \| Uncharacterized protein; afdb-uniprot50 \| AF-A0A3S0BU18-F1-MODEL\_V4 \| 1.0 \| 2.159e-05 \| 141 \| 0.144 \| 152 \| 101 \| 6 \| 5 \| 142 \| 27 \| 163 \| Uncharacterized protein \| Uncharacterized protein; afdb-uniprot50 \| AF-A0A3N1XSC8-F1-MODEL\_V4 \| 1.0 \| 2.291e-05 \| 141 \| 0.167 \| 131 \| 87 \| 10 \| 9 \| 127 \| 11 \| 131 \| Uncharacterized protein \| Uncharacterized protein; afdb-uniprot50 \| AF-A0A7K3NLH1-F1-MODEL\_V4 \| 1.0 \| 0.001712 \| 140 \| 0.141 \| 99 \| 79 \| 4 \| 6 \| 101 \| 1 \| 96 \| Uncharacterized protein \| Uncharacterized protein; afdb-uniprot50 \| AF-A0A2E3J263-F1-MODEL\_V4 \| 1.0 \| 9.461e-05 \| 140 \| 0.158 \| 126 \| 93 \| 7 \| 8 \| 125 \| 550 \| 670 \| Uncharacterized protein \| Uncharacterized protein; afdb-uniprot50 \| AF-D1AG96-F1-MODEL\_V4 \| 1.0 \| 0.0002295 \| 139 \| 0.102 \| 136 \| 110 \| 6 \| 9 \| 136 \| 21 \| 152 \| BppU\_N domain-containing protein \| BppU\_N domain-containing protein; afdb-uniprot50 \| AF-A0A090Z5B2-F1-MODEL\_V4 \| 1.0 \| 2.578e-05 \| 139 \| 0.125 \| 184 \| 126 \| 7 \| 6 \| 178 \| 15 \| 174 \| BppU\_N domain-containing protein \| BppU\_N domain-containing protein; afdb-uniprot50 \| AF-A0A2N5DZ13-F1-MODEL\_V4 \| 1.0 \| 0.002589 \| 139 \| 0.095 \| 94 \| 76 \| 6 \| 8 \| 95 \| 134 \| 224 \| Uncharacterized protein \| Uncharacterized protein; afdb-uniprot50 \| AF-C9L790-F1-MODEL\_V4 \| 1.0 \| 3.078e-05 \| 139 \| 0.148 \| 175 \| 113 \| 8 \| 8 \| 165 \| 11 \| 166 \| Uncharacterized protein \| Uncharacterized protein; afdb-uniprot50 \| AF-A0A417BVX3-F1-MODEL\_V4 \| 1.0 \| 1.919e-05 \| 139 \| 0.132 \| 174 \| 115 \| 10 \| 8 \| 166 \| 16 \| 168 \| Uncharacterized protein \| Uncharacterized protein; afdb-uniprot50 \| AF-A0A844KFR1-F1-MODEL\_V4 \| 1.0 \| 9.461e-05 \| 138 \| 0.105 \| 170 \| 118 \| 8 \| 5 \| 166 \| 16 \| 159 \| Uncharacterized protein \| Uncharacterized protein; afdb-uniprot50 \| AF-A0A3C1IA62-F1-MODEL\_V4 \| 1.0 \| 0.0001349 \| 138 \| 0.092 \| 140 \| 110 \| 10 \| 8 \| 134 \| 12 \| 147 \| Uncharacterized protein \| Uncharacterized protein; afdb-uniprot50 \| AF-A0A1N6WK55-F1-MODEL\_V4 \| 1.0 \| 0.000113 \| 138 \| 0.139 \| 165 \| 116 \| 6 \| 9 \| 165 \| 23 \| 169 \| Lysophospholipase L1 \| Lysophospholipase L1; afdb-uniprot50 \| AF-A0A350VIV5-F1-MODEL\_V4 \| 1.0 \| 5.559e-05 \| 137 \| 0.163 \| 171 \| 125 \| 8 \| 5 \| 166 \| 16 \| 177 \| Uncharacterized protein \| Uncharacterized protein; afdb-uniprot50 \| AF-A0A5P3XC75-F1-MODEL\_V4 \| 1.0 \| 2.902e-05 \| 137 \| 0.14 \| 171 \| 118 \| 7 \| 9 \| 165 \| 25 \| 180 \| DUF2479 domain-containing protein \| DUF2479 domain-containing protein; afdb-uniprot50 \| AF-A0A7X9K8L6-F1-MODEL\_V4 \| 1.0 \| 0.0008423 \| 137 \| 0.148 \| 101 \| 81 \| 3 \| 9 \| 107 \| 408 \| 505 \| Peptidase\_C25 domain-containing protein \| Peptidase\_C25 domain-containing protein; afdb-uniprot50 \| AF-A0A6N1ESY5-F1-MODEL\_V4 \| 1.0 \| 0.001006 \| 137 \| 0.121 \| 132 \| 104 \| 7 \| 7 \| 128 \| 9 \| 138 \| Pectate\_lyase\_3 domain-containing protein \| Pectate\_lyase\_3 domain-containing protein; afdb-uniprot50 \| AF-A0A7C5W505-F1-MODEL\_V4 \| 1.0 \| 0.0003907 \| 137 \| 0.178 \| 129 \| 81 \| 7 \| 7 \| 119 \| 199 \| 318 \| Uncharacterized protein \| Uncharacterized protein; afdb-uniprot50 \| AF-A0A174D4S5-F1-MODEL\_V4 \| 1.0 \| 6.256e-05 \| 137 \| 0.114 \| 174 \| 122 \| 11 \| 9 \| 165 \| 20 \| 178 \| Choline binding protein \| Choline binding protein; afdb-uniprot50 \| AF-R9N9F0-F1-MODEL\_V4 \| 1.0 \| 0.0002908 \| 137 \| 0.169 \| 124 \| 95 \| 5 \| 9 \| 128 \| 21 \| 140 \| Uncharacterized protein \| Uncharacterized protein; afdb-uniprot50 \| AF-A0A429GIY2-F1-MODEL\_V4 \| 1.0 \| 0.0003272 \| 137 \| 0.172 \| 116 \| 80 \| 6 \| 7 \| 109 \| 732 \| 844 \| CGP-CTERM sorting domain-containing protein \| CGP-CTERM sorting domain-containing protein; afdb-uniprot50 \| AF-A0A496KGP7-F1-MODEL\_V4 \| 1.0 \| 0.001613 \| 136 \| 0.12 \| 108 \| 83 \| 3 \| 9 \| 107 \| 7 \| 111 \| Uncharacterized protein \| Uncharacterized protein; afdb-uniprot50 \| AF-A0A7Y9PCR7-F1-MODEL\_V4 \| 1.0 \| 0.00244 \| 136 \| 0.193 \| 88 \| 59 \| 4 \| 26 \| 101 \| 123 \| 210 \| Uncharacterized protein \| Uncharacterized protein; afdb-uniprot50 \| AF-R8RDA6-F1-MODEL\_V4 \| 1.0 \| 3.465e-05 \| 136 \| 0.13 \| 153 \| 115 \| 6 \| 6 \| 142 \| 15 \| 165 \| BppU\_N domain-containing protein \| BppU\_N domain-containing protein; afdb-uniprot50 \| AF-A0A1H3XCM1-F1-MODEL\_V4 \| 1.0 \| 0.0001708 \| 136 \| 0.126 \| 166 \| 115 \| 7 \| 9 \| 166 \| 22 \| 165 \| BppU\_N domain-containing protein \| BppU\_N domain-containing protein; afdb-uniprot50 \| AF-A0A2A8VJ41-F1-MODEL\_V4 \| 1.0 \| 6.637e-05 \| 136 \| 0.148 \| 155 \| 110 \| 8 \| 6 \| 142 \| 15 \| 165 \| BppU\_N domain-containing protein \| BppU\_N domain-containing protein; afdb-uniprot50 \| AF-A0A2B5HPR6-F1-MODEL\_V4 \| 1.0 \| 7.041e-05 \| 136 \| 0.148 \| 135 \| 93 \| 9 \| 8 \| 127 \| 17 \| 144 \| Uncharacterized protein \| Uncharacterized protein; afdb-uniprot50 \| AF-A0A1I3QIP6-F1-MODEL\_V4 \| 1.0 \| 8.39e-06 \| 136 \| 0.121 \| 189 \| 127 \| 11 \| 9 \| 164 \| 24 \| 206 \| Uncharacterized protein \| Uncharacterized protein; afdb-uniprot50 \| AF-A0A2H3KVU9-F1-MODEL\_V4 \| 1.0 \| 0.00244 \| 135 \| 0.211 \| 90 \| 62 \| 5 \| 7 \| 95 \| 41 \| 122 \| Uncharacterized protein \| Uncharacterized protein; afdb-uniprot50 \| AF-A0A1C5U595-F1-MODEL\_V4 \| 1.0 \| 0.0002295 \| 135 \| 0.139 \| 151 \| 111 \| 10 \| 9 \| 148 \| 22 \| 164 \| Uncharacterized protein \| Uncharacterized protein; afdb-uniprot50 \| AF-A0A0P9E8S0-F1-MODEL\_V4 \| 1.0 \| 0.000113 \| 135 \| 0.155 \| 174 \| 117 \| 11 \| 8 \| 167 \| 38 \| 195 \| Uncharacterized protein \| Uncharacterized protein; afdb-uniprot50 \| AF-A0A4P6ZYX7-F1-MODEL\_V4 \| 1.0 \| 0.0002295 \| 135 \| 0.149 \| 161 \| 105 \| 9 \| 9 \| 158 \| 23 \| 162 \| Uncharacterized protein \| Uncharacterized protein; afdb-uniprot50 \| AF-A0A837IKU0-F1-MODEL\_V4 \| 1.0 \| 0.000161 \| 134 \| 0.126 \| 134 \| 105 \| 5 \| 7 \| 135 \| 31 \| 157 \| Peptidase, M23/M37 family \| Peptidase, M23/M37 family; afdb-uniprot50 \| AF-A0A1C4ERV9-F1-MODEL\_V4 \| 1.0 \| 3.078e-05 \| 134 \| 0.141 \| 156 \| 110 \| 10 \| 6 \| 142 \| 1 \| 151 \| BppU\_N domain-containing protein \| BppU\_N domain-containing protein; afdb-uniprot50 \| AF-A0A3F3S3N3-F1-MODEL\_V4 \| 1.0 \| 0.0002295 \| 134 \| 0.119 \| 142 \| 107 \| 6 \| 7 \| 135 \| 18 \| 154 \| Hep\_Hag \| Hep\_Hag; afdb-uniprot50 \| AF-A0A0E9N1Z6-F1-MODEL\_V4 \| 1.0 \| 0.003478 \| 133 \| 0.134 \| 104 \| 83 \| 5 \| 7 \| 106 \| 10 \| 110 \| Uncharacterized protein \| Uncharacterized protein; afdb-uniprot50 \| AF-A0A2H9QAH0-F1-MODEL\_V4 \| 1.0 \| 0.001201 \| 133 \| 0.162 \| 111 \| 78 \| 8 \| 5 \| 108 \| 275 \| 377 \| Uncharacterized protein \| Uncharacterized protein; afdb-uniprot50 \| AF-A0A497L8C9-F1-MODEL\_V4 \| 1.0 \| 0.007069 \| 132 \| 0.094 \| 95 \| 80 \| 3 \| 6 \| 96 \| 1 \| 93 \| Uncharacterized protein \| Uncharacterized protein; afdb-uniprot50 \| AF-A0A0W8G563-F1-MODEL\_V4 \| 1.0 \| 0.003091 \| 132 \| 0.139 \| 93 \| 71 \| 6 \| 9 \| 96 \| 12 \| 100 \| Uncharacterized protein \| Uncharacterized protein; afdb-uniprot50 \| AF-A0A848RGW3-F1-MODEL\_V4 \| 1.0 \| 5.24e-05 \| 132 \| 0.165 \| 139 \| 87 \| 9 \| 9 \| 125 \| 17 \| 148 \| BppU family phage baseplate upper protein \| BppU family phage baseplate upper protein; afdb-uniprot50 \| AF-A0A846M644-F1-MODEL\_V4 \| 1.0 \| 2.43e-05 \| 132 \| 0.12 \| 182 \| 107 \| 10 \| 8 \| 156 \| 45 \| 206 \| Uncharacterized protein \| Uncharacterized protein; afdb-uniprot50 \| AF-A0A1W7LNG8-F1-MODEL\_V4 \| 1.0 \| 0.0002295 \| 132 \| 0.083 \| 168 \| 122 \| 8 \| 9 \| 166 \| 23 \| 168 \| Uncharacterized protein \| Uncharacterized protein; afdb-uniprot50 \| AF-A0A2C4PYW6-F1-MODEL\_V4 \| 1.0 \| 8.918e-05 \| 132 \| 0.134 \| 156 \| 114 \| 9 \| 2 \| 142 \| 11 \| 160 \| DUF2479 domain-containing protein \| DUF2479 domain-containing protein; afdb-uniprot50 \| AF-A0A7J4FW38-F1-MODEL\_V4 \| 1.0 \| 0.00369 \| 132 \| 0.148 \| 94 \| 73 \| 4 \| 7 \| 94 \| 38 \| 130 \| Carboxypeptidase regulatory-like domain-containing protein \| Carboxypeptidase regulatory-like domain-containing protein; afdb-uniprot50 \| AF-A0A174X6D8-F1-MODEL\_V4 \| 1.0 \| 2.578e-05 \| 132 \| 0.137 \| 175 \| 116 \| 10 \| 8 \| 166 \| 16 \| 171 \| Uncharacterized protein \| Uncharacterized protein; afdb-uniprot50 \| AF-A0A2X3L147-F1-MODEL\_V4 \| 1.0 \| 0.0005909 \| 131 \| 0.096 \| 104 \| 83 \| 7 \| 10 \| 107 \| 2 \| 100 \| Big\_13 domain-containing protein \| Big\_13 domain-containing protein; afdb-uniprot50 \| AF-A0A3L7SAM3-F1-MODEL\_V4 \| 1.0 \| 0.0002295 \| 131 \| 0.161 \| 136 \| 74 \| 9 \| 3 \| 103 \| 287 \| 417 \| Uncharacterized protein \| Uncharacterized protein; afdb-uniprot50 \| AF-A0A3M2A537-F1-MODEL\_V4 \| 1.0 \| 0.0007484 \| 131 \| 0.236 \| 93 \| 56 \| 7 \| 7 \| 95 \| 52 \| 133 \| Uncharacterized protein \| Uncharacterized protein; afdb-uniprot50 \| AF-A0A538K330-F1-MODEL\_V4 \| 1.0 \| 0.003279 \| 131 \| 0.127 \| 94 \| 72 \| 4 \| 7 \| 95 \| 261 \| 349 \| Uncharacterized protein \| Uncharacterized protein; afdb-uniprot50 \| AF-A0A6A8DIG3-F1-MODEL\_V4 \| 1.0 \| 7.041e-05 \| 131 \| 0.145 \| 193 \| 129 \| 11 \| 8 \| 169 \| 21 \| 208 \| Uncharacterized protein \| Uncharacterized protein; afdb-uniprot50 \| AF-A0A2P9GEF6-F1-MODEL\_V4 \| 1.0 \| 9.442e-06 \| 131 \| 0.095 \| 209 \| 138 \| 10 \| 8 \| 179 \| 90 \| 284 \| Uncharacterized protein \| Uncharacterized protein; afdb-uniprot50 \| AF-A0A1L6LN89-F1-MODEL\_V4 \| 1.0 \| 0.00244 \| 130 \| 0.126 \| 111 \| 83 \| 6 \| 6 \| 107 \| 6 \| 111 \| Uncharacterized protein \| Uncharacterized protein; afdb-uniprot50 \| AF-A0A377PLH1-F1-MODEL\_V4 \| 1.0 \| 0.000113 \| 130 \| 0.134 \| 141 \| 95 \| 8 \| 8 \| 141 \| 7 \| 127 \| Big\_13 domain-containing protein \| Big\_13 domain-containing protein; afdb-uniprot50 \| AF-E3CAQ5-F1-MODEL\_V4 \| 1.0 \| 0.0003471 \| 130 \| 0.098 \| 163 \| 121 \| 6 \| 8 \| 148 \| 12 \| 170 \| Uncharacterized protein \| Uncharacterized protein; afdb-uniprot50 \| AF-A0A1G2S722-F1-MODEL\_V4 \| 1.0 \| 0.000948 \| 130 \| 0.159 \| 88 \| 68 \| 4 \| 9 \| 95 \| 430 \| 512 \| Uncharacterized protein \| Uncharacterized protein; afdb-uniprot50 \| AF-A0A354KRM3-F1-MODEL\_V4 \| 1.0 \| 2.159e-05 \| 130 \| 0.114 \| 183 \| 131 \| 9 \| 9 \| 165 \| 24 \| 201 \| Uncharacterized protein \| Uncharacterized protein; afdb-uniprot50 \| AF-A0A1I3P6U5-F1-MODEL\_V4 \| 1.0 \| 0.001351 \| 129 \| 0.153 \| 98 \| 70 \| 4 \| 9 \| 96 \| 6 \| 100 \| Uncharacterized protein \| Uncharacterized protein; afdb-uniprot50 \| AF-A0A2D8ESG9-F1-MODEL\_V4 \| 1.0 \| 0.0004949 \| 129 \| 0.108 \| 148 \| 75 \| 6 \| 6 \| 96 \| 5 \| 152 \| Uncharacterized protein \| Uncharacterized protein; afdb-uniprot50 \| AF-A0A3D5KYB9-F1-MODEL\_V4 \| 1.0 \| 2.735e-05 \| 129 \| 0.125 \| 167 \| 118 \| 7 \| 9 \| 166 \| 20 \| 167 \| BppU\_N domain-containing protein \| BppU\_N domain-containing protein; afdb-uniprot50 \| AF-A0A4S3LGE2-F1-MODEL\_V4 \| 1.0 \| 0.006663 \| 129 \| 0.172 \| 93 \| 67 \| 5 \| 8 \| 96 \| 208 \| 294 \| Ig-like domain-containing protein \| Ig-like domain-containing protein; afdb-uniprot50 \| AF-A0A3C1CTL9-F1-MODEL\_V4 \| 1.0 \| 0.0002164 \| 129 \| 0.119 \| 142 \| 100 \| 5 \| 9 \| 127 \| 19 \| 158 \| Uncharacterized protein \| Uncharacterized protein; afdb-uniprot50 \| AF-A0A366ZBU9-F1-MODEL\_V4 \| 1.0 \| 0.0001812 \| 129 \| 0.153 \| 130 \| 94 \| 9 \| 1 \| 124 \| 323 \| 442 \| Uncharacterized protein \| Uncharacterized protein; afdb-uniprot50 \| AF-A0A662KU90-F1-MODEL\_V4 \| 1.0 \| 5.897e-05 \| 128 \| 0.097 \| 154 \| 117 \| 5 \| 8 \| 160 \| 5 \| 137 \| Uncharacterized protein \| Uncharacterized protein; afdb-uniprot50 \| AF-A0A2E5D4P0-F1-MODEL\_V4 \| 1.0 \| 0.0004397 \| 128 \| 0.134 \| 104 \| 75 \| 5 \| 8 \| 100 \| 50 \| 149 \| Uncharacterized protein \| Uncharacterized protein; afdb-uniprot50 \| AF-F1TEH3-F1-MODEL\_V4 \| 1.0 \| 0.001926 \| 128 \| 0.125 \| 120 \| 84 \| 8 \| 2 \| 108 \| 13 \| 124 \| BppU\_N domain-containing protein \| BppU\_N domain-containing protein; afdb-uniprot50 \| AF-X1CLZ4-F1-MODEL\_V4 \| 1.0 \| 0.002168 \| 128 \| 0.099 \| 111 \| 85 \| 7 \| 8 \| 108 \| 132 \| 237 \| Uncharacterized protein \| Uncharacterized protein; afdb-uniprot50 \| AF-A0A7W1I8A5-F1-MODEL\_V4 \| 1.0 \| 8.406e-05 \| 128 \| 0.123 \| 146 \| 100 \| 12 \| 7 \| 134 \| 272 \| 407 \| Uncharacterized protein \| Uncharacterized protein; afdb-uniprot50 \| AF-A0A2X0V1N7-F1-MODEL\_V4 \| 1.0 \| 0.0001004 \| 128 \| 0.15 \| 173 \| 110 \| 10 \| 9 \| 164 \| 25 \| 177 \| Uncharacterized protein \| Uncharacterized protein; afdb-uniprot50 \| AF-A0A4P8S4K1-F1-MODEL\_V4 \| 1.0 \| 0.0001923 \| 128 \| 0.101 \| 177 \| 121 \| 9 \| 6 \| 164 \| 19 \| 175 \| Uncharacterized protein \| Uncharacterized protein; afdb-uniprot50 \| AF-A0A4Q7PQS7-F1-MODEL\_V4 \| 1.0 \| 0.0008936 \| 127 \| 0.125 \| 128 \| 98 \| 8 \| 9 \| 127 \| 26 \| 148 \| Uncharacterized protein \| Uncharacterized protein; afdb-uniprot50 \| AF-A5ZRU3-F1-MODEL\_V4 \| 1.0 \| 0.0001708 \| 127 \| 0.113 \| 176 \| 125 \| 8 \| 3 \| 166 \| 5 \| 161 \| BppU\_N domain-containing protein \| BppU\_N domain-containing protein; afdb-uniprot50 \| AF-A0A3E2WI43-F1-MODEL\_V4 \| 1.0 \| 4.388e-05 \| 127 \| 0.136 \| 176 \| 122 \| 6 \| 8 \| 166 \| 16 \| 178 \| Uncharacterized protein \| Uncharacterized protein; afdb-uniprot50 \| AF-A0A7X2MU43-F1-MODEL\_V4 \| 1.0 \| 0.0003683 \| 126 \| 0.095 \| 105 \| 84 \| 7 \| 9 \| 108 \| 2 \| 100 \| Big\_13 domain-containing protein \| Big\_13 domain-containing protein; afdb-uniprot50 \| AF-A0A3C0G3Z4-F1-MODEL\_V4 \| 1.0 \| 0.000794 \| 126 \| 0.079 \| 138 \| 104 \| 5 \| 9 \| 142 \| 3 \| 121 \| Uncharacterized protein \| Uncharacterized protein; afdb-uniprot50 \| AF-A0A2S4GM85-F1-MODEL\_V4 \| 1.0 \| 0.0002295 \| 126 \| 0.124 \| 169 \| 119 \| 7 \| 5 \| 166 \| 16 \| 162 \| Uncharacterized protein \| Uncharacterized protein; afdb-uniprot50 \| AF-A0A810ZRD3-F1-MODEL\_V4 \| 1.0 \| 0.001521 \| 126 \| 0.118 \| 110 \| 78 \| 7 \| 8 \| 107 \| 141 \| 241 \| Uncharacterized protein \| Uncharacterized protein; afdb-uniprot50 \| AF-R8E0D3-F1-MODEL\_V4 \| 1.0 \| 5.897e-05 \| 126 \| 0.125 \| 176 \| 128 \| 5 \| 6 \| 170 \| 15 \| 175 \| BppU\_N domain-containing protein \| BppU\_N domain-containing protein; afdb-uniprot50 \| AF-B0PAR7-F1-MODEL\_V4 \| 1.0 \| 0.0001065 \| 126 \| 0.129 \| 170 \| 122 \| 5 \| 8 \| 166 \| 159 \| 313 \| Uncharacterized protein \| Uncharacterized protein; afdb-uniprot50 \| AF-A0A349MVT2-F1-MODEL\_V4 \| 1.0 \| 0.00844 \| 126 \| 0.197 \| 91 \| 66 \| 4 \| 9 \| 95 \| 351 \| 438 \| Uncharacterized protein \| Uncharacterized protein; afdb-uniprot50 \| AF-X8HS60-F1-MODEL\_V4 \| 1.0 \| 0.0004145 \| 125 \| 0.148 \| 135 \| 96 \| 7 \| 7 \| 127 \| 20 \| 149 \| Uncharacterized protein \| Uncharacterized protein; afdb-uniprot50 \| AF-A0A417HW05-F1-MODEL\_V4 \| 1.0 \| 4.388e-05 \| 125 \| 0.189 \| 137 \| 76 \| 8 \| 15 \| 124 \| 32 \| 160 \| Uncharacterized protein \| Uncharacterized protein; afdb-uniprot50 \| AF-A0A2B9TYA6-F1-MODEL\_V4 \| 1.0 \| 0.000113 \| 125 \| 0.117 \| 153 \| 117 \| 6 \| 6 \| 142 \| 15 \| 165 \| BppU\_N domain-containing protein \| BppU\_N domain-containing protein; afdb-uniprot50 \| AF-A0A446IBB5-F1-MODEL\_V4 \| 1.0 \| 9.461e-05 \| 125 \| 0.085 \| 188 \| 137 \| 10 \| 8 \| 179 \| 16 \| 184 \| DUF2479 domain-containing protein \| DUF2479 domain-containing protein; afdb-uniprot50 \| AF-A0A2T5V3S1-F1-MODEL\_V4 \| 1.0 \| 0.0001349 \| 125 \| 0.14 \| 185 \| 122 \| 8 \| 6 \| 178 \| 15 \| 174 \| Uncharacterized protein DUF2479 \| Uncharacterized protein DUF2479; afdb-uniprot50 \| AF-R7H217-F1-MODEL\_V4 \| 1.0 \| 7.469e-05 \| 125 \| 0.129 \| 162 \| 114 \| 10 \| 6 \| 144 \| 6 \| 163 \| Uncharacterized protein \| Uncharacterized protein; afdb-uniprot50 \| AF-R9MUQ2-F1-MODEL\_V4 \| 1.0 \| 0.0001349 \| 125 \| 0.114 \| 184 \| 123 \| 8 \| 5 \| 178 \| 16 \| 169 \| Uncharacterized protein \| Uncharacterized protein; afdb-uniprot50 \| AF-A0A1T4KGE4-F1-MODEL\_V4 \| 1.0 \| 0.004406 \| 125 \| 0.145 \| 110 \| 85 \| 5 \| 4 \| 107 \| 384 \| 490 \| Uncharacterized protein \| Uncharacterized protein; afdb-uniprot50 \| AF-A0A1R7IK63-F1-MODEL\_V4 \| 1.0 \| 7.924e-05 \| 125 \| 0.091 \| 186 \| 127 \| 6 \| 3 \| 165 \| 11 \| 177 \| Phage pre-neck appendage-like protein \| Phage pre-neck appendage-like protein; afdb-uniprot50 \| AF-A0A544TAC3-F1-MODEL\_V4 \| 1.0 \| 0.000113 \| 125 \| 0.116 \| 172 \| 115 \| 11 \| 8 \| 166 \| 21 \| 168 \| NodB homology domain-containing protein \| NodB homology domain-containing protein; afdb-uniprot50 \| AF-A0A2B0JB62-F1-MODEL\_V4 \| 1.0 \| 7.041e-05 \| 124 \| 0.13 \| 138 \| 90 \| 8 \| 6 \| 124 \| 15 \| 141 \| BppU\_N domain-containing protein \| BppU\_N domain-containing protein; afdb-uniprot50 \| AF-A0A7C9H3D3-F1-MODEL\_V4 \| 1.0 \| 0.0004145 \| 124 \| 0.092 \| 108 \| 85 \| 6 \| 21 \| 124 \| 46 \| 144 \| BppU\_N domain-containing protein \| BppU\_N domain-containing protein; afdb-uniprot50 \| AF-A0A2A8VGJ7-F1-MODEL\_V4 \| 1.0 \| 0.0001349 \| 124 \| 0.135 \| 184 \| 117 \| 8 \| 6 \| 178 \| 15 \| 167 \| BppU\_N domain-containing protein \| BppU\_N domain-containing protein; afdb-uniprot50 \| AF-A0A1B1KZP8-F1-MODEL\_V4 \| 1.0 \| 0.0001708 \| 124 \| 0.146 \| 178 \| 126 \| 6 \| 4 \| 170 \| 13 \| 175 \| Phage protein \| Phage protein; afdb-uniprot50 \| AF-U2TW11-F1-MODEL\_V4 \| 1.0 \| 0.000665 \| 124 \| 0.154 \| 136 \| 99 \| 7 \| 4 \| 125 \| 17 \| 150 \| Collagen triple helix repeat protein \| Collagen triple helix repeat protein; afdb-uniprot50 \| AF-A0A396KPQ0-F1-MODEL\_V4 \| 1.0 \| 3.899e-05 \| 124 \| 0.121 \| 173 \| 124 \| 8 \| 8 \| 164 \| 16 \| 176 \| Uncharacterized protein \| Uncharacterized protein; afdb-uniprot50 \| AF-A0A3N2FNP4-F1-MODEL\_V4 \| 1.0 \| 0.0007484 \| 124 \| 0.18 \| 122 \| 90 \| 8 \| 8 \| 126 \| 602 \| 716 \| Glycine rich protein \| Glycine rich protein; afdb-uniprot50 \| AF-A0A2S4GT96-F1-MODEL\_V4 \| 1.0 \| 0.0001271 \| 123 \| 0.132 \| 166 \| 110 \| 10 \| 9 \| 165 \| 19 \| 159 \| Uncharacterized protein \| Uncharacterized protein; afdb-uniprot50 \| AF-A0A0K9FG35-F1-MODEL\_V4 \| 1.0 \| 0.0008936 \| 123 \| 0.157 \| 127 \| 84 \| 9 \| 9 \| 124 \| 22 \| 136 \| Uncharacterized protein \| Uncharacterized protein; afdb-uniprot50 \| AF-A0A506QRY5-F1-MODEL\_V4 \| 1.0 \| 0.003478 \| 123 \| 0.087 \| 91 \| 72 \| 6 \| 8 \| 93 \| 206 \| 290 \| BapA prefix-like domain-containing protein \| BapA prefix-like domain-containing protein; afdb-uniprot50 \| AF-A0A6L6H013-F1-MODEL\_V4 \| 1.0 \| 0.0001065 \| 123 \| 0.129 \| 178 \| 113 \| 9 \| 9 \| 166 \| 24 \| 179 \| Uncharacterized protein \| Uncharacterized protein; afdb-uniprot50 \| AF-A0A3R6NAE3-F1-MODEL\_V4 \| 1.0 \| 7.924e-05 \| 123 \| 0.109 \| 174 \| 122 \| 9 \| 8 \| 166 \| 16 \| 171 \| H\_lectin domain-containing protein \| H\_lectin domain-containing protein; afdb-uniprot50 \| AF-A0A658JWQ7-F1-MODEL\_V4 \| 1.0 \| 5.24e-05 \| 123 \| 0.149 \| 167 \| 100 \| 12 \| 8 \| 142 \| 16 \| 172 \| Uncharacterized protein \| Uncharacterized protein; afdb-uniprot50 \| AF-A0A2W4NIF2-F1-MODEL\_V4 \| 1.0 \| 0.00526 \| 123 \| 0.122 \| 114 \| 84 \| 6 \| 6 \| 108 \| 128 \| 236 \| Uncharacterized protein \| Uncharacterized protein; afdb-uniprot50 \| AF-A0A0L0AZV1-F1-MODEL\_V4 \| 1.0 \| 0.004406 \| 123 \| 0.139 \| 115 \| 89 \| 3 \| 3 \| 107 \| 155 \| 269 \| Uncharacterized protein \| Uncharacterized protein; afdb-uniprot50 \| AF-A0A414S2W6-F1-MODEL\_V4 \| 1.0 \| 0.0003683 \| 123 \| 0.129 \| 170 \| 129 \| 9 \| 9 \| 168 \| 21 \| 181 \| Uncharacterized protein \| Uncharacterized protein; afdb-uniprot50 \| AF-A0A3S4JY58-F1-MODEL\_V4 \| 1.0 \| 0.0004397 \| 122 \| 0.127 \| 110 \| 77 \| 8 \| 8 \| 107 \| 33 \| 133 \| Adhesin for cattle intestine colonization \| Adhesin for cattle intestine colonization; afdb-uniprot50 \| AF-A0A1C5W6G3-F1-MODEL\_V4 \| 1.0 \| 0.0002908 \| 122 \| 0.093 \| 107 \| 89 \| 4 \| 30 \| 132 \| 47 \| 149 \| BppU\_N domain-containing protein \| BppU\_N domain-containing protein; afdb-uniprot50 \| AF-A0A8B3A5B4-F1-MODEL\_V4 \| 1.0 \| 6.637e-05 \| 122 \| 0.157 \| 178 \| 116 \| 9 \| 8 \| 166 \| 16 \| 178 \| Uncharacterized protein \| Uncharacterized protein; afdb-uniprot50 \| AF-A0A7Y5ZE53-F1-MODEL\_V4 \| 1.0 \| 0.0003085 \| 122 \| 0.136 \| 132 \| 96 \| 8 \| 6 \| 128 \| 380 \| 502 \| LPXTG cell wall anchor domain-containing protein \| LPXTG cell wall anchor domain-containing protein; afdb-uniprot50 \| AF-A0A7U9RBF8-F1-MODEL\_V4 \| 1.0 \| 0.0008423 \| 122 \| 0.139 \| 136 \| 100 \| 8 \| 5 \| 128 \| 14 \| 144 \| BppU\_N domain-containing protein \| BppU\_N domain-containing protein; afdb-uniprot50 \| AF-C0VY35-F1-MODEL\_V4 \| 1.0 \| 0.0023 \| 122 \| 0.071 \| 140 \| 103 \| 6 \| 7 \| 142 \| 594 \| 710 \| LPXTG-motif cell wall anchor domain protein \| LPXTG-motif cell wall anchor domain protein; afdb-uniprot50 \| AF-A0A4R8VD28-F1-MODEL\_V4 \| 1.0 \| 0.00592 \| 122 \| 0.152 \| 105 \| 77 \| 9 \| 7 \| 107 \| 557 \| 653 \| Big\_6 domain-containing protein \| Big\_6 domain-containing protein; afdb-uniprot50 \| AF-A0A2D6WYC0-F1-MODEL\_V4 \| 1.0 \| 0.0002295 \| 121 \| 0.104 \| 153 \| 72 \| 7 \| 6 \| 96 \| 1 \| 150 \| Uncharacterized protein \| Uncharacterized protein; afdb-uniprot50 \| AF-A0A1C5KJI7-F1-MODEL\_V4 \| 1.0 \| 0.0004949 \| 121 \| 0.101 \| 128 \| 97 \| 7 \| 8 \| 124 \| 14 \| 134 \| Uncharacterized protein \| Uncharacterized protein; afdb-uniprot50 \| AF-A0A4Q9HRS1-F1-MODEL\_V4 \| 1.0 \| 0.000525 \| 121 \| 0.1 \| 139 \| 92 \| 6 \| 7 \| 142 \| 53 \| 161 \| Uncharacterized protein \| Uncharacterized protein; afdb-uniprot50 \| AF-A0A7X2SX63-F1-MODEL\_V4 \| 1.0 \| 0.0001065 \| 121 \| 0.151 \| 152 \| 102 \| 10 \| 8 \| 142 \| 60 \| 201 \| Big\_13 domain-containing protein \| Big\_13 domain-containing protein; afdb-uniprot50 \| AF-A0A1Q6MGK8-F1-MODEL\_V4 \| 1.0 \| 5.24e-05 \| 121 \| 0.138 \| 180 \| 120 \| 6 \| 8 \| 166 \| 36 \| 201 \| Uncharacterized protein \| Uncharacterized protein; afdb-uniprot50 \| AF-A0A1B1LED8-F1-MODEL\_V4 \| 1.0 \| 0.0001271 \| 121 \| 0.134 \| 171 \| 124 \| 6 \| 9 \| 170 \| 20 \| 175 \| Phage protein \| Phage protein; afdb-uniprot50 \| AF-A0A143ZVL5-F1-MODEL\_V4 \| 1.0 \| 0.0003085 \| 121 \| 0.132 \| 173 \| 119 \| 8 \| 6 \| 166 \| 18 \| 171 \| Uncharacterized protein \| Uncharacterized protein; afdb-uniprot50 \| AF-A0A7V7HH80-F1-MODEL\_V4 \| 1.0 \| 0.0004665 \| 121 \| 0.122 \| 122 \| 93 \| 7 \| 17 \| 127 \| 32 \| 150 \| DUF2479 domain-containing protein \| DUF2479 domain-containing protein; afdb-uniprot50 \| AF-A0A437UJ27-F1-MODEL\_V4 \| 1.0 \| 0.000665 \| 121 \| 0.134 \| 163 \| 112 \| 8 \| 8 \| 152 \| 24 \| 175 \| BppU\_N domain-containing protein \| BppU\_N domain-containing protein; afdb-uniprot50 \| AF-F7JWE5-F1-MODEL\_V4 \| 1.0 \| 3.899e-05 \| 120 \| 0.094 \| 148 \| 114 \| 8 \| 7 \| 141 \| 10 \| 150 \| Uncharacterized protein \| Uncharacterized protein; afdb-uniprot50 \| AF-A0A1V9QR56-F1-MODEL\_V4 \| 1.0 \| 0.0004665 \| 120 \| 0.054 \| 148 \| 120 \| 8 \| 8 \| 142 \| 9 \| 149 \| Uncharacterized protein \| Uncharacterized protein; afdb-uniprot50 \| AF-A0A3B9GR26-F1-MODEL\_V4 \| 1.0 \| 0.0002908 \| 120 \| 0.134 \| 164 \| 116 \| 8 \| 9 \| 164 \| 20 \| 165 \| Uncharacterized protein \| Uncharacterized protein; afdb-uniprot50 \| AF-A0A1C5MR39-F1-MODEL\_V4 \| 1.0 \| 0.004674 \| 120 \| 0.125 \| 104 \| 78 \| 7 \| 7 \| 106 \| 139 \| 233 \| Uncharacterized protein \| Uncharacterized protein; afdb-uniprot50 \| AF-R5RJJ3-F1-MODEL\_V4 \| 1.0 \| 0.0001708 \| 120 \| 0.126 \| 174 \| 112 \| 10 \| 8 \| 166 \| 16 \| 164 \| Uncharacterized protein \| Uncharacterized protein; afdb-uniprot50 \| AF-A0A350H2I6-F1-MODEL\_V4 \| 1.0 \| 0.0003683 \| 120 \| 0.121 \| 156 \| 108 \| 9 \| 9 \| 141 \| 20 \| 169 \| BppU\_N domain-containing protein \| BppU\_N domain-containing protein; afdb-uniprot50 \| AF-A0A6P1Q0A7-F1-MODEL\_V4 \| 1.0 \| 0.00369 \| 120 \| 0.112 \| 98 \| 77 \| 7 \| 8 \| 100 \| 62 \| 154 \| Uncharacterized protein \| Uncharacterized protein; afdb-uniprot50 \| AF-A0A0F9PSE9-F1-MODEL\_V4 \| 1.0 \| 0.005581 \| 119 \| 0.166 \| 84 \| 59 \| 4 \| 22 \| 96 \| 104 \| 185 \| Uncharacterized protein \| Uncharacterized protein; afdb-uniprot50 \| AF-A0A1C6KJ53-F1-MODEL\_V4 \| 1.0 \| 0.0001708 \| 119 \| 0.12 \| 224 \| 133 \| 14 \| 9 \| 179 \| 3 \| 215 \| Collagen triple helix repeat (20 copies) \| Collagen triple helix repeat (20 copies); afdb-uniprot50 \| AF-A0A2V2FAF0-F1-MODEL\_V4 \| 1.0 \| 6.637e-05 \| 119 \| 0.165 \| 163 \| 109 \| 7 \| 15 \| 166 \| 12 \| 158 \| Uncharacterized protein \| Uncharacterized protein; afdb-uniprot50 \| AF-A0A534IQ49-F1-MODEL\_V4 \| 1.0 \| 0.0003683 \| 119 \| 0.12 \| 149 \| 105 \| 11 \| 7 \| 142 \| 390 \| 525 \| Uncharacterized protein \| Uncharacterized protein; afdb-uniprot50 \| AF-R5C5C3-F1-MODEL\_V4 \| 1.0 \| 3.078e-05 \| 119 \| 0.132 \| 188 \| 118 \| 9 \| 6 \| 165 \| 15 \| 185 \| BppU\_N domain-containing protein \| BppU\_N domain-containing protein; afdb-uniprot50 \| AF-A0A354KQE9-F1-MODEL\_V4 \| 1.0 \| 2.902e-05 \| 119 \| 0.108 \| 203 \| 137 \| 10 \| 1 \| 163 \| 1 \| 199 \| Uncharacterized protein \| Uncharacterized protein; afdb-uniprot50 \| AF-A0A810YRC3-F1-MODEL\_V4 \| 1.0 \| 0.00526 \| 119 \| 0.138 \| 94 \| 72 \| 6 \| 8 \| 97 \| 410 \| 498 \| Uncharacterized protein \| Uncharacterized protein; afdb-uniprot50 \| AF-A0A2N1W3X1-F1-MODEL\_V4 \| 1.0 \| 0.002168 \| 119 \| 0.094 \| 95 \| 66 \| 3 \| 12 \| 95 \| 896 \| 981 \| Glyco\_hydro\_44 domain-containing protein \| Glyco\_hydro\_44 domain-containing protein; afdb-uniprot50 \| AF-A0A7J3TMW7-F1-MODEL\_V4 \| 1.0 \| 0.004153 \| 118 \| 0.112 \| 124 \| 86 \| 7 \| 7 \| 108 \| 48 \| 169 \| Uncharacterized protein \| Uncharacterized protein; afdb-uniprot50 \| AF-A0A1G4VBC8-F1-MODEL\_V4 \| 1.0 \| 0.0002164 \| 118 \| 0.139 \| 172 \| 123 \| 11 \| 3 \| 164 \| 14 \| 170 \| BppU\_N domain-containing protein \| BppU\_N domain-containing protein; afdb-uniprot50 \| AF-A0A4W6F9B7-F1-MODEL\_V4 \| 1.0 \| 6.256e-05 \| 118 \| 0.077 \| 206 \| 140 \| 9 \| 9 \| 177 \| 24 \| 216 \| Ig-like domain-containing protein \| Ig-like domain-containing protein; afdb-uniprot50 \| AF-A0A174CIG2-F1-MODEL\_V4 \| 1.0 \| 0.0008423 \| 118 \| 0.14 \| 171 \| 117 \| 8 \| 5 \| 166 \| 16 \| 165 \| Uncharacterized protein \| Uncharacterized protein; afdb-uniprot50 \| AF-A0A8B6H4W0-F1-MODEL\_V4 \| 1.0 \| 0.0004665 \| 118 \| 0.098 \| 142 \| 99 \| 7 \| 5 \| 142 \| 162 \| 278 \| Uncharacterized protein \| Uncharacterized protein; afdb-uniprot50 \| AF-A0A6N2ZND8-F1-MODEL\_V4 \| 1.0 \| 0.0001923 \| 118 \| 0.124 \| 177 \| 114 \| 9 \| 6 \| 166 \| 20 \| 171 \| BppU\_N domain-containing protein \| BppU\_N domain-containing protein; afdb-uniprot50 \| AF-A0A2H3PGQ7-F1-MODEL\_V4 \| 1.0 \| 0.0004145 \| 118 \| 0.142 \| 161 \| 114 \| 9 \| 2 \| 142 \| 11 \| 167 \| BppU\_N domain-containing protein \| BppU\_N domain-containing protein; afdb-uniprot50 \| AF-R0AR67-F1-MODEL\_V4 \| 1.0 \| 0.0001198 \| 118 \| 0.157 \| 178 \| 116 \| 9 \| 8 \| 166 \| 16 \| 178 \| Uncharacterized protein \| Uncharacterized protein; afdb-uniprot50 \| AF-R6HGZ8-F1-MODEL\_V4 \| 1.0 \| 0.0001198 \| 118 \| 0.129 \| 177 \| 110 \| 7 \| 6 \| 166 \| 16 \| 164 \| BppU\_N domain-containing protein \| BppU\_N domain-containing protein; afdb-uniprot50 \| AF-A0A1T0AWF2-F1-MODEL\_V4 \| 1.0 \| 0.009498 \| 118 \| 0.104 \| 96 \| 74 \| 5 \| 8 \| 96 \| 241 \| 331 \| Uncharacterized protein \| Uncharacterized protein; afdb-uniprot50 \| AF-A0A843BS13-F1-MODEL\_V4 \| 1.0 \| 0.0004145 \| 118 \| 0.123 \| 138 \| 100 \| 10 \| 5 \| 127 \| 441 \| 572 \| Uncharacterized protein \| Uncharacterized protein; afdb-uniprot50 \| AF-A0A1C6DXN6-F1-MODEL\_V4 \| 1.0 \| 0.001132 \| 117 \| 0.096 \| 124 \| 95 \| 6 \| 6 \| 122 \| 15 \| 128 \| Uncharacterized protein \| Uncharacterized protein; afdb-uniprot50 \| AF-A0A1Y4G926-F1-MODEL\_V4 \| 1.0 \| 0.002168 \| 117 \| 0.145 \| 131 \| 94 \| 8 \| 9 \| 125 \| 21 \| 147 \| BppU\_N domain-containing protein \| BppU\_N domain-containing protein; afdb-uniprot50 \| AF-C2VRW8-F1-MODEL\_V4 \| 1.0 \| 0.0002295 \| 117 \| 0.133 \| 150 \| 97 \| 10 \| 9 \| 142 \| 98 \| 230 \| Uncharacterized protein \| Uncharacterized protein; afdb-uniprot50 \| AF-R9JK16-F1-MODEL\_V4 \| 1.0 \| 0.0003085 \| 117 \| 0.156 \| 179 \| 118 \| 9 \| 1 \| 167 \| 18 \| 175 \| Uncharacterized protein \| Uncharacterized protein; afdb-uniprot50 \| AF-A0A7X8ZBB4-F1-MODEL\_V4 \| 1.0 \| 4.388e-05 \| 117 \| 0.173 \| 173 \| 108 \| 10 \| 9 \| 166 \| 11 \| 163 \| Uncharacterized protein \| Uncharacterized protein; afdb-uniprot50 \| AF-A0A0H5SGG3-F1-MODEL\_V4 \| 1.0 \| 0.0004145 \| 117 \| 0.14 \| 171 \| 109 \| 8 \| 9 \| 166 \| 21 \| 166 \| Uncharacterized protein \| Uncharacterized protein; afdb-uniprot50 \| AF-A0A316M9Y2-F1-MODEL\_V4 \| 1.0 \| 0.001816 \| 117 \| 0.094 \| 169 \| 125 \| 7 \| 9 \| 166 \| 22 \| 173 \| Uncharacterized protein \| Uncharacterized protein; afdb-uniprot50 \| AF-A0A411WGX7-F1-MODEL\_V4 \| 1.0 \| 0.002168 \| 116 \| 0.113 \| 106 \| 76 \| 7 \| 8 \| 104 \| 55 \| 151 \| Ig-like domain-containing protein \| Ig-like domain-containing protein; afdb-uniprot50 \| AF-A0A0F6W690-F1-MODEL\_V4 \| 1.0 \| 7.469e-05 \| 116 \| 0.156 \| 147 \| 99 \| 7 \| 7 \| 142 \| 138 \| 270 \| Uncharacterized protein \| Uncharacterized protein; afdb-uniprot50 \| AF-A0A6J7FJT6-F1-MODEL\_V4 \| 1.0 \| 0.0001923 \| 116 \| 0.178 \| 146 \| 101 \| 9 \| 9 \| 142 \| 154 \| 292 \| Unannotated protein \| Unannotated protein; afdb-uniprot50 \| AF-A0A3E2VR12-F1-MODEL\_V4 \| 1.0 \| 0.0004145 \| 116 \| 0.148 \| 175 \| 121 \| 9 \| 8 \| 166 \| 16 \| 178 \| Uncharacterized protein \| Uncharacterized protein; afdb-uniprot50 \| AF-A0A0Q9JHR9-F1-MODEL\_V4 \| 1.0 \| 0.001926 \| 116 \| 0.101 \| 108 \| 77 \| 8 \| 8 \| 107 \| 166 \| 261 \| Uncharacterized protein \| Uncharacterized protein; afdb-uniprot50 \| AF-A0A1C6ET81-F1-MODEL\_V4 \| 1.0 \| 0.001132 \| 116 \| 0.117 \| 179 \| 131 \| 9 \| 9 \| 179 \| 22 \| 181 \| Uncharacterized protein \| Uncharacterized protein; afdb-uniprot50 \| AF-A0A7G5MXH1-F1-MODEL\_V4 \| 1.0 \| 0.0002741 \| 116 \| 0.165 \| 169 \| 104 \| 9 \| 6 \| 166 \| 17 \| 156 \| Uncharacterized protein \| Uncharacterized protein; afdb-uniprot50 \| AF-A0A653VSI4-F1-MODEL\_V4 \| 1.0 \| 0.006281 \| 116 \| 0.104 \| 125 \| 99 \| 6 \| 9 \| 124 \| 23 \| 143 \| Uncharacterized protein \| Uncharacterized protein; afdb-uniprot50 \| AF-A0A7S0INU3-F1-MODEL\_V4 \| 1.0 \| 0.001067 \| 115 \| 0.122 \| 122 \| 80 \| 6 \| 7 \| 107 \| 78 \| 193 \| Hypothetical protein \| Hypothetical protein; afdb-uniprot50 \| AF-A0A6I2FDN3-F1-MODEL\_V4 \| 1.0 \| 0.0003907 \| 115 \| 0.143 \| 132 \| 92 \| 9 \| 7 \| 131 \| 38 \| 155 \| Uncharacterized protein \| Uncharacterized protein; afdb-uniprot50 \| AF-A0A843Z2H5-F1-MODEL\_V4 \| 1.0 \| 0.000794 \| 115 \| 0.133 \| 120 \| 87 \| 9 \| 17 \| 124 \| 35 \| 149 \| DUF2479 domain-containing protein \| DUF2479 domain-containing protein; afdb-uniprot50 \| AF-F8WU49-F1-MODEL\_V4 \| 1.0 \| 0.0008423 \| 115 \| 0.095 \| 168 \| 128 \| 5 \| 9 \| 169 \| 22 \| 172 \| Uncharacterized protein \| Uncharacterized protein; afdb-uniprot50 \| AF-A0A7K3XZ12-F1-MODEL\_V4 \| 1.0 \| 0.0001198 \| 115 \| 0.121 \| 181 \| 117 \| 10 \| 6 \| 166 \| 22 \| 180 \| Uncharacterized protein \| Uncharacterized protein; afdb-uniprot50 \| AF-A0A2V2FF44-F1-MODEL\_V4 \| 1.0 \| 0.0004949 \| 115 \| 0.109 \| 146 \| 106 \| 10 \| 9 \| 138 \| 27 \| 164 \| Uncharacterized protein \| Uncharacterized protein; afdb-uniprot50 \| AF-K1GH27-F1-MODEL\_V4 \| 1.0 \| 0.0002908 \| 115 \| 0.154 \| 181 \| 121 \| 8 \| 9 \| 179 \| 21 \| 179 \| Uncharacterized protein \| Uncharacterized protein; afdb-uniprot50 \| AF-A0A854ZIF3-F1-MODEL\_V4 \| 1.0 \| 0.0001812 \| 115 \| 0.139 \| 179 \| 112 \| 11 \| 8 \| 162 \| 17 \| 177 \| Uncharacterized protein \| Uncharacterized protein; afdb-uniprot50 \| AF-A0A0F5HNP9-F1-MODEL\_V4 \| 1.0 \| 0.001521 \| 115 \| 0.104 \| 163 \| 114 \| 6 \| 9 \| 164 \| 23 \| 160 \| Phage pre-neck appendage-like protein \| Phage pre-neck appendage-like protein; afdb-uniprot50 \| AF-N2B7B9-F1-MODEL\_V4 \| 1.0 \| 0.0004397 \| 114 \| 0.101 \| 158 \| 110 \| 9 \| 9 \| 142 \| 24 \| 173 \| Uncharacterized protein \| Uncharacterized protein; afdb-uniprot50 \| AF-W0U658-F1-MODEL\_V4 \| 1.0 \| 0.0007484 \| 114 \| 0.083 \| 167 \| 120 \| 7 \| 9 \| 166 \| 21 \| 163 \| BppU\_N domain-containing protein \| BppU\_N domain-containing protein; afdb-uniprot50 \| AF-A0A416RMF7-F1-MODEL\_V4 \| 1.0 \| 0.0001518 \| 114 \| 0.129 \| 177 \| 110 \| 11 \| 9 \| 166 \| 23 \| 174 \| Uncharacterized protein \| Uncharacterized protein; afdb-uniprot50 \| AF-A0A4S2EF76-F1-MODEL\_V4 \| 1.0 \| 0.000557 \| 114 \| 0.105 \| 151 \| 106 \| 9 \| 8 \| 142 \| 9 \| 146 \| Collagen-like protein \| Collagen-like protein; afdb-uniprot50 \| AF-A0A1B1LIW2-F1-MODEL\_V4 \| 1.0 \| 0.0003085 \| 114 \| 0.124 \| 177 \| 119 \| 7 \| 6 \| 166 \| 15 \| 171 \| Phage protein \| Phage protein; afdb-uniprot50 \| AF-A0A7G9G6Y2-F1-MODEL\_V4 \| 1.0 \| 0.0007484 \| 114 \| 0.13 \| 169 \| 111 \| 8 \| 9 \| 168 \| 21 \| 162 \| BppU family phage baseplate upper protein \| BppU family phage baseplate upper protein; afdb-uniprot50 \| AF-A0A3A9FK46-F1-MODEL\_V4 \| 1.0 \| 0.0001518 \| 114 \| 0.124 \| 177 \| 116 \| 8 \| 9 \| 166 \| 20 \| 176 \| Uncharacterized protein \| Uncharacterized protein; afdb-uniprot50 \| AF-A0A6N7SV12-F1-MODEL\_V4 \| 1.0 \| 0.0004665 \| 114 \| 0.081 \| 171 \| 122 \| 8 \| 9 \| 164 \| 24 \| 174 \| DUF2479 domain-containing protein \| DUF2479 domain-containing protein; afdb-uniprot50 \| AF-A0A658JQ79-F1-MODEL\_V4 \| 1.0 \| 0.001816 \| 114 \| 0.132 \| 174 \| 116 \| 9 \| 4 \| 166 \| 18 \| 167 \| DUF2479 domain-containing protein \| DUF2479 domain-containing protein; afdb-uniprot50 \| AF-A0A1Q5PL34-F1-MODEL\_V4 \| 1.0 \| 0.004153 \| 114 \| 0.131 \| 99 \| 80 \| 4 \| 10 \| 107 \| 587 \| 680 \| Fibronectin type-III domain-containing protein \| Fibronectin type-III domain-containing protein; afdb-uniprot50 \| AF-A0A177L7H1-F1-MODEL\_V4 \| 1.0 \| 0.001926 \| 114 \| 0.096 \| 165 \| 122 \| 5 \| 9 \| 166 \| 23 \| 167 \| BppU\_N domain-containing protein \| BppU\_N domain-containing protein; afdb-uniprot50 \| AF-X0SIW3-F1-MODEL\_V4 \| 1.0 \| 0.008953 \| 113 \| 0.106 \| 103 \| 84 \| 6 \| 9 \| 107 \| 2 \| 100 \| Uncharacterized protein \| Uncharacterized protein; afdb-uniprot50 \| AF-A0A2L2BRS6-F1-MODEL\_V4 \| 1.0 \| 0.001351 \| 113 \| 0.09 \| 121 \| 96 \| 8 \| 9 \| 124 \| 22 \| 133 \| Uncharacterized protein \| Uncharacterized protein; afdb-uniprot50 \| AF-A0A0M6WYK5-F1-MODEL\_V4 \| 1.0 \| 0.0002741 \| 113 \| 0.113 \| 176 \| 112 \| 11 \| 9 \| 166 \| 24 \| 173 \| BppU\_N domain-containing protein \| BppU\_N domain-containing protein; afdb-uniprot50 \| AF-A0A2H9MF23-F1-MODEL\_V4 \| 1.0 \| 0.002168 \| 113 \| 0.147 \| 102 \| 73 \| 7 \| 8 \| 104 \| 269 \| 361 \| Peptidase\_S8 domain-containing protein \| Peptidase\_S8 domain-containing protein; afdb-uniprot50 \| AF-A0A6P5KX56-F1-MODEL\_V4 \| 1.0 \| 0.0002164 \| 113 \| 0.129 \| 154 \| 104 \| 8 \| 7 \| 142 \| 225 \| 366 \| intercellular adhesion molecule 3 isoform X2 \| intercellular adhesion molecule 3 isoform X2; afdb-uniprot50 \| AF-A0A1Q9J512-F1-MODEL\_V4 \| 1.0 \| 0.0008423 \| 113 \| 0.11 \| 136 \| 95 \| 8 \| 9 \| 127 \| 22 \| 148 \| BppU\_N domain-containing protein \| BppU\_N domain-containing protein; afdb-uniprot50 \| AF-A0A033V779-F1-MODEL\_V4 \| 1.0 \| 0.0003683 \| 113 \| 0.122 \| 155 \| 104 \| 8 \| 20 \| 166 \| 48 \| 178 \| BppU\_N domain-containing protein \| BppU\_N domain-containing protein; afdb-uniprot50 \| AF-A0A6M5FW15-F1-MODEL\_V4 \| 1.0 \| 0.0005909 \| 113 \| 0.176 \| 170 \| 108 \| 9 \| 9 \| 166 \| 22 \| 171 \| Uncharacterized protein \| Uncharacterized protein; afdb-uniprot50 \| AF-A0A060YWJ5-F1-MODEL\_V4 \| 1.0 \| 0.007069 \| 112 \| 0.162 \| 111 \| 75 \| 9 \| 7 \| 104 \| 5 \| 110 \| Uncharacterized protein \| Uncharacterized protein; afdb-uniprot50 \| AF-A0A1C5W5W4-F1-MODEL\_V4 \| 1.0 \| 0.000525 \| 112 \| 0.074 \| 147 \| 105 \| 8 \| 7 \| 128 \| 11 \| 151 \| Uncharacterized protein \| Uncharacterized protein; afdb-uniprot50 \| AF-G8SD22-F1-MODEL\_V4 \| 1.0 \| 0.0006268 \| 112 \| 0.112 \| 125 \| 95 \| 9 \| 6 \| 124 \| 55 \| 169 \| Fibronectin type-III domain-containing protein \| Fibronectin type-III domain-containing protein; afdb-uniprot50 \| AF-V5GR29-F1-MODEL\_V4 \| 1.0 \| 0.0001923 \| 112 \| 0.12 \| 175 \| 123 \| 9 \| 3 \| 168 \| 6 \| 158 \| Dolichyl-diphosphooligosaccharide--protein glycosyltransferase subunit 2 \| Dolichyl-diphosphooligosaccharide--protein glycosyltransferase subunit 2; afdb-uniprot50 \| AF-W0FKU2-F1-MODEL\_V4 \| 1.0 \| 0.0003683 \| 112 \| 0.122 \| 171 \| 121 \| 5 \| 8 \| 166 \| 37 \| 190 \| Uncharacterized protein \| Uncharacterized protein; afdb-uniprot50 \| AF-A0A412T7C3-F1-MODEL\_V4 \| 1.0 \| 0.0003907 \| 112 \| 0.134 \| 171 \| 113 \| 12 \| 9 \| 164 \| 25 \| 175 \| DUF2479 domain-containing protein \| DUF2479 domain-containing protein; afdb-uniprot50 \| AF-A0A133KN76-F1-MODEL\_V4 \| 1.0 \| 0.0003085 \| 112 \| 0.176 \| 136 \| 88 \| 11 \| 9 \| 124 \| 28 \| 159 \| Uncharacterized protein \| Uncharacterized protein; afdb-uniprot50 \| AF-A0A316QDJ6-F1-MODEL\_V4 \| 1.0 \| 0.000794 \| 112 \| 0.108 \| 166 \| 121 \| 7 \| 8 \| 165 \| 18 \| 164 \| Uncharacterized protein \| Uncharacterized protein; afdb-uniprot50 \| AF-A0A348ZCH6-F1-MODEL\_V4 \| 1.0 \| 0.002913 \| 112 \| 0.09 \| 132 \| 90 \| 9 \| 9 \| 124 \| 22 \| 139 \| Uncharacterized protein \| Uncharacterized protein; afdb-uniprot50 \| AF-A0A261QKZ1-F1-MODEL\_V4 \| 1.0 \| 0.004153 \| 112 \| 0.097 \| 123 \| 96 \| 6 \| 9 \| 124 \| 23 \| 137 \| Uncharacterized protein \| Uncharacterized protein; afdb-uniprot50 \| AF-A0A4U9VLC3-F1-MODEL\_V4 \| 1.0 \| 0.005581 \| 112 \| 0.105 \| 95 \| 75 \| 6 \| 8 \| 97 \| 10 \| 99 \| HYR domain \| HYR domain; afdb-uniprot50 \| AF-A0A379AD61-F1-MODEL\_V4 \| 1.0 \| 0.004674 \| 112 \| 0.15 \| 93 \| 65 \| 8 \| 8 \| 93 \| 52 \| 137 \| Uncharacterized protein \| Uncharacterized protein; afdb-uniprot50 \| AF-A0A3D3UB27-F1-MODEL\_V4 \| 1.0 \| 0.0003471 \| 112 \| 0.142 \| 190 \| 119 \| 12 \| 6 \| 179 \| 16 \| 177 \| BppU\_N domain-containing protein \| BppU\_N domain-containing protein; afdb-uniprot50 \| AF-A0A2N2P695-F1-MODEL\_V4 \| 1.0 \| 0.007955 \| 112 \| 0.134 \| 97 \| 70 \| 8 \| 8 \| 94 \| 1066 \| 1158 \| Uncharacterized protein \| Uncharacterized protein; afdb-uniprot50 \| AF-A0A350V7B0-F1-MODEL\_V4 \| 1.0 \| 0.0004397 \| 111 \| 0.094 \| 170 \| 125 \| 9 \| 1 \| 156 \| 12 \| 166 \| Uncharacterized protein \| Uncharacterized protein; afdb-uniprot50 \| AF-A0A417GRQ1-F1-MODEL\_V4 \| 1.0 \| 4.137e-05 \| 111 \| 0.162 \| 185 \| 90 \| 11 \| 15 \| 142 \| 32 \| 208 \| Uncharacterized protein \| Uncharacterized protein; afdb-uniprot50 \| AF-T2RFS3-F1-MODEL\_V4 \| 1.0 \| 0.0006268 \| 111 \| 0.122 \| 163 \| 111 \| 5 \| 15 \| 165 \| 12 \| 154 \| BppU\_N domain-containing protein \| BppU\_N domain-containing protein; afdb-uniprot50 \| AF-A0A7C7QDX2-F1-MODEL\_V4 \| 1.0 \| 0.00244 \| 111 \| 0.174 \| 132 \| 77 \| 9 \| 7 \| 124 \| 267 \| 380 \| Uncharacterized protein \| Uncharacterized protein; afdb-uniprot50 \| AF-A0A242WLZ3-F1-MODEL\_V4 \| 1.0 \| 0.001274 \| 111 \| 0.12 \| 133 \| 97 \| 9 \| 9 \| 127 \| 26 \| 152 \| BppU\_N domain-containing protein \| BppU\_N domain-containing protein; afdb-uniprot50 \| AF-A0A1F4WKG2-F1-MODEL\_V4 \| 1.0 \| 0.004153 \| 111 \| 0.138 \| 94 \| 78 \| 2 \| 3 \| 96 \| 513 \| 603 \| Uncharacterized protein \| Uncharacterized protein; afdb-uniprot50 \| AF-A0A8B6H4W6-F1-MODEL\_V4 \| 1.0 \| 0.007955 \| 111 \| 0.094 \| 95 \| 72 \| 5 \| 7 \| 94 \| 373 \| 460 \| Uncharacterized protein \| Uncharacterized protein; afdb-uniprot50 \| AF-A0A7U9T1A0-F1-MODEL\_V4 \| 1.0 \| 0.001521 \| 111 \| 0.143 \| 167 \| 117 \| 6 \| 6 \| 166 \| 15 \| 161 \| Uncharacterized protein \| Uncharacterized protein; afdb-uniprot50 \| AF-A0A143Z024-F1-MODEL\_V4 \| 1.0 \| 0.00244 \| 111 \| 0.147 \| 170 \| 118 \| 7 \| 8 \| 166 \| 24 \| 177 \| Uncharacterized protein \| Uncharacterized protein; afdb-uniprot50 \| AF-A0A380DMD7-F1-MODEL\_V4 \| 1.0 \| 0.0007055 \| 110 \| 0.117 \| 188 \| 127 \| 10 \| 9 \| 179 \| 24 \| 189 \| Phage tail fiber protein \| Phage tail fiber protein; afdb-uniprot50 \| AF-A0A823HEB2-F1-MODEL\_V4 \| 1.0 \| 0.001816 \| 110 \| 0.082 \| 169 \| 119 \| 8 \| 9 \| 158 \| 28 \| 179 \| DUF2479 domain-containing protein \| DUF2479 domain-containing protein; afdb-uniprot50 \| AF-A0A037Z2D6-F1-MODEL\_V4 \| 1.0 \| 5.897e-05 \| 110 \| 0.175 \| 171 \| 103 \| 11 \| 9 \| 166 \| 11 \| 156 \| Uncharacterized protein \| Uncharacterized protein; afdb-uniprot50 \| AF-A0A2V2G0U0-F1-MODEL\_V4 \| 1.0 \| 0.0004145 \| 110 \| 0.127 \| 172 \| 117 \| 5 \| 7 \| 166 \| 20 \| 170 \| Uncharacterized protein \| Uncharacterized protein; afdb-uniprot50 \| AF-A0A3C2EKY6-F1-MODEL\_V4 \| 1.0 \| 0.0001518 \| 110 \| 0.195 \| 169 \| 102 \| 12 \| 9 \| 166 \| 11 \| 156 \| Uncharacterized protein \| Uncharacterized protein; afdb-uniprot50 \| AF-A0A1N7F4Z8-F1-MODEL\_V4 \| 1.0 \| 0.0001431 \| 110 \| 0.136 \| 168 \| 110 \| 10 \| 3 \| 142 \| 17 \| 177 \| BppU\_N domain-containing protein \| BppU\_N domain-containing protein; afdb-uniprot50 \| AF-A0A7M3YFZ2-F1-MODEL\_V4 \| 1.0 \| 0.0004397 \| 110 \| 0.101 \| 148 \| 113 \| 9 \| 7 \| 142 \| 472 \| 611 \| Uncharacterized protein \| Uncharacterized protein; afdb-uniprot50 \| AF-A0A523W2L9-F1-MODEL\_V4 \| 1.0 \| 0.000113 \| 110 \| 0.142 \| 176 \| 138 \| 8 \| 8 \| 179 \| 457 \| 623 \| Uncharacterized protein \| Uncharacterized protein; afdb-uniprot50 \| AF-A0A4Y8UNF9-F1-MODEL\_V4 \| 1.0 \| 0.001816 \| 110 \| 0.174 \| 126 \| 89 \| 9 \| 9 \| 126 \| 892 \| 1010 \| Fibronectin type III domain-containing protein \| Fibronectin type III domain-containing protein; afdb-uniprot50 \| AF-A0A2E1ACQ2-F1-MODEL\_V4 \| 1.0 \| 0.0008936 \| 110 \| 0.227 \| 110 \| 69 \| 5 \| 7 \| 109 \| 669 \| 769 \| Uncharacterized protein \| Uncharacterized protein; afdb-uniprot50 \| AF-A0A3B8NXB8-F1-MODEL\_V4 \| 1.0 \| 0.001613 \| 109 \| 0.135 \| 118 \| 95 \| 4 \| 14 \| 124 \| 32 \| 149 \| Uncharacterized protein \| Uncharacterized protein; afdb-uniprot50 \| AF-A0A5R9C6T5-F1-MODEL\_V4 \| 1.0 \| 0.000665 \| 109 \| 0.126 \| 190 \| 126 \| 11 \| 1 \| 170 \| 13 \| 182 \| Uncharacterized protein \| Uncharacterized protein; afdb-uniprot50 \| AF-A0A429ZSE8-F1-MODEL\_V4 \| 1.0 \| 0.001006 \| 109 \| 0.112 \| 169 \| 120 \| 9 \| 9 \| 166 \| 21 \| 170 \| BppU\_N domain-containing protein \| BppU\_N domain-containing protein; afdb-uniprot50 \| AF-A0A173Z244-F1-MODEL\_V4 \| 1.0 \| 0.0003683 \| 109 \| 0.142 \| 182 \| 116 \| 10 \| 5 \| 166 \| 23 \| 184 \| Uncharacterized protein \| Uncharacterized protein; afdb-uniprot50 \| AF-A0A6Z6TAU9-F1-MODEL\_V4 \| 1.0 \| 0.001434 \| 109 \| 0.092 \| 183 \| 122 \| 11 \| 7 \| 169 \| 25 \| 183 \| DUF2479 domain-containing protein \| DUF2479 domain-containing protein; afdb-uniprot50 \| AF-A0A8B5WFU7-F1-MODEL\_V4 \| 1.0 \| 0.002913 \| 109 \| 0.137 \| 109 \| 82 \| 5 \| 6 \| 107 \| 630 \| 733 \| Uncharacterized protein \| Uncharacterized protein; afdb-uniprot50 \| AF-A0A0R2D990-F1-MODEL\_V4 \| 1.0 \| 0.001132 \| 109 \| 0.181 \| 138 \| 84 \| 6 \| 9 \| 142 \| 340 \| 452 \| Uncharacterized protein \| Uncharacterized protein; afdb-uniprot50 \| AF-A0A1S8S9Z7-F1-MODEL\_V4 \| 1.0 \| 0.000557 \| 108 \| 0.089 \| 168 \| 124 \| 5 \| 9 \| 166 \| 20 \| 168 \| Uncharacterized protein \| Uncharacterized protein; afdb-uniprot50 \| AF-R5FZM6-F1-MODEL\_V4 \| 1.0 \| 0.0001923 \| 108 \| 0.132 \| 181 \| 114 \| 10 \| 8 \| 166 \| 16 \| 175 \| Uncharacterized protein \| Uncharacterized protein; afdb-uniprot50 \| AF-A0A1C6EHP8-F1-MODEL\_V4 \| 1.0 \| 0.0003272 \| 108 \| 0.082 \| 182 \| 126 \| 8 \| 7 \| 157 \| 18 \| 189 \| Uncharacterized protein \| Uncharacterized protein; afdb-uniprot50 \| AF-G9QBI2-F1-MODEL\_V4 \| 1.0 \| 0.0008936 \| 108 \| 0.137 \| 153 \| 99 \| 9 \| 9 \| 142 \| 235 \| 373 \| Uncharacterized protein \| Uncharacterized protein; afdb-uniprot50 \| AF-A0A1F7QTB1-F1-MODEL\_V4 \| 1.0 \| 0.003091 \| 108 \| 0.104 \| 105 \| 81 \| 5 \| 8 \| 107 \| 42 \| 138 \| Glucanase \| Glucanase; afdb-uniprot50 \| AF-R5L6I5-F1-MODEL\_V4 \| 1.0 \| 0.0006268 \| 108 \| 0.121 \| 156 \| 109 \| 9 \| 8 \| 142 \| 15 \| 163 \| Uncharacterized protein \| Uncharacterized protein; afdb-uniprot50 \| AF-R7NRH2-F1-MODEL\_V4 \| 1.0 \| 0.00369 \| 108 \| 0.083 \| 132 \| 91 \| 9 \| 9 \| 124 \| 22 \| 139 \| Uncharacterized protein \| Uncharacterized protein; afdb-uniprot50 \| AF-A0A1G7MZS4-F1-MODEL\_V4 \| 1.0 \| 0.001067 \| 108 \| 0.109 \| 128 \| 98 \| 9 \| 8 \| 128 \| 438 \| 556 \| LPXTG-motif cell wall anchor domain-containing protein \| LPXTG-motif cell wall anchor domain-containing protein; afdb-uniprot50 \| AF-A0A2H9MLH3-F1-MODEL\_V4 \| 1.0 \| 0.0003085 \| 108 \| 0.102 \| 146 \| 113 \| 7 \| 9 \| 142 \| 480 \| 619 \| Peptidase\_S8 domain-containing protein \| Peptidase\_S8 domain-containing protein; afdb-uniprot50 \| AF-A0A3A1XKV0-F1-MODEL\_V4 \| 1.0 \| 0.0002435 \| 108 \| 0.145 \| 179 \| 109 \| 13 \| 6 \| 165 \| 20 \| 173 \| Methyl-accepting transducer domain-containing protein \| Methyl-accepting transducer domain-containing protein; afdb-uniprot50 \| AF-A0A374JQW7-F1-MODEL\_V4 \| 1.0 \| 0.0008423 \| 108 \| 0.105 \| 171 \| 110 \| 10 \| 9 \| 164 \| 25 \| 167 \| DUF2479 domain-containing protein \| DUF2479 domain-containing protein; afdb-uniprot50 \| AF-A0A1C6DME9-F1-MODEL\_V4 \| 1.0 \| 0.0004397 \| 107 \| 0.122 \| 179 \| 113 \| 10 \| 9 \| 173 \| 19 \| 167 \| Domain of uncharacterized function (DUF2479) \| Domain of uncharacterized function (DUF2479); afdb-uniprot50 \| AF-B4DA57-F1-MODEL\_V4 \| 1.0 \| 0.001712 \| 107 \| 0.144 \| 138 \| 97 \| 5 \| 10 \| 128 \| 22 \| 157 \| Uncharacterized protein \| Uncharacterized protein; afdb-uniprot50 \| AF-A0A1C5T6I9-F1-MODEL\_V4 \| 1.0 \| 0.0002741 \| 107 \| 0.082 \| 182 \| 125 \| 7 \| 5 \| 166 \| 11 \| 170 \| BppU\_N domain-containing protein \| BppU\_N domain-containing protein; afdb-uniprot50 \| AF-K8DBU7-F1-MODEL\_V4 \| 1.0 \| 0.0001518 \| 107 \| 0.15 \| 146 \| 91 \| 9 \| 8 \| 142 \| 190 \| 313 \| Large repetitive protein \| Large repetitive protein; afdb-uniprot50 \| AF-A0A6N7YBU7-F1-MODEL\_V4 \| 1.0 \| 0.0001518 \| 107 \| 0.149 \| 181 \| 115 \| 8 \| 8 \| 166 \| 37 \| 200 \| Uncharacterized protein \| Uncharacterized protein; afdb-uniprot50 \| AF-A0A2D6NE59-F1-MODEL\_V4 \| 1.0 \| 0.0004665 \| 107 \| 0.136 \| 132 \| 91 \| 8 \| 8 \| 126 \| 258 \| 379 \| Uncharacterized protein \| Uncharacterized protein; afdb-uniprot50 \| AF-A0A1Q6J949-F1-MODEL\_V4 \| 1.0 \| 0.0001708 \| 107 \| 0.142 \| 183 \| 126 \| 7 \| 7 \| 166 \| 12 \| 186 \| Uncharacterized protein \| Uncharacterized protein; afdb-uniprot50 \| AF-V5MHJ2-F1-MODEL\_V4 \| 1.0 \| 0.0008936 \| 107 \| 0.118 \| 144 \| 99 \| 10 \| 3 \| 127 \| 16 \| 150 \| BppU\_N domain-containing protein \| BppU\_N domain-containing protein; afdb-uniprot50 \| AF-W4VP65-F1-MODEL\_V4 \| 1.0 \| 0.003478 \| 107 \| 0.114 \| 114 \| 83 \| 5 \| 5 \| 108 \| 608 \| 713 \| Uncharacterized protein \| Uncharacterized protein; afdb-uniprot50 \| AF-R8SYT5-F1-MODEL\_V4 \| 1.0 \| 0.001816 \| 107 \| 0.106 \| 169 \| 124 \| 10 \| 9 \| 166 \| 23 \| 175 \| SGNH\_hydro domain-containing protein \| SGNH\_hydro domain-containing protein; afdb-uniprot50 \| AF-A0A7X2RXU3-F1-MODEL\_V4 \| 1.0 \| 0.001613 \| 106 \| 0.106 \| 113 \| 82 \| 6 \| 28 \| 134 \| 49 \| 148 \| DUF2479 domain-containing protein \| DUF2479 domain-containing protein; afdb-uniprot50 \| AF-A0A1M6TYC1-F1-MODEL\_V4 \| 1.0 \| 0.0008423 \| 106 \| 0.093 \| 193 \| 140 \| 11 \| 9 \| 179 \| 26 \| 205 \| BppU\_N domain-containing protein \| BppU\_N domain-containing protein; afdb-uniprot50 \| AF-R7K9N5-F1-MODEL\_V4 \| 1.0 \| 0.0002435 \| 106 \| 0.115 \| 173 \| 117 \| 9 \| 7 \| 164 \| 14 \| 165 \| Putative antireceptor \| Putative antireceptor; afdb-uniprot50 \| AF-A0A396ZWI6-F1-MODEL\_V4 \| 1.0 \| 0.0007484 \| 106 \| 0.11 \| 172 \| 131 \| 7 \| 9 \| 168 \| 128 \| 289 \| Dolichyl-diphosphooligosaccharide--protein glycosyltransferase subunit 2 \| Dolichyl-diphosphooligosaccharide--protein glycosyltransferase subunit 2; afdb-uniprot50 \| AF-A0A395YDS6-F1-MODEL\_V4 \| 1.0 \| 0.0001708 \| 106 \| 0.112 \| 177 \| 117 \| 8 \| 8 \| 166 \| 14 \| 168 \| Uncharacterized protein \| Uncharacterized protein; afdb-uniprot50 \| AF-A0A6L8THQ1-F1-MODEL\_V4 \| 1.0 \| 0.001351 \| 106 \| 0.128 \| 171 \| 111 \| 9 \| 7 \| 163 \| 18 \| 164 \| Uncharacterized protein \| Uncharacterized protein; afdb-uniprot50 \| AF-A0A2H1FCV0-F1-MODEL\_V4 \| 1.0 \| 0.001201 \| 105 \| 0.12 \| 133 \| 93 \| 9 \| 7 \| 128 \| 47 \| 166 \| Uncharacterized protein \| Uncharacterized protein; afdb-uniprot50 \| AF-A0A376LJQ6-F1-MODEL\_V4 \| 1.0 \| 0.00592 \| 105 \| 0.092 \| 108 \| 81 \| 7 \| 8 \| 107 \| 119 \| 217 \| Adhesin for cattle intestine colonization \| Adhesin for cattle intestine colonization; afdb-uniprot50 \| AF-A0A6B1VLF8-F1-MODEL\_V4 \| 1.0 \| 0.0003272 \| 105 \| 0.127 \| 149 \| 99 \| 7 \| 6 \| 125 \| 13 \| 159 \| Uncharacterized protein \| Uncharacterized protein; afdb-uniprot50 \| AF-A0A3E3EGS6-F1-MODEL\_V4 \| 1.0 \| 0.0006268 \| 105 \| 0.116 \| 172 \| 119 \| 7 \| 7 \| 166 \| 17 \| 167 \| Uncharacterized protein \| Uncharacterized protein; afdb-uniprot50 \| AF-A0A417VHC6-F1-MODEL\_V4 \| 1.0 \| 0.001201 \| 105 \| 0.145 \| 172 \| 106 \| 12 \| 9 \| 166 \| 23 \| 167 \| DUF2479 domain-containing protein \| DUF2479 domain-containing protein; afdb-uniprot50 \| AF-A0A2D3PVV1-F1-MODEL\_V4 \| 1.0 \| 0.0006268 \| 105 \| 0.121 \| 157 \| 111 \| 6 \| 17 \| 164 \| 31 \| 169 \| Uncharacterized protein \| Uncharacterized protein; afdb-uniprot50 \| AF-A0A2W5XP03-F1-MODEL\_V4 \| 1.0 \| 0.006663 \| 105 \| 0.184 \| 130 \| 81 \| 4 \| 13 \| 124 \| 60 \| 182 \| Peptidase\_MA\_2 domain-containing protein \| Peptidase\_MA\_2 domain-containing protein; afdb-uniprot50 \| AF-A0A6J8CMK5-F1-MODEL\_V4 \| 1.0 \| 0.002589 \| 105 \| 0.15 \| 133 \| 100 \| 6 \| 15 \| 136 \| 169 \| 299 \| Opine dehydrogenase \| Opine dehydrogenase; afdb-uniprot50 \| AF-A0A1G8VNZ3-F1-MODEL\_V4 \| 1.0 \| 0.002168 \| 105 \| 0.132 \| 158 \| 107 \| 7 \| 17 \| 166 \| 35 \| 170 \| BppU\_N domain-containing protein \| BppU\_N domain-containing protein; afdb-uniprot50 \| AF-A0A6H0TDU9-F1-MODEL\_V4 \| 1.0 \| 0.001351 \| 105 \| 0.152 \| 171 \| 115 \| 10 \| 9 \| 166 \| 22 \| 175 \| BppU family phage baseplate upper protein \| BppU family phage baseplate upper protein; afdb-uniprot50 \| AF-D4M5A2-F1-MODEL\_V4 \| 1.0 \| 0.0003272 \| 105 \| 0.14 \| 178 \| 115 \| 9 \| 5 \| 166 \| 16 \| 171 \| Uncharacterized protein \| Uncharacterized protein; afdb-uniprot50 \| AF-A0A356CXZ6-F1-MODEL\_V4 \| 1.0 \| 0.001613 \| 104 \| 0.098 \| 173 \| 119 \| 8 \| 6 \| 166 \| 16 \| 163 \| BppU\_N domain-containing protein \| BppU\_N domain-containing protein; afdb-uniprot50 \| AF-A0A6P5L4Q4-F1-MODEL\_V4 \| 1.0 \| 0.0004949 \| 104 \| 0.114 \| 157 \| 103 \| 8 \| 7 \| 142 \| 138 \| 279 \| intercellular adhesion molecule 3 isoform X3 \| intercellular adhesion molecule 3 isoform X3; afdb-uniprot50 \| AF-A0A7X5CGT2-F1-MODEL\_V4 \| 1.0 \| 0.001434 \| 104 \| 0.11 \| 172 \| 116 \| 10 \| 9 \| 166 \| 59 \| 207 \| DUF2479 domain-containing protein \| DUF2479 domain-containing protein; afdb-uniprot50 \| AF-A0A376VLZ8-F1-MODEL\_V4 \| 1.0 \| 0.001006 \| 103 \| 0.097 \| 143 \| 98 \| 8 \| 8 \| 142 \| 47 \| 166 \| Adhesin for cattle intestine colonization \| Adhesin for cattle intestine colonization; afdb-uniprot50 \| AF-A0A4P7U9X2-F1-MODEL\_V4 \| 1.0 \| 0.0003907 \| 103 \| 0.164 \| 164 \| 87 \| 8 \| 8 \| 124 \| 43 \| 203 \| CopC domain-containing protein \| CopC domain-containing protein; afdb-uniprot50 \| AF-R9MX48-F1-MODEL\_V4 \| 1.0 \| 0.0003471 \| 103 \| 0.153 \| 143 \| 98 \| 5 \| 29 \| 170 \| 41 \| 161 \| Uncharacterized protein \| Uncharacterized protein; afdb-uniprot50 \| AF-A0A356J3M0-F1-MODEL\_V4 \| 1.0 \| 0.0004949 \| 103 \| 0.145 \| 172 \| 112 \| 11 \| 7 \| 166 \| 1 \| 149 \| Uncharacterized protein \| Uncharacterized protein; afdb-uniprot50 \| AF-A0A844QZW9-F1-MODEL\_V4 \| 1.0 \| 0.001351 \| 103 \| 0.138 \| 159 \| 112 \| 8 \| 27 \| 179 \| 50 \| 189 \| DUF2479 domain-containing protein \| DUF2479 domain-containing protein; afdb-uniprot50 \| AF-A0A347WIJ3-F1-MODEL\_V4 \| 1.0 \| 0.001201 \| 103 \| 0.112 \| 187 \| 107 \| 9 \| 6 \| 168 \| 21 \| 172 \| BppU\_N domain-containing protein \| BppU\_N domain-containing protein; afdb-uniprot50 \| AF-A0A822PFD8-F1-MODEL\_V4 \| 1.0 \| 0.001006 \| 103 \| 0.121 \| 165 \| 109 \| 7 \| 15 \| 165 \| 35 \| 177 \| Uncharacterized protein \| Uncharacterized protein; afdb-uniprot50 \| AF-A0A1G6RQ90-F1-MODEL\_V4 \| 1.0 \| 0.002746 \| 103 \| 0.107 \| 140 \| 98 \| 10 \| 5 \| 124 \| 23 \| 155 \| Uncharacterized protein \| Uncharacterized protein; afdb-uniprot50 \| AF-A0A1Y4LS28-F1-MODEL\_V4 \| 1.0 \| 0.001613 \| 103 \| 0.146 \| 177 \| 118 \| 7 \| 5 \| 167 \| 12 \| 169 \| Uncharacterized protein \| Uncharacterized protein; afdb-uniprot50 \| AF-A0A7V7SAY6-F1-MODEL\_V4 \| 1.0 \| 0.001521 \| 103 \| 0.147 \| 136 \| 91 \| 10 \| 9 \| 127 \| 22 \| 149 \| DUF2479 domain-containing protein \| DUF2479 domain-containing protein; afdb-uniprot50 \| AF-W7L1A8-F1-MODEL\_V4 \| 1.0 \| 0.00526 \| 103 \| 0.093 \| 171 \| 127 \| 8 \| 9 \| 166 \| 23 \| 178 \| Phage pre-neck appendage-like protein \| Phage pre-neck appendage-like protein; afdb-uniprot50 \| AF-A0A7K3W8B9-F1-MODEL\_V4 \| 1.0 \| 0.0004145 \| 103 \| 0.153 \| 130 \| 87 \| 10 \| 6 \| 124 \| 386 \| 503 \| LPXTG cell wall anchor domain-containing protein \| LPXTG cell wall anchor domain-containing protein; afdb-uniprot50 \| AF-A0A1W9QEJ3-F1-MODEL\_V4 \| 1.0 \| 0.006663 \| 103 \| 0.085 \| 128 \| 83 \| 6 \| 9 \| 108 \| 165 \| 286 \| Uncharacterized protein \| Uncharacterized protein; afdb-uniprot50 \| AF-A0A1C5VG89-F1-MODEL\_V4 \| 1.0 \| 0.007069 \| 103 \| 0.089 \| 167 \| 126 \| 9 \| 9 \| 166 \| 22 \| 171 \| Uncharacterized protein \| Uncharacterized protein; afdb-uniprot50 \| AF-W0RS98-F1-MODEL\_V4 \| 1.0 \| 0.0003471 \| 103 \| 0.161 \| 198 \| 119 \| 14 \| 6 \| 179 \| 741 \| 915 \| CHASE2 domain protein \| CHASE2 domain protein; afdb-uniprot50 \| AF-A2DAW7-F1-MODEL\_V4 \| 1.0 \| 0.002746 \| 102 \| 0.07 \| 99 \| 80 \| 7 \| 8 \| 100 \| 49 \| 141 \| Uncharacterized protein \| Uncharacterized protein; afdb-uniprot50 \| AF-A2GME7-F1-MODEL\_V4 \| 1.0 \| 0.000557 \| 102 \| 0.156 \| 128 \| 93 \| 9 \| 6 \| 124 \| 44 \| 165 \| Uncharacterized protein \| Uncharacterized protein; afdb-uniprot50 \| AF-A0A1T4K5N9-F1-MODEL\_V4 \| 1.0 \| 0.001351 \| 102 \| 0.118 \| 185 \| 124 \| 11 \| 1 \| 166 \| 15 \| 179 \| BppU\_N domain-containing protein \| BppU\_N domain-containing protein; afdb-uniprot50 \| AF-A0A1H3BKQ8-F1-MODEL\_V4 \| 1.0 \| 0.000794 \| 102 \| 0.11 \| 190 \| 119 \| 11 \| 4 \| 145 \| 21 \| 208 \| Uncharacterized protein \| Uncharacterized protein; afdb-uniprot50 \| AF-A0A662SM31-F1-MODEL\_V4 \| 1.0 \| 0.003478 \| 102 \| 0.152 \| 131 \| 84 \| 8 \| 7 \| 124 \| 269 \| 385 \| Uncharacterized protein \| Uncharacterized protein; afdb-uniprot50 \| AF-A0A1Y4DR79-F1-MODEL\_V4 \| 1.0 \| 0.007499 \| 102 \| 0.13 \| 138 \| 98 \| 8 \| 4 \| 125 \| 55 \| 186 \| BppU\_N domain-containing protein \| BppU\_N domain-containing protein; afdb-uniprot50 \| AF-A0A1S9UV81-F1-MODEL\_V4 \| 1.0 \| 0.0004949 \| 102 \| 0.164 \| 140 \| 95 \| 7 \| 19 \| 142 \| 43 \| 176 \| BppU\_N domain-containing protein \| BppU\_N domain-containing protein; afdb-uniprot50 \| AF-A0A534I7T6-F1-MODEL\_V4 \| 1.0 \| 0.009498 \| 102 \| 0.123 \| 105 \| 79 \| 6 \| 8 \| 107 \| 443 \| 539 \| Uncharacterized protein \| Uncharacterized protein; afdb-uniprot50 \| AF-A0A3E4GMK1-F1-MODEL\_V4 \| 1.0 \| 0.000204 \| 102 \| 0.106 \| 178 \| 122 \| 8 \| 7 \| 166 \| 11 \| 169 \| Uncharacterized protein \| Uncharacterized protein; afdb-uniprot50 \| AF-A0A1P8J5T2-F1-MODEL\_V4 \| 1.0 \| 0.0006268 \| 102 \| 0.129 \| 177 \| 117 \| 8 \| 6 \| 165 \| 20 \| 176 \| Methyl-accepting transducer domain-containing protein \| Methyl-accepting transducer domain-containing protein; afdb-uniprot50 \| AF-A0A522C888-F1-MODEL\_V4 \| 1.0 \| 0.001351 \| 102 \| 0.143 \| 146 \| 107 \| 6 \| 7 \| 142 \| 535 \| 672 \| DUF1929 domain-containing protein \| DUF1929 domain-containing protein; afdb-uniprot50 \| AF-A0A101XYV2-F1-MODEL\_V4 \| 1.0 \| 0.000948 \| 102 \| 0.144 \| 118 \| 82 \| 6 \| 9 \| 107 \| 671 \| 788 \| Uncharacterized protein \| Uncharacterized protein; afdb-uniprot50 \| AF-A0A2P2BRD5-F1-MODEL\_V4 \| 1.0 \| 0.001067 \| 102 \| 0.157 \| 152 \| 100 \| 5 \| 28 \| 168 \| 51 \| 185 \| BppU\_N domain-containing protein \| BppU\_N domain-containing protein; afdb-uniprot50 \| AF-A0A867I3J4-F1-MODEL\_V4 \| 1.0 \| 0.0003471 \| 102 \| 0.136 \| 176 \| 104 \| 10 \| 8 \| 163 \| 24 \| 171 \| SGNH/GDSL hydrolase family protein \| SGNH/GDSL hydrolase family protein; afdb-uniprot50 \| AF-A0A843J091-F1-MODEL\_V4 \| 1.0 \| 0.001006 \| 102 \| 0.097 \| 134 \| 98 \| 6 \| 31 \| 143 \| 51 \| 182 \| Uncharacterized protein \| Uncharacterized protein; afdb-uniprot50 \| AF-A0A2N6SD88-F1-MODEL\_V4 \| 1.0 \| 0.003279 \| 101 \| 0.098 \| 142 \| 101 \| 6 \| 9 \| 128 \| 24 \| 160 \| Uncharacterized protein \| Uncharacterized protein; afdb-uniprot50 \| AF-A0A3C0CBN8-F1-MODEL\_V4 \| 1.0 \| 0.00526 \| 101 \| 0.092 \| 130 \| 99 \| 7 \| 14 \| 135 \| 13 \| 131 \| Uncharacterized protein \| Uncharacterized protein; afdb-uniprot50 \| AF-A0A3L6US60-F1-MODEL\_V4 \| 1.0 \| 0.001274 \| 101 \| 0.133 \| 172 \| 121 \| 11 \| 9 \| 164 \| 2 \| 161 \| Dolichyl-diphosphooligosaccharide--protein glycosyltransferase subunit 2 \| Dolichyl-diphosphooligosaccharide--protein glycosyltransferase subunit 2; afdb-uniprot50 \| AF-A0A349YTI5-F1-MODEL\_V4 \| 1.0 \| 0.001067 \| 101 \| 0.116 \| 171 \| 115 \| 10 \| 9 \| 166 \| 19 \| 166 \| Uncharacterized protein \| Uncharacterized protein; afdb-uniprot50 \| AF-R6QMQ6-F1-MODEL\_V4 \| 1.0 \| 0.00244 \| 101 \| 0.108 \| 157 \| 104 \| 6 \| 29 \| 179 \| 49 \| 175 \| Hep/Hag repeat protein \| Hep/Hag repeat protein; afdb-uniprot50 \| AF-A0A3D2E0M6-F1-MODEL\_V4 \| 1.0 \| 0.0003683 \| 101 \| 0.12 \| 158 \| 110 \| 10 \| 9 \| 142 \| 18 \| 170 \| Uncharacterized protein \| Uncharacterized protein; afdb-uniprot50 \| AF-A0A822PCT9-F1-MODEL\_V4 \| 1.0 \| 0.002913 \| 101 \| 0.075 \| 173 \| 120 \| 7 \| 9 \| 166 \| 20 \| 167 \| Choline binding protein \| Choline binding protein; afdb-uniprot50 \| AF-A0A0H2PBG6-F1-MODEL\_V4 \| 1.0 \| 0.001351 \| 101 \| 0.173 \| 138 \| 88 \| 11 \| 9 \| 124 \| 26 \| 159 \| Tail fiber protein \| Tail fiber protein; afdb-uniprot50 \| AF-A0A1I0BNH9-F1-MODEL\_V4 \| 1.0 \| 0.0004949 \| 101 \| 0.095 \| 146 \| 108 \| 6 \| 22 \| 164 \| 32 \| 156 \| Uncharacterized protein \| Uncharacterized protein; afdb-uniprot50 \| AF-A0A385NWF3-F1-MODEL\_V4 \| 1.0 \| 0.006663 \| 101 \| 0.131 \| 144 \| 104 \| 7 \| 9 \| 138 \| 23 \| 159 \| Uncharacterized protein \| Uncharacterized protein; afdb-uniprot50 \| AF-A0A4Z2IX58-F1-MODEL\_V4 \| 1.0 \| 0.004153 \| 101 \| 0.111 \| 144 \| 113 \| 7 \| 7 \| 142 \| 497 \| 633 \| Deleted in malignant brain tumors 1 protein \| Deleted in malignant brain tumors 1 protein; afdb-uniprot50 \| AF-A0A0F9I0H5-F1-MODEL\_V4 \| 1.0 \| 0.008953 \| 101 \| 0.165 \| 115 \| 61 \| 8 \| 9 \| 105 \| 643 \| 740 \| Uncharacterized protein \| Uncharacterized protein; afdb-uniprot50 \| AF-A0A1I1EW75-F1-MODEL\_V4 \| 1.0 \| 0.003091 \| 101 \| 0.129 \| 170 \| 111 \| 9 \| 11 \| 166 \| 24 \| 170 \| Uncharacterized protein \| Uncharacterized protein; afdb-uniprot50 \| AF-A0A7U9X8V9-F1-MODEL\_V4 \| 1.0 \| 0.0005909 \| 101 \| 0.098 \| 193 \| 128 \| 7 \| 9 \| 179 \| 21 \| 189 \| Autotransporter adhesin EhaG \| Autotransporter adhesin EhaG; afdb-uniprot50 \| AF-A0A3M2LFC0-F1-MODEL\_V4 \| 1.0 \| 0.00369 \| 100 \| 0.136 \| 139 \| 92 \| 10 \| 9 \| 128 \| 43 \| 172 \| Uncharacterized protein \| Uncharacterized protein; afdb-uniprot50 \| AF-A0A3S2MRH1-F1-MODEL\_V4 \| 1.0 \| 0.0008936 \| 100 \| 0.106 \| 141 \| 103 \| 6 \| 9 \| 135 \| 31 \| 162 \| ig domain-containing protein \| ig domain-containing protein; afdb-uniprot50 \| AF-A0A412EM07-F1-MODEL\_V4 \| 1.0 \| 0.0007484 \| 100 \| 0.08 \| 174 \| 129 \| 6 \| 8 \| 166 \| 38 \| 195 \| Uncharacterized protein \| Uncharacterized protein; afdb-uniprot50 \| AF-A0A086ZDU3-F1-MODEL\_V4 \| 1.0 \| 0.001926 \| 100 \| 0.166 \| 150 \| 89 \| 9 \| 8 \| 127 \| 8 \| 151 \| Tail fiber protein \| Tail fiber protein; afdb-uniprot50 \| AF-A0A3B3D8Y5-F1-MODEL\_V4 \| 1.0 \| 0.002044 \| 100 \| 0.107 \| 121 \| 90 \| 6 \| 9 \| 125 \| 407 \| 513 \| Uncharacterized protein \| Uncharacterized protein; afdb-uniprot50 \| AF-A0A523YCW7-F1-MODEL\_V4 \| 1.0 \| 0.0008936 \| 100 \| 0.118 \| 144 \| 103 \| 7 \| 6 \| 142 \| 545 \| 671 \| Fibronectin type-III domain-containing protein \| Fibronectin type-III domain-containing protein; afdb-uniprot50 \| AF-A0A6B1ZRS2-F1-MODEL\_V4 \| 1.0 \| 0.001434 \| 100 \| 0.142 \| 175 \| 112 \| 9 \| 6 \| 165 \| 22 \| 173 \| Uncharacterized protein \| Uncharacterized protein; afdb-uniprot50 \| AF-A0A0N0C451-F1-MODEL\_V4 \| 1.0 \| 0.001351 \| 100 \| 0.119 \| 117 \| 85 \| 6 \| 9 \| 107 \| 854 \| 970 \| Uncharacterized protein \| Uncharacterized protein; afdb-uniprot50 \| AF-A0A1X2IR94-F1-MODEL\_V4 \| 1.0 \| 0.000948 \| 100 \| 0.168 \| 166 \| 108 \| 10 \| 9 \| 165 \| 63 \| 207 \| Uncharacterized protein \| Uncharacterized protein; afdb-uniprot50 \| AF-A0A3C0II44-F1-MODEL\_V4 \| 0.999 \| 0.004153 \| 99 \| 0.102 \| 146 \| 111 \| 9 \| 6 \| 142 \| 14 \| 148 \| Uncharacterized protein \| Uncharacterized protein; afdb-uniprot50 \| AF-A0A7V9P6P7-F1-MODEL\_V4 \| 0.999 \| 0.0007484 \| 99 \| 0.165 \| 157 \| 96 \| 12 \| 6 \| 142 \| 47 \| 188 \| Uncharacterized protein \| Uncharacterized protein; afdb-uniprot50 \| AF-A0A1D7ULP3-F1-MODEL\_V4 \| 0.999 \| 0.0008423 \| 99 \| 0.148 \| 175 \| 111 \| 10 \| 9 \| 166 \| 26 \| 179 \| Uncharacterized protein \| Uncharacterized protein; afdb-uniprot50 \| AF-A0A843IDY4-F1-MODEL\_V4 \| 0.999 \| 0.001926 \| 99 \| 0.118 \| 144 \| 97 \| 9 \| 7 \| 143 \| 106 \| 226 \| Uncharacterized protein \| Uncharacterized protein; afdb-uniprot50 \| AF-A0A6S5JXD1-F1-MODEL\_V4 \| 0.999 \| 0.00526 \| 99 \| 0.172 \| 110 \| 71 \| 8 \| 8 \| 107 \| 251 \| 350 \| Uncharacterized protein \| Uncharacterized protein; afdb-uniprot50 \| AF-A0A3R6NWS5-F1-MODEL\_V4 \| 0.999 \| 0.001712 \| 99 \| 0.135 \| 184 \| 113 \| 12 \| 8 \| 166 \| 21 \| 183 \| Uncharacterized protein \| Uncharacterized protein; afdb-uniprot50 \| AF-A0A415D505-F1-MODEL\_V4 \| 0.999 \| 0.001274 \| 99 \| 0.155 \| 116 \| 87 \| 3 \| 30 \| 138 \| 38 \| 149 \| Uncharacterized protein \| Uncharacterized protein; afdb-uniprot50 \| AF-A0A348Z913-F1-MODEL\_V4 \| 0.999 \| 0.0008936 \| 99 \| 0.114 \| 174 \| 107 \| 12 \| 9 \| 166 \| 22 \| 164 \| BppU\_N domain-containing protein \| BppU\_N domain-containing protein; afdb-uniprot50 \| AF-X1G6M4-F1-MODEL\_V4 \| 0.999 \| 0.0023 \| 99 \| 0.113 \| 141 \| 114 \| 7 \| 8 \| 145 \| 130 \| 262 \| NosD domain-containing protein \| NosD domain-containing protein; afdb-uniprot50 \| AF-C4IGZ0-F1-MODEL\_V4 \| 0.999 \| 0.001006 \| 99 \| 0.137 \| 175 \| 120 \| 8 \| 6 \| 166 \| 14 \| 171 \| Uncharacterized protein \| Uncharacterized protein; afdb-uniprot50 \| AF-A0A7D6VTN3-F1-MODEL\_V4 \| 0.999 \| 0.00526 \| 99 \| 0.109 \| 146 \| 100 \| 8 \| 8 \| 128 \| 21 \| 161 \| Uncharacterized protein \| Uncharacterized protein; afdb-uniprot50 \| AF-A0A429ZSL4-F1-MODEL\_V4 \| 0.999 \| 0.001613 \| 99 \| 0.147 \| 217 \| 119 \| 13 \| 1 \| 178 \| 1 \| 190 \| GP-PDE domain-containing protein \| GP-PDE domain-containing protein; afdb-uniprot50 \| AF-A0A6I4NWJ7-F1-MODEL\_V4 \| 0.999 \| 0.004406 \| 99 \| 0.137 \| 145 \| 90 \| 11 \| 9 \| 142 \| 606 \| 726 \| Uncharacterized protein \| Uncharacterized protein; afdb-uniprot50 \| AF-A0A4V6APB1-F1-MODEL\_V4 \| 0.999 \| 0.002589 \| 98 \| 0.138 \| 152 \| 103 \| 9 \| 7 \| 142 \| 279 \| 418 \| Deleted in malignant brain tumors 1 protein Hensin \| Deleted in malignant brain tumors 1 protein Hensin; afdb-uniprot50 \| AF-A0A3D2W3M9-F1-MODEL\_V4 \| 0.999 \| 0.001067 \| 98 \| 0.141 \| 156 \| 100 \| 9 \| 6 \| 142 \| 21 \| 161 \| BppU\_N domain-containing protein \| BppU\_N domain-containing protein; afdb-uniprot50 \| AF-A0A1D2KT36-F1-MODEL\_V4 \| 0.999 \| 0.002589 \| 98 \| 0.105 \| 142 \| 107 \| 5 \| 27 \| 163 \| 52 \| 178 \| DUF2479 domain-containing protein \| DUF2479 domain-containing protein; afdb-uniprot50 \| AF-T3DU14-F1-MODEL\_V4 \| 0.999 \| 0.001434 \| 98 \| 0.098 \| 152 \| 115 \| 9 \| 9 \| 142 \| 19 \| 166 \| PQQ-like domain protein \| PQQ-like domain protein; afdb-uniprot50 \| AF-A0A396L304-F1-MODEL\_V4 \| 0.999 \| 0.00244 \| 98 \| 0.08 \| 175 \| 127 \| 7 \| 7 \| 166 \| 11 \| 166 \| Uncharacterized protein \| Uncharacterized protein; afdb-uniprot50 \| AF-A0A162U122-F1-MODEL\_V4 \| 0.999 \| 0.001434 \| 98 \| 0.163 \| 153 \| 102 \| 7 \| 6 \| 142 \| 54 \| 196 \| Secreted sialophospho protein \| Secreted sialophospho protein; afdb-uniprot50 \| AF-A0A3Q9S137-F1-MODEL\_V4 \| 0.999 \| 0.005581 \| 98 \| 0.161 \| 105 \| 64 \| 9 \| 9 \| 98 \| 785 \| 880 \| Uncharacterized protein \| Uncharacterized protein; afdb-uniprot50 \| AF-A0A6J8ASX6-F1-MODEL\_V4 \| 0.999 \| 0.001351 \| 97 \| 0.12 \| 150 \| 110 \| 7 \| 10 \| 142 \| 9 \| 153 \| Fibronectin type-III domain-containing protein \| Fibronectin type-III domain-containing protein; afdb-uniprot50 \| AF-A0A846PSR2-F1-MODEL\_V4 \| 0.999 \| 0.003478 \| 97 \| 0.149 \| 114 \| 81 \| 7 \| 7 \| 107 \| 141 \| 251 \| Uncharacterized protein \| Uncharacterized protein; afdb-uniprot50 \| AF-A2HSA7-F1-MODEL\_V4 \| 0.999 \| 0.001067 \| 97 \| 0.179 \| 128 \| 90 \| 9 \| 6 \| 124 \| 141 \| 262 \| Uncharacterized protein \| Uncharacterized protein; afdb-uniprot50 \| AF-A0A3M6TAQ9-F1-MODEL\_V4 \| 0.999 \| 0.001006 \| 97 \| 0.083 \| 155 \| 107 \| 11 \| 4 \| 142 \| 28 \| 163 \| Uncharacterized protein \| Uncharacterized protein; afdb-uniprot50 \| AF-R0AP54-F1-MODEL\_V4 \| 0.999 \| 0.00526 \| 97 \| 0.135 \| 170 \| 112 \| 10 \| 9 \| 166 \| 23 \| 169 \| BppU\_N domain-containing protein \| BppU\_N domain-containing protein; afdb-uniprot50 \| AF-A0A397BUT5-F1-MODEL\_V4 \| 0.999 \| 0.001926 \| 97 \| 0.114 \| 184 \| 136 \| 10 \| 9 \| 178 \| 128 \| 298 \| Dolichyl-diphosphooligosaccharide--protein glycosyltransferase subunit 2 \| Dolichyl-diphosphooligosaccharide--protein glycosyltransferase subunit 2; afdb-uniprot50 \| AF-R5KP20-F1-MODEL\_V4 \| 0.999 \| 0.001434 \| 97 \| 0.119 \| 159 \| 106 \| 8 \| 22 \| 179 \| 36 \| 161 \| Uncharacterized protein \| Uncharacterized protein; afdb-uniprot50 \| AF-A0A1F1KZP2-F1-MODEL\_V4 \| 0.999 \| 0.001926 \| 97 \| 0.137 \| 174 \| 121 \| 8 \| 5 \| 166 \| 15 \| 171 \| Uncharacterized protein \| Uncharacterized protein; afdb-uniprot50 \| AF-A0A669B4J7-F1-MODEL\_V4 \| 0.999 \| 0.001521 \| 97 \| 0.188 \| 143 \| 99 \| 9 \| 9 \| 142 \| 433 \| 567 \| Uncharacterized protein \| Uncharacterized protein; afdb-uniprot50 \| AF-A0A535MAP0-F1-MODEL\_V4 \| 0.999 \| 0.0005909 \| 97 \| 0.175 \| 177 \| 109 \| 13 \| 7 \| 166 \| 34 \| 190 \| Copper resistance protein CopC \| Copper resistance protein CopC; afdb-uniprot50 \| AF-M1LNW5-F1-MODEL\_V4 \| 0.999 \| 0.00526 \| 97 \| 0.13 \| 168 \| 116 \| 8 \| 9 \| 166 \| 22 \| 169 \| Uncharacterized protein \| Uncharacterized protein; afdb-uniprot50 \| AF-A0A8B1YVX0-F1-MODEL\_V4 \| 0.999 \| 0.0004397 \| 97 \| 0.135 \| 206 \| 136 \| 11 \| 3 \| 179 \| 8 \| 200 \| Uncharacterized protein \| Uncharacterized protein; afdb-uniprot50 \| AF-A0A848B9H4-F1-MODEL\_V4 \| 0.999 \| 0.00526 \| 97 \| 0.137 \| 175 \| 117 \| 10 \| 9 \| 166 \| 22 \| 179 \| Uncharacterized protein \| Uncharacterized protein; afdb-uniprot50 \| AF-A0A6N3A7F2-F1-MODEL\_V4 \| 0.999 \| 0.0008936 \| 97 \| 0.116 \| 171 \| 119 \| 7 \| 7 \| 164 \| 34 \| 185 \| Uncharacterized protein \| Uncharacterized protein; afdb-uniprot50 \| AF-A0A7S8E8U1-F1-MODEL\_V4 \| 0.999 \| 0.001521 \| 97 \| 0.16 \| 143 \| 84 \| 7 \| 7 \| 142 \| 594 \| 707 \| PPC domain-containing protein \| PPC domain-containing protein; afdb-uniprot50 \| AF-A0A523K1L5-F1-MODEL\_V4 \| 0.999 \| 0.00526 \| 96 \| 0.107 \| 102 \| 70 \| 10 \| 8 \| 100 \| 73 \| 162 \| Uncharacterized protein \| Uncharacterized protein; afdb-uniprot50 \| AF-A0A256BLC4-F1-MODEL\_V4 \| 0.999 \| 0.002589 \| 96 \| 0.104 \| 181 \| 119 \| 10 \| 9 \| 165 \| 30 \| 191 \| BppU\_N domain-containing protein \| BppU\_N domain-containing protein; afdb-uniprot50 \| AF-A0A7G5NVU1-F1-MODEL\_V4 \| 0.999 \| 0.004959 \| 96 \| 0.114 \| 175 \| 125 \| 8 \| 5 \| 166 \| 15 \| 172 \| Uncharacterized protein \| Uncharacterized protein; afdb-uniprot50 \| AF-C0EIT1-F1-MODEL\_V4 \| 0.999 \| 0.002044 \| 96 \| 0.172 \| 145 \| 91 \| 7 \| 30 \| 166 \| 56 \| 179 \| BppU\_N domain-containing protein \| BppU\_N domain-containing protein; afdb-uniprot50 \| AF-A0A7J2R513-F1-MODEL\_V4 \| 0.999 \| 0.001201 \| 96 \| 0.126 \| 166 \| 113 \| 8 \| 15 \| 160 \| 398 \| 551 \| VWA domain-containing protein \| VWA domain-containing protein; afdb-uniprot50 \| AF-A0A2V8GIF6-F1-MODEL\_V4 \| 0.999 \| 0.001132 \| 96 \| 0.138 \| 252 \| 123 \| 14 \| 6 \| 179 \| 606 \| 841 \| Uncharacterized protein \| Uncharacterized protein; afdb-uniprot50 \| AF-A0A2G6E620-F1-MODEL\_V4 \| 0.998 \| 0.002913 \| 95 \| 0.119 \| 109 \| 80 \| 6 \| 7 \| 108 \| 55 \| 154 \| Uncharacterized protein \| Uncharacterized protein; afdb-uniprot50 \| AF-A0A543HW98-F1-MODEL\_V4 \| 0.998 \| 0.0002295 \| 95 \| 0.151 \| 172 \| 117 \| 9 \| 7 \| 166 \| 38 \| 192 \| Uncharacterized protein \| Uncharacterized protein; afdb-uniprot50 \| AF-A0A4R8ZQF4-F1-MODEL\_V4 \| 0.998 \| 0.0001431 \| 95 \| 0.165 \| 187 \| 124 \| 11 \| 9 \| 179 \| 47 \| 217 \| Copper resistance protein CopC \| Copper resistance protein CopC; afdb-uniprot50 \| AF-A0A7U9XHU5-F1-MODEL\_V4 \| 0.998 \| 0.001201 \| 95 \| 0.137 \| 138 \| 94 \| 5 \| 29 \| 166 \| 41 \| 153 \| Uncharacterized protein \| Uncharacterized protein; afdb-uniprot50 \| AF-A0A6L5YI01-F1-MODEL\_V4 \| 0.998 \| 0.001613 \| 95 \| 0.085 \| 199 \| 124 \| 13 \| 8 \| 168 \| 19 \| 197 \| Uncharacterized protein \| Uncharacterized protein; afdb-uniprot50 \| AF-A0A7V9I5K7-F1-MODEL\_V4 \| 0.998 \| 0.0007484 \| 95 \| 0.138 \| 195 \| 126 \| 12 \| 7 \| 179 \| 34 \| 208 \| Uncharacterized protein \| Uncharacterized protein; afdb-uniprot50 \| AF-A0A174XQW7-F1-MODEL\_V4 \| 0.998 \| 0.00592 \| 95 \| 0.154 \| 194 \| 133 \| 13 \| 9 \| 179 \| 21 \| 206 \| Collagen triple helix repeat-containing protein \| Collagen triple helix repeat-containing protein; afdb-uniprot50 \| AF-A0A2M7DL93-F1-MODEL\_V4 \| 0.998 \| 0.0002908 \| 95 \| 0.154 \| 149 \| 101 \| 9 \| 7 \| 136 \| 179 \| 321 \| DUF11 domain-containing protein \| DUF11 domain-containing protein; afdb-uniprot50 \| AF-W7C4F2-F1-MODEL\_V4 \| 0.998 \| 0.001132 \| 95 \| 0.147 \| 149 \| 93 \| 10 \| 28 \| 165 \| 54 \| 179 \| BppU\_N domain-containing protein \| BppU\_N domain-containing protein; afdb-uniprot50 \| AF-A0A7Y1YUI9-F1-MODEL\_V4 \| 0.998 \| 0.008953 \| 95 \| 0.163 \| 110 \| 73 \| 9 \| 8 \| 107 \| 32 \| 132 \| Uncharacterized protein \| Uncharacterized protein; afdb-uniprot50 \| AF-A0A0Q5WDI7-F1-MODEL\_V4 \| 0.998 \| 0.001926 \| 95 \| 0.157 \| 127 \| 90 \| 10 \| 8 \| 126 \| 401 \| 518 \| Uncharacterized protein \| Uncharacterized protein; afdb-uniprot50 \| AF-A0A523YH80-F1-MODEL\_V4 \| 0.998 \| 0.0008936 \| 95 \| 0.121 \| 140 \| 97 \| 6 \| 8 \| 136 \| 550 \| 674 \| Fibronectin type-III domain-containing protein \| Fibronectin type-III domain-containing protein; afdb-uniprot50 \| AF-A0A2K4ZNI6-F1-MODEL\_V4 \| 0.998 \| 0.001613 \| 95 \| 0.128 \| 140 \| 99 \| 6 \| 28 \| 164 \| 50 \| 169 \| Uncharacterized protein \| Uncharacterized protein; afdb-uniprot50 \| AF-A0A845QX82-F1-MODEL\_V4 \| 0.998 \| 0.007069 \| 95 \| 0.076 \| 171 \| 123 \| 11 \| 9 \| 166 \| 22 \| 170 \| DUF2479 domain-containing protein \| DUF2479 domain-containing protein; afdb-uniprot50 \| AF-S4B432-F1-MODEL\_V4 \| 0.998 \| 0.0023 \| 95 \| 0.125 \| 199 \| 125 \| 11 \| 6 \| 178 \| 22 \| 197 \| Peptidase\_M14 domain-containing protein \| Peptidase\_M14 domain-containing protein; afdb-uniprot50 \| AF-A0A1Z5IVA8-F1-MODEL\_V4 \| 0.998 \| 0.004153 \| 95 \| 0.089 \| 189 \| 124 \| 11 \| 6 \| 166 \| 25 \| 193 \| GDSL-like lipase/acylhydrolase \| GDSL-like lipase/acylhydrolase; afdb-uniprot50 \| AF-A0A0Q8PIW8-F1-MODEL\_V4 \| 0.998 \| 0.0001923 \| 94 \| 0.131 \| 183 \| 111 \| 10 \| 4 \| 164 \| 35 \| 191 \| CopC domain-containing protein \| CopC domain-containing protein; afdb-uniprot50 \| AF-A0A5F0FGF2-F1-MODEL\_V4 \| 0.998 \| 0.0001349 \| 94 \| 0.12 \| 174 \| 133 \| 7 \| 15 \| 179 \| 45 \| 207 \| Copper resistance protein CopC \| Copper resistance protein CopC; afdb-uniprot50 \| AF-A0A813XUR2-F1-MODEL\_V4 \| 0.998 \| 0.000665 \| 94 \| 0.097 \| 205 \| 151 \| 12 \| 3 \| 179 \| 38 \| 236 \| Hypothetical protein \| Hypothetical protein; afdb-uniprot50 \| AF-A0A3M1FDJ3-F1-MODEL\_V4 \| 0.998 \| 0.001816 \| 94 \| 0.097 \| 134 \| 93 \| 7 \| 9 \| 124 \| 245 \| 368 \| Uncharacterized protein \| Uncharacterized protein; afdb-uniprot50 \| AF-A0A2X0QIE9-F1-MODEL\_V4 \| 0.998 \| 0.002168 \| 94 \| 0.097 \| 154 \| 108 \| 9 \| 20 \| 166 \| 48 \| 177 \| BppU\_N domain-containing protein \| BppU\_N domain-containing protein; afdb-uniprot50 \| AF-A0A4Q7PK69-F1-MODEL\_V4 \| 0.998 \| 0.0023 \| 94 \| 0.154 \| 175 \| 111 \| 12 \| 9 \| 166 \| 26 \| 180 \| Uncharacterized protein \| Uncharacterized protein; afdb-uniprot50 \| AF-A0A847WF40-F1-MODEL\_V4 \| 0.998 \| 0.002746 \| 94 \| 0.129 \| 186 \| 127 \| 9 \| 9 \| 178 \| 20 \| 186 \| Uncharacterized protein \| Uncharacterized protein; afdb-uniprot50 \| AF-A0A6J8BSE7-F1-MODEL\_V4 \| 0.998 \| 0.009498 \| 94 \| 0.104 \| 143 \| 112 \| 6 \| 11 \| 142 \| 669 \| 806 \| Uncharacterized protein \| Uncharacterized protein; afdb-uniprot50 \| AF-A0A6P7P392-F1-MODEL\_V4 \| 0.998 \| 0.001274 \| 94 \| 0.127 \| 141 \| 107 \| 7 \| 9 \| 142 \| 940 \| 1071 \| scavenger receptor cysteine-rich type 1 protein M160-like isoform X2 \| scavenger receptor cysteine-rich type 1 protein M160-like isoform X2; afdb-uniprot50 \| AF-A0A328L2A5-F1-MODEL\_V4 \| 0.998 \| 0.001434 \| 93 \| 0.102 \| 176 \| 122 \| 12 \| 6 \| 166 \| 19 \| 173 \| BppU\_N domain-containing protein \| BppU\_N domain-containing protein; afdb-uniprot50 \| AF-A0A3R7YHD4-F1-MODEL\_V4 \| 0.998 \| 0.001006 \| 93 \| 0.146 \| 157 \| 87 \| 9 \| 9 \| 142 \| 185 \| 317 \| Uncharacterized protein \| Uncharacterized protein; afdb-uniprot50 \| AF-A0A1J1HAE1-F1-MODEL\_V4 \| 0.998 \| 0.002746 \| 93 \| 0.092 \| 151 \| 112 \| 6 \| 9 \| 135 \| 74 \| 223 \| Uncharacterized protein \| Uncharacterized protein; afdb-uniprot50 \| AF-A2H226-F1-MODEL\_V4 \| 0.998 \| 0.003478 \| 93 \| 0.172 \| 122 \| 92 \| 7 \| 9 \| 124 \| 230 \| 348 \| Uncharacterized protein \| Uncharacterized protein; afdb-uniprot50 \| AF-A0A4Q9XZW9-F1-MODEL\_V4 \| 0.998 \| 0.002168 \| 93 \| 0.127 \| 180 \| 114 \| 14 \| 5 \| 164 \| 16 \| 172 \| DUF2479 domain-containing protein \| DUF2479 domain-containing protein; afdb-uniprot50 \| AF-A0A420YQM6-F1-MODEL\_V4 \| 0.998 \| 0.001274 \| 93 \| 0.146 \| 218 \| 115 \| 13 \| 6 \| 162 \| 22 \| 229 \| Uncharacterized protein \| Uncharacterized protein; afdb-uniprot50 \| AF-A0A7J2ISC6-F1-MODEL\_V4 \| 0.998 \| 0.004406 \| 93 \| 0.089 \| 146 \| 117 \| 5 \| 7 \| 142 \| 306 \| 445 \| DUF1565 domain-containing protein \| DUF1565 domain-containing protein; afdb-uniprot50 \| AF-A0A1V5IGQ3-F1-MODEL\_V4 \| 0.998 \| 0.00592 \| 93 \| 0.161 \| 124 \| 77 \| 10 \| 6 \| 108 \| 522 \| 639 \| DUF3821 domain-containing protein \| DUF3821 domain-containing protein; afdb-uniprot50 \| AF-A0A0C9MY19-F1-MODEL\_V4 \| 0.998 \| 0.003091 \| 93 \| 0.127 \| 141 \| 103 \| 6 \| 9 \| 142 \| 62 \| 189 \| Uncharacterized protein \| Uncharacterized protein; afdb-uniprot50 \| AF-A0A7U9X263-F1-MODEL\_V4 \| 0.998 \| 0.007069 \| 93 \| 0.109 \| 128 \| 91 \| 6 \| 9 \| 124 \| 20 \| 136 \| Uncharacterized protein \| Uncharacterized protein; afdb-uniprot50 \| AF-A0A7T4G8C7-F1-MODEL\_V4 \| 0.998 \| 0.002746 \| 93 \| 0.109 \| 174 \| 115 \| 9 \| 6 \| 158 \| 22 \| 176 \| SGNH/GDSL hydrolase family protein \| SGNH/GDSL hydrolase family protein; afdb-uniprot50 \| AF-A0A6I7EES4-F1-MODEL\_V4 \| 0.998 \| 0.00369 \| 93 \| 0.123 \| 138 \| 97 \| 11 \| 9 \| 127 \| 21 \| 153 \| BppU family phage baseplate upper protein \| BppU family phage baseplate upper protein; afdb-uniprot50 \| AF-A0A532TIZ1-F1-MODEL\_V4 \| 0.997 \| 0.004406 \| 92 \| 0.093 \| 128 \| 98 \| 4 \| 8 \| 128 \| 47 \| 163 \| Uncharacterized protein \| Uncharacterized protein; afdb-uniprot50 \| AF-A0A495WU82-F1-MODEL\_V4 \| 0.997 \| 0.001067 \| 92 \| 0.142 \| 190 \| 123 \| 11 \| 9 \| 179 \| 55 \| 223 \| CopC domain-containing protein \| CopC domain-containing protein; afdb-uniprot50 \| AF-A0A846KDD2-F1-MODEL\_V4 \| 0.997 \| 0.000525 \| 92 \| 0.115 \| 165 \| 109 \| 12 \| 20 \| 164 \| 52 \| 199 \| BppU family phage baseplate upper protein \| BppU family phage baseplate upper protein; afdb-uniprot50 \| AF-D4FN16-F1-MODEL\_V4 \| 0.997 \| 0.001132 \| 92 \| 0.172 \| 168 \| 107 \| 7 \| 20 \| 179 \| 47 \| 190 \| BppU\_N domain-containing protein \| BppU\_N domain-containing protein; afdb-uniprot50 \| AF-A0A1V5ZC38-F1-MODEL\_V4 \| 0.997 \| 0.001006 \| 92 \| 0.118 \| 160 \| 91 \| 11 \| 6 \| 142 \| 227 \| 359 \| Uncharacterized protein \| Uncharacterized protein; afdb-uniprot50 \| AF-A0A242WL41-F1-MODEL\_V4 \| 0.997 \| 0.00526 \| 92 \| 0.139 \| 165 \| 110 \| 12 \| 8 \| 142 \| 20 \| 182 \| BppU\_N domain-containing protein \| BppU\_N domain-containing protein; afdb-uniprot50 \| AF-A0A523YCV8-F1-MODEL\_V4 \| 0.997 \| 0.000794 \| 92 \| 0.12 \| 149 \| 99 \| 7 \| 9 \| 142 \| 835 \| 966 \| Fibronectin type-III domain-containing protein \| Fibronectin type-III domain-containing protein; afdb-uniprot50 \| AF-D6SR61-F1-MODEL\_V4 \| 0.997 \| 0.002746 \| 92 \| 0.179 \| 145 \| 97 \| 9 \| 7 \| 142 \| 230 \| 361 \| Lipoprotein \| Lipoprotein; afdb-uniprot50 \| AF-A0A6I4P3U8-F1-MODEL\_V4 \| 0.997 \| 0.000557 \| 91 \| 0.147 \| 190 \| 121 \| 10 \| 9 \| 178 \| 44 \| 212 \| CopC domain-containing protein \| CopC domain-containing protein; afdb-uniprot50 \| AF-A0A6J7GNW7-F1-MODEL\_V4 \| 0.997 \| 0.004674 \| 91 \| 0.093 \| 129 \| 99 \| 10 \| 9 \| 128 \| 291 \| 410 \| Unannotated protein \| Unannotated protein; afdb-uniprot50 \| AF-A0A1C5M3U0-F1-MODEL\_V4 \| 0.997 \| 0.003915 \| 91 \| 0.155 \| 148 \| 93 \| 8 \| 28 \| 166 \| 56 \| 180 \| Uncharacterized protein \| Uncharacterized protein; afdb-uniprot50 \| AF-A0A7C4V780-F1-MODEL\_V4 \| 0.997 \| 0.003915 \| 91 \| 0.149 \| 167 \| 111 \| 14 \| 9 \| 158 \| 549 \| 701 \| Peptidase\_S8 domain-containing protein \| Peptidase\_S8 domain-containing protein; afdb-uniprot50 \| AF-A0A101XV12-F1-MODEL\_V4 \| 0.997 \| 0.001926 \| 91 \| 0.152 \| 118 \| 81 \| 7 \| 9 \| 107 \| 730 \| 847 \| Uncharacterized protein \| Uncharacterized protein; afdb-uniprot50 \| AF-N6VQ37-F1-MODEL\_V4 \| 0.997 \| 0.001351 \| 91 \| 0.099 \| 131 \| 97 \| 10 \| 5 \| 128 \| 845 \| 961 \| Uncharacterized protein \| Uncharacterized protein; afdb-uniprot50 \| AF-A0A521H572-F1-MODEL\_V4 \| 0.997 \| 0.00592 \| 91 \| 0.162 \| 135 \| 96 \| 9 \| 11 \| 136 \| 878 \| 1004 \| VWA domain-containing protein \| VWA domain-containing protein; afdb-uniprot50 \| AF-A0A3E2BTS2-F1-MODEL\_V4 \| 0.996 \| 0.003091 \| 90 \| 0.117 \| 178 \| 115 \| 9 \| 6 \| 166 \| 20 \| 172 \| BppU\_N domain-containing protein \| BppU\_N domain-containing protein; afdb-uniprot50 \| AF-A0A6A4RWM8-F1-MODEL\_V4 \| 0.996 \| 0.002589 \| 90 \| 0.113 \| 159 \| 104 \| 7 \| 6 \| 142 \| 71 \| 214 \| 40S ribosomal protein S12 \| 40S ribosomal protein S12; afdb-uniprot50 \| AF-A0A7V3ZT25-F1-MODEL\_V4 \| 0.996 \| 0.002913 \| 90 \| 0.139 \| 151 \| 99 \| 10 \| 6 \| 142 \| 334 \| 467 \| Uncharacterized protein \| Uncharacterized protein; afdb-uniprot50 \| AF-A0A844FIK1-F1-MODEL\_V4 \| 0.996 \| 0.00592 \| 90 \| 0.139 \| 186 \| 109 \| 10 \| 6 \| 166 \| 22 \| 181 \| Uncharacterized protein \| Uncharacterized protein; afdb-uniprot50 \| AF-A0A3S2MDZ9-F1-MODEL\_V4 \| 0.996 \| 0.001712 \| 90 \| 0.127 \| 157 \| 103 \| 10 \| 9 \| 142 \| 662 \| 807 \| Uncharacterized protein \| Uncharacterized protein; afdb-uniprot50 \| AF-A0A4Y5ZNF9-F1-MODEL\_V4 \| 0.996 \| 0.003915 \| 89 \| 0.123 \| 146 \| 99 \| 9 \| 8 \| 142 \| 54 \| 181 \| Ig-like domain-containing protein \| Ig-like domain-containing protein; afdb-uniprot50 \| AF-A0A3A9CCX0-F1-MODEL\_V4 \| 0.996 \| 0.0023 \| 89 \| 0.128 \| 148 \| 97 \| 7 \| 30 \| 166 \| 48 \| 174 \| DUF2479 domain-containing protein \| DUF2479 domain-containing protein; afdb-uniprot50 \| AF-A0A7J5ZBY5-F1-MODEL\_V4 \| 0.996 \| 0.001816 \| 89 \| 0.136 \| 169 \| 118 \| 11 \| 3 \| 163 \| 32 \| 180 \| Uncharacterized protein \| Uncharacterized protein; afdb-uniprot50 \| AF-L1NGG6-F1-MODEL\_V4 \| 0.996 \| 0.001006 \| 89 \| 0.137 \| 203 \| 101 \| 13 \| 9 \| 149 \| 32 \| 222 \| Uncharacterized protein \| Uncharacterized protein; afdb-uniprot50 \| AF-Q4L3Y3-F1-MODEL\_V4 \| 0.996 \| 0.004153 \| 89 \| 0.126 \| 158 \| 110 \| 7 \| 7 \| 143 \| 16 \| 166 \| Uncharacterized protein \| Uncharacterized protein; afdb-uniprot50 \| AF-A0A399UNF6-F1-MODEL\_V4 \| 0.995 \| 0.0003085 \| 88 \| 0.148 \| 202 \| 114 \| 11 \| 8 \| 179 \| 61 \| 234 \| CopC domain-containing protein \| CopC domain-containing protein; afdb-uniprot50 \| AF-A0A151MA98-F1-MODEL\_V4 \| 0.995 \| 0.000948 \| 88 \| 0.107 \| 186 \| 110 \| 12 \| 5 \| 142 \| 72 \| 249 \| Ig-like domain-containing protein \| Ig-like domain-containing protein; afdb-uniprot50 \| AF-A0A852YGZ0-F1-MODEL\_V4 \| 0.995 \| 0.001201 \| 88 \| 0.139 \| 194 \| 122 \| 13 \| 9 \| 179 \| 63 \| 234 \| Uncharacterized protein \| Uncharacterized protein; afdb-uniprot50 \| AF-A0A3M6TAX9-F1-MODEL\_V4 \| 0.995 \| 0.001816 \| 88 \| 0.132 \| 174 \| 113 \| 12 \| 7 \| 159 \| 72 \| 228 \| Uncharacterized protein \| Uncharacterized protein; afdb-uniprot50 \| AF-A0A380CHS6-F1-MODEL\_V4 \| 0.995 \| 0.007069 \| 88 \| 0.145 \| 165 \| 103 \| 13 \| 17 \| 166 \| 36 \| 177 \| Putative phage tail protein \| Putative phage tail protein; afdb-uniprot50 \| AF-A0A143Z9K5-F1-MODEL\_V4 \| 0.995 \| 0.007499 \| 88 \| 0.113 \| 176 \| 118 \| 9 \| 9 \| 164 \| 27 \| 184 \| Uncharacterized protein \| Uncharacterized protein; afdb-uniprot50 \| AF-A0A6G0I273-F1-MODEL\_V4 \| 0.995 \| 0.002746 \| 88 \| 0.198 \| 146 \| 91 \| 8 \| 9 \| 142 \| 926 \| 1057 \| Scavenger receptor cysteine-rich type 1 protein M130 Soluble CD163 \| Scavenger receptor cysteine-rich type 1 protein M130 Soluble CD163; afdb-uniprot50 \| AF-A0A524CJ65-F1-MODEL\_V4 \| 0.994 \| 0.004959 \| 87 \| 0.126 \| 126 \| 92 \| 7 \| 9 \| 126 \| 203 \| 318 \| NosD domain-containing protein \| NosD domain-containing protein; afdb-uniprot50 \| AF-J8KZ84-F1-MODEL\_V4 \| 0.994 \| 0.004153 \| 87 \| 0.14 \| 164 \| 112 \| 10 \| 8 \| 142 \| 22 \| 185 \| BppU\_N domain-containing protein \| BppU\_N domain-containing protein; afdb-uniprot50 \| AF-A0A3Q0GK91-F1-MODEL\_V4 \| 0.994 \| 0.0006268 \| 87 \| 0.106 \| 188 \| 112 \| 14 \| 4 \| 142 \| 142 \| 322 \| uncharacterized protein LOC112550110 isoform X1 \| uncharacterized protein LOC112550110 isoform X1; afdb-uniprot50 \| AF-A0A6A8SHN8-F1-MODEL\_V4 \| 0.994 \| 0.007955 \| 87 \| 0.119 \| 151 \| 103 \| 9 \| 25 \| 166 \| 46 \| 175 \| DUF2479 domain-containing protein \| DUF2479 domain-containing protein; afdb-uniprot50 \| AF-A0A351F507-F1-MODEL\_V4 \| 0.994 \| 0.004674 \| 87 \| 0.11 \| 163 \| 120 \| 7 \| 9 \| 166 \| 20 \| 162 \| Uncharacterized protein \| Uncharacterized protein; afdb-uniprot50 \| AF-A0A7M7KDJ7-F1-MODEL\_V4 \| 0.994 \| 0.008953 \| 87 \| 0.092 \| 151 \| 118 \| 10 \| 8 \| 142 \| 747 \| 894 \| Uncharacterized protein \| Uncharacterized protein; afdb-uniprot50 \| AF-A0A3E3EHK7-F1-MODEL\_V4 \| 0.993 \| 0.007955 \| 86 \| 0.075 \| 146 \| 110 \| 7 \| 22 \| 164 \| 36 \| 159 \| Uncharacterized protein \| Uncharacterized protein; afdb-uniprot50 \| AF-A0A379AD69-F1-MODEL\_V4 \| 0.993 \| 0.00592 \| 86 \| 0.139 \| 143 \| 100 \| 9 \| 8 \| 142 \| 144 \| 271 \| Big\_13 domain-containing protein \| Big\_13 domain-containing protein; afdb-uniprot50 \| AF-A0A381QTK1-F1-MODEL\_V4 \| 0.993 \| 0.003279 \| 86 \| 0.109 \| 146 \| 94 \| 8 \| 6 \| 135 \| 170 \| 295 \| Uncharacterized protein \| Uncharacterized protein; afdb-uniprot50 \| AF-R9AHF4-F1-MODEL\_V4 \| 0.993 \| 0.0004949 \| 86 \| 0.145 \| 200 \| 118 \| 15 \| 7 \| 179 \| 228 \| 401 \| Dolichyl-diphosphooligosaccharide--protein glycosyltransferase subunit WBP1 \| Dolichyl-diphosphooligosaccharide--protein glycosyltransferase subunit WBP1; afdb-uniprot50 \| AF-A0A0E0UW38-F1-MODEL\_V4 \| 0.993 \| 0.00592 \| 86 \| 0.107 \| 168 \| 121 \| 11 \| 28 \| 179 \| 53 \| 207 \| Gp19 \| Gp19; afdb-uniprot50 \| AF-A0A3B3E2G3-F1-MODEL\_V4 \| 0.993 \| 0.0023 \| 86 \| 0.098 \| 153 \| 105 \| 7 \| 9 \| 142 \| 289 \| 427 \| Uncharacterized protein \| Uncharacterized protein; afdb-uniprot50 \| AF-A0A6P7LDW9-F1-MODEL\_V4 \| 0.993 \| 0.00369 \| 86 \| 0.15 \| 146 \| 100 \| 7 \| 9 \| 142 \| 444 \| 577 \| scavenger receptor cysteine-rich type 1 protein M130-like \| scavenger receptor cysteine-rich type 1 protein M130-like; afdb-uniprot50 \| AF-A0A0B0EE87-F1-MODEL\_V4 \| 0.993 \| 0.003915 \| 86 \| 0.14 \| 164 \| 120 \| 7 \| 15 \| 166 \| 499 \| 653 \| VWFA domain-containing protein \| VWFA domain-containing protein; afdb-uniprot50 \| AF-Q32J73-F1-MODEL\_V4 \| 0.993 \| 0.006663 \| 86 \| 0.115 \| 139 \| 96 \| 9 \| 8 \| 136 \| 649 \| 770 \| Uncharacterized protein \| Uncharacterized protein; afdb-uniprot50 \| AF-A0A6H3J5S0-F1-MODEL\_V4 \| 0.992 \| 0.0005909 \| 85 \| 0.154 \| 181 \| 122 \| 9 \| 14 \| 179 \| 6 \| 170 \| Copper resistance protein CopC \| Copper resistance protein CopC; afdb-uniprot50 \| AF-A0A1H8GB40-F1-MODEL\_V4 \| 0.992 \| 0.001067 \| 85 \| 0.114 \| 157 \| 105 \| 10 \| 7 \| 136 \| 11 \| 160 \| PASTA domain, binds beta-lactams \| PASTA domain, binds beta-lactams; afdb-uniprot50 \| AF-A0A497HKX8-F1-MODEL\_V4 \| 0.992 \| 0.004153 \| 85 \| 0.097 \| 134 \| 98 \| 9 \| 8 \| 128 \| 293 \| 416 \| CARDB domain-containing protein \| CARDB domain-containing protein; afdb-uniprot50 \| AF-A0A4Y5ZTI8-F1-MODEL\_V4 \| 0.992 \| 0.007955 \| 85 \| 0.136 \| 146 \| 103 \| 10 \| 8 \| 142 \| 251 \| 384 \| Ig-like domain-containing protein \| Ig-like domain-containing protein; afdb-uniprot50 \| AF-A0A6L3ENR8-F1-MODEL\_V4 \| 0.992 \| 0.002589 \| 85 \| 0.161 \| 142 \| 102 \| 10 \| 9 \| 142 \| 315 \| 447 \| Uncharacterized protein \| Uncharacterized protein; afdb-uniprot50 \| AF-A0A354M6W3-F1-MODEL\_V4 \| 0.992 \| 0.003091 \| 85 \| 0.1 \| 160 \| 114 \| 9 \| 8 \| 142 \| 10 \| 164 \| Uncharacterized protein \| Uncharacterized protein; afdb-uniprot50 \| AF-A0A2B5I3B6-F1-MODEL\_V4 \| 0.992 \| 0.00369 \| 85 \| 0.115 \| 164 \| 106 \| 11 \| 9 \| 142 \| 23 \| 177 \| BppU\_N domain-containing protein \| BppU\_N domain-containing protein; afdb-uniprot50 \| AF-A0A368W6S3-F1-MODEL\_V4 \| 0.992 \| 0.001926 \| 85 \| 0.114 \| 148 \| 95 \| 7 \| 7 \| 125 \| 460 \| 600 \| FecR family protein \| FecR family protein; afdb-uniprot50 \| AF-A0A814IBB4-F1-MODEL\_V4 \| 0.992 \| 0.003478 \| 85 \| 0.125 \| 144 \| 112 \| 8 \| 7 \| 142 \| 213 \| 350 \| Hypothetical protein \| Hypothetical protein; afdb-uniprot50 \| AF-A0A0A6ZMX3-F1-MODEL\_V4 \| 0.992 \| 0.009498 \| 85 \| 0.11 \| 145 \| 94 \| 9 \| 8 \| 142 \| 467 \| 586 \| Membrane glycoprotein \| Membrane glycoprotein; afdb-uniprot50 \| AF-A2F0K9-F1-MODEL\_V4 \| 0.991 \| 0.0003085 \| 84 \| 0.088 \| 169 \| 110 \| 9 \| 7 \| 142 \| 48 \| 205 \| Uncharacterized protein \| Uncharacterized protein; afdb-uniprot50 \| AF-A0A7T9Z8U9-F1-MODEL\_V4 \| 0.991 \| 0.00592 \| 84 \| 0.134 \| 141 \| 97 \| 6 \| 7 \| 142 \| 51 \| 171 \| Excalibur calcium-binding domain-containing protein \| Excalibur calcium-binding domain-containing protein; afdb-uniprot50 \| AF-A0A2N6QKG2-F1-MODEL\_V4 \| 0.991 \| 0.007955 \| 84 \| 0.109 \| 182 \| 132 \| 11 \| 8 \| 166 \| 24 \| 198 \| BppU\_N domain-containing protein \| BppU\_N domain-containing protein; afdb-uniprot50 \| AF-A0A6B0AW78-F1-MODEL\_V4 \| 0.991 \| 0.00244 \| 84 \| 0.143 \| 153 \| 98 \| 9 \| 27 \| 165 \| 56 \| 189 \| DUF2479 domain-containing protein \| DUF2479 domain-containing protein; afdb-uniprot50 \| AF-S9PVW1-F1-MODEL\_V4 \| 0.991 \| 0.00844 \| 84 \| 0.125 \| 151 \| 103 \| 11 \| 7 \| 136 \| 291 \| 433 \| Dolichyl-diphosphooligosaccharide--protein glycosyltransferase subunit WBP1 \| Dolichyl-diphosphooligosaccharide--protein glycosyltransferase subunit WBP1; afdb-uniprot50 \| AF-A0A5N5MH13-F1-MODEL\_V4 \| 0.991 \| 0.005581 \| 84 \| 0.135 \| 140 \| 105 \| 8 \| 9 \| 142 \| 347 \| 476 \| Uncharacterized protein \| Uncharacterized protein; afdb-uniprot50 \| AF-D2RBM3-F1-MODEL\_V4 \| 0.991 \| 0.005581 \| 84 \| 0.122 \| 179 \| 116 \| 10 \| 6 \| 165 \| 20 \| 176 \| Methyl-accepting transducer domain-containing protein \| Methyl-accepting transducer domain-containing protein; afdb-uniprot50 \| AF-X1G5Z6-F1-MODEL\_V4 \| 0.99 \| 0.001434 \| 83 \| 0.129 \| 154 \| 101 \| 9 \| 5 \| 142 \| 27 \| 163 \| Uncharacterized protein \| Uncharacterized protein; afdb-uniprot50 \| AF-A0A3B4YJ65-F1-MODEL\_V4 \| 0.99 \| 0.00592 \| 83 \| 0.128 \| 156 \| 101 \| 7 \| 9 \| 160 \| 129 \| 253 \| Uncharacterized protein \| Uncharacterized protein; afdb-uniprot50 \| AF-A0A518GX12-F1-MODEL\_V4 \| 0.99 \| 0.003915 \| 83 \| 0.105 \| 190 \| 119 \| 9 \| 6 \| 164 \| 79 \| 248 \| Uncharacterized protein \| Uncharacterized protein; afdb-uniprot50 \| AF-A0A7J3SM07-F1-MODEL\_V4 \| 0.99 \| 0.006663 \| 83 \| 0.104 \| 162 \| 115 \| 10 \| 7 \| 158 \| 110 \| 251 \| Uncharacterized protein \| Uncharacterized protein; afdb-uniprot50 \| AF-E1RGZ0-F1-MODEL\_V4 \| 0.99 \| 0.006663 \| 83 \| 0.115 \| 130 \| 82 \| 5 \| 8 \| 115 \| 31 \| 149 \| Uncharacterized protein \| Uncharacterized protein; afdb-uniprot50 \| AF-A0A0L0SSW8-F1-MODEL\_V4 \| 0.99 \| 0.002168 \| 83 \| 0.1 \| 200 \| 133 \| 13 \| 7 \| 179 \| 294 \| 473 \| Dolichyl-diphosphooligosaccharide--protein glycosyltransferase subunit WBP1 \| Dolichyl-diphosphooligosaccharide--protein glycosyltransferase subunit WBP1; afdb-uniprot50 \| AF-A0A5M9ZIC2-F1-MODEL\_V4 \| 0.99 \| 0.002589 \| 83 \| 0.154 \| 142 \| 82 \| 12 \| 9 \| 126 \| 24 \| 151 \| Uncharacterized protein \| Uncharacterized protein; afdb-uniprot50 \| AF-A0A7J9PQ88-F1-MODEL\_V4 \| 0.99 \| 0.003091 \| 83 \| 0.134 \| 134 \| 97 \| 8 \| 9 \| 129 \| 519 \| 646 \| Uncharacterized protein \| Uncharacterized protein; afdb-uniprot50 \| AF-A0A842BP15-F1-MODEL\_V4 \| 0.988 \| 0.003478 \| 82 \| 0.156 \| 160 \| 91 \| 8 \| 9 \| 142 \| 120 \| 261 \| LPXTG cell wall anchor domain-containing protein \| LPXTG cell wall anchor domain-containing protein; afdb-uniprot50 \| AF-A0A7J9SFD5-F1-MODEL\_V4 \| 0.988 \| 0.002913 \| 82 \| 0.116 \| 129 \| 93 \| 7 \| 9 \| 124 \| 305 \| 425 \| Uncharacterized protein \| Uncharacterized protein; afdb-uniprot50 \| AF-A0A6G0ITN7-F1-MODEL\_V4 \| 0.988 \| 0.00244 \| 82 \| 0.13 \| 161 \| 99 \| 9 \| 3 \| 142 \| 296 \| 436 \| Deleted in malignant brain tumors 1 protein Glycoprotein 340 \| Deleted in malignant brain tumors 1 protein Glycoprotein 340; afdb-uniprot50 \| AF-A0A843LRD6-F1-MODEL\_V4 \| 0.988 \| 0.003478 \| 82 \| 0.106 \| 160 \| 93 \| 12 \| 6 \| 142 \| 281 \| 413 \| PGF-CTERM sorting domain-containing protein \| PGF-CTERM sorting domain-containing protein; afdb-uniprot50 \| AF-A0A5C1G595-F1-MODEL\_V4 \| 0.988 \| 0.005581 \| 82 \| 0.115 \| 173 \| 113 \| 10 \| 3 \| 142 \| 15 \| 180 \| DUF2479 domain-containing protein \| DUF2479 domain-containing protein; afdb-uniprot50 \| AF-A0A2M7AA77-F1-MODEL\_V4 \| 0.988 \| 0.007955 \| 82 \| 0.102 \| 156 \| 91 \| 12 \| 7 \| 122 \| 277 \| 423 \| Carb-bd\_dom\_fam9 domain-containing protein \| Carb-bd\_dom\_fam9 domain-containing protein; afdb-uniprot50 \| AF-A0A6P7K4J8-F1-MODEL\_V4 \| 0.986 \| 0.009498 \| 81 \| 0.144 \| 138 \| 96 \| 8 \| 9 \| 138 \| 109 \| 232 \| uncharacterized protein LOC114449958 \| uncharacterized protein LOC114449958; afdb-uniprot50 \| AF-A0A2N0G476-F1-MODEL\_V4 \| 0.986 \| 0.004959 \| 81 \| 0.109 \| 155 \| 106 \| 10 \| 8 \| 142 \| 5 \| 147 \| Uncharacterized protein \| Uncharacterized protein; afdb-uniprot50 \| AF-A0A535C1Z2-F1-MODEL\_V4 \| 0.986 \| 0.003091 \| 81 \| 0.145 \| 144 \| 100 \| 11 \| 6 \| 135 \| 168 \| 302 \| Uncharacterized protein \| Uncharacterized protein; afdb-uniprot50 \| AF-A0A1V4ZS71-F1-MODEL\_V4 \| 0.986 \| 0.006281 \| 81 \| 0.155 \| 154 \| 79 \| 12 \| 9 \| 142 \| 203 \| 325 \| Uncharacterized protein \| Uncharacterized protein; afdb-uniprot50 \| AF-A0A3M2QGJ1-F1-MODEL\_V4 \| 0.986 \| 0.003915 \| 81 \| 0.127 \| 188 \| 113 \| 14 \| 9 \| 166 \| 26 \| 192 \| Phage baseplate upper protein \| Phage baseplate upper protein; afdb-uniprot50 \| AF-A0A4V6KU45-F1-MODEL\_V4 \| 0.986 \| 0.00592 \| 81 \| 0.121 \| 189 \| 119 \| 12 \| 8 \| 178 \| 356 \| 515 \| Protein of uncharacterized function (DUF1533) \| Protein of uncharacterized function (DUF1533); afdb-uniprot50 \| AF-A0A842F260-F1-MODEL\_V4 \| 0.986 \| 0.009498 \| 81 \| 0.153 \| 117 \| 80 \| 6 \| 29 \| 142 \| 423 \| 523 \| LPXTG cell wall anchor domain-containing protein \| LPXTG cell wall anchor domain-containing protein; afdb-uniprot50 \| AF-A0A452J3I1-F1-MODEL\_V4 \| 0.984 \| 0.003478 \| 80 \| 0.071 \| 167 \| 118 \| 10 \| 9 \| 145 \| 28 \| 187 \| Ig-like domain-containing protein \| Ig-like domain-containing protein; afdb-uniprot50 \| AF-H1Z1T2-F1-MODEL\_V4 \| 0.984 \| 0.003091 \| 80 \| 0.135 \| 155 \| 87 \| 11 \| 8 \| 142 \| 193 \| 320 \| Uncharacterized protein \| Uncharacterized protein; afdb-uniprot50 \| AF-A0A1V1P2D0-F1-MODEL\_V4 \| 0.984 \| 0.003091 \| 80 \| 0.09 \| 177 \| 106 \| 11 \| 8 \| 157 \| 296 \| 444 \| Uncharacterized protein \| Uncharacterized protein; afdb-uniprot50 \| AF-A0A2R5GPP2-F1-MODEL\_V4 \| 0.984 \| 0.001434 \| 80 \| 0.141 \| 198 \| 124 \| 13 \| 15 \| 179 \| 480 \| 664 \| DnaJ protein ERDJ2A \| DnaJ protein ERDJ2A; afdb-uniprot50 \| AF-A0A1Q5P452-F1-MODEL\_V4 \| 0.984 \| 0.006663 \| 80 \| 0.12 \| 141 \| 95 \| 7 \| 31 \| 164 \| 50 \| 168 \| NodB homology domain-containing protein \| NodB homology domain-containing protein; afdb-uniprot50 \| AF-A0A3N2CWC2-F1-MODEL\_V4 \| 0.981 \| 0.007069 \| 79 \| 0.127 \| 173 \| 115 \| 9 \| 8 \| 166 \| 46 \| 196 \| CopC domain-containing protein \| CopC domain-containing protein; afdb-uniprot50 \| AF-A0A674NHX4-F1-MODEL\_V4 \| 0.981 \| 0.000948 \| 79 \| 0.091 \| 174 \| 110 \| 12 \| 7 \| 142 \| 51 \| 214 \| IGv domain-containing protein \| IGv domain-containing protein; afdb-uniprot50 \| AF-A0A556UXS5-F1-MODEL\_V4 \| 0.981 \| 0.001132 \| 79 \| 0.131 \| 160 \| 114 \| 9 \| 7 \| 149 \| 21 \| 172 \| Motile sperm domain-containing protein 1 \| Motile sperm domain-containing protein 1; afdb-uniprot50 \| AF-A0A7K4B3J4-F1-MODEL\_V4 \| 0.981 \| 0.002746 \| 79 \| 0.113 \| 158 \| 91 \| 10 \| 9 \| 142 \| 241 \| 373 \| PGF-CTERM sorting domain-containing protein \| PGF-CTERM sorting domain-containing protein; afdb-uniprot50 \| AF-A0A7D8H6E0-F1-MODEL\_V4 \| 0.981 \| 0.007069 \| 79 \| 0.134 \| 164 \| 105 \| 12 \| 8 \| 142 \| 22 \| 177 \| Uncharacterized protein \| Uncharacterized protein; afdb-uniprot50 \| AF-A0A7J5YJ57-F1-MODEL\_V4 \| 0.981 \| 0.007069 \| 79 \| 0.059 \| 202 \| 159 \| 7 \| 9 \| 179 \| 322 \| 523 \| ZP domain-containing protein \| ZP domain-containing protein; afdb-uniprot50 \| AF-A0A7J5XR45-F1-MODEL\_V4 \| 0.981 \| 0.007499 \| 79 \| 0.132 \| 151 \| 101 \| 8 \| 9 \| 142 \| 764 \| 901 \| Ig-like domain-containing protein \| Ig-like domain-containing protein; afdb-uniprot50 \| AF-A0A3M1PQZ9-F1-MODEL\_V4 \| 0.981 \| 0.002589 \| 79 \| 0.125 \| 208 \| 124 \| 12 \| 15 \| 178 \| 42 \| 235 \| Tetratricopeptide repeat protein \| Tetratricopeptide repeat protein; afdb-uniprot50 \| AF-A0A7K1J0H1-F1-MODEL\_V4 \| 0.978 \| 0.007499 \| 78 \| 0.117 \| 128 \| 95 \| 9 \| 9 \| 128 \| 102 \| 219 \| Uncharacterized protein \| Uncharacterized protein; afdb-uniprot50 \| AF-A0A3Q3ABB6-F1-MODEL\_V4 \| 0.978 \| 0.006663 \| 78 \| 0.117 \| 179 \| 111 \| 8 \| 5 \| 179 \| 304 \| 439 \| Scavenger receptor cysteine-rich type 1 protein M160-like \| Scavenger receptor cysteine-rich type 1 protein M160-like; afdb-uniprot50 \| AF-A0A834FJH2-F1-MODEL\_V4 \| 0.978 \| 0.00369 \| 78 \| 0.153 \| 196 \| 113 \| 11 \| 3 \| 178 \| 681 \| 843 \| Uncharacterized protein \| Uncharacterized protein; afdb-uniprot50 \| AF-A0A7J5XPK3-F1-MODEL\_V4 \| 0.978 \| 0.00526 \| 78 \| 0.14 \| 149 \| 97 \| 10 \| 9 \| 142 \| 839 \| 971 \| Uncharacterized protein \| Uncharacterized protein; afdb-uniprot50 \| AF-B8MP86-F1-MODEL\_V4 \| 0.975 \| 0.008953 \| 77 \| 0.096 \| 218 \| 147 \| 9 \| 3 \| 179 \| 12 \| 220 \| Uncharacterized protein \| Uncharacterized protein; afdb-uniprot50 \| AF-A0A841ZTY5-F1-MODEL\_V4 \| 0.975 \| 0.007955 \| 77 \| 0.126 \| 150 \| 92 \| 9 \| 14 \| 142 \| 286 \| 417 \| LPXTG cell wall anchor domain-containing protein \| LPXTG cell wall anchor domain-containing protein; afdb-uniprot50 \| AF-A0A537DD91-F1-MODEL\_V4 \| 0.975 \| 0.007069 \| 77 \| 0.125 \| 167 \| 128 \| 10 \| 7 \| 165 \| 642 \| 798 \| PKD domain-containing protein \| PKD domain-containing protein; afdb-uniprot50 \| AF-A0A1Y1JV51-F1-MODEL\_V4 \| 0.975 \| 0.007499 \| 77 \| 0.093 \| 193 \| 130 \| 8 \| 10 \| 164 \| 30 \| 215 \| Uncharacterized protein \| Uncharacterized protein; afdb-uniprot50 \| AF-A0A6A5FI04-F1-MODEL\_V4 \| 0.971 \| 0.00844 \| 76 \| 0.156 \| 147 \| 96 \| 10 \| 9 \| 142 \| 14 \| 145 \| Ig-like domain-containing protein \| Ig-like domain-containing protein; afdb-uniprot50 \| AF-A0A7J3DL36-F1-MODEL\_V4 \| 0.971 \| 0.0006268 \| 76 \| 0.125 \| 200 \| 134 \| 11 \| 1 \| 166 \| 17 \| 209 \| Uncharacterized protein \| Uncharacterized protein; afdb-uniprot50 \| AF-A0A1A8Q8P6-F1-MODEL\_V4 \| 0.971 \| 0.003915 \| 76 \| 0.1 \| 139 \| 105 \| 6 \| 10 \| 142 \| 103 \| 227 \| Fibronectin type-III domain-containing protein \| Fibronectin type-III domain-containing protein; afdb-uniprot50 \| AF-A0A3S1AFG5-F1-MODEL\_V4 \| 0.971 \| 0.006663 \| 76 \| 0.1 \| 149 \| 102 \| 7 \| 5 \| 142 \| 286 \| 413 \| MG3 domain-containing protein \| MG3 domain-containing protein; afdb-uniprot50 \| AF-A0A3P8Y1B5-F1-MODEL\_V4 \| 0.971 \| 0.006281 \| 76 \| 0.147 \| 149 \| 99 \| 9 \| 9 \| 142 \| 338 \| 473 \| Uncharacterized protein \| Uncharacterized protein; afdb-uniprot50 \| AF-A0A0F8AG26-F1-MODEL\_V4 \| 0.971 \| 0.003279 \| 76 \| 0.065 \| 168 \| 126 \| 7 \| 6 \| 142 \| 324 \| 491 \| Transforming growth factor beta receptor type 3 \| Transforming growth factor beta receptor type 3; afdb-uniprot50 \| AF-A0A484DDS8-F1-MODEL\_V4 \| 0.971 \| 0.009498 \| 76 \| 0.164 \| 158 \| 94 \| 9 \| 3 \| 142 \| 632 \| 769 \| Uncharacterized protein \| Uncharacterized protein; afdb-uniprot50 \| AF-A0A3P9JQK7-F1-MODEL\_V4 \| 0.967 \| 0.007499 \| 75 \| 0.114 \| 192 \| 123 \| 14 \| 8 \| 164 \| 56 \| 235 \| Ig-like domain-containing protein \| Ig-like domain-containing protein; afdb-uniprot50 \| AF-A0A517XKX4-F1-MODEL\_V4 \| 0.967 \| 0.001521 \| 75 \| 0.137 \| 175 \| 104 \| 7 \| 9 \| 141 \| 600 \| 769 \| Leupeptin-inactivating enzyme 1 \| Leupeptin-inactivating enzyme 1; afdb-uniprot50 \| AF-A0A7Y2HJU2-F1-MODEL\_V4 \| 0.967 \| 0.002913 \| 75 \| 0.158 \| 145 \| 97 \| 7 \| 8 \| 142 \| 568 \| 697 \| Uncharacterized protein \| Uncharacterized protein; afdb-uniprot50 \| AF-A0A0M0BEQ4-F1-MODEL\_V4 \| 0.967 \| 0.004674 \| 75 \| 0.074 \| 135 \| 98 \| 9 \| 5 \| 128 \| 901 \| 1019 \| Uncharacterized protein \| Uncharacterized protein; afdb-uniprot50 \| AF-A0A662BMB6-F1-MODEL\_V4 \| 0.967 \| 0.003915 \| 75 \| 0.095 \| 199 \| 127 \| 11 \| 7 \| 166 \| 957 \| 1141 \| Uncharacterized protein \| Uncharacterized protein; afdb-uniprot50 \| AF-A0A094B7D5-F1-MODEL\_V4 \| 0.961 \| 0.007499 \| 74 \| 0.137 \| 153 \| 103 \| 10 \| 8 \| 142 \| 113 \| 254 \| Uncharacterized protein \| Uncharacterized protein; afdb-uniprot50 \| AF-A0A7X0XKQ9-F1-MODEL\_V4 \| 0.961 \| 0.003279 \| 74 \| 0.125 \| 168 \| 95 \| 10 \| 9 \| 142 \| 51 \| 200 \| LPXTG cell wall anchor domain-containing protein \| LPXTG cell wall anchor domain-containing protein; afdb-uniprot50 \| AF-A0A1W9LDA3-F1-MODEL\_V4 \| 0.961 \| 0.009498 \| 74 \| 0.164 \| 158 \| 88 \| 9 \| 6 \| 129 \| 317 \| 464 \| Uncharacterized protein \| Uncharacterized protein; afdb-uniprot50 \| AF-A0A672HNP6-F1-MODEL\_V4 \| 0.956 \| 0.001434 \| 73 \| 0.112 \| 169 \| 119 \| 9 \| 7 \| 155 \| 20 \| 177 \| Motile sperm domain-containing protein 1 \| Motile sperm domain-containing protein 1; afdb-uniprot50 \| AF-A0A6I9Y835-F1-MODEL\_V4 \| 0.956 \| 0.008953 \| 73 \| 0.102 \| 185 \| 134 \| 12 \| 8 \| 179 \| 77 \| 242 \| semaphorin-3F-like \| semaphorin-3F-like; afdb-uniprot50 \| AF-A0A803VYX2-F1-MODEL\_V4 \| 0.956 \| 0.007499 \| 73 \| 0.138 \| 202 \| 111 \| 9 \| 9 \| 179 \| 740 \| 909 \| Uncharacterized protein \| Uncharacterized protein; afdb-uniprot50 \| AF-A0A4W5L7M6-F1-MODEL\_V4 \| 0.949 \| 0.001816 \| 72 \| 0.095 \| 168 \| 109 \| 10 \| 9 \| 142 \| 29 \| 187 \| Ig-like domain-containing protein \| Ig-like domain-containing protein; afdb-uniprot50 \| AF-A0A7J5XLP4-F1-MODEL\_V4 \| 0.949 \| 0.002589 \| 72 \| 0.097 \| 154 \| 113 \| 8 \| 7 \| 142 \| 47 \| 192 \| Motile sperm domain-containing protein 1 \| Motile sperm domain-containing protein 1; afdb-uniprot50 \| AF-A0A832TUV5-F1-MODEL\_V4 \| 0.949 \| 0.008953 \| 72 \| 0.111 \| 162 \| 88 \| 10 \| 6 \| 142 \| 171 \| 301 \| Uncharacterized protein \| Uncharacterized protein; afdb-uniprot50 \| AF-A0A2G9S9Q7-F1-MODEL\_V4 \| 0.949 \| 0.004959 \| 72 \| 0.152 \| 151 \| 103 \| 9 \| 2 \| 142 \| 82 \| 217 \| Ig-like domain-containing protein \| Ig-like domain-containing protein; afdb-uniprot50 \| AF-A0A6G0ITN3-F1-MODEL\_V4 \| 0.949 \| 0.00844 \| 72 \| 0.128 \| 164 \| 99 \| 10 \| 3 \| 142 \| 403 \| 546 \| Scavenger receptor cysteine-rich type 1 protein M130 \| Scavenger receptor cysteine-rich type 1 protein M130; afdb-uniprot50 \| AF-A0A843LMU9-F1-MODEL\_V4 \| 0.949 \| 0.009498 \| 72 \| 0.113 \| 159 \| 90 \| 11 \| 9 \| 142 \| 436 \| 568 \| PGF-CTERM sorting domain-containing protein \| PGF-CTERM sorting domain-containing protein; afdb-uniprot50 \| AF-A0A7J4HUN3-F1-MODEL\_V4 \| 0.941 \| 0.00526 \| 71 \| 0.112 \| 178 \| 124 \| 9 \| 7 \| 164 \| 304 \| 467 \| Uncharacterized protein \| Uncharacterized protein; afdb-uniprot50 \| AF-S8F0H2-F1-MODEL\_V4 \| 0.941 \| 0.002589 \| 71 \| 0.147 \| 197 \| 119 \| 13 \| 15 \| 179 \| 400 \| 579 \| Uncharacterized protein \| Uncharacterized protein; afdb-uniprot50 \| AF-A0A8B6DDG9-F1-MODEL\_V4 \| 0.941 \| 0.009498 \| 71 \| 0.13 \| 123 \| 95 \| 3 \| 32 \| 142 \| 159 \| 281 \| Uncharacterized protein \| Uncharacterized protein; afdb-uniprot50 \| AF-A0A2V1A8H1-F1-MODEL\_V4 \| 0.941 \| 0.00526 \| 71 \| 0.113 \| 158 \| 102 \| 10 \| 9 \| 142 \| 53 \| 196 \| SAC domain-containing protein \| SAC domain-containing protein; afdb-uniprot50 \| AF-A0A6G7Z4I0-F1-MODEL\_V4 \| 0.933 \| 0.003279 \| 70 \| 0.119 \| 151 \| 107 \| 7 \| 31 \| 166 \| 45 \| 184 \| Fibronectin type-III domain-containing protein \| Fibronectin type-III domain-containing protein; afdb-uniprot50 \| AF-A0A7J6B8F1-F1-MODEL\_V4 \| 0.923 \| 0.0023 \| 69 \| 0.109 \| 191 \| 131 \| 12 \| 7 \| 178 \| 87 \| 257 \| Motile sperm domain-containing protein 1 \| Motile sperm domain-containing protein 1; afdb-uniprot50 \| AF-A0A847I2Q6-F1-MODEL\_V4 \| 0.923 \| 0.004674 \| 69 \| 0.168 \| 172 \| 101 \| 13 \| 17 \| 163 \| 298 \| 452 \| VWA domain-containing protein \| VWA domain-containing protein; afdb-uniprot50 \| AF-A0A087TGG5-F1-MODEL\_V4 \| 0.923 \| 0.004153 \| 69 \| 0.148 \| 148 \| 95 \| 9 \| 9 \| 142 \| 585 \| 715 \| Receptor protein-tyrosine kinase \| Receptor protein-tyrosine kinase; afdb-uniprot50 \| AF-A0A2M9C4S0-F1-MODEL\_V4 \| 0.912 \| 0.008953 \| 68 \| 0.128 \| 194 \| 97 \| 11 \| 9 \| 142 \| 46 \| 227 \| Methionine-rich copper-binding protein CopC \| Methionine-rich copper-binding protein CopC; afdb-uniprot50 \| AF-A0A1J0U2V1-F1-MODEL\_V4 \| 0.912 \| 0.004959 \| 68 \| 0.154 \| 188 \| 119 \| 13 \| 9 \| 179 \| 342 \| 506 \| Cadherin\_5 domain-containing protein \| Cadherin\_5 domain-containing protein; afdb-uniprot50 \| AF-A0A3R9QQN6-F1-MODEL\_V4 \| 0.912 \| 0.004959 \| 68 \| 0.119 \| 151 \| 90 \| 9 \| 7 \| 142 \| 412 \| 534 \| Uncharacterized protein \| Uncharacterized protein; afdb-uniprot50 \| AF-A0A3Q1GFR9-F1-MODEL\_V4 \| 0.9 \| 0.004674 \| 67 \| 0.105 \| 151 \| 111 \| 7 \| 7 \| 142 \| 21 \| 162 \| Motile sperm domain-containing protein 1 \| Motile sperm domain-containing protein 1; afdb-uniprot50 \| AF-A0A3Q1J8F6-F1-MODEL\_V4 \| 0.9 \| 0.00526 \| 67 \| 0.113 \| 167 \| 121 \| 8 \| 7 \| 155 \| 112 \| 269 \| Motile sperm domain-containing protein 1 \| Motile sperm domain-containing protein 1; afdb-uniprot50 \| AF-A0A075B1J7-F1-MODEL\_V4 \| 0.9 \| 0.008953 \| 67 \| 0.101 \| 207 \| 135 \| 12 \| 8 \| 178 \| 226 \| 417 \| Uncharacterized protein \| Uncharacterized protein; afdb-uniprot50 \| AF-A0A670KDQ6-F1-MODEL\_V4 \| 0.887 \| 0.003915 \| 66 \| 0.105 \| 151 \| 108 \| 7 \| 8 \| 142 \| 24 \| 163 \| Motile sperm domain-containing protein 1 \| Motile sperm domain-containing protein 1; afdb-uniprot50 \| AF-A0A3M6TAK4-F1-MODEL\_V4 \| 0.887 \| 0.009498 \| 66 \| 0.106 \| 169 \| 113 \| 11 \| 15 \| 160 \| 25 \| 178 \| Uncharacterized protein \| Uncharacterized protein; afdb-uniprot50 \| AF-A0A0S7IV27-F1-MODEL\_V4 \| 0.872 \| 0.004959 \| 65 \| 0.126 \| 166 \| 116 \| 8 \| 8 \| 155 \| 54 \| 208 \| Motile sperm domain-containing protein 1 \| Motile sperm domain-containing protein 1; afdb-uniprot50 \| AF-A0A6F9ANK6-F1-MODEL\_V4 \| 0.872 \| 0.003478 \| 65 \| 0.101 \| 187 \| 133 \| 9 \| 8 \| 178 \| 58 \| 225 \| Motile sperm domain-containing protein 1 \| Motile sperm domain-containing protein 1; afdb-uniprot50 \| AF-A0A497MZN8-F1-MODEL\_V4 \| 0.855 \| 0.00844 \| 64 \| 0.164 \| 158 \| 94 \| 13 \| 7 \| 142 \| 126 \| 267 \| Uncharacterized protein \| Uncharacterized protein; afdb-uniprot50 \| AF-A0A2Y9RF76-F1-MODEL\_V4 \| 0.855 \| 0.00592 \| 64 \| 0.163 \| 165 \| 94 \| 9 \| 7 \| 142 \| 121 \| 270 \| T-cell-interacting, activating receptor on myeloid cells protein 1 isoform X5 \| T-cell-interacting, activating receptor on myeloid cells protein 1 isoform X5; afdb-uniprot50 \| AF-A0A7M3ZE92-F1-MODEL\_V4 \| 0.855 \| 0.007499 \| 64 \| 0.098 \| 172 \| 112 \| 8 \| 8 \| 142 \| 110 \| 275 \| Uncharacterized protein \| Uncharacterized protein; afdb-uniprot50 \| AF-A0A832Y8B9-F1-MODEL\_V4 \| 0.855 \| 0.001132 \| 64 \| 0.168 \| 184 \| 104 \| 12 \| 7 \| 164 \| 117 \| 277 \| Uncharacterized protein \| Uncharacterized protein; afdb-uniprot50 \| AF-A0A351H4X9-F1-MODEL\_V4 \| 0.837 \| 0.004153 \| 63 \| 0.087 \| 240 \| 126 \| 13 \| 5 \| 153 \| 22 \| 259 \| Uncharacterized protein \| Uncharacterized protein; afdb-uniprot50 \| AF-A0A3B5BAV1-F1-MODEL\_V4 \| 0.817 \| 0.007955 \| 62 \| 0.119 \| 167 \| 116 \| 8 \| 8 \| 155 \| 24 \| 178 \| Motile sperm domain-containing protein 1 \| Motile sperm domain-containing protein 1; afdb-uniprot50 \| AF-A0A3Q8ZFA0-F1-MODEL\_V4 \| 0.817 \| 0.00592 \| 62 \| 0.122 \| 179 \| 120 \| 8 \| 15 \| 166 \| 20 \| 188 \| Uncharacterized protein \| Uncharacterized protein; afdb-uniprot50 \| AF-A0A7X0FNP4-F1-MODEL\_V4 \| 0.817 \| 0.008953 \| 62 \| 0.191 \| 157 \| 93 \| 12 \| 6 \| 142 \| 579 \| 721 \| Uncharacterized protein \| Uncharacterized protein; afdb-uniprot50 \| AF-A0A2D5X0L3-F1-MODEL\_V4 \| 0.795 \| 0.00244 \| 61 \| 0.157 \| 210 \| 115 \| 13 \| 9 \| 179 \| 121 \| 307 \| Uncharacterized protein \| Uncharacterized protein; afdb-uniprot50 \| AF-A0A672YNI3-F1-MODEL\_V4 \| 0.747 \| 0.009498 \| 59 \| 0.113 \| 150 \| 108 \| 7 \| 8 \| 141 \| 24 \| 164 \| Motile sperm domain-containing protein 1 \| Motile sperm domain-containing protein 1; afdb-uniprot50 \| AF-A0A534JZD8-F1-MODEL\_V4 \| 0.747 \| 0.0007484 \| 59 \| 0.151 \| 172 \| 96 \| 9 \| 9 \| 142 \| 49 \| 208 \| Spt5-NGN domain-containing protein \| Spt5-NGN domain-containing protein; afdb-uniprot50 \| AF-A0A0M2LTA1-F1-MODEL\_V4 \| 0.72 \| 0.009498 \| 58 \| 0.142 \| 161 \| 90 \| 13 \| 7 \| 142 \| 541 \| 678 \| Uncharacterized protein \| Uncharacterized protein; afdb-uniprot50 \| AF-A0A075HIB3-F1-MODEL\_V4 \| 0.692 \| 0.004674 \| 57 \| 0.094 \| 179 \| 108 \| 11 \| 8 \| 142 \| 17 \| 185 \| Uncharacterized protein \| Uncharacterized protein |
| Top keywords  (threshold 1.00e-02 (evalue)) | **domain\_containing, BppU\_N, light, receptor, Fab, DUF2479, Dolichyl\_diphosphooligosaccharide\_\_protein, glycosyltransferase, Phage, immunoglobulin** |
| Output files | ../../similar\_structures/03\_FANPEZAQ\_CDS\_0003\_afdb-proteome\_foldseek.tsv ../../similar\_structures/03\_FANPEZAQ\_CDS\_0003\_afdb-uniprot50\_foldseek.tsv ../../similar\_structures/03\_FANPEZAQ\_CDS\_0003\_merged.svg ../../similar\_structures/03\_FANPEZAQ\_CDS\_0003\_pdb\_foldseek.tsv |

  
  
  

Return to summary | Go to previous | Go to next

  


---

**Sequence/structure alignments coloring**  
Each object in the alignment figures is colored according to its E-value following this color coding:

1e-100
10
