## Supplementary material for "Completing the BASEL phage collection to unlock hidden diversity for systematic exploration of phage-host interactions": Data S2: 4.html

FANPEZAQ\_CDS\_0004


Return to summary | Go to previous | Go to next

|  |  |
| --- | --- |
| FANPEZAQ\_CDS\_0004  Page creation date: 02 Sep 2024, 12:00  Project folder: n/a  Input sequences file: Escherichia\_virus\_HeidiAbel.gb | portal phage lambda fragment capsid putative duf935 head prophage and packaging domain\_containing bacteriophage minor terminase head\_tail phage\_related duf4055 spp1 preconnector hk97 gp6\_like parb protease phage\_mu\_f |

### Sequence information

|  |  |
| --- | --- |
| Name | FANPEZAQ\_CDS\_0004  04\_FANPEZAQ\_CDS\_0004 (pipeline id) |
| Imported annotations | Escherichia\_virus\_HeidiAbel Bas97 |
| Protein sequence | MQSIRAQSTGEPSLMKLWPFTRNKPAEQAQQPRTRRLPLNRLASRMFTGASVDRLGTAWG TQPLTADEVINKNQRILVARSREQAANNDYAKKFLRLCRNNIVGPKGVLLQAKSTDNTGK LDTLANEAIETAFDKWGHKSNCDVTGKLSWRAIQAACINSAAKDGEFFVRLVFGADAGPW GFALQLLDPQRCPVDMVEDRLNNGNFIRQGIEFNRYGRPVAYYFSTVDESDDNYRWSGRD FVRIPADEILHGFLDDMVGQKRGLPWMATSLFRMRHLGGMEEAALVNARAGANKMGFIEW DENSAGPEFEDDDELIIDSEPGEFQVLPQGARVKDWSPQYPNGEFAVFTKQMLRGMASGM GVPYNDLASDLEGVNFSSIRQGTLDSRENWKELQEWLIENLHQPVFEAWLPRALLAGRIT VKGRSLKPERIDRYSDIEWQPRRWAWIDPSSDVAAAERSKNNMLQSPGRIIREGGSDPDT VWKETASDVARMIEAYTQHGISREMAEKLVMQSMGMDKQLLMGQQNAEKPAE |
| Number of residues | 532 |
| Molecular weight (Da) | 60054.10 |
| Output files | ../../query\_sequences/04\_FANPEZAQ\_CDS\_0004.fasta |

### Putative domain architecture and protein family

#### Search results (HHblits)^1^

|  |  |
| --- | --- |
| Domain family databases searched | Pfam, Ncbi-cd, Cath, Phrogs |
| Results, scheme(s)  (Top layers only; threshold 1.00e-03 (evalue)) | xml version="1.0" encoding="utf-8" standalone="no"?       2024-09-02T21:08:12.560655 image/svg+xml   Matplotlib v3.7.2, https://matplotlib.org/ |
| Results, table  (E-value ≤ 1.00e-03 (evalue)) | db \| id \| prob \| evalue \| pvalue \| score \| cols \| query \| query\_len \| template \| template\_len \| name \| description; pfam \| PF06074 \| 99.8 \| 2.8e-25 \| 4.8e-29 \| 224.8 \| 351 \| (56, 479) \| 532 \| (30, 391) \| 504 \| DUF935 \| Protein of unknown function (DUF935); pfam \| PF05136 \| 99.7 \| 2.8e-23 \| 4.8e-27 \| 198.4 \| 330 \| (53, 393) \| 532 \| (2, 333) \| 334 \| Phage\_portal\_2 \| Phage portal protein, lambda family; pfam \| PF04860 \| 99.7 \| 1.3e-21 \| 2.3e-25 \| 185.4 \| 324 \| (89, 474) \| 532 \| (1, 332) \| 332 \| Phage\_portal \| Phage portal protein; pfam \| PF06381 \| 99.6 \| 3e-20 \| 5.3e-24 \| 177.5 \| 320 \| (80, 463) \| 532 \| (2, 348) \| 350 \| DUF1073 \| Protein of unknown function (DUF1073); pfam \| PF07230 \| 99.6 \| 8.3e-20 \| 1.5e-23 \| 181.3 \| 382 \| (76, 486) \| 532 \| (3, 449) \| 454 \| Portal\_Gp20 \| Bacteriophage T4-like portal protein (Gp20); pfam \| PF05133 \| 98.7 \| 8.9e-13 \| 1.6e-16 \| 128.8 \| 337 \| (87, 475) \| 532 \| (54, 422) \| 457 \| Phage\_prot\_Gp6 \| Phage portal protein, SPP1 Gp6-like; phrogs \| 21 \| 100.0 \| 3.1e-59 \| 3.7e-63 \| 486.4 \| 447 \| (40, 494) \| 532 \| (17, 502) \| 533 \| portal protein \| portal protein; Category: head and packaging; p173499 VI\_06341; phrogs \| 2662 \| 99.9 \| 6.5e-31 \| 7.7e-35 \| 272.9 \| 344 \| (81, 485) \| 532 \| (42, 391) \| 685 \| portal protein \| portal protein; Category: head and packaging; NC\_023612\_p3; phrogs \| 146 \| 99.9 \| 1.7e-29 \| 2e-33 \| 259.5 \| 365 \| (75, 492) \| 532 \| (60, 460) \| 485 \| portal protein \| portal protein; Category: head and packaging; p353614 VI\_06430; phrogs \| 12 \| 99.9 \| 3.4e-28 \| 4.2e-32 \| 248.4 \| 344 \| (76, 482) \| 532 \| (20, 369) \| 434 \| portal protein \| portal protein; Category: head and packaging; p213127 VI\_06558; phrogs \| 9232 \| 99.6 \| 5.7e-20 \| 6.6e-24 \| 173.7 \| 365 \| (79, 498) \| 532 \| (52, 425) \| 479 \| portal protein \| portal protein; Category: head and packaging; NC\_004927\_p94; phrogs \| 2271 \| 99.4 \| 6.4e-18 \| 7.2e-22 \| 152.4 \| 394 \| (57, 515) \| 532 \| (21, 418) \| 620 \| portal protein \| portal protein; Category: head and packaging; p375194 VI\_08017; phrogs \| 5929 \| 99.4 \| 9.3e-18 \| 1.1e-21 \| 145.0 \| 150 \| (310, 483) \| 532 \| (14, 167) \| 231 \| portal protein \| portal protein; Category: head and packaging; p208881 VI\_08063; phrogs \| 537 \| 99.4 \| 1.1e-17 \| 1.3e-21 \| 137.0 \| 126 \| (375, 504) \| 532 \| (2, 129) \| 154 \| portal protein \| portal protein; Category: head and packaging; p386813 VI\_08621; phrogs \| 12751 \| 99.4 \| 1.4e-17 \| 1.6e-21 \| 146.4 \| 268 \| (179, 476) \| 532 \| (12, 294) \| 361 \| NA \| NA; Category: unknown function; p305174 VI\_10042; phrogs \| 15528 \| 99.4 \| 3.8e-17 \| 4.2e-21 \| 153.8 \| 361 \| (82, 492) \| 532 \| (116, 516) \| 698 \| portal protein \| portal protein; Category: head and packaging; NC\_005263\_p18; phrogs \| 10963 \| 99.1 \| 5.1e-15 \| 5.8e-19 \| 139.4 \| 369 \| (54, 473) \| 532 \| (41, 435) \| 509 \| portal protein \| portal protein; Category: head and packaging; KU160647\_p18; phrogs \| 2739 \| 99.0 \| 4.3e-14 \| 5e-18 \| 139.1 \| 364 \| (73, 488) \| 532 \| (65, 446) \| 494 \| portal protein \| portal protein; Category: head and packaging; MF547663\_p3; phrogs \| 4660 \| 99.0 \| 7.8e-14 \| 8.8e-18 \| 132.1 \| 364 \| (80, 504) \| 532 \| (112, 501) \| 765 \| minor head protein \| minor head protein; Category: head and packaging; LC102730\_p15; phrogs \| 3753 \| 98.7 \| 1.6e-12 \| 1.8e-16 \| 124.3 \| 351 \| (77, 485) \| 532 \| (68, 448) \| 490 \| portal protein \| portal protein; Category: head and packaging; MG596799\_p2; phrogs \| 12037 \| 98.2 \| 2.7e-10 \| 3e-14 \| 90.8 \| 176 \| (43, 226) \| 532 \| (25, 205) \| 223 \| NA \| NA; Category: unknown function; p423831 VI\_07821; phrogs \| 3993 \| 98.1 \| 8e-10 \| 8.9e-14 \| 85.0 \| 131 \| (44, 175) \| 532 \| (18, 156) \| 166 \| NA \| NA; Category: unknown function; p426125 VI\_07755; phrogs \| 15615 \| 97.7 \| 1.7e-08 \| 1.9e-12 \| 81.4 \| 141 \| (74, 251) \| 532 \| (28, 179) \| 180 \| virion structural protein \| virion structural protein; Category: head and packaging; p25678 VI\_08154; phrogs \| 68 \| 97.3 \| 1.5e-07 \| 1.8e-11 \| 80.9 \| 207 \| (159, 409) \| 532 \| (2, 219) \| 243 \| portal protein \| portal protein; Category: head and packaging; p356679 VI\_02767; phrogs \| 11323 \| 96.6 \| 4.4e-06 \| 5e-10 \| 61.6 \| 69 \| (402, 479) \| 532 \| (5, 73) \| 116 \| NA \| NA; Category: unknown function; p325568 VI\_04535; phrogs \| 16240 \| 96.4 \| 1.1e-05 \| 1.2e-09 \| 76.2 \| 393 \| (73, 501) \| 532 \| (95, 515) \| 635 \| portal protein \| portal protein; Category: head and packaging; NC\_010811\_p211; phrogs \| 213 \| 96.1 \| 2.8e-05 \| 3.2e-09 \| 76.3 \| 314 \| (72, 406) \| 532 \| (46, 435) \| 580 \| portal protein \| portal protein; Category: head and packaging; NC\_019401\_p248; phrogs \| 33813 \| 96.0 \| 3.9e-05 \| 4.4e-09 \| 56.7 \| 102 \| (36, 138) \| 532 \| (22, 125) \| 160 \| NA \| NA; Category: unknown function; p223265 VI\_05666; phrogs \| 776 \| 95.6 \| 9.5e-05 \| 1.1e-08 \| 71.6 \| 354 \| (77, 485) \| 532 \| (29, 413) \| 586 \| portal protein \| portal protein; Category: head and packaging; NC\_029005\_p52; phrogs \| 27056 \| 95.6 \| 0.00012 \| 1.3e-08 \| 52.5 \| 60 \| (45, 104) \| 532 \| (22, 83) \| 135 \| NA \| NA; Category: unknown function; p391015 VI\_06918; phrogs \| 113 \| 95.4 \| 0.00015 \| 1.8e-08 \| 71.1 \| 335 \| (77, 481) \| 532 \| (52, 393) \| 474 \| portal protein \| portal protein; Category: head and packaging; p68996 VI\_00707; phrogs \| 11117 \| 95.1 \| 0.00027 \| 3.1e-08 \| 57.9 \| 137 \| (321, 478) \| 532 \| (29, 170) \| 246 \| portal protein \| portal protein; Category: head and packaging; p100773 VI\_12396; phrogs \| 11226 \| 94.9 \| 0.00041 \| 4.6e-08 \| 62.0 \| 268 \| (77, 389) \| 532 \| (33, 337) \| 387 \| portal protein \| portal protein; Category: head and packaging; KC821625\_p80 |
| Top keywords  (threshold 1.00e-03 (evalue)) | **portal, head, and, packaging, Phage, DUF935, lambda, DUF1073, Bacteriophage, T4\_like** |
| Output files | ../../domain\_architecture/04\_FANPEZAQ\_CDS\_0004\_cath.hhr ../../domain\_architecture/04\_FANPEZAQ\_CDS\_0004\_merged.svg ../../domain\_architecture/04\_FANPEZAQ\_CDS\_0004\_ncbi-cd.hhr ../../domain\_architecture/04\_FANPEZAQ\_CDS\_0004\_pfam.hhr ../../domain\_architecture/04\_FANPEZAQ\_CDS\_0004\_phrogs.hhr |

### Identical protein sequences/structures

#### Search results

|  |  |
| --- | --- |
| Protein sequence databases searched | Pdb, Swissprot, Refseq |
| Identical proteins found | -- |
| Top keywords | -- |
| Output files | -- |

### Similar protein sequences/structures

#### Sequence similarity search results (HHblits)^1^

|  |  |
| --- | --- |
| Sequence databases searched | Uniclust, Pdb70 |
| Results, scheme(s)  (Top layers only, threshold 1.00e-03 (evalue)) | xml version="1.0" encoding="utf-8" standalone="no"?       2024-09-02T21:08:29.764656 image/svg+xml   Matplotlib v3.7.2, https://matplotlib.org/ |
| Results, table(s)  (threshold 1.00e-03 (evalue)) | db \| id \| prob \| evalue \| pvalue \| score \| cols \| query \| query\_len \| template \| template\_len \| name \| description; uniclust \| UniRef100\_A0A0S2KEQ2 \| 100.0 \| 4e-104 \| 8e-110 \| 812.0 \| 482 \| (7, 519) \| 532 \| (23, 512) \| 559 \| Phage portal protein, lambda \| Phage portal protein, lambda; uniclust \| UniRef100\_A0A011VKJ3 \| 100.0 \| 2e-100 \| 3e-106 \| 810.0 \| 476 \| (12, 514) \| 532 \| (113, 598) \| 677 \| Portal protein \| Portal protein; uniclust \| UniRef100\_A0A076G8T0 \| 100.0 \| 7.7e-97 \| 2e-102 \| 758.2 \| 477 \| (13, 516) \| 532 \| (74, 562) \| 614 \| Portal protein \| Portal protein; uniclust \| UniRef100\_A0A060H8W7 \| 100.0 \| 1.1e-96 \| 2e-102 \| 783.4 \| 455 \| (43, 505) \| 532 \| (90, 547) \| 640 \| Portal protein \| Portal protein; uniclust \| UniRef100\_A0A011QUJ3 \| 100.0 \| 1.7e-95 \| 4e-101 \| 763.6 \| 450 \| (44, 512) \| 532 \| (104, 561) \| 683 \| Phage portal protein, lambda family \| Phage portal protein, lambda family; uniclust \| UniRef100\_A0A075KD33 \| 100.0 \| 2.6e-95 \| 6e-101 \| 770.2 \| 477 \| (17, 504) \| 532 \| (28, 528) \| 606 \| Phage portal protein, lambda family \| Phage portal protein, lambda family; uniclust \| UniRef100\_A0A073ITU9 \| 100.0 \| 3.8e-95 \| 8e-101 \| 761.3 \| 470 \| (17, 504) \| 532 \| (29, 507) \| 623 \| Portal protein \| Portal protein; uniclust \| UniRef100\_A0A014MMT4 \| 100.0 \| 1.6e-94 \| 3e-100 \| 765.0 \| 475 \| (15, 503) \| 532 \| (61, 567) \| 698 \| Portal protein \| Portal protein; uniclust \| UniRef100\_A0A024LS66 \| 100.0 \| 1.6e-93 \| 3.6e-99 \| 745.4 \| 447 \| (45, 510) \| 532 \| (47, 502) \| 545 \| Phage portal protein \| Phage portal protein; uniclust \| UniRef100\_A0A081NFB6 \| 100.0 \| 1.2e-86 \| 2.5e-92 \| 664.8 \| 450 \| (45, 512) \| 532 \| (55, 516) \| 588 \| Portal protein \| Portal protein; uniclust \| UniRef100\_A0A024E891 \| 100.0 \| 1.9e-86 \| 4.2e-92 \| 692.0 \| 446 \| (44, 506) \| 532 \| (49, 507) \| 598 \| Portal protein lambda family \| Portal protein lambda family; uniclust \| UniRef100\_A0A068Z022 \| 100.0 \| 3.1e-86 \| 6.6e-92 \| 702.1 \| 453 \| (46, 506) \| 532 \| (109, 600) \| 711 \| Phage portal protein \| Phage portal protein; uniclust \| UniRef100\_A0A022G2U4 \| 100.0 \| 7.1e-84 \| 1.5e-89 \| 666.9 \| 479 \| (13, 511) \| 532 \| (121, 607) \| 727 \| Portal protein \| Portal protein; uniclust \| UniRef100\_A0A0F9E9R9 \| 100.0 \| 8.5e-84 \| 1.7e-89 \| 643.4 \| 472 \| (17, 495) \| 532 \| (16, 512) \| 539 \| Phage portal protein (Fragment) \| Phage portal protein (Fragment); uniclust \| UniRef100\_A0A091FES5 \| 100.0 \| 4.7e-83 \| 9.4e-89 \| 644.0 \| 469 \| (19, 505) \| 532 \| (25, 503) \| 601 \| Phage portal protein \| Phage portal protein; uniclust \| UniRef100\_A0A0F9KBE7 \| 100.0 \| 1.4e-82 \| 2.9e-88 \| 626.8 \| 400 \| (92, 508) \| 532 \| (4, 412) \| 448 \| Phage portal protein (Fragment) \| Phage portal protein (Fragment); uniclust \| UniRef100\_A0A073IWK7 \| 100.0 \| 1.5e-82 \| 3e-88 \| 631.9 \| 462 \| (9, 495) \| 532 \| (41, 510) \| 556 \| Phage portal protein \| Phage portal protein; uniclust \| UniRef100\_A0A4Y1MVH7 \| 100.0 \| 5.7e-82 \| 1.1e-87 \| 636.3 \| 480 \| (11, 519) \| 532 \| (6, 503) \| 651 \| Portal protein \| Portal protein; uniclust \| UniRef100\_A0A060H3H7 \| 100.0 \| 9.3e-82 \| 1.9e-87 \| 628.9 \| 481 \| (9, 508) \| 532 \| (40, 532) \| 563 \| Phage portal protein \| Phage portal protein; uniclust \| UniRef100\_A0A0P1FA99 \| 100.0 \| 2.4e-81 \| 4.9e-87 \| 614.5 \| 366 \| (44, 419) \| 532 \| (34, 406) \| 437 \| Phage portal protein, lambda family \| Phage portal protein, lambda family; uniclust \| UniRef100\_A0A0S9N407 \| 100.0 \| 2.8e-79 \| 5.4e-85 \| 595.9 \| 457 \| (44, 504) \| 532 \| (7, 469) \| 499 \| Portal protein \| Portal protein; uniclust \| UniRef100\_A0A142Y555 \| 100.0 \| 2.9e-79 \| 5.6e-85 \| 600.1 \| 492 \| (12, 511) \| 532 \| (4, 512) \| 609 \| Phage portal protein, lambda family \| Phage portal protein, lambda family; uniclust \| UniRef100\_A0A090N879 \| 100.0 \| 6.1e-79 \| 1.1e-84 \| 589.6 \| 445 \| (44, 505) \| 532 \| (28, 479) \| 612 \| Phage portal protein, lambda family \| Phage portal protein, lambda family; uniclust \| UniRef100\_A0A1E3GXJ9 \| 100.0 \| 1.7e-78 \| 3.1e-84 \| 621.5 \| 474 \| (15, 516) \| 532 \| (2, 486) \| 1194 \| Phage portal protein, lambda family \| Phage portal protein, lambda family; uniclust \| UniRef100\_A0A1D8IV87 \| 100.0 \| 3.3e-77 \| 6.6e-83 \| 603.2 \| 442 \| (44, 505) \| 532 \| (43, 503) \| 669 \| Phage portal protein \| Phage portal protein; uniclust \| UniRef100\_A0A0Q6V110 \| 100.0 \| 6.7e-77 \| 1.3e-82 \| 595.0 \| 477 \| (12, 519) \| 532 \| (3, 496) \| 568 \| Portal protein \| Portal protein; uniclust \| UniRef100\_A0A1A9VKE9 \| 100.0 \| 1e-75 \| 1.9e-81 \| 563.8 \| 449 \| (45, 514) \| 532 \| (233, 692) \| 696 \| Phage portal protein \| Phage portal protein; uniclust \| UniRef100\_A0A510E8S9 \| 100.0 \| 1.3e-75 \| 2.4e-81 \| 584.0 \| 453 \| (43, 506) \| 532 \| (16, 474) \| 962 \| Phage portal protein \| Phage portal protein; uniclust \| UniRef100\_A0A066T9U0 \| 100.0 \| 1.1e-74 \| 2.2e-80 \| 600.6 \| 450 \| (50, 509) \| 532 \| (102, 588) \| 700 \| Phage portal protein \| Phage portal protein; uniclust \| UniRef100\_A0A437MJQ4 \| 100.0 \| 2.1e-73 \| 4e-79 \| 554.9 \| 465 \| (15, 511) \| 532 \| (4, 481) \| 504 \| Phage portal protein \| Phage portal protein; uniclust \| UniRef100\_A0A0Q0CI78 \| 100.0 \| 1.1e-71 \| 2.1e-77 \| 534.6 \| 346 \| (154, 513) \| 532 \| (52, 406) \| 433 \| Histidinol phosphatase \| Histidinol phosphatase; uniclust \| UniRef100\_A0A1J5EX01 \| 100.0 \| 3e-71 \| 5.9e-77 \| 545.1 \| 448 \| (47, 505) \| 532 \| (18, 492) \| 525 \| Phage portal protein \| Phage portal protein; uniclust \| UniRef100\_A0A087M4C7 \| 100.0 \| 3.6e-71 \| 7.6e-77 \| 576.0 \| 476 \| (13, 505) \| 532 \| (21, 539) \| 597 \| Capsid protein \| Capsid protein; uniclust \| UniRef100\_A0A0F9IDI9 \| 100.0 \| 4.8e-71 \| 1e-76 \| 584.0 \| 431 \| (43, 502) \| 532 \| (101, 536) \| 652 \| Portal protein \| Portal protein; uniclust \| UniRef100\_A0A0F8ZRP6 \| 100.0 \| 3.8e-70 \| 7.7e-76 \| 532.2 \| 363 \| (132, 513) \| 532 \| (3, 374) \| 405 \| Phage portal protein (Fragment) \| Phage portal protein (Fragment); uniclust \| UniRef100\_A0A159Z5V5 \| 100.0 \| 2.6e-69 \| 4.8e-75 \| 535.4 \| 446 \| (45, 507) \| 532 \| (39, 492) \| 873 \| ATP-dependent Clp protease proteolytic subunit \| ATP-dependent Clp protease proteolytic subunit; uniclust \| UniRef100\_A0A193LMF8 \| 100.0 \| 1.1e-68 \| 2.2e-74 \| 538.6 \| 431 \| (49, 493) \| 532 \| (49, 491) \| 528 \| Phage portal protein \| Phage portal protein; uniclust \| UniRef100\_A0A6G1WEZ2 \| 100.0 \| 2.1e-68 \| 4e-74 \| 527.3 \| 453 \| (44, 504) \| 532 \| (35, 525) \| 556 \| Phage portal protein (Fragment) \| Phage portal protein (Fragment); uniclust \| UniRef100\_UPI00123DC76E \| 100.0 \| 3.5e-68 \| 6.6e-74 \| 523.9 \| 472 \| (18, 503) \| 532 \| (1, 512) \| 705 \| phage portal protein \| phage portal protein; uniclust \| UniRef100\_A0A1R1LRL3 \| 100.0 \| 4e-67 \| 7.8e-73 \| 515.3 \| 457 \| (45, 514) \| 532 \| (28, 524) \| 539 \| Phage portal protein \| Phage portal protein; uniclust \| UniRef100\_A0A071LT14 \| 100.0 \| 6.4e-67 \| 1.3e-72 \| 525.3 \| 419 \| (63, 493) \| 532 \| (61, 486) \| 509 \| Portal protein \| Portal protein; uniclust \| UniRef100\_UPI002091CF7E \| 100.0 \| 7.2e-66 \| 1.3e-71 \| 506.0 \| 480 \| (16, 519) \| 532 \| (1, 500) \| 884 \| phage portal protein \| phage portal protein; uniclust \| UniRef100\_A0A2D8T9H3 \| 100.0 \| 8.7e-66 \| 1.7e-71 \| 503.2 \| 376 \| (124, 506) \| 532 \| (2, 407) \| 455 \| Phage portal protein (Fragment) \| Phage portal protein (Fragment); uniclust \| UniRef100\_A0A1H6Z348 \| 100.0 \| 1.1e-64 \| 2.2e-70 \| 495.5 \| 365 \| (132, 504) \| 532 \| (1, 393) \| 421 \| Phage portal protein, lambda family (Fragment) \| Phage portal protein, lambda family (Fragment); uniclust \| UniRef100\_A0A0C2U5P6 \| 100.0 \| 2.1e-64 \| 3.9e-70 \| 470.4 \| 441 \| (44, 505) \| 532 \| (14, 459) \| 463 \| Phage portal protein \| Phage portal protein; uniclust \| UniRef100\_A0A421BJ29 \| 100.0 \| 7.7e-64 \| 1.4e-69 \| 479.5 \| 468 \| (15, 508) \| 532 \| (150, 623) \| 644 \| Phage portal protein \| Phage portal protein; uniclust \| UniRef100\_A0A0Q6MTA0 \| 100.0 \| 5.2e-63 \| 1e-68 \| 479.1 \| 400 \| (98, 517) \| 532 \| (13, 421) \| 463 \| Phage portal protein \| Phage portal protein; uniclust \| UniRef100\_A0A2W4SHI4 \| 100.0 \| 7.9e-63 \| 1.5e-68 \| 475.8 \| 392 \| (16, 420) \| 532 \| (23, 423) \| 445 \| Phage portal protein (Fragment) \| Phage portal protein (Fragment); uniclust \| UniRef100\_A0A061KB43 \| 100.0 \| 1.5e-61 \| 2.9e-67 \| 479.2 \| 456 \| (47, 514) \| 532 \| (66, 559) \| 608 \| Phage portal protein \| Phage portal protein; uniclust \| UniRef100\_A0A1Z9JFC0 \| 100.0 \| 2.9e-61 \| 6e-67 \| 496.5 \| 421 \| (44, 502) \| 532 \| (56, 490) \| 578 \| Phage portal protein \| Phage portal protein; uniclust \| UniRef100\_A0A1G0ICK1 \| 100.0 \| 8.2e-61 \| 1.6e-66 \| 471.2 \| 488 \| (15, 516) \| 532 \| (52, 575) \| 619 \| Phage portal protein \| Phage portal protein; uniclust \| UniRef100\_A0A175R9E3 \| 100.0 \| 1e-60 \| 1.9e-66 \| 451.3 \| 435 \| (43, 505) \| 532 \| (21, 460) \| 472 \| Portal protein \| Portal protein; uniclust \| UniRef100\_A0A9E0NW25 \| 100.0 \| 1.4e-60 \| 2.7e-66 \| 462.8 \| 400 \| (92, 508) \| 532 \| (320, 728) \| 741 \| Phage portal protein \| Phage portal protein; uniclust \| UniRef100\_A0A1B4C0H5 \| 100.0 \| 1.5e-60 \| 2.7e-66 \| 450.2 \| 441 \| (45, 505) \| 532 \| (22, 480) \| 504 \| Portal protein \| Portal protein; uniclust \| UniRef100\_A0A443L3G4 \| 100.0 \| 2.9e-60 \| 5.5e-66 \| 444.8 \| 373 \| (16, 405) \| 532 \| (4, 383) \| 384 \| Phage portal protein (Fragment) \| Phage portal protein (Fragment); uniclust \| UniRef100\_A0A1V6HR06 \| 100.0 \| 4.6e-60 \| 9.2e-66 \| 482.5 \| 461 \| (16, 507) \| 532 \| (27, 500) \| 662 \| Phage portal protein, lambda family \| Phage portal protein, lambda family; uniclust \| UniRef100\_A0A380AGT5 \| 100.0 \| 3.2e-59 \| 6.2e-65 \| 458.1 \| 420 \| (63, 495) \| 532 \| (61, 489) \| 513 \| Phage portal protein, lambda family \| Phage portal protein, lambda family; uniclust \| UniRef100\_A0A353ZI73 \| 100.0 \| 3.8e-59 \| 7.3e-65 \| 457.6 \| 448 \| (45, 508) \| 532 \| (49, 523) \| 552 \| Phage portal protein \| Phage portal protein; uniclust \| UniRef100\_A0A2M9P8F9 \| 100.0 \| 8.8e-59 \| 1.7e-64 \| 419.0 \| 273 \| (16, 300) \| 532 \| (2, 280) \| 296 \| Phage portal protein (Fragment) \| Phage portal protein (Fragment); uniclust \| UniRef100\_UPI002044A97D \| 100.0 \| 1.2e-58 \| 2.2e-64 \| 450.4 \| 433 \| (43, 504) \| 532 \| (20, 459) \| 637 \| phage portal protein \| phage portal protein; uniclust \| UniRef100\_A0A1V5GTW5 \| 100.0 \| 1.3e-58 \| 2.6e-64 \| 441.0 \| 292 \| (209, 507) \| 532 \| (1, 295) \| 328 \| Phage portal protein, lambda family \| Phage portal protein, lambda family; uniclust \| UniRef100\_A0A1S6TP94 \| 100.0 \| 2.6e-58 \| 5.2e-64 \| 459.4 \| 397 \| (66, 495) \| 532 \| (49, 453) \| 486 \| Phage portal protein, lambda family \| Phage portal protein, lambda family; uniclust \| UniRef100\_A0A835Z6S7 \| 100.0 \| 3e-58 \| 5.5e-64 \| 444.6 \| 468 \| (16, 519) \| 532 \| (1, 487) \| 695 \| Phage portal protein \| Phage portal protein; uniclust \| UniRef100\_A0A349EMI8 \| 100.0 \| 1.4e-57 \| 2.9e-63 \| 442.7 \| 346 \| (15, 365) \| 532 \| (11, 385) \| 387 \| Phage portal protein \| Phage portal protein; uniclust \| UniRef100\_A0A3B8ZGJ9 \| 100.0 \| 1.6e-57 \| 3e-63 \| 442.3 \| 437 \| (43, 508) \| 532 \| (37, 479) \| 609 \| Phage portal protein \| Phage portal protein; uniclust \| UniRef100\_A0A2N9AI26 \| 100.0 \| 1.8e-57 \| 3.4e-63 \| 458.4 \| 447 \| (45, 507) \| 532 \| (767, 1241) \| 1265 \| Terminase GpA (Modular protein) \| Terminase GpA (Modular protein); uniclust \| UniRef100\_A0A0S8GDJ6 \| 100.0 \| 1.9e-57 \| 4e-63 \| 473.7 \| 428 \| (44, 494) \| 532 \| (26, 478) \| 605 \| Phage portal protein \| Phage portal protein; uniclust \| UniRef100\_A0A2E7BJT9 \| 100.0 \| 2e-57 \| 4.1e-63 \| 434.0 \| 291 \| (203, 513) \| 532 \| (12, 308) \| 350 \| Phage portal protein (Fragment) \| Phage portal protein (Fragment); uniclust \| UniRef100\_A0A0A2V4M4 \| 100.0 \| 2.8e-57 \| 5.2e-63 \| 470.7 \| 455 \| (46, 508) \| 532 \| (1183, 1679) \| 1696 \| Portal protein B \| Portal protein B; uniclust \| UniRef100\_A0A1M4S7W9 \| 100.0 \| 4.2e-57 \| 8.2e-63 \| 425.6 \| 300 \| (44, 353) \| 532 \| (28, 342) \| 342 \| Phage portal protein, lambda family \| Phage portal protein, lambda family; uniclust \| UniRef100\_A0A2X1LNZ6 \| 100.0 \| 6.6e-57 \| 1.2e-62 \| 428.1 \| 430 \| (63, 503) \| 532 \| (16, 468) \| 484 \| Lambda family phage portal protein \| Lambda family phage portal protein; uniclust \| UniRef100\_A0A1G1LMU8 \| 100.0 \| 1.1e-56 \| 2.1e-62 \| 420.6 \| 257 \| (240, 503) \| 532 \| (7, 275) \| 312 \| Phage portal protein (Fragment) \| Phage portal protein (Fragment); uniclust \| UniRef100\_A0A1V1UMH2 \| 100.0 \| 1.3e-56 \| 2.6e-62 \| 447.9 \| 434 \| (63, 523) \| 532 \| (66, 532) \| 547 \| Phage portal protein, lambda family \| Phage portal protein, lambda family; uniclust \| UniRef100\_A0A7X4AD15 \| 100.0 \| 4.5e-56 \| 8.6e-62 \| 428.6 \| 404 \| (43, 481) \| 532 \| (35, 441) \| 472 \| Phage portal protein \| Phage portal protein; uniclust \| UniRef100\_A0A9E5UZN2 \| 100.0 \| 7.8e-56 \| 1.4e-61 \| 414.8 \| 472 \| (15, 506) \| 532 \| (9, 490) \| 494 \| Phage portal protein \| Phage portal protein; uniclust \| UniRef100\_A0A011T786 \| 100.0 \| 9e-56 \| 1.8e-61 \| 450.9 \| 429 \| (61, 506) \| 532 \| (76, 550) \| 597 \| Capsid protein \| Capsid protein; uniclust \| UniRef100\_A0A354UBL5 \| 100.0 \| 1.6e-54 \| 3.1e-60 \| 416.5 \| 448 \| (44, 505) \| 532 \| (22, 479) \| 511 \| Phage portal protein \| Phage portal protein; uniclust \| UniRef100\_A0A0H4A1U0 \| 100.0 \| 1.7e-54 \| 3.3e-60 \| 426.6 \| 464 \| (14, 502) \| 532 \| (12, 482) \| 512 \| Phage portal protein \| Phage portal protein; uniclust \| UniRef100\_A0A7G9NZU8 \| 100.0 \| 4.7e-54 \| 8.7e-60 \| 413.8 \| 438 \| (44, 487) \| 532 \| (53, 527) \| 656 \| Phage portal protein \| Phage portal protein; uniclust \| UniRef100\_A0A0E2E9S4 \| 100.0 \| 6.1e-53 \| 1.3e-58 \| 434.5 \| 409 \| (71, 495) \| 532 \| (93, 543) \| 590 \| SPP1 family phage portal protein \| SPP1 family phage portal protein; uniclust \| UniRef100\_A0A2D5SAY0 \| 100.0 \| 8.9e-53 \| 1.8e-58 \| 406.2 \| 380 \| (15, 416) \| 532 \| (11, 395) \| 398 \| Phage portal protein (Fragment) \| Phage portal protein (Fragment); uniclust \| UniRef100\_A0A1F4CAA0 \| 100.0 \| 2.1e-52 \| 4e-58 \| 380.4 \| 288 \| (208, 502) \| 532 \| (1, 312) \| 338 \| Phage portal protein (Fragment) \| Phage portal protein (Fragment); uniclust \| UniRef100\_A0A368TQV2 \| 100.0 \| 2.2e-52 \| 4.1e-58 \| 410.1 \| 443 \| (43, 505) \| 532 \| (183, 647) \| 744 \| Phage portal protein \| Phage portal protein; uniclust \| UniRef100\_A0A3R8IU71 \| 100.0 \| 2.6e-52 \| 4.9e-58 \| 406.4 \| 450 \| (48, 505) \| 532 \| (57, 508) \| 748 \| Phage portal protein \| Phage portal protein; uniclust \| UniRef100\_UPI00025BA5D3 \| 100.0 \| 2.8e-52 \| 5.2e-58 \| 409.3 \| 444 \| (45, 492) \| 532 \| (29, 516) \| 823 \| phage portal protein \| phage portal protein; uniclust \| UniRef100\_A0A142WWE9 \| 100.0 \| 3.9e-52 \| 8.1e-58 \| 424.7 \| 425 \| (45, 506) \| 532 \| (29, 468) \| 513 \| Phage portal protein, lambda family \| Phage portal protein, lambda family; uniclust \| UniRef100\_A0A062V9J7 \| 100.0 \| 4.8e-52 \| 9.5e-58 \| 394.2 \| 317 \| (44, 377) \| 532 \| (24, 344) \| 344 \| Phage portal protein, lambda family (Fragment) \| Phage portal protein, lambda family (Fragment); uniclust \| UniRef100\_A0A6L6JH12 \| 100.0 \| 1.1e-51 \| 2.1e-57 \| 395.2 \| 390 \| (7, 398) \| 532 \| (8, 414) \| 425 \| Phage portal protein (Fragment) \| Phage portal protein (Fragment); uniclust \| UniRef100\_W0IYG2 \| 100.0 \| 1.8e-51 \| 3.3e-57 \| 394.3 \| 451 \| (48, 505) \| 532 \| (43, 496) \| 620 \| Portal protein \| Portal protein; uniclust \| UniRef100\_A0A0B8PGA6 \| 100.0 \| 8.5e-51 \| 1.6e-56 \| 388.4 \| 433 \| (64, 504) \| 532 \| (37, 477) \| 501 \| Phage portal protein \| Phage portal protein; uniclust \| UniRef100\_A0A345DE62 \| 100.0 \| 2.9e-50 \| 5.3e-56 \| 379.0 \| 430 \| (46, 505) \| 532 \| (30, 471) \| 509 \| Phage portal protein \| Phage portal protein; uniclust \| UniRef100\_A0A3L7RG21 \| 100.0 \| 5.4e-50 \| 1.1e-55 \| 379.8 \| 335 \| (155, 503) \| 532 \| (8, 344) \| 372 \| Phage portal protein \| Phage portal protein; uniclust \| UniRef100\_A0A2S1XL37 \| 100.0 \| 1.1e-49 \| 2.1e-55 \| 359.2 \| 253 \| (16, 280) \| 532 \| (1, 260) \| 261 \| Phage portal protein \| Phage portal protein; uniclust \| UniRef100\_A0A9E5G6J2 \| 100.0 \| 1.7e-49 \| 3.1e-55 \| 372.3 \| 466 \| (17, 505) \| 532 \| (5, 481) \| 487 \| Phage portal protein \| Phage portal protein; uniclust \| UniRef100\_A0A524IJ31 \| 100.0 \| 1.7e-49 \| 3.4e-55 \| 392.4 \| 431 \| (44, 504) \| 532 \| (34, 479) \| 542 \| Phage portal protein (Fragment) \| Phage portal protein (Fragment); uniclust \| UniRef100\_UPI001F424B1F \| 100.0 \| 2.4e-49 \| 4.4e-55 \| 395.1 \| 473 \| (19, 510) \| 532 \| (512, 996) \| 1019 \| phage portal protein \| phage portal protein; uniclust \| UniRef100\_A0A063ZY87 \| 100.0 \| 6e-49 \| 1.1e-54 \| 393.7 \| 450 \| (29, 493) \| 532 \| (574, 1056) \| 1075 \| Terminase \| Terminase; uniclust \| UniRef100\_X0U7Z2 \| 100.0 \| 9e-49 \| 1.7e-54 \| 361.5 \| 369 \| (122, 505) \| 532 \| (10, 389) \| 417 \| Phage portal protein (Fragment) \| Phage portal protein (Fragment); uniclust \| UniRef100\_UPI0009FE0364 \| 100.0 \| 1e-48 \| 1.9e-54 \| 360.4 \| 362 \| (134, 507) \| 532 \| (6, 374) \| 408 \| phage portal protein \| phage portal protein; uniclust \| UniRef100\_A0A072TET7 \| 100.0 \| 1.2e-48 \| 2.1e-54 \| 400.7 \| 459 \| (46, 512) \| 532 \| (803, 1302) \| 1593 \| Phage portal protein, lambda family protein (Fragment) \| Phage portal protein, lambda family protein (Fragment); uniclust \| UniRef100\_A0A2U3ERF1 \| 100.0 \| 1.1e-48 \| 2.2e-54 \| 382.3 \| 332 \| (46, 382) \| 532 \| (38, 413) \| 420 \| Phage portal protein (Fragment) \| Phage portal protein (Fragment); uniclust \| UniRef100\_A0A250DMY9 \| 100.0 \| 1.7e-48 \| 3.3e-54 \| 367.7 \| 259 \| (43, 301) \| 532 \| (21, 280) \| 326 \| Phage portal protein \| Phage portal protein; uniclust \| UniRef100\_A0A376NRU5 \| 100.0 \| 2.1e-48 \| 4.1e-54 \| 370.9 \| 345 \| (15, 371) \| 532 \| (22, 374) \| 391 \| Phage portal protein, lambda family \| Phage portal protein, lambda family; uniclust \| UniRef100\_A0A1T4WXT7 \| 100.0 \| 2.1e-48 \| 4.3e-54 \| 388.5 \| 447 \| (48, 504) \| 532 \| (29, 490) \| 518 \| Capsid protein \| Capsid protein; uniclust \| UniRef100\_A0A661TNZ2 \| 100.0 \| 2.6e-48 \| 4.8e-54 \| 375.0 \| 470 \| (16, 495) \| 532 \| (94, 615) \| 660 \| Phage portal protein \| Phage portal protein; uniclust \| UniRef100\_A0A3L7SGJ4 \| 100.0 \| 4.7e-48 \| 8.6e-54 \| 370.4 \| 436 \| (44, 507) \| 532 \| (33, 476) \| 605 \| Phage portal protein \| Phage portal protein; uniclust \| UniRef100\_A0A177R1N7 \| 100.0 \| 6.5e-48 \| 1.3e-53 \| 392.2 \| 402 \| (57, 495) \| 532 \| (66, 484) \| 537 \| Phage portal protein \| Phage portal protein; uniclust \| UniRef100\_A0A0S4TN73 \| 100.0 \| 7.9e-48 \| 1.5e-53 \| 355.7 \| 258 \| (16, 292) \| 532 \| (25, 287) \| 303 \| Phage portal protein (Fragment) \| Phage portal protein (Fragment); uniclust \| UniRef100\_A0A6G4KD87 \| 100.0 \| 1.1e-47 \| 2e-53 \| 359.1 \| 349 \| (87, 451) \| 532 \| (1, 358) \| 363 \| Phage portal protein (Fragment) \| Phage portal protein (Fragment); uniclust \| UniRef100\_A0A4P5XY08 \| 100.0 \| 1.1e-47 \| 2.2e-53 \| 378.9 \| 421 \| (44, 488) \| 532 \| (73, 498) \| 616 \| Phage portal protein \| Phage portal protein; uniclust \| UniRef100\_UPI000A7FFD44 \| 100.0 \| 1.2e-47 \| 2.2e-53 \| 352.7 \| 348 \| (18, 371) \| 532 \| (1, 380) \| 381 \| phage portal protein \| phage portal protein; uniclust \| UniRef100\_A0A0A1AA99 \| 100.0 \| 1.5e-47 \| 2.8e-53 \| 374.4 \| 472 \| (15, 509) \| 532 \| (22, 503) \| 669 \| Phage portal protein \| Phage portal protein; uniclust \| UniRef100\_A0A812JEV6 \| 100.0 \| 3.2e-47 \| 5.8e-53 \| 404.0 \| 419 \| (65, 495) \| 532 \| (1153, 1576) \| 4451 \| Phage portal protein \| Phage portal protein; uniclust \| UniRef100\_A0A0A0YPM0 \| 100.0 \| 3.8e-47 \| 8e-53 \| 393.2 \| 413 \| (66, 491) \| 532 \| (64, 527) \| 555 \| Portal protein \| Portal protein; uniclust \| UniRef100\_A0A0F9AB75 \| 100.0 \| 6.2e-47 \| 1.2e-52 \| 351.4 \| 258 \| (243, 505) \| 532 \| (9, 278) \| 300 \| Phage portal protein (Fragment) \| Phage portal protein (Fragment); uniclust \| UniRef100\_A0A7J6YNN5 \| 100.0 \| 6.7e-47 \| 1.2e-52 \| 363.6 \| 279 \| (217, 507) \| 532 \| (333, 616) \| 623 \| DUF1016 domain-containing protein \| DUF1016 domain-containing protein; uniclust \| UniRef100\_A0A2A2R1Z6 \| 100.0 \| 7.2e-47 \| 1.5e-52 \| 386.3 \| 428 \| (44, 503) \| 532 \| (35, 481) \| 545 \| Phage portal protein \| Phage portal protein; uniclust \| UniRef100\_A0A954DEK2 \| 100.0 \| 8.4e-47 \| 1.5e-52 \| 359.4 \| 435 \| (45, 495) \| 532 \| (60, 514) \| 560 \| Phage portal protein \| Phage portal protein; uniclust \| UniRef100\_A0A965PFI3 \| 100.0 \| 1.6e-46 \| 2.9e-52 \| 375.1 \| 463 \| (16, 510) \| 532 \| (4, 480) \| 833 \| Phage portal protein \| Phage portal protein; uniclust \| UniRef100\_A0A973DZ03 \| 100.0 \| 1.9e-46 \| 3.6e-52 \| 360.0 \| 445 \| (61, 514) \| 532 \| (18, 500) \| 522 \| Phage portal protein \| Phage portal protein; uniclust \| UniRef100\_A0A1G3M6A5 \| 100.0 \| 1.4e-45 \| 2.6e-51 \| 341.8 \| 357 \| (45, 419) \| 532 \| (58, 426) \| 429 \| Phage portal protein (Fragment) \| Phage portal protein (Fragment); uniclust \| UniRef100\_A0A2D6FNP9 \| 100.0 \| 1.6e-45 \| 3.1e-51 \| 355.4 \| 428 \| (44, 501) \| 532 \| (22, 460) \| 501 \| Phage portal protein \| Phage portal protein; uniclust \| UniRef100\_A0A389MVE2 \| 100.0 \| 2.1e-45 \| 4e-51 \| 362.7 \| 409 \| (73, 505) \| 532 \| (89, 511) \| 611 \| Phage portal protein \| Phage portal protein; uniclust \| UniRef100\_A0A1V5NFL2 \| 100.0 \| 4.9e-45 \| 9e-51 \| 367.0 \| 436 \| (43, 504) \| 532 \| (20, 482) \| 1114 \| Phage portal protein, lambda family \| Phage portal protein, lambda family; uniclust \| UniRef100\_A0A7C8EMQ6 \| 100.0 \| 9.4e-45 \| 1.7e-50 \| 342.0 \| 430 \| (46, 505) \| 532 \| (51, 492) \| 502 \| Phage portal protein \| Phage portal protein; uniclust \| UniRef100\_UPI0003A4D93C \| 100.0 \| 1.9e-44 \| 3.6e-50 \| 337.2 \| 414 \| (69, 509) \| 532 \| (42, 456) \| 464 \| phage portal protein \| phage portal protein; uniclust \| UniRef100\_A0A2X2S8A3 \| 100.0 \| 2.9e-44 \| 5.5e-50 \| 333.3 \| 317 \| (153, 472) \| 532 \| (1, 348) \| 349 \| Phage portal protein \| Phage portal protein; uniclust \| UniRef100\_A0A167I0Q7 \| 100.0 \| 3.9e-44 \| 7.6e-50 \| 342.9 \| 355 \| (141, 507) \| 532 \| (15, 400) \| 418 \| Plasmid partitioning protein ParB (Fragment) \| Plasmid partitioning protein ParB (Fragment); uniclust \| UniRef100\_A0A517Q5C2 \| 100.0 \| 4.3e-44 \| 8.1e-50 \| 343.0 \| 462 \| (15, 503) \| 532 \| (7, 481) \| 504 \| Phage portal protein, lambda family \| Phage portal protein, lambda family; uniclust \| UniRef100\_A0A3M3BPQ1 \| 100.0 \| 4.3e-44 \| 8.2e-50 \| 326.2 \| 292 \| (126, 420) \| 532 \| (2, 307) \| 314 \| Portal protein (Fragment) \| Portal protein (Fragment); uniclust \| UniRef100\_A0A564WHL9 \| 100.0 \| 6.3e-44 \| 1.2e-49 \| 335.1 \| 444 \| (47, 506) \| 532 \| (4, 466) \| 481 \| Putative Phage portal protein \| Putative Phage portal protein; uniclust \| UniRef100\_A0A120G5R1 \| 100.0 \| 6.5e-44 \| 1.3e-49 \| 334.2 \| 250 \| (248, 503) \| 532 \| (23, 290) \| 322 \| Phage portal protein, lambda family \| Phage portal protein, lambda family; uniclust \| UniRef100\_A0A0F9MU33 \| 100.0 \| 1.1e-43 \| 1.9e-49 \| 337.2 \| 459 \| (46, 523) \| 532 \| (27, 523) \| 537 \| Phage portal protein, lambda family \| Phage portal protein, lambda family; uniclust \| UniRef100\_UPI0021BB8F59 \| 100.0 \| 1.8e-43 \| 3.2e-49 \| 336.8 \| 410 \| (64, 501) \| 532 \| (62, 498) \| 555 \| phage portal protein \| phage portal protein; uniclust \| UniRef100\_UPI000AC8D934 \| 100.0 \| 7.1e-43 \| 1.3e-48 \| 344.1 \| 413 \| (65, 492) \| 532 \| (70, 500) \| 819 \| phage portal protein \| phage portal protein; uniclust \| UniRef100\_A0A965TTW4 \| 100.0 \| 8.2e-43 \| 1.5e-48 \| 337.0 \| 467 \| (17, 492) \| 532 \| (2, 507) \| 522 \| Phage portal protein \| Phage portal protein; uniclust \| UniRef100\_A0A1G0RDT4 \| 100.0 \| 8.7e-43 \| 1.6e-48 \| 331.8 \| 443 \| (43, 491) \| 532 \| (32, 497) \| 548 \| Phage portal protein \| Phage portal protein; uniclust \| UniRef100\_A0A1G3LXF9 \| 100.0 \| 1.3e-42 \| 2.5e-48 \| 320.2 \| 355 \| (146, 508) \| 532 \| (22, 389) \| 403 \| Phage portal protein \| Phage portal protein; uniclust \| UniRef100\_A0A3B9NPV5 \| 100.0 \| 1.6e-42 \| 3.2e-48 \| 317.6 \| 234 \| (17, 254) \| 532 \| (1, 237) \| 238 \| Phage portal protein (Fragment) \| Phage portal protein (Fragment); uniclust \| UniRef100\_UPI0009B970C7 \| 100.0 \| 1.6e-42 \| 3.3e-48 \| 332.1 \| 318 \| (57, 381) \| 532 \| (46, 370) \| 376 \| phage portal protein \| phage portal protein; uniclust \| UniRef100\_A0A0U1DBB2 \| 100.0 \| 2.6e-42 \| 5.1e-48 \| 311.7 \| 229 \| (271, 508) \| 532 \| (2, 238) \| 261 \| Putative phage portal protein \| Putative phage portal protein; uniclust \| UniRef100\_A0A853IJZ4 \| 100.0 \| 4.2e-42 \| 8.1e-48 \| 307.5 \| 226 \| (248, 478) \| 532 \| (6, 240) \| 240 \| Phage portal protein (Fragment) \| Phage portal protein (Fragment); uniclust \| UniRef100\_A0A0F9HAI2 \| 100.0 \| 4.2e-42 \| 8.4e-48 \| 316.5 \| 237 \| (262, 507) \| 532 \| (2, 241) \| 267 \| Phage portal protein (Fragment) \| Phage portal protein (Fragment); uniclust \| UniRef100\_A0A0G4PZK6 \| 100.0 \| 4.7e-42 \| 9.1e-48 \| 319.4 \| 283 \| (216, 511) \| 532 \| (3, 288) \| 313 \| Phage portal protein, lambda family \| Phage portal protein, lambda family; uniclust \| UniRef100\_A0A965PAQ9 \| 100.0 \| 7.4e-42 \| 1.4e-47 \| 333.5 \| 406 \| (45, 493) \| 532 \| (31, 451) \| 644 \| Phage portal protein \| Phage portal protein; uniclust \| UniRef100\_UPI0003260BA1 \| 100.0 \| 9.4e-42 \| 1.7e-47 \| 322.2 \| 347 \| (15, 373) \| 532 \| (1, 359) \| 503 \| phage portal protein \| phage portal protein; uniclust \| UniRef100\_A0A7X7YX50 \| 100.0 \| 9.8e-42 \| 1.8e-47 \| 321.4 \| 450 \| (20, 505) \| 532 \| (15, 476) \| 494 \| Phage portal protein \| Phage portal protein; uniclust \| UniRef100\_A0A965P5L4 \| 100.0 \| 9.9e-42 \| 1.8e-47 \| 340.2 \| 439 \| (43, 518) \| 532 \| (32, 476) \| 957 \| Phage portal protein \| Phage portal protein; uniclust \| UniRef100\_A0A166JYY8 \| 100.0 \| 1.5e-41 \| 2.8e-47 \| 327.7 \| 406 \| (73, 505) \| 532 \| (27, 438) \| 463 \| Portal protein \| Portal protein; uniclust \| UniRef100\_A0A349JAV8 \| 100.0 \| 2e-41 \| 3.6e-47 \| 329.5 \| 438 \| (43, 506) \| 532 \| (34, 478) \| 688 \| Phage portal protein \| Phage portal protein; uniclust \| UniRef100\_A0A2M9P8T1 \| 100.0 \| 3.4e-41 \| 6.5e-47 \| 305.5 \| 253 \| (249, 518) \| 532 \| (1, 258) \| 281 \| Phage portal protein (Fragment) \| Phage portal protein (Fragment); uniclust \| UniRef100\_A0A965I6N8 \| 100.0 \| 4.8e-41 \| 8.9e-47 \| 329.5 \| 473 \| (15, 514) \| 532 \| (1, 490) \| 667 \| Phage portal protein \| Phage portal protein; uniclust \| UniRef100\_A0A3A1Y7Y1 \| 100.0 \| 6.3e-41 \| 1.2e-46 \| 330.9 \| 430 \| (45, 504) \| 532 \| (53, 485) \| 534 \| Phage portal protein \| Phage portal protein; uniclust \| UniRef100\_A0A554XEQ4 \| 100.0 \| 6.9e-41 \| 1.3e-46 \| 316.2 \| 315 \| (43, 367) \| 532 \| (18, 336) \| 500 \| Putative esterase \| Putative esterase; uniclust \| UniRef100\_A0A7X7H5B0 \| 100.0 \| 1.2e-40 \| 2.2e-46 \| 315.6 \| 427 \| (43, 491) \| 532 \| (25, 487) \| 515 \| Phage portal protein \| Phage portal protein; uniclust \| UniRef100\_A0A7C4BYU0 \| 100.0 \| 1.2e-40 \| 2.2e-46 \| 310.9 \| 313 \| (46, 387) \| 532 \| (23, 336) \| 446 \| Phage portal protein (Fragment) \| Phage portal protein (Fragment); uniclust \| UniRef100\_C1AAN8 \| 100.0 \| 2.7e-40 \| 5e-46 \| 318.2 \| 455 \| (45, 509) \| 532 \| (41, 524) \| 608 \| Phage portal protein lambda family protein \| Phage portal protein lambda family protein; uniclust \| UniRef100\_UPI000FCA5EF8 \| 100.0 \| 3.5e-40 \| 6.5e-46 \| 290.5 \| 259 \| (141, 402) \| 532 \| (5, 271) \| 273 \| phage portal protein \| phage portal protein; uniclust \| UniRef100\_A0A062VDI3 \| 100.0 \| 3.9e-40 \| 7.9e-46 \| 299.3 \| 183 \| (317, 509) \| 532 \| (14, 199) \| 229 \| Lambda family phage portal protein \| Lambda family phage portal protein; uniclust \| UniRef100\_A0A1H9QDY8 \| 100.0 \| 5e-40 \| 9.1e-46 \| 307.0 \| 362 \| (51, 416) \| 532 \| (31, 419) \| 447 \| Phage portal protein, lambda family \| Phage portal protein, lambda family; uniclust \| UniRef100\_A0A0Q4ZHQ5 \| 100.0 \| 4.7e-40 \| 9.4e-46 \| 329.1 \| 404 \| (71, 502) \| 532 \| (65, 484) \| 530 \| Phage portal protein \| Phage portal protein; uniclust \| UniRef100\_A0A929TYB1 \| 100.0 \| 6.2e-40 \| 1.2e-45 \| 298.2 \| 280 \| (16, 301) \| 532 \| (1, 281) \| 305 \| Phage portal protein (Fragment) \| Phage portal protein (Fragment); uniclust \| UniRef100\_A0A257VIH0 \| 100.0 \| 7.9e-40 \| 1.5e-45 \| 294.0 \| 283 \| (43, 341) \| 532 \| (10, 303) \| 318 \| Phage portal protein (Fragment) \| Phage portal protein (Fragment); uniclust \| UniRef100\_A0A812INR5 \| 100.0 \| 1.4e-39 \| 2.5e-45 \| 328.9 \| 393 \| (62, 467) \| 532 \| (53, 470) \| 1131 \| B protein \| B protein; uniclust \| UniRef100\_A0A0S4U0W3 \| 100.0 \| 3.1e-39 \| 5.7e-45 \| 286.3 \| 246 \| (47, 301) \| 532 \| (8, 265) \| 285 \| Bacteriophage-like protein (Fragment) \| Bacteriophage-like protein (Fragment); uniclust \| UniRef100\_A0A0S8IF22 \| 100.0 \| 3.1e-39 \| 6e-45 \| 319.9 \| 420 \| (67, 494) \| 532 \| (63, 529) \| 552 \| Phage portal protein \| Phage portal protein; uniclust \| UniRef100\_R9L6B1 \| 100.0 \| 4.5e-39 \| 8.2e-45 \| 295.7 \| 366 \| (16, 394) \| 532 \| (3, 382) \| 383 \| Lambda family phage portal protein \| Lambda family phage portal protein; uniclust \| UniRef100\_A0A2W5C3N3 \| 100.0 \| 7e-39 \| 1.3e-44 \| 290.0 \| 311 \| (180, 503) \| 532 \| (2, 325) \| 336 \| Phage portal protein (Fragment) \| Phage portal protein (Fragment); uniclust \| UniRef100\_A0A0H2VTW3 \| 100.0 \| 8.5e-39 \| 1.6e-44 \| 297.5 \| 339 \| (165, 514) \| 532 \| (4, 372) \| 388 \| Head-tail preconnector gp5 \| Head-tail preconnector gp5; uniclust \| UniRef100\_UPI0020272EAC \| 100.0 \| 9.4e-39 \| 1.7e-44 \| 315.3 \| 357 \| (127, 490) \| 532 \| (3, 391) \| 808 \| phage portal protein \| phage portal protein; uniclust \| UniRef100\_A0A0T7A5Q8 \| 100.0 \| 9.7e-39 \| 1.8e-44 \| 302.2 \| 411 \| (52, 472) \| 532 \| (18, 488) \| 504 \| Portal protein \| Portal protein; uniclust \| UniRef100\_A0A376JWU1 \| 100.0 \| 9.8e-39 \| 1.9e-44 \| 302.3 \| 345 \| (63, 416) \| 532 \| (61, 414) \| 415 \| Phage portal protein, lambda family \| Phage portal protein, lambda family; uniclust \| UniRef100\_A0A0S4TZ33 \| 100.0 \| 1.5e-38 \| 3e-44 \| 293.7 \| 194 \| (319, 516) \| 532 \| (11, 206) \| 249 \| Bacteriophage-related protein \| Bacteriophage-related protein; uniclust \| UniRef100\_UPI0022F09839 \| 100.0 \| 3.5e-38 \| 6.5e-44 \| 292.5 \| 348 \| (126, 495) \| 532 \| (24, 378) \| 415 \| phage portal protein \| phage portal protein; uniclust \| UniRef100\_A0A352V3P3 \| 100.0 \| 4.1e-38 \| 7.6e-44 \| 287.2 \| 319 \| (159, 495) \| 532 \| (7, 331) \| 358 \| Phage portal protein \| Phage portal protein; uniclust \| UniRef100\_A0A6M3M268 \| 100.0 \| 5.4e-38 \| 9.9e-44 \| 298.2 \| 471 \| (4, 505) \| 532 \| (18, 500) \| 520 \| Putative portal protein \| Putative portal protein; uniclust \| UniRef100\_A0A0F2PJU6 \| 100.0 \| 7e-38 \| 1.3e-43 \| 305.6 \| 410 \| (79, 505) \| 532 \| (66, 509) \| 604 \| Capsid protein \| Capsid protein; uniclust \| UniRef100\_A0A165XDC1 \| 100.0 \| 8.7e-38 \| 1.6e-43 \| 306.5 \| 428 \| (75, 510) \| 532 \| (139, 727) \| 742 \| Phage portal protein, lambda family \| Phage portal protein, lambda family; uniclust \| UniRef100\_A0A356GP31 \| 100.0 \| 1e-37 \| 2e-43 \| 287.0 \| 229 \| (13, 246) \| 532 \| (11, 247) \| 260 \| Phage portal protein (Fragment) \| Phage portal protein (Fragment); uniclust \| UniRef100\_A0A7X7KP38 \| 100.0 \| 1.3e-37 \| 2.4e-43 \| 295.9 \| 422 \| (45, 487) \| 532 \| (48, 491) \| 524 \| Phage portal protein \| Phage portal protein; uniclust \| UniRef100\_A0A1V2N5C6 \| 100.0 \| 1.3e-37 \| 2.5e-43 \| 275.7 \| 185 \| (325, 519) \| 532 \| (2, 187) \| 204 \| Phage portal protein, lambda family \| Phage portal protein, lambda family; uniclust \| UniRef100\_A0A354EMX8 \| 100.0 \| 1.4e-37 \| 2.6e-43 \| 286.6 \| 291 \| (180, 477) \| 532 \| (6, 302) \| 389 \| Phage portal protein \| Phage portal protein; uniclust \| UniRef100\_A0A7W1G2Z3 \| 100.0 \| 1.4e-37 \| 2.7e-43 \| 301.8 \| 447 \| (46, 505) \| 532 \| (10, 475) \| 519 \| Phage portal protein \| Phage portal protein; uniclust \| UniRef100\_A0A1X7MMD6 \| 100.0 \| 1.7e-37 \| 3.1e-43 \| 273.0 \| 245 \| (158, 405) \| 532 \| (2, 265) \| 266 \| Phage portal protein, lambda family \| Phage portal protein, lambda family; uniclust \| UniRef100\_A0A376YBL2 \| 100.0 \| 2.6e-37 \| 5e-43 \| 286.1 \| 307 \| (78, 393) \| 532 \| (5, 347) \| 348 \| Capsid protein of prophage \| Capsid protein of prophage; uniclust \| UniRef100\_A0A8X6LDJ2 \| 100.0 \| 3.3e-37 \| 6e-43 \| 265.3 \| 205 \| (163, 372) \| 532 \| (6, 216) \| 226 \| Phage portal protein \| Phage portal protein; uniclust \| UniRef100\_A0A2V8RVC3 \| 100.0 \| 3.3e-37 \| 6.4e-43 \| 294.4 \| 336 \| (141, 506) \| 532 \| (1, 348) \| 376 \| Phage portal protein \| Phage portal protein; uniclust \| UniRef100\_A0A0F9LPE5 \| 100.0 \| 4.6e-37 \| 9e-43 \| 302.2 \| 382 \| (56, 484) \| 532 \| (42, 443) \| 478 \| Portal protein \| Portal protein; uniclust \| UniRef100\_A0A2W5DU94 \| 100.0 \| 6.5e-37 \| 1.3e-42 \| 280.5 \| 216 \| (283, 504) \| 532 \| (1, 220) \| 242 \| Phage portal protein (Fragment) \| Phage portal protein (Fragment); uniclust \| UniRef100\_A0A495YAC3 \| 100.0 \| 7.4e-37 \| 1.5e-42 \| 276.3 \| 180 \| (44, 233) \| 532 \| (24, 205) \| 226 \| Phage portal protein (Fragment) \| Phage portal protein (Fragment); uniclust \| UniRef100\_A0A0G4Q0C6 \| 100.0 \| 8.7e-37 \| 1.7e-42 \| 281.8 \| 241 \| (15, 267) \| 532 \| (22, 267) \| 267 \| Phage portal protein, lambda family \| Phage portal protein, lambda family; uniclust \| UniRef100\_A0A6J5NU14 \| 100.0 \| 1e-36 \| 2.1e-42 \| 294.5 \| 349 \| (108, 488) \| 532 \| (2, 368) \| 391 \| Phage portal protein, lambda family \| Phage portal protein, lambda family; uniclust \| UniRef100\_A0A060HCC5 \| 100.0 \| 1e-36 \| 2.1e-42 \| 275.7 \| 181 \| (318, 506) \| 532 \| (25, 205) \| 233 \| Portal protein \| Portal protein; uniclust \| UniRef100\_A0A5C7Q7S1 \| 100.0 \| 1.2e-36 \| 2.2e-42 \| 288.9 \| 424 \| (44, 490) \| 532 \| (13, 459) \| 512 \| Phage portal protein \| Phage portal protein; uniclust \| UniRef100\_A0A357ZWC2 \| 100.0 \| 2.1e-36 \| 4.1e-42 \| 305.7 \| 425 \| (48, 504) \| 532 \| (18, 465) \| 581 \| Phage portal protein \| Phage portal protein; uniclust \| UniRef100\_A0A970C2J4 \| 100.0 \| 2.4e-36 \| 4.5e-42 \| 290.6 \| 440 \| (48, 495) \| 532 \| (59, 522) \| 587 \| Phage portal protein \| Phage portal protein; uniclust \| UniRef100\_A0A9E0EC56 \| 100.0 \| 2.6e-36 \| 4.7e-42 \| 272.3 \| 315 \| (18, 361) \| 532 \| (1, 322) \| 322 \| Phage portal protein (Fragment) \| Phage portal protein (Fragment); uniclust \| UniRef100\_Q2CB91 \| 100.0 \| 2.7e-36 \| 5.1e-42 \| 266.8 \| 234 \| (249, 490) \| 532 \| (1, 237) \| 245 \| Probable bacteriophage-related protein \| Probable bacteriophage-related protein; uniclust \| UniRef100\_UPI002012B078 \| 100.0 \| 3.1e-36 \| 5.7e-42 \| 300.5 \| 334 \| (21, 360) \| 532 \| (1, 353) \| 907 \| phage portal protein \| phage portal protein; uniclust \| UniRef100\_UPI0002FED4F3 \| 100.0 \| 5.4e-36 \| 1e-41 \| 289.3 \| 433 \| (45, 503) \| 532 \| (37, 487) \| 611 \| phage portal protein \| phage portal protein; uniclust \| UniRef100\_A0A2A2RL01 \| 100.0 \| 1.1e-35 \| 2e-41 \| 289.2 \| 404 \| (66, 504) \| 532 \| (56, 469) \| 511 \| Phage portal protein \| Phage portal protein; uniclust \| UniRef100\_A0A2U1B6F7 \| 100.0 \| 1.1e-35 \| 2e-41 \| 273.0 \| 338 \| (154, 505) \| 532 \| (8, 350) \| 373 \| Lambda family phage portal protein \| Lambda family phage portal protein; uniclust \| UniRef100\_UPI000DDF7C70 \| 100.0 \| 1.1e-35 \| 2.2e-41 \| 285.6 \| 292 \| (181, 487) \| 532 \| (33, 329) \| 330 \| phage portal protein \| phage portal protein; uniclust \| UniRef100\_A0A8G2LY73 \| 100.0 \| 1.3e-35 \| 2.4e-41 \| 281.6 \| 384 \| (124, 518) \| 532 \| (3, 419) \| 430 \| Phage portal protein (Minor capsid protein) \| Phage portal protein (Minor capsid protein); uniclust \| UniRef100\_A0A0F9FZI9 \| 100.0 \| 1.6e-35 \| 3.5e-41 \| 315.0 \| 396 \| (72, 492) \| 532 \| (97, 516) \| 607 \| Phage portal protein \| Phage portal protein; uniclust \| UniRef100\_A0A142Y7Z0 \| 100.0 \| 1.8e-35 \| 3.6e-41 \| 275.3 \| 238 \| (43, 299) \| 532 \| (38, 279) \| 281 \| Phage portal protein, lambda family \| Phage portal protein, lambda family; uniclust \| UniRef100\_UPI000AC26F33 \| 100.0 \| 2.2e-35 \| 4e-41 \| 265.5 \| 270 \| (212, 490) \| 532 \| (35, 306) \| 313 \| phage portal protein \| phage portal protein; uniclust \| UniRef100\_A0A813BI55 \| 100.0 \| 2.4e-35 \| 4.4e-41 \| 305.4 \| 449 \| (46, 505) \| 532 \| (85, 552) \| 1651 \| Peptidase S49 domain-containing protein \| Peptidase S49 domain-containing protein; uniclust \| UniRef100\_UPI001BFF0A19 \| 100.0 \| 2.8e-35 \| 5.1e-41 \| 257.1 \| 233 \| (44, 285) \| 532 \| (11, 248) \| 249 \| phage portal protein \| phage portal protein; uniclust \| UniRef100\_A0A0D8NGB3 \| 100.0 \| 2.8e-35 \| 5.4e-41 \| 275.2 \| 252 \| (46, 301) \| 532 \| (33, 309) \| 360 \| Capsid protein (Fragment) \| Capsid protein (Fragment); uniclust \| UniRef100\_A0A0S8GAH5 \| 100.0 \| 3.2e-35 \| 6e-41 \| 265.0 \| 255 \| (244, 505) \| 532 \| (1, 264) \| 273 \| Phage portal protein (Fragment) \| Phage portal protein (Fragment); uniclust \| UniRef100\_A0A0F9HR86 \| 100.0 \| 5.7e-35 \| 1.1e-40 \| 285.8 \| 408 \| (72, 495) \| 532 \| (80, 528) \| 559 \| Phage portal protein \| Phage portal protein; uniclust \| UniRef100\_A0A953JK51 \| 100.0 \| 7.2e-35 \| 1.3e-40 \| 263.6 \| 294 \| (215, 523) \| 532 \| (3, 304) \| 327 \| Phage portal protein \| Phage portal protein; uniclust \| UniRef100\_A0A2E3DCB1 \| 100.0 \| 7.9e-35 \| 1.5e-40 \| 279.7 \| 448 \| (43, 503) \| 532 \| (50, 545) \| 571 \| Phage portal protein \| Phage portal protein; uniclust \| UniRef100\_A0A968GDE4 \| 100.0 \| 8.2e-35 \| 1.5e-40 \| 282.0 \| 416 \| (73, 505) \| 532 \| (57, 505) \| 524 \| Phage portal protein \| Phage portal protein; uniclust \| UniRef100\_A0A948A818 \| 100.0 \| 8.8e-35 \| 1.6e-40 \| 258.6 \| 259 \| (30, 298) \| 532 \| (24, 285) \| 285 \| Phage portal protein (Fragment) \| Phage portal protein (Fragment); uniclust \| UniRef100\_UPI002041E7F5 \| 100.0 \| 1.1e-34 \| 1.9e-40 \| 292.9 \| 381 \| (122, 514) \| 532 \| (623, 1037) \| 1052 \| phage portal protein \| phage portal protein; uniclust \| UniRef100\_A0A090G3P5 \| 100.0 \| 3.2e-34 \| 6.2e-40 \| 271.2 \| 290 \| (210, 507) \| 532 \| (4, 326) \| 349 \| Capsid protein of prophage \| Capsid protein of prophage; uniclust \| UniRef100\_UPI000AD9D3B3 \| 100.0 \| 3.4e-34 \| 6.3e-40 \| 285.3 \| 337 \| (46, 391) \| 532 \| (18, 391) \| 729 \| phage portal protein \| phage portal protein; uniclust \| UniRef100\_A0A0Q0BVW0 \| 100.0 \| 6e-34 \| 1.2e-39 \| 257.5 \| 223 \| (277, 508) \| 532 \| (1, 237) \| 259 \| Portal protein \| Portal protein; uniclust \| UniRef100\_UPI0008FB6ED6 \| 100.0 \| 1e-33 \| 1.8e-39 \| 252.3 \| 272 \| (128, 464) \| 532 \| (2, 286) \| 287 \| phage portal protein \| phage portal protein; uniclust \| UniRef100\_A0A1H2HHH0 \| 100.0 \| 1e-33 \| 2e-39 \| 275.4 \| 431 \| (45, 503) \| 532 \| (50, 496) \| 523 \| Capsid protein \| Capsid protein; uniclust \| UniRef100\_UPI00207606F1 \| 100.0 \| 1.8e-33 \| 3.3e-39 \| 264.8 \| 301 \| (205, 513) \| 532 \| (113, 442) \| 459 \| phage portal protein \| phage portal protein; uniclust \| UniRef100\_UPI00117A55C2 \| 100.0 \| 1.8e-33 \| 3.3e-39 \| 243.2 \| 215 \| (128, 348) \| 532 \| (5, 222) \| 229 \| phage portal protein \| phage portal protein; uniclust \| UniRef100\_A0A9D0S255 \| 100.0 \| 2e-33 \| 3.8e-39 \| 244.4 \| 185 \| (318, 506) \| 532 \| (12, 196) \| 208 \| Phage portal protein \| Phage portal protein; uniclust \| UniRef100\_A0A965PTT0 \| 100.0 \| 3.8e-33 \| 7.2e-39 \| 264.0 \| 287 \| (205, 505) \| 532 \| (15, 310) \| 387 \| Phage portal protein (Fragment) \| Phage portal protein (Fragment); uniclust \| UniRef100\_A0A062VID9 \| 100.0 \| 5.7e-33 \| 1.1e-38 \| 252.5 \| 231 \| (15, 284) \| 532 \| (1, 231) \| 240 \| Phage portal protein, lambda family (Fragment) \| Phage portal protein, lambda family (Fragment); uniclust \| UniRef100\_A0A955N1X2 \| 100.0 \| 7.7e-33 \| 1.4e-38 \| 254.4 \| 310 \| (186, 505) \| 532 \| (2, 327) \| 369 \| Phage portal protein (Fragment) \| Phage portal protein (Fragment); uniclust \| UniRef100\_A0A1Y1S1G3 \| 100.0 \| 8e-33 \| 1.5e-38 \| 264.0 \| 415 \| (72, 495) \| 532 \| (45, 498) \| 522 \| Phage portal protein \| Phage portal protein; uniclust \| UniRef100\_A0A1E5NHB2 \| 100.0 \| 9.4e-33 \| 1.8e-38 \| 268.6 \| 420 \| (72, 505) \| 532 \| (64, 522) \| 540 \| Phage portal protein \| Phage portal protein; uniclust \| UniRef100\_A0A5C7P2N1 \| 100.0 \| 1.4e-32 \| 2.5e-38 \| 263.0 \| 494 \| (17, 519) \| 532 \| (1, 512) \| 535 \| Phage portal protein \| Phage portal protein; uniclust \| UniRef100\_A0A5C7PT24 \| 100.0 \| 1.6e-32 \| 2.9e-38 \| 263.9 \| 480 \| (20, 521) \| 532 \| (14, 532) \| 563 \| Phage portal protein \| Phage portal protein; uniclust \| UniRef100\_UPI001C05E608 \| 100.0 \| 2.1e-32 \| 3.8e-38 \| 246.5 \| 276 \| (133, 412) \| 532 \| (2, 309) \| 310 \| phage portal protein \| phage portal protein; uniclust \| UniRef100\_UPI001D09EB93 \| 99.9 \| 2.8e-32 \| 5.2e-38 \| 272.2 \| 343 \| (48, 397) \| 532 \| (26, 402) \| 874 \| phage portal protein \| phage portal protein; uniclust \| UniRef100\_A0A3D5KD24 \| 99.9 \| 2.8e-32 \| 5.2e-38 \| 260.7 \| 451 \| (45, 516) \| 532 \| (51, 518) \| 530 \| Phage portal protein \| Phage portal protein; uniclust \| UniRef100\_A0A257JFS0 \| 99.9 \| 5e-32 \| 9.5e-38 \| 230.8 \| 168 \| (16, 192) \| 532 \| (1, 173) \| 173 \| Portal protein (Fragment) \| Portal protein (Fragment); uniclust \| UniRef100\_A0A3C1GKI0 \| 99.9 \| 5.5e-32 \| 1.1e-37 \| 249.3 \| 239 \| (249, 494) \| 532 \| (1, 241) \| 278 \| Phage portal protein \| Phage portal protein; uniclust \| UniRef100\_A0A6G1WRJ0 \| 99.9 \| 6.2e-32 \| 1.2e-37 \| 243.6 \| 187 \| (316, 506) \| 532 \| (37, 224) \| 270 \| Phage portal protein (Fragment) \| Phage portal protein (Fragment); uniclust \| UniRef100\_S2KXG2 \| 99.9 \| 1.2e-31 \| 2.2e-37 \| 240.1 \| 242 \| (253, 494) \| 532 \| (2, 255) \| 295 \| Lambda family phage portal protein \| Lambda family phage portal protein; uniclust \| UniRef100\_UPI001FA956E3 \| 99.9 \| 1.2e-31 \| 2.3e-37 \| 243.0 \| 315 \| (18, 444) \| 532 \| (1, 324) \| 325 \| phage portal protein \| phage portal protein; uniclust \| UniRef100\_A0A5I8XHP0 \| 99.9 \| 1.7e-31 \| 3.1e-37 \| 254.5 \| 303 \| (18, 331) \| 532 \| (1, 312) \| 509 \| Phage portal protein \| Phage portal protein; uniclust \| UniRef100\_A0A0F8VDT1 \| 99.9 \| 2e-31 \| 4.2e-37 \| 247.0 \| 213 \| (11, 231) \| 532 \| (8, 225) \| 234 \| Uncharacterized protein (Fragment) \| Uncharacterized protein (Fragment); uniclust \| UniRef100\_A0A7W1T9L3 \| 99.9 \| 2.4e-31 \| 4.5e-37 \| 255.2 \| 309 \| (43, 371) \| 532 \| (29, 342) \| 546 \| Phage portal protein \| Phage portal protein; uniclust \| UniRef100\_A0A357LHL5 \| 99.9 \| 3.3e-31 \| 6e-37 \| 252.6 \| 414 \| (78, 495) \| 532 \| (7, 462) \| 509 \| Phage portal protein \| Phage portal protein; uniclust \| UniRef100\_A0A068SLY6 \| 99.9 \| 4.5e-31 \| 8.3e-37 \| 254.7 \| 400 \| (72, 495) \| 532 \| (63, 475) \| 576 \| Phage portal protein, lambda family \| Phage portal protein, lambda family; uniclust \| UniRef100\_A0A0P7YIZ9 \| 99.9 \| 7e-31 \| 1.3e-36 \| 232.2 \| 202 \| (293, 505) \| 532 \| (1, 207) \| 247 \| Phage portal protein, lambda family \| Phage portal protein, lambda family; uniclust \| UniRef100\_A0A0L6CQP6 \| 99.9 \| 7.2e-31 \| 1.4e-36 \| 229.5 \| 167 \| (336, 509) \| 532 \| (2, 169) \| 209 \| Phage portal protein, lambda family \| Phage portal protein, lambda family; uniclust \| UniRef100\_A0A833G1E6 \| 99.9 \| 1.9e-30 \| 3.6e-36 \| 242.0 \| 332 \| (78, 423) \| 532 \| (29, 396) \| 412 \| Phage portal protein (Fragment) \| Phage portal protein (Fragment); uniclust \| UniRef100\_A0A0S9PWW8 \| 99.9 \| 1.8e-30 \| 3.6e-36 \| 242.6 \| 233 \| (45, 281) \| 532 \| (33, 281) \| 283 \| Phage portal protein \| Phage portal protein; uniclust \| UniRef100\_UPI00117B6109 \| 99.9 \| 2.1e-30 \| 3.9e-36 \| 236.1 \| 250 \| (47, 301) \| 532 \| (44, 309) \| 336 \| phage portal protein \| phage portal protein; uniclust \| UniRef100\_A0A7V1N7P6 \| 99.9 \| 3.9e-30 \| 7.1e-36 \| 238.5 \| 310 \| (49, 366) \| 532 \| (51, 388) \| 389 \| Phage portal protein (Fragment) \| Phage portal protein (Fragment); uniclust \| UniRef100\_A0A965PTZ3 \| 99.9 \| 4.1e-30 \| 7.6e-36 \| 234.8 \| 336 \| (82, 450) \| 532 \| (1, 341) \| 342 \| Phage portal protein (Fragment) \| Phage portal protein (Fragment); uniclust \| UniRef100\_A0A149VAP2 \| 99.9 \| 4.9e-30 \| 9.1e-36 \| 225.7 \| 187 \| (319, 509) \| 532 \| (26, 214) \| 230 \| Capsid protein (Fragment) \| Capsid protein (Fragment); uniclust \| UniRef100\_R7MYT1 \| 99.9 \| 1.4e-29 \| 2.7e-35 \| 219.6 \| 165 \| (338, 507) \| 532 \| (1, 166) \| 179 \| Phage portal protein \| Phage portal protein; uniclust \| UniRef100\_A0A965PMJ0 \| 99.9 \| 1.5e-29 \| 2.8e-35 \| 238.4 \| 304 \| (207, 524) \| 532 \| (13, 319) \| 448 \| Phage portal protein \| Phage portal protein; uniclust \| UniRef100\_UPI001E28734F \| 99.9 \| 1.7e-29 \| 3.2e-35 \| 236.2 \| 266 \| (221, 490) \| 532 \| (1, 295) \| 417 \| phage portal protein \| phage portal protein; uniclust \| UniRef100\_A0A754E6Q2 \| 99.9 \| 2.6e-29 \| 4.8e-35 \| 252.3 \| 320 \| (28, 358) \| 532 \| (1, 329) \| 907 \| Phage portal protein \| Phage portal protein; uniclust \| UniRef100\_A0A0H4TJT5 \| 99.9 \| 3e-29 \| 5.7e-35 \| 211.7 \| 158 \| (179, 341) \| 532 \| (4, 165) \| 170 \| Phage portal protein (Fragment) \| Phage portal protein (Fragment); uniclust \| UniRef100\_UPI0004956350 \| 99.9 \| 6.5e-29 \| 1.2e-34 \| 216.0 \| 155 \| (347, 505) \| 532 \| (1, 156) \| 203 \| phage portal protein \| phage portal protein; uniclust \| UniRef100\_UPI001E588232 \| 99.9 \| 6.8e-29 \| 1.3e-34 \| 239.0 \| 268 \| (218, 489) \| 532 \| (1, 297) \| 551 \| phage portal protein \| phage portal protein; uniclust \| UniRef100\_UPI00041184E5 \| 99.9 \| 7e-29 \| 1.3e-34 \| 226.4 \| 306 \| (74, 508) \| 532 \| (3, 322) \| 335 \| phage portal protein \| phage portal protein; uniclust \| UniRef100\_UPI0002D45360 \| 99.9 \| 9.2e-29 \| 1.7e-34 \| 228.5 \| 260 \| (226, 495) \| 532 \| (11, 283) \| 309 \| phage portal protein \| phage portal protein; uniclust \| UniRef100\_A0A2P7UVS4 \| 99.9 \| 1e-28 \| 2e-34 \| 223.8 \| 200 \| (178, 379) \| 532 \| (20, 256) \| 256 \| Uncharacterized protein \| Uncharacterized protein; uniclust \| UniRef100\_A0A896Z248 \| 99.9 \| 1.4e-28 \| 2.6e-34 \| 219.6 \| 206 \| (44, 251) \| 532 \| (20, 229) \| 280 \| Portal protein \| Portal protein; uniclust \| UniRef100\_A0A0F9H021 \| 99.9 \| 1.4e-28 \| 2.6e-34 \| 215.5 \| 176 \| (122, 301) \| 532 \| (11, 196) \| 217 \| Uncharacterized protein (Fragment) \| Uncharacterized protein (Fragment); uniclust \| UniRef100\_A0A2A4RG34 \| 99.9 \| 1.8e-28 \| 3.4e-34 \| 234.3 \| 416 \| (64, 504) \| 532 \| (46, 481) \| 508 \| Phage portal protein \| Phage portal protein; uniclust \| UniRef100\_A0A081B6E1 \| 99.9 \| 1.7e-28 \| 3.7e-34 \| 258.3 \| 439 \| (12, 492) \| 532 \| (19, 486) \| 532 \| Conserved protein \| Conserved protein; uniclust \| UniRef100\_UPI000B13FF44 \| 99.9 \| 2.1e-28 \| 3.9e-34 \| 238.0 \| 278 \| (122, 406) \| 532 \| (301, 609) \| 611 \| phage portal protein \| phage portal protein; uniclust \| UniRef100\_A0A0F9Q1K6 \| 99.9 \| 2.2e-28 \| 4.7e-34 \| 257.5 \| 380 \| (55, 485) \| 532 \| (64, 466) \| 510 \| Portal protein \| Portal protein; uniclust \| UniRef100\_A0A2T4JG48 \| 99.9 \| 2.8e-28 \| 5.4e-34 \| 229.4 \| 291 \| (46, 352) \| 532 \| (32, 359) \| 360 \| Phage portal protein \| Phage portal protein; uniclust \| UniRef100\_A0A1V6E2F0 \| 99.9 \| 3e-28 \| 5.5e-34 \| 233.2 \| 420 \| (46, 489) \| 532 \| (17, 456) \| 515 \| Phage portal protein, lambda family \| Phage portal protein, lambda family; uniclust \| UniRef100\_A0A812S2M1 \| 99.9 \| 3.5e-28 \| 6.4e-34 \| 244.8 \| 441 \| (45, 502) \| 532 \| (14, 478) \| 922 \| Gene 4 protein \| Gene 4 protein; uniclust \| UniRef100\_A0A7W1J8B3 \| 99.9 \| 3.6e-28 \| 6.7e-34 \| 218.4 \| 216 \| (43, 275) \| 532 \| (33, 253) \| 259 \| Phage portal protein \| Phage portal protein; uniclust \| UniRef100\_A0A827RQQ4 \| 99.9 \| 4.2e-28 \| 8.1e-34 \| 223.8 \| 230 \| (63, 299) \| 532 \| (58, 289) \| 290 \| Phage portal protein (Fragment) \| Phage portal protein (Fragment); uniclust \| UniRef100\_A0A966RAZ4 \| 99.9 \| 5.8e-28 \| 1.1e-33 \| 227.0 \| 335 \| (122, 505) \| 532 \| (3, 342) \| 431 \| Phage portal protein \| Phage portal protein; uniclust \| UniRef100\_UPI0021C2EF29 \| 99.9 \| 5.9e-28 \| 1.1e-33 \| 207.5 \| 170 \| (323, 495) \| 532 \| (1, 170) \| 191 \| phage portal protein \| phage portal protein; uniclust \| UniRef100\_UPI0018CC8607 \| 99.9 \| 7.2e-28 \| 1.3e-33 \| 218.4 \| 254 \| (43, 300) \| 532 \| (32, 312) \| 315 \| phage portal protein \| phage portal protein; uniclust \| UniRef100\_UPI0002C9D6EA \| 99.9 \| 8.2e-28 \| 1.6e-33 \| 219.8 \| 245 \| (259, 514) \| 532 \| (8, 257) \| 275 \| phage portal protein \| phage portal protein; uniclust \| UniRef100\_A0A0F9HAI8 \| 99.9 \| 8.5e-28 \| 1.6e-33 \| 224.7 \| 316 \| (183, 517) \| 532 \| (4, 341) \| 349 \| Phage portal protein (Fragment) \| Phage portal protein (Fragment); uniclust \| UniRef100\_A0A5M6I484 \| 99.9 \| 9.3e-28 \| 1.7e-33 \| 221.0 \| 220 \| (44, 267) \| 532 \| (19, 252) \| 357 \| Phage portal protein \| Phage portal protein; uniclust \| UniRef100\_A0A2W5FLC9 \| 99.9 \| 9.8e-28 \| 1.8e-33 \| 216.4 \| 245 \| (255, 503) \| 532 \| (4, 275) \| 302 \| Phage portal protein (Fragment) \| Phage portal protein (Fragment); uniclust \| UniRef100\_A0A2X3JP92 \| 99.9 \| 1.2e-27 \| 2.2e-33 \| 219.2 \| 271 \| (240, 514) \| 532 \| (18, 318) \| 333 \| Phage portal protein (Minor capsid protein) \| Phage portal protein (Minor capsid protein); uniclust \| UniRef100\_A0A376JYF4 \| 99.9 \| 1.1e-27 \| 2.3e-33 \| 231.2 \| 280 \| (216, 506) \| 532 \| (29, 339) \| 341 \| Phage portal protein (Minor capsid protein) \| Phage portal protein (Minor capsid protein); uniclust \| UniRef100\_G1V7I5 \| 99.9 \| 1.4e-27 \| 2.5e-33 \| 211.1 \| 239 \| (253, 503) \| 532 \| (2, 244) \| 258 \| Phage portal protein (Fragment) \| Phage portal protein (Fragment); uniclust \| UniRef100\_UPI00226D29AD \| 99.9 \| 2e-27 \| 3.7e-33 \| 206.8 \| 215 \| (242, 469) \| 532 \| (4, 226) \| 230 \| phage portal protein \| phage portal protein; uniclust \| UniRef100\_A0A142XEI8 \| 99.9 \| 2.1e-27 \| 3.8e-33 \| 229.9 \| 382 \| (72, 485) \| 532 \| (63, 464) \| 498 \| Phage portal protein, lambda family \| Phage portal protein, lambda family; uniclust \| UniRef100\_UPI0022710B4D \| 99.9 \| 2.2e-27 \| 4e-33 \| 243.6 \| 421 \| (68, 506) \| 532 \| (721, 1192) \| 1208 \| phage terminase large subunit family protein \| phage terminase large subunit family protein; uniclust \| UniRef100\_A0A1G0XUR8 \| 99.9 \| 2.4e-27 \| 4.5e-33 \| 234.4 \| 398 \| (73, 503) \| 532 \| (64, 498) \| 544 \| Phage portal protein \| Phage portal protein; uniclust \| UniRef100\_A0A966R5B7 \| 99.9 \| 2.9e-27 \| 5.4e-33 \| 222.3 \| 305 \| (181, 503) \| 532 \| (29, 341) \| 427 \| Phage portal protein \| Phage portal protein; uniclust \| UniRef100\_A0A0E3JS82 \| 99.9 \| 2.8e-27 \| 5.4e-33 \| 236.7 \| 406 \| (73, 511) \| 532 \| (61, 482) \| 553 \| Portal protein \| Portal protein; uniclust \| UniRef100\_A0A5E8GT99 \| 99.9 \| 3.2e-27 \| 5.8e-33 \| 228.4 \| 436 \| (57, 502) \| 532 \| (30, 531) \| 563 \| Bacteriophage capsid protein \| Bacteriophage capsid protein; uniclust \| UniRef100\_UPI001FEBC5EC \| 99.9 \| 3.3e-27 \| 6e-33 \| 211.8 \| 188 \| (261, 453) \| 532 \| (30, 225) \| 288 \| phage portal protein \| phage portal protein; uniclust \| UniRef100\_A0A6M3IP50 \| 99.9 \| 3.7e-27 \| 6.9e-33 \| 213.2 \| 261 \| (247, 511) \| 532 \| (14, 281) \| 307 \| Putative portal protein (Fragment) \| Putative portal protein (Fragment); uniclust \| UniRef100\_A0A961H371 \| 99.9 \| 4.1e-27 \| 7.6e-33 \| 217.3 \| 325 \| (45, 400) \| 532 \| (23, 362) \| 363 \| Phage portal protein (Fragment) \| Phage portal protein (Fragment); uniclust \| UniRef100\_A0A450TW67 \| 99.9 \| 5.4e-27 \| 9.8e-33 \| 217.3 \| 199 \| (99, 301) \| 532 \| (11, 222) \| 374 \| Phage portal protein, lambda family \| Phage portal protein, lambda family; uniclust \| UniRef100\_UPI0013B7ACB1 \| 99.9 \| 6.7e-27 \| 1.2e-32 \| 212.8 \| 298 \| (45, 354) \| 532 \| (14, 319) \| 321 \| phage portal protein \| phage portal protein; uniclust \| UniRef100\_A0A9D8SJC5 \| 99.9 \| 6.9e-27 \| 1.3e-32 \| 222.0 \| 424 \| (46, 488) \| 532 \| (10, 452) \| 468 \| Phage portal protein \| Phage portal protein; uniclust \| UniRef100\_A0A4Q6ELU1 \| 99.9 \| 6.9e-27 \| 1.3e-32 \| 202.0 \| 170 \| (329, 503) \| 532 \| (2, 172) \| 192 \| Phage portal protein \| Phage portal protein; uniclust \| UniRef100\_A0A3C1YTA8 \| 99.9 \| 7.3e-27 \| 1.4e-32 \| 228.3 \| 426 \| (45, 493) \| 532 \| (19, 473) \| 494 \| Phage portal protein \| Phage portal protein; uniclust \| UniRef100\_A0A6J5NYG1 \| 99.9 \| 8.1e-27 \| 1.5e-32 \| 231.0 \| 385 \| (71, 488) \| 532 \| (82, 490) \| 622 \| Portal\_lambda, phage portal protein, lambda family \| Portal\_lambda, phage portal protein, lambda family; uniclust \| UniRef100\_A0A1H5Z513 \| 99.9 \| 1.1e-26 \| 2.2e-32 \| 198.8 \| 145 \| (362, 521) \| 532 \| (1, 146) \| 167 \| Phage portal protein, lambda family \| Phage portal protein, lambda family; uniclust \| UniRef100\_X0TAA3 \| 99.9 \| 3e-26 \| 5.5e-32 \| 203.8 \| 259 \| (19, 300) \| 532 \| (4, 264) \| 267 \| Uncharacterized protein (Fragment) \| Uncharacterized protein (Fragment); uniclust \| UniRef100\_A0A2D0IJL4 \| 99.9 \| 3.3e-26 \| 6.5e-32 \| 197.4 \| 150 \| (356, 509) \| 532 \| (1, 152) \| 166 \| Capsid protein \| Capsid protein; uniclust \| UniRef100\_UPI001EE63960 \| 99.9 \| 3.6e-26 \| 6.6e-32 \| 190.1 \| 162 \| (220, 381) \| 532 \| (1, 167) \| 167 \| phage portal protein \| phage portal protein; uniclust \| UniRef100\_A0A9E3UTF1 \| 99.9 \| 4.4e-26 \| 8.1e-32 \| 219.2 \| 387 \| (73, 505) \| 532 \| (69, 480) \| 524 \| Phage portal protein \| Phage portal protein; uniclust \| UniRef100\_A0A1Z4BY38 \| 99.9 \| 4.8e-26 \| 8.8e-32 \| 192.8 \| 174 \| (44, 226) \| 532 \| (11, 186) \| 188 \| Phage portal protein \| Phage portal protein; uniclust \| UniRef100\_A0A257WRS9 \| 99.9 \| 5e-26 \| 9.9e-32 \| 218.6 \| 263 \| (207, 489) \| 532 \| (13, 290) \| 327 \| Phage portal protein \| Phage portal protein; uniclust \| UniRef100\_A0A8J3DI50 \| 99.9 \| 6.2e-26 \| 1.1e-31 \| 218.3 \| 414 \| (45, 488) \| 532 \| (30, 472) \| 527 \| Phage portal protein \| Phage portal protein; uniclust \| UniRef100\_A0A2T6DRH8 \| 99.9 \| 7e-26 \| 1.3e-31 \| 215.9 \| 400 \| (72, 505) \| 532 \| (32, 453) \| 480 \| Phage portal protein \| Phage portal protein; uniclust \| UniRef100\_A0A1G8I036 \| 99.9 \| 7.7e-26 \| 1.5e-31 \| 217.5 \| 247 \| (46, 301) \| 532 \| (41, 303) \| 400 \| Phage portal protein, lambda family (Fragment) \| Phage portal protein, lambda family (Fragment); uniclust \| UniRef100\_A0A7V2ILH1 \| 99.9 \| 9.2e-26 \| 1.7e-31 \| 207.9 \| 239 \| (44, 301) \| 532 \| (19, 261) \| 353 \| Phage portal protein \| Phage portal protein; uniclust \| UniRef100\_UPI001A7E5E67 \| 99.9 \| 1.6e-25 \| 2.9e-31 \| 224.0 \| 201 \| (319, 526) \| 532 \| (10, 215) \| 808 \| phage portal protein \| phage portal protein; uniclust \| UniRef100\_UPI000A788E98 \| 99.9 \| 1.6e-25 \| 2.9e-31 \| 189.3 \| 174 \| (157, 332) \| 532 \| (1, 185) \| 185 \| phage portal protein \| phage portal protein; uniclust \| UniRef100\_A0A2M7FBH1 \| 99.9 \| 1.8e-25 \| 3.6e-31 \| 193.0 \| 156 \| (15, 172) \| 532 \| (1, 158) \| 159 \| Phage portal protein (Fragment) \| Phage portal protein (Fragment); uniclust \| UniRef100\_A0A9E6UXJ6 \| 99.9 \| 2.6e-25 \| 5e-31 \| 185.4 \| 120 \| (351, 473) \| 532 \| (3, 123) \| 145 \| Phage portal protein \| Phage portal protein; uniclust \| UniRef100\_A0A357XXP9 \| 99.9 \| 2.7e-25 \| 5e-31 \| 211.0 \| 224 \| (45, 290) \| 532 \| (166, 393) \| 459 \| Phage portal protein (Fragment) \| Phage portal protein (Fragment); uniclust \| UniRef100\_A0A2N8HBV6 \| 99.9 \| 3.1e-25 \| 5.6e-31 \| 211.0 \| 388 \| (73, 494) \| 532 \| (49, 445) \| 466 \| Phage portal protein \| Phage portal protein; uniclust \| UniRef100\_A0A1G8HYN6 \| 99.9 \| 3.2e-25 \| 6.1e-31 \| 199.2 \| 197 \| (317, 517) \| 532 \| (26, 223) \| 244 \| Phage portal protein, lambda family (Fragment) \| Phage portal protein, lambda family (Fragment); uniclust \| UniRef100\_A0A167HQ01 \| 99.9 \| 3.3e-25 \| 6.4e-31 \| 206.3 \| 240 \| (45, 292) \| 532 \| (20, 274) \| 296 \| Phage portal protein (Fragment) \| Phage portal protein (Fragment); uniclust \| UniRef100\_UPI002265A757 \| 99.9 \| 3.6e-25 \| 6.6e-31 \| 198.8 \| 238 \| (250, 492) \| 532 \| (1, 248) \| 284 \| phage portal protein \| phage portal protein; uniclust \| UniRef100\_UPI001FD9E691 \| 99.9 \| 3.7e-25 \| 6.9e-31 \| 186.5 \| 148 \| (176, 326) \| 532 \| (11, 163) \| 165 \| phage portal protein \| phage portal protein; uniclust \| UniRef100\_A0A368KX58 \| 99.9 \| 4.1e-25 \| 7.5e-31 \| 205.1 \| 248 \| (99, 366) \| 532 \| (13, 262) \| 373 \| Phage portal protein \| Phage portal protein; uniclust \| UniRef100\_A0A2G2K5M6 \| 99.8 \| 6.6e-25 \| 1.3e-30 \| 196.7 \| 181 \| (45, 229) \| 532 \| (30, 218) \| 218 \| Phage portal protein \| Phage portal protein; uniclust \| UniRef100\_A0A3D4DT62 \| 99.8 \| 6.7e-25 \| 1.4e-30 \| 186.4 \| 134 \| (17, 173) \| 532 \| (1, 134) \| 135 \| Phage portal protein (Fragment) \| Phage portal protein (Fragment); uniclust \| UniRef100\_A0A954G1K1 \| 99.8 \| 7.5e-25 \| 1.4e-30 \| 201.4 \| 286 \| (206, 495) \| 532 \| (10, 316) \| 343 \| Phage portal protein (Fragment) \| Phage portal protein (Fragment); uniclust \| UniRef100\_A0A961U404 \| 99.8 \| 7.5e-25 \| 1.4e-30 \| 194.9 \| 217 \| (44, 269) \| 532 \| (47, 264) \| 264 \| Phage portal protein (Fragment) \| Phage portal protein (Fragment); uniclust \| UniRef100\_A0A661TPP3 \| 99.8 \| 1e-24 \| 1.9e-30 \| 193.8 \| 181 \| (319, 514) \| 532 \| (4, 185) \| 225 \| Phage portal protein (Fragment) \| Phage portal protein (Fragment); uniclust \| UniRef100\_A0A377DG37 \| 99.8 \| 1.2e-24 \| 2.3e-30 \| 196.1 \| 233 \| (57, 298) \| 532 \| (2, 248) \| 250 \| Phage portal protein (Minor capsid protein) \| Phage portal protein (Minor capsid protein); uniclust \| UniRef100\_A0A0A1DM99 \| 99.8 \| 1.5e-24 \| 3.1e-30 \| 230.9 \| 416 \| (54, 490) \| 532 \| (66, 518) \| 626 \| Phage portal protein \| Phage portal protein; uniclust \| UniRef100\_UPI00227824CA \| 99.8 \| 1.7e-24 \| 3.2e-30 \| 192.6 \| 217 \| (148, 370) \| 532 \| (2, 226) \| 263 \| phage portal protein \| phage portal protein; uniclust \| UniRef100\_A0A2E9VWF4 \| 99.8 \| 2.5e-24 \| 4.7e-30 \| 203.5 \| 305 \| (54, 385) \| 532 \| (53, 372) \| 376 \| Phage portal protein \| Phage portal protein; uniclust \| UniRef100\_UPI000FD93CDA \| 99.8 \| 3.6e-24 \| 6.6e-30 \| 192.2 \| 229 \| (44, 276) \| 532 \| (35, 278) \| 280 \| phage portal protein \| phage portal protein; uniclust \| UniRef100\_A0A2D6M8M0 \| 99.8 \| 3.8e-24 \| 7e-30 \| 208.9 \| 427 \| (57, 495) \| 532 \| (92, 556) \| 599 \| Phage portal protein \| Phage portal protein; uniclust \| UniRef100\_A0A966SJ28 \| 99.8 \| 4.4e-24 \| 8.1e-30 \| 200.5 \| 356 \| (45, 419) \| 532 \| (23, 394) \| 407 \| Phage portal protein (Fragment) \| Phage portal protein (Fragment); uniclust \| UniRef100\_A0A9E5WIJ8 \| 99.8 \| 1.1e-23 \| 2e-29 \| 186.1 \| 208 \| (122, 340) \| 532 \| (18, 229) \| 247 \| Phage portal protein \| Phage portal protein; uniclust \| UniRef100\_A0A954NMX5 \| 99.8 \| 1.2e-23 \| 2.2e-29 \| 177.0 \| 172 \| (209, 382) \| 532 \| (1, 173) \| 176 \| Phage portal protein (Fragment) \| Phage portal protein (Fragment); uniclust \| UniRef100\_A0A061P5J7 \| 99.8 \| 1.3e-23 \| 2.8e-29 \| 217.9 \| 397 \| (54, 488) \| 532 \| (51, 463) \| 498 \| Uncharacterized protein \| Uncharacterized protein; uniclust \| UniRef100\_A0A3M1R6N2 \| 99.8 \| 1.6e-23 \| 2.9e-29 \| 187.7 \| 201 \| (44, 262) \| 532 \| (39, 243) \| 243 \| Phage portal protein (Fragment) \| Phage portal protein (Fragment); uniclust \| UniRef100\_A0A7X1E5Y1 \| 99.8 \| 1.9e-23 \| 3.5e-29 \| 199.9 \| 398 \| (73, 504) \| 532 \| (46, 470) \| 480 \| Phage portal protein \| Phage portal protein; uniclust \| UniRef100\_A0A1V5U672 \| 99.8 \| 2e-23 \| 3.9e-29 \| 186.7 \| 177 \| (318, 509) \| 532 \| (5, 184) \| 207 \| Phage portal protein, lambda family \| Phage portal protein, lambda family; uniclust \| UniRef100\_UPI001E4A37DB \| 99.8 \| 2.6e-23 \| 4.7e-29 \| 172.4 \| 156 \| (287, 457) \| 532 \| (2, 159) \| 160 \| phage portal protein \| phage portal protein; uniclust \| UniRef100\_A0A376TZP2 \| 99.8 \| 2.7e-23 \| 4.9e-29 \| 194.3 \| 234 \| (277, 514) \| 532 \| (2, 258) \| 387 \| Capsid protein of prophage \| Capsid protein of prophage; uniclust \| UniRef100\_A0A0B4ZPA5 \| 99.8 \| 2.7e-23 \| 4.9e-29 \| 210.0 \| 349 \| (46, 473) \| 532 \| (88, 451) \| 881 \| Serine protease \| Serine protease; uniclust \| UniRef100\_A0A090G440 \| 99.8 \| 3.8e-23 \| 7.1e-29 \| 195.0 \| 265 \| (17, 300) \| 532 \| (5, 286) \| 346 \| Capsid protein of prophage \| Capsid protein of prophage; uniclust \| UniRef100\_A0A376NS21 \| 99.8 \| 4.5e-23 \| 8.3e-29 \| 180.7 \| 213 \| (280, 502) \| 532 \| (1, 218) \| 219 \| Head-tail preconnector protein \| Head-tail preconnector protein; uniclust \| UniRef100\_UPI002233EA83 \| 99.8 \| 4.9e-23 \| 9e-29 \| 190.8 \| 198 \| (46, 247) \| 532 \| (116, 332) \| 357 \| phage portal protein \| phage portal protein; uniclust \| UniRef100\_A0A0J1CHN1 \| 99.8 \| 5.1e-23 \| 1e-28 \| 182.2 \| 163 \| (317, 479) \| 532 \| (7, 171) \| 181 \| Plasmid partitioning protein ParB (Fragment) \| Plasmid partitioning protein ParB (Fragment); uniclust \| UniRef100\_UPI001652C7CE \| 99.8 \| 5.9e-23 \| 1.1e-28 \| 180.2 \| 225 \| (69, 299) \| 532 \| (6, 233) \| 233 \| phage portal protein \| phage portal protein; uniclust \| UniRef100\_A0A9E6BKT2 \| 99.8 \| 7.4e-23 \| 1.4e-28 \| 197.6 \| 177 \| (46, 232) \| 532 \| (21, 199) \| 521 \| Phage portal protein \| Phage portal protein; uniclust \| UniRef100\_A0A2D9C8J7 \| 99.8 \| 7.4e-23 \| 1.4e-28 \| 185.4 \| 231 \| (263, 505) \| 532 \| (1, 244) \| 271 \| Phage portal protein (Fragment) \| Phage portal protein (Fragment); uniclust \| UniRef100\_A0A497KH68 \| 99.8 \| 7.7e-23 \| 1.4e-28 \| 194.3 \| 321 \| (72, 419) \| 532 \| (80, 419) \| 444 \| Phage portal protein (Fragment) \| Phage portal protein (Fragment); uniclust \| UniRef100\_UPI001CCED0AA \| 99.8 \| 8e-23 \| 1.5e-28 \| 181.6 \| 239 \| (26, 277) \| 532 \| (2, 245) \| 255 \| phage portal protein \| phage portal protein; uniclust \| UniRef100\_A0A317F9M0 \| 99.8 \| 8.3e-23 \| 1.6e-28 \| 175.7 \| 168 \| (43, 226) \| 532 \| (15, 184) \| 185 \| Uncharacterized protein \| Uncharacterized protein; uniclust \| UniRef100\_A0A7X6FP93 \| 99.8 \| 1.2e-22 \| 2.2e-28 \| 173.4 \| 165 \| (133, 300) \| 532 \| (3, 172) \| 192 \| Phage portal protein \| Phage portal protein; uniclust \| UniRef100\_UPI000AF07CED \| 99.8 \| 1.4e-22 \| 2.7e-28 \| 175.3 \| 149 \| (348, 503) \| 532 \| (13, 162) \| 211 \| hypothetical protein \| hypothetical protein; uniclust \| UniRef100\_UPI0004A0575F \| 99.8 \| 1.4e-22 \| 2.7e-28 \| 180.2 \| 244 \| (150, 401) \| 532 \| (3, 250) \| 257 \| phage portal protein \| phage portal protein; uniclust \| UniRef100\_A0A2G1Z9F9 \| 99.8 \| 2e-22 \| 3.7e-28 \| 176.8 \| 182 \| (320, 505) \| 532 \| (30, 212) \| 231 \| Phage portal protein (Fragment) \| Phage portal protein (Fragment); uniclust \| UniRef100\_A0A2N2HIU1 \| 99.8 \| 2.5e-22 \| 4.6e-28 \| 176.6 \| 179 \| (318, 503) \| 532 \| (23, 201) \| 235 \| Phage portal protein (Fragment) \| Phage portal protein (Fragment); uniclust \| UniRef100\_A0A7W8DQU5 \| 99.8 \| 2.6e-22 \| 4.8e-28 \| 194.1 \| 423 \| (80, 524) \| 532 \| (59, 519) \| 525 \| Capsid protein \| Capsid protein; uniclust \| UniRef100\_A0A6M3JYX7 \| 99.8 \| 3.6e-22 \| 6.6e-28 \| 197.9 \| 381 \| (54, 479) \| 532 \| (47, 439) \| 678 \| Putative portal protein \| Putative portal protein; uniclust \| UniRef100\_A0A0S4U0U3 \| 99.8 \| 3.8e-22 \| 7.1e-28 \| 163.9 \| 134 \| (86, 229) \| 532 \| (2, 137) \| 137 \| Bacteriophage-related protein (Fragment) \| Bacteriophage-related protein (Fragment); uniclust \| UniRef100\_A0A250DLV1 \| 99.8 \| 3.9e-22 \| 7.1e-28 \| 179.5 \| 224 \| (280, 507) \| 532 \| (15, 244) \| 279 \| Phage portal protein \| Phage portal protein; uniclust \| UniRef100\_UPI00209508A0 \| 99.8 \| 3.9e-22 \| 7.2e-28 \| 159.5 \| 121 \| (335, 455) \| 532 \| (3, 124) \| 127 \| phage portal protein \| phage portal protein; uniclust \| UniRef100\_A0A965LHB1 \| 99.8 \| 4.7e-22 \| 8.6e-28 \| 179.7 \| 267 \| (15, 300) \| 532 \| (1, 276) \| 288 \| Phage portal protein (Fragment) \| Phage portal protein (Fragment); uniclust \| UniRef100\_UPI00054CCE67 \| 99.8 \| 4.7e-22 \| 8.7e-28 \| 172.5 \| 179 \| (240, 419) \| 532 \| (8, 193) \| 196 \| phage portal protein \| phage portal protein; uniclust \| UniRef100\_Q9JMP2 \| 99.8 \| 7e-22 \| 1.3e-27 \| 161.7 \| 127 \| (293, 419) \| 532 \| (1, 132) \| 145 \| Orf4 protein \| Orf4 protein; uniclust \| UniRef100\_A0A965PPH0 \| 99.8 \| 8.3e-22 \| 1.5e-27 \| 172.3 \| 207 \| (156, 374) \| 532 \| (3, 219) \| 224 \| Phage portal protein \| Phage portal protein; uniclust \| UniRef100\_A0A6M3JUQ8 \| 99.8 \| 8.6e-22 \| 1.6e-27 \| 197.2 \| 380 \| (73, 492) \| 532 \| (112, 539) \| 572 \| Putative portal protein \| Putative portal protein; uniclust \| UniRef100\_A0A827RRV0 \| 99.8 \| 9.6e-22 \| 1.9e-27 \| 179.3 \| 169 \| (319, 491) \| 532 \| (23, 191) \| 217 \| Phage portal protein \| Phage portal protein; uniclust \| UniRef100\_A0A381KTT9 \| 99.7 \| 1.5e-21 \| 2.8e-27 \| 171.7 \| 169 \| (248, 416) \| 532 \| (22, 208) \| 218 \| Phage portal protein \| Phage portal protein; uniclust \| UniRef100\_A0A357XSR0 \| 99.7 \| 1.8e-21 \| 3.4e-27 \| 188.2 \| 456 \| (16, 494) \| 532 \| (2, 494) \| 519 \| Phage portal protein \| Phage portal protein; uniclust \| UniRef100\_A0A2W4N7H3 \| 99.7 \| 2e-21 \| 3.6e-27 \| 188.0 \| 398 \| (54, 490) \| 532 \| (41, 474) \| 518 \| Phage portal protein \| Phage portal protein; uniclust \| UniRef100\_A0A376LQ97 \| 99.7 \| 2.1e-21 \| 3.8e-27 \| 181.4 \| 232 \| (46, 286) \| 532 \| (91, 338) \| 375 \| Phage portal protein (Minor capsid protein) \| Phage portal protein (Minor capsid protein); uniclust \| UniRef100\_A0A965N5Y9 \| 99.7 \| 2.4e-21 \| 4.3e-27 \| 158.7 \| 122 \| (317, 441) \| 532 \| (22, 144) \| 145 \| Phage portal protein (Fragment) \| Phage portal protein (Fragment); uniclust \| UniRef100\_A0A3L7VQ06 \| 99.7 \| 2.4e-21 \| 4.4e-27 \| 194.9 \| 396 \| (72, 493) \| 532 \| (76, 492) \| 707 \| Phage portal protein \| Phage portal protein; uniclust \| UniRef100\_A0A6M3J4B5 \| 99.7 \| 2.7e-21 \| 4.9e-27 \| 181.0 \| 339 \| (161, 503) \| 532 \| (6, 362) \| 380 \| Putative portal protein \| Putative portal protein; uniclust \| UniRef100\_A0A847I6D6 \| 99.7 \| 3.4e-21 \| 6.3e-27 \| 177.9 \| 325 \| (65, 417) \| 532 \| (4, 339) \| 339 \| Phage portal protein (Fragment) \| Phage portal protein (Fragment); uniclust \| UniRef100\_A0A9E1ZJH4 \| 99.7 \| 4e-21 \| 7.3e-27 \| 160.1 \| 136 \| (318, 458) \| 532 \| (2, 137) \| 161 \| Phage portal protein \| Phage portal protein; uniclust \| UniRef100\_UPI0023560E02 \| 99.7 \| 4.7e-21 \| 8.6e-27 \| 185.1 \| 408 \| (74, 519) \| 532 \| (48, 481) \| 506 \| phage portal protein \| phage portal protein; uniclust \| UniRef100\_A0A3C1GA38 \| 99.7 \| 5.2e-21 \| 9.5e-27 \| 177.4 \| 179 \| (45, 226) \| 532 \| (79, 259) \| 349 \| Phage portal protein \| Phage portal protein; uniclust \| UniRef100\_J9FFD2 \| 99.7 \| 5.4e-21 \| 9.9e-27 \| 163.0 \| 176 \| (203, 382) \| 532 \| (6, 187) \| 187 \| Phage portal protein, lambda family (Fragment) \| Phage portal protein, lambda family (Fragment); uniclust \| UniRef100\_A0A1Q6JR94 \| 99.7 \| 5.8e-21 \| 1.1e-26 \| 166.8 \| 160 \| (14, 174) \| 532 \| (3, 172) \| 178 \| Phage portal protein \| Phage portal protein; uniclust \| UniRef100\_A0A0F9FPS3 \| 99.7 \| 8.3e-21 \| 1.5e-26 \| 183.1 \| 404 \| (48, 493) \| 532 \| (45, 492) \| 498 \| Phage portal protein \| Phage portal protein; uniclust \| UniRef100\_A0A2D7VYW9 \| 99.7 \| 1e-20 \| 1.9e-26 \| 160.4 \| 133 \| (366, 507) \| 532 \| (3, 137) \| 161 \| Phage portal protein (Fragment) \| Phage portal protein (Fragment); uniclust \| UniRef100\_A0A2W5HLR3 \| 99.7 \| 1.1e-20 \| 2e-26 \| 160.4 \| 138 \| (347, 495) \| 532 \| (2, 140) \| 167 \| Phage portal protein (Fragment) \| Phage portal protein (Fragment); uniclust \| UniRef100\_A0A970YYK5 \| 99.7 \| 1.3e-20 \| 2.3e-26 \| 177.6 \| 331 \| (45, 394) \| 532 \| (52, 394) \| 397 \| Phage portal protein (Fragment) \| Phage portal protein (Fragment); uniclust \| UniRef100\_A0A965PNA5 \| 99.7 \| 1.3e-20 \| 2.4e-26 \| 185.1 \| 240 \| (259, 507) \| 532 \| (2, 250) \| 558 \| Phage portal protein (Fragment) \| Phage portal protein (Fragment); uniclust \| UniRef100\_A0A074TJP9 \| 99.7 \| 1.2e-20 \| 2.4e-26 \| 174.8 \| 189 \| (317, 515) \| 532 \| (5, 195) \| 236 \| Capsid protein (Fragment) \| Capsid protein (Fragment); uniclust \| UniRef100\_A0A661KRT0 \| 99.7 \| 1.5e-20 \| 2.8e-26 \| 178.6 \| 314 \| (130, 478) \| 532 \| (2, 328) \| 431 \| Phage portal protein \| Phage portal protein; uniclust \| UniRef100\_A0A7V3JP37 \| 99.7 \| 2.3e-20 \| 4.3e-26 \| 177.3 \| 292 \| (72, 393) \| 532 \| (60, 369) \| 382 \| Phage portal protein (Fragment) \| Phage portal protein (Fragment); uniclust \| UniRef100\_UPI002107534C \| 99.7 \| 2.9e-20 \| 5.4e-26 \| 150.2 \| 116 \| (327, 445) \| 532 \| (1, 116) \| 132 \| phage portal protein \| phage portal protein; uniclust \| UniRef100\_A0A431I932 \| 99.7 \| 3.1e-20 \| 5.8e-26 \| 180.4 \| 407 \| (63, 494) \| 532 \| (79, 511) \| 528 \| Phage portal protein \| Phage portal protein; uniclust \| UniRef100\_A0A9D8TK68 \| 99.7 \| 3.7e-20 \| 6.8e-26 \| 158.5 \| 166 \| (58, 227) \| 532 \| (7, 177) \| 190 \| Phage portal protein (Fragment) \| Phage portal protein (Fragment); uniclust \| UniRef100\_A0A7C3DW82 \| 99.7 \| 4.3e-20 \| 7.9e-26 \| 169.1 \| 249 \| (9, 301) \| 532 \| (25, 277) \| 310 \| Phage portal protein (Fragment) \| Phage portal protein (Fragment); uniclust \| UniRef100\_UPI00147320C4 \| 99.7 \| 4.3e-20 \| 7.9e-26 \| 187.8 \| 180 \| (318, 505) \| 532 \| (691, 872) \| 886 \| phage terminase large subunit family protein \| phage terminase large subunit family protein; uniclust \| UniRef100\_UPI0022B34460 \| 99.7 \| 4.5e-20 \| 8.3e-26 \| 164.7 \| 228 \| (122, 373) \| 532 \| (7, 239) \| 255 \| phage portal protein \| phage portal protein; uniclust \| UniRef100\_A0A358C6E0 \| 99.7 \| 4.8e-20 \| 8.8e-26 \| 163.7 \| 184 \| (316, 513) \| 532 \| (33, 227) \| 245 \| Phage portal protein (Fragment) \| Phage portal protein (Fragment); uniclust \| UniRef100\_A0A661H6X6 \| 99.7 \| 4.8e-20 \| 8.9e-26 \| 161.6 \| 160 \| (343, 506) \| 532 \| (14, 176) \| 224 \| Phage portal protein (Fragment) \| Phage portal protein (Fragment); uniclust \| UniRef100\_UPI001C83F08B \| 99.7 \| 5.6e-20 \| 1e-25 \| 160.7 \| 181 \| (203, 383) \| 532 \| (8, 218) \| 219 \| phage portal protein \| phage portal protein; uniclust \| UniRef100\_A0A358B2C7 \| 99.7 \| 1e-19 \| 1.8e-25 \| 175.6 \| 194 \| (44, 254) \| 532 \| (16, 209) \| 488 \| LysR substrate-binding domain-containing protein \| LysR substrate-binding domain-containing protein; uniclust \| UniRef100\_A0A0F9HE98 \| 99.7 \| 9.7e-20 \| 1.9e-25 \| 181.5 \| 375 \| (68, 495) \| 532 \| (42, 442) \| 494 \| Portal protein \| Portal protein; uniclust \| UniRef100\_UPI0006A04A95 \| 99.7 \| 1.2e-19 \| 2.2e-25 \| 168.8 \| 170 \| (340, 513) \| 532 \| (162, 332) \| 349 \| phage portal protein \| phage portal protein; uniclust \| UniRef100\_A0A2E9Y0J4 \| 99.7 \| 1.4e-19 \| 2.6e-25 \| 160.1 \| 168 \| (18, 197) \| 532 \| (1, 168) \| 219 \| Phage portal protein \| Phage portal protein; uniclust \| UniRef100\_UPI0020A414D4 \| 99.7 \| 1.9e-19 \| 3.5e-25 \| 164.6 \| 242 \| (257, 503) \| 532 \| (8, 277) \| 304 \| phage portal protein \| phage portal protein; uniclust \| UniRef100\_A0A376U7U2 \| 99.7 \| 1.9e-19 \| 3.6e-25 \| 153.3 \| 148 \| (319, 466) \| 532 \| (17, 165) \| 165 \| Capsid protein of prophage \| Capsid protein of prophage; uniclust \| UniRef100\_A0A0F9F8Y8 \| 99.7 \| 1.9e-19 \| 3.7e-25 \| 159.5 \| 152 \| (15, 173) \| 532 \| (7, 173) \| 186 \| Uncharacterized protein \| Uncharacterized protein; uniclust \| UniRef100\_A0A1Y5TZG3 \| 99.7 \| 2e-19 \| 3.7e-25 \| 157.1 \| 174 \| (209, 387) \| 532 \| (2, 177) \| 216 \| Phage portal protein, lambda family \| Phage portal protein, lambda family; uniclust \| UniRef100\_UPI0012D43280 \| 99.7 \| 2.3e-19 \| 4.2e-25 \| 142.3 \| 108 \| (158, 266) \| 532 \| (4, 117) \| 117 \| phage portal protein \| phage portal protein; uniclust \| UniRef100\_A0A4P9VMD7 \| 99.6 \| 2.5e-19 \| 4.8e-25 \| 161.3 \| 184 \| (54, 245) \| 532 \| (26, 216) \| 216 \| Phage portal protein \| Phage portal protein; uniclust \| UniRef100\_A0A4Q8RPB7 \| 99.6 \| 3.3e-19 \| 6.2e-25 \| 155.4 \| 168 \| (219, 390) \| 532 \| (2, 197) \| 197 \| Phage portal protein (Fragment) \| Phage portal protein (Fragment); uniclust \| UniRef100\_A0A1F8QWT7 \| 99.6 \| 3.4e-19 \| 6.7e-25 \| 179.2 \| 299 \| (80, 417) \| 532 \| (7, 313) \| 439 \| Uncharacterized protein \| Uncharacterized protein; uniclust \| UniRef100\_A0A941SHU7 \| 99.6 \| 3.8e-19 \| 7.1e-25 \| 172.6 \| 434 \| (45, 505) \| 532 \| (33, 484) \| 513 \| Phage portal protein \| Phage portal protein; uniclust \| UniRef100\_A0A419ENK9 \| 99.6 \| 4.4e-19 \| 8.1e-25 \| 152.9 \| 167 \| (344, 519) \| 532 \| (11, 178) \| 196 \| Phage portal protein (Fragment) \| Phage portal protein (Fragment); uniclust \| UniRef100\_UPI00210FF08D \| 99.6 \| 4.9e-19 \| 9e-25 \| 157.6 \| 168 \| (274, 447) \| 532 \| (3, 175) \| 246 \| phage portal protein \| phage portal protein; uniclust \| UniRef100\_A0A844M4D9 \| 99.6 \| 5.6e-19 \| 1.1e-24 \| 140.2 \| 99 \| (344, 453) \| 532 \| (2, 101) \| 102 \| Phage portal protein (Fragment) \| Phage portal protein (Fragment); uniclust \| UniRef100\_A0A142WUX4 \| 99.6 \| 6e-19 \| 1.1e-24 \| 170.8 \| 402 \| (46, 477) \| 532 \| (41, 456) \| 497 \| Phage portal protein, lambda family \| Phage portal protein, lambda family; uniclust \| UniRef100\_A0A6P1B243 \| 99.6 \| 6.4e-19 \| 1.2e-24 \| 141.8 \| 126 \| (96, 222) \| 532 \| (2, 127) \| 127 \| Phage portal protein (Fragment) \| Phage portal protein (Fragment); uniclust \| UniRef100\_UPI000EB38119 \| 99.6 \| 9.4e-19 \| 1.7e-24 \| 153.4 \| 168 \| (17, 198) \| 532 \| (4, 174) \| 219 \| phage portal protein \| phage portal protein; uniclust \| UniRef100\_A0A376SKF2 \| 99.6 \| 1.1e-18 \| 2.1e-24 \| 150.9 \| 167 \| (123, 300) \| 532 \| (3, 174) \| 200 \| Head-tail preconnector protein \| Head-tail preconnector protein; uniclust \| UniRef100\_UPI0004A80172 \| 99.6 \| 2e-18 \| 3.7e-24 \| 155.0 \| 177 \| (319, 502) \| 532 \| (64, 240) \| 259 \| phage portal protein \| phage portal protein; uniclust \| UniRef100\_UPI001FE83FA2 \| 99.6 \| 2.4e-18 \| 4.5e-24 \| 166.8 \| 138 \| (361, 505) \| 532 \| (322, 460) \| 500 \| ParB N-terminal domain-containing protein \| ParB N-terminal domain-containing protein; uniclust \| UniRef100\_A0A431KHF4 \| 99.6 \| 2.6e-18 \| 4.7e-24 \| 173.7 \| 387 \| (56, 488) \| 532 \| (325, 729) \| 784 \| Phage portal protein \| Phage portal protein; uniclust \| UniRef100\_A0A5C5X9Q3 \| 99.6 \| 3e-18 \| 5.6e-24 \| 172.4 \| 398 \| (74, 491) \| 532 \| (87, 525) \| 559 \| Phage portal protein, lambda family \| Phage portal protein, lambda family; uniclust \| UniRef100\_A0A175RV87 \| 99.6 \| 3.2e-18 \| 5.8e-24 \| 162.4 \| 318 \| (78, 419) \| 532 \| (68, 398) \| 406 \| Portal protein \| Portal protein; uniclust \| UniRef100\_X1FZP5 \| 99.6 \| 3.2e-18 \| 5.9e-24 \| 151.3 \| 204 \| (11, 218) \| 532 \| (16, 230) \| 231 \| Phage portal protein (Fragment) \| Phage portal protein (Fragment); uniclust \| UniRef100\_A0A3S4U6Y1 \| 99.6 \| 3.4e-18 \| 6.2e-24 \| 161.5 \| 177 \| (323, 509) \| 532 \| (1, 178) \| 390 \| ATP-dependent Clp protease proteolytic subunit \| ATP-dependent Clp protease proteolytic subunit; uniclust \| UniRef100\_A0A934HCI9 \| 99.6 \| 4.3e-18 \| 7.9e-24 \| 164.7 \| 367 \| (73, 489) \| 532 \| (96, 481) \| 484 \| Uncharacterized protein \| Uncharacterized protein; uniclust \| UniRef100\_UPI000A67C2A5 \| 99.6 \| 5e-18 \| 9.4e-24 \| 138.2 \| 118 \| (214, 335) \| 532 \| (6, 127) \| 128 \| phage portal protein \| phage portal protein; uniclust \| UniRef100\_A0A0K0VKK6 \| 99.6 \| 5.7e-18 \| 1e-23 \| 153.8 \| 187 \| (317, 507) \| 532 \| (75, 263) \| 281 \| Plasmid partitioning protein ParB \| Plasmid partitioning protein ParB; uniclust \| UniRef100\_UPI001F204F94 \| 99.6 \| 7.1e-18 \| 1.3e-23 \| 130.9 \| 95 \| (325, 419) \| 532 \| (2, 97) \| 102 \| phage portal protein \| phage portal protein; uniclust \| UniRef100\_UPI002269D2B2 \| 99.6 \| 7.1e-18 \| 1.3e-23 \| 146.9 \| 170 \| (45, 214) \| 532 \| (31, 200) \| 207 \| phage portal protein \| phage portal protein; uniclust \| UniRef100\_UPI000CEF4514 \| 99.6 \| 7.4e-18 \| 1.4e-23 \| 154.1 \| 247 \| (248, 495) \| 532 \| (5, 269) \| 296 \| phage portal protein \| phage portal protein; uniclust \| UniRef100\_F3IS71 \| 99.6 \| 8.3e-18 \| 1.5e-23 \| 136.6 \| 126 \| (127, 255) \| 532 \| (3, 131) \| 132 \| Portal protein (Fragment) \| Portal protein (Fragment); uniclust \| UniRef100\_A0A1V5NFS7 \| 99.6 \| 9.9e-18 \| 1.9e-23 \| 145.0 \| 145 \| (357, 510) \| 532 \| (1, 145) \| 168 \| Phage portal protein, lambda family \| Phage portal protein, lambda family; uniclust \| UniRef100\_A0A6J4FY55 \| 99.6 \| 1e-17 \| 1.9e-23 \| 150.7 \| 165 \| (44, 219) \| 532 \| (14, 178) \| 261 \| Phage portal protein \| Phage portal protein; uniclust \| UniRef100\_A0A517YS81 \| 99.5 \| 1.1e-17 \| 2.2e-23 \| 149.9 \| 165 \| (43, 225) \| 532 \| (29, 197) \| 212 \| Phage portal protein, lambda family \| Phage portal protein, lambda family; uniclust \| UniRef100\_A0A1U7AEE5 \| 99.5 \| 1.2e-17 \| 2.2e-23 \| 157.3 \| 238 \| (61, 301) \| 532 \| (56, 308) \| 376 \| Phage portal protein \| Phage portal protein; uniclust \| UniRef100\_A0A7H4P585 \| 99.5 \| 1.6e-17 \| 3.2e-23 \| 155.5 \| 178 \| (45, 226) \| 532 \| (37, 223) \| 253 \| Phage portal protein, lambda family \| Phage portal protein, lambda family; uniclust \| UniRef100\_A0A1H2PWF9 \| 99.5 \| 2.4e-17 \| 4.4e-23 \| 138.1 \| 137 \| (341, 488) \| 532 \| (5, 142) \| 159 \| Phage portal protein, lambda family (Fragment) \| Phage portal protein, lambda family (Fragment); uniclust \| UniRef100\_UPI00186BB8CC \| 99.5 \| 2.7e-17 \| 4.9e-23 \| 168.8 \| 170 \| (17, 198) \| 532 \| (700, 874) \| 917 \| phage terminase large subunit family protein \| phage terminase large subunit family protein; uniclust \| UniRef100\_A0A1V5SIV4 \| 99.5 \| 2.7e-17 \| 5e-23 \| 134.9 \| 104 \| (393, 503) \| 532 \| (2, 105) \| 139 \| Phage portal protein, lambda family \| Phage portal protein, lambda family; uniclust \| UniRef100\_UPI00052DD37F \| 99.5 \| 2.6e-17 \| 5e-23 \| 137.8 \| 130 \| (135, 275) \| 532 \| (2, 136) \| 138 \| phage portal protein \| phage portal protein; uniclust \| UniRef100\_A0A659UH34 \| 99.5 \| 2.9e-17 \| 5.3e-23 \| 134.7 \| 125 \| (343, 467) \| 532 \| (9, 138) \| 139 \| Phage portal protein (Fragment) \| Phage portal protein (Fragment); uniclust \| UniRef100\_UPI0020205CB6 \| 99.5 \| 3.1e-17 \| 5.7e-23 \| 144.2 \| 169 \| (319, 492) \| 532 \| (32, 201) \| 218 \| phage portal protein \| phage portal protein; uniclust \| UniRef100\_A0A662DL46 \| 99.5 \| 3.2e-17 \| 5.8e-23 \| 153.8 \| 286 \| (207, 495) \| 532 \| (22, 315) \| 359 \| Phage portal protein (Fragment) \| Phage portal protein (Fragment); uniclust \| UniRef100\_X0XGG6 \| 99.5 \| 3.3e-17 \| 6e-23 \| 146.1 \| 168 \| (319, 493) \| 532 \| (43, 213) \| 241 \| Phage portal protein (Fragment) \| Phage portal protein (Fragment); uniclust \| UniRef100\_A0A5R2MYS9 \| 99.5 \| 3.6e-17 \| 7.3e-23 \| 143.9 \| 105 \| (316, 420) \| 532 \| (36, 140) \| 155 \| Phage portal protein (Fragment) \| Phage portal protein (Fragment); uniclust \| UniRef100\_A0A3C0YQ03 \| 99.5 \| 4.9e-17 \| 9.1e-23 \| 151.1 \| 240 \| (249, 494) \| 532 \| (28, 279) \| 331 \| Phage portal protein \| Phage portal protein; uniclust \| UniRef100\_A0A377MXP9 \| 99.5 \| 5.2e-17 \| 9.5e-23 \| 144.8 \| 163 \| (347, 514) \| 532 \| (60, 225) \| 240 \| Phage portal protein \| Phage portal protein; uniclust \| UniRef100\_A0A268UA88 \| 99.5 \| 6e-17 \| 1.1e-22 \| 149.5 \| 240 \| (67, 403) \| 532 \| (50, 290) \| 313 \| Phage portal protein \| Phage portal protein; uniclust \| UniRef100\_UPI001EEF44AD \| 99.5 \| 6.7e-17 \| 1.2e-22 \| 126.6 \| 103 \| (182, 285) \| 532 \| (1, 104) \| 106 \| phage portal protein \| phage portal protein; uniclust \| UniRef100\_A0A2S1XDV8 \| 99.5 \| 7.9e-17 \| 1.5e-22 \| 146.7 \| 187 \| (32, 231) \| 532 \| (88, 280) \| 281 \| Phage portal protein \| Phage portal protein; uniclust \| UniRef100\_UPI000500391C \| 99.5 \| 8.1e-17 \| 1.5e-22 \| 135.1 \| 139 \| (88, 231) \| 532 \| (4, 147) \| 159 \| phage portal protein \| phage portal protein; uniclust \| UniRef100\_A0A966N5E4 \| 99.5 \| 8.2e-17 \| 1.5e-22 \| 152.4 \| 222 \| (66, 300) \| 532 \| (62, 302) \| 320 \| Phage portal protein \| Phage portal protein; uniclust \| UniRef100\_UPI00163E7870 \| 99.5 \| 1.2e-16 \| 2.2e-22 \| 132.1 \| 126 \| (239, 364) \| 532 \| (8, 144) \| 144 \| phage portal protein \| phage portal protein; uniclust \| UniRef100\_A0A660UHE4 \| 99.5 \| 1.2e-16 \| 2.2e-22 \| 140.8 \| 178 \| (45, 232) \| 532 \| (30, 212) \| 219 \| Phage portal protein (Fragment) \| Phage portal protein (Fragment); uniclust \| UniRef100\_A0A1H0DUF0 \| 99.5 \| 1.1e-16 \| 2.2e-22 \| 165.1 \| 371 \| (55, 469) \| 532 \| (40, 433) \| 544 \| Phage portal protein, SPP1 Gp6-like \| Phage portal protein, SPP1 Gp6-like; uniclust \| UniRef100\_A0A142XP47 \| 99.5 \| 1.3e-16 \| 2.5e-22 \| 171.3 \| 373 \| (87, 490) \| 532 \| (4, 400) \| 1882 \| Phage portal protein, lambda family \| Phage portal protein, lambda family; uniclust \| UniRef100\_A0A843Y6R3 \| 99.5 \| 1.7e-16 \| 3e-22 \| 146.2 \| 238 \| (73, 329) \| 532 \| (46, 289) \| 304 \| Phage portal protein \| Phage portal protein; uniclust \| UniRef100\_UPI001CFD494D \| 99.5 \| 1.8e-16 \| 3.3e-22 \| 138.8 \| 179 \| (317, 505) \| 532 \| (9, 189) \| 210 \| phage portal protein \| phage portal protein; uniclust \| UniRef100\_UPI0009670FCD \| 99.5 \| 1.9e-16 \| 3.6e-22 \| 135.8 \| 148 \| (317, 464) \| 532 \| (32, 181) \| 182 \| phage portal protein \| phage portal protein; uniclust \| UniRef100\_UPI0013DF9E24 \| 99.5 \| 2e-16 \| 3.6e-22 \| 138.0 \| 177 \| (326, 506) \| 532 \| (1, 178) \| 203 \| phage portal protein \| phage portal protein; uniclust \| UniRef100\_A0A961GYB0 \| 99.5 \| 2e-16 \| 3.7e-22 \| 135.7 \| 158 \| (45, 220) \| 532 \| (23, 182) \| 182 \| Phage portal protein (Fragment) \| Phage portal protein (Fragment); uniclust \| UniRef100\_A0A5C7PU34 \| 99.5 \| 2.1e-16 \| 3.9e-22 \| 148.0 \| 209 \| (269, 485) \| 532 \| (25, 234) \| 349 \| Phage portal protein \| Phage portal protein; uniclust \| UniRef100\_A0A352V3P2 \| 99.5 \| 2.4e-16 \| 4.3e-22 \| 138.2 \| 150 \| (19, 175) \| 532 \| (6, 157) \| 211 \| Phage portal protein \| Phage portal protein; uniclust \| UniRef100\_A0A379XPD9 \| 99.5 \| 2.3e-16 \| 4.4e-22 \| 137.1 \| 143 \| (356, 508) \| 532 \| (1, 144) \| 178 \| Phage portal protein, lambda family protein \| Phage portal protein, lambda family protein; uniclust \| UniRef100\_A0A661I1H8 \| 99.4 \| 3e-16 \| 5.6e-22 \| 135.8 \| 163 \| (45, 229) \| 532 \| (15, 179) \| 193 \| Phage portal protein (Fragment) \| Phage portal protein (Fragment); uniclust \| UniRef100\_F2J6P5 \| 99.4 \| 3.1e-16 \| 5.8e-22 \| 136.9 \| 128 \| (240, 367) \| 532 \| (17, 148) \| 205 \| Portal protein GPB of phage lambda-like protein \| Portal protein GPB of phage lambda-like protein; uniclust \| UniRef100\_A0A369RJ94 \| 99.4 \| 3.5e-16 \| 6.6e-22 \| 126.1 \| 105 \| (396, 508) \| 532 \| (6, 110) \| 116 \| Phage portal protein \| Phage portal protein; uniclust \| UniRef100\_B4CWU6 \| 99.4 \| 3.8e-16 \| 6.9e-22 \| 140.5 \| 177 \| (99, 301) \| 532 \| (33, 209) \| 252 \| Bacteriophage capsid protein-like protein \| Bacteriophage capsid protein-like protein; uniclust \| UniRef100\_A0A961K8S9 \| 99.4 \| 3.7e-16 \| 7e-22 \| 126.0 \| 104 \| (397, 507) \| 532 \| (2, 105) \| 115 \| Phage portal protein \| Phage portal protein; uniclust \| UniRef100\_A0A1H5ZJX6 \| 99.4 \| 4e-16 \| 7.7e-22 \| 157.7 \| 385 \| (79, 531) \| 532 \| (65, 475) \| 496 \| Portal protein \| Portal protein; uniclust \| UniRef100\_A0A076JPG9 \| 99.4 \| 3.7e-16 \| 7.9e-22 \| 167.5 \| 275 \| (148, 481) \| 532 \| (126, 404) \| 513 \| Phage portal protein \| Phage portal protein; uniclust \| UniRef100\_A0A2X1JPZ6 \| 99.4 \| 4.4e-16 \| 8.1e-22 \| 133.3 \| 151 \| (357, 517) \| 532 \| (2, 153) \| 168 \| Portal protein (Head-tail preconnector protein) from prophage \| Portal protein (Head-tail preconnector protein) from prophage; uniclust \| UniRef100\_A0A2X1K6P9 \| 99.4 \| 8e-16 \| 1.5e-21 \| 131.0 \| 158 \| (240, 398) \| 532 \| (6, 167) \| 167 \| Portal protein \| Portal protein; uniclust \| UniRef100\_UPI0018CC948E \| 99.4 \| 8.3e-16 \| 1.5e-21 \| 141.4 \| 166 \| (327, 503) \| 532 \| (1, 167) \| 298 \| phage portal protein \| phage portal protein; uniclust \| UniRef100\_A0A7W1KP99 \| 99.4 \| 1.2e-15 \| 2.2e-21 \| 149.8 \| 396 \| (54, 487) \| 532 \| (52, 477) \| 524 \| Phage portal protein \| Phage portal protein; uniclust \| UniRef100\_A0A8J2Z721 \| 99.4 \| 1.4e-15 \| 2.5e-21 \| 129.3 \| 138 \| (43, 192) \| 532 \| (27, 164) \| 169 \| Uncharacterized protein \| Uncharacterized protein; uniclust \| UniRef100\_UPI001FE2508B \| 99.4 \| 1.5e-15 \| 2.7e-21 \| 127.2 \| 116 \| (74, 198) \| 532 \| (6, 123) \| 154 \| phage portal protein \| phage portal protein; uniclust \| UniRef100\_A0A3R5RY70 \| 99.4 \| 2.2e-15 \| 4e-21 \| 139.3 \| 206 \| (73, 300) \| 532 \| (49, 257) \| 307 \| Phage portal protein \| Phage portal protein; uniclust \| UniRef100\_UPI0011BA491B \| 99.4 \| 2.2e-15 \| 4.1e-21 \| 129.0 \| 160 \| (240, 399) \| 532 \| (8, 174) \| 177 \| phage portal protein \| phage portal protein; uniclust \| UniRef100\_UPI00227E5E2A \| 99.4 \| 2.6e-15 \| 4.7e-21 \| 124.1 \| 112 \| (48, 160) \| 532 \| (21, 137) \| 141 \| phage portal protein \| phage portal protein; uniclust \| UniRef100\_B6Y8D8 \| 99.4 \| 2.6e-15 \| 4.9e-21 \| 123.2 \| 92 \| (44, 144) \| 532 \| (20, 111) \| 114 \| Putative phage portal protein \| Putative phage portal protein; uniclust \| UniRef100\_A0A939WQF3 \| 99.4 \| 2.8e-15 \| 5.2e-21 \| 147.8 \| 438 \| (45, 506) \| 532 \| (24, 505) \| 544 \| Phage portal protein \| Phage portal protein; uniclust \| UniRef100\_A0A0F8YUP6 \| 99.4 \| 2.8e-15 \| 5.3e-21 \| 119.2 \| 96 \| (16, 112) \| 532 \| (1, 99) \| 101 \| Uncharacterized protein (Fragment) \| Uncharacterized protein (Fragment); uniclust \| UniRef100\_UPI00210222F9 \| 99.4 \| 2.9e-15 \| 5.3e-21 \| 121.7 \| 117 \| (179, 300) \| 532 \| (2, 118) \| 127 \| phage portal protein \| phage portal protein; uniclust \| UniRef100\_UPI00057803E2 \| 99.4 \| 3.1e-15 \| 5.8e-21 \| 115.5 \| 80 \| (151, 231) \| 532 \| (2, 83) \| 86 \| phage portal protein \| phage portal protein; uniclust \| UniRef100\_A0A447U5T1 \| 99.3 \| 3.7e-15 \| 6.7e-21 \| 137.8 \| 178 \| (46, 227) \| 532 \| (87, 273) \| 304 \| Phage portal protein, lambda family \| Phage portal protein, lambda family; uniclust \| UniRef100\_UPI001787C57E \| 99.3 \| 4.2e-15 \| 7.7e-21 \| 129.7 \| 167 \| (253, 419) \| 532 \| (1, 189) \| 199 \| phage portal protein \| phage portal protein; uniclust \| UniRef100\_A0A8J7QZX4 \| 99.3 \| 4.9e-15 \| 9e-21 \| 124.0 \| 125 \| (369, 505) \| 532 \| (2, 126) \| 152 \| Phage portal protein, lambda family \| Phage portal protein, lambda family; uniclust \| UniRef100\_UPI0022CDD105 \| 99.3 \| 5.5e-15 \| 1e-20 \| 140.9 \| 288 \| (73, 369) \| 532 \| (65, 390) \| 392 \| phage portal protein \| phage portal protein; uniclust \| UniRef100\_A0A7X6J9Z5 \| 99.3 \| 5.7e-15 \| 1e-20 \| 120.8 \| 127 \| (319, 450) \| 532 \| (1, 130) \| 132 \| Phage portal protein \| Phage portal protein; uniclust \| UniRef100\_UPI001111D749 \| 99.3 \| 5.7e-15 \| 1.1e-20 \| 115.7 \| 95 \| (323, 417) \| 532 \| (4, 98) \| 98 \| phage portal protein \| phage portal protein; uniclust \| UniRef100\_A0A2G4JPU1 \| 99.3 \| 6.5e-15 \| 1.2e-20 \| 123.3 \| 140 \| (370, 514) \| 532 \| (2, 143) \| 152 \| Phage portal protein \| Phage portal protein; uniclust \| UniRef100\_A0A2Z3H0N2 \| 99.3 \| 6.6e-15 \| 1.3e-20 \| 156.0 \| 393 \| (72, 495) \| 532 \| (89, 505) \| 923 \| Phage portal protein \| Phage portal protein; uniclust \| UniRef100\_A0A0A7KFK9 \| 99.3 \| 7.1e-15 \| 1.4e-20 \| 151.9 \| 365 \| (72, 487) \| 532 \| (47, 443) \| 476 \| Phage portal protein \| Phage portal protein; uniclust \| UniRef100\_UPI00191C467D \| 99.3 \| 8e-15 \| 1.5e-20 \| 124.9 \| 123 \| (18, 147) \| 532 \| (1, 126) \| 169 \| phage portal protein \| phage portal protein; uniclust \| UniRef100\_A0A2M9FV85 \| 99.3 \| 8.9e-15 \| 1.6e-20 \| 124.3 \| 131 \| (374, 511) \| 532 \| (1, 132) \| 166 \| Uncharacterized protein \| Uncharacterized protein; uniclust \| UniRef100\_A0A6N7AFZ5 \| 99.3 \| 1.6e-14 \| 2.9e-20 \| 118.8 \| 97 \| (93, 198) \| 532 \| (26, 124) \| 135 \| Lambda family phage portal protein (Fragment) \| Lambda family phage portal protein (Fragment); uniclust \| UniRef100\_A0A3C1GDR3 \| 99.3 \| 1.6e-14 \| 3e-20 \| 119.9 \| 139 \| (75, 230) \| 532 \| (3, 143) \| 143 \| Phage portal protein (Fragment) \| Phage portal protein (Fragment) |
| Top keywords  (threshold 1.00e-03 (evalue)) | **portal, Phage, Fragment, lambda, Capsid, Putative, prophage, terminase, Minor, domain\_containing** |
| Output files | ../../similar\_sequences/04\_FANPEZAQ\_CDS\_0004\_merged.svg ../../similar\_sequences/04\_FANPEZAQ\_CDS\_0004\_pdb70.a3m ../../similar\_sequences/04\_FANPEZAQ\_CDS\_0004\_pdb70.hhr ../../similar\_sequences/04\_FANPEZAQ\_CDS\_0004\_uniclust.a3m ../../similar\_sequences/04\_FANPEZAQ\_CDS\_0004\_uniclust.hhr |

#### Structure prediction (AlphaFold)^2^

|  |  |
| --- | --- |
| Stats | xml version="1.0" encoding="utf-8" standalone="no"?       2024-09-02T21:09:02.438651 image/svg+xml   Matplotlib v3.7.2, https://matplotlib.org/ |
| Predicted structure | **NGL Viewer Controls:**  - Center: *Left-Click* - Rotate: *Left-Click + Drag* - Translate: *Right-Click + Drag* - Zoom: *Shift + Left-Click + Drag* |
| Output files | ../../predicted\_structures/04\_FANPEZAQ\_CDS\_0004/features.pkl ../../predicted\_structures/04\_FANPEZAQ\_CDS\_0004/ranked\_0.pdb ../../predicted\_structures/04\_FANPEZAQ\_CDS\_0004/ranked\_0\_plots.svg ../../predicted\_structures/04\_FANPEZAQ\_CDS\_0004/result\_model\_1\_ptm\_pred\_0.pkl |

#### Structure similarity search results (Foldseek)^3^

|  |  |
| --- | --- |
| Structure databases searched | Pdb, Afdb-proteome, Afdb-uniprot50 |
| Results, scheme(s)  (Top layers only, threshold 1.00e-02 (evalue)) | xml version="1.0" encoding="utf-8" standalone="no"?       2024-09-02T21:10:12.685353 image/svg+xml   Matplotlib v3.7.2, https://matplotlib.org/ |
| Results, table  (threshold 1.00e-02 (evalue)) | db \| id \| prob \| evalue \| bits \| fident \| alnlen \| mismatch \| gapopen \| qstart \| qend \| tstart \| tend \| name \| description; pdb \| 8GTD\_A \| 1.0 \| 1.243e-31 \| 1160 \| 0.328 \| 456 \| 277 \| 10 \| 47 \| 495 \| 1 \| 434 \| Portal protein \| Portal protein; pdb \| 6QJT\_A \| 1.0 \| 5.678e-09 \| 244 \| 0.124 \| 419 \| 282 \| 21 \| 68 \| 466 \| 23 \| 376 \| Portal protein \| Portal protein; pdb \| 4ZJN\_A \| 1.0 \| 3.578e-08 \| 232 \| 0.114 \| 447 \| 284 \| 21 \| 70 \| 475 \| 26 \| 401 \| Portal protein \| Portal protein; pdb \| 8FQL\_A \| 1.0 \| 1.317e-08 \| 231 \| 0.13 \| 420 \| 261 \| 26 \| 83 \| 478 \| 4 \| 343 \| Portal protein \| Portal protein; pdb \| 6TO8\_A \| 1.0 \| 4.906e-08 \| 226 \| 0.129 \| 457 \| 272 \| 22 \| 48 \| 478 \| 3 \| 359 \| Portal protein Rcc01684 \| Portal protein Rcc01684; pdb \| 5NGD\_A \| 1.0 \| 3.578e-08 \| 222 \| 0.122 \| 482 \| 289 \| 26 \| 70 \| 517 \| 21 \| 402 \| Portal protein \| Portal protein; pdb \| 6UZC\_H \| 1.0 \| 2.447e-09 \| 219 \| 0.147 \| 481 \| 312 \| 22 \| 66 \| 497 \| 58 \| 489 \| Portal protein \| Portal protein; pdb \| 6TOA\_B \| 1.0 \| 3.772e-08 \| 211 \| 0.142 \| 464 \| 264 \| 26 \| 48 \| 478 \| 3 \| 365 \| Portal protein Rcc01684 \| Portal protein Rcc01684; pdb \| 6TBA\_1A \| 1.0 \| 1.2e-07 \| 206 \| 0.13 \| 459 \| 269 \| 25 \| 48 \| 478 \| 3 \| 359 \| Portal protein Rcc01684 \| Portal protein Rcc01684; pdb \| 8CEZ\_A \| 1.0 \| 5.45e-08 \| 202 \| 0.125 \| 423 \| 260 \| 24 \| 83 \| 478 \| 24 \| 363 \| Portal protein \| Portal protein; pdb \| 3JA7\_A \| 1.0 \| 6.055e-08 \| 202 \| 0.134 \| 477 \| 300 \| 22 \| 76 \| 498 \| 5 \| 422 \| Portal protein gp20 \| Portal protein gp20; pdb \| 6TUI\_A \| 1.0 \| 1.827e-07 \| 200 \| 0.141 \| 465 \| 267 \| 29 \| 48 \| 478 \| 3 \| 369 \| Portal protein Rcc01684 \| Portal protein Rcc01684; pdb \| 8FVH\_U \| 1.0 \| 4.709e-07 \| 188 \| 0.122 \| 500 \| 296 \| 28 \| 68 \| 500 \| 5 \| 428 \| E217 portal protein gp19 \| E217 portal protein gp19; pdb \| 6QWP\_A \| 1.0 \| 7.876e-08 \| 185 \| 0.122 \| 539 \| 303 \| 31 \| 68 \| 519 \| 16 \| 471 \| Portal protein \| Portal protein; pdb \| 7BOU\_A \| 1.0 \| 8.301e-08 \| 174 \| 0.119 \| 577 \| 341 \| 34 \| 56 \| 519 \| 2 \| 524 \| Portal protein \| Portal protein; pdb \| 2JES\_M \| 1.0 \| 3.906e-05 \| 163 \| 0.091 \| 416 \| 258 \| 22 \| 90 \| 491 \| 60 \| 369 \| PORTAL PROTEIN \| PORTAL PROTEIN; pdb \| 7Z4W\_A \| 1.0 \| 4.52e-06 \| 161 \| 0.112 \| 464 \| 274 \| 29 \| 91 \| 500 \| 66 \| 445 \| Head completion protein gp15 \| Head completion protein gp15; pdb \| 6QX5\_I \| 1.0 \| 1.498e-06 \| 156 \| 0.125 \| 533 \| 315 \| 30 \| 68 \| 525 \| 14 \| 470 \| Portal protein \| Portal protein; pdb \| 7EY6\_A \| 1.0 \| 1.849e-06 \| 154 \| 0.112 \| 480 \| 306 \| 25 \| 68 \| 467 \| 13 \| 452 \| Portal protein \| Portal protein; pdb \| 7Y1C\_A \| 1.0 \| 4.709e-07 \| 153 \| 0.111 \| 576 \| 303 \| 33 \| 68 \| 518 \| 13 \| 504 \| phage connector protein \| phage connector protein; pdb \| 8DGF\_H \| 1.0 \| 2.054e-06 \| 143 \| 0.109 \| 477 \| 287 \| 27 \| 68 \| 464 \| 15 \| 433 \| Portal protein \| Portal protein; pdb \| 8CK0\_A \| 1.0 \| 3.663e-06 \| 141 \| 0.109 \| 521 \| 306 \| 28 \| 59 \| 499 \| 35 \| 477 \| Portal protein \| Portal protein; pdb \| 6QXM\_A \| 1.0 \| 9.44e-06 \| 140 \| 0.117 \| 409 \| 260 \| 24 \| 68 \| 416 \| 6 \| 373 \| Portal protein \| Portal protein; pdb \| 6R21\_A \| 1.0 \| 1.228e-05 \| 137 \| 0.118 \| 481 \| 287 \| 27 \| 68 \| 458 \| 16 \| 449 \| Tail tubular protein gp12 \| Tail tubular protein gp12; pdb \| 8PHO\_A \| 1.0 \| 1.037e-06 \| 130 \| 0.112 \| 382 \| 229 \| 24 \| 83 \| 432 \| 5 \| 308 \| Phage portal protein \| Phage portal protein; pdb \| 8PHP\_A \| 1.0 \| 7.257e-06 \| 127 \| 0.093 \| 437 \| 244 \| 23 \| 83 \| 487 \| 5 \| 321 \| Phage portal protein \| Phage portal protein; pdb \| 8PHU\_PF \| 1.0 \| 1.684e-05 \| 125 \| 0.097 \| 449 \| 238 \| 26 \| 83 \| 487 \| 5 \| 330 \| Portal protein \| Portal protein; pdb \| 8PHU\_PG \| 1.0 \| 1.684e-05 \| 125 \| 0.103 \| 446 \| 243 \| 26 \| 83 \| 487 \| 5 \| 334 \| Portal protein \| Portal protein; pdb \| 8PHU\_PH \| 1.0 \| 1.774e-05 \| 124 \| 0.098 \| 455 \| 240 \| 26 \| 83 \| 487 \| 5 \| 339 \| Portal protein \| Portal protein; pdb \| 8PHU\_PI \| 1.0 \| 1.228e-05 \| 121 \| 0.108 \| 441 \| 242 \| 24 \| 83 \| 487 \| 5 \| 330 \| Portal protein \| Portal protein; pdb \| 7Z44\_A \| 1.0 \| 6.27e-05 \| 121 \| 0.075 \| 598 \| 373 \| 33 \| 66 \| 524 \| 21 \| 577 \| Portal protein \| Portal protein; pdb \| 7KLN\_A1 \| 1.0 \| 0.001131 \| 119 \| 0.082 \| 461 \| 271 \| 26 \| 52 \| 473 \| 10 \| 357 \| Portal protein \| Portal protein; pdb \| 8PHU\_PA \| 1.0 \| 2.078e-05 \| 116 \| 0.098 \| 448 \| 238 \| 24 \| 83 \| 487 \| 5 \| 329 \| Portal protein \| Portal protein; pdb \| 8PHU\_PD \| 1.0 \| 2.078e-05 \| 116 \| 0.1 \| 449 \| 238 \| 24 \| 83 \| 487 \| 5 \| 331 \| Portal protein \| Portal protein; pdb \| 8PHU\_PE \| 1.0 \| 4.573e-05 \| 116 \| 0.096 \| 464 \| 240 \| 26 \| 83 \| 487 \| 5 \| 348 \| Portal protein \| Portal protein; pdb \| 7Y22\_B \| 1.0 \| 0.0001118 \| 113 \| 0.109 \| 422 \| 227 \| 24 \| 68 \| 418 \| 8 \| 351 \| phage connector protein \| phage connector protein; pdb \| 7Z4B\_ED \| 1.0 \| 0.0001455 \| 108 \| 0.086 \| 610 \| 373 \| 34 \| 66 \| 527 \| 21 \| 593 \| Portal protein \| Portal protein; pdb \| 8PHU\_PC \| 1.0 \| 6.27e-05 \| 107 \| 0.095 \| 460 \| 238 \| 24 \| 83 \| 487 \| 5 \| 341 \| Portal protein \| Portal protein; pdb \| 8PHU\_PJ \| 1.0 \| 6.966e-05 \| 106 \| 0.096 \| 464 \| 240 \| 24 \| 83 \| 487 \| 5 \| 348 \| Portal protein \| Portal protein; pdb \| 8PHU\_PK \| 1.0 \| 0.0002216 \| 103 \| 0.094 \| 468 \| 238 \| 26 \| 83 \| 487 \| 5 \| 349 \| Portal protein \| Portal protein; pdb \| 8PHU\_PB \| 1.0 \| 0.0001242 \| 101 \| 0.097 \| 472 \| 241 \| 24 \| 83 \| 487 \| 5 \| 358 \| Portal protein \| Portal protein; pdb \| 7WMP\_A \| 0.999 \| 7.738e-05 \| 98 \| 0.108 \| 560 \| 334 \| 30 \| 66 \| 515 \| 27 \| 530 \| Nozzle protein gp25 \| Nozzle protein gp25; pdb \| 7Z4A\_K \| 0.999 \| 0.0002335 \| 98 \| 0.09 \| 631 \| 357 \| 41 \| 39 \| 524 \| 10 \| 568 \| Portal protein \| Portal protein; pdb \| 8PHU\_PL \| 0.998 \| 0.0003375 \| 93 \| 0.093 \| 479 \| 240 \| 24 \| 83 \| 487 \| 5 \| 363 \| Portal protein \| Portal protein; pdb \| 3KDR\_C \| 0.991 \| 0.006765 \| 84 \| 0.094 \| 402 \| 208 \| 23 \| 83 \| 455 \| 10 \| 284 \| HK97 Family Phage Portal Protein \| HK97 Family Phage Portal Protein; pdb \| 1IJG\_A \| 0.988 \| 0.00249 \| 82 \| 0.132 \| 257 \| 161 \| 15 \| 155 \| 400 \| 41 \| 246 \| UPPER COLLAR PROTEIN \| UPPER COLLAR PROTEIN; pdb \| 7SYA\_H \| 0.986 \| 0.0088 \| 81 \| 0.087 \| 434 \| 279 \| 23 \| 66 \| 424 \| 18 \| 409 \| Portal protein \| Portal protein; pdb \| 7SZ6\_K \| 0.984 \| 0.004441 \| 80 \| 0.089 \| 490 \| 293 \| 28 \| 66 \| 475 \| 18 \| 434 \| Portal protein \| Portal protein; pdb \| 1H5W\_C \| 0.981 \| 0.003239 \| 79 \| 0.115 \| 260 \| 163 \| 18 \| 155 \| 399 \| 34 \| 241 \| UPPER COLLAR PROTEIN \| UPPER COLLAR PROTEIN; pdb \| 3KDR\_A \| 0.981 \| 0.009776 \| 79 \| 0.117 \| 401 \| 206 \| 25 \| 79 \| 455 \| 6 \| 282 \| HK97 Family Phage Portal Protein \| HK97 Family Phage Portal Protein; pdb \| 7SFS\_A \| 0.978 \| 0.002242 \| 78 \| 0.122 \| 627 \| 310 \| 40 \| 67 \| 522 \| 17 \| 573 \| Gene 3 protein \| Gene 3 protein; pdb \| 1H5W\_A \| 0.967 \| 0.005481 \| 75 \| 0.131 \| 258 \| 157 \| 17 \| 155 \| 399 \| 35 \| 238 \| UPPER COLLAR PROTEIN \| UPPER COLLAR PROTEIN; pdb \| 1H5W\_B \| 0.961 \| 0.006765 \| 74 \| 0.128 \| 257 \| 164 \| 15 \| 155 \| 399 \| 36 \| 244 \| UPPER COLLAR PROTEIN \| UPPER COLLAR PROTEIN; pdb \| 7SP4\_B \| 0.933 \| 0.002127 \| 70 \| 0.113 \| 626 \| 306 \| 40 \| 50 \| 522 \| 49 \| 578 \| Gene 5 protein \| Gene 5 protein; pdb \| 4V4K\_N \| 0.923 \| 0.006418 \| 69 \| 0.11 \| 615 \| 317 \| 40 \| 67 \| 524 \| 17 \| 558 \| PACKAGED DNA STABILIZATION PROTEIN GP4 \| PACKAGED DNA STABILIZATION PROTEIN GP4; pdb \| 7QOI\_FB \| 0.855 \| 0.0052 \| 64 \| 0.101 \| 631 \| 324 \| 33 \| 90 \| 523 \| 74 \| 658 \| Portal protein gp20 \| Portal protein gp20; pdb \| 7QOI\_FC \| 0.632 \| 0.00713 \| 55 \| 0.093 \| 654 \| 326 \| 36 \| 71 \| 523 \| 77 \| 664 \| Portal protein gp20 \| Portal protein gp20; afdb-proteome \| AF-Q8ZQH0-F1-MODEL\_V4 \| 1.0 \| 1.11e-31 \| 1140 \| 0.206 \| 500 \| 369 \| 12 \| 39 \| 532 \| 21 \| 498 \| Fels-1 prophage protein \| Fels-1 prophage protein; afdb-proteome \| AF-Q8ZQ95-F1-MODEL\_V4 \| 1.0 \| 1.017e-25 \| 911 \| 0.217 \| 446 \| 305 \| 14 \| 99 \| 532 \| 2 \| 415 \| Gifsy-2 prophage protein \| Gifsy-2 prophage protein; afdb-proteome \| AF-Q8ZMZ5-F1-MODEL\_V4 \| 1.0 \| 4.039e-24 \| 800 \| 0.203 \| 517 \| 345 \| 17 \| 47 \| 515 \| 21 \| 518 \| Gifsy-1 prophage protein \| Gifsy-1 prophage protein; afdb-proteome \| AF-Q2FYC7-F1-MODEL\_V4 \| 1.0 \| 3.655e-10 \| 253 \| 0.1 \| 480 \| 310 \| 18 \| 23 \| 478 \| 2 \| 383 \| Phage portal protein, HK97 family \| Phage portal protein, HK97 family; afdb-proteome \| AF-A0A0H3GWD5-F1-MODEL\_V4 \| 1.0 \| 2.729e-08 \| 220 \| 0.114 \| 490 \| 304 \| 26 \| 14 \| 478 \| 1 \| 385 \| Phage portal protein, HK97 family \| Phage portal protein, HK97 family; afdb-proteome \| AF-A0A0H3H1P8-F1-MODEL\_V4 \| 1.0 \| 6.22e-05 \| 134 \| 0.109 \| 467 \| 270 \| 27 \| 1 \| 448 \| 4 \| 343 \| Putative prophage presumed portal protein \| Putative prophage presumed portal protein; afdb-proteome \| AF-A0A0H3GQE9-F1-MODEL\_V4 \| 1.0 \| 4.537e-05 \| 127 \| 0.108 \| 478 \| 249 \| 27 \| 2 \| 446 \| 8 \| 341 \| Putative phage portal protein \| Putative phage portal protein; afdb-proteome \| AF-Q8ZMS9-F1-MODEL\_V4 \| 1.0 \| 0.0001053 \| 124 \| 0.119 \| 427 \| 247 \| 21 \| 1 \| 397 \| 4 \| 331 \| Fels-2 prophage protein \| Fels-2 prophage protein; afdb-proteome \| AF-Q5F7S3-F1-MODEL\_V4 \| 0.998 \| 0.000459 \| 94 \| 0.082 \| 695 \| 359 \| 36 \| 49 \| 516 \| 2 \| 644 \| Uncharacterized protein \| Uncharacterized protein; afdb-uniprot50 \| AF-A0A0D0QKS8-F1-MODEL\_V4 \| 1.0 \| 1.912e-69 \| 2992 \| 0.673 \| 487 \| 158 \| 1 \| 46 \| 532 \| 1 \| 486 \| Contig\_80, whole genome shotgun sequence \| Contig\_80, whole genome shotgun sequence; afdb-uniprot50 \| AF-A0A0A8TLB7-F1-MODEL\_V4 \| 1.0 \| 2.05e-63 \| 2629 \| 0.568 \| 512 \| 214 \| 3 \| 13 \| 518 \| 3 \| 513 \| Phage-related portal protein \| Phage-related portal protein; afdb-uniprot50 \| AF-A0A7Z1WDC6-F1-MODEL\_V4 \| 1.0 \| 2.698e-62 \| 2544 \| 0.537 \| 530 \| 231 \| 6 \| 4 \| 532 \| 1 \| 517 \| Phage portal protein \| Phage portal protein; afdb-uniprot50 \| AF-E4PPT4-F1-MODEL\_V4 \| 1.0 \| 5.536e-59 \| 2339 \| 0.493 \| 527 \| 250 \| 7 \| 14 \| 532 \| 1 \| 518 \| Phage portal protein, lambda family \| Phage portal protein, lambda family; afdb-uniprot50 \| AF-A0A1H9YBA9-F1-MODEL\_V4 \| 1.0 \| 1.369e-57 \| 2293 \| 0.454 \| 537 \| 264 \| 7 \| 1 \| 532 \| 13 \| 525 \| Phage portal protein, lambda family \| Phage portal protein, lambda family; afdb-uniprot50 \| AF-A0A2S0MND2-F1-MODEL\_V4 \| 1.0 \| 5.601e-58 \| 2274 \| 0.471 \| 539 \| 263 \| 8 \| 1 \| 532 \| 1 \| 524 \| Phage portal protein \| Phage portal protein; afdb-uniprot50 \| AF-A0A118DU69-F1-MODEL\_V4 \| 1.0 \| 5.733e-56 \| 2256 \| 0.498 \| 495 \| 219 \| 8 \| 46 \| 532 \| 1 \| 474 \| Portal protein \| Portal protein; afdb-uniprot50 \| AF-A0A1X7L1E7-F1-MODEL\_V4 \| 1.0 \| 9.588e-57 \| 2244 \| 0.465 \| 520 \| 255 \| 6 \| 15 \| 532 \| 3 \| 501 \| Phage portal protein, lambda family \| Phage portal protein, lambda family; afdb-uniprot50 \| AF-A0A837E1C6-F1-MODEL\_V4 \| 1.0 \| 3.613e-55 \| 2203 \| 0.463 \| 516 \| 239 \| 6 \| 27 \| 532 \| 2 \| 489 \| Portal protein \| Portal protein; afdb-uniprot50 \| AF-A0A7H8DNI9-F1-MODEL\_V4 \| 1.0 \| 2.529e-54 \| 2134 \| 0.429 \| 521 \| 281 \| 8 \| 17 \| 532 \| 1 \| 510 \| Phage portal protein \| Phage portal protein; afdb-uniprot50 \| AF-Q87CG8-F1-MODEL\_V4 \| 1.0 \| 6.257e-53 \| 2082 \| 0.449 \| 530 \| 259 \| 9 \| 1 \| 523 \| 3 \| 506 \| Phage-related portal protein \| Phage-related portal protein; afdb-uniprot50 \| AF-A0A1F9UST4-F1-MODEL\_V4 \| 1.0 \| 2.002e-47 \| 1861 \| 0.404 \| 495 \| 279 \| 6 \| 13 \| 502 \| 1 \| 484 \| Phage portal protein \| Phage portal protein; afdb-uniprot50 \| AF-A0A7J0BKD5-F1-MODEL\_V4 \| 1.0 \| 5.665e-48 \| 1833 \| 0.383 \| 524 \| 290 \| 7 \| 13 \| 532 \| 2 \| 496 \| Phage portal protein \| Phage portal protein; afdb-uniprot50 \| AF-A0A7C8HWB8-F1-MODEL\_V4 \| 1.0 \| 6.991e-48 \| 1817 \| 0.362 \| 533 \| 317 \| 7 \| 2 \| 532 \| 12 \| 523 \| Uncharacterized protein \| Uncharacterized protein; afdb-uniprot50 \| AF-A0A2G6D2V0-F1-MODEL\_V4 \| 1.0 \| 3.213e-47 \| 1809 \| 0.368 \| 535 \| 303 \| 10 \| 2 \| 528 \| 1 \| 508 \| Phage portal protein \| Phage portal protein; afdb-uniprot50 \| AF-A0A3R8IU71-F1-MODEL\_V4 \| 1.0 \| 6.293e-48 \| 1799 \| 0.401 \| 540 \| 289 \| 10 \| 1 \| 532 \| 7 \| 520 \| Phage portal protein \| Phage portal protein; afdb-uniprot50 \| AF-A0A2D7GUF0-F1-MODEL\_V4 \| 1.0 \| 1.921e-46 \| 1791 \| 0.33 \| 526 \| 318 \| 8 \| 15 \| 532 \| 1 \| 500 \| Phage portal protein \| Phage portal protein; afdb-uniprot50 \| AF-A0A661FAN3-F1-MODEL\_V4 \| 1.0 \| 1.345e-45 \| 1786 \| 0.36 \| 508 \| 300 \| 8 \| 6 \| 502 \| 7 \| 500 \| Phage portal protein \| Phage portal protein; afdb-uniprot50 \| AF-A0A1G2ZKX1-F1-MODEL\_V4 \| 1.0 \| 1.345e-45 \| 1750 \| 0.393 \| 490 \| 273 \| 6 \| 45 \| 532 \| 20 \| 487 \| Phage portal protein \| Phage portal protein; afdb-uniprot50 \| AF-A0A0U5N3Z0-F1-MODEL\_V4 \| 1.0 \| 2.529e-45 \| 1736 \| 0.385 \| 513 \| 280 \| 8 \| 29 \| 532 \| 8 \| 494 \| Putative Phage portal protein, lambda family \| Putative Phage portal protein, lambda family; afdb-uniprot50 \| AF-W0IYG2-F1-MODEL\_V4 \| 1.0 \| 4.698e-46 \| 1718 \| 0.381 \| 532 \| 293 \| 7 \| 4 \| 532 \| 10 \| 508 \| Portal protein \| Portal protein; afdb-uniprot50 \| AF-A0A348SWC9-F1-MODEL\_V4 \| 1.0 \| 3.896e-44 \| 1712 \| 0.516 \| 405 \| 176 \| 6 \| 132 \| 532 \| 2 \| 390 \| Phage portal protein \| Phage portal protein; afdb-uniprot50 \| AF-A0A542RSK1-F1-MODEL\_V4 \| 1.0 \| 1.791e-43 \| 1701 \| 0.358 \| 491 \| 289 \| 6 \| 26 \| 496 \| 13 \| 497 \| Lambda family phage portal protein \| Lambda family phage portal protein; afdb-uniprot50 \| AF-A0A3A4NUK8-F1-MODEL\_V4 \| 1.0 \| 1.512e-44 \| 1679 \| 0.35 \| 511 \| 287 \| 9 \| 39 \| 532 \| 17 \| 499 \| Phage portal protein \| Phage portal protein; afdb-uniprot50 \| AF-A0A0F9J630-F1-MODEL\_V4 \| 1.0 \| 1.548e-42 \| 1673 \| 0.372 \| 454 \| 270 \| 7 \| 46 \| 495 \| 14 \| 456 \| Uncharacterized protein \| Uncharacterized protein; afdb-uniprot50 \| AF-A0A7X4CN61-F1-MODEL\_V4 \| 1.0 \| 2.558e-44 \| 1671 \| 0.373 \| 533 \| 312 \| 7 \| 2 \| 532 \| 23 \| 535 \| Phage portal protein \| Phage portal protein; afdb-uniprot50 \| AF-A0A1Y0N464-F1-MODEL\_V4 \| 1.0 \| 7.41e-43 \| 1658 \| 0.341 \| 495 \| 302 \| 9 \| 44 \| 532 \| 26 \| 502 \| Uncharacterized protein \| Uncharacterized protein; afdb-uniprot50 \| AF-A0A2X1U6Z4-F1-MODEL\_V4 \| 1.0 \| 8.935e-45 \| 1658 \| 0.36 \| 530 \| 295 \| 11 \| 23 \| 532 \| 2 \| 507 \| Putative phage portal protein \| Putative phage portal protein; afdb-uniprot50 \| AF-A0A2A5BDL3-F1-MODEL\_V4 \| 1.0 \| 2.529e-45 \| 1658 \| 0.388 \| 522 \| 291 \| 10 \| 15 \| 532 \| 1 \| 498 \| Phage portal protein \| Phage portal protein; afdb-uniprot50 \| AF-A0A3N2E0V0-F1-MODEL\_V4 \| 1.0 \| 8.329e-42 \| 1653 \| 0.318 \| 502 \| 291 \| 10 \| 46 \| 532 \| 1 \| 466 \| Lambda family phage portal protein \| Lambda family phage portal protein; afdb-uniprot50 \| AF-A0A6M8STS1-F1-MODEL\_V4 \| 1.0 \| 2.097e-43 \| 1651 \| 0.382 \| 494 \| 267 \| 9 \| 43 \| 531 \| 19 \| 479 \| Phage portal protein \| Phage portal protein; afdb-uniprot50 \| AF-A0A5Q0TIZ8-F1-MODEL\_V4 \| 1.0 \| 2.013e-42 \| 1624 \| 0.35 \| 499 \| 292 \| 10 \| 23 \| 516 \| 1 \| 472 \| Phage portal protein \| Phage portal protein; afdb-uniprot50 \| AF-A0A1V5CVB9-F1-MODEL\_V4 \| 1.0 \| 5.935e-44 \| 1622 \| 0.319 \| 536 \| 322 \| 7 \| 15 \| 532 \| 1 \| 511 \| Phage portal protein, lambda family \| Phage portal protein, lambda family; afdb-uniprot50 \| AF-A0A5N0TED7-F1-MODEL\_V4 \| 1.0 \| 1.791e-43 \| 1621 \| 0.348 \| 522 \| 297 \| 12 \| 27 \| 531 \| 3 \| 498 \| Phage portal protein \| Phage portal protein; afdb-uniprot50 \| AF-A0A831XL27-F1-MODEL\_V4 \| 1.0 \| 4.379e-43 \| 1617 \| 0.34 \| 505 \| 294 \| 8 \| 21 \| 502 \| 1 \| 489 \| Phage portal protein \| Phage portal protein; afdb-uniprot50 \| AF-A0A6M3KNK2-F1-MODEL\_V4 \| 1.0 \| 1.239e-43 \| 1616 \| 0.34 \| 520 \| 308 \| 7 \| 23 \| 532 \| 1 \| 495 \| Putative portal protein \| Putative portal protein; afdb-uniprot50 \| AF-A0A653JFP4-F1-MODEL\_V4 \| 1.0 \| 1.129e-42 \| 1612 \| 0.333 \| 531 \| 314 \| 6 \| 15 \| 532 \| 1 \| 504 \| Phage portal protein, lambda family \| Phage portal protein, lambda family; afdb-uniprot50 \| AF-A0A1Z9Q5E6-F1-MODEL\_V4 \| 1.0 \| 3.066e-42 \| 1603 \| 0.308 \| 512 \| 321 \| 7 \| 24 \| 532 \| 4 \| 485 \| Phage portal protein \| Phage portal protein; afdb-uniprot50 \| AF-A0A6H1ZMX2-F1-MODEL\_V4 \| 1.0 \| 8.779e-42 \| 1597 \| 0.313 \| 504 \| 316 \| 10 \| 37 \| 532 \| 9 \| 490 \| Putative portal protein \| Putative portal protein; afdb-uniprot50 \| AF-A0A2E7V4F4-F1-MODEL\_V4 \| 1.0 \| 3.27e-41 \| 1594 \| 0.315 \| 514 \| 323 \| 10 \| 23 \| 532 \| 2 \| 490 \| Phage portal protein \| Phage portal protein; afdb-uniprot50 \| AF-A0A6L9M4Z4-F1-MODEL\_V4 \| 1.0 \| 8.677e-43 \| 1593 \| 0.355 \| 531 \| 307 \| 11 \| 8 \| 531 \| 1 \| 503 \| Phage portal protein \| Phage portal protein; afdb-uniprot50 \| AF-A0A2D5VNS8-F1-MODEL\_V4 \| 1.0 \| 1.071e-42 \| 1584 \| 0.324 \| 514 \| 306 \| 7 \| 39 \| 532 \| 14 \| 506 \| Phage portal protein \| Phage portal protein; afdb-uniprot50 \| AF-A0A1B3W9X7-F1-MODEL\_V4 \| 1.0 \| 2.978e-40 \| 1578 \| 0.331 \| 479 \| 288 \| 7 \| 26 \| 502 \| 30 \| 478 \| Phage portal protein \| Phage portal protein; afdb-uniprot50 \| AF-A0A1Y5TZN6-F1-MODEL\_V4 \| 1.0 \| 2.013e-42 \| 1578 \| 0.335 \| 531 \| 317 \| 11 \| 15 \| 532 \| 1 \| 508 \| Phage portal protein, lambda family \| Phage portal protein, lambda family; afdb-uniprot50 \| AF-A0A843YIQ3-F1-MODEL\_V4 \| 1.0 \| 2.147e-41 \| 1569 \| 0.287 \| 528 \| 352 \| 8 \| 5 \| 532 \| 1 \| 504 \| Phage portal protein \| Phage portal protein; afdb-uniprot50 \| AF-A0A1E4UTF6-F1-MODEL\_V4 \| 1.0 \| 1.76e-40 \| 1565 \| 0.333 \| 486 \| 304 \| 8 \| 16 \| 495 \| 1 \| 472 \| Phage portal protein \| Phage portal protein; afdb-uniprot50 \| AF-A0A7G3GB32-F1-MODEL\_V4 \| 1.0 \| 4.431e-42 \| 1562 \| 0.342 \| 529 \| 299 \| 12 \| 23 \| 532 \| 2 \| 500 \| Phage portal protein \| Phage portal protein; afdb-uniprot50 \| AF-A0A1V6GE05-F1-MODEL\_V4 \| 1.0 \| 2.122e-42 \| 1553 \| 0.339 \| 524 \| 320 \| 9 \| 18 \| 532 \| 16 \| 522 \| Phage portal protein, lambda family \| Phage portal protein, lambda family; afdb-uniprot50 \| AF-A0A1L3ZRM5-F1-MODEL\_V4 \| 1.0 \| 6.146e-41 \| 1551 \| 0.343 \| 521 \| 294 \| 7 \| 23 \| 532 \| 2 \| 485 \| Phage portal protein \| Phage portal protein; afdb-uniprot50 \| AF-A0A7G8BY97-F1-MODEL\_V4 \| 1.0 \| 3.632e-41 \| 1543 \| 0.317 \| 533 \| 322 \| 13 \| 11 \| 532 \| 1 \| 502 \| Phage portal protein \| Phage portal protein; afdb-uniprot50 \| AF-A0A7X8AGP4-F1-MODEL\_V4 \| 1.0 \| 2.172e-40 \| 1543 \| 0.36 \| 475 \| 282 \| 9 \| 39 \| 502 \| 6 \| 469 \| Phage portal protein \| Phage portal protein; afdb-uniprot50 \| AF-A0A0F9FZF8-F1-MODEL\_V4 \| 1.0 \| 4.922e-42 \| 1536 \| 0.336 \| 529 \| 294 \| 9 \| 26 \| 529 \| 10 \| 506 \| Uncharacterized protein \| Uncharacterized protein; afdb-uniprot50 \| AF-A0A4Y3TD15-F1-MODEL\_V4 \| 1.0 \| 1.584e-40 \| 1535 \| 0.364 \| 475 \| 282 \| 7 \| 29 \| 502 \| 13 \| 468 \| Phage portal protein \| Phage portal protein; afdb-uniprot50 \| AF-A0A7Y4DYP6-F1-MODEL\_V4 \| 1.0 \| 1.426e-40 \| 1535 \| 0.355 \| 475 \| 278 \| 6 \| 58 \| 532 \| 16 \| 462 \| Phage portal protein \| Phage portal protein; afdb-uniprot50 \| AF-W0E8V8-F1-MODEL\_V4 \| 1.0 \| 1.978e-39 \| 1535 \| 0.289 \| 504 \| 325 \| 7 \| 44 \| 532 \| 4 \| 489 \| Uncharacterized protein \| Uncharacterized protein; afdb-uniprot50 \| AF-A0A6L4BC33-F1-MODEL\_V4 \| 1.0 \| 7.585e-41 \| 1531 \| 0.308 \| 529 \| 333 \| 13 \| 14 \| 532 \| 1 \| 506 \| Phage portal protein \| Phage portal protein; afdb-uniprot50 \| AF-A0A1M7U252-F1-MODEL\_V4 \| 1.0 \| 1.409e-41 \| 1528 \| 0.328 \| 536 \| 316 \| 10 \| 1 \| 531 \| 7 \| 503 \| Phage portal protein, lambda family \| Phage portal protein, lambda family; afdb-uniprot50 \| AF-A0A1F8QKN1-F1-MODEL\_V4 \| 1.0 \| 2.001e-38 \| 1522 \| 0.359 \| 442 \| 266 \| 8 \| 36 \| 467 \| 16 \| 450 \| Phage portal protein \| Phage portal protein; afdb-uniprot50 \| AF-A0A1G0DQV6-F1-MODEL\_V4 \| 1.0 \| 1.353e-40 \| 1521 \| 0.3 \| 503 \| 309 \| 11 \| 44 \| 532 \| 29 \| 502 \| Phage portal protein \| Phage portal protein; afdb-uniprot50 \| AF-A0A4Q0ZSC3-F1-MODEL\_V4 \| 1.0 \| 5.249e-41 \| 1520 \| 0.34 \| 523 \| 303 \| 13 \| 14 \| 526 \| 1 \| 491 \| Phage portal protein \| Phage portal protein; afdb-uniprot50 \| AF-A0A7V8HP07-F1-MODEL\_V4 \| 1.0 \| 9.471e-40 \| 1514 \| 0.345 \| 477 \| 268 \| 8 \| 70 \| 532 \| 3 \| 449 \| Phage portal protein \| Phage portal protein; afdb-uniprot50 \| AF-Q1ILS5-F1-MODEL\_V4 \| 1.0 \| 5.249e-41 \| 1505 \| 0.317 \| 514 \| 325 \| 8 \| 1 \| 502 \| 1 \| 500 \| Phage portal protein, lambda \| Phage portal protein, lambda; afdb-uniprot50 \| AF-A0A844IP85-F1-MODEL\_V4 \| 1.0 \| 1.76e-40 \| 1498 \| 0.33 \| 509 \| 313 \| 9 \| 28 \| 518 \| 1 \| 499 \| Phage portal protein \| Phage portal protein; afdb-uniprot50 \| AF-A0A848V4U3-F1-MODEL\_V4 \| 1.0 \| 9.982e-40 \| 1496 \| 0.314 \| 487 \| 312 \| 10 \| 24 \| 502 \| 1 \| 473 \| Phage portal protein \| Phage portal protein; afdb-uniprot50 \| AF-A0A2Z6U1K4-F1-MODEL\_V4 \| 1.0 \| 6.828e-41 \| 1496 \| 0.295 \| 541 \| 350 \| 10 \| 2 \| 532 \| 1 \| 520 \| Phage-related protein \| Phage-related protein; afdb-uniprot50 \| AF-A0A2D5ZTJ2-F1-MODEL\_V4 \| 1.0 \| 7.585e-41 \| 1495 \| 0.293 \| 531 \| 341 \| 10 \| 7 \| 532 \| 8 \| 509 \| Phage portal protein \| Phage portal protein; afdb-uniprot50 \| AF-A0A227JR73-F1-MODEL\_V4 \| 1.0 \| 1.78e-39 \| 1493 \| 0.323 \| 488 \| 304 \| 11 \| 17 \| 495 \| 1 \| 471 \| Phage portal protein \| Phage portal protein; afdb-uniprot50 \| AF-A0A369R324-F1-MODEL\_V4 \| 1.0 \| 6.146e-41 \| 1489 \| 0.319 \| 539 \| 310 \| 11 \| 1 \| 531 \| 15 \| 504 \| Phage portal protein \| Phage portal protein; afdb-uniprot50 \| AF-A0A1M7FPC8-F1-MODEL\_V4 \| 1.0 \| 2.343e-38 \| 1485 \| 0.317 \| 475 \| 308 \| 8 \| 36 \| 502 \| 15 \| 481 \| Phage portal protein, lambda family \| Phage portal protein, lambda family; afdb-uniprot50 \| AF-A0A844G1T9-F1-MODEL\_V4 \| 1.0 \| 3.308e-40 \| 1466 \| 0.308 \| 534 \| 319 \| 11 \| 20 \| 532 \| 9 \| 512 \| Phage portal protein \| Phage portal protein; afdb-uniprot50 \| AF-A0A2A5DHS4-F1-MODEL\_V4 \| 1.0 \| 9.091e-39 \| 1465 \| 0.302 \| 500 \| 317 \| 9 \| 37 \| 532 \| 37 \| 508 \| Phage portal protein \| Phage portal protein; afdb-uniprot50 \| AF-A0A3S0BB35-F1-MODEL\_V4 \| 1.0 \| 3.527e-39 \| 1464 \| 0.341 \| 512 \| 304 \| 12 \| 28 \| 532 \| 14 \| 499 \| Phage portal protein \| Phage portal protein; afdb-uniprot50 \| AF-A0A3A0G2M7-F1-MODEL\_V4 \| 1.0 \| 8.183e-39 \| 1452 \| 0.287 \| 498 \| 319 \| 8 \| 45 \| 532 \| 3 \| 474 \| Phage portal protein \| Phage portal protein; afdb-uniprot50 \| AF-A0A0Q7SMQ8-F1-MODEL\_V4 \| 1.0 \| 2.47e-38 \| 1443 \| 0.307 \| 501 \| 326 \| 11 \| 39 \| 531 \| 34 \| 521 \| Uncharacterized protein \| Uncharacterized protein; afdb-uniprot50 \| AF-A0A286GNB7-F1-MODEL\_V4 \| 1.0 \| 8.986e-40 \| 1440 \| 0.318 \| 518 \| 316 \| 15 \| 15 \| 519 \| 1 \| 494 \| Phage portal protein, lambda family \| Phage portal protein, lambda family; afdb-uniprot50 \| AF-A0A0S8DLM0-F1-MODEL\_V4 \| 1.0 \| 1.183e-38 \| 1438 \| 0.304 \| 523 \| 326 \| 10 \| 14 \| 532 \| 1 \| 489 \| Uncharacterized protein \| Uncharacterized protein; afdb-uniprot50 \| AF-U5QE38-F1-MODEL\_V4 \| 1.0 \| 2.134e-37 \| 1437 \| 0.321 \| 476 \| 300 \| 10 \| 39 \| 497 \| 35 \| 504 \| Phage portal protein, lambda family \| Phage portal protein, lambda family; afdb-uniprot50 \| AF-A0A0Q2WT08-F1-MODEL\_V4 \| 1.0 \| 2.223e-38 \| 1434 \| 0.47 \| 393 \| 194 \| 8 \| 1 \| 386 \| 3 \| 388 \| Portal protein \| Portal protein; afdb-uniprot50 \| AF-A0A2A5AEH9-F1-MODEL\_V4 \| 1.0 \| 8.183e-39 \| 1434 \| 0.271 \| 508 \| 355 \| 7 \| 14 \| 515 \| 1 \| 499 \| Phage portal protein \| Phage portal protein; afdb-uniprot50 \| AF-A0A7Y5N878-F1-MODEL\_V4 \| 1.0 \| 7.764e-39 \| 1433 \| 0.3 \| 516 \| 330 \| 8 \| 26 \| 532 \| 2 \| 495 \| Phage portal protein \| Phage portal protein; afdb-uniprot50 \| AF-A0A355AHD4-F1-MODEL\_V4 \| 1.0 \| 1.898e-38 \| 1426 \| 0.295 \| 508 \| 334 \| 8 \| 30 \| 532 \| 14 \| 502 \| Phage portal protein \| Phage portal protein; afdb-uniprot50 \| AF-A0A3B9Q8L9-F1-MODEL\_V4 \| 1.0 \| 2.47e-38 \| 1415 \| 0.3 \| 515 \| 319 \| 12 \| 35 \| 532 \| 12 \| 502 \| Phage portal protein \| Phage portal protein; afdb-uniprot50 \| AF-A0A2V5PTT5-F1-MODEL\_V4 \| 1.0 \| 9.582e-39 \| 1411 \| 0.327 \| 534 \| 308 \| 13 \| 20 \| 523 \| 1 \| 513 \| Phage portal protein \| Phage portal protein; afdb-uniprot50 \| AF-A0A2M7YUE4-F1-MODEL\_V4 \| 1.0 \| 1.749e-36 \| 1410 \| 0.353 \| 456 \| 274 \| 7 \| 29 \| 475 \| 1 \| 444 \| Phage portal protein \| Phage portal protein; afdb-uniprot50 \| AF-A0A1F9VF65-F1-MODEL\_V4 \| 1.0 \| 1.689e-39 \| 1405 \| 0.292 \| 541 \| 325 \| 10 \| 15 \| 531 \| 33 \| 539 \| Phage portal protein \| Phage portal protein; afdb-uniprot50 \| AF-A0A2E7WCF7-F1-MODEL\_V4 \| 1.0 \| 1.401e-37 \| 1402 \| 0.327 \| 482 \| 300 \| 12 \| 33 \| 502 \| 12 \| 481 \| Phage portal protein \| Phage portal protein; afdb-uniprot50 \| AF-A0A166AHR0-F1-MODEL\_V4 \| 1.0 \| 4.642e-38 \| 1396 \| 0.302 \| 545 \| 315 \| 10 \| 15 \| 532 \| 1 \| 507 \| Phage portal protein, lambda family \| Phage portal protein, lambda family; afdb-uniprot50 \| AF-A0A3B9NU50-F1-MODEL\_V4 \| 1.0 \| 5.499e-37 \| 1393 \| 0.296 \| 495 \| 308 \| 9 \| 44 \| 532 \| 25 \| 485 \| Phage portal protein \| Phage portal protein; afdb-uniprot50 \| AF-A0A3S0CHF6-F1-MODEL\_V4 \| 1.0 \| 2.664e-36 \| 1382 \| 0.338 \| 482 \| 276 \| 11 \| 39 \| 490 \| 39 \| 507 \| Phage portal protein \| Phage portal protein; afdb-uniprot50 \| AF-F5R8A0-F1-MODEL\_V4 \| 1.0 \| 7.855e-38 \| 1379 \| 0.317 \| 514 \| 295 \| 10 \| 22 \| 490 \| 6 \| 508 \| Phage-related protein \| Phage-related protein; afdb-uniprot50 \| AF-A0A7Y5K1H3-F1-MODEL\_V4 \| 1.0 \| 7.071e-38 \| 1373 \| 0.282 \| 534 \| 335 \| 9 \| 20 \| 532 \| 2 \| 508 \| Phage portal protein \| Phage portal protein; afdb-uniprot50 \| AF-F4BFR2-F1-MODEL\_V4 \| 1.0 \| 1.034e-36 \| 1372 \| 0.296 \| 489 \| 301 \| 10 \| 17 \| 496 \| 1 \| 455 \| Phage-related portal protein \| Phage-related portal protein; afdb-uniprot50 \| AF-A0A3C0NVL9-F1-MODEL\_V4 \| 1.0 \| 9.415e-36 \| 1362 \| 0.339 \| 439 \| 263 \| 10 \| 85 \| 502 \| 1 \| 433 \| Phage portal protein \| Phage portal protein; afdb-uniprot50 \| AF-A0A359LSW5-F1-MODEL\_V4 \| 1.0 \| 2.37e-37 \| 1350 \| 0.281 \| 515 \| 335 \| 9 \| 45 \| 527 \| 2 \| 513 \| Phage portal protein \| Phage portal protein; afdb-uniprot50 \| AF-A0A0F9CUR0-F1-MODEL\_V4 \| 1.0 \| 2.995e-35 \| 1321 \| 0.252 \| 507 \| 346 \| 10 \| 36 \| 532 \| 31 \| 514 \| Uncharacterized protein \| Uncharacterized protein; afdb-uniprot50 \| AF-A0A7Y0NUG5-F1-MODEL\_V4 \| 1.0 \| 2.696e-35 \| 1310 \| 0.282 \| 492 \| 323 \| 11 \| 23 \| 502 \| 12 \| 485 \| Lambda family phage portal protein \| Lambda family phage portal protein; afdb-uniprot50 \| AF-A0A7T6AR04-F1-MODEL\_V4 \| 1.0 \| 8.475e-36 \| 1310 \| 0.295 \| 501 \| 312 \| 15 \| 39 \| 517 \| 26 \| 507 \| Phage portal protein \| Phage portal protein; afdb-uniprot50 \| AF-A0A0B8PGA6-F1-MODEL\_V4 \| 1.0 \| 1.058e-34 \| 1309 \| 0.303 \| 442 \| 282 \| 10 \| 99 \| 532 \| 2 \| 425 \| Phage portal protein \| Phage portal protein; afdb-uniprot50 \| AF-A0A7V3AD27-F1-MODEL\_V4 \| 1.0 \| 9.637e-34 \| 1278 \| 0.266 \| 496 \| 321 \| 12 \| 45 \| 531 \| 2 \| 463 \| Phage portal protein \| Phage portal protein; afdb-uniprot50 \| AF-A0A6J4FT41-F1-MODEL\_V4 \| 1.0 \| 4.154e-34 \| 1267 \| 0.296 \| 472 \| 304 \| 12 \| 39 \| 502 \| 5 \| 456 \| Phage portal protein, lambda family \| Phage portal protein, lambda family; afdb-uniprot50 \| AF-A0A4R2Z1J8-F1-MODEL\_V4 \| 1.0 \| 5.629e-35 \| 1265 \| 0.238 \| 544 \| 356 \| 14 \| 8 \| 532 \| 1 \| 504 \| Lambda family phage portal protein \| Lambda family phage portal protein; afdb-uniprot50 \| AF-A0A108V769-F1-MODEL\_V4 \| 1.0 \| 3.03e-34 \| 1256 \| 0.265 \| 505 \| 320 \| 11 \| 45 \| 530 \| 3 \| 475 \| Portal protein \| Portal protein; afdb-uniprot50 \| AF-A0A369T5A0-F1-MODEL\_V4 \| 1.0 \| 4.378e-34 \| 1249 \| 0.261 \| 489 \| 324 \| 11 \| 28 \| 501 \| 1 \| 467 \| Phage portal protein \| Phage portal protein; afdb-uniprot50 \| AF-A0A7C2SUL5-F1-MODEL\_V4 \| 1.0 \| 4.669e-33 \| 1246 \| 0.272 \| 470 \| 311 \| 12 \| 47 \| 502 \| 23 \| 475 \| Phage portal protein \| Phage portal protein; afdb-uniprot50 \| AF-M2ZLA5-F1-MODEL\_V4 \| 1.0 \| 5.762e-33 \| 1245 \| 0.276 \| 474 \| 288 \| 12 \| 67 \| 532 \| 5 \| 431 \| Lambda family phage portal protein \| Lambda family phage portal protein; afdb-uniprot50 \| AF-A0A427D6X7-F1-MODEL\_V4 \| 1.0 \| 2.121e-33 \| 1242 \| 0.264 \| 473 \| 317 \| 11 \| 43 \| 502 \| 5 \| 459 \| Phage portal protein \| Phage portal protein; afdb-uniprot50 \| AF-A0A6I6DRZ3-F1-MODEL\_V4 \| 1.0 \| 1.699e-34 \| 1241 \| 0.23 \| 508 \| 360 \| 13 \| 1 \| 500 \| 1 \| 485 \| Phage portal protein \| Phage portal protein; afdb-uniprot50 \| AF-A0A6N9TLF9-F1-MODEL\_V4 \| 1.0 \| 1.631e-33 \| 1238 \| 0.248 \| 491 \| 336 \| 14 \| 39 \| 515 \| 28 \| 499 \| Phage portal protein \| Phage portal protein; afdb-uniprot50 \| AF-B0KSY7-F1-MODEL\_V4 \| 1.0 \| 5.695e-34 \| 1237 \| 0.271 \| 494 \| 322 \| 11 \| 24 \| 502 \| 1 \| 471 \| Phage portal protein, lambda family \| Phage portal protein, lambda family; afdb-uniprot50 \| AF-A0A1I1UG86-F1-MODEL\_V4 \| 1.0 \| 3.405e-33 \| 1236 \| 0.237 \| 497 \| 340 \| 12 \| 17 \| 497 \| 1 \| 474 \| Phage portal protein, lambda family \| Phage portal protein, lambda family; afdb-uniprot50 \| AF-A0A2W6RMJ5-F1-MODEL\_V4 \| 1.0 \| 1.699e-34 \| 1233 \| 0.253 \| 533 \| 355 \| 11 \| 16 \| 532 \| 1 \| 506 \| Phage portal protein \| Phage portal protein; afdb-uniprot50 \| AF-A0A380N598-F1-MODEL\_V4 \| 1.0 \| 3.231e-33 \| 1227 \| 0.243 \| 513 \| 342 \| 15 \| 34 \| 532 \| 12 \| 492 \| Phage portal protein, lambda family \| Phage portal protein, lambda family; afdb-uniprot50 \| AF-A0A353UTN7-F1-MODEL\_V4 \| 1.0 \| 1.409e-32 \| 1226 \| 0.27 \| 470 \| 304 \| 13 \| 20 \| 479 \| 4 \| 444 \| Phage portal protein \| Phage portal protein; afdb-uniprot50 \| AF-A0A564WHL9-F1-MODEL\_V4 \| 1.0 \| 7.495e-33 \| 1226 \| 0.251 \| 504 \| 326 \| 12 \| 46 \| 531 \| 3 \| 473 \| Putative Phage portal protein \| Putative Phage portal protein; afdb-uniprot50 \| AF-F3Z2S4-F1-MODEL\_V4 \| 1.0 \| 6.668e-34 \| 1226 \| 0.256 \| 503 \| 339 \| 11 \| 39 \| 532 \| 30 \| 506 \| Phage portal protein, lambda family \| Phage portal protein, lambda family; afdb-uniprot50 \| AF-A0A1U7MG31-F1-MODEL\_V4 \| 1.0 \| 1.887e-34 \| 1223 \| 0.282 \| 509 \| 319 \| 12 \| 39 \| 532 \| 21 \| 498 \| Phage portal protein, lambda family \| Phage portal protein, lambda family; afdb-uniprot50 \| AF-A0A2R2IT64-F1-MODEL\_V4 \| 1.0 \| 5.467e-33 \| 1222 \| 0.252 \| 495 \| 327 \| 10 \| 46 \| 529 \| 20 \| 482 \| Portal protein \| Portal protein; afdb-uniprot50 \| AF-A0A1A9VKE9-F1-MODEL\_V4 \| 1.0 \| 2.121e-33 \| 1221 \| 0.256 \| 499 \| 326 \| 12 \| 44 \| 531 \| 232 \| 696 \| Uncharacterized protein \| Uncharacterized protein; afdb-uniprot50 \| AF-A0A0B6CWK4-F1-MODEL\_V4 \| 1.0 \| 8.326e-33 \| 1219 \| 0.242 \| 512 \| 327 \| 14 \| 20 \| 515 \| 2 \| 468 \| Phage portal protein, lambda family \| Phage portal protein, lambda family; afdb-uniprot50 \| AF-A0A550EVA9-F1-MODEL\_V4 \| 1.0 \| 1.989e-34 \| 1219 \| 0.267 \| 524 \| 323 \| 13 \| 37 \| 531 \| 2 \| 493 \| Phage portal protein \| Phage portal protein; afdb-uniprot50 \| AF-A0A255XT94-F1-MODEL\_V4 \| 1.0 \| 1.631e-33 \| 1218 \| 0.254 \| 507 \| 338 \| 12 \| 1 \| 495 \| 2 \| 480 \| Phage portal protein \| Phage portal protein; afdb-uniprot50 \| AF-A0A2N5EXY9-F1-MODEL\_V4 \| 1.0 \| 1.393e-33 \| 1216 \| 0.272 \| 488 \| 313 \| 14 \| 39 \| 502 \| 8 \| 477 \| Phage portal protein \| Phage portal protein; afdb-uniprot50 \| AF-A0A0F9M3A7-F1-MODEL\_V4 \| 1.0 \| 2.236e-33 \| 1216 \| 0.251 \| 508 \| 332 \| 13 \| 43 \| 532 \| 26 \| 503 \| Uncharacterized protein \| Uncharacterized protein; afdb-uniprot50 \| AF-A0A3N1MH20-F1-MODEL\_V4 \| 1.0 \| 3.941e-34 \| 1214 \| 0.246 \| 552 \| 349 \| 16 \| 3 \| 532 \| 1 \| 507 \| Lambda family phage portal protein \| Lambda family phage portal protein; afdb-uniprot50 \| AF-A0A2M9G2J3-F1-MODEL\_V4 \| 1.0 \| 1.547e-33 \| 1211 \| 0.258 \| 499 \| 337 \| 13 \| 15 \| 502 \| 1 \| 477 \| Phage portal protein \| Phage portal protein; afdb-uniprot50 \| AF-A0A5E6YAG0-F1-MODEL\_V4 \| 1.0 \| 9.143e-34 \| 1210 \| 0.269 \| 531 \| 325 \| 15 \| 28 \| 532 \| 1 \| 494 \| Uncharacterized protein \| Uncharacterized protein; afdb-uniprot50 \| AF-A0A024EJ49-F1-MODEL\_V4 \| 1.0 \| 1.083e-32 \| 1207 \| 0.267 \| 475 \| 313 \| 12 \| 45 \| 502 \| 8 \| 464 \| Portal protein lambda family \| Portal protein lambda family; afdb-uniprot50 \| AF-A0A3N4ABJ5-F1-MODEL\_V4 \| 1.0 \| 1.565e-32 \| 1206 \| 0.262 \| 476 \| 320 \| 11 \| 42 \| 502 \| 10 \| 469 \| Phage portal protein \| Phage portal protein; afdb-uniprot50 \| AF-A0A6I5YNG6-F1-MODEL\_V4 \| 1.0 \| 9.864e-32 \| 1204 \| 0.272 \| 463 \| 285 \| 15 \| 77 \| 531 \| 2 \| 420 \| Phage portal protein \| Phage portal protein; afdb-uniprot50 \| AF-A0A0H3K144-F1-MODEL\_V4 \| 1.0 \| 2.791e-32 \| 1204 \| 0.285 \| 470 \| 304 \| 11 \| 36 \| 502 \| 24 \| 464 \| Uncharacterized protein \| Uncharacterized protein; afdb-uniprot50 \| AF-A0A377P186-F1-MODEL\_V4 \| 1.0 \| 1.268e-32 \| 1202 \| 0.246 \| 508 \| 333 \| 13 \| 34 \| 530 \| 13 \| 481 \| Phage portal protein, lambda family \| Phage portal protein, lambda family; afdb-uniprot50 \| AF-A0A6G6IPP9-F1-MODEL\_V4 \| 1.0 \| 9.25e-33 \| 1200 \| 0.251 \| 508 \| 329 \| 12 \| 41 \| 530 \| 11 \| 485 \| Phage portal protein \| Phage portal protein; afdb-uniprot50 \| AF-A0A432ADL0-F1-MODEL\_V4 \| 1.0 \| 3.269e-32 \| 1200 \| 0.274 \| 474 \| 307 \| 12 \| 47 \| 502 \| 23 \| 477 \| Phage portal protein \| Phage portal protein; afdb-uniprot50 \| AF-A0A285IZJ2-F1-MODEL\_V4 \| 1.0 \| 7.111e-33 \| 1198 \| 0.273 \| 487 \| 314 \| 13 \| 39 \| 502 \| 8 \| 477 \| Phage portal protein, lambda family \| Phage portal protein, lambda family; afdb-uniprot50 \| AF-A0A7V4P0E4-F1-MODEL\_V4 \| 1.0 \| 3.269e-32 \| 1197 \| 0.291 \| 442 \| 285 \| 9 \| 72 \| 502 \| 2 \| 426 \| Phage portal protein \| Phage portal protein; afdb-uniprot50 \| AF-A0A3M8C9W0-F1-MODEL\_V4 \| 1.0 \| 8.326e-33 \| 1197 \| 0.238 \| 511 \| 334 \| 13 \| 34 \| 532 \| 12 \| 479 \| Phage portal protein \| Phage portal protein; afdb-uniprot50 \| AF-A0A285T1W8-F1-MODEL\_V4 \| 1.0 \| 3.445e-32 \| 1193 \| 0.248 \| 471 \| 327 \| 13 \| 35 \| 502 \| 6 \| 452 \| Phage portal protein, lambda family \| Phage portal protein, lambda family; afdb-uniprot50 \| AF-A0A1G0RDT4-F1-MODEL\_V4 \| 1.0 \| 7.408e-34 \| 1192 \| 0.235 \| 544 \| 354 \| 11 \| 23 \| 532 \| 1 \| 516 \| Phage portal protein \| Phage portal protein; afdb-uniprot50 \| AF-A0A0C6F7H6-F1-MODEL\_V4 \| 1.0 \| 6.144e-32 \| 1189 \| 0.266 \| 462 \| 313 \| 13 \| 45 \| 502 \| 15 \| 454 \| Portal protein \| Portal protein; afdb-uniprot50 \| AF-A0A075KD33-F1-MODEL\_V4 \| 1.0 \| 1.337e-32 \| 1187 \| 0.248 \| 504 \| 334 \| 11 \| 39 \| 532 \| 24 \| 492 \| Phage portal protein, lambda family \| Phage portal protein, lambda family; afdb-uniprot50 \| AF-A0A7W9SB80-F1-MODEL\_V4 \| 1.0 \| 8.879e-32 \| 1185 \| 0.224 \| 495 \| 353 \| 10 \| 23 \| 504 \| 2 \| 478 \| Lambda family phage portal protein \| Lambda family phage portal protein; afdb-uniprot50 \| AF-A0A2S6ZDS2-F1-MODEL\_V4 \| 1.0 \| 2.648e-32 \| 1185 \| 0.251 \| 474 \| 321 \| 12 \| 45 \| 502 \| 6 \| 461 \| Phage portal protein \| Phage portal protein; afdb-uniprot50 \| AF-A0A7T5UIB8-F1-MODEL\_V4 \| 1.0 \| 7.111e-33 \| 1184 \| 0.259 \| 508 \| 329 \| 13 \| 44 \| 518 \| 24 \| 517 \| Phage portal protein \| Phage portal protein; afdb-uniprot50 \| AF-A0A0F9MU33-F1-MODEL\_V4 \| 1.0 \| 5.126e-34 \| 1184 \| 0.25 \| 563 \| 338 \| 16 \| 15 \| 532 \| 1 \| 524 \| Uncharacterized protein \| Uncharacterized protein; afdb-uniprot50 \| AF-A0A2W0FY89-F1-MODEL\_V4 \| 1.0 \| 5.762e-33 \| 1183 \| 0.252 \| 518 \| 330 \| 13 \| 39 \| 532 \| 8 \| 492 \| Phage portal protein \| Phage portal protein; afdb-uniprot50 \| AF-A0A1C3WPM1-F1-MODEL\_V4 \| 1.0 \| 1.189e-33 \| 1183 \| 0.25 \| 531 \| 342 \| 14 \| 13 \| 532 \| 4 \| 489 \| Phage portal protein, lambda family \| Phage portal protein, lambda family; afdb-uniprot50 \| AF-A0A4U7J925-F1-MODEL\_V4 \| 1.0 \| 1.425e-31 \| 1183 \| 0.26 \| 500 \| 355 \| 7 \| 39 \| 532 \| 23 \| 513 \| Phage portal protein \| Phage portal protein; afdb-uniprot50 \| AF-A0A4U6RHC0-F1-MODEL\_V4 \| 1.0 \| 4.978e-32 \| 1181 \| 0.247 \| 509 \| 345 \| 12 \| 28 \| 529 \| 1 \| 478 \| Phage portal protein \| Phage portal protein; afdb-uniprot50 \| AF-D5EFB3-F1-MODEL\_V4 \| 1.0 \| 1.096e-31 \| 1181 \| 0.242 \| 474 \| 329 \| 13 \| 39 \| 502 \| 30 \| 483 \| Phage portal protein, lambda family \| Phage portal protein, lambda family; afdb-uniprot50 \| AF-A0A3M1QZI3-F1-MODEL\_V4 \| 1.0 \| 2.648e-32 \| 1181 \| 0.229 \| 501 \| 356 \| 12 \| 39 \| 532 \| 27 \| 504 \| Phage portal protein \| Phage portal protein; afdb-uniprot50 \| AF-A0A7S8C6A8-F1-MODEL\_V4 \| 1.0 \| 9.358e-32 \| 1181 \| 0.269 \| 475 \| 313 \| 13 \| 36 \| 502 \| 25 \| 473 \| Phage portal protein \| Phage portal protein; afdb-uniprot50 \| AF-A0A2E7K4K1-F1-MODEL\_V4 \| 1.0 \| 2.06e-31 \| 1181 \| 0.243 \| 480 \| 332 \| 13 \| 35 \| 502 \| 20 \| 480 \| Phage portal protein \| Phage portal protein; afdb-uniprot50 \| AF-A0A521H779-F1-MODEL\_V4 \| 1.0 \| 2.648e-32 \| 1180 \| 0.256 \| 500 \| 328 \| 11 \| 36 \| 532 \| 27 \| 485 \| Phage portal protein \| Phage portal protein; afdb-uniprot50 \| AF-A0A2S4QSC7-F1-MODEL\_V4 \| 1.0 \| 2.908e-33 \| 1180 \| 0.227 \| 541 \| 363 \| 14 \| 8 \| 532 \| 1 \| 502 \| Phage portal protein \| Phage portal protein; afdb-uniprot50 \| AF-A0A084XVZ6-F1-MODEL\_V4 \| 1.0 \| 2.384e-32 \| 1179 \| 0.238 \| 477 \| 330 \| 11 \| 39 \| 502 \| 8 \| 464 \| Phage portal protein, lambda family \| Phage portal protein, lambda family; afdb-uniprot50 \| AF-A0A7G9NZU8-F1-MODEL\_V4 \| 1.0 \| 9.358e-32 \| 1179 \| 0.249 \| 501 \| 333 \| 7 \| 39 \| 502 \| 48 \| 542 \| Phage portal protein \| Phage portal protein; afdb-uniprot50 \| AF-A0A516SAX6-F1-MODEL\_V4 \| 1.0 \| 1.932e-32 \| 1178 \| 0.266 \| 492 \| 331 \| 12 \| 23 \| 502 \| 1 \| 474 \| Phage portal protein \| Phage portal protein; afdb-uniprot50 \| AF-A0A2M9EAZ3-F1-MODEL\_V4 \| 1.0 \| 3.445e-32 \| 1177 \| 0.268 \| 470 \| 291 \| 14 \| 62 \| 502 \| 5 \| 450 \| Phage portal protein \| Phage portal protein; afdb-uniprot50 \| AF-A0A1E3G7C6-F1-MODEL\_V4 \| 1.0 \| 4.481e-32 \| 1177 \| 0.278 \| 467 \| 303 \| 11 \| 46 \| 495 \| 5 \| 454 \| Phage portal protein \| Phage portal protein; afdb-uniprot50 \| AF-A0A073ITU9-F1-MODEL\_V4 \| 1.0 \| 3.589e-33 \| 1176 \| 0.241 \| 518 \| 354 \| 12 \| 34 \| 532 \| 17 \| 514 \| Uncharacterized protein \| Uncharacterized protein; afdb-uniprot50 \| AF-A0A327Q828-F1-MODEL\_V4 \| 1.0 \| 7.495e-33 \| 1174 \| 0.264 \| 510 \| 334 \| 15 \| 36 \| 531 \| 23 \| 505 \| Lambda family phage portal protein \| Lambda family phage portal protein; afdb-uniprot50 \| AF-A0A554UE06-F1-MODEL\_V4 \| 1.0 \| 1.954e-31 \| 1174 \| 0.261 \| 486 \| 334 \| 11 \| 23 \| 502 \| 3 \| 469 \| Phage portal protein \| Phage portal protein; afdb-uniprot50 \| AF-A0A850IRK2-F1-MODEL\_V4 \| 1.0 \| 6.003e-34 \| 1174 \| 0.239 \| 527 \| 353 \| 14 \| 15 \| 530 \| 1 \| 490 \| Phage portal protein \| Phage portal protein; afdb-uniprot50 \| AF-A0A7G6PYY7-F1-MODEL\_V4 \| 1.0 \| 1.217e-31 \| 1172 \| 0.255 \| 501 \| 320 \| 17 \| 24 \| 502 \| 1 \| 470 \| Phage portal protein \| Phage portal protein; afdb-uniprot50 \| AF-F6AG24-F1-MODEL\_V4 \| 1.0 \| 3.827e-32 \| 1172 \| 0.234 \| 524 \| 348 \| 11 \| 23 \| 532 \| 1 \| 485 \| Phage portal protein, lambda family \| Phage portal protein, lambda family; afdb-uniprot50 \| AF-A0A2A4ZZY4-F1-MODEL\_V4 \| 1.0 \| 1.565e-32 \| 1172 \| 0.242 \| 511 \| 352 \| 14 \| 34 \| 532 \| 12 \| 499 \| Phage portal protein \| Phage portal protein; afdb-uniprot50 \| AF-A0A838Y2T1-F1-MODEL\_V4 \| 1.0 \| 2.679e-31 \| 1170 \| 0.272 \| 452 \| 289 \| 12 \| 71 \| 519 \| 2 \| 416 \| Phage portal protein \| Phage portal protein; afdb-uniprot50 \| AF-Q24VH0-F1-MODEL\_V4 \| 1.0 \| 1.669e-31 \| 1170 \| 0.242 \| 471 \| 331 \| 9 \| 35 \| 497 \| 18 \| 470 \| Uncharacterized protein \| Uncharacterized protein; afdb-uniprot50 \| AF-A0A6N7AIQ8-F1-MODEL\_V4 \| 1.0 \| 3.101e-32 \| 1169 \| 0.248 \| 490 \| 322 \| 14 \| 54 \| 532 \| 9 \| 463 \| Bacteriophage capsid protein \| Bacteriophage capsid protein; afdb-uniprot50 \| AF-A0A0F2S3S0-F1-MODEL\_V4 \| 1.0 \| 1.352e-31 \| 1169 \| 0.274 \| 459 \| 304 \| 12 \| 54 \| 502 \| 4 \| 443 \| Phage portal protein \| Phage portal protein; afdb-uniprot50 \| AF-A0A7C1RGC0-F1-MODEL\_V4 \| 1.0 \| 3.101e-32 \| 1168 \| 0.271 \| 509 \| 326 \| 16 \| 8 \| 502 \| 1 \| 478 \| Phage portal protein \| Phage portal protein; afdb-uniprot50 \| AF-A0A3S0HXJ2-F1-MODEL\_V4 \| 1.0 \| 1.65e-32 \| 1165 \| 0.233 \| 510 \| 348 \| 17 \| 38 \| 532 \| 19 \| 500 \| Phage portal protein \| Phage portal protein; afdb-uniprot50 \| AF-A0A839W5R8-F1-MODEL\_V4 \| 1.0 \| 2.513e-32 \| 1165 \| 0.227 \| 496 \| 350 \| 13 \| 14 \| 488 \| 1 \| 484 \| Lambda family phage portal protein \| Lambda family phage portal protein; afdb-uniprot50 \| AF-A0A159Z5V5-F1-MODEL\_V4 \| 1.0 \| 4.669e-33 \| 1164 \| 0.258 \| 523 \| 340 \| 17 \| 20 \| 532 \| 4 \| 488 \| ATP-dependent Clp protease proteolytic subunit \| ATP-dependent Clp protease proteolytic subunit; afdb-uniprot50 \| AF-A0A1I5C211-F1-MODEL\_V4 \| 1.0 \| 7.279e-31 \| 1162 \| 0.248 \| 458 \| 318 \| 12 \| 39 \| 492 \| 17 \| 452 \| Phage portal protein, lambda family \| Phage portal protein, lambda family; afdb-uniprot50 \| AF-A0A846UIP3-F1-MODEL\_V4 \| 1.0 \| 1.232e-30 \| 1162 \| 0.212 \| 471 \| 344 \| 12 \| 39 \| 496 \| 14 \| 470 \| Lambda family phage portal protein \| Lambda family phage portal protein; afdb-uniprot50 \| AF-A0A512H917-F1-MODEL\_V4 \| 1.0 \| 6.476e-32 \| 1161 \| 0.296 \| 465 \| 289 \| 15 \| 38 \| 491 \| 3 \| 440 \| Phage portal protein \| Phage portal protein; afdb-uniprot50 \| AF-A0A431KJD1-F1-MODEL\_V4 \| 1.0 \| 3.101e-32 \| 1161 \| 0.234 \| 499 \| 342 \| 13 \| 39 \| 517 \| 21 \| 499 \| Phage portal protein \| Phage portal protein; afdb-uniprot50 \| AF-A0A2N1QUE0-F1-MODEL\_V4 \| 1.0 \| 2.036e-32 \| 1159 \| 0.252 \| 500 \| 338 \| 13 \| 36 \| 531 \| 23 \| 490 \| Phage portal protein \| Phage portal protein; afdb-uniprot50 \| AF-A0A6S5K3W5-F1-MODEL\_V4 \| 1.0 \| 1.65e-32 \| 1157 \| 0.234 \| 490 \| 339 \| 13 \| 17 \| 488 \| 1 \| 472 \| Uncharacterized protein \| Uncharacterized protein; afdb-uniprot50 \| AF-A0A661TRQ9-F1-MODEL\_V4 \| 1.0 \| 4.302e-31 \| 1157 \| 0.241 \| 464 \| 322 \| 14 \| 36 \| 486 \| 19 \| 465 \| Phage portal protein \| Phage portal protein; afdb-uniprot50 \| AF-V7HLM5-F1-MODEL\_V4 \| 1.0 \| 1.854e-31 \| 1156 \| 0.264 \| 476 \| 316 \| 13 \| 38 \| 492 \| 4 \| 466 \| Phage portal protein \| Phage portal protein; afdb-uniprot50 \| AF-A0A1H0UU43-F1-MODEL\_V4 \| 1.0 \| 3.445e-32 \| 1155 \| 0.237 \| 509 \| 340 \| 14 \| 36 \| 519 \| 34 \| 519 \| Phage portal protein, lambda family \| Phage portal protein, lambda family; afdb-uniprot50 \| AF-A0A6L7FGN1-F1-MODEL\_V4 \| 1.0 \| 3.101e-32 \| 1154 \| 0.221 \| 547 \| 365 \| 18 \| 1 \| 532 \| 2 \| 502 \| Phage portal protein \| Phage portal protein; afdb-uniprot50 \| AF-R5BPP1-F1-MODEL\_V4 \| 1.0 \| 4.534e-31 \| 1150 \| 0.219 \| 483 \| 342 \| 10 \| 36 \| 502 \| 19 \| 482 \| Uncharacterized protein \| Uncharacterized protein; afdb-uniprot50 \| AF-A0A354UBL5-F1-MODEL\_V4 \| 1.0 \| 2.412e-31 \| 1149 \| 0.259 \| 494 \| 324 \| 13 \| 15 \| 502 \| 1 \| 458 \| Phage portal protein \| Phage portal protein; afdb-uniprot50 \| AF-A0A5E1A733-F1-MODEL\_V4 \| 1.0 \| 7.992e-32 \| 1149 \| 0.233 \| 488 \| 343 \| 14 \| 14 \| 487 \| 1 \| 471 \| Phage portal protein, lambda family \| Phage portal protein, lambda family; afdb-uniprot50 \| AF-A0A7U3YJH6-F1-MODEL\_V4 \| 1.0 \| 4.978e-32 \| 1148 \| 0.225 \| 514 \| 353 \| 16 \| 39 \| 532 \| 29 \| 517 \| Phage portal protein, lambda family \| Phage portal protein, lambda family; afdb-uniprot50 \| AF-A0A2A5EI97-F1-MODEL\_V4 \| 1.0 \| 7.194e-32 \| 1147 \| 0.252 \| 492 \| 335 \| 12 \| 57 \| 532 \| 2 \| 476 \| Phage portal protein \| Phage portal protein; afdb-uniprot50 \| AF-A0A661RGN5-F1-MODEL\_V4 \| 1.0 \| 4.081e-31 \| 1147 \| 0.245 \| 468 \| 322 \| 16 \| 44 \| 497 \| 12 \| 462 \| Phage portal protein \| Phage portal protein; afdb-uniprot50 \| AF-A0A654ALK5-F1-MODEL\_V4 \| 1.0 \| 1.669e-31 \| 1147 \| 0.225 \| 502 \| 345 \| 12 \| 39 \| 532 \| 21 \| 486 \| Portal protein \| Portal protein; afdb-uniprot50 \| AF-A0A439TWD0-F1-MODEL\_V4 \| 1.0 \| 8.424e-32 \| 1147 \| 0.235 \| 530 \| 347 \| 14 \| 28 \| 532 \| 1 \| 497 \| Phage portal protein \| Phage portal protein; afdb-uniprot50 \| AF-A0A1M7RIE4-F1-MODEL\_V4 \| 1.0 \| 8.879e-32 \| 1146 \| 0.244 \| 498 \| 329 \| 13 \| 39 \| 532 \| 38 \| 492 \| Phage portal protein, lambda family \| Phage portal protein, lambda family; afdb-uniprot50 \| AF-A0A1Q8G4U8-F1-MODEL\_V4 \| 1.0 \| 2.036e-32 \| 1145 \| 0.22 \| 513 \| 350 \| 14 \| 34 \| 532 \| 12 \| 488 \| Uncharacterized protein \| Uncharacterized protein; afdb-uniprot50 \| AF-A3WW86-F1-MODEL\_V4 \| 1.0 \| 3.674e-31 \| 1144 \| 0.27 \| 470 \| 311 \| 13 \| 36 \| 502 \| 5 \| 445 \| Probable bacteriophage-related protein \| Probable bacteriophage-related protein; afdb-uniprot50 \| AF-A0A2R4BQC3-F1-MODEL\_V4 \| 1.0 \| 2.572e-30 \| 1143 \| 0.27 \| 474 \| 307 \| 13 \| 46 \| 502 \| 3 \| 454 \| Phage portal protein \| Phage portal protein; afdb-uniprot50 \| AF-A0A1F3Y515-F1-MODEL\_V4 \| 1.0 \| 4.978e-32 \| 1142 \| 0.263 \| 509 \| 324 \| 15 \| 36 \| 531 \| 23 \| 493 \| Phage portal protein \| Phage portal protein; afdb-uniprot50 \| AF-A0A6S6XWA1-F1-MODEL\_V4 \| 1.0 \| 6.144e-32 \| 1141 \| 0.253 \| 508 \| 334 \| 14 \| 39 \| 532 \| 4 \| 480 \| Phage portal protein \| Phage portal protein; afdb-uniprot50 \| AF-D3NWR9-F1-MODEL\_V4 \| 1.0 \| 4.534e-31 \| 1140 \| 0.251 \| 469 \| 312 \| 14 \| 39 \| 502 \| 13 \| 447 \| Phage portal protein \| Phage portal protein; afdb-uniprot50 \| AF-A0A413FZE1-F1-MODEL\_V4 \| 1.0 \| 2.44e-30 \| 1140 \| 0.237 \| 475 \| 333 \| 7 \| 39 \| 500 \| 22 \| 480 \| Phage portal protein \| Phage portal protein; afdb-uniprot50 \| AF-H8NX02-F1-MODEL\_V4 \| 1.0 \| 1.602e-30 \| 1138 \| 0.218 \| 490 \| 336 \| 11 \| 53 \| 531 \| 1 \| 454 \| Uncharacterized protein \| Uncharacterized protein; afdb-uniprot50 \| AF-B8IET8-F1-MODEL\_V4 \| 1.0 \| 1.502e-31 \| 1138 \| 0.233 \| 510 \| 336 \| 12 \| 49 \| 532 \| 20 \| 500 \| Phage portal protein, lambda family \| Phage portal protein, lambda family; afdb-uniprot50 \| AF-A0A091FES5-F1-MODEL\_V4 \| 1.0 \| 4.534e-31 \| 1137 \| 0.224 \| 507 \| 351 \| 16 \| 37 \| 532 \| 16 \| 491 \| Uncharacterized protein \| Uncharacterized protein; afdb-uniprot50 \| AF-A0A841S0J5-F1-MODEL\_V4 \| 1.0 \| 5.898e-31 \| 1137 \| 0.225 \| 488 \| 339 \| 13 \| 34 \| 502 \| 12 \| 479 \| Lambda family phage portal protein \| Lambda family phage portal protein; afdb-uniprot50 \| AF-A0A7U0SE26-F1-MODEL\_V4 \| 1.0 \| 6.216e-31 \| 1135 \| 0.241 \| 510 \| 340 \| 11 \| 37 \| 532 \| 7 \| 483 \| Phage portal protein \| Phage portal protein; afdb-uniprot50 \| AF-A0A3L8Q0N2-F1-MODEL\_V4 \| 1.0 \| 2.857e-30 \| 1135 \| 0.219 \| 501 \| 351 \| 10 \| 39 \| 531 \| 24 \| 492 \| Phage portal protein \| Phage portal protein; afdb-uniprot50 \| AF-A0A2K4MTR6-F1-MODEL\_V4 \| 1.0 \| 5.829e-32 \| 1135 \| 0.247 \| 513 \| 326 \| 14 \| 35 \| 532 \| 18 \| 485 \| Phage portal protein \| Phage portal protein; afdb-uniprot50 \| AF-A0A439EQB9-F1-MODEL\_V4 \| 1.0 \| 5.531e-32 \| 1131 \| 0.467 \| 340 \| 165 \| 9 \| 7 \| 338 \| 9 \| 340 \| Phage portal protein \| Phage portal protein; afdb-uniprot50 \| AF-A0A1H2N8A6-F1-MODEL\_V4 \| 1.0 \| 2.679e-31 \| 1131 \| 0.208 \| 518 \| 376 \| 13 \| 32 \| 532 \| 10 \| 510 \| Phage portal protein \| Phage portal protein; afdb-uniprot50 \| AF-A0A0T5PNS0-F1-MODEL\_V4 \| 1.0 \| 6.987e-30 \| 1130 \| 0.235 \| 459 \| 325 \| 10 \| 46 \| 491 \| 22 \| 467 \| Uncharacterized protein \| Uncharacterized protein; afdb-uniprot50 \| AF-A0A810UW02-F1-MODEL\_V4 \| 1.0 \| 6.144e-32 \| 1129 \| 0.243 \| 484 \| 334 \| 12 \| 3 \| 473 \| 1 \| 465 \| Phage portal protein \| Phage portal protein; afdb-uniprot50 \| AF-A0A379VWB5-F1-MODEL\_V4 \| 1.0 \| 2.824e-31 \| 1129 \| 0.239 \| 493 \| 329 \| 14 \| 54 \| 532 \| 6 \| 466 \| Phage portal protein \| Phage portal protein; afdb-uniprot50 \| AF-A0A235H652-F1-MODEL\_V4 \| 1.0 \| 1.954e-31 \| 1128 \| 0.241 \| 502 \| 338 \| 14 \| 21 \| 502 \| 8 \| 486 \| Phage portal protein \| Phage portal protein; afdb-uniprot50 \| AF-A0A081RFJ6-F1-MODEL\_V4 \| 1.0 \| 1.217e-31 \| 1127 \| 0.252 \| 510 \| 320 \| 17 \| 35 \| 531 \| 13 \| 474 \| Phage portal protein, lambda family \| Phage portal protein, lambda family; afdb-uniprot50 \| AF-A0A843YX80-F1-MODEL\_V4 \| 1.0 \| 6.552e-31 \| 1127 \| 0.22 \| 535 \| 349 \| 17 \| 17 \| 532 \| 1 \| 486 \| Phage portal protein \| Phage portal protein; afdb-uniprot50 \| AF-A0A7K0GNI4-F1-MODEL\_V4 \| 1.0 \| 5.967e-30 \| 1127 \| 0.238 \| 462 \| 327 \| 11 \| 46 \| 497 \| 23 \| 469 \| Phage portal protein \| Phage portal protein; afdb-uniprot50 \| AF-A0A6G2BPX3-F1-MODEL\_V4 \| 1.0 \| 3.631e-32 \| 1127 \| 0.206 \| 519 \| 366 \| 15 \| 1 \| 491 \| 1 \| 501 \| Phage portal protein \| Phage portal protein; afdb-uniprot50 \| AF-A0A3S0EHX2-F1-MODEL\_V4 \| 1.0 \| 6.476e-32 \| 1126 \| 0.242 \| 527 \| 347 \| 16 \| 36 \| 532 \| 39 \| 543 \| Phage portal protein \| Phage portal protein; afdb-uniprot50 \| AF-A0A1Q4UUL1-F1-MODEL\_V4 \| 1.0 \| 1.876e-30 \| 1125 \| 0.373 \| 410 \| 219 \| 8 \| 130 \| 532 \| 2 \| 380 \| Uncharacterized protein \| Uncharacterized protein; afdb-uniprot50 \| AF-A0A1V5WQS9-F1-MODEL\_V4 \| 1.0 \| 9.468e-31 \| 1125 \| 0.264 \| 476 \| 305 \| 15 \| 39 \| 502 \| 32 \| 474 \| Phage portal protein, lambda family \| Phage portal protein, lambda family; afdb-uniprot50 \| AF-A0A7V1K4N8-F1-MODEL\_V4 \| 1.0 \| 8.326e-33 \| 1125 \| 0.218 \| 527 \| 358 \| 15 \| 1 \| 495 \| 10 \| 514 \| Phage portal protein \| Phage portal protein; afdb-uniprot50 \| AF-A0A6J4FCQ3-F1-MODEL\_V4 \| 1.0 \| 6.552e-31 \| 1123 \| 0.234 \| 507 \| 334 \| 14 \| 39 \| 531 \| 5 \| 471 \| Bacteriophage capsid protein (Modular protein) \| Bacteriophage capsid protein (Modular protein); afdb-uniprot50 \| AF-A0A258LKW1-F1-MODEL\_V4 \| 1.0 \| 1.246e-29 \| 1121 \| 0.227 \| 470 \| 326 \| 12 \| 36 \| 491 \| 6 \| 452 \| Phage portal protein \| Phage portal protein; afdb-uniprot50 \| AF-A0A560F1W8-F1-MODEL\_V4 \| 1.0 \| 5.829e-32 \| 1120 \| 0.244 \| 557 \| 350 \| 15 \| 1 \| 532 \| 6 \| 516 \| Lambda family phage portal protein \| Lambda family phage portal protein; afdb-uniprot50 \| AF-A0A3S0V4X8-F1-MODEL\_V4 \| 1.0 \| 7.279e-31 \| 1118 \| 0.234 \| 512 \| 361 \| 12 \| 1 \| 502 \| 76 \| 566 \| Phage portal protein \| Phage portal protein; afdb-uniprot50 \| AF-A0A4Y8MZU1-F1-MODEL\_V4 \| 1.0 \| 2.222e-29 \| 1117 \| 0.252 \| 472 \| 322 \| 13 \| 39 \| 502 \| 7 \| 455 \| Phage portal protein \| Phage portal protein; afdb-uniprot50 \| AF-A0A1T5K0T3-F1-MODEL\_V4 \| 1.0 \| 1.689e-30 \| 1117 \| 0.227 \| 475 \| 341 \| 14 \| 36 \| 502 \| 13 \| 469 \| Phage portal protein, lambda family \| Phage portal protein, lambda family; afdb-uniprot50 \| AF-A0A8B3PYM4-F1-MODEL\_V4 \| 1.0 \| 2.412e-31 \| 1116 \| 0.228 \| 512 \| 334 \| 15 \| 38 \| 532 \| 3 \| 470 \| Phage portal protein \| Phage portal protein; afdb-uniprot50 \| AF-A0A2N9AI26-F1-MODEL\_V4 \| 1.0 \| 1.409e-32 \| 1116 \| 0.237 \| 568 \| 357 \| 19 \| 1 \| 532 \| 726 \| 1253 \| Terminase GpA (Modular protein) \| Terminase GpA (Modular protein); afdb-uniprot50 \| AF-A0A411X2P3-F1-MODEL\_V4 \| 1.0 \| 2.572e-30 \| 1114 \| 0.225 \| 497 \| 358 \| 13 \| 35 \| 523 \| 26 \| 503 \| Phage portal protein \| Phage portal protein; afdb-uniprot50 \| AF-A0A363UL99-F1-MODEL\_V4 \| 1.0 \| 1.052e-30 \| 1113 \| 0.207 \| 506 \| 357 \| 13 \| 39 \| 532 \| 55 \| 528 \| Phage portal protein \| Phage portal protein; afdb-uniprot50 \| AF-A0A142Y555-F1-MODEL\_V4 \| 1.0 \| 8.424e-32 \| 1113 \| 0.218 \| 531 \| 366 \| 14 \| 35 \| 523 \| 18 \| 541 \| Phage portal protein, lambda family \| Phage portal protein, lambda family; afdb-uniprot50 \| AF-A0A1V5QSP9-F1-MODEL\_V4 \| 1.0 \| 1.854e-31 \| 1113 \| 0.254 \| 535 \| 340 \| 15 \| 15 \| 524 \| 1 \| 501 \| Phage portal protein, lambda family \| Phage portal protein, lambda family; afdb-uniprot50 \| AF-A0A6L6JH12-F1-MODEL\_V4 \| 1.0 \| 1.298e-30 \| 1112 \| 0.48 \| 312 \| 154 \| 4 \| 2 \| 309 \| 1 \| 308 \| Phage portal protein \| Phage portal protein; afdb-uniprot50 \| AF-A0A071LT14-F1-MODEL\_V4 \| 1.0 \| 1.442e-30 \| 1110 \| 0.227 \| 488 \| 349 \| 11 \| 17 \| 498 \| 1 \| 466 \| Uncharacterized protein \| Uncharacterized protein; afdb-uniprot50 \| AF-A0A0Q0VFZ3-F1-MODEL\_V4 \| 1.0 \| 8.623e-30 \| 1110 \| 0.237 \| 502 \| 339 \| 12 \| 37 \| 531 \| 26 \| 490 \| Uncharacterized protein \| Uncharacterized protein; afdb-uniprot50 \| AF-B3G2H0-F1-MODEL\_V4 \| 1.0 \| 8.424e-32 \| 1110 \| 0.222 \| 543 \| 378 \| 14 \| 1 \| 517 \| 92 \| 616 \| Phage portal protein \| Phage portal protein; afdb-uniprot50 \| AF-A0A7Z0SJA8-F1-MODEL\_V4 \| 1.0 \| 1.854e-31 \| 1109 \| 0.228 \| 529 \| 345 \| 16 \| 23 \| 530 \| 2 \| 488 \| Phage portal protein \| Phage portal protein; afdb-uniprot50 \| AF-A0A2M8G061-F1-MODEL\_V4 \| 1.0 \| 8.623e-30 \| 1108 \| 0.269 \| 467 \| 308 \| 14 \| 36 \| 497 \| 38 \| 476 \| Phage portal protein \| Phage portal protein; afdb-uniprot50 \| AF-A0A1F4FSB3-F1-MODEL\_V4 \| 1.0 \| 3.526e-30 \| 1107 \| 0.245 \| 473 \| 311 \| 13 \| 39 \| 502 \| 28 \| 463 \| Phage portal protein \| Phage portal protein; afdb-uniprot50 \| AF-A0A2D7VZC3-F1-MODEL\_V4 \| 1.0 \| 3.526e-30 \| 1106 \| 0.287 \| 441 \| 283 \| 9 \| 100 \| 532 \| 2 \| 419 \| Phage portal protein \| Phage portal protein; afdb-uniprot50 \| AF-A0A2E5F847-F1-MODEL\_V4 \| 1.0 \| 4.081e-31 \| 1106 \| 0.24 \| 503 \| 331 \| 15 \| 46 \| 531 \| 25 \| 493 \| Phage portal protein \| Phage portal protein; afdb-uniprot50 \| AF-A0A0G1R5P6-F1-MODEL\_V4 \| 1.0 \| 1.368e-30 \| 1106 \| 0.22 \| 509 \| 353 \| 16 \| 42 \| 532 \| 7 \| 489 \| Phage portal protein, lambda family \| Phage portal protein, lambda family; afdb-uniprot50 \| AF-E0NXK2-F1-MODEL\_V4 \| 1.0 \| 1.368e-30 \| 1106 \| 0.219 \| 505 \| 361 \| 11 \| 39 \| 532 \| 22 \| 504 \| Phage portal protein, lambda family \| Phage portal protein, lambda family; afdb-uniprot50 \| AF-A0A7W6WBD4-F1-MODEL\_V4 \| 1.0 \| 1.78e-30 \| 1103 \| 0.234 \| 512 \| 339 \| 15 \| 39 \| 532 \| 10 \| 486 \| Lambda family phage portal protein \| Lambda family phage portal protein; afdb-uniprot50 \| AF-A0A2P7S021-F1-MODEL\_V4 \| 1.0 \| 2.469e-29 \| 1100 \| 0.257 \| 462 \| 309 \| 13 \| 46 \| 502 \| 2 \| 434 \| Phage portal protein \| Phage portal protein; afdb-uniprot50 \| AF-A0A480BUD5-F1-MODEL\_V4 \| 1.0 \| 7.364e-30 \| 1100 \| 0.215 \| 477 \| 339 \| 14 \| 39 \| 502 \| 34 \| 488 \| Phage portal protein \| Phage portal protein; afdb-uniprot50 \| AF-A0A2V4FUZ4-F1-MODEL\_V4 \| 1.0 \| 6.144e-32 \| 1100 \| 0.232 \| 550 \| 359 \| 15 \| 1 \| 532 \| 2 \| 506 \| Phage portal protein \| Phage portal protein; afdb-uniprot50 \| AF-A0A858X372-F1-MODEL\_V4 \| 1.0 \| 1.689e-30 \| 1099 \| 0.271 \| 461 \| 298 \| 13 \| 44 \| 502 \| 20 \| 444 \| Phage portal protein \| Phage portal protein; afdb-uniprot50 \| AF-C6BVY9-F1-MODEL\_V4 \| 1.0 \| 4.835e-30 \| 1099 \| 0.245 \| 476 \| 326 \| 10 \| 39 \| 502 \| 28 \| 482 \| Phage portal protein, lambda family \| Phage portal protein, lambda family; afdb-uniprot50 \| AF-A0A3S2UY29-F1-MODEL\_V4 \| 1.0 \| 4.129e-30 \| 1098 \| 0.224 \| 508 \| 347 \| 15 \| 29 \| 532 \| 1 \| 465 \| Phage portal protein \| Phage portal protein; afdb-uniprot50 \| AF-A0A4R6GV31-F1-MODEL\_V4 \| 1.0 \| 4.352e-30 \| 1098 \| 0.239 \| 471 \| 315 \| 13 \| 46 \| 502 \| 10 \| 451 \| Lambda family phage portal protein \| Lambda family phage portal protein; afdb-uniprot50 \| AF-A0A3D2ITU5-F1-MODEL\_V4 \| 1.0 \| 4.129e-30 \| 1098 \| 0.23 \| 494 \| 344 \| 12 \| 35 \| 519 \| 18 \| 484 \| Phage portal protein \| Phage portal protein; afdb-uniprot50 \| AF-A0A5P9H3D4-F1-MODEL\_V4 \| 1.0 \| 8.181e-30 \| 1096 \| 0.271 \| 446 \| 288 \| 13 \| 62 \| 502 \| 3 \| 416 \| Phage portal protein, lambda family \| Phage portal protein, lambda family; afdb-uniprot50 \| AF-A0A1J5JBS4-F1-MODEL\_V4 \| 1.0 \| 6.906e-31 \| 1096 \| 0.272 \| 466 \| 289 \| 13 \| 83 \| 532 \| 2 \| 433 \| Phage portal protein \| Phage portal protein; afdb-uniprot50 \| AF-F2K1G2-F1-MODEL\_V4 \| 1.0 \| 2.315e-30 \| 1096 \| 0.208 \| 498 \| 359 \| 10 \| 38 \| 515 \| 9 \| 491 \| Phage portal protein, lambda family \| Phage portal protein, lambda family; afdb-uniprot50 \| AF-A0A348SU45-F1-MODEL\_V4 \| 1.0 \| 3.346e-30 \| 1095 \| 0.24 \| 474 \| 317 \| 13 \| 46 \| 502 \| 5 \| 452 \| Phage portal protein \| Phage portal protein; afdb-uniprot50 \| AF-A0A6N6T1F2-F1-MODEL\_V4 \| 1.0 \| 7.762e-30 \| 1095 \| 0.24 \| 495 \| 331 \| 12 \| 26 \| 502 \| 2 \| 469 \| Phage portal protein \| Phage portal protein; afdb-uniprot50 \| AF-A0A7C1N2Z9-F1-MODEL\_V4 \| 1.0 \| 2.572e-30 \| 1093 \| 0.219 \| 505 \| 349 \| 14 \| 39 \| 531 \| 39 \| 510 \| Phage portal protein \| Phage portal protein; afdb-uniprot50 \| AF-A0A7C1Y793-F1-MODEL\_V4 \| 1.0 \| 1.459e-29 \| 1093 \| 0.236 \| 478 \| 326 \| 15 \| 42 \| 496 \| 3 \| 464 \| Phage portal protein \| Phage portal protein; afdb-uniprot50 \| AF-A0A5P9JT86-F1-MODEL\_V4 \| 1.0 \| 1.109e-30 \| 1092 \| 0.247 \| 513 \| 345 \| 15 \| 1 \| 502 \| 4 \| 486 \| Phage portal protein \| Phage portal protein; afdb-uniprot50 \| AF-A0A098ARS2-F1-MODEL\_V4 \| 1.0 \| 5.727e-29 \| 1091 \| 0.269 \| 423 \| 278 \| 10 \| 37 \| 446 \| 4 \| 408 \| Portal protein, lambda family,phage portal protein, lambda family,Phage portal protein, lambda family \| Portal protein, lambda family,phage portal protein, lambda family,Phage portal protein, lambda family; afdb-uniprot50 \| AF-A0A1M7R7P3-F1-MODEL\_V4 \| 1.0 \| 2.084e-30 \| 1091 \| 0.225 \| 509 \| 339 \| 15 \| 39 \| 532 \| 29 \| 497 \| Phage portal protein, lambda family \| Phage portal protein, lambda family; afdb-uniprot50 \| AF-A0A524RVZ8-F1-MODEL\_V4 \| 1.0 \| 1.096e-31 \| 1091 \| 0.227 \| 589 \| 351 \| 17 \| 1 \| 502 \| 19 \| 590 \| Phage portal protein \| Phage portal protein; afdb-uniprot50 \| AF-A0A106BVU9-F1-MODEL\_V4 \| 1.0 \| 3.963e-29 \| 1090 \| 0.233 \| 462 \| 311 \| 13 \| 47 \| 487 \| 6 \| 445 \| Uncharacterized protein \| Uncharacterized protein; afdb-uniprot50 \| AF-A0A6N9P4E6-F1-MODEL\_V4 \| 1.0 \| 3.346e-30 \| 1088 \| 0.232 \| 500 \| 351 \| 11 \| 39 \| 532 \| 24 \| 496 \| Phage portal protein \| Phage portal protein; afdb-uniprot50 \| AF-A0A2G2DPB9-F1-MODEL\_V4 \| 1.0 \| 2.108e-29 \| 1087 \| 0.205 \| 525 \| 361 \| 15 \| 20 \| 530 \| 2 \| 484 \| Phage portal protein \| Phage portal protein; afdb-uniprot50 \| AF-A0A0H4W134-F1-MODEL\_V4 \| 1.0 \| 2.146e-32 \| 1085 \| 0.256 \| 558 \| 319 \| 15 \| 39 \| 524 \| 4 \| 537 \| Phage portal protein, lambda family \| Phage portal protein, lambda family; afdb-uniprot50 \| AF-A9C2C6-F1-MODEL\_V4 \| 1.0 \| 4.534e-31 \| 1085 \| 0.225 \| 542 \| 353 \| 17 \| 21 \| 532 \| 16 \| 520 \| Phage portal protein, lambda family \| Phage portal protein, lambda family; afdb-uniprot50 \| AF-A0A0F2NL12-F1-MODEL\_V4 \| 1.0 \| 1.538e-29 \| 1084 \| 0.238 \| 470 \| 326 \| 17 \| 46 \| 502 \| 1 \| 451 \| Uncharacterized protein \| Uncharacterized protein; afdb-uniprot50 \| AF-A0A4Q1AN27-F1-MODEL\_V4 \| 1.0 \| 5.434e-29 \| 1084 \| 0.201 \| 501 \| 359 \| 10 \| 30 \| 508 \| 1 \| 482 \| Phage portal protein \| Phage portal protein; afdb-uniprot50 \| AF-A0A151FKE0-F1-MODEL\_V4 \| 1.0 \| 7.762e-30 \| 1083 \| 0.251 \| 466 \| 305 \| 11 \| 77 \| 532 \| 2 \| 433 \| Uncharacterized protein \| Uncharacterized protein; afdb-uniprot50 \| AF-A0A7T5FFX4-F1-MODEL\_V4 \| 1.0 \| 6.289e-30 \| 1082 \| 0.224 \| 500 \| 332 \| 15 \| 46 \| 532 \| 23 \| 479 \| Phage portal protein \| Phage portal protein; afdb-uniprot50 \| AF-A0A2N0WWU8-F1-MODEL\_V4 \| 1.0 \| 1.92e-28 \| 1082 \| 0.247 \| 460 \| 322 \| 12 \| 39 \| 491 \| 21 \| 463 \| Phage portal protein \| Phage portal protein; afdb-uniprot50 \| AF-A0A1D8IV87-F1-MODEL\_V4 \| 1.0 \| 6.216e-31 \| 1082 \| 0.22 \| 504 \| 326 \| 14 \| 57 \| 532 \| 2 \| 466 \| Phage portal protein \| Phage portal protein; afdb-uniprot50 \| AF-X6D259-F1-MODEL\_V4 \| 1.0 \| 3.917e-30 \| 1080 \| 0.262 \| 483 \| 314 \| 12 \| 36 \| 502 \| 25 \| 481 \| Uncharacterized protein \| Uncharacterized protein; afdb-uniprot50 \| AF-A0A2E9SLJ8-F1-MODEL\_V4 \| 1.0 \| 2.572e-30 \| 1078 \| 0.25 \| 496 \| 309 \| 17 \| 14 \| 495 \| 1 \| 447 \| Phage portal protein \| Phage portal protein; afdb-uniprot50 \| AF-A0A059KU09-F1-MODEL\_V4 \| 1.0 \| 1.442e-30 \| 1078 \| 0.209 \| 520 \| 359 \| 15 \| 32 \| 519 \| 10 \| 509 \| Capsid protein \| Capsid protein; afdb-uniprot50 \| AF-A0A1E3G798-F1-MODEL\_V4 \| 1.0 \| 5.371e-30 \| 1077 \| 0.218 \| 504 \| 351 \| 15 \| 23 \| 502 \| 1 \| 485 \| Phage portal protein \| Phage portal protein; afdb-uniprot50 \| AF-A0A0A8K3F6-F1-MODEL\_V4 \| 1.0 \| 3.211e-29 \| 1076 \| 0.247 \| 464 \| 316 \| 16 \| 29 \| 487 \| 1 \| 436 \| Uncharacterized protein \| Uncharacterized protein; afdb-uniprot50 \| AF-A0A368TQV2-F1-MODEL\_V4 \| 1.0 \| 6.906e-31 \| 1076 \| 0.217 \| 574 \| 368 \| 16 \| 1 \| 532 \| 127 \| 661 \| Phage portal protein \| Phage portal protein; afdb-uniprot50 \| AF-A4N155-F1-MODEL\_V4 \| 1.0 \| 1.476e-28 \| 1075 \| 0.227 \| 436 \| 312 \| 12 \| 39 \| 464 \| 21 \| 441 \| Bacteriophage capsid protein \| Bacteriophage capsid protein; afdb-uniprot50 \| AF-A0A1T4W4N3-F1-MODEL\_V4 \| 1.0 \| 1.801e-29 \| 1075 \| 0.229 \| 527 \| 343 \| 19 \| 26 \| 532 \| 2 \| 485 \| Phage portal protein, lambda family \| Phage portal protein, lambda family; afdb-uniprot50 \| AF-A0A5A7N0P2-F1-MODEL\_V4 \| 1.0 \| 4.835e-30 \| 1075 \| 0.264 \| 476 \| 305 \| 15 \| 35 \| 491 \| 15 \| 464 \| Phage portal protein \| Phage portal protein; afdb-uniprot50 \| AF-A0A410P784-F1-MODEL\_V4 \| 1.0 \| 2.711e-30 \| 1074 \| 0.23 \| 521 \| 341 \| 16 \| 39 \| 532 \| 5 \| 492 \| Phage portal protein \| Phage portal protein; afdb-uniprot50 \| AF-A0A765BM50-F1-MODEL\_V4 \| 1.0 \| 4.455e-28 \| 1073 \| 0.213 \| 440 \| 325 \| 10 \| 39 \| 472 \| 21 \| 445 \| Phage portal protein \| Phage portal protein; afdb-uniprot50 \| AF-B6WS36-F1-MODEL\_V4 \| 1.0 \| 3.917e-30 \| 1073 \| 0.232 \| 538 \| 353 \| 16 \| 14 \| 529 \| 1 \| 500 \| Phage portal protein, lambda family \| Phage portal protein, lambda family; afdb-uniprot50 \| AF-A0A521CT40-F1-MODEL\_V4 \| 1.0 \| 1.246e-29 \| 1071 \| 0.22 \| 498 \| 341 \| 15 \| 39 \| 532 \| 24 \| 478 \| Phage portal protein, lambda family \| Phage portal protein, lambda family; afdb-uniprot50 \| AF-A0A6N3TG59-F1-MODEL\_V4 \| 1.0 \| 1.476e-28 \| 1071 \| 0.272 \| 463 \| 307 \| 12 \| 39 \| 497 \| 28 \| 464 \| Phage portal protein \| Phage portal protein; afdb-uniprot50 \| AF-A0A0S9N407-F1-MODEL\_V4 \| 1.0 \| 6.289e-30 \| 1070 \| 0.35 \| 428 \| 238 \| 12 \| 125 \| 532 \| 2 \| 409 \| Uncharacterized protein \| Uncharacterized protein; afdb-uniprot50 \| AF-A0A0A1W3J4-F1-MODEL\_V4 \| 1.0 \| 1.196e-28 \| 1069 \| 0.27 \| 462 \| 294 \| 14 \| 46 \| 502 \| 1 \| 424 \| Uncharacterized protein \| Uncharacterized protein; afdb-uniprot50 \| AF-A0A562C7Q2-F1-MODEL\_V4 \| 1.0 \| 9.979e-31 \| 1069 \| 0.223 \| 520 \| 348 \| 16 \| 34 \| 532 \| 12 \| 496 \| Lambda family phage portal protein \| Lambda family phage portal protein; afdb-uniprot50 \| AF-A0A1B1A8L3-F1-MODEL\_V4 \| 1.0 \| 1.077e-28 \| 1068 \| 0.235 \| 463 \| 321 \| 12 \| 36 \| 490 \| 25 \| 462 \| Uncharacterized protein \| Uncharacterized protein; afdb-uniprot50 \| AF-A0A248JRM8-F1-MODEL\_V4 \| 1.0 \| 2.222e-29 \| 1068 \| 0.237 \| 471 \| 314 \| 14 \| 46 \| 502 \| 47 \| 486 \| Phage portal protein \| Phage portal protein; afdb-uniprot50 \| AF-A0A3D1NR46-F1-MODEL\_V4 \| 1.0 \| 1.135e-28 \| 1067 \| 0.22 \| 485 \| 333 \| 13 \| 23 \| 502 \| 1 \| 445 \| Phage portal protein \| Phage portal protein; afdb-uniprot50 \| AF-E2CJU7-F1-MODEL\_V4 \| 1.0 \| 9.691e-29 \| 1066 \| 0.228 \| 472 \| 327 \| 10 \| 44 \| 497 \| 12 \| 464 \| Phage portal protein, lambda family \| Phage portal protein, lambda family; afdb-uniprot50 \| AF-A0A1M6Q0A4-F1-MODEL\_V4 \| 1.0 \| 1.232e-30 \| 1064 \| 0.247 \| 533 \| 348 \| 15 \| 2 \| 530 \| 1 \| 484 \| Phage portal protein, lambda family \| Phage portal protein, lambda family; afdb-uniprot50 \| AF-A0A640W790-F1-MODEL\_V4 \| 1.0 \| 9.579e-30 \| 1064 \| 0.186 \| 520 \| 367 \| 12 \| 36 \| 532 \| 19 \| 505 \| Phage portal protein \| Phage portal protein; afdb-uniprot50 \| AF-A0A1V5D6K6-F1-MODEL\_V4 \| 1.0 \| 1.52e-30 \| 1064 \| 0.232 \| 520 \| 349 \| 14 \| 39 \| 532 \| 33 \| 528 \| Phage portal protein, lambda family \| Phage portal protein, lambda family; afdb-uniprot50 \| AF-E6X1P5-F1-MODEL\_V4 \| 1.0 \| 7.45e-29 \| 1063 \| 0.227 \| 496 \| 333 \| 14 \| 17 \| 502 \| 1 \| 456 \| Phage portal protein, lambda family \| Phage portal protein, lambda family; afdb-uniprot50 \| AF-A0A6I6GTE6-F1-MODEL\_V4 \| 1.0 \| 7.762e-30 \| 1061 \| 0.214 \| 523 \| 371 \| 16 \| 32 \| 532 \| 10 \| 514 \| Phage portal protein \| Phage portal protein; afdb-uniprot50 \| AF-A0A255YQI4-F1-MODEL\_V4 \| 1.0 \| 6.629e-30 \| 1057 \| 0.218 \| 512 \| 353 \| 12 \| 7 \| 502 \| 1 \| 481 \| Phage portal protein \| Phage portal protein; afdb-uniprot50 \| AF-A0A5C7SHE2-F1-MODEL\_V4 \| 1.0 \| 3.917e-30 \| 1056 \| 0.22 \| 536 \| 362 \| 17 \| 20 \| 532 \| 27 \| 529 \| Phage portal protein \| Phage portal protein; afdb-uniprot50 \| AF-A0A5C7LKW9-F1-MODEL\_V4 \| 1.0 \| 4.403e-29 \| 1055 \| 0.245 \| 473 \| 327 \| 14 \| 39 \| 502 \| 21 \| 472 \| Phage portal protein \| Phage portal protein; afdb-uniprot50 \| AF-A0A085EXP1-F1-MODEL\_V4 \| 1.0 \| 6.363e-29 \| 1055 \| 0.234 \| 481 \| 324 \| 14 \| 36 \| 502 \| 28 \| 478 \| Putative Phage portal protein, lambda family \| Putative Phage portal protein, lambda family; afdb-uniprot50 \| AF-A0A4R2P5J0-F1-MODEL\_V4 \| 1.0 \| 1.261e-28 \| 1053 \| 0.252 \| 471 \| 315 \| 14 \| 42 \| 491 \| 5 \| 459 \| Lambda family phage portal protein \| Lambda family phage portal protein; afdb-uniprot50 \| AF-A0A524ITH3-F1-MODEL\_V4 \| 1.0 \| 1.801e-29 \| 1053 \| 0.233 \| 509 \| 334 \| 16 \| 35 \| 532 \| 22 \| 485 \| Phage portal protein \| Phage portal protein; afdb-uniprot50 \| AF-A0A1F9BJ96-F1-MODEL\_V4 \| 1.0 \| 1.384e-29 \| 1053 \| 0.229 \| 487 \| 327 \| 16 \| 39 \| 502 \| 33 \| 494 \| Phage portal protein \| Phage portal protein; afdb-uniprot50 \| AF-A0A1H2I371-F1-MODEL\_V4 \| 1.0 \| 2.775e-28 \| 1052 \| 0.25 \| 472 \| 320 \| 11 \| 39 \| 498 \| 19 \| 468 \| Phage portal protein, lambda family \| Phage portal protein, lambda family; afdb-uniprot50 \| AF-A0A0S2KEQ2-F1-MODEL\_V4 \| 1.0 \| 9.195e-29 \| 1048 \| 0.241 \| 501 \| 333 \| 13 \| 36 \| 532 \| 24 \| 481 \| Phage portal protein, lambda \| Phage portal protein, lambda; afdb-uniprot50 \| AF-A0A7R6ZZ48-F1-MODEL\_V4 \| 1.0 \| 6.108e-28 \| 1047 \| 0.194 \| 488 \| 333 \| 13 \| 42 \| 519 \| 5 \| 442 \| Uncharacterized protein \| Uncharacterized protein; afdb-uniprot50 \| AF-A0A3B9ZLI4-F1-MODEL\_V4 \| 1.0 \| 2.469e-29 \| 1047 \| 0.25 \| 491 \| 329 \| 15 \| 36 \| 502 \| 34 \| 509 \| Phage portal protein \| Phage portal protein; afdb-uniprot50 \| AF-B6IPY1-F1-MODEL\_V4 \| 1.0 \| 1.401e-28 \| 1045 \| 0.271 \| 476 \| 309 \| 17 \| 36 \| 502 \| 25 \| 471 \| Phage portal protein, lambda family \| Phage portal protein, lambda family; afdb-uniprot50 \| AF-A0A345DE62-F1-MODEL\_V4 \| 1.0 \| 9.579e-30 \| 1045 \| 0.195 \| 533 \| 365 \| 17 \| 13 \| 532 \| 2 \| 483 \| Uncharacterized protein \| Uncharacterized protein; afdb-uniprot50 \| AF-A0A7D7PEZ6-F1-MODEL\_V4 \| 1.0 \| 1.898e-29 \| 1043 \| 0.201 \| 552 \| 380 \| 19 \| 9 \| 532 \| 1 \| 519 \| Phage portal protein \| Phage portal protein; afdb-uniprot50 \| AF-A0A4R2R7K8-F1-MODEL\_V4 \| 1.0 \| 7.538e-28 \| 1042 \| 0.263 \| 433 \| 294 \| 11 \| 74 \| 502 \| 2 \| 413 \| Lambda family phage portal protein \| Lambda family phage portal protein; afdb-uniprot50 \| AF-A0A2E2DF85-F1-MODEL\_V4 \| 1.0 \| 1.92e-28 \| 1042 \| 0.239 \| 472 \| 321 \| 15 \| 36 \| 502 \| 23 \| 461 \| Phage portal protein \| Phage portal protein; afdb-uniprot50 \| AF-A0A494X0D8-F1-MODEL\_V4 \| 1.0 \| 2.248e-28 \| 1041 \| 0.246 \| 463 \| 310 \| 16 \| 36 \| 488 \| 23 \| 456 \| Phage portal protein \| Phage portal protein; afdb-uniprot50 \| AF-A0A2E3R5Z3-F1-MODEL\_V4 \| 1.0 \| 4.891e-29 \| 1040 \| 0.215 \| 505 \| 352 \| 14 \| 47 \| 532 \| 28 \| 507 \| Phage portal protein \| Phage portal protein; afdb-uniprot50 \| AF-A0A3T0L2W9-F1-MODEL\_V4 \| 1.0 \| 2.498e-28 \| 1038 \| 0.226 \| 477 \| 334 \| 15 \| 39 \| 497 \| 29 \| 488 \| Phage portal protein \| Phage portal protein; afdb-uniprot50 \| AF-A0A4R1Q774-F1-MODEL\_V4 \| 1.0 \| 6.289e-30 \| 1038 \| 0.228 \| 521 \| 332 \| 12 \| 37 \| 532 \| 31 \| 506 \| Lambda family phage portal protein \| Lambda family phage portal protein; afdb-uniprot50 \| AF-E4LA53-F1-MODEL\_V4 \| 1.0 \| 2.342e-29 \| 1038 \| 0.221 \| 501 \| 352 \| 11 \| 39 \| 532 \| 28 \| 497 \| Phage portal protein, lambda family \| Phage portal protein, lambda family; afdb-uniprot50 \| AF-A0A352L2C6-F1-MODEL\_V4 \| 1.0 \| 3.917e-30 \| 1038 \| 0.225 \| 515 \| 346 \| 17 \| 36 \| 532 \| 42 \| 521 \| Phage portal protein \| Phage portal protein; afdb-uniprot50 \| AF-A0A844HP12-F1-MODEL\_V4 \| 1.0 \| 5.156e-29 \| 1034 \| 0.244 \| 504 \| 335 \| 14 \| 39 \| 532 \| 30 \| 497 \| Phage portal protein \| Phage portal protein; afdb-uniprot50 \| AF-A0A5B9Y8G9-F1-MODEL\_V4 \| 1.0 \| 7.364e-30 \| 1034 \| 0.243 \| 502 \| 337 \| 16 \| 36 \| 500 \| 23 \| 518 \| Phage portal protein \| Phage portal protein; afdb-uniprot50 \| AF-A0A2S6H5A5-F1-MODEL\_V4 \| 1.0 \| 9.691e-29 \| 1033 \| 0.246 \| 466 \| 306 \| 15 \| 45 \| 502 \| 9 \| 437 \| Lambda family phage portal protein \| Lambda family phage portal protein; afdb-uniprot50 \| AF-A0A5C1B6Q3-F1-MODEL\_V4 \| 1.0 \| 3.249e-28 \| 1033 \| 0.222 \| 476 \| 333 \| 13 \| 36 \| 502 \| 27 \| 474 \| Phage portal protein \| Phage portal protein; afdb-uniprot50 \| AF-A0A1V5E5F9-F1-MODEL\_V4 \| 1.0 \| 1.021e-28 \| 1033 \| 0.231 \| 488 \| 336 \| 14 \| 39 \| 502 \| 39 \| 511 \| Phage portal protein, lambda family \| Phage portal protein, lambda family; afdb-uniprot50 \| AF-A0A268TNZ2-F1-MODEL\_V4 \| 1.0 \| 7.45e-29 \| 1032 \| 0.195 \| 531 \| 361 \| 15 \| 15 \| 530 \| 1 \| 480 \| Phage portal protein \| Phage portal protein; afdb-uniprot50 \| AF-A0A4U8YLR7-F1-MODEL\_V4 \| 1.0 \| 1.077e-28 \| 1030 \| 0.222 \| 481 \| 323 \| 12 \| 45 \| 502 \| 44 \| 496 \| Phage portal protein lambda family \| Phage portal protein lambda family; afdb-uniprot50 \| AF-A0A6P1EIH4-F1-MODEL\_V4 \| 1.0 \| 9.195e-29 \| 1030 \| 0.206 \| 465 \| 328 \| 13 \| 60 \| 502 \| 22 \| 467 \| Phage portal protein \| Phage portal protein; afdb-uniprot50 \| AF-A0A7C5BTM8-F1-MODEL\_V4 \| 1.0 \| 2.024e-28 \| 1029 \| 0.247 \| 488 \| 316 \| 17 \| 18 \| 502 \| 4 \| 443 \| Phage portal protein \| Phage portal protein; afdb-uniprot50 \| AF-A0A2E6KND4-F1-MODEL\_V4 \| 1.0 \| 1.476e-28 \| 1029 \| 0.241 \| 505 \| 343 \| 14 \| 39 \| 532 \| 25 \| 500 \| Phage portal protein \| Phage portal protein; afdb-uniprot50 \| AF-A0A2A4V1I6-F1-MODEL\_V4 \| 1.0 \| 9.195e-29 \| 1028 \| 0.217 \| 511 \| 357 \| 14 \| 35 \| 532 \| 21 \| 501 \| Phage portal protein \| Phage portal protein; afdb-uniprot50 \| AF-A0A037UQ03-F1-MODEL\_V4 \| 1.0 \| 6.363e-29 \| 1028 \| 0.236 \| 482 \| 323 \| 16 \| 37 \| 502 \| 28 \| 480 \| Portal protein \| Portal protein; afdb-uniprot50 \| AF-A0A076LEP4-F1-MODEL\_V4 \| 1.0 \| 8.374e-28 \| 1026 \| 0.215 \| 463 \| 322 \| 12 \| 78 \| 532 \| 2 \| 431 \| Phage portal protein \| Phage portal protein; afdb-uniprot50 \| AF-A0A3D1J2Q7-F1-MODEL\_V4 \| 1.0 \| 1.459e-29 \| 1025 \| 0.269 \| 490 \| 309 \| 19 \| 35 \| 502 \| 23 \| 485 \| Phage portal protein \| Phage portal protein; afdb-uniprot50 \| AF-A0A4P7EGN2-F1-MODEL\_V4 \| 1.0 \| 1.261e-28 \| 1024 \| 0.237 \| 501 \| 339 \| 16 \| 36 \| 523 \| 25 \| 495 \| Phage portal protein \| Phage portal protein; afdb-uniprot50 \| AF-A0A1L9BVE0-F1-MODEL\_V4 \| 1.0 \| 1.556e-28 \| 1023 \| 0.206 \| 509 \| 357 \| 21 \| 39 \| 532 \| 21 \| 497 \| Portal protein \| Portal protein; afdb-uniprot50 \| AF-A0A1E3ZLT6-F1-MODEL\_V4 \| 1.0 \| 2.222e-29 \| 1022 \| 0.245 \| 501 \| 318 \| 19 \| 39 \| 517 \| 18 \| 480 \| Phage portal protein \| Phage portal protein; afdb-uniprot50 \| AF-L7U545-F1-MODEL\_V4 \| 1.0 \| 9.195e-29 \| 1022 \| 0.225 \| 519 \| 344 \| 15 \| 34 \| 532 \| 13 \| 493 \| Phage portal protein \| Phage portal protein; afdb-uniprot50 \| AF-G0ER26-F1-MODEL\_V4 \| 1.0 \| 8.724e-29 \| 1021 \| 0.253 \| 477 \| 311 \| 15 \| 36 \| 502 \| 18 \| 459 \| Phage portal protein lambda family \| Phage portal protein lambda family; afdb-uniprot50 \| AF-A0A2W6W644-F1-MODEL\_V4 \| 1.0 \| 2.248e-28 \| 1019 \| 0.261 \| 463 \| 288 \| 17 \| 42 \| 502 \| 18 \| 428 \| Phage portal protein \| Phage portal protein; afdb-uniprot50 \| AF-A0A2E5AE83-F1-MODEL\_V4 \| 1.0 \| 2.275e-27 \| 1018 \| 0.25 \| 428 \| 289 \| 13 \| 78 \| 496 \| 2 \| 406 \| Phage portal protein \| Phage portal protein; afdb-uniprot50 \| AF-A0A2W5A5V5-F1-MODEL\_V4 \| 1.0 \| 1.92e-28 \| 1017 \| 0.208 \| 494 \| 350 \| 15 \| 23 \| 491 \| 3 \| 480 \| Phage portal protein \| Phage portal protein; afdb-uniprot50 \| AF-W0DY98-F1-MODEL\_V4 \| 1.0 \| 6.865e-27 \| 1016 \| 0.269 \| 430 \| 274 \| 13 \| 78 \| 502 \| 2 \| 396 \| Portal protein \| Portal protein; afdb-uniprot50 \| AF-A0A348FY34-F1-MODEL\_V4 \| 1.0 \| 9.805e-28 \| 1014 \| 0.251 \| 474 \| 315 \| 15 \| 30 \| 488 \| 1 \| 449 \| Phage portal protein \| Phage portal protein; afdb-uniprot50 \| AF-A0A7C5PYI8-F1-MODEL\_V4 \| 1.0 \| 8.277e-29 \| 1014 \| 0.248 \| 499 \| 337 \| 15 \| 34 \| 502 \| 28 \| 518 \| Phage portal protein \| Phage portal protein; afdb-uniprot50 \| AF-A0A1C3E9J0-F1-MODEL\_V4 \| 1.0 \| 4.276e-27 \| 1013 \| 0.211 \| 453 \| 326 \| 12 \| 79 \| 523 \| 2 \| 431 \| Phage portal protein \| Phage portal protein; afdb-uniprot50 \| AF-A0A353ZI73-F1-MODEL\_V4 \| 1.0 \| 2e-29 \| 1013 \| 0.191 \| 516 \| 368 \| 14 \| 45 \| 532 \| 47 \| 541 \| Phage portal protein \| Phage portal protein; afdb-uniprot50 \| AF-A0A497V0G5-F1-MODEL\_V4 \| 1.0 \| 7.151e-28 \| 1012 \| 0.202 \| 498 \| 358 \| 15 \| 35 \| 530 \| 25 \| 485 \| Lambda family phage portal protein \| Lambda family phage portal protein; afdb-uniprot50 \| AF-A0A5C7T3J8-F1-MODEL\_V4 \| 1.0 \| 7.45e-29 \| 1012 \| 0.221 \| 511 \| 349 \| 15 \| 35 \| 531 \| 22 \| 497 \| Phage portal protein \| Phage portal protein; afdb-uniprot50 \| AF-A0A841K9E4-F1-MODEL\_V4 \| 1.0 \| 7.853e-29 \| 1012 \| 0.238 \| 512 \| 337 \| 15 \| 35 \| 532 \| 18 \| 490 \| Lambda family phage portal protein \| Lambda family phage portal protein; afdb-uniprot50 \| AF-A0A437MJQ4-F1-MODEL\_V4 \| 1.0 \| 2.024e-28 \| 1011 \| 0.224 \| 482 \| 326 \| 16 \| 36 \| 502 \| 29 \| 477 \| Phage portal protein \| Phage portal protein; afdb-uniprot50 \| AF-A0A5C8P971-F1-MODEL\_V4 \| 1.0 \| 4.01e-28 \| 1010 \| 0.216 \| 504 \| 351 \| 15 \| 40 \| 532 \| 32 \| 502 \| Phage portal protein \| Phage portal protein; afdb-uniprot50 \| AF-A0A166DQ97-F1-MODEL\_V4 \| 1.0 \| 4.403e-29 \| 1009 \| 0.226 \| 507 \| 337 \| 15 \| 21 \| 502 \| 1 \| 477 \| Phage portal protein, lambda family \| Phage portal protein, lambda family; afdb-uniprot50 \| AF-A0A2E3QA51-F1-MODEL\_V4 \| 1.0 \| 3.249e-28 \| 1009 \| 0.231 \| 483 \| 326 \| 16 \| 36 \| 502 \| 32 \| 485 \| Phage portal protein \| Phage portal protein; afdb-uniprot50 \| AF-A0A2D6BG92-F1-MODEL\_V4 \| 1.0 \| 1.196e-28 \| 1008 \| 0.245 \| 465 \| 316 \| 12 \| 42 \| 502 \| 17 \| 450 \| Phage portal protein \| Phage portal protein; afdb-uniprot50 \| AF-A0A420DP32-F1-MODEL\_V4 \| 1.0 \| 2.633e-28 \| 1008 \| 0.247 \| 497 \| 318 \| 15 \| 23 \| 502 \| 2 \| 459 \| Lambda family phage portal protein \| Lambda family phage portal protein; afdb-uniprot50 \| AF-A0A6N9P582-F1-MODEL\_V4 \| 1.0 \| 4.226e-28 \| 1008 \| 0.246 \| 479 \| 322 \| 14 \| 49 \| 500 \| 10 \| 476 \| Phage portal protein \| Phage portal protein; afdb-uniprot50 \| AF-A0A6G5QFV0-F1-MODEL\_V4 \| 1.0 \| 7.235e-27 \| 1007 \| 0.239 \| 505 \| 322 \| 17 \| 15 \| 500 \| 1 \| 462 \| Phage portal protein, lambda family \| Phage portal protein, lambda family; afdb-uniprot50 \| AF-A0A840JNA6-F1-MODEL\_V4 \| 1.0 \| 1.822e-28 \| 1007 \| 0.205 \| 512 \| 363 \| 15 \| 35 \| 517 \| 21 \| 517 \| Lambda family phage portal protein \| Lambda family phage portal protein; afdb-uniprot50 \| AF-A0A518B2S3-F1-MODEL\_V4 \| 1.0 \| 3.425e-28 \| 1005 \| 0.203 \| 516 \| 375 \| 12 \| 39 \| 529 \| 9 \| 513 \| Phage portal protein, lambda family \| Phage portal protein, lambda family; afdb-uniprot50 \| AF-A0A6I2J0E3-F1-MODEL\_V4 \| 1.0 \| 3.804e-28 \| 1002 \| 0.456 \| 348 \| 175 \| 6 \| 191 \| 532 \| 1 \| 340 \| Phage portal protein \| Phage portal protein; afdb-uniprot50 \| AF-A0A1T4QCI5-F1-MODEL\_V4 \| 1.0 \| 1.033e-27 \| 1002 \| 0.211 \| 511 \| 355 \| 14 \| 17 \| 501 \| 1 \| 489 \| Phage portal protein, lambda family \| Phage portal protein, lambda family; afdb-uniprot50 \| AF-A0A075MK45-F1-MODEL\_V4 \| 1.0 \| 1.417e-27 \| 1001 \| 0.215 \| 506 \| 338 \| 16 \| 46 \| 517 \| 6 \| 486 \| Phage portal protein, lambda family \| Phage portal protein, lambda family; afdb-uniprot50 \| AF-A0A510IYQ6-F1-MODEL\_V4 \| 1.0 \| 5.007e-27 \| 1000 \| 0.228 \| 416 \| 296 \| 12 \| 35 \| 439 \| 18 \| 419 \| Phage portal protein \| Phage portal protein; afdb-uniprot50 \| AF-A0A2Z6AZ44-F1-MODEL\_V4 \| 1.0 \| 1.92e-28 \| 1000 \| 0.216 \| 512 \| 357 \| 15 \| 21 \| 500 \| 1 \| 500 \| Lambda family phage portal protein \| Lambda family phage portal protein; afdb-uniprot50 \| AF-A0A3P3D1Q4-F1-MODEL\_V4 \| 1.0 \| 1.033e-27 \| 999 \| 0.232 \| 477 \| 311 \| 15 \| 38 \| 502 \| 3 \| 436 \| Phage portal protein \| Phage portal protein; afdb-uniprot50 \| AF-A0A7D7VGU6-F1-MODEL\_V4 \| 1.0 \| 2.807e-27 \| 998 \| 0.24 \| 483 \| 325 \| 15 \| 46 \| 502 \| 11 \| 477 \| Phage portal protein \| Phage portal protein; afdb-uniprot50 \| AF-A0A6L6WAJ2-F1-MODEL\_V4 \| 1.0 \| 4.276e-27 \| 998 \| 0.202 \| 475 \| 341 \| 13 \| 36 \| 495 \| 20 \| 471 \| Phage portal protein \| Phage portal protein; afdb-uniprot50 \| AF-K0NIH4-F1-MODEL\_V4 \| 1.0 \| 9.691e-29 \| 998 \| 0.209 \| 550 \| 368 \| 16 \| 8 \| 532 \| 1 \| 508 \| Phage portal protein, lambda family \| Phage portal protein, lambda family; afdb-uniprot50 \| AF-A0A522VF55-F1-MODEL\_V4 \| 1.0 \| 2.602e-29 \| 998 \| 0.195 \| 578 \| 377 \| 16 \| 2 \| 532 \| 3 \| 539 \| Phage portal protein \| Phage portal protein; afdb-uniprot50 \| AF-A0A225NFN7-F1-MODEL\_V4 \| 1.0 \| 2.959e-27 \| 997 \| 0.245 \| 427 \| 287 \| 13 \| 77 \| 502 \| 2 \| 394 \| Uncharacterized protein \| Uncharacterized protein; afdb-uniprot50 \| AF-A0A2D8RC71-F1-MODEL\_V4 \| 1.0 \| 2.275e-27 \| 996 \| 0.248 \| 466 \| 313 \| 15 \| 39 \| 502 \| 28 \| 458 \| Phage portal protein \| Phage portal protein; afdb-uniprot50 \| AF-A5V8D4-F1-MODEL\_V4 \| 1.0 \| 1.843e-27 \| 996 \| 0.242 \| 474 \| 319 \| 17 \| 36 \| 496 \| 27 \| 473 \| Phage portal protein, lambda family \| Phage portal protein, lambda family; afdb-uniprot50 \| AF-A0A7G9GXH5-F1-MODEL\_V4 \| 1.0 \| 1.476e-28 \| 995 \| 0.23 \| 508 \| 332 \| 15 \| 39 \| 525 \| 35 \| 504 \| Phage portal protein \| Phage portal protein; afdb-uniprot50 \| AF-A0A3P3DCQ1-F1-MODEL\_V4 \| 1.0 \| 1.033e-27 \| 995 \| 0.243 \| 502 \| 349 \| 14 \| 39 \| 523 \| 21 \| 508 \| Phage portal protein \| Phage portal protein; afdb-uniprot50 \| AF-A0A1V2DSA5-F1-MODEL\_V4 \| 1.0 \| 2.664e-27 \| 994 \| 0.23 \| 485 \| 330 \| 14 \| 46 \| 502 \| 6 \| 475 \| Phage portal protein \| Phage portal protein; afdb-uniprot50 \| AF-A0A7X9FN59-F1-MODEL\_V4 \| 1.0 \| 1.92e-28 \| 994 \| 0.226 \| 547 \| 363 \| 19 \| 15 \| 532 \| 1 \| 516 \| Phage portal protein \| Phage portal protein; afdb-uniprot50 \| AF-A0A4V0XDV8-F1-MODEL\_V4 \| 1.0 \| 4.75e-27 \| 994 \| 0.214 \| 485 \| 339 \| 13 \| 34 \| 502 \| 12 \| 470 \| Phage portal protein \| Phage portal protein; afdb-uniprot50 \| AF-A0A4U8YXU7-F1-MODEL\_V4 \| 1.0 \| 1.196e-28 \| 994 \| 0.233 \| 506 \| 344 \| 15 \| 48 \| 523 \| 23 \| 514 \| Phage portal protein lambda family \| Phage portal protein lambda family; afdb-uniprot50 \| AF-A0A2W5SIJ8-F1-MODEL\_V4 \| 1.0 \| 7.853e-29 \| 993 \| 0.209 \| 564 \| 375 \| 15 \| 1 \| 532 \| 1 \| 525 \| Phage portal protein \| Phage portal protein; afdb-uniprot50 \| AF-A0A316S9H5-F1-MODEL\_V4 \| 1.0 \| 3.652e-27 \| 992 \| 0.205 \| 492 \| 357 \| 9 \| 39 \| 518 \| 34 \| 503 \| Phage portal protein \| Phage portal protein; afdb-uniprot50 \| AF-A0A225SMR3-F1-MODEL\_V4 \| 1.0 \| 3.119e-27 \| 992 \| 0.213 \| 482 \| 347 \| 15 \| 39 \| 497 \| 34 \| 506 \| Phage portal protein \| Phage portal protein; afdb-uniprot50 \| AF-A0A0F9F3I2-F1-MODEL\_V4 \| 1.0 \| 1.21e-27 \| 991 \| 0.215 \| 520 \| 353 \| 14 \| 35 \| 531 \| 37 \| 524 \| Uncharacterized protein \| Uncharacterized protein; afdb-uniprot50 \| AF-A0A0R3MA71-F1-MODEL\_V4 \| 1.0 \| 3.609e-28 \| 990 \| 0.201 \| 536 \| 377 \| 16 \| 2 \| 502 \| 1 \| 520 \| Uncharacterized protein \| Uncharacterized protein; afdb-uniprot50 \| AF-A0A4U8S355-F1-MODEL\_V4 \| 1.0 \| 3.287e-27 \| 989 \| 0.204 \| 503 \| 329 \| 15 \| 15 \| 499 \| 1 \| 450 \| Phage portal protein \| Phage portal protein; afdb-uniprot50 \| AF-A0A1I5HB10-F1-MODEL\_V4 \| 1.0 \| 1.401e-28 \| 988 \| 0.209 \| 526 \| 359 \| 16 \| 20 \| 532 \| 3 \| 484 \| Phage portal protein, lambda family \| Phage portal protein, lambda family; afdb-uniprot50 \| AF-A0A1S1TN63-F1-MODEL\_V4 \| 1.0 \| 7.151e-28 \| 988 \| 0.225 \| 488 \| 341 \| 18 \| 35 \| 498 \| 19 \| 493 \| Phage portal protein \| Phage portal protein; afdb-uniprot50 \| AF-A0A316NFS7-F1-MODEL\_V4 \| 1.0 \| 1.246e-29 \| 988 \| 0.216 \| 532 \| 344 \| 14 \| 40 \| 532 \| 36 \| 533 \| Phage portal protein \| Phage portal protein; afdb-uniprot50 \| AF-A0A2N3BA45-F1-MODEL\_V4 \| 1.0 \| 6.179e-27 \| 987 \| 0.29 \| 400 \| 261 \| 11 \| 27 \| 415 \| 1 \| 388 \| Phage portal protein \| Phage portal protein; afdb-uniprot50 \| AF-A0A7V2AKM7-F1-MODEL\_V4 \| 1.0 \| 1.344e-27 \| 985 \| 0.224 \| 539 \| 356 \| 16 \| 27 \| 532 \| 2 \| 511 \| Phage portal protein \| Phage portal protein; afdb-uniprot50 \| AF-A0A7W6NJ63-F1-MODEL\_V4 \| 1.0 \| 2e-29 \| 985 \| 0.209 \| 559 \| 380 \| 21 \| 1 \| 532 \| 2 \| 525 \| Lambda family phage portal protein \| Lambda family phage portal protein; afdb-uniprot50 \| AF-A0A7R7T6F2-F1-MODEL\_V4 \| 1.0 \| 4.276e-27 \| 984 \| 0.259 \| 485 \| 314 \| 12 \| 36 \| 502 \| 23 \| 480 \| Phage portal protein \| Phage portal protein; afdb-uniprot50 \| AF-A0A4R3AAZ5-F1-MODEL\_V4 \| 1.0 \| 8.929e-27 \| 983 \| 0.252 \| 468 \| 307 \| 12 \| 44 \| 496 \| 2 \| 441 \| Lambda family phage portal protein \| Lambda family phage portal protein; afdb-uniprot50 \| AF-A0A2E2Q7X0-F1-MODEL\_V4 \| 1.0 \| 5.498e-28 \| 983 \| 0.221 \| 514 \| 352 \| 17 \| 46 \| 532 \| 18 \| 510 \| Phage portal protein \| Phage portal protein; afdb-uniprot50 \| AF-A0A4R3HW29-F1-MODEL\_V4 \| 1.0 \| 6.513e-27 \| 983 \| 0.229 \| 480 \| 341 \| 16 \| 40 \| 497 \| 35 \| 507 \| Lambda family phage portal protein \| Lambda family phage portal protein; afdb-uniprot50 \| AF-A0A1G7C613-F1-MODEL\_V4 \| 1.0 \| 1.679e-26 \| 982 \| 0.26 \| 433 \| 286 \| 9 \| 62 \| 491 \| 1 \| 402 \| Phage portal protein, lambda family \| Phage portal protein, lambda family; afdb-uniprot50 \| AF-A0A090KJ34-F1-MODEL\_V4 \| 1.0 \| 9.92e-27 \| 982 \| 0.178 \| 504 \| 369 \| 13 \| 36 \| 532 \| 27 \| 492 \| Phage portal protein \| Phage portal protein; afdb-uniprot50 \| AF-H0T3Q1-F1-MODEL\_V4 \| 1.0 \| 1.135e-28 \| 981 \| 0.21 \| 519 \| 341 \| 15 \| 58 \| 532 \| 3 \| 496 \| Putative Phage portal protein, lambda \| Putative Phage portal protein, lambda; afdb-uniprot50 \| AF-A0A7W0HL78-F1-MODEL\_V4 \| 1.0 \| 7.945e-28 \| 981 \| 0.24 \| 520 \| 351 \| 17 \| 1 \| 498 \| 28 \| 525 \| Lambda family phage portal protein \| Lambda family phage portal protein; afdb-uniprot50 \| AF-A0A1T2XA84-F1-MODEL\_V4 \| 1.0 \| 9.195e-29 \| 980 \| 0.214 \| 551 \| 368 \| 18 \| 15 \| 515 \| 1 \| 536 \| Phage portal protein \| Phage portal protein; afdb-uniprot50 \| AF-A0A016XIN5-F1-MODEL\_V4 \| 1.0 \| 1.843e-27 \| 978 \| 0.23 \| 499 \| 336 \| 12 \| 23 \| 502 \| 2 \| 471 \| Uncharacterized protein \| Uncharacterized protein; afdb-uniprot50 \| AF-A0A117KHB7-F1-MODEL\_V4 \| 1.0 \| 4.507e-27 \| 978 \| 0.251 \| 481 \| 310 \| 19 \| 39 \| 502 \| 23 \| 470 \| Portal protein \| Portal protein; afdb-uniprot50 \| AF-A0A0Q4S2Z1-F1-MODEL\_V4 \| 1.0 \| 1.556e-28 \| 973 \| 0.243 \| 522 \| 325 \| 19 \| 35 \| 532 \| 28 \| 503 \| Uncharacterized protein \| Uncharacterized protein; afdb-uniprot50 \| AF-A0A363S2N1-F1-MODEL\_V4 \| 1.0 \| 4.949e-28 \| 973 \| 0.186 \| 510 \| 364 \| 16 \| 39 \| 517 \| 21 \| 510 \| Phage portal protein \| Phage portal protein; afdb-uniprot50 \| AF-A0A3A9B6U4-F1-MODEL\_V4 \| 1.0 \| 2.807e-27 \| 973 \| 0.202 \| 490 \| 345 \| 14 \| 43 \| 492 \| 14 \| 497 \| Phage portal protein \| Phage portal protein; afdb-uniprot50 \| AF-A0A6M3K3Y5-F1-MODEL\_V4 \| 1.0 \| 2.775e-28 \| 973 \| 0.201 \| 531 \| 372 \| 16 \| 39 \| 530 \| 52 \| 569 \| Putative portal protein \| Putative portal protein; afdb-uniprot50 \| AF-A0A5C7Q6R0-F1-MODEL\_V4 \| 1.0 \| 6.513e-27 \| 972 \| 0.23 \| 498 \| 337 \| 13 \| 40 \| 532 \| 18 \| 474 \| Phage portal protein \| Phage portal protein; afdb-uniprot50 \| AF-A0A4Y1MVH7-F1-MODEL\_V4 \| 1.0 \| 1.417e-27 \| 972 \| 0.224 \| 531 \| 331 \| 17 \| 39 \| 532 \| 27 \| 513 \| Portal protein \| Portal protein; afdb-uniprot50 \| AF-A0A238KCR7-F1-MODEL\_V4 \| 1.0 \| 2.398e-27 \| 971 \| 0.233 \| 492 \| 334 \| 17 \| 35 \| 517 \| 18 \| 475 \| Phage portal protein, lambda family \| Phage portal protein, lambda family; afdb-uniprot50 \| AF-A0A1V5N9M5-F1-MODEL\_V4 \| 1.0 \| 3.249e-28 \| 971 \| 0.213 \| 533 \| 357 \| 15 \| 17 \| 532 \| 1 \| 488 \| Phage portal protein, lambda family \| Phage portal protein, lambda family; afdb-uniprot50 \| AF-A0A0M5L0Q6-F1-MODEL\_V4 \| 1.0 \| 3.804e-28 \| 969 \| 0.21 \| 517 \| 369 \| 11 \| 39 \| 532 \| 5 \| 505 \| Phage-related portal protein \| Phage-related portal protein; afdb-uniprot50 \| AF-A0A643F376-F1-MODEL\_V4 \| 1.0 \| 3.505e-26 \| 968 \| 0.235 \| 484 \| 309 \| 16 \| 18 \| 497 \| 1 \| 427 \| Phage portal protein \| Phage portal protein; afdb-uniprot50 \| AF-A0A557QLS5-F1-MODEL\_V4 \| 1.0 \| 1.511e-26 \| 968 \| 0.211 \| 478 \| 352 \| 13 \| 39 \| 497 \| 39 \| 510 \| Phage portal protein \| Phage portal protein; afdb-uniprot50 \| AF-A0A1C1YRY8-F1-MODEL\_V4 \| 1.0 \| 2.891e-29 \| 968 \| 0.219 \| 557 \| 375 \| 21 \| 8 \| 532 \| 1 \| 529 \| Uncharacterized protein \| Uncharacterized protein; afdb-uniprot50 \| AF-A0A5C5X2S3-F1-MODEL\_V4 \| 1.0 \| 2.275e-27 \| 968 \| 0.203 \| 515 \| 357 \| 20 \| 37 \| 532 \| 35 \| 515 \| Phage portal protein, lambda family \| Phage portal protein, lambda family; afdb-uniprot50 \| AF-A0A1V6DZJ2-F1-MODEL\_V4 \| 1.0 \| 4.057e-27 \| 967 \| 0.2 \| 505 \| 355 \| 15 \| 39 \| 521 \| 18 \| 495 \| Phage portal protein, lambda family \| Phage portal protein, lambda family; afdb-uniprot50 \| AF-A0A258A0N0-F1-MODEL\_V4 \| 1.0 \| 3.465e-27 \| 967 \| 0.205 \| 526 \| 346 \| 14 \| 39 \| 532 \| 20 \| 505 \| Phage portal protein \| Phage portal protein; afdb-uniprot50 \| AF-A0A1V6E2A1-F1-MODEL\_V4 \| 1.0 \| 1.92e-28 \| 967 \| 0.246 \| 528 \| 330 \| 18 \| 33 \| 530 \| 36 \| 525 \| Phage portal protein, lambda family \| Phage portal protein, lambda family; afdb-uniprot50 \| AF-A0A1P8QQ19-F1-MODEL\_V4 \| 1.0 \| 3.652e-27 \| 967 \| 0.227 \| 484 \| 340 \| 16 \| 36 \| 496 \| 31 \| 503 \| Phage portal protein \| Phage portal protein; afdb-uniprot50 \| AF-A0A5M6HUJ4-F1-MODEL\_V4 \| 1.0 \| 5.498e-28 \| 965 \| 0.252 \| 511 \| 333 \| 15 \| 10 \| 498 \| 46 \| 529 \| Phage portal protein \| Phage portal protein; afdb-uniprot50 \| AF-A0A0Q4HVV6-F1-MODEL\_V4 \| 1.0 \| 1.046e-26 \| 964 \| 0.255 \| 443 \| 288 \| 14 \| 75 \| 502 \| 2 \| 417 \| Portal protein \| Portal protein; afdb-uniprot50 \| AF-A0A1Q6U834-F1-MODEL\_V4 \| 1.0 \| 3.465e-27 \| 964 \| 0.218 \| 480 \| 335 \| 15 \| 39 \| 488 \| 24 \| 493 \| Phage portal protein \| Phage portal protein; afdb-uniprot50 \| AF-A0A2L1U4F6-F1-MODEL\_V4 \| 1.0 \| 1.196e-28 \| 964 \| 0.215 \| 530 \| 356 \| 14 \| 34 \| 532 \| 12 \| 512 \| Phage portal protein, lambda family \| Phage portal protein, lambda family; afdb-uniprot50 \| AF-A0A259U583-F1-MODEL\_V4 \| 1.0 \| 1.728e-28 \| 964 \| 0.209 \| 553 \| 370 \| 16 \| 20 \| 532 \| 1 \| 526 \| Phage portal protein, lambda family \| Phage portal protein, lambda family; afdb-uniprot50 \| AF-A0A4Q0SRN9-F1-MODEL\_V4 \| 1.0 \| 6.513e-27 \| 963 \| 0.229 \| 471 \| 323 \| 16 \| 29 \| 484 \| 1 \| 446 \| Uncharacterized protein \| Uncharacterized protein; afdb-uniprot50 \| AF-A0A495PS99-F1-MODEL\_V4 \| 1.0 \| 2.527e-27 \| 963 \| 0.246 \| 520 \| 321 \| 17 \| 16 \| 532 \| 5 \| 456 \| Lambda family phage portal protein \| Lambda family phage portal protein; afdb-uniprot50 \| AF-A0A1X9SVH6-F1-MODEL\_V4 \| 1.0 \| 7.715e-26 \| 961 \| 0.233 \| 476 \| 287 \| 14 \| 20 \| 475 \| 2 \| 419 \| Phage portal protein, lambda family \| Phage portal protein, lambda family; afdb-uniprot50 \| AF-A0A1H9GFL6-F1-MODEL\_V4 \| 1.0 \| 6.179e-27 \| 961 \| 0.226 \| 468 \| 328 \| 17 \| 39 \| 497 \| 23 \| 465 \| Phage portal protein, lambda family \| Phage portal protein, lambda family; afdb-uniprot50 \| AF-A0A1J5EX01-F1-MODEL\_V4 \| 1.0 \| 1.556e-28 \| 961 \| 0.198 \| 520 \| 353 \| 19 \| 46 \| 532 \| 12 \| 500 \| Phage portal protein \| Phage portal protein; afdb-uniprot50 \| AF-A0A421BJ29-F1-MODEL\_V4 \| 1.0 \| 9.195e-29 \| 961 \| 0.219 \| 573 \| 356 \| 19 \| 1 \| 532 \| 111 \| 632 \| Phage portal protein \| Phage portal protein; afdb-uniprot50 \| AF-A0A1E3GXJ9-F1-MODEL\_V4 \| 1.0 \| 2.072e-26 \| 961 \| 0.25 \| 475 \| 314 \| 15 \| 39 \| 502 \| 27 \| 470 \| Phage portal protein, lambda family \| Phage portal protein, lambda family; afdb-uniprot50 \| AF-A0A2P9HNT3-F1-MODEL\_V4 \| 1.0 \| 3.849e-27 \| 960 \| 0.221 \| 461 \| 324 \| 15 \| 8 \| 456 \| 1 \| 438 \| Phage portal protein, lambda family \| Phage portal protein, lambda family; afdb-uniprot50 \| AF-A0A2U2MXN7-F1-MODEL\_V4 \| 1.0 \| 6.108e-28 \| 958 \| 0.228 \| 546 \| 353 \| 21 \| 44 \| 532 \| 21 \| 554 \| Phage portal protein \| Phage portal protein; afdb-uniprot50 \| AF-G1UXR1-F1-MODEL\_V4 \| 1.0 \| 9.303e-28 \| 957 \| 0.223 \| 515 \| 350 \| 16 \| 11 \| 490 \| 1 \| 500 \| Lambda family phage portal protein \| Lambda family phage portal protein; afdb-uniprot50 \| AF-A0A2N8HQG3-F1-MODEL\_V4 \| 1.0 \| 2.807e-27 \| 957 \| 0.187 \| 538 \| 391 \| 13 \| 27 \| 523 \| 1 \| 533 \| Phage portal protein \| Phage portal protein; afdb-uniprot50 \| AF-A0A1Y6CPS8-F1-MODEL\_V4 \| 1.0 \| 3.849e-27 \| 952 \| 0.207 \| 492 \| 340 \| 17 \| 52 \| 516 \| 11 \| 479 \| Phage portal protein, lambda family \| Phage portal protein, lambda family; afdb-uniprot50 \| AF-A0A849WA89-F1-MODEL\_V4 \| 1.0 \| 7.235e-27 \| 952 \| 0.214 \| 480 \| 333 \| 16 \| 36 \| 502 \| 31 \| 479 \| Phage portal protein \| Phage portal protein; afdb-uniprot50 \| AF-A0A1A8T9E5-F1-MODEL\_V4 \| 1.0 \| 1.943e-27 \| 952 \| 0.185 \| 523 \| 365 \| 15 \| 43 \| 531 \| 12 \| 507 \| Phage portal protein, lambda family \| Phage portal protein, lambda family; afdb-uniprot50 \| AF-A0A367FXQ0-F1-MODEL\_V4 \| 1.0 \| 3.804e-28 \| 952 \| 0.237 \| 523 \| 352 \| 16 \| 36 \| 518 \| 23 \| 538 \| Phage portal protein \| Phage portal protein; afdb-uniprot50 \| AF-A0A0P6VW69-F1-MODEL\_V4 \| 1.0 \| 1.769e-26 \| 950 \| 0.224 \| 504 \| 341 \| 14 \| 36 \| 531 \| 23 \| 484 \| Uncharacterized protein \| Uncharacterized protein; afdb-uniprot50 \| AF-A0A212IX70-F1-MODEL\_V4 \| 1.0 \| 1.769e-26 \| 950 \| 0.223 \| 488 \| 336 \| 16 \| 44 \| 495 \| 15 \| 495 \| Lambda family phage portal protein \| Lambda family phage portal protein; afdb-uniprot50 \| AF-A0A7X4AD15-F1-MODEL\_V4 \| 1.0 \| 3.652e-27 \| 949 \| 0.202 \| 519 \| 346 \| 20 \| 15 \| 530 \| 15 \| 468 \| Phage portal protein \| Phage portal protein; afdb-uniprot50 \| AF-A0A7Z3TXU7-F1-MODEL\_V4 \| 1.0 \| 1.36e-26 \| 949 \| 0.221 \| 497 \| 327 \| 18 \| 20 \| 490 \| 3 \| 465 \| Phage portal protein \| Phage portal protein; afdb-uniprot50 \| AF-A0A0K1NDB4-F1-MODEL\_V4 \| 1.0 \| 6.785e-28 \| 949 \| 0.211 \| 562 \| 342 \| 13 \| 17 \| 532 \| 8 \| 514 \| Portal protein \| Portal protein; afdb-uniprot50 \| AF-A0A2A5A4N7-F1-MODEL\_V4 \| 1.0 \| 7.715e-26 \| 948 \| 0.21 \| 471 \| 333 \| 13 \| 36 \| 497 \| 30 \| 470 \| Phage portal protein \| Phage portal protein; afdb-uniprot50 \| AF-A0A839H7H8-F1-MODEL\_V4 \| 1.0 \| 1.769e-26 \| 947 \| 0.22 \| 495 \| 330 \| 15 \| 46 \| 532 \| 31 \| 477 \| Phage portal protein \| Phage portal protein; afdb-uniprot50 \| AF-A0A7C8HT84-F1-MODEL\_V4 \| 1.0 \| 1.224e-26 \| 946 \| 0.209 \| 505 \| 332 \| 12 \| 39 \| 523 \| 34 \| 491 \| Uncharacterized protein \| Uncharacterized protein; afdb-uniprot50 \| AF-A0A1H5Z5E8-F1-MODEL\_V4 \| 1.0 \| 1.21e-27 \| 946 \| 0.222 \| 543 \| 337 \| 16 \| 5 \| 502 \| 1 \| 503 \| Phage portal protein \| Phage portal protein; afdb-uniprot50 \| AF-A3VBF2-F1-MODEL\_V4 \| 1.0 \| 2.664e-27 \| 945 \| 0.226 \| 538 \| 335 \| 20 \| 20 \| 532 \| 1 \| 482 \| Uncharacterized protein \| Uncharacterized protein; afdb-uniprot50 \| AF-E3H7C7-F1-MODEL\_V4 \| 1.0 \| 4.56e-26 \| 945 \| 0.195 \| 486 \| 349 \| 12 \| 39 \| 494 \| 29 \| 502 \| Phage portal protein, lambda family \| Phage portal protein, lambda family; afdb-uniprot50 \| AF-A0A0G1Z1U3-F1-MODEL\_V4 \| 1.0 \| 1.511e-26 \| 942 \| 0.232 \| 469 \| 320 \| 16 \| 57 \| 495 \| 2 \| 460 \| Phage portal protein, lambda family \| Phage portal protein, lambda family; afdb-uniprot50 \| AF-A0A134CAF6-F1-MODEL\_V4 \| 1.0 \| 1.21e-27 \| 942 \| 0.201 \| 521 \| 353 \| 17 \| 13 \| 488 \| 1 \| 503 \| Phage portal protein, lambda family \| Phage portal protein, lambda family; afdb-uniprot50 \| AF-M2YEI3-F1-MODEL\_V4 \| 1.0 \| 1.659e-27 \| 942 \| 0.214 \| 512 \| 349 \| 18 \| 47 \| 532 \| 51 \| 535 \| Lambda family phage portal protein \| Lambda family phage portal protein; afdb-uniprot50 \| AF-A0A2T5K540-F1-MODEL\_V4 \| 1.0 \| 3.652e-27 \| 942 \| 0.219 \| 525 \| 355 \| 18 \| 36 \| 532 \| 38 \| 535 \| Lambda family phage portal protein \| Lambda family phage portal protein; afdb-uniprot50 \| AF-A0A546X7Y9-F1-MODEL\_V4 \| 1.0 \| 2.133e-28 \| 939 \| 0.194 \| 541 \| 382 \| 20 \| 1 \| 515 \| 6 \| 518 \| Phage portal protein \| Phage portal protein; afdb-uniprot50 \| AF-A0A095YJQ5-F1-MODEL\_V4 \| 1.0 \| 5.562e-27 \| 938 \| 0.223 \| 496 \| 339 \| 17 \| 39 \| 502 \| 22 \| 503 \| Lambda family phage portal protein \| Lambda family phage portal protein; afdb-uniprot50 \| AF-A0A369BH58-F1-MODEL\_V4 \| 1.0 \| 1.401e-28 \| 938 \| 0.233 \| 530 \| 347 \| 20 \| 39 \| 523 \| 31 \| 545 \| Lambda family phage portal protein \| Lambda family phage portal protein; afdb-uniprot50 \| AF-A0A353GEE3-F1-MODEL\_V4 \| 1.0 \| 2.301e-26 \| 937 \| 0.219 \| 515 \| 360 \| 16 \| 23 \| 532 \| 1 \| 478 \| Uncharacterized protein \| Uncharacterized protein; afdb-uniprot50 \| AF-A0A2U1WJ09-F1-MODEL\_V4 \| 1.0 \| 1.046e-26 \| 937 \| 0.202 \| 500 \| 337 \| 16 \| 46 \| 502 \| 3 \| 483 \| Uncharacterized protein \| Uncharacterized protein; afdb-uniprot50 \| AF-A0A2V7CXU2-F1-MODEL\_V4 \| 1.0 \| 1.148e-27 \| 936 \| 0.207 \| 554 \| 351 \| 16 \| 9 \| 532 \| 1 \| 496 \| Phage portal protein \| Phage portal protein; afdb-uniprot50 \| AF-A0A4V3ZNW4-F1-MODEL\_V4 \| 1.0 \| 1.344e-27 \| 935 \| 0.229 \| 535 \| 345 \| 15 \| 2 \| 492 \| 82 \| 593 \| Phage portal protein \| Phage portal protein; afdb-uniprot50 \| AF-A0A7C3DQI9-F1-MODEL\_V4 \| 1.0 \| 7.715e-26 \| 934 \| 0.248 \| 479 \| 299 \| 16 \| 35 \| 502 \| 11 \| 439 \| Phage portal protein \| Phage portal protein; afdb-uniprot50 \| AF-A0A5D0RSH2-F1-MODEL\_V4 \| 1.0 \| 3.155e-26 \| 934 \| 0.226 \| 491 \| 338 \| 15 \| 40 \| 517 \| 25 \| 486 \| Phage portal protein \| Phage portal protein; afdb-uniprot50 \| AF-A0A840Z535-F1-MODEL\_V4 \| 1.0 \| 5.066e-26 \| 934 \| 0.189 \| 476 \| 342 \| 16 \| 39 \| 502 \| 27 \| 470 \| Lambda family phage portal protein \| Lambda family phage portal protein; afdb-uniprot50 \| AF-D6KLF9-F1-MODEL\_V4 \| 1.0 \| 6.437e-28 \| 934 \| 0.191 \| 564 \| 380 \| 16 \| 13 \| 532 \| 1 \| 532 \| Phage portal protein, lambda family \| Phage portal protein, lambda family; afdb-uniprot50 \| AF-A0A1Q6U9X6-F1-MODEL\_V4 \| 1.0 \| 1.224e-26 \| 933 \| 0.223 \| 483 \| 310 \| 15 \| 78 \| 532 \| 2 \| 447 \| Phage portal protein \| Phage portal protein; afdb-uniprot50 \| AF-A0A7G6RHV5-F1-MODEL\_V4 \| 1.0 \| 1.004e-25 \| 933 \| 0.229 \| 419 \| 295 \| 12 \| 85 \| 496 \| 2 \| 399 \| Phage portal protein \| Phage portal protein; afdb-uniprot50 \| AF-A0A1I3HJ62-F1-MODEL\_V4 \| 1.0 \| 3.465e-27 \| 933 \| 0.222 \| 509 \| 330 \| 16 \| 13 \| 491 \| 1 \| 473 \| Phage portal protein, lambda family \| Phage portal protein, lambda family; afdb-uniprot50 \| AF-A0A101K665-F1-MODEL\_V4 \| 1.0 \| 4.695e-28 \| 933 \| 0.213 \| 530 \| 332 \| 15 \| 39 \| 530 \| 25 \| 507 \| Phage portal protein \| Phage portal protein; afdb-uniprot50 \| AF-A0A385Q1G5-F1-MODEL\_V4 \| 1.0 \| 3.894e-26 \| 932 \| 0.224 \| 494 \| 341 \| 15 \| 36 \| 519 \| 23 \| 484 \| Phage portal protein \| Phage portal protein; afdb-uniprot50 \| AF-A0A6I4TSN8-F1-MODEL\_V4 \| 1.0 \| 1.493e-27 \| 932 \| 0.181 \| 588 \| 411 \| 19 \| 1 \| 532 \| 2 \| 574 \| Phage portal protein \| Phage portal protein; afdb-uniprot50 \| AF-A0A1E2RXZ6-F1-MODEL\_V4 \| 1.0 \| 6.945e-26 \| 930 \| 0.295 \| 399 \| 252 \| 10 \| 105 \| 496 \| 3 \| 379 \| Phage portal protein, lambda family \| Phage portal protein, lambda family; afdb-uniprot50 \| AF-A0A5S4X1N8-F1-MODEL\_V4 \| 1.0 \| 1.058e-25 \| 930 \| 0.218 \| 467 \| 334 \| 16 \| 27 \| 484 \| 1 \| 445 \| Phage portal protein \| Phage portal protein; afdb-uniprot50 \| AF-A0A4R3EC07-F1-MODEL\_V4 \| 1.0 \| 1.511e-26 \| 930 \| 0.22 \| 480 \| 324 \| 17 \| 35 \| 492 \| 18 \| 469 \| Lambda family phage portal protein \| Lambda family phage portal protein; afdb-uniprot50 \| AF-A0A356C2B9-F1-MODEL\_V4 \| 1.0 \| 4.75e-27 \| 928 \| 0.21 \| 538 \| 356 \| 12 \| 39 \| 531 \| 29 \| 542 \| Phage portal protein \| Phage portal protein; afdb-uniprot50 \| AF-A0A0Q7XTZ7-F1-MODEL\_V4 \| 1.0 \| 5.498e-28 \| 928 \| 0.197 \| 547 \| 355 \| 20 \| 20 \| 532 \| 1 \| 497 \| Uncharacterized protein \| Uncharacterized protein; afdb-uniprot50 \| AF-Q72A26-F1-MODEL\_V4 \| 1.0 \| 5.277e-27 \| 927 \| 0.203 \| 541 \| 361 \| 17 \| 23 \| 530 \| 1 \| 504 \| Portal protein \| Portal protein; afdb-uniprot50 \| AF-G1USE5-F1-MODEL\_V4 \| 1.0 \| 1.749e-27 \| 926 \| 0.209 \| 529 \| 348 \| 20 \| 39 \| 532 \| 31 \| 524 \| Lambda family phage portal protein \| Lambda family phage portal protein; afdb-uniprot50 \| AF-A0A842IW31-F1-MODEL\_V4 \| 1.0 \| 1.769e-26 \| 925 \| 0.208 \| 537 \| 362 \| 15 \| 20 \| 532 \| 1 \| 498 \| Phage portal protein \| Phage portal protein; afdb-uniprot50 \| AF-A0A1S2L7H0-F1-MODEL\_V4 \| 1.0 \| 3.287e-27 \| 924 \| 0.221 \| 523 \| 361 \| 17 \| 20 \| 515 \| 2 \| 505 \| Phage portal protein \| Phage portal protein; afdb-uniprot50 \| AF-A0A835Z6S7-F1-MODEL\_V4 \| 1.0 \| 6.945e-26 \| 924 \| 0.228 \| 482 \| 316 \| 18 \| 39 \| 502 \| 27 \| 470 \| Phage portal protein \| Phage portal protein; afdb-uniprot50 \| AF-A0A2P7V3N4-F1-MODEL\_V4 \| 1.0 \| 2.158e-27 \| 923 \| 0.212 \| 513 \| 343 \| 17 \| 36 \| 495 \| 23 \| 527 \| Phage portal protein \| Phage portal protein; afdb-uniprot50 \| AF-A0A369XXM4-F1-MODEL\_V4 \| 1.0 \| 4.104e-26 \| 922 \| 0.226 \| 468 \| 323 \| 14 \| 49 \| 497 \| 40 \| 487 \| Phage portal protein \| Phage portal protein; afdb-uniprot50 \| AF-A0A431PRZ4-F1-MODEL\_V4 \| 1.0 \| 7.715e-26 \| 922 \| 0.218 \| 476 \| 333 \| 14 \| 39 \| 502 \| 25 \| 473 \| Phage portal protein \| Phage portal protein; afdb-uniprot50 \| AF-A3K1Z6-F1-MODEL\_V4 \| 1.0 \| 5.627e-26 \| 921 \| 0.204 \| 488 \| 334 \| 15 \| 35 \| 490 \| 21 \| 486 \| Phage portal protein, lambda \| Phage portal protein, lambda; afdb-uniprot50 \| AF-A0A6L9LZ03-F1-MODEL\_V4 \| 1.0 \| 8.132e-26 \| 921 \| 0.214 \| 514 \| 356 \| 16 \| 15 \| 497 \| 1 \| 497 \| Phage portal protein \| Phage portal protein; afdb-uniprot50 \| AF-A0A7X7KP38-F1-MODEL\_V4 \| 1.0 \| 3.287e-27 \| 920 \| 0.196 \| 566 \| 360 \| 21 \| 15 \| 532 \| 1 \| 519 \| Phage portal protein \| Phage portal protein; afdb-uniprot50 \| AF-A0A259DE51-F1-MODEL\_V4 \| 1.0 \| 2.183e-26 \| 918 \| 0.233 \| 441 \| 294 \| 13 \| 98 \| 532 \| 3 \| 405 \| Phage portal protein \| Phage portal protein; afdb-uniprot50 \| AF-A0A7X7H5B0-F1-MODEL\_V4 \| 1.0 \| 6.437e-28 \| 917 \| 0.21 \| 538 \| 339 \| 18 \| 36 \| 521 \| 10 \| 513 \| Phage portal protein \| Phage portal protein; afdb-uniprot50 \| AF-A0A3L7Q8T8-F1-MODEL\_V4 \| 1.0 \| 2.301e-26 \| 917 \| 0.207 \| 512 \| 364 \| 15 \| 1 \| 502 \| 1 \| 480 \| Phage portal protein \| Phage portal protein; afdb-uniprot50 \| AF-A0A509DVN1-F1-MODEL\_V4 \| 1.0 \| 2.587e-25 \| 916 \| 0.226 \| 476 \| 327 \| 16 \| 36 \| 502 \| 26 \| 469 \| Uncharacterized protein \| Uncharacterized protein; afdb-uniprot50 \| AF-A0A5B9MAJ4-F1-MODEL\_V4 \| 1.0 \| 2.664e-27 \| 916 \| 0.206 \| 581 \| 383 \| 17 \| 12 \| 532 \| 3 \| 565 \| Phage portal protein, lambda family \| Phage portal protein, lambda family; afdb-uniprot50 \| AF-A0A3D2ICL3-F1-MODEL\_V4 \| 1.0 \| 1.102e-26 \| 915 \| 0.205 \| 521 \| 360 \| 14 \| 35 \| 515 \| 21 \| 527 \| Phage portal protein \| Phage portal protein; afdb-uniprot50 \| AF-A0A4Q7TER2-F1-MODEL\_V4 \| 1.0 \| 3.695e-26 \| 914 \| 0.222 \| 476 \| 307 \| 11 \| 41 \| 467 \| 12 \| 473 \| Lambda family phage portal protein \| Lambda family phage portal protein; afdb-uniprot50 \| AF-A0A6H1ZM60-F1-MODEL\_V4 \| 1.0 \| 5.627e-26 \| 913 \| 0.211 \| 530 \| 353 \| 18 \| 39 \| 532 \| 33 \| 533 \| Putative portal protein \| Putative portal protein; afdb-uniprot50 \| AF-A0A1M6LTT3-F1-MODEL\_V4 \| 1.0 \| 6.945e-26 \| 912 \| 0.234 \| 498 \| 335 \| 13 \| 39 \| 500 \| 17 \| 504 \| Phage portal protein, lambda family \| Phage portal protein, lambda family; afdb-uniprot50 \| AF-A0A1L3GDR9-F1-MODEL\_V4 \| 1.0 \| 1.175e-25 \| 912 \| 0.207 \| 482 \| 346 \| 15 \| 35 \| 488 \| 25 \| 498 \| Phage portal protein \| Phage portal protein; afdb-uniprot50 \| AF-A0A2U8WAV1-F1-MODEL\_V4 \| 1.0 \| 3.652e-27 \| 912 \| 0.186 \| 563 \| 385 \| 21 \| 13 \| 517 \| 1 \| 548 \| Phage portal protein \| Phage portal protein; afdb-uniprot50 \| AF-E1JR91-F1-MODEL\_V4 \| 1.0 \| 9.412e-27 \| 912 \| 0.207 \| 525 \| 348 \| 17 \| 45 \| 532 \| 18 \| 511 \| Phage portal protein, lambda family \| Phage portal protein, lambda family; afdb-uniprot50 \| AF-A0A369QQY9-F1-MODEL\_V4 \| 1.0 \| 1.511e-26 \| 911 \| 0.189 \| 548 \| 373 \| 17 \| 11 \| 532 \| 1 \| 503 \| Phage portal protein \| Phage portal protein; afdb-uniprot50 \| AF-A0A353MHD9-F1-MODEL\_V4 \| 1.0 \| 8.929e-27 \| 911 \| 0.208 \| 536 \| 368 \| 20 \| 21 \| 532 \| 9 \| 511 \| Phage portal protein \| Phage portal protein; afdb-uniprot50 \| AF-A0A854GYB8-F1-MODEL\_V4 \| 1.0 \| 2.183e-26 \| 910 \| 0.232 \| 494 \| 327 \| 16 \| 39 \| 495 \| 17 \| 495 \| Phage portal protein \| Phage portal protein; afdb-uniprot50 \| AF-A0A1H1JS74-F1-MODEL\_V4 \| 1.0 \| 1.058e-25 \| 910 \| 0.211 \| 491 \| 335 \| 13 \| 45 \| 496 \| 28 \| 505 \| Phage portal protein, lambda family \| Phage portal protein, lambda family; afdb-uniprot50 \| AF-E2CHI6-F1-MODEL\_V4 \| 1.0 \| 4.862e-25 \| 909 \| 0.215 \| 478 \| 333 \| 13 \| 39 \| 502 \| 23 \| 472 \| Phage portal protein, lambda family \| Phage portal protein, lambda family; afdb-uniprot50 \| AF-A0A5C7P2N1-F1-MODEL\_V4 \| 1.0 \| 6.513e-27 \| 909 \| 0.171 \| 549 \| 393 \| 18 \| 23 \| 532 \| 10 \| 535 \| Phage portal protein \| Phage portal protein; afdb-uniprot50 \| AF-A0A5E6U3P4-F1-MODEL\_V4 \| 1.0 \| 2.557e-26 \| 909 \| 0.217 \| 501 \| 339 \| 18 \| 39 \| 503 \| 33 \| 516 \| Uncharacterized protein \| Uncharacterized protein; afdb-uniprot50 \| AF-K2IL77-F1-MODEL\_V4 \| 1.0 \| 4.507e-27 \| 909 \| 0.183 \| 572 \| 377 \| 18 \| 15 \| 532 \| 1 \| 536 \| Portal protein \| Portal protein; afdb-uniprot50 \| AF-A0A349XPR7-F1-MODEL\_V4 \| 1.0 \| 9.634e-25 \| 908 \| 0.235 \| 428 \| 280 \| 15 \| 38 \| 452 \| 3 \| 396 \| Phage portal protein \| Phage portal protein; afdb-uniprot50 \| AF-A0A2U0YK94-F1-MODEL\_V4 \| 1.0 \| 1.376e-25 \| 908 \| 0.226 \| 451 \| 307 \| 12 \| 98 \| 532 \| 2 \| 426 \| Lambda family phage portal protein \| Lambda family phage portal protein; afdb-uniprot50 \| AF-A0A1H7TZE7-F1-MODEL\_V4 \| 1.0 \| 1.434e-26 \| 908 \| 0.187 \| 571 \| 373 \| 21 \| 20 \| 532 \| 9 \| 546 \| Phage portal protein, lambda family \| Phage portal protein, lambda family; afdb-uniprot50 \| AF-A0A7Y4VYM6-F1-MODEL\_V4 \| 1.0 \| 2.301e-26 \| 908 \| 0.197 \| 526 \| 379 \| 14 \| 39 \| 532 \| 39 \| 553 \| Phage portal protein \| Phage portal protein; afdb-uniprot50 \| AF-A0A2W4SHI4-F1-MODEL\_V4 \| 1.0 \| 2.356e-24 \| 906 \| 0.244 \| 389 \| 268 \| 12 \| 35 \| 409 \| 19 \| 395 \| Phage portal protein \| Phage portal protein; afdb-uniprot50 \| AF-H1HCQ0-F1-MODEL\_V4 \| 1.0 \| 1.434e-26 \| 904 \| 0.206 \| 509 \| 348 \| 15 \| 39 \| 517 \| 29 \| 511 \| Lambda family phage portal protein \| Lambda family phage portal protein; afdb-uniprot50 \| AF-A0A354U425-F1-MODEL\_V4 \| 1.0 \| 1.239e-25 \| 903 \| 0.216 \| 521 \| 337 \| 15 \| 20 \| 532 \| 2 \| 459 \| Uncharacterized protein \| Uncharacterized protein; afdb-uniprot50 \| AF-A0A418VVD6-F1-MODEL\_V4 \| 1.0 \| 7.235e-27 \| 903 \| 0.207 \| 525 \| 349 \| 17 \| 45 \| 519 \| 18 \| 525 \| Phage portal protein \| Phage portal protein; afdb-uniprot50 \| AF-A0A1Q6PWQ7-F1-MODEL\_V4 \| 1.0 \| 1.529e-25 \| 903 \| 0.212 \| 484 \| 342 \| 17 \| 49 \| 500 \| 51 \| 527 \| Phage portal protein \| Phage portal protein; afdb-uniprot50 \| AF-A0A367V4E8-F1-MODEL\_V4 \| 1.0 \| 5.125e-25 \| 902 \| 0.235 \| 442 \| 294 \| 15 \| 75 \| 502 \| 2 \| 413 \| Uncharacterized protein \| Uncharacterized protein; afdb-uniprot50 \| AF-A0A059KRD9-F1-MODEL\_V4 \| 1.0 \| 7.32e-26 \| 901 \| 0.204 \| 508 \| 350 \| 19 \| 39 \| 532 \| 22 \| 489 \| Uncharacterized protein \| Uncharacterized protein; afdb-uniprot50 \| AF-A0A1H2EP65-F1-MODEL\_V4 \| 1.0 \| 1.046e-26 \| 901 \| 0.211 \| 562 \| 357 \| 17 \| 20 \| 531 \| 1 \| 526 \| Phage portal protein, lambda family \| Phage portal protein, lambda family; afdb-uniprot50 \| AF-A0A521VMA3-F1-MODEL\_V4 \| 1.0 \| 2.84e-26 \| 900 \| 0.217 \| 529 \| 344 \| 15 \| 39 \| 532 \| 39 \| 532 \| Phage portal protein \| Phage portal protein; afdb-uniprot50 \| AF-A0A1H5ZJ40-F1-MODEL\_V4 \| 1.0 \| 1.253e-24 \| 899 \| 0.253 \| 442 \| 290 \| 15 \| 39 \| 466 \| 23 \| 438 \| Phage portal protein, lambda family \| Phage portal protein, lambda family; afdb-uniprot50 \| AF-A0A410NTY6-F1-MODEL\_V4 \| 1.0 \| 1.224e-26 \| 899 \| 0.217 \| 537 \| 358 \| 20 \| 33 \| 524 \| 25 \| 544 \| Phage portal protein \| Phage portal protein; afdb-uniprot50 \| AF-G5NAD5-F1-MODEL\_V4 \| 1.0 \| 2.454e-25 \| 898 \| 0.241 \| 460 \| 301 \| 15 \| 73 \| 502 \| 1 \| 442 \| Phage portal protein \| Phage portal protein; afdb-uniprot50 \| AF-A0A1W1BN33-F1-MODEL\_V4 \| 1.0 \| 1.698e-25 \| 898 \| 0.212 \| 509 \| 323 \| 19 \| 39 \| 523 \| 13 \| 467 \| Phage portal protein, lambda family \| Phage portal protein, lambda family; afdb-uniprot50 \| AF-A0A5Q3L866-F1-MODEL\_V4 \| 1.0 \| 5.627e-26 \| 898 \| 0.213 \| 535 \| 361 \| 19 \| 31 \| 532 \| 5 \| 512 \| Phage portal protein, lambda family \| Phage portal protein, lambda family; afdb-uniprot50 \| AF-A0A5A9EPH8-F1-MODEL\_V4 \| 1.0 \| 5.693e-25 \| 898 \| 0.215 \| 479 \| 327 \| 16 \| 39 \| 497 \| 28 \| 477 \| Phage portal protein \| Phage portal protein; afdb-uniprot50 \| AF-A0A5C8BL97-F1-MODEL\_V4 \| 1.0 \| 6.252e-26 \| 897 \| 0.219 \| 488 \| 334 \| 16 \| 49 \| 498 \| 46 \| 524 \| Phage portal protein \| Phage portal protein; afdb-uniprot50 \| AF-A0A1I5F286-F1-MODEL\_V4 \| 1.0 \| 8.929e-27 \| 896 \| 0.19 \| 535 \| 365 \| 17 \| 41 \| 532 \| 48 \| 557 \| Phage portal protein, lambda family \| Phage portal protein, lambda family; afdb-uniprot50 \| AF-A0A517NLJ6-F1-MODEL\_V4 \| 1.0 \| 2.183e-26 \| 895 \| 0.189 \| 550 \| 374 \| 20 \| 9 \| 531 \| 1 \| 505 \| Phage portal protein, lambda family \| Phage portal protein, lambda family; afdb-uniprot50 \| AF-A0A1V1UMH2-F1-MODEL\_V4 \| 1.0 \| 9.412e-27 \| 895 \| 0.191 \| 554 \| 374 \| 20 \| 7 \| 523 \| 1 \| 517 \| Phage portal protein, lambda family \| Phage portal protein, lambda family; afdb-uniprot50 \| AF-G4KQ83-F1-MODEL\_V4 \| 1.0 \| 1.769e-26 \| 895 \| 0.195 \| 548 \| 362 \| 17 \| 39 \| 532 \| 35 \| 557 \| Putative portal protein \| Putative portal protein; afdb-uniprot50 \| AF-A0A1B9VJM9-F1-MODEL\_V4 \| 1.0 \| 2.454e-25 \| 895 \| 0.212 \| 508 \| 333 \| 16 \| 38 \| 495 \| 13 \| 503 \| Phage portal protein, lambda family \| Phage portal protein, lambda family; afdb-uniprot50 \| AF-A0A5B8BXE4-F1-MODEL\_V4 \| 1.0 \| 8.227e-25 \| 893 \| 0.179 \| 491 \| 366 \| 15 \| 39 \| 502 \| 34 \| 514 \| Phage portal protein \| Phage portal protein; afdb-uniprot50 \| AF-A0A7Y4DJD1-F1-MODEL\_V4 \| 1.0 \| 6.325e-25 \| 892 \| 0.145 \| 488 \| 389 \| 13 \| 37 \| 517 \| 21 \| 487 \| Phage portal protein \| Phage portal protein; afdb-uniprot50 \| AF-A0A2Z5ZE91-F1-MODEL\_V4 \| 1.0 \| 1.593e-26 \| 891 \| 0.194 \| 585 \| 362 \| 20 \| 5 \| 532 \| 15 \| 546 \| Phage portal protein \| Phage portal protein; afdb-uniprot50 \| AF-A0A7C4ZHN2-F1-MODEL\_V4 \| 1.0 \| 1.434e-26 \| 890 \| 0.227 \| 480 \| 308 \| 17 \| 28 \| 491 \| 1 \| 433 \| Phage portal protein \| Phage portal protein; afdb-uniprot50 \| AF-E3HBM5-F1-MODEL\_V4 \| 1.0 \| 2.209e-25 \| 890 \| 0.204 \| 495 \| 353 \| 19 \| 31 \| 498 \| 19 \| 499 \| Phage portal protein, lambda family \| Phage portal protein, lambda family; afdb-uniprot50 \| AF-A0A349HLY6-F1-MODEL\_V4 \| 1.0 \| 8.571e-26 \| 889 \| 0.24 \| 499 \| 307 \| 19 \| 67 \| 532 \| 2 \| 461 \| Phage portal protein \| Phage portal protein; afdb-uniprot50 \| AF-A0A841G3H8-F1-MODEL\_V4 \| 1.0 \| 1.865e-26 \| 889 \| 0.204 \| 548 \| 361 \| 20 \| 1 \| 502 \| 78 \| 596 \| Lambda family phage portal protein \| Lambda family phage portal protein; afdb-uniprot50 \| AF-A0A1G5T351-F1-MODEL\_V4 \| 1.0 \| 2.183e-26 \| 888 \| 0.171 \| 555 \| 403 \| 15 \| 11 \| 532 \| 2 \| 532 \| Phage portal protein, lambda family \| Phage portal protein, lambda family; afdb-uniprot50 \| AF-A0A6N1VH77-F1-MODEL\_V4 \| 1.0 \| 1.239e-25 \| 886 \| 0.221 \| 515 \| 337 \| 18 \| 35 \| 532 \| 24 \| 491 \| Phage portal protein \| Phage portal protein; afdb-uniprot50 \| AF-A0A7J5WDZ0-F1-MODEL\_V4 \| 1.0 \| 1.887e-25 \| 886 \| 0.194 \| 525 \| 360 \| 20 \| 42 \| 532 \| 12 \| 507 \| Phage portal protein lambda \| Phage portal protein lambda; afdb-uniprot50 \| AF-A0A268THM8-F1-MODEL\_V4 \| 1.0 \| 1.467e-24 \| 885 \| 0.212 \| 480 \| 312 \| 18 \| 38 \| 500 \| 19 \| 449 \| Uncharacterized protein \| Uncharacterized protein; afdb-uniprot50 \| AF-A0A7C8EMQ6-F1-MODEL\_V4 \| 1.0 \| 1.004e-25 \| 885 \| 0.188 \| 520 \| 366 \| 15 \| 2 \| 502 \| 7 \| 489 \| Phage portal protein \| Phage portal protein; afdb-uniprot50 \| AF-A0A068SLY6-F1-MODEL\_V4 \| 1.0 \| 5.277e-27 \| 885 \| 0.195 \| 538 \| 366 \| 20 \| 17 \| 532 \| 1 \| 493 \| Phage portal protein, lambda family \| Phage portal protein, lambda family; afdb-uniprot50 \| AF-A0A1G2YTA2-F1-MODEL\_V4 \| 1.0 \| 4.377e-25 \| 884 \| 0.216 \| 495 \| 340 \| 17 \| 17 \| 502 \| 1 \| 456 \| Uncharacterized protein \| Uncharacterized protein; afdb-uniprot50 \| AF-A0A7V3NBN7-F1-MODEL\_V4 \| 1.0 \| 1.79e-25 \| 884 \| 0.208 \| 548 \| 354 \| 21 \| 4 \| 532 \| 1 \| 487 \| Phage portal protein \| Phage portal protein; afdb-uniprot50 \| AF-A0A2N5XX91-F1-MODEL\_V4 \| 1.0 \| 6.001e-25 \| 884 \| 0.228 \| 490 \| 320 \| 16 \| 36 \| 502 \| 23 \| 477 \| Phage portal protein \| Phage portal protein; afdb-uniprot50 \| AF-A0A258VI72-F1-MODEL\_V4 \| 1.0 \| 5.066e-26 \| 884 \| 0.203 \| 520 \| 352 \| 20 \| 36 \| 532 \| 21 \| 501 \| Uncharacterized protein \| Uncharacterized protein; afdb-uniprot50 \| AF-A0A389MVE2-F1-MODEL\_V4 \| 1.0 \| 4.75e-27 \| 884 \| 0.185 \| 589 \| 379 \| 18 \| 1 \| 532 \| 10 \| 554 \| Uncharacterized protein \| Uncharacterized protein; afdb-uniprot50 \| AF-A0A063ZY87-F1-MODEL\_V4 \| 1.0 \| 7.626e-27 \| 884 \| 0.211 \| 515 \| 352 \| 15 \| 39 \| 517 \| 579 \| 1075 \| Uncharacterized protein \| Uncharacterized protein; afdb-uniprot50 \| AF-A0A357BWI3-F1-MODEL\_V4 \| 1.0 \| 1.909e-24 \| 883 \| 0.23 \| 521 \| 337 \| 17 \| 17 \| 531 \| 1 \| 463 \| Uncharacterized protein \| Uncharacterized protein; afdb-uniprot50 \| AF-A0A3C1F6B3-F1-MODEL\_V4 \| 1.0 \| 5.693e-25 \| 883 \| 0.193 \| 490 \| 346 \| 16 \| 40 \| 492 \| 41 \| 518 \| Phage portal protein \| Phage portal protein; afdb-uniprot50 \| AF-A0A550KMN9-F1-MODEL\_V4 \| 1.0 \| 1.239e-25 \| 882 \| 0.192 \| 571 \| 386 \| 17 \| 6 \| 532 \| 1 \| 540 \| Phage portal protein \| Phage portal protein; afdb-uniprot50 \| AF-A0A3D5KD24-F1-MODEL\_V4 \| 1.0 \| 2.557e-26 \| 881 \| 0.2 \| 513 \| 357 \| 14 \| 37 \| 526 \| 43 \| 525 \| Uncharacterized protein \| Uncharacterized protein; afdb-uniprot50 \| AF-A0A4P8IPG7-F1-MODEL\_V4 \| 1.0 \| 5.339e-26 \| 880 \| 0.175 \| 524 \| 359 \| 16 \| 46 \| 518 \| 26 \| 527 \| Phage portal protein \| Phage portal protein; afdb-uniprot50 \| AF-A0A786U7W6-F1-MODEL\_V4 \| 1.0 \| 1.63e-24 \| 879 \| 0.219 \| 460 \| 305 \| 14 \| 70 \| 488 \| 2 \| 448 \| Phage portal protein \| Phage portal protein; afdb-uniprot50 \| AF-A0A796MCI7-F1-MODEL\_V4 \| 1.0 \| 4.104e-26 \| 879 \| 0.228 \| 534 \| 348 \| 18 \| 35 \| 532 \| 19 \| 524 \| Phage portal protein \| Phage portal protein; afdb-uniprot50 \| AF-A0A530AI76-F1-MODEL\_V4 \| 1.0 \| 3.826e-23 \| 878 \| 0.262 \| 362 \| 236 \| 10 \| 70 \| 417 \| 2 \| 346 \| Phage portal protein \| Phage portal protein; afdb-uniprot50 \| AF-A0A7X7K4M3-F1-MODEL\_V4 \| 1.0 \| 1.547e-24 \| 878 \| 0.217 \| 493 \| 323 \| 15 \| 45 \| 532 \| 4 \| 438 \| Phage portal protein \| Phage portal protein; afdb-uniprot50 \| AF-A0A7Y5S0S4-F1-MODEL\_V4 \| 1.0 \| 3.894e-26 \| 878 \| 0.209 \| 530 \| 345 \| 20 \| 22 \| 532 \| 6 \| 480 \| Phage portal protein \| Phage portal protein; afdb-uniprot50 \| AF-A0A6L8M286-F1-MODEL\_V4 \| 1.0 \| 1.058e-25 \| 878 \| 0.194 \| 510 \| 368 \| 17 \| 39 \| 532 \| 21 \| 503 \| Phage portal protein \| Phage portal protein; afdb-uniprot50 \| AF-A0A497U413-F1-MODEL\_V4 \| 1.0 \| 7.406e-25 \| 878 \| 0.217 \| 473 \| 306 \| 14 \| 73 \| 502 \| 2 \| 453 \| Phage portal protein \| Phage portal protein; afdb-uniprot50 \| AF-A0A706VWX3-F1-MODEL\_V4 \| 1.0 \| 1.679e-26 \| 878 \| 0.181 \| 585 \| 381 \| 22 \| 8 \| 532 \| 1 \| 547 \| Phage portal protein \| Phage portal protein; afdb-uniprot50 \| AF-A0A239C998-F1-MODEL\_V4 \| 1.0 \| 9.034e-26 \| 877 \| 0.206 \| 533 \| 364 \| 19 \| 27 \| 532 \| 1 \| 501 \| Phage portal protein, lambda family \| Phage portal protein, lambda family; afdb-uniprot50 \| AF-A0A661TNZ2-F1-MODEL\_V4 \| 1.0 \| 4.326e-26 \| 877 \| 0.18 \| 571 \| 376 \| 17 \| 6 \| 497 \| 59 \| 616 \| Phage portal protein \| Phage portal protein; afdb-uniprot50 \| AF-A0A4Q3P809-F1-MODEL\_V4 \| 1.0 \| 2.209e-25 \| 876 \| 0.229 \| 505 \| 331 \| 19 \| 11 \| 488 \| 1 \| 474 \| Phage portal protein \| Phage portal protein; afdb-uniprot50 \| AF-A0A5C7LHV8-F1-MODEL\_V4 \| 1.0 \| 1.058e-25 \| 876 \| 0.232 \| 521 \| 347 \| 18 \| 23 \| 532 \| 12 \| 490 \| Phage portal protein \| Phage portal protein; afdb-uniprot50 \| AF-A0A2W6XAC1-F1-MODEL\_V4 \| 1.0 \| 2.096e-25 \| 876 \| 0.225 \| 484 \| 322 \| 16 \| 39 \| 502 \| 33 \| 483 \| Phage portal protein \| Phage portal protein; afdb-uniprot50 \| AF-B6Y9F9-F1-MODEL\_V4 \| 1.0 \| 7.493e-24 \| 875 \| 0.295 \| 369 \| 239 \| 8 \| 135 \| 496 \| 4 \| 358 \| Putative phage portal protein \| Putative phage portal protein; afdb-uniprot50 \| AF-A0A1M2YW43-F1-MODEL\_V4 \| 1.0 \| 6.325e-25 \| 875 \| 0.202 \| 484 \| 324 \| 15 \| 62 \| 503 \| 3 \| 466 \| Phage portal protein \| Phage portal protein; afdb-uniprot50 \| AF-A0A2E2B1J6-F1-MODEL\_V4 \| 1.0 \| 6.666e-25 \| 875 \| 0.209 \| 507 \| 348 \| 17 \| 13 \| 502 \| 1 \| 471 \| Uncharacterized protein \| Uncharacterized protein; afdb-uniprot50 \| AF-A0A1V5QVH4-F1-MODEL\_V4 \| 1.0 \| 1.376e-25 \| 875 \| 0.204 \| 519 \| 357 \| 19 \| 40 \| 531 \| 38 \| 527 \| Phage portal protein, lambda family \| Phage portal protein, lambda family; afdb-uniprot50 \| AF-A0A5C5WDP0-F1-MODEL\_V4 \| 1.0 \| 1.529e-25 \| 874 \| 0.176 \| 527 \| 373 \| 16 \| 2 \| 497 \| 9 \| 505 \| Phage portal protein, lambda family \| Phage portal protein, lambda family; afdb-uniprot50 \| AF-A0A1B1C3C9-F1-MODEL\_V4 \| 1.0 \| 3.94e-25 \| 873 \| 0.166 \| 504 \| 373 \| 14 \| 37 \| 532 \| 26 \| 490 \| Uncharacterized protein \| Uncharacterized protein; afdb-uniprot50 \| AF-A0A2A5AG71-F1-MODEL\_V4 \| 1.0 \| 1.175e-25 \| 872 \| 0.185 \| 571 \| 370 \| 18 \| 20 \| 532 \| 1 \| 534 \| Uncharacterized protein \| Uncharacterized protein; afdb-uniprot50 \| AF-A0A2W6T5F2-F1-MODEL\_V4 \| 1.0 \| 4.862e-25 \| 871 \| 0.182 \| 509 \| 336 \| 16 \| 53 \| 503 \| 13 \| 499 \| Phage portal protein \| Phage portal protein; afdb-uniprot50 \| AF-A0A2S5TR06-F1-MODEL\_V4 \| 1.0 \| 6.001e-25 \| 871 \| 0.181 \| 501 \| 357 \| 15 \| 34 \| 495 \| 25 \| 511 \| Phage portal protein \| Phage portal protein; afdb-uniprot50 \| AF-A0A6N1VGE5-F1-MODEL\_V4 \| 1.0 \| 4.326e-26 \| 871 \| 0.204 \| 553 \| 351 \| 20 \| 36 \| 531 \| 25 \| 545 \| Phage portal protein \| Phage portal protein; afdb-uniprot50 \| AF-A0A1C7BGI4-F1-MODEL\_V4 \| 1.0 \| 4.613e-25 \| 870 \| 0.203 \| 511 \| 346 \| 19 \| 54 \| 532 \| 3 \| 484 \| Lambda family phage portal protein \| Lambda family phage portal protein; afdb-uniprot50 \| AF-A0A3B8S5Y1-F1-MODEL\_V4 \| 1.0 \| 1.45e-25 \| 870 \| 0.21 \| 512 \| 351 \| 18 \| 34 \| 499 \| 12 \| 516 \| Phage portal protein \| Phage portal protein; afdb-uniprot50 \| AF-A0A0F3IYT1-F1-MODEL\_V4 \| 1.0 \| 9.412e-27 \| 869 \| 0.243 \| 484 \| 310 \| 16 \| 36 \| 500 \| 23 \| 469 \| Uncharacterized protein \| Uncharacterized protein; afdb-uniprot50 \| AF-A0A2P2E5U7-F1-MODEL\_V4 \| 1.0 \| 8.227e-25 \| 869 \| 0.204 \| 490 \| 341 \| 18 \| 39 \| 502 \| 30 \| 496 \| Uncharacterized protein \| Uncharacterized protein; afdb-uniprot50 \| AF-A0A1I3H6N4-F1-MODEL\_V4 \| 1.0 \| 1.058e-25 \| 869 \| 0.191 \| 527 \| 360 \| 18 \| 39 \| 532 \| 27 \| 520 \| Phage portal protein, lambda family \| Phage portal protein, lambda family; afdb-uniprot50 \| AF-A0A1G3M2B9-F1-MODEL\_V4 \| 1.0 \| 1.115e-25 \| 868 \| 0.198 \| 544 \| 352 \| 22 \| 24 \| 532 \| 1 \| 495 \| Phage portal protein \| Phage portal protein; afdb-uniprot50 \| AF-A0A7C5LXR1-F1-MODEL\_V4 \| 1.0 \| 1.175e-25 \| 868 \| 0.198 \| 524 \| 362 \| 17 \| 22 \| 532 \| 28 \| 506 \| Phage portal protein \| Phage portal protein; afdb-uniprot50 \| AF-B4CXQ8-F1-MODEL\_V4 \| 1.0 \| 3.738e-25 \| 867 \| 0.199 \| 497 \| 323 \| 17 \| 38 \| 505 \| 3 \| 453 \| Portal protein lambda \| Portal protein lambda; afdb-uniprot50 \| AF-A0A6M3MCR2-F1-MODEL\_V4 \| 1.0 \| 3.505e-26 \| 867 \| 0.178 \| 515 \| 347 \| 16 \| 39 \| 529 \| 47 \| 509 \| Putative portal protein \| Putative portal protein; afdb-uniprot50 \| AF-A0A843BDF6-F1-MODEL\_V4 \| 1.0 \| 4.104e-26 \| 867 \| 0.208 \| 527 \| 349 \| 18 \| 37 \| 531 \| 34 \| 524 \| Phage portal protein \| Phage portal protein; afdb-uniprot50 \| AF-A0A2S8FUK2-F1-MODEL\_V4 \| 1.0 \| 3.029e-25 \| 866 \| 0.225 \| 509 \| 341 \| 20 \| 39 \| 527 \| 29 \| 504 \| Uncharacterized protein \| Uncharacterized protein; afdb-uniprot50 \| AF-A0A1U9NQD9-F1-MODEL\_V4 \| 1.0 \| 4.326e-26 \| 866 \| 0.198 \| 550 \| 360 \| 22 \| 14 \| 532 \| 1 \| 500 \| Phage portal protein, lambda family \| Phage portal protein, lambda family; afdb-uniprot50 \| AF-A0A2E3DCB1-F1-MODEL\_V4 \| 1.0 \| 7.32e-26 \| 866 \| 0.176 \| 550 \| 373 \| 18 \| 39 \| 532 \| 40 \| 565 \| Uncharacterized protein \| Uncharacterized protein; afdb-uniprot50 \| AF-A0A1W9HA00-F1-MODEL\_V4 \| 1.0 \| 2.907e-24 \| 865 \| 0.246 \| 486 \| 306 \| 18 \| 24 \| 502 \| 1 \| 433 \| Uncharacterized protein \| Uncharacterized protein; afdb-uniprot50 \| AF-A0A358PRF9-F1-MODEL\_V4 \| 1.0 \| 3.94e-25 \| 865 \| 0.221 \| 483 \| 323 \| 15 \| 85 \| 527 \| 3 \| 472 \| Phage portal protein \| Phage portal protein; afdb-uniprot50 \| AF-A0A3A1YQG3-F1-MODEL\_V4 \| 1.0 \| 1.306e-25 \| 865 \| 0.186 \| 542 \| 376 \| 19 \| 4 \| 532 \| 1 \| 490 \| Uncharacterized protein \| Uncharacterized protein; afdb-uniprot50 \| AF-S9S655-F1-MODEL\_V4 \| 1.0 \| 3.738e-25 \| 863 \| 0.19 \| 556 \| 367 \| 22 \| 27 \| 532 \| 13 \| 535 \| Phage portal protein, lambda family \| Phage portal protein, lambda family; afdb-uniprot50 \| AF-A0A661I4X9-F1-MODEL\_V4 \| 1.0 \| 1.63e-24 \| 862 \| 0.176 \| 493 \| 344 \| 16 \| 39 \| 521 \| 6 \| 446 \| Phage portal protein \| Phage portal protein; afdb-uniprot50 \| AF-A0A1M3AIF6-F1-MODEL\_V4 \| 1.0 \| 2.483e-24 \| 862 \| 0.176 \| 533 \| 348 \| 18 \| 52 \| 532 \| 5 \| 498 \| Phage portal protein \| Phage portal protein; afdb-uniprot50 \| AF-A0A4V0I5D1-F1-MODEL\_V4 \| 1.0 \| 3.695e-26 \| 862 \| 0.206 \| 561 \| 371 \| 21 \| 1 \| 532 \| 1 \| 516 \| Uncharacterized protein \| Uncharacterized protein; afdb-uniprot50 \| AF-A0A1Q3NUP4-F1-MODEL\_V4 \| 1.0 \| 6.666e-25 \| 862 \| 0.219 \| 506 \| 329 \| 18 \| 45 \| 503 \| 19 \| 505 \| Phage portal protein \| Phage portal protein; afdb-uniprot50 \| AF-A0A4R3MIV3-F1-MODEL\_V4 \| 1.0 \| 1.253e-24 \| 861 \| 0.273 \| 432 \| 266 \| 13 \| 15 \| 434 \| 1 \| 396 \| Lambda family phage portal protein \| Lambda family phage portal protein; afdb-uniprot50 \| AF-A0A2A5DIF0-F1-MODEL\_V4 \| 1.0 \| 4.428e-24 \| 860 \| 0.201 \| 471 \| 327 \| 14 \| 39 \| 501 \| 34 \| 463 \| Phage portal protein \| Phage portal protein; afdb-uniprot50 \| AF-A0A0F9JDA0-F1-MODEL\_V4 \| 1.0 \| 1.529e-25 \| 860 \| 0.203 \| 530 \| 351 \| 21 \| 17 \| 531 \| 1 \| 474 \| Uncharacterized protein \| Uncharacterized protein; afdb-uniprot50 \| AF-A0A2A4HXW4-F1-MODEL\_V4 \| 1.0 \| 5.465e-24 \| 860 \| 0.242 \| 471 \| 319 \| 14 \| 39 \| 502 \| 31 \| 470 \| Phage portal protein \| Phage portal protein; afdb-uniprot50 \| AF-A0A317FAF6-F1-MODEL\_V4 \| 1.0 \| 9.634e-25 \| 859 \| 0.218 \| 522 \| 325 \| 18 \| 45 \| 532 \| 12 \| 484 \| Phage portal protein \| Phage portal protein; afdb-uniprot50 \| AF-A0A5C7Q4H8-F1-MODEL\_V4 \| 1.0 \| 1.189e-24 \| 858 \| 0.217 \| 511 \| 334 \| 20 \| 27 \| 502 \| 3 \| 482 \| Phage portal protein \| Phage portal protein; afdb-uniprot50 \| AF-A0A0Q0F5Y9-F1-MODEL\_V4 \| 1.0 \| 2.483e-24 \| 857 \| 0.225 \| 453 \| 305 \| 17 \| 77 \| 496 \| 2 \| 441 \| Portal protein, lambda family \| Portal protein, lambda family; afdb-uniprot50 \| AF-A0A7X4AD17-F1-MODEL\_V4 \| 1.0 \| 3.782e-24 \| 857 \| 0.203 \| 521 \| 364 \| 16 \| 3 \| 518 \| 1 \| 475 \| Phage portal protein \| Phage portal protein; afdb-uniprot50 \| AF-A0A3L7SGJ4-F1-MODEL\_V4 \| 1.0 \| 5.066e-26 \| 857 \| 0.215 \| 544 \| 346 \| 22 \| 1 \| 532 \| 4 \| 478 \| Phage portal protein \| Phage portal protein; afdb-uniprot50 \| AF-A0A517SER7-F1-MODEL\_V4 \| 1.0 \| 1.07e-24 \| 856 \| 0.213 \| 510 \| 360 \| 19 \| 1 \| 502 \| 3 \| 479 \| Phage portal protein, lambda family \| Phage portal protein, lambda family; afdb-uniprot50 \| AF-A0A1Y0FDI5-F1-MODEL\_V4 \| 1.0 \| 3.404e-24 \| 855 \| 0.27 \| 403 \| 258 \| 14 \| 68 \| 451 \| 2 \| 387 \| Uncharacterized protein \| Uncharacterized protein; afdb-uniprot50 \| AF-A0A2T5K791-F1-MODEL\_V4 \| 1.0 \| 3.546e-25 \| 854 \| 0.232 \| 504 \| 317 \| 21 \| 15 \| 502 \| 1 \| 450 \| Lambda family phage portal protein \| Lambda family phage portal protein; afdb-uniprot50 \| AF-A0A1W1Y8S5-F1-MODEL\_V4 \| 1.0 \| 6.945e-26 \| 854 \| 0.174 \| 555 \| 377 \| 22 \| 1 \| 523 \| 4 \| 509 \| Capsid protein \| Capsid protein; afdb-uniprot50 \| AF-A0A7S6MI22-F1-MODEL\_V4 \| 1.0 \| 9.14e-25 \| 853 \| 0.192 \| 524 \| 353 \| 15 \| 42 \| 515 \| 23 \| 526 \| Phage portal protein \| Phage portal protein; afdb-uniprot50 \| AF-A0A840JB72-F1-MODEL\_V4 \| 1.0 \| 2.035e-23 \| 852 \| 0.224 \| 415 \| 291 \| 13 \| 98 \| 495 \| 2 \| 402 \| Lambda family phage portal protein \| Lambda family phage portal protein; afdb-uniprot50 \| AF-A0A3S0EMD5-F1-MODEL\_V4 \| 1.0 \| 2.235e-24 \| 852 \| 0.21 \| 508 \| 346 \| 18 \| 9 \| 502 \| 1 \| 467 \| Phage portal protein \| Phage portal protein; afdb-uniprot50 \| AF-A0A8B2PKS7-F1-MODEL\_V4 \| 1.0 \| 3.738e-25 \| 852 \| 0.21 \| 538 \| 338 \| 16 \| 46 \| 532 \| 26 \| 527 \| Uncharacterized protein \| Uncharacterized protein; afdb-uniprot50 \| AF-A0A3A1Y7Y1-F1-MODEL\_V4 \| 1.0 \| 1.015e-24 \| 852 \| 0.172 \| 505 \| 372 \| 15 \| 36 \| 532 \| 45 \| 511 \| Uncharacterized protein \| Uncharacterized protein; afdb-uniprot50 \| AF-A0A2D2C211-F1-MODEL\_V4 \| 1.0 \| 7.806e-25 \| 851 \| 0.198 \| 530 \| 352 \| 18 \| 15 \| 532 \| 1 \| 469 \| Phage portal protein \| Phage portal protein; afdb-uniprot50 \| AF-A0A7C1QAN0-F1-MODEL\_V4 \| 1.0 \| 8.227e-25 \| 851 \| 0.214 \| 517 \| 358 \| 19 \| 1 \| 502 \| 9 \| 492 \| Phage portal protein \| Phage portal protein; afdb-uniprot50 \| AF-A0A1X0SWB6-F1-MODEL\_V4 \| 1.0 \| 4.862e-25 \| 851 \| 0.205 \| 556 \| 357 \| 21 \| 9 \| 515 \| 14 \| 533 \| Phage portal protein, lambda family \| Phage portal protein, lambda family; afdb-uniprot50 \| AF-A0A6L6J9Q3-F1-MODEL\_V4 \| 1.0 \| 5.246e-23 \| 850 \| 0.516 \| 269 \| 119 \| 4 \| 266 \| 532 \| 41 \| 300 \| Phage portal protein \| Phage portal protein; afdb-uniprot50 \| AF-A0A522WDL3-F1-MODEL\_V4 \| 1.0 \| 2.261e-23 \| 850 \| 0.256 \| 393 \| 265 \| 10 \| 122 \| 502 \| 5 \| 382 \| Phage portal protein \| Phage portal protein; afdb-uniprot50 \| AF-A0A0F9KBE7-F1-MODEL\_V4 \| 1.0 \| 2.758e-24 \| 850 \| 0.274 \| 422 \| 262 \| 9 \| 127 \| 532 \| 7 \| 400 \| Uncharacterized protein \| Uncharacterized protein; afdb-uniprot50 \| AF-A0A4P5XY08-F1-MODEL\_V4 \| 1.0 \| 3.986e-24 \| 849 \| 0.199 \| 507 \| 356 \| 16 \| 34 \| 532 \| 5 \| 469 \| Uncharacterized protein \| Uncharacterized protein; afdb-uniprot50 \| AF-A0A496PDJ5-F1-MODEL\_V4 \| 1.0 \| 4.862e-25 \| 848 \| 0.202 \| 500 \| 325 \| 17 \| 81 \| 532 \| 2 \| 475 \| Phage portal protein \| Phage portal protein; afdb-uniprot50 \| AF-A0A812S2M1-F1-MODEL\_V4 \| 1.0 \| 5.76e-24 \| 847 \| 0.19 \| 483 \| 352 \| 19 \| 43 \| 502 \| 12 \| 478 \| Gene 4 protein \| Gene 4 protein; afdb-uniprot50 \| AF-A0A4Q6F5P2-F1-MODEL\_V4 \| 1.0 \| 1.336e-23 \| 846 \| 0.209 \| 415 \| 293 \| 14 \| 59 \| 449 \| 1 \| 404 \| Phage portal protein \| Phage portal protein; afdb-uniprot50 \| AF-A0A4E0QSA5-F1-MODEL\_V4 \| 1.0 \| 9.247e-24 \| 846 \| 0.232 \| 460 \| 292 \| 17 \| 39 \| 487 \| 7 \| 416 \| Phage portal protein \| Phage portal protein; afdb-uniprot50 \| AF-A0A6N8K7M5-F1-MODEL\_V4 \| 1.0 \| 5.125e-25 \| 846 \| 0.202 \| 518 \| 341 \| 19 \| 62 \| 530 \| 1 \| 495 \| Phage portal protein \| Phage portal protein; afdb-uniprot50 \| AF-A0A2T6DLK8-F1-MODEL\_V4 \| 1.0 \| 3.23e-24 \| 846 \| 0.192 \| 483 \| 332 \| 17 \| 44 \| 505 \| 23 \| 468 \| Uncharacterized protein \| Uncharacterized protein; afdb-uniprot50 \| AF-A0A518L4N8-F1-MODEL\_V4 \| 1.0 \| 6.666e-25 \| 845 \| 0.192 \| 562 \| 382 \| 16 \| 14 \| 515 \| 1 \| 550 \| Phage portal protein, lambda family \| Phage portal protein, lambda family; afdb-uniprot50 \| AF-A0A443L3G4-F1-MODEL\_V4 \| 1.0 \| 6.474e-23 \| 844 \| 0.256 \| 378 \| 256 \| 11 \| 36 \| 405 \| 28 \| 388 \| Phage portal protein \| Phage portal protein; afdb-uniprot50 \| AF-A0A2D8S0N3-F1-MODEL\_V4 \| 1.0 \| 4.377e-25 \| 844 \| 0.167 \| 538 \| 372 \| 17 \| 7 \| 502 \| 1 \| 504 \| Phage portal protein \| Phage portal protein; afdb-uniprot50 \| AF-A0A085TW01-F1-MODEL\_V4 \| 1.0 \| 2.383e-23 \| 843 \| 0.255 \| 391 \| 267 \| 11 \| 105 \| 491 \| 3 \| 373 \| Phage portal protein, lambda family \| Phage portal protein, lambda family; afdb-uniprot50 \| AF-A0A353B4R9-F1-MODEL\_V4 \| 1.0 \| 1.408e-23 \| 843 \| 0.214 \| 508 \| 329 \| 19 \| 19 \| 519 \| 11 \| 455 \| Uncharacterized protein \| Uncharacterized protein; afdb-uniprot50 \| AF-U1YH26-F1-MODEL\_V4 \| 1.0 \| 1.027e-23 \| 842 \| 0.228 \| 420 \| 283 \| 12 \| 127 \| 532 \| 3 \| 395 \| Uncharacterized protein \| Uncharacterized protein; afdb-uniprot50 \| AF-A0A1C7GVL9-F1-MODEL\_V4 \| 1.0 \| 5.76e-24 \| 842 \| 0.216 \| 435 \| 288 \| 10 \| 39 \| 468 \| 21 \| 407 \| Uncharacterized protein \| Uncharacterized protein; afdb-uniprot50 \| AF-U7GCW3-F1-MODEL\_V4 \| 1.0 \| 2.328e-25 \| 842 \| 0.193 \| 589 \| 382 \| 21 \| 1 \| 530 \| 3 \| 557 \| Portal protein \| Portal protein; afdb-uniprot50 \| AF-M5RV16-F1-MODEL\_V4 \| 1.0 \| 1.811e-24 \| 841 \| 0.201 \| 545 \| 377 \| 20 \| 1 \| 531 \| 14 \| 514 \| Phage portal protein, lambda family \| Phage portal protein, lambda family; afdb-uniprot50 \| AF-A0A1E4N739-F1-MODEL\_V4 \| 1.0 \| 1.79e-25 \| 841 \| 0.172 \| 568 \| 377 \| 21 \| 21 \| 532 \| 32 \| 562 \| Phage portal protein \| Phage portal protein; afdb-uniprot50 \| AF-A0A290MQ25-F1-MODEL\_V4 \| 1.0 \| 6.666e-25 \| 840 \| 0.189 \| 550 \| 359 \| 17 \| 39 \| 532 \| 21 \| 539 \| Phage portal protein \| Phage portal protein; afdb-uniprot50 \| AF-A0A2D5FQ67-F1-MODEL\_V4 \| 1.0 \| 1.07e-24 \| 838 \| 0.18 \| 560 \| 382 \| 21 \| 2 \| 502 \| 6 \| 547 \| Uncharacterized protein \| Uncharacterized protein; afdb-uniprot50 \| AF-A0A376ZFL5-F1-MODEL\_V4 \| 1.0 \| 4.377e-25 \| 838 \| 0.196 \| 576 \| 385 \| 21 \| 11 \| 532 \| 45 \| 596 \| Capsid protein of prophage \| Capsid protein of prophage; afdb-uniprot50 \| AF-A0A078MHH5-F1-MODEL\_V4 \| 1.0 \| 1.887e-25 \| 836 \| 0.194 \| 565 \| 365 \| 19 \| 20 \| 532 \| 1 \| 527 \| Phage portal protein, lambda family \| Phage portal protein, lambda family; afdb-uniprot50 \| AF-A0A653RBF9-F1-MODEL\_V4 \| 1.0 \| 3.738e-25 \| 835 \| 0.187 \| 556 \| 383 \| 21 \| 20 \| 532 \| 2 \| 531 \| Phage portal protein \| Phage portal protein; afdb-uniprot50 \| AF-A0A087M4C7-F1-MODEL\_V4 \| 1.0 \| 5.402e-25 \| 835 \| 0.172 \| 558 \| 400 \| 18 \| 15 \| 532 \| 1 \| 536 \| Uncharacterized protein \| Uncharacterized protein; afdb-uniprot50 \| AF-A0A4Z1R295-F1-MODEL\_V4 \| 1.0 \| 6.071e-24 \| 833 \| 0.179 \| 536 \| 352 \| 21 \| 49 \| 532 \| 4 \| 503 \| Phage portal protein \| Phage portal protein; afdb-uniprot50 \| AF-A0A4Y7RX04-F1-MODEL\_V4 \| 1.0 \| 4.153e-25 \| 833 \| 0.213 \| 548 \| 346 \| 19 \| 39 \| 532 \| 17 \| 533 \| Uncharacterized protein \| Uncharacterized protein; afdb-uniprot50 \| AF-A0A840FJN1-F1-MODEL\_V4 \| 1.0 \| 1.811e-24 \| 833 \| 0.177 \| 512 \| 366 \| 15 \| 39 \| 502 \| 29 \| 533 \| Lambda family phage portal protein \| Lambda family phage portal protein; afdb-uniprot50 \| AF-A0A5C6C1F2-F1-MODEL\_V4 \| 1.0 \| 2.758e-24 \| 832 \| 0.198 \| 530 \| 350 \| 19 \| 27 \| 532 \| 8 \| 486 \| Phage portal protein, lambda family \| Phage portal protein, lambda family; afdb-uniprot50 \| AF-A0A353BG30-F1-MODEL\_V4 \| 1.0 \| 2.012e-24 \| 832 \| 0.172 \| 528 \| 371 \| 21 \| 37 \| 532 \| 8 \| 501 \| Uncharacterized protein \| Uncharacterized protein; afdb-uniprot50 \| AF-A0A1Z9JFC0-F1-MODEL\_V4 \| 1.0 \| 2.035e-23 \| 831 \| 0.169 \| 520 \| 364 \| 19 \| 27 \| 530 \| 11 \| 478 \| Uncharacterized protein \| Uncharacterized protein; afdb-uniprot50 \| AF-A0A1Q3Z0R9-F1-MODEL\_V4 \| 1.0 \| 4.201e-24 \| 831 \| 0.211 \| 486 \| 329 \| 14 \| 58 \| 502 \| 3 \| 475 \| Phage portal protein \| Phage portal protein; afdb-uniprot50 \| AF-A0A2E2KT33-F1-MODEL\_V4 \| 1.0 \| 2.035e-23 \| 831 \| 0.194 \| 484 \| 341 \| 14 \| 46 \| 490 \| 5 \| 478 \| Phage portal protein \| Phage portal protein; afdb-uniprot50 \| AF-A0A7Y9FKF2-F1-MODEL\_V4 \| 1.0 \| 3.23e-24 \| 830 \| 0.231 \| 513 \| 322 \| 17 \| 35 \| 531 \| 24 \| 480 \| Lambda family phage portal protein \| Lambda family phage portal protein; afdb-uniprot50 \| AF-A0A285YSW8-F1-MODEL\_V4 \| 1.0 \| 1.738e-23 \| 830 \| 0.169 \| 495 \| 337 \| 15 \| 62 \| 502 \| 5 \| 479 \| Phage portal protein, lambda family \| Phage portal protein, lambda family; afdb-uniprot50 \| AF-D8JWB8-F1-MODEL\_V4 \| 1.0 \| 1.321e-24 \| 830 \| 0.194 \| 535 \| 365 \| 15 \| 18 \| 502 \| 1 \| 519 \| Phage portal protein, lambda family \| Phage portal protein, lambda family; afdb-uniprot50 \| AF-A0A270BET9-F1-MODEL\_V4 \| 1.0 \| 6.666e-25 \| 830 \| 0.182 \| 597 \| 382 \| 22 \| 9 \| 532 \| 10 \| 573 \| Phage portal protein \| Phage portal protein; afdb-uniprot50 \| AF-A0A7X3I6F0-F1-MODEL\_V4 \| 1.0 \| 6.666e-25 \| 830 \| 0.195 \| 572 \| 379 \| 16 \| 1 \| 532 \| 1 \| 531 \| Phage portal protein \| Phage portal protein; afdb-uniprot50 \| AF-A0A2E7ZPP6-F1-MODEL\_V4 \| 1.0 \| 6.745e-24 \| 829 \| 0.163 \| 515 \| 364 \| 17 \| 39 \| 532 \| 5 \| 473 \| Uncharacterized protein \| Uncharacterized protein; afdb-uniprot50 \| AF-A0A812INR5-F1-MODEL\_V4 \| 1.0 \| 1.083e-23 \| 829 \| 0.226 \| 451 \| 295 \| 18 \| 54 \| 473 \| 49 \| 476 \| B protein \| B protein; afdb-uniprot50 \| AF-A0A357XSR0-F1-MODEL\_V4 \| 1.0 \| 3.588e-24 \| 828 \| 0.169 \| 560 \| 395 \| 18 \| 1 \| 531 \| 1 \| 519 \| Uncharacterized protein \| Uncharacterized protein; afdb-uniprot50 \| AF-A0A259MIN2-F1-MODEL\_V4 \| 1.0 \| 1.07e-24 \| 828 \| 0.168 \| 557 \| 400 \| 16 \| 1 \| 532 \| 58 \| 576 \| Phage portal protein \| Phage portal protein; afdb-uniprot50 \| AF-A0A1I3ZDY8-F1-MODEL\_V4 \| 1.0 \| 2.145e-23 \| 826 \| 0.201 \| 486 \| 335 \| 18 \| 35 \| 502 \| 20 \| 470 \| Phage portal protein, lambda family \| Phage portal protein, lambda family; afdb-uniprot50 \| AF-A0A351VMH6-F1-MODEL\_V4 \| 1.0 \| 6.325e-25 \| 826 \| 0.182 \| 548 \| 373 \| 18 \| 15 \| 497 \| 1 \| 538 \| Phage portal protein \| Phage portal protein; afdb-uniprot50 \| AF-A0A6I1IVV5-F1-MODEL\_V4 \| 1.0 \| 7.493e-24 \| 825 \| 0.198 \| 515 \| 339 \| 17 \| 49 \| 522 \| 7 \| 488 \| Phage portal protein \| Phage portal protein; afdb-uniprot50 \| AF-A0A4Q3GP67-F1-MODEL\_V4 \| 1.0 \| 4.428e-24 \| 825 \| 0.189 \| 506 \| 354 \| 16 \| 23 \| 488 \| 1 \| 490 \| Phage portal protein \| Phage portal protein; afdb-uniprot50 \| AF-A0A258L750-F1-MODEL\_V4 \| 1.0 \| 1.189e-24 \| 825 \| 0.19 \| 530 \| 366 \| 19 \| 23 \| 502 \| 1 \| 517 \| Phage portal protein \| Phage portal protein; afdb-uniprot50 \| AF-A0A2U1SE48-F1-MODEL\_V4 \| 1.0 \| 2.209e-25 \| 825 \| 0.197 \| 572 \| 374 \| 19 \| 2 \| 525 \| 1 \| 535 \| Uncharacterized protein \| Uncharacterized protein; afdb-uniprot50 \| AF-A0A7X8YIN0-F1-MODEL\_V4 \| 1.0 \| 2.012e-24 \| 824 \| 0.182 \| 515 \| 363 \| 15 \| 39 \| 532 \| 20 \| 497 \| Phage portal protein \| Phage portal protein; afdb-uniprot50 \| AF-A0A518GNB7-F1-MODEL\_V4 \| 1.0 \| 1.811e-24 \| 824 \| 0.205 \| 526 \| 346 \| 22 \| 29 \| 532 \| 12 \| 487 \| Phage portal protein, lambda family \| Phage portal protein, lambda family; afdb-uniprot50 \| AF-A0A4P7EGH2-F1-MODEL\_V4 \| 1.0 \| 7.109e-24 \| 821 \| 0.194 \| 504 \| 334 \| 18 \| 51 \| 503 \| 28 \| 510 \| Phage portal protein \| Phage portal protein; afdb-uniprot50 \| AF-A0A420WDI6-F1-MODEL\_V4 \| 1.0 \| 2.454e-25 \| 821 \| 0.201 \| 562 \| 361 \| 27 \| 1 \| 502 \| 2 \| 535 \| Lambda family phage portal protein \| Lambda family phage portal protein; afdb-uniprot50 \| AF-A0A5E8P2L6-F1-MODEL\_V4 \| 1.0 \| 6.214e-22 \| 820 \| 0.222 \| 369 \| 258 \| 13 \| 93 \| 450 \| 2 \| 352 \| Phage portal protein \| Phage portal protein; afdb-uniprot50 \| AF-A0A349JAV8-F1-MODEL\_V4 \| 1.0 \| 3.63e-23 \| 820 \| 0.213 \| 478 \| 330 \| 17 \| 38 \| 502 \| 30 \| 474 \| Uncharacterized protein \| Uncharacterized protein; afdb-uniprot50 \| AF-A0A7W1L746-F1-MODEL\_V4 \| 1.0 \| 3.1e-23 \| 819 \| 0.209 \| 472 \| 325 \| 17 \| 10 \| 473 \| 14 \| 445 \| Phage portal protein \| Phage portal protein; afdb-uniprot50 \| AF-A0A530QV48-F1-MODEL\_V4 \| 1.0 \| 9.861e-23 \| 818 \| 0.303 \| 356 \| 231 \| 7 \| 148 \| 492 \| 2 \| 351 \| Phage portal protein \| Phage portal protein; afdb-uniprot50 \| AF-A0A3E0N0Y2-F1-MODEL\_V4 \| 1.0 \| 1.909e-24 \| 818 \| 0.195 \| 486 \| 326 \| 18 \| 62 \| 519 \| 1 \| 449 \| Phage portal protein \| Phage portal protein; afdb-uniprot50 \| AF-A0A7Z6W8D4-F1-MODEL\_V4 \| 1.0 \| 4.667e-24 \| 818 \| 0.18 \| 525 \| 368 \| 14 \| 11 \| 497 \| 1 \| 501 \| Phage portal protein \| Phage portal protein; afdb-uniprot50 \| AF-A0A766GFH2-F1-MODEL\_V4 \| 1.0 \| 9.747e-24 \| 817 \| 0.176 \| 522 \| 346 \| 18 \| 59 \| 523 \| 1 \| 495 \| Phage portal protein \| Phage portal protein; afdb-uniprot50 \| AF-A0A0F9R9A4-F1-MODEL\_V4 \| 1.0 \| 3.588e-24 \| 817 \| 0.183 \| 540 \| 367 \| 20 \| 23 \| 532 \| 1 \| 496 \| Uncharacterized protein \| Uncharacterized protein; afdb-uniprot50 \| AF-A0A518CTW5-F1-MODEL\_V4 \| 1.0 \| 2.907e-24 \| 817 \| 0.167 \| 543 \| 383 \| 19 \| 27 \| 532 \| 2 \| 512 \| Phage portal protein, lambda family \| Phage portal protein, lambda family; afdb-uniprot50 \| AF-A0A2D5TT87-F1-MODEL\_V4 \| 1.0 \| 3.986e-24 \| 817 \| 0.187 \| 540 \| 371 \| 22 \| 36 \| 532 \| 19 \| 533 \| Uncharacterized protein \| Uncharacterized protein; afdb-uniprot50 \| AF-A0A1F9C0U9-F1-MODEL\_V4 \| 1.0 \| 7.493e-24 \| 816 \| 0.222 \| 458 \| 303 \| 14 \| 94 \| 532 \| 2 \| 425 \| Phage portal protein \| Phage portal protein; afdb-uniprot50 \| AF-A0A518G2U8-F1-MODEL\_V4 \| 1.0 \| 1.931e-23 \| 816 \| 0.188 \| 540 \| 390 \| 19 \| 1 \| 531 \| 11 \| 511 \| Phage portal protein, lambda family \| Phage portal protein, lambda family; afdb-uniprot50 \| AF-A0A517VMM7-F1-MODEL\_V4 \| 1.0 \| 1.811e-24 \| 816 \| 0.198 \| 543 \| 370 \| 17 \| 13 \| 531 \| 1 \| 502 \| Phage portal protein, lambda family \| Phage portal protein, lambda family; afdb-uniprot50 \| AF-A0A5C5XSQ0-F1-MODEL\_V4 \| 1.0 \| 7.897e-24 \| 815 \| 0.218 \| 517 \| 334 \| 20 \| 28 \| 532 \| 14 \| 472 \| Phage portal protein, lambda family \| Phage portal protein, lambda family; afdb-uniprot50 \| AF-A0A398BXH2-F1-MODEL\_V4 \| 1.0 \| 2.12e-24 \| 815 \| 0.192 \| 583 \| 392 \| 23 \| 2 \| 532 \| 17 \| 572 \| Phage portal protein \| Phage portal protein; afdb-uniprot50 \| AF-A0A430EDN5-F1-MODEL\_V4 \| 1.0 \| 1.909e-24 \| 814 \| 0.242 \| 495 \| 312 \| 18 \| 39 \| 517 \| 32 \| 479 \| Phage portal protein \| Phage portal protein; afdb-uniprot50 \| AF-A0A1C0SI27-F1-MODEL\_V4 \| 1.0 \| 2.791e-23 \| 814 \| 0.169 \| 530 \| 361 \| 15 \| 43 \| 532 \| 7 \| 497 \| Phage portal protein \| Phage portal protein; afdb-uniprot50 \| AF-A0A7C7PZ16-F1-MODEL\_V4 \| 1.0 \| 8.672e-25 \| 814 \| 0.178 \| 533 \| 374 \| 19 \| 35 \| 532 \| 25 \| 528 \| Phage portal protein \| Phage portal protein; afdb-uniprot50 \| AF-A0A1G7NEZ5-F1-MODEL\_V4 \| 1.0 \| 2.356e-24 \| 814 \| 0.192 \| 550 \| 358 \| 25 \| 15 \| 501 \| 1 \| 527 \| Phage portal protein, lambda family \| Phage portal protein, lambda family; afdb-uniprot50 \| AF-A0A5N3PHC4-F1-MODEL\_V4 \| 1.0 \| 6.399e-24 \| 814 \| 0.169 \| 550 \| 382 \| 20 \| 24 \| 532 \| 12 \| 527 \| Phage portal protein \| Phage portal protein; afdb-uniprot50 \| AF-A0A1N7SSK9-F1-MODEL\_V4 \| 1.0 \| 1.268e-23 \| 813 \| 0.168 \| 533 \| 351 \| 17 \| 57 \| 532 \| 2 \| 499 \| Phage portal \| Phage portal; afdb-uniprot50 \| AF-A0A1M7D8F9-F1-MODEL\_V4 \| 1.0 \| 1.547e-24 \| 813 \| 0.188 \| 541 \| 351 \| 20 \| 7 \| 501 \| 1 \| 499 \| Phage portal protein, lambda family \| Phage portal protein, lambda family; afdb-uniprot50 \| AF-A0A4P7WG88-F1-MODEL\_V4 \| 1.0 \| 1.467e-24 \| 812 \| 0.193 \| 547 \| 344 \| 25 \| 47 \| 517 \| 11 \| 536 \| Phage portal protein \| Phage portal protein; afdb-uniprot50 \| AF-A0A3D4D8W6-F1-MODEL\_V4 \| 1.0 \| 9.522e-26 \| 812 \| 0.217 \| 537 \| 357 \| 17 \| 35 \| 523 \| 18 \| 539 \| Phage portal protein \| Phage portal protein; afdb-uniprot50 \| AF-A0A2W5S548-F1-MODEL\_V4 \| 1.0 \| 2.012e-24 \| 812 \| 0.183 \| 549 \| 363 \| 19 \| 23 \| 502 \| 2 \| 533 \| Phage portal protein \| Phage portal protein; afdb-uniprot50 \| AF-A0A1V5NFL2-F1-MODEL\_V4 \| 1.0 \| 2.758e-24 \| 812 \| 0.213 \| 543 \| 351 \| 18 \| 17 \| 532 \| 1 \| 494 \| Phage portal protein, lambda family \| Phage portal protein, lambda family; afdb-uniprot50 \| AF-A0A522VYK9-F1-MODEL\_V4 \| 1.0 \| 6.904e-22 \| 810 \| 0.241 \| 372 \| 254 \| 10 \| 39 \| 395 \| 2 \| 360 \| Phage portal protein \| Phage portal protein; afdb-uniprot50 \| AF-A0A7K1B1S8-F1-MODEL\_V4 \| 1.0 \| 3.404e-24 \| 810 \| 0.18 \| 550 \| 376 \| 19 \| 6 \| 532 \| 1 \| 498 \| Phage portal protein \| Phage portal protein; afdb-uniprot50 \| AF-A0A7V9JT41-F1-MODEL\_V4 \| 1.0 \| 6.142e-23 \| 808 \| 0.202 \| 480 \| 324 \| 18 \| 60 \| 532 \| 1 \| 428 \| Phage portal protein \| Phage portal protein; afdb-uniprot50 \| AF-A0A285M8B5-F1-MODEL\_V4 \| 1.0 \| 5.185e-24 \| 808 \| 0.179 \| 562 \| 375 \| 21 \| 30 \| 532 \| 14 \| 548 \| Phage portal protein, lambda family \| Phage portal protein, lambda family; afdb-uniprot50 \| AF-A0A369U8Z9-F1-MODEL\_V4 \| 1.0 \| 7.806e-25 \| 808 \| 0.172 \| 580 \| 364 \| 26 \| 30 \| 532 \| 12 \| 552 \| Phage portal protein \| Phage portal protein; afdb-uniprot50 \| AF-A0A7W6C1D9-F1-MODEL\_V4 \| 1.0 \| 3.192e-25 \| 807 \| 0.214 \| 514 \| 335 \| 17 \| 39 \| 523 \| 33 \| 506 \| Lambda family phage portal protein \| Lambda family phage portal protein; afdb-uniprot50 \| AF-A0A420DHA7-F1-MODEL\_V4 \| 1.0 \| 6.399e-24 \| 807 \| 0.182 \| 553 \| 367 \| 23 \| 2 \| 518 \| 1 \| 504 \| Capsid protein \| Capsid protein; afdb-uniprot50 \| AF-A0A1T4WXT7-F1-MODEL\_V4 \| 1.0 \| 6.142e-23 \| 805 \| 0.168 \| 499 \| 359 \| 19 \| 13 \| 491 \| 1 \| 463 \| Capsid protein \| Capsid protein; afdb-uniprot50 \| AF-A0A0B1MGE1-F1-MODEL\_V4 \| 1.0 \| 3.484e-22 \| 804 \| 0.205 \| 365 \| 260 \| 11 \| 39 \| 390 \| 21 \| 368 \| Capsid protein \| Capsid protein; afdb-uniprot50 \| AF-A0A6M1U479-F1-MODEL\_V4 \| 1.0 \| 1.268e-23 \| 804 \| 0.286 \| 409 \| 253 \| 12 \| 136 \| 532 \| 1 \| 382 \| Phage portal protein \| Phage portal protein; afdb-uniprot50 \| AF-A0A5C7Q7S1-F1-MODEL\_V4 \| 1.0 \| 8.876e-23 \| 803 \| 0.187 \| 497 \| 341 \| 17 \| 47 \| 515 \| 16 \| 477 \| Phage portal protein \| Phage portal protein; afdb-uniprot50 \| AF-A0A444KUH4-F1-MODEL\_V4 \| 1.0 \| 2.12e-24 \| 803 \| 0.209 \| 553 \| 340 \| 20 \| 46 \| 532 \| 10 \| 531 \| Phage portal protein \| Phage portal protein; afdb-uniprot50 \| AF-A0A4Z0ITW1-F1-MODEL\_V4 \| 1.0 \| 1.189e-24 \| 803 \| 0.197 \| 561 \| 358 \| 19 \| 18 \| 532 \| 1 \| 515 \| Phage portal protein \| Phage portal protein; afdb-uniprot50 \| AF-A0A1I3N5J6-F1-MODEL\_V4 \| 1.0 \| 2.356e-24 \| 802 \| 0.268 \| 402 \| 255 \| 12 \| 131 \| 516 \| 12 \| 390 \| Phage portal protein, lambda family \| Phage portal protein, lambda family; afdb-uniprot50 \| AF-A0A3G8M317-F1-MODEL\_V4 \| 1.0 \| 3.986e-24 \| 802 \| 0.181 \| 569 \| 392 \| 21 \| 1 \| 515 \| 12 \| 560 \| Phage portal protein \| Phage portal protein; afdb-uniprot50 \| AF-R9L6B1-F1-MODEL\_V4 \| 1.0 \| 9.861e-23 \| 799 \| 0.232 \| 383 \| 260 \| 9 \| 32 \| 394 \| 14 \| 382 \| Lambda family phage portal protein \| Lambda family phage portal protein; afdb-uniprot50 \| AF-A0A1V6HR06-F1-MODEL\_V4 \| 1.0 \| 3.444e-23 \| 798 \| 0.16 \| 556 \| 389 \| 19 \| 20 \| 531 \| 1 \| 522 \| Phage portal protein, lambda family \| Phage portal protein, lambda family; afdb-uniprot50 \| AF-A0A3M1LU04-F1-MODEL\_V4 \| 1.0 \| 1.039e-22 \| 797 \| 0.271 \| 394 \| 248 \| 11 \| 39 \| 421 \| 2 \| 367 \| Phage portal protein \| Phage portal protein; afdb-uniprot50 \| AF-A0A3L4XBL8-F1-MODEL\_V4 \| 1.0 \| 1.141e-23 \| 797 \| 0.221 \| 488 \| 319 \| 17 \| 86 \| 532 \| 2 \| 469 \| Phage portal protein \| Phage portal protein; afdb-uniprot50 \| AF-A0A2E5AGP1-F1-MODEL\_V4 \| 1.0 \| 4.667e-24 \| 797 \| 0.168 \| 599 \| 393 \| 20 \| 5 \| 532 \| 1 \| 565 \| Uncharacterized protein \| Uncharacterized protein; afdb-uniprot50 \| AF-I6APC7-F1-MODEL\_V4 \| 1.0 \| 1.738e-23 \| 796 \| 0.16 \| 492 \| 348 \| 16 \| 62 \| 532 \| 62 \| 509 \| Bacteriophage capsid protein \| Bacteriophage capsid protein; afdb-uniprot50 \| AF-A0A5S4WJH0-F1-MODEL\_V4 \| 1.0 \| 6.325e-25 \| 795 \| 0.179 \| 568 \| 356 \| 22 \| 40 \| 532 \| 16 \| 548 \| Phage portal protein \| Phage portal protein; afdb-uniprot50 \| AF-C1AAN8-F1-MODEL\_V4 \| 1.0 \| 1.738e-23 \| 795 \| 0.151 \| 548 \| 387 \| 17 \| 20 \| 532 \| 24 \| 528 \| Phage portal protein lambda family protein \| Phage portal protein lambda family protein; afdb-uniprot50 \| AF-I6B1B8-F1-MODEL\_V4 \| 1.0 \| 3.826e-23 \| 794 \| 0.195 \| 523 \| 354 \| 18 \| 35 \| 532 \| 25 \| 505 \| Bacteriophage capsid protein \| Bacteriophage capsid protein; afdb-uniprot50 \| AF-A0A848VGF3-F1-MODEL\_V4 \| 1.0 \| 9.634e-25 \| 793 \| 0.194 \| 541 \| 365 \| 20 \| 24 \| 532 \| 10 \| 511 \| Phage portal protein \| Phage portal protein; afdb-uniprot50 \| AF-A0A442I655-F1-MODEL\_V4 \| 1.0 \| 2.383e-23 \| 793 \| 0.172 \| 556 \| 395 \| 16 \| 14 \| 532 \| 1 \| 528 \| Phage portal protein \| Phage portal protein; afdb-uniprot50 \| AF-A0A0P7Y6W9-F1-MODEL\_V4 \| 1.0 \| 1.875e-21 \| 792 \| 0.266 \| 398 \| 252 \| 14 \| 74 \| 459 \| 3 \| 372 \| Phage portal protein, lambda family \| Phage portal protein, lambda family; afdb-uniprot50 \| AF-A0A1M7ZLP8-F1-MODEL\_V4 \| 1.0 \| 1.718e-24 \| 792 \| 0.202 \| 579 \| 353 \| 29 \| 11 \| 532 \| 1 \| 527 \| Phage portal protein, lambda family \| Phage portal protein, lambda family; afdb-uniprot50 \| AF-A0A1G3M6A5-F1-MODEL\_V4 \| 1.0 \| 1.669e-22 \| 789 \| 0.242 \| 421 \| 285 \| 14 \| 11 \| 418 \| 26 \| 425 \| Phage portal protein \| Phage portal protein; afdb-uniprot50 \| AF-A0A285D5Q3-F1-MODEL\_V4 \| 1.0 \| 3.94e-25 \| 789 \| 0.243 \| 539 \| 330 \| 17 \| 16 \| 532 \| 1 \| 483 \| Lambda family phage portal protein \| Lambda family phage portal protein; afdb-uniprot50 \| AF-T0GB94-F1-MODEL\_V4 \| 1.0 \| 1.649e-23 \| 789 \| 0.211 \| 544 \| 365 \| 21 \| 27 \| 531 \| 11 \| 529 \| Uncharacterized protein \| Uncharacterized protein; afdb-uniprot50 \| AF-A0A0F9E9R9-F1-MODEL\_V4 \| 1.0 \| 3.63e-23 \| 786 \| 0.214 \| 439 \| 293 \| 16 \| 39 \| 439 \| 24 \| 448 \| Uncharacterized protein \| Uncharacterized protein; afdb-uniprot50 \| AF-A0A236MRM7-F1-MODEL\_V4 \| 1.0 \| 4.667e-24 \| 786 \| 0.183 \| 546 \| 347 \| 20 \| 39 \| 523 \| 159 \| 666 \| Phage portal protein (Minor capsid protein) \| Phage portal protein (Minor capsid protein); afdb-uniprot50 \| AF-A0A7U9F7S2-F1-MODEL\_V4 \| 1.0 \| 1.502e-22 \| 785 \| 0.269 \| 419 \| 259 \| 14 \| 127 \| 532 \| 2 \| 386 \| Uncharacterized protein \| Uncharacterized protein; afdb-uniprot50 \| AF-A0A5C7PCJ8-F1-MODEL\_V4 \| 1.0 \| 2.758e-24 \| 785 \| 0.186 \| 537 \| 361 \| 23 \| 39 \| 530 \| 19 \| 524 \| Phage portal protein \| Phage portal protein; afdb-uniprot50 \| AF-A0A662JPW1-F1-MODEL\_V4 \| 1.0 \| 8.421e-23 \| 783 \| 0.215 \| 464 \| 308 \| 16 \| 46 \| 496 \| 14 \| 434 \| Uncharacterized protein \| Uncharacterized protein; afdb-uniprot50 \| AF-A0A3N9N4D9-F1-MODEL\_V4 \| 1.0 \| 1.039e-22 \| 783 \| 0.183 \| 534 \| 370 \| 20 \| 9 \| 517 \| 11 \| 503 \| Phage portal protein \| Phage portal protein; afdb-uniprot50 \| AF-A0A7X9G897-F1-MODEL\_V4 \| 1.0 \| 6.142e-23 \| 783 \| 0.184 \| 559 \| 358 \| 19 \| 13 \| 532 \| 1 \| 500 \| Phage portal protein \| Phage portal protein; afdb-uniprot50 \| AF-A0A1G3A9B5-F1-MODEL\_V4 \| 1.0 \| 1.141e-23 \| 782 \| 0.196 \| 530 \| 352 \| 19 \| 21 \| 532 \| 13 \| 486 \| Uncharacterized protein \| Uncharacterized protein; afdb-uniprot50 \| AF-A0A2E5UE44-F1-MODEL\_V4 \| 1.0 \| 7.192e-23 \| 782 \| 0.165 \| 539 \| 369 \| 16 \| 42 \| 532 \| 15 \| 520 \| Phage portal protein \| Phage portal protein; afdb-uniprot50 \| AF-A0A0Q4ZHQ5-F1-MODEL\_V4 \| 1.0 \| 1.485e-23 \| 781 \| 0.157 \| 559 \| 392 \| 21 \| 17 \| 532 \| 25 \| 547 \| Uncharacterized protein \| Uncharacterized protein; afdb-uniprot50 \| AF-F7X6P0-F1-MODEL\_V4 \| 1.0 \| 1.141e-23 \| 781 \| 0.205 \| 594 \| 373 \| 24 \| 1 \| 532 \| 65 \| 621 \| Lambda family phage portal protein \| Lambda family phage portal protein; afdb-uniprot50 \| AF-A0A518G4B4-F1-MODEL\_V4 \| 1.0 \| 9.356e-23 \| 780 \| 0.192 \| 505 \| 361 \| 18 \| 11 \| 495 \| 2 \| 479 \| Phage portal protein, lambda family \| Phage portal protein, lambda family; afdb-uniprot50 \| AF-A0A496YY11-F1-MODEL\_V4 \| 1.0 \| 9.356e-23 \| 779 \| 0.18 \| 525 \| 367 \| 19 \| 20 \| 532 \| 1 \| 474 \| Uncharacterized protein \| Uncharacterized protein; afdb-uniprot50 \| AF-A0A354U4Q5-F1-MODEL\_V4 \| 1.0 \| 5.529e-23 \| 779 \| 0.209 \| 530 \| 347 \| 17 \| 23 \| 532 \| 2 \| 479 \| Uncharacterized protein \| Uncharacterized protein; afdb-uniprot50 \| AF-A0A259CCE8-F1-MODEL\_V4 \| 1.0 \| 1.009e-20 \| 778 \| 0.266 \| 368 \| 247 \| 13 \| 44 \| 402 \| 3 \| 356 \| Phage portal protein \| Phage portal protein; afdb-uniprot50 \| AF-A0A3A0D2G7-F1-MODEL\_V4 \| 1.0 \| 1.779e-21 \| 778 \| 0.244 \| 401 \| 267 \| 16 \| 23 \| 415 \| 11 \| 383 \| Uncharacterized protein \| Uncharacterized protein; afdb-uniprot50 \| AF-A0A1H2HHH0-F1-MODEL\_V4 \| 1.0 \| 9.861e-23 \| 777 \| 0.16 \| 553 \| 388 \| 18 \| 13 \| 532 \| 2 \| 510 \| Capsid protein \| Capsid protein; afdb-uniprot50 \| AF-A0A0F9QX35-F1-MODEL\_V4 \| 1.0 \| 1.217e-22 \| 777 \| 0.216 \| 490 \| 340 \| 18 \| 1 \| 475 \| 9 \| 469 \| Uncharacterized protein \| Uncharacterized protein; afdb-uniprot50 \| AF-A0A6N2ZN93-F1-MODEL\_V4 \| 1.0 \| 6.071e-24 \| 775 \| 0.206 \| 513 \| 351 \| 19 \| 27 \| 495 \| 1 \| 501 \| Phage portal protein, lambda family \| Phage portal protein, lambda family; afdb-uniprot50 \| AF-A0A377PFS5-F1-MODEL\_V4 \| 1.0 \| 1.602e-21 \| 774 \| 0.236 \| 422 \| 280 \| 14 \| 122 \| 532 \| 4 \| 394 \| Phage portal protein, lambda family \| Phage portal protein, lambda family; afdb-uniprot50 \| AF-A0A4D7QII8-F1-MODEL\_V4 \| 1.0 \| 1.649e-23 \| 774 \| 0.157 \| 585 \| 406 \| 20 \| 1 \| 531 \| 7 \| 558 \| Phage portal protein \| Phage portal protein; afdb-uniprot50 \| AF-A0A480BUG0-F1-MODEL\_V4 \| 1.0 \| 9.576e-21 \| 773 \| 0.221 \| 357 \| 260 \| 8 \| 39 \| 385 \| 28 \| 376 \| Phage portal protein \| Phage portal protein; afdb-uniprot50 \| AF-A0A3M9XN02-F1-MODEL\_V4 \| 1.0 \| 4.428e-24 \| 772 \| 0.18 \| 575 \| 382 \| 25 \| 17 \| 532 \| 1 \| 545 \| Phage portal protein \| Phage portal protein; afdb-uniprot50 \| AF-Q727I3-F1-MODEL\_V4 \| 1.0 \| 1.268e-23 \| 772 \| 0.197 \| 582 \| 366 \| 23 \| 1 \| 532 \| 8 \| 538 \| Phage portal protein, lambda family \| Phage portal protein, lambda family; afdb-uniprot50 \| AF-A0A1Z5H699-F1-MODEL\_V4 \| 1.0 \| 4.128e-21 \| 770 \| 0.232 \| 404 \| 268 \| 13 \| 46 \| 418 \| 18 \| 410 \| Uncharacterized protein \| Uncharacterized protein; afdb-uniprot50 \| AF-A0A2E1P339-F1-MODEL\_V4 \| 1.0 \| 1.537e-20 \| 769 \| 0.316 \| 275 \| 181 \| 2 \| 31 \| 305 \| 12 \| 279 \| Phage portal protein \| Phage portal protein; afdb-uniprot50 \| AF-A0A7W7NWS1-F1-MODEL\_V4 \| 1.0 \| 6.071e-24 \| 769 \| 0.179 \| 557 \| 363 \| 21 \| 39 \| 532 \| 7 \| 532 \| Lambda family phage portal protein \| Lambda family phage portal protein; afdb-uniprot50 \| AF-A0A3A1YCY1-F1-MODEL\_V4 \| 1.0 \| 4.033e-23 \| 768 \| 0.18 \| 542 \| 376 \| 21 \| 16 \| 532 \| 12 \| 510 \| Uncharacterized protein \| Uncharacterized protein; afdb-uniprot50 \| AF-A0A0E3C0X6-F1-MODEL\_V4 \| 1.0 \| 1.425e-22 \| 767 \| 0.218 \| 421 \| 296 \| 12 \| 105 \| 517 \| 2 \| 397 \| Capsid protein \| Capsid protein; afdb-uniprot50 \| AF-A0A439EQ92-F1-MODEL\_V4 \| 1.0 \| 1.875e-21 \| 765 \| 0.218 \| 389 \| 278 \| 12 \| 122 \| 502 \| 1 \| 371 \| Phage portal protein \| Phage portal protein; afdb-uniprot50 \| AF-A0A7V3AHA7-F1-MODEL\_V4 \| 1.0 \| 1.537e-20 \| 764 \| 0.274 \| 331 \| 215 \| 9 \| 46 \| 366 \| 3 \| 318 \| Phage portal protein \| Phage portal protein; afdb-uniprot50 \| AF-A0A0H3X0U6-F1-MODEL\_V4 \| 1.0 \| 3.137e-22 \| 764 \| 0.178 \| 548 \| 361 \| 25 \| 44 \| 532 \| 6 \| 523 \| Phage portal protein \| Phage portal protein; afdb-uniprot50 \| AF-A0A855TGI4-F1-MODEL\_V4 \| 1.0 \| 1.217e-22 \| 763 \| 0.174 \| 482 \| 350 \| 15 \| 58 \| 531 \| 12 \| 453 \| Uncharacterized protein \| Uncharacterized protein; afdb-uniprot50 \| AF-A0A2D6XDS2-F1-MODEL\_V4 \| 1.0 \| 3.871e-22 \| 763 \| 0.187 \| 507 \| 345 \| 18 \| 39 \| 520 \| 19 \| 483 \| Uncharacterized protein \| Uncharacterized protein; afdb-uniprot50 \| AF-A0A6P2LTT1-F1-MODEL\_V4 \| 1.0 \| 4.722e-23 \| 763 \| 0.236 \| 452 \| 270 \| 17 \| 24 \| 459 \| 1 \| 393 \| Portal protein \| Portal protein; afdb-uniprot50 \| AF-A0A366G0P7-F1-MODEL\_V4 \| 1.0 \| 1.502e-22 \| 762 \| 0.191 \| 518 \| 324 \| 20 \| 74 \| 532 \| 3 \| 484 \| Lambda family phage portal protein \| Lambda family phage portal protein; afdb-uniprot50 \| AF-A0A0E2L6R3-F1-MODEL\_V4 \| 1.0 \| 6.214e-22 \| 761 \| 0.236 \| 431 \| 277 \| 14 \| 49 \| 439 \| 25 \| 443 \| Portal protein \| Portal protein; afdb-uniprot50 \| AF-A0A1Y1S1G3-F1-MODEL\_V4 \| 1.0 \| 1.832e-23 \| 760 \| 0.178 \| 504 \| 338 \| 17 \| 73 \| 532 \| 46 \| 517 \| Uncharacterized protein \| Uncharacterized protein; afdb-uniprot50 \| AF-A0A1M4WC61-F1-MODEL\_V4 \| 1.0 \| 2.17e-22 \| 759 \| 0.179 \| 536 \| 356 \| 21 \| 20 \| 502 \| 12 \| 516 \| Phage portal protein, lambda family \| Phage portal protein, lambda family; afdb-uniprot50 \| AF-A0A420Z232-F1-MODEL\_V4 \| 1.0 \| 9.861e-23 \| 758 \| 0.176 \| 561 \| 384 \| 17 \| 23 \| 532 \| 2 \| 535 \| Phage portal protein \| Phage portal protein; afdb-uniprot50 \| AF-A0A7W1G457-F1-MODEL\_V4 \| 1.0 \| 2.261e-23 \| 757 \| 0.183 \| 522 \| 352 \| 26 \| 39 \| 531 \| 20 \| 496 \| Phage portal protein \| Phage portal protein; afdb-uniprot50 \| AF-A0A1Z8KYX4-F1-MODEL\_V4 \| 1.0 \| 3.444e-23 \| 756 \| 0.194 \| 529 \| 364 \| 21 \| 39 \| 532 \| 32 \| 533 \| Phage portal protein \| Phage portal protein; afdb-uniprot50 \| AF-A0A0T7A5Q8-F1-MODEL\_V4 \| 1.0 \| 1.352e-22 \| 755 \| 0.194 \| 504 \| 318 \| 17 \| 40 \| 472 \| 2 \| 488 \| Uncharacterized protein \| Uncharacterized protein; afdb-uniprot50 \| AF-A0A154LMF9-F1-MODEL\_V4 \| 1.0 \| 2.261e-23 \| 755 \| 0.212 \| 518 \| 337 \| 19 \| 28 \| 502 \| 15 \| 504 \| Uncharacterized protein \| Uncharacterized protein; afdb-uniprot50 \| AF-A0A1A6FM45-F1-MODEL\_V4 \| 1.0 \| 3.444e-23 \| 752 \| 0.18 \| 538 \| 354 \| 23 \| 51 \| 532 \| 1 \| 507 \| Phage portal protein \| Phage portal protein; afdb-uniprot50 \| AF-A0A4R8FFK2-F1-MODEL\_V4 \| 1.0 \| 3.1e-23 \| 752 \| 0.157 \| 583 \| 381 \| 24 \| 13 \| 532 \| 2 \| 537 \| Lambda family phage portal protein \| Lambda family phage portal protein; afdb-uniprot50 \| AF-A0A3C1QZ48-F1-MODEL\_V4 \| 1.0 \| 3.345e-21 \| 750 \| 0.231 \| 437 \| 299 \| 15 \| 28 \| 456 \| 10 \| 417 \| Uncharacterized protein \| Uncharacterized protein; afdb-uniprot50 \| AF-A0A439RCC1-F1-MODEL\_V4 \| 1.0 \| 2.035e-23 \| 749 \| 0.173 \| 588 \| 374 \| 24 \| 2 \| 532 \| 7 \| 539 \| Phage portal protein \| Phage portal protein; afdb-uniprot50 \| AF-A0A4R1PSV5-F1-MODEL\_V4 \| 1.0 \| 1.231e-21 \| 747 \| 0.301 \| 358 \| 225 \| 11 \| 161 \| 502 \| 2 \| 350 \| Lambda family phage portal protein \| Lambda family phage portal protein; afdb-uniprot50 \| AF-A0A7W1IQN8-F1-MODEL\_V4 \| 1.0 \| 2.541e-22 \| 747 \| 0.192 \| 498 \| 326 \| 15 \| 46 \| 502 \| 2 \| 464 \| Phage portal protein \| Phage portal protein; afdb-uniprot50 \| AF-A0A5C5VQX8-F1-MODEL\_V4 \| 1.0 \| 4.25e-23 \| 747 \| 0.179 \| 556 \| 362 \| 22 \| 29 \| 532 \| 15 \| 528 \| Phage portal protein, lambda family \| Phage portal protein, lambda family; afdb-uniprot50 \| AF-A0A3C1YZ33-F1-MODEL\_V4 \| 1.0 \| 1.583e-22 \| 744 \| 0.167 \| 548 \| 370 \| 17 \| 37 \| 532 \| 8 \| 521 \| Uncharacterized protein \| Uncharacterized protein; afdb-uniprot50 \| AF-A0A286IFC3-F1-MODEL\_V4 \| 1.0 \| 7.192e-23 \| 744 \| 0.17 \| 521 \| 350 \| 22 \| 26 \| 503 \| 24 \| 505 \| Lambda family phage portal protein \| Lambda family phage portal protein; afdb-uniprot50 \| AF-A0A5C6BLH5-F1-MODEL\_V4 \| 1.0 \| 2.571e-21 \| 743 \| 0.184 \| 482 \| 332 \| 18 \| 39 \| 495 \| 10 \| 455 \| Phage portal protein, lambda family \| Phage portal protein, lambda family; afdb-uniprot50 \| AF-D5SMQ0-F1-MODEL\_V4 \| 1.0 \| 6.214e-22 \| 741 \| 0.18 \| 531 \| 366 \| 21 \| 24 \| 532 \| 1 \| 484 \| Bacteriophage capsid protein-like protein \| Bacteriophage capsid protein-like protein; afdb-uniprot50 \| AF-A0A537MC68-F1-MODEL\_V4 \| 1.0 \| 4.08e-22 \| 741 \| 0.16 \| 561 \| 394 \| 19 \| 1 \| 532 \| 4 \| 516 \| Phage portal protein \| Phage portal protein; afdb-uniprot50 \| AF-A0A5C8A356-F1-MODEL\_V4 \| 1.0 \| 1.352e-22 \| 737 \| 0.202 \| 558 \| 374 \| 20 \| 2 \| 532 \| 23 \| 536 \| Phage portal protein \| Phage portal protein; afdb-uniprot50 \| AF-A0A4Z0F5W2-F1-MODEL\_V4 \| 1.0 \| 3.444e-23 \| 736 \| 0.204 \| 544 \| 332 \| 27 \| 47 \| 532 \| 25 \| 525 \| Phage portal protein \| Phage portal protein; afdb-uniprot50 \| AF-A0A354TJG1-F1-MODEL\_V4 \| 1.0 \| 1.052e-21 \| 734 \| 0.17 \| 539 \| 365 \| 20 \| 20 \| 532 \| 1 \| 483 \| Uncharacterized protein \| Uncharacterized protein; afdb-uniprot50 \| AF-A0A4Q2YKT3-F1-MODEL\_V4 \| 1.0 \| 1.039e-22 \| 733 \| 0.203 \| 575 \| 371 \| 23 \| 5 \| 532 \| 3 \| 537 \| Phage portal protein \| Phage portal protein; afdb-uniprot50 \| AF-A0A2T6DRH8-F1-MODEL\_V4 \| 1.0 \| 3.525e-21 \| 730 \| 0.183 \| 517 \| 351 \| 21 \| 46 \| 532 \| 4 \| 479 \| Uncharacterized protein \| Uncharacterized protein; afdb-uniprot50 \| AF-A0A5C5Y9N3-F1-MODEL\_V4 \| 1.0 \| 1.854e-22 \| 723 \| 0.206 \| 504 \| 338 \| 21 \| 4 \| 487 \| 1 \| 462 \| Phage portal protein, lambda family \| Phage portal protein, lambda family; afdb-uniprot50 \| AF-A0A3N5X2X0-F1-MODEL\_V4 \| 1.0 \| 4.833e-21 \| 721 \| 0.175 \| 513 \| 340 \| 20 \| 44 \| 531 \| 18 \| 472 \| Phage portal protein \| Phage portal protein; afdb-uniprot50 \| AF-A0A1E5NHB2-F1-MODEL\_V4 \| 1.0 \| 1.583e-22 \| 717 \| 0.16 \| 556 \| 383 \| 21 \| 23 \| 532 \| 21 \| 538 \| Uncharacterized protein \| Uncharacterized protein; afdb-uniprot50 \| AF-A0A127CHD1-F1-MODEL\_V4 \| 1.0 \| 1.095e-22 \| 717 \| 0.195 \| 572 \| 361 \| 25 \| 35 \| 532 \| 27 \| 573 \| Portal protein \| Portal protein; afdb-uniprot50 \| AF-A0A5C7PT24-F1-MODEL\_V4 \| 1.0 \| 2.083e-21 \| 715 \| 0.177 \| 534 \| 378 \| 21 \| 22 \| 532 \| 48 \| 543 \| Phage portal protein \| Phage portal protein; afdb-uniprot50 \| AF-A0A517R7D1-F1-MODEL\_V4 \| 1.0 \| 7.362e-21 \| 714 \| 0.177 \| 523 \| 350 \| 23 \| 36 \| 532 \| 3 \| 471 \| Phage portal protein, lambda family \| Phage portal protein, lambda family; afdb-uniprot50 \| AF-A0A1W2ACJ6-F1-MODEL\_V4 \| 1.0 \| 7.277e-22 \| 714 \| 0.154 \| 549 \| 389 \| 21 \| 22 \| 532 \| 10 \| 521 \| Capsid protein \| Capsid protein; afdb-uniprot50 \| AF-A0A2E9RXI4-F1-MODEL\_V4 \| 1.0 \| 2.571e-21 \| 712 \| 0.166 \| 521 \| 354 \| 21 \| 39 \| 532 \| 4 \| 470 \| Uncharacterized protein \| Uncharacterized protein; afdb-uniprot50 \| AF-A0A2U2C460-F1-MODEL\_V4 \| 1.0 \| 1.298e-21 \| 712 \| 0.193 \| 502 \| 310 \| 20 \| 51 \| 492 \| 1 \| 467 \| Phage portal protein \| Phage portal protein; afdb-uniprot50 \| AF-A0A5C7PZR6-F1-MODEL\_V4 \| 1.0 \| 1.009e-20 \| 711 \| 0.172 \| 529 \| 359 \| 20 \| 36 \| 532 \| 6 \| 487 \| Phage portal protein \| Phage portal protein; afdb-uniprot50 \| AF-A0A258L7H2-F1-MODEL\_V4 \| 1.0 \| 1.779e-21 \| 710 \| 0.205 \| 447 \| 278 \| 17 \| 39 \| 426 \| 7 \| 435 \| Phage portal protein \| Phage portal protein; afdb-uniprot50 \| AF-A0A846QDJ1-F1-MODEL\_V4 \| 1.0 \| 6.627e-21 \| 706 \| 0.179 \| 513 \| 352 \| 17 \| 15 \| 502 \| 1 \| 469 \| Capsid protein \| Capsid protein; afdb-uniprot50 \| AF-A0A7X7IJJ8-F1-MODEL\_V4 \| 1.0 \| 3.567e-20 \| 704 \| 0.202 \| 444 \| 298 \| 14 \| 92 \| 532 \| 2 \| 392 \| Phage portal protein \| Phage portal protein; afdb-uniprot50 \| AF-A0A6M3IJ20-F1-MODEL\_V4 \| 1.0 \| 4.833e-21 \| 704 \| 0.163 \| 509 \| 365 \| 18 \| 47 \| 526 \| 7 \| 483 \| Putative portal protein \| Putative portal protein; afdb-uniprot50 \| AF-A0A2U1B6F7-F1-MODEL\_V4 \| 1.0 \| 5.154e-20 \| 703 \| 0.3 \| 340 \| 219 \| 6 \| 168 \| 502 \| 2 \| 327 \| Lambda family phage portal protein \| Lambda family phage portal protein; afdb-uniprot50 \| AF-A0A7W1RRH0-F1-MODEL\_V4 \| 1.0 \| 7.448e-20 \| 703 \| 0.157 \| 444 \| 325 \| 14 \| 75 \| 495 \| 36 \| 453 \| Phage portal protein \| Phage portal protein; afdb-uniprot50 \| AF-A0A7C5WID2-F1-MODEL\_V4 \| 1.0 \| 4.586e-21 \| 703 \| 0.155 \| 539 \| 391 \| 18 \| 19 \| 532 \| 12 \| 511 \| Phage portal protein \| Phage portal protein; afdb-uniprot50 \| AF-A0A1V5QJV4-F1-MODEL\_V4 \| 1.0 \| 1.708e-20 \| 701 \| 0.207 \| 457 \| 307 \| 14 \| 99 \| 532 \| 2 \| 426 \| Phage portal protein, lambda family \| Phage portal protein, lambda family; afdb-uniprot50 \| AF-A0A2Z6AIM3-F1-MODEL\_V4 \| 1.0 \| 2.083e-21 \| 701 \| 0.186 \| 520 \| 344 \| 24 \| 63 \| 532 \| 1 \| 491 \| Uncharacterized protein \| Uncharacterized protein; afdb-uniprot50 \| AF-A0A5C5WMX5-F1-MODEL\_V4 \| 1.0 \| 2.742e-20 \| 699 \| 0.182 \| 461 \| 324 \| 16 \| 58 \| 502 \| 3 \| 426 \| Phage portal protein, lambda family \| Phage portal protein, lambda family; afdb-uniprot50 \| AF-A0A5C5VCI3-F1-MODEL\_V4 \| 1.0 \| 1.688e-21 \| 698 \| 0.168 \| 565 \| 386 \| 21 \| 24 \| 531 \| 1 \| 538 \| Phage portal protein, lambda family \| Phage portal protein, lambda family; afdb-uniprot50 \| AF-A0A8B4NDG1-F1-MODEL\_V4 \| 1.0 \| 4.639e-20 \| 697 \| 0.236 \| 418 \| 266 \| 14 \| 49 \| 426 \| 25 \| 429 \| Capsid protein of prophage \| Capsid protein of prophage; afdb-uniprot50 \| AF-A0A3C1GAJ4-F1-MODEL\_V4 \| 1.0 \| 6.106e-19 \| 693 \| 0.383 \| 295 \| 165 \| 4 \| 234 \| 515 \| 4 \| 294 \| Phage portal protein \| Phage portal protein; afdb-uniprot50 \| AF-A0A165U8M4-F1-MODEL\_V4 \| 1.0 \| 5.369e-21 \| 691 \| 0.154 \| 589 \| 371 \| 23 \| 15 \| 532 \| 1 \| 533 \| Phage portal protein, lambda family \| Phage portal protein, lambda family; afdb-uniprot50 \| AF-A0A442NM56-F1-MODEL\_V4 \| 1.0 \| 1.121e-20 \| 690 \| 0.17 \| 492 \| 331 \| 16 \| 63 \| 503 \| 1 \| 466 \| Phage portal protein \| Phage portal protein; afdb-uniprot50 \| AF-A0A840XZF9-F1-MODEL\_V4 \| 1.0 \| 6.035e-20 \| 689 \| 0.25 \| 376 \| 236 \| 13 \| 75 \| 428 \| 4 \| 355 \| Lambda family phage portal protein \| Lambda family phage portal protein; afdb-uniprot50 \| AF-A0A0F9IQ30-F1-MODEL\_V4 \| 1.0 \| 2e-20 \| 687 \| 0.231 \| 401 \| 258 \| 14 \| 34 \| 396 \| 20 \| 408 \| Uncharacterized protein \| Uncharacterized protein; afdb-uniprot50 \| AF-G9PUH7-F1-MODEL\_V4 \| 1.0 \| 1.246e-20 \| 686 \| 0.239 \| 392 \| 268 \| 8 \| 157 \| 530 \| 2 \| 381 \| Lambda family phage portal protein \| Lambda family phage portal protein; afdb-uniprot50 \| AF-A0A833G1E6-F1-MODEL\_V4 \| 1.0 \| 1.196e-19 \| 682 \| 0.185 \| 409 \| 274 \| 16 \| 71 \| 438 \| 22 \| 412 \| Phage portal protein \| Phage portal protein; afdb-uniprot50 \| AF-A0A3D2GU98-F1-MODEL\_V4 \| 1.0 \| 1.182e-20 \| 682 \| 0.208 \| 455 \| 308 \| 19 \| 39 \| 456 \| 32 \| 471 \| Uncharacterized protein \| Uncharacterized protein; afdb-uniprot50 \| AF-A0A285MF00-F1-MODEL\_V4 \| 1.0 \| 1.52e-21 \| 680 \| 0.169 \| 566 \| 355 \| 22 \| 1 \| 498 \| 3 \| 521 \| Phage portal protein, lambda family \| Phage portal protein, lambda family; afdb-uniprot50 \| AF-X0U7Z2-F1-MODEL\_V4 \| 1.0 \| 7.448e-20 \| 679 \| 0.208 \| 437 \| 291 \| 13 \| 113 \| 532 \| 3 \| 401 \| Uncharacterized protein \| Uncharacterized protein; afdb-uniprot50 \| AF-A0A376UD89-F1-MODEL\_V4 \| 1.0 \| 1.639e-19 \| 676 \| 0.213 \| 399 \| 270 \| 13 \| 143 \| 517 \| 4 \| 382 \| Head-tail preconnector protein \| Head-tail preconnector protein; afdb-uniprot50 \| AF-A0A1V1X572-F1-MODEL\_V4 \| 1.0 \| 3.608e-19 \| 675 \| 0.169 \| 489 \| 347 \| 18 \| 37 \| 502 \| 11 \| 463 \| Uncharacterized protein \| Uncharacterized protein; afdb-uniprot50 \| AF-A0A0F9FJZ9-F1-MODEL\_V4 \| 1.0 \| 2.342e-20 \| 675 \| 0.185 \| 555 \| 362 \| 21 \| 22 \| 532 \| 43 \| 551 \| Uncharacterized protein \| Uncharacterized protein; afdb-uniprot50 \| AF-A0A1H9QDY8-F1-MODEL\_V4 \| 1.0 \| 3.046e-20 \| 672 \| 0.196 \| 453 \| 301 \| 16 \| 28 \| 433 \| 1 \| 437 \| Phage portal protein, lambda family \| Phage portal protein, lambda family; afdb-uniprot50 \| AF-A0A517QH17-F1-MODEL\_V4 \| 1.0 \| 1.196e-19 \| 670 \| 0.145 \| 496 \| 366 \| 16 \| 23 \| 502 \| 1 \| 454 \| Phage portal protein, lambda family \| Phage portal protein, lambda family; afdb-uniprot50 \| AF-A0A1G3LXF9-F1-MODEL\_V4 \| 1.0 \| 1.328e-19 \| 669 \| 0.207 \| 420 \| 287 \| 12 \| 130 \| 532 \| 2 \| 392 \| Phage portal protein \| Phage portal protein; afdb-uniprot50 \| AF-A0A6C2U4J9-F1-MODEL\_V4 \| 1.0 \| 1.384e-20 \| 667 \| 0.186 \| 493 \| 326 \| 18 \| 44 \| 497 \| 6 \| 462 \| Uncharacterized protein \| Uncharacterized protein; afdb-uniprot50 \| AF-A0A2E8E056-F1-MODEL\_V4 \| 1.0 \| 4.586e-21 \| 666 \| 0.157 \| 533 \| 361 \| 19 \| 27 \| 517 \| 3 \| 489 \| Uncharacterized protein \| Uncharacterized protein; afdb-uniprot50 \| AF-A0A5C5WP67-F1-MODEL\_V4 \| 1.0 \| 6.627e-21 \| 663 \| 0.15 \| 537 \| 377 \| 21 \| 13 \| 515 \| 2 \| 493 \| Phage portal protein, lambda family \| Phage portal protein, lambda family; afdb-uniprot50 \| AF-A0A1V6CPT6-F1-MODEL\_V4 \| 1.0 \| 6.035e-20 \| 653 \| 0.171 \| 513 \| 339 \| 25 \| 39 \| 503 \| 19 \| 493 \| Phage portal protein, lambda family \| Phage portal protein, lambda family; afdb-uniprot50 \| AF-A0A7G8VCB0-F1-MODEL\_V4 \| 1.0 \| 6.361e-20 \| 649 \| 0.174 \| 517 \| 311 \| 20 \| 89 \| 530 \| 6 \| 481 \| Phage portal protein \| Phage portal protein; afdb-uniprot50 \| AF-A0A2X1LNZ6-F1-MODEL\_V4 \| 1.0 \| 1.843e-18 \| 642 \| 0.23 \| 394 \| 275 \| 11 \| 127 \| 499 \| 11 \| 397 \| Lambda family phage portal protein \| Lambda family phage portal protein; afdb-uniprot50 \| AF-A0A2J6HXS8-F1-MODEL\_V4 \| 1.0 \| 8.569e-17 \| 638 \| 0.389 \| 244 \| 136 \| 4 \| 263 \| 502 \| 1 \| 235 \| Phage portal protein \| Phage portal protein; afdb-uniprot50 \| AF-A0A844HAF1-F1-MODEL\_V4 \| 1.0 \| 3.848e-18 \| 637 \| 0.229 \| 357 \| 226 \| 15 \| 23 \| 375 \| 9 \| 320 \| Phage portal protein \| Phage portal protein; afdb-uniprot50 \| AF-A0A410S8D1-F1-MODEL\_V4 \| 1.0 \| 8.721e-20 \| 637 \| 0.15 \| 544 \| 385 \| 22 \| 11 \| 532 \| 2 \| 490 \| Phage portal protein \| Phage portal protein; afdb-uniprot50 \| AF-A0A518CCF6-F1-MODEL\_V4 \| 1.0 \| 6.783e-19 \| 636 \| 0.223 \| 385 \| 269 \| 13 \| 17 \| 395 \| 1 \| 361 \| Phage portal protein, lambda family \| Phage portal protein, lambda family; afdb-uniprot50 \| AF-A0A3M3BPQ1-F1-MODEL\_V4 \| 1.0 \| 7.624e-18 \| 635 \| 0.323 \| 278 \| 172 \| 6 \| 160 \| 422 \| 4 \| 280 \| Portal protein \| Portal protein; afdb-uniprot50 \| AF-A0A447JGS9-F1-MODEL\_V4 \| 1.0 \| 1.275e-18 \| 634 \| 0.211 \| 436 \| 278 \| 17 \| 47 \| 433 \| 34 \| 452 \| Phage portal protein, lambda family \| Phage portal protein, lambda family; afdb-uniprot50 \| AF-A0A1H5ZK26-F1-MODEL\_V4 \| 1.0 \| 1.115e-16 \| 633 \| 0.259 \| 301 \| 208 \| 7 \| 122 \| 413 \| 13 \| 307 \| Phage portal protein, lambda family \| Phage portal protein, lambda family; afdb-uniprot50 \| AF-A0A7W1G2Z3-F1-MODEL\_V4 \| 1.0 \| 9.689e-20 \| 630 \| 0.165 \| 525 \| 346 \| 19 \| 46 \| 532 \| 10 \| 480 \| Phage portal protein \| Phage portal protein; afdb-uniprot50 \| AF-A0A827GZ56-F1-MODEL\_V4 \| 1.0 \| 3.286e-18 \| 624 \| 0.228 \| 398 \| 253 \| 14 \| 99 \| 455 \| 3 \| 387 \| Phage portal protein \| Phage portal protein; afdb-uniprot50 \| AF-A0A177R1N7-F1-MODEL\_V4 \| 1.0 \| 8.823e-19 \| 622 \| 0.198 \| 478 \| 314 \| 22 \| 35 \| 491 \| 19 \| 448 \| Uncharacterized protein \| Uncharacterized protein; afdb-uniprot50 \| AF-A0A3B9NPV5-F1-MODEL\_V4 \| 1.0 \| 2.071e-17 \| 621 \| 0.37 \| 243 \| 140 \| 6 \| 14 \| 253 \| 1 \| 233 \| Uncharacterized protein \| Uncharacterized protein; afdb-uniprot50 \| AF-A0A257JLH4-F1-MODEL\_V4 \| 1.0 \| 5.276e-18 \| 619 \| 0.301 \| 322 \| 202 \| 9 \| 185 \| 502 \| 6 \| 308 \| Phage portal protein \| Phage portal protein; afdb-uniprot50 \| AF-A0A853YYM9-F1-MODEL\_V4 \| 1.0 \| 8.469e-18 \| 614 \| 0.318 \| 336 \| 194 \| 7 \| 204 \| 532 \| 3 \| 310 \| Portal protein \| Portal protein; afdb-uniprot50 \| AF-A0A0S8GDJ6-F1-MODEL\_V4 \| 1.0 \| 4.402e-20 \| 614 \| 0.168 \| 582 \| 352 \| 23 \| 26 \| 532 \| 2 \| 526 \| Uncharacterized protein \| Uncharacterized protein; afdb-uniprot50 \| AF-A0A8B5P8M5-F1-MODEL\_V4 \| 1.0 \| 5.861e-18 \| 611 \| 0.19 \| 442 \| 283 \| 15 \| 23 \| 413 \| 2 \| 419 \| Phage portal protein \| Phage portal protein; afdb-uniprot50 \| AF-A0A439F4D3-F1-MODEL\_V4 \| 1.0 \| 1.089e-18 \| 608 \| 0.229 \| 393 \| 253 \| 14 \| 160 \| 532 \| 4 \| 366 \| Uncharacterized protein \| Uncharacterized protein; afdb-uniprot50 \| AF-A0A357LHL5-F1-MODEL\_V4 \| 1.0 \| 4.009e-19 \| 607 \| 0.167 \| 519 \| 352 \| 24 \| 73 \| 532 \| 2 \| 499 \| Phage portal protein \| Phage portal protein; afdb-uniprot50 \| AF-A0A6G4KD87-F1-MODEL\_V4 \| 1.0 \| 4.558e-17 \| 606 \| 0.215 \| 306 \| 211 \| 11 \| 93 \| 385 \| 2 \| 291 \| Phage portal protein \| Phage portal protein; afdb-uniprot50 \| AF-A0A0E3YYG8-F1-MODEL\_V4 \| 1.0 \| 1.678e-17 \| 604 \| 0.14 \| 483 \| 355 \| 15 \| 45 \| 504 \| 17 \| 462 \| Uncharacterized protein \| Uncharacterized protein; afdb-uniprot50 \| AF-A0A1M7U6E7-F1-MODEL\_V4 \| 1.0 \| 5.626e-17 \| 603 \| 0.284 \| 316 \| 214 \| 5 \| 191 \| 502 \| 11 \| 318 \| Phage portal protein, lambda family \| Phage portal protein, lambda family; afdb-uniprot50 \| AF-A0A554XEQ4-F1-MODEL\_V4 \| 1.0 \| 7.149e-19 \| 601 \| 0.246 \| 397 \| 266 \| 13 \| 36 \| 416 \| 8 \| 387 \| Putative esterase \| Putative esterase; afdb-uniprot50 \| AF-A0A5U4DB71-F1-MODEL\_V4 \| 1.0 \| 1.115e-16 \| 600 \| 0.227 \| 325 \| 224 \| 11 \| 39 \| 348 \| 7 \| 319 \| Phage portal protein \| Phage portal protein; afdb-uniprot50 \| AF-A0A380AGT5-F1-MODEL\_V4 \| 1.0 \| 1.592e-17 \| 591 \| 0.242 \| 355 \| 233 \| 8 \| 187 \| 532 \| 6 \| 333 \| Phage portal protein, lambda family \| Phage portal protein, lambda family; afdb-uniprot50 \| AF-A0A357ZWC2-F1-MODEL\_V4 \| 1.0 \| 6.511e-18 \| 588 \| 0.147 \| 507 \| 365 \| 16 \| 51 \| 532 \| 12 \| 476 \| Uncharacterized protein \| Uncharacterized protein; afdb-uniprot50 \| AF-A0A1L5KK25-F1-MODEL\_V4 \| 1.0 \| 4.749e-18 \| 585 \| 0.186 \| 476 \| 304 \| 13 \| 127 \| 531 \| 3 \| 466 \| Phage portal protein, lambda family \| Phage portal protein, lambda family; afdb-uniprot50 \| AF-A0A3W5Y8G0-F1-MODEL\_V4 \| 1.0 \| 5.123e-16 \| 584 \| 0.225 \| 319 \| 222 \| 9 \| 165 \| 474 \| 4 \| 306 \| Phage portal protein \| Phage portal protein; afdb-uniprot50 \| AF-A0A2U8HXA8-F1-MODEL\_V4 \| 1.0 \| 2.993e-17 \| 583 \| 0.268 \| 339 \| 220 \| 9 \| 207 \| 532 \| 1 \| 324 \| Phage portal protein \| Phage portal protein; afdb-uniprot50 \| AF-A0A3L7RG21-F1-MODEL\_V4 \| 1.0 \| 7.713e-17 \| 583 \| 0.217 \| 377 \| 255 \| 11 \| 160 \| 532 \| 4 \| 344 \| Phage portal protein \| Phage portal protein; afdb-uniprot50 \| AF-A0A847I6D6-F1-MODEL\_V4 \| 1.0 \| 1.988e-16 \| 577 \| 0.194 \| 371 \| 252 \| 16 \| 62 \| 417 \| 1 \| 339 \| Phage portal protein \| Phage portal protein; afdb-uniprot50 \| AF-A0A7C4BYU0-F1-MODEL\_V4 \| 1.0 \| 2.497e-19 \| 575 \| 0.243 \| 443 \| 258 \| 16 \| 37 \| 459 \| 2 \| 387 \| Phage portal protein \| Phage portal protein; afdb-uniprot50 \| AF-A0A1U7D0P3-F1-MODEL\_V4 \| 1.0 \| 5.93e-17 \| 572 \| 0.211 \| 402 \| 264 \| 14 \| 131 \| 492 \| 2 \| 390 \| Phage portal protein, lambda family \| Phage portal protein, lambda family; afdb-uniprot50 \| AF-A0A3C1YTA8-F1-MODEL\_V4 \| 1.0 \| 3.694e-17 \| 572 \| 0.133 \| 532 \| 385 \| 18 \| 20 \| 517 \| 2 \| 491 \| Uncharacterized protein \| Uncharacterized protein; afdb-uniprot50 \| AF-A0A1Y2XGJ0-F1-MODEL\_V4 \| 1.0 \| 4.558e-17 \| 569 \| 0.195 \| 409 \| 254 \| 16 \| 39 \| 396 \| 18 \| 402 \| Phage portal protein \| Phage portal protein; afdb-uniprot50 \| AF-A0A166Z3X1-F1-MODEL\_V4 \| 1.0 \| 2.839e-17 \| 568 \| 0.194 \| 427 \| 285 \| 13 \| 135 \| 519 \| 2 \| 411 \| Phage portal protein, lambda family \| Phage portal protein, lambda family; afdb-uniprot50 \| AF-B3CPN2-F1-MODEL\_V4 \| 1.0 \| 7.804e-16 \| 566 \| 0.292 \| 291 \| 193 \| 4 \| 211 \| 496 \| 112 \| 394 \| Putative phage portal protein \| Putative phage portal protein; afdb-uniprot50 \| AF-A0A5C7PV87-F1-MODEL\_V4 \| 1.0 \| 1.698e-16 \| 563 \| 0.24 \| 358 \| 237 \| 7 \| 186 \| 532 \| 17 \| 350 \| Phage portal protein \| Phage portal protein; afdb-uniprot50 \| AF-A0A0F9LPE5-F1-MODEL\_V4 \| 1.0 \| 6.863e-18 \| 555 \| 0.174 \| 516 \| 335 \| 21 \| 37 \| 531 \| 26 \| 471 \| Uncharacterized protein \| Uncharacterized protein; afdb-uniprot50 \| AF-A0A3D1IXD3-F1-MODEL\_V4 \| 1.0 \| 2.556e-17 \| 550 \| 0.127 \| 548 \| 372 \| 24 \| 51 \| 532 \| 26 \| 533 \| Uncharacterized protein \| Uncharacterized protein; afdb-uniprot50 \| AF-A0A5C7Q617-F1-MODEL\_V4 \| 1.0 \| 2.234e-15 \| 548 \| 0.245 \| 310 \| 215 \| 8 \| 187 \| 487 \| 14 \| 313 \| Phage portal protein \| Phage portal protein; afdb-uniprot50 \| AF-A0A379XPD6-F1-MODEL\_V4 \| 1.0 \| 2.453e-16 \| 548 \| 0.187 \| 347 \| 253 \| 10 \| 39 \| 374 \| 21 \| 349 \| Phage portal protein, lambda family protein \| Phage portal protein, lambda family protein; afdb-uniprot50 \| AF-A0A3S4M9N1-F1-MODEL\_V4 \| 1.0 \| 1.546e-15 \| 542 \| 0.225 \| 341 \| 228 \| 8 \| 196 \| 532 \| 1 \| 309 \| Phage portal protein, lambda family \| Phage portal protein, lambda family; afdb-uniprot50 \| AF-A0A1V5GTW5-F1-MODEL\_V4 \| 1.0 \| 2.011e-15 \| 540 \| 0.286 \| 304 \| 191 \| 5 \| 233 \| 532 \| 6 \| 287 \| Phage portal protein, lambda family \| Phage portal protein, lambda family; afdb-uniprot50 \| AF-A0A0P1FA99-F1-MODEL\_V4 \| 1.0 \| 1.305e-16 \| 534 \| 0.263 \| 319 \| 205 \| 11 \| 15 \| 328 \| 1 \| 294 \| Phage portal protein, lambda family \| Phage portal protein, lambda family; afdb-uniprot50 \| AF-A0A250DMY9-F1-MODEL\_V4 \| 1.0 \| 7.713e-17 \| 531 \| 0.3 \| 273 \| 176 \| 6 \| 67 \| 328 \| 2 \| 270 \| Uncharacterized protein \| Uncharacterized protein; afdb-uniprot50 \| AF-A0A484WP22-F1-MODEL\_V4 \| 1.0 \| 1.988e-16 \| 530 \| 0.206 \| 411 \| 269 \| 15 \| 122 \| 490 \| 3 \| 398 \| Capsid protein of prophage \| Capsid protein of prophage; afdb-uniprot50 \| AF-A0A532UBS6-F1-MODEL\_V4 \| 1.0 \| 9.917e-18 \| 530 \| 0.156 \| 497 \| 311 \| 19 \| 39 \| 490 \| 18 \| 451 \| Uncharacterized protein \| Uncharacterized protein; afdb-uniprot50 \| AF-A0A3M6EEB9-F1-MODEL\_V4 \| 1.0 \| 1.484e-14 \| 528 \| 0.291 \| 257 \| 169 \| 6 \| 58 \| 311 \| 3 \| 249 \| Portal protein \| Portal protein; afdb-uniprot50 \| AF-A0A3C1XMB6-F1-MODEL\_V4 \| 1.0 \| 7.713e-17 \| 526 \| 0.207 \| 371 \| 235 \| 13 \| 39 \| 357 \| 15 \| 378 \| Phage portal protein \| Phage portal protein; afdb-uniprot50 \| AF-A0A7W1T9L3-F1-MODEL\_V4 \| 1.0 \| 4.375e-16 \| 522 \| 0.221 \| 374 \| 246 \| 13 \| 21 \| 381 \| 15 \| 356 \| Phage portal protein \| Phage portal protein; afdb-uniprot50 \| AF-A0A7X7KMY8-F1-MODEL\_V4 \| 1.0 \| 3.587e-15 \| 519 \| 0.237 \| 341 \| 221 \| 14 \| 20 \| 352 \| 9 \| 318 \| Phage portal protein \| Phage portal protein; afdb-uniprot50 \| AF-A0A448MNG2-F1-MODEL\_V4 \| 1.0 \| 5.463e-15 \| 515 \| 0.22 \| 295 \| 208 \| 9 \| 39 \| 328 \| 21 \| 298 \| Bacteriophage capsid protein \| Bacteriophage capsid protein; afdb-uniprot50 \| AF-B2I820-F1-MODEL\_V4 \| 1.0 \| 3.825e-14 \| 513 \| 0.435 \| 216 \| 115 \| 2 \| 317 \| 532 \| 1 \| 209 \| Putative phage related protein \| Putative phage related protein; afdb-uniprot50 \| AF-A0A6C2YPG2-F1-MODEL\_V4 \| 1.0 \| 3.545e-16 \| 513 \| 0.144 \| 486 \| 349 \| 23 \| 39 \| 491 \| 58 \| 509 \| Uncharacterized protein \| Uncharacterized protein; afdb-uniprot50 \| AF-A0A827BFZ7-F1-MODEL\_V4 \| 1.0 \| 1.07e-15 \| 510 \| 0.175 \| 428 \| 284 \| 12 \| 143 \| 523 \| 6 \| 411 \| Phage portal protein \| Phage portal protein; afdb-uniprot50 \| AF-A0A7W8GZX7-F1-MODEL\_V4 \| 1.0 \| 3.229e-15 \| 504 \| 0.192 \| 400 \| 254 \| 11 \| 154 \| 502 \| 2 \| 383 \| Lambda family phage portal protein \| Lambda family phage portal protein; afdb-uniprot50 \| AF-A0A2M9P8F9-F1-MODEL\_V4 \| 1.0 \| 3.671e-13 \| 496 \| 0.267 \| 258 \| 172 \| 9 \| 39 \| 292 \| 9 \| 253 \| Phage portal protein \| Phage portal protein; afdb-uniprot50 \| AF-A0A497KH68-F1-MODEL\_V4 \| 1.0 \| 8.225e-16 \| 493 \| 0.146 \| 452 \| 302 \| 20 \| 39 \| 456 \| 42 \| 443 \| Uncharacterized protein \| Uncharacterized protein; afdb-uniprot50 \| AF-A0A7J6YNN5-F1-MODEL\_V4 \| 1.0 \| 2.906e-15 \| 491 \| 0.268 \| 316 \| 198 \| 8 \| 220 \| 530 \| 336 \| 623 \| Uncharacterized protein \| Uncharacterized protein; afdb-uniprot50 \| AF-J0PPA8-F1-MODEL\_V4 \| 1.0 \| 2.383e-14 \| 488 \| 0.232 \| 314 \| 221 \| 6 \| 201 \| 502 \| 2 \| 307 \| Lambda family phage portal protein \| Lambda family phage portal protein; afdb-uniprot50 \| AF-F0SNL3-F1-MODEL\_V4 \| 1.0 \| 6.25e-17 \| 486 \| 0.148 \| 564 \| 364 \| 25 \| 24 \| 517 \| 35 \| 551 \| Uncharacterized protein \| Uncharacterized protein; afdb-uniprot50 \| AF-A0A4C3GAR0-F1-MODEL\_V4 \| 1.0 \| 3.063e-15 \| 484 \| 0.175 \| 415 \| 285 \| 14 \| 150 \| 523 \| 376 \| 774 \| Phage terminase large subunit \| Phage terminase large subunit; afdb-uniprot50 \| AF-A0A354EMX8-F1-MODEL\_V4 \| 1.0 \| 4.079e-13 \| 482 \| 0.266 \| 300 \| 208 \| 8 \| 179 \| 473 \| 5 \| 297 \| Phage portal protein \| Phage portal protein; afdb-uniprot50 \| AF-A0A2E1P380-F1-MODEL\_V4 \| 1.0 \| 5.592e-13 \| 479 \| 0.297 \| 262 \| 162 \| 6 \| 275 \| 532 \| 2 \| 245 \| Phage portal protein \| Phage portal protein; afdb-uniprot50 \| AF-A0A7I8Z9X4-F1-MODEL\_V4 \| 1.0 \| 3.671e-13 \| 478 \| 0.271 \| 280 \| 188 \| 6 \| 205 \| 476 \| 4 \| 275 \| Hypothetical protein \| Hypothetical protein; afdb-uniprot50 \| AF-M1P158-F1-MODEL\_V4 \| 1.0 \| 5.758e-15 \| 478 \| 0.126 \| 500 \| 350 \| 20 \| 10 \| 500 \| 8 \| 429 \| Portal protein lambda \| Portal protein lambda; afdb-uniprot50 \| AF-A0A827GZ82-F1-MODEL\_V4 \| 1.0 \| 2.647e-14 \| 477 \| 0.215 \| 344 \| 232 \| 10 \| 195 \| 517 \| 13 \| 339 \| Phage portal protein \| Phage portal protein; afdb-uniprot50 \| AF-A0A6P1B243-F1-MODEL\_V4 \| 1.0 \| 1.601e-12 \| 476 \| 0.582 \| 127 \| 52 \| 1 \| 96 \| 222 \| 2 \| 127 \| Phage portal protein \| Phage portal protein; afdb-uniprot50 \| AF-A0A353UWW7-F1-MODEL\_V4 \| 1.0 \| 5.592e-13 \| 469 \| 0.266 \| 274 \| 181 \| 10 \| 39 \| 306 \| 27 \| 286 \| Phage portal protein \| Phage portal protein; afdb-uniprot50 \| AF-A0A484X983-F1-MODEL\_V4 \| 1.0 \| 1.095e-13 \| 464 \| 0.246 \| 341 \| 216 \| 10 \| 140 \| 448 \| 3 \| 334 \| Capsid protein of prophage \| Capsid protein of prophage; afdb-uniprot50 \| AF-L8M8B8-F1-MODEL\_V4 \| 1.0 \| 3.305e-13 \| 462 \| 0.28 \| 239 \| 150 \| 7 \| 23 \| 255 \| 12 \| 234 \| Bacteriophage capsid protein \| Bacteriophage capsid protein; afdb-uniprot50 \| AF-A0A0K3YI17-F1-MODEL\_V4 \| 1.0 \| 1.875e-12 \| 459 \| 0.234 \| 277 \| 199 \| 8 \| 214 \| 486 \| 4 \| 271 \| Portal protein (Head-tail preconnector protein) from prophage \| Portal protein (Head-tail preconnector protein) from prophage; afdb-uniprot50 \| AF-A0A250DLV1-F1-MODEL\_V4 \| 1.0 \| 1.601e-12 \| 457 \| 0.319 \| 263 \| 152 \| 4 \| 277 \| 532 \| 12 \| 254 \| Uncharacterized protein \| Uncharacterized protein; afdb-uniprot50 \| AF-A0A291A6H8-F1-MODEL\_V4 \| 1.0 \| 1.519e-12 \| 457 \| 0.275 \| 272 \| 176 \| 5 \| 244 \| 502 \| 3 \| 266 \| Phage portal protein \| Phage portal protein; afdb-uniprot50 \| AF-A0A6I2J5G2-F1-MODEL\_V4 \| 1.0 \| 3.136e-13 \| 453 \| 0.442 \| 190 \| 99 \| 4 \| 1 \| 189 \| 3 \| 186 \| Phage portal protein \| Phage portal protein; afdb-uniprot50 \| AF-A0A1G1LMU8-F1-MODEL\_V4 \| 1.0 \| 1.688e-12 \| 452 \| 0.288 \| 253 \| 170 \| 4 \| 253 \| 502 \| 1 \| 246 \| Phage portal protein \| Phage portal protein; afdb-uniprot50 \| AF-A0A2V8RVC3-F1-MODEL\_V4 \| 1.0 \| 1.039e-13 \| 447 \| 0.152 \| 387 \| 273 \| 14 \| 160 \| 532 \| 5 \| 350 \| Uncharacterized protein \| Uncharacterized protein; afdb-uniprot50 \| AF-A0A352V3P3-F1-MODEL\_V4 \| 1.0 \| 2.541e-13 \| 444 \| 0.225 \| 337 \| 209 \| 10 \| 210 \| 531 \| 3 \| 302 \| Uncharacterized protein \| Uncharacterized protein; afdb-uniprot50 \| AF-A0A3S4CSS4-F1-MODEL\_V4 \| 1.0 \| 5.894e-13 \| 442 \| 0.218 \| 316 \| 214 \| 8 \| 218 \| 532 \| 5 \| 288 \| Phage portal protein, lambda family \| Phage portal protein, lambda family; afdb-uniprot50 \| AF-A0A3C0I3F4-F1-MODEL\_V4 \| 1.0 \| 2.439e-12 \| 441 \| 0.244 \| 245 \| 162 \| 9 \| 34 \| 264 \| 13 \| 248 \| Uncharacterized protein \| Uncharacterized protein; afdb-uniprot50 \| AF-A0A7V6YCH4-F1-MODEL\_V4 \| 1.0 \| 1.688e-12 \| 441 \| 0.252 \| 317 \| 216 \| 6 \| 221 \| 532 \| 1 \| 301 \| Phage portal protein \| Phage portal protein; afdb-uniprot50 \| AF-A0A3M2YA61-F1-MODEL\_V4 \| 1.0 \| 1.367e-12 \| 440 \| 0.268 \| 305 \| 185 \| 5 \| 242 \| 530 \| 3 \| 285 \| Uncharacterized protein \| Uncharacterized protein; afdb-uniprot50 \| AF-A0A258REZ0-F1-MODEL\_V4 \| 1.0 \| 7.274e-13 \| 435 \| 0.273 \| 311 \| 193 \| 7 \| 234 \| 532 \| 1 \| 290 \| Phage portal protein \| Phage portal protein; afdb-uniprot50 \| AF-A0A754E6Q2-F1-MODEL\_V4 \| 1.0 \| 2.556e-17 \| 435 \| 0.157 \| 595 \| 347 \| 25 \| 39 \| 532 \| 11 \| 551 \| Phage portal protein \| Phage portal protein; afdb-uniprot50 \| AF-A0A1F4CAA0-F1-MODEL\_V4 \| 1.0 \| 3.483e-13 \| 430 \| 0.247 \| 360 \| 209 \| 12 \| 208 \| 532 \| 1 \| 333 \| Phage portal protein \| Phage portal protein; afdb-uniprot50 \| AF-A0A7C2ZHD3-F1-MODEL\_V4 \| 1.0 \| 2.975e-13 \| 429 \| 0.133 \| 493 \| 297 \| 25 \| 39 \| 515 \| 19 \| 397 \| Uncharacterized protein \| Uncharacterized protein; afdb-uniprot50 \| AF-A0A7X7IZ38-F1-MODEL\_V4 \| 1.0 \| 1.168e-12 \| 428 \| 0.216 \| 323 \| 217 \| 11 \| 24 \| 342 \| 12 \| 302 \| Phage portal protein \| Phage portal protein; afdb-uniprot50 \| AF-A0A497SVU5-F1-MODEL\_V4 \| 1.0 \| 5.244e-14 \| 425 \| 0.134 \| 467 \| 298 \| 23 \| 23 \| 470 \| 6 \| 385 \| Uncharacterized protein \| Uncharacterized protein; afdb-uniprot50 \| AF-A0A378BW42-F1-MODEL\_V4 \| 1.0 \| 9.973e-13 \| 423 \| 0.233 \| 325 \| 212 \| 14 \| 217 \| 532 \| 4 \| 300 \| Phage portal protein, lambda family \| Phage portal protein, lambda family; afdb-uniprot50 \| AF-A0A326U088-F1-MODEL\_V4 \| 1.0 \| 3.305e-13 \| 419 \| 0.141 \| 509 \| 314 \| 24 \| 27 \| 523 \| 3 \| 400 \| Uncharacterized protein \| Uncharacterized protein; afdb-uniprot50 \| AF-A0A376JWU1-F1-MODEL\_V4 \| 1.0 \| 3.715e-12 \| 418 \| 0.214 \| 261 \| 179 \| 10 \| 20 \| 264 \| 5 \| 255 \| Phage portal protein, lambda family \| Phage portal protein, lambda family; afdb-uniprot50 \| AF-A0A793I648-F1-MODEL\_V4 \| 1.0 \| 1.897e-11 \| 412 \| 0.217 \| 276 \| 201 \| 6 \| 195 \| 466 \| 13 \| 277 \| Phage portal protein \| Phage portal protein; afdb-uniprot50 \| AF-A0A2H0LJS2-F1-MODEL\_V4 \| 1.0 \| 1.537e-11 \| 412 \| 0.209 \| 282 \| 204 \| 6 \| 254 \| 532 \| 9 \| 274 \| Phage portal protein \| Phage portal protein; afdb-uniprot50 \| AF-A0A2W5H3L8-F1-MODEL\_V4 \| 1.0 \| 3.344e-12 \| 411 \| 0.211 \| 307 \| 208 \| 11 \| 36 \| 330 \| 23 \| 307 \| Phage portal protein \| Phage portal protein; afdb-uniprot50 \| AF-A0A2D8T9H3-F1-MODEL\_V4 \| 1.0 \| 2.287e-13 \| 411 \| 0.185 \| 372 \| 245 \| 11 \| 192 \| 532 \| 22 \| 366 \| Phage portal protein \| Phage portal protein; afdb-uniprot50 \| AF-K2GKL9-F1-MODEL\_V4 \| 1.0 \| 1.779e-12 \| 408 \| 0.221 \| 334 \| 203 \| 11 \| 39 \| 328 \| 27 \| 347 \| Uncharacterized protein \| Uncharacterized protein; afdb-uniprot50 \| AF-H5UYW8-F1-MODEL\_V4 \| 1.0 \| 2.107e-11 \| 406 \| 0.216 \| 310 \| 209 \| 8 \| 226 \| 532 \| 2 \| 280 \| Putative portal protein \| Putative portal protein; afdb-uniprot50 \| AF-A0A3D5T0J9-F1-MODEL\_V4 \| 1.0 \| 1.245e-11 \| 405 \| 0.238 \| 310 \| 206 \| 6 \| 226 \| 532 \| 1 \| 283 \| Phage portal protein \| Phage portal protein; afdb-uniprot50 \| AF-A0A0F9HAI2-F1-MODEL\_V4 \| 1.0 \| 5.093e-12 \| 396 \| 0.364 \| 233 \| 135 \| 4 \| 303 \| 524 \| 1 \| 231 \| Uncharacterized protein \| Uncharacterized protein; afdb-uniprot50 \| AF-A0A455T535-F1-MODEL\_V4 \| 1.0 \| 8.419e-14 \| 396 \| 0.139 \| 587 \| 348 \| 29 \| 3 \| 531 \| 6 \| 493 \| Uncharacterized protein \| Uncharacterized protein; afdb-uniprot50 \| AF-M5JSY7-F1-MODEL\_V4 \| 1.0 \| 4.832e-12 \| 395 \| 0.2 \| 374 \| 236 \| 13 \| 168 \| 502 \| 2 \| 351 \| Phage portal protein, lambda \| Phage portal protein, lambda; afdb-uniprot50 \| AF-A0A7C5MJ10-F1-MODEL\_V4 \| 1.0 \| 2.41e-13 \| 394 \| 0.113 \| 527 \| 335 \| 25 \| 39 \| 532 \| 7 \| 434 \| Phage portal protein \| Phage portal protein; afdb-uniprot50 \| AF-A0A3A4ZEQ3-F1-MODEL\_V4 \| 1.0 \| 4.249e-14 \| 394 \| 0.131 \| 556 \| 358 \| 26 \| 37 \| 518 \| 107 \| 611 \| Phage portal protein \| Phage portal protein; afdb-uniprot50 \| AF-A0A6M3XSW5-F1-MODEL\_V4 \| 1.0 \| 4.349e-12 \| 392 \| 0.166 \| 396 \| 273 \| 15 \| 160 \| 518 \| 5 \| 380 \| Putative portal protein \| Putative portal protein; afdb-uniprot50 \| AF-A0A7V4ZD36-F1-MODEL\_V4 \| 1.0 \| 1.875e-12 \| 392 \| 0.139 \| 493 \| 286 \| 31 \| 15 \| 477 \| 1 \| 385 \| Uncharacterized protein \| Uncharacterized protein; afdb-uniprot50 \| AF-A0A403NK32-F1-MODEL\_V4 \| 1.0 \| 4.127e-12 \| 391 \| 0.214 \| 345 \| 223 \| 9 \| 207 \| 515 \| 6 \| 338 \| Phage portal protein \| Phage portal protein; afdb-uniprot50 \| AF-A0A367D8L0-F1-MODEL\_V4 \| 1.0 \| 1.821e-10 \| 390 \| 0.258 \| 236 \| 165 \| 8 \| 252 \| 483 \| 3 \| 232 \| Phage portal protein \| Phage portal protein; afdb-uniprot50 \| AF-A0A6L9HC99-F1-MODEL\_V4 \| 1.0 \| 3.422e-10 \| 387 \| 0.233 \| 253 \| 182 \| 3 \| 244 \| 487 \| 1 \| 250 \| Phage portal protein \| Phage portal protein; afdb-uniprot50 \| AF-A0A6M3IP50-F1-MODEL\_V4 \| 1.0 \| 3.21e-11 \| 384 \| 0.197 \| 294 \| 212 \| 4 \| 254 \| 530 \| 21 \| 307 \| Putative portal protein \| Putative portal protein; afdb-uniprot50 \| AF-W8YFG7-F1-MODEL\_V4 \| 1.0 \| 4.584e-12 \| 384 \| 0.246 \| 329 \| 205 \| 9 \| 210 \| 502 \| 51 \| 372 \| Uncharacterized protein \| Uncharacterized protein; afdb-uniprot50 \| AF-A0A402A2E6-F1-MODEL\_V4 \| 1.0 \| 2.57e-12 \| 384 \| 0.129 \| 534 \| 353 \| 24 \| 13 \| 523 \| 2 \| 446 \| Uncharacterized protein \| Uncharacterized protein; afdb-uniprot50 \| AF-A0A0F9MZW4-F1-MODEL\_V4 \| 1.0 \| 2.383e-14 \| 384 \| 0.157 \| 579 \| 356 \| 24 \| 24 \| 532 \| 1 \| 517 \| Uncharacterized protein \| Uncharacterized protein; afdb-uniprot50 \| AF-A0A5S5MFU0-F1-MODEL\_V4 \| 1.0 \| 4.946e-10 \| 377 \| 0.244 \| 262 \| 177 \| 7 \| 252 \| 497 \| 14 \| 270 \| Phage portal protein \| Phage portal protein; afdb-uniprot50 \| AF-A0A368KX58-F1-MODEL\_V4 \| 1.0 \| 2.601e-11 \| 377 \| 0.23 \| 286 \| 186 \| 10 \| 99 \| 379 \| 13 \| 269 \| Phage portal protein \| Phage portal protein; afdb-uniprot50 \| AF-A0A6M3LXD3-F1-MODEL\_V4 \| 1.0 \| 2.822e-13 \| 377 \| 0.141 \| 515 \| 333 \| 30 \| 39 \| 525 \| 19 \| 452 \| Putative portal protein \| Putative portal protein; afdb-uniprot50 \| AF-A0A3V0YA20-F1-MODEL\_V4 \| 1.0 \| 3.422e-10 \| 375 \| 0.189 \| 275 \| 199 \| 6 \| 258 \| 530 \| 1 \| 253 \| Phage portal protein \| Phage portal protein; afdb-uniprot50 \| AF-A0A0S7BFT1-F1-MODEL\_V4 \| 1.0 \| 7.274e-13 \| 373 \| 0.158 \| 497 \| 302 \| 26 \| 28 \| 486 \| 1 \| 419 \| Phage portal protein, lambda family \| Phage portal protein, lambda family; afdb-uniprot50 \| AF-A0A402ARA1-F1-MODEL\_V4 \| 1.0 \| 1.121e-11 \| 372 \| 0.131 \| 465 \| 295 \| 23 \| 38 \| 459 \| 7 \| 405 \| Uncharacterized protein \| Uncharacterized protein; afdb-uniprot50 \| AF-A0A0F9HE98-F1-MODEL\_V4 \| 1.0 \| 6.212e-13 \| 372 \| 0.138 \| 535 \| 345 \| 23 \| 39 \| 523 \| 11 \| 479 \| Uncharacterized protein \| Uncharacterized protein; afdb-uniprot50 \| AF-A0A4P9VL88-F1-MODEL\_V4 \| 1.0 \| 1.009e-11 \| 371 \| 0.172 \| 353 \| 238 \| 10 \| 187 \| 502 \| 1 \| 336 \| Phage portal protein \| Phage portal protein; afdb-uniprot50 \| AF-A0A7C4Z3X6-F1-MODEL\_V4 \| 1.0 \| 9.083e-12 \| 370 \| 0.151 \| 529 \| 317 \| 25 \| 39 \| 532 \| 4 \| 435 \| Phage portal protein \| Phage portal protein; afdb-uniprot50 \| AF-A0A633NAI0-F1-MODEL\_V4 \| 1.0 \| 6.104e-10 \| 368 \| 0.22 \| 209 \| 146 \| 5 \| 157 \| 357 \| 2 \| 201 \| Phage portal protein \| Phage portal protein; afdb-uniprot50 \| AF-A0A357YBI7-F1-MODEL\_V4 \| 1.0 \| 3.566e-11 \| 368 \| 0.193 \| 315 \| 220 \| 8 \| 211 \| 497 \| 1 \| 309 \| Uncharacterized protein \| Uncharacterized protein; afdb-uniprot50 \| AF-A0A6J4K1M5-F1-MODEL\_V4 \| 1.0 \| 1.976e-12 \| 367 \| 0.145 \| 510 \| 312 \| 25 \| 35 \| 488 \| 7 \| 448 \| Uncharacterized protein \| Uncharacterized protein; afdb-uniprot50 \| AF-A0A7V2ILH1-F1-MODEL\_V4 \| 1.0 \| 1.707e-11 \| 364 \| 0.216 \| 323 \| 215 \| 11 \| 17 \| 328 \| 1 \| 296 \| Phage portal protein \| Phage portal protein; afdb-uniprot50 \| AF-A0A7C5LMG7-F1-MODEL\_V4 \| 1.0 \| 1.064e-11 \| 364 \| 0.163 \| 435 \| 259 \| 24 \| 39 \| 439 \| 4 \| 367 \| Uncharacterized protein \| Uncharacterized protein; afdb-uniprot50 \| AF-A0A2N2NPH9-F1-MODEL\_V4 \| 1.0 \| 6.548e-13 \| 357 \| 0.14 \| 497 \| 312 \| 28 \| 37 \| 488 \| 7 \| 433 \| Uncharacterized protein \| Uncharacterized protein; afdb-uniprot50 \| AF-A0A1R1LRL3-F1-MODEL\_V4 \| 1.0 \| 3.045e-11 \| 356 \| 0.187 \| 383 \| 233 \| 16 \| 186 \| 519 \| 7 \| 360 \| Phage portal protein \| Phage portal protein; afdb-uniprot50 \| AF-A0A0F8ZDI4-F1-MODEL\_V4 \| 1.0 \| 7.848e-11 \| 353 \| 0.133 \| 427 \| 275 \| 17 \| 10 \| 413 \| 1 \| 355 \| Uncharacterized protein \| Uncharacterized protein; afdb-uniprot50 \| AF-A0A120G5R1-F1-MODEL\_V4 \| 1.0 \| 3.422e-10 \| 349 \| 0.215 \| 301 \| 197 \| 7 \| 250 \| 532 \| 3 \| 282 \| Phage portal protein, lambda family \| Phage portal protein, lambda family; afdb-uniprot50 \| AF-A0A7A7BND2-F1-MODEL\_V4 \| 1.0 \| 1.021e-10 \| 348 \| 0.169 \| 353 \| 220 \| 12 \| 187 \| 491 \| 8 \| 335 \| Phage portal protein \| Phage portal protein; afdb-uniprot50 \| AF-A0A2T3BVF4-F1-MODEL\_V4 \| 1.0 \| 1.601e-12 \| 346 \| 0.144 \| 589 \| 340 \| 29 \| 17 \| 508 \| 1 \| 522 \| Uncharacterized protein \| Uncharacterized protein; afdb-uniprot50 \| AF-A0A383EWE0-F1-MODEL\_V4 \| 1.0 \| 1.209e-09 \| 340 \| 0.307 \| 228 \| 140 \| 5 \| 305 \| 532 \| 1 \| 210 \| Uncharacterized protein \| Uncharacterized protein; afdb-uniprot50 \| AF-A0A6M3KLI4-F1-MODEL\_V4 \| 1.0 \| 1.779e-12 \| 340 \| 0.15 \| 505 \| 326 \| 27 \| 54 \| 532 \| 21 \| 448 \| Putative portal protein \| Putative portal protein; afdb-uniprot50 \| AF-A0A662DL46-F1-MODEL\_V4 \| 1.0 \| 3.081e-10 \| 338 \| 0.168 \| 356 \| 267 \| 13 \| 191 \| 532 \| 4 \| 344 \| Uncharacterized protein \| Uncharacterized protein; afdb-uniprot50 \| AF-A0A377ECV1-F1-MODEL\_V4 \| 1.0 \| 7.94e-10 \| 336 \| 0.201 \| 307 \| 211 \| 6 \| 250 \| 532 \| 2 \| 298 \| Capsid protein of prophage \| Capsid protein of prophage; afdb-uniprot50 \| AF-A0A0F9EPK0-F1-MODEL\_V4 \| 1.0 \| 1.62e-11 \| 336 \| 0.121 \| 568 \| 355 \| 27 \| 7 \| 532 \| 1 \| 466 \| Uncharacterized protein \| Uncharacterized protein; afdb-uniprot50 \| AF-A0A0F8ZAV5-F1-MODEL\_V4 \| 1.0 \| 2.107e-11 \| 335 \| 0.148 \| 396 \| 263 \| 23 \| 39 \| 412 \| 21 \| 364 \| Uncharacterized protein \| Uncharacterized protein; afdb-uniprot50 \| AF-A0A765BQZ8-F1-MODEL\_V4 \| 1.0 \| 3.607e-10 \| 333 \| 0.204 \| 327 \| 212 \| 10 \| 233 \| 521 \| 1 \| 317 \| Phage portal protein \| Phage portal protein; afdb-uniprot50 \| AF-A0A662KLX8-F1-MODEL\_V4 \| 1.0 \| 1.121e-11 \| 330 \| 0.155 \| 528 \| 318 \| 27 \| 39 \| 523 \| 6 \| 448 \| Uncharacterized protein \| Uncharacterized protein; afdb-uniprot50 \| AF-A0A810UT01-F1-MODEL\_V4 \| 1.0 \| 7.231e-09 \| 329 \| 0.232 \| 232 \| 164 \| 5 \| 244 \| 465 \| 1 \| 228 \| Uncharacterized protein \| Uncharacterized protein; afdb-uniprot50 \| AF-A0A849R2Z3-F1-MODEL\_V4 \| 1.0 \| 1.707e-11 \| 324 \| 0.141 \| 495 \| 308 \| 25 \| 1 \| 469 \| 5 \| 408 \| Uncharacterized protein \| Uncharacterized protein; afdb-uniprot50 \| AF-A0A842MGS5-F1-MODEL\_V4 \| 1.0 \| 1.021e-10 \| 321 \| 0.132 \| 483 \| 308 \| 25 \| 54 \| 518 \| 23 \| 412 \| DUF1073 domain-containing protein \| DUF1073 domain-containing protein; afdb-uniprot50 \| AF-A0A1F9ALP8-F1-MODEL\_V4 \| 1.0 \| 1.475e-10 \| 319 \| 0.111 \| 512 \| 338 \| 23 \| 39 \| 531 \| 21 \| 434 \| Uncharacterized protein \| Uncharacterized protein; afdb-uniprot50 \| AF-A0A497GNK8-F1-MODEL\_V4 \| 1.0 \| 4.452e-10 \| 318 \| 0.126 \| 450 \| 283 \| 22 \| 78 \| 491 \| 31 \| 406 \| Uncharacterized protein \| Uncharacterized protein; afdb-uniprot50 \| AF-A0A0S8GAH5-F1-MODEL\_V4 \| 1.0 \| 6.509e-09 \| 316 \| 0.213 \| 267 \| 178 \| 7 \| 273 \| 530 \| 2 \| 245 \| Uncharacterized protein \| Uncharacterized protein; afdb-uniprot50 \| AF-A0A774PAJ4-F1-MODEL\_V4 \| 1.0 \| 2.806e-09 \| 313 \| 0.167 \| 310 \| 207 \| 7 \| 252 \| 523 \| 26 \| 322 \| Phage portal protein \| Phage portal protein; afdb-uniprot50 \| AF-A0A845CKT9-F1-MODEL\_V4 \| 1.0 \| 2.247e-10 \| 310 \| 0.14 \| 506 \| 300 \| 30 \| 50 \| 515 \| 3 \| 413 \| Uncharacterized protein \| Uncharacterized protein; afdb-uniprot50 \| AF-A0A2T0YIX7-F1-MODEL\_V4 \| 1.0 \| 8.617e-12 \| 308 \| 0.139 \| 546 \| 335 \| 33 \| 39 \| 506 \| 29 \| 517 \| Uncharacterized protein \| Uncharacterized protein; afdb-uniprot50 \| AF-A0A0A7G0D7-F1-MODEL\_V4 \| 1.0 \| 2.889e-11 \| 307 \| 0.136 \| 520 \| 325 \| 26 \| 37 \| 500 \| 7 \| 458 \| Portal domain protein \| Portal domain protein; afdb-uniprot50 \| AF-A0A6D0Y5S4-F1-MODEL\_V4 \| 1.0 \| 8.467e-09 \| 303 \| 0.18 \| 310 \| 198 \| 10 \| 220 \| 488 \| 2 \| 296 \| Phage portal protein \| Phage portal protein; afdb-uniprot50 \| AF-A0A497BG15-F1-MODEL\_V4 \| 1.0 \| 3.081e-10 \| 303 \| 0.151 \| 495 \| 291 \| 27 \| 40 \| 517 \| 17 \| 399 \| Uncharacterized protein \| Uncharacterized protein; afdb-uniprot50 \| AF-A0A520NQA9-F1-MODEL\_V4 \| 1.0 \| 1.057e-07 \| 300 \| 0.342 \| 143 \| 89 \| 1 \| 45 \| 187 \| 15 \| 152 \| Phage portal protein \| Phage portal protein; afdb-uniprot50 \| AF-A0A5C7PU34-F1-MODEL\_V4 \| 1.0 \| 9.406e-09 \| 296 \| 0.2 \| 284 \| 202 \| 6 \| 252 \| 529 \| 5 \| 269 \| Phage portal protein \| Phage portal protein; afdb-uniprot50 \| AF-A0A7M3IAY5-F1-MODEL\_V4 \| 1.0 \| 8.127e-08 \| 294 \| 0.24 \| 191 \| 134 \| 5 \| 195 \| 381 \| 13 \| 196 \| Phage portal protein \| Phage portal protein; afdb-uniprot50 \| AF-A0A496S006-F1-MODEL\_V4 \| 1.0 \| 4.452e-10 \| 291 \| 0.121 \| 484 \| 312 \| 24 \| 13 \| 474 \| 2 \| 394 \| Uncharacterized protein \| Uncharacterized protein; afdb-uniprot50 \| AF-A0A830FYU9-F1-MODEL\_V4 \| 1.0 \| 3.247e-10 \| 289 \| 0.128 \| 513 \| 326 \| 27 \| 15 \| 479 \| 1 \| 440 \| Uncharacterized protein \| Uncharacterized protein; afdb-uniprot50 \| AF-A0A1M4S7X1-F1-MODEL\_V4 \| 1.0 \| 4.102e-08 \| 287 \| 0.28 \| 207 \| 132 \| 5 \| 295 \| 490 \| 3 \| 203 \| Phage portal protein, lambda family \| Phage portal protein, lambda family; afdb-uniprot50 \| AF-A0A256WIT2-F1-MODEL\_V4 \| 1.0 \| 5.431e-11 \| 284 \| 0.119 \| 553 \| 348 \| 28 \| 15 \| 532 \| 1 \| 449 \| Uncharacterized protein \| Uncharacterized protein; afdb-uniprot50 \| AF-A0A1V5X4F2-F1-MODEL\_V4 \| 1.0 \| 5.004e-09 \| 277 \| 0.119 \| 486 \| 288 \| 21 \| 54 \| 497 \| 8 \| 395 \| Uncharacterized protein \| Uncharacterized protein; afdb-uniprot50 \| AF-A0A6L7X1R6-F1-MODEL\_V4 \| 1.0 \| 2.526e-09 \| 272 \| 0.145 \| 592 \| 362 \| 32 \| 24 \| 501 \| 1 \| 562 \| Uncharacterized protein \| Uncharacterized protein; afdb-uniprot50 \| AF-A0A7X7FQM4-F1-MODEL\_V4 \| 1.0 \| 1.697e-07 \| 271 \| 0.164 \| 262 \| 174 \| 6 \| 285 \| 532 \| 2 \| 232 \| Phage portal protein \| Phage portal protein; afdb-uniprot50 \| AF-A0A0F9M865-F1-MODEL\_V4 \| 1.0 \| 7.94e-10 \| 268 \| 0.107 \| 492 \| 325 \| 25 \| 50 \| 502 \| 42 \| 458 \| Uncharacterized protein \| Uncharacterized protein; afdb-uniprot50 \| AF-A0A2E9Y0J4-F1-MODEL\_V4 \| 1.0 \| 9.914e-09 \| 264 \| 0.247 \| 222 \| 137 \| 5 \| 45 \| 264 \| 17 \| 210 \| Uncharacterized protein \| Uncharacterized protein; afdb-uniprot50 \| AF-A0A2V4ZQX8-F1-MODEL\_V4 \| 1.0 \| 4.557e-08 \| 264 \| 0.194 \| 304 \| 186 \| 8 \| 271 \| 532 \| 2 \| 288 \| Lambda family phage portal protein \| Lambda family phage portal protein; afdb-uniprot50 \| AF-A0A1V5V193-F1-MODEL\_V4 \| 1.0 \| 8.467e-09 \| 263 \| 0.114 \| 499 \| 317 \| 27 \| 5 \| 478 \| 36 \| 434 \| Uncharacterized protein \| Uncharacterized protein; afdb-uniprot50 \| AF-A0A7C6TUL3-F1-MODEL\_V4 \| 1.0 \| 8.127e-08 \| 262 \| 0.141 \| 368 \| 219 \| 18 \| 94 \| 442 \| 2 \| 291 \| Uncharacterized protein \| Uncharacterized protein; afdb-uniprot50 \| AF-A0A258DZ30-F1-MODEL\_V4 \| 1.0 \| 1.789e-07 \| 262 \| 0.208 \| 292 \| 184 \| 5 \| 255 \| 503 \| 4 \| 291 \| Uncharacterized protein \| Uncharacterized protein; afdb-uniprot50 \| AF-A0A1V5ZM23-F1-MODEL\_V4 \| 1.0 \| 1.224e-08 \| 261 \| 0.132 \| 460 \| 278 \| 27 \| 51 \| 477 \| 10 \| 381 \| Uncharacterized protein \| Uncharacterized protein; afdb-uniprot50 \| AF-A0A2I1SLT0-F1-MODEL\_V4 \| 1.0 \| 1.033e-09 \| 260 \| 0.095 \| 461 \| 305 \| 21 \| 39 \| 478 \| 7 \| 376 \| Phage portal protein \| Phage portal protein; afdb-uniprot50 \| AF-I9B6E5-F1-MODEL\_V4 \| 1.0 \| 2.023e-10 \| 260 \| 0.132 \| 535 \| 305 \| 32 \| 67 \| 517 \| 13 \| 472 \| Bacteriophage portal protein, SPP1 Gp6-like protein \| Bacteriophage portal protein, SPP1 Gp6-like protein; afdb-uniprot50 \| AF-A0A7W3YMY9-F1-MODEL\_V4 \| 1.0 \| 4.747e-09 \| 259 \| 0.095 \| 448 \| 277 \| 20 \| 81 \| 488 \| 7 \| 366 \| Phage portal protein \| Phage portal protein; afdb-uniprot50 \| AF-A0A8B1ASQ7-F1-MODEL\_V4 \| 1.0 \| 1.101e-08 \| 258 \| 0.109 \| 448 \| 279 \| 21 \| 51 \| 478 \| 13 \| 360 \| Phage portal protein \| Phage portal protein; afdb-uniprot50 \| AF-A0A661CM99-F1-MODEL\_V4 \| 1.0 \| 3.463e-09 \| 256 \| 0.113 \| 510 \| 321 \| 26 \| 18 \| 479 \| 1 \| 427 \| Uncharacterized protein \| Uncharacterized protein; afdb-uniprot50 \| AF-A0A143HAD2-F1-MODEL\_V4 \| 1.0 \| 2.957e-09 \| 255 \| 0.105 \| 464 \| 285 \| 21 \| 39 \| 478 \| 10 \| 367 \| Portal protein \| Portal protein; afdb-uniprot50 \| AF-Q2FYC7-F1-MODEL\_V4 \| 1.0 \| 1.344e-09 \| 253 \| 0.1 \| 480 \| 310 \| 18 \| 23 \| 478 \| 2 \| 383 \| Phage portal protein, HK97 family \| Phage portal protein, HK97 family; afdb-uniprot50 \| AF-A0A822IYQ1-F1-MODEL\_V4 \| 1.0 \| 9.914e-09 \| 252 \| 0.087 \| 512 \| 356 \| 24 \| 28 \| 494 \| 1 \| 446 \| Uncharacterized protein \| Uncharacterized protein; afdb-uniprot50 \| AF-A0A7I8EK12-F1-MODEL\_V4 \| 1.0 \| 7.711e-08 \| 251 \| 0.128 \| 389 \| 262 \| 15 \| 14 \| 372 \| 1 \| 342 \| Uncharacterized protein \| Uncharacterized protein; afdb-uniprot50 \| AF-A0A4Q6DI35-F1-MODEL\_V4 \| 1.0 \| 3.027e-07 \| 249 \| 0.207 \| 222 \| 161 \| 5 \| 292 \| 502 \| 3 \| 220 \| Phage portal protein \| Phage portal protein; afdb-uniprot50 \| AF-A0A5T2E4A2-F1-MODEL\_V4 \| 1.0 \| 3.779e-06 \| 248 \| 0.246 \| 166 \| 119 \| 5 \| 270 \| 431 \| 2 \| 165 \| Phage portal protein \| Phage portal protein; afdb-uniprot50 \| AF-A0A2R6MRI0-F1-MODEL\_V4 \| 1.0 \| 4.803e-08 \| 248 \| 0.118 \| 480 \| 296 \| 28 \| 27 \| 467 \| 1 \| 392 \| Uncharacterized protein \| Uncharacterized protein; afdb-uniprot50 \| AF-A0A0F9NZU5-F1-MODEL\_V4 \| 1.0 \| 1.147e-09 \| 248 \| 0.131 \| 600 \| 340 \| 33 \| 67 \| 532 \| 24 \| 576 \| Uncharacterized protein \| Uncharacterized protein; afdb-uniprot50 \| AF-A0A432QN30-F1-MODEL\_V4 \| 1.0 \| 1.101e-08 \| 246 \| 0.105 \| 579 \| 380 \| 29 \| 49 \| 532 \| 10 \| 545 \| Uncharacterized protein \| Uncharacterized protein; afdb-uniprot50 \| AF-A0A7V1WC45-F1-MODEL\_V4 \| 1.0 \| 9.406e-09 \| 245 \| 0.125 \| 496 \| 316 \| 26 \| 11 \| 473 \| 1 \| 411 \| DUF935 family protein \| DUF935 family protein; afdb-uniprot50 \| AF-A0A6S7F1S0-F1-MODEL\_V4 \| 1.0 \| 1.209e-09 \| 245 \| 0.13 \| 512 \| 309 \| 24 \| 18 \| 464 \| 6 \| 446 \| Uncharacterized protein \| Uncharacterized protein; afdb-uniprot50 \| AF-A0A6B5FEG4-F1-MODEL\_V4 \| 1.0 \| 1.045e-08 \| 244 \| 0.105 \| 419 \| 270 \| 19 \| 83 \| 478 \| 4 \| 340 \| Phage portal protein \| Phage portal protein; afdb-uniprot50 \| AF-W2E7Z9-F1-MODEL\_V4 \| 1.0 \| 9.406e-09 \| 244 \| 0.12 \| 414 \| 256 \| 19 \| 85 \| 478 \| 41 \| 366 \| ATP-dependent Clp protease proteolytic subunit \| ATP-dependent Clp protease proteolytic subunit; afdb-uniprot50 \| AF-A0A7X7LEA5-F1-MODEL\_V4 \| 1.0 \| 5.494e-10 \| 243 \| 0.109 \| 586 \| 362 \| 32 \| 28 \| 532 \| 1 \| 507 \| Uncharacterized protein \| Uncharacterized protein; afdb-uniprot50 \| AF-A0A3A4QJC1-F1-MODEL\_V4 \| 1.0 \| 3.847e-09 \| 241 \| 0.121 \| 479 \| 285 \| 27 \| 13 \| 434 \| 2 \| 401 \| DUF935 family protein \| DUF935 family protein; afdb-uniprot50 \| AF-A0A843ANU4-F1-MODEL\_V4 \| 1.0 \| 1.864e-08 \| 240 \| 0.117 \| 527 \| 313 \| 32 \| 39 \| 532 \| 19 \| 426 \| DUF935 family protein \| DUF935 family protein; afdb-uniprot50 \| AF-A0A6G1WRJ0-F1-MODEL\_V4 \| 1.0 \| 8.667e-07 \| 239 \| 0.256 \| 218 \| 138 \| 5 \| 316 \| 532 \| 18 \| 212 \| Phage portal protein \| Phage portal protein; afdb-uniprot50 \| AF-A0A7Y9DWQ4-F1-MODEL\_V4 \| 1.0 \| 1.209e-09 \| 239 \| 0.158 \| 481 \| 300 \| 26 \| 59 \| 495 \| 46 \| 465 \| Uncharacterized protein \| Uncharacterized protein; afdb-uniprot50 \| AF-A0A2E0IZX8-F1-MODEL\_V4 \| 1.0 \| 8.467e-09 \| 238 \| 0.126 \| 484 \| 297 \| 20 \| 15 \| 478 \| 1 \| 378 \| Phage portal protein \| Phage portal protein; afdb-uniprot50 \| AF-A0A7V4AU95-F1-MODEL\_V4 \| 1.0 \| 5.062e-08 \| 237 \| 0.112 \| 427 \| 262 \| 29 \| 39 \| 426 \| 6 \| 354 \| DUF935 family protein \| DUF935 family protein; afdb-uniprot50 \| AF-A0A7V5S056-F1-MODEL\_V4 \| 1.0 \| 9.135e-07 \| 236 \| 0.106 \| 405 \| 263 \| 22 \| 24 \| 401 \| 1 \| 333 \| DUF935 family protein \| DUF935 family protein; afdb-uniprot50 \| AF-A0A662IQ42-F1-MODEL\_V4 \| 1.0 \| 1.864e-08 \| 236 \| 0.106 \| 509 \| 313 \| 26 \| 1 \| 474 \| 19 \| 420 \| Phage\_Mu\_F domain-containing protein \| Phage\_Mu\_F domain-containing protein; afdb-uniprot50 \| AF-A0A1Q9KYC6-F1-MODEL\_V4 \| 1.0 \| 1.32e-06 \| 235 \| 0.16 \| 292 \| 194 \| 7 \| 270 \| 523 \| 5 \| 283 \| Phage portal protein \| Phage portal protein; afdb-uniprot50 \| AF-A0A2I7SNJ0-F1-MODEL\_V4 \| 1.0 \| 1.592e-08 \| 235 \| 0.1 \| 586 \| 348 \| 31 \| 58 \| 532 \| 3 \| 520 \| Uncharacterized protein \| Uncharacterized protein; afdb-uniprot50 \| AF-A0A1W9LYD6-F1-MODEL\_V4 \| 1.0 \| 1.359e-08 \| 235 \| 0.13 \| 537 \| 321 \| 27 \| 39 \| 532 \| 8 \| 441 \| Topoisom\_I domain-containing protein \| Topoisom\_I domain-containing protein; afdb-uniprot50 \| AF-A0A2T0YNG5-F1-MODEL\_V4 \| 1.0 \| 4.803e-08 \| 234 \| 0.11 \| 508 \| 324 \| 29 \| 88 \| 532 \| 16 \| 458 \| SPP1 Gp6-like portal protein \| SPP1 Gp6-like portal protein; afdb-uniprot50 \| AF-A0A662KXA9-F1-MODEL\_V4 \| 1.0 \| 3.19e-07 \| 232 \| 0.11 \| 435 \| 292 \| 20 \| 26 \| 432 \| 2 \| 369 \| Uncharacterized protein \| Uncharacterized protein; afdb-uniprot50 \| AF-A0A4U3A7B6-F1-MODEL\_V4 \| 1.0 \| 3.324e-08 \| 231 \| 0.107 \| 427 \| 256 \| 18 \| 38 \| 439 \| 3 \| 329 \| Phage portal protein \| Phage portal protein; afdb-uniprot50 \| AF-A0A7C3TS71-F1-MODEL\_V4 \| 1.0 \| 2.585e-07 \| 229 \| 0.13 \| 414 \| 273 \| 21 \| 125 \| 523 \| 5 \| 346 \| Phage portal protein \| Phage portal protein; afdb-uniprot50 \| AF-A0A2K3ZHX0-F1-MODEL\_V4 \| 1.0 \| 1.964e-08 \| 229 \| 0.116 \| 506 \| 334 \| 23 \| 87 \| 531 \| 77 \| 530 \| Phage portal protein \| Phage portal protein; afdb-uniprot50 \| AF-A0A376TZP2-F1-MODEL\_V4 \| 1.0 \| 3.19e-07 \| 228 \| 0.181 \| 298 \| 198 \| 8 \| 273 \| 532 \| 2 \| 291 \| Capsid protein of prophage \| Capsid protein of prophage; afdb-uniprot50 \| AF-A0A0F9AB75-F1-MODEL\_V4 \| 1.0 \| 3.402e-06 \| 227 \| 0.205 \| 229 \| 166 \| 4 \| 285 \| 502 \| 2 \| 225 \| Uncharacterized protein \| Uncharacterized protein; afdb-uniprot50 \| AF-A0A2A9EYD0-F1-MODEL\_V4 \| 1.0 \| 7.231e-09 \| 227 \| 0.139 \| 501 \| 305 \| 30 \| 72 \| 532 \| 48 \| 462 \| SPP1 Gp6-like portal protein \| SPP1 Gp6-like portal protein; afdb-uniprot50 \| AF-A0A2H5WZA7-F1-MODEL\_V4 \| 1.0 \| 1.885e-07 \| 226 \| 0.102 \| 431 \| 272 \| 29 \| 39 \| 439 \| 11 \| 356 \| Uncharacterized protein \| Uncharacterized protein; afdb-uniprot50 \| AF-A0A2A3U1G4-F1-MODEL\_V4 \| 1.0 \| 3.503e-08 \| 226 \| 0.095 \| 419 \| 267 \| 19 \| 83 \| 478 \| 32 \| 361 \| Phage portal protein \| Phage portal protein; afdb-uniprot50 \| AF-A0A662UJK4-F1-MODEL\_V4 \| 1.0 \| 1.678e-08 \| 226 \| 0.155 \| 496 \| 304 \| 29 \| 66 \| 517 \| 20 \| 444 \| Uncharacterized protein \| Uncharacterized protein; afdb-uniprot50 \| AF-A0A7V5ZGN1-F1-MODEL\_V4 \| 1.0 \| 3.228e-06 \| 225 \| 0.107 \| 391 \| 243 \| 24 \| 85 \| 448 \| 2 \| 313 \| DUF935 family protein \| DUF935 family protein; afdb-uniprot50 \| AF-M2VUE3-F1-MODEL\_V4 \| 1.0 \| 6.175e-09 \| 224 \| 0.144 \| 506 \| 302 \| 30 \| 62 \| 502 \| 1 \| 440 \| Bacteriophage protein \| Bacteriophage protein; afdb-uniprot50 \| AF-A0A0F9JGC3-F1-MODEL\_V4 \| 1.0 \| 2.327e-07 \| 224 \| 0.113 \| 465 \| 317 \| 24 \| 40 \| 478 \| 19 \| 414 \| Uncharacterized protein \| Uncharacterized protein; afdb-uniprot50 \| AF-A0A167I0Q7-F1-MODEL\_V4 \| 1.0 \| 6.395e-06 \| 223 \| 0.234 \| 239 \| 147 \| 5 \| 252 \| 462 \| 4 \| 234 \| Plasmid partitioning protein ParB \| Plasmid partitioning protein ParB; afdb-uniprot50 \| AF-A0A1G6LT02-F1-MODEL\_V4 \| 1.0 \| 5.928e-08 \| 223 \| 0.1 \| 505 \| 321 \| 25 \| 20 \| 477 \| 1 \| 419 \| Uncharacterized protein \| Uncharacterized protein; afdb-uniprot50 \| AF-A0A2M8Q5N2-F1-MODEL\_V4 \| 1.0 \| 1.678e-08 \| 222 \| 0.121 \| 592 \| 354 \| 32 \| 43 \| 532 \| 6 \| 533 \| Uncharacterized protein \| Uncharacterized protein; afdb-uniprot50 \| AF-S2KXG2-F1-MODEL\_V4 \| 1.0 \| 9.135e-07 \| 221 \| 0.216 \| 236 \| 158 \| 5 \| 301 \| 532 \| 5 \| 217 \| Lambda family phage portal protein \| Lambda family phage portal protein; afdb-uniprot50 \| AF-A0A5C7P6L9-F1-MODEL\_V4 \| 1.0 \| 1.252e-06 \| 220 \| 0.168 \| 315 \| 215 \| 11 \| 244 \| 531 \| 4 \| 298 \| Phage portal protein \| Phage portal protein; afdb-uniprot50 \| AF-A0A844QMZ9-F1-MODEL\_V4 \| 1.0 \| 8.127e-08 \| 220 \| 0.106 \| 440 \| 276 \| 18 \| 23 \| 445 \| 2 \| 341 \| Phage portal protein \| Phage portal protein; afdb-uniprot50 \| AF-A0A099YCQ5-F1-MODEL\_V4 \| 1.0 \| 9.406e-09 \| 218 \| 0.119 \| 469 \| 277 \| 25 \| 39 \| 478 \| 6 \| 367 \| Portal protein \| Portal protein; afdb-uniprot50 \| AF-A0A7K0ZTW4-F1-MODEL\_V4 \| 1.0 \| 7.316e-08 \| 218 \| 0.114 \| 516 \| 342 \| 26 \| 1 \| 473 \| 95 \| 538 \| DUF935 family protein \| DUF935 family protein; afdb-uniprot50 \| AF-A0A0F9NYT3-F1-MODEL\_V4 \| 1.0 \| 1.29e-08 \| 217 \| 0.106 \| 547 \| 354 \| 24 \| 22 \| 532 \| 30 \| 477 \| Uncharacterized protein \| Uncharacterized protein; afdb-uniprot50 \| AF-A0A0F9J430-F1-MODEL\_V4 \| 1.0 \| 6.585e-08 \| 216 \| 0.103 \| 522 \| 320 \| 26 \| 15 \| 467 \| 1 \| 443 \| Uncharacterized protein \| Uncharacterized protein; afdb-uniprot50 \| AF-A0A6M3MEN7-F1-MODEL\_V4 \| 1.0 \| 4.374e-07 \| 214 \| 0.101 \| 423 \| 281 \| 23 \| 78 \| 478 \| 41 \| 386 \| Uncharacterized protein \| Uncharacterized protein; afdb-uniprot50 \| AF-A0A1V6GNZ4-F1-MODEL\_V4 \| 1.0 \| 2.872e-07 \| 212 \| 0.125 \| 501 \| 315 \| 28 \| 11 \| 476 \| 1 \| 413 \| Phage Mu protein F like protein \| Phage Mu protein F like protein; afdb-uniprot50 \| AF-A0A399WWF0-F1-MODEL\_V4 \| 1.0 \| 2.234e-06 \| 210 \| 0.095 \| 408 \| 255 \| 19 \| 39 \| 407 \| 6 \| 338 \| DUF935 family protein \| DUF935 family protein; afdb-uniprot50 \| AF-C5R881-F1-MODEL\_V4 \| 1.0 \| 9.516e-08 \| 210 \| 0.105 \| 473 \| 282 \| 23 \| 23 \| 476 \| 19 \| 369 \| Phage portal protein, HK97 family \| Phage portal protein, HK97 family; afdb-uniprot50 \| AF-A0A4P0YCK2-F1-MODEL\_V4 \| 1.0 \| 1.564e-05 \| 209 \| 0.201 \| 213 \| 155 \| 3 \| 321 \| 532 \| 1 \| 199 \| Capsid protein of prophage \| Capsid protein of prophage; afdb-uniprot50 \| AF-A0A4D9WRN2-F1-MODEL\_V4 \| 1.0 \| 6.585e-08 \| 209 \| 0.1 \| 498 \| 318 \| 23 \| 51 \| 515 \| 32 \| 432 \| DUF935 family protein \| DUF935 family protein; afdb-uniprot50 \| AF-A0A2H0A011-F1-MODEL\_V4 \| 1.0 \| 1.375e-07 \| 209 \| 0.105 \| 492 \| 329 \| 27 \| 27 \| 485 \| 3 \| 416 \| Uncharacterized protein \| Uncharacterized protein; afdb-uniprot50 \| AF-A0A0R2G1M7-F1-MODEL\_V4 \| 1.0 \| 7.316e-08 \| 208 \| 0.122 \| 458 \| 286 \| 26 \| 92 \| 499 \| 34 \| 425 \| Uncharacterized protein \| Uncharacterized protein; afdb-uniprot50 \| AF-A0A7U6KQJ3-F1-MODEL\_V4 \| 1.0 \| 1.003e-07 \| 207 \| 0.131 \| 473 \| 266 \| 26 \| 39 \| 467 \| 6 \| 377 \| Uncharacterized protein \| Uncharacterized protein; afdb-uniprot50 \| AF-A0A1F6GLB2-F1-MODEL\_V4 \| 1.0 \| 2.453e-07 \| 207 \| 0.102 \| 448 \| 276 \| 23 \| 58 \| 471 \| 33 \| 388 \| Uncharacterized protein \| Uncharacterized protein; afdb-uniprot50 \| AF-A0A1C3SC05-F1-MODEL\_V4 \| 1.0 \| 9.628e-07 \| 207 \| 0.106 \| 507 \| 342 \| 31 \| 67 \| 495 \| 9 \| 482 \| Phage protein \| Phage protein; afdb-uniprot50 \| AF-A0A7C6ADP8-F1-MODEL\_V4 \| 1.0 \| 1.61e-07 \| 207 \| 0.095 \| 554 \| 337 \| 32 \| 29 \| 532 \| 1 \| 440 \| DUF935 family protein \| DUF935 family protein; afdb-uniprot50 \| AF-A0A660E253-F1-MODEL\_V4 \| 1.0 \| 5.928e-08 \| 205 \| 0.095 \| 471 \| 293 \| 21 \| 38 \| 478 \| 3 \| 370 \| Phage portal protein [Lactobacillus brevis] \| Phage portal protein [Lactobacillus brevis]; afdb-uniprot50 \| AF-A0A0F9MCG4-F1-MODEL\_V4 \| 1.0 \| 5.624e-08 \| 205 \| 0.108 \| 600 \| 360 \| 31 \| 6 \| 530 \| 1 \| 500 \| Uncharacterized protein \| Uncharacterized protein; afdb-uniprot50 \| AF-A0A3C1G6A3-F1-MODEL\_V4 \| 1.0 \| 5.336e-08 \| 205 \| 0.114 \| 488 \| 324 \| 30 \| 17 \| 476 \| 1 \| 408 \| Uncharacterized protein \| Uncharacterized protein; afdb-uniprot50 \| AF-A0A7V9U6J0-F1-MODEL\_V4 \| 1.0 \| 1.885e-07 \| 202 \| 0.108 \| 535 \| 330 \| 29 \| 66 \| 532 \| 9 \| 464 \| Phage portal protein \| Phage portal protein; afdb-uniprot50 \| AF-A0A2G1Z9F9-F1-MODEL\_V4 \| 1.0 \| 6.741e-06 \| 200 \| 0.198 \| 262 \| 170 \| 8 \| 278 \| 531 \| 2 \| 231 \| Uncharacterized protein \| Uncharacterized protein; afdb-uniprot50 \| AF-A0A377Q9Y2-F1-MODEL\_V4 \| 1.0 \| 7.801e-07 \| 200 \| 0.125 \| 479 \| 288 \| 24 \| 24 \| 473 \| 9 \| 385 \| Mu-like prophage protein gp29 \| Mu-like prophage protein gp29; afdb-uniprot50 \| AF-A0A353GK31-F1-MODEL\_V4 \| 1.0 \| 3.544e-07 \| 200 \| 0.106 \| 534 \| 323 \| 27 \| 4 \| 477 \| 1 \| 440 \| Uncharacterized protein \| Uncharacterized protein; afdb-uniprot50 \| AF-A0A3D0N1B7-F1-MODEL\_V4 \| 1.0 \| 3.153e-08 \| 199 \| 0.107 \| 504 \| 321 \| 25 \| 9 \| 478 \| 32 \| 440 \| Uncharacterized protein \| Uncharacterized protein; afdb-uniprot50 \| AF-A0A1U7GKP2-F1-MODEL\_V4 \| 1.0 \| 4.61e-07 \| 198 \| 0.106 \| 547 \| 364 \| 29 \| 51 \| 532 \| 18 \| 504 \| Uncharacterized protein \| Uncharacterized protein; afdb-uniprot50 \| AF-A0A3N0ZNV0-F1-MODEL\_V4 \| 1.0 \| 2.327e-07 \| 197 \| 0.096 \| 593 \| 364 \| 30 \| 15 \| 531 \| 1 \| 497 \| DUF935 family protein \| DUF935 family protein; afdb-uniprot50 \| AF-A0A6M3L929-F1-MODEL\_V4 \| 1.0 \| 3.027e-07 \| 196 \| 0.104 \| 441 \| 289 \| 25 \| 92 \| 491 \| 74 \| 449 \| Putative portal protein \| Putative portal protein; afdb-uniprot50 \| AF-A0A4R7QR18-F1-MODEL\_V4 \| 1.0 \| 3.062e-06 \| 194 \| 0.101 \| 435 \| 295 \| 22 \| 69 \| 473 \| 34 \| 402 \| Uncharacterized protein DUF935 \| Uncharacterized protein DUF935; afdb-uniprot50 \| AF-A0A496UFM6-F1-MODEL\_V4 \| 1.0 \| 1.908e-06 \| 194 \| 0.075 \| 492 \| 295 \| 25 \| 77 \| 488 \| 8 \| 419 \| Uncharacterized protein \| Uncharacterized protein; afdb-uniprot50 \| AF-A0A3N0ZBD8-F1-MODEL\_V4 \| 1.0 \| 3.19e-07 \| 194 \| 0.091 \| 549 \| 327 \| 27 \| 51 \| 523 \| 2 \| 454 \| DUF935 family protein \| DUF935 family protein; afdb-uniprot50 \| AF-A0A5Y3MP01-F1-MODEL\_V4 \| 1.0 \| 1.987e-07 \| 191 \| 0.102 \| 584 \| 335 \| 30 \| 24 \| 521 \| 8 \| 488 \| DUF935 family protein \| DUF935 family protein; afdb-uniprot50 \| AF-A0A0F9MWD7-F1-MODEL\_V4 \| 1.0 \| 7.022e-07 \| 191 \| 0.117 \| 586 \| 373 \| 33 \| 45 \| 532 \| 10 \| 549 \| Uncharacterized protein \| Uncharacterized protein; afdb-uniprot50 \| AF-T1HRW6-F1-MODEL\_V4 \| 1.0 \| 4.374e-07 \| 190 \| 0.112 \| 567 \| 342 \| 27 \| 24 \| 532 \| 8 \| 471 \| Phage\_Mu\_F domain-containing protein \| Phage\_Mu\_F domain-containing protein; afdb-uniprot50 \| AF-A0A640WJG5-F1-MODEL\_V4 \| 1.0 \| 4.859e-07 \| 189 \| 0.096 \| 572 \| 372 \| 29 \| 67 \| 531 \| 21 \| 554 \| Phage tail protein \| Phage tail protein; afdb-uniprot50 \| AF-A0A843GSM3-F1-MODEL\_V4 \| 1.0 \| 1.127e-06 \| 188 \| 0.106 \| 546 \| 347 \| 32 \| 62 \| 532 \| 35 \| 514 \| Phage portal protein \| Phage portal protein; afdb-uniprot50 \| AF-A0A7U5HK87-F1-MODEL\_V4 \| 1.0 \| 1.885e-07 \| 187 \| 0.127 \| 432 \| 270 \| 26 \| 108 \| 498 \| 2 \| 367 \| Phage portal protein \| Phage portal protein; afdb-uniprot50 \| AF-A0A0Q4JBM3-F1-MODEL\_V4 \| 1.0 \| 1.188e-06 \| 185 \| 0.097 \| 481 \| 298 \| 22 \| 39 \| 478 \| 30 \| 415 \| Uncharacterized protein \| Uncharacterized protein; afdb-uniprot50 \| AF-A0A432QRG5-F1-MODEL\_V4 \| 1.0 \| 3.363e-07 \| 185 \| 0.108 \| 500 \| 311 \| 31 \| 87 \| 529 \| 68 \| 489 \| DUF4055 domain-containing protein \| DUF4055 domain-containing protein; afdb-uniprot50 \| AF-B8F5B1-F1-MODEL\_V4 \| 1.0 \| 7.022e-07 \| 183 \| 0.123 \| 544 \| 310 \| 33 \| 39 \| 518 \| 4 \| 444 \| Putative phage-related protein \| Putative phage-related protein; afdb-uniprot50 \| AF-D8A1C1-F1-MODEL\_V4 \| 1.0 \| 6.741e-06 \| 182 \| 0.103 \| 475 \| 293 \| 22 \| 63 \| 500 \| 1 \| 379 \| Uncharacterized protein \| Uncharacterized protein; afdb-uniprot50 \| AF-A0A7X7ERQ9-F1-MODEL\_V4 \| 1.0 \| 2.615e-06 \| 181 \| 0.09 \| 473 \| 316 \| 25 \| 28 \| 473 \| 1 \| 386 \| DUF935 family protein \| DUF935 family protein; afdb-uniprot50 \| AF-A0A090G3P5-F1-MODEL\_V4 \| 1.0 \| 1.336e-05 \| 180 \| 0.183 \| 261 \| 182 \| 8 \| 274 \| 532 \| 24 \| 255 \| Capsid protein of prophage \| Capsid protein of prophage; afdb-uniprot50 \| AF-Q30VX9-F1-MODEL\_V4 \| 1.0 \| 3.402e-06 \| 179 \| 0.104 \| 469 \| 308 \| 20 \| 38 \| 479 \| 19 \| 402 \| Uncharacterized protein \| Uncharacterized protein; afdb-uniprot50 \| AF-E9CQ96-F1-MODEL\_V4 \| 1.0 \| 6.82e-05 \| 178 \| 0.189 \| 201 \| 134 \| 6 \| 254 \| 429 \| 27 \| 223 \| Putative phage portal protein \| Putative phage portal protein; afdb-uniprot50 \| AF-A0A7X9E3K7-F1-MODEL\_V4 \| 1.0 \| 4.477e-05 \| 177 \| 0.083 \| 431 \| 287 \| 23 \| 67 \| 475 \| 2 \| 346 \| DUF935 family protein \| DUF935 family protein; afdb-uniprot50 \| AF-A0A1Y1QSY6-F1-MODEL\_V4 \| 1.0 \| 7.801e-07 \| 177 \| 0.117 \| 560 \| 305 \| 29 \| 51 \| 523 \| 16 \| 473 \| Uncharacterized protein \| Uncharacterized protein; afdb-uniprot50 \| AF-A0A1A9WW55-F1-MODEL\_V4 \| 1.0 \| 1.629e-06 \| 176 \| 0.101 \| 560 \| 339 \| 25 \| 24 \| 517 \| 8 \| 469 \| Phage\_Mu\_F domain-containing protein \| Phage\_Mu\_F domain-containing protein; afdb-uniprot50 \| AF-A0A450Z5H8-F1-MODEL\_V4 \| 1.0 \| 7.105e-06 \| 174 \| 0.103 \| 453 \| 267 \| 24 \| 45 \| 461 \| 8 \| 357 \| Uncharacterized protein \| Uncharacterized protein; afdb-uniprot50 \| AF-A0A3M0XMA1-F1-MODEL\_V4 \| 1.0 \| 1.07e-06 \| 173 \| 0.088 \| 461 \| 290 \| 23 \| 51 \| 477 \| 13 \| 377 \| DUF935 family protein \| DUF935 family protein; afdb-uniprot50 \| AF-U4T7Q4-F1-MODEL\_V4 \| 1.0 \| 1.908e-06 \| 173 \| 0.117 \| 469 \| 292 \| 22 \| 39 \| 474 \| 4 \| 383 \| Mu-like prophage FluMu protein gp29 \| Mu-like prophage FluMu protein gp29; afdb-uniprot50 \| AF-A0A149TMR3-F1-MODEL\_V4 \| 1.0 \| 4.425e-06 \| 173 \| 0.092 \| 455 \| 296 \| 21 \| 58 \| 474 \| 39 \| 414 \| Uncharacterized protein \| Uncharacterized protein; afdb-uniprot50 \| AF-A0A7W1RW72-F1-MODEL\_V4 \| 1.0 \| 2.872e-07 \| 172 \| 0.121 \| 592 \| 341 \| 33 \| 51 \| 532 \| 2 \| 524 \| DUF4055 domain-containing protein \| DUF4055 domain-containing protein; afdb-uniprot50 \| AF-A0A2A2A6L3-F1-MODEL\_V4 \| 1.0 \| 1.32e-06 \| 172 \| 0.124 \| 483 \| 289 \| 28 \| 40 \| 486 \| 21 \| 405 \| Uncharacterized protein \| Uncharacterized protein; afdb-uniprot50 \| AF-A0A840S794-F1-MODEL\_V4 \| 1.0 \| 6.395e-06 \| 171 \| 0.095 \| 472 \| 311 \| 22 \| 24 \| 472 \| 10 \| 388 \| Phage gp29-like protein \| Phage gp29-like protein; afdb-uniprot50 \| AF-A0A2V3UB21-F1-MODEL\_V4 \| 1.0 \| 1.07e-06 \| 170 \| 0.113 \| 511 \| 305 \| 32 \| 63 \| 487 \| 8 \| 456 \| Uncharacterized protein DUF4055 \| Uncharacterized protein DUF4055; afdb-uniprot50 \| AF-A0A8B3NDI1-F1-MODEL\_V4 \| 1.0 \| 1.188e-06 \| 170 \| 0.103 \| 524 \| 298 \| 31 \| 63 \| 488 \| 10 \| 459 \| DUF4055 domain-containing protein \| DUF4055 domain-containing protein; afdb-uniprot50 \| AF-A0A6G8AZX7-F1-MODEL\_V4 \| 1.0 \| 7.401e-07 \| 170 \| 0.11 \| 433 \| 265 \| 21 \| 39 \| 454 \| 4 \| 333 \| Phage portal protein \| Phage portal protein; afdb-uniprot50 \| AF-A0A7Y5LKE7-F1-MODEL\_V4 \| 1.0 \| 3.586e-06 \| 167 \| 0.11 \| 425 \| 267 \| 22 \| 39 \| 439 \| 3 \| 340 \| DUF935 family protein \| DUF935 family protein; afdb-uniprot50 \| AF-A0A849PHX0-F1-MODEL\_V4 \| 1.0 \| 8.416e-05 \| 167 \| 0.105 \| 407 \| 267 \| 21 \| 85 \| 467 \| 2 \| 335 \| DUF935 family protein \| DUF935 family protein; afdb-uniprot50 \| AF-A0A254RDH1-F1-MODEL\_V4 \| 1.0 \| 5.462e-06 \| 167 \| 0.1 \| 570 \| 353 \| 25 \| 20 \| 523 \| 1 \| 476 \| Uncharacterized protein \| Uncharacterized protein; afdb-uniprot50 \| AF-A0A7V5SQD7-F1-MODEL\_V4 \| 1.0 \| 2.51e-05 \| 166 \| 0.074 \| 415 \| 269 \| 19 \| 40 \| 420 \| 8 \| 341 \| DUF935 family protein \| DUF935 family protein; afdb-uniprot50 \| AF-A0A353QZH6-F1-MODEL\_V4 \| 1.0 \| 1.127e-06 \| 164 \| 0.094 \| 579 \| 333 \| 29 \| 90 \| 524 \| 117 \| 647 \| Uncharacterized protein \| Uncharacterized protein; afdb-uniprot50 \| AF-A0A4Y9PSX4-F1-MODEL\_V4 \| 1.0 \| 0.0001039 \| 163 \| 0.1 \| 498 \| 329 \| 27 \| 2 \| 467 \| 5 \| 415 \| Uncharacterized protein \| Uncharacterized protein; afdb-uniprot50 \| AF-A0A838BWC4-F1-MODEL\_V4 \| 1.0 \| 3.228e-06 \| 163 \| 0.097 \| 625 \| 374 \| 35 \| 18 \| 516 \| 1 \| 561 \| Head-tail connector protein \| Head-tail connector protein; afdb-uniprot50 \| AF-A0A7J4ZRG6-F1-MODEL\_V4 \| 1.0 \| 7.801e-07 \| 163 \| 0.129 \| 570 \| 337 \| 30 \| 63 \| 523 \| 3 \| 522 \| Phage tail protein \| Phage tail protein; afdb-uniprot50 \| AF-A0A6M3IKJ8-F1-MODEL\_V4 \| 1.0 \| 4.664e-06 \| 163 \| 0.094 \| 655 \| 358 \| 29 \| 66 \| 532 \| 18 \| 625 \| Uncharacterized protein \| Uncharacterized protein; afdb-uniprot50 \| AF-A0A357LVL4-F1-MODEL\_V4 \| 1.0 \| 2.481e-06 \| 163 \| 0.124 \| 596 \| 336 \| 32 \| 24 \| 532 \| 8 \| 504 \| Phage\_Mu\_F domain-containing protein \| Phage\_Mu\_F domain-containing protein; afdb-uniprot50 \| AF-A0A329VRT9-F1-MODEL\_V4 \| 1.0 \| 1.202e-05 \| 162 \| 0.094 \| 443 \| 267 \| 23 \| 51 \| 464 \| 7 \| 344 \| Uncharacterized protein \| Uncharacterized protein; afdb-uniprot50 \| AF-A0A2K2ZXI3-F1-MODEL\_V4 \| 1.0 \| 1.831e-05 \| 162 \| 0.095 \| 430 \| 284 \| 26 \| 90 \| 473 \| 86 \| 456 \| DUF4055 domain-containing protein \| DUF4055 domain-containing protein; afdb-uniprot50 \| AF-A0A0F9HG25-F1-MODEL\_V4 \| 1.0 \| 8.319e-06 \| 161 \| 0.123 \| 518 \| 284 \| 27 \| 58 \| 532 \| 14 \| 404 \| Uncharacterized protein \| Uncharacterized protein; afdb-uniprot50 \| AF-A0A840RLG2-F1-MODEL\_V4 \| 1.0 \| 2.905e-06 \| 160 \| 0.101 \| 559 \| 325 \| 28 \| 39 \| 523 \| 24 \| 478 \| Phage gp29-like protein \| Phage gp29-like protein; afdb-uniprot50 \| AF-A0A1W1XTP3-F1-MODEL\_V4 \| 1.0 \| 7.488e-06 \| 160 \| 0.091 \| 699 \| 401 \| 35 \| 7 \| 532 \| 3 \| 640 \| Uncharacterized protein \| Uncharacterized protein; afdb-uniprot50 \| AF-A0A3A8TWS2-F1-MODEL\_V4 \| 1.0 \| 1.717e-06 \| 158 \| 0.087 \| 493 \| 318 \| 31 \| 67 \| 488 \| 30 \| 461 \| DUF4055 domain-containing protein \| DUF4055 domain-containing protein; afdb-uniprot50 \| AF-A0A6L5BK53-F1-MODEL\_V4 \| 1.0 \| 0.0001039 \| 157 \| 0.106 \| 377 \| 238 \| 17 \| 49 \| 402 \| 33 \| 333 \| Uncharacterized protein \| Uncharacterized protein; afdb-uniprot50 \| AF-A0A450ZFK4-F1-MODEL\_V4 \| 1.0 \| 6.741e-06 \| 156 \| 0.119 \| 568 \| 304 \| 30 \| 27 \| 518 \| 2 \| 449 \| Mu-like prophage protein gp29 \| Mu-like prophage protein gp29; afdb-uniprot50 \| AF-A0A352Z9P4-F1-MODEL\_V4 \| 1.0 \| 1.141e-05 \| 155 \| 0.096 \| 547 \| 316 \| 30 \| 51 \| 523 \| 32 \| 474 \| DUF935 domain-containing protein \| DUF935 domain-containing protein; afdb-uniprot50 \| AF-A0A3M8QA88-F1-MODEL\_V4 \| 1.0 \| 6.741e-06 \| 155 \| 0.08 \| 635 \| 380 \| 33 \| 66 \| 532 \| 31 \| 629 \| Uncharacterized protein \| Uncharacterized protein; afdb-uniprot50 \| AF-A0A3N6V6C1-F1-MODEL\_V4 \| 1.0 \| 5.462e-06 \| 155 \| 0.118 \| 624 \| 344 \| 33 \| 68 \| 525 \| 39 \| 622 \| Uncharacterized protein \| Uncharacterized protein; afdb-uniprot50 \| AF-A0A3B1CWW8-F1-MODEL\_V4 \| 1.0 \| 0.0001095 \| 153 \| 0.088 \| 465 \| 282 \| 27 \| 58 \| 473 \| 19 \| 390 \| Uncharacterized protein \| Uncharacterized protein; afdb-uniprot50 \| AF-A0A3D9PWD8-F1-MODEL\_V4 \| 1.0 \| 2.94e-05 \| 152 \| 0.09 \| 583 \| 345 \| 29 \| 23 \| 523 \| 3 \| 481 \| Phage gp29-like protein \| Phage gp29-like protein; afdb-uniprot50 \| AF-A0A6L7L088-F1-MODEL\_V4 \| 1.0 \| 7.985e-05 \| 152 \| 0.1 \| 458 \| 288 \| 20 \| 52 \| 474 \| 35 \| 403 \| DUF935 family protein \| DUF935 family protein; afdb-uniprot50 \| AF-A0A0F9DSH9-F1-MODEL\_V4 \| 1.0 \| 8.416e-05 \| 151 \| 0.108 \| 398 \| 247 \| 18 \| 125 \| 478 \| 8 \| 341 \| Uncharacterized protein \| Uncharacterized protein; afdb-uniprot50 \| AF-A0A515ERN2-F1-MODEL\_V4 \| 1.0 \| 7.985e-05 \| 150 \| 0.105 \| 456 \| 278 \| 26 \| 56 \| 478 \| 15 \| 373 \| DUF935 family protein \| DUF935 family protein; afdb-uniprot50 \| AF-A0A7X7NIS7-F1-MODEL\_V4 \| 1.0 \| 1.027e-05 \| 148 \| 0.101 \| 564 \| 316 \| 28 \| 48 \| 523 \| 20 \| 480 \| DUF935 family protein \| DUF935 family protein; afdb-uniprot50 \| AF-A0A2A2ARI2-F1-MODEL\_V4 \| 1.0 \| 0.0002286 \| 147 \| 0.091 \| 470 \| 314 \| 21 \| 36 \| 475 \| 23 \| 409 \| Uncharacterized protein \| Uncharacterized protein; afdb-uniprot50 \| AF-A0A1V5KBQ0-F1-MODEL\_V4 \| 1.0 \| 6.395e-06 \| 146 \| 0.117 \| 586 \| 308 \| 35 \| 92 \| 532 \| 80 \| 601 \| Uncharacterized protein \| Uncharacterized protein; afdb-uniprot50 \| AF-A0A6M8VBB9-F1-MODEL\_V4 \| 1.0 \| 0.0001501 \| 145 \| 0.103 \| 387 \| 245 \| 19 \| 38 \| 398 \| 2 \| 312 \| DUF935 family protein \| DUF935 family protein; afdb-uniprot50 \| AF-A0A1V5V157-F1-MODEL\_V4 \| 1.0 \| 2.789e-05 \| 145 \| 0.094 \| 614 \| 354 \| 30 \| 24 \| 532 \| 1 \| 517 \| Phage Mu protein F like protein \| Phage Mu protein F like protein; afdb-uniprot50 \| AF-X1C5I8-F1-MODEL\_V4 \| 1.0 \| 0.0002822 \| 144 \| 0.135 \| 309 \| 213 \| 14 \| 171 \| 464 \| 2 \| 271 \| Uncharacterized protein \| Uncharacterized protein; afdb-uniprot50 \| AF-A0A2E7E6N9-F1-MODEL\_V4 \| 1.0 \| 9.35e-05 \| 143 \| 0.11 \| 453 \| 274 \| 25 \| 58 \| 480 \| 37 \| 390 \| Uncharacterized protein \| Uncharacterized protein; afdb-uniprot50 \| AF-T2GD94-F1-MODEL\_V4 \| 1.0 \| 9.242e-06 \| 142 \| 0.099 \| 575 \| 331 \| 28 \| 54 \| 523 \| 30 \| 522 \| Uncharacterized protein \| Uncharacterized protein; afdb-uniprot50 \| AF-A0A7C5UJ90-F1-MODEL\_V4 \| 1.0 \| 0.0002169 \| 141 \| 0.103 \| 318 \| 199 \| 15 \| 152 \| 446 \| 10 \| 264 \| DUF935 family protein \| DUF935 family protein; afdb-uniprot50 \| AF-A0A6M3KQJ0-F1-MODEL\_V4 \| 1.0 \| 3.098e-05 \| 141 \| 0.1 \| 546 \| 342 \| 30 \| 38 \| 523 \| 18 \| 474 \| Phage\_Mu\_F domain-containing protein \| Phage\_Mu\_F domain-containing protein; afdb-uniprot50 \| AF-A0A2S0MD11-F1-MODEL\_V4 \| 1.0 \| 4.248e-05 \| 138 \| 0.104 \| 576 \| 326 \| 28 \| 40 \| 523 \| 17 \| 494 \| DUF935 domain-containing protein \| DUF935 domain-containing protein; afdb-uniprot50 \| AF-A0A2D6X058-F1-MODEL\_V4 \| 1.0 \| 4.248e-05 \| 138 \| 0.101 \| 649 \| 364 \| 40 \| 66 \| 532 \| 20 \| 631 \| Uncharacterized protein \| Uncharacterized protein; afdb-uniprot50 \| AF-A0A0F9K8Y7-F1-MODEL\_V4 \| 1.0 \| 4.477e-05 \| 137 \| 0.11 \| 609 \| 308 \| 34 \| 66 \| 468 \| 18 \| 598 \| Uncharacterized protein \| Uncharacterized protein; afdb-uniprot50 \| AF-A0A2E7DC09-F1-MODEL\_V4 \| 1.0 \| 2.144e-05 \| 136 \| 0.09 \| 629 \| 353 \| 35 \| 53 \| 532 \| 49 \| 607 \| Uncharacterized protein \| Uncharacterized protein; afdb-uniprot50 \| AF-A0A2H0FN30-F1-MODEL\_V4 \| 1.0 \| 0.0001216 \| 135 \| 0.075 \| 464 \| 283 \| 24 \| 64 \| 468 \| 24 \| 400 \| Uncharacterized protein \| Uncharacterized protein; afdb-uniprot50 \| AF-A0A0W8G0T3-F1-MODEL\_V4 \| 1.0 \| 0.000559 \| 134 \| 0.081 \| 393 \| 264 \| 20 \| 39 \| 407 \| 6 \| 325 \| Uncharacterized protein \| Uncharacterized protein; afdb-uniprot50 \| AF-A0A847DLT8-F1-MODEL\_V4 \| 1.0 \| 5.824e-05 \| 134 \| 0.108 \| 509 \| 320 \| 35 \| 15 \| 473 \| 1 \| 425 \| DUF935 family protein \| DUF935 family protein; afdb-uniprot50 \| AF-A0A2R4BP72-F1-MODEL\_V4 \| 1.0 \| 0.0001039 \| 134 \| 0.081 \| 654 \| 358 \| 30 \| 58 \| 531 \| 7 \| 597 \| Phage protein \| Phage protein; afdb-uniprot50 \| AF-A0A357XXK9-F1-MODEL\_V4 \| 1.0 \| 0.002313 \| 131 \| 0.211 \| 142 \| 108 \| 2 \| 288 \| 427 \| 2 \| 141 \| Uncharacterized protein \| Uncharacterized protein; afdb-uniprot50 \| AF-A0A0F8YUF6-F1-MODEL\_V4 \| 1.0 \| 0.0007665 \| 130 \| 0.12 \| 341 \| 210 \| 14 \| 207 \| 519 \| 1 \| 279 \| Uncharacterized protein \| Uncharacterized protein; afdb-uniprot50 \| AF-A0A2H0Z0M0-F1-MODEL\_V4 \| 1.0 \| 0.0008975 \| 130 \| 0.084 \| 377 \| 228 \| 17 \| 62 \| 407 \| 25 \| 315 \| Uncharacterized protein \| Uncharacterized protein; afdb-uniprot50 \| AF-A0A2E0VI79-F1-MODEL\_V4 \| 1.0 \| 9.855e-05 \| 127 \| 0.09 \| 694 \| 365 \| 42 \| 66 \| 532 \| 22 \| 675 \| Uncharacterized protein \| Uncharacterized protein; afdb-uniprot50 \| AF-A0A521YNN5-F1-MODEL\_V4 \| 1.0 \| 0.0003304 \| 127 \| 0.082 \| 667 \| 375 \| 32 \| 40 \| 532 \| 6 \| 609 \| Uncharacterized protein \| Uncharacterized protein; afdb-uniprot50 \| AF-A0A4Q6ACK7-F1-MODEL\_V4 \| 1.0 \| 0.0001582 \| 126 \| 0.103 \| 414 \| 257 \| 25 \| 62 \| 397 \| 9 \| 386 \| DUF4055 domain-containing protein \| DUF4055 domain-containing protein; afdb-uniprot50 \| AF-A0A0F9CUA3-F1-MODEL\_V4 \| 1.0 \| 0.001367 \| 120 \| 0.148 \| 304 \| 218 \| 12 \| 252 \| 532 \| 17 \| 302 \| Uncharacterized protein \| Uncharacterized protein; afdb-uniprot50 \| AF-A0A3N2M4B1-F1-MODEL\_V4 \| 1.0 \| 3.628e-05 \| 120 \| 0.105 \| 453 \| 264 \| 23 \| 70 \| 407 \| 3 \| 429 \| Uncharacterized protein \| Uncharacterized protein; afdb-uniprot50 \| AF-A0A1Y1QPD1-F1-MODEL\_V4 \| 1.0 \| 0.0004298 \| 117 \| 0.097 \| 731 \| 395 \| 40 \| 21 \| 532 \| 1 \| 685 \| Uncharacterized protein \| Uncharacterized protein; afdb-uniprot50 \| AF-A0A7J0AIQ2-F1-MODEL\_V4 \| 1.0 \| 0.0001501 \| 116 \| 0.114 \| 456 \| 256 \| 23 \| 70 \| 408 \| 2 \| 426 \| Uncharacterized protein \| Uncharacterized protein; afdb-uniprot50 \| AF-A0A378VE46-F1-MODEL\_V4 \| 1.0 \| 0.0007665 \| 113 \| 0.101 \| 533 \| 340 \| 25 \| 43 \| 523 \| 21 \| 466 \| Phage protein \| Phage protein; afdb-uniprot50 \| AF-A0A2D7XHA7-F1-MODEL\_V4 \| 1.0 \| 0.0006546 \| 113 \| 0.077 \| 630 \| 324 \| 32 \| 13 \| 449 \| 2 \| 567 \| Phage portal protein \| Phage portal protein; afdb-uniprot50 \| AF-A0A4Q7AJZ6-F1-MODEL\_V4 \| 1.0 \| 0.0006546 \| 112 \| 0.101 \| 599 \| 353 \| 33 \| 66 \| 515 \| 35 \| 596 \| Portal protein p19 \| Portal protein p19; afdb-uniprot50 \| AF-A0A7Y5LIV9-F1-MODEL\_V4 \| 1.0 \| 0.002082 \| 110 \| 0.074 \| 441 \| 250 \| 25 \| 45 \| 427 \| 19 \| 359 \| DUF935 family protein \| DUF935 family protein; afdb-uniprot50 \| AF-A0A0F9BIB7-F1-MODEL\_V4 \| 1.0 \| 0.001519 \| 110 \| 0.106 \| 468 \| 241 \| 23 \| 72 \| 409 \| 7 \| 427 \| Uncharacterized protein \| Uncharacterized protein; afdb-uniprot50 \| AF-A0A4S2E2Z1-F1-MODEL\_V4 \| 1.0 \| 0.001874 \| 101 \| 0.12 \| 475 \| 255 \| 23 \| 109 \| 521 \| 19 \| 392 \| DUF935 family protein \| DUF935 family protein; afdb-uniprot50 \| AF-R7J256-F1-MODEL\_V4 \| 0.994 \| 0.000997 \| 87 \| 0.1 \| 456 \| 266 \| 26 \| 71 \| 404 \| 3 \| 436 \| Uncharacterized protein \| Uncharacterized protein |
| Top keywords  (threshold 1.00e-02 (evalue)) | **portal, Phage, lambda, DUF935, Putative, Capsid, prophage, domain\_containing, Bacteriophage, Phage\_related** |
| Output files | ../../similar\_structures/04\_FANPEZAQ\_CDS\_0004\_afdb-proteome\_foldseek.tsv ../../similar\_structures/04\_FANPEZAQ\_CDS\_0004\_afdb-uniprot50\_foldseek.tsv ../../similar\_structures/04\_FANPEZAQ\_CDS\_0004\_merged.svg ../../similar\_structures/04\_FANPEZAQ\_CDS\_0004\_pdb\_foldseek.tsv |

  
  
  

Return to summary | Go to previous | Go to next

  


---

**Sequence/structure alignments coloring**  
Each object in the alignment figures is colored according to its E-value following this color coding:

1e-100
10
