## Supplementary material for "Completing the BASEL phage collection to unlock hidden diversity for systematic exploration of phage-host interactions": Data S2: 5.html

FANPEZAQ\_CDS\_0005


Return to summary | Go to previous | Go to next

|  |  |
| --- | --- |
| FANPEZAQ\_CDS\_0005  Page creation date: 02 Sep 2024, 12:00  Project folder: n/a  Input sequences file: Escherichia\_virus\_HeidiAbel.gb | capsid phage major hk97 protease prohead head peptidase fragment putative u35 caudovirus bacteriophage and packaging domain\_containing virus gp36 gpt phage\_related capsid\_like phi\_c31 maturation tigr01554 mu predicted prophage structural mu\_like\_gpt particle mu\_like maritimacin serine gp5 virus\_like 10a encapsulin u37 hypothetical |

### Sequence information

|  |  |
| --- | --- |
| Name | FANPEZAQ\_CDS\_0005  05\_FANPEZAQ\_CDS\_0005 (pipeline id) |
| Imported annotations | Escherichia\_virus\_HeidiAbel Bas97 |
| Protein sequence | MQKNPLNSIIGAEFTRKSDSKTLPDFNKGGLVRTMEVRKFDTEKRTVELAFSSEIEVERW YGIEILDHSPESVRLDRLRDGGALLVDHDWTDQVGVVESVSIDADRRGRAVVRFGRSARA DEIFQDIVDGIRKHVSVGYRVLAAKLQETRDEWQDVYRITEWEPYEISIVSVPADASVGV GRAFDKPQEEQRGSKSDTAGTVEVPAPTNIASEGIRMEKILRDAAGNLVRAMVDEKGNIT QVLEMLERAGDDARNAEQRGREAEKARVRSITEMGEKYNARDLALKAIGEGTTAEDFQRS LLDHLNKPKDPAQPADGTRSGSKPLSEMPSSVIGMSDKDIRRYSIFNVVRALANPNDAAA QRAAAFEIECSHAAQQQYGRSAKGILIPDDVLRAFNAGGAPNTPAGAQTGSNLVGTDFMA SSFIDLLRPRTTIMRLARTMGGLVGNVEIPKQTGGATAYWIGEGDDATETTPTIGQLGLS PKTVAAYTDITRRLLMQSTPDAEAIVRGDLVAAIAQAIDYAGYYGTGTSNQPLGLANYTG INAVDFAADNPTYAELVQMETEIAADNADVNNMAYVLNARGRGAAKTTPKFASGSSVADA GVIWEPGNTLNGYRAEVTNQVQDGDVFFGNFADLVVGMWGGLDMTVDPYSLSKSGGLRIV VFQDVDFAVRRVESFALGRNAE |
| Number of residues | 682 |
| Molecular weight (Da) | 73931.80 |
| Output files | ../../query\_sequences/05\_FANPEZAQ\_CDS\_0005.fasta |

### Putative domain architecture and protein family

#### Search results (HHblits)^1^

|  |  |
| --- | --- |
| Domain family databases searched | Pfam, Ncbi-cd, Cath, Phrogs |
| Results, scheme(s)  (Top layers only; threshold 1.00e-03 (evalue)) | xml version="1.0" encoding="utf-8" standalone="no"?       2024-09-02T21:08:12.828277 image/svg+xml   Matplotlib v3.7.2, https://matplotlib.org/ |
| Results, table  (E-value ≤ 1.00e-03 (evalue)) | db \| id \| prob \| evalue \| pvalue \| score \| cols \| query \| query\_len \| template \| template\_len \| name \| description; pfam \| PF04586 \| 99.3 \| 4e-17 \| 8e-21 \| 150.7 \| 143 \| (34, 180) \| 682 \| (2, 153) \| 164 \| Peptidase\_S78 \| Caudovirus prohead serine protease; pfam \| PF05065 \| 99.1 \| 3.4e-15 \| 7.2e-19 \| 145.8 \| 255 \| (416, 680) \| 682 \| (6, 275) \| 275 \| Phage\_capsid \| Phage capsid family; pfam \| PF19821 \| 98.9 \| 1.5e-13 \| 2.9e-17 \| 140.8 \| 266 \| (416, 682) \| 682 \| (8, 342) \| 342 \| Phage\_capsid\_2 \| Phage capsid protein; pfam \| PF14550 \| 98.9 \| 1.9e-13 \| 3.7e-17 \| 119.4 \| 99 \| (36, 143) \| 682 \| (4, 121) \| 123 \| Peptidase\_S78\_2 \| Putative phage serine protease XkdF; pfam \| PF06673 \| 98.6 \| 7.3e-12 \| 1.4e-15 \| 128.4 \| 212 \| (460, 681) \| 682 \| (129, 345) \| 347 \| L\_lactis\_ph-MCP \| Lactococcus lactis bacteriophage major capsid protein; pfam \| PF05125 \| 98.4 \| 5e-11 \| 1e-14 \| 120.0 \| 261 \| (416, 680) \| 682 \| (25, 324) \| 325 \| Phage\_cap\_P2 \| Phage major capsid protein, P2 family; pfam \| PF13252 \| 97.4 \| 4.4e-08 \| 9e-12 \| 96.3 \| 262 \| (412, 681) \| 682 \| (6, 310) \| 313 \| DUF4043 \| Protein of unknown function (DUF4043); pfam \| PF20036 \| 97.3 \| 1.1e-07 \| 2.2e-11 \| 93.8 \| 218 \| (416, 641) \| 682 \| (7, 243) \| 307 \| Gp13-like \| Major capsid protein 13-like; pfam \| PF11651 \| 97.2 \| 1.4e-07 \| 2.8e-11 \| 97.3 \| 196 \| (416, 623) \| 682 \| (10, 218) \| 421 \| P22\_CoatProtein \| P22 coat protein - gene protein 5; pfam \| PF09950 \| 97.0 \| 5.7e-07 \| 1.2e-10 \| 84.9 \| 207 \| (463, 680) \| 682 \| (10, 244) \| 245 \| DUF2184 \| Uncharacterized protein conserved in bacteria (DUF2184); pfam \| PF04454 \| 96.5 \| 4.6e-06 \| 9.4e-10 \| 78.2 \| 181 \| (469, 665) \| 682 \| (62, 250) \| 251 \| Linocin\_M18 \| Encapsulating protein for peroxidase; pfam \| PF19307 \| 96.3 \| 9.2e-06 \| 1.8e-09 \| 77.1 \| 190 \| (477, 680) \| 682 \| (40, 249) \| 254 \| PCLP \| Phage capsid-like protein; cath \| 1ohgA01 \| 99.1 \| 2.6e-15 \| 5.5e-19 \| 122.4 \| 109 \| (432, 541) \| 682 \| (1, 110) \| 146 \| Major capsid protein \| CATHCODE: 3.30.2400.10 NAME: Major capsid protein. Chain: a, b, c, d, e, f, g. Synonym: gp5, head protein. Engineered: yes. Other\_details: cleaved form, residues 104-385 SOURCE: Bacteriophage hk97. Organism\_taxid: 37554. Expressed in: escherichia coli. Expression\_system\_taxid: 469008. CLASS: Alpha Beta, ARCH: 2-Layer Sandwich, TOPOL: Major capsid protein gp5 fold, HOMOL: Major capsid protein gp5; phrogs \| 617 \| 100.0 \| 9.7e-57 \| 1.2e-60 \| 480.6 \| 482 \| (51, 674) \| 682 \| (2, 551) \| 551 \| major head protein \| major head protein; Category: head and packaging; p410871 VI\_08691; phrogs \| 2263 \| 100.0 \| 2.7e-35 \| 3.4e-39 \| 317.4 \| 443 \| (37, 680) \| 682 \| (38, 509) \| 518 \| major head protein \| major head protein; Category: head and packaging; p384151 VI\_05178; phrogs \| 4523 \| 100.0 \| 6.3e-35 \| 7.5e-39 \| 315.5 \| 451 \| (29, 526) \| 682 \| (22, 501) \| 667 \| major head and protease protein \| major head and protease protein; Category: head and packaging; p99396 VI\_01094; phrogs \| 10 \| 99.8 \| 3.4e-25 \| 4.7e-29 \| 238.8 \| 263 \| (404, 681) \| 682 \| (125, 405) \| 413 \| major head protein \| major head protein; Category: head and packaging; p205037 VI\_04338; phrogs \| 80 \| 99.8 \| 1.3e-24 \| 1.6e-28 \| 208.2 \| 136 \| (42, 187) \| 682 \| (15, 161) \| 187 \| head maturation protease \| head maturation protease; Category: head and packaging; MF678788\_p6; phrogs \| 3244 \| 99.7 \| 2.9e-21 \| 3.4e-25 \| 186.8 \| 160 \| (22, 188) \| 682 \| (21, 199) \| 279 \| head maturation protease \| head maturation protease; Category: head and packaging; MF140424\_p7; phrogs \| 267 \| 99.7 \| 2.9e-21 \| 3.8e-25 \| 201.7 \| 265 \| (406, 680) \| 682 \| (18, 343) \| 347 \| major head protein \| major head protein; Category: head and packaging; NC\_031123\_p36; phrogs \| 9644 \| 99.3 \| 1e-16 \| 1.1e-20 \| 151.6 \| 152 \| (23, 186) \| 682 \| (137, 309) \| 336 \| head maturation protease \| head maturation protease; Category: head and packaging; p263721 VI\_00383; phrogs \| 13042 \| 99.2 \| 5.5e-16 \| 6.2e-20 \| 147.5 \| 143 \| (38, 185) \| 682 \| (16, 174) \| 367 \| major head protein \| major head protein; Category: head and packaging; JX486088\_p23; phrogs \| 11321 \| 99.2 \| 2.5e-15 \| 2.9e-19 \| 146.8 \| 127 \| (50, 182) \| 682 \| (7, 145) \| 353 \| NA \| NA; Category: unknown function; p210731 VI\_06219; phrogs \| 3270 \| 99.1 \| 8.6e-15 \| 1e-18 \| 150.7 \| 193 \| (471, 679) \| 682 \| (269, 473) \| 481 \| head maturation protease \| head maturation protease; Category: head and packaging; NC\_001629\_p12; phrogs \| 37668 \| 98.9 \| 1.1e-13 \| 1.2e-17 \| 125.3 \| 140 \| (39, 184) \| 682 \| (15, 167) \| 313 \| head maturation protease \| head maturation protease; Category: head and packaging; p13833 VI\_00993; phrogs \| 9427 \| 98.6 \| 4.6e-12 \| 5.3e-16 \| 122.9 \| 152 \| (27, 186) \| 682 \| (11, 171) \| 427 \| head maturation protease \| head maturation protease; Category: head and packaging; MF347638\_p47; phrogs \| 7053 \| 98.4 \| 4.3e-11 \| 4.8e-15 \| 110.9 \| 142 \| (40, 185) \| 682 \| (32, 191) \| 367 \| head maturation protease \| head maturation protease; Category: head and packaging; JF937090\_p9; phrogs \| 15928 \| 97.9 \| 5.4e-09 \| 6.3e-13 \| 88.3 \| 117 \| (552, 679) \| 682 \| (2, 130) \| 141 \| major head protein \| major head protein; Category: head and packaging; p268325 VI\_08160; phrogs \| 7015 \| 97.7 \| 2.1e-08 \| 2.4e-12 \| 88.3 \| 131 \| (40, 187) \| 682 \| (11, 143) \| 230 \| head maturation protease \| head maturation protease; Category: head and packaging; KX578043\_p18; phrogs \| 544 \| 97.3 \| 1.4e-07 \| 1.7e-11 \| 92.0 \| 247 \| (409, 678) \| 682 \| (8, 264) \| 272 \| major head protein \| major head protein; Category: head and packaging; p136878 VI\_00984; phrogs \| 1028 \| 96.8 \| 2e-06 \| 2.4e-10 \| 80.5 \| 133 \| (41, 186) \| 682 \| (24, 165) \| 257 \| head maturation protease \| head maturation protease; Category: head and packaging; MF001365\_p35; phrogs \| 13361 \| 96.5 \| 6.3e-06 \| 7.1e-10 \| 62.5 \| 70 \| (37, 114) \| 682 \| (12, 81) \| 101 \| NA \| NA; Category: unknown function; p46871 VI\_06362; phrogs \| 6538 \| 96.5 \| 6.6e-06 \| 7.3e-10 \| 69.7 \| 173 \| (495, 679) \| 682 \| (2, 203) \| 213 \| NA \| NA; Category: unknown function; p252690 VI\_08262; phrogs \| 3437 \| 96.5 \| 8e-06 \| 9.8e-10 \| 79.0 \| 257 \| (416, 679) \| 682 \| (11, 280) \| 284 \| major head protein \| major head protein; Category: head and packaging; NC\_020481\_p9; phrogs \| 17162 \| 96.1 \| 2.8e-05 \| 3.1e-09 \| 58.1 \| 73 \| (608, 681) \| 682 \| (5, 83) \| 84 \| NA \| NA; Category: unknown function; p331326 VI\_04001; phrogs \| 21715 \| 96.0 \| 3.5e-05 \| 4e-09 \| 71.0 \| 127 \| (41, 186) \| 682 \| (15, 145) \| 385 \| NA \| NA; Category: unknown function; p212986 VI\_01051; phrogs \| 1650 \| 96.0 \| 4e-05 \| 4.9e-09 \| 74.3 \| 210 \| (460, 679) \| 682 \| (78, 296) \| 298 \| major head protein \| major head protein; Category: head and packaging; NC\_026606\_p9; phrogs \| 30146 \| 95.7 \| 9.5e-05 \| 1.1e-08 \| 51.3 \| 53 \| (37, 97) \| 682 \| (12, 64) \| 65 \| NA \| NA; Category: unknown function; p428492 VI\_08052; phrogs \| 3922 \| 95.3 \| 0.00017 \| 2.1e-08 \| 70.9 \| 255 \| (408, 679) \| 682 \| (70, 337) \| 340 \| major head protein \| major head protein; Category: head and packaging; p276403 VI\_00011; phrogs \| 30620 \| 94.2 \| 0.00099 \| 1.1e-07 \| 55.4 \| 99 \| (427, 527) \| 682 \| (25, 129) \| 183 \| NA \| NA; Category: unknown function; p428682 VI\_05331 |
| Top keywords  (threshold 1.00e-03 (evalue)) | **head, and, packaging, major, protease, maturation, capsid, Phage, gp5, serine** |
| Output files | ../../domain\_architecture/05\_FANPEZAQ\_CDS\_0005\_cath.hhr ../../domain\_architecture/05\_FANPEZAQ\_CDS\_0005\_merged.svg ../../domain\_architecture/05\_FANPEZAQ\_CDS\_0005\_ncbi-cd.hhr ../../domain\_architecture/05\_FANPEZAQ\_CDS\_0005\_pfam.hhr ../../domain\_architecture/05\_FANPEZAQ\_CDS\_0005\_phrogs.hhr |

### Identical protein sequences/structures

#### Search results

|  |  |
| --- | --- |
| Protein sequence databases searched | Pdb, Swissprot, Refseq |
| Identical proteins found | -- |
| Top keywords | -- |
| Output files | -- |

### Similar protein sequences/structures

#### Sequence similarity search results (HHblits)^1^

|  |  |
| --- | --- |
| Sequence databases searched | Uniclust, Pdb70 |
| Results, scheme(s)  (Top layers only, threshold 1.00e-03 (evalue)) | xml version="1.0" encoding="utf-8" standalone="no"?       2024-09-02T21:08:30.417592 image/svg+xml   Matplotlib v3.7.2, https://matplotlib.org/ |
| Results, table(s)  (threshold 1.00e-03 (evalue)) | db \| id \| prob \| evalue \| pvalue \| score \| cols \| query \| query\_len \| template \| template\_len \| name \| description; uniclust \| UniRef100\_A0A060GZS1 \| 100.0 \| 2.5e-93 \| 5.1e-99 \| 772.2 \| 604 \| (31, 681) \| 682 \| (136, 743) \| 757 \| Peptidase U35 \| Peptidase U35; uniclust \| UniRef100\_A0A0G3G1E5 \| 100.0 \| 1.4e-88 \| 3e-94 \| 729.5 \| 580 \| (31, 681) \| 682 \| (53, 650) \| 662 \| Phage major capsid protein \| Phage major capsid protein; uniclust \| UniRef100\_A0A0K6I524 \| 100.0 \| 7.3e-88 \| 1.4e-93 \| 710.2 \| 575 \| (40, 681) \| 682 \| (118, 720) \| 733 \| Phage major capsid protein, HK97 family \| Phage major capsid protein, HK97 family; uniclust \| UniRef100\_A0A089QSP9 \| 100.0 \| 1.5e-84 \| 2.9e-90 \| 675.9 \| 557 \| (31, 681) \| 682 \| (41, 615) \| 618 \| Phage protein \| Phage protein; uniclust \| UniRef100\_A0A2E1P3K3 \| 100.0 \| 3e-84 \| 5.9e-90 \| 676.7 \| 568 \| (43, 681) \| 682 \| (273, 845) \| 848 \| Phage major capsid protein \| Phage major capsid protein; uniclust \| UniRef100\_A0A2D5VNT2 \| 100.0 \| 2.3e-57 \| 4.4e-63 \| 444.2 \| 458 \| (31, 537) \| 682 \| (33, 493) \| 499 \| Phage major capsid protein (Fragment) \| Phage major capsid protein (Fragment); uniclust \| UniRef100\_A0A164AHR0 \| 100.0 \| 3.1e-57 \| 5.8e-63 \| 440.3 \| 502 \| (32, 681) \| 682 \| (10, 534) \| 537 \| Phage capsid family protein \| Phage capsid family protein; uniclust \| UniRef100\_A0A031JPI0 \| 100.0 \| 4.5e-57 \| 8.9e-63 \| 471.0 \| 479 \| (42, 681) \| 682 \| (85, 579) \| 594 \| Phage major capsid protein, HK97 \| Phage major capsid protein, HK97; uniclust \| UniRef100\_A0A071LTJ4 \| 100.0 \| 5.3e-57 \| 1.1e-62 \| 471.4 \| 573 \| (31, 680) \| 682 \| (13, 616) \| 622 \| Phage major capsid protein \| Phage major capsid protein; uniclust \| UniRef100\_A0A193LMW5 \| 100.0 \| 8.6e-57 \| 1.7e-62 \| 465.9 \| 605 \| (31, 679) \| 682 \| (6, 663) \| 682 \| Peptidase \| Peptidase; uniclust \| UniRef100\_A0A2E7BJW1 \| 100.0 \| 4.3e-55 \| 8.2e-61 \| 440.5 \| 598 \| (32, 682) \| 682 \| (70, 711) \| 720 \| Phage major capsid protein \| Phage major capsid protein; uniclust \| UniRef100\_A0A1G0FL18 \| 100.0 \| 3.6e-53 \| 7.2e-59 \| 425.8 \| 336 \| (331, 681) \| 682 \| (88, 442) \| 445 \| Capsid protein \| Capsid protein; uniclust \| UniRef100\_A0A1G0RDU6 \| 100.0 \| 5.5e-53 \| 1e-58 \| 422.4 \| 605 \| (31, 681) \| 682 \| (18, 643) \| 644 \| Phage major capsid protein \| Phage major capsid protein; uniclust \| UniRef100\_A0A075KI09 \| 100.0 \| 1.1e-51 \| 2.1e-57 \| 438.8 \| 589 \| (31, 679) \| 682 \| (127, 757) \| 768 \| Peptidase U35 phage prohead HK97 \| Peptidase U35 phage prohead HK97; uniclust \| UniRef100\_A0A031HTD4 \| 100.0 \| 1.9e-51 \| 3.7e-57 \| 434.2 \| 574 \| (31, 682) \| 682 \| (28, 669) \| 717 \| HK97 family phage prohead protease \| HK97 family phage prohead protease; uniclust \| UniRef100\_A0A2D7GUF3 \| 100.0 \| 2.2e-51 \| 4.2e-57 \| 402.3 \| 369 \| (31, 451) \| 682 \| (25, 399) \| 399 \| Capsid protein (Fragment) \| Capsid protein (Fragment); uniclust \| UniRef100\_A0A938SX03 \| 100.0 \| 1.3e-50 \| 2.5e-56 \| 400.5 \| 580 \| (33, 679) \| 682 \| (245, 864) \| 869 \| Phage major capsid protein \| Phage major capsid protein; uniclust \| UniRef100\_A0A011TBF3 \| 100.0 \| 7.6e-50 \| 1.6e-55 \| 444.9 \| 599 \| (31, 680) \| 682 \| (119, 773) \| 862 \| Peptidase S14 \| Peptidase S14; uniclust \| UniRef100\_A0A1V4MJF0 \| 100.0 \| 1.8e-47 \| 3.3e-53 \| 382.5 \| 523 \| (31, 680) \| 682 \| (15, 571) \| 586 \| Capsid protein \| Capsid protein; uniclust \| UniRef100\_UPI001AC3FFAA \| 100.0 \| 5.3e-47 \| 9.8e-53 \| 365.5 \| 585 \| (31, 681) \| 682 \| (5, 636) \| 639 \| phage major capsid protein \| phage major capsid protein; uniclust \| UniRef100\_A0A812JEV6 \| 100.0 \| 8.6e-47 \| 1.6e-52 \| 404.4 \| 614 \| (38, 677) \| 682 \| (1617, 2266) \| 4451 \| Phage portal protein \| Phage portal protein; uniclust \| UniRef100\_A0A4Q8WY63 \| 100.0 \| 1e-46 \| 1.9e-52 \| 382.3 \| 571 \| (31, 671) \| 682 \| (10, 619) \| 635 \| HK97 family phage prohead protease (Fragment) \| HK97 family phage prohead protease (Fragment); uniclust \| UniRef100\_A0A081CWY9 \| 100.0 \| 3.5e-46 \| 7.2e-52 \| 409.6 \| 545 \| (32, 677) \| 682 \| (140, 724) \| 749 \| Peptidase U35 family protein \| Peptidase U35 family protein; uniclust \| UniRef100\_A0A0C5DX41 \| 100.0 \| 3.4e-45 \| 7e-51 \| 382.5 \| 323 \| (341, 681) \| 682 \| (97, 436) \| 446 \| Phage major capsid protein \| Phage major capsid protein; uniclust \| UniRef100\_A0A934RQD8 \| 100.0 \| 6.8e-45 \| 1.3e-50 \| 352.5 \| 576 \| (45, 681) \| 682 \| (41, 660) \| 665 \| Phage major capsid protein \| Phage major capsid protein; uniclust \| UniRef100\_A0A0F9KZ70 \| 100.0 \| 6.8e-45 \| 1.4e-50 \| 372.3 \| 331 \| (335, 681) \| 682 \| (87, 432) \| 436 \| Phage major capsid protein (Fragment) \| Phage major capsid protein (Fragment); uniclust \| UniRef100\_A0A087ND25 \| 100.0 \| 1e-44 \| 2e-50 \| 369.3 \| 315 \| (342, 681) \| 682 \| (120, 448) \| 459 \| Phage-related protein \| Phage-related protein; uniclust \| UniRef100\_A0A560FKK8 \| 100.0 \| 1.2e-43 \| 2.3e-49 \| 361.2 \| 494 \| (43, 681) \| 682 \| (278, 794) \| 796 \| HK97 family phage prohead protease/HK97 family phage major capsid protein,TIGR01554 \| HK97 family phage prohead protease/HK97 family phage major capsid protein,TIGR01554; uniclust \| UniRef100\_A0A952V6M2 \| 100.0 \| 4.1e-43 \| 7.6e-49 \| 347.4 \| 548 \| (32, 669) \| 682 \| (19, 610) \| 630 \| Phage major capsid protein \| Phage major capsid protein; uniclust \| UniRef100\_A0A7X4CLK5 \| 100.0 \| 1.4e-42 \| 2.6e-48 \| 340.4 \| 613 \| (42, 680) \| 682 \| (54, 748) \| 750 \| Phage major capsid protein \| Phage major capsid protein; uniclust \| UniRef100\_A0A0F9DVX4 \| 100.0 \| 3.4e-42 \| 6.7e-48 \| 341.9 \| 308 \| (337, 681) \| 682 \| (73, 401) \| 406 \| Phage major capsid protein \| Phage major capsid protein; uniclust \| UniRef100\_A0A0Q5PQY2 \| 100.0 \| 3.9e-42 \| 7.5e-48 \| 354.6 \| 444 \| (31, 525) \| 682 \| (20, 483) \| 649 \| Uncharacterized protein \| Uncharacterized protein; uniclust \| UniRef100\_A0A316SJM9 \| 100.0 \| 4.2e-42 \| 8.1e-48 \| 349.1 \| 574 \| (31, 678) \| 682 \| (24, 631) \| 634 \| Peptidase \| Peptidase; uniclust \| UniRef100\_A0A0F3QFF6 \| 100.0 \| 6.1e-42 \| 1.2e-47 \| 340.3 \| 465 \| (35, 679) \| 682 \| (14, 494) \| 499 \| Phage prohead protease, HK97 family \| Phage prohead protease, HK97 family; uniclust \| UniRef100\_A0A1S6TP86 \| 100.0 \| 1.3e-41 \| 2.6e-47 \| 352.2 \| 546 \| (39, 680) \| 682 \| (30, 584) \| 594 \| Phage capsid protein \| Phage capsid protein; uniclust \| UniRef100\_A0A261Q1A3 \| 100.0 \| 2.6e-41 \| 5.2e-47 \| 345.2 \| 329 \| (336, 681) \| 682 \| (134, 484) \| 492 \| Phage major capsid protein \| Phage major capsid protein; uniclust \| UniRef100\_A0A0F2RQY9 \| 100.0 \| 2.9e-41 \| 5.5e-47 \| 345.2 \| 586 \| (31, 680) \| 682 \| (18, 649) \| 652 \| Peptidase U35 \| Peptidase U35; uniclust \| UniRef100\_A0A2U1B6D9 \| 100.0 \| 1e-40 \| 2e-46 \| 327.8 \| 588 \| (31, 679) \| 682 \| (13, 611) \| 612 \| HK97 family phage prohead protease/HK97 family phage major capsid protein,TIGR01554 \| HK97 family phage prohead protease/HK97 family phage major capsid protein,TIGR01554; uniclust \| UniRef100\_A0A0A8H0T3 \| 100.0 \| 3.8e-40 \| 7.7e-46 \| 333.9 \| 270 \| (393, 680) \| 682 \| (119, 399) \| 404 \| Major capsid protein, HK97 family \| Major capsid protein, HK97 family; uniclust \| UniRef100\_A0A1I0J9W7 \| 100.0 \| 5.1e-40 \| 1e-45 \| 339.9 \| 554 \| (39, 680) \| 682 \| (25, 618) \| 620 \| Prohead serine protease \| Prohead serine protease; uniclust \| UniRef100\_A0A061QK74 \| 100.0 \| 6.9e-40 \| 1.4e-45 \| 333.0 \| 316 \| (337, 681) \| 682 \| (138, 469) \| 482 \| Phage major capsid protein, HK97 family \| Phage major capsid protein, HK97 family; uniclust \| UniRef100\_A0A0P6VLZ5 \| 100.0 \| 1.8e-39 \| 3.4e-45 \| 330.1 \| 571 \| (32, 672) \| 682 \| (38, 645) \| 654 \| Bacteriophage Mu GpT domain-containing protein \| Bacteriophage Mu GpT domain-containing protein; uniclust \| UniRef100\_A0A0F9RBZ5 \| 100.0 \| 1.5e-38 \| 3e-44 \| 339.4 \| 606 \| (31, 679) \| 682 \| (43, 713) \| 716 \| Bacteriophage Mu GpT domain-containing protein \| Bacteriophage Mu GpT domain-containing protein; uniclust \| UniRef100\_A0A087M3M1 \| 100.0 \| 2.5e-38 \| 4.8e-44 \| 332.7 \| 570 \| (47, 681) \| 682 \| (102, 730) \| 739 \| Peptidase U35 \| Peptidase U35; uniclust \| UniRef100\_A0A024P7I8 \| 100.0 \| 3.5e-38 \| 7e-44 \| 334.0 \| 489 \| (31, 680) \| 682 \| (36, 551) \| 646 \| Phage major capsid protein, HK97 family \| Phage major capsid protein, HK97 family; uniclust \| UniRef100\_A0A1G1GKX3 \| 100.0 \| 8.5e-38 \| 1.7e-43 \| 310.4 \| 324 \| (335, 681) \| 682 \| (41, 370) \| 371 \| Capsid protein \| Capsid protein; uniclust \| UniRef100\_A0A080IWL3 \| 100.0 \| 2.5e-37 \| 4.8e-43 \| 307.5 \| 502 \| (39, 679) \| 682 \| (13, 547) \| 564 \| Phage major capsid protein, HK97 family \| Phage major capsid protein, HK97 family; uniclust \| UniRef100\_A0A145VQW4 \| 100.0 \| 4.3e-37 \| 8.2e-43 \| 288.3 \| 269 \| (409, 681) \| 682 \| (23, 293) \| 298 \| Capsid protein \| Capsid protein; uniclust \| UniRef100\_A0A1E3GXJ9 \| 100.0 \| 5e-37 \| 9.3e-43 \| 322.2 \| 599 \| (31, 674) \| 682 \| (539, 1187) \| 1194 \| Phage portal protein, lambda family \| Phage portal protein, lambda family; uniclust \| UniRef100\_A0A074TEC5 \| 100.0 \| 8.2e-37 \| 1.6e-42 \| 314.2 \| 274 \| (390, 681) \| 682 \| (157, 443) \| 448 \| Capsid protein \| Capsid protein; uniclust \| UniRef100\_A0A0B6D4D0 \| 100.0 \| 9.8e-37 \| 1.8e-42 \| 293.9 \| 556 \| (43, 679) \| 682 \| (16, 591) \| 597 \| Phage major capsid protein, HK97 family \| Phage major capsid protein, HK97 family; uniclust \| UniRef100\_A0A929ZEC9 \| 100.0 \| 1e-36 \| 1.9e-42 \| 294.9 \| 376 \| (38, 475) \| 682 \| (30, 424) \| 428 \| HK97 family phage prohead protease (Fragment) \| HK97 family phage prohead protease (Fragment); uniclust \| UniRef100\_A0A354UAC1 \| 100.0 \| 2.1e-36 \| 3.9e-42 \| 293.5 \| 590 \| (43, 677) \| 682 \| (16, 633) \| 637 \| Phage major capsid protein \| Phage major capsid protein; uniclust \| UniRef100\_A0A075TQ00 \| 100.0 \| 2.7e-36 \| 5.6e-42 \| 317.4 \| 270 \| (393, 680) \| 682 \| (168, 451) \| 487 \| Capsid protein \| Capsid protein; uniclust \| UniRef100\_A0A0F7PPX3 \| 100.0 \| 5.2e-36 \| 9.9e-42 \| 301.1 \| 555 \| (44, 681) \| 682 \| (26, 640) \| 642 \| Phage prohead protease, HK97 family/phage major capsid protein, HK97 family \| Phage prohead protease, HK97 family/phage major capsid protein, HK97 family; uniclust \| UniRef100\_A0A072NGX0 \| 100.0 \| 6e-36 \| 1.2e-41 \| 310.5 \| 264 \| (406, 680) \| 682 \| (155, 434) \| 442 \| Phage major capsid protein, HK97 family \| Phage major capsid protein, HK97 family; uniclust \| UniRef100\_A0A2D2C2A3 \| 100.0 \| 9.7e-36 \| 1.9e-41 \| 308.5 \| 604 \| (31, 680) \| 682 \| (12, 692) \| 695 \| Bacteriophage Mu GpT domain-containing protein \| Bacteriophage Mu GpT domain-containing protein; uniclust \| UniRef100\_A0A3B9KKY0 \| 100.0 \| 1e-35 \| 2.1e-41 \| 311.1 \| 271 \| (393, 680) \| 682 \| (286, 569) \| 572 \| Phage major capsid protein \| Phage major capsid protein; uniclust \| UniRef100\_A0A101G6Z0 \| 100.0 \| 2.1e-35 \| 4.2e-41 \| 302.7 \| 271 \| (393, 679) \| 682 \| (109, 400) \| 426 \| HK97 family phage prohead protease \| HK97 family phage prohead protease; uniclust \| UniRef100\_A0A3N5X4U1 \| 100.0 \| 2.3e-35 \| 4.4e-41 \| 289.6 \| 326 \| (336, 681) \| 682 \| (75, 423) \| 432 \| Phage major capsid protein (Fragment) \| Phage major capsid protein (Fragment); uniclust \| UniRef100\_A0A2A8V9J7 \| 100.0 \| 2.6e-35 \| 5.2e-41 \| 293.8 \| 270 \| (393, 681) \| 682 \| (123, 403) \| 410 \| Phage major capsid protein \| Phage major capsid protein; uniclust \| UniRef100\_A0A017H7N1 \| 100.0 \| 2.7e-35 \| 5.4e-41 \| 309.7 \| 271 \| (393, 681) \| 682 \| (183, 467) \| 491 \| Phage major capsid protein \| Phage major capsid protein; uniclust \| UniRef100\_UPI0022E46B5D \| 100.0 \| 4.2e-35 \| 7.7e-41 \| 272.7 \| 399 \| (31, 475) \| 682 \| (16, 422) \| 423 \| HK97 family phage prohead protease \| HK97 family phage prohead protease; uniclust \| UniRef100\_A0A094YHB1 \| 100.0 \| 4.9e-35 \| 1e-40 \| 311.1 \| 269 \| (394, 681) \| 682 \| (203, 485) \| 497 \| Phage major capsid protein \| Phage major capsid protein; uniclust \| UniRef100\_A0A1N6XWD6 \| 100.0 \| 6.3e-35 \| 1.2e-40 \| 296.2 \| 486 \| (41, 674) \| 682 \| (41, 562) \| 581 \| Phage prohead protease, HK97 family/phage major capsid protein, HK97 family,TIGR01554 \| Phage prohead protease, HK97 family/phage major capsid protein, HK97 family,TIGR01554; uniclust \| UniRef100\_A0A0F8YXG7 \| 100.0 \| 1.3e-34 \| 2.6e-40 \| 300.0 \| 269 \| (393, 681) \| 682 \| (131, 424) \| 431 \| Phage major capsid protein (Fragment) \| Phage major capsid protein (Fragment); uniclust \| UniRef100\_A0A0D9NGH1 \| 100.0 \| 3.1e-34 \| 5.8e-40 \| 281.6 \| 550 \| (33, 680) \| 682 \| (11, 578) \| 580 \| Peptidase U35 \| Peptidase U35; uniclust \| UniRef100\_A0A6G6X8M0 \| 100.0 \| 3.5e-34 \| 6.5e-40 \| 281.0 \| 373 \| (270, 679) \| 682 \| (174, 551) \| 555 \| Phage major capsid protein \| Phage major capsid protein; uniclust \| UniRef100\_A0A076JIU2 \| 100.0 \| 3.4e-34 \| 6.9e-40 \| 311.7 \| 482 \| (31, 680) \| 682 \| (127, 636) \| 646 \| Prohead peptidase \| Prohead peptidase; uniclust \| UniRef100\_A0A0A1WC95 \| 100.0 \| 3.7e-34 \| 7.3e-40 \| 300.4 \| 565 \| (38, 675) \| 682 \| (48, 652) \| 667 \| Putative phage prohead (Fragment) \| Putative phage prohead (Fragment); uniclust \| UniRef100\_A0A074MYL0 \| 100.0 \| 3.7e-34 \| 7.5e-40 \| 299.2 \| 272 \| (393, 682) \| 682 \| (186, 471) \| 476 \| Phage major capsid protein \| Phage major capsid protein; uniclust \| UniRef100\_A0A158S6T4 \| 100.0 \| 5.3e-34 \| 1e-39 \| 278.8 \| 314 \| (341, 681) \| 682 \| (145, 470) \| 485 \| Putative phage-related protein \| Putative phage-related protein; uniclust \| UniRef100\_A0A3N2E2B1 \| 100.0 \| 8.1e-34 \| 1.5e-39 \| 284.3 \| 396 \| (261, 681) \| 682 \| (255, 656) \| 667 \| ATP-dependent Clp protease proteolytic subunit \| ATP-dependent Clp protease proteolytic subunit; uniclust \| UniRef100\_A0A164A2P9 \| 100.0 \| 1.1e-33 \| 2.1e-39 \| 287.1 \| 323 \| (339, 681) \| 682 \| (92, 429) \| 435 \| Phage major capsid protein \| Phage major capsid protein; uniclust \| UniRef100\_UPI000D2FC543 \| 100.0 \| 1.4e-33 \| 2.6e-39 \| 273.1 \| 525 \| (42, 670) \| 682 \| (29, 585) \| 608 \| phage major capsid protein \| phage major capsid protein; uniclust \| UniRef100\_A0A023WVI6 \| 100.0 \| 2.7e-33 \| 5.5e-39 \| 297.6 \| 275 \| (393, 682) \| 682 \| (186, 487) \| 580 \| HK97 family phage major capsid protein \| HK97 family phage major capsid protein; uniclust \| UniRef100\_A0A2M8B3M2 \| 100.0 \| 4.2e-33 \| 7.8e-39 \| 268.9 \| 455 \| (31, 523) \| 682 \| (10, 508) \| 516 \| Peptidase U37 (Fragment) \| Peptidase U37 (Fragment); uniclust \| UniRef100\_UPI0009EE22D6 \| 100.0 \| 4.4e-33 \| 8.1e-39 \| 265.6 \| 461 \| (41, 679) \| 682 \| (23, 515) \| 520 \| phage major capsid protein \| phage major capsid protein; uniclust \| UniRef100\_A0A7V9J303 \| 100.0 \| 5.8e-33 \| 1.1e-38 \| 267.4 \| 543 \| (31, 679) \| 682 \| (7, 570) \| 573 \| Phage major capsid protein \| Phage major capsid protein; uniclust \| UniRef100\_A0A0F5JCS1 \| 100.0 \| 6.3e-33 \| 1.2e-38 \| 274.4 \| 313 \| (341, 679) \| 682 \| (90, 407) \| 414 \| HK97 family phage major capsid protein \| HK97 family phage major capsid protein; uniclust \| UniRef100\_A0A0F3IQW8 \| 100.0 \| 9e-33 \| 1.7e-38 \| 264.5 \| 491 \| (45, 681) \| 682 \| (17, 535) \| 539 \| Phage capsid protein \| Phage capsid protein; uniclust \| UniRef100\_A0A0F9IAP4 \| 100.0 \| 1.3e-32 \| 2.6e-38 \| 305.1 \| 575 \| (45, 681) \| 682 \| (108, 738) \| 750 \| Bacteriophage Mu GpT domain-containing protein \| Bacteriophage Mu GpT domain-containing protein; uniclust \| UniRef100\_A0A1V5X326 \| 100.0 \| 1.4e-32 \| 2.7e-38 \| 292.0 \| 563 \| (45, 670) \| 682 \| (92, 717) \| 734 \| Caudovirus prohead protease \| Caudovirus prohead protease; uniclust \| UniRef100\_A0A142Y5E5 \| 100.0 \| 2.1e-32 \| 4.3e-38 \| 300.2 \| 597 \| (41, 677) \| 682 \| (79, 738) \| 745 \| Mu-like prophage major head subunit gpT \| Mu-like prophage major head subunit gpT; uniclust \| UniRef100\_A0A1C6BMQ2 \| 99.9 \| 3.5e-32 \| 6.9e-38 \| 250.2 \| 154 \| (31, 185) \| 682 \| (24, 185) \| 232 \| Phage prohead protease, HK97 family \| Phage prohead protease, HK97 family; uniclust \| UniRef100\_A0A127CGL7 \| 99.9 \| 4e-32 \| 7.8e-38 \| 282.4 \| 535 \| (38, 679) \| 682 \| (21, 592) \| 601 \| Bacteriophage Mu GpT domain-containing protein \| Bacteriophage Mu GpT domain-containing protein; uniclust \| UniRef100\_A0A965MHY9 \| 99.9 \| 4.9e-32 \| 9.1e-38 \| 252.5 \| 371 \| (271, 680) \| 682 \| (16, 388) \| 390 \| Phage major capsid protein \| Phage major capsid protein; uniclust \| UniRef100\_A0A133NSV4 \| 99.9 \| 5.4e-32 \| 1.1e-37 \| 265.9 \| 262 \| (408, 680) \| 682 \| (52, 319) \| 334 \| Phage capsid family protein \| Phage capsid family protein; uniclust \| UniRef100\_A0A290XJJ8 \| 99.9 \| 1.2e-31 \| 2.4e-37 \| 281.4 \| 535 \| (37, 680) \| 682 \| (41, 614) \| 618 \| Major capsid protein \| Major capsid protein; uniclust \| UniRef100\_A0A7I8N2L9 \| 99.9 \| 1.5e-31 \| 2.8e-37 \| 245.8 \| 340 \| (31, 454) \| 682 \| (3, 348) \| 373 \| Phage capsid and scaffold \| Phage capsid and scaffold; uniclust \| UniRef100\_A0A382Z2D1 \| 99.9 \| 1.5e-31 \| 2.9e-37 \| 242.4 \| 154 \| (31, 185) \| 682 \| (24, 179) \| 205 \| Caudovirus prohead protease (Fragment) \| Caudovirus prohead protease (Fragment); uniclust \| UniRef100\_H1CG40 \| 99.9 \| 2e-31 \| 3.7e-37 \| 261.3 \| 556 \| (43, 681) \| 682 \| (15, 611) \| 615 \| HK97 family phage prohead protease \| HK97 family phage prohead protease; uniclust \| UniRef100\_A0A965HBR8 \| 99.9 \| 2.5e-31 \| 4.8e-37 \| 244.2 \| 263 \| (40, 354) \| 682 \| (28, 297) \| 314 \| Phage major capsid protein (Fragment) \| Phage major capsid protein (Fragment); uniclust \| UniRef100\_A0A0F9CDL2 \| 99.9 \| 3.3e-31 \| 6.5e-37 \| 259.4 \| 315 \| (31, 437) \| 682 \| (12, 343) \| 371 \| Uncharacterized protein (Fragment) \| Uncharacterized protein (Fragment); uniclust \| UniRef100\_A0A6N0ZA72 \| 99.9 \| 4.4e-31 \| 8.4e-37 \| 256.5 \| 381 \| (269, 681) \| 682 \| (14, 415) \| 418 \| Phage major capsid protein \| Phage major capsid protein; uniclust \| UniRef100\_A0A167WYC7 \| 99.9 \| 7.2e-31 \| 1.5e-36 \| 277.4 \| 271 \| (393, 682) \| 682 \| (178, 463) \| 482 \| Phage major capsid protein \| Phage major capsid protein; uniclust \| UniRef100\_A0A098F6L7 \| 99.9 \| 8.9e-31 \| 1.7e-36 \| 241.6 \| 145 \| (40, 185) \| 682 \| (65, 215) \| 263 \| Caudovirus prohead protease \| Caudovirus prohead protease; uniclust \| UniRef100\_A0A3A5AR09 \| 99.9 \| 9.4e-31 \| 1.8e-36 \| 240.9 \| 216 \| (459, 681) \| 682 \| (2, 235) \| 264 \| Phage major capsid protein \| Phage major capsid protein; uniclust \| UniRef100\_A0A3M1QXF2 \| 99.9 \| 1.1e-30 \| 2.1e-36 \| 265.9 \| 578 \| (40, 675) \| 682 \| (22, 664) \| 670 \| Caudovirus prohead protease \| Caudovirus prohead protease; uniclust \| UniRef100\_A0A015WYK1 \| 99.9 \| 1.3e-30 \| 2.5e-36 \| 260.8 \| 318 \| (340, 679) \| 682 \| (130, 463) \| 482 \| Phage major capsid protein, HK97 family \| Phage major capsid protein, HK97 family; uniclust \| UniRef100\_A0A376TC30 \| 99.9 \| 1.6e-30 \| 3e-36 \| 257.5 \| 562 \| (38, 681) \| 682 \| (24, 648) \| 656 \| Putative major head protein/prohead protease \| Putative major head protein/prohead protease; uniclust \| UniRef100\_A0A0C2VPM1 \| 99.9 \| 1.7e-30 \| 3.3e-36 \| 254.7 \| 264 \| (406, 681) \| 682 \| (85, 367) \| 370 \| Phage major capsid protein \| Phage major capsid protein; uniclust \| UniRef100\_Q1ILS6 \| 99.9 \| 1.9e-30 \| 3.6e-36 \| 257.8 \| 397 \| (261, 681) \| 682 \| (353, 777) \| 781 \| Peptidase S49 \| Peptidase S49; uniclust \| UniRef100\_A0A1D8UTE4 \| 99.9 \| 2e-30 \| 3.9e-36 \| 264.6 \| 274 \| (393, 681) \| 682 \| (147, 455) \| 515 \| Major capsid protein \| Major capsid protein; uniclust \| UniRef100\_A0A6B3AAV6 \| 99.9 \| 2.2e-30 \| 4.1e-36 \| 254.5 \| 488 \| (43, 681) \| 682 \| (17, 528) \| 530 \| Phage major capsid protein \| Phage major capsid protein; uniclust \| UniRef100\_A0A0S7X9K7 \| 99.9 \| 2.3e-30 \| 4.4e-36 \| 254.1 \| 273 \| (393, 680) \| 682 \| (116, 414) \| 423 \| Phage capsid protein \| Phage capsid protein; uniclust \| UniRef100\_A0A348SWC8 \| 99.9 \| 2.6e-30 \| 4.9e-36 \| 241.2 \| 154 \| (31, 185) \| 682 \| (25, 181) \| 330 \| Phage major capsid protein (Fragment) \| Phage major capsid protein (Fragment); uniclust \| UniRef100\_A0A031HR58 \| 99.9 \| 2.5e-30 \| 5e-36 \| 264.5 \| 268 \| (393, 680) \| 682 \| (146, 442) \| 464 \| Phage major capsid protein, HK97 family \| Phage major capsid protein, HK97 family; uniclust \| UniRef100\_A0A1H7NMW7 \| 99.9 \| 3.5e-30 \| 6.8e-36 \| 269.9 \| 538 \| (44, 680) \| 682 \| (36, 612) \| 637 \| Phage prohead protease, HK97 family/phage major capsid protein, HK97 family,TIGR01554 \| Phage prohead protease, HK97 family/phage major capsid protein, HK97 family,TIGR01554; uniclust \| UniRef100\_A0A0F9ENL1 \| 99.9 \| 4.2e-30 \| 8.2e-36 \| 240.5 \| 215 \| (456, 680) \| 682 \| (9, 237) \| 243 \| Phage major capsid protein (Fragment) \| Phage major capsid protein (Fragment); uniclust \| UniRef100\_A0A021VTX6 \| 99.9 \| 1.1e-29 \| 2.2e-35 \| 262.1 \| 270 \| (393, 680) \| 682 \| (144, 427) \| 435 \| HK97 family phage major capsid protein \| HK97 family phage major capsid protein; uniclust \| UniRef100\_A0A011U0Z7 \| 99.9 \| 2.4e-29 \| 5e-35 \| 272.1 \| 268 \| (393, 680) \| 682 \| (241, 536) \| 602 \| Capsid protein \| Capsid protein; uniclust \| UniRef100\_A0A011TWP2 \| 99.9 \| 2.8e-29 \| 5.7e-35 \| 268.7 \| 272 \| (393, 680) \| 682 \| (182, 470) \| 557 \| Capsid protein \| Capsid protein; uniclust \| UniRef100\_A0A0G9K210 \| 99.9 \| 3.4e-29 \| 6.7e-35 \| 262.8 \| 571 \| (31, 678) \| 682 \| (25, 634) \| 640 \| Bacteriophage Mu GpT domain-containing protein \| Bacteriophage Mu GpT domain-containing protein; uniclust \| UniRef100\_A0A064CN03 \| 99.9 \| 7.4e-29 \| 1.5e-34 \| 249.4 \| 260 \| (412, 680) \| 682 \| (57, 322) \| 341 \| Capsid protein \| Capsid protein; uniclust \| UniRef100\_A0A0N0T0A0 \| 99.9 \| 9.7e-29 \| 1.9e-34 \| 243.7 \| 271 \| (393, 681) \| 682 \| (122, 411) \| 435 \| Phage major capsid protein \| Phage major capsid protein; uniclust \| UniRef100\_A0A022LMT4 \| 99.9 \| 1.1e-28 \| 2.1e-34 \| 242.5 \| 264 \| (406, 681) \| 682 \| (61, 336) \| 361 \| Capsid protein \| Capsid protein; uniclust \| UniRef100\_A0A2S9PYB0 \| 99.9 \| 1.2e-28 \| 2.4e-34 \| 266.3 \| 271 \| (393, 681) \| 682 \| (512, 800) \| 803 \| Phage major capsid protein \| Phage major capsid protein; uniclust \| UniRef100\_A0A258QCW1 \| 99.9 \| 1.3e-28 \| 2.5e-34 \| 242.6 \| 322 \| (342, 681) \| 682 \| (34, 375) \| 378 \| Phage capsid protein \| Phage capsid protein; uniclust \| UniRef100\_A0A0J6SLM0 \| 99.9 \| 1.7e-28 \| 3.4e-34 \| 249.6 \| 268 \| (406, 681) \| 682 \| (149, 447) \| 458 \| Capsid protein \| Capsid protein; uniclust \| UniRef100\_A0A142YPV2 \| 99.9 \| 1.9e-28 \| 3.7e-34 \| 238.0 \| 323 \| (341, 681) \| 682 \| (18, 368) \| 372 \| Phage capsid family protein \| Phage capsid family protein; uniclust \| UniRef100\_A0A142X041 \| 99.9 \| 2.2e-28 \| 4.5e-34 \| 255.9 \| 270 \| (393, 680) \| 682 \| (182, 469) \| 473 \| Phage capsid family protein \| Phage capsid family protein; uniclust \| UniRef100\_A0A0J9D0M7 \| 99.9 \| 2.8e-28 \| 5.6e-34 \| 248.3 \| 263 \| (406, 681) \| 682 \| (167, 457) \| 459 \| Phage major capsid protein \| Phage major capsid protein; uniclust \| UniRef100\_A0A2N3B9M3 \| 99.9 \| 3.7e-28 \| 7e-34 \| 235.5 \| 270 \| (31, 308) \| 682 \| (13, 307) \| 367 \| Peptidase U37 (Fragment) \| Peptidase U37 (Fragment); uniclust \| UniRef100\_A0A2E7K3Z7 \| 99.9 \| 3.9e-28 \| 7.5e-34 \| 236.1 \| 242 \| (38, 308) \| 682 \| (22, 272) \| 379 \| HK97 family phage prohead protease (Fragment) \| HK97 family phage prohead protease (Fragment); uniclust \| UniRef100\_A0A024YGR4 \| 99.9 \| 5.5e-28 \| 1.1e-33 \| 248.8 \| 271 \| (392, 680) \| 682 \| (129, 422) \| 428 \| Capsid protein \| Capsid protein; uniclust \| UniRef100\_A0A1H7ZYZ5 \| 99.9 \| 5.8e-28 \| 1.1e-33 \| 251.0 \| 605 \| (32, 674) \| 682 \| (36, 709) \| 716 \| Mu-like prophage major head subunit gpT \| Mu-like prophage major head subunit gpT; uniclust \| UniRef100\_A0A0F9KYA8 \| 99.9 \| 5.8e-28 \| 1.2e-33 \| 261.8 \| 521 \| (32, 591) \| 682 \| (75, 674) \| 788 \| Bacteriophage Mu GpT domain-containing protein \| Bacteriophage Mu GpT domain-containing protein; uniclust \| UniRef100\_A0A0K6HVN0 \| 99.9 \| 6.2e-28 \| 1.2e-33 \| 251.7 \| 268 \| (393, 680) \| 682 \| (158, 454) \| 458 \| Phage major capsid protein, HK97 family \| Phage major capsid protein, HK97 family; uniclust \| UniRef100\_A0A0G1G7G4 \| 99.9 \| 7.6e-28 \| 1.5e-33 \| 248.3 \| 267 \| (405, 680) \| 682 \| (296, 590) \| 605 \| Phage capsid family protein \| Phage capsid family protein; uniclust \| UniRef100\_UPI000B0837BE \| 99.9 \| 9.9e-28 \| 1.9e-33 \| 234.7 \| 436 \| (43, 514) \| 682 \| (21, 502) \| 519 \| HK97 family phage prohead protease \| HK97 family phage prohead protease; uniclust \| UniRef100\_A0A8T6P0T0 \| 99.9 \| 1e-27 \| 1.9e-33 \| 230.7 \| 274 \| (393, 681) \| 682 \| (240, 536) \| 538 \| Phage major capsid protein \| Phage major capsid protein; uniclust \| UniRef100\_A0A084E6F3 \| 99.9 \| 1e-27 \| 2e-33 \| 250.0 \| 314 \| (340, 681) \| 682 \| (151, 493) \| 549 \| Phage major capsid protein, HK97 family \| Phage major capsid protein, HK97 family; uniclust \| UniRef100\_A0A7W2BLG8 \| 99.9 \| 1.2e-27 \| 2.3e-33 \| 224.4 \| 271 \| (393, 681) \| 682 \| (15, 294) \| 298 \| Phage major capsid protein \| Phage major capsid protein; uniclust \| UniRef100\_A0A081BFN2 \| 99.9 \| 1.3e-27 \| 2.5e-33 \| 232.9 \| 268 \| (31, 307) \| 682 \| (31, 326) \| 328 \| Peptidase U35 phage prohead HK97 (Fragment) \| Peptidase U35 phage prohead HK97 (Fragment); uniclust \| UniRef100\_A0A132HLC0 \| 99.9 \| 1.4e-27 \| 2.6e-33 \| 242.2 \| 154 \| (31, 185) \| 682 \| (28, 184) \| 531 \| Phage major capsid protein \| Phage major capsid protein; uniclust \| UniRef100\_A0A7X8B6T2 \| 99.9 \| 1.4e-27 \| 2.8e-33 \| 250.0 \| 213 \| (455, 679) \| 682 \| (365, 583) \| 595 \| HK97 family phage prohead protease \| HK97 family phage prohead protease; uniclust \| UniRef100\_A0A0F3MPU1 \| 99.9 \| 1.5e-27 \| 2.9e-33 \| 240.5 \| 261 \| (406, 680) \| 682 \| (89, 369) \| 375 \| Phage major capsid protein, HK97 family \| Phage major capsid protein, HK97 family; uniclust \| UniRef100\_A0A399XUC0 \| 99.9 \| 1.7e-27 \| 3.2e-33 \| 234.6 \| 267 \| (407, 681) \| 682 \| (194, 480) \| 484 \| Phage major capsid protein \| Phage major capsid protein; uniclust \| UniRef100\_A0A0G1A6G8 \| 99.9 \| 1.8e-27 \| 3.3e-33 \| 229.4 \| 272 \| (391, 681) \| 682 \| (119, 401) \| 402 \| Phage major capsid protein, HK97 family \| Phage major capsid protein, HK97 family; uniclust \| UniRef100\_A0A081NWP6 \| 99.9 \| 2e-27 \| 3.9e-33 \| 248.4 \| 255 \| (406, 673) \| 682 \| (179, 457) \| 482 \| Capsid protein \| Capsid protein; uniclust \| UniRef100\_A0A662S3E0 \| 99.9 \| 2.4e-27 \| 4.5e-33 \| 232.8 \| 269 \| (406, 681) \| 682 \| (364, 654) \| 658 \| Phage major capsid protein (Fragment) \| Phage major capsid protein (Fragment); uniclust \| UniRef100\_A0A1I7EFL9 \| 99.9 \| 3.1e-27 \| 6e-33 \| 240.8 \| 468 \| (34, 677) \| 682 \| (23, 514) \| 520 \| Phage prohead protease, HK97 family \| Phage prohead protease, HK97 family; uniclust \| UniRef100\_A0A0J1FQE0 \| 99.9 \| 3.2e-27 \| 6.4e-33 \| 243.5 \| 269 \| (393, 681) \| 682 \| (141, 420) \| 431 \| Phage capsid family protein \| Phage capsid family protein; uniclust \| UniRef100\_A0A7W8FWR1 \| 99.9 \| 3.6e-27 \| 6.7e-33 \| 229.5 \| 469 \| (31, 679) \| 682 \| (6, 491) \| 495 \| HK97 family phage major capsid protein \| HK97 family phage major capsid protein; uniclust \| UniRef100\_A0A6A4R9P0 \| 99.9 \| 4.5e-27 \| 8.2e-33 \| 216.2 \| 342 \| (263, 617) \| 682 \| (3, 356) \| 358 \| Phage major capsid protein (Fragment) \| Phage major capsid protein (Fragment); uniclust \| UniRef100\_A0A1C5ZD29 \| 99.9 \| 5.1e-27 \| 9.7e-33 \| 237.2 \| 275 \| (393, 682) \| 682 \| (230, 534) \| 537 \| Predicted phage phi-C31 gp36 major capsid-like protein \| Predicted phage phi-C31 gp36 major capsid-like protein; uniclust \| UniRef100\_A0A142X159 \| 99.9 \| 6.4e-27 \| 1.3e-32 \| 244.3 \| 277 \| (404, 681) \| 682 \| (209, 512) \| 519 \| Phage capsid family protein \| Phage capsid family protein; uniclust \| UniRef100\_A0A0C7LAF9 \| 99.9 \| 7e-27 \| 1.3e-32 \| 230.6 \| 480 \| (41, 679) \| 682 \| (21, 510) \| 515 \| HK97 family phage prohead protease \| HK97 family phage prohead protease; uniclust \| UniRef100\_A0A2D6PAP5 \| 99.9 \| 7.7e-27 \| 1.4e-32 \| 197.1 \| 154 \| (31, 184) \| 682 \| (14, 169) \| 173 \| HK97 family phage prohead protease (Fragment) \| HK97 family phage prohead protease (Fragment); uniclust \| UniRef100\_A0A223I0W6 \| 99.9 \| 7.8e-27 \| 1.4e-32 \| 224.4 \| 263 \| (406, 679) \| 682 \| (120, 396) \| 529 \| Phage major capsid protein, HK97 family \| Phage major capsid protein, HK97 family; uniclust \| UniRef100\_A0A160T8V4 \| 99.9 \| 1.1e-26 \| 2.1e-32 \| 238.8 \| 254 \| (406, 671) \| 682 \| (348, 627) \| 650 \| Uncharacterized protein \| Uncharacterized protein; uniclust \| UniRef100\_A0A1G7TNP2 \| 99.9 \| 1.4e-26 \| 2.7e-32 \| 220.0 \| 262 \| (406, 680) \| 682 \| (10, 297) \| 307 \| Phage major capsid protein, HK97 family \| Phage major capsid protein, HK97 family; uniclust \| UniRef100\_A0A100Y4B7 \| 99.9 \| 1.5e-26 \| 2.9e-32 \| 232.0 \| 272 \| (392, 680) \| 682 \| (140, 422) \| 425 \| Capsid protein \| Capsid protein; uniclust \| UniRef100\_A0A6J5KKY4 \| 99.9 \| 1.8e-26 \| 3.6e-32 \| 243.5 \| 478 \| (34, 679) \| 682 \| (114, 616) \| 620 \| Phage capsid \| Phage capsid; uniclust \| UniRef100\_A0A060AEU4 \| 99.9 \| 2e-26 \| 3.9e-32 \| 230.8 \| 261 \| (408, 680) \| 682 \| (102, 375) \| 382 \| Capsid protein \| Capsid protein; uniclust \| UniRef100\_A0A2W4JAL7 \| 99.9 \| 2e-26 \| 3.9e-32 \| 219.0 \| 246 \| (429, 681) \| 682 \| (38, 310) \| 318 \| Phage major capsid protein \| Phage major capsid protein; uniclust \| UniRef100\_A0A1C5WLB0 \| 99.9 \| 2.4e-26 \| 4.7e-32 \| 207.1 \| 142 \| (42, 184) \| 682 \| (45, 192) \| 221 \| Phage prohead protease, HK97 family \| Phage prohead protease, HK97 family; uniclust \| UniRef100\_A0A0C1YAL0 \| 99.9 \| 2.7e-26 \| 5.2e-32 \| 226.1 \| 263 \| (406, 677) \| 682 \| (116, 399) \| 405 \| Capsid protein \| Capsid protein; uniclust \| UniRef100\_A0A090E9R7 \| 99.9 \| 3e-26 \| 5.7e-32 \| 228.8 \| 269 \| (406, 682) \| 682 \| (142, 438) \| 442 \| Putative Phage major capsid protein, HK97 family \| Putative Phage major capsid protein, HK97 family; uniclust \| UniRef100\_A0A075KJX9 \| 99.9 \| 3.4e-26 \| 6.8e-32 \| 236.2 \| 258 \| (408, 680) \| 682 \| (147, 419) \| 459 \| Putative major head protein \| Putative major head protein; uniclust \| UniRef100\_A0A0F9BLA7 \| 99.9 \| 3.7e-26 \| 7.4e-32 \| 241.1 \| 264 \| (393, 671) \| 682 \| (201, 487) \| 505 \| Phage major capsid protein \| Phage major capsid protein; uniclust \| UniRef100\_A0A1S9JNN0 \| 99.9 \| 4.1e-26 \| 7.9e-32 \| 228.0 \| 131 \| (36, 181) \| 682 \| (25, 164) \| 454 \| Phage major capsid protein (Fragment) \| Phage major capsid protein (Fragment); uniclust \| UniRef100\_A0A009Z9E2 \| 99.9 \| 4.1e-26 \| 8.2e-32 \| 238.1 \| 267 \| (405, 681) \| 682 \| (225, 515) \| 528 \| Phage major capsid protein, HK97 family \| Phage major capsid protein, HK97 family; uniclust \| UniRef100\_A0A3B9G3E3 \| 99.9 \| 7.6e-26 \| 1.4e-31 \| 228.5 \| 522 \| (44, 681) \| 682 \| (19, 561) \| 563 \| Phage major capsid protein (Fragment) \| Phage major capsid protein (Fragment); uniclust \| UniRef100\_A0A0B4CLV4 \| 99.9 \| 7.5e-26 \| 1.5e-31 \| 201.1 \| 184 \| (494, 681) \| 682 \| (1, 187) \| 190 \| Uncharacterized protein \| Uncharacterized protein; uniclust \| UniRef100\_UPI00047CD657 \| 99.9 \| 8.4e-26 \| 1.6e-31 \| 223.8 \| 517 \| (38, 681) \| 682 \| (12, 565) \| 569 \| phage major capsid protein \| phage major capsid protein; uniclust \| UniRef100\_A0A031I873 \| 99.9 \| 7.6e-26 \| 1.6e-31 \| 244.1 \| 266 \| (406, 680) \| 682 \| (191, 482) \| 500 \| Phage major capsid protein, HK97 family \| Phage major capsid protein, HK97 family; uniclust \| UniRef100\_A0A0B8PLS9 \| 99.9 \| 8.9e-26 \| 1.6e-31 \| 207.7 \| 279 \| (260, 568) \| 682 \| (70, 352) \| 356 \| Phage major capsid protein \| Phage major capsid protein; uniclust \| UniRef100\_A0A1A9A792 \| 99.9 \| 9.6e-26 \| 1.8e-31 \| 211.3 \| 269 \| (406, 681) \| 682 \| (18, 309) \| 312 \| Phage major capsid protein, HK97 family \| Phage major capsid protein, HK97 family; uniclust \| UniRef100\_UPI0022AE545A \| 99.9 \| 1.2e-25 \| 2.1e-31 \| 221.2 \| 578 \| (38, 677) \| 682 \| (29, 648) \| 653 \| HK97 family phage prohead protease \| HK97 family phage prohead protease; uniclust \| UniRef100\_A0A069APM8 \| 99.9 \| 1.2e-25 \| 2.3e-31 \| 230.2 \| 259 \| (409, 681) \| 682 \| (151, 410) \| 435 \| Putative phage capsid protein \| Putative phage capsid protein; uniclust \| UniRef100\_A0A0R0CNK4 \| 99.9 \| 2.1e-25 \| 4e-31 \| 224.8 \| 272 \| (393, 682) \| 682 \| (192, 479) \| 485 \| Capsid protein \| Capsid protein; uniclust \| UniRef100\_A0A0F9IZX0 \| 99.9 \| 2.1e-25 \| 4e-31 \| 218.7 \| 264 \| (406, 680) \| 682 \| (153, 437) \| 441 \| Phage major capsid protein \| Phage major capsid protein; uniclust \| UniRef100\_A0A023D045 \| 99.9 \| 2.1e-25 \| 4e-31 \| 230.0 \| 424 \| (44, 679) \| 682 \| (56, 511) \| 533 \| Uncharacterized protein \| Uncharacterized protein; uniclust \| UniRef100\_A0A099YFD8 \| 99.9 \| 2.1e-25 \| 4.3e-31 \| 242.9 \| 543 \| (44, 680) \| 682 \| (57, 627) \| 655 \| Capsid protein \| Capsid protein; uniclust \| UniRef100\_A0A316RJS4 \| 99.9 \| 2.5e-25 \| 4.7e-31 \| 208.5 \| 258 \| (406, 678) \| 682 \| (34, 293) \| 303 \| Phage major capsid protein \| Phage major capsid protein; uniclust \| UniRef100\_A0A4R0QZ35 \| 99.9 \| 3.2e-25 \| 6.1e-31 \| 212.6 \| 281 \| (381, 677) \| 682 \| (50, 335) \| 338 \| Phage major capsid protein \| Phage major capsid protein; uniclust \| UniRef100\_A0A832GWI7 \| 99.9 \| 3.5e-25 \| 6.5e-31 \| 210.3 \| 431 \| (34, 640) \| 682 \| (9, 459) \| 461 \| Phage major capsid protein (Fragment) \| Phage major capsid protein (Fragment); uniclust \| UniRef100\_A0A140E9M0 \| 99.9 \| 3.4e-25 \| 6.5e-31 \| 228.9 \| 530 \| (38, 680) \| 682 \| (19, 580) \| 584 \| Bacteriophage Mu GpT domain-containing protein \| Bacteriophage Mu GpT domain-containing protein; uniclust \| UniRef100\_A0A2D0K5M3 \| 99.9 \| 3.7e-25 \| 7e-31 \| 212.7 \| 266 \| (406, 680) \| 682 \| (38, 317) \| 324 \| Phage capsid protein \| Phage capsid protein; uniclust \| UniRef100\_A0A0D0Q3B7 \| 99.9 \| 3.8e-25 \| 7.2e-31 \| 223.9 \| 264 \| (406, 677) \| 682 \| (208, 496) \| 517 \| Phage major capsid protein \| Phage major capsid protein; uniclust \| UniRef100\_A0A0D7P7D9 \| 99.9 \| 4.1e-25 \| 7.9e-31 \| 207.7 \| 145 \| (38, 185) \| 682 \| (44, 205) \| 256 \| Peptidase U35 \| Peptidase U35; uniclust \| UniRef100\_C6M4F3 \| 99.9 \| 4.4e-25 \| 8e-31 \| 194.1 \| 148 \| (38, 185) \| 682 \| (23, 171) \| 251 \| Caudovirus prohead protease \| Caudovirus prohead protease; uniclust \| UniRef100\_A0A014CAX5 \| 99.9 \| 4.5e-25 \| 9.1e-31 \| 221.0 \| 170 \| (2, 181) \| 682 \| (33, 210) \| 319 \| Phage prohead protease, HK97 family \| Phage prohead protease, HK97 family; uniclust \| UniRef100\_A0A011UU65 \| 99.8 \| 5.2e-25 \| 1.1e-30 \| 231.9 \| 262 \| (407, 679) \| 682 \| (155, 426) \| 454 \| Capsid protein \| Capsid protein; uniclust \| UniRef100\_A0A940E8X0 \| 99.8 \| 8.8e-25 \| 1.6e-30 \| 207.0 \| 255 \| (422, 681) \| 682 \| (172, 446) \| 449 \| Phage major capsid protein \| Phage major capsid protein; uniclust \| UniRef100\_A0A087M280 \| 99.8 \| 9.6e-25 \| 1.9e-30 \| 202.2 \| 152 \| (31, 184) \| 682 \| (18, 182) \| 242 \| Peptidase U35 \| Peptidase U35; uniclust \| UniRef100\_A0A016QMV8 \| 99.8 \| 9.2e-25 \| 1.9e-30 \| 233.6 \| 262 \| (407, 679) \| 682 \| (209, 480) \| 505 \| Putative phage phi-C31 gp36 major capsid-like protein \| Putative phage phi-C31 gp36 major capsid-like protein; uniclust \| UniRef100\_A0A447LNI1 \| 99.8 \| 1.1e-24 \| 2.2e-30 \| 192.8 \| 147 \| (33, 185) \| 682 \| (13, 166) \| 199 \| Phage prohead protease, HK97 family \| Phage prohead protease, HK97 family; uniclust \| UniRef100\_A0A1Z8N5M0 \| 99.8 \| 1.2e-24 \| 2.3e-30 \| 209.5 \| 263 \| (409, 680) \| 682 \| (111, 388) \| 391 \| Phage major capsid protein \| Phage major capsid protein; uniclust \| UniRef100\_A0A7Y9W360 \| 99.8 \| 1.2e-24 \| 2.3e-30 \| 210.8 \| 345 \| (32, 398) \| 682 \| (5, 377) \| 454 \| HK97 family phage prohead protease \| HK97 family phage prohead protease; uniclust \| UniRef100\_A0A081HUR9 \| 99.8 \| 1.4e-24 \| 2.7e-30 \| 220.2 \| 268 \| (393, 680) \| 682 \| (158, 454) \| 457 \| HK97 family phage major capsid protein \| HK97 family phage major capsid protein; uniclust \| UniRef100\_UPI000A547A50 \| 99.8 \| 1.6e-24 \| 3e-30 \| 199.8 \| 302 \| (31, 371) \| 682 \| (17, 320) \| 334 \| HK97 family phage prohead protease \| HK97 family phage prohead protease; uniclust \| UniRef100\_A0A4V1UI61 \| 99.8 \| 1.6e-24 \| 3.1e-30 \| 209.1 \| 196 \| (339, 546) \| 682 \| (81, 283) \| 322 \| Phage major capsid protein (Fragment) \| Phage major capsid protein (Fragment); uniclust \| UniRef100\_A0A0H5RW03 \| 99.8 \| 1.8e-24 \| 3.6e-30 \| 210.1 \| 264 \| (408, 681) \| 682 \| (31, 298) \| 347 \| Phage major capsid protein, HK97 \| Phage major capsid protein, HK97; uniclust \| UniRef100\_A0A2N6UZ54 \| 99.8 \| 1.9e-24 \| 3.7e-30 \| 212.7 \| 202 \| (388, 607) \| 682 \| (112, 316) \| 323 \| Phage major capsid protein (Fragment) \| Phage major capsid protein (Fragment); uniclust \| UniRef100\_A0A1C5JTV2 \| 99.8 \| 2e-24 \| 3.8e-30 \| 219.1 \| 270 \| (390, 681) \| 682 \| (202, 502) \| 506 \| Phage major capsid protein, HK97 family \| Phage major capsid protein, HK97 family; uniclust \| UniRef100\_UPI0006A401C8 \| 99.8 \| 2.2e-24 \| 4e-30 \| 207.8 \| 487 \| (31, 679) \| 682 \| (5, 517) \| 521 \| phage major capsid protein \| phage major capsid protein; uniclust \| UniRef100\_A0A4Q5ADZ7 \| 99.8 \| 2.3e-24 \| 4.3e-30 \| 221.5 \| 243 \| (428, 680) \| 682 \| (424, 684) \| 694 \| Phage capsid protein \| Phage capsid protein; uniclust \| UniRef100\_A0A0D5LSQ3 \| 99.8 \| 2.2e-24 \| 4.4e-30 \| 212.7 \| 134 \| (38, 181) \| 682 \| (55, 192) \| 299 \| Peptidase U35 \| Peptidase U35; uniclust \| UniRef100\_A0A929NN34 \| 99.8 \| 2.5e-24 \| 4.5e-30 \| 200.4 \| 323 \| (342, 679) \| 682 \| (53, 381) \| 383 \| Phage major capsid protein \| Phage major capsid protein; uniclust \| UniRef100\_Q1QI78 \| 99.8 \| 2.5e-24 \| 4.6e-30 \| 222.6 \| 571 \| (48, 668) \| 682 \| (553, 1166) \| 1179 \| Phage terminase GpA \| Phage terminase GpA; uniclust \| UniRef100\_A0A263E3D0 \| 99.8 \| 3e-24 \| 5.7e-30 \| 210.0 \| 275 \| (393, 681) \| 682 \| (115, 413) \| 417 \| Phage major capsid protein \| Phage major capsid protein; uniclust \| UniRef100\_A0A1D3SNN3 \| 99.8 \| 3.8e-24 \| 7.4e-30 \| 215.9 \| 265 \| (406, 680) \| 682 \| (126, 406) \| 443 \| Phage major capsid protein \| Phage major capsid protein; uniclust \| UniRef100\_A0A1A1W0A4 \| 99.8 \| 4.6e-24 \| 8.8e-30 \| 204.2 \| 265 \| (407, 680) \| 682 \| (75, 347) \| 352 \| Phage capsid protein \| Phage capsid protein; uniclust \| UniRef100\_A0A0F9AWL4 \| 99.8 \| 4.4e-24 \| 8.8e-30 \| 216.3 \| 259 \| (406, 675) \| 682 \| (49, 330) \| 350 \| Phage major capsid protein (Fragment) \| Phage major capsid protein (Fragment); uniclust \| UniRef100\_A0A0F3N2G0 \| 99.8 \| 5.8e-24 \| 1.1e-29 \| 208.8 \| 260 \| (407, 680) \| 682 \| (84, 358) \| 365 \| Phage major capsid protein, HK97 family \| Phage major capsid protein, HK97 family; uniclust \| UniRef100\_A0A014FFI8 \| 99.8 \| 6.1e-24 \| 1.2e-29 \| 208.5 \| 263 \| (406, 679) \| 682 \| (55, 327) \| 339 \| Phage major capsid protein, HK97 family \| Phage major capsid protein, HK97 family; uniclust \| UniRef100\_A0A249MU97 \| 99.8 \| 6.3e-24 \| 1.2e-29 \| 210.1 \| 264 \| (406, 680) \| 682 \| (130, 422) \| 426 \| Phage major capsid protein \| Phage major capsid protein; uniclust \| UniRef100\_A0A3D1WNS7 \| 99.8 \| 6.9e-24 \| 1.3e-29 \| 201.8 \| 191 \| (406, 607) \| 682 \| (120, 311) \| 323 \| Phage major capsid protein (Fragment) \| Phage major capsid protein (Fragment); uniclust \| UniRef100\_A0A1V2NG87 \| 99.8 \| 7.2e-24 \| 1.4e-29 \| 206.2 \| 264 \| (393, 680) \| 682 \| (115, 392) \| 395 \| Capsid protein \| Capsid protein; uniclust \| UniRef100\_A0A011T3L4 \| 99.8 \| 7.4e-24 \| 1.4e-29 \| 197.2 \| 137 \| (31, 181) \| 682 \| (16, 162) \| 226 \| Peptidase U35 \| Peptidase U35; uniclust \| UniRef100\_A0A2D5HBK7 \| 99.8 \| 7.7e-24 \| 1.5e-29 \| 204.3 \| 258 \| (410, 680) \| 682 \| (2, 287) \| 292 \| Phage major capsid protein (Fragment) \| Phage major capsid protein (Fragment); uniclust \| UniRef100\_A0A1Y0Y2T8 \| 99.8 \| 8e-24 \| 1.6e-29 \| 218.1 \| 261 \| (410, 680) \| 682 \| (152, 442) \| 487 \| Phage major capsid protein \| Phage major capsid protein; uniclust \| UniRef100\_A0A011PIW6 \| 99.8 \| 1.2e-23 \| 2.3e-29 \| 204.7 \| 219 \| (454, 680) \| 682 \| (57, 291) \| 306 \| Phage major capsid protein, HK97 family \| Phage major capsid protein, HK97 family; uniclust \| UniRef100\_A0A061LX04 \| 99.8 \| 1.4e-23 \| 2.8e-29 \| 213.0 \| 262 \| (408, 680) \| 682 \| (143, 423) \| 444 \| Capsid protein \| Capsid protein; uniclust \| UniRef100\_A0A6J5RYQ5 \| 99.8 \| 1.7e-23 \| 3e-29 \| 199.0 \| 268 \| (406, 681) \| 682 \| (158, 450) \| 455 \| COG4653 Predicted phage phi-C31 gp36 major capsid-like protein \| COG4653 Predicted phage phi-C31 gp36 major capsid-like protein; uniclust \| UniRef100\_A0A1B0XUP8 \| 99.8 \| 1.6e-23 \| 3.1e-29 \| 217.3 \| 471 \| (43, 678) \| 682 \| (21, 515) \| 534 \| Capsid protein with prohead protease \| Capsid protein with prohead protease; uniclust \| UniRef100\_A0A3D9ZDR9 \| 99.8 \| 1.7e-23 \| 3.2e-29 \| 196.0 \| 151 \| (31, 184) \| 682 \| (7, 170) \| 273 \| HK97 family phage prohead protease \| HK97 family phage prohead protease; uniclust \| UniRef100\_A0A017RTY2 \| 99.8 \| 1.8e-23 \| 3.5e-29 \| 219.2 \| 268 \| (404, 680) \| 682 \| (194, 489) \| 504 \| Capsid protein \| Capsid protein; uniclust \| UniRef100\_A0A088FP12 \| 99.8 \| 1.8e-23 \| 3.6e-29 \| 218.7 \| 272 \| (392, 680) \| 682 \| (223, 530) \| 547 \| Major capsid protein \| Major capsid protein; uniclust \| UniRef100\_A0A0F9F1M1 \| 99.8 \| 1.9e-23 \| 3.8e-29 \| 215.3 \| 259 \| (413, 681) \| 682 \| (157, 424) \| 427 \| Phage major capsid protein \| Phage major capsid protein; uniclust \| UniRef100\_A0A1V5LY97 \| 99.8 \| 3.2e-23 \| 6.1e-29 \| 183.9 \| 171 \| (500, 680) \| 682 \| (2, 186) \| 194 \| Phage capsid family protein \| Phage capsid family protein; uniclust \| UniRef100\_A0A7D5FN44 \| 99.8 \| 3.3e-23 \| 6.2e-29 \| 194.1 \| 268 \| (406, 681) \| 682 \| (38, 334) \| 336 \| Major capsid protein (Fragment) \| Major capsid protein (Fragment); uniclust \| UniRef100\_A0A090IID5 \| 99.8 \| 3.1e-23 \| 6.3e-29 \| 202.1 \| 132 \| (41, 181) \| 682 \| (36, 173) \| 263 \| Phage prohead protease \| Phage prohead protease; uniclust \| UniRef100\_A0A017H821 \| 99.8 \| 3.4e-23 \| 6.7e-29 \| 189.7 \| 129 \| (42, 183) \| 682 \| (24, 154) \| 190 \| Peptidase U35, phage prohead HK97 \| Peptidase U35, phage prohead HK97; uniclust \| UniRef100\_A0A087DI38 \| 99.8 \| 3.9e-23 \| 7.3e-29 \| 212.1 \| 253 \| (413, 679) \| 682 \| (410, 673) \| 686 \| Capsid protein \| Capsid protein; uniclust \| UniRef100\_A0A011PJT1 \| 99.8 \| 3.8e-23 \| 7.6e-29 \| 215.9 \| 262 \| (406, 677) \| 682 \| (184, 468) \| 486 \| Phage major capsid protein, HK97 family \| Phage major capsid protein, HK97 family; uniclust \| UniRef100\_A0A015WYB3 \| 99.8 \| 4e-23 \| 7.9e-29 \| 211.1 \| 264 \| (406, 680) \| 682 \| (122, 388) \| 398 \| Phage major capsid protein, HK97 family \| Phage major capsid protein, HK97 family; uniclust \| UniRef100\_A0A1V5PUE1 \| 99.8 \| 4.2e-23 \| 8e-29 \| 205.0 \| 260 \| (393, 671) \| 682 \| (134, 416) \| 441 \| Phage capsid family protein \| Phage capsid family protein; uniclust \| UniRef100\_A0A2D7QM53 \| 99.8 \| 4.3e-23 \| 8.4e-29 \| 205.3 \| 315 \| (338, 680) \| 682 \| (84, 405) \| 408 \| Phage major capsid protein \| Phage major capsid protein; uniclust \| UniRef100\_A0A661TNK1 \| 99.8 \| 4.6e-23 \| 8.5e-29 \| 195.2 \| 397 \| (39, 508) \| 682 \| (21, 434) \| 437 \| HK97 family phage prohead protease (Fragment) \| HK97 family phage prohead protease (Fragment); uniclust \| UniRef100\_A0A0F9JCM6 \| 99.8 \| 5e-23 \| 9.9e-29 \| 218.0 \| 468 \| (45, 564) \| 682 \| (51, 555) \| 592 \| Bacteriophage Mu GpT domain-containing protein (Fragment) \| Bacteriophage Mu GpT domain-containing protein (Fragment); uniclust \| UniRef100\_A0A022MNJ2 \| 99.8 \| 4.9e-23 \| 9.9e-29 \| 215.1 \| 261 \| (407, 680) \| 682 \| (153, 431) \| 446 \| Capsid protein \| Capsid protein; uniclust \| UniRef100\_A0A938T756 \| 99.8 \| 6.1e-23 \| 1.1e-28 \| 200.6 \| 526 \| (41, 679) \| 682 \| (16, 582) \| 586 \| Phage major capsid protein \| Phage major capsid protein; uniclust \| UniRef100\_A0A229GM47 \| 99.8 \| 6.5e-23 \| 1.2e-28 \| 194.9 \| 130 \| (38, 181) \| 682 \| (38, 175) \| 295 \| Peptidase U35 \| Peptidase U35; uniclust \| UniRef100\_A0A143PPF0 \| 99.8 \| 6.7e-23 \| 1.3e-28 \| 196.7 \| 262 \| (406, 679) \| 682 \| (119, 387) \| 411 \| Phage major capsid protein, HK97 family \| Phage major capsid protein, HK97 family; uniclust \| UniRef100\_A0A7W0TDA0 \| 99.8 \| 8.1e-23 \| 1.5e-28 \| 196.5 \| 269 \| (393, 679) \| 682 \| (131, 419) \| 423 \| Phage major capsid protein \| Phage major capsid protein; uniclust \| UniRef100\_A0A011MGC8 \| 99.8 \| 7.9e-23 \| 1.6e-28 \| 213.9 \| 263 \| (406, 679) \| 682 \| (188, 460) \| 492 \| Capsid protein \| Capsid protein; uniclust \| UniRef100\_UPI000DA43CDE \| 99.8 \| 8.7e-23 \| 1.6e-28 \| 196.9 \| 375 \| (269, 680) \| 682 \| (104, 489) \| 492 \| phage major capsid protein \| phage major capsid protein; uniclust \| UniRef100\_A0A916TP50 \| 99.8 \| 9.5e-23 \| 1.7e-28 \| 193.5 \| 311 \| (38, 437) \| 682 \| (90, 419) \| 444 \| Uncharacterized protein \| Uncharacterized protein; uniclust \| UniRef100\_A0A068RC46 \| 99.8 \| 9e-23 \| 1.8e-28 \| 201.5 \| 131 \| (42, 181) \| 682 \| (76, 213) \| 291 \| Caudovirus prohead protease family protein \| Caudovirus prohead protease family protein; uniclust \| UniRef100\_A0A8S5NH81 \| 99.8 \| 1e-22 \| 1.9e-28 \| 179.7 \| 153 \| (32, 185) \| 682 \| (14, 174) \| 251 \| Major capsid protein (Fragment) \| Major capsid protein (Fragment); uniclust \| UniRef100\_A0A161IDV2 \| 99.8 \| 1e-22 \| 2.1e-28 \| 196.4 \| 136 \| (36, 181) \| 682 \| (34, 173) \| 260 \| Caudovirus prohead protease \| Caudovirus prohead protease; uniclust \| UniRef100\_A0A059IUN6 \| 99.8 \| 1.5e-22 \| 2.9e-28 \| 205.0 \| 262 \| (406, 680) \| 682 \| (149, 432) \| 515 \| HK97 family phage major capsid protein \| HK97 family phage major capsid protein; uniclust \| UniRef100\_A0A1Q3QQ17 \| 99.8 \| 1.5e-22 \| 3e-28 \| 206.9 \| 260 \| (393, 671) \| 682 \| (107, 381) \| 396 \| Capsid protein \| Capsid protein; uniclust \| UniRef100\_A0A7W1RV33 \| 99.8 \| 1.7e-22 \| 3e-28 \| 192.8 \| 268 \| (406, 681) \| 682 \| (166, 462) \| 463 \| Phage major capsid protein \| Phage major capsid protein; uniclust \| UniRef100\_A0A2D9C9C5 \| 99.8 \| 1.7e-22 \| 3.1e-28 \| 199.0 \| 571 \| (32, 678) \| 682 \| (22, 626) \| 630 \| HK97 family phage prohead protease \| HK97 family phage prohead protease; uniclust \| UniRef100\_A0A354P6A4 \| 99.8 \| 1.8e-22 \| 3.4e-28 \| 197.0 \| 277 \| (386, 681) \| 682 \| (147, 426) \| 430 \| Phage major capsid protein \| Phage major capsid protein; uniclust \| UniRef100\_UPI00131B246A \| 99.8 \| 2e-22 \| 3.6e-28 \| 196.5 \| 382 \| (262, 681) \| 682 \| (119, 561) \| 564 \| phage major capsid protein \| phage major capsid protein; uniclust \| UniRef100\_A0A432MJM0 \| 99.8 \| 3.1e-22 \| 5.6e-28 \| 189.0 \| 311 \| (342, 678) \| 682 \| (91, 419) \| 421 \| Phage major capsid protein \| Phage major capsid protein; uniclust \| UniRef100\_UPI0006943B46 \| 99.8 \| 3.1e-22 \| 5.7e-28 \| 196.8 \| 262 \| (393, 672) \| 682 \| (326, 602) \| 613 \| phage major capsid protein \| phage major capsid protein; uniclust \| UniRef100\_A0A101UZE9 \| 99.8 \| 3e-22 \| 5.8e-28 \| 202.0 \| 264 \| (406, 681) \| 682 \| (175, 456) \| 460 \| Capsid protein \| Capsid protein; uniclust \| UniRef100\_A0A2U1BFB6 \| 99.8 \| 3.3e-22 \| 6.3e-28 \| 192.3 \| 258 \| (408, 679) \| 682 \| (105, 379) \| 383 \| HK97 family phage major capsid protein \| HK97 family phage major capsid protein; uniclust \| UniRef100\_A0A8I1GCE2 \| 99.8 \| 3.4e-22 \| 6.4e-28 \| 187.3 \| 254 \| (31, 308) \| 682 \| (15, 285) \| 331 \| Peptidase U35 \| Peptidase U35; uniclust \| UniRef100\_A0A060PVD0 \| 99.8 \| 3.3e-22 \| 6.5e-28 \| 187.3 \| 131 \| (42, 181) \| 682 \| (59, 190) \| 227 \| Phage prohead protease \| Phage prohead protease; uniclust \| UniRef100\_A0A068D9Z5 \| 99.8 \| 4e-22 \| 7.8e-28 \| 203.4 \| 256 \| (406, 671) \| 682 \| (120, 402) \| 419 \| Phage major capsid protein \| Phage major capsid protein; uniclust \| UniRef100\_A0A1M3FIN7 \| 99.8 \| 4e-22 \| 8e-28 \| 206.3 \| 271 \| (393, 680) \| 682 \| (172, 478) \| 487 \| Major capsid protein \| Major capsid protein; uniclust \| UniRef100\_A0A060QM50 \| 99.8 \| 4.1e-22 \| 8.2e-28 \| 209.1 \| 262 \| (407, 679) \| 682 \| (186, 458) \| 499 \| Phage major capsid protein \| Phage major capsid protein; uniclust \| UniRef100\_D6PFS1 \| 99.8 \| 5.3e-22 \| 9.7e-28 \| 171.3 \| 207 \| (348, 559) \| 682 \| (3, 211) \| 214 \| Putative phage related protein \| Putative phage related protein; uniclust \| UniRef100\_A0A3A9EF23 \| 99.8 \| 5.3e-22 \| 9.8e-28 \| 173.3 \| 153 \| (31, 184) \| 682 \| (17, 180) \| 223 \| Caudovirus prohead protease \| Caudovirus prohead protease; uniclust \| UniRef100\_A0A084J8V6 \| 99.8 \| 5.6e-22 \| 1.1e-27 \| 195.6 \| 260 \| (409, 680) \| 682 \| (34, 316) \| 338 \| Phage major capsid protein \| Phage major capsid protein; uniclust \| UniRef100\_A0A7X8QWG9 \| 99.8 \| 6e-22 \| 1.1e-27 \| 190.5 \| 264 \| (406, 679) \| 682 \| (113, 408) \| 410 \| Phage major capsid protein \| Phage major capsid protein; uniclust \| UniRef100\_UPI000738CC35 \| 99.8 \| 6.7e-22 \| 1.3e-27 \| 169.0 \| 144 \| (31, 180) \| 682 \| (12, 161) \| 164 \| HK97 family phage prohead protease \| HK97 family phage prohead protease; uniclust \| UniRef100\_A0A013UG14 \| 99.8 \| 6.6e-22 \| 1.3e-27 \| 183.1 \| 133 \| (36, 181) \| 682 \| (27, 166) \| 229 \| Caudovirus prohead protease family protein \| Caudovirus prohead protease family protein; uniclust \| UniRef100\_A0A117JJ31 \| 99.8 \| 6.8e-22 \| 1.3e-27 \| 186.4 \| 265 \| (406, 679) \| 682 \| (11, 275) \| 278 \| PE domain-containing protein (Fragment) \| PE domain-containing protein (Fragment); uniclust \| UniRef100\_A0A0F7R9L9 \| 99.8 \| 6.8e-22 \| 1.3e-27 \| 196.5 \| 269 \| (392, 681) \| 682 \| (96, 375) \| 390 \| Phage Capsid protein \| Phage Capsid protein; uniclust \| UniRef100\_A0A4V3IRT6 \| 99.8 \| 7.2e-22 \| 1.4e-27 \| 191.6 \| 262 \| (411, 681) \| 682 \| (112, 382) \| 384 \| Phage major capsid protein \| Phage major capsid protein; uniclust \| UniRef100\_A0A083XX19 \| 99.8 \| 7.2e-22 \| 1.5e-27 \| 211.3 \| 256 \| (415, 680) \| 682 \| (153, 426) \| 468 \| Phage capsid protein \| Phage capsid protein; uniclust \| UniRef100\_A0A081I3V9 \| 99.8 \| 7.5e-22 \| 1.5e-27 \| 200.3 \| 231 \| (438, 680) \| 682 \| (66, 304) \| 363 \| HK97 family phage major capsid protein \| HK97 family phage major capsid protein; uniclust \| UniRef100\_A0A059FG22 \| 99.8 \| 7.5e-22 \| 1.5e-27 \| 187.5 \| 131 \| (41, 181) \| 682 \| (56, 190) \| 240 \| Phage prohead protease HK97 family protein \| Phage prohead protease HK97 family protein; uniclust \| UniRef100\_A0A962YQX0 \| 99.8 \| 8.6e-22 \| 1.6e-27 \| 186.2 \| 270 \| (406, 680) \| 682 \| (148, 421) \| 423 \| Phage major capsid protein \| Phage major capsid protein; uniclust \| UniRef100\_A0A060VRW9 \| 99.8 \| 7.8e-22 \| 1.6e-27 \| 195.0 \| 174 \| (1, 181) \| 682 \| (1, 182) \| 273 \| Phage prohead protease, HK97 family \| Phage prohead protease, HK97 family; uniclust \| UniRef100\_A0A1H8L3J0 \| 99.8 \| 8.9e-22 \| 1.6e-27 \| 194.2 \| 318 \| (31, 437) \| 682 \| (23, 361) \| 632 \| Prohead serine protease \| Prohead serine protease; uniclust \| UniRef100\_A0A2A9FC94 \| 99.8 \| 9.3e-22 \| 1.8e-27 \| 200.8 \| 544 \| (32, 679) \| 682 \| (17, 587) \| 595 \| HK97 family phage prohead protease/HK97 family phage major capsid protein,TIGR01554 \| HK97 family phage prohead protease/HK97 family phage major capsid protein,TIGR01554; uniclust \| UniRef100\_A0A963KA71 \| 99.8 \| 9.5e-22 \| 1.8e-27 \| 186.2 \| 324 \| (333, 681) \| 682 \| (14, 347) \| 348 \| Phage major capsid protein \| Phage major capsid protein; uniclust \| UniRef100\_A0A3S4JUU9 \| 99.8 \| 9.8e-22 \| 1.9e-27 \| 196.3 \| 258 \| (406, 680) \| 682 \| (190, 457) \| 460 \| Predicted phage phi-C31 gp36 major capsid-like protein \| Predicted phage phi-C31 gp36 major capsid-like protein; uniclust \| UniRef100\_A0A5S4ZXS2 \| 99.8 \| 1.1e-21 \| 2e-27 \| 184.0 \| 261 \| (393, 670) \| 682 \| (21, 305) \| 328 \| HK97 family phage major capsid protein \| HK97 family phage major capsid protein; uniclust \| UniRef100\_UPI00215628C7 \| 99.8 \| 1.2e-21 \| 2.2e-27 \| 192.7 \| 529 \| (43, 678) \| 682 \| (39, 605) \| 611 \| phage major capsid protein \| phage major capsid protein; uniclust \| UniRef100\_A0A084E6F4 \| 99.8 \| 1.2e-21 \| 2.3e-27 \| 181.2 \| 133 \| (36, 181) \| 682 \| (18, 157) \| 222 \| Phage prohead protease, HK97 family \| Phage prohead protease, HK97 family; uniclust \| UniRef100\_A0A086ZY60 \| 99.8 \| 1.1e-21 \| 2.3e-27 \| 196.9 \| 231 \| (440, 680) \| 682 \| (72, 310) \| 330 \| Phage major capsid protein, HK97 family \| Phage major capsid protein, HK97 family; uniclust \| UniRef100\_A0A098AU90 \| 99.8 \| 1.2e-21 \| 2.3e-27 \| 185.0 \| 136 \| (43, 183) \| 682 \| (47, 190) \| 238 \| Phage head maturation protease \| Phage head maturation protease; uniclust \| UniRef100\_A0A1V4I2J1 \| 99.7 \| 1.4e-21 \| 2.6e-27 \| 171.0 \| 145 \| (38, 184) \| 682 \| (22, 181) \| 204 \| Peptidase U35 \| Peptidase U35; uniclust \| UniRef100\_UPI000CF32F82 \| 99.7 \| 1.4e-21 \| 2.7e-27 \| 195.7 \| 421 \| (43, 680) \| 682 \| (37, 475) \| 480 \| phage major capsid protein \| phage major capsid protein; uniclust \| UniRef100\_A0A6J5S3H4 \| 99.7 \| 1.5e-21 \| 2.8e-27 \| 189.1 \| 283 \| (386, 680) \| 682 \| (123, 425) \| 429 \| COG4653 Predicted phage phi-C31 gp36 major capsid-like protein \| COG4653 Predicted phage phi-C31 gp36 major capsid-like protein; uniclust \| UniRef100\_A0A5A7MXX4 \| 99.7 \| 1.7e-21 \| 3.1e-27 \| 196.5 \| 453 \| (32, 524) \| 682 \| (24, 509) \| 801 \| Uncharacterized protein \| Uncharacterized protein; uniclust \| UniRef100\_A0A0F9HAL3 \| 99.7 \| 1.7e-21 \| 3.2e-27 \| 185.3 \| 266 \| (406, 682) \| 682 \| (84, 369) \| 370 \| Phage major capsid protein \| Phage major capsid protein; uniclust \| UniRef100\_A0A255R0K0 \| 99.7 \| 1.9e-21 \| 3.5e-27 \| 189.5 \| 265 \| (406, 682) \| 682 \| (162, 432) \| 434 \| Phage major capsid protein \| Phage major capsid protein; uniclust \| UniRef100\_A0A074U171 \| 99.7 \| 1.8e-21 \| 3.5e-27 \| 200.6 \| 260 \| (408, 679) \| 682 \| (190, 460) \| 476 \| Nucleoid-structuring protein H-NS \| Nucleoid-structuring protein H-NS; uniclust \| UniRef100\_A0A023W7M8 \| 99.7 \| 1.9e-21 \| 3.8e-27 \| 200.7 \| 259 \| (412, 681) \| 682 \| (37, 326) \| 351 \| Major capsid protein \| Major capsid protein; uniclust \| UniRef100\_A0A8S5N8Q6 \| 99.7 \| 2.3e-21 \| 4.2e-27 \| 168.9 \| 149 \| (32, 185) \| 682 \| (25, 176) \| 226 \| Major capsid protein (Fragment) \| Major capsid protein (Fragment); uniclust \| UniRef100\_A0A0M8SR27 \| 99.7 \| 2.3e-21 \| 4.4e-27 \| 195.6 \| 269 \| (393, 679) \| 682 \| (139, 436) \| 449 \| Phage head \| Phage head; uniclust \| UniRef100\_A0A0F9WTB8 \| 99.7 \| 2.3e-21 \| 4.4e-27 \| 195.1 \| 269 \| (393, 679) \| 682 \| (145, 448) \| 451 \| HK97 family phage major capsid protein \| HK97 family phage major capsid protein; uniclust \| UniRef100\_A0A1G3DZF5 \| 99.7 \| 2.3e-21 \| 4.4e-27 \| 202.3 \| 247 \| (422, 678) \| 682 \| (249, 506) \| 591 \| Capsid protein \| Capsid protein; uniclust \| UniRef100\_A0A1C3X8S9 \| 99.7 \| 2.6e-21 \| 5.2e-27 \| 199.9 \| 261 \| (405, 681) \| 682 \| (96, 387) \| 393 \| Phage major capsid protein, HK97 family \| Phage major capsid protein, HK97 family; uniclust \| UniRef100\_A0A059N3B4 \| 99.7 \| 2.8e-21 \| 5.3e-27 \| 175.0 \| 136 \| (41, 181) \| 682 \| (28, 170) \| 204 \| Phage prohead protease, HK97 family \| Phage prohead protease, HK97 family; uniclust \| UniRef100\_A0A060LW15 \| 99.7 \| 2.9e-21 \| 5.7e-27 \| 195.4 \| 271 \| (393, 681) \| 682 \| (124, 411) \| 485 \| Phage-related protein \| Phage-related protein; uniclust \| UniRef100\_A0A1E3GWI0 \| 99.7 \| 3e-21 \| 5.7e-27 \| 186.3 \| 266 \| (406, 681) \| 682 \| (51, 338) \| 344 \| Phage capsid family protein \| Phage capsid family protein; uniclust \| UniRef100\_A0A081HUR8 \| 99.7 \| 3.2e-21 \| 6.3e-27 \| 178.8 \| 148 \| (31, 184) \| 682 \| (16, 172) \| 207 \| HK97 family phage prohead protease \| HK97 family phage prohead protease; uniclust \| UniRef100\_A0A2M9PA59 \| 99.7 \| 3.4e-21 \| 6.4e-27 \| 174.1 \| 235 \| (432, 678) \| 682 \| (12, 260) \| 263 \| Phage major capsid protein (Fragment) \| Phage major capsid protein (Fragment); uniclust \| UniRef100\_A0A023DZP0 \| 99.7 \| 3.9e-21 \| 7.7e-27 \| 180.6 \| 134 \| (38, 181) \| 682 \| (29, 167) \| 237 \| Putative prohead protease \| Putative prohead protease; uniclust \| UniRef100\_A0A0F8ZWE4 \| 99.7 \| 3.9e-21 \| 7.9e-27 \| 205.3 \| 393 \| (260, 680) \| 682 \| (132, 557) \| 561 \| Bacteriophage Mu GpT domain-containing protein (Fragment) \| Bacteriophage Mu GpT domain-containing protein (Fragment); uniclust \| UniRef100\_A0A4Q3XXJ0 \| 99.7 \| 4.6e-21 \| 8.5e-27 \| 172.3 \| 152 \| (31, 183) \| 682 \| (19, 174) \| 281 \| HK97 family phage prohead protease (Fragment) \| HK97 family phage prohead protease (Fragment); uniclust \| UniRef100\_A0A1A2T1Y5 \| 99.7 \| 4.5e-21 \| 8.8e-27 \| 196.4 \| 263 \| (409, 681) \| 682 \| (155, 430) \| 438 \| Capsid protein \| Capsid protein; uniclust \| UniRef100\_A0A2W2FHR8 \| 99.7 \| 5e-21 \| 9.6e-27 \| 191.7 \| 266 \| (406, 680) \| 682 \| (133, 433) \| 445 \| Phage major capsid protein \| Phage major capsid protein; uniclust \| UniRef100\_A0A524RVZ4 \| 99.7 \| 6.4e-21 \| 1.2e-26 \| 191.1 \| 338 \| (333, 678) \| 682 \| (385, 736) \| 737 \| Phage major capsid protein \| Phage major capsid protein; uniclust \| UniRef100\_A0A652L3X9 \| 99.7 \| 6.5e-21 \| 1.2e-26 \| 185.9 \| 269 \| (393, 678) \| 682 \| (264, 549) \| 554 \| Phage major capsid protein \| Phage major capsid protein; uniclust \| UniRef100\_A0A0D6QFX9 \| 99.7 \| 7.2e-21 \| 1.4e-26 \| 188.7 \| 253 \| (406, 670) \| 682 \| (125, 401) \| 424 \| Phage capsid family protein \| Phage capsid family protein; uniclust \| UniRef100\_A0A518EKY1 \| 99.7 \| 7.9e-21 \| 1.5e-26 \| 183.2 \| 432 \| (38, 506) \| 682 \| (22, 495) \| 497 \| Caudovirus prohead protease \| Caudovirus prohead protease; uniclust \| UniRef100\_A0A068QRW5 \| 99.7 \| 7.7e-21 \| 1.5e-26 \| 190.7 \| 267 \| (393, 679) \| 682 \| (110, 400) \| 501 \| Putative major capsid protein \| Putative major capsid protein; uniclust \| UniRef100\_A0A0H3A784 \| 99.7 \| 8.7e-21 \| 1.7e-26 \| 184.4 \| 133 \| (42, 181) \| 682 \| (60, 203) \| 287 \| Phage prohead protease, HK97 family \| Phage prohead protease, HK97 family; uniclust \| UniRef100\_A0A1I1GTH7 \| 99.7 \| 9.3e-21 \| 1.8e-26 \| 187.0 \| 260 \| (406, 679) \| 682 \| (118, 399) \| 422 \| Phage major capsid protein, HK97 family \| Phage major capsid protein, HK97 family; uniclust \| UniRef100\_A0A0J5GQW0 \| 99.7 \| 9.2e-21 \| 1.8e-26 \| 188.7 \| 260 \| (408, 681) \| 682 \| (140, 408) \| 420 \| Phage capsid protein (Fragment) \| Phage capsid protein (Fragment); uniclust \| UniRef100\_A0A1V5D5J1 \| 99.7 \| 9.6e-21 \| 1.8e-26 \| 180.6 \| 150 \| (29, 182) \| 682 \| (6, 165) \| 322 \| Caudovirus prohead protease \| Caudovirus prohead protease; uniclust \| UniRef100\_UPI001E304D27 \| 99.7 \| 1.4e-20 \| 2.5e-26 \| 171.9 \| 269 \| (31, 307) \| 682 \| (23, 297) \| 313 \| HK97 family phage prohead protease \| HK97 family phage prohead protease; uniclust \| UniRef100\_A0A0C5E3N1 \| 99.7 \| 1.5e-20 \| 2.9e-26 \| 173.3 \| 137 \| (36, 181) \| 682 \| (22, 161) \| 227 \| HK97 family phage prohead protease \| HK97 family phage prohead protease; uniclust \| UniRef100\_A0A3A8JJQ6 \| 99.7 \| 1.6e-20 \| 3.1e-26 \| 187.5 \| 265 \| (408, 682) \| 682 \| (136, 413) \| 416 \| Phage major capsid protein \| Phage major capsid protein; uniclust \| UniRef100\_A0A843H9F1 \| 99.7 \| 1.8e-20 \| 3.3e-26 \| 173.6 \| 268 \| (406, 680) \| 682 \| (70, 349) \| 350 \| Phage major capsid protein \| Phage major capsid protein; uniclust \| UniRef100\_A0A0F9MYQ3 \| 99.7 \| 2e-20 \| 3.7e-26 \| 186.9 \| 589 \| (41, 676) \| 682 \| (33, 692) \| 702 \| Phage major capsid protein \| Phage major capsid protein; uniclust \| UniRef100\_A0A640YC84 \| 99.7 \| 2.1e-20 \| 3.8e-26 \| 180.6 \| 473 \| (31, 680) \| 682 \| (13, 500) \| 502 \| HK97 family phage prohead protease \| HK97 family phage prohead protease; uniclust \| UniRef100\_A0A150IJM8 \| 99.7 \| 2.1e-20 \| 3.9e-26 \| 177.6 \| 258 \| (412, 680) \| 682 \| (71, 342) \| 345 \| Phage capsid family protein \| Phage capsid family protein; uniclust \| UniRef100\_A0A212I573 \| 99.7 \| 2.3e-20 \| 4.3e-26 \| 175.7 \| 249 \| (421, 681) \| 682 \| (14, 295) \| 305 \| Structural protein \| Structural protein; uniclust \| UniRef100\_UPI001FCE965E \| 99.7 \| 2.5e-20 \| 4.6e-26 \| 172.3 \| 281 \| (385, 679) \| 682 \| (44, 328) \| 331 \| phage major capsid protein \| phage major capsid protein; uniclust \| UniRef100\_A0A259YXH9 \| 99.7 \| 2.7e-20 \| 5.1e-26 \| 185.7 \| 468 \| (41, 681) \| 682 \| (12, 514) \| 515 \| Phage major capsid protein \| Phage major capsid protein; uniclust \| UniRef100\_A0A1V5LQQ8 \| 99.7 \| 2.8e-20 \| 5.2e-26 \| 168.7 \| 262 \| (406, 680) \| 682 \| (10, 295) \| 297 \| Phage capsid family protein \| Phage capsid family protein; uniclust \| UniRef100\_A0A011UW27 \| 99.7 \| 2.7e-20 \| 5.4e-26 \| 183.5 \| 129 \| (44, 181) \| 682 \| (80, 212) \| 293 \| Peptidase U35 \| Peptidase U35; uniclust \| UniRef100\_UPI0009B17BBC \| 99.7 \| 3.4e-20 \| 6.4e-26 \| 177.3 \| 146 \| (31, 184) \| 682 \| (8, 159) \| 355 \| HK97 family phage prohead protease \| HK97 family phage prohead protease; uniclust \| UniRef100\_A0A7Z2S8T5 \| 99.7 \| 3.5e-20 \| 6.5e-26 \| 163.2 \| 151 \| (32, 185) \| 682 \| (23, 177) \| 240 \| Caudovirus prohead protease \| Caudovirus prohead protease; uniclust \| UniRef100\_A0A084J9C1 \| 99.7 \| 3.3e-20 \| 6.5e-26 \| 191.2 \| 259 \| (406, 680) \| 682 \| (144, 421) \| 434 \| Capsid protein \| Capsid protein; uniclust \| UniRef100\_A0A0Q4Y1E9 \| 99.7 \| 3.5e-20 \| 6.6e-26 \| 168.4 \| 128 \| (41, 181) \| 682 \| (28, 162) \| 218 \| Peptidase U35 \| Peptidase U35; uniclust \| UniRef100\_F5YBI3 \| 99.7 \| 3.6e-20 \| 6.7e-26 \| 168.6 \| 253 \| (412, 678) \| 682 \| (33, 302) \| 304 \| Phage capsid family \| Phage capsid family; uniclust \| UniRef100\_A0A285D3S6 \| 99.7 \| 3.6e-20 \| 6.7e-26 \| 170.8 \| 214 \| (31, 308) \| 682 \| (23, 246) \| 292 \| Prohead serine protease \| Prohead serine protease; uniclust \| UniRef100\_A0A164A2N1 \| 99.7 \| 3.5e-20 \| 6.9e-26 \| 174.1 \| 135 \| (43, 185) \| 682 \| (25, 168) \| 217 \| Uncharacterized protein \| Uncharacterized protein; uniclust \| UniRef100\_A0A0Q6BU62 \| 99.7 \| 4.3e-20 \| 8e-26 \| 172.8 \| 152 \| (31, 184) \| 682 \| (13, 177) \| 290 \| Peptidase U35 \| Peptidase U35; uniclust \| UniRef100\_A0A0G1JMI3 \| 99.7 \| 4.5e-20 \| 8.4e-26 \| 183.3 \| 265 \| (407, 681) \| 682 \| (174, 471) \| 475 \| HK97 family major capsid protein \| HK97 family major capsid protein; uniclust \| UniRef100\_A0A1C3WL59 \| 99.7 \| 5.3e-20 \| 1e-25 \| 180.0 \| 301 \| (337, 673) \| 682 \| (65, 380) \| 393 \| Phage major capsid protein, HK97 family \| Phage major capsid protein, HK97 family; uniclust \| UniRef100\_A0A3C1GA37 \| 99.7 \| 5.8e-20 \| 1.1e-25 \| 170.3 \| 228 \| (32, 307) \| 682 \| (84, 313) \| 348 \| Uncharacterized protein \| Uncharacterized protein; uniclust \| UniRef100\_UPI0022B09E7B \| 99.7 \| 5.8e-20 \| 1.1e-25 \| 181.5 \| 271 \| (393, 681) \| 682 \| (319, 615) \| 616 \| phage major capsid protein \| phage major capsid protein; uniclust \| UniRef100\_A0A2V5Y3W0 \| 99.7 \| 5.9e-20 \| 1.1e-25 \| 171.2 \| 327 \| (337, 679) \| 682 \| (20, 359) \| 364 \| Phage major capsid protein \| Phage major capsid protein; uniclust \| UniRef100\_A0A1I0EDA5 \| 99.7 \| 5.9e-20 \| 1.2e-25 \| 195.2 \| 263 \| (406, 680) \| 682 \| (239, 522) \| 526 \| Phage major capsid protein, HK97 family \| Phage major capsid protein, HK97 family; uniclust \| UniRef100\_A0A246JDV9 \| 99.7 \| 6.7e-20 \| 1.3e-25 \| 185.9 \| 272 \| (403, 680) \| 682 \| (182, 480) \| 495 \| Phage major capsid protein \| Phage major capsid protein; uniclust \| UniRef100\_A0A0Q7IVE6 \| 99.7 \| 7.2e-20 \| 1.4e-25 \| 164.2 \| 133 \| (41, 182) \| 682 \| (21, 159) \| 197 \| Peptidase U35 \| Peptidase U35; uniclust \| UniRef100\_A0A136PUC6 \| 99.7 \| 7.5e-20 \| 1.5e-25 \| 205.3 \| 523 \| (39, 679) \| 682 \| (576, 1129) \| 1146 \| Uncharacterized protein \| Uncharacterized protein; uniclust \| UniRef100\_A0A083XX18 \| 99.7 \| 7.9e-20 \| 1.5e-25 \| 172.7 \| 142 \| (31, 181) \| 682 \| (7, 164) \| 255 \| Peptidase \| Peptidase; uniclust \| UniRef100\_A0A0F9BNT3 \| 99.7 \| 8.1e-20 \| 1.5e-25 \| 181.0 \| 261 \| (409, 681) \| 682 \| (143, 424) \| 427 \| Phage major capsid protein \| Phage major capsid protein; uniclust \| UniRef100\_UPI0014703FA7 \| 99.7 \| 8.5e-20 \| 1.6e-25 \| 175.9 \| 366 \| (269, 679) \| 682 \| (40, 407) \| 410 \| phage major capsid protein \| phage major capsid protein; uniclust \| UniRef100\_A0A0C9QCT1 \| 99.7 \| 8.2e-20 \| 1.6e-25 \| 179.2 \| 137 \| (43, 183) \| 682 \| (61, 204) \| 286 \| HK97 family phage prohead protease \| HK97 family phage prohead protease; uniclust \| UniRef100\_A0A015W478 \| 99.7 \| 8.3e-20 \| 1.7e-25 \| 183.9 \| 171 \| (1, 181) \| 682 \| (2, 181) \| 274 \| Phage prohead protease, HK97 family \| Phage prohead protease, HK97 family; uniclust \| UniRef100\_A0A062VEV4 \| 99.7 \| 9.7e-20 \| 1.8e-25 \| 156.3 \| 147 \| (31, 183) \| 682 \| (7, 158) \| 201 \| Peptidase U35, phage prohead HK97 (Fragment) \| Peptidase U35, phage prohead HK97 (Fragment); uniclust \| UniRef100\_A0A1V5ID83 \| 99.7 \| 1e-19 \| 2e-25 \| 173.9 \| 263 \| (406, 680) \| 682 \| (95, 369) \| 372 \| Phage capsid family protein \| Phage capsid family protein; uniclust \| UniRef100\_A0A087DDT0 \| 99.7 \| 9.8e-20 \| 2e-25 \| 196.4 \| 247 \| (422, 680) \| 682 \| (211, 465) \| 484 \| Phage prohead protease, HK97 family protein \| Phage prohead protease, HK97 family protein; uniclust \| UniRef100\_A0A840YDI0 \| 99.7 \| 1.1e-19 \| 2.1e-25 \| 177.7 \| 150 \| (32, 184) \| 682 \| (24, 184) \| 418 \| Peptidase U35 (Fragment) \| Peptidase U35 (Fragment); uniclust \| UniRef100\_UPI001CFB8F9B \| 99.7 \| 1.2e-19 \| 2.2e-25 \| 178.0 \| 461 \| (35, 678) \| 682 \| (8, 503) \| 512 \| HK97 family phage prohead protease \| HK97 family phage prohead protease; uniclust \| UniRef100\_A0A178MUR1 \| 99.7 \| 1.3e-19 \| 2.4e-25 \| 170.5 \| 137 \| (39, 181) \| 682 \| (42, 188) \| 256 \| Peptidase U35 \| Peptidase U35; uniclust \| UniRef100\_A0A149TLR9 \| 99.7 \| 1.3e-19 \| 2.5e-25 \| 170.5 \| 138 \| (33, 181) \| 682 \| (29, 173) \| 233 \| Uncharacterized protein \| Uncharacterized protein; uniclust \| UniRef100\_A0A1V5WQJ7 \| 99.7 \| 1.4e-19 \| 2.5e-25 \| 178.4 \| 529 \| (40, 678) \| 682 \| (12, 588) \| 594 \| Phage capsid family protein \| Phage capsid family protein; uniclust \| UniRef100\_A0A0F7L8Q8 \| 99.7 \| 1.3e-19 \| 2.5e-25 \| 185.7 \| 256 \| (411, 679) \| 682 \| (133, 403) \| 418 \| Phage major capsid protein HK97 \| Phage major capsid protein HK97; uniclust \| UniRef100\_A0A0W1GEU5 \| 99.7 \| 1.4e-19 \| 2.6e-25 \| 168.9 \| 132 \| (43, 181) \| 682 \| (17, 152) \| 248 \| Peptidase U35 \| Peptidase U35; uniclust \| UniRef100\_A0A3E1K5B0 \| 99.7 \| 1.5e-19 \| 2.7e-25 \| 174.3 \| 265 \| (406, 681) \| 682 \| (213, 479) \| 483 \| Phage major capsid protein \| Phage major capsid protein; uniclust \| UniRef100\_A0A3A6NR20 \| 99.7 \| 1.5e-19 \| 2.7e-25 \| 174.6 \| 267 \| (406, 681) \| 682 \| (103, 405) \| 408 \| Phage major capsid protein \| Phage major capsid protein; uniclust \| UniRef100\_A0A2D8XER8 \| 99.7 \| 1.5e-19 \| 2.8e-25 \| 178.5 \| 539 \| (63, 678) \| 682 \| (44, 605) \| 611 \| Phage major capsid protein \| Phage major capsid protein; uniclust \| UniRef100\_A0A846L7P7 \| 99.7 \| 1.6e-19 \| 2.9e-25 \| 176.5 \| 263 \| (406, 681) \| 682 \| (238, 526) \| 529 \| Phage major capsid protein \| Phage major capsid protein; uniclust \| UniRef100\_A0A9E3PA23 \| 99.7 \| 1.6e-19 \| 3e-25 \| 170.5 \| 279 \| (393, 681) \| 682 \| (98, 400) \| 403 \| Phage major capsid protein \| Phage major capsid protein; uniclust \| UniRef100\_A0A031HUL5 \| 99.7 \| 1.6e-19 \| 3.2e-25 \| 166.9 \| 134 \| (38, 181) \| 682 \| (21, 158) \| 233 \| HK97 family phage prohead protease \| HK97 family phage prohead protease; uniclust \| UniRef100\_A0A359LVB0 \| 99.7 \| 1.9e-19 \| 3.4e-25 \| 177.2 \| 293 \| (42, 380) \| 682 \| (27, 322) \| 588 \| Tripartite tricarboxylate transporter substrate binding protein \| Tripartite tricarboxylate transporter substrate binding protein; uniclust \| UniRef100\_A0A1H7U3C7 \| 99.7 \| 2e-19 \| 3.7e-25 \| 172.0 \| 262 \| (408, 680) \| 682 \| (92, 361) \| 379 \| Phage major capsid protein, HK97 family \| Phage major capsid protein, HK97 family; uniclust \| UniRef100\_A0A1I6URX3 \| 99.7 \| 2e-19 \| 4e-25 \| 168.8 \| 149 \| (31, 184) \| 682 \| (24, 189) \| 239 \| Phage prohead protease, HK97 family \| Phage prohead protease, HK97 family; uniclust \| UniRef100\_A0A0F9FEI0 \| 99.7 \| 2.1e-19 \| 4.1e-25 \| 169.3 \| 225 \| (447, 681) \| 682 \| (42, 277) \| 280 \| Phage major capsid protein (Fragment) \| Phage major capsid protein (Fragment); uniclust \| UniRef100\_A0A0D1C1F0 \| 99.7 \| 2.2e-19 \| 4.3e-25 \| 166.5 \| 138 \| (43, 184) \| 682 \| (35, 181) \| 216 \| Phage prohead protein \| Phage prohead protein; uniclust \| UniRef100\_A0A1C5S2N3 \| 99.7 \| 2.2e-19 \| 4.3e-25 \| 178.7 \| 252 \| (407, 680) \| 682 \| (105, 365) \| 378 \| Predicted phage phi-C31 gp36 major capsid-like protein \| Predicted phage phi-C31 gp36 major capsid-like protein; uniclust \| UniRef100\_A0A0C5CQK6 \| 99.6 \| 2.5e-19 \| 4.5e-25 \| 166.9 \| 221 \| (406, 639) \| 682 \| (29, 256) \| 358 \| Phage major capsid protein \| Phage major capsid protein; uniclust \| UniRef100\_A0A098AP33 \| 99.6 \| 2.5e-19 \| 5e-25 \| 193.7 \| 261 \| (410, 679) \| 682 \| (163, 448) \| 564 \| Putative prophage protein \| Putative prophage protein; uniclust \| UniRef100\_A0A7V7PNP3 \| 99.6 \| 2.8e-19 \| 5.1e-25 \| 174.9 \| 251 \| (406, 666) \| 682 \| (102, 368) \| 550 \| Phage major capsid protein \| Phage major capsid protein; uniclust \| UniRef100\_A0A7K2K343 \| 99.6 \| 2.7e-19 \| 5.1e-25 \| 186.0 \| 506 \| (45, 680) \| 682 \| (58, 599) \| 613 \| Phage major capsid protein \| Phage major capsid protein; uniclust \| UniRef100\_A0A2K8PMM0 \| 99.6 \| 2.7e-19 \| 5.2e-25 \| 184.9 \| 430 \| (31, 678) \| 682 \| (43, 499) \| 575 \| Caudovirus prohead protease \| Caudovirus prohead protease; uniclust \| UniRef100\_A0A3R9V132 \| 99.6 \| 2.9e-19 \| 5.4e-25 \| 175.7 \| 267 \| (406, 680) \| 682 \| (148, 424) \| 426 \| Phage major capsid protein \| Phage major capsid protein; uniclust \| UniRef100\_A0A5D0XCB7 \| 99.6 \| 3.1e-19 \| 5.7e-25 \| 169.0 \| 266 \| (407, 682) \| 682 \| (52, 338) \| 340 \| Phage major capsid protein \| Phage major capsid protein; uniclust \| UniRef100\_A0A0F9PEJ2 \| 99.6 \| 3.2e-19 \| 6.2e-25 \| 183.1 \| 133 \| (43, 181) \| 682 \| (22, 158) \| 449 \| Phage-like element PBSX protein XkdF domain-containing protein \| Phage-like element PBSX protein XkdF domain-containing protein; uniclust \| UniRef100\_A0A170SMX2 \| 99.6 \| 3.8e-19 \| 7.1e-25 \| 169.5 \| 262 \| (406, 680) \| 682 \| (100, 386) \| 390 \| HK97 family phage major capsid protein (Fragment) \| HK97 family phage major capsid protein (Fragment); uniclust \| UniRef100\_A0A0Q5UI83 \| 99.6 \| 3.8e-19 \| 7.6e-25 \| 195.0 \| 385 \| (259, 680) \| 682 \| (314, 729) \| 733 \| ATP-dependent Clp protease proteolytic subunit \| ATP-dependent Clp protease proteolytic subunit; uniclust \| UniRef100\_A0A165XDA1 \| 99.6 \| 4.3e-19 \| 7.9e-25 \| 159.9 \| 144 \| (39, 184) \| 682 \| (22, 169) \| 277 \| Caudovirus prohead protease \| Caudovirus prohead protease; uniclust \| UniRef100\_A0A061LZA8 \| 99.6 \| 4e-19 \| 7.9e-25 \| 172.5 \| 132 \| (44, 184) \| 682 \| (51, 189) \| 263 \| Peptidase U35 \| Peptidase U35; uniclust \| UniRef100\_A0A084UCG9 \| 99.6 \| 4.3e-19 \| 8.3e-25 \| 167.1 \| 130 \| (43, 181) \| 682 \| (36, 169) \| 256 \| HK97 family phage prohead protease \| HK97 family phage prohead protease; uniclust \| UniRef100\_A0A1A2PL73 \| 99.6 \| 4.3e-19 \| 8.5e-25 \| 182.6 \| 261 \| (411, 681) \| 682 \| (146, 418) \| 424 \| Major capsid protein \| Major capsid protein; uniclust \| UniRef100\_A0A0H1RDA6 \| 99.6 \| 4.9e-19 \| 9.5e-25 \| 162.2 \| 130 \| (43, 182) \| 682 \| (38, 171) \| 206 \| Peptidase U35 \| Peptidase U35; uniclust \| UniRef100\_A0A031HP15 \| 99.6 \| 4.7e-19 \| 9.6e-25 \| 182.8 \| 241 \| (421, 681) \| 682 \| (89, 357) \| 382 \| Phage major capsid protein, HK97 family \| Phage major capsid protein, HK97 family; uniclust \| UniRef100\_A0A6I2TYG7 \| 99.6 \| 5.3e-19 \| 9.7e-25 \| 168.5 \| 282 \| (376, 679) \| 682 \| (146, 430) \| 432 \| Phage major capsid protein \| Phage major capsid protein; uniclust \| UniRef100\_A0A0H5CWQ9 \| 99.6 \| 5.2e-19 \| 9.9e-25 \| 161.3 \| 129 \| (43, 181) \| 682 \| (39, 171) \| 220 \| Phage prohead protease, HK97 family \| Phage prohead protease, HK97 family; uniclust \| UniRef100\_A0A7C4PRQ6 \| 99.6 \| 5.4e-19 \| 1e-24 \| 178.2 \| 278 \| (386, 681) \| 682 \| (170, 470) \| 472 \| Phage major capsid protein \| Phage major capsid protein; uniclust \| UniRef100\_A0A091B3N3 \| 99.6 \| 5.3e-19 \| 1.1e-24 \| 184.8 \| 256 \| (415, 679) \| 682 \| (143, 405) \| 414 \| Capsid protein \| Capsid protein; uniclust \| UniRef100\_UPI0009494E27 \| 99.6 \| 6.5e-19 \| 1.2e-24 \| 164.7 \| 149 \| (31, 185) \| 682 \| (4, 165) \| 295 \| HK97 family phage prohead protease \| HK97 family phage prohead protease; uniclust \| UniRef100\_A0A0G4KA63 \| 99.6 \| 6.7e-19 \| 1.3e-24 \| 171.6 \| 264 \| (406, 679) \| 682 \| (130, 419) \| 439 \| Phage major capsid protein, HK97 family \| Phage major capsid protein, HK97 family; uniclust \| UniRef100\_A0A0K2JIG5 \| 99.6 \| 7.6e-19 \| 1.4e-24 \| 177.2 \| 257 \| (407, 678) \| 682 \| (322, 592) \| 595 \| Phage major capsid protein \| Phage major capsid protein; uniclust \| UniRef100\_A0A6J5MIS0 \| 99.6 \| 7.8e-19 \| 1.5e-24 \| 171.8 \| 262 \| (406, 679) \| 682 \| (95, 369) \| 371 \| COG4653 Predicted phage phi-C31 gp36 major capsid-like protein \| COG4653 Predicted phage phi-C31 gp36 major capsid-like protein; uniclust \| UniRef100\_A0A174DQP5 \| 99.6 \| 8e-19 \| 1.6e-24 \| 177.2 \| 258 \| (409, 681) \| 682 \| (94, 361) \| 369 \| Phage major capsid protein, HK97 family \| Phage major capsid protein, HK97 family; uniclust \| UniRef100\_A0A016QNE7 \| 99.6 \| 8.9e-19 \| 1.7e-24 \| 173.5 \| 137 \| (35, 181) \| 682 \| (118, 257) \| 377 \| Phage prohead protease, HK97 family \| Phage prohead protease, HK97 family; uniclust \| UniRef100\_A0A1Q3QR31 \| 99.6 \| 9.1e-19 \| 1.8e-24 \| 167.8 \| 139 \| (41, 184) \| 682 \| (27, 174) \| 250 \| HK97 family phage prohead protease \| HK97 family phage prohead protease; uniclust \| UniRef100\_A0A062XI24 \| 99.6 \| 9.1e-19 \| 1.8e-24 \| 165.2 \| 128 \| (47, 181) \| 682 \| (44, 180) \| 232 \| Prohead protease \| Prohead protease; uniclust \| UniRef100\_A0A011URX1 \| 99.6 \| 9.8e-19 \| 1.9e-24 \| 166.1 \| 142 \| (38, 185) \| 682 \| (33, 182) \| 234 \| Peptidase U35 \| Peptidase U35; uniclust \| UniRef100\_A0A059G8L6 \| 99.6 \| 1e-18 \| 2e-24 \| 180.3 \| 316 \| (341, 681) \| 682 \| (113, 450) \| 519 \| Phage major capsid protein \| Phage major capsid protein; uniclust \| UniRef100\_A0A142XTA2 \| 99.6 \| 1.1e-18 \| 2.1e-24 \| 175.3 \| 259 \| (406, 671) \| 682 \| (168, 447) \| 468 \| Phage capsid family protein \| Phage capsid family protein; uniclust \| UniRef100\_A0A0F9LI03 \| 99.6 \| 1.1e-18 \| 2.1e-24 \| 174.8 \| 265 \| (407, 681) \| 682 \| (163, 435) \| 435 \| Phage major capsid protein \| Phage major capsid protein; uniclust \| UniRef100\_A0A2G2LGM9 \| 99.6 \| 1.1e-18 \| 2.1e-24 \| 168.7 \| 138 \| (35, 181) \| 682 \| (22, 165) \| 294 \| HK97 family phage prohead protease \| HK97 family phage prohead protease; uniclust \| UniRef100\_A0A2W7AVB5 \| 99.6 \| 1.1e-18 \| 2.1e-24 \| 167.4 \| 146 \| (40, 185) \| 682 \| (35, 188) \| 405 \| HK97 family phage prohead protease \| HK97 family phage prohead protease; uniclust \| UniRef100\_A0A023W708 \| 99.6 \| 1.1e-18 \| 2.2e-24 \| 180.5 \| 253 \| (411, 673) \| 682 \| (41, 319) \| 417 \| Major capsid protein \| Major capsid protein; uniclust \| UniRef100\_A0A6N7GTQ9 \| 99.6 \| 1.2e-18 \| 2.2e-24 \| 166.6 \| 135 \| (32, 181) \| 682 \| (5, 150) \| 296 \| Uncharacterized protein \| Uncharacterized protein; uniclust \| UniRef100\_A0A0B5CU19 \| 99.6 \| 1.1e-18 \| 2.2e-24 \| 176.2 \| 254 \| (413, 680) \| 682 \| (38, 316) \| 412 \| Main capsid protein \| Main capsid protein; uniclust \| UniRef100\_A0A1M6LHF3 \| 99.6 \| 1.2e-18 \| 2.3e-24 \| 173.3 \| 131 \| (41, 181) \| 682 \| (53, 192) \| 410 \| Phage prohead protease, HK97 family \| Phage prohead protease, HK97 family; uniclust \| UniRef100\_A0A956YZ53 \| 99.6 \| 1.3e-18 \| 2.4e-24 \| 168.4 \| 259 \| (406, 676) \| 682 \| (100, 385) \| 404 \| Phage major capsid protein \| Phage major capsid protein; uniclust \| UniRef100\_A0A965HA95 \| 99.6 \| 1.3e-18 \| 2.4e-24 \| 154.1 \| 214 \| (466, 681) \| 682 \| (2, 242) \| 244 \| Phage major capsid protein \| Phage major capsid protein; uniclust \| UniRef100\_A0A0X3XWX8 \| 99.6 \| 1.3e-18 \| 2.5e-24 \| 181.2 \| 236 \| (433, 679) \| 682 \| (198, 446) \| 459 \| Phage capsid protein \| Phage capsid protein; uniclust \| UniRef100\_A0A6M3KXC0 \| 99.6 \| 1.3e-18 \| 2.6e-24 \| 156.7 \| 132 \| (38, 181) \| 682 \| (12, 147) \| 202 \| Putative prohead protease \| Putative prohead protease; uniclust \| UniRef100\_A0A0D0KY24 \| 99.6 \| 1.3e-18 \| 2.6e-24 \| 180.9 \| 379 \| (260, 670) \| 682 \| (138, 544) \| 559 \| Bacteriophage Mu GpT domain-containing protein \| Bacteriophage Mu GpT domain-containing protein; uniclust \| UniRef100\_A0A015V7E1 \| 99.6 \| 1.3e-18 \| 2.6e-24 \| 172.5 \| 149 \| (31, 184) \| 682 \| (57, 219) \| 270 \| Caudovirus prohead protease family protein \| Caudovirus prohead protease family protein; uniclust \| UniRef100\_A0A081I8G1 \| 99.6 \| 1.3e-18 \| 2.7e-24 \| 182.0 \| 260 \| (413, 681) \| 682 \| (191, 462) \| 470 \| HK97 family phage major capsid protein \| HK97 family phage major capsid protein; uniclust \| UniRef100\_A0A0J5GP20 \| 99.6 \| 1.3e-18 \| 2.7e-24 \| 164.7 \| 145 \| (31, 183) \| 682 \| (18, 176) \| 216 \| Peptidase U35 \| Peptidase U35; uniclust \| UniRef100\_A0A045IH31 \| 99.6 \| 1.5e-18 \| 2.9e-24 \| 163.2 \| 142 \| (31, 181) \| 682 \| (18, 169) \| 208 \| PhiRv2 phage protease \| PhiRv2 phage protease; uniclust \| UniRef100\_A0A812Y0T0 \| 99.6 \| 1.6e-18 \| 3e-24 \| 171.3 \| 269 \| (393, 680) \| 682 \| (147, 425) \| 603 \| Phage major capsid protein (Fragment) \| Phage major capsid protein (Fragment); uniclust \| UniRef100\_A0A0P7D8F3 \| 99.6 \| 1.6e-18 \| 3e-24 \| 156.3 \| 136 \| (36, 181) \| 682 \| (18, 160) \| 230 \| Primosome assembly protein PriA \| Primosome assembly protein PriA; uniclust \| UniRef100\_A0A077M1Z5 \| 99.6 \| 1.5e-18 \| 3e-24 \| 187.9 \| 503 \| (39, 679) \| 682 \| (42, 578) \| 585 \| Uncharacterized protein \| Uncharacterized protein; uniclust \| UniRef100\_A0A7C4QW79 \| 99.6 \| 1.7e-18 \| 3.1e-24 \| 172.6 \| 259 \| (413, 680) \| 682 \| (373, 651) \| 654 \| Phage major capsid protein \| Phage major capsid protein; uniclust \| UniRef100\_A0A0F9FAM1 \| 99.6 \| 1.6e-18 \| 3.1e-24 \| 164.6 \| 139 \| (38, 184) \| 682 \| (20, 161) \| 290 \| HK97 family phage prohead protease \| HK97 family phage prohead protease; uniclust \| UniRef100\_A0A1H3NUF2 \| 99.6 \| 1.7e-18 \| 3.3e-24 \| 159.3 \| 210 \| (462, 680) \| 682 \| (3, 218) \| 222 \| Phage major capsid protein, HK97 family \| Phage major capsid protein, HK97 family; uniclust \| UniRef100\_A0A0D5XUH0 \| 99.6 \| 1.7e-18 \| 3.3e-24 \| 181.9 \| 261 \| (406, 679) \| 682 \| (164, 433) \| 451 \| Capsid protein \| Capsid protein; uniclust \| UniRef100\_A0A0U4IJH3 \| 99.6 \| 1.7e-18 \| 3.4e-24 \| 190.3 \| 478 \| (39, 679) \| 682 \| (52, 544) \| 576 \| Major capsid and protease fusion protein \| Major capsid and protease fusion protein; uniclust \| UniRef100\_A0A068D609 \| 99.6 \| 1.8e-18 \| 3.4e-24 \| 153.1 \| 135 \| (42, 183) \| 682 \| (27, 165) \| 198 \| HK97 family phage prohead protease \| HK97 family phage prohead protease; uniclust \| UniRef100\_A0A6J5PI70 \| 99.6 \| 1.9e-18 \| 3.4e-24 \| 172.4 \| 513 \| (31, 679) \| 682 \| (5, 580) \| 584 \| Major\_cap\_HK97, phage major capsid protein, HK97 family \| Major\_cap\_HK97, phage major capsid protein, HK97 family; uniclust \| UniRef100\_A0A0Q7DEB2 \| 99.6 \| 1.8e-18 \| 3.5e-24 \| 156.0 \| 129 \| (38, 181) \| 682 \| (44, 173) \| 205 \| Peptidase U35 \| Peptidase U35; uniclust \| UniRef100\_A0A0F9JEK1 \| 99.6 \| 1.8e-18 \| 3.6e-24 \| 180.2 \| 263 \| (406, 676) \| 682 \| (188, 475) \| 493 \| Phage major capsid protein \| Phage major capsid protein; uniclust \| UniRef100\_A0A7W7ZFU5 \| 99.6 \| 2e-18 \| 3.8e-24 \| 169.6 \| 264 \| (406, 680) \| 682 \| (120, 399) \| 402 \| HK97 family phage major capsid protein \| HK97 family phage major capsid protein; uniclust \| UniRef100\_A0A2U3NJA2 \| 99.6 \| 2e-18 \| 3.8e-24 \| 156.5 \| 209 \| (462, 681) \| 682 \| (2, 219) \| 227 \| Capsid protein (Fragment) \| Capsid protein (Fragment); uniclust \| UniRef100\_A0A6J5LD92 \| 99.6 \| 2.1e-18 \| 4e-24 \| 154.4 \| 133 \| (41, 180) \| 682 \| (15, 150) \| 205 \| COG3740 Phage head maturation protease \| COG3740 Phage head maturation protease; uniclust \| UniRef100\_A0A0A0YV29 \| 99.6 \| 2.2e-18 \| 4.1e-24 \| 153.6 \| 129 \| (43, 181) \| 682 \| (22, 155) \| 195 \| Prohead maturation protease \| Prohead maturation protease; uniclust \| UniRef100\_A0A059NYL5 \| 99.6 \| 2.6e-18 \| 5.1e-24 \| 173.8 \| 253 \| (413, 680) \| 682 \| (45, 318) \| 362 \| Phage major capsid protein, HK97 family \| Phage major capsid protein, HK97 family; uniclust \| UniRef100\_A0A2U2DX69 \| 99.6 \| 2.8e-18 \| 5.2e-24 \| 167.4 \| 262 \| (406, 680) \| 682 \| (139, 436) \| 440 \| Phage major capsid protein \| Phage major capsid protein; uniclust \| UniRef100\_A0A0K1JGP8 \| 99.6 \| 2.8e-18 \| 5.4e-24 \| 168.2 \| 260 \| (408, 679) \| 682 \| (127, 408) \| 409 \| Capsid protein \| Capsid protein; uniclust \| UniRef100\_A0A0S7CWH9 \| 99.6 \| 2.8e-18 \| 5.4e-24 \| 166.5 \| 137 \| (33, 180) \| 682 \| (12, 154) \| 306 \| Prohead protease \| Prohead protease; uniclust \| UniRef100\_A0A497JQ78 \| 99.6 \| 2.8e-18 \| 5.4e-24 \| 168.5 \| 250 \| (414, 680) \| 682 \| (70, 323) \| 324 \| Phage major capsid protein \| Phage major capsid protein; uniclust \| UniRef100\_A0A0F9W0Z4 \| 99.6 \| 3e-18 \| 5.5e-24 \| 162.8 \| 271 \| (393, 681) \| 682 \| (133, 412) \| 414 \| Phage major capsid protein \| Phage major capsid protein; uniclust \| UniRef100\_A0A7I9ZGE4 \| 99.6 \| 3.2e-18 \| 5.9e-24 \| 169.4 \| 520 \| (31, 681) \| 682 \| (17, 601) \| 606 \| Major capsid protein \| Major capsid protein; uniclust \| UniRef100\_A0A2W7RG09 \| 99.6 \| 3.3e-18 \| 6.1e-24 \| 161.5 \| 319 \| (43, 509) \| 682 \| (15, 340) \| 343 \| HK97 family phage prohead protease/HK97 family phage major capsid protein,TIGR01554 (Fragment) \| HK97 family phage prohead protease/HK97 family phage major capsid protein,TIGR01554 (Fragment); uniclust \| UniRef100\_A0A484YZ81 \| 99.6 \| 3.3e-18 \| 6.1e-24 \| 154.4 \| 212 \| (462, 680) \| 682 \| (2, 226) \| 232 \| HK97 family phage major capsid protein \| HK97 family phage major capsid protein; uniclust \| UniRef100\_A0A966TPN7 \| 99.6 \| 3.6e-18 \| 6.7e-24 \| 157.0 \| 260 \| (411, 681) \| 682 \| (41, 317) \| 318 \| Phage major capsid protein \| Phage major capsid protein; uniclust \| UniRef100\_A0A6I4E5J7 \| 99.6 \| 3.6e-18 \| 7e-24 \| 165.9 \| 133 \| (41, 181) \| 682 \| (89, 228) \| 290 \| HK97 family phage prohead protease \| HK97 family phage prohead protease; uniclust \| UniRef100\_A0A101WGC7 \| 99.6 \| 3.7e-18 \| 7e-24 \| 173.6 \| 264 \| (406, 677) \| 682 \| (186, 469) \| 488 \| Phage capsid protein \| Phage capsid protein; uniclust \| UniRef100\_A0A839V847 \| 99.6 \| 4.1e-18 \| 7.5e-24 \| 158.2 \| 253 \| (344, 614) \| 682 \| (83, 343) \| 343 \| HK97 family phage major capsid protein \| HK97 family phage major capsid protein; uniclust \| UniRef100\_A0A443IQP8 \| 99.6 \| 4.1e-18 \| 7.7e-24 \| 163.7 \| 251 \| (422, 682) \| 682 \| (40, 326) \| 336 \| Phage major capsid protein \| Phage major capsid protein; uniclust \| UniRef100\_A0A0A8K3H1 \| 99.6 \| 4.5e-18 \| 8.6e-24 \| 161.6 \| 131 \| (41, 181) \| 682 \| (49, 182) \| 299 \| Gene transfer agent prohead protease \| Gene transfer agent prohead protease; uniclust \| UniRef100\_UPI001FCE8D07 \| 99.6 \| 4.8e-18 \| 8.8e-24 \| 157.2 \| 149 \| (31, 185) \| 682 \| (12, 168) \| 333 \| HK97 family phage prohead protease \| HK97 family phage prohead protease; uniclust \| UniRef100\_A0A6N8YYY6 \| 99.6 \| 4.7e-18 \| 9.1e-24 \| 157.5 \| 132 \| (45, 183) \| 682 \| (56, 189) \| 231 \| HK97 family phage prohead protease \| HK97 family phage prohead protease; uniclust \| UniRef100\_A0A090KKN6 \| 99.6 \| 4.8e-18 \| 9.4e-24 \| 158.5 \| 138 \| (41, 184) \| 682 \| (27, 172) \| 217 \| Uncultured bacterium genome assembly Metasoil\_fosmids\_resub \| Uncultured bacterium genome assembly Metasoil\_fosmids\_resub; uniclust \| UniRef100\_A0A1G5SX48 \| 99.6 \| 5.1e-18 \| 9.5e-24 \| 166.3 \| 151 \| (32, 184) \| 682 \| (7, 170) \| 429 \| Phage prohead protease, HK97 family \| Phage prohead protease, HK97 family; uniclust \| UniRef100\_A0A1V6D5P9 \| 99.6 \| 5.3e-18 \| 9.9e-24 \| 166.4 \| 269 \| (404, 681) \| 682 \| (161, 452) \| 454 \| Phage capsid family protein \| Phage capsid family protein; uniclust \| UniRef100\_A0A3D3LX94 \| 99.6 \| 5.5e-18 \| 1e-23 \| 163.4 \| 263 \| (406, 680) \| 682 \| (108, 397) \| 399 \| Phage major capsid protein \| Phage major capsid protein; uniclust \| UniRef100\_A0A090C747 \| 99.6 \| 5.2e-18 \| 1e-23 \| 177.2 \| 263 \| (406, 680) \| 682 \| (132, 415) \| 436 \| Putative capsid protein \| Putative capsid protein; uniclust \| UniRef100\_A0A068QTX5 \| 99.6 \| 5.5e-18 \| 1e-23 \| 159.6 \| 261 \| (406, 679) \| 682 \| (15, 294) \| 301 \| Putative major capsid protein \| Putative major capsid protein; uniclust \| UniRef100\_A0A7V2IQW9 \| 99.6 \| 6e-18 \| 1.1e-23 \| 157.8 \| 220 \| (445, 672) \| 682 \| (7, 251) \| 269 \| Phage major capsid protein \| Phage major capsid protein; uniclust \| UniRef100\_A0A1H0N3V1 \| 99.6 \| 6.1e-18 \| 1.2e-23 \| 165.3 \| 257 \| (408, 680) \| 682 \| (26, 286) \| 297 \| Phage major capsid protein, HK97 family \| Phage major capsid protein, HK97 family; uniclust \| UniRef100\_A0A2S6MR39 \| 99.6 \| 6.3e-18 \| 1.2e-23 \| 158.9 \| 131 \| (44, 181) \| 682 \| (70, 204) \| 242 \| HK97 family phage prohead protease \| HK97 family phage prohead protease; uniclust \| UniRef100\_A0A0A7RXR2 \| 99.6 \| 6.4e-18 \| 1.2e-23 \| 176.0 \| 263 \| (406, 680) \| 682 \| (247, 531) \| 553 \| Major capsid protein \| Major capsid protein; uniclust \| UniRef100\_A0A2A7MT55 \| 99.6 \| 8e-18 \| 1.5e-23 \| 157.4 \| 199 \| (470, 678) \| 682 \| (25, 248) \| 255 \| Phage major capsid protein \| Phage major capsid protein; uniclust \| UniRef100\_A0A0C1X3C7 \| 99.6 \| 7.6e-18 \| 1.5e-23 \| 179.2 \| 260 \| (410, 680) \| 682 \| (176, 474) \| 487 \| Phage major capsid protein \| Phage major capsid protein; uniclust \| UniRef100\_A0A068SKZ3 \| 99.6 \| 7.9e-18 \| 1.5e-23 \| 169.9 \| 146 \| (31, 184) \| 682 \| (26, 193) \| 376 \| Uncharacterized protein \| Uncharacterized protein; uniclust \| UniRef100\_A0A3D2THT3 \| 99.6 \| 8.5e-18 \| 1.6e-23 \| 166.5 \| 261 \| (406, 672) \| 682 \| (175, 456) \| 473 \| Phage major capsid protein \| Phage major capsid protein; uniclust \| UniRef100\_A0A0F9NU21 \| 99.6 \| 8e-18 \| 1.6e-23 \| 163.9 \| 134 \| (46, 183) \| 682 \| (47, 187) \| 271 \| HK97 family phage prohead protease \| HK97 family phage prohead protease; uniclust \| UniRef100\_A0A3M0WW28 \| 99.6 \| 8.2e-18 \| 1.6e-23 \| 159.3 \| 137 \| (44, 184) \| 682 \| (30, 174) \| 224 \| HK97 family phage prohead protease \| HK97 family phage prohead protease; uniclust \| UniRef100\_A0A6J7WJ50 \| 99.6 \| 8.7e-18 \| 1.6e-23 \| 165.8 \| 301 \| (336, 671) \| 682 \| (86, 412) \| 436 \| Major\_cap\_HK97, phage major capsid protein, HK97 family \| Major\_cap\_HK97, phage major capsid protein, HK97 family; uniclust \| UniRef100\_A0A0H3LJ84 \| 99.6 \| 8.2e-18 \| 1.7e-23 \| 183.7 \| 263 \| (406, 680) \| 682 \| (269, 553) \| 574 \| PhiRV1 phage protein \| PhiRV1 phage protein; uniclust \| UniRef100\_A0A060PI05 \| 99.6 \| 8.9e-18 \| 1.8e-23 \| 181.1 \| 248 \| (406, 679) \| 682 \| (161, 427) \| 492 \| Phage major capsid protein \| Phage major capsid protein; uniclust \| UniRef100\_A0A0S3PZ93 \| 99.6 \| 9.2e-18 \| 1.8e-23 \| 170.3 \| 197 \| (393, 607) \| 682 \| (137, 349) \| 355 \| Phage capsid family protein \| Phage capsid family protein; uniclust \| UniRef100\_A0A1U7CX52 \| 99.6 \| 1e-17 \| 1.8e-23 \| 160.3 \| 308 \| (363, 680) \| 682 \| (98, 430) \| 434 \| Phage capsid family protein \| Phage capsid family protein; uniclust \| UniRef100\_A0A0X3Y2E8 \| 99.6 \| 9.6e-18 \| 1.9e-23 \| 176.8 \| 259 \| (408, 680) \| 682 \| (310, 593) \| 595 \| Phage capsid protein \| Phage capsid protein; uniclust \| UniRef100\_UPI00158CF282 \| 99.6 \| 1e-17 \| 1.9e-23 \| 140.8 \| 129 \| (40, 169) \| 682 \| (4, 132) \| 171 \| hypothetical protein \| hypothetical protein; uniclust \| UniRef100\_A0A1V2ZD93 \| 99.5 \| 1.1e-17 \| 2.1e-23 \| 159.8 \| 206 \| (463, 679) \| 682 \| (128, 340) \| 345 \| Major capsid protein \| Major capsid protein; uniclust \| UniRef100\_A0A660TAH7 \| 99.5 \| 1.3e-17 \| 2.4e-23 \| 157.9 \| 263 \| (406, 679) \| 682 \| (114, 395) \| 397 \| Phage major capsid protein \| Phage major capsid protein; uniclust \| UniRef100\_A0A212I3J2 \| 99.5 \| 1.2e-17 \| 2.4e-23 \| 161.4 \| 214 \| (456, 679) \| 682 \| (72, 294) \| 315 \| Major capsid protein \| Major capsid protein; uniclust \| UniRef100\_A0A1Y0H480 \| 99.5 \| 1.3e-17 \| 2.4e-23 \| 164.1 \| 234 \| (438, 680) \| 682 \| (320, 564) \| 565 \| Phage major capsid protein \| Phage major capsid protein; uniclust \| UniRef100\_UPI0010AE5AB6 \| 99.5 \| 1.4e-17 \| 2.5e-23 \| 145.9 \| 149 \| (31, 185) \| 682 \| (7, 163) \| 222 \| hypothetical protein \| hypothetical protein; uniclust \| UniRef100\_A0A2D5TTA0 \| 99.5 \| 1.7e-17 \| 3.2e-23 \| 154.0 \| 136 \| (40, 183) \| 682 \| (41, 182) \| 257 \| HK97 family phage prohead protease \| HK97 family phage prohead protease; uniclust \| UniRef100\_A0A0A0RPP6 \| 99.5 \| 1.7e-17 \| 3.3e-23 \| 151.5 \| 128 \| (43, 181) \| 682 \| (32, 163) \| 212 \| Prohead protease \| Prohead protease; uniclust \| UniRef100\_A0A2M7EMM4 \| 99.5 \| 1.8e-17 \| 3.4e-23 \| 154.5 \| 128 \| (44, 181) \| 682 \| (46, 179) \| 252 \| HK97 family phage prohead protease \| HK97 family phage prohead protease; uniclust \| UniRef100\_A0A158FYM0 \| 99.5 \| 1.9e-17 \| 3.7e-23 \| 151.1 \| 130 \| (43, 181) \| 682 \| (27, 161) \| 226 \| Phage-like protein \| Phage-like protein; uniclust \| UniRef100\_A0A0L0KLK6 \| 99.5 \| 1.9e-17 \| 3.8e-23 \| 167.5 \| 261 \| (411, 680) \| 682 \| (59, 355) \| 364 \| Phage capsid family protein \| Phage capsid family protein; uniclust \| UniRef100\_A0A2S3W008 \| 99.5 \| 2e-17 \| 3.8e-23 \| 156.3 \| 134 \| (42, 181) \| 682 \| (78, 221) \| 282 \| Caudovirus prohead protease \| Caudovirus prohead protease; uniclust \| UniRef100\_A0A9E3QNN9 \| 99.5 \| 2.3e-17 \| 4.2e-23 \| 158.1 \| 270 \| (393, 680) \| 682 \| (134, 435) \| 438 \| Phage major capsid protein \| Phage major capsid protein; uniclust \| UniRef100\_A0A0F0GPG3 \| 99.5 \| 2.3e-17 \| 4.3e-23 \| 164.5 \| 251 \| (408, 670) \| 682 \| (151, 427) \| 514 \| Phage major capsid protein \| Phage major capsid protein; uniclust \| UniRef100\_UPI000C9B78D3 \| 99.5 \| 2.3e-17 \| 4.3e-23 \| 159.8 \| 250 \| (413, 681) \| 682 \| (145, 404) \| 405 \| phage major capsid protein \| phage major capsid protein; uniclust \| UniRef100\_A0A5Q2RKZ8 \| 99.5 \| 2.4e-17 \| 4.4e-23 \| 159.4 \| 265 \| (409, 681) \| 682 \| (194, 472) \| 475 \| Phage major capsid protein \| Phage major capsid protein; uniclust \| UniRef100\_A0A136KYT3 \| 99.5 \| 2.3e-17 \| 4.4e-23 \| 174.1 \| 269 \| (406, 680) \| 682 \| (301, 621) \| 624 \| Phage capsid family protein \| Phage capsid family protein; uniclust \| UniRef100\_A0A2S5XKA8 \| 99.5 \| 2.7e-17 \| 5.3e-23 \| 173.6 \| 245 \| (423, 679) \| 682 \| (168, 435) \| 477 \| Phage major capsid protein \| Phage major capsid protein; uniclust \| UniRef100\_A0A0K9T4G1 \| 99.5 \| 2.8e-17 \| 5.3e-23 \| 161.2 \| 265 \| (405, 681) \| 682 \| (113, 410) \| 418 \| Phage major capsid protein (Fragment) \| Phage major capsid protein (Fragment); uniclust \| UniRef100\_A0A239MKE3 \| 99.5 \| 2.9e-17 \| 5.3e-23 \| 159.9 \| 266 \| (406, 679) \| 682 \| (146, 438) \| 448 \| Phage major capsid protein, HK97 family \| Phage major capsid protein, HK97 family; uniclust \| UniRef100\_A0A960HG97 \| 99.5 \| 2.9e-17 \| 5.5e-23 \| 163.9 \| 261 \| (409, 680) \| 682 \| (164, 433) \| 439 \| Phage major capsid protein \| Phage major capsid protein; uniclust \| UniRef100\_A0A6B9T8U5 \| 99.5 \| 3e-17 \| 5.6e-23 \| 158.9 \| 263 \| (406, 679) \| 682 \| (102, 382) \| 386 \| Phage major capsid protein \| Phage major capsid protein; uniclust \| UniRef100\_A0A2X2J2H8 \| 99.5 \| 3.1e-17 \| 5.9e-23 \| 147.4 \| 136 \| (32, 183) \| 682 \| (15, 162) \| 202 \| Caudovirus prohead protease \| Caudovirus prohead protease; uniclust \| UniRef100\_A0A3C1LU60 \| 99.5 \| 3.2e-17 \| 5.9e-23 \| 154.2 \| 256 \| (410, 678) \| 682 \| (96, 368) \| 374 \| Phage major capsid protein \| Phage major capsid protein; uniclust \| UniRef100\_G7QC57 \| 99.5 \| 3.3e-17 \| 6.1e-23 \| 146.4 \| 137 \| (38, 181) \| 682 \| (29, 170) \| 254 \| Phage prohead protease, HK97 family \| Phage prohead protease, HK97 family; uniclust \| UniRef100\_A0A069AVH7 \| 99.5 \| 3.2e-17 \| 6.6e-23 \| 174.7 \| 248 \| (406, 679) \| 682 \| (126, 395) \| 426 \| Phage major capsid protein, HK97 family \| Phage major capsid protein, HK97 family; uniclust \| UniRef100\_A0A0F7PH54 \| 99.5 \| 3.2e-17 \| 6.6e-23 \| 171.0 \| 252 \| (406, 681) \| 682 \| (83, 368) \| 374 \| Phage capsid family protein \| Phage capsid family protein; pdb70 \| 3QPR\_A \| 99.2 \| 5.4e-16 \| 5.7e-20 \| 163.2 \| 262 \| (409, 681) \| 682 \| (111, 382) \| 385 \| Major capsid protein \| 3QPR\_A Major capsid protein Virus Procapsid particles, VIRUS; pdb70 \| 3QPR\_C \| 99.2 \| 5.4e-16 \| 5.7e-20 \| 163.2 \| 262 \| (409, 681) \| 682 \| (111, 382) \| 385 \| Major capsid protein \| 3QPR\_C Major capsid protein Virus Procapsid particles, VIRUS; pdb70 \| 1OHG\_G \| 99.2 \| 2.7e-15 \| 3.1e-19 \| 147.1 \| 263 \| (408, 681) \| 682 \| (7, 279) \| 282 \| MAJOR CAPSID PROTEIN \| 1OHG\_G MAJOR CAPSID PROTEIN VIRUS, VIRUS COAT PROTEIN, VIRUS/VIRAL; pdb70 \| 6B0X\_F \| 99.2 \| 3.4e-15 \| 3.7e-19 \| 152.4 \| 261 \| (410, 681) \| 682 \| (34, 313) \| 324 \| Major head protein, Scaffold protein \| 6B0X\_F Major head protein, Scaffold protein major capsid protein, HK97-like fold; pdb70 \| 6B23\_C \| 99.2 \| 3.4e-15 \| 3.7e-19 \| 152.4 \| 261 \| (410, 681) \| 682 \| (34, 313) \| 324 \| Major head protein, Capsid morphogenesis \| 6B23\_C Major head protein, Capsid morphogenesis major capsid protein, HK97-like fold; pdb70 \| 3JB5\_D \| 99.1 \| 4.8e-15 \| 5.2e-19 \| 150.1 \| 272 \| (408, 681) \| 682 \| (6, 304) \| 315 \| major capsid protein \| 3JB5\_D major capsid protein acne, bacteriophage, HK97-like, VIRUS; pdb70 \| 1IF0\_G \| 99.0 \| 2.3e-14 \| 2.6e-18 \| 138.6 \| 242 \| (430, 681) \| 682 \| (4, 255) \| 256 \| MAJOR CAPSID PROTEIN GP5 \| 1IF0\_G MAJOR CAPSID PROTEIN GP5 Bacteriophage, Virus, Capsid, cryoEM, Pseudo-atomic; pdb70 \| 6OMA\_F \| 98.8 \| 3.6e-13 \| 4e-17 \| 134.2 \| 268 \| (406, 681) \| 682 \| (6, 297) \| 299 \| Major capsid protein \| 6OMA\_F Major capsid protein capsid, HK97-fold, dsDNA-phage, icosahedral, VIRUS; pdb70 \| 6OMC\_B \| 98.8 \| 3.6e-13 \| 4e-17 \| 134.2 \| 268 \| (406, 681) \| 682 \| (6, 297) \| 299 \| Major capsid protein \| 6OMC\_B Major capsid protein capsid, HK97-fold, dsDNA-phage, icosahedral, VIRUS; pdb70 \| 4PT2\_B \| 98.7 \| 2.3e-12 \| 2.5e-16 \| 127.3 \| 250 \| (418, 681) \| 682 \| (19, 281) \| 287 \| Encapsulin protein \| 4PT2\_B Encapsulin protein HK97 fold, Shell protein, VIRUS; pdb70 \| 6NJ8\_A \| 98.7 \| 3e-12 \| 3.3e-16 \| 126.3 \| 251 \| (418, 682) \| 682 \| (19, 281) \| 282 \| Quasibacillus thermotolerans encapsulin, targeting peptide \| 6NJ8\_A Quasibacillus thermotolerans encapsulin, targeting peptide encapsulin, iron storage, IMEF, icosahedral; pdb70 \| 3J7V\_E \| 98.7 \| 4.4e-12 \| 4.7e-16 \| 130.1 \| 275 \| (400, 681) \| 682 \| (13, 343) \| 345 \| Major capsid protein 10A \| 3J7V\_E Major capsid protein 10A maturation, DNA packaging, Procapsid, Non-covalent; pdb70 \| 3J7W\_D \| 98.7 \| 4.4e-12 \| 4.7e-16 \| 130.1 \| 275 \| (400, 681) \| 682 \| (13, 343) \| 345 \| Major capsid protein 10A \| 3J7W\_D Major capsid protein 10A maturation, DNA packaging, Procapsid, Non-covalent; pdb70 \| 3J1A\_G \| 98.6 \| 1.2e-11 \| 1.4e-15 \| 114.6 \| 193 \| (481, 679) \| 682 \| (2, 199) \| 199 \| capsid protein \| 3J1A\_G capsid protein halophage, bacteriophage HK97, bacteriophage T7; pdb70 \| 3J1A\_B \| 98.5 \| 1.7e-11 \| 1.9e-15 \| 113.5 \| 193 \| (481, 679) \| 682 \| (2, 199) \| 199 \| capsid protein \| 3J1A\_B capsid protein halophage, bacteriophage HK97, bacteriophage T7; pdb70 \| 4BML\_B \| 98.4 \| 9e-11 \| 9.9e-15 \| 118.3 \| 266 \| (410, 680) \| 682 \| (24, 331) \| 332 \| MAJOR CAPSID PROTEIN \| 4BML\_B MAJOR CAPSID PROTEIN VIRUS, MARINE VIRUS, OUTER CAPSID; pdb70 \| 3DKT\_I \| 98.1 \| 7e-10 \| 7.7e-14 \| 108.0 \| 235 \| (419, 679) \| 682 \| (19, 264) \| 265 \| Maritimacin, Putative uncharacterized protein \| 3DKT\_I Maritimacin, Putative uncharacterized protein enzyme encapsulation, nanocompartment, oxidative stress; pdb70 \| 2XD8\_B \| 98.1 \| 1e-09 \| 1.1e-13 \| 113.8 \| 277 \| (401, 681) \| 682 \| (16, 368) \| 375 \| T7-LIKE CAPSID PROTEIN \| 2XD8\_B T7-LIKE CAPSID PROTEIN MARINE PODOVIRUS, T7-LIKE VIRUS, VIRUS; pdb70 \| 6I9G\_N \| 97.9 \| 4.2e-09 \| 4.6e-13 \| 102.8 \| 235 \| (419, 680) \| 682 \| (19, 265) \| 265 \| Uncharacterized protein \| 6I9G\_N Uncharacterized protein Nanocompartment, nanocage, packaging of biocatalysts HET: GOL, SO4; pdb70 \| 3J40\_B \| 97.4 \| 1.4e-07 \| 1.4e-11 \| 96.6 \| 112 \| (416, 528) \| 682 \| (20, 135) \| 335 \| gp10, gp7 \| 3J40\_B gp10, gp7 capsid, accessory protein, VIRUS; pdb70 \| 6R3A\_A \| 96.9 \| 1.3e-06 \| 1.4e-10 \| 87.6 \| 206 \| (410, 624) \| 682 \| (5, 224) \| 323 \| Major capsid protein \| 6R3A\_A Major capsid protein Bacteriophage, Capsid protein, image processing; pdb70 \| 6RTL\_E \| 96.9 \| 1.3e-06 \| 1.4e-10 \| 87.6 \| 206 \| (410, 624) \| 682 \| (5, 224) \| 323 \| Major capsid protein \| 6RTL\_E Major capsid protein Bacteriophage, maturation process, cryo electron; pdb70 \| 2E0Z\_B \| 96.9 \| 1.7e-06 \| 1.7e-10 \| 88.5 \| 184 \| (475, 679) \| 682 \| (159, 345) \| 345 \| Virus-like particle \| 2E0Z\_B Virus-like particle virus-like particle, virus, bacteriophage, HK97; pdb70 \| 2E0Z\_C \| 96.9 \| 1.7e-06 \| 1.7e-10 \| 88.5 \| 184 \| (475, 679) \| 682 \| (159, 345) \| 345 \| Virus-like particle \| 2E0Z\_C Virus-like particle virus-like particle, virus, bacteriophage, HK97; pdb70 \| 6J3Q\_U \| 96.8 \| 2.6e-06 \| 2.7e-10 \| 86.2 \| 263 \| (415, 681) \| 682 \| (9, 345) \| 348 \| the major capsid protein, the \| 6J3Q\_U the major capsid protein, the cyanophage, Siphoviridae, capsid, VIRUS; pdb70 \| 3BJQ\_F \| 96.4 \| 1.2e-05 \| 1.3e-09 \| 79.7 \| 261 \| (414, 681) \| 682 \| (12, 311) \| 316 \| Phage-related protein \| 3BJQ\_F Phage-related protein PHAGE-RELATED PROTEIN, STRUCTURAL GENOMICS, JOINT HET: PG4, MSE; pdb70 \| 3BJQ\_E \| 96.4 \| 1.2e-05 \| 1.3e-09 \| 79.7 \| 261 \| (414, 681) \| 682 \| (12, 311) \| 316 \| Phage-related protein \| 3BJQ\_E Phage-related protein PHAGE-RELATED PROTEIN, STRUCTURAL GENOMICS, JOINT HET: PG4; pdb70 \| 5WK1\_F \| 96.3 \| 1.6e-05 \| 1.6e-09 \| 81.5 \| 249 \| (418, 673) \| 682 \| (68, 342) \| 352 \| Capsid Stabilizing Protein, Major Capsid \| 5WK1\_F Capsid Stabilizing Protein, Major Capsid Major Capsid Protein, Capsid Stabilizing; pdb70 \| 5LII\_P \| 95.4 \| 0.00018 \| 1.8e-08 \| 76.2 \| 206 \| (374, 588) \| 682 \| (10, 230) \| 463 \| Major capsid protein \| 5LII\_P Major capsid protein polyvalent staphylococcal bactoriophage, Myoviridae, tail |
| Top keywords  (threshold 1.00e-03 (evalue)) | **Phage, capsid, major, HK97, prohead, protease, Fragment, Peptidase, U35, Bacteriophage** |
| Output files | ../../similar\_sequences/05\_FANPEZAQ\_CDS\_0005\_merged.svg ../../similar\_sequences/05\_FANPEZAQ\_CDS\_0005\_pdb70.a3m ../../similar\_sequences/05\_FANPEZAQ\_CDS\_0005\_pdb70.hhr ../../similar\_sequences/05\_FANPEZAQ\_CDS\_0005\_uniclust.a3m ../../similar\_sequences/05\_FANPEZAQ\_CDS\_0005\_uniclust.hhr |

#### Structure prediction (AlphaFold)^2^

|  |  |
| --- | --- |
| Stats | xml version="1.0" encoding="utf-8" standalone="no"?       2024-09-02T21:09:04.565873 image/svg+xml   Matplotlib v3.7.2, https://matplotlib.org/ |
| Predicted structure | **NGL Viewer Controls:**  - Center: *Left-Click* - Rotate: *Left-Click + Drag* - Translate: *Right-Click + Drag* - Zoom: *Shift + Left-Click + Drag* |
| Output files | ../../predicted\_structures/05\_FANPEZAQ\_CDS\_0005/features.pkl ../../predicted\_structures/05\_FANPEZAQ\_CDS\_0005/ranked\_0.pdb ../../predicted\_structures/05\_FANPEZAQ\_CDS\_0005/ranked\_0\_plots.svg ../../predicted\_structures/05\_FANPEZAQ\_CDS\_0005/result\_model\_1\_ptm\_pred\_0.pkl |

#### Structure similarity search results (Foldseek)^3^

|  |  |
| --- | --- |
| Structure databases searched | Pdb, Afdb-proteome, Afdb-uniprot50 |
| Results, scheme(s)  (Top layers only, threshold 1.00e-02 (evalue)) | xml version="1.0" encoding="utf-8" standalone="no"?       2024-09-02T21:10:16.128244 image/svg+xml   Matplotlib v3.7.2, https://matplotlib.org/ |
| Results, table  (threshold 1.00e-02 (evalue)) | db \| id \| prob \| evalue \| bits \| fident \| alnlen \| mismatch \| gapopen \| qstart \| qend \| tstart \| tend \| name \| description; pdb \| 8GTA\_G \| 1.0 \| 2.164e-26 \| 869 \| 0.407 \| 275 \| 153 \| 6 \| 410 \| 681 \| 1 \| 268 \| Major capsid protein \| Major capsid protein; pdb \| 6TB9\_C5 \| 1.0 \| 7.413e-16 \| 479 \| 0.193 \| 315 \| 204 \| 17 \| 400 \| 682 \| 1 \| 297 \| Major capsid protein Rcc01687 \| Major capsid protein Rcc01687; pdb \| 6TSU\_U4 \| 1.0 \| 1.102e-15 \| 473 \| 0.196 \| 285 \| 189 \| 14 \| 423 \| 682 \| 22 \| 291 \| Major capsid protein Rcc01687 \| Major capsid protein Rcc01687; pdb \| 6TUI\_B5 \| 1.0 \| 2.352e-14 \| 446 \| 0.182 \| 291 \| 187 \| 17 \| 422 \| 682 \| 1 \| 270 \| Phage major capsid protein, HK97 family \| Phage major capsid protein, HK97 family; pdb \| 1OHG\_A \| 1.0 \| 1.087e-13 \| 444 \| 0.175 \| 274 \| 198 \| 13 \| 423 \| 682 \| 21 \| 280 \| MAJOR CAPSID PROTEIN \| MAJOR CAPSID PROTEIN; pdb \| 6TUI\_A5 \| 1.0 \| 6.388e-15 \| 440 \| 0.187 \| 315 \| 203 \| 17 \| 398 \| 682 \| 3 \| 294 \| Phage major capsid protein, HK97 family \| Phage major capsid protein, HK97 family; pdb \| 1IF0\_D \| 1.0 \| 7.307e-14 \| 431 \| 0.176 \| 266 \| 193 \| 12 \| 430 \| 682 \| 4 \| 256 \| PROTEIN (MAJOR CAPSID PROTEIN GP5) \| PROTEIN (MAJOR CAPSID PROTEIN GP5); pdb \| 2FS3\_F \| 1.0 \| 1.175e-12 \| 416 \| 0.172 \| 273 \| 197 \| 14 \| 424 \| 682 \| 1 \| 258 \| Major capsid protein \| Major capsid protein; pdb \| 2FS3\_B \| 1.0 \| 1.616e-13 \| 414 \| 0.175 \| 285 \| 200 \| 14 \| 420 \| 682 \| 9 \| 280 \| Major capsid protein \| Major capsid protein; pdb \| 6TB9\_B5 \| 1.0 \| 6.905e-14 \| 412 \| 0.158 \| 291 \| 195 \| 15 \| 422 \| 682 \| 1 \| 271 \| Major capsid protein Rcc01687 \| Major capsid protein Rcc01687; pdb \| 8CFA\_D \| 1.0 \| 1.746e-12 \| 397 \| 0.171 \| 263 \| 181 \| 11 \| 430 \| 681 \| 2 \| 238 \| Major capsid subunit \| Major capsid subunit; pdb \| 8EDU\_G \| 1.0 \| 3.19e-13 \| 395 \| 0.158 \| 315 \| 224 \| 12 \| 398 \| 681 \| 1 \| 305 \| Capsid \| Capsid; pdb \| 8CFA\_F \| 1.0 \| 1.956e-12 \| 394 \| 0.16 \| 261 \| 181 \| 12 \| 430 \| 679 \| 3 \| 236 \| Major capsid subunit \| Major capsid subunit; pdb \| 2FRP\_G \| 1.0 \| 1.956e-12 \| 392 \| 0.165 \| 266 \| 183 \| 14 \| 430 \| 682 \| 4 \| 243 \| Major capsid protein \| Major capsid protein; pdb \| 8CFA\_A \| 1.0 \| 1.392e-12 \| 388 \| 0.154 \| 265 \| 180 \| 13 \| 430 \| 681 \| 1 \| 234 \| Major capsid subunit \| Major capsid subunit; pdb \| 8ECK\_E \| 1.0 \| 1.315e-12 \| 388 \| 0.135 \| 303 \| 230 \| 14 \| 406 \| 682 \| 18 \| 314 \| Major capsid protein \| Major capsid protein; pdb \| 2FTE\_G \| 1.0 \| 4.577e-12 \| 380 \| 0.161 \| 266 \| 183 \| 13 \| 430 \| 682 \| 4 \| 242 \| major capsid protein \| major capsid protein; pdb \| 3JB5\_B \| 1.0 \| 7.464e-13 \| 374 \| 0.181 \| 298 \| 195 \| 14 \| 422 \| 680 \| 13 \| 300 \| major capsid protein \| major capsid protein; pdb \| 8CFA\_E \| 1.0 \| 1.593e-11 \| 366 \| 0.166 \| 265 \| 184 \| 11 \| 427 \| 681 \| 2 \| 239 \| Major capsid subunit \| Major capsid subunit; pdb \| 3QPR\_G \| 1.0 \| 6.431e-12 \| 366 \| 0.166 \| 265 \| 188 \| 11 \| 430 \| 682 \| 4 \| 247 \| Major capsid protein \| Major capsid protein; pdb \| 3E8K\_G \| 1.0 \| 5.127e-12 \| 361 \| 0.157 \| 260 \| 186 \| 11 \| 435 \| 682 \| 9 \| 247 \| Major capsid protein \| Major capsid protein; pdb \| 6NJ8\_B \| 1.0 \| 1.998e-11 \| 355 \| 0.132 \| 264 \| 201 \| 10 \| 432 \| 682 \| 28 \| 276 \| Encapsulating protein for a DyP-type peroxidase \| Encapsulating protein for a DyP-type peroxidase; pdb \| 8EDU\_D \| 1.0 \| 4.577e-12 \| 353 \| 0.152 \| 321 \| 230 \| 14 \| 398 \| 681 \| 1 \| 316 \| Capsid \| Capsid; pdb \| 8ECN\_F \| 1.0 \| 7.203e-12 \| 352 \| 0.143 \| 300 \| 207 \| 14 \| 423 \| 681 \| 18 \| 308 \| Major capsid protein \| Major capsid protein; pdb \| 8FQK\_E \| 1.0 \| 1.505e-11 \| 347 \| 0.155 \| 263 \| 183 \| 10 \| 432 \| 682 \| 33 \| 268 \| Scaffolding domain delta \| Scaffolding domain delta; pdb \| 7MH2\_D \| 1.0 \| 1.27e-11 \| 343 \| 0.14 \| 263 \| 192 \| 12 \| 435 \| 682 \| 35 \| 278 \| T4GALA Engineered Protein Nanocage \| T4GALA Engineered Protein Nanocage; pdb \| 8EB4\_G \| 1.0 \| 1.344e-11 \| 340 \| 0.137 \| 305 \| 206 \| 15 \| 421 \| 680 \| 18 \| 310 \| Major capsid protein \| Major capsid protein; pdb \| 3JB5\_G \| 1.0 \| 1.784e-11 \| 339 \| 0.168 \| 291 \| 183 \| 15 \| 422 \| 680 \| 13 \| 276 \| major capsid protein \| major capsid protein; pdb \| 2GP1\_A \| 1.0 \| 1.452e-10 \| 333 \| 0.171 \| 274 \| 187 \| 16 \| 423 \| 682 \| 7 \| 254 \| Major capsid protein \| Major capsid protein; pdb \| 8FQK\_D \| 1.0 \| 3.144e-11 \| 332 \| 0.161 \| 266 \| 178 \| 13 \| 432 \| 682 \| 30 \| 265 \| Scaffolding domain delta \| Scaffolding domain delta; pdb \| 8E16\_C \| 1.0 \| 6.568e-11 \| 326 \| 0.124 \| 282 \| 193 \| 16 \| 423 \| 678 \| 14 \| 267 \| Major capsid protein, gp6 \| Major capsid protein, gp6; pdb \| 8EB4\_H \| 1.0 \| 3.944e-11 \| 313 \| 0.13 \| 314 \| 209 \| 14 \| 422 \| 682 \| 19 \| 321 \| Major capsid protein \| Major capsid protein; pdb \| 7MH2\_C \| 1.0 \| 1.372e-10 \| 311 \| 0.13 \| 260 \| 193 \| 12 \| 435 \| 680 \| 35 \| 275 \| T4GALA Engineered Protein Nanocage \| T4GALA Engineered Protein Nanocage; pdb \| 8ECJ\_E \| 1.0 \| 1.372e-10 \| 308 \| 0.105 \| 304 \| 224 \| 13 \| 413 \| 681 \| 11 \| 301 \| Major capsid protein \| Major capsid protein; pdb \| 8EC2\_C \| 1.0 \| 2.419e-10 \| 307 \| 0.1 \| 299 \| 211 \| 15 \| 426 \| 682 \| 22 \| 304 \| Major capsid protein \| Major capsid protein; pdb \| 8ECO\_F \| 1.0 \| 6.568e-11 \| 306 \| 0.14 \| 299 \| 204 \| 14 \| 423 \| 681 \| 17 \| 302 \| Major capsid protein \| Major capsid protein; pdb \| 7S2T\_B \| 1.0 \| 3.211e-10 \| 303 \| 0.126 \| 260 \| 194 \| 11 \| 434 \| 677 \| 27 \| 269 \| EncA \| EncA; pdb \| 7KMX\_F \| 1.0 \| 2.368e-11 \| 301 \| 0.155 \| 308 \| 209 \| 21 \| 410 \| 681 \| 2 \| 294 \| Major capsid protein \| Major capsid protein; pdb \| 8ECI\_C \| 1.0 \| 3.327e-11 \| 297 \| 0.14 \| 348 \| 215 \| 19 \| 382 \| 679 \| 1 \| 314 \| Major capsid protein \| Major capsid protein; pdb \| 8EC8\_B \| 1.0 \| 4.512e-10 \| 295 \| 0.111 \| 297 \| 211 \| 15 \| 421 \| 680 \| 18 \| 298 \| Major capsid protein \| Major capsid protein; pdb \| 7OZ4\_C \| 1.0 \| 3.398e-10 \| 295 \| 0.124 \| 297 \| 218 \| 13 \| 423 \| 680 \| 8 \| 301 \| p2 family phage major capsid protein \| p2 family phage major capsid protein; pdb \| 2E0Z\_C \| 1.0 \| 1.861e-09 \| 287 \| 0.087 \| 252 \| 183 \| 12 \| 432 \| 679 \| 20 \| 228 \| Virus-like particle \| Virus-like particle; pdb \| 2E0Z\_B \| 1.0 \| 1.402e-09 \| 284 \| 0.087 \| 251 \| 190 \| 10 \| 433 \| 679 \| 21 \| 236 \| Virus-like particle \| Virus-like particle; pdb \| 7Y23\_F \| 1.0 \| 5.865e-11 \| 280 \| 0.105 \| 351 \| 227 \| 21 \| 402 \| 681 \| 6 \| 340 \| phage capsid protein \| phage capsid protein; pdb \| 5TJT\_C \| 1.0 \| 4.512e-10 \| 269 \| 0.129 \| 285 \| 206 \| 14 \| 423 \| 678 \| 19 \| 290 \| Major capsid protein \| Major capsid protein; pdb \| 7S21\_A \| 1.0 \| 2.084e-09 \| 267 \| 0.117 \| 256 \| 187 \| 15 \| 435 \| 677 \| 35 \| 264 \| EncA \| EncA; pdb \| 7KQ5\_A \| 1.0 \| 5.348e-10 \| 263 \| 0.135 \| 258 \| 184 \| 16 \| 434 \| 679 \| 34 \| 264 \| Maritimacin \| Maritimacin; pdb \| 7Y23\_G \| 1.0 \| 2.506e-11 \| 263 \| 0.118 \| 338 \| 230 \| 19 \| 402 \| 682 \| 1 \| 327 \| phage capsid protein \| phage capsid protein; pdb \| 3J7X\_F \| 1.0 \| 6.709e-10 \| 263 \| 0.115 \| 345 \| 228 \| 20 \| 402 \| 682 \| 12 \| 343 \| Major capsid protein 10A \| Major capsid protein 10A; pdb \| 7RWZ\_D \| 1.0 \| 1.969e-09 \| 261 \| 0.136 \| 272 \| 187 \| 18 \| 426 \| 681 \| 14 \| 253 \| Major capsid protein \| Major capsid protein; pdb \| 7Y3T\_C \| 1.0 \| 7.1e-10 \| 256 \| 0.104 \| 343 \| 235 \| 21 \| 399 \| 682 \| 1 \| 330 \| phage major capsid protein \| phage major capsid protein; pdb \| 6WKV\_A \| 1.0 \| 1.056e-09 \| 250 \| 0.123 \| 252 \| 185 \| 13 \| 435 \| 679 \| 34 \| 256 \| Encapsulin \| Encapsulin; pdb \| 7BCV\_A \| 1.0 \| 2.767e-09 \| 250 \| 0.12 \| 257 \| 193 \| 14 \| 434 \| 681 \| 34 \| 266 \| Linocin-M18 \| Linocin-M18; pdb \| 7Z4B\_AY \| 1.0 \| 3.597e-10 \| 243 \| 0.1 \| 360 \| 231 \| 16 \| 399 \| 681 \| 1 \| 344 \| Major head protein \| Major head protein; pdb \| 8HDT\_A \| 1.0 \| 1.661e-09 \| 238 \| 0.116 \| 310 \| 227 \| 17 \| 407 \| 681 \| 3 \| 300 \| Major capsid \| Major capsid; pdb \| 6J3Q\_A \| 1.0 \| 2.253e-08 \| 236 \| 0.115 \| 354 \| 223 \| 19 \| 399 \| 682 \| 1 \| 334 \| major capsid protein \| major capsid protein; pdb \| 7OE2\_A \| 1.0 \| 4.447e-08 \| 235 \| 0.122 \| 262 \| 193 \| 13 \| 431 \| 681 \| 31 \| 266 \| Linocin\_M18 bacteriocin protein \| Linocin\_M18 bacteriocin protein; pdb \| 7LIJ\_A \| 1.0 \| 2.614e-09 \| 231 \| 0.123 \| 260 \| 191 \| 14 \| 431 \| 679 \| 31 \| 264 \| Maritimacin \| Maritimacin; pdb \| 5WK1\_B \| 1.0 \| 2.47e-09 \| 231 \| 0.128 \| 272 \| 202 \| 13 \| 433 \| 678 \| 31 \| 293 \| Major Capsid Protein \| Major Capsid Protein; pdb \| 7LIS\_A \| 1.0 \| 4.608e-09 \| 228 \| 0.138 \| 260 \| 177 \| 17 \| 434 \| 679 \| 34 \| 260 \| Maritimacin \| Maritimacin; pdb \| 8GIU\_G \| 1.0 \| 1.208e-08 \| 227 \| 0.098 \| 305 \| 213 \| 14 \| 433 \| 681 \| 45 \| 343 \| Capsid protein \| Capsid protein; pdb \| 8IKA\_A1 \| 1.0 \| 1.603e-08 \| 225 \| 0.096 \| 259 \| 193 \| 16 \| 435 \| 680 \| 35 \| 265 \| Type 1 encapsulin shell protein \| Type 1 encapsulin shell protein; pdb \| 7JW1\_A \| 1.0 \| 9.976e-10 \| 225 \| 0.089 \| 292 \| 192 \| 15 \| 432 \| 679 \| 14 \| 275 \| Major capsid protein gpN \| Major capsid protein gpN; pdb \| 6I9G\_B \| 1.0 \| 3.165e-08 \| 224 \| 0.124 \| 258 \| 185 \| 18 \| 435 \| 679 \| 35 \| 264 \| Linocin-M18 \| Linocin-M18; pdb \| 6WKK\_F \| 1.0 \| 2.991e-08 \| 223 \| 0.096 \| 290 \| 243 \| 15 \| 402 \| 682 \| 1 \| 280 \| Gp27 major capsid protein \| Gp27 major capsid protein; pdb \| 7LII\_A \| 1.0 \| 1.278e-08 \| 221 \| 0.119 \| 251 \| 187 \| 12 \| 435 \| 679 \| 35 \| 257 \| Maritimacin \| Maritimacin; pdb \| 8FRS\_E \| 1.0 \| 9.097e-09 \| 219 \| 0.12 \| 332 \| 225 \| 22 \| 402 \| 681 \| 1 \| 317 \| Major structural protein \| Major structural protein; pdb \| 7LIM\_A \| 1.0 \| 2.826e-08 \| 217 \| 0.126 \| 252 \| 186 \| 12 \| 434 \| 679 \| 34 \| 257 \| Maritimacin \| Maritimacin; pdb \| 7LIL\_A \| 1.0 \| 6.852e-09 \| 217 \| 0.126 \| 260 \| 190 \| 16 \| 431 \| 679 \| 31 \| 264 \| Maritimacin \| Maritimacin; pdb \| 7LIK\_A \| 1.0 \| 9.627e-09 \| 217 \| 0.126 \| 260 \| 184 \| 14 \| 434 \| 679 \| 34 \| 264 \| Maritimacin \| Maritimacin; pdb \| 6X8T\_A \| 1.0 \| 3.752e-08 \| 216 \| 0.118 \| 329 \| 201 \| 20 \| 385 \| 682 \| 1 \| 271 \| Protein SrpI \| Protein SrpI; pdb \| 7F38\_I \| 1.0 \| 3.752e-08 \| 214 \| 0.118 \| 362 \| 223 \| 21 \| 401 \| 680 \| 3 \| 350 \| Putative major capsid protein \| Putative major capsid protein; pdb \| 8DN9\_A \| 1.0 \| 6.613e-08 \| 213 \| 0.114 \| 263 \| 188 \| 14 \| 430 \| 677 \| 30 \| 262 \| 29 kDa antigen cfp29 \| 29 kDa antigen cfp29; pdb \| 7BOJ\_A \| 1.0 \| 6.248e-08 \| 213 \| 0.136 \| 257 \| 183 \| 16 \| 435 \| 679 \| 35 \| 264 \| 29 kDa antigen Cfp29 \| 29 kDa antigen Cfp29; pdb \| 6B23\_D \| 1.0 \| 4.202e-08 \| 209 \| 0.131 \| 281 \| 186 \| 18 \| 431 \| 681 \| 30 \| 282 \| Capsid morphogenesis B protein \| Capsid morphogenesis B protein; pdb \| 6J3Q\_U \| 1.0 \| 4.447e-08 \| 209 \| 0.107 \| 334 \| 220 \| 14 \| 423 \| 680 \| 12 \| 343 \| major capsid protein \| major capsid protein; pdb \| 7JW1\_C \| 1.0 \| 2.991e-08 \| 208 \| 0.089 \| 290 \| 202 \| 15 \| 432 \| 682 \| 14 \| 280 \| Major capsid protein gpN \| Major capsid protein gpN; pdb \| 7JW1\_B \| 1.0 \| 1.353e-08 \| 208 \| 0.082 \| 303 \| 208 \| 15 \| 432 \| 682 \| 38 \| 322 \| Major capsid protein gpN \| Major capsid protein gpN; pdb \| 7RWZ\_A \| 1.0 \| 3.421e-07 \| 205 \| 0.148 \| 215 \| 147 \| 15 \| 480 \| 681 \| 69 \| 260 \| Cos capsid morphogenesis protein (Ccm) \| Cos capsid morphogenesis protein (Ccm); pdb \| 8GIU\_C \| 1.0 \| 2.253e-08 \| 202 \| 0.094 \| 317 \| 215 \| 17 \| 435 \| 681 \| 48 \| 362 \| Capsid protein \| Capsid protein; pdb \| 7F38\_D \| 1.0 \| 1.547e-07 \| 200 \| 0.142 \| 358 \| 216 \| 18 \| 412 \| 679 \| 3 \| 359 \| Putative major capsid protein \| Putative major capsid protein; pdb \| 6XGQ\_E \| 1.0 \| 1.796e-08 \| 195 \| 0.108 \| 368 \| 223 \| 24 \| 405 \| 682 \| 1 \| 353 \| YSD1\_16 \| YSD1\_16; pdb \| 6OKB\_F \| 1.0 \| 8.473e-07 \| 194 \| 0.14 \| 284 \| 187 \| 15 \| 436 \| 682 \| 26 \| 289 \| Major capsid protein \| Major capsid protein; pdb \| 2XVR\_C \| 1.0 \| 1.77e-06 \| 184 \| 0.123 \| 260 \| 196 \| 13 \| 427 \| 669 \| 1 \| 245 \| MAJOR CAPSID PROTEIN 10A \| MAJOR CAPSID PROTEIN 10A; pdb \| 7Y3T\_G \| 1.0 \| 6.248e-08 \| 180 \| 0.118 \| 338 \| 226 \| 20 \| 399 \| 682 \| 1 \| 320 \| phage major capsid protein \| phage major capsid protein; pdb \| 7EEL\_A \| 1.0 \| 8.296e-08 \| 178 \| 0.102 \| 372 \| 227 \| 21 \| 407 \| 681 \| 2 \| 363 \| Major capsid proteins \| Major capsid proteins; pdb \| 8H89\_B \| 1.0 \| 6.613e-08 \| 175 \| 0.125 \| 312 \| 196 \| 19 \| 444 \| 682 \| 54 \| 361 \| Major capsid protein \| Major capsid protein; pdb \| 8PHQ\_AC \| 1.0 \| 8.296e-08 \| 173 \| 0.109 \| 301 \| 195 \| 23 \| 435 \| 681 \| 30 \| 311 \| Major capsid protein \| Major capsid protein; pdb \| 8PKH\_AG \| 1.0 \| 9.291e-08 \| 172 \| 0.103 \| 308 \| 185 \| 20 \| 435 \| 681 \| 30 \| 307 \| Major capsid protein \| Major capsid protein; pdb \| 7JW1\_D \| 1.0 \| 5.384e-07 \| 171 \| 0.095 \| 294 \| 192 \| 16 \| 433 \| 682 \| 39 \| 302 \| Major capsid protein gpN \| Major capsid protein gpN; pdb \| 8PHS\_CD \| 1.0 \| 2.054e-07 \| 170 \| 0.104 \| 297 \| 201 \| 25 \| 435 \| 681 \| 14 \| 295 \| Major capsid protein \| Major capsid protein; pdb \| 7Z46\_H \| 1.0 \| 8.779e-08 \| 170 \| 0.113 \| 371 \| 209 \| 19 \| 400 \| 681 \| 1 \| 340 \| Major head protein \| Major head protein; pdb \| 6XGQ\_G \| 1.0 \| 3.832e-07 \| 168 \| 0.097 \| 328 \| 208 \| 20 \| 430 \| 682 \| 18 \| 332 \| YSD1\_16 \| YSD1\_16; pdb \| 7Z4A\_C \| 1.0 \| 6.754e-07 \| 167 \| 0.114 \| 341 \| 192 \| 19 \| 422 \| 681 \| 2 \| 313 \| Major head protein \| Major head protein; pdb \| 7Z48\_C \| 1.0 \| 8.473e-07 \| 164 \| 0.119 \| 344 \| 187 \| 20 \| 423 \| 681 \| 5 \| 317 \| Major head protein \| Major head protein; pdb \| 7Z48\_D \| 1.0 \| 2.487e-06 \| 162 \| 0.097 \| 337 \| 224 \| 18 \| 411 \| 681 \| 1 \| 323 \| Major head protein \| Major head protein; pdb \| 3BJQ\_C \| 1.0 \| 8.177e-06 \| 158 \| 0.151 \| 238 \| 154 \| 13 \| 478 \| 681 \| 67 \| 290 \| Phage-related protein \| Phage-related protein; pdb \| 5L35\_A \| 1.0 \| 1.063e-06 \| 155 \| 0.083 \| 443 \| 208 \| 19 \| 410 \| 682 \| 8 \| 422 \| Gene 5 protein \| Gene 5 protein; pdb \| 6QVK\_1B \| 1.0 \| 3.055e-07 \| 154 \| 0.111 \| 405 \| 240 \| 22 \| 330 \| 682 \| 6 \| 342 \| Major capsid protein \| Major capsid protein; pdb \| 7Z4A\_G \| 1.0 \| 5.82e-06 \| 148 \| 0.106 \| 329 \| 210 \| 18 \| 411 \| 681 \| 1 \| 303 \| Major head protein \| Major head protein; pdb \| 8CJZ\_G \| 1.0 \| 1.149e-05 \| 147 \| 0.156 \| 287 \| 149 \| 15 \| 421 \| 680 \| 24 \| 244 \| Major Capsid Protein \| Major Capsid Protein; pdb \| 3BJQ\_G \| 1.0 \| 3.997e-05 \| 146 \| 0.159 \| 238 \| 153 \| 11 \| 478 \| 681 \| 66 \| 290 \| Phage-related protein \| Phage-related protein; pdb \| 3J4U\_G \| 1.0 \| 5.196e-06 \| 145 \| 0.127 \| 275 \| 160 \| 15 \| 423 \| 680 \| 24 \| 235 \| major capsid protein \| major capsid protein; pdb \| 7F2P\_I \| 1.0 \| 2.487e-06 \| 145 \| 0.09 \| 388 \| 228 \| 24 \| 395 \| 681 \| 4 \| 367 \| KHP40 MCP \| KHP40 MCP; pdb \| 8GMO\_AD \| 1.0 \| 2.221e-06 \| 144 \| 0.107 \| 280 \| 176 \| 13 \| 467 \| 680 \| 165 \| 436 \| Mature major capsid protein \| Mature major capsid protein; pdb \| 3BJQ\_H \| 1.0 \| 1.441e-05 \| 143 \| 0.145 \| 310 \| 183 \| 19 \| 420 \| 681 \| 13 \| 288 \| Phage-related protein \| Phage-related protein; pdb \| 3BJQ\_F \| 1.0 \| 1.216e-05 \| 143 \| 0.145 \| 309 \| 187 \| 22 \| 420 \| 681 \| 13 \| 291 \| Phage-related protein \| Phage-related protein; pdb \| 8CJZ\_H \| 1.0 \| 4.639e-06 \| 143 \| 0.148 \| 290 \| 150 \| 14 \| 421 \| 681 \| 24 \| 245 \| Spike Base Protein \| Spike Base Protein; pdb \| 3J4U\_C \| 1.0 \| 3.569e-05 \| 141 \| 0.125 \| 278 \| 159 \| 16 \| 422 \| 680 \| 23 \| 235 \| major capsid protein \| major capsid protein; pdb \| 7F2P\_A \| 1.0 \| 6.159e-06 \| 139 \| 0.081 \| 392 \| 222 \| 25 \| 398 \| 680 \| 2 \| 364 \| KHP40 MCP \| KHP40 MCP; pdb \| 7DN2\_C \| 1.0 \| 4.142e-06 \| 139 \| 0.071 \| 390 \| 232 \| 17 \| 398 \| 682 \| 4 \| 368 \| Major structural protein ORF14 \| Major structural protein ORF14; pdb \| 7F2P\_B \| 1.0 \| 9.692e-06 \| 135 \| 0.09 \| 396 \| 210 \| 26 \| 402 \| 682 \| 3 \| 363 \| KHP40 MCP \| KHP40 MCP; pdb \| 3BJQ\_E \| 1.0 \| 5.307e-05 \| 134 \| 0.143 \| 313 \| 192 \| 22 \| 420 \| 681 \| 13 \| 300 \| Phage-related protein \| Phage-related protein; pdb \| 6Q3G\_AD \| 1.0 \| 7.301e-06 \| 134 \| 0.086 \| 371 \| 231 \| 20 \| 402 \| 681 \| 1 \| 354 \| Major head protein \| Major head protein; pdb \| 8I1T\_G \| 1.0 \| 2.221e-06 \| 133 \| 0.075 \| 439 \| 227 \| 17 \| 407 \| 680 \| 1 \| 425 \| Major capsid protein \| Major capsid protein; pdb \| 3BJQ\_B \| 1.0 \| 2.845e-05 \| 132 \| 0.144 \| 298 \| 177 \| 21 \| 438 \| 681 \| 26 \| 299 \| Phage-related protein \| Phage-related protein; pdb \| 3BJQ\_I \| 1.0 \| 7.046e-05 \| 130 \| 0.14 \| 291 \| 177 \| 20 \| 433 \| 678 \| 23 \| 285 \| Phage-related protein \| Phage-related protein; pdb \| 3BJQ\_A \| 1.0 \| 4.738e-05 \| 130 \| 0.143 \| 314 \| 183 \| 22 \| 420 \| 681 \| 13 \| 292 \| Phage-related protein \| Phage-related protein; pdb \| 7DN2\_I \| 1.0 \| 7.726e-06 \| 130 \| 0.073 \| 382 \| 219 \| 25 \| 408 \| 679 \| 5 \| 361 \| Major structural protein ORF14 \| Major structural protein ORF14; pdb \| 5LII\_P \| 1.0 \| 3.187e-05 \| 130 \| 0.098 \| 437 \| 226 \| 20 \| 409 \| 682 \| 1 \| 432 \| Major capsid protein \| Major capsid protein; pdb \| 7VS5\_AA \| 1.0 \| 5.82e-06 \| 130 \| 0.114 \| 296 \| 186 \| 16 \| 452 \| 681 \| 171 \| 456 \| Major capsid protein \| Major capsid protein; pdb \| 3BJQ\_J \| 1.0 \| 7.457e-05 \| 129 \| 0.153 \| 306 \| 184 \| 20 \| 420 \| 678 \| 13 \| 290 \| Phage-related protein \| Phage-related protein; pdb \| 7DN2\_H \| 1.0 \| 2.143e-05 \| 129 \| 0.083 \| 393 \| 213 \| 23 \| 402 \| 679 \| 1 \| 361 \| Major structural protein ORF14 \| Major structural protein ORF14; pdb \| 6OKB\_E \| 1.0 \| 0.0001472 \| 127 \| 0.142 \| 302 \| 188 \| 19 \| 417 \| 681 \| 20 \| 287 \| Major capsid protein \| Major capsid protein; pdb \| 8EGT\_A \| 1.0 \| 2.948e-06 \| 127 \| 0.098 \| 406 \| 221 \| 21 \| 392 \| 682 \| 13 \| 388 \| Major capsid protein \| Major capsid protein; pdb \| 3BJQ\_D \| 1.0 \| 4.477e-05 \| 126 \| 0.15 \| 305 \| 174 \| 20 \| 433 \| 681 \| 23 \| 298 \| Phage-related protein \| Phage-related protein; pdb \| 7F2P\_G \| 1.0 \| 1.362e-05 \| 126 \| 0.075 \| 397 \| 212 \| 24 \| 402 \| 679 \| 8 \| 368 \| KHP40 MCP \| KHP40 MCP; pdb \| 6IAW\_O \| 1.0 \| 1.086e-05 \| 124 \| 0.097 \| 399 \| 227 \| 21 \| 392 \| 681 \| 12 \| 386 \| Major head protein \| Major head protein; pdb \| 3J7V\_B \| 1.0 \| 7.046e-05 \| 122 \| 0.08 \| 297 \| 212 \| 17 \| 434 \| 681 \| 25 \| 309 \| Major capsid protein 10A \| Major capsid protein 10A; pdb \| 8GMO\_6 \| 1.0 \| 1.614e-05 \| 122 \| 0.109 \| 274 \| 174 \| 13 \| 471 \| 680 \| 195 \| 462 \| Mature major capsid protein \| Mature major capsid protein; pdb \| 7VRT\_FB \| 1.0 \| 1.441e-05 \| 121 \| 0.123 \| 275 \| 162 \| 18 \| 468 \| 681 \| 115 \| 371 \| Major capsid protein \| Major capsid protein; pdb \| 6I9E\_A \| 1.0 \| 2.098e-06 \| 121 \| 0.098 \| 394 \| 240 \| 20 \| 399 \| 682 \| 6 \| 394 \| Major head protein \| Major head protein; pdb \| 1YUE\_A \| 1.0 \| 6.518e-06 \| 120 \| 0.128 \| 289 \| 161 \| 17 \| 465 \| 682 \| 122 \| 390 \| Head vertex protein Gp24 \| Head vertex protein Gp24; pdb \| 6Q3G\_KD \| 1.0 \| 5.499e-06 \| 117 \| 0.093 \| 415 \| 229 \| 25 \| 378 \| 681 \| 4 \| 382 \| Major head protein \| Major head protein; pdb \| 7VS5\_DW \| 1.0 \| 8.352e-05 \| 117 \| 0.108 \| 287 \| 168 \| 16 \| 469 \| 681 \| 158 \| 430 \| Major capsid protein \| Major capsid protein; pdb \| 7VS5\_HG \| 1.0 \| 5.015e-05 \| 116 \| 0.144 \| 305 \| 140 \| 21 \| 463 \| 682 \| 142 \| 410 \| Capsid vertex protein \| Capsid vertex protein; pdb \| 5VF3\_L \| 1.0 \| 2.688e-05 \| 114 \| 0.093 \| 277 \| 171 \| 16 \| 474 \| 681 \| 189 \| 454 \| Major capsid protein \| Major capsid protein; pdb \| 7QOF\_A \| 1.0 \| 7.891e-05 \| 114 \| 0.106 \| 291 \| 177 \| 15 \| 465 \| 681 \| 187 \| 468 \| Auxiliary capsid protein gp36 \| Auxiliary capsid protein gp36; pdb \| 6XGP\_B \| 1.0 \| 0.0001048 \| 111 \| 0.098 \| 275 \| 164 \| 17 \| 478 \| 682 \| 70 \| 330 \| YSD1\_17 major capsid protein \| YSD1\_17 major capsid protein; pdb \| 7DN2\_E \| 1.0 \| 5.944e-05 \| 111 \| 0.09 \| 397 \| 203 \| 29 \| 406 \| 681 \| 11 \| 370 \| Major structural protein ORF14 \| Major structural protein ORF14; pdb \| 4AN5\_C \| 1.0 \| 0.0009027 \| 110 \| 0.11 \| 272 \| 173 \| 18 \| 433 \| 676 \| 3 \| 233 \| COAT PROTEIN \| COAT PROTEIN; pdb \| 6XGP\_A \| 1.0 \| 9.354e-05 \| 106 \| 0.099 \| 272 \| 164 \| 19 \| 478 \| 682 \| 70 \| 327 \| YSD1\_17 major capsid protein \| YSD1\_17 major capsid protein; pdb \| 7VI9\_G \| 1.0 \| 0.0005122 \| 104 \| 0.111 \| 341 \| 219 \| 20 \| 418 \| 681 \| 2 \| 335 \| Major capsid protein \| Major capsid protein; pdb \| 7QOF\_B \| 1.0 \| 0.0001109 \| 102 \| 0.089 \| 323 \| 178 \| 19 \| 465 \| 681 \| 187 \| 499 \| Auxiliary capsid protein gp36 \| Auxiliary capsid protein gp36; pdb \| 3BQW\_A \| 1.0 \| 0.0001109 \| 100 \| 0.105 \| 295 \| 158 \| 18 \| 474 \| 681 \| 72 \| 347 \| Putative capsid protein of prophage \| Putative capsid protein of prophage; pdb \| 7SJ5\_A \| 0.999 \| 7.891e-05 \| 97 \| 0.12 \| 365 \| 172 \| 26 \| 424 \| 681 \| 18 \| 340 \| Major capsid protein \| Major capsid protein; pdb \| 6LGN\_Z \| 0.999 \| 0.0001472 \| 97 \| 0.084 \| 413 \| 204 \| 22 \| 422 \| 681 \| 6 \| 397 \| Small capsomere-interacting protein \| Small capsomere-interacting protein; pdb \| 5JBL\_A \| 0.997 \| 0.002504 \| 92 \| 0.096 \| 176 \| 107 \| 17 \| 30 \| 170 \| 5 \| 163 \| Prohead core protein protease \| Prohead core protein protease; pdb \| 3J7V\_G \| 0.996 \| 0.0005122 \| 90 \| 0.11 \| 300 \| 200 \| 20 \| 437 \| 681 \| 30 \| 317 \| Major capsid protein 10A \| Major capsid protein 10A; pdb \| 7QOI\_CD \| 0.996 \| 0.0007196 \| 90 \| 0.102 \| 294 \| 167 \| 18 \| 466 \| 681 \| 173 \| 447 \| Auxiliary capsid protein gp36 \| Auxiliary capsid protein gp36; pdb \| 5JBL\_B \| 0.996 \| 0.005859 \| 89 \| 0.107 \| 177 \| 108 \| 16 \| 30 \| 173 \| 5 \| 164 \| Prohead core protein protease \| Prohead core protein protease; pdb \| 3J40\_E \| 0.996 \| 0.0005737 \| 89 \| 0.102 \| 311 \| 182 \| 16 \| 391 \| 682 \| 11 \| 243 \| gp7 \| gp7; pdb \| 8I1V\_D \| 0.993 \| 0.0003255 \| 86 \| 0.106 \| 404 \| 189 \| 19 \| 431 \| 680 \| 23 \| 408 \| Major capsid protein \| Major capsid protein; pdb \| 7QOI\_AD \| 0.993 \| 0.001011 \| 86 \| 0.1 \| 298 \| 168 \| 14 \| 466 \| 681 \| 173 \| 452 \| Auxiliary capsid protein gp36 \| Auxiliary capsid protein gp36; pdb \| 8I1V\_G \| 0.99 \| 0.0004321 \| 83 \| 0.103 \| 416 \| 185 \| 23 \| 431 \| 680 \| 23 \| 416 \| Major capsid protein \| Major capsid protein; pdb \| 6ODM\_W \| 0.99 \| 0.005536 \| 83 \| 0.155 \| 58 \| 43 \| 3 \| 625 \| 681 \| 1055 \| 1107 \| Major capsid protein \| Major capsid protein; pdb \| 8I1V\_A \| 0.984 \| 0.0009554 \| 80 \| 0.102 \| 409 \| 189 \| 21 \| 431 \| 680 \| 22 \| 411 \| Major capsid protein \| Major capsid protein; pdb \| 8I1V\_F \| 0.984 \| 0.0009027 \| 80 \| 0.084 \| 428 \| 182 \| 19 \| 431 \| 680 \| 23 \| 418 \| Major capsid protein \| Major capsid protein; pdb \| 6ODM\_S \| 0.984 \| 0.004943 \| 80 \| 0.161 \| 62 \| 45 \| 4 \| 621 \| 681 \| 1013 \| 1068 \| Major capsid protein \| Major capsid protein; pdb \| 8I1V\_B \| 0.855 \| 0.001684 \| 64 \| 0.096 \| 406 \| 193 \| 18 \| 431 \| 680 \| 23 \| 410 \| Major capsid protein \| Major capsid protein; afdb-proteome \| AF-Q2FX56-F1-MODEL\_V4 \| 1.0 \| 1.607e-14 \| 429 \| 0.132 \| 324 \| 221 \| 12 \| 402 \| 682 \| 8 \| 314 \| Phage head protein, putative \| Phage head protein, putative; afdb-proteome \| AF-A0A0H3GTX7-F1-MODEL\_V4 \| 1.0 \| 2.34e-15 \| 396 \| 0.133 \| 463 \| 276 \| 24 \| 258 \| 681 \| 8 \| 384 \| Phage major capsid protein, HK97 family \| Phage major capsid protein, HK97 family; afdb-proteome \| AF-A0A0H3GZA0-F1-MODEL\_V4 \| 1.0 \| 9.717e-12 \| 348 \| 0.138 \| 295 \| 215 \| 15 \| 410 \| 681 \| 12 \| 290 \| Uncharacterized protein \| Uncharacterized protein; afdb-proteome \| AF-A0A133CKM2-F1-MODEL\_V4 \| 1.0 \| 1.813e-11 \| 251 \| 0.131 \| 455 \| 256 \| 23 \| 268 \| 679 \| 6 \| 364 \| Major capsid protein \| Major capsid protein; afdb-proteome \| AF-A0A242FSQ3-F1-MODEL\_V4 \| 1.0 \| 3.527e-09 \| 215 \| 0.109 \| 393 \| 244 \| 20 \| 338 \| 681 \| 46 \| 381 \| Phage capsid protein \| Phage capsid protein; afdb-uniprot50 \| AF-B2I5F3-F1-MODEL\_V4 \| 1.0 \| 8.792e-81 \| 2210 \| 0.472 \| 656 \| 297 \| 11 \| 29 \| 681 \| 15 \| 624 \| Peptidase U35 phage prohead HK97 \| Peptidase U35 phage prohead HK97; afdb-uniprot50 \| AF-A0A4Q3Z182-F1-MODEL\_V4 \| 1.0 \| 5.592e-89 \| 2171 \| 0.513 \| 676 \| 305 \| 10 \| 19 \| 682 \| 1 \| 664 \| Phage major capsid protein \| Phage major capsid protein; afdb-uniprot50 \| AF-A0A420WVK6-F1-MODEL\_V4 \| 1.0 \| 4.038e-91 \| 2168 \| 0.569 \| 717 \| 253 \| 11 \| 1 \| 682 \| 37 \| 732 \| HK97 family phage major capsid protein \| HK97 family phage major capsid protein; afdb-uniprot50 \| AF-A0A1H2ZH10-F1-MODEL\_V4 \| 1.0 \| 2.527e-81 \| 2004 \| 0.5 \| 662 \| 294 \| 9 \| 36 \| 682 \| 6 \| 645 \| Phage prohead protease, HK97 family/phage major capsid protein, HK97 family,TIGR01554 \| Phage prohead protease, HK97 family/phage major capsid protein, HK97 family,TIGR01554; afdb-uniprot50 \| AF-A0A7H8DLL2-F1-MODEL\_V4 \| 1.0 \| 5.789e-82 \| 1964 \| 0.471 \| 679 \| 320 \| 15 \| 19 \| 682 \| 2 \| 656 \| Phage major capsid protein \| Phage major capsid protein; afdb-uniprot50 \| AF-A0A7C8LYK9-F1-MODEL\_V4 \| 1.0 \| 3.647e-69 \| 1916 \| 0.392 \| 675 \| 347 \| 14 \| 11 \| 681 \| 9 \| 624 \| Uncharacterized protein \| Uncharacterized protein; afdb-uniprot50 \| AF-A0A4Q3L4Z9-F1-MODEL\_V4 \| 1.0 \| 3.571e-70 \| 1901 \| 0.351 \| 672 \| 381 \| 10 \| 30 \| 682 \| 10 \| 645 \| Phage major capsid protein \| Phage major capsid protein; afdb-uniprot50 \| AF-A0A0C1ILN4-F1-MODEL\_V4 \| 1.0 \| 2.491e-79 \| 1875 \| 0.473 \| 708 \| 311 \| 13 \| 1 \| 682 \| 1 \| 672 \| Peptidase U35 \| Peptidase U35; afdb-uniprot50 \| AF-A0A149SVI9-F1-MODEL\_V4 \| 1.0 \| 3.808e-75 \| 1838 \| 0.477 \| 677 \| 280 \| 11 \| 28 \| 682 \| 14 \| 638 \| Uncharacterized protein \| Uncharacterized protein; afdb-uniprot50 \| AF-A0A0Q7SGT0-F1-MODEL\_V4 \| 1.0 \| 2.159e-67 \| 1838 \| 0.375 \| 709 \| 334 \| 19 \| 19 \| 681 \| 6 \| 651 \| Uncharacterized protein \| Uncharacterized protein; afdb-uniprot50 \| AF-A0A166AHQ0-F1-MODEL\_V4 \| 1.0 \| 7.782e-68 \| 1761 \| 0.38 \| 662 \| 341 \| 13 \| 30 \| 681 \| 11 \| 613 \| Phage capsid family protein \| Phage capsid family protein; afdb-uniprot50 \| AF-A0A7J0BZB5-F1-MODEL\_V4 \| 1.0 \| 7.2e-69 \| 1755 \| 0.449 \| 647 \| 282 \| 11 \| 48 \| 681 \| 2 \| 587 \| Uncharacterized protein \| Uncharacterized protein; afdb-uniprot50 \| AF-A0A501X0T1-F1-MODEL\_V4 \| 1.0 \| 7.403e-65 \| 1746 \| 0.397 \| 675 \| 329 \| 16 \| 19 \| 682 \| 32 \| 639 \| Phage major capsid protein \| Phage major capsid protein; afdb-uniprot50 \| AF-A0A143DDT8-F1-MODEL\_V4 \| 1.0 \| 7.41e-73 \| 1738 \| 0.445 \| 674 \| 307 \| 15 \| 18 \| 679 \| 2 \| 620 \| Uncharacterized protein \| Uncharacterized protein; afdb-uniprot50 \| AF-A0A1S1CXE5-F1-MODEL\_V4 \| 1.0 \| 1.078e-65 \| 1736 \| 0.372 \| 685 \| 345 \| 21 \| 19 \| 680 \| 1 \| 623 \| Major capsid protein \| Major capsid protein; afdb-uniprot50 \| AF-A0A2E9II77-F1-MODEL\_V4 \| 1.0 \| 5.696e-64 \| 1729 \| 0.351 \| 677 \| 351 \| 19 \| 12 \| 678 \| 13 \| 611 \| Phage major capsid protein \| Phage major capsid protein; afdb-uniprot50 \| AF-A0A2E1P3K3-F1-MODEL\_V4 \| 1.0 \| 1.547e-64 \| 1670 \| 0.364 \| 674 \| 338 \| 19 \| 17 \| 678 \| 269 \| 863 \| Phage major capsid protein \| Phage major capsid protein; afdb-uniprot50 \| AF-A0A3A4XAN1-F1-MODEL\_V4 \| 1.0 \| 2.237e-68 \| 1668 \| 0.396 \| 663 \| 320 \| 14 \| 30 \| 682 \| 40 \| 632 \| Phage major capsid protein \| Phage major capsid protein; afdb-uniprot50 \| AF-A0A2E0ENB5-F1-MODEL\_V4 \| 1.0 \| 2.505e-68 \| 1663 \| 0.39 \| 682 \| 344 \| 15 \| 19 \| 681 \| 9 \| 637 \| Phage major capsid protein \| Phage major capsid protein; afdb-uniprot50 \| AF-A0A5P9J3N0-F1-MODEL\_V4 \| 1.0 \| 2.847e-70 \| 1662 \| 0.409 \| 669 \| 337 \| 17 \| 39 \| 682 \| 44 \| 679 \| Phage capsid family protein \| Phage capsid family protein; afdb-uniprot50 \| AF-A0A2V4KHZ2-F1-MODEL\_V4 \| 1.0 \| 1.093e-67 \| 1652 \| 0.366 \| 676 \| 337 \| 15 \| 30 \| 682 \| 11 \| 617 \| Phage major capsid protein \| Phage major capsid protein; afdb-uniprot50 \| AF-A0A1X7L142-F1-MODEL\_V4 \| 1.0 \| 5.305e-62 \| 1632 \| 0.443 \| 586 \| 278 \| 8 \| 99 \| 682 \| 4 \| 543 \| Phage major capsid protein, HK97 family \| Phage major capsid protein, HK97 family; afdb-uniprot50 \| AF-A0A1Y0N5R1-F1-MODEL\_V4 \| 1.0 \| 7.248e-66 \| 1615 \| 0.396 \| 689 \| 314 \| 22 \| 19 \| 680 \| 2 \| 615 \| Uncharacterized protein \| Uncharacterized protein; afdb-uniprot50 \| AF-Q3RBW7-F1-MODEL\_V4 \| 1.0 \| 4.282e-48 \| 1608 \| 0.527 \| 370 \| 164 \| 5 \| 313 \| 681 \| 1 \| 360 \| Uncharacterized protein \| Uncharacterized protein; afdb-uniprot50 \| AF-A0A5C7PYX8-F1-MODEL\_V4 \| 1.0 \| 2.284e-67 \| 1598 \| 0.416 \| 668 \| 321 \| 12 \| 19 \| 678 \| 2 \| 608 \| Phage major capsid protein \| Phage major capsid protein; afdb-uniprot50 \| AF-A0A7X8AGV1-F1-MODEL\_V4 \| 1.0 \| 2.927e-66 \| 1589 \| 0.394 \| 679 \| 320 \| 17 \| 29 \| 682 \| 19 \| 631 \| Phage major capsid protein \| Phage major capsid protein; afdb-uniprot50 \| AF-F4BFR1-F1-MODEL\_V4 \| 1.0 \| 1.696e-65 \| 1585 \| 0.356 \| 676 \| 327 \| 18 \| 25 \| 681 \| 14 \| 600 \| Phage protein \| Phage protein; afdb-uniprot50 \| AF-A0A0F9KU40-F1-MODEL\_V4 \| 1.0 \| 1.071e-68 \| 1575 \| 0.391 \| 722 \| 347 \| 19 \| 16 \| 682 \| 33 \| 716 \| Uncharacterized protein \| Uncharacterized protein; afdb-uniprot50 \| AF-A0A8B3RDF4-F1-MODEL\_V4 \| 1.0 \| 2.885e-64 \| 1520 \| 0.351 \| 680 \| 349 \| 21 \| 19 \| 682 \| 1 \| 604 \| Phage major capsid protein \| Phage major capsid protein; afdb-uniprot50 \| AF-A0A3A4P4D4-F1-MODEL\_V4 \| 1.0 \| 7.403e-65 \| 1511 \| 0.377 \| 696 \| 340 \| 22 \| 16 \| 680 \| 3 \| 635 \| Phage major capsid protein \| Phage major capsid protein; afdb-uniprot50 \| AF-A0A355AHA6-F1-MODEL\_V4 \| 1.0 \| 1.018e-65 \| 1511 \| 0.335 \| 665 \| 389 \| 14 \| 20 \| 678 \| 215 \| 832 \| Phage major capsid protein \| Phage major capsid protein; afdb-uniprot50 \| AF-A0A1W9SXD3-F1-MODEL\_V4 \| 1.0 \| 1.873e-63 \| 1495 \| 0.384 \| 692 \| 306 \| 16 \| 11 \| 681 \| 24 \| 616 \| Uncharacterized protein \| Uncharacterized protein; afdb-uniprot50 \| AF-A0A6M3XKV2-F1-MODEL\_V4 \| 1.0 \| 6.604e-57 \| 1459 \| 0.294 \| 687 \| 364 \| 20 \| 26 \| 682 \| 11 \| 606 \| Putative capsid protein \| Putative capsid protein; afdb-uniprot50 \| AF-A0A1G2ZKV7-F1-MODEL\_V4 \| 1.0 \| 2.885e-64 \| 1451 \| 0.365 \| 686 \| 336 \| 22 \| 29 \| 681 \| 11 \| 629 \| Uncharacterized protein \| Uncharacterized protein; afdb-uniprot50 \| AF-A0A5C9BFK7-F1-MODEL\_V4 \| 1.0 \| 1.844e-53 \| 1420 \| 0.32 \| 670 \| 343 \| 23 \| 39 \| 680 \| 12 \| 596 \| Putative phage major capsid protein \| Putative phage major capsid protein; afdb-uniprot50 \| AF-A0A2H6GL21-F1-MODEL\_V4 \| 1.0 \| 7.775e-60 \| 1419 \| 0.33 \| 672 \| 342 \| 19 \| 24 \| 678 \| 4 \| 584 \| Phage capsid family protein \| Phage capsid family protein; afdb-uniprot50 \| AF-A0A3A0FXT4-F1-MODEL\_V4 \| 1.0 \| 5.817e-63 \| 1416 \| 0.366 \| 688 \| 357 \| 19 \| 30 \| 681 \| 33 \| 677 \| Phage major capsid protein \| Phage major capsid protein; afdb-uniprot50 \| AF-A0A448XYU4-F1-MODEL\_V4 \| 1.0 \| 3.323e-60 \| 1414 \| 0.348 \| 680 \| 378 \| 18 \| 14 \| 679 \| 3 \| 631 \| HK97 family phage major capsid protein \| HK97 family phage major capsid protein; afdb-uniprot50 \| AF-A0A2D3WU31-F1-MODEL\_V4 \| 1.0 \| 6.422e-61 \| 1413 \| 0.339 \| 662 \| 348 \| 14 \| 30 \| 682 \| 14 \| 594 \| Phage major capsid protein \| Phage major capsid protein; afdb-uniprot50 \| AF-A0A661FCZ5-F1-MODEL\_V4 \| 1.0 \| 3.644e-61 \| 1407 \| 0.368 \| 664 \| 326 \| 13 \| 30 \| 678 \| 14 \| 598 \| Phage major capsid protein \| Phage major capsid protein; afdb-uniprot50 \| AF-A0A0H3ZS84-F1-MODEL\_V4 \| 1.0 \| 7.775e-60 \| 1390 \| 0.376 \| 664 \| 345 \| 14 \| 29 \| 682 \| 10 \| 614 \| Phage major capsid protein \| Phage major capsid protein; afdb-uniprot50 \| AF-A0A254T9Q6-F1-MODEL\_V4 \| 1.0 \| 2.882e-56 \| 1378 \| 0.341 \| 689 \| 340 \| 23 \| 29 \| 681 \| 9 \| 619 \| Uncharacterized protein \| Uncharacterized protein; afdb-uniprot50 \| AF-A0A661QEN8-F1-MODEL\_V4 \| 1.0 \| 1.659e-58 \| 1360 \| 0.37 \| 664 \| 311 \| 19 \| 38 \| 681 \| 21 \| 597 \| Phage major capsid protein \| Phage major capsid protein; afdb-uniprot50 \| AF-A0A2E7V2Q0-F1-MODEL\_V4 \| 1.0 \| 5.455e-58 \| 1359 \| 0.331 \| 660 \| 346 \| 20 \| 40 \| 680 \| 41 \| 623 \| Phage major capsid protein \| Phage major capsid protein; afdb-uniprot50 \| AF-A0A6M8SY00-F1-MODEL\_V4 \| 1.0 \| 1.858e-58 \| 1356 \| 0.373 \| 672 \| 327 \| 24 \| 40 \| 681 \| 20 \| 627 \| Phage major capsid protein \| Phage major capsid protein; afdb-uniprot50 \| AF-A0A2A5AEG6-F1-MODEL\_V4 \| 1.0 \| 2.157e-59 \| 1350 \| 0.328 \| 675 \| 371 \| 16 \| 26 \| 682 \| 12 \| 621 \| Phage major capsid protein \| Phage major capsid protein; afdb-uniprot50 \| AF-A0A3C0NVR1-F1-MODEL\_V4 \| 1.0 \| 1.285e-54 \| 1347 \| 0.3 \| 708 \| 383 \| 24 \| 10 \| 681 \| 17 \| 647 \| Phage major capsid protein \| Phage major capsid protein; afdb-uniprot50 \| AF-A0A238KH43-F1-MODEL\_V4 \| 1.0 \| 2.574e-56 \| 1315 \| 0.319 \| 685 \| 399 \| 20 \| 19 \| 680 \| 1 \| 641 \| Phage capsid family protein \| Phage capsid family protein; afdb-uniprot50 \| AF-A0A3M1CK54-F1-MODEL\_V4 \| 1.0 \| 3.051e-56 \| 1314 \| 0.32 \| 668 \| 347 \| 20 \| 30 \| 679 \| 5 \| 583 \| Phage major capsid protein \| Phage major capsid protein; afdb-uniprot50 \| AF-A0A2J8GXS4-F1-MODEL\_V4 \| 1.0 \| 2.611e-58 \| 1301 \| 0.367 \| 669 \| 333 \| 19 \| 30 \| 682 \| 12 \| 606 \| Phage major capsid protein \| Phage major capsid protein; afdb-uniprot50 \| AF-A0A227JQC9-F1-MODEL\_V4 \| 1.0 \| 3.965e-57 \| 1300 \| 0.296 \| 699 \| 378 \| 21 \| 31 \| 680 \| 3 \| 636 \| Phage major capsid protein \| Phage major capsid protein; afdb-uniprot50 \| AF-A0A0G3G1E5-F1-MODEL\_V4 \| 1.0 \| 4.164e-52 \| 1287 \| 0.309 \| 646 \| 373 \| 16 \| 46 \| 681 \| 2 \| 584 \| Uncharacterized protein \| Uncharacterized protein; afdb-uniprot50 \| AF-A0A2A5DI76-F1-MODEL\_V4 \| 1.0 \| 4.903e-55 \| 1281 \| 0.302 \| 672 \| 364 \| 21 \| 41 \| 682 \| 33 \| 629 \| Phage major capsid protein \| Phage major capsid protein; afdb-uniprot50 \| AF-A0A5N0TEK3-F1-MODEL\_V4 \| 1.0 \| 1.793e-57 \| 1270 \| 0.353 \| 685 \| 353 \| 19 \| 16 \| 681 \| 5 \| 618 \| Phage major capsid protein \| Phage major capsid protein; afdb-uniprot50 \| AF-A0A0F2NXV9-F1-MODEL\_V4 \| 1.0 \| 4.315e-53 \| 1268 \| 0.305 \| 678 \| 355 \| 20 \| 18 \| 682 \| 10 \| 584 \| Uncharacterized protein \| Uncharacterized protein; afdb-uniprot50 \| AF-A0A3B9Q986-F1-MODEL\_V4 \| 1.0 \| 8.461e-56 \| 1268 \| 0.3 \| 698 \| 392 \| 24 \| 19 \| 680 \| 1 \| 637 \| Phage major capsid protein \| Phage major capsid protein; afdb-uniprot50 \| AF-A0A7Y5NBG9-F1-MODEL\_V4 \| 1.0 \| 7.995e-56 \| 1254 \| 0.306 \| 707 \| 378 \| 24 \| 20 \| 681 \| 1 \| 639 \| Phage major capsid protein \| Phage major capsid protein; afdb-uniprot50 \| AF-A0A1Q3WRH7-F1-MODEL\_V4 \| 1.0 \| 1.206e-57 \| 1252 \| 0.336 \| 734 \| 353 \| 30 \| 30 \| 681 \| 18 \| 699 \| Uncharacterized protein \| Uncharacterized protein; afdb-uniprot50 \| AF-A0A369R3L5-F1-MODEL\_V4 \| 1.0 \| 8.34e-54 \| 1245 \| 0.301 \| 714 \| 364 \| 29 \| 29 \| 682 \| 16 \| 654 \| Phage major capsid protein \| Phage major capsid protein; afdb-uniprot50 \| AF-A0A831U076-F1-MODEL\_V4 \| 1.0 \| 6.373e-56 \| 1228 \| 0.331 \| 697 \| 366 \| 27 \| 29 \| 681 \| 16 \| 656 \| Phage major capsid protein \| Phage major capsid protein; afdb-uniprot50 \| AF-A0A3N2E2B1-F1-MODEL\_V4 \| 1.0 \| 4.759e-43 \| 1222 \| 0.393 \| 435 \| 237 \| 11 \| 252 \| 682 \| 246 \| 657 \| ATP-dependent Clp protease proteolytic subunit \| ATP-dependent Clp protease proteolytic subunit; afdb-uniprot50 \| AF-A0A259F2V0-F1-MODEL\_V4 \| 1.0 \| 3.908e-55 \| 1222 \| 0.329 \| 695 \| 349 \| 22 \| 12 \| 682 \| 135 \| 736 \| Uncharacterized protein \| Uncharacterized protein; afdb-uniprot50 \| AF-A0A5K7YQH8-F1-MODEL\_V4 \| 1.0 \| 2.958e-36 \| 1210 \| 0.426 \| 359 \| 194 \| 6 \| 331 \| 682 \| 10 \| 363 \| Uncharacterized protein \| Uncharacterized protein; afdb-uniprot50 \| AF-A0A6M1TYU1-F1-MODEL\_V4 \| 1.0 \| 6.46e-50 \| 1209 \| 0.283 \| 659 \| 355 \| 21 \| 42 \| 681 \| 21 \| 580 \| Phage major capsid protein \| Phage major capsid protein; afdb-uniprot50 \| AF-A0A259FKU1-F1-MODEL\_V4 \| 1.0 \| 9.007e-45 \| 1203 \| 0.348 \| 542 \| 263 \| 19 \| 164 \| 681 \| 2 \| 477 \| Uncharacterized protein \| Uncharacterized protein; afdb-uniprot50 \| AF-A0A7C4G5Z6-F1-MODEL\_V4 \| 1.0 \| 2.981e-41 \| 1202 \| 0.393 \| 455 \| 221 \| 13 \| 252 \| 681 \| 2 \| 426 \| Phage major capsid protein \| Phage major capsid protein; afdb-uniprot50 \| AF-A0A2V5PK61-F1-MODEL\_V4 \| 1.0 \| 6.79e-53 \| 1201 \| 0.303 \| 701 \| 352 \| 24 \| 30 \| 681 \| 10 \| 622 \| Phage major capsid protein \| Phage major capsid protein; afdb-uniprot50 \| AF-A0A7Y0IMP7-F1-MODEL\_V4 \| 1.0 \| 1.706e-54 \| 1197 \| 0.262 \| 717 \| 409 \| 20 \| 17 \| 682 \| 7 \| 654 \| HK97 family phage major capsid protein/HK97 family phage prohead protease \| HK97 family phage major capsid protein/HK97 family phage prohead protease; afdb-uniprot50 \| AF-A0A7G8BY98-F1-MODEL\_V4 \| 1.0 \| 7.881e-54 \| 1190 \| 0.302 \| 711 \| 399 \| 24 \| 19 \| 682 \| 2 \| 662 \| Phage major capsid protein \| Phage major capsid protein; afdb-uniprot50 \| AF-A0A3S0D8V4-F1-MODEL\_V4 \| 1.0 \| 1.046e-53 \| 1189 \| 0.288 \| 708 \| 400 \| 21 \| 12 \| 682 \| 27 \| 667 \| Phage major capsid protein \| Phage major capsid protein; afdb-uniprot50 \| AF-A0A4P5QRR4-F1-MODEL\_V4 \| 1.0 \| 2.413e-51 \| 1187 \| 0.29 \| 684 \| 391 \| 22 \| 30 \| 680 \| 11 \| 632 \| Uncharacterized protein \| Uncharacterized protein; afdb-uniprot50 \| AF-A0A081RAC8-F1-MODEL\_V4 \| 1.0 \| 4.598e-50 \| 1185 \| 0.288 \| 682 \| 379 \| 20 \| 17 \| 681 \| 3 \| 594 \| Phage major capsid protein, HK97 family \| Phage major capsid protein, HK97 family; afdb-uniprot50 \| AF-A0A1M7GV20-F1-MODEL\_V4 \| 1.0 \| 1.818e-51 \| 1185 \| 0.297 \| 678 \| 363 \| 21 \| 41 \| 682 \| 15 \| 614 \| Phage prohead protease, HK97 family/phage major capsid protein, HK97 family,TIGR01554 \| Phage prohead protease, HK97 family/phage major capsid protein, HK97 family,TIGR01554; afdb-uniprot50 \| AF-A0A2W7GS67-F1-MODEL\_V4 \| 1.0 \| 1.205e-49 \| 1183 \| 0.288 \| 662 \| 369 \| 18 \| 38 \| 681 \| 13 \| 590 \| HK97 family phage major capsid protein \| HK97 family phage major capsid protein; afdb-uniprot50 \| AF-A0A1H0VBY7-F1-MODEL\_V4 \| 1.0 \| 4.501e-51 \| 1171 \| 0.279 \| 683 \| 374 \| 21 \| 29 \| 681 \| 10 \| 604 \| Phage major capsid protein, HK97 family \| Phage major capsid protein, HK97 family; afdb-uniprot50 \| AF-A0A0P8D6G1-F1-MODEL\_V4 \| 1.0 \| 9.137e-47 \| 1159 \| 0.341 \| 620 \| 306 \| 20 \| 89 \| 681 \| 1 \| 544 \| Phage major capsid protein, HK97 family \| Phage major capsid protein, HK97 family; afdb-uniprot50 \| AF-A0A259FDJ6-F1-MODEL\_V4 \| 1.0 \| 1.669e-47 \| 1159 \| 0.298 \| 681 \| 335 \| 19 \| 23 \| 682 \| 1 \| 559 \| Uncharacterized protein \| Uncharacterized protein; afdb-uniprot50 \| AF-A0A0S9N309-F1-MODEL\_V4 \| 1.0 \| 4.16e-44 \| 1157 \| 0.335 \| 567 \| 314 \| 16 \| 124 \| 679 \| 2 \| 516 \| Uncharacterized protein \| Uncharacterized protein; afdb-uniprot50 \| AF-A0A1G0RDU6-F1-MODEL\_V4 \| 1.0 \| 2.608e-50 \| 1154 \| 0.258 \| 731 \| 431 \| 26 \| 1 \| 682 \| 1 \| 669 \| Uncharacterized protein \| Uncharacterized protein; afdb-uniprot50 \| AF-A0A6H9I617-F1-MODEL\_V4 \| 1.0 \| 1.459e-48 \| 1145 \| 0.296 \| 679 \| 355 \| 26 \| 38 \| 681 \| 11 \| 601 \| Phage major capsid protein \| Phage major capsid protein; afdb-uniprot50 \| AF-U5QHN1-F1-MODEL\_V4 \| 1.0 \| 1.883e-52 \| 1129 \| 0.272 \| 744 \| 420 \| 25 \| 1 \| 682 \| 26 \| 709 \| Uncharacterized protein \| Uncharacterized protein; afdb-uniprot50 \| AF-A0A660NJM1-F1-MODEL\_V4 \| 1.0 \| 4.407e-52 \| 1113 \| 0.473 \| 526 \| 235 \| 9 \| 14 \| 534 \| 13 \| 501 \| Phage major capsid protein \| Phage major capsid protein; afdb-uniprot50 \| AF-C6M4F4-F1-MODEL\_V4 \| 1.0 \| 5.636e-35 \| 1112 \| 0.406 \| 364 \| 184 \| 10 \| 333 \| 680 \| 15 \| 362 \| Putative phage capsid family \| Putative phage capsid family; afdb-uniprot50 \| AF-A0A2M7QL94-F1-MODEL\_V4 \| 1.0 \| 3.665e-50 \| 1111 \| 0.29 \| 686 \| 385 \| 22 \| 19 \| 681 \| 16 \| 622 \| Phage major capsid protein \| Phage major capsid protein; afdb-uniprot50 \| AF-A0A419ENJ7-F1-MODEL\_V4 \| 1.0 \| 4.437e-49 \| 1111 \| 0.248 \| 712 \| 434 \| 22 \| 1 \| 682 \| 3 \| 643 \| Phage major capsid protein \| Phage major capsid protein; afdb-uniprot50 \| AF-A0A2W5L8E5-F1-MODEL\_V4 \| 1.0 \| 3.463e-50 \| 1104 \| 0.282 \| 730 \| 403 \| 23 \| 30 \| 682 \| 2 \| 687 \| Phage major capsid protein \| Phage major capsid protein; afdb-uniprot50 \| AF-A0A844G5I7-F1-MODEL\_V4 \| 1.0 \| 1.609e-38 \| 1102 \| 0.345 \| 452 \| 237 \| 14 \| 261 \| 682 \| 43 \| 465 \| Phage major capsid protein \| Phage major capsid protein; afdb-uniprot50 \| AF-A0A2E7WAE6-F1-MODEL\_V4 \| 1.0 \| 3.513e-52 \| 1102 \| 0.282 \| 694 \| 385 \| 21 \| 30 \| 680 \| 215 \| 838 \| Phage major capsid protein \| Phage major capsid protein; afdb-uniprot50 \| AF-A0A3B9NV55-F1-MODEL\_V4 \| 1.0 \| 2.429e-48 \| 1090 \| 0.282 \| 709 \| 367 \| 24 \| 40 \| 680 \| 14 \| 648 \| Phage major capsid protein \| Phage major capsid protein; afdb-uniprot50 \| AF-A0A1V5CVG4-F1-MODEL\_V4 \| 1.0 \| 8.886e-51 \| 1084 \| 0.304 \| 706 \| 378 \| 26 \| 30 \| 681 \| 22 \| 668 \| Phage capsid family protein \| Phage capsid family protein; afdb-uniprot50 \| AF-A0A2N1Z9S1-F1-MODEL\_V4 \| 1.0 \| 2.293e-40 \| 1082 \| 0.348 \| 490 \| 235 \| 14 \| 249 \| 682 \| 7 \| 468 \| Phage major capsid protein \| Phage major capsid protein; afdb-uniprot50 \| AF-A0A7X4CLK5-F1-MODEL\_V4 \| 1.0 \| 1.693e-49 \| 1073 \| 0.268 \| 776 \| 418 \| 25 \| 19 \| 682 \| 13 \| 750 \| Phage major capsid protein \| Phage major capsid protein; afdb-uniprot50 \| AF-A0A2E0R7B6-F1-MODEL\_V4 \| 1.0 \| 3.962e-49 \| 1072 \| 0.253 \| 695 \| 425 \| 26 \| 25 \| 680 \| 14 \| 653 \| Phage major capsid protein \| Phage major capsid protein; afdb-uniprot50 \| AF-A0A259RGV4-F1-MODEL\_V4 \| 1.0 \| 1.83e-48 \| 1067 \| 0.276 \| 730 \| 382 \| 25 \| 6 \| 681 \| 9 \| 645 \| Uncharacterized protein \| Uncharacterized protein; afdb-uniprot50 \| AF-A0A2U1B6D9-F1-MODEL\_V4 \| 1.0 \| 3.179e-46 \| 1056 \| 0.221 \| 667 \| 434 \| 25 \| 30 \| 678 \| 4 \| 602 \| HK97 family phage prohead protease/HK97 family phage major capsid protein,TIGR01554 \| HK97 family phage prohead protease/HK97 family phage major capsid protein,TIGR01554; afdb-uniprot50 \| AF-A0A1G0FL18-F1-MODEL\_V4 \| 1.0 \| 4.307e-37 \| 1054 \| 0.335 \| 453 \| 252 \| 16 \| 258 \| 681 \| 7 \| 439 \| Uncharacterized protein \| Uncharacterized protein; afdb-uniprot50 \| AF-A0A285D6R0-F1-MODEL\_V4 \| 1.0 \| 3.849e-45 \| 1049 \| 0.302 \| 672 \| 357 \| 25 \| 41 \| 682 \| 31 \| 620 \| HK97 family phage major capsid protein \| HK97 family phage major capsid protein; afdb-uniprot50 \| AF-A0A2A2GC02-F1-MODEL\_V4 \| 1.0 \| 1.136e-33 \| 1036 \| 0.299 \| 441 \| 251 \| 15 \| 258 \| 680 \| 20 \| 420 \| Phage major capsid protein \| Phage major capsid protein; afdb-uniprot50 \| AF-A0A3S0EPF2-F1-MODEL\_V4 \| 1.0 \| 1.284e-46 \| 1031 \| 0.33 \| 674 \| 330 \| 25 \| 78 \| 681 \| 2 \| 623 \| Phage major capsid protein \| Phage major capsid protein; afdb-uniprot50 \| AF-N6XHI8-F1-MODEL\_V4 \| 1.0 \| 2.755e-34 \| 1028 \| 0.351 \| 395 \| 217 \| 12 \| 308 \| 681 \| 11 \| 387 \| Caudovirus prohead protease family protein \| Caudovirus prohead protease family protein; afdb-uniprot50 \| AF-A0A7V8PSM8-F1-MODEL\_V4 \| 1.0 \| 1.816e-43 \| 1021 \| 0.222 \| 684 \| 380 \| 22 \| 30 \| 681 \| 7 \| 570 \| Phage major capsid protein \| Phage major capsid protein; afdb-uniprot50 \| AF-A0A4R2YZE6-F1-MODEL\_V4 \| 1.0 \| 3.068e-45 \| 1018 \| 0.235 \| 710 \| 414 \| 29 \| 30 \| 681 \| 3 \| 641 \| HK97 family phage prohead protease/HK97 family phage major capsid protein,TIGR01554 \| HK97 family phage prohead protease/HK97 family phage major capsid protein,TIGR01554; afdb-uniprot50 \| AF-A0A7G8GUS2-F1-MODEL\_V4 \| 1.0 \| 1.097e-32 \| 1002 \| 0.351 \| 367 \| 199 \| 10 \| 338 \| 682 \| 18 \| 367 \| Phage major capsid protein/ HK97 family \| Phage major capsid protein/ HK97 family; afdb-uniprot50 \| AF-A0A1F9VF43-F1-MODEL\_V4 \| 1.0 \| 8.818e-46 \| 987 \| 0.298 \| 708 \| 403 \| 27 \| 17 \| 682 \| 7 \| 662 \| Uncharacterized protein \| Uncharacterized protein; afdb-uniprot50 \| AF-A0A7G8DXJ8-F1-MODEL\_V4 \| 1.0 \| 6.046e-29 \| 985 \| 0.378 \| 309 \| 160 \| 9 \| 398 \| 682 \| 15 \| 315 \| Phage major capsid protein/ HK97 family \| Phage major capsid protein/ HK97 family; afdb-uniprot50 \| AF-A0A2A7U7A2-F1-MODEL\_V4 \| 1.0 \| 2.311e-45 \| 985 \| 0.225 \| 700 \| 441 \| 29 \| 30 \| 682 \| 4 \| 649 \| Major capsid protein \| Major capsid protein; afdb-uniprot50 \| AF-Q1ILS6-F1-MODEL\_V4 \| 1.0 \| 1.467e-37 \| 985 \| 0.271 \| 523 \| 315 \| 15 \| 183 \| 682 \| 299 \| 778 \| Peptidase S49 \| Peptidase S49; afdb-uniprot50 \| AF-A0A286GN99-F1-MODEL\_V4 \| 1.0 \| 6.134e-31 \| 977 \| 0.282 \| 425 \| 252 \| 13 \| 272 \| 682 \| 7 \| 392 \| Phage major capsid protein, HK97 family \| Phage major capsid protein, HK97 family; afdb-uniprot50 \| AF-A0A1G1GKX3-F1-MODEL\_V4 \| 1.0 \| 1.211e-30 \| 967 \| 0.312 \| 371 \| 222 \| 13 \| 325 \| 682 \| 22 \| 372 \| Uncharacterized protein \| Uncharacterized protein; afdb-uniprot50 \| AF-A0A2N3B852-F1-MODEL\_V4 \| 1.0 \| 5.295e-46 \| 962 \| 0.383 \| 615 \| 293 \| 15 \| 30 \| 620 \| 9 \| 561 \| Phage major capsid protein \| Phage major capsid protein; afdb-uniprot50 \| AF-A0A7G9P270-F1-MODEL\_V4 \| 1.0 \| 1.69e-33 \| 955 \| 0.316 \| 442 \| 235 \| 16 \| 261 \| 681 \| 8 \| 403 \| Phage major capsid protein \| Phage major capsid protein; afdb-uniprot50 \| AF-A0A382CXB3-F1-MODEL\_V4 \| 1.0 \| 6.355e-32 \| 952 \| 0.335 \| 388 \| 207 \| 13 \| 315 \| 680 \| 1 \| 359 \| Uncharacterized protein \| Uncharacterized protein; afdb-uniprot50 \| AF-A0A2E9VJ27-F1-MODEL\_V4 \| 1.0 \| 1.477e-34 \| 952 \| 0.334 \| 467 \| 242 \| 14 \| 258 \| 682 \| 7 \| 446 \| Phage major capsid protein \| Phage major capsid protein; afdb-uniprot50 \| AF-A0A6G6X8M0-F1-MODEL\_V4 \| 1.0 \| 7.701e-39 \| 946 \| 0.221 \| 683 \| 372 \| 25 \| 6 \| 681 \| 24 \| 553 \| Phage major capsid protein \| Phage major capsid protein; afdb-uniprot50 \| AF-A4ACG5-F1-MODEL\_V4 \| 1.0 \| 1.68e-44 \| 946 \| 0.268 \| 716 \| 370 \| 22 \| 18 \| 682 \| 26 \| 638 \| Phage major capsid protein, HK97 family \| Phage major capsid protein, HK97 family; afdb-uniprot50 \| AF-A0A5C7LIE0-F1-MODEL\_V4 \| 1.0 \| 1.643e-37 \| 938 \| 0.283 \| 582 \| 345 \| 20 \| 115 \| 680 \| 1 \| 526 \| Phage major capsid protein \| Phage major capsid protein; afdb-uniprot50 \| AF-A0A0T9LH47-F1-MODEL\_V4 \| 1.0 \| 9.596e-42 \| 938 \| 0.21 \| 735 \| 385 \| 28 \| 16 \| 682 \| 2 \| 608 \| Phage major capsid protein, HK97 family \| Phage major capsid protein, HK97 family; afdb-uniprot50 \| AF-A0A1F5QTK4-F1-MODEL\_V4 \| 1.0 \| 3.924e-28 \| 932 \| 0.325 \| 338 \| 197 \| 12 \| 367 \| 681 \| 7 \| 336 \| Uncharacterized protein \| Uncharacterized protein; afdb-uniprot50 \| AF-A0A318GPG6-F1-MODEL\_V4 \| 1.0 \| 4.723e-38 \| 926 \| 0.232 \| 693 \| 367 \| 24 \| 1 \| 682 \| 13 \| 551 \| HK97 family phage major capsid protein \| HK97 family phage major capsid protein; afdb-uniprot50 \| AF-A0A7U8KI29-F1-MODEL\_V4 \| 1.0 \| 1.007e-28 \| 925 \| 0.247 \| 351 \| 236 \| 11 \| 338 \| 682 \| 9 \| 337 \| Putative bacteriochlorophyll 4-vinyl reductase \| Putative bacteriochlorophyll 4-vinyl reductase; afdb-uniprot50 \| AF-A0A258RTP1-F1-MODEL\_V4 \| 1.0 \| 1.328e-31 \| 924 \| 0.239 \| 451 \| 285 \| 20 \| 253 \| 682 \| 4 \| 417 \| Uncharacterized protein \| Uncharacterized protein; afdb-uniprot50 \| AF-A0A3U4W9B0-F1-MODEL\_V4 \| 1.0 \| 4.528e-40 \| 924 \| 0.239 \| 667 \| 405 \| 28 \| 40 \| 682 \| 21 \| 608 \| Phage major capsid protein \| Phage major capsid protein; afdb-uniprot50 \| AF-A0A1N6KUG4-F1-MODEL\_V4 \| 1.0 \| 1.015e-33 \| 917 \| 0.295 \| 481 \| 262 \| 18 \| 252 \| 682 \| 3 \| 456 \| Phage major capsid protein, HK97 family \| Phage major capsid protein, HK97 family; afdb-uniprot50 \| AF-A0A0F9KZ70-F1-MODEL\_V4 \| 1.0 \| 3.43e-29 \| 915 \| 0.318 \| 383 \| 225 \| 11 \| 313 \| 682 \| 1 \| 360 \| Uncharacterized protein \| Uncharacterized protein; afdb-uniprot50 \| AF-A0A7W4FCS3-F1-MODEL\_V4 \| 1.0 \| 5.674e-32 \| 913 \| 0.266 \| 435 \| 270 \| 15 \| 262 \| 682 \| 11 \| 410 \| Phage major capsid protein \| Phage major capsid protein; afdb-uniprot50 \| AF-A0A071LTJ4-F1-MODEL\_V4 \| 1.0 \| 1.059e-39 \| 912 \| 0.222 \| 687 \| 380 \| 26 \| 41 \| 682 \| 18 \| 595 \| Uncharacterized protein \| Uncharacterized protein; afdb-uniprot50 \| AF-A0A5T3MXQ8-F1-MODEL\_V4 \| 1.0 \| 1.691e-41 \| 910 \| 0.23 \| 715 \| 411 \| 31 \| 42 \| 682 \| 19 \| 667 \| Phage major capsid protein \| Phage major capsid protein; afdb-uniprot50 \| AF-A0A1F4EFY0-F1-MODEL\_V4 \| 1.0 \| 3.606e-32 \| 906 \| 0.278 \| 460 \| 265 \| 18 \| 255 \| 682 \| 4 \| 428 \| Uncharacterized protein \| Uncharacterized protein; afdb-uniprot50 \| AF-A0A6N8Q9L6-F1-MODEL\_V4 \| 1.0 \| 2.981e-41 \| 905 \| 0.239 \| 709 \| 393 \| 28 \| 17 \| 681 \| 6 \| 611 \| Phage major capsid protein \| Phage major capsid protein; afdb-uniprot50 \| AF-A0A7C7SWM6-F1-MODEL\_V4 \| 1.0 \| 1.479e-42 \| 894 \| 0.35 \| 594 \| 291 \| 19 \| 40 \| 622 \| 39 \| 548 \| Phage major capsid protein \| Phage major capsid protein; afdb-uniprot50 \| AF-A0A5E9ZST6-F1-MODEL\_V4 \| 1.0 \| 2.376e-41 \| 888 \| 0.207 \| 737 \| 445 \| 31 \| 13 \| 681 \| 3 \| 668 \| Major capsid protein \| Major capsid protein; afdb-uniprot50 \| AF-A0A1F4EF80-F1-MODEL\_V4 \| 1.0 \| 1.328e-31 \| 886 \| 0.275 \| 458 \| 270 \| 18 \| 254 \| 682 \| 4 \| 428 \| Uncharacterized protein \| Uncharacterized protein; afdb-uniprot50 \| AF-A0A558BSV8-F1-MODEL\_V4 \| 1.0 \| 4.274e-32 \| 874 \| 0.254 \| 459 \| 279 \| 17 \| 255 \| 682 \| 5 \| 431 \| Phage major capsid protein \| Phage major capsid protein; afdb-uniprot50 \| AF-A0A8B5U5C5-F1-MODEL\_V4 \| 1.0 \| 2.259e-30 \| 867 \| 0.244 \| 441 \| 264 \| 17 \| 271 \| 682 \| 3 \| 403 \| Phage major capsid protein \| Phage major capsid protein; afdb-uniprot50 \| AF-A0A3D1JKW7-F1-MODEL\_V4 \| 1.0 \| 1.497e-28 \| 864 \| 0.252 \| 444 \| 265 \| 21 \| 258 \| 682 \| 7 \| 402 \| Phage major capsid protein \| Phage major capsid protein; afdb-uniprot50 \| AF-A0A439EPL4-F1-MODEL\_V4 \| 1.0 \| 6.128e-23 \| 858 \| 0.566 \| 217 \| 90 \| 3 \| 420 \| 635 \| 2 \| 215 \| Phage major capsid protein \| Phage major capsid protein; afdb-uniprot50 \| AF-A0A0Q5UAE4-F1-MODEL\_V4 \| 1.0 \| 2.477e-31 \| 858 \| 0.288 \| 465 \| 259 \| 19 \| 253 \| 682 \| 5 \| 432 \| Uncharacterized protein \| Uncharacterized protein; afdb-uniprot50 \| AF-A0A855TD60-F1-MODEL\_V4 \| 1.0 \| 6.636e-38 \| 858 \| 0.211 \| 689 \| 400 \| 28 \| 16 \| 681 \| 7 \| 574 \| Phage major capsid protein \| Phage major capsid protein; afdb-uniprot50 \| AF-A0A7W9N1B8-F1-MODEL\_V4 \| 1.0 \| 1.738e-29 \| 856 \| 0.254 \| 433 \| 264 \| 18 \| 258 \| 681 \| 7 \| 389 \| HK97 family phage major capsid protein \| HK97 family phage major capsid protein; afdb-uniprot50 \| AF-A0A439EPB1-F1-MODEL\_V4 \| 1.0 \| 5.707e-21 \| 853 \| 0.566 \| 157 \| 66 \| 2 \| 29 \| 184 \| 15 \| 170 \| Phage major capsid protein \| Phage major capsid protein; afdb-uniprot50 \| AF-A0A6B3SWS1-F1-MODEL\_V4 \| 1.0 \| 2.134e-30 \| 848 \| 0.248 \| 454 \| 265 \| 20 \| 255 \| 681 \| 4 \| 408 \| Phage major capsid protein \| Phage major capsid protein; afdb-uniprot50 \| AF-A0A2D7GUF3-F1-MODEL\_V4 \| 1.0 \| 1.961e-34 \| 839 \| 0.398 \| 402 \| 173 \| 8 \| 26 \| 422 \| 11 \| 348 \| Capsid protein \| Capsid protein; afdb-uniprot50 \| AF-A0A7C2Y7C3-F1-MODEL\_V4 \| 1.0 \| 1.356e-30 \| 838 \| 0.262 \| 449 \| 280 \| 15 \| 253 \| 680 \| 4 \| 422 \| Phage major capsid protein \| Phage major capsid protein; afdb-uniprot50 \| AF-A0A5B9W0Y3-F1-MODEL\_V4 \| 1.0 \| 5.028e-27 \| 837 \| 0.275 \| 377 \| 223 \| 15 \| 335 \| 682 \| 1 \| 356 \| Phage capsid family protein \| Phage capsid family protein; afdb-uniprot50 \| AF-A0A838LXB9-F1-MODEL\_V4 \| 1.0 \| 1.839e-29 \| 837 \| 0.255 \| 447 \| 270 \| 17 \| 264 \| 680 \| 7 \| 420 \| Phage major capsid protein \| Phage major capsid protein; afdb-uniprot50 \| AF-A0A268U1K3-F1-MODEL\_V4 \| 1.0 \| 6.636e-38 \| 837 \| 0.201 \| 686 \| 412 \| 27 \| 19 \| 682 \| 10 \| 581 \| Uncharacterized protein \| Uncharacterized protein; afdb-uniprot50 \| AF-A0A5V0EAD3-F1-MODEL\_V4 \| 1.0 \| 6.497e-39 \| 834 \| 0.205 \| 701 \| 419 \| 25 \| 40 \| 682 \| 26 \| 646 \| Major capsid protein \| Major capsid protein; afdb-uniprot50 \| AF-A0A2K8V901-F1-MODEL\_V4 \| 1.0 \| 2.39e-30 \| 827 \| 0.251 \| 461 \| 258 \| 19 \| 271 \| 682 \| 11 \| 433 \| Uncharacterized protein \| Uncharacterized protein; afdb-uniprot50 \| AF-A0A838RT86-F1-MODEL\_V4 \| 1.0 \| 5.598e-38 \| 825 \| 0.225 \| 670 \| 382 \| 26 \| 38 \| 682 \| 16 \| 573 \| Phage major capsid protein \| Phage major capsid protein; afdb-uniprot50 \| AF-A0A2M9PA59-F1-MODEL\_V4 \| 1.0 \| 1.08e-22 \| 821 \| 0.295 \| 274 \| 167 \| 10 \| 422 \| 681 \| 2 \| 263 \| Phage major capsid protein \| Phage major capsid protein; afdb-uniprot50 \| AF-A0A2N3DXN2-F1-MODEL\_V4 \| 1.0 \| 6.916e-28 \| 819 \| 0.229 \| 458 \| 280 \| 22 \| 254 \| 681 \| 4 \| 418 \| Phage major capsid protein \| Phage major capsid protein; afdb-uniprot50 \| AF-A0A2E9GN38-F1-MODEL\_V4 \| 1.0 \| 2.677e-30 \| 818 \| 0.295 \| 440 \| 252 \| 16 \| 271 \| 678 \| 7 \| 420 \| Phage major capsid protein \| Phage major capsid protein; afdb-uniprot50 \| AF-A0A839W6R8-F1-MODEL\_V4 \| 1.0 \| 2.032e-35 \| 815 \| 0.2 \| 687 \| 413 \| 27 \| 38 \| 680 \| 15 \| 608 \| HK97 family phage major capsid protein/HK97 family phage prohead protease \| HK97 family phage major capsid protein/HK97 family phage prohead protease; afdb-uniprot50 \| AF-A0A7W4YX49-F1-MODEL\_V4 \| 1.0 \| 6.535e-28 \| 814 \| 0.222 \| 431 \| 259 \| 16 \| 282 \| 678 \| 3 \| 391 \| HK97 family phage major capsid protein \| HK97 family phage major capsid protein; afdb-uniprot50 \| AF-A0A261Q1A3-F1-MODEL\_V4 \| 1.0 \| 1.058e-31 \| 812 \| 0.263 \| 486 \| 274 \| 19 \| 252 \| 682 \| 10 \| 466 \| Phage major capsid protein \| Phage major capsid protein; afdb-uniprot50 \| AF-C6M4F3-F1-MODEL\_V4 \| 1.0 \| 1.355e-22 \| 807 \| 0.465 \| 202 \| 101 \| 5 \| 19 \| 215 \| 1 \| 200 \| Caudovirus prohead protease \| Caudovirus prohead protease; afdb-uniprot50 \| AF-A0A382Z2D1-F1-MODEL\_V4 \| 1.0 \| 1.167e-21 \| 786 \| 0.444 \| 189 \| 96 \| 4 \| 30 \| 215 \| 5 \| 187 \| Uncharacterized protein \| Uncharacterized protein; afdb-uniprot50 \| AF-A0A348SWC8-F1-MODEL\_V4 \| 1.0 \| 2.356e-28 \| 786 \| 0.501 \| 269 \| 120 \| 3 \| 16 \| 279 \| 2 \| 261 \| Phage major capsid protein \| Phage major capsid protein; afdb-uniprot50 \| AF-A0A0F9DVX4-F1-MODEL\_V4 \| 1.0 \| 1.219e-27 \| 786 \| 0.252 \| 456 \| 246 \| 17 \| 252 \| 682 \| 13 \| 398 \| Uncharacterized protein \| Uncharacterized protein; afdb-uniprot50 \| AF-W0E3R9-F1-MODEL\_V4 \| 1.0 \| 1.377e-40 \| 786 \| 0.251 \| 728 \| 381 \| 31 \| 12 \| 681 \| 7 \| 628 \| Uncharacterized protein \| Uncharacterized protein; afdb-uniprot50 \| AF-A0A6N0ZIS5-F1-MODEL\_V4 \| 1.0 \| 4.489e-27 \| 785 \| 0.242 \| 437 \| 281 \| 12 \| 264 \| 681 \| 7 \| 412 \| Phage major capsid protein \| Phage major capsid protein; afdb-uniprot50 \| AF-A0A2D5VNT2-F1-MODEL\_V4 \| 1.0 \| 1.894e-41 \| 785 \| 0.366 \| 527 \| 282 \| 12 \| 1 \| 511 \| 8 \| 498 \| Phage major capsid protein \| Phage major capsid protein; afdb-uniprot50 \| AF-A0A418M3V2-F1-MODEL\_V4 \| 1.0 \| 1.497e-28 \| 781 \| 0.217 \| 473 \| 286 \| 18 \| 263 \| 682 \| 3 \| 444 \| Phage major capsid protein \| Phage major capsid protein; afdb-uniprot50 \| AF-A0A4U8TCV4-F1-MODEL\_V4 \| 1.0 \| 4.097e-34 \| 777 \| 0.184 \| 667 \| 393 \| 25 \| 40 \| 680 \| 15 \| 556 \| Phage major capsid protein \| Phage major capsid protein; afdb-uniprot50 \| AF-A0A558C376-F1-MODEL\_V4 \| 1.0 \| 2.894e-29 \| 774 \| 0.229 \| 457 \| 286 \| 14 \| 258 \| 680 \| 5 \| 429 \| Phage major capsid protein \| Phage major capsid protein; afdb-uniprot50 \| AF-A0A837Q3L9-F1-MODEL\_V4 \| 1.0 \| 7.98e-40 \| 774 \| 0.351 \| 512 \| 266 \| 14 \| 33 \| 532 \| 9 \| 466 \| Peptidase \| Peptidase; afdb-uniprot50 \| AF-A0A6N8DIV7-F1-MODEL\_V4 \| 1.0 \| 1.128e-28 \| 772 \| 0.263 \| 444 \| 260 \| 19 \| 264 \| 682 \| 14 \| 415 \| Phage major capsid protein \| Phage major capsid protein; afdb-uniprot50 \| AF-A0A180F243-F1-MODEL\_V4 \| 1.0 \| 1.128e-28 \| 771 \| 0.248 \| 455 \| 266 \| 17 \| 264 \| 682 \| 14 \| 428 \| Putative phage phi-C31 gp36 major capsid-like protein \| Putative phage phi-C31 gp36 major capsid-like protein; afdb-uniprot50 \| AF-A0A8A8EP07-F1-MODEL\_V4 \| 1.0 \| 1.219e-27 \| 765 \| 0.256 \| 445 \| 275 \| 17 \| 253 \| 681 \| 2 \| 406 \| Phage major capsid protein \| Phage major capsid protein; afdb-uniprot50 \| AF-A0A2V5Y3W0-F1-MODEL\_V4 \| 1.0 \| 9.783e-25 \| 764 \| 0.215 \| 362 \| 251 \| 12 \| 335 \| 681 \| 18 \| 361 \| Phage major capsid protein \| Phage major capsid protein; afdb-uniprot50 \| AF-A0A239HKB7-F1-MODEL\_V4 \| 1.0 \| 4.554e-29 \| 760 \| 0.248 \| 471 \| 262 \| 20 \| 261 \| 682 \| 8 \| 435 \| Phage major capsid protein, HK97 family \| Phage major capsid protein, HK97 family; afdb-uniprot50 \| AF-A0A1G3FQ64-F1-MODEL\_V4 \| 1.0 \| 1.088e-27 \| 757 \| 0.259 \| 455 \| 265 \| 20 \| 255 \| 681 \| 5 \| 415 \| Uncharacterized protein \| Uncharacterized protein; afdb-uniprot50 \| AF-A0A371WU99-F1-MODEL\_V4 \| 1.0 \| 6.051e-37 \| 749 \| 0.209 \| 727 \| 427 \| 31 \| 14 \| 681 \| 34 \| 671 \| Phage major capsid protein \| Phage major capsid protein; afdb-uniprot50 \| AF-A0A1U7GEM9-F1-MODEL\_V4 \| 1.0 \| 8.373e-27 \| 744 \| 0.235 \| 428 \| 260 \| 18 \| 279 \| 682 \| 43 \| 427 \| Uncharacterized protein \| Uncharacterized protein; afdb-uniprot50 \| AF-A0A6A4R9W0-F1-MODEL\_V4 \| 1.0 \| 1.042e-21 \| 741 \| 0.443 \| 203 \| 93 \| 4 \| 18 \| 201 \| 2 \| 203 \| Phage major capsid protein \| Phage major capsid protein; afdb-uniprot50 \| AF-A0A7C5ALH4-F1-MODEL\_V4 \| 1.0 \| 1.851e-26 \| 739 \| 0.25 \| 428 \| 239 \| 17 \| 285 \| 682 \| 11 \| 386 \| Phage major capsid protein \| Phage major capsid protein; afdb-uniprot50 \| AF-A0A5P9CVW4-F1-MODEL\_V4 \| 1.0 \| 4.489e-27 \| 738 \| 0.221 \| 446 \| 277 \| 18 \| 261 \| 679 \| 8 \| 410 \| Phage capsid family protein \| Phage capsid family protein; afdb-uniprot50 \| AF-A0A432MJM0-F1-MODEL\_V4 \| 1.0 \| 1.749e-26 \| 735 \| 0.242 \| 457 \| 260 \| 21 \| 264 \| 678 \| 7 \| 419 \| Phage major capsid protein \| Phage major capsid protein; afdb-uniprot50 \| AF-A0A6L8UIY2-F1-MODEL\_V4 \| 1.0 \| 9.925e-27 \| 734 \| 0.248 \| 446 \| 270 \| 17 \| 251 \| 681 \| 20 \| 415 \| Phage major capsid protein \| Phage major capsid protein; afdb-uniprot50 \| AF-A0A1U7CX52-F1-MODEL\_V4 \| 1.0 \| 2.753e-26 \| 732 \| 0.223 \| 460 \| 288 \| 18 \| 258 \| 682 \| 7 \| 432 \| Uncharacterized protein \| Uncharacterized protein; afdb-uniprot50 \| AF-A0A7G7KL91-F1-MODEL\_V4 \| 1.0 \| 1.065e-20 \| 731 \| 0.264 \| 261 \| 176 \| 9 \| 428 \| 679 \| 6 \| 259 \| Phage major capsid protein \| Phage major capsid protein; afdb-uniprot50 \| AF-A0A6L8U335-F1-MODEL\_V4 \| 1.0 \| 6.675e-27 \| 731 \| 0.245 \| 457 \| 271 \| 20 \| 258 \| 681 \| 8 \| 423 \| Phage major capsid protein \| Phage major capsid protein; afdb-uniprot50 \| AF-A0A268TJ56-F1-MODEL\_V4 \| 1.0 \| 3.086e-34 \| 731 \| 0.184 \| 677 \| 429 \| 29 \| 40 \| 682 \| 15 \| 602 \| Uncharacterized protein \| Uncharacterized protein; afdb-uniprot50 \| AF-A0A2D8LFG7-F1-MODEL\_V4 \| 1.0 \| 2.458e-26 \| 723 \| 0.213 \| 458 \| 296 \| 20 \| 258 \| 681 \| 8 \| 435 \| Phage major capsid protein \| Phage major capsid protein; afdb-uniprot50 \| AF-A0A1U7CNJ6-F1-MODEL\_V4 \| 1.0 \| 4.241e-27 \| 723 \| 0.237 \| 472 \| 287 \| 20 \| 256 \| 682 \| 4 \| 447 \| Uncharacterized protein \| Uncharacterized protein; afdb-uniprot50 \| AF-A0A3D1J480-F1-MODEL\_V4 \| 1.0 \| 1.776e-36 \| 723 \| 0.203 \| 737 \| 432 \| 32 \| 13 \| 681 \| 50 \| 699 \| Phage major capsid protein \| Phage major capsid protein; afdb-uniprot50 \| AF-A0A2D9C9C5-F1-MODEL\_V4 \| 1.0 \| 5.439e-34 \| 720 \| 0.185 \| 705 \| 437 \| 32 \| 26 \| 682 \| 16 \| 630 \| HK97 family phage prohead protease \| HK97 family phage prohead protease; afdb-uniprot50 \| AF-A0A2W5AUZ1-F1-MODEL\_V4 \| 1.0 \| 8.081e-26 \| 714 \| 0.251 \| 397 \| 242 \| 16 \| 306 \| 682 \| 36 \| 397 \| Phage major capsid protein \| Phage major capsid protein; afdb-uniprot50 \| AF-A0A519K9R5-F1-MODEL\_V4 \| 1.0 \| 1.299e-24 \| 711 \| 0.22 \| 458 \| 294 \| 17 \| 249 \| 678 \| 6 \| 428 \| Phage major capsid protein \| Phage major capsid protein; afdb-uniprot50 \| AF-A0A7G8CD93-F1-MODEL\_V4 \| 1.0 \| 1.904e-22 \| 710 \| 0.244 \| 348 \| 229 \| 14 \| 343 \| 679 \| 11 \| 335 \| Phage major capsid protein/ HK97 family \| Phage major capsid protein/ HK97 family; afdb-uniprot50 \| AF-A0A1V6GVR6-F1-MODEL\_V4 \| 1.0 \| 5.959e-27 \| 705 \| 0.254 \| 472 \| 253 \| 19 \| 271 \| 681 \| 4 \| 437 \| Phage capsid family protein \| Phage capsid family protein; afdb-uniprot50 \| AF-A0A7W6QQW4-F1-MODEL\_V4 \| 1.0 \| 2.601e-26 \| 703 \| 0.221 \| 457 \| 271 \| 20 \| 258 \| 682 \| 8 \| 411 \| HK97 family phage major capsid protein \| HK97 family phage major capsid protein; afdb-uniprot50 \| AF-A0A5C7R6U5-F1-MODEL\_V4 \| 1.0 \| 1.12e-31 \| 688 \| 0.196 \| 693 \| 312 \| 19 \| 26 \| 677 \| 17 \| 505 \| Uncharacterized protein \| Uncharacterized protein; afdb-uniprot50 \| AF-A0A3D0WRI2-F1-MODEL\_V4 \| 1.0 \| 5.17e-23 \| 687 \| 0.203 \| 438 \| 257 \| 17 \| 265 \| 682 \| 4 \| 369 \| Phage major capsid protein \| Phage major capsid protein; afdb-uniprot50 \| AF-A0A7G8FKV0-F1-MODEL\_V4 \| 1.0 \| 1.7e-22 \| 686 \| 0.228 \| 355 \| 237 \| 17 \| 343 \| 681 \| 7 \| 340 \| Phage major capsid protein/ HK97 family \| Phage major capsid protein/ HK97 family; afdb-uniprot50 \| AF-A0A2D9FXS3-F1-MODEL\_V4 \| 1.0 \| 2.074e-26 \| 686 \| 0.197 \| 487 \| 302 \| 25 \| 250 \| 682 \| 8 \| 459 \| Phage major capsid protein \| Phage major capsid protein; afdb-uniprot50 \| AF-A0A7Y7NKU6-F1-MODEL\_V4 \| 1.0 \| 2.812e-25 \| 685 \| 0.237 \| 458 \| 260 \| 21 \| 260 \| 681 \| 9 \| 413 \| Phage major capsid protein \| Phage major capsid protein; afdb-uniprot50 \| AF-A0A2Z4IND8-F1-MODEL\_V4 \| 1.0 \| 3.733e-25 \| 682 \| 0.221 \| 466 \| 288 \| 16 \| 263 \| 682 \| 4 \| 440 \| Phage major capsid protein \| Phage major capsid protein; afdb-uniprot50 \| AF-A0A3M1UTX0-F1-MODEL\_V4 \| 1.0 \| 2.032e-35 \| 682 \| 0.328 \| 600 \| 308 \| 18 \| 19 \| 611 \| 13 \| 524 \| Phage major capsid protein \| Phage major capsid protein; afdb-uniprot50 \| AF-A0A843JDH5-F1-MODEL\_V4 \| 1.0 \| 4.035e-24 \| 679 \| 0.203 \| 443 \| 266 \| 17 \| 258 \| 682 \| 7 \| 380 \| Phage major capsid protein \| Phage major capsid protein; afdb-uniprot50 \| AF-A0A1C6BMQ2-F1-MODEL\_V4 \| 1.0 \| 2.456e-18 \| 672 \| 0.418 \| 165 \| 90 \| 3 \| 28 \| 187 \| 7 \| 170 \| Phage prohead protease, HK97 family \| Phage prohead protease, HK97 family; afdb-uniprot50 \| AF-Q08QD1-F1-MODEL\_V4 \| 1.0 \| 2.272e-19 \| 670 \| 0.223 \| 295 \| 196 \| 14 \| 402 \| 679 \| 1 \| 279 \| Phage capsid family \| Phage capsid family; afdb-uniprot50 \| AF-A0A368NXR3-F1-MODEL\_V4 \| 1.0 \| 9.378e-27 \| 667 \| 0.214 \| 526 \| 319 \| 24 \| 191 \| 679 \| 5 \| 473 \| Phage major capsid protein \| Phage major capsid protein; afdb-uniprot50 \| AF-A0A4R8I8I6-F1-MODEL\_V4 \| 1.0 \| 5.062e-24 \| 665 \| 0.207 \| 457 \| 283 \| 21 \| 255 \| 682 \| 4 \| 410 \| HK97 family phage major capsid protein \| HK97 family phage major capsid protein; afdb-uniprot50 \| AF-U5BPA1-F1-MODEL\_V4 \| 1.0 \| 6.963e-25 \| 664 \| 0.212 \| 457 \| 275 \| 21 \| 258 \| 682 \| 8 \| 411 \| Uncharacterized protein \| Uncharacterized protein; afdb-uniprot50 \| AF-A0A212KXM3-F1-MODEL\_V4 \| 1.0 \| 2.104e-28 \| 664 \| 0.157 \| 661 \| 396 \| 31 \| 38 \| 682 \| 11 \| 526 \| Phage major capsid protein, HK97 family \| Phage major capsid protein, HK97 family; afdb-uniprot50 \| AF-A0A226I7W7-F1-MODEL\_V4 \| 1.0 \| 8.253e-25 \| 659 \| 0.206 \| 451 \| 302 \| 21 \| 258 \| 680 \| 8 \| 430 \| Uncharacterized protein \| Uncharacterized protein; afdb-uniprot50 \| AF-A0A2T4JHM7-F1-MODEL\_V4 \| 1.0 \| 9.916e-19 \| 656 \| 0.238 \| 298 \| 194 \| 13 \| 402 \| 682 \| 1 \| 282 \| Phage major capsid protein \| Phage major capsid protein; afdb-uniprot50 \| AF-A0A4R0QZ35-F1-MODEL\_V4 \| 1.0 \| 7.419e-22 \| 654 \| 0.222 \| 359 \| 226 \| 15 \| 343 \| 679 \| 11 \| 338 \| Phage major capsid protein \| Phage major capsid protein; afdb-uniprot50 \| AF-A0A1G7G3W8-F1-MODEL\_V4 \| 1.0 \| 2.976e-25 \| 654 \| 0.205 \| 457 \| 300 \| 17 \| 258 \| 682 \| 10 \| 435 \| Phage major capsid protein, HK97 family \| Phage major capsid protein, HK97 family; afdb-uniprot50 \| AF-A0A0F9MYQ3-F1-MODEL\_V4 \| 1.0 \| 3.606e-32 \| 649 \| 0.174 \| 747 \| 462 \| 35 \| 26 \| 680 \| 13 \| 696 \| Uncharacterized protein \| Uncharacterized protein; afdb-uniprot50 \| AF-A0A4Q7LWN4-F1-MODEL\_V4 \| 1.0 \| 5.13e-18 \| 643 \| 0.198 \| 262 \| 192 \| 8 \| 431 \| 682 \| 29 \| 282 \| HK97 family phage major capsid protein \| HK97 family phage major capsid protein; afdb-uniprot50 \| AF-A0A7I9V3A8-F1-MODEL\_V4 \| 1.0 \| 1.423e-17 \| 640 \| 0.171 \| 256 \| 202 \| 4 \| 431 \| 681 \| 28 \| 278 \| Uncharacterized protein \| Uncharacterized protein; afdb-uniprot50 \| AF-A0A378RQE6-F1-MODEL\_V4 \| 1.0 \| 5.357e-24 \| 636 \| 0.244 \| 459 \| 269 \| 20 \| 258 \| 681 \| 8 \| 423 \| Predicted phage phi-C31 gp36 major capsid-like protein \| Predicted phage phi-C31 gp36 major capsid-like protein; afdb-uniprot50 \| AF-A0A506U2R2-F1-MODEL\_V4 \| 1.0 \| 4.149e-20 \| 635 \| 0.219 \| 324 \| 212 \| 11 \| 371 \| 679 \| 18 \| 315 \| Phage major capsid protein \| Phage major capsid protein; afdb-uniprot50 \| AF-F3Z7W5-F1-MODEL\_V4 \| 1.0 \| 9.042e-18 \| 632 \| 0.16 \| 274 \| 204 \| 11 \| 423 \| 682 \| 6 \| 267 \| Uncharacterized protein \| Uncharacterized protein; afdb-uniprot50 \| AF-A0A849MU29-F1-MODEL\_V4 \| 1.0 \| 1.227e-24 \| 632 \| 0.205 \| 463 \| 288 \| 18 \| 264 \| 678 \| 10 \| 440 \| Phage major capsid protein \| Phage major capsid protein; afdb-uniprot50 \| AF-A0A3A9H2E6-F1-MODEL\_V4 \| 1.0 \| 4.519e-24 \| 632 \| 0.207 \| 448 \| 261 \| 19 \| 296 \| 679 \| 81 \| 498 \| Phage major capsid protein \| Phage major capsid protein; afdb-uniprot50 \| AF-A0A1A9A792-F1-MODEL\_V4 \| 1.0 \| 1.651e-18 \| 631 \| 0.202 \| 277 \| 191 \| 8 \| 430 \| 682 \| 40 \| 310 \| Phage major capsid protein, HK97 family \| Phage major capsid protein, HK97 family; afdb-uniprot50 \| AF-A0A7L8AFB1-F1-MODEL\_V4 \| 1.0 \| 6.35e-24 \| 631 \| 0.23 \| 456 \| 268 \| 20 \| 263 \| 679 \| 3 \| 414 \| Phage major capsid protein \| Phage major capsid protein; afdb-uniprot50 \| AF-A0A5C7P846-F1-MODEL\_V4 \| 1.0 \| 1.748e-18 \| 630 \| 0.221 \| 271 \| 178 \| 10 \| 444 \| 682 \| 1 \| 270 \| Phage major capsid protein \| Phage major capsid protein; afdb-uniprot50 \| AF-A0A6I2TYG7-F1-MODEL\_V4 \| 1.0 \| 1.096e-24 \| 630 \| 0.238 \| 453 \| 262 \| 18 \| 282 \| 681 \| 10 \| 432 \| Phage major capsid protein \| Phage major capsid protein; afdb-uniprot50 \| AF-A0A1A0TGW3-F1-MODEL\_V4 \| 1.0 \| 3.524e-17 \| 628 \| 0.195 \| 256 \| 193 \| 6 \| 430 \| 682 \| 28 \| 273 \| Capsid protein \| Capsid protein; afdb-uniprot50 \| AF-A0A5F0FN25-F1-MODEL\_V4 \| 1.0 \| 2.809e-17 \| 628 \| 0.17 \| 258 \| 203 \| 8 \| 431 \| 682 \| 28 \| 280 \| Phage major capsid protein \| Phage major capsid protein; afdb-uniprot50 \| AF-A0A7J9VRD2-F1-MODEL\_V4 \| 1.0 \| 1.345e-17 \| 627 \| 0.196 \| 259 \| 192 \| 7 \| 431 \| 682 \| 29 \| 278 \| Phage major capsid protein \| Phage major capsid protein; afdb-uniprot50 \| AF-B5IP42-F1-MODEL\_V4 \| 1.0 \| 7.419e-22 \| 626 \| 0.213 \| 412 \| 272 \| 16 \| 293 \| 682 \| 1 \| 382 \| Uncharacterized protein \| Uncharacterized protein; afdb-uniprot50 \| AF-A0A6S6N9V6-F1-MODEL\_V4 \| 1.0 \| 9.051e-26 \| 625 \| 0.14 \| 684 \| 392 \| 31 \| 19 \| 681 \| 3 \| 511 \| Peptidase U35 \| Peptidase U35; afdb-uniprot50 \| AF-A0A1J4P7K5-F1-MODEL\_V4 \| 1.0 \| 1.785e-17 \| 623 \| 0.191 \| 251 \| 197 \| 5 \| 433 \| 681 \| 30 \| 276 \| Uncharacterized protein \| Uncharacterized protein; afdb-uniprot50 \| AF-A0A553GPR9-F1-MODEL\_V4 \| 1.0 \| 3.977e-22 \| 622 \| 0.2 \| 434 \| 273 \| 19 \| 268 \| 679 \| 11 \| 392 \| Phage major capsid protein \| Phage major capsid protein; afdb-uniprot50 \| AF-A0A496WP50-F1-MODEL\_V4 \| 1.0 \| 3.17e-22 \| 621 \| 0.192 \| 415 \| 268 \| 18 \| 295 \| 681 \| 13 \| 388 \| Phage major capsid protein \| Phage major capsid protein; afdb-uniprot50 \| AF-A0A367A8R2-F1-MODEL\_V4 \| 1.0 \| 7.469e-19 \| 620 \| 0.186 \| 316 \| 216 \| 10 \| 381 \| 682 \| 2 \| 290 \| Phage major capsid protein \| Phage major capsid protein; afdb-uniprot50 \| AF-E3FHX8-F1-MODEL\_V4 \| 1.0 \| 4.918e-20 \| 620 \| 0.209 \| 358 \| 227 \| 18 \| 335 \| 679 \| 3 \| 317 \| Phage major capsid protein, HK97 \| Phage major capsid protein, HK97; afdb-uniprot50 \| AF-A0A2V3ZYP9-F1-MODEL\_V4 \| 1.0 \| 1.385e-29 \| 618 \| 0.149 \| 691 \| 424 \| 33 \| 28 \| 680 \| 3 \| 567 \| Phage major capsid protein \| Phage major capsid protein; afdb-uniprot50 \| AF-A0A4V2Y1S1-F1-MODEL\_V4 \| 1.0 \| 2.654e-17 \| 617 \| 0.217 \| 262 \| 184 \| 8 \| 431 \| 682 \| 28 \| 278 \| Phage major capsid protein \| Phage major capsid protein; afdb-uniprot50 \| AF-X6KKD3-F1-MODEL\_V4 \| 1.0 \| 2.852e-27 \| 617 \| 0.171 \| 687 \| 382 \| 29 \| 19 \| 682 \| 4 \| 525 \| Capsid protein \| Capsid protein; afdb-uniprot50 \| AF-A0A7J9XPI0-F1-MODEL\_V4 \| 1.0 \| 1.889e-17 \| 615 \| 0.189 \| 259 \| 194 \| 7 \| 431 \| 682 \| 34 \| 283 \| Phage major capsid protein \| Phage major capsid protein; afdb-uniprot50 \| AF-A0A0B5E1B7-F1-MODEL\_V4 \| 1.0 \| 1.813e-27 \| 615 \| 0.165 \| 687 \| 363 \| 34 \| 17 \| 679 \| 3 \| 502 \| HK97 family phage major capsid protein \| HK97 family phage major capsid protein; afdb-uniprot50 \| AF-A0A3C1F4S7-F1-MODEL\_V4 \| 1.0 \| 1.917e-19 \| 614 \| 0.21 \| 318 \| 198 \| 13 \| 398 \| 682 \| 14 \| 311 \| Phage major capsid protein \| Phage major capsid protein; afdb-uniprot50 \| AF-A0A5J4QQM2-F1-MODEL\_V4 \| 1.0 \| 4.09e-18 \| 613 \| 0.177 \| 276 \| 197 \| 8 \| 409 \| 680 \| 2 \| 251 \| Uncharacterized protein \| Uncharacterized protein; afdb-uniprot50 \| AF-A0A841KJN3-F1-MODEL\_V4 \| 1.0 \| 1.136e-33 \| 612 \| 0.204 \| 744 \| 447 \| 38 \| 16 \| 682 \| 11 \| 686 \| Uncharacterized protein \| Uncharacterized protein; afdb-uniprot50 \| AF-A0A1Q4SYY1-F1-MODEL\_V4 \| 1.0 \| 2.37e-17 \| 611 \| 0.223 \| 259 \| 183 \| 9 \| 433 \| 682 \| 31 \| 280 \| Capsid protein \| Capsid protein; afdb-uniprot50 \| AF-A0A251YMX4-F1-MODEL\_V4 \| 1.0 \| 5.869e-17 \| 610 \| 0.188 \| 260 \| 196 \| 7 \| 431 \| 682 \| 30 \| 282 \| Phage capsid family protein \| Phage capsid family protein; afdb-uniprot50 \| AF-A0A436RU37-F1-MODEL\_V4 \| 1.0 \| 1.006e-20 \| 610 \| 0.206 \| 407 \| 238 \| 20 \| 295 \| 682 \| 4 \| 344 \| Phage major capsid protein \| Phage major capsid protein; afdb-uniprot50 \| AF-A0A1I5E198-F1-MODEL\_V4 \| 1.0 \| 4.885e-23 \| 610 \| 0.215 \| 463 \| 284 \| 21 \| 261 \| 681 \| 9 \| 434 \| Phage major capsid protein, HK97 family \| Phage major capsid protein, HK97 family; afdb-uniprot50 \| AF-A0A5C7UGA7-F1-MODEL\_V4 \| 1.0 \| 6.72e-24 \| 610 \| 0.198 \| 469 \| 292 \| 13 \| 295 \| 682 \| 13 \| 478 \| Phage major capsid protein \| Phage major capsid protein; afdb-uniprot50 \| AF-A0A3E3J0E7-F1-MODEL\_V4 \| 1.0 \| 9.244e-25 \| 610 \| 0.206 \| 493 \| 286 \| 22 \| 266 \| 682 \| 65 \| 528 \| Phage major capsid protein \| Phage major capsid protein; afdb-uniprot50 \| AF-A0A2T0R256-F1-MODEL\_V4 \| 1.0 \| 1.423e-17 \| 609 \| 0.182 \| 263 \| 196 \| 6 \| 431 \| 682 \| 29 \| 283 \| HK97 family phage major capsid protein \| HK97 family phage major capsid protein; afdb-uniprot50 \| AF-A0A087M199-F1-MODEL\_V4 \| 1.0 \| 4.956e-25 \| 607 \| 0.135 \| 663 \| 383 \| 25 \| 39 \| 682 \| 11 \| 502 \| Capsid protein \| Capsid protein; afdb-uniprot50 \| AF-A0A0F9CUT8-F1-MODEL\_V4 \| 1.0 \| 9.57e-18 \| 606 \| 0.233 \| 300 \| 203 \| 9 \| 398 \| 682 \| 30 \| 317 \| Uncharacterized protein \| Uncharacterized protein; afdb-uniprot50 \| AF-A0A2T0QNI1-F1-MODEL\_V4 \| 1.0 \| 1.999e-17 \| 605 \| 0.183 \| 262 \| 191 \| 9 \| 431 \| 680 \| 32 \| 282 \| HK97 family phage major capsid protein \| HK97 family phage major capsid protein; afdb-uniprot50 \| AF-A0A2W4JAL7-F1-MODEL\_V4 \| 1.0 \| 2.599e-18 \| 604 \| 0.242 \| 272 \| 168 \| 11 \| 438 \| 681 \| 4 \| 265 \| Phage major capsid protein \| Phage major capsid protein; afdb-uniprot50 \| AF-A0A1I1CFH0-F1-MODEL\_V4 \| 1.0 \| 3.33e-17 \| 604 \| 0.196 \| 260 \| 188 \| 6 \| 431 \| 682 \| 29 \| 275 \| Phage major capsid protein, HK97 family \| Phage major capsid protein, HK97 family; afdb-uniprot50 \| AF-A0A7W8JSS7-F1-MODEL\_V4 \| 1.0 \| 1.143e-22 \| 604 \| 0.22 \| 468 \| 286 \| 21 \| 245 \| 680 \| 1 \| 421 \| HK97 family phage major capsid protein \| HK97 family phage major capsid protein; afdb-uniprot50 \| AF-A0A662S3E0-F1-MODEL\_V4 \| 1.0 \| 1.642e-29 \| 604 \| 0.194 \| 698 \| 385 \| 30 \| 15 \| 681 \| 104 \| 654 \| Phage major capsid protein \| Phage major capsid protein; afdb-uniprot50 \| AF-A0A142X159-F1-MODEL\_V4 \| 1.0 \| 4.121e-23 \| 603 \| 0.204 \| 464 \| 296 \| 21 \| 279 \| 682 \| 20 \| 470 \| Phage capsid family protein \| Phage capsid family protein; afdb-uniprot50 \| AF-A0A2D9CCN7-F1-MODEL\_V4 \| 1.0 \| 1.799e-22 \| 602 \| 0.189 \| 444 \| 278 \| 18 \| 264 \| 681 \| 14 \| 401 \| Phage major capsid protein \| Phage major capsid protein; afdb-uniprot50 \| AF-A0A378YEU3-F1-MODEL\_V4 \| 1.0 \| 5.869e-17 \| 600 \| 0.194 \| 262 \| 194 \| 7 \| 431 \| 682 \| 29 \| 283 \| Predicted phage phi-C31 gp36 major capsid-like protein \| Predicted phage phi-C31 gp36 major capsid-like protein; afdb-uniprot50 \| AF-A0A1H7U3C7-F1-MODEL\_V4 \| 1.0 \| 2.693e-19 \| 600 \| 0.206 \| 348 \| 219 \| 20 \| 346 \| 679 \| 24 \| 328 \| Phage major capsid protein, HK97 family \| Phage major capsid protein, HK97 family; afdb-uniprot50 \| AF-A0A1G3BD02-F1-MODEL\_V4 \| 1.0 \| 1.042e-21 \| 600 \| 0.192 \| 457 \| 250 \| 20 \| 273 \| 682 \| 4 \| 388 \| Uncharacterized protein \| Uncharacterized protein; afdb-uniprot50 \| AF-A0A372GN26-F1-MODEL\_V4 \| 1.0 \| 2.404e-19 \| 598 \| 0.183 \| 338 \| 214 \| 11 \| 381 \| 682 \| 2 \| 313 \| Phage major capsid protein \| Phage major capsid protein; afdb-uniprot50 \| AF-A0A263DU06-F1-MODEL\_V4 \| 1.0 \| 3.947e-17 \| 597 \| 0.19 \| 263 \| 192 \| 8 \| 431 \| 682 \| 30 \| 282 \| Phage major capsid protein \| Phage major capsid protein; afdb-uniprot50 \| AF-A0A4Q0M7Y1-F1-MODEL\_V4 \| 1.0 \| 1.227e-24 \| 597 \| 0.15 \| 683 \| 389 \| 30 \| 19 \| 682 \| 4 \| 513 \| Phage major capsid protein \| Phage major capsid protein; afdb-uniprot50 \| AF-A0A098BVT6-F1-MODEL\_V4 \| 1.0 \| 3.146e-17 \| 595 \| 0.187 \| 262 \| 191 \| 7 \| 431 \| 682 \| 28 \| 277 \| Phage major capsid protein, HK97 family \| Phage major capsid protein, HK97 family; afdb-uniprot50 \| AF-U6RGH7-F1-MODEL\_V4 \| 1.0 \| 4.062e-21 \| 595 \| 0.206 \| 432 \| 271 \| 18 \| 267 \| 682 \| 3 \| 378 \| HK97 family phage major capsid protein \| HK97 family phage major capsid protein; afdb-uniprot50 \| AF-A0A7U9F631-F1-MODEL\_V4 \| 1.0 \| 3.599e-16 \| 594 \| 0.348 \| 175 \| 101 \| 5 \| 30 \| 194 \| 13 \| 184 \| Uncharacterized protein \| Uncharacterized protein; afdb-uniprot50 \| AF-A0A839FEI7-F1-MODEL\_V4 \| 1.0 \| 6.956e-17 \| 593 \| 0.177 \| 248 \| 191 \| 7 \| 436 \| 681 \| 35 \| 271 \| HK97 family phage major capsid protein \| HK97 family phage major capsid protein; afdb-uniprot50 \| AF-A0A2S2D0Z2-F1-MODEL\_V4 \| 1.0 \| 7.419e-22 \| 593 \| 0.172 \| 458 \| 288 \| 22 \| 265 \| 681 \| 8 \| 415 \| Phage major capsid protein \| Phage major capsid protein; afdb-uniprot50 \| AF-A0A511HNN2-F1-MODEL\_V4 \| 1.0 \| 6.485e-23 \| 593 \| 0.199 \| 441 \| 281 \| 20 \| 279 \| 680 \| 16 \| 423 \| Uncharacterized protein \| Uncharacterized protein; afdb-uniprot50 \| AF-A0A0J6SLM0-F1-MODEL\_V4 \| 1.0 \| 1.664e-23 \| 593 \| 0.172 \| 486 \| 304 \| 25 \| 256 \| 681 \| 4 \| 451 \| Uncharacterized protein \| Uncharacterized protein; afdb-uniprot50 \| AF-A0A6C2YSM9-F1-MODEL\_V4 \| 1.0 \| 1.606e-22 \| 593 \| 0.2 \| 465 \| 293 \| 18 \| 285 \| 682 \| 15 \| 467 \| Uncharacterized protein \| Uncharacterized protein; afdb-uniprot50 \| AF-A0A2G2J0Z9-F1-MODEL\_V4 \| 1.0 \| 2.933e-23 \| 593 \| 0.192 \| 493 \| 291 \| 19 \| 252 \| 681 \| 36 \| 484 \| Phage major capsid protein \| Phage major capsid protein; afdb-uniprot50 \| AF-A0A3D4V181-F1-MODEL\_V4 \| 1.0 \| 3.401e-16 \| 592 \| 0.162 \| 265 \| 205 \| 7 \| 431 \| 682 \| 4 \| 264 \| Phage major capsid protein \| Phage major capsid protein; afdb-uniprot50 \| AF-A0A0F9FEI0-F1-MODEL\_V4 \| 1.0 \| 1.226e-16 \| 590 \| 0.146 \| 280 \| 215 \| 9 \| 410 \| 681 \| 1 \| 264 \| Uncharacterized protein \| Uncharacterized protein; afdb-uniprot50 \| AF-A0A5C1R7K0-F1-MODEL\_V4 \| 1.0 \| 3.809e-16 \| 589 \| 0.206 \| 261 \| 189 \| 9 \| 431 \| 680 \| 29 \| 282 \| Phage major capsid protein \| Phage major capsid protein; afdb-uniprot50 \| AF-A0A7Y4JL20-F1-MODEL\_V4 \| 1.0 \| 5.096e-21 \| 588 \| 0.211 \| 420 \| 257 \| 19 \| 286 \| 680 \| 6 \| 376 \| Phage major capsid protein \| Phage major capsid protein; afdb-uniprot50 \| AF-A0A1X2FVP9-F1-MODEL\_V4 \| 1.0 \| 5.096e-21 \| 587 \| 0.216 \| 420 \| 229 \| 19 \| 294 \| 681 \| 15 \| 366 \| Uncharacterized protein \| Uncharacterized protein; afdb-uniprot50 \| AF-A0A0A2G587-F1-MODEL\_V4 \| 1.0 \| 1.675e-20 \| 587 \| 0.203 \| 442 \| 255 \| 17 \| 268 \| 681 \| 7 \| 379 \| Uncharacterized protein \| Uncharacterized protein; afdb-uniprot50 \| AF-A0A1B7LDB2-F1-MODEL\_V4 \| 1.0 \| 1.496e-20 \| 587 \| 0.202 \| 414 \| 242 \| 17 \| 296 \| 681 \| 23 \| 376 \| Uncharacterized protein \| Uncharacterized protein; afdb-uniprot50 \| AF-I9WTH1-F1-MODEL\_V4 \| 1.0 \| 1.486e-23 \| 587 \| 0.204 \| 484 \| 298 \| 22 \| 261 \| 681 \| 7 \| 466 \| Phage major capsid protein, HK97 family \| Phage major capsid protein, HK97 family; afdb-uniprot50 \| AF-A0A7B4HVD7-F1-MODEL\_V4 \| 1.0 \| 1.999e-17 \| 586 \| 0.197 \| 284 \| 186 \| 11 \| 430 \| 681 \| 2 \| 275 \| Phage major capsid protein \| Phage major capsid protein; afdb-uniprot50 \| AF-A0A090E9R7-F1-MODEL\_V4 \| 1.0 \| 7.264e-23 \| 586 \| 0.2 \| 464 \| 288 \| 21 \| 264 \| 681 \| 5 \| 431 \| Putative Phage major capsid protein, HK97 family \| Putative Phage major capsid protein, HK97 family; afdb-uniprot50 \| AF-A0A1Q9VAW5-F1-MODEL\_V4 \| 1.0 \| 4.266e-16 \| 585 \| 0.185 \| 259 \| 194 \| 7 \| 431 \| 681 \| 28 \| 277 \| Capsid protein \| Capsid protein; afdb-uniprot50 \| AF-A0A3S9X822-F1-MODEL\_V4 \| 1.0 \| 4.951e-17 \| 585 \| 0.171 \| 292 \| 219 \| 10 \| 402 \| 682 \| 2 \| 281 \| Uncharacterized protein \| Uncharacterized protein; afdb-uniprot50 \| AF-A0A5A9ZIC2-F1-MODEL\_V4 \| 1.0 \| 2.287e-16 \| 585 \| 0.169 \| 259 \| 201 \| 5 \| 431 \| 682 \| 34 \| 285 \| Phage major capsid protein \| Phage major capsid protein; afdb-uniprot50 \| AF-A0A543KX21-F1-MODEL\_V4 \| 1.0 \| 9.774e-17 \| 584 \| 0.186 \| 257 \| 192 \| 9 \| 431 \| 681 \| 27 \| 272 \| HK97 family phage major capsid protein \| HK97 family phage major capsid protein; afdb-uniprot50 \| AF-A0A429N598-F1-MODEL\_V4 \| 1.0 \| 3.401e-16 \| 583 \| 0.16 \| 262 \| 200 \| 7 \| 430 \| 681 \| 29 \| 280 \| Phage major capsid protein \| Phage major capsid protein; afdb-uniprot50 \| AF-A0A3A8RP75-F1-MODEL\_V4 \| 1.0 \| 1.042e-21 \| 583 \| 0.184 \| 445 \| 283 \| 23 \| 271 \| 682 \| 6 \| 403 \| Phage major capsid protein \| Phage major capsid protein; afdb-uniprot50 \| AF-A0A4V2RW77-F1-MODEL\_V4 \| 1.0 \| 1.218e-19 \| 582 \| 0.183 \| 403 \| 246 \| 15 \| 295 \| 681 \| 27 \| 362 \| HK97 family phage major capsid protein \| HK97 family phage major capsid protein; afdb-uniprot50 \| AF-A0A518I8W7-F1-MODEL\_V4 \| 1.0 \| 4.519e-24 \| 582 \| 0.172 \| 584 \| 342 \| 28 \| 180 \| 682 \| 7 \| 529 \| Phage capsid family protein \| Phage capsid family protein; afdb-uniprot50 \| AF-A0A2U1FA90-F1-MODEL\_V4 \| 1.0 \| 2.287e-16 \| 580 \| 0.178 \| 257 \| 199 \| 7 \| 431 \| 682 \| 31 \| 280 \| HK97 family phage major capsid protein \| HK97 family phage major capsid protein; afdb-uniprot50 \| AF-A0A7I9ZGE4-F1-MODEL\_V4 \| 1.0 \| 1.618e-27 \| 580 \| 0.147 \| 723 \| 432 \| 33 \| 15 \| 678 \| 2 \| 598 \| Uncharacterized protein \| Uncharacterized protein; afdb-uniprot50 \| AF-A0A3A8JWE7-F1-MODEL\_V4 \| 1.0 \| 1.192e-20 \| 578 \| 0.193 \| 408 \| 262 \| 22 \| 298 \| 682 \| 35 \| 398 \| Phage major capsid protein \| Phage major capsid protein; afdb-uniprot50 \| AF-A0A3A4UX11-F1-MODEL\_V4 \| 1.0 \| 1.904e-22 \| 578 \| 0.18 \| 494 \| 308 \| 16 \| 270 \| 682 \| 2 \| 479 \| Phage major capsid protein \| Phage major capsid protein; afdb-uniprot50 \| AF-A0A5C5AK74-F1-MODEL\_V4 \| 1.0 \| 9.925e-27 \| 578 \| 0.128 \| 708 \| 432 \| 37 \| 23 \| 682 \| 1 \| 571 \| Phage major capsid protein \| Phage major capsid protein; afdb-uniprot50 \| AF-A0A7Y9DIE2-F1-MODEL\_V4 \| 1.0 \| 1.484e-15 \| 574 \| 0.154 \| 259 \| 204 \| 5 \| 431 \| 682 \| 29 \| 279 \| HK97 family phage major capsid protein \| HK97 family phage major capsid protein; afdb-uniprot50 \| AF-A0A5C8ZHA4-F1-MODEL\_V4 \| 1.0 \| 3.401e-16 \| 574 \| 0.211 \| 260 \| 189 \| 8 \| 431 \| 682 \| 28 \| 279 \| Phage major capsid protein \| Phage major capsid protein; afdb-uniprot50 \| AF-A0A436KKN4-F1-MODEL\_V4 \| 1.0 \| 6.624e-22 \| 574 \| 0.192 \| 468 \| 291 \| 19 \| 264 \| 681 \| 5 \| 435 \| Phage major capsid protein \| Phage major capsid protein; afdb-uniprot50 \| AF-A0A0Q7IS85-F1-MODEL\_V4 \| 1.0 \| 1.583e-20 \| 573 \| 0.212 \| 457 \| 266 \| 16 \| 238 \| 681 \| 5 \| 380 \| Capsid protein \| Capsid protein; afdb-uniprot50 \| AF-A0A6I2R3H1-F1-MODEL\_V4 \| 1.0 \| 9.112e-23 \| 573 \| 0.215 \| 488 \| 275 \| 18 \| 271 \| 680 \| 46 \| 503 \| Phage major capsid protein \| Phage major capsid protein; afdb-uniprot50 \| AF-A0A2K1Q3K7-F1-MODEL\_V4 \| 1.0 \| 8.495e-29 \| 573 \| 0.201 \| 724 \| 336 \| 29 \| 18 \| 681 \| 9 \| 550 \| Caudovirus prohead protease \| Caudovirus prohead protease; afdb-uniprot50 \| AF-A0A3N1D5M0-F1-MODEL\_V4 \| 1.0 \| 3.599e-16 \| 572 \| 0.163 \| 257 \| 205 \| 5 \| 431 \| 682 \| 32 \| 283 \| HK97 family phage major capsid protein \| HK97 family phage major capsid protein; afdb-uniprot50 \| AF-A0A7S7QRL3-F1-MODEL\_V4 \| 1.0 \| 1.675e-20 \| 572 \| 0.193 \| 398 \| 266 \| 16 \| 313 \| 680 \| 1 \| 373 \| Phage major capsid protein \| Phage major capsid protein; afdb-uniprot50 \| AF-A0A843HXH6-F1-MODEL\_V4 \| 1.0 \| 1.944e-21 \| 572 \| 0.18 \| 477 \| 300 \| 17 \| 249 \| 681 \| 3 \| 432 \| Phage major capsid protein \| Phage major capsid protein; afdb-uniprot50 \| AF-A0A643EYU7-F1-MODEL\_V4 \| 1.0 \| 5.321e-27 \| 572 \| 0.159 \| 701 \| 402 \| 31 \| 13 \| 682 \| 3 \| 546 \| Phage major capsid protein \| Phage major capsid protein; afdb-uniprot50 \| AF-A0A5A9GUV5-F1-MODEL\_V4 \| 1.0 \| 6.535e-28 \| 572 \| 0.187 \| 709 \| 377 \| 34 \| 5 \| 681 \| 63 \| 604 \| Phage major capsid protein \| Phage major capsid protein; afdb-uniprot50 \| AF-A0A356UJ62-F1-MODEL\_V4 \| 1.0 \| 8.081e-26 \| 571 \| 0.152 \| 693 \| 377 \| 32 \| 38 \| 682 \| 6 \| 535 \| Uncharacterized protein \| Uncharacterized protein; afdb-uniprot50 \| AF-A0A1B3ZE75-F1-MODEL\_V4 \| 1.0 \| 1.042e-21 \| 570 \| 0.201 \| 471 \| 291 \| 22 \| 255 \| 681 \| 4 \| 433 \| Uncharacterized protein \| Uncharacterized protein; afdb-uniprot50 \| AF-A0A255XX39-F1-MODEL\_V4 \| 1.0 \| 6.817e-26 \| 569 \| 0.179 \| 691 \| 354 \| 33 \| 38 \| 681 \| 11 \| 535 \| Phage major capsid protein \| Phage major capsid protein; afdb-uniprot50 \| AF-A0A1M7RID9-F1-MODEL\_V4 \| 1.0 \| 4.523e-32 \| 569 \| 0.19 \| 735 \| 442 \| 38 \| 28 \| 681 \| 2 \| 664 \| Prohead serine protease \| Prohead serine protease; afdb-uniprot50 \| AF-A0A7C9H8Y0-F1-MODEL\_V4 \| 1.0 \| 5.707e-21 \| 566 \| 0.185 \| 437 \| 256 \| 17 \| 295 \| 680 \| 22 \| 409 \| Phage major capsid protein \| Phage major capsid protein; afdb-uniprot50 \| AF-A0A5B9R260-F1-MODEL\_V4 \| 1.0 \| 1.021e-22 \| 565 \| 0.196 \| 508 \| 284 \| 18 \| 252 \| 681 \| 16 \| 477 \| Phage capsid family protein \| Phage capsid family protein; afdb-uniprot50 \| AF-A0A1V5LQQ8-F1-MODEL\_V4 \| 1.0 \| 1.056e-15 \| 563 \| 0.169 \| 277 \| 192 \| 8 \| 431 \| 682 \| 34 \| 297 \| Phage capsid family protein \| Phage capsid family protein; afdb-uniprot50 \| AF-A0A521CQZ1-F1-MODEL\_V4 \| 1.0 \| 1.763e-31 \| 562 \| 0.157 \| 731 \| 460 \| 34 \| 12 \| 681 \| 111 \| 746 \| Phage prohead protease, HK97 family \| Phage prohead protease, HK97 family; afdb-uniprot50 \| AF-A0A2E5AE13-F1-MODEL\_V4 \| 1.0 \| 3.898e-31 \| 560 \| 0.182 \| 729 \| 437 \| 37 \| 30 \| 681 \| 13 \| 659 \| Uncharacterized protein \| Uncharacterized protein; afdb-uniprot50 \| AF-C0W521-F1-MODEL\_V4 \| 1.0 \| 1.298e-16 \| 557 \| 0.191 \| 266 \| 190 \| 8 \| 433 \| 681 \| 27 \| 284 \| Putative phage capsid family \| Putative phage capsid family; afdb-uniprot50 \| AF-A0A2E2XBH4-F1-MODEL\_V4 \| 1.0 \| 1.236e-21 \| 557 \| 0.183 \| 468 \| 302 \| 19 \| 258 \| 681 \| 3 \| 434 \| Phage major capsid protein \| Phage major capsid protein; afdb-uniprot50 \| AF-A0A1W1VX51-F1-MODEL\_V4 \| 1.0 \| 6.91e-20 \| 556 \| 0.209 \| 458 \| 253 \| 20 \| 251 \| 681 \| 7 \| 382 \| Phage major capsid protein, HK97 family \| Phage major capsid protein, HK97 family; afdb-uniprot50 \| AF-A0A1V6D5P9-F1-MODEL\_V4 \| 1.0 \| 1.289e-19 \| 556 \| 0.202 \| 376 \| 236 \| 19 \| 336 \| 681 \| 109 \| 450 \| Phage capsid family protein \| Phage capsid family protein; afdb-uniprot50 \| AF-A0A3P6KQL2-F1-MODEL\_V4 \| 1.0 \| 2.259e-30 \| 556 \| 0.175 \| 731 \| 430 \| 36 \| 19 \| 681 \| 1 \| 626 \| Mu-like prophage major head subunit gpT \| Mu-like prophage major head subunit gpT; afdb-uniprot50 \| AF-A0A8B2NHU5-F1-MODEL\_V4 \| 1.0 \| 9.783e-25 \| 555 \| 0.16 \| 673 \| 372 \| 29 \| 37 \| 681 \| 11 \| 518 \| Phage major capsid protein \| Phage major capsid protein; afdb-uniprot50 \| AF-A0A512HAW7-F1-MODEL\_V4 \| 1.0 \| 4.852e-26 \| 555 \| 0.165 \| 694 \| 397 \| 32 \| 19 \| 682 \| 4 \| 545 \| Peptidase U35 \| Peptidase U35; afdb-uniprot50 \| AF-A0A498R0I8-F1-MODEL\_V4 \| 1.0 \| 1.252e-15 \| 554 \| 0.188 \| 260 \| 192 \| 5 \| 436 \| 682 \| 36 \| 289 \| Uncharacterized protein \| Uncharacterized protein; afdb-uniprot50 \| AF-A0A2U1YTD1-F1-MODEL\_V4 \| 1.0 \| 4.062e-21 \| 554 \| 0.183 \| 484 \| 286 \| 21 \| 266 \| 682 \| 4 \| 445 \| Uncharacterized protein \| Uncharacterized protein; afdb-uniprot50 \| AF-A0A318RLQ4-F1-MODEL\_V4 \| 1.0 \| 7.105e-16 \| 553 \| 0.186 \| 258 \| 190 \| 9 \| 435 \| 682 \| 34 \| 281 \| HK97 family phage major capsid protein \| HK97 family phage major capsid protein; afdb-uniprot50 \| AF-A0A132HLC0-F1-MODEL\_V4 \| 1.0 \| 1.8e-30 \| 553 \| 0.214 \| 680 \| 311 \| 27 \| 27 \| 682 \| 15 \| 495 \| Uncharacterized protein \| Uncharacterized protein; afdb-uniprot50 \| AF-A0A7V3IXB5-F1-MODEL\_V4 \| 1.0 \| 3.101e-15 \| 552 \| 0.195 \| 266 \| 191 \| 8 \| 431 \| 682 \| 29 \| 285 \| Phage major capsid protein \| Phage major capsid protein; afdb-uniprot50 \| AF-A0A7U0UNX5-F1-MODEL\_V4 \| 1.0 \| 2.869e-16 \| 552 \| 0.221 \| 262 \| 172 \| 9 \| 437 \| 678 \| 35 \| 284 \| Phage major capsid protein \| Phage major capsid protein; afdb-uniprot50 \| AF-A0A177VZ34-F1-MODEL\_V4 \| 1.0 \| 1.628e-16 \| 552 \| 0.168 \| 308 \| 210 \| 11 \| 402 \| 681 \| 3 \| 292 \| Phage capsid family protein \| Phage capsid family protein; afdb-uniprot50 \| AF-A0A4R6RGM9-F1-MODEL\_V4 \| 1.0 \| 6.529e-20 \| 551 \| 0.175 \| 508 \| 289 \| 21 \| 191 \| 681 \| 2 \| 396 \| HK97 family phage major capsid protein \| HK97 family phage major capsid protein; afdb-uniprot50 \| AF-A0A442WPD5-F1-MODEL\_V4 \| 1.0 \| 1.736e-21 \| 551 \| 0.174 \| 511 \| 307 \| 20 \| 204 \| 682 \| 1 \| 428 \| Phage major capsid protein \| Phage major capsid protein; afdb-uniprot50 \| AF-A0A6I0BNG2-F1-MODEL\_V4 \| 1.0 \| 2.074e-26 \| 551 \| 0.144 \| 707 \| 398 \| 33 \| 32 \| 681 \| 5 \| 561 \| Phage major capsid protein \| Phage major capsid protein; afdb-uniprot50 \| AF-A0A5C4Y014-F1-MODEL\_V4 \| 1.0 \| 1.663e-15 \| 549 \| 0.158 \| 265 \| 200 \| 8 \| 431 \| 681 \| 38 \| 293 \| Phage major capsid protein \| Phage major capsid protein; afdb-uniprot50 \| AF-A0A416XLM7-F1-MODEL\_V4 \| 1.0 \| 6.301e-19 \| 549 \| 0.197 \| 364 \| 249 \| 13 \| 334 \| 682 \| 28 \| 363 \| Phage major capsid protein \| Phage major capsid protein; afdb-uniprot50 \| AF-A0A4Q7UYC6-F1-MODEL\_V4 \| 1.0 \| 5.664e-16 \| 548 \| 0.158 \| 259 \| 195 \| 10 \| 436 \| 682 \| 39 \| 286 \| HK97 family phage major capsid protein \| HK97 family phage major capsid protein; afdb-uniprot50 \| AF-A0A7K3LU50-F1-MODEL\_V4 \| 1.0 \| 5.24e-17 \| 548 \| 0.167 \| 317 \| 235 \| 13 \| 374 \| 681 \| 29 \| 325 \| Phage major capsid protein \| Phage major capsid protein; afdb-uniprot50 \| AF-A0A1E3GZU2-F1-MODEL\_V4 \| 1.0 \| 3.307e-20 \| 548 \| 0.191 \| 507 \| 280 \| 22 \| 191 \| 681 \| 1 \| 393 \| Phage capsid family protein \| Phage capsid family protein; afdb-uniprot50 \| AF-A0A3D2INS8-F1-MODEL\_V4 \| 1.0 \| 1.183e-15 \| 547 \| 0.322 \| 177 \| 110 \| 5 \| 19 \| 186 \| 11 \| 186 \| Caudovirus prohead protease \| Caudovirus prohead protease; afdb-uniprot50 \| AF-A0A3N0BSI3-F1-MODEL\_V4 \| 1.0 \| 1.862e-15 \| 547 \| 0.179 \| 256 \| 192 \| 8 \| 436 \| 682 \| 37 \| 283 \| Phage major capsid protein \| Phage major capsid protein; afdb-uniprot50 \| AF-A0A329BHT4-F1-MODEL\_V4 \| 1.0 \| 1.701e-30 \| 547 \| 0.197 \| 730 \| 288 \| 31 \| 15 \| 681 \| 9 \| 503 \| HK97 family phage prohead protease \| HK97 family phage prohead protease; afdb-uniprot50 \| AF-V7KIQ7-F1-MODEL\_V4 \| 1.0 \| 7.105e-16 \| 545 \| 0.169 \| 260 \| 190 \| 9 \| 435 \| 681 \| 33 \| 279 \| Capsid protein \| Capsid protein; afdb-uniprot50 \| AF-A0A642F139-F1-MODEL\_V4 \| 1.0 \| 5.023e-19 \| 545 \| 0.182 \| 378 \| 267 \| 18 \| 338 \| 682 \| 8 \| 376 \| Phage major capsid protein \| Phage major capsid protein; afdb-uniprot50 \| AF-A0A1I2GJW8-F1-MODEL\_V4 \| 1.0 \| 2.564e-24 \| 545 \| 0.119 \| 696 \| 380 \| 29 \| 32 \| 682 \| 4 \| 511 \| Phage prohead protease, HK97 family/phage major capsid protein, HK97 family,TIGR01554 \| Phage prohead protease, HK97 family/phage major capsid protein, HK97 family,TIGR01554; afdb-uniprot50 \| AF-A0A7U6ITT4-F1-MODEL\_V4 \| 1.0 \| 3.838e-21 \| 544 \| 0.147 \| 522 \| 332 \| 24 \| 191 \| 681 \| 3 \| 442 \| Uncharacterized protein \| Uncharacterized protein; afdb-uniprot50 \| AF-A0A259MPF3-F1-MODEL\_V4 \| 1.0 \| 1.128e-28 \| 544 \| 0.182 \| 736 \| 430 \| 34 \| 37 \| 682 \| 11 \| 664 \| Uncharacterized protein \| Uncharacterized protein; afdb-uniprot50 \| AF-A0A2P9HI32-F1-MODEL\_V4 \| 1.0 \| 4.683e-25 \| 543 \| 0.142 \| 701 \| 384 \| 31 \| 17 \| 681 \| 3 \| 522 \| Phage major capsid protein \| Phage major capsid protein; afdb-uniprot50 \| AF-A0A348AC18-F1-MODEL\_V4 \| 1.0 \| 1.571e-15 \| 542 \| 0.159 \| 269 \| 197 \| 10 \| 431 \| 681 \| 89 \| 346 \| Phage major capsid protein, HK97 family \| Phage major capsid protein, HK97 family; afdb-uniprot50 \| AF-A0A0F9LI03-F1-MODEL\_V4 \| 1.0 \| 4.815e-21 \| 542 \| 0.159 \| 463 \| 326 \| 19 \| 249 \| 679 \| 5 \| 436 \| Uncharacterized protein \| Uncharacterized protein; afdb-uniprot50 \| AF-A0A2W6Y6L7-F1-MODEL\_V4 \| 1.0 \| 2.872e-24 \| 541 \| 0.138 \| 677 \| 380 \| 30 \| 43 \| 681 \| 17 \| 527 \| Phage major capsid protein \| Phage major capsid protein; afdb-uniprot50 \| AF-A0A380N679-F1-MODEL\_V4 \| 1.0 \| 4.554e-29 \| 540 \| 0.204 \| 715 \| 395 \| 39 \| 39 \| 681 \| 21 \| 633 \| Mu-like prophage major head subunit gpT \| Mu-like prophage major head subunit gpT; afdb-uniprot50 \| AF-A0A4Q3XXJ0-F1-MODEL\_V4 \| 1.0 \| 2.072e-18 \| 539 \| 0.434 \| 235 \| 106 \| 7 \| 14 \| 235 \| 14 \| 234 \| Uncharacterized protein \| Uncharacterized protein; afdb-uniprot50 \| AF-A0A1D3R7K8-F1-MODEL\_V4 \| 1.0 \| 4.683e-25 \| 539 \| 0.124 \| 689 \| 398 \| 31 \| 32 \| 682 \| 7 \| 527 \| Putative phage capsid protein \| Putative phage capsid protein; afdb-uniprot50 \| AF-A0A0Q9R783-F1-MODEL\_V4 \| 1.0 \| 1.403e-15 \| 538 \| 0.161 \| 260 \| 195 \| 9 \| 431 \| 677 \| 4 \| 253 \| Uncharacterized protein \| Uncharacterized protein; afdb-uniprot50 \| AF-A0A852TM38-F1-MODEL\_V4 \| 1.0 \| 1.571e-15 \| 538 \| 0.156 \| 262 \| 196 \| 8 \| 433 \| 682 \| 29 \| 277 \| HK97 family phage major capsid protein \| HK97 family phage major capsid protein; afdb-uniprot50 \| AF-A0A1Z3VNT3-F1-MODEL\_V4 \| 1.0 \| 1.056e-15 \| 537 \| 0.211 \| 265 \| 185 \| 8 \| 430 \| 682 \| 32 \| 284 \| Phage major capsid protein \| Phage major capsid protein; afdb-uniprot50 \| AF-A0A1X2DT10-F1-MODEL\_V4 \| 1.0 \| 5.785e-15 \| 537 \| 0.162 \| 258 \| 197 \| 6 \| 436 \| 682 \| 36 \| 285 \| Capsid protein \| Capsid protein; afdb-uniprot50 \| AF-A0A4Q4AXM0-F1-MODEL\_V4 \| 1.0 \| 1.56e-18 \| 537 \| 0.184 \| 384 \| 238 \| 17 \| 313 \| 679 \| 35 \| 360 \| Phage major capsid protein \| Phage major capsid protein; afdb-uniprot50 \| AF-A0A661DS55-F1-MODEL\_V4 \| 1.0 \| 2.058e-21 \| 536 \| 0.159 \| 489 \| 326 \| 20 \| 252 \| 680 \| 6 \| 469 \| Phage major capsid protein \| Phage major capsid protein; afdb-uniprot50 \| AF-G2I0U6-F1-MODEL\_V4 \| 1.0 \| 7.419e-22 \| 536 \| 0.215 \| 482 \| 285 \| 21 \| 249 \| 680 \| 20 \| 458 \| Bacteriophage protein \| Bacteriophage protein; afdb-uniprot50 \| AF-A0A843LCP9-F1-MODEL\_V4 \| 1.0 \| 8.913e-16 \| 535 \| 0.171 \| 251 \| 189 \| 9 \| 433 \| 681 \| 54 \| 287 \| Phage major capsid protein \| Phage major capsid protein; afdb-uniprot50 \| AF-A0A0Q5VUD7-F1-MODEL\_V4 \| 1.0 \| 5.508e-20 \| 535 \| 0.17 \| 499 \| 283 \| 20 \| 203 \| 681 \| 1 \| 388 \| Capsid protein \| Capsid protein; afdb-uniprot50 \| AF-A0A3C1INU1-F1-MODEL\_V4 \| 1.0 \| 3.762e-30 \| 535 \| 0.188 \| 722 \| 407 \| 45 \| 26 \| 681 \| 15 \| 623 \| Mu-like\_gpT domain-containing protein \| Mu-like\_gpT domain-containing protein; afdb-uniprot50 \| AF-U2DD62-F1-MODEL\_V4 \| 1.0 \| 7.958e-16 \| 534 \| 0.353 \| 184 \| 100 \| 7 \| 30 \| 203 \| 17 \| 191 \| Caudovirus prohead protease \| Caudovirus prohead protease; afdb-uniprot50 \| AF-A0A3R6SVE9-F1-MODEL\_V4 \| 1.0 \| 1.929e-16 \| 533 \| 0.357 \| 196 \| 107 \| 7 \| 13 \| 198 \| 8 \| 194 \| Caudovirus prohead protease \| Caudovirus prohead protease; afdb-uniprot50 \| AF-A0A2M7SYD1-F1-MODEL\_V4 \| 1.0 \| 2.456e-18 \| 533 \| 0.166 \| 361 \| 277 \| 12 \| 334 \| 681 \| 33 \| 382 \| Phage major capsid protein \| Phage major capsid protein; afdb-uniprot50 \| AF-A4TBM3-F1-MODEL\_V4 \| 1.0 \| 9.992e-24 \| 533 \| 0.134 \| 700 \| 435 \| 33 \| 31 \| 682 \| 2 \| 578 \| Phage prohead protease, HK97 family \| Phage prohead protease, HK97 family; afdb-uniprot50 \| AF-A0A3N5EJT9-F1-MODEL\_V4 \| 1.0 \| 2.149e-27 \| 533 \| 0.19 \| 702 \| 427 \| 37 \| 37 \| 681 \| 10 \| 626 \| Phage major capsid protein \| Phage major capsid protein; afdb-uniprot50 \| AF-A0A8A6JDC6-F1-MODEL\_V4 \| 1.0 \| 1.971e-15 \| 532 \| 0.172 \| 261 \| 193 \| 11 \| 431 \| 681 \| 28 \| 275 \| Phage major capsid protein \| Phage major capsid protein; afdb-uniprot50 \| AF-I4EHG9-F1-MODEL\_V4 \| 1.0 \| 1.496e-20 \| 532 \| 0.178 \| 459 \| 291 \| 20 \| 258 \| 682 \| 8 \| 414 \| Putative Phage major capsid protein, HK97 family \| Putative Phage major capsid protein, HK97 family; afdb-uniprot50 \| AF-A0A640YC84-F1-MODEL\_V4 \| 1.0 \| 1.404e-23 \| 532 \| 0.118 \| 691 \| 393 \| 30 \| 19 \| 682 \| 1 \| 502 \| HK97 family phage prohead protease \| HK97 family phage prohead protease; afdb-uniprot50 \| AF-A0A3M1QXF2-F1-MODEL\_V4 \| 1.0 \| 3.43e-29 \| 532 \| 0.175 \| 741 \| 434 \| 43 \| 30 \| 681 \| 9 \| 661 \| Uncharacterized protein \| Uncharacterized protein; afdb-uniprot50 \| AF-A0A1A7QM62-F1-MODEL\_V4 \| 1.0 \| 5.24e-17 \| 531 \| 0.176 \| 329 \| 226 \| 15 \| 376 \| 682 \| 57 \| 362 \| Uncharacterized protein \| Uncharacterized protein; afdb-uniprot50 \| AF-A0A1F8NTD0-F1-MODEL\_V4 \| 1.0 \| 1.087e-19 \| 530 \| 0.178 \| 454 \| 286 \| 18 \| 249 \| 682 \| 11 \| 397 \| Uncharacterized protein \| Uncharacterized protein; afdb-uniprot50 \| AF-A0A0S7X9K7-F1-MODEL\_V4 \| 1.0 \| 3.192e-19 \| 529 \| 0.203 \| 428 \| 245 \| 22 \| 295 \| 681 \| 27 \| 399 \| Uncharacterized protein \| Uncharacterized protein; afdb-uniprot50 \| AF-A0A316S7Y7-F1-MODEL\_V4 \| 1.0 \| 8.727e-17 \| 528 \| 0.32 \| 206 \| 119 \| 7 \| 5 \| 199 \| 6 \| 201 \| Caudovirus prohead protease \| Caudovirus prohead protease; afdb-uniprot50 \| AF-A0A1H4QQ99-F1-MODEL\_V4 \| 1.0 \| 4.117e-15 \| 528 \| 0.191 \| 256 \| 190 \| 8 \| 431 \| 680 \| 29 \| 273 \| Phage major capsid protein, HK97 family \| Phage major capsid protein, HK97 family; afdb-uniprot50 \| AF-A0A2S6AID0-F1-MODEL\_V4 \| 1.0 \| 7.362e-17 \| 528 \| 0.19 \| 304 \| 209 \| 12 \| 398 \| 682 \| 34 \| 319 \| Phage major capsid protein \| Phage major capsid protein; afdb-uniprot50 \| AF-A0A060QKT2-F1-MODEL\_V4 \| 1.0 \| 6.573e-17 \| 528 \| 0.174 \| 316 \| 231 \| 15 \| 381 \| 682 \| 12 \| 311 \| Phage major capsid protein \| Phage major capsid protein; afdb-uniprot50 \| AF-B5ZDC2-F1-MODEL\_V4 \| 1.0 \| 7.16e-21 \| 527 \| 0.177 \| 478 \| 284 \| 20 \| 271 \| 680 \| 3 \| 439 \| Uncharacterized protein \| Uncharacterized protein; afdb-uniprot50 \| AF-A0A443W106-F1-MODEL\_V4 \| 1.0 \| 2.695e-27 \| 527 \| 0.221 \| 632 \| 353 \| 25 \| 40 \| 631 \| 20 \| 552 \| Major capsid protein \| Major capsid protein; afdb-uniprot50 \| AF-A0A0Q9ES63-F1-MODEL\_V4 \| 1.0 \| 9.514e-29 \| 527 \| 0.174 \| 738 \| 435 \| 42 \| 35 \| 682 \| 10 \| 662 \| Uncharacterized protein \| Uncharacterized protein; afdb-uniprot50 \| AF-A0A543NSK2-F1-MODEL\_V4 \| 1.0 \| 8.913e-16 \| 526 \| 0.176 \| 289 \| 213 \| 10 \| 409 \| 682 \| 3 \| 281 \| HK97 family phage major capsid protein \| HK97 family phage major capsid protein; afdb-uniprot50 \| AF-A0A564G6H4-F1-MODEL\_V4 \| 1.0 \| 1.167e-21 \| 525 \| 0.191 \| 481 \| 306 \| 20 \| 258 \| 680 \| 5 \| 460 \| Uncharacterized protein \| Uncharacterized protein; afdb-uniprot50 \| AF-A0A6M3L763-F1-MODEL\_V4 \| 1.0 \| 2.891e-21 \| 525 \| 0.18 \| 521 \| 314 \| 22 \| 249 \| 682 \| 12 \| 506 \| Putative capsid protein \| Putative capsid protein; afdb-uniprot50 \| AF-A0A165XGL2-F1-MODEL\_V4 \| 1.0 \| 2.999e-30 \| 525 \| 0.186 \| 745 \| 431 \| 38 \| 30 \| 677 \| 13 \| 679 \| Caudovirus prohead protease \| Caudovirus prohead protease; afdb-uniprot50 \| AF-C0ZMF4-F1-MODEL\_V4 \| 1.0 \| 7.257e-15 \| 524 \| 0.175 \| 251 \| 194 \| 6 \| 436 \| 681 \| 41 \| 283 \| Uncharacterized protein \| Uncharacterized protein; afdb-uniprot50 \| AF-A0A3A4XE90-F1-MODEL\_V4 \| 1.0 \| 5.626e-19 \| 524 \| 0.179 \| 401 \| 262 \| 17 \| 309 \| 682 \| 2 \| 362 \| Phage major capsid protein \| Phage major capsid protein; afdb-uniprot50 \| AF-A0A537KSG3-F1-MODEL\_V4 \| 1.0 \| 1.496e-20 \| 523 \| 0.165 \| 490 \| 300 \| 21 \| 262 \| 682 \| 7 \| 456 \| Phage major capsid protein \| Phage major capsid protein; afdb-uniprot50 \| AF-A0A7V6QN74-F1-MODEL\_V4 \| 1.0 \| 2.472e-15 \| 522 \| 0.17 \| 264 \| 190 \| 12 \| 433 \| 682 \| 56 \| 304 \| Phage major capsid protein \| Phage major capsid protein; afdb-uniprot50 \| AF-A0A3S4V390-F1-MODEL\_V4 \| 1.0 \| 1.354e-14 \| 521 \| 0.183 \| 261 \| 192 \| 9 \| 431 \| 681 \| 29 \| 278 \| Predicted phage phi-C31 gp36 major capsid-like protein \| Predicted phage phi-C31 gp36 major capsid-like protein; afdb-uniprot50 \| AF-A0A419VU13-F1-MODEL\_V4 \| 1.0 \| 1.079e-14 \| 521 \| 0.155 \| 258 \| 197 \| 9 \| 433 \| 679 \| 57 \| 304 \| HK97 family phage major capsid protein \| HK97 family phage major capsid protein; afdb-uniprot50 \| AF-A0A0N8W5Z7-F1-MODEL\_V4 \| 1.0 \| 1.007e-28 \| 521 \| 0.172 \| 729 \| 457 \| 38 \| 19 \| 681 \| 3 \| 650 \| Mu-like\_gpT domain-containing protein \| Mu-like\_gpT domain-containing protein; afdb-uniprot50 \| AF-A0A7R7RKY8-F1-MODEL\_V4 \| 1.0 \| 1.079e-14 \| 520 \| 0.159 \| 264 \| 198 \| 7 \| 430 \| 677 \| 29 \| 284 \| Uncharacterized protein \| Uncharacterized protein; afdb-uniprot50 \| AF-A0A7X7WBZ3-F1-MODEL\_V4 \| 1.0 \| 5.954e-19 \| 520 \| 0.193 \| 433 \| 255 \| 18 \| 271 \| 681 \| 3 \| 363 \| Phage major capsid protein \| Phage major capsid protein; afdb-uniprot50 \| AF-L7U6P8-F1-MODEL\_V4 \| 1.0 \| 2.894e-29 \| 519 \| 0.185 \| 726 \| 442 \| 36 \| 40 \| 681 \| 10 \| 669 \| ClpP family peptidase \| ClpP family peptidase; afdb-uniprot50 \| AF-A0A4S5BFZ4-F1-MODEL\_V4 \| 1.0 \| 3.019e-27 \| 519 \| 0.178 \| 712 \| 450 \| 33 \| 37 \| 679 \| 12 \| 657 \| Phage major capsid protein \| Phage major capsid protein; afdb-uniprot50 \| AF-A0A827RVM8-F1-MODEL\_V4 \| 1.0 \| 7.844e-14 \| 518 \| 0.37 \| 162 \| 90 \| 2 \| 40 \| 195 \| 18 \| 173 \| Uncharacterized protein \| Uncharacterized protein; afdb-uniprot50 \| AF-A0A7W0ZDQ7-F1-MODEL\_V4 \| 1.0 \| 6.857e-15 \| 518 \| 0.182 \| 279 \| 194 \| 11 \| 418 \| 681 \| 2 \| 261 \| Phage major capsid protein \| Phage major capsid protein; afdb-uniprot50 \| AF-A0A4V3IRT6-F1-MODEL\_V4 \| 1.0 \| 3.378e-19 \| 518 \| 0.189 \| 385 \| 247 \| 12 \| 334 \| 679 \| 31 \| 389 \| Phage major capsid protein \| Phage major capsid protein; afdb-uniprot50 \| AF-A0A7C1UH41-F1-MODEL\_V4 \| 1.0 \| 9.505e-21 \| 518 \| 0.156 \| 499 \| 283 \| 24 \| 258 \| 681 \| 3 \| 438 \| Phage major capsid protein \| Phage major capsid protein; afdb-uniprot50 \| AF-Q31QA3-F1-MODEL\_V4 \| 1.0 \| 3.63e-29 \| 518 \| 0.195 \| 733 \| 416 \| 41 \| 13 \| 682 \| 2 \| 623 \| Mu-like\_gpT domain-containing protein \| Mu-like\_gpT domain-containing protein; afdb-uniprot50 \| AF-A0A157ZYR5-F1-MODEL\_V4 \| 1.0 \| 1.112e-26 \| 517 \| 0.185 \| 711 \| 428 \| 34 \| 19 \| 679 \| 15 \| 624 \| Phage-like protein \| Phage-like protein; afdb-uniprot50 \| AF-A0A442RKY7-F1-MODEL\_V4 \| 1.0 \| 4.746e-19 \| 516 \| 0.206 \| 407 \| 258 \| 18 \| 319 \| 681 \| 2 \| 387 \| Phage major capsid protein \| Phage major capsid protein; afdb-uniprot50 \| AF-A0A7Y3PR89-F1-MODEL\_V4 \| 1.0 \| 2.404e-19 \| 516 \| 0.183 \| 409 \| 266 \| 18 \| 313 \| 681 \| 19 \| 399 \| Phage major capsid protein \| Phage major capsid protein; afdb-uniprot50 \| AF-A0A367AXK8-F1-MODEL\_V4 \| 1.0 \| 2.178e-21 \| 516 \| 0.207 \| 486 \| 278 \| 24 \| 250 \| 682 \| 20 \| 451 \| Phage major capsid protein \| Phage major capsid protein; afdb-uniprot50 \| AF-A0A0B0DDJ1-F1-MODEL\_V4 \| 1.0 \| 1.698e-14 \| 515 \| 0.184 \| 249 \| 184 \| 7 \| 443 \| 681 \| 1 \| 240 \| Uncharacterized protein \| Uncharacterized protein; afdb-uniprot50 \| AF-A0A084GIL3-F1-MODEL\_V4 \| 1.0 \| 1.434e-22 \| 515 \| 0.13 \| 691 \| 364 \| 30 \| 16 \| 681 \| 4 \| 482 \| Uncharacterized protein \| Uncharacterized protein; afdb-uniprot50 \| AF-A0A1S1WT94-F1-MODEL\_V4 \| 1.0 \| 1.701e-30 \| 515 \| 0.198 \| 761 \| 431 \| 41 \| 16 \| 682 \| 2 \| 677 \| Uncharacterized protein \| Uncharacterized protein; afdb-uniprot50 \| AF-A0A7W4NV52-F1-MODEL\_V4 \| 1.0 \| 1.006e-20 \| 514 \| 0.2 \| 478 \| 279 \| 22 \| 264 \| 679 \| 8 \| 444 \| Phage major capsid protein \| Phage major capsid protein; afdb-uniprot50 \| AF-A0A1N6U3E7-F1-MODEL\_V4 \| 1.0 \| 4.651e-28 \| 514 \| 0.175 \| 722 \| 443 \| 31 \| 12 \| 682 \| 6 \| 625 \| Phage prohead protease, HK97 family \| Phage prohead protease, HK97 family; afdb-uniprot50 \| AF-A0A2V2E9I6-F1-MODEL\_V4 \| 1.0 \| 8.668e-20 \| 513 \| 0.185 \| 436 \| 265 \| 20 \| 294 \| 680 \| 13 \| 407 \| Phage major capsid protein \| Phage major capsid protein; afdb-uniprot50 \| AF-A0A839NAF1-F1-MODEL\_V4 \| 1.0 \| 6.668e-19 \| 512 \| 0.166 \| 390 \| 277 \| 16 \| 334 \| 682 \| 1 \| 383 \| HK97 family phage major capsid protein \| HK97 family phage major capsid protein; afdb-uniprot50 \| AF-A0A6A7UM03-F1-MODEL\_V4 \| 1.0 \| 6.811e-18 \| 512 \| 0.187 \| 379 \| 264 \| 14 \| 337 \| 681 \| 16 \| 384 \| Phage major capsid protein \| Phage major capsid protein; afdb-uniprot50 \| AF-A0A073IUD9-F1-MODEL\_V4 \| 1.0 \| 1.946e-29 \| 512 \| 0.168 \| 794 \| 432 \| 38 \| 15 \| 681 \| 4 \| 695 \| Uncharacterized protein \| Uncharacterized protein; afdb-uniprot50 \| AF-A0A1G7D256-F1-MODEL\_V4 \| 1.0 \| 9.983e-16 \| 511 \| 0.187 \| 293 \| 204 \| 9 \| 410 \| 682 \| 1 \| 279 \| Phage major capsid protein, HK97 family \| Phage major capsid protein, HK97 family; afdb-uniprot50 \| AF-A0A0N1BIW0-F1-MODEL\_V4 \| 1.0 \| 1.134e-17 \| 510 \| 0.161 \| 366 \| 259 \| 13 \| 346 \| 679 \| 9 \| 358 \| Uncharacterized protein \| Uncharacterized protein; afdb-uniprot50 \| AF-A0A857LMJ1-F1-MODEL\_V4 \| 1.0 \| 9.783e-25 \| 510 \| 0.138 \| 752 \| 420 \| 35 \| 30 \| 682 \| 3 \| 625 \| HK97 family phage prohead protease \| HK97 family phage prohead protease; afdb-uniprot50 \| AF-A0A1B1C378-F1-MODEL\_V4 \| 1.0 \| 4.395e-28 \| 510 \| 0.173 \| 733 \| 460 \| 36 \| 31 \| 681 \| 2 \| 670 \| Uncharacterized protein \| Uncharacterized protein; afdb-uniprot50 \| AF-A0A1X0BKY4-F1-MODEL\_V4 \| 1.0 \| 1.244e-18 \| 509 \| 0.152 \| 459 \| 294 \| 14 \| 249 \| 682 \| 2 \| 390 \| Uncharacterized protein \| Uncharacterized protein; afdb-uniprot50 \| AF-M7A7R2-F1-MODEL\_V4 \| 1.0 \| 5.994e-16 \| 508 \| 0.166 \| 318 \| 239 \| 12 \| 381 \| 682 \| 44 \| 351 \| Phage capsid protein \| Phage capsid protein; afdb-uniprot50 \| AF-A0A2H0LLW2-F1-MODEL\_V4 \| 1.0 \| 4.651e-28 \| 508 \| 0.192 \| 713 \| 410 \| 37 \| 40 \| 681 \| 1 \| 618 \| Uncharacterized protein \| Uncharacterized protein; afdb-uniprot50 \| AF-A0A560FKK8-F1-MODEL\_V4 \| 1.0 \| 1.245e-26 \| 508 \| 0.16 \| 717 \| 393 \| 34 \| 5 \| 681 \| 269 \| 816 \| HK97 family phage prohead protease/HK97 family phage major capsid protein,TIGR01554 \| HK97 family phage prohead protease/HK97 family phage major capsid protein,TIGR01554; afdb-uniprot50 \| AF-A0A7C6P388-F1-MODEL\_V4 \| 1.0 \| 1.325e-15 \| 507 \| 0.147 \| 258 \| 194 \| 11 \| 433 \| 682 \| 51 \| 290 \| Phage major capsid protein \| Phage major capsid protein; afdb-uniprot50 \| AF-A0A250JC89-F1-MODEL\_V4 \| 1.0 \| 1.316e-18 \| 507 \| 0.214 \| 420 \| 246 \| 22 \| 284 \| 680 \| 6 \| 364 \| Major capsid protein \| Major capsid protein; afdb-uniprot50 \| AF-A0A1Q2M5H8-F1-MODEL\_V4 \| 1.0 \| 3.146e-17 \| 507 \| 0.155 \| 367 \| 275 \| 14 \| 338 \| 681 \| 22 \| 376 \| Uncharacterized protein \| Uncharacterized protein; afdb-uniprot50 \| AF-A0A1I3MTS0-F1-MODEL\_V4 \| 1.0 \| 9.182e-28 \| 507 \| 0.189 \| 746 \| 430 \| 40 \| 1 \| 681 \| 1 \| 636 \| Uncharacterized protein \| Uncharacterized protein; afdb-uniprot50 \| AF-A0A7G6VRU1-F1-MODEL\_V4 \| 1.0 \| 1.072e-17 \| 506 \| 0.164 \| 444 \| 279 \| 17 \| 251 \| 682 \| 3 \| 366 \| Phage major capsid protein \| Phage major capsid protein; afdb-uniprot50 \| AF-A0A515KLR5-F1-MODEL\_V4 \| 1.0 \| 4.58e-18 \| 506 \| 0.182 \| 399 \| 270 \| 17 \| 313 \| 679 \| 4 \| 378 \| Phage major capsid protein \| Phage major capsid protein; afdb-uniprot50 \| AF-A0A263E3D0-F1-MODEL\_V4 \| 1.0 \| 5.023e-19 \| 506 \| 0.19 \| 399 \| 245 \| 14 \| 338 \| 682 \| 12 \| 386 \| Phage major capsid protein \| Phage major capsid protein; afdb-uniprot50 \| AF-A0A3F3ITH6-F1-MODEL\_V4 \| 1.0 \| 7.16e-21 \| 506 \| 0.182 \| 477 \| 291 \| 22 \| 252 \| 681 \| 2 \| 426 \| Capsid protein \| Capsid protein; afdb-uniprot50 \| AF-A0A160NYS6-F1-MODEL\_V4 \| 1.0 \| 7.739e-20 \| 505 \| 0.161 \| 457 \| 302 \| 20 \| 268 \| 681 \| 4 \| 422 \| Phage major capsid protein \| Phage major capsid protein; afdb-uniprot50 \| AF-A0A143PU17-F1-MODEL\_V4 \| 1.0 \| 2.147e-19 \| 504 \| 0.16 \| 450 \| 275 \| 17 \| 264 \| 680 \| 10 \| 389 \| Phage major capsid protein, HK97 family \| Phage major capsid protein, HK97 family; afdb-uniprot50 \| AF-A0A059MSK3-F1-MODEL\_V4 \| 1.0 \| 2.192e-18 \| 504 \| 0.173 \| 460 \| 279 \| 19 \| 265 \| 681 \| 2 \| 403 \| Uncharacterized protein \| Uncharacterized protein; afdb-uniprot50 \| AF-S9Q4D5-F1-MODEL\_V4 \| 1.0 \| 1.444e-19 \| 504 \| 0.208 \| 446 \| 267 \| 24 \| 276 \| 680 \| 8 \| 408 \| Uncharacterized protein \| Uncharacterized protein; afdb-uniprot50 \| AF-A0A1X0JZU7-F1-MODEL\_V4 \| 1.0 \| 2.013e-14 \| 503 \| 0.166 \| 295 \| 207 \| 11 \| 407 \| 682 \| 2 \| 276 \| Capsid protein \| Capsid protein; afdb-uniprot50 \| AF-A0A5C7FZT0-F1-MODEL\_V4 \| 1.0 \| 1.175e-18 \| 503 \| 0.141 \| 396 \| 264 \| 16 \| 338 \| 680 \| 12 \| 384 \| Phage major capsid protein \| Phage major capsid protein; afdb-uniprot50 \| AF-A0A4P1JU19-F1-MODEL\_V4 \| 1.0 \| 5.398e-29 \| 503 \| 0.173 \| 738 \| 457 \| 35 \| 23 \| 679 \| 20 \| 685 \| Phage prohead protease, HK97 family \| Phage prohead protease, HK97 family; afdb-uniprot50 \| AF-A0A418L4Y0-F1-MODEL\_V4 \| 1.0 \| 2.192e-18 \| 502 \| 0.155 \| 412 \| 293 \| 16 \| 313 \| 682 \| 4 \| 402 \| Phage major capsid protein \| Phage major capsid protein; afdb-uniprot50 \| AF-A0A7S8C5X4-F1-MODEL\_V4 \| 1.0 \| 4.066e-29 \| 502 \| 0.191 \| 738 \| 441 \| 39 \| 30 \| 681 \| 15 \| 682 \| Mu-like prophage major head subunit gpT family protein \| Mu-like prophage major head subunit gpT family protein; afdb-uniprot50 \| AF-A0A0S9DAB5-F1-MODEL\_V4 \| 1.0 \| 4.421e-17 \| 501 \| 0.141 \| 367 \| 286 \| 11 \| 335 \| 681 \| 1 \| 358 \| Uncharacterized protein \| Uncharacterized protein; afdb-uniprot50 \| AF-A0A2S0LYN6-F1-MODEL\_V4 \| 1.0 \| 2.086e-15 \| 500 \| 0.161 \| 267 \| 204 \| 9 \| 431 \| 682 \| 31 \| 292 \| Phage major capsid protein \| Phage major capsid protein; afdb-uniprot50 \| AF-A0A810PMP6-F1-MODEL\_V4 \| 1.0 \| 2.616e-15 \| 500 \| 0.191 \| 297 \| 192 \| 13 \| 419 \| 678 \| 1 \| 286 \| Uncharacterized protein \| Uncharacterized protein; afdb-uniprot50 \| AF-A0A3N5X4U1-F1-MODEL\_V4 \| 1.0 \| 2.404e-19 \| 500 \| 0.221 \| 420 \| 243 \| 20 \| 282 \| 666 \| 35 \| 405 \| Phage major capsid protein \| Phage major capsid protein; afdb-uniprot50 \| AF-A0A0F9IZX0-F1-MODEL\_V4 \| 1.0 \| 1.316e-18 \| 500 \| 0.145 \| 453 \| 302 \| 17 \| 253 \| 680 \| 9 \| 401 \| Uncharacterized protein \| Uncharacterized protein; afdb-uniprot50 \| AF-A0A5C7LSW7-F1-MODEL\_V4 \| 1.0 \| 3.5e-20 \| 500 \| 0.169 \| 484 \| 314 \| 19 \| 238 \| 681 \| 5 \| 440 \| Phage major capsid protein \| Phage major capsid protein; afdb-uniprot50 \| AF-A0A4P5VJG7-F1-MODEL\_V4 \| 1.0 \| 5.513e-28 \| 500 \| 0.193 \| 732 \| 419 \| 39 \| 18 \| 681 \| 5 \| 632 \| Uncharacterized protein \| Uncharacterized protein; afdb-uniprot50 \| AF-A0A3P5XMZ8-F1-MODEL\_V4 \| 1.0 \| 1.506e-17 \| 499 \| 0.169 \| 442 \| 282 \| 15 \| 261 \| 682 \| 4 \| 380 \| Phage capsid family protein \| Phage capsid family protein; afdb-uniprot50 \| AF-A0A0Q7QKI6-F1-MODEL\_V4 \| 1.0 \| 6.765e-21 \| 499 \| 0.191 \| 474 \| 290 \| 20 \| 247 \| 682 \| 42 \| 460 \| Uncharacterized protein \| Uncharacterized protein; afdb-uniprot50 \| AF-A0A2E6KNG2-F1-MODEL\_V4 \| 1.0 \| 2.406e-27 \| 499 \| 0.177 \| 726 \| 446 \| 36 \| 16 \| 681 \| 18 \| 652 \| Peptidase U37 \| Peptidase U37; afdb-uniprot50 \| AF-H5TTQ1-F1-MODEL\_V4 \| 1.0 \| 1.889e-17 \| 498 \| 0.151 \| 442 \| 295 \| 12 \| 262 \| 682 \| 6 \| 388 \| Uncharacterized protein \| Uncharacterized protein; afdb-uniprot50 \| AF-A0A1A3PAA7-F1-MODEL\_V4 \| 1.0 \| 1.013e-17 \| 498 \| 0.158 \| 460 \| 284 \| 22 \| 251 \| 682 \| 7 \| 391 \| Uncharacterized protein \| Uncharacterized protein; afdb-uniprot50 \| AF-D3NSU8-F1-MODEL\_V4 \| 1.0 \| 1.316e-18 \| 498 \| 0.166 \| 456 \| 288 \| 20 \| 251 \| 682 \| 22 \| 409 \| Phage major capsid protein \| Phage major capsid protein; afdb-uniprot50 \| AF-A0A1X1V399-F1-MODEL\_V4 \| 1.0 \| 5.626e-19 \| 498 \| 0.154 \| 454 \| 325 \| 18 \| 263 \| 682 \| 5 \| 433 \| Uncharacterized protein \| Uncharacterized protein; afdb-uniprot50 \| AF-A0A1E3GXJ9-F1-MODEL\_V4 \| 1.0 \| 8.495e-29 \| 498 \| 0.175 \| 718 \| 459 \| 39 \| 41 \| 681 \| 554 \| 1215 \| Phage portal protein, lambda family \| Phage portal protein, lambda family; afdb-uniprot50 \| AF-A0A0F2L3X1-F1-MODEL\_V4 \| 1.0 \| 3.651e-18 \| 497 \| 0.194 \| 407 \| 263 \| 16 \| 314 \| 681 \| 1 \| 381 \| Uncharacterized protein \| Uncharacterized protein; afdb-uniprot50 \| AF-A0A1Q8I8P9-F1-MODEL\_V4 \| 1.0 \| 1.134e-17 \| 497 \| 0.153 \| 462 \| 290 \| 21 \| 251 \| 681 \| 5 \| 396 \| Uncharacterized protein \| Uncharacterized protein; afdb-uniprot50 \| AF-A0A7C6NXR1-F1-MODEL\_V4 \| 1.0 \| 2.013e-14 \| 496 \| 0.113 \| 264 \| 203 \| 12 \| 433 \| 681 \| 58 \| 305 \| Phage major capsid protein \| Phage major capsid protein; afdb-uniprot50 \| AF-A0A4R8Y418-F1-MODEL\_V4 \| 1.0 \| 6.529e-20 \| 496 \| 0.139 \| 539 \| 330 \| 19 \| 178 \| 680 \| 3 \| 443 \| Phage major capsid protein \| Phage major capsid protein; afdb-uniprot50 \| AF-A0A626M3G7-F1-MODEL\_V4 \| 1.0 \| 1.118e-15 \| 495 \| 0.312 \| 221 \| 114 \| 5 \| 19 \| 213 \| 2 \| 210 \| Major capsid protein \| Major capsid protein; afdb-uniprot50 \| AF-A0A251ZWI0-F1-MODEL\_V4 \| 1.0 \| 2.239e-17 \| 495 \| 0.152 \| 393 \| 258 \| 17 \| 327 \| 681 \| 5 \| 360 \| Uncharacterized protein \| Uncharacterized protein; afdb-uniprot50 \| AF-A0A0B2K2J4-F1-MODEL\_V4 \| 1.0 \| 1.931e-24 \| 495 \| 0.217 \| 501 \| 278 \| 20 \| 18 \| 480 \| 2 \| 426 \| Uncharacterized protein \| Uncharacterized protein; afdb-uniprot50 \| AF-W4A1A5-F1-MODEL\_V4 \| 1.0 \| 4.09e-18 \| 494 \| 0.168 \| 434 \| 257 \| 22 \| 282 \| 682 \| 7 \| 369 \| Phage major capsid protein, HK97 family \| Phage major capsid protein, HK97 family; afdb-uniprot50 \| AF-A0A7J9XQM0-F1-MODEL\_V4 \| 1.0 \| 1.474e-18 \| 494 \| 0.151 \| 475 \| 281 \| 20 \| 249 \| 680 \| 5 \| 400 \| Phage major capsid protein \| Phage major capsid protein; afdb-uniprot50 \| AF-G7QC56-F1-MODEL\_V4 \| 1.0 \| 1.087e-19 \| 494 \| 0.19 \| 478 \| 283 \| 26 \| 256 \| 681 \| 6 \| 431 \| HK97 family phage major capsid protein \| HK97 family phage major capsid protein; afdb-uniprot50 \| AF-C0ZW70-F1-MODEL\_V4 \| 1.0 \| 4.27e-24 \| 494 \| 0.13 \| 713 \| 429 \| 36 \| 16 \| 682 \| 3 \| 570 \| Uncharacterized protein \| Uncharacterized protein; afdb-uniprot50 \| AF-A0A292QWT5-F1-MODEL\_V4 \| 1.0 \| 1.497e-28 \| 494 \| 0.158 \| 795 \| 427 \| 37 \| 17 \| 681 \| 26 \| 708 \| Peptidase \| Peptidase; afdb-uniprot50 \| AF-A0A5J6GE42-F1-MODEL\_V4 \| 1.0 \| 3.45e-18 \| 493 \| 0.176 \| 448 \| 275 \| 18 \| 259 \| 682 \| 2 \| 379 \| Phage major capsid protein \| Phage major capsid protein; afdb-uniprot50 \| AF-A0A7Z0GMJ1-F1-MODEL\_V4 \| 1.0 \| 1.474e-18 \| 493 \| 0.145 \| 467 \| 288 \| 17 \| 258 \| 682 \| 8 \| 405 \| HK97 family phage major capsid protein \| HK97 family phage major capsid protein; afdb-uniprot50 \| AF-A0A6V8U7K8-F1-MODEL\_V4 \| 1.0 \| 7.057e-19 \| 493 \| 0.169 \| 449 \| 296 \| 20 \| 251 \| 682 \| 61 \| 449 \| Major capsid protein \| Major capsid protein; afdb-uniprot50 \| AF-A0A7W1RV33-F1-MODEL\_V4 \| 1.0 \| 9.505e-21 \| 493 \| 0.184 \| 510 \| 281 \| 19 \| 258 \| 682 \| 4 \| 463 \| Phage major capsid protein \| Phage major capsid protein; afdb-uniprot50 \| AF-A0A7C2HM19-F1-MODEL\_V4 \| 1.0 \| 1.959e-26 \| 493 \| 0.167 \| 712 \| 414 \| 36 \| 43 \| 681 \| 21 \| 626 \| Peptidase U37 \| Peptidase U37; afdb-uniprot50 \| AF-R7JPW2-F1-MODEL\_V4 \| 1.0 \| 2.956e-28 \| 493 \| 0.181 \| 745 \| 457 \| 34 \| 40 \| 681 \| 15 \| 709 \| Peptidase U35 phage prohead HK97 \| Peptidase U35 phage prohead HK97; afdb-uniprot50 \| AF-A0A317CYD6-F1-MODEL\_V4 \| 1.0 \| 1.85e-18 \| 492 \| 0.164 \| 463 \| 293 \| 17 \| 250 \| 682 \| 4 \| 402 \| Phage major capsid protein \| Phage major capsid protein; afdb-uniprot50 \| AF-A0A2D9CEY6-F1-MODEL\_V4 \| 1.0 \| 2.85e-19 \| 492 \| 0.19 \| 457 \| 281 \| 21 \| 255 \| 680 \| 5 \| 403 \| Phage major capsid protein \| Phage major capsid protein; afdb-uniprot50 \| AF-A0A1U7GJ79-F1-MODEL\_V4 \| 1.0 \| 1.049e-18 \| 492 \| 0.2 \| 474 \| 270 \| 26 \| 264 \| 682 \| 3 \| 422 \| Uncharacterized protein \| Uncharacterized protein; afdb-uniprot50 \| AF-A8SB58-F1-MODEL\_V4 \| 1.0 \| 5.785e-15 \| 491 \| 0.366 \| 183 \| 100 \| 6 \| 23 \| 194 \| 2 \| 179 \| Caudovirus prohead protease \| Caudovirus prohead protease; afdb-uniprot50 \| AF-A0A7Z0GNW8-F1-MODEL\_V4 \| 1.0 \| 6.714e-16 \| 490 \| 0.178 \| 320 \| 206 \| 14 \| 381 \| 682 \| 9 \| 289 \| HK97 family phage major capsid protein \| HK97 family phage major capsid protein; afdb-uniprot50 \| AF-A0A1F9FW66-F1-MODEL\_V4 \| 1.0 \| 3.787e-27 \| 490 \| 0.18 \| 708 \| 420 \| 32 \| 40 \| 680 \| 31 \| 644 \| Uncharacterized protein \| Uncharacterized protein; afdb-uniprot50 \| AF-A1T829-F1-MODEL\_V4 \| 1.0 \| 2.828e-14 \| 489 \| 0.141 \| 262 \| 206 \| 12 \| 431 \| 682 \| 29 \| 281 \| Phage major capsid protein, HK97 \| Phage major capsid protein, HK97; afdb-uniprot50 \| AF-A0A3S0DN68-F1-MODEL\_V4 \| 1.0 \| 6.573e-17 \| 489 \| 0.161 \| 372 \| 267 \| 17 \| 335 \| 681 \| 1 \| 352 \| Phage major capsid protein \| Phage major capsid protein; afdb-uniprot50 \| AF-A0A3N2KEJ4-F1-MODEL\_V4 \| 1.0 \| 1.27e-17 \| 489 \| 0.177 \| 383 \| 261 \| 13 \| 338 \| 682 \| 6 \| 372 \| Phage major capsid protein \| Phage major capsid protein; afdb-uniprot50 \| AF-A0A502HT88-F1-MODEL\_V4 \| 1.0 \| 6.811e-18 \| 489 \| 0.169 \| 453 \| 278 \| 21 \| 252 \| 682 \| 20 \| 396 \| Phage major capsid protein \| Phage major capsid protein; afdb-uniprot50 \| AF-A0A0P0D9A9-F1-MODEL\_V4 \| 1.0 \| 2.072e-18 \| 489 \| 0.14 \| 426 \| 293 \| 19 \| 279 \| 681 \| 63 \| 438 \| Capsid protein \| Capsid protein; afdb-uniprot50 \| AF-A0A2P8NKG7-F1-MODEL\_V4 \| 1.0 \| 1.176e-26 \| 489 \| 0.163 \| 729 \| 440 \| 33 \| 12 \| 682 \| 6 \| 622 \| Mu-like\_gpT domain-containing protein \| Mu-like\_gpT domain-containing protein; afdb-uniprot50 \| AF-A0A2I1R5G1-F1-MODEL\_V4 \| 1.0 \| 2.386e-14 \| 488 \| 0.145 \| 261 \| 204 \| 11 \| 433 \| 681 \| 31 \| 284 \| Phage major capsid protein \| Phage major capsid protein; afdb-uniprot50 \| AF-A0A7C6WF68-F1-MODEL\_V4 \| 1.0 \| 9.433e-16 \| 488 \| 0.138 \| 339 \| 252 \| 13 \| 369 \| 679 \| 2 \| 328 \| Phage major capsid protein \| Phage major capsid protein; afdb-uniprot50 \| AF-A0A1V2NG87-F1-MODEL\_V4 \| 1.0 \| 5.746e-18 \| 488 \| 0.155 \| 456 \| 300 \| 19 \| 252 \| 682 \| 10 \| 405 \| Uncharacterized protein \| Uncharacterized protein; afdb-uniprot50 \| AF-A0A6I2J2E1-F1-MODEL\_V4 \| 1.0 \| 7.469e-19 \| 487 \| 0.445 \| 249 \| 121 \| 4 \| 282 \| 529 \| 3 \| 235 \| Phage major capsid protein \| Phage major capsid protein; afdb-uniprot50 \| AF-A0A2E0EK67-F1-MODEL\_V4 \| 1.0 \| 2.809e-17 \| 487 \| 0.173 \| 374 \| 265 \| 13 \| 338 \| 679 \| 29 \| 390 \| Phage major capsid protein \| Phage major capsid protein; afdb-uniprot50 \| AF-W7D9N4-F1-MODEL\_V4 \| 1.0 \| 6.668e-19 \| 487 \| 0.15 \| 452 \| 300 \| 21 \| 261 \| 682 \| 4 \| 401 \| Uncharacterized protein \| Uncharacterized protein; afdb-uniprot50 \| AF-A0A7Y5LZT5-F1-MODEL\_V4 \| 1.0 \| 2.072e-18 \| 487 \| 0.178 \| 470 \| 281 \| 24 \| 261 \| 682 \| 8 \| 420 \| Phage major capsid protein \| Phage major capsid protein; afdb-uniprot50 \| AF-A0A7M2XU64-F1-MODEL\_V4 \| 1.0 \| 9.235e-17 \| 486 \| 0.149 \| 448 \| 282 \| 20 \| 264 \| 682 \| 3 \| 380 \| Phage major capsid protein \| Phage major capsid protein; afdb-uniprot50 \| AF-A0A0N0T0A0-F1-MODEL\_V4 \| 1.0 \| 1.85e-18 \| 486 \| 0.17 \| 386 \| 262 \| 10 \| 338 \| 681 \| 11 \| 380 \| Uncharacterized protein \| Uncharacterized protein; afdb-uniprot50 \| AF-A0A0F9KP71-F1-MODEL\_V4 \| 1.0 \| 1.175e-18 \| 486 \| 0.143 \| 466 \| 306 \| 18 \| 234 \| 681 \| 31 \| 421 \| Uncharacterized protein \| Uncharacterized protein; afdb-uniprot50 \| AF-A0A147IZI7-F1-MODEL\_V4 \| 1.0 \| 4.004e-19 \| 486 \| 0.189 \| 475 \| 290 \| 20 \| 255 \| 681 \| 6 \| 433 \| Uncharacterized protein \| Uncharacterized protein; afdb-uniprot50 \| AF-A0A2V8NCC7-F1-MODEL\_V4 \| 1.0 \| 5.626e-19 \| 486 \| 0.149 \| 428 \| 295 \| 19 \| 278 \| 679 \| 59 \| 443 \| Phage major capsid protein \| Phage major capsid protein; afdb-uniprot50 \| AF-A0A841QKT1-F1-MODEL\_V4 \| 1.0 \| 8.365e-19 \| 486 \| 0.162 \| 462 \| 311 \| 16 \| 249 \| 680 \| 35 \| 450 \| HK97 family phage major capsid protein \| HK97 family phage major capsid protein; afdb-uniprot50 \| AF-A0A4V2DDR1-F1-MODEL\_V4 \| 1.0 \| 2.356e-28 \| 486 \| 0.149 \| 742 \| 469 \| 35 \| 18 \| 681 \| 3 \| 660 \| Mu-like\_gpT domain-containing protein \| Mu-like\_gpT domain-containing protein; afdb-uniprot50 \| AF-A0A143PPF0-F1-MODEL\_V4 \| 1.0 \| 5.954e-19 \| 485 \| 0.177 \| 444 \| 277 \| 17 \| 271 \| 682 \| 3 \| 390 \| Phage major capsid protein, HK97 family \| Phage major capsid protein, HK97 family; afdb-uniprot50 \| AF-A0A2T5W6J4-F1-MODEL\_V4 \| 1.0 \| 5.626e-19 \| 484 \| 0.173 \| 467 \| 294 \| 18 \| 263 \| 682 \| 3 \| 424 \| HK97 family phage major capsid protein \| HK97 family phage major capsid protein; afdb-uniprot50 \| AF-A0A1S7QBP3-F1-MODEL\_V4 \| 1.0 \| 3.019e-27 \| 484 \| 0.166 \| 716 \| 466 \| 34 \| 17 \| 681 \| 9 \| 644 \| Peptidase U35 phage prohead HK97 \| Peptidase U35 phage prohead HK97; afdb-uniprot50 \| AF-A0A3M1YKP9-F1-MODEL\_V4 \| 1.0 \| 2.657e-25 \| 484 \| 0.145 \| 798 \| 458 \| 40 \| 15 \| 679 \| 3 \| 709 \| Phage major capsid protein \| Phage major capsid protein; afdb-uniprot50 \| AF-A0A7R8N8Q9-F1-MODEL\_V4 \| 1.0 \| 2.437e-13 \| 483 \| 0.234 \| 230 \| 155 \| 9 \| 457 \| 682 \| 1 \| 213 \| Uncharacterized protein \| Uncharacterized protein; afdb-uniprot50 \| AF-A0A329M2M9-F1-MODEL\_V4 \| 1.0 \| 1.134e-17 \| 483 \| 0.15 \| 433 \| 313 \| 11 \| 267 \| 682 \| 3 \| 397 \| Phage major capsid protein \| Phage major capsid protein; afdb-uniprot50 \| AF-A0A1G3YGU2-F1-MODEL\_V4 \| 1.0 \| 1.2e-17 \| 483 \| 0.162 \| 474 \| 276 \| 24 \| 258 \| 679 \| 5 \| 409 \| Uncharacterized protein \| Uncharacterized protein; afdb-uniprot50 \| AF-M5JSC1-F1-MODEL\_V4 \| 1.0 \| 1.529e-27 \| 483 \| 0.177 \| 742 \| 478 \| 37 \| 31 \| 682 \| 1 \| 699 \| Peptidase U35 phage prohead HK97 \| Peptidase U35 phage prohead HK97; afdb-uniprot50 \| AF-A0A753E0J0-F1-MODEL\_V4 \| 1.0 \| 3.548e-14 \| 482 \| 0.143 \| 292 \| 226 \| 11 \| 401 \| 679 \| 13 \| 293 \| Phage major capsid protein \| Phage major capsid protein; afdb-uniprot50 \| AF-A0A7K2HUA6-F1-MODEL\_V4 \| 1.0 \| 8.246e-17 \| 482 \| 0.147 \| 460 \| 289 \| 21 \| 251 \| 681 \| 3 \| 388 \| Phage major capsid protein \| Phage major capsid protein; afdb-uniprot50 \| AF-A0A196LBE7-F1-MODEL\_V4 \| 1.0 \| 8.544e-18 \| 482 \| 0.133 \| 433 \| 307 \| 16 \| 264 \| 682 \| 6 \| 384 \| Uncharacterized protein \| Uncharacterized protein; afdb-uniprot50 \| AF-A0A5S3Y9T7-F1-MODEL\_V4 \| 1.0 \| 2.544e-19 \| 482 \| 0.192 \| 463 \| 285 \| 19 \| 268 \| 681 \| 4 \| 426 \| Phage major capsid protein \| Phage major capsid protein; afdb-uniprot50 \| AF-A0A7X4X9J4-F1-MODEL\_V4 \| 1.0 \| 3.378e-19 \| 482 \| 0.184 \| 476 \| 282 \| 21 \| 258 \| 680 \| 7 \| 429 \| Phage major capsid protein \| Phage major capsid protein; afdb-uniprot50 \| AF-A0A1H7NMW7-F1-MODEL\_V4 \| 1.0 \| 3.104e-23 \| 482 \| 0.124 \| 697 \| 420 \| 30 \| 39 \| 682 \| 12 \| 570 \| Phage prohead protease, HK97 family/phage major capsid protein, HK97 family,TIGR01554 \| Phage prohead protease, HK97 family/phage major capsid protein, HK97 family,TIGR01554; afdb-uniprot50 \| AF-A0A2T0Q9X2-F1-MODEL\_V4 \| 1.0 \| 9.916e-19 \| 481 \| 0.155 \| 462 \| 291 \| 21 \| 262 \| 678 \| 7 \| 414 \| HK97 family phage major capsid protein \| HK97 family phage major capsid protein; afdb-uniprot50 \| AF-A0A521H793-F1-MODEL\_V4 \| 1.0 \| 6.307e-27 \| 481 \| 0.17 \| 749 \| 463 \| 39 \| 17 \| 677 \| 5 \| 682 \| Peptidase U37 \| Peptidase U37; afdb-uniprot50 \| AF-T0U3G2-F1-MODEL\_V4 \| 1.0 \| 9.84e-14 \| 480 \| 0.199 \| 246 \| 172 \| 9 \| 448 \| 681 \| 9 \| 241 \| Phage major capsid protein, HK97 family \| Phage major capsid protein, HK97 family; afdb-uniprot50 \| AF-A0A857MKH2-F1-MODEL\_V4 \| 1.0 \| 4.611e-15 \| 480 \| 0.161 \| 298 \| 216 \| 10 \| 398 \| 681 \| 5 \| 282 \| Phage major capsid protein \| Phage major capsid protein; afdb-uniprot50 \| AF-A0A6L7WRA2-F1-MODEL\_V4 \| 1.0 \| 4.951e-17 \| 480 \| 0.19 \| 362 \| 250 \| 16 \| 338 \| 680 \| 14 \| 351 \| Phage major capsid protein \| Phage major capsid protein; afdb-uniprot50 \| AF-A0A5F0E144-F1-MODEL\_V4 \| 1.0 \| 1.889e-17 \| 480 \| 0.13 \| 391 \| 273 \| 15 \| 338 \| 682 \| 7 \| 376 \| Phage major capsid protein \| Phage major capsid protein; afdb-uniprot50 \| AF-A0A1T4SDY1-F1-MODEL\_V4 \| 1.0 \| 3.651e-18 \| 480 \| 0.195 \| 465 \| 260 \| 20 \| 238 \| 679 \| 2 \| 375 \| Phage major capsid protein, HK97 family \| Phage major capsid protein, HK97 family; afdb-uniprot50 \| AF-A0A6L6JA71-F1-MODEL\_V4 \| 1.0 \| 1.134e-17 \| 480 \| 0.162 \| 413 \| 284 \| 18 \| 298 \| 682 \| 8 \| 386 \| Phage major capsid protein \| Phage major capsid protein; afdb-uniprot50 \| AF-K9D228-F1-MODEL\_V4 \| 1.0 \| 3.578e-27 \| 480 \| 0.164 \| 728 \| 442 \| 40 \| 16 \| 682 \| 8 \| 629 \| HK97 family phage prohead protease \| HK97 family phage prohead protease; afdb-uniprot50 \| AF-A0A177JN28-F1-MODEL\_V4 \| 1.0 \| 9.634e-15 \| 478 \| 0.147 \| 264 \| 205 \| 7 \| 433 \| 681 \| 31 \| 289 \| Uncharacterized protein \| Uncharacterized protein; afdb-uniprot50 \| AF-A0A2N3QVS0-F1-MODEL\_V4 \| 1.0 \| 5.429e-18 \| 478 \| 0.156 \| 460 \| 290 \| 20 \| 269 \| 681 \| 5 \| 413 \| Phage major capsid protein \| Phage major capsid protein; afdb-uniprot50 \| AF-A0A6A7LQA8-F1-MODEL\_V4 \| 1.0 \| 8.544e-18 \| 478 \| 0.134 \| 477 \| 304 \| 21 \| 253 \| 680 \| 2 \| 418 \| Phage major capsid protein \| Phage major capsid protein; afdb-uniprot50 \| AF-A0A7I7UUS7-F1-MODEL\_V4 \| 1.0 \| 1.095e-16 \| 477 \| 0.175 \| 404 \| 254 \| 19 \| 313 \| 678 \| 1 \| 363 \| Uncharacterized protein \| Uncharacterized protein; afdb-uniprot50 \| AF-A0A839ID07-F1-MODEL\_V4 \| 1.0 \| 1.628e-16 \| 476 \| 0.135 \| 391 \| 283 \| 14 \| 335 \| 682 \| 1 \| 379 \| Phage major capsid protein \| Phage major capsid protein; afdb-uniprot50 \| AF-A0A542VQW9-F1-MODEL\_V4 \| 1.0 \| 1.85e-18 \| 476 \| 0.15 \| 458 \| 277 \| 19 \| 268 \| 682 \| 5 \| 393 \| HK97 family phage major capsid protein \| HK97 family phage major capsid protein; afdb-uniprot50 \| AF-A0A0F9R051-F1-MODEL\_V4 \| 1.0 \| 5.746e-18 \| 476 \| 0.143 \| 480 \| 307 \| 20 \| 251 \| 680 \| 2 \| 427 \| Uncharacterized protein \| Uncharacterized protein; afdb-uniprot50 \| AF-A0A5C7M207-F1-MODEL\_V4 \| 1.0 \| 1.56e-18 \| 476 \| 0.164 \| 468 \| 293 \| 22 \| 251 \| 681 \| 28 \| 434 \| Phage major capsid protein \| Phage major capsid protein; afdb-uniprot50 \| AF-I2GKQ2-F1-MODEL\_V4 \| 1.0 \| 8.365e-19 \| 475 \| 0.183 \| 473 \| 282 \| 27 \| 251 \| 681 \| 3 \| 413 \| Phage major capsid protein, HK97 family \| Phage major capsid protein, HK97 family; afdb-uniprot50 \| AF-A0A7J9XKN3-F1-MODEL\_V4 \| 1.0 \| 4.678e-17 \| 474 \| 0.133 \| 473 \| 280 \| 22 \| 264 \| 682 \| 3 \| 399 \| Phage major capsid protein \| Phage major capsid protein; afdb-uniprot50 \| AF-A0A3M2B028-F1-MODEL\_V4 \| 1.0 \| 4.848e-18 \| 474 \| 0.15 \| 479 \| 288 \| 20 \| 250 \| 681 \| 9 \| 415 \| Phage major capsid protein \| Phage major capsid protein; afdb-uniprot50 \| AF-A0A1V6C830-F1-MODEL\_V4 \| 1.0 \| 3.524e-17 \| 473 \| 0.161 \| 416 \| 256 \| 19 \| 296 \| 681 \| 25 \| 377 \| Phage capsid family protein \| Phage capsid family protein; afdb-uniprot50 \| AF-A0A7H5ET99-F1-MODEL\_V4 \| 1.0 \| 3.45e-18 \| 473 \| 0.155 \| 449 \| 284 \| 19 \| 258 \| 682 \| 13 \| 390 \| Phage major capsid protein \| Phage major capsid protein; afdb-uniprot50 \| AF-A0A848HSK9-F1-MODEL\_V4 \| 1.0 \| 1.202e-25 \| 473 \| 0.196 \| 704 \| 438 \| 41 \| 50 \| 681 \| 2 \| 649 \| Uncharacterized protein \| Uncharacterized protein; afdb-uniprot50 \| AF-A0A644ZYJ0-F1-MODEL\_V4 \| 1.0 \| 4.778e-16 \| 472 \| 0.162 \| 394 \| 242 \| 19 \| 313 \| 682 \| 1 \| 330 \| Uncharacterized protein \| Uncharacterized protein; afdb-uniprot50 \| AF-A0A856N7Q3-F1-MODEL\_V4 \| 1.0 \| 1.316e-18 \| 472 \| 0.155 \| 481 \| 323 \| 23 \| 252 \| 680 \| 9 \| 458 \| Phage major capsid protein \| Phage major capsid protein; afdb-uniprot50 \| AF-A0A7W3IZJ2-F1-MODEL\_V4 \| 1.0 \| 4.004e-19 \| 472 \| 0.177 \| 496 \| 300 \| 20 \| 262 \| 681 \| 7 \| 470 \| HK97 family phage major capsid protein \| HK97 family phage major capsid protein; afdb-uniprot50 \| AF-A4L2R1-F1-MODEL\_V4 \| 1.0 \| 4.177e-17 \| 471 \| 0.149 \| 415 \| 244 \| 16 \| 295 \| 681 \| 41 \| 374 \| Phage major capsid protein \| Phage major capsid protein; afdb-uniprot50 \| AF-A0A5C8IZR3-F1-MODEL\_V4 \| 1.0 \| 5.24e-17 \| 471 \| 0.161 \| 378 \| 269 \| 17 \| 338 \| 679 \| 82 \| 447 \| Phage major capsid protein \| Phage major capsid protein; afdb-uniprot50 \| AF-A0A3E2T6Y0-F1-MODEL\_V4 \| 1.0 \| 5.702e-13 \| 470 \| 0.372 \| 153 \| 89 \| 3 \| 42 \| 188 \| 12 \| 163 \| Caudovirus prohead protease \| Caudovirus prohead protease; afdb-uniprot50 \| AF-A0A7W4VP00-F1-MODEL\_V4 \| 1.0 \| 8.128e-15 \| 470 \| 0.175 \| 279 \| 179 \| 13 \| 438 \| 680 \| 16 \| 279 \| HK97 family phage major capsid protein \| HK97 family phage major capsid protein; afdb-uniprot50 \| AF-A0A3G6WSW8-F1-MODEL\_V4 \| 1.0 \| 3.146e-17 \| 470 \| 0.168 \| 427 \| 265 \| 19 \| 294 \| 681 \| 9 \| 384 \| Phage major capsid protein \| Phage major capsid protein; afdb-uniprot50 \| AF-A0A7C7DF73-F1-MODEL\_V4 \| 1.0 \| 1.748e-18 \| 470 \| 0.163 \| 465 \| 293 \| 21 \| 258 \| 680 \| 7 \| 417 \| Phage major capsid protein \| Phage major capsid protein; afdb-uniprot50 \| AF-A0A0P0Z4E5-F1-MODEL\_V4 \| 1.0 \| 4.951e-17 \| 469 \| 0.182 \| 456 \| 275 \| 18 \| 243 \| 682 \| 8 \| 381 \| Phage major capsid protein, HK97 family \| Phage major capsid protein, HK97 family; afdb-uniprot50 \| AF-A0A1R0KE28-F1-MODEL\_V4 \| 1.0 \| 2.654e-17 \| 469 \| 0.144 \| 463 \| 289 \| 21 \| 250 \| 682 \| 15 \| 400 \| Uncharacterized protein \| Uncharacterized protein; afdb-uniprot50 \| AF-A0A3N5Z1L7-F1-MODEL\_V4 \| 1.0 \| 7.105e-16 \| 469 \| 0.152 \| 367 \| 271 \| 15 \| 338 \| 682 \| 34 \| 382 \| Phage major capsid protein \| Phage major capsid protein; afdb-uniprot50 \| AF-A0A6I4SN09-F1-MODEL\_V4 \| 1.0 \| 1.628e-16 \| 468 \| 0.138 \| 376 \| 275 \| 15 \| 335 \| 679 \| 1 \| 358 \| Phage major capsid protein \| Phage major capsid protein; afdb-uniprot50 \| AF-A0A1Y3LZN4-F1-MODEL\_V4 \| 1.0 \| 7.791e-17 \| 468 \| 0.156 \| 448 \| 287 \| 18 \| 258 \| 682 \| 4 \| 383 \| Phage major capsid protein \| Phage major capsid protein; afdb-uniprot50 \| AF-A0A7V3I586-F1-MODEL\_V4 \| 1.0 \| 2.711e-16 \| 468 \| 0.186 \| 370 \| 248 \| 18 \| 341 \| 681 \| 53 \| 398 \| Phage major capsid protein \| Phage major capsid protein; afdb-uniprot50 \| AF-A0A431JK96-F1-MODEL\_V4 \| 1.0 \| 3.45e-18 \| 468 \| 0.126 \| 467 \| 317 \| 22 \| 234 \| 681 \| 40 \| 434 \| Phage major capsid protein \| Phage major capsid protein; afdb-uniprot50 \| AF-A0A1G8RXE6-F1-MODEL\_V4 \| 1.0 \| 7.469e-19 \| 468 \| 0.183 \| 480 \| 294 \| 24 \| 251 \| 681 \| 5 \| 435 \| Phage major capsid protein, HK97 family \| Phage major capsid protein, HK97 family; afdb-uniprot50 \| AF-A0A5D4JM19-F1-MODEL\_V4 \| 1.0 \| 7.904e-19 \| 468 \| 0.176 \| 487 \| 287 \| 19 \| 246 \| 678 \| 14 \| 440 \| Phage major capsid protein \| Phage major capsid protein; afdb-uniprot50 \| AF-A0A136PD24-F1-MODEL\_V4 \| 1.0 \| 3.401e-16 \| 468 \| 0.166 \| 372 \| 237 \| 14 \| 338 \| 680 \| 134 \| 461 \| Phage capsid family protein \| Phage capsid family protein; afdb-uniprot50 \| AF-K2JM81-F1-MODEL\_V4 \| 1.0 \| 1.629e-24 \| 468 \| 0.181 \| 700 \| 372 \| 33 \| 16 \| 681 \| 227 \| 759 \| Phage major capsid protein, HK97 family \| Phage major capsid protein, HK97 family; afdb-uniprot50 \| AF-A0A8B3A685-F1-MODEL\_V4 \| 1.0 \| 2.176e-13 \| 467 \| 0.341 \| 158 \| 97 \| 3 \| 38 \| 189 \| 6 \| 162 \| Caudovirus prohead protease \| Caudovirus prohead protease; afdb-uniprot50 \| AF-A0A3A9D626-F1-MODEL\_V4 \| 1.0 \| 6.618e-14 \| 467 \| 0.337 \| 181 \| 109 \| 4 \| 34 \| 208 \| 25 \| 200 \| Caudovirus prohead protease \| Caudovirus prohead protease; afdb-uniprot50 \| AF-A0A315TC63-F1-MODEL\_V4 \| 1.0 \| 6.211e-17 \| 467 \| 0.153 \| 450 \| 289 \| 21 \| 258 \| 681 \| 5 \| 388 \| HK97 family phage major capsid protein \| HK97 family phage major capsid protein; afdb-uniprot50 \| AF-A0A5F0LGR9-F1-MODEL\_V4 \| 1.0 \| 2.072e-18 \| 467 \| 0.176 \| 475 \| 284 \| 24 \| 255 \| 681 \| 4 \| 419 \| Phage major capsid protein \| Phage major capsid protein; afdb-uniprot50 \| AF-A0A238WSY7-F1-MODEL\_V4 \| 1.0 \| 3.864e-18 \| 467 \| 0.181 \| 469 \| 289 \| 23 \| 252 \| 680 \| 17 \| 430 \| Phage major capsid protein, HK97 family \| Phage major capsid protein, HK97 family; afdb-uniprot50 \| AF-A0A441D124-F1-MODEL\_V4 \| 1.0 \| 6.169e-20 \| 467 \| 0.173 \| 589 \| 319 \| 25 \| 116 \| 679 \| 21 \| 466 \| Phage major capsid protein \| Phage major capsid protein; afdb-uniprot50 \| AF-A0A7T7CAQ5-F1-MODEL\_V4 \| 1.0 \| 3.26e-18 \| 466 \| 0.16 \| 462 \| 279 \| 21 \| 271 \| 682 \| 3 \| 405 \| Phage major capsid protein \| Phage major capsid protein; afdb-uniprot50 \| AF-M7NVL3-F1-MODEL\_V4 \| 1.0 \| 3.651e-18 \| 466 \| 0.191 \| 474 \| 275 \| 23 \| 252 \| 680 \| 9 \| 419 \| Putative phage phi-C31 gp36 major capsid-like protein \| Putative phage phi-C31 gp36 major capsid-like protein; afdb-uniprot50 \| AF-R9LVI7-F1-MODEL\_V4 \| 1.0 \| 4.985e-14 \| 465 \| 0.361 \| 177 \| 106 \| 3 \| 19 \| 189 \| 5 \| 180 \| Uncharacterized protein \| Uncharacterized protein; afdb-uniprot50 \| AF-A0A2G2G701-F1-MODEL\_V4 \| 1.0 \| 7.208e-18 \| 465 \| 0.18 \| 448 \| 261 \| 19 \| 262 \| 676 \| 5 \| 379 \| Phage major capsid protein \| Phage major capsid protein; afdb-uniprot50 \| AF-A0A497HDL0-F1-MODEL\_V4 \| 1.0 \| 4.951e-17 \| 465 \| 0.171 \| 374 \| 255 \| 14 \| 345 \| 681 \| 31 \| 386 \| Phage major capsid protein \| Phage major capsid protein; afdb-uniprot50 \| AF-A0A177PC93-F1-MODEL\_V4 \| 1.0 \| 9.57e-18 \| 465 \| 0.179 \| 469 \| 266 \| 24 \| 243 \| 682 \| 12 \| 390 \| Uncharacterized protein \| Uncharacterized protein; afdb-uniprot50 \| AF-A0A2L0TWR2-F1-MODEL\_V4 \| 1.0 \| 1.298e-16 \| 465 \| 0.146 \| 456 \| 296 \| 22 \| 258 \| 682 \| 7 \| 400 \| Phage major capsid protein \| Phage major capsid protein; afdb-uniprot50 \| AF-A0A1Y3SJ18-F1-MODEL\_V4 \| 1.0 \| 1.864e-23 \| 465 \| 0.135 \| 746 \| 389 \| 34 \| 27 \| 681 \| 2 \| 582 \| Phage major capsid protein \| Phage major capsid protein; afdb-uniprot50 \| AF-A0A150IJM8-F1-MODEL\_V4 \| 1.0 \| 1.538e-16 \| 464 \| 0.169 \| 371 \| 257 \| 16 \| 336 \| 681 \| 1 \| 345 \| Phage capsid family protein \| Phage capsid family protein; afdb-uniprot50 \| AF-A0A7C7C6E2-F1-MODEL\_V4 \| 1.0 \| 1.538e-16 \| 464 \| 0.15 \| 385 \| 278 \| 16 \| 336 \| 682 \| 3 \| 376 \| Phage major capsid protein \| Phage major capsid protein; afdb-uniprot50 \| AF-A0A2S5UU29-F1-MODEL\_V4 \| 1.0 \| 2.192e-18 \| 464 \| 0.139 \| 459 \| 293 \| 22 \| 263 \| 681 \| 9 \| 405 \| Phage major capsid protein \| Phage major capsid protein; afdb-uniprot50 \| AF-A0A3C1GGD5-F1-MODEL\_V4 \| 1.0 \| 2.809e-17 \| 464 \| 0.144 \| 463 \| 306 \| 22 \| 252 \| 682 \| 4 \| 408 \| Phage major capsid protein \| Phage major capsid protein; afdb-uniprot50 \| AF-A0A352HSX6-F1-MODEL\_V4 \| 1.0 \| 1.999e-17 \| 463 \| 0.189 \| 432 \| 272 \| 16 \| 271 \| 681 \| 4 \| 378 \| Phage major capsid protein \| Phage major capsid protein; afdb-uniprot50 \| AF-A0A433WXB3-F1-MODEL\_V4 \| 1.0 \| 4.177e-17 \| 463 \| 0.167 \| 449 \| 288 \| 19 \| 253 \| 680 \| 8 \| 391 \| Phage major capsid protein \| Phage major capsid protein; afdb-uniprot50 \| AF-A0A3A0CKV3-F1-MODEL\_V4 \| 1.0 \| 5.316e-19 \| 463 \| 0.169 \| 483 \| 288 \| 23 \| 258 \| 682 \| 9 \| 436 \| Phage major capsid protein \| Phage major capsid protein; afdb-uniprot50 \| AF-A0A504UPN0-F1-MODEL\_V4 \| 1.0 \| 2.118e-25 \| 463 \| 0.17 \| 729 \| 406 \| 38 \| 19 \| 682 \| 6 \| 600 \| Peptidase U35 \| Peptidase U35; afdb-uniprot50 \| AF-A0A133ZYY2-F1-MODEL\_V4 \| 1.0 \| 2.13e-14 \| 462 \| 0.314 \| 191 \| 115 \| 5 \| 11 \| 193 \| 14 \| 196 \| Caudovirus prohead protease \| Caudovirus prohead protease; afdb-uniprot50 \| AF-A0A0K9YYM7-F1-MODEL\_V4 \| 1.0 \| 3.33e-17 \| 462 \| 0.133 \| 449 \| 313 \| 21 \| 256 \| 681 \| 4 \| 399 \| Uncharacterized protein \| Uncharacterized protein; afdb-uniprot50 \| AF-A0A7C1KK58-F1-MODEL\_V4 \| 1.0 \| 2.599e-18 \| 462 \| 0.188 \| 477 \| 297 \| 23 \| 251 \| 682 \| 2 \| 433 \| Phage major capsid protein \| Phage major capsid protein; afdb-uniprot50 \| AF-A0A4R4A8U2-F1-MODEL\_V4 \| 1.0 \| 9.369e-19 \| 462 \| 0.175 \| 468 \| 272 \| 20 \| 252 \| 682 \| 56 \| 446 \| HK97 family phage major capsid protein \| HK97 family phage major capsid protein; afdb-uniprot50 \| AF-F8KPN2-F1-MODEL\_V4 \| 1.0 \| 4.27e-24 \| 462 \| 0.189 \| 619 \| 392 \| 21 \| 40 \| 622 \| 27 \| 571 \| Uncharacterized protein \| Uncharacterized protein; afdb-uniprot50 \| AF-A0A1V5WQJ7-F1-MODEL\_V4 \| 1.0 \| 6.259e-22 \| 462 \| 0.127 \| 719 \| 422 \| 39 \| 35 \| 682 \| 9 \| 592 \| Phage capsid family protein \| Phage capsid family protein; afdb-uniprot50 \| AF-A0A8A4FL43-F1-MODEL\_V4 \| 1.0 \| 7.064e-27 \| 462 \| 0.181 \| 765 \| 432 \| 47 \| 19 \| 681 \| 1 \| 672 \| Peptidase U35 \| Peptidase U35; afdb-uniprot50 \| AF-A0A4R2H804-F1-MODEL\_V4 \| 1.0 \| 2.056e-13 \| 461 \| 0.144 \| 256 \| 201 \| 8 \| 431 \| 681 \| 29 \| 271 \| HK97 family phage major capsid protein \| HK97 family phage major capsid protein; afdb-uniprot50 \| AF-A0A7G5NYF7-F1-MODEL\_V4 \| 1.0 \| 9.84e-14 \| 461 \| 0.161 \| 260 \| 192 \| 7 \| 433 \| 682 \| 60 \| 303 \| Phage major capsid protein \| Phage major capsid protein; afdb-uniprot50 \| AF-A0A432G3A4-F1-MODEL\_V4 \| 1.0 \| 2.562e-16 \| 461 \| 0.15 \| 440 \| 296 \| 15 \| 252 \| 682 \| 5 \| 375 \| Phage major capsid protein \| Phage major capsid protein; afdb-uniprot50 \| AF-A0A259XLU9-F1-MODEL\_V4 \| 1.0 \| 2.042e-16 \| 461 \| 0.137 \| 421 \| 295 \| 14 \| 271 \| 682 \| 29 \| 390 \| Phage major capsid protein \| Phage major capsid protein; afdb-uniprot50 \| AF-A0A260BZ92-F1-MODEL\_V4 \| 1.0 \| 7.105e-16 \| 461 \| 0.146 \| 375 \| 263 \| 13 \| 346 \| 677 \| 36 \| 396 \| Phage major capsid protein \| Phage major capsid protein; afdb-uniprot50 \| AF-A0A0F9JLY2-F1-MODEL\_V4 \| 1.0 \| 1.785e-17 \| 461 \| 0.219 \| 388 \| 214 \| 17 \| 331 \| 678 \| 59 \| 397 \| Uncharacterized protein \| Uncharacterized protein; afdb-uniprot50 \| AF-A0A561ECB5-F1-MODEL\_V4 \| 1.0 \| 4.678e-17 \| 461 \| 0.137 \| 445 \| 321 \| 13 \| 258 \| 682 \| 8 \| 409 \| HK97 family phage major capsid protein \| HK97 family phage major capsid protein; afdb-uniprot50 \| AF-A0A494SQV9-F1-MODEL\_V4 \| 1.0 \| 1.56e-18 \| 461 \| 0.153 \| 475 \| 282 \| 19 \| 264 \| 682 \| 11 \| 421 \| Phage major capsid protein \| Phage major capsid protein; afdb-uniprot50 \| AF-A0A142IR20-F1-MODEL\_V4 \| 1.0 \| 3.708e-28 \| 461 \| 0.18 \| 731 \| 422 \| 34 \| 38 \| 681 \| 11 \| 651 \| Mu-like\_gpT domain-containing protein \| Mu-like\_gpT domain-containing protein; afdb-uniprot50 \| AF-A0A2S5N3Y2-F1-MODEL\_V4 \| 1.0 \| 2.074e-26 \| 461 \| 0.161 \| 742 \| 486 \| 36 \| 11 \| 681 \| 14 \| 689 \| Uncharacterized protein \| Uncharacterized protein; afdb-uniprot50 \| AF-A0A151FZE6-F1-MODEL\_V4 \| 1.0 \| 3.216e-24 \| 461 \| 0.16 \| 749 \| 452 \| 33 \| 14 \| 679 \| 82 \| 736 \| Uncharacterized protein \| Uncharacterized protein; afdb-uniprot50 \| AF-A0A084J8V6-F1-MODEL\_V4 \| 1.0 \| 1.166e-13 \| 460 \| 0.163 \| 269 \| 193 \| 9 \| 431 \| 682 \| 32 \| 285 \| Uncharacterized protein \| Uncharacterized protein; afdb-uniprot50 \| AF-A0A1H9KBY5-F1-MODEL\_V4 \| 1.0 \| 1.595e-25 \| 460 \| 0.152 \| 714 \| 418 \| 38 \| 40 \| 681 \| 16 \| 614 \| Uncharacterized protein \| Uncharacterized protein; afdb-uniprot50 \| AF-A0A520YI95-F1-MODEL\_V4 \| 1.0 \| 2.601e-26 \| 460 \| 0.182 \| 761 \| 440 \| 43 \| 18 \| 675 \| 11 \| 692 \| Uncharacterized protein \| Uncharacterized protein; afdb-uniprot50 \| AF-A0A2E7K3Z7-F1-MODEL\_V4 \| 1.0 \| 2.072e-18 \| 459 \| 0.278 \| 355 \| 185 \| 14 \| 17 \| 358 \| 3 \| 299 \| Uncharacterized protein \| Uncharacterized protein; afdb-uniprot50 \| AF-A0A2I1VRC1-F1-MODEL\_V4 \| 1.0 \| 5.275e-14 \| 459 \| 0.145 \| 274 \| 207 \| 7 \| 431 \| 681 \| 34 \| 303 \| Phage major capsid protein \| Phage major capsid protein; afdb-uniprot50 \| AF-A0A0F9RQJ3-F1-MODEL\_V4 \| 1.0 \| 1.594e-17 \| 459 \| 0.164 \| 469 \| 292 \| 19 \| 243 \| 682 \| 19 \| 416 \| Uncharacterized protein \| Uncharacterized protein; afdb-uniprot50 \| AF-A0A7V5XDB4-F1-MODEL\_V4 \| 1.0 \| 7.369e-25 \| 459 \| 0.169 \| 703 \| 414 \| 33 \| 41 \| 681 \| 16 \| 610 \| Peptidase U35 \| Peptidase U35; afdb-uniprot50 \| AF-A0A1E5HBV2-F1-MODEL\_V4 \| 1.0 \| 3.783e-19 \| 459 \| 0.183 \| 479 \| 284 \| 21 \| 252 \| 682 \| 207 \| 626 \| ATP-dependent Clp protease proteolytic subunit \| ATP-dependent Clp protease proteolytic subunit; afdb-uniprot50 \| AF-A0A0N8GFX9-F1-MODEL\_V4 \| 1.0 \| 1.688e-25 \| 459 \| 0.179 \| 743 \| 453 \| 39 \| 11 \| 681 \| 21 \| 678 \| Mu-like\_gpT domain-containing protein \| Mu-like\_gpT domain-containing protein; afdb-uniprot50 \| AF-H9UJE2-F1-MODEL\_V4 \| 1.0 \| 7.362e-17 \| 458 \| 0.128 \| 466 \| 276 \| 22 \| 255 \| 682 \| 2 \| 375 \| Putative phage phi-C31 gp36 major capsid-like protein \| Putative phage phi-C31 gp36 major capsid-like protein; afdb-uniprot50 \| AF-A0A3D5RUL1-F1-MODEL\_V4 \| 1.0 \| 8.544e-18 \| 458 \| 0.162 \| 467 \| 284 \| 22 \| 252 \| 680 \| 7 \| 404 \| Phage major capsid protein \| Phage major capsid protein; afdb-uniprot50 \| AF-A0A7Z2M7Y8-F1-MODEL\_V4 \| 1.0 \| 1.889e-17 \| 458 \| 0.148 \| 464 \| 307 \| 20 \| 249 \| 682 \| 8 \| 413 \| Capsid protein \| Capsid protein; afdb-uniprot50 \| AF-A0A7Y5QI34-F1-MODEL\_V4 \| 1.0 \| 8.544e-18 \| 458 \| 0.144 \| 469 \| 292 \| 22 \| 264 \| 678 \| 13 \| 426 \| Phage major capsid protein \| Phage major capsid protein; afdb-uniprot50 \| AF-A0A1Y4L4F4-F1-MODEL\_V4 \| 1.0 \| 8.301e-14 \| 457 \| 0.178 \| 269 \| 186 \| 14 \| 436 \| 682 \| 2 \| 257 \| Phage major capsid protein \| Phage major capsid protein; afdb-uniprot50 \| AF-A0A1H0Y9H5-F1-MODEL\_V4 \| 1.0 \| 4.177e-17 \| 457 \| 0.12 \| 447 \| 306 \| 16 \| 258 \| 682 \| 4 \| 385 \| Phage major capsid protein, HK97 family \| Phage major capsid protein, HK97 family; afdb-uniprot50 \| AF-F1YJE6-F1-MODEL\_V4 \| 1.0 \| 3.213e-16 \| 457 \| 0.154 \| 408 \| 293 \| 13 \| 315 \| 681 \| 1 \| 397 \| Uncharacterized protein \| Uncharacterized protein; afdb-uniprot50 \| AF-A0A1R0X157-F1-MODEL\_V4 \| 1.0 \| 1.244e-18 \| 457 \| 0.162 \| 481 \| 289 \| 20 \| 255 \| 682 \| 10 \| 429 \| Uncharacterized protein \| Uncharacterized protein; afdb-uniprot50 \| AF-Q14M91-F1-MODEL\_V4 \| 1.0 \| 1.761e-23 \| 457 \| 0.133 \| 747 \| 426 \| 44 \| 1 \| 681 \| 1 \| 591 \| Hypothetical prophage major capsid protein \| Hypothetical prophage major capsid protein; afdb-uniprot50 \| AF-X0XIA7-F1-MODEL\_V4 \| 1.0 \| 4.295e-13 \| 456 \| 0.227 \| 251 \| 161 \| 7 \| 443 \| 670 \| 1 \| 241 \| Uncharacterized protein \| Uncharacterized protein; afdb-uniprot50 \| AF-A0A5C7M8D0-F1-MODEL\_V4 \| 1.0 \| 3.551e-22 \| 456 \| 0.145 \| 721 \| 423 \| 40 \| 31 \| 680 \| 1 \| 599 \| Phage major capsid protein \| Phage major capsid protein; afdb-uniprot50 \| AF-A0A7W2BLG8-F1-MODEL\_V4 \| 1.0 \| 7.844e-14 \| 455 \| 0.211 \| 217 \| 148 \| 9 \| 478 \| 682 \| 20 \| 225 \| Phage major capsid protein \| Phage major capsid protein; afdb-uniprot50 \| AF-A0A7H8ZSI3-F1-MODEL\_V4 \| 1.0 \| 1.687e-17 \| 455 \| 0.168 \| 470 \| 275 \| 19 \| 251 \| 681 \| 3 \| 395 \| Phage major capsid protein \| Phage major capsid protein; afdb-uniprot50 \| AF-A0A6B1I8G0-F1-MODEL\_V4 \| 1.0 \| 1.244e-18 \| 455 \| 0.181 \| 468 \| 296 \| 21 \| 253 \| 680 \| 40 \| 460 \| Phage major capsid protein \| Phage major capsid protein; afdb-uniprot50 \| AF-A0A399XUC0-F1-MODEL\_V4 \| 1.0 \| 2.272e-19 \| 455 \| 0.165 \| 561 \| 321 \| 22 \| 172 \| 682 \| 12 \| 475 \| Phage major capsid protein \| Phage major capsid protein; afdb-uniprot50 \| AF-A0A5C7YF14-F1-MODEL\_V4 \| 1.0 \| 2.888e-13 \| 454 \| 0.188 \| 260 \| 192 \| 11 \| 431 \| 681 \| 29 \| 278 \| Phage major capsid protein \| Phage major capsid protein; afdb-uniprot50 \| AF-A0A0F9IVS0-F1-MODEL\_V4 \| 1.0 \| 1.252e-15 \| 454 \| 0.146 \| 389 \| 279 \| 18 \| 305 \| 681 \| 6 \| 353 \| Uncharacterized protein \| Uncharacterized protein; afdb-uniprot50 \| AF-A0A443TLW2-F1-MODEL\_V4 \| 1.0 \| 3.809e-16 \| 454 \| 0.143 \| 445 \| 276 \| 20 \| 262 \| 681 \| 9 \| 373 \| Phage major capsid protein \| Phage major capsid protein; afdb-uniprot50 \| AF-A0A7W8J095-F1-MODEL\_V4 \| 1.0 \| 3.036e-16 \| 454 \| 0.134 \| 447 \| 309 \| 19 \| 251 \| 682 \| 2 \| 385 \| HK97 family phage major capsid protein \| HK97 family phage major capsid protein; afdb-uniprot50 \| AF-A0A2A8V9J7-F1-MODEL\_V4 \| 1.0 \| 7.208e-18 \| 454 \| 0.154 \| 461 \| 284 \| 21 \| 264 \| 680 \| 23 \| 421 \| Phage major capsid protein \| Phage major capsid protein; afdb-uniprot50 \| AF-A0A6L5R7M5-F1-MODEL\_V4 \| 1.0 \| 4.421e-17 \| 453 \| 0.135 \| 450 \| 293 \| 21 \| 260 \| 682 \| 7 \| 387 \| Phage major capsid protein \| Phage major capsid protein; afdb-uniprot50 \| AF-A0A6N8I321-F1-MODEL\_V4 \| 1.0 \| 4.951e-17 \| 453 \| 0.161 \| 452 \| 293 \| 23 \| 256 \| 681 \| 4 \| 395 \| Phage capsid family protein \| Phage capsid family protein; afdb-uniprot50 \| AF-J5E045-F1-MODEL\_V4 \| 1.0 \| 2.239e-17 \| 453 \| 0.126 \| 473 \| 298 \| 19 \| 252 \| 682 \| 3 \| 402 \| Uncharacterized protein \| Uncharacterized protein; afdb-uniprot50 \| AF-A0A562NCU5-F1-MODEL\_V4 \| 1.0 \| 3.33e-17 \| 453 \| 0.141 \| 465 \| 305 \| 22 \| 245 \| 681 \| 1 \| 399 \| HK97 family phage major capsid protein \| HK97 family phage major capsid protein; afdb-uniprot50 \| AF-A0A198AI12-F1-MODEL\_V4 \| 1.0 \| 3.401e-16 \| 453 \| 0.162 \| 388 \| 228 \| 21 \| 337 \| 682 \| 72 \| 404 \| Uncharacterized protein \| Uncharacterized protein; afdb-uniprot50 \| AF-A0A1E2SJE2-F1-MODEL\_V4 \| 1.0 \| 2.809e-17 \| 453 \| 0.152 \| 447 \| 279 \| 20 \| 275 \| 681 \| 40 \| 426 \| Uncharacterized protein \| Uncharacterized protein; afdb-uniprot50 \| AF-A0A7V7PNP3-F1-MODEL\_V4 \| 1.0 \| 2.911e-18 \| 451 \| 0.187 \| 458 \| 269 \| 22 \| 243 \| 682 \| 12 \| 384 \| Phage major capsid protein \| Phage major capsid protein; afdb-uniprot50 \| AF-A0A542J6Z7-F1-MODEL\_V4 \| 1.0 \| 6.714e-16 \| 450 \| 0.139 \| 446 \| 289 \| 18 \| 261 \| 682 \| 5 \| 379 \| HK97 family phage major capsid protein \| HK97 family phage major capsid protein; afdb-uniprot50 \| AF-A0A1H2XPJ1-F1-MODEL\_V4 \| 1.0 \| 6.344e-16 \| 450 \| 0.138 \| 367 \| 280 \| 14 \| 338 \| 682 \| 29 \| 381 \| Phage major capsid protein, HK97 family \| Phage major capsid protein, HK97 family; afdb-uniprot50 \| AF-A0A4Y8RT87-F1-MODEL\_V4 \| 1.0 \| 1.226e-16 \| 450 \| 0.177 \| 411 \| 271 \| 19 \| 309 \| 682 \| 1 \| 381 \| Phage major capsid protein \| Phage major capsid protein; afdb-uniprot50 \| AF-A0A1A0S8B5-F1-MODEL\_V4 \| 1.0 \| 1.183e-15 \| 450 \| 0.14 \| 457 \| 297 \| 18 \| 251 \| 682 \| 3 \| 388 \| Uncharacterized protein \| Uncharacterized protein; afdb-uniprot50 \| AF-A0A7W6CYU8-F1-MODEL\_V4 \| 1.0 \| 2.161e-16 \| 450 \| 0.13 \| 466 \| 303 \| 21 \| 264 \| 682 \| 6 \| 416 \| HK97 family phage major capsid protein \| HK97 family phage major capsid protein; afdb-uniprot50 \| AF-A0A7X4F2D3-F1-MODEL\_V4 \| 1.0 \| 1.687e-17 \| 450 \| 0.147 \| 460 \| 302 \| 21 \| 264 \| 681 \| 7 \| 418 \| Phage major capsid protein \| Phage major capsid protein; afdb-uniprot50 \| AF-A0A379E0G1-F1-MODEL\_V4 \| 1.0 \| 1.27e-17 \| 449 \| 0.193 \| 428 \| 243 \| 19 \| 295 \| 681 \| 7 \| 373 \| Predicted phage phi-C31 gp36 major capsid-like protein \| Predicted phage phi-C31 gp36 major capsid-like protein; afdb-uniprot50 \| AF-A0A2G2L0U3-F1-MODEL\_V4 \| 1.0 \| 6.344e-16 \| 449 \| 0.141 \| 425 \| 294 \| 14 \| 272 \| 681 \| 3 \| 371 \| Phage major capsid protein \| Phage major capsid protein; afdb-uniprot50 \| AF-A0A349YIG3-F1-MODEL\_V4 \| 1.0 \| 5.352e-16 \| 449 \| 0.128 \| 444 \| 298 \| 20 \| 259 \| 681 \| 6 \| 381 \| Phage major capsid protein \| Phage major capsid protein; afdb-uniprot50 \| AF-A0A3D3Z4A9-F1-MODEL\_V4 \| 1.0 \| 5.746e-18 \| 449 \| 0.161 \| 472 \| 286 \| 20 \| 264 \| 678 \| 7 \| 425 \| Phage major capsid protein \| Phage major capsid protein; afdb-uniprot50 \| AF-A0A2D8K5D9-F1-MODEL\_V4 \| 1.0 \| 2.241e-25 \| 449 \| 0.182 \| 733 \| 447 \| 39 \| 17 \| 681 \| 28 \| 675 \| Uncharacterized protein \| Uncharacterized protein; afdb-uniprot50 \| AF-A0A4Y1N435-F1-MODEL\_V4 \| 1.0 \| 8.544e-18 \| 448 \| 0.177 \| 497 \| 264 \| 24 \| 261 \| 680 \| 3 \| 431 \| Uncharacterized protein \| Uncharacterized protein; afdb-uniprot50 \| AF-A0A3N4SE98-F1-MODEL\_V4 \| 1.0 \| 3.192e-19 \| 448 \| 0.169 \| 556 \| 349 \| 25 \| 196 \| 681 \| 1 \| 513 \| HK97 family phage major capsid protein \| HK97 family phage major capsid protein; afdb-uniprot50 \| AF-A0A2N3F2A1-F1-MODEL\_V4 \| 1.0 \| 4.295e-13 \| 447 \| 0.164 \| 274 \| 189 \| 15 \| 434 \| 681 \| 2 \| 261 \| Phage major capsid protein \| Phage major capsid protein; afdb-uniprot50 \| AF-A0A6B9T8U5-F1-MODEL\_V4 \| 1.0 \| 1.76e-15 \| 447 \| 0.14 \| 371 \| 279 \| 17 \| 338 \| 681 \| 14 \| 371 \| Phage major capsid protein \| Phage major capsid protein; afdb-uniprot50 \| AF-A0A0F7RMS0-F1-MODEL\_V4 \| 1.0 \| 6.956e-17 \| 447 \| 0.142 \| 429 \| 281 \| 19 \| 279 \| 682 \| 32 \| 398 \| Phage Capsid protein \| Phage Capsid protein; afdb-uniprot50 \| AF-A0A1V5NDX4-F1-MODEL\_V4 \| 1.0 \| 1.095e-16 \| 447 \| 0.126 \| 465 \| 327 \| 16 \| 243 \| 681 \| 6 \| 417 \| Phage capsid family protein \| Phage capsid family protein; afdb-uniprot50 \| AF-A0A845RNZ3-F1-MODEL\_V4 \| 1.0 \| 9.103e-15 \| 446 \| 0.15 \| 358 \| 227 \| 16 \| 344 \| 682 \| 5 \| 304 \| Phage major capsid protein \| Phage major capsid protein; afdb-uniprot50 \| AF-A0A2M9M606-F1-MODEL\_V4 \| 1.0 \| 5.24e-17 \| 446 \| 0.163 \| 433 \| 278 \| 22 \| 279 \| 682 \| 3 \| 380 \| Phage major capsid protein \| Phage major capsid protein; afdb-uniprot50 \| AF-A0A7W1RVR0-F1-MODEL\_V4 \| 1.0 \| 1.628e-16 \| 446 \| 0.137 \| 458 \| 292 \| 19 \| 258 \| 681 \| 21 \| 409 \| Phage major capsid protein \| Phage major capsid protein; afdb-uniprot50 \| AF-A0A1A0PLW6-F1-MODEL\_V4 \| 1.0 \| 1.835e-13 \| 445 \| 0.171 \| 285 \| 180 \| 12 \| 431 \| 681 \| 30 \| 292 \| Uncharacterized protein \| Uncharacterized protein; afdb-uniprot50 \| AF-A0A7V9S9R7-F1-MODEL\_V4 \| 1.0 \| 1.862e-15 \| 445 \| 0.17 \| 357 \| 265 \| 13 \| 346 \| 677 \| 17 \| 367 \| Phage major capsid protein \| Phage major capsid protein; afdb-uniprot50 \| AF-B2KDP2-F1-MODEL\_V4 \| 1.0 \| 8.246e-17 \| 445 \| 0.162 \| 455 \| 275 \| 19 \| 274 \| 681 \| 5 \| 400 \| Phage major capsid protein \| Phage major capsid protein; afdb-uniprot50 \| AF-A0A521ZM45-F1-MODEL\_V4 \| 1.0 \| 1.785e-17 \| 445 \| 0.145 \| 487 \| 310 \| 23 \| 215 \| 682 \| 1 \| 400 \| Phage major capsid protein \| Phage major capsid protein; afdb-uniprot50 \| AF-A0A2E9VXI6-F1-MODEL\_V4 \| 1.0 \| 5.24e-17 \| 445 \| 0.161 \| 482 \| 283 \| 28 \| 254 \| 682 \| 5 \| 418 \| Phage major capsid protein \| Phage major capsid protein; afdb-uniprot50 \| AF-A0A1Y0Y2T8-F1-MODEL\_V4 \| 1.0 \| 4.848e-18 \| 445 \| 0.163 \| 489 \| 298 \| 21 \| 262 \| 680 \| 5 \| 452 \| Uncharacterized protein \| Uncharacterized protein; afdb-uniprot50 \| AF-A0A1X2EPP6-F1-MODEL\_V4 \| 1.0 \| 2.888e-13 \| 444 \| 0.16 \| 261 \| 194 \| 11 \| 433 \| 681 \| 32 \| 279 \| Major capsid protein \| Major capsid protein; afdb-uniprot50 \| AF-A0A4Q3PQP0-F1-MODEL\_V4 \| 1.0 \| 4.266e-16 \| 444 \| 0.164 \| 376 \| 275 \| 14 \| 338 \| 682 \| 8 \| 375 \| Phage major capsid protein \| Phage major capsid protein; afdb-uniprot50 \| AF-A0A1P8YNF5-F1-MODEL\_V4 \| 1.0 \| 1.298e-16 \| 444 \| 0.141 \| 452 \| 305 \| 20 \| 258 \| 682 \| 6 \| 401 \| Phage major capsid protein, HK97 family \| Phage major capsid protein, HK97 family; afdb-uniprot50 \| AF-A0A7X0D7X9-F1-MODEL\_V4 \| 1.0 \| 2.711e-16 \| 444 \| 0.125 \| 407 \| 276 \| 21 \| 338 \| 682 \| 15 \| 403 \| HK97 family phage major capsid protein \| HK97 family phage major capsid protein; afdb-uniprot50 \| AF-A0A0H4KS24-F1-MODEL\_V4 \| 1.0 \| 3.08e-18 \| 444 \| 0.194 \| 468 \| 276 \| 26 \| 258 \| 681 \| 5 \| 415 \| Phage major capsid protein, HK97 family \| Phage major capsid protein, HK97 family; afdb-uniprot50 \| AF-A0A5J6G9W9-F1-MODEL\_V4 \| 1.0 \| 3.08e-18 \| 444 \| 0.146 \| 477 \| 282 \| 22 \| 262 \| 678 \| 9 \| 420 \| Phage major capsid protein \| Phage major capsid protein; afdb-uniprot50 \| AF-A0A2U3L962-F1-MODEL\_V4 \| 1.0 \| 1.27e-17 \| 444 \| 0.141 \| 481 \| 315 \| 23 \| 253 \| 682 \| 3 \| 436 \| Putative Phage major capsid protein, HK97 family \| Putative Phage major capsid protein, HK97 family; afdb-uniprot50 \| AF-A0A2E0J2G0-F1-MODEL\_V4 \| 1.0 \| 3.45e-18 \| 444 \| 0.198 \| 468 \| 294 \| 21 \| 256 \| 681 \| 4 \| 432 \| Phage major capsid protein \| Phage major capsid protein; afdb-uniprot50 \| AF-A0A840ZNF0-F1-MODEL\_V4 \| 1.0 \| 8.544e-18 \| 444 \| 0.153 \| 521 \| 334 \| 26 \| 195 \| 682 \| 2 \| 448 \| HK97 family phage major capsid protein \| HK97 family phage major capsid protein; afdb-uniprot50 \| AF-A0A7X7T206-F1-MODEL\_V4 \| 1.0 \| 3.26e-18 \| 444 \| 0.144 \| 492 \| 303 \| 22 \| 251 \| 681 \| 5 \| 439 \| Phage major capsid protein \| Phage major capsid protein; afdb-uniprot50 \| AF-A0A100WUZ1-F1-MODEL\_V4 \| 1.0 \| 2.508e-17 \| 444 \| 0.164 \| 492 \| 294 \| 23 \| 258 \| 678 \| 8 \| 453 \| Phage-related major capsid protein \| Phage-related major capsid protein; afdb-uniprot50 \| AF-A0A5F2I8T1-F1-MODEL\_V4 \| 1.0 \| 4.683e-25 \| 444 \| 0.147 \| 698 \| 456 \| 35 \| 40 \| 681 \| 50 \| 664 \| Mu-like\_gpT domain-containing protein \| Mu-like\_gpT domain-containing protein; afdb-uniprot50 \| AF-A0A177P7V0-F1-MODEL\_V4 \| 1.0 \| 3.527e-25 \| 444 \| 0.148 \| 748 \| 463 \| 33 \| 16 \| 681 \| 18 \| 673 \| Uncharacterized protein \| Uncharacterized protein; afdb-uniprot50 \| AF-A0A2U1BFB6-F1-MODEL\_V4 \| 1.0 \| 1.971e-15 \| 443 \| 0.161 \| 340 \| 226 \| 16 \| 375 \| 682 \| 20 \| 332 \| HK97 family phage major capsid protein \| HK97 family phage major capsid protein; afdb-uniprot50 \| AF-F5RN15-F1-MODEL\_V4 \| 1.0 \| 2.508e-17 \| 443 \| 0.159 \| 469 \| 277 \| 24 \| 252 \| 682 \| 2 \| 391 \| HK97 family major capsid protein \| HK97 family major capsid protein; afdb-uniprot50 \| AF-A0A317LFV3-F1-MODEL\_V4 \| 1.0 \| 2.287e-16 \| 443 \| 0.157 \| 450 \| 309 \| 20 \| 253 \| 682 \| 2 \| 401 \| Phage major capsid protein \| Phage major capsid protein; afdb-uniprot50 \| AF-A0A4R7IC04-F1-MODEL\_V4 \| 1.0 \| 2.116e-17 \| 443 \| 0.155 \| 501 \| 298 \| 20 \| 252 \| 681 \| 10 \| 456 \| HK97 family phage major capsid protein \| HK97 family phage major capsid protein; afdb-uniprot50 \| AF-A0A7C4QW79-F1-MODEL\_V4 \| 1.0 \| 1.7e-22 \| 443 \| 0.146 \| 702 \| 438 \| 32 \| 16 \| 681 \| 77 \| 652 \| Phage major capsid protein \| Phage major capsid protein; afdb-uniprot50 \| AF-A0A430EDT7-F1-MODEL\_V4 \| 1.0 \| 5.551e-25 \| 443 \| 0.165 \| 750 \| 434 \| 37 \| 17 \| 681 \| 78 \| 720 \| Mu-like\_gpT domain-containing protein \| Mu-like\_gpT domain-containing protein; afdb-uniprot50 \| AF-A0A229VZ15-F1-MODEL\_V4 \| 1.0 \| 1.639e-13 \| 442 \| 0.131 \| 282 \| 208 \| 10 \| 425 \| 682 \| 11 \| 279 \| Major capsid protein \| Major capsid protein; afdb-uniprot50 \| AF-A0A542HV74-F1-MODEL\_V4 \| 1.0 \| 1.823e-16 \| 442 \| 0.132 \| 446 \| 270 \| 18 \| 279 \| 682 \| 36 \| 406 \| HK97 family phage major capsid protein \| HK97 family phage major capsid protein; afdb-uniprot50 \| AF-A0A2K8LT81-F1-MODEL\_V4 \| 1.0 \| 5.583e-14 \| 440 \| 0.182 \| 280 \| 189 \| 9 \| 431 \| 681 \| 26 \| 294 \| Phage major capsid protein \| Phage major capsid protein; afdb-uniprot50 \| AF-A0A5C8L259-F1-MODEL\_V4 \| 1.0 \| 1.538e-16 \| 440 \| 0.149 \| 447 \| 283 \| 22 \| 268 \| 682 \| 4 \| 385 \| Phage major capsid protein \| Phage major capsid protein; afdb-uniprot50 \| AF-A0A5J5J5J3-F1-MODEL\_V4 \| 1.0 \| 5.545e-17 \| 440 \| 0.165 \| 471 \| 284 \| 20 \| 257 \| 682 \| 5 \| 411 \| Phage major capsid protein \| Phage major capsid protein; afdb-uniprot50 \| AF-A0A3D8NG00-F1-MODEL\_V4 \| 1.0 \| 1.785e-17 \| 440 \| 0.155 \| 451 \| 298 \| 21 \| 267 \| 681 \| 11 \| 414 \| Phage major capsid protein \| Phage major capsid protein; afdb-uniprot50 \| AF-G6XIV7-F1-MODEL\_V4 \| 1.0 \| 1.345e-17 \| 440 \| 0.139 \| 502 \| 304 \| 22 \| 261 \| 681 \| 8 \| 462 \| Uncharacterized protein \| Uncharacterized protein; afdb-uniprot50 \| AF-A0A6L9G9S4-F1-MODEL\_V4 \| 1.0 \| 4.299e-21 \| 440 \| 0.11 \| 671 \| 397 \| 29 \| 46 \| 682 \| 3 \| 507 \| Uncharacterized protein \| Uncharacterized protein; afdb-uniprot50 \| AF-A0A653VEB3-F1-MODEL\_V4 \| 1.0 \| 4.266e-16 \| 439 \| 0.169 \| 442 \| 270 \| 18 \| 265 \| 682 \| 7 \| 375 \| Phage capsid family protein \| Phage capsid family protein; afdb-uniprot50 \| AF-A0A355J1L3-F1-MODEL\_V4 \| 1.0 \| 1.785e-17 \| 439 \| 0.142 \| 456 \| 295 \| 24 \| 258 \| 680 \| 8 \| 400 \| Uncharacterized protein \| Uncharacterized protein; afdb-uniprot50 \| AF-A0A6N9BAL6-F1-MODEL\_V4 \| 1.0 \| 1.594e-17 \| 439 \| 0.174 \| 458 \| 282 \| 21 \| 258 \| 680 \| 5 \| 401 \| Uncharacterized protein \| Uncharacterized protein; afdb-uniprot50 \| AF-A0A1I6GEC2-F1-MODEL\_V4 \| 1.0 \| 7.791e-17 \| 439 \| 0.146 \| 479 \| 303 \| 22 \| 243 \| 682 \| 26 \| 437 \| Phage major capsid protein, HK97 family \| Phage major capsid protein, HK97 family; afdb-uniprot50 \| AF-A0A7W6FWA4-F1-MODEL\_V4 \| 1.0 \| 3.146e-17 \| 439 \| 0.158 \| 461 \| 299 \| 22 \| 249 \| 682 \| 75 \| 473 \| HK97 family phage major capsid protein \| HK97 family phage major capsid protein; afdb-uniprot50 \| AF-A0A2Z5ZI90-F1-MODEL\_V4 \| 1.0 \| 7.904e-19 \| 439 \| 0.143 \| 501 \| 308 \| 21 \| 252 \| 681 \| 7 \| 457 \| Major capsid protein \| Major capsid protein; afdb-uniprot50 \| AF-A0A7T5K4D8-F1-MODEL\_V4 \| 1.0 \| 2.639e-28 \| 439 \| 0.197 \| 739 \| 404 \| 39 \| 38 \| 681 \| 11 \| 655 \| Mu-like prophage major head subunit gpT family protein \| Mu-like prophage major head subunit gpT family protein; afdb-uniprot50 \| AF-A0A5N8X7L1-F1-MODEL\_V4 \| 1.0 \| 3.477e-23 \| 439 \| 0.14 \| 719 \| 435 \| 41 \| 12 \| 682 \| 318 \| 901 \| Phage portal protein \| Phage portal protein; afdb-uniprot50 \| AF-A0A2U3NJA2-F1-MODEL\_V4 \| 1.0 \| 6.759e-13 \| 438 \| 0.198 \| 237 \| 166 \| 8 \| 453 \| 682 \| 2 \| 221 \| Capsid protein \| Capsid protein; afdb-uniprot50 \| AF-A0A416AP22-F1-MODEL\_V4 \| 1.0 \| 1.102e-13 \| 438 \| 0.152 \| 268 \| 193 \| 15 \| 431 \| 681 \| 54 \| 304 \| Phage major capsid protein \| Phage major capsid protein; afdb-uniprot50 \| AF-A0A6N0LFZ4-F1-MODEL\_V4 \| 1.0 \| 2.336e-15 \| 438 \| 0.15 \| 372 \| 270 \| 12 \| 345 \| 682 \| 7 \| 366 \| Phage major capsid protein \| Phage major capsid protein; afdb-uniprot50 \| AF-A0A2C9WSZ8-F1-MODEL\_V4 \| 1.0 \| 5.057e-16 \| 438 \| 0.132 \| 452 \| 298 \| 20 \| 252 \| 682 \| 13 \| 391 \| Phage major capsid protein \| Phage major capsid protein; afdb-uniprot50 \| AF-M4NDD1-F1-MODEL\_V4 \| 1.0 \| 3.213e-16 \| 438 \| 0.151 \| 415 \| 293 \| 16 \| 306 \| 679 \| 7 \| 403 \| Phage capsid family protein \| Phage capsid family protein; afdb-uniprot50 \| AF-A0A149TLS8-F1-MODEL\_V4 \| 1.0 \| 3.651e-18 \| 438 \| 0.17 \| 480 \| 304 \| 20 \| 264 \| 682 \| 2 \| 448 \| Uncharacterized protein \| Uncharacterized protein; afdb-uniprot50 \| AF-A0A737BVU5-F1-MODEL\_V4 \| 1.0 \| 1.875e-12 \| 437 \| 0.136 \| 256 \| 198 \| 10 \| 436 \| 679 \| 2 \| 246 \| Phage major capsid protein \| Phage major capsid protein; afdb-uniprot50 \| AF-A0A2A9ENW6-F1-MODEL\_V4 \| 1.0 \| 5.352e-16 \| 437 \| 0.149 \| 369 \| 280 \| 14 \| 338 \| 681 \| 9 \| 368 \| HK97 family phage major capsid protein \| HK97 family phage major capsid protein; afdb-uniprot50 \| AF-A0A1V5ID83-F1-MODEL\_V4 \| 1.0 \| 6.714e-16 \| 437 \| 0.161 \| 409 \| 254 \| 15 \| 295 \| 681 \| 29 \| 370 \| Phage capsid family protein \| Phage capsid family protein; afdb-uniprot50 \| AF-A0A7G6Z6C8-F1-MODEL\_V4 \| 1.0 \| 1.226e-16 \| 437 \| 0.142 \| 448 \| 278 \| 20 \| 264 \| 677 \| 3 \| 378 \| Phage major capsid protein \| Phage major capsid protein; afdb-uniprot50 \| AF-E1QM51-F1-MODEL\_V4 \| 1.0 \| 1.929e-16 \| 437 \| 0.17 \| 423 \| 255 \| 17 \| 335 \| 681 \| 1 \| 403 \| Major capsid protein HK97 \| Major capsid protein HK97; afdb-uniprot50 \| AF-A0A1Q6K3Y8-F1-MODEL\_V4 \| 1.0 \| 5.869e-17 \| 437 \| 0.149 \| 449 \| 294 \| 22 \| 251 \| 682 \| 27 \| 404 \| Uncharacterized protein \| Uncharacterized protein; afdb-uniprot50 \| AF-A0A7Z1AWA5-F1-MODEL\_V4 \| 1.0 \| 7.965e-24 \| 437 \| 0.136 \| 709 \| 424 \| 37 \| 39 \| 679 \| 426 \| 1013 \| Uncharacterized protein \| Uncharacterized protein; afdb-uniprot50 \| AF-A0A1M5PQD6-F1-MODEL\_V4 \| 1.0 \| 7.519e-16 \| 436 \| 0.143 \| 459 \| 277 \| 20 \| 244 \| 682 \| 12 \| 374 \| Phage major capsid protein, HK97 family \| Phage major capsid protein, HK97 family; afdb-uniprot50 \| AF-A0A1C6UT23-F1-MODEL\_V4 \| 1.0 \| 7.519e-16 \| 436 \| 0.12 \| 425 \| 290 \| 14 \| 281 \| 681 \| 15 \| 379 \| Phage major capsid protein, HK97 family \| Phage major capsid protein, HK97 family; afdb-uniprot50 \| AF-A0A7W3Y5V0-F1-MODEL\_V4 \| 1.0 \| 3.809e-16 \| 436 \| 0.138 \| 418 \| 301 \| 18 \| 284 \| 679 \| 52 \| 432 \| Phage major capsid protein \| Phage major capsid protein; afdb-uniprot50 \| AF-A0A2M9MMU3-F1-MODEL\_V4 \| 1.0 \| 2.088e-23 \| 436 \| 0.13 \| 721 \| 389 \| 30 \| 26 \| 680 \| 2 \| 550 \| Uncharacterized protein \| Uncharacterized protein; afdb-uniprot50 \| AF-A0A2W4ZXB8-F1-MODEL\_V4 \| 1.0 \| 3.216e-24 \| 436 \| 0.157 \| 717 \| 393 \| 35 \| 40 \| 677 \| 1 \| 585 \| Uncharacterized protein \| Uncharacterized protein; afdb-uniprot50 \| AF-A0A4V2BF65-F1-MODEL\_V4 \| 1.0 \| 2.074e-26 \| 436 \| 0.141 \| 728 \| 487 \| 44 \| 17 \| 679 \| 3 \| 657 \| Uncharacterized protein \| Uncharacterized protein; afdb-uniprot50 \| AF-A0A1Y3TL93-F1-MODEL\_V4 \| 1.0 \| 3.497e-12 \| 435 \| 0.383 \| 159 \| 88 \| 5 \| 38 \| 190 \| 9 \| 163 \| Caudovirus prohead protease \| Caudovirus prohead protease; afdb-uniprot50 \| AF-A0A1B1LIJ3-F1-MODEL\_V4 \| 1.0 \| 1.639e-13 \| 435 \| 0.172 \| 267 \| 193 \| 13 \| 431 \| 681 \| 17 \| 271 \| Major capsid protein b \| Major capsid protein b; afdb-uniprot50 \| AF-A0A7W8J549-F1-MODEL\_V4 \| 1.0 \| 4.611e-15 \| 435 \| 0.124 \| 353 \| 276 \| 15 \| 349 \| 681 \| 34 \| 373 \| HK97 family phage major capsid protein \| HK97 family phage major capsid protein; afdb-uniprot50 \| AF-A0A497AYE2-F1-MODEL\_V4 \| 1.0 \| 5.096e-21 \| 435 \| 0.158 \| 708 \| 390 \| 43 \| 19 \| 681 \| 2 \| 548 \| Phage major capsid protein \| Phage major capsid protein; afdb-uniprot50 \| AF-A0A0C1KQP5-F1-MODEL\_V4 \| 1.0 \| 1.862e-15 \| 434 \| 0.156 \| 370 \| 270 \| 14 \| 345 \| 682 \| 4 \| 363 \| Uncharacterized protein \| Uncharacterized protein; afdb-uniprot50 \| AF-U7UE74-F1-MODEL\_V4 \| 1.0 \| 4.177e-17 \| 434 \| 0.144 \| 465 \| 284 \| 20 \| 265 \| 682 \| 4 \| 401 \| Phage major capsid protein, HK97 family \| Phage major capsid protein, HK97 family; afdb-uniprot50 \| AF-A0A164R867-F1-MODEL\_V4 \| 1.0 \| 1.76e-15 \| 434 \| 0.156 \| 447 \| 287 \| 18 \| 261 \| 677 \| 9 \| 395 \| Uncharacterized protein \| Uncharacterized protein; afdb-uniprot50 \| AF-A0A5C6UM44-F1-MODEL\_V4 \| 1.0 \| 3.401e-16 \| 434 \| 0.117 \| 424 \| 306 \| 15 \| 304 \| 679 \| 42 \| 445 \| Phage major capsid protein \| Phage major capsid protein; afdb-uniprot50 \| AF-A0A6M3ZTZ3-F1-MODEL\_V4 \| 1.0 \| 2.711e-16 \| 433 \| 0.146 \| 423 \| 290 \| 17 \| 285 \| 682 \| 53 \| 429 \| Phage major capsid protein \| Phage major capsid protein; afdb-uniprot50 \| AF-A0A5C7M8X5-F1-MODEL\_V4 \| 1.0 \| 6.386e-13 \| 432 \| 0.167 \| 263 \| 194 \| 12 \| 431 \| 681 \| 29 \| 278 \| Phage major capsid protein \| Phage major capsid protein; afdb-uniprot50 \| AF-A0A3B0IS29-F1-MODEL\_V4 \| 1.0 \| 2.056e-13 \| 432 \| 0.136 \| 308 \| 231 \| 14 \| 392 \| 677 \| 43 \| 337 \| Uncharacterized protein \| Uncharacterized protein; afdb-uniprot50 \| AF-A0A359AJM0-F1-MODEL\_V4 \| 1.0 \| 8.246e-17 \| 432 \| 0.173 \| 491 \| 283 \| 26 \| 217 \| 680 \| 1 \| 395 \| Phage major capsid protein \| Phage major capsid protein; afdb-uniprot50 \| AF-A0A5C7QAV4-F1-MODEL\_V4 \| 1.0 \| 1.929e-16 \| 432 \| 0.123 \| 477 \| 309 \| 24 \| 249 \| 679 \| 10 \| 423 \| Phage major capsid protein \| Phage major capsid protein; afdb-uniprot50 \| AF-A0A7C3X2B0-F1-MODEL\_V4 \| 1.0 \| 1.723e-16 \| 432 \| 0.131 \| 470 \| 317 \| 16 \| 246 \| 680 \| 18 \| 431 \| Phage major capsid protein \| Phage major capsid protein; afdb-uniprot50 \| AF-A0A2A4EDF3-F1-MODEL\_V4 \| 1.0 \| 1.453e-16 \| 431 \| 0.146 \| 457 \| 279 \| 20 \| 271 \| 682 \| 3 \| 393 \| Uncharacterized protein \| Uncharacterized protein; afdb-uniprot50 \| AF-A0A3D3LX94-F1-MODEL\_V4 \| 1.0 \| 2.161e-16 \| 431 \| 0.163 \| 433 \| 268 \| 20 \| 305 \| 681 \| 4 \| 398 \| Phage major capsid protein \| Phage major capsid protein; afdb-uniprot50 \| AF-A0A239A9U8-F1-MODEL\_V4 \| 1.0 \| 1.298e-16 \| 431 \| 0.143 \| 461 \| 294 \| 19 \| 250 \| 679 \| 5 \| 395 \| Phage major capsid protein, HK97 family \| Phage major capsid protein, HK97 family; afdb-uniprot50 \| AF-A0A4R8JD56-F1-MODEL\_V4 \| 1.0 \| 2.42e-16 \| 431 \| 0.141 \| 458 \| 307 \| 16 \| 243 \| 682 \| 40 \| 429 \| HK97 family phage major capsid protein \| HK97 family phage major capsid protein; afdb-uniprot50 \| AF-A0A2G6IFW5-F1-MODEL\_V4 \| 1.0 \| 7.362e-17 \| 431 \| 0.161 \| 488 \| 297 \| 22 \| 251 \| 681 \| 5 \| 437 \| Phage major capsid protein \| Phage major capsid protein; afdb-uniprot50 \| AF-A0A1G3WMU9-F1-MODEL\_V4 \| 1.0 \| 2.086e-15 \| 430 \| 0.132 \| 391 \| 275 \| 14 \| 335 \| 680 \| 1 \| 372 \| Uncharacterized protein \| Uncharacterized protein; afdb-uniprot50 \| AF-A0A3P6KYD8-F1-MODEL\_V4 \| 1.0 \| 7.687e-23 \| 430 \| 0.207 \| 540 \| 295 \| 23 \| 19 \| 519 \| 1 \| 446 \| Mu-like prophage major head subunit gpT \| Mu-like prophage major head subunit gpT; afdb-uniprot50 \| AF-S5DTM3-F1-MODEL\_V4 \| 1.0 \| 6.344e-16 \| 429 \| 0.146 \| 450 \| 267 \| 19 \| 271 \| 680 \| 3 \| 375 \| Phage major capsid protein \| Phage major capsid protein; afdb-uniprot50 \| AF-B9XDE7-F1-MODEL\_V4 \| 1.0 \| 4.031e-16 \| 429 \| 0.16 \| 455 \| 295 \| 22 \| 252 \| 681 \| 11 \| 403 \| Major capsid protein HK97 \| Major capsid protein HK97; afdb-uniprot50 \| AF-A0A0F9W0Z4-F1-MODEL\_V4 \| 1.0 \| 1.594e-17 \| 429 \| 0.173 \| 466 \| 296 \| 26 \| 252 \| 681 \| 2 \| 414 \| Uncharacterized protein \| Uncharacterized protein; afdb-uniprot50 \| AF-A0A6P1YHM1-F1-MODEL\_V4 \| 1.0 \| 4.678e-17 \| 429 \| 0.137 \| 503 \| 330 \| 23 \| 204 \| 679 \| 1 \| 426 \| Phage major capsid protein \| Phage major capsid protein; afdb-uniprot50 \| AF-A0A522ASE0-F1-MODEL\_V4 \| 1.0 \| 6.811e-18 \| 429 \| 0.14 \| 477 \| 307 \| 21 \| 237 \| 680 \| 37 \| 443 \| Phage major capsid protein \| Phage major capsid protein; afdb-uniprot50 \| AF-A0A4R5TM46-F1-MODEL\_V4 \| 1.0 \| 7.958e-16 \| 428 \| 0.142 \| 435 \| 302 \| 14 \| 252 \| 679 \| 15 \| 385 \| Phage major capsid protein \| Phage major capsid protein; afdb-uniprot50 \| AF-A0A516NU19-F1-MODEL\_V4 \| 1.0 \| 5.057e-16 \| 428 \| 0.142 \| 450 \| 293 \| 20 \| 261 \| 681 \| 11 \| 396 \| Phage major capsid protein \| Phage major capsid protein; afdb-uniprot50 \| AF-A0A2A3ETC9-F1-MODEL\_V4 \| 1.0 \| 1.095e-16 \| 428 \| 0.153 \| 468 \| 284 \| 17 \| 263 \| 678 \| 21 \| 428 \| Phage major capsid protein \| Phage major capsid protein; afdb-uniprot50 \| AF-R5UUT8-F1-MODEL\_V4 \| 1.0 \| 9.983e-16 \| 428 \| 0.144 \| 380 \| 266 \| 16 \| 349 \| 679 \| 66 \| 435 \| Uncharacterized protein \| Uncharacterized protein; afdb-uniprot50 \| AF-A0A2D6F059-F1-MODEL\_V4 \| 1.0 \| 3.146e-17 \| 428 \| 0.152 \| 480 \| 286 \| 20 \| 282 \| 678 \| 3 \| 444 \| Phage major capsid protein \| Phage major capsid protein; afdb-uniprot50 \| AF-A0A4Q5NL72-F1-MODEL\_V4 \| 1.0 \| 3.524e-17 \| 428 \| 0.15 \| 498 \| 311 \| 27 \| 249 \| 682 \| 3 \| 452 \| Phage major capsid protein \| Phage major capsid protein; afdb-uniprot50 \| AF-A0A7X7FKI2-F1-MODEL\_V4 \| 1.0 \| 4.177e-17 \| 428 \| 0.166 \| 487 \| 324 \| 20 \| 243 \| 681 \| 19 \| 471 \| Phage major capsid protein \| Phage major capsid protein; afdb-uniprot50 \| AF-A0A286DYE7-F1-MODEL\_V4 \| 1.0 \| 1.785e-17 \| 428 \| 0.138 \| 498 \| 310 \| 25 \| 268 \| 677 \| 9 \| 475 \| Phage major capsid protein, HK97 family \| Phage major capsid protein, HK97 family; afdb-uniprot50 \| AF-A0A6L9HU12-F1-MODEL\_V4 \| 1.0 \| 1.787e-25 \| 428 \| 0.161 \| 815 \| 430 \| 37 \| 16 \| 682 \| 5 \| 713 \| Uncharacterized protein \| Uncharacterized protein; afdb-uniprot50 \| AF-A0A5S3XG18-F1-MODEL\_V4 \| 1.0 \| 1.423e-17 \| 427 \| 0.173 \| 473 \| 286 \| 23 \| 261 \| 681 \| 7 \| 426 \| Phage major capsid protein \| Phage major capsid protein; afdb-uniprot50 \| AF-A0A429P0Y7-F1-MODEL\_V4 \| 1.0 \| 6.436e-18 \| 427 \| 0.14 \| 550 \| 356 \| 23 \| 178 \| 682 \| 3 \| 480 \| Phage major capsid protein \| Phage major capsid protein; afdb-uniprot50 \| AF-A0A433J4X0-F1-MODEL\_V4 \| 1.0 \| 1.506e-17 \| 427 \| 0.169 \| 455 \| 275 \| 22 \| 279 \| 682 \| 97 \| 499 \| Phage major capsid protein \| Phage major capsid protein; afdb-uniprot50 \| AF-A0A7Y3T6V6-F1-MODEL\_V4 \| 1.0 \| 3.039e-24 \| 427 \| 0.155 \| 790 \| 428 \| 43 \| 40 \| 681 \| 6 \| 703 \| Uncharacterized protein \| Uncharacterized protein; afdb-uniprot50 \| AF-A0A1H0SZG1-F1-MODEL\_V4 \| 1.0 \| 3.676e-15 \| 426 \| 0.145 \| 448 \| 280 \| 22 \| 253 \| 682 \| 2 \| 364 \| Phage major capsid protein, HK97 family \| Phage major capsid protein, HK97 family; afdb-uniprot50 \| AF-A0A316QDD0-F1-MODEL\_V4 \| 1.0 \| 4.117e-15 \| 426 \| 0.144 \| 374 \| 276 \| 13 \| 338 \| 680 \| 14 \| 374 \| Phage major capsid protein \| Phage major capsid protein; afdb-uniprot50 \| AF-A0A3M1Y489-F1-MODEL\_V4 \| 1.0 \| 9.433e-16 \| 426 \| 0.154 \| 452 \| 306 \| 16 \| 251 \| 679 \| 3 \| 401 \| Phage major capsid protein \| Phage major capsid protein; afdb-uniprot50 \| AF-A0A7C2D4S3-F1-MODEL\_V4 \| 1.0 \| 4.515e-16 \| 426 \| 0.143 \| 425 \| 298 \| 21 \| 294 \| 682 \| 6 \| 400 \| Phage major capsid protein \| Phage major capsid protein; afdb-uniprot50 \| AF-A0A6N6T4Y3-F1-MODEL\_V4 \| 1.0 \| 3.809e-16 \| 426 \| 0.13 \| 483 \| 297 \| 21 \| 244 \| 681 \| 5 \| 409 \| Phage major capsid protein \| Phage major capsid protein; afdb-uniprot50 \| AF-A0A223I0W6-F1-MODEL\_V4 \| 1.0 \| 9.57e-18 \| 426 \| 0.163 \| 459 \| 291 \| 21 \| 252 \| 682 \| 6 \| 399 \| Phage major capsid protein, HK97 family \| Phage major capsid protein, HK97 family; afdb-uniprot50 \| AF-A0A5N0UV25-F1-MODEL\_V4 \| 1.0 \| 1.041e-13 \| 425 \| 0.143 \| 278 \| 202 \| 12 \| 431 \| 682 \| 27 \| 294 \| Phage major capsid protein \| Phage major capsid protein; afdb-uniprot50 \| AF-A0A1Y2QCN7-F1-MODEL\_V4 \| 1.0 \| 1.862e-15 \| 425 \| 0.172 \| 376 \| 262 \| 15 \| 343 \| 679 \| 5 \| 370 \| Phage major capsid protein \| Phage major capsid protein; afdb-uniprot50 \| AF-A0A7C0VZ44-F1-MODEL\_V4 \| 1.0 \| 3.599e-16 \| 425 \| 0.154 \| 435 \| 273 \| 18 \| 295 \| 681 \| 9 \| 396 \| Phage major capsid protein \| Phage major capsid protein; afdb-uniprot50 \| AF-A0A5C5XQ98-F1-MODEL\_V4 \| 1.0 \| 3.213e-16 \| 425 \| 0.153 \| 464 \| 271 \| 23 \| 261 \| 682 \| 10 \| 393 \| Phage capsid family protein \| Phage capsid family protein; afdb-uniprot50 \| AF-A0A5M6HTD8-F1-MODEL\_V4 \| 1.0 \| 3.602e-24 \| 425 \| 0.149 \| 734 \| 449 \| 36 \| 19 \| 681 \| 3 \| 631 \| Peptidase U35 \| Peptidase U35; afdb-uniprot50 \| AF-A0A4Q7ZQ54-F1-MODEL\_V4 \| 1.0 \| 8.727e-17 \| 424 \| 0.155 \| 476 \| 270 \| 25 \| 264 \| 682 \| 8 \| 408 \| HK97 family phage major capsid protein \| HK97 family phage major capsid protein; afdb-uniprot50 \| AF-A0A1G7YG34-F1-MODEL\_V4 \| 1.0 \| 1.2e-17 \| 424 \| 0.176 \| 505 \| 307 \| 23 \| 252 \| 682 \| 8 \| 477 \| Phage major capsid protein, HK97 family \| Phage major capsid protein, HK97 family; afdb-uniprot50 \| AF-A0A0G1VYU8-F1-MODEL\_V4 \| 1.0 \| 7.257e-15 \| 423 \| 0.118 \| 381 \| 273 \| 16 \| 335 \| 682 \| 56 \| 406 \| Uncharacterized protein \| Uncharacterized protein; afdb-uniprot50 \| AF-A0A127R0F9-F1-MODEL\_V4 \| 1.0 \| 8.727e-17 \| 423 \| 0.152 \| 510 \| 333 \| 20 \| 193 \| 679 \| 22 \| 455 \| Phage major capsid protein, HK97 family \| Phage major capsid protein, HK97 family; afdb-uniprot50 \| AF-A0A0Q6RSL7-F1-MODEL\_V4 \| 1.0 \| 2.088e-23 \| 423 \| 0.17 \| 763 \| 424 \| 40 \| 1 \| 681 \| 1 \| 636 \| Uncharacterized protein \| Uncharacterized protein; afdb-uniprot50 \| AF-A0A4Q6IAB5-F1-MODEL\_V4 \| 1.0 \| 2.352e-12 \| 422 \| 0.15 \| 273 \| 192 \| 13 \| 431 \| 682 \| 129 \| 382 \| Phage major capsid protein \| Phage major capsid protein; afdb-uniprot50 \| AF-V8CV52-F1-MODEL\_V4 \| 1.0 \| 2.769e-15 \| 422 \| 0.115 \| 460 \| 308 \| 16 \| 267 \| 681 \| 2 \| 407 \| Head protein \| Head protein; afdb-uniprot50 \| AF-A0A6M3JT93-F1-MODEL\_V4 \| 1.0 \| 5.352e-16 \| 422 \| 0.145 \| 482 \| 282 \| 25 \| 258 \| 680 \| 8 \| 418 \| Putative capsid protein \| Putative capsid protein; afdb-uniprot50 \| AF-A0A6G6Z5J1-F1-MODEL\_V4 \| 1.0 \| 9.496e-13 \| 421 \| 0.156 \| 287 \| 194 \| 10 \| 433 \| 680 \| 90 \| 367 \| Phage major capsid protein \| Phage major capsid protein; afdb-uniprot50 \| AF-A0A0S8JG41-F1-MODEL\_V4 \| 1.0 \| 1.183e-15 \| 421 \| 0.155 \| 458 \| 295 \| 22 \| 252 \| 681 \| 16 \| 409 \| Uncharacterized protein \| Uncharacterized protein; afdb-uniprot50 \| AF-A0A812Y0T0-F1-MODEL\_V4 \| 1.0 \| 6.811e-18 \| 421 \| 0.181 \| 479 \| 293 \| 22 \| 246 \| 682 \| 8 \| 429 \| Hypothetical protein \| Hypothetical protein; afdb-uniprot50 \| AF-A0A0L8EZU3-F1-MODEL\_V4 \| 1.0 \| 4.357e-15 \| 420 \| 0.128 \| 443 \| 297 \| 20 \| 264 \| 682 \| 6 \| 383 \| Uncharacterized protein \| Uncharacterized protein; afdb-uniprot50 \| AF-C2PQK7-F1-MODEL\_V4 \| 1.0 \| 1.354e-14 \| 420 \| 0.133 \| 404 \| 274 \| 14 \| 335 \| 682 \| 1 \| 384 \| Phage major capsid protein \| Phage major capsid protein; afdb-uniprot50 \| AF-A0A1I0J9W7-F1-MODEL\_V4 \| 1.0 \| 3.477e-23 \| 420 \| 0.17 \| 716 \| 411 \| 38 \| 40 \| 682 \| 24 \| 629 \| Prohead serine protease \| Prohead serine protease; afdb-uniprot50 \| AF-A0A0F9MYG6-F1-MODEL\_V4 \| 1.0 \| 5.24e-17 \| 419 \| 0.163 \| 446 \| 262 \| 22 \| 284 \| 681 \| 19 \| 401 \| Uncharacterized protein \| Uncharacterized protein; afdb-uniprot50 \| AF-A0A419HBW7-F1-MODEL\_V4 \| 1.0 \| 6.573e-17 \| 419 \| 0.164 \| 505 \| 313 \| 25 \| 250 \| 681 \| 2 \| 470 \| Phage major capsid protein \| Phage major capsid protein; afdb-uniprot50 \| AF-A0A840CSE2-F1-MODEL\_V4 \| 1.0 \| 3.524e-17 \| 419 \| 0.13 \| 460 \| 300 \| 18 \| 249 \| 679 \| 97 \| 485 \| HK97 family phage major capsid protein \| HK97 family phage major capsid protein; afdb-uniprot50 \| AF-A0A654D1K5-F1-MODEL\_V4 \| 1.0 \| 5.664e-16 \| 418 \| 0.159 \| 445 \| 306 \| 17 \| 250 \| 679 \| 14 \| 405 \| Putative Phage major capsid protein, HK97 family \| Putative Phage major capsid protein, HK97 family; afdb-uniprot50 \| AF-A0A810K9P5-F1-MODEL\_V4 \| 1.0 \| 3.401e-16 \| 418 \| 0.16 \| 435 \| 281 \| 20 \| 294 \| 678 \| 19 \| 419 \| Uncharacterized protein \| Uncharacterized protein; afdb-uniprot50 \| AF-A0A437LLE3-F1-MODEL\_V4 \| 1.0 \| 2.207e-15 \| 417 \| 0.14 \| 450 \| 297 \| 19 \| 251 \| 679 \| 3 \| 383 \| Phage major capsid protein \| Phage major capsid protein; afdb-uniprot50 \| AF-A0A2E7HPZ8-F1-MODEL\_V4 \| 1.0 \| 1.298e-16 \| 417 \| 0.147 \| 455 \| 285 \| 20 \| 279 \| 679 \| 18 \| 423 \| Phage major capsid protein \| Phage major capsid protein; afdb-uniprot50 \| AF-A0A2E0X6G4-F1-MODEL\_V4 \| 1.0 \| 7.791e-17 \| 417 \| 0.134 \| 470 \| 304 \| 23 \| 280 \| 680 \| 26 \| 461 \| Phage major capsid protein \| Phage major capsid protein; afdb-uniprot50 \| AF-A0A1X1W1I0-F1-MODEL\_V4 \| 1.0 \| 1.875e-12 \| 416 \| 0.151 \| 264 \| 199 \| 9 \| 431 \| 682 \| 29 \| 279 \| Major capsid protein \| Major capsid protein; afdb-uniprot50 \| AF-A0A5X1QJ83-F1-MODEL\_V4 \| 1.0 \| 1.76e-15 \| 416 \| 0.152 \| 445 \| 296 \| 17 \| 265 \| 682 \| 9 \| 399 \| Phage major capsid protein \| Phage major capsid protein; afdb-uniprot50 \| AF-A0A7X4F2B2-F1-MODEL\_V4 \| 1.0 \| 4.678e-17 \| 416 \| 0.149 \| 481 \| 289 \| 24 \| 264 \| 682 \| 5 \| 427 \| Phage major capsid protein \| Phage major capsid protein; afdb-uniprot50 \| AF-A0A519P9Z1-F1-MODEL\_V4 \| 1.0 \| 7.791e-17 \| 416 \| 0.148 \| 437 \| 299 \| 18 \| 306 \| 679 \| 4 \| 430 \| Phage major capsid protein \| Phage major capsid protein; afdb-uniprot50 \| AF-A0A431MZP7-F1-MODEL\_V4 \| 1.0 \| 2.161e-16 \| 415 \| 0.156 \| 461 \| 295 \| 20 \| 249 \| 682 \| 2 \| 395 \| Phage major capsid protein \| Phage major capsid protein; afdb-uniprot50 \| AF-J2G792-F1-MODEL\_V4 \| 1.0 \| 2.42e-16 \| 415 \| 0.144 \| 513 \| 323 \| 27 \| 198 \| 679 \| 2 \| 429 \| Phage major capsid protein, HK97 family \| Phage major capsid protein, HK97 family; afdb-uniprot50 \| AF-A0A239JPQ6-F1-MODEL\_V4 \| 1.0 \| 9.042e-18 \| 415 \| 0.201 \| 481 \| 323 \| 17 \| 182 \| 648 \| 3 \| 436 \| Phage major capsid protein, HK97 family \| Phage major capsid protein, HK97 family; afdb-uniprot50 \| AF-A0A554S931-F1-MODEL\_V4 \| 1.0 \| 8.422e-16 \| 414 \| 0.141 \| 459 \| 297 \| 18 \| 252 \| 682 \| 35 \| 424 \| Phage major capsid protein \| Phage major capsid protein; afdb-uniprot50 \| AF-A0A0F9WTB8-F1-MODEL\_V4 \| 1.0 \| 4.678e-17 \| 414 \| 0.156 \| 485 \| 314 \| 24 \| 256 \| 682 \| 2 \| 449 \| Uncharacterized protein \| Uncharacterized protein; afdb-uniprot50 \| AF-A0A353J4Q1-F1-MODEL\_V4 \| 1.0 \| 7.628e-18 \| 414 \| 0.146 \| 465 \| 295 \| 19 \| 252 \| 680 \| 19 \| 417 \| Phage major capsid protein \| Phage major capsid protein; afdb-uniprot50 \| AF-A0A7V7XCR5-F1-MODEL\_V4 \| 1.0 \| 1.298e-16 \| 413 \| 0.146 \| 457 \| 295 \| 20 \| 285 \| 682 \| 20 \| 440 \| Phage major capsid protein \| Phage major capsid protein; afdb-uniprot50 \| AF-A0A0F9MYU0-F1-MODEL\_V4 \| 1.0 \| 1.902e-14 \| 412 \| 0.137 \| 343 \| 248 \| 13 \| 375 \| 682 \| 16 \| 345 \| Uncharacterized protein \| Uncharacterized protein; afdb-uniprot50 \| AF-C7N4P4-F1-MODEL\_V4 \| 1.0 \| 3.036e-16 \| 411 \| 0.169 \| 460 \| 277 \| 24 \| 262 \| 682 \| 9 \| 402 \| Phage major capsid protein, HK97 family \| Phage major capsid protein, HK97 family; afdb-uniprot50 \| AF-A0A0F8Z7I4-F1-MODEL\_V4 \| 1.0 \| 6.759e-13 \| 410 \| 0.166 \| 301 \| 209 \| 15 \| 412 \| 680 \| 1 \| 291 \| Uncharacterized protein \| Uncharacterized protein; afdb-uniprot50 \| AF-A0A1C4B7A3-F1-MODEL\_V4 \| 1.0 \| 2.93e-15 \| 410 \| 0.155 \| 456 \| 278 \| 22 \| 258 \| 682 \| 4 \| 383 \| Phage major capsid protein, HK97 family \| Phage major capsid protein, HK97 family; afdb-uniprot50 \| AF-A0A1V5PUE1-F1-MODEL\_V4 \| 1.0 \| 5.466e-15 \| 410 \| 0.155 \| 374 \| 235 \| 14 \| 379 \| 682 \| 21 \| 383 \| Phage capsid family protein \| Phage capsid family protein; afdb-uniprot50 \| AF-A0A644VRL6-F1-MODEL\_V4 \| 1.0 \| 7.57e-13 \| 410 \| 0.19 \| 278 \| 184 \| 10 \| 431 \| 681 \| 33 \| 296 \| S-l\_SbsC\_C domain-containing protein \| S-l\_SbsC\_C domain-containing protein; afdb-uniprot50 \| AF-A0A5Q0DKA8-F1-MODEL\_V4 \| 1.0 \| 1.079e-14 \| 410 \| 0.144 \| 380 \| 279 \| 13 \| 338 \| 681 \| 27 \| 396 \| Phage major capsid protein \| Phage major capsid protein; afdb-uniprot50 \| AF-A0A1M3EVZ0-F1-MODEL\_V4 \| 1.0 \| 4.177e-17 \| 410 \| 0.143 \| 508 \| 322 \| 27 \| 252 \| 677 \| 4 \| 480 \| Uncharacterized protein \| Uncharacterized protein; afdb-uniprot50 \| AF-A0A327QB84-F1-MODEL\_V4 \| 1.0 \| 2.872e-24 \| 410 \| 0.168 \| 724 \| 396 \| 43 \| 33 \| 681 \| 8 \| 600 \| Mu-like prophage major head subunit gpT \| Mu-like prophage major head subunit gpT; afdb-uniprot50 \| AF-A0A5C7T964-F1-MODEL\_V4 \| 1.0 \| 2.42e-16 \| 409 \| 0.139 \| 460 \| 318 \| 17 \| 234 \| 681 \| 9 \| 402 \| Phage major capsid protein \| Phage major capsid protein; afdb-uniprot50 \| AF-A0A0Q6UYN3-F1-MODEL\_V4 \| 1.0 \| 1.572e-23 \| 409 \| 0.145 \| 790 \| 474 \| 33 \| 17 \| 681 \| 14 \| 727 \| Uncharacterized protein \| Uncharacterized protein; afdb-uniprot50 \| AF-A0A143YDP5-F1-MODEL\_V4 \| 1.0 \| 5.466e-15 \| 408 \| 0.166 \| 373 \| 246 \| 13 \| 335 \| 682 \| 1 \| 333 \| Phage capsid \| Phage capsid; afdb-uniprot50 \| AF-A0A1X0K6U6-F1-MODEL\_V4 \| 1.0 \| 3.473e-15 \| 408 \| 0.131 \| 411 \| 296 \| 16 \| 295 \| 682 \| 19 \| 391 \| Uncharacterized protein \| Uncharacterized protein; afdb-uniprot50 \| AF-A0A7J5TYE1-F1-MODEL\_V4 \| 1.0 \| 4.177e-17 \| 408 \| 0.13 \| 474 \| 301 \| 21 \| 252 \| 679 \| 46 \| 454 \| Phage major capsid protein \| Phage major capsid protein; afdb-uniprot50 \| AF-A0A645AX28-F1-MODEL\_V4 \| 1.0 \| 2.254e-14 \| 407 \| 0.16 \| 386 \| 268 \| 16 \| 336 \| 681 \| 1 \| 370 \| Uncharacterized protein \| Uncharacterized protein; afdb-uniprot50 \| AF-A0A2M8TG61-F1-MODEL\_V4 \| 1.0 \| 6.714e-16 \| 407 \| 0.109 \| 482 \| 309 \| 25 \| 233 \| 681 \| 22 \| 416 \| Phage major capsid protein \| Phage major capsid protein; afdb-uniprot50 \| AF-A0A6L6ES98-F1-MODEL\_V4 \| 1.0 \| 3.575e-19 \| 407 \| 0.173 \| 628 \| 362 \| 28 \| 171 \| 681 \| 61 \| 648 \| Phage major capsid protein \| Phage major capsid protein; afdb-uniprot50 \| AF-A0A7V8XZY4-F1-MODEL\_V4 \| 1.0 \| 5.018e-11 \| 406 \| 0.214 \| 210 \| 149 \| 7 \| 480 \| 682 \| 7 \| 207 \| Phage major capsid protein \| Phage major capsid protein; afdb-uniprot50 \| AF-A0A4T0UXI4-F1-MODEL\_V4 \| 1.0 \| 1.915e-11 \| 406 \| 0.109 \| 256 \| 207 \| 9 \| 430 \| 679 \| 13 \| 253 \| Phage major capsid protein \| Phage major capsid protein; afdb-uniprot50 \| AF-A0A327SK31-F1-MODEL\_V4 \| 1.0 \| 7.519e-16 \| 406 \| 0.151 \| 450 \| 301 \| 21 \| 249 \| 679 \| 21 \| 408 \| HK97 family phage major capsid protein \| HK97 family phage major capsid protein; afdb-uniprot50 \| AF-A0A7X3Y810-F1-MODEL\_V4 \| 1.0 \| 1.034e-16 \| 406 \| 0.168 \| 463 \| 290 \| 25 \| 267 \| 682 \| 4 \| 418 \| Uncharacterized protein \| Uncharacterized protein; afdb-uniprot50 \| AF-A0A858WUY8-F1-MODEL\_V4 \| 1.0 \| 4.815e-21 \| 406 \| 0.128 \| 701 \| 444 \| 40 \| 40 \| 682 \| 19 \| 610 \| Peptidase \| Peptidase; afdb-uniprot50 \| AF-R4KAR4-F1-MODEL\_V4 \| 1.0 \| 7.105e-16 \| 405 \| 0.148 \| 459 \| 298 \| 23 \| 262 \| 682 \| 11 \| 414 \| Putative phage phi-C31 gp36 major capsid-like protein \| Putative phage phi-C31 gp36 major capsid-like protein; afdb-uniprot50 \| AF-A0A3D4JJQ9-F1-MODEL\_V4 \| 1.0 \| 8.012e-13 \| 404 \| 0.132 \| 257 \| 192 \| 12 \| 435 \| 681 \| 61 \| 296 \| Phage major capsid protein \| Phage major capsid protein; afdb-uniprot50 \| AF-A0A5R8Y4J3-F1-MODEL\_V4 \| 1.0 \| 1.971e-15 \| 404 \| 0.145 \| 454 \| 270 \| 22 \| 264 \| 681 \| 13 \| 384 \| Phage major capsid protein \| Phage major capsid protein; afdb-uniprot50 \| AF-A0A401HW27-F1-MODEL\_V4 \| 1.0 \| 4.357e-15 \| 404 \| 0.148 \| 405 \| 273 \| 19 \| 335 \| 682 \| 1 \| 390 \| Putative phage major capsid protein \| Putative phage major capsid protein; afdb-uniprot50 \| AF-A0A496WAE6-F1-MODEL\_V4 \| 1.0 \| 1.663e-15 \| 404 \| 0.141 \| 481 \| 287 \| 22 \| 249 \| 681 \| 2 \| 404 \| Phage major capsid protein \| Phage major capsid protein; afdb-uniprot50 \| AF-A0A527P893-F1-MODEL\_V4 \| 1.0 \| 4.357e-15 \| 404 \| 0.158 \| 453 \| 306 \| 17 \| 246 \| 679 \| 12 \| 408 \| Phage major capsid protein \| Phage major capsid protein; afdb-uniprot50 \| AF-A0A1Y3RFP0-F1-MODEL\_V4 \| 1.0 \| 8.913e-16 \| 403 \| 0.142 \| 442 \| 284 \| 21 \| 264 \| 682 \| 26 \| 395 \| Phage major capsid protein \| Phage major capsid protein; afdb-uniprot50 \| AF-A0A1H7A3R3-F1-MODEL\_V4 \| 1.0 \| 1.929e-16 \| 403 \| 0.146 \| 485 \| 299 \| 23 \| 251 \| 682 \| 2 \| 424 \| Phage major capsid protein, HK97 family \| Phage major capsid protein, HK97 family; afdb-uniprot50 \| AF-A0A1Q8DAV2-F1-MODEL\_V4 \| 1.0 \| 4.88e-15 \| 402 \| 0.138 \| 412 \| 277 \| 19 \| 295 \| 681 \| 5 \| 363 \| Phage capsid protein \| Phage capsid protein; afdb-uniprot50 \| AF-A0A522EXF1-F1-MODEL\_V4 \| 1.0 \| 1.209e-14 \| 402 \| 0.141 \| 480 \| 268 \| 22 \| 258 \| 679 \| 2 \| 395 \| Phage major capsid protein \| Phage major capsid protein; afdb-uniprot50 \| AF-A0A4R4I9T1-F1-MODEL\_V4 \| 1.0 \| 2.711e-16 \| 402 \| 0.149 \| 476 \| 301 \| 21 \| 238 \| 679 \| 57 \| 462 \| Phage major capsid protein \| Phage major capsid protein; afdb-uniprot50 \| AF-A0A256C0Q1-F1-MODEL\_V4 \| 1.0 \| 3.89e-15 \| 401 \| 0.137 \| 459 \| 304 \| 18 \| 252 \| 681 \| 6 \| 401 \| Phage major capsid protein \| Phage major capsid protein; afdb-uniprot50 \| AF-A0A6L5FY65-F1-MODEL\_V4 \| 1.0 \| 4.515e-16 \| 401 \| 0.162 \| 486 \| 263 \| 26 \| 271 \| 682 \| 4 \| 419 \| Phage major capsid protein \| Phage major capsid protein; afdb-uniprot50 \| AF-A0A7C1VPJ3-F1-MODEL\_V4 \| 1.0 \| 3.036e-16 \| 401 \| 0.144 \| 490 \| 291 \| 25 \| 252 \| 681 \| 2 \| 423 \| Phage major capsid protein \| Phage major capsid protein; afdb-uniprot50 \| AF-A0A5K7Z4E0-F1-MODEL\_V4 \| 1.0 \| 1.929e-16 \| 401 \| 0.139 \| 489 \| 286 \| 30 \| 264 \| 681 \| 5 \| 429 \| Phage capsid protein \| Phage capsid protein; afdb-uniprot50 \| AF-A0A1F5QST1-F1-MODEL\_V4 \| 1.0 \| 6.956e-17 \| 401 \| 0.148 \| 490 \| 305 \| 21 \| 262 \| 682 \| 10 \| 456 \| Uncharacterized protein \| Uncharacterized protein; afdb-uniprot50 \| AF-A0A2W6SDJ2-F1-MODEL\_V4 \| 1.0 \| 5.062e-24 \| 401 \| 0.147 \| 774 \| 446 \| 42 \| 18 \| 681 \| 30 \| 699 \| Uncharacterized protein \| Uncharacterized protein; afdb-uniprot50 \| AF-A0A370KRT5-F1-MODEL\_V4 \| 1.0 \| 5.091e-13 \| 400 \| 0.173 \| 283 \| 180 \| 10 \| 433 \| 682 \| 9 \| 270 \| Uncharacterized protein \| Uncharacterized protein; afdb-uniprot50 \| AF-A0A1P8PSS7-F1-MODEL\_V4 \| 1.0 \| 2.769e-15 \| 400 \| 0.145 \| 461 \| 278 \| 20 \| 262 \| 681 \| 5 \| 390 \| Uncharacterized protein \| Uncharacterized protein; afdb-uniprot50 \| AF-A0A518EKY1-F1-MODEL\_V4 \| 1.0 \| 3.307e-20 \| 400 \| 0.186 \| 537 \| 333 \| 26 \| 30 \| 516 \| 12 \| 494 \| Caudovirus prohead protease \| Caudovirus prohead protease; afdb-uniprot50 \| AF-A0A7I8Z983-F1-MODEL\_V4 \| 1.0 \| 2.636e-20 \| 399 \| 0.226 \| 516 \| 275 \| 24 \| 19 \| 482 \| 1 \| 444 \| Hypothetical protein \| Hypothetical protein; afdb-uniprot50 \| AF-A0A2G8MBD4-F1-MODEL\_V4 \| 1.0 \| 2.95e-12 \| 398 \| 0.274 \| 204 \| 144 \| 4 \| 12 \| 213 \| 17 \| 218 \| Uncharacterized protein \| Uncharacterized protein; afdb-uniprot50 \| AF-A0A0F8Y0I6-F1-MODEL\_V4 \| 1.0 \| 1.412e-12 \| 398 \| 0.116 \| 308 \| 242 \| 14 \| 389 \| 679 \| 28 \| 322 \| Uncharacterized protein \| Uncharacterized protein; afdb-uniprot50 \| AF-R6SUA7-F1-MODEL\_V4 \| 1.0 \| 2.93e-15 \| 398 \| 0.126 \| 458 \| 281 \| 24 \| 264 \| 681 \| 2 \| 380 \| Phage major capsid protein HK97 family \| Phage major capsid protein HK97 family; afdb-uniprot50 \| AF-A9NFU9-F1-MODEL\_V4 \| 1.0 \| 3.167e-14 \| 397 \| 0.138 \| 397 \| 261 \| 15 \| 338 \| 682 \| 8 \| 375 \| Uncharacterized protein \| Uncharacterized protein; afdb-uniprot50 \| AF-A0A1Q3QQ17-F1-MODEL\_V4 \| 1.0 \| 5.466e-15 \| 397 \| 0.147 \| 467 \| 275 \| 26 \| 271 \| 682 \| 3 \| 401 \| Uncharacterized protein \| Uncharacterized protein; afdb-uniprot50 \| AF-A0A502HSQ5-F1-MODEL\_V4 \| 1.0 \| 4.811e-13 \| 397 \| 0.122 \| 318 \| 232 \| 13 \| 392 \| 677 \| 106 \| 408 \| Phage major capsid protein \| Phage major capsid protein; afdb-uniprot50 \| AF-A0A3R9P8L0-F1-MODEL\_V4 \| 1.0 \| 7.257e-15 \| 397 \| 0.141 \| 423 \| 304 \| 15 \| 307 \| 682 \| 9 \| 419 \| Phage major capsid protein \| Phage major capsid protein; afdb-uniprot50 \| AF-A0A0Q6CCC3-F1-MODEL\_V4 \| 1.0 \| 5.466e-15 \| 397 \| 0.151 \| 501 \| 322 \| 21 \| 196 \| 679 \| 1 \| 415 \| Uncharacterized protein \| Uncharacterized protein; afdb-uniprot50 \| AF-A0A7C6A307-F1-MODEL\_V4 \| 1.0 \| 1.736e-21 \| 397 \| 0.137 \| 725 \| 394 \| 39 \| 41 \| 681 \| 22 \| 598 \| Peptidase U35 \| Peptidase U35; afdb-uniprot50 \| AF-A0A2D6X839-F1-MODEL\_V4 \| 1.0 \| 2.993e-14 \| 396 \| 0.167 \| 334 \| 218 \| 17 \| 400 \| 681 \| 15 \| 340 \| Phage major capsid protein \| Phage major capsid protein; afdb-uniprot50 \| AF-A0A3B8XY37-F1-MODEL\_V4 \| 1.0 \| 2.013e-14 \| 396 \| 0.124 \| 379 \| 267 \| 19 \| 338 \| 681 \| 9 \| 357 \| Uncharacterized protein \| Uncharacterized protein; afdb-uniprot50 \| AF-A0A7C4KPZ2-F1-MODEL\_V4 \| 1.0 \| 4.611e-15 \| 396 \| 0.129 \| 472 \| 292 \| 22 \| 251 \| 681 \| 2 \| 395 \| Phage major capsid protein \| Phage major capsid protein; afdb-uniprot50 \| AF-A0A6L9XYJ8-F1-MODEL\_V4 \| 1.0 \| 8.246e-17 \| 396 \| 0.161 \| 490 \| 324 \| 26 \| 249 \| 681 \| 28 \| 487 \| Phage major capsid protein \| Phage major capsid protein; afdb-uniprot50 \| AF-A0A1G3FT84-F1-MODEL\_V4 \| 1.0 \| 9.433e-16 \| 395 \| 0.245 \| 407 \| 237 \| 19 \| 255 \| 635 \| 5 \| 367 \| Uncharacterized protein \| Uncharacterized protein; afdb-uniprot50 \| AF-A0A2D9H4D9-F1-MODEL\_V4 \| 1.0 \| 6.122e-15 \| 395 \| 0.136 \| 389 \| 265 \| 16 \| 338 \| 682 \| 21 \| 382 \| Phage major capsid protein \| Phage major capsid protein; afdb-uniprot50 \| AF-A0A3D5CRX3-F1-MODEL\_V4 \| 1.0 \| 7.958e-16 \| 395 \| 0.137 \| 464 \| 298 \| 21 \| 252 \| 681 \| 23 \| 418 \| Phage major capsid protein \| Phage major capsid protein; afdb-uniprot50 \| AF-A0A1M4Y9R8-F1-MODEL\_V4 \| 1.0 \| 8.246e-17 \| 395 \| 0.168 \| 492 \| 290 \| 24 \| 252 \| 682 \| 48 \| 481 \| Phage major capsid protein, HK97 family \| Phage major capsid protein, HK97 family; afdb-uniprot50 \| AF-A0A497LYT2-F1-MODEL\_V4 \| 1.0 \| 3.304e-12 \| 394 \| 0.092 \| 293 \| 235 \| 7 \| 398 \| 682 \| 3 \| 272 \| Uncharacterized protein \| Uncharacterized protein; afdb-uniprot50 \| AF-A0A1F8UQS0-F1-MODEL\_V4 \| 1.0 \| 1.823e-16 \| 394 \| 0.136 \| 463 \| 315 \| 17 \| 252 \| 680 \| 23 \| 434 \| Uncharacterized protein \| Uncharacterized protein; afdb-uniprot50 \| AF-A0A3S3AHQ4-F1-MODEL\_V4 \| 1.0 \| 3.33e-17 \| 394 \| 0.152 \| 532 \| 347 \| 20 \| 171 \| 681 \| 67 \| 515 \| Phage major capsid protein \| Phage major capsid protein; afdb-uniprot50 \| AF-A0A653F5R1-F1-MODEL\_V4 \| 1.0 \| 4.145e-12 \| 393 \| 0.154 \| 266 \| 196 \| 13 \| 433 \| 682 \| 32 \| 284 \| Phage capsid family protein \| Phage capsid family protein; afdb-uniprot50 \| AF-G4I9Y4-F1-MODEL\_V4 \| 1.0 \| 1.734e-13 \| 392 \| 0.167 \| 341 \| 213 \| 14 \| 398 \| 682 \| 6 \| 331 \| Uncharacterized protein \| Uncharacterized protein; afdb-uniprot50 \| AF-A0A522Q9N9-F1-MODEL\_V4 \| 1.0 \| 1.373e-16 \| 392 \| 0.172 \| 555 \| 327 \| 32 \| 191 \| 681 \| 1 \| 486 \| Phage major capsid protein \| Phage major capsid protein; afdb-uniprot50 \| AF-H2JIC2-F1-MODEL\_V4 \| 1.0 \| 4.811e-13 \| 391 \| 0.117 \| 340 \| 241 \| 15 \| 349 \| 681 \| 10 \| 297 \| Phage capsid family protein \| Phage capsid family protein; afdb-uniprot50 \| AF-A0A7C3UHG4-F1-MODEL\_V4 \| 1.0 \| 5.165e-15 \| 391 \| 0.154 \| 383 \| 259 \| 20 \| 338 \| 681 \| 13 \| 369 \| Phage major capsid protein \| Phage major capsid protein; afdb-uniprot50 \| AF-A0A3M2A8H0-F1-MODEL\_V4 \| 1.0 \| 2.93e-15 \| 391 \| 0.165 \| 464 \| 283 \| 25 \| 252 \| 682 \| 2 \| 394 \| Phage major capsid protein \| Phage major capsid protein; afdb-uniprot50 \| AF-I0GRZ0-F1-MODEL\_V4 \| 1.0 \| 4.88e-15 \| 391 \| 0.134 \| 460 \| 297 \| 18 \| 252 \| 682 \| 48 \| 435 \| Putative phage major capsid protein \| Putative phage major capsid protein; afdb-uniprot50 \| AF-A0A522RY69-F1-MODEL\_V4 \| 1.0 \| 6.211e-17 \| 391 \| 0.156 \| 498 \| 303 \| 26 \| 244 \| 681 \| 19 \| 459 \| Phage major capsid protein \| Phage major capsid protein; afdb-uniprot50 \| AF-A0A352DNU6-F1-MODEL\_V4 \| 1.0 \| 9.433e-16 \| 390 \| 0.138 \| 476 \| 282 \| 23 \| 251 \| 680 \| 35 \| 428 \| Phage major capsid protein \| Phage major capsid protein; afdb-uniprot50 \| AF-A0A0S8GFL0-F1-MODEL\_V4 \| 1.0 \| 1.118e-15 \| 390 \| 0.137 \| 494 \| 295 \| 25 \| 266 \| 682 \| 2 \| 441 \| Uncharacterized protein \| Uncharacterized protein; afdb-uniprot50 \| AF-A0A6F9E8C9-F1-MODEL\_V4 \| 1.0 \| 1.118e-15 \| 389 \| 0.156 \| 447 \| 292 \| 20 \| 262 \| 682 \| 19 \| 406 \| Phage major capsid protein, HK97 family \| Phage major capsid protein, HK97 family; afdb-uniprot50 \| AF-A0A2B0JB50-F1-MODEL\_V4 \| 1.0 \| 1.412e-12 \| 388 \| 0.127 \| 267 \| 199 \| 11 \| 433 \| 679 \| 43 \| 295 \| Phage major capsid protein \| Phage major capsid protein; afdb-uniprot50 \| AF-A0A1Y4UTW1-F1-MODEL\_V4 \| 1.0 \| 5.275e-14 \| 388 \| 0.122 \| 473 \| 295 \| 24 \| 252 \| 681 \| 3 \| 398 \| Phage major capsid protein \| Phage major capsid protein; afdb-uniprot50 \| AF-A0A7X0H6V8-F1-MODEL\_V4 \| 1.0 \| 2.386e-14 \| 388 \| 0.173 \| 387 \| 243 \| 19 \| 332 \| 682 \| 86 \| 431 \| HK97 family phage major capsid protein \| HK97 family phage major capsid protein; afdb-uniprot50 \| AF-A0A1I5HWK7-F1-MODEL\_V4 \| 1.0 \| 5.664e-16 \| 386 \| 0.185 \| 470 \| 267 \| 24 \| 268 \| 681 \| 6 \| 415 \| Phage major capsid protein, HK97 family \| Phage major capsid protein, HK97 family; afdb-uniprot50 \| AF-A0A6N9Q8R2-F1-MODEL\_V4 \| 1.0 \| 6.857e-15 \| 386 \| 0.117 \| 468 \| 320 \| 22 \| 262 \| 681 \| 4 \| 426 \| Phage major capsid protein \| Phage major capsid protein; afdb-uniprot50 \| AF-A0A436GWX9-F1-MODEL\_V4 \| 1.0 \| 5.785e-15 \| 386 \| 0.131 \| 455 \| 305 \| 20 \| 252 \| 682 \| 66 \| 454 \| Phage major capsid protein \| Phage major capsid protein; afdb-uniprot50 \| AF-A0A7C3UYK7-F1-MODEL\_V4 \| 1.0 \| 1.02e-14 \| 385 \| 0.259 \| 385 \| 199 \| 16 \| 282 \| 643 \| 40 \| 361 \| Phage major capsid protein \| Phage major capsid protein; afdb-uniprot50 \| AF-A0A4Q3H3Y4-F1-MODEL\_V4 \| 1.0 \| 1.748e-18 \| 384 \| 0.3 \| 390 \| 178 \| 9 \| 30 \| 397 \| 2 \| 318 \| Uncharacterized protein \| Uncharacterized protein; afdb-uniprot50 \| AF-A0A2A5S652-F1-MODEL\_V4 \| 1.0 \| 1.142e-14 \| 384 \| 0.146 \| 390 \| 248 \| 20 \| 312 \| 681 \| 2 \| 326 \| Uncharacterized protein \| Uncharacterized protein; afdb-uniprot50 \| AF-A0A6M1WCW4-F1-MODEL\_V4 \| 1.0 \| 1.209e-14 \| 384 \| 0.127 \| 461 \| 278 \| 23 \| 264 \| 681 \| 2 \| 381 \| Phage major capsid protein \| Phage major capsid protein; afdb-uniprot50 \| AF-A0A1Q9JGH3-F1-MODEL\_V4 \| 1.0 \| 2.254e-14 \| 384 \| 0.119 \| 420 \| 268 \| 19 \| 335 \| 681 \| 1 \| 391 \| Uncharacterized protein \| Uncharacterized protein; afdb-uniprot50 \| AF-A0A2S3VZY0-F1-MODEL\_V4 \| 1.0 \| 2.809e-17 \| 384 \| 0.16 \| 505 \| 307 \| 20 \| 252 \| 678 \| 6 \| 471 \| Phage capsid family protein \| Phage capsid family protein; afdb-uniprot50 \| AF-A0A1G8R0Y5-F1-MODEL\_V4 \| 1.0 \| 9.103e-15 \| 383 \| 0.124 \| 450 \| 312 \| 20 \| 249 \| 682 \| 2 \| 385 \| Phage major capsid protein, HK97 family \| Phage major capsid protein, HK97 family; afdb-uniprot50 \| AF-A0A2A8XVM2-F1-MODEL\_V4 \| 1.0 \| 9.298e-14 \| 383 \| 0.131 \| 387 \| 280 \| 15 \| 338 \| 681 \| 12 \| 385 \| Phage major capsid protein \| Phage major capsid protein; afdb-uniprot50 \| AF-A0A859DMT2-F1-MODEL\_V4 \| 1.0 \| 2.769e-15 \| 383 \| 0.162 \| 444 \| 292 \| 19 \| 262 \| 682 \| 19 \| 405 \| Phage major capsid protein \| Phage major capsid protein; afdb-uniprot50 \| AF-A0A3S0UJX5-F1-MODEL\_V4 \| 1.0 \| 1.403e-15 \| 381 \| 0.248 \| 390 \| 235 \| 18 \| 252 \| 620 \| 2 \| 354 \| Phage major capsid protein \| Phage major capsid protein; afdb-uniprot50 \| AF-A0A1H4BNJ7-F1-MODEL\_V4 \| 1.0 \| 2.254e-14 \| 381 \| 0.149 \| 454 \| 280 \| 22 \| 268 \| 681 \| 3 \| 390 \| Phage major capsid protein, HK97 family \| Phage major capsid protein, HK97 family; afdb-uniprot50 \| AF-A0A644XRB4-F1-MODEL\_V4 \| 1.0 \| 1.279e-14 \| 380 \| 0.114 \| 461 \| 315 \| 23 \| 251 \| 682 \| 5 \| 401 \| Uncharacterized protein \| Uncharacterized protein; afdb-uniprot50 \| AF-A0A7K1ZMJ6-F1-MODEL\_V4 \| 1.0 \| 3.676e-15 \| 380 \| 0.139 \| 458 \| 305 \| 25 \| 264 \| 681 \| 7 \| 415 \| Phage major capsid protein \| Phage major capsid protein; afdb-uniprot50 \| AF-A0A077L3E3-F1-MODEL\_V4 \| 1.0 \| 1.663e-15 \| 380 \| 0.146 \| 484 \| 301 \| 25 \| 256 \| 679 \| 4 \| 435 \| Uncharacterized protein \| Uncharacterized protein; afdb-uniprot50 \| AF-A0A7X4ACZ8-F1-MODEL\_V4 \| 1.0 \| 1.889e-17 \| 380 \| 0.134 \| 567 \| 357 \| 24 \| 172 \| 680 \| 72 \| 562 \| Phage major capsid protein \| Phage major capsid protein; afdb-uniprot50 \| AF-A0A7Z9KVM6-F1-MODEL\_V4 \| 1.0 \| 1.442e-11 \| 379 \| 0.09 \| 287 \| 245 \| 6 \| 407 \| 681 \| 2 \| 284 \| Uncharacterized protein \| Uncharacterized protein; afdb-uniprot50 \| AF-A0A0G1JMI3-F1-MODEL\_V4 \| 1.0 \| 3.473e-15 \| 379 \| 0.139 \| 493 \| 300 \| 28 \| 267 \| 682 \| 3 \| 448 \| HK97 family major capsid protein \| HK97 family major capsid protein; afdb-uniprot50 \| AF-A0A843HQ06-F1-MODEL\_V4 \| 1.0 \| 1.209e-14 \| 378 \| 0.139 \| 467 \| 293 \| 24 \| 254 \| 682 \| 2 \| 397 \| Phage major capsid protein \| Phage major capsid protein; afdb-uniprot50 \| AF-A0A7G1IRW9-F1-MODEL\_V4 \| 1.0 \| 3.497e-12 \| 377 \| 0.128 \| 358 \| 184 \| 11 \| 438 \| 679 \| 36 \| 381 \| Uncharacterized protein \| Uncharacterized protein; afdb-uniprot50 \| AF-A0A3L6ZMY8-F1-MODEL\_V4 \| 1.0 \| 1.433e-14 \| 377 \| 0.149 \| 462 \| 282 \| 22 \| 264 \| 682 \| 7 \| 400 \| Phage major capsid protein \| Phage major capsid protein; afdb-uniprot50 \| AF-A0A0U1L1V2-F1-MODEL\_V4 \| 1.0 \| 1.142e-14 \| 377 \| 0.128 \| 467 \| 297 \| 20 \| 261 \| 680 \| 10 \| 413 \| Phage major capsid protein \| Phage major capsid protein; afdb-uniprot50 \| AF-A0A1X7A4I7-F1-MODEL\_V4 \| 1.0 \| 5.466e-15 \| 377 \| 0.158 \| 472 \| 293 \| 22 \| 264 \| 682 \| 8 \| 428 \| Phage capsid family protein \| Phage capsid family protein; afdb-uniprot50 \| AF-A0A2V0MXJ8-F1-MODEL\_V4 \| 1.0 \| 7.68e-15 \| 377 \| 0.134 \| 475 \| 311 \| 22 \| 251 \| 680 \| 3 \| 422 \| Uncharacterized protein \| Uncharacterized protein; afdb-uniprot50 \| AF-A0A0G1YGY1-F1-MODEL\_V4 \| 1.0 \| 1.484e-15 \| 377 \| 0.142 \| 492 \| 315 \| 25 \| 263 \| 681 \| 5 \| 462 \| Phage major capsid protein, HK97 family \| Phage major capsid protein, HK97 family; afdb-uniprot50 \| AF-A0A1L8I4U9-F1-MODEL\_V4 \| 1.0 \| 1.005e-12 \| 376 \| 0.151 \| 390 \| 248 \| 16 \| 313 \| 682 \| 1 \| 327 \| Uncharacterized protein \| Uncharacterized protein; afdb-uniprot50 \| AF-A0A2V5TLL7-F1-MODEL\_V4 \| 1.0 \| 1.639e-13 \| 376 \| 0.158 \| 397 \| 272 \| 18 \| 313 \| 681 \| 2 \| 364 \| Phage major capsid protein \| Phage major capsid protein; afdb-uniprot50 \| AF-A0A0Q7I3W7-F1-MODEL\_V4 \| 1.0 \| 1.538e-16 \| 376 \| 0.184 \| 494 \| 293 \| 28 \| 252 \| 682 \| 6 \| 452 \| Uncharacterized protein \| Uncharacterized protein; afdb-uniprot50 \| AF-A0A4R5WCP3-F1-MODEL\_V4 \| 1.0 \| 1.956e-10 \| 375 \| 0.146 \| 212 \| 159 \| 8 \| 481 \| 682 \| 5 \| 204 \| Phage major capsid protein \| Phage major capsid protein; afdb-uniprot50 \| AF-A0A6I5WKB9-F1-MODEL\_V4 \| 1.0 \| 5.908e-14 \| 375 \| 0.137 \| 408 \| 251 \| 21 \| 338 \| 682 \| 13 \| 382 \| Phage major capsid protein \| Phage major capsid protein; afdb-uniprot50 \| AF-C5R879-F1-MODEL\_V4 \| 1.0 \| 4.205e-14 \| 375 \| 0.158 \| 455 \| 287 \| 24 \| 252 \| 681 \| 5 \| 388 \| Dyp-type peroxidase family protein \| Dyp-type peroxidase family protein; afdb-uniprot50 \| AF-A0A174XIH6-F1-MODEL\_V4 \| 1.0 \| 3.101e-15 \| 375 \| 0.153 \| 464 \| 281 \| 25 \| 261 \| 682 \| 4 \| 397 \| Phage major capsid protein, HK97 family \| Phage major capsid protein, HK97 family; afdb-uniprot50 \| AF-A0A7X3RYI9-F1-MODEL\_V4 \| 1.0 \| 3.101e-15 \| 374 \| 0.135 \| 487 \| 312 \| 26 \| 249 \| 681 \| 3 \| 434 \| Uncharacterized protein \| Uncharacterized protein; afdb-uniprot50 \| AF-A0A402D5R7-F1-MODEL\_V4 \| 1.0 \| 5.352e-16 \| 374 \| 0.16 \| 492 \| 304 \| 23 \| 263 \| 682 \| 3 \| 457 \| Uncharacterized protein \| Uncharacterized protein; afdb-uniprot50 \| AF-A0A1E5XQL2-F1-MODEL\_V4 \| 1.0 \| 4.88e-15 \| 373 \| 0.17 \| 433 \| 248 \| 18 \| 300 \| 681 \| 21 \| 393 \| Phage capsid protein \| Phage capsid protein; afdb-uniprot50 \| AF-A0A829PFN5-F1-MODEL\_V4 \| 1.0 \| 1.026e-11 \| 372 \| 0.157 \| 292 \| 213 \| 13 \| 410 \| 681 \| 10 \| 288 \| Phage major capsid protein, HK97 family \| Phage major capsid protein, HK97 family; afdb-uniprot50 \| AF-A0A0R1GSB6-F1-MODEL\_V4 \| 1.0 \| 3.189e-11 \| 372 \| 0.148 \| 282 \| 187 \| 15 \| 433 \| 681 \| 39 \| 300 \| Phage capsid family protein \| Phage capsid family protein; afdb-uniprot50 \| AF-A0A6M5YN86-F1-MODEL\_V4 \| 1.0 \| 1.571e-15 \| 372 \| 0.14 \| 483 \| 311 \| 28 \| 256 \| 681 \| 9 \| 444 \| Phage major capsid protein \| Phage major capsid protein; afdb-uniprot50 \| AF-E4V9K9-F1-MODEL\_V4 \| 1.0 \| 2.634e-12 \| 371 \| 0.161 \| 322 \| 216 \| 13 \| 398 \| 681 \| 86 \| 391 \| Putative phage major capsid protein, HK97 family \| Putative phage major capsid protein, HK97 family; afdb-uniprot50 \| AF-A0A349PNZ6-F1-MODEL\_V4 \| 1.0 \| 1.041e-13 \| 370 \| 0.15 \| 371 \| 262 \| 21 \| 334 \| 680 \| 85 \| 426 \| Phage major capsid protein \| Phage major capsid protein; afdb-uniprot50 \| AF-A0A6M1X6G5-F1-MODEL\_V4 \| 1.0 \| 1.494e-12 \| 369 \| 0.142 \| 281 \| 184 \| 14 \| 433 \| 679 \| 50 \| 307 \| Phage major capsid protein \| Phage major capsid protein; afdb-uniprot50 \| AF-A0A437MNA2-F1-MODEL\_V4 \| 1.0 \| 3.676e-15 \| 369 \| 0.162 \| 485 \| 296 \| 24 \| 252 \| 681 \| 13 \| 442 \| Phage major capsid protein \| Phage major capsid protein; afdb-uniprot50 \| AF-A0A1E8RGG7-F1-MODEL\_V4 \| 1.0 \| 1.298e-16 \| 369 \| 0.195 \| 456 \| 278 \| 25 \| 252 \| 681 \| 53 \| 445 \| Uncharacterized protein \| Uncharacterized protein; afdb-uniprot50 \| AF-A0A0R2CK44-F1-MODEL\_V4 \| 1.0 \| 5.954e-19 \| 369 \| 0.136 \| 705 \| 418 \| 37 \| 18 \| 682 \| 4 \| 557 \| Phage capsid protein \| Phage capsid protein; afdb-uniprot50 \| AF-A0A7X3TZU9-F1-MODEL\_V4 \| 1.0 \| 1.005e-12 \| 368 \| 0.156 \| 338 \| 226 \| 17 \| 386 \| 677 \| 1 \| 325 \| Phage major capsid protein \| Phage major capsid protein; afdb-uniprot50 \| AF-A0A0F9QUI3-F1-MODEL\_V4 \| 1.0 \| 1.209e-14 \| 368 \| 0.144 \| 469 \| 273 \| 19 \| 271 \| 682 \| 4 \| 401 \| Uncharacterized protein \| Uncharacterized protein; afdb-uniprot50 \| AF-A0A7Z1WZG9-F1-MODEL\_V4 \| 1.0 \| 6.479e-15 \| 368 \| 0.129 \| 479 \| 304 \| 26 \| 246 \| 679 \| 1 \| 411 \| Phage major capsid protein \| Phage major capsid protein; afdb-uniprot50 \| AF-A0A0F9FJ22-F1-MODEL\_V4 \| 1.0 \| 4.742e-11 \| 367 \| 0.12 \| 299 \| 214 \| 11 \| 411 \| 681 \| 1 \| 278 \| Uncharacterized protein \| Uncharacterized protein; afdb-uniprot50 \| AF-A0A3B0FX46-F1-MODEL\_V4 \| 1.0 \| 3.101e-15 \| 367 \| 0.163 \| 484 \| 300 \| 22 \| 253 \| 680 \| 7 \| 441 \| Phage major capsid protein \| Phage major capsid protein; afdb-uniprot50 \| AF-A0A3D1VAZ0-F1-MODEL\_V4 \| 1.0 \| 2.93e-15 \| 367 \| 0.148 \| 485 \| 303 \| 17 \| 251 \| 680 \| 24 \| 453 \| Phage major capsid protein \| Phage major capsid protein; afdb-uniprot50 \| AF-A0A7G9R3H2-F1-MODEL\_V4 \| 1.0 \| 4.611e-15 \| 366 \| 0.139 \| 446 \| 285 \| 16 \| 263 \| 679 \| 7 \| 382 \| Uncharacterized protein \| Uncharacterized protein; afdb-uniprot50 \| AF-A0A1W1ZBJ4-F1-MODEL\_V4 \| 1.0 \| 8.301e-14 \| 366 \| 0.14 \| 455 \| 297 \| 20 \| 246 \| 681 \| 21 \| 400 \| Phage major capsid protein, HK97 family \| Phage major capsid protein, HK97 family; afdb-uniprot50 \| AF-A0A2W7AVB5-F1-MODEL\_V4 \| 1.0 \| 7.958e-16 \| 365 \| 0.219 \| 424 \| 223 \| 15 \| 18 \| 432 \| 4 \| 328 \| Uncharacterized protein \| Uncharacterized protein; afdb-uniprot50 \| AF-A0A0M8X8L3-F1-MODEL\_V4 \| 1.0 \| 7.003e-14 \| 365 \| 0.133 \| 450 \| 271 \| 21 \| 262 \| 682 \| 12 \| 371 \| Uncharacterized protein \| Uncharacterized protein; afdb-uniprot50 \| AF-A0A3D2THT3-F1-MODEL\_V4 \| 1.0 \| 5.664e-16 \| 365 \| 0.151 \| 523 \| 290 \| 26 \| 251 \| 682 \| 10 \| 469 \| Phage major capsid protein \| Phage major capsid protein; afdb-uniprot50 \| AF-D8JVB9-F1-MODEL\_V4 \| 1.0 \| 7.958e-16 \| 365 \| 0.136 \| 600 \| 319 \| 28 \| 175 \| 679 \| 2 \| 497 \| Phage major capsid protein, HK97 family \| Phage major capsid protein, HK97 family; afdb-uniprot50 \| AF-X1AHU6-F1-MODEL\_V4 \| 1.0 \| 9.361e-11 \| 364 \| 0.383 \| 193 \| 111 \| 4 \| 394 \| 583 \| 4 \| 191 \| Uncharacterized protein \| Uncharacterized protein; afdb-uniprot50 \| AF-A0A1T4ZWM4-F1-MODEL\_V4 \| 1.0 \| 6.714e-16 \| 364 \| 0.172 \| 509 \| 285 \| 23 \| 264 \| 681 \| 3 \| 466 \| Phage major capsid protein, HK97 family \| Phage major capsid protein, HK97 family; afdb-uniprot50 \| AF-A0A5Q2P6R5-F1-MODEL\_V4 \| 1.0 \| 2.847e-11 \| 363 \| 0.159 \| 282 \| 184 \| 15 \| 433 \| 681 \| 39 \| 300 \| Phage major capsid protein \| Phage major capsid protein; afdb-uniprot50 \| AF-A0A7X7E744-F1-MODEL\_V4 \| 1.0 \| 1.118e-15 \| 363 \| 0.131 \| 493 \| 323 \| 19 \| 251 \| 680 \| 3 \| 453 \| Phage major capsid protein \| Phage major capsid protein; afdb-uniprot50 \| AF-A0A6M3ISE2-F1-MODEL\_V4 \| 1.0 \| 9.165e-12 \| 362 \| 0.149 \| 308 \| 219 \| 16 \| 400 \| 681 \| 2 \| 292 \| Putative capsid protein \| Putative capsid protein; afdb-uniprot50 \| AF-A0A2B9DNW7-F1-MODEL\_V4 \| 1.0 \| 1.134e-17 \| 362 \| 0.114 \| 653 \| 369 \| 30 \| 81 \| 681 \| 8 \| 502 \| Uncharacterized protein \| Uncharacterized protein; afdb-uniprot50 \| AF-X6C314-F1-MODEL\_V4 \| 1.0 \| 3.623e-13 \| 361 \| 0.153 \| 403 \| 252 \| 20 \| 320 \| 681 \| 1 \| 355 \| Uncharacterized protein \| Uncharacterized protein; afdb-uniprot50 \| AF-A0A7C2FJD4-F1-MODEL\_V4 \| 1.0 \| 9.433e-16 \| 361 \| 0.146 \| 498 \| 296 \| 24 \| 264 \| 682 \| 6 \| 453 \| Phage major capsid protein \| Phage major capsid protein; afdb-uniprot50 \| AF-A0A7R8N572-F1-MODEL\_V4 \| 1.0 \| 2.769e-15 \| 360 \| 0.231 \| 415 \| 195 \| 14 \| 12 \| 356 \| 3 \| 363 \| Uncharacterized protein \| Uncharacterized protein; afdb-uniprot50 \| AF-A0A0F9PBS1-F1-MODEL\_V4 \| 1.0 \| 3.548e-14 \| 360 \| 0.159 \| 433 \| 263 \| 24 \| 295 \| 682 \| 15 \| 391 \| Uncharacterized protein \| Uncharacterized protein; afdb-uniprot50 \| AF-A0A2A4XWW1-F1-MODEL\_V4 \| 1.0 \| 6.618e-14 \| 360 \| 0.137 \| 464 \| 268 \| 26 \| 251 \| 682 \| 49 \| 412 \| Phage major capsid protein \| Phage major capsid protein; afdb-uniprot50 \| AF-A0A6I5U7Z2-F1-MODEL\_V4 \| 1.0 \| 1.226e-16 \| 360 \| 0.153 \| 594 \| 343 \| 27 \| 175 \| 680 \| 3 \| 524 \| Phage major capsid protein \| Phage major capsid protein; afdb-uniprot50 \| AF-A0A3B8RSS5-F1-MODEL\_V4 \| 1.0 \| 2.672e-14 \| 359 \| 0.144 \| 395 \| 265 \| 19 \| 335 \| 678 \| 5 \| 377 \| Phage major capsid protein \| Phage major capsid protein; afdb-uniprot50 \| AF-A0A5C7M720-F1-MODEL\_V4 \| 1.0 \| 3.282e-15 \| 358 \| 0.167 \| 501 \| 324 \| 22 \| 249 \| 681 \| 3 \| 478 \| Phage major capsid protein \| Phage major capsid protein; afdb-uniprot50 \| AF-A0A7K0EIY7-F1-MODEL\_V4 \| 1.0 \| 6.479e-15 \| 358 \| 0.131 \| 480 \| 328 \| 21 \| 243 \| 679 \| 35 \| 468 \| Phage major capsid protein \| Phage major capsid protein; afdb-uniprot50 \| AF-A0A660ZTR1-F1-MODEL\_V4 \| 1.0 \| 1.02e-14 \| 357 \| 0.142 \| 490 \| 280 \| 30 \| 251 \| 681 \| 3 \| 411 \| Phage major capsid protein \| Phage major capsid protein; afdb-uniprot50 \| AF-A0A7C3V1U7-F1-MODEL\_V4 \| 1.0 \| 1.956e-10 \| 356 \| 0.12 \| 273 \| 209 \| 10 \| 425 \| 682 \| 37 \| 293 \| Uncharacterized protein \| Uncharacterized protein; afdb-uniprot50 \| AF-A0A7X2FSL1-F1-MODEL\_V4 \| 1.0 \| 5.018e-11 \| 356 \| 0.094 \| 296 \| 232 \| 10 \| 413 \| 682 \| 8 \| 293 \| Uncharacterized protein \| Uncharacterized protein; afdb-uniprot50 \| AF-A0A386WCQ4-F1-MODEL\_V4 \| 1.0 \| 7.958e-16 \| 355 \| 0.154 \| 517 \| 292 \| 25 \| 252 \| 680 \| 40 \| 499 \| Phage major capsid protein \| Phage major capsid protein; afdb-uniprot50 \| AF-R6UVK3-F1-MODEL\_V4 \| 1.0 \| 2.42e-16 \| 355 \| 0.137 \| 487 \| 305 \| 22 \| 249 \| 681 \| 24 \| 449 \| Uncharacterized protein \| Uncharacterized protein; afdb-uniprot50 \| AF-A0A0F9ML94-F1-MODEL\_V4 \| 1.0 \| 4.234e-11 \| 354 \| 0.099 \| 302 \| 233 \| 11 \| 404 \| 680 \| 1 \| 288 \| Uncharacterized protein \| Uncharacterized protein; afdb-uniprot50 \| AF-A0A0X8H1D3-F1-MODEL\_V4 \| 1.0 \| 2.888e-13 \| 354 \| 0.13 \| 430 \| 260 \| 22 \| 279 \| 682 \| 42 \| 383 \| Uncharacterized protein \| Uncharacterized protein; afdb-uniprot50 \| AF-A0A2N2DQ71-F1-MODEL\_V4 \| 1.0 \| 3.973e-14 \| 354 \| 0.134 \| 477 \| 305 \| 25 \| 262 \| 682 \| 4 \| 428 \| Phage major capsid protein \| Phage major capsid protein; afdb-uniprot50 \| AF-A0A7Y7IFX1-F1-MODEL\_V4 \| 1.0 \| 3.282e-15 \| 354 \| 0.151 \| 529 \| 277 \| 26 \| 258 \| 679 \| 8 \| 471 \| Phage major capsid protein \| Phage major capsid protein; afdb-uniprot50 \| AF-A0A840YDI0-F1-MODEL\_V4 \| 1.0 \| 6.714e-16 \| 353 \| 0.236 \| 368 \| 218 \| 14 \| 19 \| 358 \| 15 \| 347 \| Uncharacterized protein \| Uncharacterized protein; afdb-uniprot50 \| AF-A0A496M1E1-F1-MODEL\_V4 \| 1.0 \| 2.847e-11 \| 352 \| 0.126 \| 268 \| 196 \| 11 \| 442 \| 682 \| 26 \| 282 \| Phage major capsid protein \| Phage major capsid protein; afdb-uniprot50 \| AF-C3EDP4-F1-MODEL\_V4 \| 1.0 \| 4.71e-14 \| 352 \| 0.161 \| 366 \| 237 \| 23 \| 338 \| 681 \| 11 \| 328 \| Major capsid protein b \| Major capsid protein b; afdb-uniprot50 \| AF-A0A3D5PUN7-F1-MODEL\_V4 \| 1.0 \| 2.908e-10 \| 351 \| 0.154 \| 233 \| 163 \| 11 \| 466 \| 682 \| 3 \| 217 \| Phage major capsid protein \| Phage major capsid protein; afdb-uniprot50 \| AF-A0A0F9Q956-F1-MODEL\_V4 \| 1.0 \| 1.809e-11 \| 351 \| 0.133 \| 300 \| 213 \| 13 \| 413 \| 681 \| 8 \| 291 \| Uncharacterized protein \| Uncharacterized protein; afdb-uniprot50 \| AF-A0A5Q0Q946-F1-MODEL\_V4 \| 1.0 \| 1.166e-13 \| 351 \| 0.141 \| 467 \| 314 \| 19 \| 245 \| 679 \| 1 \| 412 \| Phage major capsid protein \| Phage major capsid protein; afdb-uniprot50 \| AF-A0A7M2A7V1-F1-MODEL\_V4 \| 1.0 \| 2.207e-15 \| 350 \| 0.163 \| 477 \| 297 \| 25 \| 251 \| 682 \| 24 \| 443 \| Phage major capsid protein \| Phage major capsid protein; afdb-uniprot50 \| AF-I0KCY2-F1-MODEL\_V4 \| 1.0 \| 1.484e-15 \| 348 \| 0.123 \| 501 \| 316 \| 21 \| 238 \| 679 \| 30 \| 466 \| Major capsid protein Gp5 \| Major capsid protein Gp5; afdb-uniprot50 \| AF-A0A523WV01-F1-MODEL\_V4 \| 1.0 \| 1.15e-11 \| 347 \| 0.124 \| 305 \| 218 \| 10 \| 399 \| 682 \| 18 \| 294 \| Uncharacterized protein \| Uncharacterized protein; afdb-uniprot50 \| AF-A0A084XUA4-F1-MODEL\_V4 \| 1.0 \| 3.572e-11 \| 347 \| 0.107 \| 299 \| 230 \| 12 \| 407 \| 681 \| 8 \| 293 \| Uncharacterized protein \| Uncharacterized protein; afdb-uniprot50 \| AF-A0A7G5H2I5-F1-MODEL\_V4 \| 1.0 \| 8.301e-14 \| 347 \| 0.131 \| 464 \| 311 \| 19 \| 252 \| 679 \| 5 \| 412 \| Phage major capsid protein \| Phage major capsid protein; afdb-uniprot50 \| AF-A0A0D7F6Q0-F1-MODEL\_V4 \| 1.0 \| 7.462e-11 \| 344 \| 0.092 \| 303 \| 223 \| 15 \| 402 \| 680 \| 2 \| 276 \| Uncharacterized protein \| Uncharacterized protein; afdb-uniprot50 \| AF-A0A839V847-F1-MODEL\_V4 \| 1.0 \| 8.301e-14 \| 343 \| 0.24 \| 378 \| 217 \| 15 \| 264 \| 613 \| 7 \| 342 \| HK97 family phage major capsid protein \| HK97 family phage major capsid protein; afdb-uniprot50 \| AF-A0A1B0Z1T9-F1-MODEL\_V4 \| 1.0 \| 7.051e-11 \| 342 \| 0.121 \| 296 \| 228 \| 13 \| 410 \| 682 \| 1 \| 287 \| Putative major capsid protein \| Putative major capsid protein; afdb-uniprot50 \| AF-A0A2D6EYL3-F1-MODEL\_V4 \| 1.0 \| 1.442e-11 \| 342 \| 0.111 \| 305 \| 239 \| 10 \| 389 \| 682 \| 61 \| 344 \| Uncharacterized protein \| Uncharacterized protein; afdb-uniprot50 \| AF-A0A6M7TFY0-F1-MODEL\_V4 \| 1.0 \| 1.548e-13 \| 342 \| 0.16 \| 441 \| 236 \| 20 \| 308 \| 681 \| 17 \| 390 \| Phage major capsid protein \| Phage major capsid protein; afdb-uniprot50 \| AF-A0A426QMC7-F1-MODEL\_V4 \| 1.0 \| 9.983e-16 \| 342 \| 0.148 \| 598 \| 349 \| 35 \| 172 \| 682 \| 2 \| 526 \| Phage major capsid protein \| Phage major capsid protein; afdb-uniprot50 \| AF-A0A3G8M216-F1-MODEL\_V4 \| 1.0 \| 9.361e-11 \| 341 \| 0.294 \| 214 \| 119 \| 9 \| 31 \| 213 \| 1 \| 213 \| Uncharacterized protein \| Uncharacterized protein; afdb-uniprot50 \| AF-A0A5C7UJM2-F1-MODEL\_V4 \| 1.0 \| 2.672e-14 \| 340 \| 0.129 \| 502 \| 361 \| 23 \| 195 \| 679 \| 1 \| 443 \| Phage major capsid protein \| Phage major capsid protein; afdb-uniprot50 \| AF-A0A1N6YF10-F1-MODEL\_V4 \| 1.0 \| 2.013e-14 \| 340 \| 0.14 \| 521 \| 302 \| 32 \| 261 \| 681 \| 3 \| 477 \| Phage major capsid protein, HK97 family \| Phage major capsid protein, HK97 family; afdb-uniprot50 \| AF-A0A1A3P540-F1-MODEL\_V4 \| 1.0 \| 4.742e-11 \| 339 \| 0.15 \| 293 \| 215 \| 17 \| 410 \| 681 \| 2 \| 281 \| Uncharacterized protein \| Uncharacterized protein; afdb-uniprot50 \| AF-A0A7J9XIJ9-F1-MODEL\_V4 \| 1.0 \| 8.786e-14 \| 339 \| 0.136 \| 497 \| 295 \| 24 \| 251 \| 681 \| 3 \| 431 \| Phage major capsid protein \| Phage major capsid protein; afdb-uniprot50 \| AF-A0A1Y4R494-F1-MODEL\_V4 \| 1.0 \| 5.165e-15 \| 339 \| 0.139 \| 474 \| 300 \| 21 \| 246 \| 682 \| 43 \| 445 \| Phage major capsid protein \| Phage major capsid protein; afdb-uniprot50 \| AF-A0A6G6YPV7-F1-MODEL\_V4 \| 1.0 \| 4.811e-13 \| 337 \| 0.178 \| 421 \| 235 \| 23 \| 309 \| 681 \| 2 \| 359 \| Phage major capsid protein \| Phage major capsid protein; afdb-uniprot50 \| AF-A0A2N1R2F5-F1-MODEL\_V4 \| 1.0 \| 6.122e-15 \| 336 \| 0.143 \| 473 \| 271 \| 18 \| 249 \| 681 \| 227 \| 605 \| Phage major capsid protein \| Phage major capsid protein; afdb-uniprot50 \| AF-A0A6P2DJV8-F1-MODEL\_V4 \| 1.0 \| 5.466e-15 \| 335 \| 0.173 \| 513 \| 287 \| 28 \| 256 \| 682 \| 4 \| 465 \| Uncharacterized protein \| Uncharacterized protein; afdb-uniprot50 \| AF-A0A7K1SKS3-F1-MODEL\_V4 \| 1.0 \| 2.437e-13 \| 335 \| 0.121 \| 478 \| 322 \| 22 \| 246 \| 679 \| 33 \| 456 \| Phage major capsid protein \| Phage major capsid protein; afdb-uniprot50 \| AF-A0A381E722-F1-MODEL\_V4 \| 1.0 \| 1.639e-13 \| 333 \| 0.146 \| 458 \| 277 \| 23 \| 271 \| 681 \| 3 \| 393 \| HK97 family major capsid protein \| HK97 family major capsid protein; afdb-uniprot50 \| AF-A0A545TU80-F1-MODEL\_V4 \| 1.0 \| 2.318e-10 \| 332 \| 0.098 \| 304 \| 229 \| 12 \| 405 \| 682 \| 8 \| 292 \| Uncharacterized protein \| Uncharacterized protein; afdb-uniprot50 \| AF-A0A3C0YSV1-F1-MODEL\_V4 \| 1.0 \| 2.1e-12 \| 332 \| 0.14 \| 470 \| 317 \| 20 \| 251 \| 682 \| 32 \| 452 \| Phage major capsid protein \| Phage major capsid protein; afdb-uniprot50 \| AF-A0A5K1IT80-F1-MODEL\_V4 \| 1.0 \| 6.386e-13 \| 330 \| 0.123 \| 446 \| 288 \| 26 \| 269 \| 680 \| 16 \| 392 \| Phage capsid family protein \| Phage capsid family protein; afdb-uniprot50 \| AF-A0A1Y4LTW7-F1-MODEL\_V4 \| 1.0 \| 1.516e-14 \| 330 \| 0.15 \| 471 \| 301 \| 17 \| 244 \| 679 \| 42 \| 448 \| Phage major capsid protein \| Phage major capsid protein; afdb-uniprot50 \| AF-X0SBZ6-F1-MODEL\_V4 \| 1.0 \| 4.811e-13 \| 328 \| 0.193 \| 419 \| 230 \| 20 \| 294 \| 662 \| 10 \| 370 \| Uncharacterized protein \| Uncharacterized protein; afdb-uniprot50 \| AF-A0A0D6DVL7-F1-MODEL\_V4 \| 1.0 \| 1.433e-14 \| 328 \| 0.144 \| 478 \| 285 \| 27 \| 252 \| 682 \| 13 \| 413 \| Major capsid protein \| Major capsid protein; afdb-uniprot50 \| AF-A0A832B3P5-F1-MODEL\_V4 \| 1.0 \| 5.664e-16 \| 327 \| 0.138 \| 598 \| 361 \| 30 \| 171 \| 682 \| 250 \| 779 \| Phage major capsid protein \| Phage major capsid protein; afdb-uniprot50 \| AF-A0A0F9K1J2-F1-MODEL\_V4 \| 1.0 \| 1.956e-10 \| 325 \| 0.125 \| 312 \| 229 \| 17 \| 398 \| 682 \| 1 \| 295 \| Uncharacterized protein \| Uncharacterized protein; afdb-uniprot50 \| AF-A0A6C2CA91-F1-MODEL\_V4 \| 1.0 \| 3.835e-13 \| 325 \| 0.142 \| 366 \| 243 \| 18 \| 356 \| 681 \| 7 \| 341 \| Phage major capsid protein \| Phage major capsid protein; afdb-uniprot50 \| AF-A0A5C5VFR5-F1-MODEL\_V4 \| 1.0 \| 9.298e-14 \| 325 \| 0.173 \| 495 \| 291 \| 26 \| 253 \| 681 \| 9 \| 451 \| Phage capsid family protein \| Phage capsid family protein; afdb-uniprot50 \| AF-A0A1S9YK65-F1-MODEL\_V4 \| 1.0 \| 1.559e-10 \| 324 \| 0.144 \| 277 \| 190 \| 13 \| 434 \| 680 \| 8 \| 267 \| Capsid protein \| Capsid protein; afdb-uniprot50 \| AF-A0A661D5X9-F1-MODEL\_V4 \| 1.0 \| 2.318e-10 \| 324 \| 0.126 \| 284 \| 191 \| 12 \| 431 \| 680 \| 29 \| 289 \| Uncharacterized protein \| Uncharacterized protein; afdb-uniprot50 \| AF-A0A2S6XCT0-F1-MODEL\_V4 \| 1.0 \| 1.15e-11 \| 324 \| 0.117 \| 393 \| 259 \| 19 \| 313 \| 681 \| 1 \| 329 \| Uncharacterized protein \| Uncharacterized protein; afdb-uniprot50 \| AF-A0A2A7GD25-F1-MODEL\_V4 \| 1.0 \| 2.056e-13 \| 324 \| 0.154 \| 394 \| 249 \| 21 \| 336 \| 681 \| 1 \| 358 \| Phage major capsid protein \| Phage major capsid protein; afdb-uniprot50 \| AF-A0A0F8ZHG2-F1-MODEL\_V4 \| 1.0 \| 9.361e-11 \| 323 \| 0.142 \| 281 \| 193 \| 14 \| 420 \| 679 \| 35 \| 288 \| Uncharacterized protein \| Uncharacterized protein; afdb-uniprot50 \| AF-A0A7J4ZR60-F1-MODEL\_V4 \| 1.0 \| 3.375e-11 \| 322 \| 0.115 \| 329 \| 244 \| 15 \| 391 \| 682 \| 14 \| 332 \| Uncharacterized protein \| Uncharacterized protein; afdb-uniprot50 \| AF-A0A2V4VIH6-F1-MODEL\_V4 \| 1.0 \| 1.771e-12 \| 322 \| 0.154 \| 381 \| 223 \| 18 \| 333 \| 680 \| 6 \| 320 \| HK97 family phage major capsid protein \| HK97 family phage major capsid protein; afdb-uniprot50 \| AF-A0A661TNK1-F1-MODEL\_V4 \| 1.0 \| 5.545e-17 \| 322 \| 0.194 \| 514 \| 276 \| 22 \| 39 \| 515 \| 21 \| 433 \| Uncharacterized protein \| Uncharacterized protein; afdb-uniprot50 \| AF-A0A371IJQ6-F1-MODEL\_V4 \| 1.0 \| 2.056e-13 \| 320 \| 0.121 \| 378 \| 245 \| 16 \| 338 \| 681 \| 42 \| 366 \| Phage major capsid protein \| Phage major capsid protein; afdb-uniprot50 \| AF-A0A4V1UI61-F1-MODEL\_V4 \| 1.0 \| 1.306e-13 \| 318 \| 0.258 \| 306 \| 177 \| 9 \| 255 \| 541 \| 4 \| 278 \| Phage major capsid protein \| Phage major capsid protein; afdb-uniprot50 \| AF-A0A6B5FRH0-F1-MODEL\_V4 \| 1.0 \| 1.048e-10 \| 318 \| 0.155 \| 225 \| 152 \| 10 \| 481 \| 680 \| 56 \| 267 \| Phage major capsid protein \| Phage major capsid protein; afdb-uniprot50 \| AF-A0A7J3JAD0-F1-MODEL\_V4 \| 1.0 \| 8.358e-11 \| 318 \| 0.112 \| 302 \| 223 \| 12 \| 398 \| 679 \| 30 \| 306 \| Uncharacterized protein \| Uncharacterized protein; afdb-uniprot50 \| AF-A0A841A8H5-F1-MODEL\_V4 \| 1.0 \| 4.985e-14 \| 318 \| 0.206 \| 383 \| 210 \| 10 \| 35 \| 324 \| 12 \| 393 \| Uncharacterized protein \| Uncharacterized protein; afdb-uniprot50 \| AF-W0JUC6-F1-MODEL\_V4 \| 1.0 \| 5.702e-13 \| 317 \| 0.146 \| 430 \| 271 \| 21 \| 292 \| 681 \| 47 \| 420 \| Uncharacterized protein \| Uncharacterized protein; afdb-uniprot50 \| AF-A0A068D9Z5-F1-MODEL\_V4 \| 1.0 \| 4.811e-13 \| 317 \| 0.158 \| 436 \| 251 \| 23 \| 300 \| 681 \| 60 \| 433 \| Phage major capsid protein \| Phage major capsid protein; afdb-uniprot50 \| AF-A0A0F9Q477-F1-MODEL\_V4 \| 1.0 \| 9.561e-10 \| 316 \| 0.099 \| 343 \| 247 \| 13 \| 392 \| 682 \| 54 \| 386 \| Uncharacterized protein \| Uncharacterized protein; afdb-uniprot50 \| AF-A0A836P8B0-F1-MODEL\_V4 \| 1.0 \| 1.848e-10 \| 315 \| 0.143 \| 307 \| 201 \| 13 \| 405 \| 674 \| 15 \| 296 \| Major capsid subunit gp9 \| Major capsid subunit gp9; afdb-uniprot50 \| AF-A0A842E264-F1-MODEL\_V4 \| 1.0 \| 1.166e-13 \| 315 \| 0.137 \| 421 \| 246 \| 23 \| 324 \| 679 \| 2 \| 370 \| Phage major capsid protein \| Phage major capsid protein; afdb-uniprot50 \| AF-A0A0F9LNJ4-F1-MODEL\_V4 \| 1.0 \| 8.845e-11 \| 313 \| 0.116 \| 310 \| 225 \| 18 \| 402 \| 682 \| 2 \| 291 \| Uncharacterized protein \| Uncharacterized protein; afdb-uniprot50 \| AF-D3NXQ3-F1-MODEL\_V4 \| 1.0 \| 2.596e-10 \| 312 \| 0.092 \| 315 \| 236 \| 15 \| 400 \| 682 \| 1 \| 297 \| Uncharacterized protein \| Uncharacterized protein; afdb-uniprot50 \| AF-A0A810UUB9-F1-MODEL\_V4 \| 1.0 \| 1.298e-16 \| 312 \| 0.221 \| 496 \| 248 \| 21 \| 17 \| 432 \| 2 \| 439 \| Uncharacterized protein \| Uncharacterized protein; afdb-uniprot50 \| AF-A0A139CW77-F1-MODEL\_V4 \| 1.0 \| 5.388e-13 \| 312 \| 0.145 \| 399 \| 255 \| 22 \| 335 \| 681 \| 4 \| 368 \| Major capsid protein \| Major capsid protein; afdb-uniprot50 \| AF-A0A7W1JBE0-F1-MODEL\_V4 \| 1.0 \| 7.153e-13 \| 312 \| 0.142 \| 464 \| 311 \| 25 \| 272 \| 681 \| 40 \| 470 \| Phage major capsid protein \| Phage major capsid protein; afdb-uniprot50 \| AF-A0A4S2EV56-F1-MODEL\_V4 \| 1.0 \| 5.388e-13 \| 311 \| 0.124 \| 386 \| 253 \| 17 \| 338 \| 679 \| 37 \| 381 \| Phage major capsid protein \| Phage major capsid protein; afdb-uniprot50 \| AF-A0A6M3XUP1-F1-MODEL\_V4 \| 1.0 \| 3.189e-11 \| 309 \| 0.111 \| 429 \| 290 \| 16 \| 313 \| 680 \| 1 \| 399 \| Putative structural protein \| Putative structural protein; afdb-uniprot50 \| AF-A8SKN8-F1-MODEL\_V4 \| 1.0 \| 3.917e-12 \| 306 \| 0.099 \| 421 \| 285 \| 18 \| 279 \| 682 \| 31 \| 374 \| Phage major capsid protein, HK97 family \| Phage major capsid protein, HK97 family; afdb-uniprot50 \| AF-A0A2E7LMR5-F1-MODEL\_V4 \| 1.0 \| 9.034e-10 \| 304 \| 0.096 \| 270 \| 214 \| 13 \| 428 \| 681 \| 24 \| 279 \| Uncharacterized protein \| Uncharacterized protein; afdb-uniprot50 \| AF-F9RFV9-F1-MODEL\_V4 \| 1.0 \| 2.07e-10 \| 304 \| 0.16 \| 268 \| 194 \| 18 \| 431 \| 682 \| 33 \| 285 \| Uncharacterized protein \| Uncharacterized protein; afdb-uniprot50 \| AF-A0A255DW94-F1-MODEL\_V4 \| 1.0 \| 2.908e-10 \| 303 \| 0.12 \| 348 \| 180 \| 13 \| 449 \| 681 \| 2 \| 338 \| Phage major capsid protein \| Phage major capsid protein; afdb-uniprot50 \| AF-A0A5C7M0U3-F1-MODEL\_V4 \| 1.0 \| 9.57e-18 \| 303 \| 0.12 \| 688 \| 434 \| 36 \| 67 \| 682 \| 16 \| 604 \| Phage major capsid protein \| Phage major capsid protein; afdb-uniprot50 \| AF-A0A0F9EE26-F1-MODEL\_V4 \| 1.0 \| 2.07e-10 \| 301 \| 0.087 \| 333 \| 239 \| 16 \| 402 \| 681 \| 3 \| 323 \| Uncharacterized protein \| Uncharacterized protein; afdb-uniprot50 \| AF-A0A2A5FHH9-F1-MODEL\_V4 \| 1.0 \| 1.412e-12 \| 301 \| 0.157 \| 501 \| 287 \| 26 \| 249 \| 680 \| 5 \| 439 \| Phage major capsid protein \| Phage major capsid protein; afdb-uniprot50 \| AF-A0A0R2JSR1-F1-MODEL\_V4 \| 1.0 \| 5.311e-11 \| 300 \| 0.161 \| 278 \| 176 \| 14 \| 439 \| 681 \| 83 \| 338 \| Major head protein Cps \| Major head protein Cps; afdb-uniprot50 \| AF-A0A2D6X1N3-F1-MODEL\_V4 \| 1.0 \| 1.11e-10 \| 300 \| 0.103 \| 356 \| 233 \| 18 \| 402 \| 681 \| 13 \| 358 \| Uncharacterized protein \| Uncharacterized protein; afdb-uniprot50 \| AF-A0A0F9GE20-F1-MODEL\_V4 \| 1.0 \| 6.075e-10 \| 296 \| 0.092 \| 282 \| 229 \| 12 \| 422 \| 680 \| 15 \| 292 \| Uncharacterized protein \| Uncharacterized protein; afdb-uniprot50 \| AF-A0A1V4ZC61-F1-MODEL\_V4 \| 1.0 \| 1.217e-11 \| 296 \| 0.131 \| 434 \| 265 \| 20 \| 309 \| 682 \| 18 \| 399 \| Uncharacterized protein \| Uncharacterized protein; afdb-uniprot50 \| AF-A0A3M2BE71-F1-MODEL\_V4 \| 1.0 \| 3.036e-16 \| 293 \| 0.136 \| 666 \| 409 \| 36 \| 107 \| 682 \| 83 \| 672 \| Uncharacterized protein \| Uncharacterized protein; afdb-uniprot50 \| AF-A0A6A8SHP3-F1-MODEL\_V4 \| 1.0 \| 4.643e-12 \| 292 \| 0.118 \| 397 \| 263 \| 16 \| 334 \| 679 \| 2 \| 362 \| Phage major capsid protein \| Phage major capsid protein; afdb-uniprot50 \| AF-A0A1M3CGP7-F1-MODEL\_V4 \| 1.0 \| 9.561e-10 \| 290 \| 0.111 \| 314 \| 225 \| 21 \| 404 \| 680 \| 1 \| 297 \| Uncharacterized protein \| Uncharacterized protein; afdb-uniprot50 \| AF-A0A1H4H424-F1-MODEL\_V4 \| 1.0 \| 1.771e-12 \| 290 \| 0.13 \| 430 \| 258 \| 20 \| 316 \| 679 \| 1 \| 380 \| Phage major capsid protein, HK97 family \| Phage major capsid protein, HK97 family; afdb-uniprot50 \| AF-X0S7V5-F1-MODEL\_V4 \| 1.0 \| 4.742e-11 \| 287 \| 0.129 \| 463 \| 283 \| 24 \| 252 \| 666 \| 11 \| 401 \| Uncharacterized protein \| Uncharacterized protein; afdb-uniprot50 \| AF-A0A7X2TNW2-F1-MODEL\_V4 \| 1.0 \| 1.582e-12 \| 282 \| 0.134 \| 431 \| 259 \| 23 \| 334 \| 681 \| 3 \| 402 \| Phage major capsid protein \| Phage major capsid protein; afdb-uniprot50 \| AF-A0A380JMN2-F1-MODEL\_V4 \| 1.0 \| 2.402e-11 \| 281 \| 0.136 \| 404 \| 263 \| 19 \| 334 \| 681 \| 2 \| 375 \| Predicted phage phi-C31 gp36 major capsid-like protein \| Predicted phage phi-C31 gp36 major capsid-like protein; afdb-uniprot50 \| AF-A0A0F9EU19-F1-MODEL\_V4 \| 1.0 \| 5.311e-11 \| 281 \| 0.092 \| 380 \| 267 \| 17 \| 337 \| 680 \| 48 \| 385 \| Uncharacterized protein \| Uncharacterized protein; afdb-uniprot50 \| AF-A0A0M5M622-F1-MODEL\_V4 \| 1.0 \| 1.592e-09 \| 278 \| 0.083 \| 276 \| 203 \| 13 \| 435 \| 682 \| 30 \| 283 \| Uncharacterized protein \| Uncharacterized protein; afdb-uniprot50 \| AF-A0A2V2RBB4-F1-MODEL\_V4 \| 1.0 \| 1.086e-11 \| 277 \| 0.29 \| 279 \| 170 \| 7 \| 257 \| 521 \| 4 \| 268 \| Phage major capsid protein \| Phage major capsid protein; afdb-uniprot50 \| AF-A0A3M2C237-F1-MODEL\_V4 \| 1.0 \| 6.386e-13 \| 277 \| 0.122 \| 434 \| 275 \| 22 \| 334 \| 680 \| 3 \| 417 \| Phage major capsid protein \| Phage major capsid protein; afdb-uniprot50 \| AF-A0A0E0US10-F1-MODEL\_V4 \| 1.0 \| 3.375e-11 \| 276 \| 0.14 \| 399 \| 255 \| 22 \| 335 \| 680 \| 1 \| 364 \| Major capsid protein \| Major capsid protein; afdb-uniprot50 \| AF-A0A841X480-F1-MODEL\_V4 \| 1.0 \| 1.086e-11 \| 275 \| 0.121 \| 451 \| 270 \| 25 \| 264 \| 679 \| 6 \| 365 \| Phage major capsid protein \| Phage major capsid protein; afdb-uniprot50 \| AF-A0A540WHX5-F1-MODEL\_V4 \| 1.0 \| 5.741e-10 \| 274 \| 0.178 \| 365 \| 242 \| 20 \| 295 \| 641 \| 27 \| 351 \| Phage major capsid protein \| Phage major capsid protein; afdb-uniprot50 \| AF-A0A2E8F3P1-F1-MODEL\_V4 \| 1.0 \| 1.126e-12 \| 272 \| 0.132 \| 573 \| 334 \| 28 \| 194 \| 678 \| 6 \| 503 \| Phage major capsid protein \| Phage major capsid protein; afdb-uniprot50 \| AF-A0A0F9VRZ0-F1-MODEL\_V4 \| 1.0 \| 3.189e-11 \| 271 \| 0.136 \| 411 \| 271 \| 18 \| 308 \| 682 \| 34 \| 396 \| Uncharacterized protein \| Uncharacterized protein; afdb-uniprot50 \| AF-A0A202F7U9-F1-MODEL\_V4 \| 1.0 \| 1.363e-11 \| 267 \| 0.129 \| 402 \| 256 \| 23 \| 335 \| 679 \| 1 \| 365 \| Uncharacterized protein \| Uncharacterized protein; afdb-uniprot50 \| AF-A0A2T4M4G5-F1-MODEL\_V4 \| 1.0 \| 1.809e-11 \| 266 \| 0.139 \| 394 \| 264 \| 18 \| 338 \| 681 \| 18 \| 386 \| Phage major capsid protein \| Phage major capsid protein; afdb-uniprot50 \| AF-A0A5R9FB27-F1-MODEL\_V4 \| 1.0 \| 1.11e-10 \| 264 \| 0.112 \| 372 \| 254 \| 19 \| 338 \| 679 \| 66 \| 391 \| Phage major capsid protein \| Phage major capsid protein; afdb-uniprot50 \| AF-A0A3D9YKZ9-F1-MODEL\_V4 \| 1.0 \| 2.614e-07 \| 264 \| 0.224 \| 147 \| 96 \| 6 \| 42 \| 178 \| 19 \| 157 \| HK97 family phage prohead protease \| HK97 family phage prohead protease; afdb-uniprot50 \| AF-A0A5C7Q7W6-F1-MODEL\_V4 \| 1.0 \| 3.398e-08 \| 260 \| 0.177 \| 180 \| 125 \| 9 \| 19 \| 188 \| 6 \| 172 \| HK97 family phage prohead protease \| HK97 family phage prohead protease; afdb-uniprot50 \| AF-X0SVC7-F1-MODEL\_V4 \| 1.0 \| 5.235e-09 \| 260 \| 0.089 \| 279 \| 198 \| 13 \| 430 \| 676 \| 1 \| 255 \| Uncharacterized protein \| Uncharacterized protein; afdb-uniprot50 \| AF-L1QG40-F1-MODEL\_V4 \| 1.0 \| 1.559e-10 \| 259 \| 0.102 \| 441 \| 293 \| 20 \| 269 \| 682 \| 6 \| 370 \| Phage major capsid protein, HK97 family \| Phage major capsid protein, HK97 family; afdb-uniprot50 \| AF-A0A380F728-F1-MODEL\_V4 \| 1.0 \| 6.662e-11 \| 258 \| 0.121 \| 412 \| 269 \| 20 \| 334 \| 680 \| 2 \| 385 \| Phage protein \| Phage protein; afdb-uniprot50 \| AF-A0A497US25-F1-MODEL\_V4 \| 1.0 \| 3.375e-11 \| 257 \| 0.122 \| 505 \| 285 \| 24 \| 252 \| 679 \| 21 \| 444 \| Uncharacterized protein \| Uncharacterized protein; afdb-uniprot50 \| AF-A0A0F9ITE6-F1-MODEL\_V4 \| 1.0 \| 9.907e-11 \| 255 \| 0.263 \| 334 \| 186 \| 12 \| 261 \| 568 \| 4 \| 303 \| Uncharacterized protein \| Uncharacterized protein; afdb-uniprot50 \| AF-A0A7K0J5W9-F1-MODEL\_V4 \| 1.0 \| 7.051e-11 \| 255 \| 0.136 \| 461 \| 239 \| 19 \| 233 \| 673 \| 25 \| 346 \| Phage major capsid protein \| Phage major capsid protein; afdb-uniprot50 \| AF-A0A0F9T8B6-F1-MODEL\_V4 \| 1.0 \| 2.453e-10 \| 255 \| 0.123 \| 365 \| 254 \| 18 \| 338 \| 680 \| 88 \| 408 \| Uncharacterized protein \| Uncharacterized protein; afdb-uniprot50 \| AF-A0A1I4ZM31-F1-MODEL\_V4 \| 1.0 \| 1.11e-10 \| 254 \| 0.112 \| 416 \| 250 \| 24 \| 335 \| 681 \| 1 \| 366 \| Phage major capsid protein, HK97 family \| Phage major capsid protein, HK97 family; afdb-uniprot50 \| AF-A0A1G7NZP4-F1-MODEL\_V4 \| 1.0 \| 3.375e-11 \| 251 \| 0.121 \| 421 \| 254 \| 24 \| 323 \| 681 \| 2 \| 368 \| Phage major capsid protein, HK97 family \| Phage major capsid protein, HK97 family; afdb-uniprot50 \| AF-A0A497X2P2-F1-MODEL\_V4 \| 1.0 \| 5.989e-08 \| 246 \| 0.175 \| 188 \| 123 \| 13 \| 19 \| 192 \| 2 \| 171 \| Uncharacterized protein \| Uncharacterized protein; afdb-uniprot50 \| AF-A0A512JIM5-F1-MODEL\_V4 \| 1.0 \| 1.217e-11 \| 246 \| 0.186 \| 402 \| 219 \| 12 \| 51 \| 353 \| 2 \| 394 \| Uncharacterized protein \| Uncharacterized protein; afdb-uniprot50 \| AF-A0A135IJY6-F1-MODEL\_V4 \| 1.0 \| 1.661e-07 \| 234 \| 0.174 \| 195 \| 134 \| 9 \| 27 \| 208 \| 4 \| 184 \| Peptidase U35 \| Peptidase U35; afdb-uniprot50 \| AF-A0A3A6MU59-F1-MODEL\_V4 \| 1.0 \| 5.347e-08 \| 231 \| 0.207 \| 183 \| 117 \| 10 \| 6 \| 173 \| 98 \| 267 \| HK97 family phage prohead protease \| HK97 family phage prohead protease; afdb-uniprot50 \| AF-A0A2D9VGK2-F1-MODEL\_V4 \| 1.0 \| 3.034e-08 \| 221 \| 0.194 \| 216 \| 128 \| 14 \| 18 \| 211 \| 2 \| 193 \| HK97 family phage prohead protease \| HK97 family phage prohead protease; afdb-uniprot50 \| AF-A0A7W7LNM0-F1-MODEL\_V4 \| 1.0 \| 1.296e-08 \| 217 \| 0.112 \| 454 \| 224 \| 20 \| 400 \| 681 \| 1 \| 447 \| Uncharacterized protein \| Uncharacterized protein; afdb-uniprot50 \| AF-A0A073IXT7-F1-MODEL\_V4 \| 1.0 \| 1.483e-07 \| 216 \| 0.188 \| 186 \| 121 \| 8 \| 17 \| 185 \| 2 \| 174 \| Primosome assembly protein PriA \| Primosome assembly protein PriA; afdb-uniprot50 \| AF-A0A7W3UAB7-F1-MODEL\_V4 \| 1.0 \| 2.748e-10 \| 215 \| 0.158 \| 441 \| 241 \| 21 \| 40 \| 426 \| 25 \| 389 \| HK97 family phage prohead protease \| HK97 family phage prohead protease; afdb-uniprot50 \| AF-A0A4R5EYX0-F1-MODEL\_V4 \| 1.0 \| 8.905e-08 \| 204 \| 0.216 \| 203 \| 116 \| 12 \| 11 \| 193 \| 14 \| 193 \| HK97 family phage prohead protease \| HK97 family phage prohead protease; afdb-uniprot50 \| AF-A0A3G8GRJ2-F1-MODEL\_V4 \| 1.0 \| 4.353e-07 \| 204 \| 0.19 \| 221 \| 134 \| 12 \| 43 \| 258 \| 28 \| 208 \| Uncharacterized protein \| Uncharacterized protein; afdb-uniprot50 \| AF-A0A2E6W8T6-F1-MODEL\_V4 \| 1.0 \| 4.876e-07 \| 195 \| 0.127 \| 291 \| 197 \| 18 \| 420 \| 682 \| 28 \| 289 \| Uncharacterized protein \| Uncharacterized protein; afdb-uniprot50 \| AF-A0A0F8WVU2-F1-MODEL\_V4 \| 1.0 \| 1.9e-06 \| 194 \| 0.134 \| 215 \| 158 \| 11 \| 481 \| 682 \| 15 \| 214 \| Uncharacterized protein \| Uncharacterized protein; afdb-uniprot50 \| AF-B9XA48-F1-MODEL\_V4 \| 1.0 \| 8.294e-06 \| 191 \| 0.285 \| 189 \| 114 \| 4 \| 367 \| 534 \| 5 \| 193 \| Putative phage-related protein \| Putative phage-related protein; afdb-uniprot50 \| AF-A0A1E4IEB7-F1-MODEL\_V4 \| 1.0 \| 1.86e-07 \| 188 \| 0.106 \| 338 \| 215 \| 22 \| 410 \| 681 \| 1 \| 317 \| Uncharacterized protein \| Uncharacterized protein; afdb-uniprot50 \| AF-A0A2P8DWF2-F1-MODEL\_V4 \| 1.0 \| 4.262e-08 \| 186 \| 0.135 \| 369 \| 219 \| 17 \| 39 \| 379 \| 6 \| 302 \| HK97 family phage prohead protease \| HK97 family phage prohead protease; afdb-uniprot50 \| AF-A0A443KJC0-F1-MODEL\_V4 \| 1.0 \| 1.696e-06 \| 180 \| 0.181 \| 176 \| 112 \| 10 \| 31 \| 192 \| 4 \| 161 \| HK97 family phage prohead protease \| HK97 family phage prohead protease; afdb-uniprot50 \| AF-A0A517R7E1-F1-MODEL\_V4 \| 1.0 \| 5.903e-06 \| 175 \| 0.122 \| 277 \| 159 \| 13 \| 418 \| 680 \| 16 \| 222 \| Uncharacterized protein \| Uncharacterized protein; afdb-uniprot50 \| AF-A0A3R5RK36-F1-MODEL\_V4 \| 1.0 \| 3.887e-07 \| 173 \| 0.133 \| 321 \| 178 \| 17 \| 382 \| 682 \| 3 \| 243 \| Uncharacterized protein \| Uncharacterized protein; afdb-uniprot50 \| AF-A0A368YH54-F1-MODEL\_V4 \| 1.0 \| 7.95e-08 \| 164 \| 0.141 \| 453 \| 222 \| 21 \| 18 \| 330 \| 7 \| 432 \| HK97 family phage prohead protease \| HK97 family phage prohead protease; afdb-uniprot50 \| AF-A0A6B1C0U0-F1-MODEL\_V4 \| 1.0 \| 3.887e-07 \| 149 \| 0.149 \| 489 \| 260 \| 26 \| 261 \| 671 \| 8 \| 418 \| Uncharacterized protein \| Uncharacterized protein; afdb-uniprot50 \| AF-A0A0F9U9R9-F1-MODEL\_V4 \| 1.0 \| 2.04e-08 \| 140 \| 0.128 \| 481 \| 255 \| 23 \| 12 \| 396 \| 121 \| 533 \| Uncharacterized protein \| Uncharacterized protein; afdb-uniprot50 \| AF-A0A0F9N5R3-F1-MODEL\_V4 \| 1.0 \| 1.056e-07 \| 136 \| 0.136 \| 447 \| 259 \| 27 \| 38 \| 421 \| 19 \| 401 \| Uncharacterized protein \| Uncharacterized protein |
| Top keywords  (threshold 1.00e-02 (evalue)) | **capsid, major, Phage, HK97, prohead, protease, Putative, Peptidase, head, Caudovirus** |
| Output files | ../../similar\_structures/05\_FANPEZAQ\_CDS\_0005\_afdb-proteome\_foldseek.tsv ../../similar\_structures/05\_FANPEZAQ\_CDS\_0005\_afdb-uniprot50\_foldseek.tsv ../../similar\_structures/05\_FANPEZAQ\_CDS\_0005\_merged.svg ../../similar\_structures/05\_FANPEZAQ\_CDS\_0005\_pdb\_foldseek.tsv |

  
  
  

Return to summary | Go to previous | Go to next

  


---

**Sequence/structure alignments coloring**  
Each object in the alignment figures is colored according to its E-value following this color coding:

1e-100
10
