## Supplementary material for "Completing the BASEL phage collection to unlock hidden diversity for systematic exploration of phage-host interactions": Data S2: 6.html

FANPEZAQ\_CDS\_0006


Return to summary | Go to previous | Go to next

|  |  |
| --- | --- |
| FANPEZAQ\_CDS\_0006  Page creation date: 02 Sep 2024, 12:00  Project folder: n/a  Input sequences file: Escherichia\_virus\_HeidiAbel.gb | domain\_containing rho termination factor n\_terminal heh lem fragment transcription hypothetical sap phage rho\_n ribosomal 50s endonuclease vii mu\_like prophage flumu and a t4 recombination l21 in dimerisation tail factor\_like alpha yes rna\_binding isoform putative rna\_bd |

### Sequence information

|  |  |
| --- | --- |
| Name | FANPEZAQ\_CDS\_0006  06\_FANPEZAQ\_CDS\_0006 (pipeline id) |
| Imported annotations | Escherichia\_virus\_HeidiAbel Bas97 |
| Protein sequence | MSGTYVLKLTSAIAISGEVMKAGSLVEVSELEAKNLLSRGKAEMHDEQPARDDDVEITKM NKDQLIAIAEQMEIEGADKMTKAQLIEAIEAAGETKEGDE |
| Number of residues | 100 |
| Molecular weight (Da) | 10809.12 |
| Output files | ../../query\_sequences/06\_FANPEZAQ\_CDS\_0006.fasta |

### Putative domain architecture and protein family

#### Search results (HHblits)^1^

|  |  |
| --- | --- |
| Domain family databases searched | Pfam, Ncbi-cd, Cath, Phrogs |
| Results, scheme(s)  (Top layers only; threshold 1.00e-03 (evalue)) | xml version="1.0" encoding="utf-8" standalone="no"?       2024-09-02T21:08:13.120903 image/svg+xml   Matplotlib v3.7.2, https://matplotlib.org/ |
| Results, table  (E-value ≤ 1.00e-03 (evalue)) | db \| id \| prob \| evalue \| pvalue \| score \| cols \| query \| query\_len \| template \| template\_len \| name \| description; pfam \| PF09124 \| 97.4 \| 6.3e-08 \| 1.3e-11 \| 54.1 \| 48 \| (48, 95) \| 100 \| (4, 51) \| 54 \| Endonuc-dimeris \| T4 recombination endonuclease VII, dimerisation; pfam \| PF10281 \| 97.3 \| 1e-07 \| 2.2e-11 \| 49.0 \| 35 \| (57, 91) \| 100 \| (1, 36) \| 37 \| Ish1 \| Putative nuclear envelope organisation protein; pfam \| PF12949 \| 95.4 \| 7.9e-05 \| 1.9e-08 \| 34.0 \| 33 \| (58, 90) \| 100 \| (2, 34) \| 35 \| HeH \| HeH/LEM domain; ncbi-cd \| cd12935 \| 94.6 \| 0.00027 \| 7e-08 \| 31.8 \| 34 \| (58, 91) \| 100 \| (2, 35) \| 36 \| LEM\_like \| cd12935 LEM\_like; LEM-like domain of lamina-associated polypeptide 2 (LAP2) and similar proteins. LAP2, also termed thymopoietin (TP), or thymopoietin-related peptide (TPRP), is composed of isoform alpha and isoforms beta/gamma and may be involved in chromatin organization and postmitotic reassembly.; cath \| 1e7lA02 \| 98.2 \| 1.8e-10 \| 2.9e-14 \| 67.4 \| 47 \| (48, 94) \| 100 \| (5, 51) \| 55 \| Recombination endonuclease vii \| CATHCODE: 1.10.720.10 NAME: Recombination endonuclease vii. Chain: a, b. Synonym: gp49. Engineered: yes. Mutation: yes. Other\_details: one zn bound to cys 23,26,58,61 of each chain SOURCE: Bacteriophage t4. Organism\_taxid: 10665. Gene: gp49. Expressed in: escherichia coli. Expression\_system\_taxid: 511693. CLASS: Mainly Alpha, ARCH: Orthogonal Bundle, TOPOL: Transcription Termination Factor Rho, Rna-binding Domain; Chain A, Domain 1, HOMOL: Transcription Termination Factor Rho, Rna-binding Domain; Chain A, Domain 1; cath \| 2outA02 \| 96.2 \| 1.2e-05 \| 2e-09 \| 41.3 \| 39 \| (59, 97) \| 100 \| (1, 39) \| 41 \| Mu-like prophage flumu protein gp35, protein hi1507 in mu-like prophage flumu region \| CATHCODE: 1.10.720.30 NAME: Mu-like prophage flumu protein gp35, protein hi1507 in mu-like prophage flumu region. Chain: a. Engineered: yes SOURCE: Haemophilus influenzae. Organism\_taxid: 727. Strain: dsm 11121, kw20, rd. Atcc: 51907, 51907. Gene: hi1506, hi1507. Expressed in: escherichia coli. Expression\_system\_taxid: 562. CLASS: Mainly Alpha, ARCH: Orthogonal Bundle, TOPOL: Transcription Termination Factor Rho, Rna-binding Domain; Chain A, Domain 1, HOMOL: SAP domain; phrogs \| 3281 \| 99.7 \| 1.7e-21 \| 2.1e-25 \| 134.3 \| 94 \| (1, 98) \| 100 \| (6, 130) \| 145 \| endonuclease VII \| endonuclease VII; Category: DNA, RNA and nucleotide metabolism; MG711467\_p48; phrogs \| 390 \| 98.8 \| 5.9e-13 \| 7.5e-17 \| 85.1 \| 63 \| (23, 92) \| 100 \| (30, 94) \| 95 \| Arc-like repressor \| Arc-like repressor; Category: transcription regulation; p204416 VI\_04755; phrogs \| 2724 \| 98.4 \| 4e-11 \| 4.9e-15 \| 74.0 \| 67 \| (26, 95) \| 100 \| (10, 78) \| 79 \| NA \| NA; Category: unknown function; p122757 VI\_05802; phrogs \| 2250 \| 98.2 \| 3.4e-10 \| 4.2e-14 \| 67.7 \| 44 \| (52, 95) \| 100 \| (21, 64) \| 66 \| NA \| NA; Category: unknown function; p428684 VI\_05331; phrogs \| 1539 \| 97.9 \| 5.1e-09 \| 6.3e-13 \| 62.4 \| 36 \| (56, 91) \| 100 \| (31, 66) \| 66 \| Arc-like repressor \| Arc-like repressor; Category: transcription regulation; NC\_004586\_p34; phrogs \| 10541 \| 97.0 \| 9.6e-07 \| 1.2e-10 \| 55.8 \| 44 \| (51, 94) \| 100 \| (55, 98) \| 100 \| NA \| NA; Category: unknown function; NC\_029029\_p8; phrogs \| 1772 \| 95.7 \| 9e-05 \| 1e-08 \| 46.4 \| 38 \| (53, 90) \| 100 \| (72, 109) \| 111 \| NA \| NA; Category: unknown function; p424151 VI\_10171; phrogs \| 38661 \| 95.6 \| 0.00011 \| 1.3e-08 \| 42.7 \| 40 \| (53, 92) \| 100 \| (31, 70) \| 71 \| NA \| NA; Category: unknown function; p193317 VI\_04290; phrogs \| 19159 \| 95.4 \| 0.00014 \| 1.6e-08 \| 44.2 \| 36 \| (58, 93) \| 100 \| (52, 87) \| 89 \| NA \| NA; Category: unknown function; p84411 VI\_10196; phrogs \| 9487 \| 95.4 \| 0.00014 \| 1.7e-08 \| 47.2 \| 42 \| (56, 97) \| 100 \| (2, 43) \| 133 \| NA \| NA; Category: unknown function; MF347638\_p49; phrogs \| 17694 \| 95.1 \| 0.00027 \| 3e-08 \| 43.2 \| 39 \| (53, 91) \| 100 \| (59, 97) \| 98 \| NA \| NA; Category: unknown function; p62152 VI\_08379; phrogs \| 8953 \| 94.5 \| 0.00063 \| 7.5e-08 \| 44.2 \| 40 \| (58, 97) \| 100 \| (90, 129) \| 134 \| NA \| NA; Category: unknown function; p258246 VI\_06608 |
| Top keywords  (threshold 1.00e-03 (evalue)) | **and, A, Transcription, in, endonuclease, VII, Alpha, yes, Termination, Factor** |
| Output files | ../../domain\_architecture/06\_FANPEZAQ\_CDS\_0006\_cath.hhr ../../domain\_architecture/06\_FANPEZAQ\_CDS\_0006\_merged.svg ../../domain\_architecture/06\_FANPEZAQ\_CDS\_0006\_ncbi-cd.hhr ../../domain\_architecture/06\_FANPEZAQ\_CDS\_0006\_pfam.hhr ../../domain\_architecture/06\_FANPEZAQ\_CDS\_0006\_phrogs.hhr |

### Identical protein sequences/structures

#### Search results

|  |  |
| --- | --- |
| Protein sequence databases searched | Pdb, Swissprot, Refseq |
| Identical proteins found | -- |
| Top keywords | -- |
| Output files | -- |

### Similar protein sequences/structures

#### Sequence similarity search results (HHblits)^1^

|  |  |
| --- | --- |
| Sequence databases searched | Uniclust, Pdb70 |
| Results, scheme(s)  (Top layers only, threshold 1.00e-03 (evalue)) | xml version="1.0" encoding="utf-8" standalone="no"?       2024-09-02T21:08:31.054110 image/svg+xml   Matplotlib v3.7.2, https://matplotlib.org/ |
| Results, table(s)  (threshold 1.00e-03 (evalue)) | db \| id \| prob \| evalue \| pvalue \| score \| cols \| query \| query\_len \| template \| template\_len \| name \| description; uniclust \| UniRef100\_A0A098MGF8 \| 99.6 \| 2.9e-19 \| 6.2e-25 \| 115.8 \| 96 \| (2, 97) \| 100 \| (6, 108) \| 122 \| Rho termination factor N-terminal domain-containing protein \| Rho termination factor N-terminal domain-containing protein; uniclust \| UniRef100\_A0A088FQV7 \| 99.6 \| 5e-19 \| 9.8e-25 \| 112.9 \| 97 \| (1, 97) \| 100 \| (16, 113) \| 121 \| Rho termination factor N-terminal domain-containing protein \| Rho termination factor N-terminal domain-containing protein; uniclust \| UniRef100\_A0A2N1AP72 \| 99.6 \| 2.3e-18 \| 4.5e-24 \| 107.5 \| 94 \| (3, 96) \| 100 \| (6, 104) \| 106 \| Rho termination factor N-terminal domain-containing protein \| Rho termination factor N-terminal domain-containing protein; uniclust \| UniRef100\_A0A168CE49 \| 99.5 \| 2.6e-17 \| 5.5e-23 \| 105.3 \| 90 \| (4, 97) \| 100 \| (1, 91) \| 112 \| Phage protein \| Phage protein; uniclust \| UniRef100\_A0A0P8YT19 \| 99.5 \| 2.8e-17 \| 5.7e-23 \| 104.3 \| 95 \| (2, 97) \| 100 \| (18, 112) \| 113 \| Rho termination factor N-terminal domain-containing protein \| Rho termination factor N-terminal domain-containing protein; uniclust \| UniRef100\_A0A089LLN8 \| 99.5 \| 8.7e-17 \| 1.8e-22 \| 99.6 \| 88 \| (6, 98) \| 100 \| (3, 90) \| 94 \| Rho termination factor N-terminal domain-containing protein \| Rho termination factor N-terminal domain-containing protein; uniclust \| UniRef100\_A0A024YGT7 \| 99.4 \| 2.2e-16 \| 4.6e-22 \| 102.4 \| 93 \| (4, 96) \| 100 \| (1, 105) \| 121 \| Rho termination factor N-terminal domain-containing protein \| Rho termination factor N-terminal domain-containing protein; uniclust \| UniRef100\_A0A1C7DPF7 \| 99.3 \| 2.9e-15 \| 6.1e-21 \| 94.3 \| 82 \| (6, 96) \| 100 \| (3, 84) \| 101 \| Rho termination factor N-terminal domain-containing protein \| Rho termination factor N-terminal domain-containing protein; uniclust \| UniRef100\_A0A0E3TAL3 \| 99.2 \| 9.8e-14 \| 2e-19 \| 88.5 \| 93 \| (1, 93) \| 100 \| (1, 105) \| 107 \| Rho termination factor domain protein \| Rho termination factor domain protein; uniclust \| UniRef100\_A0A7Y8VF85 \| 99.1 \| 3.9e-13 \| 7.5e-19 \| 83.7 \| 94 \| (2, 95) \| 100 \| (3, 97) \| 98 \| Rho termination factor N-terminal domain-containing protein \| Rho termination factor N-terminal domain-containing protein; uniclust \| UniRef100\_A0A1F1ZVW8 \| 99.1 \| 3.7e-13 \| 7.7e-19 \| 87.7 \| 90 \| (2, 92) \| 100 \| (3, 115) \| 116 \| Rho termination factor N-terminal domain-containing protein \| Rho termination factor N-terminal domain-containing protein; uniclust \| UniRef100\_A0A0Q3TXV1 \| 99.1 \| 5e-13 \| 9.7e-19 \| 84.8 \| 94 \| (2, 96) \| 100 \| (1, 94) \| 111 \| Rho termination factor N-terminal domain-containing protein \| Rho termination factor N-terminal domain-containing protein; uniclust \| UniRef100\_A0A645IWH2 \| 99.1 \| 5.1e-13 \| 1e-18 \| 86.2 \| 94 \| (4, 97) \| 100 \| (1, 102) \| 119 \| Rho termination factor N-terminal domain-containing protein \| Rho termination factor N-terminal domain-containing protein; uniclust \| UniRef100\_A0A0Q4FWF8 \| 99.1 \| 6.2e-13 \| 1.3e-18 \| 84.9 \| 97 \| (1, 97) \| 100 \| (1, 103) \| 105 \| Rho termination factor N-terminal domain-containing protein \| Rho termination factor N-terminal domain-containing protein; uniclust \| UniRef100\_A0A150KSC3 \| 99.1 \| 7.8e-13 \| 1.6e-18 \| 84.2 \| 95 \| (1, 95) \| 100 \| (1, 97) \| 106 \| Uncharacterized protein \| Uncharacterized protein; uniclust \| UniRef100\_A0A1Y4BC71 \| 99.0 \| 1.1e-12 \| 2.1e-18 \| 84.0 \| 93 \| (4, 97) \| 100 \| (1, 110) \| 117 \| Rho termination factor N-terminal domain-containing protein \| Rho termination factor N-terminal domain-containing protein; uniclust \| UniRef100\_A0A1I5MMY0 \| 99.0 \| 2.3e-12 \| 4.6e-18 \| 85.8 \| 92 \| (3, 94) \| 100 \| (1, 110) \| 136 \| Heat induced stress protein YflT \| Heat induced stress protein YflT; uniclust \| UniRef100\_A0A290GHN0 \| 99.0 \| 2.9e-12 \| 5.8e-18 \| 80.1 \| 90 \| (2, 93) \| 100 \| (1, 91) \| 96 \| HeH/LEM domain-containing protein \| HeH/LEM domain-containing protein; uniclust \| UniRef100\_A0A2P8DHJ0 \| 98.9 \| 8.9e-12 \| 1.8e-17 \| 81.1 \| 76 \| (22, 97) \| 100 \| (10, 88) \| 114 \| Rho termination factor-like protein \| Rho termination factor-like protein; uniclust \| UniRef100\_A0A0Q4LHG8 \| 98.9 \| 1.6e-11 \| 3.1e-17 \| 77.4 \| 91 \| (6, 97) \| 100 \| (3, 94) \| 99 \| Rho termination factor N-terminal domain-containing protein \| Rho termination factor N-terminal domain-containing protein; uniclust \| UniRef100\_A0A0N9YLG6 \| 98.8 \| 2.3e-11 \| 4.8e-17 \| 79.2 \| 96 \| (2, 97) \| 100 \| (4, 105) \| 114 \| HeH/LEM domain-containing protein \| HeH/LEM domain-containing protein; uniclust \| UniRef100\_A0A553SNK1 \| 98.8 \| 2.7e-11 \| 5.3e-17 \| 80.4 \| 92 \| (4, 95) \| 100 \| (1, 121) \| 133 \| Uncharacterized protein \| Uncharacterized protein; uniclust \| UniRef100\_A0A1Q9PIP6 \| 98.8 \| 3.3e-11 \| 6.8e-17 \| 80.2 \| 92 \| (3, 94) \| 100 \| (5, 115) \| 131 \| Rho termination factor N-terminal domain-containing protein \| Rho termination factor N-terminal domain-containing protein; uniclust \| UniRef100\_A0A1M4Y9M8 \| 98.8 \| 4.4e-11 \| 8.9e-17 \| 78.7 \| 96 \| (1, 96) \| 100 \| (10, 120) \| 122 \| Mu-like prophage FluMu N-terminal domain-containing protein \| Mu-like prophage FluMu N-terminal domain-containing protein; uniclust \| UniRef100\_A0A0M3B2K8 \| 98.8 \| 4.3e-11 \| 9e-17 \| 73.1 \| 72 \| (6, 96) \| 100 \| (3, 74) \| 77 \| Rho termination factor N-terminal domain-containing protein \| Rho termination factor N-terminal domain-containing protein; uniclust \| UniRef100\_A0A0Z8CJH6 \| 98.8 \| 6.8e-11 \| 1.3e-16 \| 75.2 \| 82 \| (13, 94) \| 100 \| (12, 101) \| 102 \| Phage protein \| Phage protein; uniclust \| UniRef100\_A0A1H8QNZ5 \| 98.7 \| 8.5e-11 \| 1.7e-16 \| 74.3 \| 92 \| (2, 93) \| 100 \| (4, 96) \| 99 \| Rho termination factor, N-terminal domain \| Rho termination factor, N-terminal domain; uniclust \| UniRef100\_A0A174TMF5 \| 98.7 \| 9.7e-11 \| 2e-16 \| 81.0 \| 93 \| (2, 94) \| 100 \| (19, 150) \| 169 \| Rho termination factor N-terminal domain-containing protein \| Rho termination factor N-terminal domain-containing protein; uniclust \| UniRef100\_A0A089JTJ5 \| 98.7 \| 1.1e-10 \| 2.3e-16 \| 74.4 \| 78 \| (12, 94) \| 100 \| (12, 90) \| 104 \| Rho termination factor N-terminal domain-containing protein \| Rho termination factor N-terminal domain-containing protein; uniclust \| UniRef100\_A0A090HXG2 \| 98.7 \| 1.2e-10 \| 2.5e-16 \| 72.5 \| 76 \| (11, 91) \| 100 \| (9, 85) \| 87 \| HeH/LEM domain-containing protein \| HeH/LEM domain-containing protein; uniclust \| UniRef100\_A0A972MRU9 \| 98.7 \| 1.5e-10 \| 2.8e-16 \| 70.4 \| 87 \| (7, 95) \| 100 \| (4, 90) \| 92 \| Rho termination factor N-terminal domain-containing protein \| Rho termination factor N-terminal domain-containing protein; uniclust \| UniRef100\_A0A1Y4VDP2 \| 98.7 \| 1.4e-10 \| 2.9e-16 \| 78.4 \| 94 \| (3, 96) \| 100 \| (1, 124) \| 138 \| Rho termination factor N-terminal domain-containing protein \| Rho termination factor N-terminal domain-containing protein; uniclust \| UniRef100\_A0A173R5M0 \| 98.7 \| 1.6e-10 \| 3.2e-16 \| 78.2 \| 75 \| (22, 96) \| 100 \| (30, 124) \| 147 \| Rho termination factor N-terminal domain-containing protein \| Rho termination factor N-terminal domain-containing protein; uniclust \| UniRef100\_A0A0N1AA76 \| 98.7 \| 1.7e-10 \| 3.6e-16 \| 74.4 \| 94 \| (1, 96) \| 100 \| (1, 101) \| 103 \| Rho termination factor N-terminal domain-containing protein \| Rho termination factor N-terminal domain-containing protein; uniclust \| UniRef100\_A0A0F0C809 \| 98.7 \| 2e-10 \| 4.2e-16 \| 71.5 \| 66 \| (28, 93) \| 100 \| (13, 80) \| 83 \| Rho termination factor N-terminal domain-containing protein \| Rho termination factor N-terminal domain-containing protein; uniclust \| UniRef100\_A0A099WPM7 \| 98.7 \| 2.2e-10 \| 4.4e-16 \| 72.5 \| 87 \| (6, 95) \| 100 \| (7, 96) \| 98 \| Uncharacterized protein \| Uncharacterized protein; uniclust \| UniRef100\_A0A1C5LBF0 \| 98.6 \| 2.4e-10 \| 5e-16 \| 71.4 \| 66 \| (29, 94) \| 100 \| (14, 83) \| 86 \| Rho termination factor, N-terminal domain \| Rho termination factor, N-terminal domain; uniclust \| UniRef100\_A0A101WGP7 \| 98.6 \| 3.4e-10 \| 7e-16 \| 76.4 \| 93 \| (4, 96) \| 100 \| (1, 120) \| 142 \| Rho termination factor N-terminal domain-containing protein \| Rho termination factor N-terminal domain-containing protein; uniclust \| UniRef100\_A0A0A8IK76 \| 98.6 \| 4.1e-10 \| 8.2e-16 \| 69.0 \| 74 \| (9, 97) \| 100 \| (4, 79) \| 80 \| Rho termination factor N-terminal domain-containing protein \| Rho termination factor N-terminal domain-containing protein; uniclust \| UniRef100\_A0A0V8D0U7 \| 98.6 \| 5.3e-10 \| 1.1e-15 \| 73.4 \| 91 \| (4, 94) \| 100 \| (1, 94) \| 117 \| Conjugal transfer protein \| Conjugal transfer protein; uniclust \| UniRef100\_A0A0S7Z717 \| 98.6 \| 5.9e-10 \| 1.1e-15 \| 72.0 \| 88 \| (5, 93) \| 100 \| (1, 97) \| 120 \| Rho termination factor N-terminal domain-containing protein \| Rho termination factor N-terminal domain-containing protein; uniclust \| UniRef100\_A0A2G1LY20 \| 98.6 \| 6.3e-10 \| 1.2e-15 \| 69.5 \| 90 \| (6, 95) \| 100 \| (6, 96) \| 99 \| Rho termination factor N-terminal domain-containing protein \| Rho termination factor N-terminal domain-containing protein; uniclust \| UniRef100\_A0A1G9TZJ8 \| 98.6 \| 7.2e-10 \| 1.4e-15 \| 64.3 \| 37 \| (53, 89) \| 100 \| (24, 60) \| 61 \| Rho termination factor, N-terminal domain \| Rho termination factor, N-terminal domain; uniclust \| UniRef100\_A0A1M6EXN4 \| 98.5 \| 8.1e-10 \| 1.7e-15 \| 73.3 \| 95 \| (1, 95) \| 100 \| (5, 104) \| 122 \| Rho termination factor, N-terminal domain \| Rho termination factor, N-terminal domain; uniclust \| UniRef100\_A0A5K7YYK9 \| 98.5 \| 1e-09 \| 1.9e-15 \| 68.6 \| 90 \| (6, 95) \| 100 \| (4, 95) \| 99 \| HeH/LEM domain-containing protein \| HeH/LEM domain-containing protein; uniclust \| UniRef100\_A0A1W7ABQ3 \| 98.5 \| 9.3e-10 \| 2e-15 \| 70.7 \| 84 \| (11, 94) \| 100 \| (9, 98) \| 99 \| Phage protein \| Phage protein; uniclust \| UniRef100\_A0A1C6JH25 \| 98.5 \| 1.2e-09 \| 2.5e-15 \| 74.5 \| 94 \| (4, 97) \| 100 \| (1, 128) \| 150 \| Rho termination factor N-terminal domain-containing protein \| Rho termination factor N-terminal domain-containing protein; uniclust \| UniRef100\_A0A845NDA8 \| 98.5 \| 1.8e-09 \| 3.4e-15 \| 66.6 \| 93 \| (2, 96) \| 100 \| (1, 93) \| 97 \| Rho termination factor N-terminal domain-containing protein \| Rho termination factor N-terminal domain-containing protein; uniclust \| UniRef100\_A0A0U9HBT6 \| 98.5 \| 2.1e-09 \| 3.8e-15 \| 66.3 \| 89 \| (5, 93) \| 100 \| (1, 91) \| 96 \| Uncharacterized protein \| Uncharacterized protein; uniclust \| UniRef100\_A0A349GRC7 \| 98.4 \| 2.4e-09 \| 4.5e-15 \| 67.4 \| 91 \| (2, 94) \| 100 \| (1, 101) \| 103 \| Rho termination factor N-terminal domain-containing protein \| Rho termination factor N-terminal domain-containing protein; uniclust \| UniRef100\_A0A1W6JNK7 \| 98.4 \| 2.3e-09 \| 4.7e-15 \| 68.1 \| 89 \| (1, 93) \| 100 \| (1, 91) \| 96 \| Transcriptional terminator \| Transcriptional terminator; uniclust \| UniRef100\_A0A0Z8DCB5 \| 98.4 \| 2.5e-09 \| 4.9e-15 \| 67.5 \| 91 \| (1, 93) \| 100 \| (1, 97) \| 98 \| HeH/LEM domain \| HeH/LEM domain; uniclust \| UniRef100\_A0A0A8JM68 \| 98.4 \| 2.7e-09 \| 5.3e-15 \| 71.1 \| 92 \| (4, 95) \| 100 \| (2, 114) \| 130 \| Uncharacterized protein \| Uncharacterized protein; uniclust \| UniRef100\_A0A1C5YVF9 \| 98.4 \| 2.9e-09 \| 5.8e-15 \| 64.0 \| 38 \| (54, 91) \| 100 \| (32, 69) \| 70 \| HeH/LEM domain \| HeH/LEM domain; uniclust \| UniRef100\_A0A1R0X056 \| 98.4 \| 4.2e-09 \| 7.7e-15 \| 67.9 \| 91 \| (7, 97) \| 100 \| (12, 106) \| 124 \| Rho termination factor N-terminal domain-containing protein \| Rho termination factor N-terminal domain-containing protein; uniclust \| UniRef100\_A0A1M6AHL9 \| 98.4 \| 3.8e-09 \| 7.8e-15 \| 67.0 \| 65 \| (31, 95) \| 100 \| (18, 88) \| 91 \| HeH/LEM domain-containing protein \| HeH/LEM domain-containing protein; uniclust \| UniRef100\_A0A0U3P188 \| 98.4 \| 4.3e-09 \| 8.3e-15 \| 65.4 \| 85 \| (3, 92) \| 100 \| (2, 88) \| 89 \| T4 recombination endonuclease VII dimerisation domain-containing protein \| T4 recombination endonuclease VII dimerisation domain-containing protein; uniclust \| UniRef100\_A0A1I4PMR5 \| 98.4 \| 4.3e-09 \| 8.8e-15 \| 70.3 \| 88 \| (3, 90) \| 100 \| (1, 99) \| 125 \| Heat induced stress protein YflT \| Heat induced stress protein YflT; uniclust \| UniRef100\_A0A1Y3TP18 \| 98.4 \| 4.5e-09 \| 9.2e-15 \| 72.7 \| 94 \| (2, 95) \| 100 \| (1, 147) \| 160 \| Rho termination factor N-terminal domain-containing protein \| Rho termination factor N-terminal domain-containing protein; uniclust \| UniRef100\_A0A0T6DTS4 \| 98.4 \| 5e-09 \| 9.6e-15 \| 66.1 \| 70 \| (20, 89) \| 100 \| (16, 93) \| 96 \| T4 recombination endonuclease VII dimerisation domain-containing protein \| T4 recombination endonuclease VII dimerisation domain-containing protein; uniclust \| UniRef100\_A0A069A8G7 \| 98.4 \| 4.8e-09 \| 9.8e-15 \| 65.4 \| 60 \| (31, 95) \| 100 \| (19, 78) \| 84 \| Rho termination factor N-terminal domain-containing protein \| Rho termination factor N-terminal domain-containing protein; uniclust \| UniRef100\_UPI0006231A7A \| 98.3 \| 6.5e-09 \| 1.2e-14 \| 65.8 \| 72 \| (23, 94) \| 100 \| (25, 99) \| 111 \| Rho termination factor N-terminal domain-containing protein \| Rho termination factor N-terminal domain-containing protein; uniclust \| UniRef100\_A0A0J7Y8Q0 \| 98.3 \| 6.2e-09 \| 1.3e-14 \| 67.3 \| 73 \| (19, 96) \| 100 \| (27, 101) \| 105 \| Rho termination factor N-terminal domain-containing protein \| Rho termination factor N-terminal domain-containing protein; uniclust \| UniRef100\_A0A943MCU2 \| 98.3 \| 7.1e-09 \| 1.3e-14 \| 67.2 \| 91 \| (5, 95) \| 100 \| (1, 110) \| 117 \| Uncharacterized protein \| Uncharacterized protein; uniclust \| UniRef100\_A0A0D7X2K7 \| 98.3 \| 7.1e-09 \| 1.5e-14 \| 69.9 \| 85 \| (2, 88) \| 100 \| (1, 106) \| 127 \| Uncharacterized protein \| Uncharacterized protein; uniclust \| UniRef100\_A0A7V7HMI4 \| 98.3 \| 8.9e-09 \| 1.7e-14 \| 62.5 \| 67 \| (22, 90) \| 100 \| (9, 75) \| 77 \| Rho termination factor N-terminal domain-containing protein \| Rho termination factor N-terminal domain-containing protein; uniclust \| UniRef100\_A0A0X1RV11 \| 98.3 \| 1.2e-08 \| 2.4e-14 \| 63.7 \| 45 \| (53, 97) \| 100 \| (30, 74) \| 85 \| HeH/LEM domain protein \| HeH/LEM domain protein; uniclust \| UniRef100\_A0A845R2Z6 \| 98.3 \| 1.4e-08 \| 2.5e-14 \| 62.1 \| 83 \| (2, 91) \| 100 \| (1, 88) \| 89 \| Rho termination protein \| Rho termination protein; uniclust \| UniRef100\_A0A0T7K7C9 \| 98.2 \| 1.4e-08 \| 2.7e-14 \| 60.1 \| 41 \| (55, 95) \| 100 \| (17, 57) \| 65 \| Transcription termination factor \| Transcription termination factor; uniclust \| UniRef100\_A0A142KCR9 \| 98.2 \| 1.3e-08 \| 2.8e-14 \| 70.7 \| 93 \| (1, 93) \| 100 \| (9, 149) \| 151 \| Tail assembly chaperone \| Tail assembly chaperone; uniclust \| UniRef100\_A0A1B1G6D0 \| 98.2 \| 1.5e-08 \| 2.9e-14 \| 63.1 \| 92 \| (1, 93) \| 100 \| (1, 94) \| 95 \| Rho termination factor N-terminal domain-containing protein \| Rho termination factor N-terminal domain-containing protein; uniclust \| UniRef100\_A0A383RBW5 \| 98.2 \| 1.5e-08 \| 3e-14 \| 63.3 \| 42 \| (53, 94) \| 100 \| (38, 79) \| 82 \| Rho termination factor N-terminal domain-containing protein \| Rho termination factor N-terminal domain-containing protein; uniclust \| UniRef100\_A0A2D4DNJ5 \| 98.2 \| 1.6e-08 \| 3.3e-14 \| 63.6 \| 75 \| (14, 96) \| 100 \| (12, 86) \| 89 \| Phage protein \| Phage protein; uniclust \| UniRef100\_A0A0M2LYI1 \| 98.2 \| 1.9e-08 \| 3.7e-14 \| 64.3 \| 74 \| (23, 99) \| 100 \| (10, 83) \| 96 \| Rho termination factor N-terminal domain-containing protein \| Rho termination factor N-terminal domain-containing protein; uniclust \| UniRef100\_A0A1G8IAC0 \| 98.2 \| 1.9e-08 \| 3.7e-14 \| 62.6 \| 39 \| (53, 91) \| 100 \| (42, 80) \| 82 \| Rho termination factor, N-terminal domain \| Rho termination factor, N-terminal domain; uniclust \| UniRef100\_A0A0L0W633 \| 98.2 \| 2.4e-08 \| 4.5e-14 \| 55.5 \| 42 \| (55, 96) \| 100 \| (8, 49) \| 52 \| Transcription termination factor \| Transcription termination factor; uniclust \| UniRef100\_UPI002033AFD5 \| 98.2 \| 2.4e-08 \| 4.6e-14 \| 63.1 \| 80 \| (13, 92) \| 100 \| (11, 101) \| 103 \| Rho termination factor N-terminal domain-containing protein \| Rho termination factor N-terminal domain-containing protein; uniclust \| UniRef100\_A0A352NNW6 \| 98.2 \| 2.2e-08 \| 4.6e-14 \| 62.9 \| 39 \| (55, 93) \| 100 \| (45, 83) \| 85 \| Rho termination factor N-terminal domain-containing protein \| Rho termination factor N-terminal domain-containing protein; uniclust \| UniRef100\_A0A2W4IJI6 \| 98.1 \| 3.9e-08 \| 7.4e-14 \| 63.6 \| 85 \| (3, 91) \| 100 \| (4, 92) \| 115 \| Rho termination factor N-terminal domain-containing protein \| Rho termination factor N-terminal domain-containing protein; uniclust \| UniRef100\_A0A1H4QZW6 \| 98.1 \| 3.8e-08 \| 7.9e-14 \| 63.6 \| 89 \| (1, 99) \| 100 \| (1, 89) \| 97 \| HeH/LEM domain-containing protein \| HeH/LEM domain-containing protein; uniclust \| UniRef100\_A0A089MTA0 \| 98.1 \| 4.4e-08 \| 8.7e-14 \| 64.8 \| 92 \| (2, 93) \| 100 \| (1, 121) \| 123 \| Rho termination factor N-terminal domain-containing protein \| Rho termination factor N-terminal domain-containing protein; uniclust \| UniRef100\_A0A0F6Z7H9 \| 98.1 \| 4.2e-08 \| 9.2e-14 \| 67.5 \| 94 \| (1, 94) \| 100 \| (1, 138) \| 140 \| Uncharacterized protein \| Uncharacterized protein; uniclust \| UniRef100\_E6X1P2 \| 98.1 \| 5e-08 \| 9.3e-14 \| 59.4 \| 81 \| (7, 93) \| 100 \| (4, 84) \| 85 \| Rho termination factor domain protein \| Rho termination factor domain protein; uniclust \| UniRef100\_A0A1E2SK37 \| 98.1 \| 5.4e-08 \| 1.1e-13 \| 64.6 \| 73 \| (22, 94) \| 100 \| (33, 111) \| 127 \| Rho termination factor N-terminal domain-containing protein \| Rho termination factor N-terminal domain-containing protein; uniclust \| UniRef100\_C4L0Q0 \| 98.1 \| 5.6e-08 \| 1.1e-13 \| 62.7 \| 93 \| (3, 95) \| 100 \| (4, 106) \| 112 \| Rho termination factor N-terminal domain-containing protein \| Rho termination factor N-terminal domain-containing protein; uniclust \| UniRef100\_A0A1J0TTB7 \| 98.1 \| 5.5e-08 \| 1.1e-13 \| 64.4 \| 37 \| (54, 90) \| 100 \| (63, 99) \| 117 \| Rho termination factor N-terminal domain-containing protein \| Rho termination factor N-terminal domain-containing protein; uniclust \| UniRef100\_A0A414PRA9 \| 98.1 \| 6.3e-08 \| 1.2e-13 \| 60.4 \| 86 \| (5, 91) \| 100 \| (1, 93) \| 98 \| HeH/LEM domain-containing protein \| HeH/LEM domain-containing protein; uniclust \| UniRef100\_A0A1M2ZYQ4 \| 98.1 \| 6.2e-08 \| 1.2e-13 \| 62.4 \| 90 \| (6, 95) \| 100 \| (4, 105) \| 106 \| Rho termination factor N-terminal domain-containing protein \| Rho termination factor N-terminal domain-containing protein; uniclust \| UniRef100\_A0A0H3J9M1 \| 98.1 \| 5.7e-08 \| 1.2e-13 \| 65.2 \| 92 \| (2, 94) \| 100 \| (1, 97) \| 122 \| Rho termination factor N-terminal domain-containing protein \| Rho termination factor N-terminal domain-containing protein; uniclust \| UniRef100\_A0A2D9NX61 \| 98.0 \| 7.6e-08 \| 1.5e-13 \| 60.6 \| 76 \| (13, 98) \| 100 \| (14, 91) \| 93 \| Rho termination factor N-terminal domain-containing protein \| Rho termination factor N-terminal domain-containing protein; uniclust \| UniRef100\_UPI0003493A31 \| 98.0 \| 8.7e-08 \| 1.6e-13 \| 60.5 \| 82 \| (12, 93) \| 100 \| (11, 93) \| 105 \| Rho termination factor N-terminal domain-containing protein \| Rho termination factor N-terminal domain-containing protein; uniclust \| UniRef100\_A0A173R5B0 \| 98.0 \| 8.6e-08 \| 1.7e-13 \| 61.9 \| 87 \| (10, 98) \| 100 \| (8, 97) \| 108 \| Rho termination factor N-terminal domain-containing protein \| Rho termination factor N-terminal domain-containing protein; uniclust \| UniRef100\_A0A373L6M1 \| 98.0 \| 8.5e-08 \| 1.7e-13 \| 58.4 \| 42 \| (53, 94) \| 100 \| (23, 64) \| 74 \| Rho termination factor N-terminal domain-containing protein \| Rho termination factor N-terminal domain-containing protein; uniclust \| UniRef100\_A0A951JXR5 \| 98.0 \| 1e-07 \| 1.9e-13 \| 48.6 \| 32 \| (60, 91) \| 100 \| (1, 32) \| 33 \| Rho termination factor N-terminal domain-containing protein \| Rho termination factor N-terminal domain-containing protein; uniclust \| UniRef100\_A0A1C5TAF6 \| 98.0 \| 9.5e-08 \| 1.9e-13 \| 62.0 \| 43 \| (52, 94) \| 100 \| (59, 101) \| 102 \| Rho termination factor, N-terminal domain \| Rho termination factor, N-terminal domain; uniclust \| UniRef100\_A0A3D1GE26 \| 98.0 \| 1e-07 \| 1.9e-13 \| 60.1 \| 39 \| (53, 91) \| 100 \| (55, 93) \| 94 \| 50S ribosomal protein L21 (Fragment) \| 50S ribosomal protein L21 (Fragment); uniclust \| UniRef100\_A0A022LHQ7 \| 98.0 \| 9.9e-08 \| 2e-13 \| 63.5 \| 66 \| (29, 94) \| 100 \| (33, 112) \| 116 \| Rho termination factor \| Rho termination factor; uniclust \| UniRef100\_A0A6G4H4V0 \| 98.0 \| 1.1e-07 \| 2.1e-13 \| 53.3 \| 38 \| (54, 91) \| 100 \| (14, 51) \| 52 \| Termination factor Rho \| Termination factor Rho; uniclust \| UniRef100\_A0A1D9GT57 \| 98.0 \| 1e-07 \| 2.2e-13 \| 65.3 \| 82 \| (15, 96) \| 100 \| (24, 131) \| 142 \| Phage tail protein \| Phage tail protein; uniclust \| UniRef100\_A0A1G6NU17 \| 98.0 \| 1.1e-07 \| 2.4e-13 \| 66.5 \| 78 \| (16, 93) \| 100 \| (17, 101) \| 163 \| Rho termination factor N-terminal domain-containing protein \| Rho termination factor N-terminal domain-containing protein; uniclust \| UniRef100\_A0A1W6RR01 \| 98.0 \| 1.2e-07 \| 2.4e-13 \| 60.6 \| 66 \| (27, 92) \| 100 \| (28, 98) \| 99 \| HeH/LEM domain-containing protein \| HeH/LEM domain-containing protein; uniclust \| UniRef100\_A0A1M4Z686 \| 98.0 \| 1.3e-07 \| 2.5e-13 \| 52.2 \| 39 \| (54, 92) \| 100 \| (6, 44) \| 46 \| Rho termination factor, N-terminal domain \| Rho termination factor, N-terminal domain; uniclust \| UniRef100\_A0A359B3E5 \| 98.0 \| 1.5e-07 \| 2.8e-13 \| 61.2 \| 94 \| (5, 98) \| 100 \| (1, 102) \| 116 \| Rho termination factor N-terminal domain-containing protein \| Rho termination factor N-terminal domain-containing protein; uniclust \| UniRef100\_A0A0L6JGZ7 \| 98.0 \| 1.4e-07 \| 2.8e-13 \| 61.3 \| 69 \| (20, 94) \| 100 \| (22, 90) \| 100 \| Rho termination factor N-terminal domain-containing protein \| Rho termination factor N-terminal domain-containing protein; uniclust \| UniRef100\_A0A0B5Q9F7 \| 98.0 \| 1.4e-07 \| 2.8e-13 \| 59.5 \| 64 \| (30, 93) \| 100 \| (19, 86) \| 89 \| Rho termination factor N-terminal domain-containing protein \| Rho termination factor N-terminal domain-containing protein; uniclust \| UniRef100\_A0A074L0V4 \| 98.0 \| 1.5e-07 \| 3e-13 \| 61.8 \| 67 \| (29, 95) \| 100 \| (27, 101) \| 109 \| Rho termination factor \| Rho termination factor; uniclust \| UniRef100\_A0A076NJ41 \| 97.9 \| 1.6e-07 \| 3.3e-13 \| 62.6 \| 91 \| (4, 94) \| 100 \| (1, 117) \| 120 \| Phage protein \| Phage protein; uniclust \| UniRef100\_A0A921G371 \| 97.9 \| 1.8e-07 \| 3.4e-13 \| 60.3 \| 88 \| (7, 94) \| 100 \| (5, 101) \| 108 \| Uncharacterized protein \| Uncharacterized protein; uniclust \| UniRef100\_A0A2I8AAC4 \| 97.9 \| 1.8e-07 \| 3.5e-13 \| 48.2 \| 32 \| (60, 91) \| 100 \| (1, 32) \| 33 \| Rho termination factor N-terminal domain-containing protein \| Rho termination factor N-terminal domain-containing protein; uniclust \| UniRef100\_A0A1C5WIW1 \| 97.9 \| 1.7e-07 \| 3.5e-13 \| 59.1 \| 77 \| (9, 91) \| 100 \| (3, 79) \| 83 \| Rho termination factor N-terminal domain-containing protein \| Rho termination factor N-terminal domain-containing protein; uniclust \| UniRef100\_A0A1A0LLB8 \| 97.9 \| 2.1e-07 \| 3.8e-13 \| 58.7 \| 86 \| (4, 89) \| 100 \| (3, 98) \| 99 \| Uncharacterized protein \| Uncharacterized protein; uniclust \| UniRef100\_A5EUZ2 \| 97.9 \| 2.2e-07 \| 4.1e-13 \| 55.6 \| 74 \| (5, 90) \| 100 \| (1, 74) \| 75 \| T4 recombination endonuclease VII dimerisation domain-containing protein \| T4 recombination endonuclease VII dimerisation domain-containing protein; uniclust \| UniRef100\_A0A0A1W8H8 \| 97.9 \| 2.1e-07 \| 4.2e-13 \| 60.7 \| 85 \| (10, 95) \| 100 \| (12, 99) \| 103 \| Rho termination factor N-terminal domain-containing protein \| Rho termination factor N-terminal domain-containing protein; uniclust \| UniRef100\_A0A8J6SEE7 \| 97.9 \| 2.3e-07 \| 4.3e-13 \| 49.0 \| 36 \| (60, 95) \| 100 \| (1, 36) \| 38 \| Rho termination factor N-terminal domain-containing protein \| Rho termination factor N-terminal domain-containing protein; uniclust \| UniRef100\_A0A3G3BWG3 \| 97.9 \| 2.3e-07 \| 4.4e-13 \| 61.5 \| 92 \| (4, 95) \| 100 \| (1, 111) \| 124 \| Tail assembly chaperone \| Tail assembly chaperone; uniclust \| UniRef100\_A0A1I1D1B8 \| 97.9 \| 2.4e-07 \| 4.8e-13 \| 60.5 \| 95 \| (4, 98) \| 100 \| (2, 103) \| 105 \| HeH/LEM domain-containing protein \| HeH/LEM domain-containing protein; uniclust \| UniRef100\_UPI000490AEB4 \| 97.9 \| 2.9e-07 \| 5.3e-13 \| 56.9 \| 86 \| (7, 95) \| 100 \| (3, 88) \| 91 \| HeH/LEM domain-containing protein \| HeH/LEM domain-containing protein; uniclust \| UniRef100\_A0A849D8D2 \| 97.9 \| 2.8e-07 \| 5.3e-13 \| 53.1 \| 37 \| (54, 90) \| 100 \| (19, 55) \| 56 \| 50S ribosomal protein L21 \| 50S ribosomal protein L21; uniclust \| UniRef100\_A0A175R7S3 \| 97.9 \| 2.8e-07 \| 5.4e-13 \| 56.1 \| 68 \| (9, 92) \| 100 \| (4, 71) \| 72 \| Rho termination factor N-terminal domain-containing protein \| Rho termination factor N-terminal domain-containing protein; uniclust \| UniRef100\_A0A1C5THF0 \| 97.9 \| 2.8e-07 \| 5.5e-13 \| 59.6 \| 63 \| (32, 94) \| 100 \| (40, 102) \| 104 \| Rho termination factor N-terminal domain-containing protein \| Rho termination factor N-terminal domain-containing protein; uniclust \| UniRef100\_UPI000D3A7A5A \| 97.9 \| 3.3e-07 \| 6.1e-13 \| 58.2 \| 89 \| (5, 93) \| 100 \| (1, 105) \| 107 \| Rho termination factor N-terminal domain-containing protein \| Rho termination factor N-terminal domain-containing protein; uniclust \| UniRef100\_A0A957ZEC5 \| 97.8 \| 3.8e-07 \| 7.1e-13 \| 57.8 \| 76 \| (19, 94) \| 100 \| (20, 102) \| 105 \| Rho termination factor N-terminal domain-containing protein \| Rho termination factor N-terminal domain-containing protein; uniclust \| UniRef100\_A0A049DS68 \| 97.8 \| 3.5e-07 \| 7.5e-13 \| 62.4 \| 96 \| (1, 96) \| 100 \| (4, 124) \| 130 \| Uncharacterized protein \| Uncharacterized protein; uniclust \| UniRef100\_A0A1C6BBU0 \| 97.8 \| 3.9e-07 \| 7.5e-13 \| 55.4 \| 37 \| (54, 90) \| 100 \| (39, 75) \| 76 \| Transcription termination factor Rho \| Transcription termination factor Rho; uniclust \| UniRef100\_UPI00203B3015 \| 97.8 \| 4.1e-07 \| 7.5e-13 \| 56.4 \| 83 \| (1, 92) \| 100 \| (1, 90) \| 92 \| Rho termination factor N-terminal domain-containing protein \| Rho termination factor N-terminal domain-containing protein; uniclust \| UniRef100\_UPI001CCEC438 \| 97.8 \| 4.1e-07 \| 7.6e-13 \| 57.7 \| 80 \| (15, 94) \| 100 \| (20, 105) \| 106 \| hypothetical protein \| hypothetical protein; uniclust \| UniRef100\_G9X2L6 \| 97.8 \| 4.2e-07 \| 8.1e-13 \| 56.2 \| 57 \| (33, 93) \| 100 \| (23, 79) \| 82 \| Rho termination factor N-terminal domain-containing protein \| Rho termination factor N-terminal domain-containing protein; uniclust \| UniRef100\_A0A4R8FYN6 \| 97.8 \| 4.7e-07 \| 8.6e-13 \| 57.6 \| 41 \| (53, 93) \| 100 \| (58, 98) \| 107 \| Rho termination factor-like protein \| Rho termination factor-like protein; uniclust \| UniRef100\_UPI000D0298A2 \| 97.8 \| 4.7e-07 \| 8.6e-13 \| 56.4 \| 85 \| (2, 91) \| 100 \| (1, 92) \| 94 \| hypothetical protein \| hypothetical protein; uniclust \| UniRef100\_A0A150NVU6 \| 97.8 \| 4.7e-07 \| 9.1e-13 \| 56.1 \| 40 \| (54, 93) \| 100 \| (37, 76) \| 80 \| Phage transcriptional terminator \| Phage transcriptional terminator; uniclust \| UniRef100\_A0A0C6FQZ8 \| 97.8 \| 4.6e-07 \| 9.2e-13 \| 58.0 \| 72 \| (22, 93) \| 100 \| (13, 93) \| 95 \| Uncharacterized protein \| Uncharacterized protein; uniclust \| UniRef100\_A0A1C6AJB8 \| 97.8 \| 4.7e-07 \| 9.3e-13 \| 57.4 \| 48 \| (52, 99) \| 100 \| (42, 89) \| 89 \| Rho termination factor N-terminal domain-containing protein \| Rho termination factor N-terminal domain-containing protein; uniclust \| UniRef100\_A0A073J6G9 \| 97.8 \| 4.9e-07 \| 9.8e-13 \| 61.4 \| 68 \| (29, 96) \| 100 \| (48, 128) \| 134 \| Rho termination factor \| Rho termination factor; uniclust \| UniRef100\_A0A8S5MRJ0 \| 97.8 \| 5.6e-07 \| 1.1e-12 \| 47.8 \| 35 \| (60, 94) \| 100 \| (1, 36) \| 37 \| HeH/LEM domain \| HeH/LEM domain; uniclust \| UniRef100\_A0A4Q8R804 \| 97.8 \| 6.6e-07 \| 1.3e-12 \| 63.6 \| 75 \| (18, 92) \| 100 \| (32, 144) \| 198 \| Rho termination factor N-terminal domain-containing protein \| Rho termination factor N-terminal domain-containing protein; uniclust \| UniRef100\_A0A017RV20 \| 97.8 \| 6.3e-07 \| 1.3e-12 \| 59.5 \| 79 \| (17, 95) \| 100 \| (17, 97) \| 112 \| Rho termination factor N-terminal domain-containing protein \| Rho termination factor N-terminal domain-containing protein; uniclust \| UniRef100\_A0A1Q9PJD2 \| 97.7 \| 6.8e-07 \| 1.3e-12 \| 57.4 \| 45 \| (53, 97) \| 100 \| (51, 95) \| 99 \| Rho termination factor N-terminal domain-containing protein \| Rho termination factor N-terminal domain-containing protein; uniclust \| UniRef100\_UPI0022FF5F71 \| 97.7 \| 7.8e-07 \| 1.4e-12 \| 57.3 \| 81 \| (14, 95) \| 100 \| (16, 96) \| 115 \| Rho termination factor N-terminal domain-containing protein \| Rho termination factor N-terminal domain-containing protein; uniclust \| UniRef100\_A0A7X9A1P3 \| 97.7 \| 8e-07 \| 1.5e-12 \| 56.9 \| 74 \| (17, 90) \| 100 \| (19, 111) \| 111 \| Rho termination factor N-terminal domain-containing protein \| Rho termination factor N-terminal domain-containing protein; uniclust \| UniRef100\_A0A1L6BZF2 \| 97.7 \| 7.4e-07 \| 1.5e-12 \| 60.0 \| 88 \| (4, 92) \| 100 \| (1, 121) \| 126 \| Uncharacterized protein \| Uncharacterized protein; uniclust \| UniRef100\_UPI00214DC62F \| 97.7 \| 9.4e-07 \| 1.7e-12 \| 52.2 \| 41 \| (54, 94) \| 100 \| (27, 67) \| 68 \| Rho termination factor N-terminal domain-containing protein \| Rho termination factor N-terminal domain-containing protein; uniclust \| UniRef100\_UPI00073A2B65 \| 97.7 \| 9.5e-07 \| 1.8e-12 \| 54.7 \| 88 \| (5, 93) \| 100 \| (1, 88) \| 90 \| hypothetical protein \| hypothetical protein; uniclust \| UniRef100\_A0A7X9AD68 \| 97.7 \| 1.1e-06 \| 2e-12 \| 53.4 \| 74 \| (13, 89) \| 100 \| (5, 78) \| 80 \| Rho termination factor N-terminal domain-containing protein \| Rho termination factor N-terminal domain-containing protein; uniclust \| UniRef100\_A0A385TUN3 \| 97.7 \| 1.2e-06 \| 2.1e-12 \| 53.6 \| 70 \| (11, 91) \| 100 \| (8, 79) \| 81 \| Rho termination factor N-terminal domain-containing protein \| Rho termination factor N-terminal domain-containing protein; uniclust \| UniRef100\_A0A4Y8UXJ0 \| 97.7 \| 1.1e-06 \| 2.2e-12 \| 57.0 \| 86 \| (5, 92) \| 100 \| (2, 90) \| 110 \| SAP domain-containing protein \| SAP domain-containing protein; uniclust \| UniRef100\_A0A0U3F3Q7 \| 97.7 \| 1.2e-06 \| 2.4e-12 \| 58.6 \| 92 \| (5, 96) \| 100 \| (3, 106) \| 115 \| HeH/LEM domain-containing protein \| HeH/LEM domain-containing protein; uniclust \| UniRef100\_A0A8I0I2U5 \| 97.7 \| 1.2e-06 \| 2.5e-12 \| 58.4 \| 87 \| (13, 99) \| 100 \| (18, 117) \| 117 \| Uncharacterized protein \| Uncharacterized protein; uniclust \| UniRef100\_A0A1U9WR62 \| 97.6 \| 1.4e-06 \| 2.6e-12 \| 56.6 \| 88 \| (6, 93) \| 100 \| (3, 109) \| 112 \| Rho termination factor N-terminal domain-containing protein \| Rho termination factor N-terminal domain-containing protein; uniclust \| UniRef100\_A0A3N9UVK7 \| 97.6 \| 1.5e-06 \| 2.9e-12 \| 56.1 \| 76 \| (14, 97) \| 100 \| (13, 91) \| 97 \| Rho termination factor N-terminal domain-containing protein \| Rho termination factor N-terminal domain-containing protein; uniclust \| UniRef100\_E3PRZ2 \| 97.6 \| 1.6e-06 \| 2.9e-12 \| 54.3 \| 79 \| (13, 91) \| 100 \| (13, 93) \| 95 \| Rho termination factor N-terminal domain-containing protein \| Rho termination factor N-terminal domain-containing protein; uniclust \| UniRef100\_A0A173SN79 \| 97.6 \| 1.5e-06 \| 3e-12 \| 60.1 \| 88 \| (6, 93) \| 100 \| (3, 126) \| 150 \| Rho termination factor N-terminal domain-containing protein \| Rho termination factor N-terminal domain-containing protein; uniclust \| UniRef100\_A0A1Y5NZV1 \| 97.6 \| 1.6e-06 \| 3.1e-12 \| 51.6 \| 37 \| (55, 91) \| 100 \| (28, 64) \| 65 \| Rho termination factor N-terminal domain-containing protein \| Rho termination factor N-terminal domain-containing protein; uniclust \| UniRef100\_A0A1V4VSQ7 \| 97.6 \| 1.7e-06 \| 3.2e-12 \| 54.9 \| 73 \| (18, 91) \| 100 \| (14, 101) \| 103 \| Rho termination factor N-terminal domain-containing protein \| Rho termination factor N-terminal domain-containing protein; uniclust \| UniRef100\_A0A8B4Q926 \| 97.6 \| 1.7e-06 \| 3.2e-12 \| 54.8 \| 33 \| (54, 86) \| 100 \| (49, 81) \| 92 \| Rho termination factor, N-terminal domain \| Rho termination factor, N-terminal domain; uniclust \| UniRef100\_A0A0Q4GFT6 \| 97.6 \| 1.7e-06 \| 3.2e-12 \| 54.0 \| 79 \| (6, 94) \| 100 \| (3, 82) \| 85 \| Rho termination factor N-terminal domain-containing protein \| Rho termination factor N-terminal domain-containing protein; uniclust \| UniRef100\_UPI001FE188BC \| 97.6 \| 1.9e-06 \| 3.6e-12 \| 53.5 \| 38 \| (53, 90) \| 100 \| (46, 83) \| 84 \| Rho termination factor N-terminal domain-containing protein \| Rho termination factor N-terminal domain-containing protein; uniclust \| UniRef100\_A0A1X1IFA2 \| 97.6 \| 2e-06 \| 3.7e-12 \| 55.2 \| 85 \| (5, 89) \| 100 \| (3, 100) \| 102 \| HeH/LEM domain-containing protein \| HeH/LEM domain-containing protein; uniclust \| UniRef100\_A0A3E0KHC7 \| 97.6 \| 2e-06 \| 3.7e-12 \| 46.0 \| 37 \| (55, 91) \| 100 \| (3, 39) \| 40 \| Rho termination factor N-terminal domain-containing protein \| Rho termination factor N-terminal domain-containing protein; uniclust \| UniRef100\_A0A059N3B0 \| 97.6 \| 1.9e-06 \| 3.8e-12 \| 59.8 \| 47 \| (53, 99) \| 100 \| (92, 139) \| 146 \| Rho termination factor protein \| Rho termination factor protein; uniclust \| UniRef100\_A0A011PZG2 \| 97.6 \| 1.8e-06 \| 3.8e-12 \| 54.0 \| 44 \| (1, 44) \| 100 \| (8, 51) \| 77 \| Uncharacterized protein \| Uncharacterized protein; uniclust \| UniRef100\_A0A926I9L6 \| 97.6 \| 2.1e-06 \| 3.8e-12 \| 47.0 \| 37 \| (55, 91) \| 100 \| (6, 42) \| 45 \| Rho termination factor N-terminal domain-containing protein \| Rho termination factor N-terminal domain-containing protein; uniclust \| UniRef100\_A0A249P9B4 \| 97.6 \| 2.2e-06 \| 4.3e-12 \| 55.5 \| 74 \| (21, 94) \| 100 \| (15, 95) \| 98 \| Phage protein \| Phage protein; uniclust \| UniRef100\_UPI001F25D97B \| 97.6 \| 2.3e-06 \| 4.3e-12 \| 52.8 \| 78 \| (9, 92) \| 100 \| (9, 86) \| 87 \| hypothetical protein \| hypothetical protein; uniclust \| UniRef100\_A0A3A6W6V7 \| 97.6 \| 2.3e-06 \| 4.4e-12 \| 57.1 \| 88 \| (7, 94) \| 100 \| (3, 95) \| 133 \| Rho termination factor N-terminal domain-containing protein \| Rho termination factor N-terminal domain-containing protein; uniclust \| UniRef100\_A0A1H9BTI4 \| 97.6 \| 2.4e-06 \| 4.7e-12 \| 55.3 \| 40 \| (53, 92) \| 100 \| (55, 94) \| 97 \| Rho termination factor, N-terminal domain \| Rho termination factor, N-terminal domain; uniclust \| UniRef100\_A0A7D7WFP1 \| 97.6 \| 2.4e-06 \| 4.7e-12 \| 55.4 \| 42 \| (55, 96) \| 100 \| (40, 81) \| 100 \| Rho termination factor N-terminal domain-containing protein \| Rho termination factor N-terminal domain-containing protein; uniclust \| UniRef100\_A0A2E0V1P7 \| 97.6 \| 2.6e-06 \| 4.8e-12 \| 52.2 \| 39 \| (54, 92) \| 100 \| (42, 80) \| 83 \| Rho termination factor N-terminal domain-containing protein \| Rho termination factor N-terminal domain-containing protein; uniclust \| UniRef100\_A0A0C2HKT8 \| 97.5 \| 2.6e-06 \| 5.3e-12 \| 53.5 \| 36 \| (60, 95) \| 100 \| (2, 37) \| 78 \| SAP domain-containing protein \| SAP domain-containing protein; uniclust \| UniRef100\_A0A1Y3ULW9 \| 97.5 \| 2.9e-06 \| 5.8e-12 \| 54.9 \| 71 \| (22, 92) \| 100 \| (9, 97) \| 99 \| HeH/LEM domain-containing protein \| HeH/LEM domain-containing protein; uniclust \| UniRef100\_A0A1C5TV19 \| 97.5 \| 3.1e-06 \| 5.9e-12 \| 52.8 \| 67 \| (28, 94) \| 100 \| (13, 81) \| 84 \| Transcription termination factor Rho \| Transcription termination factor Rho; uniclust \| UniRef100\_UPI0008A26671 \| 97.5 \| 3.2e-06 \| 5.9e-12 \| 53.4 \| 41 \| (53, 93) \| 100 \| (58, 98) \| 99 \| Rho termination factor N-terminal domain-containing protein \| Rho termination factor N-terminal domain-containing protein; uniclust \| UniRef100\_A0A3Q7F3B7 \| 97.5 \| 3.3e-06 \| 6.2e-12 \| 48.2 \| 38 \| (54, 91) \| 100 \| (13, 50) \| 52 \| Rho termination factor N-terminal domain-containing protein \| Rho termination factor N-terminal domain-containing protein; uniclust \| UniRef100\_UPI000419D3AE \| 97.5 \| 3.7e-06 \| 6.8e-12 \| 53.4 \| 62 \| (31, 92) \| 100 \| (39, 101) \| 102 \| hypothetical protein \| hypothetical protein; uniclust \| UniRef100\_A0A417YJN1 \| 97.5 \| 3.5e-06 \| 7e-12 \| 54.5 \| 85 \| (4, 89) \| 100 \| (1, 92) \| 98 \| Rho termination factor N-terminal domain-containing protein \| Rho termination factor N-terminal domain-containing protein; uniclust \| UniRef100\_UPI001EFFA8A0 \| 97.5 \| 4e-06 \| 7.3e-12 \| 52.9 \| 44 \| (54, 97) \| 100 \| (38, 81) \| 98 \| Rho termination factor N-terminal domain-containing protein \| Rho termination factor N-terminal domain-containing protein; uniclust \| UniRef100\_A0A0U2WI47 \| 97.5 \| 4.1e-06 \| 7.6e-12 \| 54.7 \| 76 \| (18, 93) \| 100 \| (35, 118) \| 121 \| Rho termination factor N-terminal domain-containing protein \| Rho termination factor N-terminal domain-containing protein; uniclust \| UniRef100\_A0A0H4TI36 \| 97.5 \| 3.9e-06 \| 7.8e-12 \| 55.6 \| 81 \| (13, 93) \| 100 \| (12, 97) \| 109 \| Rho termination factor \| Rho termination factor; uniclust \| UniRef100\_A0A0M0F5X6 \| 97.5 \| 4.1e-06 \| 8.6e-12 \| 59.3 \| 64 \| (31, 94) \| 100 \| (32, 96) \| 160 \| Rho termination factor N-terminal domain-containing protein \| Rho termination factor N-terminal domain-containing protein; uniclust \| UniRef100\_A0A4P8S989 \| 97.4 \| 5.7e-06 \| 1.1e-11 \| 52.3 \| 90 \| (5, 94) \| 100 \| (3, 94) \| 98 \| HeH/LEM domain-containing protein \| HeH/LEM domain-containing protein; uniclust \| UniRef100\_A0A088T1Y5 \| 97.4 \| 5.9e-06 \| 1.1e-11 \| 53.8 \| 36 \| (55, 90) \| 100 \| (71, 106) \| 107 \| Rho termination factor N-terminal domain-containing protein \| Rho termination factor N-terminal domain-containing protein; uniclust \| UniRef100\_B1BPQ6 \| 97.4 \| 6.2e-06 \| 1.1e-11 \| 48.7 \| 40 \| (53, 92) \| 100 \| (24, 63) \| 64 \| 50S ribosomal protein L21 \| 50S ribosomal protein L21; uniclust \| UniRef100\_A0A3T0L2P0 \| 97.4 \| 6.3e-06 \| 1.2e-11 \| 54.4 \| 90 \| (3, 93) \| 100 \| (2, 105) \| 129 \| Rho termination factor N-terminal domain-containing protein \| Rho termination factor N-terminal domain-containing protein; uniclust \| UniRef100\_UPI001F28AE23 \| 97.4 \| 6.4e-06 \| 1.2e-11 \| 52.4 \| 75 \| (16, 90) \| 100 \| (19, 101) \| 102 \| hypothetical protein \| hypothetical protein; uniclust \| UniRef100\_A0A2D3WNW5 \| 97.4 \| 6.5e-06 \| 1.2e-11 \| 53.0 \| 87 \| (7, 93) \| 100 \| (4, 107) \| 109 \| Rho termination factor N-terminal domain-containing protein \| Rho termination factor N-terminal domain-containing protein; uniclust \| UniRef100\_A0A9D0ZY95 \| 97.4 \| 6.6e-06 \| 1.2e-11 \| 42.4 \| 31 \| (60, 90) \| 100 \| (1, 31) \| 33 \| Rho termination factor N-terminal domain-containing protein \| Rho termination factor N-terminal domain-containing protein; uniclust \| UniRef100\_A0A7W3VUZ5 \| 97.4 \| 6.4e-06 \| 1.2e-11 \| 53.5 \| 75 \| (21, 97) \| 100 \| (14, 96) \| 102 \| SAP domain-containing protein \| SAP domain-containing protein; uniclust \| UniRef100\_A0A844J7S0 \| 97.4 \| 7e-06 \| 1.3e-11 \| 48.2 \| 38 \| (54, 91) \| 100 \| (25, 62) \| 64 \| Rho termination factor N-terminal domain-containing protein \| Rho termination factor N-terminal domain-containing protein; uniclust \| UniRef100\_A0A161WMN4 \| 97.4 \| 6.8e-06 \| 1.3e-11 \| 57.5 \| 43 \| (53, 95) \| 100 \| (94, 136) \| 166 \| Termination factor Rho \| Termination factor Rho; uniclust \| UniRef100\_A0A0C2V9W8 \| 97.4 \| 6.7e-06 \| 1.4e-11 \| 53.0 \| 44 \| (53, 96) \| 100 \| (34, 77) \| 88 \| Uncharacterized protein \| Uncharacterized protein; uniclust \| UniRef100\_UPI0018F8AC73 \| 97.4 \| 7.8e-06 \| 1.4e-11 \| 49.1 \| 39 \| (53, 91) \| 100 \| (31, 69) \| 72 \| Rho termination factor N-terminal domain-containing protein \| Rho termination factor N-terminal domain-containing protein; uniclust \| UniRef100\_A0A2D7UWP3 \| 97.4 \| 8.4e-06 \| 1.5e-11 \| 54.1 \| 39 \| (53, 91) \| 100 \| (93, 131) \| 133 \| Rho termination factor N-terminal domain-containing protein \| Rho termination factor N-terminal domain-containing protein; uniclust \| UniRef100\_K4JW68 \| 97.4 \| 8.2e-06 \| 1.6e-11 \| 48.5 \| 42 \| (54, 95) \| 100 \| (16, 57) \| 61 \| Rho termination factor N-terminal domain-containing protein \| Rho termination factor N-terminal domain-containing protein; uniclust \| UniRef100\_A0A967ZS15 \| 97.3 \| 8.7e-06 \| 1.6e-11 \| 47.0 \| 38 \| (53, 90) \| 100 \| (20, 57) \| 58 \| Rho termination factor N-terminal domain-containing protein \| Rho termination factor N-terminal domain-containing protein; uniclust \| UniRef100\_A0A352RPP2 \| 97.3 \| 8.7e-06 \| 1.6e-11 \| 50.5 \| 80 \| (10, 93) \| 100 \| (4, 85) \| 87 \| Rho termination factor N-terminal domain-containing protein \| Rho termination factor N-terminal domain-containing protein; uniclust \| UniRef100\_A0A4P9THH5 \| 97.3 \| 9.1e-06 \| 1.7e-11 \| 51.4 \| 90 \| (1, 90) \| 100 \| (1, 96) \| 97 \| Rho termination factor N-terminal domain-containing protein \| Rho termination factor N-terminal domain-containing protein; uniclust \| UniRef100\_A0A1Q8EFU1 \| 97.3 \| 9.4e-06 \| 1.7e-11 \| 41.9 \| 32 \| (60, 91) \| 100 \| (1, 32) \| 33 \| Rho termination factor N-terminal domain-containing protein \| Rho termination factor N-terminal domain-containing protein; uniclust \| UniRef100\_A0A1G6NTL8 \| 97.3 \| 9e-06 \| 1.7e-11 \| 55.5 \| 92 \| (1, 92) \| 100 \| (1, 101) \| 139 \| Uncharacterized protein \| Uncharacterized protein; uniclust \| UniRef100\_A0A024Q8V2 \| 97.3 \| 9e-06 \| 1.8e-11 \| 48.2 \| 46 \| (1, 46) \| 100 \| (1, 46) \| 58 \| Phage protein \| Phage protein; uniclust \| UniRef100\_A0A7Y0HZA0 \| 97.3 \| 9.5e-06 \| 1.8e-11 \| 51.3 \| 44 \| (54, 97) \| 100 \| (4, 47) \| 91 \| Transcription termination factor Rho (Fragment) \| Transcription termination factor Rho (Fragment); uniclust \| UniRef100\_A0A0D0Q5G0 \| 97.3 \| 9.1e-06 \| 1.9e-11 \| 55.9 \| 42 \| (55, 96) \| 100 \| (85, 127) \| 132 \| Phage protein \| Phage protein; uniclust \| UniRef100\_UPI001D192B90 \| 97.3 \| 1.1e-05 \| 2e-11 \| 50.0 \| 73 \| (15, 91) \| 100 \| (11, 84) \| 86 \| hypothetical protein \| hypothetical protein; uniclust \| UniRef100\_A0A2E6JK78 \| 97.3 \| 1.1e-05 \| 2.1e-11 \| 50.8 \| 38 \| (53, 90) \| 100 \| (55, 92) \| 93 \| Rho termination factor \| Rho termination factor; uniclust \| UniRef100\_UPI00040F1258 \| 97.3 \| 1.1e-05 \| 2.1e-11 \| 50.2 \| 42 \| (54, 95) \| 100 \| (34, 75) \| 81 \| hypothetical protein \| hypothetical protein; uniclust \| UniRef100\_A0A2N2BUP2 \| 97.3 \| 1.1e-05 \| 2.1e-11 \| 45.8 \| 39 \| (54, 92) \| 100 \| (13, 51) \| 53 \| Termination factor Rho \| Termination factor Rho; uniclust \| UniRef100\_UPI001147A498 \| 97.3 \| 1.1e-05 \| 2.1e-11 \| 53.7 \| 89 \| (3, 91) \| 100 \| (1, 106) \| 120 \| hypothetical protein \| hypothetical protein; uniclust \| UniRef100\_A0A0B0HXU4 \| 97.3 \| 1e-05 \| 2.1e-11 \| 48.5 \| 41 \| (4, 44) \| 100 \| (1, 42) \| 59 \| Uncharacterized protein \| Uncharacterized protein; uniclust \| UniRef100\_R9BU95 \| 97.3 \| 1.2e-05 \| 2.3e-11 \| 45.3 \| 40 \| (53, 92) \| 100 \| (10, 49) \| 51 \| Rho termination factor N-terminal domain-containing protein \| Rho termination factor N-terminal domain-containing protein; uniclust \| UniRef100\_A0A352P5R9 \| 97.3 \| 1.2e-05 \| 2.3e-11 \| 47.1 \| 42 \| (56, 97) \| 100 \| (4, 45) \| 63 \| Transcription termination factor Rho (Fragment) \| Transcription termination factor Rho (Fragment); uniclust \| UniRef100\_A0A559IX56 \| 97.3 \| 1.2e-05 \| 2.3e-11 \| 54.6 \| 89 \| (7, 95) \| 100 \| (5, 121) \| 126 \| Uncharacterized protein \| Uncharacterized protein; uniclust \| UniRef100\_UPI0004939A68 \| 97.3 \| 1.3e-05 \| 2.5e-11 \| 51.9 \| 75 \| (16, 90) \| 100 \| (15, 100) \| 101 \| hypothetical protein \| hypothetical protein; uniclust \| UniRef100\_A0A6L9MLT1 \| 97.3 \| 1.3e-05 \| 2.5e-11 \| 51.6 \| 40 \| (53, 92) \| 100 \| (53, 92) \| 93 \| Rho termination factor N-terminal domain-containing protein \| Rho termination factor N-terminal domain-containing protein; uniclust \| UniRef100\_A0A6N9YN10 \| 97.3 \| 1.4e-05 \| 2.6e-11 \| 51.7 \| 77 \| (23, 99) \| 100 \| (23, 102) \| 112 \| Rho termination factor N-terminal domain-containing protein \| Rho termination factor N-terminal domain-containing protein; uniclust \| UniRef100\_A0A1T0A0C1 \| 97.3 \| 1.3e-05 \| 2.6e-11 \| 53.6 \| 48 \| (52, 99) \| 100 \| (62, 109) \| 114 \| HeH/LEM domain \| HeH/LEM domain; uniclust \| UniRef100\_A0A1G6LUK0 \| 97.2 \| 1.5e-05 \| 2.8e-11 \| 50.1 \| 86 \| (5, 91) \| 100 \| (1, 91) \| 93 \| Rho termination factor N-terminal domain-containing protein \| Rho termination factor N-terminal domain-containing protein; uniclust \| UniRef100\_A0A2R3MVX2 \| 97.2 \| 1.5e-05 \| 2.8e-11 \| 51.3 \| 83 \| (11, 93) \| 100 \| (8, 101) \| 104 \| Rho termination factor N-terminal domain-containing protein \| Rho termination factor N-terminal domain-containing protein; uniclust \| UniRef100\_A0A2Z4WB78 \| 97.2 \| 1.6e-05 \| 2.9e-11 \| 47.2 \| 39 \| (53, 91) \| 100 \| (23, 61) \| 64 \| Rho termination factor N-terminal domain-containing protein \| Rho termination factor N-terminal domain-containing protein; uniclust \| UniRef100\_A0A4V2NXY4 \| 97.2 \| 1.6e-05 \| 3e-11 \| 52.3 \| 38 \| (54, 91) \| 100 \| (85, 122) \| 124 \| Rho termination factor N-terminal domain-containing protein \| Rho termination factor N-terminal domain-containing protein; uniclust \| UniRef100\_A0A965B2Z3 \| 97.2 \| 1.6e-05 \| 3e-11 \| 44.5 \| 40 \| (60, 99) \| 100 \| (1, 40) \| 49 \| Rho termination factor N-terminal domain-containing protein (Fragment) \| Rho termination factor N-terminal domain-containing protein (Fragment); uniclust \| UniRef100\_A0A2S8GSH5 \| 97.2 \| 1.6e-05 \| 3e-11 \| 49.8 \| 71 \| (19, 92) \| 100 \| (17, 87) \| 91 \| Rho termination factor N-terminal domain-containing protein \| Rho termination factor N-terminal domain-containing protein; uniclust \| UniRef100\_A0A3T1D355 \| 97.2 \| 1.6e-05 \| 3.1e-11 \| 45.4 \| 41 \| (54, 94) \| 100 \| (9, 50) \| 52 \| Rho termination factor N-terminal domain-containing protein \| Rho termination factor N-terminal domain-containing protein; uniclust \| UniRef100\_A0A961E4S7 \| 97.2 \| 1.6e-05 \| 3.1e-11 \| 51.5 \| 45 \| (54, 98) \| 100 \| (58, 102) \| 106 \| Rho termination factor N-terminal domain-containing protein (Fragment) \| Rho termination factor N-terminal domain-containing protein (Fragment); uniclust \| UniRef100\_A0A3Q9S8J1 \| 97.2 \| 1.7e-05 \| 3.1e-11 \| 41.8 \| 34 \| (60, 93) \| 100 \| (1, 34) \| 36 \| Rho termination factor N-terminal domain-containing protein \| Rho termination factor N-terminal domain-containing protein; uniclust \| UniRef100\_A0A838E6T1 \| 97.2 \| 1.7e-05 \| 3.2e-11 \| 42.0 \| 35 \| (58, 92) \| 100 \| (2, 36) \| 37 \| Rho termination factor N-terminal domain-containing protein \| Rho termination factor N-terminal domain-containing protein; uniclust \| UniRef100\_A0A6I1ZKL4 \| 97.2 \| 1.7e-05 \| 3.2e-11 \| 45.1 \| 44 \| (55, 98) \| 100 \| (5, 48) \| 51 \| Transcription termination factor Rho (Fragment) \| Transcription termination factor Rho (Fragment); uniclust \| UniRef100\_UPI002245C791 \| 97.2 \| 1.8e-05 \| 3.4e-11 \| 51.1 \| 90 \| (6, 95) \| 100 \| (3, 105) \| 109 \| HeH/LEM domain-containing protein \| HeH/LEM domain-containing protein; uniclust \| UniRef100\_A0A0C2VDB3 \| 97.2 \| 1.7e-05 \| 3.4e-11 \| 54.2 \| 89 \| (3, 91) \| 100 \| (14, 123) \| 127 \| Uncharacterized protein \| Uncharacterized protein; uniclust \| UniRef100\_UPI001BEABB8A \| 97.2 \| 1.9e-05 \| 3.6e-11 \| 53.1 \| 91 \| (4, 94) \| 100 \| (1, 113) \| 127 \| hypothetical protein \| hypothetical protein; uniclust \| UniRef100\_A0A022QNQ7 \| 97.2 \| 1.7e-05 \| 3.6e-11 \| 63.1 \| 39 \| (56, 94) \| 100 \| (284, 322) \| 371 \| Rho termination factor N-terminal domain-containing protein \| Rho termination factor N-terminal domain-containing protein; uniclust \| UniRef100\_A0A1C7WI36 \| 97.2 \| 1.8e-05 \| 3.6e-11 \| 49.1 \| 44 \| (2, 45) \| 100 \| (4, 47) \| 73 \| DUF2635 domain-containing protein \| DUF2635 domain-containing protein; uniclust \| UniRef100\_A0A357D622 \| 97.2 \| 2e-05 \| 3.7e-11 \| 49.9 \| 41 \| (54, 94) \| 100 \| (54, 94) \| 97 \| Rho termination factor N-terminal domain-containing protein \| Rho termination factor N-terminal domain-containing protein; uniclust \| UniRef100\_A0A1V4PGT5 \| 97.2 \| 1.9e-05 \| 3.7e-11 \| 54.8 \| 44 \| (54, 97) \| 100 \| (47, 90) \| 155 \| Rho termination factor N-terminal domain-containing protein \| Rho termination factor N-terminal domain-containing protein; uniclust \| UniRef100\_UPI0004717B1F \| 97.2 \| 2.1e-05 \| 3.8e-11 \| 49.6 \| 69 \| (25, 95) \| 100 \| (11, 80) \| 94 \| SAP domain-containing protein \| SAP domain-containing protein; uniclust \| UniRef100\_A0A3A9CS94 \| 97.2 \| 2.1e-05 \| 3.8e-11 \| 52.2 \| 86 \| (10, 95) \| 100 \| (5, 124) \| 130 \| Rho termination factor N-terminal domain-containing protein \| Rho termination factor N-terminal domain-containing protein; uniclust \| UniRef100\_A0A9D1GLC9 \| 97.2 \| 2.1e-05 \| 3.9e-11 \| 45.1 \| 38 \| (60, 97) \| 100 \| (1, 38) \| 55 \| Rho termination factor N-terminal domain-containing protein \| Rho termination factor N-terminal domain-containing protein; uniclust \| UniRef100\_A0A533S1U8 \| 97.2 \| 2.2e-05 \| 4.1e-11 \| 51.1 \| 90 \| (5, 95) \| 100 \| (11, 109) \| 114 \| Rho termination factor N-terminal domain-containing protein \| Rho termination factor N-terminal domain-containing protein; uniclust \| UniRef100\_A0A2H0LEZ5 \| 97.2 \| 2.2e-05 \| 4.4e-11 \| 53.2 \| 44 \| (55, 98) \| 100 \| (10, 53) \| 126 \| Transcription termination factor Rho (Fragment) \| Transcription termination factor Rho (Fragment); uniclust \| UniRef100\_A0A161XAR9 \| 97.2 \| 2.3e-05 \| 4.4e-11 \| 53.5 \| 88 \| (1, 88) \| 100 \| (1, 101) \| 143 \| Rho termination factor N-terminal domain-containing protein \| Rho termination factor N-terminal domain-containing protein; uniclust \| UniRef100\_A0A094YJ18 \| 97.2 \| 2e-05 \| 4.4e-11 \| 51.7 \| 37 \| (58, 95) \| 100 \| (13, 49) \| 94 \| SAP domain-containing protein \| SAP domain-containing protein; uniclust \| UniRef100\_A0A0Q1AB16 \| 97.2 \| 2.1e-05 \| 4.5e-11 \| 54.9 \| 45 \| (53, 97) \| 100 \| (20, 65) \| 140 \| Rho termination protein \| Rho termination protein; uniclust \| UniRef100\_C7RHA8 \| 97.2 \| 2.4e-05 \| 4.6e-11 \| 49.1 \| 69 \| (22, 90) \| 100 \| (9, 84) \| 85 \| Rho termination factor N-terminal domain-containing protein \| Rho termination factor N-terminal domain-containing protein; uniclust \| UniRef100\_UPI00047A0816 \| 97.2 \| 2.4e-05 \| 4.6e-11 \| 46.7 \| 42 \| (54, 95) \| 100 \| (20, 61) \| 63 \| Rho termination factor N-terminal domain-containing protein \| Rho termination factor N-terminal domain-containing protein; uniclust \| UniRef100\_A0A142K9H1 \| 97.2 \| 2.2e-05 \| 4.6e-11 \| 55.9 \| 92 \| (6, 97) \| 100 \| (14, 154) \| 158 \| Head-to-tail connector protein \| Head-to-tail connector protein; uniclust \| UniRef100\_A0A942AZ65 \| 97.2 \| 2.5e-05 \| 4.7e-11 \| 49.2 \| 38 \| (53, 90) \| 100 \| (55, 92) \| 93 \| Rho termination factor N-terminal domain-containing protein \| Rho termination factor N-terminal domain-containing protein; uniclust \| UniRef100\_A0A117DZC3 \| 97.2 \| 2.4e-05 \| 4.7e-11 \| 51.1 \| 39 \| (53, 91) \| 100 \| (63, 101) \| 102 \| Rho termination factor \| Rho termination factor; uniclust \| UniRef100\_X0S7Z5 \| 97.1 \| 2.6e-05 \| 4.8e-11 \| 50.0 \| 93 \| (5, 97) \| 100 \| (1, 97) \| 104 \| Rho termination factor N-terminal domain-containing protein \| Rho termination factor N-terminal domain-containing protein; uniclust \| UniRef100\_A0A7J9WAB8 \| 97.1 \| 2.7e-05 \| 4.9e-11 \| 41.1 \| 35 \| (57, 91) \| 100 \| (1, 35) \| 36 \| Rho termination factor \| Rho termination factor; uniclust \| UniRef100\_A0A8F4FSC2 \| 97.1 \| 2.5e-05 \| 4.9e-11 \| 56.7 \| 78 \| (16, 93) \| 100 \| (110, 191) \| 209 \| Rho termination factor N-terminal domain-containing protein \| Rho termination factor N-terminal domain-containing protein; uniclust \| UniRef100\_A0A2W1W9H2 \| 97.1 \| 2.7e-05 \| 4.9e-11 \| 49.4 \| 90 \| (4, 94) \| 100 \| (1, 93) \| 97 \| Rho termination factor N-terminal domain-containing protein \| Rho termination factor N-terminal domain-containing protein; uniclust \| UniRef100\_A0A5B8NRI3 \| 97.1 \| 2.7e-05 \| 5e-11 \| 41.8 \| 35 \| (60, 94) \| 100 \| (1, 35) \| 39 \| Rho termination factor N-terminal domain-containing protein \| Rho termination factor N-terminal domain-containing protein; uniclust \| UniRef100\_A0A496N535 \| 97.1 \| 2.8e-05 \| 5.1e-11 \| 43.7 \| 35 \| (56, 90) \| 100 \| (3, 37) \| 49 \| Rho termination factor N-terminal domain-containing protein (Fragment) \| Rho termination factor N-terminal domain-containing protein (Fragment); uniclust \| UniRef100\_A0A6G7WWK3 \| 97.1 \| 2.7e-05 \| 5.1e-11 \| 41.1 \| 33 \| (59, 91) \| 100 \| (2, 34) \| 35 \| Rho termination factor N-terminal domain-containing protein \| Rho termination factor N-terminal domain-containing protein; uniclust \| UniRef100\_A0A0A2XDX3 \| 97.1 \| 2.7e-05 \| 5.3e-11 \| 56.1 \| 85 \| (13, 97) \| 100 \| (45, 154) \| 183 \| Mu-like prophage FluMu N-terminal domain-containing protein \| Mu-like prophage FluMu N-terminal domain-containing protein; uniclust \| UniRef100\_A0A7T4KSI9 \| 97.1 \| 2.9e-05 \| 5.4e-11 \| 42.3 \| 36 \| (55, 90) \| 100 \| (4, 39) \| 40 \| HeH/LEM domain-containing protein \| HeH/LEM domain-containing protein; uniclust \| UniRef100\_A0A1B7LYA6 \| 97.1 \| 2.8e-05 \| 5.5e-11 \| 53.2 \| 42 \| (54, 95) \| 100 \| (86, 127) \| 132 \| Rho termination factor N-terminal domain-containing protein \| Rho termination factor N-terminal domain-containing protein; uniclust \| UniRef100\_A0A1H7RLC7 \| 97.1 \| 2.9e-05 \| 5.7e-11 \| 48.6 \| 58 \| (32, 89) \| 100 \| (18, 78) \| 80 \| Rho termination factor, N-terminal domain \| Rho termination factor, N-terminal domain; uniclust \| UniRef100\_A0A6C0APZ7 \| 97.1 \| 2.7e-05 \| 5.7e-11 \| 59.2 \| 41 \| (55, 95) \| 100 \| (177, 217) \| 251 \| Rho termination factor N-terminal domain-containing protein \| Rho termination factor N-terminal domain-containing protein; uniclust \| UniRef100\_A0A699P7Y3 \| 97.1 \| 3.2e-05 \| 5.8e-11 \| 49.6 \| 37 \| (54, 90) \| 100 \| (65, 101) \| 103 \| SAP-like protein BP-73 (Fragment) \| SAP-like protein BP-73 (Fragment); uniclust \| UniRef100\_A0A3B8T676 \| 97.1 \| 3.3e-05 \| 6e-11 \| 42.5 \| 36 \| (54, 89) \| 100 \| (8, 43) \| 44 \| 30S ribosomal protein S2 (Fragment) \| 30S ribosomal protein S2 (Fragment); uniclust \| UniRef100\_UPI001CF4ECA6 \| 97.1 \| 3.3e-05 \| 6e-11 \| 51.9 \| 78 \| (15, 92) \| 100 \| (17, 94) \| 139 \| hypothetical protein \| hypothetical protein; uniclust \| UniRef100\_A0A415MTC6 \| 97.1 \| 3.3e-05 \| 6.2e-11 \| 52.4 \| 89 \| (5, 95) \| 100 \| (3, 128) \| 133 \| SAP domain-containing protein \| SAP domain-containing protein; uniclust \| UniRef100\_A0A3E4QZK6 \| 97.1 \| 3.2e-05 \| 6.2e-11 \| 57.3 \| 46 \| (54, 99) \| 100 \| (198, 243) \| 245 \| Phage tail protein \| Phage tail protein; uniclust \| UniRef100\_A0A0S3U001 \| 97.1 \| 3e-05 \| 6.3e-11 \| 59.1 \| 42 \| (54, 95) \| 100 \| (193, 234) \| 261 \| Uncharacterized protein \| Uncharacterized protein; uniclust \| UniRef100\_A0A7X6FP84 \| 97.1 \| 3.3e-05 \| 6.5e-11 \| 51.9 \| 91 \| (4, 94) \| 100 \| (1, 115) \| 118 \| HeH/LEM domain-containing protein \| HeH/LEM domain-containing protein; uniclust \| UniRef100\_J2YWN5 \| 97.1 \| 3.4e-05 \| 6.6e-11 \| 46.7 \| 38 \| (54, 91) \| 100 \| (11, 48) \| 66 \| Rho termination factor, N-terminal domain protein (Fragment) \| Rho termination factor, N-terminal domain protein (Fragment); uniclust \| UniRef100\_A0A100W6R9 \| 97.1 \| 3.5e-05 \| 6.7e-11 \| 51.2 \| 71 \| (19, 89) \| 100 \| (19, 110) \| 111 \| Uncharacterized protein \| Uncharacterized protein; uniclust \| UniRef100\_A0A501VFZ7 \| 97.1 \| 3.5e-05 \| 6.9e-11 \| 48.8 \| 55 \| (34, 91) \| 100 \| (21, 75) \| 82 \| HeH/LEM domain-containing protein \| HeH/LEM domain-containing protein; uniclust \| UniRef100\_A0A4Z0ISG8 \| 97.1 \| 3.8e-05 \| 7.1e-11 \| 52.5 \| 82 \| (13, 94) \| 100 \| (12, 123) \| 147 \| Rho termination factor N-terminal domain-containing protein \| Rho termination factor N-terminal domain-containing protein; uniclust \| UniRef100\_UPI001020ADB0 \| 97.1 \| 3.8e-05 \| 7.2e-11 \| 52.2 \| 62 \| (33, 94) \| 100 \| (36, 99) \| 137 \| hypothetical protein \| hypothetical protein; uniclust \| UniRef100\_A0A075WTM6 \| 97.1 \| 3.5e-05 \| 7.3e-11 \| 54.5 \| 44 \| (53, 96) \| 100 \| (39, 83) \| 154 \| Rho termination factor N-terminal domain-containing protein \| Rho termination factor N-terminal domain-containing protein; uniclust \| UniRef100\_UPI001D057D6D \| 97.1 \| 3.9e-05 \| 7.3e-11 \| 46.8 \| 39 \| (53, 91) \| 100 \| (32, 70) \| 71 \| Rho termination factor N-terminal domain-containing protein \| Rho termination factor N-terminal domain-containing protein; uniclust \| UniRef100\_A0A1X3P896 \| 97.1 \| 3.9e-05 \| 7.3e-11 \| 52.2 \| 85 \| (5, 91) \| 100 \| (1, 136) \| 137 \| SAP domain-containing protein \| SAP domain-containing protein; uniclust \| UniRef100\_A0A1H3FGL4 \| 97.1 \| 3.8e-05 \| 7.3e-11 \| 47.5 \| 40 \| (54, 93) \| 100 \| (31, 70) \| 73 \| HeH/LEM domain-containing protein \| HeH/LEM domain-containing protein; uniclust \| UniRef100\_UPI000744C40B \| 97.1 \| 3.9e-05 \| 7.3e-11 \| 49.4 \| 80 \| (13, 92) \| 100 \| (10, 93) \| 95 \| HeH/LEM domain-containing protein \| HeH/LEM domain-containing protein; uniclust \| UniRef100\_A0A096KKD3 \| 97.1 \| 3.6e-05 \| 7.4e-11 \| 53.0 \| 40 \| (54, 93) \| 100 \| (89, 128) \| 131 \| Rho termination factor N-terminal domain-containing protein \| Rho termination factor N-terminal domain-containing protein; uniclust \| UniRef100\_UPI0021E05E95 \| 97.1 \| 4.2e-05 \| 7.7e-11 \| 53.6 \| 80 \| (17, 96) \| 100 \| (17, 106) \| 185 \| hypothetical protein \| hypothetical protein; uniclust \| UniRef100\_A0A119CYA6 \| 97.0 \| 4e-05 \| 8.5e-11 \| 47.4 \| 45 \| (2, 46) \| 100 \| (1, 45) \| 67 \| Uncharacterized protein \| Uncharacterized protein; uniclust \| UniRef100\_A0A418KHN6 \| 97.0 \| 4.3e-05 \| 8.6e-11 \| 47.4 \| 36 \| (60, 95) \| 100 \| (1, 36) \| 72 \| Rho termination factor N-terminal domain-containing protein (Fragment) \| Rho termination factor N-terminal domain-containing protein (Fragment); uniclust \| UniRef100\_UPI0013DFA039 \| 97.0 \| 4.8e-05 \| 8.8e-11 \| 52.2 \| 86 \| (5, 94) \| 100 \| (1, 90) \| 158 \| hypothetical protein \| hypothetical protein; uniclust \| UniRef100\_A0A2W5QPU5 \| 97.0 \| 4.5e-05 \| 8.9e-11 \| 54.8 \| 44 \| (1, 44) \| 100 \| (1, 46) \| 182 \| Rho termination factor N-terminal domain-containing protein \| Rho termination factor N-terminal domain-containing protein; uniclust \| UniRef100\_A0A7L7KTL4 \| 97.0 \| 4.9e-05 \| 9.1e-11 \| 49.6 \| 36 \| (54, 89) \| 100 \| (78, 113) \| 114 \| Rho termination factor N-terminal domain-containing protein \| Rho termination factor N-terminal domain-containing protein; uniclust \| UniRef100\_A0A1L6BXY7 \| 97.0 \| 5e-05 \| 9.2e-11 \| 42.2 \| 39 \| (54, 92) \| 100 \| (5, 43) \| 46 \| Rho termination factor N-terminal domain-containing protein \| Rho termination factor N-terminal domain-containing protein; uniclust \| UniRef100\_A0A3M1CYN9 \| 97.0 \| 5e-05 \| 9.2e-11 \| 43.8 \| 35 \| (57, 91) \| 100 \| (21, 55) \| 56 \| 50S ribosomal protein L21 (Fragment) \| 50S ribosomal protein L21 (Fragment); uniclust \| UniRef100\_A0A0U2VH56 \| 97.0 \| 4.9e-05 \| 9.4e-11 \| 43.1 \| 38 \| (54, 91) \| 100 \| (9, 46) \| 47 \| HeH/LEM domain-containing protein \| HeH/LEM domain-containing protein; uniclust \| UniRef100\_UPI00096A3CC9 \| 97.0 \| 5.1e-05 \| 9.6e-11 \| 46.0 \| 42 \| (54, 95) \| 100 \| (23, 64) \| 68 \| Rho termination factor N-terminal domain-containing protein \| Rho termination factor N-terminal domain-containing protein; uniclust \| UniRef100\_A0A1G5CMI0 \| 97.0 \| 5.3e-05 \| 9.8e-11 \| 50.0 \| 40 \| (53, 92) \| 100 \| (83, 122) \| 123 \| Rho termination factor, N-terminal domain \| Rho termination factor, N-terminal domain; uniclust \| UniRef100\_A0A8S0V1Y1 \| 97.0 \| 5.4e-05 \| 9.9e-11 \| 50.6 \| 35 \| (55, 89) \| 100 \| (97, 131) \| 132 \| Rho-N domain-containing 1, chloroplastic \| Rho-N domain-containing 1, chloroplastic; uniclust \| UniRef100\_UPI000EAE16BA \| 97.0 \| 5.4e-05 \| 9.9e-11 \| 48.6 \| 69 \| (20, 97) \| 100 \| (18, 87) \| 103 \| Rho termination factor N-terminal domain-containing protein \| Rho termination factor N-terminal domain-containing protein; uniclust \| UniRef100\_A0A0Q6VKW7 \| 97.0 \| 4.9e-05 \| 9.9e-11 \| 54.4 \| 36 \| (55, 90) \| 100 \| (134, 169) \| 170 \| Rho termination factor N-terminal domain-containing protein \| Rho termination factor N-terminal domain-containing protein; uniclust \| UniRef100\_A0A523V6Q3 \| 97.0 \| 5.2e-05 \| 1e-10 \| 45.9 \| 44 \| (55, 98) \| 100 \| (8, 51) \| 65 \| Transcription termination factor Rho (Fragment) \| Transcription termination factor Rho (Fragment); uniclust \| UniRef100\_A0A3C0G1P5 \| 97.0 \| 5.6e-05 \| 1e-10 \| 40.5 \| 33 \| (59, 91) \| 100 \| (3, 35) \| 38 \| 50S ribosomal protein L21 (Fragment) \| 50S ribosomal protein L21 (Fragment); uniclust \| UniRef100\_A0A0Q6AJX2 \| 97.0 \| 5.3e-05 \| 1.1e-10 \| 48.1 \| 47 \| (1, 47) \| 100 \| (1, 47) \| 80 \| Uncharacterized protein \| Uncharacterized protein; uniclust \| UniRef100\_A0A0B5CYN2 \| 97.0 \| 5.9e-05 \| 1.1e-10 \| 43.5 \| 39 \| (54, 92) \| 100 \| (14, 52) \| 53 \| HeH/LEM domain-containing protein \| HeH/LEM domain-containing protein; uniclust \| UniRef100\_A0A0N0K113 \| 97.0 \| 6e-05 \| 1.1e-10 \| 49.8 \| 73 \| (21, 93) \| 100 \| (20, 97) \| 118 \| Rho termination factor N-terminal domain-containing protein \| Rho termination factor N-terminal domain-containing protein; uniclust \| UniRef100\_A0A2D7WBG9 \| 97.0 \| 6.1e-05 \| 1.1e-10 \| 52.4 \| 38 \| (53, 90) \| 100 \| (131, 168) \| 169 \| Rho termination factor N-terminal domain-containing protein \| Rho termination factor N-terminal domain-containing protein; uniclust \| UniRef100\_A0A6C0DL56 \| 97.0 \| 6.1e-05 \| 1.1e-10 \| 39.2 \| 32 \| (60, 91) \| 100 \| (1, 32) \| 33 \| Rho termination factor N-terminal domain-containing protein \| Rho termination factor N-terminal domain-containing protein; uniclust \| UniRef100\_A0A1V2UCW6 \| 97.0 \| 6.2e-05 \| 1.1e-10 \| 39.5 \| 33 \| (60, 92) \| 100 \| (1, 33) \| 34 \| HeH/LEM domain-containing protein \| HeH/LEM domain-containing protein; uniclust \| UniRef100\_A0A022PR37 \| 97.0 \| 5.6e-05 \| 1.2e-10 \| 62.8 \| 40 \| (54, 93) \| 100 \| (438, 477) \| 542 \| Uncharacterized protein (Fragment) \| Uncharacterized protein (Fragment); uniclust \| UniRef100\_A0A5B9Y7U7 \| 97.0 \| 6.5e-05 \| 1.2e-10 \| 49.2 \| 88 \| (11, 98) \| 100 \| (9, 112) \| 115 \| Rho termination factor N-terminal domain-containing protein \| Rho termination factor N-terminal domain-containing protein; uniclust \| UniRef100\_A0A960CIP6 \| 96.9 \| 6.9e-05 \| 1.3e-10 \| 44.5 \| 37 \| (55, 91) \| 100 \| (26, 62) \| 63 \| Rho termination factor N-terminal domain-containing protein \| Rho termination factor N-terminal domain-containing protein; uniclust \| UniRef100\_A0A142KC13 \| 96.9 \| 6.5e-05 \| 1.3e-10 \| 53.1 \| 74 \| (18, 91) \| 100 \| (41, 155) \| 157 \| Head-to-tail connector protein \| Head-to-tail connector protein; uniclust \| UniRef100\_A0A3N5SVS3 \| 96.9 \| 7e-05 \| 1.3e-10 \| 47.0 \| 43 \| (54, 96) \| 100 \| (6, 49) \| 83 \| Transcription termination factor Rho (Fragment) \| Transcription termination factor Rho (Fragment); uniclust \| UniRef100\_A0A081PN78 \| 96.9 \| 6.7e-05 \| 1.3e-10 \| 47.1 \| 42 \| (56, 97) \| 100 \| (31, 73) \| 76 \| HeH/LEM domain protein \| HeH/LEM domain protein; uniclust \| UniRef100\_A0A7L5ABR4 \| 96.9 \| 7.5e-05 \| 1.4e-10 \| 48.3 \| 86 \| (9, 94) \| 100 \| (13, 102) \| 106 \| Uncharacterized protein \| Uncharacterized protein; uniclust \| UniRef100\_A0A6L6TN84 \| 96.9 \| 7.7e-05 \| 1.4e-10 \| 45.6 \| 58 \| (32, 90) \| 100 \| (18, 75) \| 76 \| Rho termination factor N-terminal domain-containing protein \| Rho termination factor N-terminal domain-containing protein; uniclust \| UniRef100\_A0A3N5Q9R5 \| 96.9 \| 7.9e-05 \| 1.5e-10 \| 46.3 \| 40 \| (54, 93) \| 100 \| (37, 76) \| 78 \| Addiction module toxin RelE (Fragment) \| Addiction module toxin RelE (Fragment); uniclust \| UniRef100\_A0A165PJJ3 \| 96.9 \| 7.4e-05 \| 1.5e-10 \| 51.7 \| 41 \| (5, 45) \| 100 \| (9, 49) \| 129 \| Mu-like prophage FluMu N-terminal domain-containing protein \| Mu-like prophage FluMu N-terminal domain-containing protein; uniclust \| UniRef100\_A0A1V3XH09 \| 96.9 \| 8.4e-05 \| 1.6e-10 \| 47.1 \| 44 \| (55, 98) \| 100 \| (15, 58) \| 94 \| Rho termination factor, N-terminal domain protein \| Rho termination factor, N-terminal domain protein; uniclust \| UniRef100\_UPI0009886066 \| 96.9 \| 8.6e-05 \| 1.6e-10 \| 46.5 \| 37 \| (54, 90) \| 100 \| (50, 86) \| 87 \| Rho termination factor N-terminal domain-containing protein \| Rho termination factor N-terminal domain-containing protein; uniclust \| UniRef100\_A0A2N3FBN1 \| 96.9 \| 8.8e-05 \| 1.6e-10 \| 39.1 \| 32 \| (60, 91) \| 100 \| (1, 32) \| 35 \| Rho termination factor N-terminal domain-containing protein (Fragment) \| Rho termination factor N-terminal domain-containing protein (Fragment); uniclust \| UniRef100\_A0A0U1DLB0 \| 96.9 \| 8.1e-05 \| 1.6e-10 \| 59.3 \| 45 \| (54, 98) \| 100 \| (43, 87) \| 398 \| Rho RNA-BD domain-containing protein \| Rho RNA-BD domain-containing protein; uniclust \| UniRef100\_A0A838H6R9 \| 96.9 \| 8.3e-05 \| 1.6e-10 \| 45.0 \| 46 \| (1, 46) \| 100 \| (1, 48) \| 62 \| Uncharacterized protein \| Uncharacterized protein; uniclust \| UniRef100\_A0A355UCP4 \| 96.9 \| 9.2e-05 \| 1.7e-10 \| 50.4 \| 37 \| (54, 90) \| 100 \| (111, 147) \| 148 \| Rho termination factor N-terminal domain-containing protein (Fragment) \| Rho termination factor N-terminal domain-containing protein (Fragment); uniclust \| UniRef100\_A0A255GPY9 \| 96.9 \| 9.3e-05 \| 1.7e-10 \| 45.5 \| 38 \| (54, 91) \| 100 \| (40, 77) \| 78 \| Rho termination factor N-terminal domain-containing protein \| Rho termination factor N-terminal domain-containing protein; uniclust \| UniRef100\_UPI001C8CC8D9 \| 96.9 \| 0.0001 \| 1.8e-10 \| 43.1 \| 39 \| (54, 92) \| 100 \| (19, 57) \| 59 \| Rho termination factor N-terminal domain-containing protein \| Rho termination factor N-terminal domain-containing protein; uniclust \| UniRef100\_A0A926IAT8 \| 96.9 \| 9.8e-05 \| 1.8e-10 \| 48.5 \| 84 \| (13, 96) \| 100 \| (12, 105) \| 110 \| Rho termination factor N-terminal domain-containing protein \| Rho termination factor N-terminal domain-containing protein; uniclust \| UniRef100\_A0A399NTV9 \| 96.9 \| 9.9e-05 \| 2e-10 \| 49.4 \| 43 \| (55, 97) \| 100 \| (11, 53) \| 109 \| Transcription termination factor Rho (Fragment) \| Transcription termination factor Rho (Fragment); uniclust \| UniRef100\_UPI00209F097A \| 96.9 \| 0.00011 \| 2e-10 \| 48.4 \| 85 \| (8, 92) \| 100 \| (3, 112) \| 118 \| hypothetical protein \| hypothetical protein; uniclust \| UniRef100\_A0A8S0HYC0 \| 96.9 \| 0.00011 \| 2e-10 \| 47.5 \| 81 \| (16, 96) \| 100 \| (3, 98) \| 101 \| Rho termination factor N-terminal domain-containing protein \| Rho termination factor N-terminal domain-containing protein; uniclust \| UniRef100\_A0A349AY30 \| 96.9 \| 0.0001 \| 2e-10 \| 50.7 \| 42 \| (55, 96) \| 100 \| (10, 51) \| 126 \| Transcription termination factor Rho (Fragment) \| Transcription termination factor Rho (Fragment); uniclust \| UniRef100\_A0A0Q5HJK0 \| 96.8 \| 0.0001 \| 2e-10 \| 53.7 \| 37 \| (54, 90) \| 100 \| (154, 190) \| 191 \| Rho termination factor N-terminal domain-containing protein \| Rho termination factor N-terminal domain-containing protein; uniclust \| UniRef100\_A0A067ZJ05 \| 96.8 \| 0.00011 \| 2e-10 \| 47.4 \| 83 \| (13, 95) \| 100 \| (10, 101) \| 104 \| Rho termination factor N-terminal domain-containing protein \| Rho termination factor N-terminal domain-containing protein; uniclust \| UniRef100\_A0A6L6JB59 \| 96.8 \| 0.00011 \| 2.1e-10 \| 47.2 \| 45 \| (1, 45) \| 100 \| (1, 45) \| 95 \| Uncharacterized protein \| Uncharacterized protein; uniclust \| UniRef100\_A0A078GBV8 \| 96.8 \| 0.00011 \| 2.2e-10 \| 60.3 \| 38 \| (55, 92) \| 100 \| (422, 459) \| 475 \| BnaC03g17360D protein \| BnaC03g17360D protein; uniclust \| UniRef100\_A0A143HCV3 \| 96.8 \| 0.00012 \| 2.2e-10 \| 46.9 \| 80 \| (13, 92) \| 100 \| (12, 92) \| 99 \| Rho termination factor N-terminal domain-containing protein \| Rho termination factor N-terminal domain-containing protein; uniclust \| UniRef100\_A0A446I8R2 \| 96.8 \| 0.00012 \| 2.3e-10 \| 43.6 \| 37 \| (53, 89) \| 100 \| (26, 62) \| 63 \| Rho termination factor N-terminal domain-containing protein \| Rho termination factor N-terminal domain-containing protein; uniclust \| UniRef100\_A0A163MUE0 \| 96.8 \| 0.00011 \| 2.3e-10 \| 49.5 \| 91 \| (2, 92) \| 100 \| (1, 112) \| 114 \| HeH/LEM domain-containing protein \| HeH/LEM domain-containing protein; uniclust \| UniRef100\_A0A2S4YIP2 \| 96.8 \| 0.00012 \| 2.3e-10 \| 46.0 \| 41 \| (54, 94) \| 100 \| (46, 86) \| 89 \| Rho termination factor N-terminal domain-containing protein \| Rho termination factor N-terminal domain-containing protein; uniclust \| UniRef100\_A0A1X0IMZ6 \| 96.8 \| 0.00013 \| 2.4e-10 \| 45.5 \| 37 \| (55, 91) \| 100 \| (47, 83) \| 84 \| Rho termination factor \| Rho termination factor; uniclust \| UniRef100\_UPI00156691C1 \| 96.8 \| 0.00013 \| 2.5e-10 \| 46.9 \| 69 \| (15, 93) \| 100 \| (17, 85) \| 101 \| hypothetical protein \| hypothetical protein; uniclust \| UniRef100\_A0A085LDJ2 \| 96.8 \| 0.00013 \| 2.5e-10 \| 50.5 \| 40 \| (55, 94) \| 100 \| (39, 78) \| 131 \| Rho termination factor N-terminal domain-containing protein (Fragment) \| Rho termination factor N-terminal domain-containing protein (Fragment); uniclust \| UniRef100\_A0A7X9A4X6 \| 96.8 \| 0.00014 \| 2.6e-10 \| 50.3 \| 39 \| (54, 92) \| 100 \| (122, 160) \| 162 \| Rho termination factor N-terminal domain-containing protein \| Rho termination factor N-terminal domain-containing protein; uniclust \| UniRef100\_A0A3L8PIB7 \| 96.8 \| 0.00014 \| 2.6e-10 \| 51.6 \| 38 \| (54, 91) \| 100 \| (143, 180) \| 181 \| Rho termination factor N-terminal domain-containing protein \| Rho termination factor N-terminal domain-containing protein; uniclust \| UniRef100\_K0V9V4 \| 96.8 \| 0.00014 \| 2.6e-10 \| 52.4 \| 45 \| (54, 98) \| 100 \| (26, 70) \| 201 \| Transcription termination factor Rho (Fragment) \| Transcription termination factor Rho (Fragment); uniclust \| UniRef100\_A0A2A7MKH2 \| 96.8 \| 0.00014 \| 2.6e-10 \| 44.5 \| 41 \| (54, 94) \| 100 \| (25, 65) \| 69 \| Rho termination factor N-terminal domain-containing protein \| Rho termination factor N-terminal domain-containing protein; uniclust \| UniRef100\_A0A2Z6ZZW1 \| 96.8 \| 0.00013 \| 2.6e-10 \| 55.3 \| 40 \| (55, 94) \| 100 \| (218, 257) \| 267 \| SAP-like protein BP-73 (Fragment) \| SAP-like protein BP-73 (Fragment); uniclust \| UniRef100\_A0A022N6K0 \| 96.8 \| 0.00014 \| 2.7e-10 \| 42.4 \| 39 \| (54, 92) \| 100 \| (16, 54) \| 55 \| HeH/LEM domain-containing protein \| HeH/LEM domain-containing protein; uniclust \| UniRef100\_A0A1N7MP98 \| 96.8 \| 0.00014 \| 2.7e-10 \| 50.8 \| 43 \| (54, 96) \| 100 \| (98, 140) \| 146 \| Rho termination factor, N-terminal domain \| Rho termination factor, N-terminal domain; uniclust \| UniRef100\_UPI00065B46D6 \| 96.8 \| 0.00014 \| 2.7e-10 \| 47.5 \| 42 \| (55, 96) \| 100 \| (21, 62) \| 95 \| Rho termination factor N-terminal domain-containing protein \| Rho termination factor N-terminal domain-containing protein; uniclust \| UniRef100\_A0A0A2SJL7 \| 96.8 \| 0.00014 \| 2.8e-10 \| 50.9 \| 82 \| (11, 92) \| 100 \| (6, 105) \| 155 \| Rho termination factor N-terminal domain-containing protein \| Rho termination factor N-terminal domain-containing protein; uniclust \| UniRef100\_A0A3G6ISY3 \| 96.8 \| 0.00015 \| 2.8e-10 \| 48.8 \| 80 \| (16, 95) \| 100 \| (18, 121) \| 124 \| Uncharacterized protein \| Uncharacterized protein; uniclust \| UniRef100\_UPI00195E8D1B \| 96.8 \| 0.00015 \| 2.8e-10 \| 54.6 \| 37 \| (54, 90) \| 100 \| (226, 262) \| 263 \| Rho termination factor N-terminal domain-containing protein \| Rho termination factor N-terminal domain-containing protein; uniclust \| UniRef100\_A0A1I5II67 \| 96.8 \| 0.00015 \| 2.9e-10 \| 48.5 \| 74 \| (20, 93) \| 100 \| (27, 122) \| 123 \| Uncharacterized protein \| Uncharacterized protein; uniclust \| UniRef100\_A0A1C5GJ85 \| 96.8 \| 0.00015 \| 2.9e-10 \| 49.6 \| 41 \| (54, 94) \| 100 \| (14, 54) \| 131 \| Rho termination factor, N-terminal domain (Fragment) \| Rho termination factor, N-terminal domain (Fragment); uniclust \| UniRef100\_A0A8I1NIZ1 \| 96.8 \| 0.00016 \| 3e-10 \| 48.8 \| 35 \| (58, 92) \| 100 \| (84, 118) \| 119 \| Rho termination factor N-terminal domain-containing protein \| Rho termination factor N-terminal domain-containing protein; uniclust \| UniRef100\_F0SXG1 \| 96.8 \| 0.00016 \| 3e-10 \| 44.7 \| 66 \| (27, 94) \| 100 \| (11, 76) \| 80 \| Rho termination factor N-terminal domain-containing protein \| Rho termination factor N-terminal domain-containing protein; uniclust \| UniRef100\_A0A8T7B3C8 \| 96.8 \| 0.00016 \| 3e-10 \| 48.4 \| 39 \| (53, 91) \| 100 \| (78, 116) \| 118 \| Rho termination factor N-terminal domain-containing protein \| Rho termination factor N-terminal domain-containing protein; uniclust \| UniRef100\_UPI0008A58C56 \| 96.8 \| 0.00017 \| 3.1e-10 \| 47.0 \| 84 \| (5, 89) \| 100 \| (1, 105) \| 108 \| hypothetical protein \| hypothetical protein; uniclust \| UniRef100\_A0A0U4K5G7 \| 96.8 \| 0.00016 \| 3.1e-10 \| 43.5 \| 40 \| (5, 44) \| 100 \| (2, 41) \| 62 \| Uncharacterized protein \| Uncharacterized protein; uniclust \| UniRef100\_A0A943ECP9 \| 96.8 \| 0.00017 \| 3.1e-10 \| 37.9 \| 32 \| (60, 91) \| 100 \| (1, 32) \| 34 \| SAP domain-containing protein \| SAP domain-containing protein; uniclust \| UniRef100\_A0A4U7J7L9 \| 96.7 \| 0.00017 \| 3.2e-10 \| 45.4 \| 85 \| (7, 92) \| 100 \| (2, 88) \| 89 \| Rho termination factor N-terminal domain-containing protein \| Rho termination factor N-terminal domain-containing protein; uniclust \| UniRef100\_A0A842EMJ7 \| 96.7 \| 0.00017 \| 3.2e-10 \| 38.9 \| 36 \| (60, 95) \| 100 \| (1, 36) \| 38 \| HeH/LEM domain-containing protein \| HeH/LEM domain-containing protein; uniclust \| UniRef100\_A0A3R6KE31 \| 96.7 \| 0.00017 \| 3.2e-10 \| 51.2 \| 41 \| (56, 96) \| 100 \| (12, 52) \| 163 \| Rho termination factor N-terminal domain-containing protein \| Rho termination factor N-terminal domain-containing protein; uniclust \| UniRef100\_A0A1C2DD79 \| 96.7 \| 0.00016 \| 3.2e-10 \| 54.1 \| 40 \| (55, 94) \| 100 \| (192, 231) \| 233 \| Rho termination factor N-terminal domain-containing protein \| Rho termination factor N-terminal domain-containing protein; uniclust \| UniRef100\_A0A399G9E4 \| 96.7 \| 0.00017 \| 3.3e-10 \| 43.7 \| 38 \| (54, 91) \| 100 \| (26, 63) \| 66 \| Rho termination factor N-terminal domain-containing protein \| Rho termination factor N-terminal domain-containing protein; uniclust \| UniRef100\_A0A0A1VZC2 \| 96.7 \| 0.00017 \| 3.3e-10 \| 56.1 \| 42 \| (54, 95) \| 100 \| (190, 231) \| 314 \| Rho termination factor N-terminal domain-containing protein \| Rho termination factor N-terminal domain-containing protein; uniclust \| UniRef100\_UPI00191E8781 \| 96.7 \| 0.00018 \| 3.3e-10 \| 48.3 \| 72 \| (20, 91) \| 100 \| (34, 128) \| 131 \| hypothetical protein \| hypothetical protein; uniclust \| UniRef100\_A0A0A1MNV9 \| 96.7 \| 0.00016 \| 3.3e-10 \| 44.2 \| 40 \| (4, 43) \| 100 \| (1, 40) \| 62 \| Uncharacterized protein \| Uncharacterized protein; uniclust \| UniRef100\_A0A4Q5N7H8 \| 96.7 \| 0.00018 \| 3.3e-10 \| 44.2 \| 39 \| (53, 91) \| 100 \| (31, 69) \| 71 \| Rho termination factor \| Rho termination factor; uniclust \| UniRef100\_A0A0M2H758 \| 96.7 \| 0.00016 \| 3.3e-10 \| 52.4 \| 43 \| (53, 97) \| 100 \| (80, 122) \| 173 \| SAP domain protein \| SAP domain protein; uniclust \| UniRef100\_A0A0N9NCV3 \| 96.7 \| 0.00017 \| 3.3e-10 \| 52.2 \| 33 \| (57, 89) \| 100 \| (144, 176) \| 177 \| Rho termination factor N-terminal domain-containing protein \| Rho termination factor N-terminal domain-containing protein; uniclust \| UniRef100\_A0A0N0Z495 \| 96.7 \| 0.00017 \| 3.4e-10 \| 53.2 \| 45 \| (54, 98) \| 100 \| (21, 65) \| 202 \| Rho RNA-BD domain-containing protein \| Rho RNA-BD domain-containing protein; uniclust \| UniRef100\_A0A1X1X990 \| 96.7 \| 0.00017 \| 3.4e-10 \| 48.7 \| 80 \| (15, 94) \| 100 \| (16, 104) \| 116 \| Rho termination factor N-terminal domain-containing protein \| Rho termination factor N-terminal domain-containing protein; uniclust \| UniRef100\_A0A936ZSD3 \| 96.7 \| 0.00017 \| 3.4e-10 \| 43.9 \| 43 \| (3, 45) \| 100 \| (1, 43) \| 65 \| Uncharacterized protein \| Uncharacterized protein; uniclust \| UniRef100\_A0A336N748 \| 96.7 \| 0.00017 \| 3.4e-10 \| 43.9 \| 43 \| (2, 44) \| 100 \| (1, 43) \| 61 \| Uncharacterized protein \| Uncharacterized protein; uniclust \| UniRef100\_A0A959YTN7 \| 96.7 \| 0.00019 \| 3.4e-10 \| 38.6 \| 35 \| (57, 91) \| 100 \| (3, 37) \| 38 \| Rho termination factor N-terminal domain-containing protein \| Rho termination factor N-terminal domain-containing protein; uniclust \| UniRef100\_A0A1E4VLJ2 \| 96.7 \| 0.00018 \| 3.4e-10 \| 50.8 \| 41 \| (54, 94) \| 100 \| (106, 146) \| 169 \| Rho termination factor N-terminal domain-containing protein \| Rho termination factor N-terminal domain-containing protein; uniclust \| UniRef100\_A0A960ZEV2 \| 96.7 \| 0.00019 \| 3.5e-10 \| 43.3 \| 41 \| (54, 94) \| 100 \| (27, 67) \| 69 \| Rho termination factor N-terminal domain-containing protein (Fragment) \| Rho termination factor N-terminal domain-containing protein (Fragment); uniclust \| UniRef100\_A0A2Z4PZN3 \| 96.7 \| 0.00019 \| 3.6e-10 \| 45.7 \| 72 \| (23, 95) \| 100 \| (9, 80) \| 83 \| HeH/LEM domain-containing protein \| HeH/LEM domain-containing protein; uniclust \| UniRef100\_A0A7V7QJT4 \| 96.7 \| 0.0002 \| 3.7e-10 \| 43.5 \| 39 \| (54, 92) \| 100 \| (33, 71) \| 72 \| Rho termination factor N-terminal domain-containing protein \| Rho termination factor N-terminal domain-containing protein; uniclust \| UniRef100\_UPI001E3C94D1 \| 96.7 \| 0.0002 \| 3.7e-10 \| 41.4 \| 37 \| (55, 91) \| 100 \| (18, 54) \| 55 \| Rho termination factor N-terminal domain-containing protein \| Rho termination factor N-terminal domain-containing protein; uniclust \| UniRef100\_A0A088T2U9 \| 96.7 \| 0.00018 \| 3.8e-10 \| 55.5 \| 41 \| (54, 94) \| 100 \| (188, 228) \| 273 \| 50S ribosomal protein L20 \| 50S ribosomal protein L20; uniclust \| UniRef100\_A0A161I2L0 \| 96.7 \| 0.00018 \| 3.9e-10 \| 57.5 \| 42 \| (54, 95) \| 100 \| (261, 302) \| 360 \| Rho termination factor N-terminal domain-containing protein \| Rho termination factor N-terminal domain-containing protein; uniclust \| UniRef100\_A0A2G1CVH0 \| 96.7 \| 0.00021 \| 3.9e-10 \| 47.4 \| 89 \| (6, 95) \| 100 \| (8, 110) \| 114 \| Uncharacterized protein \| Uncharacterized protein; uniclust \| UniRef100\_UPI0011A4A6DC \| 96.7 \| 0.00021 \| 3.9e-10 \| 46.2 \| 39 \| (54, 92) \| 100 \| (64, 102) \| 103 \| Rho termination factor N-terminal domain-containing protein \| Rho termination factor N-terminal domain-containing protein; uniclust \| UniRef100\_A0A1A0IIT4 \| 96.7 \| 0.0002 \| 4e-10 \| 51.6 \| 33 \| (59, 91) \| 100 \| (139, 171) \| 173 \| Rho termination factor N-terminal domain-containing protein \| Rho termination factor N-terminal domain-containing protein; uniclust \| UniRef100\_A0A1Q9T603 \| 96.7 \| 0.0002 \| 4.1e-10 \| 52.0 \| 40 \| (55, 94) \| 100 \| (19, 58) \| 182 \| Rho termination factor N-terminal domain-containing protein \| Rho termination factor N-terminal domain-containing protein; uniclust \| UniRef100\_A0A0Q8PJF9 \| 96.7 \| 0.00023 \| 4.1e-10 \| 42.9 \| 41 \| (5, 45) \| 100 \| (1, 41) \| 68 \| Mu-like prophage FluMu N-terminal domain-containing protein \| Mu-like prophage FluMu N-terminal domain-containing protein; uniclust \| UniRef100\_UPI0015585255 \| 96.7 \| 0.00023 \| 4.2e-10 \| 42.4 \| 42 \| (51, 92) \| 100 \| (21, 63) \| 64 \| Rho termination factor N-terminal domain-containing protein \| Rho termination factor N-terminal domain-containing protein; uniclust \| UniRef100\_A0A0J1C4L7 \| 96.7 \| 0.00022 \| 4.3e-10 \| 44.6 \| 44 \| (53, 96) \| 100 \| (23, 66) \| 73 \| HeH/LEM domain-containing protein \| HeH/LEM domain-containing protein; uniclust \| UniRef100\_A0A845R497 \| 96.7 \| 0.00023 \| 4.3e-10 \| 46.6 \| 67 \| (26, 94) \| 100 \| (29, 96) \| 107 \| HeH/LEM domain-containing protein \| HeH/LEM domain-containing protein; uniclust \| UniRef100\_A0A5C7NE73 \| 96.7 \| 0.00024 \| 4.4e-10 \| 46.8 \| 77 \| (14, 91) \| 100 \| (21, 112) \| 115 \| Rho termination factor N-terminal domain-containing protein \| Rho termination factor N-terminal domain-containing protein; uniclust \| UniRef100\_A0A3D2M3Y4 \| 96.7 \| 0.00024 \| 4.4e-10 \| 44.5 \| 76 \| (19, 96) \| 100 \| (5, 81) \| 85 \| HeH/LEM domain-containing protein (Fragment) \| HeH/LEM domain-containing protein (Fragment); uniclust \| UniRef100\_A0A1G4FHX2 \| 96.7 \| 0.00023 \| 4.4e-10 \| 44.5 \| 39 \| (54, 92) \| 100 \| (35, 73) \| 74 \| HeH/LEM domain protein \| HeH/LEM domain protein; uniclust \| UniRef100\_A0A4Y9NA30 \| 96.7 \| 0.00022 \| 4.4e-10 \| 48.4 \| 41 \| (55, 95) \| 100 \| (35, 75) \| 119 \| Transcription termination factor Rho (Fragment) \| Transcription termination factor Rho (Fragment); uniclust \| UniRef100\_A0A847LPG8 \| 96.7 \| 0.00024 \| 4.5e-10 \| 48.0 \| 41 \| (55, 95) \| 100 \| (4, 44) \| 136 \| Starch-binding protein \| Starch-binding protein; uniclust \| UniRef100\_A0A3G9A8C3 \| 96.7 \| 0.00022 \| 4.5e-10 \| 48.6 \| 90 \| (1, 90) \| 100 \| (1, 112) \| 115 \| Uncharacterized protein \| Uncharacterized protein; uniclust \| UniRef100\_H3NPD2 \| 96.7 \| 0.00024 \| 4.5e-10 \| 45.0 \| 67 \| (22, 90) \| 100 \| (20, 86) \| 88 \| HeH/LEM domain-containing protein \| HeH/LEM domain-containing protein; uniclust \| UniRef100\_A0A0J8DFR9 \| 96.7 \| 0.00024 \| 4.5e-10 \| 43.8 \| 42 \| (5, 46) \| 100 \| (1, 43) \| 72 \| Uncharacterized protein \| Uncharacterized protein; uniclust \| UniRef100\_A0A0M3SMX1 \| 96.7 \| 0.00023 \| 4.7e-10 \| 44.9 \| 39 \| (57, 95) \| 100 \| (35, 73) \| 75 \| HeH/LEM domain-containing protein \| HeH/LEM domain-containing protein; uniclust \| UniRef100\_A0A1B1U2N6 \| 96.7 \| 0.00025 \| 4.7e-10 \| 45.4 \| 36 \| (54, 89) \| 100 \| (53, 88) \| 89 \| Rho termination factor N-terminal domain-containing protein \| Rho termination factor N-terminal domain-containing protein; uniclust \| UniRef100\_A0A6B0BAF2 \| 96.6 \| 0.00025 \| 4.7e-10 \| 47.0 \| 88 \| (1, 93) \| 100 \| (1, 94) \| 108 \| Phage protein \| Phage protein; uniclust \| UniRef100\_A0A3S4QE94 \| 96.6 \| 0.00026 \| 4.8e-10 \| 45.0 \| 87 \| (3, 90) \| 100 \| (2, 88) \| 89 \| HeH/LEM domain \| HeH/LEM domain; uniclust \| UniRef100\_UPI0006915928 \| 96.6 \| 0.00026 \| 4.8e-10 \| 44.9 \| 38 \| (54, 91) \| 100 \| (54, 91) \| 92 \| Rho termination factor N-terminal domain-containing protein \| Rho termination factor N-terminal domain-containing protein; uniclust \| UniRef100\_A0A8S9SLN2 \| 96.6 \| 0.00027 \| 5.1e-10 \| 45.6 \| 39 \| (53, 91) \| 100 \| (55, 93) \| 94 \| Rho termination factor N-terminal domain-containing protein \| Rho termination factor N-terminal domain-containing protein; uniclust \| UniRef100\_A0A0G0RI45 \| 96.6 \| 0.00027 \| 5.1e-10 \| 48.0 \| 71 \| (23, 93) \| 100 \| (41, 119) \| 121 \| HeH/LEM domain-containing protein \| HeH/LEM domain-containing protein; uniclust \| UniRef100\_Q1ILS8 \| 96.6 \| 0.00028 \| 5.1e-10 \| 47.0 \| 76 \| (19, 94) \| 100 \| (31, 119) \| 123 \| Rho termination factor N-terminal domain-containing protein \| Rho termination factor N-terminal domain-containing protein; uniclust \| UniRef100\_A0A2S8MDY3 \| 96.6 \| 0.00027 \| 5.2e-10 \| 42.0 \| 37 \| (54, 90) \| 100 \| (21, 57) \| 59 \| Rho termination factor (Fragment) \| Rho termination factor (Fragment); uniclust \| UniRef100\_A0A7C6FK88 \| 96.6 \| 0.00026 \| 5.3e-10 \| 60.0 \| 44 \| (54, 97) \| 100 \| (27, 70) \| 649 \| Transcription termination factor Rho \| Transcription termination factor Rho; uniclust \| UniRef100\_A0A164DND9 \| 96.6 \| 0.00028 \| 5.3e-10 \| 51.1 \| 37 \| (54, 90) \| 100 \| (156, 192) \| 193 \| Rho termination factor N-terminal domain-containing protein \| Rho termination factor N-terminal domain-containing protein; uniclust \| UniRef100\_A0A399NFB3 \| 96.6 \| 0.00027 \| 5.3e-10 \| 45.6 \| 42 \| (54, 95) \| 100 \| (10, 51) \| 89 \| Transcription termination factor Rho (Fragment) \| Transcription termination factor Rho (Fragment); uniclust \| UniRef100\_A0A512CPT3 \| 96.6 \| 0.00029 \| 5.4e-10 \| 46.4 \| 76 \| (8, 90) \| 100 \| (4, 81) \| 115 \| Rho termination factor N-terminal domain-containing protein \| Rho termination factor N-terminal domain-containing protein; uniclust \| UniRef100\_A0A2A2M454 \| 96.6 \| 0.00028 \| 5.4e-10 \| 50.2 \| 45 \| (54, 98) \| 100 \| (15, 59) \| 159 \| Rho RNA-BD domain-containing protein \| Rho RNA-BD domain-containing protein; uniclust \| UniRef100\_A0A2J6WPY3 \| 96.6 \| 0.00027 \| 5.4e-10 \| 54.6 \| 40 \| (55, 94) \| 100 \| (21, 60) \| 278 \| Rho termination factor N-terminal domain-containing protein \| Rho termination factor N-terminal domain-containing protein; uniclust \| UniRef100\_A0A353RPN2 \| 96.6 \| 0.00026 \| 5.5e-10 \| 48.2 \| 44 \| (55, 98) \| 100 \| (11, 54) \| 113 \| Transcription termination factor Rho (Fragment) \| Transcription termination factor Rho (Fragment); uniclust \| UniRef100\_A0A363D603 \| 96.6 \| 0.00029 \| 5.5e-10 \| 46.4 \| 41 \| (55, 95) \| 100 \| (5, 45) \| 106 \| Rho termination factor N-terminal domain-containing protein \| Rho termination factor N-terminal domain-containing protein; uniclust \| UniRef100\_A0A1C6JYU0 \| 96.6 \| 0.00027 \| 5.5e-10 \| 48.3 \| 41 \| (56, 96) \| 100 \| (66, 106) \| 115 \| Uncharacterized protein \| Uncharacterized protein; uniclust \| UniRef100\_A0A097PBF3 \| 96.6 \| 0.00028 \| 5.8e-10 \| 51.1 \| 95 \| (1, 95) \| 100 \| (5, 125) \| 170 \| HeH/LEM domain-containing protein \| HeH/LEM domain-containing protein; uniclust \| UniRef100\_A0A661IZL4 \| 96.6 \| 0.00032 \| 5.9e-10 \| 44.5 \| 41 \| (54, 94) \| 100 \| (33, 73) \| 91 \| Rho termination factor N-terminal domain-containing protein (Fragment) \| Rho termination factor N-terminal domain-containing protein (Fragment); uniclust \| UniRef100\_A0A1Y1RN50 \| 96.6 \| 0.00029 \| 5.9e-10 \| 55.2 \| 40 \| (54, 93) \| 100 \| (274, 313) \| 314 \| Rho termination factor N-terminal domain-containing protein \| Rho termination factor N-terminal domain-containing protein; uniclust \| UniRef100\_A0A651FJL6 \| 96.6 \| 0.00033 \| 6.1e-10 \| 45.0 \| 89 \| (1, 89) \| 100 \| (1, 97) \| 99 \| Rho termination factor N-terminal domain-containing protein \| Rho termination factor N-terminal domain-containing protein; uniclust \| UniRef100\_A0A3C1A1A0 \| 96.6 \| 0.00034 \| 6.2e-10 \| 40.0 \| 32 \| (54, 85) \| 100 \| (20, 51) \| 51 \| Rho termination factor N-terminal domain-containing protein (Fragment) \| Rho termination factor N-terminal domain-containing protein (Fragment); uniclust \| UniRef100\_UPI0021069BFE \| 96.6 \| 0.00034 \| 6.2e-10 \| 41.6 \| 39 \| (56, 94) \| 100 \| (23, 61) \| 63 \| Rho termination factor N-terminal domain-containing protein \| Rho termination factor N-terminal domain-containing protein; uniclust \| UniRef100\_A0A171NWN3 \| 96.6 \| 0.00033 \| 6.4e-10 \| 48.8 \| 38 \| (54, 91) \| 100 \| (108, 145) \| 148 \| 50S ribosomal protein (Fragment) \| 50S ribosomal protein (Fragment); uniclust \| UniRef100\_A0A6N7EVP9 \| 96.6 \| 0.00034 \| 6.5e-10 \| 43.8 \| 37 \| (60, 96) \| 100 \| (1, 37) \| 80 \| Rho termination factor N-terminal domain-containing protein (Fragment) \| Rho termination factor N-terminal domain-containing protein (Fragment); uniclust \| UniRef100\_UPI000D1A72FA \| 96.6 \| 0.00036 \| 6.6e-10 \| 45.1 \| 46 \| (54, 99) \| 100 \| (48, 93) \| 102 \| Rho termination factor N-terminal domain-containing protein \| Rho termination factor N-terminal domain-containing protein; uniclust \| UniRef100\_A0A3G9JYC5 \| 96.6 \| 0.00035 \| 6.7e-10 \| 50.1 \| 38 \| (54, 91) \| 100 \| (135, 172) \| 174 \| Rho termination factor N-terminal domain-containing protein \| Rho termination factor N-terminal domain-containing protein; uniclust \| UniRef100\_A0A2R5F593 \| 96.6 \| 0.00037 \| 6.7e-10 \| 44.3 \| 41 \| (54, 94) \| 100 \| (7, 47) \| 91 \| Transcription termination factor Rho \| Transcription termination factor Rho; uniclust \| UniRef100\_A0A1C2XPW0 \| 96.6 \| 0.00035 \| 6.8e-10 \| 44.8 \| 39 \| (54, 93) \| 100 \| (46, 84) \| 85 \| Rho termination factor N-terminal domain-containing protein \| Rho termination factor N-terminal domain-containing protein; uniclust \| UniRef100\_A0A061GFL4 \| 96.6 \| 0.00035 \| 6.8e-10 \| 54.8 \| 36 \| (55, 90) \| 100 \| (306, 341) \| 342 \| Rho termination factor, putative isoform 1 \| Rho termination factor, putative isoform 1; uniclust \| UniRef100\_A0A5C6ULJ2 \| 96.6 \| 0.00036 \| 6.9e-10 \| 45.5 \| 84 \| (4, 95) \| 100 \| (1, 96) \| 100 \| Peripheral subunit-binding (PSBD) domain-containing protein \| Peripheral subunit-binding (PSBD) domain-containing protein; uniclust \| UniRef100\_A0A6C0DS71 \| 96.5 \| 0.00037 \| 7e-10 \| 51.4 \| 39 \| (55, 93) \| 100 \| (152, 190) \| 215 \| Rho termination factor N-terminal domain-containing protein \| Rho termination factor N-terminal domain-containing protein; uniclust \| UniRef100\_UPI0020195133 \| 96.5 \| 0.00039 \| 7.2e-10 \| 47.1 \| 76 \| (20, 98) \| 100 \| (21, 97) \| 137 \| hypothetical protein \| hypothetical protein; uniclust \| UniRef100\_A0A950YI39 \| 96.5 \| 0.00041 \| 7.5e-10 \| 37.6 \| 33 \| (60, 92) \| 100 \| (1, 33) \| 39 \| Rho termination factor N-terminal domain-containing protein \| Rho termination factor N-terminal domain-containing protein; uniclust \| UniRef100\_A0A1B1ISD2 \| 96.5 \| 0.00036 \| 7.5e-10 \| 53.3 \| 39 \| (54, 92) \| 100 \| (202, 240) \| 241 \| Sarcoplasmic reticulum histidine-rich calcium-binding protein-like \| Sarcoplasmic reticulum histidine-rich calcium-binding protein-like; uniclust \| UniRef100\_UPI0020561702 \| 96.5 \| 0.00041 \| 7.5e-10 \| 44.8 \| 80 \| (14, 93) \| 100 \| (13, 100) \| 101 \| hypothetical protein \| hypothetical protein; uniclust \| UniRef100\_A0A9D4E6J0 \| 96.5 \| 0.00041 \| 7.6e-10 \| 42.6 \| 39 \| (59, 97) \| 100 \| (3, 41) \| 75 \| Uncharacterized protein \| Uncharacterized protein; uniclust \| UniRef100\_A0A1V2VNK8 \| 96.5 \| 0.00037 \| 7.6e-10 \| 46.8 \| 84 \| (2, 95) \| 100 \| (1, 88) \| 102 \| HeH/LEM domain-containing protein \| HeH/LEM domain-containing protein; uniclust \| UniRef100\_A0A2D6X3A0 \| 96.5 \| 0.00042 \| 7.6e-10 \| 40.9 \| 42 \| (5, 46) \| 100 \| (1, 42) \| 60 \| Uncharacterized protein \| Uncharacterized protein; uniclust \| UniRef100\_A0A0A7PDZ3 \| 96.5 \| 0.00038 \| 7.7e-10 \| 45.5 \| 85 \| (5, 95) \| 100 \| (1, 87) \| 90 \| Secreted protein \| Secreted protein; uniclust \| UniRef100\_A0A2N0VMG3 \| 96.5 \| 0.00041 \| 7.9e-10 \| 48.3 \| 40 \| (54, 93) \| 100 \| (103, 142) \| 145 \| Rho termination factor N-terminal domain-containing protein \| Rho termination factor N-terminal domain-containing protein; uniclust \| UniRef100\_A0A498IVK1 \| 96.5 \| 0.0004 \| 8e-10 \| 58.3 \| 39 \| (53, 91) \| 100 \| (566, 604) \| 607 \| Rho termination factor N-terminal domain-containing protein \| Rho termination factor N-terminal domain-containing protein; uniclust \| UniRef100\_A0A382MFM9 \| 96.5 \| 0.00044 \| 8e-10 \| 39.9 \| 41 \| (5, 45) \| 100 \| (1, 41) \| 53 \| Uncharacterized protein (Fragment) \| Uncharacterized protein (Fragment); uniclust \| UniRef100\_A0A844Q625 \| 96.5 \| 0.00044 \| 8.1e-10 \| 45.6 \| 83 \| (11, 93) \| 100 \| (7, 106) \| 114 \| Rho termination factor N-terminal domain-containing protein \| Rho termination factor N-terminal domain-containing protein; uniclust \| UniRef100\_A0A1H0FZA0 \| 96.5 \| 0.00044 \| 8.2e-10 \| 46.0 \| 69 \| (22, 91) \| 100 \| (33, 115) \| 116 \| Rho termination factor, N-terminal domain \| Rho termination factor, N-terminal domain; uniclust \| UniRef100\_A0A1C3H478 \| 96.5 \| 0.00044 \| 8.3e-10 \| 41.3 \| 42 \| (5, 46) \| 100 \| (1, 42) \| 58 \| Uncharacterized protein \| Uncharacterized protein; uniclust \| UniRef100\_UPI001382ED58 \| 96.5 \| 0.00044 \| 8.4e-10 \| 45.4 \| 41 \| (55, 95) \| 100 \| (10, 50) \| 104 \| Rho termination factor N-terminal domain-containing protein \| Rho termination factor N-terminal domain-containing protein; uniclust \| UniRef100\_A0A3D2JCI4 \| 96.5 \| 0.00042 \| 8.5e-10 \| 48.5 \| 42 \| (54, 95) \| 100 \| (82, 123) \| 134 \| Rho termination factor N-terminal domain-containing protein (Fragment) \| Rho termination factor N-terminal domain-containing protein (Fragment); uniclust \| UniRef100\_UPI00178A52FA \| 96.5 \| 0.00047 \| 8.7e-10 \| 46.9 \| 92 \| (3, 95) \| 100 \| (2, 104) \| 139 \| Rho termination factor N-terminal domain-containing protein \| Rho termination factor N-terminal domain-containing protein; uniclust \| UniRef100\_A0A6V8CT67 \| 96.5 \| 0.00046 \| 8.7e-10 \| 41.8 \| 38 \| (56, 95) \| 100 \| (3, 40) \| 63 \| SAP domain-containing protein (Fragment) \| SAP domain-containing protein (Fragment); uniclust \| UniRef100\_A0A8S5M0J8 \| 96.5 \| 0.00046 \| 8.8e-10 \| 37.3 \| 33 \| (60, 92) \| 100 \| (1, 34) \| 36 \| Transcription regulator-like protein \| Transcription regulator-like protein; uniclust \| UniRef100\_A0A0B4EK77 \| 96.5 \| 0.00046 \| 8.8e-10 \| 52.8 \| 43 \| (54, 96) \| 100 \| (106, 148) \| 293 \| Rho termination factor N-terminal domain-containing protein \| Rho termination factor N-terminal domain-containing protein; uniclust \| UniRef100\_A0A0R3MJG7 \| 96.5 \| 0.00048 \| 8.9e-10 \| 44.9 \| 76 \| (16, 91) \| 100 \| (25, 106) \| 107 \| Rho termination factor N-terminal domain-containing protein \| Rho termination factor N-terminal domain-containing protein; uniclust \| UniRef100\_A0A0C2ZSS3 \| 96.5 \| 0.00048 \| 8.9e-10 \| 46.2 \| 41 \| (55, 95) \| 100 \| (45, 85) \| 119 \| Rho termination factor N-terminal domain-containing protein (Fragment) \| Rho termination factor N-terminal domain-containing protein (Fragment); uniclust \| UniRef100\_A0A482MWG0 \| 96.5 \| 0.00049 \| 9e-10 \| 43.7 \| 41 \| (53, 93) \| 100 \| (46, 86) \| 88 \| Rho termination factor N-terminal domain-containing protein \| Rho termination factor N-terminal domain-containing protein; uniclust \| UniRef100\_A0A411YG54 \| 96.5 \| 0.00047 \| 9.1e-10 \| 52.6 \| 39 \| (54, 92) \| 100 \| (228, 266) \| 267 \| Rho termination factor N-terminal domain-containing protein \| Rho termination factor N-terminal domain-containing protein; uniclust \| UniRef100\_A0A6I2UUF3 \| 96.5 \| 0.0005 \| 9.2e-10 \| 42.9 \| 44 \| (1, 44) \| 100 \| (1, 44) \| 82 \| Uncharacterized protein \| Uncharacterized protein; uniclust \| UniRef100\_UPI002356B0F1 \| 96.5 \| 0.0005 \| 9.2e-10 \| 47.3 \| 88 \| (7, 94) \| 100 \| (3, 113) \| 151 \| hypothetical protein \| hypothetical protein; uniclust \| UniRef100\_A0A061AAU7 \| 96.5 \| 0.00046 \| 9.2e-10 \| 50.9 \| 43 \| (55, 97) \| 100 \| (150, 192) \| 196 \| 50S ribosomal protein L20 \| 50S ribosomal protein L20; uniclust \| UniRef100\_A0A7W6RF31 \| 96.5 \| 0.00049 \| 9.3e-10 \| 41.7 \| 35 \| (12, 46) \| 100 \| (14, 48) \| 63 \| Uncharacterized protein \| Uncharacterized protein; uniclust \| UniRef100\_UPI001FE300D8 \| 96.5 \| 0.00051 \| 9.4e-10 \| 40.2 \| 38 \| (59, 96) \| 100 \| (3, 40) \| 57 \| Rho termination factor N-terminal domain-containing protein \| Rho termination factor N-terminal domain-containing protein; uniclust \| UniRef100\_UPI001FD29EAF \| 96.5 \| 0.00051 \| 9.4e-10 \| 47.1 \| 77 \| (19, 95) \| 100 \| (29, 123) \| 147 \| hypothetical protein \| hypothetical protein; uniclust \| UniRef100\_UPI0002F4E584 \| 96.5 \| 0.0005 \| 9.5e-10 \| 46.6 \| 45 \| (54, 98) \| 100 \| (10, 54) \| 119 \| Rho termination factor N-terminal domain-containing protein \| Rho termination factor N-terminal domain-containing protein; uniclust \| UniRef100\_UPI001F182A02 \| 96.5 \| 0.00051 \| 9.5e-10 \| 43.6 \| 38 \| (53, 90) \| 100 \| (47, 84) \| 84 \| Rho termination factor N-terminal domain-containing protein \| Rho termination factor N-terminal domain-containing protein; uniclust \| UniRef100\_UPI001BB11BDD \| 96.5 \| 0.00052 \| 9.6e-10 \| 40.9 \| 37 \| (60, 96) \| 100 \| (1, 37) \| 63 \| Rho termination factor N-terminal domain-containing protein \| Rho termination factor N-terminal domain-containing protein; uniclust \| UniRef100\_UPI001918AB4A \| 96.5 \| 0.00051 \| 9.8e-10 \| 49.9 \| 41 \| (56, 96) \| 100 \| (143, 183) \| 189 \| hypothetical protein \| hypothetical protein; uniclust \| UniRef100\_A0A372JKS1 \| 96.5 \| 0.00051 \| 9.8e-10 \| 51.8 \| 41 \| (55, 95) \| 100 \| (36, 76) \| 251 \| Transcription termination factor Rho (Fragment) \| Transcription termination factor Rho (Fragment); uniclust \| UniRef100\_A0A1V6HXP6 \| 96.5 \| 0.00052 \| 9.9e-10 \| 52.6 \| 41 \| (56, 96) \| 100 \| (3, 43) \| 296 \| Rho termination factor N-terminal domain-containing protein \| Rho termination factor N-terminal domain-containing protein; uniclust \| UniRef100\_A0A0F7L7M2 \| 96.5 \| 0.0005 \| 1e-09 \| 44.4 \| 42 \| (4, 45) \| 100 \| (14, 55) \| 84 \| Uncharacterized protein \| Uncharacterized protein; uniclust \| UniRef100\_A0A1H8W1H1 \| 96.5 \| 0.00053 \| 1e-09 \| 47.7 \| 71 \| (22, 92) \| 100 \| (63, 145) \| 146 \| Rho termination factor, N-terminal domain \| Rho termination factor, N-terminal domain; uniclust \| UniRef100\_A0A1H1TF00 \| 96.5 \| 0.00051 \| 1e-09 \| 46.5 \| 40 \| (54, 93) \| 100 \| (69, 108) \| 113 \| Rho termination factor, N-terminal domain \| Rho termination factor, N-terminal domain; uniclust \| UniRef100\_A0A094P6K3 \| 96.4 \| 0.00055 \| 1.1e-09 \| 48.1 \| 36 \| (55, 90) \| 100 \| (114, 149) \| 151 \| Rho termination factor N-terminal domain-containing protein \| Rho termination factor N-terminal domain-containing protein; uniclust \| UniRef100\_A0A7C6WBF3 \| 96.4 \| 0.00058 \| 1.1e-09 \| 46.8 \| 35 \| (57, 91) \| 100 \| (112, 146) \| 147 \| 30S ribosomal protein S20 \| 30S ribosomal protein S20; uniclust \| UniRef100\_A0A1H7ZYP4 \| 96.4 \| 0.00057 \| 1.1e-09 \| 46.2 \| 47 \| (53, 99) \| 100 \| (65, 111) \| 113 \| HeH/LEM domain-containing protein \| HeH/LEM domain-containing protein; uniclust \| UniRef100\_A0A0M2LXZ2 \| 96.4 \| 0.00056 \| 1.1e-09 \| 43.5 \| 46 \| (1, 46) \| 100 \| (1, 46) \| 78 \| Uncharacterized protein \| Uncharacterized protein; uniclust \| UniRef100\_A0A1S1TMF4 \| 96.4 \| 0.00061 \| 1.1e-09 \| 45.1 \| 77 \| (18, 94) \| 100 \| (19, 107) \| 110 \| Uncharacterized protein \| Uncharacterized protein; uniclust \| UniRef100\_X0WVX3 \| 96.4 \| 0.00063 \| 1.2e-09 \| 38.5 \| 35 \| (55, 90) \| 100 \| (14, 48) \| 48 \| Rho termination factor N-terminal domain-containing protein (Fragment) \| Rho termination factor N-terminal domain-containing protein (Fragment); uniclust \| UniRef100\_A0A0S8E8A4 \| 96.4 \| 0.00057 \| 1.2e-09 \| 43.5 \| 35 \| (59, 94) \| 100 \| (6, 40) \| 75 \| SAP domain-containing protein \| SAP domain-containing protein; uniclust \| UniRef100\_UPI000375EB70 \| 96.4 \| 0.00061 \| 1.2e-09 \| 47.6 \| 42 \| (54, 95) \| 100 \| (7, 49) \| 151 \| hypothetical protein \| hypothetical protein; uniclust \| UniRef100\_A0A7X0ZKM8 \| 96.4 \| 0.00062 \| 1.2e-09 \| 39.7 \| 39 \| (53, 91) \| 100 \| (12, 50) \| 52 \| HeH/LEM domain-containing protein \| HeH/LEM domain-containing protein; uniclust \| UniRef100\_A0A2E5TCS1 \| 96.4 \| 0.0006 \| 1.2e-09 \| 43.4 \| 43 \| (3, 45) \| 100 \| (7, 49) \| 78 \| DUF2635 domain-containing protein \| DUF2635 domain-containing protein; uniclust \| UniRef100\_UPI0021E5D87B \| 96.4 \| 0.00064 \| 1.2e-09 \| 43.1 \| 40 \| (54, 93) \| 100 \| (49, 88) \| 89 \| Rho termination factor N-terminal domain-containing protein \| Rho termination factor N-terminal domain-containing protein; uniclust \| UniRef100\_A0A7J9VEG8 \| 96.4 \| 0.0006 \| 1.2e-09 \| 54.4 \| 41 \| (55, 95) \| 100 \| (34, 74) \| 372 \| Transcription termination factor Rho (Fragment) \| Transcription termination factor Rho (Fragment); uniclust \| UniRef100\_A0A519NL56 \| 96.4 \| 0.00063 \| 1.2e-09 \| 40.1 \| 43 \| (55, 97) \| 100 \| (5, 47) \| 55 \| Rho termination factor N-terminal domain-containing protein (Fragment) \| Rho termination factor N-terminal domain-containing protein (Fragment); uniclust \| UniRef100\_A0A076JPF1 \| 96.4 \| 0.00059 \| 1.2e-09 \| 59.1 \| 43 \| (54, 96) \| 100 \| (59, 101) \| 780 \| Transcription termination factor Rho \| Transcription termination factor Rho; uniclust \| UniRef100\_A0A0A9Q8Y3 \| 96.4 \| 0.00066 \| 1.2e-09 \| 35.9 \| 32 \| (60, 91) \| 100 \| (1, 32) \| 34 \| Rho termination factor N-terminal domain-containing protein \| Rho termination factor N-terminal domain-containing protein; uniclust \| UniRef100\_UPI00210A2392 \| 96.4 \| 0.00068 \| 1.2e-09 \| 38.9 \| 41 \| (54, 94) \| 100 \| (8, 48) \| 51 \| HeH/LEM domain-containing protein \| HeH/LEM domain-containing protein; uniclust \| UniRef100\_A0A2H6HI62 \| 96.4 \| 0.00064 \| 1.3e-09 \| 54.8 \| 43 \| (54, 96) \| 100 \| (6, 48) \| 398 \| 2-acyl-glycerophospho-ethanolamine acyltransferase \| 2-acyl-glycerophospho-ethanolamine acyltransferase; uniclust \| UniRef100\_A0A6L7B069 \| 96.4 \| 0.0007 \| 1.3e-09 \| 37.9 \| 40 \| (60, 99) \| 100 \| (1, 40) \| 45 \| HeH/LEM domain-containing protein \| HeH/LEM domain-containing protein; uniclust \| UniRef100\_A0A1T2CHS8 \| 96.4 \| 0.00066 \| 1.3e-09 \| 48.0 \| 43 \| (4, 46) \| 100 \| (13, 55) \| 145 \| Uncharacterized protein \| Uncharacterized protein; uniclust \| UniRef100\_A0A379FQN9 \| 96.4 \| 0.00064 \| 1.3e-09 \| 46.1 \| 41 \| (4, 44) \| 100 \| (1, 41) \| 108 \| Procyclic acidic repetitive protein (PARP) \| Procyclic acidic repetitive protein (PARP); uniclust \| UniRef100\_A0A2S8FHG4 \| 96.4 \| 0.00069 \| 1.3e-09 \| 41.6 \| 38 \| (54, 91) \| 100 \| (25, 63) \| 65 \| Rho termination factor N-terminal domain-containing protein \| Rho termination factor N-terminal domain-containing protein; uniclust \| UniRef100\_A0A954APN1 \| 96.4 \| 0.00069 \| 1.4e-09 \| 46.0 \| 44 \| (3, 46) \| 100 \| (13, 56) \| 114 \| Uncharacterized protein \| Uncharacterized protein; uniclust \| UniRef100\_A0A9D4B561 \| 96.4 \| 0.00074 \| 1.4e-09 \| 41.1 \| 38 \| (59, 96) \| 100 \| (3, 40) \| 66 \| Uncharacterized protein \| Uncharacterized protein; uniclust \| UniRef100\_A0A3A0ET32 \| 96.4 \| 0.00075 \| 1.4e-09 \| 39.8 \| 37 \| (55, 91) \| 100 \| (22, 58) \| 59 \| Rho termination factor N-terminal domain-containing protein \| Rho termination factor N-terminal domain-containing protein; uniclust \| UniRef100\_A0A7W9N0T6 \| 96.4 \| 0.00077 \| 1.4e-09 \| 46.4 \| 81 \| (15, 95) \| 100 \| (8, 102) \| 149 \| Rho termination factor \| Rho termination factor; uniclust \| UniRef100\_UPI00218C784C \| 96.4 \| 0.00078 \| 1.4e-09 \| 39.1 \| 39 \| (6, 44) \| 100 \| (2, 40) \| 54 \| hypothetical protein \| hypothetical protein; uniclust \| UniRef100\_X7Y4V5 \| 96.4 \| 0.00076 \| 1.4e-09 \| 49.9 \| 45 \| (54, 98) \| 100 \| (27, 71) \| 232 \| Rho termination factor, N-terminal domain protein \| Rho termination factor, N-terminal domain protein; uniclust \| UniRef100\_A0A848L2M3 \| 96.4 \| 0.00078 \| 1.4e-09 \| 41.1 \| 40 \| (54, 93) \| 100 \| (31, 70) \| 71 \| Rho termination factor N-terminal domain-containing protein \| Rho termination factor N-terminal domain-containing protein; uniclust \| UniRef100\_A0A2E0K4L4 \| 96.4 \| 0.00076 \| 1.4e-09 \| 37.7 \| 35 \| (54, 90) \| 100 \| (7, 41) \| 42 \| SAP domain-containing protein (Fragment) \| SAP domain-containing protein (Fragment); uniclust \| UniRef100\_A0A0M8Y5M5 \| 96.4 \| 0.00074 \| 1.4e-09 \| 51.9 \| 42 \| (53, 94) \| 100 \| (31, 72) \| 278 \| Rho termination factor N-terminal domain-containing protein \| Rho termination factor N-terminal domain-containing protein; uniclust \| UniRef100\_A0A2E0E7I7 \| 96.3 \| 0.00075 \| 1.5e-09 \| 51.2 \| 39 \| (54, 92) \| 100 \| (214, 252) \| 253 \| Rho termination factor N-terminal domain-containing protein \| Rho termination factor N-terminal domain-containing protein; uniclust \| UniRef100\_A0A009IA11 \| 96.3 \| 0.00075 \| 1.5e-09 \| 47.4 \| 41 \| (53, 93) \| 100 \| (98, 138) \| 145 \| HeH/LEM domain protein \| HeH/LEM domain protein; uniclust \| UniRef100\_A0A973TXR3 \| 96.3 \| 0.0008 \| 1.5e-09 \| 42.6 \| 43 \| (55, 97) \| 100 \| (42, 84) \| 88 \| Transcription termination factor Rho (Fragment) \| Transcription termination factor Rho (Fragment); uniclust \| UniRef100\_A0A0B5AVK8 \| 96.3 \| 0.00068 \| 1.5e-09 \| 47.1 \| 43 \| (1, 43) \| 100 \| (1, 46) \| 121 \| Uncharacterized protein \| Uncharacterized protein; uniclust \| UniRef100\_UPI001F2021B5 \| 96.3 \| 0.00079 \| 1.5e-09 \| 47.3 \| 40 \| (54, 93) \| 100 \| (93, 132) \| 156 \| Rho termination factor N-terminal domain-containing protein \| Rho termination factor N-terminal domain-containing protein; uniclust \| UniRef100\_UPI0005C4E073 \| 96.3 \| 0.00081 \| 1.5e-09 \| 42.9 \| 41 \| (55, 95) \| 100 \| (6, 46) \| 92 \| Rho termination factor N-terminal domain-containing protein \| Rho termination factor N-terminal domain-containing protein; uniclust \| UniRef100\_A0A4R2IZA5 \| 96.3 \| 0.00076 \| 1.5e-09 \| 44.1 \| 36 \| (57, 92) \| 100 \| (53, 88) \| 90 \| Rho termination factor-like protein \| Rho termination factor-like protein; uniclust \| UniRef100\_A0A971BR62 \| 96.3 \| 0.00082 \| 1.5e-09 \| 42.7 \| 44 \| (53, 96) \| 100 \| (7, 50) \| 90 \| Transcription termination factor Rho (Fragment) \| Transcription termination factor Rho (Fragment); uniclust \| UniRef100\_A0A2G5NW73 \| 96.3 \| 0.0008 \| 1.5e-09 \| 45.0 \| 37 \| (54, 90) \| 100 \| (74, 110) \| 111 \| Rho termination protein \| Rho termination protein; uniclust \| UniRef100\_A0A178V4Q9 \| 96.3 \| 0.00083 \| 1.5e-09 \| 44.7 \| 37 \| (55, 91) \| 100 \| (82, 118) \| 119 \| Rho termination factor N-terminal domain-containing protein \| Rho termination factor N-terminal domain-containing protein |
| Top keywords  (threshold 1.00e-03 (evalue)) | **Rho, termination, domain\_containing, factor, N\_terminal, HeH, LEM, Fragment, hypothetical, Transcription** |
| Output files | ../../similar\_sequences/06\_FANPEZAQ\_CDS\_0006\_merged.svg ../../similar\_sequences/06\_FANPEZAQ\_CDS\_0006\_pdb70.a3m ../../similar\_sequences/06\_FANPEZAQ\_CDS\_0006\_pdb70.hhr ../../similar\_sequences/06\_FANPEZAQ\_CDS\_0006\_uniclust.a3m ../../similar\_sequences/06\_FANPEZAQ\_CDS\_0006\_uniclust.hhr |

#### Structure prediction (AlphaFold)^2^

|  |  |
| --- | --- |
| Stats | xml version="1.0" encoding="utf-8" standalone="no"?       2024-09-02T21:09:05.778690 image/svg+xml   Matplotlib v3.7.2, https://matplotlib.org/ |
| Predicted structure | **NGL Viewer Controls:**  - Center: *Left-Click* - Rotate: *Left-Click + Drag* - Translate: *Right-Click + Drag* - Zoom: *Shift + Left-Click + Drag* |
| Output files | ../../predicted\_structures/06\_FANPEZAQ\_CDS\_0006/features.pkl ../../predicted\_structures/06\_FANPEZAQ\_CDS\_0006/ranked\_0.pdb ../../predicted\_structures/06\_FANPEZAQ\_CDS\_0006/ranked\_0\_plots.svg ../../predicted\_structures/06\_FANPEZAQ\_CDS\_0006/result\_model\_1\_ptm\_pred\_0.pkl |

#### Structure similarity search results (Foldseek)^3^

|  |  |
| --- | --- |
| Structure databases searched | Pdb, Afdb-proteome, Afdb-uniprot50 |
| Results, scheme(s)  (Top layers only, threshold 1.00e-02 (evalue)) | xml version="1.0" encoding="utf-8" standalone="no"?       2024-09-02T21:10:16.399809 image/svg+xml   Matplotlib v3.7.2, https://matplotlib.org/ |
| Results, table  (threshold 1.00e-02 (evalue)) | db \| id \| prob \| evalue \| bits \| fident \| alnlen \| mismatch \| gapopen \| qstart \| qend \| tstart \| tend \| name \| description; afdb-uniprot50 \| AF-A0A2Z3IK27-F1-MODEL\_V4 \| 1.0 \| 6.197e-10 \| 263 \| 0.531 \| 96 \| 40 \| 3 \| 1 \| 93 \| 1 \| 94 \| Rho\_N domain-containing protein \| Rho\_N domain-containing protein; afdb-uniprot50 \| AF-A0A1H2R4E9-F1-MODEL\_V4 \| 1.0 \| 6.471e-09 \| 255 \| 0.431 \| 95 \| 51 \| 2 \| 1 \| 94 \| 2 \| 94 \| Rho termination factor, N-terminal domain \| Rho termination factor, N-terminal domain; afdb-uniprot50 \| AF-A0A515BE84-F1-MODEL\_V4 \| 1.0 \| 2.091e-08 \| 234 \| 0.406 \| 96 \| 55 \| 2 \| 1 \| 95 \| 1 \| 95 \| Uncharacterized protein \| Uncharacterized protein; afdb-uniprot50 \| AF-A0A238JAW9-F1-MODEL\_V4 \| 1.0 \| 4.878e-08 \| 226 \| 0.391 \| 92 \| 53 \| 1 \| 2 \| 93 \| 5 \| 93 \| Uncharacterized protein \| Uncharacterized protein; afdb-uniprot50 \| AF-A0A0P8YT19-F1-MODEL\_V4 \| 1.0 \| 8.768e-08 \| 225 \| 0.45 \| 91 \| 45 \| 1 \| 3 \| 93 \| 19 \| 104 \| Rho\_N domain-containing protein \| Rho\_N domain-containing protein; afdb-uniprot50 \| AF-A0A420WVL7-F1-MODEL\_V4 \| 1.0 \| 5.931e-08 \| 219 \| 0.371 \| 105 \| 59 \| 3 \| 1 \| 100 \| 3 \| 105 \| Rho termination factor-like protein \| Rho termination factor-like protein; afdb-uniprot50 \| AF-A0A291LZ12-F1-MODEL\_V4 \| 1.0 \| 6.33e-08 \| 218 \| 0.355 \| 107 \| 55 \| 1 \| 3 \| 95 \| 2 \| 108 \| Rho\_N domain-containing protein \| Rho\_N domain-containing protein; afdb-uniprot50 \| AF-A0A0A8TH08-F1-MODEL\_V4 \| 1.0 \| 3.024e-07 \| 209 \| 0.333 \| 93 \| 58 \| 2 \| 3 \| 94 \| 2 \| 91 \| Uncharacterized protein \| Uncharacterized protein; afdb-uniprot50 \| AF-A0A7V7G2K9-F1-MODEL\_V4 \| 1.0 \| 8.215e-08 \| 204 \| 0.392 \| 107 \| 55 \| 4 \| 2 \| 100 \| 11 \| 115 \| Rho\_N domain-containing protein \| Rho\_N domain-containing protein; afdb-uniprot50 \| AF-A0A845NDA8-F1-MODEL\_V4 \| 1.0 \| 5.675e-06 \| 193 \| 0.285 \| 98 \| 68 \| 1 \| 3 \| 100 \| 2 \| 97 \| Uncharacterized protein \| Uncharacterized protein; afdb-uniprot50 \| AF-E4PPT2-F1-MODEL\_V4 \| 1.0 \| 8.577e-07 \| 192 \| 0.333 \| 99 \| 59 \| 3 \| 2 \| 96 \| 3 \| 98 \| Protein containing Rho termination factor, N-terminal domain \| Protein containing Rho termination factor, N-terminal domain; afdb-uniprot50 \| AF-A0A0D0QEA8-F1-MODEL\_V4 \| 1.0 \| 4.47e-07 \| 191 \| 0.459 \| 98 \| 40 \| 2 \| 3 \| 91 \| 2 \| 95 \| Contig\_80, whole genome shotgun sequence \| Contig\_80, whole genome shotgun sequence; afdb-uniprot50 \| AF-A0A103EIE9-F1-MODEL\_V4 \| 1.0 \| 0.0001575 \| 190 \| 0.465 \| 58 \| 28 \| 1 \| 2 \| 56 \| 6 \| 63 \| Uncharacterized protein \| Uncharacterized protein; afdb-uniprot50 \| AF-Q24VH2-F1-MODEL\_V4 \| 1.0 \| 2.597e-06 \| 182 \| 0.268 \| 97 \| 68 \| 2 \| 1 \| 96 \| 1 \| 95 \| Rho\_N domain-containing protein \| Rho\_N domain-containing protein; afdb-uniprot50 \| AF-A0A8A6KC69-F1-MODEL\_V4 \| 1.0 \| 0.000857 \| 181 \| 0.375 \| 56 \| 35 \| 0 \| 1 \| 56 \| 1 \| 56 \| Uncharacterized protein \| Uncharacterized protein; afdb-uniprot50 \| AF-A0A6I2UUF3-F1-MODEL\_V4 \| 1.0 \| 0.0003921 \| 178 \| 0.436 \| 55 \| 26 \| 1 \| 1 \| 55 \| 1 \| 50 \| Uncharacterized protein \| Uncharacterized protein; afdb-uniprot50 \| AF-A0A6I2JDE3-F1-MODEL\_V4 \| 1.0 \| 0.001267 \| 175 \| 0.272 \| 55 \| 40 \| 0 \| 1 \| 55 \| 2 \| 56 \| Uncharacterized protein \| Uncharacterized protein; afdb-uniprot50 \| AF-A0A2S0MNH5-F1-MODEL\_V4 \| 1.0 \| 0.0001475 \| 171 \| 0.4 \| 65 \| 35 \| 1 \| 1 \| 61 \| 2 \| 66 \| Uncharacterized protein \| Uncharacterized protein; afdb-uniprot50 \| AF-A0A315BS07-F1-MODEL\_V4 \| 1.0 \| 0.0001794 \| 163 \| 0.393 \| 66 \| 30 \| 1 \| 1 \| 56 \| 1 \| 66 \| Uncharacterized protein \| Uncharacterized protein; afdb-uniprot50 \| AF-A0A177H989-F1-MODEL\_V4 \| 1.0 \| 0.006053 \| 158 \| 0.365 \| 52 \| 32 \| 1 \| 1 \| 52 \| 126 \| 176 \| Uncharacterized protein \| Uncharacterized protein; afdb-uniprot50 \| AF-A0A143DF48-F1-MODEL\_V4 \| 1.0 \| 0.003155 \| 157 \| 0.296 \| 54 \| 38 \| 0 \| 3 \| 56 \| 10 \| 63 \| Uncharacterized protein \| Uncharacterized protein; afdb-uniprot50 \| AF-A0A844C2Q1-F1-MODEL\_V4 \| 1.0 \| 1.089e-05 \| 157 \| 0.326 \| 95 \| 62 \| 2 \| 3 \| 95 \| 2 \| 96 \| Uncharacterized protein \| Uncharacterized protein; afdb-uniprot50 \| AF-A0A7C8GRC8-F1-MODEL\_V4 \| 1.0 \| 7.69e-05 \| 156 \| 0.221 \| 95 \| 72 \| 2 \| 5 \| 97 \| 2 \| 96 \| Uncharacterized protein \| Uncharacterized protein; afdb-uniprot50 \| AF-J2IAJ1-F1-MODEL\_V4 \| 1.0 \| 0.002278 \| 155 \| 0.307 \| 52 \| 33 \| 1 \| 1 \| 52 \| 1 \| 49 \| Uncharacterized protein \| Uncharacterized protein; afdb-uniprot50 \| AF-A0A371P075-F1-MODEL\_V4 \| 1.0 \| 3.755e-05 \| 155 \| 0.268 \| 97 \| 67 \| 3 \| 6 \| 100 \| 2 \| 96 \| Uncharacterized protein \| Uncharacterized protein; afdb-uniprot50 \| AF-A0A844HNJ4-F1-MODEL\_V4 \| 1.0 \| 0.0001915 \| 155 \| 0.352 \| 71 \| 36 \| 1 \| 1 \| 61 \| 1 \| 71 \| Uncharacterized protein \| Uncharacterized protein; afdb-uniprot50 \| AF-A0A2T5NTL6-F1-MODEL\_V4 \| 1.0 \| 0.006053 \| 154 \| 0.277 \| 54 \| 38 \| 1 \| 2 \| 55 \| 3 \| 55 \| Uncharacterized protein \| Uncharacterized protein; afdb-uniprot50 \| AF-A0A1X7L195-F1-MODEL\_V4 \| 1.0 \| 3.297e-05 \| 154 \| 0.255 \| 98 \| 69 \| 2 \| 1 \| 97 \| 1 \| 95 \| Uncharacterized protein \| Uncharacterized protein; afdb-uniprot50 \| AF-A0A845TX33-F1-MODEL\_V4 \| 1.0 \| 0.007855 \| 150 \| 0.215 \| 51 \| 40 \| 0 \| 3 \| 53 \| 5 \| 55 \| Uncharacterized protein \| Uncharacterized protein; afdb-uniprot50 \| AF-F4BFR0-F1-MODEL\_V4 \| 1.0 \| 0.003367 \| 150 \| 0.266 \| 60 \| 43 \| 1 \| 3 \| 61 \| 2 \| 61 \| Uncharacterized protein \| Uncharacterized protein; afdb-uniprot50 \| AF-A0A2A3MN15-F1-MODEL\_V4 \| 1.0 \| 0.003836 \| 148 \| 0.339 \| 53 \| 33 \| 1 \| 1 \| 53 \| 1 \| 51 \| Uncharacterized protein \| Uncharacterized protein; afdb-uniprot50 \| AF-A0A656JJH8-F1-MODEL\_V4 \| 1.0 \| 0.004094 \| 147 \| 0.218 \| 55 \| 43 \| 0 \| 2 \| 56 \| 30 \| 84 \| Uncharacterized protein \| Uncharacterized protein; afdb-uniprot50 \| AF-A0A4U7J7L9-F1-MODEL\_V4 \| 1.0 \| 3.519e-05 \| 145 \| 0.329 \| 88 \| 56 \| 3 \| 7 \| 92 \| 2 \| 88 \| Uncharacterized protein \| Uncharacterized protein; afdb-uniprot50 \| AF-A0A2G1LY20-F1-MODEL\_V4 \| 1.0 \| 4.874e-05 \| 145 \| 0.282 \| 99 \| 66 \| 2 \| 1 \| 96 \| 1 \| 97 \| Uncharacterized protein \| Uncharacterized protein; afdb-uniprot50 \| AF-A0A838H6R9-F1-MODEL\_V4 \| 1.0 \| 0.009551 \| 141 \| 0.254 \| 55 \| 39 \| 1 \| 1 \| 53 \| 1 \| 55 \| Uncharacterized protein \| Uncharacterized protein; afdb-uniprot50 \| AF-A0A336N748-F1-MODEL\_V4 \| 1.0 \| 0.006053 \| 141 \| 0.22 \| 59 \| 44 \| 1 \| 3 \| 61 \| 2 \| 58 \| Uncharacterized protein \| Uncharacterized protein; afdb-uniprot50 \| AF-A0A7H1MI15-F1-MODEL\_V4 \| 1.0 \| 0.003836 \| 141 \| 0.228 \| 57 \| 44 \| 0 \| 4 \| 60 \| 5 \| 61 \| Uncharacterized protein \| Uncharacterized protein; afdb-uniprot50 \| AF-A0A6L6JB59-F1-MODEL\_V4 \| 1.0 \| 7.69e-05 \| 140 \| 0.408 \| 93 \| 41 \| 3 \| 1 \| 93 \| 1 \| 79 \| Uncharacterized protein \| Uncharacterized protein; afdb-uniprot50 \| AF-A0A511XP19-F1-MODEL\_V4 \| 1.0 \| 0.008384 \| 139 \| 0.147 \| 61 \| 50 \| 1 \| 1 \| 61 \| 1 \| 59 \| Uncharacterized protein \| Uncharacterized protein; afdb-uniprot50 \| AF-A0A378R0P4-F1-MODEL\_V4 \| 1.0 \| 0.00646 \| 138 \| 0.206 \| 58 \| 45 \| 1 \| 4 \| 61 \| 10 \| 66 \| Uncharacterized protein \| Uncharacterized protein; afdb-uniprot50 \| AF-A0A024YGT7-F1-MODEL\_V4 \| 1.0 \| 5.926e-05 \| 138 \| 0.262 \| 99 \| 68 \| 1 \| 5 \| 98 \| 2 \| 100 \| Uncharacterized protein \| Uncharacterized protein; afdb-uniprot50 \| AF-A0A831RBJ2-F1-MODEL\_V4 \| 1.0 \| 0.006053 \| 137 \| 0.203 \| 59 \| 47 \| 0 \| 3 \| 61 \| 1 \| 59 \| Uncharacterized protein \| Uncharacterized protein; afdb-uniprot50 \| AF-A0A3N4VST1-F1-MODEL\_V4 \| 1.0 \| 0.004094 \| 136 \| 0.209 \| 62 \| 48 \| 1 \| 1 \| 61 \| 1 \| 62 \| Uncharacterized protein \| Uncharacterized protein; afdb-uniprot50 \| AF-A0A3T0L2P0-F1-MODEL\_V4 \| 1.0 \| 0.0002652 \| 136 \| 0.254 \| 110 \| 67 \| 2 \| 3 \| 98 \| 2 \| 110 \| Uncharacterized protein \| Uncharacterized protein; afdb-uniprot50 \| AF-A0A4Q0ZTB4-F1-MODEL\_V4 \| 1.0 \| 0.0001794 \| 134 \| 0.24 \| 104 \| 61 \| 3 \| 4 \| 92 \| 5 \| 105 \| Uncharacterized protein \| Uncharacterized protein; afdb-uniprot50 \| AF-A0A1H3BTJ3-F1-MODEL\_V4 \| 1.0 \| 0.00646 \| 133 \| 0.229 \| 61 \| 47 \| 0 \| 1 \| 61 \| 1 \| 61 \| Uncharacterized protein \| Uncharacterized protein; afdb-uniprot50 \| AF-A0A3C0F2T7-F1-MODEL\_V4 \| 1.0 \| 0.005313 \| 133 \| 0.2 \| 60 \| 46 \| 2 \| 2 \| 61 \| 16 \| 73 \| Uncharacterized protein \| Uncharacterized protein; afdb-uniprot50 \| AF-A0A2D7VZD1-F1-MODEL\_V4 \| 1.0 \| 0.005313 \| 133 \| 0.237 \| 59 \| 39 \| 1 \| 4 \| 56 \| 12 \| 70 \| Uncharacterized protein \| Uncharacterized protein; afdb-uniprot50 \| AF-A0A7V8FKE8-F1-MODEL\_V4 \| 1.0 \| 0.006053 \| 133 \| 0.311 \| 61 \| 39 \| 1 \| 3 \| 60 \| 2 \| 62 \| Uncharacterized protein \| Uncharacterized protein; afdb-uniprot50 \| AF-A0A165R271-F1-MODEL\_V4 \| 1.0 \| 0.008384 \| 132 \| 0.245 \| 61 \| 46 \| 0 \| 1 \| 61 \| 1 \| 61 \| Uncharacterized protein \| Uncharacterized protein; afdb-uniprot50 \| AF-A0A0G4Q915-F1-MODEL\_V4 \| 1.0 \| 0.00646 \| 132 \| 0.218 \| 64 \| 47 \| 1 \| 1 \| 61 \| 6 \| 69 \| Uncharacterized protein \| Uncharacterized protein; afdb-uniprot50 \| AF-A0A2W1W9H2-F1-MODEL\_V4 \| 1.0 \| 0.0004185 \| 132 \| 0.278 \| 97 \| 64 \| 4 \| 5 \| 97 \| 2 \| 96 \| Uncharacterized protein \| Uncharacterized protein; afdb-uniprot50 \| AF-A0A2D7TS63-F1-MODEL\_V4 \| 1.0 \| 0.00437 \| 132 \| 0.215 \| 65 \| 43 \| 2 \| 1 \| 65 \| 103 \| 159 \| Uncharacterized protein \| Uncharacterized protein; afdb-uniprot50 \| AF-A0A0C1INV0-F1-MODEL\_V4 \| 1.0 \| 0.008948 \| 131 \| 0.25 \| 56 \| 41 \| 1 \| 2 \| 56 \| 7 \| 62 \| Uncharacterized protein \| Uncharacterized protein; afdb-uniprot50 \| AF-A0A119CYA6-F1-MODEL\_V4 \| 1.0 \| 0.008384 \| 131 \| 0.258 \| 62 \| 43 \| 1 \| 3 \| 61 \| 2 \| 63 \| Uncharacterized protein \| Uncharacterized protein; afdb-uniprot50 \| AF-A0A1L3I567-F1-MODEL\_V4 \| 1.0 \| 0.005313 \| 131 \| 0.275 \| 58 \| 42 \| 0 \| 4 \| 61 \| 9 \| 66 \| Uncharacterized protein \| Uncharacterized protein; afdb-uniprot50 \| AF-A0A1Q8Q264-F1-MODEL\_V4 \| 1.0 \| 0.0002652 \| 131 \| 0.229 \| 96 \| 64 \| 3 \| 5 \| 94 \| 2 \| 93 \| Uncharacterized protein \| Uncharacterized protein; afdb-uniprot50 \| AF-A0A2N8HQH7-F1-MODEL\_V4 \| 1.0 \| 0.0001295 \| 131 \| 0.226 \| 97 \| 67 \| 3 \| 5 \| 93 \| 2 \| 98 \| Rho\_N domain-containing protein \| Rho\_N domain-containing protein; afdb-uniprot50 \| AF-A0A496KBK1-F1-MODEL\_V4 \| 1.0 \| 5.202e-05 \| 131 \| 0.213 \| 117 \| 67 \| 5 \| 5 \| 99 \| 2 \| 115 \| Uncharacterized protein \| Uncharacterized protein; afdb-uniprot50 \| AF-A0A4P8S989-F1-MODEL\_V4 \| 1.0 \| 0.0001137 \| 130 \| 0.225 \| 102 \| 70 \| 4 \| 3 \| 99 \| 1 \| 98 \| Uncharacterized protein \| Uncharacterized protein; afdb-uniprot50 \| AF-A0A1Q3R4D4-F1-MODEL\_V4 \| 1.0 \| 0.008948 \| 129 \| 0.169 \| 59 \| 49 \| 0 \| 3 \| 61 \| 5 \| 63 \| Uncharacterized protein \| Uncharacterized protein; afdb-uniprot50 \| AF-A0A175RVJ5-F1-MODEL\_V4 \| 1.0 \| 0.001443 \| 127 \| 0.166 \| 96 \| 78 \| 2 \| 3 \| 96 \| 1 \| 96 \| Rho\_N domain-containing protein \| Rho\_N domain-containing protein; afdb-uniprot50 \| AF-A0A841TSU2-F1-MODEL\_V4 \| 1.0 \| 0.0002328 \| 126 \| 0.237 \| 101 \| 63 \| 3 \| 6 \| 94 \| 2 \| 100 \| Uncharacterized protein \| Uncharacterized protein; afdb-uniprot50 \| AF-A0A4Y8UXJ0-F1-MODEL\_V4 \| 1.0 \| 0.0005431 \| 125 \| 0.247 \| 97 \| 65 \| 3 \| 6 \| 98 \| 2 \| 94 \| SAP domain-containing protein \| SAP domain-containing protein; afdb-uniprot50 \| AF-D5SZE1-F1-MODEL\_V4 \| 1.0 \| 0.005313 \| 125 \| 0.241 \| 58 \| 38 \| 1 \| 4 \| 55 \| 6 \| 63 \| Uncharacterized protein \| Uncharacterized protein; afdb-uniprot50 \| AF-A0A0K9MD59-F1-MODEL\_V4 \| 1.0 \| 0.0009763 \| 125 \| 0.272 \| 99 \| 68 \| 2 \| 6 \| 100 \| 2 \| 100 \| Rho\_N domain-containing protein \| Rho\_N domain-containing protein; afdb-uniprot50 \| AF-A0A2K1EF22-F1-MODEL\_V4 \| 1.0 \| 0.0005089 \| 122 \| 0.292 \| 89 \| 63 \| 0 \| 6 \| 94 \| 2 \| 90 \| Uncharacterized protein \| Uncharacterized protein; afdb-uniprot50 \| AF-C4L0Q0-F1-MODEL\_V4 \| 1.0 \| 0.0006187 \| 121 \| 0.292 \| 99 \| 58 \| 3 \| 6 \| 93 \| 2 \| 99 \| Uncharacterized protein \| Uncharacterized protein; afdb-uniprot50 \| AF-A0A098MGF8-F1-MODEL\_V4 \| 1.0 \| 0.0006187 \| 119 \| 0.214 \| 98 \| 76 \| 1 \| 4 \| 100 \| 8 \| 105 \| Uncharacterized protein \| Uncharacterized protein; afdb-uniprot50 \| AF-A0A1H2ZIW5-F1-MODEL\_V4 \| 1.0 \| 0.007359 \| 118 \| 0.253 \| 63 \| 42 \| 1 \| 4 \| 61 \| 9 \| 71 \| Uncharacterized protein \| Uncharacterized protein; afdb-uniprot50 \| AF-A0A2N1AP72-F1-MODEL\_V4 \| 1.0 \| 0.001644 \| 117 \| 0.376 \| 85 \| 48 \| 2 \| 14 \| 93 \| 1 \| 85 \| Rho\_N domain-containing protein \| Rho\_N domain-containing protein; afdb-uniprot50 \| AF-A0A654DQC9-F1-MODEL\_V4 \| 1.0 \| 0.002278 \| 117 \| 0.213 \| 89 \| 66 \| 2 \| 5 \| 93 \| 2 \| 86 \| Uncharacterized protein \| Uncharacterized protein; afdb-uniprot50 \| AF-A0A559IZJ5-F1-MODEL\_V4 \| 1.0 \| 0.002769 \| 117 \| 0.163 \| 104 \| 77 \| 4 \| 3 \| 100 \| 2 \| 101 \| Uncharacterized protein \| Uncharacterized protein; afdb-uniprot50 \| AF-A0A1A9BP13-F1-MODEL\_V4 \| 1.0 \| 0.0009763 \| 117 \| 0.283 \| 106 \| 63 \| 3 \| 5 \| 100 \| 2 \| 104 \| Rho termination factor, N-terminal domain \| Rho termination factor, N-terminal domain; afdb-uniprot50 \| AF-A0A5C0ZYA2-F1-MODEL\_V4 \| 1.0 \| 0.0003021 \| 117 \| 0.237 \| 101 \| 68 \| 4 \| 6 \| 100 \| 2 \| 99 \| Uncharacterized protein \| Uncharacterized protein; afdb-uniprot50 \| AF-A0A2W4IJI6-F1-MODEL\_V4 \| 1.0 \| 0.000803 \| 116 \| 0.257 \| 97 \| 67 \| 2 \| 1 \| 97 \| 1 \| 92 \| Rho\_N domain-containing protein \| Rho\_N domain-containing protein; afdb-uniprot50 \| AF-A0A6I2MAN7-F1-MODEL\_V4 \| 1.0 \| 0.001644 \| 110 \| 0.21 \| 100 \| 67 \| 3 \| 1 \| 92 \| 1 \| 96 \| Uncharacterized protein \| Uncharacterized protein; afdb-uniprot50 \| AF-A0A2N0EX80-F1-MODEL\_V4 \| 1.0 \| 0.001755 \| 109 \| 0.148 \| 101 \| 80 \| 3 \| 1 \| 99 \| 1 \| 97 \| Uncharacterized protein \| Uncharacterized protein; afdb-uniprot50 \| AF-A0A6A2ERZ4-F1-MODEL\_V4 \| 1.0 \| 0.001267 \| 108 \| 0.23 \| 113 \| 66 \| 3 \| 5 \| 96 \| 2 \| 114 \| Uncharacterized protein \| Uncharacterized protein; afdb-uniprot50 \| AF-A0A1R1C583-F1-MODEL\_V4 \| 1.0 \| 0.001443 \| 107 \| 0.262 \| 99 \| 68 \| 1 \| 6 \| 99 \| 2 \| 100 \| Uncharacterized protein \| Uncharacterized protein; afdb-uniprot50 \| AF-A0A1R0YQ56-F1-MODEL\_V4 \| 1.0 \| 0.000857 \| 107 \| 0.176 \| 102 \| 78 \| 2 \| 4 \| 100 \| 8 \| 108 \| Uncharacterized protein \| Uncharacterized protein; afdb-uniprot50 \| AF-A0A150KSC3-F1-MODEL\_V4 \| 1.0 \| 0.003367 \| 106 \| 0.24 \| 100 \| 68 \| 2 \| 6 \| 97 \| 2 \| 101 \| Uncharacterized protein \| Uncharacterized protein; afdb-uniprot50 \| AF-A0A1N7MP98-F1-MODEL\_V4 \| 1.0 \| 0.002956 \| 104 \| 0.208 \| 120 \| 73 \| 6 \| 2 \| 99 \| 7 \| 126 \| Rho termination factor, N-terminal domain \| Rho termination factor, N-terminal domain; afdb-uniprot50 \| AF-A0A101JYR9-F1-MODEL\_V4 \| 1.0 \| 0.004978 \| 104 \| 0.146 \| 130 \| 71 \| 4 \| 3 \| 98 \| 1 \| 124 \| Uncharacterized protein \| Uncharacterized protein; afdb-uniprot50 \| AF-A0A4R6MWN5-F1-MODEL\_V4 \| 1.0 \| 0.005313 \| 102 \| 0.215 \| 93 \| 68 \| 3 \| 3 \| 92 \| 1 \| 91 \| Uncharacterized protein \| Uncharacterized protein; afdb-uniprot50 \| AF-A0A7D8AP78-F1-MODEL\_V4 \| 1.0 \| 0.006053 \| 102 \| 0.18 \| 100 \| 74 \| 4 \| 3 \| 94 \| 1 \| 100 \| Uncharacterized protein \| Uncharacterized protein; afdb-uniprot50 \| AF-A0A838B6P4-F1-MODEL\_V4 \| 1.0 \| 0.003836 \| 102 \| 0.13 \| 115 \| 82 \| 3 \| 1 \| 97 \| 1 \| 115 \| Uncharacterized protein \| Uncharacterized protein; afdb-uniprot50 \| AF-A0A2E2Q9J7-F1-MODEL\_V4 \| 1.0 \| 0.007359 \| 102 \| 0.225 \| 93 \| 63 \| 4 \| 6 \| 92 \| 2 \| 91 \| Uncharacterized protein \| Uncharacterized protein; afdb-uniprot50 \| AF-K2KD66-F1-MODEL\_V4 \| 1.0 \| 0.002134 \| 102 \| 0.176 \| 147 \| 74 \| 8 \| 1 \| 100 \| 1 \| 147 \| Rho termination factor domain protein \| Rho termination factor domain protein; afdb-uniprot50 \| AF-A0A656YVZ8-F1-MODEL\_V4 \| 1.0 \| 0.006053 \| 102 \| 0.17 \| 82 \| 65 \| 2 \| 2 \| 81 \| 27 \| 107 \| Uncharacterized protein \| Uncharacterized protein; afdb-uniprot50 \| AF-A0A7U1GLP5-F1-MODEL\_V4 \| 1.0 \| 0.006895 \| 101 \| 0.15 \| 100 \| 82 \| 2 \| 4 \| 100 \| 3 \| 102 \| SAP domain-containing protein \| SAP domain-containing protein; afdb-uniprot50 \| AF-A0A1V2IRA7-F1-MODEL\_V4 \| 1.0 \| 0.004094 \| 101 \| 0.19 \| 110 \| 71 \| 2 \| 6 \| 99 \| 2 \| 109 \| Uncharacterized protein \| Uncharacterized protein; afdb-uniprot50 \| AF-A0A7C3DC32-F1-MODEL\_V4 \| 1.0 \| 0.003836 \| 101 \| 0.15 \| 113 \| 72 \| 5 \| 1 \| 92 \| 1 \| 110 \| Uncharacterized protein \| Uncharacterized protein; afdb-uniprot50 \| AF-A0A5B0BSY5-F1-MODEL\_V4 \| 0.999 \| 0.009551 \| 98 \| 0.122 \| 98 \| 82 \| 3 \| 4 \| 100 \| 3 \| 97 \| Uncharacterized protein \| Uncharacterized protein; afdb-uniprot50 \| AF-A0A7C3CLX3-F1-MODEL\_V4 \| 0.999 \| 0.002134 \| 98 \| 0.148 \| 128 \| 70 \| 5 \| 1 \| 94 \| 26 \| 148 \| Helix-hairpin-helix domain-containing protein \| Helix-hairpin-helix domain-containing protein; afdb-uniprot50 \| AF-A0A7Z8B2W0-F1-MODEL\_V4 \| 0.999 \| 0.002769 \| 98 \| 0.21 \| 138 \| 64 \| 6 \| 1 \| 93 \| 5 \| 142 \| Rho\_N domain-containing protein \| Rho\_N domain-containing protein; afdb-uniprot50 \| AF-A0A271M6K9-F1-MODEL\_V4 \| 0.999 \| 0.008384 \| 97 \| 0.198 \| 116 \| 70 \| 4 \| 6 \| 100 \| 2 \| 115 \| Uncharacterized protein \| Uncharacterized protein; afdb-uniprot50 \| AF-A0A0Q3TXV1-F1-MODEL\_V4 \| 0.999 \| 0.008948 \| 96 \| 0.23 \| 100 \| 72 \| 3 \| 4 \| 100 \| 3 \| 100 \| Uncharacterized protein \| Uncharacterized protein; afdb-uniprot50 \| AF-D2TCD3-F1-MODEL\_V4 \| 0.999 \| 0.007855 \| 96 \| 0.226 \| 97 \| 60 \| 1 \| 2 \| 98 \| 12 \| 93 \| Putative membrane protein ycf1 \| Putative membrane protein ycf1; afdb-uniprot50 \| AF-A0A5E9UEY1-F1-MODEL\_V4 \| 0.998 \| 0.005313 \| 95 \| 0.226 \| 97 \| 73 \| 1 \| 6 \| 100 \| 2 \| 98 \| Uncharacterized protein \| Uncharacterized protein; afdb-uniprot50 \| AF-S3A3S2-F1-MODEL\_V4 \| 0.998 \| 0.007855 \| 95 \| 0.179 \| 106 \| 66 \| 4 \| 6 \| 91 \| 2 \| 106 \| Uncharacterized protein \| Uncharacterized protein; afdb-uniprot50 \| AF-A0A0N1AA76-F1-MODEL\_V4 \| 0.998 \| 0.006053 \| 93 \| 0.257 \| 97 \| 60 \| 5 \| 6 \| 93 \| 2 \| 95 \| Uncharacterized protein \| Uncharacterized protein; afdb-uniprot50 \| AF-A0A851HHB7-F1-MODEL\_V4 \| 0.997 \| 0.00437 \| 92 \| 0.173 \| 121 \| 61 \| 5 \| 6 \| 92 \| 2 \| 117 \| Uncharacterized protein \| Uncharacterized protein; afdb-uniprot50 \| AF-A0A564G4X5-F1-MODEL\_V4 \| 0.997 \| 0.007855 \| 92 \| 0.159 \| 119 \| 79 \| 3 \| 1 \| 98 \| 1 \| 119 \| Uncharacterized protein \| Uncharacterized protein; afdb-uniprot50 \| AF-A0A497KAL2-F1-MODEL\_V4 \| 0.997 \| 0.003594 \| 92 \| 0.218 \| 110 \| 67 \| 6 \| 2 \| 95 \| 209 \| 315 \| Uncharacterized protein \| Uncharacterized protein; afdb-uniprot50 \| AF-A0A2X4RUV3-F1-MODEL\_V4 \| 0.997 \| 0.001873 \| 91 \| 0.22 \| 109 \| 62 \| 6 \| 1 \| 88 \| 1 \| 107 \| Uncharacterized protein \| Uncharacterized protein; afdb-uniprot50 \| AF-A0A165T6X8-F1-MODEL\_V4 \| 0.997 \| 0.006895 \| 91 \| 0.21 \| 114 \| 67 \| 4 \| 2 \| 96 \| 3 \| 112 \| Uncharacterized protein \| Uncharacterized protein; afdb-uniprot50 \| AF-A0A1A9G202-F1-MODEL\_V4 \| 0.997 \| 0.003836 \| 91 \| 0.221 \| 113 \| 73 \| 3 \| 1 \| 98 \| 1 \| 113 \| Uncharacterized protein \| Uncharacterized protein; afdb-uniprot50 \| AF-A0A512J8Y4-F1-MODEL\_V4 \| 0.996 \| 0.00646 \| 89 \| 0.214 \| 98 \| 65 \| 4 \| 3 \| 89 \| 1 \| 97 \| Uncharacterized protein \| Uncharacterized protein; afdb-uniprot50 \| AF-A0A1H2EPI0-F1-MODEL\_V4 \| 0.993 \| 0.005671 \| 86 \| 0.23 \| 100 \| 63 \| 7 \| 1 \| 92 \| 3 \| 96 \| Uncharacterized protein \| Uncharacterized protein; afdb-uniprot50 \| AF-A0A1C6VDX7-F1-MODEL\_V4 \| 0.992 \| 0.006895 \| 85 \| 0.181 \| 127 \| 74 \| 5 \| 3 \| 100 \| 2 \| 127 \| Uncharacterized protein \| Uncharacterized protein; afdb-uniprot50 \| AF-A0A1G8GDC6-F1-MODEL\_V4 \| 0.991 \| 0.003594 \| 84 \| 0.214 \| 107 \| 59 \| 5 \| 1 \| 92 \| 1 \| 97 \| Uncharacterized protein \| Uncharacterized protein |
| Top keywords  (threshold 1.00e-02 (evalue)) | **domain\_containing, Rho\_N, Rho, termination, factor, N\_terminal, SAP, factor\_like, containing, Contig\_80** |
| Output files | ../../similar\_structures/06\_FANPEZAQ\_CDS\_0006\_afdb-proteome\_foldseek.tsv ../../similar\_structures/06\_FANPEZAQ\_CDS\_0006\_afdb-uniprot50\_foldseek.tsv ../../similar\_structures/06\_FANPEZAQ\_CDS\_0006\_merged.svg ../../similar\_structures/06\_FANPEZAQ\_CDS\_0006\_pdb\_foldseek.tsv |

  
  
  

Return to summary | Go to previous | Go to next

  


---

**Sequence/structure alignments coloring**  
Each object in the alignment figures is colored according to its E-value following this color coding:

1e-100
10
