## Supplementary material for "Completing the BASEL phage collection to unlock hidden diversity for systematic exploration of phage-host interactions": Data S2: 9.html

FANPEZAQ\_CDS\_0009


Return to summary | Go to previous | Go to next

|  |  |
| --- | --- |
| FANPEZAQ\_CDS\_0009  Page creation date: 02 Sep 2024, 12:00  Project folder: n/a  Input sequences file: Escherichia\_virus\_HeidiAbel.gb | phage hypothetical tail portal fragment |

### Sequence information

|  |  |
| --- | --- |
| Name | FANPEZAQ\_CDS\_0009  09\_FANPEZAQ\_CDS\_0009 (pipeline id) |
| Imported annotations | Escherichia\_virus\_HeidiAbel Bas97 |
| Protein sequence | MPIKDKAEVFIEDEIFTQVEEIFWKHFERDIKARVKYFDTSDRWA |
| Number of residues | 45 |
| Molecular weight (Da) | 5608.30 |
| Output files | ../../query\_sequences/09\_FANPEZAQ\_CDS\_0009.fasta |

### Putative domain architecture and protein family

#### Search results (HHblits)^1^

|  |  |
| --- | --- |
| Domain family databases searched | Pfam, Ncbi-cd, Cath, Phrogs |
| Results, scheme(s)  (Top layers only; threshold 1.00e-03 (evalue)) | xml version="1.0" encoding="utf-8" standalone="no"?       2024-09-02T21:08:13.707031 image/svg+xml   Matplotlib v3.7.2, https://matplotlib.org/ |
| Results, table  (E-value ≤ 1.00e-03 (evalue)) | -- |
| Top keywords  (threshold 1.00e-03 (evalue)) | -- |
| Output files | ../../domain\_architecture/09\_FANPEZAQ\_CDS\_0009\_cath.hhr ../../domain\_architecture/09\_FANPEZAQ\_CDS\_0009\_merged.svg ../../domain\_architecture/09\_FANPEZAQ\_CDS\_0009\_ncbi-cd.hhr ../../domain\_architecture/09\_FANPEZAQ\_CDS\_0009\_pfam.hhr ../../domain\_architecture/09\_FANPEZAQ\_CDS\_0009\_phrogs.hhr |

### Identical protein sequences/structures

#### Search results

|  |  |
| --- | --- |
| Protein sequence databases searched | Pdb, Swissprot, Refseq |
| Identical proteins found | -- |
| Top keywords | -- |
| Output files | -- |

### Similar protein sequences/structures

#### Sequence similarity search results (HHblits)^1^

|  |  |
| --- | --- |
| Sequence databases searched | Uniclust, Pdb70 |
| Results, scheme(s)  (Top layers only, threshold 1.00e-03 (evalue)) | xml version="1.0" encoding="utf-8" standalone="no"?       2024-09-02T21:08:31.522522 image/svg+xml   Matplotlib v3.7.2, https://matplotlib.org/ |
| Results, table(s)  (threshold 1.00e-03 (evalue)) | db \| id \| prob \| evalue \| pvalue \| score \| cols \| query \| query\_len \| template \| template\_len \| name \| description; uniclust \| UniRef100\_A0A061YFQ4 \| 99.5 \| 3e-17 \| 8.1e-23 \| 111.6 \| 43 \| (1, 43) \| 45 \| (164, 206) \| 234 \| Uncharacterized protein \| Uncharacterized protein; uniclust \| UniRef100\_A0A085ASD5 \| 99.5 \| 1.2e-16 \| 2.8e-22 \| 110.8 \| 44 \| (1, 44) \| 45 \| (218, 261) \| 272 \| Phage protein \| Phage protein; uniclust \| UniRef100\_A0A4V6JHM3 \| 99.5 \| 1.8e-16 \| 3.5e-22 \| 92.4 \| 44 \| (1, 44) \| 45 \| (27, 70) \| 72 \| Uncharacterized protein \| Uncharacterized protein; uniclust \| UniRef100\_A0A088FVG9 \| 99.4 \| 9e-16 \| 2.2e-21 \| 103.4 \| 44 \| (1, 44) \| 45 \| (153, 196) \| 205 \| Phage tail protein \| Phage tail protein; uniclust \| UniRef100\_A0A6L3XNY4 \| 99.1 \| 6.6e-13 \| 1.2e-18 \| 87.1 \| 40 \| (1, 40) \| 45 \| (125, 164) \| 165 \| Phage portal protein (Fragment) \| Phage portal protein (Fragment); uniclust \| UniRef100\_A0A0B4BLP4 \| 98.8 \| 2.7e-11 \| 5.6e-17 \| 82.6 \| 45 \| (1, 45) \| 45 \| (156, 200) \| 205 \| Tail protein \| Tail protein; uniclust \| UniRef100\_UPI001E3A7150 \| 98.1 \| 5.5e-08 \| 1e-13 \| 62.0 \| 41 \| (1, 41) \| 45 \| (71, 111) \| 112 \| hypothetical protein \| hypothetical protein; uniclust \| UniRef100\_UPI0006B67FA6 \| 98.0 \| 8.3e-08 \| 1.5e-13 \| 63.7 \| 38 \| (1, 38) \| 45 \| (113, 150) \| 152 \| hypothetical protein \| hypothetical protein; uniclust \| UniRef100\_A0A8S7U9Y6 \| 97.2 \| 2.1e-05 \| 3.9e-11 \| 54.6 \| 39 \| (1, 39) \| 45 \| (153, 191) \| 192 \| Uncharacterized protein \| Uncharacterized protein; uniclust \| UniRef100\_UPI00235E9506 \| 97.0 \| 6.5e-05 \| 1.2e-10 \| 43.3 \| 33 \| (12, 44) \| 45 \| (13, 45) \| 50 \| hypothetical protein \| hypothetical protein |
| Top keywords  (threshold 1.00e-03 (evalue)) | **Phage, hypothetical, tail, portal, Fragment** |
| Output files | ../../similar\_sequences/09\_FANPEZAQ\_CDS\_0009\_merged.svg ../../similar\_sequences/09\_FANPEZAQ\_CDS\_0009\_pdb70.a3m ../../similar\_sequences/09\_FANPEZAQ\_CDS\_0009\_pdb70.hhr ../../similar\_sequences/09\_FANPEZAQ\_CDS\_0009\_uniclust.a3m ../../similar\_sequences/09\_FANPEZAQ\_CDS\_0009\_uniclust.hhr |

#### Structure prediction (AlphaFold)^2^

|  |  |
| --- | --- |
| Stats | xml version="1.0" encoding="utf-8" standalone="no"?       2024-09-02T21:09:08.159697 image/svg+xml   Matplotlib v3.7.2, https://matplotlib.org/ |
| Predicted structure | **NGL Viewer Controls:**  - Center: *Left-Click* - Rotate: *Left-Click + Drag* - Translate: *Right-Click + Drag* - Zoom: *Shift + Left-Click + Drag* |
| Output files | ../../predicted\_structures/09\_FANPEZAQ\_CDS\_0009/features.pkl ../../predicted\_structures/09\_FANPEZAQ\_CDS\_0009/ranked\_0.pdb ../../predicted\_structures/09\_FANPEZAQ\_CDS\_0009/ranked\_0\_plots.svg ../../predicted\_structures/09\_FANPEZAQ\_CDS\_0009/result\_model\_1\_ptm\_pred\_0.pkl |

#### Structure similarity search results (Foldseek)^3^

|  |  |
| --- | --- |
| Structure databases searched | Pdb, Afdb-proteome, Afdb-uniprot50 |
| Results, scheme(s)  (Top layers only, threshold 1.00e-02 (evalue)) | xml version="1.0" encoding="utf-8" standalone="no"?       2024-09-02T21:10:16.750126 image/svg+xml   Matplotlib v3.7.2, https://matplotlib.org/ |
| Results, table  (threshold 1.00e-02 (evalue)) | -- |
| Top keywords  (threshold 1.00e-02 (evalue)) | -- |
| Output files | ../../similar\_structures/09\_FANPEZAQ\_CDS\_0009\_afdb-proteome\_foldseek.tsv ../../similar\_structures/09\_FANPEZAQ\_CDS\_0009\_afdb-uniprot50\_foldseek.tsv ../../similar\_structures/09\_FANPEZAQ\_CDS\_0009\_merged.svg ../../similar\_structures/09\_FANPEZAQ\_CDS\_0009\_pdb\_foldseek.tsv |

  
  
  

Return to summary | Go to previous | Go to next

  


---

**Sequence/structure alignments coloring**  
Each object in the alignment figures is colored according to its E-value following this color coding:

1e-100
10
