## Supplementary material for "Completing the BASEL phage collection to unlock hidden diversity for systematic exploration of phage-host interactions": Data S2: 10.html

FANPEZAQ\_CDS\_0010


Return to summary | Go to previous | Go to next

|  |  |
| --- | --- |
| FANPEZAQ\_CDS\_0010  Page creation date: 02 Sep 2024, 12:00  Project folder: n/a  Input sequences file: Escherichia\_virus\_HeidiAbel.gb | hypothetical gp37 phage duf1834 tail prophage domain\_containing mu\_like minor u fragment putative duf3168 ankyrin phage\_related duf4128 associated gp37\_like duf806 atp\_binding |

### Sequence information

|  |  |
| --- | --- |
| Name | FANPEZAQ\_CDS\_0010  10\_FANPEZAQ\_CDS\_0010 (pipeline id) |
| Imported annotations | Escherichia\_virus\_HeidiAbel Bas97 |
| Protein sequence | MNADTHIDLEVLHNAIVADIKAAFPQLVTVEFYQDDPEARKTLPIPACLLTISELEADNE IDPGTEQLAVVATFEAHFIINSIRTPRAALAIRTLAAAFMAWLRKRRWTNPADPARKLPT GPANVVGGFPDDFTPELDKYEVWRVEWQQVIHLGESVWNDDGITPSEVYFSWSPEIGFGH EQDYEKVTP |
| Number of residues | 189 |
| Molecular weight (Da) | 21268.69 |
| Output files | ../../query\_sequences/10\_FANPEZAQ\_CDS\_0010.fasta |

### Putative domain architecture and protein family

#### Search results (HHblits)^1^

|  |  |
| --- | --- |
| Domain family databases searched | Pfam, Ncbi-cd, Cath, Phrogs |
| Results, scheme(s)  (Top layers only; threshold 1.00e-03 (evalue)) | xml version="1.0" encoding="utf-8" standalone="no"?       2024-09-02T21:08:13.914589 image/svg+xml   Matplotlib v3.7.2, https://matplotlib.org/ |
| Results, table  (E-value ≤ 1.00e-03 (evalue)) | db \| id \| prob \| evalue \| pvalue \| score \| cols \| query \| query\_len \| template \| template\_len \| name \| description; phrogs \| 1072 \| 100.0 \| 1.7e-39 \| 2.1e-43 \| 254.3 \| 166 \| (9, 188) \| 189 \| (1, 177) \| 183 \| NA \| NA; Category: unknown function; p140225 VI\_01698; phrogs \| 490 \| 99.9 \| 5.2e-27 \| 6.8e-31 \| 183.3 \| 138 \| (9, 157) \| 189 \| (2, 145) \| 183 \| portal protein \| portal protein; Category: head and packaging; p377723 VI\_06234; phrogs \| 16426 \| 99.0 \| 4e-14 \| 4.6e-18 \| 106.3 \| 144 \| (8, 159) \| 189 \| (6, 161) \| 188 \| NA \| NA; Category: unknown function; p190981 VI\_07687; phrogs \| 6911 \| 96.5 \| 6.8e-06 \| 8.4e-10 \| 59.5 \| 130 \| (10, 153) \| 189 \| (2, 144) \| 149 \| NA \| NA; Category: unknown function; p352299 VI\_01044; phrogs \| 20428 \| 95.3 \| 0.0002 \| 2.3e-08 \| 51.0 \| 161 \| (11, 188) \| 189 \| (6, 171) \| 175 \| tail terminator \| tail terminator; Category: connector; p329161 VI\_04355; phrogs \| 209 \| 95.3 \| 0.00018 \| 2.3e-08 \| 50.2 \| 94 \| (12, 109) \| 189 \| (2, 106) \| 131 \| portal protein \| portal protein; Category: head and packaging; p338775 VI\_10901 |
| Top keywords  (threshold 1.00e-03 (evalue)) | **portal, head, and, packaging, p140225, VI\_01698, p377723, VI\_06234, p190981, VI\_07687** |
| Output files | ../../domain\_architecture/10\_FANPEZAQ\_CDS\_0010\_cath.hhr ../../domain\_architecture/10\_FANPEZAQ\_CDS\_0010\_merged.svg ../../domain\_architecture/10\_FANPEZAQ\_CDS\_0010\_ncbi-cd.hhr ../../domain\_architecture/10\_FANPEZAQ\_CDS\_0010\_pfam.hhr ../../domain\_architecture/10\_FANPEZAQ\_CDS\_0010\_phrogs.hhr |

### Identical protein sequences/structures

#### Search results

|  |  |
| --- | --- |
| Protein sequence databases searched | Pdb, Swissprot, Refseq |
| Identical proteins found | -- |
| Top keywords | -- |
| Output files | -- |

### Similar protein sequences/structures

#### Sequence similarity search results (HHblits)^1^

|  |  |
| --- | --- |
| Sequence databases searched | Uniclust, Pdb70 |
| Results, scheme(s)  (Top layers only, threshold 1.00e-03 (evalue)) | xml version="1.0" encoding="utf-8" standalone="no"?       2024-09-02T21:08:31.783454 image/svg+xml   Matplotlib v3.7.2, https://matplotlib.org/ |
| Results, table(s)  (threshold 1.00e-03 (evalue)) | db \| id \| prob \| evalue \| pvalue \| score \| cols \| query \| query\_len \| template \| template\_len \| name \| description; uniclust \| UniRef100\_A0A024E9X0 \| 100.0 \| 1.8e-45 \| 3.8e-51 \| 283.3 \| 173 \| (2, 188) \| 189 \| (11, 183) \| 201 \| Phage protein \| Phage protein; uniclust \| UniRef100\_A0A088FRT4 \| 100.0 \| 2.1e-45 \| 4.5e-51 \| 290.1 \| 179 \| (1, 189) \| 189 \| (17, 195) \| 231 \| Uncharacterized protein \| Uncharacterized protein; uniclust \| UniRef100\_A0A142BA55 \| 100.0 \| 3.4e-42 \| 7.5e-48 \| 267.1 \| 175 \| (1, 189) \| 189 \| (3, 178) \| 196 \| Phage protein \| Phage protein; uniclust \| UniRef100\_A0A024HA76 \| 100.0 \| 4.3e-40 \| 9.1e-46 \| 255.0 \| 171 \| (4, 188) \| 189 \| (29, 200) \| 213 \| Phage protein \| Phage protein; uniclust \| UniRef100\_A0A081MYL3 \| 100.0 \| 5.7e-38 \| 1.2e-43 \| 238.1 \| 176 \| (3, 188) \| 189 \| (8, 184) \| 191 \| Phage protein \| Phage protein; uniclust \| UniRef100\_A0A066T1B0 \| 100.0 \| 1.2e-36 \| 2.5e-42 \| 236.8 \| 173 \| (2, 189) \| 189 \| (5, 181) \| 205 \| Uncharacterized protein \| Uncharacterized protein; uniclust \| UniRef100\_A0A167H3U2 \| 100.0 \| 1.3e-36 \| 2.8e-42 \| 242.1 \| 175 \| (2, 188) \| 189 \| (42, 222) \| 243 \| Uncharacterized protein \| Uncharacterized protein; uniclust \| UniRef100\_A0A066RR18 \| 100.0 \| 3.8e-36 \| 7.6e-42 \| 224.6 \| 174 \| (3, 189) \| 189 \| (1, 174) \| 181 \| Phage protein \| Phage protein; uniclust \| UniRef100\_A0A059IUK2 \| 100.0 \| 3.1e-34 \| 6e-40 \| 215.7 \| 172 \| (4, 188) \| 189 \| (42, 213) \| 217 \| Phage related protein \| Phage related protein; uniclust \| UniRef100\_A0A0Q2U909 \| 99.9 \| 3.9e-32 \| 8.2e-38 \| 209.2 \| 177 \| (2, 189) \| 189 \| (12, 193) \| 198 \| Uncharacterized protein \| Uncharacterized protein; uniclust \| UniRef100\_A0A031FSS6 \| 99.9 \| 8e-31 \| 1.6e-36 \| 198.2 \| 172 \| (1, 189) \| 189 \| (1, 173) \| 188 \| Putative phage-like protein \| Putative phage-like protein; uniclust \| UniRef100\_A0A853IM63 \| 99.9 \| 1.3e-30 \| 2.5e-36 \| 192.8 \| 174 \| (3, 186) \| 189 \| (5, 182) \| 185 \| Uncharacterized protein (Fragment) \| Uncharacterized protein (Fragment); uniclust \| UniRef100\_A0A0J7J7S1 \| 99.9 \| 1.7e-30 \| 3.6e-36 \| 201.6 \| 144 \| (2, 156) \| 189 \| (3, 153) \| 192 \| Mu-like prophage protein gp37 \| Mu-like prophage protein gp37; uniclust \| UniRef100\_A0A104N6L4 \| 99.9 \| 3.5e-30 \| 6.9e-36 \| 185.8 \| 123 \| (55, 188) \| 189 \| (5, 127) \| 139 \| Phage tail protein (Fragment) \| Phage tail protein (Fragment); uniclust \| UniRef100\_A0A219Y8Y8 \| 99.9 \| 7.1e-30 \| 1.4e-35 \| 189.2 \| 156 \| (3, 170) \| 189 \| (1, 156) \| 180 \| Uncharacterized protein \| Uncharacterized protein; uniclust \| UniRef100\_A0A1C2K2M7 \| 99.9 \| 5.2e-29 \| 1.1e-34 \| 190.5 \| 176 \| (1, 188) \| 189 \| (10, 186) \| 191 \| Uncharacterized protein \| Uncharacterized protein; uniclust \| UniRef100\_A0A142BH88 \| 99.9 \| 3.1e-28 \| 6.3e-34 \| 198.9 \| 165 \| (7, 189) \| 189 \| (136, 300) \| 305 \| Uncharacterized protein \| Uncharacterized protein; uniclust \| UniRef100\_A0A2J8GXZ0 \| 99.9 \| 8.3e-28 \| 1.6e-33 \| 179.9 \| 171 \| (5, 188) \| 189 \| (1, 175) \| 180 \| Phage tail protein \| Phage tail protein; uniclust \| UniRef100\_A0A060H8T0 \| 99.9 \| 2e-27 \| 3.7e-33 \| 173.1 \| 176 \| (4, 189) \| 189 \| (5, 180) \| 180 \| Tail protein \| Tail protein; uniclust \| UniRef100\_A0A0J5JHL4 \| 99.9 \| 2.4e-27 \| 4.8e-33 \| 177.7 \| 134 \| (8, 156) \| 189 \| (4, 139) \| 167 \| Gp37 protein \| Gp37 protein; uniclust \| UniRef100\_A0A1A9VKF9 \| 99.9 \| 3.4e-26 \| 6.5e-32 \| 185.1 \| 160 \| (4, 176) \| 189 \| (52, 211) \| 378 \| Phage\_base\_V domain-containing protein \| Phage\_base\_V domain-containing protein; uniclust \| UniRef100\_UPI0021CE98C7 \| 99.9 \| 7.7e-26 \| 1.4e-31 \| 165.2 \| 175 \| (4, 187) \| 189 \| (4, 184) \| 185 \| hypothetical protein \| hypothetical protein; uniclust \| UniRef100\_A0A1T2CHW9 \| 99.9 \| 1.4e-25 \| 2.8e-31 \| 167.5 \| 154 \| (5, 174) \| 189 \| (1, 154) \| 170 \| Phage protein \| Phage protein; uniclust \| UniRef100\_A0A9D8J6R2 \| 99.8 \| 1.2e-23 \| 2.2e-29 \| 147.6 \| 126 \| (6, 146) \| 189 \| (1, 126) \| 126 \| Uncharacterized protein (Fragment) \| Uncharacterized protein (Fragment); uniclust \| UniRef100\_Q9PFF2 \| 99.8 \| 7.2e-23 \| 1.3e-28 \| 143.6 \| 137 \| (45, 189) \| 189 \| (1, 137) \| 137 \| Phage-related protein \| Phage-related protein; uniclust \| UniRef100\_A0A654AL97 \| 99.8 \| 2.2e-22 \| 4.1e-28 \| 148.1 \| 138 \| (7, 155) \| 189 \| (13, 150) \| 189 \| Phage protein \| Phage protein; uniclust \| UniRef100\_A0A0W8JFJ1 \| 99.8 \| 3.2e-22 \| 6.1e-28 \| 150.8 \| 171 \| (6, 188) \| 189 \| (8, 178) \| 185 \| Phage protein \| Phage protein; uniclust \| UniRef100\_A0A085AFN8 \| 99.8 \| 4.1e-22 \| 7.9e-28 \| 151.5 \| 175 \| (2, 189) \| 189 \| (1, 176) \| 197 \| Phage protein \| Phage protein; uniclust \| UniRef100\_UPI002263F815 \| 99.7 \| 1.7e-21 \| 3.2e-27 \| 157.8 \| 153 \| (3, 168) \| 189 \| (2, 154) \| 411 \| DUF4113 domain-containing protein \| DUF4113 domain-containing protein; uniclust \| UniRef100\_A0A380T1F7 \| 99.7 \| 2.1e-21 \| 4e-27 \| 127.5 \| 79 \| (101, 187) \| 189 \| (3, 81) \| 85 \| Uncharacterized protein \| Uncharacterized protein; uniclust \| UniRef100\_UPI000B195A94 \| 99.7 \| 2.3e-21 \| 4.5e-27 \| 140.5 \| 129 \| (49, 188) \| 189 \| (2, 133) \| 143 \| hypothetical protein \| hypothetical protein; uniclust \| UniRef100\_A0A5E4NQZ0 \| 99.7 \| 4.8e-21 \| 8.7e-27 \| 163.1 \| 148 \| (10, 171) \| 189 \| (282, 429) \| 691 \| Bacteriophage lambda, GpZ, minor tail,Ankyrin repeat-containing domain,Ankyrin repeat \| Bacteriophage lambda, GpZ, minor tail,Ankyrin repeat-containing domain,Ankyrin repeat; uniclust \| UniRef100\_A0A1G8I0M9 \| 99.7 \| 6.6e-21 \| 1.3e-26 \| 144.3 \| 171 \| (3, 188) \| 189 \| (6, 181) \| 192 \| Gp37 protein \| Gp37 protein; uniclust \| UniRef100\_A0A1H2FMJ7 \| 99.7 \| 7.6e-21 \| 1.5e-26 \| 145.5 \| 157 \| (2, 171) \| 189 \| (13, 176) \| 192 \| Phage protein Gp37/Gp68 \| Phage protein Gp37/Gp68; uniclust \| UniRef100\_A0A0H2M1V4 \| 99.7 \| 8.3e-21 \| 1.7e-26 \| 148.4 \| 144 \| (1, 154) \| 189 \| (1, 152) \| 206 \| DUF1834 family protein \| DUF1834 family protein; uniclust \| UniRef100\_A0A1Z3U7W0 \| 99.7 \| 4.9e-20 \| 9.3e-26 \| 139.5 \| 171 \| (6, 186) \| 189 \| (1, 181) \| 195 \| Uncharacterized protein \| Uncharacterized protein; uniclust \| UniRef100\_F2K221 \| 99.7 \| 4.9e-20 \| 9.5e-26 \| 139.6 \| 173 \| (2, 188) \| 189 \| (1, 177) \| 184 \| Gp37 protein \| Gp37 protein; uniclust \| UniRef100\_A0A839IZ53 \| 99.7 \| 7.9e-20 \| 1.5e-25 \| 137.7 \| 173 \| (2, 188) \| 189 \| (5, 185) \| 197 \| Uncharacterized protein \| Uncharacterized protein; uniclust \| UniRef100\_A0A853IIQ5 \| 99.7 \| 1e-19 \| 1.9e-25 \| 126.8 \| 118 \| (63, 187) \| 189 \| (5, 125) \| 127 \| Uncharacterized protein \| Uncharacterized protein; uniclust \| UniRef100\_A0A1E7HY00 \| 99.6 \| 3.4e-19 \| 6.5e-25 \| 134.7 \| 161 \| (13, 188) \| 189 \| (6, 179) \| 187 \| Uncharacterized protein \| Uncharacterized protein; uniclust \| UniRef100\_A0A059IW70 \| 99.6 \| 3.4e-19 \| 6.5e-25 \| 124.1 \| 110 \| (53, 172) \| 189 \| (2, 111) \| 115 \| Phage related protein \| Phage related protein; uniclust \| UniRef100\_A0A1N6I1G9 \| 99.6 \| 3.7e-19 \| 7.1e-25 \| 135.0 \| 170 \| (7, 188) \| 189 \| (15, 190) \| 195 \| Uncharacterized protein \| Uncharacterized protein; uniclust \| UniRef100\_A0A7J6YLT3 \| 99.6 \| 3.9e-19 \| 7.2e-25 \| 147.8 \| 144 \| (20, 176) \| 189 \| (2, 145) \| 508 \| Baseplate protein J-like domain-containing protein \| Baseplate protein J-like domain-containing protein; uniclust \| UniRef100\_A0A090SU28 \| 99.6 \| 4.8e-19 \| 8.9e-25 \| 131.9 \| 172 \| (9, 188) \| 189 \| (3, 183) \| 197 \| Phage protein \| Phage protein; uniclust \| UniRef100\_A0A071M5B9 \| 99.6 \| 6.4e-19 \| 1.3e-24 \| 141.3 \| 154 \| (12, 177) \| 189 \| (26, 194) \| 234 \| Uncharacterized protein \| Uncharacterized protein; uniclust \| UniRef100\_UPI00037FEC58 \| 99.6 \| 1.4e-18 \| 2.5e-24 \| 127.7 \| 169 \| (6, 187) \| 189 \| (9, 177) \| 177 \| hypothetical protein \| hypothetical protein; uniclust \| UniRef100\_A0A1I5KYE0 \| 99.6 \| 2.1e-18 \| 3.9e-24 \| 126.4 \| 155 \| (6, 174) \| 189 \| (2, 158) \| 170 \| Gp37 protein \| Gp37 protein; uniclust \| UniRef100\_A0A068T9Q2 \| 99.6 \| 2e-18 \| 4.1e-24 \| 138.4 \| 141 \| (8, 159) \| 189 \| (20, 177) \| 223 \| Uncharacterized protein \| Uncharacterized protein; uniclust \| UniRef100\_UPI001565E1E4 \| 99.6 \| 3.6e-18 \| 6.6e-24 \| 125.9 \| 172 \| (3, 188) \| 189 \| (1, 174) \| 181 \| hypothetical protein \| hypothetical protein; uniclust \| UniRef100\_A0A8S4QQF2 \| 99.5 \| 1.1e-17 \| 2.1e-23 \| 138.2 \| 131 \| (28, 176) \| 189 \| (152, 282) \| 409 \| Jg15362 protein \| Jg15362 protein; uniclust \| UniRef100\_A0A0P7IJV4 \| 99.5 \| 1.3e-17 \| 2.6e-23 \| 128.4 \| 153 \| (4, 168) \| 189 \| (1, 161) \| 181 \| Uncharacterized protein \| Uncharacterized protein; uniclust \| UniRef100\_A0A1G5ACN2 \| 99.5 \| 3.7e-17 \| 6.8e-23 \| 120.9 \| 169 \| (7, 188) \| 189 \| (1, 174) \| 177 \| Uncharacterized protein \| Uncharacterized protein; uniclust \| UniRef100\_UPI000F02A975 \| 99.5 \| 4.5e-17 \| 8.5e-23 \| 121.6 \| 145 \| (1, 157) \| 189 \| (1, 145) \| 170 \| hypothetical protein \| hypothetical protein; uniclust \| UniRef100\_A0A0A1Z018 \| 99.5 \| 5.8e-17 \| 1.1e-22 \| 120.6 \| 155 \| (3, 171) \| 189 \| (1, 157) \| 185 \| Uncharacterized protein \| Uncharacterized protein; uniclust \| UniRef100\_A0A1D2X525 \| 99.5 \| 9.8e-17 \| 1.9e-22 \| 118.9 \| 140 \| (5, 152) \| 189 \| (7, 153) \| 156 \| Uncharacterized protein \| Uncharacterized protein; uniclust \| UniRef100\_A0A4P0TMC0 \| 99.5 \| 2.1e-16 \| 3.9e-22 \| 104.3 \| 71 \| (117, 187) \| 189 \| (8, 78) \| 82 \| Phage protein \| Phage protein; uniclust \| UniRef100\_A0A4V2PRI1 \| 99.4 \| 3.2e-16 \| 5.8e-22 \| 115.9 \| 165 \| (7, 189) \| 189 \| (10, 174) \| 179 \| Gp37 protein \| Gp37 protein; uniclust \| UniRef100\_A0A0C5VSK4 \| 99.4 \| 3.6e-16 \| 7.3e-22 \| 123.8 \| 143 \| (6, 157) \| 189 \| (1, 175) \| 214 \| Uncharacterized protein \| Uncharacterized protein; uniclust \| UniRef100\_A0A4P9VGP3 \| 99.4 \| 4.4e-16 \| 8.2e-22 \| 116.4 \| 133 \| (41, 186) \| 189 \| (55, 187) \| 193 \| DUF1834 family protein \| DUF1834 family protein; uniclust \| UniRef100\_A0A327JK07 \| 99.4 \| 4.8e-16 \| 9.3e-22 \| 118.7 \| 158 \| (1, 171) \| 189 \| (8, 172) \| 184 \| Uncharacterized protein \| Uncharacterized protein; uniclust \| UniRef100\_UPI000B7E93C5 \| 99.4 \| 3e-15 \| 5.6e-21 \| 109.7 \| 132 \| (3, 149) \| 189 \| (1, 137) \| 154 \| hypothetical protein \| hypothetical protein; uniclust \| UniRef100\_A0A0X8JJA6 \| 99.2 \| 5.3e-14 \| 1e-19 \| 107.6 \| 138 \| (7, 157) \| 189 \| (1, 146) \| 185 \| Osmotically inducible protein OsmC \| Osmotically inducible protein OsmC; uniclust \| UniRef100\_A0A1F9BJG0 \| 99.2 \| 4.9e-14 \| 1e-19 \| 108.4 \| 136 \| (3, 152) \| 189 \| (7, 150) \| 155 \| DUF1834 domain-containing protein \| DUF1834 domain-containing protein; uniclust \| UniRef100\_A0A450Z1H3 \| 99.2 \| 6e-14 \| 1.1e-19 \| 99.9 \| 128 \| (6, 145) \| 189 \| (1, 132) \| 133 \| Uncharacterized protein \| Uncharacterized protein; uniclust \| UniRef100\_A0A5E4XFY4 \| 99.2 \| 1e-13 \| 1.9e-19 \| 105.4 \| 143 \| (7, 160) \| 189 \| (1, 151) \| 188 \| Uncharacterized protein \| Uncharacterized protein; uniclust \| UniRef100\_UPI001E3A84FA \| 99.1 \| 2.6e-13 \| 4.8e-19 \| 85.4 \| 64 \| (126, 189) \| 189 \| (2, 65) \| 65 \| hypothetical protein \| hypothetical protein; uniclust \| UniRef100\_UPI0004168F22 \| 99.1 \| 2.6e-13 \| 4.9e-19 \| 104.6 \| 167 \| (7, 188) \| 189 \| (8, 192) \| 195 \| hypothetical protein \| hypothetical protein; uniclust \| UniRef100\_A0A450TJH1 \| 99.1 \| 2.7e-13 \| 5e-19 \| 89.3 \| 61 \| (128, 188) \| 189 \| (16, 76) \| 79 \| Uncharacterized protein \| Uncharacterized protein; uniclust \| UniRef100\_A0A1E3G659 \| 99.1 \| 3.6e-13 \| 6.6e-19 \| 100.6 \| 163 \| (9, 187) \| 189 \| (2, 164) \| 181 \| Uncharacterized protein \| Uncharacterized protein; uniclust \| UniRef100\_A0A0W1G855 \| 99.1 \| 3.7e-13 \| 7.8e-19 \| 111.8 \| 144 \| (2, 155) \| 189 \| (8, 171) \| 256 \| DUF1834 family protein \| DUF1834 family protein; uniclust \| UniRef100\_A0A3N2E1B1 \| 99.1 \| 4.6e-13 \| 8.5e-19 \| 100.1 \| 173 \| (3, 188) \| 189 \| (1, 175) \| 181 \| Gp37 protein \| Gp37 protein; uniclust \| UniRef100\_UPI0013D3F786 \| 99.1 \| 9.3e-13 \| 1.7e-18 \| 97.3 \| 146 \| (2, 156) \| 189 \| (1, 147) \| 165 \| hypothetical protein \| hypothetical protein; uniclust \| UniRef100\_A0A5S9P2S6 \| 99.1 \| 9.9e-13 \| 1.8e-18 \| 99.9 \| 172 \| (9, 189) \| 189 \| (10, 198) \| 201 \| Uncharacterized protein \| Uncharacterized protein; uniclust \| UniRef100\_A0A059ZVM0 \| 99.0 \| 1.3e-12 \| 2.7e-18 \| 106.4 \| 166 \| (1, 188) \| 189 \| (37, 216) \| 234 \| Uncharacterized protein \| Uncharacterized protein; uniclust \| UniRef100\_A0A0E4BWM4 \| 99.0 \| 1.5e-12 \| 3e-18 \| 101.9 \| 139 \| (6, 156) \| 189 \| (1, 146) \| 194 \| Uncharacterized protein \| Uncharacterized protein; uniclust \| UniRef100\_UPI001E56994E \| 99.0 \| 1.8e-12 \| 3.3e-18 \| 87.2 \| 86 \| (1, 87) \| 189 \| (1, 88) \| 92 \| hypothetical protein \| hypothetical protein; uniclust \| UniRef100\_A0A8H8X087 \| 99.0 \| 2e-12 \| 3.8e-18 \| 100.5 \| 160 \| (5, 174) \| 189 \| (1, 179) \| 208 \| Uncharacterized protein \| Uncharacterized protein; uniclust \| UniRef100\_A0A9E3PK95 \| 99.0 \| 3.1e-12 \| 6e-18 \| 99.5 \| 146 \| (10, 165) \| 189 \| (19, 180) \| 204 \| Uncharacterized protein \| Uncharacterized protein; uniclust \| UniRef100\_A0A1C3EBP2 \| 99.0 \| 3.4e-12 \| 6.2e-18 \| 96.2 \| 165 \| (3, 177) \| 189 \| (1, 181) \| 187 \| Phage tail protein \| Phage tail protein; uniclust \| UniRef100\_A0A2A4XUH6 \| 99.0 \| 3.5e-12 \| 6.4e-18 \| 94.7 \| 140 \| (6, 157) \| 189 \| (1, 147) \| 169 \| Uncharacterized protein \| Uncharacterized protein; uniclust \| UniRef100\_A0A009YIC9 \| 98.9 \| 5.7e-12 \| 1.2e-17 \| 103.4 \| 143 \| (2, 155) \| 189 \| (17, 176) \| 224 \| DUF1834 family protein \| DUF1834 family protein; uniclust \| UniRef100\_A0A650ENI9 \| 98.9 \| 1e-11 \| 2.1e-17 \| 97.4 \| 135 \| (3, 155) \| 189 \| (1, 146) \| 184 \| Uncharacterized protein \| Uncharacterized protein; uniclust \| UniRef100\_A0A2K8U790 \| 98.9 \| 1.2e-11 \| 2.2e-17 \| 90.8 \| 138 \| (8, 153) \| 189 \| (7, 151) \| 154 \| Uncharacterized protein \| Uncharacterized protein; uniclust \| UniRef100\_UPI002152489E \| 98.8 \| 2.4e-11 \| 4.3e-17 \| 82.6 \| 85 \| (93, 187) \| 189 \| (2, 87) \| 95 \| hypothetical protein \| hypothetical protein; uniclust \| UniRef100\_A0A2D8QD62 \| 98.8 \| 2.8e-11 \| 5.1e-17 \| 92.8 \| 177 \| (7, 187) \| 189 \| (4, 194) \| 207 \| Uncharacterized protein \| Uncharacterized protein; uniclust \| UniRef100\_A0A4P0TLN7 \| 98.8 \| 3.2e-11 \| 5.9e-17 \| 85.5 \| 107 \| (4, 129) \| 189 \| (3, 109) \| 117 \| DUF3168 domain-containing protein \| DUF3168 domain-containing protein; uniclust \| UniRef100\_UPI001EDABC2D \| 98.8 \| 3.2e-11 \| 5.9e-17 \| 87.3 \| 130 \| (46, 186) \| 189 \| (1, 135) \| 139 \| hypothetical protein \| hypothetical protein; uniclust \| UniRef100\_A0A4U9HEQ6 \| 98.8 \| 3.8e-11 \| 7.1e-17 \| 80.0 \| 70 \| (119, 188) \| 189 \| (11, 80) \| 85 \| Uncharacterized protein \| Uncharacterized protein; uniclust \| UniRef100\_A0A8J2Z5Y1 \| 98.8 \| 3.6e-11 \| 7.1e-17 \| 92.7 \| 161 \| (2, 177) \| 189 \| (1, 161) \| 175 \| Uncharacterized protein \| Uncharacterized protein; uniclust \| UniRef100\_A0A022PGL8 \| 98.8 \| 3.7e-11 \| 7.8e-17 \| 97.3 \| 135 \| (2, 153) \| 189 \| (14, 157) \| 209 \| Mu-like prophage protein gp37 \| Mu-like prophage protein gp37; uniclust \| UniRef100\_A0A0H3ZQZ5 \| 98.8 \| 5.2e-11 \| 1.1e-16 \| 93.9 \| 158 \| (2, 174) \| 189 \| (4, 161) \| 183 \| Uncharacterized protein \| Uncharacterized protein; uniclust \| UniRef100\_A0A7X3ZHV1 \| 98.8 \| 6.6e-11 \| 1.2e-16 \| 91.8 \| 154 \| (24, 187) \| 189 \| (47, 216) \| 223 \| Uncharacterized protein \| Uncharacterized protein; uniclust \| UniRef100\_A0A2N8GRJ2 \| 98.7 \| 9e-11 \| 1.7e-16 \| 78.4 \| 77 \| (40, 127) \| 189 \| (8, 84) \| 85 \| Uncharacterized protein (Fragment) \| Uncharacterized protein (Fragment); uniclust \| UniRef100\_A0A1H1G3G9 \| 98.7 \| 8.7e-11 \| 1.7e-16 \| 92.2 \| 139 \| (7, 155) \| 189 \| (11, 158) \| 189 \| Uncharacterized protein \| Uncharacterized protein; uniclust \| UniRef100\_A0A846QLX3 \| 98.7 \| 9.4e-11 \| 1.7e-16 \| 79.7 \| 86 \| (92, 188) \| 189 \| (3, 88) \| 94 \| Uncharacterized protein \| Uncharacterized protein; uniclust \| UniRef100\_UPI001E4FC916 \| 98.7 \| 9.6e-11 \| 1.8e-16 \| 84.6 \| 98 \| (6, 109) \| 189 \| (1, 98) \| 135 \| hypothetical protein \| hypothetical protein; uniclust \| UniRef100\_A0A1S8YKZ2 \| 98.7 \| 1.4e-10 \| 2.5e-16 \| 87.7 \| 166 \| (9, 188) \| 189 \| (7, 176) \| 182 \| Phage tail protein \| Phage tail protein; uniclust \| UniRef100\_UPI0018F1F2B8 \| 98.7 \| 1.5e-10 \| 2.7e-16 \| 87.8 \| 168 \| (5, 188) \| 189 \| (2, 170) \| 175 \| hypothetical protein \| hypothetical protein; uniclust \| UniRef100\_B7RNM1 \| 98.7 \| 2.1e-10 \| 3.9e-16 \| 77.3 \| 70 \| (117, 186) \| 189 \| (15, 84) \| 89 \| Uncharacterized protein \| Uncharacterized protein; uniclust \| UniRef100\_A0A9D9F620 \| 98.6 \| 2.7e-10 \| 4.9e-16 \| 87.2 \| 161 \| (7, 176) \| 189 \| (13, 183) \| 197 \| Uncharacterized protein \| Uncharacterized protein; uniclust \| UniRef100\_A0A3N0V7T1 \| 98.6 \| 2.7e-10 \| 5.2e-16 \| 87.6 \| 137 \| (7, 156) \| 189 \| (1, 143) \| 182 \| DUF1834 family protein \| DUF1834 family protein; uniclust \| UniRef100\_UPI001596FEA7 \| 98.6 \| 3.3e-10 \| 6.1e-16 \| 86.3 \| 112 \| (42, 161) \| 189 \| (41, 155) \| 191 \| hypothetical protein \| hypothetical protein; uniclust \| UniRef100\_A0A962EV07 \| 98.6 \| 5e-10 \| 9.2e-16 \| 83.7 \| 143 \| (3, 157) \| 189 \| (1, 150) \| 166 \| Uncharacterized protein \| Uncharacterized protein; uniclust \| UniRef100\_UPI001C25C53F \| 98.6 \| 6.1e-10 \| 1.1e-15 \| 75.8 \| 70 \| (5, 79) \| 189 \| (1, 70) \| 93 \| hypothetical protein \| hypothetical protein; uniclust \| UniRef100\_UPI0004983960 \| 98.6 \| 7.1e-10 \| 1.3e-15 \| 85.9 \| 136 \| (7, 153) \| 189 \| (1, 148) \| 213 \| hypothetical protein \| hypothetical protein; uniclust \| UniRef100\_A0A3R8IUE9 \| 98.6 \| 7.2e-10 \| 1.3e-15 \| 84.8 \| 159 \| (4, 176) \| 189 \| (6, 176) \| 187 \| DUF1834 family protein \| DUF1834 family protein; uniclust \| UniRef100\_UPI001F146AA5 \| 98.6 \| 7.8e-10 \| 1.4e-15 \| 82.6 \| 113 \| (13, 145) \| 189 \| (3, 121) \| 164 \| hypothetical protein \| hypothetical protein; uniclust \| UniRef100\_A0A2V3UBZ9 \| 98.5 \| 8.5e-10 \| 1.6e-15 \| 83.9 \| 142 \| (6, 155) \| 189 \| (1, 148) \| 185 \| Uncharacterized protein DUF1834 \| Uncharacterized protein DUF1834; uniclust \| UniRef100\_A0A1E3UZA7 \| 98.5 \| 8.8e-10 \| 1.6e-15 \| 84.1 \| 153 \| (15, 181) \| 189 \| (16, 183) \| 191 \| Uncharacterized protein \| Uncharacterized protein; uniclust \| UniRef100\_A0A063BJE5 \| 98.5 \| 7.9e-10 \| 1.7e-15 \| 92.3 \| 162 \| (1, 173) \| 189 \| (8, 192) \| 243 \| Mu-like prophage protein gp37 \| Mu-like prophage protein gp37; uniclust \| UniRef100\_A0A0J7LV93 \| 98.5 \| 9.2e-10 \| 1.9e-15 \| 89.4 \| 155 \| (4, 176) \| 189 \| (8, 176) \| 201 \| Mu-like prophage protein gp37 \| Mu-like prophage protein gp37; uniclust \| UniRef100\_A0A136Q971 \| 98.5 \| 1.5e-09 \| 2.7e-15 \| 72.6 \| 70 \| (6, 80) \| 189 \| (13, 82) \| 83 \| DUF3168 domain-containing protein (Fragment) \| DUF3168 domain-containing protein (Fragment); uniclust \| UniRef100\_A0A5P9F0H4 \| 98.5 \| 1.8e-09 \| 3.5e-15 \| 83.4 \| 179 \| (6, 189) \| 189 \| (1, 186) \| 190 \| Uncharacterized protein \| Uncharacterized protein; uniclust \| UniRef100\_A0A8T5Z499 \| 98.4 \| 2.3e-09 \| 4.3e-15 \| 73.6 \| 83 \| (99, 188) \| 189 \| (2, 85) \| 96 \| Phage tail protein \| Phage tail protein; uniclust \| UniRef100\_UPI0020A2AFB5 \| 98.4 \| 3.3e-09 \| 6.1e-15 \| 88.5 \| 152 \| (13, 177) \| 189 \| (16, 182) \| 379 \| hypothetical protein \| hypothetical protein; uniclust \| UniRef100\_A0A439U2R2 \| 98.4 \| 3.5e-09 \| 6.7e-15 \| 82.8 \| 172 \| (2, 188) \| 189 \| (1, 182) \| 188 \| Uncharacterized protein \| Uncharacterized protein; uniclust \| UniRef100\_UPI000B8EB34D \| 98.4 \| 4.2e-09 \| 7.8e-15 \| 74.8 \| 95 \| (87, 188) \| 189 \| (13, 111) \| 116 \| hypothetical protein \| hypothetical protein; uniclust \| UniRef100\_UPI00227F3D8E \| 98.4 \| 4.5e-09 \| 8.2e-15 \| 79.8 \| 155 \| (8, 175) \| 189 \| (7, 164) \| 179 \| hypothetical protein \| hypothetical protein; uniclust \| UniRef100\_A0A8S5QI65 \| 98.4 \| 5.2e-09 \| 1e-14 \| 79.7 \| 140 \| (6, 155) \| 189 \| (1, 148) \| 157 \| Uncharacterized protein \| Uncharacterized protein; uniclust \| UniRef100\_UPI0020C04467 \| 98.4 \| 5.6e-09 \| 1e-14 \| 72.0 \| 95 \| (5, 108) \| 189 \| (1, 95) \| 97 \| hypothetical protein \| hypothetical protein; uniclust \| UniRef100\_A0A4U8YKI5 \| 98.4 \| 5.7e-09 \| 1.1e-14 \| 78.7 \| 154 \| (6, 170) \| 189 \| (1, 160) \| 171 \| Uncharacterized protein \| Uncharacterized protein; uniclust \| UniRef100\_UPI001C99755B \| 98.3 \| 6.6e-09 \| 1.2e-14 \| 72.6 \| 94 \| (3, 98) \| 189 \| (1, 102) \| 105 \| DUF1834 family protein \| DUF1834 family protein; uniclust \| UniRef100\_A0A0X8JK81 \| 98.3 \| 6.5e-09 \| 1.3e-14 \| 85.7 \| 141 \| (3, 153) \| 189 \| (1, 156) \| 231 \| Uncharacterized protein \| Uncharacterized protein; uniclust \| UniRef100\_F2J640 \| 98.3 \| 9.5e-09 \| 1.8e-14 \| 88.3 \| 136 \| (6, 152) \| 189 \| (1, 142) \| 489 \| Uncharacterized protein \| Uncharacterized protein; uniclust \| UniRef100\_UPI00234BD1FE \| 98.3 \| 1.1e-08 \| 2.1e-14 \| 72.7 \| 78 \| (5, 85) \| 189 \| (6, 83) \| 115 \| hypothetical protein \| hypothetical protein; uniclust \| UniRef100\_UPI0020BFFED6 \| 98.3 \| 1.2e-08 \| 2.1e-14 \| 71.2 \| 80 \| (88, 174) \| 189 \| (9, 88) \| 102 \| hypothetical protein \| hypothetical protein; uniclust \| UniRef100\_UPI001903E94F \| 98.2 \| 1.5e-08 \| 2.7e-14 \| 76.9 \| 149 \| (1, 156) \| 189 \| (1, 166) \| 175 \| hypothetical protein \| hypothetical protein; uniclust \| UniRef100\_A0A1D2QS75 \| 98.2 \| 2.3e-08 \| 4.3e-14 \| 84.6 \| 153 \| (6, 162) \| 189 \| (1, 182) \| 411 \| Uncharacterized protein \| Uncharacterized protein; uniclust \| UniRef100\_A0A1G8G2T2 \| 98.2 \| 3e-08 \| 5.9e-14 \| 79.8 \| 141 \| (4, 155) \| 189 \| (3, 162) \| 206 \| Mu-like prophage protein gp37 (Fragment) \| Mu-like prophage protein gp37 (Fragment); uniclust \| UniRef100\_UPI000AB590E9 \| 98.2 \| 3.2e-08 \| 6e-14 \| 71.5 \| 104 \| (64, 177) \| 189 \| (7, 112) \| 126 \| hypothetical protein \| hypothetical protein; uniclust \| UniRef100\_A0A345DE57 \| 98.1 \| 3.5e-08 \| 6.5e-14 \| 75.6 \| 140 \| (7, 161) \| 189 \| (1, 141) \| 184 \| Uncharacterized protein \| Uncharacterized protein; uniclust \| UniRef100\_A0A432QS63 \| 98.1 \| 3.9e-08 \| 7.1e-14 \| 75.5 \| 144 \| (9, 162) \| 189 \| (8, 157) \| 186 \| DUF1834 domain-containing protein \| DUF1834 domain-containing protein; uniclust \| UniRef100\_UPI00217D2FA9 \| 98.1 \| 4.9e-08 \| 9e-14 \| 69.5 \| 92 \| (5, 102) \| 189 \| (4, 99) \| 114 \| hypothetical protein \| hypothetical protein; uniclust \| UniRef100\_A0A142BHP0 \| 98.1 \| 5.7e-08 \| 1e-13 \| 68.3 \| 84 \| (64, 154) \| 189 \| (15, 101) \| 105 \| Phage protein \| Phage protein; uniclust \| UniRef100\_Q4E9T1 \| 98.0 \| 7.8e-08 \| 1.5e-13 \| 61.2 \| 52 \| (119, 170) \| 189 \| (3, 54) \| 58 \| Uncharacterized protein \| Uncharacterized protein; uniclust \| UniRef100\_UPI002271FB79 \| 98.0 \| 8.3e-08 \| 1.5e-13 \| 73.9 \| 138 \| (7, 155) \| 189 \| (1, 151) \| 188 \| hypothetical protein \| hypothetical protein; uniclust \| UniRef100\_UPI001F062539 \| 98.0 \| 1e-07 \| 1.9e-13 \| 65.4 \| 79 \| (2, 85) \| 189 \| (3, 82) \| 91 \| hypothetical protein \| hypothetical protein; uniclust \| UniRef100\_A0A0A1HZ01 \| 98.0 \| 1.2e-07 \| 2.2e-13 \| 64.9 \| 78 \| (103, 188) \| 189 \| (10, 88) \| 89 \| Uncharacterized protein \| Uncharacterized protein; uniclust \| UniRef100\_A0A431IBE3 \| 98.0 \| 1.2e-07 \| 2.2e-13 \| 76.6 \| 129 \| (26, 171) \| 189 \| (110, 245) \| 266 \| DUF1834 family protein \| DUF1834 family protein; uniclust \| UniRef100\_Q4EDG4 \| 98.0 \| 1.3e-07 \| 2.5e-13 \| 62.8 \| 50 \| (7, 59) \| 189 \| (1, 50) \| 69 \| Uncharacterized protein \| Uncharacterized protein; uniclust \| UniRef100\_A0A7X6JH33 \| 98.0 \| 1.7e-07 \| 3e-13 \| 60.3 \| 39 \| (148, 186) \| 189 \| (19, 57) \| 63 \| Uncharacterized protein \| Uncharacterized protein; uniclust \| UniRef100\_A0A0A2Y1K5 \| 97.9 \| 1.7e-07 \| 3.5e-13 \| 78.3 \| 141 \| (3, 151) \| 189 \| (20, 174) \| 244 \| Mu-like prophage protein gp37 \| Mu-like prophage protein gp37; uniclust \| UniRef100\_A0A059IWS3 \| 97.9 \| 2.3e-07 \| 4.3e-13 \| 61.1 \| 59 \| (8, 71) \| 189 \| (2, 60) \| 67 \| Uncharacterized protein \| Uncharacterized protein; uniclust \| UniRef100\_A0A193QHJ5 \| 97.8 \| 3.3e-07 \| 6.6e-13 \| 75.2 \| 139 \| (4, 151) \| 189 \| (1, 153) \| 219 \| Uncharacterized protein \| Uncharacterized protein; uniclust \| UniRef100\_UPI001931093B \| 97.8 \| 3.7e-07 \| 6.9e-13 \| 60.9 \| 69 \| (75, 152) \| 189 \| (1, 74) \| 76 \| hypothetical protein \| hypothetical protein; uniclust \| UniRef100\_UPI001FC8A895 \| 97.8 \| 3.9e-07 \| 7.2e-13 \| 60.4 \| 59 \| (127, 187) \| 189 \| (9, 67) \| 73 \| hypothetical protein \| hypothetical protein; uniclust \| UniRef100\_A0A8J3HUP9 \| 97.8 \| 4.5e-07 \| 8.3e-13 \| 57.9 \| 56 \| (83, 145) \| 189 \| (5, 60) \| 60 \| Uncharacterized protein \| Uncharacterized protein; uniclust \| UniRef100\_UPI0013D70C2A \| 97.8 \| 4.6e-07 \| 8.5e-13 \| 62.8 \| 89 \| (46, 148) \| 189 \| (2, 92) \| 94 \| hypothetical protein \| hypothetical protein; uniclust \| UniRef100\_A0A962YS06 \| 97.8 \| 5.2e-07 \| 9.5e-13 \| 67.6 \| 137 \| (6, 152) \| 189 \| (3, 147) \| 150 \| Uncharacterized protein \| Uncharacterized protein; uniclust \| UniRef100\_A0A0Q8AN87 \| 97.8 \| 5.4e-07 \| 1e-12 \| 70.9 \| 146 \| (6, 159) \| 189 \| (1, 160) \| 182 \| Suppressor of fused-like domain-containing protein \| Suppressor of fused-like domain-containing protein; uniclust \| UniRef100\_A0A962B4M7 \| 97.7 \| 7.8e-07 \| 1.4e-12 \| 69.5 \| 151 \| (2, 160) \| 189 \| (16, 172) \| 198 \| Uncharacterized protein \| Uncharacterized protein; uniclust \| UniRef100\_A0A969Q5I0 \| 97.7 \| 8.7e-07 \| 1.6e-12 \| 58.4 \| 44 \| (142, 188) \| 189 \| (24, 67) \| 70 \| Uncharacterized protein \| Uncharacterized protein; uniclust \| UniRef100\_A0A085NVH9 \| 97.7 \| 1.1e-06 \| 2e-12 \| 70.0 \| 126 \| (6, 148) \| 189 \| (1, 135) \| 190 \| DUF1834 family protein \| DUF1834 family protein; uniclust \| UniRef100\_A0A1X4N607 \| 97.7 \| 1.1e-06 \| 2.2e-12 \| 71.4 \| 163 \| (7, 182) \| 189 \| (1, 181) \| 198 \| Uncharacterized protein \| Uncharacterized protein; uniclust \| UniRef100\_A0A933SHU2 \| 97.6 \| 1.5e-06 \| 2.9e-12 \| 67.5 \| 135 \| (5, 153) \| 189 \| (1, 145) \| 148 \| DUF1834 family protein \| DUF1834 family protein; uniclust \| UniRef100\_A0A800BWL1 \| 97.6 \| 2.3e-06 \| 4.1e-12 \| 58.4 \| 75 \| (6, 87) \| 189 \| (1, 76) \| 83 \| Uncharacterized protein (Fragment) \| Uncharacterized protein (Fragment); uniclust \| UniRef100\_Z9JKS2 \| 97.6 \| 2.4e-06 \| 4.4e-12 \| 67.9 \| 76 \| (4, 81) \| 189 \| (5, 80) \| 217 \| Uncharacterized protein \| Uncharacterized protein; uniclust \| UniRef100\_A0A090KJ28 \| 97.5 \| 3.1e-06 \| 6.3e-12 \| 68.2 \| 137 \| (8, 157) \| 189 \| (15, 151) \| 180 \| Uncharacterized phage protein \| Uncharacterized phage protein; uniclust \| UniRef100\_UPI00082BE2D3 \| 97.5 \| 3.9e-06 \| 7.1e-12 \| 64.0 \| 137 \| (7, 157) \| 189 \| (5, 146) \| 162 \| hypothetical protein \| hypothetical protein; uniclust \| UniRef100\_UPI0004263E9E \| 97.5 \| 4.4e-06 \| 8e-12 \| 65.0 \| 138 \| (8, 157) \| 189 \| (4, 147) \| 185 \| hypothetical protein \| hypothetical protein; uniclust \| UniRef100\_UPI001C04E967 \| 97.4 \| 5.6e-06 \| 1e-11 \| 61.0 \| 98 \| (5, 109) \| 189 \| (3, 104) \| 129 \| hypothetical protein \| hypothetical protein; uniclust \| UniRef100\_A0A1G3LVL6 \| 97.4 \| 7.1e-06 \| 1.4e-11 \| 64.9 \| 130 \| (5, 149) \| 189 \| (1, 144) \| 152 \| Uncharacterized protein \| Uncharacterized protein; uniclust \| UniRef100\_UPI001A9E6345 \| 97.3 \| 1.2e-05 \| 2.1e-11 \| 55.4 \| 70 \| (5, 81) \| 189 \| (4, 73) \| 83 \| hypothetical protein \| hypothetical protein; uniclust \| UniRef100\_UPI001BAEFF64 \| 97.3 \| 1.2e-05 \| 2.2e-11 \| 67.1 \| 146 \| (7, 161) \| 189 \| (4, 184) \| 307 \| hypothetical protein \| hypothetical protein; uniclust \| UniRef100\_A0A853I2M1 \| 97.3 \| 1.2e-05 \| 2.2e-11 \| 64.0 \| 170 \| (7, 187) \| 189 \| (1, 177) \| 182 \| Uncharacterized protein \| Uncharacterized protein; uniclust \| UniRef100\_UPI001C98E712 \| 97.3 \| 1.2e-05 \| 2.3e-11 \| 57.6 \| 67 \| (83, 157) \| 189 \| (7, 75) \| 107 \| hypothetical protein \| hypothetical protein; uniclust \| UniRef100\_A0A0Q0ZM37 \| 97.3 \| 1.2e-05 \| 2.5e-11 \| 66.8 \| 148 \| (8, 173) \| 189 \| (23, 173) \| 206 \| Uncharacterized protein \| Uncharacterized protein; uniclust \| UniRef100\_A0A0F4VEU8 \| 97.2 \| 1.5e-05 \| 2.9e-11 \| 62.4 \| 137 \| (7, 154) \| 189 \| (1, 148) \| 148 \| Uncharacterized protein \| Uncharacterized protein; uniclust \| UniRef100\_A0A6L8GFP2 \| 97.2 \| 2.3e-05 \| 4.2e-11 \| 63.4 \| 148 \| (28, 187) \| 189 \| (40, 227) \| 234 \| Uncharacterized protein \| Uncharacterized protein; uniclust \| UniRef100\_A0A145VQF5 \| 97.2 \| 2.5e-05 \| 4.6e-11 \| 62.8 \| 109 \| (44, 163) \| 189 \| (23, 133) \| 212 \| Uncharacterized protein \| Uncharacterized protein; uniclust \| UniRef100\_A0A376K4X6 \| 97.1 \| 2.6e-05 \| 4.8e-11 \| 56.6 \| 86 \| (83, 177) \| 189 \| (5, 93) \| 113 \| Uncharacterized protein \| Uncharacterized protein; uniclust \| UniRef100\_A0A127MKR6 \| 97.1 \| 2.7e-05 \| 5.1e-11 \| 60.7 \| 135 \| (8, 151) \| 189 \| (5, 149) \| 155 \| DUF1834 domain-containing protein \| DUF1834 domain-containing protein; uniclust \| UniRef100\_A0A956AIV7 \| 97.1 \| 3e-05 \| 5.5e-11 \| 60.8 \| 137 \| (8, 155) \| 189 \| (2, 149) \| 186 \| Uncharacterized protein \| Uncharacterized protein; uniclust \| UniRef100\_A0A072T9J4 \| 97.1 \| 2.8e-05 \| 6.1e-11 \| 65.5 \| 144 \| (1, 156) \| 189 \| (26, 178) \| 198 \| DUF1834 family protein \| DUF1834 family protein; uniclust \| UniRef100\_A0A839HEF9 \| 97.1 \| 3.6e-05 \| 6.5e-11 \| 59.7 \| 147 \| (1, 155) \| 189 \| (1, 166) \| 171 \| Uncharacterized protein \| Uncharacterized protein; uniclust \| UniRef100\_A0A1E3G7H1 \| 97.1 \| 3.7e-05 \| 6.9e-11 \| 60.0 \| 169 \| (7, 187) \| 189 \| (3, 178) \| 178 \| Uncharacterized protein \| Uncharacterized protein; uniclust \| UniRef100\_A0A0K0VKS5 \| 97.0 \| 5.1e-05 \| 9.4e-11 \| 57.1 \| 76 \| (4, 85) \| 189 \| (3, 79) \| 139 \| Uncharacterized protein \| Uncharacterized protein; uniclust \| UniRef100\_UPI0010CB9643 \| 97.0 \| 5.1e-05 \| 9.4e-11 \| 49.4 \| 46 \| (143, 188) \| 189 \| (2, 48) \| 59 \| hypothetical protein \| hypothetical protein; uniclust \| UniRef100\_A0A8D9PAR5 \| 96.9 \| 7.3e-05 \| 1.3e-10 \| 56.4 \| 135 \| (4, 153) \| 189 \| (2, 136) \| 140 \| Uncharacterized protein \| Uncharacterized protein; uniclust \| UniRef100\_A0A066RQJ8 \| 96.9 \| 7.3e-05 \| 1.5e-10 \| 61.5 \| 166 \| (2, 188) \| 189 \| (4, 178) \| 192 \| Phage protein \| Phage protein; uniclust \| UniRef100\_UPI001F4243D2 \| 96.9 \| 9.5e-05 \| 1.7e-10 \| 54.6 \| 112 \| (7, 133) \| 189 \| (5, 117) \| 121 \| hypothetical protein \| hypothetical protein; uniclust \| UniRef100\_A0A1F4PEF5 \| 96.8 \| 0.00012 \| 2.5e-10 \| 58.8 \| 122 \| (3, 136) \| 189 \| (1, 135) \| 154 \| DUF1834 domain-containing protein \| DUF1834 domain-containing protein; uniclust \| UniRef100\_A0A0G0CA56 \| 96.8 \| 0.00013 \| 2.7e-10 \| 59.7 \| 125 \| (5, 148) \| 189 \| (1, 129) \| 167 \| Gp37 protein \| Gp37 protein; uniclust \| UniRef100\_A0A1A8TCK2 \| 96.7 \| 0.00017 \| 3.4e-10 \| 58.4 \| 125 \| (12, 151) \| 189 \| (5, 134) \| 176 \| Uncharacterized protein \| Uncharacterized protein; uniclust \| UniRef100\_A0A0X6ZPF0 \| 96.7 \| 0.00021 \| 4.5e-10 \| 61.0 \| 159 \| (4, 176) \| 189 \| (9, 181) \| 208 \| Uncharacterized protein \| Uncharacterized protein; uniclust \| UniRef100\_A0A8J3HUL9 \| 96.7 \| 0.00025 \| 4.7e-10 \| 43.1 \| 40 \| (25, 69) \| 189 \| (2, 41) \| 42 \| Uncharacterized protein \| Uncharacterized protein; uniclust \| UniRef100\_A0A5P9CRF5 \| 96.6 \| 0.00029 \| 5.5e-10 \| 56.1 \| 106 \| (41, 157) \| 189 \| (43, 148) \| 180 \| Uncharacterized protein \| Uncharacterized protein; uniclust \| UniRef100\_A0A0T7E1W9 \| 96.5 \| 0.00037 \| 7.1e-10 \| 55.9 \| 128 \| (15, 156) \| 189 \| (16, 144) \| 173 \| Uncharacterized protein \| Uncharacterized protein; uniclust \| UniRef100\_A0A8T7D0H9 \| 96.5 \| 0.00041 \| 7.5e-10 \| 63.6 \| 101 \| (7, 109) \| 189 \| (8, 113) \| 624 \| Uncharacterized protein \| Uncharacterized protein; uniclust \| UniRef100\_A0A522X2S5 \| 96.5 \| 0.00041 \| 7.8e-10 \| 57.5 \| 135 \| (7, 153) \| 189 \| (1, 159) \| 210 \| DUF1834 family protein \| DUF1834 family protein; uniclust \| UniRef100\_A0A833L7J3 \| 96.5 \| 0.00044 \| 8e-10 \| 55.3 \| 135 \| (6, 151) \| 189 \| (1, 145) \| 195 \| DUF1834 family protein \| DUF1834 family protein; uniclust \| UniRef100\_A0A3A5AI19 \| 96.5 \| 0.00041 \| 8.4e-10 \| 56.1 \| 109 \| (5, 124) \| 189 \| (1, 120) \| 155 \| DUF1834 family protein \| DUF1834 family protein; uniclust \| UniRef100\_A0A074LMC7 \| 96.5 \| 0.0004 \| 8.7e-10 \| 57.7 \| 98 \| (6, 108) \| 189 \| (1, 98) \| 172 \| Uncharacterized protein \| Uncharacterized protein; uniclust \| UniRef100\_A0A095ZCL4 \| 96.5 \| 0.00042 \| 9.2e-10 \| 57.7 \| 135 \| (5, 152) \| 189 \| (2, 149) \| 172 \| Uncharacterized protein \| Uncharacterized protein; uniclust \| UniRef100\_A0A134CJH4 \| 96.4 \| 0.0007 \| 1.4e-09 \| 54.4 \| 100 \| (2, 108) \| 189 \| (4, 103) \| 143 \| Uncharacterized protein \| Uncharacterized protein; uniclust \| UniRef100\_A0A0C5E1K8 \| 96.3 \| 0.00078 \| 1.5e-09 \| 55.8 \| 111 \| (8, 128) \| 189 \| (1, 128) \| 189 \| DUF1834 family protein \| DUF1834 family protein; uniclust \| UniRef100\_A0A1V5IG09 \| 96.3 \| 0.00088 \| 1.7e-09 \| 53.2 \| 136 \| (4, 154) \| 189 \| (1, 141) \| 146 \| Uncharacterized protein \| Uncharacterized protein; uniclust \| UniRef100\_A0A1B8Q520 \| 96.3 \| 0.00084 \| 1.7e-09 \| 55.1 \| 141 \| (1, 157) \| 189 \| (4, 146) \| 165 \| Gp37 protein \| Gp37 protein; uniclust \| UniRef100\_A0A5M8P6C2 \| 96.3 \| 0.00092 \| 1.8e-09 \| 55.4 \| 136 \| (5, 152) \| 189 \| (20, 161) \| 187 \| DUF3168 domain-containing protein \| DUF3168 domain-containing protein |
| Top keywords  (threshold 1.00e-03 (evalue)) | **hypothetical, Phage, DUF1834, Gp37, domain\_containing, Fragment, tail, Mu\_like, prophage, DUF3168** |
| Output files | ../../similar\_sequences/10\_FANPEZAQ\_CDS\_0010\_merged.svg ../../similar\_sequences/10\_FANPEZAQ\_CDS\_0010\_pdb70.a3m ../../similar\_sequences/10\_FANPEZAQ\_CDS\_0010\_pdb70.hhr ../../similar\_sequences/10\_FANPEZAQ\_CDS\_0010\_uniclust.a3m ../../similar\_sequences/10\_FANPEZAQ\_CDS\_0010\_uniclust.hhr |

#### Structure prediction (AlphaFold)^2^

|  |  |
| --- | --- |
| Stats | xml version="1.0" encoding="utf-8" standalone="no"?       2024-09-02T21:09:09.106565 image/svg+xml   Matplotlib v3.7.2, https://matplotlib.org/ |
| Predicted structure | **NGL Viewer Controls:**  - Center: *Left-Click* - Rotate: *Left-Click + Drag* - Translate: *Right-Click + Drag* - Zoom: *Shift + Left-Click + Drag* |
| Output files | ../../predicted\_structures/10\_FANPEZAQ\_CDS\_0010/features.pkl ../../predicted\_structures/10\_FANPEZAQ\_CDS\_0010/ranked\_0.pdb ../../predicted\_structures/10\_FANPEZAQ\_CDS\_0010/ranked\_0\_plots.svg ../../predicted\_structures/10\_FANPEZAQ\_CDS\_0010/result\_model\_1\_ptm\_pred\_0.pkl |

#### Structure similarity search results (Foldseek)^3^

|  |  |
| --- | --- |
| Structure databases searched | Pdb, Afdb-proteome, Afdb-uniprot50 |
| Results, scheme(s)  (Top layers only, threshold 1.00e-02 (evalue)) | xml version="1.0" encoding="utf-8" standalone="no"?       2024-09-02T21:10:18.111435 image/svg+xml   Matplotlib v3.7.2, https://matplotlib.org/ |
| Results, table  (threshold 1.00e-02 (evalue)) | db \| id \| prob \| evalue \| bits \| fident \| alnlen \| mismatch \| gapopen \| qstart \| qend \| tstart \| tend \| name \| description; pdb \| 6U5F\_A \| 1.0 \| 4.707e-09 \| 294 \| 0.279 \| 179 \| 110 \| 4 \| 6 \| 184 \| 2 \| 161 \| Tube PA0623 \| Tube PA0623; pdb \| 2GJV\_E \| 1.0 \| 0.001048 \| 134 \| 0.094 \| 158 \| 111 \| 10 \| 7 \| 156 \| 3 \| 136 \| putative cytoplasmic protein \| putative cytoplasmic protein; pdb \| 3FZB\_A \| 1.0 \| 0.001768 \| 112 \| 0.102 \| 156 \| 106 \| 9 \| 6 \| 153 \| 1 \| 130 \| Minor tail protein U \| Minor tail protein U; pdb \| 4ACV\_B \| 1.0 \| 0.00223 \| 103 \| 0.077 \| 154 \| 108 \| 10 \| 1 \| 152 \| 1 \| 122 \| PROPHAGE LAMBDALM01, ANTIGEN B \| PROPHAGE LAMBDALM01, ANTIGEN B; pdb \| 3FZ2\_B \| 1.0 \| 0.004224 \| 103 \| 0.109 \| 155 \| 105 \| 9 \| 6 \| 153 \| 1 \| 129 \| Minor tail protein U \| Minor tail protein U; pdb \| 3FZB\_F \| 1.0 \| 0.005028 \| 103 \| 0.111 \| 153 \| 108 \| 8 \| 6 \| 153 \| 1 \| 130 \| Minor tail protein U \| Minor tail protein U; pdb \| 3FZB\_G \| 1.0 \| 0.008002 \| 101 \| 0.091 \| 153 \| 112 \| 8 \| 6 \| 153 \| 2 \| 132 \| Minor tail protein U \| Minor tail protein U; pdb \| 3FZB\_B \| 0.999 \| 0.003348 \| 99 \| 0.108 \| 147 \| 105 \| 7 \| 13 \| 153 \| 4 \| 130 \| Minor tail protein U \| Minor tail protein U; pdb \| 2L25\_A \| 0.999 \| 0.001247 \| 99 \| 0.125 \| 168 \| 101 \| 13 \| 7 \| 156 \| 2 \| 141 \| Uncharacterized protein \| Uncharacterized protein; pdb \| 3FZB\_H \| 0.999 \| 0.004477 \| 98 \| 0.095 \| 147 \| 107 \| 7 \| 13 \| 153 \| 3 \| 129 \| Minor tail protein U \| Minor tail protein U; pdb \| 3FZ2\_F \| 0.999 \| 0.004477 \| 98 \| 0.115 \| 156 \| 106 \| 9 \| 6 \| 153 \| 2 \| 133 \| Minor tail protein U \| Minor tail protein U; pdb \| 3FZB\_I \| 0.996 \| 0.006722 \| 90 \| 0.108 \| 148 \| 102 \| 9 \| 13 \| 153 \| 3 \| 127 \| Minor tail protein U \| Minor tail protein U; pdb \| 8GTF\_K \| 0.978 \| 0.006343 \| 78 \| 0.168 \| 154 \| 90 \| 12 \| 18 \| 153 \| 4 \| 137 \| Terminator protein \| Terminator protein; afdb-proteome \| AF-G3XD82-F1-MODEL\_V4 \| 1.0 \| 8.093e-14 \| 465 \| 0.277 \| 180 \| 111 \| 4 \| 6 \| 185 \| 5 \| 165 \| Uncharacterized protein \| Uncharacterized protein; afdb-proteome \| AF-P44231-F1-MODEL\_V4 \| 1.0 \| 1.18e-06 \| 195 \| 0.144 \| 201 \| 142 \| 9 \| 1 \| 187 \| 1 \| 185 \| Mu-like prophage FluMu protein gp37 \| Mu-like prophage FluMu protein gp37; afdb-proteome \| AF-Q8ZN03-F1-MODEL\_V4 \| 1.0 \| 0.0007022 \| 140 \| 0.096 \| 145 \| 112 \| 7 \| 10 \| 152 \| 5 \| 132 \| Gifsy-1 prophage protein \| Gifsy-1 prophage protein; afdb-proteome \| AF-Q8ZKJ0-F1-MODEL\_V4 \| 1.0 \| 0.0005899 \| 133 \| 0.102 \| 166 \| 119 \| 9 \| 7 \| 165 \| 2 \| 144 \| Putative cytoplasmic protein \| Putative cytoplasmic protein; afdb-proteome \| AF-Q8ZQG6-F1-MODEL\_V4 \| 1.0 \| 0.00133 \| 129 \| 0.104 \| 144 \| 108 \| 8 \| 12 \| 152 \| 7 \| 132 \| Fels-1 prophage protein \| Fels-1 prophage protein; afdb-proteome \| AF-Q5F997-F1-MODEL\_V4 \| 1.0 \| 0.0005252 \| 119 \| 0.123 \| 154 \| 111 \| 11 \| 7 \| 150 \| 1 \| 140 \| Phage associated protein \| Phage associated protein; afdb-proteome \| AF-Q8ZQ90-F1-MODEL\_V4 \| 1.0 \| 0.006022 \| 112 \| 0.124 \| 145 \| 106 \| 8 \| 12 \| 153 \| 7 \| 133 \| Gifsy-2 prophage probable minor tail protein \| Gifsy-2 prophage probable minor tail protein; afdb-proteome \| AF-Q9I547-F1-MODEL\_V4 \| 0.442 \| 0.007597 \| 49 \| 0.152 \| 190 \| 93 \| 13 \| 13 \| 166 \| 10 \| 167 \| Uncharacterized protein \| Uncharacterized protein; afdb-uniprot50 \| AF-A0A6M4YGW3-F1-MODEL\_V4 \| 1.0 \| 2.154e-23 \| 889 \| 0.56 \| 189 \| 69 \| 3 \| 1 \| 178 \| 1 \| 186 \| Uncharacterized protein \| Uncharacterized protein; afdb-uniprot50 \| AF-A0A124PC40-F1-MODEL\_V4 \| 1.0 \| 3.371e-21 \| 784 \| 0.541 \| 181 \| 71 \| 5 \| 12 \| 189 \| 2 \| 173 \| Uncharacterized protein \| Uncharacterized protein; afdb-uniprot50 \| AF-A0A430HF56-F1-MODEL\_V4 \| 1.0 \| 1.141e-20 \| 780 \| 0.492 \| 189 \| 85 \| 4 \| 1 \| 189 \| 1 \| 178 \| Uncharacterized protein \| Uncharacterized protein; afdb-uniprot50 \| AF-A0A410UF72-F1-MODEL\_V4 \| 1.0 \| 1.1e-19 \| 762 \| 0.377 \| 188 \| 107 \| 3 \| 2 \| 189 \| 3 \| 180 \| Uncharacterized protein \| Uncharacterized protein; afdb-uniprot50 \| AF-A0A291LZL3-F1-MODEL\_V4 \| 1.0 \| 6.519e-20 \| 743 \| 0.478 \| 190 \| 86 \| 4 \| 1 \| 189 \| 1 \| 178 \| Uncharacterized protein \| Uncharacterized protein; afdb-uniprot50 \| AF-A0A315BPV6-F1-MODEL\_V4 \| 1.0 \| 6.655e-19 \| 702 \| 0.508 \| 187 \| 79 \| 4 \| 2 \| 187 \| 3 \| 177 \| Uncharacterized protein \| Uncharacterized protein; afdb-uniprot50 \| AF-A0A0B4BHC6-F1-MODEL\_V4 \| 1.0 \| 2.683e-18 \| 697 \| 0.394 \| 185 \| 100 \| 3 \| 5 \| 189 \| 3 \| 175 \| Uncharacterized protein \| Uncharacterized protein; afdb-uniprot50 \| AF-A0A143DEE7-F1-MODEL\_V4 \| 1.0 \| 1.59e-18 \| 688 \| 0.468 \| 188 \| 88 \| 4 \| 1 \| 188 \| 1 \| 176 \| Uncharacterized protein \| Uncharacterized protein; afdb-uniprot50 \| AF-A0A844HSY5-F1-MODEL\_V4 \| 1.0 \| 3.076e-17 \| 662 \| 0.404 \| 188 \| 98 \| 4 \| 2 \| 189 \| 3 \| 176 \| Uncharacterized protein \| Uncharacterized protein; afdb-uniprot50 \| AF-A0A2N1AP90-F1-MODEL\_V4 \| 1.0 \| 7.631e-18 \| 661 \| 0.423 \| 189 \| 97 \| 3 \| 1 \| 189 \| 1 \| 177 \| Uncharacterized protein \| Uncharacterized protein; afdb-uniprot50 \| AF-A0A7V8FKG6-F1-MODEL\_V4 \| 1.0 \| 2.902e-17 \| 660 \| 0.38 \| 189 \| 105 \| 4 \| 1 \| 188 \| 1 \| 178 \| Uncharacterized protein \| Uncharacterized protein; afdb-uniprot50 \| AF-A0A7J0BX95-F1-MODEL\_V4 \| 1.0 \| 2.301e-17 \| 653 \| 0.349 \| 189 \| 108 \| 6 \| 1 \| 189 \| 1 \| 174 \| Uncharacterized protein \| Uncharacterized protein; afdb-uniprot50 \| AF-A0A1H9YBP8-F1-MODEL\_V4 \| 1.0 \| 4.895e-17 \| 605 \| 0.356 \| 188 \| 108 \| 5 \| 1 \| 187 \| 1 \| 176 \| Uncharacterized protein \| Uncharacterized protein; afdb-uniprot50 \| AF-A0A238J9A3-F1-MODEL\_V4 \| 1.0 \| 8.428e-16 \| 600 \| 0.358 \| 184 \| 106 \| 3 \| 6 \| 189 \| 4 \| 175 \| Uncharacterized protein \| Uncharacterized protein; afdb-uniprot50 \| AF-A0A846VUR4-F1-MODEL\_V4 \| 1.0 \| 2.963e-16 \| 599 \| 0.354 \| 186 \| 106 \| 5 \| 4 \| 188 \| 2 \| 174 \| Uncharacterized protein \| Uncharacterized protein; afdb-uniprot50 \| AF-A0A1X7L1R0-F1-MODEL\_V4 \| 1.0 \| 4.449e-16 \| 597 \| 0.302 \| 182 \| 113 \| 4 \| 6 \| 187 \| 3 \| 170 \| Uncharacterized protein \| Uncharacterized protein; afdb-uniprot50 \| AF-A0A158E8C2-F1-MODEL\_V4 \| 1.0 \| 2.091e-16 \| 594 \| 0.357 \| 190 \| 110 \| 7 \| 2 \| 187 \| 1 \| 182 \| Uncharacterized protein \| Uncharacterized protein; afdb-uniprot50 \| AF-K5Z1V0-F1-MODEL\_V4 \| 1.0 \| 6.545e-17 \| 585 \| 0.321 \| 190 \| 113 \| 4 \| 2 \| 189 \| 3 \| 178 \| Uncharacterized protein \| Uncharacterized protein; afdb-uniprot50 \| AF-A0A7W4WZS5-F1-MODEL\_V4 \| 1.0 \| 1.314e-16 \| 583 \| 0.338 \| 201 \| 110 \| 8 \| 2 \| 189 \| 1 \| 191 \| Uncharacterized protein \| Uncharacterized protein; afdb-uniprot50 \| AF-A0A149SVH3-F1-MODEL\_V4 \| 1.0 \| 1.597e-15 \| 577 \| 0.366 \| 191 \| 104 \| 6 \| 2 \| 188 \| 4 \| 181 \| Uncharacterized protein \| Uncharacterized protein; afdb-uniprot50 \| AF-A0A1X7CH01-F1-MODEL\_V4 \| 1.0 \| 1.597e-15 \| 576 \| 0.331 \| 187 \| 110 \| 6 \| 1 \| 187 \| 1 \| 172 \| Uncharacterized protein \| Uncharacterized protein; afdb-uniprot50 \| AF-A0A3A8EIU2-F1-MODEL\_V4 \| 1.0 \| 4.449e-16 \| 569 \| 0.404 \| 193 \| 102 \| 8 \| 2 \| 188 \| 1 \| 186 \| Uncharacterized protein \| Uncharacterized protein; afdb-uniprot50 \| AF-Q9PFF2-F1-MODEL\_V4 \| 1.0 \| 1.292e-14 \| 546 \| 0.4 \| 145 \| 79 \| 2 \| 45 \| 189 \| 1 \| 137 \| Uncharacterized protein \| Uncharacterized protein; afdb-uniprot50 \| AF-A0A7H8NU11-F1-MODEL\_V4 \| 1.0 \| 7.504e-16 \| 541 \| 0.357 \| 196 \| 107 \| 9 \| 1 \| 189 \| 1 \| 184 \| Uncharacterized protein \| Uncharacterized protein; afdb-uniprot50 \| AF-A0A2E3M0I4-F1-MODEL\_V4 \| 1.0 \| 2.693e-15 \| 532 \| 0.342 \| 181 \| 105 \| 4 \| 9 \| 189 \| 4 \| 170 \| Uncharacterized protein \| Uncharacterized protein; afdb-uniprot50 \| AF-A0A7W2B841-F1-MODEL\_V4 \| 1.0 \| 3.206e-15 \| 528 \| 0.372 \| 188 \| 99 \| 5 \| 1 \| 188 \| 1 \| 169 \| Uncharacterized protein \| Uncharacterized protein; afdb-uniprot50 \| AF-A0A4T2ACD2-F1-MODEL\_V4 \| 1.0 \| 5.73e-15 \| 524 \| 0.317 \| 195 \| 114 \| 8 \| 2 \| 189 \| 1 \| 183 \| Uncharacterized protein \| Uncharacterized protein; afdb-uniprot50 \| AF-A0A2Z6AZ36-F1-MODEL\_V4 \| 1.0 \| 1.369e-14 \| 518 \| 0.281 \| 181 \| 116 \| 3 \| 8 \| 188 \| 1 \| 167 \| Uncharacterized protein \| Uncharacterized protein; afdb-uniprot50 \| AF-A0A4U1KS61-F1-MODEL\_V4 \| 1.0 \| 3.468e-14 \| 505 \| 0.338 \| 189 \| 111 \| 4 \| 1 \| 189 \| 1 \| 175 \| Uncharacterized protein \| Uncharacterized protein; afdb-uniprot50 \| AF-A0A081MYL3-F1-MODEL\_V4 \| 1.0 \| 7.821e-14 \| 495 \| 0.278 \| 190 \| 124 \| 6 \| 1 \| 189 \| 1 \| 178 \| Uncharacterized protein \| Uncharacterized protein; afdb-uniprot50 \| AF-A0A446AZW9-F1-MODEL\_V4 \| 1.0 \| 2.807e-13 \| 492 \| 0.35 \| 180 \| 103 \| 5 \| 8 \| 187 \| 3 \| 168 \| Phage protein \| Phage protein; afdb-uniprot50 \| AF-A0A1H6X2H3-F1-MODEL\_V4 \| 1.0 \| 4.734e-13 \| 475 \| 0.304 \| 184 \| 112 \| 5 \| 5 \| 187 \| 2 \| 170 \| Uncharacterized protein \| Uncharacterized protein; afdb-uniprot50 \| AF-A0A3M3CQE2-F1-MODEL\_V4 \| 1.0 \| 5.317e-13 \| 473 \| 0.288 \| 184 \| 117 \| 4 \| 2 \| 185 \| 1 \| 170 \| Uncharacterized protein \| Uncharacterized protein; afdb-uniprot50 \| AF-A0A450ZAR3-F1-MODEL\_V4 \| 1.0 \| 4.215e-13 \| 473 \| 0.245 \| 187 \| 125 \| 7 \| 7 \| 189 \| 2 \| 176 \| Uncharacterized protein \| Uncharacterized protein; afdb-uniprot50 \| AF-A0A2S6H5L1-F1-MODEL\_V4 \| 1.0 \| 2.099e-13 \| 472 \| 0.312 \| 189 \| 117 \| 5 \| 5 \| 189 \| 2 \| 181 \| Uncharacterized protein \| Uncharacterized protein; afdb-uniprot50 \| AF-S2EUJ0-F1-MODEL\_V4 \| 1.0 \| 8.461e-13 \| 470 \| 0.312 \| 176 \| 108 \| 5 \| 12 \| 187 \| 2 \| 164 \| Uncharacterized protein \| Uncharacterized protein; afdb-uniprot50 \| AF-A6UZB9-F1-MODEL\_V4 \| 1.0 \| 2.099e-13 \| 468 \| 0.263 \| 186 \| 118 \| 4 \| 2 \| 187 \| 1 \| 167 \| Uncharacterized protein \| Uncharacterized protein; afdb-uniprot50 \| AF-A0A061JGS2-F1-MODEL\_V4 \| 1.0 \| 1.57e-13 \| 464 \| 0.378 \| 161 \| 86 \| 4 \| 27 \| 187 \| 3 \| 149 \| Uncharacterized protein \| Uncharacterized protein; afdb-uniprot50 \| AF-A0A0Q8D856-F1-MODEL\_V4 \| 1.0 \| 9.504e-13 \| 464 \| 0.301 \| 186 \| 116 \| 4 \| 2 \| 187 \| 1 \| 172 \| Uncharacterized protein \| Uncharacterized protein; afdb-uniprot50 \| AF-A0A7S6S605-F1-MODEL\_V4 \| 1.0 \| 1.869e-13 \| 463 \| 0.296 \| 182 \| 114 \| 4 \| 8 \| 189 \| 4 \| 171 \| Uncharacterized protein \| Uncharacterized protein; afdb-uniprot50 \| AF-A0A7V7NR68-F1-MODEL\_V4 \| 1.0 \| 7.533e-13 \| 463 \| 0.273 \| 183 \| 120 \| 4 \| 7 \| 189 \| 5 \| 174 \| Uncharacterized protein \| Uncharacterized protein; afdb-uniprot50 \| AF-S2KFP5-F1-MODEL\_V4 \| 1.0 \| 6.707e-13 \| 463 \| 0.327 \| 186 \| 112 \| 4 \| 2 \| 187 \| 1 \| 173 \| Uncharacterized protein \| Uncharacterized protein; afdb-uniprot50 \| AF-A0A7W5B8Q7-F1-MODEL\_V4 \| 1.0 \| 5.635e-13 \| 461 \| 0.263 \| 186 \| 123 \| 4 \| 4 \| 189 \| 2 \| 173 \| Uncharacterized protein \| Uncharacterized protein; afdb-uniprot50 \| AF-Q73HC3-F1-MODEL\_V4 \| 1.0 \| 7.108e-13 \| 460 \| 0.325 \| 172 \| 102 \| 5 \| 7 \| 178 \| 1 \| 158 \| Uncharacterized protein \| Uncharacterized protein; afdb-uniprot50 \| AF-A0A7C8M1D4-F1-MODEL\_V4 \| 1.0 \| 2.974e-13 \| 456 \| 0.319 \| 191 \| 114 \| 5 \| 2 \| 188 \| 5 \| 183 \| Uncharacterized protein \| Uncharacterized protein; afdb-uniprot50 \| AF-A0A1Y3KWD3-F1-MODEL\_V4 \| 1.0 \| 1.271e-12 \| 449 \| 0.289 \| 183 \| 109 \| 5 \| 9 \| 187 \| 4 \| 169 \| Uncharacterized protein \| Uncharacterized protein; afdb-uniprot50 \| AF-A0A142BA55-F1-MODEL\_V4 \| 1.0 \| 1.271e-12 \| 445 \| 0.268 \| 186 \| 119 \| 9 \| 7 \| 188 \| 3 \| 175 \| Uncharacterized protein \| Uncharacterized protein; afdb-uniprot50 \| AF-H8L2J7-F1-MODEL\_V4 \| 1.0 \| 2.807e-13 \| 441 \| 0.297 \| 195 \| 115 \| 6 \| 1 \| 187 \| 1 \| 181 \| Uncharacterized protein \| Uncharacterized protein; afdb-uniprot50 \| AF-A0A5E4NQZ0-F1-MODEL\_V4 \| 1.0 \| 1.427e-12 \| 438 \| 0.337 \| 166 \| 96 \| 4 \| 9 \| 174 \| 281 \| 432 \| Bacteriophage lambda, GpZ, minor tail,Ankyrin repeat-containing domain,Ankyrin repeat \| Bacteriophage lambda, GpZ, minor tail,Ankyrin repeat-containing domain,Ankyrin repeat; afdb-uniprot50 \| AF-A0A7U3R1S8-F1-MODEL\_V4 \| 1.0 \| 1.603e-12 \| 436 \| 0.313 \| 182 \| 109 \| 5 \| 8 \| 189 \| 4 \| 169 \| Uncharacterized protein \| Uncharacterized protein; afdb-uniprot50 \| AF-A0A258LKP1-F1-MODEL\_V4 \| 1.0 \| 6.847e-12 \| 435 \| 0.274 \| 182 \| 117 \| 7 \| 7 \| 188 \| 2 \| 168 \| Uncharacterized protein \| Uncharacterized protein; afdb-uniprot50 \| AF-A0A240TSR2-F1-MODEL\_V4 \| 1.0 \| 1.427e-12 \| 431 \| 0.318 \| 182 \| 108 \| 5 \| 8 \| 189 \| 4 \| 169 \| Uncharacterized protein \| Uncharacterized protein; afdb-uniprot50 \| AF-A0A1C3HH22-F1-MODEL\_V4 \| 1.0 \| 3.037e-12 \| 430 \| 0.234 \| 188 \| 122 \| 6 \| 6 \| 188 \| 4 \| 174 \| Uncharacterized protein \| Uncharacterized protein; afdb-uniprot50 \| AF-A0A379YFI5-F1-MODEL\_V4 \| 1.0 \| 9.155e-12 \| 423 \| 0.229 \| 192 \| 127 \| 11 \| 1 \| 188 \| 1 \| 175 \| Uncharacterized protein \| Uncharacterized protein; afdb-uniprot50 \| AF-A0A0D5Y351-F1-MODEL\_V4 \| 1.0 \| 1.375e-11 \| 417 \| 0.28 \| 171 \| 110 \| 5 \| 17 \| 187 \| 2 \| 159 \| Uncharacterized protein \| Uncharacterized protein; afdb-uniprot50 \| AF-A0A059PCW0-F1-MODEL\_V4 \| 1.0 \| 1.457e-11 \| 416 \| 0.333 \| 159 \| 93 \| 4 \| 7 \| 165 \| 1 \| 146 \| Uncharacterized protein \| Uncharacterized protein; afdb-uniprot50 \| AF-A0A1X7MHZ6-F1-MODEL\_V4 \| 1.0 \| 3.411e-12 \| 415 \| 0.274 \| 182 \| 110 \| 6 \| 8 \| 189 \| 3 \| 162 \| Uncharacterized protein \| Uncharacterized protein; afdb-uniprot50 \| AF-A0A3N2E1B1-F1-MODEL\_V4 \| 1.0 \| 3.831e-12 \| 414 \| 0.209 \| 186 \| 132 \| 7 \| 4 \| 187 \| 2 \| 174 \| Uncharacterized protein \| Uncharacterized protein; afdb-uniprot50 \| AF-A0A7Y3Z3N8-F1-MODEL\_V4 \| 1.0 \| 3.615e-12 \| 414 \| 0.216 \| 189 \| 130 \| 6 \| 6 \| 189 \| 4 \| 179 \| Uncharacterized protein \| Uncharacterized protein; afdb-uniprot50 \| AF-A0A089PYZ4-F1-MODEL\_V4 \| 1.0 \| 6.096e-12 \| 414 \| 0.209 \| 186 \| 131 \| 5 \| 6 \| 188 \| 4 \| 176 \| Uncharacterized protein \| Uncharacterized protein; afdb-uniprot50 \| AF-A0A3G6ZVQ7-F1-MODEL\_V4 \| 1.0 \| 6.461e-12 \| 414 \| 0.304 \| 187 \| 117 \| 4 \| 1 \| 187 \| 42 \| 215 \| Uncharacterized protein \| Uncharacterized protein; afdb-uniprot50 \| AF-A0A7Z0MZG6-F1-MODEL\_V4 \| 1.0 \| 1.603e-12 \| 413 \| 0.287 \| 188 \| 109 \| 7 \| 7 \| 188 \| 2 \| 170 \| Uncharacterized protein \| Uncharacterized protein; afdb-uniprot50 \| AF-A0A6L9FI07-F1-MODEL\_V4 \| 1.0 \| 1.09e-11 \| 411 \| 0.229 \| 183 \| 128 \| 4 \| 6 \| 188 \| 3 \| 172 \| Uncharacterized protein \| Uncharacterized protein; afdb-uniprot50 \| AF-A0A509JCF5-F1-MODEL\_V4 \| 1.0 \| 9.702e-12 \| 410 \| 0.254 \| 185 \| 124 \| 4 \| 1 \| 185 \| 1 \| 171 \| Uncharacterized protein \| Uncharacterized protein; afdb-uniprot50 \| AF-A0A1A9VKF9-F1-MODEL\_V4 \| 1.0 \| 3.285e-11 \| 410 \| 0.3 \| 160 \| 99 \| 4 \| 1 \| 160 \| 44 \| 190 \| Phage\_base\_V domain-containing protein \| Phage\_base\_V domain-containing protein; afdb-uniprot50 \| AF-A0A1H1EGK5-F1-MODEL\_V4 \| 1.0 \| 5.228e-11 \| 404 \| 0.236 \| 186 \| 124 \| 5 \| 2 \| 187 \| 1 \| 168 \| Uncharacterized protein \| Uncharacterized protein; afdb-uniprot50 \| AF-A0A450W6V5-F1-MODEL\_V4 \| 1.0 \| 1.838e-11 \| 403 \| 0.245 \| 187 \| 126 \| 7 \| 6 \| 189 \| 2 \| 176 \| Uncharacterized protein \| Uncharacterized protein; afdb-uniprot50 \| AF-A0A7K0GNI0-F1-MODEL\_V4 \| 1.0 \| 5.121e-12 \| 401 \| 0.252 \| 182 \| 124 \| 6 \| 7 \| 187 \| 4 \| 174 \| Uncharacterized protein \| Uncharacterized protein; afdb-uniprot50 \| AF-A0A4V2PRI1-F1-MODEL\_V4 \| 1.0 \| 1.224e-11 \| 395 \| 0.209 \| 191 \| 129 \| 8 \| 1 \| 189 \| 4 \| 174 \| Uncharacterized protein \| Uncharacterized protein; afdb-uniprot50 \| AF-A0A2S9B8M0-F1-MODEL\_V4 \| 1.0 \| 2.064e-11 \| 394 \| 0.281 \| 185 \| 108 \| 6 \| 2 \| 177 \| 1 \| 169 \| Uncharacterized protein \| Uncharacterized protein; afdb-uniprot50 \| AF-B8GS13-F1-MODEL\_V4 \| 1.0 \| 3.911e-11 \| 393 \| 0.267 \| 191 \| 120 \| 9 \| 7 \| 189 \| 1 \| 179 \| Uncharacterized protein \| Uncharacterized protein; afdb-uniprot50 \| AF-A0A3M5CXK4-F1-MODEL\_V4 \| 1.0 \| 3.482e-11 \| 390 \| 0.322 \| 152 \| 92 \| 3 \| 36 \| 187 \| 1 \| 141 \| Uncharacterized protein \| Uncharacterized protein; afdb-uniprot50 \| AF-A0A3M6E3B6-F1-MODEL\_V4 \| 1.0 \| 2.509e-10 \| 390 \| 0.313 \| 166 \| 98 \| 5 \| 14 \| 178 \| 2 \| 152 \| Uncharacterized protein \| Uncharacterized protein; afdb-uniprot50 \| AF-A0A1I1UB35-F1-MODEL\_V4 \| 1.0 \| 1.324e-10 \| 388 \| 0.293 \| 177 \| 110 \| 5 \| 7 \| 183 \| 2 \| 163 \| Uncharacterized protein \| Uncharacterized protein; afdb-uniprot50 \| AF-A0A826WKL5-F1-MODEL\_V4 \| 1.0 \| 2.925e-11 \| 388 \| 0.236 \| 186 \| 123 \| 7 \| 8 \| 188 \| 1 \| 172 \| Uncharacterized protein \| Uncharacterized protein; afdb-uniprot50 \| AF-A0A2N0CX69-F1-MODEL\_V4 \| 1.0 \| 5.872e-11 \| 387 \| 0.234 \| 188 \| 125 \| 7 \| 6 \| 189 \| 3 \| 175 \| Uncharacterized protein \| Uncharacterized protein; afdb-uniprot50 \| AF-A0A0H3ZM50-F1-MODEL\_V4 \| 1.0 \| 2.188e-11 \| 384 \| 0.286 \| 157 \| 102 \| 3 \| 33 \| 189 \| 1 \| 147 \| Uncharacterized protein \| Uncharacterized protein; afdb-uniprot50 \| AF-A0A6S5X6Y7-F1-MODEL\_V4 \| 1.0 \| 7.408e-11 \| 384 \| 0.279 \| 172 \| 112 \| 5 \| 1 \| 172 \| 1 \| 160 \| Uncharacterized protein \| Uncharacterized protein; afdb-uniprot50 \| AF-A0A6H2NT10-F1-MODEL\_V4 \| 1.0 \| 1.324e-10 \| 382 \| 0.368 \| 144 \| 78 \| 4 \| 30 \| 173 \| 2 \| 132 \| Uncharacterized protein \| Uncharacterized protein; afdb-uniprot50 \| AF-A0A212KN54-F1-MODEL\_V4 \| 1.0 \| 6.461e-12 \| 381 \| 0.221 \| 194 \| 123 \| 8 \| 7 \| 187 \| 2 \| 180 \| Uncharacterized protein \| Uncharacterized protein; afdb-uniprot50 \| AF-A0A7J6YLT3-F1-MODEL\_V4 \| 1.0 \| 1.576e-10 \| 375 \| 0.34 \| 147 \| 84 \| 4 \| 20 \| 166 \| 2 \| 135 \| Uncharacterized protein \| Uncharacterized protein; afdb-uniprot50 \| AF-A0A3G2IKE0-F1-MODEL\_V4 \| 1.0 \| 2.064e-11 \| 365 \| 0.189 \| 201 \| 133 \| 11 \| 1 \| 188 \| 1 \| 184 \| Uncharacterized protein \| Uncharacterized protein; afdb-uniprot50 \| AF-A0A059KU41-F1-MODEL\_V4 \| 1.0 \| 5.657e-10 \| 364 \| 0.226 \| 181 \| 122 \| 8 \| 8 \| 187 \| 3 \| 166 \| Uncharacterized protein \| Uncharacterized protein; afdb-uniprot50 \| AF-A0A1V2DSI8-F1-MODEL\_V4 \| 1.0 \| 5.541e-11 \| 363 \| 0.305 \| 170 \| 102 \| 7 \| 9 \| 178 \| 4 \| 157 \| Uncharacterized protein \| Uncharacterized protein; afdb-uniprot50 \| AF-A0A1E3G659-F1-MODEL\_V4 \| 1.0 \| 1.179e-10 \| 361 \| 0.187 \| 181 \| 127 \| 7 \| 8 \| 186 \| 1 \| 163 \| Uncharacterized protein \| Uncharacterized protein; afdb-uniprot50 \| AF-A0A5E7KJV8-F1-MODEL\_V4 \| 1.0 \| 1.989e-10 \| 359 \| 0.267 \| 198 \| 117 \| 6 \| 1 \| 189 \| 4 \| 182 \| Uncharacterized protein \| Uncharacterized protein; afdb-uniprot50 \| AF-Q31HU1-F1-MODEL\_V4 \| 1.0 \| 2.818e-10 \| 357 \| 0.232 \| 172 \| 117 \| 6 \| 8 \| 178 \| 3 \| 160 \| Uncharacterized protein \| Uncharacterized protein; afdb-uniprot50 \| AF-A0A1E3LHB7-F1-MODEL\_V4 \| 1.0 \| 3.767e-10 \| 352 \| 0.319 \| 166 \| 95 \| 5 \| 8 \| 173 \| 4 \| 151 \| Uncharacterized protein \| Uncharacterized protein; afdb-uniprot50 \| AF-A0A7X3ZHV1-F1-MODEL\_V4 \| 1.0 \| 2.76e-11 \| 350 \| 0.23 \| 226 \| 123 \| 9 \| 1 \| 188 \| 5 \| 217 \| Uncharacterized protein \| Uncharacterized protein; afdb-uniprot50 \| AF-A0A4P9VGK5-F1-MODEL\_V4 \| 1.0 \| 3.767e-10 \| 345 \| 0.291 \| 175 \| 107 \| 7 \| 2 \| 170 \| 1 \| 164 \| Uncharacterized protein \| Uncharacterized protein; afdb-uniprot50 \| AF-A0A266LMM3-F1-MODEL\_V4 \| 1.0 \| 2.561e-09 \| 342 \| 0.277 \| 166 \| 107 \| 5 \| 12 \| 177 \| 2 \| 154 \| Uncharacterized protein \| Uncharacterized protein; afdb-uniprot50 \| AF-A0A6I2UNP3-F1-MODEL\_V4 \| 1.0 \| 5.995e-10 \| 340 \| 0.197 \| 197 \| 138 \| 11 \| 2 \| 189 \| 1 \| 186 \| Uncharacterized protein \| Uncharacterized protein; afdb-uniprot50 \| AF-A0A1S8YKZ2-F1-MODEL\_V4 \| 1.0 \| 1.249e-10 \| 339 \| 0.207 \| 183 \| 129 \| 6 \| 9 \| 188 \| 7 \| 176 \| Uncharacterized protein \| Uncharacterized protein; afdb-uniprot50 \| AF-A0A1W0CDL2-F1-MODEL\_V4 \| 1.0 \| 4.484e-10 \| 337 \| 0.279 \| 172 \| 103 \| 8 \| 8 \| 179 \| 1 \| 151 \| Uncharacterized protein \| Uncharacterized protein; afdb-uniprot50 \| AF-A0A1T2CHW9-F1-MODEL\_V4 \| 1.0 \| 2.509e-10 \| 337 \| 0.227 \| 185 \| 118 \| 9 \| 2 \| 181 \| 1 \| 165 \| Uncharacterized protein \| Uncharacterized protein; afdb-uniprot50 \| AF-A0A377QAJ9-F1-MODEL\_V4 \| 1.0 \| 5.657e-10 \| 336 \| 0.258 \| 178 \| 103 \| 7 \| 7 \| 171 \| 1 \| 162 \| Uncharacterized protein \| Uncharacterized protein; afdb-uniprot50 \| AF-A0A1T4W4R6-F1-MODEL\_V4 \| 1.0 \| 6.354e-10 \| 335 \| 0.218 \| 197 \| 131 \| 11 \| 1 \| 189 \| 1 \| 182 \| Uncharacterized protein \| Uncharacterized protein; afdb-uniprot50 \| AF-A0A654AL97-F1-MODEL\_V4 \| 1.0 \| 5.036e-10 \| 334 \| 0.248 \| 177 \| 113 \| 6 \| 1 \| 168 \| 1 \| 166 \| Uncharacterized protein \| Uncharacterized protein; afdb-uniprot50 \| AF-F3LHZ3-F1-MODEL\_V4 \| 1.0 \| 3.554e-10 \| 331 \| 0.218 \| 192 \| 123 \| 10 \| 7 \| 188 \| 1 \| 175 \| Uncharacterized protein \| Uncharacterized protein; afdb-uniprot50 \| AF-A0A090SU28-F1-MODEL\_V4 \| 1.0 \| 2.151e-09 \| 325 \| 0.235 \| 191 \| 129 \| 5 \| 7 \| 188 \| 1 \| 183 \| Uncharacterized protein \| Uncharacterized protein; afdb-uniprot50 \| AF-A0A348HHI8-F1-MODEL\_V4 \| 1.0 \| 2.03e-09 \| 324 \| 0.216 \| 189 \| 134 \| 9 \| 1 \| 188 \| 1 \| 176 \| Aspartatecarbamoyltransferase, catalytic chain \| Aspartatecarbamoyltransferase, catalytic chain; afdb-uniprot50 \| AF-A0A837E8U7-F1-MODEL\_V4 \| 1.0 \| 5.995e-10 \| 323 \| 0.197 \| 187 \| 129 \| 8 \| 9 \| 188 \| 8 \| 180 \| Uncharacterized protein \| Uncharacterized protein; afdb-uniprot50 \| AF-A0A1G5ACN2-F1-MODEL\_V4 \| 1.0 \| 2.03e-09 \| 319 \| 0.235 \| 195 \| 119 \| 9 \| 7 \| 189 \| 1 \| 177 \| Uncharacterized protein \| Uncharacterized protein; afdb-uniprot50 \| AF-A0A0Q2U909-F1-MODEL\_V4 \| 1.0 \| 2.416e-09 \| 316 \| 0.209 \| 196 \| 129 \| 12 \| 1 \| 188 \| 1 \| 178 \| Uncharacterized protein \| Uncharacterized protein; afdb-uniprot50 \| AF-A0A853IEN6-F1-MODEL\_V4 \| 1.0 \| 3.231e-09 \| 314 \| 0.21 \| 190 \| 131 \| 9 \| 7 \| 189 \| 1 \| 178 \| Uncharacterized protein \| Uncharacterized protein; afdb-uniprot50 \| AF-A0A1M6B7W3-F1-MODEL\_V4 \| 1.0 \| 4.076e-09 \| 314 \| 0.194 \| 185 \| 135 \| 6 \| 5 \| 188 \| 2 \| 173 \| Uncharacterized protein \| Uncharacterized protein; afdb-uniprot50 \| AF-A0A145VQF5-F1-MODEL\_V4 \| 1.0 \| 5.895e-08 \| 313 \| 0.262 \| 118 \| 78 \| 2 \| 47 \| 164 \| 26 \| 134 \| Uncharacterized protein \| Uncharacterized protein; afdb-uniprot50 \| AF-A0A7Z7VMF7-F1-MODEL\_V4 \| 1.0 \| 2.876e-09 \| 308 \| 0.226 \| 199 \| 121 \| 11 \| 5 \| 178 \| 2 \| 192 \| Uncharacterized protein \| Uncharacterized protein; afdb-uniprot50 \| AF-A0A1I5W3T1-F1-MODEL\_V4 \| 1.0 \| 2.876e-09 \| 298 \| 0.203 \| 187 \| 129 \| 11 \| 9 \| 188 \| 7 \| 180 \| Uncharacterized protein \| Uncharacterized protein; afdb-uniprot50 \| AF-A0A1S2TBK1-F1-MODEL\_V4 \| 1.0 \| 2.614e-08 \| 297 \| 0.248 \| 165 \| 104 \| 5 \| 25 \| 189 \| 2 \| 146 \| Uncharacterized protein \| Uncharacterized protein; afdb-uniprot50 \| AF-A0A839IZ53-F1-MODEL\_V4 \| 1.0 \| 1.741e-08 \| 292 \| 0.184 \| 201 \| 135 \| 13 \| 1 \| 188 \| 1 \| 185 \| Uncharacterized protein \| Uncharacterized protein; afdb-uniprot50 \| AF-F4BFQ7-F1-MODEL\_V4 \| 1.0 \| 2.328e-08 \| 285 \| 0.191 \| 178 \| 129 \| 7 \| 1 \| 178 \| 1 \| 163 \| Uncharacterized protein \| Uncharacterized protein; afdb-uniprot50 \| AF-A0A1C3EBP2-F1-MODEL\_V4 \| 1.0 \| 8.672e-09 \| 284 \| 0.203 \| 187 \| 131 \| 9 \| 5 \| 179 \| 3 \| 183 \| Uncharacterized protein \| Uncharacterized protein; afdb-uniprot50 \| AF-A0A2N0CKM3-F1-MODEL\_V4 \| 1.0 \| 1.054e-07 \| 283 \| 0.206 \| 179 \| 118 \| 7 \| 6 \| 180 \| 3 \| 161 \| Uncharacterized protein \| Uncharacterized protein; afdb-uniprot50 \| AF-A0A447INZ1-F1-MODEL\_V4 \| 1.0 \| 1.094e-08 \| 278 \| 0.195 \| 189 \| 128 \| 10 \| 9 \| 188 \| 8 \| 181 \| Uncharacterized protein \| Uncharacterized protein; afdb-uniprot50 \| AF-A0A4P9VGP3-F1-MODEL\_V4 \| 1.0 \| 9.19e-09 \| 276 \| 0.2 \| 190 \| 130 \| 7 \| 1 \| 189 \| 23 \| 191 \| Uncharacterized protein \| Uncharacterized protein; afdb-uniprot50 \| AF-A0A345DE57-F1-MODEL\_V4 \| 1.0 \| 1.409e-07 \| 271 \| 0.243 \| 164 \| 102 \| 8 \| 7 \| 166 \| 1 \| 146 \| Uncharacterized protein \| Uncharacterized protein; afdb-uniprot50 \| AF-A0A839HEF9-F1-MODEL\_V4 \| 1.0 \| 1.184e-07 \| 269 \| 0.166 \| 168 \| 125 \| 7 \| 1 \| 155 \| 1 \| 166 \| Uncharacterized protein \| Uncharacterized protein; afdb-uniprot50 \| AF-A0A4U8YKI5-F1-MODEL\_V4 \| 1.0 \| 4.953e-08 \| 265 \| 0.215 \| 181 \| 117 \| 8 \| 7 \| 179 \| 2 \| 165 \| Uncharacterized protein \| Uncharacterized protein; afdb-uniprot50 \| AF-A0A6A4R943-F1-MODEL\_V4 \| 1.0 \| 4.673e-08 \| 262 \| 0.182 \| 197 \| 140 \| 10 \| 1 \| 189 \| 1 \| 184 \| Uncharacterized protein \| Uncharacterized protein; afdb-uniprot50 \| AF-A0A7D7X3U7-F1-MODEL\_V4 \| 1.0 \| 1.884e-07 \| 260 \| 0.151 \| 192 \| 139 \| 11 \| 1 \| 185 \| 1 \| 175 \| Uncharacterized protein \| Uncharacterized protein; afdb-uniprot50 \| AF-A0A853IIQ5-F1-MODEL\_V4 \| 1.0 \| 7.592e-07 \| 259 \| 0.295 \| 132 \| 85 \| 3 \| 60 \| 189 \| 2 \| 127 \| Uncharacterized protein \| Uncharacterized protein; afdb-uniprot50 \| AF-A7MWA0-F1-MODEL\_V4 \| 1.0 \| 1.677e-07 \| 256 \| 0.157 \| 197 \| 139 \| 11 \| 1 \| 189 \| 1 \| 178 \| Uncharacterized protein \| Uncharacterized protein; afdb-uniprot50 \| AF-A0A379CNZ4-F1-MODEL\_V4 \| 1.0 \| 5.358e-07 \| 255 \| 0.165 \| 193 \| 137 \| 11 \| 2 \| 188 \| 1 \| 175 \| Uncharacterized protein \| Uncharacterized protein; afdb-uniprot50 \| AF-A0A432QS63-F1-MODEL\_V4 \| 1.0 \| 1.184e-07 \| 255 \| 0.166 \| 168 \| 120 \| 6 \| 2 \| 161 \| 1 \| 156 \| Uncharacterized protein \| Uncharacterized protein; afdb-uniprot50 \| AF-A0A7W6WLT8-F1-MODEL\_V4 \| 1.0 \| 3.177e-07 \| 254 \| 0.185 \| 194 \| 131 \| 10 \| 1 \| 186 \| 1 \| 175 \| Uncharacterized protein \| Uncharacterized protein; afdb-uniprot50 \| AF-A0A4P9VH08-F1-MODEL\_V4 \| 1.0 \| 3.112e-08 \| 254 \| 0.198 \| 186 \| 133 \| 4 \| 2 \| 187 \| 120 \| 289 \| Uncharacterized protein \| Uncharacterized protein; afdb-uniprot50 \| AF-A0A3S0H8F4-F1-MODEL\_V4 \| 1.0 \| 2.998e-07 \| 253 \| 0.164 \| 189 \| 133 \| 10 \| 7 \| 189 \| 2 \| 171 \| Uncharacterized protein \| Uncharacterized protein; afdb-uniprot50 \| AF-A0A7T8NTX8-F1-MODEL\_V4 \| 1.0 \| 7.437e-08 \| 250 \| 0.186 \| 188 \| 132 \| 10 \| 8 \| 185 \| 3 \| 179 \| Uncharacterized protein \| Uncharacterized protein; afdb-uniprot50 \| AF-A0A6I2KE43-F1-MODEL\_V4 \| 1.0 \| 2.669e-07 \| 248 \| 0.405 \| 106 \| 56 \| 1 \| 84 \| 189 \| 3 \| 101 \| Uncharacterized protein \| Uncharacterized protein; afdb-uniprot50 \| AF-A0A292QWT2-F1-MODEL\_V4 \| 1.0 \| 2.426e-06 \| 248 \| 0.177 \| 152 \| 103 \| 11 \| 9 \| 156 \| 7 \| 140 \| Uncharacterized protein \| Uncharacterized protein; afdb-uniprot50 \| AF-A0A7Y6Z4J4-F1-MODEL\_V4 \| 1.0 \| 6.018e-07 \| 248 \| 0.192 \| 177 \| 128 \| 7 \| 1 \| 172 \| 1 \| 167 \| Uncharacterized protein \| Uncharacterized protein; afdb-uniprot50 \| AF-A0A085AFN8-F1-MODEL\_V4 \| 1.0 \| 6.248e-08 \| 248 \| 0.169 \| 165 \| 123 \| 8 \| 27 \| 189 \| 1 \| 153 \| Phage protein \| Phage protein; afdb-uniprot50 \| AF-A0A5S9P2S6-F1-MODEL\_V4 \| 1.0 \| 6.248e-08 \| 245 \| 0.202 \| 202 \| 127 \| 12 \| 9 \| 189 \| 10 \| 198 \| Uncharacterized protein \| Uncharacterized protein; afdb-uniprot50 \| AF-A0A5P9F0H4-F1-MODEL\_V4 \| 1.0 \| 1.777e-07 \| 243 \| 0.17 \| 194 \| 135 \| 9 \| 9 \| 188 \| 4 \| 185 \| Uncharacterized protein \| Uncharacterized protein; afdb-uniprot50 \| AF-A0A522WEL2-F1-MODEL\_V4 \| 1.0 \| 2.998e-07 \| 241 \| 0.234 \| 175 \| 103 \| 10 \| 9 \| 169 \| 3 \| 160 \| Uncharacterized protein \| Uncharacterized protein; afdb-uniprot50 \| AF-A0A1V5V144-F1-MODEL\_V4 \| 1.0 \| 1.076e-06 \| 240 \| 0.215 \| 158 \| 103 \| 9 \| 1 \| 154 \| 1 \| 141 \| Uncharacterized protein \| Uncharacterized protein; afdb-uniprot50 \| AF-A0A849VN83-F1-MODEL\_V4 \| 1.0 \| 3.782e-07 \| 239 \| 0.17 \| 193 \| 133 \| 13 \| 7 \| 189 \| 1 \| 176 \| Uncharacterized protein \| Uncharacterized protein; afdb-uniprot50 \| AF-A0A3B0MY09-F1-MODEL\_V4 \| 1.0 \| 7.592e-07 \| 238 \| 0.156 \| 172 \| 126 \| 8 \| 1 \| 165 \| 1 \| 160 \| Uncharacterized protein \| Uncharacterized protein; afdb-uniprot50 \| AF-A0A853I879-F1-MODEL\_V4 \| 1.0 \| 4.008e-07 \| 238 \| 0.206 \| 189 \| 123 \| 9 \| 1 \| 188 \| 123 \| 285 \| Uncharacterized protein \| Uncharacterized protein; afdb-uniprot50 \| AF-A0A515ERM5-F1-MODEL\_V4 \| 1.0 \| 1.281e-06 \| 233 \| 0.185 \| 151 \| 104 \| 7 \| 7 \| 152 \| 1 \| 137 \| Uncharacterized protein \| Uncharacterized protein; afdb-uniprot50 \| AF-A0A2D8ACY1-F1-MODEL\_V4 \| 1.0 \| 2.998e-07 \| 233 \| 0.196 \| 188 \| 128 \| 10 \| 7 \| 187 \| 1 \| 172 \| Uncharacterized protein \| Uncharacterized protein; afdb-uniprot50 \| AF-A0A7X4AMR2-F1-MODEL\_V4 \| 1.0 \| 1.615e-06 \| 227 \| 0.182 \| 192 \| 128 \| 11 \| 7 \| 189 \| 2 \| 173 \| Uncharacterized protein \| Uncharacterized protein; afdb-uniprot50 \| AF-I3TTD7-F1-MODEL\_V4 \| 1.0 \| 4.502e-07 \| 227 \| 0.203 \| 201 \| 130 \| 10 \| 9 \| 189 \| 6 \| 196 \| Uncharacterized protein \| Uncharacterized protein; afdb-uniprot50 \| AF-A0A365XR84-F1-MODEL\_V4 \| 1.0 \| 4.336e-06 \| 226 \| 0.128 \| 171 \| 127 \| 6 \| 9 \| 173 \| 5 \| 159 \| Uncharacterized protein \| Uncharacterized protein; afdb-uniprot50 \| AF-A0A6S7D6J5-F1-MODEL\_V4 \| 1.0 \| 6.018e-07 \| 225 \| 0.17 \| 200 \| 137 \| 10 \| 1 \| 183 \| 1 \| 188 \| Uncharacterized protein \| Uncharacterized protein; afdb-uniprot50 \| AF-A0A833IRE7-F1-MODEL\_V4 \| 1.0 \| 2.571e-06 \| 224 \| 0.174 \| 189 \| 138 \| 7 \| 7 \| 189 \| 1 \| 177 \| Uncharacterized protein \| Uncharacterized protein; afdb-uniprot50 \| AF-A0A640W7I2-F1-MODEL\_V4 \| 1.0 \| 4.87e-06 \| 223 \| 0.153 \| 156 \| 109 \| 7 \| 1 \| 154 \| 1 \| 135 \| Uncharacterized protein \| Uncharacterized protein; afdb-uniprot50 \| AF-A0A1G9UBD2-F1-MODEL\_V4 \| 1.0 \| 1.036e-05 \| 223 \| 0.182 \| 148 \| 100 \| 5 \| 13 \| 152 \| 3 \| 137 \| Uncharacterized protein \| Uncharacterized protein; afdb-uniprot50 \| AF-A0A496VZB9-F1-MODEL\_V4 \| 1.0 \| 5.162e-06 \| 223 \| 0.158 \| 164 \| 112 \| 7 \| 1 \| 159 \| 1 \| 143 \| Uncharacterized protein \| Uncharacterized protein; afdb-uniprot50 \| AF-A0A3D4RUC7-F1-MODEL\_V4 \| 1.0 \| 2.038e-06 \| 223 \| 0.126 \| 221 \| 135 \| 10 \| 7 \| 189 \| 2 \| 202 \| Uncharacterized protein \| Uncharacterized protein; afdb-uniprot50 \| AF-A0A4S8HNS9-F1-MODEL\_V4 \| 1.0 \| 2.725e-06 \| 222 \| 0.136 \| 176 \| 128 \| 7 \| 9 \| 177 \| 6 \| 164 \| Uncharacterized protein \| Uncharacterized protein; afdb-uniprot50 \| AF-A0A7X2NZB9-F1-MODEL\_V4 \| 1.0 \| 9.226e-06 \| 221 \| 0.43 \| 93 \| 45 \| 1 \| 96 \| 188 \| 2 \| 86 \| Uncharacterized protein \| Uncharacterized protein; afdb-uniprot50 \| AF-A0A2N1TYS7-F1-MODEL\_V4 \| 1.0 \| 8.214e-06 \| 221 \| 0.152 \| 151 \| 110 \| 7 \| 7 \| 154 \| 2 \| 137 \| Uncharacterized protein \| Uncharacterized protein; afdb-uniprot50 \| AF-A0A2I6S9D2-F1-MODEL\_V4 \| 1.0 \| 7.751e-06 \| 220 \| 0.177 \| 158 \| 113 \| 7 \| 2 \| 155 \| 1 \| 145 \| Uncharacterized protein \| Uncharacterized protein; afdb-uniprot50 \| AF-A0A348FYH7-F1-MODEL\_V4 \| 1.0 \| 9.038e-07 \| 220 \| 0.178 \| 174 \| 118 \| 11 \| 1 \| 164 \| 1 \| 159 \| Uncharacterized protein \| Uncharacterized protein; afdb-uniprot50 \| AF-A0A1X7NDV7-F1-MODEL\_V4 \| 1.0 \| 3.643e-06 \| 219 \| 0.165 \| 175 \| 126 \| 8 \| 1 \| 165 \| 1 \| 165 \| Uncharacterized protein \| Uncharacterized protein; afdb-uniprot50 \| AF-A0A1M3PZ62-F1-MODEL\_V4 \| 1.0 \| 7.751e-06 \| 215 \| 0.177 \| 152 \| 112 \| 5 \| 10 \| 159 \| 7 \| 147 \| Uncharacterized protein \| Uncharacterized protein; afdb-uniprot50 \| AF-A0A7C1ULK3-F1-MODEL\_V4 \| 1.0 \| 9.778e-06 \| 214 \| 0.125 \| 152 \| 103 \| 8 \| 13 \| 152 \| 2 \| 135 \| Uncharacterized protein \| Uncharacterized protein; afdb-uniprot50 \| AF-A0A2P5N8U8-F1-MODEL\_V4 \| 1.0 \| 8.706e-06 \| 214 \| 0.104 \| 153 \| 119 \| 7 \| 12 \| 159 \| 5 \| 144 \| Uncharacterized protein \| Uncharacterized protein; afdb-uniprot50 \| AF-A0A2W7JFE3-F1-MODEL\_V4 \| 1.0 \| 4.336e-06 \| 214 \| 0.148 \| 155 \| 117 \| 5 \| 15 \| 166 \| 5 \| 147 \| Uncharacterized protein \| Uncharacterized protein; afdb-uniprot50 \| AF-A0A1F4PEF5-F1-MODEL\_V4 \| 1.0 \| 7.751e-06 \| 213 \| 0.117 \| 162 \| 112 \| 9 \| 6 \| 152 \| 2 \| 147 \| Uncharacterized protein \| Uncharacterized protein; afdb-uniprot50 \| AF-A0A134B7B8-F1-MODEL\_V4 \| 1.0 \| 2.571e-06 \| 212 \| 0.128 \| 179 \| 127 \| 7 \| 10 \| 172 \| 2 \| 167 \| Uncharacterized protein \| Uncharacterized protein; afdb-uniprot50 \| AF-A0A109C5L1-F1-MODEL\_V4 \| 1.0 \| 9.778e-06 \| 209 \| 0.165 \| 163 \| 113 \| 9 \| 7 \| 165 \| 2 \| 145 \| Uncharacterized protein \| Uncharacterized protein; afdb-uniprot50 \| AF-A0A3S1EBT0-F1-MODEL\_V4 \| 1.0 \| 2.725e-06 \| 209 \| 0.197 \| 192 \| 128 \| 12 \| 1 \| 178 \| 1 \| 180 \| Uncharacterized protein \| Uncharacterized protein; afdb-uniprot50 \| AF-A0A7U5AJB0-F1-MODEL\_V4 \| 1.0 \| 2.571e-06 \| 209 \| 0.208 \| 182 \| 126 \| 9 \| 4 \| 177 \| 2 \| 173 \| Uncharacterized protein \| Uncharacterized protein; afdb-uniprot50 \| AF-A0A1Q3ZHY0-F1-MODEL\_V4 \| 1.0 \| 5.797e-06 \| 208 \| 0.127 \| 172 \| 126 \| 11 \| 9 \| 173 \| 4 \| 158 \| Uncharacterized protein \| Uncharacterized protein; afdb-uniprot50 \| AF-A0A2W7QSY0-F1-MODEL\_V4 \| 1.0 \| 2.337e-05 \| 207 \| 0.196 \| 183 \| 126 \| 11 \| 1 \| 174 \| 1 \| 171 \| Uncharacterized protein \| Uncharacterized protein; afdb-uniprot50 \| AF-A0A6L8MMN6-F1-MODEL\_V4 \| 1.0 \| 2.081e-05 \| 205 \| 0.141 \| 148 \| 105 \| 5 \| 13 \| 152 \| 2 \| 135 \| Uncharacterized protein \| Uncharacterized protein; afdb-uniprot50 \| AF-A0A1H6CBN8-F1-MODEL\_V4 \| 1.0 \| 7.314e-06 \| 205 \| 0.133 \| 157 \| 112 \| 7 \| 7 \| 156 \| 5 \| 144 \| Uncharacterized protein \| Uncharacterized protein; afdb-uniprot50 \| AF-R6VPC3-F1-MODEL\_V4 \| 1.0 \| 8.528e-07 \| 205 \| 0.179 \| 178 \| 114 \| 9 \| 10 \| 172 \| 2 \| 162 \| Uncharacterized protein \| Uncharacterized protein; afdb-uniprot50 \| AF-T2G8C6-F1-MODEL\_V4 \| 1.0 \| 3.311e-05 \| 204 \| 0.186 \| 161 \| 116 \| 9 \| 1 \| 155 \| 1 \| 152 \| Uncharacterized protein \| Uncharacterized protein; afdb-uniprot50 \| AF-A0A349PRD1-F1-MODEL\_V4 \| 1.0 \| 1.468e-05 \| 204 \| 0.123 \| 170 \| 126 \| 12 \| 8 \| 168 \| 1 \| 156 \| Uncharacterized protein \| Uncharacterized protein; afdb-uniprot50 \| AF-A0A849MY44-F1-MODEL\_V4 \| 1.0 \| 1.712e-06 \| 204 \| 0.169 \| 171 \| 119 \| 7 \| 14 \| 179 \| 4 \| 156 \| Uncharacterized protein \| Uncharacterized protein; afdb-uniprot50 \| AF-A0A844H9D9-F1-MODEL\_V4 \| 1.0 \| 3.243e-06 \| 204 \| 0.132 \| 204 \| 138 \| 9 \| 2 \| 179 \| 9 \| 199 \| Uncharacterized protein \| Uncharacterized protein; afdb-uniprot50 \| AF-A0A7J5VW47-F1-MODEL\_V4 \| 1.0 \| 7.751e-06 \| 202 \| 0.131 \| 183 \| 129 \| 9 \| 8 \| 178 \| 1 \| 165 \| Uncharacterized protein \| Uncharacterized protein; afdb-uniprot50 \| AF-A0A1G5ZJI4-F1-MODEL\_V4 \| 1.0 \| 1.357e-06 \| 202 \| 0.132 \| 204 \| 134 \| 9 \| 15 \| 189 \| 5 \| 194 \| Uncharacterized protein \| Uncharacterized protein; afdb-uniprot50 \| AF-A0A446AZ98-F1-MODEL\_V4 \| 1.0 \| 7.164e-07 \| 202 \| 0.161 \| 217 \| 128 \| 15 \| 7 \| 189 \| 1 \| 197 \| Uncharacterized protein \| Uncharacterized protein; afdb-uniprot50 \| AF-A0A450ZWN7-F1-MODEL\_V4 \| 1.0 \| 3.124e-05 \| 201 \| 0.151 \| 158 \| 107 \| 7 \| 1 \| 153 \| 1 \| 136 \| Uncharacterized protein \| Uncharacterized protein; afdb-uniprot50 \| AF-I3TN74-F1-MODEL\_V4 \| 1.0 \| 1.556e-05 \| 200 \| 0.198 \| 151 \| 104 \| 8 \| 10 \| 153 \| 2 \| 142 \| Uncharacterized protein \| Uncharacterized protein; afdb-uniprot50 \| AF-A0A1X3RM65-F1-MODEL\_V4 \| 1.0 \| 1.963e-05 \| 200 \| 0.177 \| 169 \| 123 \| 6 \| 7 \| 173 \| 1 \| 155 \| Uncharacterized protein \| Uncharacterized protein; afdb-uniprot50 \| AF-A0A4R3UQ50-F1-MODEL\_V4 \| 1.0 \| 8.706e-06 \| 200 \| 0.169 \| 159 \| 113 \| 7 \| 14 \| 165 \| 4 \| 150 \| Uncharacterized protein \| Uncharacterized protein; afdb-uniprot50 \| AF-W0HQ32-F1-MODEL\_V4 \| 1.0 \| 1.468e-05 \| 200 \| 0.164 \| 164 \| 117 \| 9 \| 8 \| 165 \| 1 \| 150 \| Uncharacterized protein \| Uncharacterized protein; afdb-uniprot50 \| AF-A0A349MJN4-F1-MODEL\_V4 \| 1.0 \| 2.081e-05 \| 199 \| 0.147 \| 149 \| 106 \| 7 \| 12 \| 153 \| 5 \| 139 \| Uncharacterized protein \| Uncharacterized protein; afdb-uniprot50 \| AF-A0A2A4XUH6-F1-MODEL\_V4 \| 1.0 \| 1.615e-06 \| 199 \| 0.21 \| 176 \| 111 \| 10 \| 8 \| 168 \| 3 \| 165 \| Uncharacterized protein \| Uncharacterized protein; afdb-uniprot50 \| AF-A0A4Z0Q1Z6-F1-MODEL\_V4 \| 1.0 \| 4.092e-06 \| 198 \| 0.138 \| 181 \| 122 \| 8 \| 8 \| 168 \| 1 \| 167 \| Uncharacterized protein \| Uncharacterized protein; afdb-uniprot50 \| AF-A0A3D5BS75-F1-MODEL\_V4 \| 1.0 \| 2.477e-05 \| 196 \| 0.112 \| 169 \| 117 \| 10 \| 8 \| 166 \| 1 \| 146 \| Uncharacterized protein \| Uncharacterized protein; afdb-uniprot50 \| AF-A0A8B3TEH3-F1-MODEL\_V4 \| 1.0 \| 8.214e-06 \| 196 \| 0.112 \| 187 \| 142 \| 9 \| 9 \| 187 \| 4 \| 174 \| DUF1834 family protein \| DUF1834 family protein; afdb-uniprot50 \| AF-A0A6A7YJM2-F1-MODEL\_V4 \| 1.0 \| 2.038e-06 \| 196 \| 0.158 \| 189 \| 124 \| 10 \| 14 \| 182 \| 4 \| 177 \| Uncharacterized protein \| Uncharacterized protein; afdb-uniprot50 \| AF-A0A3N2E265-F1-MODEL\_V4 \| 1.0 \| 4.336e-06 \| 196 \| 0.141 \| 191 \| 120 \| 9 \| 7 \| 165 \| 2 \| 180 \| Uncharacterized protein \| Uncharacterized protein; afdb-uniprot50 \| AF-A0A0Q4UF98-F1-MODEL\_V4 \| 1.0 \| 4.972e-05 \| 195 \| 0.18 \| 150 \| 101 \| 8 \| 7 \| 153 \| 2 \| 132 \| Uncharacterized protein \| Uncharacterized protein; afdb-uniprot50 \| AF-A0A3M1MUF4-F1-MODEL\_V4 \| 1.0 \| 0.0001684 \| 195 \| 0.093 \| 149 \| 118 \| 5 \| 9 \| 154 \| 3 \| 137 \| Uncharacterized protein \| Uncharacterized protein; afdb-uniprot50 \| AF-A0A840MMI4-F1-MODEL\_V4 \| 1.0 \| 5.269e-05 \| 195 \| 0.131 \| 167 \| 128 \| 6 \| 7 \| 172 \| 1 \| 151 \| Uncharacterized protein \| Uncharacterized protein; afdb-uniprot50 \| AF-A0A450YW35-F1-MODEL\_V4 \| 1.0 \| 1.649e-05 \| 195 \| 0.174 \| 172 \| 123 \| 7 \| 1 \| 171 \| 1 \| 154 \| Gp37 protein \| Gp37 protein; afdb-uniprot50 \| AF-V9HKZ2-F1-MODEL\_V4 \| 1.0 \| 2.948e-05 \| 194 \| 0.146 \| 157 \| 108 \| 8 \| 8 \| 153 \| 3 \| 144 \| Uncharacterized protein \| Uncharacterized protein; afdb-uniprot50 \| AF-A0A6N7B6E2-F1-MODEL\_V4 \| 1.0 \| 2.337e-05 \| 194 \| 0.155 \| 167 \| 120 \| 7 \| 1 \| 166 \| 1 \| 147 \| Uncharacterized protein \| Uncharacterized protein; afdb-uniprot50 \| AF-A0A450WGT7-F1-MODEL\_V4 \| 1.0 \| 2.625e-05 \| 194 \| 0.169 \| 171 \| 124 \| 7 \| 1 \| 170 \| 1 \| 154 \| Gp37 protein \| Gp37 protein; afdb-uniprot50 \| AF-A0A5C7J5X6-F1-MODEL\_V4 \| 1.0 \| 6.144e-06 \| 194 \| 0.157 \| 210 \| 143 \| 12 \| 2 \| 189 \| 17 \| 214 \| DUF1834 family protein \| DUF1834 family protein; afdb-uniprot50 \| AF-A0A1G8LDR9-F1-MODEL\_V4 \| 1.0 \| 3.311e-05 \| 193 \| 0.175 \| 148 \| 105 \| 9 \| 13 \| 155 \| 2 \| 137 \| Uncharacterized protein \| Uncharacterized protein; afdb-uniprot50 \| AF-A0A177N1G7-F1-MODEL\_V4 \| 1.0 \| 9.419e-05 \| 193 \| 0.146 \| 150 \| 105 \| 7 \| 12 \| 153 \| 4 \| 138 \| Uncharacterized protein \| Uncharacterized protein; afdb-uniprot50 \| AF-A0A5I6PT12-F1-MODEL\_V4 \| 1.0 \| 1.556e-05 \| 193 \| 0.142 \| 175 \| 120 \| 9 \| 9 \| 166 \| 3 \| 164 \| DUF1834 family protein \| DUF1834 family protein; afdb-uniprot50 \| AF-V4NRC4-F1-MODEL\_V4 \| 1.0 \| 6.901e-06 \| 193 \| 0.152 \| 203 \| 130 \| 13 \| 12 \| 186 \| 5 \| 193 \| Uncharacterized protein \| Uncharacterized protein; afdb-uniprot50 \| AF-A0A4R3LGT3-F1-MODEL\_V4 \| 1.0 \| 4.092e-06 \| 192 \| 0.19 \| 184 \| 129 \| 8 \| 10 \| 189 \| 5 \| 172 \| Uncharacterized protein \| Uncharacterized protein; afdb-uniprot50 \| AF-A0A285P0D4-F1-MODEL\_V4 \| 1.0 \| 2.571e-06 \| 192 \| 0.136 \| 205 \| 132 \| 11 \| 15 \| 188 \| 5 \| 195 \| Uncharacterized protein \| Uncharacterized protein; afdb-uniprot50 \| AF-A0A2C8YIV8-F1-MODEL\_V4 \| 1.0 \| 2.625e-05 \| 191 \| 0.184 \| 163 \| 111 \| 8 \| 1 \| 156 \| 9 \| 156 \| Uncharacterized protein \| Uncharacterized protein; afdb-uniprot50 \| AF-G2H4N2-F1-MODEL\_V4 \| 1.0 \| 5.797e-06 \| 191 \| 0.154 \| 207 \| 140 \| 13 \| 8 \| 189 \| 1 \| 197 \| Uncharacterized protein \| Uncharacterized protein; afdb-uniprot50 \| AF-A0A2S6N2V3-F1-MODEL\_V4 \| 1.0 \| 1.164e-05 \| 191 \| 0.134 \| 178 \| 124 \| 9 \| 8 \| 166 \| 1 \| 167 \| Uncharacterized protein \| Uncharacterized protein; afdb-uniprot50 \| AF-A0A345DDG3-F1-MODEL\_V4 \| 1.0 \| 8.214e-06 \| 191 \| 0.152 \| 217 \| 125 \| 13 \| 12 \| 180 \| 4 \| 209 \| Uncharacterized protein \| Uncharacterized protein; afdb-uniprot50 \| AF-A0A5C7TSP4-F1-MODEL\_V4 \| 1.0 \| 5.269e-05 \| 190 \| 0.15 \| 166 \| 114 \| 7 \| 1 \| 163 \| 1 \| 142 \| Uncharacterized protein \| Uncharacterized protein; afdb-uniprot50 \| AF-A0A2S0MD53-F1-MODEL\_V4 \| 1.0 \| 4.427e-05 \| 190 \| 0.173 \| 161 \| 112 \| 9 \| 1 \| 153 \| 1 \| 148 \| Uncharacterized protein \| Uncharacterized protein; afdb-uniprot50 \| AF-A0A6M3ZW34-F1-MODEL\_V4 \| 1.0 \| 3.06e-06 \| 190 \| 0.125 \| 207 \| 135 \| 11 \| 15 \| 189 \| 5 \| 197 \| Uncharacterized protein \| Uncharacterized protein; afdb-uniprot50 \| AF-A0A0F9FP65-F1-MODEL\_V4 \| 1.0 \| 8.214e-06 \| 190 \| 0.145 \| 179 \| 121 \| 8 \| 10 \| 166 \| 6 \| 174 \| Uncharacterized protein \| Uncharacterized protein; afdb-uniprot50 \| AF-A0A104N6L4-F1-MODEL\_V4 \| 1.0 \| 2.782e-05 \| 189 \| 0.286 \| 115 \| 71 \| 3 \| 71 \| 185 \| 3 \| 106 \| Uncharacterized protein \| Uncharacterized protein; afdb-uniprot50 \| AF-A0A1G0I709-F1-MODEL\_V4 \| 1.0 \| 5.918e-05 \| 189 \| 0.18 \| 150 \| 106 \| 6 \| 7 \| 152 \| 1 \| 137 \| Uncharacterized protein \| Uncharacterized protein; afdb-uniprot50 \| AF-I3YEI7-F1-MODEL\_V4 \| 1.0 \| 1.307e-05 \| 189 \| 0.204 \| 176 \| 107 \| 11 \| 1 \| 166 \| 1 \| 153 \| Uncharacterized protein \| Uncharacterized protein; afdb-uniprot50 \| AF-A0A249STX7-F1-MODEL\_V4 \| 1.0 \| 1.556e-05 \| 189 \| 0.155 \| 180 \| 109 \| 11 \| 10 \| 165 \| 2 \| 162 \| Uncharacterized protein \| Uncharacterized protein; afdb-uniprot50 \| AF-A0A2A2JXH5-F1-MODEL\_V4 \| 1.0 \| 3.719e-05 \| 188 \| 0.163 \| 159 \| 103 \| 8 \| 6 \| 156 \| 2 \| 138 \| Uncharacterized protein \| Uncharacterized protein; afdb-uniprot50 \| AF-A0A194AGD2-F1-MODEL\_V4 \| 1.0 \| 0.0002528 \| 188 \| 0.13 \| 153 \| 117 \| 6 \| 7 \| 155 \| 2 \| 142 \| Uncharacterized protein \| Uncharacterized protein; afdb-uniprot50 \| AF-W8F0Q4-F1-MODEL\_V4 \| 1.0 \| 1.098e-05 \| 188 \| 0.136 \| 191 \| 125 \| 11 \| 8 \| 189 \| 1 \| 160 \| Uncharacterized protein \| Uncharacterized protein; afdb-uniprot50 \| AF-A0A379S6I8-F1-MODEL\_V4 \| 1.0 \| 3.861e-06 \| 188 \| 0.129 \| 208 \| 144 \| 15 \| 1 \| 189 \| 165 \| 354 \| Phage protein \| Phage protein; afdb-uniprot50 \| AF-A0A059IW70-F1-MODEL\_V4 \| 1.0 \| 8.386e-05 \| 187 \| 0.315 \| 95 \| 57 \| 2 \| 71 \| 165 \| 3 \| 89 \| Uncharacterized protein \| Uncharacterized protein; afdb-uniprot50 \| AF-A0A286GYR4-F1-MODEL\_V4 \| 1.0 \| 7.913e-05 \| 187 \| 0.168 \| 148 \| 107 \| 6 \| 13 \| 155 \| 4 \| 140 \| Uncharacterized protein \| Uncharacterized protein; afdb-uniprot50 \| AF-A0A833GRL2-F1-MODEL\_V4 \| 1.0 \| 3.941e-05 \| 187 \| 0.158 \| 164 \| 113 \| 9 \| 8 \| 164 \| 1 \| 146 \| Uncharacterized protein \| Uncharacterized protein; afdb-uniprot50 \| AF-B6IMF8-F1-MODEL\_V4 \| 1.0 \| 2.337e-05 \| 187 \| 0.177 \| 169 \| 121 \| 6 \| 7 \| 173 \| 1 \| 153 \| Uncharacterized protein \| Uncharacterized protein; afdb-uniprot50 \| AF-A0A386HSH2-F1-MODEL\_V4 \| 1.0 \| 2.948e-05 \| 187 \| 0.134 \| 186 \| 134 \| 7 \| 1 \| 166 \| 7 \| 185 \| Uncharacterized protein \| Uncharacterized protein; afdb-uniprot50 \| AF-A0A1M7B4C0-F1-MODEL\_V4 \| 1.0 \| 9.419e-05 \| 185 \| 0.142 \| 161 \| 104 \| 7 \| 1 \| 156 \| 2 \| 133 \| Uncharacterized protein \| Uncharacterized protein; afdb-uniprot50 \| AF-A0A661DES6-F1-MODEL\_V4 \| 1.0 \| 1.556e-05 \| 185 \| 0.212 \| 155 \| 100 \| 12 \| 7 \| 153 \| 1 \| 141 \| Uncharacterized protein \| Uncharacterized protein; afdb-uniprot50 \| AF-A0A7S8C7I5-F1-MODEL\_V4 \| 1.0 \| 0.0001499 \| 185 \| 0.169 \| 165 \| 117 \| 9 \| 7 \| 165 \| 2 \| 152 \| Uncharacterized protein \| Uncharacterized protein; afdb-uniprot50 \| AF-A0A0Q0A4D7-F1-MODEL\_V4 \| 1.0 \| 6.901e-06 \| 185 \| 0.13 \| 222 \| 144 \| 13 \| 1 \| 189 \| 65 \| 270 \| Uncharacterized protein \| Uncharacterized protein; afdb-uniprot50 \| AF-A0A2E1RHS9-F1-MODEL\_V4 \| 1.0 \| 9.982e-05 \| 183 \| 0.172 \| 145 \| 101 \| 6 \| 13 \| 156 \| 2 \| 128 \| Uncharacterized protein \| Uncharacterized protein; afdb-uniprot50 \| AF-A0A3A0EP56-F1-MODEL\_V4 \| 1.0 \| 5.918e-05 \| 183 \| 0.153 \| 150 \| 108 \| 6 \| 7 \| 153 \| 1 \| 134 \| Uncharacterized protein \| Uncharacterized protein; afdb-uniprot50 \| AF-A0A7C1A7F0-F1-MODEL\_V4 \| 1.0 \| 0.000284 \| 183 \| 0.131 \| 152 \| 114 \| 7 \| 8 \| 154 \| 1 \| 139 \| DUF1834 family protein \| DUF1834 family protein; afdb-uniprot50 \| AF-A0A355TT27-F1-MODEL\_V4 \| 1.0 \| 0.0001335 \| 183 \| 0.135 \| 162 \| 119 \| 7 \| 7 \| 165 \| 1 \| 144 \| Uncharacterized protein \| Uncharacterized protein; afdb-uniprot50 \| AF-A0A109BYV0-F1-MODEL\_V4 \| 1.0 \| 4.691e-05 \| 183 \| 0.137 \| 153 \| 118 \| 9 \| 7 \| 153 \| 2 \| 146 \| Uncharacterized protein \| Uncharacterized protein; afdb-uniprot50 \| AF-A0A0F9PV67-F1-MODEL\_V4 \| 1.0 \| 2.081e-05 \| 183 \| 0.146 \| 177 \| 118 \| 14 \| 1 \| 155 \| 1 \| 166 \| Uncharacterized protein \| Uncharacterized protein; afdb-uniprot50 \| AF-A0A2K4MRJ3-F1-MODEL\_V4 \| 1.0 \| 2.782e-05 \| 183 \| 0.128 \| 179 \| 126 \| 11 \| 2 \| 174 \| 34 \| 188 \| Uncharacterized protein \| Uncharacterized protein; afdb-uniprot50 \| AF-A0A1B9LF31-F1-MODEL\_V4 \| 1.0 \| 5.584e-05 \| 183 \| 0.133 \| 150 \| 109 \| 8 \| 14 \| 155 \| 4 \| 140 \| Uncharacterized protein \| Uncharacterized protein; afdb-uniprot50 \| AF-A0A3B9IJU4-F1-MODEL\_V4 \| 1.0 \| 7.913e-05 \| 182 \| 0.153 \| 150 \| 103 \| 7 \| 13 \| 153 \| 2 \| 136 \| Uncharacterized protein \| Uncharacterized protein; afdb-uniprot50 \| AF-A0A853YIQ0-F1-MODEL\_V4 \| 1.0 \| 0.0001058 \| 182 \| 0.163 \| 153 \| 103 \| 6 \| 13 \| 156 \| 2 \| 138 \| Uncharacterized protein \| Uncharacterized protein; afdb-uniprot50 \| AF-A0A6A8RIJ7-F1-MODEL\_V4 \| 1.0 \| 0.0002251 \| 182 \| 0.124 \| 161 \| 117 \| 7 \| 7 \| 155 \| 3 \| 151 \| Uncharacterized protein \| Uncharacterized protein; afdb-uniprot50 \| AF-A0A4R1AH59-F1-MODEL\_V4 \| 1.0 \| 7.045e-05 \| 182 \| 0.114 \| 166 \| 118 \| 11 \| 7 \| 159 \| 1 \| 150 \| Uncharacterized protein \| Uncharacterized protein; afdb-uniprot50 \| AF-A0A423PRY6-F1-MODEL\_V4 \| 1.0 \| 3.643e-06 \| 182 \| 0.159 \| 194 \| 120 \| 11 \| 1 \| 188 \| 1 \| 157 \| Uncharacterized protein \| Uncharacterized protein; afdb-uniprot50 \| AF-V7HLN1-F1-MODEL\_V4 \| 1.0 \| 1.438e-06 \| 182 \| 0.179 \| 201 \| 131 \| 11 \| 1 \| 188 \| 1 \| 180 \| Uncharacterized protein \| Uncharacterized protein; afdb-uniprot50 \| AF-A0A3N7BFR2-F1-MODEL\_V4 \| 1.0 \| 1.357e-06 \| 182 \| 0.176 \| 210 \| 120 \| 17 \| 15 \| 189 \| 5 \| 196 \| Uncharacterized protein \| Uncharacterized protein; afdb-uniprot50 \| AF-A0A1B4FYB9-F1-MODEL\_V4 \| 1.0 \| 1.234e-05 \| 182 \| 0.138 \| 216 \| 134 \| 14 \| 6 \| 179 \| 2 \| 207 \| Uncharacterized protein \| Uncharacterized protein; afdb-uniprot50 \| AF-A0A2P8EAQ1-F1-MODEL\_V4 \| 1.0 \| 3.124e-05 \| 181 \| 0.175 \| 160 \| 100 \| 10 \| 7 \| 152 \| 1 \| 142 \| Uncharacterized protein \| Uncharacterized protein; afdb-uniprot50 \| AF-A0A2P5GLV5-F1-MODEL\_V4 \| 1.0 \| 5.584e-05 \| 181 \| 0.147 \| 170 \| 117 \| 8 \| 7 \| 168 \| 1 \| 150 \| Uncharacterized protein \| Uncharacterized protein; afdb-uniprot50 \| AF-A0A7I0KP35-F1-MODEL\_V4 \| 1.0 \| 6.272e-05 \| 181 \| 0.155 \| 167 \| 127 \| 5 \| 7 \| 173 \| 1 \| 153 \| Uncharacterized protein \| Uncharacterized protein; afdb-uniprot50 \| AF-A0A482IRV5-F1-MODEL\_V4 \| 1.0 \| 3.509e-05 \| 181 \| 0.173 \| 173 \| 125 \| 6 \| 8 \| 179 \| 1 \| 156 \| Uncharacterized protein \| Uncharacterized protein; afdb-uniprot50 \| AF-A0A747D8U5-F1-MODEL\_V4 \| 1.0 \| 0.0001335 \| 181 \| 0.133 \| 165 \| 119 \| 8 \| 7 \| 157 \| 1 \| 155 \| DUF1834 family protein \| DUF1834 family protein; afdb-uniprot50 \| AF-A0A0Q4UDE4-F1-MODEL\_V4 \| 1.0 \| 0.0001589 \| 180 \| 0.197 \| 147 \| 99 \| 7 \| 13 \| 152 \| 2 \| 136 \| Uncharacterized protein \| Uncharacterized protein; afdb-uniprot50 \| AF-A0A844GC83-F1-MODEL\_V4 \| 1.0 \| 0.0003189 \| 180 \| 0.113 \| 150 \| 117 \| 7 \| 6 \| 152 \| 3 \| 139 \| Uncharacterized protein \| Uncharacterized protein; afdb-uniprot50 \| AF-A0A418W4C9-F1-MODEL\_V4 \| 1.0 \| 5.584e-05 \| 180 \| 0.209 \| 153 \| 96 \| 8 \| 14 \| 154 \| 5 \| 144 \| Uncharacterized protein \| Uncharacterized protein; afdb-uniprot50 \| AF-A0A0Q4TYV2-F1-MODEL\_V4 \| 1.0 \| 7.913e-05 \| 180 \| 0.182 \| 148 \| 101 \| 9 \| 9 \| 150 \| 5 \| 138 \| Uncharacterized protein \| Uncharacterized protein; afdb-uniprot50 \| AF-A0A1B4SJZ4-F1-MODEL\_V4 \| 1.0 \| 4.691e-05 \| 180 \| 0.204 \| 171 \| 115 \| 8 \| 7 \| 174 \| 2 \| 154 \| Uncharacterized protein \| Uncharacterized protein; afdb-uniprot50 \| AF-A0A450VPX4-F1-MODEL\_V4 \| 1.0 \| 2.205e-05 \| 179 \| 0.107 \| 195 \| 138 \| 10 \| 1 \| 188 \| 1 \| 166 \| Gp37 protein \| Gp37 protein; afdb-uniprot50 \| AF-A0A4Y6VZN6-F1-MODEL\_V4 \| 1.0 \| 9.226e-06 \| 179 \| 0.19 \| 194 \| 125 \| 14 \| 10 \| 183 \| 7 \| 188 \| DUF1834 family protein \| DUF1834 family protein; afdb-uniprot50 \| AF-A0A089YFT4-F1-MODEL\_V4 \| 1.0 \| 7.751e-06 \| 178 \| 0.128 \| 203 \| 138 \| 11 \| 7 \| 187 \| 1 \| 186 \| Uncharacterized protein \| Uncharacterized protein; afdb-uniprot50 \| AF-A0A7W8DYF7-F1-MODEL\_V4 \| 1.0 \| 3.311e-05 \| 178 \| 0.175 \| 194 \| 118 \| 12 \| 1 \| 168 \| 1 \| 178 \| Uncharacterized protein \| Uncharacterized protein; afdb-uniprot50 \| AF-A0A5M6I4T3-F1-MODEL\_V4 \| 1.0 \| 1.748e-05 \| 178 \| 0.212 \| 188 \| 117 \| 10 \| 13 \| 187 \| 6 \| 175 \| Uncharacterized protein \| Uncharacterized protein; afdb-uniprot50 \| AF-A0A1E8PKI9-F1-MODEL\_V4 \| 1.0 \| 3.719e-05 \| 178 \| 0.156 \| 211 \| 135 \| 11 \| 8 \| 189 \| 1 \| 197 \| Uncharacterized protein \| Uncharacterized protein; afdb-uniprot50 \| AF-A0A1T1AP14-F1-MODEL\_V4 \| 1.0 \| 5.584e-05 \| 178 \| 0.141 \| 191 \| 136 \| 10 \| 7 \| 172 \| 1 \| 188 \| Uncharacterized protein \| Uncharacterized protein; afdb-uniprot50 \| AF-X6QBR1-F1-MODEL\_V4 \| 1.0 \| 8.887e-05 \| 178 \| 0.147 \| 176 \| 121 \| 10 \| 8 \| 165 \| 1 \| 165 \| PF08873 domain protein \| PF08873 domain protein; afdb-uniprot50 \| AF-A0A4V2V2Q1-F1-MODEL\_V4 \| 1.0 \| 0.0001891 \| 177 \| 0.177 \| 158 \| 98 \| 7 \| 7 \| 154 \| 1 \| 136 \| Uncharacterized protein \| Uncharacterized protein; afdb-uniprot50 \| AF-A0A1D2X525-F1-MODEL\_V4 \| 1.0 \| 8.386e-05 \| 177 \| 0.119 \| 159 \| 111 \| 9 \| 8 \| 152 \| 1 \| 144 \| Uncharacterized protein \| Uncharacterized protein; afdb-uniprot50 \| AF-A0A7X7S173-F1-MODEL\_V4 \| 1.0 \| 5.269e-05 \| 177 \| 0.168 \| 172 \| 125 \| 7 \| 7 \| 177 \| 1 \| 155 \| Uncharacterized protein \| Uncharacterized protein; afdb-uniprot50 \| AF-A0A847H596-F1-MODEL\_V4 \| 1.0 \| 0.0002528 \| 176 \| 0.113 \| 150 \| 110 \| 6 \| 7 \| 152 \| 1 \| 131 \| Uncharacterized protein \| Uncharacterized protein; afdb-uniprot50 \| AF-A0A1C0SPE6-F1-MODEL\_V4 \| 1.0 \| 0.0001589 \| 176 \| 0.161 \| 149 \| 106 \| 8 \| 14 \| 156 \| 3 \| 138 \| Uncharacterized protein \| Uncharacterized protein; afdb-uniprot50 \| AF-A0A2W6T3K4-F1-MODEL\_V4 \| 1.0 \| 0.0001188 \| 176 \| 0.131 \| 160 \| 112 \| 10 \| 14 \| 166 \| 9 \| 148 \| Uncharacterized protein \| Uncharacterized protein; afdb-uniprot50 \| AF-A0A4V3WAW4-F1-MODEL\_V4 \| 1.0 \| 0.0001891 \| 176 \| 0.159 \| 163 \| 110 \| 10 \| 5 \| 156 \| 2 \| 148 \| Uncharacterized protein \| Uncharacterized protein; afdb-uniprot50 \| AF-A0A1R1JMQ1-F1-MODEL\_V4 \| 1.0 \| 8.706e-06 \| 176 \| 0.118 \| 203 \| 134 \| 10 \| 13 \| 185 \| 3 \| 190 \| Uncharacterized protein \| Uncharacterized protein; afdb-uniprot50 \| AF-A0A2M9WHN8-F1-MODEL\_V4 \| 1.0 \| 7.045e-05 \| 176 \| 0.137 \| 175 \| 119 \| 10 \| 12 \| 166 \| 5 \| 167 \| Uncharacterized protein \| Uncharacterized protein; afdb-uniprot50 \| AF-A0A559Q8J5-F1-MODEL\_V4 \| 1.0 \| 9.419e-05 \| 175 \| 0.166 \| 168 \| 110 \| 8 \| 7 \| 165 \| 4 \| 150 \| Uncharacterized protein \| Uncharacterized protein; afdb-uniprot50 \| AF-A0A4R1FMA6-F1-MODEL\_V4 \| 1.0 \| 0.0001589 \| 175 \| 0.113 \| 168 \| 130 \| 6 \| 7 \| 172 \| 2 \| 152 \| Gp37 protein \| Gp37 protein; afdb-uniprot50 \| AF-A0A1Q3QXE1-F1-MODEL\_V4 \| 1.0 \| 3.509e-05 \| 175 \| 0.139 \| 172 \| 129 \| 10 \| 5 \| 165 \| 3 \| 166 \| Uncharacterized protein \| Uncharacterized protein; afdb-uniprot50 \| AF-Q1MNR2-F1-MODEL\_V4 \| 1.0 \| 2.782e-05 \| 175 \| 0.174 \| 189 \| 125 \| 11 \| 8 \| 177 \| 1 \| 177 \| Uncharacterized protein \| Uncharacterized protein; afdb-uniprot50 \| AF-A0A2V4BZS6-F1-MODEL\_V4 \| 1.0 \| 0.0001058 \| 174 \| 0.096 \| 186 \| 130 \| 11 \| 1 \| 166 \| 2 \| 169 \| Uncharacterized protein \| Uncharacterized protein; afdb-uniprot50 \| AF-A0A0M4M4N7-F1-MODEL\_V4 \| 1.0 \| 4.427e-05 \| 174 \| 0.123 \| 186 \| 136 \| 7 \| 10 \| 187 \| 5 \| 171 \| Putative prophage protein \| Putative prophage protein; afdb-uniprot50 \| AF-A0A376HZ26-F1-MODEL\_V4 \| 1.0 \| 3.124e-05 \| 174 \| 0.095 \| 189 \| 141 \| 10 \| 8 \| 187 \| 1 \| 168 \| Mu-like prophage protein gp37 \| Mu-like prophage protein gp37; afdb-uniprot50 \| AF-A0A6I1NPD5-F1-MODEL\_V4 \| 1.0 \| 7.314e-06 \| 174 \| 0.154 \| 188 \| 117 \| 13 \| 15 \| 177 \| 5 \| 175 \| Uncharacterized protein \| Uncharacterized protein; afdb-uniprot50 \| AF-W0BVV9-F1-MODEL\_V4 \| 1.0 \| 5.918e-05 \| 174 \| 0.146 \| 177 \| 123 \| 9 \| 8 \| 166 \| 1 \| 167 \| Uncharacterized protein \| Uncharacterized protein; afdb-uniprot50 \| AF-A0A1F4QWC5-F1-MODEL\_V4 \| 1.0 \| 0.0001784 \| 173 \| 0.196 \| 153 \| 102 \| 7 \| 7 \| 155 \| 1 \| 136 \| Uncharacterized protein \| Uncharacterized protein; afdb-uniprot50 \| AF-A0A318KU15-F1-MODEL\_V4 \| 1.0 \| 8.887e-05 \| 173 \| 0.111 \| 170 \| 123 \| 8 \| 7 \| 165 \| 2 \| 154 \| Gp37 protein \| Gp37 protein; afdb-uniprot50 \| AF-G4CJF9-F1-MODEL\_V4 \| 1.0 \| 0.0001891 \| 173 \| 0.111 \| 170 \| 130 \| 7 \| 8 \| 174 \| 3 \| 154 \| Phage Gp37 protein \| Phage Gp37 protein; afdb-uniprot50 \| AF-A0A3G7TJT5-F1-MODEL\_V4 \| 1.0 \| 0.0001188 \| 173 \| 0.142 \| 175 \| 133 \| 6 \| 2 \| 174 \| 1 \| 160 \| Uncharacterized protein \| Uncharacterized protein; afdb-uniprot50 \| AF-A0A6L8L6D7-F1-MODEL\_V4 \| 1.0 \| 4.177e-05 \| 173 \| 0.144 \| 194 \| 137 \| 10 \| 8 \| 187 \| 1 \| 179 \| DUF1834 family protein \| DUF1834 family protein; afdb-uniprot50 \| AF-A0A143DC32-F1-MODEL\_V4 \| 1.0 \| 5.584e-05 \| 173 \| 0.14 \| 178 \| 120 \| 9 \| 13 \| 172 \| 2 \| 164 \| Uncharacterized protein \| Uncharacterized protein; afdb-uniprot50 \| AF-A0A2Z6GCD5-F1-MODEL\_V4 \| 1.0 \| 0.0002004 \| 172 \| 0.141 \| 148 \| 108 \| 8 \| 7 \| 152 \| 1 \| 131 \| Uncharacterized protein \| Uncharacterized protein; afdb-uniprot50 \| AF-A0A1G3JTJ3-F1-MODEL\_V4 \| 1.0 \| 0.0002386 \| 172 \| 0.16 \| 150 \| 106 \| 7 \| 7 \| 152 \| 2 \| 135 \| Uncharacterized protein \| Uncharacterized protein; afdb-uniprot50 \| AF-A0A143XXV9-F1-MODEL\_V4 \| 1.0 \| 0.0001784 \| 172 \| 0.144 \| 159 \| 112 \| 9 \| 10 \| 155 \| 2 \| 149 \| Uncharacterized protein \| Uncharacterized protein; afdb-uniprot50 \| AF-A0A1J5D4A8-F1-MODEL\_V4 \| 1.0 \| 0.0001121 \| 172 \| 0.118 \| 169 \| 122 \| 9 \| 6 \| 163 \| 2 \| 154 \| Uncharacterized protein \| Uncharacterized protein; afdb-uniprot50 \| AF-A0A7L5ABA4-F1-MODEL\_V4 \| 1.0 \| 0.0001684 \| 172 \| 0.137 \| 174 \| 138 \| 6 \| 1 \| 172 \| 2 \| 165 \| Uncharacterized protein \| Uncharacterized protein; afdb-uniprot50 \| AF-A0A5N0TG56-F1-MODEL\_V4 \| 1.0 \| 0.0002679 \| 171 \| 0.151 \| 145 \| 101 \| 9 \| 13 \| 155 \| 4 \| 128 \| Uncharacterized protein \| Uncharacterized protein; afdb-uniprot50 \| AF-A0A5A8F1R4-F1-MODEL\_V4 \| 1.0 \| 0.0002251 \| 171 \| 0.142 \| 154 \| 105 \| 8 \| 8 \| 153 \| 1 \| 135 \| Uncharacterized protein \| Uncharacterized protein; afdb-uniprot50 \| AF-A0A5E8D1X6-F1-MODEL\_V4 \| 1.0 \| 0.0002386 \| 171 \| 0.134 \| 163 \| 106 \| 12 \| 10 \| 159 \| 2 \| 142 \| Uncharacterized protein \| Uncharacterized protein; afdb-uniprot50 \| AF-C1FA28-F1-MODEL\_V4 \| 1.0 \| 5.918e-05 \| 171 \| 0.168 \| 178 \| 112 \| 11 \| 1 \| 162 \| 1 \| 158 \| Uncharacterized protein \| Uncharacterized protein; afdb-uniprot50 \| AF-A0A1B7KXZ2-F1-MODEL\_V4 \| 1.0 \| 7.314e-06 \| 171 \| 0.13 \| 200 \| 141 \| 12 \| 7 \| 189 \| 1 \| 184 \| Uncharacterized protein \| Uncharacterized protein; afdb-uniprot50 \| AF-A0A3M3KWV5-F1-MODEL\_V4 \| 1.0 \| 9.226e-06 \| 171 \| 0.134 \| 208 \| 131 \| 14 \| 15 \| 189 \| 5 \| 196 \| Uncharacterized protein \| Uncharacterized protein; afdb-uniprot50 \| AF-A0A3M5JCG4-F1-MODEL\_V4 \| 1.0 \| 9.226e-06 \| 171 \| 0.134 \| 208 \| 131 \| 14 \| 15 \| 189 \| 5 \| 196 \| Uncharacterized protein \| Uncharacterized protein; afdb-uniprot50 \| AF-A0A1Z8PY23-F1-MODEL\_V4 \| 1.0 \| 9.778e-06 \| 171 \| 0.163 \| 202 \| 117 \| 16 \| 8 \| 178 \| 1 \| 181 \| Uncharacterized protein \| Uncharacterized protein; afdb-uniprot50 \| AF-A0A2D0JLA8-F1-MODEL\_V4 \| 1.0 \| 6.272e-05 \| 171 \| 0.144 \| 173 \| 116 \| 10 \| 8 \| 162 \| 1 \| 159 \| Uncharacterized protein \| Uncharacterized protein; afdb-uniprot50 \| AF-A0A2D9DW63-F1-MODEL\_V4 \| 1.0 \| 0.0001784 \| 170 \| 0.135 \| 148 \| 103 \| 7 \| 14 \| 155 \| 3 \| 131 \| Uncharacterized protein \| Uncharacterized protein; afdb-uniprot50 \| AF-A0A560II09-F1-MODEL\_V4 \| 1.0 \| 0.0002004 \| 170 \| 0.164 \| 152 \| 103 \| 10 \| 11 \| 152 \| 2 \| 139 \| Uncharacterized protein \| Uncharacterized protein; afdb-uniprot50 \| AF-A0A7V1ZNY4-F1-MODEL\_V4 \| 1.0 \| 0.000284 \| 170 \| 0.125 \| 168 \| 117 \| 8 \| 8 \| 165 \| 1 \| 148 \| DUF1834 family protein \| DUF1834 family protein; afdb-uniprot50 \| AF-A0A329W903-F1-MODEL\_V4 \| 1.0 \| 0.0001259 \| 170 \| 0.173 \| 167 \| 124 \| 4 \| 7 \| 173 \| 1 \| 153 \| Uncharacterized protein \| Uncharacterized protein; afdb-uniprot50 \| AF-A0A6M0FVW8-F1-MODEL\_V4 \| 1.0 \| 0.0001058 \| 170 \| 0.116 \| 171 \| 128 \| 8 \| 7 \| 166 \| 3 \| 161 \| Uncharacterized protein \| Uncharacterized protein; afdb-uniprot50 \| AF-A0A6B2KN64-F1-MODEL\_V4 \| 1.0 \| 2.948e-05 \| 170 \| 0.123 \| 203 \| 134 \| 12 \| 15 \| 189 \| 5 \| 191 \| Uncharacterized protein \| Uncharacterized protein; afdb-uniprot50 \| AF-A0A7T8TA87-F1-MODEL\_V4 \| 1.0 \| 1.234e-05 \| 170 \| 0.123 \| 211 \| 130 \| 14 \| 15 \| 189 \| 5 \| 196 \| Uncharacterized protein \| Uncharacterized protein; afdb-uniprot50 \| AF-B9Z305-F1-MODEL\_V4 \| 1.0 \| 3.941e-05 \| 170 \| 0.155 \| 174 \| 120 \| 10 \| 9 \| 166 \| 7 \| 169 \| Uncharacterized protein \| Uncharacterized protein; afdb-uniprot50 \| AF-A0A6M8SQE1-F1-MODEL\_V4 \| 1.0 \| 0.0005379 \| 169 \| 0.132 \| 158 \| 113 \| 8 \| 9 \| 165 \| 4 \| 138 \| Uncharacterized protein \| Uncharacterized protein; afdb-uniprot50 \| AF-A0A376YHH1-F1-MODEL\_V4 \| 1.0 \| 4.972e-05 \| 169 \| 0.131 \| 152 \| 120 \| 5 \| 7 \| 155 \| 1 \| 143 \| Phage-related protein Gp37-like protein \| Phage-related protein Gp37-like protein; afdb-uniprot50 \| AF-C1D844-F1-MODEL\_V4 \| 1.0 \| 0.0003797 \| 169 \| 0.124 \| 161 \| 117 \| 9 \| 8 \| 164 \| 1 \| 141 \| Gp37 domain containing protein \| Gp37 domain containing protein; afdb-uniprot50 \| AF-A0A763CFX9-F1-MODEL\_V4 \| 1.0 \| 0.0001589 \| 169 \| 0.162 \| 179 \| 124 \| 8 \| 7 \| 177 \| 1 \| 161 \| Uncharacterized protein \| Uncharacterized protein; afdb-uniprot50 \| AF-A0A653JWK4-F1-MODEL\_V4 \| 1.0 \| 5.584e-05 \| 169 \| 0.164 \| 194 \| 125 \| 10 \| 12 \| 179 \| 5 \| 187 \| Phage gp37-like protein \| Phage gp37-like protein; afdb-uniprot50 \| AF-A0A812IPU7-F1-MODEL\_V4 \| 1.0 \| 0.0001188 \| 169 \| 0.127 \| 157 \| 117 \| 7 \| 2 \| 156 \| 569 \| 707 \| Hypothetical protein \| Hypothetical protein; afdb-uniprot50 \| AF-A0A6B0W596-F1-MODEL\_V4 \| 1.0 \| 0.000338 \| 168 \| 0.118 \| 152 \| 124 \| 5 \| 7 \| 156 \| 1 \| 144 \| Uncharacterized protein \| Uncharacterized protein; afdb-uniprot50 \| AF-A0A7C6JL55-F1-MODEL\_V4 \| 1.0 \| 0.0001684 \| 168 \| 0.16 \| 156 \| 101 \| 8 \| 10 \| 152 \| 2 \| 140 \| Uncharacterized protein \| Uncharacterized protein; afdb-uniprot50 \| AF-A0A7C9K240-F1-MODEL\_V4 \| 1.0 \| 2.948e-05 \| 168 \| 0.105 \| 208 \| 148 \| 13 \| 1 \| 186 \| 4 \| 195 \| Uncharacterized protein \| Uncharacterized protein; afdb-uniprot50 \| AF-A0A2X2DE00-F1-MODEL\_V4 \| 1.0 \| 0.0001259 \| 168 \| 0.125 \| 184 \| 131 \| 10 \| 8 \| 176 \| 1 \| 169 \| Mu-like prophage protein gp37 \| Mu-like prophage protein gp37; afdb-uniprot50 \| AF-A0A7T9FPJ0-F1-MODEL\_V4 \| 1.0 \| 3.311e-05 \| 168 \| 0.162 \| 191 \| 128 \| 11 \| 1 \| 165 \| 1 \| 185 \| DUF1834 family protein \| DUF1834 family protein; afdb-uniprot50 \| AF-A0A221KCS7-F1-MODEL\_V4 \| 1.0 \| 0.0002528 \| 167 \| 0.166 \| 156 \| 104 \| 12 \| 8 \| 153 \| 13 \| 152 \| Uncharacterized protein \| Uncharacterized protein; afdb-uniprot50 \| AF-A0A4R2GW33-F1-MODEL\_V4 \| 1.0 \| 0.0002251 \| 167 \| 0.109 \| 165 \| 128 \| 8 \| 2 \| 163 \| 16 \| 164 \| Gp37 protein \| Gp37 protein; afdb-uniprot50 \| AF-Q126A0-F1-MODEL\_V4 \| 1.0 \| 5.269e-05 \| 167 \| 0.138 \| 195 \| 133 \| 12 \| 7 \| 182 \| 1 \| 179 \| Uncharacterized protein \| Uncharacterized protein; afdb-uniprot50 \| AF-A0A2E2ISM1-F1-MODEL\_V4 \| 1.0 \| 3.941e-05 \| 167 \| 0.188 \| 196 \| 121 \| 13 \| 12 \| 182 \| 5 \| 187 \| Uncharacterized protein \| Uncharacterized protein; afdb-uniprot50 \| AF-A0A085ELB4-F1-MODEL\_V4 \| 1.0 \| 0.0002004 \| 166 \| 0.128 \| 171 \| 116 \| 12 \| 7 \| 163 \| 3 \| 154 \| Uncharacterized protein \| Uncharacterized protein; afdb-uniprot50 \| AF-A0A840LZ35-F1-MODEL\_V4 \| 1.0 \| 8.386e-05 \| 166 \| 0.138 \| 180 \| 122 \| 9 \| 1 \| 165 \| 1 \| 162 \| Uncharacterized protein \| Uncharacterized protein; afdb-uniprot50 \| AF-A0A3S0EG19-F1-MODEL\_V4 \| 1.0 \| 5.269e-05 \| 166 \| 0.146 \| 177 \| 122 \| 9 \| 7 \| 166 \| 1 \| 165 \| Uncharacterized protein \| Uncharacterized protein; afdb-uniprot50 \| AF-A0A0L0GQ01-F1-MODEL\_V4 \| 1.0 \| 9.419e-05 \| 166 \| 0.124 \| 225 \| 137 \| 12 \| 8 \| 189 \| 1 \| 208 \| Uncharacterized protein \| Uncharacterized protein; afdb-uniprot50 \| AF-A0A327J9N9-F1-MODEL\_V4 \| 1.0 \| 0.0007192 \| 165 \| 0.089 \| 146 \| 113 \| 7 \| 9 \| 152 \| 5 \| 132 \| Uncharacterized protein \| Uncharacterized protein; afdb-uniprot50 \| AF-A0A0C1G7G9-F1-MODEL\_V4 \| 1.0 \| 5.584e-05 \| 165 \| 0.144 \| 180 \| 121 \| 12 \| 2 \| 173 \| 3 \| 157 \| Uncharacterized protein \| Uncharacterized protein; afdb-uniprot50 \| AF-A0A2D3T3I7-F1-MODEL\_V4 \| 1.0 \| 7.466e-05 \| 165 \| 0.122 \| 171 \| 120 \| 6 \| 15 \| 166 \| 5 \| 164 \| Uncharacterized protein \| Uncharacterized protein; afdb-uniprot50 \| AF-A0A2A2H0E7-F1-MODEL\_V4 \| 1.0 \| 4.972e-05 \| 165 \| 0.106 \| 187 \| 141 \| 11 \| 15 \| 189 \| 5 \| 177 \| Uncharacterized protein \| Uncharacterized protein; afdb-uniprot50 \| AF-A0A3R6C1V5-F1-MODEL\_V4 \| 1.0 \| 0.0005379 \| 164 \| 0.09 \| 154 \| 107 \| 7 \| 18 \| 156 \| 2 \| 137 \| Uncharacterized protein \| Uncharacterized protein; afdb-uniprot50 \| AF-A0A0F5PGA2-F1-MODEL\_V4 \| 1.0 \| 0.0001589 \| 164 \| 0.173 \| 150 \| 94 \| 10 \| 17 \| 153 \| 3 \| 135 \| Uncharacterized protein \| Uncharacterized protein; afdb-uniprot50 \| AF-A0A2N8NK38-F1-MODEL\_V4 \| 1.0 \| 0.0001684 \| 164 \| 0.12 \| 166 \| 111 \| 10 \| 1 \| 152 \| 1 \| 145 \| Uncharacterized protein \| Uncharacterized protein; afdb-uniprot50 \| AF-A0A7U6QP29-F1-MODEL\_V4 \| 1.0 \| 0.0002679 \| 164 \| 0.117 \| 170 \| 123 \| 8 \| 1 \| 165 \| 1 \| 148 \| Uncharacterized protein \| Uncharacterized protein; afdb-uniprot50 \| AF-F3PKG9-F1-MODEL\_V4 \| 1.0 \| 0.0002251 \| 164 \| 0.151 \| 165 \| 109 \| 10 \| 8 \| 157 \| 1 \| 149 \| Uncharacterized protein \| Uncharacterized protein; afdb-uniprot50 \| AF-A0A1M7RI50-F1-MODEL\_V4 \| 1.0 \| 0.0001414 \| 164 \| 0.151 \| 165 \| 121 \| 7 \| 7 \| 166 \| 1 \| 151 \| Gp37 protein \| Gp37 protein; afdb-uniprot50 \| AF-A0A225E4L8-F1-MODEL\_V4 \| 1.0 \| 5.584e-05 \| 164 \| 0.156 \| 179 \| 120 \| 14 \| 6 \| 163 \| 7 \| 175 \| Uncharacterized protein \| Uncharacterized protein; afdb-uniprot50 \| AF-D2TJ30-F1-MODEL\_V4 \| 1.0 \| 7.913e-05 \| 164 \| 0.11 \| 209 \| 131 \| 11 \| 8 \| 174 \| 1 \| 196 \| Hypothetical prophage protein \| Hypothetical prophage protein; afdb-uniprot50 \| AF-A0A0G0CA56-F1-MODEL\_V4 \| 1.0 \| 0.001213 \| 163 \| 0.117 \| 153 \| 116 \| 7 \| 7 \| 157 \| 1 \| 136 \| Uncharacterized protein \| Uncharacterized protein; afdb-uniprot50 \| AF-A0A509KZR1-F1-MODEL\_V4 \| 1.0 \| 0.0003009 \| 163 \| 0.092 \| 152 \| 112 \| 7 \| 13 \| 157 \| 9 \| 141 \| Uncharacterized protein \| Uncharacterized protein; afdb-uniprot50 \| AF-L1P217-F1-MODEL\_V4 \| 1.0 \| 0.0001414 \| 163 \| 0.12 \| 182 \| 134 \| 8 \| 1 \| 179 \| 1 \| 159 \| Gp37 protein \| Gp37 protein; afdb-uniprot50 \| AF-A0A4Q3RNJ9-F1-MODEL\_V4 \| 1.0 \| 9.419e-05 \| 163 \| 0.127 \| 181 \| 130 \| 9 \| 7 \| 174 \| 2 \| 167 \| Uncharacterized protein \| Uncharacterized protein; afdb-uniprot50 \| AF-A0A5C9D9K9-F1-MODEL\_V4 \| 1.0 \| 0.0001891 \| 163 \| 0.151 \| 172 \| 117 \| 12 \| 9 \| 164 \| 14 \| 172 \| Uncharacterized protein \| Uncharacterized protein; afdb-uniprot50 \| AF-A0A6A2FMX2-F1-MODEL\_V4 \| 1.0 \| 8.386e-05 \| 163 \| 0.142 \| 182 \| 118 \| 9 \| 7 \| 166 \| 2 \| 167 \| Uncharacterized protein \| Uncharacterized protein; afdb-uniprot50 \| AF-A0A837QHE5-F1-MODEL\_V4 \| 1.0 \| 3.124e-05 \| 163 \| 0.117 \| 178 \| 135 \| 7 \| 25 \| 187 \| 7 \| 177 \| Uncharacterized protein \| Uncharacterized protein; afdb-uniprot50 \| AF-A0A2V3UBZ9-F1-MODEL\_V4 \| 1.0 \| 5.918e-05 \| 163 \| 0.164 \| 182 \| 125 \| 12 \| 8 \| 179 \| 3 \| 167 \| Uncharacterized protein DUF1834 \| Uncharacterized protein DUF1834; afdb-uniprot50 \| AF-A0A437T2J4-F1-MODEL\_V4 \| 1.0 \| 3.311e-05 \| 163 \| 0.105 \| 200 \| 136 \| 9 \| 15 \| 189 \| 5 \| 186 \| Uncharacterized protein \| Uncharacterized protein; afdb-uniprot50 \| AF-A0A7C4DI82-F1-MODEL\_V4 \| 1.0 \| 0.001286 \| 162 \| 0.093 \| 150 \| 117 \| 7 \| 8 \| 154 \| 1 \| 134 \| DUF1834 family protein \| DUF1834 family protein; afdb-uniprot50 \| AF-A0A2D5PCB8-F1-MODEL\_V4 \| 1.0 \| 0.0001891 \| 162 \| 0.132 \| 158 \| 111 \| 11 \| 7 \| 159 \| 2 \| 138 \| Uncharacterized protein \| Uncharacterized protein; afdb-uniprot50 \| AF-A0A3C0TZ33-F1-MODEL\_V4 \| 1.0 \| 0.0003009 \| 162 \| 0.136 \| 147 \| 110 \| 7 \| 15 \| 153 \| 5 \| 142 \| Uncharacterized protein \| Uncharacterized protein; afdb-uniprot50 \| AF-A0A5P9CRF5-F1-MODEL\_V4 \| 1.0 \| 0.0001684 \| 162 \| 0.162 \| 160 \| 118 \| 7 \| 9 \| 166 \| 11 \| 156 \| Uncharacterized protein \| Uncharacterized protein; afdb-uniprot50 \| AF-A0A011NCS5-F1-MODEL\_V4 \| 1.0 \| 4.972e-05 \| 162 \| 0.152 \| 184 \| 127 \| 11 \| 1 \| 177 \| 1 \| 162 \| Uncharacterized protein \| Uncharacterized protein; afdb-uniprot50 \| AF-B3QTJ8-F1-MODEL\_V4 \| 1.0 \| 0.0001058 \| 162 \| 0.102 \| 176 \| 131 \| 8 \| 9 \| 168 \| 8 \| 172 \| Uncharacterized protein \| Uncharacterized protein; afdb-uniprot50 \| AF-A0A0F9GHP4-F1-MODEL\_V4 \| 1.0 \| 0.0001414 \| 162 \| 0.116 \| 163 \| 125 \| 8 \| 1 \| 154 \| 20 \| 172 \| Uncharacterized protein \| Uncharacterized protein; afdb-uniprot50 \| AF-A0A4V3AUT9-F1-MODEL\_V4 \| 1.0 \| 0.0002386 \| 162 \| 0.122 \| 172 \| 137 \| 4 \| 7 \| 178 \| 1 \| 158 \| Uncharacterized protein \| Uncharacterized protein; afdb-uniprot50 \| AF-A0A1Y1Q6H6-F1-MODEL\_V4 \| 1.0 \| 7.466e-05 \| 162 \| 0.102 \| 176 \| 125 \| 10 \| 12 \| 176 \| 3 \| 156 \| Uncharacterized protein \| Uncharacterized protein; afdb-uniprot50 \| AF-A0A7C1A4S1-F1-MODEL\_V4 \| 1.0 \| 0.000284 \| 161 \| 0.12 \| 166 \| 118 \| 9 \| 6 \| 155 \| 3 \| 156 \| Uncharacterized protein \| Uncharacterized protein; afdb-uniprot50 \| AF-A0A7I6PS33-F1-MODEL\_V4 \| 1.0 \| 8.887e-05 \| 161 \| 0.137 \| 167 \| 122 \| 10 \| 14 \| 166 \| 4 \| 162 \| Uncharacterized protein \| Uncharacterized protein; afdb-uniprot50 \| AF-A0A7Y7NQN0-F1-MODEL\_V4 \| 1.0 \| 0.0001335 \| 160 \| 0.14 \| 150 \| 111 \| 8 \| 8 \| 148 \| 1 \| 141 \| Uncharacterized protein \| Uncharacterized protein; afdb-uniprot50 \| AF-A0A381E3W2-F1-MODEL\_V4 \| 1.0 \| 0.0005379 \| 160 \| 0.12 \| 158 \| 123 \| 10 \| 7 \| 156 \| 2 \| 151 \| Uncharacterized protein \| Uncharacterized protein; afdb-uniprot50 \| AF-R6T089-F1-MODEL\_V4 \| 1.0 \| 0.0001589 \| 160 \| 0.134 \| 171 \| 103 \| 10 \| 10 \| 154 \| 2 \| 153 \| Uncharacterized protein \| Uncharacterized protein; afdb-uniprot50 \| AF-A0A1C3HHN1-F1-MODEL\_V4 \| 1.0 \| 0.0002124 \| 160 \| 0.156 \| 172 \| 129 \| 6 \| 7 \| 177 \| 1 \| 157 \| Gp37 protein \| Gp37 protein; afdb-uniprot50 \| AF-A0A515EKE4-F1-MODEL\_V4 \| 1.0 \| 3.941e-05 \| 160 \| 0.137 \| 204 \| 132 \| 11 \| 10 \| 178 \| 3 \| 197 \| Uncharacterized protein \| Uncharacterized protein; afdb-uniprot50 \| AF-A0A845BQV7-F1-MODEL\_V4 \| 1.0 \| 0.0001891 \| 159 \| 0.15 \| 153 \| 110 \| 6 \| 7 \| 156 \| 3 \| 138 \| Uncharacterized protein \| Uncharacterized protein; afdb-uniprot50 \| AF-A0A249DY56-F1-MODEL\_V4 \| 1.0 \| 0.0001335 \| 159 \| 0.146 \| 157 \| 106 \| 7 \| 15 \| 159 \| 5 \| 145 \| Uncharacterized protein \| Uncharacterized protein; afdb-uniprot50 \| AF-U6Q2Q7-F1-MODEL\_V4 \| 1.0 \| 0.00108 \| 159 \| 0.168 \| 160 \| 114 \| 11 \| 7 \| 156 \| 1 \| 151 \| Uncharacterized protein \| Uncharacterized protein; afdb-uniprot50 \| AF-A0A1N6M5S0-F1-MODEL\_V4 \| 1.0 \| 3.509e-05 \| 159 \| 0.128 \| 194 \| 136 \| 12 \| 1 \| 173 \| 3 \| 184 \| Uncharacterized protein \| Uncharacterized protein; afdb-uniprot50 \| AF-A0A0Q8STR6-F1-MODEL\_V4 \| 1.0 \| 0.0005701 \| 158 \| 0.132 \| 159 \| 106 \| 8 \| 7 \| 156 \| 1 \| 136 \| Uncharacterized protein \| Uncharacterized protein; afdb-uniprot50 \| AF-A0A0T5PNX3-F1-MODEL\_V4 \| 1.0 \| 0.00108 \| 158 \| 0.131 \| 152 \| 118 \| 8 \| 10 \| 156 \| 6 \| 148 \| Uncharacterized protein \| Uncharacterized protein; afdb-uniprot50 \| AF-G2IX39-F1-MODEL\_V4 \| 1.0 \| 0.0002386 \| 158 \| 0.127 \| 172 \| 125 \| 9 \| 8 \| 174 \| 1 \| 152 \| Gp37 protein \| Gp37 protein; afdb-uniprot50 \| AF-A0A2A2GYG9-F1-MODEL\_V4 \| 1.0 \| 0.0007192 \| 157 \| 0.138 \| 152 \| 105 \| 8 \| 8 \| 153 \| 1 \| 132 \| Uncharacterized protein \| Uncharacterized protein; afdb-uniprot50 \| AF-A0A1G0MRP2-F1-MODEL\_V4 \| 1.0 \| 0.0003797 \| 157 \| 0.082 \| 157 \| 128 \| 5 \| 8 \| 154 \| 1 \| 151 \| Uncharacterized protein \| Uncharacterized protein; afdb-uniprot50 \| AF-A0A1J0WHQ0-F1-MODEL\_V4 \| 1.0 \| 0.0002124 \| 157 \| 0.159 \| 169 \| 104 \| 12 \| 1 \| 153 \| 1 \| 147 \| Uncharacterized protein \| Uncharacterized protein; afdb-uniprot50 \| AF-A0A178HLQ9-F1-MODEL\_V4 \| 1.0 \| 0.0001784 \| 157 \| 0.123 \| 178 \| 127 \| 8 \| 8 \| 179 \| 1 \| 155 \| Uncharacterized protein \| Uncharacterized protein; afdb-uniprot50 \| AF-A0A3M1IM86-F1-MODEL\_V4 \| 1.0 \| 0.0002251 \| 157 \| 0.143 \| 174 \| 113 \| 9 \| 10 \| 172 \| 2 \| 150 \| Uncharacterized protein \| Uncharacterized protein; afdb-uniprot50 \| AF-A0A0J7L1A0-F1-MODEL\_V4 \| 1.0 \| 0.0002528 \| 157 \| 0.105 \| 189 \| 131 \| 10 \| 4 \| 166 \| 2 \| 178 \| Uncharacterized protein \| Uncharacterized protein; afdb-uniprot50 \| AF-A0A1G6LS75-F1-MODEL\_V4 \| 1.0 \| 0.0002386 \| 157 \| 0.089 \| 179 \| 142 \| 4 \| 1 \| 166 \| 1 \| 171 \| Uncharacterized protein \| Uncharacterized protein; afdb-uniprot50 \| AF-A0A7W4IJG5-F1-MODEL\_V4 \| 1.0 \| 0.0001121 \| 157 \| 0.129 \| 193 \| 141 \| 11 \| 1 \| 179 \| 1 \| 180 \| Uncharacterized protein \| Uncharacterized protein; afdb-uniprot50 \| AF-A0A2D6E5V7-F1-MODEL\_V4 \| 1.0 \| 4.972e-05 \| 157 \| 0.104 \| 172 \| 128 \| 11 \| 7 \| 165 \| 5 \| 163 \| Uncharacterized protein \| Uncharacterized protein; afdb-uniprot50 \| AF-A0A1V5PMC3-F1-MODEL\_V4 \| 1.0 \| 0.00108 \| 156 \| 0.104 \| 153 \| 115 \| 8 \| 7 \| 155 \| 1 \| 135 \| Gp37 protein \| Gp37 protein; afdb-uniprot50 \| AF-A0A4Q3M822-F1-MODEL\_V4 \| 1.0 \| 0.0001684 \| 156 \| 0.15 \| 159 \| 102 \| 9 \| 8 \| 152 \| 1 \| 140 \| Uncharacterized protein \| Uncharacterized protein; afdb-uniprot50 \| AF-X2HAW8-F1-MODEL\_V4 \| 1.0 \| 0.0002124 \| 156 \| 0.093 \| 171 \| 127 \| 10 \| 1 \| 166 \| 1 \| 148 \| Uncharacterized protein \| Uncharacterized protein; afdb-uniprot50 \| AF-A0A4Y9VRF7-F1-MODEL\_V4 \| 1.0 \| 6.272e-05 \| 156 \| 0.133 \| 203 \| 133 \| 13 \| 13 \| 189 \| 2 \| 187 \| Uncharacterized protein \| Uncharacterized protein; afdb-uniprot50 \| AF-A0A2S5TZJ7-F1-MODEL\_V4 \| 1.0 \| 0.0001589 \| 156 \| 0.132 \| 204 \| 143 \| 13 \| 1 \| 189 \| 1 \| 185 \| Uncharacterized protein \| Uncharacterized protein; afdb-uniprot50 \| AF-A0A826W9J1-F1-MODEL\_V4 \| 1.0 \| 0.0002386 \| 156 \| 0.142 \| 169 \| 111 \| 11 \| 8 \| 155 \| 1 \| 156 \| DUF1834 family protein \| DUF1834 family protein; afdb-uniprot50 \| AF-A0A545T5R9-F1-MODEL\_V4 \| 1.0 \| 0.0001589 \| 156 \| 0.122 \| 221 \| 141 \| 9 \| 1 \| 182 \| 1 \| 207 \| Uncharacterized protein \| Uncharacterized protein; afdb-uniprot50 \| AF-A0A7V3JM37-F1-MODEL\_V4 \| 1.0 \| 0.0004519 \| 155 \| 0.12 \| 158 \| 114 \| 10 \| 8 \| 155 \| 1 \| 143 \| Uncharacterized protein \| Uncharacterized protein; afdb-uniprot50 \| AF-A0A2W0B4M6-F1-MODEL\_V4 \| 1.0 \| 0.0009616 \| 155 \| 0.131 \| 152 \| 115 \| 8 \| 8 \| 151 \| 4 \| 146 \| Uncharacterized protein \| Uncharacterized protein; afdb-uniprot50 \| AF-A0A411WIP5-F1-MODEL\_V4 \| 1.0 \| 0.0006786 \| 155 \| 0.118 \| 160 \| 123 \| 6 \| 7 \| 165 \| 4 \| 146 \| Uncharacterized protein \| Uncharacterized protein; afdb-uniprot50 \| AF-A0A380MYP2-F1-MODEL\_V4 \| 1.0 \| 0.001444 \| 155 \| 0.1 \| 160 \| 126 \| 9 \| 8 \| 156 \| 3 \| 155 \| Uncharacterized protein \| Uncharacterized protein; afdb-uniprot50 \| AF-A0A0Q3ETN6-F1-MODEL\_V4 \| 1.0 \| 9.982e-05 \| 155 \| 0.138 \| 180 \| 120 \| 9 \| 2 \| 166 \| 1 \| 160 \| Uncharacterized protein \| Uncharacterized protein; afdb-uniprot50 \| AF-A0A7K1U6C2-F1-MODEL\_V4 \| 1.0 \| 0.000338 \| 155 \| 0.127 \| 173 \| 124 \| 9 \| 9 \| 168 \| 5 \| 163 \| Uncharacterized protein \| Uncharacterized protein; afdb-uniprot50 \| AF-A0A2N1XYT5-F1-MODEL\_V4 \| 1.0 \| 9.982e-05 \| 155 \| 0.109 \| 191 \| 129 \| 13 \| 7 \| 177 \| 1 \| 170 \| Uncharacterized protein \| Uncharacterized protein; afdb-uniprot50 \| AF-A0A3G9GEK2-F1-MODEL\_V4 \| 1.0 \| 9.982e-05 \| 155 \| 0.139 \| 229 \| 128 \| 15 \| 12 \| 188 \| 6 \| 217 \| Mu-like prophage FluMu protein gp37 \| Mu-like prophage FluMu protein gp37; afdb-uniprot50 \| AF-A0A2N9YI19-F1-MODEL\_V4 \| 1.0 \| 0.0008078 \| 155 \| 0.153 \| 163 \| 118 \| 9 \| 1 \| 156 \| 147 \| 296 \| Uncharacterized protein \| Uncharacterized protein; afdb-uniprot50 \| AF-A0A661DU83-F1-MODEL\_V4 \| 1.0 \| 0.0004789 \| 154 \| 0.107 \| 149 \| 119 \| 6 \| 7 \| 155 \| 1 \| 135 \| Uncharacterized protein \| Uncharacterized protein; afdb-uniprot50 \| AF-A0A2E4SSH5-F1-MODEL\_V4 \| 1.0 \| 0.0004789 \| 154 \| 0.101 \| 157 \| 119 \| 7 \| 6 \| 154 \| 2 \| 144 \| Uncharacterized protein \| Uncharacterized protein; afdb-uniprot50 \| AF-A0A0R2LKL6-F1-MODEL\_V4 \| 1.0 \| 0.0005701 \| 154 \| 0.12 \| 166 \| 120 \| 13 \| 6 \| 165 \| 2 \| 147 \| Uncharacterized protein \| Uncharacterized protein; afdb-uniprot50 \| AF-A0A2J9QIZ5-F1-MODEL\_V4 \| 1.0 \| 0.0003582 \| 154 \| 0.115 \| 173 \| 122 \| 11 \| 1 \| 166 \| 1 \| 149 \| Uncharacterized protein \| Uncharacterized protein; afdb-uniprot50 \| AF-A5EUZ0-F1-MODEL\_V4 \| 1.0 \| 0.0002528 \| 154 \| 0.167 \| 161 \| 113 \| 11 \| 7 \| 157 \| 6 \| 155 \| Uncharacterized protein \| Uncharacterized protein; afdb-uniprot50 \| AF-A0A4R6U5E4-F1-MODEL\_V4 \| 1.0 \| 6.647e-05 \| 154 \| 0.143 \| 174 \| 127 \| 8 \| 7 \| 174 \| 2 \| 159 \| Gp37 protein \| Gp37 protein; afdb-uniprot50 \| AF-A0A4R8IH74-F1-MODEL\_V4 \| 1.0 \| 0.000284 \| 154 \| 0.089 \| 190 \| 133 \| 9 \| 7 \| 178 \| 3 \| 170 \| Uncharacterized protein \| Uncharacterized protein; afdb-uniprot50 \| AF-A0A3B0JB18-F1-MODEL\_V4 \| 1.0 \| 7.466e-05 \| 154 \| 0.096 \| 176 \| 139 \| 6 \| 22 \| 188 \| 9 \| 173 \| Uncharacterized protein \| Uncharacterized protein; afdb-uniprot50 \| AF-Q602Y7-F1-MODEL\_V4 \| 1.0 \| 0.0001414 \| 154 \| 0.189 \| 174 \| 117 \| 9 \| 1 \| 170 \| 1 \| 154 \| Uncharacterized protein \| Uncharacterized protein; afdb-uniprot50 \| AF-A0A6M8HNB9-F1-MODEL\_V4 \| 1.0 \| 0.0001058 \| 154 \| 0.146 \| 198 \| 126 \| 14 \| 7 \| 188 \| 1 \| 171 \| Uncharacterized protein \| Uncharacterized protein; afdb-uniprot50 \| AF-A0A1X0TIU2-F1-MODEL\_V4 \| 1.0 \| 0.0002251 \| 154 \| 0.081 \| 172 \| 134 \| 9 \| 1 \| 166 \| 31 \| 184 \| Uncharacterized protein \| Uncharacterized protein; afdb-uniprot50 \| AF-A0A0A4ABV7-F1-MODEL\_V4 \| 1.0 \| 0.0002124 \| 154 \| 0.137 \| 182 \| 119 \| 10 \| 8 \| 166 \| 1 \| 167 \| Uncharacterized protein \| Uncharacterized protein; afdb-uniprot50 \| AF-A0A518HBN8-F1-MODEL\_V4 \| 1.0 \| 0.0003009 \| 154 \| 0.141 \| 177 \| 135 \| 10 \| 7 \| 178 \| 3 \| 167 \| Uncharacterized protein \| Uncharacterized protein; afdb-uniprot50 \| AF-A0A1E4AHP0-F1-MODEL\_V4 \| 1.0 \| 0.0001784 \| 153 \| 0.152 \| 170 \| 118 \| 9 \| 10 \| 171 \| 4 \| 155 \| Uncharacterized protein \| Uncharacterized protein; afdb-uniprot50 \| AF-A0A166FAH8-F1-MODEL\_V4 \| 1.0 \| 0.0003797 \| 153 \| 0.117 \| 170 \| 123 \| 7 \| 9 \| 166 \| 9 \| 163 \| Uncharacterized protein \| Uncharacterized protein; afdb-uniprot50 \| AF-E2CGF9-F1-MODEL\_V4 \| 1.0 \| 8.887e-05 \| 153 \| 0.109 \| 183 \| 134 \| 10 \| 3 \| 177 \| 14 \| 175 \| Uncharacterized protein \| Uncharacterized protein; afdb-uniprot50 \| AF-A0A857FM52-F1-MODEL\_V4 \| 1.0 \| 7.045e-05 \| 153 \| 0.139 \| 193 \| 134 \| 12 \| 7 \| 188 \| 1 \| 172 \| Uncharacterized protein \| Uncharacterized protein; afdb-uniprot50 \| AF-A0A420WGS9-F1-MODEL\_V4 \| 1.0 \| 0.0003582 \| 152 \| 0.111 \| 153 \| 118 \| 6 \| 7 \| 157 \| 1 \| 137 \| Uncharacterized protein DUF4128 \| Uncharacterized protein DUF4128; afdb-uniprot50 \| AF-A0A5C7P964-F1-MODEL\_V4 \| 1.0 \| 0.0008078 \| 152 \| 0.114 \| 157 \| 114 \| 8 \| 9 \| 155 \| 5 \| 146 \| Uncharacterized protein \| Uncharacterized protein; afdb-uniprot50 \| AF-A0A1B1G143-F1-MODEL\_V4 \| 1.0 \| 0.0004789 \| 152 \| 0.109 \| 173 \| 111 \| 11 \| 10 \| 165 \| 2 \| 148 \| Uncharacterized protein \| Uncharacterized protein; afdb-uniprot50 \| AF-A0A1C3FDG3-F1-MODEL\_V4 \| 1.0 \| 0.0006403 \| 152 \| 0.126 \| 174 \| 133 \| 6 \| 7 \| 178 \| 1 \| 157 \| Uncharacterized protein \| Uncharacterized protein; afdb-uniprot50 \| AF-A0A1Q3Q8M2-F1-MODEL\_V4 \| 1.0 \| 0.0001121 \| 152 \| 0.17 \| 199 \| 121 \| 8 \| 1 \| 185 \| 1 \| 169 \| Uncharacterized protein \| Uncharacterized protein; afdb-uniprot50 \| AF-A0A1G7SE70-F1-MODEL\_V4 \| 1.0 \| 6.272e-05 \| 152 \| 0.183 \| 191 \| 119 \| 12 \| 8 \| 179 \| 1 \| 173 \| Uncharacterized protein \| Uncharacterized protein; afdb-uniprot50 \| AF-A0A2D3TE26-F1-MODEL\_V4 \| 1.0 \| 0.0001058 \| 152 \| 0.145 \| 185 \| 126 \| 7 \| 15 \| 178 \| 5 \| 178 \| Uncharacterized protein \| Uncharacterized protein; afdb-uniprot50 \| AF-A0A0J6T1Z2-F1-MODEL\_V4 \| 1.0 \| 0.0008078 \| 151 \| 0.178 \| 151 \| 97 \| 9 \| 14 \| 153 \| 4 \| 138 \| Uncharacterized protein \| Uncharacterized protein; afdb-uniprot50 \| AF-A0A5U3CNK9-F1-MODEL\_V4 \| 1.0 \| 0.0007622 \| 151 \| 0.142 \| 161 \| 117 \| 9 \| 7 \| 163 \| 1 \| 144 \| Uncharacterized protein \| Uncharacterized protein; afdb-uniprot50 \| AF-A0A2J8HSC1-F1-MODEL\_V4 \| 1.0 \| 0.0004519 \| 151 \| 0.106 \| 178 \| 125 \| 11 \| 3 \| 166 \| 5 \| 162 \| Uncharacterized protein \| Uncharacterized protein; afdb-uniprot50 \| AF-A0A2A2HG93-F1-MODEL\_V4 \| 1.0 \| 0.0004519 \| 151 \| 0.149 \| 161 \| 116 \| 9 \| 8 \| 164 \| 1 \| 144 \| Uncharacterized protein \| Uncharacterized protein; afdb-uniprot50 \| AF-A0A444HEH6-F1-MODEL\_V4 \| 1.0 \| 0.0004519 \| 151 \| 0.13 \| 184 \| 127 \| 9 \| 9 \| 178 \| 4 \| 168 \| Uncharacterized protein \| Uncharacterized protein; afdb-uniprot50 \| AF-A0A5B7ZK92-F1-MODEL\_V4 \| 1.0 \| 6.647e-05 \| 151 \| 0.135 \| 199 \| 138 \| 10 \| 7 \| 188 \| 1 \| 182 \| Uncharacterized protein \| Uncharacterized protein; afdb-uniprot50 \| AF-A0A7Y1CQT2-F1-MODEL\_V4 \| 1.0 \| 5.584e-05 \| 151 \| 0.123 \| 211 \| 145 \| 9 \| 7 \| 189 \| 1 \| 199 \| Uncharacterized protein \| Uncharacterized protein; afdb-uniprot50 \| AF-A0A1E3WMG0-F1-MODEL\_V4 \| 1.0 \| 8.386e-05 \| 151 \| 0.1 \| 209 \| 147 \| 13 \| 1 \| 187 \| 11 \| 200 \| Uncharacterized protein \| Uncharacterized protein; afdb-uniprot50 \| AF-A0A1Q4T414-F1-MODEL\_V4 \| 1.0 \| 0.00153 \| 150 \| 0.125 \| 159 \| 114 \| 10 \| 1 \| 156 \| 3 \| 139 \| Uncharacterized protein \| Uncharacterized protein; afdb-uniprot50 \| AF-A0A1G1N144-F1-MODEL\_V4 \| 1.0 \| 0.0004789 \| 150 \| 0.141 \| 156 \| 106 \| 13 \| 8 \| 151 \| 1 \| 140 \| Uncharacterized protein \| Uncharacterized protein; afdb-uniprot50 \| AF-A0A3C2D818-F1-MODEL\_V4 \| 1.0 \| 0.000338 \| 150 \| 0.119 \| 167 \| 108 \| 12 \| 8 \| 158 \| 1 \| 144 \| Uncharacterized protein \| Uncharacterized protein; afdb-uniprot50 \| AF-A0A4R1C4H1-F1-MODEL\_V4 \| 1.0 \| 0.001213 \| 150 \| 0.105 \| 152 \| 120 \| 7 \| 7 \| 152 \| 2 \| 143 \| Uncharacterized protein \| Uncharacterized protein; afdb-uniprot50 \| AF-A0A075WUA5-F1-MODEL\_V4 \| 1.0 \| 0.00153 \| 150 \| 0.098 \| 162 \| 123 \| 8 \| 8 \| 166 \| 1 \| 142 \| Uncharacterized protein \| Uncharacterized protein; afdb-uniprot50 \| AF-A0A806Z7M4-F1-MODEL\_V4 \| 1.0 \| 0.0001684 \| 150 \| 0.156 \| 160 \| 106 \| 9 \| 27 \| 168 \| 3 \| 151 \| Uncharacterized protein \| Uncharacterized protein; afdb-uniprot50 \| AF-A0A2N7L8B1-F1-MODEL\_V4 \| 1.0 \| 0.0001335 \| 150 \| 0.128 \| 194 \| 139 \| 12 \| 1 \| 174 \| 1 \| 184 \| Uncharacterized protein \| Uncharacterized protein; afdb-uniprot50 \| AF-A0A4P2QPQ9-F1-MODEL\_V4 \| 1.0 \| 0.0005701 \| 150 \| 0.148 \| 148 \| 104 \| 9 \| 27 \| 166 \| 124 \| 257 \| Uncharacterized protein \| Uncharacterized protein; afdb-uniprot50 \| AF-A0A1Q3WT67-F1-MODEL\_V4 \| 1.0 \| 0.0006042 \| 149 \| 0.1 \| 180 \| 128 \| 11 \| 6 \| 166 \| 2 \| 166 \| Uncharacterized protein \| Uncharacterized protein; afdb-uniprot50 \| AF-A0A0H3ZYQ3-F1-MODEL\_V4 \| 1.0 \| 0.0002528 \| 149 \| 0.094 \| 190 \| 139 \| 11 \| 8 \| 187 \| 6 \| 172 \| Uncharacterized protein \| Uncharacterized protein; afdb-uniprot50 \| AF-B6WSR7-F1-MODEL\_V4 \| 1.0 \| 2.948e-05 \| 149 \| 0.164 \| 201 \| 133 \| 11 \| 10 \| 189 \| 5 \| 191 \| Uncharacterized protein \| Uncharacterized protein; afdb-uniprot50 \| AF-A0A7U2KPY8-F1-MODEL\_V4 \| 1.0 \| 0.00153 \| 148 \| 0.102 \| 156 \| 111 \| 7 \| 7 \| 154 \| 1 \| 135 \| Uncharacterized protein \| Uncharacterized protein; afdb-uniprot50 \| AF-A0A1G7TC54-F1-MODEL\_V4 \| 1.0 \| 0.0004024 \| 148 \| 0.095 \| 157 \| 101 \| 11 \| 18 \| 156 \| 9 \| 142 \| Uncharacterized protein \| Uncharacterized protein; afdb-uniprot50 \| AF-A0A365QUT4-F1-MODEL\_V4 \| 1.0 \| 0.001444 \| 148 \| 0.185 \| 156 \| 106 \| 12 \| 7 \| 152 \| 3 \| 147 \| Uncharacterized protein \| Uncharacterized protein; afdb-uniprot50 \| AF-A0A2D7VXQ5-F1-MODEL\_V4 \| 1.0 \| 0.003451 \| 148 \| 0.111 \| 153 \| 124 \| 6 \| 5 \| 153 \| 2 \| 146 \| Uncharacterized protein \| Uncharacterized protein; afdb-uniprot50 \| AF-A0A5C7Q4P6-F1-MODEL\_V4 \| 1.0 \| 0.0006042 \| 148 \| 0.186 \| 161 \| 112 \| 8 \| 10 \| 162 \| 5 \| 154 \| Uncharacterized protein \| Uncharacterized protein; afdb-uniprot50 \| AF-A0A7Y4DJI8-F1-MODEL\_V4 \| 1.0 \| 0.0003189 \| 148 \| 0.094 \| 201 \| 137 \| 16 \| 5 \| 189 \| 3 \| 174 \| Uncharacterized protein \| Uncharacterized protein; afdb-uniprot50 \| AF-A0A377HPJ9-F1-MODEL\_V4 \| 1.0 \| 7.466e-05 \| 148 \| 0.135 \| 206 \| 143 \| 12 \| 1 \| 187 \| 1 \| 190 \| Uncharacterized protein \| Uncharacterized protein; afdb-uniprot50 \| AF-A0A4V2W9H1-F1-MODEL\_V4 \| 1.0 \| 0.0009073 \| 147 \| 0.107 \| 149 \| 117 \| 6 \| 8 \| 155 \| 3 \| 136 \| Uncharacterized protein \| Uncharacterized protein; afdb-uniprot50 \| AF-A0A5D9DCM1-F1-MODEL\_V4 \| 1.0 \| 0.0006403 \| 147 \| 0.137 \| 153 \| 114 \| 6 \| 7 \| 156 \| 2 \| 139 \| Uncharacterized protein \| Uncharacterized protein; afdb-uniprot50 \| AF-A0A1G6A6F1-F1-MODEL\_V4 \| 1.0 \| 0.004353 \| 147 \| 0.067 \| 148 \| 122 \| 5 \| 10 \| 153 \| 5 \| 140 \| Gp37 protein \| Gp37 protein; afdb-uniprot50 \| AF-T0YPM2-F1-MODEL\_V4 \| 1.0 \| 0.00153 \| 147 \| 0.114 \| 157 \| 120 \| 9 \| 6 \| 154 \| 3 \| 148 \| Phage associated protein \| Phage associated protein; afdb-uniprot50 \| AF-A0A1U7GIF2-F1-MODEL\_V4 \| 1.0 \| 0.00153 \| 147 \| 0.122 \| 163 \| 120 \| 10 \| 9 \| 165 \| 6 \| 151 \| Uncharacterized protein \| Uncharacterized protein; afdb-uniprot50 \| AF-A0A2G5KHB3-F1-MODEL\_V4 \| 1.0 \| 0.0005701 \| 147 \| 0.138 \| 173 \| 114 \| 12 \| 7 \| 166 \| 1 \| 151 \| Uncharacterized protein \| Uncharacterized protein; afdb-uniprot50 \| AF-A0A2D6EYR9-F1-MODEL\_V4 \| 1.0 \| 0.0007192 \| 147 \| 0.13 \| 168 \| 124 \| 6 \| 5 \| 154 \| 3 \| 166 \| Uncharacterized protein \| Uncharacterized protein; afdb-uniprot50 \| AF-A0A7W4PA56-F1-MODEL\_V4 \| 1.0 \| 0.0002528 \| 147 \| 0.122 \| 179 \| 134 \| 10 \| 13 \| 179 \| 8 \| 175 \| Uncharacterized protein \| Uncharacterized protein; afdb-uniprot50 \| AF-A0A1J0EKE0-F1-MODEL\_V4 \| 1.0 \| 0.0001414 \| 147 \| 0.153 \| 182 \| 125 \| 14 \| 8 \| 174 \| 1 \| 168 \| Uncharacterized protein \| Uncharacterized protein; afdb-uniprot50 \| AF-X5MMY5-F1-MODEL\_V4 \| 1.0 \| 0.001444 \| 146 \| 0.131 \| 152 \| 108 \| 8 \| 8 \| 152 \| 6 \| 140 \| Uncharacterized protein \| Uncharacterized protein; afdb-uniprot50 \| AF-A0A1Y3QT12-F1-MODEL\_V4 \| 1.0 \| 0.0006042 \| 146 \| 0.124 \| 161 \| 119 \| 8 \| 7 \| 155 \| 5 \| 155 \| Uncharacterized protein \| Uncharacterized protein; afdb-uniprot50 \| AF-A0A2R4VQW9-F1-MODEL\_V4 \| 1.0 \| 0.00153 \| 146 \| 0.136 \| 176 \| 122 \| 10 \| 1 \| 165 \| 1 \| 157 \| Uncharacterized protein \| Uncharacterized protein; afdb-uniprot50 \| AF-A0A318NGV1-F1-MODEL\_V4 \| 1.0 \| 0.0006042 \| 146 \| 0.112 \| 177 \| 112 \| 10 \| 8 \| 165 \| 1 \| 151 \| Uncharacterized protein \| Uncharacterized protein; afdb-uniprot50 \| AF-A0A4S2GWA7-F1-MODEL\_V4 \| 1.0 \| 0.002298 \| 145 \| 0.163 \| 153 \| 104 \| 9 \| 9 \| 154 \| 11 \| 146 \| DUF3168 domain-containing protein \| DUF3168 domain-containing protein; afdb-uniprot50 \| AF-A0A840EF97-F1-MODEL\_V4 \| 1.0 \| 0.001213 \| 145 \| 0.096 \| 176 \| 120 \| 11 \| 1 \| 166 \| 1 \| 147 \| Uncharacterized protein \| Uncharacterized protein; afdb-uniprot50 \| AF-A0A4P7L3J1-F1-MODEL\_V4 \| 1.0 \| 0.0001259 \| 145 \| 0.119 \| 176 \| 125 \| 9 \| 7 \| 166 \| 1 \| 162 \| Uncharacterized protein \| Uncharacterized protein; afdb-uniprot50 \| AF-A0A1D9BCM1-F1-MODEL\_V4 \| 1.0 \| 0.0001058 \| 145 \| 0.129 \| 193 \| 129 \| 12 \| 15 \| 184 \| 5 \| 181 \| Uncharacterized protein \| Uncharacterized protein; afdb-uniprot50 \| AF-A0A7C5WMU7-F1-MODEL\_V4 \| 1.0 \| 0.006168 \| 144 \| 0.097 \| 154 \| 120 \| 7 \| 7 \| 155 \| 2 \| 141 \| Uncharacterized protein \| Uncharacterized protein; afdb-uniprot50 \| AF-A0A4R1PSF6-F1-MODEL\_V4 \| 1.0 \| 0.00153 \| 144 \| 0.132 \| 159 \| 107 \| 10 \| 7 \| 152 \| 1 \| 141 \| Uncharacterized protein \| Uncharacterized protein; afdb-uniprot50 \| AF-A0A1T2L9Q7-F1-MODEL\_V4 \| 1.0 \| 0.001444 \| 144 \| 0.116 \| 154 \| 111 \| 10 \| 13 \| 154 \| 4 \| 144 \| Uncharacterized protein \| Uncharacterized protein; afdb-uniprot50 \| AF-A0A5S9PHL1-F1-MODEL\_V4 \| 1.0 \| 0.001286 \| 144 \| 0.115 \| 156 \| 114 \| 7 \| 12 \| 156 \| 2 \| 144 \| Uncharacterized protein \| Uncharacterized protein; afdb-uniprot50 \| AF-A0A7W6P0F7-F1-MODEL\_V4 \| 1.0 \| 0.0008078 \| 144 \| 0.116 \| 163 \| 133 \| 6 \| 7 \| 164 \| 1 \| 157 \| Uncharacterized protein \| Uncharacterized protein; afdb-uniprot50 \| AF-L7U9W8-F1-MODEL\_V4 \| 1.0 \| 0.0008078 \| 144 \| 0.141 \| 177 \| 125 \| 12 \| 10 \| 176 \| 5 \| 164 \| Uncharacterized protein \| Uncharacterized protein; afdb-uniprot50 \| AF-A0A2E3P790-F1-MODEL\_V4 \| 1.0 \| 0.0002386 \| 144 \| 0.162 \| 203 \| 129 \| 14 \| 8 \| 185 \| 1 \| 187 \| Uncharacterized protein \| Uncharacterized protein; afdb-uniprot50 \| AF-A0A0D8CN10-F1-MODEL\_V4 \| 1.0 \| 0.0001589 \| 144 \| 0.136 \| 205 \| 143 \| 11 \| 6 \| 188 \| 2 \| 194 \| Uncharacterized protein \| Uncharacterized protein; afdb-uniprot50 \| AF-G6XIS4-F1-MODEL\_V4 \| 1.0 \| 0.0001499 \| 144 \| 0.12 \| 183 \| 127 \| 12 \| 7 \| 173 \| 1 \| 165 \| Uncharacterized protein \| Uncharacterized protein; afdb-uniprot50 \| AF-A0A5D0CT21-F1-MODEL\_V4 \| 1.0 \| 0.0006403 \| 144 \| 0.164 \| 146 \| 109 \| 6 \| 8 \| 150 \| 1 \| 136 \| Uncharacterized protein \| Uncharacterized protein; afdb-uniprot50 \| AF-A0A6M3IP32-F1-MODEL\_V4 \| 1.0 \| 0.000338 \| 143 \| 0.131 \| 190 \| 143 \| 11 \| 1 \| 179 \| 1 \| 179 \| Uncharacterized protein \| Uncharacterized protein; afdb-uniprot50 \| AF-D3A6M2-F1-MODEL\_V4 \| 1.0 \| 0.0006786 \| 143 \| 0.15 \| 173 \| 114 \| 11 \| 8 \| 161 \| 1 \| 159 \| Uncharacterized protein \| Uncharacterized protein; afdb-uniprot50 \| AF-A0A2E9DC73-F1-MODEL\_V4 \| 1.0 \| 0.0009616 \| 142 \| 0.133 \| 157 \| 107 \| 11 \| 7 \| 156 \| 4 \| 138 \| Uncharacterized protein \| Uncharacterized protein; afdb-uniprot50 \| AF-A0A644VLE7-F1-MODEL\_V4 \| 1.0 \| 0.001622 \| 142 \| 0.099 \| 161 \| 115 \| 8 \| 10 \| 154 \| 4 \| 150 \| Uncharacterized protein \| Uncharacterized protein; afdb-uniprot50 \| AF-A0A096AIF8-F1-MODEL\_V4 \| 1.0 \| 0.001822 \| 142 \| 0.118 \| 169 \| 131 \| 7 \| 7 \| 173 \| 1 \| 153 \| Uncharacterized protein \| Uncharacterized protein; afdb-uniprot50 \| AF-A0A223D2E0-F1-MODEL\_V4 \| 1.0 \| 0.0003189 \| 142 \| 0.112 \| 187 \| 133 \| 14 \| 5 \| 166 \| 2 \| 180 \| Uncharacterized protein \| Uncharacterized protein; afdb-uniprot50 \| AF-A0A2S7SPC8-F1-MODEL\_V4 \| 1.0 \| 0.0001684 \| 142 \| 0.118 \| 194 \| 133 \| 10 \| 2 \| 178 \| 23 \| 195 \| Uncharacterized protein \| Uncharacterized protein; afdb-uniprot50 \| AF-A0A7X6EJH3-F1-MODEL\_V4 \| 1.0 \| 6.647e-05 \| 142 \| 0.141 \| 212 \| 131 \| 13 \| 15 \| 189 \| 5 \| 202 \| Uncharacterized protein \| Uncharacterized protein; afdb-uniprot50 \| AF-H0EA80-F1-MODEL\_V4 \| 1.0 \| 0.0001589 \| 142 \| 0.139 \| 215 \| 136 \| 11 \| 1 \| 182 \| 1 \| 199 \| Uncharacterized protein \| Uncharacterized protein; afdb-uniprot50 \| AF-A0A5A9EMM4-F1-MODEL\_V4 \| 1.0 \| 0.0003797 \| 142 \| 0.147 \| 190 \| 136 \| 11 \| 1 \| 178 \| 1 \| 176 \| Uncharacterized protein \| Uncharacterized protein; afdb-uniprot50 \| AF-A0A2D6AI45-F1-MODEL\_V4 \| 1.0 \| 0.004613 \| 141 \| 0.098 \| 162 \| 121 \| 11 \| 1 \| 156 \| 1 \| 143 \| Uncharacterized protein \| Uncharacterized protein; afdb-uniprot50 \| AF-A0A521BNQ1-F1-MODEL\_V4 \| 1.0 \| 0.000338 \| 141 \| 0.131 \| 167 \| 99 \| 12 \| 9 \| 157 \| 6 \| 144 \| Uncharacterized protein \| Uncharacterized protein; afdb-uniprot50 \| AF-C9MV92-F1-MODEL\_V4 \| 1.0 \| 0.0002679 \| 140 \| 0.125 \| 167 \| 114 \| 11 \| 10 \| 158 \| 2 \| 154 \| Uncharacterized protein \| Uncharacterized protein; afdb-uniprot50 \| AF-A0A5C7QCW3-F1-MODEL\_V4 \| 1.0 \| 0.001719 \| 140 \| 0.134 \| 171 \| 121 \| 10 \| 9 \| 164 \| 5 \| 163 \| Uncharacterized protein \| Uncharacterized protein; afdb-uniprot50 \| AF-A0A7D4FTT1-F1-MODEL\_V4 \| 1.0 \| 0.001286 \| 139 \| 0.089 \| 179 \| 117 \| 11 \| 10 \| 166 \| 2 \| 156 \| Uncharacterized protein \| Uncharacterized protein; afdb-uniprot50 \| AF-A0A1D8UTD4-F1-MODEL\_V4 \| 1.0 \| 0.0002528 \| 139 \| 0.128 \| 195 \| 136 \| 12 \| 7 \| 187 \| 1 \| 175 \| Uncharacterized protein \| Uncharacterized protein; afdb-uniprot50 \| AF-A0A1S1TJW6-F1-MODEL\_V4 \| 1.0 \| 5.584e-05 \| 139 \| 0.162 \| 209 \| 127 \| 12 \| 1 \| 189 \| 1 \| 181 \| Uncharacterized protein \| Uncharacterized protein; afdb-uniprot50 \| AF-A0A644WAV0-F1-MODEL\_V4 \| 1.0 \| 0.0002124 \| 139 \| 0.149 \| 201 \| 122 \| 13 \| 8 \| 189 \| 1 \| 171 \| Uncharacterized protein \| Uncharacterized protein; afdb-uniprot50 \| AF-A0A0N9Y5D3-F1-MODEL\_V4 \| 1.0 \| 0.002581 \| 138 \| 0.118 \| 144 \| 102 \| 7 \| 12 \| 153 \| 3 \| 123 \| Uncharacterized protein \| Uncharacterized protein; afdb-uniprot50 \| AF-A0A3D5AF28-F1-MODEL\_V4 \| 1.0 \| 0.0008078 \| 138 \| 0.094 \| 159 \| 113 \| 10 \| 12 \| 157 \| 2 \| 142 \| Uncharacterized protein \| Uncharacterized protein; afdb-uniprot50 \| AF-A0A1F8X037-F1-MODEL\_V4 \| 1.0 \| 0.003657 \| 138 \| 0.12 \| 158 \| 113 \| 9 \| 7 \| 154 \| 1 \| 142 \| Uncharacterized protein \| Uncharacterized protein; afdb-uniprot50 \| AF-A0A7K3ZIE1-F1-MODEL\_V4 \| 1.0 \| 0.0008078 \| 138 \| 0.105 \| 170 \| 115 \| 10 \| 7 \| 165 \| 2 \| 145 \| Uncharacterized protein \| Uncharacterized protein; afdb-uniprot50 \| AF-A0A1V4QIE0-F1-MODEL\_V4 \| 1.0 \| 0.0008561 \| 138 \| 0.129 \| 154 \| 116 \| 4 \| 7 \| 153 \| 1 \| 143 \| Uncharacterized protein \| Uncharacterized protein; afdb-uniprot50 \| AF-A0A428MSH7-F1-MODEL\_V4 \| 1.0 \| 0.0004519 \| 138 \| 0.104 \| 173 \| 131 \| 12 \| 2 \| 166 \| 4 \| 160 \| Uncharacterized protein \| Uncharacterized protein; afdb-uniprot50 \| AF-A0A1V6HT77-F1-MODEL\_V4 \| 1.0 \| 0.00193 \| 138 \| 0.104 \| 173 \| 127 \| 12 \| 2 \| 153 \| 1 \| 166 \| Uncharacterized protein \| Uncharacterized protein; afdb-uniprot50 \| AF-A0A1Y0Y7R5-F1-MODEL\_V4 \| 1.0 \| 0.0002251 \| 138 \| 0.137 \| 175 \| 130 \| 9 \| 1 \| 166 \| 34 \| 196 \| Uncharacterized protein \| Uncharacterized protein; afdb-uniprot50 \| AF-A0A317DUZ0-F1-MODEL\_V4 \| 1.0 \| 0.006537 \| 138 \| 0.201 \| 154 \| 106 \| 10 \| 9 \| 156 \| 170 \| 312 \| Uncharacterized protein \| Uncharacterized protein; afdb-uniprot50 \| AF-A0A4R5VXB1-F1-MODEL\_V4 \| 1.0 \| 0.001286 \| 137 \| 0.138 \| 152 \| 106 \| 12 \| 7 \| 152 \| 1 \| 133 \| Uncharacterized protein \| Uncharacterized protein; afdb-uniprot50 \| AF-A0A7G2T4T1-F1-MODEL\_V4 \| 1.0 \| 0.001213 \| 137 \| 0.136 \| 169 \| 106 \| 12 \| 10 \| 162 \| 2 \| 146 \| Uncharacterized protein \| Uncharacterized protein; afdb-uniprot50 \| AF-A0A1L4D144-F1-MODEL\_V4 \| 1.0 \| 0.002168 \| 137 \| 0.121 \| 165 \| 125 \| 6 \| 9 \| 168 \| 5 \| 154 \| Uncharacterized protein \| Uncharacterized protein; afdb-uniprot50 \| AF-B7RNM1-F1-MODEL\_V4 \| 1.0 \| 0.001213 \| 136 \| 0.442 \| 70 \| 39 \| 0 \| 120 \| 189 \| 18 \| 87 \| Uncharacterized protein \| Uncharacterized protein; afdb-uniprot50 \| AF-A0A7W9S927-F1-MODEL\_V4 \| 1.0 \| 0.001286 \| 136 \| 0.111 \| 152 \| 110 \| 7 \| 7 \| 152 \| 2 \| 134 \| Uncharacterized protein \| Uncharacterized protein; afdb-uniprot50 \| AF-A0A4Q7TEQ2-F1-MODEL\_V4 \| 1.0 \| 0.004353 \| 136 \| 0.107 \| 149 \| 111 \| 6 \| 10 \| 153 \| 4 \| 135 \| Uncharacterized protein \| Uncharacterized protein; afdb-uniprot50 \| AF-A0A7J2K9N9-F1-MODEL\_V4 \| 1.0 \| 0.009262 \| 136 \| 0.12 \| 149 \| 117 \| 7 \| 7 \| 152 \| 1 \| 138 \| Uncharacterized protein \| Uncharacterized protein; afdb-uniprot50 \| AF-A0A6M3IGG6-F1-MODEL\_V4 \| 1.0 \| 0.002168 \| 136 \| 0.134 \| 163 \| 116 \| 10 \| 9 \| 162 \| 6 \| 152 \| Uncharacterized protein \| Uncharacterized protein; afdb-uniprot50 \| AF-R7JPZ3-F1-MODEL\_V4 \| 1.0 \| 0.0007622 \| 136 \| 0.109 \| 173 \| 130 \| 7 \| 1 \| 168 \| 3 \| 156 \| Phage-related protein Gp37-like protein \| Phage-related protein Gp37-like protein; afdb-uniprot50 \| AF-A0A6C2YKQ7-F1-MODEL\_V4 \| 1.0 \| 0.001622 \| 136 \| 0.09 \| 165 \| 129 \| 7 \| 6 \| 156 \| 8 \| 165 \| Uncharacterized protein \| Uncharacterized protein; afdb-uniprot50 \| AF-A0A6G9AQR9-F1-MODEL\_V4 \| 1.0 \| 0.0005701 \| 136 \| 0.166 \| 174 \| 118 \| 12 \| 7 \| 162 \| 1 \| 165 \| Uncharacterized protein \| Uncharacterized protein; afdb-uniprot50 \| AF-A0A0R1RDJ7-F1-MODEL\_V4 \| 1.0 \| 0.002298 \| 135 \| 0.13 \| 153 \| 104 \| 7 \| 1 \| 153 \| 1 \| 124 \| Uncharacterized protein \| Uncharacterized protein; afdb-uniprot50 \| AF-A0A3V2JIY3-F1-MODEL\_V4 \| 1.0 \| 0.003072 \| 135 \| 0.087 \| 171 \| 129 \| 7 \| 7 \| 172 \| 1 \| 149 \| Uncharacterized protein \| Uncharacterized protein; afdb-uniprot50 \| AF-A0A6M0S828-F1-MODEL\_V4 \| 1.0 \| 0.001213 \| 135 \| 0.139 \| 158 \| 109 \| 10 \| 10 \| 153 \| 2 \| 146 \| Uncharacterized protein \| Uncharacterized protein; afdb-uniprot50 \| AF-A0A4U0GUD8-F1-MODEL\_V4 \| 1.0 \| 0.001622 \| 135 \| 0.12 \| 166 \| 119 \| 10 \| 7 \| 154 \| 1 \| 157 \| Uncharacterized protein \| Uncharacterized protein; afdb-uniprot50 \| AF-A0A7V7SWA7-F1-MODEL\_V4 \| 1.0 \| 0.000338 \| 135 \| 0.152 \| 184 \| 114 \| 13 \| 1 \| 154 \| 1 \| 172 \| Uncharacterized protein \| Uncharacterized protein; afdb-uniprot50 \| AF-A0A3G9IET3-F1-MODEL\_V4 \| 1.0 \| 0.000284 \| 135 \| 0.1 \| 180 \| 129 \| 8 \| 8 \| 166 \| 7 \| 174 \| Uncharacterized protein \| Uncharacterized protein; afdb-uniprot50 \| AF-A0A0F9TI92-F1-MODEL\_V4 \| 1.0 \| 0.0002124 \| 135 \| 0.111 \| 161 \| 125 \| 8 \| 9 \| 159 \| 53 \| 205 \| Uncharacterized protein \| Uncharacterized protein; afdb-uniprot50 \| AF-A0A4Y9J0T5-F1-MODEL\_V4 \| 1.0 \| 0.003451 \| 134 \| 0.088 \| 136 \| 97 \| 8 \| 29 \| 156 \| 3 \| 119 \| Uncharacterized protein \| Uncharacterized protein; afdb-uniprot50 \| AF-A0A0X8JIR5-F1-MODEL\_V4 \| 1.0 \| 0.00193 \| 134 \| 0.12 \| 149 \| 113 \| 7 \| 7 \| 152 \| 1 \| 134 \| Uncharacterized protein \| Uncharacterized protein; afdb-uniprot50 \| AF-A0A3F3PDR2-F1-MODEL\_V4 \| 1.0 \| 0.003657 \| 134 \| 0.103 \| 164 \| 122 \| 12 \| 1 \| 155 \| 1 \| 148 \| Uncharacterized protein \| Uncharacterized protein; afdb-uniprot50 \| AF-A0A3A4ZKA5-F1-MODEL\_V4 \| 1.0 \| 0.0009073 \| 134 \| 0.137 \| 160 \| 118 \| 9 \| 5 \| 156 \| 2 \| 149 \| Uncharacterized protein \| Uncharacterized protein; afdb-uniprot50 \| AF-A0A838MQ51-F1-MODEL\_V4 \| 1.0 \| 0.001719 \| 134 \| 0.153 \| 163 \| 116 \| 10 \| 9 \| 165 \| 9 \| 155 \| Uncharacterized protein \| Uncharacterized protein; afdb-uniprot50 \| AF-A0A1E4HY13-F1-MODEL\_V4 \| 1.0 \| 0.0006786 \| 134 \| 0.137 \| 174 \| 123 \| 8 \| 1 \| 165 \| 1 \| 156 \| Uncharacterized protein \| Uncharacterized protein; afdb-uniprot50 \| AF-A0A5P8MWM5-F1-MODEL\_V4 \| 1.0 \| 0.0006042 \| 134 \| 0.158 \| 196 \| 128 \| 18 \| 9 \| 188 \| 14 \| 188 \| Uncharacterized protein \| Uncharacterized protein; afdb-uniprot50 \| AF-A0A7W8JWA4-F1-MODEL\_V4 \| 1.0 \| 0.001822 \| 133 \| 0.111 \| 161 \| 129 \| 5 \| 7 \| 165 \| 1 \| 149 \| Uncharacterized protein \| Uncharacterized protein; afdb-uniprot50 \| AF-A0A661KVG1-F1-MODEL\_V4 \| 1.0 \| 0.002581 \| 133 \| 0.105 \| 161 \| 121 \| 10 \| 9 \| 159 \| 13 \| 160 \| Uncharacterized protein \| Uncharacterized protein; afdb-uniprot50 \| AF-A0A179CYM1-F1-MODEL\_V4 \| 1.0 \| 0.0001891 \| 133 \| 0.148 \| 188 \| 127 \| 12 \| 1 \| 178 \| 1 \| 165 \| Uncharacterized protein \| Uncharacterized protein; afdb-uniprot50 \| AF-A0A524QZ15-F1-MODEL\_V4 \| 1.0 \| 0.001822 \| 133 \| 0.118 \| 152 \| 105 \| 11 \| 9 \| 153 \| 48 \| 177 \| Uncharacterized protein \| Uncharacterized protein; afdb-uniprot50 \| AF-A0A850KD82-F1-MODEL\_V4 \| 1.0 \| 0.0008561 \| 133 \| 0.106 \| 179 \| 129 \| 10 \| 10 \| 169 \| 7 \| 173 \| DUF1834 family protein \| DUF1834 family protein; afdb-uniprot50 \| AF-A0A2A2HEB9-F1-MODEL\_V4 \| 1.0 \| 0.000338 \| 133 \| 0.125 \| 191 \| 125 \| 12 \| 1 \| 166 \| 1 \| 174 \| Uncharacterized protein \| Uncharacterized protein; afdb-uniprot50 \| AF-A0A4U9HEQ6-F1-MODEL\_V4 \| 1.0 \| 0.00108 \| 132 \| 0.333 \| 69 \| 46 \| 0 \| 120 \| 188 \| 12 \| 80 \| Uncharacterized protein \| Uncharacterized protein; afdb-uniprot50 \| AF-A0A4Y6CYD7-F1-MODEL\_V4 \| 1.0 \| 0.005492 \| 132 \| 0.147 \| 156 \| 108 \| 4 \| 1 \| 152 \| 1 \| 135 \| Uncharacterized protein \| Uncharacterized protein; afdb-uniprot50 \| AF-A0A7C1U118-F1-MODEL\_V4 \| 1.0 \| 0.006928 \| 132 \| 0.128 \| 156 \| 114 \| 7 \| 8 \| 154 \| 1 \| 143 \| Uncharacterized protein \| Uncharacterized protein; afdb-uniprot50 \| AF-A0A7Z7NIL0-F1-MODEL\_V4 \| 1.0 \| 0.0007192 \| 132 \| 0.111 \| 170 \| 116 \| 10 \| 1 \| 157 \| 1 \| 148 \| Uncharacterized protein \| Uncharacterized protein; afdb-uniprot50 \| AF-A8TYE4-F1-MODEL\_V4 \| 1.0 \| 0.0002004 \| 132 \| 0.165 \| 163 \| 107 \| 9 \| 33 \| 187 \| 2 \| 143 \| Uncharacterized protein \| Uncharacterized protein; afdb-uniprot50 \| AF-A0A2A5A3B0-F1-MODEL\_V4 \| 1.0 \| 0.001362 \| 132 \| 0.136 \| 176 \| 129 \| 12 \| 9 \| 173 \| 5 \| 168 \| Uncharacterized protein \| Uncharacterized protein; afdb-uniprot50 \| AF-A0A4R2USQ7-F1-MODEL\_V4 \| 1.0 \| 0.004108 \| 131 \| 0.121 \| 156 \| 107 \| 11 \| 5 \| 154 \| 2 \| 133 \| Uncharacterized protein \| Uncharacterized protein; afdb-uniprot50 \| AF-F5R8A5-F1-MODEL\_V4 \| 1.0 \| 0.003657 \| 131 \| 0.164 \| 146 \| 106 \| 8 \| 10 \| 153 \| 6 \| 137 \| Uncharacterized protein \| Uncharacterized protein; afdb-uniprot50 \| AF-A0A2N5XX45-F1-MODEL\_V4 \| 1.0 \| 0.001213 \| 131 \| 0.118 \| 160 \| 120 \| 6 \| 8 \| 153 \| 1 \| 153 \| Uncharacterized protein \| Uncharacterized protein; afdb-uniprot50 \| AF-A0A7K1B1R7-F1-MODEL\_V4 \| 1.0 \| 0.002581 \| 131 \| 0.122 \| 171 \| 121 \| 9 \| 7 \| 161 \| 1 \| 158 \| Uncharacterized protein \| Uncharacterized protein; afdb-uniprot50 \| AF-A0A327K8K4-F1-MODEL\_V4 \| 1.0 \| 0.0003189 \| 131 \| 0.172 \| 191 \| 129 \| 11 \| 1 \| 177 \| 1 \| 176 \| Uncharacterized protein \| Uncharacterized protein; afdb-uniprot50 \| AF-A0A6P2CLK3-F1-MODEL\_V4 \| 1.0 \| 0.005492 \| 130 \| 0.093 \| 150 \| 109 \| 5 \| 7 \| 155 \| 1 \| 124 \| DUF806 family protein \| DUF806 family protein; afdb-uniprot50 \| AF-A0A084XW78-F1-MODEL\_V4 \| 1.0 \| 0.001213 \| 130 \| 0.084 \| 153 \| 114 \| 9 \| 6 \| 152 \| 2 \| 134 \| Uncharacterized protein \| Uncharacterized protein; afdb-uniprot50 \| AF-A0A543NXI2-F1-MODEL\_V4 \| 1.0 \| 0.002735 \| 130 \| 0.154 \| 162 \| 107 \| 13 \| 1 \| 154 \| 1 \| 140 \| Uncharacterized protein DUF3168 \| Uncharacterized protein DUF3168; afdb-uniprot50 \| AF-A0A1V4QT86-F1-MODEL\_V4 \| 1.0 \| 0.000338 \| 130 \| 0.15 \| 146 \| 106 \| 8 \| 15 \| 154 \| 8 \| 141 \| Uncharacterized protein \| Uncharacterized protein; afdb-uniprot50 \| AF-A0A518CIC7-F1-MODEL\_V4 \| 1.0 \| 0.002168 \| 130 \| 0.108 \| 157 \| 122 \| 10 \| 7 \| 154 \| 1 \| 148 \| Uncharacterized protein \| Uncharacterized protein; afdb-uniprot50 \| AF-A0A2E8E032-F1-MODEL\_V4 \| 1.0 \| 0.00582 \| 130 \| 0.097 \| 154 \| 121 \| 8 \| 9 \| 155 \| 5 \| 147 \| Uncharacterized protein \| Uncharacterized protein; afdb-uniprot50 \| AF-A0A0G3XEA1-F1-MODEL\_V4 \| 1.0 \| 0.001622 \| 130 \| 0.11 \| 163 \| 121 \| 9 \| 9 \| 163 \| 9 \| 155 \| Uncharacterized protein \| Uncharacterized protein; afdb-uniprot50 \| AF-D3DZY8-F1-MODEL\_V4 \| 1.0 \| 0.002046 \| 130 \| 0.122 \| 171 \| 126 \| 9 \| 9 \| 163 \| 7 \| 169 \| Uncharacterized protein \| Uncharacterized protein; afdb-uniprot50 \| AF-A0A1Y2LFX2-F1-MODEL\_V4 \| 1.0 \| 0.0004024 \| 130 \| 0.144 \| 201 \| 133 \| 13 \| 8 \| 187 \| 1 \| 183 \| Uncharacterized protein \| Uncharacterized protein; afdb-uniprot50 \| AF-A0A2G6EXL3-F1-MODEL\_V4 \| 1.0 \| 0.006168 \| 129 \| 0.127 \| 110 \| 86 \| 4 \| 46 \| 152 \| 5 \| 107 \| Uncharacterized protein \| Uncharacterized protein; afdb-uniprot50 \| AF-A0A7T8RK97-F1-MODEL\_V4 \| 1.0 \| 0.006168 \| 129 \| 0.115 \| 147 \| 107 \| 5 \| 7 \| 153 \| 1 \| 124 \| DUF806 family protein \| DUF806 family protein; afdb-uniprot50 \| AF-A0A6L7Y352-F1-MODEL\_V4 \| 1.0 \| 0.007342 \| 129 \| 0.129 \| 155 \| 113 \| 7 \| 7 \| 156 \| 1 \| 138 \| Uncharacterized protein \| Uncharacterized protein; afdb-uniprot50 \| AF-A0A524GQA9-F1-MODEL\_V4 \| 1.0 \| 0.00193 \| 129 \| 0.13 \| 168 \| 122 \| 13 \| 1 \| 155 \| 1 \| 157 \| Uncharacterized protein \| Uncharacterized protein; afdb-uniprot50 \| AF-A0A0T9QYM9-F1-MODEL\_V4 \| 1.0 \| 0.001362 \| 129 \| 0.107 \| 158 \| 112 \| 9 \| 27 \| 166 \| 2 \| 148 \| Mu-like prophage protein gp37 \| Mu-like prophage protein gp37; afdb-uniprot50 \| AF-A0A2W5QQ02-F1-MODEL\_V4 \| 1.0 \| 0.003451 \| 128 \| 0.136 \| 147 \| 100 \| 10 \| 14 \| 153 \| 2 \| 128 \| Uncharacterized protein \| Uncharacterized protein; afdb-uniprot50 \| AF-A0A246S189-F1-MODEL\_V4 \| 1.0 \| 0.001719 \| 128 \| 0.124 \| 153 \| 106 \| 9 \| 7 \| 152 \| 3 \| 134 \| Uncharacterized protein \| Uncharacterized protein; afdb-uniprot50 \| AF-A0A4Q4GUU5-F1-MODEL\_V4 \| 1.0 \| 0.001286 \| 128 \| 0.114 \| 166 \| 123 \| 10 \| 1 \| 158 \| 2 \| 151 \| Uncharacterized protein \| Uncharacterized protein; afdb-uniprot50 \| AF-A0A2I0CTH9-F1-MODEL\_V4 \| 1.0 \| 0.004889 \| 128 \| 0.104 \| 163 \| 117 \| 13 \| 9 \| 155 \| 5 \| 154 \| Uncharacterized protein \| Uncharacterized protein; afdb-uniprot50 \| AF-A0A016XM23-F1-MODEL\_V4 \| 1.0 \| 0.003072 \| 128 \| 0.109 \| 183 \| 132 \| 9 \| 9 \| 179 \| 4 \| 167 \| Uncharacterized protein \| Uncharacterized protein; afdb-uniprot50 \| AF-A0A3N1J5Z6-F1-MODEL\_V4 \| 1.0 \| 0.001822 \| 128 \| 0.098 \| 183 \| 135 \| 10 \| 7 \| 176 \| 1 \| 166 \| Uncharacterized protein \| Uncharacterized protein; afdb-uniprot50 \| AF-A0A1M3E549-F1-MODEL\_V4 \| 1.0 \| 0.001622 \| 128 \| 0.12 \| 208 \| 141 \| 13 \| 1 \| 186 \| 2 \| 189 \| Uncharacterized protein \| Uncharacterized protein; afdb-uniprot50 \| AF-A0A179C349-F1-MODEL\_V4 \| 1.0 \| 0.003657 \| 127 \| 0.116 \| 146 \| 100 \| 7 \| 11 \| 154 \| 4 \| 122 \| Uncharacterized protein \| Uncharacterized protein; afdb-uniprot50 \| AF-A0A0R2K2G6-F1-MODEL\_V4 \| 1.0 \| 0.002298 \| 127 \| 0.135 \| 148 \| 98 \| 8 \| 10 \| 152 \| 2 \| 124 \| Uncharacterized protein \| Uncharacterized protein; afdb-uniprot50 \| AF-A0A1G3BXB4-F1-MODEL\_V4 \| 1.0 \| 0.003876 \| 127 \| 0.089 \| 167 \| 126 \| 10 \| 7 \| 165 \| 2 \| 150 \| Uncharacterized protein \| Uncharacterized protein; afdb-uniprot50 \| AF-A0A7X9RUM6-F1-MODEL\_V4 \| 1.0 \| 0.002046 \| 127 \| 0.119 \| 167 \| 125 \| 10 \| 12 \| 166 \| 5 \| 161 \| Uncharacterized protein \| Uncharacterized protein; afdb-uniprot50 \| AF-A0A6H1ZMY2-F1-MODEL\_V4 \| 1.0 \| 0.0004789 \| 127 \| 0.113 \| 176 \| 132 \| 10 \| 4 \| 163 \| 2 \| 169 \| Uncharacterized protein \| Uncharacterized protein; afdb-uniprot50 \| AF-A0A841QIH0-F1-MODEL\_V4 \| 1.0 \| 0.0006786 \| 127 \| 0.126 \| 182 \| 121 \| 13 \| 1 \| 165 \| 4 \| 164 \| Uncharacterized protein \| Uncharacterized protein; afdb-uniprot50 \| AF-A0A3A6MU72-F1-MODEL\_V4 \| 1.0 \| 0.0006042 \| 127 \| 0.101 \| 167 \| 125 \| 8 \| 7 \| 166 \| 1 \| 149 \| Uncharacterized protein \| Uncharacterized protein; afdb-uniprot50 \| AF-A0A177PHJ4-F1-MODEL\_V4 \| 1.0 \| 0.003876 \| 126 \| 0.162 \| 154 \| 101 \| 10 \| 7 \| 152 \| 2 \| 135 \| Uncharacterized protein \| Uncharacterized protein; afdb-uniprot50 \| AF-A0A2U2DWH8-F1-MODEL\_V4 \| 1.0 \| 0.002046 \| 126 \| 0.097 \| 164 \| 122 \| 9 \| 7 \| 156 \| 2 \| 153 \| Uncharacterized protein \| Uncharacterized protein; afdb-uniprot50 \| AF-A0A3N5LHM6-F1-MODEL\_V4 \| 1.0 \| 0.001145 \| 126 \| 0.13 \| 161 \| 114 \| 11 \| 9 \| 152 \| 6 \| 157 \| Uncharacterized protein \| Uncharacterized protein; afdb-uniprot50 \| AF-A0A089ZCH0-F1-MODEL\_V4 \| 1.0 \| 0.00108 \| 126 \| 0.124 \| 161 \| 117 \| 9 \| 9 \| 153 \| 9 \| 161 \| Phage-related protein \| Phage-related protein; afdb-uniprot50 \| AF-X0VBA1-F1-MODEL\_V4 \| 1.0 \| 6.647e-05 \| 126 \| 0.141 \| 184 \| 125 \| 10 \| 30 \| 187 \| 14 \| 190 \| Uncharacterized protein \| Uncharacterized protein; afdb-uniprot50 \| AF-A0A3N7CE51-F1-MODEL\_V4 \| 1.0 \| 0.001822 \| 126 \| 0.149 \| 201 \| 134 \| 12 \| 1 \| 182 \| 1 \| 183 \| Uncharacterized protein \| Uncharacterized protein; afdb-uniprot50 \| AF-E6YRT6-F1-MODEL\_V4 \| 1.0 \| 0.001145 \| 126 \| 0.111 \| 188 \| 140 \| 8 \| 10 \| 188 \| 46 \| 215 \| Uncharacterized protein \| Uncharacterized protein; afdb-uniprot50 \| AF-A0A2N6QWT3-F1-MODEL\_V4 \| 1.0 \| 0.004353 \| 125 \| 0.119 \| 159 \| 111 \| 7 \| 7 \| 162 \| 1 \| 133 \| Phage tail protein \| Phage tail protein; afdb-uniprot50 \| AF-A0A0N1BU45-F1-MODEL\_V4 \| 1.0 \| 0.0006403 \| 125 \| 0.144 \| 159 \| 101 \| 9 \| 7 \| 152 \| 1 \| 137 \| Uncharacterized protein \| Uncharacterized protein; afdb-uniprot50 \| AF-A0A255YQH8-F1-MODEL\_V4 \| 1.0 \| 0.002435 \| 125 \| 0.173 \| 156 \| 111 \| 7 \| 1 \| 152 \| 1 \| 142 \| Uncharacterized protein \| Uncharacterized protein; afdb-uniprot50 \| AF-A0A1H3TG38-F1-MODEL\_V4 \| 1.0 \| 0.002298 \| 125 \| 0.156 \| 172 \| 109 \| 11 \| 9 \| 165 \| 6 \| 156 \| Uncharacterized protein \| Uncharacterized protein; afdb-uniprot50 \| AF-A0A6I7MP73-F1-MODEL\_V4 \| 1.0 \| 0.002046 \| 125 \| 0.14 \| 171 \| 115 \| 10 \| 10 \| 162 \| 2 \| 158 \| Uncharacterized protein \| Uncharacterized protein; afdb-uniprot50 \| AF-A0A1V6DHH5-F1-MODEL\_V4 \| 1.0 \| 0.001286 \| 125 \| 0.125 \| 176 \| 123 \| 12 \| 8 \| 166 \| 1 \| 162 \| Uncharacterized protein \| Uncharacterized protein; afdb-uniprot50 \| AF-C1D6N8-F1-MODEL\_V4 \| 1.0 \| 0.001286 \| 125 \| 0.137 \| 232 \| 139 \| 14 \| 1 \| 189 \| 1 \| 214 \| DUF1834 domain containing protein \| DUF1834 domain containing protein; afdb-uniprot50 \| AF-A0A844GBQ3-F1-MODEL\_V4 \| 1.0 \| 0.009262 \| 124 \| 0.1 \| 129 \| 94 \| 10 \| 29 \| 152 \| 5 \| 116 \| Uncharacterized protein \| Uncharacterized protein; afdb-uniprot50 \| AF-A0A0R2BTK5-F1-MODEL\_V4 \| 1.0 \| 0.006928 \| 124 \| 0.099 \| 151 \| 108 \| 7 \| 7 \| 155 \| 2 \| 126 \| Uncharacterized protein \| Uncharacterized protein; afdb-uniprot50 \| AF-A0A366FMK0-F1-MODEL\_V4 \| 1.0 \| 0.002168 \| 124 \| 0.201 \| 154 \| 103 \| 11 \| 7 \| 151 \| 1 \| 143 \| Uncharacterized protein \| Uncharacterized protein; afdb-uniprot50 \| AF-W0RPE6-F1-MODEL\_V4 \| 1.0 \| 0.0009616 \| 124 \| 0.16 \| 175 \| 111 \| 12 \| 1 \| 157 \| 1 \| 157 \| Uncharacterized protein \| Uncharacterized protein; afdb-uniprot50 \| AF-A0A126R8T4-F1-MODEL\_V4 \| 1.0 \| 0.0008078 \| 124 \| 0.174 \| 166 \| 111 \| 10 \| 9 \| 165 \| 9 \| 157 \| Uncharacterized protein \| Uncharacterized protein; afdb-uniprot50 \| AF-A0A142XV75-F1-MODEL\_V4 \| 1.0 \| 0.00582 \| 124 \| 0.116 \| 171 \| 121 \| 10 \| 1 \| 155 \| 1 \| 157 \| Uncharacterized protein \| Uncharacterized protein; afdb-uniprot50 \| AF-A0A7V3W3S9-F1-MODEL\_V4 \| 1.0 \| 0.001019 \| 124 \| 0.13 \| 184 \| 125 \| 13 \| 1 \| 157 \| 1 \| 176 \| Uncharacterized protein \| Uncharacterized protein; afdb-uniprot50 \| AF-R7I5X1-F1-MODEL\_V4 \| 1.0 \| 0.001622 \| 124 \| 0.09 \| 199 \| 149 \| 11 \| 7 \| 189 \| 1 \| 183 \| Uncharacterized protein \| Uncharacterized protein; afdb-uniprot50 \| AF-A0A502HRN3-F1-MODEL\_V4 \| 1.0 \| 0.0002124 \| 124 \| 0.135 \| 192 \| 118 \| 13 \| 30 \| 189 \| 108 \| 283 \| Uncharacterized protein \| Uncharacterized protein; afdb-uniprot50 \| AF-A0A0F2IYW9-F1-MODEL\_V4 \| 1.0 \| 0.009262 \| 123 \| 0.134 \| 119 \| 90 \| 5 \| 40 \| 157 \| 2 \| 108 \| Uncharacterized protein \| Uncharacterized protein; afdb-uniprot50 \| AF-A0A2E7W9M7-F1-MODEL\_V4 \| 1.0 \| 0.002735 \| 123 \| 0.129 \| 154 \| 110 \| 7 \| 7 \| 154 \| 1 \| 136 \| Uncharacterized protein \| Uncharacterized protein; afdb-uniprot50 \| AF-A0A4R7BTN2-F1-MODEL\_V4 \| 1.0 \| 0.004108 \| 123 \| 0.129 \| 154 \| 108 \| 9 \| 7 \| 153 \| 1 \| 135 \| Uncharacterized protein DUF4128 \| Uncharacterized protein DUF4128; afdb-uniprot50 \| AF-A0A2D6XI96-F1-MODEL\_V4 \| 1.0 \| 0.006537 \| 123 \| 0.147 \| 149 \| 115 \| 5 \| 8 \| 153 \| 5 \| 144 \| Uncharacterized protein \| Uncharacterized protein; afdb-uniprot50 \| AF-A0A7X9JN73-F1-MODEL\_V4 \| 1.0 \| 0.003072 \| 123 \| 0.111 \| 152 \| 107 \| 8 \| 9 \| 152 \| 19 \| 150 \| Uncharacterized protein \| Uncharacterized protein; afdb-uniprot50 \| AF-A0A496QXT6-F1-MODEL\_V4 \| 1.0 \| 0.001286 \| 123 \| 0.118 \| 152 \| 117 \| 9 \| 9 \| 154 \| 12 \| 152 \| Uncharacterized protein \| Uncharacterized protein; afdb-uniprot50 \| AF-A0A4Z0IPR0-F1-MODEL\_V4 \| 1.0 \| 0.006537 \| 123 \| 0.156 \| 160 \| 111 \| 8 \| 9 \| 155 \| 4 \| 152 \| Uncharacterized protein \| Uncharacterized protein; afdb-uniprot50 \| AF-A0A2D6MCC7-F1-MODEL\_V4 \| 1.0 \| 0.002581 \| 123 \| 0.141 \| 170 \| 118 \| 12 \| 7 \| 164 \| 3 \| 156 \| Uncharacterized protein \| Uncharacterized protein; afdb-uniprot50 \| AF-A0A4U6D146-F1-MODEL\_V4 \| 1.0 \| 0.002899 \| 123 \| 0.089 \| 178 \| 130 \| 8 \| 12 \| 178 \| 4 \| 160 \| Uncharacterized protein \| Uncharacterized protein; afdb-uniprot50 \| AF-A0A521H788-F1-MODEL\_V4 \| 1.0 \| 0.003256 \| 123 \| 0.09 \| 188 \| 141 \| 9 \| 12 \| 189 \| 8 \| 175 \| Uncharacterized protein \| Uncharacterized protein; afdb-uniprot50 \| AF-A0A4Z0SC48-F1-MODEL\_V4 \| 1.0 \| 0.006168 \| 122 \| 0.132 \| 151 \| 102 \| 8 \| 7 \| 155 \| 2 \| 125 \| Uncharacterized protein \| Uncharacterized protein; afdb-uniprot50 \| AF-A0A069CSJ6-F1-MODEL\_V4 \| 1.0 \| 0.00582 \| 122 \| 0.111 \| 152 \| 107 \| 7 \| 7 \| 153 \| 1 \| 129 \| Phage tail protein \| Phage tail protein; afdb-uniprot50 \| AF-A0A2P8FP87-F1-MODEL\_V4 \| 1.0 \| 0.004353 \| 122 \| 0.103 \| 155 \| 108 \| 7 \| 7 \| 153 \| 1 \| 132 \| Uncharacterized protein DUF4128 \| Uncharacterized protein DUF4128; afdb-uniprot50 \| AF-A0A661DGF6-F1-MODEL\_V4 \| 1.0 \| 0.001822 \| 122 \| 0.128 \| 148 \| 107 \| 12 \| 9 \| 153 \| 8 \| 136 \| Uncharacterized protein \| Uncharacterized protein; afdb-uniprot50 \| AF-A0A2S5CGG2-F1-MODEL\_V4 \| 1.0 \| 0.00153 \| 122 \| 0.153 \| 163 \| 106 \| 10 \| 6 \| 155 \| 3 \| 146 \| Uncharacterized protein \| Uncharacterized protein; afdb-uniprot50 \| AF-A0A1E7Q8E4-F1-MODEL\_V4 \| 1.0 \| 0.002435 \| 122 \| 0.115 \| 164 \| 109 \| 12 \| 9 \| 154 \| 4 \| 149 \| Uncharacterized protein \| Uncharacterized protein; afdb-uniprot50 \| AF-K9RS62-F1-MODEL\_V4 \| 1.0 \| 0.005182 \| 122 \| 0.106 \| 169 \| 125 \| 8 \| 1 \| 163 \| 1 \| 149 \| Uncharacterized protein \| Uncharacterized protein; afdb-uniprot50 \| AF-A0A1V5IG05-F1-MODEL\_V4 \| 1.0 \| 0.003072 \| 122 \| 0.122 \| 171 \| 117 \| 9 \| 1 \| 155 \| 1 \| 154 \| Uncharacterized protein \| Uncharacterized protein; afdb-uniprot50 \| AF-A0A5C8UQH2-F1-MODEL\_V4 \| 1.0 \| 0.004353 \| 122 \| 0.119 \| 167 \| 125 \| 9 \| 10 \| 166 \| 6 \| 160 \| Uncharacterized protein \| Uncharacterized protein; afdb-uniprot50 \| AF-A0A3D5AU90-F1-MODEL\_V4 \| 1.0 \| 0.006928 \| 121 \| 0.15 \| 113 \| 82 \| 4 \| 48 \| 157 \| 5 \| 106 \| Uncharacterized protein \| Uncharacterized protein; afdb-uniprot50 \| AF-A0A2G2PNR6-F1-MODEL\_V4 \| 1.0 \| 0.003072 \| 121 \| 0.12 \| 150 \| 100 \| 10 \| 9 \| 152 \| 8 \| 131 \| Uncharacterized protein \| Uncharacterized protein; afdb-uniprot50 \| AF-A0A175RB75-F1-MODEL\_V4 \| 1.0 \| 0.004613 \| 121 \| 0.136 \| 147 \| 103 \| 10 \| 16 \| 155 \| 9 \| 138 \| Uncharacterized protein \| Uncharacterized protein; afdb-uniprot50 \| AF-A0A0F9QW21-F1-MODEL\_V4 \| 1.0 \| 0.003256 \| 121 \| 0.116 \| 163 \| 121 \| 11 \| 1 \| 152 \| 2 \| 152 \| Uncharacterized protein \| Uncharacterized protein; afdb-uniprot50 \| AF-A0A830FZ37-F1-MODEL\_V4 \| 1.0 \| 0.001362 \| 121 \| 0.135 \| 200 \| 120 \| 12 \| 14 \| 173 \| 4 \| 190 \| Uncharacterized protein \| Uncharacterized protein; afdb-uniprot50 \| AF-A0A1C0V8E1-F1-MODEL\_V4 \| 1.0 \| 0.001362 \| 121 \| 0.125 \| 191 \| 137 \| 11 \| 9 \| 189 \| 5 \| 175 \| Uncharacterized protein \| Uncharacterized protein; afdb-uniprot50 \| AF-A0A645AWW2-F1-MODEL\_V4 \| 1.0 \| 0.006168 \| 120 \| 0.1 \| 159 \| 107 \| 10 \| 8 \| 154 \| 1 \| 135 \| Uncharacterized protein \| Uncharacterized protein; afdb-uniprot50 \| AF-A0A848WCE3-F1-MODEL\_V4 \| 1.0 \| 0.007781 \| 120 \| 0.086 \| 151 \| 109 \| 11 \| 9 \| 153 \| 7 \| 134 \| Uncharacterized protein \| Uncharacterized protein; afdb-uniprot50 \| AF-A0A0A8K5T5-F1-MODEL\_V4 \| 1.0 \| 0.004353 \| 120 \| 0.177 \| 158 \| 98 \| 11 \| 7 \| 154 \| 2 \| 137 \| Uncharacterized protein \| Uncharacterized protein; afdb-uniprot50 \| AF-A0A1W9Z0J1-F1-MODEL\_V4 \| 1.0 \| 0.005182 \| 120 \| 0.103 \| 165 \| 116 \| 13 \| 1 \| 156 \| 2 \| 143 \| Uncharacterized protein \| Uncharacterized protein; afdb-uniprot50 \| AF-A0A2S3W4U1-F1-MODEL\_V4 \| 1.0 \| 0.00153 \| 120 \| 0.109 \| 155 \| 111 \| 8 \| 43 \| 188 \| 1 \| 137 \| Uncharacterized protein \| Uncharacterized protein; afdb-uniprot50 \| AF-A0A7W6K7J7-F1-MODEL\_V4 \| 1.0 \| 0.00193 \| 120 \| 0.12 \| 166 \| 109 \| 8 \| 1 \| 152 \| 13 \| 155 \| Uncharacterized protein \| Uncharacterized protein; afdb-uniprot50 \| AF-A0A7C3USK7-F1-MODEL\_V4 \| 1.0 \| 0.001213 \| 120 \| 0.142 \| 176 \| 115 \| 12 \| 1 \| 156 \| 2 \| 161 \| Uncharacterized protein \| Uncharacterized protein; afdb-uniprot50 \| AF-Q141R8-F1-MODEL\_V4 \| 1.0 \| 0.001145 \| 120 \| 0.169 \| 165 \| 121 \| 7 \| 10 \| 172 \| 8 \| 158 \| Uncharacterized protein \| Uncharacterized protein; afdb-uniprot50 \| AF-A0A7C6E6J6-F1-MODEL\_V4 \| 1.0 \| 0.007342 \| 119 \| 0.102 \| 136 \| 96 \| 7 \| 30 \| 154 \| 9 \| 129 \| Uncharacterized protein \| Uncharacterized protein; afdb-uniprot50 \| AF-Q5F997-F1-MODEL\_V4 \| 1.0 \| 0.00193 \| 119 \| 0.123 \| 154 \| 111 \| 11 \| 7 \| 150 \| 1 \| 140 \| Phage associated protein \| Phage associated protein; afdb-uniprot50 \| AF-A0A1B7X3R8-F1-MODEL\_V4 \| 1.0 \| 0.002581 \| 119 \| 0.098 \| 152 \| 117 \| 9 \| 7 \| 152 \| 2 \| 139 \| Uncharacterized protein \| Uncharacterized protein; afdb-uniprot50 \| AF-A0A7V0NNN3-F1-MODEL\_V4 \| 1.0 \| 0.008247 \| 119 \| 0.144 \| 166 \| 112 \| 11 \| 1 \| 154 \| 1 \| 148 \| Uncharacterized protein \| Uncharacterized protein; afdb-uniprot50 \| AF-A0A6I1J9G2-F1-MODEL\_V4 \| 1.0 \| 0.002046 \| 119 \| 0.168 \| 154 \| 112 \| 10 \| 7 \| 152 \| 3 \| 148 \| Uncharacterized protein \| Uncharacterized protein; afdb-uniprot50 \| AF-A0A661TVI5-F1-MODEL\_V4 \| 1.0 \| 0.005492 \| 119 \| 0.118 \| 160 \| 114 \| 7 \| 9 \| 154 \| 4 \| 150 \| Uncharacterized protein \| Uncharacterized protein; afdb-uniprot50 \| AF-A0A402A527-F1-MODEL\_V4 \| 1.0 \| 0.002581 \| 119 \| 0.129 \| 178 \| 114 \| 10 \| 5 \| 165 \| 3 \| 156 \| Uncharacterized protein \| Uncharacterized protein; afdb-uniprot50 \| AF-A0A0F9BJT1-F1-MODEL\_V4 \| 1.0 \| 0.006928 \| 119 \| 0.09 \| 177 \| 140 \| 10 \| 1 \| 165 \| 2 \| 169 \| Uncharacterized protein \| Uncharacterized protein; afdb-uniprot50 \| AF-A0A2D6XLF9-F1-MODEL\_V4 \| 1.0 \| 0.00193 \| 119 \| 0.145 \| 165 \| 121 \| 11 \| 1 \| 154 \| 17 \| 172 \| Uncharacterized protein \| Uncharacterized protein; afdb-uniprot50 \| AF-A0A1H9Q9R0-F1-MODEL\_V4 \| 1.0 \| 0.0004789 \| 119 \| 0.173 \| 184 \| 121 \| 11 \| 1 \| 175 \| 1 \| 162 \| Uncharacterized protein \| Uncharacterized protein; afdb-uniprot50 \| AF-A0A1I6VDK5-F1-MODEL\_V4 \| 1.0 \| 0.0005701 \| 119 \| 0.13 \| 207 \| 142 \| 15 \| 1 \| 187 \| 1 \| 189 \| Uncharacterized protein \| Uncharacterized protein; afdb-uniprot50 \| AF-A0A3N5UZL3-F1-MODEL\_V4 \| 1.0 \| 0.004108 \| 118 \| 0.111 \| 162 \| 113 \| 11 \| 5 \| 155 \| 2 \| 143 \| Uncharacterized protein \| Uncharacterized protein; afdb-uniprot50 \| AF-A0A427UIF4-F1-MODEL\_V4 \| 1.0 \| 0.004889 \| 118 \| 0.132 \| 166 \| 117 \| 11 \| 1 \| 160 \| 2 \| 146 \| Uncharacterized protein \| Uncharacterized protein; afdb-uniprot50 \| AF-A0A1E4HKH2-F1-MODEL\_V4 \| 1.0 \| 0.009262 \| 118 \| 0.143 \| 153 \| 109 \| 11 \| 9 \| 152 \| 10 \| 149 \| Uncharacterized protein \| Uncharacterized protein; afdb-uniprot50 \| AF-A0A497AEL7-F1-MODEL\_V4 \| 1.0 \| 0.004613 \| 118 \| 0.086 \| 162 \| 126 \| 11 \| 6 \| 152 \| 2 \| 156 \| Uncharacterized protein \| Uncharacterized protein; afdb-uniprot50 \| AF-A0A6M3LF59-F1-MODEL\_V4 \| 1.0 \| 0.003072 \| 118 \| 0.104 \| 172 \| 126 \| 13 \| 1 \| 156 \| 3 \| 162 \| Uncharacterized protein \| Uncharacterized protein; afdb-uniprot50 \| AF-A0A843ACS1-F1-MODEL\_V4 \| 1.0 \| 0.0004789 \| 118 \| 0.104 \| 182 \| 126 \| 12 \| 1 \| 164 \| 1 \| 163 \| Uncharacterized protein \| Uncharacterized protein; afdb-uniprot50 \| AF-A0A6M3IUD6-F1-MODEL\_V4 \| 1.0 \| 0.001822 \| 118 \| 0.136 \| 169 \| 121 \| 7 \| 7 \| 168 \| 1 \| 151 \| Uncharacterized protein \| Uncharacterized protein; afdb-uniprot50 \| AF-A0A661HGP8-F1-MODEL\_V4 \| 1.0 \| 0.003876 \| 118 \| 0.094 \| 180 \| 142 \| 10 \| 11 \| 181 \| 6 \| 173 \| Uncharacterized protein \| Uncharacterized protein; afdb-uniprot50 \| AF-A0A506TZ73-F1-MODEL\_V4 \| 1.0 \| 0.001145 \| 117 \| 0.165 \| 163 \| 105 \| 13 \| 1 \| 152 \| 3 \| 145 \| Uncharacterized protein \| Uncharacterized protein; afdb-uniprot50 \| AF-A0A2I0FU84-F1-MODEL\_V4 \| 1.0 \| 0.002581 \| 117 \| 0.123 \| 194 \| 131 \| 12 \| 15 \| 186 \| 5 \| 181 \| Uncharacterized protein \| Uncharacterized protein; afdb-uniprot50 \| AF-A0A1H8IRD0-F1-MODEL\_V4 \| 1.0 \| 0.006928 \| 116 \| 0.1 \| 169 \| 130 \| 7 \| 1 \| 164 \| 1 \| 152 \| Uncharacterized protein \| Uncharacterized protein; afdb-uniprot50 \| AF-A0A5C5VW99-F1-MODEL\_V4 \| 1.0 \| 0.002046 \| 116 \| 0.107 \| 167 \| 116 \| 14 \| 14 \| 165 \| 4 \| 152 \| Uncharacterized protein \| Uncharacterized protein; afdb-uniprot50 \| AF-D2U1L2-F1-MODEL\_V4 \| 1.0 \| 0.005492 \| 116 \| 0.094 \| 137 \| 99 \| 5 \| 43 \| 165 \| 16 \| 141 \| Conserved hypothetical phage protein \| Conserved hypothetical phage protein; afdb-uniprot50 \| AF-A0A2D6F2E1-F1-MODEL\_V4 \| 1.0 \| 0.001622 \| 116 \| 0.165 \| 181 \| 112 \| 14 \| 7 \| 166 \| 2 \| 164 \| Uncharacterized protein \| Uncharacterized protein; afdb-uniprot50 \| AF-A0A3D3SH89-F1-MODEL\_V4 \| 1.0 \| 0.002581 \| 116 \| 0.084 \| 190 \| 142 \| 12 \| 7 \| 183 \| 1 \| 171 \| Uncharacterized protein \| Uncharacterized protein; afdb-uniprot50 \| AF-A0A1B7X0V1-F1-MODEL\_V4 \| 1.0 \| 0.000284 \| 116 \| 0.148 \| 209 \| 137 \| 13 \| 7 \| 189 \| 3 \| 196 \| Uncharacterized protein \| Uncharacterized protein; afdb-uniprot50 \| AF-A0A841LE00-F1-MODEL\_V4 \| 1.0 \| 0.006537 \| 116 \| 0.11 \| 227 \| 140 \| 15 \| 1 \| 189 \| 2 \| 204 \| Uncharacterized protein \| Uncharacterized protein; afdb-uniprot50 \| AF-A0A1I3HI47-F1-MODEL\_V4 \| 1.0 \| 0.008247 \| 115 \| 0.14 \| 150 \| 103 \| 10 \| 10 \| 153 \| 6 \| 135 \| Uncharacterized protein \| Uncharacterized protein; afdb-uniprot50 \| AF-A0A1E4ZIU0-F1-MODEL\_V4 \| 1.0 \| 0.005182 \| 115 \| 0.093 \| 161 \| 117 \| 12 \| 9 \| 164 \| 6 \| 142 \| Uncharacterized protein \| Uncharacterized protein; afdb-uniprot50 \| AF-A0A7W9VZR3-F1-MODEL\_V4 \| 1.0 \| 0.006537 \| 115 \| 0.136 \| 161 \| 108 \| 12 \| 7 \| 156 \| 1 \| 141 \| Uncharacterized protein \| Uncharacterized protein; afdb-uniprot50 \| AF-A0A3S0BCC7-F1-MODEL\_V4 \| 1.0 \| 0.00874 \| 115 \| 0.124 \| 169 \| 115 \| 12 \| 9 \| 159 \| 4 \| 157 \| Uncharacterized protein \| Uncharacterized protein; afdb-uniprot50 \| AF-A0A5C7MB17-F1-MODEL\_V4 \| 1.0 \| 0.004889 \| 115 \| 0.167 \| 161 \| 110 \| 10 \| 9 \| 154 \| 16 \| 167 \| Uncharacterized protein \| Uncharacterized protein; afdb-uniprot50 \| AF-A0A523Z9A7-F1-MODEL\_V4 \| 1.0 \| 0.009262 \| 115 \| 0.133 \| 165 \| 125 \| 8 \| 6 \| 156 \| 7 \| 167 \| Uncharacterized protein \| Uncharacterized protein; afdb-uniprot50 \| AF-A0A0U1K6V6-F1-MODEL\_V4 \| 1.0 \| 0.00153 \| 115 \| 0.117 \| 205 \| 133 \| 14 \| 1 \| 187 \| 26 \| 200 \| Putative ATP-binding protein \| Putative ATP-binding protein; afdb-uniprot50 \| AF-A0A0Q5ZKT3-F1-MODEL\_V4 \| 1.0 \| 0.007342 \| 114 \| 0.113 \| 150 \| 113 \| 9 \| 7 \| 152 \| 1 \| 134 \| Uncharacterized protein \| Uncharacterized protein; afdb-uniprot50 \| AF-A0A1H9QD02-F1-MODEL\_V4 \| 1.0 \| 0.002581 \| 114 \| 0.14 \| 157 \| 104 \| 9 \| 7 \| 154 \| 2 \| 136 \| Uncharacterized protein \| Uncharacterized protein; afdb-uniprot50 \| AF-A0A257ZLI7-F1-MODEL\_V4 \| 1.0 \| 0.004353 \| 114 \| 0.104 \| 162 \| 113 \| 7 \| 7 \| 152 \| 2 \| 147 \| Uncharacterized protein \| Uncharacterized protein; afdb-uniprot50 \| AF-A0A1H8A9M5-F1-MODEL\_V4 \| 1.0 \| 0.001719 \| 114 \| 0.131 \| 182 \| 127 \| 9 \| 7 \| 165 \| 2 \| 175 \| Uncharacterized protein \| Uncharacterized protein; afdb-uniprot50 \| AF-I4E5S8-F1-MODEL\_V4 \| 1.0 \| 0.004108 \| 114 \| 0.133 \| 203 \| 119 \| 11 \| 27 \| 187 \| 2 \| 189 \| Uncharacterized protein \| Uncharacterized protein; afdb-uniprot50 \| AF-A0A3A4VTU4-F1-MODEL\_V4 \| 1.0 \| 0.006928 \| 113 \| 0.118 \| 152 \| 110 \| 10 \| 12 \| 154 \| 3 \| 139 \| Uncharacterized protein \| Uncharacterized protein; afdb-uniprot50 \| AF-A0A0F9N372-F1-MODEL\_V4 \| 1.0 \| 0.00874 \| 113 \| 0.115 \| 165 \| 123 \| 12 \| 9 \| 165 \| 5 \| 154 \| Uncharacterized protein \| Uncharacterized protein; afdb-uniprot50 \| AF-A0A3D4ZXF2-F1-MODEL\_V4 \| 1.0 \| 0.004613 \| 113 \| 0.174 \| 183 \| 107 \| 15 \| 1 \| 153 \| 1 \| 169 \| Uncharacterized protein \| Uncharacterized protein; afdb-uniprot50 \| AF-A0A249PA02-F1-MODEL\_V4 \| 1.0 \| 0.001213 \| 112 \| 0.118 \| 161 \| 112 \| 10 \| 1 \| 154 \| 1 \| 138 \| Uncharacterized protein \| Uncharacterized protein; afdb-uniprot50 \| AF-A0A6B0W5K2-F1-MODEL\_V4 \| 1.0 \| 0.004613 \| 112 \| 0.144 \| 159 \| 109 \| 10 \| 11 \| 156 \| 8 \| 152 \| Uncharacterized protein \| Uncharacterized protein; afdb-uniprot50 \| AF-A0A1S2VDH5-F1-MODEL\_V4 \| 1.0 \| 0.003072 \| 112 \| 0.141 \| 184 \| 107 \| 14 \| 13 \| 165 \| 3 \| 166 \| Uncharacterized protein \| Uncharacterized protein; afdb-uniprot50 \| AF-A0A0F9X9T6-F1-MODEL\_V4 \| 1.0 \| 0.003256 \| 112 \| 0.111 \| 179 \| 116 \| 12 \| 1 \| 153 \| 1 \| 162 \| Uncharacterized protein \| Uncharacterized protein; afdb-uniprot50 \| AF-A0A1B7LDB7-F1-MODEL\_V4 \| 1.0 \| 0.004108 \| 112 \| 0.142 \| 169 \| 115 \| 9 \| 9 \| 166 \| 10 \| 159 \| Uncharacterized protein \| Uncharacterized protein; afdb-uniprot50 \| AF-A0A0F9UL68-F1-MODEL\_V4 \| 1.0 \| 0.001622 \| 112 \| 0.15 \| 233 \| 142 \| 17 \| 1 \| 188 \| 1 \| 222 \| Uncharacterized protein \| Uncharacterized protein; afdb-uniprot50 \| AF-A0A2E8NYT8-F1-MODEL\_V4 \| 1.0 \| 0.001213 \| 112 \| 0.113 \| 238 \| 129 \| 14 \| 1 \| 166 \| 1 \| 228 \| Uncharacterized protein \| Uncharacterized protein; afdb-uniprot50 \| AF-I9N4S3-F1-MODEL\_V4 \| 1.0 \| 0.005492 \| 111 \| 0.128 \| 148 \| 106 \| 7 \| 14 \| 155 \| 8 \| 138 \| Uncharacterized protein \| Uncharacterized protein; afdb-uniprot50 \| AF-A0A1M4Y9T1-F1-MODEL\_V4 \| 1.0 \| 0.004613 \| 111 \| 0.138 \| 152 \| 106 \| 9 \| 14 \| 152 \| 4 \| 143 \| Uncharacterized protein \| Uncharacterized protein; afdb-uniprot50 \| AF-A0A1V5WR40-F1-MODEL\_V4 \| 1.0 \| 0.006537 \| 111 \| 0.114 \| 183 \| 136 \| 13 \| 1 \| 174 \| 1 \| 166 \| Uncharacterized protein \| Uncharacterized protein; afdb-uniprot50 \| AF-A0A5C7QH98-F1-MODEL\_V4 \| 1.0 \| 0.00193 \| 111 \| 0.142 \| 190 \| 120 \| 15 \| 10 \| 168 \| 22 \| 199 \| Uncharacterized protein \| Uncharacterized protein; afdb-uniprot50 \| AF-A0A660MBK1-F1-MODEL\_V4 \| 1.0 \| 0.006928 \| 110 \| 0.169 \| 159 \| 108 \| 9 \| 12 \| 156 \| 5 \| 153 \| Uncharacterized protein \| Uncharacterized protein; afdb-uniprot50 \| AF-A0A0F9BKL4-F1-MODEL\_V4 \| 1.0 \| 0.005182 \| 110 \| 0.118 \| 186 \| 136 \| 10 \| 7 \| 179 \| 1 \| 171 \| Uncharacterized protein \| Uncharacterized protein; afdb-uniprot50 \| AF-A0A212KK97-F1-MODEL\_V4 \| 1.0 \| 0.005492 \| 110 \| 0.149 \| 167 \| 116 \| 6 \| 9 \| 161 \| 4 \| 158 \| Uncharacterized protein \| Uncharacterized protein; afdb-uniprot50 \| AF-A0A1E3H1X1-F1-MODEL\_V4 \| 1.0 \| 0.007342 \| 110 \| 0.116 \| 155 \| 121 \| 5 \| 1 \| 150 \| 110 \| 253 \| Uncharacterized protein \| Uncharacterized protein; afdb-uniprot50 \| AF-A0A2D7VXM3-F1-MODEL\_V4 \| 1.0 \| 0.004353 \| 110 \| 0.11 \| 136 \| 91 \| 7 \| 30 \| 153 \| 227 \| 344 \| Uncharacterized protein \| Uncharacterized protein; afdb-uniprot50 \| AF-A0A7C1PIS6-F1-MODEL\_V4 \| 1.0 \| 0.003451 \| 110 \| 0.147 \| 156 \| 104 \| 11 \| 29 \| 166 \| 265 \| 409 \| Uncharacterized protein \| Uncharacterized protein; afdb-uniprot50 \| AF-A0A3D9E5J7-F1-MODEL\_V4 \| 1.0 \| 0.007342 \| 109 \| 0.099 \| 151 \| 122 \| 8 \| 10 \| 153 \| 11 \| 154 \| Uncharacterized protein \| Uncharacterized protein; afdb-uniprot50 \| AF-A0A371JLB5-F1-MODEL\_V4 \| 1.0 \| 0.002168 \| 109 \| 0.126 \| 214 \| 136 \| 14 \| 1 \| 178 \| 13 \| 211 \| Uncharacterized protein \| Uncharacterized protein; afdb-uniprot50 \| AF-A0A523RTI9-F1-MODEL\_V4 \| 1.0 \| 0.003256 \| 108 \| 0.108 \| 157 \| 115 \| 9 \| 9 \| 157 \| 10 \| 149 \| Uncharacterized protein \| Uncharacterized protein; afdb-uniprot50 \| AF-A0A1M7AHU1-F1-MODEL\_V4 \| 1.0 \| 0.003876 \| 108 \| 0.102 \| 185 \| 132 \| 14 \| 7 \| 172 \| 1 \| 170 \| Uncharacterized protein \| Uncharacterized protein; afdb-uniprot50 \| AF-A0A559IZI9-F1-MODEL\_V4 \| 1.0 \| 0.006537 \| 108 \| 0.091 \| 186 \| 123 \| 10 \| 10 \| 165 \| 4 \| 173 \| Uncharacterized protein \| Uncharacterized protein; afdb-uniprot50 \| AF-A0A2J9S7X7-F1-MODEL\_V4 \| 1.0 \| 0.004889 \| 108 \| 0.116 \| 198 \| 130 \| 12 \| 7 \| 187 \| 1 \| 170 \| ATP-binding protein \| ATP-binding protein; afdb-uniprot50 \| AF-A0A2U8WAI8-F1-MODEL\_V4 \| 1.0 \| 0.003451 \| 108 \| 0.14 \| 192 \| 141 \| 10 \| 1 \| 187 \| 1 \| 173 \| Uncharacterized protein \| Uncharacterized protein; afdb-uniprot50 \| AF-A0A2E7BJZ9-F1-MODEL\_V4 \| 1.0 \| 0.002899 \| 107 \| 0.13 \| 168 \| 121 \| 9 \| 9 \| 164 \| 5 \| 159 \| Uncharacterized protein \| Uncharacterized protein; afdb-uniprot50 \| AF-A0A5C7JA81-F1-MODEL\_V4 \| 1.0 \| 0.003451 \| 107 \| 0.105 \| 209 \| 118 \| 10 \| 1 \| 154 \| 1 \| 195 \| Uncharacterized protein \| Uncharacterized protein; afdb-uniprot50 \| AF-A0A1Z9SNZ4-F1-MODEL\_V4 \| 1.0 \| 0.004353 \| 107 \| 0.092 \| 226 \| 135 \| 17 \| 2 \| 172 \| 1 \| 211 \| Uncharacterized protein \| Uncharacterized protein; afdb-uniprot50 \| AF-A0A7U7G943-F1-MODEL\_V4 \| 1.0 \| 0.006168 \| 106 \| 0.125 \| 159 \| 102 \| 13 \| 11 \| 154 \| 4 \| 140 \| Uncharacterized protein \| Uncharacterized protein; afdb-uniprot50 \| AF-A0A2S5N3V3-F1-MODEL\_V4 \| 1.0 \| 0.00874 \| 106 \| 0.167 \| 155 \| 106 \| 12 \| 7 \| 153 \| 1 \| 140 \| Uncharacterized protein \| Uncharacterized protein; afdb-uniprot50 \| AF-A0A1V5V8V5-F1-MODEL\_V4 \| 1.0 \| 0.005182 \| 106 \| 0.098 \| 172 \| 123 \| 13 \| 6 \| 163 \| 3 \| 156 \| Uncharacterized protein \| Uncharacterized protein; afdb-uniprot50 \| AF-A0A5M9HA02-F1-MODEL\_V4 \| 1.0 \| 0.00874 \| 106 \| 0.128 \| 187 \| 117 \| 13 \| 7 \| 165 \| 1 \| 169 \| Uncharacterized protein \| Uncharacterized protein; afdb-uniprot50 \| AF-A0A1A9RL24-F1-MODEL\_V4 \| 1.0 \| 0.002899 \| 106 \| 0.128 \| 195 \| 120 \| 11 \| 7 \| 171 \| 1 \| 175 \| Uncharacterized protein \| Uncharacterized protein; afdb-uniprot50 \| AF-A0A0S8AR84-F1-MODEL\_V4 \| 1.0 \| 0.005492 \| 106 \| 0.097 \| 184 \| 123 \| 12 \| 7 \| 178 \| 1 \| 153 \| Uncharacterized protein \| Uncharacterized protein; afdb-uniprot50 \| AF-A0A0R2MKJ7-F1-MODEL\_V4 \| 1.0 \| 0.003072 \| 105 \| 0.129 \| 155 \| 106 \| 8 \| 7 \| 159 \| 1 \| 128 \| Uncharacterized protein \| Uncharacterized protein; afdb-uniprot50 \| AF-A0A7J2R9I8-F1-MODEL\_V4 \| 1.0 \| 0.00582 \| 105 \| 0.142 \| 161 \| 112 \| 11 \| 5 \| 154 \| 2 \| 147 \| Uncharacterized protein \| Uncharacterized protein; afdb-uniprot50 \| AF-A0A348V8F1-F1-MODEL\_V4 \| 1.0 \| 0.009262 \| 105 \| 0.129 \| 177 \| 114 \| 12 \| 12 \| 163 \| 4 \| 165 \| Uncharacterized protein \| Uncharacterized protein; afdb-uniprot50 \| AF-A0A239ABH2-F1-MODEL\_V4 \| 1.0 \| 0.004889 \| 105 \| 0.129 \| 178 \| 128 \| 11 \| 13 \| 178 \| 4 \| 166 \| Uncharacterized protein \| Uncharacterized protein; afdb-uniprot50 \| AF-A0A7Y6T7P5-F1-MODEL\_V4 \| 1.0 \| 0.002168 \| 105 \| 0.107 \| 196 \| 148 \| 11 \| 1 \| 187 \| 1 \| 178 \| Uncharacterized protein \| Uncharacterized protein; afdb-uniprot50 \| AF-N2AZG9-F1-MODEL\_V4 \| 1.0 \| 0.009262 \| 105 \| 0.109 \| 183 \| 127 \| 15 \| 7 \| 166 \| 1 \| 170 \| Uncharacterized protein \| Uncharacterized protein; afdb-uniprot50 \| AF-A0A1Y4RYL6-F1-MODEL\_V4 \| 1.0 \| 0.006168 \| 104 \| 0.15 \| 179 \| 110 \| 11 \| 13 \| 164 \| 2 \| 165 \| Uncharacterized protein \| Uncharacterized protein; afdb-uniprot50 \| AF-A0A2S2F818-F1-MODEL\_V4 \| 1.0 \| 0.001719 \| 104 \| 0.152 \| 242 \| 122 \| 16 \| 4 \| 173 \| 2 \| 232 \| Uncharacterized protein \| Uncharacterized protein; afdb-uniprot50 \| AF-A0A661VEU5-F1-MODEL\_V4 \| 1.0 \| 0.004108 \| 104 \| 0.119 \| 159 \| 105 \| 11 \| 29 \| 166 \| 181 \| 325 \| Uncharacterized protein \| Uncharacterized protein; afdb-uniprot50 \| AF-A0A833LAQ3-F1-MODEL\_V4 \| 1.0 \| 0.007342 \| 103 \| 0.131 \| 145 \| 96 \| 7 \| 15 \| 153 \| 6 \| 126 \| DUF806 family protein \| DUF806 family protein; afdb-uniprot50 \| AF-A0A6M3IDV3-F1-MODEL\_V4 \| 1.0 \| 0.003256 \| 103 \| 0.1 \| 169 \| 128 \| 10 \| 1 \| 156 \| 3 \| 160 \| Uncharacterized protein \| Uncharacterized protein; afdb-uniprot50 \| AF-A0A7R7T6R5-F1-MODEL\_V4 \| 1.0 \| 0.002435 \| 103 \| 0.123 \| 194 \| 145 \| 13 \| 2 \| 187 \| 1 \| 177 \| Uncharacterized protein \| Uncharacterized protein; afdb-uniprot50 \| AF-A0A437MN64-F1-MODEL\_V4 \| 1.0 \| 0.008247 \| 102 \| 0.161 \| 149 \| 96 \| 10 \| 14 \| 152 \| 4 \| 133 \| Uncharacterized protein \| Uncharacterized protein; afdb-uniprot50 \| AF-A0A7W8VCX7-F1-MODEL\_V4 \| 1.0 \| 0.002435 \| 102 \| 0.126 \| 166 \| 112 \| 12 \| 2 \| 155 \| 1 \| 145 \| Uncharacterized protein \| Uncharacterized protein; afdb-uniprot50 \| AF-A0A0L7MPP8-F1-MODEL\_V4 \| 1.0 \| 0.00153 \| 102 \| 0.168 \| 178 \| 114 \| 12 \| 1 \| 165 \| 1 \| 157 \| Uncharacterized protein \| Uncharacterized protein; afdb-uniprot50 \| AF-A0A7X6FP76-F1-MODEL\_V4 \| 1.0 \| 0.002899 \| 102 \| 0.14 \| 164 \| 118 \| 5 \| 30 \| 189 \| 21 \| 165 \| Uncharacterized protein \| Uncharacterized protein; afdb-uniprot50 \| AF-A0A2D7IIR0-F1-MODEL\_V4 \| 1.0 \| 0.006928 \| 102 \| 0.11 \| 200 \| 110 \| 10 \| 28 \| 166 \| 40 \| 232 \| Uncharacterized protein \| Uncharacterized protein; afdb-uniprot50 \| AF-A0A662EAC9-F1-MODEL\_V4 \| 1.0 \| 0.008247 \| 101 \| 0.13 \| 153 \| 104 \| 11 \| 6 \| 152 \| 2 \| 131 \| Uncharacterized protein \| Uncharacterized protein; afdb-uniprot50 \| AF-A0A3N1M918-F1-MODEL\_V4 \| 1.0 \| 0.00582 \| 101 \| 0.133 \| 150 \| 101 \| 11 \| 9 \| 152 \| 7 \| 133 \| Uncharacterized protein DUF4128 \| Uncharacterized protein DUF4128; afdb-uniprot50 \| AF-A0A838TS93-F1-MODEL\_V4 \| 1.0 \| 0.007342 \| 101 \| 0.111 \| 152 \| 114 \| 6 \| 9 \| 154 \| 9 \| 145 \| Uncharacterized protein \| Uncharacterized protein; afdb-uniprot50 \| AF-A0A7G3E4R6-F1-MODEL\_V4 \| 1.0 \| 0.009816 \| 101 \| 0.129 \| 170 \| 118 \| 9 \| 2 \| 156 \| 1 \| 155 \| Uncharacterized protein \| Uncharacterized protein; afdb-uniprot50 \| AF-A0A1U7GKN0-F1-MODEL\_V4 \| 1.0 \| 0.002735 \| 101 \| 0.125 \| 175 \| 118 \| 12 \| 1 \| 153 \| 7 \| 168 \| Uncharacterized protein \| Uncharacterized protein; afdb-uniprot50 \| AF-A0A1E4N746-F1-MODEL\_V4 \| 1.0 \| 0.001444 \| 101 \| 0.092 \| 195 \| 135 \| 9 \| 1 \| 174 \| 4 \| 177 \| Uncharacterized protein \| Uncharacterized protein; afdb-uniprot50 \| AF-A0A1P9XTF9-F1-MODEL\_V4 \| 1.0 \| 0.00193 \| 101 \| 0.165 \| 169 \| 105 \| 7 \| 16 \| 172 \| 4 \| 148 \| Uncharacterized protein \| Uncharacterized protein; afdb-uniprot50 \| AF-A0A0F9HRR7-F1-MODEL\_V4 \| 1.0 \| 0.002581 \| 101 \| 0.113 \| 194 \| 139 \| 9 \| 10 \| 178 \| 18 \| 203 \| Uncharacterized protein \| Uncharacterized protein; afdb-uniprot50 \| AF-A0A0R2A4Y7-F1-MODEL\_V4 \| 1.0 \| 0.00874 \| 100 \| 0.098 \| 152 \| 110 \| 7 \| 7 \| 157 \| 1 \| 126 \| Phage tail protein \| Phage tail protein; afdb-uniprot50 \| AF-A0A7Y8VDE0-F1-MODEL\_V4 \| 1.0 \| 0.007781 \| 100 \| 0.135 \| 155 \| 102 \| 12 \| 7 \| 152 \| 2 \| 133 \| Uncharacterized protein \| Uncharacterized protein; afdb-uniprot50 \| AF-A0A1U7CX60-F1-MODEL\_V4 \| 1.0 \| 0.008247 \| 100 \| 0.094 \| 158 \| 121 \| 7 \| 1 \| 152 \| 13 \| 154 \| Uncharacterized protein \| Uncharacterized protein; afdb-uniprot50 \| AF-A0A7C6KND4-F1-MODEL\_V4 \| 1.0 \| 0.005492 \| 100 \| 0.113 \| 194 \| 119 \| 15 \| 9 \| 165 \| 7 \| 184 \| Uncharacterized protein \| Uncharacterized protein; afdb-uniprot50 \| AF-A0A0F9UER8-F1-MODEL\_V4 \| 1.0 \| 0.002581 \| 100 \| 0.112 \| 186 \| 123 \| 12 \| 29 \| 187 \| 187 \| 357 \| Uncharacterized protein \| Uncharacterized protein; afdb-uniprot50 \| AF-A0A2E4IAJ2-F1-MODEL\_V4 \| 1.0 \| 0.009262 \| 100 \| 0.074 \| 201 \| 118 \| 9 \| 28 \| 166 \| 16 \| 210 \| Uncharacterized protein \| Uncharacterized protein; afdb-uniprot50 \| AF-A0A380Q6K4-F1-MODEL\_V4 \| 0.999 \| 0.002735 \| 99 \| 0.095 \| 189 \| 136 \| 11 \| 1 \| 172 \| 12 \| 182 \| Putative ATP-binding protein \| Putative ATP-binding protein; afdb-uniprot50 \| AF-A0A7Z0QH09-F1-MODEL\_V4 \| 0.999 \| 0.004108 \| 99 \| 0.121 \| 198 \| 144 \| 14 \| 1 \| 186 \| 1 \| 180 \| Uncharacterized protein \| Uncharacterized protein; afdb-uniprot50 \| AF-A0A7Y6T8L2-F1-MODEL\_V4 \| 0.999 \| 0.004889 \| 98 \| 0.158 \| 158 \| 104 \| 10 \| 13 \| 152 \| 4 \| 150 \| Uncharacterized protein \| Uncharacterized protein; afdb-uniprot50 \| AF-G0ER30-F1-MODEL\_V4 \| 0.999 \| 0.006168 \| 98 \| 0.135 \| 185 \| 124 \| 11 \| 1 \| 171 \| 1 \| 163 \| Uncharacterized protein \| Uncharacterized protein; afdb-uniprot50 \| AF-A0A2D0AHM2-F1-MODEL\_V4 \| 0.999 \| 0.00874 \| 98 \| 0.098 \| 173 \| 118 \| 15 \| 9 \| 166 \| 11 \| 160 \| Uncharacterized protein \| Uncharacterized protein; afdb-uniprot50 \| AF-A0A6H1ZM14-F1-MODEL\_V4 \| 0.999 \| 0.003657 \| 97 \| 0.126 \| 214 \| 127 \| 15 \| 2 \| 189 \| 1 \| 180 \| Uncharacterized protein \| Uncharacterized protein; afdb-uniprot50 \| AF-A0A4R6IIY1-F1-MODEL\_V4 \| 0.999 \| 0.003451 \| 96 \| 0.151 \| 165 \| 112 \| 11 \| 1 \| 154 \| 1 \| 148 \| Uncharacterized protein \| Uncharacterized protein; afdb-uniprot50 \| AF-A0A4R8FHZ6-F1-MODEL\_V4 \| 0.999 \| 0.004889 \| 96 \| 0.154 \| 168 \| 113 \| 8 \| 9 \| 164 \| 4 \| 154 \| Uncharacterized protein \| Uncharacterized protein; afdb-uniprot50 \| AF-A0A3E0N2N0-F1-MODEL\_V4 \| 0.998 \| 0.006537 \| 95 \| 0.132 \| 166 \| 106 \| 10 \| 1 \| 153 \| 1 \| 141 \| Uncharacterized protein \| Uncharacterized protein; afdb-uniprot50 \| AF-A0A2D6AID0-F1-MODEL\_V4 \| 0.998 \| 0.008247 \| 95 \| 0.09 \| 166 \| 107 \| 13 \| 7 \| 153 \| 2 \| 142 \| Uncharacterized protein \| Uncharacterized protein; afdb-uniprot50 \| AF-A0A382XXG6-F1-MODEL\_V4 \| 0.998 \| 0.001362 \| 95 \| 0.117 \| 179 \| 124 \| 9 \| 9 \| 166 \| 19 \| 184 \| Uncharacterized protein \| Uncharacterized protein; afdb-uniprot50 \| AF-A0A1H3W4X8-F1-MODEL\_V4 \| 0.998 \| 0.007342 \| 94 \| 0.12 \| 166 \| 119 \| 9 \| 1 \| 156 \| 2 \| 150 \| Uncharacterized protein \| Uncharacterized protein; afdb-uniprot50 \| AF-A0A7X7Z267-F1-MODEL\_V4 \| 0.998 \| 0.005492 \| 94 \| 0.168 \| 178 \| 109 \| 12 \| 1 \| 156 \| 1 \| 161 \| Uncharacterized protein \| Uncharacterized protein; afdb-uniprot50 \| AF-A0A564G526-F1-MODEL\_V4 \| 0.998 \| 0.004889 \| 94 \| 0.149 \| 214 \| 132 \| 17 \| 1 \| 189 \| 1 \| 189 \| Uncharacterized protein \| Uncharacterized protein; afdb-uniprot50 \| AF-A0A0F9B3J1-F1-MODEL\_V4 \| 0.998 \| 0.005182 \| 94 \| 0.137 \| 197 \| 115 \| 12 \| 7 \| 175 \| 4 \| 173 \| Uncharacterized protein \| Uncharacterized protein; afdb-uniprot50 \| AF-A0A2E2JTZ9-F1-MODEL\_V4 \| 0.998 \| 0.008247 \| 93 \| 0.18 \| 166 \| 96 \| 14 \| 7 \| 155 \| 2 \| 144 \| Uncharacterized protein \| Uncharacterized protein; afdb-uniprot50 \| AF-A0A1C3WP40-F1-MODEL\_V4 \| 0.998 \| 0.001719 \| 93 \| 0.158 \| 196 \| 118 \| 15 \| 13 \| 186 \| 5 \| 175 \| Uncharacterized protein \| Uncharacterized protein; afdb-uniprot50 \| AF-H1XTE3-F1-MODEL\_V4 \| 0.997 \| 0.007342 \| 91 \| 0.128 \| 164 \| 105 \| 12 \| 7 \| 151 \| 2 \| 146 \| Uncharacterized protein \| Uncharacterized protein; afdb-uniprot50 \| AF-A0A5C7TWN4-F1-MODEL\_V4 \| 0.996 \| 0.004353 \| 89 \| 0.184 \| 168 \| 100 \| 15 \| 9 \| 154 \| 5 \| 157 \| Uncharacterized protein \| Uncharacterized protein; afdb-uniprot50 \| AF-A0A1I5MX81-F1-MODEL\_V4 \| 0.996 \| 0.007342 \| 89 \| 0.137 \| 174 \| 113 \| 11 \| 8 \| 168 \| 1 \| 150 \| Uncharacterized protein \| Uncharacterized protein; afdb-uniprot50 \| AF-A0A2D6AIL0-F1-MODEL\_V4 \| 0.993 \| 0.007781 \| 86 \| 0.143 \| 146 \| 99 \| 10 \| 21 \| 154 \| 10 \| 141 \| Uncharacterized protein \| Uncharacterized protein; afdb-uniprot50 \| AF-A0A102DFT8-F1-MODEL\_V4 \| 0.993 \| 0.002735 \| 86 \| 0.111 \| 206 \| 129 \| 14 \| 3 \| 179 \| 11 \| 191 \| Uncharacterized protein \| Uncharacterized protein; afdb-uniprot50 \| AF-A0A7C9TIT4-F1-MODEL\_V4 \| 0.986 \| 0.006928 \| 81 \| 0.134 \| 208 \| 121 \| 13 \| 4 \| 166 \| 23 \| 216 \| Uncharacterized protein \| Uncharacterized protein; afdb-uniprot50 \| AF-A0A838DQ39-F1-MODEL\_V4 \| 0.984 \| 0.00582 \| 80 \| 0.137 \| 197 \| 113 \| 13 \| 1 \| 165 \| 1 \| 172 \| Uncharacterized protein \| Uncharacterized protein; afdb-uniprot50 \| AF-A0A6J5FR96-F1-MODEL\_V4 \| 0.984 \| 0.006537 \| 80 \| 0.096 \| 207 \| 131 \| 14 \| 7 \| 187 \| 2 \| 178 \| Uncharacterized protein \| Uncharacterized protein; afdb-uniprot50 \| AF-A0A7C1ZHV5-F1-MODEL\_V4 \| 0.981 \| 0.008247 \| 79 \| 0.12 \| 183 \| 107 \| 13 \| 11 \| 152 \| 10 \| 179 \| Uncharacterized protein \| Uncharacterized protein; afdb-uniprot50 \| AF-A0A146H6I1-F1-MODEL\_V4 \| 0.981 \| 0.004108 \| 79 \| 0.134 \| 193 \| 130 \| 10 \| 12 \| 185 \| 408 \| 582 \| Uncharacterized protein \| Uncharacterized protein |
| Top keywords  (threshold 1.00e-02 (evalue)) | **Gp37, DUF1834, tail, Phage, prophage, Minor, U, Putative, Mu\_like, DUF4128** |
| Output files | ../../similar\_structures/10\_FANPEZAQ\_CDS\_0010\_afdb-proteome\_foldseek.tsv ../../similar\_structures/10\_FANPEZAQ\_CDS\_0010\_afdb-uniprot50\_foldseek.tsv ../../similar\_structures/10\_FANPEZAQ\_CDS\_0010\_merged.svg ../../similar\_structures/10\_FANPEZAQ\_CDS\_0010\_pdb\_foldseek.tsv |

  
  
  

Return to summary | Go to previous | Go to next

  


---

**Sequence/structure alignments coloring**  
Each object in the alignment figures is colored according to its E-value following this color coding:

1e-100
10
