## Supplementary material for "Completing the BASEL phage collection to unlock hidden diversity for systematic exploration of phage-host interactions": Data S2: 11.html

FANPEZAQ\_CDS\_0011


Return to summary | Go to previous | Go to next

|  |  |
| --- | --- |
| FANPEZAQ\_CDS\_0011  Page creation date: 02 Sep 2024, 12:00  Project folder: n/a  Input sequences file: Escherichia\_virus\_HeidiAbel.gb | baseplate assembly phage v type secretion domain\_containing vi system vgr ob\_fold gp5 phage\_base\_v vgrg tip phage\_related fragment rhs gpv p2 putative element in beta\_helix spike bacteriophage vgrg1 n\_terminal conserved iv gp138 gp45 tail prophage gene bacteria mu secreted beta ob structural so4 yes fold d t4 and a engineered escherichia |

### Sequence information

|  |  |
| --- | --- |
| Name | FANPEZAQ\_CDS\_0011  11\_FANPEZAQ\_CDS\_0011 (pipeline id) |
| Imported annotations | Escherichia\_virus\_HeidiAbel Bas97 |
| Protein sequence | MSADVSELNRQLNNVVRIGTIKQLDLANARAKVSVAGCTTDWLPWGTNRAGKRRDWSPPV IGEQVVLFAPYGDLGQAVIGPSIFQEDHAAPAASADQETTVYPDGTTVDYNSASNTLTIT VAGAGNVIINCKQANVNTETATVTASTSVTLDTPDTFCTGNLTVAKSLTMGQEGGSATMK GAVSIEGPSLTHNGKNVGSDHSHTGVQPGGGNTGAPA |
| Number of residues | 217 |
| Molecular weight (Da) | 22267.36 |
| Output files | ../../query\_sequences/11\_FANPEZAQ\_CDS\_0011.fasta |

### Putative domain architecture and protein family

#### Search results (HHblits)^1^

|  |  |
| --- | --- |
| Domain family databases searched | Pfam, Ncbi-cd, Cath, Phrogs |
| Results, scheme(s)  (Top layers only; threshold 1.00e-03 (evalue)) | xml version="1.0" encoding="utf-8" standalone="no"?       2024-09-02T21:08:14.150912 image/svg+xml   Matplotlib v3.7.2, https://matplotlib.org/ |
| Results, table  (E-value ≤ 1.00e-03 (evalue)) | db \| id \| prob \| evalue \| pvalue \| score \| cols \| query \| query\_len \| template \| template\_len \| name \| description; pfam \| PF04717 \| 96.3 \| 6.2e-06 \| 1.6e-09 \| 48.8 \| 43 \| (42, 84) \| 217 \| (32, 75) \| 75 \| Phage\_base\_V \| Type VI secretion system/phage-baseplate injector OB domain; pfam \| PF18715 \| 93.9 \| 0.00069 \| 1.6e-07 \| 38.8 \| 39 \| (139, 182) \| 217 \| (3, 41) \| 47 \| Phage\_spike \| Phage spike trimer; cath \| 3qr8A02 \| 98.4 \| 3.9e-11 \| 6e-15 \| 78.5 \| 71 \| (90, 162) \| 217 \| (2, 73) \| 73 \| Baseplate assembly protein v \| CATHCODE: 6.20.150.10 NAME: Baseplate assembly protein v. Chain: a. Synonym: gpv. Engineered: yes SOURCE: Enterobacteria phage p2. Bacteriophage p2. Organism\_taxid: 10679. Gene: v. Expressed in: escherichia coli. Expression\_system\_taxid: 562 CLASS: Special, ARCH: Other non-globular, TOPOL: Chondroitinase Ac; Chain A, domain 3, HOMOL: Chondroitinase Ac; Chain A, domain 3; cath \| 3qr8A01 \| 98.2 \| 1.8e-10 \| 2.9e-14 \| 76.2 \| 87 \| (1, 87) \| 217 \| (3, 89) \| 90 \| Baseplate assembly protein v \| CATHCODE: 2.40.50.230 NAME: Baseplate assembly protein v. Chain: a. Synonym: gpv. Engineered: yes SOURCE: Enterobacteria phage p2. Bacteriophage p2. Organism\_taxid: 10679. Gene: v. Expressed in: escherichia coli. Expression\_system\_taxid: 562 CLASS: Mainly Beta, ARCH: Beta Barrel, TOPOL: OB fold (Dihydrolipoamide Acetyltransferase, E2P), HOMOL: Gp5 N-terminal domain; cath \| 1wthA01 \| 97.6 \| 2.1e-08 \| 3.3e-12 \| 72.1 \| 81 \| (14, 94) \| 217 \| (8, 111) \| 128 \| Tail-associated lysozyme \| CATHCODE: 2.40.50.260 NAME: Tail-associated lysozyme. Chain: a. Synonym: protein gp5. Engineered: yes. Mutation: yes. Baseplate structural protein gp27. Chain: d. Synonym: hub protein 27. Engineered: yes SOURCE: Enterobacteria phage t4. Organism\_taxid: 10665. Gene: 5. Expressed in: escherichia coli bl21(de3). Expression\_system\_taxid: 469008. CLASS: Mainly Beta, ARCH: Beta Barrel, TOPOL: OB fold (Dihydrolipoamide Acetyltransferase, E2P), HOMOL: Nucleic acid-binding protein domain; cath \| 4uhvA04 \| 96.2 \| 1.3e-05 \| 2.2e-09 \| 50.2 \| 74 \| (14, 87) \| 217 \| (14, 102) \| 103 \| Vgrg1, valine-glycine repeat protein g1 \| CATHCODE: 2.40.50.230 NAME: Vgrg1, valine-glycine repeat protein g1. Chain: a, b. Engineered: yes SOURCE: Pseudomonas aeruginosa. Organism\_taxid: 208964. Strain: pao1. Expressed in: escherichia coli. Expression\_system\_taxid: 469008. CLASS: Mainly Beta, ARCH: Beta Barrel, TOPOL: OB fold (Dihydrolipoamide Acetyltransferase, E2P), HOMOL: Gp5 N-terminal domain; cath \| 2p5zX04 \| 96.2 \| 1.5e-05 \| 2.5e-09 \| 48.4 \| 55 \| (29, 83) \| 217 \| (22, 85) \| 85 \| Type vi secretion system component \| CATHCODE: 2.40.50.230 NAME: Type vi secretion system component. Chain: x. Fragment: n-terminal domain. Engineered: yes SOURCE: Escherichia coli o6. Organism\_taxid: 217992. Strain: o6:h1, cft073, upec. Atcc: 700928. Gene: c3393. Expressed in: escherichia coli bl21(de3). Expression\_system\_taxid: 469008. CLASS: Mainly Beta, ARCH: Beta Barrel, TOPOL: OB fold (Dihydrolipoamide Acetyltransferase, E2P), HOMOL: Gp5 N-terminal domain; phrogs \| 9992 \| 99.8 \| 2.8e-25 \| 3.2e-29 \| 164.5 \| 127 \| (1, 128) \| 217 \| (4, 136) \| 223 \| baseplate assembly protein \| baseplate assembly protein; Category: tail; p200092 VI\_07688; phrogs \| 11 \| 99.7 \| 1.8e-22 \| 2.2e-26 \| 160.8 \| 180 \| (7, 213) \| 217 \| (19, 255) \| 257 \| baseplate spike \| baseplate spike; Category: tail; KY883640\_p56; phrogs \| 6142 \| 99.5 \| 1.9e-18 \| 2.2e-22 \| 115.7 \| 84 \| (1, 85) \| 217 \| (3, 87) \| 107 \| baseplate assembly protein \| baseplate assembly protein; Category: tail; p403853 VI\_11032 |
| Top keywords  (threshold 1.00e-03 (evalue)) | **Beta, yes, Baseplate, a, Engineered, escherichia, coli, OB, Organism\_taxid, Expressed** |
| Output files | ../../domain\_architecture/11\_FANPEZAQ\_CDS\_0011\_cath.hhr ../../domain\_architecture/11\_FANPEZAQ\_CDS\_0011\_merged.svg ../../domain\_architecture/11\_FANPEZAQ\_CDS\_0011\_ncbi-cd.hhr ../../domain\_architecture/11\_FANPEZAQ\_CDS\_0011\_pfam.hhr ../../domain\_architecture/11\_FANPEZAQ\_CDS\_0011\_phrogs.hhr |

### Identical protein sequences/structures

#### Search results

|  |  |
| --- | --- |
| Protein sequence databases searched | Pdb, Swissprot, Refseq |
| Identical proteins found | -- |
| Top keywords | -- |
| Output files | -- |

### Similar protein sequences/structures

#### Sequence similarity search results (HHblits)^1^

|  |  |
| --- | --- |
| Sequence databases searched | Uniclust, Pdb70 |
| Results, scheme(s)  (Top layers only, threshold 1.00e-03 (evalue)) | xml version="1.0" encoding="utf-8" standalone="no"?       2024-09-02T21:08:32.411119 image/svg+xml   Matplotlib v3.7.2, https://matplotlib.org/ |
| Results, table(s)  (threshold 1.00e-03 (evalue)) | db \| id \| prob \| evalue \| pvalue \| score \| cols \| query \| query\_len \| template \| template\_len \| name \| description; uniclust \| UniRef100\_A0A023Q1D6 \| 100.0 \| 1.1e-45 \| 2.1e-51 \| 295.6 \| 214 \| (1, 216) \| 217 \| (54, 274) \| 306 \| Gp5/Type VI secretion system Vgr protein OB-fold domain-containing protein \| Gp5/Type VI secretion system Vgr protein OB-fold domain-containing protein; uniclust \| UniRef100\_A0A068Z7F4 \| 100.0 \| 6e-44 \| 1.2e-49 \| 283.6 \| 209 \| (1, 217) \| 217 \| (54, 264) \| 282 \| Phage baseplate assembly protein V \| Phage baseplate assembly protein V; uniclust \| UniRef100\_A0A0L1KF84 \| 100.0 \| 4.3e-43 \| 8.5e-49 \| 262.2 \| 184 \| (1, 217) \| 217 \| (15, 198) \| 209 \| Phage baseplate assembly protein V \| Phage baseplate assembly protein V; uniclust \| UniRef100\_A0A031GY36 \| 100.0 \| 4.9e-43 \| 1e-48 \| 280.1 \| 201 \| (1, 217) \| 217 \| (38, 246) \| 291 \| Phage P2 baseplate assembly gpV protein \| Phage P2 baseplate assembly gpV protein; uniclust \| UniRef100\_A0A3A9HX61 \| 100.0 \| 5.9e-43 \| 1.1e-48 \| 256.5 \| 204 \| (1, 217) \| 217 \| (1, 207) \| 208 \| Phage baseplate assembly protein V \| Phage baseplate assembly protein V; uniclust \| UniRef100\_A0A016XIR2 \| 100.0 \| 1e-42 \| 2e-48 \| 264.3 \| 207 \| (3, 217) \| 217 \| (6, 225) \| 255 \| Baseplate assembly protein \| Baseplate assembly protein; uniclust \| UniRef100\_A0A011MGN7 \| 100.0 \| 1.8e-42 \| 3.4e-48 \| 259.7 \| 197 \| (1, 217) \| 217 \| (30, 229) \| 232 \| Baseplate assembly protein \| Baseplate assembly protein; uniclust \| UniRef100\_A0A061JPC8 \| 100.0 \| 2.8e-42 \| 5.6e-48 \| 269.4 \| 133 \| (1, 133) \| 217 \| (30, 162) \| 252 \| Baseplate assembly protein \| Baseplate assembly protein; uniclust \| UniRef100\_A0A0Y5J9M2 \| 100.0 \| 1.8e-40 \| 3.5e-46 \| 255.0 \| 194 \| (1, 217) \| 217 \| (42, 237) \| 267 \| Phage baseplate assembly protein V \| Phage baseplate assembly protein V; uniclust \| UniRef100\_A0A0P6WI63 \| 100.0 \| 5.1e-40 \| 9.9e-46 \| 242.2 \| 182 \| (2, 217) \| 217 \| (12, 193) \| 195 \| Gp5/Type VI secretion system Vgr protein OB-fold domain-containing protein \| Gp5/Type VI secretion system Vgr protein OB-fold domain-containing protein; uniclust \| UniRef100\_A0A077L0E8 \| 100.0 \| 7.3e-40 \| 1.5e-45 \| 247.5 \| 190 \| (1, 217) \| 217 \| (10, 199) \| 207 \| Gp5/Type VI secretion system Vgr protein OB-fold domain-containing protein \| Gp5/Type VI secretion system Vgr protein OB-fold domain-containing protein; uniclust \| UniRef100\_A0A064AG27 \| 100.0 \| 2.1e-39 \| 4.2e-45 \| 246.4 \| 205 \| (1, 217) \| 217 \| (2, 217) \| 220 \| Baseplate assembly protein \| Baseplate assembly protein; uniclust \| UniRef100\_A0A014N999 \| 100.0 \| 1.2e-38 \| 2.2e-44 \| 238.7 \| 192 \| (1, 217) \| 217 \| (54, 245) \| 245 \| Baseplate assembly protein \| Baseplate assembly protein; uniclust \| UniRef100\_A0A0D8L226 \| 100.0 \| 2e-38 \| 3.8e-44 \| 234.6 \| 205 \| (3, 217) \| 217 \| (1, 207) \| 210 \| Baseplate assembly protein (Fragment) \| Baseplate assembly protein (Fragment); uniclust \| UniRef100\_A0A011VGA0 \| 100.0 \| 4.5e-37 \| 9e-43 \| 232.7 \| 186 \| (1, 217) \| 217 \| (17, 203) \| 207 \| Phage baseplate assembly protein V \| Phage baseplate assembly protein V; uniclust \| UniRef100\_A0A1M7R7X5 \| 100.0 \| 5.3e-37 \| 1.1e-42 \| 237.6 \| 212 \| (2, 217) \| 217 \| (11, 247) \| 249 \| Phage baseplate assembly protein V \| Phage baseplate assembly protein V; uniclust \| UniRef100\_A0A059IUA8 \| 100.0 \| 5.6e-37 \| 1.1e-42 \| 232.9 \| 131 \| (1, 133) \| 217 \| (23, 153) \| 209 \| Prophage LambdaW5, baseplate assembly protein V \| Prophage LambdaW5, baseplate assembly protein V; uniclust \| UniRef100\_A0A149SVM5 \| 100.0 \| 1e-36 \| 2.1e-42 \| 234.8 \| 182 \| (2, 203) \| 217 \| (10, 197) \| 223 \| Gp5/Type VI secretion system Vgr protein OB-fold domain-containing protein \| Gp5/Type VI secretion system Vgr protein OB-fold domain-containing protein; uniclust \| UniRef100\_A0A076PVW1 \| 100.0 \| 3.8e-36 \| 7.2e-42 \| 228.1 \| 198 \| (4, 217) \| 217 \| (42, 253) \| 283 \| Baseplate assembly protein \| Baseplate assembly protein; uniclust \| UniRef100\_A0A239C9W6 \| 100.0 \| 4.8e-36 \| 9.5e-42 \| 228.2 \| 194 \| (1, 217) \| 217 \| (20, 219) \| 227 \| Phage baseplate assembly protein V \| Phage baseplate assembly protein V; uniclust \| UniRef100\_A0A011LYC8 \| 100.0 \| 5.7e-36 \| 1.1e-41 \| 233.5 \| 203 \| (2, 216) \| 217 \| (24, 240) \| 242 \| Phage baseplate protein \| Phage baseplate protein; uniclust \| UniRef100\_A0A1E3G706 \| 100.0 \| 7.2e-36 \| 1.4e-41 \| 221.3 \| 124 \| (1, 124) \| 217 \| (3, 126) \| 198 \| Gp5/Type VI secretion system Vgr protein OB-fold domain-containing protein \| Gp5/Type VI secretion system Vgr protein OB-fold domain-containing protein; uniclust \| UniRef100\_A0A0L8A9N2 \| 100.0 \| 1.5e-35 \| 2.9e-41 \| 218.9 \| 195 \| (1, 216) \| 217 \| (1, 197) \| 199 \| Baseplate assembly protein \| Baseplate assembly protein; uniclust \| UniRef100\_A0A5S9Q467 \| 100.0 \| 5.1e-35 \| 9.7e-41 \| 214.2 \| 205 \| (1, 217) \| 217 \| (3, 213) \| 213 \| Gp5/Type VI secretion system Vgr protein OB-fold domain-containing protein \| Gp5/Type VI secretion system Vgr protein OB-fold domain-containing protein; uniclust \| UniRef100\_A0A136H344 \| 100.0 \| 6.9e-34 \| 1.4e-39 \| 213.8 \| 184 \| (1, 217) \| 217 \| (29, 212) \| 214 \| Gp5/Type VI secretion system Vgr protein OB-fold domain-containing protein \| Gp5/Type VI secretion system Vgr protein OB-fold domain-containing protein; uniclust \| UniRef100\_A0A066RUV3 \| 100.0 \| 9.5e-34 \| 1.9e-39 \| 217.8 \| 152 \| (3, 161) \| 217 \| (37, 190) \| 244 \| Phage baseplate assembly protein V \| Phage baseplate assembly protein V; uniclust \| UniRef100\_A0A0B4CLV9 \| 100.0 \| 8.6e-33 \| 1.6e-38 \| 199.0 \| 182 \| (1, 217) \| 217 \| (6, 189) \| 189 \| Baseplate assembly protein \| Baseplate assembly protein; uniclust \| UniRef100\_A0A098AYV7 \| 100.0 \| 1e-32 \| 2e-38 \| 203.8 \| 162 \| (3, 217) \| 217 \| (12, 177) \| 186 \| Phage baseplate assembly protein V \| Phage baseplate assembly protein V; uniclust \| UniRef100\_A0A009PIX5 \| 100.0 \| 1.9e-32 \| 3.6e-38 \| 195.2 \| 183 \| (9, 217) \| 217 \| (5, 187) \| 188 \| Phage baseplate assembly V family protein \| Phage baseplate assembly V family protein; uniclust \| UniRef100\_A0A101WGP2 \| 99.9 \| 1.8e-31 \| 3.5e-37 \| 193.5 \| 162 \| (1, 217) \| 217 \| (4, 169) \| 172 \| Baseplate assembly protein \| Baseplate assembly protein; uniclust \| UniRef100\_A0A4R2MLJ2 \| 99.9 \| 4.2e-31 \| 8.1e-37 \| 186.4 \| 129 \| (3, 133) \| 217 \| (7, 135) \| 156 \| Phage baseplate assembly protein V \| Phage baseplate assembly protein V; uniclust \| UniRef100\_A0A085ASD7 \| 99.9 \| 4.9e-31 \| 9.5e-37 \| 199.4 \| 133 \| (1, 133) \| 217 \| (19, 153) \| 231 \| Phage-related baseplate protein \| Phage-related baseplate protein; uniclust \| UniRef100\_A0A024E8J9 \| 99.9 \| 5e-31 \| 9.6e-37 \| 199.5 \| 198 \| (1, 210) \| 217 \| (31, 228) \| 244 \| Pyocin R2\_PP \| Pyocin R2\_PP; uniclust \| UniRef100\_A0A0F7XZI7 \| 99.9 \| 2.4e-30 \| 4.7e-36 \| 184.7 \| 124 \| (1, 124) \| 217 \| (11, 134) \| 157 \| Phage baseplate assembly protein V \| Phage baseplate assembly protein V; uniclust \| UniRef100\_A0A4V2RT09 \| 99.9 \| 4.2e-30 \| 8e-36 \| 181.9 \| 118 \| (4, 121) \| 217 \| (9, 126) \| 150 \| Phage baseplate assembly protein V \| Phage baseplate assembly protein V; uniclust \| UniRef100\_V5ZC64 \| 99.9 \| 6.3e-30 \| 1.2e-35 \| 189.1 \| 191 \| (1, 216) \| 217 \| (84, 274) \| 275 \| Baseplate assembly protein V GpV \| Baseplate assembly protein V GpV; uniclust \| UniRef100\_A0A0A1AFI6 \| 99.9 \| 1.4e-29 \| 2.6e-35 \| 192.5 \| 194 \| (16, 216) \| 217 \| (10, 243) \| 257 \| Phage baseplate assembly protein V \| Phage baseplate assembly protein V; uniclust \| UniRef100\_A0A0M5KYZ3 \| 99.9 \| 2.8e-29 \| 5.3e-35 \| 185.3 \| 214 \| (3, 217) \| 217 \| (10, 232) \| 232 \| Phage-related baseplate assembly protein \| Phage-related baseplate assembly protein; uniclust \| UniRef100\_A0A0U2B1K8 \| 99.9 \| 4.5e-29 \| 8.5e-35 \| 179.5 \| 172 \| (10, 216) \| 217 \| (6, 181) \| 181 \| Phage baseplate assembly protein V \| Phage baseplate assembly protein V; uniclust \| UniRef100\_A0A1I3LN83 \| 99.9 \| 6.8e-29 \| 1.3e-34 \| 195.7 \| 210 \| (1, 217) \| 217 \| (1, 211) \| 473 \| Phage baseplate assembly protein V \| Phage baseplate assembly protein V; uniclust \| UniRef100\_A0A156SSS7 \| 99.9 \| 1.5e-28 \| 2.8e-34 \| 173.6 \| 130 \| (1, 132) \| 217 \| (3, 132) \| 159 \| Phage baseplate assembly-like protein \| Phage baseplate assembly-like protein; uniclust \| UniRef100\_A0A0M5MIR9 \| 99.9 \| 1.6e-28 \| 3.1e-34 \| 185.5 \| 195 \| (5, 207) \| 217 \| (3, 208) \| 216 \| Phage baseplate assembly protein V \| Phage baseplate assembly protein V; uniclust \| UniRef100\_A0A1G5B4F2 \| 99.9 \| 5.7e-28 \| 1.1e-33 \| 181.0 \| 129 \| (3, 133) \| 217 \| (10, 141) \| 206 \| Phage baseplate assembly protein V \| Phage baseplate assembly protein V; uniclust \| UniRef100\_A0A376DH93 \| 99.9 \| 1.1e-27 \| 2.1e-33 \| 186.6 \| 191 \| (4, 215) \| 217 \| (215, 405) \| 418 \| Putative phage tail completion protein S \| Putative phage tail completion protein S; uniclust \| UniRef100\_A0A0G0BNF7 \| 99.9 \| 2.1e-27 \| 4.1e-33 \| 175.9 \| 110 \| (9, 119) \| 217 \| (6, 118) \| 180 \| Gp5/Type VI secretion system Vgr protein OB-fold domain-containing protein \| Gp5/Type VI secretion system Vgr protein OB-fold domain-containing protein; uniclust \| UniRef100\_A0A0Q7A745 \| 99.9 \| 3.5e-27 \| 6.9e-33 \| 178.9 \| 169 \| (2, 217) \| 217 \| (22, 195) \| 206 \| Gp5/Type VI secretion system Vgr protein OB-fold domain-containing protein \| Gp5/Type VI secretion system Vgr protein OB-fold domain-containing protein; uniclust \| UniRef100\_A0A2C9EHC3 \| 99.9 \| 3.7e-27 \| 7e-33 \| 178.3 \| 193 \| (3, 217) \| 217 \| (48, 241) \| 243 \| Baseplate assembly protein V \| Baseplate assembly protein V; uniclust \| UniRef100\_A0A1C2K2I7 \| 99.9 \| 4.9e-27 \| 9.4e-33 \| 177.6 \| 207 \| (2, 216) \| 217 \| (24, 242) \| 243 \| Gp5/Type VI secretion system Vgr protein OB-fold domain-containing protein \| Gp5/Type VI secretion system Vgr protein OB-fold domain-containing protein; uniclust \| UniRef100\_A0A074TD51 \| 99.9 \| 8.8e-27 \| 1.7e-32 \| 174.5 \| 163 \| (2, 217) \| 217 \| (36, 205) \| 215 \| Baseplate assembly protein \| Baseplate assembly protein; uniclust \| UniRef100\_A0A0A6D4K9 \| 99.9 \| 9.1e-27 \| 1.7e-32 \| 170.8 \| 186 \| (10, 217) \| 217 \| (3, 195) \| 195 \| Baseplate assembly protein \| Baseplate assembly protein; uniclust \| UniRef100\_A0A7X4AMQ9 \| 99.9 \| 1.1e-26 \| 2e-32 \| 174.9 \| 131 \| (3, 133) \| 217 \| (87, 219) \| 308 \| Phage baseplate assembly protein V \| Phage baseplate assembly protein V; uniclust \| UniRef100\_A0A017HCC4 \| 99.9 \| 1.2e-26 \| 2.3e-32 \| 182.6 \| 160 \| (1, 217) \| 217 \| (73, 237) \| 266 \| Phage-related baseplate assembly protein V \| Phage-related baseplate assembly protein V; uniclust \| UniRef100\_A0A074MGB6 \| 99.9 \| 2e-26 \| 3.9e-32 \| 174.3 \| 117 \| (3, 121) \| 217 \| (15, 134) \| 190 \| Phage baseplate assembly protein V \| Phage baseplate assembly protein V; uniclust \| UniRef100\_A0A075KC29 \| 99.9 \| 2.8e-26 \| 5.6e-32 \| 172.2 \| 161 \| (1, 217) \| 217 \| (6, 173) \| 181 \| Phage baseplate assembly protein V \| Phage baseplate assembly protein V; uniclust \| UniRef100\_A0A010S5G8 \| 99.9 \| 3.3e-26 \| 6.6e-32 \| 180.0 \| 181 \| (17, 217) \| 217 \| (53, 240) \| 255 \| Baseplate assembly protein \| Baseplate assembly protein; uniclust \| UniRef100\_A0A177RV31 \| 99.9 \| 4.7e-26 \| 8.8e-32 \| 163.5 \| 123 \| (1, 123) \| 217 \| (21, 144) \| 176 \| Baseplate assembly protein (Fragment) \| Baseplate assembly protein (Fragment); uniclust \| UniRef100\_A0A377KA20 \| 99.9 \| 5.8e-26 \| 1.1e-31 \| 156.2 \| 121 \| (1, 121) \| 217 \| (3, 123) \| 139 \| Phage baseplate assembly protein V \| Phage baseplate assembly protein V; uniclust \| UniRef100\_A0A378AZS7 \| 99.9 \| 7e-26 \| 1.3e-31 \| 155.4 \| 120 \| (3, 122) \| 217 \| (1, 120) \| 142 \| Baseplate assembly protein V \| Baseplate assembly protein V; uniclust \| UniRef100\_A0A081MYL2 \| 99.9 \| 7.3e-26 \| 1.4e-31 \| 169.1 \| 130 \| (3, 133) \| 217 \| (11, 144) \| 213 \| Gp5/Type VI secretion system Vgr protein OB-fold domain-containing protein \| Gp5/Type VI secretion system Vgr protein OB-fold domain-containing protein; uniclust \| UniRef100\_A0A1N7LRL3 \| 99.9 \| 7.4e-26 \| 1.4e-31 \| 169.5 \| 119 \| (3, 121) \| 217 \| (20, 138) \| 215 \| Phage baseplate assembly protein V \| Phage baseplate assembly protein V; uniclust \| UniRef100\_A0A5M7Q9F8 \| 99.9 \| 7.9e-26 \| 1.4e-31 \| 166.9 \| 193 \| (1, 216) \| 217 \| (1, 261) \| 262 \| Phage baseplate assembly protein V \| Phage baseplate assembly protein V; uniclust \| UniRef100\_A0A286C7H0 \| 99.9 \| 7.9e-26 \| 1.5e-31 \| 168.8 \| 196 \| (1, 216) \| 217 \| (6, 215) \| 217 \| Phage baseplate assembly protein V \| Phage baseplate assembly protein V; uniclust \| UniRef100\_A0A1Y6CPE2 \| 99.9 \| 8.8e-26 \| 1.7e-31 \| 165.8 \| 122 \| (3, 124) \| 217 \| (5, 127) \| 211 \| Phage baseplate assembly protein V \| Phage baseplate assembly protein V; uniclust \| UniRef100\_A0A2J4XY04 \| 99.9 \| 1.3e-25 \| 2.4e-31 \| 148.9 \| 113 \| (3, 115) \| 217 \| (1, 113) \| 114 \| Phage baseplate assembly protein V (Fragment) \| Phage baseplate assembly protein V (Fragment); uniclust \| UniRef100\_A0A0A1AKB5 \| 99.9 \| 1.5e-25 \| 2.8e-31 \| 162.3 \| 185 \| (4, 205) \| 217 \| (6, 200) \| 212 \| Phage baseplate assembly V family protein \| Phage baseplate assembly V family protein; uniclust \| UniRef100\_A0A0A8H9Z0 \| 99.9 \| 2.2e-25 \| 4.4e-31 \| 169.5 \| 146 \| (1, 159) \| 217 \| (3, 148) \| 187 \| Phage baseplate assembly protein V \| Phage baseplate assembly protein V; uniclust \| UniRef100\_A0A085AFN7 \| 99.9 \| 2.3e-25 \| 4.5e-31 \| 174.1 \| 126 \| (2, 131) \| 217 \| (37, 164) \| 250 \| Phage-related baseplate protein \| Phage-related baseplate protein; uniclust \| UniRef100\_UPI001E4BCFD7 \| 99.9 \| 2.8e-25 \| 5.2e-31 \| 156.5 \| 152 \| (48, 217) \| 217 \| (3, 157) \| 160 \| phage baseplate assembly protein V \| phage baseplate assembly protein V; uniclust \| UniRef100\_A0A381EC13 \| 99.9 \| 3.7e-25 \| 7e-31 \| 157.5 \| 166 \| (1, 217) \| 217 \| (1, 170) \| 170 \| Phage P2 baseplate assembly protein gpV \| Phage P2 baseplate assembly protein gpV; uniclust \| UniRef100\_UPI0021A5E1A5 \| 99.9 \| 4.2e-25 \| 7.7e-31 \| 167.4 \| 184 \| (6, 217) \| 217 \| (110, 310) \| 310 \| phage baseplate assembly protein V \| phage baseplate assembly protein V; uniclust \| UniRef100\_A0A167H3T4 \| 99.9 \| 4.2e-25 \| 8e-31 \| 163.6 \| 122 \| (1, 122) \| 217 \| (5, 126) \| 191 \| Gp5/Type VI secretion system Vgr protein OB-fold domain-containing protein \| Gp5/Type VI secretion system Vgr protein OB-fold domain-containing protein; uniclust \| UniRef100\_A0A078MK26 \| 99.9 \| 4.1e-25 \| 8.1e-31 \| 162.1 \| 129 \| (3, 133) \| 217 \| (4, 135) \| 175 \| Phage-related baseplate assembly protein \| Phage-related baseplate assembly protein; uniclust \| UniRef100\_A0A0B1R3F1 \| 99.8 \| 5.2e-25 \| 1e-30 \| 170.0 \| 174 \| (16, 216) \| 217 \| (25, 209) \| 226 \| Baseplate assembly protein \| Baseplate assembly protein; uniclust \| UniRef100\_A0A013SVT5 \| 99.8 \| 6.9e-25 \| 1.4e-30 \| 178.8 \| 175 \| (17, 217) \| 217 \| (65, 278) \| 304 \| Putative transmembrane phage protein \| Putative transmembrane phage protein; uniclust \| UniRef100\_A0A0M4DFV7 \| 99.8 \| 7.7e-25 \| 1.5e-30 \| 170.6 \| 195 \| (1, 217) \| 217 \| (13, 225) \| 238 \| Mu-like prophage protein gp45 \| Mu-like prophage protein gp45; uniclust \| UniRef100\_A0A0B2K158 \| 99.8 \| 9.4e-25 \| 1.8e-30 \| 155.5 \| 117 \| (7, 123) \| 217 \| (4, 123) \| 150 \| Gp5/Type VI secretion system Vgr protein OB-fold domain-containing protein \| Gp5/Type VI secretion system Vgr protein OB-fold domain-containing protein; uniclust \| UniRef100\_A0A927DLG2 \| 99.8 \| 1.2e-24 \| 2.3e-30 \| 166.6 \| 180 \| (1, 205) \| 217 \| (1, 180) \| 310 \| Phage baseplate assembly protein V \| Phage baseplate assembly protein V; uniclust \| UniRef100\_UPI001EFC9B91 \| 99.8 \| 3e-24 \| 5.5e-30 \| 161.2 \| 162 \| (1, 172) \| 217 \| (1, 162) \| 294 \| phage baseplate assembly protein V \| phage baseplate assembly protein V; uniclust \| UniRef100\_A0A099PAQ9 \| 99.8 \| 2.9e-24 \| 5.9e-30 \| 161.7 \| 141 \| (14, 161) \| 217 \| (21, 174) \| 187 \| Gp5/Type VI secretion system Vgr protein OB-fold domain-containing protein \| Gp5/Type VI secretion system Vgr protein OB-fold domain-containing protein; uniclust \| UniRef100\_A0A060GZS5 \| 99.8 \| 9.3e-24 \| 1.7e-29 \| 153.5 \| 181 \| (4, 208) \| 217 \| (33, 215) \| 224 \| Baseplate assembly protein \| Baseplate assembly protein; uniclust \| UniRef100\_UPI0020131D42 \| 99.8 \| 1.3e-23 \| 2.3e-29 \| 163.7 \| 148 \| (1, 164) \| 217 \| (1, 148) \| 396 \| phage baseplate assembly protein V \| phage baseplate assembly protein V; uniclust \| UniRef100\_A0A0J6K5M1 \| 99.8 \| 1.7e-23 \| 3.2e-29 \| 160.0 \| 177 \| (2, 216) \| 217 \| (15, 197) \| 221 \| Baseplate assembly protein \| Baseplate assembly protein; uniclust \| UniRef100\_A0A066PI41 \| 99.8 \| 2e-23 \| 4e-29 \| 160.9 \| 99 \| (15, 113) \| 217 \| (26, 127) \| 212 \| Phage baseplate assembly protein V \| Phage baseplate assembly protein V; uniclust \| UniRef100\_UPI0004DCBFF1 \| 99.8 \| 4.4e-23 \| 8e-29 \| 154.1 \| 117 \| (4, 122) \| 217 \| (8, 124) \| 278 \| phage baseplate assembly protein V \| phage baseplate assembly protein V; uniclust \| UniRef100\_A0A1H4BQX8 \| 99.8 \| 5.5e-23 \| 1e-28 \| 150.6 \| 166 \| (49, 216) \| 217 \| (2, 192) \| 194 \| Phage baseplate assembly protein V \| Phage baseplate assembly protein V; uniclust \| UniRef100\_A0A063BCW0 \| 99.8 \| 7.8e-23 \| 1.6e-28 \| 163.3 \| 187 \| (1, 205) \| 217 \| (36, 243) \| 278 \| Phage baseplate assembly protein V \| Phage baseplate assembly protein V; uniclust \| UniRef100\_A0A0S4L9Z0 \| 99.8 \| 1.2e-22 \| 2.4e-28 \| 161.3 \| 144 \| (16, 159) \| 217 \| (25, 193) \| 294 \| Gp5/Type VI secretion system Vgr protein OB-fold domain-containing protein \| Gp5/Type VI secretion system Vgr protein OB-fold domain-containing protein; uniclust \| UniRef100\_A0A0T7DUY3 \| 99.8 \| 1.4e-22 \| 2.7e-28 \| 155.3 \| 118 \| (4, 121) \| 217 \| (11, 134) \| 265 \| Putative Baseplate \| Putative Baseplate; uniclust \| UniRef100\_A0A6L7TC77 \| 99.8 \| 2.1e-22 \| 3.9e-28 \| 158.5 \| 127 \| (3, 129) \| 217 \| (236, 368) \| 428 \| Phage baseplate assembly protein V \| Phage baseplate assembly protein V; uniclust \| UniRef100\_A0A1M6LDL7 \| 99.8 \| 2.1e-22 \| 4.2e-28 \| 148.7 \| 163 \| (1, 217) \| 217 \| (6, 172) \| 174 \| Bacteriophage Mu Gp45 protein \| Bacteriophage Mu Gp45 protein; uniclust \| UniRef100\_A0A1I5MP97 \| 99.8 \| 2.7e-22 \| 5.2e-28 \| 149.8 \| 123 \| (6, 129) \| 217 \| (7, 132) \| 205 \| Phage P2 baseplate assembly protein gpV \| Phage P2 baseplate assembly protein gpV; uniclust \| UniRef100\_A0A0X1U7T7 \| 99.8 \| 3.3e-22 \| 6.5e-28 \| 154.0 \| 97 \| (12, 114) \| 217 \| (20, 122) \| 227 \| Phage-related baseplate assembly protein \| Phage-related baseplate assembly protein; uniclust \| UniRef100\_A0A0Q2UFL2 \| 99.8 \| 3.9e-22 \| 7.4e-28 \| 144.8 \| 157 \| (8, 217) \| 217 \| (7, 166) \| 172 \| Gp5/Type VI secretion system Vgr protein OB-fold domain-containing protein \| Gp5/Type VI secretion system Vgr protein OB-fold domain-containing protein; uniclust \| UniRef100\_A0A2G1CUF1 \| 99.8 \| 4.1e-22 \| 7.8e-28 \| 151.4 \| 129 \| (2, 132) \| 217 \| (1, 137) \| 248 \| Baseplate assembly protein \| Baseplate assembly protein; uniclust \| UniRef100\_A0A022PBR6 \| 99.8 \| 4e-22 \| 8.1e-28 \| 156.8 \| 175 \| (1, 210) \| 217 \| (19, 197) \| 238 \| Phage baseplate assembly protein V \| Phage baseplate assembly protein V; uniclust \| UniRef100\_A0A1J5E6Z0 \| 99.8 \| 5.4e-22 \| 1.1e-27 \| 155.4 \| 143 \| (18, 160) \| 217 \| (30, 188) \| 253 \| Gp5/Type VI secretion system Vgr protein OB-fold domain-containing protein \| Gp5/Type VI secretion system Vgr protein OB-fold domain-containing protein; uniclust \| UniRef100\_A0A7W6WMQ5 \| 99.8 \| 6.8e-22 \| 1.3e-27 \| 141.7 \| 128 \| (4, 133) \| 217 \| (12, 139) \| 203 \| Phage baseplate assembly protein V \| Phage baseplate assembly protein V; uniclust \| UniRef100\_A0A219Y8Y7 \| 99.8 \| 6.7e-22 \| 1.3e-27 \| 147.7 \| 122 \| (2, 124) \| 217 \| (4, 126) \| 233 \| Baseplate assembly protein V \| Baseplate assembly protein V; uniclust \| UniRef100\_A0A1U9VIH8 \| 99.8 \| 6.9e-22 \| 1.3e-27 \| 147.1 \| 180 \| (2, 215) \| 217 \| (7, 197) \| 202 \| Phage baseplate protein \| Phage baseplate protein; uniclust \| UniRef100\_A0A094ZPX1 \| 99.8 \| 6.9e-22 \| 1.4e-27 \| 150.4 \| 178 \| (1, 217) \| 217 \| (7, 191) \| 200 \| Phage baseplate assembly protein V \| Phage baseplate assembly protein V; uniclust \| UniRef100\_A0A077QIF1 \| 99.8 \| 7.2e-22 \| 1.4e-27 \| 151.3 \| 174 \| (17, 217) \| 217 \| (29, 239) \| 240 \| Putative bacteriophage protein \| Putative bacteriophage protein; uniclust \| UniRef100\_A0A0Q8B2S5 \| 99.8 \| 7.3e-22 \| 1.4e-27 \| 149.7 \| 162 \| (2, 217) \| 217 \| (26, 187) \| 200 \| Gp5/Type VI secretion system Vgr protein OB-fold domain-containing protein \| Gp5/Type VI secretion system Vgr protein OB-fold domain-containing protein; uniclust \| UniRef100\_A0A062V419 \| 99.8 \| 8.8e-22 \| 1.8e-27 \| 158.9 \| 116 \| (16, 131) \| 217 \| (47, 179) \| 284 \| Gp5/Type VI secretion system Vgr protein OB-fold domain-containing protein \| Gp5/Type VI secretion system Vgr protein OB-fold domain-containing protein; uniclust \| UniRef100\_A0A014LI06 \| 99.8 \| 1e-21 \| 1.9e-27 \| 147.5 \| 143 \| (18, 160) \| 217 \| (22, 180) \| 234 \| Type IV secretion protein Rhs \| Type IV secretion protein Rhs; uniclust \| UniRef100\_A0A2G3L892 \| 99.8 \| 1.2e-21 \| 2.3e-27 \| 147.0 \| 129 \| (1, 132) \| 217 \| (1, 130) \| 284 \| Oxidoreductase \| Oxidoreductase; uniclust \| UniRef100\_A0A376X398 \| 99.7 \| 1.5e-21 \| 2.8e-27 \| 141.5 \| 149 \| (52, 213) \| 217 \| (8, 164) \| 168 \| Phage baseplate assembly protein V \| Phage baseplate assembly protein V; uniclust \| UniRef100\_A0A016XKF3 \| 99.7 \| 1.6e-21 \| 3.2e-27 \| 154.2 \| 138 \| (2, 162) \| 217 \| (22, 162) \| 231 \| Phage baseplate assembly protein V \| Phage baseplate assembly protein V; uniclust \| UniRef100\_A0A238TH44 \| 99.7 \| 1.8e-21 \| 3.3e-27 \| 141.5 \| 180 \| (12, 213) \| 217 \| (17, 200) \| 203 \| Phage-related baseplate assembly protein \| Phage-related baseplate assembly protein; uniclust \| UniRef100\_A0A0Q6VK61 \| 99.7 \| 2.1e-21 \| 4.1e-27 \| 154.0 \| 145 \| (17, 161) \| 217 \| (52, 217) \| 281 \| Gp5/Type VI secretion system Vgr protein OB-fold domain-containing protein \| Gp5/Type VI secretion system Vgr protein OB-fold domain-containing protein; uniclust \| UniRef100\_A0A1T2XAL1 \| 99.7 \| 2.4e-21 \| 4.7e-27 \| 138.6 \| 107 \| (14, 123) \| 217 \| (3, 112) \| 140 \| Phage baseplate assembly protein V \| Phage baseplate assembly protein V; uniclust \| UniRef100\_UPI000C8213C5 \| 99.7 \| 3.4e-21 \| 6.2e-27 \| 149.4 \| 159 \| (1, 168) \| 217 \| (1, 159) \| 368 \| phage baseplate assembly protein V \| phage baseplate assembly protein V; uniclust \| UniRef100\_UPI001403B0E1 \| 99.7 \| 4e-21 \| 7.4e-27 \| 143.3 \| 187 \| (18, 215) \| 217 \| (79, 267) \| 269 \| phage baseplate assembly protein V \| phage baseplate assembly protein V; uniclust \| UniRef100\_A0A7S8EHN4 \| 99.7 \| 4.4e-21 \| 8.3e-27 \| 141.7 \| 88 \| (4, 92) \| 217 \| (29, 116) \| 198 \| Phage baseplate assembly protein V \| Phage baseplate assembly protein V; uniclust \| UniRef100\_A0A1Z5H6B0 \| 99.7 \| 4.6e-21 \| 8.8e-27 \| 135.7 \| 133 \| (47, 210) \| 217 \| (3, 135) \| 145 \| Gp5/Type VI secretion system Vgr protein OB-fold domain-containing protein \| Gp5/Type VI secretion system Vgr protein OB-fold domain-containing protein; uniclust \| UniRef100\_A0A256CB18 \| 99.7 \| 5.4e-21 \| 1e-26 \| 143.6 \| 206 \| (2, 215) \| 217 \| (1, 218) \| 220 \| Uncharacterized protein \| Uncharacterized protein; uniclust \| UniRef100\_A0A0A2WM62 \| 99.7 \| 5.4e-21 \| 1.1e-26 \| 146.8 \| 121 \| (12, 132) \| 217 \| (23, 158) \| 221 \| Rhs element Vgr protein \| Rhs element Vgr protein; uniclust \| UniRef100\_A0A4Q6DAY4 \| 99.7 \| 6.8e-21 \| 1.2e-26 \| 136.6 \| 123 \| (1, 123) \| 217 \| (1, 123) \| 195 \| Phage baseplate assembly protein V (Fragment) \| Phage baseplate assembly protein V (Fragment); uniclust \| UniRef100\_A0A547PW88 \| 99.7 \| 7.2e-21 \| 1.3e-26 \| 136.4 \| 149 \| (1, 217) \| 217 \| (51, 199) \| 200 \| Phage baseplate assembly protein V \| Phage baseplate assembly protein V; uniclust \| UniRef100\_UPI001558E326 \| 99.7 \| 8.8e-21 \| 1.6e-26 \| 131.3 \| 126 \| (6, 133) \| 217 \| (4, 129) \| 159 \| phage baseplate assembly protein V \| phage baseplate assembly protein V; uniclust \| UniRef100\_A0A1Q7BLJ5 \| 99.7 \| 8.6e-21 \| 1.7e-26 \| 147.3 \| 147 \| (16, 166) \| 217 \| (25, 189) \| 251 \| Gp5/Type VI secretion system Vgr protein OB-fold domain-containing protein \| Gp5/Type VI secretion system Vgr protein OB-fold domain-containing protein; uniclust \| UniRef100\_A0A074LQM8 \| 99.7 \| 1.2e-20 \| 2.3e-26 \| 141.4 \| 152 \| (17, 217) \| 217 \| (27, 181) \| 182 \| Gp5/Type VI secretion system Vgr protein OB-fold domain-containing protein \| Gp5/Type VI secretion system Vgr protein OB-fold domain-containing protein; uniclust \| UniRef100\_A0A8X6Y3B4 \| 99.7 \| 1.5e-20 \| 2.8e-26 \| 127.9 \| 118 \| (3, 120) \| 217 \| (7, 124) \| 135 \| Spike protein \| Spike protein; uniclust \| UniRef100\_A0A0H3WP77 \| 99.7 \| 1.5e-20 \| 3e-26 \| 144.8 \| 150 \| (3, 168) \| 217 \| (20, 175) \| 219 \| Baseplate assembly protein \| Baseplate assembly protein; uniclust \| UniRef100\_A0A1X7MG63 \| 99.7 \| 2e-20 \| 3.8e-26 \| 122.0 \| 86 \| (3, 88) \| 217 \| (8, 95) \| 98 \| Phage-related baseplate assembly protein V (Fragment) \| Phage-related baseplate assembly protein V (Fragment); uniclust \| UniRef100\_A0A011NHQ0 \| 99.7 \| 2.9e-20 \| 5.8e-26 \| 148.1 \| 116 \| (16, 131) \| 217 \| (60, 191) \| 271 \| Rhs element Vgr protein \| Rhs element Vgr protein; uniclust \| UniRef100\_UPI001BAEB46D \| 99.7 \| 5.7e-20 \| 1.1e-25 \| 118.4 \| 92 \| (3, 94) \| 217 \| (8, 99) \| 102 \| phage baseplate assembly protein V \| phage baseplate assembly protein V; uniclust \| UniRef100\_A0A0Q4KYF8 \| 99.7 \| 6e-20 \| 1.2e-25 \| 138.8 \| 141 \| (15, 155) \| 217 \| (18, 172) \| 213 \| Gp5/Type VI secretion system Vgr protein OB-fold domain-containing protein \| Gp5/Type VI secretion system Vgr protein OB-fold domain-containing protein; uniclust \| UniRef100\_A0A1Q7MAI9 \| 99.7 \| 6e-20 \| 1.2e-25 \| 141.1 \| 145 \| (16, 160) \| 217 \| (26, 188) \| 237 \| Gp5/Type VI secretion system Vgr protein OB-fold domain-containing protein \| Gp5/Type VI secretion system Vgr protein OB-fold domain-containing protein; uniclust \| UniRef100\_UPI00156D5AAA \| 99.7 \| 8.6e-20 \| 1.6e-25 \| 132.1 \| 117 \| (4, 120) \| 217 \| (2, 119) \| 212 \| phage baseplate assembly protein V \| phage baseplate assembly protein V; uniclust \| UniRef100\_G1UXN9 \| 99.7 \| 9.8e-20 \| 1.8e-25 \| 146.7 \| 199 \| (14, 216) \| 217 \| (22, 230) \| 512 \| Gp5/Type VI secretion system Vgr protein OB-fold domain-containing protein \| Gp5/Type VI secretion system Vgr protein OB-fold domain-containing protein; uniclust \| UniRef100\_A0A376FH76 \| 99.7 \| 1e-19 \| 1.9e-25 \| 127.5 \| 137 \| (63, 207) \| 217 \| (1, 139) \| 143 \| Phage Baseplate Assembly protein V \| Phage Baseplate Assembly protein V; uniclust \| UniRef100\_A0A1E3G5Z4 \| 99.7 \| 1.2e-19 \| 2.2e-25 \| 129.2 \| 130 \| (2, 133) \| 217 \| (20, 153) \| 188 \| Gp5/Type VI secretion system Vgr protein OB-fold domain-containing protein \| Gp5/Type VI secretion system Vgr protein OB-fold domain-containing protein; uniclust \| UniRef100\_A0A165RMS4 \| 99.7 \| 1.3e-19 \| 2.5e-25 \| 133.7 \| 168 \| (3, 217) \| 217 \| (12, 179) \| 180 \| Phage-related baseplate assembly protein \| Phage-related baseplate assembly protein; uniclust \| UniRef100\_A0A0C4WPL1 \| 99.7 \| 1.4e-19 \| 2.7e-25 \| 133.4 \| 168 \| (1, 217) \| 217 \| (3, 173) \| 174 \| Phage baseplate assembly V family protein \| Phage baseplate assembly V family protein; uniclust \| UniRef100\_A0A1H2R423 \| 99.7 \| 1.5e-19 \| 2.8e-25 \| 126.7 \| 84 \| (2, 86) \| 217 \| (13, 97) \| 140 \| Phage baseplate assembly protein V \| Phage baseplate assembly protein V; uniclust \| UniRef100\_UPI0008A6FAC2 \| 99.7 \| 1.7e-19 \| 3.1e-25 \| 128.9 \| 152 \| (4, 171) \| 217 \| (2, 154) \| 193 \| phage baseplate assembly protein V \| phage baseplate assembly protein V; uniclust \| UniRef100\_UPI00135400ED \| 99.7 \| 1.8e-19 \| 3.3e-25 \| 126.3 \| 130 \| (3, 132) \| 217 \| (5, 136) \| 150 \| phage baseplate assembly protein V \| phage baseplate assembly protein V; uniclust \| UniRef100\_A0A0A8WRA5 \| 99.7 \| 1.8e-19 \| 3.4e-25 \| 141.2 \| 78 \| (16, 93) \| 217 \| (20, 107) \| 290 \| Putative type VI secretion system protein VgrGA \| Putative type VI secretion system protein VgrGA; uniclust \| UniRef100\_A0A087KRL2 \| 99.7 \| 2.1e-19 \| 4.2e-25 \| 139.4 \| 167 \| (1, 217) \| 217 \| (7, 181) \| 211 \| Prophage baseplate assembly protein V \| Prophage baseplate assembly protein V; uniclust \| UniRef100\_A0A1D2QMX7 \| 99.6 \| 2.9e-19 \| 5.4e-25 \| 130.3 \| 120 \| (3, 122) \| 217 \| (11, 135) \| 199 \| Gp5/Type VI secretion system Vgr protein OB-fold domain-containing protein \| Gp5/Type VI secretion system Vgr protein OB-fold domain-containing protein; uniclust \| UniRef100\_UPI0013D0D801 \| 99.6 \| 3e-19 \| 5.6e-25 \| 116.8 \| 91 \| (30, 120) \| 217 \| (2, 92) \| 98 \| phage baseplate assembly protein V \| phage baseplate assembly protein V; uniclust \| UniRef100\_A0A061YJY3 \| 99.6 \| 3.6e-19 \| 6.7e-25 \| 137.8 \| 127 \| (3, 129) \| 217 \| (17, 144) \| 316 \| Phage baseplate assembly protein V \| Phage baseplate assembly protein V; uniclust \| UniRef100\_A0A955WDS1 \| 99.6 \| 3.9e-19 \| 7.6e-25 \| 145.5 \| 104 \| (16, 119) \| 217 \| (23, 138) \| 356 \| Gp5/Type VI secretion system Vgr protein OB-fold domain-containing protein \| Gp5/Type VI secretion system Vgr protein OB-fold domain-containing protein; uniclust \| UniRef100\_A0A021X9L8 \| 99.6 \| 4.9e-19 \| 9.7e-25 \| 139.4 \| 103 \| (2, 104) \| 217 \| (48, 155) \| 235 \| Phage P2 baseplate assembly protein GpV \| Phage P2 baseplate assembly protein GpV; uniclust \| UniRef100\_A0A015YA37 \| 99.6 \| 5.2e-19 \| 1e-24 \| 158.1 \| 119 \| (15, 133) \| 217 \| (391, 522) \| 651 \| Phage late control D family protein \| Phage late control D family protein; uniclust \| UniRef100\_A0A1I7E546 \| 99.6 \| 7.1e-19 \| 1.3e-24 \| 130.8 \| 131 \| (1, 133) \| 217 \| (1, 132) \| 254 \| Phage baseplate assembly protein V \| Phage baseplate assembly protein V; uniclust \| UniRef100\_A0A2N2D1G0 \| 99.6 \| 8.1e-19 \| 1.6e-24 \| 136.1 \| 108 \| (16, 123) \| 217 \| (21, 142) \| 248 \| Phage tail protein \| Phage tail protein; uniclust \| UniRef100\_UPI000DFC22FD \| 99.6 \| 9.1e-19 \| 1.7e-24 \| 123.0 \| 137 \| (52, 205) \| 217 \| (15, 158) \| 170 \| phage baseplate assembly protein V \| phage baseplate assembly protein V; uniclust \| UniRef100\_A0A8S4QQF2 \| 99.6 \| 1e-18 \| 1.9e-24 \| 139.6 \| 126 \| (6, 133) \| 217 \| (264, 389) \| 409 \| Jg15362 protein \| Jg15362 protein; uniclust \| UniRef100\_A0A080M624 \| 99.6 \| 9.4e-19 \| 1.9e-24 \| 143.8 \| 117 \| (17, 133) \| 217 \| (39, 170) \| 304 \| Phage-related baseplate assembly protein \| Phage-related baseplate assembly protein; uniclust \| UniRef100\_A0A1A9VKF9 \| 99.6 \| 1.4e-18 \| 2.6e-24 \| 138.1 \| 119 \| (13, 133) \| 217 \| (200, 318) \| 378 \| Phage\_base\_V domain-containing protein \| Phage\_base\_V domain-containing protein; uniclust \| UniRef100\_A0A085ZBA7 \| 99.6 \| 1.5e-18 \| 3e-24 \| 160.8 \| 132 \| (16, 153) \| 217 \| (475, 616) \| 858 \| Rhs element Vgr protein \| Rhs element Vgr protein; uniclust \| UniRef100\_A0A7X2CFJ2 \| 99.6 \| 1.7e-18 \| 3.2e-24 \| 120.2 \| 141 \| (55, 217) \| 217 \| (2, 142) \| 142 \| Phage baseplate assembly protein V \| Phage baseplate assembly protein V; uniclust \| UniRef100\_A0A077NCM3 \| 99.6 \| 1.8e-18 \| 3.6e-24 \| 138.3 \| 169 \| (17, 217) \| 217 \| (47, 249) \| 257 \| Putative Similar to bacteriophage protein \| Putative Similar to bacteriophage protein; uniclust \| UniRef100\_A0A512ISE8 \| 99.6 \| 2e-18 \| 3.9e-24 \| 132.1 \| 155 \| (15, 171) \| 217 \| (25, 187) \| 221 \| Gp5/Type VI secretion system Vgr protein OB-fold domain-containing protein \| Gp5/Type VI secretion system Vgr protein OB-fold domain-containing protein; uniclust \| UniRef100\_A0A285CU56 \| 99.6 \| 2.1e-18 \| 4.1e-24 \| 145.6 \| 118 \| (16, 133) \| 217 \| (236, 364) \| 456 \| Gp5/Type VI secretion system Vgr protein OB-fold domain-containing protein \| Gp5/Type VI secretion system Vgr protein OB-fold domain-containing protein; uniclust \| UniRef100\_A0A1A1YQV4 \| 99.6 \| 2.3e-18 \| 4.4e-24 \| 136.0 \| 117 \| (16, 132) \| 217 \| (12, 139) \| 275 \| Gp5/Type VI secretion system Vgr protein OB-fold domain-containing protein \| Gp5/Type VI secretion system Vgr protein OB-fold domain-containing protein; uniclust \| UniRef100\_A0A2N1PDD4 \| 99.6 \| 2.4e-18 \| 4.8e-24 \| 153.2 \| 139 \| (14, 152) \| 217 \| (412, 566) \| 648 \| Gp5/Type VI secretion system Vgr protein OB-fold domain-containing protein \| Gp5/Type VI secretion system Vgr protein OB-fold domain-containing protein; uniclust \| UniRef100\_A0A014LZB8 \| 99.6 \| 2.5e-18 \| 5e-24 \| 155.3 \| 118 \| (16, 133) \| 217 \| (395, 524) \| 640 \| Type VI secretion protein VgrG \| Type VI secretion protein VgrG; uniclust \| UniRef100\_A0A730JXJ6 \| 99.6 \| 3e-18 \| 5.6e-24 \| 110.2 \| 90 \| (22, 111) \| 217 \| (2, 91) \| 91 \| Phage baseplate assembly protein V (Fragment) \| Phage baseplate assembly protein V (Fragment); uniclust \| UniRef100\_A0A1M7YBP6 \| 99.6 \| 3.1e-18 \| 5.9e-24 \| 122.9 \| 106 \| (10, 121) \| 217 \| (4, 112) \| 159 \| Phage baseplate assembly protein V \| Phage baseplate assembly protein V; uniclust \| UniRef100\_A0A0A1IUD8 \| 99.6 \| 3.3e-18 \| 6.5e-24 \| 135.0 \| 187 \| (17, 217) \| 217 \| (31, 257) \| 259 \| Phage protein Gp138 N-terminal domain-containing protein \| Phage protein Gp138 N-terminal domain-containing protein; uniclust \| UniRef100\_A0A1M6PFI3 \| 99.6 \| 3.8e-18 \| 7.2e-24 \| 131.5 \| 141 \| (15, 155) \| 217 \| (20, 176) \| 251 \| Gp5/Type VI secretion system Vgr protein OB-fold domain-containing protein (Fragment) \| Gp5/Type VI secretion system Vgr protein OB-fold domain-containing protein (Fragment); uniclust \| UniRef100\_J1JE15 \| 99.6 \| 4.8e-18 \| 8.8e-24 \| 116.2 \| 119 \| (3, 122) \| 217 \| (7, 125) \| 142 \| Phage baseplate assembly protein V (Fragment) \| Phage baseplate assembly protein V (Fragment); uniclust \| UniRef100\_A0A090IQB5 \| 99.6 \| 4.8e-18 \| 9e-24 \| 130.3 \| 198 \| (3, 207) \| 217 \| (53, 257) \| 268 \| Phage baseplate assembly protein V \| Phage baseplate assembly protein V; uniclust \| UniRef100\_A0A6I1HQM0 \| 99.6 \| 5.5e-18 \| 1e-23 \| 129.8 \| 119 \| (1, 121) \| 217 \| (4, 123) \| 314 \| Phage baseplate assembly protein V \| Phage baseplate assembly protein V; uniclust \| UniRef100\_A0A190XCB5 \| 99.6 \| 5.5e-18 \| 1.1e-23 \| 132.6 \| 160 \| (16, 217) \| 217 \| (43, 217) \| 222 \| Putative baseplate assembly protein \| Putative baseplate assembly protein; uniclust \| UniRef100\_A0A0H3ZTQ1 \| 99.6 \| 5.8e-18 \| 1.1e-23 \| 131.7 \| 199 \| (3, 216) \| 217 \| (35, 263) \| 266 \| Baseplate assembly protein V \| Baseplate assembly protein V; uniclust \| UniRef100\_A0A0V0Q6U3 \| 99.6 \| 5.8e-18 \| 1.1e-23 \| 131.6 \| 119 \| (2, 120) \| 217 \| (17, 169) \| 251 \| Gp5/Type VI secretion system Vgr protein OB-fold domain-containing protein \| Gp5/Type VI secretion system Vgr protein OB-fold domain-containing protein; uniclust \| UniRef100\_A0A516S9U8 \| 99.6 \| 6.7e-18 \| 1.2e-23 \| 122.5 \| 130 \| (3, 132) \| 217 \| (56, 186) \| 209 \| Phage baseplate assembly protein V \| Phage baseplate assembly protein V; uniclust \| UniRef100\_A0A0C1ZK93 \| 99.6 \| 6.8e-18 \| 1.3e-23 \| 129.7 \| 131 \| (3, 133) \| 217 \| (5, 149) \| 209 \| VgrG protein \| VgrG protein; uniclust \| UniRef100\_A0A0R9Q039 \| 99.6 \| 7.3e-18 \| 1.3e-23 \| 125.1 \| 104 \| (17, 121) \| 217 \| (7, 113) \| 240 \| Baseplate protein \| Baseplate protein; uniclust \| UniRef100\_A0A139D6P1 \| 99.6 \| 7.7e-18 \| 1.4e-23 \| 121.9 \| 120 \| (5, 133) \| 217 \| (5, 127) \| 184 \| Phage baseplate assembly protein V \| Phage baseplate assembly protein V; uniclust \| UniRef100\_A0A0B1XX70 \| 99.6 \| 7.4e-18 \| 1.5e-23 \| 149.1 \| 119 \| (15, 133) \| 217 \| (375, 504) \| 589 \| Type VI secretion protein VgrG \| Type VI secretion protein VgrG; uniclust \| UniRef100\_A0A0X8C3M9 \| 99.6 \| 8.6e-18 \| 1.7e-23 \| 135.6 \| 131 \| (17, 153) \| 217 \| (72, 211) \| 322 \| Gp5/Type VI secretion system Vgr protein OB-fold domain-containing protein \| Gp5/Type VI secretion system Vgr protein OB-fold domain-containing protein; uniclust \| UniRef100\_UPI000C036BF4 \| 99.6 \| 9.8e-18 \| 1.8e-23 \| 110.2 \| 89 \| (35, 123) \| 217 \| (3, 91) \| 101 \| phage baseplate assembly protein V \| phage baseplate assembly protein V; uniclust \| UniRef100\_A0A016A3P0 \| 99.6 \| 9e-18 \| 1.8e-23 \| 152.7 \| 134 \| (16, 155) \| 217 \| (406, 553) \| 721 \| Phage late control D family protein \| Phage late control D family protein; uniclust \| UniRef100\_A0A068R1Y7 \| 99.6 \| 9.2e-18 \| 1.8e-23 \| 132.9 \| 178 \| (17, 217) \| 217 \| (40, 237) \| 243 \| Phage protein Gp138 N-terminal domain-containing protein \| Phage protein Gp138 N-terminal domain-containing protein; uniclust \| UniRef100\_A0A0J9EDT8 \| 99.6 \| 1e-17 \| 2e-23 \| 119.1 \| 151 \| (4, 217) \| 217 \| (8, 159) \| 159 \| Phage baseplate assembly protein V \| Phage baseplate assembly protein V; uniclust \| UniRef100\_A0A927HQP9 \| 99.5 \| 1.4e-17 \| 2.6e-23 \| 122.7 \| 171 \| (10, 205) \| 217 \| (2, 172) \| 232 \| Phage baseplate assembly protein V \| Phage baseplate assembly protein V; uniclust \| UniRef100\_A0A0K9NC70 \| 99.5 \| 1.3e-17 \| 2.7e-23 \| 135.8 \| 138 \| (17, 154) \| 217 \| (38, 208) \| 279 \| Gp5/Type VI secretion system Vgr protein OB-fold domain-containing protein \| Gp5/Type VI secretion system Vgr protein OB-fold domain-containing protein; uniclust \| UniRef100\_A0A0M0SQ28 \| 99.5 \| 1.4e-17 \| 2.7e-23 \| 142.5 \| 132 \| (17, 153) \| 217 \| (377, 525) \| 572 \| Gp5/Type VI secretion system Vgr protein OB-fold domain-containing protein \| Gp5/Type VI secretion system Vgr protein OB-fold domain-containing protein; uniclust \| UniRef100\_A0A4Q4AGX4 \| 99.5 \| 1.5e-17 \| 2.8e-23 \| 106.8 \| 83 \| (4, 86) \| 217 \| (10, 95) \| 98 \| Phage baseplate assembly protein V (Fragment) \| Phage baseplate assembly protein V (Fragment); uniclust \| UniRef100\_A0A069QGK5 \| 99.5 \| 1.5e-17 \| 3e-23 \| 135.1 \| 117 \| (17, 133) \| 217 \| (20, 150) \| 280 \| Gp5/Type VI secretion system Vgr protein OB-fold domain-containing protein \| Gp5/Type VI secretion system Vgr protein OB-fold domain-containing protein; uniclust \| UniRef100\_A0A4P9VER7 \| 99.5 \| 1.7e-17 \| 3.2e-23 \| 119.3 \| 130 \| (3, 133) \| 217 \| (9, 142) \| 191 \| Phage baseplate assembly protein V \| Phage baseplate assembly protein V; uniclust \| UniRef100\_A0A264Y5K0 \| 99.5 \| 1.7e-17 \| 3.3e-23 \| 132.2 \| 118 \| (16, 133) \| 217 \| (64, 192) \| 285 \| Gp5/Type VI secretion system Vgr protein OB-fold domain-containing protein \| Gp5/Type VI secretion system Vgr protein OB-fold domain-containing protein; uniclust \| UniRef100\_UPI000488E995 \| 99.5 \| 2.1e-17 \| 3.8e-23 \| 122.3 \| 128 \| (3, 132) \| 217 \| (28, 155) \| 239 \| phage baseplate assembly protein V \| phage baseplate assembly protein V; uniclust \| UniRef100\_A0A016QM46 \| 99.5 \| 2.2e-17 \| 4.5e-23 \| 153.1 \| 117 \| (15, 131) \| 217 \| (552, 686) \| 899 \| Rhs element Vgr protein \| Rhs element Vgr protein; uniclust \| UniRef100\_A0A0D8L1Y3 \| 99.5 \| 2.5e-17 \| 4.8e-23 \| 114.4 \| 117 \| (80, 217) \| 217 \| (2, 118) \| 119 \| Baseplate assembly protein (Fragment) \| Baseplate assembly protein (Fragment); uniclust \| UniRef100\_A0A0D2JZP5 \| 99.5 \| 2.9e-17 \| 5.4e-23 \| 118.9 \| 126 \| (5, 132) \| 217 \| (12, 140) \| 181 \| Baseplate protein \| Baseplate protein; uniclust \| UniRef100\_A0A353GTI7 \| 99.5 \| 3.1e-17 \| 5.7e-23 \| 120.5 \| 177 \| (11, 217) \| 217 \| (45, 225) \| 225 \| Phage baseplate assembly protein V \| Phage baseplate assembly protein V; uniclust \| UniRef100\_A0A6I5RL13 \| 99.5 \| 3.1e-17 \| 5.9e-23 \| 108.3 \| 85 \| (1, 85) \| 217 \| (5, 89) \| 91 \| Phage baseplate assembly protein V (Fragment) \| Phage baseplate assembly protein V (Fragment); uniclust \| UniRef100\_F4QG87 \| 99.5 \| 3.1e-17 \| 6e-23 \| 122.7 \| 186 \| (1, 216) \| 217 \| (1, 189) \| 199 \| Phage baseplate assembly protein V family protein \| Phage baseplate assembly protein V family protein; uniclust \| UniRef100\_A0A1A7KHP9 \| 99.5 \| 3.1e-17 \| 6.1e-23 \| 149.6 \| 133 \| (15, 153) \| 217 \| (146, 286) \| 854 \| Uncharacterized protein \| Uncharacterized protein; uniclust \| UniRef100\_A0A0Q1FEE3 \| 99.5 \| 3.4e-17 \| 6.5e-23 \| 126.4 \| 191 \| (3, 217) \| 217 \| (9, 216) \| 232 \| Baseplate assembly protein \| Baseplate assembly protein; uniclust \| UniRef100\_A0A023WVH6 \| 99.5 \| 3.6e-17 \| 7.2e-23 \| 126.3 \| 115 \| (1, 132) \| 217 \| (12, 129) \| 205 \| Phage baseplate assembly protein V \| Phage baseplate assembly protein V; uniclust \| UniRef100\_A0A3N5MAF8 \| 99.5 \| 4.1e-17 \| 8e-23 \| 136.0 \| 119 \| (15, 133) \| 217 \| (187, 322) \| 402 \| Gp5/Type VI secretion system Vgr protein OB-fold domain-containing protein \| Gp5/Type VI secretion system Vgr protein OB-fold domain-containing protein; uniclust \| UniRef100\_A0A840REX7 \| 99.5 \| 4.6e-17 \| 8.4e-23 \| 115.6 \| 118 \| (4, 121) \| 217 \| (3, 122) \| 178 \| Phage baseplate assembly protein V \| Phage baseplate assembly protein V; uniclust \| UniRef100\_UPI001F2EC826 \| 99.5 \| 5.4e-17 \| 1e-22 \| 109.9 \| 114 \| (8, 121) \| 217 \| (2, 115) \| 127 \| phage baseplate assembly protein V \| phage baseplate assembly protein V; uniclust \| UniRef100\_A0A257KYV8 \| 99.5 \| 5.4e-17 \| 1e-22 \| 124.2 \| 118 \| (16, 133) \| 217 \| (27, 155) \| 232 \| Type IV secretion protein Rhs (Fragment) \| Type IV secretion protein Rhs (Fragment); uniclust \| UniRef100\_A0A143XMY6 \| 99.5 \| 5.8e-17 \| 1.1e-22 \| 122.1 \| 107 \| (9, 121) \| 217 \| (11, 123) \| 177 \| Phage-related baseplate assembly protein \| Phage-related baseplate assembly protein; uniclust \| UniRef100\_A0A0S8CC80 \| 99.5 \| 6.1e-17 \| 1.2e-22 \| 142.4 \| 140 \| (17, 156) \| 217 \| (503, 663) \| 728 \| Gp5/Type VI secretion system Vgr protein OB-fold domain-containing protein \| Gp5/Type VI secretion system Vgr protein OB-fold domain-containing protein; uniclust \| UniRef100\_A0A4Q1SII8 \| 99.5 \| 6.1e-17 \| 1.2e-22 \| 140.0 \| 116 \| (9, 124) \| 217 \| (392, 523) \| 596 \| Type VI secretion system tip protein VgrG \| Type VI secretion system tip protein VgrG; uniclust \| UniRef100\_A0A149TN28 \| 99.5 \| 6.2e-17 \| 1.2e-22 \| 121.1 \| 178 \| (1, 217) \| 217 \| (10, 192) \| 193 \| Baseplate assembly protein \| Baseplate assembly protein; uniclust \| UniRef100\_A0A8S5LES5 \| 99.5 \| 6.6e-17 \| 1.2e-22 \| 118.2 \| 165 \| (15, 185) \| 217 \| (6, 174) \| 208 \| Baseplate assembly protein \| Baseplate assembly protein; uniclust \| UniRef100\_A0A0G3A0D9 \| 99.5 \| 6.4e-17 \| 1.3e-22 \| 143.4 \| 135 \| (13, 154) \| 217 \| (379, 523) \| 568 \| Phage baseplate assembly protein V \| Phage baseplate assembly protein V; uniclust \| UniRef100\_A0A4U0Z1D4 \| 99.5 \| 7.5e-17 \| 1.4e-22 \| 122.9 \| 150 \| (1, 217) \| 217 \| (146, 295) \| 296 \| Phage baseplate assembly protein V \| Phage baseplate assembly protein V; uniclust \| UniRef100\_A0A085HGF5 \| 99.5 \| 7.9e-17 \| 1.5e-22 \| 122.8 \| 181 \| (1, 217) \| 217 \| (1, 224) \| 225 \| Putative phage protein \| Putative phage protein; uniclust \| UniRef100\_A0A538PGR1 \| 99.5 \| 8.9e-17 \| 1.7e-22 \| 134.7 \| 100 \| (16, 115) \| 217 \| (22, 134) \| 455 \| OmpA-like domain-containing protein (Fragment) \| OmpA-like domain-containing protein (Fragment); uniclust \| UniRef100\_A0A248UJ04 \| 99.5 \| 1.4e-16 \| 2.6e-22 \| 114.3 \| 94 \| (1, 94) \| 217 \| (15, 113) \| 149 \| Phage-related baseplate assembly family protein \| Phage-related baseplate assembly family protein; uniclust \| UniRef100\_A0A0J7KLX9 \| 99.5 \| 1.4e-16 \| 2.7e-22 \| 138.0 \| 132 \| (16, 153) \| 217 \| (366, 506) \| 608 \| Gp5/Type VI secretion system Vgr protein OB-fold domain-containing protein (Fragment) \| Gp5/Type VI secretion system Vgr protein OB-fold domain-containing protein (Fragment); uniclust \| UniRef100\_A0A1H4BCX2 \| 99.5 \| 1.5e-16 \| 2.9e-22 \| 139.3 \| 117 \| (16, 132) \| 217 \| (378, 505) \| 583 \| Rhs element Vgr protein \| Rhs element Vgr protein; uniclust \| UniRef100\_UPI0005E17221 \| 99.5 \| 1.8e-16 \| 3.5e-22 \| 101.7 \| 74 \| (1, 74) \| 217 \| (2, 75) \| 86 \| phage baseplate assembly protein V \| phage baseplate assembly protein V; uniclust \| UniRef100\_A0A0J7IFW6 \| 99.5 \| 1.8e-16 \| 3.5e-22 \| 123.5 \| 119 \| (29, 153) \| 217 \| (1, 126) \| 227 \| Gp5/Type VI secretion system Vgr protein OB-fold domain-containing protein \| Gp5/Type VI secretion system Vgr protein OB-fold domain-containing protein; uniclust \| UniRef100\_A0A519RXB4 \| 99.5 \| 1.9e-16 \| 3.6e-22 \| 126.8 \| 118 \| (15, 132) \| 217 \| (100, 229) \| 307 \| Gp5/Type VI secretion system Vgr protein OB-fold domain-containing protein \| Gp5/Type VI secretion system Vgr protein OB-fold domain-containing protein; uniclust \| UniRef100\_A0A0X3APK5 \| 99.5 \| 2.1e-16 \| 4.2e-22 \| 143.2 \| 136 \| (16, 156) \| 217 \| (412, 559) \| 813 \| Uncharacterized conserved protein, implicated in type VI secretion and phage assembly \| Uncharacterized conserved protein, implicated in type VI secretion and phage assembly; uniclust \| UniRef100\_A0A1V5WGM0 \| 99.5 \| 2.2e-16 \| 4.3e-22 \| 120.3 \| 145 \| (15, 159) \| 217 \| (7, 176) \| 214 \| Phage-related baseplate assembly protein \| Phage-related baseplate assembly protein; uniclust \| UniRef100\_A0A0M3AAX9 \| 99.4 \| 2.4e-16 \| 4.6e-22 \| 117.4 \| 166 \| (7, 217) \| 217 \| (8, 179) \| 184 \| Uncharacterized protein \| Uncharacterized protein; uniclust \| UniRef100\_A0A077KP81 \| 99.4 \| 2.6e-16 \| 5.2e-22 \| 148.4 \| 132 \| (16, 153) \| 217 \| (439, 579) \| 1149 \| Gp5/Type VI secretion system Vgr protein OB-fold domain-containing protein \| Gp5/Type VI secretion system Vgr protein OB-fold domain-containing protein; uniclust \| UniRef100\_UPI000D607F75 \| 99.4 \| 3.4e-16 \| 6.3e-22 \| 116.3 \| 187 \| (11, 217) \| 217 \| (5, 212) \| 216 \| phage baseplate assembly protein \| phage baseplate assembly protein; uniclust \| UniRef100\_E8KKS1 \| 99.4 \| 3.5e-16 \| 6.6e-22 \| 107.0 \| 110 \| (11, 121) \| 217 \| (9, 121) \| 125 \| Gp5/Type VI secretion system Vgr protein OB-fold domain-containing protein (Fragment) \| Gp5/Type VI secretion system Vgr protein OB-fold domain-containing protein (Fragment); uniclust \| UniRef100\_A0A2V3UB47 \| 99.4 \| 3.8e-16 \| 7.3e-22 \| 113.3 \| 93 \| (2, 95) \| 217 \| (20, 112) \| 158 \| Phage baseplate assembly protein V \| Phage baseplate assembly protein V; uniclust \| UniRef100\_A0A3D3TAS6 \| 99.4 \| 4e-16 \| 7.6e-22 \| 123.1 \| 120 \| (14, 133) \| 217 \| (19, 154) \| 273 \| DUF2345 domain-containing protein (Fragment) \| DUF2345 domain-containing protein (Fragment); uniclust \| UniRef100\_UPI002155723C \| 99.4 \| 4.3e-16 \| 7.9e-22 \| 120.3 \| 148 \| (15, 170) \| 217 \| (1, 150) \| 324 \| phage baseplate assembly protein V \| phage baseplate assembly protein V; uniclust \| UniRef100\_A0A538CIT5 \| 99.4 \| 4.4e-16 \| 8.1e-22 \| 110.9 \| 142 \| (15, 161) \| 217 \| (16, 166) \| 179 \| Gp5/Type VI secretion system Vgr protein OB-fold domain-containing protein \| Gp5/Type VI secretion system Vgr protein OB-fold domain-containing protein; uniclust \| UniRef100\_A0A958AIE9 \| 99.4 \| 4.5e-16 \| 8.4e-22 \| 112.5 \| 114 \| (17, 130) \| 217 \| (22, 151) \| 176 \| Gp5/Type VI secretion system Vgr protein OB-fold domain-containing protein \| Gp5/Type VI secretion system Vgr protein OB-fold domain-containing protein; uniclust \| UniRef100\_A0A0A2ECB3 \| 99.4 \| 4.3e-16 \| 8.5e-22 \| 141.5 \| 120 \| (14, 133) \| 217 \| (463, 597) \| 733 \| Gp5/Type VI secretion system Vgr protein OB-fold domain-containing protein \| Gp5/Type VI secretion system Vgr protein OB-fold domain-containing protein; uniclust \| UniRef100\_A0A1Q3KMT8 \| 99.4 \| 4.6e-16 \| 8.7e-22 \| 115.6 \| 139 \| (16, 160) \| 217 \| (19, 172) \| 193 \| Gp5/Type VI secretion system Vgr protein OB-fold domain-containing protein \| Gp5/Type VI secretion system Vgr protein OB-fold domain-containing protein; uniclust \| UniRef100\_A0A239WJT3 \| 99.4 \| 4.6e-16 \| 8.9e-22 \| 131.6 \| 117 \| (17, 133) \| 217 \| (210, 340) \| 473 \| Uncharacterized protein conserved in bacteria \| Uncharacterized protein conserved in bacteria; uniclust \| UniRef100\_A0A0D8FQF9 \| 99.4 \| 4.6e-16 \| 9.2e-22 \| 139.1 \| 97 \| (16, 112) \| 217 \| (420, 528) \| 645 \| Phage-like baseplate assembly protein \| Phage-like baseplate assembly protein; uniclust \| UniRef100\_A0A016CPP4 \| 99.4 \| 5.4e-16 \| 1.1e-21 \| 139.1 \| 118 \| (16, 133) \| 217 \| (390, 521) \| 745 \| Phage-related baseplate assembly family protein \| Phage-related baseplate assembly family protein; uniclust \| UniRef100\_A0A0X8C270 \| 99.4 \| 5.4e-16 \| 1.1e-21 \| 144.6 \| 120 \| (14, 133) \| 217 \| (401, 530) \| 1104 \| Gp5/Type VI secretion system Vgr protein OB-fold domain-containing protein \| Gp5/Type VI secretion system Vgr protein OB-fold domain-containing protein; uniclust \| UniRef100\_F0IFV9 \| 99.4 \| 6e-16 \| 1.1e-21 \| 120.0 \| 116 \| (17, 133) \| 217 \| (60, 185) \| 294 \| Gp5/Type VI secretion system Vgr protein OB-fold domain-containing protein \| Gp5/Type VI secretion system Vgr protein OB-fold domain-containing protein; uniclust \| UniRef100\_A0A174LDR4 \| 99.4 \| 6.3e-16 \| 1.2e-21 \| 121.9 \| 144 \| (15, 158) \| 217 \| (33, 205) \| 269 \| Uncharacterized protein conserved in bacteria \| Uncharacterized protein conserved in bacteria; uniclust \| UniRef100\_A0A0C3MRL4 \| 99.4 \| 6.3e-16 \| 1.2e-21 \| 118.8 \| 144 \| (15, 160) \| 217 \| (17, 180) \| 219 \| Phage-related baseplate assembly family protein \| Phage-related baseplate assembly family protein; uniclust \| UniRef100\_A0A1Z3U8S6 \| 99.4 \| 6.6e-16 \| 1.2e-21 \| 115.0 \| 160 \| (5, 164) \| 217 \| (26, 192) \| 222 \| Phage baseplate assembly protein V \| Phage baseplate assembly protein V; uniclust \| UniRef100\_A0A1E7J785 \| 99.4 \| 6.3e-16 \| 1.3e-21 \| 125.4 \| 107 \| (15, 121) \| 217 \| (18, 143) \| 279 \| Gp5/Type VI secretion system Vgr protein OB-fold domain-containing protein \| Gp5/Type VI secretion system Vgr protein OB-fold domain-containing protein; uniclust \| UniRef100\_A0A2A4XT07 \| 99.4 \| 6.8e-16 \| 1.3e-21 \| 112.8 \| 143 \| (6, 166) \| 217 \| (25, 167) \| 190 \| Gp5/Type VI secretion system Vgr protein OB-fold domain-containing protein \| Gp5/Type VI secretion system Vgr protein OB-fold domain-containing protein; uniclust \| UniRef100\_UPI00217DEE8E \| 99.4 \| 6.9e-16 \| 1.3e-21 \| 112.6 \| 130 \| (3, 132) \| 217 \| (11, 141) \| 211 \| phage baseplate assembly protein V \| phage baseplate assembly protein V; uniclust \| UniRef100\_A0A2V8SPQ8 \| 99.4 \| 7.2e-16 \| 1.4e-21 \| 112.5 \| 99 \| (17, 115) \| 217 \| (24, 133) \| 164 \| Gp5/Type VI secretion system Vgr protein OB-fold domain-containing protein (Fragment) \| Gp5/Type VI secretion system Vgr protein OB-fold domain-containing protein (Fragment); uniclust \| UniRef100\_A0A378VX02 \| 99.4 \| 8e-16 \| 1.5e-21 \| 104.9 \| 132 \| (48, 216) \| 217 \| (3, 134) \| 135 \| Putative phage associated protein \| Putative phage associated protein; uniclust \| UniRef100\_A0A011QIW0 \| 99.4 \| 7.6e-16 \| 1.5e-21 \| 124.4 \| 103 \| (1, 106) \| 217 \| (4, 125) \| 252 \| Rhs element Vgr protein \| Rhs element Vgr protein; uniclust \| UniRef100\_A0A2D9F2J4 \| 99.4 \| 8e-16 \| 1.6e-21 \| 118.6 \| 132 \| (2, 164) \| 217 \| (12, 146) \| 198 \| Phage baseplate assembly protein V \| Phage baseplate assembly protein V; uniclust \| UniRef100\_A0A376KHX7 \| 99.4 \| 8.8e-16 \| 1.6e-21 \| 97.6 \| 83 \| (4, 86) \| 217 \| (1, 83) \| 85 \| Baseplate assembly protein V \| Baseplate assembly protein V; uniclust \| UniRef100\_A0A261QLM5 \| 99.4 \| 8.8e-16 \| 1.6e-21 \| 103.9 \| 104 \| (17, 121) \| 217 \| (7, 113) \| 118 \| Baseplate protein (Fragment) \| Baseplate protein (Fragment); uniclust \| UniRef100\_A0A1G9EXB4 \| 99.4 \| 9.5e-16 \| 1.8e-21 \| 118.4 \| 119 \| (15, 133) \| 217 \| (22, 153) \| 226 \| Gp5/Type VI secretion system Vgr protein OB-fold domain-containing protein \| Gp5/Type VI secretion system Vgr protein OB-fold domain-containing protein; uniclust \| UniRef100\_A0A1M6M230 \| 99.4 \| 1.2e-15 \| 2.2e-21 \| 124.0 \| 139 \| (15, 153) \| 217 \| (135, 285) \| 378 \| Gp5/Type VI secretion system Vgr protein OB-fold domain-containing protein (Fragment) \| Gp5/Type VI secretion system Vgr protein OB-fold domain-containing protein (Fragment); uniclust \| UniRef100\_A0A9E8D7C5 \| 99.4 \| 1.3e-15 \| 2.4e-21 \| 102.5 \| 95 \| (50, 152) \| 217 \| (9, 103) \| 113 \| Phage baseplate assembly protein V \| Phage baseplate assembly protein V; uniclust \| UniRef100\_A0A090V6M7 \| 99.4 \| 1.3e-15 \| 2.4e-21 \| 117.2 \| 100 \| (16, 115) \| 217 \| (35, 171) \| 223 \| Phage protein Gp138 N-terminal domain-containing protein \| Phage protein Gp138 N-terminal domain-containing protein; uniclust \| UniRef100\_A0A7Y6Z4G0 \| 99.4 \| 1.4e-15 \| 2.5e-21 \| 110.7 \| 129 \| (3, 133) \| 217 \| (9, 138) \| 205 \| Phage baseplate assembly protein V \| Phage baseplate assembly protein V; uniclust \| UniRef100\_A0A090G3N9 \| 99.4 \| 1.4e-15 \| 2.7e-21 \| 112.4 \| 110 \| (2, 113) \| 217 \| (17, 132) \| 184 \| Gp5/Type VI secretion system Vgr protein OB-fold domain-containing protein \| Gp5/Type VI secretion system Vgr protein OB-fold domain-containing protein; uniclust \| UniRef100\_A0A3D3M4P4 \| 99.4 \| 1.5e-15 \| 2.8e-21 \| 106.8 \| 116 \| (50, 172) \| 217 \| (2, 117) \| 137 \| Phage baseplate assembly protein V (Fragment) \| Phage baseplate assembly protein V (Fragment); uniclust \| UniRef100\_A0A0T9QI62 \| 99.4 \| 1.5e-15 \| 2.8e-21 \| 99.2 \| 67 \| (3, 69) \| 217 \| (17, 83) \| 84 \| Phage-related baseplate assembly protein \| Phage-related baseplate assembly protein; uniclust \| UniRef100\_A0A031FUD1 \| 99.4 \| 1.5e-15 \| 2.9e-21 \| 113.3 \| 122 \| (1, 122) \| 217 \| (20, 143) \| 203 \| Phage baseplate assembly protein V \| Phage baseplate assembly protein V; uniclust \| UniRef100\_A0A0E3V6K5 \| 99.4 \| 1.5e-15 \| 3e-21 \| 136.4 \| 150 \| (8, 159) \| 217 \| (361, 540) \| 610 \| Gp5/Type VI secretion system Vgr protein OB-fold domain-containing protein \| Gp5/Type VI secretion system Vgr protein OB-fold domain-containing protein; uniclust \| UniRef100\_A0A538Q0J0 \| 99.4 \| 1.6e-15 \| 3e-21 \| 120.1 \| 100 \| (16, 115) \| 217 \| (22, 134) \| 307 \| Gp5/Type VI secretion system Vgr protein OB-fold domain-containing protein (Fragment) \| Gp5/Type VI secretion system Vgr protein OB-fold domain-containing protein (Fragment); uniclust \| UniRef100\_Q31HU0 \| 99.4 \| 2.1e-15 \| 3.8e-21 \| 111.0 \| 191 \| (15, 209) \| 217 \| (1, 215) \| 221 \| Phage-related baseplate assembly protein V \| Phage-related baseplate assembly protein V; uniclust \| UniRef100\_A0A0E3KUE2 \| 99.4 \| 2.2e-15 \| 4.1e-21 \| 128.2 \| 117 \| (15, 131) \| 217 \| (376, 505) \| 574 \| VgrG protein \| VgrG protein; uniclust \| UniRef100\_A0A1H2FMJ1 \| 99.4 \| 2.1e-15 \| 4.2e-21 \| 116.8 \| 101 \| (3, 104) \| 217 \| (23, 131) \| 207 \| Phage baseplate assembly protein V \| Phage baseplate assembly protein V; uniclust \| UniRef100\_A0A0S2ZDV9 \| 99.4 \| 2.3e-15 \| 4.4e-21 \| 114.7 \| 160 \| (17, 209) \| 217 \| (21, 206) \| 218 \| Phage protein Gp138 N-terminal domain-containing protein \| Phage protein Gp138 N-terminal domain-containing protein; uniclust \| UniRef100\_A0A134C1J7 \| 99.4 \| 2.4e-15 \| 4.4e-21 \| 102.7 \| 111 \| (4, 114) \| 217 \| (2, 115) \| 120 \| Uncharacterized protein (Fragment) \| Uncharacterized protein (Fragment); uniclust \| UniRef100\_A0A350LX97 \| 99.4 \| 2.9e-15 \| 5.4e-21 \| 117.0 \| 84 \| (5, 88) \| 217 \| (206, 289) \| 349 \| Gp5/Type VI secretion system Vgr protein OB-fold domain-containing protein \| Gp5/Type VI secretion system Vgr protein OB-fold domain-containing protein; uniclust \| UniRef100\_A0A2D4SLA2 \| 99.4 \| 2.9e-15 \| 5.7e-21 \| 116.4 \| 109 \| (12, 120) \| 217 \| (8, 128) \| 219 \| Gp5/Type VI secretion system Vgr protein OB-fold domain-containing protein \| Gp5/Type VI secretion system Vgr protein OB-fold domain-containing protein; uniclust \| UniRef100\_A0A1D8UTH1 \| 99.3 \| 3.5e-15 \| 6.6e-21 \| 109.4 \| 159 \| (1, 217) \| 217 \| (1, 164) \| 166 \| Baseplate assembly protein \| Baseplate assembly protein; uniclust \| UniRef100\_A0A0N1CB70 \| 99.3 \| 3.4e-15 \| 6.7e-21 \| 117.3 \| 103 \| (2, 104) \| 217 \| (20, 129) \| 231 \| Gp5/Type VI secretion system Vgr protein OB-fold domain-containing protein \| Gp5/Type VI secretion system Vgr protein OB-fold domain-containing protein; uniclust \| UniRef100\_A0A069QD16 \| 99.3 \| 4.4e-15 \| 8.4e-21 \| 121.0 \| 139 \| (17, 155) \| 217 \| (105, 262) \| 354 \| Gp5/Type VI secretion system Vgr protein OB-fold domain-containing protein (Fragment) \| Gp5/Type VI secretion system Vgr protein OB-fold domain-containing protein (Fragment); uniclust \| UniRef100\_A0A543G2B2 \| 99.3 \| 4.9e-15 \| 9.3e-21 \| 113.8 \| 125 \| (28, 153) \| 217 \| (10, 144) \| 239 \| Gp5/Type VI secretion system Vgr protein OB-fold domain-containing protein \| Gp5/Type VI secretion system Vgr protein OB-fold domain-containing protein; uniclust \| UniRef100\_A0A096D7L8 \| 99.3 \| 4.6e-15 \| 9.5e-21 \| 131.3 \| 143 \| (7, 153) \| 217 \| (300, 452) \| 493 \| Gp5/Type VI secretion system Vgr protein OB-fold domain-containing protein \| Gp5/Type VI secretion system Vgr protein OB-fold domain-containing protein; uniclust \| UniRef100\_A0A0T5NZY3 \| 99.3 \| 4.8e-15 \| 9.5e-21 \| 131.6 \| 119 \| (14, 132) \| 217 \| (380, 513) \| 591 \| Gp5/Type VI secretion system Vgr protein OB-fold domain-containing protein \| Gp5/Type VI secretion system Vgr protein OB-fold domain-containing protein; uniclust \| UniRef100\_A0A3D9EV90 \| 99.3 \| 5.6e-15 \| 1e-20 \| 103.1 \| 114 \| (2, 115) \| 217 \| (1, 120) \| 154 \| Phage baseplate assembly protein V \| Phage baseplate assembly protein V; uniclust \| UniRef100\_A0A381EFJ1 \| 99.3 \| 5.9e-15 \| 1.1e-20 \| 106.7 \| 139 \| (18, 217) \| 217 \| (22, 163) \| 164 \| Mu-like prophage protein gp45 \| Mu-like prophage protein gp45; uniclust \| UniRef100\_A0A023D2H7 \| 99.3 \| 5.7e-15 \| 1.1e-20 \| 115.8 \| 76 \| (17, 92) \| 217 \| (29, 106) \| 228 \| Phage related baseplate assembly protein \| Phage related baseplate assembly protein; uniclust \| UniRef100\_A0A0Q8LLQ7 \| 99.3 \| 5.8e-15 \| 1.1e-20 \| 128.5 \| 118 \| (15, 132) \| 217 \| (376, 505) \| 601 \| Gp5/Type VI secretion system Vgr protein OB-fold domain-containing protein \| Gp5/Type VI secretion system Vgr protein OB-fold domain-containing protein; uniclust \| UniRef100\_UPI0020135E1D \| 99.3 \| 6.1e-15 \| 1.1e-20 \| 104.7 \| 134 \| (69, 205) \| 217 \| (14, 158) \| 172 \| phage baseplate assembly protein V \| phage baseplate assembly protein V; uniclust \| UniRef100\_A0A0D6TN34 \| 99.3 \| 6.2e-15 \| 1.2e-20 \| 116.0 \| 124 \| (29, 153) \| 217 \| (28, 161) \| 254 \| Uncharacterized protein \| Uncharacterized protein; uniclust \| UniRef100\_A0A0C5AEP6 \| 99.3 \| 8.2e-15 \| 1.5e-20 \| 103.4 \| 110 \| (10, 120) \| 217 \| (20, 131) \| 146 \| Baseplate assembly protein \| Baseplate assembly protein; uniclust \| UniRef100\_A0A2C7A8I5 \| 99.3 \| 8.5e-15 \| 1.6e-20 \| 106.6 \| 185 \| (3, 217) \| 217 \| (12, 202) \| 202 \| Gp5/Type VI secretion system Vgr protein OB-fold domain-containing protein \| Gp5/Type VI secretion system Vgr protein OB-fold domain-containing protein; uniclust \| UniRef100\_A0A150S1K0 \| 99.3 \| 8.9e-15 \| 1.7e-20 \| 108.3 \| 104 \| (28, 131) \| 217 \| (21, 135) \| 187 \| Gp5/Type VI secretion system Vgr protein OB-fold domain-containing protein \| Gp5/Type VI secretion system Vgr protein OB-fold domain-containing protein; uniclust \| UniRef100\_A0A1H4H7U1 \| 99.3 \| 1e-14 \| 2e-20 \| 129.9 \| 133 \| (16, 154) \| 217 \| (399, 550) \| 643 \| Uncharacterized conserved protein, implicated in type VI secretion and phage assembly \| Uncharacterized conserved protein, implicated in type VI secretion and phage assembly; uniclust \| UniRef100\_A0A0D9NFY0 \| 99.3 \| 1e-14 \| 2e-20 \| 121.6 \| 117 \| (17, 133) \| 217 \| (152, 282) \| 424 \| Type IV secretion protein Rhs \| Type IV secretion protein Rhs; uniclust \| UniRef100\_A0A2G6EWU3 \| 99.3 \| 1.1e-14 \| 2.1e-20 \| 102.3 \| 108 \| (5, 112) \| 217 \| (6, 123) \| 160 \| Gp5/Type VI secretion system Vgr protein OB-fold domain-containing protein \| Gp5/Type VI secretion system Vgr protein OB-fold domain-containing protein; uniclust \| UniRef100\_A0A7J6YLT3 \| 99.3 \| 1.1e-14 \| 2.1e-20 \| 118.7 \| 109 \| (15, 123) \| 217 \| (136, 244) \| 508 \| Baseplate protein J-like domain-containing protein \| Baseplate protein J-like domain-containing protein; uniclust \| UniRef100\_A0A345DE56 \| 99.3 \| 1.2e-14 \| 2.2e-20 \| 103.5 \| 141 \| (4, 162) \| 217 \| (7, 147) \| 174 \| Gp5/Type VI secretion system Vgr protein OB-fold domain-containing protein \| Gp5/Type VI secretion system Vgr protein OB-fold domain-containing protein; uniclust \| UniRef100\_A0A1D2QM78 \| 99.3 \| 1.3e-14 \| 2.5e-20 \| 115.5 \| 120 \| (2, 121) \| 217 \| (20, 169) \| 303 \| Gp5/Type VI secretion system Vgr protein OB-fold domain-containing protein \| Gp5/Type VI secretion system Vgr protein OB-fold domain-containing protein; uniclust \| UniRef100\_A0A3S4GGV8 \| 99.3 \| 1.3e-14 \| 2.5e-20 \| 94.5 \| 70 \| (54, 123) \| 217 \| (3, 72) \| 93 \| Phage baseplate assembly protein V \| Phage baseplate assembly protein V; uniclust \| UniRef100\_A0A376DPX4 \| 99.3 \| 1.5e-14 \| 2.9e-20 \| 116.9 \| 134 \| (17, 156) \| 217 \| (81, 227) \| 325 \| Uncharacterized protein conserved in bacteria \| Uncharacterized protein conserved in bacteria; uniclust \| UniRef100\_A0A1Y0Y2S8 \| 99.3 \| 1.6e-14 \| 3.1e-20 \| 110.5 \| 189 \| (3, 217) \| 217 \| (15, 210) \| 210 \| Baseplate assembly protein \| Baseplate assembly protein; uniclust \| UniRef100\_A0A7W0QEQ5 \| 99.3 \| 1.7e-14 \| 3.1e-20 \| 118.6 \| 116 \| (18, 133) \| 217 \| (30, 156) \| 454 \| Gp5/Type VI secretion system Vgr protein OB-fold domain-containing protein \| Gp5/Type VI secretion system Vgr protein OB-fold domain-containing protein; uniclust \| UniRef100\_A0A0E3ZX09 \| 99.3 \| 1.8e-14 \| 3.5e-20 \| 124.4 \| 153 \| (16, 168) \| 217 \| (351, 526) \| 576 \| Gp5/Type VI secretion system Vgr protein OB-fold domain-containing protein \| Gp5/Type VI secretion system Vgr protein OB-fold domain-containing protein; uniclust \| UniRef100\_A0A2S8ACA2 \| 99.3 \| 2e-14 \| 3.9e-20 \| 117.1 \| 140 \| (14, 154) \| 217 \| (133, 287) \| 372 \| Type IV secretion protein Rhs (Fragment) \| Type IV secretion protein Rhs (Fragment); uniclust \| UniRef100\_A0A0W8FUT7 \| 99.3 \| 2.1e-14 \| 4.1e-20 \| 111.5 \| 138 \| (7, 168) \| 217 \| (34, 173) \| 223 \| Prophage baseplate assembly protein v \| Prophage baseplate assembly protein v; uniclust \| UniRef100\_UPI000F060CC4 \| 99.3 \| 2.3e-14 \| 4.2e-20 \| 101.6 \| 121 \| (1, 121) \| 217 \| (21, 143) \| 148 \| phage baseplate assembly protein V \| phage baseplate assembly protein V; uniclust \| UniRef100\_A0A349GR45 \| 99.3 \| 2.4e-14 \| 4.5e-20 \| 106.4 \| 117 \| (14, 133) \| 217 \| (4, 123) \| 199 \| Baseplate assembly protein \| Baseplate assembly protein; uniclust \| UniRef100\_A0A077KAY0 \| 99.3 \| 2.3e-14 \| 4.5e-20 \| 117.7 \| 172 \| (15, 217) \| 217 \| (65, 260) \| 313 \| Putative baseplate protein \| Putative baseplate protein; uniclust \| UniRef100\_A0A1J0MF38 \| 99.3 \| 2.4e-14 \| 4.6e-20 \| 113.4 \| 195 \| (13, 210) \| 217 \| (13, 252) \| 263 \| Baseplate central spike \| Baseplate central spike; uniclust \| UniRef100\_A0A0Q0D424 \| 99.3 \| 2.7e-14 \| 5e-20 \| 96.5 \| 110 \| (2, 111) \| 217 \| (3, 113) \| 113 \| Baseplate assembly protein V \| Baseplate assembly protein V; uniclust \| UniRef100\_A0A0F0H101 \| 99.3 \| 2.5e-14 \| 5e-20 \| 128.8 \| 98 \| (16, 113) \| 217 \| (424, 539) \| 653 \| Gp5/Type VI secretion system Vgr protein OB-fold domain-containing protein \| Gp5/Type VI secretion system Vgr protein OB-fold domain-containing protein; uniclust \| UniRef100\_A0A450Z2A7 \| 99.2 \| 3e-14 \| 5.6e-20 \| 96.2 \| 100 \| (32, 133) \| 217 \| (2, 101) \| 114 \| Phage baseplate assembly protein V \| Phage baseplate assembly protein V; uniclust \| UniRef100\_A0A075MMT0 \| 99.2 \| 3.1e-14 \| 6e-20 \| 108.3 \| 117 \| (17, 133) \| 217 \| (26, 153) \| 185 \| Gp5/Type VI secretion system Vgr protein OB-fold domain-containing protein \| Gp5/Type VI secretion system Vgr protein OB-fold domain-containing protein; uniclust \| UniRef100\_A0A094QB96 \| 99.2 \| 3.4e-14 \| 6.5e-20 \| 121.2 \| 97 \| (14, 110) \| 217 \| (367, 476) \| 596 \| Gp5/Type VI secretion system Vgr protein OB-fold domain-containing protein \| Gp5/Type VI secretion system Vgr protein OB-fold domain-containing protein; uniclust \| UniRef100\_A0A376U5W8 \| 99.2 \| 3.5e-14 \| 6.7e-20 \| 102.1 \| 132 \| (61, 217) \| 217 \| (15, 149) \| 149 \| Putative phage related protein \| Putative phage related protein; uniclust \| UniRef100\_A0A0D1BTC3 \| 99.2 \| 3.4e-14 \| 6.7e-20 \| 112.7 \| 90 \| (17, 106) \| 217 \| (43, 168) \| 232 \| Phage protein Gp138 N-terminal domain-containing protein \| Phage protein Gp138 N-terminal domain-containing protein; uniclust \| UniRef100\_A0A2M6UU70 \| 99.2 \| 3.6e-14 \| 6.8e-20 \| 105.5 \| 173 \| (1, 216) \| 217 \| (14, 196) \| 203 \| Baseplate assembly protein \| Baseplate assembly protein; uniclust \| UniRef100\_A0A3G4V723 \| 99.2 \| 3.8e-14 \| 7.2e-20 \| 104.8 \| 116 \| (5, 121) \| 217 \| (23, 139) \| 203 \| Phage baseplate assembly protein V \| Phage baseplate assembly protein V; uniclust \| UniRef100\_A0A1C6T0T6 \| 99.2 \| 4.1e-14 \| 8.1e-20 \| 110.7 \| 117 \| (17, 133) \| 217 \| (25, 161) \| 220 \| Phage-related baseplate assembly protein \| Phage-related baseplate assembly protein; uniclust \| UniRef100\_UPI002355B909 \| 99.2 \| 4.8e-14 \| 8.8e-20 \| 104.8 \| 128 \| (4, 131) \| 217 \| (22, 151) \| 230 \| phage baseplate assembly protein V \| phage baseplate assembly protein V; uniclust \| UniRef100\_A0A0C5V150 \| 99.2 \| 4.7e-14 \| 8.8e-20 \| 109.7 \| 114 \| (3, 116) \| 217 \| (24, 169) \| 263 \| Phage P2 baseplate assembly protein gpV \| Phage P2 baseplate assembly protein gpV; uniclust \| UniRef100\_UPI001EE5592A \| 99.2 \| 4.9e-14 \| 8.9e-20 \| 100.6 \| 91 \| (6, 96) \| 217 \| (8, 101) \| 174 \| phage baseplate assembly protein V \| phage baseplate assembly protein V; uniclust \| UniRef100\_A0A0Q4L4M4 \| 99.2 \| 4.6e-14 \| 9e-20 \| 113.5 \| 129 \| (1, 133) \| 217 \| (6, 147) \| 276 \| Gp5/Type VI secretion system Vgr protein OB-fold domain-containing protein \| Gp5/Type VI secretion system Vgr protein OB-fold domain-containing protein; uniclust \| UniRef100\_A0A1G6Z7R3 \| 99.2 \| 4.9e-14 \| 9.6e-20 \| 125.5 \| 104 \| (17, 120) \| 217 \| (354, 466) \| 627 \| Uncharacterized conserved protein, implicated in type VI secretion and phage assembly \| Uncharacterized conserved protein, implicated in type VI secretion and phage assembly; uniclust \| UniRef100\_A0A8S5QP55 \| 99.2 \| 5.1e-14 \| 9.6e-20 \| 111.3 \| 166 \| (41, 217) \| 217 \| (88, 287) \| 287 \| Baseplate protein \| Baseplate protein; uniclust \| UniRef100\_A0A7C2HLH3 \| 99.2 \| 5.4e-14 \| 1e-19 \| 105.8 \| 91 \| (24, 114) \| 217 \| (96, 199) \| 251 \| Gp5/Type VI secretion system Vgr protein OB-fold domain-containing protein \| Gp5/Type VI secretion system Vgr protein OB-fold domain-containing protein; uniclust \| UniRef100\_A0A1R1MK78 \| 99.2 \| 5.4e-14 \| 1e-19 \| 106.1 \| 141 \| (7, 153) \| 217 \| (4, 160) \| 194 \| Gp5/Type VI secretion system Vgr protein OB-fold domain-containing protein \| Gp5/Type VI secretion system Vgr protein OB-fold domain-containing protein; uniclust \| UniRef100\_A0A0N0UY75 \| 99.2 \| 6e-14 \| 1.1e-19 \| 121.7 \| 114 \| (16, 129) \| 217 \| (376, 497) \| 595 \| Gp5/Type VI secretion system Vgr protein OB-fold domain-containing protein \| Gp5/Type VI secretion system Vgr protein OB-fold domain-containing protein; uniclust \| UniRef100\_A0A7V8ABK9 \| 99.2 \| 6.3e-14 \| 1.2e-19 \| 96.6 \| 119 \| (9, 131) \| 217 \| (11, 129) \| 139 \| Baseplate assembly protein V \| Baseplate assembly protein V; uniclust \| UniRef100\_A0A6L4A988 \| 99.2 \| 6.5e-14 \| 1.2e-19 \| 117.5 \| 117 \| (17, 133) \| 217 \| (255, 393) \| 487 \| VgrG-related protein (Fragment) \| VgrG-related protein (Fragment); uniclust \| UniRef100\_A0A2C9EHI6 \| 99.2 \| 7.1e-14 \| 1.3e-19 \| 105.2 \| 161 \| (11, 217) \| 217 \| (71, 234) \| 237 \| Prophage MuSo2, baseplate assembly protein V \| Prophage MuSo2, baseplate assembly protein V; uniclust \| UniRef100\_A0A9D2KBF5 \| 99.2 \| 7.5e-14 \| 1.4e-19 \| 104.0 \| 142 \| (10, 154) \| 217 \| (4, 157) \| 234 \| Phage baseplate assembly protein V \| Phage baseplate assembly protein V; uniclust \| UniRef100\_A0A014MQT2 \| 99.2 \| 7.2e-14 \| 1.4e-19 \| 110.8 \| 107 \| (17, 123) \| 217 \| (28, 144) \| 220 \| Baseplate assembly protein \| Baseplate assembly protein; uniclust \| UniRef100\_A0A3M8AFK2 \| 99.2 \| 7.7e-14 \| 1.5e-19 \| 115.6 \| 143 \| (8, 154) \| 217 \| (103, 257) \| 328 \| Gp5/Type VI secretion system Vgr protein OB-fold domain-containing protein (Fragment) \| Gp5/Type VI secretion system Vgr protein OB-fold domain-containing protein (Fragment); uniclust \| UniRef100\_A0A066RIQ8 \| 99.2 \| 7.9e-14 \| 1.6e-19 \| 125.1 \| 81 \| (16, 96) \| 217 \| (366, 461) \| 666 \| Type IV secretion protein Rhs \| Type IV secretion protein Rhs; uniclust \| UniRef100\_A0A812QV09 \| 99.2 \| 9.4e-14 \| 1.7e-19 \| 122.0 \| 115 \| (17, 133) \| 217 \| (944, 1058) \| 1173 \| XkdF protein \| XkdF protein; uniclust \| UniRef100\_A0A2P8AQI4 \| 99.2 \| 9e-14 \| 1.7e-19 \| 110.8 \| 138 \| (10, 154) \| 217 \| (95, 242) \| 283 \| Gp5/Type VI secretion system Vgr protein OB-fold domain-containing protein \| Gp5/Type VI secretion system Vgr protein OB-fold domain-containing protein; uniclust \| UniRef100\_A0A0B6S1Y0 \| 99.2 \| 9e-14 \| 1.7e-19 \| 108.5 \| 169 \| (16, 210) \| 217 \| (34, 222) \| 233 \| Phage baseplate assembly protein V \| Phage baseplate assembly protein V; uniclust \| UniRef100\_A0A1E3WN49 \| 99.2 \| 1.1e-13 \| 2.1e-19 \| 104.2 \| 182 \| (2, 210) \| 217 \| (9, 193) \| 205 \| Baseplate puncturing device gp45 \| Baseplate puncturing device gp45; uniclust \| UniRef100\_A0A0A2ZWF3 \| 99.2 \| 1.2e-13 \| 2.4e-19 \| 111.0 \| 71 \| (1, 72) \| 217 \| (4, 98) \| 231 \| Phage baseplate protein \| Phage baseplate protein; uniclust \| UniRef100\_A0A1Y0UW95 \| 99.2 \| 1.3e-13 \| 2.4e-19 \| 101.8 \| 137 \| (39, 217) \| 217 \| (16, 155) \| 158 \| Baseplate assembly protein \| Baseplate assembly protein; uniclust \| UniRef100\_A0A075R0S9 \| 99.2 \| 1.3e-13 \| 2.6e-19 \| 126.9 \| 112 \| (9, 122) \| 217 \| (384, 504) \| 753 \| Putative phage protein \| Putative phage protein; uniclust \| UniRef100\_A0A3D3TEQ1 \| 99.2 \| 1.4e-13 \| 2.6e-19 \| 119.4 \| 130 \| (17, 153) \| 217 \| (372, 516) \| 620 \| Gp5/Type VI secretion system Vgr protein OB-fold domain-containing protein \| Gp5/Type VI secretion system Vgr protein OB-fold domain-containing protein; uniclust \| UniRef100\_A0A067Z827 \| 99.2 \| 1.3e-13 \| 2.6e-19 \| 114.5 \| 175 \| (17, 217) \| 217 \| (85, 279) \| 299 \| Phage protein Gp138 N-terminal domain-containing protein \| Phage protein Gp138 N-terminal domain-containing protein; uniclust \| UniRef100\_A0A066ZM10 \| 99.2 \| 1.5e-13 \| 2.9e-19 \| 104.7 \| 148 \| (3, 172) \| 217 \| (9, 160) \| 217 \| Baseplate assembly protein \| Baseplate assembly protein; uniclust \| UniRef100\_A0A376EWE4 \| 99.2 \| 1.7e-13 \| 3.2e-19 \| 113.1 \| 142 \| (17, 167) \| 217 \| (397, 541) \| 550 \| Gene D protein \| Gene D protein; uniclust \| UniRef100\_A0A1N7HSV3 \| 99.2 \| 1.8e-13 \| 3.4e-19 \| 113.2 \| 136 \| (17, 158) \| 217 \| (84, 228) \| 424 \| Gp5/Type VI secretion system Vgr protein OB-fold domain-containing protein (Fragment) \| Gp5/Type VI secretion system Vgr protein OB-fold domain-containing protein (Fragment); uniclust \| UniRef100\_A0A6N6MEQ4 \| 99.2 \| 1.9e-13 \| 3.5e-19 \| 103.1 \| 95 \| (3, 99) \| 217 \| (16, 117) \| 217 \| Phage baseplate assembly protein V \| Phage baseplate assembly protein V; uniclust \| UniRef100\_A0A3N5RDY0 \| 99.2 \| 1.9e-13 \| 3.6e-19 \| 108.6 \| 105 \| (17, 121) \| 217 \| (61, 181) \| 284 \| Phage tail protein \| Phage tail protein; uniclust \| UniRef100\_A0A5C8AZF4 \| 99.2 \| 2e-13 \| 3.7e-19 \| 100.8 \| 179 \| (13, 205) \| 217 \| (4, 196) \| 216 \| Phage baseplate assembly protein V \| Phage baseplate assembly protein V; uniclust \| UniRef100\_A0A0X3XDX9 \| 99.2 \| 1.9e-13 \| 3.7e-19 \| 119.0 \| 75 \| (16, 90) \| 217 \| (249, 329) \| 505 \| Gp5/Type VI secretion system Vgr protein OB-fold domain-containing protein \| Gp5/Type VI secretion system Vgr protein OB-fold domain-containing protein; uniclust \| UniRef100\_A0A193QLB5 \| 99.1 \| 2e-13 \| 3.8e-19 \| 94.2 \| 75 \| (140, 217) \| 217 \| (30, 106) \| 106 \| Uncharacterized protein \| Uncharacterized protein; uniclust \| UniRef100\_A0A0A8IL55 \| 99.1 \| 2.2e-13 \| 4.2e-19 \| 102.8 \| 91 \| (1, 96) \| 217 \| (12, 107) \| 183 \| Baseplate assembly protein \| Baseplate assembly protein; uniclust \| UniRef100\_A0A158D125 \| 99.1 \| 2.8e-13 \| 5.2e-19 \| 99.1 \| 134 \| (43, 216) \| 217 \| (12, 145) \| 168 \| Bacteriophage Mu Gp45 protein \| Bacteriophage Mu Gp45 protein; uniclust \| UniRef100\_A0A1H8YYE7 \| 99.1 \| 2.7e-13 \| 5.3e-19 \| 106.1 \| 159 \| (17, 216) \| 217 \| (34, 223) \| 226 \| Phage protein Gp138 N-terminal domain-containing protein \| Phage protein Gp138 N-terminal domain-containing protein; uniclust \| UniRef100\_G8X7R7 \| 99.1 \| 3.1e-13 \| 5.9e-19 \| 101.6 \| 119 \| (29, 153) \| 217 \| (31, 156) \| 209 \| Rhs element Vgr protein \| Rhs element Vgr protein; uniclust \| UniRef100\_A0A0D0H1Q9 \| 99.1 \| 3.3e-13 \| 6.2e-19 \| 100.3 \| 185 \| (6, 217) \| 217 \| (2, 194) \| 198 \| Bacteriophage Mu Gp45 family protein \| Bacteriophage Mu Gp45 family protein; uniclust \| UniRef100\_UPI00201700A5 \| 99.1 \| 3.7e-13 \| 6.7e-19 \| 91.1 \| 86 \| (36, 121) \| 217 \| (4, 89) \| 123 \| phage baseplate assembly protein V \| phage baseplate assembly protein V; uniclust \| UniRef100\_A0A0A2W1E5 \| 99.1 \| 3.9e-13 \| 7.1e-19 \| 121.8 \| 160 \| (16, 216) \| 217 \| (1052, 1248) \| 1885 \| Uncharacterized protein \| Uncharacterized protein; uniclust \| UniRef100\_A0A088C4R9 \| 99.1 \| 4.4e-13 \| 8.6e-19 \| 108.2 \| 162 \| (17, 203) \| 217 \| (49, 231) \| 264 \| p2 gpV-like protein \| p2 gpV-like protein; uniclust \| UniRef100\_A0A2E3N1P5 \| 99.1 \| 5.4e-13 \| 9.9e-19 \| 111.3 \| 163 \| (3, 169) \| 217 \| (18, 202) \| 601 \| Gp5/Type VI secretion system Vgr protein OB-fold domain-containing protein \| Gp5/Type VI secretion system Vgr protein OB-fold domain-containing protein; uniclust \| UniRef100\_UPI00187624BB \| 99.1 \| 5.9e-13 \| 1.1e-18 \| 103.6 \| 115 \| (16, 131) \| 217 \| (18, 139) \| 317 \| phage baseplate assembly protein V \| phage baseplate assembly protein V; uniclust \| UniRef100\_A0A155I046 \| 99.1 \| 6.4e-13 \| 1.2e-18 \| 100.9 \| 173 \| (2, 216) \| 217 \| (41, 216) \| 227 \| Phage baseplate assembly protein V \| Phage baseplate assembly protein V; uniclust \| UniRef100\_A0A1E8PJ79 \| 99.1 \| 6.5e-13 \| 1.2e-18 \| 101.0 \| 176 \| (17, 217) \| 217 \| (33, 220) \| 221 \| Phage baseplate assembly protein V \| Phage baseplate assembly protein V; uniclust \| UniRef100\_A0A369KV77 \| 99.1 \| 6.8e-13 \| 1.3e-18 \| 95.4 \| 133 \| (3, 158) \| 217 \| (9, 144) \| 146 \| Phage baseplate assembly protein V (Fragment) \| Phage baseplate assembly protein V (Fragment); uniclust \| UniRef100\_UPI0022B86340 \| 99.1 \| 7.2e-13 \| 1.3e-18 \| 104.8 \| 150 \| (41, 215) \| 217 \| (184, 361) \| 363 \| Gp138 family membrane-puncturing spike protein \| Gp138 family membrane-puncturing spike protein; uniclust \| UniRef100\_A0A562GH31 \| 99.1 \| 7.6e-13 \| 1.4e-18 \| 92.0 \| 100 \| (12, 113) \| 217 \| (8, 111) \| 132 \| Phage baseplate assembly protein V \| Phage baseplate assembly protein V; uniclust \| UniRef100\_A0A2V8QTC4 \| 99.1 \| 9e-13 \| 1.7e-18 \| 98.1 \| 104 \| (28, 131) \| 217 \| (7, 118) \| 225 \| Phage tail protein \| Phage tail protein; uniclust \| UniRef100\_A0A126QN32 \| 99.1 \| 8.8e-13 \| 1.7e-18 \| 104.5 \| 190 \| (14, 216) \| 217 \| (37, 250) \| 262 \| Baseplate assembly protein \| Baseplate assembly protein; uniclust \| UniRef100\_A0A9E3GUV8 \| 99.1 \| 9.1e-13 \| 1.7e-18 \| 99.2 \| 186 \| (6, 217) \| 217 \| (7, 195) \| 201 \| Uncharacterized protein \| Uncharacterized protein; uniclust \| UniRef100\_UPI00222FD497 \| 99.1 \| 1e-12 \| 1.8e-18 \| 90.2 \| 111 \| (17, 127) \| 217 \| (2, 122) \| 132 \| phage baseplate assembly protein V \| phage baseplate assembly protein V; uniclust \| UniRef100\_A0A318ELH1 \| 99.1 \| 9.8e-13 \| 1.9e-18 \| 102.6 \| 141 \| (13, 160) \| 217 \| (3, 150) \| 236 \| Tail fiber protein \| Tail fiber protein; uniclust \| UniRef100\_A0A2Z5T171 \| 99.1 \| 1e-12 \| 1.9e-18 \| 101.8 \| 171 \| (41, 216) \| 217 \| (65, 272) \| 276 \| Baseplate assembly protein \| Baseplate assembly protein; uniclust \| UniRef100\_UPI001CDCD071 \| 99.1 \| 1e-12 \| 1.9e-18 \| 91.1 \| 109 \| (15, 123) \| 217 \| (1, 112) \| 141 \| phage baseplate assembly protein V \| phage baseplate assembly protein V; uniclust \| UniRef100\_A0A2I7R3I7 \| 99.1 \| 1e-12 \| 1.9e-18 \| 98.0 \| 123 \| (4, 132) \| 217 \| (9, 135) \| 228 \| Baseplate assembly protein V \| Baseplate assembly protein V; uniclust \| UniRef100\_E0WUR2 \| 99.1 \| 1e-12 \| 1.9e-18 \| 88.8 \| 104 \| (5, 108) \| 217 \| (3, 107) \| 121 \| Putative phage baseplate protein \| Putative phage baseplate protein; uniclust \| UniRef100\_A0A250KKR6 \| 99.1 \| 1e-12 \| 1.9e-18 \| 113.7 \| 117 \| (17, 133) \| 217 \| (359, 489) \| 593 \| Type IV secretion protein Rhs \| Type IV secretion protein Rhs; uniclust \| UniRef100\_A0A285JXF0 \| 99.1 \| 1e-12 \| 2e-18 \| 109.3 \| 153 \| (4, 159) \| 217 \| (6, 195) \| 494 \| Gp5/Type VI secretion system Vgr protein OB-fold domain-containing protein \| Gp5/Type VI secretion system Vgr protein OB-fold domain-containing protein; uniclust \| UniRef100\_A0A956GQI0 \| 99.0 \| 1.1e-12 \| 2.1e-18 \| 115.2 \| 119 \| (18, 136) \| 217 \| (28, 158) \| 1114 \| Gp5/Type VI secretion system Vgr protein OB-fold domain-containing protein \| Gp5/Type VI secretion system Vgr protein OB-fold domain-containing protein; uniclust \| UniRef100\_A0A2D9B6H3 \| 99.0 \| 1.3e-12 \| 2.5e-18 \| 102.9 \| 105 \| (16, 120) \| 217 \| (28, 141) \| 251 \| Gp5/Type VI secretion system Vgr protein OB-fold domain-containing protein \| Gp5/Type VI secretion system Vgr protein OB-fold domain-containing protein; uniclust \| UniRef100\_A0A061PST6 \| 99.0 \| 1.3e-12 \| 2.6e-18 \| 118.6 \| 67 \| (27, 93) \| 217 \| (399, 472) \| 676 \| VgrG protein \| VgrG protein; uniclust \| UniRef100\_A0A0A7S6V5 \| 99.0 \| 1.4e-12 \| 2.7e-18 \| 104.3 \| 70 \| (2, 71) \| 217 \| (4, 92) \| 266 \| Phage protein Gp138 N-terminal domain-containing protein \| Phage protein Gp138 N-terminal domain-containing protein; uniclust \| UniRef100\_A0A0C4YA02 \| 99.0 \| 1.5e-12 \| 2.7e-18 \| 95.4 \| 95 \| (16, 110) \| 217 \| (25, 122) \| 167 \| Phage-related baseplate assembly protein \| Phage-related baseplate assembly protein; uniclust \| UniRef100\_UPI000A7B1761 \| 99.0 \| 1.8e-12 \| 3.4e-18 \| 91.2 \| 102 \| (29, 130) \| 217 \| (11, 124) \| 154 \| phage baseplate assembly protein V \| phage baseplate assembly protein V; uniclust \| UniRef100\_A0A1A8ZH78 \| 99.0 \| 1.7e-12 \| 3.4e-18 \| 103.2 \| 117 \| (17, 133) \| 217 \| (29, 153) \| 246 \| Gp5/Type VI secretion system Vgr protein OB-fold domain-containing protein \| Gp5/Type VI secretion system Vgr protein OB-fold domain-containing protein; uniclust \| UniRef100\_A0A5C4P976 \| 99.0 \| 2.4e-12 \| 4.4e-18 \| 90.3 \| 89 \| (2, 90) \| 217 \| (1, 89) \| 150 \| Phage baseplate assembly protein V \| Phage baseplate assembly protein V; uniclust \| UniRef100\_A0A077EH50 \| 99.0 \| 2.3e-12 \| 4.5e-18 \| 117.2 \| 117 \| (17, 133) \| 217 \| (490, 619) \| 842 \| VgrG protein \| VgrG protein; uniclust \| UniRef100\_A0A7V9PH58 \| 99.0 \| 2.3e-12 \| 4.5e-18 \| 112.6 \| 105 \| (17, 124) \| 217 \| (354, 466) \| 627 \| Gp5/Type VI secretion system Vgr protein OB-fold domain-containing protein \| Gp5/Type VI secretion system Vgr protein OB-fold domain-containing protein; uniclust \| UniRef100\_A0A8I2CH63 \| 99.0 \| 2.5e-12 \| 4.6e-18 \| 90.7 \| 85 \| (2, 86) \| 217 \| (1, 85) \| 155 \| Phage baseplate assembly protein V \| Phage baseplate assembly protein V; uniclust \| UniRef100\_A0A009YIF6 \| 99.0 \| 2.3e-12 \| 4.6e-18 \| 99.1 \| 107 \| (1, 122) \| 217 \| (14, 122) \| 181 \| Bacteriophage Mu Gp45 family protein \| Bacteriophage Mu Gp45 family protein; uniclust \| UniRef100\_A0A348FYH6 \| 99.0 \| 2.7e-12 \| 5e-18 \| 96.2 \| 93 \| (2, 96) \| 217 \| (9, 101) \| 200 \| Gp5/Type VI secretion system Vgr protein OB-fold domain-containing protein \| Gp5/Type VI secretion system Vgr protein OB-fold domain-containing protein; uniclust \| UniRef100\_A0A5C8XT18 \| 99.0 \| 2.9e-12 \| 5.3e-18 \| 84.3 \| 88 \| (58, 161) \| 217 \| (2, 89) \| 102 \| Phage baseplate assembly protein V (Fragment) \| Phage baseplate assembly protein V (Fragment); uniclust \| UniRef100\_A0A0H2Y5Z8 \| 99.0 \| 2.9e-12 \| 5.3e-18 \| 86.5 \| 73 \| (1, 73) \| 217 \| (36, 108) \| 119 \| Gp5/Type VI secretion system Vgr protein OB-fold domain-containing protein \| Gp5/Type VI secretion system Vgr protein OB-fold domain-containing protein; uniclust \| UniRef100\_UPI00209A7807 \| 99.0 \| 3.1e-12 \| 5.8e-18 \| 91.0 \| 78 \| (1, 78) \| 217 \| (1, 78) \| 164 \| phage baseplate assembly protein V \| phage baseplate assembly protein V; uniclust \| UniRef100\_A0A950GBI3 \| 99.0 \| 3.1e-12 \| 5.8e-18 \| 88.4 \| 68 \| (40, 107) \| 217 \| (2, 70) \| 137 \| Baseplate assembly protein \| Baseplate assembly protein; uniclust \| UniRef100\_A0A3N9EW37 \| 99.0 \| 3.1e-12 \| 5.9e-18 \| 91.5 \| 135 \| (66, 216) \| 217 \| (3, 142) \| 147 \| Phage baseplate assembly protein V (Fragment) \| Phage baseplate assembly protein V (Fragment); uniclust \| UniRef100\_A0A072TF05 \| 99.0 \| 3.3e-12 \| 6.1e-18 \| 116.0 \| 159 \| (6, 173) \| 217 \| (470, 629) \| 1809 \| Bacteriophage regulatory protein, putative (Fragment) \| Bacteriophage regulatory protein, putative (Fragment); uniclust \| UniRef100\_A0A3D4Z671 \| 99.0 \| 3.3e-12 \| 6.1e-18 \| 92.9 \| 136 \| (3, 169) \| 217 \| (44, 182) \| 190 \| Phage baseplate assembly protein V \| Phage baseplate assembly protein V; uniclust \| UniRef100\_A0A1Y4GV21 \| 99.0 \| 3.6e-12 \| 6.9e-18 \| 113.1 \| 105 \| (17, 121) \| 217 \| (500, 623) \| 776 \| Gp5/Type VI secretion system Vgr protein OB-fold domain-containing protein \| Gp5/Type VI secretion system Vgr protein OB-fold domain-containing protein; uniclust \| UniRef100\_A0A8T7CZW0 \| 99.0 \| 3.8e-12 \| 7e-18 \| 99.0 \| 111 \| (4, 114) \| 217 \| (7, 151) \| 308 \| Phage baseplate assembly protein V \| Phage baseplate assembly protein V; uniclust \| UniRef100\_UPI002021AE60 \| 99.0 \| 4e-12 \| 7.4e-18 \| 91.7 \| 72 \| (21, 92) \| 217 \| (2, 73) \| 179 \| phage baseplate assembly protein V \| phage baseplate assembly protein V; uniclust \| UniRef100\_A0A193LS04 \| 99.0 \| 4.2e-12 \| 7.8e-18 \| 94.4 \| 173 \| (3, 216) \| 217 \| (11, 187) \| 188 \| Phage baseplate protein \| Phage baseplate protein; uniclust \| UniRef100\_A0A0B1M335 \| 99.0 \| 4e-12 \| 7.9e-18 \| 103.9 \| 192 \| (14, 216) \| 217 \| (42, 245) \| 291 \| Phage baseplate protein \| Phage baseplate protein; uniclust \| UniRef100\_A0A7Y5WP02 \| 99.0 \| 4.3e-12 \| 8e-18 \| 99.0 \| 115 \| (15, 130) \| 217 \| (26, 152) \| 275 \| Gp5/Type VI secretion system Vgr protein OB-fold domain-containing protein \| Gp5/Type VI secretion system Vgr protein OB-fold domain-containing protein; uniclust \| UniRef100\_UPI00077A5E31 \| 99.0 \| 4.5e-12 \| 8.4e-18 \| 90.5 \| 116 \| (1, 133) \| 217 \| (1, 119) \| 136 \| phage baseplate assembly protein V \| phage baseplate assembly protein V; uniclust \| UniRef100\_A0A344TT64 \| 99.0 \| 4.7e-12 \| 8.7e-18 \| 106.4 \| 142 \| (17, 159) \| 217 \| (242, 398) \| 483 \| Gp5/Type VI secretion system Vgr protein OB-fold domain-containing protein \| Gp5/Type VI secretion system Vgr protein OB-fold domain-containing protein; uniclust \| UniRef100\_A0A0R3L044 \| 99.0 \| 4.9e-12 \| 9e-18 \| 94.8 \| 100 \| (4, 104) \| 217 \| (13, 118) \| 233 \| Gp5/Type VI secretion system Vgr protein OB-fold domain-containing protein \| Gp5/Type VI secretion system Vgr protein OB-fold domain-containing protein; uniclust \| UniRef100\_A0A1Q6LL85 \| 98.9 \| 5.1e-12 \| 9.9e-18 \| 92.3 \| 104 \| (13, 122) \| 217 \| (2, 110) \| 142 \| Uncharacterized protein \| Uncharacterized protein; uniclust \| UniRef100\_A0A0X8JVG9 \| 98.9 \| 5.4e-12 \| 1e-17 \| 97.5 \| 163 \| (14, 217) \| 217 \| (26, 207) \| 208 \| Phage protein Gp138 N-terminal domain-containing protein \| Phage protein Gp138 N-terminal domain-containing protein; uniclust \| UniRef100\_A0A7H8YU12 \| 98.9 \| 5.7e-12 \| 1.1e-17 \| 113.2 \| 137 \| (16, 153) \| 217 \| (397, 546) \| 1055 \| Rhs element Vgr protein \| Rhs element Vgr protein; uniclust \| UniRef100\_A0A827UUR8 \| 98.9 \| 5.6e-12 \| 1.1e-17 \| 90.0 \| 107 \| (2, 122) \| 217 \| (5, 114) \| 137 \| Baseplate assembly protein (Fragment) \| Baseplate assembly protein (Fragment); uniclust \| UniRef100\_UPI001BC86C81 \| 98.9 \| 6.3e-12 \| 1.2e-17 \| 99.8 \| 115 \| (17, 131) \| 217 \| (41, 165) \| 296 \| phage baseplate assembly protein V \| phage baseplate assembly protein V; uniclust \| UniRef100\_UPI000C7FE3C4 \| 98.9 \| 6.7e-12 \| 1.2e-17 \| 89.7 \| 58 \| (48, 105) \| 217 \| (4, 61) \| 167 \| phage baseplate assembly protein V \| phage baseplate assembly protein V; uniclust \| UniRef100\_A0A3S4S5U6 \| 98.9 \| 6.9e-12 \| 1.3e-17 \| 86.6 \| 107 \| (15, 123) \| 217 \| (1, 108) \| 135 \| Baseplate assembly protein V \| Baseplate assembly protein V; uniclust \| UniRef100\_A0A1Q1PVM9 \| 98.9 \| 7.2e-12 \| 1.4e-17 \| 98.7 \| 129 \| (16, 152) \| 217 \| (24, 160) \| 240 \| D-like protein \| D-like protein; uniclust \| UniRef100\_A0A653V686 \| 98.9 \| 7.8e-12 \| 1.5e-17 \| 108.2 \| 118 \| (16, 133) \| 217 \| (399, 526) \| 636 \| Gp5/Type VI secretion system Vgr protein OB-fold domain-containing protein \| Gp5/Type VI secretion system Vgr protein OB-fold domain-containing protein; uniclust \| UniRef100\_C9MS63 \| 98.9 \| 8.2e-12 \| 1.5e-17 \| 113.3 \| 117 \| (17, 133) \| 217 \| (361, 491) \| 1047 \| Gp5/Type VI secretion system Vgr protein OB-fold domain-containing protein \| Gp5/Type VI secretion system Vgr protein OB-fold domain-containing protein; uniclust \| UniRef100\_A0A022PDU6 \| 98.9 \| 7.9e-12 \| 1.6e-17 \| 112.7 \| 107 \| (16, 122) \| 217 \| (371, 491) \| 575 \| Gp5/Type VI secretion system Vgr protein OB-fold domain-containing protein \| Gp5/Type VI secretion system Vgr protein OB-fold domain-containing protein; uniclust \| UniRef100\_A0A7C1CYC4 \| 98.9 \| 1e-11 \| 1.9e-17 \| 98.6 \| 117 \| (16, 132) \| 217 \| (141, 268) \| 316 \| Type IV secretion protein Rhs \| Type IV secretion protein Rhs; uniclust \| UniRef100\_UPI000B280816 \| 98.9 \| 1e-11 \| 2e-17 \| 91.1 \| 147 \| (11, 169) \| 217 \| (3, 156) \| 176 \| phage baseplate assembly protein V \| phage baseplate assembly protein V; uniclust \| UniRef100\_UPI001C8C52A2 \| 98.9 \| 1.1e-11 \| 2e-17 \| 95.5 \| 80 \| (16, 95) \| 217 \| (103, 189) \| 283 \| phage baseplate assembly protein V \| phage baseplate assembly protein V; uniclust \| UniRef100\_A0A0D9NAU1 \| 98.9 \| 1.1e-11 \| 2.2e-17 \| 98.0 \| 108 \| (24, 131) \| 217 \| (5, 125) \| 248 \| Type IV secretion protein Rhs \| Type IV secretion protein Rhs; uniclust \| UniRef100\_A0A951W2Z1 \| 98.9 \| 1.3e-11 \| 2.3e-17 \| 93.1 \| 114 \| (16, 132) \| 217 \| (2, 118) \| 240 \| Phage baseplate assembly protein V \| Phage baseplate assembly protein V; uniclust \| UniRef100\_UPI00202674E5 \| 98.9 \| 1.3e-11 \| 2.3e-17 \| 104.0 \| 133 \| (27, 159) \| 217 \| (424, 571) \| 626 \| phage baseplate assembly protein V \| phage baseplate assembly protein V; uniclust \| UniRef100\_A0A2D6I496 \| 98.9 \| 1.3e-11 \| 2.5e-17 \| 105.4 \| 75 \| (17, 91) \| 217 \| (347, 428) \| 570 \| Gp5/Type VI secretion system Vgr protein OB-fold domain-containing protein \| Gp5/Type VI secretion system Vgr protein OB-fold domain-containing protein; uniclust \| UniRef100\_R9B5T6 \| 98.9 \| 1.4e-11 \| 2.5e-17 \| 85.5 \| 98 \| (6, 103) \| 217 \| (2, 99) \| 137 \| Gp5/Type VI secretion system Vgr protein OB-fold domain-containing protein \| Gp5/Type VI secretion system Vgr protein OB-fold domain-containing protein; uniclust \| UniRef100\_A0A2V2GM48 \| 98.9 \| 1.4e-11 \| 2.5e-17 \| 95.7 \| 97 \| (12, 113) \| 217 \| (87, 189) \| 300 \| Gp5/Type VI secretion system Vgr protein OB-fold domain-containing protein \| Gp5/Type VI secretion system Vgr protein OB-fold domain-containing protein; uniclust \| UniRef100\_A0A096CE48 \| 98.9 \| 1.6e-11 \| 3.1e-17 \| 107.6 \| 97 \| (12, 114) \| 217 \| (295, 400) \| 482 \| Gp5/Type VI secretion system Vgr protein OB-fold domain-containing protein \| Gp5/Type VI secretion system Vgr protein OB-fold domain-containing protein; uniclust \| UniRef100\_A0A534P4W5 \| 98.9 \| 1.8e-11 \| 3.2e-17 \| 97.8 \| 105 \| (16, 120) \| 217 \| (26, 143) \| 374 \| Gp5/Type VI secretion system Vgr protein OB-fold domain-containing protein (Fragment) \| Gp5/Type VI secretion system Vgr protein OB-fold domain-containing protein (Fragment); uniclust \| UniRef100\_A0A2S6HRN7 \| 98.9 \| 1.7e-11 \| 3.4e-17 \| 105.8 \| 110 \| (11, 124) \| 217 \| (267, 384) \| 452 \| Late control gene D protein (GPD) \| Late control gene D protein (GPD); uniclust \| UniRef100\_A0A2S0MND7 \| 98.9 \| 1.9e-11 \| 3.5e-17 \| 88.1 \| 77 \| (5, 83) \| 217 \| (52, 129) \| 175 \| Gp5/Type VI secretion system Vgr protein OB-fold domain-containing protein \| Gp5/Type VI secretion system Vgr protein OB-fold domain-containing protein; uniclust \| UniRef100\_A0A0B5KGS8 \| 98.9 \| 1.9e-11 \| 3.6e-17 \| 111.5 \| 67 \| (27, 93) \| 217 \| (594, 669) \| 848 \| ImpA family type VI secretion-associated protein \| ImpA family type VI secretion-associated protein; uniclust \| UniRef100\_A0A1D2R0D3 \| 98.9 \| 1.9e-11 \| 3.7e-17 \| 100.2 \| 94 \| (15, 111) \| 217 \| (52, 156) \| 314 \| Gp5/Type VI secretion system Vgr protein OB-fold domain-containing protein \| Gp5/Type VI secretion system Vgr protein OB-fold domain-containing protein; uniclust \| UniRef100\_A0A1H3JED8 \| 98.9 \| 2e-11 \| 4e-17 \| 108.5 \| 113 \| (16, 129) \| 217 \| (378, 503) \| 567 \| Rhs element Vgr protein \| Rhs element Vgr protein; uniclust \| UniRef100\_A0A023XLV0 \| 98.8 \| 2.1e-11 \| 4.2e-17 \| 96.6 \| 117 \| (17, 133) \| 217 \| (39, 171) \| 217 \| Baseplate assembly protein V \| Baseplate assembly protein V; uniclust \| UniRef100\_A0A1H9NJ96 \| 98.8 \| 2.2e-11 \| 4.3e-17 \| 92.9 \| 69 \| (14, 82) \| 217 \| (8, 82) \| 180 \| Baseplate assembly protein \| Baseplate assembly protein; uniclust \| UniRef100\_UPI0011A2738F \| 98.8 \| 2.3e-11 \| 4.4e-17 \| 99.5 \| 109 \| (1, 116) \| 217 \| (1, 109) \| 363 \| contractile injection system protein, VgrG/Pvc8 family \| contractile injection system protein, VgrG/Pvc8 family; uniclust \| UniRef100\_A0A0F9W262 \| 98.8 \| 2.3e-11 \| 4.4e-17 \| 95.4 \| 143 \| (16, 160) \| 217 \| (20, 185) \| 228 \| Gp5/Type VI secretion system Vgr protein OB-fold domain-containing protein (Fragment) \| Gp5/Type VI secretion system Vgr protein OB-fold domain-containing protein (Fragment); uniclust \| UniRef100\_A0A173SAA9 \| 98.8 \| 2.4e-11 \| 4.6e-17 \| 87.1 \| 76 \| (9, 84) \| 217 \| (9, 114) \| 123 \| Uncharacterized protein \| Uncharacterized protein; uniclust \| UniRef100\_G0A9X5 \| 98.8 \| 2.7e-11 \| 4.9e-17 \| 92.0 \| 76 \| (2, 77) \| 217 \| (39, 114) \| 249 \| Site-specific recombinase \| Site-specific recombinase; uniclust \| UniRef100\_A0A064AML4 \| 98.8 \| 2.6e-11 \| 5.2e-17 \| 93.1 \| 86 \| (17, 103) \| 217 \| (8, 96) \| 183 \| Phage baseplate protein \| Phage baseplate protein; uniclust \| UniRef100\_A0A6P0Q5Z6 \| 98.8 \| 2.8e-11 \| 5.2e-17 \| 83.0 \| 67 \| (28, 94) \| 217 \| (37, 108) \| 120 \| Phage tail protein (Fragment) \| Phage tail protein (Fragment); uniclust \| UniRef100\_UPI001F3F1939 \| 98.8 \| 2.8e-11 \| 5.2e-17 \| 80.4 \| 89 \| (1, 91) \| 217 \| (1, 89) \| 101 \| phage baseplate assembly protein V \| phage baseplate assembly protein V; uniclust \| UniRef100\_A0A1F8VKX0 \| 98.8 \| 2.9e-11 \| 5.6e-17 \| 92.1 \| 166 \| (13, 217) \| 217 \| (6, 173) \| 187 \| Baseplate assembly protein \| Baseplate assembly protein; uniclust \| UniRef100\_A0A1P8ENB8 \| 98.8 \| 3.3e-11 \| 6.3e-17 \| 91.3 \| 162 \| (16, 215) \| 217 \| (8, 176) \| 179 \| Uncharacterized protein \| Uncharacterized protein; uniclust \| UniRef100\_UPI001EE4E6FE \| 98.8 \| 3.6e-11 \| 6.6e-17 \| 80.5 \| 73 \| (16, 88) \| 217 \| (18, 97) \| 110 \| phage baseplate assembly protein V \| phage baseplate assembly protein V; uniclust \| UniRef100\_A0A0T9PA02 \| 98.8 \| 3.8e-11 \| 7.2e-17 \| 94.3 \| 184 \| (17, 216) \| 217 \| (32, 245) \| 254 \| Phage protein Gp138 N-terminal domain-containing protein \| Phage protein Gp138 N-terminal domain-containing protein; uniclust \| UniRef100\_UPI00201ED154 \| 98.8 \| 3.9e-11 \| 7.2e-17 \| 89.2 \| 122 \| (1, 122) \| 217 \| (1, 130) \| 214 \| hypothetical protein \| hypothetical protein; uniclust \| UniRef100\_UPI001F403EA4 \| 98.8 \| 4.2e-11 \| 7.8e-17 \| 95.2 \| 108 \| (17, 124) \| 217 \| (174, 301) \| 357 \| phage baseplate assembly protein V \| phage baseplate assembly protein V; uniclust \| UniRef100\_UPI00164A91CE \| 98.8 \| 4.3e-11 \| 7.9e-17 \| 79.5 \| 76 \| (52, 127) \| 217 \| (2, 77) \| 105 \| phage baseplate assembly protein V \| phage baseplate assembly protein V; uniclust \| UniRef100\_A0A845R707 \| 98.8 \| 4.3e-11 \| 7.9e-17 \| 87.9 \| 118 \| (10, 131) \| 217 \| (60, 183) \| 197 \| Gp5/Type VI secretion system Vgr protein OB-fold domain-containing protein \| Gp5/Type VI secretion system Vgr protein OB-fold domain-containing protein; uniclust \| UniRef100\_A0A0X3AMS8 \| 98.8 \| 4.4e-11 \| 8.3e-17 \| 100.5 \| 117 \| (17, 133) \| 217 \| (167, 293) \| 414 \| Phage-related baseplate assembly protein (Fragment) \| Phage-related baseplate assembly protein (Fragment); uniclust \| UniRef100\_A0A0C1ET35 \| 98.8 \| 4.6e-11 \| 8.9e-17 \| 93.2 \| 99 \| (53, 157) \| 217 \| (16, 115) \| 222 \| Uncharacterized protein \| Uncharacterized protein; uniclust \| UniRef100\_A0A7S8C6X6 \| 98.8 \| 5.8e-11 \| 1.1e-16 \| 98.4 \| 96 \| (3, 99) \| 217 \| (330, 431) \| 515 \| Phage virion morphogenesis protein \| Phage virion morphogenesis protein; uniclust \| UniRef100\_A0A377I2R7 \| 98.8 \| 5.9e-11 \| 1.1e-16 \| 92.0 \| 169 \| (18, 205) \| 217 \| (24, 208) \| 289 \| Phage baseplate assembly protein V \| Phage baseplate assembly protein V; uniclust \| UniRef100\_A0A379VYI9 \| 98.8 \| 6.3e-11 \| 1.2e-16 \| 83.7 \| 103 \| (17, 120) \| 217 \| (7, 112) \| 129 \| Baseplate protein \| Baseplate protein; uniclust \| UniRef100\_A0A919LS40 \| 98.8 \| 6.5e-11 \| 1.2e-16 \| 73.8 \| 71 \| (4, 74) \| 217 \| (1, 71) \| 72 \| Uncharacterized protein \| Uncharacterized protein; uniclust \| UniRef100\_UPI0002DB02C0 \| 98.8 \| 6.9e-11 \| 1.3e-16 \| 87.2 \| 133 \| (17, 154) \| 217 \| (34, 182) \| 201 \| phage baseplate assembly protein V \| phage baseplate assembly protein V; uniclust \| UniRef100\_UPI001A19B5C2 \| 98.8 \| 6.9e-11 \| 1.3e-16 \| 103.6 \| 117 \| (17, 133) \| 217 \| (137, 263) \| 963 \| phage baseplate assembly protein V \| phage baseplate assembly protein V; uniclust \| UniRef100\_A0A088C368 \| 98.8 \| 6.5e-11 \| 1.3e-16 \| 97.1 \| 192 \| (17, 216) \| 217 \| (48, 281) \| 283 \| p2 gpV-like protein \| p2 gpV-like protein; uniclust \| UniRef100\_UPI002115C689 \| 98.8 \| 7.1e-11 \| 1.3e-16 \| 74.5 \| 69 \| (23, 91) \| 217 \| (3, 71) \| 79 \| phage baseplate assembly protein V \| phage baseplate assembly protein V; uniclust \| UniRef100\_UPI000371D434 \| 98.8 \| 7.2e-11 \| 1.3e-16 \| 87.2 \| 102 \| (3, 104) \| 217 \| (13, 119) \| 181 \| phage baseplate assembly protein V \| phage baseplate assembly protein V; uniclust \| UniRef100\_A0A977FHW3 \| 98.8 \| 7.4e-11 \| 1.4e-16 \| 82.0 \| 127 \| (41, 216) \| 217 \| (10, 136) \| 137 \| Phage baseplate assembly protein \| Phage baseplate assembly protein; uniclust \| UniRef100\_A0A928JDR0 \| 98.7 \| 8e-11 \| 1.5e-16 \| 90.3 \| 74 \| (17, 90) \| 217 \| (24, 128) \| 265 \| Phage protein Gp138 N-terminal domain-containing protein \| Phage protein Gp138 N-terminal domain-containing protein; uniclust \| UniRef100\_A0A7V1ZNU7 \| 98.7 \| 9.6e-11 \| 1.8e-16 \| 85.2 \| 90 \| (5, 94) \| 217 \| (12, 104) \| 181 \| Gp5/Type VI secretion system Vgr protein OB-fold domain-containing protein \| Gp5/Type VI secretion system Vgr protein OB-fold domain-containing protein; uniclust \| UniRef100\_A0A0Q7SRW7 \| 98.7 \| 9.4e-11 \| 1.9e-16 \| 111.1 \| 140 \| (11, 153) \| 217 \| (292, 450) \| 1001 \| Uncharacterized protein \| Uncharacterized protein; uniclust \| UniRef100\_A0A161W0R2 \| 98.7 \| 1e-10 \| 2e-16 \| 79.1 \| 71 \| (7, 83) \| 217 \| (4, 77) \| 84 \| Uncharacterized protein \| Uncharacterized protein; uniclust \| UniRef100\_A0A9E3FXI7 \| 98.7 \| 1.1e-10 \| 2e-16 \| 85.1 \| 139 \| (8, 158) \| 217 \| (19, 160) \| 184 \| Uncharacterized protein \| Uncharacterized protein; uniclust \| UniRef100\_A0A0U5B8F1 \| 98.7 \| 1.1e-10 \| 2.1e-16 \| 101.0 \| 141 \| (11, 154) \| 217 \| (285, 438) \| 476 \| Gp5/Type VI secretion system Vgr protein OB-fold domain-containing protein \| Gp5/Type VI secretion system Vgr protein OB-fold domain-containing protein; uniclust \| UniRef100\_UPI0004B6FA99 \| 98.7 \| 1.1e-10 \| 2.1e-16 \| 78.1 \| 84 \| (30, 113) \| 217 \| (15, 99) \| 103 \| phage baseplate assembly protein V \| phage baseplate assembly protein V; uniclust \| UniRef100\_A0A1Y4DVJ7 \| 98.7 \| 1.2e-10 \| 2.2e-16 \| 104.3 \| 99 \| (17, 115) \| 217 \| (821, 940) \| 1023 \| Gp5/Type VI secretion system Vgr protein OB-fold domain-containing protein \| Gp5/Type VI secretion system Vgr protein OB-fold domain-containing protein; uniclust \| UniRef100\_A0A2N5KH98 \| 98.7 \| 1.2e-10 \| 2.3e-16 \| 96.7 \| 108 \| (17, 124) \| 217 \| (194, 317) \| 435 \| Gp5/Type VI secretion system Vgr protein OB-fold domain-containing protein \| Gp5/Type VI secretion system Vgr protein OB-fold domain-containing protein; uniclust \| UniRef100\_A0A162L1Y1 \| 98.7 \| 1.2e-10 \| 2.3e-16 \| 106.4 \| 77 \| (17, 93) \| 217 \| (387, 478) \| 702 \| Gp5/Type VI secretion system Vgr protein OB-fold domain-containing protein \| Gp5/Type VI secretion system Vgr protein OB-fold domain-containing protein; uniclust \| UniRef100\_A0A2S0JRF5 \| 98.7 \| 1.3e-10 \| 2.4e-16 \| 83.2 \| 115 \| (15, 133) \| 217 \| (30, 147) \| 163 \| Uncharacterized protein \| Uncharacterized protein; uniclust \| UniRef100\_A0A1H7UA08 \| 98.7 \| 1.3e-10 \| 2.5e-16 \| 89.8 \| 88 \| (16, 106) \| 217 \| (23, 117) \| 219 \| Gp5/Type VI secretion system Vgr protein OB-fold domain-containing protein \| Gp5/Type VI secretion system Vgr protein OB-fold domain-containing protein; uniclust \| UniRef100\_A0A430BCY6 \| 98.7 \| 1.4e-10 \| 2.6e-16 \| 94.4 \| 174 \| (8, 216) \| 217 \| (175, 359) \| 364 \| Baseplate assembly protein \| Baseplate assembly protein; uniclust \| UniRef100\_A0A1N6X9L1 \| 98.7 \| 1.5e-10 \| 2.7e-16 \| 80.6 \| 82 \| (12, 93) \| 217 \| (4, 92) \| 137 \| Type VI secretion system, phage-baseplate injector (Fragment) \| Type VI secretion system, phage-baseplate injector (Fragment); uniclust \| UniRef100\_A0A1A6FJR2 \| 98.7 \| 1.6e-10 \| 2.9e-16 \| 90.9 \| 87 \| (6, 94) \| 217 \| (22, 114) \| 290 \| Gp5/Type VI secretion system Vgr protein OB-fold domain-containing protein \| Gp5/Type VI secretion system Vgr protein OB-fold domain-containing protein; uniclust \| UniRef100\_A0A023D7D4 \| 98.7 \| 1.5e-10 \| 2.9e-16 \| 105.2 \| 67 \| (27, 93) \| 217 \| (407, 482) \| 675 \| Secretion system Type VI Rhs element Vgr \| Secretion system Type VI Rhs element Vgr; uniclust \| UniRef100\_A0A068CE12 \| 98.7 \| 1.5e-10 \| 3e-16 \| 87.8 \| 98 \| (42, 169) \| 217 \| (47, 144) \| 170 \| Baseplate assembly protein V \| Baseplate assembly protein V; uniclust \| UniRef100\_A0A0D2GHS9 \| 98.7 \| 1.6e-10 \| 3e-16 \| 91.1 \| 114 \| (4, 133) \| 217 \| (13, 131) \| 216 \| Uncharacterized protein \| Uncharacterized protein; uniclust \| UniRef100\_UPI001F5DA9EC \| 98.7 \| 1.9e-10 \| 3.5e-16 \| 79.5 \| 93 \| (39, 133) \| 217 \| (2, 99) \| 131 \| phage baseplate assembly protein V \| phage baseplate assembly protein V; uniclust \| UniRef100\_A0A3M1ATA3 \| 98.7 \| 1.9e-10 \| 3.5e-16 \| 80.3 \| 75 \| (18, 92) \| 217 \| (24, 105) \| 139 \| Gp5/Type VI secretion system Vgr protein OB-fold domain-containing protein \| Gp5/Type VI secretion system Vgr protein OB-fold domain-containing protein; uniclust \| UniRef100\_A0A968SXI1 \| 98.7 \| 1.9e-10 \| 3.5e-16 \| 97.1 \| 110 \| (6, 115) \| 217 \| (352, 468) \| 605 \| Type VI secretion system tip protein VgrG \| Type VI secretion system tip protein VgrG; uniclust \| UniRef100\_UPI0004064B79 \| 98.7 \| 2e-10 \| 3.6e-16 \| 84.7 \| 117 \| (3, 121) \| 217 \| (40, 159) \| 197 \| phage baseplate assembly protein V \| phage baseplate assembly protein V; uniclust \| UniRef100\_UPI002012D1F8 \| 98.7 \| 2e-10 \| 3.6e-16 \| 94.0 \| 117 \| (17, 133) \| 217 \| (24, 154) \| 447 \| phage baseplate assembly protein V \| phage baseplate assembly protein V; uniclust \| UniRef100\_A0A327VQM2 \| 98.7 \| 2.1e-10 \| 4e-16 \| 98.6 \| 115 \| (17, 131) \| 217 \| (350, 474) \| 583 \| Uncharacterized protein involved in type VI secretion and phage assembly \| Uncharacterized protein involved in type VI secretion and phage assembly; uniclust \| UniRef100\_A0A1C5P7Q4 \| 98.7 \| 2.3e-10 \| 4.5e-16 \| 98.4 \| 117 \| (14, 133) \| 217 \| (285, 412) \| 474 \| Phage protein D \| Phage protein D; uniclust \| UniRef100\_UPI00068998DA \| 98.7 \| 2.5e-10 \| 4.5e-16 \| 92.3 \| 116 \| (17, 132) \| 217 \| (115, 248) \| 368 \| phage baseplate assembly protein V \| phage baseplate assembly protein V; uniclust \| UniRef100\_A0A1I4TS69 \| 98.6 \| 2.4e-10 \| 4.8e-16 \| 106.1 \| 77 \| (17, 93) \| 217 \| (382, 473) \| 713 \| Type VI secretion system secreted protein VgrG \| Type VI secretion system secreted protein VgrG; uniclust \| UniRef100\_A0A1M4ZE13 \| 98.6 \| 2.6e-10 \| 4.9e-16 \| 97.0 \| 120 \| (13, 132) \| 217 \| (364, 494) \| 556 \| Uncharacterized conserved protein, implicated in type VI secretion and phage assembly \| Uncharacterized conserved protein, implicated in type VI secretion and phage assembly; uniclust \| UniRef100\_A0A377BG29 \| 98.6 \| 2.8e-10 \| 5.2e-16 \| 83.1 \| 115 \| (2, 133) \| 217 \| (17, 132) \| 157 \| Bacteriophage Mu Gp45 protein \| Bacteriophage Mu Gp45 protein; uniclust \| UniRef100\_UPI0021AB61CB \| 98.6 \| 2.9e-10 \| 5.4e-16 \| 96.3 \| 139 \| (18, 161) \| 217 \| (78, 234) \| 621 \| phage baseplate assembly protein V \| phage baseplate assembly protein V; uniclust \| UniRef100\_A0A917BFD9 \| 98.6 \| 2.9e-10 \| 5.6e-16 \| 84.1 \| 97 \| (15, 122) \| 217 \| (7, 108) \| 162 \| Gp5/Type VI secretion system Vgr protein OB-fold domain-containing protein \| Gp5/Type VI secretion system Vgr protein OB-fold domain-containing protein; uniclust \| UniRef100\_A0A2W6ZDC9 \| 98.6 \| 3e-10 \| 5.6e-16 \| 86.7 \| 83 \| (3, 88) \| 217 \| (13, 100) \| 220 \| Gp5/Type VI secretion system Vgr protein OB-fold domain-containing protein \| Gp5/Type VI secretion system Vgr protein OB-fold domain-containing protein; uniclust \| UniRef100\_A0A812U007 \| 98.6 \| 3.2e-10 \| 5.8e-16 \| 102.0 \| 136 \| (25, 160) \| 217 \| (1119, 1274) \| 1306 \| VgrG5 protein (Fragment) \| VgrG5 protein (Fragment); uniclust \| UniRef100\_A0A094RX34 \| 98.6 \| 3.2e-10 \| 6e-16 \| 88.2 \| 57 \| (17, 73) \| 217 \| (44, 112) \| 225 \| Phage protein Gp138 N-terminal domain-containing protein (Fragment) \| Phage protein Gp138 N-terminal domain-containing protein (Fragment); uniclust \| UniRef100\_A0A1W1Z4L7 \| 98.6 \| 3.2e-10 \| 6.1e-16 \| 83.8 \| 105 \| (3, 107) \| 217 \| (17, 128) \| 164 \| Type VI secretion system, phage-baseplate injector \| Type VI secretion system, phage-baseplate injector; uniclust \| UniRef100\_A0A7Y1K0F9 \| 98.6 \| 3.3e-10 \| 6.4e-16 \| 98.0 \| 76 \| (17, 92) \| 217 \| (205, 295) \| 514 \| Type VI secretion system tip protein VgrG \| Type VI secretion system tip protein VgrG; uniclust \| UniRef100\_A0A0S3U5G4 \| 98.6 \| 3.4e-10 \| 6.6e-16 \| 94.1 \| 76 \| (17, 92) \| 217 \| (36, 118) \| 314 \| Rhs element Vgr protein \| Rhs element Vgr protein; uniclust \| UniRef100\_A0A075K3F1 \| 98.6 \| 3.4e-10 \| 6.7e-16 \| 106.6 \| 79 \| (17, 95) \| 217 \| (505, 598) \| 918 \| Type IV secretion protein Rhs \| Type IV secretion protein Rhs; uniclust \| UniRef100\_A0A061Q576 \| 98.6 \| 3.6e-10 \| 7e-16 \| 105.1 \| 78 \| (16, 93) \| 217 \| (384, 476) \| 848 \| VgrG protein \| VgrG protein; uniclust \| UniRef100\_UPI001569D794 \| 98.6 \| 3.9e-10 \| 7.3e-16 \| 77.1 \| 104 \| (81, 208) \| 217 \| (2, 106) \| 115 \| phage baseplate assembly protein V \| phage baseplate assembly protein V; uniclust \| UniRef100\_A0A959CMX5 \| 98.6 \| 4.5e-10 \| 8.3e-16 \| 88.2 \| 100 \| (16, 115) \| 217 \| (29, 139) \| 311 \| Gp5/Type VI secretion system Vgr protein OB-fold domain-containing protein \| Gp5/Type VI secretion system Vgr protein OB-fold domain-containing protein; uniclust \| UniRef100\_A0A371WU32 \| 98.6 \| 4.9e-10 \| 8.9e-16 \| 82.6 \| 92 \| (3, 95) \| 217 \| (20, 117) \| 195 \| Gp5/Type VI secretion system Vgr protein OB-fold domain-containing protein \| Gp5/Type VI secretion system Vgr protein OB-fold domain-containing protein; uniclust \| UniRef100\_A0A6B8KG73 \| 98.6 \| 5e-10 \| 9.2e-16 \| 82.1 \| 106 \| (2, 111) \| 217 \| (14, 120) \| 188 \| Gp5/Type VI secretion system Vgr protein OB-fold domain-containing protein \| Gp5/Type VI secretion system Vgr protein OB-fold domain-containing protein; uniclust \| UniRef100\_A0A1A3AFQ8 \| 98.6 \| 5.3e-10 \| 9.8e-16 \| 82.3 \| 111 \| (16, 133) \| 217 \| (14, 129) \| 175 \| Gp5/Type VI secretion system Vgr protein OB-fold domain-containing protein \| Gp5/Type VI secretion system Vgr protein OB-fold domain-containing protein; uniclust \| UniRef100\_A0A023Q190 \| 98.6 \| 5e-10 \| 1e-15 \| 104.7 \| 97 \| (17, 113) \| 217 \| (439, 545) \| 766 \| Type IV secretion protein Rhs \| Type IV secretion protein Rhs; uniclust \| UniRef100\_A0A8J6YCD9 \| 98.6 \| 5.5e-10 \| 1e-15 \| 73.2 \| 86 \| (44, 131) \| 217 \| (2, 88) \| 95 \| Phage baseplate assembly protein V \| Phage baseplate assembly protein V; uniclust \| UniRef100\_UPI000E2E4993 \| 98.6 \| 5.7e-10 \| 1.1e-15 \| 68.1 \| 62 \| (1, 62) \| 217 \| (1, 62) \| 62 \| phage baseplate assembly protein V \| phage baseplate assembly protein V; uniclust \| UniRef100\_A0A3C1JKA1 \| 98.6 \| 5.8e-10 \| 1.1e-15 \| 91.6 \| 95 \| (15, 109) \| 217 \| (225, 332) \| 453 \| Gp5/Type VI secretion system Vgr protein OB-fold domain-containing protein \| Gp5/Type VI secretion system Vgr protein OB-fold domain-containing protein; uniclust \| UniRef100\_V5ZAQ6 \| 98.6 \| 5.9e-10 \| 1.1e-15 \| 74.0 \| 72 \| (1, 72) \| 217 \| (1, 72) \| 102 \| Baseplate assembly protein V GpV \| Baseplate assembly protein V GpV; uniclust \| UniRef100\_A0A0P8YXD2 \| 98.6 \| 6.5e-10 \| 1.2e-15 \| 74.2 \| 51 \| (32, 82) \| 217 \| (7, 58) \| 105 \| Gp5/Type VI secretion system Vgr protein OB-fold domain-containing protein (Fragment) \| Gp5/Type VI secretion system Vgr protein OB-fold domain-containing protein (Fragment); uniclust \| UniRef100\_A0A925MG30 \| 98.6 \| 6.6e-10 \| 1.2e-15 \| 83.5 \| 117 \| (16, 133) \| 217 \| (18, 148) \| 192 \| Rhs element Vgr protein \| Rhs element Vgr protein; pdb70 \| 3QR8\_A \| 99.9 \| 2.6e-33 \| 2e-37 \| 207.3 \| 207 \| (2, 217) \| 217 \| (4, 211) \| 211 \| Baseplate assembly protein V \| 3QR8\_A Baseplate assembly protein V Beta-Helix, OB-Fold, Phage Baseplate, Iron-Binding; pdb70 \| 4S37\_K \| 99.9 \| 2.8e-32 \| 2.1e-36 \| 197.8 \| 184 \| (1, 203) \| 217 \| (1, 185) \| 185 \| Phage baseplate protein \| 4S37\_K Phage baseplate protein Cell Puncturing Device, Trimer, beta-helix HET: SO4, PGE, CL, EDO; pdb70 \| 4S37\_L \| 99.9 \| 2.8e-32 \| 2.1e-36 \| 197.8 \| 184 \| (1, 203) \| 217 \| (1, 185) \| 185 \| Phage baseplate protein \| 4S37\_L Phage baseplate protein Cell Puncturing Device, Trimer, beta-helix HET: PGE, CL, SO4, EDO; pdb70 \| 3PQI\_A \| 99.7 \| 1.3e-22 \| 9.7e-27 \| 152.7 \| 176 \| (13, 213) \| 217 \| (23, 239) \| 247 \| gene product 138 \| 3PQI\_A gene product 138 Beta-Helix, OB-Fold, Phage Baseplate, Iron-Binding; pdb70 \| 4RU3\_A \| 99.7 \| 4.7e-21 \| 3.5e-25 \| 141.8 \| 155 \| (13, 216) \| 217 \| (32, 218) \| 221 \| puncturing protein gp41 \| 4RU3\_A puncturing protein gp41 gp41, membrane translocation, phage baseplate; pdb70 \| 6J0M\_C \| 99.4 \| 1.3e-17 \| 1e-21 \| 137.7 \| 140 \| (13, 159) \| 217 \| (361, 514) \| 538 \| Pvc8 \| 6J0M\_C Pvc8 assembly, Photorhabdus asymbiotica, PVC, contractile; pdb70 \| 6RBK\_C \| 99.4 \| 4.7e-17 \| 3.6e-21 \| 134.2 \| 139 \| (13, 158) \| 217 \| (352, 504) \| 529 \| Afp1, Afp2, Afp3, Afp16 \| 6RBK\_C Afp1, Afp2, Afp3, Afp16 Anti-feeding prophage, secretion system, AFP; pdb70 \| 4MTK\_A \| 99.2 \| 1.5e-15 \| 1.1e-19 \| 128.2 \| 109 \| (13, 121) \| 217 \| (364, 500) \| 643 \| VgrG1 \| 4MTK\_A VgrG1 Beta-barrel, OB-fold, Beta-helix, Type VI HET: SO4, TAM; pdb70 \| 4MTK\_B \| 99.2 \| 1.5e-15 \| 1.1e-19 \| 128.2 \| 109 \| (13, 121) \| 217 \| (364, 500) \| 643 \| VgrG1 \| 4MTK\_B VgrG1 Beta-barrel, OB-fold, Beta-helix, Type VI HET: TAM, SO4; pdb70 \| 4MTK\_C \| 99.2 \| 1.5e-15 \| 1.1e-19 \| 128.2 \| 109 \| (13, 121) \| 217 \| (364, 500) \| 643 \| VgrG1 \| 4MTK\_C VgrG1 Beta-barrel, OB-fold, Beta-helix, Type VI HET: TAM, SO4; pdb70 \| 4MTK\_D \| 99.2 \| 1.5e-15 \| 1.1e-19 \| 128.2 \| 109 \| (13, 121) \| 217 \| (364, 500) \| 643 \| VgrG1 \| 4MTK\_D VgrG1 Beta-barrel, OB-fold, Beta-helix, Type VI HET: SO4, TAM; pdb70 \| 4MTK\_E \| 99.2 \| 1.5e-15 \| 1.1e-19 \| 128.2 \| 109 \| (13, 121) \| 217 \| (364, 500) \| 643 \| VgrG1 \| 4MTK\_E VgrG1 Beta-barrel, OB-fold, Beta-helix, Type VI HET: SO4, TAM; pdb70 \| 4MTK\_F \| 99.2 \| 1.5e-15 \| 1.1e-19 \| 128.2 \| 109 \| (13, 121) \| 217 \| (364, 500) \| 643 \| VgrG1 \| 4MTK\_F VgrG1 Beta-barrel, OB-fold, Beta-helix, Type VI HET: SO4, TAM; pdb70 \| 4UHV\_A \| 99.2 \| 1.5e-15 \| 1.1e-19 \| 128.3 \| 109 \| (13, 121) \| 217 \| (364, 500) \| 651 \| VGRG1, VALINE-GLYCINE REPEAT PROTEIN G1 \| 4UHV\_A VGRG1, VALINE-GLYCINE REPEAT PROTEIN G1 STRUCTURAL PROTEIN, VGRG1, VIRULENCE, TOXIN HET: NA, MSE, CL; pdb70 \| 4UHV\_B \| 99.2 \| 1.5e-15 \| 1.1e-19 \| 128.3 \| 109 \| (13, 121) \| 217 \| (364, 500) \| 651 \| VGRG1, VALINE-GLYCINE REPEAT PROTEIN G1 \| 4UHV\_B VGRG1, VALINE-GLYCINE REPEAT PROTEIN G1 STRUCTURAL PROTEIN, VGRG1, VIRULENCE, TOXIN HET: CL, NA, MSE; pdb70 \| 3AQJ\_A \| 99.0 \| 8.5e-14 \| 6.3e-18 \| 94.0 \| 122 \| (87, 217) \| 217 \| (12, 134) \| 134 \| Baseplate assembly protein V \| 3AQJ\_A Baseplate assembly protein V bacteriophage, tail spike, iron binding; pdb70 \| 3AQJ\_Q \| 99.0 \| 8.5e-14 \| 6.3e-18 \| 94.0 \| 122 \| (87, 217) \| 217 \| (12, 134) \| 134 \| Baseplate assembly protein V \| 3AQJ\_Q Baseplate assembly protein V bacteriophage, tail spike, iron binding; pdb70 \| 3QR7\_B \| 98.9 \| 4.4e-13 \| 3.3e-17 \| 88.2 \| 112 \| (97, 217) \| 217 \| (3, 115) \| 115 \| Baseplate assembly protein V \| 3QR7\_B Baseplate assembly protein V Beta-Helix, OB-Fold, Phage Baseplate, Iron-Binding; pdb70 \| 2P5Z\_X \| 98.8 \| 1.3e-12 \| 9.8e-17 \| 106.5 \| 81 \| (14, 94) \| 217 \| (384, 481) \| 491 \| Type VI secretion system component \| 2P5Z\_X Type VI secretion system component structural genomics, UNKNOWN FUNCTION, PSI-2; pdb70 \| 2Z6B\_A \| 98.7 \| 4.5e-12 \| 3.3e-16 \| 105.0 \| 81 \| (12, 92) \| 217 \| (6, 109) \| 584 \| Tail-associated lysozyme (E.C.3.2.1.17), Baseplate structural \| 2Z6B\_A Tail-associated lysozyme (E.C.3.2.1.17), Baseplate structural Protein containing metal complexes, Antimicrobial; pdb70 \| 3VTO\_B \| 98.7 \| 6.2e-12 \| 4.6e-16 \| 82.4 \| 98 \| (99, 211) \| 217 \| (6, 103) \| 115 \| Protein gp45 \| 3VTO\_B Protein gp45 beta-helix, central spike, Mu phage; pdb70 \| 3VTO\_Q \| 98.7 \| 6.2e-12 \| 4.6e-16 \| 82.4 \| 98 \| (99, 211) \| 217 \| (6, 103) \| 115 \| Protein gp45 \| 3VTO\_Q Protein gp45 beta-helix, central spike, Mu phage; pdb70 \| 3VTN\_A \| 98.6 \| 8.8e-12 \| 6.6e-16 \| 79.8 \| 98 \| (102, 216) \| 217 \| (3, 100) \| 101 \| Protein gp45 \| 3VTN\_A Protein gp45 beta-helix, central spike, Mu phage; pdb70 \| 1K28\_A \| 98.6 \| 1.7e-11 \| 1.3e-15 \| 101.5 \| 78 \| (14, 91) \| 217 \| (8, 108) \| 584 \| TAIL-ASSOCIATED LYSOZYME, BASEPLATE STRUCTURAL PROTEIN \| 1K28\_A TAIL-ASSOCIATED LYSOZYME, BASEPLATE STRUCTURAL PROTEIN Triple-stranded beta-helix, OB fold, pseudohexamer; pdb70 \| 5IV5\_YA \| 98.1 \| 1.1e-09 \| 8e-14 \| 90.6 \| 75 \| (15, 89) \| 217 \| (9, 106) \| 575 \| Baseplate wedge protein gp6, Baseplate \| 5IV5\_YA Baseplate wedge protein gp6, Baseplate T4, baseplate-tail tube complex, pre-attachment; pdb70 \| 5IV5\_YB \| 98.1 \| 1.1e-09 \| 8e-14 \| 90.6 \| 75 \| (15, 89) \| 217 \| (9, 106) \| 575 \| Baseplate wedge protein gp6, Baseplate \| 5IV5\_YB Baseplate wedge protein gp6, Baseplate T4, baseplate-tail tube complex, pre-attachment; pdb70 \| 5IV5\_YC \| 98.1 \| 1.1e-09 \| 8e-14 \| 90.6 \| 75 \| (15, 89) \| 217 \| (9, 106) \| 575 \| Baseplate wedge protein gp6, Baseplate \| 5IV5\_YC Baseplate wedge protein gp6, Baseplate T4, baseplate-tail tube complex, pre-attachment; pdb70 \| 1WTH\_A \| 98.1 \| 1.3e-09 \| 9.4e-14 \| 90.2 \| 73 \| (14, 86) \| 217 \| (8, 103) \| 584 \| Tail-associated lysozyme (E.C.3.2.1.17)/Baseplate structural protein \| 1WTH\_A Tail-associated lysozyme (E.C.3.2.1.17)/Baseplate structural protein Triple-stranded beta-helix, OB fold, pseudohexamer; pdb70 \| 4S36\_A \| 98.1 \| 1.4e-09 \| 1e-13 \| 67.7 \| 88 \| (98, 203) \| 217 \| (3, 90) \| 90 \| Phage baseplate protein \| 4S36\_A Phage baseplate protein Cell Puncturing Device, Trimer, beta-helix; pdb70 \| 3PQH\_A \| 97.4 \| 2.1e-07 \| 1.6e-11 \| 60.9 \| 75 \| (101, 207) \| 217 \| (34, 111) \| 127 \| gene product 138 C-terminus \| 3PQH\_A gene product 138 C-terminus Beta-Helix, OB-Fold, Phage Baseplate, Iron-Binding; pdb70 \| 5IV5\_YA \| 97.3 \| 3.4e-07 \| 2.6e-11 \| 75.3 \| 82 \| (40, 122) \| 217 \| (325, 417) \| 575 \| Baseplate wedge protein gp6, Baseplate \| 5IV5\_YA Baseplate wedge protein gp6, Baseplate T4, baseplate-tail tube complex, pre-attachment; pdb70 \| 5IV5\_YB \| 97.3 \| 3.4e-07 \| 2.6e-11 \| 75.3 \| 82 \| (40, 122) \| 217 \| (325, 417) \| 575 \| Baseplate wedge protein gp6, Baseplate \| 5IV5\_YB Baseplate wedge protein gp6, Baseplate T4, baseplate-tail tube complex, pre-attachment; pdb70 \| 5IV5\_YC \| 97.3 \| 3.4e-07 \| 2.6e-11 \| 75.3 \| 82 \| (40, 122) \| 217 \| (325, 417) \| 575 \| Baseplate wedge protein gp6, Baseplate \| 5IV5\_YC Baseplate wedge protein gp6, Baseplate T4, baseplate-tail tube complex, pre-attachment; pdb70 \| 1K28\_A \| 97.1 \| 9e-07 \| 6.7e-11 \| 72.8 \| 81 \| (40, 121) \| 217 \| (314, 416) \| 584 \| TAIL-ASSOCIATED LYSOZYME, BASEPLATE STRUCTURAL PROTEIN \| 1K28\_A TAIL-ASSOCIATED LYSOZYME, BASEPLATE STRUCTURAL PROTEIN Triple-stranded beta-helix, OB fold, pseudohexamer |
| Top keywords  (threshold 1.00e-03 (evalue)) | **baseplate, assembly, Phage, V, Type, secretion, VI, domain\_containing, Vgr, system** |
| Output files | ../../similar\_sequences/11\_FANPEZAQ\_CDS\_0011\_merged.svg ../../similar\_sequences/11\_FANPEZAQ\_CDS\_0011\_pdb70.a3m ../../similar\_sequences/11\_FANPEZAQ\_CDS\_0011\_pdb70.hhr ../../similar\_sequences/11\_FANPEZAQ\_CDS\_0011\_uniclust.a3m ../../similar\_sequences/11\_FANPEZAQ\_CDS\_0011\_uniclust.hhr |

#### Structure prediction (AlphaFold)^2^

|  |  |
| --- | --- |
| Stats | xml version="1.0" encoding="utf-8" standalone="no"?       2024-09-02T21:09:09.981572 image/svg+xml   Matplotlib v3.7.2, https://matplotlib.org/ |
| Predicted structure | **NGL Viewer Controls:**  - Center: *Left-Click* - Rotate: *Left-Click + Drag* - Translate: *Right-Click + Drag* - Zoom: *Shift + Left-Click + Drag* |
| Output files | ../../predicted\_structures/11\_FANPEZAQ\_CDS\_0011/features.pkl ../../predicted\_structures/11\_FANPEZAQ\_CDS\_0011/ranked\_0.pdb ../../predicted\_structures/11\_FANPEZAQ\_CDS\_0011/ranked\_0\_plots.svg ../../predicted\_structures/11\_FANPEZAQ\_CDS\_0011/result\_model\_1\_ptm\_pred\_0.pkl |

#### Structure similarity search results (Foldseek)^3^

|  |  |
| --- | --- |
| Structure databases searched | Pdb, Afdb-proteome, Afdb-uniprot50 |
| Results, scheme(s)  (Top layers only, threshold 1.00e-02 (evalue)) | xml version="1.0" encoding="utf-8" standalone="no"?       2024-09-02T21:10:19.348302 image/svg+xml   Matplotlib v3.7.2, https://matplotlib.org/ |
| Results, table  (threshold 1.00e-02 (evalue)) | db \| id \| prob \| evalue \| bits \| fident \| alnlen \| mismatch \| gapopen \| qstart \| qend \| tstart \| tend \| name \| description; pdb \| 3QR8\_A \| 1.0 \| 4.718e-16 \| 601 \| 0.382 \| 157 \| 96 \| 1 \| 6 \| 162 \| 2 \| 157 \| Baseplate assembly protein V \| Baseplate assembly protein V; pdb \| 4S37\_G \| 1.0 \| 1.524e-13 \| 488 \| 0.274 \| 186 \| 123 \| 3 \| 2 \| 186 \| 1 \| 175 \| Phage baseplate protein \| Phage baseplate protein; pdb \| 4S37\_Q \| 1.0 \| 8.936e-13 \| 449 \| 0.268 \| 175 \| 116 \| 3 \| 14 \| 187 \| 3 \| 166 \| Phage baseplate protein \| Phage baseplate protein; pdb \| 3AQJ\_R \| 1.0 \| 4.55e-05 \| 198 \| 0.326 \| 104 \| 65 \| 3 \| 93 \| 196 \| 1 \| 99 \| Baseplate assembly protein V \| Baseplate assembly protein V; pdb \| 7AEF\_Q \| 1.0 \| 2.54e-07 \| 192 \| 0.13 \| 223 \| 131 \| 7 \| 16 \| 193 \| 376 \| 580 \| Phosphoserine phosphatase SerB \| Phosphoserine phosphatase SerB; pdb \| 8GRA\_G \| 1.0 \| 6.943e-08 \| 191 \| 0.195 \| 245 \| 146 \| 12 \| 15 \| 216 \| 363 \| 599 \| Type VI secretion system spike protein VgrG \| Type VI secretion system spike protein VgrG; pdb \| 6H3N\_B \| 1.0 \| 5.448e-06 \| 180 \| 0.141 \| 247 \| 162 \| 11 \| 16 \| 217 \| 354 \| 595 \| VgrG1 \| VgrG1; pdb \| 6J0M\_A \| 1.0 \| 3.021e-06 \| 176 \| 0.102 \| 176 \| 124 \| 9 \| 16 \| 159 \| 362 \| 535 \| Pvc8 \| Pvc8; pdb \| 7B5H\_AJ \| 1.0 \| 2.54e-07 \| 174 \| 0.156 \| 223 \| 134 \| 9 \| 16 \| 193 \| 372 \| 585 \| All3320 protein \| All3320 protein; pdb \| 4UHV\_A \| 1.0 \| 8.731e-06 \| 162 \| 0.151 \| 244 \| 159 \| 14 \| 16 \| 215 \| 367 \| 606 \| VGRG1, VALINE-GLYCINE REPEAT PROTEIN G1 \| VGRG1, VALINE-GLYCINE REPEAT PROTEIN G1; pdb \| 7Q5P\_B \| 1.0 \| 3.594e-05 \| 159 \| 0.135 \| 244 \| 167 \| 11 \| 16 \| 217 \| 354 \| 595 \| Type VI secretion protein VgrG \| Type VI secretion protein VgrG; pdb \| 8EON\_P \| 1.0 \| 3.388e-05 \| 158 \| 0.146 \| 177 \| 108 \| 10 \| 4 \| 144 \| 20 \| 189 \| Baseplate spike gp43 \| Baseplate spike gp43; pdb \| 6RBK\_C \| 1.0 \| 4.55e-05 \| 153 \| 0.127 \| 172 \| 126 \| 9 \| 16 \| 164 \| 332 \| 502 \| Afp8 \| Afp8; pdb \| 4RU3\_A \| 1.0 \| 5.119e-05 \| 126 \| 0.141 \| 233 \| 107 \| 12 \| 4 \| 197 \| 6 \| 184 \| puncturing protein gp41 \| puncturing protein gp41; pdb \| 3I4O\_B \| 1.0 \| 0.005085 \| 114 \| 0.197 \| 76 \| 43 \| 8 \| 15 \| 83 \| 2 \| 66 \| Translation initiation factor IF-1 \| Translation initiation factor IF-1; pdb \| 2OQK\_A \| 1.0 \| 0.006069 \| 108 \| 0.146 \| 75 \| 56 \| 6 \| 15 \| 88 \| 11 \| 78 \| Putative translation initiation factor eIF-1A \| Putative translation initiation factor eIF-1A; pdb \| 3PQI\_A \| 1.0 \| 0.006829 \| 104 \| 0.154 \| 110 \| 79 \| 3 \| 16 \| 111 \| 3 \| 112 \| gene product 138 \| gene product 138; pdb \| 7YFZ\_Y \| 0.997 \| 0.006829 \| 92 \| 0.115 \| 156 \| 96 \| 7 \| 1 \| 117 \| 11 \| 163 \| Pam3 spike gp20 \| Pam3 spike gp20; pdb \| 3FP9\_K \| 0.992 \| 0.004261 \| 85 \| 0.121 \| 107 \| 71 \| 8 \| 16 \| 119 \| 2 \| 88 \| Proteasome-associated ATPase \| Proteasome-associated ATPase; pdb \| 3FP9\_G \| 0.991 \| 0.005085 \| 84 \| 0.121 \| 107 \| 71 \| 8 \| 16 \| 119 \| 2 \| 88 \| Proteasome-associated ATPase \| Proteasome-associated ATPase; pdb \| 5KWA\_A \| 0.986 \| 0.008645 \| 81 \| 0.121 \| 107 \| 71 \| 8 \| 16 \| 119 \| 3 \| 89 \| Proteasome-associated ATPase \| Proteasome-associated ATPase; afdb-proteome \| AF-A0A0H3GU25-F1-MODEL\_V4 \| 1.0 \| 2.964e-18 \| 692 \| 0.356 \| 202 \| 124 \| 4 \| 16 \| 217 \| 2 \| 197 \| Putative prophage baseplate assembly protein \| Putative prophage baseplate assembly protein; afdb-proteome \| AF-Q8ZMU1-F1-MODEL\_V4 \| 1.0 \| 3.98e-18 \| 558 \| 0.269 \| 223 \| 126 \| 3 \| 1 \| 217 \| 1 \| 192 \| Fels-2 prophage protein \| Fels-2 prophage protein; afdb-proteome \| AF-A0A0H3GM22-F1-MODEL\_V4 \| 1.0 \| 3.954e-16 \| 517 \| 0.265 \| 196 \| 120 \| 3 \| 1 \| 196 \| 1 \| 172 \| Baseplate assembly protein V \| Baseplate assembly protein V; afdb-proteome \| AF-G3XCT9-F1-MODEL\_V4 \| 1.0 \| 5.954e-15 \| 504 \| 0.275 \| 203 \| 129 \| 5 \| 1 \| 203 \| 1 \| 185 \| Phage\_base\_V domain-containing protein \| Phage\_base\_V domain-containing protein; afdb-proteome \| AF-Q9I0F3-F1-MODEL\_V4 \| 1.0 \| 5.137e-06 \| 170 \| 0.142 \| 246 \| 163 \| 10 \| 16 \| 217 \| 407 \| 648 \| Type VI secretion system spike protein VgrG1c \| Type VI secretion system spike protein VgrG1c; afdb-proteome \| AF-Q9I1A6-F1-MODEL\_V4 \| 1.0 \| 1.772e-05 \| 169 \| 0.137 \| 232 \| 169 \| 11 \| 16 \| 217 \| 352 \| 582 \| Phage\_base\_V domain-containing protein \| Phage\_base\_V domain-containing protein; afdb-proteome \| AF-Q9HTT3-F1-MODEL\_V4 \| 1.0 \| 4.057e-06 \| 152 \| 0.177 \| 265 \| 154 \| 16 \| 16 \| 217 \| 372 \| 635 \| Phage\_base\_V domain-containing protein \| Phage\_base\_V domain-containing protein; afdb-proteome \| AF-Q9HU93-F1-MODEL\_V4 \| 1.0 \| 8.232e-06 \| 147 \| 0.162 \| 264 \| 158 \| 14 \| 16 \| 217 \| 377 \| 639 \| Type VI secretion system spike protein VgrG5 \| Type VI secretion system spike protein VgrG5; afdb-proteome \| AF-Q9I737-F1-MODEL\_V4 \| 1.0 \| 0.0004031 \| 139 \| 0.142 \| 232 \| 167 \| 10 \| 16 \| 217 \| 370 \| 599 \| Type VI secretion system spike protein VgrG1b \| Type VI secretion system spike protein VgrG1b; afdb-proteome \| AF-Q9HYC3-F1-MODEL\_V4 \| 1.0 \| 3.012e-05 \| 137 \| 0.153 \| 234 \| 155 \| 12 \| 16 \| 210 \| 377 \| 606 \| Type VI secretion system spike protein VgrG4b \| Type VI secretion system spike protein VgrG4b; afdb-uniprot50 \| AF-A0A8A6KD57-F1-MODEL\_V4 \| 1.0 \| 1.044e-22 \| 799 \| 0.471 \| 214 \| 104 \| 3 \| 4 \| 217 \| 5 \| 209 \| Phage baseplate assembly protein V \| Phage baseplate assembly protein V; afdb-uniprot50 \| AF-A0A2Z3IDQ9-F1-MODEL\_V4 \| 1.0 \| 2.691e-23 \| 782 \| 0.536 \| 218 \| 93 \| 2 \| 1 \| 217 \| 1 \| 211 \| Phage baseplate assembly protein V \| Phage baseplate assembly protein V; afdb-uniprot50 \| AF-A0A1I3LN83-F1-MODEL\_V4 \| 1.0 \| 3.577e-20 \| 780 \| 0.364 \| 217 \| 132 \| 4 \| 1 \| 217 \| 1 \| 211 \| Phage baseplate assembly protein V \| Phage baseplate assembly protein V; afdb-uniprot50 \| AF-A0A2D8NW13-F1-MODEL\_V4 \| 1.0 \| 1.163e-19 \| 774 \| 0.359 \| 217 \| 135 \| 4 \| 1 \| 217 \| 1 \| 213 \| Baseplate assembly protein \| Baseplate assembly protein; afdb-uniprot50 \| AF-A0A486S9U3-F1-MODEL\_V4 \| 1.0 \| 5.405e-20 \| 769 \| 0.376 \| 215 \| 128 \| 5 \| 3 \| 217 \| 39 \| 247 \| Baseplate assembly protein V \| Baseplate assembly protein V; afdb-uniprot50 \| AF-A0A081RRN8-F1-MODEL\_V4 \| 1.0 \| 1.388e-19 \| 764 \| 0.367 \| 215 \| 128 \| 5 \| 4 \| 217 \| 32 \| 239 \| Phage baseplate assembly protein V \| Phage baseplate assembly protein V; afdb-uniprot50 \| AF-A0A068QXN4-F1-MODEL\_V4 \| 1.0 \| 1.657e-19 \| 759 \| 0.331 \| 214 \| 137 \| 4 \| 4 \| 217 \| 29 \| 236 \| Baseplate assembly protein V \| Baseplate assembly protein V; afdb-uniprot50 \| AF-A0A0W2DNN1-F1-MODEL\_V4 \| 1.0 \| 7.258e-20 \| 758 \| 0.388 \| 211 \| 123 \| 4 \| 3 \| 213 \| 5 \| 209 \| Baseplate assembly protein \| Baseplate assembly protein; afdb-uniprot50 \| AF-A0A8B3QQT6-F1-MODEL\_V4 \| 1.0 \| 4.256e-19 \| 753 \| 0.36 \| 194 \| 119 \| 3 \| 3 \| 196 \| 5 \| 193 \| Phage baseplate assembly protein V \| Phage baseplate assembly protein V; afdb-uniprot50 \| AF-A0A370T1W2-F1-MODEL\_V4 \| 1.0 \| 3.169e-19 \| 749 \| 0.358 \| 215 \| 132 \| 4 \| 3 \| 217 \| 21 \| 229 \| Phage baseplate assembly protein V \| Phage baseplate assembly protein V; afdb-uniprot50 \| AF-E3G776-F1-MODEL\_V4 \| 1.0 \| 1.097e-19 \| 741 \| 0.398 \| 221 \| 119 \| 8 \| 1 \| 217 \| 1 \| 211 \| Phage baseplate assembly protein V \| Phage baseplate assembly protein V; afdb-uniprot50 \| AF-A0A7Y1SGM7-F1-MODEL\_V4 \| 1.0 \| 1.752e-18 \| 740 \| 0.336 \| 217 \| 141 \| 3 \| 1 \| 217 \| 1 \| 214 \| Phage baseplate assembly protein V \| Phage baseplate assembly protein V; afdb-uniprot50 \| AF-A0A7T9YBU7-F1-MODEL\_V4 \| 1.0 \| 5.387e-19 \| 733 \| 0.333 \| 213 \| 131 \| 5 \| 1 \| 213 \| 1 \| 202 \| Phage baseplate assembly protein V \| Phage baseplate assembly protein V; afdb-uniprot50 \| AF-A0A0F3UMW4-F1-MODEL\_V4 \| 1.0 \| 1.159e-18 \| 728 \| 0.365 \| 194 \| 118 \| 3 \| 3 \| 196 \| 5 \| 193 \| Baseplate assembly protein \| Baseplate assembly protein; afdb-uniprot50 \| AF-A0A376PNH2-F1-MODEL\_V4 \| 1.0 \| 1.159e-18 \| 721 \| 0.366 \| 199 \| 119 \| 4 \| 1 \| 197 \| 1 \| 194 \| Baseplate assembly protein V (GpV) \| Baseplate assembly protein V (GpV); afdb-uniprot50 \| AF-A0A6M8RDL6-F1-MODEL\_V4 \| 1.0 \| 1.971e-18 \| 720 \| 0.377 \| 217 \| 126 \| 7 \| 1 \| 217 \| 1 \| 208 \| Phage baseplate assembly protein V \| Phage baseplate assembly protein V; afdb-uniprot50 \| AF-A0A0T9KKH2-F1-MODEL\_V4 \| 1.0 \| 4.514e-19 \| 719 \| 0.391 \| 217 \| 120 \| 7 \| 3 \| 217 \| 5 \| 211 \| Baseplate assembly protein V \| Baseplate assembly protein V; afdb-uniprot50 \| AF-A0A2W5L0U8-F1-MODEL\_V4 \| 1.0 \| 1.769e-21 \| 717 \| 0.425 \| 214 \| 99 \| 3 \| 4 \| 217 \| 6 \| 195 \| Phage baseplate assembly protein V \| Phage baseplate assembly protein V; afdb-uniprot50 \| AF-A0A2A7T2L7-F1-MODEL\_V4 \| 1.0 \| 6.43e-19 \| 714 \| 0.382 \| 217 \| 122 \| 7 \| 3 \| 217 \| 5 \| 211 \| Phage baseplate assembly protein V \| Phage baseplate assembly protein V; afdb-uniprot50 \| AF-A0A7U3BDQ0-F1-MODEL\_V4 \| 1.0 \| 7.234e-19 \| 706 \| 0.364 \| 217 \| 126 \| 6 \| 3 \| 217 \| 5 \| 211 \| Baseplate assembly protein \| Baseplate assembly protein; afdb-uniprot50 \| AF-A0A2S8HW35-F1-MODEL\_V4 \| 1.0 \| 7.234e-19 \| 705 \| 0.375 \| 221 \| 124 \| 7 \| 1 \| 217 \| 1 \| 211 \| Baseplate assembly protein \| Baseplate assembly protein; afdb-uniprot50 \| AF-A0A377CCX9-F1-MODEL\_V4 \| 1.0 \| 8.14e-19 \| 705 \| 0.373 \| 217 \| 124 \| 6 \| 3 \| 217 \| 5 \| 211 \| Baseplate assembly protein V (GpV) \| Baseplate assembly protein V (GpV); afdb-uniprot50 \| AF-A0A8A7TZ35-F1-MODEL\_V4 \| 1.0 \| 9.158e-19 \| 704 \| 0.387 \| 217 \| 121 \| 6 \| 3 \| 217 \| 5 \| 211 \| Phage baseplate assembly protein V \| Phage baseplate assembly protein V; afdb-uniprot50 \| AF-A0A810U4Z0-F1-MODEL\_V4 \| 1.0 \| 4.514e-19 \| 704 \| 0.373 \| 217 \| 124 \| 6 \| 3 \| 217 \| 17 \| 223 \| Phage baseplate assembly protein \| Phage baseplate assembly protein; afdb-uniprot50 \| AF-A0A761QNR2-F1-MODEL\_V4 \| 1.0 \| 2.638e-17 \| 702 \| 0.371 \| 164 \| 102 \| 1 \| 3 \| 166 \| 5 \| 167 \| Phage baseplate assembly protein V \| Phage baseplate assembly protein V; afdb-uniprot50 \| AF-A0A484X0R0-F1-MODEL\_V4 \| 1.0 \| 6.041e-18 \| 700 \| 0.369 \| 195 \| 116 \| 4 \| 1 \| 193 \| 1 \| 190 \| Baseplate assembly protein V (GpV) \| Baseplate assembly protein V (GpV); afdb-uniprot50 \| AF-F4N734-F1-MODEL\_V4 \| 1.0 \| 6.022e-17 \| 691 \| 0.38 \| 176 \| 106 \| 2 \| 1 \| 176 \| 1 \| 173 \| Baseplate assembly protein V \| Baseplate assembly protein V; afdb-uniprot50 \| AF-A0A7Z1A3H5-F1-MODEL\_V4 \| 1.0 \| 1.379e-17 \| 688 \| 0.353 \| 215 \| 131 \| 7 \| 6 \| 217 \| 2 \| 211 \| Phage baseplate assembly protein V \| Phage baseplate assembly protein V; afdb-uniprot50 \| AF-A0A376TE63-F1-MODEL\_V4 \| 1.0 \| 1.971e-18 \| 683 \| 0.364 \| 217 \| 126 \| 6 \| 3 \| 217 \| 28 \| 234 \| Baseplate assembly protein V (GpV) \| Baseplate assembly protein V (GpV); afdb-uniprot50 \| AF-A0A1T4W1B8-F1-MODEL\_V4 \| 1.0 \| 3.35e-18 \| 680 \| 0.36 \| 222 \| 135 \| 3 \| 1 \| 217 \| 1 \| 220 \| Phage baseplate assembly protein V \| Phage baseplate assembly protein V; afdb-uniprot50 \| AF-A0A660NMC7-F1-MODEL\_V4 \| 1.0 \| 2.36e-19 \| 678 \| 0.486 \| 189 \| 83 \| 5 \| 1 \| 187 \| 1 \| 177 \| Phage baseplate assembly protein V \| Phage baseplate assembly protein V; afdb-uniprot50 \| AF-A0A826YSP9-F1-MODEL\_V4 \| 1.0 \| 1.846e-16 \| 670 \| 0.375 \| 149 \| 92 \| 1 \| 3 \| 151 \| 5 \| 152 \| Phage baseplate assembly protein V \| Phage baseplate assembly protein V; afdb-uniprot50 \| AF-A0A2P5N6T8-F1-MODEL\_V4 \| 1.0 \| 1.082e-15 \| 664 \| 0.344 \| 145 \| 95 \| 0 \| 1 \| 145 \| 1 \| 145 \| Baseplate assembly protein \| Baseplate assembly protein; afdb-uniprot50 \| AF-A0A118DV18-F1-MODEL\_V4 \| 1.0 \| 1.037e-20 \| 663 \| 0.42 \| 221 \| 116 \| 4 \| 1 \| 217 \| 1 \| 213 \| Uncharacterized protein \| Uncharacterized protein; afdb-uniprot50 \| AF-A0A3G9FLY0-F1-MODEL\_V4 \| 1.0 \| 7.674e-19 \| 662 \| 0.367 \| 223 \| 126 \| 6 \| 1 \| 217 \| 1 \| 214 \| Phage baseplate assembly protein \| Phage baseplate assembly protein; afdb-uniprot50 \| AF-A0A212KMZ7-F1-MODEL\_V4 \| 1.0 \| 1.37e-15 \| 660 \| 0.411 \| 136 \| 79 \| 1 \| 4 \| 138 \| 7 \| 142 \| Phage baseplate assembly protein V (Modular protein) \| Phage baseplate assembly protein V (Modular protein); afdb-uniprot50 \| AF-A0A554X0S7-F1-MODEL\_V4 \| 1.0 \| 8.059e-16 \| 658 \| 0.379 \| 137 \| 80 \| 1 \| 1 \| 132 \| 1 \| 137 \| Phage baseplate assembly protein V \| Phage baseplate assembly protein V; afdb-uniprot50 \| AF-A0A516WAE0-F1-MODEL\_V4 \| 1.0 \| 5.677e-17 \| 657 \| 0.292 \| 219 \| 147 \| 6 \| 1 \| 217 \| 1 \| 213 \| Phage baseplate assembly protein V \| Phage baseplate assembly protein V; afdb-uniprot50 \| AF-A0A369RHM3-F1-MODEL\_V4 \| 1.0 \| 3.733e-15 \| 656 \| 0.423 \| 118 \| 68 \| 0 \| 4 \| 121 \| 7 \| 124 \| Phage P2 baseplate assembly protein gpV \| Phage P2 baseplate assembly protein gpV; afdb-uniprot50 \| AF-A0A7V8FPR0-F1-MODEL\_V4 \| 1.0 \| 1.458e-16 \| 655 \| 0.317 \| 227 \| 138 \| 8 \| 3 \| 216 \| 5 \| 227 \| Phage\_base\_V domain-containing protein \| Phage\_base\_V domain-containing protein; afdb-uniprot50 \| AF-A0A6N7JJ19-F1-MODEL\_V4 \| 1.0 \| 1.375e-16 \| 654 \| 0.316 \| 202 \| 129 \| 6 \| 4 \| 203 \| 7 \| 201 \| Phage baseplate assembly protein V \| Phage baseplate assembly protein V; afdb-uniprot50 \| AF-A0A7X2BHN0-F1-MODEL\_V4 \| 1.0 \| 5.352e-17 \| 654 \| 0.298 \| 218 \| 146 \| 6 \| 1 \| 215 \| 1 \| 214 \| Phage baseplate assembly protein V \| Phage baseplate assembly protein V; afdb-uniprot50 \| AF-N6USL5-F1-MODEL\_V4 \| 1.0 \| 5.045e-17 \| 647 \| 0.322 \| 223 \| 141 \| 6 \| 4 \| 217 \| 8 \| 229 \| Phage baseplate assembly protein V \| Phage baseplate assembly protein V; afdb-uniprot50 \| AF-A0A430HF75-F1-MODEL\_V4 \| 1.0 \| 2.998e-20 \| 645 \| 0.492 \| 207 \| 89 \| 3 \| 5 \| 197 \| 2 \| 206 \| Phage baseplate assembly protein V \| Phage baseplate assembly protein V; afdb-uniprot50 \| AF-A0A4R2MLJ2-F1-MODEL\_V4 \| 1.0 \| 5.982e-15 \| 644 \| 0.406 \| 128 \| 76 \| 0 \| 5 \| 132 \| 2 \| 129 \| Phage baseplate assembly protein V \| Phage baseplate assembly protein V; afdb-uniprot50 \| AF-A0A3N9U432-F1-MODEL\_V4 \| 1.0 \| 2.503e-19 \| 644 \| 0.345 \| 217 \| 108 \| 4 \| 1 \| 217 \| 1 \| 183 \| Phage baseplate assembly protein V \| Phage baseplate assembly protein V; afdb-uniprot50 \| AF-A0A7W8YF54-F1-MODEL\_V4 \| 1.0 \| 1.757e-19 \| 641 \| 0.385 \| 218 \| 110 \| 7 \| 1 \| 217 \| 1 \| 195 \| Phage baseplate assembly protein V \| Phage baseplate assembly protein V; afdb-uniprot50 \| AF-A0A1M7KGY4-F1-MODEL\_V4 \| 1.0 \| 9.038e-15 \| 637 \| 0.391 \| 120 \| 73 \| 0 \| 2 \| 121 \| 1 \| 120 \| Phage baseplate assembly protein V \| Phage baseplate assembly protein V; afdb-uniprot50 \| AF-A0A4Q6FFT3-F1-MODEL\_V4 \| 1.0 \| 3.959e-15 \| 636 \| 0.307 \| 143 \| 99 \| 0 \| 1 \| 143 \| 1 \| 143 \| Phage baseplate assembly protein V \| Phage baseplate assembly protein V; afdb-uniprot50 \| AF-A0A418HAA9-F1-MODEL\_V4 \| 1.0 \| 2.337e-16 \| 636 \| 0.375 \| 181 \| 108 \| 3 \| 16 \| 196 \| 2 \| 177 \| Phage baseplate assembly protein V \| Phage baseplate assembly protein V; afdb-uniprot50 \| AF-A0A258LJQ0-F1-MODEL\_V4 \| 1.0 \| 5.029e-16 \| 635 \| 0.335 \| 194 \| 118 \| 6 \| 1 \| 194 \| 1 \| 183 \| Phage\_base\_V domain-containing protein \| Phage\_base\_V domain-containing protein; afdb-uniprot50 \| AF-A0A447N5E7-F1-MODEL\_V4 \| 1.0 \| 1.214e-14 \| 634 \| 0.344 \| 125 \| 82 \| 0 \| 1 \| 125 \| 1 \| 125 \| Phage baseplate assembly protein V \| Phage baseplate assembly protein V; afdb-uniprot50 \| AF-A0A1Y2K0V2-F1-MODEL\_V4 \| 1.0 \| 2.188e-14 \| 628 \| 0.366 \| 120 \| 76 \| 0 \| 1 \| 120 \| 1 \| 120 \| Putative baseplate assembly protein V \| Putative baseplate assembly protein V; afdb-uniprot50 \| AF-A1VSH0-F1-MODEL\_V4 \| 1.0 \| 4.241e-18 \| 627 \| 0.349 \| 229 \| 123 \| 7 \| 1 \| 217 \| 3 \| 217 \| Phage baseplate assembly protein V \| Phage baseplate assembly protein V; afdb-uniprot50 \| AF-A0A376WR96-F1-MODEL\_V4 \| 1.0 \| 8.113e-18 \| 627 \| 0.301 \| 219 \| 147 \| 4 \| 1 \| 217 \| 1 \| 215 \| Baseplate assembly protein V (GpV) \| Baseplate assembly protein V (GpV); afdb-uniprot50 \| AF-A0A6C1U1Y1-F1-MODEL\_V4 \| 1.0 \| 5.639e-15 \| 626 \| 0.372 \| 129 \| 81 \| 0 \| 4 \| 132 \| 7 \| 135 \| Phage baseplate assembly protein V \| Phage baseplate assembly protein V; afdb-uniprot50 \| AF-I3TTD6-F1-MODEL\_V4 \| 1.0 \| 2.196e-15 \| 624 \| 0.405 \| 148 \| 81 \| 3 \| 1 \| 143 \| 1 \| 146 \| Phage baseplate assembly protein V \| Phage baseplate assembly protein V; afdb-uniprot50 \| AF-A0A2E3M0K1-F1-MODEL\_V4 \| 1.0 \| 1.079e-14 \| 623 \| 0.345 \| 133 \| 85 \| 1 \| 1 \| 131 \| 1 \| 133 \| Baseplate assembly protein V \| Baseplate assembly protein V; afdb-uniprot50 \| AF-A0A1Y6CPE2-F1-MODEL\_V4 \| 1.0 \| 7.573e-15 \| 623 \| 0.315 \| 133 \| 88 \| 1 \| 4 \| 136 \| 6 \| 135 \| Phage baseplate assembly protein V \| Phage baseplate assembly protein V; afdb-uniprot50 \| AF-A0A1C3EL99-F1-MODEL\_V4 \| 1.0 \| 2.62e-15 \| 623 \| 0.322 \| 152 \| 103 \| 0 \| 4 \| 155 \| 8 \| 159 \| Baseplate assembly protein \| Baseplate assembly protein; afdb-uniprot50 \| AF-A0A376YK40-F1-MODEL\_V4 \| 1.0 \| 9.618e-16 \| 622 \| 0.368 \| 171 \| 103 \| 3 \| 16 \| 186 \| 2 \| 167 \| Baseplate assembly protein V (GpV) \| Baseplate assembly protein V (GpV); afdb-uniprot50 \| AF-A0A1N6UJB4-F1-MODEL\_V4 \| 1.0 \| 1.453e-15 \| 619 \| 0.229 \| 218 \| 156 \| 6 \| 1 \| 217 \| 1 \| 207 \| Phage baseplate assembly protein V \| Phage baseplate assembly protein V; afdb-uniprot50 \| AF-A0A2E5BTY2-F1-MODEL\_V4 \| 1.0 \| 1.834e-14 \| 618 \| 0.372 \| 129 \| 81 \| 0 \| 4 \| 132 \| 15 \| 143 \| Baseplate assembly protein V \| Baseplate assembly protein V; afdb-uniprot50 \| AF-A0A6P1S8D3-F1-MODEL\_V4 \| 1.0 \| 2.503e-19 \| 616 \| 0.368 \| 217 \| 114 \| 7 \| 1 \| 217 \| 1 \| 194 \| Phage baseplate assembly protein V \| Phage baseplate assembly protein V; afdb-uniprot50 \| AF-A0A4P9VEW0-F1-MODEL\_V4 \| 1.0 \| 1.536e-14 \| 615 \| 0.341 \| 129 \| 85 \| 0 \| 9 \| 137 \| 2 \| 130 \| Phage baseplate assembly protein V \| Phage baseplate assembly protein V; afdb-uniprot50 \| AF-A0A3B0LVV6-F1-MODEL\_V4 \| 1.0 \| 5.334e-16 \| 613 \| 0.294 \| 217 \| 143 \| 6 \| 1 \| 217 \| 33 \| 239 \| Uncharacterized protein \| Uncharacterized protein; afdb-uniprot50 \| AF-A0A2J9GZ67-F1-MODEL\_V4 \| 1.0 \| 3.18e-20 \| 612 \| 0.36 \| 233 \| 133 \| 3 \| 1 \| 217 \| 7 \| 239 \| Phage baseplate assembly protein V \| Phage baseplate assembly protein V; afdb-uniprot50 \| AF-A0A2S8HYS2-F1-MODEL\_V4 \| 1.0 \| 3.339e-17 \| 611 \| 0.292 \| 226 \| 144 \| 4 \| 2 \| 213 \| 1 \| 224 \| Phage baseplate assembly protein V \| Phage baseplate assembly protein V; afdb-uniprot50 \| AF-A0A377KA20-F1-MODEL\_V4 \| 1.0 \| 2.462e-14 \| 609 \| 0.33 \| 127 \| 85 \| 0 \| 1 \| 127 \| 1 \| 127 \| Phage baseplate assembly protein V \| Phage baseplate assembly protein V; afdb-uniprot50 \| AF-A0A4S3LTU7-F1-MODEL\_V4 \| 1.0 \| 2.817e-19 \| 609 \| 0.375 \| 213 \| 118 \| 5 \| 1 \| 213 \| 1 \| 198 \| Phage baseplate assembly protein V \| Phage baseplate assembly protein V; afdb-uniprot50 \| AF-A0A143DE80-F1-MODEL\_V4 \| 1.0 \| 3.757e-17 \| 608 \| 0.367 \| 215 \| 123 \| 6 \| 1 \| 205 \| 1 \| 212 \| Uncharacterized protein \| Uncharacterized protein; afdb-uniprot50 \| AF-A0A2U2DBJ1-F1-MODEL\_V4 \| 1.0 \| 2.638e-17 \| 606 \| 0.281 \| 220 \| 148 \| 6 \| 2 \| 217 \| 1 \| 214 \| Phage baseplate assembly protein V \| Phage baseplate assembly protein V; afdb-uniprot50 \| AF-A0A4U8YLR0-F1-MODEL\_V4 \| 1.0 \| 6.73e-15 \| 605 \| 0.378 \| 148 \| 92 \| 0 \| 4 \| 151 \| 7 \| 154 \| Phage baseplate assembly protein v/gp45 \| Phage baseplate assembly protein v/gp45; afdb-uniprot50 \| AF-W1J949-F1-MODEL\_V4 \| 1.0 \| 9.587e-15 \| 601 \| 0.377 \| 127 \| 75 \| 2 \| 1 \| 127 \| 1 \| 123 \| Baseplate assembly protein V \| Baseplate assembly protein V; afdb-uniprot50 \| AF-A0A2E3Q547-F1-MODEL\_V4 \| 1.0 \| 5.299e-14 \| 601 \| 0.31 \| 132 \| 91 \| 0 \| 1 \| 132 \| 1 \| 132 \| Phage baseplate protein \| Phage baseplate protein; afdb-uniprot50 \| AF-C3X1Y6-F1-MODEL\_V4 \| 1.0 \| 1.468e-18 \| 601 \| 0.334 \| 206 \| 122 \| 5 \| 1 \| 203 \| 1 \| 194 \| Phage baseplate assembly protein V \| Phage baseplate assembly protein V; afdb-uniprot50 \| AF-A0A771BAN4-F1-MODEL\_V4 \| 1.0 \| 2.62e-15 \| 600 \| 0.366 \| 180 \| 109 \| 3 \| 17 \| 196 \| 1 \| 175 \| Phage baseplate assembly protein V \| Phage baseplate assembly protein V; afdb-uniprot50 \| AF-A0A741LDN8-F1-MODEL\_V4 \| 1.0 \| 2.47e-15 \| 599 \| 0.372 \| 180 \| 108 \| 3 \| 17 \| 196 \| 1 \| 175 \| Phage baseplate assembly protein V \| Phage baseplate assembly protein V; afdb-uniprot50 \| AF-Q31Q99-F1-MODEL\_V4 \| 1.0 \| 1.079e-14 \| 598 \| 0.272 \| 169 \| 113 \| 4 \| 2 \| 160 \| 18 \| 186 \| Phage baseplate assembly protein V \| Phage baseplate assembly protein V; afdb-uniprot50 \| AF-A0A5A9ENI4-F1-MODEL\_V4 \| 1.0 \| 3.117e-14 \| 596 \| 0.361 \| 130 \| 82 \| 1 \| 4 \| 132 \| 11 \| 140 \| Phage baseplate assembly protein V \| Phage baseplate assembly protein V; afdb-uniprot50 \| AF-A0A7Z1R6V5-F1-MODEL\_V4 \| 1.0 \| 3.317e-15 \| 595 \| 0.25 \| 204 \| 146 \| 4 \| 1 \| 203 \| 1 \| 198 \| Phage baseplate protein \| Phage baseplate protein; afdb-uniprot50 \| AF-B4EF38-F1-MODEL\_V4 \| 1.0 \| 4.227e-17 \| 594 \| 0.368 \| 220 \| 117 \| 8 \| 1 \| 203 \| 42 \| 256 \| Phage baseplate assembly protein \| Phage baseplate assembly protein; afdb-uniprot50 \| AF-A0A5W3AAX1-F1-MODEL\_V4 \| 1.0 \| 2.07e-15 \| 593 \| 0.365 \| 156 \| 81 \| 2 \| 1 \| 154 \| 1 \| 140 \| Phage baseplate assembly protein V \| Phage baseplate assembly protein V; afdb-uniprot50 \| AF-A0A4R7NQH3-F1-MODEL\_V4 \| 1.0 \| 9.747e-20 \| 591 \| 0.373 \| 222 \| 128 \| 5 \| 4 \| 217 \| 34 \| 252 \| Phage baseplate assembly protein V \| Phage baseplate assembly protein V; afdb-uniprot50 \| AF-A0A7W8K684-F1-MODEL\_V4 \| 1.0 \| 2.07e-15 \| 590 \| 0.386 \| 168 \| 84 \| 4 \| 1 \| 167 \| 1 \| 150 \| Phage baseplate assembly protein V \| Phage baseplate assembly protein V; afdb-uniprot50 \| AF-A0A396RYI9-F1-MODEL\_V4 \| 1.0 \| 6.022e-17 \| 588 \| 0.259 \| 216 \| 122 \| 4 \| 2 \| 217 \| 1 \| 178 \| Phage baseplate assembly protein V \| Phage baseplate assembly protein V; afdb-uniprot50 \| AF-A0A7V8FKD6-F1-MODEL\_V4 \| 1.0 \| 3.159e-18 \| 588 \| 0.372 \| 220 \| 114 \| 6 \| 1 \| 217 \| 1 \| 199 \| Phage\_base\_V domain-containing protein \| Phage\_base\_V domain-containing protein; afdb-uniprot50 \| AF-A0A142JGU2-F1-MODEL\_V4 \| 1.0 \| 1.226e-17 \| 587 \| 0.327 \| 220 \| 110 \| 5 \| 1 \| 217 \| 1 \| 185 \| Uncharacterized protein \| Uncharacterized protein; afdb-uniprot50 \| AF-A0A3E0X1A7-F1-MODEL\_V4 \| 1.0 \| 6.002e-16 \| 586 \| 0.393 \| 150 \| 77 \| 1 \| 2 \| 137 \| 1 \| 150 \| Phage\_base\_V domain-containing protein \| Phage\_base\_V domain-containing protein; afdb-uniprot50 \| AF-A0A741LDS5-F1-MODEL\_V4 \| 1.0 \| 1.84e-15 \| 586 \| 0.389 \| 203 \| 112 \| 6 \| 17 \| 217 \| 1 \| 193 \| Phage baseplate assembly protein V \| Phage baseplate assembly protein V; afdb-uniprot50 \| AF-A0A6H2NSY1-F1-MODEL\_V4 \| 1.0 \| 1.828e-13 \| 584 \| 0.418 \| 110 \| 64 \| 0 \| 4 \| 113 \| 7 \| 116 \| Phage baseplate assembly protein V \| Phage baseplate assembly protein V; afdb-uniprot50 \| AF-A0A1B8QD27-F1-MODEL\_V4 \| 1.0 \| 4.469e-16 \| 581 \| 0.289 \| 214 \| 129 \| 5 \| 1 \| 214 \| 1 \| 191 \| Phage\_base\_V domain-containing protein \| Phage\_base\_V domain-containing protein; afdb-uniprot50 \| AF-A0A376DH93-F1-MODEL\_V4 \| 1.0 \| 4.012e-19 \| 580 \| 0.347 \| 216 \| 120 \| 4 \| 2 \| 217 \| 213 \| 407 \| Putative phage tail completion protein S \| Putative phage tail completion protein S; afdb-uniprot50 \| AF-A0A5C8XUI7-F1-MODEL\_V4 \| 1.0 \| 3.53e-16 \| 578 \| 0.329 \| 176 \| 97 \| 3 \| 1 \| 176 \| 1 \| 155 \| Phage baseplate assembly protein V \| Phage baseplate assembly protein V; afdb-uniprot50 \| AF-A0A235EWV9-F1-MODEL\_V4 \| 1.0 \| 4.788e-19 \| 577 \| 0.355 \| 225 \| 120 \| 4 \| 1 \| 217 \| 1 \| 208 \| Uncharacterized protein \| Uncharacterized protein; afdb-uniprot50 \| AF-A0A6L2ZPM3-F1-MODEL\_V4 \| 1.0 \| 5.37e-18 \| 576 \| 0.351 \| 216 \| 119 \| 4 \| 2 \| 217 \| 3 \| 197 \| Baseplate assembly protein V (GpV) \| Baseplate assembly protein V (GpV); afdb-uniprot50 \| AF-A0A6N4PBL9-F1-MODEL\_V4 \| 1.0 \| 2.495e-18 \| 575 \| 0.333 \| 213 \| 121 \| 4 \| 5 \| 217 \| 2 \| 193 \| Phage baseplate assembly protein V \| Phage baseplate assembly protein V; afdb-uniprot50 \| AF-A0A081B6D4-F1-MODEL\_V4 \| 1.0 \| 6.022e-17 \| 575 \| 0.311 \| 212 \| 139 \| 3 \| 4 \| 215 \| 7 \| 211 \| Phage baseplate assembly protein V \| Phage baseplate assembly protein V; afdb-uniprot50 \| AF-A0A7X4JXC7-F1-MODEL\_V4 \| 1.0 \| 2.344e-17 \| 574 \| 0.3 \| 216 \| 113 \| 2 \| 2 \| 217 \| 1 \| 178 \| Phage baseplate assembly protein V \| Phage baseplate assembly protein V; afdb-uniprot50 \| AF-A0A5E5AEH0-F1-MODEL\_V4 \| 1.0 \| 1.082e-15 \| 574 \| 0.299 \| 224 \| 136 \| 5 \| 1 \| 217 \| 1 \| 210 \| Baseplate assembly protein \| Baseplate assembly protein; afdb-uniprot50 \| AF-A0A7H4NMM9-F1-MODEL\_V4 \| 1.0 \| 5.982e-15 \| 571 \| 0.307 \| 140 \| 93 \| 1 \| 1 \| 136 \| 1 \| 140 \| Baseplate assembly protein V \| Baseplate assembly protein V; afdb-uniprot50 \| AF-A0A178GUC9-F1-MODEL\_V4 \| 1.0 \| 1.366e-14 \| 571 \| 0.331 \| 148 \| 85 \| 1 \| 4 \| 137 \| 7 \| 154 \| Baseplate assembly protein V \| Baseplate assembly protein V; afdb-uniprot50 \| AF-A0A502CNP6-F1-MODEL\_V4 \| 1.0 \| 2.798e-17 \| 571 \| 0.296 \| 216 \| 114 \| 4 \| 2 \| 217 \| 1 \| 178 \| Phage baseplate assembly protein V \| Phage baseplate assembly protein V; afdb-uniprot50 \| AF-A0A2U9LGE2-F1-MODEL\_V4 \| 1.0 \| 5.37e-18 \| 571 \| 0.342 \| 213 \| 119 \| 4 \| 5 \| 217 \| 2 \| 193 \| Phage baseplate assembly protein V \| Phage baseplate assembly protein V; afdb-uniprot50 \| AF-A0A244BW49-F1-MODEL\_V4 \| 1.0 \| 5.696e-18 \| 570 \| 0.338 \| 213 \| 120 \| 4 \| 5 \| 217 \| 2 \| 193 \| Baseplate assembly protein \| Baseplate assembly protein; afdb-uniprot50 \| AF-A0A379ZSB9-F1-MODEL\_V4 \| 1.0 \| 1.63e-14 \| 569 \| 0.304 \| 138 \| 90 \| 1 \| 1 \| 132 \| 1 \| 138 \| Phage baseplate assembly protein V \| Phage baseplate assembly protein V; afdb-uniprot50 \| AF-A0A1V2N3K0-F1-MODEL\_V4 \| 1.0 \| 6.73e-15 \| 568 \| 0.351 \| 148 \| 82 \| 2 \| 4 \| 137 \| 7 \| 154 \| Phage baseplate assembly protein V family protein \| Phage baseplate assembly protein V family protein; afdb-uniprot50 \| AF-A0A6N4T373-F1-MODEL\_V4 \| 1.0 \| 2.929e-13 \| 568 \| 0.365 \| 126 \| 79 \| 1 \| 1 \| 125 \| 1 \| 126 \| Phage\_base\_V domain-containing protein \| Phage\_base\_V domain-containing protein; afdb-uniprot50 \| AF-A0A1C7W7X8-F1-MODEL\_V4 \| 1.0 \| 1.958e-16 \| 567 \| 0.292 \| 222 \| 144 \| 5 \| 2 \| 217 \| 1 \| 215 \| Phage\_base\_V domain-containing protein \| Phage\_base\_V domain-containing protein; afdb-uniprot50 \| AF-A0A653KXX6-F1-MODEL\_V4 \| 1.0 \| 2.487e-17 \| 566 \| 0.337 \| 237 \| 87 \| 6 \| 1 \| 217 \| 1 \| 187 \| Contig\_41, whole genome shotgun sequence \| Contig\_41, whole genome shotgun sequence; afdb-uniprot50 \| AF-A0A4Y9VQX4-F1-MODEL\_V4 \| 1.0 \| 2.091e-18 \| 566 \| 0.315 \| 219 \| 127 \| 6 \| 1 \| 217 \| 1 \| 198 \| Phage baseplate assembly protein V \| Phage baseplate assembly protein V; afdb-uniprot50 \| AF-A0A2S6H5F0-F1-MODEL\_V4 \| 1.0 \| 1.531e-13 \| 564 \| 0.335 \| 140 \| 85 \| 2 \| 1 \| 132 \| 1 \| 140 \| Phage baseplate assembly protein V \| Phage baseplate assembly protein V; afdb-uniprot50 \| AF-A0A1G6JDU4-F1-MODEL\_V4 \| 1.0 \| 3.35e-18 \| 563 \| 0.362 \| 232 \| 125 \| 6 \| 1 \| 217 \| 1 \| 224 \| Phage baseplate assembly protein V \| Phage baseplate assembly protein V; afdb-uniprot50 \| AF-A0A853SGL8-F1-MODEL\_V4 \| 1.0 \| 2.612e-14 \| 562 \| 0.41 \| 139 \| 78 \| 3 \| 3 \| 139 \| 6 \| 142 \| Phage-related baseplate assembly protein \| Phage-related baseplate assembly protein; afdb-uniprot50 \| AF-A0A2S8QRV5-F1-MODEL\_V4 \| 1.0 \| 8.059e-16 \| 562 \| 0.242 \| 194 \| 142 \| 3 \| 1 \| 193 \| 1 \| 190 \| Phage baseplate assembly protein V \| Phage baseplate assembly protein V; afdb-uniprot50 \| AF-A0A3S5B410-F1-MODEL\_V4 \| 1.0 \| 7.648e-18 \| 561 \| 0.386 \| 212 \| 118 \| 4 \| 6 \| 217 \| 28 \| 227 \| Phage P2 baseplate assembly protein gpV \| Phage P2 baseplate assembly protein gpV; afdb-uniprot50 \| AF-A0A7W6RFW2-F1-MODEL\_V4 \| 1.0 \| 5.352e-17 \| 560 \| 0.324 \| 216 \| 103 \| 3 \| 2 \| 217 \| 6 \| 178 \| Phage baseplate assembly protein V \| Phage baseplate assembly protein V; afdb-uniprot50 \| AF-A0A774TGT6-F1-MODEL\_V4 \| 1.0 \| 5.696e-18 \| 560 \| 0.345 \| 217 \| 118 \| 5 \| 1 \| 217 \| 1 \| 193 \| Phage baseplate assembly protein V \| Phage baseplate assembly protein V; afdb-uniprot50 \| AF-A0A6N8Q3A4-F1-MODEL\_V4 \| 1.0 \| 1.734e-15 \| 559 \| 0.287 \| 181 \| 118 \| 3 \| 1 \| 181 \| 1 \| 170 \| Phage baseplate assembly protein V \| Phage baseplate assembly protein V; afdb-uniprot50 \| AF-A0A829E378-F1-MODEL\_V4 \| 1.0 \| 6.022e-17 \| 559 \| 0.313 \| 217 \| 115 \| 6 \| 1 \| 217 \| 1 \| 183 \| Uncharacterized protein \| Uncharacterized protein; afdb-uniprot50 \| AF-A0A6N4T205-F1-MODEL\_V4 \| 1.0 \| 1.939e-13 \| 558 \| 0.359 \| 128 \| 81 \| 1 \| 1 \| 127 \| 1 \| 128 \| Phage\_base\_V domain-containing protein \| Phage\_base\_V domain-containing protein; afdb-uniprot50 \| AF-Q8ZMU1-F1-MODEL\_V4 \| 1.0 \| 1.463e-17 \| 558 \| 0.269 \| 223 \| 126 \| 3 \| 1 \| 217 \| 1 \| 192 \| Fels-2 prophage protein \| Fels-2 prophage protein; afdb-uniprot50 \| AF-A0A376HNC6-F1-MODEL\_V4 \| 1.0 \| 7.648e-18 \| 558 \| 0.341 \| 217 \| 119 \| 5 \| 1 \| 217 \| 1 \| 193 \| Baseplate assembly protein V \| Baseplate assembly protein V; afdb-uniprot50 \| AF-A0A1H0NN53-F1-MODEL\_V4 \| 1.0 \| 2.647e-18 \| 558 \| 0.36 \| 225 \| 116 \| 6 \| 1 \| 217 \| 1 \| 205 \| Phage baseplate assembly protein V \| Phage baseplate assembly protein V; afdb-uniprot50 \| AF-A0A1A9VKF9-F1-MODEL\_V4 \| 1.0 \| 2.063e-14 \| 558 \| 0.376 \| 146 \| 77 \| 1 \| 6 \| 137 \| 191 \| 336 \| Phage\_base\_V domain-containing protein \| Phage\_base\_V domain-containing protein; afdb-uniprot50 \| AF-J1JE15-F1-MODEL\_V4 \| 1.0 \| 4.172e-13 \| 557 \| 0.333 \| 135 \| 88 \| 2 \| 4 \| 137 \| 8 \| 141 \| Phage baseplate assembly protein V \| Phage baseplate assembly protein V; afdb-uniprot50 \| AF-A0A7Z8DPV0-F1-MODEL\_V4 \| 1.0 \| 7.21e-18 \| 556 \| 0.35 \| 217 \| 117 \| 5 \| 1 \| 217 \| 1 \| 193 \| Phage baseplate assembly protein V \| Phage baseplate assembly protein V; afdb-uniprot50 \| AF-C0R4U2-F1-MODEL\_V4 \| 1.0 \| 1.63e-14 \| 555 \| 0.348 \| 149 \| 81 \| 3 \| 4 \| 137 \| 7 \| 154 \| Baseplate assembly protein V \| Baseplate assembly protein V; afdb-uniprot50 \| AF-A0A3S5YL67-F1-MODEL\_V4 \| 1.0 \| 9.682e-18 \| 555 \| 0.341 \| 217 \| 119 \| 5 \| 1 \| 217 \| 1 \| 193 \| Baseplate assembly protein \| Baseplate assembly protein; afdb-uniprot50 \| AF-A0A2X2ZC27-F1-MODEL\_V4 \| 1.0 \| 9.682e-18 \| 555 \| 0.341 \| 217 \| 119 \| 5 \| 1 \| 217 \| 1 \| 193 \| Baseplate assembly protein V \| Baseplate assembly protein V; afdb-uniprot50 \| AF-A0A376SKQ2-F1-MODEL\_V4 \| 1.0 \| 7.163e-16 \| 553 \| 0.333 \| 177 \| 95 \| 2 \| 3 \| 179 \| 5 \| 158 \| Baseplate assembly protein V (GpV) \| Baseplate assembly protein V (GpV); afdb-uniprot50 \| AF-A0A6D0PZJ4-F1-MODEL\_V4 \| 1.0 \| 1.089e-17 \| 553 \| 0.341 \| 217 \| 119 \| 5 \| 1 \| 217 \| 1 \| 193 \| Phage baseplate assembly protein V \| Phage baseplate assembly protein V; afdb-uniprot50 \| AF-C1DBK4-F1-MODEL\_V4 \| 1.0 \| 1.214e-14 \| 552 \| 0.342 \| 143 \| 86 \| 2 \| 1 \| 137 \| 1 \| 141 \| Phage baseplate assembly protein V \| Phage baseplate assembly protein V; afdb-uniprot50 \| AF-A0A2Z6GCH3-F1-MODEL\_V4 \| 1.0 \| 4.484e-17 \| 551 \| 0.327 \| 214 \| 103 \| 6 \| 6 \| 217 \| 3 \| 177 \| Baseplate assembly protein \| Baseplate assembly protein; afdb-uniprot50 \| AF-A0A1X7L1D4-F1-MODEL\_V4 \| 1.0 \| 5.299e-14 \| 550 \| 0.291 \| 151 \| 101 \| 2 \| 1 \| 145 \| 4 \| 154 \| Phage baseplate assembly protein V \| Phage baseplate assembly protein V; afdb-uniprot50 \| AF-A0A2W5AAE8-F1-MODEL\_V4 \| 1.0 \| 2.077e-16 \| 550 \| 0.284 \| 218 \| 130 \| 6 \| 1 \| 217 \| 1 \| 193 \| Phage baseplate assembly protein V \| Phage baseplate assembly protein V; afdb-uniprot50 \| AF-A0A166Z6Q3-F1-MODEL\_V4 \| 1.0 \| 1.024e-16 \| 550 \| 0.288 \| 215 \| 130 \| 6 \| 3 \| 217 \| 5 \| 196 \| Baseplate assembly protein \| Baseplate assembly protein; afdb-uniprot50 \| AF-A0A0T9TB75-F1-MODEL\_V4 \| 1.0 \| 1.082e-15 \| 549 \| 0.278 \| 219 \| 99 \| 4 \| 1 \| 217 \| 1 \| 162 \| Baseplate assembly protein \| Baseplate assembly protein; afdb-uniprot50 \| AF-A0A6S6Y3K5-F1-MODEL\_V4 \| 1.0 \| 1.746e-17 \| 549 \| 0.337 \| 216 \| 136 \| 5 \| 4 \| 217 \| 9 \| 219 \| Phage baseplate assembly protein V (Modular protein) \| Phage baseplate assembly protein V (Modular protein); afdb-uniprot50 \| AF-W0L460-F1-MODEL\_V4 \| 1.0 \| 2.337e-16 \| 548 \| 0.31 \| 216 \| 117 \| 4 \| 2 \| 217 \| 1 \| 184 \| Baseplate assembly protein \| Baseplate assembly protein; afdb-uniprot50 \| AF-A0A482PH41-F1-MODEL\_V4 \| 1.0 \| 1.379e-17 \| 548 \| 0.331 \| 217 \| 121 \| 5 \| 1 \| 217 \| 1 \| 193 \| Phage baseplate assembly protein V \| Phage baseplate assembly protein V; afdb-uniprot50 \| AF-A0A559QNI7-F1-MODEL\_V4 \| 1.0 \| 4.214e-16 \| 548 \| 0.331 \| 193 \| 111 \| 2 \| 2 \| 194 \| 1 \| 175 \| Phage baseplate assembly protein V \| Phage baseplate assembly protein V; afdb-uniprot50 \| AF-A0A519FIN0-F1-MODEL\_V4 \| 1.0 \| 1.01e-12 \| 546 \| 0.31 \| 132 \| 89 \| 2 \| 1 \| 131 \| 1 \| 131 \| Phage baseplate assembly protein V \| Phage baseplate assembly protein V; afdb-uniprot50 \| AF-A0A843NDI5-F1-MODEL\_V4 \| 1.0 \| 1.448e-14 \| 545 \| 0.331 \| 160 \| 91 \| 1 \| 5 \| 164 \| 2 \| 145 \| Phage baseplate assembly protein V \| Phage baseplate assembly protein V; afdb-uniprot50 \| AF-A0A1E3G706-F1-MODEL\_V4 \| 1.0 \| 1.283e-13 \| 545 \| 0.331 \| 145 \| 95 \| 1 \| 1 \| 143 \| 1 \| 145 \| Phage\_base\_V domain-containing protein \| Phage\_base\_V domain-containing protein; afdb-uniprot50 \| AF-A0A1X0W485-F1-MODEL\_V4 \| 1.0 \| 4.757e-17 \| 545 \| 0.269 \| 223 \| 118 \| 4 \| 1 \| 217 \| 1 \| 184 \| Baseplate assembly protein \| Baseplate assembly protein; afdb-uniprot50 \| AF-A0A1E7PZX9-F1-MODEL\_V4 \| 1.0 \| 8.549e-16 \| 545 \| 0.336 \| 193 \| 91 \| 4 \| 1 \| 193 \| 1 \| 156 \| Phage\_base\_V domain-containing protein \| Phage\_base\_V domain-containing protein; afdb-uniprot50 \| AF-A0A158E8E3-F1-MODEL\_V4 \| 1.0 \| 1.226e-17 \| 545 \| 0.288 \| 225 \| 136 \| 4 \| 1 \| 208 \| 1 \| 218 \| Phage baseplate assembly protein V \| Phage baseplate assembly protein V; afdb-uniprot50 \| AF-W1HV66-F1-MODEL\_V4 \| 1.0 \| 1.932e-12 \| 544 \| 0.371 \| 105 \| 66 \| 0 \| 16 \| 120 \| 2 \| 106 \| Baseplate assembly protein V \| Baseplate assembly protein V; afdb-uniprot50 \| AF-A0A149QJX7-F1-MODEL\_V4 \| 1.0 \| 3.542e-17 \| 544 \| 0.304 \| 217 \| 122 \| 6 \| 1 \| 217 \| 1 \| 188 \| Baseplate assembly protein \| Baseplate assembly protein; afdb-uniprot50 \| AF-A0A0M5KYZ3-F1-MODEL\_V4 \| 1.0 \| 2.968e-17 \| 544 \| 0.308 \| 217 \| 136 \| 4 \| 4 \| 217 \| 8 \| 213 \| Phage-related baseplate assembly protein \| Phage-related baseplate assembly protein; afdb-uniprot50 \| AF-A0A366IM94-F1-MODEL\_V4 \| 1.0 \| 1.027e-17 \| 542 \| 0.331 \| 220 \| 120 \| 5 \| 3 \| 217 \| 44 \| 241 \| Phage baseplate assembly protein V \| Phage baseplate assembly protein V; afdb-uniprot50 \| AF-A0A1B3E8F5-F1-MODEL\_V4 \| 1.0 \| 5.029e-16 \| 541 \| 0.328 \| 195 \| 106 \| 4 \| 2 \| 196 \| 1 \| 170 \| Phage baseplate assembly protein V \| Phage baseplate assembly protein V; afdb-uniprot50 \| AF-A0A7Z0MZS5-F1-MODEL\_V4 \| 1.0 \| 9.618e-16 \| 540 \| 0.331 \| 223 \| 124 \| 8 \| 4 \| 217 \| 10 \| 216 \| Phage baseplate assembly protein V \| Phage baseplate assembly protein V; afdb-uniprot50 \| AF-A0A6D0J6U3-F1-MODEL\_V4 \| 1.0 \| 4.694e-13 \| 539 \| 0.346 \| 130 \| 84 \| 1 \| 16 \| 145 \| 2 \| 130 \| Phage baseplate assembly protein V \| Phage baseplate assembly protein V; afdb-uniprot50 \| AF-A0A4V2RT09-F1-MODEL\_V4 \| 1.0 \| 3.496e-13 \| 539 \| 0.282 \| 131 \| 90 \| 2 \| 5 \| 135 \| 9 \| 135 \| Phage baseplate assembly protein V \| Phage baseplate assembly protein V; afdb-uniprot50 \| AF-A0A5X8YW87-F1-MODEL\_V4 \| 1.0 \| 4.469e-16 \| 538 \| 0.322 \| 192 \| 107 \| 4 \| 1 \| 192 \| 1 \| 169 \| Phage baseplate assembly protein V \| Phage baseplate assembly protein V; afdb-uniprot50 \| AF-A0A853I7W8-F1-MODEL\_V4 \| 1.0 \| 2.929e-13 \| 538 \| 0.307 \| 140 \| 97 \| 0 \| 16 \| 155 \| 2 \| 141 \| Phage baseplate assembly protein V \| Phage baseplate assembly protein V; afdb-uniprot50 \| AF-A0A7W5K784-F1-MODEL\_V4 \| 1.0 \| 9.65e-17 \| 538 \| 0.304 \| 217 \| 129 \| 7 \| 1 \| 217 \| 1 \| 195 \| Phage baseplate assembly protein V \| Phage baseplate assembly protein V; afdb-uniprot50 \| AF-A0A7W6WMQ5-F1-MODEL\_V4 \| 1.0 \| 7.187e-17 \| 538 \| 0.352 \| 190 \| 104 \| 4 \| 6 \| 193 \| 14 \| 186 \| Phage baseplate assembly protein V \| Phage baseplate assembly protein V; afdb-uniprot50 \| AF-A0A7U9IYM4-F1-MODEL\_V4 \| 1.0 \| 3.757e-17 \| 537 \| 0.341 \| 217 \| 119 \| 5 \| 1 \| 217 \| 1 \| 193 \| Phage protein gp15 \| Phage protein gp15; afdb-uniprot50 \| AF-Q877J8-F1-MODEL\_V4 \| 1.0 \| 3.53e-16 \| 537 \| 0.319 \| 219 \| 123 \| 6 \| 1 \| 217 \| 1 \| 195 \| Phage-related baseplate assembly protein \| Phage-related baseplate assembly protein; afdb-uniprot50 \| AF-A0A485AXI1-F1-MODEL\_V4 \| 1.0 \| 1.148e-15 \| 537 \| 0.273 \| 194 \| 117 \| 3 \| 3 \| 196 \| 22 \| 191 \| Phage P2 baseplate assembly protein gpV \| Phage P2 baseplate assembly protein gpV; afdb-uniprot50 \| AF-A0A7W4KR34-F1-MODEL\_V4 \| 1.0 \| 1.822e-12 \| 536 \| 0.327 \| 113 \| 76 \| 0 \| 10 \| 122 \| 2 \| 114 \| Phage baseplate assembly protein V \| Phage baseplate assembly protein V; afdb-uniprot50 \| AF-A0A4R3Y8K0-F1-MODEL\_V4 \| 1.0 \| 2.329e-15 \| 536 \| 0.289 \| 214 \| 110 \| 5 \| 4 \| 217 \| 6 \| 177 \| Phage baseplate assembly protein V \| Phage baseplate assembly protein V; afdb-uniprot50 \| AF-A0A2U1XZ31-F1-MODEL\_V4 \| 1.0 \| 1.296e-16 \| 536 \| 0.304 \| 230 \| 136 \| 8 \| 4 \| 217 \| 18 \| 239 \| Phage\_base\_V domain-containing protein \| Phage\_base\_V domain-containing protein; afdb-uniprot50 \| AF-A0A8B4LYP3-F1-MODEL\_V4 \| 1.0 \| 2.798e-17 \| 534 \| 0.36 \| 219 \| 110 \| 7 \| 1 \| 217 \| 1 \| 191 \| Baseplate assembly protein V \| Baseplate assembly protein V; afdb-uniprot50 \| AF-A0A7X5KA07-F1-MODEL\_V4 \| 1.0 \| 6.041e-18 \| 534 \| 0.262 \| 248 \| 151 \| 5 \| 1 \| 217 \| 1 \| 247 \| Phage baseplate assembly protein V \| Phage baseplate assembly protein V; afdb-uniprot50 \| AF-V5ZC64-F1-MODEL\_V4 \| 1.0 \| 5.677e-17 \| 534 \| 0.288 \| 215 \| 128 \| 4 \| 3 \| 217 \| 86 \| 275 \| Baseplate assembly protein V GpV \| Baseplate assembly protein V GpV; afdb-uniprot50 \| AF-A0A061JPC8-F1-MODEL\_V4 \| 1.0 \| 4.725e-15 \| 532 \| 0.337 \| 166 \| 93 \| 3 \| 1 \| 166 \| 1 \| 149 \| Baseplate assembly protein \| Baseplate assembly protein; afdb-uniprot50 \| AF-A0A1F2LPI2-F1-MODEL\_V4 \| 1.0 \| 9.068e-16 \| 532 \| 0.28 \| 196 \| 117 \| 3 \| 1 \| 196 \| 1 \| 172 \| Baseplate assembly protein \| Baseplate assembly protein; afdb-uniprot50 \| AF-A0A3N8PDL7-F1-MODEL\_V4 \| 1.0 \| 3.127e-15 \| 532 \| 0.272 \| 224 \| 143 \| 5 \| 3 \| 215 \| 4 \| 218 \| Phage baseplate assembly protein V \| Phage baseplate assembly protein V; afdb-uniprot50 \| AF-A0A369YJU0-F1-MODEL\_V4 \| 1.0 \| 3.985e-17 \| 531 \| 0.327 \| 217 \| 106 \| 7 \| 1 \| 217 \| 1 \| 177 \| Phage baseplate assembly protein V \| Phage baseplate assembly protein V; afdb-uniprot50 \| AF-A0A2A5BPR3-F1-MODEL\_V4 \| 1.0 \| 1.292e-15 \| 531 \| 0.24 \| 212 \| 142 \| 4 \| 2 \| 213 \| 1 \| 193 \| Uncharacterized protein \| Uncharacterized protein; afdb-uniprot50 \| AF-A0A4P9VFF4-F1-MODEL\_V4 \| 1.0 \| 8.059e-16 \| 531 \| 0.262 \| 202 \| 107 \| 4 \| 1 \| 197 \| 9 \| 173 \| Phage baseplate assembly protein V \| Phage baseplate assembly protein V; afdb-uniprot50 \| AF-A0A2E4CSJ4-F1-MODEL\_V4 \| 1.0 \| 2.337e-16 \| 531 \| 0.299 \| 204 \| 127 \| 5 \| 14 \| 217 \| 22 \| 209 \| Baseplate assembly protein \| Baseplate assembly protein; afdb-uniprot50 \| AF-A0A797M4Y5-F1-MODEL\_V4 \| 1.0 \| 1.723e-13 \| 529 \| 0.3 \| 153 \| 88 \| 2 \| 1 \| 153 \| 1 \| 134 \| Phage baseplate assembly protein V \| Phage baseplate assembly protein V; afdb-uniprot50 \| AF-A0A420WVL0-F1-MODEL\_V4 \| 1.0 \| 3.328e-16 \| 529 \| 0.305 \| 193 \| 110 \| 4 \| 4 \| 196 \| 6 \| 174 \| Phage baseplate assembly protein V \| Phage baseplate assembly protein V; afdb-uniprot50 \| AF-A0A2K4JEB7-F1-MODEL\_V4 \| 1.0 \| 5.029e-16 \| 529 \| 0.298 \| 218 \| 130 \| 2 \| 1 \| 217 \| 1 \| 196 \| Phage baseplate assembly protein V \| Phage baseplate assembly protein V; afdb-uniprot50 \| AF-A0A833FRJ1-F1-MODEL\_V4 \| 1.0 \| 2.78e-15 \| 528 \| 0.273 \| 219 \| 117 \| 4 \| 4 \| 217 \| 8 \| 189 \| Phage baseplate assembly protein V \| Phage baseplate assembly protein V; afdb-uniprot50 \| AF-A0A7X3ZHR6-F1-MODEL\_V4 \| 1.0 \| 1.137e-12 \| 527 \| 0.266 \| 135 \| 98 \| 1 \| 4 \| 137 \| 11 \| 145 \| Phage baseplate assembly protein V \| Phage baseplate assembly protein V; afdb-uniprot50 \| AF-U2B529-F1-MODEL\_V4 \| 1.0 \| 4.425e-13 \| 526 \| 0.307 \| 143 \| 90 \| 4 \| 1 \| 137 \| 1 \| 140 \| Baseplate assembly protein \| Baseplate assembly protein; afdb-uniprot50 \| AF-A0A516SJA1-F1-MODEL\_V4 \| 1.0 \| 1.137e-12 \| 526 \| 0.288 \| 135 \| 93 \| 2 \| 1 \| 132 \| 1 \| 135 \| Phage baseplate assembly protein V \| Phage baseplate assembly protein V; afdb-uniprot50 \| AF-A0A4T2A4P9-F1-MODEL\_V4 \| 1.0 \| 2.62e-15 \| 526 \| 0.312 \| 192 \| 114 \| 4 \| 2 \| 193 \| 1 \| 174 \| Phage baseplate assembly protein V \| Phage baseplate assembly protein V; afdb-uniprot50 \| AF-B0UTN4-F1-MODEL\_V4 \| 1.0 \| 1.296e-16 \| 526 \| 0.289 \| 204 \| 126 \| 3 \| 3 \| 206 \| 5 \| 189 \| Phage baseplate assembly protein V \| Phage baseplate assembly protein V; afdb-uniprot50 \| AF-A0A318KM99-F1-MODEL\_V4 \| 1.0 \| 6.022e-17 \| 526 \| 0.331 \| 229 \| 137 \| 7 \| 1 \| 217 \| 1 \| 225 \| Phage baseplate assembly protein V \| Phage baseplate assembly protein V; afdb-uniprot50 \| AF-A0A516SHS7-F1-MODEL\_V4 \| 1.0 \| 7.21e-18 \| 526 \| 0.328 \| 225 \| 113 \| 4 \| 4 \| 217 \| 41 \| 238 \| Phage baseplate assembly protein V \| Phage baseplate assembly protein V; afdb-uniprot50 \| AF-A0A2G3L892-F1-MODEL\_V4 \| 1.0 \| 3.757e-17 \| 526 \| 0.32 \| 243 \| 121 \| 9 \| 1 \| 217 \| 1 \| 225 \| Oxidoreductase \| Oxidoreductase; afdb-uniprot50 \| AF-A0A7Z7RJ47-F1-MODEL\_V4 \| 1.0 \| 3.554e-18 \| 525 \| 0.337 \| 228 \| 130 \| 5 \| 4 \| 217 \| 35 \| 255 \| Phage P2 baseplate assembly protein gpV \| Phage P2 baseplate assembly protein gpV; afdb-uniprot50 \| AF-A0A7X0AGS3-F1-MODEL\_V4 \| 1.0 \| 4.214e-16 \| 522 \| 0.294 \| 214 \| 127 \| 5 \| 2 \| 209 \| 1 \| 196 \| Phage baseplate assembly protein V \| Phage baseplate assembly protein V; afdb-uniprot50 \| AF-A0A2W5FLQ5-F1-MODEL\_V4 \| 1.0 \| 1.547e-16 \| 522 \| 0.255 \| 239 \| 120 \| 6 \| 2 \| 217 \| 3 \| 206 \| Phage baseplate assembly protein V \| Phage baseplate assembly protein V; afdb-uniprot50 \| AF-A0A1V2N5A2-F1-MODEL\_V4 \| 1.0 \| 7.548e-14 \| 521 \| 0.352 \| 139 \| 76 \| 2 \| 13 \| 137 \| 2 \| 140 \| Phage baseplate assembly protein V family protein \| Phage baseplate assembly protein V family protein; afdb-uniprot50 \| AF-A0A5J6LDB1-F1-MODEL\_V4 \| 1.0 \| 3.317e-15 \| 521 \| 0.271 \| 217 \| 110 \| 4 \| 1 \| 217 \| 1 \| 169 \| Phage baseplate assembly protein V \| Phage baseplate assembly protein V; afdb-uniprot50 \| AF-A0A542V4K0-F1-MODEL\_V4 \| 1.0 \| 9.068e-16 \| 521 \| 0.282 \| 195 \| 121 \| 4 \| 2 \| 196 \| 1 \| 176 \| Phage baseplate assembly protein V \| Phage baseplate assembly protein V; afdb-uniprot50 \| AF-A0A377A641-F1-MODEL\_V4 \| 1.0 \| 1.536e-14 \| 520 \| 0.335 \| 170 \| 94 \| 2 \| 1 \| 170 \| 1 \| 151 \| Baseplate assembly protein V \| Baseplate assembly protein V; afdb-uniprot50 \| AF-E2CJU1-F1-MODEL\_V4 \| 1.0 \| 9.068e-16 \| 520 \| 0.276 \| 217 \| 132 \| 3 \| 3 \| 217 \| 9 \| 202 \| Phage-related baseplate assembly protein V \| Phage-related baseplate assembly protein V; afdb-uniprot50 \| AF-A0A5S9Q467-F1-MODEL\_V4 \| 1.0 \| 3.733e-15 \| 520 \| 0.262 \| 221 \| 143 \| 5 \| 1 \| 217 \| 1 \| 205 \| Phage\_base\_V domain-containing protein \| Phage\_base\_V domain-containing protein; afdb-uniprot50 \| AF-A0A375AA54-F1-MODEL\_V4 \| 1.0 \| 3.53e-16 \| 520 \| 0.357 \| 210 \| 95 \| 3 \| 7 \| 216 \| 3 \| 172 \| Baseplate assembly protein V \| Baseplate assembly protein V; afdb-uniprot50 \| AF-A0A840REX7-F1-MODEL\_V4 \| 1.0 \| 1.723e-13 \| 519 \| 0.367 \| 136 \| 78 \| 2 \| 2 \| 129 \| 1 \| 136 \| Phage baseplate assembly protein V \| Phage baseplate assembly protein V; afdb-uniprot50 \| AF-A0A430BCB8-F1-MODEL\_V4 \| 1.0 \| 4.214e-16 \| 519 \| 0.267 \| 217 \| 132 \| 6 \| 1 \| 217 \| 1 \| 190 \| Phage baseplate assembly protein V \| Phage baseplate assembly protein V; afdb-uniprot50 \| AF-A0A516SJG1-F1-MODEL\_V4 \| 1.0 \| 3.92e-12 \| 519 \| 0.276 \| 130 \| 93 \| 1 \| 4 \| 132 \| 44 \| 173 \| Phage baseplate assembly protein V \| Phage baseplate assembly protein V; afdb-uniprot50 \| AF-A0A212KBU0-F1-MODEL\_V4 \| 1.0 \| 4.455e-15 \| 518 \| 0.265 \| 207 \| 144 \| 4 \| 4 \| 208 \| 13 \| 213 \| Phage P2 baseplate assembly gpV-like protein (Modular protein) \| Phage P2 baseplate assembly gpV-like protein (Modular protein); afdb-uniprot50 \| AF-A0A410UF52-F1-MODEL\_V4 \| 1.0 \| 1.086e-16 \| 518 \| 0.279 \| 215 \| 135 \| 5 \| 4 \| 217 \| 20 \| 215 \| Phage baseplate assembly protein V \| Phage baseplate assembly protein V; afdb-uniprot50 \| AF-A0A0H3GM22-F1-MODEL\_V4 \| 1.0 \| 1.453e-15 \| 517 \| 0.265 \| 196 \| 120 \| 3 \| 1 \| 196 \| 1 \| 172 \| Baseplate assembly protein V \| Baseplate assembly protein V; afdb-uniprot50 \| AF-A0A377AJR1-F1-MODEL\_V4 \| 1.0 \| 4.996e-14 \| 516 \| 0.341 \| 155 \| 83 \| 2 \| 1 \| 155 \| 1 \| 136 \| Baseplate assembly protein V \| Baseplate assembly protein V; afdb-uniprot50 \| AF-A0A5M6I3L2-F1-MODEL\_V4 \| 1.0 \| 3.097e-12 \| 516 \| 0.3 \| 123 \| 86 \| 0 \| 4 \| 126 \| 8 \| 130 \| Phage baseplate assembly protein V \| Phage baseplate assembly protein V; afdb-uniprot50 \| AF-A0A1B3Z862-F1-MODEL\_V4 \| 1.0 \| 2.78e-15 \| 516 \| 0.262 \| 217 \| 126 \| 6 \| 1 \| 217 \| 1 \| 183 \| Baseplate assembly protein \| Baseplate assembly protein; afdb-uniprot50 \| AF-A0A4Q6F5K6-F1-MODEL\_V4 \| 1.0 \| 8.949e-12 \| 515 \| 0.321 \| 112 \| 76 \| 0 \| 2 \| 113 \| 1 \| 112 \| Phage baseplate assembly protein V \| Phage baseplate assembly protein V; afdb-uniprot50 \| AF-A0A653NAD7-F1-MODEL\_V4 \| 1.0 \| 2.063e-14 \| 515 \| 0.314 \| 162 \| 95 \| 4 \| 4 \| 157 \| 29 \| 182 \| Phage\_base\_V domain-containing protein \| Phage\_base\_V domain-containing protein; afdb-uniprot50 \| AF-A0A5C7C8J3-F1-MODEL\_V4 \| 1.0 \| 2.203e-16 \| 515 \| 0.301 \| 219 \| 127 \| 6 \| 2 \| 217 \| 1 \| 196 \| Phage baseplate assembly protein V \| Phage baseplate assembly protein V; afdb-uniprot50 \| AF-A0A177RV31-F1-MODEL\_V4 \| 1.0 \| 2.919e-12 \| 514 \| 0.385 \| 114 \| 67 \| 2 \| 1 \| 113 \| 1 \| 112 \| Baseplate assembly protein \| Baseplate assembly protein; afdb-uniprot50 \| AF-A0A1M3PG72-F1-MODEL\_V4 \| 1.0 \| 3.328e-16 \| 514 \| 0.328 \| 216 \| 116 \| 7 \| 2 \| 217 \| 1 \| 187 \| Phage\_base\_V domain-containing protein \| Phage\_base\_V domain-containing protein; afdb-uniprot50 \| AF-A0A1N7B0F6-F1-MODEL\_V4 \| 1.0 \| 9.618e-16 \| 514 \| 0.263 \| 216 \| 119 \| 5 \| 4 \| 217 \| 15 \| 192 \| Phage baseplate assembly protein V \| Phage baseplate assembly protein V; afdb-uniprot50 \| AF-A0A1E1F5H0-F1-MODEL\_V4 \| 1.0 \| 1.292e-15 \| 513 \| 0.259 \| 208 \| 118 \| 6 \| 1 \| 208 \| 1 \| 172 \| Baseplate assembly protein \| Baseplate assembly protein; afdb-uniprot50 \| AF-A0A5A7M6L4-F1-MODEL\_V4 \| 1.0 \| 1.024e-16 \| 513 \| 0.334 \| 227 \| 123 \| 6 \| 4 \| 217 \| 42 \| 253 \| Baseplate assembly protein \| Baseplate assembly protein; afdb-uniprot50 \| AF-A0A250DVL7-F1-MODEL\_V4 \| 1.0 \| 5.029e-16 \| 513 \| 0.288 \| 260 \| 141 \| 10 \| 1 \| 217 \| 1 \| 259 \| Phage baseplate protein \| Phage baseplate protein; afdb-uniprot50 \| AF-A0A515BE25-F1-MODEL\_V4 \| 1.0 \| 6.708e-14 \| 511 \| 0.244 \| 217 \| 117 \| 4 \| 1 \| 217 \| 1 \| 170 \| Phage baseplate assembly protein V \| Phage baseplate assembly protein V; afdb-uniprot50 \| AF-G2HX98-F1-MODEL\_V4 \| 1.0 \| 1.541e-15 \| 511 \| 0.327 \| 226 \| 125 \| 11 \| 1 \| 209 \| 1 \| 216 \| Phage baseplate assembly protein \| Phage baseplate assembly protein; afdb-uniprot50 \| AF-A0A3S8YQ07-F1-MODEL\_V4 \| 1.0 \| 5.299e-14 \| 510 \| 0.323 \| 164 \| 95 \| 4 \| 2 \| 157 \| 1 \| 156 \| Phage baseplate assembly protein V \| Phage baseplate assembly protein V; afdb-uniprot50 \| AF-A0A2G3K3I8-F1-MODEL\_V4 \| 1.0 \| 2.798e-17 \| 510 \| 0.303 \| 214 \| 133 \| 3 \| 2 \| 199 \| 1 \| 214 \| Baseplate assembly protein \| Baseplate assembly protein; afdb-uniprot50 \| AF-A0A6L6JFU3-F1-MODEL\_V4 \| 1.0 \| 7.548e-14 \| 509 \| 0.286 \| 213 \| 89 \| 5 \| 6 \| 217 \| 2 \| 152 \| Phage baseplate assembly protein V \| Phage baseplate assembly protein V; afdb-uniprot50 \| AF-A0A8A8P9Z4-F1-MODEL\_V4 \| 1.0 \| 2.05e-12 \| 508 \| 0.336 \| 125 \| 81 \| 1 \| 1 \| 123 \| 1 \| 125 \| Phage baseplate assembly protein V \| Phage baseplate assembly protein V; afdb-uniprot50 \| AF-A0A329VPT4-F1-MODEL\_V4 \| 1.0 \| 1.852e-17 \| 508 \| 0.266 \| 259 \| 147 \| 6 \| 1 \| 217 \| 1 \| 258 \| Phage baseplate protein \| Phage baseplate protein; afdb-uniprot50 \| AF-A0A1V2DS20-F1-MODEL\_V4 \| 1.0 \| 3.519e-15 \| 507 \| 0.273 \| 227 \| 116 \| 6 \| 1 \| 216 \| 1 \| 189 \| Phage\_base\_V domain-containing protein \| Phage\_base\_V domain-containing protein; afdb-uniprot50 \| AF-A0A2E9SIQ4-F1-MODEL\_V4 \| 1.0 \| 1.287e-14 \| 507 \| 0.231 \| 229 \| 148 \| 9 \| 1 \| 217 \| 1 \| 213 \| Baseplate assembly protein \| Baseplate assembly protein; afdb-uniprot50 \| AF-A0A6B3LC16-F1-MODEL\_V4 \| 1.0 \| 1.082e-15 \| 505 \| 0.285 \| 228 \| 133 \| 8 \| 1 \| 213 \| 1 \| 213 \| Phage baseplate assembly protein V \| Phage baseplate assembly protein V; afdb-uniprot50 \| AF-A0A378AZS7-F1-MODEL\_V4 \| 1.0 \| 3.72e-14 \| 503 \| 0.313 \| 150 \| 96 \| 2 \| 1 \| 145 \| 1 \| 148 \| Baseplate assembly protein V \| Baseplate assembly protein V; afdb-uniprot50 \| AF-Q3RF60-F1-MODEL\_V4 \| 1.0 \| 4.741e-16 \| 502 \| 0.314 \| 216 \| 122 \| 6 \| 4 \| 217 \| 47 \| 238 \| Phage baseplate assembly protein V \| Phage baseplate assembly protein V; afdb-uniprot50 \| AF-A0A516SJL7-F1-MODEL\_V4 \| 1.0 \| 1.816e-11 \| 501 \| 0.392 \| 102 \| 62 \| 0 \| 2 \| 103 \| 1 \| 102 \| Phage baseplate assembly protein V \| Phage baseplate assembly protein V; afdb-uniprot50 \| AF-A0A1T0AWZ3-F1-MODEL\_V4 \| 1.0 \| 4.469e-16 \| 500 \| 0.336 \| 196 \| 105 \| 6 \| 1 \| 196 \| 1 \| 171 \| Baseplate assembly protein \| Baseplate assembly protein; afdb-uniprot50 \| AF-A0A849VJT8-F1-MODEL\_V4 \| 1.0 \| 6.753e-16 \| 500 \| 0.275 \| 218 \| 150 \| 5 \| 1 \| 217 \| 1 \| 211 \| Phage baseplate assembly protein V \| Phage baseplate assembly protein V; afdb-uniprot50 \| AF-A0A7U9AYN8-F1-MODEL\_V4 \| 1.0 \| 7.573e-15 \| 499 \| 0.344 \| 174 \| 94 \| 3 \| 1 \| 174 \| 1 \| 154 \| Baseplate assembly protein V \| Baseplate assembly protein V; afdb-uniprot50 \| AF-A0A6I3XBP6-F1-MODEL\_V4 \| 1.0 \| 5.982e-15 \| 499 \| 0.252 \| 202 \| 123 \| 5 \| 1 \| 198 \| 1 \| 178 \| Phage baseplate assembly protein V \| Phage baseplate assembly protein V; afdb-uniprot50 \| AF-A0A1W9H429-F1-MODEL\_V4 \| 1.0 \| 2.47e-15 \| 498 \| 0.271 \| 217 \| 117 \| 6 \| 1 \| 217 \| 1 \| 176 \| Phage\_base\_V domain-containing protein \| Phage\_base\_V domain-containing protein; afdb-uniprot50 \| AF-A0A1X3DKD5-F1-MODEL\_V4 \| 1.0 \| 7.163e-16 \| 498 \| 0.326 \| 233 \| 116 \| 6 \| 5 \| 217 \| 2 \| 213 \| Uncharacterized protein \| Uncharacterized protein; afdb-uniprot50 \| AF-W0ITY9-F1-MODEL\_V4 \| 1.0 \| 1.079e-14 \| 497 \| 0.276 \| 224 \| 112 \| 7 \| 1 \| 217 \| 1 \| 181 \| Baseplate assembly protein \| Baseplate assembly protein; afdb-uniprot50 \| AF-A0A4Y6U991-F1-MODEL\_V4 \| 1.0 \| 2.62e-15 \| 497 \| 0.303 \| 214 \| 114 \| 7 \| 4 \| 217 \| 14 \| 192 \| Phage baseplate assembly protein V \| Phage baseplate assembly protein V; afdb-uniprot50 \| AF-A0A418VVD7-F1-MODEL\_V4 \| 1.0 \| 9.587e-15 \| 497 \| 0.297 \| 225 \| 118 \| 5 \| 4 \| 203 \| 8 \| 217 \| Phage baseplate assembly protein V \| Phage baseplate assembly protein V; afdb-uniprot50 \| AF-F3LHZ4-F1-MODEL\_V4 \| 1.0 \| 5.299e-14 \| 496 \| 0.254 \| 212 \| 115 \| 4 \| 4 \| 213 \| 8 \| 178 \| Phage\_base\_V domain-containing protein \| Phage\_base\_V domain-containing protein; afdb-uniprot50 \| AF-A0A1I1UBL3-F1-MODEL\_V4 \| 1.0 \| 1.214e-14 \| 495 \| 0.261 \| 191 \| 123 \| 2 \| 4 \| 194 \| 6 \| 178 \| Phage baseplate assembly protein V \| Phage baseplate assembly protein V; afdb-uniprot50 \| AF-A0A2D3WVI8-F1-MODEL\_V4 \| 1.0 \| 7.548e-14 \| 494 \| 0.3 \| 180 \| 109 \| 6 \| 1 \| 177 \| 1 \| 166 \| Phage\_base\_V domain-containing protein \| Phage\_base\_V domain-containing protein; afdb-uniprot50 \| AF-A0A379C9Y4-F1-MODEL\_V4 \| 1.0 \| 8.549e-16 \| 494 \| 0.293 \| 218 \| 123 \| 7 \| 2 \| 217 \| 1 \| 189 \| Phage P2 baseplate assembly protein gpV \| Phage P2 baseplate assembly protein gpV; afdb-uniprot50 \| AF-A0A7R7V081-F1-MODEL\_V4 \| 1.0 \| 1.375e-16 \| 494 \| 0.3 \| 220 \| 121 \| 5 \| 1 \| 217 \| 23 \| 212 \| Baseplate assembly protein \| Baseplate assembly protein; afdb-uniprot50 \| AF-A0A2X2BK34-F1-MODEL\_V4 \| 1.0 \| 1.439e-12 \| 493 \| 0.371 \| 194 \| 116 \| 5 \| 24 \| 217 \| 1 \| 188 \| Phage baseplate assembly protein \| Phage baseplate assembly protein; afdb-uniprot50 \| AF-A0A7G8VI14-F1-MODEL\_V4 \| 1.0 \| 5.352e-17 \| 493 \| 0.289 \| 242 \| 146 \| 4 \| 1 \| 217 \| 11 \| 251 \| Phage baseplate assembly protein V \| Phage baseplate assembly protein V; afdb-uniprot50 \| AF-A0A1B8YNH2-F1-MODEL\_V4 \| 1.0 \| 9.65e-17 \| 493 \| 0.254 \| 259 \| 150 \| 6 \| 1 \| 217 \| 45 \| 302 \| Phage-related baseplate assembly protein \| Phage-related baseplate assembly protein; afdb-uniprot50 \| AF-A0A016XIR2-F1-MODEL\_V4 \| 1.0 \| 1.02e-15 \| 492 \| 0.306 \| 225 \| 119 \| 7 \| 1 \| 217 \| 1 \| 196 \| Baseplate assembly protein \| Baseplate assembly protein; afdb-uniprot50 \| AF-F3YY65-F1-MODEL\_V4 \| 1.0 \| 1.932e-12 \| 492 \| 0.237 \| 164 \| 119 \| 1 \| 4 \| 161 \| 8 \| 171 \| Phage baseplate assembly protein V \| Phage baseplate assembly protein V; afdb-uniprot50 \| AF-A0A7X4GDK7-F1-MODEL\_V4 \| 1.0 \| 2.063e-14 \| 491 \| 0.227 \| 207 \| 119 \| 5 \| 12 \| 217 \| 11 \| 177 \| Phage baseplate assembly protein V \| Phage baseplate assembly protein V; afdb-uniprot50 \| AF-A0A7W6REL9-F1-MODEL\_V4 \| 1.0 \| 1.635e-15 \| 491 \| 0.317 \| 189 \| 87 \| 3 \| 4 \| 153 \| 12 \| 197 \| Phage baseplate assembly protein V \| Phage baseplate assembly protein V; afdb-uniprot50 \| AF-F1Z9C2-F1-MODEL\_V4 \| 1.0 \| 2.919e-12 \| 489 \| 0.294 \| 136 \| 89 \| 5 \| 13 \| 146 \| 12 \| 142 \| Phage P2 baseplate assembly gpV-like protein \| Phage P2 baseplate assembly gpV-like protein; afdb-uniprot50 \| AF-A0A524RVZ0-F1-MODEL\_V4 \| 1.0 \| 5.264e-12 \| 488 \| 0.284 \| 151 \| 98 \| 4 \| 3 \| 145 \| 18 \| 166 \| Phage baseplate assembly protein V \| Phage baseplate assembly protein V; afdb-uniprot50 \| AF-A0A1I7E546-F1-MODEL\_V4 \| 1.0 \| 3.117e-14 \| 488 \| 0.258 \| 197 \| 127 \| 5 \| 1 \| 196 \| 1 \| 179 \| Phage baseplate assembly protein V \| Phage baseplate assembly protein V; afdb-uniprot50 \| AF-A0A0T9QWS2-F1-MODEL\_V4 \| 1.0 \| 5.299e-14 \| 487 \| 0.279 \| 168 \| 108 \| 2 \| 1 \| 155 \| 1 \| 168 \| Baseplate assembly protein V \| Baseplate assembly protein V; afdb-uniprot50 \| AF-A0A6L8GG55-F1-MODEL\_V4 \| 1.0 \| 1.453e-15 \| 487 \| 0.277 \| 227 \| 131 \| 9 \| 4 \| 217 \| 9 \| 215 \| Phage baseplate assembly protein V \| Phage baseplate assembly protein V; afdb-uniprot50 \| AF-A0A167H3T4-F1-MODEL\_V4 \| 1.0 \| 1.079e-14 \| 486 \| 0.283 \| 194 \| 105 \| 3 \| 1 \| 194 \| 1 \| 160 \| Phage\_base\_V domain-containing protein \| Phage\_base\_V domain-containing protein; afdb-uniprot50 \| AF-U3U4P9-F1-MODEL\_V4 \| 1.0 \| 3.708e-13 \| 486 \| 0.308 \| 198 \| 121 \| 5 \| 16 \| 209 \| 6 \| 191 \| Uncharacterized protein \| Uncharacterized protein; afdb-uniprot50 \| AF-A0A2C9EHC3-F1-MODEL\_V4 \| 1.0 \| 1.541e-15 \| 486 \| 0.304 \| 223 \| 122 \| 6 \| 1 \| 217 \| 46 \| 241 \| Baseplate assembly protein V \| Baseplate assembly protein V; afdb-uniprot50 \| AF-A0A450W6W7-F1-MODEL\_V4 \| 1.0 \| 2.056e-13 \| 485 \| 0.306 \| 150 \| 98 \| 1 \| 1 \| 144 \| 1 \| 150 \| Phage baseplate assembly protein V \| Phage baseplate assembly protein V; afdb-uniprot50 \| AF-A0A2U0SJV9-F1-MODEL\_V4 \| 1.0 \| 2.063e-14 \| 485 \| 0.308 \| 201 \| 106 \| 5 \| 17 \| 217 \| 2 \| 169 \| Phage baseplate assembly protein V \| Phage baseplate assembly protein V; afdb-uniprot50 \| AF-D1P8A9-F1-MODEL\_V4 \| 1.0 \| 1.202e-11 \| 484 \| 0.31 \| 145 \| 98 \| 2 \| 17 \| 161 \| 1 \| 143 \| Phage-related baseplate assembly protein \| Phage-related baseplate assembly protein; afdb-uniprot50 \| AF-A0A5E7YKZ5-F1-MODEL\_V4 \| 1.0 \| 9.618e-16 \| 484 \| 0.274 \| 222 \| 134 \| 7 \| 2 \| 217 \| 6 \| 206 \| Phage baseplate assembly protein V \| Phage baseplate assembly protein V; afdb-uniprot50 \| AF-A0A521PR02-F1-MODEL\_V4 \| 1.0 \| 1.017e-14 \| 483 \| 0.281 \| 213 \| 113 \| 7 \| 12 \| 217 \| 5 \| 184 \| Phage baseplate assembly protein V \| Phage baseplate assembly protein V; afdb-uniprot50 \| AF-A0A1G5ACL8-F1-MODEL\_V4 \| 1.0 \| 5.012e-15 \| 480 \| 0.284 \| 204 \| 103 \| 4 \| 4 \| 206 \| 8 \| 169 \| Phage baseplate assembly protein V \| Phage baseplate assembly protein V; afdb-uniprot50 \| AF-A0A1M6B7Z3-F1-MODEL\_V4 \| 1.0 \| 8.033e-15 \| 480 \| 0.267 \| 191 \| 116 \| 3 \| 4 \| 194 \| 5 \| 171 \| Phage-related baseplate assembly protein \| Phage-related baseplate assembly protein; afdb-uniprot50 \| AF-A0A4V0XDV9-F1-MODEL\_V4 \| 1.0 \| 3.117e-14 \| 480 \| 0.293 \| 201 \| 120 \| 7 \| 1 \| 193 \| 15 \| 201 \| Phage\_base\_V domain-containing protein \| Phage\_base\_V domain-containing protein; afdb-uniprot50 \| AF-V4N894-F1-MODEL\_V4 \| 1.0 \| 5.658e-16 \| 480 \| 0.328 \| 210 \| 118 \| 5 \| 4 \| 213 \| 32 \| 218 \| Baseplate assembly protein \| Baseplate assembly protein; afdb-uniprot50 \| AF-A0A381EC13-F1-MODEL\_V4 \| 1.0 \| 3.72e-14 \| 479 \| 0.271 \| 221 \| 106 \| 3 \| 1 \| 217 \| 1 \| 170 \| Phage P2 baseplate assembly protein gpV \| Phage P2 baseplate assembly protein gpV; afdb-uniprot50 \| AF-A0A0P6WI63-F1-MODEL\_V4 \| 1.0 \| 3.306e-14 \| 479 \| 0.285 \| 207 \| 127 \| 7 \| 16 \| 217 \| 2 \| 192 \| Phage\_base\_V domain-containing protein \| Phage\_base\_V domain-containing protein; afdb-uniprot50 \| AF-A0A7X4GVY2-F1-MODEL\_V4 \| 1.0 \| 1.74e-16 \| 478 \| 0.33 \| 230 \| 117 \| 6 \| 1 \| 196 \| 1 \| 227 \| Phage baseplate assembly protein V \| Phage baseplate assembly protein V; afdb-uniprot50 \| AF-A0A1X7CH99-F1-MODEL\_V4 \| 1.0 \| 1.02e-15 \| 478 \| 0.284 \| 253 \| 143 \| 7 \| 1 \| 216 \| 1 \| 252 \| Phage baseplate assembly protein V \| Phage baseplate assembly protein V; afdb-uniprot50 \| AF-A0A285JXF0-F1-MODEL\_V4 \| 1.0 \| 1.635e-15 \| 477 \| 0.237 \| 236 \| 135 \| 3 \| 1 \| 192 \| 3 \| 237 \| Phage\_base\_V domain-containing protein \| Phage\_base\_V domain-containing protein; afdb-uniprot50 \| AF-A0A326LB63-F1-MODEL\_V4 \| 1.0 \| 1.287e-14 \| 476 \| 0.24 \| 216 \| 133 \| 8 \| 3 \| 217 \| 8 \| 193 \| Phage baseplate assembly protein V \| Phage baseplate assembly protein V; afdb-uniprot50 \| AF-A0A5E4XG36-F1-MODEL\_V4 \| 1.0 \| 2.196e-15 \| 476 \| 0.294 \| 224 \| 142 \| 4 \| 1 \| 217 \| 1 \| 215 \| Baseplate protein \| Baseplate protein; afdb-uniprot50 \| AF-A0A2N2S1D1-F1-MODEL\_V4 \| 1.0 \| 1.958e-16 \| 476 \| 0.305 \| 236 \| 127 \| 7 \| 1 \| 217 \| 1 \| 218 \| Phage baseplate assembly protein V \| Phage baseplate assembly protein V; afdb-uniprot50 \| AF-A0A212J3Z5-F1-MODEL\_V4 \| 1.0 \| 3.117e-14 \| 476 \| 0.26 \| 219 \| 153 \| 5 \| 1 \| 215 \| 1 \| 214 \| Putative Phage baseplate assembly protein \| Putative Phage baseplate assembly protein; afdb-uniprot50 \| AF-A0A1E3G5Z4-F1-MODEL\_V4 \| 1.0 \| 3.306e-14 \| 474 \| 0.276 \| 195 \| 106 \| 6 \| 4 \| 192 \| 15 \| 180 \| Phage\_base\_V domain-containing protein \| Phage\_base\_V domain-containing protein; afdb-uniprot50 \| AF-A0A0M4TG07-F1-MODEL\_V4 \| 1.0 \| 5.317e-15 \| 474 \| 0.267 \| 217 \| 128 \| 5 \| 1 \| 217 \| 1 \| 186 \| Phage\_base\_V domain-containing protein \| Phage\_base\_V domain-containing protein; afdb-uniprot50 \| AF-A0A7Y7HZL4-F1-MODEL\_V4 \| 1.0 \| 3.53e-16 \| 474 \| 0.323 \| 210 \| 124 \| 5 \| 1 \| 198 \| 1 \| 204 \| Phage baseplate assembly protein V \| Phage baseplate assembly protein V; afdb-uniprot50 \| AF-A0A397NIY9-F1-MODEL\_V4 \| 1.0 \| 1.214e-14 \| 472 \| 0.354 \| 189 \| 99 \| 7 \| 16 \| 197 \| 2 \| 174 \| Phage baseplate assembly protein V \| Phage baseplate assembly protein V; afdb-uniprot50 \| AF-A0A1G5B4F2-F1-MODEL\_V4 \| 1.0 \| 2.586e-11 \| 471 \| 0.231 \| 138 \| 103 \| 2 \| 4 \| 141 \| 7 \| 141 \| Phage baseplate assembly protein V \| Phage baseplate assembly protein V; afdb-uniprot50 \| AF-K5XX39-F1-MODEL\_V4 \| 1.0 \| 1.828e-13 \| 471 \| 0.276 \| 224 \| 110 \| 7 \| 1 \| 217 \| 1 \| 179 \| Phage P2 baseplate assembly gpV-like protein \| Phage P2 baseplate assembly gpV-like protein; afdb-uniprot50 \| AF-A4P0H7-F1-MODEL\_V4 \| 1.0 \| 5.012e-15 \| 471 \| 0.318 \| 201 \| 112 \| 5 \| 1 \| 201 \| 1 \| 176 \| Predicted phage P2-like baseplate assembly protein \| Predicted phage P2-like baseplate assembly protein; afdb-uniprot50 \| AF-A0A2W4T7L3-F1-MODEL\_V4 \| 1.0 \| 1.712e-11 \| 471 \| 0.301 \| 136 \| 92 \| 2 \| 1 \| 133 \| 1 \| 136 \| Phage baseplate assembly protein V \| Phage baseplate assembly protein V; afdb-uniprot50 \| AF-A0A3Q8U0Q9-F1-MODEL\_V4 \| 1.0 \| 5.299e-14 \| 471 \| 0.24 \| 220 \| 144 \| 7 \| 1 \| 203 \| 1 \| 214 \| Phage baseplate assembly protein V \| Phage baseplate assembly protein V; afdb-uniprot50 \| AF-A0A0Q7ECU9-F1-MODEL\_V4 \| 1.0 \| 9.038e-15 \| 469 \| 0.266 \| 210 \| 115 \| 6 \| 1 \| 203 \| 1 \| 178 \| Phage\_base\_V domain-containing protein \| Phage\_base\_V domain-containing protein; afdb-uniprot50 \| AF-A0A1H3ZWK9-F1-MODEL\_V4 \| 1.0 \| 1.84e-15 \| 469 \| 0.276 \| 217 \| 143 \| 6 \| 1 \| 217 \| 1 \| 203 \| Phage baseplate assembly protein V \| Phage baseplate assembly protein V; afdb-uniprot50 \| AF-A0A345DE56-F1-MODEL\_V4 \| 1.0 \| 5.962e-14 \| 468 \| 0.225 \| 200 \| 123 \| 6 \| 1 \| 196 \| 1 \| 172 \| Phage\_base\_V domain-containing protein \| Phage\_base\_V domain-containing protein; afdb-uniprot50 \| AF-G1UXN9-F1-MODEL\_V4 \| 1.0 \| 4.172e-13 \| 468 \| 0.25 \| 236 \| 150 \| 10 \| 2 \| 217 \| 3 \| 231 \| Uncharacterized protein \| Uncharacterized protein; afdb-uniprot50 \| AF-S6GRC2-F1-MODEL\_V4 \| 1.0 \| 3.933e-13 \| 467 \| 0.272 \| 143 \| 96 \| 2 \| 1 \| 137 \| 1 \| 141 \| Phage\_base\_V domain-containing protein \| Phage\_base\_V domain-containing protein; afdb-uniprot50 \| AF-A0A0J9EDT8-F1-MODEL\_V4 \| 1.0 \| 1.01e-12 \| 467 \| 0.237 \| 219 \| 102 \| 7 \| 1 \| 217 \| 1 \| 156 \| Phage baseplate assembly protein V \| Phage baseplate assembly protein V; afdb-uniprot50 \| AF-H8L2J8-F1-MODEL\_V4 \| 1.0 \| 2.761e-13 \| 467 \| 0.244 \| 209 \| 136 \| 3 \| 1 \| 203 \| 1 \| 193 \| Phage baseplate assembly protein V \| Phage baseplate assembly protein V; afdb-uniprot50 \| AF-A0A1E7HY32-F1-MODEL\_V4 \| 1.0 \| 4.214e-16 \| 467 \| 0.278 \| 269 \| 142 \| 7 \| 1 \| 217 \| 2 \| 270 \| Uncharacterized protein \| Uncharacterized protein; afdb-uniprot50 \| AF-A0A7J6YLT3-F1-MODEL\_V4 \| 1.0 \| 4.425e-13 \| 466 \| 0.287 \| 153 \| 105 \| 2 \| 15 \| 167 \| 136 \| 284 \| Uncharacterized protein \| Uncharacterized protein; afdb-uniprot50 \| AF-A0A367WWC3-F1-MODEL\_V4 \| 1.0 \| 1.932e-12 \| 465 \| 0.237 \| 160 \| 114 \| 2 \| 2 \| 161 \| 1 \| 152 \| Phage\_base\_V domain-containing protein \| Phage\_base\_V domain-containing protein; afdb-uniprot50 \| AF-A0A6G8F2D8-F1-MODEL\_V4 \| 1.0 \| 5.317e-15 \| 465 \| 0.283 \| 219 \| 136 \| 8 \| 1 \| 213 \| 1 \| 204 \| Bacteriophage baseplate assembly protein V \| Bacteriophage baseplate assembly protein V; afdb-uniprot50 \| AF-A0A4V0IE64-F1-MODEL\_V4 \| 1.0 \| 3.274e-11 \| 463 \| 0.457 \| 107 \| 53 \| 2 \| 1 \| 107 \| 1 \| 102 \| Phage P2 baseplate assembly protein gpV \| Phage P2 baseplate assembly protein gpV; afdb-uniprot50 \| AF-A0A7W5BZS0-F1-MODEL\_V4 \| 1.0 \| 1.361e-13 \| 463 \| 0.288 \| 201 \| 106 \| 5 \| 1 \| 193 \| 1 \| 172 \| Phage baseplate assembly protein V \| Phage baseplate assembly protein V; afdb-uniprot50 \| AF-A0A7U2VHG3-F1-MODEL\_V4 \| 1.0 \| 4.996e-14 \| 462 \| 0.287 \| 209 \| 112 \| 5 \| 9 \| 217 \| 3 \| 174 \| Phage baseplate assembly protein V \| Phage baseplate assembly protein V; afdb-uniprot50 \| AF-A0A4R7Q4T9-F1-MODEL\_V4 \| 1.0 \| 5.658e-16 \| 462 \| 0.204 \| 293 \| 156 \| 6 \| 1 \| 217 \| 1 \| 292 \| Phage baseplate assembly protein V \| Phage baseplate assembly protein V; afdb-uniprot50 \| AF-E1PLB0-F1-MODEL\_V4 \| 1.0 \| 2.9e-10 \| 461 \| 0.361 \| 94 \| 60 \| 0 \| 1 \| 94 \| 1 \| 94 \| Baseplate assembly protein V \| Baseplate assembly protein V; afdb-uniprot50 \| AF-A0A4R1K4T2-F1-MODEL\_V4 \| 1.0 \| 3.296e-13 \| 461 \| 0.304 \| 174 \| 102 \| 7 \| 1 \| 166 \| 1 \| 163 \| Phage baseplate assembly protein V \| Phage baseplate assembly protein V; afdb-uniprot50 \| AF-A0A623T421-F1-MODEL\_V4 \| 1.0 \| 1.075e-13 \| 461 \| 0.256 \| 199 \| 125 \| 6 \| 1 \| 197 \| 1 \| 178 \| Phage baseplate assembly protein V \| Phage baseplate assembly protein V; afdb-uniprot50 \| AF-A0A373FPN0-F1-MODEL\_V4 \| 1.0 \| 5.012e-15 \| 461 \| 0.295 \| 227 \| 121 \| 5 \| 1 \| 203 \| 1 \| 212 \| Phage baseplate assembly protein V \| Phage baseplate assembly protein V; afdb-uniprot50 \| AF-A0A1G8I115-F1-MODEL\_V4 \| 1.0 \| 1.945e-14 \| 460 \| 0.264 \| 193 \| 114 \| 4 \| 4 \| 194 \| 26 \| 192 \| Phage baseplate assembly protein V \| Phage baseplate assembly protein V; afdb-uniprot50 \| AF-A0A692CV12-F1-MODEL\_V4 \| 1.0 \| 2.167e-11 \| 456 \| 0.302 \| 142 \| 93 \| 4 \| 16 \| 156 \| 2 \| 138 \| Phage baseplate assembly protein V \| Phage baseplate assembly protein V; afdb-uniprot50 \| AF-A0A5Q0TIR4-F1-MODEL\_V4 \| 1.0 \| 2.752e-12 \| 456 \| 0.326 \| 144 \| 87 \| 4 \| 4 \| 137 \| 6 \| 149 \| Phage baseplate assembly protein V \| Phage baseplate assembly protein V; afdb-uniprot50 \| AF-A0A812QV09-F1-MODEL\_V4 \| 1.0 \| 1.608e-10 \| 456 \| 0.322 \| 124 \| 82 \| 1 \| 15 \| 138 \| 942 \| 1063 \| XkdF protein \| XkdF protein; afdb-uniprot50 \| AF-A0A7S7WDY0-F1-MODEL\_V4 \| 1.0 \| 3.959e-15 \| 455 \| 0.294 \| 224 \| 141 \| 6 \| 1 \| 217 \| 1 \| 214 \| Phage baseplate assembly protein V \| Phage baseplate assembly protein V; afdb-uniprot50 \| AF-A0A1R1MK78-F1-MODEL\_V4 \| 1.0 \| 4.144e-11 \| 452 \| 0.258 \| 143 \| 97 \| 6 \| 15 \| 151 \| 4 \| 143 \| Phage\_base\_V domain-containing protein \| Phage\_base\_V domain-containing protein; afdb-uniprot50 \| AF-A0A6G5QFQ9-F1-MODEL\_V4 \| 1.0 \| 8.978e-13 \| 452 \| 0.263 \| 209 \| 133 \| 9 \| 16 \| 209 \| 2 \| 204 \| Phage baseplate assembly protein V \| Phage baseplate assembly protein V; afdb-uniprot50 \| AF-A7MWA1-F1-MODEL\_V4 \| 1.0 \| 2.603e-13 \| 452 \| 0.284 \| 172 \| 104 \| 4 \| 4 \| 172 \| 26 \| 181 \| Phage\_base\_V domain-containing protein \| Phage\_base\_V domain-containing protein; afdb-uniprot50 \| AF-A0A5E7A8F9-F1-MODEL\_V4 \| 1.0 \| 2.036e-10 \| 450 \| 0.289 \| 121 \| 85 \| 1 \| 12 \| 132 \| 2 \| 121 \| Uncharacterized protein \| Uncharacterized protein; afdb-uniprot50 \| AF-A0A2D3W2I8-F1-MODEL\_V4 \| 1.0 \| 1.939e-13 \| 450 \| 0.257 \| 206 \| 119 \| 8 \| 1 \| 203 \| 1 \| 175 \| Uncharacterized protein \| Uncharacterized protein; afdb-uniprot50 \| AF-A0A2T5J1G2-F1-MODEL\_V4 \| 1.0 \| 8.492e-14 \| 450 \| 0.231 \| 225 \| 128 \| 7 \| 1 \| 213 \| 1 \| 192 \| Phage baseplate assembly protein V \| Phage baseplate assembly protein V; afdb-uniprot50 \| AF-A0A2E2N1A4-F1-MODEL\_V4 \| 1.0 \| 5.903e-11 \| 450 \| 0.239 \| 138 \| 100 \| 3 \| 9 \| 141 \| 2 \| 139 \| Uncharacterized protein \| Uncharacterized protein; afdb-uniprot50 \| AF-A0A837JGD3-F1-MODEL\_V4 \| 1.0 \| 1.357e-12 \| 450 \| 0.254 \| 224 \| 146 \| 8 \| 5 \| 214 \| 4 \| 220 \| Baseplate assembly protein \| Baseplate assembly protein; afdb-uniprot50 \| AF-X1P9Y8-F1-MODEL\_V4 \| 1.0 \| 2.056e-13 \| 449 \| 0.247 \| 194 \| 133 \| 6 \| 1 \| 185 \| 1 \| 190 \| Phage\_base\_V domain-containing protein \| Phage\_base\_V domain-containing protein; afdb-uniprot50 \| AF-A0A381C5Y2-F1-MODEL\_V4 \| 1.0 \| 1.723e-13 \| 449 \| 0.254 \| 236 \| 141 \| 8 \| 1 \| 216 \| 1 \| 221 \| Phage P2 baseplate assembly protein gpV \| Phage P2 baseplate assembly protein gpV; afdb-uniprot50 \| AF-A0A5S3S1K6-F1-MODEL\_V4 \| 1.0 \| 5.621e-14 \| 449 \| 0.271 \| 199 \| 130 \| 4 \| 4 \| 195 \| 11 \| 201 \| Phage baseplate assembly protein V \| Phage baseplate assembly protein V; afdb-uniprot50 \| AF-A0A2G6EI12-F1-MODEL\_V4 \| 1.0 \| 2.438e-11 \| 448 \| 0.236 \| 161 \| 116 \| 5 \| 1 \| 158 \| 1 \| 157 \| Phage\_base\_V domain-containing protein \| Phage\_base\_V domain-containing protein; afdb-uniprot50 \| AF-A0A7U8ZG09-F1-MODEL\_V4 \| 1.0 \| 5.012e-15 \| 448 \| 0.287 \| 216 \| 116 \| 6 \| 3 \| 217 \| 36 \| 214 \| Uncharacterized protein \| Uncharacterized protein; afdb-uniprot50 \| AF-A0A4Q0ZNI2-F1-MODEL\_V4 \| 1.0 \| 3.484e-12 \| 447 \| 0.303 \| 158 \| 102 \| 5 \| 1 \| 155 \| 1 \| 153 \| Uncharacterized protein \| Uncharacterized protein; afdb-uniprot50 \| AF-A0A136H344-F1-MODEL\_V4 \| 1.0 \| 1.017e-14 \| 447 \| 0.301 \| 202 \| 118 \| 3 \| 3 \| 197 \| 8 \| 193 \| Uncharacterized protein \| Uncharacterized protein; afdb-uniprot50 \| AF-A0A6L2ZS80-F1-MODEL\_V4 \| 1.0 \| 2.188e-14 \| 446 \| 0.307 \| 218 \| 121 \| 7 \| 1 \| 215 \| 1 \| 191 \| Putative phage baseplate protein \| Putative phage baseplate protein; afdb-uniprot50 \| AF-A0A3B6Y4E8-F1-MODEL\_V4 \| 1.0 \| 7.116e-14 \| 445 \| 0.246 \| 195 \| 133 \| 3 \| 4 \| 192 \| 11 \| 197 \| Phage baseplate assembly protein V \| Phage baseplate assembly protein V; afdb-uniprot50 \| AF-A0A6N3IEF8-F1-MODEL\_V4 \| 1.0 \| 3.285e-12 \| 445 \| 0.213 \| 234 \| 159 \| 9 \| 1 \| 216 \| 2 \| 228 \| Uncharacterized protein \| Uncharacterized protein; afdb-uniprot50 \| AF-A0A423PRV3-F1-MODEL\_V4 \| 1.0 \| 2.948e-15 \| 444 \| 0.294 \| 221 \| 128 \| 5 \| 4 \| 196 \| 7 \| 227 \| Phage\_base\_V domain-containing protein \| Phage\_base\_V domain-containing protein; afdb-uniprot50 \| AF-E4PPS8-F1-MODEL\_V4 \| 1.0 \| 3.972e-16 \| 443 \| 0.266 \| 251 \| 121 \| 6 \| 1 \| 196 \| 1 \| 243 \| Phage-related baseplate assembly protein V \| Phage-related baseplate assembly protein V; afdb-uniprot50 \| AF-V4PX29-F1-MODEL\_V4 \| 1.0 \| 2.454e-13 \| 442 \| 0.239 \| 217 \| 150 \| 4 \| 1 \| 217 \| 1 \| 202 \| Uncharacterized protein \| Uncharacterized protein; afdb-uniprot50 \| AF-A0A853IBN9-F1-MODEL\_V4 \| 1.0 \| 5.942e-13 \| 440 \| 0.283 \| 201 \| 102 \| 6 \| 1 \| 190 \| 1 \| 170 \| Phage baseplate assembly protein V \| Phage baseplate assembly protein V; afdb-uniprot50 \| AF-A0A5C8XTJ4-F1-MODEL\_V4 \| 1.0 \| 9.43e-10 \| 439 \| 0.383 \| 86 \| 50 \| 1 \| 1 \| 86 \| 1 \| 83 \| Phage baseplate assembly protein V \| Phage baseplate assembly protein V; afdb-uniprot50 \| AF-A0A6L9FGD0-F1-MODEL\_V4 \| 1.0 \| 8.006e-14 \| 439 \| 0.256 \| 230 \| 134 \| 8 \| 1 \| 214 \| 1 \| 209 \| Phage baseplate assembly protein V \| Phage baseplate assembly protein V; afdb-uniprot50 \| AF-A0A2E6BPD3-F1-MODEL\_V4 \| 1.0 \| 1.828e-13 \| 437 \| 0.294 \| 204 \| 106 \| 5 \| 14 \| 217 \| 8 \| 173 \| Baseplate assembly protein \| Baseplate assembly protein; afdb-uniprot50 \| AF-A0A258A0W3-F1-MODEL\_V4 \| 1.0 \| 8.006e-14 \| 436 \| 0.278 \| 219 \| 107 \| 8 \| 4 \| 217 \| 18 \| 190 \| Phage\_base\_V domain-containing protein \| Phage\_base\_V domain-containing protein; afdb-uniprot50 \| AF-A0A5H2Y3Q4-F1-MODEL\_V4 \| 1.0 \| 7.573e-15 \| 435 \| 0.251 \| 235 \| 146 \| 8 \| 4 \| 217 \| 11 \| 236 \| Bacteriophage baseplate assembly protein V \| Bacteriophage baseplate assembly protein V; afdb-uniprot50 \| AF-A0A1B7XDP0-F1-MODEL\_V4 \| 1.0 \| 1.287e-14 \| 435 \| 0.283 \| 219 \| 129 \| 6 \| 1 \| 193 \| 1 \| 217 \| Phage\_base\_V domain-containing protein \| Phage\_base\_V domain-containing protein; afdb-uniprot50 \| AF-A0A8B2NVK9-F1-MODEL\_V4 \| 1.0 \| 1.516e-10 \| 434 \| 0.265 \| 143 \| 94 \| 3 \| 4 \| 140 \| 11 \| 148 \| Baseplate assembly protein \| Baseplate assembly protein; afdb-uniprot50 \| AF-A0A1S6U8Y2-F1-MODEL\_V4 \| 1.0 \| 3.92e-12 \| 434 \| 0.267 \| 202 \| 123 \| 7 \| 16 \| 217 \| 4 \| 180 \| Phage baseplate assembly protein V \| Phage baseplate assembly protein V; afdb-uniprot50 \| AF-A0A824EIG8-F1-MODEL\_V4 \| 1.0 \| 2.291e-10 \| 433 \| 0.318 \| 113 \| 74 \| 2 \| 16 \| 127 \| 2 \| 112 \| Phage baseplate assembly protein V \| Phage baseplate assembly protein V; afdb-uniprot50 \| AF-A0A4R1J4B1-F1-MODEL\_V4 \| 1.0 \| 1.075e-13 \| 433 \| 0.268 \| 227 \| 154 \| 6 \| 1 \| 217 \| 1 \| 225 \| Phage baseplate assembly protein V \| Phage baseplate assembly protein V; afdb-uniprot50 \| AF-A0A0U9HBQ1-F1-MODEL\_V4 \| 1.0 \| 6.642e-11 \| 432 \| 0.246 \| 138 \| 98 \| 3 \| 1 \| 136 \| 1 \| 134 \| Phage baseplate assembly protein V \| Phage baseplate assembly protein V; afdb-uniprot50 \| AF-A0A4R0DU02-F1-MODEL\_V4 \| 1.0 \| 1.624e-13 \| 432 \| 0.221 \| 217 \| 122 \| 6 \| 1 \| 217 \| 1 \| 170 \| Phage baseplate assembly protein V \| Phage baseplate assembly protein V; afdb-uniprot50 \| AF-F4QJB9-F1-MODEL\_V4 \| 1.0 \| 1.828e-13 \| 432 \| 0.267 \| 202 \| 125 \| 7 \| 16 \| 217 \| 2 \| 180 \| Baseplate assembly protein V \| Baseplate assembly protein V; afdb-uniprot50 \| AF-A0A149SVM5-F1-MODEL\_V4 \| 1.0 \| 1.624e-13 \| 432 \| 0.253 \| 201 \| 118 \| 7 \| 1 \| 194 \| 1 \| 176 \| Phage\_base\_V domain-containing protein \| Phage\_base\_V domain-containing protein; afdb-uniprot50 \| AF-N6VFT2-F1-MODEL\_V4 \| 1.0 \| 5.962e-14 \| 432 \| 0.263 \| 224 \| 134 \| 7 \| 4 \| 208 \| 8 \| 219 \| Phage baseplate assembly protein V \| Phage baseplate assembly protein V; afdb-uniprot50 \| AF-A0A6S5X6Y8-F1-MODEL\_V4 \| 1.0 \| 1.129e-10 \| 430 \| 0.278 \| 133 \| 94 \| 2 \| 15 \| 146 \| 4 \| 135 \| Phage\_base\_V domain-containing protein \| Phage\_base\_V domain-containing protein; afdb-uniprot50 \| AF-S5MPB7-F1-MODEL\_V4 \| 1.0 \| 1.511e-09 \| 429 \| 0.361 \| 94 \| 60 \| 0 \| 1 \| 94 \| 1 \| 94 \| Baseplate assembly protein V \| Baseplate assembly protein V; afdb-uniprot50 \| AF-A0A3D9EV90-F1-MODEL\_V4 \| 1.0 \| 2.05e-12 \| 429 \| 0.272 \| 154 \| 102 \| 3 \| 2 \| 145 \| 1 \| 154 \| Phage baseplate assembly protein V \| Phage baseplate assembly protein V; afdb-uniprot50 \| AF-A0A380AJE4-F1-MODEL\_V4 \| 1.0 \| 7.927e-11 \| 428 \| 0.193 \| 145 \| 108 \| 3 \| 2 \| 137 \| 1 \| 145 \| Phage P2 baseplate assembly protein gpV \| Phage P2 baseplate assembly protein gpV; afdb-uniprot50 \| AF-A0A2D9W381-F1-MODEL\_V4 \| 1.0 \| 1.279e-12 \| 426 \| 0.236 \| 233 \| 112 \| 4 \| 2 \| 217 \| 1 \| 184 \| Baseplate assembly protein \| Baseplate assembly protein; afdb-uniprot50 \| AF-A0A269PJ60-F1-MODEL\_V4 \| 1.0 \| 5.942e-13 \| 424 \| 0.248 \| 221 \| 149 \| 4 \| 4 \| 217 \| 11 \| 221 \| Phage\_base\_V domain-containing protein \| Phage\_base\_V domain-containing protein; afdb-uniprot50 \| AF-A0A6G3I739-F1-MODEL\_V4 \| 1.0 \| 4.632e-09 \| 423 \| 0.349 \| 83 \| 54 \| 0 \| 1 \| 83 \| 1 \| 83 \| Phage baseplate assembly protein V \| Phage baseplate assembly protein V; afdb-uniprot50 \| AF-A0A4Q0YIW9-F1-MODEL\_V4 \| 1.0 \| 2.603e-13 \| 423 \| 0.238 \| 197 \| 120 \| 2 \| 1 \| 197 \| 1 \| 167 \| Phage\_base\_V domain-containing protein \| Phage\_base\_V domain-containing protein; afdb-uniprot50 \| AF-A0A7Z7KMS9-F1-MODEL\_V4 \| 1.0 \| 2.306e-12 \| 422 \| 0.242 \| 206 \| 132 \| 6 \| 1 \| 203 \| 1 \| 185 \| Phage P2 baseplate assembly protein gpV \| Phage P2 baseplate assembly protein gpV; afdb-uniprot50 \| AF-A0A7W4RHJ5-F1-MODEL\_V4 \| 1.0 \| 1.075e-13 \| 421 \| 0.31 \| 235 \| 133 \| 8 \| 1 \| 217 \| 7 \| 230 \| Phage baseplate assembly protein V \| Phage baseplate assembly protein V; afdb-uniprot50 \| AF-A0A024LSX6-F1-MODEL\_V4 \| 1.0 \| 2.89e-09 \| 420 \| 0.344 \| 90 \| 59 \| 0 \| 3 \| 92 \| 7 \| 96 \| Phage baseplate assembly protein V1 \| Phage baseplate assembly protein V1; afdb-uniprot50 \| AF-A0A2P7V3Q8-F1-MODEL\_V4 \| 1.0 \| 1.198e-10 \| 420 \| 0.189 \| 148 \| 117 \| 1 \| 2 \| 146 \| 1 \| 148 \| Baseplate assembly protein \| Baseplate assembly protein; afdb-uniprot50 \| AF-A0A1N7LRL3-F1-MODEL\_V4 \| 1.0 \| 1.816e-11 \| 418 \| 0.285 \| 147 \| 93 \| 5 \| 1 \| 138 \| 1 \| 144 \| Phage baseplate assembly protein V \| Phage baseplate assembly protein V; afdb-uniprot50 \| AF-A0A6I1J2X0-F1-MODEL\_V4 \| 1.0 \| 1.357e-12 \| 418 \| 0.228 \| 201 \| 130 \| 7 \| 1 \| 197 \| 1 \| 180 \| Phage baseplate assembly protein V \| Phage baseplate assembly protein V; afdb-uniprot50 \| AF-A0A3S4JW88-F1-MODEL\_V4 \| 1.0 \| 6.241e-10 \| 416 \| 0.368 \| 95 \| 60 \| 0 \| 1 \| 95 \| 1 \| 95 \| Phage P2 baseplate assembly protein gpV \| Phage P2 baseplate assembly protein gpV; afdb-uniprot50 \| AF-A0A081MYL2-F1-MODEL\_V4 \| 1.0 \| 6.686e-13 \| 416 \| 0.229 \| 222 \| 127 \| 8 \| 4 \| 213 \| 10 \| 199 \| Phage\_base\_V domain-containing protein \| Phage\_base\_V domain-containing protein; afdb-uniprot50 \| AF-A0A7J5WFZ4-F1-MODEL\_V4 \| 1.0 \| 1.202e-11 \| 416 \| 0.217 \| 221 \| 162 \| 5 \| 4 \| 213 \| 10 \| 230 \| Phage baseplate assembly protein \| Phage baseplate assembly protein; afdb-uniprot50 \| AF-A0A5T0IXE4-F1-MODEL\_V4 \| 1.0 \| 4.663e-11 \| 415 \| 0.312 \| 144 \| 86 \| 5 \| 16 \| 151 \| 2 \| 140 \| Phage baseplate assembly protein V \| Phage baseplate assembly protein V; afdb-uniprot50 \| AF-A0A1H9YBJ7-F1-MODEL\_V4 \| 1.0 \| 1.007e-11 \| 414 \| 0.238 \| 222 \| 108 \| 7 \| 1 \| 217 \| 2 \| 167 \| Phage baseplate assembly protein V \| Phage baseplate assembly protein V; afdb-uniprot50 \| AF-A0A2W5KZA5-F1-MODEL\_V4 \| 1.0 \| 7.98e-13 \| 413 \| 0.312 \| 195 \| 108 \| 5 \| 23 \| 217 \| 2 \| 170 \| Phage baseplate assembly protein V \| Phage baseplate assembly protein V; afdb-uniprot50 \| AF-A0A839IVK7-F1-MODEL\_V4 \| 1.0 \| 4.382e-10 \| 412 \| 0.198 \| 136 \| 103 \| 3 \| 3 \| 135 \| 29 \| 161 \| Phage baseplate assembly protein V \| Phage baseplate assembly protein V; afdb-uniprot50 \| AF-A0A6M8SWQ4-F1-MODEL\_V4 \| 1.0 \| 7.069e-12 \| 412 \| 0.227 \| 207 \| 141 \| 8 \| 16 \| 217 \| 8 \| 200 \| Phage baseplate assembly protein V \| Phage baseplate assembly protein V; afdb-uniprot50 \| AF-A0A809B5H8-F1-MODEL\_V4 \| 1.0 \| 2.056e-13 \| 412 \| 0.242 \| 243 \| 156 \| 5 \| 1 \| 215 \| 1 \| 243 \| Phage baseplate assembly protein V \| Phage baseplate assembly protein V; afdb-uniprot50 \| AF-A0A8B4NZR2-F1-MODEL\_V4 \| 1.0 \| 1.425e-09 \| 411 \| 0.389 \| 95 \| 55 \| 1 \| 1 \| 95 \| 1 \| 92 \| Baseplate assembly protein V \| Baseplate assembly protein V; afdb-uniprot50 \| AF-A0A1G3UAP2-F1-MODEL\_V4 \| 1.0 \| 4.425e-13 \| 411 \| 0.234 \| 209 \| 123 \| 7 \| 1 \| 208 \| 1 \| 173 \| Uncharacterized protein \| Uncharacterized protein; afdb-uniprot50 \| AF-A0A7J0BHU1-F1-MODEL\_V4 \| 1.0 \| 9.492e-12 \| 411 \| 0.202 \| 222 \| 163 \| 7 \| 4 \| 217 \| 8 \| 223 \| Baseplate protein \| Baseplate protein; afdb-uniprot50 \| AF-J4UQU3-F1-MODEL\_V4 \| 1.0 \| 2.291e-10 \| 408 \| 0.228 \| 140 \| 98 \| 3 \| 1 \| 137 \| 1 \| 133 \| Phage-like baseplate assembly protein \| Phage-like baseplate assembly protein; afdb-uniprot50 \| AF-A0A7V7G2T5-F1-MODEL\_V4 \| 1.0 \| 5.281e-13 \| 407 \| 0.257 \| 241 \| 99 \| 6 \| 1 \| 217 \| 1 \| 185 \| Phage baseplate assembly protein V \| Phage baseplate assembly protein V; afdb-uniprot50 \| AF-G2ZT35-F1-MODEL\_V4 \| 1.0 \| 2.446e-12 \| 403 \| 0.22 \| 227 \| 159 \| 9 \| 4 \| 217 \| 7 \| 228 \| Putative Phage baseplate assembly protein V \| Putative Phage baseplate assembly protein V; afdb-uniprot50 \| AF-A0A4V3M376-F1-MODEL\_V4 \| 1.0 \| 2.569e-09 \| 402 \| 0.393 \| 89 \| 54 \| 0 \| 8 \| 96 \| 5 \| 93 \| Phage baseplate assembly protein V \| Phage baseplate assembly protein V; afdb-uniprot50 \| AF-A0A3G6WBA3-F1-MODEL\_V4 \| 1.0 \| 7.98e-13 \| 402 \| 0.231 \| 233 \| 90 \| 6 \| 1 \| 214 \| 1 \| 163 \| Phage baseplate assembly protein V \| Phage baseplate assembly protein V; afdb-uniprot50 \| AF-A0A6L7TC77-F1-MODEL\_V4 \| 1.0 \| 2.603e-13 \| 402 \| 0.324 \| 225 \| 102 \| 8 \| 4 \| 216 \| 237 \| 423 \| Phage baseplate assembly protein V \| Phage baseplate assembly protein V; afdb-uniprot50 \| AF-A0A3G4V723-F1-MODEL\_V4 \| 1.0 \| 1.926e-11 \| 401 \| 0.247 \| 194 \| 116 \| 6 \| 1 \| 193 \| 4 \| 168 \| Phage baseplate assembly protein V \| Phage baseplate assembly protein V; afdb-uniprot50 \| AF-A0A2N1AP79-F1-MODEL\_V4 \| 1.0 \| 2.174e-12 \| 401 \| 0.288 \| 177 \| 92 \| 2 \| 18 \| 194 \| 2 \| 144 \| Phage baseplate assembly protein V \| Phage baseplate assembly protein V; afdb-uniprot50 \| AF-D5EFA0-F1-MODEL\_V4 \| 1.0 \| 1.129e-10 \| 400 \| 0.168 \| 160 \| 114 \| 3 \| 4 \| 160 \| 7 \| 150 \| Phage baseplate assembly protein V \| Phage baseplate assembly protein V; afdb-uniprot50 \| AF-E2CGG9-F1-MODEL\_V4 \| 1.0 \| 1.279e-12 \| 400 \| 0.278 \| 205 \| 104 \| 6 \| 16 \| 217 \| 14 \| 177 \| Phage baseplate component \| Phage baseplate component; afdb-uniprot50 \| AF-A0A6A4R6B7-F1-MODEL\_V4 \| 1.0 \| 1.439e-12 \| 399 \| 0.229 \| 231 \| 112 \| 3 \| 4 \| 217 \| 26 \| 207 \| Phage baseplate assembly protein V \| Phage baseplate assembly protein V; afdb-uniprot50 \| AF-E2CN36-F1-MODEL\_V4 \| 1.0 \| 1.614e-11 \| 398 \| 0.231 \| 216 \| 107 \| 4 \| 3 \| 217 \| 12 \| 169 \| Putative baseplate assembly protein V \| Putative baseplate assembly protein V; afdb-uniprot50 \| AF-A0A1C3EE72-F1-MODEL\_V4 \| 1.0 \| 3.107e-13 \| 397 \| 0.239 \| 205 \| 123 \| 3 \| 4 \| 208 \| 31 \| 202 \| Phage\_base\_V domain-containing protein \| Phage\_base\_V domain-containing protein; afdb-uniprot50 \| AF-A0A0K0VKK2-F1-MODEL\_V4 \| 1.0 \| 1.194e-09 \| 396 \| 0.171 \| 134 \| 107 \| 3 \| 3 \| 134 \| 22 \| 153 \| Uncharacterized protein \| Uncharacterized protein; afdb-uniprot50 \| AF-A0A1Y1QXP2-F1-MODEL\_V4 \| 1.0 \| 2.743e-11 \| 395 \| 0.195 \| 215 \| 116 \| 5 \| 4 \| 217 \| 8 \| 166 \| Uncharacterized protein \| Uncharacterized protein; afdb-uniprot50 \| AF-A0A827JR60-F1-MODEL\_V4 \| 1.0 \| 5.584e-12 \| 394 \| 0.279 \| 197 \| 117 \| 4 \| 21 \| 217 \| 2 \| 173 \| Phage baseplate assembly protein V \| Phage baseplate assembly protein V; afdb-uniprot50 \| AF-A0A349GR45-F1-MODEL\_V4 \| 1.0 \| 4.648e-10 \| 394 \| 0.219 \| 155 \| 114 \| 5 \| 15 \| 165 \| 5 \| 156 \| Baseplate assembly protein \| Baseplate assembly protein; afdb-uniprot50 \| AF-G2JAY8-F1-MODEL\_V4 \| 1.0 \| 7.069e-12 \| 394 \| 0.216 \| 222 \| 162 \| 8 \| 4 \| 217 \| 10 \| 227 \| Putative phage-related baseplate assembly protein V \| Putative phage-related baseplate assembly protein V; afdb-uniprot50 \| AF-A0A6P1LVS1-F1-MODEL\_V4 \| 1.0 \| 7.069e-12 \| 393 \| 0.216 \| 217 \| 125 \| 6 \| 1 \| 217 \| 1 \| 172 \| Phage baseplate assembly protein V \| Phage baseplate assembly protein V; afdb-uniprot50 \| AF-A0A828D047-F1-MODEL\_V4 \| 1.0 \| 2.595e-12 \| 392 \| 0.286 \| 185 \| 120 \| 5 \| 16 \| 191 \| 2 \| 183 \| Phage baseplate assembly protein V \| Phage baseplate assembly protein V; afdb-uniprot50 \| AF-A0A806CKJ7-F1-MODEL\_V4 \| 1.0 \| 5.923e-12 \| 391 \| 0.23 \| 200 \| 121 \| 6 \| 16 \| 215 \| 2 \| 168 \| Phage baseplate assembly protein V \| Phage baseplate assembly protein V; afdb-uniprot50 \| AF-A0A1I5RPU5-F1-MODEL\_V4 \| 1.0 \| 4.131e-10 \| 390 \| 0.24 \| 137 \| 90 \| 6 \| 1 \| 129 \| 1 \| 131 \| Phage P2 baseplate assembly protein gpV \| Phage P2 baseplate assembly protein gpV; afdb-uniprot50 \| AF-A0A4R2GWQ4-F1-MODEL\_V4 \| 1.0 \| 1.275e-11 \| 390 \| 0.24 \| 208 \| 109 \| 7 \| 16 \| 217 \| 7 \| 171 \| Phage baseplate assembly protein V \| Phage baseplate assembly protein V; afdb-uniprot50 \| AF-E1QHA2-F1-MODEL\_V4 \| 1.0 \| 1.27e-10 \| 390 \| 0.204 \| 147 \| 108 \| 2 \| 4 \| 141 \| 11 \| 157 \| Phage baseplate assembly protein V \| Phage baseplate assembly protein V; afdb-uniprot50 \| AF-A0A2T3NIK5-F1-MODEL\_V4 \| 1.0 \| 2.422e-09 \| 390 \| 0.246 \| 134 \| 95 \| 4 \| 4 \| 135 \| 22 \| 151 \| Phage baseplate assembly protein V \| Phage baseplate assembly protein V; afdb-uniprot50 \| AF-A0A708DHR8-F1-MODEL\_V4 \| 1.0 \| 1.907e-08 \| 389 \| 0.447 \| 76 \| 42 \| 0 \| 3 \| 78 \| 5 \| 80 \| Phage baseplate assembly protein V \| Phage baseplate assembly protein V; afdb-uniprot50 \| AF-A0A259UCZ3-F1-MODEL\_V4 \| 1.0 \| 2.743e-11 \| 389 \| 0.204 \| 220 \| 113 \| 5 \| 1 \| 217 \| 1 \| 161 \| Phage-related baseplate assembly protein \| Phage-related baseplate assembly protein; afdb-uniprot50 \| AF-A0A3N2E0Q5-F1-MODEL\_V4 \| 1.0 \| 5.903e-11 \| 388 \| 0.31 \| 148 \| 92 \| 6 \| 5 \| 148 \| 6 \| 147 \| Phage baseplate assembly protein V \| Phage baseplate assembly protein V; afdb-uniprot50 \| AF-A0A1C3EBJ8-F1-MODEL\_V4 \| 1.0 \| 1.202e-11 \| 387 \| 0.223 \| 224 \| 148 \| 10 \| 4 \| 213 \| 7 \| 218 \| Phage\_base\_V domain-containing protein \| Phage\_base\_V domain-containing protein; afdb-uniprot50 \| AF-A0A2J4XY04-F1-MODEL\_V4 \| 1.0 \| 1.506e-08 \| 386 \| 0.383 \| 86 \| 51 \| 1 \| 1 \| 86 \| 1 \| 84 \| Phage baseplate assembly protein V \| Phage baseplate assembly protein V; afdb-uniprot50 \| AF-A0A2T5HH03-F1-MODEL\_V4 \| 1.0 \| 3.907e-11 \| 386 \| 0.29 \| 141 \| 83 \| 5 \| 1 \| 130 \| 1 \| 135 \| Phage baseplate assembly protein V \| Phage baseplate assembly protein V; afdb-uniprot50 \| AF-A0A239C9W6-F1-MODEL\_V4 \| 1.0 \| 6.664e-12 \| 386 \| 0.222 \| 220 \| 153 \| 4 \| 4 \| 217 \| 8 \| 215 \| Phage baseplate assembly protein V \| Phage baseplate assembly protein V; afdb-uniprot50 \| AF-A0A366D029-F1-MODEL\_V4 \| 1.0 \| 3.696e-12 \| 386 \| 0.217 \| 193 \| 123 \| 5 \| 4 \| 194 \| 26 \| 192 \| Phage baseplate assembly protein V \| Phage baseplate assembly protein V; afdb-uniprot50 \| AF-A0A0A6D4K9-F1-MODEL\_V4 \| 1.0 \| 2.16e-10 \| 385 \| 0.235 \| 178 \| 120 \| 7 \| 16 \| 187 \| 5 \| 172 \| Baseplate assembly protein \| Baseplate assembly protein; afdb-uniprot50 \| AF-A0A5S9P2X6-F1-MODEL\_V4 \| 1.0 \| 7.498e-12 \| 384 \| 0.23 \| 230 \| 128 \| 13 \| 1 \| 217 \| 1 \| 194 \| Phage\_base\_V domain-containing protein \| Phage\_base\_V domain-containing protein; afdb-uniprot50 \| AF-A0A5Y3B0Q3-F1-MODEL\_V4 \| 1.0 \| 7.523e-13 \| 384 \| 0.217 \| 234 \| 155 \| 8 \| 1 \| 215 \| 1 \| 225 \| Phage baseplate assembly protein V \| Phage baseplate assembly protein V; afdb-uniprot50 \| AF-A0A7M3MAY2-F1-MODEL\_V4 \| 1.0 \| 1.275e-11 \| 383 \| 0.198 \| 237 \| 163 \| 9 \| 4 \| 217 \| 8 \| 240 \| Uncharacterized protein \| Uncharacterized protein; afdb-uniprot50 \| AF-A0A825DZB9-F1-MODEL\_V4 \| 1.0 \| 3.696e-12 \| 382 \| 0.293 \| 191 \| 118 \| 7 \| 16 \| 195 \| 2 \| 186 \| Phage baseplate assembly protein V \| Phage baseplate assembly protein V; afdb-uniprot50 \| AF-A0A1L6I2B1-F1-MODEL\_V4 \| 1.0 \| 6.642e-11 \| 381 \| 0.256 \| 218 \| 146 \| 9 \| 4 \| 217 \| 8 \| 213 \| Uncharacterized protein \| Uncharacterized protein; afdb-uniprot50 \| AF-A0A3Q0L1B6-F1-MODEL\_V4 \| 1.0 \| 2.446e-12 \| 380 \| 0.25 \| 208 \| 140 \| 7 \| 1 \| 193 \| 1 \| 207 \| Phage P2 baseplate assembly protein gpV \| Phage P2 baseplate assembly protein gpV; afdb-uniprot50 \| AF-A0A0U2B1K8-F1-MODEL\_V4 \| 1.0 \| 1.822e-12 \| 379 \| 0.241 \| 211 \| 118 \| 8 \| 13 \| 216 \| 6 \| 181 \| Uncharacterized protein \| Uncharacterized protein; afdb-uniprot50 \| AF-A0A6I1HQM0-F1-MODEL\_V4 \| 1.0 \| 1.624e-13 \| 379 \| 0.234 \| 315 \| 128 \| 6 \| 3 \| 213 \| 6 \| 311 \| Phage baseplate assembly protein V \| Phage baseplate assembly protein V; afdb-uniprot50 \| AF-A0A6H0T7G9-F1-MODEL\_V4 \| 1.0 \| 2.752e-12 \| 376 \| 0.221 \| 208 \| 128 \| 8 \| 16 \| 217 \| 14 \| 193 \| Phage baseplate protein \| Phage baseplate protein; afdb-uniprot50 \| AF-A0A1E4AHK4-F1-MODEL\_V4 \| 1.0 \| 1.348e-10 \| 376 \| 0.197 \| 208 \| 149 \| 11 \| 16 \| 213 \| 2 \| 201 \| Uncharacterized protein \| Uncharacterized protein; afdb-uniprot50 \| AF-A0A0J9DZ44-F1-MODEL\_V4 \| 1.0 \| 5.923e-12 \| 376 \| 0.259 \| 227 \| 144 \| 10 \| 4 \| 217 \| 7 \| 222 \| Phage baseplate assembly protein V \| Phage baseplate assembly protein V; afdb-uniprot50 \| AF-A0A0P0ADA7-F1-MODEL\_V4 \| 1.0 \| 2.036e-10 \| 375 \| 0.231 \| 194 \| 122 \| 9 \| 16 \| 203 \| 4 \| 176 \| Uncharacterized protein \| Uncharacterized protein; afdb-uniprot50 \| AF-A0A2N0CXD6-F1-MODEL\_V4 \| 1.0 \| 5.923e-12 \| 375 \| 0.263 \| 220 \| 124 \| 6 \| 1 \| 206 \| 1 \| 196 \| Phage baseplate assembly protein V \| Phage baseplate assembly protein V; afdb-uniprot50 \| AF-A0A602TUF8-F1-MODEL\_V4 \| 1.0 \| 5.565e-11 \| 374 \| 0.223 \| 192 \| 123 \| 6 \| 16 \| 203 \| 6 \| 175 \| Phage baseplate assembly protein V \| Phage baseplate assembly protein V; afdb-uniprot50 \| AF-A0A2M7G5P7-F1-MODEL\_V4 \| 1.0 \| 3.671e-10 \| 374 \| 0.204 \| 181 \| 125 \| 7 \| 16 \| 191 \| 12 \| 178 \| Phage baseplate assembly protein V \| Phage baseplate assembly protein V; afdb-uniprot50 \| AF-A0A2S2E577-F1-MODEL\_V4 \| 1.0 \| 1.434e-11 \| 374 \| 0.214 \| 214 \| 144 \| 7 \| 5 \| 217 \| 2 \| 192 \| Phage\_base\_V domain-containing protein \| Phage\_base\_V domain-containing protein; afdb-uniprot50 \| AF-A0A6D2GFI2-F1-MODEL\_V4 \| 1.0 \| 6.241e-10 \| 370 \| 0.244 \| 172 \| 118 \| 5 \| 16 \| 182 \| 6 \| 170 \| Baseplate protein \| Baseplate protein; afdb-uniprot50 \| AF-A0A376EWE4-F1-MODEL\_V4 \| 1.0 \| 6.642e-11 \| 370 \| 0.235 \| 174 \| 110 \| 5 \| 16 \| 185 \| 396 \| 550 \| Gene D protein \| Gene D protein; afdb-uniprot50 \| AF-A0A6C1BQR0-F1-MODEL\_V4 \| 1.0 \| 8.381e-10 \| 369 \| 0.206 \| 131 \| 95 \| 2 \| 16 \| 137 \| 2 \| 132 \| Phage baseplate assembly protein V \| Phage baseplate assembly protein V; afdb-uniprot50 \| AF-A0A7L6UL61-F1-MODEL\_V4 \| 1.0 \| 9.43e-10 \| 369 \| 0.323 \| 167 \| 105 \| 5 \| 32 \| 196 \| 16 \| 176 \| Phage baseplate assembly protein V \| Phage baseplate assembly protein V; afdb-uniprot50 \| AF-A0A7L5ZYB0-F1-MODEL\_V4 \| 1.0 \| 5.529e-09 \| 368 \| 0.192 \| 140 \| 105 \| 2 \| 15 \| 149 \| 3 \| 139 \| Phage baseplate assembly protein V \| Phage baseplate assembly protein V; afdb-uniprot50 \| AF-A0A1G0LH00-F1-MODEL\_V4 \| 1.0 \| 2.725e-09 \| 368 \| 0.254 \| 153 \| 104 \| 6 \| 17 \| 163 \| 2 \| 150 \| Phage\_base\_V domain-containing protein \| Phage\_base\_V domain-containing protein; afdb-uniprot50 \| AF-A0A541BHN8-F1-MODEL\_V4 \| 1.0 \| 6.303e-13 \| 367 \| 0.227 \| 264 \| 150 \| 9 \| 4 \| 217 \| 11 \| 270 \| Phage baseplate assembly protein V \| Phage baseplate assembly protein V; afdb-uniprot50 \| AF-R9B5T6-F1-MODEL\_V4 \| 1.0 \| 1.122e-08 \| 366 \| 0.27 \| 111 \| 76 \| 2 \| 1 \| 111 \| 1 \| 106 \| Phage\_base\_V domain-containing protein \| Phage\_base\_V domain-containing protein; afdb-uniprot50 \| AF-A0A285NJ63-F1-MODEL\_V4 \| 1.0 \| 3.683e-11 \| 366 \| 0.18 \| 222 \| 125 \| 4 \| 4 \| 211 \| 11 \| 189 \| Phage baseplate assembly protein V \| Phage baseplate assembly protein V; afdb-uniprot50 \| AF-A0A2T6F922-F1-MODEL\_V4 \| 1.0 \| 2.586e-11 \| 366 \| 0.212 \| 226 \| 155 \| 9 \| 1 \| 217 \| 1 \| 212 \| Phage baseplate assembly protein V \| Phage baseplate assembly protein V; afdb-uniprot50 \| AF-A0A6B3ITL1-F1-MODEL\_V4 \| 1.0 \| 8.436e-12 \| 365 \| 0.223 \| 233 \| 124 \| 7 \| 1 \| 216 \| 1 \| 193 \| Phage baseplate assembly protein V \| Phage baseplate assembly protein V; afdb-uniprot50 \| AF-A0A2N7G847-F1-MODEL\_V4 \| 1.0 \| 3.274e-11 \| 365 \| 0.231 \| 207 \| 151 \| 6 \| 1 \| 203 \| 1 \| 203 \| Uncharacterized protein \| Uncharacterized protein; afdb-uniprot50 \| AF-A0A838Y7A3-F1-MODEL\_V4 \| 1.0 \| 7.927e-11 \| 364 \| 0.202 \| 202 \| 143 \| 10 \| 3 \| 196 \| 7 \| 198 \| Phage baseplate assembly protein V \| Phage baseplate assembly protein V; afdb-uniprot50 \| AF-A0A4P6G5E4-F1-MODEL\_V4 \| 1.0 \| 4.963e-12 \| 362 \| 0.212 \| 226 \| 137 \| 8 \| 1 \| 217 \| 1 \| 194 \| Phage baseplate protein \| Phage baseplate protein; afdb-uniprot50 \| AF-A0A602MX29-F1-MODEL\_V4 \| 1.0 \| 1.007e-11 \| 362 \| 0.248 \| 217 \| 133 \| 7 \| 1 \| 207 \| 1 \| 197 \| Phage baseplate assembly protein V \| Phage baseplate assembly protein V; afdb-uniprot50 \| AF-A0A2K4IH05-F1-MODEL\_V4 \| 1.0 \| 1.926e-11 \| 362 \| 0.232 \| 211 \| 134 \| 9 \| 2 \| 203 \| 27 \| 218 \| Phage baseplate assembly protein V \| Phage baseplate assembly protein V; afdb-uniprot50 \| AF-A0A4R5HEX7-F1-MODEL\_V4 \| 1.0 \| 5.903e-11 \| 361 \| 0.191 \| 214 \| 137 \| 6 \| 3 \| 208 \| 25 \| 210 \| Phage baseplate assembly protein V \| Phage baseplate assembly protein V; afdb-uniprot50 \| AF-D1AFB2-F1-MODEL\_V4 \| 1.0 \| 2.9e-10 \| 360 \| 0.163 \| 208 \| 138 \| 6 \| 12 \| 216 \| 3 \| 177 \| Phage baseplate assembly protein V \| Phage baseplate assembly protein V; afdb-uniprot50 \| AF-A0A327QB94-F1-MODEL\_V4 \| 1.0 \| 9.461e-11 \| 360 \| 0.231 \| 207 \| 128 \| 11 \| 16 \| 217 \| 3 \| 183 \| Phage baseplate assembly protein V \| Phage baseplate assembly protein V; afdb-uniprot50 \| AF-A0A165THH2-F1-MODEL\_V4 \| 1.0 \| 3.086e-11 \| 359 \| 0.237 \| 232 \| 117 \| 4 \| 4 \| 217 \| 15 \| 204 \| Phage-related baseplate assembly protein \| Phage-related baseplate assembly protein; afdb-uniprot50 \| AF-G8PUM2-F1-MODEL\_V4 \| 1.0 \| 2.043e-11 \| 359 \| 0.241 \| 232 \| 116 \| 4 \| 4 \| 217 \| 15 \| 204 \| Phage-related baseplate assembly protein V \| Phage-related baseplate assembly protein V; afdb-uniprot50 \| AF-A0A2S7JRA1-F1-MODEL\_V4 \| 1.0 \| 5.903e-11 \| 359 \| 0.237 \| 215 \| 136 \| 7 \| 16 \| 217 \| 8 \| 207 \| Phage baseplate assembly protein V \| Phage baseplate assembly protein V; afdb-uniprot50 \| AF-A0A411WI65-F1-MODEL\_V4 \| 1.0 \| 1.926e-11 \| 358 \| 0.219 \| 205 \| 135 \| 8 \| 16 \| 217 \| 3 \| 185 \| Phage baseplate assembly protein V \| Phage baseplate assembly protein V; afdb-uniprot50 \| AF-A0A1N6M5J8-F1-MODEL\_V4 \| 1.0 \| 3.473e-11 \| 358 \| 0.219 \| 223 \| 141 \| 9 \| 1 \| 216 \| 4 \| 200 \| Phage-related baseplate assembly protein \| Phage-related baseplate assembly protein; afdb-uniprot50 \| AF-A0A075KC29-F1-MODEL\_V4 \| 1.0 \| 2.029e-09 \| 357 \| 0.211 \| 142 \| 100 \| 3 \| 1 \| 139 \| 1 \| 133 \| Phage baseplate assembly protein V \| Phage baseplate assembly protein V; afdb-uniprot50 \| AF-A0A0C1EUH5-F1-MODEL\_V4 \| 1.0 \| 1.129e-10 \| 357 \| 0.25 \| 188 \| 120 \| 8 \| 17 \| 197 \| 2 \| 175 \| Baseplate protein \| Baseplate protein; afdb-uniprot50 \| AF-A0A2G6EWU3-F1-MODEL\_V4 \| 1.0 \| 7.901e-10 \| 355 \| 0.246 \| 162 \| 102 \| 6 \| 7 \| 158 \| 8 \| 159 \| Phage\_base\_V domain-containing protein \| Phage\_base\_V domain-containing protein; afdb-uniprot50 \| AF-A0A660N066-F1-MODEL\_V4 \| 1.0 \| 4.158e-12 \| 355 \| 0.248 \| 205 \| 122 \| 7 \| 16 \| 209 \| 12 \| 195 \| Phage baseplate assembly protein V \| Phage baseplate assembly protein V; afdb-uniprot50 \| AF-A0A4U7BE93-F1-MODEL\_V4 \| 1.0 \| 2.167e-11 \| 354 \| 0.222 \| 189 \| 125 \| 7 \| 16 \| 203 \| 2 \| 169 \| Phage baseplate assembly protein V \| Phage baseplate assembly protein V; afdb-uniprot50 \| AF-A0A7D5LN82-F1-MODEL\_V4 \| 1.0 \| 7.901e-10 \| 353 \| 0.21 \| 176 \| 122 \| 6 \| 16 \| 186 \| 5 \| 168 \| Phage baseplate assembly protein V \| Phage baseplate assembly protein V; afdb-uniprot50 \| AF-A0A5C4P976-F1-MODEL\_V4 \| 1.0 \| 1.064e-10 \| 351 \| 0.238 \| 168 \| 110 \| 2 \| 2 \| 169 \| 1 \| 150 \| Phage baseplate assembly protein V \| Phage baseplate assembly protein V; afdb-uniprot50 \| AF-A0A1H1G342-F1-MODEL\_V4 \| 1.0 \| 2.276e-08 \| 351 \| 0.19 \| 126 \| 97 \| 3 \| 4 \| 128 \| 8 \| 129 \| Uncharacterized protein \| Uncharacterized protein; afdb-uniprot50 \| AF-A0A840ML35-F1-MODEL\_V4 \| 1.0 \| 1.712e-11 \| 351 \| 0.28 \| 200 \| 116 \| 8 \| 16 \| 197 \| 12 \| 201 \| Phage baseplate assembly protein V \| Phage baseplate assembly protein V; afdb-uniprot50 \| AF-A0A0D8L226-F1-MODEL\_V4 \| 1.0 \| 1.339e-08 \| 350 \| 0.329 \| 170 \| 106 \| 5 \| 49 \| 217 \| 2 \| 164 \| Baseplate assembly protein \| Baseplate assembly protein; afdb-uniprot50 \| AF-A0A4R5W1B5-F1-MODEL\_V4 \| 1.0 \| 1.521e-11 \| 348 \| 0.218 \| 224 \| 128 \| 11 \| 1 \| 216 \| 1 \| 185 \| Phage baseplate assembly protein V \| Phage baseplate assembly protein V; afdb-uniprot50 \| AF-A0A7C3GU98-F1-MODEL\_V4 \| 1.0 \| 4.144e-11 \| 348 \| 0.165 \| 217 \| 155 \| 5 \| 4 \| 217 \| 8 \| 201 \| Phage baseplate assembly protein V \| Phage baseplate assembly protein V; afdb-uniprot50 \| AF-A0A5A8F6F2-F1-MODEL\_V4 \| 1.0 \| 6.642e-11 \| 348 \| 0.147 \| 217 \| 151 \| 7 \| 4 \| 208 \| 11 \| 205 \| Phage baseplate assembly protein V \| Phage baseplate assembly protein V; afdb-uniprot50 \| AF-A0A257PQI7-F1-MODEL\_V4 \| 1.0 \| 2.036e-10 \| 347 \| 0.2 \| 220 \| 120 \| 7 \| 4 \| 217 \| 5 \| 174 \| Phage\_base\_V domain-containing protein \| Phage\_base\_V domain-containing protein; afdb-uniprot50 \| AF-A0A6L8M2W9-F1-MODEL\_V4 \| 1.0 \| 2.43e-10 \| 346 \| 0.216 \| 222 \| 152 \| 10 \| 4 \| 217 \| 10 \| 217 \| Phage baseplate assembly protein V \| Phage baseplate assembly protein V; afdb-uniprot50 \| AF-A0A5B9Y7V9-F1-MODEL\_V4 \| 1.0 \| 1.343e-09 \| 345 \| 0.207 \| 183 \| 122 \| 4 \| 12 \| 191 \| 9 \| 171 \| Phage baseplate assembly protein V \| Phage baseplate assembly protein V; afdb-uniprot50 \| AF-A0A1D2QMX7-F1-MODEL\_V4 \| 1.0 \| 1.348e-10 \| 345 \| 0.244 \| 209 \| 122 \| 6 \| 4 \| 208 \| 12 \| 188 \| Phage\_base\_V domain-containing protein \| Phage\_base\_V domain-containing protein; afdb-uniprot50 \| AF-A0A5C8BDY7-F1-MODEL\_V4 \| 1.0 \| 3.647e-08 \| 344 \| 0.238 \| 126 \| 91 \| 3 \| 14 \| 137 \| 5 \| 127 \| Phage baseplate assembly protein V \| Phage baseplate assembly protein V; afdb-uniprot50 \| AF-A0A345J3I8-F1-MODEL\_V4 \| 1.0 \| 9.492e-12 \| 344 \| 0.237 \| 219 \| 145 \| 8 \| 1 \| 211 \| 2 \| 206 \| Phage baseplate assembly protein V \| Phage baseplate assembly protein V; afdb-uniprot50 \| AF-A0A4Q8MEL0-F1-MODEL\_V4 \| 1.0 \| 3.907e-11 \| 343 \| 0.197 \| 228 \| 100 \| 3 \| 4 \| 216 \| 20 \| 179 \| Uncharacterized protein \| Uncharacterized protein; afdb-uniprot50 \| AF-A0A3G2EA58-F1-MODEL\_V4 \| 1.0 \| 7.927e-11 \| 342 \| 0.186 \| 220 \| 143 \| 8 \| 3 \| 217 \| 4 \| 192 \| Phage\_base\_V domain-containing protein \| Phage\_base\_V domain-containing protein; afdb-uniprot50 \| AF-A0A2S0JB32-F1-MODEL\_V4 \| 1.0 \| 2.43e-10 \| 341 \| 0.206 \| 189 \| 108 \| 5 \| 16 \| 203 \| 5 \| 152 \| Phage baseplate assembly protein V \| Phage baseplate assembly protein V; afdb-uniprot50 \| AF-A0A1M7RIP4-F1-MODEL\_V4 \| 1.0 \| 1.198e-10 \| 341 \| 0.224 \| 218 \| 138 \| 10 \| 16 \| 217 \| 5 \| 207 \| Phage baseplate assembly protein V \| Phage baseplate assembly protein V; afdb-uniprot50 \| AF-A0A2V2AA94-F1-MODEL\_V4 \| 1.0 \| 3.461e-10 \| 341 \| 0.216 \| 254 \| 127 \| 8 \| 1 \| 217 \| 1 \| 219 \| Phage baseplate assembly protein V \| Phage baseplate assembly protein V; afdb-uniprot50 \| AF-A0A4Y7RWH3-F1-MODEL\_V4 \| 1.0 \| 4.396e-11 \| 340 \| 0.242 \| 194 \| 123 \| 5 \| 11 \| 186 \| 4 \| 191 \| Uncharacterized protein \| Uncharacterized protein; afdb-uniprot50 \| AF-A0A7T5ENC6-F1-MODEL\_V4 \| 1.0 \| 1.348e-10 \| 339 \| 0.21 \| 195 \| 131 \| 8 \| 12 \| 202 \| 2 \| 177 \| Phage baseplate assembly protein V \| Phage baseplate assembly protein V; afdb-uniprot50 \| AF-A0A1V4IV34-F1-MODEL\_V4 \| 1.0 \| 4.632e-09 \| 336 \| 0.228 \| 149 \| 99 \| 4 \| 1 \| 145 \| 1 \| 137 \| Phage-related baseplate assembly protein \| Phage-related baseplate assembly protein; afdb-uniprot50 \| AF-A0A1W1Z4L7-F1-MODEL\_V4 \| 1.0 \| 3.881e-09 \| 336 \| 0.189 \| 148 \| 106 \| 4 \| 4 \| 138 \| 18 \| 164 \| Type VI secretion system, phage-baseplate injector \| Type VI secretion system, phage-baseplate injector; afdb-uniprot50 \| AF-A0A6G5QN06-F1-MODEL\_V4 \| 1.0 \| 1.81e-10 \| 335 \| 0.259 \| 189 \| 103 \| 7 \| 16 \| 203 \| 2 \| 154 \| Phage baseplate assembly protein V \| Phage baseplate assembly protein V; afdb-uniprot50 \| AF-A0A1H4QYN2-F1-MODEL\_V4 \| 1.0 \| 1.133e-11 \| 335 \| 0.216 \| 226 \| 143 \| 9 \| 1 \| 217 \| 1 \| 201 \| Phage\_base\_V domain-containing protein \| Phage\_base\_V domain-containing protein; afdb-uniprot50 \| AF-A0A2X3AA95-F1-MODEL\_V4 \| 1.0 \| 5.903e-11 \| 333 \| 0.333 \| 186 \| 103 \| 4 \| 32 \| 217 \| 2 \| 166 \| Baseplate assembly protein V \| Baseplate assembly protein V; afdb-uniprot50 \| AF-B3QTI6-F1-MODEL\_V4 \| 1.0 \| 7.022e-10 \| 333 \| 0.201 \| 208 \| 129 \| 7 \| 16 \| 214 \| 2 \| 181 \| Phage baseplate assembly protein V \| Phage baseplate assembly protein V; afdb-uniprot50 \| AF-A0A2X0V6P6-F1-MODEL\_V4 \| 1.0 \| 1.27e-10 \| 333 \| 0.238 \| 210 \| 124 \| 6 \| 1 \| 193 \| 1 \| 191 \| Phage P2 baseplate assembly protein gpV \| Phage P2 baseplate assembly protein gpV; afdb-uniprot50 \| AF-Q602Z7-F1-MODEL\_V4 \| 1.0 \| 7.901e-10 \| 333 \| 0.219 \| 196 \| 135 \| 8 \| 16 \| 203 \| 13 \| 198 \| Prophage MuMc02, baseplate assembly protein V \| Prophage MuMc02, baseplate assembly protein V; afdb-uniprot50 \| AF-K1K036-F1-MODEL\_V4 \| 1.0 \| 5.565e-11 \| 332 \| 0.264 \| 204 \| 128 \| 4 \| 15 \| 196 \| 12 \| 215 \| Phage baseplate assembly protein V \| Phage baseplate assembly protein V; afdb-uniprot50 \| AF-A0A2E2NAE7-F1-MODEL\_V4 \| 1.0 \| 1.19e-08 \| 330 \| 0.253 \| 134 \| 78 \| 3 \| 1 \| 134 \| 1 \| 112 \| Phage\_base\_V domain-containing protein \| Phage\_base\_V domain-containing protein; afdb-uniprot50 \| AF-A0A827QZX9-F1-MODEL\_V4 \| 1.0 \| 7.449e-10 \| 329 \| 0.218 \| 192 \| 112 \| 8 \| 16 \| 203 \| 2 \| 159 \| Phage baseplate assembly protein V \| Phage baseplate assembly protein V; afdb-uniprot50 \| AF-A0A375J5W5-F1-MODEL\_V4 \| 1.0 \| 4.396e-11 \| 329 \| 0.357 \| 193 \| 99 \| 5 \| 32 \| 217 \| 2 \| 176 \| Bacteriophage P2 Baseplate assembly protein GPV \| Bacteriophage P2 Baseplate assembly protein GPV; afdb-uniprot50 \| AF-A0A4Q6DAY4-F1-MODEL\_V4 \| 1.0 \| 1.792e-07 \| 328 \| 0.289 \| 107 \| 76 \| 0 \| 40 \| 146 \| 1 \| 107 \| Phage baseplate assembly protein V \| Phage baseplate assembly protein V; afdb-uniprot50 \| AF-A0A348AJ13-F1-MODEL\_V4 \| 1.0 \| 1.343e-09 \| 327 \| 0.205 \| 195 \| 111 \| 6 \| 12 \| 203 \| 4 \| 157 \| Phage-related baseplate assembly protein \| Phage-related baseplate assembly protein; afdb-uniprot50 \| AF-G2IX47-F1-MODEL\_V4 \| 1.0 \| 3.076e-10 \| 327 \| 0.185 \| 199 \| 144 \| 9 \| 16 \| 208 \| 10 \| 196 \| Phage baseplate assembly protein V \| Phage baseplate assembly protein V; afdb-uniprot50 \| AF-A0A7J4VI64-F1-MODEL\_V4 \| 1.0 \| 1.81e-10 \| 326 \| 0.207 \| 227 \| 120 \| 6 \| 1 \| 217 \| 1 \| 177 \| Phage\_base\_V domain-containing protein \| Phage\_base\_V domain-containing protein; afdb-uniprot50 \| AF-A0A139D6P1-F1-MODEL\_V4 \| 1.0 \| 4.648e-10 \| 326 \| 0.203 \| 211 \| 133 \| 7 \| 1 \| 203 \| 1 \| 184 \| Phage baseplate assembly protein V \| Phage baseplate assembly protein V; afdb-uniprot50 \| AF-A0A081NV30-F1-MODEL\_V4 \| 1.0 \| 3.252e-09 \| 323 \| 0.233 \| 171 \| 103 \| 6 \| 12 \| 179 \| 4 \| 149 \| Uncharacterized protein \| Uncharacterized protein; afdb-uniprot50 \| AF-A0A5E5BJ14-F1-MODEL\_V4 \| 1.0 \| 2.881e-08 \| 320 \| 0.204 \| 166 \| 124 \| 4 \| 3 \| 161 \| 7 \| 171 \| Phage-related baseplate assembly protein \| Phage-related baseplate assembly protein; afdb-uniprot50 \| AF-A0A096AIG4-F1-MODEL\_V4 \| 1.0 \| 7.022e-10 \| 319 \| 0.229 \| 192 \| 115 \| 7 \| 15 \| 203 \| 10 \| 171 \| Baseplate protein \| Baseplate protein; afdb-uniprot50 \| AF-A0A0G3W8Q4-F1-MODEL\_V4 \| 1.0 \| 3.45e-09 \| 317 \| 0.198 \| 166 \| 117 \| 4 \| 1 \| 157 \| 3 \| 161 \| Phage baseplate assembly protein V \| Phage baseplate assembly protein V; afdb-uniprot50 \| AF-R7JNY1-F1-MODEL\_V4 \| 1.0 \| 6.598e-09 \| 317 \| 0.24 \| 162 \| 106 \| 6 \| 16 \| 172 \| 2 \| 151 \| Putative bacteriophage baseplate protein \| Putative bacteriophage baseplate protein; afdb-uniprot50 \| AF-A0A380Z445-F1-MODEL\_V4 \| 1.0 \| 2.036e-10 \| 316 \| 0.221 \| 190 \| 118 \| 6 \| 16 \| 193 \| 10 \| 181 \| Phage P2 baseplate assembly protein gpV \| Phage P2 baseplate assembly protein gpV; afdb-uniprot50 \| AF-W4AM65-F1-MODEL\_V4 \| 1.0 \| 8.353e-09 \| 313 \| 0.196 \| 183 \| 113 \| 3 \| 11 \| 190 \| 3 \| 154 \| Phage baseplate assembly protein V \| Phage baseplate assembly protein V; afdb-uniprot50 \| AF-A0A1B9NZR2-F1-MODEL\_V4 \| 1.0 \| 1.343e-09 \| 312 \| 0.168 \| 196 \| 136 \| 5 \| 4 \| 197 \| 22 \| 192 \| Uncharacterized protein \| Uncharacterized protein; afdb-uniprot50 \| AF-A0A662Z9P3-F1-MODEL\_V4 \| 1.0 \| 2.569e-09 \| 310 \| 0.198 \| 166 \| 104 \| 4 \| 1 \| 137 \| 1 \| 166 \| Phage baseplate assembly protein V \| Phage baseplate assembly protein V; afdb-uniprot50 \| AF-A0A285MD33-F1-MODEL\_V4 \| 1.0 \| 1.706e-10 \| 310 \| 0.207 \| 236 \| 112 \| 5 \| 4 \| 217 \| 13 \| 195 \| Phage baseplate assembly protein V \| Phage baseplate assembly protein V; afdb-uniprot50 \| AF-A0A6S7B3J0-F1-MODEL\_V4 \| 1.0 \| 5.884e-10 \| 308 \| 0.202 \| 207 \| 121 \| 7 \| 1 \| 191 \| 1 \| 179 \| Phage\_base\_V domain-containing protein \| Phage\_base\_V domain-containing protein; afdb-uniprot50 \| AF-A0A561IRQ8-F1-MODEL\_V4 \| 1.0 \| 7.045e-11 \| 307 \| 0.213 \| 220 \| 150 \| 10 \| 15 \| 216 \| 14 \| 228 \| Phage baseplate assembly protein V \| Phage baseplate assembly protein V; afdb-uniprot50 \| AF-A0A209AE83-F1-MODEL\_V4 \| 1.0 \| 3.076e-10 \| 306 \| 0.223 \| 179 \| 83 \| 6 \| 1 \| 125 \| 1 \| 177 \| Phage\_base\_V domain-containing protein \| Phage\_base\_V domain-containing protein; afdb-uniprot50 \| AF-Q31HU0-F1-MODEL\_V4 \| 1.0 \| 2.291e-10 \| 306 \| 0.244 \| 225 \| 142 \| 10 \| 16 \| 217 \| 2 \| 221 \| Phage-related baseplate assembly protein V \| Phage-related baseplate assembly protein V; afdb-uniprot50 \| AF-A0A074MGB6-F1-MODEL\_V4 \| 1.0 \| 7.399e-08 \| 303 \| 0.15 \| 160 \| 117 \| 3 \| 12 \| 168 \| 3 \| 146 \| Uncharacterized protein \| Uncharacterized protein; afdb-uniprot50 \| AF-A0A3C0RRX5-F1-MODEL\_V4 \| 1.0 \| 6.598e-09 \| 303 \| 0.18 \| 161 \| 115 \| 6 \| 4 \| 158 \| 8 \| 157 \| Uncharacterized protein \| Uncharacterized protein; afdb-uniprot50 \| AF-D1Y2B5-F1-MODEL\_V4 \| 1.0 \| 1.122e-08 \| 302 \| 0.189 \| 148 \| 101 \| 4 \| 16 \| 145 \| 9 \| 155 \| Phage-related baseplate assembly protein \| Phage-related baseplate assembly protein; afdb-uniprot50 \| AF-A0A561QKN0-F1-MODEL\_V4 \| 1.0 \| 1.689e-07 \| 300 \| 0.18 \| 133 \| 101 \| 3 \| 16 \| 145 \| 2 \| 129 \| Phage baseplate assembly protein V \| Phage baseplate assembly protein V; afdb-uniprot50 \| AF-A0A1C3NRR4-F1-MODEL\_V4 \| 1.0 \| 1.061e-09 \| 300 \| 0.292 \| 178 \| 99 \| 5 \| 40 \| 217 \| 3 \| 153 \| Phage-related baseplate protein \| Phage-related baseplate protein; afdb-uniprot50 \| AF-A0A149PP11-F1-MODEL\_V4 \| 1.0 \| 1.603e-09 \| 299 \| 0.368 \| 179 \| 93 \| 6 \| 40 \| 217 \| 1 \| 160 \| Baseplate assembly protein \| Baseplate assembly protein; afdb-uniprot50 \| AF-A0A239EJ28-F1-MODEL\_V4 \| 1.0 \| 1.913e-09 \| 297 \| 0.301 \| 146 \| 77 \| 5 \| 4 \| 149 \| 5 \| 125 \| Phage baseplate assembly protein V \| Phage baseplate assembly protein V; afdb-uniprot50 \| AF-A0A285M2J1-F1-MODEL\_V4 \| 1.0 \| 2.414e-08 \| 297 \| 0.182 \| 159 \| 109 \| 7 \| 1 \| 145 \| 1 \| 152 \| Uncharacterized protein \| Uncharacterized protein; afdb-uniprot50 \| AF-A0A8A8M598-F1-MODEL\_V4 \| 1.0 \| 3.683e-11 \| 293 \| 0.238 \| 252 \| 143 \| 11 \| 1 \| 217 \| 1 \| 238 \| Uncharacterized protein \| Uncharacterized protein; afdb-uniprot50 \| AF-A0A7Y3Z3N5-F1-MODEL\_V4 \| 1.0 \| 2.577e-10 \| 293 \| 0.199 \| 241 \| 158 \| 9 \| 4 \| 213 \| 11 \| 247 \| Phage baseplate assembly protein V \| Phage baseplate assembly protein V; afdb-uniprot50 \| AF-G9PUJ1-F1-MODEL\_V4 \| 1.0 \| 6.576e-08 \| 291 \| 0.172 \| 145 \| 102 \| 2 \| 13 \| 139 \| 5 \| 149 \| Uncharacterized protein \| Uncharacterized protein; afdb-uniprot50 \| AF-A0A843SF22-F1-MODEL\_V4 \| 1.0 \| 1.27e-10 \| 291 \| 0.205 \| 238 \| 145 \| 10 \| 4 \| 217 \| 7 \| 224 \| Phage baseplate assembly protein V \| Phage baseplate assembly protein V; afdb-uniprot50 \| AF-A0A547PW88-F1-MODEL\_V4 \| 1.0 \| 6.62e-10 \| 287 \| 0.224 \| 214 \| 98 \| 6 \| 4 \| 217 \| 54 \| 199 \| Phage baseplate assembly protein V \| Phage baseplate assembly protein V; afdb-uniprot50 \| AF-D8IV20-F1-MODEL\_V4 \| 1.0 \| 4.367e-09 \| 284 \| 0.252 \| 206 \| 133 \| 10 \| 16 \| 217 \| 8 \| 196 \| Bacteriophage baseplate assembly (GpV) protein \| Bacteriophage baseplate assembly (GpV) protein; afdb-uniprot50 \| AF-A0A1I5W3T5-F1-MODEL\_V4 \| 1.0 \| 8.353e-09 \| 283 \| 0.201 \| 218 \| 103 \| 8 \| 1 \| 217 \| 1 \| 148 \| Phage baseplate assembly protein V \| Phage baseplate assembly protein V; afdb-uniprot50 \| AF-A0A181WXG7-F1-MODEL\_V4 \| 1.0 \| 1.81e-10 \| 280 \| 0.259 \| 231 \| 125 \| 8 \| 16 \| 203 \| 6 \| 233 \| Baseplate assembly protein V \| Baseplate assembly protein V; afdb-uniprot50 \| AF-A0A1I5MP97-F1-MODEL\_V4 \| 1.0 \| 1.334e-07 \| 279 \| 0.165 \| 151 \| 109 \| 5 \| 1 \| 141 \| 1 \| 144 \| Phage P2 baseplate assembly protein gpV \| Phage P2 baseplate assembly protein gpV; afdb-uniprot50 \| AF-A0A377Z5C8-F1-MODEL\_V4 \| 1.0 \| 1.894e-06 \| 273 \| 0.346 \| 130 \| 80 \| 3 \| 67 \| 196 \| 2 \| 126 \| Putative prophage baseplate assembly protein \| Putative prophage baseplate assembly protein; afdb-uniprot50 \| AF-A0A847G8C6-F1-MODEL\_V4 \| 1.0 \| 3.659e-09 \| 271 \| 0.204 \| 244 \| 128 \| 9 \| 16 \| 198 \| 2 \| 240 \| Phage baseplate assembly protein V \| Phage baseplate assembly protein V; afdb-uniprot50 \| AF-A0A842J1J9-F1-MODEL\_V4 \| 1.0 \| 2.406e-07 \| 270 \| 0.15 \| 180 \| 137 \| 7 \| 16 \| 192 \| 7 \| 173 \| Uncharacterized protein \| Uncharacterized protein; afdb-uniprot50 \| AF-A0A5E5P1X8-F1-MODEL\_V4 \| 1.0 \| 3.241e-08 \| 270 \| 0.174 \| 223 \| 156 \| 10 \| 3 \| 209 \| 7 \| 217 \| Baseplate assembly protein \| Baseplate assembly protein; afdb-uniprot50 \| AF-A0A5C6Y6D1-F1-MODEL\_V4 \| 1.0 \| 4.632e-09 \| 267 \| 0.148 \| 236 \| 144 \| 10 \| 3 \| 216 \| 5 \| 205 \| Baseplate assembly protein \| Baseplate assembly protein; afdb-uniprot50 \| AF-A0A447M7I0-F1-MODEL\_V4 \| 1.0 \| 3.231e-07 \| 266 \| 0.293 \| 143 \| 90 \| 3 \| 40 \| 182 \| 6 \| 137 \| Phage baseplate assembly protein V \| Phage baseplate assembly protein V; afdb-uniprot50 \| AF-A0A1S2FCU4-F1-MODEL\_V4 \| 1.0 \| 5.195e-08 \| 265 \| 0.186 \| 198 \| 120 \| 9 \| 1 \| 195 \| 1 \| 160 \| Uncharacterized protein \| Uncharacterized protein; afdb-uniprot50 \| AF-A0A6G7AI25-F1-MODEL\_V4 \| 1.0 \| 1.913e-09 \| 264 \| 0.245 \| 224 \| 112 \| 7 \| 14 \| 217 \| 11 \| 197 \| Phage baseplate assembly protein V \| Phage baseplate assembly protein V; afdb-uniprot50 \| AF-A0A3A9CSA3-F1-MODEL\_V4 \| 1.0 \| 2.406e-07 \| 262 \| 0.177 \| 192 \| 136 \| 8 \| 14 \| 202 \| 3 \| 175 \| Uncharacterized protein \| Uncharacterized protein; afdb-uniprot50 \| AF-A0A826UHE7-F1-MODEL\_V4 \| 1.0 \| 7.424e-09 \| 261 \| 0.337 \| 169 \| 91 \| 4 \| 49 \| 217 \| 1 \| 148 \| Phage baseplate assembly protein V \| Phage baseplate assembly protein V; afdb-uniprot50 \| AF-U2F4C9-F1-MODEL\_V4 \| 1.0 \| 2.552e-07 \| 260 \| 0.258 \| 186 \| 116 \| 8 \| 40 \| 209 \| 3 \| 182 \| Baseplate assembly protein V, putative \| Baseplate assembly protein V, putative; afdb-uniprot50 \| AF-A0A2J9HGD4-F1-MODEL\_V4 \| 1.0 \| 6.22e-09 \| 258 \| 0.32 \| 184 \| 109 \| 3 \| 48 \| 215 \| 7 \| 190 \| Uncharacterized protein \| Uncharacterized protein; afdb-uniprot50 \| AF-A0A2A2HCZ4-F1-MODEL\_V4 \| 1.0 \| 5.178e-07 \| 256 \| 0.282 \| 156 \| 101 \| 4 \| 49 \| 196 \| 1 \| 153 \| Baseplate protein \| Baseplate protein; afdb-uniprot50 \| AF-A0A353GTI7-F1-MODEL\_V4 \| 1.0 \| 1.262e-08 \| 256 \| 0.206 \| 213 \| 125 \| 8 \| 1 \| 196 \| 23 \| 208 \| Uncharacterized protein \| Uncharacterized protein; afdb-uniprot50 \| AF-A0A1H4BQX8-F1-MODEL\_V4 \| 1.0 \| 5.884e-10 \| 254 \| 0.319 \| 194 \| 105 \| 5 \| 49 \| 217 \| 2 \| 193 \| Phage baseplate assembly protein V \| Phage baseplate assembly protein V; afdb-uniprot50 \| AF-A0A133NE44-F1-MODEL\_V4 \| 1.0 \| 5.474e-06 \| 251 \| 0.163 \| 183 \| 136 \| 8 \| 16 \| 195 \| 7 \| 175 \| Uncharacterized protein \| Uncharacterized protein; afdb-uniprot50 \| AF-A0A1M6IZQ5-F1-MODEL\_V4 \| 1.0 \| 5.492e-07 \| 247 \| 0.15 \| 179 \| 143 \| 5 \| 5 \| 182 \| 6 \| 176 \| Bacteriophage Mu Gp45 protein \| Bacteriophage Mu Gp45 protein; afdb-uniprot50 \| AF-A0A367FZE3-F1-MODEL\_V4 \| 1.0 \| 1.415e-07 \| 242 \| 0.201 \| 169 \| 122 \| 4 \| 15 \| 179 \| 4 \| 163 \| Uncharacterized protein \| Uncharacterized protein; afdb-uniprot50 \| AF-A0A6C8C1W7-F1-MODEL\_V4 \| 1.0 \| 1.786e-06 \| 239 \| 0.314 \| 127 \| 76 \| 4 \| 40 \| 158 \| 4 \| 127 \| Phage baseplate assembly protein V \| Phage baseplate assembly protein V; afdb-uniprot50 \| AF-A0A198XEV1-F1-MODEL\_V4 \| 1.0 \| 2.707e-07 \| 234 \| 0.235 \| 178 \| 100 \| 6 \| 40 \| 216 \| 3 \| 145 \| Baseplate assembly protein V \| Baseplate assembly protein V; afdb-uniprot50 \| AF-A0A2S9WYZ5-F1-MODEL\_V4 \| 1.0 \| 9.398e-09 \| 233 \| 0.191 \| 224 \| 140 \| 9 \| 16 \| 217 \| 3 \| 207 \| Phage\_base\_V domain-containing protein \| Phage\_base\_V domain-containing protein; afdb-uniprot50 \| AF-R0KQF0-F1-MODEL\_V4 \| 1.0 \| 2.852e-05 \| 231 \| 0.32 \| 128 \| 82 \| 3 \| 67 \| 194 \| 2 \| 124 \| Baseplate assembly protein V \| Baseplate assembly protein V; afdb-uniprot50 \| AF-A0A4P0TM83-F1-MODEL\_V4 \| 1.0 \| 2.871e-07 \| 231 \| 0.244 \| 180 \| 102 \| 5 \| 32 \| 203 \| 2 \| 155 \| Putative baseplate assembly protein V \| Putative baseplate assembly protein V; afdb-uniprot50 \| AF-A0A064AML4-F1-MODEL\_V4 \| 1.0 \| 1.587e-06 \| 224 \| 0.189 \| 179 \| 117 \| 7 \| 19 \| 190 \| 2 \| 159 \| Phage baseplate protein \| Phage baseplate protein; afdb-uniprot50 \| AF-A0A376FH76-F1-MODEL\_V4 \| 1.0 \| 4.572e-05 \| 223 \| 0.317 \| 126 \| 81 \| 4 \| 67 \| 192 \| 3 \| 123 \| Phage Baseplate Assembly protein V \| Phage Baseplate Assembly protein V; afdb-uniprot50 \| AF-D0LVW0-F1-MODEL\_V4 \| 1.0 \| 1.901e-07 \| 223 \| 0.179 \| 217 \| 154 \| 10 \| 4 \| 203 \| 7 \| 216 \| Phage\_base\_V domain-containing protein \| Phage\_base\_V domain-containing protein; afdb-uniprot50 \| AF-A0A3N2DJS8-F1-MODEL\_V4 \| 1.0 \| 4.338e-07 \| 223 \| 0.197 \| 218 \| 155 \| 10 \| 16 \| 215 \| 377 \| 592 \| Type VI secretion system secreted protein VgrG \| Type VI secretion system secreted protein VgrG; afdb-uniprot50 \| AF-A0A210A7I4-F1-MODEL\_V4 \| 1.0 \| 2.268e-07 \| 221 \| 0.337 \| 163 \| 87 \| 4 \| 55 \| 217 \| 1 \| 142 \| Baseplate assembly protein \| Baseplate assembly protein; afdb-uniprot50 \| AF-A0A2X3JT57-F1-MODEL\_V4 \| 1.0 \| 8.831e-08 \| 220 \| 0.323 \| 170 \| 94 \| 4 \| 48 \| 217 \| 20 \| 168 \| Baseplate assembly protein V \| Baseplate assembly protein V; afdb-uniprot50 \| AF-A0A828EP35-F1-MODEL\_V4 \| 1.0 \| 2.016e-07 \| 218 \| 0.327 \| 162 \| 88 \| 4 \| 56 \| 217 \| 1 \| 141 \| Phage baseplate assembly protein V \| Phage baseplate assembly protein V; afdb-uniprot50 \| AF-A0A1Z5H6B0-F1-MODEL\_V4 \| 1.0 \| 1.496e-06 \| 218 \| 0.217 \| 175 \| 94 \| 4 \| 48 \| 216 \| 4 \| 141 \| Phage\_base\_V domain-containing protein \| Phage\_base\_V domain-containing protein; afdb-uniprot50 \| AF-A0A6F9JMD1-F1-MODEL\_V4 \| 1.0 \| 3.856e-07 \| 214 \| 0.26 \| 173 \| 112 \| 6 \| 45 \| 203 \| 2 \| 172 \| Phage baseplate assembly protein V \| Phage baseplate assembly protein V; afdb-uniprot50 \| AF-A0A847TFE5-F1-MODEL\_V4 \| 1.0 \| 3.231e-07 \| 213 \| 0.176 \| 238 \| 160 \| 8 \| 16 \| 217 \| 365 \| 602 \| Uncharacterized protein \| Uncharacterized protein; afdb-uniprot50 \| AF-A0A4U0Z1D4-F1-MODEL\_V4 \| 1.0 \| 3.046e-07 \| 212 \| 0.23 \| 217 \| 100 \| 7 \| 1 \| 217 \| 146 \| 295 \| Phage baseplate assembly protein V \| Phage baseplate assembly protein V; afdb-uniprot50 \| AF-Q1H2H9-F1-MODEL\_V4 \| 1.0 \| 3.856e-07 \| 212 \| 0.148 \| 222 \| 126 \| 8 \| 15 \| 193 \| 344 \| 545 \| Rhs element Vgr protein \| Rhs element Vgr protein; afdb-uniprot50 \| AF-A0A545SPM8-F1-MODEL\_V4 \| 1.0 \| 1.182e-06 \| 212 \| 0.156 \| 243 \| 164 \| 12 \| 16 \| 217 \| 374 \| 616 \| Type VI secretion system tip protein VgrG \| Type VI secretion system tip protein VgrG; afdb-uniprot50 \| AF-A0A2Z5T171-F1-MODEL\_V4 \| 1.0 \| 3.869e-08 \| 210 \| 0.212 \| 250 \| 142 \| 12 \| 1 \| 197 \| 2 \| 249 \| Uncharacterized protein \| Uncharacterized protein; afdb-uniprot50 \| AF-A0A350LX97-F1-MODEL\_V4 \| 1.0 \| 5.492e-07 \| 208 \| 0.212 \| 216 \| 98 \| 6 \| 2 \| 217 \| 203 \| 346 \| Phage\_base\_V domain-containing protein \| Phage\_base\_V domain-containing protein; afdb-uniprot50 \| AF-A0A736MH22-F1-MODEL\_V4 \| 1.0 \| 4.338e-07 \| 207 \| 0.333 \| 156 \| 83 \| 4 \| 58 \| 213 \| 1 \| 135 \| Phage baseplate assembly protein V \| Phage baseplate assembly protein V; afdb-uniprot50 \| AF-A0A2K9LLB3-F1-MODEL\_V4 \| 1.0 \| 1.33e-06 \| 206 \| 0.171 \| 233 \| 162 \| 13 \| 16 \| 217 \| 375 \| 607 \| Phage\_base\_V domain-containing protein \| Phage\_base\_V domain-containing protein; afdb-uniprot50 \| AF-G6XKW3-F1-MODEL\_V4 \| 1.0 \| 1.411e-06 \| 205 \| 0.171 \| 222 \| 140 \| 8 \| 2 \| 217 \| 1 \| 184 \| Uncharacterized protein \| Uncharacterized protein; afdb-uniprot50 \| AF-A0A358LCR7-F1-MODEL\_V4 \| 1.0 \| 1.689e-07 \| 202 \| 0.196 \| 250 \| 158 \| 12 \| 1 \| 211 \| 1 \| 246 \| Baseplate assembly protein \| Baseplate assembly protein; afdb-uniprot50 \| AF-A5L163-F1-MODEL\_V4 \| 1.0 \| 3.036e-06 \| 201 \| 0.147 \| 231 \| 154 \| 10 \| 16 \| 217 \| 266 \| 482 \| Rhs element Vgr protein:Gp5-like protein \| Rhs element Vgr protein:Gp5-like protein; afdb-uniprot50 \| AF-A0A376X398-F1-MODEL\_V4 \| 1.0 \| 9.306e-06 \| 197 \| 0.251 \| 171 \| 106 \| 5 \| 48 \| 209 \| 4 \| 161 \| Phage baseplate assembly protein V \| Phage baseplate assembly protein V; afdb-uniprot50 \| AF-A0A6P1T758-F1-MODEL\_V4 \| 1.0 \| 2.268e-07 \| 197 \| 0.168 \| 219 \| 147 \| 11 \| 16 \| 213 \| 374 \| 578 \| Type VI secretion system tip protein VgrG \| Type VI secretion system tip protein VgrG; afdb-uniprot50 \| AF-A0A1H7LDT2-F1-MODEL\_V4 \| 1.0 \| 5.178e-07 \| 197 \| 0.15 \| 252 \| 164 \| 11 \| 16 \| 217 \| 366 \| 617 \| Uncharacterized conserved protein, implicated in type VI secretion and phage assembly \| Uncharacterized conserved protein, implicated in type VI secretion and phage assembly; afdb-uniprot50 \| AF-A0A847KZN7-F1-MODEL\_V4 \| 1.0 \| 7.351e-06 \| 196 \| 0.134 \| 223 \| 147 \| 10 \| 2 \| 217 \| 1 \| 184 \| Uncharacterized protein \| Uncharacterized protein; afdb-uniprot50 \| AF-A0A150RNN9-F1-MODEL\_V4 \| 1.0 \| 1.406e-05 \| 195 \| 0.17 \| 246 \| 160 \| 12 \| 15 \| 217 \| 377 \| 621 \| Uncharacterized protein \| Uncharacterized protein; afdb-uniprot50 \| AF-A0A1Y1RHY7-F1-MODEL\_V4 \| 1.0 \| 1.047e-05 \| 194 \| 0.132 \| 227 \| 170 \| 9 \| 14 \| 217 \| 372 \| 594 \| Uncharacterized protein \| Uncharacterized protein; afdb-uniprot50 \| AF-A0A2R7QZ58-F1-MODEL\_V4 \| 1.0 \| 5.144e-05 \| 193 \| 0.24 \| 154 \| 95 \| 6 \| 51 \| 203 \| 6 \| 138 \| Phage baseplate assembly protein V \| Phage baseplate assembly protein V; afdb-uniprot50 \| AF-A0A4Q7HXN1-F1-MODEL\_V4 \| 1.0 \| 1.491e-05 \| 193 \| 0.149 \| 247 \| 171 \| 13 \| 1 \| 217 \| 1 \| 238 \| Uncharacterized protein \| Uncharacterized protein; afdb-uniprot50 \| AF-A0A2D5EKJ4-F1-MODEL\_V4 \| 1.0 \| 1.047e-05 \| 193 \| 0.147 \| 258 \| 163 \| 12 \| 16 \| 217 \| 24 \| 280 \| Type VI secretion system tip protein VgrG \| Type VI secretion system tip protein VgrG; afdb-uniprot50 \| AF-A0A2L0ESL0-F1-MODEL\_V4 \| 1.0 \| 1.491e-05 \| 192 \| 0.16 \| 237 \| 163 \| 12 \| 15 \| 216 \| 422 \| 657 \| Uncharacterized protein \| Uncharacterized protein; afdb-uniprot50 \| AF-A0A5M9RDN8-F1-MODEL\_V4 \| 1.0 \| 6.533e-06 \| 191 \| 0.146 \| 218 \| 158 \| 6 \| 14 \| 207 \| 352 \| 565 \| Type VI secretion system tip protein VgrG \| Type VI secretion system tip protein VgrG; afdb-uniprot50 \| AF-A0A510UF59-F1-MODEL\_V4 \| 1.0 \| 3.209e-05 \| 190 \| 0.12 \| 248 \| 179 \| 11 \| 1 \| 215 \| 1 \| 242 \| Uncharacterized protein \| Uncharacterized protein; afdb-uniprot50 \| AF-A0A1D7W984-F1-MODEL\_V4 \| 1.0 \| 1.111e-05 \| 188 \| 0.117 \| 229 \| 171 \| 10 \| 15 \| 217 \| 320 \| 543 \| Uncharacterized protein \| Uncharacterized protein; afdb-uniprot50 \| AF-A0A3A6UBA1-F1-MODEL\_V4 \| 1.0 \| 1.684e-06 \| 188 \| 0.175 \| 256 \| 157 \| 12 \| 16 \| 217 \| 362 \| 617 \| Type VI secretion system tip protein VgrG \| Type VI secretion system tip protein VgrG; afdb-uniprot50 \| AF-A0A1C3EDR2-F1-MODEL\_V4 \| 1.0 \| 8.773e-06 \| 187 \| 0.174 \| 223 \| 147 \| 10 \| 16 \| 209 \| 362 \| 576 \| Type IV secretion protein Rhs \| Type IV secretion protein Rhs; afdb-uniprot50 \| AF-A0A7H0GLJ8-F1-MODEL\_V4 \| 1.0 \| 4.063e-05 \| 186 \| 0.176 \| 238 \| 160 \| 11 \| 16 \| 217 \| 426 \| 663 \| Type VI secretion system tip protein VgrG \| Type VI secretion system tip protein VgrG; afdb-uniprot50 \| AF-A0A1Y1R6F0-F1-MODEL\_V4 \| 1.0 \| 1.25e-05 \| 183 \| 0.15 \| 226 \| 167 \| 6 \| 16 \| 217 \| 368 \| 592 \| Phage\_base\_V domain-containing protein \| Phage\_base\_V domain-containing protein; afdb-uniprot50 \| AF-A0A607GYE7-F1-MODEL\_V4 \| 1.0 \| 9.871e-06 \| 182 \| 0.262 \| 175 \| 106 \| 5 \| 40 \| 207 \| 3 \| 161 \| Phage baseplate assembly protein V \| Phage baseplate assembly protein V; afdb-uniprot50 \| AF-A0A5C6RYB4-F1-MODEL\_V4 \| 1.0 \| 2.852e-05 \| 182 \| 0.139 \| 223 \| 165 \| 10 \| 16 \| 213 \| 59 \| 279 \| Type VI secretion system tip protein VgrG \| Type VI secretion system tip protein VgrG; afdb-uniprot50 \| AF-A0A7U7EJN7-F1-MODEL\_V4 \| 1.0 \| 1.178e-05 \| 181 \| 0.139 \| 236 \| 168 \| 11 \| 16 \| 217 \| 22 \| 256 \| Actin cross-linking toxin VgrG1 \| Actin cross-linking toxin VgrG1; afdb-uniprot50 \| AF-A0A3L8C1Y2-F1-MODEL\_V4 \| 1.0 \| 1.491e-05 \| 180 \| 0.114 \| 228 \| 175 \| 10 \| 16 \| 217 \| 339 \| 565 \| Type VI secretion system tip protein VgrG \| Type VI secretion system tip protein VgrG; afdb-uniprot50 \| AF-A0A4P2QEK0-F1-MODEL\_V4 \| 1.0 \| 1.78e-05 \| 180 \| 0.142 \| 232 \| 165 \| 10 \| 15 \| 216 \| 373 \| 600 \| Uncharacterized protein \| Uncharacterized protein; afdb-uniprot50 \| AF-A0A4U9UU72-F1-MODEL\_V4 \| 1.0 \| 3.83e-05 \| 179 \| 0.115 \| 233 \| 169 \| 9 \| 16 \| 217 \| 123 \| 349 \| Uncharacterized protein conserved in bacteria \| Uncharacterized protein conserved in bacteria; afdb-uniprot50 \| AF-A0A448MIP6-F1-MODEL\_V4 \| 1.0 \| 4.063e-05 \| 178 \| 0.131 \| 244 \| 169 \| 9 \| 16 \| 217 \| 179 \| 421 \| Uncharacterized protein conserved in bacteria \| Uncharacterized protein conserved in bacteria; afdb-uniprot50 \| AF-A0A509DVS8-F1-MODEL\_V4 \| 1.0 \| 4.063e-05 \| 178 \| 0.157 \| 228 \| 161 \| 10 \| 16 \| 217 \| 371 \| 593 \| Actin cross-linking toxin VgrG1 \| Actin cross-linking toxin VgrG1; afdb-uniprot50 \| AF-A0A3S4W3R1-F1-MODEL\_V4 \| 1.0 \| 3.026e-05 \| 177 \| 0.131 \| 244 \| 169 \| 9 \| 16 \| 217 \| 65 \| 307 \| Uncharacterized protein conserved in bacteria \| Uncharacterized protein conserved in bacteria; afdb-uniprot50 \| AF-A0A7Y5KHH1-F1-MODEL\_V4 \| 1.0 \| 3.611e-05 \| 177 \| 0.156 \| 230 \| 166 \| 9 \| 16 \| 217 \| 97 \| 326 \| Uncharacterized protein \| Uncharacterized protein; afdb-uniprot50 \| AF-A0A1N7GD70-F1-MODEL\_V4 \| 1.0 \| 3.83e-05 \| 177 \| 0.131 \| 229 \| 163 \| 11 \| 16 \| 216 \| 363 \| 583 \| Type VI secretion system secreted protein VgrG \| Type VI secretion system secreted protein VgrG; afdb-uniprot50 \| AF-A0A3B0ZXQ5-F1-MODEL\_V4 \| 1.0 \| 2.689e-05 \| 177 \| 0.129 \| 239 \| 164 \| 12 \| 16 \| 215 \| 364 \| 597 \| VgrG protein \| VgrG protein; afdb-uniprot50 \| AF-A0A368X9H0-F1-MODEL\_V4 \| 1.0 \| 5.144e-05 \| 176 \| 0.151 \| 258 \| 162 \| 12 \| 16 \| 217 \| 4 \| 260 \| Type VI secretion system secreted protein VgrG \| Type VI secretion system secreted protein VgrG; afdb-uniprot50 \| AF-A0A2N1IN93-F1-MODEL\_V4 \| 1.0 \| 1.888e-05 \| 176 \| 0.144 \| 236 \| 167 \| 10 \| 16 \| 217 \| 77 \| 311 \| Type VI secretion system tip protein VgrG \| Type VI secretion system tip protein VgrG; afdb-uniprot50 \| AF-A0A509MIG7-F1-MODEL\_V4 \| 1.0 \| 2.689e-05 \| 175 \| 0.147 \| 223 \| 159 \| 10 \| 16 \| 210 \| 167 \| 386 \| Type VI secretion system tip protein VgrG \| Type VI secretion system tip protein VgrG; afdb-uniprot50 \| AF-A0A6M0CR85-F1-MODEL\_V4 \| 1.0 \| 4.849e-05 \| 175 \| 0.126 \| 229 \| 171 \| 9 \| 16 \| 217 \| 201 \| 427 \| Type VI secretion system tip protein VgrG \| Type VI secretion system tip protein VgrG; afdb-uniprot50 \| AF-A0A3A0D0H8-F1-MODEL\_V4 \| 1.0 \| 6.139e-05 \| 174 \| 0.147 \| 231 \| 167 \| 8 \| 16 \| 217 \| 374 \| 603 \| Uncharacterized protein \| Uncharacterized protein; afdb-uniprot50 \| AF-A0A1B8YCK0-F1-MODEL\_V4 \| 1.0 \| 2.253e-05 \| 173 \| 0.126 \| 229 \| 163 \| 13 \| 16 \| 217 \| 57 \| 275 \| Phage-related baseplate assembly protein \| Phage-related baseplate assembly protein; afdb-uniprot50 \| AF-A0A7X4WP42-F1-MODEL\_V4 \| 1.0 \| 5.787e-05 \| 173 \| 0.142 \| 231 \| 165 \| 10 \| 16 \| 217 \| 61 \| 287 \| Type VI secretion system tip protein VgrG \| Type VI secretion system tip protein VgrG; afdb-uniprot50 \| AF-A0A167ADJ3-F1-MODEL\_V4 \| 1.0 \| 2.535e-05 \| 173 \| 0.148 \| 236 \| 165 \| 11 \| 14 \| 217 \| 663 \| 894 \| Uncharacterized protein \| Uncharacterized protein; afdb-uniprot50 \| AF-A0A1I6HDB9-F1-MODEL\_V4 \| 1.0 \| 3.026e-05 \| 172 \| 0.13 \| 230 \| 171 \| 10 \| 16 \| 217 \| 378 \| 606 \| Type VI secretion system secreted protein VgrG \| Type VI secretion system secreted protein VgrG; afdb-uniprot50 \| AF-A0A0F7PK02-F1-MODEL\_V4 \| 1.0 \| 3.209e-05 \| 171 \| 0.155 \| 232 \| 165 \| 12 \| 16 \| 217 \| 12 \| 242 \| Rhs element Vgr protein \| Rhs element Vgr protein; afdb-uniprot50 \| AF-A0A0F7LRQ6-F1-MODEL\_V4 \| 1.0 \| 1.582e-05 \| 171 \| 0.12 \| 225 \| 162 \| 13 \| 16 \| 213 \| 283 \| 498 \| Vrg family protein \| Vrg family protein; afdb-uniprot50 \| AF-A0A178KKM7-F1-MODEL\_V4 \| 1.0 \| 8.773e-06 \| 171 \| 0.157 \| 267 \| 156 \| 13 \| 16 \| 217 \| 368 \| 630 \| Type IV secretion protein Rhs \| Type IV secretion protein Rhs; afdb-uniprot50 \| AF-A0A1B1A454-F1-MODEL\_V4 \| 1.0 \| 5.144e-05 \| 171 \| 0.145 \| 241 \| 166 \| 12 \| 16 \| 217 \| 374 \| 613 \| Type IV secretion protein Rhs \| Type IV secretion protein Rhs; afdb-uniprot50 \| AF-A0A6B2IKF0-F1-MODEL\_V4 \| 1.0 \| 0.0001044 \| 171 \| 0.144 \| 236 \| 167 \| 12 \| 16 \| 217 \| 376 \| 610 \| Type VI secretion system tip protein VgrG \| Type VI secretion system tip protein VgrG; afdb-uniprot50 \| AF-A0A5P8P3V2-F1-MODEL\_V4 \| 1.0 \| 8.243e-05 \| 171 \| 0.12 \| 233 \| 176 \| 8 \| 14 \| 217 \| 379 \| 611 \| Type VI secretion system tip protein VgrG \| Type VI secretion system tip protein VgrG; afdb-uniprot50 \| AF-A0A4U1H893-F1-MODEL\_V4 \| 1.0 \| 7.771e-05 \| 171 \| 0.152 \| 236 \| 156 \| 13 \| 16 \| 216 \| 381 \| 607 \| Type VI secretion system tip protein VgrG \| Type VI secretion system tip protein VgrG; afdb-uniprot50 \| AF-A0A763CHS0-F1-MODEL\_V4 \| 1.0 \| 3.611e-05 \| 170 \| 0.148 \| 250 \| 163 \| 13 \| 1 \| 217 \| 93 \| 325 \| Uncharacterized protein \| Uncharacterized protein; afdb-uniprot50 \| AF-A0A848EAM8-F1-MODEL\_V4 \| 1.0 \| 1.582e-05 \| 170 \| 0.164 \| 225 \| 161 \| 11 \| 16 \| 217 \| 388 \| 608 \| Type VI secretion system tip protein VgrG \| Type VI secretion system tip protein VgrG; afdb-uniprot50 \| AF-A0A128EUM3-F1-MODEL\_V4 \| 1.0 \| 3.611e-05 \| 169 \| 0.157 \| 222 \| 156 \| 7 \| 16 \| 210 \| 362 \| 579 \| Phage-related baseplate assembly protein \| Phage-related baseplate assembly protein; afdb-uniprot50 \| AF-A0A521TV16-F1-MODEL\_V4 \| 1.0 \| 9.306e-06 \| 168 \| 0.138 \| 231 \| 155 \| 11 \| 14 \| 217 \| 387 \| 600 \| Type VI secretion system tip protein VgrG \| Type VI secretion system tip protein VgrG; afdb-uniprot50 \| AF-A0A561C8P1-F1-MODEL\_V4 \| 1.0 \| 6.512e-05 \| 168 \| 0.165 \| 229 \| 159 \| 10 \| 16 \| 217 \| 417 \| 640 \| Type VI secretion system secreted protein VgrG \| Type VI secretion system secreted protein VgrG; afdb-uniprot50 \| AF-A0A4U1IUQ7-F1-MODEL\_V4 \| 1.0 \| 3.22e-06 \| 168 \| 0.165 \| 260 \| 157 \| 11 \| 15 \| 217 \| 467 \| 723 \| Type VI secretion system tip protein VgrG \| Type VI secretion system tip protein VgrG; afdb-uniprot50 \| AF-A0A655SXW0-F1-MODEL\_V4 \| 1.0 \| 9.306e-06 \| 167 \| 0.126 \| 246 \| 171 \| 12 \| 1 \| 208 \| 10 \| 249 \| Uncharacterized protein \| Uncharacterized protein; afdb-uniprot50 \| AF-U7NGG0-F1-MODEL\_V4 \| 1.0 \| 4.324e-06 \| 167 \| 0.178 \| 230 \| 142 \| 12 \| 16 \| 217 \| 52 \| 262 \| Phage\_base\_V domain-containing protein \| Phage\_base\_V domain-containing protein; afdb-uniprot50 \| AF-A0A2S8Q0N4-F1-MODEL\_V4 \| 1.0 \| 4.849e-05 \| 167 \| 0.122 \| 229 \| 164 \| 13 \| 16 \| 217 \| 229 \| 447 \| Type VI secretion system tip protein VgrG \| Type VI secretion system tip protein VgrG; afdb-uniprot50 \| AF-A0A017T821-F1-MODEL\_V4 \| 1.0 \| 4.849e-05 \| 167 \| 0.128 \| 241 \| 167 \| 11 \| 16 \| 217 \| 402 \| 638 \| VgrG protein \| VgrG protein; afdb-uniprot50 \| AF-A0A2E4NX28-F1-MODEL\_V4 \| 1.0 \| 1.111e-05 \| 166 \| 0.154 \| 226 \| 153 \| 12 \| 16 \| 213 \| 5 \| 220 \| Type VI secretion system tip protein VgrG \| Type VI secretion system tip protein VgrG; afdb-uniprot50 \| AF-A0A366AC90-F1-MODEL\_V4 \| 1.0 \| 3.404e-05 \| 166 \| 0.142 \| 225 \| 157 \| 11 \| 16 \| 213 \| 76 \| 291 \| Type VI secretion system tip protein VgrG \| Type VI secretion system tip protein VgrG; afdb-uniprot50 \| AF-A0A0E4FZN6-F1-MODEL\_V4 \| 1.0 \| 6.953e-07 \| 166 \| 0.188 \| 228 \| 146 \| 8 \| 9 \| 214 \| 2 \| 212 \| Uncharacterized protein \| Uncharacterized protein; afdb-uniprot50 \| AF-A0A291HKX6-F1-MODEL\_V4 \| 1.0 \| 8.744e-05 \| 166 \| 0.12 \| 248 \| 179 \| 11 \| 1 \| 217 \| 230 \| 469 \| Uncharacterized protein \| Uncharacterized protein; afdb-uniprot50 \| AF-A0A2G6PY94-F1-MODEL\_V4 \| 1.0 \| 4.849e-05 \| 166 \| 0.122 \| 220 \| 166 \| 9 \| 16 \| 213 \| 368 \| 582 \| Phage\_base\_V domain-containing protein \| Phage\_base\_V domain-containing protein; afdb-uniprot50 \| AF-A0A557PFK3-F1-MODEL\_V4 \| 1.0 \| 3.611e-05 \| 166 \| 0.15 \| 232 \| 155 \| 12 \| 16 \| 217 \| 375 \| 594 \| Type VI secretion system tip protein VgrG \| Type VI secretion system tip protein VgrG; afdb-uniprot50 \| AF-A0A7Y5FUS3-F1-MODEL\_V4 \| 1.0 \| 0.0001486 \| 165 \| 0.116 \| 231 \| 174 \| 8 \| 16 \| 217 \| 246 \| 475 \| Type VI secretion system tip protein VgrG \| Type VI secretion system tip protein VgrG; afdb-uniprot50 \| AF-A0A0N0IAF2-F1-MODEL\_V4 \| 1.0 \| 8.744e-05 \| 165 \| 0.139 \| 223 \| 161 \| 9 \| 16 \| 210 \| 357 \| 576 \| VgrG family protein \| VgrG family protein; afdb-uniprot50 \| AF-A0A2X1VG03-F1-MODEL\_V4 \| 1.0 \| 9.275e-05 \| 165 \| 0.142 \| 239 \| 164 \| 11 \| 16 \| 217 \| 368 \| 602 \| Uncharacterized protein conserved in bacteria \| Uncharacterized protein conserved in bacteria; afdb-uniprot50 \| AF-A0A844CPB5-F1-MODEL\_V4 \| 1.0 \| 5.787e-05 \| 165 \| 0.144 \| 243 \| 163 \| 12 \| 16 \| 217 \| 406 \| 644 \| Type VI secretion system tip protein VgrG \| Type VI secretion system tip protein VgrG; afdb-uniprot50 \| AF-A0A2P1VB00-F1-MODEL\_V4 \| 1.0 \| 6.512e-05 \| 165 \| 0.171 \| 228 \| 159 \| 10 \| 16 \| 217 \| 416 \| 639 \| Type VI secretion system tip protein VgrG \| Type VI secretion system tip protein VgrG; afdb-uniprot50 \| AF-U7HMH7-F1-MODEL\_V4 \| 1.0 \| 2.253e-05 \| 164 \| 0.147 \| 258 \| 163 \| 9 \| 16 \| 217 \| 55 \| 311 \| Phage\_base\_V domain-containing protein \| Phage\_base\_V domain-containing protein; afdb-uniprot50 \| AF-A0A1C4BKP2-F1-MODEL\_V4 \| 1.0 \| 0.0001486 \| 164 \| 0.13 \| 223 \| 163 \| 10 \| 16 \| 210 \| 16 \| 235 \| Rhs element Vgr protein \| Rhs element Vgr protein; afdb-uniprot50 \| AF-A0A6N7B6D2-F1-MODEL\_V4 \| 1.0 \| 6.93e-06 \| 163 \| 0.227 \| 229 \| 134 \| 11 \| 29 \| 217 \| 7 \| 232 \| Phage baseplate \| Phage baseplate; afdb-uniprot50 \| AF-A0A349J5T5-F1-MODEL\_V4 \| 1.0 \| 1.178e-05 \| 163 \| 0.155 \| 232 \| 165 \| 8 \| 16 \| 217 \| 9 \| 239 \| Phage\_base\_V domain-containing protein \| Phage\_base\_V domain-containing protein; afdb-uniprot50 \| AF-Q49TP6-F1-MODEL\_V4 \| 1.0 \| 2.852e-05 \| 163 \| 0.134 \| 230 \| 154 \| 11 \| 16 \| 210 \| 42 \| 261 \| VgrG \| VgrG; afdb-uniprot50 \| AF-A0A2D5EBA9-F1-MODEL\_V4 \| 1.0 \| 1.111e-05 \| 163 \| 0.175 \| 233 \| 139 \| 12 \| 16 \| 217 \| 39 \| 249 \| Lysozyme \| Lysozyme; afdb-uniprot50 \| AF-A0A7W1IND0-F1-MODEL\_V4 \| 1.0 \| 0.0001577 \| 163 \| 0.12 \| 224 \| 169 \| 10 \| 16 \| 217 \| 371 \| 588 \| Type VI secretion system tip protein VgrG \| Type VI secretion system tip protein VgrG; afdb-uniprot50 \| AF-A0A7Y3ZDT9-F1-MODEL\_V4 \| 1.0 \| 6.512e-05 \| 162 \| 0.127 \| 228 \| 170 \| 10 \| 16 \| 217 \| 3 \| 227 \| Type VI secretion system tip protein VgrG \| Type VI secretion system tip protein VgrG; afdb-uniprot50 \| AF-A0A2T1A7F3-F1-MODEL\_V4 \| 1.0 \| 1.25e-05 \| 162 \| 0.147 \| 223 \| 159 \| 11 \| 16 \| 217 \| 257 \| 469 \| Type VI secretion system secreted protein VgrG \| Type VI secretion system secreted protein VgrG; afdb-uniprot50 \| AF-A0A2G6HUW3-F1-MODEL\_V4 \| 1.0 \| 0.000268 \| 162 \| 0.166 \| 228 \| 160 \| 10 \| 16 \| 217 \| 369 \| 592 \| Phage\_base\_V domain-containing protein \| Phage\_base\_V domain-containing protein; afdb-uniprot50 \| AF-A0A3N9U6C8-F1-MODEL\_V4 \| 1.0 \| 7.326e-05 \| 161 \| 0.151 \| 225 \| 155 \| 11 \| 16 \| 213 \| 376 \| 591 \| Type VI secretion system tip protein VgrG \| Type VI secretion system tip protein VgrG; afdb-uniprot50 \| AF-A0A4Q6DCT9-F1-MODEL\_V4 \| 1.0 \| 7.771e-05 \| 160 \| 0.181 \| 231 \| 156 \| 10 \| 16 \| 217 \| 64 \| 290 \| Type VI secretion system tip protein VgrG \| Type VI secretion system tip protein VgrG; afdb-uniprot50 \| AF-A0A5C7CBJ7-F1-MODEL\_V4 \| 1.0 \| 0.0002843 \| 160 \| 0.116 \| 250 \| 184 \| 12 \| 1 \| 217 \| 237 \| 482 \| Uncharacterized protein \| Uncharacterized protein; afdb-uniprot50 \| AF-A0A5E6WXA5-F1-MODEL\_V4 \| 1.0 \| 8.243e-05 \| 160 \| 0.139 \| 237 \| 168 \| 10 \| 16 \| 217 \| 266 \| 501 \| Actin cross-linking toxin VgrG1 \| Actin cross-linking toxin VgrG1; afdb-uniprot50 \| AF-A0A5J6LH20-F1-MODEL\_V4 \| 1.0 \| 2.689e-05 \| 160 \| 0.152 \| 256 \| 162 \| 13 \| 16 \| 217 \| 375 \| 629 \| Type VI secretion system tip protein VgrG \| Type VI secretion system tip protein VgrG; afdb-uniprot50 \| AF-A0A2S5FHP1-F1-MODEL\_V4 \| 1.0 \| 1.78e-05 \| 159 \| 0.137 \| 226 \| 160 \| 11 \| 16 \| 217 \| 220 \| 434 \| Type VI secretion system tip protein VgrG \| Type VI secretion system tip protein VgrG; afdb-uniprot50 \| AF-F8G2Q8-F1-MODEL\_V4 \| 1.0 \| 0.0003393 \| 159 \| 0.143 \| 237 \| 164 \| 12 \| 16 \| 217 \| 334 \| 566 \| Type VI secretion system Vgr family protein \| Type VI secretion system Vgr family protein; afdb-uniprot50 \| AF-A0A1D2X8R5-F1-MODEL\_V4 \| 1.0 \| 0.0002382 \| 159 \| 0.152 \| 242 \| 158 \| 14 \| 16 \| 217 \| 372 \| 606 \| Phage\_base\_V domain-containing protein \| Phage\_base\_V domain-containing protein; afdb-uniprot50 \| AF-A0A1I5SLG1-F1-MODEL\_V4 \| 1.0 \| 2.689e-05 \| 158 \| 0.14 \| 228 \| 161 \| 11 \| 16 \| 217 \| 110 \| 328 \| Rhs element Vgr protein \| Rhs element Vgr protein; afdb-uniprot50 \| AF-A0A175VFY3-F1-MODEL\_V4 \| 1.0 \| 1.325e-05 \| 158 \| 0.146 \| 226 \| 158 \| 11 \| 16 \| 217 \| 156 \| 370 \| Type IV secretion protein Rhs \| Type IV secretion protein Rhs; afdb-uniprot50 \| AF-A0A379DP27-F1-MODEL\_V4 \| 1.0 \| 1.582e-05 \| 158 \| 0.127 \| 244 \| 170 \| 11 \| 16 \| 217 \| 358 \| 600 \| Uncharacterized protein conserved in bacteria \| Uncharacterized protein conserved in bacteria; afdb-uniprot50 \| AF-A0A5C6S553-F1-MODEL\_V4 \| 1.0 \| 2.535e-05 \| 157 \| 0.173 \| 230 \| 156 \| 11 \| 16 \| 217 \| 102 \| 325 \| Type VI secretion system tip protein VgrG \| Type VI secretion system tip protein VgrG; afdb-uniprot50 \| AF-A0A6N7YBH3-F1-MODEL\_V4 \| 1.0 \| 9.871e-06 \| 157 \| 0.192 \| 234 \| 153 \| 11 \| 2 \| 217 \| 347 \| 562 \| VgrG-related protein \| VgrG-related protein; afdb-uniprot50 \| AF-A0A5S9QB60-F1-MODEL\_V4 \| 1.0 \| 3.611e-05 \| 157 \| 0.162 \| 227 \| 155 \| 9 \| 16 \| 217 \| 367 \| 583 \| Actin cross-linking toxin VgrG1 \| Actin cross-linking toxin VgrG1; afdb-uniprot50 \| AF-U7QPZ3-F1-MODEL\_V4 \| 1.0 \| 0.0001577 \| 156 \| 0.1 \| 218 \| 170 \| 8 \| 16 \| 209 \| 39 \| 254 \| Phage\_base\_V domain-containing protein \| Phage\_base\_V domain-containing protein; afdb-uniprot50 \| AF-A0A0X8XCU6-F1-MODEL\_V4 \| 1.0 \| 8.744e-05 \| 156 \| 0.151 \| 231 \| 161 \| 12 \| 16 \| 217 \| 403 \| 627 \| VgrG protein \| VgrG protein; afdb-uniprot50 \| AF-R1INV5-F1-MODEL\_V4 \| 1.0 \| 2.852e-05 \| 155 \| 0.135 \| 228 \| 162 \| 11 \| 16 \| 217 \| 122 \| 340 \| VgrG protein \| VgrG protein; afdb-uniprot50 \| AF-D0GK50-F1-MODEL\_V4 \| 1.0 \| 0.0001486 \| 155 \| 0.124 \| 217 \| 151 \| 8 \| 16 \| 204 \| 291 \| 496 \| Uncharacterized protein \| Uncharacterized protein; afdb-uniprot50 \| AF-A0A379GKP7-F1-MODEL\_V4 \| 1.0 \| 2.253e-05 \| 155 \| 0.126 \| 261 \| 166 \| 13 \| 16 \| 217 \| 261 \| 518 \| Uncharacterized protein conserved in bacteria \| Uncharacterized protein conserved in bacteria; afdb-uniprot50 \| AF-A0A447QXA9-F1-MODEL\_V4 \| 1.0 \| 2.689e-05 \| 155 \| 0.147 \| 264 \| 175 \| 13 \| 1 \| 217 \| 233 \| 493 \| Phage protein \| Phage protein; afdb-uniprot50 \| AF-A0A1H5U3E3-F1-MODEL\_V4 \| 1.0 \| 5.144e-05 \| 155 \| 0.131 \| 273 \| 165 \| 11 \| 16 \| 217 \| 389 \| 660 \| Rhs element Vgr protein \| Rhs element Vgr protein; afdb-uniprot50 \| AF-E6V5I3-F1-MODEL\_V4 \| 1.0 \| 5.787e-05 \| 154 \| 0.17 \| 252 \| 158 \| 10 \| 16 \| 217 \| 417 \| 667 \| Type VI secretion system Vgr family protein \| Type VI secretion system Vgr family protein; afdb-uniprot50 \| AF-A0A8B3VQM5-F1-MODEL\_V4 \| 1.0 \| 0.0001245 \| 153 \| 0.24 \| 166 \| 87 \| 6 \| 53 \| 217 \| 1 \| 128 \| Putative phage associated protein \| Putative phage associated protein; afdb-uniprot50 \| AF-A0A2S8PXI7-F1-MODEL\_V4 \| 1.0 \| 6.139e-05 \| 153 \| 0.114 \| 227 \| 161 \| 11 \| 16 \| 213 \| 286 \| 501 \| Type VI secretion system tip protein VgrG \| Type VI secretion system tip protein VgrG; afdb-uniprot50 \| AF-A0A1E7PYR3-F1-MODEL\_V4 \| 1.0 \| 3.404e-05 \| 152 \| 0.153 \| 235 \| 154 \| 13 \| 16 \| 217 \| 129 \| 351 \| Phage\_base\_V domain-containing protein \| Phage\_base\_V domain-containing protein; afdb-uniprot50 \| AF-A0A7H1J8Z0-F1-MODEL\_V4 \| 1.0 \| 4.572e-05 \| 152 \| 0.129 \| 263 \| 167 \| 11 \| 16 \| 217 \| 376 \| 637 \| Type VI secretion system tip protein VgrG \| Type VI secretion system tip protein VgrG; afdb-uniprot50 \| AF-A0A1X4N450-F1-MODEL\_V4 \| 1.0 \| 0.0002843 \| 152 \| 0.154 \| 233 \| 165 \| 10 \| 16 \| 217 \| 362 \| 593 \| Phage\_base\_V domain-containing protein \| Phage\_base\_V domain-containing protein; afdb-uniprot50 \| AF-M1FL72-F1-MODEL\_V4 \| 1.0 \| 2.689e-05 \| 151 \| 0.157 \| 241 \| 143 \| 11 \| 16 \| 217 \| 67 \| 286 \| Phage\_base\_V domain-containing protein \| Phage\_base\_V domain-containing protein; afdb-uniprot50 \| AF-A0A2X1TPE1-F1-MODEL\_V4 \| 1.0 \| 3.209e-05 \| 151 \| 0.132 \| 227 \| 164 \| 9 \| 16 \| 217 \| 55 \| 273 \| Uncharacterized protein conserved in bacteria \| Uncharacterized protein conserved in bacteria; afdb-uniprot50 \| AF-A6FCK4-F1-MODEL\_V4 \| 1.0 \| 5.787e-05 \| 150 \| 0.094 \| 243 \| 183 \| 9 \| 4 \| 213 \| 13 \| 251 \| Uncharacterized protein \| Uncharacterized protein; afdb-uniprot50 \| AF-A0A624E833-F1-MODEL\_V4 \| 1.0 \| 0.0001401 \| 150 \| 0.104 \| 248 \| 182 \| 12 \| 1 \| 213 \| 43 \| 285 \| Uncharacterized protein \| Uncharacterized protein; afdb-uniprot50 \| AF-A0A2C6DHW8-F1-MODEL\_V4 \| 1.0 \| 4.572e-05 \| 150 \| 0.107 \| 241 \| 178 \| 10 \| 4 \| 211 \| 13 \| 249 \| Uncharacterized protein \| Uncharacterized protein; afdb-uniprot50 \| AF-A0A1G5PCP2-F1-MODEL\_V4 \| 1.0 \| 0.0002246 \| 150 \| 0.108 \| 230 \| 167 \| 11 \| 16 \| 217 \| 2 \| 221 \| Rhs element Vgr protein \| Rhs element Vgr protein; afdb-uniprot50 \| AF-A0A1R4B2C8-F1-MODEL\_V4 \| 1.0 \| 0.0002117 \| 150 \| 0.138 \| 224 \| 159 \| 10 \| 16 \| 213 \| 375 \| 590 \| Phage-related baseplate assembly protein \| Phage-related baseplate assembly protein; afdb-uniprot50 \| AF-A0A6L9FHW8-F1-MODEL\_V4 \| 1.0 \| 0.0001107 \| 149 \| 0.108 \| 249 \| 172 \| 12 \| 1 \| 217 \| 1 \| 231 \| Uncharacterized protein \| Uncharacterized protein; afdb-uniprot50 \| AF-A0A855LDI3-F1-MODEL\_V4 \| 1.0 \| 0.0001882 \| 149 \| 0.129 \| 231 \| 168 \| 9 \| 16 \| 217 \| 13 \| 239 \| Type VI secretion system tip protein VgrG \| Type VI secretion system tip protein VgrG; afdb-uniprot50 \| AF-A0A128FL67-F1-MODEL\_V4 \| 1.0 \| 6.139e-05 \| 149 \| 0.136 \| 227 \| 163 \| 10 \| 16 \| 217 \| 274 \| 492 \| Phage-related baseplate assembly protein \| Phage-related baseplate assembly protein; afdb-uniprot50 \| AF-A0A246WKE1-F1-MODEL\_V4 \| 1.0 \| 9.838e-05 \| 149 \| 0.142 \| 231 \| 159 \| 8 \| 16 \| 213 \| 398 \| 622 \| Phage\_base\_V domain-containing protein \| Phage\_base\_V domain-containing protein; afdb-uniprot50 \| AF-F3BJN5-F1-MODEL\_V4 \| 1.0 \| 3.209e-05 \| 148 \| 0.162 \| 246 \| 159 \| 14 \| 4 \| 211 \| 13 \| 249 \| Uncharacterized protein \| Uncharacterized protein; afdb-uniprot50 \| AF-A0A840G0V0-F1-MODEL\_V4 \| 1.0 \| 0.0001774 \| 147 \| 0.165 \| 248 \| 160 \| 12 \| 16 \| 217 \| 416 \| 662 \| Type VI secretion system secreted protein VgrG \| Type VI secretion system secreted protein VgrG; afdb-uniprot50 \| AF-A0A1Z2SEM0-F1-MODEL\_V4 \| 1.0 \| 4.063e-05 \| 147 \| 0.154 \| 266 \| 160 \| 12 \| 16 \| 217 \| 374 \| 638 \| Phage\_base\_V domain-containing protein \| Phage\_base\_V domain-containing protein; afdb-uniprot50 \| AF-Q9KN42-F1-MODEL\_V4 \| 1.0 \| 0.0001774 \| 147 \| 0.15 \| 266 \| 158 \| 14 \| 16 \| 217 \| 375 \| 636 \| Type VI secretion system spike protein VgrG3 \| Type VI secretion system spike protein VgrG3; afdb-uniprot50 \| AF-B2PU61-F1-MODEL\_V4 \| 1.0 \| 0.0004556 \| 146 \| 0.13 \| 238 \| 167 \| 11 \| 16 \| 216 \| 227 \| 461 \| Rhs element Vgr protein \| Rhs element Vgr protein; afdb-uniprot50 \| AF-A0A158C7T3-F1-MODEL\_V4 \| 1.0 \| 0.0004296 \| 146 \| 0.135 \| 236 \| 169 \| 10 \| 16 \| 217 \| 370 \| 604 \| Rhs element Vgr protein \| Rhs element Vgr protein; afdb-uniprot50 \| AF-A0A3G5KE21-F1-MODEL\_V4 \| 1.0 \| 0.0002117 \| 145 \| 0.121 \| 231 \| 165 \| 12 \| 16 \| 217 \| 472 \| 693 \| Type VI secretion system tip protein VgrG \| Type VI secretion system tip protein VgrG; afdb-uniprot50 \| AF-A0A161UY31-F1-MODEL\_V4 \| 1.0 \| 0.0004556 \| 143 \| 0.144 \| 236 \| 167 \| 11 \| 16 \| 217 \| 381 \| 615 \| Phage-related baseplate assembly protein \| Phage-related baseplate assembly protein; afdb-uniprot50 \| AF-A0A271JCW4-F1-MODEL\_V4 \| 1.0 \| 0.0009244 \| 143 \| 0.165 \| 236 \| 161 \| 11 \| 16 \| 217 \| 375 \| 608 \| Phage\_base\_V domain-containing protein \| Phage\_base\_V domain-containing protein; afdb-uniprot50 \| AF-A0A553JLK0-F1-MODEL\_V4 \| 1.0 \| 0.0001321 \| 142 \| 0.155 \| 232 \| 156 \| 11 \| 16 \| 217 \| 372 \| 593 \| Type VI secretion system tip protein VgrG \| Type VI secretion system tip protein VgrG; afdb-uniprot50 \| AF-A0A833IVZ1-F1-MODEL\_V4 \| 1.0 \| 0.0001107 \| 142 \| 0.147 \| 264 \| 162 \| 13 \| 16 \| 217 \| 368 \| 630 \| Type VI secretion system tip protein VgrG \| Type VI secretion system tip protein VgrG; afdb-uniprot50 \| AF-C5CK96-F1-MODEL\_V4 \| 1.0 \| 0.0001107 \| 142 \| 0.142 \| 253 \| 164 \| 11 \| 16 \| 217 \| 417 \| 667 \| Type VI secretion system Vgr family protein \| Type VI secretion system Vgr family protein; afdb-uniprot50 \| AF-A0A7Y3Y1U2-F1-MODEL\_V4 \| 1.0 \| 0.0001486 \| 142 \| 0.145 \| 268 \| 161 \| 11 \| 14 \| 217 \| 374 \| 637 \| Type VI secretion system tip protein VgrG \| Type VI secretion system tip protein VgrG; afdb-uniprot50 \| AF-A0A840FVA2-F1-MODEL\_V4 \| 1.0 \| 3.209e-05 \| 141 \| 0.292 \| 140 \| 80 \| 3 \| 81 \| 201 \| 9 \| 148 \| Phage baseplate assembly protein V \| Phage baseplate assembly protein V; afdb-uniprot50 \| AF-A0A2W7GN04-F1-MODEL\_V4 \| 1.0 \| 6.139e-05 \| 141 \| 0.162 \| 252 \| 160 \| 10 \| 16 \| 217 \| 315 \| 565 \| Rhs element Vgr protein \| Rhs element Vgr protein; afdb-uniprot50 \| AF-N6WVX3-F1-MODEL\_V4 \| 1.0 \| 0.0001774 \| 140 \| 0.14 \| 263 \| 164 \| 12 \| 16 \| 217 \| 369 \| 630 \| Type VI secretion system tip protein VgrG \| Type VI secretion system tip protein VgrG; afdb-uniprot50 \| AF-A0A6L9PNP9-F1-MODEL\_V4 \| 1.0 \| 8.243e-05 \| 140 \| 0.15 \| 252 \| 163 \| 10 \| 16 \| 217 \| 417 \| 667 \| Type VI secretion system tip protein VgrG \| Type VI secretion system tip protein VgrG; afdb-uniprot50 \| AF-A0A6B3LAR9-F1-MODEL\_V4 \| 1.0 \| 0.0001321 \| 139 \| 0.172 \| 226 \| 152 \| 10 \| 16 \| 217 \| 375 \| 589 \| Type VI secretion system tip protein VgrG \| Type VI secretion system tip protein VgrG; afdb-uniprot50 \| AF-A0A0F5V8P1-F1-MODEL\_V4 \| 1.0 \| 0.0002117 \| 139 \| 0.143 \| 230 \| 159 \| 11 \| 16 \| 213 \| 380 \| 603 \| Type VI secretion protein VgrG \| Type VI secretion protein VgrG; afdb-uniprot50 \| AF-A0A6G7LP93-F1-MODEL\_V4 \| 1.0 \| 0.0001882 \| 138 \| 0.147 \| 244 \| 164 \| 9 \| 16 \| 216 \| 376 \| 618 \| Type VI secretion system tip protein VgrG \| Type VI secretion system tip protein VgrG; afdb-uniprot50 \| AF-A0A3A0E6C2-F1-MODEL\_V4 \| 1.0 \| 7.326e-05 \| 137 \| 0.093 \| 266 \| 176 \| 8 \| 16 \| 217 \| 374 \| 638 \| Phage\_base\_V domain-containing protein \| Phage\_base\_V domain-containing protein; afdb-uniprot50 \| AF-D0XBY7-F1-MODEL\_V4 \| 1.0 \| 0.0001996 \| 133 \| 0.138 \| 260 \| 165 \| 11 \| 16 \| 217 \| 9 \| 267 \| Phage\_base\_V domain-containing protein \| Phage\_base\_V domain-containing protein; afdb-uniprot50 \| AF-A0A0K2E151-F1-MODEL\_V4 \| 1.0 \| 0.0002246 \| 133 \| 0.144 \| 221 \| 157 \| 10 \| 16 \| 208 \| 372 \| 588 \| Rhs element Vgr family protein \| Rhs element Vgr family protein; afdb-uniprot50 \| AF-D1AH50-F1-MODEL\_V4 \| 1.0 \| 0.0002246 \| 132 \| 0.143 \| 209 \| 136 \| 10 \| 16 \| 192 \| 308 \| 505 \| Uncharacterized protein \| Uncharacterized protein; afdb-uniprot50 \| AF-A0A2D0J1V8-F1-MODEL\_V4 \| 1.0 \| 0.0001044 \| 132 \| 0.12 \| 266 \| 166 \| 12 \| 16 \| 217 \| 357 \| 618 \| Rhs element Vgr family protein \| Rhs element Vgr family protein; afdb-uniprot50 \| AF-A0A2V5E8F6-F1-MODEL\_V4 \| 1.0 \| 0.0004556 \| 131 \| 0.156 \| 236 \| 164 \| 10 \| 16 \| 217 \| 392 \| 626 \| Type VI secretion system secreted protein VgrG \| Type VI secretion system secreted protein VgrG; afdb-uniprot50 \| AF-A0A3M8FG17-F1-MODEL\_V4 \| 1.0 \| 0.0002843 \| 129 \| 0.134 \| 215 \| 165 \| 9 \| 16 \| 217 \| 227 \| 433 \| Type VI secretion system tip protein VgrG \| Type VI secretion system tip protein VgrG; afdb-uniprot50 \| AF-A0A063Y972-F1-MODEL\_V4 \| 1.0 \| 0.0004556 \| 129 \| 0.139 \| 244 \| 164 \| 13 \| 16 \| 217 \| 354 \| 593 \| VgrG protein \| VgrG protein; afdb-uniprot50 \| AF-A0A455UDC9-F1-MODEL\_V4 \| 1.0 \| 0.0001672 \| 125 \| 0.126 \| 261 \| 160 \| 12 \| 16 \| 217 \| 7 \| 258 \| Phage\_base\_V domain-containing protein \| Phage\_base\_V domain-containing protein; afdb-uniprot50 \| AF-A0A660LA17-F1-MODEL\_V4 \| 1.0 \| 6.512e-05 \| 123 \| 0.151 \| 218 \| 145 \| 10 \| 16 \| 214 \| 372 \| 568 \| Uncharacterized protein involved in type VI secretion and phage assembly \| Uncharacterized protein involved in type VI secretion and phage assembly; afdb-uniprot50 \| AF-A0A6J4RIX7-F1-MODEL\_V4 \| 1.0 \| 0.0001245 \| 121 \| 0.165 \| 218 \| 144 \| 9 \| 16 \| 209 \| 366 \| 569 \| Phage\_base\_V domain-containing protein \| Phage\_base\_V domain-containing protein; afdb-uniprot50 \| AF-A0A7Y2C7P2-F1-MODEL\_V4 \| 1.0 \| 0.001768 \| 121 \| 0.134 \| 246 \| 168 \| 9 \| 16 \| 217 \| 325 \| 569 \| Type VI secretion system tip protein VgrG \| Type VI secretion system tip protein VgrG; afdb-uniprot50 \| AF-A0A6L7D310-F1-MODEL\_V4 \| 1.0 \| 0.004281 \| 119 \| 0.13 \| 192 \| 151 \| 6 \| 39 \| 217 \| 547 \| 735 \| Phage\_base\_V domain-containing protein \| Phage\_base\_V domain-containing protein; afdb-uniprot50 \| AF-A0A377B6W3-F1-MODEL\_V4 \| 1.0 \| 9.838e-05 \| 117 \| 0.284 \| 183 \| 88 \| 8 \| 61 \| 203 \| 1 \| 180 \| Putative phage baseplate component \| Putative phage baseplate component; afdb-uniprot50 \| AF-A0A496WMG0-F1-MODEL\_V4 \| 1.0 \| 0.00117 \| 115 \| 0.117 \| 263 \| 164 \| 10 \| 16 \| 217 \| 407 \| 662 \| Uncharacterized protein \| Uncharacterized protein; afdb-uniprot50 \| AF-A0A134A0Y4-F1-MODEL\_V4 \| 1.0 \| 0.008686 \| 114 \| 0.097 \| 205 \| 146 \| 10 \| 30 \| 210 \| 366 \| 555 \| Uncharacterized protein \| Uncharacterized protein; afdb-uniprot50 \| AF-A0A3A9A3H4-F1-MODEL\_V4 \| 1.0 \| 6.512e-05 \| 114 \| 0.161 \| 248 \| 153 \| 11 \| 16 \| 216 \| 288 \| 527 \| Uncharacterized protein \| Uncharacterized protein; afdb-uniprot50 \| AF-A0A177W637-F1-MODEL\_V4 \| 1.0 \| 0.002374 \| 111 \| 0.151 \| 224 \| 166 \| 9 \| 16 \| 217 \| 372 \| 593 \| Phage-related baseplate assembly protein \| Phage-related baseplate assembly protein; afdb-uniprot50 \| AF-A0A376MVV2-F1-MODEL\_V4 \| 1.0 \| 0.0007302 \| 107 \| 0.272 \| 154 \| 89 \| 6 \| 65 \| 217 \| 3 \| 134 \| Baseplate assembly protein V \| Baseplate assembly protein V; afdb-uniprot50 \| AF-A0A813EA03-F1-MODEL\_V4 \| 0.872 \| 0.009773 \| 65 \| 0.189 \| 232 \| 155 \| 13 \| 1 \| 217 \| 450 \| 663 \| Hypothetical protein \| Hypothetical protein |
| Top keywords  (threshold 1.00e-02 (evalue)) | **baseplate, assembly, V, Phage, Phage\_base\_V, domain\_containing, Type, secretion, VI, system** |
| Output files | ../../similar\_structures/11\_FANPEZAQ\_CDS\_0011\_afdb-proteome\_foldseek.tsv ../../similar\_structures/11\_FANPEZAQ\_CDS\_0011\_afdb-uniprot50\_foldseek.tsv ../../similar\_structures/11\_FANPEZAQ\_CDS\_0011\_merged.svg ../../similar\_structures/11\_FANPEZAQ\_CDS\_0011\_pdb\_foldseek.tsv |

  
  
  

Return to summary | Go to previous | Go to next

  


---

**Sequence/structure alignments coloring**  
Each object in the alignment figures is colored according to its E-value following this color coding:

1e-100
10
