## Supplementary material for "Completing the BASEL phage collection to unlock hidden diversity for systematic exploration of phage-host interactions": Data S2: 12.html

FANPEZAQ\_CDS\_0012


Return to summary | Go to previous | Go to next

|  |  |
| --- | --- |
| FANPEZAQ\_CDS\_0012  Page creation date: 02 Sep 2024, 12:00  Project folder: n/a  Input sequences file: Escherichia\_virus\_HeidiAbel.gb | domain\_containing gpw\_gp25 baseplate gp25 gpw assembly irad gp25\_like phage lysozyme w putative 25\_like duf2634 gene wedge secretion type fragment vi system tail tsse hypothetical bacteriophage integrase lysozyme\_like phage\_like prophage a sheath t4 complex phage\_related element pbsx gp46 gp6 baseplate\_tail tube pre\_attachment xkds oxidoreductase |

### Sequence information

|  |  |
| --- | --- |
| Name | FANPEZAQ\_CDS\_0012  12\_FANPEZAQ\_CDS\_0012 (pipeline id) |
| Imported annotations | Escherichia\_virus\_HeidiAbel Bas97 |
| Protein sequence | MKGTNAATGAPLDGMDHLRQSIRDILTTPIGTRVMRRDYGSRLPYLVDAPMNRSTLLDLY AATAEALETWEPRISVQQVTATAAEPGRVELSISGEYLPDGKPITIDGIVVT |
| Number of residues | 112 |
| Molecular weight (Da) | 12140.70 |
| Output files | ../../query\_sequences/12\_FANPEZAQ\_CDS\_0012.fasta |

### Putative domain architecture and protein family

#### Search results (HHblits)^1^

|  |  |
| --- | --- |
| Domain family databases searched | Pfam, Ncbi-cd, Cath, Phrogs |
| Results, scheme(s)  (Top layers only; threshold 1.00e-03 (evalue)) | xml version="1.0" encoding="utf-8" standalone="no"?       2024-09-02T21:08:14.430277 image/svg+xml   Matplotlib v3.7.2, https://matplotlib.org/ |
| Results, table  (E-value ≤ 1.00e-03 (evalue)) | db \| id \| prob \| evalue \| pvalue \| score \| cols \| query \| query\_len \| template \| template\_len \| name \| description; pfam \| PF04965 \| 99.1 \| 2.4e-15 \| 4.5e-19 \| 85.0 \| 90 \| (12, 101) \| 112 \| (1, 92) \| 93 \| GPW\_gp25 \| Baseplate wedge protein gp25; pfam \| PF10761 \| 99.0 \| 3.6e-14 \| 6.4e-18 \| 82.3 \| 81 \| (9, 102) \| 112 \| (17, 99) \| 102 \| DUF2590 \| Protein of unknown function (DUF2590); pfam \| PF10934 \| 98.9 \| 1.8e-13 \| 3.3e-17 \| 78.3 \| 73 \| (10, 83) \| 112 \| (19, 93) \| 104 \| DUF2634 \| Protein of unknown function (DUF2634); pfam \| PF07409 \| 97.7 \| 8e-09 \| 1.5e-12 \| 59.4 \| 62 \| (38, 100) \| 112 \| (44, 108) \| 110 \| GP46 \| Phage protein GP46; cath \| 2ia7A00 \| 99.4 \| 4.5e-18 \| 6.1e-22 \| 100.4 \| 97 \| (11, 107) \| 112 \| (27, 125) \| 134 \| Tail lysozyme, putative \| CATHCODE: 3.10.450.40 NAME: Tail lysozyme, putative. Chain: a. Engineered: yes SOURCE: Geobacter sulfurreducens. Organism\_taxid: 35554. Gene: np\_952040.1. Expressed in: escherichia coli. Expression\_system\_taxid: 562. Expression\_system\_vector\_type: plasmid CLASS: Alpha Beta, ARCH: Roll, TOPOL: Nuclear Transport Factor 2; Chain: A, , HOMOL: Nuclear Transport Factor 2; Chain: A,; cath \| 4hrzB00 \| 99.4 \| 1.3e-17 \| 1.8e-21 \| 98.1 \| 94 \| (10, 103) \| 112 \| (26, 120) \| 134 \| Tail lysozyme \| CATHCODE: 3.10.450.40 NAME: Tail lysozyme. Chain: a, b. Synonym: sheath polymerization initiator protein gp25, outer wedge of baseplate protein, protein gp25. Engineered: yes SOURCE: Enterobacteria phage t4. Organism\_taxid: 10665. Gene: 25. Expressed in: escherichia coli. Expression\_system\_taxid: 562. CLASS: Alpha Beta, ARCH: Roll, TOPOL: Nuclear Transport Factor 2; Chain: A, , HOMOL: Nuclear Transport Factor 2; Chain: A,; phrogs \| 261 \| 99.9 \| 3.2e-27 \| 4e-31 \| 152.7 \| 99 \| (5, 104) \| 112 \| (27, 127) \| 139 \| baseplate wedge subunit \| baseplate wedge subunit; Category: tail; KU686208\_p77; phrogs \| 43 \| 99.8 \| 2.8e-26 \| 3.6e-30 \| 146.3 \| 107 \| (1, 107) \| 112 \| (4, 111) \| 127 \| baseplate wedge subunit \| baseplate wedge subunit; Category: tail; p108457 VI\_04298; phrogs \| 975 \| 99.5 \| 2e-18 \| 2.4e-22 \| 118.2 \| 96 \| (10, 110) \| 112 \| (127, 223) \| 236 \| baseplate protein \| baseplate protein; Category: tail; NC\_020871\_p93; phrogs \| 120 \| 99.2 \| 1.2e-15 \| 1.4e-19 \| 95.1 \| 70 \| (10, 83) \| 112 \| (14, 84) \| 118 \| baseplate wedge subunit \| baseplate wedge subunit; Category: tail; p345492 VI\_12445; phrogs \| 37990 \| 98.6 \| 1.2e-11 \| 1.3e-15 \| 67.8 \| 74 \| (36, 110) \| 112 \| (1, 74) \| 80 \| baseplate wedge subunit \| baseplate wedge subunit; Category: tail; NC\_028820\_p187; phrogs \| 2669 \| 98.0 \| 2.1e-09 \| 2.4e-13 \| 66.1 \| 59 \| (10, 68) \| 112 \| (20, 78) \| 138 \| baseplate protein \| baseplate protein; Category: tail; JN204348\_p134; phrogs \| 1896 \| 97.7 \| 1.3e-08 \| 1.4e-12 \| 61.2 \| 95 \| (7, 101) \| 112 \| (15, 113) \| 118 \| NA \| NA; Category: unknown function; p6628 VI\_08092; phrogs \| 4233 \| 97.3 \| 1.7e-07 \| 1.9e-11 \| 61.1 \| 80 \| (12, 96) \| 112 \| (26, 107) \| 208 \| NA \| NA; Category: unknown function; KX552041\_p157; phrogs \| 15729 \| 96.8 \| 2.5e-06 \| 2.8e-10 \| 48.1 \| 62 \| (19, 84) \| 112 \| (10, 71) \| 94 \| baseplate protein \| baseplate protein; Category: tail; p127195 VI\_01263; phrogs \| 5026 \| 96.4 \| 9e-06 \| 1e-09 \| 47.8 \| 74 \| (16, 99) \| 112 \| (31, 106) \| 117 \| baseplate protein \| baseplate protein; Category: tail; MF185718\_p38; phrogs \| 33750 \| 96.1 \| 3.1e-05 \| 3.5e-09 \| 42.7 \| 62 \| (21, 84) \| 112 \| (16, 77) \| 91 \| NA \| NA; Category: unknown function; p321351 VI\_06335; phrogs \| 4033 \| 96.0 \| 3.3e-05 \| 3.8e-09 \| 52.5 \| 87 \| (10, 102) \| 112 \| (226, 313) \| 320 \| NA \| NA; Category: unknown function; p55048 VI\_12418; phrogs \| 180 \| 94.6 \| 0.00054 \| 6.7e-08 \| 40.8 \| 75 \| (28, 103) \| 112 \| (34, 108) \| 120 \| baseplate wedge subunit \| baseplate wedge subunit; Category: tail; p189333 VI\_09191; phrogs \| 33571 \| 94.4 \| 0.00076 \| 8.5e-08 \| 40.2 \| 71 \| (10, 83) \| 112 \| (59, 130) \| 161 \| NA \| NA; Category: unknown function; NC\_008584\_p54 |
| Top keywords  (threshold 1.00e-03 (evalue)) | **baseplate, tail, wedge, A, Nuclear, Transport, Factor, gp25, Phage, lysozyme** |
| Output files | ../../domain\_architecture/12\_FANPEZAQ\_CDS\_0012\_cath.hhr ../../domain\_architecture/12\_FANPEZAQ\_CDS\_0012\_merged.svg ../../domain\_architecture/12\_FANPEZAQ\_CDS\_0012\_ncbi-cd.hhr ../../domain\_architecture/12\_FANPEZAQ\_CDS\_0012\_pfam.hhr ../../domain\_architecture/12\_FANPEZAQ\_CDS\_0012\_phrogs.hhr |

### Identical protein sequences/structures

#### Search results

|  |  |
| --- | --- |
| Protein sequence databases searched | Pdb, Swissprot, Refseq |
| Identical proteins found | -- |
| Top keywords | -- |
| Output files | -- |

### Similar protein sequences/structures

#### Sequence similarity search results (HHblits)^1^

|  |  |
| --- | --- |
| Sequence databases searched | Uniclust, Pdb70 |
| Results, scheme(s)  (Top layers only, threshold 1.00e-03 (evalue)) | xml version="1.0" encoding="utf-8" standalone="no"?       2024-09-02T21:08:33.068464 image/svg+xml   Matplotlib v3.7.2, https://matplotlib.org/ |
| Results, table(s)  (threshold 1.00e-03 (evalue)) | db \| id \| prob \| evalue \| pvalue \| score \| cols \| query \| query\_len \| template \| template\_len \| name \| description; uniclust \| UniRef100\_A0A011NBA4 \| 99.9 \| 2.1e-30 \| 4.2e-36 \| 164.1 \| 110 \| (1, 110) \| 112 \| (27, 136) \| 154 \| IraD/Gp25-like domain-containing protein \| IraD/Gp25-like domain-containing protein; uniclust \| UniRef100\_A0A143DG48 \| 99.9 \| 4.2e-30 \| 8.1e-36 \| 158.6 \| 110 \| (1, 110) \| 112 \| (23, 132) \| 140 \| IraD/Gp25-like domain-containing protein \| IraD/Gp25-like domain-containing protein; uniclust \| UniRef100\_A0A062INA0 \| 99.9 \| 2.5e-29 \| 4.9e-35 \| 157.1 \| 111 \| (1, 111) \| 112 \| (5, 115) \| 144 \| Lysozyme family protein \| Lysozyme family protein; uniclust \| UniRef100\_A0A024E981 \| 99.9 \| 1.4e-28 \| 2.8e-34 \| 155.9 \| 110 \| (1, 111) \| 112 \| (31, 140) \| 153 \| Baseplate assembly protein W \| Baseplate assembly protein W; uniclust \| UniRef100\_A0A016XHY4 \| 99.9 \| 1.7e-28 \| 3.5e-34 \| 155.6 \| 111 \| (1, 111) \| 112 \| (24, 135) \| 151 \| Baseplate assembly protein \| Baseplate assembly protein; uniclust \| UniRef100\_A0A011NH43 \| 99.9 \| 2.6e-28 \| 5.2e-34 \| 164.3 \| 106 \| (4, 110) \| 112 \| (51, 160) \| 227 \| Baseplate wedge subunit \| Baseplate wedge subunit; uniclust \| UniRef100\_A0A014PYG1 \| 99.9 \| 3.1e-28 \| 6e-34 \| 148.5 \| 111 \| (1, 111) \| 112 \| (3, 113) \| 124 \| Baseplate assembly protein (Fragment) \| Baseplate assembly protein (Fragment); uniclust \| UniRef100\_A0A069PL87 \| 99.9 \| 6.8e-28 \| 1.3e-33 \| 144.9 \| 107 \| (1, 107) \| 112 \| (7, 113) \| 121 \| IraD/Gp25-like domain-containing protein \| IraD/Gp25-like domain-containing protein; uniclust \| UniRef100\_A0A062UZG7 \| 99.9 \| 9.7e-28 \| 1.9e-33 \| 148.6 \| 100 \| (11, 110) \| 112 \| (26, 127) \| 137 \| Phage baseplate assembly protein W \| Phage baseplate assembly protein W; uniclust \| UniRef100\_A0A1H3KA32 \| 99.9 \| 1.3e-27 \| 2.6e-33 \| 149.2 \| 101 \| (11, 111) \| 112 \| (34, 135) \| 144 \| IraD/Gp25-like domain-containing protein \| IraD/Gp25-like domain-containing protein; uniclust \| UniRef100\_A0A061KN01 \| 99.9 \| 5.7e-27 \| 1.1e-32 \| 146.9 \| 110 \| (1, 110) \| 112 \| (5, 114) \| 144 \| Baseplate assembly protein \| Baseplate assembly protein; uniclust \| UniRef100\_A0A066RM20 \| 99.9 \| 6.5e-27 \| 1.3e-32 \| 143.9 \| 107 \| (1, 107) \| 112 \| (15, 121) \| 129 \| dTDP-glucose pyrophosphorylase \| dTDP-glucose pyrophosphorylase; uniclust \| UniRef100\_A0A1M5Y4I2 \| 99.9 \| 7e-27 \| 1.4e-32 \| 146.3 \| 103 \| (8, 110) \| 112 \| (19, 125) \| 149 \| Gene 25-like lysozyme \| Gene 25-like lysozyme; uniclust \| UniRef100\_A0A0Q4L1R6 \| 99.9 \| 1.5e-26 \| 2.9e-32 \| 146.0 \| 108 \| (3, 110) \| 112 \| (27, 138) \| 145 \| IraD/Gp25-like domain-containing protein \| IraD/Gp25-like domain-containing protein; uniclust \| UniRef100\_A0A0Q8XJN5 \| 99.9 \| 2.1e-26 \| 4.1e-32 \| 145.0 \| 100 \| (11, 110) \| 112 \| (31, 132) \| 150 \| IraD/Gp25-like domain-containing protein \| IraD/Gp25-like domain-containing protein; uniclust \| UniRef100\_A0A0C5AE75 \| 99.9 \| 2.4e-26 \| 4.6e-32 \| 141.7 \| 107 \| (4, 110) \| 112 \| (25, 135) \| 139 \| Baseplate wedge subunit \| Baseplate wedge subunit; uniclust \| UniRef100\_A0A0E3EPQ9 \| 99.9 \| 2.5e-26 \| 5.2e-32 \| 146.3 \| 108 \| (3, 110) \| 112 \| (32, 142) \| 148 \| Base plate wedge subunit \| Base plate wedge subunit; uniclust \| UniRef100\_A0A084SKL2 \| 99.9 \| 6.5e-26 \| 1.3e-31 \| 145.6 \| 103 \| (8, 110) \| 112 \| (37, 143) \| 182 \| IraD/Gp25-like domain-containing protein \| IraD/Gp25-like domain-containing protein; uniclust \| UniRef100\_A0A0F9DR72 \| 99.9 \| 9.8e-26 \| 1.9e-31 \| 143.6 \| 97 \| (15, 111) \| 112 \| (38, 138) \| 171 \| IraD/Gp25-like domain-containing protein \| IraD/Gp25-like domain-containing protein; uniclust \| UniRef100\_A0A1D7SKY7 \| 99.9 \| 2.1e-25 \| 4.2e-31 \| 144.2 \| 109 \| (2, 110) \| 112 \| (40, 151) \| 167 \| Baseplate wedge subunit \| Baseplate wedge subunit; uniclust \| UniRef100\_A0A1I2FX92 \| 99.9 \| 3.5e-25 \| 7e-31 \| 135.6 \| 100 \| (12, 111) \| 112 \| (19, 118) \| 123 \| Gene 25-like lysozyme \| Gene 25-like lysozyme; uniclust \| UniRef100\_A0A022PCH8 \| 99.9 \| 3.8e-25 \| 7.6e-31 \| 141.9 \| 105 \| (4, 108) \| 112 \| (40, 147) \| 169 \| Phage baseplate assembly protein W \| Phage baseplate assembly protein W; uniclust \| UniRef100\_A0A060AMS0 \| 99.9 \| 4.1e-25 \| 8.1e-31 \| 139.2 \| 109 \| (2, 110) \| 112 \| (27, 139) \| 147 \| Putative baseplate wedge subunit \| Putative baseplate wedge subunit; uniclust \| UniRef100\_A0A011NTL3 \| 99.9 \| 4.2e-25 \| 8.2e-31 \| 137.5 \| 102 \| (8, 109) \| 112 \| (29, 134) \| 144 \| Baseplate wedge subunit \| Baseplate wedge subunit; uniclust \| UniRef100\_A0A067XRJ3 \| 99.9 \| 4.2e-25 \| 8.4e-31 \| 140.5 \| 107 \| (4, 110) \| 112 \| (37, 146) \| 151 \| IraD/Gp25-like domain-containing protein \| IraD/Gp25-like domain-containing protein; uniclust \| UniRef100\_A0A103EKP3 \| 99.8 \| 8e-25 \| 1.5e-30 \| 127.6 \| 84 \| (1, 84) \| 112 \| (9, 92) \| 101 \| Phage baseplate protein (Fragment) \| Phage baseplate protein (Fragment); uniclust \| UniRef100\_A0A023Q0J9 \| 99.8 \| 1.3e-24 \| 2.5e-30 \| 142.0 \| 100 \| (11, 110) \| 112 \| (60, 161) \| 179 \| IraD/Gp25-like domain-containing protein \| IraD/Gp25-like domain-containing protein; uniclust \| UniRef100\_A0A1H8VXY1 \| 99.8 \| 8.6e-24 \| 1.7e-29 \| 130.6 \| 111 \| (1, 111) \| 112 \| (17, 129) \| 134 \| IraD/Gp25-like domain-containing protein \| IraD/Gp25-like domain-containing protein; uniclust \| UniRef100\_A0A348FYH4 \| 99.8 \| 1.8e-23 \| 3.6e-29 \| 125.8 \| 103 \| (1, 103) \| 112 \| (9, 112) \| 119 \| IraD/Gp25-like domain-containing protein \| IraD/Gp25-like domain-containing protein; uniclust \| UniRef100\_A0A011UMM6 \| 99.8 \| 4.1e-23 \| 8.2e-29 \| 133.9 \| 102 \| (1, 102) \| 112 \| (34, 136) \| 168 \| IraD/Gp25-like domain-containing protein \| IraD/Gp25-like domain-containing protein; uniclust \| UniRef100\_A0A4R2C4F1 \| 99.8 \| 7.3e-23 \| 1.4e-28 \| 128.0 \| 103 \| (8, 110) \| 112 \| (20, 126) \| 169 \| IraD/Gp25-like domain-containing protein \| IraD/Gp25-like domain-containing protein; uniclust \| UniRef100\_A0A0Q7E412 \| 99.8 \| 1.3e-22 \| 2.4e-28 \| 126.0 \| 103 \| (8, 110) \| 112 \| (32, 140) \| 148 \| IraD/Gp25-like domain-containing protein \| IraD/Gp25-like domain-containing protein; uniclust \| UniRef100\_A0A2E4ZEV5 \| 99.8 \| 2.9e-22 \| 5.6e-28 \| 122.4 \| 100 \| (11, 110) \| 112 \| (25, 125) \| 130 \| IraD/Gp25-like domain-containing protein \| IraD/Gp25-like domain-containing protein; uniclust \| UniRef100\_A0A5S3XXE0 \| 99.8 \| 4.1e-22 \| 7.9e-28 \| 122.0 \| 100 \| (11, 110) \| 112 \| (20, 123) \| 129 \| IraD/Gp25-like domain-containing protein \| IraD/Gp25-like domain-containing protein; uniclust \| UniRef100\_A0A0D0KR61 \| 99.8 \| 7.7e-22 \| 1.5e-27 \| 124.1 \| 106 \| (2, 107) \| 112 \| (5, 119) \| 144 \| Baseplate assembly protein W (GpW) \| Baseplate assembly protein W (GpW); uniclust \| UniRef100\_A0A291FP80 \| 99.8 \| 8.8e-22 \| 1.6e-27 \| 124.3 \| 107 \| (1, 107) \| 112 \| (73, 179) \| 184 \| Putative baseplate assembly protein W \| Putative baseplate assembly protein W; uniclust \| UniRef100\_A0A1Z8UZN9 \| 99.7 \| 1.5e-21 \| 2.8e-27 \| 123.7 \| 107 \| (4, 110) \| 112 \| (44, 153) \| 159 \| IraD/Gp25-like domain-containing protein \| IraD/Gp25-like domain-containing protein; uniclust \| UniRef100\_A0A0C1GGW4 \| 99.7 \| 1.5e-21 \| 2.9e-27 \| 123.7 \| 94 \| (8, 102) \| 112 \| (34, 127) \| 149 \| IraD/Gp25-like domain-containing protein \| IraD/Gp25-like domain-containing protein; uniclust \| UniRef100\_A0A0E3G425 \| 99.7 \| 1.5e-21 \| 3e-27 \| 123.6 \| 107 \| (4, 110) \| 112 \| (32, 142) \| 147 \| Base plate wedge subunit \| Base plate wedge subunit; uniclust \| UniRef100\_A0A376JCH0 \| 99.7 \| 1.7e-21 \| 3.3e-27 \| 114.8 \| 81 \| (2, 82) \| 112 \| (6, 86) \| 98 \| Phage baseplate protein \| Phage baseplate protein; uniclust \| UniRef100\_D4HTV2 \| 99.7 \| 1.8e-21 \| 3.3e-27 \| 118.2 \| 105 \| (1, 105) \| 112 \| (43, 147) \| 154 \| Gp13 protein \| Gp13 protein; uniclust \| UniRef100\_A0A059IUA3 \| 99.7 \| 2.4e-21 \| 4.6e-27 \| 115.1 \| 107 \| (1, 107) \| 112 \| (1, 107) \| 119 \| Putative prophage LambdaW5, baseplate assembly protein W,putative \| Putative prophage LambdaW5, baseplate assembly protein W,putative; uniclust \| UniRef100\_A0A1R1MKK7 \| 99.7 \| 3.6e-21 \| 7e-27 \| 118.1 \| 86 \| (11, 98) \| 112 \| (20, 107) \| 125 \| IraD/Gp25-like domain-containing protein \| IraD/Gp25-like domain-containing protein; uniclust \| UniRef100\_A0A410T6G4 \| 99.7 \| 6.3e-21 \| 1.2e-26 \| 114.7 \| 91 \| (11, 101) \| 112 \| (15, 106) \| 120 \| Baseplate wedge subunit \| Baseplate wedge subunit; uniclust \| UniRef100\_A0A1V5RRZ4 \| 99.7 \| 7.2e-21 \| 1.4e-26 \| 116.7 \| 100 \| (11, 110) \| 112 \| (38, 138) \| 139 \| Gene 25-like lysozyme \| Gene 25-like lysozyme; uniclust \| UniRef100\_A0A011P705 \| 99.7 \| 8e-21 \| 1.6e-26 \| 121.3 \| 91 \| (9, 101) \| 112 \| (35, 125) \| 151 \| Phage baseplate protein \| Phage baseplate protein; uniclust \| UniRef100\_A0A7X5DY26 \| 99.7 \| 1.4e-20 \| 2.5e-26 \| 111.7 \| 100 \| (11, 110) \| 112 \| (19, 119) \| 122 \| Baseplate protein \| Baseplate protein; uniclust \| UniRef100\_A0A0A0YS61 \| 99.7 \| 1.9e-20 \| 3.7e-26 \| 117.5 \| 108 \| (3, 110) \| 112 \| (25, 135) \| 139 \| Base plate wedge subunit \| Base plate wedge subunit; uniclust \| UniRef100\_A0A8I0FE23 \| 99.7 \| 2.2e-20 \| 4e-26 \| 111.0 \| 98 \| (1, 98) \| 112 \| (12, 109) \| 127 \| GPW/gp25 family protein \| GPW/gp25 family protein; uniclust \| UniRef100\_A0A023W5U8 \| 99.7 \| 2.2e-20 \| 4.3e-26 \| 114.7 \| 92 \| (11, 102) \| 112 \| (26, 118) \| 137 \| Baseplate wedge subunit \| Baseplate wedge subunit; uniclust \| UniRef100\_A0A191W8N1 \| 99.7 \| 2.9e-20 \| 5.6e-26 \| 114.1 \| 100 \| (8, 109) \| 112 \| (14, 115) \| 126 \| IraD/Gp25-like domain-containing protein \| IraD/Gp25-like domain-containing protein; uniclust \| UniRef100\_A0A0A8HAU6 \| 99.7 \| 3.4e-20 \| 6.6e-26 \| 109.5 \| 82 \| (16, 97) \| 112 \| (7, 88) \| 100 \| Phage baseplate assembly protein W \| Phage baseplate assembly protein W; uniclust \| UniRef100\_A0A178HLN5 \| 99.7 \| 3.9e-20 \| 7.6e-26 \| 118.2 \| 94 \| (9, 103) \| 112 \| (47, 141) \| 157 \| IraD/Gp25-like domain-containing protein \| IraD/Gp25-like domain-containing protein; uniclust \| UniRef100\_A0A031JP30 \| 99.7 \| 5.9e-20 \| 1.2e-25 \| 118.5 \| 105 \| (4, 110) \| 112 \| (36, 143) \| 167 \| GPW/gp25 family protein \| GPW/gp25 family protein; uniclust \| UniRef100\_A0A1D2QY07 \| 99.7 \| 7.7e-20 \| 1.5e-25 \| 113.9 \| 102 \| (8, 110) \| 112 \| (29, 137) \| 142 \| IraD/Gp25-like domain-containing protein \| IraD/Gp25-like domain-containing protein; uniclust \| UniRef100\_A0A6G3XJJ0 \| 99.7 \| 1.1e-19 \| 2e-25 \| 105.9 \| 90 \| (4, 93) \| 112 \| (16, 108) \| 109 \| GPW/gp25 family protein (Fragment) \| GPW/gp25 family protein (Fragment); uniclust \| UniRef100\_A0A3C1ALA6 \| 99.7 \| 1.6e-19 \| 3e-25 \| 104.1 \| 92 \| (12, 103) \| 112 \| (1, 92) \| 97 \| Phage baseplate protein (Fragment) \| Phage baseplate protein (Fragment); uniclust \| UniRef100\_A0A1Z9HXH3 \| 99.6 \| 2.3e-19 \| 4.7e-25 \| 114.3 \| 98 \| (12, 109) \| 112 \| (37, 136) \| 144 \| IraD/Gp25-like domain-containing protein \| IraD/Gp25-like domain-containing protein; uniclust \| UniRef100\_A0A0F9S7G6 \| 99.6 \| 3.7e-19 \| 7.4e-25 \| 115.9 \| 98 \| (11, 110) \| 112 \| (43, 140) \| 170 \| IraD/Gp25-like domain-containing protein \| IraD/Gp25-like domain-containing protein; uniclust \| UniRef100\_A0A091APV5 \| 99.6 \| 5e-19 \| 9.8e-25 \| 106.4 \| 76 \| (8, 83) \| 112 \| (26, 103) \| 106 \| IraD/Gp25-like domain-containing protein (Fragment) \| IraD/Gp25-like domain-containing protein (Fragment); uniclust \| UniRef100\_UPI001573BC7E \| 99.6 \| 6e-19 \| 1.1e-24 \| 108.3 \| 101 \| (11, 111) \| 112 \| (40, 142) \| 150 \| type VI secretion system baseplate subunit TssE \| type VI secretion system baseplate subunit TssE; uniclust \| UniRef100\_A0A1M6M9N6 \| 99.6 \| 1e-18 \| 2e-24 \| 109.6 \| 100 \| (8, 108) \| 112 \| (34, 136) \| 144 \| Gene 25-like lysozyme \| Gene 25-like lysozyme; uniclust \| UniRef100\_A0A0Q1ADI1 \| 99.6 \| 1.1e-18 \| 2.2e-24 \| 124.4 \| 99 \| (11, 110) \| 112 \| (33, 131) \| 348 \| IraD/Gp25-like domain-containing protein \| IraD/Gp25-like domain-containing protein; uniclust \| UniRef100\_A0A927MQK2 \| 99.6 \| 1.4e-18 \| 2.5e-24 \| 109.2 \| 103 \| (8, 110) \| 112 \| (33, 139) \| 189 \| Phage baseplate assembly protein W \| Phage baseplate assembly protein W; uniclust \| UniRef100\_A0A067ZJ13 \| 99.6 \| 1.6e-18 \| 3.2e-24 \| 108.9 \| 88 \| (13, 102) \| 112 \| (27, 116) \| 125 \| IraD/Gp25-like domain-containing protein \| IraD/Gp25-like domain-containing protein; uniclust \| UniRef100\_D5SL83 \| 99.6 \| 2.2e-18 \| 4e-24 \| 105.2 \| 108 \| (3, 110) \| 112 \| (15, 125) \| 149 \| GPW/gp25 family protein \| GPW/gp25 family protein; uniclust \| UniRef100\_A0A2E2FJP9 \| 99.6 \| 2.4e-18 \| 4.5e-24 \| 105.4 \| 103 \| (8, 110) \| 112 \| (21, 126) \| 132 \| Phage baseplate protein \| Phage baseplate protein; uniclust \| UniRef100\_A8TYF6 \| 99.6 \| 2.6e-18 \| 4.7e-24 \| 104.8 \| 95 \| (4, 98) \| 112 \| (1, 95) \| 153 \| IraD/Gp25-like domain-containing protein \| IraD/Gp25-like domain-containing protein; uniclust \| UniRef100\_A0A1V3IM03 \| 99.6 \| 3.1e-18 \| 5.8e-24 \| 100.6 \| 97 \| (13, 110) \| 112 \| (3, 99) \| 105 \| Baseplate assembly protein W (Fragment) \| Baseplate assembly protein W (Fragment); uniclust \| UniRef100\_A0A023XMK4 \| 99.6 \| 3e-18 \| 6.1e-24 \| 111.8 \| 99 \| (11, 110) \| 112 \| (42, 141) \| 164 \| IraD/Gp25-like domain-containing protein \| IraD/Gp25-like domain-containing protein; uniclust \| UniRef100\_A0A0Q2YY05 \| 99.6 \| 3.1e-18 \| 6.1e-24 \| 108.9 \| 96 \| (4, 100) \| 112 \| (28, 123) \| 147 \| IraD/Gp25-like domain-containing protein \| IraD/Gp25-like domain-containing protein; uniclust \| UniRef100\_A0A0F9FRJ8 \| 99.6 \| 3.8e-18 \| 7.5e-24 \| 106.9 \| 97 \| (11, 108) \| 112 \| (24, 121) \| 133 \| IraD/Gp25-like domain-containing protein \| IraD/Gp25-like domain-containing protein; uniclust \| UniRef100\_A0A0S4XLR8 \| 99.6 \| 6.6e-18 \| 1.3e-23 \| 104.5 \| 93 \| (15, 107) \| 112 \| (36, 129) \| 135 \| Putative phage-related baseplate assembly protein (GPW-like) \| Putative phage-related baseplate assembly protein (GPW-like); uniclust \| UniRef100\_A0A011PP45 \| 99.6 \| 7.3e-18 \| 1.5e-23 \| 110.4 \| 100 \| (11, 111) \| 112 \| (57, 162) \| 173 \| Uncharacterized protein \| Uncharacterized protein; uniclust \| UniRef100\_UPI001BAAC7D5 \| 99.6 \| 9.3e-18 \| 1.7e-23 \| 108.1 \| 106 \| (5, 110) \| 112 \| (108, 216) \| 224 \| GPW/gp25 family protein \| GPW/gp25 family protein; uniclust \| UniRef100\_A0A1X7MCL8 \| 99.6 \| 9.6e-18 \| 1.8e-23 \| 97.8 \| 91 \| (20, 111) \| 112 \| (3, 93) \| 98 \| GPW/gp25 family protein \| GPW/gp25 family protein; uniclust \| UniRef100\_A0A081C215 \| 99.6 \| 9.9e-18 \| 2e-23 \| 109.7 \| 99 \| (11, 110) \| 112 \| (57, 160) \| 174 \| Conserved domain protein \| Conserved domain protein; uniclust \| UniRef100\_A0A0Q8LUA8 \| 99.5 \| 1.6e-17 \| 3e-23 \| 104.9 \| 100 \| (11, 110) \| 112 \| (40, 141) \| 150 \| IraD/Gp25-like domain-containing protein \| IraD/Gp25-like domain-containing protein; uniclust \| UniRef100\_A0A1B1IUU0 \| 99.5 \| 1.7e-17 \| 3.3e-23 \| 106.2 \| 106 \| (5, 110) \| 112 \| (21, 130) \| 156 \| Baseplate protein \| Baseplate protein; uniclust \| UniRef100\_A0A074LW06 \| 99.5 \| 2e-17 \| 4e-23 \| 102.6 \| 86 \| (13, 100) \| 112 \| (22, 108) \| 119 \| IraD/Gp25-like domain-containing protein \| IraD/Gp25-like domain-containing protein; uniclust \| UniRef100\_A0A127AW49 \| 99.5 \| 2.3e-17 \| 4.2e-23 \| 102.6 \| 107 \| (4, 110) \| 112 \| (37, 147) \| 156 \| Sheath initiation protein \| Sheath initiation protein; uniclust \| UniRef100\_UPI001F313682 \| 99.5 \| 2.5e-17 \| 4.6e-23 \| 100.6 \| 105 \| (1, 106) \| 112 \| (1, 107) \| 152 \| GPW/gp25 family protein \| GPW/gp25 family protein; uniclust \| UniRef100\_A0A927DLG2 \| 99.5 \| 3.7e-17 \| 6.9e-23 \| 111.0 \| 108 \| (2, 110) \| 112 \| (195, 303) \| 310 \| Phage baseplate assembly protein V \| Phage baseplate assembly protein V; uniclust \| UniRef100\_A0A014LIJ6 \| 99.5 \| 5.2e-17 \| 1e-22 \| 104.3 \| 96 \| (8, 104) \| 112 \| (55, 153) \| 170 \| IraD/Gp25-like domain-containing protein \| IraD/Gp25-like domain-containing protein; uniclust \| UniRef100\_A0A061NG15 \| 99.5 \| 5.3e-17 \| 1.1e-22 \| 102.8 \| 96 \| (8, 108) \| 112 \| (18, 117) \| 128 \| DUF2634 domain-containing protein \| DUF2634 domain-containing protein; uniclust \| UniRef100\_A0A2E3HQV9 \| 99.5 \| 5.8e-17 \| 1.1e-22 \| 99.7 \| 100 \| (5, 104) \| 112 \| (18, 120) \| 131 \| IraD/Gp25-like domain-containing protein \| IraD/Gp25-like domain-containing protein; uniclust \| UniRef100\_A0A0N1C7H2 \| 99.5 \| 7.1e-17 \| 1.4e-22 \| 103.8 \| 101 \| (2, 102) \| 112 \| (21, 122) \| 161 \| Integrase \| Integrase; uniclust \| UniRef100\_Q1QI83 \| 99.5 \| 7.5e-17 \| 1.4e-22 \| 99.8 \| 100 \| (2, 101) \| 112 \| (43, 143) \| 165 \| GPW/gp25 \| GPW/gp25; uniclust \| UniRef100\_A0A064AK59 \| 99.5 \| 7e-17 \| 1.4e-22 \| 102.1 \| 87 \| (12, 100) \| 112 \| (35, 122) \| 132 \| Isoleucyl-tRNA synthetase \| Isoleucyl-tRNA synthetase; uniclust \| UniRef100\_A0A1T4WV80 \| 99.5 \| 8.2e-17 \| 1.5e-22 \| 101.8 \| 82 \| (2, 83) \| 112 \| (81, 162) \| 192 \| Phage baseplate assembly protein W \| Phage baseplate assembly protein W; uniclust \| UniRef100\_A0A0F9T6N1 \| 99.5 \| 7.9e-17 \| 1.5e-22 \| 104.4 \| 89 \| (11, 99) \| 112 \| (38, 128) \| 171 \| IraD/Gp25-like domain-containing protein \| IraD/Gp25-like domain-containing protein; uniclust \| UniRef100\_UPI000291EF94 \| 99.5 \| 8.6e-17 \| 1.6e-22 \| 94.4 \| 108 \| (2, 109) \| 112 \| (6, 113) \| 116 \| GPW/gp25 family protein \| GPW/gp25 family protein; uniclust \| UniRef100\_A0A3D2C224 \| 99.5 \| 9e-17 \| 1.7e-22 \| 95.7 \| 78 \| (3, 80) \| 112 \| (46, 125) \| 127 \| IraD/Gp25-like domain-containing protein (Fragment) \| IraD/Gp25-like domain-containing protein (Fragment); uniclust \| UniRef100\_UPI0022B1D31B \| 99.5 \| 9.5e-17 \| 1.7e-22 \| 94.9 \| 80 \| (4, 83) \| 112 \| (1, 80) \| 121 \| GPW/gp25 family protein \| GPW/gp25 family protein; uniclust \| UniRef100\_A0A086ML21 \| 99.5 \| 9.8e-17 \| 1.9e-22 \| 99.9 \| 100 \| (8, 108) \| 112 \| (20, 122) \| 131 \| IraD/Gp25-like domain-containing protein \| IraD/Gp25-like domain-containing protein; uniclust \| UniRef100\_UPI0003640FF3 \| 99.5 \| 1.1e-16 \| 2.1e-22 \| 97.1 \| 110 \| (2, 111) \| 112 \| (14, 127) \| 134 \| GPW/gp25 family protein \| GPW/gp25 family protein; uniclust \| UniRef100\_A0A0X1U7T5 \| 99.5 \| 1.8e-16 \| 3.7e-22 \| 99.7 \| 87 \| (12, 100) \| 112 \| (31, 117) \| 126 \| Protein 25-like lysozyme \| Protein 25-like lysozyme; uniclust \| UniRef100\_A0A117MMK7 \| 99.4 \| 3.1e-16 \| 6e-22 \| 97.7 \| 99 \| (11, 110) \| 112 \| (28, 127) \| 135 \| IraD/Gp25-like domain-containing protein \| IraD/Gp25-like domain-containing protein; uniclust \| UniRef100\_A0A2I7QP25 \| 99.4 \| 3.4e-16 \| 6.7e-22 \| 98.8 \| 102 \| (1, 102) \| 112 \| (19, 126) \| 143 \| Uncharacterized protein \| Uncharacterized protein; uniclust \| UniRef100\_A0A1H1G2U4 \| 99.4 \| 4e-16 \| 7.5e-22 \| 94.4 \| 108 \| (1, 109) \| 112 \| (1, 120) \| 122 \| IraD/Gp25-like domain-containing protein \| IraD/Gp25-like domain-containing protein; uniclust \| UniRef100\_A0A2D5K3P5 \| 99.4 \| 4.3e-16 \| 8.1e-22 \| 89.9 \| 63 \| (9, 71) \| 112 \| (23, 87) \| 87 \| IraD/Gp25-like domain-containing protein (Fragment) \| IraD/Gp25-like domain-containing protein (Fragment); uniclust \| UniRef100\_A0A069F3H4 \| 99.4 \| 4.9e-16 \| 1e-21 \| 98.3 \| 93 \| (11, 108) \| 112 \| (24, 118) \| 125 \| Uncharacterized protein \| Uncharacterized protein; uniclust \| UniRef100\_A0A0E4BWU6 \| 99.4 \| 5.7e-16 \| 1.1e-21 \| 94.4 \| 99 \| (3, 101) \| 112 \| (12, 111) \| 121 \| IraD/Gp25-like domain-containing protein \| IraD/Gp25-like domain-containing protein; uniclust \| UniRef100\_A0A031FQW9 \| 99.4 \| 5.6e-16 \| 1.1e-21 \| 96.8 \| 107 \| (1, 110) \| 112 \| (5, 114) \| 124 \| Phage baseplate assembly protein W \| Phage baseplate assembly protein W; uniclust \| UniRef100\_A0A024B013 \| 99.4 \| 7.8e-16 \| 1.6e-21 \| 106.9 \| 98 \| (11, 110) \| 112 \| (129, 227) \| 246 \| Baseplate protein \| Baseplate protein; uniclust \| UniRef100\_A0A0C2DUF9 \| 99.4 \| 8.4e-16 \| 1.6e-21 \| 99.5 \| 104 \| (6, 110) \| 112 \| (61, 173) \| 182 \| IraD/Gp25-like domain-containing protein \| IraD/Gp25-like domain-containing protein; uniclust \| UniRef100\_A0A069QGW3 \| 99.4 \| 8.3e-16 \| 1.7e-21 \| 101.4 \| 97 \| (11, 107) \| 112 \| (48, 158) \| 177 \| Putative lysozyme \| Putative lysozyme; uniclust \| UniRef100\_A0A0A7K9I0 \| 99.4 \| 9.4e-16 \| 1.9e-21 \| 101.5 \| 98 \| (11, 108) \| 112 \| (49, 161) \| 175 \| IraD/Gp25-like domain-containing protein \| IraD/Gp25-like domain-containing protein; uniclust \| UniRef100\_A0A2E0GD41 \| 99.4 \| 1.1e-15 \| 2.1e-21 \| 97.7 \| 98 \| (11, 108) \| 112 \| (34, 136) \| 156 \| IraD/Gp25-like domain-containing protein \| IraD/Gp25-like domain-containing protein; uniclust \| UniRef100\_A0A6J5L6U0 \| 99.4 \| 1.5e-15 \| 2.9e-21 \| 95.6 \| 107 \| (3, 110) \| 112 \| (17, 127) \| 142 \| GpW/Gp25/anti-adapter protein IraD \| GpW/Gp25/anti-adapter protein IraD; uniclust \| UniRef100\_A0A370LRT3 \| 99.4 \| 1.7e-15 \| 3.1e-21 \| 93.9 \| 73 \| (11, 83) \| 112 \| (22, 94) \| 143 \| IraD/Gp25-like domain-containing protein \| IraD/Gp25-like domain-containing protein; uniclust \| UniRef100\_A0A085ZBD9 \| 99.4 \| 1.8e-15 \| 3.7e-21 \| 99.2 \| 98 \| (11, 108) \| 112 \| (43, 153) \| 163 \| IraD/Gp25-like domain-containing protein \| IraD/Gp25-like domain-containing protein; uniclust \| UniRef100\_A0A1F9LNZ8 \| 99.4 \| 2.6e-15 \| 4.9e-21 \| 94.0 \| 96 \| (4, 99) \| 112 \| (24, 123) \| 150 \| IraD/Gp25-like domain-containing protein (Fragment) \| IraD/Gp25-like domain-containing protein (Fragment); uniclust \| UniRef100\_A0A1Z9PTU6 \| 99.4 \| 2.8e-15 \| 5.4e-21 \| 95.6 \| 94 \| (11, 106) \| 112 \| (42, 135) \| 156 \| IraD/Gp25-like domain-containing protein \| IraD/Gp25-like domain-containing protein; uniclust \| UniRef100\_A0A6P0Q329 \| 99.3 \| 3.3e-15 \| 6.1e-21 \| 84.7 \| 73 \| (4, 76) \| 112 \| (18, 92) \| 92 \| GPW/gp25 family protein (Fragment) \| GPW/gp25 family protein (Fragment); uniclust \| UniRef100\_A0A1V5LTW0 \| 99.3 \| 3.4e-15 \| 6.3e-21 \| 87.2 \| 79 \| (12, 91) \| 112 \| (29, 107) \| 110 \| Gene 25-like lysozyme \| Gene 25-like lysozyme; uniclust \| UniRef100\_A0A068T9E9 \| 99.3 \| 3.5e-15 \| 6.7e-21 \| 92.6 \| 102 \| (2, 103) \| 112 \| (9, 111) \| 137 \| Putative phage gp25 protein \| Putative phage gp25 protein; uniclust \| UniRef100\_A0A1J0EEV2 \| 99.3 \| 3.9e-15 \| 7.4e-21 \| 90.8 \| 90 \| (8, 99) \| 112 \| (22, 111) \| 125 \| Baseplate protein \| Baseplate protein; uniclust \| UniRef100\_A0A090HXB8 \| 99.3 \| 4.1e-15 \| 8.2e-21 \| 94.1 \| 97 \| (8, 109) \| 112 \| (25, 125) \| 130 \| Conserved domain protein \| Conserved domain protein; uniclust \| UniRef100\_A0A562GHU8 \| 99.3 \| 4.4e-15 \| 8.3e-21 \| 91.8 \| 100 \| (7, 108) \| 112 \| (31, 133) \| 144 \| Phage baseplate assembly protein W \| Phage baseplate assembly protein W; uniclust \| UniRef100\_A0A081NYB0 \| 99.3 \| 5e-15 \| 9.9e-21 \| 94.2 \| 88 \| (8, 96) \| 112 \| (23, 114) \| 142 \| IraD/Gp25-like domain-containing protein \| IraD/Gp25-like domain-containing protein; uniclust \| UniRef100\_A0A318L0U5 \| 99.3 \| 5.6e-15 \| 1e-20 \| 88.1 \| 102 \| (2, 103) \| 112 \| (9, 111) \| 125 \| IraD/Gp25-like domain-containing protein \| IraD/Gp25-like domain-containing protein; uniclust \| UniRef100\_A0A3C1NZW4 \| 99.3 \| 6.5e-15 \| 1.2e-20 \| 77.2 \| 57 \| (20, 76) \| 112 \| (3, 59) \| 59 \| Baseplate protein (Fragment) \| Baseplate protein (Fragment); uniclust \| UniRef100\_A0A0B5FRN4 \| 99.3 \| 7.4e-15 \| 1.4e-20 \| 99.6 \| 97 \| (13, 109) \| 112 \| (31, 131) \| 305 \| IraD/Gp25-like domain-containing protein \| IraD/Gp25-like domain-containing protein; uniclust \| UniRef100\_A0A381WMD8 \| 99.3 \| 7.5e-15 \| 1.4e-20 \| 84.9 \| 73 \| (4, 76) \| 112 \| (18, 92) \| 94 \| IraD/Gp25-like domain-containing protein \| IraD/Gp25-like domain-containing protein; uniclust \| UniRef100\_A0A2T6NFS1 \| 99.3 \| 8.8e-15 \| 1.6e-20 \| 92.5 \| 100 \| (11, 110) \| 112 \| (77, 178) \| 185 \| IraD/Gp25-like domain-containing protein \| IraD/Gp25-like domain-containing protein; uniclust \| UniRef100\_Q7MB49 \| 99.3 \| 9.1e-15 \| 1.7e-20 \| 95.7 \| 96 \| (6, 101) \| 112 \| (121, 219) \| 239 \| Photorhabdus luminescens subsp. laumondii TTO1 complete genome; segment 6/17 \| Photorhabdus luminescens subsp. laumondii TTO1 complete genome; segment 6/17; uniclust \| UniRef100\_A0A550H8T3 \| 99.3 \| 9.8e-15 \| 1.8e-20 \| 90.6 \| 98 \| (12, 109) \| 112 \| (39, 140) \| 146 \| IraD/Gp25-like domain-containing protein \| IraD/Gp25-like domain-containing protein; uniclust \| UniRef100\_A0A6I7PSS5 \| 99.3 \| 1.9e-14 \| 3.6e-20 \| 86.2 \| 98 \| (13, 110) \| 112 \| (26, 125) \| 127 \| IraD/Gp25-like domain-containing protein \| IraD/Gp25-like domain-containing protein; uniclust \| UniRef100\_A0A853IK11 \| 99.3 \| 2e-14 \| 3.7e-20 \| 77.5 \| 52 \| (1, 52) \| 112 \| (9, 60) \| 63 \| Baseplate assembly protein W (Fragment) \| Baseplate assembly protein W (Fragment); uniclust \| UniRef100\_A0A1T2XCA8 \| 99.3 \| 1.9e-14 \| 3.7e-20 \| 87.3 \| 85 \| (16, 103) \| 112 \| (17, 101) \| 108 \| Phage baseplate assembly protein W \| Phage baseplate assembly protein W; uniclust \| UniRef100\_A0A023D1J8 \| 99.3 \| 1.9e-14 \| 3.7e-20 \| 91.2 \| 100 \| (8, 111) \| 112 \| (21, 125) \| 130 \| Phage tail protein \| Phage tail protein; uniclust \| UniRef100\_A0A4U0FCL7 \| 99.3 \| 2.1e-14 \| 3.9e-20 \| 89.8 \| 106 \| (4, 110) \| 112 \| (24, 136) \| 153 \| DUF2634 domain-containing protein \| DUF2634 domain-containing protein; uniclust \| UniRef100\_A0A080M8X4 \| 99.3 \| 2e-14 \| 4e-20 \| 91.1 \| 98 \| (11, 110) \| 112 \| (38, 140) \| 145 \| DUF2634 domain-containing protein \| DUF2634 domain-containing protein; uniclust \| UniRef100\_UPI0022B9D4FA \| 99.3 \| 2.2e-14 \| 4e-20 \| 93.4 \| 97 \| (13, 109) \| 112 \| (31, 131) \| 228 \| GPW/gp25 family protein \| GPW/gp25 family protein; uniclust \| UniRef100\_A0A2D6F2M5 \| 99.3 \| 2.1e-14 \| 4.1e-20 \| 92.3 \| 87 \| (13, 99) \| 112 \| (53, 140) \| 161 \| IraD/Gp25-like domain-containing protein \| IraD/Gp25-like domain-containing protein; uniclust \| UniRef100\_A0A0X3VIR3 \| 99.3 \| 2.2e-14 \| 4.4e-20 \| 91.6 \| 92 \| (8, 99) \| 112 \| (18, 114) \| 142 \| IraD/Gp25-like domain-containing protein \| IraD/Gp25-like domain-containing protein; uniclust \| UniRef100\_A0A1W9VM06 \| 99.3 \| 2.6e-14 \| 4.8e-20 \| 85.2 \| 95 \| (12, 107) \| 112 \| (15, 110) \| 115 \| IraD/Gp25-like domain-containing protein \| IraD/Gp25-like domain-containing protein; uniclust \| UniRef100\_A0A090WBV0 \| 99.2 \| 2.8e-14 \| 5.6e-20 \| 93.3 \| 96 \| (11, 106) \| 112 \| (28, 137) \| 154 \| Gene 25-like lysozyme \| Gene 25-like lysozyme; uniclust \| UniRef100\_A0A1J5PQ93 \| 99.2 \| 3.2e-14 \| 5.8e-20 \| 89.5 \| 74 \| (9, 83) \| 112 \| (19, 92) \| 176 \| Protein 25-like lysozyme \| Protein 25-like lysozyme; uniclust \| UniRef100\_A0A4Q3KFV4 \| 99.2 \| 5.7e-14 \| 1.1e-19 \| 96.8 \| 107 \| (4, 110) \| 112 \| (1, 113) \| 365 \| Baseplate assembly protein \| Baseplate assembly protein; uniclust \| UniRef100\_A0A0H4WV92 \| 99.2 \| 5.4e-14 \| 1.1e-19 \| 89.4 \| 96 \| (11, 108) \| 112 \| (16, 112) \| 133 \| IraD/Gp25-like domain-containing protein \| IraD/Gp25-like domain-containing protein; uniclust \| UniRef100\_A0A936U591 \| 99.2 \| 6.1e-14 \| 1.1e-19 \| 90.1 \| 72 \| (12, 83) \| 112 \| (4, 75) \| 203 \| GPW/gp25 family protein \| GPW/gp25 family protein; uniclust \| UniRef100\_A0A0J8DAG7 \| 99.2 \| 7.3e-14 \| 1.4e-19 \| 86.4 \| 87 \| (11, 99) \| 112 \| (19, 106) \| 114 \| Putative lysozyme \| Putative lysozyme; uniclust \| UniRef100\_A0A0N7CEU8 \| 99.2 \| 8.2e-14 \| 1.5e-19 \| 85.5 \| 106 \| (2, 107) \| 112 \| (22, 130) \| 137 \| Putative baseplate wedge subunit \| Putative baseplate wedge subunit; uniclust \| UniRef100\_A0A7X5D8Q1 \| 99.2 \| 8.7e-14 \| 1.6e-19 \| 86.6 \| 101 \| (3, 103) \| 112 \| (45, 149) \| 162 \| IraD/Gp25-like domain-containing protein \| IraD/Gp25-like domain-containing protein; uniclust \| UniRef100\_A0A7L6A4Z7 \| 99.2 \| 1e-13 \| 1.8e-19 \| 89.1 \| 103 \| (8, 110) \| 112 \| (25, 130) \| 202 \| GPW/gp25 family protein \| GPW/gp25 family protein; uniclust \| UniRef100\_A0A084SY44 \| 99.2 \| 9.6e-14 \| 1.9e-19 \| 89.1 \| 90 \| (11, 102) \| 112 \| (38, 129) \| 150 \| IraD/Gp25-like domain-containing protein \| IraD/Gp25-like domain-containing protein; uniclust \| UniRef100\_A0A158DVL2 \| 99.2 \| 9.8e-14 \| 1.9e-19 \| 96.3 \| 89 \| (8, 98) \| 112 \| (109, 200) \| 248 \| Bacteriophage protein \| Bacteriophage protein; uniclust \| UniRef100\_A0A074LKB3 \| 99.2 \| 1e-13 \| 2e-19 \| 87.4 \| 93 \| (8, 101) \| 112 \| (21, 116) \| 133 \| Phage baseplate protein \| Phage baseplate protein; uniclust \| UniRef100\_A0A0E2HGZ0 \| 99.2 \| 1e-13 \| 2e-19 \| 89.2 \| 103 \| (7, 110) \| 112 \| (25, 130) \| 142 \| Phage protein \| Phage protein; uniclust \| UniRef100\_A0A0F9Q0T0 \| 99.2 \| 1.2e-13 \| 2.2e-19 \| 88.7 \| 93 \| (11, 103) \| 112 \| (48, 151) \| 164 \| IraD/Gp25-like domain-containing protein \| IraD/Gp25-like domain-containing protein; uniclust \| UniRef100\_UPI0011A0CDE0 \| 99.2 \| 1.3e-13 \| 2.3e-19 \| 95.3 \| 99 \| (1, 99) \| 112 \| (6, 104) \| 368 \| contractile injection system protein, VgrG/Pvc8 family \| contractile injection system protein, VgrG/Pvc8 family; uniclust \| UniRef100\_A0A2N1PDC8 \| 99.2 \| 1.2e-13 \| 2.4e-19 \| 85.3 \| 94 \| (11, 104) \| 112 \| (17, 115) \| 124 \| IraD/Gp25-like domain-containing protein \| IraD/Gp25-like domain-containing protein; uniclust \| UniRef100\_A0A017T4E9 \| 99.2 \| 1.3e-13 \| 2.6e-19 \| 92.6 \| 88 \| (13, 102) \| 112 \| (46, 136) \| 178 \| IraD/Gp25-like domain-containing protein \| IraD/Gp25-like domain-containing protein; uniclust \| UniRef100\_A0A379WI79 \| 99.2 \| 1.4e-13 \| 2.7e-19 \| 81.1 \| 74 \| (9, 83) \| 112 \| (20, 93) \| 101 \| Baseplate protein \| Baseplate protein; uniclust \| UniRef100\_A0A061NU27 \| 99.2 \| 1.4e-13 \| 2.7e-19 \| 84.2 \| 85 \| (13, 99) \| 112 \| (19, 104) \| 107 \| IraD/Gp25-like domain-containing protein \| IraD/Gp25-like domain-containing protein; uniclust \| UniRef100\_A0A1E5XW96 \| 99.2 \| 1.4e-13 \| 2.8e-19 \| 89.8 \| 99 \| (11, 110) \| 112 \| (62, 165) \| 170 \| IraD/Gp25-like domain-containing protein \| IraD/Gp25-like domain-containing protein; uniclust \| UniRef100\_A0A150Q962 \| 99.2 \| 1.7e-13 \| 3.2e-19 \| 86.9 \| 99 \| (8, 109) \| 112 \| (23, 123) \| 141 \| IraD/Gp25-like domain-containing protein \| IraD/Gp25-like domain-containing protein; uniclust \| UniRef100\_A0A323U6E9 \| 99.2 \| 1.7e-13 \| 3.4e-19 \| 87.7 \| 99 \| (9, 110) \| 112 \| (21, 119) \| 138 \| IraD/Gp25-like domain-containing protein \| IraD/Gp25-like domain-containing protein; uniclust \| UniRef100\_A0A812QV09 \| 99.2 \| 1.9e-13 \| 3.5e-19 \| 104.4 \| 97 \| (11, 107) \| 112 \| (1069, 1169) \| 1173 \| XkdF protein \| XkdF protein; uniclust \| UniRef100\_A0A135ZED6 \| 99.2 \| 1.8e-13 \| 3.6e-19 \| 86.2 \| 96 \| (3, 101) \| 112 \| (16, 115) \| 133 \| DUF2634 domain-containing protein \| DUF2634 domain-containing protein; uniclust \| UniRef100\_A0A1Y3QHP6 \| 99.2 \| 1.9e-13 \| 3.6e-19 \| 87.4 \| 101 \| (8, 109) \| 112 \| (34, 138) \| 153 \| Uncharacterized protein (Fragment) \| Uncharacterized protein (Fragment); uniclust \| UniRef100\_A0A174Z8M2 \| 99.1 \| 2e-13 \| 4e-19 \| 86.1 \| 88 \| (11, 100) \| 112 \| (36, 124) \| 131 \| Gene 25-like lysozyme \| Gene 25-like lysozyme; uniclust \| UniRef100\_A0A381ZQG4 \| 99.1 \| 2.6e-13 \| 4.9e-19 \| 79.3 \| 84 \| (19, 102) \| 112 \| (5, 89) \| 99 \| IraD/Gp25-like domain-containing protein \| IraD/Gp25-like domain-containing protein; uniclust \| UniRef100\_A0A662G9R2 \| 99.1 \| 3.7e-13 \| 7e-19 \| 82.8 \| 97 \| (8, 105) \| 112 \| (24, 127) \| 137 \| Uncharacterized protein (Fragment) \| Uncharacterized protein (Fragment); uniclust \| UniRef100\_A0A010SSS2 \| 99.1 \| 4e-13 \| 7.7e-19 \| 86.1 \| 99 \| (8, 111) \| 112 \| (25, 128) \| 146 \| Phage tail protein \| Phage tail protein; uniclust \| UniRef100\_A0A1V5ZPA0 \| 99.1 \| 4.5e-13 \| 8.4e-19 \| 83.5 \| 101 \| (8, 108) \| 112 \| (30, 133) \| 145 \| Gene 25-like lysozyme \| Gene 25-like lysozyme; uniclust \| UniRef100\_A0A2D5F9D6 \| 99.1 \| 5.1e-13 \| 9.5e-19 \| 83.8 \| 96 \| (5, 100) \| 112 \| (39, 136) \| 147 \| IraD/Gp25-like domain-containing protein \| IraD/Gp25-like domain-containing protein; uniclust \| UniRef100\_A0A2D4SL70 \| 99.1 \| 5.8e-13 \| 1.1e-18 \| 85.9 \| 72 \| (11, 82) \| 112 \| (20, 91) \| 151 \| IraD/Gp25-like domain-containing protein \| IraD/Gp25-like domain-containing protein; uniclust \| UniRef100\_A0A0C1PQ31 \| 99.1 \| 6.6e-13 \| 1.3e-18 \| 84.2 \| 86 \| (11, 99) \| 112 \| (22, 109) \| 126 \| DUF2634 domain-containing protein \| DUF2634 domain-containing protein; uniclust \| UniRef100\_A0A060QLL0 \| 99.1 \| 6.8e-13 \| 1.3e-18 \| 84.0 \| 100 \| (8, 111) \| 112 \| (23, 127) \| 130 \| Putative phage tail protein \| Putative phage tail protein; uniclust \| UniRef100\_A0A7Z0LWZ0 \| 99.1 \| 7.3e-13 \| 1.4e-18 \| 96.3 \| 79 \| (1, 79) \| 112 \| (1, 79) \| 598 \| GPW/gp25 family protein \| GPW/gp25 family protein; uniclust \| UniRef100\_A0A069AW33 \| 99.1 \| 6.9e-13 \| 1.4e-18 \| 88.9 \| 92 \| (4, 98) \| 112 \| (45, 141) \| 178 \| Putative phage protein \| Putative phage protein; uniclust \| UniRef100\_A0A4Y7RJW3 \| 99.1 \| 7.6e-13 \| 1.5e-18 \| 85.1 \| 102 \| (8, 110) \| 112 \| (25, 130) \| 152 \| IraD/Gp25-like domain-containing protein \| IraD/Gp25-like domain-containing protein; uniclust \| UniRef100\_A0A3C2A343 \| 99.1 \| 8.4e-13 \| 1.5e-18 \| 83.5 \| 99 \| (11, 109) \| 112 \| (69, 169) \| 176 \| IraD/Gp25-like domain-containing protein \| IraD/Gp25-like domain-containing protein; uniclust \| UniRef100\_A0A2G6EIV6 \| 99.1 \| 8.8e-13 \| 1.6e-18 \| 87.5 \| 75 \| (2, 76) \| 112 \| (11, 85) \| 253 \| IraD/Gp25-like domain-containing protein \| IraD/Gp25-like domain-containing protein; uniclust \| UniRef100\_A0A6L3Z3G6 \| 99.1 \| 9e-13 \| 1.6e-18 \| 81.8 \| 102 \| (1, 102) \| 112 \| (43, 144) \| 154 \| IraD/Gp25-like domain-containing protein \| IraD/Gp25-like domain-containing protein; uniclust \| UniRef100\_A0A2S6HUW9 \| 99.1 \| 8.9e-13 \| 1.7e-18 \| 81.6 \| 84 \| (13, 99) \| 112 \| (20, 103) \| 123 \| IraD/Gp25-like domain-containing protein \| IraD/Gp25-like domain-containing protein; uniclust \| UniRef100\_A0A0H3ZTY4 \| 99.1 \| 8.9e-13 \| 1.7e-18 \| 81.1 \| 100 \| (1, 102) \| 112 \| (3, 105) \| 112 \| Phage protein \| Phage protein; uniclust \| UniRef100\_A0A161YFA5 \| 99.1 \| 9.1e-13 \| 1.7e-18 \| 78.8 \| 80 \| (17, 98) \| 112 \| (16, 95) \| 100 \| Gene 25-like lysozyme \| Gene 25-like lysozyme; uniclust \| UniRef100\_A0A0G3AVB6 \| 99.1 \| 9.6e-13 \| 1.8e-18 \| 81.6 \| 96 \| (13, 108) \| 112 \| (22, 119) \| 124 \| IraD/Gp25-like domain-containing protein \| IraD/Gp25-like domain-containing protein; uniclust \| UniRef100\_UPI0011E4B62D \| 99.1 \| 1e-12 \| 1.9e-18 \| 72.5 \| 71 \| (11, 82) \| 112 \| (1, 71) \| 75 \| GPW/gp25 family protein \| GPW/gp25 family protein; uniclust \| UniRef100\_A0A126QLP9 \| 99.0 \| 1.1e-12 \| 2.2e-18 \| 86.6 \| 104 \| (7, 111) \| 112 \| (51, 158) \| 183 \| Baseplate assembly protein \| Baseplate assembly protein; uniclust \| UniRef100\_UPI001E5F099E \| 99.0 \| 1.4e-12 \| 2.6e-18 \| 76.0 \| 70 \| (15, 84) \| 112 \| (5, 74) \| 103 \| GPW/gp25 family protein \| GPW/gp25 family protein; uniclust \| UniRef100\_A0A2V8VC79 \| 99.0 \| 1.3e-12 \| 2.6e-18 \| 85.1 \| 88 \| (13, 102) \| 112 \| (34, 124) \| 146 \| Type VI secretion system baseplate subunit TssE \| Type VI secretion system baseplate subunit TssE; uniclust \| UniRef100\_A0A061Q550 \| 99.0 \| 1.3e-12 \| 2.7e-18 \| 85.2 \| 94 \| (13, 108) \| 112 \| (46, 142) \| 157 \| IraD/Gp25-like domain-containing protein \| IraD/Gp25-like domain-containing protein; uniclust \| UniRef100\_A0A081NWR0 \| 99.0 \| 1.5e-12 \| 2.8e-18 \| 79.9 \| 84 \| (11, 98) \| 112 \| (21, 106) \| 120 \| Phage protein \| Phage protein; uniclust \| UniRef100\_A0A7C2B4J0 \| 99.0 \| 1.5e-12 \| 2.8e-18 \| 79.1 \| 86 \| (11, 97) \| 112 \| (25, 110) \| 134 \| IraD/Gp25-like domain-containing protein \| IraD/Gp25-like domain-containing protein; uniclust \| UniRef100\_A0A0R2JEB0 \| 99.0 \| 1.6e-12 \| 3e-18 \| 79.4 \| 84 \| (11, 98) \| 112 \| (17, 102) \| 115 \| IraD/Gp25-like domain-containing protein \| IraD/Gp25-like domain-containing protein; uniclust \| UniRef100\_A0A5C7JCT6 \| 99.0 \| 1.7e-12 \| 3.2e-18 \| 83.1 \| 79 \| (5, 83) \| 112 \| (22, 104) \| 159 \| IraD/Gp25-like domain-containing protein \| IraD/Gp25-like domain-containing protein; uniclust \| UniRef100\_A0A037YM48 \| 99.0 \| 1.6e-12 \| 3.2e-18 \| 86.1 \| 96 \| (12, 109) \| 112 \| (46, 145) \| 172 \| Type VI secretion protein \| Type VI secretion protein; uniclust \| UniRef100\_A0A0F9TG53 \| 99.0 \| 1.8e-12 \| 3.4e-18 \| 80.8 \| 98 \| (11, 109) \| 112 \| (40, 139) \| 159 \| IraD/Gp25-like domain-containing protein \| IraD/Gp25-like domain-containing protein; uniclust \| UniRef100\_A0A1C5ZER0 \| 99.0 \| 2e-12 \| 3.8e-18 \| 81.0 \| 104 \| (3, 110) \| 112 \| (10, 124) \| 126 \| Protein of uncharacterized function (DUF2634) \| Protein of uncharacterized function (DUF2634); uniclust \| UniRef100\_A0A1Q1PVL8 \| 99.0 \| 2.2e-12 \| 4.3e-18 \| 82.8 \| 94 \| (12, 106) \| 112 \| (37, 130) \| 145 \| Type IV secretion protein \| Type IV secretion protein; uniclust \| UniRef100\_A0A061NKR2 \| 99.0 \| 2.2e-12 \| 4.5e-18 \| 86.6 \| 93 \| (3, 98) \| 112 \| (50, 147) \| 177 \| Phage-like element PBSX protein XkdS \| Phage-like element PBSX protein XkdS; uniclust \| UniRef100\_A0A968VPK2 \| 99.0 \| 2.7e-12 \| 5e-18 \| 82.9 \| 73 \| (11, 83) \| 112 \| (27, 99) \| 203 \| GPW/gp25 family protein \| GPW/gp25 family protein; uniclust \| UniRef100\_D6H7Q3 \| 99.0 \| 2.8e-12 \| 5.2e-18 \| 87.0 \| 98 \| (5, 102) \| 112 \| (195, 294) \| 300 \| IraD/Gp25-like domain-containing protein \| IraD/Gp25-like domain-containing protein; uniclust \| UniRef100\_A0A4Q6D2R8 \| 99.0 \| 2.9e-12 \| 5.4e-18 \| 75.3 \| 98 \| (1, 99) \| 112 \| (1, 98) \| 106 \| Uncharacterized protein \| Uncharacterized protein; uniclust \| UniRef100\_A0A973HYV5 \| 99.0 \| 3e-12 \| 5.5e-18 \| 76.5 \| 104 \| (5, 108) \| 112 \| (9, 116) \| 119 \| GPW/gp25 family protein \| GPW/gp25 family protein; uniclust \| UniRef100\_B3QTI5 \| 99.0 \| 3.1e-12 \| 5.7e-18 \| 94.4 \| 90 \| (11, 102) \| 112 \| (22, 111) \| 687 \| GPW/gp25 family protein \| GPW/gp25 family protein; uniclust \| UniRef100\_A0A023PTA0 \| 99.0 \| 3.1e-12 \| 6.4e-18 \| 85.3 \| 89 \| (11, 101) \| 112 \| (50, 141) \| 166 \| Type VI secretion system protein \| Type VI secretion system protein; uniclust \| UniRef100\_A0A084JID6 \| 99.0 \| 3.6e-12 \| 7.1e-18 \| 81.6 \| 89 \| (7, 98) \| 112 \| (29, 122) \| 138 \| Phage protein \| Phage protein; uniclust \| UniRef100\_UPI000B2BD334 \| 99.0 \| 4e-12 \| 7.3e-18 \| 78.3 \| 74 \| (9, 83) \| 112 \| (37, 110) \| 144 \| GPW/gp25 family protein \| GPW/gp25 family protein; uniclust \| UniRef100\_A0A377AWY0 \| 99.0 \| 4.4e-12 \| 8.2e-18 \| 74.0 \| 86 \| (25, 110) \| 112 \| (1, 86) \| 94 \| Baseplate assembly protein W \| Baseplate assembly protein W; uniclust \| UniRef100\_A0A6J5KRL7 \| 99.0 \| 4.4e-12 \| 8.7e-18 \| 81.9 \| 86 \| (13, 99) \| 112 \| (31, 117) \| 143 \| Baseplate wedge subunit \| Baseplate wedge subunit; uniclust \| UniRef100\_A0A936LRJ0 \| 99.0 \| 4.8e-12 \| 8.9e-18 \| 78.2 \| 95 \| (14, 108) \| 112 \| (27, 124) \| 135 \| GPW/gp25 family protein \| GPW/gp25 family protein; uniclust \| UniRef100\_A0A958S4Z7 \| 99.0 \| 5e-12 \| 9.2e-18 \| 75.8 \| 100 \| (11, 110) \| 112 \| (18, 119) \| 121 \| GPW/gp25 family protein \| GPW/gp25 family protein; uniclust \| UniRef100\_A0A5J4RAP1 \| 98.9 \| 5.1e-12 \| 9.8e-18 \| 78.9 \| 91 \| (14, 104) \| 112 \| (29, 121) \| 129 \| IraD/Gp25-like domain-containing protein \| IraD/Gp25-like domain-containing protein; uniclust \| UniRef100\_A0A3D4L1F4 \| 98.9 \| 5.5e-12 \| 1e-17 \| 75.3 \| 85 \| (13, 99) \| 112 \| (19, 103) \| 107 \| IraD/Gp25-like domain-containing protein \| IraD/Gp25-like domain-containing protein; uniclust \| UniRef100\_A0A1Z9GYA1 \| 98.9 \| 5.6e-12 \| 1e-17 \| 81.4 \| 98 \| (11, 109) \| 112 \| (52, 150) \| 179 \| IraD/Gp25-like domain-containing protein \| IraD/Gp25-like domain-containing protein; uniclust \| UniRef100\_E2CFL3 \| 98.9 \| 5.7e-12 \| 1.1e-17 \| 76.5 \| 100 \| (2, 102) \| 112 \| (3, 103) \| 120 \| GpW/gp25 family protein \| GpW/gp25 family protein; uniclust \| UniRef100\_A0A645HFP9 \| 98.9 \| 5.9e-12 \| 1.1e-17 \| 78.1 \| 100 \| (11, 111) \| 112 \| (44, 148) \| 151 \| IraD/Gp25-like domain-containing protein \| IraD/Gp25-like domain-containing protein; uniclust \| UniRef100\_A0A0M0SE81 \| 98.9 \| 6.6e-12 \| 1.3e-17 \| 78.2 \| 93 \| (3, 102) \| 112 \| (12, 108) \| 117 \| IraD/Gp25-like domain-containing protein \| IraD/Gp25-like domain-containing protein; uniclust \| UniRef100\_UPI00106CED4F \| 98.9 \| 7.2e-12 \| 1.3e-17 \| 77.0 \| 100 \| (8, 107) \| 112 \| (23, 126) \| 133 \| GPW/gp25 family protein \| GPW/gp25 family protein; uniclust \| UniRef100\_A0A2X1LMF2 \| 98.9 \| 8.7e-12 \| 1.6e-17 \| 66.5 \| 51 \| (2, 52) \| 112 \| (6, 56) \| 58 \| Baseplate assembly protein W \| Baseplate assembly protein W; uniclust \| UniRef100\_A0A952LGA6 \| 98.9 \| 8.8e-12 \| 1.7e-17 \| 72.4 \| 66 \| (34, 99) \| 112 \| (2, 68) \| 86 \| GPW/gp25 family protein \| GPW/gp25 family protein; uniclust \| UniRef100\_A0A090SWE0 \| 98.9 \| 8.9e-12 \| 1.7e-17 \| 76.3 \| 108 \| (2, 109) \| 112 \| (5, 115) \| 124 \| Uncharacterized protein \| Uncharacterized protein; uniclust \| UniRef100\_A0A2U3KMN3 \| 98.9 \| 9.6e-12 \| 1.8e-17 \| 76.5 \| 95 \| (13, 108) \| 112 \| (26, 121) \| 126 \| IraD/Gp25-like domain-containing protein \| IraD/Gp25-like domain-containing protein; uniclust \| UniRef100\_A0A8S5UG22 \| 98.9 \| 1.1e-11 \| 2e-17 \| 77.0 \| 100 \| (11, 110) \| 112 \| (24, 126) \| 131 \| Baseplate wedge protein \| Baseplate wedge protein; uniclust \| UniRef100\_A0A015WIV6 \| 98.9 \| 1.2e-11 \| 2.4e-17 \| 79.5 \| 96 \| (12, 107) \| 112 \| (24, 133) \| 144 \| Lysozyme family protein \| Lysozyme family protein; uniclust \| UniRef100\_A0A0W0VMK7 \| 98.9 \| 1.3e-11 \| 2.5e-17 \| 78.4 \| 102 \| (4, 108) \| 112 \| (26, 129) \| 144 \| IraD/Gp25-like domain-containing protein \| IraD/Gp25-like domain-containing protein; uniclust \| UniRef100\_A0A1G8FCC5 \| 98.9 \| 1.3e-11 \| 2.5e-17 \| 78.0 \| 80 \| (11, 92) \| 112 \| (33, 113) \| 138 \| Phage baseplate assembly protein W \| Phage baseplate assembly protein W; uniclust \| UniRef100\_A0A3A9EVM2 \| 98.9 \| 1.7e-11 \| 3.1e-17 \| 74.5 \| 87 \| (4, 93) \| 112 \| (40, 126) \| 131 \| IraD/Gp25-like domain-containing protein \| IraD/Gp25-like domain-containing protein; uniclust \| UniRef100\_A0A096B5Z2 \| 98.9 \| 1.7e-11 \| 3.4e-17 \| 79.2 \| 88 \| (11, 101) \| 112 \| (21, 108) \| 133 \| Uncharacterized protein \| Uncharacterized protein; uniclust \| UniRef100\_A0A0D0QWP6 \| 98.9 \| 1.8e-11 \| 3.5e-17 \| 79.5 \| 93 \| (11, 105) \| 112 \| (43, 138) \| 146 \| Phage-like element PBSX protein xkdS \| Phage-like element PBSX protein xkdS; uniclust \| UniRef100\_A0A497PPN4 \| 98.9 \| 1.9e-11 \| 3.5e-17 \| 75.1 \| 85 \| (11, 96) \| 112 \| (30, 116) \| 131 \| IraD/Gp25-like domain-containing protein \| IraD/Gp25-like domain-containing protein; uniclust \| UniRef100\_UPI0020131D42 \| 98.9 \| 1.9e-11 \| 3.6e-17 \| 85.8 \| 75 \| (1, 75) \| 112 \| (151, 225) \| 396 \| phage baseplate assembly protein V \| phage baseplate assembly protein V; uniclust \| UniRef100\_A0A1V5IBU2 \| 98.8 \| 2.2e-11 \| 4.1e-17 \| 75.8 \| 70 \| (13, 82) \| 112 \| (26, 96) \| 153 \| Gene 25-like lysozyme \| Gene 25-like lysozyme; uniclust \| UniRef100\_A0A0F7L855 \| 98.8 \| 2.6e-11 \| 4.9e-17 \| 76.3 \| 91 \| (12, 102) \| 112 \| (37, 131) \| 144 \| IraD/Gp25-like domain-containing protein \| IraD/Gp25-like domain-containing protein; uniclust \| UniRef100\_A0A353BET6 \| 98.8 \| 2.8e-11 \| 5.1e-17 \| 76.0 \| 86 \| (13, 98) \| 112 \| (44, 137) \| 161 \| IraD/Gp25-like domain-containing protein \| IraD/Gp25-like domain-containing protein; uniclust \| UniRef100\_A0A2K8LN18 \| 98.8 \| 3e-11 \| 5.5e-17 \| 77.2 \| 88 \| (15, 103) \| 112 \| (84, 172) \| 182 \| IraD/Gp25-like domain-containing protein \| IraD/Gp25-like domain-containing protein; uniclust \| UniRef100\_A0A0B6CV24 \| 98.8 \| 3.4e-11 \| 6.2e-17 \| 71.5 \| 96 \| (3, 99) \| 112 \| (2, 98) \| 112 \| 25-like lysozyme family protein \| 25-like lysozyme family protein; uniclust \| UniRef100\_A0A2D6XAB7 \| 98.8 \| 3.8e-11 \| 7.2e-17 \| 79.9 \| 93 \| (13, 105) \| 112 \| (91, 189) \| 219 \| IraD/Gp25-like domain-containing protein \| IraD/Gp25-like domain-containing protein; uniclust \| UniRef100\_A0A023Q087 \| 98.8 \| 3.7e-11 \| 7.4e-17 \| 81.5 \| 90 \| (15, 106) \| 112 \| (47, 138) \| 190 \| Type VI secretion protein \| Type VI secretion protein; uniclust \| UniRef100\_A0A7X6JWS6 \| 98.8 \| 4e-11 \| 7.5e-17 \| 71.9 \| 74 \| (33, 106) \| 112 \| (1, 74) \| 109 \| IraD/Gp25-like domain-containing protein \| IraD/Gp25-like domain-containing protein; uniclust \| UniRef100\_A0A8S5MEP5 \| 98.8 \| 4.1e-11 \| 7.6e-17 \| 73.3 \| 89 \| (11, 99) \| 112 \| (25, 116) \| 135 \| Baseplate wedge protein \| Baseplate wedge protein; uniclust \| UniRef100\_A0A1M7CV50 \| 98.8 \| 4.1e-11 \| 7.7e-17 \| 74.9 \| 99 \| (11, 109) \| 112 \| (23, 123) \| 133 \| Gene 25-like lysozyme \| Gene 25-like lysozyme; uniclust \| UniRef100\_A0A0B0SDG5 \| 98.8 \| 3.9e-11 \| 7.7e-17 \| 81.9 \| 99 \| (11, 111) \| 112 \| (110, 208) \| 214 \| LysM domain-containing protein \| LysM domain-containing protein; uniclust \| UniRef100\_A0A0R1NHW6 \| 98.8 \| 4.4e-11 \| 8.3e-17 \| 74.1 \| 85 \| (11, 100) \| 112 \| (23, 108) \| 121 \| Phage protein \| Phage protein; uniclust \| UniRef100\_A0A1I1MPJ1 \| 98.8 \| 4.7e-11 \| 8.9e-17 \| 75.2 \| 73 \| (11, 83) \| 112 \| (23, 96) \| 140 \| Phage baseplate assembly protein W \| Phage baseplate assembly protein W; uniclust \| UniRef100\_A0A2D3VZ43 \| 98.8 \| 5e-11 \| 9.1e-17 \| 69.9 \| 87 \| (18, 106) \| 112 \| (7, 93) \| 103 \| Baseplate assembly protein \| Baseplate assembly protein; uniclust \| UniRef100\_A0A1Q6RI12 \| 98.8 \| 4.7e-11 \| 9.2e-17 \| 75.1 \| 88 \| (11, 100) \| 112 \| (21, 108) \| 123 \| IraD/Gp25-like domain-containing protein \| IraD/Gp25-like domain-containing protein; uniclust \| UniRef100\_A0A1R4HML8 \| 98.8 \| 5.1e-11 \| 9.6e-17 \| 74.1 \| 84 \| (11, 99) \| 112 \| (19, 105) \| 127 \| Uncharacterized protein similar to VCA0109 \| Uncharacterized protein similar to VCA0109; uniclust \| UniRef100\_A0A268THM0 \| 98.8 \| 5.4e-11 \| 1e-16 \| 70.9 \| 67 \| (17, 84) \| 112 \| (5, 72) \| 91 \| IraD/Gp25-like domain-containing protein \| IraD/Gp25-like domain-containing protein; uniclust \| UniRef100\_A0A0F7FGC4 \| 98.8 \| 6.1e-11 \| 1.1e-16 \| 71.2 \| 97 \| (11, 108) \| 112 \| (17, 113) \| 119 \| IraD/Gp25-like domain-containing protein \| IraD/Gp25-like domain-containing protein; uniclust \| UniRef100\_A0A3S4FEP7 \| 98.8 \| 6e-11 \| 1.1e-16 \| 70.2 \| 71 \| (1, 71) \| 112 \| (4, 74) \| 102 \| Phage baseplate assembly protein W \| Phage baseplate assembly protein W; uniclust \| UniRef100\_A0A482MLI0 \| 98.8 \| 6.1e-11 \| 1.1e-16 \| 76.2 \| 99 \| (10, 108) \| 112 \| (43, 146) \| 156 \| Baseplate assembly protein \| Baseplate assembly protein; uniclust \| UniRef100\_X0WMB6 \| 98.8 \| 7.4e-11 \| 1.4e-16 \| 69.1 \| 76 \| (24, 99) \| 112 \| (2, 78) \| 102 \| IraD/Gp25-like domain-containing protein (Fragment) \| IraD/Gp25-like domain-containing protein (Fragment); uniclust \| UniRef100\_A0A0M1LJ08 \| 98.8 \| 7.3e-11 \| 1.4e-16 \| 71.9 \| 71 \| (12, 84) \| 112 \| (16, 86) \| 104 \| IraD/Gp25-like domain-containing protein \| IraD/Gp25-like domain-containing protein; uniclust \| UniRef100\_A0A1Y6CPE4 \| 98.7 \| 8.3e-11 \| 1.5e-16 \| 69.4 \| 97 \| (1, 98) \| 112 \| (1, 97) \| 106 \| IraD/Gp25-like domain-containing protein \| IraD/Gp25-like domain-containing protein; uniclust \| UniRef100\_A0A0A3Z8U7 \| 98.7 \| 7.9e-11 \| 1.6e-16 \| 74.5 \| 105 \| (4, 110) \| 112 \| (6, 119) \| 122 \| Phage GP46 family protein \| Phage GP46 family protein; uniclust \| UniRef100\_A0A023WLJ8 \| 98.7 \| 8.1e-11 \| 1.6e-16 \| 78.4 \| 92 \| (13, 106) \| 112 \| (32, 126) \| 173 \| GPW/gp25 family protein \| GPW/gp25 family protein; uniclust \| UniRef100\_A0A010SNZ9 \| 98.7 \| 8.1e-11 \| 1.6e-16 \| 77.4 \| 92 \| (14, 106) \| 112 \| (36, 134) \| 154 \| Type VI secretion protein \| Type VI secretion protein; uniclust \| UniRef100\_A0A0A7GG36 \| 98.7 \| 8.6e-11 \| 1.6e-16 \| 72.1 \| 86 \| (11, 99) \| 112 \| (24, 110) \| 119 \| IraD/Gp25-like domain-containing protein \| IraD/Gp25-like domain-containing protein; uniclust \| UniRef100\_A0A399IS01 \| 98.7 \| 8.8e-11 \| 1.6e-16 \| 71.4 \| 97 \| (8, 105) \| 112 \| (16, 115) \| 123 \| DUF2634 domain-containing protein \| DUF2634 domain-containing protein; uniclust \| UniRef100\_A0A1B7X0G4 \| 98.7 \| 9e-11 \| 1.8e-16 \| 74.5 \| 89 \| (7, 102) \| 112 \| (22, 114) \| 127 \| Uncharacterized protein \| Uncharacterized protein; uniclust \| UniRef100\_A0A126R1Q4 \| 98.7 \| 9.5e-11 \| 1.8e-16 \| 73.2 \| 91 \| (7, 99) \| 112 \| (20, 114) \| 131 \| Phage-related protein \| Phage-related protein; uniclust \| UniRef100\_A0A0F4NK88 \| 98.7 \| 1e-10 \| 1.8e-16 \| 70.3 \| 70 \| (15, 84) \| 112 \| (35, 104) \| 119 \| IraD/Gp25-like domain-containing protein \| IraD/Gp25-like domain-containing protein; uniclust \| UniRef100\_UPI00211A9560 \| 98.7 \| 1e-10 \| 1.9e-16 \| 72.5 \| 70 \| (13, 82) \| 112 \| (74, 143) \| 145 \| GPW/gp25 family protein \| GPW/gp25 family protein; uniclust \| UniRef100\_A0A0F3RXN9 \| 98.7 \| 1e-10 \| 1.9e-16 \| 73.8 \| 91 \| (5, 99) \| 112 \| (22, 115) \| 135 \| IraD/Gp25-like domain-containing protein \| IraD/Gp25-like domain-containing protein; uniclust \| UniRef100\_A0A833ALP0 \| 98.7 \| 1e-10 \| 1.9e-16 \| 65.3 \| 46 \| (8, 53) \| 112 \| (24, 71) \| 71 \| Phage tail protein (Fragment) \| Phage tail protein (Fragment); uniclust \| UniRef100\_UPI00114C9B31 \| 98.7 \| 1.3e-10 \| 2.3e-16 \| 69.1 \| 79 \| (1, 79) \| 112 \| (1, 79) \| 103 \| hypothetical protein \| hypothetical protein; uniclust \| UniRef100\_A0A9D2RLV6 \| 98.7 \| 1.3e-10 \| 2.5e-16 \| 69.0 \| 84 \| (14, 99) \| 112 \| (10, 94) \| 97 \| Uncharacterized protein \| Uncharacterized protein; uniclust \| UniRef100\_A0A2A4Y9P5 \| 98.7 \| 1.3e-10 \| 2.6e-16 \| 74.5 \| 92 \| (14, 105) \| 112 \| (26, 119) \| 138 \| Type VI secretion system baseplate subunit TssE \| Type VI secretion system baseplate subunit TssE; uniclust \| UniRef100\_A0A1Q7YH61 \| 98.7 \| 1.4e-10 \| 2.7e-16 \| 73.3 \| 98 \| (11, 110) \| 112 \| (22, 131) \| 144 \| IraD/Gp25-like domain-containing protein \| IraD/Gp25-like domain-containing protein; uniclust \| UniRef100\_A0A074VBM7 \| 98.7 \| 1.6e-10 \| 2.9e-16 \| 72.7 \| 87 \| (11, 99) \| 112 \| (57, 143) \| 159 \| Phage baseplate assembly protein W \| Phage baseplate assembly protein W; uniclust \| UniRef100\_A0A3C0GL75 \| 98.7 \| 1.6e-10 \| 3e-16 \| 66.0 \| 70 \| (14, 83) \| 112 \| (3, 72) \| 87 \| IraD/Gp25-like domain-containing protein (Fragment) \| IraD/Gp25-like domain-containing protein (Fragment); uniclust \| UniRef100\_UPI001F535E75 \| 98.7 \| 1.8e-10 \| 3.4e-16 \| 73.7 \| 72 \| (2, 73) \| 112 \| (4, 75) \| 179 \| GPW/gp25 family protein \| GPW/gp25 family protein; uniclust \| UniRef100\_A0A352A2I0 \| 98.7 \| 2e-10 \| 4e-16 \| 75.6 \| 87 \| (11, 99) \| 112 \| (46, 135) \| 154 \| DUF2634 domain-containing protein \| DUF2634 domain-containing protein; uniclust \| UniRef100\_A0A075R4E3 \| 98.7 \| 2.2e-10 \| 4.3e-16 \| 73.2 \| 103 \| (8, 110) \| 112 \| (18, 128) \| 134 \| DUF2634 domain-containing protein \| DUF2634 domain-containing protein; uniclust \| UniRef100\_A0A0C1PSI5 \| 98.7 \| 2.3e-10 \| 4.6e-16 \| 73.8 \| 99 \| (3, 106) \| 112 \| (14, 118) \| 134 \| DUF2634 domain-containing protein \| DUF2634 domain-containing protein; uniclust \| UniRef100\_A0A843GHX7 \| 98.7 \| 2.5e-10 \| 4.6e-16 \| 70.0 \| 104 \| (4, 107) \| 112 \| (13, 123) \| 134 \| GPW/gp25 family protein \| GPW/gp25 family protein; uniclust \| UniRef100\_A0A0W0SAI8 \| 98.6 \| 2.6e-10 \| 4.8e-16 \| 73.1 \| 89 \| (13, 102) \| 112 \| (39, 130) \| 150 \| IraD/Gp25-like domain-containing protein \| IraD/Gp25-like domain-containing protein; uniclust \| UniRef100\_A0A8S5NX59 \| 98.6 \| 2.9e-10 \| 5.2e-16 \| 65.3 \| 84 \| (16, 100) \| 112 \| (2, 86) \| 89 \| Baseplate assembly protein \| Baseplate assembly protein; uniclust \| UniRef100\_A0A350PF63 \| 98.6 \| 2.9e-10 \| 5.4e-16 \| 67.8 \| 69 \| (13, 82) \| 112 \| (26, 94) \| 97 \| IraD/Gp25-like domain-containing protein (Fragment) \| IraD/Gp25-like domain-containing protein (Fragment); uniclust \| UniRef100\_A0A014M2C2 \| 98.6 \| 2.9e-10 \| 5.5e-16 \| 60.8 \| 49 \| (2, 50) \| 112 \| (7, 55) \| 55 \| Baseplate assembly protein (Fragment) \| Baseplate assembly protein (Fragment); uniclust \| UniRef100\_A0A084J9B0 \| 98.6 \| 2.8e-10 \| 5.6e-16 \| 77.9 \| 91 \| (8, 100) \| 112 \| (52, 148) \| 191 \| DUF2634 domain-containing protein \| DUF2634 domain-containing protein; uniclust \| UniRef100\_A0A1F5AKW5 \| 98.6 \| 3.2e-10 \| 6.1e-16 \| 73.9 \| 85 \| (15, 101) \| 112 \| (39, 126) \| 156 \| IraD/Gp25-like domain-containing protein \| IraD/Gp25-like domain-containing protein; uniclust \| UniRef100\_A0A150QKC3 \| 98.6 \| 3.4e-10 \| 6.8e-16 \| 73.5 \| 105 \| (1, 109) \| 112 \| (9, 115) \| 139 \| IraD/Gp25-like domain-containing protein \| IraD/Gp25-like domain-containing protein; uniclust \| UniRef100\_A0A7U3VTF8 \| 98.6 \| 4e-10 \| 7.3e-16 \| 67.2 \| 106 \| (1, 107) \| 112 \| (1, 106) \| 111 \| Baseplate assembly protein GpW \| Baseplate assembly protein GpW; uniclust \| UniRef100\_A0A8S4QTC3 \| 98.6 \| 4e-10 \| 7.4e-16 \| 87.2 \| 90 \| (11, 100) \| 112 \| (641, 730) \| 1131 \| Jg18748 protein \| Jg18748 protein; uniclust \| UniRef100\_A0A066TKD3 \| 98.6 \| 3.8e-10 \| 7.4e-16 \| 75.0 \| 83 \| (15, 98) \| 112 \| (43, 127) \| 175 \| IraD/Gp25-like domain-containing protein \| IraD/Gp25-like domain-containing protein; uniclust \| UniRef100\_UPI00035FE005 \| 98.6 \| 3.9e-10 \| 7.4e-16 \| 79.3 \| 97 \| (8, 108) \| 112 \| (194, 295) \| 304 \| DUF2634 domain-containing protein \| DUF2634 domain-containing protein; uniclust \| UniRef100\_UPI001F3F1559 \| 98.6 \| 4.2e-10 \| 7.8e-16 \| 70.6 \| 94 \| (11, 109) \| 112 \| (56, 151) \| 155 \| DUF2634 domain-containing protein \| DUF2634 domain-containing protein; uniclust \| UniRef100\_UPI0020B36D29 \| 98.6 \| 4.7e-10 \| 8.6e-16 \| 62.1 \| 62 \| (1, 62) \| 112 \| (2, 63) \| 72 \| hypothetical protein \| hypothetical protein; uniclust \| UniRef100\_A0A1F9MHG2 \| 98.6 \| 4.5e-10 \| 8.7e-16 \| 71.7 \| 86 \| (11, 97) \| 112 \| (25, 116) \| 138 \| IraD/Gp25-like domain-containing protein \| IraD/Gp25-like domain-containing protein; uniclust \| UniRef100\_A0A1S6L331 \| 98.6 \| 4.7e-10 \| 8.8e-16 \| 71.1 \| 99 \| (10, 108) \| 112 \| (31, 135) \| 141 \| IraD/Gp25-like domain-containing protein \| IraD/Gp25-like domain-containing protein; uniclust \| UniRef100\_UPI00211E9846 \| 98.6 \| 4.9e-10 \| 9e-16 \| 66.2 \| 71 \| (5, 75) \| 112 \| (19, 90) \| 105 \| GPW/gp25 family protein \| GPW/gp25 family protein; uniclust \| UniRef100\_A0A014N481 \| 98.6 \| 4.8e-10 \| 9.2e-16 \| 76.3 \| 85 \| (16, 101) \| 112 \| (52, 137) \| 204 \| IraD/Gp25-like domain-containing protein \| IraD/Gp25-like domain-containing protein; uniclust \| UniRef100\_S6ANJ8 \| 98.6 \| 5.2e-10 \| 9.6e-16 \| 70.3 \| 84 \| (11, 97) \| 112 \| (41, 126) \| 156 \| Uncharacterized protein \| Uncharacterized protein; uniclust \| UniRef100\_A0A4P9VI48 \| 98.6 \| 5.3e-10 \| 9.7e-16 \| 64.6 \| 78 \| (2, 81) \| 112 \| (3, 80) \| 92 \| Baseplate assembly protein W \| Baseplate assembly protein W; uniclust \| UniRef100\_A0A1F2QFX3 \| 98.6 \| 5.3e-10 \| 1e-15 \| 70.8 \| 105 \| (5, 110) \| 112 \| (15, 125) \| 138 \| IraD/Gp25-like domain-containing protein \| IraD/Gp25-like domain-containing protein; uniclust \| UniRef100\_UPI001FEB107A \| 98.6 \| 5.5e-10 \| 1e-15 \| 63.8 \| 81 \| (27, 108) \| 112 \| (1, 82) \| 86 \| GPW/gp25 family protein \| GPW/gp25 family protein; uniclust \| UniRef100\_A0A0B4ZXG6 \| 98.6 \| 5.7e-10 \| 1e-15 \| 63.0 \| 66 \| (37, 102) \| 112 \| (2, 68) \| 80 \| Baseplate wedge subunit \| Baseplate wedge subunit; uniclust \| UniRef100\_A0A7U6KDW8 \| 98.6 \| 5.5e-10 \| 1e-15 \| 68.0 \| 87 \| (11, 99) \| 112 \| (15, 101) \| 105 \| IraD/Gp25-like domain-containing protein \| IraD/Gp25-like domain-containing protein; uniclust \| UniRef100\_A0A317FGN1 \| 98.6 \| 5.8e-10 \| 1.1e-15 \| 75.2 \| 95 \| (11, 105) \| 112 \| (155, 251) \| 262 \| IraD/Gp25-like domain-containing protein \| IraD/Gp25-like domain-containing protein; uniclust \| UniRef100\_A0A0N9ML22 \| 98.6 \| 6.2e-10 \| 1.2e-15 \| 70.1 \| 83 \| (11, 99) \| 112 \| (27, 111) \| 123 \| Uncharacterized protein \| Uncharacterized protein; uniclust \| UniRef100\_A0A7V2KVL8 \| 98.6 \| 6.9e-10 \| 1.3e-15 \| 68.6 \| 88 \| (14, 101) \| 112 \| (15, 104) \| 139 \| GPW/gp25 family protein \| GPW/gp25 family protein; uniclust \| UniRef100\_A0A9D7PFK3 \| 98.6 \| 7.7e-10 \| 1.4e-15 \| 63.6 \| 74 \| (26, 99) \| 112 \| (1, 75) \| 89 \| GPW/gp25 family protein \| GPW/gp25 family protein; uniclust \| UniRef100\_A0A0M9DJQ4 \| 98.6 \| 7.5e-10 \| 1.4e-15 \| 68.2 \| 84 \| (11, 98) \| 112 \| (17, 102) \| 118 \| Phage portal protein \| Phage portal protein; uniclust \| UniRef100\_A0A074M821 \| 98.5 \| 7.8e-10 \| 1.5e-15 \| 70.6 \| 97 \| (11, 109) \| 112 \| (36, 133) \| 140 \| Phage portal protein \| Phage portal protein; uniclust \| UniRef100\_A0A8J3EAG9 \| 98.5 \| 9.2e-10 \| 1.7e-15 \| 60.0 \| 52 \| (1, 52) \| 112 \| (1, 52) \| 66 \| Uncharacterized protein \| Uncharacterized protein; uniclust \| UniRef100\_A0A1C3NFC8 \| 98.5 \| 9.7e-10 \| 1.8e-15 \| 69.8 \| 89 \| (10, 98) \| 112 \| (58, 150) \| 166 \| IraD/Gp25-like domain-containing protein \| IraD/Gp25-like domain-containing protein; uniclust \| UniRef100\_A0A350PFI1 \| 98.5 \| 9.8e-10 \| 1.8e-15 \| 66.6 \| 90 \| (11, 102) \| 112 \| (22, 111) \| 122 \| IraD/Gp25-like domain-containing protein \| IraD/Gp25-like domain-containing protein; uniclust \| UniRef100\_A0A947D6A6 \| 98.5 \| 9.7e-10 \| 1.8e-15 \| 71.9 \| 80 \| (2, 81) \| 112 \| (36, 115) \| 170 \| Uncharacterized protein \| Uncharacterized protein; uniclust \| UniRef100\_A0A1S1YVC0 \| 98.5 \| 1e-09 \| 1.9e-15 \| 66.4 \| 85 \| (13, 98) \| 112 \| (24, 108) \| 120 \| IraD/Gp25-like domain-containing protein \| IraD/Gp25-like domain-containing protein; uniclust \| UniRef100\_A0A063BG98 \| 98.5 \| 1e-09 \| 2e-15 \| 72.2 \| 107 \| (3, 110) \| 112 \| (37, 146) \| 164 \| GP46 family protein \| GP46 family protein; uniclust \| UniRef100\_F4BFR9 \| 98.5 \| 1.2e-09 \| 2.3e-15 \| 61.6 \| 73 \| (35, 107) \| 112 \| (1, 75) \| 79 \| IraD/Gp25-like domain-containing protein \| IraD/Gp25-like domain-containing protein; uniclust \| UniRef100\_UPI000586A777 \| 98.5 \| 1.3e-09 \| 2.3e-15 \| 63.1 \| 73 \| (31, 103) \| 112 \| (1, 75) \| 91 \| GPW/gp25 family protein \| GPW/gp25 family protein; uniclust \| UniRef100\_A0A0F6SF01 \| 98.5 \| 1.2e-09 \| 2.3e-15 \| 71.2 \| 99 \| (11, 110) \| 112 \| (45, 146) \| 159 \| IraD/Gp25-like domain-containing protein \| IraD/Gp25-like domain-containing protein; uniclust \| UniRef100\_UPI001475626A \| 98.5 \| 1.3e-09 \| 2.5e-15 \| 57.2 \| 45 \| (14, 58) \| 112 \| (4, 48) \| 54 \| GPW/gp25 family protein \| GPW/gp25 family protein; uniclust \| UniRef100\_A0A8S0HU87 \| 98.5 \| 1.4e-09 \| 2.6e-15 \| 59.5 \| 57 \| (27, 83) \| 112 \| (1, 57) \| 67 \| IraD/Gp25-like domain-containing protein \| IraD/Gp25-like domain-containing protein; uniclust \| UniRef100\_A0A2V8SEE1 \| 98.5 \| 1.5e-09 \| 2.8e-15 \| 61.4 \| 57 \| (54, 110) \| 112 \| (5, 62) \| 69 \| Baseplate protein (Fragment) \| Baseplate protein (Fragment); uniclust \| UniRef100\_A0A1Z8WSX8 \| 98.5 \| 1.6e-09 \| 3.1e-15 \| 73.5 \| 87 \| (13, 99) \| 112 \| (67, 159) \| 175 \| IraD/Gp25-like domain-containing protein \| IraD/Gp25-like domain-containing protein; uniclust \| UniRef100\_G8LQ32 \| 98.5 \| 1.8e-09 \| 3.3e-15 \| 64.7 \| 65 \| (2, 66) \| 112 \| (6, 70) \| 112 \| GPW/Gp25 Family Protein \| GPW/Gp25 Family Protein; uniclust \| UniRef100\_UPI00058EE6FE \| 98.5 \| 1.8e-09 \| 3.4e-15 \| 63.3 \| 81 \| (13, 95) \| 112 \| (16, 96) \| 99 \| hypothetical protein \| hypothetical protein; uniclust \| UniRef100\_A0A398BF99 \| 98.5 \| 1.8e-09 \| 3.5e-15 \| 69.8 \| 102 \| (5, 108) \| 112 \| (35, 142) \| 147 \| DUF2634 domain-containing protein \| DUF2634 domain-containing protein; uniclust \| UniRef100\_UPI001D00D725 \| 98.5 \| 2.1e-09 \| 3.8e-15 \| 71.7 \| 89 \| (11, 99) \| 112 \| (127, 215) \| 234 \| GPW/gp25 family protein \| GPW/gp25 family protein; uniclust \| UniRef100\_A0A0N1JQ98 \| 98.5 \| 2.1e-09 \| 3.9e-15 \| 69.5 \| 68 \| (15, 83) \| 112 \| (38, 105) \| 145 \| ImpF \| ImpF; uniclust \| UniRef100\_A0A2D8ERB6 \| 98.4 \| 2.2e-09 \| 4.1e-15 \| 66.2 \| 99 \| (8, 106) \| 112 \| (17, 124) \| 134 \| IraD/Gp25-like domain-containing protein \| IraD/Gp25-like domain-containing protein; uniclust \| UniRef100\_A0A174T7Q2 \| 98.4 \| 2.1e-09 \| 4.2e-15 \| 66.0 \| 85 \| (12, 98) \| 112 \| (7, 93) \| 99 \| Early E1A protein \| Early E1A protein; uniclust \| UniRef100\_A0A7C4DJH8 \| 98.4 \| 2.5e-09 \| 4.6e-15 \| 69.2 \| 65 \| (11, 75) \| 112 \| (27, 91) \| 186 \| IraD/Gp25-like domain-containing protein \| IraD/Gp25-like domain-containing protein; uniclust \| UniRef100\_A0A968I5W3 \| 98.4 \| 2.6e-09 \| 4.7e-15 \| 66.4 \| 90 \| (11, 100) \| 112 \| (20, 114) \| 141 \| GPW/gp25 family protein \| GPW/gp25 family protein; uniclust \| UniRef100\_A0A0K1N0J6 \| 98.4 \| 2.6e-09 \| 4.9e-15 \| 67.9 \| 83 \| (13, 97) \| 112 \| (28, 113) \| 138 \| Type VI secretion system baseplate subunit TssE \| Type VI secretion system baseplate subunit TssE; uniclust \| UniRef100\_A0A509B7Z6 \| 98.4 \| 3.2e-09 \| 5.9e-15 \| 64.9 \| 86 \| (14, 101) \| 112 \| (4, 93) \| 113 \| Type VI secretion system lysozyme-like protein \| Type VI secretion system lysozyme-like protein; uniclust \| UniRef100\_A0A962Q7G8 \| 98.4 \| 3.6e-09 \| 6.6e-15 \| 64.6 \| 87 \| (11, 98) \| 112 \| (21, 108) \| 125 \| GPW/gp25 family protein \| GPW/gp25 family protein; uniclust \| UniRef100\_UPI001EF54EB6 \| 98.4 \| 3.6e-09 \| 6.6e-15 \| 60.7 \| 48 \| (8, 55) \| 112 \| (30, 79) \| 86 \| GPW/gp25 family protein \| GPW/gp25 family protein; uniclust \| UniRef100\_A0A327JB01 \| 98.4 \| 3.9e-09 \| 7.2e-15 \| 65.1 \| 91 \| (8, 100) \| 112 \| (33, 124) \| 134 \| IraD/Gp25-like domain-containing protein \| IraD/Gp25-like domain-containing protein; uniclust \| UniRef100\_A0A7C3ETG1 \| 98.4 \| 4e-09 \| 7.4e-15 \| 66.2 \| 84 \| (15, 98) \| 112 \| (36, 127) \| 141 \| IraD/Gp25-like domain-containing protein \| IraD/Gp25-like domain-containing protein; uniclust \| UniRef100\_A0A8J7EID1 \| 98.4 \| 4.8e-09 \| 8.7e-15 \| 62.6 \| 71 \| (13, 84) \| 112 \| (2, 75) \| 108 \| GPW/gp25 family protein \| GPW/gp25 family protein; uniclust \| UniRef100\_A0A9D2GTM2 \| 98.4 \| 5e-09 \| 9.1e-15 \| 60.4 \| 78 \| (12, 92) \| 112 \| (6, 83) \| 88 \| GPW/gp25 family protein \| GPW/gp25 family protein; uniclust \| UniRef100\_A0A248SJW8 \| 98.4 \| 5.1e-09 \| 9.3e-15 \| 63.9 \| 98 \| (11, 109) \| 112 \| (23, 121) \| 124 \| IraD/Gp25-like domain-containing protein \| IraD/Gp25-like domain-containing protein; uniclust \| UniRef100\_A0A4R6LWW8 \| 98.4 \| 4.9e-09 \| 9.4e-15 \| 65.5 \| 85 \| (11, 99) \| 112 \| (22, 108) \| 120 \| Uncharacterized protein \| Uncharacterized protein; uniclust \| UniRef100\_A0A662Q525 \| 98.4 \| 5.3e-09 \| 9.8e-15 \| 64.8 \| 86 \| (14, 99) \| 112 \| (38, 123) \| 137 \| IraD/Gp25-like domain-containing protein \| IraD/Gp25-like domain-containing protein; uniclust \| UniRef100\_A0A509B7P4 \| 98.4 \| 5.5e-09 \| 1e-14 \| 58.7 \| 42 \| (11, 52) \| 112 \| (23, 64) \| 71 \| Putative bacteriophage baseplate protein \| Putative bacteriophage baseplate protein; uniclust \| UniRef100\_UPI001CA5D6B8 \| 98.3 \| 6e-09 \| 1.1e-14 \| 63.2 \| 93 \| (12, 107) \| 112 \| (22, 115) \| 119 \| hypothetical protein \| hypothetical protein; uniclust \| UniRef100\_A0A6J5RXQ6 \| 98.3 \| 6.2e-09 \| 1.1e-14 \| 63.4 \| 100 \| (11, 110) \| 112 \| (20, 119) \| 123 \| COG3628 Phage baseplate assembly protein W \| COG3628 Phage baseplate assembly protein W; uniclust \| UniRef100\_U2F0H1 \| 98.3 \| 6.4e-09 \| 1.2e-14 \| 58.9 \| 64 \| (35, 98) \| 112 \| (1, 64) \| 73 \| Baseplate assembly protein W, putative \| Baseplate assembly protein W, putative; uniclust \| UniRef100\_A0A1C5WXX2 \| 98.3 \| 6e-09 \| 1.2e-14 \| 67.5 \| 82 \| (11, 97) \| 112 \| (31, 112) \| 131 \| Uncharacterized protein \| Uncharacterized protein; uniclust \| UniRef100\_A0A173R5E2 \| 98.3 \| 6.6e-09 \| 1.3e-14 \| 64.6 \| 83 \| (13, 98) \| 112 \| (7, 91) \| 106 \| Early E1A protein \| Early E1A protein; uniclust \| UniRef100\_A0A0F6TGM0 \| 98.3 \| 7.2e-09 \| 1.4e-14 \| 65.0 \| 100 \| (5, 105) \| 112 \| (18, 120) \| 129 \| Putative base plate wedge subunit \| Putative base plate wedge subunit; uniclust \| UniRef100\_A0A021X862 \| 98.3 \| 7.6e-09 \| 1.5e-14 \| 68.2 \| 88 \| (7, 98) \| 112 \| (36, 128) \| 147 \| Putative structural protein \| Putative structural protein; uniclust \| UniRef100\_A0A1E3G6B2 \| 98.3 \| 8.5e-09 \| 1.6e-14 \| 63.1 \| 104 \| (2, 105) \| 112 \| (9, 120) \| 125 \| IraD/Gp25-like domain-containing protein \| IraD/Gp25-like domain-containing protein; uniclust \| UniRef100\_A0A0C2UZ28 \| 98.3 \| 8.7e-09 \| 1.6e-14 \| 72.4 \| 92 \| (7, 100) \| 112 \| (184, 278) \| 295 \| LysM domain-containing protein \| LysM domain-containing protein; uniclust \| UniRef100\_A0A069DI41 \| 98.3 \| 8.7e-09 \| 1.7e-14 \| 66.7 \| 75 \| (8, 83) \| 112 \| (26, 102) \| 141 \| Uncharacterized protein \| Uncharacterized protein; uniclust \| UniRef100\_UPI0011682DE8 \| 98.3 \| 9.2e-09 \| 1.7e-14 \| 58.1 \| 66 \| (10, 77) \| 112 \| (12, 77) \| 78 \| GPW/gp25 family protein \| GPW/gp25 family protein; uniclust \| UniRef100\_A0A9E7CRK2 \| 98.3 \| 9.7e-09 \| 1.8e-14 \| 64.7 \| 85 \| (11, 96) \| 112 \| (36, 121) \| 142 \| DUF2634 domain-containing protein \| DUF2634 domain-containing protein; uniclust \| UniRef100\_R7UZ90 \| 98.3 \| 1e-08 \| 1.9e-14 \| 73.8 \| 67 \| (11, 77) \| 112 \| (366, 432) \| 433 \| Gp5/Type VI secretion system Vgr protein OB-fold domain-containing protein (Fragment) \| Gp5/Type VI secretion system Vgr protein OB-fold domain-containing protein (Fragment); uniclust \| UniRef100\_A0A1Y3AZE0 \| 98.3 \| 1e-08 \| 1.9e-14 \| 84.6 \| 87 \| (11, 97) \| 112 \| (3006, 3094) \| 3230 \| Chitinase-like protein \| Chitinase-like protein; uniclust \| UniRef100\_A8V442 \| 98.3 \| 1.1e-08 \| 2e-14 \| 59.9 \| 67 \| (8, 75) \| 112 \| (19, 87) \| 95 \| Uncharacterized protein \| Uncharacterized protein; uniclust \| UniRef100\_A0A017RUG6 \| 98.3 \| 9.6e-09 \| 2e-14 \| 69.1 \| 87 \| (11, 99) \| 112 \| (51, 140) \| 156 \| Phage protein \| Phage protein; uniclust \| UniRef100\_A0A932TVE7 \| 98.3 \| 1.2e-08 \| 2.2e-14 \| 68.5 \| 88 \| (11, 99) \| 112 \| (131, 223) \| 239 \| DUF2634 domain-containing protein \| DUF2634 domain-containing protein; uniclust \| UniRef100\_A0A933KI50 \| 98.3 \| 1.2e-08 \| 2.2e-14 \| 73.2 \| 100 \| (8, 109) \| 112 \| (317, 419) \| 423 \| DUF2634 domain-containing protein \| DUF2634 domain-containing protein; uniclust \| UniRef100\_A0A3N7GF44 \| 98.3 \| 1.3e-08 \| 2.4e-14 \| 65.0 \| 93 \| (15, 108) \| 112 \| (69, 162) \| 166 \| IraD/Gp25-like domain-containing protein \| IraD/Gp25-like domain-containing protein; uniclust \| UniRef100\_A0A920HWM8 \| 98.3 \| 1.4e-08 \| 2.6e-14 \| 60.2 \| 78 \| (5, 82) \| 112 \| (20, 99) \| 103 \| IraD/Gp25-like domain-containing protein \| IraD/Gp25-like domain-containing protein; uniclust \| UniRef100\_A0A8I0H272 \| 98.2 \| 1.6e-08 \| 2.9e-14 \| 55.0 \| 44 \| (10, 53) \| 112 \| (6, 49) \| 63 \| Baseplate protein (Fragment) \| Baseplate protein (Fragment); uniclust \| UniRef100\_A0A1V3RQV3 \| 98.2 \| 1.8e-08 \| 3.4e-14 \| 63.6 \| 72 \| (11, 83) \| 112 \| (38, 109) \| 153 \| IraD/Gp25-like domain-containing protein \| IraD/Gp25-like domain-containing protein; uniclust \| UniRef100\_UPI001C6791EA \| 98.2 \| 1.9e-08 \| 3.4e-14 \| 59.1 \| 84 \| (21, 105) \| 112 \| (4, 88) \| 96 \| GPW/gp25 family protein \| GPW/gp25 family protein; uniclust \| UniRef100\_A0A166CB13 \| 98.2 \| 1.8e-08 \| 3.5e-14 \| 63.4 \| 73 \| (8, 81) \| 112 \| (17, 92) \| 124 \| DUF2634 domain-containing protein \| DUF2634 domain-containing protein; uniclust \| UniRef100\_UPI001FCA211C \| 98.2 \| 2.1e-08 \| 3.8e-14 \| 61.0 \| 52 \| (11, 62) \| 112 \| (24, 75) \| 119 \| GPW/gp25 family protein \| GPW/gp25 family protein; uniclust \| UniRef100\_UPI00132FC3C4 \| 98.2 \| 2.3e-08 \| 4.2e-14 \| 57.9 \| 72 \| (5, 81) \| 112 \| (8, 79) \| 88 \| type VI secretion system baseplate subunit TssE \| type VI secretion system baseplate subunit TssE; uniclust \| UniRef100\_A0A1X1NFH7 \| 98.2 \| 2.4e-08 \| 4.6e-14 \| 72.8 \| 92 \| (8, 99) \| 112 \| (246, 341) \| 366 \| Baseplate protein J-like domain-containing protein \| Baseplate protein J-like domain-containing protein; uniclust \| UniRef100\_A0A379FGB5 \| 98.2 \| 2.5e-08 \| 4.8e-14 \| 61.7 \| 68 \| (11, 80) \| 112 \| (33, 101) \| 108 \| TssC \| TssC; uniclust \| UniRef100\_A0A061P919 \| 98.2 \| 2.6e-08 \| 5.1e-14 \| 66.9 \| 92 \| (4, 97) \| 112 \| (60, 159) \| 178 \| Phage-like element PBSX protein XkdS \| Phage-like element PBSX protein XkdS; uniclust \| UniRef100\_A0A1C5PJ69 \| 98.2 \| 2.7e-08 \| 5.4e-14 \| 66.1 \| 89 \| (8, 98) \| 112 \| (48, 141) \| 156 \| Protein of uncharacterized function (DUF2634) \| Protein of uncharacterized function (DUF2634); uniclust \| UniRef100\_A0A521S4V9 \| 98.2 \| 2.9e-08 \| 5.4e-14 \| 70.2 \| 87 \| (14, 101) \| 112 \| (200, 287) \| 300 \| IraD/Gp25-like domain-containing protein \| IraD/Gp25-like domain-containing protein; uniclust \| UniRef100\_A0A3S0EL96 \| 98.2 \| 3.2e-08 \| 5.8e-14 \| 64.5 \| 87 \| (15, 102) \| 112 \| (89, 176) \| 187 \| IraD/Gp25-like domain-containing protein \| IraD/Gp25-like domain-containing protein; uniclust \| UniRef100\_UPI001F4C0C0A \| 98.2 \| 3.2e-08 \| 5.9e-14 \| 62.4 \| 91 \| (11, 102) \| 112 \| (44, 135) \| 149 \| GPW/gp25 family protein \| GPW/gp25 family protein; uniclust \| UniRef100\_UPI001EF48225 \| 98.1 \| 3.5e-08 \| 6.5e-14 \| 63.1 \| 87 \| (13, 100) \| 112 \| (62, 149) \| 164 \| GPW/gp25 family protein \| GPW/gp25 family protein; uniclust \| UniRef100\_A0A694VIP3 \| 98.1 \| 3.6e-08 \| 6.7e-14 \| 54.1 \| 43 \| (11, 53) \| 112 \| (15, 57) \| 65 \| Uncharacterized protein (Fragment) \| Uncharacterized protein (Fragment); uniclust \| UniRef100\_B1T4E9 \| 98.1 \| 3.7e-08 \| 6.8e-14 \| 57.2 \| 73 \| (35, 107) \| 112 \| (1, 75) \| 89 \| GPW/gp25 family protein \| GPW/gp25 family protein; uniclust \| UniRef100\_A0A0S2ZE39 \| 98.1 \| 3.5e-08 \| 6.9e-14 \| 60.1 \| 75 \| (20, 97) \| 112 \| (15, 89) \| 93 \| Uncharacterized protein \| Uncharacterized protein; uniclust \| UniRef100\_A0A961Y5S4 \| 98.1 \| 3.8e-08 \| 7e-14 \| 59.9 \| 105 \| (2, 108) \| 112 \| (4, 117) \| 119 \| Uncharacterized protein \| Uncharacterized protein; uniclust \| UniRef100\_UPI0021D421C0 \| 98.1 \| 4e-08 \| 7.3e-14 \| 57.3 \| 75 \| (36, 110) \| 112 \| (9, 85) \| 91 \| GPW/gp25 family protein \| GPW/gp25 family protein; uniclust \| UniRef100\_UPI000A5AA828 \| 98.1 \| 4e-08 \| 7.4e-14 \| 73.0 \| 71 \| (13, 83) \| 112 \| (34, 105) \| 579 \| GPW/gp25 family protein \| GPW/gp25 family protein; uniclust \| UniRef100\_A0A084SH40 \| 98.1 \| 4e-08 \| 7.7e-14 \| 62.7 \| 94 \| (12, 108) \| 112 \| (20, 113) \| 131 \| IraD/Gp25-like domain-containing protein \| IraD/Gp25-like domain-containing protein; uniclust \| UniRef100\_A0A4Z0GYW9 \| 98.1 \| 4.3e-08 \| 8e-14 \| 63.1 \| 107 \| (4, 111) \| 112 \| (57, 165) \| 171 \| IraD/Gp25-like domain-containing protein \| IraD/Gp25-like domain-containing protein; uniclust \| UniRef100\_A0A017T1I7 \| 98.1 \| 4.2e-08 \| 8.4e-14 \| 68.1 \| 96 \| (14, 110) \| 112 \| (74, 179) \| 197 \| IraD/Gp25-like domain-containing protein \| IraD/Gp25-like domain-containing protein; uniclust \| UniRef100\_UPI0008F12106 \| 98.1 \| 4.7e-08 \| 8.6e-14 \| 56.2 \| 78 \| (2, 110) \| 112 \| (6, 83) \| 84 \| baseplate assembly protein \| baseplate assembly protein; uniclust \| UniRef100\_A0A661Z4X6 \| 98.1 \| 4.7e-08 \| 8.6e-14 \| 60.8 \| 94 \| (5, 98) \| 112 \| (12, 107) \| 135 \| IraD/Gp25-like domain-containing protein \| IraD/Gp25-like domain-containing protein; uniclust \| UniRef100\_A0A428MS47 \| 98.1 \| 5e-08 \| 9.3e-14 \| 62.6 \| 89 \| (4, 95) \| 112 \| (33, 127) \| 148 \| DUF2634 domain-containing protein \| DUF2634 domain-containing protein; uniclust \| UniRef100\_A0A0A1IWV2 \| 98.1 \| 4.8e-08 \| 9.7e-14 \| 64.5 \| 88 \| (8, 99) \| 112 \| (24, 116) \| 140 \| Uncharacterized protein \| Uncharacterized protein; uniclust \| UniRef100\_A0A7C7FT58 \| 98.1 \| 5.3e-08 \| 9.8e-14 \| 57.9 \| 81 \| (18, 98) \| 112 \| (2, 88) \| 102 \| IraD/Gp25-like domain-containing protein \| IraD/Gp25-like domain-containing protein; uniclust \| UniRef100\_UPI00053BFFF5 \| 98.1 \| 5.9e-08 \| 1.1e-13 \| 54.2 \| 65 \| (17, 83) \| 112 \| (5, 69) \| 71 \| GPW/gp25 family protein \| GPW/gp25 family protein; uniclust \| UniRef100\_A0A959FP97 \| 98.1 \| 6e-08 \| 1.1e-13 \| 55.5 \| 44 \| (11, 54) \| 112 \| (30, 73) \| 82 \| GPW/gp25 family protein (Fragment) \| GPW/gp25 family protein (Fragment); uniclust \| UniRef100\_UPI0009E8DA5C \| 98.1 \| 6.4e-08 \| 1.2e-13 \| 61.7 \| 66 \| (16, 82) \| 112 \| (31, 96) \| 149 \| type VI secretion system baseplate subunit TssE \| type VI secretion system baseplate subunit TssE; uniclust \| UniRef100\_A0A382IDM1 \| 98.1 \| 6.3e-08 \| 1.2e-13 \| 58.1 \| 49 \| (4, 52) \| 112 \| (43, 93) \| 94 \| Uncharacterized protein (Fragment) \| Uncharacterized protein (Fragment); uniclust \| UniRef100\_A0A3N7GKM9 \| 98.1 \| 6.6e-08 \| 1.2e-13 \| 54.3 \| 39 \| (4, 42) \| 112 \| (19, 59) \| 73 \| IraD/Gp25-like domain-containing protein (Fragment) \| IraD/Gp25-like domain-containing protein (Fragment); uniclust \| UniRef100\_A0A7K4MRV6 \| 98.1 \| 6.8e-08 \| 1.3e-13 \| 54.2 \| 52 \| (4, 55) \| 112 \| (17, 70) \| 73 \| Baseplate wedge subunit (Fragment) \| Baseplate wedge subunit (Fragment); uniclust \| UniRef100\_A0A143XKD4 \| 98.1 \| 6.8e-08 \| 1.4e-13 \| 60.8 \| 86 \| (12, 99) \| 112 \| (17, 102) \| 106 \| Gene 25-like lysozyme \| Gene 25-like lysozyme; uniclust \| UniRef100\_A0A1S2W8T3 \| 98.0 \| 7.7e-08 \| 1.4e-13 \| 62.1 \| 80 \| (3, 83) \| 112 \| (35, 120) \| 147 \| DUF2634 domain-containing protein \| DUF2634 domain-containing protein; uniclust \| UniRef100\_A0A6J5KJ56 \| 98.0 \| 8e-08 \| 1.5e-13 \| 60.6 \| 80 \| (18, 98) \| 112 \| (27, 107) \| 132 \| GpW/Gp25/anti-adapter protein IraD \| GpW/Gp25/anti-adapter protein IraD; uniclust \| UniRef100\_A0A1I1H4X3 \| 98.0 \| 8e-08 \| 1.5e-13 \| 61.5 \| 72 \| (11, 82) \| 112 \| (14, 85) \| 132 \| Predicted component of the type VI protein secretion system \| Predicted component of the type VI protein secretion system; uniclust \| UniRef100\_A0A1V5M2G2 \| 98.0 \| 8.8e-08 \| 1.6e-13 \| 55.5 \| 40 \| (13, 52) \| 112 \| (30, 69) \| 87 \| Gene 25-like lysozyme \| Gene 25-like lysozyme; uniclust \| UniRef100\_A0A1W9S296 \| 98.0 \| 9.1e-08 \| 1.7e-13 \| 61.1 \| 99 \| (7, 109) \| 112 \| (25, 125) \| 131 \| IraD/Gp25-like domain-containing protein \| IraD/Gp25-like domain-containing protein; uniclust \| UniRef100\_A0A0K0NKR9 \| 98.0 \| 8.7e-08 \| 1.7e-13 \| 62.0 \| 87 \| (4, 100) \| 112 \| (12, 102) \| 125 \| Putative lysozyme \| Putative lysozyme; uniclust \| UniRef100\_D8NFC7 \| 98.0 \| 9.9e-08 \| 1.8e-13 \| 56.8 \| 51 \| (1, 51) \| 112 \| (1, 51) \| 102 \| Putative bacteriophage baseplate assembly-like protein \| Putative bacteriophage baseplate assembly-like protein; uniclust \| UniRef100\_UPI0010AB85DE \| 98.0 \| 1e-07 \| 1.8e-13 \| 62.1 \| 86 \| (11, 97) \| 112 \| (75, 165) \| 181 \| GPW/gp25 family protein \| GPW/gp25 family protein; uniclust \| UniRef100\_A0A076G5J9 \| 98.0 \| 9.7e-08 \| 1.9e-13 \| 62.1 \| 84 \| (11, 98) \| 112 \| (22, 108) \| 128 \| Baseplate component \| Baseplate component; uniclust \| UniRef100\_A0A218MMN8 \| 98.0 \| 1.1e-07 \| 2e-13 \| 59.3 \| 84 \| (15, 100) \| 112 \| (40, 124) \| 135 \| Putative baseplate wedge protein \| Putative baseplate wedge protein; uniclust \| UniRef100\_A0A164MCD3 \| 98.0 \| 1.1e-07 \| 2.1e-13 \| 62.7 \| 95 \| (11, 107) \| 112 \| (40, 140) \| 147 \| IraD/Gp25-like domain-containing protein \| IraD/Gp25-like domain-containing protein; uniclust \| UniRef100\_A0A6N2TUW9 \| 98.0 \| 1.1e-07 \| 2.1e-13 \| 60.6 \| 91 \| (11, 103) \| 112 \| (20, 110) \| 122 \| Histidine kinase \| Histidine kinase; uniclust \| UniRef100\_A0A450X8I7 \| 98.0 \| 1.2e-07 \| 2.2e-13 \| 55.0 \| 40 \| (11, 50) \| 112 \| (47, 86) \| 87 \| Uncharacterized protein (Fragment) \| Uncharacterized protein (Fragment); uniclust \| UniRef100\_A0A3B8IUD1 \| 98.0 \| 1.4e-07 \| 2.5e-13 \| 54.2 \| 42 \| (11, 52) \| 112 \| (32, 75) \| 82 \| Lysozyme (Fragment) \| Lysozyme (Fragment); uniclust \| UniRef100\_A0A0X8WQV4 \| 98.0 \| 1.5e-07 \| 2.8e-13 \| 56.3 \| 93 \| (13, 105) \| 112 \| (3, 96) \| 105 \| IraD/Gp25-like domain-containing protein \| IraD/Gp25-like domain-containing protein; uniclust \| UniRef100\_A0A1F2Z7U0 \| 98.0 \| 1.5e-07 \| 3e-13 \| 63.4 \| 94 \| (15, 110) \| 112 \| (43, 147) \| 167 \| IraD/Gp25-like domain-containing protein \| IraD/Gp25-like domain-containing protein; uniclust \| UniRef100\_A0A0P7GKB2 \| 98.0 \| 1.6e-07 \| 3e-13 \| 57.7 \| 43 \| (1, 43) \| 112 \| (79, 121) \| 123 \| Baseplate wedge subunit \| Baseplate wedge subunit; uniclust \| UniRef100\_A0A1H4EGE0 \| 98.0 \| 1.6e-07 \| 3e-13 \| 62.1 \| 94 \| (14, 107) \| 112 \| (21, 144) \| 152 \| IraD/Gp25-like domain-containing protein \| IraD/Gp25-like domain-containing protein; uniclust \| UniRef100\_UPI000BBDD989 \| 98.0 \| 1.7e-07 \| 3.1e-13 \| 56.4 \| 68 \| (15, 83) \| 112 \| (13, 80) \| 107 \| type VI secretion system baseplate subunit TssE \| type VI secretion system baseplate subunit TssE; uniclust \| UniRef100\_A0A7C6EBT5 \| 97.9 \| 1.7e-07 \| 3.1e-13 \| 59.0 \| 101 \| (8, 110) \| 112 \| (21, 124) \| 130 \| IraD/Gp25-like domain-containing protein \| IraD/Gp25-like domain-containing protein; uniclust \| UniRef100\_K8GN81 \| 97.9 \| 2e-07 \| 3.7e-13 \| 59.3 \| 102 \| (8, 110) \| 112 \| (35, 150) \| 153 \| Uncharacterized protein \| Uncharacterized protein; uniclust \| UniRef100\_A0A509B4A1 \| 97.9 \| 2e-07 \| 3.8e-13 \| 54.0 \| 65 \| (36, 102) \| 112 \| (2, 66) \| 77 \| Putative bacteriophage baseplate protein \| Putative bacteriophage baseplate protein; uniclust \| UniRef100\_A0A4S2ABK4 \| 97.9 \| 2e-07 \| 3.8e-13 \| 58.4 \| 83 \| (13, 97) \| 112 \| (18, 100) \| 118 \| Uncharacterized protein \| Uncharacterized protein; uniclust \| UniRef100\_A0A8S5V9K4 \| 97.9 \| 2.1e-07 \| 3.8e-13 \| 57.1 \| 87 \| (11, 97) \| 112 \| (27, 113) \| 120 \| Baseplate wedge protein \| Baseplate wedge protein; uniclust \| UniRef100\_UPI001D1663D5 \| 97.9 \| 2.1e-07 \| 3.9e-13 \| 52.5 \| 69 \| (4, 72) \| 112 \| (1, 70) \| 74 \| hypothetical protein \| hypothetical protein; uniclust \| UniRef100\_A0A2E6PSE7 \| 97.9 \| 2.1e-07 \| 3.9e-13 \| 60.7 \| 97 \| (13, 110) \| 112 \| (57, 158) \| 182 \| IraD/Gp25-like domain-containing protein \| IraD/Gp25-like domain-containing protein; uniclust \| UniRef100\_A0A1J0MHE5 \| 97.9 \| 2.2e-07 \| 4.1e-13 \| 52.2 \| 42 \| (13, 54) \| 112 \| (27, 68) \| 72 \| Baseplate wedge subunit \| Baseplate wedge subunit; uniclust \| UniRef100\_A0A7C3UU77 \| 97.9 \| 2.3e-07 \| 4.2e-13 \| 57.6 \| 85 \| (15, 99) \| 112 \| (30, 116) \| 130 \| IraD/Gp25-like domain-containing protein \| IraD/Gp25-like domain-containing protein; uniclust \| UniRef100\_A0A090QXN9 \| 97.9 \| 2e-07 \| 4.2e-13 \| 61.5 \| 74 \| (11, 84) \| 112 \| (23, 97) \| 134 \| Uncharacterized protein \| Uncharacterized protein; uniclust \| UniRef100\_A0A0G9K5Z4 \| 97.9 \| 2.3e-07 \| 4.2e-13 \| 56.6 \| 79 \| (19, 97) \| 112 \| (8, 93) \| 105 \| Baseplate assembly protein \| Baseplate assembly protein; uniclust \| UniRef100\_A0A068NSI2 \| 97.9 \| 2.2e-07 \| 4.3e-13 \| 63.8 \| 87 \| (11, 98) \| 112 \| (57, 154) \| 190 \| Bacteriophage protein \| Bacteriophage protein; uniclust \| UniRef100\_A0A3C0GCP4 \| 97.9 \| 2.4e-07 \| 4.4e-13 \| 55.1 \| 79 \| (25, 103) \| 112 \| (2, 93) \| 100 \| IraD/Gp25-like domain-containing protein \| IraD/Gp25-like domain-containing protein; uniclust \| UniRef100\_A0A060H4I9 \| 97.9 \| 2.6e-07 \| 5.1e-13 \| 61.4 \| 86 \| (11, 99) \| 112 \| (42, 128) \| 147 \| Phage-related protein \| Phage-related protein; uniclust \| UniRef100\_A0A1C5REL3 \| 97.9 \| 2.7e-07 \| 5.3e-13 \| 62.7 \| 97 \| (11, 110) \| 112 \| (67, 167) \| 173 \| Protein of uncharacterized function (DUF2634) \| Protein of uncharacterized function (DUF2634); uniclust \| UniRef100\_A0A381SUY6 \| 97.9 \| 3.2e-07 \| 5.8e-13 \| 58.3 \| 96 \| (4, 99) \| 112 \| (37, 136) \| 150 \| IraD/Gp25-like domain-containing protein \| IraD/Gp25-like domain-containing protein; uniclust \| UniRef100\_A0A2W4L8B5 \| 97.9 \| 3.4e-07 \| 6.2e-13 \| 57.6 \| 95 \| (12, 109) \| 112 \| (12, 107) \| 128 \| IraD/Gp25-like domain-containing protein \| IraD/Gp25-like domain-containing protein; uniclust \| UniRef100\_A0A7C7EU41 \| 97.9 \| 3.2e-07 \| 6.3e-13 \| 58.0 \| 83 \| (11, 96) \| 112 \| (20, 103) \| 113 \| IraD/Gp25-like domain-containing protein \| IraD/Gp25-like domain-containing protein; uniclust \| UniRef100\_A0A377AHZ9 \| 97.9 \| 3.4e-07 \| 6.4e-13 \| 55.3 \| 67 \| (16, 83) \| 112 \| (16, 82) \| 98 \| Type VI secretion system lysozyme-like protein \| Type VI secretion system lysozyme-like protein; uniclust \| UniRef100\_A0A6J4HR49 \| 97.8 \| 3.7e-07 \| 6.9e-13 \| 59.1 \| 72 \| (11, 82) \| 112 \| (22, 102) \| 145 \| IraD/Gp25-like domain-containing protein \| IraD/Gp25-like domain-containing protein; uniclust \| UniRef100\_UPI0013DCF42C \| 97.8 \| 3.8e-07 \| 7.1e-13 \| 53.3 \| 51 \| (8, 58) \| 112 \| (19, 71) \| 83 \| GPW/gp25 family protein \| GPW/gp25 family protein; uniclust \| UniRef100\_A0A450Z097 \| 97.8 \| 4e-07 \| 7.4e-13 \| 53.0 \| 42 \| (1, 42) \| 112 \| (1, 42) \| 87 \| Uncharacterized protein \| Uncharacterized protein; uniclust \| UniRef100\_A0A1T4W4S5 \| 97.8 \| 4.1e-07 \| 7.5e-13 \| 59.6 \| 98 \| (2, 99) \| 112 \| (78, 178) \| 185 \| Phage baseplate assembly protein W \| Phage baseplate assembly protein W; uniclust \| UniRef100\_A0A843FHN1 \| 97.8 \| 4.2e-07 \| 8e-13 \| 58.4 \| 87 \| (11, 99) \| 112 \| (33, 121) \| 134 \| IraD/Gp25-like domain-containing protein \| IraD/Gp25-like domain-containing protein; uniclust \| UniRef100\_A0A521U4A4 \| 97.8 \| 4.3e-07 \| 8.1e-13 \| 58.1 \| 98 \| (11, 110) \| 112 \| (25, 124) \| 139 \| IraD/Gp25-like domain-containing protein \| IraD/Gp25-like domain-containing protein; uniclust \| UniRef100\_A0A1C6BMP3 \| 97.8 \| 4.3e-07 \| 8.1e-13 \| 54.8 \| 79 \| (13, 94) \| 112 \| (16, 94) \| 95 \| Uncharacterized protein \| Uncharacterized protein; uniclust \| UniRef100\_A0A085W9S1 \| 97.8 \| 4.1e-07 \| 8.2e-13 \| 63.9 \| 95 \| (15, 110) \| 112 \| (71, 175) \| 203 \| IraD/Gp25-like domain-containing protein \| IraD/Gp25-like domain-containing protein; uniclust \| UniRef100\_A0A2D6E551 \| 97.8 \| 4.7e-07 \| 8.6e-13 \| 58.5 \| 93 \| (6, 98) \| 112 \| (13, 111) \| 166 \| IraD/Gp25-like domain-containing protein \| IraD/Gp25-like domain-containing protein; uniclust \| UniRef100\_A0A1V6N1Y4 \| 97.8 \| 5e-07 \| 9.4e-13 \| 57.5 \| 103 \| (4, 109) \| 112 \| (18, 125) \| 131 \| DUF2634 domain-containing protein \| DUF2634 domain-containing protein; uniclust \| UniRef100\_A0A0B6RSR4 \| 97.8 \| 5.6e-07 \| 1e-12 \| 52.8 \| 59 \| (1, 59) \| 112 \| (1, 59) \| 91 \| Putative phage baseplate assembly protein W \| Putative phage baseplate assembly protein W; uniclust \| UniRef100\_A0A0X8JJW6 \| 97.8 \| 5.9e-07 \| 1.1e-12 \| 56.7 \| 69 \| (11, 81) \| 112 \| (17, 86) \| 129 \| Type VI secretion protein \| Type VI secretion protein; uniclust \| UniRef100\_A0A1Q6PX45 \| 97.8 \| 5.9e-07 \| 1.1e-12 \| 56.1 \| 81 \| (15, 98) \| 112 \| (21, 101) \| 109 \| IraD/Gp25-like domain-containing protein \| IraD/Gp25-like domain-containing protein; uniclust \| UniRef100\_A0A090D3B0 \| 97.8 \| 5.9e-07 \| 1.2e-12 \| 60.2 \| 82 \| (11, 95) \| 112 \| (30, 113) \| 147 \| Phage protein \| Phage protein; uniclust \| UniRef100\_A0A6J5N592 \| 97.8 \| 6.3e-07 \| 1.2e-12 \| 57.9 \| 85 \| (15, 99) \| 112 \| (22, 113) \| 138 \| Baseplate wedge subunit \| Baseplate wedge subunit; uniclust \| UniRef100\_A0A1V3NED6 \| 97.8 \| 6.8e-07 \| 1.3e-12 \| 56.0 \| 87 \| (11, 98) \| 112 \| (26, 114) \| 135 \| IraD/Gp25-like domain-containing protein \| IraD/Gp25-like domain-containing protein; uniclust \| UniRef100\_A0A7W2XXK3 \| 97.8 \| 7e-07 \| 1.3e-12 \| 56.3 \| 81 \| (15, 97) \| 112 \| (30, 114) \| 141 \| Type VI secretion system baseplate subunit TssE \| Type VI secretion system baseplate subunit TssE; uniclust \| UniRef100\_A0A1C0B348 \| 97.7 \| 7.3e-07 \| 1.4e-12 \| 58.1 \| 88 \| (15, 103) \| 112 \| (29, 118) \| 141 \| 25-like lysozyme \| 25-like lysozyme; uniclust \| UniRef100\_UPI001F083AAE \| 97.7 \| 8.5e-07 \| 1.6e-12 \| 58.1 \| 100 \| (8, 107) \| 112 \| (56, 171) \| 181 \| GPW/gp25 family protein \| GPW/gp25 family protein; uniclust \| UniRef100\_A0A0G1YBV7 \| 97.7 \| 8.7e-07 \| 1.6e-12 \| 60.6 \| 97 \| (14, 111) \| 112 \| (151, 247) \| 250 \| Uncharacterized protein \| Uncharacterized protein; uniclust \| UniRef100\_A0A2V2E4P3 \| 97.7 \| 8.8e-07 \| 1.7e-12 \| 56.1 \| 97 \| (8, 109) \| 112 \| (20, 119) \| 122 \| DUF2634 domain-containing protein \| DUF2634 domain-containing protein; uniclust \| UniRef100\_A0A1I5RRA4 \| 97.7 \| 9.2e-07 \| 1.7e-12 \| 50.7 \| 62 \| (35, 97) \| 112 \| (1, 64) \| 72 \| Uncharacterized protein \| Uncharacterized protein; uniclust \| UniRef100\_A0A430VBK1 \| 97.7 \| 1e-06 \| 1.8e-12 \| 55.4 \| 89 \| (14, 104) \| 112 \| (37, 126) \| 137 \| IraD/Gp25-like domain-containing protein \| IraD/Gp25-like domain-containing protein; uniclust \| UniRef100\_E3HBK7 \| 97.7 \| 1e-06 \| 1.9e-12 \| 52.5 \| 82 \| (12, 96) \| 112 \| (14, 95) \| 98 \| GPW/gp25 family protein \| GPW/gp25 family protein; uniclust \| UniRef100\_A0A344TT62 \| 97.7 \| 1e-06 \| 1.9e-12 \| 59.7 \| 73 \| (11, 83) \| 112 \| (90, 163) \| 207 \| IraD/Gp25-like domain-containing protein \| IraD/Gp25-like domain-containing protein; uniclust \| UniRef100\_A0A0F4LD00 \| 97.7 \| 1.1e-06 \| 2e-12 \| 59.8 \| 90 \| (7, 98) \| 112 \| (62, 156) \| 178 \| DUF2634 domain-containing protein \| DUF2634 domain-containing protein; uniclust \| UniRef100\_A0A8S1IKS8 \| 97.7 \| 1.1e-06 \| 2.1e-12 \| 68.0 \| 92 \| (8, 103) \| 112 \| (310, 405) \| 822 \| Baseplate protein J-like domain-containing protein \| Baseplate protein J-like domain-containing protein; uniclust \| UniRef100\_A0A0P9KCS3 \| 97.7 \| 1.2e-06 \| 2.2e-12 \| 56.7 \| 81 \| (16, 98) \| 112 \| (16, 99) \| 137 \| Type VI secretion system lysozyme-related protein \| Type VI secretion system lysozyme-related protein; uniclust \| UniRef100\_A0A8B3NLL6 \| 97.7 \| 1.2e-06 \| 2.2e-12 \| 63.7 \| 80 \| (2, 81) \| 112 \| (4, 83) \| 419 \| Baseplate protein J-like domain-containing protein \| Baseplate protein J-like domain-containing protein; uniclust \| UniRef100\_A0A068ZX24 \| 97.7 \| 1.1e-06 \| 2.3e-12 \| 60.6 \| 88 \| (8, 97) \| 112 \| (55, 147) \| 171 \| DUF2634 domain-containing protein \| DUF2634 domain-containing protein; uniclust \| UniRef100\_UPI001C89A3A3 \| 97.7 \| 1.2e-06 \| 2.3e-12 \| 53.5 \| 58 \| (13, 79) \| 112 \| (23, 80) \| 103 \| GPW/gp25 family protein \| GPW/gp25 family protein; uniclust \| UniRef100\_A0A7C7BV91 \| 97.7 \| 1.3e-06 \| 2.3e-12 \| 52.3 \| 49 \| (11, 59) \| 112 \| (46, 94) \| 100 \| IraD/Gp25-like domain-containing protein \| IraD/Gp25-like domain-containing protein; uniclust \| UniRef100\_A0A2D6MFM3 \| 97.7 \| 1.3e-06 \| 2.4e-12 \| 58.1 \| 67 \| (15, 81) \| 112 \| (91, 157) \| 201 \| IraD/Gp25-like domain-containing protein \| IraD/Gp25-like domain-containing protein; uniclust \| UniRef100\_A0A3M2C415 \| 97.6 \| 1.5e-06 \| 2.8e-12 \| 52.8 \| 85 \| (13, 99) \| 112 \| (10, 94) \| 109 \| IraD/Gp25-like domain-containing protein (Fragment) \| IraD/Gp25-like domain-containing protein (Fragment); uniclust \| UniRef100\_A0A081P4G2 \| 97.6 \| 1.4e-06 \| 2.8e-12 \| 60.3 \| 90 \| (8, 99) \| 112 \| (51, 145) \| 180 \| DUF2634 domain-containing protein \| DUF2634 domain-containing protein; uniclust \| UniRef100\_A0A9E3QY51 \| 97.6 \| 1.6e-06 \| 3e-12 \| 54.3 \| 87 \| (11, 99) \| 112 \| (35, 122) \| 133 \| Uncharacterized protein \| Uncharacterized protein; uniclust \| UniRef100\_A0A021XFJ7 \| 97.6 \| 1.4e-06 \| 3e-12 \| 62.3 \| 88 \| (13, 102) \| 112 \| (82, 183) \| 216 \| Type VI secretion system lysozyme-like protein \| Type VI secretion system lysozyme-like protein; uniclust \| UniRef100\_A0A0H5M096 \| 97.6 \| 1.6e-06 \| 3e-12 \| 57.6 \| 85 \| (16, 101) \| 112 \| (35, 120) \| 178 \| Type VI secretion system lysozyme-like protein \| Type VI secretion system lysozyme-like protein; uniclust \| UniRef100\_A0A0F0HIV6 \| 97.6 \| 1.7e-06 \| 3.2e-12 \| 54.2 \| 86 \| (13, 99) \| 112 \| (21, 107) \| 132 \| IraD/Gp25-like domain-containing protein \| IraD/Gp25-like domain-containing protein; uniclust \| UniRef100\_E1RFI1 \| 97.6 \| 1.7e-06 \| 3.2e-12 \| 54.3 \| 90 \| (11, 107) \| 112 \| (26, 116) \| 119 \| GPW/gp25 family protein \| GPW/gp25 family protein; uniclust \| UniRef100\_UPI000BB729C8 \| 97.6 \| 1.8e-06 \| 3.3e-12 \| 56.4 \| 89 \| (5, 96) \| 112 \| (42, 135) \| 154 \| DUF2634 domain-containing protein \| DUF2634 domain-containing protein; uniclust \| UniRef100\_UPI002020CE7C \| 97.6 \| 1.8e-06 \| 3.3e-12 \| 50.3 \| 60 \| (1, 60) \| 112 \| (1, 60) \| 85 \| hypothetical protein \| hypothetical protein; uniclust \| UniRef100\_A0A2I7R3K5 \| 97.6 \| 1.8e-06 \| 3.3e-12 \| 52.7 \| 95 \| (2, 98) \| 112 \| (3, 100) \| 112 \| Lysozyme \| Lysozyme; uniclust \| UniRef100\_A0A378EEK6 \| 97.6 \| 1.9e-06 \| 3.5e-12 \| 51.9 \| 64 \| (37, 101) \| 112 \| (2, 66) \| 92 \| Type VI secretion system lysozyme-like protein \| Type VI secretion system lysozyme-like protein; uniclust \| UniRef100\_A0A1T1HD64 \| 97.6 \| 2e-06 \| 3.7e-12 \| 56.6 \| 84 \| (11, 94) \| 112 \| (19, 102) \| 150 \| IraD/Gp25-like domain-containing protein \| IraD/Gp25-like domain-containing protein; uniclust \| UniRef100\_A0A1I1DKW3 \| 97.6 \| 2.1e-06 \| 3.9e-12 \| 52.9 \| 91 \| (11, 102) \| 112 \| (20, 111) \| 119 \| Phage baseplate assembly protein W \| Phage baseplate assembly protein W; uniclust \| UniRef100\_UPI0014204378 \| 97.6 \| 2.2e-06 \| 4e-12 \| 50.3 \| 69 \| (4, 74) \| 112 \| (15, 87) \| 88 \| DUF2634 domain-containing protein \| DUF2634 domain-containing protein; uniclust \| UniRef100\_UPI0001816813 \| 97.6 \| 2.3e-06 \| 4.3e-12 \| 47.3 \| 52 \| (4, 55) \| 112 \| (1, 52) \| 64 \| hypothetical protein \| hypothetical protein; uniclust \| UniRef100\_A0A4R5N831 \| 97.6 \| 2.4e-06 \| 4.4e-12 \| 55.4 \| 99 \| (7, 107) \| 112 \| (51, 154) \| 157 \| DUF2634 domain-containing protein \| DUF2634 domain-containing protein; uniclust \| UniRef100\_A0A023Y2J8 \| 97.6 \| 2.2e-06 \| 4.5e-12 \| 60.6 \| 92 \| (14, 107) \| 112 \| (55, 162) \| 209 \| Type VI secretion protein \| Type VI secretion protein; uniclust \| UniRef100\_A0A075DXX6 \| 97.6 \| 2.4e-06 \| 4.6e-12 \| 55.8 \| 84 \| (13, 99) \| 112 \| (24, 108) \| 131 \| Putative baseplate component \| Putative baseplate component; uniclust \| UniRef100\_A0A257Q2E2 \| 97.5 \| 2.7e-06 \| 5e-12 \| 59.5 \| 99 \| (8, 110) \| 112 \| (185, 288) \| 289 \| Gp5/Type VI secretion system Vgr protein OB-fold domain-containing protein \| Gp5/Type VI secretion system Vgr protein OB-fold domain-containing protein; uniclust \| UniRef100\_H6WG16 \| 97.5 \| 2.8e-06 \| 5.1e-12 \| 55.7 \| 85 \| (11, 100) \| 112 \| (81, 165) \| 177 \| Virion structural protein \| Virion structural protein; uniclust \| UniRef100\_UPI000A200276 \| 97.5 \| 2.9e-06 \| 5.3e-12 \| 50.9 \| 66 \| (35, 100) \| 112 \| (1, 67) \| 100 \| hypothetical protein \| hypothetical protein; uniclust \| UniRef100\_A0A0F9MK12 \| 97.5 \| 2.8e-06 \| 5.4e-12 \| 54.9 \| 74 \| (11, 84) \| 112 \| (26, 100) \| 131 \| Uncharacterized protein (Fragment) \| Uncharacterized protein (Fragment); uniclust \| UniRef100\_A0A099VCP5 \| 97.5 \| 3.1e-06 \| 6e-12 \| 55.3 \| 85 \| (13, 99) \| 112 \| (23, 109) \| 130 \| Type VI secretion system baseplate subunit TssE \| Type VI secretion system baseplate subunit TssE; uniclust \| UniRef100\_A0A0Q8PDY9 \| 97.5 \| 3e-06 \| 6.1e-12 \| 57.7 \| 94 \| (15, 110) \| 112 \| (35, 139) \| 160 \| Type VI secretion protein, lysozyme-like protein \| Type VI secretion protein, lysozyme-like protein; uniclust \| UniRef100\_A0A0Q8ANH2 \| 97.5 \| 3.4e-06 \| 6.3e-12 \| 53.4 \| 65 \| (2, 66) \| 112 \| (5, 69) \| 126 \| IraD/Gp25-like domain-containing protein \| IraD/Gp25-like domain-containing protein; uniclust \| UniRef100\_A0A1H8JI52 \| 97.5 \| 3.3e-06 \| 6.3e-12 \| 55.3 \| 87 \| (11, 99) \| 112 \| (40, 128) \| 143 \| DUF2634 domain-containing protein \| DUF2634 domain-containing protein; uniclust \| UniRef100\_A0A009YPJ6 \| 97.5 \| 3.5e-06 \| 6.7e-12 \| 53.6 \| 108 \| (1, 109) \| 112 \| (1, 115) \| 120 \| Phage GP46 family protein \| Phage GP46 family protein; uniclust \| UniRef100\_A0A1Q6LL95 \| 97.5 \| 3.6e-06 \| 6.9e-12 \| 51.1 \| 77 \| (14, 93) \| 112 \| (11, 87) \| 93 \| Uncharacterized protein \| Uncharacterized protein; uniclust \| UniRef100\_A0A0K8MDE3 \| 97.5 \| 3.6e-06 \| 7.1e-12 \| 58.8 \| 95 \| (15, 110) \| 112 \| (65, 173) \| 201 \| Gene 25-like lysozyme \| Gene 25-like lysozyme; uniclust \| UniRef100\_A0A497N9T7 \| 97.5 \| 3.9e-06 \| 7.2e-12 \| 52.8 \| 89 \| (11, 99) \| 112 \| (26, 116) \| 133 \| IraD/Gp25-like domain-containing protein \| IraD/Gp25-like domain-containing protein; uniclust \| UniRef100\_A0A349NQ61 \| 97.5 \| 4e-06 \| 7.3e-12 \| 49.1 \| 69 \| (32, 100) \| 112 \| (1, 73) \| 86 \| IraD/Gp25-like domain-containing protein (Fragment) \| IraD/Gp25-like domain-containing protein (Fragment); uniclust \| UniRef100\_A0A022FQR5 \| 97.5 \| 3.7e-06 \| 7.3e-12 \| 58.7 \| 95 \| (15, 110) \| 112 \| (58, 162) \| 186 \| Type VI secretion system lysozyme \| Type VI secretion system lysozyme; uniclust \| UniRef100\_A0A843GVE6 \| 97.5 \| 3.9e-06 \| 7.4e-12 \| 53.7 \| 87 \| (11, 97) \| 112 \| (19, 106) \| 127 \| IraD/Gp25-like domain-containing protein \| IraD/Gp25-like domain-containing protein; uniclust \| UniRef100\_A0A352JSI9 \| 97.5 \| 4.1e-06 \| 7.5e-12 \| 53.6 \| 97 \| (11, 108) \| 112 \| (19, 123) \| 139 \| IraD/Gp25-like domain-containing protein \| IraD/Gp25-like domain-containing protein; uniclust \| UniRef100\_A0A943Z1H4 \| 97.5 \| 4.2e-06 \| 7.8e-12 \| 50.4 \| 81 \| (13, 95) \| 112 \| (19, 99) \| 102 \| Uncharacterized protein \| Uncharacterized protein; uniclust \| UniRef100\_A0A354D7Q5 \| 97.5 \| 4.1e-06 \| 7.9e-12 \| 68.9 \| 99 \| (8, 110) \| 112 \| (759, 860) \| 863 \| Baseplate protein J-like domain-containing protein \| Baseplate protein J-like domain-containing protein; uniclust \| UniRef100\_X1ADF5 \| 97.5 \| 4.4e-06 \| 8e-12 \| 49.5 \| 44 \| (11, 54) \| 112 \| (34, 77) \| 92 \| Uncharacterized protein \| Uncharacterized protein; uniclust \| UniRef100\_A0A1G3MBB2 \| 97.5 \| 4.4e-06 \| 8.1e-12 \| 53.1 \| 70 \| (14, 83) \| 112 \| (39, 108) \| 143 \| IraD/Gp25-like domain-containing protein \| IraD/Gp25-like domain-containing protein; pdb70 \| 2IA7\_A \| 99.6 \| 5.4e-20 \| 4.3e-24 \| 111.2 \| 101 \| (8, 108) \| 112 \| (24, 126) \| 134 \| Crystal structure of putative tail \| 2IA7\_A Crystal structure of putative tail NP\_952040.1, putative tail lysozyme, Structural; pdb70 \| 5IV5\_BD \| 99.6 \| 6.3e-20 \| 5e-24 \| 110.7 \| 102 \| (7, 108) \| 112 \| (21, 124) \| 132 \| Baseplate wedge protein gp6, Baseplate \| 5IV5\_BD Baseplate wedge protein gp6, Baseplate T4, baseplate-tail tube complex, pre-attachment; pdb70 \| 5IV5\_DG \| 99.6 \| 6.3e-20 \| 5e-24 \| 110.7 \| 102 \| (7, 108) \| 112 \| (21, 124) \| 132 \| Baseplate wedge protein gp6, Baseplate \| 5IV5\_DG Baseplate wedge protein gp6, Baseplate T4, baseplate-tail tube complex, pre-attachment; pdb70 \| 5IV5\_FJ \| 99.6 \| 6.3e-20 \| 5e-24 \| 110.7 \| 102 \| (7, 108) \| 112 \| (21, 124) \| 132 \| Baseplate wedge protein gp6, Baseplate \| 5IV5\_FJ Baseplate wedge protein gp6, Baseplate T4, baseplate-tail tube complex, pre-attachment; pdb70 \| 5IV5\_IC \| 99.6 \| 6.3e-20 \| 5e-24 \| 110.7 \| 102 \| (7, 108) \| 112 \| (21, 124) \| 132 \| Baseplate wedge protein gp6, Baseplate \| 5IV5\_IC Baseplate wedge protein gp6, Baseplate T4, baseplate-tail tube complex, pre-attachment; pdb70 \| 5IV5\_T \| 99.6 \| 6.3e-20 \| 5e-24 \| 110.7 \| 102 \| (7, 108) \| 112 \| (21, 124) \| 132 \| Baseplate wedge protein gp6, Baseplate \| 5IV5\_T Baseplate wedge protein gp6, Baseplate T4, baseplate-tail tube complex, pre-attachment; pdb70 \| 5IW9\_A \| 99.6 \| 6.6e-20 \| 5.3e-24 \| 110.5 \| 103 \| (7, 109) \| 112 \| (20, 124) \| 131 \| Baseplate wedge protein gp25 \| 5IW9\_A Baseplate wedge protein gp25 contractile sheath, baseplate, wedge, sheath HET: MSE; pdb70 \| 6OD1\_B \| 99.6 \| 1.5e-19 \| 1.2e-23 \| 106.9 \| 97 \| (11, 109) \| 112 \| (3, 100) \| 115 \| Regulator of RpoS, Anti-adapter protein \| 6OD1\_B Regulator of RpoS, Anti-adapter protein response regulator, ClpXP adaptor, anti-adaptor; pdb70 \| 6RAO\_H \| 99.6 \| 1.8e-19 \| 1.4e-23 \| 109.7 \| 102 \| (7, 108) \| 112 \| (24, 129) \| 140 \| Afp1, Afp2, Afp3, Afp5, Afp9 \| 6RAO\_H Afp1, Afp2, Afp3, Afp5, Afp9 Anti-feeding prophage, secretion system, AFP; pdb70 \| 6GJ1\_D \| 99.6 \| 2.3e-19 \| 1.8e-23 \| 109.9 \| 100 \| (8, 109) \| 112 \| (23, 123) \| 143 \| Putative type VI secretion protein \| 6GJ1\_D Putative type VI secretion protein Secretion, baseplate, complex, STRUCTURAL PROTEIN; pdb70 \| 6J0N\_H \| 99.5 \| 3.5e-19 \| 2.8e-23 \| 108.5 \| 103 \| (7, 109) \| 112 \| (24, 130) \| 140 \| Pvc1, Pvc9, Pvc11, Pvc12, Pvc4 \| 6J0N\_H Pvc1, Pvc9, Pvc11, Pvc12, Pvc4 assembly, Photorhabdus asymbiotica, PVC, contractile |
| Top keywords  (threshold 1.00e-03 (evalue)) | **domain\_containing, IraD, Gp25\_like, Baseplate, GPW, gp25, assembly, Phage, Fragment, DUF2634** |
| Output files | ../../similar\_sequences/12\_FANPEZAQ\_CDS\_0012\_merged.svg ../../similar\_sequences/12\_FANPEZAQ\_CDS\_0012\_pdb70.a3m ../../similar\_sequences/12\_FANPEZAQ\_CDS\_0012\_pdb70.hhr ../../similar\_sequences/12\_FANPEZAQ\_CDS\_0012\_uniclust.a3m ../../similar\_sequences/12\_FANPEZAQ\_CDS\_0012\_uniclust.hhr |

#### Structure prediction (AlphaFold)^2^

|  |  |
| --- | --- |
| Stats | xml version="1.0" encoding="utf-8" standalone="no"?       2024-09-02T21:09:10.763989 image/svg+xml   Matplotlib v3.7.2, https://matplotlib.org/ |
| Predicted structure | **NGL Viewer Controls:**  - Center: *Left-Click* - Rotate: *Left-Click + Drag* - Translate: *Right-Click + Drag* - Zoom: *Shift + Left-Click + Drag* |
| Output files | ../../predicted\_structures/12\_FANPEZAQ\_CDS\_0012/features.pkl ../../predicted\_structures/12\_FANPEZAQ\_CDS\_0012/ranked\_0.pdb ../../predicted\_structures/12\_FANPEZAQ\_CDS\_0012/ranked\_0\_plots.svg ../../predicted\_structures/12\_FANPEZAQ\_CDS\_0012/result\_model\_1\_ptm\_pred\_0.pkl |

#### Structure similarity search results (Foldseek)^3^

|  |  |
| --- | --- |
| Structure databases searched | Pdb, Afdb-proteome, Afdb-uniprot50 |
| Results, scheme(s)  (Top layers only, threshold 1.00e-02 (evalue)) | xml version="1.0" encoding="utf-8" standalone="no"?       2024-09-02T21:10:23.559066 image/svg+xml   Matplotlib v3.7.2, https://matplotlib.org/ |
| Results, table  (threshold 1.00e-02 (evalue)) | db \| id \| prob \| evalue \| bits \| fident \| alnlen \| mismatch \| gapopen \| qstart \| qend \| tstart \| tend \| name \| description; pdb \| 6U5B\_A \| 1.0 \| 2.212e-08 \| 302 \| 0.443 \| 97 \| 54 \| 0 \| 1 \| 97 \| 1 \| 97 \| Sheath Initiator PA0617 \| Sheath Initiator PA0617; pdb \| 6U5K\_B \| 1.0 \| 2.18e-07 \| 255 \| 0.443 \| 97 \| 54 \| 0 \| 1 \| 97 \| 1 \| 97 \| Sheath Initiator PA0617 \| Sheath Initiator PA0617; pdb \| 2IA7\_A \| 1.0 \| 5.181e-07 \| 243 \| 0.34 \| 97 \| 61 \| 2 \| 12 \| 105 \| 4 \| 100 \| Tail lysozyme, putative \| Tail lysozyme, putative; pdb \| 5IW9\_B \| 1.0 \| 0.0001057 \| 166 \| 0.202 \| 94 \| 69 \| 4 \| 17 \| 105 \| 27 \| 119 \| Baseplate wedge protein gp25 \| Baseplate wedge protein gp25; pdb \| 6RAO\_H \| 1.0 \| 0.0009203 \| 138 \| 0.22 \| 100 \| 75 \| 2 \| 2 \| 98 \| 15 \| 114 \| Afp9 \| Afp9; pdb \| 7B5H\_AI \| 1.0 \| 9.934e-05 \| 130 \| 0.187 \| 112 \| 73 \| 4 \| 2 \| 107 \| 16 \| 115 \| All3318 protein \| All3318 protein; pdb \| 7AEB\_S \| 1.0 \| 0.0002512 \| 127 \| 0.231 \| 108 \| 73 \| 2 \| 2 \| 107 \| 14 \| 113 \| Putative tail lysozyme \| Putative tail lysozyme; pdb \| 6J0N\_D \| 1.0 \| 0.002056 \| 126 \| 0.163 \| 98 \| 79 \| 2 \| 2 \| 96 \| 17 \| 114 \| Pvc7 \| Pvc7; afdb-proteome \| AF-G3XD42-F1-MODEL\_V4 \| 1.0 \| 4.826e-12 \| 463 \| 0.416 \| 108 \| 61 \| 1 \| 1 \| 108 \| 1 \| 106 \| Probable bacteriophage protein \| Probable bacteriophage protein; afdb-proteome \| AF-Q8ZMU2-F1-MODEL\_V4 \| 1.0 \| 3.714e-11 \| 424 \| 0.38 \| 105 \| 64 \| 1 \| 1 \| 105 \| 4 \| 107 \| Fels-2 prophage protein \| Fels-2 prophage protein; afdb-proteome \| AF-A0A0H3H1Q9-F1-MODEL\_V4 \| 1.0 \| 3.085e-11 \| 412 \| 0.37 \| 108 \| 66 \| 1 \| 2 \| 107 \| 6 \| 113 \| Putative prophage baseplate protein \| Putative prophage baseplate protein; afdb-proteome \| AF-A0A0H3GUQ6-F1-MODEL\_V4 \| 1.0 \| 1.063e-10 \| 400 \| 0.39 \| 105 \| 62 \| 2 \| 2 \| 105 \| 5 \| 108 \| Baseplate assembly protein W \| Baseplate assembly protein W; afdb-proteome \| AF-A0A0H4IV38-F1-MODEL\_V4 \| 1.0 \| 5.228e-09 \| 341 \| 0.38 \| 105 \| 62 \| 2 \| 3 \| 105 \| 1 \| 104 \| Phage baseplate assembly protein \| Phage baseplate assembly protein; afdb-proteome \| AF-Q8ZKK2-F1-MODEL\_V4 \| 1.0 \| 3.965e-07 \| 271 \| 0.25 \| 108 \| 77 \| 4 \| 1 \| 107 \| 11 \| 115 \| Putative phage baseplate protein \| Putative phage baseplate protein; afdb-proteome \| AF-A0A0H3GZN2-F1-MODEL\_V4 \| 1.0 \| 0.0001922 \| 145 \| 0.189 \| 95 \| 68 \| 5 \| 15 \| 105 \| 30 \| 119 \| Type VI secretion system lysozyme-related protein \| Type VI secretion system lysozyme-related protein; afdb-proteome \| AF-P44239-F1-MODEL\_V4 \| 1.0 \| 0.001674 \| 123 \| 0.221 \| 131 \| 80 \| 10 \| 1 \| 112 \| 5 \| 132 \| Mu-like prophage FluMu protein gp46 \| Mu-like prophage FluMu protein gp46; afdb-proteome \| AF-Q9I366-F1-MODEL\_V4 \| 1.0 \| 0.002745 \| 118 \| 0.244 \| 98 \| 64 \| 6 \| 16 \| 106 \| 26 \| 120 \| GPW\_gp25 domain-containing protein \| GPW\_gp25 domain-containing protein; afdb-proteome \| AF-Q9I1B1-F1-MODEL\_V4 \| 0.998 \| 0.00479 \| 95 \| 0.178 \| 123 \| 81 \| 6 \| 1 \| 105 \| 1 \| 121 \| GPW\_gp25 domain-containing protein \| GPW\_gp25 domain-containing protein; afdb-uniprot50 \| AF-A0A6M4YGB1-F1-MODEL\_V4 \| 1.0 \| 4.39e-18 \| 740 \| 0.675 \| 111 \| 36 \| 0 \| 1 \| 111 \| 1 \| 111 \| Phage baseplate protein \| Phage baseplate protein; afdb-uniprot50 \| AF-A0A066RRA4-F1-MODEL\_V4 \| 1.0 \| 8.029e-17 \| 695 \| 0.598 \| 112 \| 45 \| 0 \| 1 \| 112 \| 1 \| 112 \| Phage baseplate protein \| Phage baseplate protein; afdb-uniprot50 \| AF-A0A2S0MNS1-F1-MODEL\_V4 \| 1.0 \| 3.329e-16 \| 655 \| 0.589 \| 112 \| 46 \| 0 \| 1 \| 112 \| 1 \| 112 \| Phage baseplate protein \| Phage baseplate protein; afdb-uniprot50 \| AF-I7DR34-F1-MODEL\_V4 \| 1.0 \| 4.825e-16 \| 650 \| 0.669 \| 112 \| 37 \| 0 \| 1 \| 112 \| 39 \| 150 \| Putative baseplate assembly protein W \| Putative baseplate assembly protein W; afdb-uniprot50 \| AF-Q3RBV9-F1-MODEL\_V4 \| 1.0 \| 3.767e-16 \| 649 \| 0.598 \| 112 \| 45 \| 0 \| 1 \| 112 \| 1 \| 112 \| GPW/gp25 \| GPW/gp25; afdb-uniprot50 \| AF-A0A2U1XZ29-F1-MODEL\_V4 \| 1.0 \| 2.264e-15 \| 633 \| 0.594 \| 111 \| 45 \| 0 \| 1 \| 111 \| 1 \| 111 \| GPW\_gp25 domain-containing protein \| GPW\_gp25 domain-containing protein; afdb-uniprot50 \| AF-A0A212KBR9-F1-MODEL\_V4 \| 1.0 \| 3.281e-15 \| 631 \| 0.473 \| 112 \| 59 \| 0 \| 1 \| 112 \| 1 \| 112 \| Putative Baseplate assembly protein W \| Putative Baseplate assembly protein W; afdb-uniprot50 \| AF-A0A1D2QM84-F1-MODEL\_V4 \| 1.0 \| 6.478e-15 \| 614 \| 0.504 \| 111 \| 55 \| 0 \| 1 \| 111 \| 1 \| 111 \| GPW\_gp25 domain-containing protein \| GPW\_gp25 domain-containing protein; afdb-uniprot50 \| AF-A0A7U9KG42-F1-MODEL\_V4 \| 1.0 \| 1.279e-14 \| 602 \| 0.5 \| 112 \| 56 \| 0 \| 1 \| 112 \| 1 \| 112 \| Putative lysozyme \| Putative lysozyme; afdb-uniprot50 \| AF-A0A3N2E0S6-F1-MODEL\_V4 \| 1.0 \| 1.972e-14 \| 585 \| 0.526 \| 112 \| 53 \| 0 \| 1 \| 112 \| 1 \| 112 \| GPW\_gp25 domain-containing protein \| GPW\_gp25 domain-containing protein; afdb-uniprot50 \| AF-A0A6G8F2Q6-F1-MODEL\_V4 \| 1.0 \| 6.384e-14 \| 569 \| 0.383 \| 112 \| 69 \| 0 \| 1 \| 112 \| 1 \| 112 \| Bacteriophage baseplate assembly protein W \| Bacteriophage baseplate assembly protein W; afdb-uniprot50 \| AF-N6VCT6-F1-MODEL\_V4 \| 1.0 \| 3.187e-13 \| 548 \| 0.565 \| 99 \| 43 \| 0 \| 14 \| 112 \| 3 \| 101 \| Phage baseplate assembly protein GpW \| Phage baseplate assembly protein GpW; afdb-uniprot50 \| AF-A0A3G2IL39-F1-MODEL\_V4 \| 1.0 \| 2.199e-13 \| 545 \| 0.446 \| 112 \| 62 \| 0 \| 1 \| 112 \| 1 \| 112 \| Phage baseplate protein \| Phage baseplate protein; afdb-uniprot50 \| AF-A0A6L2ZTB2-F1-MODEL\_V4 \| 1.0 \| 6.384e-14 \| 543 \| 0.489 \| 139 \| 42 \| 2 \| 1 \| 110 \| 1 \| 139 \| Baseplate assembly protein \| Baseplate assembly protein; afdb-uniprot50 \| AF-A0A0F4Q7B2-F1-MODEL\_V4 \| 1.0 \| 3.44e-14 \| 542 \| 0.518 \| 110 \| 53 \| 0 \| 1 \| 110 \| 1 \| 110 \| Baseplate assembly protein W \| Baseplate assembly protein W; afdb-uniprot50 \| AF-A0A430DQS5-F1-MODEL\_V4 \| 1.0 \| 1.614e-13 \| 541 \| 0.513 \| 115 \| 51 \| 3 \| 1 \| 111 \| 1 \| 114 \| Oxidoreductase \| Oxidoreductase; afdb-uniprot50 \| AF-A0A149SVH9-F1-MODEL\_V4 \| 1.0 \| 1.517e-13 \| 535 \| 0.517 \| 112 \| 53 \| 1 \| 1 \| 112 \| 1 \| 111 \| GPW\_gp25 domain-containing protein \| GPW\_gp25 domain-containing protein; afdb-uniprot50 \| AF-A0A2Z4UFI3-F1-MODEL\_V4 \| 1.0 \| 2.647e-13 \| 528 \| 0.495 \| 113 \| 56 \| 1 \| 1 \| 112 \| 1 \| 113 \| Gene 25-like lysozyme \| Gene 25-like lysozyme; afdb-uniprot50 \| AF-A0A3N2E0Q2-F1-MODEL\_V4 \| 1.0 \| 4.618e-13 \| 527 \| 0.414 \| 111 \| 65 \| 0 \| 1 \| 111 \| 1 \| 111 \| GPW\_gp25 domain-containing protein \| GPW\_gp25 domain-containing protein; afdb-uniprot50 \| AF-A0A7X3ZHR5-F1-MODEL\_V4 \| 1.0 \| 2.816e-13 \| 526 \| 0.464 \| 112 \| 58 \| 1 \| 1 \| 112 \| 1 \| 110 \| Phage baseplate protein \| Phage baseplate protein; afdb-uniprot50 \| AF-A0A432WB53-F1-MODEL\_V4 \| 1.0 \| 2.199e-13 \| 526 \| 0.526 \| 112 \| 53 \| 0 \| 1 \| 112 \| 3 \| 114 \| GPW\_gp25 domain-containing protein \| GPW\_gp25 domain-containing protein; afdb-uniprot50 \| AF-A0A5P9F0H1-F1-MODEL\_V4 \| 1.0 \| 6.291e-13 \| 521 \| 0.427 \| 110 \| 63 \| 0 \| 1 \| 110 \| 3 \| 112 \| Gene 25-like lysozyme \| Gene 25-like lysozyme; afdb-uniprot50 \| AF-A0A4U8YRG6-F1-MODEL\_V4 \| 1.0 \| 4.081e-13 \| 519 \| 0.509 \| 110 \| 53 \| 1 \| 1 \| 110 \| 2 \| 110 \| Gpw/gp25/anti-adapter protein irad \| Gpw/gp25/anti-adapter protein irad; afdb-uniprot50 \| AF-A0A2D8QD63-F1-MODEL\_V4 \| 1.0 \| 5.914e-13 \| 519 \| 0.518 \| 110 \| 51 \| 1 \| 3 \| 110 \| 17 \| 126 \| Baseplate assembly protein \| Baseplate assembly protein; afdb-uniprot50 \| AF-A0A192C892-F1-MODEL\_V4 \| 1.0 \| 1.242e-12 \| 516 \| 0.372 \| 110 \| 69 \| 0 \| 1 \| 110 \| 1 \| 110 \| Baseplate assembly protein W \| Baseplate assembly protein W; afdb-uniprot50 \| AF-W8ZTA4-F1-MODEL\_V4 \| 1.0 \| 1.321e-12 \| 514 \| 0.372 \| 110 \| 69 \| 0 \| 1 \| 110 \| 17 \| 126 \| Putative phage baseplate assembly protein \| Putative phage baseplate assembly protein; afdb-uniprot50 \| AF-A0A238UMY5-F1-MODEL\_V4 \| 1.0 \| 9.699e-13 \| 503 \| 0.513 \| 111 \| 49 \| 2 \| 1 \| 111 \| 1 \| 106 \| GPW\_gp25 domain-containing protein \| GPW\_gp25 domain-containing protein; afdb-uniprot50 \| AF-A0A6B8QMM4-F1-MODEL\_V4 \| 1.0 \| 8.057e-13 \| 498 \| 0.527 \| 108 \| 49 \| 1 \| 1 \| 106 \| 1 \| 108 \| Baseplate assembly protein \| Baseplate assembly protein; afdb-uniprot50 \| AF-A0A1I5W3Y1-F1-MODEL\_V4 \| 1.0 \| 5.15e-12 \| 488 \| 0.495 \| 107 \| 53 \| 1 \| 1 \| 107 \| 1 \| 106 \| GPW\_gp25 domain-containing protein \| GPW\_gp25 domain-containing protein; afdb-uniprot50 \| AF-A0A7T8CNB6-F1-MODEL\_V4 \| 1.0 \| 2.775e-12 \| 484 \| 0.447 \| 123 \| 57 \| 1 \| 1 \| 112 \| 1 \| 123 \| Phage baseplate protein \| Phage baseplate protein; afdb-uniprot50 \| AF-A0A2W4T7K0-F1-MODEL\_V4 \| 1.0 \| 2.775e-12 \| 483 \| 0.513 \| 109 \| 51 \| 1 \| 1 \| 109 \| 10 \| 116 \| Phage baseplate protein \| Phage baseplate protein; afdb-uniprot50 \| AF-A0A7Y0FCZ2-F1-MODEL\_V4 \| 1.0 \| 6.2e-12 \| 473 \| 0.5 \| 108 \| 51 \| 2 \| 1 \| 106 \| 1 \| 107 \| GPW\_gp25 domain-containing protein \| GPW\_gp25 domain-containing protein; afdb-uniprot50 \| AF-A0A0Q2Y5K4-F1-MODEL\_V4 \| 1.0 \| 4.841e-12 \| 471 \| 0.447 \| 105 \| 56 \| 1 \| 1 \| 105 \| 1 \| 103 \| GPW\_gp25 domain-containing protein \| GPW\_gp25 domain-containing protein; afdb-uniprot50 \| AF-A0A2N7UDN8-F1-MODEL\_V4 \| 1.0 \| 2.167e-12 \| 471 \| 0.509 \| 108 \| 50 \| 2 \| 1 \| 106 \| 1 \| 107 \| Baseplate assembly protein \| Baseplate assembly protein; afdb-uniprot50 \| AF-A0A6H2NTS6-F1-MODEL\_V4 \| 1.0 \| 7.94e-12 \| 469 \| 0.547 \| 95 \| 43 \| 0 \| 17 \| 111 \| 12 \| 106 \| Baseplate assembly protein W \| Baseplate assembly protein W; afdb-uniprot50 \| AF-A0A2K4MJA3-F1-MODEL\_V4 \| 1.0 \| 7.94e-12 \| 467 \| 0.417 \| 115 \| 63 \| 2 \| 1 \| 112 \| 3 \| 116 \| Baseplate assembly protein \| Baseplate assembly protein; afdb-uniprot50 \| AF-A0A369VRM6-F1-MODEL\_V4 \| 1.0 \| 3.341e-12 \| 460 \| 0.403 \| 114 \| 62 \| 3 \| 1 \| 111 \| 1 \| 111 \| Oxidoreductase \| Oxidoreductase; afdb-uniprot50 \| AF-A0A2Z6GBZ8-F1-MODEL\_V4 \| 1.0 \| 1.474e-11 \| 458 \| 0.414 \| 111 \| 61 \| 2 \| 1 \| 107 \| 3 \| 113 \| Baseplate assembly protein \| Baseplate assembly protein; afdb-uniprot50 \| AF-A0A1G5TK42-F1-MODEL\_V4 \| 1.0 \| 7.94e-12 \| 458 \| 0.4 \| 115 \| 62 \| 3 \| 1 \| 111 \| 1 \| 112 \| GPW\_gp25 domain-containing protein \| GPW\_gp25 domain-containing protein; afdb-uniprot50 \| AF-B4F0J8-F1-MODEL\_V4 \| 1.0 \| 2.136e-11 \| 457 \| 0.401 \| 112 \| 65 \| 2 \| 1 \| 112 \| 3 \| 112 \| Putative phage baseplate assembly protein \| Putative phage baseplate assembly protein; afdb-uniprot50 \| AF-A0A8A8MM88-F1-MODEL\_V4 \| 1.0 \| 1.474e-11 \| 457 \| 0.428 \| 105 \| 60 \| 0 \| 1 \| 105 \| 5 \| 109 \| GPW/gp25 family protein \| GPW/gp25 family protein; afdb-uniprot50 \| AF-A0A4R7GGA6-F1-MODEL\_V4 \| 1.0 \| 1.668e-11 \| 456 \| 0.467 \| 107 \| 57 \| 0 \| 1 \| 107 \| 4 \| 110 \| GPW\_gp25 domain-containing protein \| GPW\_gp25 domain-containing protein; afdb-uniprot50 \| AF-A0A1H9HC27-F1-MODEL\_V4 \| 1.0 \| 2.735e-11 \| 454 \| 0.428 \| 112 \| 63 \| 1 \| 1 \| 112 \| 1 \| 111 \| GPW\_gp25 domain-containing protein \| GPW\_gp25 domain-containing protein; afdb-uniprot50 \| AF-A0A345RIW3-F1-MODEL\_V4 \| 1.0 \| 7.016e-12 \| 454 \| 0.482 \| 112 \| 56 \| 2 \| 1 \| 111 \| 1 \| 111 \| Baseplate assembly protein \| Baseplate assembly protein; afdb-uniprot50 \| AF-A0A6I4T4J5-F1-MODEL\_V4 \| 1.0 \| 5.15e-12 \| 453 \| 0.459 \| 111 \| 55 \| 2 \| 1 \| 106 \| 4 \| 114 \| Oxidoreductase \| Oxidoreductase; afdb-uniprot50 \| AF-A0A631DZ78-F1-MODEL\_V4 \| 1.0 \| 1.668e-11 \| 452 \| 0.367 \| 106 \| 67 \| 0 \| 1 \| 106 \| 5 \| 110 \| Baseplate assembly protein \| Baseplate assembly protein; afdb-uniprot50 \| AF-C3X1Y5-F1-MODEL\_V4 \| 1.0 \| 2.136e-11 \| 452 \| 0.385 \| 109 \| 62 \| 2 \| 1 \| 106 \| 1 \| 107 \| GPW\_gp25 domain-containing protein \| GPW\_gp25 domain-containing protein; afdb-uniprot50 \| AF-A0A7A6VT71-F1-MODEL\_V4 \| 1.0 \| 2.136e-11 \| 447 \| 0.358 \| 106 \| 68 \| 0 \| 1 \| 106 \| 5 \| 110 \| Baseplate assembly protein \| Baseplate assembly protein; afdb-uniprot50 \| AF-A0A158S0X6-F1-MODEL\_V4 \| 1.0 \| 2.909e-11 \| 447 \| 0.459 \| 111 \| 55 \| 3 \| 1 \| 107 \| 6 \| 115 \| Baseplate assembly protein W \| Baseplate assembly protein W; afdb-uniprot50 \| AF-A0A7G7WLN4-F1-MODEL\_V4 \| 1.0 \| 3.726e-11 \| 446 \| 0.387 \| 111 \| 66 \| 2 \| 2 \| 112 \| 4 \| 112 \| GPW/gp25 family protein \| GPW/gp25 family protein; afdb-uniprot50 \| AF-A0A812QV09-F1-MODEL\_V4 \| 1.0 \| 5.479e-12 \| 445 \| 0.495 \| 113 \| 53 \| 2 \| 3 \| 111 \| 1061 \| 1173 \| XkdF protein \| XkdF protein; afdb-uniprot50 \| AF-A0A4Z0W627-F1-MODEL\_V4 \| 1.0 \| 1.002e-10 \| 444 \| 0.37 \| 108 \| 68 \| 0 \| 1 \| 108 \| 1 \| 108 \| Baseplate assembly protein W \| Baseplate assembly protein W; afdb-uniprot50 \| AF-A0A829QVE9-F1-MODEL\_V4 \| 1.0 \| 5.744e-11 \| 444 \| 0.351 \| 108 \| 70 \| 0 \| 1 \| 108 \| 3 \| 110 \| Putative lysozyme \| Putative lysozyme; afdb-uniprot50 \| AF-A0A854BCP0-F1-MODEL\_V4 \| 1.0 \| 1.302e-11 \| 442 \| 0.428 \| 112 \| 62 \| 2 \| 1 \| 111 \| 1 \| 111 \| Baseplate assembly protein \| Baseplate assembly protein; afdb-uniprot50 \| AF-A0A6H1ZRT2-F1-MODEL\_V4 \| 1.0 \| 8.324e-11 \| 440 \| 0.345 \| 113 \| 72 \| 2 \| 1 \| 112 \| 8 \| 119 \| Putative baseplate wedge subunit \| Putative baseplate wedge subunit; afdb-uniprot50 \| AF-A0A1R3WEX9-F1-MODEL\_V4 \| 1.0 \| 2.909e-11 \| 440 \| 0.441 \| 111 \| 56 \| 2 \| 1 \| 106 \| 1 \| 110 \| GPW\_gp25 domain-containing protein \| GPW\_gp25 domain-containing protein; afdb-uniprot50 \| AF-A0A2U3F139-F1-MODEL\_V4 \| 1.0 \| 3.292e-11 \| 439 \| 0.439 \| 107 \| 58 \| 2 \| 2 \| 108 \| 6 \| 110 \| Baseplate assembly protein \| Baseplate assembly protein; afdb-uniprot50 \| AF-A0A3Y9JE76-F1-MODEL\_V4 \| 1.0 \| 3.963e-11 \| 439 \| 0.38 \| 105 \| 65 \| 0 \| 1 \| 105 \| 5 \| 109 \| Baseplate assembly protein \| Baseplate assembly protein; afdb-uniprot50 \| AF-A0A142JGU1-F1-MODEL\_V4 \| 1.0 \| 1.302e-11 \| 437 \| 0.45 \| 111 \| 58 \| 2 \| 1 \| 108 \| 1 \| 111 \| GPW\_gp25 domain-containing protein \| GPW\_gp25 domain-containing protein; afdb-uniprot50 \| AF-A0A8B2NXK0-F1-MODEL\_V4 \| 1.0 \| 1.887e-11 \| 433 \| 0.486 \| 113 \| 50 \| 2 \| 2 \| 106 \| 5 \| 117 \| Baseplate assembly protein \| Baseplate assembly protein; afdb-uniprot50 \| AF-A0A1T4WV80-F1-MODEL\_V4 \| 1.0 \| 6.596e-12 \| 433 \| 0.477 \| 111 \| 53 \| 1 \| 1 \| 106 \| 80 \| 190 \| Phage baseplate assembly protein W \| Phage baseplate assembly protein W; afdb-uniprot50 \| AF-A0A2L0ACC8-F1-MODEL\_V4 \| 1.0 \| 2.909e-11 \| 432 \| 0.444 \| 108 \| 57 \| 2 \| 1 \| 105 \| 1 \| 108 \| Oxidoreductase \| Oxidoreductase; afdb-uniprot50 \| AF-A0A7V8SF12-F1-MODEL\_V4 \| 1.0 \| 4.771e-11 \| 432 \| 0.422 \| 109 \| 59 \| 3 \| 1 \| 105 \| 1 \| 109 \| GPW\_gp25 domain-containing protein \| GPW\_gp25 domain-containing protein; afdb-uniprot50 \| AF-A0A6L2ZQK7-F1-MODEL\_V4 \| 1.0 \| 6.915e-11 \| 431 \| 0.413 \| 104 \| 61 \| 0 \| 2 \| 105 \| 4 \| 107 \| Phage baseplate assembly protein \| Phage baseplate assembly protein; afdb-uniprot50 \| AF-A0A854E0F0-F1-MODEL\_V4 \| 1.0 \| 3.502e-11 \| 431 \| 0.443 \| 106 \| 58 \| 1 \| 1 \| 105 \| 1 \| 106 \| Baseplate assembly protein \| Baseplate assembly protein; afdb-uniprot50 \| AF-A0A2V3UCJ0-F1-MODEL\_V4 \| 1.0 \| 9.559e-12 \| 431 \| 0.474 \| 116 \| 51 \| 1 \| 1 \| 106 \| 3 \| 118 \| GPW\_gp25 domain-containing protein \| GPW\_gp25 domain-containing protein; afdb-uniprot50 \| AF-A0A326GHB1-F1-MODEL\_V4 \| 1.0 \| 2.909e-11 \| 429 \| 0.462 \| 108 \| 58 \| 0 \| 1 \| 108 \| 1 \| 108 \| Oxidoreductase \| Oxidoreductase; afdb-uniprot50 \| AF-A0A2A4XT86-F1-MODEL\_V4 \| 1.0 \| 6.5e-11 \| 428 \| 0.419 \| 105 \| 61 \| 0 \| 1 \| 105 \| 1 \| 105 \| GPW\_gp25 domain-containing protein \| GPW\_gp25 domain-containing protein; afdb-uniprot50 \| AF-A0A5E4SXB6-F1-MODEL\_V4 \| 1.0 \| 5.744e-11 \| 428 \| 0.364 \| 118 \| 68 \| 3 \| 1 \| 112 \| 3 \| 119 \| Phage baseplate protein \| Phage baseplate protein; afdb-uniprot50 \| AF-A0A1V0LL55-F1-MODEL\_V4 \| 1.0 \| 6.5e-11 \| 427 \| 0.371 \| 105 \| 65 \| 1 \| 1 \| 105 \| 5 \| 108 \| Baseplate assembly protein \| Baseplate assembly protein; afdb-uniprot50 \| AF-A0A328XFA0-F1-MODEL\_V4 \| 1.0 \| 8.855e-11 \| 427 \| 0.449 \| 109 \| 56 \| 1 \| 1 \| 105 \| 1 \| 109 \| GPW\_gp25 domain-containing protein \| GPW\_gp25 domain-containing protein; afdb-uniprot50 \| AF-A0A367WUC5-F1-MODEL\_V4 \| 1.0 \| 3.502e-11 \| 426 \| 0.41 \| 112 \| 62 \| 2 \| 1 \| 111 \| 1 \| 109 \| GPW\_gp25 domain-containing protein \| GPW\_gp25 domain-containing protein; afdb-uniprot50 \| AF-A0A7Z0LWZ0-F1-MODEL\_V4 \| 1.0 \| 2.735e-11 \| 426 \| 0.467 \| 107 \| 54 \| 2 \| 1 \| 105 \| 1 \| 106 \| GPW/gp25 family protein \| GPW/gp25 family protein; afdb-uniprot50 \| AF-A0A828IES2-F1-MODEL\_V4 \| 1.0 \| 9.42e-11 \| 425 \| 0.419 \| 105 \| 60 \| 1 \| 2 \| 105 \| 7 \| 111 \| Baseplate assembly protein \| Baseplate assembly protein; afdb-uniprot50 \| AF-A0A3R7GMZ6-F1-MODEL\_V4 \| 1.0 \| 7.825e-11 \| 424 \| 0.416 \| 108 \| 60 \| 1 \| 1 \| 105 \| 1 \| 108 \| Baseplate assembly protein \| Baseplate assembly protein; afdb-uniprot50 \| AF-A0A6P1LA29-F1-MODEL\_V4 \| 1.0 \| 3.726e-11 \| 423 \| 0.467 \| 109 \| 54 \| 2 \| 1 \| 105 \| 1 \| 109 \| Baseplate assembly protein \| Baseplate assembly protein; afdb-uniprot50 \| AF-A0A3E0X3W0-F1-MODEL\_V4 \| 1.0 \| 7.464e-12 \| 423 \| 0.465 \| 116 \| 54 \| 2 \| 1 \| 108 \| 1 \| 116 \| Baseplate assembly protein \| Baseplate assembly protein; afdb-uniprot50 \| AF-Q1QI83-F1-MODEL\_V4 \| 1.0 \| 2.136e-11 \| 422 \| 0.426 \| 122 \| 56 \| 3 \| 1 \| 108 \| 42 \| 163 \| GPW/gp25 \| GPW/gp25; afdb-uniprot50 \| AF-A0A286B0L7-F1-MODEL\_V4 \| 1.0 \| 7.016e-12 \| 420 \| 0.5 \| 112 \| 50 \| 2 \| 1 \| 106 \| 1 \| 112 \| GPW\_gp25 domain-containing protein \| GPW\_gp25 domain-containing protein; afdb-uniprot50 \| AF-Q9P9R9-F1-MODEL\_V4 \| 1.0 \| 2.417e-11 \| 419 \| 0.566 \| 90 \| 39 \| 0 \| 23 \| 112 \| 2 \| 91 \| Phage-related baseplate assembly protein \| Phage-related baseplate assembly protein; afdb-uniprot50 \| AF-A0A2S4LWD4-F1-MODEL\_V4 \| 1.0 \| 7.356e-11 \| 419 \| 0.436 \| 110 \| 56 \| 2 \| 1 \| 105 \| 1 \| 109 \| GPW\_gp25 domain-containing protein \| GPW\_gp25 domain-containing protein; afdb-uniprot50 \| AF-A0A1N7LRK4-F1-MODEL\_V4 \| 1.0 \| 1.86e-10 \| 418 \| 0.33 \| 112 \| 75 \| 0 \| 1 \| 112 \| 1 \| 112 \| GPW\_gp25 domain-containing protein \| GPW\_gp25 domain-containing protein; afdb-uniprot50 \| AF-A0A6I3XA50-F1-MODEL\_V4 \| 1.0 \| 1.002e-10 \| 418 \| 0.398 \| 113 \| 63 \| 2 \| 1 \| 111 \| 1 \| 110 \| Baseplate assembly protein \| Baseplate assembly protein; afdb-uniprot50 \| AF-A0A1Y1QY83-F1-MODEL\_V4 \| 1.0 \| 2.695e-10 \| 417 \| 0.504 \| 105 \| 50 \| 2 \| 4 \| 107 \| 1 \| 104 \| GPW\_gp25 domain-containing protein \| GPW\_gp25 domain-containing protein; afdb-uniprot50 \| AF-A0A812RF50-F1-MODEL\_V4 \| 1.0 \| 2.037e-12 \| 417 \| 0.46 \| 126 \| 52 \| 1 \| 1 \| 110 \| 63 \| 188 \| W protein \| W protein; afdb-uniprot50 \| AF-A0A1B4ETV1-F1-MODEL\_V4 \| 1.0 \| 4.155e-10 \| 415 \| 0.37 \| 108 \| 66 \| 1 \| 1 \| 108 \| 4 \| 109 \| Phage baseplate protein \| Phage baseplate protein; afdb-uniprot50 \| AF-A0A806CDY2-F1-MODEL\_V4 \| 1.0 \| 1.206e-10 \| 415 \| 0.4 \| 105 \| 63 \| 0 \| 1 \| 105 \| 1 \| 105 \| GPW/gp25 family protein \| GPW/gp25 family protein; afdb-uniprot50 \| AF-A8TYF6-F1-MODEL\_V4 \| 1.0 \| 2.534e-10 \| 414 \| 0.583 \| 96 \| 40 \| 0 \| 4 \| 99 \| 1 \| 96 \| GPW\_gp25 domain-containing protein \| GPW\_gp25 domain-containing protein; afdb-uniprot50 \| AF-A0A4Q0Y939-F1-MODEL\_V4 \| 1.0 \| 4.155e-10 \| 412 \| 0.593 \| 91 \| 35 \| 1 \| 20 \| 110 \| 1 \| 89 \| Phage baseplate protein \| Phage baseplate protein; afdb-uniprot50 \| AF-A0A0H3H1Q9-F1-MODEL\_V4 \| 1.0 \| 1.134e-10 \| 412 \| 0.37 \| 108 \| 66 \| 1 \| 2 \| 107 \| 6 \| 113 \| Putative prophage baseplate protein \| Putative prophage baseplate protein; afdb-uniprot50 \| AF-A0A5M6I4S2-F1-MODEL\_V4 \| 1.0 \| 1.134e-10 \| 412 \| 0.373 \| 107 \| 66 \| 1 \| 3 \| 108 \| 4 \| 110 \| GPW\_gp25 domain-containing protein \| GPW\_gp25 domain-containing protein; afdb-uniprot50 \| AF-A0A431QP25-F1-MODEL\_V4 \| 1.0 \| 5.399e-11 \| 412 \| 0.486 \| 109 \| 50 \| 2 \| 1 \| 103 \| 1 \| 109 \| Baseplate assembly protein \| Baseplate assembly protein; afdb-uniprot50 \| AF-A0A829R2Z7-F1-MODEL\_V4 \| 1.0 \| 2.105e-10 \| 411 \| 0.375 \| 104 \| 65 \| 0 \| 2 \| 105 \| 4 \| 107 \| Putative lysozyme \| Putative lysozyme; afdb-uniprot50 \| AF-A0A7Z1A3K0-F1-MODEL\_V4 \| 1.0 \| 2.867e-10 \| 411 \| 0.432 \| 104 \| 59 \| 0 \| 3 \| 106 \| 1 \| 104 \| Phage baseplate assembly protein \| Phage baseplate assembly protein; afdb-uniprot50 \| AF-A0A1M4WDA9-F1-MODEL\_V4 \| 1.0 \| 1.643e-10 \| 410 \| 0.485 \| 103 \| 50 \| 2 \| 4 \| 105 \| 1 \| 101 \| GPW\_gp25 domain-containing protein \| GPW\_gp25 domain-containing protein; afdb-uniprot50 \| AF-A0A250DSI4-F1-MODEL\_V4 \| 1.0 \| 2.105e-10 \| 410 \| 0.428 \| 105 \| 60 \| 0 \| 1 \| 105 \| 1 \| 105 \| GPW\_gp25 domain-containing protein \| GPW\_gp25 domain-containing protein; afdb-uniprot50 \| AF-A0A2G6IRN8-F1-MODEL\_V4 \| 1.0 \| 3.906e-10 \| 409 \| 0.361 \| 108 \| 68 \| 1 \| 1 \| 107 \| 1 \| 108 \| GPW\_gp25 domain-containing protein \| GPW\_gp25 domain-containing protein; afdb-uniprot50 \| AF-N9R592-F1-MODEL\_V4 \| 1.0 \| 2.239e-10 \| 409 \| 0.367 \| 106 \| 66 \| 1 \| 3 \| 107 \| 1 \| 106 \| GPW\_gp25 domain-containing protein \| GPW\_gp25 domain-containing protein; afdb-uniprot50 \| AF-A0A6L9JNR0-F1-MODEL\_V4 \| 1.0 \| 7.825e-11 \| 408 \| 0.412 \| 109 \| 63 \| 1 \| 1 \| 108 \| 3 \| 111 \| Baseplate assembly protein \| Baseplate assembly protein; afdb-uniprot50 \| AF-A0A1C3K3F0-F1-MODEL\_V4 \| 1.0 \| 1.002e-10 \| 408 \| 0.417 \| 115 \| 60 \| 3 \| 1 \| 111 \| 4 \| 115 \| Phage baseplate assembly protein \| Phage baseplate assembly protein; afdb-uniprot50 \| AF-A0A2W6T5C9-F1-MODEL\_V4 \| 1.0 \| 2.909e-11 \| 408 \| 0.413 \| 116 \| 62 \| 4 \| 1 \| 111 \| 4 \| 118 \| Baseplate assembly protein \| Baseplate assembly protein; afdb-uniprot50 \| AF-A0A7X2UKL5-F1-MODEL\_V4 \| 1.0 \| 3.05e-10 \| 407 \| 0.419 \| 105 \| 60 \| 1 \| 1 \| 105 \| 3 \| 106 \| Baseplate assembly protein \| Baseplate assembly protein; afdb-uniprot50 \| AF-A0A4P7L9H6-F1-MODEL\_V4 \| 1.0 \| 2.867e-10 \| 407 \| 0.368 \| 103 \| 64 \| 1 \| 3 \| 105 \| 5 \| 106 \| Gene 25-like lysozyme \| Gene 25-like lysozyme; afdb-uniprot50 \| AF-A0A7V8DBK9-F1-MODEL\_V4 \| 1.0 \| 3.05e-10 \| 406 \| 0.4 \| 105 \| 63 \| 0 \| 1 \| 105 \| 1 \| 105 \| GPW\_gp25 domain-containing protein \| GPW\_gp25 domain-containing protein; afdb-uniprot50 \| AF-A0A8B4MIR0-F1-MODEL\_V4 \| 1.0 \| 3.906e-10 \| 406 \| 0.342 \| 105 \| 69 \| 0 \| 3 \| 107 \| 1 \| 105 \| Phage-related baseplate assembly protein (GPW-like) \| Phage-related baseplate assembly protein (GPW-like); afdb-uniprot50 \| AF-A0A1Z9Z2R2-F1-MODEL\_V4 \| 1.0 \| 4.485e-11 \| 405 \| 0.39 \| 110 \| 63 \| 1 \| 3 \| 108 \| 1 \| 110 \| Baseplate assembly protein \| Baseplate assembly protein; afdb-uniprot50 \| AF-R9V0R8-F1-MODEL\_V4 \| 1.0 \| 2.382e-10 \| 405 \| 0.451 \| 104 \| 56 \| 1 \| 4 \| 106 \| 1 \| 104 \| Baseplate assembly protein \| Baseplate assembly protein; afdb-uniprot50 \| AF-A0A0M1HPW6-F1-MODEL\_V4 \| 1.0 \| 5.002e-10 \| 403 \| 0.355 \| 104 \| 67 \| 0 \| 3 \| 106 \| 1 \| 104 \| Baseplate assembly protein \| Baseplate assembly protein; afdb-uniprot50 \| AF-A0A1G7SDT3-F1-MODEL\_V4 \| 1.0 \| 3.672e-10 \| 402 \| 0.407 \| 108 \| 63 \| 1 \| 1 \| 107 \| 1 \| 108 \| GPW\_gp25 domain-containing protein \| GPW\_gp25 domain-containing protein; afdb-uniprot50 \| AF-A0A5P2STB1-F1-MODEL\_V4 \| 1.0 \| 2.105e-10 \| 402 \| 0.359 \| 103 \| 65 \| 1 \| 4 \| 105 \| 1 \| 103 \| Baseplate assembly protein \| Baseplate assembly protein; afdb-uniprot50 \| AF-A0A0E4BWU6-F1-MODEL\_V4 \| 1.0 \| 7.712e-10 \| 402 \| 0.292 \| 106 \| 74 \| 1 \| 1 \| 105 \| 8 \| 113 \| GPW\_gp25 domain-containing protein \| GPW\_gp25 domain-containing protein; afdb-uniprot50 \| AF-A0A2S8VCL2-F1-MODEL\_V4 \| 1.0 \| 1.978e-10 \| 401 \| 0.464 \| 112 \| 55 \| 3 \| 4 \| 111 \| 1 \| 111 \| Baseplate assembly protein \| Baseplate assembly protein; afdb-uniprot50 \| AF-A0A348HI76-F1-MODEL\_V4 \| 1.0 \| 4.155e-10 \| 401 \| 0.359 \| 103 \| 66 \| 0 \| 3 \| 105 \| 2 \| 104 \| Phage baseplate assemblyprotein W \| Phage baseplate assemblyprotein W; afdb-uniprot50 \| AF-A0A1B9JGD1-F1-MODEL\_V4 \| 1.0 \| 6.406e-10 \| 401 \| 0.4 \| 105 \| 61 \| 2 \| 2 \| 105 \| 4 \| 107 \| GPW\_gp25 domain-containing protein \| GPW\_gp25 domain-containing protein; afdb-uniprot50 \| AF-A0A0C4YEA8-F1-MODEL\_V4 \| 1.0 \| 5.744e-11 \| 401 \| 0.4 \| 115 \| 63 \| 1 \| 1 \| 109 \| 4 \| 118 \| Phage baseplate assembly protein \| Phage baseplate assembly protein; afdb-uniprot50 \| AF-Q1QKS3-F1-MODEL\_V4 \| 1.0 \| 1.134e-10 \| 401 \| 0.396 \| 121 \| 60 \| 3 \| 1 \| 108 \| 3 \| 123 \| GPW/gp25 \| GPW/gp25; afdb-uniprot50 \| AF-A0A143I3T2-F1-MODEL\_V4 \| 1.0 \| 3.906e-10 \| 399 \| 0.413 \| 104 \| 60 \| 1 \| 4 \| 106 \| 1 \| 104 \| GPW\_gp25 domain-containing protein \| GPW\_gp25 domain-containing protein; afdb-uniprot50 \| AF-A0A2V4DZL0-F1-MODEL\_V4 \| 1.0 \| 4.155e-10 \| 399 \| 0.378 \| 111 \| 64 \| 3 \| 2 \| 111 \| 4 \| 110 \| GPW\_gp25 domain-containing protein \| GPW\_gp25 domain-containing protein; afdb-uniprot50 \| AF-A0A350LX95-F1-MODEL\_V4 \| 1.0 \| 1.283e-10 \| 398 \| 0.42 \| 107 \| 62 \| 0 \| 1 \| 107 \| 1 \| 107 \| Phage baseplate protein \| Phage baseplate protein; afdb-uniprot50 \| AF-A0A376D4F9-F1-MODEL\_V4 \| 1.0 \| 9.284e-10 \| 397 \| 0.384 \| 104 \| 63 \| 1 \| 2 \| 105 \| 32 \| 134 \| GPW/gp25 family protein \| GPW/gp25 family protein; afdb-uniprot50 \| AF-A0A5Y3W5M6-F1-MODEL\_V4 \| 1.0 \| 4.155e-10 \| 396 \| 0.409 \| 110 \| 63 \| 2 \| 1 \| 109 \| 3 \| 111 \| Baseplate assembly protein \| Baseplate assembly protein; afdb-uniprot50 \| AF-G8PUM4-F1-MODEL\_V4 \| 1.0 \| 5.076e-11 \| 396 \| 0.396 \| 121 \| 60 \| 2 \| 1 \| 108 \| 3 \| 123 \| GPW/gp25 \| GPW/gp25; afdb-uniprot50 \| AF-E2CJU0-F1-MODEL\_V4 \| 1.0 \| 1.365e-10 \| 396 \| 0.378 \| 119 \| 61 \| 3 \| 1 \| 107 \| 3 \| 120 \| GPW/gp25 \| GPW/gp25; afdb-uniprot50 \| AF-A0A7X9X5F1-F1-MODEL\_V4 \| 1.0 \| 2.239e-10 \| 394 \| 0.417 \| 115 \| 60 \| 1 \| 1 \| 108 \| 1 \| 115 \| Baseplate assembly protein \| Baseplate assembly protein; afdb-uniprot50 \| AF-A0A318L0U5-F1-MODEL\_V4 \| 1.0 \| 3.906e-10 \| 394 \| 0.371 \| 105 \| 65 \| 1 \| 2 \| 105 \| 9 \| 113 \| GPW\_gp25 domain-containing protein \| GPW\_gp25 domain-containing protein; afdb-uniprot50 \| AF-A0A1I1UBH6-F1-MODEL\_V4 \| 1.0 \| 6.815e-10 \| 393 \| 0.419 \| 105 \| 59 \| 2 \| 3 \| 106 \| 2 \| 105 \| GPW\_gp25 domain-containing protein \| GPW\_gp25 domain-containing protein; afdb-uniprot50 \| AF-V4PNX9-F1-MODEL\_V4 \| 1.0 \| 4.155e-10 \| 393 \| 0.439 \| 107 \| 58 \| 1 \| 1 \| 105 \| 1 \| 107 \| GPW\_gp25 domain-containing protein \| GPW\_gp25 domain-containing protein; afdb-uniprot50 \| AF-A0A2E6BQ95-F1-MODEL\_V4 \| 1.0 \| 2.239e-10 \| 393 \| 0.44 \| 109 \| 55 \| 2 \| 4 \| 108 \| 1 \| 107 \| GPW\_gp25 domain-containing protein \| GPW\_gp25 domain-containing protein; afdb-uniprot50 \| AF-A0A2N7P6U4-F1-MODEL\_V4 \| 1.0 \| 6.022e-10 \| 393 \| 0.327 \| 113 \| 73 \| 1 \| 1 \| 110 \| 2 \| 114 \| Uncharacterized protein \| Uncharacterized protein; afdb-uniprot50 \| AF-A0A212J4Q2-F1-MODEL\_V4 \| 1.0 \| 7.249e-10 \| 392 \| 0.388 \| 108 \| 66 \| 0 \| 1 \| 108 \| 6 \| 113 \| Putative Baseplate assembly protein W \| Putative Baseplate assembly protein W; afdb-uniprot50 \| AF-D4M9N2-F1-MODEL\_V4 \| 1.0 \| 2.534e-10 \| 390 \| 0.443 \| 115 \| 56 \| 4 \| 1 \| 111 \| 4 \| 114 \| Phage baseplate assembly protein W \| Phage baseplate assembly protein W; afdb-uniprot50 \| AF-A0A2W5X821-F1-MODEL\_V4 \| 1.0 \| 3.906e-10 \| 390 \| 0.372 \| 110 \| 65 \| 2 \| 1 \| 106 \| 1 \| 110 \| Uncharacterized protein \| Uncharacterized protein; afdb-uniprot50 \| AF-A0A5J6LCN8-F1-MODEL\_V4 \| 1.0 \| 2.382e-10 \| 389 \| 0.436 \| 110 \| 56 \| 3 \| 3 \| 111 \| 5 \| 109 \| GPW\_gp25 domain-containing protein \| GPW\_gp25 domain-containing protein; afdb-uniprot50 \| AF-A0A556S966-F1-MODEL\_V4 \| 1.0 \| 3.05e-10 \| 387 \| 0.38 \| 113 \| 64 \| 2 \| 1 \| 108 \| 3 \| 114 \| Baseplate assembly protein \| Baseplate assembly protein; afdb-uniprot50 \| AF-A0A6L3Y821-F1-MODEL\_V4 \| 1.0 \| 1.86e-10 \| 387 \| 0.404 \| 121 \| 58 \| 2 \| 1 \| 107 \| 3 \| 123 \| Baseplate assembly protein \| Baseplate assembly protein; afdb-uniprot50 \| AF-A0A4Q3KFV4-F1-MODEL\_V4 \| 1.0 \| 5.321e-10 \| 387 \| 0.432 \| 111 \| 57 \| 2 \| 4 \| 108 \| 1 \| 111 \| Uncharacterized protein \| Uncharacterized protein; afdb-uniprot50 \| AF-A0A1G8TJ30-F1-MODEL\_V4 \| 1.0 \| 2.867e-10 \| 386 \| 0.514 \| 107 \| 49 \| 2 \| 4 \| 108 \| 1 \| 106 \| GPW\_gp25 domain-containing protein \| GPW\_gp25 domain-containing protein; afdb-uniprot50 \| AF-A0A856Y7R7-F1-MODEL\_V4 \| 1.0 \| 1.118e-09 \| 386 \| 0.5 \| 98 \| 48 \| 1 \| 1 \| 97 \| 1 \| 98 \| Baseplate assembly protein \| Baseplate assembly protein; afdb-uniprot50 \| AF-A0A2S9RND6-F1-MODEL\_V4 \| 1.0 \| 9.284e-10 \| 386 \| 0.358 \| 106 \| 66 \| 2 \| 4 \| 108 \| 1 \| 105 \| Gene 25-like lysozyme \| Gene 25-like lysozyme; afdb-uniprot50 \| AF-A0A348FYH4-F1-MODEL\_V4 \| 1.0 \| 3.906e-10 \| 385 \| 0.419 \| 105 \| 60 \| 1 \| 3 \| 106 \| 2 \| 106 \| GPW\_gp25 domain-containing protein \| GPW\_gp25 domain-containing protein; afdb-uniprot50 \| AF-A0A2T5J1F8-F1-MODEL\_V4 \| 1.0 \| 2.695e-10 \| 385 \| 0.377 \| 106 \| 66 \| 0 \| 1 \| 106 \| 3 \| 108 \| GPW\_gp25 domain-containing protein \| GPW\_gp25 domain-containing protein; afdb-uniprot50 \| AF-A0A2W7IMC6-F1-MODEL\_V4 \| 1.0 \| 4.42e-10 \| 384 \| 0.403 \| 109 \| 61 \| 2 \| 4 \| 111 \| 1 \| 106 \| GPW\_gp25 domain-containing protein \| GPW\_gp25 domain-containing protein; afdb-uniprot50 \| AF-A0A5E4XG18-F1-MODEL\_V4 \| 1.0 \| 6.815e-10 \| 384 \| 0.382 \| 115 \| 64 \| 2 \| 1 \| 111 \| 1 \| 112 \| Phage baseplate protein \| Phage baseplate protein; afdb-uniprot50 \| AF-Q31HT9-F1-MODEL\_V4 \| 1.0 \| 8.727e-10 \| 383 \| 0.299 \| 117 \| 74 \| 4 \| 1 \| 112 \| 1 \| 114 \| Phage baseplate assembly protein \| Phage baseplate assembly protein; afdb-uniprot50 \| AF-A0A554XC26-F1-MODEL\_V4 \| 1.0 \| 8.727e-10 \| 382 \| 0.457 \| 105 \| 57 \| 0 \| 1 \| 105 \| 23 \| 127 \| Lysozyme \| Lysozyme; afdb-uniprot50 \| AF-A0A5B9ACT0-F1-MODEL\_V4 \| 1.0 \| 6.022e-10 \| 381 \| 0.402 \| 92 \| 55 \| 0 \| 14 \| 105 \| 1 \| 92 \| Baseplate assembly protein \| Baseplate assembly protein; afdb-uniprot50 \| AF-A0A256CAS9-F1-MODEL\_V4 \| 1.0 \| 2.206e-09 \| 380 \| 0.4 \| 105 \| 62 \| 1 \| 4 \| 107 \| 1 \| 105 \| GPW\_gp25 domain-containing protein \| GPW\_gp25 domain-containing protein; afdb-uniprot50 \| AF-A0A0T9QIN0-F1-MODEL\_V4 \| 1.0 \| 9.876e-10 \| 380 \| 0.39 \| 105 \| 64 \| 0 \| 1 \| 105 \| 5 \| 109 \| Phage baseplate assembly protein W \| Phage baseplate assembly protein W; afdb-uniprot50 \| AF-A0A542V4K3-F1-MODEL\_V4 \| 1.0 \| 1.523e-09 \| 378 \| 0.415 \| 106 \| 61 \| 1 \| 4 \| 108 \| 1 \| 106 \| GPW\_gp25 domain-containing protein \| GPW\_gp25 domain-containing protein; afdb-uniprot50 \| AF-A0A2S2E6D5-F1-MODEL\_V4 \| 1.0 \| 8.727e-10 \| 377 \| 0.3 \| 113 \| 74 \| 2 \| 1 \| 109 \| 2 \| 113 \| Uncharacterized protein \| Uncharacterized protein; afdb-uniprot50 \| AF-A0A423PQJ1-F1-MODEL\_V4 \| 1.0 \| 5.579e-09 \| 375 \| 0.417 \| 103 \| 60 \| 0 \| 4 \| 106 \| 1 \| 103 \| Baseplate assembly protein \| Baseplate assembly protein; afdb-uniprot50 \| AF-A0A516W9Y9-F1-MODEL\_V4 \| 1.0 \| 6.022e-10 \| 375 \| 0.476 \| 105 \| 53 \| 1 \| 4 \| 106 \| 1 \| 105 \| Baseplate assembly protein \| Baseplate assembly protein; afdb-uniprot50 \| AF-S6GUS5-F1-MODEL\_V4 \| 1.0 \| 1.189e-09 \| 375 \| 0.401 \| 107 \| 60 \| 3 \| 1 \| 106 \| 8 \| 111 \| W protein \| W protein; afdb-uniprot50 \| AF-A0A1Y3CLD4-F1-MODEL\_V4 \| 1.0 \| 1.833e-09 \| 375 \| 0.38 \| 105 \| 63 \| 2 \| 3 \| 105 \| 1 \| 105 \| Baseplate assembly protein \| Baseplate assembly protein; afdb-uniprot50 \| AF-A0A0U3ICT5-F1-MODEL\_V4 \| 1.0 \| 2.206e-09 \| 374 \| 0.388 \| 103 \| 63 \| 0 \| 3 \| 105 \| 1 \| 103 \| Baseplate assembly protein \| Baseplate assembly protein; afdb-uniprot50 \| AF-A0A7T9Q7H3-F1-MODEL\_V4 \| 1.0 \| 2.074e-09 \| 374 \| 0.388 \| 103 \| 63 \| 0 \| 3 \| 105 \| 1 \| 103 \| GPW/gp25 family protein \| GPW/gp25 family protein; afdb-uniprot50 \| AF-A0A166Z6R0-F1-MODEL\_V4 \| 1.0 \| 5.661e-10 \| 373 \| 0.412 \| 109 \| 60 \| 2 \| 1 \| 108 \| 3 \| 108 \| Baseplate assembly protein \| Baseplate assembly protein; afdb-uniprot50 \| AF-A0A2N3KSL6-F1-MODEL\_V4 \| 1.0 \| 1.431e-09 \| 373 \| 0.364 \| 107 \| 66 \| 1 \| 1 \| 105 \| 1 \| 107 \| Baseplate assembly protein W \| Baseplate assembly protein W; afdb-uniprot50 \| AF-A0A1M3HHS7-F1-MODEL\_V4 \| 1.0 \| 1.345e-09 \| 372 \| 0.416 \| 108 \| 60 \| 2 \| 1 \| 105 \| 1 \| 108 \| GPW\_gp25 domain-containing protein \| GPW\_gp25 domain-containing protein; afdb-uniprot50 \| AF-A0A4R0EZX5-F1-MODEL\_V4 \| 1.0 \| 2.347e-09 \| 372 \| 0.378 \| 103 \| 64 \| 0 \| 3 \| 105 \| 1 \| 103 \| Baseplate assembly protein \| Baseplate assembly protein; afdb-uniprot50 \| AF-A0A7S8HE74-F1-MODEL\_V4 \| 1.0 \| 5.321e-10 \| 371 \| 0.388 \| 108 \| 55 \| 2 \| 12 \| 111 \| 1 \| 105 \| GPW/gp25 family protein \| GPW/gp25 family protein; afdb-uniprot50 \| AF-A0A8B0TIL9-F1-MODEL\_V4 \| 1.0 \| 2.656e-09 \| 371 \| 0.401 \| 102 \| 61 \| 0 \| 4 \| 105 \| 1 \| 102 \| GPW/gp25 family protein \| GPW/gp25 family protein; afdb-uniprot50 \| AF-A0A4R0FND4-F1-MODEL\_V4 \| 1.0 \| 2.826e-09 \| 369 \| 0.368 \| 103 \| 65 \| 0 \| 3 \| 105 \| 1 \| 103 \| Baseplate assembly protein \| Baseplate assembly protein; afdb-uniprot50 \| AF-C8N768-F1-MODEL\_V4 \| 1.0 \| 1.95e-09 \| 369 \| 0.361 \| 108 \| 67 \| 2 \| 2 \| 108 \| 19 \| 125 \| Putative lysozyme \| Putative lysozyme; afdb-uniprot50 \| AF-A0A103EKP3-F1-MODEL\_V4 \| 1.0 \| 4.095e-09 \| 368 \| 0.675 \| 80 \| 26 \| 0 \| 1 \| 80 \| 1 \| 80 \| Phage baseplate protein \| Phage baseplate protein; afdb-uniprot50 \| AF-A0A074TCW7-F1-MODEL\_V4 \| 1.0 \| 1.118e-09 \| 368 \| 0.371 \| 113 \| 61 \| 1 \| 3 \| 105 \| 2 \| 114 \| GPW\_gp25 domain-containing protein \| GPW\_gp25 domain-containing protein; afdb-uniprot50 \| AF-A0A1I1UZQ7-F1-MODEL\_V4 \| 1.0 \| 1.95e-09 \| 368 \| 0.359 \| 114 \| 64 \| 2 \| 1 \| 105 \| 2 \| 115 \| GPW\_gp25 domain-containing protein \| GPW\_gp25 domain-containing protein; afdb-uniprot50 \| AF-A0A7H2V930-F1-MODEL\_V4 \| 1.0 \| 3.198e-09 \| 367 \| 0.378 \| 103 \| 64 \| 0 \| 3 \| 105 \| 1 \| 103 \| GPW/gp25 family protein \| GPW/gp25 family protein; afdb-uniprot50 \| AF-A0A239RWY0-F1-MODEL\_V4 \| 1.0 \| 3.619e-09 \| 367 \| 0.388 \| 103 \| 63 \| 0 \| 3 \| 105 \| 1 \| 103 \| Baseplate assembly protein \| Baseplate assembly protein; afdb-uniprot50 \| AF-A0A258L738-F1-MODEL\_V4 \| 1.0 \| 1.265e-09 \| 367 \| 0.368 \| 114 \| 63 \| 2 \| 1 \| 105 \| 4 \| 117 \| GPW\_gp25 domain-containing protein \| GPW\_gp25 domain-containing protein; afdb-uniprot50 \| AF-A0A5N4WTB4-F1-MODEL\_V4 \| 1.0 \| 3.198e-09 \| 366 \| 0.378 \| 103 \| 64 \| 0 \| 3 \| 105 \| 1 \| 103 \| Baseplate assembly protein \| Baseplate assembly protein; afdb-uniprot50 \| AF-A0A2A2B5A6-F1-MODEL\_V4 \| 1.0 \| 9.876e-10 \| 366 \| 0.471 \| 106 \| 52 \| 3 \| 3 \| 105 \| 7 \| 111 \| Baseplate assembly protein \| Baseplate assembly protein; afdb-uniprot50 \| AF-A0A1W1XJR9-F1-MODEL\_V4 \| 1.0 \| 1.345e-09 \| 365 \| 0.368 \| 103 \| 64 \| 1 \| 4 \| 105 \| 1 \| 103 \| GPW\_gp25 domain-containing protein \| GPW\_gp25 domain-containing protein; afdb-uniprot50 \| AF-A0A846VGN2-F1-MODEL\_V4 \| 1.0 \| 4.356e-09 \| 365 \| 0.28 \| 114 \| 73 \| 2 \| 1 \| 105 \| 6 \| 119 \| Uncharacterized protein \| Uncharacterized protein; afdb-uniprot50 \| AF-A0A345DE47-F1-MODEL\_V4 \| 1.0 \| 1.265e-09 \| 364 \| 0.467 \| 107 \| 54 \| 2 \| 4 \| 107 \| 1 \| 107 \| GPW\_gp25 domain-containing protein \| GPW\_gp25 domain-containing protein; afdb-uniprot50 \| AF-A0A359KBY6-F1-MODEL\_V4 \| 1.0 \| 2.206e-09 \| 362 \| 0.373 \| 107 \| 64 \| 2 \| 1 \| 105 \| 1 \| 106 \| Uncharacterized protein \| Uncharacterized protein; afdb-uniprot50 \| AF-A0A1B8QCY8-F1-MODEL\_V4 \| 1.0 \| 7.144e-09 \| 361 \| 0.352 \| 105 \| 68 \| 0 \| 4 \| 108 \| 1 \| 105 \| GPW\_gp25 domain-containing protein \| GPW\_gp25 domain-containing protein; afdb-uniprot50 \| AF-A0A547PWD4-F1-MODEL\_V4 \| 1.0 \| 2.074e-09 \| 361 \| 0.495 \| 103 \| 49 \| 1 \| 1 \| 103 \| 1 \| 100 \| Phage baseplate protein \| Phage baseplate protein; afdb-uniprot50 \| AF-A0A2G6E136-F1-MODEL\_V4 \| 1.0 \| 9.876e-10 \| 360 \| 0.419 \| 112 \| 54 \| 2 \| 4 \| 106 \| 1 \| 110 \| GPW\_gp25 domain-containing protein \| GPW\_gp25 domain-containing protein; afdb-uniprot50 \| AF-A0A2G6EWW6-F1-MODEL\_V4 \| 1.0 \| 1.95e-09 \| 359 \| 0.446 \| 103 \| 57 \| 0 \| 3 \| 105 \| 10 \| 112 \| GPW\_gp25 domain-containing protein \| GPW\_gp25 domain-containing protein; afdb-uniprot50 \| AF-A0A377AI41-F1-MODEL\_V4 \| 1.0 \| 6.313e-09 \| 357 \| 0.422 \| 97 \| 56 \| 0 \| 1 \| 97 \| 5 \| 101 \| Baseplate assembly protein W \| Baseplate assembly protein W; afdb-uniprot50 \| AF-D6H7Q3-F1-MODEL\_V4 \| 1.0 \| 6.716e-09 \| 357 \| 0.377 \| 106 \| 63 \| 2 \| 2 \| 105 \| 192 \| 296 \| Uncharacterized protein \| Uncharacterized protein; afdb-uniprot50 \| AF-C4GGH2-F1-MODEL\_V4 \| 1.0 \| 4.095e-09 \| 355 \| 0.386 \| 106 \| 60 \| 3 \| 3 \| 105 \| 1 \| 104 \| Putative lysozyme \| Putative lysozyme; afdb-uniprot50 \| AF-A0A7U3VTF8-F1-MODEL\_V4 \| 1.0 \| 6.716e-09 \| 355 \| 0.294 \| 112 \| 78 \| 1 \| 1 \| 112 \| 1 \| 111 \| Baseplate assembly protein GpW \| Baseplate assembly protein GpW; afdb-uniprot50 \| AF-C1DBK2-F1-MODEL\_V4 \| 1.0 \| 7.144e-09 \| 354 \| 0.411 \| 107 \| 59 \| 3 \| 4 \| 106 \| 1 \| 107 \| GpW/GP25 family protein \| GpW/GP25 family protein; afdb-uniprot50 \| AF-A0A2M9VR53-F1-MODEL\_V4 \| 1.0 \| 4.095e-09 \| 353 \| 0.445 \| 92 \| 51 \| 0 \| 15 \| 106 \| 1 \| 92 \| Phage baseplate protein \| Phage baseplate protein; afdb-uniprot50 \| AF-A0A0A2YRZ1-F1-MODEL\_V4 \| 1.0 \| 5.579e-09 \| 351 \| 0.388 \| 103 \| 61 \| 2 \| 4 \| 105 \| 1 \| 102 \| Baseplate wedge subunit \| Baseplate wedge subunit; afdb-uniprot50 \| AF-A0A371WUQ6-F1-MODEL\_V4 \| 1.0 \| 3.619e-09 \| 349 \| 0.35 \| 120 \| 63 \| 2 \| 1 \| 105 \| 20 \| 139 \| Baseplate assembly protein \| Baseplate assembly protein; afdb-uniprot50 \| AF-A0A241WJJ5-F1-MODEL\_V4 \| 1.0 \| 1.265e-09 \| 348 \| 0.408 \| 120 \| 59 \| 4 \| 1 \| 108 \| 1 \| 120 \| GPW\_gp25 domain-containing protein \| GPW\_gp25 domain-containing protein; afdb-uniprot50 \| AF-A0A6B8KGJ0-F1-MODEL\_V4 \| 1.0 \| 2.074e-09 \| 348 \| 0.33 \| 115 \| 72 \| 3 \| 1 \| 110 \| 2 \| 116 \| Uncharacterized protein \| Uncharacterized protein; afdb-uniprot50 \| AF-A0A7W6RF26-F1-MODEL\_V4 \| 1.0 \| 9.149e-09 \| 347 \| 0.352 \| 125 \| 64 \| 4 \| 1 \| 108 \| 1 \| 125 \| Uncharacterized protein \| Uncharacterized protein; afdb-uniprot50 \| AF-A0A6D0J2V1-F1-MODEL\_V4 \| 1.0 \| 5.579e-09 \| 346 \| 0.382 \| 89 \| 55 \| 0 \| 17 \| 105 \| 6 \| 94 \| Baseplate assembly protein \| Baseplate assembly protein; afdb-uniprot50 \| AF-A0A4R3Y632-F1-MODEL\_V4 \| 1.0 \| 4.634e-09 \| 345 \| 0.35 \| 100 \| 60 \| 3 \| 17 \| 112 \| 7 \| 105 \| GPW\_gp25 domain-containing protein \| GPW\_gp25 domain-containing protein; afdb-uniprot50 \| AF-A0A326L1I8-F1-MODEL\_V4 \| 1.0 \| 5.579e-09 \| 345 \| 0.371 \| 113 \| 64 \| 3 \| 1 \| 106 \| 3 \| 115 \| GPW\_gp25 domain-containing protein \| GPW\_gp25 domain-containing protein; afdb-uniprot50 \| AF-A0A809X5W8-F1-MODEL\_V4 \| 1.0 \| 1.41e-08 \| 344 \| 0.317 \| 104 \| 64 \| 2 \| 1 \| 100 \| 8 \| 108 \| Uncharacterized protein \| Uncharacterized protein; afdb-uniprot50 \| AF-A0A516SJB4-F1-MODEL\_V4 \| 1.0 \| 3.198e-09 \| 344 \| 0.37 \| 116 \| 63 \| 3 \| 1 \| 108 \| 6 \| 119 \| Phage baseplate protein \| Phage baseplate protein; afdb-uniprot50 \| AF-A0A535HZP7-F1-MODEL\_V4 \| 1.0 \| 6.716e-09 \| 343 \| 0.408 \| 98 \| 56 \| 1 \| 10 \| 105 \| 2 \| 99 \| GPW/gp25 family protein \| GPW/gp25 family protein; afdb-uniprot50 \| AF-A0A7C6PU32-F1-MODEL\_V4 \| 1.0 \| 6.716e-09 \| 343 \| 0.279 \| 111 \| 73 \| 2 \| 1 \| 105 \| 7 \| 116 \| GPW/gp25 family protein \| GPW/gp25 family protein; afdb-uniprot50 \| AF-A0A7W8YF20-F1-MODEL\_V4 \| 1.0 \| 1.806e-08 \| 342 \| 0.558 \| 86 \| 38 \| 0 \| 1 \| 86 \| 1 \| 86 \| GPW\_gp25 domain-containing protein \| GPW\_gp25 domain-containing protein; afdb-uniprot50 \| AF-A0A037UUA6-F1-MODEL\_V4 \| 1.0 \| 5.244e-09 \| 342 \| 0.415 \| 106 \| 57 \| 1 \| 1 \| 101 \| 1 \| 106 \| GPW\_gp25 domain-containing protein \| GPW\_gp25 domain-containing protein; afdb-uniprot50 \| AF-A0A0H4IV38-F1-MODEL\_V4 \| 1.0 \| 1.922e-08 \| 341 \| 0.38 \| 105 \| 62 \| 2 \| 3 \| 105 \| 1 \| 104 \| Phage baseplate assembly protein \| Phage baseplate assembly protein; afdb-uniprot50 \| AF-A0A6L7EAJ2-F1-MODEL\_V4 \| 1.0 \| 1.172e-08 \| 340 \| 0.389 \| 95 \| 54 \| 2 \| 17 \| 111 \| 1 \| 91 \| Baseplate assembly protein \| Baseplate assembly protein; afdb-uniprot50 \| AF-A0A3L7AJZ0-F1-MODEL\_V4 \| 1.0 \| 4.93e-09 \| 340 \| 0.312 \| 112 \| 68 \| 2 \| 1 \| 103 \| 1 \| 112 \| Baseplate assembly protein \| Baseplate assembly protein; afdb-uniprot50 \| AF-A0A7W8AJL1-F1-MODEL\_V4 \| 1.0 \| 6.313e-09 \| 340 \| 0.324 \| 114 \| 68 \| 3 \| 1 \| 105 \| 8 \| 121 \| GPW\_gp25 domain-containing protein \| GPW\_gp25 domain-containing protein; afdb-uniprot50 \| AF-A0A2A5BNQ3-F1-MODEL\_V4 \| 1.0 \| 1.118e-09 \| 338 \| 0.342 \| 114 \| 65 \| 3 \| 1 \| 105 \| 1 \| 113 \| GPW\_gp25 domain-containing protein \| GPW\_gp25 domain-containing protein; afdb-uniprot50 \| AF-A0A839IXR3-F1-MODEL\_V4 \| 1.0 \| 8.601e-09 \| 338 \| 0.293 \| 116 \| 72 \| 4 \| 1 \| 112 \| 1 \| 110 \| Phage baseplate protein \| Phage baseplate protein; afdb-uniprot50 \| AF-A0A1L6M2Z8-F1-MODEL\_V4 \| 1.0 \| 3.619e-09 \| 338 \| 0.324 \| 117 \| 67 \| 2 \| 1 \| 105 \| 9 \| 125 \| GPW\_gp25 domain-containing protein \| GPW\_gp25 domain-containing protein; afdb-uniprot50 \| AF-B7RNM3-F1-MODEL\_V4 \| 1.0 \| 3.151e-08 \| 337 \| 0.576 \| 78 \| 33 \| 0 \| 35 \| 112 \| 1 \| 78 \| GPW/gp25 \| GPW/gp25; afdb-uniprot50 \| AF-A0A1Y6CPE4-F1-MODEL\_V4 \| 1.0 \| 1.596e-08 \| 337 \| 0.314 \| 108 \| 71 \| 2 \| 1 \| 108 \| 1 \| 105 \| Uncharacterized protein \| Uncharacterized protein; afdb-uniprot50 \| AF-A0A5E8NU52-F1-MODEL\_V4 \| 1.0 \| 3.151e-08 \| 334 \| 0.387 \| 93 \| 57 \| 0 \| 1 \| 93 \| 5 \| 97 \| Baseplate assembly protein \| Baseplate assembly protein; afdb-uniprot50 \| AF-E2CFL3-F1-MODEL\_V4 \| 1.0 \| 5.935e-09 \| 334 \| 0.275 \| 116 \| 79 \| 3 \| 1 \| 112 \| 2 \| 116 \| GpW/gp25 family protein \| GpW/gp25 family protein; afdb-uniprot50 \| AF-Q4E960-F1-MODEL\_V4 \| 1.0 \| 3.352e-08 \| 333 \| 0.521 \| 94 \| 45 \| 0 \| 4 \| 97 \| 1 \| 94 \| GPW\_gp25 domain-containing protein \| GPW\_gp25 domain-containing protein; afdb-uniprot50 \| AF-A0A7Y2L5A3-F1-MODEL\_V4 \| 1.0 \| 1.922e-08 \| 330 \| 0.5 \| 84 \| 41 \| 1 \| 23 \| 105 \| 2 \| 85 \| Baseplate assembly protein \| Baseplate assembly protein; afdb-uniprot50 \| AF-A0A5C7PFA8-F1-MODEL\_V4 \| 1.0 \| 3.619e-09 \| 329 \| 0.363 \| 121 \| 64 \| 3 \| 1 \| 108 \| 6 \| 126 \| Baseplate assembly protein \| Baseplate assembly protein; afdb-uniprot50 \| AF-A0A7W2BN75-F1-MODEL\_V4 \| 1.0 \| 1.172e-08 \| 329 \| 0.321 \| 112 \| 69 \| 2 \| 1 \| 105 \| 2 \| 113 \| Baseplate assembly protein \| Baseplate assembly protein; afdb-uniprot50 \| AF-A0A6N7C1I5-F1-MODEL\_V4 \| 1.0 \| 1.596e-08 \| 328 \| 0.44 \| 100 \| 53 \| 1 \| 1 \| 97 \| 13 \| 112 \| Phage baseplate assembly protein \| Phage baseplate assembly protein; afdb-uniprot50 \| AF-A0A1E3G6B2-F1-MODEL\_V4 \| 1.0 \| 2.044e-08 \| 327 \| 0.362 \| 113 \| 64 \| 1 \| 1 \| 105 \| 8 \| 120 \| Uncharacterized protein \| Uncharacterized protein; afdb-uniprot50 \| AF-A0A6N8TE09-F1-MODEL\_V4 \| 1.0 \| 9.149e-09 \| 327 \| 0.313 \| 115 \| 72 \| 2 \| 1 \| 108 \| 8 \| 122 \| Integrase \| Integrase; afdb-uniprot50 \| AF-A0A4V1LS71-F1-MODEL\_V4 \| 1.0 \| 2.461e-08 \| 326 \| 0.449 \| 89 \| 48 \| 1 \| 17 \| 105 \| 14 \| 101 \| GPW\_gp25 domain-containing protein \| GPW\_gp25 domain-containing protein; afdb-uniprot50 \| AF-A0A6A4R9J3-F1-MODEL\_V4 \| 1.0 \| 2.175e-08 \| 324 \| 0.256 \| 117 \| 74 \| 5 \| 1 \| 112 \| 1 \| 109 \| Phage baseplate protein \| Phage baseplate protein; afdb-uniprot50 \| AF-A0A258KUE2-F1-MODEL\_V4 \| 1.0 \| 8.085e-09 \| 323 \| 0.37 \| 108 \| 62 \| 2 \| 2 \| 103 \| 10 \| 117 \| GPW\_gp25 domain-containing protein \| GPW\_gp25 domain-containing protein; afdb-uniprot50 \| AF-A0A484YKT9-F1-MODEL\_V4 \| 1.0 \| 3.151e-08 \| 322 \| 0.416 \| 84 \| 48 \| 1 \| 23 \| 105 \| 2 \| 85 \| Baseplate Assembly protein W \| Baseplate Assembly protein W; afdb-uniprot50 \| AF-A0A379CBH3-F1-MODEL\_V4 \| 1.0 \| 3.352e-08 \| 320 \| 0.411 \| 107 \| 57 \| 3 \| 4 \| 105 \| 1 \| 106 \| Gene 25-like lysozyme \| Gene 25-like lysozyme; afdb-uniprot50 \| AF-B7KB51-F1-MODEL\_V4 \| 1.0 \| 9.733e-09 \| 320 \| 0.339 \| 109 \| 68 \| 3 \| 2 \| 107 \| 19 \| 126 \| GPW/gp25 family protein \| GPW/gp25 family protein; afdb-uniprot50 \| AF-A0A1M3AHU0-F1-MODEL\_V4 \| 1.0 \| 1.596e-08 \| 320 \| 0.351 \| 108 \| 64 \| 2 \| 2 \| 103 \| 7 \| 114 \| GPW\_gp25 domain-containing protein \| GPW\_gp25 domain-containing protein; afdb-uniprot50 \| AF-A0A7W6N996-F1-MODEL\_V4 \| 1.0 \| 5.168e-08 \| 319 \| 0.373 \| 123 \| 61 \| 3 \| 1 \| 107 \| 1 \| 123 \| GPW\_gp25 domain-containing protein \| GPW\_gp25 domain-containing protein; afdb-uniprot50 \| AF-A0A2H0PCR0-F1-MODEL\_V4 \| 1.0 \| 8.085e-09 \| 319 \| 0.296 \| 108 \| 71 \| 3 \| 2 \| 105 \| 13 \| 119 \| Baseplate protein \| Baseplate protein; afdb-uniprot50 \| AF-A0A3G4W5G7-F1-MODEL\_V4 \| 1.0 \| 2.044e-08 \| 318 \| 0.327 \| 107 \| 71 \| 1 \| 2 \| 107 \| 14 \| 120 \| Gene 25-like lysozyme \| Gene 25-like lysozyme; afdb-uniprot50 \| AF-A0A4Q8MC05-F1-MODEL\_V4 \| 1.0 \| 1.246e-08 \| 318 \| 0.361 \| 119 \| 62 \| 3 \| 1 \| 105 \| 3 \| 121 \| Uncharacterized protein \| Uncharacterized protein; afdb-uniprot50 \| AF-A0A376ZFF8-F1-MODEL\_V4 \| 1.0 \| 1.155e-07 \| 316 \| 0.5 \| 78 \| 39 \| 0 \| 1 \| 78 \| 5 \| 82 \| Baseplate assembly protein W \| Baseplate assembly protein W; afdb-uniprot50 \| AF-A0A0R3KWV8-F1-MODEL\_V4 \| 1.0 \| 3.352e-08 \| 316 \| 0.33 \| 115 \| 71 \| 2 \| 1 \| 109 \| 1 \| 115 \| GPW\_gp25 domain-containing protein \| GPW\_gp25 domain-containing protein; afdb-uniprot50 \| AF-A0A836PEE7-F1-MODEL\_V4 \| 1.0 \| 7.49e-08 \| 315 \| 0.283 \| 113 \| 76 \| 2 \| 1 \| 109 \| 2 \| 113 \| dTDP-glucose pyrophosphorylase \| dTDP-glucose pyrophosphorylase; afdb-uniprot50 \| AF-A0A2R4K2P8-F1-MODEL\_V4 \| 1.0 \| 1.806e-08 \| 315 \| 0.33 \| 106 \| 69 \| 2 \| 2 \| 105 \| 15 \| 120 \| Phage baseplate assembly protein W \| Phage baseplate assembly protein W; afdb-uniprot50 \| AF-A0A367QB00-F1-MODEL\_V4 \| 1.0 \| 2.461e-08 \| 315 \| 0.348 \| 109 \| 67 \| 3 \| 2 \| 107 \| 18 \| 125 \| Baseplate protein \| Baseplate protein; afdb-uniprot50 \| AF-A0A4Q6D2R8-F1-MODEL\_V4 \| 1.0 \| 3.794e-08 \| 313 \| 0.259 \| 108 \| 77 \| 2 \| 1 \| 108 \| 1 \| 105 \| Uncharacterized protein \| Uncharacterized protein; afdb-uniprot50 \| AF-A0A327JMQ1-F1-MODEL\_V4 \| 1.0 \| 9.592e-08 \| 312 \| 0.341 \| 123 \| 70 \| 5 \| 1 \| 112 \| 3 \| 125 \| Baseplate assembly protein \| Baseplate assembly protein; afdb-uniprot50 \| AF-A0A1Z4UHS5-F1-MODEL\_V4 \| 1.0 \| 2.785e-08 \| 312 \| 0.348 \| 109 \| 67 \| 3 \| 2 \| 107 \| 20 \| 127 \| GPW/gp25 family protein \| GPW/gp25 family protein; afdb-uniprot50 \| AF-A0A2C9D702-F1-MODEL\_V4 \| 1.0 \| 5.498e-08 \| 312 \| 0.268 \| 123 \| 72 \| 2 \| 1 \| 105 \| 1 \| 123 \| Baseplate wedge subunit \| Baseplate wedge subunit; afdb-uniprot50 \| AF-A0A3B8IJI9-F1-MODEL\_V4 \| 1.0 \| 1.035e-08 \| 312 \| 0.247 \| 109 \| 78 \| 2 \| 2 \| 106 \| 15 \| 123 \| GPW\_gp25 domain-containing protein \| GPW\_gp25 domain-containing protein; afdb-uniprot50 \| AF-N2IFU2-F1-MODEL\_V4 \| 1.0 \| 6.222e-08 \| 310 \| 0.36 \| 86 \| 54 \| 1 \| 21 \| 105 \| 1 \| 86 \| GPW\_gp25 domain-containing protein \| GPW\_gp25 domain-containing protein; afdb-uniprot50 \| AF-A0A7Y3X776-F1-MODEL\_V4 \| 1.0 \| 2.785e-08 \| 310 \| 0.348 \| 109 \| 67 \| 3 \| 2 \| 107 \| 13 \| 120 \| GPW/gp25 family protein \| GPW/gp25 family protein; afdb-uniprot50 \| AF-E2CI67-F1-MODEL\_V4 \| 1.0 \| 7.041e-08 \| 310 \| 0.271 \| 114 \| 74 \| 2 \| 1 \| 105 \| 8 \| 121 \| Putative phage gp25 protein \| Putative phage gp25 protein; afdb-uniprot50 \| AF-A0A845ZX24-F1-MODEL\_V4 \| 1.0 \| 3.352e-08 \| 309 \| 0.345 \| 110 \| 66 \| 4 \| 2 \| 107 \| 21 \| 128 \| GPW/gp25 family protein \| GPW/gp25 family protein; afdb-uniprot50 \| AF-W5WMR1-F1-MODEL\_V4 \| 1.0 \| 3.151e-08 \| 308 \| 0.357 \| 112 \| 64 \| 3 \| 2 \| 105 \| 9 \| 120 \| GPW\_gp25 domain-containing protein \| GPW\_gp25 domain-containing protein; afdb-uniprot50 \| AF-Q7NKR4-F1-MODEL\_V4 \| 1.0 \| 3.794e-08 \| 308 \| 0.357 \| 109 \| 66 \| 3 \| 2 \| 107 \| 24 \| 131 \| Gll1413 protein \| Gll1413 protein; afdb-uniprot50 \| AF-A0A7X6FTX0-F1-MODEL\_V4 \| 1.0 \| 1.894e-07 \| 307 \| 0.33 \| 100 \| 66 \| 1 \| 1 \| 99 \| 8 \| 107 \| GPW\_gp25 domain-containing protein \| GPW\_gp25 domain-containing protein; afdb-uniprot50 \| AF-A0A521CK88-F1-MODEL\_V4 \| 1.0 \| 1.228e-07 \| 307 \| 0.201 \| 114 \| 84 \| 1 \| 1 \| 107 \| 4 \| 117 \| Phage baseplate assembly protein W \| Phage baseplate assembly protein W; afdb-uniprot50 \| AF-A0A6I5NLR7-F1-MODEL\_V4 \| 1.0 \| 2.313e-08 \| 307 \| 0.342 \| 108 \| 67 \| 3 \| 2 \| 106 \| 22 \| 128 \| GPW/gp25 family protein \| GPW/gp25 family protein; afdb-uniprot50 \| AF-A0A0Q8ANH2-F1-MODEL\_V4 \| 1.0 \| 2.175e-08 \| 306 \| 0.318 \| 113 \| 70 \| 2 \| 2 \| 107 \| 5 \| 117 \| Uncharacterized protein \| Uncharacterized protein; afdb-uniprot50 \| AF-A0A7C5ASQ0-F1-MODEL\_V4 \| 1.0 \| 2.785e-08 \| 306 \| 0.327 \| 107 \| 69 \| 2 \| 2 \| 105 \| 14 \| 120 \| Phage baseplate protein \| Phage baseplate protein; afdb-uniprot50 \| AF-A0A3B8JYJ5-F1-MODEL\_V4 \| 1.0 \| 3.352e-08 \| 306 \| 0.345 \| 110 \| 66 \| 4 \| 2 \| 107 \| 21 \| 128 \| Baseplate protein \| Baseplate protein; afdb-uniprot50 \| AF-A0A5S9QTJ7-F1-MODEL\_V4 \| 1.0 \| 5.498e-08 \| 305 \| 0.236 \| 114 \| 79 \| 4 \| 1 \| 111 \| 2 \| 110 \| Uncharacterized protein \| Uncharacterized protein; afdb-uniprot50 \| AF-A0A1C3F4Y7-F1-MODEL\_V4 \| 1.0 \| 3.352e-08 \| 305 \| 0.268 \| 108 \| 75 \| 2 \| 2 \| 105 \| 14 \| 121 \| GPW/gp25 family protein \| GPW/gp25 family protein; afdb-uniprot50 \| AF-A0A2E2NRP7-F1-MODEL\_V4 \| 1.0 \| 4.567e-08 \| 305 \| 0.273 \| 106 \| 75 \| 1 \| 2 \| 105 \| 16 \| 121 \| Baseplate protein \| Baseplate protein; afdb-uniprot50 \| AF-A0A7X6CDB5-F1-MODEL\_V4 \| 1.0 \| 4.567e-08 \| 305 \| 0.357 \| 109 \| 66 \| 3 \| 2 \| 107 \| 22 \| 129 \| GPW/gp25 family protein \| GPW/gp25 family protein; afdb-uniprot50 \| AF-A0A369RGC9-F1-MODEL\_V4 \| 1.0 \| 1.307e-07 \| 304 \| 0.519 \| 77 \| 37 \| 0 \| 35 \| 111 \| 1 \| 77 \| Baseplate assembly protein W \| Baseplate assembly protein W; afdb-uniprot50 \| AF-A0A5M8P8S7-F1-MODEL\_V4 \| 1.0 \| 1.228e-07 \| 304 \| 0.299 \| 117 \| 70 \| 3 \| 1 \| 105 \| 2 \| 118 \| Baseplate assembly protein \| Baseplate assembly protein; afdb-uniprot50 \| AF-A0A2T4UGD6-F1-MODEL\_V4 \| 1.0 \| 4.036e-08 \| 304 \| 0.289 \| 107 \| 73 \| 2 \| 2 \| 105 \| 13 \| 119 \| Baseplate protein \| Baseplate protein; afdb-uniprot50 \| AF-A0A533SQR7-F1-MODEL\_V4 \| 1.0 \| 1.02e-07 \| 304 \| 0.278 \| 115 \| 74 \| 3 \| 1 \| 106 \| 11 \| 125 \| GPW/gp25 family protein \| GPW/gp25 family protein; afdb-uniprot50 \| AF-A0A6I1JFW3-F1-MODEL\_V4 \| 1.0 \| 3.794e-08 \| 304 \| 0.315 \| 114 \| 68 \| 2 \| 1 \| 104 \| 7 \| 120 \| GPW\_gp25 domain-containing protein \| GPW\_gp25 domain-containing protein; afdb-uniprot50 \| AF-A0A836SDZ9-F1-MODEL\_V4 \| 1.0 \| 3.566e-08 \| 304 \| 0.3 \| 103 \| 69 \| 2 \| 5 \| 105 \| 30 \| 131 \| Baseplate protein \| Baseplate protein; afdb-uniprot50 \| AF-A0A4R5ICH3-F1-MODEL\_V4 \| 1.0 \| 2.313e-08 \| 304 \| 0.287 \| 108 \| 72 \| 3 \| 2 \| 105 \| 14 \| 120 \| Phage baseplate protein \| Phage baseplate protein; afdb-uniprot50 \| AF-A0A6G3ZD83-F1-MODEL\_V4 \| 1.0 \| 4.036e-08 \| 304 \| 0.317 \| 107 \| 69 \| 3 \| 2 \| 105 \| 22 \| 127 \| GPW/gp25 family protein \| GPW/gp25 family protein; afdb-uniprot50 \| AF-A0A4C2ENV8-F1-MODEL\_V4 \| 1.0 \| 5.849e-08 \| 304 \| 0.327 \| 107 \| 69 \| 2 \| 2 \| 105 \| 46 \| 152 \| Baseplate protein \| Baseplate protein; afdb-uniprot50 \| AF-A0A2U0SVH0-F1-MODEL\_V4 \| 1.0 \| 4.501e-07 \| 303 \| 0.605 \| 71 \| 28 \| 0 \| 1 \| 71 \| 1 \| 71 \| Gene 25-like lysozyme \| Gene 25-like lysozyme; afdb-uniprot50 \| AF-A0A1A9AYR2-F1-MODEL\_V4 \| 1.0 \| 1.573e-07 \| 303 \| 0.245 \| 114 \| 77 \| 4 \| 1 \| 111 \| 1 \| 108 \| Gene 25-like lysozyme \| Gene 25-like lysozyme; afdb-uniprot50 \| AF-H8GWR0-F1-MODEL\_V4 \| 1.0 \| 1.02e-07 \| 303 \| 0.354 \| 96 \| 60 \| 1 \| 12 \| 105 \| 2 \| 97 \| GPW/gp25 family protein \| GPW/gp25 family protein; afdb-uniprot50 \| AF-A0A090SWE0-F1-MODEL\_V4 \| 1.0 \| 1.228e-07 \| 303 \| 0.229 \| 109 \| 80 \| 2 \| 1 \| 106 \| 4 \| 111 \| Uncharacterized protein \| Uncharacterized protein; afdb-uniprot50 \| AF-E2CHJ3-F1-MODEL\_V4 \| 1.0 \| 2.785e-08 \| 303 \| 0.376 \| 117 \| 60 \| 3 \| 1 \| 104 \| 3 \| 119 \| GPW/gp25 \| GPW/gp25; afdb-uniprot50 \| AF-A0A485AJM6-F1-MODEL\_V4 \| 1.0 \| 3.977e-07 \| 302 \| 0.445 \| 83 \| 46 \| 0 \| 1 \| 83 \| 5 \| 87 \| Gene 25-like lysozyme \| Gene 25-like lysozyme; afdb-uniprot50 \| AF-A0A327JS28-F1-MODEL\_V4 \| 1.0 \| 1.307e-07 \| 302 \| 0.252 \| 119 \| 77 \| 4 \| 1 \| 108 \| 20 \| 137 \| Uncharacterized protein \| Uncharacterized protein; afdb-uniprot50 \| AF-A0A484HJL0-F1-MODEL\_V4 \| 1.0 \| 2.313e-08 \| 302 \| 0.272 \| 110 \| 76 \| 2 \| 2 \| 107 \| 37 \| 146 \| Putative phage baseplate protein \| Putative phage baseplate protein; afdb-uniprot50 \| AF-A0A286EJF8-F1-MODEL\_V4 \| 1.0 \| 5.849e-08 \| 301 \| 0.283 \| 113 \| 72 \| 3 \| 2 \| 105 \| 22 \| 134 \| GPW\_gp25 domain-containing protein \| GPW\_gp25 domain-containing protein; afdb-uniprot50 \| AF-A0A0U3ESH1-F1-MODEL\_V4 \| 1.0 \| 4.036e-08 \| 300 \| 0.229 \| 109 \| 81 \| 2 \| 2 \| 107 \| 12 \| 120 \| GPW\_gp25 domain-containing protein \| GPW\_gp25 domain-containing protein; afdb-uniprot50 \| AF-A0A0N1NB24-F1-MODEL\_V4 \| 1.0 \| 3.794e-08 \| 300 \| 0.345 \| 113 \| 66 \| 3 \| 1 \| 105 \| 28 \| 140 \| GPW/gp25 family protein \| GPW/gp25 family protein; afdb-uniprot50 \| AF-A0A5P8MV09-F1-MODEL\_V4 \| 1.0 \| 9.017e-08 \| 299 \| 0.306 \| 124 \| 73 \| 3 \| 1 \| 111 \| 6 \| 129 \| Integrase \| Integrase; afdb-uniprot50 \| AF-A0A521DJN8-F1-MODEL\_V4 \| 1.0 \| 5.849e-08 \| 299 \| 0.254 \| 106 \| 76 \| 2 \| 2 \| 105 \| 23 \| 127 \| GPW\_gp25 domain-containing protein \| GPW\_gp25 domain-containing protein; afdb-uniprot50 \| AF-A0A6P0YQM2-F1-MODEL\_V4 \| 1.0 \| 3.794e-08 \| 299 \| 0.229 \| 109 \| 79 \| 3 \| 2 \| 106 \| 21 \| 128 \| GPW\_gp25 domain-containing protein \| GPW\_gp25 domain-containing protein; afdb-uniprot50 \| AF-B9NUV2-F1-MODEL\_V4 \| 1.0 \| 7.041e-08 \| 299 \| 0.259 \| 108 \| 76 \| 2 \| 2 \| 105 \| 25 \| 132 \| Phage baseplate assembly protein W \| Phage baseplate assembly protein W; afdb-uniprot50 \| AF-A0A528VJI5-F1-MODEL\_V4 \| 1.0 \| 1.78e-07 \| 298 \| 0.343 \| 102 \| 66 \| 1 \| 1 \| 101 \| 1 \| 102 \| GPW\_gp25 domain-containing protein \| GPW\_gp25 domain-containing protein; afdb-uniprot50 \| AF-A0A1H2U8Y5-F1-MODEL\_V4 \| 1.0 \| 1.228e-07 \| 298 \| 0.305 \| 108 \| 72 \| 2 \| 2 \| 107 \| 14 \| 120 \| GPW\_gp25 domain-containing protein \| GPW\_gp25 domain-containing protein; afdb-uniprot50 \| AF-A0A428YAU4-F1-MODEL\_V4 \| 1.0 \| 3.352e-08 \| 297 \| 0.311 \| 106 \| 70 \| 2 \| 2 \| 105 \| 14 \| 118 \| Phage baseplate protein \| Phage baseplate protein; afdb-uniprot50 \| AF-A0A4R7QFY3-F1-MODEL\_V4 \| 1.0 \| 4.293e-08 \| 297 \| 0.259 \| 108 \| 77 \| 2 \| 2 \| 106 \| 21 \| 128 \| GPW\_gp25 domain-containing protein \| GPW\_gp25 domain-containing protein; afdb-uniprot50 \| AF-A0A2D9B6I8-F1-MODEL\_V4 \| 1.0 \| 1.228e-07 \| 296 \| 0.245 \| 114 \| 79 \| 2 \| 1 \| 107 \| 8 \| 121 \| GPW\_gp25 domain-containing protein \| GPW\_gp25 domain-containing protein; afdb-uniprot50 \| AF-A0A2G9WV35-F1-MODEL\_V4 \| 1.0 \| 8.476e-08 \| 296 \| 0.31 \| 116 \| 71 \| 3 \| 1 \| 108 \| 7 \| 121 \| Integrase \| Integrase; afdb-uniprot50 \| AF-A0A428Z2I7-F1-MODEL\_V4 \| 1.0 \| 1.02e-07 \| 296 \| 0.268 \| 108 \| 77 \| 2 \| 2 \| 107 \| 18 \| 125 \| Baseplate protein \| Baseplate protein; afdb-uniprot50 \| AF-A0A442I641-F1-MODEL\_V4 \| 1.0 \| 6.619e-08 \| 296 \| 0.305 \| 118 \| 73 \| 3 \| 1 \| 109 \| 27 \| 144 \| GPW\_gp25 domain-containing protein \| GPW\_gp25 domain-containing protein; afdb-uniprot50 \| AF-A0A2A2HU34-F1-MODEL\_V4 \| 1.0 \| 9.592e-08 \| 295 \| 0.293 \| 109 \| 73 \| 3 \| 2 \| 107 \| 14 \| 121 \| Baseplate protein \| Baseplate protein; afdb-uniprot50 \| AF-A0A1K2HLI9-F1-MODEL\_V4 \| 1.0 \| 1.155e-07 \| 295 \| 0.292 \| 113 \| 72 \| 3 \| 1 \| 105 \| 9 \| 121 \| GPW\_gp25 domain-containing protein \| GPW\_gp25 domain-containing protein; afdb-uniprot50 \| AF-M0DEF1-F1-MODEL\_V4 \| 1.0 \| 1.155e-07 \| 295 \| 0.299 \| 107 \| 72 \| 2 \| 2 \| 105 \| 14 \| 120 \| GPW/gp25 family protein \| GPW/gp25 family protein; afdb-uniprot50 \| AF-A0A6G3R8A8-F1-MODEL\_V4 \| 1.0 \| 7.968e-08 \| 295 \| 0.289 \| 107 \| 73 \| 2 \| 2 \| 105 \| 14 \| 120 \| GPW/gp25 family protein \| GPW/gp25 family protein; afdb-uniprot50 \| AF-A0A7H9VBC7-F1-MODEL\_V4 \| 1.0 \| 1.573e-07 \| 295 \| 0.267 \| 112 \| 75 \| 3 \| 1 \| 105 \| 35 \| 146 \| Baseplate assembly protein \| Baseplate assembly protein; afdb-uniprot50 \| AF-A0A3N5Q1L3-F1-MODEL\_V4 \| 1.0 \| 3.977e-07 \| 294 \| 0.315 \| 95 \| 63 \| 1 \| 14 \| 106 \| 1 \| 95 \| Phage baseplate protein \| Phage baseplate protein; afdb-uniprot50 \| AF-R7JS48-F1-MODEL\_V4 \| 1.0 \| 4.788e-07 \| 294 \| 0.271 \| 107 \| 76 \| 2 \| 1 \| 106 \| 9 \| 114 \| Baseplate \| Baseplate; afdb-uniprot50 \| AF-A0A661RQ76-F1-MODEL\_V4 \| 1.0 \| 9.017e-08 \| 294 \| 0.305 \| 108 \| 70 \| 3 \| 2 \| 105 \| 14 \| 120 \| Baseplate protein \| Baseplate protein; afdb-uniprot50 \| AF-A0A3N0FWT9-F1-MODEL\_V4 \| 1.0 \| 4.036e-08 \| 294 \| 0.25 \| 108 \| 77 \| 2 \| 2 \| 105 \| 16 \| 123 \| GPW\_gp25 domain-containing protein \| GPW\_gp25 domain-containing protein; afdb-uniprot50 \| AF-A0A840YD88-F1-MODEL\_V4 \| 1.0 \| 1.228e-07 \| 293 \| 0.291 \| 96 \| 66 \| 1 \| 12 \| 105 \| 2 \| 97 \| Uncharacterized protein \| Uncharacterized protein; afdb-uniprot50 \| AF-A0A084T291-F1-MODEL\_V4 \| 1.0 \| 7.968e-08 \| 293 \| 0.293 \| 109 \| 73 \| 2 \| 2 \| 106 \| 14 \| 122 \| Baseplate protein \| Baseplate protein; afdb-uniprot50 \| AF-A0A1W2F8A0-F1-MODEL\_V4 \| 1.0 \| 1.479e-07 \| 293 \| 0.333 \| 105 \| 66 \| 1 \| 1 \| 101 \| 1 \| 105 \| GPW\_gp25 domain-containing protein \| GPW\_gp25 domain-containing protein; afdb-uniprot50 \| AF-A0A1P8FGT4-F1-MODEL\_V4 \| 1.0 \| 9.592e-08 \| 293 \| 0.307 \| 114 \| 70 \| 3 \| 1 \| 105 \| 11 \| 124 \| Baseplate assembly protein \| Baseplate assembly protein; afdb-uniprot50 \| AF-A0A4S2RPJ9-F1-MODEL\_V4 \| 1.0 \| 1.894e-07 \| 293 \| 0.309 \| 113 \| 70 \| 3 \| 1 \| 105 \| 7 \| 119 \| Baseplate protein \| Baseplate protein; afdb-uniprot50 \| AF-A0A561IRH5-F1-MODEL\_V4 \| 1.0 \| 1.673e-07 \| 293 \| 0.283 \| 106 \| 74 \| 2 \| 1 \| 105 \| 11 \| 115 \| Gene 25-like lysozyme \| Gene 25-like lysozyme; afdb-uniprot50 \| AF-A0A2P2GC78-F1-MODEL\_V4 \| 1.0 \| 1.085e-07 \| 292 \| 0.289 \| 107 \| 73 \| 2 \| 2 \| 105 \| 14 \| 120 \| Baseplate protein \| Baseplate protein; afdb-uniprot50 \| AF-A0A016QL71-F1-MODEL\_V4 \| 1.0 \| 6.222e-08 \| 292 \| 0.379 \| 108 \| 62 \| 3 \| 2 \| 105 \| 13 \| 119 \| GPW/gp25 family protein \| GPW/gp25 family protein; afdb-uniprot50 \| AF-A0A3N5W7B2-F1-MODEL\_V4 \| 1.0 \| 4.858e-08 \| 292 \| 0.333 \| 108 \| 67 \| 3 \| 2 \| 105 \| 22 \| 128 \| Baseplate protein \| Baseplate protein; afdb-uniprot50 \| AF-A0A2G6EIV6-F1-MODEL\_V4 \| 1.0 \| 3.977e-07 \| 292 \| 0.431 \| 95 \| 54 \| 0 \| 3 \| 97 \| 12 \| 106 \| GPW\_gp25 domain-containing protein \| GPW\_gp25 domain-containing protein; afdb-uniprot50 \| AF-A0A5P9CRI3-F1-MODEL\_V4 \| 1.0 \| 4.501e-07 \| 291 \| 0.254 \| 110 \| 76 \| 3 \| 1 \| 107 \| 1 \| 107 \| Uncharacterized protein \| Uncharacterized protein; afdb-uniprot50 \| AF-A0A7Y2SMU2-F1-MODEL\_V4 \| 1.0 \| 1.39e-07 \| 291 \| 0.261 \| 111 \| 75 \| 4 \| 2 \| 107 \| 13 \| 121 \| GPW/gp25 family protein \| GPW/gp25 family protein; afdb-uniprot50 \| AF-A0A7X5QSP2-F1-MODEL\_V4 \| 1.0 \| 7.041e-08 \| 291 \| 0.203 \| 108 \| 82 \| 2 \| 2 \| 105 \| 13 \| 120 \| GPW/gp25 family protein \| GPW/gp25 family protein; afdb-uniprot50 \| AF-A0A6C7EF79-F1-MODEL\_V4 \| 1.0 \| 1.02e-07 \| 291 \| 0.296 \| 108 \| 71 \| 3 \| 2 \| 105 \| 20 \| 126 \| GPW\_gp25 domain-containing protein \| GPW\_gp25 domain-containing protein; afdb-uniprot50 \| AF-A0A833LM40-F1-MODEL\_V4 \| 1.0 \| 1.894e-07 \| 290 \| 0.312 \| 112 \| 70 \| 3 \| 1 \| 105 \| 8 \| 119 \| Baseplate assembly protein \| Baseplate assembly protein; afdb-uniprot50 \| AF-A0A150P6E2-F1-MODEL\_V4 \| 1.0 \| 1.39e-07 \| 290 \| 0.28 \| 107 \| 74 \| 2 \| 2 \| 105 \| 17 \| 123 \| GPW\_gp25 domain-containing protein \| GPW\_gp25 domain-containing protein; afdb-uniprot50 \| AF-A0A1T3NP84-F1-MODEL\_V4 \| 1.0 \| 1.894e-07 \| 290 \| 0.298 \| 114 \| 71 \| 3 \| 1 \| 105 \| 17 \| 130 \| GPW\_gp25 domain-containing protein \| GPW\_gp25 domain-containing protein; afdb-uniprot50 \| AF-A0A842NAI3-F1-MODEL\_V4 \| 1.0 \| 5.418e-07 \| 289 \| 0.264 \| 102 \| 73 \| 2 \| 12 \| 112 \| 4 \| 104 \| GPW/gp25 family protein \| GPW/gp25 family protein; afdb-uniprot50 \| AF-A0A212S7U4-F1-MODEL\_V4 \| 1.0 \| 1.479e-07 \| 289 \| 0.325 \| 123 \| 69 \| 4 \| 1 \| 110 \| 2 \| 123 \| GPW\_gp25 domain-containing protein \| GPW\_gp25 domain-containing protein; afdb-uniprot50 \| AF-A0A7V9GIA2-F1-MODEL\_V4 \| 1.0 \| 9.592e-08 \| 289 \| 0.297 \| 111 \| 73 \| 3 \| 2 \| 108 \| 14 \| 123 \| GPW/gp25 family protein \| GPW/gp25 family protein; afdb-uniprot50 \| AF-A0A4U7JDR4-F1-MODEL\_V4 \| 1.0 \| 1.78e-07 \| 289 \| 0.311 \| 106 \| 70 \| 2 \| 2 \| 105 \| 15 \| 119 \| GPW/gp25 family protein \| GPW/gp25 family protein; afdb-uniprot50 \| AF-A0A4P8HN78-F1-MODEL\_V4 \| 1.0 \| 1.228e-07 \| 289 \| 0.272 \| 110 \| 75 \| 3 \| 2 \| 107 \| 16 \| 124 \| Uncharacterized protein \| Uncharacterized protein; afdb-uniprot50 \| AF-A0A0N0JDW9-F1-MODEL\_V4 \| 1.0 \| 1.307e-07 \| 289 \| 0.283 \| 113 \| 73 \| 3 \| 2 \| 106 \| 14 \| 126 \| GPW\_gp25 domain-containing protein \| GPW\_gp25 domain-containing protein; afdb-uniprot50 \| AF-A0A0R3MPT4-F1-MODEL\_V4 \| 1.0 \| 5.498e-08 \| 289 \| 0.35 \| 120 \| 63 \| 4 \| 1 \| 105 \| 1 \| 120 \| Phage baseplate protein \| Phage baseplate protein; afdb-uniprot50 \| AF-B8IDQ3-F1-MODEL\_V4 \| 1.0 \| 9.592e-08 \| 289 \| 0.28 \| 114 \| 72 \| 2 \| 1 \| 104 \| 3 \| 116 \| GPW/gp25 family protein \| GPW/gp25 family protein; afdb-uniprot50 \| AF-A0A2Z3Z422-F1-MODEL\_V4 \| 1.0 \| 3.106e-07 \| 288 \| 0.354 \| 96 \| 60 \| 1 \| 12 \| 105 \| 2 \| 97 \| Phage baseplate protein \| Phage baseplate protein; afdb-uniprot50 \| AF-A0A3S0WS72-F1-MODEL\_V4 \| 1.0 \| 2.015e-07 \| 288 \| 0.22 \| 109 \| 79 \| 3 \| 1 \| 106 \| 1 \| 106 \| Uncharacterized protein \| Uncharacterized protein; afdb-uniprot50 \| AF-A0A371K5T5-F1-MODEL\_V4 \| 1.0 \| 1.155e-07 \| 288 \| 0.259 \| 104 \| 76 \| 1 \| 2 \| 105 \| 11 \| 113 \| Baseplate assembly protein \| Baseplate assembly protein; afdb-uniprot50 \| AF-A0A6N9YFS0-F1-MODEL\_V4 \| 1.0 \| 1.307e-07 \| 288 \| 0.292 \| 106 \| 72 \| 2 \| 3 \| 105 \| 2 \| 107 \| GPW/gp25 family protein \| GPW/gp25 family protein; afdb-uniprot50 \| AF-A0A1I6IQR2-F1-MODEL\_V4 \| 1.0 \| 1.307e-07 \| 288 \| 0.308 \| 107 \| 71 \| 2 \| 2 \| 105 \| 14 \| 120 \| GPW\_gp25 domain-containing protein \| GPW\_gp25 domain-containing protein; afdb-uniprot50 \| AF-A0A1F3S636-F1-MODEL\_V4 \| 1.0 \| 7.49e-08 \| 288 \| 0.259 \| 108 \| 76 \| 2 \| 2 \| 105 \| 15 \| 122 \| GPW\_gp25 domain-containing protein \| GPW\_gp25 domain-containing protein; afdb-uniprot50 \| AF-A0A4Y9PT98-F1-MODEL\_V4 \| 1.0 \| 2.015e-07 \| 288 \| 0.289 \| 114 \| 72 \| 3 \| 1 \| 105 \| 7 \| 120 \| Baseplate protein \| Baseplate protein; afdb-uniprot50 \| AF-A0A515KQR3-F1-MODEL\_V4 \| 1.0 \| 7.49e-08 \| 288 \| 0.218 \| 110 \| 82 \| 2 \| 2 \| 107 \| 17 \| 126 \| GPW/gp25 family protein \| GPW/gp25 family protein; afdb-uniprot50 \| AF-A0A2M9J5R0-F1-MODEL\_V4 \| 1.0 \| 1.39e-07 \| 288 \| 0.299 \| 107 \| 72 \| 2 \| 2 \| 105 \| 14 \| 120 \| Baseplate protein \| Baseplate protein; afdb-uniprot50 \| AF-A0A6P0NDR6-F1-MODEL\_V4 \| 1.0 \| 2.744e-07 \| 287 \| 0.276 \| 112 \| 74 \| 2 \| 1 \| 105 \| 7 \| 118 \| GPW/gp25 family protein \| GPW/gp25 family protein; afdb-uniprot50 \| AF-A0A6V8KJ56-F1-MODEL\_V4 \| 1.0 \| 5.498e-08 \| 287 \| 0.297 \| 111 \| 75 \| 2 \| 1 \| 108 \| 9 \| 119 \| GPW\_gp25 domain-containing protein \| GPW\_gp25 domain-containing protein; afdb-uniprot50 \| AF-A0A1H7YK45-F1-MODEL\_V4 \| 1.0 \| 9.017e-08 \| 287 \| 0.203 \| 108 \| 82 \| 2 \| 2 \| 105 \| 17 \| 124 \| GPW\_gp25 domain-containing protein \| GPW\_gp25 domain-containing protein; afdb-uniprot50 \| AF-A0A7T7I7G9-F1-MODEL\_V4 \| 1.0 \| 2.58e-07 \| 287 \| 0.278 \| 115 \| 73 \| 3 \| 1 \| 105 \| 7 \| 121 \| GPW/gp25 family protein \| GPW/gp25 family protein; afdb-uniprot50 \| AF-A0A1Z4QPU1-F1-MODEL\_V4 \| 1.0 \| 5.498e-08 \| 287 \| 0.35 \| 114 \| 64 \| 4 \| 1 \| 105 \| 42 \| 154 \| GPW / gp25 family protein \| GPW / gp25 family protein; afdb-uniprot50 \| AF-A0A3C2A343-F1-MODEL\_V4 \| 1.0 \| 5.849e-08 \| 287 \| 0.222 \| 108 \| 80 \| 2 \| 2 \| 105 \| 58 \| 165 \| GPW\_gp25 domain-containing protein \| GPW\_gp25 domain-containing protein; afdb-uniprot50 \| AF-A0A3M1SPN9-F1-MODEL\_V4 \| 1.0 \| 3.977e-07 \| 286 \| 0.351 \| 91 \| 57 \| 1 \| 17 \| 105 \| 7 \| 97 \| Baseplate protein \| Baseplate protein; afdb-uniprot50 \| AF-A0A1N7S7K8-F1-MODEL\_V4 \| 1.0 \| 1.894e-07 \| 286 \| 0.311 \| 106 \| 71 \| 1 \| 2 \| 105 \| 14 \| 119 \| GPW/gp25 family protein \| GPW/gp25 family protein; afdb-uniprot50 \| AF-A0A1V2QXE0-F1-MODEL\_V4 \| 1.0 \| 1.307e-07 \| 285 \| 0.273 \| 106 \| 74 \| 2 \| 2 \| 105 \| 11 \| 115 \| GPW\_gp25 domain-containing protein \| GPW\_gp25 domain-containing protein; afdb-uniprot50 \| AF-A0A6M3K799-F1-MODEL\_V4 \| 1.0 \| 3.515e-07 \| 285 \| 0.173 \| 115 \| 87 \| 3 \| 1 \| 107 \| 7 \| 121 \| Putative baseplate wedge subunit \| Putative baseplate wedge subunit; afdb-uniprot50 \| AF-E2SQJ7-F1-MODEL\_V4 \| 1.0 \| 7.968e-08 \| 285 \| 0.271 \| 107 \| 73 \| 4 \| 2 \| 105 \| 13 \| 117 \| Putative lysozyme \| Putative lysozyme; afdb-uniprot50 \| AF-A0A3N5XQ15-F1-MODEL\_V4 \| 1.0 \| 7.968e-08 \| 285 \| 0.296 \| 108 \| 71 \| 3 \| 2 \| 105 \| 12 \| 118 \| Baseplate protein \| Baseplate protein; afdb-uniprot50 \| AF-A0A1F8LUG1-F1-MODEL\_V4 \| 1.0 \| 1.78e-07 \| 285 \| 0.261 \| 107 \| 76 \| 2 \| 2 \| 105 \| 18 \| 124 \| Phage baseplate protein \| Phage baseplate protein; afdb-uniprot50 \| AF-A0A371PUU5-F1-MODEL\_V4 \| 1.0 \| 1.307e-07 \| 285 \| 0.296 \| 108 \| 71 \| 3 \| 2 \| 105 \| 37 \| 143 \| Baseplate protein \| Baseplate protein; afdb-uniprot50 \| AF-F5XJ49-F1-MODEL\_V4 \| 1.0 \| 1.39e-07 \| 285 \| 0.333 \| 108 \| 67 \| 3 \| 2 \| 105 \| 15 \| 121 \| GPW\_gp25 domain-containing protein \| GPW\_gp25 domain-containing protein; afdb-uniprot50 \| AF-A0A7S8IJW2-F1-MODEL\_V4 \| 1.0 \| 1.479e-07 \| 284 \| 0.349 \| 123 \| 61 \| 2 \| 4 \| 107 \| 1 \| 123 \| Baseplate assembly protein \| Baseplate assembly protein; afdb-uniprot50 \| AF-A0A6N6KES2-F1-MODEL\_V4 \| 1.0 \| 1.78e-07 \| 284 \| 0.252 \| 107 \| 78 \| 1 \| 2 \| 106 \| 20 \| 126 \| GPW\_gp25 domain-containing protein \| GPW\_gp25 domain-containing protein; afdb-uniprot50 \| AF-A0A1V0Q088-F1-MODEL\_V4 \| 1.0 \| 3.515e-07 \| 284 \| 0.25 \| 116 \| 78 \| 3 \| 1 \| 107 \| 6 \| 121 \| GPW\_gp25 domain-containing protein \| GPW\_gp25 domain-containing protein; afdb-uniprot50 \| AF-A0A6I2SHR9-F1-MODEL\_V4 \| 1.0 \| 1.573e-07 \| 284 \| 0.324 \| 108 \| 68 \| 3 \| 2 \| 105 \| 14 \| 120 \| Baseplate protein \| Baseplate protein; afdb-uniprot50 \| AF-A0A437RHL5-F1-MODEL\_V4 \| 1.0 \| 1.78e-07 \| 284 \| 0.283 \| 113 \| 73 \| 3 \| 2 \| 106 \| 14 \| 126 \| Baseplate assembly protein \| Baseplate assembly protein; afdb-uniprot50 \| AF-A0A356KNI6-F1-MODEL\_V4 \| 1.0 \| 1.673e-07 \| 284 \| 0.308 \| 107 \| 71 \| 2 \| 2 \| 105 \| 29 \| 135 \| Baseplate protein \| Baseplate protein; afdb-uniprot50 \| AF-A0A7K0JP37-F1-MODEL\_V4 \| 1.0 \| 1.228e-07 \| 284 \| 0.318 \| 110 \| 68 \| 4 \| 2 \| 105 \| 28 \| 136 \| Baseplate protein \| Baseplate protein; afdb-uniprot50 \| AF-A0A0U5IN79-F1-MODEL\_V4 \| 1.0 \| 3.566e-08 \| 284 \| 0.269 \| 115 \| 75 \| 4 \| 1 \| 107 \| 33 \| 146 \| GPW/gp25 family protein \| GPW/gp25 family protein; afdb-uniprot50 \| AF-A0A828DTS9-F1-MODEL\_V4 \| 1.0 \| 1.457e-06 \| 283 \| 0.433 \| 83 \| 47 \| 0 \| 1 \| 83 \| 4 \| 86 \| Baseplate assembly protein \| Baseplate assembly protein; afdb-uniprot50 \| AF-E8U3P0-F1-MODEL\_V4 \| 1.0 \| 1.228e-07 \| 283 \| 0.361 \| 108 \| 64 \| 3 \| 2 \| 105 \| 13 \| 119 \| GPW/gp25 family protein \| GPW/gp25 family protein; afdb-uniprot50 \| AF-A0A239FMN3-F1-MODEL\_V4 \| 1.0 \| 1.085e-07 \| 283 \| 0.33 \| 115 \| 67 \| 3 \| 1 \| 105 \| 3 \| 117 \| GPW\_gp25 domain-containing protein \| GPW\_gp25 domain-containing protein; afdb-uniprot50 \| AF-X5L3Y3-F1-MODEL\_V4 \| 1.0 \| 2.58e-07 \| 283 \| 0.263 \| 114 \| 75 \| 3 \| 1 \| 105 \| 7 \| 120 \| GPW/gp25 family protein \| GPW/gp25 family protein; afdb-uniprot50 \| AF-A0A2M8MC75-F1-MODEL\_V4 \| 1.0 \| 1.673e-07 \| 283 \| 0.299 \| 107 \| 72 \| 2 \| 2 \| 105 \| 14 \| 120 \| Baseplate protein \| Baseplate protein; afdb-uniprot50 \| AF-A0A1H6Z5N9-F1-MODEL\_V4 \| 1.0 \| 7.041e-08 \| 283 \| 0.339 \| 109 \| 69 \| 2 \| 1 \| 106 \| 18 \| 126 \| GPW\_gp25 domain-containing protein \| GPW\_gp25 domain-containing protein; afdb-uniprot50 \| AF-A0A2U0SFJ6-F1-MODEL\_V4 \| 1.0 \| 1.155e-07 \| 283 \| 0.242 \| 107 \| 78 \| 2 \| 2 \| 105 \| 36 \| 142 \| GPW\_gp25 domain-containing protein \| GPW\_gp25 domain-containing protein; afdb-uniprot50 \| AF-A0A2M8Q5J2-F1-MODEL\_V4 \| 1.0 \| 1.573e-07 \| 282 \| 0.305 \| 108 \| 70 \| 3 \| 2 \| 105 \| 13 \| 119 \| Baseplate protein \| Baseplate protein; afdb-uniprot50 \| AF-A0A329LW89-F1-MODEL\_V4 \| 1.0 \| 1.39e-07 \| 282 \| 0.297 \| 111 \| 70 \| 3 \| 2 \| 105 \| 14 \| 123 \| Baseplate protein \| Baseplate protein; afdb-uniprot50 \| AF-A0A2V8RLG8-F1-MODEL\_V4 \| 1.0 \| 1.573e-07 \| 282 \| 0.256 \| 117 \| 80 \| 4 \| 2 \| 112 \| 18 \| 133 \| Baseplate protein \| Baseplate protein; afdb-uniprot50 \| AF-A0A561BKI6-F1-MODEL\_V4 \| 1.0 \| 1.479e-07 \| 282 \| 0.345 \| 107 \| 66 \| 2 \| 2 \| 105 \| 13 \| 118 \| GPW\_gp25 domain-containing protein \| GPW\_gp25 domain-containing protein; afdb-uniprot50 \| AF-A0A2P2CID1-F1-MODEL\_V4 \| 1.0 \| 1.573e-07 \| 282 \| 0.287 \| 108 \| 72 \| 3 \| 2 \| 105 \| 23 \| 129 \| Putative GPW / gp25 family \| Putative GPW / gp25 family; afdb-uniprot50 \| AF-A0A1B6AS33-F1-MODEL\_V4 \| 1.0 \| 1.39e-07 \| 282 \| 0.261 \| 107 \| 76 \| 2 \| 2 \| 105 \| 14 \| 120 \| GPW\_gp25 domain-containing protein \| GPW\_gp25 domain-containing protein; afdb-uniprot50 \| AF-A0A7Y4FP63-F1-MODEL\_V4 \| 1.0 \| 9.453e-07 \| 281 \| 0.243 \| 111 \| 78 \| 3 \| 1 \| 108 \| 1 \| 108 \| Phage baseplate protein \| Phage baseplate protein; afdb-uniprot50 \| AF-A0A2S5N393-F1-MODEL\_V4 \| 1.0 \| 8.886e-07 \| 281 \| 0.239 \| 117 \| 80 \| 2 \| 1 \| 108 \| 5 \| 121 \| Integrase \| Integrase; afdb-uniprot50 \| AF-A0A412YNF3-F1-MODEL\_V4 \| 1.0 \| 1.894e-07 \| 281 \| 0.236 \| 110 \| 80 \| 2 \| 2 \| 107 \| 18 \| 127 \| GPW\_gp25 domain-containing protein \| GPW\_gp25 domain-containing protein; afdb-uniprot50 \| AF-A0A497W433-F1-MODEL\_V4 \| 1.0 \| 9.017e-08 \| 281 \| 0.333 \| 108 \| 67 \| 3 \| 2 \| 105 \| 17 \| 123 \| GPW\_gp25 domain-containing protein \| GPW\_gp25 domain-containing protein; afdb-uniprot50 \| AF-A0A1V5CGG0-F1-MODEL\_V4 \| 1.0 \| 1.39e-07 \| 281 \| 0.214 \| 107 \| 81 \| 3 \| 2 \| 105 \| 39 \| 145 \| Gene 25-like lysozyme \| Gene 25-like lysozyme; afdb-uniprot50 \| AF-A0A376CQH6-F1-MODEL\_V4 \| 1.0 \| 1.866e-06 \| 280 \| 0.454 \| 77 \| 42 \| 0 \| 1 \| 77 \| 4 \| 80 \| GPW/gp25 family protein \| GPW/gp25 family protein; afdb-uniprot50 \| AF-A0A3S6F5U6-F1-MODEL\_V4 \| 1.0 \| 9.453e-07 \| 280 \| 0.459 \| 87 \| 47 \| 0 \| 1 \| 87 \| 3 \| 89 \| GPW\_gp25 domain-containing protein \| GPW\_gp25 domain-containing protein; afdb-uniprot50 \| AF-A0A0N8GF33-F1-MODEL\_V4 \| 1.0 \| 1.211e-06 \| 280 \| 0.238 \| 113 \| 77 \| 2 \| 4 \| 107 \| 1 \| 113 \| Uncharacterized protein \| Uncharacterized protein; afdb-uniprot50 \| AF-A0A136LCY3-F1-MODEL\_V4 \| 1.0 \| 2.744e-07 \| 280 \| 0.28 \| 107 \| 73 \| 3 \| 2 \| 105 \| 12 \| 117 \| GPW/gp25 family protein \| GPW/gp25 family protein; afdb-uniprot50 \| AF-A0A2D5YH56-F1-MODEL\_V4 \| 1.0 \| 2.425e-07 \| 280 \| 0.268 \| 108 \| 76 \| 2 \| 3 \| 107 \| 24 \| 131 \| GPW\_gp25 domain-containing protein \| GPW\_gp25 domain-containing protein; afdb-uniprot50 \| AF-A0A2E5AW95-F1-MODEL\_V4 \| 1.0 \| 3.106e-07 \| 280 \| 0.212 \| 108 \| 82 \| 2 \| 3 \| 107 \| 25 \| 132 \| GPW\_gp25 domain-containing protein \| GPW\_gp25 domain-containing protein; afdb-uniprot50 \| AF-A0A2D6MEG6-F1-MODEL\_V4 \| 1.0 \| 3.739e-07 \| 280 \| 0.203 \| 108 \| 83 \| 2 \| 3 \| 107 \| 26 \| 133 \| GPW\_gp25 domain-containing protein \| GPW\_gp25 domain-containing protein; afdb-uniprot50 \| AF-A0A2T0T080-F1-MODEL\_V4 \| 1.0 \| 5.498e-08 \| 279 \| 0.307 \| 114 \| 72 \| 3 \| 2 \| 108 \| 14 \| 127 \| GPW\_gp25 domain-containing protein \| GPW\_gp25 domain-containing protein; afdb-uniprot50 \| AF-A0A1R3VJW4-F1-MODEL\_V4 \| 1.0 \| 2.28e-07 \| 279 \| 0.238 \| 109 \| 81 \| 1 \| 2 \| 108 \| 17 \| 125 \| GPW\_gp25 domain-containing protein \| GPW\_gp25 domain-containing protein; afdb-uniprot50 \| AF-A0A410CPI4-F1-MODEL\_V4 \| 1.0 \| 1.479e-07 \| 279 \| 0.24 \| 108 \| 78 \| 2 \| 2 \| 105 \| 16 \| 123 \| GPW\_gp25 domain-containing protein \| GPW\_gp25 domain-containing protein; afdb-uniprot50 \| AF-A0A542Q250-F1-MODEL\_V4 \| 1.0 \| 3.515e-07 \| 279 \| 0.266 \| 124 \| 79 \| 4 \| 1 \| 112 \| 7 \| 130 \| GPW\_gp25 domain-containing protein \| GPW\_gp25 domain-containing protein; afdb-uniprot50 \| AF-A0A5B8ASK0-F1-MODEL\_V4 \| 1.0 \| 2.015e-07 \| 279 \| 0.298 \| 114 \| 71 \| 3 \| 1 \| 105 \| 19 \| 132 \| GPW/gp25 family protein \| GPW/gp25 family protein; afdb-uniprot50 \| AF-A0A4P8T4C7-F1-MODEL\_V4 \| 1.0 \| 3.106e-07 \| 279 \| 0.254 \| 106 \| 77 \| 2 \| 2 \| 105 \| 14 \| 119 \| Baseplate protein \| Baseplate protein; afdb-uniprot50 \| AF-A0A376KHY4-F1-MODEL\_V4 \| 1.0 \| 1.754e-06 \| 278 \| 0.513 \| 72 \| 35 \| 0 \| 1 \| 72 \| 5 \| 76 \| Baseplate assembly protein W \| Baseplate assembly protein W; afdb-uniprot50 \| AF-A0A1M4ZEM6-F1-MODEL\_V4 \| 1.0 \| 1.155e-07 \| 278 \| 0.229 \| 109 \| 80 \| 2 \| 2 \| 106 \| 16 \| 124 \| GPW\_gp25 domain-containing protein \| GPW\_gp25 domain-containing protein; afdb-uniprot50 \| AF-A0A7W7H4Y6-F1-MODEL\_V4 \| 1.0 \| 2.919e-07 \| 278 \| 0.271 \| 107 \| 75 \| 2 \| 2 \| 105 \| 14 \| 120 \| Phage baseplate assembly protein W \| Phage baseplate assembly protein W; afdb-uniprot50 \| AF-A0A3A0A4P4-F1-MODEL\_V4 \| 1.0 \| 3.106e-07 \| 278 \| 0.194 \| 108 \| 84 \| 2 \| 2 \| 106 \| 16 \| 123 \| GPW\_gp25 domain-containing protein \| GPW\_gp25 domain-containing protein; afdb-uniprot50 \| AF-A0A1C5YWG8-F1-MODEL\_V4 \| 1.0 \| 3.515e-07 \| 278 \| 0.269 \| 115 \| 73 \| 5 \| 1 \| 105 \| 8 \| 121 \| Type VI secretion system lysozyme-like protein \| Type VI secretion system lysozyme-like protein; afdb-uniprot50 \| AF-A0A3D1NSF4-F1-MODEL\_V4 \| 1.0 \| 5.093e-07 \| 277 \| 0.288 \| 104 \| 71 \| 2 \| 3 \| 106 \| 1 \| 101 \| Baseplate protein \| Baseplate protein; afdb-uniprot50 \| AF-A0A1E5G2K2-F1-MODEL\_V4 \| 1.0 \| 1.155e-07 \| 277 \| 0.304 \| 115 \| 70 \| 3 \| 1 \| 105 \| 2 \| 116 \| GPW\_gp25 domain-containing protein \| GPW\_gp25 domain-containing protein; afdb-uniprot50 \| AF-A0A3B8I8Y8-F1-MODEL\_V4 \| 1.0 \| 1.479e-07 \| 277 \| 0.222 \| 108 \| 80 \| 2 \| 2 \| 105 \| 15 \| 122 \| GPW\_gp25 domain-containing protein \| GPW\_gp25 domain-containing protein; afdb-uniprot50 \| AF-A0A1Q9LK30-F1-MODEL\_V4 \| 1.0 \| 2.015e-07 \| 277 \| 0.261 \| 107 \| 76 \| 2 \| 2 \| 105 \| 14 \| 120 \| Baseplate protein \| Baseplate protein; afdb-uniprot50 \| AF-A0A1C5CGJ3-F1-MODEL\_V4 \| 1.0 \| 2.744e-07 \| 277 \| 0.324 \| 108 \| 68 \| 3 \| 2 \| 105 \| 14 \| 120 \| GPW\_gp25 domain-containing protein \| GPW\_gp25 domain-containing protein; afdb-uniprot50 \| AF-A0A7W3I8F7-F1-MODEL\_V4 \| 1.0 \| 2.247e-06 \| 276 \| 0.426 \| 82 \| 47 \| 0 \| 2 \| 83 \| 5 \| 86 \| GPW/gp25 family protein \| GPW/gp25 family protein; afdb-uniprot50 \| AF-A0A7W7Z0T9-F1-MODEL\_V4 \| 1.0 \| 1.78e-07 \| 276 \| 0.304 \| 115 \| 62 \| 4 \| 1 \| 106 \| 1 \| 106 \| GPW\_gp25 domain-containing protein \| GPW\_gp25 domain-containing protein; afdb-uniprot50 \| AF-A0A085AFN6-F1-MODEL\_V4 \| 1.0 \| 6.523e-07 \| 276 \| 0.23 \| 117 \| 77 \| 5 \| 1 \| 112 \| 1 \| 109 \| Phage baseplate assembly protein \| Phage baseplate assembly protein; afdb-uniprot50 \| AF-A0A662AHS8-F1-MODEL\_V4 \| 1.0 \| 2.744e-07 \| 276 \| 0.216 \| 111 \| 81 \| 3 \| 1 \| 106 \| 1 \| 110 \| GPW\_gp25 domain-containing protein \| GPW\_gp25 domain-containing protein; afdb-uniprot50 \| AF-A0A3A4NZZ6-F1-MODEL\_V4 \| 1.0 \| 1.02e-07 \| 276 \| 0.258 \| 112 \| 77 \| 3 \| 2 \| 108 \| 14 \| 124 \| Baseplate protein \| Baseplate protein; afdb-uniprot50 \| AF-A0A2N7W3T6-F1-MODEL\_V4 \| 1.0 \| 5.764e-07 \| 276 \| 0.286 \| 115 \| 73 \| 3 \| 2 \| 107 \| 10 \| 124 \| Baseplate assembly protein \| Baseplate assembly protein; afdb-uniprot50 \| AF-A0A1Q7WS59-F1-MODEL\_V4 \| 1.0 \| 2.28e-07 \| 276 \| 0.261 \| 107 \| 76 \| 2 \| 2 \| 105 \| 14 \| 120 \| Baseplate protein \| Baseplate protein; afdb-uniprot50 \| AF-A0A7X0CIC5-F1-MODEL\_V4 \| 1.0 \| 2.919e-07 \| 276 \| 0.242 \| 107 \| 78 \| 2 \| 2 \| 105 \| 16 \| 122 \| GPW\_gp25 domain-containing protein \| GPW\_gp25 domain-containing protein; afdb-uniprot50 \| AF-A0A2U0TQZ5-F1-MODEL\_V4 \| 1.0 \| 1.479e-07 \| 276 \| 0.24 \| 108 \| 78 \| 2 \| 2 \| 105 \| 15 \| 122 \| GPW\_gp25 domain-containing protein \| GPW\_gp25 domain-containing protein; afdb-uniprot50 \| AF-A0A542J108-F1-MODEL\_V4 \| 1.0 \| 5.418e-07 \| 276 \| 0.289 \| 114 \| 72 \| 3 \| 1 \| 105 \| 7 \| 120 \| GPW\_gp25 domain-containing protein \| GPW\_gp25 domain-containing protein; afdb-uniprot50 \| AF-A0A1J5PQ93-F1-MODEL\_V4 \| 1.0 \| 4.231e-07 \| 276 \| 0.247 \| 109 \| 78 \| 3 \| 1 \| 106 \| 10 \| 117 \| Protein 25-like lysozyme \| Protein 25-like lysozyme; afdb-uniprot50 \| AF-A0A5E4XAT6-F1-MODEL\_V4 \| 1.0 \| 9.453e-07 \| 275 \| 0.263 \| 91 \| 65 \| 1 \| 17 \| 105 \| 7 \| 97 \| Baseplate protein \| Baseplate protein; afdb-uniprot50 \| AF-A0A0F9DR72-F1-MODEL\_V4 \| 1.0 \| 3.106e-07 \| 275 \| 0.226 \| 115 \| 79 \| 4 \| 1 \| 106 \| 2 \| 115 \| GPW\_gp25 domain-containing protein \| GPW\_gp25 domain-containing protein; afdb-uniprot50 \| AF-A0A2A3D7N8-F1-MODEL\_V4 \| 1.0 \| 3.977e-07 \| 275 \| 0.281 \| 110 \| 72 \| 3 \| 4 \| 106 \| 1 \| 110 \| Baseplate assembly protein \| Baseplate assembly protein; afdb-uniprot50 \| AF-A0A6P0KR28-F1-MODEL\_V4 \| 1.0 \| 3.304e-07 \| 275 \| 0.266 \| 109 \| 77 \| 2 \| 2 \| 107 \| 14 \| 122 \| GPW/gp25 family protein \| GPW/gp25 family protein; afdb-uniprot50 \| AF-A0A1Q8A383-F1-MODEL\_V4 \| 1.0 \| 3.515e-07 \| 275 \| 0.301 \| 106 \| 72 \| 1 \| 2 \| 105 \| 16 \| 121 \| Baseplate protein \| Baseplate protein; afdb-uniprot50 \| AF-A0A1M7YV13-F1-MODEL\_V4 \| 1.0 \| 5.764e-07 \| 275 \| 0.247 \| 117 \| 78 \| 3 \| 1 \| 107 \| 7 \| 123 \| Gene 25-like lysozyme \| Gene 25-like lysozyme; afdb-uniprot50 \| AF-I5B7H2-F1-MODEL\_V4 \| 1.0 \| 6.132e-07 \| 275 \| 0.263 \| 114 \| 75 \| 3 \| 1 \| 105 \| 8 \| 121 \| Phage baseplate assembly protein W \| Phage baseplate assembly protein W; afdb-uniprot50 \| AF-A0A1I7EL93-F1-MODEL\_V4 \| 1.0 \| 2.785e-08 \| 275 \| 0.403 \| 109 \| 61 \| 2 \| 1 \| 105 \| 24 \| 132 \| GPW\_gp25 domain-containing protein \| GPW\_gp25 domain-containing protein; afdb-uniprot50 \| AF-A0A386WG07-F1-MODEL\_V4 \| 1.0 \| 1.39e-07 \| 275 \| 0.333 \| 108 \| 69 \| 2 \| 1 \| 105 \| 18 \| 125 \| GPW\_gp25 domain-containing protein \| GPW\_gp25 domain-containing protein; afdb-uniprot50 \| AF-A0A4Q5SKL8-F1-MODEL\_V4 \| 1.0 \| 2.425e-07 \| 275 \| 0.28 \| 107 \| 74 \| 2 \| 2 \| 105 \| 26 \| 132 \| GPW\_gp25 domain-containing protein \| GPW\_gp25 domain-containing protein; afdb-uniprot50 \| AF-A0A4Z0H7I9-F1-MODEL\_V4 \| 1.0 \| 4.501e-07 \| 275 \| 0.289 \| 114 \| 72 \| 3 \| 1 \| 105 \| 5 \| 118 \| Baseplate protein \| Baseplate protein; afdb-uniprot50 \| AF-A0A850GIX7-F1-MODEL\_V4 \| 1.0 \| 6.523e-07 \| 274 \| 0.204 \| 98 \| 76 \| 1 \| 12 \| 107 \| 2 \| 99 \| GPW/gp25 family protein \| GPW/gp25 family protein; afdb-uniprot50 \| AF-A0A3M1B2Y6-F1-MODEL\_V4 \| 1.0 \| 1.307e-07 \| 274 \| 0.252 \| 107 \| 77 \| 2 \| 2 \| 105 \| 11 \| 117 \| GPW\_gp25 domain-containing protein \| GPW\_gp25 domain-containing protein; afdb-uniprot50 \| AF-A0A382EFG4-F1-MODEL\_V4 \| 1.0 \| 3.739e-07 \| 274 \| 0.25 \| 108 \| 78 \| 2 \| 3 \| 107 \| 16 \| 123 \| GPW\_gp25 domain-containing protein \| GPW\_gp25 domain-containing protein; afdb-uniprot50 \| AF-A0A6M0FMQ1-F1-MODEL\_V4 \| 1.0 \| 2.744e-07 \| 274 \| 0.22 \| 109 \| 82 \| 2 \| 2 \| 107 \| 16 \| 124 \| GPW/gp25 family protein \| GPW/gp25 family protein; afdb-uniprot50 \| AF-A0A4Y3L060-F1-MODEL\_V4 \| 1.0 \| 7.381e-07 \| 274 \| 0.272 \| 125 \| 75 \| 5 \| 1 \| 112 \| 7 \| 128 \| GPW\_gp25 domain-containing protein \| GPW\_gp25 domain-containing protein; afdb-uniprot50 \| AF-A0A6S6SM83-F1-MODEL\_V4 \| 1.0 \| 3.106e-07 \| 274 \| 0.275 \| 109 \| 76 \| 2 \| 2 \| 107 \| 23 \| 131 \| GPW\_gp25 domain-containing protein \| GPW\_gp25 domain-containing protein; afdb-uniprot50 \| AF-A0A0F9TG53-F1-MODEL\_V4 \| 1.0 \| 2.919e-07 \| 274 \| 0.261 \| 107 \| 75 \| 3 \| 2 \| 105 \| 30 \| 135 \| GPW\_gp25 domain-containing protein \| GPW\_gp25 domain-containing protein; afdb-uniprot50 \| AF-A0A1T4W4S5-F1-MODEL\_V4 \| 1.0 \| 1.07e-06 \| 274 \| 0.268 \| 108 \| 74 \| 2 \| 1 \| 105 \| 77 \| 182 \| Phage baseplate assembly protein W \| Phage baseplate assembly protein W; afdb-uniprot50 \| AF-A0A810Z3I0-F1-MODEL\_V4 \| 1.0 \| 1.07e-06 \| 273 \| 0.228 \| 92 \| 68 \| 2 \| 17 \| 106 \| 7 \| 97 \| Baseplate protein \| Baseplate protein; afdb-uniprot50 \| AF-A0A3B0VNW2-F1-MODEL\_V4 \| 1.0 \| 4.231e-07 \| 273 \| 0.273 \| 106 \| 74 \| 2 \| 3 \| 105 \| 16 \| 121 \| GPW\_gp25 domain-containing protein \| GPW\_gp25 domain-containing protein; afdb-uniprot50 \| AF-A0A4Q9HLM9-F1-MODEL\_V4 \| 1.0 \| 4.501e-07 \| 273 \| 0.271 \| 107 \| 75 \| 2 \| 2 \| 105 \| 14 \| 120 \| Baseplate protein \| Baseplate protein; afdb-uniprot50 \| AF-A0A0D8FVT8-F1-MODEL\_V4 \| 1.0 \| 3.739e-07 \| 273 \| 0.289 \| 107 \| 73 \| 2 \| 2 \| 105 \| 42 \| 148 \| 25-like lysozyme \| 25-like lysozyme; afdb-uniprot50 \| AF-A0A2P8QYN7-F1-MODEL\_V4 \| 1.0 \| 2.744e-07 \| 272 \| 0.347 \| 95 \| 56 \| 2 \| 12 \| 106 \| 4 \| 92 \| Baseplate assembly protein \| Baseplate assembly protein; afdb-uniprot50 \| AF-A0A4P8G0E4-F1-MODEL\_V4 \| 1.0 \| 4.231e-07 \| 272 \| 0.223 \| 112 \| 80 \| 4 \| 1 \| 107 \| 2 \| 111 \| Phage baseplate protein \| Phage baseplate protein; afdb-uniprot50 \| AF-C1D831-F1-MODEL\_V4 \| 1.0 \| 3.106e-07 \| 272 \| 0.219 \| 114 \| 82 \| 4 \| 1 \| 111 \| 8 \| 117 \| Putative phage baseplate protein \| Putative phage baseplate protein; afdb-uniprot50 \| AF-A0A1W1X795-F1-MODEL\_V4 \| 1.0 \| 5.093e-07 \| 272 \| 0.269 \| 115 \| 78 \| 4 \| 2 \| 112 \| 12 \| 124 \| GPW\_gp25 domain-containing protein \| GPW\_gp25 domain-containing protein; afdb-uniprot50 \| AF-A0A7X5EZA1-F1-MODEL\_V4 \| 1.0 \| 7.852e-07 \| 272 \| 0.241 \| 116 \| 81 \| 2 \| 1 \| 109 \| 8 \| 123 \| Integrase \| Integrase; afdb-uniprot50 \| AF-A0A1M7AAS8-F1-MODEL\_V4 \| 1.0 \| 2.143e-07 \| 272 \| 0.201 \| 109 \| 83 \| 2 \| 2 \| 106 \| 13 \| 121 \| GPW\_gp25 domain-containing protein \| GPW\_gp25 domain-containing protein; afdb-uniprot50 \| AF-A0A2W6DBF0-F1-MODEL\_V4 \| 1.0 \| 6.939e-07 \| 272 \| 0.28 \| 114 \| 73 \| 3 \| 1 \| 105 \| 7 \| 120 \| Baseplate protein \| Baseplate protein; afdb-uniprot50 \| AF-A0A3S2YJR9-F1-MODEL\_V4 \| 1.0 \| 7.381e-07 \| 272 \| 0.235 \| 106 \| 76 \| 1 \| 1 \| 101 \| 3 \| 108 \| Uncharacterized protein \| Uncharacterized protein; afdb-uniprot50 \| AF-A0A6J4TRM4-F1-MODEL\_V4 \| 1.0 \| 3.304e-07 \| 272 \| 0.324 \| 108 \| 68 \| 3 \| 2 \| 105 \| 27 \| 133 \| GPW\_gp25 domain-containing protein \| GPW\_gp25 domain-containing protein; afdb-uniprot50 \| AF-A0A1A9HJ71-F1-MODEL\_V4 \| 1.0 \| 3.106e-07 \| 272 \| 0.229 \| 109 \| 80 \| 2 \| 2 \| 106 \| 16 \| 124 \| GPW\_gp25 domain-containing protein \| GPW\_gp25 domain-containing protein; afdb-uniprot50 \| AF-A0A1G3UAL2-F1-MODEL\_V4 \| 1.0 \| 9.017e-08 \| 271 \| 0.37 \| 100 \| 54 \| 3 \| 12 \| 108 \| 3 \| 96 \| Baseplate assembly protein \| Baseplate assembly protein; afdb-uniprot50 \| AF-A0A285NMD3-F1-MODEL\_V4 \| 1.0 \| 3.061e-06 \| 271 \| 0.276 \| 94 \| 64 \| 2 \| 17 \| 108 \| 13 \| 104 \| GPW\_gp25 domain-containing protein \| GPW\_gp25 domain-containing protein; afdb-uniprot50 \| AF-A0A0Q7TJ69-F1-MODEL\_V4 \| 1.0 \| 6.222e-08 \| 271 \| 0.321 \| 109 \| 69 \| 3 \| 1 \| 105 \| 13 \| 120 \| GPW\_gp25 domain-containing protein \| GPW\_gp25 domain-containing protein; afdb-uniprot50 \| AF-A0A2S6I8X0-F1-MODEL\_V4 \| 1.0 \| 2.015e-07 \| 271 \| 0.275 \| 109 \| 75 \| 2 \| 2 \| 106 \| 14 \| 122 \| GPW\_gp25 domain-containing protein \| GPW\_gp25 domain-containing protein; afdb-uniprot50 \| AF-A0A1Q4UKA2-F1-MODEL\_V4 \| 1.0 \| 2.58e-07 \| 271 \| 0.293 \| 109 \| 72 \| 3 \| 2 \| 106 \| 23 \| 130 \| GPW\_gp25 domain-containing protein \| GPW\_gp25 domain-containing protein; afdb-uniprot50 \| AF-A0A7Y6XH67-F1-MODEL\_V4 \| 1.0 \| 2.919e-07 \| 271 \| 0.25 \| 108 \| 77 \| 2 \| 2 \| 105 \| 17 \| 124 \| GPW/gp25 family protein \| GPW/gp25 family protein; afdb-uniprot50 \| AF-A0A2G1XB05-F1-MODEL\_V4 \| 1.0 \| 1.07e-06 \| 271 \| 0.263 \| 114 \| 75 \| 3 \| 1 \| 105 \| 14 \| 127 \| Baseplate protein \| Baseplate protein; afdb-uniprot50 \| AF-A0A1N6M9S4-F1-MODEL\_V4 \| 1.0 \| 2.143e-07 \| 271 \| 0.203 \| 108 \| 82 \| 2 \| 2 \| 105 \| 15 \| 122 \| Baseplate wedge subunit \| Baseplate wedge subunit; afdb-uniprot50 \| AF-A0A0M4DWU2-F1-MODEL\_V4 \| 1.0 \| 4.231e-07 \| 271 \| 0.342 \| 108 \| 66 \| 3 \| 2 \| 105 \| 14 \| 120 \| Baseplate protein \| Baseplate protein; afdb-uniprot50 \| AF-A0A7Y7B5C7-F1-MODEL\_V4 \| 1.0 \| 4.231e-07 \| 271 \| 0.271 \| 107 \| 75 \| 2 \| 2 \| 105 \| 14 \| 120 \| GPW/gp25 family protein \| GPW/gp25 family protein; afdb-uniprot50 \| AF-A0A376J3V4-F1-MODEL\_V4 \| 1.0 \| 3.256e-06 \| 270 \| 0.513 \| 72 \| 35 \| 0 \| 1 \| 72 \| 5 \| 76 \| Baseplate assembly protein W \| Baseplate assembly protein W; afdb-uniprot50 \| AF-A0A535E2Y7-F1-MODEL\_V4 \| 1.0 \| 2.015e-07 \| 270 \| 0.292 \| 106 \| 69 \| 3 \| 12 \| 112 \| 2 \| 106 \| GPW/gp25 family protein \| GPW/gp25 family protein; afdb-uniprot50 \| AF-A0A1D2QMW4-F1-MODEL\_V4 \| 1.0 \| 5.418e-07 \| 270 \| 0.225 \| 111 \| 75 \| 5 \| 1 \| 105 \| 2 \| 107 \| Uncharacterized protein \| Uncharacterized protein; afdb-uniprot50 \| AF-A0A2D6WZT7-F1-MODEL\_V4 \| 1.0 \| 4.788e-07 \| 270 \| 0.24 \| 108 \| 79 \| 2 \| 3 \| 107 \| 16 \| 123 \| GPW\_gp25 domain-containing protein \| GPW\_gp25 domain-containing protein; afdb-uniprot50 \| AF-A0A1S2N8L0-F1-MODEL\_V4 \| 1.0 \| 5.764e-07 \| 270 \| 0.207 \| 111 \| 83 \| 3 \| 2 \| 108 \| 19 \| 128 \| Putative tail lysozyme \| Putative tail lysozyme; afdb-uniprot50 \| AF-A0A4R7ZWG3-F1-MODEL\_V4 \| 1.0 \| 9.453e-07 \| 269 \| 0.23 \| 113 \| 78 \| 3 \| 1 \| 105 \| 1 \| 112 \| GPW\_gp25 domain-containing protein \| GPW\_gp25 domain-containing protein; afdb-uniprot50 \| AF-A0A0K1JCX4-F1-MODEL\_V4 \| 1.0 \| 5.764e-07 \| 269 \| 0.277 \| 108 \| 75 \| 2 \| 3 \| 107 \| 16 \| 123 \| GPW/gp25 family protein \| GPW/gp25 family protein; afdb-uniprot50 \| AF-A0A841BZN7-F1-MODEL\_V4 \| 1.0 \| 3.106e-07 \| 269 \| 0.33 \| 109 \| 68 \| 3 \| 3 \| 107 \| 15 \| 122 \| Phage baseplate assembly protein W \| Phage baseplate assembly protein W; afdb-uniprot50 \| AF-A0A3A9FX35-F1-MODEL\_V4 \| 1.0 \| 2.015e-07 \| 269 \| 0.275 \| 116 \| 72 \| 4 \| 1 \| 106 \| 10 \| 123 \| Baseplate protein \| Baseplate protein; afdb-uniprot50 \| AF-A0A6I3M8T7-F1-MODEL\_V4 \| 1.0 \| 4.788e-07 \| 269 \| 0.287 \| 108 \| 72 \| 3 \| 2 \| 105 \| 14 \| 120 \| Baseplate protein \| Baseplate protein; afdb-uniprot50 \| AF-A0A3N6FWS9-F1-MODEL\_V4 \| 1.0 \| 3.106e-07 \| 269 \| 0.293 \| 109 \| 74 \| 2 \| 2 \| 107 \| 14 \| 122 \| Gene 25-like lysozyme \| Gene 25-like lysozyme; afdb-uniprot50 \| AF-A0A350PEH1-F1-MODEL\_V4 \| 1.0 \| 3.304e-07 \| 268 \| 0.198 \| 111 \| 83 \| 3 \| 3 \| 107 \| 19 \| 129 \| GPW\_gp25 domain-containing protein \| GPW\_gp25 domain-containing protein; afdb-uniprot50 \| AF-A0A1Q4FZT8-F1-MODEL\_V4 \| 1.0 \| 2.919e-07 \| 268 \| 0.22 \| 109 \| 83 \| 1 \| 1 \| 107 \| 17 \| 125 \| GPW\_gp25 domain-containing protein \| GPW\_gp25 domain-containing protein; afdb-uniprot50 \| AF-A0A2E0M339-F1-MODEL\_V4 \| 1.0 \| 7.852e-07 \| 268 \| 0.162 \| 111 \| 89 \| 3 \| 3 \| 110 \| 25 \| 134 \| GPW\_gp25 domain-containing protein \| GPW\_gp25 domain-containing protein; afdb-uniprot50 \| AF-F9U9K7-F1-MODEL\_V4 \| 1.0 \| 2.744e-07 \| 268 \| 0.212 \| 108 \| 81 \| 2 \| 2 \| 105 \| 17 \| 124 \| GPW/gp25 family protein \| GPW/gp25 family protein; afdb-uniprot50 \| AF-A0A518BPS1-F1-MODEL\_V4 \| 1.0 \| 6.619e-08 \| 268 \| 0.297 \| 111 \| 72 \| 3 \| 1 \| 105 \| 30 \| 140 \| Gene 25-like lysozyme \| Gene 25-like lysozyme; afdb-uniprot50 \| AF-R5P778-F1-MODEL\_V4 \| 1.0 \| 1.288e-06 \| 267 \| 0.208 \| 96 \| 74 \| 1 \| 12 \| 105 \| 2 \| 97 \| GPW/gp25 family protein \| GPW/gp25 family protein; afdb-uniprot50 \| AF-A0A6M0B625-F1-MODEL\_V4 \| 1.0 \| 5.418e-07 \| 267 \| 0.192 \| 109 \| 85 \| 2 \| 2 \| 107 \| 16 \| 124 \| GPW/gp25 family protein \| GPW/gp25 family protein; afdb-uniprot50 \| AF-D8IV19-F1-MODEL\_V4 \| 1.0 \| 7.381e-07 \| 267 \| 0.3 \| 103 \| 69 \| 2 \| 5 \| 105 \| 31 \| 132 \| Bacteriophage baseplate assembly protein W \| Bacteriophage baseplate assembly protein W; afdb-uniprot50 \| AF-A0A0M8SZZ8-F1-MODEL\_V4 \| 1.0 \| 4.501e-07 \| 267 \| 0.292 \| 106 \| 72 \| 2 \| 3 \| 105 \| 2 \| 107 \| Baseplate protein \| Baseplate protein; afdb-uniprot50 \| AF-A0A7C2B4J0-F1-MODEL\_V4 \| 1.0 \| 3.977e-07 \| 266 \| 0.315 \| 111 \| 68 \| 4 \| 1 \| 105 \| 9 \| 117 \| GPW\_gp25 domain-containing protein \| GPW\_gp25 domain-containing protein; afdb-uniprot50 \| AF-A0A1Z4U7A0-F1-MODEL\_V4 \| 1.0 \| 4.788e-07 \| 266 \| 0.183 \| 109 \| 85 \| 2 \| 2 \| 106 \| 17 \| 125 \| GPW\_gp25 domain-containing protein \| GPW\_gp25 domain-containing protein; afdb-uniprot50 \| AF-A0A0S4XLR8-F1-MODEL\_V4 \| 1.0 \| 1.288e-06 \| 265 \| 0.309 \| 97 \| 63 \| 2 \| 12 \| 107 \| 4 \| 97 \| Putative phage-related baseplate assembly protein (GPW-like) \| Putative phage-related baseplate assembly protein (GPW-like); afdb-uniprot50 \| AF-A0A327JTI1-F1-MODEL\_V4 \| 1.0 \| 5.093e-07 \| 265 \| 0.252 \| 115 \| 70 \| 4 \| 1 \| 107 \| 1 \| 107 \| GPW\_gp25 domain-containing protein \| GPW\_gp25 domain-containing protein; afdb-uniprot50 \| AF-A0A3D4RXG2-F1-MODEL\_V4 \| 1.0 \| 6.132e-07 \| 265 \| 0.25 \| 108 \| 76 \| 3 \| 2 \| 105 \| 16 \| 122 \| Phage tail protein \| Phage tail protein; afdb-uniprot50 \| AF-A0A3S0SRP4-F1-MODEL\_V4 \| 1.0 \| 3.739e-07 \| 265 \| 0.268 \| 108 \| 75 \| 3 \| 1 \| 106 \| 16 \| 121 \| GPW\_gp25 domain-containing protein \| GPW\_gp25 domain-containing protein; afdb-uniprot50 \| AF-A0A254S6U8-F1-MODEL\_V4 \| 1.0 \| 7.381e-07 \| 265 \| 0.168 \| 107 \| 88 \| 1 \| 2 \| 107 \| 15 \| 121 \| GPW\_gp25 domain-containing protein \| GPW\_gp25 domain-containing protein; afdb-uniprot50 \| AF-A0A451AWS4-F1-MODEL\_V4 \| 1.0 \| 5.418e-07 \| 265 \| 0.209 \| 110 \| 83 \| 2 \| 2 \| 107 \| 16 \| 125 \| GPW\_gp25 domain-containing protein \| GPW\_gp25 domain-containing protein; afdb-uniprot50 \| AF-A0A3S0JB46-F1-MODEL\_V4 \| 1.0 \| 4.231e-07 \| 265 \| 0.276 \| 105 \| 74 \| 1 \| 3 \| 105 \| 29 \| 133 \| GPW\_gp25 domain-containing protein \| GPW\_gp25 domain-containing protein; afdb-uniprot50 \| AF-A0A2E5QDB5-F1-MODEL\_V4 \| 1.0 \| 6.132e-07 \| 265 \| 0.242 \| 107 \| 76 \| 3 \| 3 \| 105 \| 31 \| 136 \| GPW\_gp25 domain-containing protein \| GPW\_gp25 domain-containing protein; afdb-uniprot50 \| AF-A0A1W9WDK2-F1-MODEL\_V4 \| 1.0 \| 6.132e-07 \| 265 \| 0.275 \| 109 \| 74 \| 3 \| 2 \| 106 \| 20 \| 127 \| GPW\_gp25 domain-containing protein \| GPW\_gp25 domain-containing protein; afdb-uniprot50 \| AF-A0A2S2DR56-F1-MODEL\_V4 \| 1.0 \| 1.985e-06 \| 264 \| 0.234 \| 94 \| 70 \| 1 \| 17 \| 108 \| 7 \| 100 \| GPW\_gp25 domain-containing protein \| GPW\_gp25 domain-containing protein; afdb-uniprot50 \| AF-A0A6P0Y0Z1-F1-MODEL\_V4 \| 1.0 \| 6.939e-07 \| 264 \| 0.237 \| 97 \| 70 \| 2 \| 17 \| 109 \| 16 \| 112 \| GPW/gp25 family protein \| GPW/gp25 family protein; afdb-uniprot50 \| AF-A0A6M8STK6-F1-MODEL\_V4 \| 1.0 \| 1.55e-06 \| 264 \| 0.224 \| 107 \| 81 \| 2 \| 2 \| 107 \| 11 \| 116 \| GPW/gp25 family protein \| GPW/gp25 family protein; afdb-uniprot50 \| AF-A0A6N8GMG1-F1-MODEL\_V4 \| 1.0 \| 6.939e-07 \| 264 \| 0.28 \| 107 \| 74 \| 2 \| 2 \| 105 \| 14 \| 120 \| Baseplate protein \| Baseplate protein; afdb-uniprot50 \| AF-A0A6I7NUI7-F1-MODEL\_V4 \| 1.0 \| 5.093e-07 \| 264 \| 0.231 \| 108 \| 79 \| 2 \| 2 \| 105 \| 16 \| 123 \| GPW\_gp25 domain-containing protein \| GPW\_gp25 domain-containing protein; afdb-uniprot50 \| AF-A0A6D1IH24-F1-MODEL\_V4 \| 1.0 \| 7.852e-07 \| 264 \| 0.311 \| 109 \| 68 \| 4 \| 2 \| 105 \| 14 \| 120 \| Baseplate protein \| Baseplate protein; afdb-uniprot50 \| AF-A0A1S6FJH3-F1-MODEL\_V4 \| 1.0 \| 3.106e-07 \| 264 \| 0.296 \| 108 \| 72 \| 2 \| 2 \| 105 \| 17 \| 124 \| GPW\_gp25 domain-containing protein \| GPW\_gp25 domain-containing protein; afdb-uniprot50 \| AF-A0A1I0ZTA3-F1-MODEL\_V4 \| 1.0 \| 4.501e-07 \| 264 \| 0.203 \| 108 \| 82 \| 2 \| 2 \| 105 \| 19 \| 126 \| GPW\_gp25 domain-containing protein \| GPW\_gp25 domain-containing protein; afdb-uniprot50 \| AF-A0A2E0S336-F1-MODEL\_V4 \| 1.0 \| 6.939e-07 \| 264 \| 0.226 \| 106 \| 79 \| 2 \| 3 \| 105 \| 42 \| 147 \| GPW\_gp25 domain-containing protein \| GPW\_gp25 domain-containing protein; afdb-uniprot50 \| AF-N1UP64-F1-MODEL\_V4 \| 1.0 \| 3.106e-07 \| 264 \| 0.324 \| 114 \| 73 \| 3 \| 1 \| 111 \| 24 \| 136 \| GPW\_gp25 domain-containing protein \| GPW\_gp25 domain-containing protein; afdb-uniprot50 \| AF-A0A4R7JV64-F1-MODEL\_V4 \| 1.0 \| 3.304e-07 \| 264 \| 0.282 \| 117 \| 75 \| 3 \| 2 \| 109 \| 29 \| 145 \| GPW\_gp25 domain-containing protein \| GPW\_gp25 domain-containing protein; afdb-uniprot50 \| AF-A0A3C0WX89-F1-MODEL\_V4 \| 1.0 \| 1.006e-06 \| 263 \| 0.238 \| 109 \| 80 \| 2 \| 2 \| 107 \| 12 \| 120 \| Baseplate protein \| Baseplate protein; afdb-uniprot50 \| AF-A0A2M8BUW0-F1-MODEL\_V4 \| 1.0 \| 5.093e-07 \| 263 \| 0.275 \| 109 \| 72 \| 5 \| 2 \| 105 \| 13 \| 119 \| Baseplate protein \| Baseplate protein; afdb-uniprot50 \| AF-A0A7W8JUU5-F1-MODEL\_V4 \| 1.0 \| 2.425e-07 \| 263 \| 0.231 \| 108 \| 80 \| 2 \| 1 \| 105 \| 26 \| 133 \| GPW\_gp25 domain-containing protein \| GPW\_gp25 domain-containing protein; afdb-uniprot50 \| AF-A0A440VHY5-F1-MODEL\_V4 \| 1.0 \| 7.381e-07 \| 262 \| 0.289 \| 107 \| 69 \| 3 \| 6 \| 105 \| 1 \| 107 \| Baseplate assembly protein \| Baseplate assembly protein; afdb-uniprot50 \| AF-A0A1Z8SB79-F1-MODEL\_V4 \| 1.0 \| 5.764e-07 \| 262 \| 0.226 \| 106 \| 79 \| 2 \| 3 \| 105 \| 18 \| 123 \| GPW\_gp25 domain-containing protein \| GPW\_gp25 domain-containing protein; afdb-uniprot50 \| AF-R5L6F4-F1-MODEL\_V4 \| 1.0 \| 5.418e-07 \| 262 \| 0.25 \| 108 \| 76 \| 4 \| 2 \| 105 \| 16 \| 122 \| GPW\_gp25 domain-containing protein \| GPW\_gp25 domain-containing protein; afdb-uniprot50 \| AF-A0A368JP81-F1-MODEL\_V4 \| 1.0 \| 6.132e-07 \| 262 \| 0.198 \| 111 \| 83 \| 2 \| 2 \| 106 \| 16 \| 126 \| GPW\_gp25 domain-containing protein \| GPW\_gp25 domain-containing protein; afdb-uniprot50 \| AF-A0A0U3LKX5-F1-MODEL\_V4 \| 1.0 \| 4.788e-07 \| 262 \| 0.194 \| 108 \| 83 \| 2 \| 2 \| 105 \| 16 \| 123 \| GPW/gp25 family protein \| GPW/gp25 family protein; afdb-uniprot50 \| AF-A0A2A4UTL9-F1-MODEL\_V4 \| 1.0 \| 3.061e-06 \| 261 \| 0.25 \| 96 \| 71 \| 1 \| 12 \| 106 \| 2 \| 97 \| GPW\_gp25 domain-containing protein \| GPW\_gp25 domain-containing protein; afdb-uniprot50 \| AF-A0A416TLR2-F1-MODEL\_V4 \| 1.0 \| 5.764e-07 \| 261 \| 0.212 \| 108 \| 80 \| 4 \| 2 \| 105 \| 15 \| 121 \| Baseplate assembly protein \| Baseplate assembly protein; afdb-uniprot50 \| AF-A0A1V5RRZ4-F1-MODEL\_V4 \| 1.0 \| 6.939e-07 \| 261 \| 0.279 \| 111 \| 78 \| 2 \| 3 \| 112 \| 30 \| 139 \| Gene 25-like lysozyme \| Gene 25-like lysozyme; afdb-uniprot50 \| AF-A0A285QW58-F1-MODEL\_V4 \| 1.0 \| 3.106e-07 \| 260 \| 0.268 \| 108 \| 75 \| 3 \| 2 \| 106 \| 9 \| 115 \| GPW\_gp25 domain-containing protein \| GPW\_gp25 domain-containing protein; afdb-uniprot50 \| AF-A0A4V2PNK1-F1-MODEL\_V4 \| 1.0 \| 9.453e-07 \| 260 \| 0.226 \| 106 \| 80 \| 1 \| 2 \| 105 \| 14 \| 119 \| GPW\_gp25 domain-containing protein \| GPW\_gp25 domain-containing protein; afdb-uniprot50 \| AF-A0A0Q7WE07-F1-MODEL\_V4 \| 1.0 \| 2.143e-07 \| 260 \| 0.305 \| 108 \| 70 \| 3 \| 3 \| 105 \| 13 \| 120 \| GPW\_gp25 domain-containing protein \| GPW\_gp25 domain-containing protein; afdb-uniprot50 \| AF-A0A0K3BU55-F1-MODEL\_V4 \| 1.0 \| 1.02e-07 \| 260 \| 0.28 \| 114 \| 73 \| 2 \| 1 \| 105 \| 5 \| 118 \| GPW\_gp25 domain-containing protein \| GPW\_gp25 domain-containing protein; afdb-uniprot50 \| AF-W7IHM5-F1-MODEL\_V4 \| 1.0 \| 7.852e-07 \| 260 \| 0.194 \| 108 \| 82 \| 3 \| 2 \| 105 \| 15 \| 121 \| GPW\_gp25 domain-containing protein \| GPW\_gp25 domain-containing protein; afdb-uniprot50 \| AF-A0A1G7A6G8-F1-MODEL\_V4 \| 1.0 \| 6.523e-07 \| 260 \| 0.194 \| 108 \| 84 \| 2 \| 2 \| 106 \| 17 \| 124 \| GPW\_gp25 domain-containing protein \| GPW\_gp25 domain-containing protein; afdb-uniprot50 \| AF-A0A7W0L027-F1-MODEL\_V4 \| 1.0 \| 8.353e-07 \| 259 \| 0.287 \| 108 \| 74 \| 2 \| 2 \| 106 \| 15 \| 122 \| GPW/gp25 family protein \| GPW/gp25 family protein; afdb-uniprot50 \| AF-A0A1T5EG95-F1-MODEL\_V4 \| 1.0 \| 7.852e-07 \| 259 \| 0.179 \| 106 \| 85 \| 1 \| 2 \| 105 \| 16 \| 121 \| GPW\_gp25 domain-containing protein \| GPW\_gp25 domain-containing protein; afdb-uniprot50 \| AF-A0A5D0Q715-F1-MODEL\_V4 \| 1.0 \| 1.155e-07 \| 259 \| 0.357 \| 112 \| 66 \| 3 \| 3 \| 108 \| 19 \| 130 \| GPW/gp25 family protein \| GPW/gp25 family protein; afdb-uniprot50 \| AF-A0A1Z9DJT3-F1-MODEL\_V4 \| 1.0 \| 1.754e-06 \| 259 \| 0.203 \| 108 \| 83 \| 2 \| 3 \| 107 \| 24 \| 131 \| GPW\_gp25 domain-containing protein \| GPW\_gp25 domain-containing protein; afdb-uniprot50 \| AF-A0A2P7TGY9-F1-MODEL\_V4 \| 1.0 \| 1.006e-06 \| 259 \| 0.228 \| 105 \| 79 \| 1 \| 3 \| 105 \| 37 \| 141 \| GPW\_gp25 domain-containing protein \| GPW\_gp25 domain-containing protein; afdb-uniprot50 \| AF-A0A081KAP0-F1-MODEL\_V4 \| 1.0 \| 6.523e-07 \| 258 \| 0.245 \| 106 \| 77 \| 2 \| 3 \| 105 \| 17 \| 122 \| GPW\_gp25 domain-containing protein \| GPW\_gp25 domain-containing protein; afdb-uniprot50 \| AF-A0A2W4RIS3-F1-MODEL\_V4 \| 1.0 \| 1.138e-06 \| 258 \| 0.25 \| 108 \| 77 \| 3 \| 2 \| 105 \| 17 \| 124 \| GPW\_gp25 domain-containing protein \| GPW\_gp25 domain-containing protein; afdb-uniprot50 \| AF-A0A3A8AUY7-F1-MODEL\_V4 \| 1.0 \| 1.649e-06 \| 258 \| 0.207 \| 106 \| 82 \| 2 \| 2 \| 105 \| 16 \| 121 \| Baseplate protein \| Baseplate protein; afdb-uniprot50 \| AF-A0A2W2BBX1-F1-MODEL\_V4 \| 1.0 \| 7.852e-07 \| 258 \| 0.252 \| 107 \| 77 \| 2 \| 2 \| 105 \| 18 \| 124 \| GPW\_gp25 domain-containing protein \| GPW\_gp25 domain-containing protein; afdb-uniprot50 \| AF-A0A7Y4WZF6-F1-MODEL\_V4 \| 1.0 \| 7.381e-07 \| 258 \| 0.207 \| 111 \| 82 \| 2 \| 2 \| 106 \| 22 \| 132 \| GPW/gp25 family protein \| GPW/gp25 family protein; afdb-uniprot50 \| AF-A0A521BNH3-F1-MODEL\_V4 \| 1.0 \| 3.256e-06 \| 257 \| 0.186 \| 102 \| 81 \| 1 \| 8 \| 107 \| 3 \| 104 \| GPW\_gp25 domain-containing protein \| GPW\_gp25 domain-containing protein; afdb-uniprot50 \| AF-A0A2Z3LGG4-F1-MODEL\_V4 \| 1.0 \| 5.418e-07 \| 257 \| 0.231 \| 108 \| 79 \| 2 \| 2 \| 105 \| 15 \| 122 \| GPW\_gp25 domain-containing protein \| GPW\_gp25 domain-containing protein; afdb-uniprot50 \| AF-A0A1C6SZV0-F1-MODEL\_V4 \| 1.0 \| 9.453e-07 \| 257 \| 0.287 \| 108 \| 72 \| 3 \| 2 \| 105 \| 14 \| 120 \| GPW\_gp25 domain-containing protein \| GPW\_gp25 domain-containing protein; afdb-uniprot50 \| AF-A0A4Y7B9B6-F1-MODEL\_V4 \| 1.0 \| 7.852e-07 \| 257 \| 0.241 \| 116 \| 78 \| 2 \| 1 \| 106 \| 30 \| 145 \| GPW\_gp25 domain-containing protein \| GPW\_gp25 domain-containing protein; afdb-uniprot50 \| AF-A0A0A7KPC6-F1-MODEL\_V4 \| 1.0 \| 3.739e-07 \| 256 \| 0.271 \| 107 \| 74 \| 2 \| 3 \| 105 \| 12 \| 118 \| GPW\_gp25 domain-containing protein \| GPW\_gp25 domain-containing protein; afdb-uniprot50 \| AF-A0A1Z9Q4U1-F1-MODEL\_V4 \| 1.0 \| 6.523e-07 \| 256 \| 0.232 \| 112 \| 78 \| 4 \| 3 \| 107 \| 15 \| 125 \| GPW\_gp25 domain-containing protein \| GPW\_gp25 domain-containing protein; afdb-uniprot50 \| AF-A0A7W1N5U2-F1-MODEL\_V4 \| 1.0 \| 4.788e-07 \| 256 \| 0.214 \| 107 \| 81 \| 2 \| 2 \| 105 \| 13 \| 119 \| GPW/gp25 family protein \| GPW/gp25 family protein; afdb-uniprot50 \| AF-A0A0S3UC95-F1-MODEL\_V4 \| 1.0 \| 1.006e-06 \| 256 \| 0.362 \| 113 \| 62 \| 4 \| 1 \| 104 \| 15 \| 126 \| GPW / gp25 family protein \| GPW / gp25 family protein; afdb-uniprot50 \| AF-A0A6I5PFW1-F1-MODEL\_V4 \| 1.0 \| 1.288e-06 \| 255 \| 0.236 \| 93 \| 69 \| 1 \| 17 \| 107 \| 7 \| 99 \| GPW/gp25 family protein \| GPW/gp25 family protein; afdb-uniprot50 \| AF-A0A0M8QQ48-F1-MODEL\_V4 \| 1.0 \| 1.138e-06 \| 255 \| 0.283 \| 106 \| 73 \| 2 \| 3 \| 105 \| 2 \| 107 \| Baseplate protein \| Baseplate protein; afdb-uniprot50 \| AF-A0A7W7MKI6-F1-MODEL\_V4 \| 1.0 \| 8.886e-07 \| 255 \| 0.299 \| 107 \| 71 \| 3 \| 2 \| 105 \| 16 \| 121 \| Phage baseplate assembly protein W \| Phage baseplate assembly protein W; afdb-uniprot50 \| AF-A0A1I3WUF2-F1-MODEL\_V4 \| 1.0 \| 4.231e-07 \| 255 \| 0.205 \| 107 \| 82 \| 2 \| 2 \| 105 \| 17 \| 123 \| GPW\_gp25 domain-containing protein \| GPW\_gp25 domain-containing protein; afdb-uniprot50 \| AF-A0A3E5B1A9-F1-MODEL\_V4 \| 1.0 \| 2.247e-06 \| 255 \| 0.196 \| 107 \| 85 \| 1 \| 2 \| 107 \| 22 \| 128 \| GPW\_gp25 domain-containing protein \| GPW\_gp25 domain-containing protein; afdb-uniprot50 \| AF-A0A2E4W5C2-F1-MODEL\_V4 \| 1.0 \| 1.649e-06 \| 254 \| 0.243 \| 111 \| 79 \| 2 \| 1 \| 107 \| 5 \| 114 \| GPW\_gp25 domain-containing protein \| GPW\_gp25 domain-containing protein; afdb-uniprot50 \| AF-E7G9C5-F1-MODEL\_V4 \| 1.0 \| 5.418e-07 \| 254 \| 0.263 \| 114 \| 76 \| 4 \| 2 \| 108 \| 13 \| 125 \| GPW\_gp25 domain-containing protein \| GPW\_gp25 domain-containing protein; afdb-uniprot50 \| AF-A0A177NGE0-F1-MODEL\_V4 \| 1.0 \| 9.453e-07 \| 254 \| 0.256 \| 109 \| 77 \| 2 \| 3 \| 107 \| 14 \| 122 \| GPW\_gp25 domain-containing protein \| GPW\_gp25 domain-containing protein; afdb-uniprot50 \| AF-A0A3B1A623-F1-MODEL\_V4 \| 1.0 \| 9.453e-07 \| 254 \| 0.172 \| 116 \| 86 \| 3 \| 2 \| 107 \| 12 \| 127 \| GPW\_gp25 domain-containing protein \| GPW\_gp25 domain-containing protein; afdb-uniprot50 \| AF-A0A1I0XDA2-F1-MODEL\_V4 \| 1.0 \| 1.211e-06 \| 254 \| 0.231 \| 108 \| 78 \| 3 \| 2 \| 105 \| 16 \| 122 \| GPW\_gp25 domain-containing protein \| GPW\_gp25 domain-containing protein; afdb-uniprot50 \| AF-A0A424SVR1-F1-MODEL\_V4 \| 1.0 \| 1.754e-06 \| 254 \| 0.231 \| 108 \| 80 \| 2 \| 3 \| 107 \| 25 \| 132 \| GPW\_gp25 domain-containing protein \| GPW\_gp25 domain-containing protein; afdb-uniprot50 \| AF-A0A291IN73-F1-MODEL\_V4 \| 1.0 \| 6.939e-07 \| 254 \| 0.27 \| 111 \| 78 \| 2 \| 1 \| 108 \| 25 \| 135 \| GPW\_gp25 domain-containing protein \| GPW\_gp25 domain-containing protein; afdb-uniprot50 \| AF-A0A1E5BWS4-F1-MODEL\_V4 \| 1.0 \| 4.436e-06 \| 253 \| 0.254 \| 118 \| 78 \| 5 \| 1 \| 110 \| 1 \| 116 \| GPW\_gp25 domain-containing protein \| GPW\_gp25 domain-containing protein; afdb-uniprot50 \| AF-X5QYV2-F1-MODEL\_V4 \| 1.0 \| 1.138e-06 \| 253 \| 0.234 \| 111 \| 78 \| 2 \| 2 \| 105 \| 16 \| 126 \| Baseplate protein \| Baseplate protein; afdb-uniprot50 \| AF-A0A554VPK4-F1-MODEL\_V4 \| 1.0 \| 1.55e-06 \| 253 \| 0.165 \| 109 \| 87 \| 2 \| 2 \| 106 \| 18 \| 126 \| GPW/gp25 family protein \| GPW/gp25 family protein; afdb-uniprot50 \| AF-A0A327QS96-F1-MODEL\_V4 \| 1.0 \| 8.886e-07 \| 253 \| 0.194 \| 113 \| 83 \| 2 \| 1 \| 105 \| 17 \| 129 \| GPW\_gp25 domain-containing protein \| GPW\_gp25 domain-containing protein; afdb-uniprot50 \| AF-A0A5K7YHV8-F1-MODEL\_V4 \| 1.0 \| 1.07e-06 \| 253 \| 0.206 \| 116 \| 82 \| 2 \| 2 \| 107 \| 15 \| 130 \| GPW\_gp25 domain-containing protein \| GPW\_gp25 domain-containing protein; afdb-uniprot50 \| AF-A0A701NJT1-F1-MODEL\_V4 \| 1.0 \| 3.061e-06 \| 252 \| 0.404 \| 84 \| 50 \| 0 \| 12 \| 95 \| 3 \| 86 \| Baseplate assembly protein \| Baseplate assembly protein; afdb-uniprot50 \| AF-A0A1I2FX92-F1-MODEL\_V4 \| 1.0 \| 1.866e-06 \| 252 \| 0.311 \| 106 \| 70 \| 2 \| 2 \| 105 \| 12 \| 116 \| Gene 25-like lysozyme \| Gene 25-like lysozyme; afdb-uniprot50 \| AF-A0A3P1UU22-F1-MODEL\_V4 \| 1.0 \| 7.381e-07 \| 252 \| 0.3 \| 120 \| 62 \| 8 \| 1 \| 108 \| 17 \| 126 \| Baseplate assembly protein \| Baseplate assembly protein; afdb-uniprot50 \| AF-A0A2V7X7B0-F1-MODEL\_V4 \| 1.0 \| 6.838e-06 \| 251 \| 0.258 \| 93 \| 67 \| 1 \| 17 \| 107 \| 9 \| 101 \| Baseplate protein \| Baseplate protein; afdb-uniprot50 \| AF-A0A2E6EKB3-F1-MODEL\_V4 \| 1.0 \| 1.866e-06 \| 251 \| 0.256 \| 109 \| 75 \| 4 \| 2 \| 105 \| 14 \| 121 \| Baseplate protein \| Baseplate protein; afdb-uniprot50 \| AF-A0A4Q5Y1B4-F1-MODEL\_V4 \| 1.0 \| 1.37e-06 \| 251 \| 0.203 \| 108 \| 82 \| 2 \| 2 \| 105 \| 11 \| 118 \| GPW\_gp25 domain-containing protein \| GPW\_gp25 domain-containing protein; afdb-uniprot50 \| AF-A0A7X7DI45-F1-MODEL\_V4 \| 1.0 \| 1.288e-06 \| 251 \| 0.266 \| 109 \| 77 \| 2 \| 2 \| 107 \| 22 \| 130 \| GPW/gp25 family protein \| GPW/gp25 family protein; afdb-uniprot50 \| AF-A0A353GUZ2-F1-MODEL\_V4 \| 1.0 \| 1.37e-06 \| 250 \| 0.13 \| 115 \| 95 \| 3 \| 2 \| 112 \| 15 \| 128 \| GPW\_gp25 domain-containing protein \| GPW\_gp25 domain-containing protein; afdb-uniprot50 \| AF-A0A165U8T9-F1-MODEL\_V4 \| 1.0 \| 4.719e-06 \| 250 \| 0.218 \| 119 \| 79 \| 2 \| 1 \| 105 \| 1 \| 119 \| Gene 25-like lysozyme \| Gene 25-like lysozyme; afdb-uniprot50 \| AF-A9DX78-F1-MODEL\_V4 \| 1.0 \| 1.866e-06 \| 250 \| 0.183 \| 109 \| 85 \| 2 \| 2 \| 106 \| 17 \| 125 \| Tail lysozyme, putative \| Tail lysozyme, putative; afdb-uniprot50 \| AF-A0A4Y8P3N7-F1-MODEL\_V4 \| 1.0 \| 1.985e-06 \| 250 \| 0.283 \| 106 \| 74 \| 1 \| 2 \| 105 \| 23 \| 128 \| Baseplate protein \| Baseplate protein; afdb-uniprot50 \| AF-A0A0S2FBB8-F1-MODEL\_V4 \| 1.0 \| 5.093e-07 \| 249 \| 0.271 \| 114 \| 74 \| 4 \| 2 \| 107 \| 13 \| 125 \| Gene 25-like lysozyme family protein \| Gene 25-like lysozyme family protein; afdb-uniprot50 \| AF-U2HWE8-F1-MODEL\_V4 \| 1.0 \| 1.985e-06 \| 249 \| 0.188 \| 106 \| 84 \| 1 \| 2 \| 105 \| 16 \| 121 \| GPW\_gp25 domain-containing protein \| GPW\_gp25 domain-containing protein; afdb-uniprot50 \| AF-A0A1Z8QEK9-F1-MODEL\_V4 \| 1.0 \| 1.55e-06 \| 249 \| 0.185 \| 108 \| 84 \| 3 \| 3 \| 106 \| 18 \| 125 \| GPW\_gp25 domain-containing protein \| GPW\_gp25 domain-containing protein; afdb-uniprot50 \| AF-A0A7Y6XND0-F1-MODEL\_V4 \| 1.0 \| 2.877e-06 \| 249 \| 0.181 \| 116 \| 85 \| 3 \| 1 \| 106 \| 11 \| 126 \| GPW/gp25 family protein \| GPW/gp25 family protein; afdb-uniprot50 \| AF-K1RIV8-F1-MODEL\_V4 \| 1.0 \| 3.977e-07 \| 249 \| 0.218 \| 110 \| 79 \| 4 \| 1 \| 105 \| 22 \| 129 \| GPW/gp25 family protein \| GPW/gp25 family protein; afdb-uniprot50 \| AF-A0A497CQ18-F1-MODEL\_V4 \| 1.0 \| 1.55e-06 \| 249 \| 0.206 \| 116 \| 82 \| 3 \| 2 \| 107 \| 11 \| 126 \| GPW\_gp25 domain-containing protein \| GPW\_gp25 domain-containing protein; afdb-uniprot50 \| AF-A0A2T7L1T8-F1-MODEL\_V4 \| 1.0 \| 3.685e-06 \| 249 \| 0.263 \| 114 \| 75 \| 3 \| 1 \| 105 \| 19 \| 132 \| Baseplate protein \| Baseplate protein; afdb-uniprot50 \| AF-A0A252DTP5-F1-MODEL\_V4 \| 1.0 \| 2.112e-06 \| 249 \| 0.201 \| 109 \| 83 \| 2 \| 2 \| 106 \| 26 \| 134 \| GPW\_gp25 domain-containing protein \| GPW\_gp25 domain-containing protein; afdb-uniprot50 \| AF-A0A4U8UBX3-F1-MODEL\_V4 \| 1.0 \| 3.106e-07 \| 248 \| 0.285 \| 105 \| 62 \| 2 \| 1 \| 105 \| 1 \| 92 \| Baseplate assembly protein \| Baseplate assembly protein; afdb-uniprot50 \| AF-A0A380Q839-F1-MODEL\_V4 \| 1.0 \| 3.061e-06 \| 248 \| 0.482 \| 87 \| 41 \| 1 \| 1 \| 87 \| 6 \| 88 \| Phage-like protein \| Phage-like protein; afdb-uniprot50 \| AF-A0A6G3XXE1-F1-MODEL\_V4 \| 1.0 \| 2.39e-06 \| 248 \| 0.266 \| 105 \| 74 \| 2 \| 4 \| 105 \| 1 \| 105 \| GPW/gp25 family protein \| GPW/gp25 family protein; afdb-uniprot50 \| AF-A0A2G2LH18-F1-MODEL\_V4 \| 1.0 \| 2.247e-06 \| 248 \| 0.196 \| 107 \| 83 \| 2 \| 3 \| 106 \| 21 \| 127 \| GPW\_gp25 domain-containing protein \| GPW\_gp25 domain-containing protein; afdb-uniprot50 \| AF-A1ZC12-F1-MODEL\_V4 \| 1.0 \| 1.138e-06 \| 248 \| 0.201 \| 109 \| 81 \| 3 \| 2 \| 105 \| 17 \| 124 \| Phage baseplate assembly protein W \| Phage baseplate assembly protein W; afdb-uniprot50 \| AF-C5BJK3-F1-MODEL\_V4 \| 1.0 \| 1.754e-06 \| 248 \| 0.179 \| 117 \| 85 \| 3 \| 2 \| 107 \| 18 \| 134 \| Phage GpW/Gp25 family protein \| Phage GpW/Gp25 family protein; afdb-uniprot50 \| AF-A0A7C4AKA1-F1-MODEL\_V4 \| 1.0 \| 3.061e-06 \| 247 \| 0.247 \| 109 \| 75 \| 4 \| 3 \| 108 \| 6 \| 110 \| Baseplate protein \| Baseplate protein; afdb-uniprot50 \| AF-A0A2D5ICS9-F1-MODEL\_V4 \| 1.0 \| 2.877e-06 \| 247 \| 0.234 \| 111 \| 81 \| 3 \| 3 \| 110 \| 18 \| 127 \| GPW\_gp25 domain-containing protein \| GPW\_gp25 domain-containing protein; afdb-uniprot50 \| AF-A0A1H6RDH1-F1-MODEL\_V4 \| 1.0 \| 1.457e-06 \| 247 \| 0.224 \| 107 \| 80 \| 2 \| 2 \| 105 \| 17 \| 123 \| GPW\_gp25 domain-containing protein \| GPW\_gp25 domain-containing protein; afdb-uniprot50 \| AF-A0A6S5Y5N2-F1-MODEL\_V4 \| 1.0 \| 1.07e-06 \| 247 \| 0.25 \| 120 \| 79 \| 3 \| 1 \| 111 \| 18 \| 135 \| Baseplate protein \| Baseplate protein; afdb-uniprot50 \| AF-A0A1J4Z8F3-F1-MODEL\_V4 \| 1.0 \| 1.211e-06 \| 247 \| 0.188 \| 106 \| 84 \| 1 \| 2 \| 105 \| 20 \| 125 \| GPW\_gp25 domain-containing protein \| GPW\_gp25 domain-containing protein; afdb-uniprot50 \| AF-A0A7W1U792-F1-MODEL\_V4 \| 1.0 \| 4.788e-07 \| 247 \| 0.212 \| 108 \| 81 \| 2 \| 2 \| 105 \| 22 \| 129 \| GPW/gp25 family protein \| GPW/gp25 family protein; afdb-uniprot50 \| AF-U3AL81-F1-MODEL\_V4 \| 1.0 \| 8.757e-06 \| 246 \| 0.172 \| 110 \| 84 \| 4 \| 1 \| 107 \| 2 \| 107 \| Uncharacterized protein \| Uncharacterized protein; afdb-uniprot50 \| AF-A0A8A3HHQ9-F1-MODEL\_V4 \| 1.0 \| 2.705e-06 \| 246 \| 0.265 \| 113 \| 74 \| 4 \| 1 \| 105 \| 1 \| 112 \| GPW/gp25 family protein \| GPW/gp25 family protein; afdb-uniprot50 \| AF-A0A1C4VFP2-F1-MODEL\_V4 \| 1.0 \| 6.428e-06 \| 246 \| 0.201 \| 114 \| 82 \| 3 \| 1 \| 106 \| 1 \| 113 \| GPW\_gp25 domain-containing protein \| GPW\_gp25 domain-containing protein; afdb-uniprot50 \| AF-A0A246RHT3-F1-MODEL\_V4 \| 1.0 \| 3.464e-06 \| 246 \| 0.205 \| 107 \| 82 \| 3 \| 2 \| 106 \| 8 \| 113 \| GPW\_gp25 domain-containing protein \| GPW\_gp25 domain-containing protein; afdb-uniprot50 \| AF-A0A426VQD0-F1-MODEL\_V4 \| 1.0 \| 3.464e-06 \| 246 \| 0.162 \| 111 \| 86 \| 4 \| 1 \| 105 \| 4 \| 113 \| GPW\_gp25 domain-containing protein \| GPW\_gp25 domain-containing protein; afdb-uniprot50 \| AF-A0A6I7PSS5-F1-MODEL\_V4 \| 1.0 \| 2.705e-06 \| 246 \| 0.279 \| 93 \| 61 \| 2 \| 17 \| 105 \| 30 \| 120 \| GPW\_gp25 domain-containing protein \| GPW\_gp25 domain-containing protein; afdb-uniprot50 \| AF-A0A7G2RQA6-F1-MODEL\_V4 \| 1.0 \| 2.112e-06 \| 246 \| 0.149 \| 107 \| 88 \| 2 \| 3 \| 106 \| 19 \| 125 \| GPW\_gp25 domain-containing protein \| GPW\_gp25 domain-containing protein; afdb-uniprot50 \| AF-A0A3D1U7G8-F1-MODEL\_V4 \| 1.0 \| 5.764e-07 \| 246 \| 0.179 \| 117 \| 82 \| 5 \| 1 \| 105 \| 5 \| 119 \| Baseplate assembly protein \| Baseplate assembly protein; afdb-uniprot50 \| AF-A0A7K3ZDA3-F1-MODEL\_V4 \| 1.0 \| 2.247e-06 \| 246 \| 0.211 \| 109 \| 82 \| 3 \| 5 \| 112 \| 34 \| 139 \| GPW/gp25 family protein \| GPW/gp25 family protein; afdb-uniprot50 \| AF-A0A0B6CV24-F1-MODEL\_V4 \| 1.0 \| 7.852e-07 \| 245 \| 0.272 \| 110 \| 76 \| 3 \| 3 \| 109 \| 2 \| 110 \| 25-like lysozyme family protein \| 25-like lysozyme family protein; afdb-uniprot50 \| AF-A0A519PID9-F1-MODEL\_V4 \| 1.0 \| 1.649e-06 \| 245 \| 0.212 \| 108 \| 81 \| 3 \| 2 \| 105 \| 16 \| 123 \| GPW\_gp25 domain-containing protein \| GPW\_gp25 domain-containing protein; afdb-uniprot50 \| AF-A0A0F9S7G6-F1-MODEL\_V4 \| 1.0 \| 1.37e-06 \| 245 \| 0.219 \| 114 \| 79 \| 4 \| 2 \| 107 \| 13 \| 124 \| GPW\_gp25 domain-containing protein \| GPW\_gp25 domain-containing protein; afdb-uniprot50 \| AF-A0A1D9F2U1-F1-MODEL\_V4 \| 1.0 \| 9.453e-07 \| 245 \| 0.272 \| 110 \| 75 \| 3 \| 1 \| 106 \| 24 \| 132 \| GPW\_gp25 domain-containing protein \| GPW\_gp25 domain-containing protein; afdb-uniprot50 \| AF-H0FAR8-F1-MODEL\_V4 \| 1.0 \| 2.112e-06 \| 244 \| 0.242 \| 107 \| 76 \| 4 \| 2 \| 105 \| 8 \| 112 \| GPW/gp25 family protein \| GPW/gp25 family protein; afdb-uniprot50 \| AF-A0A0C5V444-F1-MODEL\_V4 \| 1.0 \| 3.464e-06 \| 244 \| 0.207 \| 106 \| 81 \| 2 \| 3 \| 105 \| 17 \| 122 \| Phage baseplate assembly protein W \| Phage baseplate assembly protein W; afdb-uniprot50 \| AF-A0A4Q9KAQ7-F1-MODEL\_V4 \| 1.0 \| 1.07e-06 \| 244 \| 0.265 \| 113 \| 76 \| 3 \| 2 \| 107 \| 19 \| 131 \| Phage tail protein \| Phage tail protein; afdb-uniprot50 \| AF-A0A6C0JT84-F1-MODEL\_V4 \| 1.0 \| 3.061e-06 \| 244 \| 0.165 \| 109 \| 86 \| 3 \| 3 \| 107 \| 21 \| 128 \| GPW\_gp25 domain-containing protein \| GPW\_gp25 domain-containing protein; afdb-uniprot50 \| AF-A0A2T0HIB9-F1-MODEL\_V4 \| 1.0 \| 1.528e-05 \| 243 \| 0.519 \| 77 \| 37 \| 0 \| 7 \| 83 \| 2 \| 78 \| Baseplate assembly protein \| Baseplate assembly protein; afdb-uniprot50 \| AF-A0A1H3JDF8-F1-MODEL\_V4 \| 1.0 \| 2.877e-06 \| 243 \| 0.244 \| 98 \| 69 \| 2 \| 17 \| 109 \| 7 \| 104 \| GPW\_gp25 domain-containing protein \| GPW\_gp25 domain-containing protein; afdb-uniprot50 \| AF-A0A1W9VM06-F1-MODEL\_V4 \| 1.0 \| 2.247e-06 \| 243 \| 0.25 \| 112 \| 77 \| 4 \| 1 \| 107 \| 1 \| 110 \| GPW\_gp25 domain-containing protein \| GPW\_gp25 domain-containing protein; afdb-uniprot50 \| AF-A0A1F4G5G2-F1-MODEL\_V4 \| 1.0 \| 7.381e-07 \| 243 \| 0.226 \| 115 \| 80 \| 4 \| 2 \| 108 \| 15 \| 128 \| Phage tail protein \| Phage tail protein; afdb-uniprot50 \| AF-L7UKW4-F1-MODEL\_V4 \| 1.0 \| 7.739e-06 \| 242 \| 0.254 \| 114 \| 76 \| 4 \| 1 \| 106 \| 1 \| 113 \| GPW\_gp25 domain-containing protein \| GPW\_gp25 domain-containing protein; afdb-uniprot50 \| AF-A0A172UEQ1-F1-MODEL\_V4 \| 1.0 \| 7.381e-07 \| 242 \| 0.262 \| 118 \| 81 \| 3 \| 1 \| 112 \| 8 \| 125 \| GPW\_gp25 domain-containing protein \| GPW\_gp25 domain-containing protein; afdb-uniprot50 \| AF-A0A4Y4CYZ4-F1-MODEL\_V4 \| 1.0 \| 3.685e-06 \| 242 \| 0.273 \| 106 \| 75 \| 1 \| 4 \| 107 \| 15 \| 120 \| GPW\_gp25 domain-containing protein \| GPW\_gp25 domain-containing protein; afdb-uniprot50 \| AF-A0A0Q6WZD8-F1-MODEL\_V4 \| 1.0 \| 2.705e-06 \| 242 \| 0.21 \| 114 \| 85 \| 4 \| 3 \| 112 \| 19 \| 131 \| GPW\_gp25 domain-containing protein \| GPW\_gp25 domain-containing protein; afdb-uniprot50 \| AF-A0A2W5H172-F1-MODEL\_V4 \| 1.0 \| 4.17e-06 \| 242 \| 0.256 \| 113 \| 77 \| 2 \| 1 \| 106 \| 16 \| 128 \| Integrase \| Integrase; afdb-uniprot50 \| AF-A0A1W0CS24-F1-MODEL\_V4 \| 1.0 \| 7.381e-07 \| 242 \| 0.238 \| 109 \| 79 \| 2 \| 1 \| 105 \| 18 \| 126 \| GPW\_gp25 domain-containing protein \| GPW\_gp25 domain-containing protein; afdb-uniprot50 \| AF-G5IB78-F1-MODEL\_V4 \| 1.0 \| 1.211e-06 \| 242 \| 0.212 \| 108 \| 80 \| 4 \| 2 \| 105 \| 15 \| 121 \| GPW\_gp25 domain-containing protein \| GPW\_gp25 domain-containing protein; afdb-uniprot50 \| AF-A0A3N9GJI3-F1-MODEL\_V4 \| 1.0 \| 6.939e-07 \| 242 \| 0.216 \| 111 \| 81 \| 2 \| 1 \| 105 \| 25 \| 135 \| Phage baseplate protein \| Phage baseplate protein; afdb-uniprot50 \| AF-A0A349ZQF9-F1-MODEL\_V4 \| 1.0 \| 2.112e-06 \| 242 \| 0.2 \| 110 \| 83 \| 4 \| 2 \| 107 \| 16 \| 124 \| Baseplate protein \| Baseplate protein; afdb-uniprot50 \| AF-A0A7V4G7W4-F1-MODEL\_V4 \| 1.0 \| 2.081e-05 \| 241 \| 0.266 \| 90 \| 64 \| 2 \| 17 \| 105 \| 2 \| 90 \| GPW\_gp25 domain-containing protein \| GPW\_gp25 domain-containing protein; afdb-uniprot50 \| AF-A0A5B1BHT0-F1-MODEL\_V4 \| 1.0 \| 7.275e-06 \| 241 \| 0.244 \| 98 \| 71 \| 2 \| 12 \| 106 \| 1 \| 98 \| GPW/gp25 family protein \| GPW/gp25 family protein; afdb-uniprot50 \| AF-A0A7K3XK95-F1-MODEL\_V4 \| 1.0 \| 9.91e-06 \| 241 \| 0.229 \| 96 \| 72 \| 1 \| 12 \| 105 \| 2 \| 97 \| GPW/gp25 family protein \| GPW/gp25 family protein; afdb-uniprot50 \| AF-A0A511QWX5-F1-MODEL\_V4 \| 1.0 \| 5.34e-06 \| 241 \| 0.264 \| 106 \| 73 \| 4 \| 1 \| 105 \| 5 \| 106 \| Uncharacterized protein \| Uncharacterized protein; afdb-uniprot50 \| AF-A0A2E6YZN4-F1-MODEL\_V4 \| 1.0 \| 5.02e-06 \| 241 \| 0.222 \| 108 \| 81 \| 2 \| 3 \| 107 \| 17 \| 124 \| GPW\_gp25 domain-containing protein \| GPW\_gp25 domain-containing protein; afdb-uniprot50 \| AF-A0A318TME3-F1-MODEL\_V4 \| 1.0 \| 3.256e-06 \| 241 \| 0.309 \| 110 \| 66 \| 2 \| 4 \| 103 \| 1 \| 110 \| GPW\_gp25 domain-containing protein \| GPW\_gp25 domain-containing protein; afdb-uniprot50 \| AF-A0A2R7JXZ8-F1-MODEL\_V4 \| 1.0 \| 9.453e-07 \| 241 \| 0.278 \| 122 \| 71 \| 4 \| 2 \| 106 \| 12 \| 133 \| Baseplate protein \| Baseplate protein; afdb-uniprot50 \| AF-A0A2N8BBF2-F1-MODEL\_V4 \| 1.0 \| 6.939e-07 \| 241 \| 0.233 \| 107 \| 77 \| 3 \| 2 \| 105 \| 17 \| 121 \| Phage baseplate protein \| Phage baseplate protein; afdb-uniprot50 \| AF-A0A1I5C9M8-F1-MODEL\_V4 \| 1.0 \| 9.453e-07 \| 241 \| 0.194 \| 108 \| 82 \| 4 \| 2 \| 105 \| 15 \| 121 \| GPW\_gp25 domain-containing protein \| GPW\_gp25 domain-containing protein; afdb-uniprot50 \| AF-A0A2W4YAP4-F1-MODEL\_V4 \| 1.0 \| 4.501e-07 \| 241 \| 0.276 \| 112 \| 73 \| 3 \| 4 \| 107 \| 26 \| 137 \| Baseplate assembly protein \| Baseplate assembly protein; afdb-uniprot50 \| AF-A0A5E4SAR3-F1-MODEL\_V4 \| 1.0 \| 2.112e-06 \| 241 \| 0.266 \| 109 \| 76 \| 2 \| 1 \| 105 \| 17 \| 125 \| Baseplate protein \| Baseplate protein; afdb-uniprot50 \| AF-A0A652LXD1-F1-MODEL\_V4 \| 1.0 \| 5.68e-06 \| 241 \| 0.224 \| 107 \| 80 \| 2 \| 2 \| 105 \| 15 \| 121 \| Baseplate protein \| Baseplate protein; afdb-uniprot50 \| AF-B9M7I0-F1-MODEL\_V4 \| 1.0 \| 1.649e-06 \| 241 \| 0.194 \| 108 \| 83 \| 2 \| 2 \| 105 \| 20 \| 127 \| Phage baseplate outer wedge protein (Acidic lysozyme), putative \| Phage baseplate outer wedge protein (Acidic lysozyme), putative; afdb-uniprot50 \| AF-A0A2D3VZ43-F1-MODEL\_V4 \| 1.0 \| 1.866e-06 \| 240 \| 0.379 \| 87 \| 50 \| 1 \| 19 \| 105 \| 8 \| 90 \| Baseplate assembly protein \| Baseplate assembly protein; afdb-uniprot50 \| AF-A0A7W7AUQ8-F1-MODEL\_V4 \| 1.0 \| 9.91e-06 \| 240 \| 0.252 \| 91 \| 65 \| 3 \| 17 \| 105 \| 7 \| 96 \| GPW\_gp25 domain-containing protein \| GPW\_gp25 domain-containing protein; afdb-uniprot50 \| AF-A0A235IXZ5-F1-MODEL\_V4 \| 1.0 \| 4.436e-06 \| 240 \| 0.207 \| 106 \| 81 \| 3 \| 2 \| 105 \| 21 \| 125 \| GPW\_gp25 domain-containing protein \| GPW\_gp25 domain-containing protein; afdb-uniprot50 \| AF-A0A2D5SX85-F1-MODEL\_V4 \| 1.0 \| 4.17e-06 \| 240 \| 0.179 \| 106 \| 85 \| 1 \| 3 \| 106 \| 29 \| 134 \| GPW\_gp25 domain-containing protein \| GPW\_gp25 domain-containing protein; afdb-uniprot50 \| AF-A0A4R7CKT7-F1-MODEL\_V4 \| 1.0 \| 1.985e-06 \| 240 \| 0.203 \| 108 \| 83 \| 2 \| 2 \| 106 \| 24 \| 131 \| GPW\_gp25 domain-containing protein \| GPW\_gp25 domain-containing protein; afdb-uniprot50 \| AF-A0A2E0HYC8-F1-MODEL\_V4 \| 1.0 \| 2.705e-06 \| 240 \| 0.174 \| 109 \| 86 \| 3 \| 3 \| 107 \| 31 \| 139 \| GPW\_gp25 domain-containing protein \| GPW\_gp25 domain-containing protein; afdb-uniprot50 \| AF-A0A369UQV3-F1-MODEL\_V4 \| 1.0 \| 5.68e-06 \| 239 \| 0.216 \| 106 \| 80 \| 3 \| 2 \| 105 \| 9 \| 113 \| GPW\_gp25 domain-containing protein \| GPW\_gp25 domain-containing protein; afdb-uniprot50 \| AF-A0A4Q3AGJ7-F1-MODEL\_V4 \| 1.0 \| 8.757e-06 \| 239 \| 0.2 \| 110 \| 81 \| 3 \| 1 \| 105 \| 7 \| 114 \| GPW\_gp25 domain-containing protein \| GPW\_gp25 domain-containing protein; afdb-uniprot50 \| AF-A0A661I6D3-F1-MODEL\_V4 \| 1.0 \| 6.838e-06 \| 239 \| 0.269 \| 104 \| 69 \| 2 \| 1 \| 97 \| 8 \| 111 \| Baseplate protein \| Baseplate protein; afdb-uniprot50 \| AF-A0A2S8QG71-F1-MODEL\_V4 \| 1.0 \| 1.985e-06 \| 239 \| 0.214 \| 112 \| 84 \| 3 \| 2 \| 110 \| 18 \| 128 \| Phage baseplate protein \| Phage baseplate protein; afdb-uniprot50 \| AF-A0A7R7DSV2-F1-MODEL\_V4 \| 1.0 \| 8.886e-07 \| 239 \| 0.294 \| 112 \| 72 \| 4 \| 2 \| 107 \| 14 \| 124 \| GPW\_gp25 domain-containing protein \| GPW\_gp25 domain-containing protein; afdb-uniprot50 \| AF-A0A1V5KBP1-F1-MODEL\_V4 \| 1.0 \| 2.39e-06 \| 239 \| 0.223 \| 121 \| 80 \| 4 \| 1 \| 107 \| 8 \| 128 \| Uncharacterized protein \| Uncharacterized protein; afdb-uniprot50 \| AF-A0A4R8CYG7-F1-MODEL\_V4 \| 1.0 \| 4.17e-06 \| 239 \| 0.242 \| 107 \| 78 \| 2 \| 2 \| 105 \| 24 \| 130 \| GPW\_gp25 domain-containing protein \| GPW\_gp25 domain-containing protein; afdb-uniprot50 \| AF-A0A1Z9KI13-F1-MODEL\_V4 \| 1.0 \| 6.043e-06 \| 239 \| 0.166 \| 108 \| 87 \| 2 \| 3 \| 107 \| 34 \| 141 \| GPW\_gp25 domain-containing protein \| GPW\_gp25 domain-containing protein; afdb-uniprot50 \| AF-A0A136LBR9-F1-MODEL\_V4 \| 1.0 \| 2.112e-06 \| 239 \| 0.201 \| 109 \| 83 \| 2 \| 2 \| 106 \| 26 \| 134 \| Phage baseplate assembly protein W \| Phage baseplate assembly protein W; afdb-uniprot50 \| AF-A0A4P2Q2Y6-F1-MODEL\_V4 \| 1.0 \| 7.739e-06 \| 238 \| 0.28 \| 114 \| 73 \| 4 \| 1 \| 106 \| 1 \| 113 \| GPW\_gp25 domain-containing protein \| GPW\_gp25 domain-containing protein; afdb-uniprot50 \| AF-A0A1I2LH68-F1-MODEL\_V4 \| 1.0 \| 5.02e-06 \| 238 \| 0.216 \| 120 \| 82 \| 5 \| 1 \| 109 \| 1 \| 119 \| GPW\_gp25 domain-containing protein \| GPW\_gp25 domain-containing protein; afdb-uniprot50 \| AF-A0A7Y5LM71-F1-MODEL\_V4 \| 1.0 \| 3.256e-06 \| 238 \| 0.302 \| 109 \| 69 \| 5 \| 2 \| 105 \| 14 \| 120 \| GPW/gp25 family protein \| GPW/gp25 family protein; afdb-uniprot50 \| AF-A0A1X0D4T4-F1-MODEL\_V4 \| 1.0 \| 3.685e-06 \| 238 \| 0.241 \| 112 \| 81 \| 4 \| 2 \| 111 \| 20 \| 129 \| GPW\_gp25 domain-containing protein \| GPW\_gp25 domain-containing protein; afdb-uniprot50 \| AF-W3ZZ38-F1-MODEL\_V4 \| 1.0 \| 1.07e-06 \| 238 \| 0.267 \| 116 \| 74 \| 5 \| 2 \| 108 \| 18 \| 131 \| GPW/gp25 family protein \| GPW/gp25 family protein; afdb-uniprot50 \| AF-A0A1G5HAB2-F1-MODEL\_V4 \| 1.0 \| 1.457e-06 \| 238 \| 0.231 \| 108 \| 78 \| 4 \| 2 \| 105 \| 14 \| 120 \| GPW\_gp25 domain-containing protein \| GPW\_gp25 domain-containing protein; afdb-uniprot50 \| AF-A0A2W6NE62-F1-MODEL\_V4 \| 1.0 \| 2.705e-06 \| 237 \| 0.301 \| 93 \| 59 \| 2 \| 12 \| 104 \| 4 \| 90 \| Baseplate assembly protein \| Baseplate assembly protein; afdb-uniprot50 \| AF-A0A399IS01-F1-MODEL\_V4 \| 1.0 \| 3.061e-06 \| 237 \| 0.192 \| 109 \| 83 \| 4 \| 2 \| 107 \| 11 \| 117 \| DUF2634 domain-containing protein \| DUF2634 domain-containing protein; afdb-uniprot50 \| AF-A0A2D8ESE3-F1-MODEL\_V4 \| 1.0 \| 6.428e-06 \| 237 \| 0.216 \| 106 \| 80 \| 2 \| 3 \| 105 \| 17 \| 122 \| GPW\_gp25 domain-containing protein \| GPW\_gp25 domain-containing protein; afdb-uniprot50 \| AF-A0A1Q3WGE0-F1-MODEL\_V4 \| 1.0 \| 2.542e-06 \| 237 \| 0.212 \| 108 \| 81 \| 3 \| 2 \| 105 \| 16 \| 123 \| GPW\_gp25 domain-containing protein \| GPW\_gp25 domain-containing protein; afdb-uniprot50 \| AF-A0A1H7PQT0-F1-MODEL\_V4 \| 1.0 \| 3.061e-06 \| 237 \| 0.194 \| 108 \| 83 \| 2 \| 2 \| 105 \| 16 \| 123 \| GPW\_gp25 domain-containing protein \| GPW\_gp25 domain-containing protein; afdb-uniprot50 \| AF-A0A2L2XJP6-F1-MODEL\_V4 \| 1.0 \| 3.256e-06 \| 236 \| 0.212 \| 108 \| 80 \| 4 \| 2 \| 106 \| 16 \| 121 \| GPW\_gp25 domain-containing protein \| GPW\_gp25 domain-containing protein; afdb-uniprot50 \| AF-C0BXW5-F1-MODEL\_V4 \| 1.0 \| 1.866e-06 \| 236 \| 0.268 \| 108 \| 74 \| 4 \| 2 \| 105 \| 15 \| 121 \| Putative lysozyme \| Putative lysozyme; afdb-uniprot50 \| AF-A0A7W1PHH8-F1-MODEL\_V4 \| 1.0 \| 1.288e-06 \| 236 \| 0.219 \| 114 \| 80 \| 4 \| 2 \| 107 \| 15 \| 127 \| GPW/gp25 family protein \| GPW/gp25 family protein; afdb-uniprot50 \| AF-A0A353G6I0-F1-MODEL\_V4 \| 1.0 \| 1.649e-06 \| 236 \| 0.25 \| 120 \| 77 \| 4 \| 1 \| 111 \| 16 \| 131 \| Baseplate \| Baseplate; afdb-uniprot50 \| AF-A0A661CBH9-F1-MODEL\_V4 \| 1.0 \| 3.685e-06 \| 236 \| 0.225 \| 111 \| 81 \| 2 \| 4 \| 109 \| 27 \| 137 \| GPW\_gp25 domain-containing protein \| GPW\_gp25 domain-containing protein; afdb-uniprot50 \| AF-A0A2E6C478-F1-MODEL\_V4 \| 1.0 \| 5.68e-06 \| 236 \| 0.242 \| 107 \| 78 \| 2 \| 3 \| 106 \| 27 \| 133 \| GPW\_gp25 domain-containing protein \| GPW\_gp25 domain-containing protein; afdb-uniprot50 \| AF-A0A372IVD6-F1-MODEL\_V4 \| 1.0 \| 5.02e-06 \| 236 \| 0.323 \| 99 \| 64 \| 2 \| 2 \| 97 \| 14 \| 112 \| Baseplate protein \| Baseplate protein; afdb-uniprot50 \| AF-A0A6L3Z3G6-F1-MODEL\_V4 \| 1.0 \| 2.247e-06 \| 236 \| 0.294 \| 119 \| 64 \| 8 \| 1 \| 108 \| 43 \| 152 \| GPW\_gp25 domain-containing protein \| GPW\_gp25 domain-containing protein; afdb-uniprot50 \| AF-A0A2E8DY83-F1-MODEL\_V4 \| 1.0 \| 4.719e-06 \| 236 \| 0.18 \| 105 \| 84 \| 2 \| 3 \| 105 \| 40 \| 144 \| GPW\_gp25 domain-containing protein \| GPW\_gp25 domain-containing protein; afdb-uniprot50 \| AF-A0A3S3QZP6-F1-MODEL\_V4 \| 1.0 \| 4.371e-05 \| 235 \| 0.27 \| 85 \| 62 \| 0 \| 1 \| 85 \| 3 \| 87 \| GPW\_gp25 domain-containing protein \| GPW\_gp25 domain-containing protein; afdb-uniprot50 \| AF-A0A832Z8E4-F1-MODEL\_V4 \| 1.0 \| 4.17e-06 \| 235 \| 0.222 \| 108 \| 81 \| 2 \| 1 \| 107 \| 11 \| 116 \| Uncharacterized protein \| Uncharacterized protein; afdb-uniprot50 \| AF-A0A7Y5NVN2-F1-MODEL\_V4 \| 1.0 \| 3.685e-06 \| 235 \| 0.191 \| 120 \| 87 \| 5 \| 1 \| 111 \| 10 \| 128 \| Uncharacterized protein \| Uncharacterized protein; afdb-uniprot50 \| AF-A0A553JK30-F1-MODEL\_V4 \| 1.0 \| 2.705e-06 \| 235 \| 0.182 \| 115 \| 86 \| 4 \| 2 \| 109 \| 11 \| 124 \| GPW/gp25 family protein \| GPW/gp25 family protein; afdb-uniprot50 \| AF-A0A6I2YZG1-F1-MODEL\_V4 \| 1.0 \| 8.232e-06 \| 235 \| 0.245 \| 106 \| 78 \| 2 \| 2 \| 105 \| 15 \| 120 \| Baseplate protein \| Baseplate protein; afdb-uniprot50 \| AF-A0A7D7VPT3-F1-MODEL\_V4 \| 1.0 \| 3.464e-06 \| 235 \| 0.222 \| 108 \| 80 \| 2 \| 2 \| 105 \| 16 \| 123 \| GPW\_gp25 domain-containing protein \| GPW\_gp25 domain-containing protein; afdb-uniprot50 \| AF-A0A661HHD5-F1-MODEL\_V4 \| 1.0 \| 1.985e-06 \| 235 \| 0.19 \| 110 \| 83 \| 3 \| 1 \| 106 \| 19 \| 126 \| GPW\_gp25 domain-containing protein \| GPW\_gp25 domain-containing protein; afdb-uniprot50 \| AF-A0A832UUD0-F1-MODEL\_V4 \| 1.0 \| 5.68e-06 \| 235 \| 0.175 \| 108 \| 84 \| 4 \| 3 \| 106 \| 31 \| 137 \| Uncharacterized protein \| Uncharacterized protein; afdb-uniprot50 \| AF-A0A2E9JHW6-F1-MODEL\_V4 \| 1.0 \| 5.02e-06 \| 235 \| 0.201 \| 109 \| 83 \| 3 \| 3 \| 107 \| 32 \| 140 \| GPW\_gp25 domain-containing protein \| GPW\_gp25 domain-containing protein; afdb-uniprot50 \| AF-A0A4Y8PTA1-F1-MODEL\_V4 \| 1.0 \| 3.464e-06 \| 235 \| 0.33 \| 106 \| 67 \| 2 \| 4 \| 105 \| 32 \| 137 \| GPW\_gp25 domain-containing protein \| GPW\_gp25 domain-containing protein; afdb-uniprot50 \| AF-Q7MB49-F1-MODEL\_V4 \| 1.0 \| 6.043e-06 \| 235 \| 0.214 \| 98 \| 75 \| 1 \| 10 \| 105 \| 126 \| 223 \| GPW\_gp25 domain-containing protein \| GPW\_gp25 domain-containing protein; afdb-uniprot50 \| AF-M7XD14-F1-MODEL\_V4 \| 1.0 \| 8.757e-06 \| 234 \| 0.263 \| 91 \| 65 \| 1 \| 17 \| 105 \| 7 \| 97 \| GPW/gp25 family protein \| GPW/gp25 family protein; afdb-uniprot50 \| AF-A0A419A4P0-F1-MODEL\_V4 \| 1.0 \| 2.247e-06 \| 234 \| 0.271 \| 107 \| 73 \| 4 \| 2 \| 105 \| 8 \| 112 \| GPW\_gp25 domain-containing protein \| GPW\_gp25 domain-containing protein; afdb-uniprot50 \| AF-A0A1Y3QHP6-F1-MODEL\_V4 \| 1.0 \| 2.39e-06 \| 234 \| 0.207 \| 106 \| 80 \| 3 \| 3 \| 105 \| 2 \| 106 \| Uncharacterized protein \| Uncharacterized protein; afdb-uniprot50 \| AF-A0A7T7L1C1-F1-MODEL\_V4 \| 1.0 \| 3.739e-07 \| 234 \| 0.263 \| 110 \| 77 \| 2 \| 2 \| 107 \| 15 \| 124 \| GPW/gp25 family protein \| GPW/gp25 family protein; afdb-uniprot50 \| AF-A0A1J0VCH8-F1-MODEL\_V4 \| 1.0 \| 5.34e-06 \| 234 \| 0.228 \| 114 \| 78 \| 4 \| 1 \| 105 \| 8 \| 120 \| GPW\_gp25 domain-containing protein \| GPW\_gp25 domain-containing protein; afdb-uniprot50 \| AF-A0A7Y9DPZ0-F1-MODEL\_V4 \| 1.0 \| 4.436e-06 \| 234 \| 0.24 \| 108 \| 77 \| 4 \| 2 \| 106 \| 22 \| 127 \| Uncharacterized protein \| Uncharacterized protein; afdb-uniprot50 \| AF-A0A4V3QZC8-F1-MODEL\_V4 \| 1.0 \| 2.705e-06 \| 234 \| 0.256 \| 109 \| 77 \| 3 \| 1 \| 106 \| 23 \| 130 \| Baseplate assembly protein \| Baseplate assembly protein; afdb-uniprot50 \| AF-A0A420YZC3-F1-MODEL\_V4 \| 1.0 \| 3.92e-06 \| 234 \| 0.16 \| 112 \| 86 \| 3 \| 2 \| 105 \| 11 \| 122 \| Uncharacterized protein \| Uncharacterized protein; afdb-uniprot50 \| AF-A0A3C1ALA6-F1-MODEL\_V4 \| 1.0 \| 1.193e-05 \| 233 \| 0.363 \| 77 \| 47 \| 1 \| 31 \| 107 \| 1 \| 75 \| Phage baseplate protein \| Phage baseplate protein; afdb-uniprot50 \| AF-A0A7C3GT18-F1-MODEL\_V4 \| 1.0 \| 6.838e-06 \| 233 \| 0.261 \| 107 \| 76 \| 3 \| 2 \| 106 \| 7 \| 112 \| GPW\_gp25 domain-containing protein \| GPW\_gp25 domain-containing protein; afdb-uniprot50 \| AF-A0A3S4U5L3-F1-MODEL\_V4 \| 1.0 \| 4.501e-07 \| 233 \| 0.3 \| 113 \| 70 \| 2 \| 1 \| 107 \| 8 \| 117 \| Baseplate assembly protein W \| Baseplate assembly protein W; afdb-uniprot50 \| AF-A0A1Z9B2C4-F1-MODEL\_V4 \| 1.0 \| 6.043e-06 \| 233 \| 0.188 \| 101 \| 81 \| 1 \| 5 \| 105 \| 12 \| 111 \| Uncharacterized protein \| Uncharacterized protein; afdb-uniprot50 \| AF-A0A1B1G1R7-F1-MODEL\_V4 \| 1.0 \| 9.316e-06 \| 233 \| 0.162 \| 111 \| 88 \| 2 \| 2 \| 107 \| 11 \| 121 \| GPW/gp25 family protein \| GPW/gp25 family protein; afdb-uniprot50 \| AF-A0A2S3UMZ3-F1-MODEL\_V4 \| 1.0 \| 1.193e-05 \| 233 \| 0.201 \| 114 \| 83 \| 4 \| 1 \| 107 \| 12 \| 124 \| Phage baseplate assembly protein W \| Phage baseplate assembly protein W; afdb-uniprot50 \| AF-A0A2E5YV38-F1-MODEL\_V4 \| 1.0 \| 4.719e-06 \| 233 \| 0.205 \| 112 \| 79 \| 4 \| 3 \| 107 \| 18 \| 126 \| GPW\_gp25 domain-containing protein \| GPW\_gp25 domain-containing protein; afdb-uniprot50 \| AF-A0A7Y6PTF8-F1-MODEL\_V4 \| 1.0 \| 8.886e-07 \| 233 \| 0.266 \| 109 \| 73 \| 3 \| 2 \| 105 \| 16 \| 122 \| GPW/gp25 family protein \| GPW/gp25 family protein; afdb-uniprot50 \| AF-A0A383EU63-F1-MODEL\_V4 \| 1.0 \| 8.757e-06 \| 233 \| 0.175 \| 108 \| 86 \| 2 \| 3 \| 107 \| 23 \| 130 \| GPW\_gp25 domain-containing protein \| GPW\_gp25 domain-containing protein; afdb-uniprot50 \| AF-A0A7X5H4C3-F1-MODEL\_V4 \| 1.0 \| 2.877e-06 \| 233 \| 0.186 \| 107 \| 84 \| 2 \| 2 \| 105 \| 18 \| 124 \| Phage baseplate protein \| Phage baseplate protein; afdb-uniprot50 \| AF-A0A562GHU8-F1-MODEL\_V4 \| 1.0 \| 2.877e-06 \| 233 \| 0.252 \| 107 \| 74 \| 5 \| 2 \| 105 \| 27 \| 130 \| Phage baseplate assembly protein W \| Phage baseplate assembly protein W; afdb-uniprot50 \| AF-A0A2D8I5K2-F1-MODEL\_V4 \| 1.0 \| 1.269e-05 \| 233 \| 0.173 \| 115 \| 87 \| 2 \| 1 \| 107 \| 24 \| 138 \| GPW\_gp25 domain-containing protein \| GPW\_gp25 domain-containing protein; afdb-uniprot50 \| AF-A0A4P5VFE5-F1-MODEL\_V4 \| 1.0 \| 1.288e-06 \| 233 \| 0.245 \| 114 \| 79 \| 3 \| 1 \| 107 \| 34 \| 147 \| GPW\_gp25 domain-containing protein \| GPW\_gp25 domain-containing protein; afdb-uniprot50 \| AF-A0A833DBQ1-F1-MODEL\_V4 \| 1.0 \| 5.68e-06 \| 232 \| 0.181 \| 116 \| 83 \| 4 \| 3 \| 108 \| 33 \| 146 \| Uncharacterized protein \| Uncharacterized protein; afdb-uniprot50 \| AF-A0A2D7BZP7-F1-MODEL\_V4 \| 1.0 \| 3.685e-06 \| 232 \| 0.148 \| 108 \| 87 \| 3 \| 3 \| 105 \| 36 \| 143 \| GPW\_gp25 domain-containing protein \| GPW\_gp25 domain-containing protein; afdb-uniprot50 \| AF-A0A2T6NFS1-F1-MODEL\_V4 \| 1.0 \| 9.316e-06 \| 232 \| 0.252 \| 111 \| 76 \| 3 \| 2 \| 105 \| 63 \| 173 \| GPW\_gp25 domain-containing protein \| GPW\_gp25 domain-containing protein; afdb-uniprot50 \| AF-A0A7L6A4Z7-F1-MODEL\_V4 \| 1.0 \| 3.256e-06 \| 232 \| 0.198 \| 106 \| 83 \| 1 \| 2 \| 105 \| 20 \| 125 \| GPW/gp25 family protein \| GPW/gp25 family protein; afdb-uniprot50 \| AF-A0A4R4TT05-F1-MODEL\_V4 \| 1.0 \| 7.275e-06 \| 231 \| 0.216 \| 111 \| 84 \| 3 \| 2 \| 110 \| 18 \| 127 \| GPW\_gp25 domain-containing protein \| GPW\_gp25 domain-containing protein; afdb-uniprot50 \| AF-A0A7Y4XEU9-F1-MODEL\_V4 \| 1.0 \| 1.866e-06 \| 231 \| 0.22 \| 118 \| 75 \| 4 \| 5 \| 107 \| 14 \| 129 \| GPW\_gp25 domain-containing protein \| GPW\_gp25 domain-containing protein; afdb-uniprot50 \| AF-A0A1Y4IQ66-F1-MODEL\_V4 \| 1.0 \| 6.428e-06 \| 231 \| 0.247 \| 109 \| 76 \| 4 \| 2 \| 105 \| 22 \| 129 \| GPW\_gp25 domain-containing protein \| GPW\_gp25 domain-containing protein; afdb-uniprot50 \| AF-A6FZ69-F1-MODEL\_V4 \| 1.0 \| 5.34e-06 \| 231 \| 0.203 \| 108 \| 82 \| 2 \| 2 \| 105 \| 24 \| 131 \| GPW/gp25 \| GPW/gp25; afdb-uniprot50 \| AF-A0A1R3WBS2-F1-MODEL\_V4 \| 1.0 \| 1.866e-06 \| 231 \| 0.177 \| 107 \| 84 \| 2 \| 3 \| 105 \| 23 \| 129 \| GPW\_gp25 domain-containing protein \| GPW\_gp25 domain-containing protein; afdb-uniprot50 \| AF-A0A3A8PSW3-F1-MODEL\_V4 \| 1.0 \| 9.316e-06 \| 231 \| 0.214 \| 107 \| 81 \| 2 \| 2 \| 105 \| 26 \| 132 \| GPW\_gp25 domain-containing protein \| GPW\_gp25 domain-containing protein; afdb-uniprot50 \| AF-A0A3N7GF44-F1-MODEL\_V4 \| 1.0 \| 1.625e-05 \| 231 \| 0.226 \| 119 \| 81 \| 5 \| 1 \| 110 \| 47 \| 163 \| GPW\_gp25 domain-containing protein \| GPW\_gp25 domain-containing protein; afdb-uniprot50 \| AF-A0A825MSY2-F1-MODEL\_V4 \| 1.0 \| 5.34e-06 \| 230 \| 0.272 \| 99 \| 63 \| 3 \| 17 \| 107 \| 21 \| 118 \| GPW/gp25 family protein \| GPW/gp25 family protein; afdb-uniprot50 \| AF-A0A2D7G6P2-F1-MODEL\_V4 \| 1.0 \| 1.35e-05 \| 230 \| 0.181 \| 110 \| 87 \| 2 \| 5 \| 112 \| 25 \| 133 \| GPW\_gp25 domain-containing protein \| GPW\_gp25 domain-containing protein; afdb-uniprot50 \| AF-A0A4P7CYP9-F1-MODEL\_V4 \| 1.0 \| 9.316e-06 \| 230 \| 0.261 \| 107 \| 74 \| 4 \| 2 \| 105 \| 27 \| 131 \| GPW\_gp25 domain-containing protein \| GPW\_gp25 domain-containing protein; afdb-uniprot50 \| AF-A0A848G1D1-F1-MODEL\_V4 \| 1.0 \| 6.428e-06 \| 230 \| 0.16 \| 106 \| 87 \| 1 \| 2 \| 105 \| 16 \| 121 \| GPW/gp25 family protein \| GPW/gp25 family protein; afdb-uniprot50 \| AF-A0A2D8EMQ8-F1-MODEL\_V4 \| 1.0 \| 2.705e-06 \| 230 \| 0.166 \| 120 \| 87 \| 4 \| 1 \| 107 \| 28 \| 147 \| GPW\_gp25 domain-containing protein \| GPW\_gp25 domain-containing protein; afdb-uniprot50 \| AF-A0A6I6N486-F1-MODEL\_V4 \| 1.0 \| 6.043e-06 \| 230 \| 0.28 \| 107 \| 72 \| 4 \| 2 \| 105 \| 43 \| 147 \| GPW\_gp25 domain-containing protein \| GPW\_gp25 domain-containing protein; afdb-uniprot50 \| AF-A0A2E6W7H5-F1-MODEL\_V4 \| 1.0 \| 7.275e-06 \| 229 \| 0.194 \| 113 \| 81 \| 3 \| 7 \| 110 \| 2 \| 113 \| GPW\_gp25 domain-containing protein \| GPW\_gp25 domain-containing protein; afdb-uniprot50 \| AF-A4YW46-F1-MODEL\_V4 \| 1.0 \| 1.121e-05 \| 229 \| 0.214 \| 107 \| 81 \| 3 \| 2 \| 106 \| 8 \| 113 \| GPW\_gp25 domain-containing protein \| GPW\_gp25 domain-containing protein; afdb-uniprot50 \| AF-A0A7X5DY26-F1-MODEL\_V4 \| 1.0 \| 3.061e-06 \| 229 \| 0.222 \| 108 \| 79 \| 4 \| 2 \| 105 \| 8 \| 114 \| Baseplate protein \| Baseplate protein; afdb-uniprot50 \| AF-A0A7W3LYA6-F1-MODEL\_V4 \| 1.0 \| 3.92e-06 \| 229 \| 0.213 \| 117 \| 80 \| 6 \| 2 \| 108 \| 7 \| 121 \| GPW\_gp25 domain-containing protein \| GPW\_gp25 domain-containing protein; afdb-uniprot50 \| AF-A0A535AQ84-F1-MODEL\_V4 \| 1.0 \| 5.68e-06 \| 229 \| 0.3 \| 100 \| 65 \| 3 \| 2 \| 97 \| 4 \| 102 \| GPW/gp25 family protein \| GPW/gp25 family protein; afdb-uniprot50 \| AF-A0A524M6P3-F1-MODEL\_V4 \| 1.0 \| 7.275e-06 \| 229 \| 0.203 \| 113 \| 83 \| 4 \| 1 \| 107 \| 9 \| 120 \| Uncharacterized protein \| Uncharacterized protein; afdb-uniprot50 \| AF-N1ZFS0-F1-MODEL\_V4 \| 1.0 \| 1.138e-06 \| 229 \| 0.218 \| 110 \| 78 \| 3 \| 2 \| 105 \| 11 \| 118 \| GPW\_gp25 domain-containing protein \| GPW\_gp25 domain-containing protein; afdb-uniprot50 \| AF-A0A2N8ML17-F1-MODEL\_V4 \| 1.0 \| 1.138e-06 \| 229 \| 0.232 \| 125 \| 82 \| 4 \| 1 \| 111 \| 1 \| 125 \| GPW\_gp25 domain-containing protein \| GPW\_gp25 domain-containing protein; afdb-uniprot50 \| AF-A0A0Q9UE51-F1-MODEL\_V4 \| 1.0 \| 1.457e-06 \| 229 \| 0.228 \| 114 \| 79 \| 4 \| 3 \| 108 \| 14 \| 126 \| GPW\_gp25 domain-containing protein \| GPW\_gp25 domain-containing protein; afdb-uniprot50 \| AF-A0A2V7RA86-F1-MODEL\_V4 \| 1.0 \| 3.256e-06 \| 229 \| 0.266 \| 109 \| 75 \| 4 \| 1 \| 105 \| 16 \| 123 \| Baseplate assembly protein \| Baseplate assembly protein; afdb-uniprot50 \| AF-A0A2D8UQ71-F1-MODEL\_V4 \| 1.0 \| 8.757e-06 \| 229 \| 0.24 \| 108 \| 79 \| 2 \| 2 \| 106 \| 24 \| 131 \| GPW\_gp25 domain-containing protein \| GPW\_gp25 domain-containing protein; afdb-uniprot50 \| AF-A0A1H5RLH3-F1-MODEL\_V4 \| 1.0 \| 4.17e-06 \| 229 \| 0.185 \| 108 \| 83 \| 4 \| 2 \| 105 \| 16 \| 122 \| GPW\_gp25 domain-containing protein \| GPW\_gp25 domain-containing protein; afdb-uniprot50 \| AF-A0A1G8B3G6-F1-MODEL\_V4 \| 1.0 \| 3.256e-06 \| 229 \| 0.2 \| 110 \| 81 \| 6 \| 1 \| 107 \| 97 \| 202 \| Phage baseplate assembly protein W \| Phage baseplate assembly protein W; afdb-uniprot50 \| AF-A0A7H4NMM7-F1-MODEL\_V4 \| 1.0 \| 4.109e-05 \| 228 \| 0.478 \| 69 \| 36 \| 0 \| 3 \| 71 \| 6 \| 74 \| Baseplate assembly protein W \| Baseplate assembly protein W; afdb-uniprot50 \| AF-A0A1M6UMM3-F1-MODEL\_V4 \| 1.0 \| 2.081e-05 \| 228 \| 0.211 \| 90 \| 70 \| 1 \| 17 \| 105 \| 7 \| 96 \| GPW\_gp25 domain-containing protein \| GPW\_gp25 domain-containing protein; afdb-uniprot50 \| AF-A0A4Q7FYR9-F1-MODEL\_V4 \| 1.0 \| 1.649e-06 \| 228 \| 0.252 \| 119 \| 67 \| 3 \| 12 \| 108 \| 2 \| 120 \| Baseplate assembly protein \| Baseplate assembly protein; afdb-uniprot50 \| AF-A0A497N9T7-F1-MODEL\_V4 \| 1.0 \| 1.193e-05 \| 228 \| 0.225 \| 111 \| 82 \| 4 \| 3 \| 110 \| 18 \| 127 \| Uncharacterized protein \| Uncharacterized protein; afdb-uniprot50 \| AF-A0A2E2ZB32-F1-MODEL\_V4 \| 1.0 \| 1.436e-05 \| 228 \| 0.155 \| 109 \| 86 \| 3 \| 1 \| 105 \| 29 \| 135 \| GPW\_gp25 domain-containing protein \| GPW\_gp25 domain-containing protein; afdb-uniprot50 \| AF-A0A7J5DNJ6-F1-MODEL\_V4 \| 1.0 \| 2.081e-05 \| 227 \| 0.282 \| 85 \| 59 \| 2 \| 22 \| 105 \| 2 \| 85 \| GPW/gp25 family protein \| GPW/gp25 family protein; afdb-uniprot50 \| AF-A0A1I3ATS5-F1-MODEL\_V4 \| 1.0 \| 1.625e-05 \| 227 \| 0.254 \| 114 \| 75 \| 5 \| 1 \| 105 \| 1 \| 113 \| GPW\_gp25 domain-containing protein \| GPW\_gp25 domain-containing protein; afdb-uniprot50 \| AF-A0A094QGT3-F1-MODEL\_V4 \| 1.0 \| 1.121e-05 \| 227 \| 0.275 \| 98 \| 69 \| 2 \| 2 \| 97 \| 14 \| 111 \| GPW/gp25 family protein \| GPW/gp25 family protein; afdb-uniprot50 \| AF-A0A3A9G8M7-F1-MODEL\_V4 \| 1.0 \| 1.457e-06 \| 227 \| 0.247 \| 109 \| 75 \| 3 \| 2 \| 105 \| 15 \| 121 \| Baseplate protein \| Baseplate protein; afdb-uniprot50 \| AF-A0A7X5D8Q1-F1-MODEL\_V4 \| 1.0 \| 1.649e-06 \| 227 \| 0.211 \| 109 \| 80 \| 3 \| 2 \| 105 \| 44 \| 151 \| GPW\_gp25 domain-containing protein \| GPW\_gp25 domain-containing protein; afdb-uniprot50 \| AF-A0A5D4S1J9-F1-MODEL\_V4 \| 1.0 \| 3.92e-06 \| 226 \| 0.163 \| 116 \| 83 \| 5 \| 2 \| 107 \| 3 \| 114 \| DUF2634 domain-containing protein \| DUF2634 domain-containing protein; afdb-uniprot50 \| AF-A0A1Q4ZX73-F1-MODEL\_V4 \| 1.0 \| 7.739e-06 \| 226 \| 0.242 \| 107 \| 76 \| 4 \| 2 \| 105 \| 9 \| 113 \| GPW\_gp25 domain-containing protein \| GPW\_gp25 domain-containing protein; afdb-uniprot50 \| AF-A0A1V5V4U5-F1-MODEL\_V4 \| 1.0 \| 2.112e-06 \| 226 \| 0.23 \| 117 \| 79 \| 4 \| 2 \| 108 \| 12 \| 127 \| Gene 25-like lysozyme \| Gene 25-like lysozyme; afdb-uniprot50 \| AF-A0A0F5JZG5-F1-MODEL\_V4 \| 1.0 \| 5.02e-06 \| 226 \| 0.203 \| 108 \| 83 \| 2 \| 2 \| 106 \| 19 \| 126 \| GPW\_gp25 domain-containing protein \| GPW\_gp25 domain-containing protein; afdb-uniprot50 \| AF-A0A3R7S1T8-F1-MODEL\_V4 \| 1.0 \| 5.68e-06 \| 226 \| 0.203 \| 108 \| 82 \| 3 \| 3 \| 107 \| 20 \| 126 \| GPW\_gp25 domain-containing protein \| GPW\_gp25 domain-containing protein; afdb-uniprot50 \| AF-A0A1Z9QID3-F1-MODEL\_V4 \| 1.0 \| 1.054e-05 \| 225 \| 0.18 \| 105 \| 82 \| 3 \| 5 \| 107 \| 12 \| 114 \| Uncharacterized protein \| Uncharacterized protein; afdb-uniprot50 \| AF-A0A3B1EJD1-F1-MODEL\_V4 \| 1.0 \| 8.757e-06 \| 225 \| 0.263 \| 110 \| 76 \| 4 \| 2 \| 107 \| 12 \| 120 \| Mlr6561 protein \| Mlr6561 protein; afdb-uniprot50 \| AF-A0A7J4UPF6-F1-MODEL\_V4 \| 1.0 \| 8.232e-06 \| 225 \| 0.169 \| 106 \| 85 \| 2 \| 3 \| 105 \| 14 \| 119 \| Uncharacterized protein \| Uncharacterized protein; afdb-uniprot50 \| AF-A0A328FDP2-F1-MODEL\_V4 \| 1.0 \| 1.121e-05 \| 225 \| 0.252 \| 107 \| 78 \| 1 \| 1 \| 105 \| 23 \| 129 \| Baseplate assembly protein \| Baseplate assembly protein; afdb-uniprot50 \| AF-A0A2S8QIP7-F1-MODEL\_V4 \| 1.0 \| 4.436e-06 \| 225 \| 0.214 \| 107 \| 81 \| 2 \| 2 \| 105 \| 18 \| 124 \| Phage baseplate protein \| Phage baseplate protein; afdb-uniprot50 \| AF-A0A1Y3WKR7-F1-MODEL\_V4 \| 1.0 \| 6.428e-06 \| 225 \| 0.247 \| 109 \| 76 \| 4 \| 2 \| 105 \| 17 \| 124 \| GPW\_gp25 domain-containing protein \| GPW\_gp25 domain-containing protein; afdb-uniprot50 \| AF-A0A1Y4HZE3-F1-MODEL\_V4 \| 1.0 \| 6.838e-06 \| 225 \| 0.222 \| 108 \| 79 \| 4 \| 2 \| 105 \| 38 \| 144 \| GPW\_gp25 domain-containing protein \| GPW\_gp25 domain-containing protein; afdb-uniprot50 \| AF-F2K224-F1-MODEL\_V4 \| 1.0 \| 8.757e-06 \| 224 \| 0.247 \| 109 \| 76 \| 3 \| 3 \| 108 \| 2 \| 107 \| Uncharacterized protein \| Uncharacterized protein; afdb-uniprot50 \| AF-A0A158EG26-F1-MODEL\_V4 \| 1.0 \| 1.729e-05 \| 224 \| 0.209 \| 105 \| 81 \| 2 \| 2 \| 105 \| 8 \| 111 \| Gene 25-like lysozyme \| Gene 25-like lysozyme; afdb-uniprot50 \| AF-A0A4T0V9V5-F1-MODEL\_V4 \| 1.0 \| 5.68e-06 \| 224 \| 0.233 \| 107 \| 77 \| 4 \| 2 \| 105 \| 10 \| 114 \| GPW\_gp25 domain-containing protein \| GPW\_gp25 domain-containing protein; afdb-uniprot50 \| AF-A0A2E6W7U8-F1-MODEL\_V4 \| 1.0 \| 2.355e-05 \| 224 \| 0.155 \| 109 \| 86 \| 3 \| 1 \| 105 \| 6 \| 112 \| GPW\_gp25 domain-containing protein \| GPW\_gp25 domain-containing protein; afdb-uniprot50 \| AF-A0A2R4SX79-F1-MODEL\_V4 \| 1.0 \| 1.193e-05 \| 224 \| 0.235 \| 106 \| 78 \| 3 \| 2 \| 105 \| 15 \| 119 \| GPW\_gp25 domain-containing protein \| GPW\_gp25 domain-containing protein; afdb-uniprot50 \| AF-A0A2D6E1F5-F1-MODEL\_V4 \| 1.0 \| 6.428e-06 \| 224 \| 0.212 \| 113 \| 82 \| 2 \| 2 \| 107 \| 10 \| 122 \| GPW\_gp25 domain-containing protein \| GPW\_gp25 domain-containing protein; afdb-uniprot50 \| AF-A0A2D8ESJ7-F1-MODEL\_V4 \| 1.0 \| 1.436e-05 \| 224 \| 0.224 \| 107 \| 80 \| 2 \| 3 \| 106 \| 27 \| 133 \| GPW\_gp25 domain-containing protein \| GPW\_gp25 domain-containing protein; afdb-uniprot50 \| AF-A0A2E8NRW1-F1-MODEL\_V4 \| 1.0 \| 1.121e-05 \| 224 \| 0.192 \| 109 \| 82 \| 3 \| 1 \| 105 \| 11 \| 117 \| GPW\_gp25 domain-containing protein \| GPW\_gp25 domain-containing protein; afdb-uniprot50 \| AF-A0A1Q4RQU7-F1-MODEL\_V4 \| 1.0 \| 3.515e-07 \| 224 \| 0.342 \| 108 \| 63 \| 2 \| 2 \| 108 \| 21 \| 121 \| Baseplate protein \| Baseplate protein; afdb-uniprot50 \| AF-A0A376ZFR1-F1-MODEL\_V4 \| 1.0 \| 1.839e-05 \| 223 \| 0.366 \| 71 \| 45 \| 0 \| 35 \| 105 \| 1 \| 71 \| Baseplate assembly protein W \| Baseplate assembly protein W; afdb-uniprot50 \| AF-A0A1M7S3A9-F1-MODEL\_V4 \| 1.0 \| 1.625e-05 \| 223 \| 0.245 \| 106 \| 77 \| 3 \| 2 \| 105 \| 8 \| 112 \| GPW\_gp25 domain-containing protein \| GPW\_gp25 domain-containing protein; afdb-uniprot50 \| AF-A0A2R6EXZ2-F1-MODEL\_V4 \| 1.0 \| 1.193e-05 \| 223 \| 0.33 \| 100 \| 62 \| 3 \| 2 \| 97 \| 14 \| 112 \| Baseplate protein \| Baseplate protein; afdb-uniprot50 \| AF-A0A1M7TQV3-F1-MODEL\_V4 \| 1.0 \| 3.256e-06 \| 223 \| 0.234 \| 115 \| 77 \| 4 \| 1 \| 105 \| 11 \| 124 \| GPW\_gp25 domain-containing protein \| GPW\_gp25 domain-containing protein; afdb-uniprot50 \| AF-A0A6B2YTY6-F1-MODEL\_V4 \| 1.0 \| 1.528e-05 \| 223 \| 0.273 \| 106 \| 68 \| 3 \| 1 \| 97 \| 8 \| 113 \| GPW/gp25 family protein \| GPW/gp25 family protein; afdb-uniprot50 \| AF-A0A212K6S4-F1-MODEL\_V4 \| 1.0 \| 2.705e-06 \| 223 \| 0.205 \| 117 \| 79 \| 5 \| 1 \| 105 \| 13 \| 127 \| Uncharacterized protein \| Uncharacterized protein; afdb-uniprot50 \| AF-A0A437M149-F1-MODEL\_V4 \| 1.0 \| 1.729e-05 \| 223 \| 0.173 \| 121 \| 85 \| 5 \| 1 \| 111 \| 94 \| 209 \| Uncharacterized protein \| Uncharacterized protein; afdb-uniprot50 \| AF-A0A521S4V9-F1-MODEL\_V4 \| 1.0 \| 1.436e-05 \| 223 \| 0.245 \| 106 \| 77 \| 3 \| 2 \| 105 \| 188 \| 292 \| GPW\_gp25 domain-containing protein \| GPW\_gp25 domain-containing protein; afdb-uniprot50 \| AF-A0A511N0N2-F1-MODEL\_V4 \| 1.0 \| 6.428e-06 \| 222 \| 0.223 \| 112 \| 81 \| 2 \| 2 \| 107 \| 14 \| 125 \| GPW\_gp25 domain-containing protein \| GPW\_gp25 domain-containing protein; afdb-uniprot50 \| AF-A0A7Z9QJ22-F1-MODEL\_V4 \| 1.0 \| 1.436e-05 \| 222 \| 0.169 \| 106 \| 85 \| 2 \| 3 \| 105 \| 17 \| 122 \| GPW\_gp25 domain-containing protein \| GPW\_gp25 domain-containing protein; afdb-uniprot50 \| AF-A0A2D8C9G0-F1-MODEL\_V4 \| 1.0 \| 1.839e-05 \| 222 \| 0.165 \| 109 \| 86 \| 3 \| 3 \| 107 \| 20 \| 127 \| GPW\_gp25 domain-containing protein \| GPW\_gp25 domain-containing protein; afdb-uniprot50 \| AF-A0A7Y6UVS1-F1-MODEL\_V4 \| 1.0 \| 1.625e-05 \| 222 \| 0.306 \| 98 \| 66 \| 1 \| 2 \| 97 \| 20 \| 117 \| GPW/gp25 family protein \| GPW/gp25 family protein; afdb-uniprot50 \| AF-A0A5S9NBS3-F1-MODEL\_V4 \| 1.0 \| 1.054e-05 \| 222 \| 0.227 \| 110 \| 79 \| 3 \| 1 \| 105 \| 13 \| 121 \| Uncharacterized protein \| Uncharacterized protein; afdb-uniprot50 \| AF-A0A1Q4P0G2-F1-MODEL\_V4 \| 1.0 \| 9.316e-06 \| 222 \| 0.188 \| 106 \| 84 \| 1 \| 2 \| 105 \| 18 \| 123 \| GPW\_gp25 domain-containing protein \| GPW\_gp25 domain-containing protein; afdb-uniprot50 \| AF-A0A4R0P0Z5-F1-MODEL\_V4 \| 1.0 \| 8.757e-06 \| 222 \| 0.189 \| 111 \| 85 \| 2 \| 1 \| 106 \| 17 \| 127 \| GPW\_gp25 domain-containing protein \| GPW\_gp25 domain-containing protein; afdb-uniprot50 \| AF-A0A1W9Z4R0-F1-MODEL\_V4 \| 1.0 \| 3.016e-05 \| 221 \| 0.277 \| 90 \| 63 \| 2 \| 17 \| 105 \| 17 \| 105 \| GPW\_gp25 domain-containing protein \| GPW\_gp25 domain-containing protein; afdb-uniprot50 \| AF-A0A525HH40-F1-MODEL\_V4 \| 1.0 \| 1.35e-05 \| 221 \| 0.259 \| 108 \| 75 \| 4 \| 2 \| 106 \| 9 \| 114 \| GPW\_gp25 domain-containing protein \| GPW\_gp25 domain-containing protein; afdb-uniprot50 \| AF-A0A6J4Y8V7-F1-MODEL\_V4 \| 1.0 \| 1.436e-05 \| 221 \| 0.229 \| 109 \| 80 \| 4 \| 2 \| 107 \| 10 \| 117 \| GPW\_gp25 domain-containing protein \| GPW\_gp25 domain-containing protein; afdb-uniprot50 \| AF-A0A2D5MRJ6-F1-MODEL\_V4 \| 1.0 \| 2.836e-05 \| 221 \| 0.166 \| 108 \| 85 \| 3 \| 1 \| 105 \| 27 \| 132 \| GPW\_gp25 domain-containing protein \| GPW\_gp25 domain-containing protein; afdb-uniprot50 \| AF-A0A3C1NZC1-F1-MODEL\_V4 \| 1.0 \| 2.665e-05 \| 221 \| 0.219 \| 105 \| 74 \| 3 \| 1 \| 97 \| 8 \| 112 \| Baseplate protein \| Baseplate protein; afdb-uniprot50 \| AF-A0A846DAZ9-F1-MODEL\_V4 \| 1.0 \| 1.957e-05 \| 220 \| 0.302 \| 96 \| 66 \| 1 \| 2 \| 96 \| 14 \| 109 \| GPW/gp25 family protein \| GPW/gp25 family protein; afdb-uniprot50 \| AF-A0A800JQS2-F1-MODEL\_V4 \| 1.0 \| 4.719e-06 \| 220 \| 0.214 \| 107 \| 81 \| 1 \| 2 \| 105 \| 10 \| 116 \| GPW\_gp25 domain-containing protein \| GPW\_gp25 domain-containing protein; afdb-uniprot50 \| AF-A0A2A5GV64-F1-MODEL\_V4 \| 1.0 \| 2.214e-05 \| 220 \| 0.168 \| 113 \| 90 \| 3 \| 3 \| 112 \| 14 \| 125 \| GPW\_gp25 domain-containing protein \| GPW\_gp25 domain-containing protein; afdb-uniprot50 \| AF-A0A1V5QG84-F1-MODEL\_V4 \| 1.0 \| 8.232e-06 \| 220 \| 0.257 \| 105 \| 66 \| 3 \| 2 \| 97 \| 15 \| 116 \| Gene 25-like lysozyme \| Gene 25-like lysozyme; afdb-uniprot50 \| AF-F6DTE8-F1-MODEL\_V4 \| 1.0 \| 4.719e-06 \| 220 \| 0.284 \| 109 \| 73 \| 4 \| 1 \| 106 \| 30 \| 136 \| GPW/gp25 family protein \| GPW/gp25 family protein; afdb-uniprot50 \| AF-A0A1J5BI56-F1-MODEL\_V4 \| 1.0 \| 2.836e-05 \| 219 \| 0.268 \| 93 \| 63 \| 3 \| 16 \| 105 \| 9 \| 99 \| GPW\_gp25 domain-containing protein \| GPW\_gp25 domain-containing protein; afdb-uniprot50 \| AF-A0A382YZB8-F1-MODEL\_V4 \| 1.0 \| 1.121e-05 \| 219 \| 0.186 \| 107 \| 83 \| 3 \| 3 \| 105 \| 4 \| 110 \| GPW\_gp25 domain-containing protein \| GPW\_gp25 domain-containing protein; afdb-uniprot50 \| AF-A0A1I6DPB9-F1-MODEL\_V4 \| 1.0 \| 6.428e-06 \| 219 \| 0.212 \| 113 \| 82 \| 4 \| 2 \| 108 \| 12 \| 123 \| GPW\_gp25 domain-containing protein \| GPW\_gp25 domain-containing protein; afdb-uniprot50 \| AF-A0A2D6MIZ8-F1-MODEL\_V4 \| 1.0 \| 8.757e-06 \| 219 \| 0.188 \| 106 \| 83 \| 3 \| 3 \| 105 \| 14 \| 119 \| GPW\_gp25 domain-containing protein \| GPW\_gp25 domain-containing protein; afdb-uniprot50 \| AF-A0A2K3J5A4-F1-MODEL\_V4 \| 1.0 \| 2.214e-05 \| 219 \| 0.205 \| 107 \| 80 \| 3 \| 4 \| 106 \| 20 \| 125 \| GPW\_gp25 domain-containing protein \| GPW\_gp25 domain-containing protein; afdb-uniprot50 \| AF-U1KRE2-F1-MODEL\_V4 \| 1.0 \| 2.112e-06 \| 219 \| 0.182 \| 115 \| 82 \| 2 \| 3 \| 105 \| 8 \| 122 \| GPW/gp25 family protein \| GPW/gp25 family protein; afdb-uniprot50 \| AF-A0A2D4ZE95-F1-MODEL\_V4 \| 1.0 \| 1.957e-05 \| 219 \| 0.168 \| 107 \| 86 \| 2 \| 3 \| 106 \| 27 \| 133 \| GPW\_gp25 domain-containing protein \| GPW\_gp25 domain-containing protein; afdb-uniprot50 \| AF-A0A5F1HX77-F1-MODEL\_V4 \| 1.0 \| 1.121e-05 \| 219 \| 0.185 \| 108 \| 85 \| 2 \| 2 \| 106 \| 17 \| 124 \| GPW/gp25 family protein \| GPW/gp25 family protein; afdb-uniprot50 \| AF-A0A3R9V7X8-F1-MODEL\_V4 \| 1.0 \| 1.193e-05 \| 219 \| 0.31 \| 100 \| 64 \| 3 \| 2 \| 97 \| 32 \| 130 \| Baseplate protein \| Baseplate protein; afdb-uniprot50 \| AF-A0A661PMX4-F1-MODEL\_V4 \| 1.0 \| 1.729e-05 \| 218 \| 0.189 \| 116 \| 87 \| 4 \| 1 \| 112 \| 5 \| 117 \| GPW\_gp25 domain-containing protein \| GPW\_gp25 domain-containing protein; afdb-uniprot50 \| AF-A0A1V5BCJ8-F1-MODEL\_V4 \| 1.0 \| 2.081e-05 \| 218 \| 0.221 \| 113 \| 84 \| 4 \| 2 \| 112 \| 8 \| 118 \| Gene 25-like lysozyme \| Gene 25-like lysozyme; afdb-uniprot50 \| AF-B0T4J0-F1-MODEL\_V4 \| 1.0 \| 1.436e-05 \| 218 \| 0.176 \| 113 \| 87 \| 4 \| 2 \| 109 \| 9 \| 120 \| GPW\_gp25 domain-containing protein \| GPW\_gp25 domain-containing protein; afdb-uniprot50 \| AF-A0A117MMK7-F1-MODEL\_V4 \| 1.0 \| 2.355e-05 \| 218 \| 0.245 \| 106 \| 78 \| 2 \| 2 \| 107 \| 12 \| 115 \| GPW\_gp25 domain-containing protein \| GPW\_gp25 domain-containing protein; afdb-uniprot50 \| AF-A0A0K9NA34-F1-MODEL\_V4 \| 1.0 \| 1.006e-06 \| 218 \| 0.275 \| 109 \| 72 \| 3 \| 2 \| 105 \| 15 \| 121 \| Putative tail lysozyme \| Putative tail lysozyme; afdb-uniprot50 \| AF-A0A550H8T3-F1-MODEL\_V4 \| 1.0 \| 6.838e-06 \| 218 \| 0.206 \| 116 \| 80 \| 4 \| 4 \| 107 \| 16 \| 131 \| GPW\_gp25 domain-containing protein \| GPW\_gp25 domain-containing protein; afdb-uniprot50 \| AF-A0A7W7CIY7-F1-MODEL\_V4 \| 1.0 \| 1.436e-05 \| 218 \| 0.261 \| 107 \| 74 \| 4 \| 2 \| 105 \| 28 \| 132 \| Phage baseplate assembly protein W \| Phage baseplate assembly protein W; afdb-uniprot50 \| AF-A0A1Z8S7X6-F1-MODEL\_V4 \| 1.0 \| 6.838e-06 \| 218 \| 0.212 \| 108 \| 80 \| 3 \| 1 \| 105 \| 27 \| 132 \| GPW\_gp25 domain-containing protein \| GPW\_gp25 domain-containing protein; afdb-uniprot50 \| AF-A0A4R1YZ54-F1-MODEL\_V4 \| 1.0 \| 1.121e-05 \| 218 \| 0.252 \| 103 \| 67 \| 3 \| 2 \| 97 \| 17 \| 116 \| GPW\_gp25 domain-containing protein \| GPW\_gp25 domain-containing protein; afdb-uniprot50 \| AF-A0A1V5ZPA0-F1-MODEL\_V4 \| 1.0 \| 1.35e-05 \| 218 \| 0.214 \| 112 \| 80 \| 4 \| 1 \| 106 \| 20 \| 129 \| Gene 25-like lysozyme \| Gene 25-like lysozyme; afdb-uniprot50 \| AF-A0A4D7BC41-F1-MODEL\_V4 \| 1.0 \| 2.355e-05 \| 218 \| 0.215 \| 116 \| 84 \| 3 \| 1 \| 111 \| 14 \| 127 \| Uncharacterized protein \| Uncharacterized protein; afdb-uniprot50 \| AF-A0A381SJG5-F1-MODEL\_V4 \| 1.0 \| 1.729e-05 \| 218 \| 0.119 \| 109 \| 92 \| 3 \| 3 \| 107 \| 41 \| 149 \| GPW\_gp25 domain-containing protein \| GPW\_gp25 domain-containing protein; afdb-uniprot50 \| AF-A0A3S0EL96-F1-MODEL\_V4 \| 1.0 \| 1.957e-05 \| 218 \| 0.245 \| 118 \| 79 \| 5 \| 1 \| 110 \| 68 \| 183 \| GPW\_gp25 domain-containing protein \| GPW\_gp25 domain-containing protein; afdb-uniprot50 \| AF-A0A5M6ITN1-F1-MODEL\_V4 \| 1.0 \| 6.428e-06 \| 218 \| 0.272 \| 110 \| 75 \| 5 \| 1 \| 108 \| 104 \| 210 \| Uncharacterized protein \| Uncharacterized protein; afdb-uniprot50 \| AF-A0A4Q3KH27-F1-MODEL\_V4 \| 1.0 \| 2.506e-05 \| 217 \| 0.189 \| 116 \| 83 \| 4 \| 1 \| 107 \| 6 \| 119 \| GPW\_gp25 domain-containing protein \| GPW\_gp25 domain-containing protein; afdb-uniprot50 \| AF-A0A2S9XIG4-F1-MODEL\_V4 \| 1.0 \| 3.209e-05 \| 217 \| 0.172 \| 110 \| 86 \| 3 \| 2 \| 107 \| 12 \| 120 \| Gene 25-like lysozyme \| Gene 25-like lysozyme; afdb-uniprot50 \| AF-A0A2N8PEC8-F1-MODEL\_V4 \| 1.0 \| 1.839e-05 \| 217 \| 0.242 \| 107 \| 76 \| 4 \| 2 \| 105 \| 17 \| 121 \| GPW\_gp25 domain-containing protein \| GPW\_gp25 domain-containing protein; afdb-uniprot50 \| AF-A0A561S9T7-F1-MODEL\_V4 \| 1.0 \| 1.269e-05 \| 217 \| 0.16 \| 143 \| 83 \| 7 \| 1 \| 111 \| 1 \| 138 \| Uncharacterized protein \| Uncharacterized protein; afdb-uniprot50 \| AF-A0A1R3VQV6-F1-MODEL\_V4 \| 1.0 \| 8.232e-06 \| 217 \| 0.203 \| 113 \| 82 \| 3 \| 2 \| 106 \| 19 \| 131 \| GPW\_gp25 domain-containing protein \| GPW\_gp25 domain-containing protein; afdb-uniprot50 \| AF-A0A3D8NEX0-F1-MODEL\_V4 \| 1.0 \| 2.355e-05 \| 217 \| 0.283 \| 106 \| 67 \| 3 \| 1 \| 97 \| 19 \| 124 \| Baseplate protein \| Baseplate protein; afdb-uniprot50 \| AF-A0A1Z9I0E6-F1-MODEL\_V4 \| 1.0 \| 1.528e-05 \| 217 \| 0.177 \| 107 \| 83 \| 4 \| 4 \| 105 \| 24 \| 130 \| GPW\_gp25 domain-containing protein \| GPW\_gp25 domain-containing protein; afdb-uniprot50 \| AF-A0A1K1LB84-F1-MODEL\_V4 \| 1.0 \| 5.68e-06 \| 217 \| 0.181 \| 116 \| 83 \| 4 \| 1 \| 105 \| 34 \| 148 \| Baseplate assembly protein, putative \| Baseplate assembly protein, putative; afdb-uniprot50 \| AF-A8ZRG2-F1-MODEL\_V4 \| 1.0 \| 5.68e-06 \| 217 \| 0.243 \| 115 \| 74 \| 4 \| 1 \| 105 \| 79 \| 190 \| GPW/gp25 family protein \| GPW/gp25 family protein; afdb-uniprot50 \| AF-A0A844ZL85-F1-MODEL\_V4 \| 1.0 \| 1.269e-05 \| 216 \| 0.259 \| 108 \| 75 \| 4 \| 2 \| 106 \| 9 \| 114 \| Uncharacterized protein \| Uncharacterized protein; afdb-uniprot50 \| AF-A0A285M3X1-F1-MODEL\_V4 \| 1.0 \| 2.081e-05 \| 216 \| 0.238 \| 109 \| 81 \| 2 \| 3 \| 111 \| 16 \| 122 \| Uncharacterized protein \| Uncharacterized protein; afdb-uniprot50 \| AF-A0A0E3Z1I2-F1-MODEL\_V4 \| 1.0 \| 2.506e-05 \| 216 \| 0.224 \| 107 \| 80 \| 3 \| 2 \| 106 \| 11 \| 116 \| GPW\_gp25 domain-containing protein \| GPW\_gp25 domain-containing protein; afdb-uniprot50 \| AF-A0A2P5KAU3-F1-MODEL\_V4 \| 1.0 \| 5.34e-06 \| 216 \| 0.166 \| 114 \| 89 \| 3 \| 1 \| 108 \| 18 \| 131 \| GPW\_gp25 domain-containing protein \| GPW\_gp25 domain-containing protein; afdb-uniprot50 \| AF-A0A327Q7K0-F1-MODEL\_V4 \| 1.0 \| 1.729e-05 \| 216 \| 0.288 \| 104 \| 70 \| 2 \| 5 \| 105 \| 11 \| 113 \| Gene 25-like lysozyme \| Gene 25-like lysozyme; afdb-uniprot50 \| AF-A0A4Q3TBW4-F1-MODEL\_V4 \| 1.0 \| 2.506e-05 \| 215 \| 0.24 \| 104 \| 73 \| 3 \| 3 \| 105 \| 2 \| 100 \| GPW\_gp25 domain-containing protein \| GPW\_gp25 domain-containing protein; afdb-uniprot50 \| AF-A0A1M6K9Z9-F1-MODEL\_V4 \| 1.0 \| 3.209e-05 \| 215 \| 0.242 \| 107 \| 78 \| 3 \| 2 \| 106 \| 7 \| 112 \| GPW\_gp25 domain-containing protein \| GPW\_gp25 domain-containing protein; afdb-uniprot50 \| AF-A0A5Q4DGZ4-F1-MODEL\_V4 \| 1.0 \| 1.121e-05 \| 215 \| 0.283 \| 113 \| 71 \| 5 \| 1 \| 105 \| 1 \| 111 \| GPW\_gp25 domain-containing protein \| GPW\_gp25 domain-containing protein; afdb-uniprot50 \| AF-A0A238LBG6-F1-MODEL\_V4 \| 1.0 \| 2.355e-05 \| 215 \| 0.194 \| 108 \| 81 \| 4 \| 2 \| 105 \| 8 \| 113 \| Gene 25-like lysozyme \| Gene 25-like lysozyme; afdb-uniprot50 \| AF-A0A2D5MUE6-F1-MODEL\_V4 \| 1.0 \| 3.631e-05 \| 215 \| 0.188 \| 117 \| 85 \| 4 \| 1 \| 107 \| 8 \| 124 \| GPW\_gp25 domain-containing protein \| GPW\_gp25 domain-containing protein; afdb-uniprot50 \| AF-F2NYC6-F1-MODEL\_V4 \| 1.0 \| 1.957e-05 \| 215 \| 0.262 \| 99 \| 70 \| 2 \| 2 \| 97 \| 13 \| 111 \| GPW/gp25 family protein \| GPW/gp25 family protein; afdb-uniprot50 \| AF-A0A369WU76-F1-MODEL\_V4 \| 1.0 \| 4.719e-06 \| 215 \| 0.212 \| 108 \| 80 \| 2 \| 3 \| 105 \| 15 \| 122 \| GPW\_gp25 domain-containing protein \| GPW\_gp25 domain-containing protein; afdb-uniprot50 \| AF-A0A2K8LN18-F1-MODEL\_V4 \| 1.0 \| 2.665e-05 \| 215 \| 0.276 \| 112 \| 75 \| 5 \| 2 \| 110 \| 70 \| 178 \| GPW\_gp25 domain-containing protein \| GPW\_gp25 domain-containing protein; afdb-uniprot50 \| AF-A0A4P9VI48-F1-MODEL\_V4 \| 1.0 \| 9.767e-05 \| 214 \| 0.402 \| 67 \| 40 \| 0 \| 1 \| 67 \| 2 \| 68 \| Baseplate assembly protein W \| Baseplate assembly protein W; afdb-uniprot50 \| AF-H5SBU9-F1-MODEL\_V4 \| 1.0 \| 4.371e-05 \| 214 \| 0.322 \| 93 \| 62 \| 1 \| 13 \| 105 \| 8 \| 99 \| Prophage MuMc02, baseplate assembly protein W \| Prophage MuMc02, baseplate assembly protein W; afdb-uniprot50 \| AF-A0A1I1DKW3-F1-MODEL\_V4 \| 1.0 \| 2.506e-05 \| 214 \| 0.179 \| 117 \| 81 \| 5 \| 1 \| 105 \| 1 \| 114 \| Uncharacterized protein \| Uncharacterized protein; afdb-uniprot50 \| AF-A0A4R4UB04-F1-MODEL\_V4 \| 1.0 \| 1.193e-05 \| 214 \| 0.225 \| 111 \| 81 \| 4 \| 2 \| 108 \| 10 \| 119 \| GPW\_gp25 domain-containing protein \| GPW\_gp25 domain-containing protein; afdb-uniprot50 \| AF-A0A1K1LCB6-F1-MODEL\_V4 \| 1.0 \| 1.269e-05 \| 214 \| 0.242 \| 95 \| 69 \| 2 \| 13 \| 105 \| 3 \| 96 \| Baseplate assembly protein, putative \| Baseplate assembly protein, putative; afdb-uniprot50 \| AF-A0A075MN45-F1-MODEL\_V4 \| 1.0 \| 3.413e-05 \| 214 \| 0.198 \| 106 \| 82 \| 3 \| 2 \| 105 \| 17 \| 121 \| Phage baseplate assembly protein W \| Phage baseplate assembly protein W; afdb-uniprot50 \| AF-A0A3M1VGJ9-F1-MODEL\_V4 \| 1.0 \| 2.665e-05 \| 214 \| 0.149 \| 107 \| 88 \| 2 \| 3 \| 106 \| 17 \| 123 \| GPW\_gp25 domain-containing protein \| GPW\_gp25 domain-containing protein; afdb-uniprot50 \| AF-A0A2E8NZT2-F1-MODEL\_V4 \| 1.0 \| 4.109e-05 \| 214 \| 0.196 \| 112 \| 84 \| 4 \| 3 \| 110 \| 17 \| 126 \| GPW\_gp25 domain-containing protein \| GPW\_gp25 domain-containing protein; afdb-uniprot50 \| AF-A0A0A6VNM1-F1-MODEL\_V4 \| 1.0 \| 2.214e-05 \| 214 \| 0.262 \| 103 \| 66 \| 3 \| 2 \| 97 \| 20 \| 119 \| Baseplate protein \| Baseplate protein; afdb-uniprot50 \| AF-A0A3S0DBJ3-F1-MODEL\_V4 \| 1.0 \| 6.838e-06 \| 214 \| 0.204 \| 127 \| 81 \| 5 \| 2 \| 109 \| 12 \| 137 \| GPW\_gp25 domain-containing protein \| GPW\_gp25 domain-containing protein; afdb-uniprot50 \| AF-A0A317I630-F1-MODEL\_V4 \| 1.0 \| 1.957e-05 \| 214 \| 0.233 \| 124 \| 75 \| 4 \| 2 \| 106 \| 17 \| 139 \| Baseplate protein \| Baseplate protein; afdb-uniprot50 \| AF-A0A2D6RI59-F1-MODEL\_V4 \| 1.0 \| 6.428e-06 \| 213 \| 0.162 \| 111 \| 88 \| 3 \| 2 \| 107 \| 11 \| 121 \| GPW\_gp25 domain-containing protein \| GPW\_gp25 domain-containing protein; afdb-uniprot50 \| AF-A0A2D5MQ93-F1-MODEL\_V4 \| 1.0 \| 2.355e-05 \| 213 \| 0.185 \| 108 \| 85 \| 2 \| 3 \| 107 \| 17 \| 124 \| GPW\_gp25 domain-containing protein \| GPW\_gp25 domain-containing protein; afdb-uniprot50 \| AF-A0A0F9QCS5-F1-MODEL\_V4 \| 1.0 \| 2.836e-05 \| 213 \| 0.186 \| 102 \| 82 \| 1 \| 5 \| 105 \| 21 \| 122 \| Uncharacterized protein \| Uncharacterized protein; afdb-uniprot50 \| AF-A0A5P2UWR7-F1-MODEL\_V4 \| 1.0 \| 3.016e-05 \| 213 \| 0.282 \| 99 \| 68 \| 2 \| 2 \| 97 \| 14 \| 112 \| Baseplate protein \| Baseplate protein; afdb-uniprot50 \| AF-A0A6V8KGV1-F1-MODEL\_V4 \| 1.0 \| 2.081e-05 \| 213 \| 0.337 \| 83 \| 54 \| 1 \| 3 \| 84 \| 15 \| 97 \| Baseplate protein \| Baseplate protein; afdb-uniprot50 \| AF-A0A7X6JAI8-F1-MODEL\_V4 \| 1.0 \| 4.65e-05 \| 212 \| 0.474 \| 78 \| 36 \| 1 \| 1 \| 78 \| 3 \| 75 \| Baseplate assembly protein \| Baseplate assembly protein; afdb-uniprot50 \| AF-A0A399WMQ1-F1-MODEL\_V4 \| 1.0 \| 4.947e-05 \| 212 \| 0.196 \| 107 \| 83 \| 3 \| 2 \| 106 \| 11 \| 116 \| GPW\_gp25 domain-containing protein \| GPW\_gp25 domain-containing protein; afdb-uniprot50 \| AF-A0A1I0F144-F1-MODEL\_V4 \| 1.0 \| 4.371e-05 \| 212 \| 0.205 \| 107 \| 80 \| 4 \| 2 \| 105 \| 12 \| 116 \| GPW\_gp25 domain-containing protein \| GPW\_gp25 domain-containing protein; afdb-uniprot50 \| AF-A0A5C4QRE3-F1-MODEL\_V4 \| 1.0 \| 5.262e-05 \| 212 \| 0.168 \| 113 \| 86 \| 4 \| 1 \| 106 \| 4 \| 115 \| GPW/gp25 family protein \| GPW/gp25 family protein; afdb-uniprot50 \| AF-A0A7C7GT66-F1-MODEL\_V4 \| 1.0 \| 1.957e-05 \| 212 \| 0.15 \| 120 \| 92 \| 3 \| 2 \| 112 \| 11 \| 129 \| GPW\_gp25 domain-containing protein \| GPW\_gp25 domain-containing protein; afdb-uniprot50 \| AF-A0A535DVR5-F1-MODEL\_V4 \| 1.0 \| 1.957e-05 \| 212 \| 0.301 \| 106 \| 69 \| 3 \| 2 \| 103 \| 14 \| 118 \| GPW/gp25 family protein \| GPW/gp25 family protein; afdb-uniprot50 \| AF-A0A6L9JRF5-F1-MODEL\_V4 \| 1.0 \| 9.316e-06 \| 212 \| 0.205 \| 107 \| 83 \| 1 \| 1 \| 105 \| 21 \| 127 \| Phage baseplate protein \| Phage baseplate protein; afdb-uniprot50 \| AF-A0A2D0KA99-F1-MODEL\_V4 \| 1.0 \| 9.91e-06 \| 212 \| 0.171 \| 111 \| 86 \| 2 \| 1 \| 105 \| 19 \| 129 \| Phage baseplate protein \| Phage baseplate protein; afdb-uniprot50 \| AF-A0A2M7WDH6-F1-MODEL\_V4 \| 1.0 \| 1.957e-05 \| 212 \| 0.201 \| 109 \| 81 \| 5 \| 3 \| 108 \| 94 \| 199 \| Uncharacterized protein \| Uncharacterized protein; afdb-uniprot50 \| AF-A0A6B3IQI9-F1-MODEL\_V4 \| 1.0 \| 1.625e-05 \| 211 \| 0.282 \| 99 \| 68 \| 2 \| 2 \| 97 \| 15 \| 113 \| GPW/gp25 family protein \| GPW/gp25 family protein; afdb-uniprot50 \| AF-A0A0F9FRJ8-F1-MODEL\_V4 \| 1.0 \| 9.91e-06 \| 211 \| 0.188 \| 117 \| 84 \| 5 \| 1 \| 107 \| 4 \| 119 \| GPW\_gp25 domain-containing protein \| GPW\_gp25 domain-containing protein; afdb-uniprot50 \| AF-A0A382SJV7-F1-MODEL\_V4 \| 1.0 \| 2.506e-05 \| 211 \| 0.158 \| 120 \| 85 \| 7 \| 1 \| 107 \| 5 \| 121 \| GPW\_gp25 domain-containing protein \| GPW\_gp25 domain-containing protein; afdb-uniprot50 \| AF-A0A842WRC3-F1-MODEL\_V4 \| 1.0 \| 3.209e-05 \| 211 \| 0.221 \| 113 \| 83 \| 5 \| 1 \| 111 \| 20 \| 129 \| DUF2634 domain-containing protein \| DUF2634 domain-containing protein; afdb-uniprot50 \| AF-A0A5M8P5N3-F1-MODEL\_V4 \| 1.0 \| 2.355e-05 \| 211 \| 0.205 \| 112 \| 83 \| 3 \| 1 \| 107 \| 14 \| 124 \| Phage baseplate protein \| Phage baseplate protein; afdb-uniprot50 \| AF-A0A3D5E673-F1-MODEL\_V4 \| 1.0 \| 2.506e-05 \| 211 \| 0.159 \| 119 \| 84 \| 7 \| 1 \| 105 \| 12 \| 128 \| GPW\_gp25 domain-containing protein \| GPW\_gp25 domain-containing protein; afdb-uniprot50 \| AF-A0A7W5F3D6-F1-MODEL\_V4 \| 1.0 \| 1.193e-05 \| 211 \| 0.252 \| 111 \| 78 \| 4 \| 2 \| 108 \| 29 \| 138 \| GPW\_gp25 domain-containing protein \| GPW\_gp25 domain-containing protein; afdb-uniprot50 \| AF-A0A2T6G6G7-F1-MODEL\_V4 \| 1.0 \| 1.528e-05 \| 210 \| 0.304 \| 92 \| 60 \| 3 \| 17 \| 105 \| 7 \| 97 \| Baseplate protein \| Baseplate protein; afdb-uniprot50 \| AF-A0A7Z6PT91-F1-MODEL\_V4 \| 1.0 \| 2.836e-05 \| 210 \| 0.254 \| 102 \| 69 \| 2 \| 12 \| 106 \| 2 \| 103 \| Integrase \| Integrase; afdb-uniprot50 \| AF-A0A5S4YIV1-F1-MODEL\_V4 \| 1.0 \| 4.109e-05 \| 210 \| 0.224 \| 107 \| 80 \| 3 \| 2 \| 106 \| 7 \| 112 \| GPW/gp25 family protein \| GPW/gp25 family protein; afdb-uniprot50 \| AF-A0A838AGQ5-F1-MODEL\_V4 \| 1.0 \| 1.957e-05 \| 210 \| 0.148 \| 108 \| 87 \| 3 \| 1 \| 105 \| 4 \| 109 \| GPW/gp25 family protein \| GPW/gp25 family protein; afdb-uniprot50 \| AF-A0A286DXR1-F1-MODEL\_V4 \| 1.0 \| 1.528e-05 \| 210 \| 0.214 \| 107 \| 79 \| 4 \| 2 \| 105 \| 15 \| 119 \| GPW\_gp25 domain-containing protein \| GPW\_gp25 domain-containing protein; afdb-uniprot50 \| AF-A0A2E8FZA1-F1-MODEL\_V4 \| 1.0 \| 1.957e-05 \| 210 \| 0.201 \| 109 \| 81 \| 3 \| 4 \| 106 \| 28 \| 136 \| GPW\_gp25 domain-containing protein \| GPW\_gp25 domain-containing protein; afdb-uniprot50 \| AF-A0A853ICS2-F1-MODEL\_V4 \| 1.0 \| 1.35e-05 \| 210 \| 0.254 \| 106 \| 70 \| 3 \| 1 \| 97 \| 20 \| 125 \| GPW/gp25 family protein \| GPW/gp25 family protein; afdb-uniprot50 \| AF-A0A8B3NLL6-F1-MODEL\_V4 \| 1.0 \| 0.0002469 \| 210 \| 0.27 \| 85 \| 62 \| 0 \| 1 \| 85 \| 3 \| 87 \| Uncharacterized protein \| Uncharacterized protein; afdb-uniprot50 \| AF-A0A4Q0ZJN2-F1-MODEL\_V4 \| 1.0 \| 1.436e-05 \| 209 \| 0.304 \| 105 \| 61 \| 5 \| 4 \| 106 \| 1 \| 95 \| GPW\_gp25 domain-containing protein \| GPW\_gp25 domain-containing protein; afdb-uniprot50 \| AF-A0A1M6M9N6-F1-MODEL\_V4 \| 1.0 \| 2.081e-05 \| 209 \| 0.223 \| 103 \| 76 \| 3 \| 6 \| 105 \| 13 \| 114 \| Gene 25-like lysozyme \| Gene 25-like lysozyme; afdb-uniprot50 \| AF-A0A0Q4KZB0-F1-MODEL\_V4 \| 1.0 \| 2.355e-05 \| 209 \| 0.276 \| 105 \| 74 \| 1 \| 2 \| 104 \| 18 \| 122 \| Baseplate assembly protein \| Baseplate assembly protein; afdb-uniprot50 \| AF-A0A2T2YA09-F1-MODEL\_V4 \| 1.0 \| 1.35e-05 \| 209 \| 0.31 \| 100 \| 65 \| 2 \| 2 \| 97 \| 17 \| 116 \| GPW\_gp25 domain-containing protein \| GPW\_gp25 domain-containing protein; afdb-uniprot50 \| AF-A0A7X5QIW4-F1-MODEL\_V4 \| 1.0 \| 5.955e-05 \| 209 \| 0.163 \| 98 \| 80 \| 1 \| 10 \| 105 \| 40 \| 137 \| Phage baseplate protein \| Phage baseplate protein; afdb-uniprot50 \| AF-A0A238JEQ2-F1-MODEL\_V4 \| 1.0 \| 7.739e-06 \| 209 \| 0.241 \| 116 \| 76 \| 3 \| 1 \| 105 \| 31 \| 145 \| GPW\_gp25 domain-containing protein \| GPW\_gp25 domain-containing protein; afdb-uniprot50 \| AF-A0A6I7P0B8-F1-MODEL\_V4 \| 1.0 \| 2.081e-05 \| 209 \| 0.28 \| 100 \| 68 \| 2 \| 2 \| 97 \| 38 \| 137 \| GPW\_gp25 domain-containing protein \| GPW\_gp25 domain-containing protein; afdb-uniprot50 \| AF-A0A1Y6CTE9-F1-MODEL\_V4 \| 1.0 \| 2.355e-05 \| 209 \| 0.247 \| 113 \| 77 \| 7 \| 1 \| 110 \| 100 \| 207 \| Phage baseplate assembly protein W \| Phage baseplate assembly protein W; afdb-uniprot50 \| AF-A0A6L4A963-F1-MODEL\_V4 \| 1.0 \| 3.209e-05 \| 208 \| 0.33 \| 100 \| 62 \| 3 \| 2 \| 97 \| 15 \| 113 \| GPW/gp25 family protein \| GPW/gp25 family protein; afdb-uniprot50 \| AF-A0A523UWB7-F1-MODEL\_V4 \| 1.0 \| 4.109e-05 \| 208 \| 0.215 \| 116 \| 80 \| 4 \| 1 \| 106 \| 1 \| 115 \| GPW\_gp25 domain-containing protein \| GPW\_gp25 domain-containing protein; afdb-uniprot50 \| AF-X5KSN9-F1-MODEL\_V4 \| 1.0 \| 5.598e-05 \| 208 \| 0.169 \| 106 \| 85 \| 3 \| 2 \| 105 \| 9 \| 113 \| GPW/gp25 family protein \| GPW/gp25 family protein; afdb-uniprot50 \| AF-A0A1D2WBQ4-F1-MODEL\_V4 \| 1.0 \| 1.729e-05 \| 208 \| 0.258 \| 112 \| 74 \| 6 \| 3 \| 111 \| 14 \| 119 \| Uncharacterized protein \| Uncharacterized protein; afdb-uniprot50 \| AF-A0A0X3VMP2-F1-MODEL\_V4 \| 1.0 \| 1.35e-05 \| 208 \| 0.258 \| 112 \| 76 \| 5 \| 2 \| 108 \| 17 \| 126 \| GPW\_gp25 domain-containing protein \| GPW\_gp25 domain-containing protein; afdb-uniprot50 \| AF-A0A074M821-F1-MODEL\_V4 \| 1.0 \| 2.836e-05 \| 208 \| 0.194 \| 108 \| 83 \| 4 \| 1 \| 106 \| 22 \| 127 \| Uncharacterized protein \| Uncharacterized protein; afdb-uniprot50 \| AF-A0A4R2GXL5-F1-MODEL\_V4 \| 1.0 \| 3.631e-05 \| 208 \| 0.191 \| 115 \| 85 \| 3 \| 1 \| 108 \| 18 \| 131 \| Uncharacterized protein \| Uncharacterized protein; afdb-uniprot50 \| AF-A0A4U8YRK1-F1-MODEL\_V4 \| 1.0 \| 2.542e-06 \| 208 \| 0.172 \| 133 \| 84 \| 5 \| 1 \| 108 \| 8 \| 139 \| Uncharacterized protein \| Uncharacterized protein; afdb-uniprot50 \| AF-A0A4Y3VBU6-F1-MODEL\_V4 \| 1.0 \| 1.625e-05 \| 208 \| 0.234 \| 111 \| 80 \| 4 \| 2 \| 108 \| 35 \| 144 \| GPW\_gp25 domain-containing protein \| GPW\_gp25 domain-containing protein; afdb-uniprot50 \| AF-A0A1V5AF29-F1-MODEL\_V4 \| 1.0 \| 3.631e-05 \| 207 \| 0.261 \| 107 \| 74 \| 4 \| 2 \| 105 \| 34 \| 138 \| Gene 25-like lysozyme \| Gene 25-like lysozyme; afdb-uniprot50 \| AF-A0A1G4U040-F1-MODEL\_V4 \| 1.0 \| 3.256e-06 \| 206 \| 0.275 \| 116 \| 73 \| 7 \| 2 \| 112 \| 15 \| 124 \| Uncharacterized protein \| Uncharacterized protein; afdb-uniprot50 \| AF-A0A2T4TTD4-F1-MODEL\_V4 \| 1.0 \| 1.957e-05 \| 206 \| 0.174 \| 109 \| 85 \| 2 \| 2 \| 105 \| 13 \| 121 \| GPW\_gp25 domain-containing protein \| GPW\_gp25 domain-containing protein; afdb-uniprot50 \| AF-A0A229UMK7-F1-MODEL\_V4 \| 1.0 \| 5.955e-05 \| 206 \| 0.132 \| 113 \| 91 \| 3 \| 2 \| 107 \| 14 \| 126 \| Uncharacterized protein \| Uncharacterized protein; afdb-uniprot50 \| AF-A0A0Q1D4W1-F1-MODEL\_V4 \| 1.0 \| 2.214e-05 \| 206 \| 0.242 \| 107 \| 76 \| 4 \| 2 \| 105 \| 19 \| 123 \| GPW\_gp25 domain-containing protein \| GPW\_gp25 domain-containing protein; afdb-uniprot50 \| AF-A8T9K2-F1-MODEL\_V4 \| 1.0 \| 2.506e-05 \| 206 \| 0.13 \| 107 \| 90 \| 2 \| 2 \| 105 \| 13 \| 119 \| GPW\_gp25 domain-containing protein \| GPW\_gp25 domain-containing protein; afdb-uniprot50 \| AF-A0A1Z9ULM3-F1-MODEL\_V4 \| 1.0 \| 2.355e-05 \| 205 \| 0.2 \| 100 \| 73 \| 3 \| 14 \| 107 \| 36 \| 134 \| GPW\_gp25 domain-containing protein \| GPW\_gp25 domain-containing protein; afdb-uniprot50 \| AF-A0A3N7HI14-F1-MODEL\_V4 \| 1.0 \| 8.757e-06 \| 205 \| 0.241 \| 120 \| 78 \| 6 \| 1 \| 108 \| 19 \| 137 \| Baseplate assembly protein \| Baseplate assembly protein; afdb-uniprot50 \| AF-A0A843F7L3-F1-MODEL\_V4 \| 1.0 \| 3.863e-05 \| 205 \| 0.163 \| 110 \| 86 \| 4 \| 3 \| 108 \| 35 \| 142 \| Uncharacterized protein \| Uncharacterized protein; afdb-uniprot50 \| AF-A0A381NWC8-F1-MODEL\_V4 \| 1.0 \| 4.65e-05 \| 205 \| 0.138 \| 123 \| 87 \| 2 \| 3 \| 106 \| 24 \| 146 \| GPW\_gp25 domain-containing protein \| GPW\_gp25 domain-containing protein; afdb-uniprot50 \| AF-A0A381TRD1-F1-MODEL\_V4 \| 1.0 \| 2.355e-05 \| 205 \| 0.176 \| 125 \| 82 \| 3 \| 3 \| 106 \| 34 \| 158 \| GPW\_gp25 domain-containing protein \| GPW\_gp25 domain-containing protein; afdb-uniprot50 \| AF-V4NPI9-F1-MODEL\_V4 \| 1.0 \| 0.0001928 \| 204 \| 0.476 \| 63 \| 33 \| 0 \| 1 \| 63 \| 6 \| 68 \| Baseplate assembly protein \| Baseplate assembly protein; afdb-uniprot50 \| AF-G8LQ32-F1-MODEL\_V4 \| 1.0 \| 0.0001176 \| 204 \| 0.469 \| 66 \| 35 \| 0 \| 1 \| 66 \| 5 \| 70 \| GPW/Gp25 Family Protein \| GPW/Gp25 Family Protein; afdb-uniprot50 \| AF-A0A2P5MMF8-F1-MODEL\_V4 \| 1.0 \| 4.371e-05 \| 204 \| 0.245 \| 106 \| 77 \| 3 \| 2 \| 105 \| 7 \| 111 \| GPW\_gp25 domain-containing protein \| GPW\_gp25 domain-containing protein; afdb-uniprot50 \| AF-A0A7W5FEZ8-F1-MODEL\_V4 \| 1.0 \| 2.836e-05 \| 204 \| 0.252 \| 107 \| 75 \| 4 \| 2 \| 105 \| 9 \| 113 \| GPW\_gp25 domain-containing protein \| GPW\_gp25 domain-containing protein; afdb-uniprot50 \| AF-A0A7W7HJ55-F1-MODEL\_V4 \| 1.0 \| 4.65e-05 \| 204 \| 0.242 \| 107 \| 76 \| 4 \| 2 \| 105 \| 9 \| 113 \| Phage baseplate assembly protein W \| Phage baseplate assembly protein W; afdb-uniprot50 \| AF-A0A2E3DKA2-F1-MODEL\_V4 \| 1.0 \| 5.955e-05 \| 204 \| 0.196 \| 122 \| 78 \| 6 \| 1 \| 107 \| 7 \| 123 \| GPW\_gp25 domain-containing protein \| GPW\_gp25 domain-containing protein; afdb-uniprot50 \| AF-D3CW99-F1-MODEL\_V4 \| 1.0 \| 4.371e-05 \| 204 \| 0.267 \| 101 \| 66 \| 3 \| 2 \| 97 \| 22 \| 119 \| GPW/gp25 family protein \| GPW/gp25 family protein; afdb-uniprot50 \| AF-A0A7X6AE11-F1-MODEL\_V4 \| 1.0 \| 3.863e-05 \| 204 \| 0.372 \| 86 \| 51 \| 2 \| 2 \| 85 \| 14 \| 98 \| Baseplate protein \| Baseplate protein; afdb-uniprot50 \| AF-A0A317FGN1-F1-MODEL\_V4 \| 1.0 \| 2.665e-05 \| 204 \| 0.194 \| 118 \| 84 \| 3 \| 1 \| 107 \| 140 \| 257 \| GPW\_gp25 domain-containing protein \| GPW\_gp25 domain-containing protein; afdb-uniprot50 \| AF-F2K1R6-F1-MODEL\_V4 \| 1.0 \| 2.665e-05 \| 203 \| 0.302 \| 96 \| 63 \| 2 \| 3 \| 97 \| 10 \| 102 \| GPW/gp25 family protein \| GPW/gp25 family protein; afdb-uniprot50 \| AF-A0A4Q5X4U8-F1-MODEL\_V4 \| 1.0 \| 5.262e-05 \| 203 \| 0.233 \| 107 \| 77 \| 4 \| 2 \| 105 \| 7 \| 111 \| GPW\_gp25 domain-containing protein \| GPW\_gp25 domain-containing protein; afdb-uniprot50 \| AF-A0A4R5QHB6-F1-MODEL\_V4 \| 1.0 \| 3.631e-05 \| 203 \| 0.261 \| 107 \| 74 \| 5 \| 2 \| 105 \| 7 \| 111 \| Uncharacterized protein \| Uncharacterized protein; afdb-uniprot50 \| AF-A0A1Z8QW37-F1-MODEL\_V4 \| 1.0 \| 1.528e-05 \| 203 \| 0.26 \| 100 \| 70 \| 2 \| 2 \| 97 \| 17 \| 116 \| GPW\_gp25 domain-containing protein \| GPW\_gp25 domain-containing protein; afdb-uniprot50 \| AF-A0A432PWH9-F1-MODEL\_V4 \| 1.0 \| 2.836e-05 \| 202 \| 0.257 \| 97 \| 68 \| 4 \| 10 \| 105 \| 5 \| 98 \| GPW\_gp25 domain-containing protein \| GPW\_gp25 domain-containing protein; afdb-uniprot50 \| AF-A0A0Q6XWC6-F1-MODEL\_V4 \| 1.0 \| 5.262e-05 \| 202 \| 0.23 \| 113 \| 80 \| 5 \| 2 \| 110 \| 7 \| 116 \| GPW\_gp25 domain-containing protein \| GPW\_gp25 domain-containing protein; afdb-uniprot50 \| AF-A0A0H3ZLU4-F1-MODEL\_V4 \| 1.0 \| 1.193e-05 \| 202 \| 0.171 \| 105 \| 83 \| 1 \| 1 \| 105 \| 13 \| 113 \| Uncharacterized protein \| Uncharacterized protein; afdb-uniprot50 \| AF-A0A2E4W7B1-F1-MODEL\_V4 \| 1.0 \| 1.625e-05 \| 202 \| 0.203 \| 118 \| 81 \| 5 \| 3 \| 108 \| 17 \| 133 \| GPW\_gp25 domain-containing protein \| GPW\_gp25 domain-containing protein; afdb-uniprot50 \| AF-A0A1G3KNN2-F1-MODEL\_V4 \| 1.0 \| 2.506e-05 \| 201 \| 0.211 \| 104 \| 77 \| 3 \| 5 \| 105 \| 14 \| 115 \| Uncharacterized protein \| Uncharacterized protein; afdb-uniprot50 \| AF-A0A2D6MIS0-F1-MODEL\_V4 \| 1.0 \| 1.193e-05 \| 201 \| 0.188 \| 117 \| 84 \| 4 \| 2 \| 108 \| 20 \| 135 \| GPW\_gp25 domain-containing protein \| GPW\_gp25 domain-containing protein; afdb-uniprot50 \| AF-A0A2E6ZKV4-F1-MODEL\_V4 \| 1.0 \| 3.863e-05 \| 201 \| 0.13 \| 107 \| 88 \| 3 \| 3 \| 105 \| 36 \| 141 \| GPW\_gp25 domain-containing protein \| GPW\_gp25 domain-containing protein; afdb-uniprot50 \| AF-A0A447N0I4-F1-MODEL\_V4 \| 1.0 \| 0.0001039 \| 200 \| 0.408 \| 71 \| 42 \| 0 \| 1 \| 71 \| 5 \| 75 \| Baseplate assembly protein \| Baseplate assembly protein; afdb-uniprot50 \| AF-A0A3D2XG26-F1-MODEL\_V4 \| 1.0 \| 3.92e-06 \| 200 \| 0.207 \| 106 \| 70 \| 2 \| 2 \| 105 \| 17 \| 110 \| GPW\_gp25 domain-containing protein \| GPW\_gp25 domain-containing protein; afdb-uniprot50 \| AF-A0A832CQ52-F1-MODEL\_V4 \| 1.0 \| 1.729e-05 \| 200 \| 0.25 \| 108 \| 74 \| 5 \| 1 \| 105 \| 13 \| 116 \| Uncharacterized protein \| Uncharacterized protein; afdb-uniprot50 \| AF-A0A661Z4X6-F1-MODEL\_V4 \| 1.0 \| 2.836e-05 \| 200 \| 0.175 \| 108 \| 86 \| 2 \| 1 \| 105 \| 7 \| 114 \| GPW\_gp25 domain-containing protein \| GPW\_gp25 domain-containing protein; afdb-uniprot50 \| AF-A0A662Q525-F1-MODEL\_V4 \| 1.0 \| 2.214e-05 \| 200 \| 0.161 \| 124 \| 87 \| 4 \| 1 \| 107 \| 12 \| 135 \| GPW\_gp25 domain-containing protein \| GPW\_gp25 domain-containing protein; afdb-uniprot50 \| AF-A0A6N8MVK6-F1-MODEL\_V4 \| 1.0 \| 0.0002627 \| 199 \| 0.428 \| 63 \| 36 \| 0 \| 1 \| 63 \| 5 \| 67 \| GPW\_gp25 domain-containing protein \| GPW\_gp25 domain-containing protein; afdb-uniprot50 \| AF-A0A843GVE6-F1-MODEL\_V4 \| 1.0 \| 0.0001415 \| 199 \| 0.141 \| 113 \| 88 \| 3 \| 1 \| 105 \| 1 \| 112 \| Uncharacterized protein \| Uncharacterized protein; afdb-uniprot50 \| AF-A0A2D8EMY5-F1-MODEL\_V4 \| 1.0 \| 7.169e-05 \| 199 \| 0.168 \| 107 \| 86 \| 2 \| 3 \| 106 \| 12 \| 118 \| GPW\_gp25 domain-containing protein \| GPW\_gp25 domain-containing protein; afdb-uniprot50 \| AF-A0A2E7WDX9-F1-MODEL\_V4 \| 1.0 \| 8.63e-05 \| 199 \| 0.137 \| 109 \| 89 \| 3 \| 3 \| 107 \| 15 \| 122 \| Uncharacterized protein \| Uncharacterized protein; afdb-uniprot50 \| AF-A0A1G8FCC5-F1-MODEL\_V4 \| 1.0 \| 3.631e-05 \| 199 \| 0.14 \| 107 \| 88 \| 3 \| 1 \| 106 \| 23 \| 126 \| Phage baseplate assembly protein W \| Phage baseplate assembly protein W; afdb-uniprot50 \| AF-A0A2E0GD41-F1-MODEL\_V4 \| 1.0 \| 3.413e-05 \| 199 \| 0.182 \| 115 \| 85 \| 4 \| 2 \| 108 \| 20 \| 133 \| GPW\_gp25 domain-containing protein \| GPW\_gp25 domain-containing protein; afdb-uniprot50 \| AF-A0A2E5T0R5-F1-MODEL\_V4 \| 1.0 \| 0.0001602 \| 199 \| 0.139 \| 136 \| 88 \| 5 \| 4 \| 112 \| 21 \| 154 \| Uncharacterized protein \| Uncharacterized protein; afdb-uniprot50 \| AF-A0A3C0G267-F1-MODEL\_V4 \| 1.0 \| 3.016e-05 \| 199 \| 0.186 \| 107 \| 80 \| 2 \| 6 \| 105 \| 83 \| 189 \| GPW\_gp25 domain-containing protein \| GPW\_gp25 domain-containing protein; afdb-uniprot50 \| AF-A0A3G8C6Y1-F1-MODEL\_V4 \| 1.0 \| 0.0001704 \| 198 \| 0.362 \| 69 \| 42 \| 1 \| 39 \| 107 \| 3 \| 69 \| Phage baseplate assembly protein \| Phage baseplate assembly protein; afdb-uniprot50 \| AF-A0A1H6APR4-F1-MODEL\_V4 \| 1.0 \| 3.631e-05 \| 198 \| 0.223 \| 112 \| 80 \| 5 \| 2 \| 108 \| 10 \| 119 \| GPW\_gp25 domain-containing protein \| GPW\_gp25 domain-containing protein; afdb-uniprot50 \| AF-A0A3C0G5D2-F1-MODEL\_V4 \| 1.0 \| 3.631e-05 \| 198 \| 0.187 \| 112 \| 85 \| 2 \| 2 \| 107 \| 13 \| 124 \| GPW\_gp25 domain-containing protein \| GPW\_gp25 domain-containing protein; afdb-uniprot50 \| AF-A0A101NH81-F1-MODEL\_V4 \| 1.0 \| 3.863e-05 \| 198 \| 0.225 \| 111 \| 81 \| 4 \| 2 \| 108 \| 13 \| 122 \| GPW\_gp25 domain-containing protein \| GPW\_gp25 domain-containing protein; afdb-uniprot50 \| AF-A0A6M3ZUD3-F1-MODEL\_V4 \| 1.0 \| 4.371e-05 \| 198 \| 0.242 \| 99 \| 72 \| 2 \| 2 \| 97 \| 16 \| 114 \| GPW\_gp25 domain-containing protein \| GPW\_gp25 domain-containing protein; afdb-uniprot50 \| AF-A0A2D7BZB0-F1-MODEL\_V4 \| 1.0 \| 8.113e-05 \| 198 \| 0.141 \| 106 \| 88 \| 2 \| 3 \| 105 \| 35 \| 140 \| GPW\_gp25 domain-containing protein \| GPW\_gp25 domain-containing protein; afdb-uniprot50 \| AF-A0A5C7PYR2-F1-MODEL\_V4 \| 1.0 \| 4.371e-05 \| 198 \| 0.21 \| 114 \| 84 \| 5 \| 1 \| 112 \| 91 \| 200 \| Uncharacterized protein \| Uncharacterized protein; afdb-uniprot50 \| AF-A0A009Q2W6-F1-MODEL\_V4 \| 1.0 \| 6.739e-05 \| 197 \| 0.366 \| 71 \| 45 \| 0 \| 35 \| 105 \| 1 \| 71 \| Lysozyme family protein \| Lysozyme family protein; afdb-uniprot50 \| AF-A0A429AGL9-F1-MODEL\_V4 \| 1.0 \| 0.0001331 \| 197 \| 0.228 \| 105 \| 76 \| 3 \| 3 \| 106 \| 6 \| 106 \| GPW\_gp25 domain-containing protein \| GPW\_gp25 domain-containing protein; afdb-uniprot50 \| AF-A0A1G3K418-F1-MODEL\_V4 \| 1.0 \| 8.113e-05 \| 197 \| 0.24 \| 108 \| 77 \| 4 \| 2 \| 106 \| 8 \| 113 \| GPW\_gp25 domain-containing protein \| GPW\_gp25 domain-containing protein; afdb-uniprot50 \| AF-A0A1L3FAG4-F1-MODEL\_V4 \| 1.0 \| 3.631e-05 \| 197 \| 0.254 \| 114 \| 76 \| 5 \| 2 \| 108 \| 9 \| 120 \| GPW\_gp25 domain-containing protein \| GPW\_gp25 domain-containing protein; afdb-uniprot50 \| AF-A0A1I6PRV0-F1-MODEL\_V4 \| 1.0 \| 0.0001039 \| 197 \| 0.224 \| 107 \| 78 \| 4 \| 2 \| 105 \| 12 \| 116 \| GPW\_gp25 domain-containing protein \| GPW\_gp25 domain-containing protein; afdb-uniprot50 \| AF-A0A1Z9UJC8-F1-MODEL\_V4 \| 1.0 \| 7.626e-05 \| 197 \| 0.159 \| 119 \| 91 \| 5 \| 2 \| 112 \| 27 \| 144 \| GPW\_gp25 domain-containing protein \| GPW\_gp25 domain-containing protein; afdb-uniprot50 \| AF-A0A4R2Q0H0-F1-MODEL\_V4 \| 1.0 \| 0.0001704 \| 196 \| 0.216 \| 106 \| 80 \| 3 \| 2 \| 105 \| 7 \| 111 \| GPW\_gp25 domain-containing protein \| GPW\_gp25 domain-containing protein; afdb-uniprot50 \| AF-A0A831ZHY3-F1-MODEL\_V4 \| 1.0 \| 2.836e-05 \| 196 \| 0.191 \| 115 \| 84 \| 5 \| 2 \| 108 \| 12 \| 125 \| Uncharacterized protein \| Uncharacterized protein; afdb-uniprot50 \| AF-A0A4Z0GYW9-F1-MODEL\_V4 \| 1.0 \| 2.665e-05 \| 196 \| 0.178 \| 112 \| 85 \| 4 \| 2 \| 107 \| 55 \| 165 \| GPW\_gp25 domain-containing protein \| GPW\_gp25 domain-containing protein; afdb-uniprot50 \| AF-A0A3D2D4U8-F1-MODEL\_V4 \| 1.0 \| 0.0003807 \| 195 \| 0.359 \| 64 \| 41 \| 0 \| 1 \| 64 \| 2 \| 65 \| Baseplate assembly protein \| Baseplate assembly protein; afdb-uniprot50 \| AF-A0A3T0L2L2-F1-MODEL\_V4 \| 1.0 \| 0.0001415 \| 195 \| 0.303 \| 89 \| 58 \| 2 \| 6 \| 93 \| 8 \| 93 \| Baseplate assembly protein \| Baseplate assembly protein; afdb-uniprot50 \| AF-A0A1S1YVC0-F1-MODEL\_V4 \| 1.0 \| 0.0001176 \| 195 \| 0.198 \| 106 \| 82 \| 3 \| 3 \| 107 \| 14 \| 117 \| GPW\_gp25 domain-containing protein \| GPW\_gp25 domain-containing protein; afdb-uniprot50 \| AF-A0A0G1YBV7-F1-MODEL\_V4 \| 1.0 \| 3.631e-05 \| 195 \| 0.182 \| 126 \| 84 \| 3 \| 1 \| 111 \| 123 \| 244 \| Uncharacterized protein \| Uncharacterized protein; afdb-uniprot50 \| AF-A0A1G8I0U2-F1-MODEL\_V4 \| 1.0 \| 0.0005186 \| 194 \| 0.298 \| 67 \| 47 \| 0 \| 1 \| 67 \| 1 \| 67 \| Uncharacterized protein \| Uncharacterized protein; afdb-uniprot50 \| AF-A0A0Q2Z3G8-F1-MODEL\_V4 \| 1.0 \| 5.955e-05 \| 194 \| 0.247 \| 109 \| 77 \| 2 \| 3 \| 107 \| 7 \| 114 \| GPW\_gp25 domain-containing protein \| GPW\_gp25 domain-containing protein; afdb-uniprot50 \| AF-A0A7C6EBT5-F1-MODEL\_V4 \| 1.0 \| 5.262e-05 \| 194 \| 0.189 \| 111 \| 84 \| 5 \| 2 \| 111 \| 17 \| 122 \| Uncharacterized protein \| Uncharacterized protein; afdb-uniprot50 \| AF-A0A2D8EIK4-F1-MODEL\_V4 \| 1.0 \| 0.0001331 \| 194 \| 0.205 \| 107 \| 82 \| 2 \| 3 \| 106 \| 27 \| 133 \| GPW\_gp25 domain-containing protein \| GPW\_gp25 domain-containing protein; afdb-uniprot50 \| AF-A0A0B6RSR4-F1-MODEL\_V4 \| 1.0 \| 8.113e-05 \| 193 \| 0.435 \| 78 \| 41 \| 1 \| 1 \| 78 \| 1 \| 75 \| Putative phage baseplate assembly protein W \| Putative phage baseplate assembly protein W; afdb-uniprot50 \| AF-A0A543PFP7-F1-MODEL\_V4 \| 1.0 \| 2.214e-05 \| 193 \| 0.205 \| 112 \| 82 \| 5 \| 2 \| 108 \| 7 \| 116 \| GPW\_gp25 domain-containing protein \| GPW\_gp25 domain-containing protein; afdb-uniprot50 \| AF-A0A424JID7-F1-MODEL\_V4 \| 1.0 \| 1.625e-05 \| 193 \| 0.2 \| 120 \| 83 \| 3 \| 1 \| 107 \| 1 \| 120 \| Baseplate protein \| Baseplate protein; afdb-uniprot50 \| AF-A0A0J1FSM7-F1-MODEL\_V4 \| 1.0 \| 3.016e-05 \| 193 \| 0.22 \| 109 \| 77 \| 7 \| 1 \| 105 \| 16 \| 120 \| 25-like lysozyme \| 25-like lysozyme; afdb-uniprot50 \| AF-A0A350IEC1-F1-MODEL\_V4 \| 1.0 \| 0.0001105 \| 193 \| 0.132 \| 113 \| 90 \| 3 \| 1 \| 105 \| 27 \| 139 \| GPW\_gp25 domain-containing protein \| GPW\_gp25 domain-containing protein; afdb-uniprot50 \| AF-A0A2D7XHE7-F1-MODEL\_V4 \| 1.0 \| 2.665e-05 \| 193 \| 0.203 \| 113 \| 82 \| 4 \| 1 \| 105 \| 33 \| 145 \| GPW\_gp25 domain-containing protein \| GPW\_gp25 domain-containing protein; afdb-uniprot50 \| AF-A0A7C1ZK85-F1-MODEL\_V4 \| 1.0 \| 5.598e-05 \| 192 \| 0.26 \| 119 \| 73 \| 5 \| 3 \| 108 \| 6 \| 122 \| GPW\_gp25 domain-containing protein \| GPW\_gp25 domain-containing protein; afdb-uniprot50 \| AF-A0A350PIZ7-F1-MODEL\_V4 \| 1.0 \| 2.355e-05 \| 192 \| 0.159 \| 113 \| 87 \| 3 \| 3 \| 107 \| 13 \| 125 \| GPW\_gp25 domain-containing protein \| GPW\_gp25 domain-containing protein; afdb-uniprot50 \| AF-A0A1I3W1X3-F1-MODEL\_V4 \| 1.0 \| 7.626e-05 \| 192 \| 0.157 \| 114 \| 86 \| 4 \| 1 \| 105 \| 2 \| 114 \| GPW\_gp25 domain-containing protein \| GPW\_gp25 domain-containing protein; afdb-uniprot50 \| AF-A0A7Y4R463-F1-MODEL\_V4 \| 1.0 \| 0.0001176 \| 192 \| 0.18 \| 100 \| 78 \| 2 \| 2 \| 97 \| 16 \| 115 \| GPW/gp25 family protein \| GPW/gp25 family protein; afdb-uniprot50 \| AF-A0A2D9N2U0-F1-MODEL\_V4 \| 1.0 \| 5.262e-05 \| 192 \| 0.137 \| 109 \| 89 \| 2 \| 3 \| 106 \| 37 \| 145 \| GPW\_gp25 domain-containing protein \| GPW\_gp25 domain-containing protein; afdb-uniprot50 \| AF-A0A353BET6-F1-MODEL\_V4 \| 1.0 \| 7.739e-06 \| 192 \| 0.2 \| 135 \| 77 \| 4 \| 3 \| 106 \| 27 \| 161 \| GPW\_gp25 domain-containing protein \| GPW\_gp25 domain-containing protein; afdb-uniprot50 \| AF-A0A519DRW4-F1-MODEL\_V4 \| 1.0 \| 0.0002051 \| 191 \| 0.255 \| 90 \| 64 \| 3 \| 24 \| 112 \| 3 \| 90 \| GPW\_gp25 domain-containing protein \| GPW\_gp25 domain-containing protein; afdb-uniprot50 \| AF-A0A7V8YDK1-F1-MODEL\_V4 \| 1.0 \| 4.65e-05 \| 191 \| 0.3 \| 90 \| 63 \| 0 \| 17 \| 106 \| 7 \| 96 \| GPW/gp25 family protein \| GPW/gp25 family protein; afdb-uniprot50 \| AF-A0A7C7GCM5-F1-MODEL\_V4 \| 1.0 \| 7.626e-05 \| 191 \| 0.154 \| 110 \| 89 \| 3 \| 2 \| 107 \| 10 \| 119 \| GPW\_gp25 domain-containing protein \| GPW\_gp25 domain-containing protein; afdb-uniprot50 \| AF-R5P4K2-F1-MODEL\_V4 \| 1.0 \| 4.371e-05 \| 191 \| 0.156 \| 115 \| 90 \| 4 \| 2 \| 110 \| 13 \| 126 \| GPW/gp25 family protein \| GPW/gp25 family protein; afdb-uniprot50 \| AF-A0A0Q9UAP1-F1-MODEL\_V4 \| 1.0 \| 3.016e-05 \| 191 \| 0.3 \| 113 \| 70 \| 3 \| 1 \| 104 \| 20 \| 132 \| GPW\_gp25 domain-containing protein \| GPW\_gp25 domain-containing protein; afdb-uniprot50 \| AF-A0A158DVL2-F1-MODEL\_V4 \| 1.0 \| 0.0001251 \| 191 \| 0.238 \| 109 \| 76 \| 6 \| 1 \| 106 \| 97 \| 201 \| Uncharacterized protein \| Uncharacterized protein; afdb-uniprot50 \| AF-A0A7R7TBH0-F1-MODEL\_V4 \| 1.0 \| 8.63e-05 \| 191 \| 0.227 \| 110 \| 78 \| 6 \| 1 \| 107 \| 98 \| 203 \| Uncharacterized protein \| Uncharacterized protein; afdb-uniprot50 \| AF-A0A328S0R0-F1-MODEL\_V4 \| 1.0 \| 0.0002051 \| 191 \| 0.198 \| 106 \| 80 \| 5 \| 3 \| 105 \| 16 \| 119 \| Uncharacterized protein \| Uncharacterized protein; afdb-uniprot50 \| AF-A0A1E4AXV8-F1-MODEL\_V4 \| 1.0 \| 0.0002182 \| 190 \| 0.247 \| 109 \| 77 \| 4 \| 5 \| 110 \| 1 \| 107 \| GPW\_gp25 domain-containing protein \| GPW\_gp25 domain-containing protein; afdb-uniprot50 \| AF-A0A0F9T6N1-F1-MODEL\_V4 \| 1.0 \| 8.63e-05 \| 190 \| 0.203 \| 108 \| 79 \| 4 \| 6 \| 106 \| 15 \| 122 \| GPW\_gp25 domain-containing protein \| GPW\_gp25 domain-containing protein; afdb-uniprot50 \| AF-A0A3A8KT25-F1-MODEL\_V4 \| 1.0 \| 0.0002469 \| 190 \| 0.169 \| 106 \| 85 \| 3 \| 2 \| 105 \| 10 \| 114 \| GPW\_gp25 domain-containing protein \| GPW\_gp25 domain-containing protein; afdb-uniprot50 \| AF-A0A0X3VIR3-F1-MODEL\_V4 \| 1.0 \| 3.92e-06 \| 190 \| 0.22 \| 118 \| 79 \| 6 \| 2 \| 107 \| 9 \| 125 \| GPW\_gp25 domain-containing protein \| GPW\_gp25 domain-containing protein; afdb-uniprot50 \| AF-A0A1G3L9L2-F1-MODEL\_V4 \| 1.0 \| 3.863e-05 \| 190 \| 0.175 \| 108 \| 84 \| 3 \| 1 \| 105 \| 15 \| 120 \| Uncharacterized protein \| Uncharacterized protein; afdb-uniprot50 \| AF-A0A1Z9M5W6-F1-MODEL\_V4 \| 1.0 \| 3.631e-05 \| 190 \| 0.165 \| 115 \| 86 \| 4 \| 4 \| 108 \| 22 \| 136 \| GPW\_gp25 domain-containing protein \| GPW\_gp25 domain-containing protein; afdb-uniprot50 \| AF-A0A6F8YYN1-F1-MODEL\_V4 \| 1.0 \| 9.767e-05 \| 190 \| 0.29 \| 100 \| 66 \| 3 \| 2 \| 97 \| 26 \| 124 \| Baseplate protein \| Baseplate protein; afdb-uniprot50 \| AF-A0A2S6WCH7-F1-MODEL\_V4 \| 1.0 \| 4.109e-05 \| 189 \| 0.241 \| 116 \| 77 \| 5 \| 2 \| 108 \| 13 \| 126 \| GPW\_gp25 domain-containing protein \| GPW\_gp25 domain-containing protein; afdb-uniprot50 \| AF-G8R6Y2-F1-MODEL\_V4 \| 1.0 \| 8.63e-05 \| 189 \| 0.257 \| 101 \| 70 \| 3 \| 2 \| 97 \| 16 \| 116 \| Phage baseplate assembly protein W \| Phage baseplate assembly protein W; afdb-uniprot50 \| AF-A0A6G3XJJ0-F1-MODEL\_V4 \| 1.0 \| 0.0003364 \| 188 \| 0.379 \| 79 \| 47 \| 1 \| 17 \| 93 \| 10 \| 88 \| GPW/gp25 family protein \| GPW/gp25 family protein; afdb-uniprot50 \| AF-A0A1V5LTW0-F1-MODEL\_V4 \| 1.0 \| 6.335e-05 \| 188 \| 0.265 \| 98 \| 65 \| 2 \| 2 \| 93 \| 13 \| 109 \| Gene 25-like lysozyme \| Gene 25-like lysozyme; afdb-uniprot50 \| AF-A0A1M7CV50-F1-MODEL\_V4 \| 1.0 \| 0.0002321 \| 188 \| 0.178 \| 101 \| 81 \| 1 \| 9 \| 107 \| 21 \| 121 \| Gene 25-like lysozyme \| Gene 25-like lysozyme; afdb-uniprot50 \| AF-A0A2P2GFT5-F1-MODEL\_V4 \| 1.0 \| 4.65e-05 \| 188 \| 0.226 \| 115 \| 79 \| 7 \| 1 \| 111 \| 20 \| 128 \| Uncharacterized protein \| Uncharacterized protein; afdb-uniprot50 \| AF-A0A656XVQ3-F1-MODEL\_V4 \| 1.0 \| 5.262e-05 \| 188 \| 0.263 \| 110 \| 75 \| 2 \| 1 \| 104 \| 18 \| 127 \| GPW\_gp25 domain-containing protein \| GPW\_gp25 domain-containing protein; afdb-uniprot50 \| AF-A0A1Z8QIV7-F1-MODEL\_V4 \| 1.0 \| 1.193e-05 \| 188 \| 0.254 \| 106 \| 65 \| 2 \| 2 \| 105 \| 17 \| 110 \| GPW\_gp25 domain-containing protein \| GPW\_gp25 domain-containing protein; afdb-uniprot50 \| AF-A0A258B3I1-F1-MODEL\_V4 \| 1.0 \| 9.767e-05 \| 188 \| 0.288 \| 104 \| 67 \| 3 \| 1 \| 97 \| 18 \| 121 \| GPW\_gp25 domain-containing protein \| GPW\_gp25 domain-containing protein; afdb-uniprot50 \| AF-A0A5B0W7Y0-F1-MODEL\_V4 \| 1.0 \| 5.955e-05 \| 188 \| 0.163 \| 110 \| 85 \| 3 \| 3 \| 105 \| 43 \| 152 \| GPW/gp25 family protein \| GPW/gp25 family protein; afdb-uniprot50 \| AF-A0A5C7Q1Z4-F1-MODEL\_V4 \| 1.0 \| 0.0001176 \| 188 \| 0.212 \| 108 \| 78 \| 6 \| 1 \| 105 \| 98 \| 201 \| Uncharacterized protein \| Uncharacterized protein; afdb-uniprot50 \| AF-T0C511-F1-MODEL\_V4 \| 1.0 \| 3.016e-05 \| 187 \| 0.228 \| 114 \| 80 \| 5 \| 1 \| 108 \| 1 \| 112 \| Uncharacterized protein \| Uncharacterized protein; afdb-uniprot50 \| AF-A0A2A5M1Q7-F1-MODEL\_V4 \| 1.0 \| 0.0001704 \| 187 \| 0.194 \| 113 \| 82 \| 4 \| 1 \| 105 \| 1 \| 112 \| GPW\_gp25 domain-containing protein \| GPW\_gp25 domain-containing protein; afdb-uniprot50 \| AF-A0A2D7NVZ0-F1-MODEL\_V4 \| 1.0 \| 2.081e-05 \| 187 \| 0.241 \| 112 \| 71 \| 3 \| 2 \| 109 \| 17 \| 118 \| GPW\_gp25 domain-containing protein \| GPW\_gp25 domain-containing protein; afdb-uniprot50 \| AF-A0A1P8YPY5-F1-MODEL\_V4 \| 1.0 \| 0.0005186 \| 186 \| 0.274 \| 91 \| 64 \| 2 \| 16 \| 105 \| 8 \| 97 \| Lysozyme family protein \| Lysozyme family protein; afdb-uniprot50 \| AF-A0A2E7W0M6-F1-MODEL\_V4 \| 1.0 \| 0.0001176 \| 186 \| 0.14 \| 114 \| 90 \| 3 \| 2 \| 107 \| 10 \| 123 \| GPW\_gp25 domain-containing protein \| GPW\_gp25 domain-containing protein; afdb-uniprot50 \| AF-R5PAT8-F1-MODEL\_V4 \| 1.0 \| 0.0001039 \| 186 \| 0.18 \| 105 \| 82 \| 2 \| 5 \| 105 \| 17 \| 121 \| GPW\_gp25 domain-containing protein \| GPW\_gp25 domain-containing protein; afdb-uniprot50 \| AF-A0A511V854-F1-MODEL\_V4 \| 1.0 \| 4.371e-05 \| 186 \| 0.2 \| 110 \| 80 \| 6 \| 1 \| 105 \| 15 \| 121 \| Uncharacterized protein \| Uncharacterized protein; afdb-uniprot50 \| AF-A0A5C1YDB3-F1-MODEL\_V4 \| 1.0 \| 1.528e-05 \| 186 \| 0.321 \| 109 \| 59 \| 3 \| 2 \| 108 \| 14 \| 109 \| GPW/gp25 family protein \| GPW/gp25 family protein; afdb-uniprot50 \| AF-A0A1I0VPW8-F1-MODEL\_V4 \| 1.0 \| 0.0001039 \| 185 \| 0.146 \| 116 \| 87 \| 4 \| 1 \| 105 \| 1 \| 115 \| GPW\_gp25 domain-containing protein \| GPW\_gp25 domain-containing protein; afdb-uniprot50 \| AF-A0A502HL99-F1-MODEL\_V4 \| 1.0 \| 0.0001331 \| 185 \| 0.18 \| 105 \| 78 \| 5 \| 3 \| 106 \| 18 \| 115 \| Uncharacterized protein \| Uncharacterized protein; afdb-uniprot50 \| AF-A0A2E9V569-F1-MODEL\_V4 \| 1.0 \| 9.767e-05 \| 185 \| 0.163 \| 110 \| 88 \| 3 \| 2 \| 107 \| 10 \| 119 \| GPW\_gp25 domain-containing protein \| GPW\_gp25 domain-containing protein; afdb-uniprot50 \| AF-A0A126QLP9-F1-MODEL\_V4 \| 1.0 \| 3.016e-05 \| 185 \| 0.165 \| 133 \| 83 \| 6 \| 1 \| 107 \| 8 \| 138 \| Baseplate assembly protein \| Baseplate assembly protein; afdb-uniprot50 \| AF-A0A417HJ04-F1-MODEL\_V4 \| 1.0 \| 4.371e-05 \| 184 \| 0.189 \| 111 \| 81 \| 4 \| 2 \| 105 \| 11 \| 119 \| GPW\_gp25 domain-containing protein \| GPW\_gp25 domain-containing protein; afdb-uniprot50 \| AF-A0A843GHX7-F1-MODEL\_V4 \| 1.0 \| 0.0001928 \| 184 \| 0.126 \| 119 \| 91 \| 3 \| 2 \| 107 \| 5 \| 123 \| GPW/gp25 family protein \| GPW/gp25 family protein; afdb-uniprot50 \| AF-A0A2U1W193-F1-MODEL\_V4 \| 1.0 \| 0.0001105 \| 184 \| 0.185 \| 108 \| 83 \| 5 \| 1 \| 106 \| 96 \| 200 \| Uncharacterized protein \| Uncharacterized protein; afdb-uniprot50 \| AF-A0A7Y5UCD9-F1-MODEL\_V4 \| 1.0 \| 0.0003162 \| 183 \| 0.325 \| 83 \| 54 \| 1 \| 25 \| 105 \| 2 \| 84 \| GPW/gp25 family protein \| GPW/gp25 family protein; afdb-uniprot50 \| AF-A0A2T5J3U2-F1-MODEL\_V4 \| 1.0 \| 6.739e-05 \| 183 \| 0.215 \| 116 \| 80 \| 3 \| 1 \| 106 \| 1 \| 115 \| Phage gp46-like protein \| Phage gp46-like protein; afdb-uniprot50 \| AF-A0A502IRR1-F1-MODEL\_V4 \| 1.0 \| 0.0001176 \| 183 \| 0.132 \| 113 \| 92 \| 4 \| 1 \| 107 \| 14 \| 126 \| DUF2634 domain-containing protein \| DUF2634 domain-containing protein; afdb-uniprot50 \| AF-A0A1Y4WCG8-F1-MODEL\_V4 \| 1.0 \| 0.0003579 \| 182 \| 0.282 \| 92 \| 62 \| 3 \| 17 \| 105 \| 7 \| 97 \| GPW\_gp25 domain-containing protein \| GPW\_gp25 domain-containing protein; afdb-uniprot50 \| AF-A0A1M7ZBF7-F1-MODEL\_V4 \| 1.0 \| 0.0002627 \| 182 \| 0.23 \| 100 \| 74 \| 2 \| 2 \| 98 \| 16 \| 115 \| GPW\_gp25 domain-containing protein \| GPW\_gp25 domain-containing protein; afdb-uniprot50 \| AF-A0A2E5AVQ0-F1-MODEL\_V4 \| 1.0 \| 9.767e-05 \| 182 \| 0.207 \| 111 \| 80 \| 4 \| 3 \| 107 \| 22 \| 130 \| GPW\_gp25 domain-containing protein \| GPW\_gp25 domain-containing protein; afdb-uniprot50 \| AF-E3PRX9-F1-MODEL\_V4 \| 1.0 \| 0.0001105 \| 181 \| 0.203 \| 108 \| 79 \| 5 \| 1 \| 106 \| 12 \| 114 \| Uncharacterized protein \| Uncharacterized protein; afdb-uniprot50 \| AF-A0A327JB01-F1-MODEL\_V4 \| 1.0 \| 0.0005186 \| 181 \| 0.148 \| 101 \| 85 \| 1 \| 5 \| 105 \| 30 \| 129 \| Uncharacterized protein \| Uncharacterized protein; afdb-uniprot50 \| AF-A0A2W6A5B1-F1-MODEL\_V4 \| 1.0 \| 4.371e-05 \| 181 \| 0.26 \| 115 \| 67 \| 5 \| 2 \| 109 \| 14 \| 117 \| Baseplate protein \| Baseplate protein; afdb-uniprot50 \| AF-A0A0F7FGC4-F1-MODEL\_V4 \| 1.0 \| 0.0001251 \| 180 \| 0.186 \| 107 \| 80 \| 4 \| 5 \| 107 \| 9 \| 112 \| GPW\_gp25 domain-containing protein \| GPW\_gp25 domain-containing protein; afdb-uniprot50 \| AF-A0A2D6AX80-F1-MODEL\_V4 \| 1.0 \| 0.0001039 \| 180 \| 0.15 \| 113 \| 88 \| 4 \| 2 \| 107 \| 28 \| 139 \| GPW\_gp25 domain-containing protein \| GPW\_gp25 domain-containing protein; afdb-uniprot50 \| AF-A0A2E2ZDB2-F1-MODEL\_V4 \| 1.0 \| 9.767e-05 \| 180 \| 0.185 \| 113 \| 84 \| 4 \| 3 \| 107 \| 35 \| 147 \| GPW\_gp25 domain-containing protein \| GPW\_gp25 domain-containing protein; afdb-uniprot50 \| AF-A0A1Z8W2C4-F1-MODEL\_V4 \| 1.0 \| 9.181e-05 \| 180 \| 0.193 \| 119 \| 83 \| 4 \| 2 \| 107 \| 27 \| 145 \| GPW\_gp25 domain-containing protein \| GPW\_gp25 domain-containing protein; afdb-uniprot50 \| AF-A0A6P0S081-F1-MODEL\_V4 \| 1.0 \| 0.0001506 \| 179 \| 0.255 \| 86 \| 62 \| 1 \| 2 \| 85 \| 14 \| 99 \| GPW/gp25 family protein \| GPW/gp25 family protein; afdb-uniprot50 \| AF-A0A7C3UU77-F1-MODEL\_V4 \| 1.0 \| 0.0001251 \| 178 \| 0.154 \| 110 \| 87 \| 3 \| 2 \| 105 \| 13 \| 122 \| GPW\_gp25 domain-containing protein \| GPW\_gp25 domain-containing protein; afdb-uniprot50 \| AF-A0A223D2Z4-F1-MODEL\_V4 \| 1.0 \| 0.0001506 \| 178 \| 0.198 \| 106 \| 80 \| 4 \| 1 \| 105 \| 15 \| 116 \| Uncharacterized protein \| Uncharacterized protein; afdb-uniprot50 \| AF-A0A1Z4H0Z3-F1-MODEL\_V4 \| 1.0 \| 0.0002469 \| 178 \| 0.24 \| 100 \| 72 \| 2 \| 2 \| 97 \| 17 \| 116 \| GPW\_gp25 domain-containing protein \| GPW\_gp25 domain-containing protein; afdb-uniprot50 \| AF-A0A382CDC4-F1-MODEL\_V4 \| 1.0 \| 0.0004583 \| 177 \| 0.244 \| 98 \| 71 \| 2 \| 3 \| 97 \| 6 \| 103 \| GPW\_gp25 domain-containing protein \| GPW\_gp25 domain-containing protein; afdb-uniprot50 \| AF-A0A6I5ZQ21-F1-MODEL\_V4 \| 1.0 \| 0.0001105 \| 177 \| 0.153 \| 111 \| 86 \| 5 \| 1 \| 105 \| 14 \| 122 \| Uncharacterized protein \| Uncharacterized protein; afdb-uniprot50 \| AF-A0A2D6X0Z9-F1-MODEL\_V4 \| 1.0 \| 9.181e-05 \| 177 \| 0.118 \| 118 \| 92 \| 5 \| 2 \| 107 \| 27 \| 144 \| GPW\_gp25 domain-containing protein \| GPW\_gp25 domain-containing protein; afdb-uniprot50 \| AF-A0A1V5IBU2-F1-MODEL\_V4 \| 1.0 \| 0.0003162 \| 177 \| 0.157 \| 133 \| 85 \| 3 \| 2 \| 107 \| 15 \| 147 \| Gene 25-like lysozyme \| Gene 25-like lysozyme; afdb-uniprot50 \| AF-A0A7X6JWS6-F1-MODEL\_V4 \| 1.0 \| 0.0001039 \| 176 \| 0.415 \| 77 \| 40 \| 1 \| 33 \| 104 \| 1 \| 77 \| GPW\_gp25 domain-containing protein \| GPW\_gp25 domain-containing protein; afdb-uniprot50 \| AF-A0A239C2Q9-F1-MODEL\_V4 \| 1.0 \| 7.169e-05 \| 176 \| 0.201 \| 114 \| 82 \| 5 \| 2 \| 108 \| 7 \| 118 \| GPW\_gp25 domain-containing protein \| GPW\_gp25 domain-containing protein; afdb-uniprot50 \| AF-J1FA57-F1-MODEL\_V4 \| 1.0 \| 0.0002794 \| 176 \| 0.169 \| 118 \| 87 \| 4 \| 1 \| 108 \| 2 \| 118 \| Phage baseplate assembly protein W \| Phage baseplate assembly protein W; afdb-uniprot50 \| AF-A0A2M9X3I2-F1-MODEL\_V4 \| 1.0 \| 0.0001602 \| 176 \| 0.176 \| 102 \| 76 \| 3 \| 10 \| 105 \| 24 \| 123 \| Baseplate wedge subunit \| Baseplate wedge subunit; afdb-uniprot50 \| AF-A0A7C5ISD4-F1-MODEL\_V4 \| 1.0 \| 4.947e-05 \| 176 \| 0.188 \| 106 \| 71 \| 2 \| 2 \| 105 \| 17 \| 109 \| GPW\_gp25 domain-containing protein \| GPW\_gp25 domain-containing protein; afdb-uniprot50 \| AF-A0A0P0QIR3-F1-MODEL\_V4 \| 1.0 \| 3.863e-05 \| 176 \| 0.203 \| 113 \| 79 \| 4 \| 1 \| 105 \| 21 \| 130 \| GPW\_gp25 domain-containing protein \| GPW\_gp25 domain-containing protein; afdb-uniprot50 \| AF-A0A2E7N873-F1-MODEL\_V4 \| 1.0 \| 0.0001813 \| 176 \| 0.163 \| 110 \| 83 \| 4 \| 3 \| 105 \| 33 \| 140 \| GPW\_gp25 domain-containing protein \| GPW\_gp25 domain-containing protein; afdb-uniprot50 \| AF-V4QXD0-F1-MODEL\_V4 \| 1.0 \| 0.0003364 \| 175 \| 0.309 \| 71 \| 49 \| 0 \| 35 \| 105 \| 2 \| 72 \| Baseplate assembly protein \| Baseplate assembly protein; afdb-uniprot50 \| AF-A0A661CXS2-F1-MODEL\_V4 \| 1.0 \| 0.0002051 \| 175 \| 0.131 \| 114 \| 85 \| 4 \| 3 \| 106 \| 16 \| 125 \| GPW\_gp25 domain-containing protein \| GPW\_gp25 domain-containing protein; afdb-uniprot50 \| AF-A0A3B8IBV9-F1-MODEL\_V4 \| 1.0 \| 0.0002794 \| 175 \| 0.242 \| 99 \| 73 \| 1 \| 1 \| 97 \| 17 \| 115 \| GPW\_gp25 domain-containing protein \| GPW\_gp25 domain-containing protein; afdb-uniprot50 \| AF-A0A1Q4EDA0-F1-MODEL\_V4 \| 1.0 \| 0.0001415 \| 175 \| 0.176 \| 113 \| 84 \| 5 \| 2 \| 107 \| 9 \| 119 \| GPW\_gp25 domain-containing protein \| GPW\_gp25 domain-containing protein; afdb-uniprot50 \| AF-A0A0F6SF01-F1-MODEL\_V4 \| 1.0 \| 0.0003364 \| 175 \| 0.18 \| 105 \| 79 \| 6 \| 6 \| 105 \| 40 \| 142 \| Uncharacterized protein \| Uncharacterized protein; afdb-uniprot50 \| AF-A0A2D6E562-F1-MODEL\_V4 \| 1.0 \| 9.181e-05 \| 175 \| 0.195 \| 123 \| 83 \| 5 \| 2 \| 111 \| 48 \| 167 \| Uncharacterized protein \| Uncharacterized protein; afdb-uniprot50 \| AF-E8WUR4-F1-MODEL\_V4 \| 1.0 \| 2.506e-05 \| 175 \| 0.221 \| 113 \| 72 \| 4 \| 2 \| 108 \| 14 \| 116 \| GPW/gp25 family protein \| GPW/gp25 family protein; afdb-uniprot50 \| AF-A0A2A2B515-F1-MODEL\_V4 \| 1.0 \| 3.413e-05 \| 174 \| 0.203 \| 113 \| 82 \| 4 \| 3 \| 108 \| 10 \| 121 \| Uncharacterized protein \| Uncharacterized protein; afdb-uniprot50 \| AF-A0A2E4W4D6-F1-MODEL\_V4 \| 1.0 \| 0.0001704 \| 174 \| 0.203 \| 113 \| 74 \| 2 \| 5 \| 106 \| 1 \| 108 \| Uncharacterized protein \| Uncharacterized protein; afdb-uniprot50 \| AF-A0A1J0LUS2-F1-MODEL\_V4 \| 1.0 \| 0.0001176 \| 174 \| 0.228 \| 114 \| 76 \| 7 \| 1 \| 107 \| 5 \| 113 \| Uncharacterized protein \| Uncharacterized protein; afdb-uniprot50 \| AF-A0A3P6JCI1-F1-MODEL\_V4 \| 1.0 \| 4.65e-05 \| 174 \| 0.236 \| 114 \| 75 \| 3 \| 3 \| 105 \| 10 \| 122 \| Phage baseplate assembly protein W \| Phage baseplate assembly protein W; afdb-uniprot50 \| AF-A0A0C2UZ28-F1-MODEL\_V4 \| 1.0 \| 0.0001176 \| 174 \| 0.175 \| 114 \| 86 \| 5 \| 1 \| 111 \| 170 \| 278 \| Uncharacterized protein \| Uncharacterized protein; afdb-uniprot50 \| AF-A0A7C7FT58-F1-MODEL\_V4 \| 1.0 \| 0.0002627 \| 173 \| 0.154 \| 97 \| 75 \| 4 \| 18 \| 107 \| 2 \| 98 \| GPW\_gp25 domain-containing protein \| GPW\_gp25 domain-containing protein; afdb-uniprot50 \| AF-S4XZU3-F1-MODEL\_V4 \| 1.0 \| 0.0001415 \| 173 \| 0.194 \| 113 \| 86 \| 5 \| 2 \| 111 \| 12 \| 122 \| GPW\_gp25 domain-containing protein \| GPW\_gp25 domain-containing protein; afdb-uniprot50 \| AF-A0A2E0MEK5-F1-MODEL\_V4 \| 1.0 \| 0.0003162 \| 173 \| 0.169 \| 112 \| 86 \| 3 \| 3 \| 107 \| 20 \| 131 \| GPW\_gp25 domain-containing protein \| GPW\_gp25 domain-containing protein; afdb-uniprot50 \| AF-A0A3D1IZH3-F1-MODEL\_V4 \| 1.0 \| 0.0005186 \| 172 \| 0.163 \| 110 \| 87 \| 5 \| 1 \| 108 \| 76 \| 182 \| Uncharacterized protein \| Uncharacterized protein; afdb-uniprot50 \| AF-A0A6I6E3I6-F1-MODEL\_V4 \| 1.0 \| 6.739e-05 \| 171 \| 0.256 \| 113 \| 75 \| 3 \| 1 \| 105 \| 1 \| 112 \| GPW\_gp25 domain-containing protein \| GPW\_gp25 domain-containing protein; afdb-uniprot50 \| AF-A0A382V1Y7-F1-MODEL\_V4 \| 1.0 \| 7.169e-05 \| 171 \| 0.176 \| 113 \| 82 \| 6 \| 3 \| 107 \| 26 \| 135 \| GPW\_gp25 domain-containing protein \| GPW\_gp25 domain-containing protein; afdb-uniprot50 \| AF-A0A1G3MBB2-F1-MODEL\_V4 \| 1.0 \| 0.0001813 \| 171 \| 0.223 \| 112 \| 82 \| 3 \| 5 \| 112 \| 28 \| 138 \| GPW\_gp25 domain-containing protein \| GPW\_gp25 domain-containing protein; afdb-uniprot50 \| AF-A0A1Q7Z6P2-F1-MODEL\_V4 \| 1.0 \| 0.0006243 \| 170 \| 0.305 \| 85 \| 57 \| 2 \| 2 \| 85 \| 14 \| 97 \| GPW\_gp25 domain-containing protein \| GPW\_gp25 domain-containing protein; afdb-uniprot50 \| AF-A0A2T5P176-F1-MODEL\_V4 \| 1.0 \| 0.0002469 \| 170 \| 0.25 \| 108 \| 75 \| 5 \| 2 \| 105 \| 3 \| 108 \| Uncharacterized protein \| Uncharacterized protein; afdb-uniprot50 \| AF-A0A0F4NK88-F1-MODEL\_V4 \| 1.0 \| 0.0006642 \| 170 \| 0.342 \| 76 \| 49 \| 1 \| 8 \| 82 \| 27 \| 102 \| GPW\_gp25 domain-containing protein \| GPW\_gp25 domain-containing protein; afdb-uniprot50 \| AF-A0A3D4UYF4-F1-MODEL\_V4 \| 1.0 \| 0.0001704 \| 170 \| 0.185 \| 113 \| 84 \| 3 \| 3 \| 107 \| 15 \| 127 \| Uncharacterized protein \| Uncharacterized protein; afdb-uniprot50 \| AF-A0A7C3ETG1-F1-MODEL\_V4 \| 1.0 \| 0.0001928 \| 170 \| 0.155 \| 116 \| 86 \| 4 \| 2 \| 105 \| 10 \| 125 \| GPW\_gp25 domain-containing protein \| GPW\_gp25 domain-containing protein; afdb-uniprot50 \| AF-A0A1Z4C3B0-F1-MODEL\_V4 \| 1.0 \| 0.0004308 \| 170 \| 0.165 \| 115 \| 89 \| 6 \| 1 \| 112 \| 85 \| 195 \| Uncharacterized protein \| Uncharacterized protein; afdb-uniprot50 \| AF-A0A7M2RZX2-F1-MODEL\_V4 \| 1.0 \| 0.0002051 \| 169 \| 0.171 \| 105 \| 87 \| 0 \| 3 \| 107 \| 10 \| 114 \| Uncharacterized protein \| Uncharacterized protein; afdb-uniprot50 \| AF-A0A350PIY9-F1-MODEL\_V4 \| 1.0 \| 0.0003364 \| 169 \| 0.119 \| 109 \| 91 \| 2 \| 2 \| 105 \| 18 \| 126 \| GPW\_gp25 domain-containing protein \| GPW\_gp25 domain-containing protein; afdb-uniprot50 \| AF-A0A511N764-F1-MODEL\_V4 \| 1.0 \| 0.0006243 \| 169 \| 0.163 \| 116 \| 87 \| 2 \| 1 \| 107 \| 82 \| 196 \| Uncharacterized protein \| Uncharacterized protein; afdb-uniprot50 \| AF-A0A5S9R4I1-F1-MODEL\_V4 \| 1.0 \| 0.0002973 \| 168 \| 0.257 \| 97 \| 60 \| 2 \| 23 \| 107 \| 2 \| 98 \| Uncharacterized protein \| Uncharacterized protein; afdb-uniprot50 \| AF-A0A2D0N0Z6-F1-MODEL\_V4 \| 1.0 \| 9.181e-05 \| 168 \| 0.196 \| 112 \| 76 \| 3 \| 2 \| 109 \| 21 \| 122 \| GPW\_gp25 domain-containing protein \| GPW\_gp25 domain-containing protein; afdb-uniprot50 \| AF-A0A209AE83-F1-MODEL\_V4 \| 1.0 \| 0.0001928 \| 168 \| 0.303 \| 79 \| 53 \| 1 \| 27 \| 105 \| 92 \| 168 \| Phage\_base\_V domain-containing protein \| Phage\_base\_V domain-containing protein; afdb-uniprot50 \| AF-A0A3C0GIN9-F1-MODEL\_V4 \| 1.0 \| 0.0001251 \| 167 \| 0.162 \| 123 \| 82 \| 6 \| 2 \| 106 \| 10 \| 129 \| GPW\_gp25 domain-containing protein \| GPW\_gp25 domain-containing protein; afdb-uniprot50 \| AF-A0A2K9P038-F1-MODEL\_V4 \| 1.0 \| 0.0002627 \| 167 \| 0.228 \| 114 \| 76 \| 8 \| 1 \| 107 \| 24 \| 132 \| Phage protein \| Phage protein; afdb-uniprot50 \| AF-A0A8A6B8Z7-F1-MODEL\_V4 \| 1.0 \| 0.0002321 \| 167 \| 0.196 \| 112 \| 81 \| 6 \| 2 \| 108 \| 26 \| 133 \| DUF2634 domain-containing protein \| DUF2634 domain-containing protein; afdb-uniprot50 \| AF-A0A7U9RXB1-F1-MODEL\_V4 \| 1.0 \| 0.0001176 \| 167 \| 0.247 \| 113 \| 78 \| 5 \| 1 \| 108 \| 36 \| 146 \| Uncharacterized protein \| Uncharacterized protein; afdb-uniprot50 \| AF-A0A841VB05-F1-MODEL\_V4 \| 1.0 \| 0.0005869 \| 166 \| 0.325 \| 86 \| 54 \| 2 \| 2 \| 86 \| 14 \| 96 \| GPW/gp25 family protein \| GPW/gp25 family protein; afdb-uniprot50 \| AF-A0A6M0RCZ2-F1-MODEL\_V4 \| 1.0 \| 0.0005186 \| 166 \| 0.166 \| 126 \| 87 \| 6 \| 1 \| 112 \| 1 \| 122 \| Uncharacterized protein \| Uncharacterized protein; afdb-uniprot50 \| AF-A0A517VBS1-F1-MODEL\_V4 \| 1.0 \| 0.0005186 \| 166 \| 0.15 \| 120 \| 89 \| 5 \| 1 \| 108 \| 1 \| 119 \| Gene 25-like lysozyme \| Gene 25-like lysozyme; afdb-uniprot50 \| AF-N1ZEV8-F1-MODEL\_V4 \| 1.0 \| 8.113e-05 \| 166 \| 0.196 \| 122 \| 82 \| 5 \| 1 \| 107 \| 1 \| 121 \| Uncharacterized protein \| Uncharacterized protein; afdb-uniprot50 \| AF-V7ZNH0-F1-MODEL\_V4 \| 1.0 \| 0.0002321 \| 166 \| 0.187 \| 112 \| 84 \| 5 \| 2 \| 108 \| 16 \| 125 \| Uncharacterized protein \| Uncharacterized protein; afdb-uniprot50 \| AF-A0A1W9S296-F1-MODEL\_V4 \| 1.0 \| 0.0004308 \| 166 \| 0.179 \| 106 \| 85 \| 2 \| 1 \| 105 \| 15 \| 119 \| Uncharacterized protein \| Uncharacterized protein; afdb-uniprot50 \| AF-A0A1I0WHG9-F1-MODEL\_V4 \| 1.0 \| 0.0005186 \| 166 \| 0.221 \| 113 \| 77 \| 6 \| 2 \| 107 \| 23 \| 131 \| Uncharacterized protein \| Uncharacterized protein; afdb-uniprot50 \| AF-A0A4R7BFE6-F1-MODEL\_V4 \| 1.0 \| 0.0001176 \| 166 \| 0.214 \| 112 \| 74 \| 3 \| 2 \| 109 \| 13 \| 114 \| GPW\_gp25 domain-containing protein \| GPW\_gp25 domain-containing protein; afdb-uniprot50 \| AF-A0A1X1NFH7-F1-MODEL\_V4 \| 1.0 \| 0.0001251 \| 166 \| 0.201 \| 124 \| 79 \| 4 \| 2 \| 105 \| 247 \| 370 \| Uncharacterized protein \| Uncharacterized protein; afdb-uniprot50 \| AF-A0A7W5K3U6-F1-MODEL\_V4 \| 1.0 \| 8.113e-05 \| 165 \| 0.209 \| 110 \| 77 \| 4 \| 2 \| 105 \| 14 \| 119 \| GPW\_gp25 domain-containing protein \| GPW\_gp25 domain-containing protein; afdb-uniprot50 \| AF-A0A539E6I3-F1-MODEL\_V4 \| 1.0 \| 0.0007995 \| 165 \| 0.205 \| 102 \| 75 \| 2 \| 2 \| 97 \| 19 \| 120 \| GPW/gp25 \| GPW/gp25; afdb-uniprot50 \| AF-A0A2K8Z4R5-F1-MODEL\_V4 \| 1.0 \| 5.598e-05 \| 165 \| 0.203 \| 108 \| 72 \| 2 \| 2 \| 107 \| 14 \| 109 \| GPW\_gp25 domain-containing protein \| GPW\_gp25 domain-containing protein; afdb-uniprot50 \| AF-A0A0L6JGL6-F1-MODEL\_V4 \| 1.0 \| 0.0005869 \| 165 \| 0.215 \| 116 \| 84 \| 4 \| 1 \| 112 \| 24 \| 136 \| Phage-like element PBSX protein, XkdS \| Phage-like element PBSX protein, XkdS; afdb-uniprot50 \| AF-C6HYF2-F1-MODEL\_V4 \| 1.0 \| 0.0002051 \| 164 \| 0.276 \| 94 \| 62 \| 5 \| 17 \| 106 \| 32 \| 123 \| GPW/gp25 family protein \| GPW/gp25 family protein; afdb-uniprot50 \| AF-A0A1H9YIU9-F1-MODEL\_V4 \| 1.0 \| 0.0004875 \| 164 \| 0.222 \| 108 \| 78 \| 6 \| 1 \| 105 \| 27 \| 131 \| Uncharacterized protein \| Uncharacterized protein; afdb-uniprot50 \| AF-A0A2D6F2M5-F1-MODEL\_V4 \| 1.0 \| 0.0002469 \| 164 \| 0.229 \| 109 \| 74 \| 8 \| 3 \| 105 \| 54 \| 158 \| Uncharacterized protein \| Uncharacterized protein; afdb-uniprot50 \| AF-A0A2A5HV32-F1-MODEL\_V4 \| 1.0 \| 0.0001039 \| 164 \| 0.226 \| 106 \| 68 \| 2 \| 2 \| 105 \| 49 \| 142 \| GPW\_gp25 domain-containing protein \| GPW\_gp25 domain-containing protein; afdb-uniprot50 \| AF-A0A2D5F9D6-F1-MODEL\_V4 \| 1.0 \| 0.0009625 \| 163 \| 0.132 \| 113 \| 92 \| 3 \| 2 \| 111 \| 36 \| 145 \| GPW\_gp25 domain-containing protein \| GPW\_gp25 domain-containing protein; afdb-uniprot50 \| AF-A0A252EC99-F1-MODEL\_V4 \| 1.0 \| 0.001024 \| 163 \| 0.205 \| 107 \| 81 \| 2 \| 2 \| 104 \| 33 \| 139 \| GPW\_gp25 domain-containing protein \| GPW\_gp25 domain-containing protein; afdb-uniprot50 \| AF-A0A5J4RAP1-F1-MODEL\_V4 \| 1.0 \| 0.0007516 \| 162 \| 0.13 \| 100 \| 82 \| 3 \| 2 \| 97 \| 13 \| 111 \| GPW\_gp25 domain-containing protein \| GPW\_gp25 domain-containing protein; afdb-uniprot50 \| AF-A0A7C7C1N4-F1-MODEL\_V4 \| 1.0 \| 0.0006642 \| 162 \| 0.155 \| 109 \| 88 \| 3 \| 2 \| 106 \| 10 \| 118 \| GPW\_gp25 domain-containing protein \| GPW\_gp25 domain-containing protein; afdb-uniprot50 \| AF-A0A4Q7YHD4-F1-MODEL\_V4 \| 1.0 \| 0.0003364 \| 162 \| 0.245 \| 106 \| 74 \| 4 \| 2 \| 102 \| 9 \| 113 \| GPW\_gp25 domain-containing protein \| GPW\_gp25 domain-containing protein; afdb-uniprot50 \| AF-A0A1M6M4V6-F1-MODEL\_V4 \| 1.0 \| 0.000405 \| 162 \| 0.176 \| 113 \| 82 \| 7 \| 1 \| 107 \| 36 \| 143 \| Uncharacterized protein \| Uncharacterized protein; afdb-uniprot50 \| AF-C3X352-F1-MODEL\_V4 \| 1.0 \| 0.0006243 \| 162 \| 0.171 \| 111 \| 86 \| 5 \| 1 \| 110 \| 94 \| 199 \| Uncharacterized protein \| Uncharacterized protein; afdb-uniprot50 \| AF-A0A5E4PHX7-F1-MODEL\_V4 \| 1.0 \| 0.0002973 \| 161 \| 0.184 \| 114 \| 81 \| 5 \| 1 \| 107 \| 10 \| 118 \| Uncharacterized protein \| Uncharacterized protein; afdb-uniprot50 \| AF-A0A074LKB3-F1-MODEL\_V4 \| 1.0 \| 0.0001176 \| 161 \| 0.175 \| 114 \| 83 \| 8 \| 2 \| 110 \| 16 \| 123 \| Uncharacterized protein \| Uncharacterized protein; afdb-uniprot50 \| AF-A0A842TDK5-F1-MODEL\_V4 \| 1.0 \| 0.0003364 \| 161 \| 0.224 \| 107 \| 79 \| 3 \| 3 \| 106 \| 17 \| 122 \| Uncharacterized protein \| Uncharacterized protein; afdb-uniprot50 \| AF-L0R441-F1-MODEL\_V4 \| 1.0 \| 0.0007995 \| 161 \| 0.193 \| 93 \| 70 \| 4 \| 17 \| 105 \| 31 \| 122 \| Putative Type VI secretion system lysozyme-related protein \| Putative Type VI secretion system lysozyme-related protein; afdb-uniprot50 \| AF-A0A2A5HX32-F1-MODEL\_V4 \| 1.0 \| 0.0001602 \| 161 \| 0.221 \| 113 \| 72 \| 4 \| 2 \| 109 \| 16 \| 117 \| GPW\_gp25 domain-containing protein \| GPW\_gp25 domain-containing protein; afdb-uniprot50 \| AF-A0A3C0G474-F1-MODEL\_V4 \| 1.0 \| 0.0002469 \| 161 \| 0.121 \| 123 \| 87 \| 6 \| 1 \| 105 \| 22 \| 141 \| GPW\_gp25 domain-containing protein \| GPW\_gp25 domain-containing protein; afdb-uniprot50 \| AF-A0A1J0LXL0-F1-MODEL\_V4 \| 1.0 \| 7.626e-05 \| 160 \| 0.192 \| 109 \| 77 \| 6 \| 1 \| 107 \| 11 \| 110 \| Uncharacterized protein \| Uncharacterized protein; afdb-uniprot50 \| AF-A0A4Z0PFZ4-F1-MODEL\_V4 \| 1.0 \| 0.0001506 \| 160 \| 0.216 \| 106 \| 69 \| 2 \| 2 \| 105 \| 26 \| 119 \| GPW\_gp25 domain-containing protein \| GPW\_gp25 domain-containing protein; afdb-uniprot50 \| AF-A0A090QXN9-F1-MODEL\_V4 \| 1.0 \| 0.0004875 \| 159 \| 0.213 \| 103 \| 77 \| 2 \| 3 \| 105 \| 1 \| 99 \| Uncharacterized protein \| Uncharacterized protein; afdb-uniprot50 \| AF-A0A7J5EX88-F1-MODEL\_V4 \| 1.0 \| 0.0006243 \| 159 \| 0.18 \| 105 \| 80 \| 3 \| 2 \| 106 \| 9 \| 107 \| Uncharacterized protein \| Uncharacterized protein; afdb-uniprot50 \| AF-C3X496-F1-MODEL\_V4 \| 1.0 \| 0.0001176 \| 159 \| 0.188 \| 122 \| 82 \| 6 \| 3 \| 108 \| 1 \| 121 \| GPW\_gp25 domain-containing protein \| GPW\_gp25 domain-containing protein; afdb-uniprot50 \| AF-R5H3R4-F1-MODEL\_V4 \| 1.0 \| 0.0007516 \| 159 \| 0.177 \| 118 \| 85 \| 8 \| 1 \| 112 \| 20 \| 131 \| Uncharacterized protein \| Uncharacterized protein; afdb-uniprot50 \| AF-A0A522CGL7-F1-MODEL\_V4 \| 1.0 \| 0.001395 \| 159 \| 0.16 \| 112 \| 84 \| 6 \| 1 \| 105 \| 10 \| 118 \| Uncharacterized protein \| Uncharacterized protein; afdb-uniprot50 \| AF-A0A7M1PWR8-F1-MODEL\_V4 \| 1.0 \| 0.0008505 \| 158 \| 0.177 \| 118 \| 85 \| 8 \| 1 \| 112 \| 23 \| 134 \| DUF2634 domain-containing protein \| DUF2634 domain-containing protein; afdb-uniprot50 \| AF-A0A1I3XYL5-F1-MODEL\_V4 \| 1.0 \| 0.0001039 \| 158 \| 0.208 \| 115 \| 73 \| 5 \| 2 \| 109 \| 16 \| 119 \| GPW\_gp25 domain-containing protein \| GPW\_gp25 domain-containing protein; afdb-uniprot50 \| AF-A0A7C0UDU3-F1-MODEL\_V4 \| 1.0 \| 0.000405 \| 157 \| 0.18 \| 111 \| 79 \| 6 \| 2 \| 105 \| 13 \| 118 \| GPW\_gp25 domain-containing protein \| GPW\_gp25 domain-containing protein; afdb-uniprot50 \| AF-A0A6N4GXB3-F1-MODEL\_V4 \| 1.0 \| 0.0002794 \| 157 \| 0.216 \| 106 \| 68 \| 2 \| 2 \| 105 \| 16 \| 108 \| GPW\_gp25 domain-containing protein \| GPW\_gp25 domain-containing protein; afdb-uniprot50 \| AF-A0A2D0MZD6-F1-MODEL\_V4 \| 1.0 \| 0.0001506 \| 157 \| 0.216 \| 106 \| 69 \| 2 \| 2 \| 105 \| 16 \| 109 \| GPW\_gp25 domain-containing protein \| GPW\_gp25 domain-containing protein; afdb-uniprot50 \| AF-A0A1C6BPB7-F1-MODEL\_V4 \| 1.0 \| 0.0002469 \| 156 \| 0.229 \| 109 \| 79 \| 4 \| 1 \| 106 \| 9 \| 115 \| Uncharacterized protein \| Uncharacterized protein; afdb-uniprot50 \| AF-H6NDT0-F1-MODEL\_V4 \| 1.0 \| 0.0002627 \| 156 \| 0.196 \| 117 \| 79 \| 8 \| 1 \| 108 \| 11 \| 121 \| Uncharacterized protein \| Uncharacterized protein; afdb-uniprot50 \| AF-A0A1F6G8V8-F1-MODEL\_V4 \| 1.0 \| 0.0001813 \| 156 \| 0.188 \| 106 \| 71 \| 2 \| 2 \| 105 \| 15 \| 107 \| GPW\_gp25 domain-containing protein \| GPW\_gp25 domain-containing protein; afdb-uniprot50 \| AF-A0A2E3HR82-F1-MODEL\_V4 \| 1.0 \| 0.0007995 \| 156 \| 0.162 \| 111 \| 85 \| 5 \| 3 \| 107 \| 22 \| 130 \| GPW\_gp25 domain-containing protein \| GPW\_gp25 domain-containing protein; afdb-uniprot50 \| AF-A0A2A6Z7N1-F1-MODEL\_V4 \| 1.0 \| 0.0005186 \| 156 \| 0.155 \| 109 \| 88 \| 4 \| 2 \| 107 \| 41 \| 148 \| Uncharacterized protein \| Uncharacterized protein; afdb-uniprot50 \| AF-A0A2D6MFM3-F1-MODEL\_V4 \| 1.0 \| 0.0003579 \| 156 \| 0.131 \| 122 \| 88 \| 5 \| 4 \| 107 \| 76 \| 197 \| GPW\_gp25 domain-containing protein \| GPW\_gp25 domain-containing protein; afdb-uniprot50 \| AF-A0A0R1ZJK1-F1-MODEL\_V4 \| 1.0 \| 0.0005186 \| 155 \| 0.169 \| 118 \| 84 \| 8 \| 2 \| 112 \| 15 \| 125 \| Uncharacterized protein \| Uncharacterized protein; afdb-uniprot50 \| AF-A0A5C7JCT6-F1-MODEL\_V4 \| 1.0 \| 0.001484 \| 155 \| 0.11 \| 109 \| 92 \| 4 \| 2 \| 105 \| 13 \| 121 \| GPW\_gp25 domain-containing protein \| GPW\_gp25 domain-containing protein; afdb-uniprot50 \| AF-A0A1I3LYC4-F1-MODEL\_V4 \| 1.0 \| 0.0005186 \| 155 \| 0.175 \| 108 \| 78 \| 5 \| 6 \| 105 \| 14 \| 118 \| Uncharacterized protein \| Uncharacterized protein; afdb-uniprot50 \| AF-A0A822PRL6-F1-MODEL\_V4 \| 1.0 \| 0.001395 \| 155 \| 0.179 \| 117 \| 86 \| 7 \| 1 \| 112 \| 21 \| 132 \| Phage-like element PBSX protein \| Phage-like element PBSX protein; afdb-uniprot50 \| AF-A0A1C6I381-F1-MODEL\_V4 \| 1.0 \| 0.0003579 \| 155 \| 0.154 \| 123 \| 90 \| 7 \| 1 \| 112 \| 16 \| 135 \| Uncharacterized protein \| Uncharacterized protein; afdb-uniprot50 \| AF-A0A7X7JM68-F1-MODEL\_V4 \| 1.0 \| 0.0001813 \| 154 \| 0.174 \| 109 \| 79 \| 6 \| 3 \| 106 \| 16 \| 118 \| Type VI secretion system baseplate subunit TssE \| Type VI secretion system baseplate subunit TssE; afdb-uniprot50 \| AF-A0A1V5B845-F1-MODEL\_V4 \| 1.0 \| 0.0002794 \| 154 \| 0.173 \| 121 \| 88 \| 4 \| 1 \| 109 \| 23 \| 143 \| Uncharacterized protein \| Uncharacterized protein; afdb-uniprot50 \| AF-A0A7X9FIB8-F1-MODEL\_V4 \| 1.0 \| 0.0001602 \| 152 \| 0.184 \| 125 \| 79 \| 6 \| 1 \| 108 \| 1 \| 119 \| GPW/gp25 family protein \| GPW/gp25 family protein; afdb-uniprot50 \| AF-X0PT30-F1-MODEL\_V4 \| 1.0 \| 0.001024 \| 152 \| 0.178 \| 112 \| 83 \| 6 \| 2 \| 108 \| 9 \| 116 \| Uncharacterized protein \| Uncharacterized protein; afdb-uniprot50 \| AF-A0A7X8EVE5-F1-MODEL\_V4 \| 1.0 \| 0.000405 \| 152 \| 0.173 \| 115 \| 84 \| 7 \| 1 \| 107 \| 22 \| 133 \| DUF2634 domain-containing protein \| DUF2634 domain-containing protein; afdb-uniprot50 \| AF-A0A1I4QKV8-F1-MODEL\_V4 \| 1.0 \| 0.0009625 \| 152 \| 0.196 \| 117 \| 84 \| 7 \| 1 \| 112 \| 23 \| 134 \| Uncharacterized protein \| Uncharacterized protein; afdb-uniprot50 \| AF-A0A4R5N831-F1-MODEL\_V4 \| 1.0 \| 0.0005869 \| 152 \| 0.219 \| 114 \| 78 \| 7 \| 2 \| 107 \| 46 \| 156 \| Uncharacterized protein \| Uncharacterized protein; afdb-uniprot50 \| AF-A0A3Q8US91-F1-MODEL\_V4 \| 1.0 \| 0.0002973 \| 152 \| 0.207 \| 135 \| 78 \| 7 \| 1 \| 108 \| 232 \| 364 \| GPW\_gp25 domain-containing protein \| GPW\_gp25 domain-containing protein; afdb-uniprot50 \| AF-A0A431IHC8-F1-MODEL\_V4 \| 1.0 \| 0.0007065 \| 151 \| 0.137 \| 116 \| 88 \| 4 \| 3 \| 107 \| 10 \| 124 \| Uncharacterized protein \| Uncharacterized protein; afdb-uniprot50 \| AF-A0A7X8C4E8-F1-MODEL\_V4 \| 1.0 \| 0.0009048 \| 151 \| 0.169 \| 112 \| 84 \| 6 \| 1 \| 107 \| 21 \| 128 \| DUF2634 domain-containing protein \| DUF2634 domain-containing protein; afdb-uniprot50 \| AF-A0A2E1V253-F1-MODEL\_V4 \| 1.0 \| 0.0002182 \| 151 \| 0.198 \| 106 \| 71 \| 2 \| 2 \| 105 \| 17 \| 110 \| GPW\_gp25 domain-containing protein \| GPW\_gp25 domain-containing protein; afdb-uniprot50 \| AF-A0A5C8AT77-F1-MODEL\_V4 \| 1.0 \| 0.001395 \| 151 \| 0.156 \| 134 \| 82 \| 4 \| 2 \| 105 \| 20 \| 152 \| GPW\_gp25 domain-containing protein \| GPW\_gp25 domain-containing protein; afdb-uniprot50 \| AF-A0A2D5PHG3-F1-MODEL\_V4 \| 1.0 \| 0.0008505 \| 150 \| 0.169 \| 106 \| 82 \| 2 \| 2 \| 105 \| 12 \| 113 \| Uncharacterized protein \| Uncharacterized protein; afdb-uniprot50 \| AF-V8I9F2-F1-MODEL\_V4 \| 1.0 \| 0.001786 \| 150 \| 0.207 \| 111 \| 79 \| 6 \| 2 \| 107 \| 15 \| 121 \| Uncharacterized protein \| Uncharacterized protein; afdb-uniprot50 \| AF-G4Q4I2-F1-MODEL\_V4 \| 1.0 \| 0.002022 \| 150 \| 0.168 \| 119 \| 85 \| 6 \| 1 \| 106 \| 25 \| 142 \| Phage protein \| Phage protein; afdb-uniprot50 \| AF-A0A4S2BQZ7-F1-MODEL\_V4 \| 1.0 \| 0.0002794 \| 150 \| 0.219 \| 114 \| 74 \| 8 \| 2 \| 107 \| 49 \| 155 \| DUF2634 domain-containing protein \| DUF2634 domain-containing protein; afdb-uniprot50 \| AF-A0A4Q6BJX5-F1-MODEL\_V4 \| 1.0 \| 0.001395 \| 149 \| 0.178 \| 112 \| 81 \| 6 \| 1 \| 105 \| 6 \| 113 \| Uncharacterized protein \| Uncharacterized protein; afdb-uniprot50 \| AF-A0A1H8JI52-F1-MODEL\_V4 \| 1.0 \| 0.002754 \| 149 \| 0.135 \| 118 \| 90 \| 8 \| 2 \| 112 \| 29 \| 141 \| Uncharacterized protein \| Uncharacterized protein; afdb-uniprot50 \| AF-A0A7U4Y7W0-F1-MODEL\_V4 \| 1.0 \| 0.001089 \| 149 \| 0.139 \| 93 \| 74 \| 3 \| 17 \| 105 \| 41 \| 131 \| Type VI secretion protein \| Type VI secretion protein; afdb-uniprot50 \| AF-A0A254RDD5-F1-MODEL\_V4 \| 1.0 \| 0.0008505 \| 148 \| 0.189 \| 95 \| 74 \| 2 \| 13 \| 105 \| 1 \| 94 \| Uncharacterized protein \| Uncharacterized protein; afdb-uniprot50 \| AF-A0A559J3Y0-F1-MODEL\_V4 \| 1.0 \| 0.001484 \| 148 \| 0.149 \| 107 \| 85 \| 5 \| 3 \| 107 \| 1 \| 103 \| DUF2634 domain-containing protein \| DUF2634 domain-containing protein; afdb-uniprot50 \| AF-A0A3F3H3F3-F1-MODEL\_V4 \| 1.0 \| 0.001311 \| 148 \| 0.154 \| 110 \| 84 \| 6 \| 2 \| 106 \| 14 \| 119 \| Uncharacterized protein \| Uncharacterized protein; afdb-uniprot50 \| AF-A0A844ECA9-F1-MODEL\_V4 \| 1.0 \| 0.0003807 \| 148 \| 0.191 \| 115 \| 80 \| 7 \| 2 \| 107 \| 29 \| 139 \| DUF2634 domain-containing protein \| DUF2634 domain-containing protein; afdb-uniprot50 \| AF-A0A1Y4UHE5-F1-MODEL\_V4 \| 1.0 \| 0.001089 \| 147 \| 0.194 \| 113 \| 84 \| 5 \| 1 \| 108 \| 20 \| 130 \| Uncharacterized protein \| Uncharacterized protein; afdb-uniprot50 \| AF-A0A0S2W0M2-F1-MODEL\_V4 \| 1.0 \| 0.0019 \| 147 \| 0.215 \| 116 \| 81 \| 7 \| 1 \| 111 \| 17 \| 127 \| Phage-like element PBSX protein xkdS \| Phage-like element PBSX protein xkdS; afdb-uniprot50 \| AF-J5GPF1-F1-MODEL\_V4 \| 1.0 \| 0.001395 \| 147 \| 0.188 \| 117 \| 85 \| 7 \| 1 \| 112 \| 23 \| 134 \| PF10934 family protein \| PF10934 family protein; afdb-uniprot50 \| AF-A0A5B1BKF6-F1-MODEL\_V4 \| 1.0 \| 0.0003807 \| 147 \| 0.186 \| 107 \| 72 \| 3 \| 2 \| 105 \| 16 \| 110 \| GPW/gp25 family protein \| GPW/gp25 family protein; afdb-uniprot50 \| AF-A0A1V6IG61-F1-MODEL\_V4 \| 1.0 \| 0.002754 \| 146 \| 0.106 \| 113 \| 92 \| 4 \| 1 \| 105 \| 2 \| 113 \| Uncharacterized protein \| Uncharacterized protein; afdb-uniprot50 \| AF-A0A4U3FI15-F1-MODEL\_V4 \| 1.0 \| 0.001395 \| 146 \| 0.169 \| 112 \| 84 \| 6 \| 1 \| 107 \| 25 \| 132 \| DUF2634 domain-containing protein \| DUF2634 domain-containing protein; afdb-uniprot50 \| AF-A0A1Y4U156-F1-MODEL\_V4 \| 1.0 \| 0.000405 \| 146 \| 0.225 \| 120 \| 78 \| 8 \| 2 \| 112 \| 22 \| 135 \| Uncharacterized protein \| Uncharacterized protein; afdb-uniprot50 \| AF-A0A2T0WPJ3-F1-MODEL\_V4 \| 1.0 \| 0.0002794 \| 146 \| 0.245 \| 106 \| 66 \| 2 \| 2 \| 105 \| 16 \| 109 \| GPW\_gp25 domain-containing protein \| GPW\_gp25 domain-containing protein; afdb-uniprot50 \| AF-A0A1P8Q4D0-F1-MODEL\_V4 \| 1.0 \| 0.002022 \| 145 \| 0.178 \| 112 \| 83 \| 6 \| 2 \| 108 \| 5 \| 112 \| Uncharacterized protein \| Uncharacterized protein; afdb-uniprot50 \| AF-A0A2V2E0J3-F1-MODEL\_V4 \| 1.0 \| 0.001233 \| 145 \| 0.245 \| 110 \| 74 \| 6 \| 1 \| 105 \| 21 \| 126 \| DUF2634 domain-containing protein \| DUF2634 domain-containing protein; afdb-uniprot50 \| AF-A0A396SP14-F1-MODEL\_V4 \| 1.0 \| 0.001159 \| 144 \| 0.177 \| 118 \| 83 \| 8 \| 2 \| 112 \| 22 \| 132 \| DUF2634 domain-containing protein \| DUF2634 domain-containing protein; afdb-uniprot50 \| AF-A0A826VXZ4-F1-MODEL\_V4 \| 1.0 \| 0.001311 \| 143 \| 0.314 \| 70 \| 48 \| 0 \| 41 \| 110 \| 1 \| 70 \| Baseplate assembly protein \| Baseplate assembly protein; afdb-uniprot50 \| AF-A0A3E3IVZ8-F1-MODEL\_V4 \| 1.0 \| 0.001311 \| 143 \| 0.155 \| 116 \| 86 \| 6 \| 1 \| 107 \| 11 \| 123 \| DUF2634 domain-containing protein \| DUF2634 domain-containing protein; afdb-uniprot50 \| AF-A0A1T4UK52-F1-MODEL\_V4 \| 1.0 \| 0.001395 \| 142 \| 0.112 \| 107 \| 89 \| 3 \| 4 \| 107 \| 8 \| 111 \| Uncharacterized protein \| Uncharacterized protein; afdb-uniprot50 \| AF-A0A7Y7HXH9-F1-MODEL\_V4 \| 1.0 \| 0.0007516 \| 142 \| 0.243 \| 111 \| 74 \| 7 \| 4 \| 106 \| 12 \| 120 \| Phage gp46-like protein \| Phage gp46-like protein; afdb-uniprot50 \| AF-A8LSA1-F1-MODEL\_V4 \| 1.0 \| 0.000405 \| 142 \| 0.219 \| 114 \| 68 \| 3 \| 1 \| 105 \| 18 \| 119 \| GPW\_gp25 domain-containing protein \| GPW\_gp25 domain-containing protein; afdb-uniprot50 \| AF-A0A3M0N0X0-F1-MODEL\_V4 \| 1.0 \| 0.001395 \| 142 \| 0.157 \| 114 \| 83 \| 7 \| 2 \| 108 \| 40 \| 147 \| DUF2634 domain-containing protein \| DUF2634 domain-containing protein; afdb-uniprot50 \| AF-A0A2E9PUI9-F1-MODEL\_V4 \| 1.0 \| 0.001786 \| 141 \| 0.169 \| 112 \| 82 \| 4 \| 3 \| 108 \| 16 \| 122 \| GPW\_gp25 domain-containing protein \| GPW\_gp25 domain-containing protein; afdb-uniprot50 \| AF-A0A3M0NK27-F1-MODEL\_V4 \| 1.0 \| 0.001311 \| 141 \| 0.157 \| 114 \| 83 \| 7 \| 2 \| 108 \| 55 \| 162 \| DUF2634 domain-containing protein \| DUF2634 domain-containing protein; afdb-uniprot50 \| AF-E6X1N5-F1-MODEL\_V4 \| 1.0 \| 0.006153 \| 139 \| 0.228 \| 92 \| 65 \| 1 \| 2 \| 93 \| 8 \| 93 \| Uncharacterized protein \| Uncharacterized protein; afdb-uniprot50 \| AF-R5SUR6-F1-MODEL\_V4 \| 1.0 \| 0.001311 \| 139 \| 0.194 \| 118 \| 87 \| 5 \| 1 \| 112 \| 24 \| 139 \| Phage protein \| Phage protein; afdb-uniprot50 \| AF-A0A4R5NMZ8-F1-MODEL\_V4 \| 1.0 \| 0.0019 \| 139 \| 0.203 \| 113 \| 77 \| 7 \| 2 \| 107 \| 55 \| 161 \| Uncharacterized protein \| Uncharacterized protein; afdb-uniprot50 \| AF-A0A1V5B2H8-F1-MODEL\_V4 \| 1.0 \| 0.00293 \| 138 \| 0.185 \| 108 \| 81 \| 3 \| 1 \| 105 \| 23 \| 126 \| Uncharacterized protein \| Uncharacterized protein; afdb-uniprot50 \| AF-F4GSE7-F1-MODEL\_V4 \| 1.0 \| 0.001395 \| 137 \| 0.215 \| 93 \| 67 \| 3 \| 17 \| 105 \| 13 \| 103 \| GPW\_gp25 domain-containing protein \| GPW\_gp25 domain-containing protein; afdb-uniprot50 \| AF-A0A1C6AKL0-F1-MODEL\_V4 \| 1.0 \| 0.00788 \| 136 \| 0.175 \| 114 \| 83 \| 6 \| 2 \| 108 \| 25 \| 134 \| Protein of uncharacterized function (DUF2634) \| Protein of uncharacterized function (DUF2634); afdb-uniprot50 \| AF-C4GJQ7-F1-MODEL\_V4 \| 1.0 \| 0.001089 \| 136 \| 0.241 \| 116 \| 75 \| 5 \| 1 \| 107 \| 40 \| 151 \| Uncharacterized protein \| Uncharacterized protein; afdb-uniprot50 \| AF-A0A173S783-F1-MODEL\_V4 \| 1.0 \| 0.001395 \| 135 \| 0.17 \| 123 \| 82 \| 8 \| 1 \| 107 \| 25 \| 143 \| Protein of uncharacterized function (DUF2634) \| Protein of uncharacterized function (DUF2634); afdb-uniprot50 \| AF-A0A2D6XAB7-F1-MODEL\_V4 \| 1.0 \| 0.003991 \| 134 \| 0.174 \| 126 \| 79 \| 5 \| 5 \| 108 \| 52 \| 174 \| GPW\_gp25 domain-containing protein \| GPW\_gp25 domain-containing protein; afdb-uniprot50 \| AF-A0A7C4DJH8-F1-MODEL\_V4 \| 1.0 \| 0.006153 \| 133 \| 0.25 \| 84 \| 57 \| 2 \| 5 \| 86 \| 21 \| 100 \| GPW\_gp25 domain-containing protein \| GPW\_gp25 domain-containing protein; afdb-uniprot50 \| AF-A0A2E7CJE8-F1-MODEL\_V4 \| 1.0 \| 0.005111 \| 132 \| 0.159 \| 88 \| 66 \| 2 \| 3 \| 90 \| 8 \| 87 \| Uncharacterized protein \| Uncharacterized protein; afdb-uniprot50 \| AF-A0A6H1Z7N3-F1-MODEL\_V4 \| 1.0 \| 0.006153 \| 132 \| 0.123 \| 113 \| 90 \| 3 \| 2 \| 105 \| 24 \| 136 \| Uncharacterized protein \| Uncharacterized protein; afdb-uniprot50 \| AF-A0A2E5G1A5-F1-MODEL\_V4 \| 1.0 \| 0.003752 \| 131 \| 0.155 \| 103 \| 84 \| 2 \| 3 \| 105 \| 10 \| 109 \| Uncharacterized protein \| Uncharacterized protein; afdb-uniprot50 \| AF-A0A1H8IY64-F1-MODEL\_V4 \| 1.0 \| 0.007407 \| 131 \| 0.159 \| 113 \| 86 \| 7 \| 2 \| 108 \| 19 \| 128 \| Uncharacterized protein \| Uncharacterized protein; afdb-uniprot50 \| AF-A0A243EHU0-F1-MODEL\_V4 \| 1.0 \| 0.002288 \| 131 \| 0.25 \| 104 \| 70 \| 5 \| 3 \| 101 \| 133 \| 233 \| Uncharacterized protein \| Uncharacterized protein; afdb-uniprot50 \| AF-A0A098CX13-F1-MODEL\_V4 \| 1.0 \| 0.004805 \| 129 \| 0.176 \| 113 \| 80 \| 7 \| 2 \| 107 \| 27 \| 133 \| Uncharacterized protein \| Uncharacterized protein; afdb-uniprot50 \| AF-A0A3C0F5D4-F1-MODEL\_V4 \| 1.0 \| 0.008382 \| 127 \| 0.166 \| 102 \| 77 \| 3 \| 2 \| 95 \| 10 \| 111 \| GPW\_gp25 domain-containing protein \| GPW\_gp25 domain-containing protein; afdb-uniprot50 \| AF-A0A431IP84-F1-MODEL\_V4 \| 1.0 \| 0.004246 \| 127 \| 0.178 \| 112 \| 85 \| 5 \| 3 \| 110 \| 16 \| 124 \| Uncharacterized protein \| Uncharacterized protein; afdb-uniprot50 \| AF-A0A1Y4D1W7-F1-MODEL\_V4 \| 1.0 \| 0.001311 \| 126 \| 0.186 \| 129 \| 75 \| 8 \| 3 \| 107 \| 18 \| 140 \| GPW\_gp25 domain-containing protein \| GPW\_gp25 domain-containing protein; afdb-uniprot50 \| AF-A0A3A9F4C1-F1-MODEL\_V4 \| 1.0 \| 0.00788 \| 120 \| 0.168 \| 113 \| 83 \| 6 \| 1 \| 105 \| 35 \| 144 \| DUF2634 domain-containing protein \| DUF2634 domain-containing protein; afdb-uniprot50 \| AF-A0A1Y4GIV1-F1-MODEL\_V4 \| 1.0 \| 0.002754 \| 116 \| 0.209 \| 124 \| 78 \| 8 \| 5 \| 111 \| 16 \| 136 \| GPW\_gp25 domain-containing protein \| GPW\_gp25 domain-containing protein; afdb-uniprot50 \| AF-F2Q843-F1-MODEL\_V4 \| 1.0 \| 0.006545 \| 111 \| 0.205 \| 102 \| 59 \| 8 \| 18 \| 105 \| 30 \| 123 \| GPW\_gp25 domain-containing protein \| GPW\_gp25 domain-containing protein |
| Top keywords  (threshold 1.00e-02 (evalue)) | **domain\_containing, GPW\_gp25, Baseplate, assembly, gp25, GPW, Phage, lysozyme, W, 25\_like** |
| Output files | ../../similar\_structures/12\_FANPEZAQ\_CDS\_0012\_afdb-proteome\_foldseek.tsv ../../similar\_structures/12\_FANPEZAQ\_CDS\_0012\_afdb-uniprot50\_foldseek.tsv ../../similar\_structures/12\_FANPEZAQ\_CDS\_0012\_merged.svg ../../similar\_structures/12\_FANPEZAQ\_CDS\_0012\_pdb\_foldseek.tsv |

  
  
  

Return to summary | Go to previous | Go to next

  


---

**Sequence/structure alignments coloring**  
Each object in the alignment figures is colored according to its E-value following this color coding:

1e-100
10
