## Supplementary material for "Completing the BASEL phage collection to unlock hidden diversity for systematic exploration of phage-host interactions": Data S2: 13.html

FANPEZAQ\_CDS\_0013


Return to summary | Go to previous | Go to next

|  |  |
| --- | --- |
| FANPEZAQ\_CDS\_0013  Page creation date: 02 Sep 2024, 12:00  Project folder: n/a  Input sequences file: Escherichia\_virus\_HeidiAbel.gb | baseplate domain\_containing assembly baseplate\_j j\_like j gp47 phage putative phage\_related tail fragment jaye wedge mu bacteriophage homolog gp6 gpj tube t4 complex baseplate\_tail pre\_attachment structural prophage |

### Sequence information

|  |  |
| --- | --- |
| Name | FANPEZAQ\_CDS\_0013  13\_FANPEZAQ\_CDS\_0013 (pipeline id) |
| Imported annotations | Escherichia\_virus\_HeidiAbel Bas97 |
| Protein sequence | MAGSYTVIDLSQLPAPTIVEQLSFETILAEMIADLQARDSTFTALVESDPAYKILEVCAF RELLVRQRANEEAKAVMLAYASGTDLDQLGANVGVKRLVIRPADPNAVPPVPAVMESDNE FRARIQLAPEGYTTAGSEGSYAFHALGADADVKDVQPTSPTPGHVVVYVLSRTGDGAAPQ PLLDKVNATLTKEEVRPLTDNVTVQSAQIITYTISAELTLLPGPDSEVVRQAAFDAVAAY AEGQRRIGYDVTMSGLYRALHQPGVQNVKLTAPTVSLVLGDGQASYCTGITVTVAGETDV |
| Number of residues | 300 |
| Molecular weight (Da) | 31848.61 |
| Output files | ../../query\_sequences/13\_FANPEZAQ\_CDS\_0013.fasta |

### Putative domain architecture and protein family

#### Search results (HHblits)^1^

|  |  |
| --- | --- |
| Domain family databases searched | Pfam, Ncbi-cd, Cath, Phrogs |
| Results, scheme(s)  (Top layers only; threshold 1.00e-03 (evalue)) | xml version="1.0" encoding="utf-8" standalone="no"?       2024-09-02T21:08:14.706197 image/svg+xml   Matplotlib v3.7.2, https://matplotlib.org/ |
| Results, table  (E-value ≤ 1.00e-03 (evalue)) | db \| id \| prob \| evalue \| pvalue \| score \| cols \| query \| query\_len \| template \| template\_len \| name \| description; pfam \| PF03434 \| 99.4 \| 3e-18 \| 6.1e-22 \| 152.2 \| 155 \| (115, 274) \| 300 \| (66, 247) \| 289 \| DUF276 \| DUF276; pfam \| PF04865 \| 99.3 \| 6.6e-17 \| 1.4e-20 \| 138.5 \| 143 \| (109, 259) \| 300 \| (104, 250) \| 250 \| Baseplate\_J \| Baseplate J-like protein; pfam \| PF11041 \| 98.6 \| 3e-12 \| 6.1e-16 \| 104.4 \| 103 \| (69, 174) \| 300 \| (32, 156) \| 181 \| DUF2612 \| Protein of unknown function (DUF2612); pfam \| PF09684 \| 94.2 \| 0.00057 \| 1.1e-07 \| 50.0 \| 68 \| (75, 159) \| 300 \| (23, 90) \| 138 \| Tail\_P2\_I \| Phage tail protein (Tail\_P2\_I); phrogs \| 219 \| 100.0 \| 1.7e-36 \| 2e-40 \| 285.5 \| 244 \| (14, 262) \| 300 \| (1, 449) \| 676 \| baseplate wedge subunit \| baseplate wedge subunit; Category: tail; KU686208\_p78; phrogs \| 4226 \| 99.8 \| 3.9e-26 \| 4.5e-30 \| 182.8 \| 230 \| (70, 299) \| 300 \| (2, 231) \| 231 \| baseplate protein \| baseplate protein; Category: tail; p428496 VI\_02892; phrogs \| 6 \| 99.8 \| 8.9e-25 \| 1.1e-28 \| 199.5 \| 283 \| (8, 296) \| 300 \| (2, 471) \| 472 \| baseplate wedge subunit \| baseplate wedge subunit; Category: tail; p248291 VI\_09499; phrogs \| 1608 \| 99.7 \| 1.3e-21 \| 1.5e-25 \| 156.9 \| 179 \| (110, 292) \| 300 \| (34, 232) \| 232 \| tail protein \| tail protein; Category: tail; p256756 VI\_05872; phrogs \| 2103 \| 99.2 \| 1.3e-15 \| 1.5e-19 \| 119.2 \| 157 \| (14, 172) \| 300 \| (1, 234) \| 245 \| baseplate protein \| baseplate protein; Category: tail; p226045 VI\_12003; phrogs \| 1160 \| 98.3 \| 2.2e-10 \| 2.6e-14 \| 92.3 \| 106 \| (30, 157) \| 300 \| (4, 111) \| 209 \| baseplate protein \| baseplate protein; Category: tail; p105495 VI\_00644; phrogs \| 7378 \| 98.2 \| 3.3e-10 \| 3.8e-14 \| 96.8 \| 98 \| (47, 162) \| 300 \| (18, 117) \| 462 \| baseplate protein \| baseplate protein; Category: tail; NC\_024792\_p167; phrogs \| 4056 \| 98.1 \| 9.1e-10 \| 1e-13 \| 85.7 \| 117 \| (20, 159) \| 300 \| (10, 127) \| 228 \| baseplate protein \| baseplate protein; Category: tail; NC\_024134\_p34; phrogs \| 6277 \| 98.0 \| 2e-09 \| 2.2e-13 \| 87.1 \| 105 \| (49, 170) \| 300 \| (21, 125) \| 283 \| NA \| NA; Category: unknown function; KY883640\_p59; phrogs \| 4411 \| 97.8 \| 6.3e-09 \| 7.1e-13 \| 83.6 \| 97 \| (47, 161) \| 300 \| (18, 116) \| 340 \| virion structural protein \| virion structural protein; Category: head and packaging; JN797798\_p181; phrogs \| 2386 \| 97.8 \| 1.1e-08 \| 1.2e-12 \| 94.3 \| 99 \| (45, 161) \| 300 \| (19, 119) \| 1309 \| virion structural protein \| virion structural protein; Category: head and packaging; NC\_020871\_p95; phrogs \| 31189 \| 97.8 \| 1.1e-08 \| 1.3e-12 \| 85.3 \| 143 \| (110, 262) \| 300 \| (263, 406) \| 662 \| baseplate protein \| baseplate protein; Category: tail; NC\_008584\_p150; phrogs \| 22472 \| 97.5 \| 4.5e-08 \| 5e-12 \| 77.2 \| 107 \| (45, 168) \| 300 \| (20, 127) \| 304 \| NA \| NA; Category: unknown function; MF370964\_p74; phrogs \| 34452 \| 97.3 \| 2.3e-07 \| 2.5e-11 \| 65.7 \| 86 \| (12, 97) \| 300 \| (2, 87) \| 175 \| NA \| NA; Category: unknown function; p43846 VI\_12438; phrogs \| 13726 \| 97.1 \| 6e-07 \| 6.8e-11 \| 79.5 \| 96 \| (48, 161) \| 300 \| (22, 119) \| 803 \| NA \| NA; Category: unknown function; NC\_027335\_p71; phrogs \| 1369 \| 96.9 \| 1.7e-06 \| 2e-10 \| 75.3 \| 128 \| (110, 243) \| 300 \| (268, 410) \| 477 \| baseplate protein \| baseplate protein; Category: tail; p133507 VI\_09775; phrogs \| 23130 \| 96.4 \| 9.7e-06 \| 1.1e-09 \| 62.9 \| 94 \| (51, 161) \| 300 \| (19, 114) \| 362 \| tail protein \| tail protein; Category: tail; NC\_022761\_p93; phrogs \| 26379 \| 96.2 \| 1.9e-05 \| 2.1e-09 \| 58.9 \| 105 \| (24, 151) \| 300 \| (6, 114) \| 220 \| NA \| NA; Category: unknown function; p152986 VI\_08595; phrogs \| 521 \| 96.2 \| 1.8e-05 \| 2.3e-09 \| 61.1 \| 107 \| (47, 172) \| 300 \| (14, 123) \| 180 \| tail protein \| tail protein; Category: tail; p222013 VI\_01195; phrogs \| 5502 \| 96.2 \| 2e-05 \| 2.3e-09 \| 71.5 \| 85 \| (60, 161) \| 300 \| (31, 116) \| 1585 \| NA \| NA; Category: unknown function; NC\_031037\_p116; phrogs \| 10646 \| 96.1 \| 3e-05 \| 3.5e-09 \| 70.5 \| 85 \| (14, 98) \| 300 \| (23, 109) \| 1154 \| baseplate wedge subunit \| baseplate wedge subunit; Category: tail; NC\_019526\_p71; phrogs \| 14699 \| 95.9 \| 5.2e-05 \| 5.8e-09 \| 46.1 \| 63 \| (237, 299) \| 300 \| (4, 66) \| 66 \| NA \| NA; Category: unknown function; p301057 VI\_10846; phrogs \| 587 \| 95.7 \| 7.4e-05 \| 9e-09 \| 60.1 \| 124 \| (47, 171) \| 300 \| (35, 181) \| 237 \| structural protein \| structural protein; Category: head and packaging; MF663761\_p45; phrogs \| 21670 \| 95.6 \| 0.00011 \| 1.3e-08 \| 51.4 \| 81 \| (16, 98) \| 300 \| (5, 85) \| 164 \| NA \| NA; Category: unknown function; p402686 VI\_09540; phrogs \| 12386 \| 95.3 \| 0.00019 \| 2.1e-08 \| 42.9 \| 62 \| (20, 85) \| 300 \| (2, 63) \| 65 \| NA \| NA; Category: unknown function; p419451 VI\_05447; phrogs \| 30339 \| 94.8 \| 0.00044 \| 4.8e-08 \| 47.1 \| 95 \| (4, 98) \| 300 \| (6, 100) \| 150 \| NA \| NA; Category: unknown function; p424932 VI\_08726; phrogs \| 4017 \| 94.8 \| 0.00046 \| 5.2e-08 \| 53.0 \| 109 \| (48, 172) \| 300 \| (24, 135) \| 285 \| baseplate protein \| baseplate protein; Category: tail; KY626162\_p27; phrogs \| 5857 \| 94.3 \| 0.0009 \| 1e-07 \| 49.7 \| 68 \| (47, 129) \| 300 \| (26, 93) \| 221 \| baseplate wedge subunit \| baseplate wedge subunit; Category: tail; p149245 VI\_12398 |
| Top keywords  (threshold 1.00e-03 (evalue)) | **tail, baseplate, wedge, structural, head, and, packaging, virion, DUF276, J\_like** |
| Output files | ../../domain\_architecture/13\_FANPEZAQ\_CDS\_0013\_cath.hhr ../../domain\_architecture/13\_FANPEZAQ\_CDS\_0013\_merged.svg ../../domain\_architecture/13\_FANPEZAQ\_CDS\_0013\_ncbi-cd.hhr ../../domain\_architecture/13\_FANPEZAQ\_CDS\_0013\_pfam.hhr ../../domain\_architecture/13\_FANPEZAQ\_CDS\_0013\_phrogs.hhr |

### Identical protein sequences/structures

#### Search results

|  |  |
| --- | --- |
| Protein sequence databases searched | Pdb, Swissprot, Refseq |
| Identical proteins found | -- |
| Top keywords | -- |
| Output files | -- |

### Similar protein sequences/structures

#### Sequence similarity search results (HHblits)^1^

|  |  |
| --- | --- |
| Sequence databases searched | Uniclust, Pdb70 |
| Results, scheme(s)  (Top layers only, threshold 1.00e-03 (evalue)) | xml version="1.0" encoding="utf-8" standalone="no"?       2024-09-02T21:08:33.705521 image/svg+xml   Matplotlib v3.7.2, https://matplotlib.org/ |
| Results, table(s)  (threshold 1.00e-03 (evalue)) | db \| id \| prob \| evalue \| pvalue \| score \| cols \| query \| query\_len \| template \| template\_len \| name \| description; uniclust \| UniRef100\_A0A016XHS1 \| 100.0 \| 3.9e-55 \| 7.7e-61 \| 373.6 \| 298 \| (1, 298) \| 300 \| (26, 325) \| 337 \| Baseplate assembly protein \| Baseplate assembly protein; uniclust \| UniRef100\_A0A024E877 \| 100.0 \| 4.5e-54 \| 9e-60 \| 370.1 \| 298 \| (1, 298) \| 300 \| (32, 330) \| 343 \| Baseplate J-like protein \| Baseplate J-like protein; uniclust \| UniRef100\_A0A009PKZ2 \| 100.0 \| 3.5e-51 \| 7e-57 \| 359.1 \| 296 \| (1, 297) \| 300 \| (57, 359) \| 398 \| Baseplate J-like family protein \| Baseplate J-like family protein; uniclust \| UniRef100\_A0A0P9B5C1 \| 100.0 \| 4.2e-51 \| 8e-57 \| 332.2 \| 298 \| (1, 298) \| 300 \| (6, 310) \| 312 \| Baseplate protein J-like domain-containing protein (Fragment) \| Baseplate protein J-like domain-containing protein (Fragment); uniclust \| UniRef100\_A0A059IU93 \| 100.0 \| 1.4e-49 \| 2.8e-55 \| 329.6 \| 282 \| (2, 299) \| 300 \| (1, 282) \| 301 \| Putative prophage LambdaW1, baseplate assembly protein J,putative \| Putative prophage LambdaW1, baseplate assembly protein J,putative; uniclust \| UniRef100\_A0A096AN02 \| 100.0 \| 1.9e-48 \| 3.9e-54 \| 343.3 \| 288 \| (2, 297) \| 300 \| (1, 369) \| 376 \| Baseplate protein J-like domain-containing protein \| Baseplate protein J-like domain-containing protein; uniclust \| UniRef100\_A0A066RVH9 \| 100.0 \| 7.5e-48 \| 1.5e-53 \| 337.2 \| 294 \| (2, 299) \| 300 \| (35, 329) \| 381 \| Baseplate assembly protein \| Baseplate assembly protein; uniclust \| UniRef100\_A0A011NCT2 \| 100.0 \| 1.8e-46 \| 3.5e-52 \| 339.1 \| 293 \| (1, 298) \| 300 \| (67, 442) \| 523 \| Phage baseplate protein \| Phage baseplate protein; uniclust \| UniRef100\_A0A077Q3X4 \| 100.0 \| 8.2e-46 \| 1.6e-51 \| 303.5 \| 295 \| (5, 299) \| 300 \| (1, 302) \| 307 \| Baseplate assembly protein J (GpJ) \| Baseplate assembly protein J (GpJ); uniclust \| UniRef100\_A0A061P986 \| 100.0 \| 9.7e-46 \| 2e-51 \| 330.2 \| 280 \| (12, 297) \| 300 \| (38, 400) \| 438 \| Phage baseplate \| Phage baseplate; uniclust \| UniRef100\_A0A096DBD2 \| 100.0 \| 1.3e-45 \| 2.6e-51 \| 326.0 \| 269 \| (21, 297) \| 300 \| (37, 389) \| 400 \| Baseplate protein J-like domain-containing protein \| Baseplate protein J-like domain-containing protein; uniclust \| UniRef100\_A0A0A8HCY4 \| 100.0 \| 1.8e-44 \| 3.6e-50 \| 310.7 \| 284 \| (3, 296) \| 300 \| (12, 376) \| 386 \| Phage baseplate assembly protein J, putative \| Phage baseplate assembly protein J, putative; uniclust \| UniRef100\_A0A068ZW73 \| 100.0 \| 2.3e-44 \| 4.7e-50 \| 320.5 \| 279 \| (10, 296) \| 300 \| (16, 386) \| 407 \| Baseplate protein J-like domain-containing protein \| Baseplate protein J-like domain-containing protein; uniclust \| UniRef100\_A0A061NGS0 \| 100.0 \| 2.5e-44 \| 4.8e-50 \| 303.4 \| 270 \| (19, 296) \| 300 \| (7, 358) \| 370 \| Phage-like element PBSX protein XkdT \| Phage-like element PBSX protein XkdT; uniclust \| UniRef100\_A0A081NYB1 \| 100.0 \| 3.5e-44 \| 6.9e-50 \| 315.2 \| 271 \| (19, 297) \| 300 \| (18, 394) \| 418 \| Baseplate protein J-like domain-containing protein \| Baseplate protein J-like domain-containing protein; uniclust \| UniRef100\_A0A0B5QQX8 \| 100.0 \| 3.7e-43 \| 7.2e-49 \| 300.9 \| 272 \| (18, 296) \| 300 \| (12, 370) \| 373 \| Baseplate J protein \| Baseplate J protein; uniclust \| UniRef100\_A0A1S9ZY08 \| 100.0 \| 1.3e-42 \| 2.6e-48 \| 282.4 \| 273 \| (6, 298) \| 300 \| (1, 274) \| 278 \| Baseplate protein J-like domain-containing protein \| Baseplate protein J-like domain-containing protein; uniclust \| UniRef100\_A0A063BJF3 \| 100.0 \| 3e-42 \| 6.2e-48 \| 310.2 \| 269 \| (20, 296) \| 300 \| (64, 412) \| 455 \| Baseplate J family protein \| Baseplate J family protein; uniclust \| UniRef100\_A0A090C6U6 \| 100.0 \| 1.5e-41 \| 2.9e-47 \| 289.4 \| 269 \| (18, 297) \| 300 \| (8, 356) \| 360 \| Putative baseplate-J protein \| Putative baseplate-J protein; uniclust \| UniRef100\_A0A0B0S945 \| 100.0 \| 1.9e-41 \| 3.7e-47 \| 296.6 \| 280 \| (4, 296) \| 300 \| (7, 376) \| 380 \| Baseplate protein J-like domain-containing protein \| Baseplate protein J-like domain-containing protein; uniclust \| UniRef100\_A0A0H3ZT21 \| 100.0 \| 5.8e-41 \| 1.1e-46 \| 275.4 \| 275 \| (2, 299) \| 300 \| (1, 282) \| 294 \| Phage-related baseplate assembly protein \| Phage-related baseplate assembly protein; uniclust \| UniRef100\_A0A017RWE9 \| 100.0 \| 5.8e-41 \| 1.2e-46 \| 295.2 \| 278 \| (16, 299) \| 300 \| (26, 387) \| 391 \| Baseplate protein J-like domain-containing protein \| Baseplate protein J-like domain-containing protein; uniclust \| UniRef100\_A0A072YE02 \| 100.0 \| 1.8e-40 \| 3.5e-46 \| 290.2 \| 274 \| (17, 296) \| 300 \| (19, 390) \| 393 \| Phage tail protein \| Phage tail protein; uniclust \| UniRef100\_A0A016XIM7 \| 100.0 \| 2.9e-40 \| 5.8e-46 \| 297.4 \| 270 \| (20, 296) \| 300 \| (38, 387) \| 451 \| Baseplate J protein \| Baseplate J protein; uniclust \| UniRef100\_UPI00082B4CAF \| 100.0 \| 3.5e-40 \| 6.7e-46 \| 275.4 \| 280 \| (5, 294) \| 300 \| (3, 362) \| 363 \| baseplate J/gp47 family protein \| baseplate J/gp47 family protein; uniclust \| UniRef100\_A0A0B1YBX4 \| 100.0 \| 3.7e-40 \| 7.4e-46 \| 288.0 \| 270 \| (21, 296) \| 300 \| (14, 368) \| 379 \| Baseplate protein J-like domain-containing protein \| Baseplate protein J-like domain-containing protein; uniclust \| UniRef100\_A0A2W1AVR7 \| 100.0 \| 4.2e-40 \| 8.1e-46 \| 277.4 \| 266 \| (21, 294) \| 300 \| (4, 355) \| 357 \| Baseplate protein J-like domain-containing protein \| Baseplate protein J-like domain-containing protein; uniclust \| UniRef100\_A0A8T0C4P9 \| 100.0 \| 2.2e-39 \| 4.3e-45 \| 263.0 \| 246 \| (53, 298) \| 300 \| (3, 252) \| 255 \| Baseplate protein J-like domain-containing protein \| Baseplate protein J-like domain-containing protein; uniclust \| UniRef100\_A0A2D9T280 \| 100.0 \| 5.1e-39 \| 9.7e-45 \| 266.8 \| 266 \| (20, 293) \| 300 \| (5, 346) \| 346 \| Baseplate protein J-like domain-containing protein \| Baseplate protein J-like domain-containing protein; uniclust \| UniRef100\_A0A074LIS1 \| 100.0 \| 5.3e-39 \| 1e-44 \| 275.4 \| 269 \| (22, 295) \| 300 \| (7, 365) \| 367 \| Baseplate protein J-like domain-containing protein \| Baseplate protein J-like domain-containing protein; uniclust \| UniRef100\_A0A059N0R3 \| 100.0 \| 8.9e-39 \| 1.8e-44 \| 282.4 \| 279 \| (2, 295) \| 300 \| (4, 387) \| 393 \| Baseplate protein J-like domain-containing protein \| Baseplate protein J-like domain-containing protein; uniclust \| UniRef100\_A0A2P9HMM4 \| 100.0 \| 5.3e-38 \| 1e-43 \| 261.6 \| 276 \| (2, 281) \| 300 \| (1, 288) \| 339 \| Baseplate assembly protein J \| Baseplate assembly protein J; uniclust \| UniRef100\_A0A0A1AFQ4 \| 100.0 \| 6.2e-38 \| 1.2e-43 \| 266.1 \| 291 \| (4, 299) \| 300 \| (1, 375) \| 391 \| Baseplate J/gp47 family protein \| Baseplate J/gp47 family protein; uniclust \| UniRef100\_A0A021XBA6 \| 100.0 \| 1.9e-37 \| 3.8e-43 \| 260.0 \| 295 \| (1, 299) \| 300 \| (4, 306) \| 307 \| Baseplate J family protein \| Baseplate J family protein; uniclust \| UniRef100\_A0A061N9X7 \| 100.0 \| 1.9e-37 \| 3.9e-43 \| 278.9 \| 270 \| (16, 295) \| 300 \| (26, 407) \| 429 \| Putative bacteriophage protein \| Putative bacteriophage protein; uniclust \| UniRef100\_A0A0M0T6B2 \| 100.0 \| 2.8e-37 \| 5.3e-43 \| 257.0 \| 267 \| (22, 296) \| 300 \| (6, 333) \| 339 \| Tail protein (Fragment) \| Tail protein (Fragment); uniclust \| UniRef100\_A0A031FRM4 \| 100.0 \| 5.3e-37 \| 1e-42 \| 263.6 \| 297 \| (1, 297) \| 300 \| (1, 335) \| 346 \| Baseplate J-like protein \| Baseplate J-like protein; uniclust \| UniRef100\_A0A5C0SDS7 \| 100.0 \| 9.9e-37 \| 1.9e-42 \| 252.9 \| 267 \| (21, 294) \| 300 \| (6, 360) \| 361 \| Baseplate J/gp47 family protein \| Baseplate J/gp47 family protein; uniclust \| UniRef100\_A0A1I7ZUF6 \| 100.0 \| 1.4e-36 \| 2.6e-42 \| 242.0 \| 220 \| (2, 221) \| 300 \| (3, 223) \| 229 \| Baseplate\_J domain-containing protein \| Baseplate\_J domain-containing protein; uniclust \| UniRef100\_A0A0C5VGB9 \| 100.0 \| 1.4e-36 \| 2.7e-42 \| 256.5 \| 289 \| (2, 292) \| 300 \| (1, 293) \| 465 \| Phage-related baseplate assembly protein \| Phage-related baseplate assembly protein; uniclust \| UniRef100\_A0A161YLM2 \| 100.0 \| 2.8e-36 \| 5.5e-42 \| 248.6 \| 278 \| (4, 297) \| 300 \| (1, 284) \| 286 \| Baseplate protein J-like domain-containing protein \| Baseplate protein J-like domain-containing protein; uniclust \| UniRef100\_A0A0C7NEI0 \| 100.0 \| 4.9e-36 \| 9.4e-42 \| 262.0 \| 272 \| (16, 293) \| 300 \| (8, 527) \| 527 \| Baseplate assembly protein J-like, predicted \| Baseplate assembly protein J-like, predicted; uniclust \| UniRef100\_A0A068Z4R0 \| 100.0 \| 1.5e-35 \| 2.9e-41 \| 262.1 \| 271 \| (19, 295) \| 300 \| (55, 424) \| 427 \| Baseplate J/gp47 family protein \| Baseplate J/gp47 family protein; uniclust \| UniRef100\_A0A0M0SY60 \| 100.0 \| 2e-35 \| 3.7e-41 \| 247.8 \| 268 \| (20, 293) \| 300 \| (14, 373) \| 373 \| Baseplate protein J-like domain-containing protein \| Baseplate protein J-like domain-containing protein; uniclust \| UniRef100\_A0A0C1G7H8 \| 100.0 \| 5e-35 \| 9.9e-41 \| 254.9 \| 283 \| (9, 299) \| 300 \| (14, 379) \| 382 \| Baseplate protein J-like domain-containing protein \| Baseplate protein J-like domain-containing protein; uniclust \| UniRef100\_A0A173RD27 \| 100.0 \| 5.2e-35 \| 9.9e-41 \| 247.7 \| 285 \| (6, 297) \| 300 \| (1, 381) \| 389 \| Uncharacterized homolog of phage Mu protein gp47 \| Uncharacterized homolog of phage Mu protein gp47; uniclust \| UniRef100\_A0A074LIV6 \| 100.0 \| 7.1e-35 \| 1.3e-40 \| 244.2 \| 269 \| (22, 296) \| 300 \| (4, 368) \| 369 \| Baseplate protein J-like domain-containing protein \| Baseplate protein J-like domain-containing protein; uniclust \| UniRef100\_A0A0T9LR98 \| 100.0 \| 9.4e-35 \| 1.8e-40 \| 236.6 \| 231 \| (1, 231) \| 300 \| (19, 249) \| 293 \| Baseplate assembly protein \| Baseplate assembly protein; uniclust \| UniRef100\_A0A0F4NJK8 \| 100.0 \| 2.1e-34 \| 3.9e-40 \| 230.2 \| 268 \| (6, 296) \| 300 \| (1, 268) \| 274 \| Baseplate protein J-like domain-containing protein \| Baseplate protein J-like domain-containing protein; uniclust \| UniRef100\_A0A375ABY7 \| 100.0 \| 5.7e-34 \| 1.1e-39 \| 237.3 \| 290 \| (5, 294) \| 300 \| (1, 297) \| 504 \| Phage baseplate assembly protein J \| Phage baseplate assembly protein J; uniclust \| UniRef100\_UPI000B7DA598 \| 100.0 \| 1.2e-33 \| 2.2e-39 \| 222.4 \| 224 \| (1, 224) \| 300 \| (1, 228) \| 233 \| baseplate J/gp47 family protein \| baseplate J/gp47 family protein; uniclust \| UniRef100\_A0A085EXM5 \| 100.0 \| 1.3e-33 \| 2.6e-39 \| 242.4 \| 269 \| (19, 294) \| 300 \| (10, 376) \| 379 \| Putative bacteriophage protein \| Putative bacteriophage protein; uniclust \| UniRef100\_A0A8J2Z5N4 \| 100.0 \| 1.6e-33 \| 3e-39 \| 225.1 \| 277 \| (4, 295) \| 300 \| (1, 281) \| 283 \| Bacteriophage protein \| Bacteriophage protein; uniclust \| UniRef100\_A0A081MYF5 \| 100.0 \| 1.7e-33 \| 3.3e-39 \| 223.5 \| 231 \| (65, 297) \| 300 \| (2, 232) \| 234 \| Baseplate protein J-like domain-containing protein (Fragment) \| Baseplate protein J-like domain-containing protein (Fragment); uniclust \| UniRef100\_A0A0E2H3N7 \| 100.0 \| 5.8e-33 \| 1.1e-38 \| 239.5 \| 272 \| (21, 297) \| 300 \| (14, 377) \| 385 \| Baseplate protein J-like domain-containing protein \| Baseplate protein J-like domain-containing protein; uniclust \| UniRef100\_A0A075RF08 \| 100.0 \| 6.9e-33 \| 1.3e-38 \| 227.5 \| 265 \| (3, 292) \| 300 \| (9, 283) \| 286 \| Baseplate assembly protein \| Baseplate assembly protein; uniclust \| UniRef100\_A0A090BV61 \| 100.0 \| 1.2e-32 \| 2.4e-38 \| 253.9 \| 277 \| (14, 295) \| 300 \| (64, 553) \| 580 \| Baseplate protein J-like domain-containing protein \| Baseplate protein J-like domain-containing protein; uniclust \| UniRef100\_A0A0F2QX06 \| 100.0 \| 1.5e-32 \| 2.8e-38 \| 221.7 \| 250 \| (21, 295) \| 300 \| (5, 259) \| 260 \| Baseplate protein J-like domain-containing protein \| Baseplate protein J-like domain-containing protein; uniclust \| UniRef100\_A0A081BI31 \| 100.0 \| 1.7e-32 \| 3.2e-38 \| 237.8 \| 271 \| (16, 295) \| 300 \| (13, 389) \| 400 \| Phage protein \| Phage protein; uniclust \| UniRef100\_A0A1H8P6M7 \| 99.9 \| 2.6e-32 \| 5e-38 \| 213.7 \| 205 \| (3, 207) \| 300 \| (6, 217) \| 218 \| Baseplate assembly protein \| Baseplate assembly protein; uniclust \| UniRef100\_A0A1C5WAX4 \| 99.9 \| 6e-32 \| 1.2e-37 \| 229.2 \| 270 \| (18, 293) \| 300 \| (1, 352) \| 354 \| Uncharacterized homolog of phage Mu protein gp47 \| Uncharacterized homolog of phage Mu protein gp47; uniclust \| UniRef100\_A0A1I2BHZ6 \| 99.9 \| 8.7e-32 \| 1.7e-37 \| 225.3 \| 246 \| (22, 285) \| 300 \| (9, 332) \| 341 \| Uncharacterized phage protein gp47/JayE \| Uncharacterized phage protein gp47/JayE; uniclust \| UniRef100\_A0A010RU55 \| 99.9 \| 3.4e-31 \| 6.8e-37 \| 232.9 \| 270 \| (21, 296) \| 300 \| (15, 387) \| 415 \| Baseplate protein \| Baseplate protein; uniclust \| UniRef100\_A0A0T7DUX5 \| 99.9 \| 7.3e-31 \| 1.4e-36 \| 225.3 \| 288 \| (3, 294) \| 300 \| (33, 414) \| 427 \| Baseplate assembly protein J \| Baseplate assembly protein J; uniclust \| UniRef100\_A0A926IEU8 \| 99.9 \| 8.8e-31 \| 1.7e-36 \| 210.6 \| 193 \| (17, 217) \| 300 \| (3, 264) \| 264 \| Baseplate J/gp47 family protein (Fragment) \| Baseplate J/gp47 family protein (Fragment); uniclust \| UniRef100\_A0A812QW15 \| 99.9 \| 9.1e-31 \| 1.7e-36 \| 233.4 \| 290 \| (2, 291) \| 300 \| (1, 297) \| 1062 \| J protein \| J protein; uniclust \| UniRef100\_A0A064AIK0 \| 99.9 \| 1.4e-30 \| 2.7e-36 \| 222.6 \| 271 \| (22, 298) \| 300 \| (12, 366) \| 372 \| Baseplate assembly protein \| Baseplate assembly protein; uniclust \| UniRef100\_A0A081J876 \| 99.9 \| 2e-30 \| 4e-36 \| 239.9 \| 273 \| (16, 297) \| 300 \| (52, 523) \| 529 \| Baseplate protein J-like domain-containing protein \| Baseplate protein J-like domain-containing protein; uniclust \| UniRef100\_A0A174TPL6 \| 99.9 \| 3e-30 \| 5.8e-36 \| 218.7 \| 279 \| (16, 299) \| 300 \| (27, 397) \| 398 \| Uncharacterized homolog of phage Mu protein gp47 \| Uncharacterized homolog of phage Mu protein gp47; uniclust \| UniRef100\_A0A0F9B3X2 \| 99.9 \| 3.8e-30 \| 7.5e-36 \| 228.6 \| 267 \| (21, 293) \| 300 \| (6, 438) \| 440 \| Baseplate protein J-like domain-containing protein (Fragment) \| Baseplate protein J-like domain-containing protein (Fragment); uniclust \| UniRef100\_A0A166EHE8 \| 99.9 \| 1.3e-29 \| 2.4e-35 \| 215.6 \| 269 \| (21, 294) \| 300 \| (24, 393) \| 395 \| Baseplate J-like protein \| Baseplate J-like protein; uniclust \| UniRef100\_A0A1I3REQ3 \| 99.9 \| 2.5e-29 \| 4.9e-35 \| 217.3 \| 267 \| (19, 295) \| 300 \| (12, 394) \| 397 \| Baseplate J-like protein \| Baseplate J-like protein; uniclust \| UniRef100\_A0A1F1SBS6 \| 99.9 \| 2.5e-29 \| 4.9e-35 \| 210.5 \| 193 \| (20, 218) \| 300 \| (4, 273) \| 280 \| Baseplate protein J-like domain-containing protein (Fragment) \| Baseplate protein J-like domain-containing protein (Fragment); uniclust \| UniRef100\_A0A4Q7FYT9 \| 99.9 \| 2.8e-29 \| 5.3e-35 \| 205.0 \| 284 \| (1, 299) \| 300 \| (4, 302) \| 310 \| Baseplate protein J-like domain-containing protein \| Baseplate protein J-like domain-containing protein; uniclust \| UniRef100\_A0A1D2QY08 \| 99.9 \| 3.7e-29 \| 7.2e-35 \| 222.6 \| 242 \| (42, 297) \| 300 \| (106, 526) \| 531 \| Baseplate protein J-like domain-containing protein \| Baseplate protein J-like domain-containing protein; uniclust \| UniRef100\_A0A0F6YHW2 \| 99.9 \| 4.5e-29 \| 8.9e-35 \| 219.8 \| 273 \| (19, 296) \| 300 \| (11, 389) \| 398 \| Baseplate assembly protein, putative \| Baseplate assembly protein, putative; uniclust \| UniRef100\_A0A0D1VV23 \| 99.9 \| 6e-29 \| 1.1e-34 \| 209.2 \| 270 \| (16, 294) \| 300 \| (8, 383) \| 386 \| Uncharacterized phage protein gp47/JayE \| Uncharacterized phage protein gp47/JayE; uniclust \| UniRef100\_A0A0M1HGJ4 \| 99.9 \| 6.2e-29 \| 1.2e-34 \| 193.8 \| 199 \| (45, 243) \| 300 \| (15, 213) \| 215 \| Baseplate assembly protein (Fragment) \| Baseplate assembly protein (Fragment); uniclust \| UniRef100\_A0A2X1W9Z3 \| 99.9 \| 9.9e-29 \| 1.8e-34 \| 195.7 \| 279 \| (7, 287) \| 300 \| (3, 288) \| 295 \| Baseplate protein phage associated protein \| Baseplate protein phage associated protein; uniclust \| UniRef100\_A0A0J1FTG2 \| 99.9 \| 3.9e-28 \| 7.3e-34 \| 193.0 \| 231 \| (17, 283) \| 300 \| (7, 239) \| 248 \| Baseplate J-like protein \| Baseplate J-like protein; uniclust \| UniRef100\_A0A348Z5F5 \| 99.9 \| 4e-28 \| 7.6e-34 \| 194.4 \| 180 \| (111, 294) \| 300 \| (58, 243) \| 244 \| Baseplate protein J-like domain-containing protein (Fragment) \| Baseplate protein J-like domain-containing protein (Fragment); uniclust \| UniRef100\_A0A0C2VDI9 \| 99.9 \| 4e-28 \| 8e-34 \| 213.4 \| 268 \| (19, 298) \| 300 \| (4, 374) \| 378 \| Baseplate protein J-like domain-containing protein \| Baseplate protein J-like domain-containing protein; uniclust \| UniRef100\_A0A135YQ64 \| 99.9 \| 4.4e-28 \| 8.3e-34 \| 202.1 \| 262 \| (21, 296) \| 300 \| (6, 345) \| 347 \| Baseplate J-like protein \| Baseplate J-like protein; uniclust \| UniRef100\_A0A061NPP2 \| 99.9 \| 4.4e-28 \| 8.6e-34 \| 205.1 \| 238 \| (6, 298) \| 300 \| (8, 326) \| 328 \| Phage-related baseplate assembly protein \| Phage-related baseplate assembly protein; uniclust \| UniRef100\_A0A1C6CKU7 \| 99.9 \| 7.6e-28 \| 1.4e-33 \| 189.6 \| 247 \| (30, 295) \| 300 \| (5, 258) \| 260 \| Uncharacterized homolog of phage Mu protein gp47 \| Uncharacterized homolog of phage Mu protein gp47; uniclust \| UniRef100\_A0A0B7GXK9 \| 99.9 \| 1.2e-27 \| 2.4e-33 \| 207.8 \| 266 \| (20, 298) \| 300 \| (8, 379) \| 387 \| Baseplate J/gp47 family protein \| Baseplate J/gp47 family protein; uniclust \| UniRef100\_A0A193QL68 \| 99.9 \| 1.5e-27 \| 2.7e-33 \| 193.7 \| 291 \| (4, 298) \| 300 \| (79, 377) \| 378 \| Baseplate J-like protein \| Baseplate J-like protein; uniclust \| UniRef100\_A0A258L4A6 \| 99.9 \| 1.8e-27 \| 3.5e-33 \| 186.1 \| 193 \| (1, 193) \| 300 \| (15, 209) \| 213 \| Baseplate protein J-like domain-containing protein \| Baseplate protein J-like domain-containing protein; uniclust \| UniRef100\_A0A1S6TP77 \| 99.9 \| 2.1e-27 \| 4.2e-33 \| 203.8 \| 266 \| (16, 295) \| 300 \| (7, 343) \| 358 \| Phage baseplate assembly protein J, putative \| Phage baseplate assembly protein J, putative; uniclust \| UniRef100\_A0A2I0CWU0 \| 99.9 \| 2.5e-27 \| 4.6e-33 \| 198.0 \| 263 \| (17, 294) \| 300 \| (9, 364) \| 365 \| Baseplate protein J-like domain-containing protein \| Baseplate protein J-like domain-containing protein; uniclust \| UniRef100\_A0A7C6EBI1 \| 99.9 \| 3e-27 \| 5.4e-33 \| 191.2 \| 266 \| (24, 293) \| 300 \| (4, 354) \| 364 \| Baseplate protein J-like domain-containing protein \| Baseplate protein J-like domain-containing protein; uniclust \| UniRef100\_A0A024AZH8 \| 99.9 \| 2.8e-27 \| 5.5e-33 \| 205.1 \| 261 \| (21, 297) \| 300 \| (7, 354) \| 362 \| Baseplate j protein \| Baseplate j protein; uniclust \| UniRef100\_B6G011 \| 99.9 \| 3.7e-27 \| 6.9e-33 \| 192.8 \| 267 \| (20, 294) \| 300 \| (3, 350) \| 354 \| Baseplate J-like protein \| Baseplate J-like protein; uniclust \| UniRef100\_A0A440V0E4 \| 99.9 \| 3.7e-27 \| 7.1e-33 \| 185.0 \| 205 \| (2, 223) \| 300 \| (1, 214) \| 218 \| Baseplate J protein \| Baseplate J protein; uniclust \| UniRef100\_A9HP66 \| 99.9 \| 4.2e-27 \| 8.1e-33 \| 195.4 \| 174 \| (112, 293) \| 300 \| (109, 285) \| 288 \| Baseplate protein J-like domain-containing protein \| Baseplate protein J-like domain-containing protein; uniclust \| UniRef100\_A0A6L2ZSF9 \| 99.9 \| 7.4e-27 \| 1.4e-32 \| 196.1 \| 277 \| (6, 282) \| 300 \| (6, 472) \| 507 \| Baseplate assembly protein \| Baseplate assembly protein; uniclust \| UniRef100\_UPI0022DF30D1 \| 99.9 \| 8.2e-27 \| 1.5e-32 \| 193.2 \| 240 \| (19, 266) \| 300 \| (3, 318) \| 446 \| baseplate J/gp47 family protein \| baseplate J/gp47 family protein; uniclust \| UniRef100\_A0A013SL59 \| 99.9 \| 1e-26 \| 2.1e-32 \| 209.9 \| 273 \| (14, 295) \| 300 \| (86, 467) \| 490 \| Baseplate J-like family protein \| Baseplate J-like family protein; uniclust \| UniRef100\_A0A128FC77 \| 99.9 \| 1.2e-26 \| 2.3e-32 \| 199.4 \| 274 \| (19, 298) \| 300 \| (3, 363) \| 365 \| Baseplate J-like protein \| Baseplate J-like protein; uniclust \| UniRef100\_A0A261QSM0 \| 99.9 \| 1.3e-26 \| 2.4e-32 \| 196.6 \| 268 \| (19, 292) \| 300 \| (6, 360) \| 361 \| Baseplate protein J-like domain-containing protein \| Baseplate protein J-like domain-containing protein; uniclust \| UniRef100\_UPI00080E3582 \| 99.9 \| 1.3e-26 \| 2.4e-32 \| 191.4 \| 284 \| (4, 295) \| 300 \| (2, 380) \| 382 \| baseplate J/gp47 family protein \| baseplate J/gp47 family protein; uniclust \| UniRef100\_A0A1F5C406 \| 99.9 \| 1.7e-26 \| 3.3e-32 \| 204.9 \| 273 \| (19, 294) \| 300 \| (4, 463) \| 464 \| Baseplate protein J-like domain-containing protein \| Baseplate protein J-like domain-containing protein; uniclust \| UniRef100\_A0A376EX36 \| 99.9 \| 3.1e-26 \| 5.9e-32 \| 187.4 \| 187 \| (109, 298) \| 300 \| (85, 276) \| 278 \| Baseplate assembly protein J \| Baseplate assembly protein J; uniclust \| UniRef100\_A0A8S4QUR0 \| 99.9 \| 4.7e-26 \| 8.6e-32 \| 197.4 \| 249 \| (14, 284) \| 300 \| (2, 252) \| 611 \| Jg15363 protein \| Jg15363 protein; uniclust \| UniRef100\_A0A0W8FUR1 \| 99.9 \| 4.7e-26 \| 8.8e-32 \| 195.8 \| 248 \| (21, 293) \| 300 \| (53, 448) \| 449 \| Phage flumu protein gp47 \| Phage flumu protein gp47; uniclust \| UniRef100\_A0A6B8KJI2 \| 99.9 \| 4.9e-26 \| 9.3e-32 \| 186.6 \| 280 \| (1, 295) \| 300 \| (5, 312) \| 315 \| Baseplate protein J-like domain-containing protein \| Baseplate protein J-like domain-containing protein; uniclust \| UniRef100\_A0A0F0HJ75 \| 99.9 \| 6.5e-26 \| 1.2e-31 \| 200.9 \| 285 \| (7, 296) \| 300 \| (2, 490) \| 494 \| Baseplate protein J-like domain-containing protein \| Baseplate protein J-like domain-containing protein; uniclust \| UniRef100\_A0A8I1NZ44 \| 99.9 \| 6.7e-26 \| 1.3e-31 \| 187.0 \| 287 \| (7, 295) \| 300 \| (14, 336) \| 338 \| Baseplate J/gp47 family protein \| Baseplate J/gp47 family protein; uniclust \| UniRef100\_A0A089K1A9 \| 99.9 \| 8.8e-26 \| 1.7e-31 \| 196.1 \| 258 \| (13, 295) \| 300 \| (16, 321) \| 339 \| Baseplate protein J-like domain-containing protein \| Baseplate protein J-like domain-containing protein; uniclust \| UniRef100\_A0A0E2E9P5 \| 99.9 \| 9.7e-26 \| 1.9e-31 \| 198.0 \| 274 \| (21, 297) \| 300 \| (6, 394) \| 396 \| Baseplate protein J-like domain-containing protein \| Baseplate protein J-like domain-containing protein; uniclust \| UniRef100\_A0A1U7N2A8 \| 99.9 \| 1.3e-25 \| 2.4e-31 \| 207.7 \| 241 \| (19, 268) \| 300 \| (3, 360) \| 748 \| Baseplate protein J-like domain-containing protein \| Baseplate protein J-like domain-containing protein; uniclust \| UniRef100\_A0A3A9BP88 \| 99.9 \| 1.3e-25 \| 2.5e-31 \| 186.1 \| 266 \| (22, 295) \| 300 \| (3, 355) \| 362 \| Baseplate protein J-like domain-containing protein \| Baseplate protein J-like domain-containing protein; uniclust \| UniRef100\_UPI0001AF5308 \| 99.9 \| 1.7e-25 \| 3.1e-31 \| 180.6 \| 257 \| (8, 267) \| 300 \| (4, 267) \| 351 \| baseplate J/gp47 family protein \| baseplate J/gp47 family protein; uniclust \| UniRef100\_A0A3N6RV94 \| 99.9 \| 1.9e-25 \| 3.5e-31 \| 185.6 \| 235 \| (5, 239) \| 300 \| (1, 242) \| 418 \| Phage tail protein I \| Phage tail protein I; uniclust \| UniRef100\_A0A2R3MZM3 \| 99.9 \| 2.3e-25 \| 4.4e-31 \| 188.8 \| 266 \| (18, 291) \| 300 \| (10, 380) \| 388 \| Baseplate protein J-like domain-containing protein \| Baseplate protein J-like domain-containing protein; uniclust \| UniRef100\_A0A328CA52 \| 99.9 \| 2.3e-25 \| 4.5e-31 \| 191.2 \| 271 \| (19, 294) \| 300 \| (13, 387) \| 390 \| Baseplate protein J-like domain-containing protein \| Baseplate protein J-like domain-containing protein; uniclust \| UniRef100\_A0A3A9EFF6 \| 99.9 \| 2.5e-25 \| 4.8e-31 \| 175.3 \| 187 \| (111, 299) \| 300 \| (17, 206) \| 207 \| Phage baseplate protein (Fragment) \| Phage baseplate protein (Fragment); uniclust \| UniRef100\_A0A080LY68 \| 99.9 \| 2.5e-25 \| 4.9e-31 \| 200.1 \| 237 \| (21, 269) \| 300 \| (14, 351) \| 523 \| Baseplate protein J-like domain-containing protein \| Baseplate protein J-like domain-containing protein; uniclust \| UniRef100\_A0A0A2WKW8 \| 99.9 \| 2.7e-25 \| 5.4e-31 \| 210.1 \| 181 \| (112, 295) \| 300 \| (439, 648) \| 679 \| Baseplate J family protein \| Baseplate J family protein; uniclust \| UniRef100\_A0A1M6UTJ7 \| 99.8 \| 5.7e-25 \| 1.1e-30 \| 186.2 \| 234 \| (17, 271) \| 300 \| (17, 275) \| 311 \| Baseplate J-like protein \| Baseplate J-like protein; uniclust \| UniRef100\_A0A022PMN6 \| 99.8 \| 7.1e-25 \| 1.4e-30 \| 197.3 \| 281 \| (10, 297) \| 300 \| (92, 476) \| 493 \| Putative phage Mu protein gp47-like protein \| Putative phage Mu protein gp47-like protein; uniclust \| UniRef100\_A0A0R3MV37 \| 99.8 \| 7.7e-25 \| 1.4e-30 \| 177.2 \| 277 \| (4, 297) \| 300 \| (4, 288) \| 290 \| Baseplate assembly protein \| Baseplate assembly protein; uniclust \| UniRef100\_A0A0A7HBT3 \| 99.8 \| 7.6e-25 \| 1.5e-30 \| 191.9 \| 276 \| (8, 298) \| 300 \| (14, 389) \| 391 \| Baseplate J-like protein \| Baseplate J-like protein; uniclust \| UniRef100\_UPI00188A944F \| 99.8 \| 1e-24 \| 1.9e-30 \| 175.7 \| 222 \| (74, 295) \| 300 \| (1, 249) \| 251 \| baseplate J/gp47 family protein \| baseplate J/gp47 family protein; uniclust \| UniRef100\_A0A0F9SMT4 \| 99.8 \| 1.2e-24 \| 2.3e-30 \| 191.6 \| 287 \| (8, 296) \| 300 \| (36, 535) \| 540 \| Baseplate protein J-like domain-containing protein \| Baseplate protein J-like domain-containing protein; uniclust \| UniRef100\_A0A068NXX0 \| 99.8 \| 1.4e-24 \| 2.7e-30 \| 195.3 \| 264 \| (16, 288) \| 300 \| (71, 484) \| 498 \| Baseplate assembly protein, putative \| Baseplate assembly protein, putative; uniclust \| UniRef100\_A0A0A7G316 \| 99.8 \| 1.5e-24 \| 2.9e-30 \| 178.6 \| 247 \| (21, 292) \| 300 \| (6, 267) \| 272 \| Baseplate J-like family protein \| Baseplate J-like family protein; uniclust \| UniRef100\_A0A164ZAU4 \| 99.8 \| 2.1e-24 \| 3.9e-30 \| 167.9 \| 181 \| (112, 294) \| 300 \| (8, 206) \| 209 \| Phage baseplate \| Phage baseplate; uniclust \| UniRef100\_A0A062V567 \| 99.8 \| 2.4e-24 \| 4.7e-30 \| 181.8 \| 261 \| (13, 297) \| 300 \| (7, 311) \| 324 \| Baseplate J-like protein \| Baseplate J-like protein; uniclust \| UniRef100\_A0A1I7E360 \| 99.8 \| 3e-24 \| 5.6e-30 \| 187.3 \| 269 \| (5, 273) \| 300 \| (1, 276) \| 557 \| Phage tail tape measure protein, TP901 family, core region \| Phage tail tape measure protein, TP901 family, core region; uniclust \| UniRef100\_A0A0E3NKG6 \| 99.8 \| 2.9e-24 \| 5.8e-30 \| 206.6 \| 183 \| (112, 297) \| 300 \| (502, 716) \| 756 \| Baseplate protein J-like domain-containing protein \| Baseplate protein J-like domain-containing protein; uniclust \| UniRef100\_A0A2A2HC74 \| 99.8 \| 3.3e-24 \| 6.2e-30 \| 167.0 \| 185 \| (111, 298) \| 300 \| (20, 205) \| 209 \| Baseplate protein J-like domain-containing protein \| Baseplate protein J-like domain-containing protein; uniclust \| UniRef100\_A0A8I2FQ02 \| 99.8 \| 3.4e-24 \| 6.5e-30 \| 177.9 \| 256 \| (13, 295) \| 300 \| (3, 290) \| 299 \| Baseplate protein J-like domain-containing protein \| Baseplate protein J-like domain-containing protein; uniclust \| UniRef100\_A0A258KUT3 \| 99.8 \| 3.8e-24 \| 7.2e-30 \| 165.6 \| 190 \| (109, 298) \| 300 \| (2, 191) \| 193 \| Baseplate protein J-like domain-containing protein \| Baseplate protein J-like domain-containing protein; uniclust \| UniRef100\_UPI000469242B \| 99.8 \| 4e-24 \| 7.4e-30 \| 177.0 \| 265 \| (16, 292) \| 300 \| (14, 366) \| 367 \| baseplate J/gp47 family protein \| baseplate J/gp47 family protein; uniclust \| UniRef100\_A0A0A8WTQ2 \| 99.8 \| 3.8e-24 \| 7.5e-30 \| 186.8 \| 248 \| (20, 289) \| 300 \| (48, 331) \| 374 \| Baseplate protein J-like domain-containing protein \| Baseplate protein J-like domain-containing protein; uniclust \| UniRef100\_A0A3N9ESU6 \| 99.8 \| 4.6e-24 \| 8.8e-30 \| 163.7 \| 170 \| (115, 291) \| 300 \| (3, 176) \| 183 \| Phage tail protein (Fragment) \| Phage tail protein (Fragment); uniclust \| UniRef100\_A0A1H5ZLM5 \| 99.8 \| 5.4e-24 \| 1e-29 \| 184.2 \| 268 \| (19, 296) \| 300 \| (10, 409) \| 413 \| Baseplate J family protein \| Baseplate J family protein; uniclust \| UniRef100\_A0A060H3Z0 \| 99.8 \| 6e-24 \| 1.2e-29 \| 187.1 \| 276 \| (9, 296) \| 300 \| (12, 391) \| 423 \| Baseplate protein J-like domain-containing protein \| Baseplate protein J-like domain-containing protein; uniclust \| UniRef100\_A0A0A0GWF9 \| 99.8 \| 6.2e-24 \| 1.2e-29 \| 164.2 \| 163 \| (1, 163) \| 300 \| (7, 176) \| 178 \| Baseplate assembly protein J \| Baseplate assembly protein J; uniclust \| UniRef100\_A0A7G5CAE3 \| 99.8 \| 7.8e-24 \| 1.4e-29 \| 159.4 \| 193 \| (14, 222) \| 300 \| (2, 194) \| 206 \| Baseplate assembly protein J \| Baseplate assembly protein J; uniclust \| UniRef100\_A0A1E5NH39 \| 99.8 \| 8.5e-24 \| 1.6e-29 \| 180.8 \| 268 \| (21, 293) \| 300 \| (4, 366) \| 367 \| Baseplate protein J-like domain-containing protein \| Baseplate protein J-like domain-containing protein; uniclust \| UniRef100\_A0A3D4RUD6 \| 99.8 \| 9.2e-24 \| 1.7e-29 \| 172.4 \| 268 \| (2, 285) \| 300 \| (1, 275) \| 325 \| Baseplate protein J-like domain-containing protein \| Baseplate protein J-like domain-containing protein; uniclust \| UniRef100\_A0A8S5PDI9 \| 99.8 \| 9.5e-24 \| 1.7e-29 \| 171.0 \| 261 \| (18, 287) \| 300 \| (1, 339) \| 354 \| Baseplate J like protein \| Baseplate J like protein; uniclust \| UniRef100\_A0A2S8P413 \| 99.8 \| 1.3e-23 \| 2.5e-29 \| 179.6 \| 271 \| (15, 292) \| 300 \| (4, 377) \| 538 \| Baseplate J family protein \| Baseplate J family protein; uniclust \| UniRef100\_UPI000A8775AC \| 99.8 \| 1.7e-23 \| 3.2e-29 \| 169.7 \| 211 \| (9, 221) \| 300 \| (5, 300) \| 303 \| baseplate J/gp47 family protein \| baseplate J/gp47 family protein; uniclust \| UniRef100\_A0A7V6NHE1 \| 99.8 \| 1.8e-23 \| 3.5e-29 \| 169.4 \| 177 \| (112, 295) \| 300 \| (88, 270) \| 273 \| Baseplate J protein \| Baseplate J protein; uniclust \| UniRef100\_A0A212JYZ3 \| 99.8 \| 1.9e-23 \| 3.6e-29 \| 172.8 \| 178 \| (111, 295) \| 300 \| (100, 284) \| 287 \| Baseplate J family protein \| Baseplate J family protein; uniclust \| UniRef100\_A0A3G2R557 \| 99.8 \| 2.1e-23 \| 3.8e-29 \| 174.5 \| 268 \| (22, 294) \| 300 \| (3, 466) \| 467 \| Baseplate protein J-like domain-containing protein \| Baseplate protein J-like domain-containing protein; uniclust \| UniRef100\_A0A098MR91 \| 99.8 \| 2.2e-23 \| 4.2e-29 \| 178.6 \| 272 \| (15, 294) \| 300 \| (7, 391) \| 395 \| Baseplate protein J \| Baseplate protein J; uniclust \| UniRef100\_A0A022MMA0 \| 99.8 \| 2.2e-23 \| 4.4e-29 \| 201.1 \| 156 \| (112, 269) \| 300 \| (564, 742) \| 876 \| Baseplate J family protein \| Baseplate J family protein; uniclust \| UniRef100\_A0A0P9WA78 \| 99.8 \| 3e-23 \| 5.7e-29 \| 172.1 \| 177 \| (111, 294) \| 300 \| (97, 297) \| 298 \| Baseplate J-like protein \| Baseplate J-like protein; uniclust \| UniRef100\_A0A2D6NHB6 \| 99.8 \| 3.2e-23 \| 6.1e-29 \| 177.6 \| 260 \| (21, 288) \| 300 \| (5, 368) \| 381 \| Baseplate protein J-like domain-containing protein \| Baseplate protein J-like domain-containing protein; uniclust \| UniRef100\_A0A2N9MST4 \| 99.8 \| 3.3e-23 \| 6.3e-29 \| 172.8 \| 238 \| (13, 269) \| 300 \| (3, 265) \| 327 \| Baseplate protein J-like domain-containing protein \| Baseplate protein J-like domain-containing protein; uniclust \| UniRef100\_A0A0R1YCE5 \| 99.8 \| 3.5e-23 \| 6.7e-29 \| 170.2 \| 170 \| (17, 192) \| 300 \| (8, 249) \| 264 \| Baseplate J-like protein \| Baseplate J-like protein; uniclust \| UniRef100\_A0A4P0TK43 \| 99.8 \| 4.5e-23 \| 8.3e-29 \| 156.1 \| 205 \| (94, 298) \| 300 \| (2, 206) \| 207 \| Putative phage baseplate assembly protein \| Putative phage baseplate assembly protein; uniclust \| UniRef100\_A0A084ZEL5 \| 99.8 \| 5.6e-23 \| 1.1e-28 \| 170.1 \| 175 \| (21, 201) \| 300 \| (6, 258) \| 265 \| Phage baseplate assembly protein J (Fragment) \| Phage baseplate assembly protein J (Fragment); uniclust \| UniRef100\_A0A3D4RUG0 \| 99.8 \| 6.2e-23 \| 1.2e-28 \| 175.3 \| 270 \| (8, 295) \| 300 \| (15, 305) \| 416 \| Baseplate protein J-like domain-containing protein \| Baseplate protein J-like domain-containing protein; uniclust \| UniRef100\_A0A009ZC54 \| 99.8 \| 8e-23 \| 1.5e-28 \| 177.0 \| 264 \| (18, 294) \| 300 \| (89, 434) \| 437 \| Baseplate J-like family protein \| Baseplate J-like family protein; uniclust \| UniRef100\_A0A0B6CQ96 \| 99.8 \| 1e-22 \| 1.9e-28 \| 159.6 \| 256 \| (19, 294) \| 300 \| (12, 267) \| 270 \| Baseplate J-like family protein \| Baseplate J-like family protein; uniclust \| UniRef100\_A0A3B8INQ5 \| 99.8 \| 1.1e-22 \| 2.2e-28 \| 185.5 \| 237 \| (21, 268) \| 300 \| (5, 335) \| 804 \| Baseplate protein J-like domain-containing protein \| Baseplate protein J-like domain-containing protein; uniclust \| UniRef100\_A0A074MBI6 \| 99.8 \| 1.1e-22 \| 2.2e-28 \| 192.6 \| 181 \| (112, 297) \| 300 \| (530, 746) \| 751 \| Baseplate protein J-like domain-containing protein \| Baseplate protein J-like domain-containing protein; uniclust \| UniRef100\_A0A8S5MVE0 \| 99.8 \| 1.2e-22 \| 2.2e-28 \| 170.3 \| 266 \| (22, 293) \| 300 \| (107, 470) \| 472 \| Baseplate J like protein \| Baseplate J like protein; uniclust \| UniRef100\_A0A2W4SAT6 \| 99.8 \| 1.2e-22 \| 2.2e-28 \| 169.2 \| 173 \| (16, 191) \| 300 \| (7, 264) \| 296 \| Baseplate protein J-like domain-containing protein (Fragment) \| Baseplate protein J-like domain-containing protein (Fragment); uniclust \| UniRef100\_A0A258DW01 \| 99.8 \| 1.6e-22 \| 3.1e-28 \| 169.7 \| 269 \| (21, 296) \| 300 \| (5, 363) \| 367 \| Baseplate protein J-like domain-containing protein \| Baseplate protein J-like domain-containing protein; uniclust \| UniRef100\_A0A0B0SG84 \| 99.8 \| 1.6e-22 \| 3.1e-28 \| 173.2 \| 256 \| (21, 295) \| 300 \| (7, 354) \| 356 \| Baseplate protein J-like domain-containing protein \| Baseplate protein J-like domain-containing protein; uniclust \| UniRef100\_A0A0C1PPW9 \| 99.8 \| 2e-22 \| 3.8e-28 \| 173.4 \| 270 \| (18, 295) \| 300 \| (10, 410) \| 414 \| Putative bacteriophage protein \| Putative bacteriophage protein; uniclust \| UniRef100\_A0A0A0RK69 \| 99.8 \| 2.7e-22 \| 5.1e-28 \| 178.2 \| 245 \| (17, 270) \| 300 \| (9, 449) \| 496 \| Baseplate component \| Baseplate component; uniclust \| UniRef100\_A0A8J4SQD0 \| 99.8 \| 2.8e-22 \| 5.1e-28 \| 179.1 \| 267 \| (21, 294) \| 300 \| (430, 779) \| 935 \| Baseplate protein J-like domain-containing protein \| Baseplate protein J-like domain-containing protein; uniclust \| UniRef100\_A0A0P6XAW3 \| 99.8 \| 4.6e-22 \| 9e-28 \| 186.2 \| 155 \| (112, 268) \| 300 \| (490, 680) \| 758 \| Baseplate protein J-like domain-containing protein \| Baseplate protein J-like domain-containing protein; uniclust \| UniRef100\_A0A2A2HFP1 \| 99.8 \| 4.9e-22 \| 9.3e-28 \| 174.2 \| 256 \| (7, 268) \| 300 \| (16, 385) \| 481 \| Baseplate J-like protein \| Baseplate J-like protein; uniclust \| UniRef100\_A0A1Y3ZYD9 \| 99.8 \| 7.4e-22 \| 1.4e-27 \| 156.6 \| 180 \| (111, 294) \| 300 \| (26, 224) \| 262 \| Baseplate protein J-like domain-containing protein \| Baseplate protein J-like domain-containing protein; uniclust \| UniRef100\_A0A662W1X1 \| 99.8 \| 8.3e-22 \| 1.5e-27 \| 159.1 \| 218 \| (21, 268) \| 300 \| (13, 232) \| 337 \| Baseplate protein J-like domain-containing protein \| Baseplate protein J-like domain-containing protein; uniclust \| UniRef100\_A0A0K6HHF2 \| 99.8 \| 9e-22 \| 1.7e-27 \| 163.3 \| 221 \| (21, 245) \| 300 \| (5, 302) \| 317 \| Uncharacterized phage protein gp47/JayE (Fragment) \| Uncharacterized phage protein gp47/JayE (Fragment); uniclust \| UniRef100\_A0A0F9SSY1 \| 99.8 \| 9.3e-22 \| 1.8e-27 \| 170.6 \| 240 \| (43, 289) \| 300 \| (30, 362) \| 385 \| Baseplate protein J-like domain-containing protein \| Baseplate protein J-like domain-containing protein; uniclust \| UniRef100\_A0A024QHR7 \| 99.8 \| 9.8e-22 \| 1.9e-27 \| 163.9 \| 261 \| (21, 291) \| 300 \| (6, 281) \| 283 \| Baseplate protein J-like domain-containing protein \| Baseplate protein J-like domain-containing protein; uniclust \| UniRef100\_A0A1G6XUY3 \| 99.8 \| 1e-21 \| 2e-27 \| 183.7 \| 268 \| (21, 296) \| 300 \| (72, 492) \| 608 \| Uncharacterized phage protein gp47/JayE \| Uncharacterized phage protein gp47/JayE; uniclust \| UniRef100\_A0A6J5KNL2 \| 99.8 \| 1.2e-21 \| 2.4e-27 \| 178.7 \| 288 \| (5, 295) \| 300 \| (1, 518) \| 623 \| Baseplate protein J-like \| Baseplate protein J-like; uniclust \| UniRef100\_A0A1J5EX15 \| 99.7 \| 1.3e-21 \| 2.6e-27 \| 171.3 \| 245 \| (20, 270) \| 300 \| (14, 351) \| 398 \| Baseplate protein J-like domain-containing protein \| Baseplate protein J-like domain-containing protein; uniclust \| UniRef100\_A0A068R2C5 \| 99.7 \| 1.4e-21 \| 2.8e-27 \| 176.4 \| 269 \| (17, 295) \| 300 \| (31, 415) \| 421 \| Phage protein \| Phage protein; uniclust \| UniRef100\_A0A011MDS5 \| 99.7 \| 1.5e-21 \| 3e-27 \| 188.7 \| 182 \| (112, 296) \| 300 \| (512, 729) \| 808 \| Baseplate assembly protein \| Baseplate assembly protein; uniclust \| UniRef100\_UPI00233ECD7D \| 99.7 \| 1.8e-21 \| 3.2e-27 \| 147.7 \| 199 \| (5, 203) \| 300 \| (1, 210) \| 211 \| baseplate J/gp47 family protein \| baseplate J/gp47 family protein; uniclust \| UniRef100\_A0A220MKU4 \| 99.7 \| 1.8e-21 \| 3.3e-27 \| 153.1 \| 182 \| (111, 294) \| 300 \| (46, 239) \| 240 \| Baseplate protein J-like domain-containing protein \| Baseplate protein J-like domain-containing protein; uniclust \| UniRef100\_A0A545T5R2 \| 99.7 \| 1.8e-21 \| 3.3e-27 \| 156.2 \| 280 \| (2, 289) \| 300 \| (1, 298) \| 320 \| Baseplate protein J-like domain-containing protein \| Baseplate protein J-like domain-containing protein; uniclust \| UniRef100\_A0A068C8J3 \| 99.7 \| 1.8e-21 \| 3.6e-27 \| 178.7 \| 272 \| (17, 296) \| 300 \| (52, 440) \| 510 \| Putative baseplate J-like protein \| Putative baseplate J-like protein; uniclust \| UniRef100\_D1P8B4 \| 99.7 \| 1.9e-21 \| 3.6e-27 \| 148.5 \| 180 \| (13, 192) \| 300 \| (2, 188) \| 191 \| Baseplate J-like protein (Fragment) \| Baseplate J-like protein (Fragment); uniclust \| UniRef100\_A0A133ZRX4 \| 99.7 \| 2.2e-21 \| 4.3e-27 \| 175.7 \| 269 \| (16, 293) \| 300 \| (22, 494) \| 504 \| Baseplate protein J-like domain-containing protein \| Baseplate protein J-like domain-containing protein; uniclust \| UniRef100\_A0A0H5RM30 \| 99.7 \| 2.6e-21 \| 5.1e-27 \| 178.3 \| 238 \| (42, 288) \| 300 \| (181, 510) \| 635 \| Baseplate J-like protein \| Baseplate J-like protein; uniclust \| UniRef100\_UPI0005E01794 \| 99.7 \| 3.3e-21 \| 6.1e-27 \| 153.0 \| 199 \| (5, 203) \| 300 \| (1, 206) \| 292 \| baseplate J/gp47 family protein \| baseplate J/gp47 family protein; uniclust \| UniRef100\_A0A101WDK8 \| 99.7 \| 3.7e-21 \| 7.2e-27 \| 179.5 \| 180 \| (112, 298) \| 300 \| (488, 694) \| 698 \| Baseplate protein J-like domain-containing protein \| Baseplate protein J-like domain-containing protein; uniclust \| UniRef100\_A0A3P8KWB0 \| 99.7 \| 4.2e-21 \| 8e-27 \| 146.3 \| 147 \| (154, 300) \| 300 \| (8, 154) \| 154 \| Uncharacterized homolog of phage Mu protein gp47 \| Uncharacterized homolog of phage Mu protein gp47; uniclust \| UniRef100\_A0A084SKL3 \| 99.7 \| 4.8e-21 \| 9.5e-27 \| 185.9 \| 154 \| (112, 268) \| 300 \| (303, 486) \| 886 \| Baseplate protein J-like domain-containing protein \| Baseplate protein J-like domain-containing protein; uniclust \| UniRef100\_A0A0K1RXW7 \| 99.7 \| 5.2e-21 \| 1e-26 \| 168.8 \| 231 \| (12, 268) \| 300 \| (15, 271) \| 457 \| Baseplate protein J-like domain-containing protein \| Baseplate protein J-like domain-containing protein; uniclust \| UniRef100\_A0A6P1Y489 \| 99.7 \| 5.4e-21 \| 1e-26 \| 158.5 \| 264 \| (16, 293) \| 300 \| (4, 348) \| 352 \| Baseplate protein J-like domain-containing protein \| Baseplate protein J-like domain-containing protein; uniclust \| UniRef100\_A0A376CQY9 \| 99.7 \| 6.8e-21 \| 1.3e-26 \| 144.5 \| 173 \| (127, 299) \| 300 \| (10, 182) \| 190 \| Baseplate assembly protein GpJ \| Baseplate assembly protein GpJ; uniclust \| UniRef100\_A0A835Z0B9 \| 99.7 \| 7.1e-21 \| 1.3e-26 \| 177.8 \| 274 \| (7, 296) \| 300 \| (5, 284) \| 1777 \| Baseplate protein J-like domain-containing protein \| Baseplate protein J-like domain-containing protein; uniclust \| UniRef100\_A0A021X854 \| 99.7 \| 8.8e-21 \| 1.7e-26 \| 162.2 \| 258 \| (14, 295) \| 300 \| (9, 300) \| 310 \| Baseplate protein J-like domain-containing protein \| Baseplate protein J-like domain-containing protein; uniclust \| UniRef100\_A0A101G5B9 \| 99.7 \| 9.8e-21 \| 1.8e-26 \| 151.8 \| 183 \| (9, 204) \| 300 \| (2, 273) \| 275 \| Baseplate J-like protein \| Baseplate J-like protein; uniclust \| UniRef100\_A0A1B1IUR1 \| 99.7 \| 1.1e-20 \| 2.1e-26 \| 179.0 \| 155 \| (112, 268) \| 300 \| (399, 564) \| 626 \| Baseplate wedge subunit \| Baseplate wedge subunit; uniclust \| UniRef100\_UPI000D012F51 \| 99.7 \| 1.3e-20 \| 2.3e-26 \| 158.5 \| 245 \| (18, 271) \| 300 \| (8, 353) \| 390 \| baseplate J/gp47 family protein \| baseplate J/gp47 family protein; uniclust \| UniRef100\_A0A166CB08 \| 99.7 \| 1.2e-20 \| 2.4e-26 \| 162.8 \| 273 \| (8, 295) \| 300 \| (5, 368) \| 374 \| Baseplate J-like protein \| Baseplate J-like protein; uniclust \| UniRef100\_A0A973IBV9 \| 99.7 \| 1.3e-20 \| 2.5e-26 \| 150.4 \| 250 \| (21, 292) \| 300 \| (5, 265) \| 266 \| Baseplate J/gp47 family protein \| Baseplate J/gp47 family protein; uniclust \| UniRef100\_A0A134ARI0 \| 99.7 \| 1.5e-20 \| 2.9e-26 \| 160.5 \| 246 \| (43, 293) \| 300 \| (34, 366) \| 369 \| Baseplate protein J-like domain-containing protein \| Baseplate protein J-like domain-containing protein; uniclust \| UniRef100\_A0A7J6YK41 \| 99.7 \| 1.7e-20 \| 3.1e-26 \| 152.7 \| 192 \| (12, 222) \| 300 \| (138, 329) \| 353 \| Baseplate protein J-like domain-containing protein \| Baseplate protein J-like domain-containing protein; uniclust \| UniRef100\_A0A353REU8 \| 99.7 \| 1.8e-20 \| 3.3e-26 \| 151.6 \| 179 \| (111, 295) \| 300 \| (75, 256) \| 258 \| Baseplate protein J-like domain-containing protein \| Baseplate protein J-like domain-containing protein; uniclust \| UniRef100\_UPI0020BFF1D4 \| 99.7 \| 1.8e-20 \| 3.4e-26 \| 157.8 \| 202 \| (43, 244) \| 300 \| (18, 223) \| 475 \| baseplate J/gp47 family protein \| baseplate J/gp47 family protein; uniclust \| UniRef100\_A0A088C3H8 \| 99.7 \| 1.8e-20 \| 3.5e-26 \| 168.1 \| 243 \| (15, 266) \| 300 \| (6, 433) \| 476 \| p2 gpJ-like protein \| p2 gpJ-like protein; uniclust \| UniRef100\_A0A1N7LRY2 \| 99.7 \| 1.9e-20 \| 3.5e-26 \| 150.2 \| 274 \| (14, 292) \| 300 \| (10, 289) \| 299 \| Phage-related baseplate assembly protein \| Phage-related baseplate assembly protein; uniclust \| UniRef100\_A0A857ED91 \| 99.7 \| 2e-20 \| 3.6e-26 \| 146.2 \| 185 \| (111, 295) \| 300 \| (9, 199) \| 257 \| Baseplate protein J-like domain-containing protein \| Baseplate protein J-like domain-containing protein; uniclust \| UniRef100\_A0A261QMI5 \| 99.7 \| 2.1e-20 \| 3.9e-26 \| 140.6 \| 192 \| (13, 211) \| 300 \| (5, 196) \| 197 \| Phage tail protein (Fragment) \| Phage tail protein (Fragment); uniclust \| UniRef100\_A0A7T7D8U4 \| 99.7 \| 2.1e-20 \| 3.9e-26 \| 161.1 \| 263 \| (10, 273) \| 300 \| (4, 354) \| 596 \| Phage tail protein I \| Phage tail protein I; uniclust \| UniRef100\_A0A8D8XD97 \| 99.7 \| 2.3e-20 \| 4.2e-26 \| 138.9 \| 168 \| (27, 209) \| 300 \| (1, 170) \| 177 \| Baseplate protein J (Fragment) \| Baseplate protein J (Fragment); uniclust \| UniRef100\_UPI0012CD38E9 \| 99.7 \| 2.4e-20 \| 4.4e-26 \| 153.3 \| 217 \| (62, 293) \| 300 \| (159, 379) \| 382 \| prophage tail fiber N-terminal domain-containing protein \| prophage tail fiber N-terminal domain-containing protein; uniclust \| UniRef100\_A0A1Q1PVP2 \| 99.7 \| 2.3e-20 \| 4.5e-26 \| 171.5 \| 247 \| (21, 271) \| 300 \| (12, 441) \| 573 \| Baseplate \| Baseplate; uniclust \| UniRef100\_A0A1L8I3J9 \| 99.7 \| 2.7e-20 \| 5e-26 \| 150.4 \| 155 \| (16, 172) \| 300 \| (8, 247) \| 257 \| Baseplate protein J-like domain-containing protein \| Baseplate protein J-like domain-containing protein; uniclust \| UniRef100\_A0A284VS02 \| 99.7 \| 2.9e-20 \| 5.5e-26 \| 158.2 \| 249 \| (13, 268) \| 300 \| (3, 274) \| 313 \| Baseplate protein J-like domain-containing protein \| Baseplate protein J-like domain-containing protein; uniclust \| UniRef100\_A0A7G8VI16 \| 99.7 \| 3.3e-20 \| 6e-26 \| 138.5 \| 144 \| (6, 149) \| 300 \| (1, 144) \| 187 \| Baseplate J/gp47 family protein \| Baseplate J/gp47 family protein; uniclust \| UniRef100\_UPI000327E1DD \| 99.7 \| 3.5e-20 \| 6.3e-26 \| 154.2 \| 198 \| (5, 202) \| 300 \| (1, 205) \| 422 \| contractile injection system protein, VgrG/Pvc8 family \| contractile injection system protein, VgrG/Pvc8 family; uniclust \| UniRef100\_A0A2N2GRU7 \| 99.7 \| 3.8e-20 \| 7.2e-26 \| 152.2 \| 178 \| (112, 296) \| 300 \| (107, 295) \| 299 \| Baseplate assembly protein (Fragment) \| Baseplate assembly protein (Fragment); uniclust \| UniRef100\_A0A2D9F577 \| 99.7 \| 3.9e-20 \| 7.3e-26 \| 153.3 \| 262 \| (21, 289) \| 300 \| (5, 348) \| 362 \| Baseplate protein J-like domain-containing protein (Fragment) \| Baseplate protein J-like domain-containing protein (Fragment); uniclust \| UniRef100\_A0A8X6FB47 \| 99.7 \| 4.2e-20 \| 7.8e-26 \| 155.9 \| 220 \| (39, 280) \| 300 \| (9, 230) \| 480 \| Baseplate protein J \| Baseplate protein J; uniclust \| UniRef100\_A0A8B3NLL6 \| 99.7 \| 4.5e-20 \| 8.2e-26 \| 153.4 \| 276 \| (7, 298) \| 300 \| (136, 418) \| 419 \| Baseplate protein J-like domain-containing protein \| Baseplate protein J-like domain-containing protein; uniclust \| UniRef100\_E2CFL2 \| 99.7 \| 4.5e-20 \| 8.3e-26 \| 149.2 \| 273 \| (6, 295) \| 300 \| (1, 280) \| 293 \| Putative baseplate assembly protein J \| Putative baseplate assembly protein J; uniclust \| UniRef100\_A0A644Z6C5 \| 99.7 \| 4.8e-20 \| 8.9e-26 \| 141.6 \| 180 \| (111, 296) \| 300 \| (19, 201) \| 204 \| Baseplate protein J-like domain-containing protein \| Baseplate protein J-like domain-containing protein; uniclust \| UniRef100\_A0A011N873 \| 99.7 \| 5.6e-20 \| 1.1e-25 \| 170.9 \| 215 \| (42, 268) \| 300 \| (115, 484) \| 644 \| Baseplate protein J-like domain-containing protein \| Baseplate protein J-like domain-containing protein; uniclust \| UniRef100\_A0A238HJ41 \| 99.7 \| 6.1e-20 \| 1.1e-25 \| 142.6 \| 178 \| (116, 296) \| 300 \| (17, 202) \| 207 \| Baseplate J-like protein \| Baseplate J-like protein; uniclust \| UniRef100\_A0A094SKS8 \| 99.7 \| 6.3e-20 \| 1.2e-25 \| 164.0 \| 271 \| (16, 295) \| 300 \| (38, 433) \| 471 \| Phage baseplate protein (Fragment) \| Phage baseplate protein (Fragment); uniclust \| UniRef100\_A0A0M0SE30 \| 99.7 \| 6.6e-20 \| 1.3e-25 \| 160.9 \| 271 \| (13, 298) \| 300 \| (9, 376) \| 396 \| Baseplate protein J-like domain-containing protein \| Baseplate protein J-like domain-containing protein; uniclust \| UniRef100\_A0A840NW02 \| 99.7 \| 7e-20 \| 1.3e-25 \| 141.2 \| 204 \| (78, 297) \| 300 \| (2, 206) \| 208 \| Phage-related baseplate assembly protein \| Phage-related baseplate assembly protein; uniclust \| UniRef100\_A0A0N9N791 \| 99.7 \| 6.7e-20 \| 1.3e-25 \| 163.0 \| 273 \| (19, 294) \| 300 \| (28, 440) \| 446 \| Baseplate protein J-like domain-containing protein \| Baseplate protein J-like domain-containing protein; uniclust \| UniRef100\_A0A7J4ETU4 \| 99.7 \| 8.2e-20 \| 1.5e-25 \| 152.5 \| 220 \| (37, 270) \| 300 \| (5, 391) \| 433 \| Baseplate protein J-like domain-containing protein \| Baseplate protein J-like domain-containing protein; uniclust \| UniRef100\_A0A0Q7JQC0 \| 99.7 \| 7.7e-20 \| 1.5e-25 \| 174.1 \| 183 \| (112, 297) \| 300 \| (478, 694) \| 748 \| Baseplate protein J-like domain-containing protein \| Baseplate protein J-like domain-containing protein; uniclust \| UniRef100\_UPI001F2080EB \| 99.7 \| 8.4e-20 \| 1.5e-25 \| 141.5 \| 228 \| (66, 293) \| 300 \| (3, 235) \| 240 \| baseplate J/gp47 family protein \| baseplate J/gp47 family protein; uniclust \| UniRef100\_A0A2S6N2W1 \| 99.7 \| 1.1e-19 \| 2.1e-25 \| 150.6 \| 277 \| (4, 295) \| 300 \| (112, 403) \| 407 \| Baseplate protein J-like domain-containing protein \| Baseplate protein J-like domain-containing protein; uniclust \| UniRef100\_A0A1C5PVS5 \| 99.7 \| 1.1e-19 \| 2.1e-25 \| 173.8 \| 180 \| (112, 296) \| 300 \| (503, 708) \| 714 \| Baseplate protein J-like domain-containing protein \| Baseplate protein J-like domain-containing protein; uniclust \| UniRef100\_A0A3D8VNC0 \| 99.7 \| 1.2e-19 \| 2.2e-25 \| 144.8 \| 151 \| (11, 166) \| 300 \| (2, 234) \| 235 \| Baseplate J family protein (Fragment) \| Baseplate J family protein (Fragment); uniclust \| UniRef100\_A0A5C7QHU8 \| 99.7 \| 1.2e-19 \| 2.3e-25 \| 157.7 \| 269 \| (18, 294) \| 300 \| (11, 442) \| 452 \| Baseplate protein J-like domain-containing protein \| Baseplate protein J-like domain-containing protein; uniclust \| UniRef100\_A0A423XTX1 \| 99.7 \| 1.6e-19 \| 3e-25 \| 152.0 \| 240 \| (53, 299) \| 300 \| (39, 359) \| 467 \| Phage baseplate protein \| Phage baseplate protein; uniclust \| UniRef100\_A0A521CK67 \| 99.7 \| 1.6e-19 \| 3e-25 \| 151.3 \| 271 \| (21, 294) \| 300 \| (6, 446) \| 447 \| Baseplate J-like protein \| Baseplate J-like protein; uniclust \| UniRef100\_A0A951P3G3 \| 99.7 \| 1.6e-19 \| 3e-25 \| 148.5 \| 260 \| (19, 287) \| 300 \| (3, 363) \| 380 \| Baseplate J/gp47 family protein \| Baseplate J/gp47 family protein; uniclust \| UniRef100\_A0A6V7D981 \| 99.7 \| 1.7e-19 \| 3.4e-25 \| 142.1 \| 130 \| (2, 131) \| 300 \| (13, 142) \| 169 \| Baseplate assembly protein J \| Baseplate assembly protein J; uniclust \| UniRef100\_UPI001AFE5812 \| 99.7 \| 1.9e-19 \| 3.5e-25 \| 155.4 \| 269 \| (18, 294) \| 300 \| (6, 382) \| 593 \| baseplate J/gp47 family protein \| baseplate J/gp47 family protein; uniclust \| UniRef100\_A0A7J6YLT3 \| 99.6 \| 2.4e-19 \| 4.5e-25 \| 152.4 \| 192 \| (12, 222) \| 300 \| (293, 484) \| 508 \| Baseplate protein J-like domain-containing protein \| Baseplate protein J-like domain-containing protein; uniclust \| UniRef100\_A0A327JML8 \| 99.6 \| 3.1e-19 \| 5.8e-25 \| 145.7 \| 284 \| (6, 297) \| 300 \| (7, 298) \| 300 \| Baseplate protein J-like domain-containing protein \| Baseplate protein J-like domain-containing protein; uniclust \| UniRef100\_UPI0002DE2F33 \| 99.6 \| 3.2e-19 \| 5.9e-25 \| 150.6 \| 221 \| (19, 245) \| 300 \| (2, 293) \| 474 \| baseplate J/gp47 family protein \| baseplate J/gp47 family protein; uniclust \| UniRef100\_A0A076G4G9 \| 99.6 \| 3.1e-19 \| 6e-25 \| 156.8 \| 269 \| (18, 295) \| 300 \| (12, 382) \| 388 \| Baseplate protein J-like domain-containing protein \| Baseplate protein J-like domain-containing protein; uniclust \| UniRef100\_A0A0R3AT35 \| 99.6 \| 3.6e-19 \| 6.8e-25 \| 153.4 \| 287 \| (2, 295) \| 300 \| (1, 391) \| 394 \| Phage protein \| Phage protein; uniclust \| UniRef100\_UPI000BB9B8A1 \| 99.6 \| 3.9e-19 \| 7.3e-25 \| 153.9 \| 265 \| (16, 287) \| 300 \| (7, 482) \| 488 \| baseplate J/gp47 family protein \| baseplate J/gp47 family protein; uniclust \| UniRef100\_A0A8X6KGD0 \| 99.6 \| 4.1e-19 \| 7.5e-25 \| 153.0 \| 218 \| (43, 282) \| 300 \| (13, 232) \| 573 \| Baseplate protein J \| Baseplate protein J; uniclust \| UniRef100\_J9PV89 \| 99.6 \| 4.2e-19 \| 7.8e-25 \| 150.0 \| 272 \| (21, 296) \| 300 \| (13, 470) \| 476 \| Putative baseplate J family protein \| Putative baseplate J family protein; uniclust \| UniRef100\_A0A3S0DD06 \| 99.6 \| 4.4e-19 \| 8.2e-25 \| 156.7 \| 178 \| (112, 295) \| 300 \| (428, 616) \| 621 \| Baseplate protein J-like domain-containing protein \| Baseplate protein J-like domain-containing protein; uniclust \| UniRef100\_A0A4P1QFN8 \| 99.6 \| 4.6e-19 \| 8.6e-25 \| 142.7 \| 215 \| (3, 221) \| 300 \| (1, 222) \| 278 \| Baseplate J-like family protein \| Baseplate J-like family protein; uniclust \| UniRef100\_A0A081C216 \| 99.6 \| 4.8e-19 \| 9.3e-25 \| 161.2 \| 220 \| (42, 272) \| 300 \| (109, 415) \| 588 \| Baseplate protein J-like domain-containing protein \| Baseplate protein J-like domain-containing protein; uniclust \| UniRef100\_A0A2M8G072 \| 99.6 \| 5.1e-19 \| 9.3e-25 \| 150.0 \| 174 \| (113, 293) \| 300 \| (301, 489) \| 491 \| Baseplate protein J-like domain-containing protein \| Baseplate protein J-like domain-containing protein; uniclust \| UniRef100\_A0A8D9LH44 \| 99.6 \| 5.9e-19 \| 1.1e-24 \| 139.2 \| 177 \| (111, 294) \| 300 \| (84, 267) \| 270 \| Baseplate J-like protein \| Baseplate J-like protein; uniclust \| UniRef100\_F5SA56 \| 99.6 \| 6.2e-19 \| 1.2e-24 \| 142.8 \| 186 \| (108, 296) \| 300 \| (95, 288) \| 293 \| Phage baseplate protein (Fragment) \| Phage baseplate protein (Fragment); uniclust \| UniRef100\_UPI001BAF66FD \| 99.6 \| 6.9e-19 \| 1.3e-24 \| 141.3 \| 261 \| (16, 291) \| 300 \| (14, 292) \| 309 \| baseplate J/gp47 family protein \| baseplate J/gp47 family protein; uniclust \| UniRef100\_A0A7C2BDY4 \| 99.6 \| 7.2e-19 \| 1.3e-24 \| 143.7 \| 258 \| (21, 292) \| 300 \| (11, 351) \| 354 \| Baseplate protein J-like domain-containing protein \| Baseplate protein J-like domain-containing protein; uniclust \| UniRef100\_A0A1V5Z4C1 \| 99.6 \| 9.3e-19 \| 1.8e-24 \| 149.1 \| 157 \| (111, 270) \| 300 \| (106, 278) \| 322 \| Baseplate J-like protein \| Baseplate J-like protein; uniclust \| UniRef100\_A0A4Q3KFV4 \| 99.6 \| 1e-18 \| 1.9e-24 \| 143.3 \| 237 \| (45, 296) \| 300 \| (124, 363) \| 365 \| Baseplate assembly protein \| Baseplate assembly protein; uniclust \| UniRef100\_A0A4V2AQW5 \| 99.6 \| 1e-18 \| 2e-24 \| 152.7 \| 261 \| (21, 294) \| 300 \| (20, 387) \| 396 \| Baseplate protein J-like domain-containing protein \| Baseplate protein J-like domain-containing protein; uniclust \| UniRef100\_A0A0M0T5G9 \| 99.6 \| 1.3e-18 \| 2.5e-24 \| 132.4 \| 147 \| (141, 294) \| 300 \| (1, 155) \| 157 \| Tail protein (Fragment) \| Tail protein (Fragment); uniclust \| UniRef100\_A0A318KRB1 \| 99.6 \| 1.5e-18 \| 2.7e-24 \| 139.3 \| 284 \| (5, 292) \| 300 \| (1, 300) \| 305 \| Phage-related baseplate assembly protein \| Phage-related baseplate assembly protein; uniclust \| UniRef100\_A0A2A5BN42 \| 99.6 \| 1.5e-18 \| 2.7e-24 \| 139.8 \| 275 \| (7, 296) \| 300 \| (11, 311) \| 314 \| Baseplate protein J-like domain-containing protein \| Baseplate protein J-like domain-containing protein; uniclust \| UniRef100\_A0A060AG40 \| 99.6 \| 1.8e-18 \| 3.6e-24 \| 158.4 \| 276 \| (10, 295) \| 300 \| (21, 512) \| 516 \| Baseplate component \| Baseplate component; uniclust \| UniRef100\_A0A6J5N751 \| 99.6 \| 2.1e-18 \| 4e-24 \| 152.2 \| 269 \| (21, 293) \| 300 \| (12, 478) \| 488 \| Baseplate wedge subunit \| Baseplate wedge subunit; uniclust \| UniRef100\_A0A963HKG0 \| 99.6 \| 2.3e-18 \| 4.4e-24 \| 157.3 \| 156 \| (112, 269) \| 300 \| (304, 490) \| 699 \| Baseplate J/gp47 family protein (Fragment) \| Baseplate J/gp47 family protein (Fragment); uniclust \| UniRef100\_A0A0D8TUY8 \| 99.6 \| 2.8e-18 \| 5.3e-24 \| 139.3 \| 154 \| (19, 173) \| 300 \| (3, 244) \| 264 \| Phage baseplate J-like protein (Fragment) \| Phage baseplate J-like protein (Fragment); uniclust \| UniRef100\_A0A1A9WW65 \| 99.6 \| 3e-18 \| 5.6e-24 \| 132.9 \| 170 \| (126, 298) \| 300 \| (15, 189) \| 191 \| Baseplate\_J domain-containing protein \| Baseplate\_J domain-containing protein; uniclust \| UniRef100\_H1D5E3 \| 99.6 \| 3.5e-18 \| 6.6e-24 \| 141.7 \| 230 \| (65, 296) \| 300 \| (9, 342) \| 344 \| Baseplate protein J-like domain-containing protein \| Baseplate protein J-like domain-containing protein; uniclust \| UniRef100\_A0A016QLU4 \| 99.6 \| 3.6e-18 \| 7e-24 \| 146.6 \| 183 \| (111, 295) \| 300 \| (81, 300) \| 308 \| Baseplate protein J-like domain-containing protein \| Baseplate protein J-like domain-containing protein; uniclust \| UniRef100\_A0A149VI46 \| 99.6 \| 3.6e-18 \| 7.1e-24 \| 147.8 \| 169 \| (17, 192) \| 300 \| (26, 278) \| 326 \| Baseplate protein J-like domain-containing protein (Fragment) \| Baseplate protein J-like domain-containing protein (Fragment); uniclust \| UniRef100\_A0A0A0YUY5 \| 99.6 \| 3.9e-18 \| 7.6e-24 \| 150.1 \| 267 \| (21, 292) \| 300 \| (5, 391) \| 393 \| Baseplate protein J-like domain-containing protein \| Baseplate protein J-like domain-containing protein; uniclust \| UniRef100\_A0A2G2PNA8 \| 99.6 \| 5.4e-18 \| 1.1e-23 \| 153.8 \| 231 \| (18, 260) \| 300 \| (21, 360) \| 486 \| Baseplate protein J-like domain-containing protein \| Baseplate protein J-like domain-containing protein; uniclust \| UniRef100\_A0A069PT37 \| 99.6 \| 5.7e-18 \| 1.1e-23 \| 150.2 \| 232 \| (19, 268) \| 300 \| (18, 286) \| 390 \| Baseplate protein J-like domain-containing protein \| Baseplate protein J-like domain-containing protein; uniclust \| UniRef100\_A0A1C5KYZ3 \| 99.6 \| 6.1e-18 \| 1.2e-23 \| 156.7 \| 179 \| (112, 295) \| 300 \| (410, 618) \| 619 \| Uncharacterized homolog of phage Mu protein gp47 \| Uncharacterized homolog of phage Mu protein gp47; uniclust \| UniRef100\_UPI001C11CD17 \| 99.6 \| 7.4e-18 \| 1.4e-23 \| 131.7 \| 171 \| (19, 197) \| 300 \| (2, 247) \| 247 \| baseplate J/gp47 family protein \| baseplate J/gp47 family protein; uniclust \| UniRef100\_A0A9E5G387 \| 99.6 \| 8.3e-18 \| 1.6e-23 \| 136.0 \| 162 \| (23, 189) \| 300 \| (2, 245) \| 257 \| Uncharacterized protein \| Uncharacterized protein; uniclust \| UniRef100\_A0A075DXJ3 \| 99.6 \| 8.2e-18 \| 1.6e-23 \| 150.6 \| 268 \| (19, 295) \| 300 \| (10, 480) \| 482 \| Putative baseplate component \| Putative baseplate component; uniclust \| UniRef100\_A0A383RGV8 \| 99.6 \| 8.8e-18 \| 1.6e-23 \| 137.3 \| 179 \| (112, 292) \| 300 \| (82, 278) \| 283 \| Baseplate J family protein \| Baseplate J family protein; uniclust \| UniRef100\_UPI001EFEDC3E \| 99.6 \| 9.2e-18 \| 1.7e-23 \| 140.4 \| 231 \| (59, 296) \| 300 \| (2, 313) \| 424 \| baseplate J/gp47 family protein \| baseplate J/gp47 family protein; uniclust \| UniRef100\_A0A8T3QP22 \| 99.6 \| 9.5e-18 \| 1.8e-23 \| 146.9 \| 198 \| (19, 221) \| 300 \| (3, 285) \| 460 \| Baseplate J/gp47 family protein \| Baseplate J/gp47 family protein; uniclust \| UniRef100\_A0A929SH31 \| 99.6 \| 9.7e-18 \| 1.8e-23 \| 138.7 \| 168 \| (111, 287) \| 300 \| (102, 269) \| 311 \| Baseplate J/gp47 family protein \| Baseplate J/gp47 family protein; uniclust \| UniRef100\_A0A5S9MGE0 \| 99.5 \| 1.1e-17 \| 2.1e-23 \| 134.8 \| 180 \| (112, 293) \| 300 \| (84, 274) \| 278 \| Baseplate protein J-like domain-containing protein \| Baseplate protein J-like domain-containing protein; uniclust \| UniRef100\_A0A0A7NU29 \| 99.5 \| 1.5e-17 \| 2.9e-23 \| 147.4 \| 235 \| (21, 270) \| 300 \| (21, 341) \| 505 \| Baseplate J family protein \| Baseplate J family protein; uniclust \| UniRef100\_A0A069PV97 \| 99.5 \| 1.5e-17 \| 3e-23 \| 168.7 \| 180 \| (111, 293) \| 300 \| (987, 1198) \| 1219 \| Baseplate protein J-like domain-containing protein \| Baseplate protein J-like domain-containing protein; uniclust \| UniRef100\_A0A0S4I2J3 \| 99.5 \| 1.7e-17 \| 3.2e-23 \| 127.0 \| 175 \| (121, 296) \| 300 \| (1, 176) \| 178 \| Baseplate J-like protein \| Baseplate J-like protein; uniclust \| UniRef100\_A0A072TF05 \| 99.5 \| 1.8e-17 \| 3.4e-23 \| 156.2 \| 242 \| (12, 258) \| 300 \| (652, 992) \| 1809 \| Bacteriophage regulatory protein, putative (Fragment) \| Bacteriophage regulatory protein, putative (Fragment); uniclust \| UniRef100\_A0A0H7L4M1 \| 99.5 \| 1.8e-17 \| 3.4e-23 \| 127.9 \| 152 \| (138, 296) \| 300 \| (2, 158) \| 163 \| Tail protein \| Tail protein; uniclust \| UniRef100\_A0A1V5RRS7 \| 99.5 \| 1.9e-17 \| 3.5e-23 \| 149.9 \| 244 \| (21, 268) \| 300 \| (15, 442) \| 787 \| Baseplate protein J-like domain-containing protein \| Baseplate protein J-like domain-containing protein; uniclust \| UniRef100\_B8IDQ2 \| 99.5 \| 2.5e-17 \| 4.5e-23 \| 130.0 \| 231 \| (4, 237) \| 300 \| (6, 257) \| 263 \| Baseplate J family protein \| Baseplate J family protein; uniclust \| UniRef100\_A0A1L4D176 \| 99.5 \| 2.5e-17 \| 4.7e-23 \| 137.4 \| 269 \| (20, 295) \| 300 \| (4, 357) \| 366 \| Baseplate protein J-like domain-containing protein \| Baseplate protein J-like domain-containing protein; uniclust \| UniRef100\_A0A080M2Y7 \| 99.5 \| 2.6e-17 \| 5e-23 \| 153.2 \| 156 \| (111, 268) \| 300 \| (356, 536) \| 604 \| Baseplate protein J-like domain-containing protein \| Baseplate protein J-like domain-containing protein; uniclust \| UniRef100\_A0A0K9NAJ6 \| 99.5 \| 2.8e-17 \| 5.5e-23 \| 157.2 \| 182 \| (112, 297) \| 300 \| (497, 697) \| 703 \| Baseplate protein J-like domain-containing protein \| Baseplate protein J-like domain-containing protein; uniclust \| UniRef100\_A0A0T9RK09 \| 99.5 \| 3.3e-17 \| 6e-23 \| 140.9 \| 267 \| (21, 291) \| 300 \| (166, 537) \| 539 \| Baseplate J family protein \| Baseplate J family protein; uniclust \| UniRef100\_A0A812RFY7 \| 99.5 \| 4.3e-17 \| 7.9e-23 \| 145.0 \| 221 \| (44, 264) \| 300 \| (11, 259) \| 772 \| J protein \| J protein; uniclust \| UniRef100\_A0A1E3G789 \| 99.5 \| 4.7e-17 \| 8.7e-23 \| 130.9 \| 275 \| (4, 293) \| 300 \| (1, 296) \| 302 \| Baseplate protein J-like domain-containing protein \| Baseplate protein J-like domain-containing protein; uniclust \| UniRef100\_A0A430RXZ0 \| 99.5 \| 4.8e-17 \| 8.9e-23 \| 130.2 \| 173 \| (112, 293) \| 300 \| (98, 274) \| 276 \| Baseplate protein J-like domain-containing protein \| Baseplate protein J-like domain-containing protein; uniclust \| UniRef100\_A0A0F9WMS5 \| 99.5 \| 5e-17 \| 9.6e-23 \| 145.7 \| 284 \| (9, 294) \| 300 \| (5, 513) \| 521 \| Baseplate protein J-like domain-containing protein \| Baseplate protein J-like domain-containing protein; uniclust \| UniRef100\_UPI001EFE5B4D \| 99.5 \| 5.3e-17 \| 9.8e-23 \| 126.1 \| 159 \| (2, 160) \| 300 \| (1, 159) \| 203 \| hypothetical protein \| hypothetical protein; uniclust \| UniRef100\_A0A353T659 \| 99.5 \| 5.2e-17 \| 9.9e-23 \| 142.2 \| 273 \| (16, 295) \| 300 \| (10, 410) \| 413 \| Baseplate protein J-like domain-containing protein \| Baseplate protein J-like domain-containing protein; uniclust \| UniRef100\_A0A2G8MJ99 \| 99.5 \| 5.5e-17 \| 1e-22 \| 142.3 \| 185 \| (111, 297) \| 300 \| (178, 391) \| 394 \| Putative baseplate assembly protein (Fragment) \| Putative baseplate assembly protein (Fragment); uniclust \| UniRef100\_UPI00187D24B2 \| 99.5 \| 5.8e-17 \| 1.1e-22 \| 119.6 \| 127 \| (114, 244) \| 300 \| (2, 130) \| 147 \| baseplate J/gp47 family protein \| baseplate J/gp47 family protein; uniclust \| UniRef100\_UPI001591BDFE \| 99.5 \| 6.4e-17 \| 1.2e-22 \| 123.1 \| 179 \| (54, 232) \| 300 \| (2, 180) \| 202 \| baseplate J/gp47 family protein \| baseplate J/gp47 family protein; uniclust \| UniRef100\_A0A0R2FQN8 \| 99.5 \| 6.6e-17 \| 1.3e-22 \| 135.0 \| 177 \| (111, 295) \| 300 \| (78, 284) \| 288 \| Phage Mu gp47 related protein \| Phage Mu gp47 related protein; uniclust \| UniRef100\_A0A2K9P040 \| 99.5 \| 7e-17 \| 1.3e-22 \| 141.1 \| 242 \| (21, 268) \| 300 \| (5, 336) \| 574 \| Baseplate J family protein \| Baseplate J family protein; uniclust \| UniRef100\_A0A2N8K8B6 \| 99.5 \| 7e-17 \| 1.3e-22 \| 133.5 \| 154 \| (19, 173) \| 300 \| (3, 233) \| 259 \| Baseplate protein J-like domain-containing protein \| Baseplate protein J-like domain-containing protein; uniclust \| UniRef100\_UPI001177B6DF \| 99.5 \| 7.5e-17 \| 1.4e-22 \| 133.6 \| 182 \| (110, 293) \| 300 \| (34, 381) \| 381 \| baseplate J/gp47 family protein \| baseplate J/gp47 family protein; uniclust \| UniRef100\_A0A929JGZ9 \| 99.5 \| 7.6e-17 \| 1.4e-22 \| 136.2 \| 173 \| (111, 287) \| 300 \| (164, 350) \| 373 \| Baseplate J/gp47 family protein \| Baseplate J/gp47 family protein; uniclust \| UniRef100\_A0A1H8RZN7 \| 99.5 \| 8.4e-17 \| 1.6e-22 \| 141.2 \| 243 \| (16, 268) \| 300 \| (6, 381) \| 428 \| Uncharacterized phage protein gp47/JayE \| Uncharacterized phage protein gp47/JayE; uniclust \| UniRef100\_A0A1V6ET53 \| 99.5 \| 8.5e-17 \| 1.6e-22 \| 140.6 \| 232 \| (17, 260) \| 300 \| (8, 344) \| 395 \| Baseplate J-like protein \| Baseplate J-like protein; uniclust \| UniRef100\_A0A6J5M9M0 \| 99.5 \| 1e-16 \| 1.9e-22 \| 133.1 \| 242 \| (21, 270) \| 300 \| (6, 348) \| 389 \| XkdT Uncharacterized homolog of phage Mu protein gp47 \| XkdT Uncharacterized homolog of phage Mu protein gp47; uniclust \| UniRef100\_A0A2E9U4S9 \| 99.5 \| 1e-16 \| 1.9e-22 \| 131.1 \| 228 \| (49, 285) \| 300 \| (28, 339) \| 343 \| Baseplate protein J-like domain-containing protein \| Baseplate protein J-like domain-containing protein; uniclust \| UniRef100\_A0A1I0R9Q5 \| 99.5 \| 1e-16 \| 2e-22 \| 140.1 \| 264 \| (24, 289) \| 300 \| (9, 360) \| 399 \| Baseplate J-like protein \| Baseplate J-like protein; uniclust \| UniRef100\_A0A943UN36 \| 99.5 \| 1.1e-16 \| 2e-22 \| 141.7 \| 185 \| (112, 298) \| 300 \| (525, 713) \| 715 \| Baseplate J/gp47 family protein \| Baseplate J/gp47 family protein; uniclust \| UniRef100\_A0A090AGM4 \| 99.5 \| 1.1e-16 \| 2e-22 \| 140.1 \| 156 \| (112, 271) \| 300 \| (318, 475) \| 638 \| Baseplate protein J-like domain-containing protein \| Baseplate protein J-like domain-containing protein; uniclust \| UniRef100\_A0A136KPX1 \| 99.5 \| 1.1e-16 \| 2.2e-22 \| 147.3 \| 183 \| (111, 295) \| 300 \| (268, 485) \| 501 \| Baseplate J-like protein \| Baseplate J-like protein; uniclust \| UniRef100\_A0A511MW83 \| 99.5 \| 1.6e-16 \| 2.9e-22 \| 131.0 \| 260 \| (24, 293) \| 300 \| (13, 361) \| 363 \| Baseplate protein J-like domain-containing protein \| Baseplate protein J-like domain-containing protein; uniclust \| UniRef100\_A0A098MIJ1 \| 99.5 \| 1.5e-16 \| 2.9e-22 \| 139.6 \| 257 \| (19, 288) \| 300 \| (35, 409) \| 426 \| Phage baseplate protein \| Phage baseplate protein; uniclust \| UniRef100\_UPI00016A6C05 \| 99.5 \| 1.8e-16 \| 3.4e-22 \| 123.5 \| 215 \| (80, 298) \| 300 \| (1, 215) \| 217 \| baseplate J/gp47 family protein \| baseplate J/gp47 family protein; uniclust \| UniRef100\_A0A0B0HE24 \| 99.5 \| 2e-16 \| 3.8e-22 \| 121.2 \| 146 \| (151, 297) \| 300 \| (3, 159) \| 163 \| Baseplate J-like protein \| Baseplate J-like protein; uniclust \| UniRef100\_A0A1V0Q0D1 \| 99.5 \| 2.1e-16 \| 4e-22 \| 151.9 \| 178 \| (112, 292) \| 300 \| (574, 788) \| 943 \| Baseplate protein J-like domain-containing protein \| Baseplate protein J-like domain-containing protein; uniclust \| UniRef100\_A0A094RWT6 \| 99.5 \| 2.1e-16 \| 4.1e-22 \| 135.0 \| 209 \| (3, 216) \| 300 \| (2, 302) \| 303 \| Phage protein (Fragment) \| Phage protein (Fragment); uniclust \| UniRef100\_UPI0009B77170 \| 99.5 \| 2.2e-16 \| 4.1e-22 \| 134.6 \| 170 \| (19, 192) \| 300 \| (2, 245) \| 487 \| baseplate J/gp47 family protein \| baseplate J/gp47 family protein; uniclust \| UniRef100\_A0A5Y3ZZF9 \| 99.5 \| 2.3e-16 \| 4.2e-22 \| 138.3 \| 234 \| (55, 295) \| 300 \| (321, 634) \| 639 \| Uncharacterized protein \| Uncharacterized protein; uniclust \| UniRef100\_A0A1S9JEE3 \| 99.5 \| 2.3e-16 \| 4.3e-22 \| 125.9 \| 179 \| (111, 296) \| 300 \| (54, 237) \| 242 \| Phage tail protein \| Phage tail protein; uniclust \| UniRef100\_A0A0Q7YYN4 \| 99.5 \| 2.2e-16 \| 4.3e-22 \| 138.1 \| 174 \| (111, 288) \| 300 \| (147, 346) \| 388 \| Baseplate protein J-like domain-containing protein \| Baseplate protein J-like domain-containing protein; uniclust \| UniRef100\_A0A0K4XJP0 \| 99.5 \| 2.4e-16 \| 4.4e-22 \| 130.9 \| 234 \| (53, 293) \| 300 \| (39, 352) \| 352 \| Putative phage baseplate protein \| Putative phage baseplate protein; uniclust \| UniRef100\_A0A1G0RA06 \| 99.4 \| 2.5e-16 \| 4.6e-22 \| 130.7 \| 269 \| (22, 294) \| 300 \| (3, 366) \| 383 \| Baseplate protein J-like domain-containing protein \| Baseplate protein J-like domain-containing protein; uniclust \| UniRef100\_A0A0S2ZD70 \| 99.4 \| 2.7e-16 \| 5.2e-22 \| 135.7 \| 266 \| (19, 294) \| 300 \| (9, 377) \| 383 \| Baseplate protein J-like domain-containing protein \| Baseplate protein J-like domain-containing protein; uniclust \| UniRef100\_UPI001E545E66 \| 99.4 \| 3.3e-16 \| 6.1e-22 \| 135.4 \| 234 \| (20, 260) \| 300 \| (4, 314) \| 556 \| baseplate J/gp47 family protein \| baseplate J/gp47 family protein; uniclust \| UniRef100\_A0A0F9SDC1 \| 99.4 \| 3.4e-16 \| 6.4e-22 \| 136.1 \| 271 \| (21, 295) \| 300 \| (4, 473) \| 475 \| Baseplate protein J-like domain-containing protein \| Baseplate protein J-like domain-containing protein; uniclust \| UniRef100\_A0A165ZZT9 \| 99.4 \| 3.5e-16 \| 6.6e-22 \| 135.5 \| 264 \| (20, 293) \| 300 \| (11, 386) \| 388 \| Baseplate J-like protein \| Baseplate J-like protein; uniclust \| UniRef100\_UPI001C0FE55C \| 99.4 \| 3.8e-16 \| 7e-22 \| 131.6 \| 198 \| (23, 243) \| 300 \| (2, 202) \| 435 \| baseplate J/gp47 family protein \| baseplate J/gp47 family protein; uniclust \| UniRef100\_A0A067XR41 \| 99.4 \| 4.6e-16 \| 9.2e-22 \| 152.5 \| 244 \| (20, 267) \| 300 \| (13, 461) \| 794 \| Baseplate wedge \| Baseplate wedge; uniclust \| UniRef100\_A0A0C2HQB0 \| 99.4 \| 5.2e-16 \| 1e-21 \| 141.0 \| 215 \| (42, 270) \| 300 \| (110, 419) \| 533 \| Baseplate protein J-like domain-containing protein \| Baseplate protein J-like domain-containing protein; uniclust \| UniRef100\_A0A356K030 \| 99.4 \| 5.3e-16 \| 1e-21 \| 147.4 \| 187 \| (111, 299) \| 300 \| (624, 837) \| 840 \| Putative baseplate assembly protein (Fragment) \| Putative baseplate assembly protein (Fragment); uniclust \| UniRef100\_A0A109WD85 \| 99.4 \| 5.8e-16 \| 1.1e-21 \| 134.4 \| 246 \| (16, 270) \| 300 \| (19, 367) \| 405 \| Baseplate protein J-like domain-containing protein \| Baseplate protein J-like domain-containing protein; uniclust \| UniRef100\_UPI00161E54F1 \| 99.4 \| 6.1e-16 \| 1.1e-21 \| 112.2 \| 132 \| (114, 252) \| 300 \| (3, 137) \| 140 \| baseplate J/gp47 family protein \| baseplate J/gp47 family protein; uniclust \| UniRef100\_A0A0E3L2V3 \| 99.4 \| 6.6e-16 \| 1.3e-21 \| 152.2 \| 183 \| (111, 295) \| 300 \| (761, 971) \| 979 \| Baseplate protein J-like domain-containing protein \| Baseplate protein J-like domain-containing protein; uniclust \| UniRef100\_A0A5C8CQS2 \| 99.4 \| 8.1e-16 \| 1.5e-21 \| 130.5 \| 261 \| (21, 292) \| 300 \| (5, 350) \| 351 \| Acetyltransferase \| Acetyltransferase; uniclust \| UniRef100\_X1GT58 \| 99.4 \| 9e-16 \| 1.7e-21 \| 113.8 \| 151 \| (139, 293) \| 300 \| (1, 157) \| 168 \| Baseplate protein J-like domain-containing protein (Fragment) \| Baseplate protein J-like domain-containing protein (Fragment); uniclust \| UniRef100\_A0A830FUX9 \| 99.4 \| 1e-15 \| 2e-21 \| 130.7 \| 259 \| (20, 294) \| 300 \| (9, 389) \| 390 \| Putative phage protein gp47/JayE \| Putative phage protein gp47/JayE; uniclust \| UniRef100\_A0A285J5J6 \| 99.4 \| 1.1e-15 \| 2e-21 \| 115.6 \| 160 \| (136, 296) \| 300 \| (1, 161) \| 163 \| Phage-related baseplate assembly protein (Fragment) \| Phage-related baseplate assembly protein (Fragment); uniclust \| UniRef100\_UPI00037164A3 \| 99.4 \| 1.2e-15 \| 2.2e-21 \| 122.7 \| 190 \| (23, 217) \| 300 \| (6, 286) \| 293 \| baseplate J/gp47 family protein \| baseplate J/gp47 family protein; uniclust \| UniRef100\_A0A842IZ09 \| 99.4 \| 1.3e-15 \| 2.3e-21 \| 123.8 \| 230 \| (64, 297) \| 300 \| (5, 317) \| 318 \| Baseplate J/gp47 family protein \| Baseplate J/gp47 family protein; uniclust \| UniRef100\_A0A3D4RY72 \| 99.4 \| 1.2e-15 \| 2.4e-21 \| 141.2 \| 181 \| (112, 295) \| 300 \| (494, 706) \| 741 \| Baseplate protein J-like domain-containing protein \| Baseplate protein J-like domain-containing protein; uniclust \| UniRef100\_A0A086ML20 \| 99.4 \| 1.4e-15 \| 2.7e-21 \| 146.7 \| 174 \| (111, 288) \| 300 \| (636, 835) \| 854 \| Baseplate protein J-like domain-containing protein \| Baseplate protein J-like domain-containing protein; uniclust \| UniRef100\_A0A098N215 \| 99.4 \| 1.4e-15 \| 2.7e-21 \| 132.7 \| 263 \| (22, 293) \| 300 \| (5, 371) \| 379 \| Phage baseplate protein \| Phage baseplate protein; uniclust \| UniRef100\_A0A0C2QJG0 \| 99.4 \| 1.5e-15 \| 2.8e-21 \| 134.1 \| 259 \| (21, 295) \| 300 \| (9, 362) \| 385 \| Baseplate protein J-like domain-containing protein \| Baseplate protein J-like domain-containing protein; uniclust \| UniRef100\_A0A7X6FRN3 \| 99.4 \| 1.6e-15 \| 3e-21 \| 116.3 \| 153 \| (5, 172) \| 300 \| (9, 161) \| 208 \| Baseplate protein J-like domain-containing protein \| Baseplate protein J-like domain-containing protein; uniclust \| UniRef100\_A0A257P1Y5 \| 99.4 \| 1.6e-15 \| 3e-21 \| 126.4 \| 179 \| (111, 295) \| 300 \| (81, 270) \| 272 \| Baseplate protein J-like domain-containing protein \| Baseplate protein J-like domain-containing protein; uniclust \| UniRef100\_UPI0015897FEC \| 99.4 \| 1.6e-15 \| 3e-21 \| 125.3 \| 261 \| (26, 294) \| 300 \| (14, 361) \| 363 \| baseplate J/gp47 family protein \| baseplate J/gp47 family protein; uniclust \| UniRef100\_A0A077NCN1 \| 99.4 \| 1.6e-15 \| 3e-21 \| 130.5 \| 274 \| (10, 293) \| 300 \| (10, 394) \| 405 \| Bacteriophage protein \| Bacteriophage protein; uniclust \| UniRef100\_A0A014N6D0 \| 99.4 \| 1.6e-15 \| 3.3e-21 \| 152.3 \| 172 \| (112, 287) \| 300 \| (761, 959) \| 1002 \| Baseplate protein J-like domain-containing protein \| Baseplate protein J-like domain-containing protein; uniclust \| UniRef100\_UPI000A1870BF \| 99.4 \| 1.8e-15 \| 3.5e-21 \| 113.0 \| 147 \| (115, 264) \| 300 \| (4, 150) \| 150 \| baseplate J/gp47 family protein \| baseplate J/gp47 family protein; uniclust \| UniRef100\_A0A242CHC8 \| 99.4 \| 2e-15 \| 3.6e-21 \| 120.2 \| 250 \| (21, 292) \| 300 \| (4, 268) \| 269 \| Baseplate protein J-like domain-containing protein \| Baseplate protein J-like domain-containing protein; uniclust \| UniRef100\_A0A0S6WQZ6 \| 99.4 \| 1.9e-15 \| 3.7e-21 \| 139.2 \| 175 \| (111, 289) \| 300 \| (309, 509) \| 528 \| Baseplate protein J-like domain-containing protein \| Baseplate protein J-like domain-containing protein; uniclust \| UniRef100\_A0A6M0A0U9 \| 99.4 \| 2.1e-15 \| 3.9e-21 \| 130.7 \| 219 \| (21, 270) \| 300 \| (10, 386) \| 449 \| Baseplate protein J-like domain-containing protein \| Baseplate protein J-like domain-containing protein; uniclust \| UniRef100\_A0A3N5LBL1 \| 99.4 \| 2.2e-15 \| 4e-21 \| 128.8 \| 245 \| (23, 271) \| 300 \| (5, 348) \| 487 \| Baseplate protein J-like domain-containing protein \| Baseplate protein J-like domain-containing protein; uniclust \| UniRef100\_A0A2W2DLG7 \| 99.4 \| 2.5e-15 \| 4.8e-21 \| 138.3 \| 155 \| (112, 270) \| 300 \| (421, 589) \| 636 \| Baseplate assembly protein \| Baseplate assembly protein; uniclust \| UniRef100\_UPI001FAC64A8 \| 99.4 \| 2.7e-15 \| 4.9e-21 \| 123.6 \| 223 \| (70, 294) \| 300 \| (2, 348) \| 350 \| baseplate J/gp47 family protein \| baseplate J/gp47 family protein; uniclust \| UniRef100\_A0A2E2FJ34 \| 99.4 \| 2.9e-15 \| 5.5e-21 \| 134.6 \| 248 \| (20, 271) \| 300 \| (12, 439) \| 555 \| Baseplate protein J-like domain-containing protein \| Baseplate protein J-like domain-containing protein; uniclust \| UniRef100\_A0A3M4ZB42 \| 99.4 \| 3.1e-15 \| 5.8e-21 \| 108.9 \| 128 \| (117, 244) \| 300 \| (1, 128) \| 131 \| Baseplate assembly protein J (Fragment) \| Baseplate assembly protein J (Fragment); uniclust \| UniRef100\_A0A0U1DT04 \| 99.3 \| 3.3e-15 \| 6.3e-21 \| 131.8 \| 194 \| (42, 244) \| 300 \| (129, 406) \| 420 \| Baseplate J-like protein \| Baseplate J-like protein; uniclust \| UniRef100\_K8GRQ2 \| 99.3 \| 3.5e-15 \| 6.5e-21 \| 123.3 \| 267 \| (20, 294) \| 300 \| (3, 358) \| 361 \| Putative phage Mu protein gp47-like protein \| Putative phage Mu protein gp47-like protein; uniclust \| UniRef100\_A0A524PY89 \| 99.3 \| 3.7e-15 \| 7.1e-21 \| 136.2 \| 187 \| (111, 299) \| 300 \| (382, 592) \| 594 \| Putative baseplate assembly protein (Fragment) \| Putative baseplate assembly protein (Fragment); uniclust \| UniRef100\_A0A5C7PKD9 \| 99.3 \| 4.6e-15 \| 8.5e-21 \| 124.0 \| 261 \| (23, 287) \| 300 \| (16, 387) \| 396 \| Baseplate protein J-like domain-containing protein \| Baseplate protein J-like domain-containing protein; uniclust \| UniRef100\_A0A1M6CQV3 \| 99.3 \| 5.5e-15 \| 1e-20 \| 104.9 \| 107 \| (134, 244) \| 300 \| (3, 111) \| 113 \| Baseplate J-like protein \| Baseplate J-like protein; uniclust \| UniRef100\_A0A7W6DEC2 \| 99.3 \| 6.2e-15 \| 1.1e-20 \| 126.0 \| 219 \| (21, 245) \| 300 \| (5, 305) \| 482 \| Putative phage protein gp47/JayE \| Putative phage protein gp47/JayE; uniclust \| UniRef100\_I8RM93 \| 99.3 \| 6.6e-15 \| 1.2e-20 \| 127.1 \| 262 \| (24, 292) \| 300 \| (15, 526) \| 526 \| Baseplate J family protein \| Baseplate J family protein; uniclust \| UniRef100\_A0A8S5ST71 \| 99.3 \| 6.6e-15 \| 1.2e-20 \| 126.8 \| 267 \| (19, 293) \| 300 \| (12, 395) \| 398 \| Baseplate wedge protein \| Baseplate wedge protein; uniclust \| UniRef100\_A0A1D9FVB1 \| 99.3 \| 7e-15 \| 1.4e-20 \| 135.3 \| 174 \| (112, 289) \| 300 \| (320, 519) \| 537 \| Baseplate assembly protein \| Baseplate assembly protein; uniclust \| UniRef100\_A0A150SA95 \| 99.3 \| 7.6e-15 \| 1.5e-20 \| 132.9 \| 199 \| (20, 220) \| 300 \| (10, 291) \| 498 \| Baseplate protein J-like domain-containing protein \| Baseplate protein J-like domain-containing protein; uniclust \| UniRef100\_A0A0Q6V809 \| 99.3 \| 7.6e-15 \| 1.5e-20 \| 147.3 \| 184 \| (111, 296) \| 300 \| (891, 1101) \| 1139 \| Baseplate protein J-like domain-containing protein \| Baseplate protein J-like domain-containing protein; uniclust \| UniRef100\_A0A843G9W3 \| 99.3 \| 8.4e-15 \| 1.6e-20 \| 126.8 \| 178 \| (112, 295) \| 300 \| (263, 455) \| 457 \| Baseplate J/gp47 family protein \| Baseplate J/gp47 family protein; uniclust \| UniRef100\_A0A1G6LTE0 \| 99.3 \| 8.5e-15 \| 1.7e-20 \| 130.2 \| 253 \| (22, 288) \| 300 \| (10, 369) \| 380 \| Baseplate J-like protein \| Baseplate J-like protein; uniclust \| UniRef100\_A0A8S5UKQ2 \| 99.3 \| 9.2e-15 \| 1.7e-20 \| 124.8 \| 280 \| (12, 294) \| 300 \| (3, 473) \| 474 \| Baseplate wedge protein \| Baseplate wedge protein; uniclust \| UniRef100\_A0A261E6U1 \| 99.3 \| 1.1e-14 \| 2.1e-20 \| 117.5 \| 162 \| (108, 270) \| 300 \| (27, 192) \| 237 \| Baseplate protein J-like domain-containing protein \| Baseplate protein J-like domain-containing protein; uniclust \| UniRef100\_A0A2E5KBW0 \| 99.3 \| 1.2e-14 \| 2.1e-20 \| 124.9 \| 267 \| (17, 293) \| 300 \| (7, 492) \| 496 \| Baseplate protein J-like domain-containing protein \| Baseplate protein J-like domain-containing protein; uniclust \| UniRef100\_A0A7X8LQY5 \| 99.3 \| 1.3e-14 \| 2.4e-20 \| 104.8 \| 132 \| (21, 170) \| 300 \| (3, 139) \| 140 \| Baseplate protein J-like domain-containing protein (Fragment) \| Baseplate protein J-like domain-containing protein (Fragment); uniclust \| UniRef100\_A0A3B8HH03 \| 99.3 \| 1.4e-14 \| 2.6e-20 \| 115.1 \| 164 \| (129, 295) \| 300 \| (9, 202) \| 209 \| Baseplate protein J-like domain-containing protein (Fragment) \| Baseplate protein J-like domain-containing protein (Fragment); uniclust \| UniRef100\_A0A5T0ECW8 \| 99.3 \| 1.6e-14 \| 2.9e-20 \| 119.3 \| 218 \| (42, 270) \| 300 \| (49, 344) \| 353 \| Baseplate assembly protein (Fragment) \| Baseplate assembly protein (Fragment); uniclust \| UniRef100\_A0A522CNZ0 \| 99.3 \| 1.6e-14 \| 3e-20 \| 120.3 \| 247 \| (42, 295) \| 300 \| (34, 374) \| 379 \| Baseplate protein J-like domain-containing protein \| Baseplate protein J-like domain-containing protein; uniclust \| UniRef100\_A0A284VRY0 \| 99.3 \| 1.8e-14 \| 3.4e-20 \| 140.3 \| 185 \| (111, 295) \| 300 \| (793, 1004) \| 1005 \| Baseplate protein J-like domain-containing protein \| Baseplate protein J-like domain-containing protein; uniclust \| UniRef100\_A0A2V8S3H9 \| 99.3 \| 1.7e-14 \| 3.4e-20 \| 138.4 \| 181 \| (112, 295) \| 300 \| (535, 774) \| 777 \| Putative baseplate assembly protein \| Putative baseplate assembly protein; uniclust \| UniRef100\_A0A098LJS2 \| 99.3 \| 1.7e-14 \| 3.4e-20 \| 146.3 \| 229 \| (5, 269) \| 300 \| (14, 312) \| 1032 \| Uncharacterized protein \| Uncharacterized protein; uniclust \| UniRef100\_UPI00038D138D \| 99.3 \| 1.9e-14 \| 3.5e-20 \| 114.5 \| 158 \| (21, 184) \| 300 \| (4, 238) \| 243 \| baseplate J/gp47 family protein \| baseplate J/gp47 family protein; uniclust \| UniRef100\_A0A350Y2A6 \| 99.3 \| 2e-14 \| 3.6e-20 \| 123.2 \| 146 \| (19, 169) \| 300 \| (3, 238) \| 482 \| Baseplate protein J-like domain-containing protein \| Baseplate protein J-like domain-containing protein; uniclust \| UniRef100\_UPI000F5FDBD6 \| 99.3 \| 2e-14 \| 3.6e-20 \| 124.7 \| 176 \| (112, 293) \| 300 \| (337, 540) \| 541 \| baseplate J/gp47 family protein \| baseplate J/gp47 family protein; uniclust \| UniRef100\_A0A3L7X434 \| 99.3 \| 2.1e-14 \| 3.9e-20 \| 124.0 \| 156 \| (111, 268) \| 300 \| (124, 315) \| 403 \| Baseplate protein J-like domain-containing protein (Fragment) \| Baseplate protein J-like domain-containing protein (Fragment); uniclust \| UniRef100\_A0A136JVI8 \| 99.3 \| 2.2e-14 \| 4.1e-20 \| 136.4 \| 175 \| (111, 289) \| 300 \| (628, 828) \| 846 \| Baseplate protein J-like domain-containing protein \| Baseplate protein J-like domain-containing protein; uniclust \| UniRef100\_A0A1F8MNF7 \| 99.3 \| 2.2e-14 \| 4.2e-20 \| 136.9 \| 251 \| (1, 287) \| 300 \| (1, 328) \| 730 \| Uncharacterized protein \| Uncharacterized protein; uniclust \| UniRef100\_A0A5U6MH76 \| 99.3 \| 2.3e-14 \| 4.2e-20 \| 106.6 \| 140 \| (111, 256) \| 300 \| (22, 164) \| 167 \| Phage tail protein (Fragment) \| Phage tail protein (Fragment); uniclust \| UniRef100\_A0A1Q6LBP8 \| 99.3 \| 2.3e-14 \| 4.2e-20 \| 119.5 \| 236 \| (22, 265) \| 300 \| (3, 321) \| 356 \| Baseplate protein J-like domain-containing protein \| Baseplate protein J-like domain-containing protein; uniclust \| UniRef100\_A0A842LGU8 \| 99.3 \| 2.5e-14 \| 4.5e-20 \| 119.5 \| 235 \| (52, 295) \| 300 \| (45, 381) \| 383 \| Baseplate J/gp47 family protein \| Baseplate J/gp47 family protein; uniclust \| UniRef100\_A0A6V7F7S0 \| 99.3 \| 2.5e-14 \| 4.6e-20 \| 110.4 \| 155 \| (2, 158) \| 300 \| (1, 155) \| 212 \| Phage-related baseplate assembly protein \| Phage-related baseplate assembly protein; uniclust \| UniRef100\_UPI0013748030 \| 99.3 \| 2.8e-14 \| 5.2e-20 \| 110.1 \| 116 \| (111, 230) \| 300 \| (82, 199) \| 203 \| baseplate J/gp47 family protein \| baseplate J/gp47 family protein; uniclust \| UniRef100\_A0A4Q0YJP7 \| 99.2 \| 3.1e-14 \| 5.7e-20 \| 112.2 \| 198 \| (6, 203) \| 300 \| (5, 239) \| 245 \| Baseplate assembly protein \| Baseplate assembly protein; uniclust \| UniRef100\_A0A1H4QY22 \| 99.2 \| 3.2e-14 \| 5.9e-20 \| 108.2 \| 145 \| (130, 280) \| 300 \| (2, 149) \| 169 \| Baseplate J-like protein \| Baseplate J-like protein; uniclust \| UniRef100\_A0A1Y1QPN6 \| 99.2 \| 3.5e-14 \| 6.4e-20 \| 116.9 \| 259 \| (21, 290) \| 300 \| (8, 338) \| 341 \| Baseplate protein J-like domain-containing protein \| Baseplate protein J-like domain-containing protein; uniclust \| UniRef100\_A0A058ZQ54 \| 99.2 \| 3.4e-14 \| 6.7e-20 \| 141.3 \| 174 \| (111, 288) \| 300 \| (838, 1037) \| 1176 \| Putative phage tail region protein \| Putative phage tail region protein; uniclust \| UniRef100\_A0A3S0BRQ0 \| 99.2 \| 3.7e-14 \| 6.9e-20 \| 123.2 \| 258 \| (22, 287) \| 300 \| (13, 468) \| 478 \| Baseplate protein J-like domain-containing protein \| Baseplate protein J-like domain-containing protein; uniclust \| UniRef100\_A0A2W4YN20 \| 99.2 \| 4.5e-14 \| 8.5e-20 \| 128.5 \| 157 \| (111, 269) \| 300 \| (322, 516) \| 568 \| Putative baseplate assembly protein \| Putative baseplate assembly protein; uniclust \| UniRef100\_A0A3P1VZS5 \| 99.2 \| 4.7e-14 \| 9.1e-20 \| 124.6 \| 226 \| (19, 256) \| 300 \| (4, 326) \| 374 \| Baseplate protein J-like domain-containing protein \| Baseplate protein J-like domain-containing protein; uniclust \| UniRef100\_A0A0E2HC00 \| 99.2 \| 5.1e-14 \| 9.6e-20 \| 113.5 \| 219 \| (22, 295) \| 300 \| (5, 236) \| 237 \| Baseplate protein J-like domain-containing protein \| Baseplate protein J-like domain-containing protein; uniclust \| UniRef100\_A0A1Q7YH65 \| 99.2 \| 5.4e-14 \| 1e-19 \| 120.9 \| 242 \| (40, 294) \| 300 \| (60, 400) \| 402 \| Baseplate protein J-like domain-containing protein \| Baseplate protein J-like domain-containing protein; uniclust \| UniRef100\_A0A074LTY1 \| 99.2 \| 5.2e-14 \| 1e-19 \| 140.1 \| 156 \| (111, 268) \| 300 \| (828, 1005) \| 1071 \| Baseplate protein J-like domain-containing protein \| Baseplate protein J-like domain-containing protein; uniclust \| UniRef100\_A0A398CJB0 \| 99.2 \| 5.9e-14 \| 1.1e-19 \| 119.5 \| 237 \| (22, 269) \| 300 \| (8, 363) \| 451 \| Baseplate protein J-like domain-containing protein \| Baseplate protein J-like domain-containing protein; uniclust \| UniRef100\_UPI0017498F18 \| 99.2 \| 6e-14 \| 1.1e-19 \| 108.9 \| 105 \| (112, 220) \| 300 \| (78, 184) \| 186 \| baseplate J/gp47 family protein \| baseplate J/gp47 family protein; uniclust \| UniRef100\_A0A080LYP8 \| 99.2 \| 6e-14 \| 1.2e-19 \| 126.1 \| 267 \| (21, 290) \| 300 \| (40, 379) \| 398 \| Uncharacterized protein \| Uncharacterized protein; uniclust \| UniRef100\_A0A1V5QEC7 \| 99.2 \| 6.4e-14 \| 1.2e-19 \| 108.3 \| 181 \| (111, 295) \| 300 \| (20, 208) \| 213 \| Baseplate J-like protein \| Baseplate J-like protein; uniclust \| UniRef100\_A0A150RG01 \| 99.2 \| 6.4e-14 \| 1.2e-19 \| 131.2 \| 155 \| (112, 269) \| 300 \| (499, 690) \| 756 \| Baseplate protein J-like domain-containing protein \| Baseplate protein J-like domain-containing protein; uniclust \| UniRef100\_A0A336NF78 \| 99.2 \| 6.5e-14 \| 1.2e-19 \| 100.3 \| 111 \| (2, 127) \| 300 \| (1, 116) \| 118 \| Phage related protein \| Phage related protein; uniclust \| UniRef100\_A0A2E6X3G2 \| 99.2 \| 6.9e-14 \| 1.3e-19 \| 121.0 \| 273 \| (20, 300) \| 300 \| (6, 416) \| 416 \| Baseplate protein J-like domain-containing protein \| Baseplate protein J-like domain-containing protein; uniclust \| UniRef100\_A0A6M0L2F8 \| 99.2 \| 7.2e-14 \| 1.4e-19 \| 126.9 \| 152 \| (112, 267) \| 300 \| (409, 577) \| 625 \| Baseplate protein J-like domain-containing protein \| Baseplate protein J-like domain-containing protein; uniclust \| UniRef100\_A0A7C1ESV1 \| 99.2 \| 7.6e-14 \| 1.4e-19 \| 96.9 \| 106 \| (22, 143) \| 300 \| (5, 110) \| 110 \| Baseplate protein J-like domain-containing protein (Fragment) \| Baseplate protein J-like domain-containing protein (Fragment); uniclust \| UniRef100\_A0A1B8SLD9 \| 99.2 \| 7.8e-14 \| 1.5e-19 \| 127.6 \| 269 \| (21, 293) \| 300 \| (3, 576) \| 588 \| Baseplate protein J-like domain-containing protein \| Baseplate protein J-like domain-containing protein; uniclust \| UniRef100\_A0A928YGD7 \| 99.2 \| 8.4e-14 \| 1.5e-19 \| 118.6 \| 244 \| (21, 270) \| 300 \| (13, 356) \| 451 \| Baseplate protein J-like domain-containing protein \| Baseplate protein J-like domain-containing protein; uniclust \| UniRef100\_A0A0U1DD70 \| 99.2 \| 8.2e-14 \| 1.6e-19 \| 105.8 \| 133 \| (165, 298) \| 300 \| (7, 139) \| 140 \| Baseplate assembly protein GpJ \| Baseplate assembly protein GpJ; uniclust \| UniRef100\_A0A3A8I9C4 \| 99.2 \| 8.1e-14 \| 1.6e-19 \| 123.9 \| 245 \| (21, 267) \| 300 \| (12, 332) \| 405 \| Baseplate protein J-like domain-containing protein \| Baseplate protein J-like domain-containing protein; uniclust \| UniRef100\_A0A2A5EI74 \| 99.2 \| 8.6e-14 \| 1.6e-19 \| 121.2 \| 150 \| (21, 170) \| 300 \| (9, 237) \| 552 \| Baseplate protein J-like domain-containing protein \| Baseplate protein J-like domain-containing protein; uniclust \| UniRef100\_A0A2X3LXX5 \| 99.2 \| 8.6e-14 \| 1.6e-19 \| 103.3 \| 105 \| (194, 298) \| 300 \| (14, 118) \| 123 \| Baseplate assembly protein J \| Baseplate assembly protein J; uniclust \| UniRef100\_A0A9D0GYQ9 \| 99.2 \| 9.1e-14 \| 1.7e-19 \| 122.4 \| 180 \| (112, 294) \| 300 \| (404, 606) \| 617 \| Uncharacterized protein \| Uncharacterized protein; uniclust \| UniRef100\_A0A521U4G1 \| 99.2 \| 9.5e-14 \| 1.8e-19 \| 119.4 \| 241 \| (21, 266) \| 300 \| (12, 356) \| 396 \| Baseplate protein J-like domain-containing protein \| Baseplate protein J-like domain-containing protein; uniclust \| UniRef100\_UPI000374DC89 \| 99.2 \| 9.7e-14 \| 1.8e-19 \| 120.6 \| 236 \| (20, 265) \| 300 \| (10, 339) \| 540 \| baseplate J/gp47 family protein \| baseplate J/gp47 family protein; uniclust \| UniRef100\_A0A645G9Q1 \| 99.2 \| 1e-13 \| 1.9e-19 \| 106.8 \| 184 \| (111, 296) \| 300 \| (15, 202) \| 206 \| Baseplate protein J-like domain-containing protein \| Baseplate protein J-like domain-containing protein; uniclust \| UniRef100\_A0A953PSF3 \| 99.2 \| 1.1e-13 \| 2e-19 \| 115.3 \| 224 \| (20, 269) \| 300 \| (9, 335) \| 369 \| Baseplate J/gp47 family protein \| Baseplate J/gp47 family protein; uniclust \| UniRef100\_A0A2X3CRG2 \| 99.2 \| 1.1e-13 \| 2.1e-19 \| 117.9 \| 183 \| (2, 191) \| 300 \| (2, 267) \| 305 \| Putative bacteriophage protein \| Putative bacteriophage protein; uniclust \| UniRef100\_A0A7Z2GCV1 \| 99.2 \| 1.1e-13 \| 2.1e-19 \| 135.2 \| 174 \| (111, 288) \| 300 \| (880, 1079) \| 1098 \| Putative baseplate assembly protein \| Putative baseplate assembly protein; uniclust \| UniRef100\_A0A398CKB0 \| 99.2 \| 1.3e-13 \| 2.4e-19 \| 122.9 \| 182 \| (112, 298) \| 300 \| (299, 516) \| 518 \| Putative baseplate assembly protein \| Putative baseplate assembly protein; uniclust \| UniRef100\_A0A967LZV4 \| 99.2 \| 1.4e-13 \| 2.6e-19 \| 120.7 \| 252 \| (18, 277) \| 300 \| (9, 463) \| 587 \| Baseplate protein J-like domain-containing protein \| Baseplate protein J-like domain-containing protein; uniclust \| UniRef100\_UPI0006C89834 \| 99.2 \| 1.7e-13 \| 3.2e-19 \| 107.7 \| 183 \| (113, 296) \| 300 \| (17, 233) \| 236 \| baseplate J/gp47 family protein \| baseplate J/gp47 family protein; uniclust \| UniRef100\_A0A523TX46 \| 99.2 \| 1.8e-13 \| 3.4e-19 \| 109.7 \| 163 \| (131, 296) \| 300 \| (42, 237) \| 242 \| Baseplate protein J-like domain-containing protein \| Baseplate protein J-like domain-containing protein; uniclust \| UniRef100\_A0A3P1VXU5 \| 99.2 \| 1.9e-13 \| 3.6e-19 \| 118.2 \| 260 \| (21, 289) \| 300 \| (19, 385) \| 394 \| Baseplate protein J-like domain-containing protein \| Baseplate protein J-like domain-containing protein; uniclust \| UniRef100\_A0A524BHK3 \| 99.1 \| 2.2e-13 \| 4e-19 \| 114.0 \| 271 \| (19, 294) \| 300 \| (17, 376) \| 381 \| Baseplate protein J-like domain-containing protein (Fragment) \| Baseplate protein J-like domain-containing protein (Fragment); uniclust \| UniRef100\_A0A1H4BJM8 \| 99.1 \| 2.4e-13 \| 4.8e-19 \| 140.0 \| 229 \| (21, 288) \| 300 \| (20, 327) \| 1375 \| Baseplate J-like protein \| Baseplate J-like protein; uniclust \| UniRef100\_A0A376TVP9 \| 99.1 \| 2.5e-13 \| 5e-19 \| 103.9 \| 104 \| (73, 176) \| 300 \| (16, 119) \| 125 \| Baseplate assembly protein GpJ \| Baseplate assembly protein GpJ; uniclust \| UniRef100\_A0A8S5LX07 \| 99.1 \| 3.1e-13 \| 5.8e-19 \| 119.3 \| 277 \| (21, 299) \| 300 \| (14, 492) \| 496 \| Baseplate wedge protein \| Baseplate wedge protein; uniclust \| UniRef100\_A0A0T9TB71 \| 99.1 \| 3.3e-13 \| 6e-19 \| 111.1 \| 145 \| (5, 149) \| 300 \| (1, 152) \| 332 \| Regulator of late gene expression \| Regulator of late gene expression; uniclust \| UniRef100\_A0A6L6YM94 \| 99.1 \| 3.3e-13 \| 6.1e-19 \| 105.0 \| 102 \| (70, 172) \| 300 \| (2, 180) \| 192 \| Baseplate J protein (Fragment) \| Baseplate J protein (Fragment); uniclust \| UniRef100\_A0A7U6KQX5 \| 99.1 \| 3.4e-13 \| 6.2e-19 \| 99.0 \| 129 \| (30, 159) \| 300 \| (7, 137) \| 150 \| Baseplate protein J-like domain-containing protein \| Baseplate protein J-like domain-containing protein; uniclust \| UniRef100\_A0A524PSP0 \| 99.1 \| 3.3e-13 \| 6.3e-19 \| 110.4 \| 185 \| (111, 296) \| 300 \| (52, 251) \| 255 \| Baseplate protein J-like domain-containing protein (Fragment) \| Baseplate protein J-like domain-containing protein (Fragment); uniclust \| UniRef100\_A0A0M0T7J9 \| 99.1 \| 3.4e-13 \| 6.4e-19 \| 109.7 \| 123 \| (20, 143) \| 300 \| (4, 200) \| 219 \| Tail protein (Fragment) \| Tail protein (Fragment); uniclust \| UniRef100\_A0A062UU35 \| 99.1 \| 3.6e-13 \| 6.9e-19 \| 134.1 \| 158 \| (111, 268) \| 300 \| (820, 998) \| 1288 \| Baseplate protein J-like domain-containing protein \| Baseplate protein J-like domain-containing protein; uniclust \| UniRef100\_UPI0018D02FB2 \| 99.1 \| 3.8e-13 \| 7.2e-19 \| 106.9 \| 176 \| (111, 292) \| 300 \| (7, 215) \| 217 \| baseplate J/gp47 family protein \| baseplate J/gp47 family protein; uniclust \| UniRef100\_A0A0A2VPK4 \| 99.1 \| 4e-13 \| 7.4e-19 \| 127.2 \| 251 \| (23, 279) \| 300 \| (958, 1317) \| 1510 \| Protein gp47 \| Protein gp47; uniclust \| UniRef100\_A0A072TEA8 \| 99.1 \| 4.3e-13 \| 8e-19 \| 131.1 \| 245 \| (27, 277) \| 300 \| (1133, 1491) \| 2834 \| Phage late control protein \| Phage late control protein; uniclust \| UniRef100\_A0A1E4QYI6 \| 99.1 \| 4.7e-13 \| 9e-19 \| 113.1 \| 229 \| (19, 270) \| 300 \| (4, 235) \| 270 \| Baseplate protein J-like domain-containing protein \| Baseplate protein J-like domain-containing protein; uniclust \| UniRef100\_A0A328TDR2 \| 99.1 \| 5.2e-13 \| 9.7e-19 \| 97.8 \| 87 \| (111, 202) \| 300 \| (32, 120) \| 130 \| Baseplate J-like family protein (Fragment) \| Baseplate J-like family protein (Fragment); uniclust \| UniRef100\_A0A931K983 \| 99.1 \| 5.6e-13 \| 1e-18 \| 113.4 \| 251 \| (31, 288) \| 300 \| (85, 426) \| 433 \| Baseplate protein J-like domain-containing protein \| Baseplate protein J-like domain-containing protein; uniclust \| UniRef100\_A0A5B8AU34 \| 99.1 \| 5.8e-13 \| 1.1e-18 \| 109.8 \| 264 \| (20, 296) \| 300 \| (9, 326) \| 332 \| Baseplate protein J-like domain-containing protein \| Baseplate protein J-like domain-containing protein; uniclust \| UniRef100\_A0A8S5M026 \| 99.1 \| 6e-13 \| 1.1e-18 \| 111.8 \| 269 \| (18, 293) \| 300 \| (7, 381) \| 389 \| Baseplate J like protein \| Baseplate J like protein; uniclust \| UniRef100\_UPI00082E9BF2 \| 99.1 \| 6.2e-13 \| 1.1e-18 \| 106.8 \| 166 \| (23, 192) \| 300 \| (4, 248) \| 272 \| baseplate J/gp47 family protein \| baseplate J/gp47 family protein; uniclust \| UniRef100\_J8V6S9 \| 99.1 \| 6.3e-13 \| 1.2e-18 \| 103.1 \| 154 \| (86, 239) \| 300 \| (1, 155) \| 212 \| Baseplate J family protein (Fragment) \| Baseplate J family protein (Fragment); uniclust \| UniRef100\_A0A0A6VQK1 \| 99.1 \| 5.9e-13 \| 1.2e-18 \| 129.2 \| 182 \| (112, 295) \| 300 \| (418, 627) \| 643 \| Baseplate protein J-like domain-containing protein \| Baseplate protein J-like domain-containing protein; uniclust \| UniRef100\_A0A3D4L1P0 \| 99.1 \| 6.4e-13 \| 1.2e-18 \| 121.3 \| 181 \| (112, 296) \| 300 \| (488, 700) \| 701 \| Baseplate protein J-like domain-containing protein \| Baseplate protein J-like domain-containing protein; uniclust \| UniRef100\_A0A6S6TBW8 \| 99.1 \| 6.8e-13 \| 1.3e-18 \| 132.1 \| 248 \| (2, 288) \| 300 \| (1, 304) \| 937 \| Uncharacterized protein \| Uncharacterized protein; uniclust \| UniRef100\_A0A2X3LRF3 \| 99.1 \| 7.4e-13 \| 1.4e-18 \| 112.0 \| 154 \| (138, 293) \| 300 \| (5, 161) \| 362 \| Tail protein I \| Tail protein I; uniclust \| UniRef100\_A0A3M2G4S3 \| 99.1 \| 7.1e-13 \| 1.4e-18 \| 121.5 \| 180 \| (111, 293) \| 300 \| (268, 481) \| 515 \| Putative baseplate assembly protein (Fragment) \| Putative baseplate assembly protein (Fragment); uniclust \| UniRef100\_M1PWF4 \| 99.1 \| 7.3e-13 \| 1.4e-18 \| 121.0 \| 85 \| (21, 105) \| 300 \| (10, 95) \| 609 \| Uncharacterized protein \| Uncharacterized protein; uniclust \| UniRef100\_UPI0018CE8811 \| 99.1 \| 8e-13 \| 1.5e-18 \| 105.2 \| 176 \| (112, 292) \| 300 \| (60, 252) \| 253 \| baseplate J/gp47 family protein \| baseplate J/gp47 family protein; uniclust \| UniRef100\_A0A951FB51 \| 99.1 \| 8.1e-13 \| 1.5e-18 \| 110.7 \| 157 \| (111, 270) \| 300 \| (98, 257) \| 322 \| Baseplate J/gp47 family protein \| Baseplate J/gp47 family protein; uniclust \| UniRef100\_A9H6K2 \| 99.1 \| 8.5e-13 \| 1.6e-18 \| 107.7 \| 152 \| (19, 171) \| 300 \| (3, 232) \| 267 \| Putative bacteriophage related protein \| Putative bacteriophage related protein; uniclust \| UniRef100\_A0A059V4B9 \| 99.1 \| 8.3e-13 \| 1.6e-18 \| 97.4 \| 110 \| (188, 297) \| 300 \| (3, 113) \| 117 \| Phage-related baseplate assembly protein \| Phage-related baseplate assembly protein; uniclust \| UniRef100\_A0A6P0SM54 \| 99.1 \| 9.1e-13 \| 1.7e-18 \| 101.8 \| 174 \| (20, 220) \| 300 \| (9, 205) \| 206 \| Baseplate protein J-like domain-containing protein (Fragment) \| Baseplate protein J-like domain-containing protein (Fragment); uniclust \| UniRef100\_UPI0002EC6E08 \| 99.1 \| 9.3e-13 \| 1.7e-18 \| 97.3 \| 138 \| (150, 289) \| 300 \| (12, 149) \| 154 \| baseplate J/gp47 family protein \| baseplate J/gp47 family protein; uniclust \| UniRef100\_UPI00167ACCF4 \| 99.1 \| 9.3e-13 \| 1.7e-18 \| 108.7 \| 285 \| (1, 297) \| 300 \| (1, 329) \| 334 \| baseplate J/gp47 family protein \| baseplate J/gp47 family protein; uniclust \| UniRef100\_A0A661PQS9 \| 99.1 \| 1.1e-12 \| 2e-18 \| 115.1 \| 240 \| (42, 288) \| 300 \| (117, 449) \| 570 \| Baseplate protein J-like domain-containing protein \| Baseplate protein J-like domain-containing protein; uniclust \| UniRef100\_A0A6L9YMA4 \| 99.1 \| 1.1e-12 \| 2e-18 \| 116.1 \| 226 \| (21, 287) \| 300 \| (18, 319) \| 387 \| Uncharacterized protein \| Uncharacterized protein; uniclust \| UniRef100\_A0A1D8TTT2 \| 99.1 \| 1.1e-12 \| 2e-18 \| 123.1 \| 156 \| (112, 271) \| 300 \| (609, 778) \| 826 \| Baseplate assembly protein \| Baseplate assembly protein; uniclust \| UniRef100\_UPI001FA8C027 \| 99.0 \| 1.1e-12 \| 2.1e-18 \| 99.5 \| 132 \| (112, 245) \| 300 \| (29, 179) \| 183 \| baseplate J/gp47 family protein \| baseplate J/gp47 family protein; uniclust \| UniRef100\_A0A1V1PGV0 \| 99.0 \| 1.3e-12 \| 2.3e-18 \| 106.1 \| 246 \| (19, 285) \| 300 \| (7, 280) \| 292 \| Baseplate protein J-like domain-containing protein \| Baseplate protein J-like domain-containing protein; uniclust \| UniRef100\_A0A081P6S8 \| 99.0 \| 1.2e-12 \| 2.4e-18 \| 134.6 \| 157 \| (111, 269) \| 300 \| (825, 1003) \| 1055 \| Baseplate protein J-like domain-containing protein \| Baseplate protein J-like domain-containing protein; uniclust \| UniRef100\_A0A0J7K9G9 \| 99.0 \| 1.4e-12 \| 2.5e-18 \| 105.9 \| 162 \| (54, 218) \| 300 \| (4, 273) \| 290 \| Baseplate j family protein (Fragment) \| Baseplate j family protein (Fragment); uniclust \| UniRef100\_A0A084Z0M7 \| 99.0 \| 1.3e-12 \| 2.5e-18 \| 102.4 \| 155 \| (133, 293) \| 300 \| (8, 175) \| 177 \| Gp47 family phage protein \| Gp47 family phage protein; uniclust \| UniRef100\_A0A0M2LYF3 \| 99.0 \| 1.3e-12 \| 2.5e-18 \| 108.4 \| 135 \| (12, 149) \| 300 \| (7, 223) \| 240 \| Baseplate protein (Fragment) \| Baseplate protein (Fragment); uniclust \| UniRef100\_A0A844GAV7 \| 99.0 \| 1.4e-12 \| 2.6e-18 \| 107.7 \| 170 \| (19, 191) \| 300 \| (3, 255) \| 290 \| Baseplate protein J-like domain-containing protein \| Baseplate protein J-like domain-containing protein; uniclust \| UniRef100\_UPI002272A788 \| 99.0 \| 1.5e-12 \| 2.7e-18 \| 110.9 \| 270 \| (22, 295) \| 300 \| (24, 395) \| 405 \| baseplate J/gp47 family protein \| baseplate J/gp47 family protein; uniclust \| UniRef100\_A0A0T9P9X2 \| 99.0 \| 1.5e-12 \| 2.8e-18 \| 116.2 \| 268 \| (21, 296) \| 300 \| (13, 472) \| 478 \| Uncharacterized homolog of phage Mu protein gp47 \| Uncharacterized homolog of phage Mu protein gp47; uniclust \| UniRef100\_A0A2G1CU44 \| 99.0 \| 1.7e-12 \| 3.1e-18 \| 108.5 \| 267 \| (10, 285) \| 300 \| (8, 353) \| 367 \| Baseplate protein J-like domain-containing protein \| Baseplate protein J-like domain-containing protein; uniclust \| UniRef100\_A0A1X3ISQ9 \| 99.0 \| 1.7e-12 \| 3.2e-18 \| 94.4 \| 124 \| (175, 298) \| 300 \| (2, 125) \| 126 \| Phage baseplate assembly protein \| Phage baseplate assembly protein; uniclust \| UniRef100\_A0A2G6CJI5 \| 99.0 \| 1.8e-12 \| 3.5e-18 \| 109.4 \| 182 \| (111, 296) \| 300 \| (96, 313) \| 318 \| T4-like baseplate wedge (Fragment) \| T4-like baseplate wedge (Fragment); uniclust \| UniRef100\_UPI000B20AC30 \| 99.0 \| 1.9e-12 \| 3.5e-18 \| 97.0 \| 140 \| (5, 160) \| 300 \| (1, 140) \| 167 \| baseplate J/gp47 family protein \| baseplate J/gp47 family protein; uniclust \| UniRef100\_A0A0A0DRH3 \| 99.0 \| 1.8e-12 \| 3.6e-18 \| 131.5 \| 155 \| (112, 268) \| 300 \| (808, 985) \| 1055 \| Baseplate protein J-like domain-containing protein \| Baseplate protein J-like domain-containing protein; uniclust \| UniRef100\_UPI0016896A8C \| 99.0 \| 2e-12 \| 3.6e-18 \| 107.2 \| 230 \| (43, 286) \| 300 \| (13, 328) \| 340 \| baseplate J/gp47 family protein \| baseplate J/gp47 family protein; uniclust \| UniRef100\_A0A2Z7A2S1 \| 99.0 \| 2e-12 \| 3.7e-18 \| 125.5 \| 227 \| (56, 289) \| 300 \| (1585, 1929) \| 2224 \| Protein gp47 \| Protein gp47; uniclust \| UniRef100\_A0A8I2K3Q2 \| 99.0 \| 2e-12 \| 3.7e-18 \| 112.3 \| 184 \| (111, 296) \| 300 \| (312, 513) \| 515 \| Putative baseplate assembly protein \| Putative baseplate assembly protein; uniclust \| UniRef100\_UPI001F4C956E \| 99.0 \| 2.1e-12 \| 3.9e-18 \| 96.6 \| 136 \| (8, 158) \| 300 \| (2, 142) \| 166 \| hypothetical protein \| hypothetical protein; uniclust \| UniRef100\_A0A0B0EQK2 \| 99.0 \| 2.2e-12 \| 4.4e-18 \| 129.3 \| 157 \| (111, 271) \| 300 \| (623, 784) \| 852 \| Baseplate protein J-like domain-containing protein \| Baseplate protein J-like domain-containing protein; uniclust \| UniRef100\_A0A963MCN6 \| 99.0 \| 2.6e-12 \| 4.8e-18 \| 118.6 \| 158 \| (111, 270) \| 300 \| (447, 633) \| 675 \| Baseplate assembly protein \| Baseplate assembly protein; uniclust \| UniRef100\_A0A146H551 \| 99.0 \| 2.7e-12 \| 5e-18 \| 116.6 \| 151 \| (21, 172) \| 300 \| (5, 245) \| 818 \| Baseplate protein J-like domain-containing protein (Fragment) \| Baseplate protein J-like domain-containing protein (Fragment); uniclust \| UniRef100\_A0A7V7B0G8 \| 99.0 \| 3e-12 \| 5.4e-18 \| 103.3 \| 217 \| (21, 260) \| 300 \| (12, 233) \| 276 \| Baseplate protein J-like domain-containing protein \| Baseplate protein J-like domain-containing protein; uniclust \| UniRef100\_A0A1W9S1W4 \| 99.0 \| 3.1e-12 \| 5.8e-18 \| 100.7 \| 171 \| (22, 210) \| 300 \| (6, 179) \| 184 \| Uncharacterized protein \| Uncharacterized protein; uniclust \| UniRef100\_UPI0020C2AC94 \| 99.0 \| 3.3e-12 \| 6e-18 \| 96.1 \| 107 \| (112, 223) \| 300 \| (53, 162) \| 171 \| baseplate J/gp47 family protein \| baseplate J/gp47 family protein; uniclust \| UniRef100\_UPI00064655BD \| 99.0 \| 3.3e-12 \| 6.1e-18 \| 100.0 \| 176 \| (113, 294) \| 300 \| (3, 198) \| 200 \| baseplate J/gp47 family protein \| baseplate J/gp47 family protein; uniclust \| UniRef100\_A0A5E4HLP5 \| 99.0 \| 3.3e-12 \| 6.1e-18 \| 109.2 \| 251 \| (21, 276) \| 300 \| (5, 413) \| 442 \| Baseplate J-like protein \| Baseplate J-like protein; uniclust \| UniRef100\_UPI0011BCFB36 \| 99.0 \| 3.4e-12 \| 6.3e-18 \| 98.1 \| 85 \| (111, 200) \| 300 \| (80, 166) \| 167 \| baseplate J/gp47 family protein \| baseplate J/gp47 family protein; uniclust \| UniRef100\_A0A6G5YFG6 \| 99.0 \| 3.5e-12 \| 6.5e-18 \| 117.8 \| 237 \| (19, 271) \| 300 \| (7, 329) \| 752 \| Baseplate protein J-like domain-containing protein \| Baseplate protein J-like domain-containing protein; uniclust \| UniRef100\_X5F9B2 \| 99.0 \| 3.6e-12 \| 6.7e-18 \| 90.6 \| 110 \| (130, 245) \| 300 \| (5, 117) \| 121 \| Phage-like element PBSX protein xkdT \| Phage-like element PBSX protein xkdT; pdb70 \| 6RAO\_J \| 99.9 \| 1.4e-32 \| 1.1e-36 \| 279.9 \| 219 \| (16, 271) \| 300 \| (9, 320) \| 963 \| Afp1, Afp2, Afp3, Afp5, Afp9 \| 6RAO\_J Afp1, Afp2, Afp3, Afp5, Afp9 Anti-feeding prophage, secretion system, AFP; pdb70 \| 6J0N\_S \| 99.9 \| 1.9e-32 \| 1.6e-36 \| 278.2 \| 218 \| (16, 267) \| 300 \| (9, 314) \| 950 \| Pvc1, Pvc9, Pvc11, Pvc12, Pvc4 \| 6J0N\_S Pvc1, Pvc9, Pvc11, Pvc12, Pvc4 assembly, Photorhabdus asymbiotica, PVC, contractile; pdb70 \| 6J0N\_O \| 99.7 \| 2.4e-21 \| 2.1e-25 \| 194.3 \| 185 \| (107, 293) \| 300 \| (500, 718) \| 728 \| Pvc1, Pvc9, Pvc11, Pvc12, Pvc4 \| 6J0N\_O Pvc1, Pvc9, Pvc11, Pvc12, Pvc4 assembly, Photorhabdus asymbiotica, PVC, contractile; pdb70 \| 6RAO\_I \| 99.7 \| 3.2e-21 \| 2.8e-25 \| 189.9 \| 181 \| (107, 289) \| 300 \| (382, 601) \| 607 \| Afp1, Afp2, Afp3, Afp5, Afp9 \| 6RAO\_I Afp1, Afp2, Afp3, Afp5, Afp9 Anti-feeding prophage, secretion system, AFP; pdb70 \| 5IV5\_BH \| 99.3 \| 1.5e-16 \| 1.3e-20 \| 157.1 \| 158 \| (107, 267) \| 300 \| (311, 477) \| 660 \| Baseplate wedge protein gp6, Baseplate \| 5IV5\_BH Baseplate wedge protein gp6, Baseplate T4, baseplate-tail tube complex, pre-attachment; pdb70 \| 5IV5\_BI \| 99.3 \| 1.5e-16 \| 1.3e-20 \| 157.1 \| 158 \| (107, 267) \| 300 \| (311, 477) \| 660 \| Baseplate wedge protein gp6, Baseplate \| 5IV5\_BI Baseplate wedge protein gp6, Baseplate T4, baseplate-tail tube complex, pre-attachment; pdb70 \| 5IV5\_EA \| 99.3 \| 1.5e-16 \| 1.3e-20 \| 157.1 \| 158 \| (107, 267) \| 300 \| (311, 477) \| 660 \| Baseplate wedge protein gp6, Baseplate \| 5IV5\_EA Baseplate wedge protein gp6, Baseplate T4, baseplate-tail tube complex, pre-attachment; pdb70 \| 5IV5\_EB \| 99.3 \| 1.5e-16 \| 1.3e-20 \| 157.1 \| 158 \| (107, 267) \| 300 \| (311, 477) \| 660 \| Baseplate wedge protein gp6, Baseplate \| 5IV5\_EB Baseplate wedge protein gp6, Baseplate T4, baseplate-tail tube complex, pre-attachment; pdb70 \| 5IV5\_GD \| 99.3 \| 1.5e-16 \| 1.3e-20 \| 157.1 \| 158 \| (107, 267) \| 300 \| (311, 477) \| 660 \| Baseplate wedge protein gp6, Baseplate \| 5IV5\_GD Baseplate wedge protein gp6, Baseplate T4, baseplate-tail tube complex, pre-attachment; pdb70 \| 5IV5\_GE \| 99.3 \| 1.5e-16 \| 1.3e-20 \| 157.1 \| 158 \| (107, 267) \| 300 \| (311, 477) \| 660 \| Baseplate wedge protein gp6, Baseplate \| 5IV5\_GE Baseplate wedge protein gp6, Baseplate T4, baseplate-tail tube complex, pre-attachment; pdb70 \| 5HX2\_E \| 99.3 \| 1.5e-16 \| 1.3e-20 \| 157.1 \| 158 \| (107, 267) \| 300 \| (311, 477) \| 660 \| Baseplate wedge protein gp7, Baseplate \| 5HX2\_E Baseplate wedge protein gp7, Baseplate T4, baseplate, complex, VIRAL PROTEIN; pdb70 \| 5IV5\_A \| 99.3 \| 1.5e-16 \| 1.3e-20 \| 157.1 \| 158 \| (107, 267) \| 300 \| (311, 477) \| 660 \| Baseplate wedge protein gp6, Baseplate \| 5IV5\_A Baseplate wedge protein gp6, Baseplate T4, baseplate-tail tube complex, pre-attachment; pdb70 \| 5IV5\_B \| 99.3 \| 1.5e-16 \| 1.3e-20 \| 157.1 \| 158 \| (107, 267) \| 300 \| (311, 477) \| 660 \| Baseplate wedge protein gp6, Baseplate \| 5IV5\_B Baseplate wedge protein gp6, Baseplate T4, baseplate-tail tube complex, pre-attachment; pdb70 \| 5IV5\_X \| 99.3 \| 1.5e-16 \| 1.3e-20 \| 157.1 \| 158 \| (107, 267) \| 300 \| (311, 477) \| 660 \| Baseplate wedge protein gp6, Baseplate \| 5IV5\_X Baseplate wedge protein gp6, Baseplate T4, baseplate-tail tube complex, pre-attachment; pdb70 \| 5IV5\_Y \| 99.3 \| 1.5e-16 \| 1.3e-20 \| 157.1 \| 158 \| (107, 267) \| 300 \| (311, 477) \| 660 \| Baseplate wedge protein gp6, Baseplate \| 5IV5\_Y Baseplate wedge protein gp6, Baseplate T4, baseplate-tail tube complex, pre-attachment; pdb70 \| 3H2T\_A \| 99.3 \| 1.7e-16 \| 1.4e-20 \| 144.5 \| 132 \| (131, 265) \| 300 \| (2, 141) \| 335 \| Baseplate structural protein Gp6 \| 3H2T\_A Baseplate structural protein Gp6 viral protein, Virion; pdb70 \| 3H3Y\_I \| 99.3 \| 1.7e-16 \| 1.4e-20 \| 144.5 \| 132 \| (131, 265) \| 300 \| (2, 141) \| 335 \| Baseplate structural protein Gp6 \| 3H3Y\_I Baseplate structural protein Gp6 viral structural protein, Virion, VIRAL; pdb70 \| 5CES\_A \| 97.9 \| 5.4e-09 \| 4.8e-13 \| 77.7 \| 92 \| (207, 298) \| 300 \| (2, 93) \| 102 \| PA0618 \| 5CES\_A PA0618 gpJ, gp6, structural protein |
| Top keywords  (threshold 1.00e-03 (evalue)) | **Baseplate, J\_like, domain\_containing, J, gp47, assembly, Phage, Fragment, Putative, wedge** |
| Output files | ../../similar\_sequences/13\_FANPEZAQ\_CDS\_0013\_merged.svg ../../similar\_sequences/13\_FANPEZAQ\_CDS\_0013\_pdb70.a3m ../../similar\_sequences/13\_FANPEZAQ\_CDS\_0013\_pdb70.hhr ../../similar\_sequences/13\_FANPEZAQ\_CDS\_0013\_uniclust.a3m ../../similar\_sequences/13\_FANPEZAQ\_CDS\_0013\_uniclust.hhr |

#### Structure prediction (AlphaFold)^2^

|  |  |
| --- | --- |
| Stats | xml version="1.0" encoding="utf-8" standalone="no"?       2024-09-02T21:09:11.794890 image/svg+xml   Matplotlib v3.7.2, https://matplotlib.org/ |
| Predicted structure | **NGL Viewer Controls:**  - Center: *Left-Click* - Rotate: *Left-Click + Drag* - Translate: *Right-Click + Drag* - Zoom: *Shift + Left-Click + Drag* |
| Output files | ../../predicted\_structures/13\_FANPEZAQ\_CDS\_0013/features.pkl ../../predicted\_structures/13\_FANPEZAQ\_CDS\_0013/ranked\_0.pdb ../../predicted\_structures/13\_FANPEZAQ\_CDS\_0013/ranked\_0\_plots.svg ../../predicted\_structures/13\_FANPEZAQ\_CDS\_0013/result\_model\_1\_ptm\_pred\_0.pkl |

#### Structure similarity search results (Foldseek)^3^

|  |  |
| --- | --- |
| Structure databases searched | Pdb, Afdb-proteome, Afdb-uniprot50 |
| Results, scheme(s)  (Top layers only, threshold 1.00e-02 (evalue)) | xml version="1.0" encoding="utf-8" standalone="no"?       2024-09-02T21:10:27.755946 image/svg+xml   Matplotlib v3.7.2, https://matplotlib.org/ |
| Results, table  (threshold 1.00e-02 (evalue)) | db \| id \| prob \| evalue \| bits \| fident \| alnlen \| mismatch \| gapopen \| qstart \| qend \| tstart \| tend \| name \| description; pdb \| 6U5B\_M \| 1.0 \| 7.017e-27 \| 787 \| 0.482 \| 294 \| 143 \| 5 \| 8 \| 296 \| 2 \| 291 \| Sheath PA0622 \| Sheath PA0622; pdb \| 6U5B\_S \| 1.0 \| 5.705e-28 \| 683 \| 0.498 \| 291 \| 143 \| 3 \| 8 \| 296 \| 3 \| 292 \| Tube PA0623 \| Tube PA0623; pdb \| 5CES\_B \| 1.0 \| 3.943e-09 \| 354 \| 0.439 \| 91 \| 51 \| 0 \| 206 \| 296 \| 1 \| 91 \| PA0618 \| PA0618; pdb \| 7YFZ\_E \| 1.0 \| 5.489e-10 \| 217 \| 0.17 \| 317 \| 191 \| 17 \| 21 \| 295 \| 12 \| 298 \| Pam3 tube initiator gp17 \| Pam3 tube initiator gp17; pdb \| 5HX2\_D \| 1.0 \| 5.777e-06 \| 159 \| 0.12 \| 183 \| 140 \| 10 \| 102 \| 270 \| 292 \| 467 \| Baseplate wedge protein gp6 \| Baseplate wedge protein gp6; pdb \| 7AEB\_A \| 1.0 \| 7.136e-07 \| 149 \| 0.107 \| 372 \| 205 \| 19 \| 13 \| 296 \| 6 \| 338 \| Phospholipid/glycerol acyltransferase \| Phospholipid/glycerol acyltransferase; pdb \| 3H2T\_A \| 1.0 \| 2.573e-05 \| 144 \| 0.115 \| 130 \| 102 \| 7 \| 151 \| 271 \| 17 \| 142 \| Baseplate structural protein Gp6 \| Baseplate structural protein Gp6; pdb \| 7B5H\_AH \| 1.0 \| 1.183e-05 \| 137 \| 0.147 \| 231 \| 141 \| 14 \| 114 \| 294 \| 974 \| 1198 \| All3317 protein \| All3317 protein; pdb \| 5IV7\_B \| 1.0 \| 5.94e-05 \| 129 \| 0.127 \| 181 \| 139 \| 9 \| 104 \| 271 \| 296 \| 470 \| Baseplate wedge protein gp11 \| Baseplate wedge protein gp11; pdb \| 8EON\_Q \| 1.0 \| 2.573e-05 \| 125 \| 0.15 \| 413 \| 192 \| 27 \| 27 \| 296 \| 7 \| 403 \| Triplex gp44-b \| Triplex gp44-b; pdb \| 6J0N\_J \| 1.0 \| 3.909e-05 \| 121 \| 0.16 \| 218 \| 136 \| 15 \| 110 \| 289 \| 393 \| 601 \| Pvc2 \| Pvc2; pdb \| 7AEB\_G \| 1.0 \| 0.0001292 \| 113 \| 0.131 \| 221 \| 144 \| 12 \| 110 \| 284 \| 789 \| 1007 \| Putative phage tail sheath protein FI \| Putative phage tail sheath protein FI; pdb \| 7KH1\_A5 \| 1.0 \| 4.285e-06 \| 111 \| 0.128 \| 411 \| 202 \| 25 \| 20 \| 294 \| 11 \| 401 \| baseplate wedge protein, gp16 \| baseplate wedge protein, gp16; pdb \| 7B5H\_AD \| 1.0 \| 4.406e-05 \| 103 \| 0.134 \| 401 \| 172 \| 21 \| 15 \| 295 \| 13 \| 358 \| All3315 protein \| All3315 protein; pdb \| 6RAO\_I \| 0.999 \| 0.0007755 \| 98 \| 0.169 \| 206 \| 140 \| 12 \| 110 \| 293 \| 246 \| 442 \| Afp11 \| Afp11; pdb \| 7YFZ\_G \| 0.998 \| 0.001046 \| 93 \| 0.128 \| 225 \| 130 \| 13 \| 21 \| 204 \| 5 \| 204 \| Pam3 baseplate wedge gp23 \| Pam3 baseplate wedge gp23; pdb \| 6J0N\_P \| 0.996 \| 0.0005752 \| 89 \| 0.129 \| 387 \| 191 \| 22 \| 17 \| 296 \| 5 \| 352 \| Pvc5 \| Pvc5; pdb \| 6RAO\_J \| 0.967 \| 0.000874 \| 75 \| 0.109 \| 348 \| 182 \| 22 \| 16 \| 287 \| 6 \| 301 \| Afp12 \| Afp12; afdb-proteome \| AF-G3XCX5-F1-MODEL\_V4 \| 1.0 \| 3.625e-42 \| 1612 \| 0.498 \| 293 \| 146 \| 1 \| 8 \| 299 \| 3 \| 295 \| Probable bacteriophage protein \| Probable bacteriophage protein; afdb-proteome \| AF-A0A0H3GRZ1-F1-MODEL\_V4 \| 1.0 \| 2.604e-41 \| 1554 \| 0.485 \| 301 \| 148 \| 1 \| 6 \| 299 \| 2 \| 302 \| Putative prophage baseplate assembly protein \| Putative prophage baseplate assembly protein; afdb-proteome \| AF-A0A0H3GKN7-F1-MODEL\_V4 \| 1.0 \| 7.192e-41 \| 1524 \| 0.471 \| 301 \| 152 \| 1 \| 6 \| 299 \| 2 \| 302 \| Baseplate assembly protein J \| Baseplate assembly protein J; afdb-proteome \| AF-Q8ZMU3-F1-MODEL\_V4 \| 1.0 \| 2.593e-39 \| 1471 \| 0.485 \| 301 \| 148 \| 2 \| 6 \| 299 \| 2 \| 302 \| Fels-2 prophage protein \| Fels-2 prophage protein; afdb-proteome \| AF-Q8ZKK3-F1-MODEL\_V4 \| 1.0 \| 2.758e-26 \| 843 \| 0.26 \| 376 \| 184 \| 10 \| 14 \| 300 \| 1 \| 371 \| Putative phage baseplate protein \| Putative phage baseplate protein; afdb-proteome \| AF-Q5F8P1-F1-MODEL\_V4 \| 1.0 \| 1.197e-13 \| 459 \| 0.431 \| 169 \| 79 \| 5 \| 1 \| 162 \| 1 \| 159 \| Baseplate protein \| Baseplate protein; afdb-proteome \| AF-P44240-F1-MODEL\_V4 \| 1.0 \| 6.587e-14 \| 384 \| 0.13 \| 367 \| 219 \| 15 \| 13 \| 292 \| 1 \| 354 \| Mu-like prophage FluMu protein gp47 \| Mu-like prophage FluMu protein gp47; afdb-proteome \| AF-P75981-F1-MODEL\_V4 \| 1.0 \| 5.795e-10 \| 294 \| 0.145 \| 199 \| 147 \| 9 \| 110 \| 293 \| 73 \| 263 \| Putative protein JayE \| Putative protein JayE; afdb-proteome \| AF-A0A0H3GMA9-F1-MODEL\_V4 \| 1.0 \| 1.118e-09 \| 242 \| 0.107 \| 390 \| 223 \| 18 \| 20 \| 295 \| 20 \| 398 \| Putative bacteriophage protein \| Putative bacteriophage protein; afdb-proteome \| AF-A0A0H3GKS8-F1-MODEL\_V4 \| 1.0 \| 7.128e-09 \| 214 \| 0.13 \| 407 \| 208 \| 18 \| 20 \| 296 \| 19 \| 409 \| Baseplate\_J domain-containing protein \| Baseplate\_J domain-containing protein; afdb-proteome \| AF-A0A0H4ISQ6-F1-MODEL\_V4 \| 1.0 \| 0.002261 \| 134 \| 0.271 \| 107 \| 76 \| 1 \| 158 \| 264 \| 1 \| 105 \| Baseplate protein \| Baseplate protein; afdb-proteome \| AF-Q32CZ0-F1-MODEL\_V4 \| 1.0 \| 0.0001838 \| 107 \| 0.131 \| 198 \| 131 \| 12 \| 40 \| 206 \| 6 \| 193 \| Hypothetical bacteriophage protein \| Hypothetical bacteriophage protein; afdb-proteome \| AF-A0A0H3GT66-F1-MODEL\_V4 \| 0.991 \| 0.002548 \| 84 \| 0.093 \| 203 \| 137 \| 12 \| 38 \| 206 \| 4 \| 193 \| Uncharacterized protein \| Uncharacterized protein; afdb-proteome \| AF-A0A0H3GSD8-F1-MODEL\_V4 \| 0.941 \| 0.004363 \| 71 \| 0.094 \| 191 \| 116 \| 10 \| 74 \| 208 \| 6 \| 195 \| Putative bacteriophage protein \| Putative bacteriophage protein; afdb-proteome \| AF-A0A077ZHG9-F1-MODEL\_V4 \| 0.941 \| 0.005219 \| 71 \| 0.145 \| 193 \| 125 \| 13 \| 25 \| 206 \| 1966 \| 2129 \| DUF2184 and DUF2612 domain containing protein \| DUF2184 and DUF2612 domain containing protein; afdb-uniprot50 \| AF-A0A7W8U3F8-F1-MODEL\_V4 \| 1.0 \| 2.582e-43 \| 1697 \| 0.579 \| 295 \| 124 \| 0 \| 5 \| 299 \| 3 \| 297 \| Phage-related baseplate assembly protein \| Phage-related baseplate assembly protein; afdb-uniprot50 \| AF-A0A066RR24-F1-MODEL\_V4 \| 1.0 \| 2.501e-42 \| 1654 \| 0.518 \| 291 \| 140 \| 0 \| 9 \| 299 \| 5 \| 295 \| Baseplate assembly protein \| Baseplate assembly protein; afdb-uniprot50 \| AF-A0A0V8T620-F1-MODEL\_V4 \| 1.0 \| 2.501e-42 \| 1631 \| 0.529 \| 287 \| 134 \| 1 \| 15 \| 300 \| 1 \| 287 \| Baseplate assembly protein \| Baseplate assembly protein; afdb-uniprot50 \| AF-A0A2T6F1A4-F1-MODEL\_V4 \| 1.0 \| 4.282e-42 \| 1629 \| 0.568 \| 292 \| 124 \| 1 \| 8 \| 299 \| 6 \| 295 \| Baseplate assembly protein \| Baseplate assembly protein; afdb-uniprot50 \| AF-G3XCX5-F1-MODEL\_V4 \| 1.0 \| 1.332e-41 \| 1612 \| 0.498 \| 293 \| 146 \| 1 \| 8 \| 299 \| 3 \| 295 \| Probable bacteriophage protein \| Probable bacteriophage protein; afdb-uniprot50 \| AF-A0A149SVY4-F1-MODEL\_V4 \| 1.0 \| 2.729e-41 \| 1587 \| 0.513 \| 300 \| 146 \| 0 \| 1 \| 300 \| 2 \| 301 \| Baseplate\_J domain-containing protein \| Baseplate\_J domain-containing protein; afdb-uniprot50 \| AF-A0A5N7XEQ3-F1-MODEL\_V4 \| 1.0 \| 2.729e-41 \| 1585 \| 0.518 \| 305 \| 139 \| 2 \| 1 \| 298 \| 1 \| 304 \| Baseplate assembly protein \| Baseplate assembly protein; afdb-uniprot50 \| AF-A0A2S6H5I2-F1-MODEL\_V4 \| 1.0 \| 6.327e-43 \| 1583 \| 0.535 \| 314 \| 123 \| 3 \| 8 \| 298 \| 6 \| 319 \| Phage-related baseplate assembly protein \| Phage-related baseplate assembly protein; afdb-uniprot50 \| AF-A0A410SH26-F1-MODEL\_V4 \| 1.0 \| 4.961e-41 \| 1572 \| 0.478 \| 303 \| 150 \| 1 \| 5 \| 299 \| 2 \| 304 \| Baseplate assembly protein \| Baseplate assembly protein; afdb-uniprot50 \| AF-A0A0H3GRZ1-F1-MODEL\_V4 \| 1.0 \| 9.572e-41 \| 1554 \| 0.485 \| 301 \| 148 \| 1 \| 6 \| 299 \| 2 \| 302 \| Putative prophage baseplate assembly protein \| Putative prophage baseplate assembly protein; afdb-uniprot50 \| AF-B5JY66-F1-MODEL\_V4 \| 1.0 \| 2.21e-40 \| 1551 \| 0.449 \| 334 \| 148 \| 4 \| 1 \| 299 \| 1 \| 333 \| Baseplate J-like protein \| Baseplate J-like protein; afdb-uniprot50 \| AF-A0A0Q8D2W6-F1-MODEL\_V4 \| 1.0 \| 5.414e-40 \| 1546 \| 0.46 \| 332 \| 143 \| 2 \| 2 \| 299 \| 3 \| 332 \| Baseplate assembly protein \| Baseplate assembly protein; afdb-uniprot50 \| AF-A0A812QW15-F1-MODEL\_V4 \| 1.0 \| 1.961e-40 \| 1545 \| 0.514 \| 305 \| 140 \| 2 \| 3 \| 300 \| 2 \| 305 \| J protein \| J protein; afdb-uniprot50 \| AF-A0A291N2G8-F1-MODEL\_V4 \| 1.0 \| 2.081e-40 \| 1544 \| 0.49 \| 300 \| 149 \| 3 \| 1 \| 299 \| 1 \| 297 \| Baseplate assembly protein \| Baseplate assembly protein; afdb-uniprot50 \| AF-A0A2T5J1G0-F1-MODEL\_V4 \| 1.0 \| 1.216e-40 \| 1538 \| 0.479 \| 298 \| 153 \| 1 \| 3 \| 298 \| 2 \| 299 \| Phage-related baseplate assembly protein \| Phage-related baseplate assembly protein; afdb-uniprot50 \| AF-A0A4Z0E2N6-F1-MODEL\_V4 \| 1.0 \| 1.454e-40 \| 1532 \| 0.481 \| 299 \| 148 \| 1 \| 8 \| 299 \| 4 \| 302 \| Baseplate assembly protein \| Baseplate assembly protein; afdb-uniprot50 \| AF-A0A0A1FE10-F1-MODEL\_V4 \| 1.0 \| 6.477e-40 \| 1531 \| 0.478 \| 297 \| 154 \| 1 \| 4 \| 299 \| 2 \| 298 \| Baseplate assembly protein J \| Baseplate assembly protein J; afdb-uniprot50 \| AF-E1SFX6-F1-MODEL\_V4 \| 1.0 \| 1.74e-40 \| 1528 \| 0.5 \| 306 \| 142 \| 2 \| 1 \| 299 \| 1 \| 302 \| Baseplate assembly protein J (GpJ) \| Baseplate assembly protein J (GpJ); afdb-uniprot50 \| AF-A0A0H3GKN7-F1-MODEL\_V4 \| 1.0 \| 2.643e-40 \| 1524 \| 0.471 \| 301 \| 152 \| 1 \| 6 \| 299 \| 2 \| 302 \| Baseplate assembly protein J \| Baseplate assembly protein J; afdb-uniprot50 \| AF-G0AIQ8-F1-MODEL\_V4 \| 1.0 \| 9.27e-40 \| 1509 \| 0.506 \| 300 \| 143 \| 2 \| 1 \| 300 \| 1 \| 295 \| Baseplate J-like protein \| Baseplate J-like protein; afdb-uniprot50 \| AF-A0A3R8WZD4-F1-MODEL\_V4 \| 1.0 \| 1.495e-39 \| 1506 \| 0.46 \| 302 \| 156 \| 1 \| 5 \| 299 \| 2 \| 303 \| Baseplate assembly protein \| Baseplate assembly protein; afdb-uniprot50 \| AF-A0A806D8P7-F1-MODEL\_V4 \| 1.0 \| 6.477e-40 \| 1505 \| 0.447 \| 302 \| 161 \| 3 \| 2 \| 299 \| 6 \| 305 \| Baseplate J family protein \| Baseplate J family protein; afdb-uniprot50 \| AF-A0A397NE13-F1-MODEL\_V4 \| 1.0 \| 2.14e-39 \| 1501 \| 0.481 \| 297 \| 152 \| 1 \| 3 \| 299 \| 2 \| 296 \| Phage-related baseplate assembly protein \| Phage-related baseplate assembly protein; afdb-uniprot50 \| AF-A0A094SSD9-F1-MODEL\_V4 \| 1.0 \| 8.226e-40 \| 1500 \| 0.473 \| 306 \| 151 \| 2 \| 1 \| 299 \| 1 \| 303 \| Baseplate assembly protein \| Baseplate assembly protein; afdb-uniprot50 \| AF-A0A2Z6GCU5-F1-MODEL\_V4 \| 1.0 \| 2.016e-39 \| 1494 \| 0.458 \| 301 \| 157 \| 3 \| 1 \| 298 \| 1 \| 298 \| Baseplate Assembly protein \| Baseplate Assembly protein; afdb-uniprot50 \| AF-A0A1S1NVP6-F1-MODEL\_V4 \| 1.0 \| 1.685e-39 \| 1492 \| 0.478 \| 299 \| 149 \| 1 \| 8 \| 299 \| 5 \| 303 \| Baseplate assembly protein \| Baseplate assembly protein; afdb-uniprot50 \| AF-A4WEL5-F1-MODEL\_V4 \| 1.0 \| 3.251e-39 \| 1489 \| 0.478 \| 299 \| 149 \| 1 \| 8 \| 299 \| 4 \| 302 \| Baseplate J family protein \| Baseplate J family protein; afdb-uniprot50 \| AF-A0A064DPN7-F1-MODEL\_V4 \| 1.0 \| 2.718e-39 \| 1487 \| 0.481 \| 299 \| 148 \| 1 \| 8 \| 299 \| 4 \| 302 \| Baseplate\_J domain-containing protein \| Baseplate\_J domain-containing protein; afdb-uniprot50 \| AF-A0A3N1XDJ3-F1-MODEL\_V4 \| 1.0 \| 3.063e-39 \| 1486 \| 0.489 \| 298 \| 145 \| 1 \| 8 \| 298 \| 2 \| 299 \| Phage-related baseplate assembly protein \| Phage-related baseplate assembly protein; afdb-uniprot50 \| AF-A0A1H9HC55-F1-MODEL\_V4 \| 1.0 \| 5.244e-39 \| 1485 \| 0.469 \| 296 \| 155 \| 1 \| 5 \| 298 \| 4 \| 299 \| Phage-related baseplate assembly protein \| Phage-related baseplate assembly protein; afdb-uniprot50 \| AF-A0A561GUB6-F1-MODEL\_V4 \| 1.0 \| 4.129e-39 \| 1484 \| 0.508 \| 299 \| 141 \| 2 \| 1 \| 299 \| 1 \| 293 \| Phage-related baseplate assembly protein \| Phage-related baseplate assembly protein; afdb-uniprot50 \| AF-A0A483GRP4-F1-MODEL\_V4 \| 1.0 \| 3.063e-39 \| 1483 \| 0.471 \| 299 \| 151 \| 1 \| 8 \| 299 \| 4 \| 302 \| Baseplate assembly protein \| Baseplate assembly protein; afdb-uniprot50 \| AF-A0A1D2QS87-F1-MODEL\_V4 \| 1.0 \| 7.069e-39 \| 1483 \| 0.459 \| 298 \| 159 \| 1 \| 5 \| 300 \| 4 \| 301 \| Baseplate\_J domain-containing protein \| Baseplate\_J domain-containing protein; afdb-uniprot50 \| AF-A0A375ABY7-F1-MODEL\_V4 \| 1.0 \| 3.664e-39 \| 1482 \| 0.511 \| 297 \| 138 \| 2 \| 6 \| 295 \| 2 \| 298 \| Phage baseplate assembly protein J \| Phage baseplate assembly protein J; afdb-uniprot50 \| AF-A0A0Q4N0X4-F1-MODEL\_V4 \| 1.0 \| 2.412e-39 \| 1472 \| 0.498 \| 301 \| 144 \| 1 \| 6 \| 299 \| 2 \| 302 \| Baseplate assembly protein \| Baseplate assembly protein; afdb-uniprot50 \| AF-C6BVX8-F1-MODEL\_V4 \| 1.0 \| 3.548e-38 \| 1469 \| 0.452 \| 292 \| 159 \| 1 \| 4 \| 295 \| 2 \| 292 \| Baseplate J family protein \| Baseplate J family protein; afdb-uniprot50 \| AF-A0A1Q8DQX3-F1-MODEL\_V4 \| 1.0 \| 5.244e-39 \| 1469 \| 0.464 \| 297 \| 155 \| 3 \| 6 \| 300 \| 2 \| 296 \| Baseplate\_J domain-containing protein \| Baseplate\_J domain-containing protein; afdb-uniprot50 \| AF-A0A4Q9EIG5-F1-MODEL\_V4 \| 1.0 \| 3.063e-39 \| 1467 \| 0.493 \| 302 \| 140 \| 2 \| 8 \| 299 \| 4 \| 302 \| Baseplate assembly protein \| Baseplate assembly protein; afdb-uniprot50 \| AF-A0A845BNS7-F1-MODEL\_V4 \| 1.0 \| 1.14e-38 \| 1461 \| 0.464 \| 299 \| 153 \| 2 \| 8 \| 299 \| 4 \| 302 \| Baseplate assembly protein \| Baseplate assembly protein; afdb-uniprot50 \| AF-A0A0D8CM68-F1-MODEL\_V4 \| 1.0 \| 2.072e-38 \| 1460 \| 0.438 \| 308 \| 160 \| 2 \| 4 \| 298 \| 3 \| 310 \| Baseplate J protein \| Baseplate J protein; afdb-uniprot50 \| AF-A0A6B3J9L7-F1-MODEL\_V4 \| 1.0 \| 1.21e-38 \| 1457 \| 0.48 \| 275 \| 139 \| 2 \| 28 \| 299 \| 2 \| 275 \| Baseplate assembly protein \| Baseplate assembly protein; afdb-uniprot50 \| AF-A0A447R011-F1-MODEL\_V4 \| 1.0 \| 1.012e-38 \| 1456 \| 0.465 \| 288 \| 147 \| 1 \| 19 \| 299 \| 2 \| 289 \| Baseplate assembly protein \| Baseplate assembly protein; afdb-uniprot50 \| AF-A0A1I1UBY4-F1-MODEL\_V4 \| 1.0 \| 2.072e-38 \| 1454 \| 0.438 \| 303 \| 159 \| 1 \| 8 \| 299 \| 4 \| 306 \| Phage-related baseplate assembly protein \| Phage-related baseplate assembly protein; afdb-uniprot50 \| AF-A0A269PHY2-F1-MODEL\_V4 \| 1.0 \| 4.506e-38 \| 1442 \| 0.47 \| 291 \| 151 \| 2 \| 5 \| 295 \| 2 \| 289 \| Baseplate assembly protein \| Baseplate assembly protein; afdb-uniprot50 \| AF-A0A2N2S1G7-F1-MODEL\_V4 \| 1.0 \| 7.716e-38 \| 1442 \| 0.433 \| 295 \| 164 \| 2 \| 8 \| 299 \| 3 \| 297 \| Baseplate assembly protein \| Baseplate assembly protein; afdb-uniprot50 \| AF-A0A2S9S4Q7-F1-MODEL\_V4 \| 1.0 \| 3.767e-38 \| 1441 \| 0.426 \| 300 \| 164 \| 2 \| 8 \| 300 \| 5 \| 303 \| Baseplate assembly protein \| Baseplate assembly protein; afdb-uniprot50 \| AF-A0A636GAS7-F1-MODEL\_V4 \| 1.0 \| 3.548e-38 \| 1440 \| 0.448 \| 299 \| 158 \| 1 \| 8 \| 299 \| 4 \| 302 \| Baseplate assembly protein \| Baseplate assembly protein; afdb-uniprot50 \| AF-A0A6I1IU11-F1-MODEL\_V4 \| 1.0 \| 4.245e-38 \| 1438 \| 0.45 \| 284 \| 152 \| 1 \| 21 \| 300 \| 1 \| 284 \| Baseplate assembly protein \| Baseplate assembly protein; afdb-uniprot50 \| AF-A0A1E7RC48-F1-MODEL\_V4 \| 1.0 \| 5.567e-39 \| 1437 \| 0.488 \| 293 \| 142 \| 2 \| 8 \| 296 \| 4 \| 292 \| Baseplate J family protein \| Baseplate J family protein; afdb-uniprot50 \| AF-A0A4Y9VR74-F1-MODEL\_V4 \| 1.0 \| 6.847e-38 \| 1437 \| 0.436 \| 298 \| 163 \| 3 \| 5 \| 298 \| 2 \| 298 \| Baseplate assembly protein \| Baseplate assembly protein; afdb-uniprot50 \| AF-A0A1E7HY03-F1-MODEL\_V4 \| 1.0 \| 4.245e-38 \| 1435 \| 0.43 \| 297 \| 167 \| 2 \| 1 \| 296 \| 1 \| 296 \| Baseplate\_J domain-containing protein \| Baseplate\_J domain-containing protein; afdb-uniprot50 \| AF-A0A7X6WY54-F1-MODEL\_V4 \| 1.0 \| 2.2e-38 \| 1435 \| 0.43 \| 300 \| 163 \| 2 \| 4 \| 295 \| 2 \| 301 \| Baseplate assembly protein \| Baseplate assembly protein; afdb-uniprot50 \| AF-H8L2K1-F1-MODEL\_V4 \| 1.0 \| 6.847e-38 \| 1430 \| 0.427 \| 295 \| 166 \| 2 \| 6 \| 299 \| 2 \| 294 \| Phage-related baseplate assembly protein \| Phage-related baseplate assembly protein; afdb-uniprot50 \| AF-A0A3F3J766-F1-MODEL\_V4 \| 1.0 \| 6.45e-38 \| 1429 \| 0.453 \| 298 \| 156 \| 1 \| 8 \| 298 \| 4 \| 301 \| Baseplate assembly protein \| Baseplate assembly protein; afdb-uniprot50 \| AF-A0A541BB49-F1-MODEL\_V4 \| 1.0 \| 2.632e-38 \| 1429 \| 0.426 \| 326 \| 154 \| 2 \| 6 \| 298 \| 7 \| 332 \| Baseplate protein \| Baseplate protein; afdb-uniprot50 \| AF-A0A162BXM4-F1-MODEL\_V4 \| 1.0 \| 2.966e-38 \| 1427 \| 0.416 \| 324 \| 156 \| 2 \| 8 \| 298 \| 9 \| 332 \| Baseplate assembly protein \| Baseplate assembly protein; afdb-uniprot50 \| AF-Q31HT8-F1-MODEL\_V4 \| 1.0 \| 5.078e-38 \| 1424 \| 0.452 \| 296 \| 156 \| 3 \| 8 \| 298 \| 3 \| 297 \| Phage baseplate J-like protein \| Phage baseplate J-like protein; afdb-uniprot50 \| AF-A0A454TSQ4-F1-MODEL\_V4 \| 1.0 \| 5.078e-38 \| 1417 \| 0.438 \| 301 \| 162 \| 2 \| 6 \| 299 \| 2 \| 302 \| Baseplate assembly protein \| Baseplate assembly protein; afdb-uniprot50 \| AF-A0A4Z0WEF0-F1-MODEL\_V4 \| 1.0 \| 6.075e-38 \| 1417 \| 0.415 \| 330 \| 158 \| 3 \| 1 \| 295 \| 1 \| 330 \| Baseplate assembly protein \| Baseplate assembly protein; afdb-uniprot50 \| AF-A0A376S4C0-F1-MODEL\_V4 \| 1.0 \| 1.104e-37 \| 1414 \| 0.489 \| 282 \| 137 \| 1 \| 25 \| 299 \| 2 \| 283 \| Baseplate assembly protein \| Baseplate assembly protein; afdb-uniprot50 \| AF-A0A199YL26-F1-MODEL\_V4 \| 1.0 \| 6.847e-38 \| 1410 \| 0.449 \| 309 \| 159 \| 2 \| 1 \| 298 \| 1 \| 309 \| Baseplate\_J domain-containing protein \| Baseplate\_J domain-containing protein; afdb-uniprot50 \| AF-A0A7W6RGM7-F1-MODEL\_V4 \| 1.0 \| 2.401e-37 \| 1409 \| 0.385 \| 366 \| 154 \| 5 \| 4 \| 298 \| 3 \| 368 \| Phage-related baseplate assembly protein \| Phage-related baseplate assembly protein; afdb-uniprot50 \| AF-A0A482ISR2-F1-MODEL\_V4 \| 1.0 \| 2.131e-37 \| 1404 \| 0.44 \| 300 \| 160 \| 2 \| 8 \| 299 \| 5 \| 304 \| Baseplate assembly protein \| Baseplate assembly protein; afdb-uniprot50 \| AF-A0A6L8GFW9-F1-MODEL\_V4 \| 1.0 \| 8.421e-37 \| 1403 \| 0.413 \| 300 \| 171 \| 1 \| 1 \| 295 \| 1 \| 300 \| Baseplate assembly protein \| Baseplate assembly protein; afdb-uniprot50 \| AF-A0A4R1ZHR1-F1-MODEL\_V4 \| 1.0 \| 2.706e-37 \| 1402 \| 0.451 \| 297 \| 158 \| 2 \| 8 \| 300 \| 3 \| 298 \| Phage-related baseplate assembly protein \| Phage-related baseplate assembly protein; afdb-uniprot50 \| AF-A0A7V7NR69-F1-MODEL\_V4 \| 1.0 \| 5.078e-38 \| 1399 \| 0.444 \| 299 \| 163 \| 2 \| 1 \| 298 \| 2 \| 298 \| Baseplate assembly protein \| Baseplate assembly protein; afdb-uniprot50 \| AF-B8GS00-F1-MODEL\_V4 \| 1.0 \| 8.94e-37 \| 1398 \| 0.457 \| 291 \| 154 \| 4 \| 6 \| 295 \| 2 \| 289 \| Baseplate J-like protein \| Baseplate J-like protein; afdb-uniprot50 \| AF-A0A2W6Y6Y1-F1-MODEL\_V4 \| 1.0 \| 7.039e-37 \| 1395 \| 0.433 \| 300 \| 164 \| 3 \| 1 \| 298 \| 1 \| 296 \| Baseplate assembly protein \| Baseplate assembly protein; afdb-uniprot50 \| AF-A0A776QGR6-F1-MODEL\_V4 \| 1.0 \| 1.725e-36 \| 1393 \| 0.503 \| 254 \| 126 \| 0 \| 46 \| 299 \| 6 \| 259 \| Baseplate assembly protein \| Baseplate assembly protein; afdb-uniprot50 \| AF-A0A4P9VSJ3-F1-MODEL\_V4 \| 1.0 \| 6.659e-39 \| 1391 \| 0.476 \| 296 \| 146 \| 3 \| 3 \| 294 \| 2 \| 292 \| Baseplate assembly protein \| Baseplate assembly protein; afdb-uniprot50 \| AF-A0A2A9KD31-F1-MODEL\_V4 \| 1.0 \| 1.678e-37 \| 1388 \| 0.464 \| 299 \| 155 \| 2 \| 1 \| 299 \| 1 \| 294 \| Phage-related baseplate assembly protein \| Phage-related baseplate assembly protein; afdb-uniprot50 \| AF-A0A1M4WDH6-F1-MODEL\_V4 \| 1.0 \| 8.421e-37 \| 1386 \| 0.445 \| 292 \| 158 \| 2 \| 5 \| 295 \| 2 \| 290 \| Phage-related baseplate assembly protein \| Phage-related baseplate assembly protein; afdb-uniprot50 \| AF-A0A4P7J5J5-F1-MODEL\_V4 \| 1.0 \| 9.799e-38 \| 1386 \| 0.461 \| 310 \| 148 \| 3 \| 1 \| 299 \| 1 \| 302 \| Baseplate assembly protein \| Baseplate assembly protein; afdb-uniprot50 \| AF-A0A7H8UVW2-F1-MODEL\_V4 \| 1.0 \| 4.364e-37 \| 1386 \| 0.404 \| 304 \| 167 \| 2 \| 8 \| 300 \| 5 \| 305 \| Baseplate J/gp47 family protein \| Baseplate J/gp47 family protein; afdb-uniprot50 \| AF-A0A380TV19-F1-MODEL\_V4 \| 1.0 \| 4.918e-37 \| 1384 \| 0.416 \| 300 \| 167 \| 3 \| 8 \| 300 \| 5 \| 303 \| Uncharacterized homolog of phage Mu protein gp47 \| Uncharacterized homolog of phage Mu protein gp47; afdb-uniprot50 \| AF-S6HS98-F1-MODEL\_V4 \| 1.0 \| 3.135e-36 \| 1383 \| 0.392 \| 303 \| 179 \| 3 \| 1 \| 299 \| 1 \| 302 \| Baseplate\_J domain-containing protein \| Baseplate\_J domain-containing protein; afdb-uniprot50 \| AF-A0A1G7S895-F1-MODEL\_V4 \| 1.0 \| 4.364e-37 \| 1374 \| 0.456 \| 298 \| 157 \| 3 \| 4 \| 298 \| 2 \| 297 \| Phage-related baseplate assembly protein \| Phage-related baseplate assembly protein; afdb-uniprot50 \| AF-A0A285VRF8-F1-MODEL\_V4 \| 1.0 \| 6.049e-36 \| 1370 \| 0.429 \| 298 \| 162 \| 3 \| 5 \| 294 \| 2 \| 299 \| Phage-related baseplate assembly protein \| Phage-related baseplate assembly protein; afdb-uniprot50 \| AF-A0A0T7PAJ5-F1-MODEL\_V4 \| 1.0 \| 6.422e-36 \| 1369 \| 0.527 \| 254 \| 120 \| 0 \| 46 \| 299 \| 7 \| 260 \| Baseplate assembly protein J \| Baseplate assembly protein J; afdb-uniprot50 \| AF-A0A1I3U816-F1-MODEL\_V4 \| 1.0 \| 3.982e-36 \| 1369 \| 0.411 \| 294 \| 171 \| 1 \| 8 \| 299 \| 4 \| 297 \| Phage-related baseplate assembly protein \| Phage-related baseplate assembly protein; afdb-uniprot50 \| AF-A0A423PQH8-F1-MODEL\_V4 \| 1.0 \| 1.725e-36 \| 1364 \| 0.426 \| 326 \| 153 \| 4 \| 8 \| 299 \| 4 \| 329 \| Baseplate assembly protein \| Baseplate assembly protein; afdb-uniprot50 \| AF-A0A1B3E8E6-F1-MODEL\_V4 \| 1.0 \| 1.358e-36 \| 1362 \| 0.423 \| 333 \| 151 \| 4 \| 8 \| 300 \| 3 \| 334 \| Baseplate assembly protein \| Baseplate assembly protein; afdb-uniprot50 \| AF-A0A0E3BVE1-F1-MODEL\_V4 \| 1.0 \| 8.658e-36 \| 1360 \| 0.427 \| 295 \| 152 \| 3 \| 8 \| 300 \| 3 \| 282 \| Baseplate assembly protein \| Baseplate assembly protein; afdb-uniprot50 \| AF-A0A1H0SWN7-F1-MODEL\_V4 \| 1.0 \| 3.135e-36 \| 1359 \| 0.425 \| 301 \| 165 \| 2 \| 6 \| 298 \| 3 \| 303 \| Phage-related baseplate assembly protein \| Phage-related baseplate assembly protein; afdb-uniprot50 \| AF-A0A853STM6-F1-MODEL\_V4 \| 1.0 \| 2.469e-36 \| 1357 \| 0.459 \| 296 \| 153 \| 2 \| 8 \| 296 \| 5 \| 300 \| Baseplate J-like protein \| Baseplate J-like protein; afdb-uniprot50 \| AF-A0A4V6X7E1-F1-MODEL\_V4 \| 1.0 \| 2.621e-36 \| 1350 \| 0.475 \| 303 \| 136 \| 4 \| 5 \| 300 \| 2 \| 288 \| Baseplate assembly protein \| Baseplate assembly protein; afdb-uniprot50 \| AF-A0A285V0V5-F1-MODEL\_V4 \| 1.0 \| 6.422e-36 \| 1350 \| 0.385 \| 301 \| 176 \| 4 \| 1 \| 299 \| 1 \| 294 \| Phage-related baseplate assembly protein \| Phage-related baseplate assembly protein; afdb-uniprot50 \| AF-A0A5Y3B1N0-F1-MODEL\_V4 \| 1.0 \| 4.227e-36 \| 1348 \| 0.435 \| 280 \| 152 \| 1 \| 21 \| 294 \| 1 \| 280 \| Baseplate assembly protein \| Baseplate assembly protein; afdb-uniprot50 \| AF-A0A7Y7D7U6-F1-MODEL\_V4 \| 1.0 \| 1.725e-36 \| 1348 \| 0.402 \| 363 \| 146 \| 6 \| 5 \| 299 \| 4 \| 363 \| Baseplate J/gp47 family protein \| Baseplate J/gp47 family protein; afdb-uniprot50 \| AF-A0A2N1AP80-F1-MODEL\_V4 \| 1.0 \| 3.751e-36 \| 1346 \| 0.411 \| 299 \| 168 \| 2 \| 6 \| 296 \| 3 \| 301 \| Baseplate assembly protein \| Baseplate assembly protein; afdb-uniprot50 \| AF-A0A3T0W3K4-F1-MODEL\_V4 \| 1.0 \| 2.191e-36 \| 1344 \| 0.417 \| 304 \| 164 \| 2 \| 5 \| 295 \| 2 \| 305 \| Baseplate J protein \| Baseplate J protein; afdb-uniprot50 \| AF-A0A377LB58-F1-MODEL\_V4 \| 1.0 \| 3.223e-35 \| 1342 \| 0.46 \| 256 \| 138 \| 0 \| 44 \| 299 \| 5 \| 260 \| Baseplate assembly protein \| Baseplate assembly protein; afdb-uniprot50 \| AF-A0A4U8YL31-F1-MODEL\_V4 \| 1.0 \| 1.531e-36 \| 1339 \| 0.424 \| 332 \| 149 \| 4 \| 6 \| 295 \| 2 \| 333 \| Baseplate protein j-like \| Baseplate protein j-like; afdb-uniprot50 \| AF-A0A1E3G8B9-F1-MODEL\_V4 \| 1.0 \| 4.613e-35 \| 1332 \| 0.404 \| 287 \| 167 \| 1 \| 10 \| 296 \| 8 \| 290 \| Baseplate\_J domain-containing protein \| Baseplate\_J domain-containing protein; afdb-uniprot50 \| AF-A0A1H2M4W7-F1-MODEL\_V4 \| 1.0 \| 1.036e-35 \| 1329 \| 0.453 \| 300 \| 155 \| 4 \| 4 \| 295 \| 3 \| 301 \| Phage-related baseplate assembly protein \| Phage-related baseplate assembly protein; afdb-uniprot50 \| AF-A0A1D7ZGU3-F1-MODEL\_V4 \| 1.0 \| 4.897e-35 \| 1326 \| 0.356 \| 297 \| 188 \| 2 \| 6 \| 300 \| 2 \| 297 \| Baseplate J protein \| Baseplate J protein; afdb-uniprot50 \| AF-A0A3N2E0Q3-F1-MODEL\_V4 \| 1.0 \| 9.757e-36 \| 1325 \| 0.389 \| 298 \| 175 \| 2 \| 5 \| 295 \| 4 \| 301 \| Phage-related baseplate assembly protein \| Phage-related baseplate assembly protein; afdb-uniprot50 \| AF-A0A377U0P4-F1-MODEL\_V4 \| 1.0 \| 1.524e-34 \| 1323 \| 0.488 \| 254 \| 130 \| 0 \| 46 \| 299 \| 7 \| 260 \| Baseplate assembly protein J \| Baseplate assembly protein J; afdb-uniprot50 \| AF-A0A3N2E200-F1-MODEL\_V4 \| 1.0 \| 2.782e-36 \| 1323 \| 0.447 \| 286 \| 156 \| 1 \| 14 \| 299 \| 3 \| 286 \| Phage-related baseplate assembly protein \| Phage-related baseplate assembly protein; afdb-uniprot50 \| AF-A0A6C8X2I9-F1-MODEL\_V4 \| 1.0 \| 7.009e-35 \| 1321 \| 0.513 \| 255 \| 122 \| 2 \| 45 \| 299 \| 1 \| 253 \| Baseplate assembly protein \| Baseplate assembly protein; afdb-uniprot50 \| AF-A0A212KBX8-F1-MODEL\_V4 \| 1.0 \| 1.773e-35 \| 1321 \| 0.389 \| 298 \| 172 \| 2 \| 8 \| 295 \| 7 \| 304 \| Baseplate assembly protein J (GPJ) \| Baseplate assembly protein J (GPJ); afdb-uniprot50 \| AF-A0A3S0YC89-F1-MODEL\_V4 \| 1.0 \| 1.315e-35 \| 1316 \| 0.466 \| 296 \| 148 \| 4 \| 8 \| 296 \| 5 \| 297 \| Baseplate assembly protein \| Baseplate assembly protein; afdb-uniprot50 \| AF-A0A837E5S2-F1-MODEL\_V4 \| 1.0 \| 1.999e-35 \| 1316 \| 0.382 \| 335 \| 169 \| 5 \| 1 \| 298 \| 1 \| 334 \| Uncharacterized protein \| Uncharacterized protein; afdb-uniprot50 \| AF-A0A7W6RE52-F1-MODEL\_V4 \| 1.0 \| 1.936e-34 \| 1313 \| 0.416 \| 298 \| 157 \| 4 \| 3 \| 300 \| 2 \| 282 \| Phage-related baseplate assembly protein \| Phage-related baseplate assembly protein; afdb-uniprot50 \| AF-A0A5N9VV02-F1-MODEL\_V4 \| 1.0 \| 2.694e-35 \| 1312 \| 0.481 \| 266 \| 138 \| 0 \| 34 \| 299 \| 2 \| 267 \| Baseplate assembly protein \| Baseplate assembly protein; afdb-uniprot50 \| AF-A0A0C2IH57-F1-MODEL\_V4 \| 1.0 \| 2.86e-35 \| 1312 \| 0.423 \| 300 \| 164 \| 3 \| 4 \| 295 \| 3 \| 301 \| Phage tail fiber protein \| Phage tail fiber protein; afdb-uniprot50 \| AF-A0A2S9M0X9-F1-MODEL\_V4 \| 1.0 \| 4.743e-34 \| 1311 \| 0.387 \| 297 \| 179 \| 2 \| 6 \| 300 \| 2 \| 297 \| Baseplate J protein \| Baseplate J protein; afdb-uniprot50 \| AF-A0A1F0H4T9-F1-MODEL\_V4 \| 1.0 \| 2.064e-36 \| 1307 \| 0.385 \| 306 \| 174 \| 2 \| 5 \| 299 \| 2 \| 304 \| Baseplate\_J domain-containing protein \| Baseplate\_J domain-containing protein; afdb-uniprot50 \| AF-F4QJC2-F1-MODEL\_V4 \| 1.0 \| 8.658e-36 \| 1302 \| 0.447 \| 306 \| 146 \| 5 \| 1 \| 299 \| 1 \| 290 \| Baseplate J-like family protein \| Baseplate J-like family protein; afdb-uniprot50 \| AF-A0A016XHS1-F1-MODEL\_V4 \| 1.0 \| 1.574e-35 \| 1301 \| 0.383 \| 297 \| 181 \| 1 \| 5 \| 299 \| 2 \| 298 \| Baseplate assembly protein \| Baseplate assembly protein; afdb-uniprot50 \| AF-A0A2E5BTS9-F1-MODEL\_V4 \| 1.0 \| 7.933e-37 \| 1300 \| 0.423 \| 321 \| 156 \| 2 \| 8 \| 299 \| 7 \| 327 \| Baseplate assembly protein \| Baseplate assembly protein; afdb-uniprot50 \| AF-A0A6P3BRR2-F1-MODEL\_V4 \| 1.0 \| 7.441e-35 \| 1298 \| 0.382 \| 306 \| 179 \| 3 \| 1 \| 299 \| 1 \| 303 \| Baseplate assembly protein \| Baseplate assembly protein; afdb-uniprot50 \| AF-A0A514B9U7-F1-MODEL\_V4 \| 1.0 \| 4.094e-35 \| 1297 \| 0.434 \| 304 \| 155 \| 4 \| 4 \| 295 \| 3 \| 301 \| Baseplate assembly protein \| Baseplate assembly protein; afdb-uniprot50 \| AF-A0A1E7WJG0-F1-MODEL\_V4 \| 1.0 \| 1.574e-35 \| 1296 \| 0.462 \| 296 \| 149 \| 3 \| 8 \| 296 \| 4 \| 296 \| Baseplate J-like protein \| Baseplate J-like protein; afdb-uniprot50 \| AF-A0A1X7L2D2-F1-MODEL\_V4 \| 1.0 \| 1.352e-34 \| 1295 \| 0.447 \| 293 \| 146 \| 3 \| 8 \| 300 \| 2 \| 278 \| Phage-related baseplate assembly protein \| Phage-related baseplate assembly protein; afdb-uniprot50 \| AF-A0A5E4XFI6-F1-MODEL\_V4 \| 1.0 \| 3.633e-35 \| 1295 \| 0.389 \| 311 \| 172 \| 4 \| 1 \| 300 \| 1 \| 304 \| Baseplate assembly protein \| Baseplate assembly protein; afdb-uniprot50 \| AF-Q9KW05-F1-MODEL\_V4 \| 1.0 \| 2.527e-33 \| 1291 \| 0.495 \| 230 \| 116 \| 0 \| 70 \| 299 \| 3 \| 232 \| Baseplate/ tail fiber protein \| Baseplate/ tail fiber protein; afdb-uniprot50 \| AF-A0A1C3EE52-F1-MODEL\_V4 \| 1.0 \| 1.065e-34 \| 1285 \| 0.371 \| 326 \| 168 \| 4 \| 10 \| 298 \| 9 \| 334 \| Baseplate\_J domain-containing protein \| Baseplate\_J domain-containing protein; afdb-uniprot50 \| AF-A0A7Y1LY14-F1-MODEL\_V4 \| 1.0 \| 5.519e-35 \| 1283 \| 0.438 \| 303 \| 155 \| 4 \| 4 \| 295 \| 3 \| 301 \| Baseplate assembly protein \| Baseplate assembly protein; afdb-uniprot50 \| AF-A0A1X4NS19-F1-MODEL\_V4 \| 1.0 \| 1.2e-34 \| 1282 \| 0.381 \| 299 \| 177 \| 3 \| 4 \| 300 \| 3 \| 295 \| Baseplate\_J domain-containing protein \| Baseplate\_J domain-containing protein; afdb-uniprot50 \| AF-A0A8A8LZL8-F1-MODEL\_V4 \| 1.0 \| 3.422e-35 \| 1277 \| 0.417 \| 302 \| 165 \| 3 \| 4 \| 298 \| 2 \| 299 \| Uncharacterized protein \| Uncharacterized protein; afdb-uniprot50 \| AF-A0A496L501-F1-MODEL\_V4 \| 1.0 \| 1.936e-34 \| 1276 \| 0.372 \| 295 \| 178 \| 1 \| 8 \| 295 \| 4 \| 298 \| Baseplate protein \| Baseplate protein; afdb-uniprot50 \| AF-A0A1V0B6E1-F1-MODEL\_V4 \| 1.0 \| 1.352e-34 \| 1275 \| 0.438 \| 301 \| 152 \| 4 \| 8 \| 296 \| 6 \| 301 \| Baseplate assembly protein \| Baseplate assembly protein; afdb-uniprot50 \| AF-A0A2J8GXT0-F1-MODEL\_V4 \| 1.0 \| 1.875e-33 \| 1269 \| 0.39 \| 287 \| 174 \| 1 \| 8 \| 294 \| 3 \| 288 \| Phage baseplate protein \| Phage baseplate protein; afdb-uniprot50 \| AF-A0A6S4USR9-F1-MODEL\_V4 \| 1.0 \| 1.065e-34 \| 1269 \| 0.442 \| 296 \| 156 \| 3 \| 8 \| 295 \| 4 \| 298 \| Baseplate assembly protein \| Baseplate assembly protein; afdb-uniprot50 \| AF-A0A6S4ZT36-F1-MODEL\_V4 \| 1.0 \| 5.035e-34 \| 1258 \| 0.428 \| 292 \| 157 \| 3 \| 8 \| 292 \| 4 \| 292 \| Baseplate assembly protein \| Baseplate assembly protein; afdb-uniprot50 \| AF-A0A853I3D0-F1-MODEL\_V4 \| 1.0 \| 6.049e-36 \| 1257 \| 0.465 \| 307 \| 148 \| 5 \| 5 \| 298 \| 2 \| 305 \| Baseplate J/gp47 family protein \| Baseplate J/gp47 family protein; afdb-uniprot50 \| AF-A0A345DE46-F1-MODEL\_V4 \| 1.0 \| 2.609e-34 \| 1253 \| 0.394 \| 307 \| 175 \| 5 \| 1 \| 298 \| 7 \| 311 \| Baseplate\_J domain-containing protein \| Baseplate\_J domain-containing protein; afdb-uniprot50 \| AF-A0A348FYH3-F1-MODEL\_V4 \| 1.0 \| 5.035e-34 \| 1250 \| 0.417 \| 297 \| 165 \| 4 \| 1 \| 296 \| 1 \| 290 \| Baseplate assembly protein \| Baseplate assembly protein; afdb-uniprot50 \| AF-A0A3L7AM87-F1-MODEL\_V4 \| 1.0 \| 8.121e-34 \| 1250 \| 0.381 \| 299 \| 170 \| 2 \| 9 \| 296 \| 7 \| 301 \| Baseplate assembly protein \| Baseplate assembly protein; afdb-uniprot50 \| AF-A0A4R0DR47-F1-MODEL\_V4 \| 1.0 \| 3.518e-34 \| 1250 \| 0.347 \| 322 \| 179 \| 3 \| 6 \| 296 \| 2 \| 323 \| Baseplate\_J domain-containing protein \| Baseplate\_J domain-containing protein; afdb-uniprot50 \| AF-A0A2C9D6M2-F1-MODEL\_V4 \| 1.0 \| 2.243e-33 \| 1248 \| 0.352 \| 360 \| 170 \| 7 \| 3 \| 300 \| 2 \| 360 \| Baseplate assembly protein \| Baseplate assembly protein; afdb-uniprot50 \| AF-A0A8A8M399-F1-MODEL\_V4 \| 1.0 \| 1.618e-34 \| 1247 \| 0.42 \| 300 \| 163 \| 3 \| 8 \| 300 \| 6 \| 301 \| Uncharacterized protein \| Uncharacterized protein; afdb-uniprot50 \| AF-A0A258L8U5-F1-MODEL\_V4 \| 1.0 \| 1.352e-34 \| 1247 \| 0.379 \| 306 \| 176 \| 3 \| 3 \| 298 \| 2 \| 303 \| Baseplate\_J domain-containing protein \| Baseplate\_J domain-containing protein; afdb-uniprot50 \| AF-A0A3E0X3Z4-F1-MODEL\_V4 \| 1.0 \| 4.209e-34 \| 1246 \| 0.385 \| 301 \| 180 \| 3 \| 1 \| 298 \| 1 \| 299 \| Baseplate\_J domain-containing protein \| Baseplate\_J domain-containing protein; afdb-uniprot50 \| AF-A0A1X7MCB0-F1-MODEL\_V4 \| 1.0 \| 5.013e-32 \| 1244 \| 0.511 \| 223 \| 109 \| 0 \| 77 \| 299 \| 1 \| 223 \| Phage baseplate assembly protein \| Phage baseplate assembly protein; afdb-uniprot50 \| AF-A0A212J3W2-F1-MODEL\_V4 \| 1.0 \| 4.743e-34 \| 1244 \| 0.402 \| 323 \| 161 \| 3 \| 5 \| 295 \| 4 \| 326 \| Baseplate assembly protein J \| Baseplate assembly protein J; afdb-uniprot50 \| AF-A0A2S2E5D8-F1-MODEL\_V4 \| 1.0 \| 8.121e-34 \| 1240 \| 0.409 \| 288 \| 154 \| 3 \| 8 \| 295 \| 3 \| 274 \| Baseplate protein J \| Baseplate protein J; afdb-uniprot50 \| AF-A0A7Y3Z3Q6-F1-MODEL\_V4 \| 1.0 \| 2.046e-32 \| 1236 \| 0.324 \| 290 \| 195 \| 1 \| 6 \| 295 \| 2 \| 290 \| Phage baseplate protein \| Phage baseplate protein; afdb-uniprot50 \| AF-A0A7V8DKV5-F1-MODEL\_V4 \| 1.0 \| 3.314e-34 \| 1235 \| 0.408 \| 299 \| 159 \| 5 \| 1 \| 299 \| 1 \| 281 \| Putative phage baseplate assembly protein \| Putative phage baseplate assembly protein; afdb-uniprot50 \| AF-A0A3N8B0H1-F1-MODEL\_V4 \| 1.0 \| 2.941e-34 \| 1232 \| 0.421 \| 304 \| 165 \| 3 \| 4 \| 300 \| 2 \| 301 \| Baseplate assembly protein \| Baseplate assembly protein; afdb-uniprot50 \| AF-A0A4Z1QXZ8-F1-MODEL\_V4 \| 1.0 \| 1.815e-32 \| 1223 \| 0.391 \| 294 \| 171 \| 4 \| 8 \| 298 \| 3 \| 291 \| Baseplate assembly protein \| Baseplate assembly protein; afdb-uniprot50 \| AF-A0A0K6GTQ4-F1-MODEL\_V4 \| 1.0 \| 1.095e-33 \| 1221 \| 0.417 \| 297 \| 159 \| 4 \| 8 \| 293 \| 3 \| 296 \| Phage-related baseplate assembly protein \| Phage-related baseplate assembly protein; afdb-uniprot50 \| AF-A0A3G9FXX2-F1-MODEL\_V4 \| 1.0 \| 3.84e-33 \| 1220 \| 0.361 \| 304 \| 178 \| 4 \| 1 \| 296 \| 1 \| 296 \| Baseplate assembly protein J \| Baseplate assembly protein J; afdb-uniprot50 \| AF-A0A484X2Q2-F1-MODEL\_V4 \| 1.0 \| 3.314e-34 \| 1220 \| 0.469 \| 296 \| 142 \| 4 \| 8 \| 294 \| 4 \| 293 \| Baseplate assembly protein \| Baseplate assembly protein; afdb-uniprot50 \| AF-A0A840X390-F1-MODEL\_V4 \| 1.0 \| 4.209e-34 \| 1219 \| 0.413 \| 334 \| 146 \| 4 \| 1 \| 299 \| 1 \| 319 \| Phage-related baseplate assembly protein \| Phage-related baseplate assembly protein; afdb-uniprot50 \| AF-Q7NAB1-F1-MODEL\_V4 \| 1.0 \| 1.031e-33 \| 1217 \| 0.429 \| 328 \| 136 \| 5 \| 8 \| 300 \| 4 \| 315 \| Baseplate\_J domain-containing protein \| Baseplate\_J domain-containing protein; afdb-uniprot50 \| AF-A0A318TPV5-F1-MODEL\_V4 \| 1.0 \| 1.823e-34 \| 1216 \| 0.379 \| 316 \| 174 \| 6 \| 1 \| 296 \| 3 \| 316 \| Phage-related baseplate assembly protein \| Phage-related baseplate assembly protein; afdb-uniprot50 \| AF-A0A2K2G607-F1-MODEL\_V4 \| 1.0 \| 6.979e-33 \| 1216 \| 0.352 \| 329 \| 174 \| 3 \| 4 \| 298 \| 7 \| 330 \| Baseplate\_J domain-containing protein \| Baseplate\_J domain-containing protein; afdb-uniprot50 \| AF-A0A258UWD9-F1-MODEL\_V4 \| 1.0 \| 1.162e-33 \| 1213 \| 0.408 \| 301 \| 159 \| 4 \| 1 \| 298 \| 1 \| 285 \| Baseplate\_J domain-containing protein \| Baseplate\_J domain-containing protein; afdb-uniprot50 \| AF-A0A6I4T7M6-F1-MODEL\_V4 \| 1.0 \| 6.979e-33 \| 1209 \| 0.4 \| 290 \| 167 \| 3 \| 8 \| 296 \| 6 \| 289 \| Baseplate assembly protein \| Baseplate assembly protein; afdb-uniprot50 \| AF-A0A7S8ACS8-F1-MODEL\_V4 \| 1.0 \| 3.617e-33 \| 1207 \| 0.365 \| 328 \| 166 \| 5 \| 8 \| 295 \| 2 \| 327 \| Phage tail protein \| Phage tail protein; afdb-uniprot50 \| AF-A0A510U3L2-F1-MODEL\_V4 \| 1.0 \| 1.06e-32 \| 1204 \| 0.372 \| 303 \| 165 \| 4 \| 4 \| 296 \| 2 \| 289 \| Baseplate assembly protein \| Baseplate assembly protein; afdb-uniprot50 \| AF-A0A5S9Q3R8-F1-MODEL\_V4 \| 1.0 \| 1.99e-33 \| 1203 \| 0.356 \| 362 \| 160 \| 6 \| 9 \| 300 \| 7 \| 365 \| Baseplate\_J domain-containing protein \| Baseplate\_J domain-containing protein; afdb-uniprot50 \| AF-A0A256CB14-F1-MODEL\_V4 \| 1.0 \| 1.766e-33 \| 1202 \| 0.314 \| 296 \| 200 \| 2 \| 8 \| 300 \| 2 \| 297 \| Baseplate\_J domain-containing protein \| Baseplate\_J domain-containing protein; afdb-uniprot50 \| AF-A0A1Q6U844-F1-MODEL\_V4 \| 1.0 \| 8.863e-33 \| 1199 \| 0.352 \| 372 \| 152 \| 5 \| 1 \| 299 \| 1 \| 356 \| Baseplate\_J domain-containing protein \| Baseplate\_J domain-containing protein; afdb-uniprot50 \| AF-A0A2A4XTY4-F1-MODEL\_V4 \| 1.0 \| 3.407e-33 \| 1198 \| 0.37 \| 294 \| 178 \| 3 \| 5 \| 296 \| 4 \| 292 \| Baseplate assembly protein \| Baseplate assembly protein; afdb-uniprot50 \| AF-A0A7X9WRR9-F1-MODEL\_V4 \| 1.0 \| 4.327e-33 \| 1195 \| 0.424 \| 299 \| 149 \| 5 \| 1 \| 295 \| 1 \| 280 \| Baseplate assembly protein \| Baseplate assembly protein; afdb-uniprot50 \| AF-A0A193QL68-F1-MODEL\_V4 \| 1.0 \| 1.31e-33 \| 1194 \| 0.4 \| 302 \| 169 \| 4 \| 6 \| 299 \| 81 \| 378 \| Baseplate J-like protein \| Baseplate J-like protein; afdb-uniprot50 \| AF-A0A495BKE9-F1-MODEL\_V4 \| 1.0 \| 1.567e-33 \| 1192 \| 0.379 \| 324 \| 162 \| 4 \| 8 \| 295 \| 2 \| 322 \| Phage-related baseplate assembly protein \| Phage-related baseplate assembly protein; afdb-uniprot50 \| AF-A0A6L2ZSF9-F1-MODEL\_V4 \| 1.0 \| 6.367e-32 \| 1192 \| 0.289 \| 491 \| 152 \| 3 \| 1 \| 294 \| 1 \| 491 \| Baseplate assembly protein \| Baseplate assembly protein; afdb-uniprot50 \| AF-A0A7H2V929-F1-MODEL\_V4 \| 1.0 \| 4.191e-32 \| 1191 \| 0.332 \| 298 \| 187 \| 3 \| 7 \| 296 \| 2 \| 295 \| Baseplate J/gp47 family protein \| Baseplate J/gp47 family protein; afdb-uniprot50 \| AF-A0A1A6FKN6-F1-MODEL\_V4 \| 1.0 \| 1.567e-33 \| 1191 \| 0.395 \| 301 \| 169 \| 3 \| 8 \| 299 \| 9 \| 305 \| Baseplate\_J domain-containing protein \| Baseplate\_J domain-containing protein; afdb-uniprot50 \| AF-A0A0N9WAW7-F1-MODEL\_V4 \| 1.0 \| 2.448e-32 \| 1189 \| 0.416 \| 300 \| 158 \| 4 \| 8 \| 295 \| 1 \| 295 \| Baseplate assembly protein \| Baseplate assembly protein; afdb-uniprot50 \| AF-A0A2T5UR67-F1-MODEL\_V4 \| 1.0 \| 2.37e-31 \| 1188 \| 0.37 \| 289 \| 166 \| 2 \| 9 \| 296 \| 8 \| 281 \| Phage-related baseplate assembly protein \| Phage-related baseplate assembly protein; afdb-uniprot50 \| AF-A0A8A8MKX7-F1-MODEL\_V4 \| 1.0 \| 8.863e-33 \| 1185 \| 0.399 \| 313 \| 166 \| 6 \| 1 \| 299 \| 1 \| 305 \| Baseplate J/gp47 family protein \| Baseplate J/gp47 family protein; afdb-uniprot50 \| AF-A0A5P9F1N3-F1-MODEL\_V4 \| 1.0 \| 3.023e-33 \| 1185 \| 0.37 \| 329 \| 155 \| 5 \| 4 \| 295 \| 3 \| 316 \| Baseplate J-like protein \| Baseplate J-like protein; afdb-uniprot50 \| AF-A0A1H2N8N7-F1-MODEL\_V4 \| 1.0 \| 8.584e-32 \| 1182 \| 0.351 \| 293 \| 174 \| 3 \| 8 \| 299 \| 3 \| 280 \| Phage-related baseplate assembly protein \| Phage-related baseplate assembly protein; afdb-uniprot50 \| AF-G8PUM5-F1-MODEL\_V4 \| 1.0 \| 7.175e-32 \| 1182 \| 0.338 \| 319 \| 185 \| 4 \| 3 \| 300 \| 2 \| 315 \| Baseplate assembly protein J, predicted \| Baseplate assembly protein J, predicted; afdb-uniprot50 \| AF-A0A0L0ETV6-F1-MODEL\_V4 \| 1.0 \| 4.191e-32 \| 1180 \| 0.347 \| 305 \| 174 \| 4 \| 5 \| 299 \| 4 \| 293 \| Baseplate\_J domain-containing protein \| Baseplate\_J domain-containing protein; afdb-uniprot50 \| AF-A0A5S9HNH2-F1-MODEL\_V4 \| 1.0 \| 4.876e-33 \| 1172 \| 0.387 \| 307 \| 162 \| 6 \| 8 \| 293 \| 2 \| 303 \| Baseplate assembly protein \| Baseplate assembly protein; afdb-uniprot50 \| AF-A0A2U1XZ04-F1-MODEL\_V4 \| 1.0 \| 1.611e-32 \| 1167 \| 0.41 \| 292 \| 156 \| 3 \| 7 \| 298 \| 2 \| 277 \| Baseplate\_J domain-containing protein \| Baseplate\_J domain-containing protein; afdb-uniprot50 \| AF-A0A6N7JGM8-F1-MODEL\_V4 \| 1.0 \| 5.035e-34 \| 1163 \| 0.411 \| 289 \| 155 \| 2 \| 8 \| 296 \| 3 \| 276 \| Baseplate assembly protein \| Baseplate assembly protein; afdb-uniprot50 \| AF-A0A2W5GCC1-F1-MODEL\_V4 \| 1.0 \| 5.65e-32 \| 1163 \| 0.371 \| 304 \| 172 \| 4 \| 8 \| 296 \| 9 \| 308 \| Baseplate assembly protein \| Baseplate assembly protein; afdb-uniprot50 \| AF-A0A611ETK6-F1-MODEL\_V4 \| 1.0 \| 1.085e-29 \| 1160 \| 0.461 \| 208 \| 112 \| 0 \| 92 \| 299 \| 1 \| 208 \| Baseplate assembly protein \| Baseplate assembly protein; afdb-uniprot50 \| AF-A0A3S0TV89-F1-MODEL\_V4 \| 1.0 \| 5.322e-32 \| 1160 \| 0.372 \| 290 \| 166 \| 3 \| 9 \| 298 \| 3 \| 276 \| Baseplate assembly protein J \| Baseplate assembly protein J; afdb-uniprot50 \| AF-A0A3A9I1H5-F1-MODEL\_V4 \| 1.0 \| 7.175e-32 \| 1160 \| 0.36 \| 372 \| 155 \| 5 \| 8 \| 296 \| 2 \| 373 \| Baseplate\_J domain-containing protein \| Baseplate\_J domain-containing protein; afdb-uniprot50 \| AF-A0A359KE54-F1-MODEL\_V4 \| 1.0 \| 3.823e-31 \| 1153 \| 0.376 \| 292 \| 164 \| 4 \| 7 \| 296 \| 4 \| 279 \| Baseplate assembly protein \| Baseplate assembly protein; afdb-uniprot50 \| AF-A0A1T4WV84-F1-MODEL\_V4 \| 1.0 \| 8.584e-32 \| 1153 \| 0.369 \| 330 \| 170 \| 5 \| 1 \| 296 \| 2 \| 327 \| Phage-related baseplate assembly protein \| Phage-related baseplate assembly protein; afdb-uniprot50 \| AF-A0A166WKK1-F1-MODEL\_V4 \| 1.0 \| 2.928e-32 \| 1145 \| 0.348 \| 310 \| 174 \| 7 \| 1 \| 296 \| 1 \| 296 \| Baseplate\_J domain-containing protein \| Baseplate\_J domain-containing protein; afdb-uniprot50 \| AF-A0A0T9RLX7-F1-MODEL\_V4 \| 1.0 \| 1.031e-33 \| 1144 \| 0.468 \| 301 \| 142 \| 6 \| 6 \| 295 \| 2 \| 295 \| Bacteriophage P2-like protein \| Bacteriophage P2-like protein; afdb-uniprot50 \| AF-A0A1S1U393-F1-MODEL\_V4 \| 1.0 \| 9.674e-32 \| 1137 \| 0.403 \| 300 \| 155 \| 5 \| 5 \| 295 \| 2 \| 286 \| Baseplate\_J domain-containing protein \| Baseplate\_J domain-containing protein; afdb-uniprot50 \| AF-A0A1S1TKX1-F1-MODEL\_V4 \| 1.0 \| 1.384e-31 \| 1137 \| 0.339 \| 327 \| 172 \| 5 \| 1 \| 298 \| 5 \| 316 \| Baseplate\_J domain-containing protein \| Baseplate\_J domain-containing protein; afdb-uniprot50 \| AF-A0A6N6MK70-F1-MODEL\_V4 \| 1.0 \| 1.758e-31 \| 1137 \| 0.325 \| 320 \| 186 \| 6 \| 5 \| 299 \| 4 \| 318 \| Baseplate J protein \| Baseplate J protein; afdb-uniprot50 \| AF-I3TTD3-F1-MODEL\_V4 \| 1.0 \| 6.759e-32 \| 1132 \| 0.4 \| 307 \| 157 \| 6 \| 1 \| 299 \| 1 \| 288 \| Phage baseplate J-like protein \| Phage baseplate J-like protein; afdb-uniprot50 \| AF-A0A2P9HMM4-F1-MODEL\_V4 \| 1.0 \| 9.674e-32 \| 1127 \| 0.368 \| 301 \| 173 \| 3 \| 7 \| 295 \| 2 \| 297 \| Baseplate assembly protein J \| Baseplate assembly protein J; afdb-uniprot50 \| AF-A0A4P5VKK1-F1-MODEL\_V4 \| 1.0 \| 1.703e-30 \| 1126 \| 0.419 \| 291 \| 153 \| 3 \| 8 \| 298 \| 4 \| 278 \| Baseplate assembly protein \| Baseplate assembly protein; afdb-uniprot50 \| AF-A0A7Z0MZG7-F1-MODEL\_V4 \| 1.0 \| 1.604e-30 \| 1124 \| 0.363 \| 305 \| 163 \| 6 \| 8 \| 296 \| 3 \| 292 \| Baseplate J/gp47 family protein \| Baseplate J/gp47 family protein; afdb-uniprot50 \| AF-A0A4Q6CUH1-F1-MODEL\_V4 \| 1.0 \| 3.931e-30 \| 1123 \| 0.44 \| 236 \| 130 \| 2 \| 61 \| 296 \| 2 \| 235 \| Baseplate\_J domain-containing protein \| Baseplate\_J domain-containing protein; afdb-uniprot50 \| AF-A0A833KQV3-F1-MODEL\_V4 \| 1.0 \| 2.824e-29 \| 1122 \| 0.418 \| 232 \| 131 \| 1 \| 68 \| 299 \| 2 \| 229 \| Baseplate assembly protein \| Baseplate assembly protein; afdb-uniprot50 \| AF-A0A5S9P369-F1-MODEL\_V4 \| 1.0 \| 5.472e-31 \| 1119 \| 0.402 \| 291 \| 156 \| 5 \| 7 \| 296 \| 4 \| 277 \| Baseplate\_J domain-containing protein \| Baseplate\_J domain-containing protein; afdb-uniprot50 \| AF-A0A1C3CVC7-F1-MODEL\_V4 \| 1.0 \| 4.059e-31 \| 1117 \| 0.303 \| 303 \| 200 \| 5 \| 8 \| 299 \| 4 \| 306 \| Phage baseplate protein \| Phage baseplate protein; afdb-uniprot50 \| AF-A0A7W6S3E7-F1-MODEL\_V4 \| 1.0 \| 1.751e-29 \| 1113 \| 0.484 \| 250 \| 129 \| 0 \| 3 \| 252 \| 2 \| 251 \| Phage-related baseplate assembly protein \| Phage-related baseplate assembly protein; afdb-uniprot50 \| AF-A0A5E8GZJ4-F1-MODEL\_V4 \| 1.0 \| 1.703e-30 \| 1112 \| 0.31 \| 328 \| 187 \| 6 \| 8 \| 296 \| 7 \| 334 \| Phage-related baseplate assembly protein \| Phage-related baseplate assembly protein; afdb-uniprot50 \| AF-A0A1I1WB92-F1-MODEL\_V4 \| 1.0 \| 5.472e-31 \| 1108 \| 0.42 \| 300 \| 159 \| 6 \| 5 \| 295 \| 4 \| 297 \| Phage-related baseplate assembly protein \| Phage-related baseplate assembly protein; afdb-uniprot50 \| AF-E2CN33-F1-MODEL\_V4 \| 1.0 \| 7.831e-31 \| 1108 \| 0.356 \| 306 \| 187 \| 4 \| 1 \| 300 \| 1 \| 302 \| Baseplate assembly protein J \| Baseplate assembly protein J; afdb-uniprot50 \| AF-V0X2S6-F1-MODEL\_V4 \| 1.0 \| 3.272e-28 \| 1105 \| 0.485 \| 202 \| 104 \| 0 \| 98 \| 299 \| 2 \| 203 \| Baseplate J-like protein \| Baseplate J-like protein; afdb-uniprot50 \| AF-A0A6I6R2A9-F1-MODEL\_V4 \| 1.0 \| 8.051e-30 \| 1104 \| 0.354 \| 293 \| 171 \| 4 \| 7 \| 296 \| 2 \| 279 \| Baseplate assembly protein \| Baseplate assembly protein; afdb-uniprot50 \| AF-A0A5S3XLF5-F1-MODEL\_V4 \| 1.0 \| 1.19e-30 \| 1097 \| 0.304 \| 309 \| 194 \| 4 \| 4 \| 299 \| 3 \| 303 \| Baseplate J protein \| Baseplate J protein; afdb-uniprot50 \| AF-F4BFR8-F1-MODEL\_V4 \| 1.0 \| 1.299e-29 \| 1093 \| 0.318 \| 295 \| 182 \| 4 \| 5 \| 295 \| 2 \| 281 \| Phage-related baseplate assembly protein \| Phage-related baseplate assembly protein; afdb-uniprot50 \| AF-A0A4R7SRP0-F1-MODEL\_V4 \| 1.0 \| 4.173e-30 \| 1092 \| 0.31 \| 296 \| 202 \| 2 \| 1 \| 296 \| 1 \| 294 \| Phage-related baseplate assembly protein \| Phage-related baseplate assembly protein; afdb-uniprot50 \| AF-A0A447M7C9-F1-MODEL\_V4 \| 1.0 \| 7.145e-30 \| 1085 \| 0.481 \| 245 \| 127 \| 0 \| 36 \| 280 \| 9 \| 253 \| Phage baseplate assembly protein J \| Phage baseplate assembly protein J; afdb-uniprot50 \| AF-A0A1H1G315-F1-MODEL\_V4 \| 1.0 \| 2.36e-29 \| 1083 \| 0.29 \| 300 \| 206 \| 4 \| 1 \| 300 \| 1 \| 293 \| Phage-related baseplate assembly protein \| Phage-related baseplate assembly protein; afdb-uniprot50 \| AF-A0A6S5X4A4-F1-MODEL\_V4 \| 1.0 \| 1.604e-30 \| 1079 \| 0.38 \| 302 \| 159 \| 8 \| 1 \| 298 \| 1 \| 278 \| Bacteriophage protein \| Bacteriophage protein; afdb-uniprot50 \| AF-A0A0E4BVF6-F1-MODEL\_V4 \| 1.0 \| 3.378e-29 \| 1075 \| 0.309 \| 304 \| 188 \| 6 \| 2 \| 300 \| 3 \| 289 \| Baseplate\_J domain-containing protein \| Baseplate\_J domain-containing protein; afdb-uniprot50 \| AF-A0A6L7TC79-F1-MODEL\_V4 \| 1.0 \| 5.626e-30 \| 1072 \| 0.475 \| 282 \| 116 \| 4 \| 31 \| 295 \| 1 \| 267 \| Baseplate assembly protein \| Baseplate assembly protein; afdb-uniprot50 \| AF-A0A2N3KSP3-F1-MODEL\_V4 \| 1.0 \| 4.29e-29 \| 1067 \| 0.337 \| 290 \| 182 \| 4 \| 13 \| 295 \| 1 \| 287 \| Baseplate\_J domain-containing protein \| Baseplate\_J domain-containing protein; afdb-uniprot50 \| AF-A0A7X3ZHS2-F1-MODEL\_V4 \| 1.0 \| 1.703e-30 \| 1064 \| 0.328 \| 283 \| 187 \| 2 \| 7 \| 287 \| 2 \| 283 \| Baseplate\_J domain-containing protein \| Baseplate\_J domain-containing protein; afdb-uniprot50 \| AF-A0A285M4J8-F1-MODEL\_V4 \| 1.0 \| 2.094e-29 \| 1062 \| 0.341 \| 293 \| 182 \| 5 \| 8 \| 300 \| 8 \| 289 \| Phage-related baseplate assembly protein \| Phage-related baseplate assembly protein; afdb-uniprot50 \| AF-A0A1D2QMV6-F1-MODEL\_V4 \| 1.0 \| 7.345e-29 \| 1061 \| 0.346 \| 297 \| 176 \| 5 \| 5 \| 300 \| 2 \| 281 \| Baseplate\_J domain-containing protein \| Baseplate\_J domain-containing protein; afdb-uniprot50 \| AF-A0A0M3V9B1-F1-MODEL\_V4 \| 1.0 \| 1.649e-29 \| 1061 \| 0.316 \| 313 \| 193 \| 5 \| 5 \| 299 \| 5 \| 314 \| Baseplate assembly protein \| Baseplate assembly protein; afdb-uniprot50 \| AF-A0A833LKM6-F1-MODEL\_V4 \| 1.0 \| 2.506e-29 \| 1059 \| 0.337 \| 299 \| 177 \| 4 \| 3 \| 296 \| 2 \| 284 \| Baseplate assembly protein \| Baseplate assembly protein; afdb-uniprot50 \| AF-A0A1S1NBP5-F1-MODEL\_V4 \| 1.0 \| 3.807e-29 \| 1053 \| 0.305 \| 304 \| 184 \| 4 \| 5 \| 296 \| 4 \| 292 \| Baseplate\_J domain-containing protein \| Baseplate\_J domain-containing protein; afdb-uniprot50 \| AF-A0A0P7JS55-F1-MODEL\_V4 \| 1.0 \| 2.998e-29 \| 1051 \| 0.303 \| 300 \| 188 \| 6 \| 3 \| 298 \| 2 \| 284 \| Baseplate\_J domain-containing protein \| Baseplate\_J domain-containing protein; afdb-uniprot50 \| AF-A0A0R3L0F6-F1-MODEL\_V4 \| 1.0 \| 3.931e-30 \| 1051 \| 0.344 \| 308 \| 185 \| 6 \| 1 \| 296 \| 1 \| 303 \| Baseplate\_J domain-containing protein \| Baseplate\_J domain-containing protein; afdb-uniprot50 \| AF-A0A1Y1QY37-F1-MODEL\_V4 \| 1.0 \| 8.313e-31 \| 1051 \| 0.334 \| 359 \| 153 \| 7 \| 8 \| 298 \| 5 \| 345 \| Baseplate\_J domain-containing protein \| Baseplate\_J domain-containing protein; afdb-uniprot50 \| AF-A0A847G9J0-F1-MODEL\_V4 \| 1.0 \| 1.696e-28 \| 1049 \| 0.319 \| 294 \| 179 \| 6 \| 8 \| 300 \| 3 \| 276 \| Phage tail protein \| Phage tail protein; afdb-uniprot50 \| AF-A0A0P6VRH8-F1-MODEL\_V4 \| 1.0 \| 8.825e-31 \| 1042 \| 0.322 \| 301 \| 189 \| 7 \| 8 \| 300 \| 9 \| 302 \| Baseplate\_J domain-containing protein \| Baseplate\_J domain-containing protein; afdb-uniprot50 \| AF-A0A4U9HA10-F1-MODEL\_V4 \| 1.0 \| 2.36e-29 \| 1036 \| 0.483 \| 271 \| 130 \| 3 \| 6 \| 267 \| 2 \| 271 \| Uncharacterized homolog of phage Mu protein gp47 \| Uncharacterized homolog of phage Mu protein gp47; afdb-uniprot50 \| AF-A0A6M4Y9L2-F1-MODEL\_V4 \| 1.0 \| 2.286e-28 \| 1034 \| 0.592 \| 228 \| 93 \| 0 \| 1 \| 228 \| 1 \| 228 \| Baseplate assembly protein \| Baseplate assembly protein; afdb-uniprot50 \| AF-A0A2A5BN42-F1-MODEL\_V4 \| 1.0 \| 3.807e-29 \| 1031 \| 0.294 \| 323 \| 187 \| 5 \| 3 \| 299 \| 7 \| 314 \| Baseplate\_J domain-containing protein \| Baseplate\_J domain-containing protein; afdb-uniprot50 \| AF-A0A5S3R796-F1-MODEL\_V4 \| 1.0 \| 4.041e-29 \| 1026 \| 0.311 \| 321 \| 180 \| 9 \| 1 \| 300 \| 1 \| 301 \| Baseplate J protein \| Baseplate J protein; afdb-uniprot50 \| AF-A0A2V4KD55-F1-MODEL\_V4 \| 1.0 \| 4.059e-31 \| 1024 \| 0.319 \| 322 \| 181 \| 4 \| 12 \| 296 \| 10 \| 330 \| Baseplate J protein \| Baseplate J protein; afdb-uniprot50 \| AF-A0A4P9VEN9-F1-MODEL\_V4 \| 1.0 \| 1.185e-28 \| 1016 \| 0.363 \| 297 \| 164 \| 7 \| 5 \| 299 \| 4 \| 277 \| Baseplate\_J domain-containing protein \| Baseplate\_J domain-containing protein; afdb-uniprot50 \| AF-A0A8B2NS25-F1-MODEL\_V4 \| 1.0 \| 5.784e-29 \| 1016 \| 0.321 \| 302 \| 177 \| 5 \| 8 \| 296 \| 8 \| 294 \| Baseplate assembly protein \| Baseplate assembly protein; afdb-uniprot50 \| AF-A0A090SXL0-F1-MODEL\_V4 \| 1.0 \| 6.14e-29 \| 1016 \| 0.324 \| 296 \| 184 \| 3 \| 5 \| 299 \| 2 \| 282 \| Phage-related baseplate assembly protein \| Phage-related baseplate assembly protein; afdb-uniprot50 \| AF-A0A4R6U4A6-F1-MODEL\_V4 \| 1.0 \| 2.085e-27 \| 1014 \| 0.332 \| 289 \| 173 \| 5 \| 13 \| 300 \| 1 \| 270 \| Phage-related baseplate assembly protein \| Phage-related baseplate assembly protein; afdb-uniprot50 \| AF-A0A212LCZ5-F1-MODEL\_V4 \| 1.0 \| 4.29e-29 \| 1014 \| 0.311 \| 305 \| 187 \| 7 \| 8 \| 296 \| 3 \| 300 \| Putative phage-related baseplate assembly protein \| Putative phage-related baseplate assembly protein; afdb-uniprot50 \| AF-A0A085ASA5-F1-MODEL\_V4 \| 1.0 \| 4.43e-30 \| 1012 \| 0.37 \| 294 \| 158 \| 5 \| 31 \| 300 \| 2 \| 292 \| Baseplate assembly protein J \| Baseplate assembly protein J; afdb-uniprot50 \| AF-A0A838CFD0-F1-MODEL\_V4 \| 1.0 \| 2.416e-26 \| 1008 \| 0.382 \| 235 \| 143 \| 1 \| 68 \| 300 \| 2 \| 236 \| Baseplate J protein \| Baseplate J protein; afdb-uniprot50 \| AF-A0A7U5UIH5-F1-MODEL\_V4 \| 1.0 \| 3.364e-27 \| 1008 \| 0.446 \| 262 \| 138 \| 1 \| 8 \| 262 \| 4 \| 265 \| Baseplate assembly protein \| Baseplate assembly protein; afdb-uniprot50 \| AF-A0A1H3ZWS2-F1-MODEL\_V4 \| 1.0 \| 4.041e-29 \| 1008 \| 0.31 \| 328 \| 177 \| 5 \| 6 \| 299 \| 2 \| 314 \| Phage-related baseplate assembly protein \| Phage-related baseplate assembly protein; afdb-uniprot50 \| AF-A0A2E3N1J6-F1-MODEL\_V4 \| 1.0 \| 1.696e-28 \| 1007 \| 0.33 \| 342 \| 163 \| 4 \| 7 \| 298 \| 2 \| 327 \| Baseplate\_J domain-containing protein \| Baseplate\_J domain-containing protein; afdb-uniprot50 \| AF-A0A5A9EMT3-F1-MODEL\_V4 \| 1.0 \| 6.919e-29 \| 1004 \| 0.375 \| 288 \| 162 \| 4 \| 12 \| 298 \| 8 \| 278 \| Baseplate assembly protein J \| Baseplate assembly protein J; afdb-uniprot50 \| AF-A0A142JGU0-F1-MODEL\_V4 \| 1.0 \| 1.973e-29 \| 1004 \| 0.346 \| 323 \| 164 \| 6 \| 8 \| 298 \| 4 \| 311 \| Baseplate\_J domain-containing protein \| Baseplate\_J domain-containing protein; afdb-uniprot50 \| AF-A0A318KRB1-F1-MODEL\_V4 \| 1.0 \| 1.505e-28 \| 1001 \| 0.351 \| 304 \| 177 \| 8 \| 9 \| 296 \| 5 \| 304 \| Phage-related baseplate assembly protein \| Phage-related baseplate assembly protein; afdb-uniprot50 \| AF-A0A554XFX2-F1-MODEL\_V4 \| 1.0 \| 6.14e-29 \| 993 \| 0.331 \| 329 \| 169 \| 7 \| 6 \| 299 \| 2 \| 314 \| Baseplate J-like protein \| Baseplate J-like protein; afdb-uniprot50 \| AF-A0A2V3UAW4-F1-MODEL\_V4 \| 1.0 \| 7.345e-29 \| 990 \| 0.361 \| 310 \| 174 \| 9 \| 4 \| 296 \| 3 \| 305 \| Phage-related baseplate assembly protein \| Phage-related baseplate assembly protein; afdb-uniprot50 \| AF-A0A1N7LRY2-F1-MODEL\_V4 \| 1.0 \| 1.218e-27 \| 988 \| 0.321 \| 302 \| 188 \| 7 \| 8 \| 300 \| 1 \| 294 \| Phage-related baseplate assembly protein \| Phage-related baseplate assembly protein; afdb-uniprot50 \| AF-A0A1S0V677-F1-MODEL\_V4 \| 1.0 \| 6.087e-25 \| 987 \| 0.491 \| 185 \| 94 \| 0 \| 115 \| 299 \| 1 \| 185 \| Baseplate assembly protein \| Baseplate assembly protein; afdb-uniprot50 \| AF-A0A7W8XVJ7-F1-MODEL\_V4 \| 1.0 \| 3.082e-28 \| 986 \| 0.313 \| 300 \| 189 \| 8 \| 6 \| 298 \| 7 \| 296 \| Phage-related baseplate assembly protein \| Phage-related baseplate assembly protein; afdb-uniprot50 \| AF-A0A433LI32-F1-MODEL\_V4 \| 1.0 \| 1.051e-28 \| 984 \| 0.269 \| 338 \| 203 \| 8 \| 1 \| 296 \| 1 \| 336 \| Baseplate\_J domain-containing protein \| Baseplate\_J domain-containing protein; afdb-uniprot50 \| AF-A0A4D7AUY0-F1-MODEL\_V4 \| 1.0 \| 1.417e-28 \| 983 \| 0.32 \| 303 \| 195 \| 6 \| 1 \| 296 \| 4 \| 302 \| Baseplate\_J domain-containing protein \| Baseplate\_J domain-containing protein; afdb-uniprot50 \| AF-A0A0F4QK12-F1-MODEL\_V4 \| 1.0 \| 1.751e-29 \| 981 \| 0.37 \| 294 \| 160 \| 7 \| 5 \| 296 \| 2 \| 272 \| Baseplate protein \| Baseplate protein; afdb-uniprot50 \| AF-A0A1Y6ITR8-F1-MODEL\_V4 \| 1.0 \| 7.552e-28 \| 980 \| 0.337 \| 302 \| 172 \| 5 \| 8 \| 300 \| 4 \| 286 \| Baseplate J-like protein \| Baseplate J-like protein; afdb-uniprot50 \| AF-A0A317PF16-F1-MODEL\_V4 \| 1.0 \| 6.313e-28 \| 979 \| 0.28 \| 314 \| 202 \| 8 \| 1 \| 298 \| 1 \| 306 \| Phage-related baseplate assembly protein \| Phage-related baseplate assembly protein; afdb-uniprot50 \| AF-A0A6N8K8T8-F1-MODEL\_V4 \| 1.0 \| 3.571e-27 \| 978 \| 0.465 \| 247 \| 128 \| 1 \| 1 \| 243 \| 1 \| 247 \| Baseplate assembly protein \| Baseplate assembly protein; afdb-uniprot50 \| AF-A0A1C3EBK2-F1-MODEL\_V4 \| 1.0 \| 1.051e-28 \| 976 \| 0.341 \| 302 \| 168 \| 9 \| 8 \| 295 \| 4 \| 288 \| Baseplate\_J domain-containing protein \| Baseplate\_J domain-containing protein; afdb-uniprot50 \| AF-A0A839IVB1-F1-MODEL\_V4 \| 1.0 \| 7.345e-29 \| 976 \| 0.304 \| 322 \| 186 \| 4 \| 12 \| 296 \| 7 \| 327 \| Baseplate J/gp47 family protein \| Baseplate J/gp47 family protein; afdb-uniprot50 \| AF-A0A1N6M6H6-F1-MODEL\_V4 \| 1.0 \| 2.735e-28 \| 976 \| 0.257 \| 326 \| 203 \| 4 \| 12 \| 298 \| 7 \| 332 \| Baseplate J-like protein \| Baseplate J-like protein; afdb-uniprot50 \| AF-A0A5P0JGH1-F1-MODEL\_V4 \| 1.0 \| 3.671e-26 \| 973 \| 0.522 \| 222 \| 99 \| 1 \| 8 \| 222 \| 4 \| 225 \| Baseplate assembly protein \| Baseplate assembly protein; afdb-uniprot50 \| AF-A0A1W2EJW9-F1-MODEL\_V4 \| 1.0 \| 1.218e-27 \| 972 \| 0.247 \| 315 \| 212 \| 9 \| 1 \| 298 \| 1 \| 307 \| Phage-related baseplate assembly protein \| Phage-related baseplate assembly protein; afdb-uniprot50 \| AF-A0A371WU65-F1-MODEL\_V4 \| 1.0 \| 1.121e-30 \| 962 \| 0.336 \| 306 \| 192 \| 2 \| 1 \| 296 \| 2 \| 306 \| Baseplate\_J domain-containing protein \| Baseplate\_J domain-containing protein; afdb-uniprot50 \| AF-B0UK53-F1-MODEL\_V4 \| 1.0 \| 2.812e-27 \| 961 \| 0.349 \| 335 \| 165 \| 12 \| 5 \| 296 \| 4 \| 328 \| Baseplate J family protein \| Baseplate J family protein; afdb-uniprot50 \| AF-A0A1R1MK94-F1-MODEL\_V4 \| 1.0 \| 1.147e-27 \| 956 \| 0.279 \| 379 \| 183 \| 10 \| 8 \| 300 \| 2 \| 376 \| Baseplate\_J domain-containing protein \| Baseplate\_J domain-containing protein; afdb-uniprot50 \| AF-A0A7X3VHX5-F1-MODEL\_V4 \| 1.0 \| 5.734e-25 \| 955 \| 0.518 \| 216 \| 100 \| 1 \| 4 \| 219 \| 2 \| 213 \| Baseplate assembly protein \| Baseplate assembly protein; afdb-uniprot50 \| AF-A0A2D2D5N1-F1-MODEL\_V4 \| 1.0 \| 1.792e-26 \| 955 \| 0.39 \| 238 \| 137 \| 2 \| 66 \| 299 \| 2 \| 235 \| Baseplate assembly protein \| Baseplate assembly protein; afdb-uniprot50 \| AF-A0A450W779-F1-MODEL\_V4 \| 1.0 \| 2.649e-27 \| 954 \| 0.311 \| 295 \| 181 \| 6 \| 6 \| 298 \| 2 \| 276 \| Phage-related baseplate assembly protein \| Phage-related baseplate assembly protein; afdb-uniprot50 \| AF-A0A7T8CNP9-F1-MODEL\_V4 \| 1.0 \| 2.812e-27 \| 951 \| 0.308 \| 318 \| 186 \| 7 \| 8 \| 296 \| 2 \| 314 \| Baseplate\_J domain-containing protein \| Baseplate\_J domain-containing protein; afdb-uniprot50 \| AF-Q3RBT0-F1-MODEL\_V4 \| 1.0 \| 4.137e-26 \| 943 \| 0.517 \| 224 \| 106 \| 1 \| 1 \| 224 \| 1 \| 222 \| Baseplate J-like protein \| Baseplate J-like protein; afdb-uniprot50 \| AF-A0A1G8HUM0-F1-MODEL\_V4 \| 1.0 \| 1.018e-27 \| 942 \| 0.251 \| 330 \| 204 \| 6 \| 10 \| 296 \| 7 \| 336 \| Phage-related baseplate assembly protein \| Phage-related baseplate assembly protein; afdb-uniprot50 \| AF-A0A7K1PHL9-F1-MODEL\_V4 \| 1.0 \| 8.242e-27 \| 937 \| 0.276 \| 300 \| 193 \| 4 \| 8 \| 299 \| 2 \| 285 \| Baseplate J protein \| Baseplate J protein; afdb-uniprot50 \| AF-A0A545T5R2-F1-MODEL\_V4 \| 1.0 \| 7.552e-28 \| 937 \| 0.317 \| 318 \| 183 \| 6 \| 4 \| 295 \| 3 \| 312 \| Baseplate\_J domain-containing protein \| Baseplate\_J domain-containing protein; afdb-uniprot50 \| AF-A0A383SE97-F1-MODEL\_V4 \| 1.0 \| 8.996e-26 \| 935 \| 0.518 \| 239 \| 110 \| 3 \| 1 \| 239 \| 1 \| 234 \| Phage-related baseplate assembly protein \| Phage-related baseplate assembly protein; afdb-uniprot50 \| AF-A0A2N7P721-F1-MODEL\_V4 \| 1.0 \| 1.018e-27 \| 935 \| 0.343 \| 300 \| 170 \| 8 \| 1 \| 296 \| 1 \| 277 \| Baseplate\_J domain-containing protein \| Baseplate\_J domain-containing protein; afdb-uniprot50 \| AF-A0A6B8KJI2-F1-MODEL\_V4 \| 1.0 \| 1.965e-27 \| 930 \| 0.33 \| 324 \| 169 \| 11 \| 8 \| 298 \| 7 \| 315 \| Baseplate\_J domain-containing protein \| Baseplate\_J domain-containing protein; afdb-uniprot50 \| AF-A0A2N6FYW7-F1-MODEL\_V4 \| 1.0 \| 1.965e-27 \| 930 \| 0.237 \| 366 \| 196 \| 6 \| 8 \| 295 \| 2 \| 362 \| Baseplate assembly protein \| Baseplate assembly protein; afdb-uniprot50 \| AF-A0A7W2BN55-F1-MODEL\_V4 \| 1.0 \| 7.764e-27 \| 929 \| 0.309 \| 320 \| 189 \| 7 \| 4 \| 296 \| 3 \| 317 \| Baseplate J/gp47 family protein \| Baseplate J/gp47 family protein; afdb-uniprot50 \| AF-A0A853IJD9-F1-MODEL\_V4 \| 1.0 \| 1.451e-25 \| 927 \| 0.403 \| 208 \| 120 \| 2 \| 92 \| 295 \| 20 \| 227 \| Baseplate J/gp47 family protein \| Baseplate J/gp47 family protein; afdb-uniprot50 \| AF-A0A7U2YWJ9-F1-MODEL\_V4 \| 1.0 \| 1.252e-26 \| 927 \| 0.357 \| 277 \| 160 \| 4 \| 10 \| 275 \| 6 \| 275 \| Baseplate protein \| Baseplate protein; afdb-uniprot50 \| AF-A0A1D2QRG1-F1-MODEL\_V4 \| 1.0 \| 1.142e-25 \| 921 \| 0.266 \| 293 \| 192 \| 7 \| 8 \| 299 \| 2 \| 272 \| Baseplate\_J domain-containing protein \| Baseplate\_J domain-containing protein; afdb-uniprot50 \| AF-A0A651FVY8-F1-MODEL\_V4 \| 1.0 \| 1.903e-26 \| 921 \| 0.298 \| 302 \| 190 \| 5 \| 3 \| 298 \| 2 \| 287 \| Baseplate\_J domain-containing protein \| Baseplate\_J domain-containing protein; afdb-uniprot50 \| AF-A0A379CNV1-F1-MODEL\_V4 \| 1.0 \| 2.812e-27 \| 920 \| 0.262 \| 331 \| 201 \| 6 \| 7 \| 296 \| 5 \| 333 \| Baseplate J-like protein \| Baseplate J-like protein; afdb-uniprot50 \| AF-A0A2S6N2W1-F1-MODEL\_V4 \| 1.0 \| 5.759e-27 \| 919 \| 0.279 \| 311 \| 194 \| 9 \| 1 \| 296 \| 109 \| 404 \| Baseplate\_J domain-containing protein \| Baseplate\_J domain-containing protein; afdb-uniprot50 \| AF-A0A524RVY7-F1-MODEL\_V4 \| 1.0 \| 1.293e-27 \| 918 \| 0.331 \| 293 \| 174 \| 5 \| 4 \| 294 \| 2 \| 274 \| Baseplate\_J domain-containing protein \| Baseplate\_J domain-containing protein; afdb-uniprot50 \| AF-A0A0Q8AUB0-F1-MODEL\_V4 \| 1.0 \| 3.671e-26 \| 916 \| 0.31 \| 296 \| 181 \| 6 \| 8 \| 296 \| 3 \| 282 \| Baseplate\_J domain-containing protein \| Baseplate\_J domain-containing protein; afdb-uniprot50 \| AF-A0A0N0KIE2-F1-MODEL\_V4 \| 1.0 \| 3.169e-27 \| 916 \| 0.273 \| 318 \| 201 \| 9 \| 1 \| 296 \| 1 \| 310 \| Baseplate\_J domain-containing protein \| Baseplate\_J domain-containing protein; afdb-uniprot50 \| AF-A0A6L2YE65-F1-MODEL\_V4 \| 1.0 \| 2.723e-26 \| 914 \| 0.361 \| 271 \| 155 \| 4 \| 10 \| 269 \| 6 \| 269 \| Baseplate protein \| Baseplate protein; afdb-uniprot50 \| AF-A0A0J9BHG4-F1-MODEL\_V4 \| 1.0 \| 1.445e-23 \| 911 \| 0.462 \| 173 \| 93 \| 0 \| 127 \| 299 \| 3 \| 175 \| Baseplate\_J domain-containing protein \| Baseplate\_J domain-containing protein; afdb-uniprot50 \| AF-A0A5P9BN28-F1-MODEL\_V4 \| 1.0 \| 2.276e-26 \| 911 \| 0.292 \| 297 \| 188 \| 6 \| 8 \| 298 \| 2 \| 282 \| Baseplate J-like protein \| Baseplate J-like protein; afdb-uniprot50 \| AF-A0A654BHD8-F1-MODEL\_V4 \| 1.0 \| 4.773e-23 \| 910 \| 0.472 \| 165 \| 87 \| 0 \| 135 \| 299 \| 2 \| 166 \| Phage-related baseplate assembly protein \| Phage-related baseplate assembly protein; afdb-uniprot50 \| AF-A0A2W6Z6F0-F1-MODEL\_V4 \| 1.0 \| 2.565e-26 \| 909 \| 0.287 \| 351 \| 186 \| 11 \| 4 \| 296 \| 3 \| 347 \| Baseplate\_J domain-containing protein \| Baseplate\_J domain-containing protein; afdb-uniprot50 \| AF-A0A1S9ZY08-F1-MODEL\_V4 \| 1.0 \| 5.402e-25 \| 905 \| 0.263 \| 292 \| 193 \| 6 \| 12 \| 300 \| 3 \| 275 \| Baseplate\_J domain-containing protein \| Baseplate\_J domain-containing protein; afdb-uniprot50 \| AF-A0A381GJS6-F1-MODEL\_V4 \| 1.0 \| 1.785e-24 \| 903 \| 0.473 \| 226 \| 112 \| 1 \| 8 \| 226 \| 4 \| 229 \| Phage baseplate assembly protein J \| Phage baseplate assembly protein J; afdb-uniprot50 \| AF-A0A377VBV5-F1-MODEL\_V4 \| 1.0 \| 2.34e-25 \| 903 \| 0.46 \| 250 \| 128 \| 1 \| 6 \| 248 \| 2 \| 251 \| Baseplate assembly protein J \| Baseplate assembly protein J; afdb-uniprot50 \| AF-Q4EB01-F1-MODEL\_V4 \| 1.0 \| 1.635e-25 \| 902 \| 0.378 \| 251 \| 140 \| 3 \| 42 \| 292 \| 1 \| 235 \| Baseplate J-like protein \| Baseplate J-like protein; afdb-uniprot50 \| AF-A0A3D1NTG9-F1-MODEL\_V4 \| 1.0 \| 1.047e-26 \| 894 \| 0.243 \| 370 \| 191 \| 9 \| 14 \| 300 \| 6 \| 369 \| Baseplate protein \| Baseplate protein; afdb-uniprot50 \| AF-A0A645F3X1-F1-MODEL\_V4 \| 1.0 \| 8.957e-24 \| 892 \| 0.317 \| 208 \| 139 \| 2 \| 94 \| 298 \| 7 \| 214 \| Baseplate\_J domain-containing protein \| Baseplate\_J domain-containing protein; afdb-uniprot50 \| AF-A0A327JML8-F1-MODEL\_V4 \| 1.0 \| 4.662e-26 \| 892 \| 0.262 \| 308 \| 211 \| 8 \| 1 \| 300 \| 1 \| 300 \| Baseplate\_J domain-containing protein \| Baseplate\_J domain-containing protein; afdb-uniprot50 \| AF-A0A0S9F1A4-F1-MODEL\_V4 \| 1.0 \| 3.349e-25 \| 891 \| 0.322 \| 298 \| 179 \| 5 \| 6 \| 298 \| 2 \| 281 \| Baseplate J protein \| Baseplate J protein; afdb-uniprot50 \| AF-F2J637-F1-MODEL\_V4 \| 1.0 \| 1.014e-25 \| 889 \| 0.304 \| 292 \| 181 \| 5 \| 13 \| 298 \| 6 \| 281 \| Phage protein gp26 \| Phage protein gp26; afdb-uniprot50 \| AF-A0A0M4LII5-F1-MODEL\_V4 \| 1.0 \| 8.474e-26 \| 888 \| 0.294 \| 289 \| 186 \| 5 \| 11 \| 298 \| 3 \| 274 \| Baseplate assembly protein J \| Baseplate assembly protein J; afdb-uniprot50 \| AF-A0A0L6KDG2-F1-MODEL\_V4 \| 1.0 \| 1.956e-25 \| 884 \| 0.257 \| 315 \| 202 \| 11 \| 8 \| 298 \| 6 \| 312 \| Baseplate\_J domain-containing protein \| Baseplate\_J domain-containing protein; afdb-uniprot50 \| AF-A0A1M7YYV2-F1-MODEL\_V4 \| 1.0 \| 1.498e-26 \| 881 \| 0.304 \| 296 \| 185 \| 6 \| 6 \| 298 \| 3 \| 280 \| Baseplate J-like protein \| Baseplate J-like protein; afdb-uniprot50 \| AF-E2CI68-F1-MODEL\_V4 \| 1.0 \| 1.014e-25 \| 881 \| 0.309 \| 297 \| 183 \| 5 \| 8 \| 298 \| 2 \| 282 \| Baseplate assembly protein J \| Baseplate assembly protein J; afdb-uniprot50 \| AF-A0A5M8P9P5-F1-MODEL\_V4 \| 1.0 \| 6.286e-26 \| 880 \| 0.279 \| 293 \| 189 \| 5 \| 12 \| 298 \| 4 \| 280 \| Baseplate assembly protein \| Baseplate assembly protein; afdb-uniprot50 \| AF-A0A2W5VFF5-F1-MODEL\_V4 \| 1.0 \| 7.983e-26 \| 880 \| 0.273 \| 326 \| 187 \| 7 \| 5 \| 296 \| 3 \| 312 \| Uncharacterized protein \| Uncharacterized protein; afdb-uniprot50 \| AF-G2HXA0-F1-MODEL\_V4 \| 1.0 \| 7.983e-26 \| 879 \| 0.205 \| 370 \| 209 \| 8 \| 8 \| 296 \| 2 \| 367 \| Phage baseplate assembly protein \| Phage baseplate assembly protein; afdb-uniprot50 \| AF-A0A1I1V0V6-F1-MODEL\_V4 \| 1.0 \| 1.843e-25 \| 877 \| 0.25 \| 316 \| 204 \| 9 \| 1 \| 296 \| 1 \| 303 \| Phage-related baseplate assembly protein \| Phage-related baseplate assembly protein; afdb-uniprot50 \| AF-A0A5B9DJ78-F1-MODEL\_V4 \| 1.0 \| 1.213e-25 \| 874 \| 0.299 \| 297 \| 186 \| 6 \| 8 \| 298 \| 5 \| 285 \| Baseplate\_J domain-containing protein \| Baseplate\_J domain-containing protein; afdb-uniprot50 \| AF-A0A7X4WDI4-F1-MODEL\_V4 \| 1.0 \| 8.75e-27 \| 870 \| 0.25 \| 328 \| 198 \| 8 \| 13 \| 296 \| 1 \| 324 \| Baseplate J protein \| Baseplate J protein; afdb-uniprot50 \| AF-A0A4R6M7P6-F1-MODEL\_V4 \| 1.0 \| 1.635e-25 \| 869 \| 0.29 \| 296 \| 183 \| 6 \| 1 \| 296 \| 1 \| 269 \| Phage-related baseplate assembly protein \| Phage-related baseplate assembly protein; afdb-uniprot50 \| AF-A0A1G3UAG2-F1-MODEL\_V4 \| 1.0 \| 8.511e-28 \| 868 \| 0.264 \| 367 \| 184 \| 9 \| 8 \| 296 \| 1 \| 359 \| Baseplate\_J domain-containing protein \| Baseplate\_J domain-containing protein; afdb-uniprot50 \| AF-A0A0F4NJK8-F1-MODEL\_V4 \| 1.0 \| 1.213e-25 \| 867 \| 0.283 \| 293 \| 188 \| 5 \| 8 \| 300 \| 2 \| 272 \| Baseplate\_J domain-containing protein \| Baseplate\_J domain-containing protein; afdb-uniprot50 \| AF-A0A4Q7FYT9-F1-MODEL\_V4 \| 1.0 \| 4.392e-26 \| 865 \| 0.259 \| 316 \| 202 \| 8 \| 1 \| 299 \| 1 \| 301 \| Baseplate\_J domain-containing protein \| Baseplate\_J domain-containing protein; afdb-uniprot50 \| AF-A0A1S7MRG2-F1-MODEL\_V4 \| 1.0 \| 8.207e-25 \| 861 \| 0.273 \| 318 \| 201 \| 10 \| 4 \| 298 \| 2 \| 312 \| Phage-related baseplate assembly protein \| Phage-related baseplate assembly protein; afdb-uniprot50 \| AF-A0A376BTH5-F1-MODEL\_V4 \| 1.0 \| 2.8e-25 \| 861 \| 0.237 \| 383 \| 197 \| 11 \| 8 \| 300 \| 1 \| 378 \| Uncharacterized homolog of phage Mu protein gp47 \| Uncharacterized homolog of phage Mu protein gp47; afdb-uniprot50 \| AF-A0A136Q9J0-F1-MODEL\_V4 \| 1.0 \| 4.007e-25 \| 857 \| 0.392 \| 285 \| 137 \| 5 \| 8 \| 257 \| 2 \| 285 \| Baseplate assembly protein \| Baseplate assembly protein; afdb-uniprot50 \| AF-A0A1I5RPD0-F1-MODEL\_V4 \| 1.0 \| 1.843e-25 \| 856 \| 0.248 \| 370 \| 188 \| 11 \| 8 \| 295 \| 2 \| 363 \| Baseplate J-like protein \| Baseplate J-like protein; afdb-uniprot50 \| AF-A0A239CA17-F1-MODEL\_V4 \| 1.0 \| 4.007e-25 \| 854 \| 0.248 \| 382 \| 198 \| 10 \| 1 \| 298 \| 1 \| 377 \| Phage-related baseplate assembly protein \| Phage-related baseplate assembly protein; afdb-uniprot50 \| AF-A0A833KP28-F1-MODEL\_V4 \| 1.0 \| 5.71e-23 \| 853 \| 0.438 \| 203 \| 110 \| 1 \| 68 \| 270 \| 2 \| 200 \| Baseplate assembly protein \| Baseplate assembly protein; afdb-uniprot50 \| AF-U3A3E9-F1-MODEL\_V4 \| 1.0 \| 5.578e-26 \| 851 \| 0.238 \| 331 \| 202 \| 7 \| 12 \| 296 \| 3 \| 329 \| Putative phage baseplate assembly protein \| Putative phage baseplate assembly protein; afdb-uniprot50 \| AF-A0A857ED91-F1-MODEL\_V4 \| 1.0 \| 3.54e-23 \| 849 \| 0.442 \| 199 \| 105 \| 1 \| 104 \| 296 \| 2 \| 200 \| Uncharacterized protein \| Uncharacterized protein; afdb-uniprot50 \| AF-A0A2D3VSP7-F1-MODEL\_V4 \| 1.0 \| 3.671e-26 \| 847 \| 0.272 \| 370 \| 182 \| 11 \| 8 \| 296 \| 1 \| 364 \| Baseplate\_J domain-containing protein \| Baseplate\_J domain-containing protein; afdb-uniprot50 \| AF-B4RLZ3-F1-MODEL\_V4 \| 1.0 \| 5.896e-24 \| 845 \| 0.369 \| 271 \| 157 \| 5 \| 1 \| 264 \| 1 \| 264 \| Putative baseplate protein, putative phage associated protein \| Putative baseplate protein, putative phage associated protein; afdb-uniprot50 \| AF-C8N767-F1-MODEL\_V4 \| 1.0 \| 8.242e-27 \| 845 \| 0.314 \| 302 \| 177 \| 6 \| 4 \| 298 \| 2 \| 280 \| Baseplate J-like protein \| Baseplate J-like protein; afdb-uniprot50 \| AF-A0A0F6A5M1-F1-MODEL\_V4 \| 1.0 \| 1.736e-25 \| 844 \| 0.309 \| 297 \| 182 \| 7 \| 1 \| 294 \| 1 \| 277 \| Baseplate\_J domain-containing protein \| Baseplate\_J domain-containing protein; afdb-uniprot50 \| AF-A0A841GJI4-F1-MODEL\_V4 \| 1.0 \| 1.681e-24 \| 842 \| 0.286 \| 342 \| 188 \| 13 \| 5 \| 300 \| 4 \| 335 \| Phage-related baseplate assembly protein \| Phage-related baseplate assembly protein; afdb-uniprot50 \| AF-A0A659JW46-F1-MODEL\_V4 \| 1.0 \| 3.897e-26 \| 842 \| 0.257 \| 377 \| 183 \| 10 \| 13 \| 300 \| 3 \| 371 \| Baseplate protein \| Baseplate protein; afdb-uniprot50 \| AF-A0A7W8AJB6-F1-MODEL\_V4 \| 1.0 \| 2.34e-25 \| 840 \| 0.322 \| 295 \| 179 \| 7 \| 8 \| 298 \| 3 \| 280 \| Phage-related baseplate assembly protein \| Phage-related baseplate assembly protein; afdb-uniprot50 \| AF-A0A061JGS7-F1-MODEL\_V4 \| 1.0 \| 3.335e-23 \| 839 \| 0.327 \| 226 \| 136 \| 3 \| 73 \| 298 \| 2 \| 211 \| Baseplate assembly protein J \| Baseplate assembly protein J; afdb-uniprot50 \| AF-A0A411WHR4-F1-MODEL\_V4 \| 1.0 \| 7.487e-24 \| 839 \| 0.222 \| 378 \| 199 \| 9 \| 12 \| 300 \| 3 \| 374 \| Baseplate protein \| Baseplate protein; afdb-uniprot50 \| AF-A0A6M8SXS4-F1-MODEL\_V4 \| 1.0 \| 1.175e-24 \| 838 \| 0.245 \| 379 \| 187 \| 12 \| 12 \| 300 \| 5 \| 374 \| Baseplate J/gp47 family protein \| Baseplate J/gp47 family protein; afdb-uniprot50 \| AF-A0A1X3ITE5-F1-MODEL\_V4 \| 1.0 \| 3.758e-23 \| 837 \| 0.43 \| 244 \| 128 \| 3 \| 8 \| 244 \| 4 \| 243 \| Baseplate assembly protein J (GpJ) \| Baseplate assembly protein J (GpJ); afdb-uniprot50 \| AF-A0A6L8LZX8-F1-MODEL\_V4 \| 1.0 \| 5.088e-25 \| 835 \| 0.235 \| 327 \| 201 \| 6 \| 13 \| 294 \| 1 \| 323 \| Baseplate\_J domain-containing protein \| Baseplate\_J domain-containing protein; afdb-uniprot50 \| AF-A0A837JBU7-F1-MODEL\_V4 \| 1.0 \| 2.554e-24 \| 834 \| 0.219 \| 396 \| 194 \| 8 \| 8 \| 295 \| 2 \| 390 \| Baseplate assembly protein \| Baseplate assembly protein; afdb-uniprot50 \| AF-L0RFG6-F1-MODEL\_V4 \| 1.0 \| 1.202e-21 \| 831 \| 0.384 \| 182 \| 111 \| 1 \| 114 \| 295 \| 6 \| 186 \| Baseplate J family protein \| Baseplate J family protein; afdb-uniprot50 \| AF-A0A7U6KQJ6-F1-MODEL\_V4 \| 1.0 \| 5.379e-23 \| 829 \| 0.336 \| 247 \| 146 \| 4 \| 55 \| 300 \| 2 \| 231 \| Baseplate assembly protein \| Baseplate assembly protein; afdb-uniprot50 \| AF-A0A3S4DC92-F1-MODEL\_V4 \| 1.0 \| 4.354e-22 \| 828 \| 0.414 \| 181 \| 104 \| 1 \| 116 \| 296 \| 3 \| 181 \| Baseplate J-like protein \| Baseplate J-like protein; afdb-uniprot50 \| AF-A0A6I5SY72-F1-MODEL\_V4 \| 1.0 \| 5.554e-24 \| 827 \| 0.382 \| 259 \| 139 \| 5 \| 18 \| 274 \| 3 \| 242 \| Baseplate assembly protein J \| Baseplate assembly protein J; afdb-uniprot50 \| AF-A0A327Q7V2-F1-MODEL\_V4 \| 1.0 \| 1.405e-24 \| 825 \| 0.219 \| 369 \| 201 \| 9 \| 13 \| 298 \| 1 \| 365 \| Phage-related baseplate assembly protein \| Phage-related baseplate assembly protein; afdb-uniprot50 \| AF-A0A7W5FTX4-F1-MODEL\_V4 \| 1.0 \| 1.681e-24 \| 823 \| 0.297 \| 296 \| 181 \| 7 \| 7 \| 298 \| 3 \| 275 \| Phage-related baseplate assembly protein \| Phage-related baseplate assembly protein; afdb-uniprot50 \| AF-A0A8B3NLL6-F1-MODEL\_V4 \| 1.0 \| 3.897e-26 \| 822 \| 0.318 \| 308 \| 184 \| 7 \| 1 \| 298 \| 127 \| 418 \| Uncharacterized protein \| Uncharacterized protein; afdb-uniprot50 \| AF-A0A7X7RZR2-F1-MODEL\_V4 \| 1.0 \| 2.406e-24 \| 819 \| 0.253 \| 386 \| 183 \| 12 \| 12 \| 300 \| 3 \| 380 \| Baseplate J/gp47 family protein \| Baseplate J/gp47 family protein; afdb-uniprot50 \| AF-A0A0B6CQ96-F1-MODEL\_V4 \| 1.0 \| 6.86e-25 \| 818 \| 0.233 \| 296 \| 200 \| 7 \| 1 \| 296 \| 1 \| 269 \| Baseplate J-like family protein \| Baseplate J-like family protein; afdb-uniprot50 \| AF-A0A3A9CZV9-F1-MODEL\_V4 \| 1.0 \| 3.443e-24 \| 818 \| 0.223 \| 376 \| 202 \| 9 \| 13 \| 300 \| 6 \| 379 \| Baseplate J/gp47 family protein \| Baseplate J/gp47 family protein; afdb-uniprot50 \| AF-A0A849V9Q1-F1-MODEL\_V4 \| 1.0 \| 6.644e-24 \| 814 \| 0.232 \| 382 \| 195 \| 12 \| 8 \| 300 \| 1 \| 373 \| Baseplate J/gp47 family protein \| Baseplate J/gp47 family protein; afdb-uniprot50 \| AF-A0A7S8C8N1-F1-MODEL\_V4 \| 1.0 \| 1.142e-25 \| 813 \| 0.336 \| 291 \| 172 \| 7 \| 12 \| 298 \| 3 \| 276 \| Baseplate J/gp47 family protein \| Baseplate J/gp47 family protein; afdb-uniprot50 \| AF-A0A1M3AMW9-F1-MODEL\_V4 \| 1.0 \| 1.728e-23 \| 811 \| 0.257 \| 318 \| 197 \| 7 \| 13 \| 296 \| 3 \| 315 \| Baseplate\_J domain-containing protein \| Baseplate\_J domain-containing protein; afdb-uniprot50 \| AF-A0A165U8U7-F1-MODEL\_V4 \| 1.0 \| 2.068e-23 \| 809 \| 0.25 \| 304 \| 199 \| 6 \| 1 \| 298 \| 1 \| 281 \| Baseplate J-like protein \| Baseplate J-like protein; afdb-uniprot50 \| AF-A0A1C0V8C6-F1-MODEL\_V4 \| 1.0 \| 1.628e-23 \| 807 \| 0.229 \| 379 \| 191 \| 10 \| 13 \| 295 \| 1 \| 374 \| Baseplate\_J domain-containing protein \| Baseplate\_J domain-containing protein; afdb-uniprot50 \| AF-A0A021XBA6-F1-MODEL\_V4 \| 1.0 \| 3.897e-26 \| 806 \| 0.298 \| 312 \| 184 \| 9 \| 1 \| 294 \| 1 \| 295 \| Baseplate J family protein \| Baseplate J family protein; afdb-uniprot50 \| AF-A0A4U8YSZ3-F1-MODEL\_V4 \| 1.0 \| 1.138e-23 \| 806 \| 0.227 \| 378 \| 201 \| 12 \| 8 \| 300 \| 1 \| 372 \| Baseplate protein j-like \| Baseplate protein j-like; afdb-uniprot50 \| AF-A0A659UR57-F1-MODEL\_V4 \| 1.0 \| 1.736e-25 \| 805 \| 0.333 \| 300 \| 174 \| 7 \| 9 \| 298 \| 3 \| 286 \| Baseplate J protein \| Baseplate J protein; afdb-uniprot50 \| AF-A0A327KHD7-F1-MODEL\_V4 \| 1.0 \| 3.774e-25 \| 805 \| 0.293 \| 300 \| 184 \| 7 \| 8 \| 295 \| 4 \| 287 \| Baseplate\_J domain-containing protein \| Baseplate\_J domain-containing protein; afdb-uniprot50 \| AF-A0A521DS17-F1-MODEL\_V4 \| 1.0 \| 6.259e-24 \| 804 \| 0.221 \| 366 \| 194 \| 11 \| 22 \| 300 \| 5 \| 366 \| Phage-related baseplate assembly protein \| Phage-related baseplate assembly protein; afdb-uniprot50 \| AF-A0A3T0L2T8-F1-MODEL\_V4 \| 1.0 \| 3.774e-25 \| 802 \| 0.24 \| 366 \| 194 \| 9 \| 13 \| 300 \| 1 \| 360 \| Baseplate\_J domain-containing protein \| Baseplate\_J domain-containing protein; afdb-uniprot50 \| AF-E6X1N4-F1-MODEL\_V4 \| 1.0 \| 1.009e-23 \| 802 \| 0.237 \| 366 \| 195 \| 9 \| 13 \| 300 \| 1 \| 360 \| Baseplate J family protein \| Baseplate J family protein; afdb-uniprot50 \| AF-A0A1M7R7L6-F1-MODEL\_V4 \| 1.0 \| 7.283e-25 \| 802 \| 0.237 \| 371 \| 196 \| 9 \| 13 \| 300 \| 1 \| 367 \| Phage-related baseplate assembly protein \| Phage-related baseplate assembly protein; afdb-uniprot50 \| AF-A0A0P9B5C1-F1-MODEL\_V4 \| 1.0 \| 6.035e-21 \| 801 \| 0.401 \| 214 \| 124 \| 2 \| 3 \| 214 \| 2 \| 213 \| Baseplate\_J domain-containing protein \| Baseplate\_J domain-containing protein; afdb-uniprot50 \| AF-A0A2N9Y980-F1-MODEL\_V4 \| 1.0 \| 1.956e-25 \| 801 \| 0.308 \| 305 \| 186 \| 8 \| 1 \| 298 \| 1 \| 287 \| Baseplate assembly protein \| Baseplate assembly protein; afdb-uniprot50 \| AF-Q2NRT7-F1-MODEL\_V4 \| 1.0 \| 8.675e-23 \| 800 \| 0.48 \| 227 \| 104 \| 3 \| 8 \| 227 \| 4 \| 223 \| Baseplate J-like protein \| Baseplate J-like protein; afdb-uniprot50 \| AF-A0A5S9R4F2-F1-MODEL\_V4 \| 1.0 \| 8.957e-24 \| 799 \| 0.245 \| 310 \| 204 \| 7 \| 1 \| 298 \| 1 \| 292 \| Baseplate\_J domain-containing protein \| Baseplate\_J domain-containing protein; afdb-uniprot50 \| AF-A0A5C7WA37-F1-MODEL\_V4 \| 1.0 \| 1.208e-23 \| 795 \| 0.245 \| 379 \| 185 \| 11 \| 13 \| 300 \| 4 \| 372 \| Baseplate protein \| Baseplate protein; afdb-uniprot50 \| AF-A0A8A9QTJ7-F1-MODEL\_V4 \| 1.0 \| 7.698e-23 \| 791 \| 0.259 \| 293 \| 191 \| 7 \| 13 \| 298 \| 3 \| 276 \| Baseplate J/gp47 family protein \| Baseplate J/gp47 family protein; afdb-uniprot50 \| AF-A0A7X6G3F7-F1-MODEL\_V4 \| 1.0 \| 1.57e-20 \| 789 \| 0.54 \| 174 \| 80 \| 0 \| 1 \| 174 \| 1 \| 174 \| Baseplate\_J domain-containing protein \| Baseplate\_J domain-containing protein; afdb-uniprot50 \| AF-A0A5C1DKC4-F1-MODEL\_V4 \| 1.0 \| 1.242e-22 \| 788 \| 0.241 \| 381 \| 190 \| 12 \| 11 \| 300 \| 61 \| 433 \| Baseplate protein \| Baseplate protein; afdb-uniprot50 \| AF-A0A1T4W4N2-F1-MODEL\_V4 \| 1.0 \| 7.487e-24 \| 785 \| 0.269 \| 301 \| 188 \| 8 \| 12 \| 298 \| 5 \| 287 \| Phage-related baseplate assembly protein \| Phage-related baseplate assembly protein; afdb-uniprot50 \| AF-J6H0S6-F1-MODEL\_V4 \| 1.0 \| 1.102e-22 \| 785 \| 0.205 \| 370 \| 207 \| 9 \| 14 \| 300 \| 1 \| 366 \| Baseplate J-like protein \| Baseplate J-like protein; afdb-uniprot50 \| AF-A0A5B9YAA6-F1-MODEL\_V4 \| 1.0 \| 1.102e-22 \| 785 \| 0.232 \| 374 \| 199 \| 10 \| 10 \| 300 \| 3 \| 371 \| Baseplate J/gp47 family protein \| Baseplate J/gp47 family protein; afdb-uniprot50 \| AF-A0A2A7AQP1-F1-MODEL\_V4 \| 1.0 \| 2.474e-23 \| 785 \| 0.205 \| 379 \| 211 \| 8 \| 10 \| 300 \| 8 \| 384 \| Phage tail protein \| Phage tail protein; afdb-uniprot50 \| AF-A0A485AH88-F1-MODEL\_V4 \| 1.0 \| 9.734e-21 \| 783 \| 0.472 \| 197 \| 97 \| 1 \| 8 \| 197 \| 4 \| 200 \| Baseplate J-like protein \| Baseplate J-like protein; afdb-uniprot50 \| AF-A0A779QIS1-F1-MODEL\_V4 \| 1.0 \| 9.17e-21 \| 782 \| 0.449 \| 207 \| 103 \| 3 \| 8 \| 207 \| 4 \| 206 \| Baseplate assembly protein \| Baseplate assembly protein; afdb-uniprot50 \| AF-A0A3L7JDR9-F1-MODEL\_V4 \| 1.0 \| 1.175e-24 \| 782 \| 0.299 \| 307 \| 183 \| 9 \| 8 \| 298 \| 7 \| 297 \| Baseplate\_J domain-containing protein \| Baseplate\_J domain-containing protein; afdb-uniprot50 \| AF-A0A4V3CHL3-F1-MODEL\_V4 \| 1.0 \| 2.33e-23 \| 782 \| 0.236 \| 377 \| 196 \| 10 \| 12 \| 300 \| 4 \| 376 \| Phage-related baseplate assembly protein \| Phage-related baseplate assembly protein; afdb-uniprot50 \| AF-A0A497UL74-F1-MODEL\_V4 \| 1.0 \| 3.897e-26 \| 781 \| 0.258 \| 371 \| 188 \| 8 \| 13 \| 300 \| 1 \| 367 \| Phage-related baseplate assembly protein \| Phage-related baseplate assembly protein; afdb-uniprot50 \| AF-A0A380Z425-F1-MODEL\_V4 \| 1.0 \| 4.119e-24 \| 781 \| 0.239 \| 364 \| 186 \| 10 \| 20 \| 300 \| 9 \| 364 \| Uncharacterized homolog of phage Mu protein gp47 \| Uncharacterized homolog of phage Mu protein gp47; afdb-uniprot50 \| AF-A0A3D4RUG0-F1-MODEL\_V4 \| 1.0 \| 1.843e-25 \| 779 \| 0.299 \| 321 \| 180 \| 10 \| 1 \| 294 \| 1 \| 303 \| Baseplate\_J domain-containing protein \| Baseplate\_J domain-containing protein; afdb-uniprot50 \| AF-A0A0K1NFZ3-F1-MODEL\_V4 \| 1.0 \| 5.209e-22 \| 778 \| 0.253 \| 292 \| 192 \| 8 \| 11 \| 295 \| 2 \| 274 \| Baseplate\_J domain-containing protein \| Baseplate\_J domain-containing protein; afdb-uniprot50 \| AF-A0A075WT46-F1-MODEL\_V4 \| 1.0 \| 5.896e-24 \| 777 \| 0.218 \| 362 \| 195 \| 9 \| 23 \| 300 \| 8 \| 365 \| Baseplate\_J domain-containing protein \| Baseplate\_J domain-containing protein; afdb-uniprot50 \| AF-M5JP61-F1-MODEL\_V4 \| 1.0 \| 1.009e-23 \| 776 \| 0.272 \| 297 \| 192 \| 7 \| 9 \| 298 \| 6 \| 285 \| Baseplate J family protein \| Baseplate J family protein; afdb-uniprot50 \| AF-A0A098MGN3-F1-MODEL\_V4 \| 1.0 \| 1.242e-22 \| 776 \| 0.244 \| 380 \| 195 \| 12 \| 8 \| 300 \| 2 \| 376 \| Baseplate J protein \| Baseplate J protein; afdb-uniprot50 \| AF-A0A450VII4-F1-MODEL\_V4 \| 1.0 \| 8.675e-23 \| 771 \| 0.3 \| 260 \| 162 \| 5 \| 39 \| 298 \| 3 \| 242 \| Phage-related baseplate assembly protein \| Phage-related baseplate assembly protein; afdb-uniprot50 \| AF-D1P879-F1-MODEL\_V4 \| 1.0 \| 1.931e-19 \| 766 \| 0.424 \| 153 \| 88 \| 0 \| 147 \| 299 \| 3 \| 155 \| Baseplate J-like protein \| Baseplate J-like protein; afdb-uniprot50 \| AF-A0A1E3G789-F1-MODEL\_V4 \| 1.0 \| 4.928e-24 \| 765 \| 0.247 \| 319 \| 199 \| 10 \| 1 \| 296 \| 1 \| 301 \| Baseplate\_J domain-containing protein \| Baseplate\_J domain-containing protein; afdb-uniprot50 \| AF-A0A0D6ARM2-F1-MODEL\_V4 \| 1.0 \| 4.102e-22 \| 760 \| 0.173 \| 421 \| 211 \| 8 \| 12 \| 300 \| 3 \| 418 \| Phage-related baseplate assembly protein \| Phage-related baseplate assembly protein; afdb-uniprot50 \| AF-E0LW08-F1-MODEL\_V4 \| 1.0 \| 2.385e-20 \| 759 \| 0.437 \| 167 \| 88 \| 2 \| 138 \| 299 \| 5 \| 170 \| Baseplate J family protein \| Baseplate J family protein; afdb-uniprot50 \| AF-Q4EBT1-F1-MODEL\_V4 \| 1.0 \| 1.827e-21 \| 759 \| 0.396 \| 227 \| 122 \| 2 \| 13 \| 239 \| 1 \| 212 \| Baseplate J-like protein \| Baseplate J-like protein; afdb-uniprot50 \| AF-A0A6G5QY03-F1-MODEL\_V4 \| 1.0 \| 1.282e-23 \| 756 \| 0.256 \| 374 \| 183 \| 15 \| 8 \| 300 \| 2 \| 361 \| Phage baseplate assembly protein J, putative \| Phage baseplate assembly protein J, putative; afdb-uniprot50 \| AF-A0A512JPE9-F1-MODEL\_V4 \| 1.0 \| 9.468e-22 \| 756 \| 0.234 \| 366 \| 202 \| 15 \| 2 \| 296 \| 10 \| 368 \| Baseplate\_J domain-containing protein \| Baseplate\_J domain-containing protein; afdb-uniprot50 \| AF-A0A367DB17-F1-MODEL\_V4 \| 1.0 \| 1.674e-22 \| 754 \| 0.193 \| 373 \| 212 \| 12 \| 12 \| 300 \| 7 \| 374 \| Baseplate J protein \| Baseplate J protein; afdb-uniprot50 \| AF-A0A1V4GFY9-F1-MODEL\_V4 \| 1.0 \| 4.622e-22 \| 746 \| 0.227 \| 378 \| 198 \| 13 \| 12 \| 300 \| 6 \| 378 \| Baseplate\_J domain-containing protein \| Baseplate\_J domain-containing protein; afdb-uniprot50 \| AF-A0A4D9IGF7-F1-MODEL\_V4 \| 1.0 \| 9.209e-23 \| 745 \| 0.271 \| 331 \| 155 \| 6 \| 53 \| 300 \| 2 \| 329 \| Baseplate protein \| Baseplate protein; afdb-uniprot50 \| AF-D1AFB5-F1-MODEL\_V4 \| 1.0 \| 3.64e-22 \| 744 \| 0.191 \| 365 \| 204 \| 11 \| 20 \| 298 \| 9 \| 368 \| Baseplate J family protein \| Baseplate J family protein; afdb-uniprot50 \| AF-A0A0R3MV37-F1-MODEL\_V4 \| 1.0 \| 6.232e-22 \| 743 \| 0.216 \| 296 \| 205 \| 9 \| 12 \| 298 \| 7 \| 284 \| Baseplate assembly protein \| Baseplate assembly protein; afdb-uniprot50 \| AF-N6VUC8-F1-MODEL\_V4 \| 1.0 \| 1.728e-23 \| 742 \| 0.296 \| 277 \| 175 \| 7 \| 25 \| 298 \| 2 \| 261 \| Phage baseplate assembly protein GpJ \| Phage baseplate assembly protein GpJ; afdb-uniprot50 \| AF-A0A1D9LNN0-F1-MODEL\_V4 \| 1.0 \| 6.061e-23 \| 742 \| 0.231 \| 380 \| 194 \| 11 \| 12 \| 298 \| 6 \| 380 \| Baseplate\_J domain-containing protein \| Baseplate\_J domain-containing protein; afdb-uniprot50 \| AF-A0A823V5N8-F1-MODEL\_V4 \| 1.0 \| 2.959e-23 \| 738 \| 0.225 \| 395 \| 191 \| 13 \| 4 \| 300 \| 10 \| 387 \| Baseplate assembly protein \| Baseplate assembly protein; afdb-uniprot50 \| AF-A0A4Y7RWI7-F1-MODEL\_V4 \| 1.0 \| 2.7e-22 \| 736 \| 0.219 \| 369 \| 201 \| 10 \| 12 \| 298 \| 5 \| 368 \| Baseplate\_J domain-containing protein \| Baseplate\_J domain-containing protein; afdb-uniprot50 \| AF-A0A2N1QGA1-F1-MODEL\_V4 \| 1.0 \| 7.455e-22 \| 736 \| 0.191 \| 381 \| 217 \| 11 \| 8 \| 300 \| 3 \| 380 \| Baseplate\_J domain-containing protein \| Baseplate\_J domain-containing protein; afdb-uniprot50 \| AF-A0A6I2URJ4-F1-MODEL\_V4 \| 1.0 \| 3.955e-19 \| 735 \| 0.494 \| 186 \| 92 \| 1 \| 1 \| 184 \| 1 \| 186 \| Baseplate assembly protein \| Baseplate assembly protein; afdb-uniprot50 \| AF-A0A7X5KSN5-F1-MODEL\_V4 \| 1.0 \| 2.543e-22 \| 735 \| 0.162 \| 382 \| 227 \| 10 \| 8 \| 300 \| 2 \| 379 \| Phage tail protein \| Phage tail protein; afdb-uniprot50 \| AF-A0A4Q8MDG2-F1-MODEL\_V4 \| 1.0 \| 2.406e-24 \| 732 \| 0.273 \| 303 \| 191 \| 9 \| 13 \| 298 \| 5 \| 295 \| Baseplate\_J domain-containing protein \| Baseplate\_J domain-containing protein; afdb-uniprot50 \| AF-N9KZE1-F1-MODEL\_V4 \| 1.0 \| 4.732e-19 \| 728 \| 0.278 \| 183 \| 129 \| 2 \| 119 \| 298 \| 3 \| 185 \| Baseplate\_J domain-containing protein \| Baseplate\_J domain-containing protein; afdb-uniprot50 \| AF-A0A4Q8MFD0-F1-MODEL\_V4 \| 1.0 \| 5.209e-22 \| 728 \| 0.223 \| 331 \| 208 \| 12 \| 1 \| 300 \| 1 \| 313 \| Baseplate\_J domain-containing protein \| Baseplate\_J domain-containing protein; afdb-uniprot50 \| AF-A0A4Q6D773-F1-MODEL\_V4 \| 1.0 \| 1.318e-22 \| 723 \| 0.26 \| 300 \| 184 \| 9 \| 14 \| 300 \| 3 \| 277 \| Baseplate\_J domain-containing protein \| Baseplate\_J domain-containing protein; afdb-uniprot50 \| AF-F3YY52-F1-MODEL\_V4 \| 1.0 \| 2.002e-22 \| 722 \| 0.222 \| 382 \| 200 \| 12 \| 8 \| 300 \| 1 \| 374 \| Baseplate J family protein \| Baseplate J family protein; afdb-uniprot50 \| AF-A0A812RFY7-F1-MODEL\_V4 \| 1.0 \| 8.919e-22 \| 720 \| 0.437 \| 258 \| 116 \| 2 \| 36 \| 264 \| 2 \| 259 \| J protein \| J protein; afdb-uniprot50 \| AF-A0A840NW02-F1-MODEL\_V4 \| 1.0 \| 5.506e-20 \| 717 \| 0.312 \| 224 \| 135 \| 6 \| 77 \| 298 \| 1 \| 207 \| Phage-related baseplate assembly protein \| Phage-related baseplate assembly protein; afdb-uniprot50 \| AF-A0A2B7YKZ8-F1-MODEL\_V4 \| 1.0 \| 1.005e-21 \| 717 \| 0.199 \| 376 \| 206 \| 11 \| 13 \| 300 \| 2 \| 370 \| Bacteriocin \| Bacteriocin; afdb-uniprot50 \| AF-A0A2H1SQX0-F1-MODEL\_V4 \| 1.0 \| 4.732e-19 \| 716 \| 0.531 \| 173 \| 79 \| 1 \| 2 \| 174 \| 9 \| 179 \| Baseplate assembly protein J \| Baseplate assembly protein J; afdb-uniprot50 \| AF-A0A2G6EWY7-F1-MODEL\_V4 \| 1.0 \| 9.17e-21 \| 714 \| 0.26 \| 292 \| 186 \| 6 \| 13 \| 298 \| 1 \| 268 \| Baseplate\_J domain-containing protein \| Baseplate\_J domain-containing protein; afdb-uniprot50 \| AF-E5Y5X0-F1-MODEL\_V4 \| 1.0 \| 1.485e-22 \| 714 \| 0.234 \| 379 \| 196 \| 10 \| 9 \| 300 \| 5 \| 376 \| Baseplate\_J domain-containing protein \| Baseplate\_J domain-containing protein; afdb-uniprot50 \| AF-A0A7X4ZSW6-F1-MODEL\_V4 \| 1.0 \| 8.402e-22 \| 714 \| 0.208 \| 378 \| 207 \| 10 \| 10 \| 298 \| 6 \| 380 \| Baseplate J/gp47 family protein \| Baseplate J/gp47 family protein; afdb-uniprot50 \| AF-R7JDJ8-F1-MODEL\_V4 \| 1.0 \| 1.242e-22 \| 713 \| 0.22 \| 377 \| 196 \| 11 \| 12 \| 300 \| 3 \| 369 \| Probable bacteriophage tail fiber protein \| Probable bacteriophage tail fiber protein; afdb-uniprot50 \| AF-A0A4Q2XB46-F1-MODEL\_V4 \| 1.0 \| 2.237e-18 \| 712 \| 0.383 \| 154 \| 95 \| 0 \| 146 \| 299 \| 2 \| 155 \| Baseplate assembly protein \| Baseplate assembly protein; afdb-uniprot50 \| AF-A0A0S4XLP9-F1-MODEL\_V4 \| 1.0 \| 1.527e-21 \| 708 \| 0.234 \| 282 \| 186 \| 6 \| 13 \| 294 \| 1 \| 252 \| Putative Baseplate J family protein \| Putative Baseplate J family protein; afdb-uniprot50 \| AF-A0A6M3J3B0-F1-MODEL\_V4 \| 1.0 \| 2.565e-26 \| 707 \| 0.276 \| 322 \| 196 \| 4 \| 12 \| 296 \| 8 \| 329 \| Putative baseplate protein \| Putative baseplate protein; afdb-uniprot50 \| AF-A0A4Q3KFV4-F1-MODEL\_V4 \| 1.0 \| 2.7e-22 \| 704 \| 0.302 \| 271 \| 163 \| 6 \| 31 \| 298 \| 118 \| 365 \| Uncharacterized protein \| Uncharacterized protein; afdb-uniprot50 \| AF-A0A3Q9PYZ0-F1-MODEL\_V4 \| 1.0 \| 4.865e-18 \| 700 \| 0.465 \| 146 \| 78 \| 0 \| 154 \| 299 \| 2 \| 147 \| Baseplate assembly protein \| Baseplate assembly protein; afdb-uniprot50 \| AF-A0A2S5F8Y6-F1-MODEL\_V4 \| 1.0 \| 1.621e-21 \| 700 \| 0.207 \| 385 \| 208 \| 13 \| 1 \| 298 \| 1 \| 375 \| Phage baseplate protein \| Phage baseplate protein; afdb-uniprot50 \| AF-A0A836P245-F1-MODEL\_V4 \| 1.0 \| 6.38e-19 \| 698 \| 0.53 \| 179 \| 81 \| 2 \| 1 \| 179 \| 1 \| 176 \| Baseplate assembly protein \| Baseplate assembly protein; afdb-uniprot50 \| AF-A0A0Q1F5V0-F1-MODEL\_V4 \| 1.0 \| 8.996e-26 \| 696 \| 0.275 \| 370 \| 182 \| 8 \| 13 \| 300 \| 3 \| 368 \| Baseplate\_J domain-containing protein \| Baseplate\_J domain-containing protein; afdb-uniprot50 \| AF-E2CJT9-F1-MODEL\_V4 \| 1.0 \| 3.847e-20 \| 694 \| 0.216 \| 291 \| 205 \| 7 \| 13 \| 296 \| 1 \| 275 \| Baseplate J family protein \| Baseplate J family protein; afdb-uniprot50 \| AF-A0A4U7JBE4-F1-MODEL\_V4 \| 1.0 \| 1.276e-21 \| 694 \| 0.217 \| 373 \| 200 \| 10 \| 14 \| 299 \| 1 \| 368 \| Baseplate J/gp47 family protein \| Baseplate J/gp47 family protein; afdb-uniprot50 \| AF-K1JKP8-F1-MODEL\_V4 \| 1.0 \| 7.053e-24 \| 694 \| 0.235 \| 374 \| 197 \| 10 \| 12 \| 300 \| 9 \| 378 \| Baseplate\_J domain-containing protein \| Baseplate\_J domain-containing protein; afdb-uniprot50 \| AF-Q0FZ02-F1-MODEL\_V4 \| 1.0 \| 3.23e-22 \| 689 \| 0.254 \| 295 \| 196 \| 9 \| 12 \| 296 \| 8 \| 288 \| Baseplate J-like protein \| Baseplate J-like protein; afdb-uniprot50 \| AF-A8TTZ7-F1-MODEL\_V4 \| 1.0 \| 6.963e-18 \| 688 \| 0.448 \| 145 \| 80 \| 0 \| 152 \| 296 \| 2 \| 146 \| Baseplate J-like protein \| Baseplate J-like protein; afdb-uniprot50 \| AF-A0A6G8CYG2-F1-MODEL\_V4 \| 1.0 \| 7.665e-21 \| 686 \| 0.307 \| 270 \| 158 \| 6 \| 32 \| 298 \| 8 \| 251 \| Baseplate J protein \| Baseplate J protein; afdb-uniprot50 \| AF-A0A378XFF5-F1-MODEL\_V4 \| 1.0 \| 3.016e-18 \| 683 \| 0.339 \| 174 \| 111 \| 3 \| 128 \| 300 \| 28 \| 198 \| Baseplate J-like protein \| Baseplate J-like protein; afdb-uniprot50 \| AF-A0A2J4P2N7-F1-MODEL\_V4 \| 1.0 \| 4.317e-18 \| 682 \| 0.506 \| 164 \| 81 \| 0 \| 77 \| 240 \| 3 \| 166 \| Baseplate assembly protein \| Baseplate assembly protein; afdb-uniprot50 \| AF-A0A827ZIF5-F1-MODEL\_V4 \| 1.0 \| 8.601e-19 \| 681 \| 0.34 \| 200 \| 117 \| 4 \| 113 \| 300 \| 22 \| 218 \| Baseplate protein \| Baseplate protein; afdb-uniprot50 \| AF-A0A1T4QDZ6-F1-MODEL\_V4 \| 1.0 \| 7.665e-21 \| 680 \| 0.177 \| 360 \| 208 \| 9 \| 23 \| 298 \| 10 \| 365 \| Phage-related baseplate assembly protein \| Phage-related baseplate assembly protein; afdb-uniprot50 \| AF-A0A376EX36-F1-MODEL\_V4 \| 1.0 \| 4.732e-19 \| 679 \| 0.315 \| 200 \| 122 \| 4 \| 113 \| 300 \| 74 \| 270 \| Baseplate assembly protein J \| Baseplate assembly protein J; afdb-uniprot50 \| AF-A0A1H1FF82-F1-MODEL\_V4 \| 1.0 \| 4.752e-21 \| 679 \| 0.244 \| 295 \| 192 \| 11 \| 13 \| 296 \| 9 \| 283 \| Phage-related baseplate assembly protein \| Phage-related baseplate assembly protein; afdb-uniprot50 \| AF-A0A545T5S2-F1-MODEL\_V4 \| 1.0 \| 1.827e-21 \| 678 \| 0.262 \| 373 \| 173 \| 12 \| 7 \| 300 \| 8 \| 357 \| Uncharacterized protein \| Uncharacterized protein; afdb-uniprot50 \| AF-A0A7D5QQY9-F1-MODEL\_V4 \| 1.0 \| 6.616e-22 \| 676 \| 0.237 \| 316 \| 152 \| 6 \| 72 \| 300 \| 6 \| 319 \| Baseplate J/gp47 family protein \| Baseplate J/gp47 family protein; afdb-uniprot50 \| AF-A0A6M6IDN5-F1-MODEL\_V4 \| 1.0 \| 1.521e-19 \| 674 \| 0.383 \| 237 \| 128 \| 3 \| 6 \| 239 \| 3 \| 224 \| Baseplate assembly protein J \| Baseplate assembly protein J; afdb-uniprot50 \| AF-D1P8B4-F1-MODEL\_V4 \| 1.0 \| 4.583e-18 \| 673 \| 0.477 \| 180 \| 87 \| 1 \| 13 \| 185 \| 2 \| 181 \| Baseplate J-like protein \| Baseplate J-like protein; afdb-uniprot50 \| AF-A0A828HR86-F1-MODEL\_V4 \| 1.0 \| 7.847e-18 \| 673 \| 0.476 \| 191 \| 89 \| 3 \| 8 \| 191 \| 4 \| 190 \| Baseplate assembly protein \| Baseplate assembly protein; afdb-uniprot50 \| AF-A0A238HJ41-F1-MODEL\_V4 \| 1.0 \| 5.661e-19 \| 672 \| 0.34 \| 191 \| 115 \| 4 \| 118 \| 300 \| 11 \| 198 \| Baseplate J-like protein \| Baseplate J-like protein; afdb-uniprot50 \| AF-A0A064AIK0-F1-MODEL\_V4 \| 1.0 \| 1.393e-20 \| 671 \| 0.181 \| 374 \| 212 \| 13 \| 13 \| 299 \| 1 \| 367 \| Baseplate assembly protein \| Baseplate assembly protein; afdb-uniprot50 \| AF-G2FT51-F1-MODEL\_V4 \| 1.0 \| 2.059e-21 \| 671 \| 0.237 \| 396 \| 183 \| 15 \| 14 \| 298 \| 1 \| 388 \| Baseplate J-like family protein \| Baseplate J-like family protein; afdb-uniprot50 \| AF-A0A081MYF5-F1-MODEL\_V4 \| 1.0 \| 2.592e-17 \| 670 \| 0.41 \| 156 \| 89 \| 2 \| 144 \| 299 \| 2 \| 154 \| Baseplate\_J domain-containing protein \| Baseplate\_J domain-containing protein; afdb-uniprot50 \| AF-A0A1V3RQM2-F1-MODEL\_V4 \| 1.0 \| 2.077e-25 \| 670 \| 0.246 \| 378 \| 188 \| 9 \| 13 \| 300 \| 8 \| 378 \| Phage tail protein \| Phage tail protein; afdb-uniprot50 \| AF-D8IV11-F1-MODEL\_V4 \| 1.0 \| 8.207e-25 \| 669 \| 0.261 \| 379 \| 179 \| 10 \| 12 \| 300 \| 7 \| 374 \| Bacteriophage tail fiber protein \| Bacteriophage tail fiber protein; afdb-uniprot50 \| AF-A0A2G6F072-F1-MODEL\_V4 \| 1.0 \| 1.312e-20 \| 668 \| 0.172 \| 371 \| 216 \| 10 \| 14 \| 300 \| 1 \| 364 \| Baseplate assembly protein \| Baseplate assembly protein; afdb-uniprot50 \| AF-G2IX29-F1-MODEL\_V4 \| 1.0 \| 5.187e-20 \| 668 \| 0.218 \| 370 \| 194 \| 11 \| 13 \| 294 \| 1 \| 363 \| Phage baseplate J-like protein \| Phage baseplate J-like protein; afdb-uniprot50 \| AF-A0A2X1N0C4-F1-MODEL\_V4 \| 1.0 \| 2.452e-19 \| 667 \| 0.396 \| 212 \| 121 \| 2 \| 31 \| 239 \| 1 \| 208 \| Baseplate assembly protein J \| Baseplate assembly protein J; afdb-uniprot50 \| AF-J8V6S9-F1-MODEL\_V4 \| 1.0 \| 4.886e-20 \| 666 \| 0.34 \| 220 \| 124 \| 5 \| 87 \| 295 \| 2 \| 211 \| Baseplate J family protein \| Baseplate J family protein; afdb-uniprot50 \| AF-A0A258KUT3-F1-MODEL\_V4 \| 1.0 \| 5.82e-18 \| 658 \| 0.352 \| 170 \| 110 \| 0 \| 131 \| 300 \| 5 \| 174 \| Baseplate\_J domain-containing protein \| Baseplate\_J domain-containing protein; afdb-uniprot50 \| AF-F5SA56-F1-MODEL\_V4 \| 1.0 \| 1.762e-18 \| 655 \| 0.319 \| 197 \| 122 \| 5 \| 113 \| 300 \| 94 \| 287 \| Phage baseplate protein \| Phage baseplate protein; afdb-uniprot50 \| AF-A0A1W0C8U1-F1-MODEL\_V4 \| 1.0 \| 4.928e-24 \| 651 \| 0.254 \| 385 \| 185 \| 10 \| 14 \| 300 \| 1 \| 381 \| Baseplate protein \| Baseplate protein; afdb-uniprot50 \| AF-D8IV18-F1-MODEL\_V4 \| 1.0 \| 6.462e-25 \| 651 \| 0.251 \| 386 \| 186 \| 10 \| 13 \| 300 \| 8 \| 388 \| Bacteriophage tail fiber protein \| Bacteriophage tail fiber protein; afdb-uniprot50 \| AF-A0A095Z753-F1-MODEL\_V4 \| 1.0 \| 1.894e-24 \| 649 \| 0.26 \| 372 \| 181 \| 13 \| 13 \| 298 \| 1 \| 364 \| Baseplate\_J domain-containing protein \| Baseplate\_J domain-containing protein; afdb-uniprot50 \| AF-A0A2W5J717-F1-MODEL\_V4 \| 1.0 \| 9.091e-17 \| 647 \| 0.534 \| 159 \| 72 \| 1 \| 1 \| 159 \| 2 \| 158 \| Baseplate assembly protein \| Baseplate assembly protein; afdb-uniprot50 \| AF-A0A1W1Z473-F1-MODEL\_V4 \| 1.0 \| 3.655e-24 \| 647 \| 0.283 \| 293 \| 186 \| 6 \| 11 \| 296 \| 12 \| 287 \| Phage-related baseplate assembly protein \| Phage-related baseplate assembly protein; afdb-uniprot50 \| AF-E2CFL2-F1-MODEL\_V4 \| 1.0 \| 1.621e-21 \| 645 \| 0.278 \| 294 \| 186 \| 10 \| 10 \| 295 \| 5 \| 280 \| Putative baseplate assembly protein J \| Putative baseplate assembly protein J; afdb-uniprot50 \| AF-A0A2M7G5N1-F1-MODEL\_V4 \| 1.0 \| 6.587e-20 \| 645 \| 0.19 \| 373 \| 206 \| 11 \| 13 \| 300 \| 1 \| 362 \| Baseplate protein \| Baseplate protein; afdb-uniprot50 \| AF-A0A4Q0YJP7-F1-MODEL\_V4 \| 1.0 \| 6.38e-19 \| 644 \| 0.35 \| 237 \| 117 \| 4 \| 6 \| 205 \| 5 \| 241 \| Baseplate assembly protein \| Baseplate assembly protein; afdb-uniprot50 \| AF-A0A832CMI4-F1-MODEL\_V4 \| 1.0 \| 2.934e-19 \| 637 \| 0.163 \| 373 \| 210 \| 11 \| 18 \| 298 \| 2 \| 364 \| Baseplate J/gp47 family protein \| Baseplate J/gp47 family protein; afdb-uniprot50 \| AF-A0A1Y6CV72-F1-MODEL\_V4 \| 1.0 \| 5.187e-20 \| 636 \| 0.244 \| 299 \| 186 \| 10 \| 13 \| 298 \| 1 \| 272 \| Phage-related baseplate assembly protein \| Phage-related baseplate assembly protein; afdb-uniprot50 \| AF-A0A827RX96-F1-MODEL\_V4 \| 1.0 \| 9.091e-17 \| 632 \| 0.459 \| 159 \| 82 \| 1 \| 68 \| 226 \| 2 \| 156 \| Baseplate assembly protein \| Baseplate assembly protein; afdb-uniprot50 \| AF-A0A845A584-F1-MODEL\_V4 \| 1.0 \| 1.226e-16 \| 629 \| 0.515 \| 165 \| 78 \| 1 \| 1 \| 165 \| 4 \| 166 \| Uncharacterized protein \| Uncharacterized protein; afdb-uniprot50 \| AF-A0A410X4X2-F1-MODEL\_V4 \| 1.0 \| 6.587e-20 \| 628 \| 0.24 \| 329 \| 159 \| 9 \| 61 \| 300 \| 4 \| 330 \| Baseplate J protein \| Baseplate J protein; afdb-uniprot50 \| AF-A0A1M7RIH8-F1-MODEL\_V4 \| 1.0 \| 2.959e-23 \| 628 \| 0.218 \| 371 \| 201 \| 9 \| 13 \| 300 \| 5 \| 369 \| Phage-related baseplate assembly protein \| Phage-related baseplate assembly protein; afdb-uniprot50 \| AF-A0A7Z1M1I0-F1-MODEL\_V4 \| 1.0 \| 1.282e-23 \| 626 \| 0.23 \| 373 \| 192 \| 10 \| 19 \| 300 \| 5 \| 373 \| Phage-related baseplate assembly protein \| Phage-related baseplate assembly protein; afdb-uniprot50 \| AF-A0A482ISU8-F1-MODEL\_V4 \| 1.0 \| 2.788e-23 \| 626 \| 0.241 \| 377 \| 192 \| 7 \| 13 \| 300 \| 7 \| 378 \| Baseplate protein \| Baseplate protein; afdb-uniprot50 \| AF-R7HY31-F1-MODEL\_V4 \| 1.0 \| 1.948e-23 \| 625 \| 0.244 \| 380 \| 198 \| 10 \| 8 \| 300 \| 3 \| 380 \| Baseplate J-like protein \| Baseplate J-like protein; afdb-uniprot50 \| AF-A0A7J6YLT3-F1-MODEL\_V4 \| 1.0 \| 1.001e-19 \| 625 \| 0.373 \| 249 \| 132 \| 7 \| 1 \| 249 \| 283 \| 507 \| Uncharacterized protein \| Uncharacterized protein; afdb-uniprot50 \| AF-A0A6A2F414-F1-MODEL\_V4 \| 1.0 \| 7.487e-24 \| 624 \| 0.237 \| 370 \| 197 \| 10 \| 12 \| 300 \| 5 \| 370 \| Baseplate J/gp47 family protein \| Baseplate J/gp47 family protein; afdb-uniprot50 \| AF-A0A0X1U7S3-F1-MODEL\_V4 \| 1.0 \| 6.259e-24 \| 624 \| 0.23 \| 378 \| 202 \| 9 \| 10 \| 300 \| 7 \| 382 \| Baseplate J-like protein \| Baseplate J-like protein; afdb-uniprot50 \| AF-A0A7I8DIE6-F1-MODEL\_V4 \| 1.0 \| 7.423e-20 \| 622 \| 0.189 \| 364 \| 205 \| 11 \| 20 \| 298 \| 5 \| 363 \| Baseplate\_J domain-containing protein \| Baseplate\_J domain-containing protein; afdb-uniprot50 \| AF-A0A7U6QPG4-F1-MODEL\_V4 \| 1.0 \| 6.743e-17 \| 621 \| 0.421 \| 147 \| 83 \| 2 \| 155 \| 300 \| 6 \| 151 \| Baseplate\_J domain-containing protein \| Baseplate\_J domain-containing protein; afdb-uniprot50 \| AF-A0A285J5J6-F1-MODEL\_V4 \| 1.0 \| 3.188e-16 \| 621 \| 0.377 \| 159 \| 97 \| 2 \| 138 \| 295 \| 3 \| 160 \| Phage-related baseplate assembly protein \| Phage-related baseplate assembly protein; afdb-uniprot50 \| AF-A0A716MJC6-F1-MODEL\_V4 \| 1.0 \| 2.452e-19 \| 619 \| 0.257 \| 345 \| 167 \| 7 \| 14 \| 273 \| 1 \| 341 \| Baseplate J/gp47 family protein \| Baseplate J/gp47 family protein; afdb-uniprot50 \| AF-A0A5S9NBD4-F1-MODEL\_V4 \| 1.0 \| 3.115e-19 \| 616 \| 0.179 \| 356 \| 201 \| 10 \| 23 \| 294 \| 26 \| 374 \| Baseplate\_J domain-containing protein \| Baseplate\_J domain-containing protein; afdb-uniprot50 \| AF-A0A3A6CL84-F1-MODEL\_V4 \| 1.0 \| 5.165e-18 \| 616 \| 0.187 \| 383 \| 219 \| 11 \| 6 \| 300 \| 3 \| 381 \| Baseplate J/gp47 family protein \| Baseplate J/gp47 family protein; afdb-uniprot50 \| AF-A0A078MEN5-F1-MODEL\_V4 \| 1.0 \| 6.435e-23 \| 615 \| 0.215 \| 372 \| 204 \| 10 \| 12 \| 300 \| 3 \| 369 \| Baseplate J-like protein \| Baseplate J-like protein; afdb-uniprot50 \| AF-A0A0C1G7H8-F1-MODEL\_V4 \| 1.0 \| 2.108e-18 \| 615 \| 0.188 \| 376 \| 208 \| 13 \| 6 \| 296 \| 7 \| 370 \| Baseplate\_J domain-containing protein \| Baseplate\_J domain-containing protein; afdb-uniprot50 \| AF-A0A4Q2QS73-F1-MODEL\_V4 \| 1.0 \| 2.365e-16 \| 614 \| 0.539 \| 141 \| 65 \| 0 \| 86 \| 226 \| 2 \| 142 \| Baseplate assembly protein \| Baseplate assembly protein; afdb-uniprot50 \| AF-A0A2G2BGS6-F1-MODEL\_V4 \| 1.0 \| 3.51e-19 \| 614 \| 0.157 \| 382 \| 231 \| 11 \| 1 \| 298 \| 4 \| 378 \| Phage baseplate assembly protein \| Phage baseplate assembly protein; afdb-uniprot50 \| AF-A0A327JA39-F1-MODEL\_V4 \| 1.0 \| 1.87e-18 \| 613 \| 0.16 \| 418 \| 211 \| 11 \| 13 \| 296 \| 1 \| 412 \| Baseplate\_J domain-containing protein \| Baseplate\_J domain-containing protein; afdb-uniprot50 \| AF-A0A447T5K9-F1-MODEL\_V4 \| 1.0 \| 1.426e-17 \| 612 \| 0.364 \| 195 \| 104 \| 4 \| 116 \| 292 \| 18 \| 210 \| Uncharacterized homolog of phage Mu protein gp47 \| Uncharacterized homolog of phage Mu protein gp47; afdb-uniprot50 \| AF-A0A7C3HFY9-F1-MODEL\_V4 \| 1.0 \| 2.764e-19 \| 612 \| 0.196 \| 367 \| 202 \| 13 \| 13 \| 296 \| 1 \| 357 \| Baseplate J protein \| Baseplate J protein; afdb-uniprot50 \| AF-A0A827JCJ0-F1-MODEL\_V4 \| 1.0 \| 4.844e-16 \| 611 \| 0.537 \| 160 \| 67 \| 1 \| 8 \| 160 \| 4 \| 163 \| Baseplate assembly protein \| Baseplate assembly protein; afdb-uniprot50 \| AF-A0A0B0HE83-F1-MODEL\_V4 \| 1.0 \| 5.31e-17 \| 611 \| 0.281 \| 192 \| 133 \| 4 \| 112 \| 300 \| 88 \| 277 \| Baseplate J-like protein \| Baseplate J-like protein; afdb-uniprot50 \| AF-A0A0T7DUX5-F1-MODEL\_V4 \| 1.0 \| 7.189e-19 \| 611 \| 0.178 \| 381 \| 210 \| 15 \| 12 \| 296 \| 7 \| 380 \| Baseplate assembly protein J \| Baseplate assembly protein J; afdb-uniprot50 \| AF-A0A6M0STE5-F1-MODEL\_V4 \| 1.0 \| 1.001e-19 \| 609 \| 0.194 \| 386 \| 197 \| 15 \| 20 \| 298 \| 5 \| 383 \| Baseplate\_J domain-containing protein \| Baseplate\_J domain-containing protein; afdb-uniprot50 \| AF-A0A3A9B522-F1-MODEL\_V4 \| 1.0 \| 2.615e-21 \| 609 \| 0.187 \| 379 \| 218 \| 10 \| 10 \| 300 \| 9 \| 385 \| Baseplate J/gp47 family protein \| Baseplate J/gp47 family protein; afdb-uniprot50 \| AF-A0A450WH23-F1-MODEL\_V4 \| 1.0 \| 1.674e-22 \| 608 \| 0.246 \| 377 \| 189 \| 12 \| 12 \| 300 \| 3 \| 372 \| Phage-related baseplate assembly protein \| Phage-related baseplate assembly protein; afdb-uniprot50 \| AF-A0A7U7FD77-F1-MODEL\_V4 \| 1.0 \| 5.53e-22 \| 607 \| 0.337 \| 361 \| 139 \| 8 \| 6 \| 294 \| 2 \| 334 \| Putative phage-related protein \| Putative phage-related protein; afdb-uniprot50 \| AF-A0A5M8P6R0-F1-MODEL\_V4 \| 1.0 \| 6.772e-19 \| 606 \| 0.185 \| 383 \| 219 \| 13 \| 1 \| 300 \| 5 \| 377 \| Baseplate\_J domain-containing protein \| Baseplate\_J domain-containing protein; afdb-uniprot50 \| AF-A0A377AJ05-F1-MODEL\_V4 \| 1.0 \| 6.743e-17 \| 603 \| 0.276 \| 188 \| 131 \| 4 \| 116 \| 300 \| 94 \| 279 \| Phage-related baseplate assembly protein J \| Phage-related baseplate assembly protein J; afdb-uniprot50 \| AF-A0A2U2DG57-F1-MODEL\_V4 \| 1.0 \| 1.226e-16 \| 602 \| 0.44 \| 186 \| 95 \| 2 \| 8 \| 188 \| 8 \| 189 \| Baseplate assembly protein \| Baseplate assembly protein; afdb-uniprot50 \| AF-A0A654C1I4-F1-MODEL\_V4 \| 1.0 \| 1.66e-18 \| 602 \| 0.379 \| 232 \| 136 \| 4 \| 1 \| 231 \| 3 \| 227 \| Uncharacterized protein \| Uncharacterized protein; afdb-uniprot50 \| AF-A0A729S941-F1-MODEL\_V4 \| 1.0 \| 4.163e-15 \| 601 \| 0.503 \| 129 \| 64 \| 0 \| 87 \| 215 \| 1 \| 129 \| Baseplate assembly protein \| Baseplate assembly protein; afdb-uniprot50 \| AF-A0A0T9KKG0-F1-MODEL\_V4 \| 1.0 \| 3.814e-16 \| 598 \| 0.471 \| 142 \| 75 \| 0 \| 158 \| 299 \| 2 \| 143 \| Baseplate assembly protein \| Baseplate assembly protein; afdb-uniprot50 \| AF-A0A844Q921-F1-MODEL\_V4 \| 1.0 \| 7.698e-23 \| 598 \| 0.213 \| 365 \| 203 \| 10 \| 12 \| 298 \| 7 \| 365 \| Baseplate J protein \| Baseplate J protein; afdb-uniprot50 \| AF-D6H7Q2-F1-MODEL\_V4 \| 1.0 \| 7.813e-16 \| 597 \| 0.414 \| 181 \| 96 \| 4 \| 10 \| 183 \| 6 \| 183 \| Baseplate\_J domain-containing protein \| Baseplate\_J domain-containing protein; afdb-uniprot50 \| AF-A0A553SNF2-F1-MODEL\_V4 \| 1.0 \| 1.777e-22 \| 595 \| 0.18 \| 372 \| 218 \| 10 \| 12 \| 300 \| 5 \| 372 \| Baseplate\_J domain-containing protein \| Baseplate\_J domain-containing protein; afdb-uniprot50 \| AF-A0A133NQP8-F1-MODEL\_V4 \| 1.0 \| 1.762e-18 \| 592 \| 0.163 \| 361 \| 211 \| 14 \| 23 \| 298 \| 10 \| 364 \| Baseplate J-like protein \| Baseplate J-like protein; afdb-uniprot50 \| AF-R7Z8V1-F1-MODEL\_V4 \| 1.0 \| 8.402e-22 \| 592 \| 0.22 \| 372 \| 198 \| 11 \| 12 \| 298 \| 5 \| 369 \| Baseplate\_J domain-containing protein \| Baseplate\_J domain-containing protein; afdb-uniprot50 \| AF-A0A285NE15-F1-MODEL\_V4 \| 1.0 \| 2.126e-22 \| 591 \| 0.258 \| 364 \| 178 \| 12 \| 19 \| 295 \| 2 \| 360 \| Phage-related baseplate assembly protein \| Phage-related baseplate assembly protein; afdb-uniprot50 \| AF-A0A7G8VI16-F1-MODEL\_V4 \| 1.0 \| 2.665e-16 \| 589 \| 0.535 \| 181 \| 79 \| 3 \| 8 \| 184 \| 3 \| 182 \| Baseplate J/gp47 family protein \| Baseplate J/gp47 family protein; afdb-uniprot50 \| AF-R6IMP8-F1-MODEL\_V4 \| 1.0 \| 2.7e-22 \| 588 \| 0.217 \| 377 \| 206 \| 10 \| 8 \| 300 \| 1 \| 372 \| Putative bacteriophage baseplate assembly protein \| Putative bacteriophage baseplate assembly protein; afdb-uniprot50 \| AF-A0A7U9RBG4-F1-MODEL\_V4 \| 1.0 \| 6.743e-17 \| 587 \| 0.288 \| 194 \| 130 \| 6 \| 111 \| 300 \| 41 \| 230 \| Baseplate\_J domain-containing protein \| Baseplate\_J domain-containing protein; afdb-uniprot50 \| AF-A0A098F587-F1-MODEL\_V4 \| 1.0 \| 2.126e-22 \| 586 \| 0.203 \| 373 \| 208 \| 10 \| 12 \| 300 \| 5 \| 372 \| Baseplate J-like protein \| Baseplate J-like protein; afdb-uniprot50 \| AF-A0A143HC25-F1-MODEL\_V4 \| 1.0 \| 2.866e-22 \| 580 \| 0.192 \| 369 \| 215 \| 9 \| 12 \| 300 \| 5 \| 370 \| Baseplate\_J domain-containing protein \| Baseplate\_J domain-containing protein; afdb-uniprot50 \| AF-A0A1A9G8H2-F1-MODEL\_V4 \| 1.0 \| 3.831e-18 \| 580 \| 0.158 \| 378 \| 228 \| 11 \| 6 \| 300 \| 3 \| 373 \| Phage baseplate assembly protein \| Phage baseplate assembly protein; afdb-uniprot50 \| AF-A0A162QMQ9-F1-MODEL\_V4 \| 1.0 \| 3.864e-22 \| 580 \| 0.215 \| 381 \| 208 \| 10 \| 8 \| 300 \| 1 \| 378 \| Baseplate J-like protein \| Baseplate J-like protein; afdb-uniprot50 \| AF-A0A511Z870-F1-MODEL\_V4 \| 1.0 \| 2.002e-22 \| 578 \| 0.233 \| 381 \| 200 \| 12 \| 8 \| 300 \| 1 \| 377 \| Baseplate\_J domain-containing protein \| Baseplate\_J domain-containing protein; afdb-uniprot50 \| AF-A0A2S0JKS5-F1-MODEL\_V4 \| 1.0 \| 2.543e-22 \| 577 \| 0.218 \| 376 \| 200 \| 14 \| 12 \| 300 \| 4 \| 372 \| Baseplate J protein \| Baseplate J protein; afdb-uniprot50 \| AF-G9PUJ5-F1-MODEL\_V4 \| 1.0 \| 1.577e-22 \| 577 \| 0.242 \| 391 \| 187 \| 14 \| 10 \| 300 \| 4 \| 385 \| Baseplate\_J domain-containing protein \| Baseplate\_J domain-containing protein; afdb-uniprot50 \| AF-A0A2U2DFB6-F1-MODEL\_V4 \| 1.0 \| 9.923e-16 \| 576 \| 0.355 \| 149 \| 96 \| 0 \| 148 \| 296 \| 1 \| 149 \| Baseplate\_J domain-containing protein \| Baseplate\_J domain-containing protein; afdb-uniprot50 \| AF-A0A6L9AAM5-F1-MODEL\_V4 \| 1.0 \| 9.052e-15 \| 576 \| 0.329 \| 161 \| 104 \| 3 \| 141 \| 300 \| 2 \| 159 \| Baseplate protein \| Baseplate protein; afdb-uniprot50 \| AF-A0A830KCY6-F1-MODEL\_V4 \| 1.0 \| 2.603e-19 \| 574 \| 0.236 \| 355 \| 175 \| 9 \| 1 \| 270 \| 1 \| 344 \| Baseplate assembly protein \| Baseplate assembly protein; afdb-uniprot50 \| AF-A0A7X2NKK3-F1-MODEL\_V4 \| 1.0 \| 7.455e-22 \| 572 \| 0.214 \| 383 \| 212 \| 8 \| 4 \| 300 \| 2 \| 381 \| Baseplate J/gp47 family protein \| Baseplate J/gp47 family protein; afdb-uniprot50 \| AF-A0A645G9Q1-F1-MODEL\_V4 \| 1.0 \| 2.829e-16 \| 571 \| 0.291 \| 192 \| 130 \| 5 \| 113 \| 300 \| 17 \| 206 \| Baseplate\_J domain-containing protein \| Baseplate\_J domain-containing protein; afdb-uniprot50 \| AF-A0A5Z4ZU36-F1-MODEL\_V4 \| 1.0 \| 7.813e-16 \| 571 \| 0.271 \| 188 \| 132 \| 4 \| 116 \| 300 \| 25 \| 210 \| Baseplate J protein \| Baseplate J protein; afdb-uniprot50 \| AF-A0A7J6YK41-F1-MODEL\_V4 \| 1.0 \| 1.058e-17 \| 571 \| 0.379 \| 237 \| 124 \| 6 \| 13 \| 249 \| 139 \| 352 \| Baseplate\_J domain-containing protein \| Baseplate\_J domain-containing protein; afdb-uniprot50 \| AF-A0A3A9EFF6-F1-MODEL\_V4 \| 1.0 \| 3.385e-16 \| 568 \| 0.291 \| 192 \| 128 \| 4 \| 116 \| 300 \| 22 \| 212 \| Phage baseplate protein \| Phage baseplate protein; afdb-uniprot50 \| AF-A0A542AK95-F1-MODEL\_V4 \| 1.0 \| 7.023e-22 \| 568 \| 0.216 \| 365 \| 196 \| 10 \| 20 \| 298 \| 3 \| 363 \| Phage-related baseplate assembly protein \| Phage-related baseplate assembly protein; afdb-uniprot50 \| AF-A0A7T2U7T2-F1-MODEL\_V4 \| 1.0 \| 1.854e-14 \| 567 \| 0.437 \| 135 \| 74 \| 1 \| 166 \| 300 \| 2 \| 134 \| Baseplate J/gp47 family protein \| Baseplate J/gp47 family protein; afdb-uniprot50 \| AF-B8IDQ2-F1-MODEL\_V4 \| 1.0 \| 5.637e-17 \| 566 \| 0.353 \| 269 \| 138 \| 11 \| 1 \| 241 \| 1 \| 261 \| Baseplate J family protein \| Baseplate J family protein; afdb-uniprot50 \| AF-A0A4U8UDU1-F1-MODEL\_V4 \| 1.0 \| 6.061e-23 \| 565 \| 0.225 \| 368 \| 193 \| 11 \| 15 \| 300 \| 1 \| 358 \| Baseplate\_J domain-containing protein \| Baseplate\_J domain-containing protein; afdb-uniprot50 \| AF-A0A6L9HNT2-F1-MODEL\_V4 \| 1.0 \| 1.133e-21 \| 564 \| 0.212 \| 382 \| 210 \| 9 \| 8 \| 300 \| 3 \| 382 \| Baseplate J/gp47 family protein \| Baseplate J/gp47 family protein; afdb-uniprot50 \| AF-A0A7J0BHX2-F1-MODEL\_V4 \| 1.0 \| 9.17e-21 \| 563 \| 0.228 \| 372 \| 197 \| 10 \| 13 \| 300 \| 1 \| 366 \| Baseplate protein \| Baseplate protein; afdb-uniprot50 \| AF-Q4E9A8-F1-MODEL\_V4 \| 1.0 \| 1.155e-16 \| 562 \| 0.285 \| 210 \| 130 \| 5 \| 85 \| 294 \| 94 \| 283 \| Baseplate\_J domain-containing protein \| Baseplate\_J domain-containing protein; afdb-uniprot50 \| AF-A0A496P7D6-F1-MODEL\_V4 \| 1.0 \| 3.23e-22 \| 562 \| 0.232 \| 365 \| 192 \| 10 \| 10 \| 290 \| 3 \| 363 \| Baseplate J/gp47 family protein \| Baseplate J/gp47 family protein; afdb-uniprot50 \| AF-A0A7J5WEF0-F1-MODEL\_V4 \| 1.0 \| 1.202e-21 \| 562 \| 0.237 \| 379 \| 192 \| 11 \| 10 \| 300 \| 5 \| 374 \| Baseplate J family \| Baseplate J family; afdb-uniprot50 \| AF-A0A1Y4WAU9-F1-MODEL\_V4 \| 1.0 \| 3.742e-21 \| 562 \| 0.221 \| 383 \| 207 \| 10 \| 7 \| 300 \| 5 \| 385 \| Phage tail protein \| Phage tail protein; afdb-uniprot50 \| AF-A0A1M7LUN9-F1-MODEL\_V4 \| 1.0 \| 3.525e-21 \| 561 \| 0.195 \| 378 \| 215 \| 8 \| 10 \| 300 \| 8 \| 383 \| Phage-related baseplate assembly protein \| Phage-related baseplate assembly protein; afdb-uniprot50 \| AF-V7ZEQ2-F1-MODEL\_V4 \| 1.0 \| 2.431e-15 \| 560 \| 0.436 \| 174 \| 94 \| 3 \| 1 \| 174 \| 1 \| 170 \| Phage-related baseplate assembly protein \| Phage-related baseplate assembly protein; afdb-uniprot50 \| AF-A0A150KS38-F1-MODEL\_V4 \| 1.0 \| 3.128e-21 \| 559 \| 0.193 \| 373 \| 212 \| 9 \| 12 \| 300 \| 5 \| 372 \| Baseplate\_J domain-containing protein \| Baseplate\_J domain-containing protein; afdb-uniprot50 \| AF-A0A6N7B8U0-F1-MODEL\_V4 \| 1.0 \| 1.621e-21 \| 555 \| 0.204 \| 396 \| 200 \| 13 \| 13 \| 300 \| 4 \| 392 \| Baseplate\_J domain-containing protein \| Baseplate\_J domain-containing protein; afdb-uniprot50 \| AF-A0A1X7M9Q2-F1-MODEL\_V4 \| 1.0 \| 2.817e-14 \| 553 \| 0.454 \| 121 \| 66 \| 0 \| 179 \| 299 \| 1 \| 121 \| Baseplate assembly protein J \| Baseplate assembly protein J; afdb-uniprot50 \| AF-A0A4U8S342-F1-MODEL\_V4 \| 1.0 \| 1.058e-17 \| 550 \| 0.16 \| 356 \| 211 \| 12 \| 13 \| 294 \| 1 \| 342 \| Baseplate\_J domain-containing protein \| Baseplate\_J domain-containing protein; afdb-uniprot50 \| AF-A0A0M1HFG5-F1-MODEL\_V4 \| 1.0 \| 2.909e-15 \| 549 \| 0.381 \| 173 \| 98 \| 3 \| 55 \| 226 \| 2 \| 166 \| Baseplate assembly protein \| Baseplate assembly protein; afdb-uniprot50 \| AF-A0A7X5CJX7-F1-MODEL\_V4 \| 1.0 \| 5.685e-21 \| 547 \| 0.214 \| 383 \| 209 \| 10 \| 7 \| 300 \| 5 \| 384 \| Baseplate J/gp47 family protein \| Baseplate J/gp47 family protein; afdb-uniprot50 \| AF-F0CB87-F1-MODEL\_V4 \| 1.0 \| 1.854e-14 \| 547 \| 0.503 \| 163 \| 77 \| 3 \| 1 \| 163 \| 1 \| 159 \| Alpha-L-arabinofuranosidase \| Alpha-L-arabinofuranosidase; afdb-uniprot50 \| AF-A0A714TMI2-F1-MODEL\_V4 \| 1.0 \| 7.128e-15 \| 546 \| 0.518 \| 164 \| 72 \| 2 \| 6 \| 162 \| 2 \| 165 \| Baseplate assembly protein \| Baseplate assembly protein; afdb-uniprot50 \| AF-V7IAG1-F1-MODEL\_V4 \| 1.0 \| 3.694e-15 \| 546 \| 0.331 \| 172 \| 104 \| 4 \| 132 \| 295 \| 3 \| 171 \| Baseplate\_J domain-containing protein \| Baseplate\_J domain-containing protein; afdb-uniprot50 \| AF-A0A744CQV0-F1-MODEL\_V4 \| 1.0 \| 7.392e-18 \| 546 \| 0.204 \| 313 \| 163 \| 9 \| 72 \| 300 \| 4 \| 314 \| Baseplate J/gp47 family protein \| Baseplate J/gp47 family protein; afdb-uniprot50 \| AF-A0A5C8BGL4-F1-MODEL\_V4 \| 1.0 \| 4.084e-20 \| 546 \| 0.181 \| 374 \| 215 \| 12 \| 10 \| 298 \| 5 \| 372 \| Baseplate\_J domain-containing protein \| Baseplate\_J domain-containing protein; afdb-uniprot50 \| AF-E5VKI5-F1-MODEL\_V4 \| 1.0 \| 9.734e-21 \| 546 \| 0.196 \| 382 \| 217 \| 10 \| 7 \| 300 \| 5 \| 384 \| Baseplate J-like protein \| Baseplate J-like protein; afdb-uniprot50 \| AF-A0A0T9LR98-F1-MODEL\_V4 \| 1.0 \| 1.053e-15 \| 545 \| 0.378 \| 193 \| 109 \| 3 \| 6 \| 190 \| 2 \| 191 \| Baseplate assembly protein \| Baseplate assembly protein; afdb-uniprot50 \| AF-F8KPM6-F1-MODEL\_V4 \| 1.0 \| 1.473e-18 \| 544 \| 0.193 \| 361 \| 201 \| 14 \| 14 \| 298 \| 1 \| 347 \| Baseplate assembly protein J, putative \| Baseplate assembly protein J, putative; afdb-uniprot50 \| AF-A0A3R6RHW9-F1-MODEL\_V4 \| 1.0 \| 1.557e-16 \| 544 \| 0.143 \| 382 \| 234 \| 11 \| 10 \| 300 \| 8 \| 387 \| Baseplate\_J domain-containing protein \| Baseplate\_J domain-containing protein; afdb-uniprot50 \| AF-A0A261QMI5-F1-MODEL\_V4 \| 1.0 \| 2.158e-15 \| 543 \| 0.336 \| 196 \| 117 \| 6 \| 12 \| 204 \| 4 \| 189 \| Phage tail protein \| Phage tail protein; afdb-uniprot50 \| AF-A0A6N3FJC1-F1-MODEL\_V4 \| 1.0 \| 2.947e-21 \| 543 \| 0.205 \| 390 \| 211 \| 12 \| 4 \| 300 \| 11 \| 394 \| Baseplate J-like protein \| Baseplate J-like protein; afdb-uniprot50 \| AF-A0A385Q5A3-F1-MODEL\_V4 \| 1.0 \| 4.217e-21 \| 542 \| 0.174 \| 379 \| 223 \| 10 \| 10 \| 300 \| 2 \| 378 \| Baseplate J/gp47 family protein \| Baseplate J/gp47 family protein; afdb-uniprot50 \| AF-A0A416B7S2-F1-MODEL\_V4 \| 1.0 \| 2.032e-15 \| 541 \| 0.313 \| 172 \| 113 \| 4 \| 132 \| 300 \| 5 \| 174 \| Baseplate\_J domain-containing protein \| Baseplate\_J domain-containing protein; afdb-uniprot50 \| AF-A0A377BF30-F1-MODEL\_V4 \| 1.0 \| 3.188e-16 \| 541 \| 0.265 \| 211 \| 137 \| 6 \| 106 \| 300 \| 39 \| 247 \| Phage-related baseplate assembly protein J \| Phage-related baseplate assembly protein J; afdb-uniprot50 \| AF-A0A379YH24-F1-MODEL\_V4 \| 1.0 \| 2.218e-14 \| 540 \| 0.555 \| 144 \| 57 \| 1 \| 8 \| 144 \| 4 \| 147 \| Baseplate J-like protein \| Baseplate J-like protein; afdb-uniprot50 \| AF-A0A369Y1S1-F1-MODEL\_V4 \| 1.0 \| 2.688e-20 \| 538 \| 0.19 \| 363 \| 208 \| 8 \| 20 \| 300 \| 7 \| 365 \| Baseplate\_J domain-containing protein \| Baseplate\_J domain-containing protein; afdb-uniprot50 \| AF-A0A0Q3ETP3-F1-MODEL\_V4 \| 1.0 \| 3.71e-17 \| 537 \| 0.186 \| 381 \| 211 \| 15 \| 1 \| 294 \| 1 \| 369 \| Baseplate\_J domain-containing protein \| Baseplate\_J domain-containing protein; afdb-uniprot50 \| AF-A0A285M2R0-F1-MODEL\_V4 \| 1.0 \| 9.387e-18 \| 537 \| 0.172 \| 388 \| 222 \| 13 \| 1 \| 300 \| 5 \| 381 \| Phage-related baseplate assembly protein \| Phage-related baseplate assembly protein; afdb-uniprot50 \| AF-A0A2N3D694-F1-MODEL\_V4 \| 1.0 \| 1.46e-14 \| 536 \| 0.537 \| 162 \| 71 \| 1 \| 1 \| 162 \| 7 \| 164 \| Baseplate assembly protein \| Baseplate assembly protein; afdb-uniprot50 \| AF-U2EPS9-F1-MODEL\_V4 \| 1.0 \| 4.98e-15 \| 530 \| 0.276 \| 181 \| 116 \| 8 \| 122 \| 300 \| 2 \| 169 \| Baseplate assembly protein J, putative \| Baseplate assembly protein J, putative; afdb-uniprot50 \| AF-A0A810Q3S1-F1-MODEL\_V4 \| 1.0 \| 4.084e-20 \| 530 \| 0.212 \| 381 \| 207 \| 11 \| 10 \| 300 \| 7 \| 384 \| Uncharacterized protein \| Uncharacterized protein; afdb-uniprot50 \| AF-B3QTJ3-F1-MODEL\_V4 \| 1.0 \| 2.776e-21 \| 529 \| 0.22 \| 381 \| 193 \| 13 \| 12 \| 294 \| 3 \| 377 \| Baseplate J family protein \| Baseplate J family protein; afdb-uniprot50 \| AF-A0A6V7E4B2-F1-MODEL\_V4 \| 1.0 \| 3.694e-15 \| 529 \| 0.456 \| 186 \| 88 \| 4 \| 1 \| 179 \| 1 \| 180 \| Alpha-L-AF\_C domain-containing protein \| Alpha-L-AF\_C domain-containing protein; afdb-uniprot50 \| AF-E3HBK6-F1-MODEL\_V4 \| 1.0 \| 4.199e-19 \| 528 \| 0.153 \| 371 \| 220 \| 11 \| 20 \| 300 \| 7 \| 373 \| Baseplate J family protein \| Baseplate J family protein; afdb-uniprot50 \| AF-A0A2X3LRF3-F1-MODEL\_V4 \| 1.0 \| 7.567e-15 \| 527 \| 0.327 \| 162 \| 105 \| 4 \| 133 \| 292 \| 1 \| 160 \| Tail protein I \| Tail protein I; afdb-uniprot50 \| AF-A0A2E3R244-F1-MODEL\_V4 \| 1.0 \| 6.587e-20 \| 527 \| 0.224 \| 370 \| 195 \| 13 \| 13 \| 298 \| 1 \| 362 \| Baseplate\_J domain-containing protein \| Baseplate\_J domain-containing protein; afdb-uniprot50 \| AF-A0A1K1LB82-F1-MODEL\_V4 \| 1.0 \| 2.167e-17 \| 527 \| 0.176 \| 391 \| 208 \| 20 \| 15 \| 299 \| 1 \| 383 \| Baseplate assembly protein, putative \| Baseplate assembly protein, putative; afdb-uniprot50 \| AF-A0A7U9S6B3-F1-MODEL\_V4 \| 1.0 \| 2.355e-14 \| 526 \| 0.269 \| 171 \| 118 \| 5 \| 133 \| 298 \| 5 \| 173 \| Baseplate\_J domain-containing protein \| Baseplate\_J domain-containing protein; afdb-uniprot50 \| AF-A0A416HFK7-F1-MODEL\_V4 \| 1.0 \| 2.921e-17 \| 526 \| 0.137 \| 378 \| 223 \| 12 \| 17 \| 300 \| 9 \| 377 \| Baseplate\_J domain-containing protein \| Baseplate\_J domain-containing protein; afdb-uniprot50 \| AF-A0A2X2UEX9-F1-MODEL\_V4 \| 1.0 \| 7.423e-20 \| 526 \| 0.195 \| 378 \| 213 \| 10 \| 12 \| 300 \| 4 \| 379 \| Baseplate J family protein \| Baseplate J family protein; afdb-uniprot50 \| AF-A0A1G3LAY8-F1-MODEL\_V4 \| 1.0 \| 1.301e-16 \| 525 \| 0.159 \| 377 \| 216 \| 14 \| 18 \| 300 \| 2 \| 371 \| Baseplate\_J domain-containing protein \| Baseplate\_J domain-containing protein; afdb-uniprot50 \| AF-A0A1W2CW87-F1-MODEL\_V4 \| 1.0 \| 4.865e-18 \| 525 \| 0.171 \| 403 \| 216 \| 14 \| 6 \| 300 \| 2 \| 394 \| Phage-related baseplate assembly protein \| Phage-related baseplate assembly protein; afdb-uniprot50 \| AF-A0A1Q6JPW8-F1-MODEL\_V4 \| 1.0 \| 2.688e-20 \| 524 \| 0.218 \| 366 \| 197 \| 8 \| 22 \| 300 \| 1 \| 364 \| Baseplate\_J domain-containing protein \| Baseplate\_J domain-containing protein; afdb-uniprot50 \| AF-A0A7U9RWZ8-F1-MODEL\_V4 \| 1.0 \| 2.375e-18 \| 523 \| 0.19 \| 325 \| 171 \| 11 \| 64 \| 300 \| 2 \| 322 \| Baseplate\_J domain-containing protein \| Baseplate\_J domain-containing protein; afdb-uniprot50 \| AF-A0A1H8P6M7-F1-MODEL\_V4 \| 1.0 \| 1.692e-13 \| 520 \| 0.46 \| 150 \| 74 \| 1 \| 8 \| 150 \| 4 \| 153 \| Uncharacterized protein \| Uncharacterized protein; afdb-uniprot50 \| AF-A0A4R2GWK6-F1-MODEL\_V4 \| 1.0 \| 9.651e-17 \| 515 \| 0.17 \| 393 \| 217 \| 14 \| 1 \| 295 \| 6 \| 387 \| Phage-related baseplate assembly protein \| Phage-related baseplate assembly protein; afdb-uniprot50 \| AF-A0A4P0TK43-F1-MODEL\_V4 \| 1.0 \| 8.767e-14 \| 514 \| 0.452 \| 146 \| 80 \| 0 \| 116 \| 261 \| 6 \| 151 \| Putative phage baseplate assembly protein \| Putative phage baseplate assembly protein; afdb-uniprot50 \| AF-A0A841I6H3-F1-MODEL\_V4 \| 1.0 \| 2.728e-13 \| 512 \| 0.5 \| 148 \| 67 \| 1 \| 8 \| 148 \| 4 \| 151 \| Phage-related baseplate assembly protein \| Phage-related baseplate assembly protein; afdb-uniprot50 \| AF-A0A2V2GNU0-F1-MODEL\_V4 \| 1.0 \| 1.128e-19 \| 510 \| 0.195 \| 378 \| 212 \| 8 \| 12 \| 300 \| 9 \| 383 \| Phage tail protein \| Phage tail protein; afdb-uniprot50 \| AF-A0A7C7ANJ6-F1-MODEL\_V4 \| 1.0 \| 1.301e-16 \| 508 \| 0.16 \| 362 \| 210 \| 13 \| 21 \| 296 \| 1 \| 354 \| Baseplate J/gp47 family protein \| Baseplate J/gp47 family protein; afdb-uniprot50 \| AF-A0A6I5ZPJ9-F1-MODEL\_V4 \| 1.0 \| 8.564e-17 \| 508 \| 0.165 \| 381 \| 211 \| 16 \| 20 \| 300 \| 8 \| 381 \| Baseplate J-like protein \| Baseplate J-like protein; afdb-uniprot50 \| AF-A0A3A9GRB5-F1-MODEL\_V4 \| 1.0 \| 1.479e-20 \| 508 \| 0.196 \| 377 \| 208 \| 12 \| 12 \| 300 \| 12 \| 381 \| Baseplate J/gp47 family protein \| Baseplate J/gp47 family protein; afdb-uniprot50 \| AF-A0A1B1NTC2-F1-MODEL\_V4 \| 1.0 \| 1.57e-20 \| 506 \| 0.2 \| 379 \| 207 \| 14 \| 10 \| 300 \| 5 \| 375 \| Baseplate\_J domain-containing protein \| Baseplate\_J domain-containing protein; afdb-uniprot50 \| AF-A0A7C9M632-F1-MODEL\_V4 \| 1.0 \| 1.667e-20 \| 505 \| 0.212 \| 372 \| 194 \| 12 \| 20 \| 298 \| 5 \| 370 \| Baseplate J/gp47 family protein \| Baseplate J/gp47 family protein; afdb-uniprot50 \| AF-A0A3N6RV94-F1-MODEL\_V4 \| 1.0 \| 4.336e-20 \| 505 \| 0.302 \| 367 \| 154 \| 12 \| 6 \| 300 \| 2 \| 338 \| Phage tail protein I \| Phage tail protein I; afdb-uniprot50 \| AF-D1P8F2-F1-MODEL\_V4 \| 1.0 \| 2.5e-14 \| 501 \| 0.481 \| 158 \| 79 \| 1 \| 25 \| 179 \| 2 \| 159 \| Baseplate J-like protein \| Baseplate J-like protein; afdb-uniprot50 \| AF-A0A2V2FW36-F1-MODEL\_V4 \| 1.0 \| 2.041e-17 \| 501 \| 0.169 \| 461 \| 198 \| 16 \| 15 \| 300 \| 10 \| 460 \| Uncharacterized protein \| Uncharacterized protein; afdb-uniprot50 \| AF-A0A3G2Q4N8-F1-MODEL\_V4 \| 1.0 \| 1.349e-19 \| 499 \| 0.188 \| 381 \| 213 \| 13 \| 12 \| 300 \| 8 \| 384 \| Phage baseplate protein \| Phage baseplate protein; afdb-uniprot50 \| AF-A0A4V3WD86-F1-MODEL\_V4 \| 1.0 \| 1.923e-17 \| 494 \| 0.191 \| 376 \| 202 \| 16 \| 12 \| 296 \| 8 \| 372 \| Baseplate J protein \| Baseplate J protein; afdb-uniprot50 \| AF-A0A2C6MEH4-F1-MODEL\_V4 \| 1.0 \| 1.652e-16 \| 493 \| 0.134 \| 373 \| 224 \| 12 \| 17 \| 296 \| 9 \| 375 \| Baseplate J protein \| Baseplate J protein; afdb-uniprot50 \| AF-A0A2A7AEU6-F1-MODEL\_V4 \| 1.0 \| 1.652e-16 \| 493 \| 0.151 \| 384 \| 227 \| 12 \| 13 \| 300 \| 1 \| 381 \| Phage baseplate protein \| Phage baseplate protein; afdb-uniprot50 \| AF-A0A323U4H9-F1-MODEL\_V4 \| 1.0 \| 1.26e-15 \| 492 \| 0.175 \| 377 \| 211 \| 15 \| 13 \| 300 \| 1 \| 366 \| Baseplate J protein \| Baseplate J protein; afdb-uniprot50 \| AF-A0A373C6W2-F1-MODEL\_V4 \| 1.0 \| 5.023e-19 \| 492 \| 0.212 \| 382 \| 206 \| 13 \| 11 \| 300 \| 2 \| 380 \| Baseplate J/gp47 family protein \| Baseplate J/gp47 family protein; afdb-uniprot50 \| AF-A0A2N8HQF9-F1-MODEL\_V4 \| 1.0 \| 5.023e-19 \| 489 \| 0.204 \| 391 \| 199 \| 17 \| 12 \| 298 \| 2 \| 384 \| Baseplate\_J domain-containing protein \| Baseplate\_J domain-containing protein; afdb-uniprot50 \| AF-A0A413J6B9-F1-MODEL\_V4 \| 1.0 \| 9.131e-19 \| 489 \| 0.198 \| 378 \| 212 \| 11 \| 10 \| 298 \| 8 \| 383 \| Phage tail protein \| Phage tail protein; afdb-uniprot50 \| AF-A0A0T9TB71-F1-MODEL\_V4 \| 1.0 \| 7.328e-14 \| 488 \| 0.464 \| 168 \| 75 \| 2 \| 6 \| 158 \| 2 \| 169 \| Regulator of late gene expression \| Regulator of late gene expression; afdb-uniprot50 \| AF-A0A3D9PVP8-F1-MODEL\_V4 \| 1.0 \| 1.6e-15 \| 488 \| 0.162 \| 364 \| 211 \| 11 \| 13 \| 292 \| 1 \| 354 \| Putative phage protein gp47/JayE \| Putative phage protein gp47/JayE; afdb-uniprot50 \| AF-A0A0K6A654-F1-MODEL\_V4 \| 1.0 \| 6.298e-13 \| 487 \| 0.395 \| 134 \| 81 \| 0 \| 166 \| 299 \| 2 \| 135 \| Baseplate assembly protein J \| Baseplate assembly protein J; afdb-uniprot50 \| AF-A0A824DTY0-F1-MODEL\_V4 \| 1.0 \| 3.922e-15 \| 487 \| 0.246 \| 203 \| 119 \| 5 \| 116 \| 300 \| 16 \| 202 \| Baseplate assembly protein \| Baseplate assembly protein; afdb-uniprot50 \| AF-A0A3S5XQX9-F1-MODEL\_V4 \| 1.0 \| 2.489e-12 \| 484 \| 0.554 \| 128 \| 50 \| 1 \| 6 \| 126 \| 3 \| 130 \| Baseplate J-like protein \| Baseplate J-like protein; afdb-uniprot50 \| AF-A0A366IJ99-F1-MODEL\_V4 \| 1.0 \| 6.475e-12 \| 482 \| 0.484 \| 99 \| 51 \| 0 \| 201 \| 299 \| 1 \| 99 \| Uncharacterized protein \| Uncharacterized protein; afdb-uniprot50 \| AF-A0A352X1L5-F1-MODEL\_V4 \| 1.0 \| 5.795e-16 \| 481 \| 0.163 \| 380 \| 214 \| 14 \| 1 \| 295 \| 1 \| 361 \| Baseplate\_J domain-containing protein \| Baseplate\_J domain-containing protein; afdb-uniprot50 \| AF-A0A418GPV5-F1-MODEL\_V4 \| 1.0 \| 9.88e-14 \| 480 \| 0.348 \| 175 \| 100 \| 3 \| 113 \| 275 \| 68 \| 240 \| Baseplate J/gp47 family protein \| Baseplate J/gp47 family protein; afdb-uniprot50 \| AF-A0A1H8DLC4-F1-MODEL\_V4 \| 1.0 \| 1.301e-16 \| 478 \| 0.176 \| 368 \| 204 \| 15 \| 21 \| 300 \| 1 \| 357 \| Uncharacterized phage protein gp47/JayE \| Uncharacterized phage protein gp47/JayE; afdb-uniprot50 \| AF-A0A080HTY9-F1-MODEL\_V4 \| 1.0 \| 1.016e-12 \| 477 \| 0.355 \| 135 \| 87 \| 0 \| 161 \| 295 \| 1 \| 135 \| Baseplate J-like family protein \| Baseplate J-like family protein; afdb-uniprot50 \| AF-A0A2G2D437-F1-MODEL\_V4 \| 1.0 \| 2.237e-18 \| 477 \| 0.164 \| 383 \| 231 \| 11 \| 1 \| 300 \| 4 \| 380 \| Phage baseplate assembly protein \| Phage baseplate assembly protein; afdb-uniprot50 \| AF-W4RXX9-F1-MODEL\_V4 \| 1.0 \| 6.686e-13 \| 475 \| 0.532 \| 139 \| 62 \| 2 \| 1 \| 139 \| 1 \| 136 \| Phage-related baseplate assembly protein \| Phage-related baseplate assembly protein; afdb-uniprot50 \| AF-A0A7I9SRD3-F1-MODEL\_V4 \| 1.0 \| 7.159e-17 \| 474 \| 0.205 \| 331 \| 149 \| 12 \| 67 \| 300 \| 2 \| 315 \| Baseplate assembly protein \| Baseplate assembly protein; afdb-uniprot50 \| AF-A0A268TIH4-F1-MODEL\_V4 \| 1.0 \| 2.099e-16 \| 474 \| 0.171 \| 356 \| 200 \| 14 \| 16 \| 294 \| 7 \| 344 \| Baseplate\_J domain-containing protein \| Baseplate\_J domain-containing protein; afdb-uniprot50 \| AF-A0A7V5GUY7-F1-MODEL\_V4 \| 1.0 \| 7.78e-14 \| 473 \| 0.254 \| 185 \| 128 \| 6 \| 116 \| 296 \| 4 \| 182 \| Baseplate\_J domain-containing protein \| Baseplate\_J domain-containing protein; afdb-uniprot50 \| AF-A0A842IZ09-F1-MODEL\_V4 \| 1.0 \| 5.264e-13 \| 472 \| 0.225 \| 186 \| 138 \| 5 \| 115 \| 298 \| 137 \| 318 \| Baseplate J/gp47 family protein \| Baseplate J/gp47 family protein; afdb-uniprot50 \| AF-W4RXF7-F1-MODEL\_V4 \| 1.0 \| 3.678e-13 \| 471 \| 0.484 \| 163 \| 78 \| 3 \| 1 \| 163 \| 1 \| 157 \| Phage-related baseplate assembly protein \| Phage-related baseplate assembly protein; afdb-uniprot50 \| AF-W4SEG4-F1-MODEL\_V4 \| 1.0 \| 4.015e-12 \| 470 \| 0.514 \| 138 \| 64 \| 2 \| 1 \| 138 \| 1 \| 135 \| Uncharacterized protein \| Uncharacterized protein; afdb-uniprot50 \| AF-A0A5U3E105-F1-MODEL\_V4 \| 1.0 \| 5.933e-13 \| 469 \| 0.51 \| 145 \| 64 \| 1 \| 8 \| 145 \| 1 \| 145 \| Baseplate assembly protein \| Baseplate assembly protein; afdb-uniprot50 \| AF-A0A511NB39-F1-MODEL\_V4 \| 1.0 \| 6.715e-15 \| 468 \| 0.155 \| 372 \| 213 \| 15 \| 14 \| 295 \| 3 \| 363 \| Baseplate\_J domain-containing protein \| Baseplate\_J domain-containing protein; afdb-uniprot50 \| AF-A0A767SN15-F1-MODEL\_V4 \| 1.0 \| 4.651e-11 \| 466 \| 0.44 \| 93 \| 52 \| 0 \| 207 \| 299 \| 1 \| 93 \| Baseplate assembly protein \| Baseplate assembly protein; afdb-uniprot50 \| AF-A0A4D7AZ74-F1-MODEL\_V4 \| 1.0 \| 9.88e-14 \| 464 \| 0.214 \| 196 \| 140 \| 7 \| 110 \| 296 \| 100 \| 290 \| Baseplate\_J domain-containing protein \| Baseplate\_J domain-containing protein; afdb-uniprot50 \| AF-A0A4S0JS99-F1-MODEL\_V4 \| 1.0 \| 3.356e-12 \| 463 \| 0.532 \| 122 \| 55 \| 1 \| 66 \| 187 \| 2 \| 121 \| Baseplate assembly protein \| Baseplate assembly protein; afdb-uniprot50 \| AF-D8A2F7-F1-MODEL\_V4 \| 1.0 \| 1.699e-15 \| 463 \| 0.265 \| 286 \| 122 \| 6 \| 14 \| 215 \| 1 \| 282 \| Baseplate J-like protein \| Baseplate J-like protein; afdb-uniprot50 \| AF-A0A3S4VYP8-F1-MODEL\_V4 \| 1.0 \| 5.795e-16 \| 463 \| 0.254 \| 299 \| 139 \| 7 \| 25 \| 243 \| 2 \| 296 \| Uncharacterized homolog of phage Mu protein gp47 \| Uncharacterized homolog of phage Mu protein gp47; afdb-uniprot50 \| AF-A0A1T5KWL5-F1-MODEL\_V4 \| 1.0 \| 1.26e-15 \| 462 \| 0.134 \| 379 \| 224 \| 13 \| 17 \| 299 \| 8 \| 378 \| Uncharacterized phage protein gp47/JayE \| Uncharacterized phage protein gp47/JayE; afdb-uniprot50 \| AF-W4RY38-F1-MODEL\_V4 \| 1.0 \| 1.906e-13 \| 461 \| 0.469 \| 162 \| 81 \| 3 \| 1 \| 162 \| 1 \| 157 \| Phage-related baseplate assembly protein \| Phage-related baseplate assembly protein; afdb-uniprot50 \| AF-Q5F8P1-F1-MODEL\_V4 \| 1.0 \| 4.401e-13 \| 459 \| 0.431 \| 169 \| 79 \| 5 \| 1 \| 162 \| 1 \| 159 \| Baseplate protein \| Baseplate protein; afdb-uniprot50 \| AF-A0A842J386-F1-MODEL\_V4 \| 1.0 \| 1.369e-12 \| 457 \| 0.227 \| 185 \| 137 \| 4 \| 116 \| 298 \| 87 \| 267 \| Baseplate J/gp47 family protein \| Baseplate J/gp47 family protein; afdb-uniprot50 \| AF-A0A846JQB4-F1-MODEL\_V4 \| 1.0 \| 7.813e-16 \| 456 \| 0.148 \| 371 \| 216 \| 18 \| 12 \| 294 \| 3 \| 361 \| Baseplate J/gp47 family protein \| Baseplate J/gp47 family protein; afdb-uniprot50 \| AF-A0A2T4JNF5-F1-MODEL\_V4 \| 1.0 \| 1.326e-11 \| 455 \| 0.43 \| 123 \| 70 \| 0 \| 173 \| 295 \| 1 \| 123 \| Baseplate assembly protein \| Baseplate assembly protein; afdb-uniprot50 \| AF-A0A3G2R4D9-F1-MODEL\_V4 \| 1.0 \| 1.426e-17 \| 455 \| 0.221 \| 325 \| 164 \| 9 \| 61 \| 298 \| 6 \| 328 \| Baseplate J/gp47 family protein \| Baseplate J/gp47 family protein; afdb-uniprot50 \| AF-A0A1L5KT79-F1-MODEL\_V4 \| 1.0 \| 2.805e-12 \| 454 \| 0.309 \| 152 \| 97 \| 4 \| 154 \| 300 \| 1 \| 149 \| Baseplate J-like protein \| Baseplate J-like protein; afdb-uniprot50 \| AF-A0A3G4W4D3-F1-MODEL\_V4 \| 1.0 \| 5.958e-15 \| 454 \| 0.185 \| 431 \| 187 \| 16 \| 22 \| 296 \| 13 \| 435 \| Baseplate J-like protein \| Baseplate J-like protein; afdb-uniprot50 \| AF-A0A0J9B6B6-F1-MODEL\_V4 \| 1.0 \| 1.408e-11 \| 453 \| 0.534 \| 131 \| 50 \| 2 \| 7 \| 128 \| 3 \| 131 \| Uncharacterized protein \| Uncharacterized protein; afdb-uniprot50 \| AF-A0A1M3AIG6-F1-MODEL\_V4 \| 1.0 \| 7.998e-13 \| 453 \| 0.216 \| 194 \| 136 \| 7 \| 115 \| 296 \| 1 \| 190 \| Uncharacterized protein \| Uncharacterized protein; afdb-uniprot50 \| AF-A0A695HWH1-F1-MODEL\_V4 \| 1.0 \| 2.218e-14 \| 452 \| 0.241 \| 203 \| 120 \| 5 \| 116 \| 300 \| 68 \| 254 \| Baseplate assembly protein \| Baseplate assembly protein; afdb-uniprot50 \| AF-E1SEF6-F1-MODEL\_V4 \| 1.0 \| 8.767e-14 \| 447 \| 0.493 \| 166 \| 70 \| 4 \| 6 \| 163 \| 2 \| 161 \| Baseplate assembly protein J (GpJ) \| Baseplate assembly protein J (GpJ); afdb-uniprot50 \| AF-A0A1M6M9S1-F1-MODEL\_V4 \| 1.0 \| 5.483e-18 \| 447 \| 0.173 \| 357 \| 202 \| 12 \| 21 \| 292 \| 4 \| 352 \| Uncharacterized phage protein gp47/JayE \| Uncharacterized phage protein gp47/JayE; afdb-uniprot50 \| AF-A0A5C7PTQ6-F1-MODEL\_V4 \| 1.0 \| 3.922e-15 \| 446 \| 0.156 \| 389 \| 217 \| 19 \| 12 \| 300 \| 4 \| 381 \| Baseplate\_J domain-containing protein \| Baseplate\_J domain-containing protein; afdb-uniprot50 \| AF-A0A3D3Y478-F1-MODEL\_V4 \| 1.0 \| 3.593e-16 \| 445 \| 0.163 \| 361 \| 205 \| 13 \| 22 \| 295 \| 1 \| 351 \| Baseplate protein J \| Baseplate protein J; afdb-uniprot50 \| AF-A0A7Y7LP48-F1-MODEL\_V4 \| 1.0 \| 1.466e-16 \| 445 \| 0.175 \| 364 \| 203 \| 14 \| 13 \| 292 \| 1 \| 351 \| Putative phage protein gp47/JayE \| Putative phage protein gp47/JayE; afdb-uniprot50 \| AF-A0A5D4RZE4-F1-MODEL\_V4 \| 1.0 \| 1.053e-15 \| 445 \| 0.132 \| 376 \| 213 \| 16 \| 20 \| 295 \| 4 \| 366 \| Baseplate\_J domain-containing protein \| Baseplate\_J domain-containing protein; afdb-uniprot50 \| AF-A0A345J3J0-F1-MODEL\_V4 \| 1.0 \| 1.563e-18 \| 444 \| 0.201 \| 377 \| 209 \| 15 \| 4 \| 300 \| 2 \| 366 \| Phage baseplate assembly protein J \| Phage baseplate assembly protein J; afdb-uniprot50 \| AF-A0A1E3UGI8-F1-MODEL\_V4 \| 1.0 \| 3.593e-16 \| 444 \| 0.163 \| 372 \| 212 \| 11 \| 22 \| 299 \| 14 \| 380 \| Baseplate J protein \| Baseplate J protein; afdb-uniprot50 \| AF-A0A074LIS1-F1-MODEL\_V4 \| 1.0 \| 3.016e-18 \| 442 \| 0.187 \| 357 \| 196 \| 14 \| 22 \| 293 \| 3 \| 350 \| Baseplate\_J domain-containing protein \| Baseplate\_J domain-containing protein; afdb-uniprot50 \| AF-A0A7U6KQX5-F1-MODEL\_V4 \| 1.0 \| 1.543e-12 \| 441 \| 0.445 \| 157 \| 77 \| 3 \| 25 \| 179 \| 2 \| 150 \| Baseplate\_J domain-containing protein \| Baseplate\_J domain-containing protein; afdb-uniprot50 \| AF-A0A521CK67-F1-MODEL\_V4 \| 1.0 \| 2.654e-14 \| 440 \| 0.137 \| 465 \| 213 \| 14 \| 1 \| 295 \| 1 \| 447 \| Baseplate J-like protein \| Baseplate J-like protein; afdb-uniprot50 \| AF-A0A7X6FRN3-F1-MODEL\_V4 \| 1.0 \| 2.281e-13 \| 438 \| 0.346 \| 202 \| 112 \| 5 \| 1 \| 198 \| 4 \| 189 \| Baseplate\_J domain-containing protein \| Baseplate\_J domain-containing protein; afdb-uniprot50 \| AF-A0A376J4L9-F1-MODEL\_V4 \| 1.0 \| 2.642e-12 \| 437 \| 0.412 \| 155 \| 84 \| 2 \| 31 \| 182 \| 1 \| 151 \| Baseplate assembly protein J \| Baseplate assembly protein J; afdb-uniprot50 \| AF-A0A316MUY0-F1-MODEL\_V4 \| 1.0 \| 4.438e-17 \| 437 \| 0.152 \| 360 \| 208 \| 14 \| 22 \| 295 \| 1 \| 349 \| Baseplate protein J \| Baseplate protein J; afdb-uniprot50 \| AF-A0A7W6S2L4-F1-MODEL\_V4 \| 1.0 \| 5.099e-12 \| 436 \| 0.467 \| 122 \| 64 \| 1 \| 1 \| 122 \| 1 \| 121 \| Phage-related baseplate assembly protein \| Phage-related baseplate assembly protein; afdb-uniprot50 \| AF-A0A379ZTU5-F1-MODEL\_V4 \| 1.0 \| 3.356e-12 \| 436 \| 0.514 \| 138 \| 60 \| 1 \| 7 \| 137 \| 3 \| 140 \| Phage baseplate assembly protein J \| Phage baseplate assembly protein J; afdb-uniprot50 \| AF-E3PRX8-F1-MODEL\_V4 \| 1.0 \| 9.347e-16 \| 434 \| 0.129 \| 364 \| 222 \| 11 \| 19 \| 294 \| 3 \| 359 \| Baseplate\_J domain-containing protein \| Baseplate\_J domain-containing protein; afdb-uniprot50 \| AF-A0A1G3KNM2-F1-MODEL\_V4 \| 1.0 \| 5.483e-18 \| 434 \| 0.172 \| 376 \| 207 \| 13 \| 18 \| 300 \| 2 \| 366 \| Baseplate\_J domain-containing protein \| Baseplate\_J domain-containing protein; afdb-uniprot50 \| AF-R6BFZ4-F1-MODEL\_V4 \| 1.0 \| 2.581e-15 \| 433 \| 0.126 \| 355 \| 216 \| 11 \| 23 \| 293 \| 1 \| 345 \| Baseplate J-like protein \| Baseplate J-like protein; afdb-uniprot50 \| AF-A0A357B384-F1-MODEL\_V4 \| 1.0 \| 9.387e-18 \| 432 \| 0.194 \| 359 \| 192 \| 12 \| 21 \| 293 \| 1 \| 348 \| Phage tail protein \| Phage tail protein; afdb-uniprot50 \| AF-A0A7Z8LKE5-F1-MODEL\_V4 \| 1.0 \| 1.02e-14 \| 432 \| 0.166 \| 378 \| 204 \| 16 \| 20 \| 295 \| 3 \| 371 \| Putative phage protein gp47/JayE \| Putative phage protein gp47/JayE; afdb-uniprot50 \| AF-A0A843AG30-F1-MODEL\_V4 \| 1.0 \| 9.651e-17 \| 431 \| 0.158 \| 373 \| 211 \| 11 \| 22 \| 295 \| 13 \| 381 \| Baseplate J/gp47 family protein \| Baseplate J/gp47 family protein; afdb-uniprot50 \| AF-A0A1A9WW65-F1-MODEL\_V4 \| 1.0 \| 1.788e-11 \| 427 \| 0.323 \| 142 \| 92 \| 3 \| 158 \| 298 \| 1 \| 139 \| Baseplate\_J domain-containing protein \| Baseplate\_J domain-containing protein; afdb-uniprot50 \| AF-A0A2A2K3F7-F1-MODEL\_V4 \| 1.0 \| 9.528e-11 \| 427 \| 0.504 \| 123 \| 60 \| 1 \| 8 \| 129 \| 4 \| 126 \| Uncharacterized protein \| Uncharacterized protein; afdb-uniprot50 \| AF-A0A6H1Z9D6-F1-MODEL\_V4 \| 1.0 \| 9.88e-14 \| 427 \| 0.152 \| 375 \| 204 \| 16 \| 23 \| 296 \| 6 \| 367 \| Putative baseplate protein \| Putative baseplate protein; afdb-uniprot50 \| AF-Q9K5D9-F1-MODEL\_V4 \| 1.0 \| 1.375e-14 \| 425 \| 0.24 \| 325 \| 154 \| 7 \| 1 \| 239 \| 1 \| 318 \| Putative base plate assembly protein J \| Putative base plate assembly protein J; afdb-uniprot50 \| AF-J5WM03-F1-MODEL\_V4 \| 1.0 \| 2.281e-13 \| 424 \| 0.113 \| 388 \| 218 \| 14 \| 20 \| 295 \| 10 \| 383 \| Baseplate J-like protein \| Baseplate J-like protein; afdb-uniprot50 \| AF-A0A7X6FPD5-F1-MODEL\_V4 \| 1.0 \| 1.495e-11 \| 422 \| 0.376 \| 117 \| 72 \| 1 \| 184 \| 300 \| 2 \| 117 \| Baseplate\_J domain-containing protein \| Baseplate\_J domain-containing protein; afdb-uniprot50 \| AF-A0A0A0GWF9-F1-MODEL\_V4 \| 1.0 \| 1.074e-10 \| 421 \| 0.495 \| 123 \| 55 \| 1 \| 8 \| 123 \| 4 \| 126 \| Baseplate assembly protein J \| Baseplate assembly protein J; afdb-uniprot50 \| AF-A6NQ14-F1-MODEL\_V4 \| 1.0 \| 9.651e-17 \| 421 \| 0.17 \| 364 \| 206 \| 13 \| 21 \| 300 \| 1 \| 352 \| Baseplate J-like protein \| Baseplate J-like protein; afdb-uniprot50 \| AF-A0A1E8PKJ1-F1-MODEL\_V4 \| 1.0 \| 9.923e-16 \| 421 \| 0.164 \| 364 \| 209 \| 14 \| 12 \| 293 \| 2 \| 352 \| Baseplate\_J domain-containing protein \| Baseplate\_J domain-containing protein; afdb-uniprot50 \| AF-A0A6L5X9Z4-F1-MODEL\_V4 \| 1.0 \| 2.167e-17 \| 420 \| 0.189 \| 374 \| 197 \| 12 \| 20 \| 298 \| 9 \| 371 \| Baseplate\_J domain-containing protein \| Baseplate\_J domain-containing protein; afdb-uniprot50 \| AF-A0A3D4CFS7-F1-MODEL\_V4 \| 1.0 \| 2.909e-15 \| 419 \| 0.149 \| 369 \| 208 \| 14 \| 20 \| 296 \| 1 \| 355 \| Baseplate\_J domain-containing protein \| Baseplate\_J domain-containing protein; afdb-uniprot50 \| AF-A0A376YPQ4-F1-MODEL\_V4 \| 1.0 \| 6.073e-10 \| 418 \| 0.468 \| 94 \| 50 \| 0 \| 206 \| 299 \| 3 \| 96 \| Baseplate assembly protein J \| Baseplate assembly protein J; afdb-uniprot50 \| AF-A0A5T1QS83-F1-MODEL\_V4 \| 1.0 \| 9.61e-15 \| 418 \| 0.231 \| 319 \| 154 \| 9 \| 30 \| 270 \| 2 \| 307 \| Baseplate assembly protein \| Baseplate assembly protein; afdb-uniprot50 \| AF-A0A268TWY5-F1-MODEL\_V4 \| 1.0 \| 7.813e-16 \| 418 \| 0.158 \| 354 \| 213 \| 13 \| 13 \| 296 \| 1 \| 339 \| Baseplate\_J domain-containing protein \| Baseplate\_J domain-containing protein; afdb-uniprot50 \| AF-Q0F1S6-F1-MODEL\_V4 \| 1.0 \| 7.36e-16 \| 418 \| 0.151 \| 364 \| 215 \| 15 \| 13 \| 293 \| 1 \| 353 \| Tail protein, putative \| Tail protein, putative; afdb-uniprot50 \| AF-A0A378VWY1-F1-MODEL\_V4 \| 1.0 \| 4.128e-11 \| 416 \| 0.444 \| 144 \| 69 \| 3 \| 1 \| 137 \| 1 \| 140 \| Baseplate protein, phage associated protein \| Baseplate protein, phage associated protein; afdb-uniprot50 \| AF-A0A1Y4NG73-F1-MODEL\_V4 \| 1.0 \| 1.754e-16 \| 416 \| 0.157 \| 363 \| 209 \| 13 \| 20 \| 296 \| 1 \| 352 \| Baseplate\_J domain-containing protein \| Baseplate\_J domain-containing protein; afdb-uniprot50 \| AF-A0A6B4SDA4-F1-MODEL\_V4 \| 1.0 \| 2.5e-14 \| 415 \| 0.146 \| 361 \| 209 \| 16 \| 21 \| 294 \| 3 \| 351 \| Baseplate J/gp47 family protein \| Baseplate J/gp47 family protein; afdb-uniprot50 \| AF-A0A3D4I4D2-F1-MODEL\_V4 \| 1.0 \| 4.844e-16 \| 414 \| 0.16 \| 361 \| 208 \| 12 \| 21 \| 296 \| 1 \| 351 \| Baseplate protein J \| Baseplate protein J; afdb-uniprot50 \| AF-A0A177XJU5-F1-MODEL\_V4 \| 1.0 \| 5.143e-16 \| 413 \| 0.182 \| 291 \| 209 \| 9 \| 12 \| 294 \| 5 \| 274 \| Baseplate\_J domain-containing protein \| Baseplate\_J domain-containing protein; afdb-uniprot50 \| AF-A0A6N3FBS6-F1-MODEL\_V4 \| 1.0 \| 8.564e-17 \| 412 \| 0.213 \| 384 \| 196 \| 18 \| 14 \| 300 \| 1 \| 375 \| Baseplate J-like protein \| Baseplate J-like protein; afdb-uniprot50 \| AF-A0A485AEW3-F1-MODEL\_V4 \| 1.0 \| 1.14e-10 \| 411 \| 0.46 \| 100 \| 54 \| 0 \| 196 \| 295 \| 3 \| 102 \| Bacteriophage P2-related tail formation protein \| Bacteriophage P2-related tail formation protein; afdb-uniprot50 \| AF-A0A417N973-F1-MODEL\_V4 \| 1.0 \| 2.829e-16 \| 411 \| 0.189 \| 358 \| 199 \| 12 \| 21 \| 295 \| 1 \| 350 \| Phage tail protein \| Phage tail protein; afdb-uniprot50 \| AF-A0A380C6G5-F1-MODEL\_V4 \| 1.0 \| 3.25e-11 \| 410 \| 0.504 \| 105 \| 52 \| 0 \| 191 \| 295 \| 11 \| 115 \| Baseplate J-like protein \| Baseplate J-like protein; afdb-uniprot50 \| AF-A0A0L0W6R4-F1-MODEL\_V4 \| 1.0 \| 1.508e-15 \| 409 \| 0.166 \| 354 \| 204 \| 16 \| 23 \| 294 \| 6 \| 350 \| Baseplate\_J domain-containing protein \| Baseplate\_J domain-containing protein; afdb-uniprot50 \| AF-A0A6V6Y4B3-F1-MODEL\_V4 \| 1.0 \| 1.22e-14 \| 409 \| 0.16 \| 362 \| 208 \| 16 \| 16 \| 292 \| 1 \| 351 \| Baseplate\_J domain-containing protein \| Baseplate\_J domain-containing protein; afdb-uniprot50 \| AF-R6GTF5-F1-MODEL\_V4 \| 1.0 \| 3.814e-16 \| 409 \| 0.141 \| 361 \| 214 \| 13 \| 21 \| 294 \| 1 \| 352 \| Baseplate J-like protein \| Baseplate J-like protein; afdb-uniprot50 \| AF-A0A502HRM3-F1-MODEL\_V4 \| 1.0 \| 5.121e-14 \| 409 \| 0.127 \| 438 \| 211 \| 14 \| 20 \| 292 \| 4 \| 435 \| Baseplate\_J domain-containing protein \| Baseplate\_J domain-containing protein; afdb-uniprot50 \| AF-A0A5C0SDS7-F1-MODEL\_V4 \| 1.0 \| 5.31e-17 \| 408 \| 0.157 \| 362 \| 209 \| 12 \| 20 \| 293 \| 2 \| 355 \| Baseplate J/gp47 family protein \| Baseplate J/gp47 family protein; afdb-uniprot50 \| AF-K8GRQ2-F1-MODEL\_V4 \| 1.0 \| 1.854e-14 \| 408 \| 0.144 \| 368 \| 216 \| 12 \| 19 \| 296 \| 2 \| 360 \| Putative phage Mu protein gp47-like protein \| Putative phage Mu protein gp47-like protein; afdb-uniprot50 \| AF-A0A108C9J6-F1-MODEL\_V4 \| 1.0 \| 7.067e-11 \| 407 \| 0.343 \| 137 \| 87 \| 2 \| 150 \| 284 \| 5 \| 140 \| Baseplate\_J domain-containing protein \| Baseplate\_J domain-containing protein; afdb-uniprot50 \| AF-A0A1G4VD53-F1-MODEL\_V4 \| 1.0 \| 4.049e-16 \| 407 \| 0.166 \| 361 \| 208 \| 11 \| 20 \| 295 \| 1 \| 353 \| Uncharacterized phage protein gp47/JayE \| Uncharacterized phage protein gp47/JayE; afdb-uniprot50 \| AF-Q03RF8-F1-MODEL\_V4 \| 1.0 \| 4.824e-14 \| 407 \| 0.142 \| 400 \| 215 \| 17 \| 16 \| 300 \| 1 \| 387 \| Phage Mu gp47 related protein \| Phage Mu gp47 related protein; afdb-uniprot50 \| AF-A0A1A9S102-F1-MODEL\_V4 \| 1.0 \| 6.933e-16 \| 406 \| 0.157 \| 356 \| 212 \| 15 \| 20 \| 294 \| 2 \| 350 \| Baseplate\_J domain-containing protein \| Baseplate\_J domain-containing protein; afdb-uniprot50 \| AF-A0A848RMJ7-F1-MODEL\_V4 \| 1.0 \| 6.503e-14 \| 406 \| 0.138 \| 361 \| 215 \| 13 \| 20 \| 295 \| 2 \| 351 \| Baseplate J/gp47 family protein \| Baseplate J/gp47 family protein; afdb-uniprot50 \| AF-A0A374UL92-F1-MODEL\_V4 \| 1.0 \| 9.347e-16 \| 406 \| 0.136 \| 360 \| 216 \| 13 \| 21 \| 294 \| 1 \| 351 \| Baseplate\_J domain-containing protein \| Baseplate\_J domain-containing protein; afdb-uniprot50 \| AF-A0A0E1UFA6-F1-MODEL\_V4 \| 1.0 \| 2.13e-09 \| 405 \| 0.361 \| 94 \| 60 \| 0 \| 207 \| 300 \| 6 \| 99 \| Phage baseplate assembly protein \| Phage baseplate assembly protein; afdb-uniprot50 \| AF-A0A8B4NGU9-F1-MODEL\_V4 \| 1.0 \| 1.587e-11 \| 405 \| 0.354 \| 155 \| 97 \| 2 \| 116 \| 269 \| 14 \| 166 \| Phage-related baseplate assembly protein J \| Phage-related baseplate assembly protein J; afdb-uniprot50 \| AF-A0A3D0Z2Q3-F1-MODEL\_V4 \| 1.0 \| 8.805e-16 \| 405 \| 0.164 \| 358 \| 207 \| 11 \| 21 \| 294 \| 1 \| 350 \| Baseplate\_J domain-containing protein \| Baseplate\_J domain-containing protein; afdb-uniprot50 \| AF-A0A644UZQ0-F1-MODEL\_V4 \| 1.0 \| 4.299e-16 \| 405 \| 0.147 \| 380 \| 215 \| 14 \| 18 \| 294 \| 8 \| 381 \| Baseplate\_J domain-containing protein \| Baseplate\_J domain-containing protein; afdb-uniprot50 \| AF-G9YSH5-F1-MODEL\_V4 \| 1.0 \| 2.345e-12 \| 404 \| 0.252 \| 182 \| 127 \| 6 \| 116 \| 294 \| 24 \| 199 \| Baseplate J-like protein \| Baseplate J-like protein; afdb-uniprot50 \| AF-A0A3D0NPG6-F1-MODEL\_V4 \| 1.0 \| 1.862e-16 \| 404 \| 0.146 \| 362 \| 214 \| 15 \| 20 \| 295 \| 1 \| 353 \| Baseplate\_J domain-containing protein \| Baseplate\_J domain-containing protein; afdb-uniprot50 \| AF-A0A2S8I2A0-F1-MODEL\_V4 \| 1.0 \| 1.631e-10 \| 403 \| 0.431 \| 139 \| 69 \| 3 \| 1 \| 132 \| 1 \| 136 \| Baseplate assembly protein \| Baseplate assembly protein; afdb-uniprot50 \| AF-R5AB96-F1-MODEL\_V4 \| 1.0 \| 2.581e-15 \| 403 \| 0.175 \| 360 \| 202 \| 12 \| 21 \| 294 \| 1 \| 351 \| Baseplate J-like protein \| Baseplate J-like protein; afdb-uniprot50 \| AF-A0A5I0MME8-F1-MODEL\_V4 \| 1.0 \| 3.922e-15 \| 402 \| 0.141 \| 368 \| 219 \| 16 \| 13 \| 296 \| 1 \| 355 \| Baseplate J/gp47 family protein \| Baseplate J/gp47 family protein; afdb-uniprot50 \| AF-A0A7M1Q5P2-F1-MODEL\_V4 \| 1.0 \| 1.55e-14 \| 401 \| 0.147 \| 365 \| 205 \| 17 \| 23 \| 293 \| 7 \| 359 \| Baseplate J/gp47 family protein \| Baseplate J/gp47 family protein; afdb-uniprot50 \| AF-B6WSP9-F1-MODEL\_V4 \| 1.0 \| 4.419e-15 \| 398 \| 0.151 \| 363 \| 214 \| 12 \| 13 \| 294 \| 1 \| 350 \| Baseplate J-like protein \| Baseplate J-like protein; afdb-uniprot50 \| AF-A0A3S0DD06-F1-MODEL\_V4 \| 1.0 \| 3.264e-13 \| 398 \| 0.127 \| 519 \| 217 \| 19 \| 1 \| 296 \| 1 \| 506 \| Uncharacterized protein \| Uncharacterized protein; afdb-uniprot50 \| AF-A0A1C7FSD0-F1-MODEL\_V4 \| 1.0 \| 7.998e-13 \| 397 \| 0.243 \| 193 \| 132 \| 7 \| 110 \| 294 \| 100 \| 286 \| Baseplate J protein \| Baseplate J protein; afdb-uniprot50 \| AF-C0DSG2-F1-MODEL\_V4 \| 1.0 \| 2.909e-15 \| 397 \| 0.156 \| 358 \| 210 \| 14 \| 20 \| 294 \| 2 \| 350 \| Baseplate J-like protein \| Baseplate J-like protein; afdb-uniprot50 \| AF-A0A412AW35-F1-MODEL\_V4 \| 1.0 \| 9.091e-17 \| 397 \| 0.163 \| 367 \| 209 \| 14 \| 20 \| 299 \| 1 \| 356 \| Baseplate protein J \| Baseplate protein J; afdb-uniprot50 \| AF-R6QTQ5-F1-MODEL\_V4 \| 1.0 \| 9.61e-15 \| 396 \| 0.142 \| 364 \| 213 \| 13 \| 20 \| 295 \| 1 \| 353 \| Baseplate J-like protein \| Baseplate J-like protein; afdb-uniprot50 \| AF-A0A4Q3L8H7-F1-MODEL\_V4 \| 1.0 \| 3.578e-14 \| 395 \| 0.146 \| 472 \| 199 \| 19 \| 20 \| 293 \| 4 \| 469 \| Baseplate\_J domain-containing protein \| Baseplate\_J domain-containing protein; afdb-uniprot50 \| AF-U7UJ67-F1-MODEL\_V4 \| 1.0 \| 1.747e-14 \| 394 \| 0.151 \| 362 \| 206 \| 15 \| 19 \| 294 \| 2 \| 348 \| Baseplate J-like protein \| Baseplate J-like protein; afdb-uniprot50 \| AF-A0A4R2XGE2-F1-MODEL\_V4 \| 1.0 \| 2.896e-13 \| 394 \| 0.15 \| 366 \| 217 \| 15 \| 14 \| 293 \| 1 \| 358 \| Putative phage protein gp47/JayE \| Putative phage protein gp47/JayE; afdb-uniprot50 \| AF-A0A1I1ENG9-F1-MODEL\_V4 \| 1.0 \| 3.814e-16 \| 394 \| 0.149 \| 368 \| 211 \| 14 \| 20 \| 296 \| 3 \| 359 \| Uncharacterized phage protein gp47/JayE \| Uncharacterized phage protein gp47/JayE; afdb-uniprot50 \| AF-A0A7C3YSU9-F1-MODEL\_V4 \| 1.0 \| 2.024e-13 \| 394 \| 0.124 \| 482 \| 219 \| 16 \| 18 \| 299 \| 2 \| 480 \| Baseplate\_J domain-containing protein \| Baseplate\_J domain-containing protein; afdb-uniprot50 \| AF-A0A826UG88-F1-MODEL\_V4 \| 1.0 \| 1.104e-09 \| 393 \| 0.435 \| 108 \| 57 \| 1 \| 98 \| 205 \| 1 \| 104 \| Baseplate assembly protein \| Baseplate assembly protein; afdb-uniprot50 \| AF-X1XQF5-F1-MODEL\_V4 \| 1.0 \| 1.364e-10 \| 392 \| 0.426 \| 122 \| 70 \| 0 \| 179 \| 300 \| 1 \| 122 \| Baseplate\_J domain-containing protein \| Baseplate\_J domain-containing protein; afdb-uniprot50 \| AF-A0A4Q9TJX4-F1-MODEL\_V4 \| 1.0 \| 2.29e-15 \| 391 \| 0.156 \| 357 \| 213 \| 9 \| 18 \| 293 \| 2 \| 351 \| Baseplate J/gp47 family protein \| Baseplate J/gp47 family protein; afdb-uniprot50 \| AF-A0A2S6QXZ3-F1-MODEL\_V4 \| 1.0 \| 4.98e-15 \| 391 \| 0.129 \| 362 \| 225 \| 10 \| 13 \| 293 \| 1 \| 353 \| Baseplate\_J domain-containing protein \| Baseplate\_J domain-containing protein; afdb-uniprot50 \| AF-A0A0F6LF31-F1-MODEL\_V4 \| 1.0 \| 4.782e-10 \| 390 \| 0.381 \| 110 \| 68 \| 0 \| 191 \| 300 \| 5 \| 114 \| Baseplate J-like family protein \| Baseplate J-like family protein; afdb-uniprot50 \| AF-A0A1H0EKE3-F1-MODEL\_V4 \| 1.0 \| 2.355e-14 \| 390 \| 0.137 \| 364 \| 222 \| 12 \| 21 \| 300 \| 1 \| 356 \| Uncharacterized phage protein gp47/JayE \| Uncharacterized phage protein gp47/JayE; afdb-uniprot50 \| AF-A0A7C6EBI1-F1-MODEL\_V4 \| 1.0 \| 1.053e-15 \| 390 \| 0.169 \| 360 \| 210 \| 11 \| 21 \| 295 \| 1 \| 356 \| Baseplate\_J domain-containing protein \| Baseplate\_J domain-containing protein; afdb-uniprot50 \| AF-A0A090ZAN9-F1-MODEL\_V4 \| 1.0 \| 3.593e-16 \| 389 \| 0.188 \| 376 \| 198 \| 13 \| 14 \| 295 \| 3 \| 365 \| Baseplate J-like family protein \| Baseplate J-like family protein; afdb-uniprot50 \| AF-B6WS15-F1-MODEL\_V4 \| 1.0 \| 1.26e-15 \| 388 \| 0.148 \| 363 \| 218 \| 12 \| 20 \| 300 \| 4 \| 357 \| Baseplate J-like protein \| Baseplate J-like protein; afdb-uniprot50 \| AF-C0XGU3-F1-MODEL\_V4 \| 1.0 \| 2.421e-13 \| 388 \| 0.138 \| 384 \| 214 \| 17 \| 20 \| 295 \| 8 \| 382 \| Baseplate\_J domain-containing protein \| Baseplate\_J domain-containing protein; afdb-uniprot50 \| AF-A0A1Q4ZD92-F1-MODEL\_V4 \| 1.0 \| 9.88e-14 \| 388 \| 0.185 \| 437 \| 187 \| 19 \| 22 \| 296 \| 13 \| 442 \| Baseplate\_J domain-containing protein \| Baseplate\_J domain-containing protein; afdb-uniprot50 \| AF-A0A1F8WPI2-F1-MODEL\_V4 \| 1.0 \| 1.118e-15 \| 387 \| 0.149 \| 354 \| 214 \| 10 \| 20 \| 293 \| 4 \| 350 \| Baseplate\_J domain-containing protein \| Baseplate\_J domain-containing protein; afdb-uniprot50 \| AF-A0A3D5NIG5-F1-MODEL\_V4 \| 1.0 \| 5.287e-15 \| 387 \| 0.146 \| 363 \| 209 \| 14 \| 21 \| 294 \| 1 \| 351 \| Baseplate\_J domain-containing protein \| Baseplate\_J domain-containing protein; afdb-uniprot50 \| AF-A0A7G3F0Q5-F1-MODEL\_V4 \| 1.0 \| 8.692e-10 \| 386 \| 0.526 \| 112 \| 46 \| 1 \| 15 \| 119 \| 7 \| 118 \| Baseplate J-like protein \| Baseplate J-like protein; afdb-uniprot50 \| AF-A0A2A7WUL9-F1-MODEL\_V4 \| 1.0 \| 1.55e-14 \| 386 \| 0.164 \| 353 \| 206 \| 14 \| 21 \| 293 \| 1 \| 344 \| Baseplate j family protein \| Baseplate j family protein; afdb-uniprot50 \| AF-A0A223D2X1-F1-MODEL\_V4 \| 1.0 \| 7.36e-16 \| 386 \| 0.175 \| 364 \| 206 \| 13 \| 20 \| 295 \| 4 \| 361 \| Baseplate\_J domain-containing protein \| Baseplate\_J domain-containing protein; afdb-uniprot50 \| AF-A0A6I6XZA3-F1-MODEL\_V4 \| 1.0 \| 8.033e-15 \| 384 \| 0.128 \| 358 \| 226 \| 12 \| 18 \| 296 \| 2 \| 352 \| Baseplate J/gp47 family protein \| Baseplate J/gp47 family protein; afdb-uniprot50 \| AF-A0A6Y1Z4C1-F1-MODEL\_V4 \| 1.0 \| 7.93e-09 \| 383 \| 0.459 \| 87 \| 47 \| 0 \| 213 \| 299 \| 1 \| 87 \| Baseplate assembly protein \| Baseplate assembly protein; afdb-uniprot50 \| AF-A0A844UU35-F1-MODEL\_V4 \| 1.0 \| 3.148e-10 \| 383 \| 0.482 \| 116 \| 60 \| 0 \| 1 \| 116 \| 1 \| 116 \| Baseplate assembly protein \| Baseplate assembly protein; afdb-uniprot50 \| AF-A0A7X5DX78-F1-MODEL\_V4 \| 1.0 \| 1.684e-11 \| 383 \| 0.165 \| 206 \| 149 \| 6 \| 113 \| 300 \| 77 \| 277 \| Baseplate\_J domain-containing protein \| Baseplate\_J domain-containing protein; afdb-uniprot50 \| AF-C6LG69-F1-MODEL\_V4 \| 1.0 \| 9.052e-15 \| 383 \| 0.143 \| 384 \| 223 \| 17 \| 8 \| 295 \| 2 \| 375 \| Baseplate J-like protein \| Baseplate J-like protein; afdb-uniprot50 \| AF-U2QTS5-F1-MODEL\_V4 \| 1.0 \| 3.264e-13 \| 383 \| 0.16 \| 392 \| 198 \| 17 \| 20 \| 298 \| 9 \| 382 \| Baseplate\_J domain-containing protein \| Baseplate\_J domain-containing protein; afdb-uniprot50 \| AF-S7V8X9-F1-MODEL\_V4 \| 1.0 \| 6.126e-14 \| 383 \| 0.151 \| 377 \| 210 \| 13 \| 23 \| 295 \| 22 \| 392 \| Baseplate J family protein \| Baseplate J family protein; afdb-uniprot50 \| AF-Q727K4-F1-MODEL\_V4 \| 1.0 \| 3.48e-15 \| 382 \| 0.189 \| 364 \| 203 \| 11 \| 13 \| 295 \| 1 \| 353 \| Tail protein, putative \| Tail protein, putative; afdb-uniprot50 \| AF-A0A3C1LVA0-F1-MODEL\_V4 \| 1.0 \| 2.431e-15 \| 382 \| 0.16 \| 367 \| 204 \| 15 \| 23 \| 300 \| 3 \| 354 \| Baseplate\_J domain-containing protein \| Baseplate\_J domain-containing protein; afdb-uniprot50 \| AF-R6U9B8-F1-MODEL\_V4 \| 1.0 \| 2.355e-14 \| 381 \| 0.138 \| 361 \| 216 \| 12 \| 21 \| 295 \| 1 \| 352 \| Baseplate J-like protein \| Baseplate J-like protein; afdb-uniprot50 \| AF-A0A2T5J3V8-F1-MODEL\_V4 \| 1.0 \| 4.98e-15 \| 380 \| 0.137 \| 363 \| 220 \| 14 \| 13 \| 294 \| 1 \| 351 \| Putative phage protein gp47/JayE \| Putative phage protein gp47/JayE; afdb-uniprot50 \| AF-A0A176L6Z2-F1-MODEL\_V4 \| 1.0 \| 1.049e-13 \| 380 \| 0.153 \| 442 \| 206 \| 15 \| 14 \| 298 \| 1 \| 431 \| Baseplate\_J domain-containing protein \| Baseplate\_J domain-containing protein; afdb-uniprot50 \| AF-A0A0D2GH44-F1-MODEL\_V4 \| 1.0 \| 1.915e-15 \| 379 \| 0.155 \| 353 \| 214 \| 9 \| 20 \| 292 \| 4 \| 352 \| Baseplate J protein \| Baseplate J protein; afdb-uniprot50 \| AF-A0A7C6PX97-F1-MODEL\_V4 \| 1.0 \| 1.638e-12 \| 379 \| 0.12 \| 481 \| 215 \| 16 \| 13 \| 295 \| 3 \| 473 \| Baseplate\_J domain-containing protein \| Baseplate\_J domain-containing protein; afdb-uniprot50 \| AF-S5MVA2-F1-MODEL\_V4 \| 1.0 \| 5.219e-09 \| 378 \| 0.411 \| 90 \| 53 \| 0 \| 206 \| 295 \| 3 \| 92 \| Baseplate assembly protein J \| Baseplate assembly protein J; afdb-uniprot50 \| AF-A0A413G370-F1-MODEL\_V4 \| 1.0 \| 4.98e-15 \| 378 \| 0.162 \| 363 \| 212 \| 14 \| 20 \| 300 \| 1 \| 353 \| Baseplate\_J domain-containing protein \| Baseplate\_J domain-containing protein; afdb-uniprot50 \| AF-A0A7V2IPE5-F1-MODEL\_V4 \| 1.0 \| 2.139e-11 \| 378 \| 0.186 \| 198 \| 145 \| 9 \| 110 \| 296 \| 247 \| 439 \| Baseplate J protein \| Baseplate J protein; afdb-uniprot50 \| AF-A0A4P9VHM3-F1-MODEL\_V4 \| 1.0 \| 2.199e-10 \| 377 \| 0.336 \| 116 \| 77 \| 0 \| 180 \| 295 \| 4 \| 119 \| Baseplate assembly protein \| Baseplate assembly protein; afdb-uniprot50 \| AF-W1L2P9-F1-MODEL\_V4 \| 1.0 \| 5.565e-11 \| 377 \| 0.323 \| 167 \| 95 \| 5 \| 1 \| 154 \| 2 \| 163 \| Uncharacterized protein \| Uncharacterized protein; afdb-uniprot50 \| AF-A0A2H1SQY8-F1-MODEL\_V4 \| 1.0 \| 7.713e-10 \| 375 \| 0.432 \| 104 \| 59 \| 0 \| 196 \| 299 \| 2 \| 105 \| Baseplate assembly protein J \| Baseplate assembly protein J; afdb-uniprot50 \| AF-A0A329KAH2-F1-MODEL\_V4 \| 1.0 \| 9.923e-16 \| 375 \| 0.176 \| 363 \| 200 \| 13 \| 19 \| 295 \| 2 \| 351 \| Baseplate j family protein \| Baseplate j family protein; afdb-uniprot50 \| AF-R5KQQ2-F1-MODEL\_V4 \| 1.0 \| 1.02e-14 \| 375 \| 0.142 \| 365 \| 213 \| 14 \| 20 \| 294 \| 1 \| 355 \| Baseplate J-like protein \| Baseplate J-like protein; afdb-uniprot50 \| AF-A0A328UJU0-F1-MODEL\_V4 \| 1.0 \| 1.747e-14 \| 375 \| 0.162 \| 363 \| 213 \| 12 \| 20 \| 300 \| 7 \| 360 \| Baseplate protein J \| Baseplate protein J; afdb-uniprot50 \| AF-A0A371WT16-F1-MODEL\_V4 \| 1.0 \| 4.824e-14 \| 375 \| 0.153 \| 379 \| 208 \| 13 \| 20 \| 293 \| 5 \| 375 \| Baseplate protein \| Baseplate protein; afdb-uniprot50 \| AF-A0A077Q3X4-F1-MODEL\_V4 \| 1.0 \| 8.692e-10 \| 373 \| 0.432 \| 104 \| 59 \| 0 \| 196 \| 299 \| 2 \| 105 \| Baseplate assembly protein J (GpJ) \| Baseplate assembly protein J (GpJ); afdb-uniprot50 \| AF-A0A243TCZ4-F1-MODEL\_V4 \| 1.0 \| 7.266e-10 \| 373 \| 0.388 \| 103 \| 60 \| 1 \| 198 \| 300 \| 119 \| 218 \| Uncharacterized protein \| Uncharacterized protein; afdb-uniprot50 \| AF-A0A1T2XKG9-F1-MODEL\_V4 \| 1.0 \| 2.431e-15 \| 373 \| 0.158 \| 360 \| 205 \| 14 \| 20 \| 293 \| 3 \| 350 \| Baseplate\_J domain-containing protein \| Baseplate\_J domain-containing protein; afdb-uniprot50 \| AF-A0A1Y0G5P4-F1-MODEL\_V4 \| 1.0 \| 4.544e-14 \| 372 \| 0.15 \| 360 \| 215 \| 15 \| 20 \| 295 \| 4 \| 356 \| Phage tail protein \| Phage tail protein; afdb-uniprot50 \| AF-A0A1G8FC75-F1-MODEL\_V4 \| 1.0 \| 1.02e-14 \| 372 \| 0.155 \| 372 \| 211 \| 14 \| 20 \| 295 \| 4 \| 368 \| Uncharacterized phage protein gp47/JayE \| Uncharacterized phage protein gp47/JayE; afdb-uniprot50 \| AF-A0A379YIU1-F1-MODEL\_V4 \| 1.0 \| 5.541e-09 \| 371 \| 0.344 \| 93 \| 61 \| 0 \| 207 \| 299 \| 17 \| 109 \| Baseplate J-like protein \| Baseplate J-like protein; afdb-uniprot50 \| AF-A0A522WFF2-F1-MODEL\_V4 \| 1.0 \| 2.355e-14 \| 370 \| 0.147 \| 360 \| 219 \| 13 \| 20 \| 296 \| 4 \| 358 \| Baseplate J/gp47 family protein \| Baseplate J/gp47 family protein; afdb-uniprot50 \| AF-I0GWS4-F1-MODEL\_V4 \| 1.0 \| 5.287e-15 \| 367 \| 0.16 \| 361 \| 207 \| 14 \| 19 \| 294 \| 3 \| 352 \| Putative baseplate protein \| Putative baseplate protein; afdb-uniprot50 \| AF-A0A348Z8Z5-F1-MODEL\_V4 \| 1.0 \| 1.854e-14 \| 367 \| 0.136 \| 359 \| 214 \| 13 \| 22 \| 293 \| 4 \| 353 \| Baseplate\_J domain-containing protein \| Baseplate\_J domain-containing protein; afdb-uniprot50 \| AF-A0A6N2T8J3-F1-MODEL\_V4 \| 1.0 \| 4.98e-15 \| 366 \| 0.181 \| 364 \| 196 \| 18 \| 22 \| 295 \| 1 \| 352 \| Baseplate J-like protein \| Baseplate J-like protein; afdb-uniprot50 \| AF-A0A1S2WAC6-F1-MODEL\_V4 \| 1.0 \| 2.829e-16 \| 366 \| 0.193 \| 373 \| 198 \| 15 \| 12 \| 293 \| 2 \| 362 \| Baseplate\_J domain-containing protein \| Baseplate\_J domain-containing protein; afdb-uniprot50 \| AF-A0A1V2H5A7-F1-MODEL\_V4 \| 1.0 \| 3.694e-15 \| 366 \| 0.185 \| 389 \| 196 \| 17 \| 11 \| 293 \| 2 \| 375 \| Baseplate\_J domain-containing protein \| Baseplate\_J domain-containing protein; afdb-uniprot50 \| AF-A0A3M4Z394-F1-MODEL\_V4 \| 1.0 \| 1.321e-09 \| 365 \| 0.433 \| 120 \| 67 \| 1 \| 117 \| 235 \| 1 \| 120 \| Baseplate assembly protein J \| Baseplate assembly protein J; afdb-uniprot50 \| AF-A0A2A7HD50-F1-MODEL\_V4 \| 1.0 \| 5.589e-13 \| 364 \| 0.129 \| 354 \| 214 \| 17 \| 21 \| 290 \| 1 \| 344 \| Baseplate J protein \| Baseplate J protein; afdb-uniprot50 \| AF-A0A3D0XAJ1-F1-MODEL\_V4 \| 1.0 \| 1.249e-11 \| 363 \| 0.225 \| 186 \| 135 \| 6 \| 112 \| 294 \| 76 \| 255 \| Baseplate\_J domain-containing protein \| Baseplate\_J domain-containing protein; afdb-uniprot50 \| AF-A0A845SWX3-F1-MODEL\_V4 \| 1.0 \| 8.527e-15 \| 363 \| 0.175 \| 360 \| 206 \| 14 \| 23 \| 300 \| 5 \| 355 \| Uncharacterized protein \| Uncharacterized protein; afdb-uniprot50 \| AF-A0A081BI31-F1-MODEL\_V4 \| 1.0 \| 5.933e-13 \| 363 \| 0.152 \| 381 \| 202 \| 13 \| 22 \| 294 \| 16 \| 383 \| Phage protein \| Phage protein; afdb-uniprot50 \| AF-A0A7Y4S5C2-F1-MODEL\_V4 \| 1.0 \| 1.182e-13 \| 362 \| 0.137 \| 357 \| 219 \| 13 \| 20 \| 293 \| 4 \| 354 \| Baseplate J/gp47 family protein \| Baseplate J/gp47 family protein; afdb-uniprot50 \| AF-B9Z2Z3-F1-MODEL\_V4 \| 1.0 \| 1.255e-13 \| 362 \| 0.135 \| 361 \| 217 \| 14 \| 20 \| 294 \| 4 \| 355 \| Baseplate J family protein \| Baseplate J family protein; afdb-uniprot50 \| AF-J1H457-F1-MODEL\_V4 \| 1.0 \| 3.175e-14 \| 361 \| 0.166 \| 367 \| 209 \| 14 \| 21 \| 300 \| 1 \| 357 \| Baseplate J-like protein \| Baseplate J-like protein; afdb-uniprot50 \| AF-A0A1S8PIF2-F1-MODEL\_V4 \| 1.0 \| 1.332e-13 \| 361 \| 0.14 \| 370 \| 210 \| 17 \| 22 \| 295 \| 6 \| 363 \| Baseplate J-like protein \| Baseplate J-like protein; afdb-uniprot50 \| AF-A0A662W1X1-F1-MODEL\_V4 \| 1.0 \| 9.307e-14 \| 360 \| 0.169 \| 355 \| 183 \| 13 \| 22 \| 295 \| 14 \| 337 \| Baseplate\_J domain-containing protein \| Baseplate\_J domain-containing protein; afdb-uniprot50 \| AF-M3AAC0-F1-MODEL\_V4 \| 1.0 \| 8.767e-14 \| 360 \| 0.136 \| 367 \| 216 \| 12 \| 13 \| 293 \| 1 \| 352 \| Tail protein \| Tail protein; afdb-uniprot50 \| AF-A0A1C6CHI6-F1-MODEL\_V4 \| 1.0 \| 5.436e-14 \| 359 \| 0.145 \| 386 \| 207 \| 15 \| 20 \| 294 \| 8 \| 381 \| Uncharacterized homolog of phage Mu protein gp47 \| Uncharacterized homolog of phage Mu protein gp47; afdb-uniprot50 \| AF-G9XBC3-F1-MODEL\_V4 \| 1.0 \| 9.052e-15 \| 358 \| 0.154 \| 357 \| 207 \| 14 \| 21 \| 295 \| 4 \| 347 \| Baseplate\_J domain-containing protein \| Baseplate\_J domain-containing protein; afdb-uniprot50 \| AF-A0A358PSA7-F1-MODEL\_V4 \| 1.0 \| 3.088e-15 \| 358 \| 0.175 \| 377 \| 206 \| 16 \| 1 \| 294 \| 11 \| 365 \| Baseplate\_J domain-containing protein \| Baseplate\_J domain-containing protein; afdb-uniprot50 \| AF-A0A1J4X5M8-F1-MODEL\_V4 \| 1.0 \| 1.215e-12 \| 358 \| 0.136 \| 381 \| 211 \| 15 \| 19 \| 293 \| 3 \| 371 \| Baseplate\_J domain-containing protein \| Baseplate\_J domain-containing protein; afdb-uniprot50 \| AF-D7MYP8-F1-MODEL\_V4 \| 1.0 \| 7.502e-11 \| 357 \| 0.216 \| 189 \| 133 \| 10 \| 116 \| 295 \| 8 \| 190 \| Baseplate J-like protein \| Baseplate J-like protein; afdb-uniprot50 \| AF-A0A3Q9S8Y2-F1-MODEL\_V4 \| 1.0 \| 4.28e-14 \| 356 \| 0.143 \| 363 \| 216 \| 14 \| 18 \| 293 \| 2 \| 356 \| Baseplate J/gp47 family protein \| Baseplate J/gp47 family protein; afdb-uniprot50 \| AF-A0A502HGK3-F1-MODEL\_V4 \| 1.0 \| 2.158e-15 \| 355 \| 0.163 \| 372 \| 202 \| 15 \| 20 \| 295 \| 4 \| 362 \| Baseplate\_J domain-containing protein \| Baseplate\_J domain-containing protein; afdb-uniprot50 \| AF-A0A7C3WLZ2-F1-MODEL\_V4 \| 1.0 \| 2.642e-12 \| 354 \| 0.117 \| 468 \| 216 \| 16 \| 20 \| 294 \| 1 \| 464 \| Baseplate\_J domain-containing protein \| Baseplate\_J domain-containing protein; afdb-uniprot50 \| AF-A0A7G9WGX3-F1-MODEL\_V4 \| 1.0 \| 3.37e-14 \| 353 \| 0.169 \| 360 \| 203 \| 15 \| 21 \| 293 \| 1 \| 351 \| Baseplate J/gp47 family protein \| Baseplate J/gp47 family protein; afdb-uniprot50 \| AF-S0FX69-F1-MODEL\_V4 \| 1.0 \| 1.645e-14 \| 353 \| 0.154 \| 362 \| 217 \| 13 \| 23 \| 300 \| 1 \| 357 \| Baseplate J-like protein \| Baseplate J-like protein; afdb-uniprot50 \| AF-A0A5C8XTI4-F1-MODEL\_V4 \| 1.0 \| 1.89e-09 \| 352 \| 0.479 \| 98 \| 51 \| 0 \| 195 \| 292 \| 3 \| 100 \| Baseplate assembly protein \| Baseplate assembly protein; afdb-uniprot50 \| AF-A0A1X3ISQ9-F1-MODEL\_V4 \| 1.0 \| 1.279e-08 \| 352 \| 0.354 \| 93 \| 60 \| 0 \| 207 \| 299 \| 17 \| 109 \| Phage baseplate assembly protein \| Phage baseplate assembly protein; afdb-uniprot50 \| AF-A0A3D0E1W7-F1-MODEL\_V4 \| 1.0 \| 1.508e-15 \| 351 \| 0.155 \| 372 \| 215 \| 15 \| 13 \| 293 \| 6 \| 369 \| Baseplate\_J domain-containing protein \| Baseplate\_J domain-containing protein; afdb-uniprot50 \| AF-A0A258L253-F1-MODEL\_V4 \| 1.0 \| 1.58e-09 \| 350 \| 0.411 \| 107 \| 63 \| 0 \| 190 \| 296 \| 12 \| 118 \| Baseplate\_J domain-containing protein \| Baseplate\_J domain-containing protein; afdb-uniprot50 \| AF-X1GT58-F1-MODEL\_V4 \| 1.0 \| 5.077e-10 \| 350 \| 0.209 \| 162 \| 118 \| 6 \| 140 \| 295 \| 2 \| 159 \| Baseplate\_J domain-containing protein \| Baseplate\_J domain-containing protein; afdb-uniprot50 \| AF-A0A418H6F8-F1-MODEL\_V4 \| 1.0 \| 3.048e-09 \| 348 \| 0.416 \| 108 \| 63 \| 0 \| 192 \| 299 \| 1 \| 108 \| Baseplate assembly protein \| Baseplate assembly protein; afdb-uniprot50 \| AF-A0A1V5QEC7-F1-MODEL\_V4 \| 1.0 \| 8.692e-10 \| 348 \| 0.183 \| 174 \| 130 \| 6 \| 131 \| 296 \| 40 \| 209 \| Baseplate J-like protein \| Baseplate J-like protein; afdb-uniprot50 \| AF-A0A1M5REH3-F1-MODEL\_V4 \| 1.0 \| 1.645e-14 \| 348 \| 0.139 \| 386 \| 220 \| 18 \| 20 \| 300 \| 4 \| 382 \| Uncharacterized phage protein gp47/JayE \| Uncharacterized phage protein gp47/JayE; afdb-uniprot50 \| AF-A0A3D0KMD1-F1-MODEL\_V4 \| 1.0 \| 3.905e-13 \| 347 \| 0.139 \| 351 \| 209 \| 11 \| 31 \| 296 \| 1 \| 343 \| Baseplate\_J domain-containing protein \| Baseplate\_J domain-containing protein; afdb-uniprot50 \| AF-A0A3N0V7H0-F1-MODEL\_V4 \| 1.0 \| 6.126e-14 \| 347 \| 0.161 \| 354 \| 210 \| 10 \| 20 \| 293 \| 4 \| 350 \| Baseplate J/gp47 family protein \| Baseplate J/gp47 family protein; afdb-uniprot50 \| AF-A0A376TU70-F1-MODEL\_V4 \| 1.0 \| 1.04e-09 \| 346 \| 0.407 \| 113 \| 67 \| 0 \| 187 \| 299 \| 13 \| 125 \| Baseplate assembly protein GpJ \| Baseplate assembly protein GpJ; afdb-uniprot50 \| AF-W4BPJ9-F1-MODEL\_V4 \| 1.0 \| 3.264e-13 \| 346 \| 0.158 \| 303 \| 213 \| 13 \| 7 \| 295 \| 2 \| 276 \| Baseplate\_J domain-containing protein \| Baseplate\_J domain-containing protein; afdb-uniprot50 \| AF-A0A2G1Z9I9-F1-MODEL\_V4 \| 1.0 \| 7.328e-14 \| 346 \| 0.128 \| 366 \| 228 \| 11 \| 13 \| 293 \| 1 \| 360 \| Baseplate\_J domain-containing protein \| Baseplate\_J domain-containing protein; afdb-uniprot50 \| AF-Q15XZ6-F1-MODEL\_V4 \| 1.0 \| 1.215e-12 \| 346 \| 0.132 \| 400 \| 227 \| 19 \| 1 \| 294 \| 1 \| 386 \| Uncharacterized phage Mu protein GP47-like protein \| Uncharacterized phage Mu protein GP47-like protein; afdb-uniprot50 \| AF-A0A291RK52-F1-MODEL\_V4 \| 1.0 \| 6.903e-14 \| 346 \| 0.148 \| 438 \| 194 \| 19 \| 22 \| 296 \| 14 \| 435 \| Baseplate\_J domain-containing protein \| Baseplate\_J domain-containing protein; afdb-uniprot50 \| AF-A0A644Z6C5-F1-MODEL\_V4 \| 1.0 \| 3.766e-10 \| 345 \| 0.217 \| 170 \| 120 \| 7 \| 129 \| 295 \| 9 \| 168 \| Baseplate\_J domain-containing protein \| Baseplate\_J domain-containing protein; afdb-uniprot50 \| AF-A0A2W1CQX1-F1-MODEL\_V4 \| 1.0 \| 2.199e-10 \| 345 \| 0.209 \| 191 \| 139 \| 6 \| 112 \| 293 \| 92 \| 279 \| TehA, Tellurite resistance protein \| TehA, Tellurite resistance protein; afdb-uniprot50 \| AF-A0A0Q4XSB4-F1-MODEL\_V4 \| 1.0 \| 8.767e-14 \| 345 \| 0.152 \| 367 \| 215 \| 15 \| 19 \| 295 \| 4 \| 364 \| Baseplate\_J domain-containing protein \| Baseplate\_J domain-containing protein; afdb-uniprot50 \| AF-A0A2U1W182-F1-MODEL\_V4 \| 1.0 \| 1.96e-12 \| 345 \| 0.135 \| 376 \| 212 \| 16 \| 20 \| 295 \| 4 \| 366 \| Baseplate\_J domain-containing protein \| Baseplate\_J domain-containing protein; afdb-uniprot50 \| AF-A0A2V2YWP1-F1-MODEL\_V4 \| 1.0 \| 9.838e-12 \| 345 \| 0.206 \| 208 \| 140 \| 7 \| 112 \| 296 \| 430 \| 635 \| Putative phage baseplate assembly protein \| Putative phage baseplate assembly protein; afdb-uniprot50 \| AF-B1TCV6-F1-MODEL\_V4 \| 1.0 \| 4.782e-10 \| 344 \| 0.387 \| 124 \| 69 \| 3 \| 101 \| 224 \| 3 \| 119 \| Baseplate J family protein \| Baseplate J family protein; afdb-uniprot50 \| AF-A0A3A9FCY9-F1-MODEL\_V4 \| 1.0 \| 9.307e-14 \| 344 \| 0.12 \| 372 \| 230 \| 11 \| 17 \| 296 \| 9 \| 375 \| Baseplate\_J domain-containing protein \| Baseplate\_J domain-containing protein; afdb-uniprot50 \| AF-A0A4V2EGN7-F1-MODEL\_V4 \| 1.0 \| 1.369e-12 \| 344 \| 0.133 \| 382 \| 216 \| 16 \| 21 \| 295 \| 19 \| 392 \| Baseplate\_J domain-containing protein \| Baseplate\_J domain-containing protein; afdb-uniprot50 \| AF-A0A3D1LC94-F1-MODEL\_V4 \| 1.0 \| 5.121e-14 \| 343 \| 0.196 \| 311 \| 156 \| 13 \| 74 \| 300 \| 2 \| 302 \| Phage tail protein \| Phage tail protein; afdb-uniprot50 \| AF-A0A5S3YLY6-F1-MODEL\_V4 \| 1.0 \| 3.048e-09 \| 342 \| 0.413 \| 116 \| 60 \| 3 \| 74 \| 189 \| 2 \| 109 \| Baseplate J protein \| Baseplate J protein; afdb-uniprot50 \| AF-A0A3S4J4W8-F1-MODEL\_V4 \| 1.0 \| 2.705e-09 \| 342 \| 0.538 \| 117 \| 47 \| 1 \| 8 \| 117 \| 4 \| 120 \| Baseplate assembly protein \| Baseplate assembly protein; afdb-uniprot50 \| AF-A0A7C1VC68-F1-MODEL\_V4 \| 1.0 \| 9.88e-14 \| 342 \| 0.139 \| 359 \| 217 \| 12 \| 20 \| 295 \| 4 \| 353 \| Baseplate J/gp47 family protein \| Baseplate J/gp47 family protein; afdb-uniprot50 \| AF-A0A4P8QBU6-F1-MODEL\_V4 \| 1.0 \| 2.148e-13 \| 342 \| 0.15 \| 378 \| 210 \| 11 \| 20 \| 293 \| 4 \| 374 \| Baseplate protein \| Baseplate protein; afdb-uniprot50 \| AF-A0A4R3HUV0-F1-MODEL\_V4 \| 1.0 \| 1.113e-13 \| 341 \| 0.15 \| 358 \| 215 \| 11 \| 20 \| 296 \| 4 \| 353 \| Putative phage protein gp47/JayE \| Putative phage protein gp47/JayE; afdb-uniprot50 \| AF-A0A5Z1E065-F1-MODEL\_V4 \| 1.0 \| 1.537e-10 \| 340 \| 0.282 \| 163 \| 101 \| 4 \| 113 \| 270 \| 28 \| 179 \| Baseplate assembly protein \| Baseplate assembly protein; afdb-uniprot50 \| AF-A0A7J5WLS9-F1-MODEL\_V4 \| 1.0 \| 1.638e-12 \| 339 \| 0.159 \| 364 \| 204 \| 16 \| 20 \| 294 \| 3 \| 353 \| Baseplate J family \| Baseplate J family; afdb-uniprot50 \| AF-A0A2E7U201-F1-MODEL\_V4 \| 1.0 \| 1.501e-13 \| 339 \| 0.139 \| 459 \| 210 \| 21 \| 14 \| 293 \| 1 \| 453 \| Baseplate\_J domain-containing protein \| Baseplate\_J domain-containing protein; afdb-uniprot50 \| AF-A0A826JR68-F1-MODEL\_V4 \| 1.0 \| 3.766e-10 \| 338 \| 0.366 \| 139 \| 75 \| 2 \| 113 \| 240 \| 61 \| 197 \| Baseplate J/gp47 family protein \| Baseplate J/gp47 family protein; afdb-uniprot50 \| AF-A0A1Y1S2G0-F1-MODEL\_V4 \| 1.0 \| 5.121e-14 \| 338 \| 0.133 \| 368 \| 223 \| 12 \| 15 \| 293 \| 1 \| 361 \| Baseplate\_J domain-containing protein \| Baseplate\_J domain-containing protein; afdb-uniprot50 \| AF-A0A0C2VDI9-F1-MODEL\_V4 \| 1.0 \| 2.345e-12 \| 338 \| 0.156 \| 389 \| 202 \| 19 \| 13 \| 293 \| 1 \| 371 \| Baseplate\_J domain-containing protein \| Baseplate\_J domain-containing protein; afdb-uniprot50 \| AF-A0A5C4QQ94-F1-MODEL\_V4 \| 1.0 \| 1.587e-11 \| 338 \| 0.124 \| 500 \| 220 \| 15 \| 11 \| 298 \| 2 \| 495 \| Baseplate\_J domain-containing protein \| Baseplate\_J domain-containing protein; afdb-uniprot50 \| AF-A0A1Y4L6I3-F1-MODEL\_V4 \| 1.0 \| 9.88e-14 \| 337 \| 0.155 \| 359 \| 209 \| 15 \| 23 \| 295 \| 1 \| 351 \| Baseplate\_J domain-containing protein \| Baseplate\_J domain-containing protein; afdb-uniprot50 \| AF-A0A1M6ZB03-F1-MODEL\_V4 \| 1.0 \| 2.896e-13 \| 337 \| 0.13 \| 397 \| 219 \| 19 \| 20 \| 296 \| 2 \| 392 \| Uncharacterized phage protein gp47/JayE \| Uncharacterized phage protein gp47/JayE; afdb-uniprot50 \| AF-A0A2N1TNI5-F1-MODEL\_V4 \| 1.0 \| 5.121e-14 \| 336 \| 0.152 \| 388 \| 216 \| 18 \| 13 \| 295 \| 1 \| 380 \| Baseplate\_J domain-containing protein \| Baseplate\_J domain-containing protein; afdb-uniprot50 \| AF-A0A3E2SY13-F1-MODEL\_V4 \| 1.0 \| 7.998e-13 \| 335 \| 0.148 \| 349 \| 205 \| 13 \| 34 \| 300 \| 4 \| 342 \| Baseplate protein J \| Baseplate protein J; afdb-uniprot50 \| AF-C8NCB0-F1-MODEL\_V4 \| 1.0 \| 3.647e-09 \| 333 \| 0.289 \| 121 \| 85 \| 1 \| 178 \| 298 \| 1 \| 120 \| Uncharacterized protein \| Uncharacterized protein; afdb-uniprot50 \| AF-A0A3A9HYN4-F1-MODEL\_V4 \| 1.0 \| 2.148e-13 \| 333 \| 0.172 \| 377 \| 210 \| 17 \| 12 \| 293 \| 3 \| 372 \| Baseplate J/gp47 family protein \| Baseplate J/gp47 family protein; afdb-uniprot50 \| AF-R4YKQ9-F1-MODEL\_V4 \| 1.0 \| 3.161e-12 \| 333 \| 0.115 \| 382 \| 225 \| 14 \| 21 \| 296 \| 22 \| 396 \| Baseplate\_J domain-containing protein \| Baseplate\_J domain-containing protein; afdb-uniprot50 \| AF-D1JAN0-F1-MODEL\_V4 \| 1.0 \| 9.267e-12 \| 333 \| 0.175 \| 216 \| 150 \| 10 \| 110 \| 300 \| 410 \| 622 \| Baseplate\_J domain-containing protein \| Baseplate\_J domain-containing protein; afdb-uniprot50 \| AF-A0A379YC97-F1-MODEL\_V4 \| 1.0 \| 1.285e-10 \| 332 \| 0.438 \| 130 \| 73 \| 0 \| 170 \| 299 \| 1 \| 130 \| Baseplate J-like protein \| Baseplate J-like protein; afdb-uniprot50 \| AF-A0A1B9ZS99-F1-MODEL\_V4 \| 1.0 \| 3.905e-13 \| 332 \| 0.211 \| 298 \| 141 \| 9 \| 77 \| 285 \| 1 \| 293 \| Baseplate\_J domain-containing protein \| Baseplate\_J domain-containing protein; afdb-uniprot50 \| AF-A0A3Q9SAV9-F1-MODEL\_V4 \| 1.0 \| 3.798e-14 \| 332 \| 0.175 \| 337 \| 188 \| 13 \| 44 \| 293 \| 10 \| 343 \| Baseplate J/gp47 family protein \| Baseplate J/gp47 family protein; afdb-uniprot50 \| AF-A0A3G2R557-F1-MODEL\_V4 \| 1.0 \| 8.491e-13 \| 332 \| 0.123 \| 470 \| 215 \| 12 \| 20 \| 295 \| 1 \| 467 \| Baseplate\_J domain-containing protein \| Baseplate\_J domain-containing protein; afdb-uniprot50 \| AF-A0A4Q3L733-F1-MODEL\_V4 \| 1.0 \| 3.465e-13 \| 332 \| 0.138 \| 476 \| 209 \| 19 \| 13 \| 293 \| 1 \| 470 \| Baseplate\_J domain-containing protein \| Baseplate\_J domain-containing protein; afdb-uniprot50 \| AF-A0A7C2U0W7-F1-MODEL\_V4 \| 1.0 \| 8.73e-12 \| 331 \| 0.125 \| 446 \| 215 \| 15 \| 14 \| 296 \| 1 \| 434 \| Baseplate\_J domain-containing protein \| Baseplate\_J domain-containing protein; afdb-uniprot50 \| AF-A0A6G4XH46-F1-MODEL\_V4 \| 1.0 \| 3.45e-11 \| 331 \| 0.214 \| 224 \| 140 \| 13 \| 110 \| 300 \| 416 \| 636 \| Putative baseplate assembly protein \| Putative baseplate assembly protein; afdb-uniprot50 \| AF-R6P8N6-F1-MODEL\_V4 \| 1.0 \| 2.631e-10 \| 330 \| 0.226 \| 172 \| 117 \| 9 \| 129 \| 295 \| 9 \| 169 \| Baseplate J-like protein \| Baseplate J-like protein; afdb-uniprot50 \| AF-A0A0T9QWQ1-F1-MODEL\_V4 \| 1.0 \| 1.724e-08 \| 329 \| 0.532 \| 109 \| 51 \| 0 \| 42 \| 150 \| 2 \| 110 \| Baseplate assembly protein \| Baseplate assembly protein; afdb-uniprot50 \| AF-Q1MNP3-F1-MODEL\_V4 \| 1.0 \| 1.732e-10 \| 329 \| 0.163 \| 196 \| 143 \| 9 \| 113 \| 294 \| 49 \| 237 \| Baseplate\_J domain-containing protein \| Baseplate\_J domain-containing protein; afdb-uniprot50 \| AF-A0A1A7PWG3-F1-MODEL\_V4 \| 1.0 \| 5.242e-11 \| 329 \| 0.211 \| 274 \| 126 \| 7 \| 14 \| 202 \| 4 \| 272 \| Phage baseplate protein \| Phage baseplate protein; afdb-uniprot50 \| AF-A0A2W7M1F1-F1-MODEL\_V4 \| 1.0 \| 2.642e-12 \| 329 \| 0.125 \| 397 \| 218 \| 17 \| 20 \| 295 \| 2 \| 390 \| Putative phage protein gp47/JayE \| Putative phage protein gp47/JayE; afdb-uniprot50 \| AF-A0A2D5TJV2-F1-MODEL\_V4 \| 1.0 \| 3.161e-12 \| 328 \| 0.142 \| 457 \| 206 \| 21 \| 20 \| 293 \| 5 \| 458 \| Baseplate\_J domain-containing protein \| Baseplate\_J domain-containing protein; afdb-uniprot50 \| AF-A0A7Y7LSY4-F1-MODEL\_V4 \| 1.0 \| 5.589e-13 \| 327 \| 0.12 \| 358 \| 225 \| 10 \| 20 \| 293 \| 4 \| 355 \| Putative phage protein gp47/JayE \| Putative phage protein gp47/JayE; afdb-uniprot50 \| AF-E1RFI2-F1-MODEL\_V4 \| 1.0 \| 8.767e-14 \| 327 \| 0.154 \| 370 \| 203 \| 15 \| 20 \| 295 \| 12 \| 365 \| Baseplate J family protein \| Baseplate J family protein; afdb-uniprot50 \| AF-A0A1Y6CTB1-F1-MODEL\_V4 \| 1.0 \| 2.642e-12 \| 327 \| 0.166 \| 385 \| 200 \| 18 \| 20 \| 294 \| 4 \| 377 \| Uncharacterized phage protein gp47/JayE \| Uncharacterized phage protein gp47/JayE; afdb-uniprot50 \| AF-A0A524BHK3-F1-MODEL\_V4 \| 1.0 \| 2.654e-14 \| 327 \| 0.129 \| 377 \| 216 \| 15 \| 17 \| 295 \| 15 \| 377 \| Baseplate\_J domain-containing protein \| Baseplate\_J domain-containing protein; afdb-uniprot50 \| AF-A0A2V4HQX5-F1-MODEL\_V4 \| 1.0 \| 4.524e-12 \| 327 \| 0.126 \| 396 \| 219 \| 18 \| 20 \| 295 \| 2 \| 390 \| Baseplate\_J domain-containing protein \| Baseplate\_J domain-containing protein; afdb-uniprot50 \| AF-A0A2E2FJ34-F1-MODEL\_V4 \| 1.0 \| 1.402e-09 \| 327 \| 0.18 \| 161 \| 126 \| 5 \| 113 \| 271 \| 275 \| 431 \| Baseplate\_J domain-containing protein \| Baseplate\_J domain-containing protein; afdb-uniprot50 \| AF-A0A017RWE9-F1-MODEL\_V4 \| 1.0 \| 1.078e-12 \| 326 \| 0.168 \| 363 \| 207 \| 13 \| 21 \| 295 \| 1 \| 356 \| Baseplate\_J domain-containing protein \| Baseplate\_J domain-containing protein; afdb-uniprot50 \| AF-A0A510IHW7-F1-MODEL\_V4 \| 1.0 \| 1.684e-11 \| 326 \| 0.151 \| 402 \| 220 \| 17 \| 14 \| 300 \| 5 \| 400 \| Baseplate\_J domain-containing protein \| Baseplate\_J domain-containing protein; afdb-uniprot50 \| AF-A0A2S3UN17-F1-MODEL\_V4 \| 1.0 \| 2.411e-11 \| 326 \| 0.111 \| 510 \| 220 \| 19 \| 12 \| 300 \| 5 \| 502 \| Putative phage baseplate assembly protein \| Putative phage baseplate assembly protein; afdb-uniprot50 \| AF-A0A1Q6UET1-F1-MODEL\_V4 \| 1.0 \| 9.88e-14 \| 325 \| 0.171 \| 362 \| 199 \| 15 \| 13 \| 292 \| 4 \| 346 \| Baseplate\_J domain-containing protein \| Baseplate\_J domain-containing protein; afdb-uniprot50 \| AF-A0A109QIW4-F1-MODEL\_V4 \| 1.0 \| 5.589e-13 \| 325 \| 0.142 \| 372 \| 215 \| 15 \| 20 \| 294 \| 3 \| 367 \| Baseplate\_J domain-containing protein \| Baseplate\_J domain-containing protein; afdb-uniprot50 \| AF-A0A431JII9-F1-MODEL\_V4 \| 1.0 \| 9.569e-13 \| 325 \| 0.132 \| 468 \| 209 \| 14 \| 20 \| 293 \| 4 \| 468 \| Baseplate\_J domain-containing protein \| Baseplate\_J domain-containing protein; afdb-uniprot50 \| AF-A0A522CGM2-F1-MODEL\_V4 \| 1.0 \| 4.128e-11 \| 325 \| 0.118 \| 496 \| 219 \| 18 \| 1 \| 294 \| 1 \| 480 \| Baseplate\_J domain-containing protein \| Baseplate\_J domain-containing protein; afdb-uniprot50 \| AF-A0A7X6ADI4-F1-MODEL\_V4 \| 1.0 \| 7.067e-11 \| 325 \| 0.178 \| 207 \| 148 \| 9 \| 110 \| 296 \| 309 \| 513 \| Putative baseplate assembly protein \| Putative baseplate assembly protein; afdb-uniprot50 \| AF-A0A255Y0J9-F1-MODEL\_V4 \| 1.0 \| 1.96e-12 \| 324 \| 0.167 \| 347 \| 203 \| 11 \| 29 \| 293 \| 2 \| 344 \| Baseplate\_J domain-containing protein \| Baseplate\_J domain-containing protein; afdb-uniprot50 \| AF-Q5E379-F1-MODEL\_V4 \| 1.0 \| 1.332e-13 \| 324 \| 0.136 \| 359 \| 211 \| 12 \| 21 \| 294 \| 4 \| 348 \| Phage protein \| Phage protein; afdb-uniprot50 \| AF-A0A7X6JCW7-F1-MODEL\_V4 \| 1.0 \| 3.134e-08 \| 323 \| 0.3 \| 113 \| 78 \| 1 \| 184 \| 295 \| 3 \| 115 \| Baseplate\_J domain-containing protein \| Baseplate\_J domain-containing protein; afdb-uniprot50 \| AF-A0A412G6U6-F1-MODEL\_V4 \| 1.0 \| 2.281e-13 \| 323 \| 0.17 \| 363 \| 202 \| 19 \| 23 \| 296 \| 5 \| 357 \| Baseplate protein J \| Baseplate protein J; afdb-uniprot50 \| AF-A0A3A3GJ63-F1-MODEL\_V4 \| 1.0 \| 2.57e-13 \| 323 \| 0.141 \| 395 \| 212 \| 17 \| 20 \| 296 \| 8 \| 393 \| Baseplate\_J domain-containing protein \| Baseplate\_J domain-containing protein; afdb-uniprot50 \| AF-A0A1I0CD50-F1-MODEL\_V4 \| 1.0 \| 6.271e-11 \| 321 \| 0.117 \| 390 \| 209 \| 17 \| 20 \| 294 \| 10 \| 379 \| Uncharacterized phage protein gp47/JayE \| Uncharacterized phage protein gp47/JayE; afdb-uniprot50 \| AF-A0A853IQN6-F1-MODEL\_V4 \| 1.0 \| 1.773e-07 \| 320 \| 0.285 \| 91 \| 64 \| 1 \| 209 \| 299 \| 8 \| 97 \| Uncharacterized protein \| Uncharacterized protein; afdb-uniprot50 \| AF-A0A5B8AU34-F1-MODEL\_V4 \| 1.0 \| 7.535e-13 \| 320 \| 0.161 \| 352 \| 201 \| 16 \| 13 \| 296 \| 1 \| 326 \| Baseplate\_J domain-containing protein \| Baseplate\_J domain-containing protein; afdb-uniprot50 \| AF-A0A510KG83-F1-MODEL\_V4 \| 1.0 \| 3.264e-13 \| 320 \| 0.128 \| 359 \| 221 \| 14 \| 22 \| 294 \| 1 \| 353 \| Baseplate J-like protein \| Baseplate J-like protein; afdb-uniprot50 \| AF-A0A0F9LBF1-F1-MODEL\_V4 \| 1.0 \| 1.692e-13 \| 320 \| 0.161 \| 389 \| 209 \| 18 \| 13 \| 295 \| 1 \| 378 \| Baseplate\_J domain-containing protein \| Baseplate\_J domain-containing protein; afdb-uniprot50 \| AF-A0A6I1HEM5-F1-MODEL\_V4 \| 1.0 \| 3.663e-11 \| 319 \| 0.405 \| 185 \| 90 \| 4 \| 8 \| 180 \| 5 \| 181 \| Uncharacterized protein \| Uncharacterized protein; afdb-uniprot50 \| AF-A0A412CI32-F1-MODEL\_V4 \| 1.0 \| 1.182e-13 \| 319 \| 0.151 \| 356 \| 213 \| 12 \| 19 \| 292 \| 2 \| 350 \| Baseplate\_J domain-containing protein \| Baseplate\_J domain-containing protein; afdb-uniprot50 \| AF-A0A258NW90-F1-MODEL\_V4 \| 1.0 \| 1.796e-13 \| 319 \| 0.157 \| 369 \| 214 \| 15 \| 11 \| 295 \| 2 \| 357 \| Baseplate\_J domain-containing protein \| Baseplate\_J domain-containing protein; afdb-uniprot50 \| AF-A0A0F9F7E7-F1-MODEL\_V4 \| 1.0 \| 4.015e-12 \| 319 \| 0.137 \| 372 \| 210 \| 16 \| 20 \| 295 \| 1 \| 357 \| Baseplate\_J domain-containing protein \| Baseplate\_J domain-containing protein; afdb-uniprot50 \| AF-A0A2G2PW46-F1-MODEL\_V4 \| 1.0 \| 9.569e-13 \| 319 \| 0.142 \| 386 \| 210 \| 16 \| 23 \| 298 \| 28 \| 402 \| Baseplate\_J domain-containing protein \| Baseplate\_J domain-containing protein; afdb-uniprot50 \| AF-A0A653R9L3-F1-MODEL\_V4 \| 1.0 \| 4.672e-13 \| 318 \| 0.161 \| 359 \| 210 \| 14 \| 20 \| 293 \| 4 \| 356 \| Baseplate\_J domain-containing protein \| Baseplate\_J domain-containing protein; afdb-uniprot50 \| AF-A0A1G9EXV1-F1-MODEL\_V4 \| 1.0 \| 1.011e-10 \| 318 \| 0.204 \| 215 \| 144 \| 9 \| 110 \| 299 \| 421 \| 633 \| Putative baseplate assembly protein \| Putative baseplate assembly protein; afdb-uniprot50 \| AF-A0A450RV24-F1-MODEL\_V4 \| 1.0 \| 4.803e-12 \| 317 \| 0.143 \| 397 \| 216 \| 20 \| 14 \| 295 \| 6 \| 393 \| Uncharacterized phage protein gp47/JayE \| Uncharacterized phage protein gp47/JayE; afdb-uniprot50 \| AF-A0A3N5XVA3-F1-MODEL\_V4 \| 1.0 \| 1.074e-10 \| 317 \| 0.185 \| 216 \| 143 \| 10 \| 111 \| 296 \| 813 \| 1025 \| Putative baseplate assembly protein \| Putative baseplate assembly protein; afdb-uniprot50 \| AF-A0A6I2SNH4-F1-MODEL\_V4 \| 1.0 \| 1.255e-13 \| 316 \| 0.193 \| 336 \| 195 \| 16 \| 1 \| 295 \| 34 \| 334 \| Baseplate\_J domain-containing protein \| Baseplate\_J domain-containing protein; afdb-uniprot50 \| AF-A0A2N2BUM8-F1-MODEL\_V4 \| 1.0 \| 5.907e-11 \| 316 \| 0.121 \| 396 \| 213 \| 17 \| 20 \| 295 \| 12 \| 392 \| Baseplate\_J domain-containing protein \| Baseplate\_J domain-containing protein; afdb-uniprot50 \| AF-A0A4Q3NS83-F1-MODEL\_V4 \| 1.0 \| 1.998e-07 \| 315 \| 0.538 \| 91 \| 41 \| 1 \| 1 \| 91 \| 1 \| 90 \| Baseplate assembly protein \| Baseplate assembly protein; afdb-uniprot50 \| AF-A0A166EHE8-F1-MODEL\_V4 \| 1.0 \| 2.57e-13 \| 315 \| 0.142 \| 365 \| 212 \| 16 \| 20 \| 293 \| 23 \| 377 \| Baseplate J-like protein \| Baseplate J-like protein; afdb-uniprot50 \| AF-A0A7X9CZY0-F1-MODEL\_V4 \| 1.0 \| 4.959e-13 \| 315 \| 0.136 \| 397 \| 208 \| 17 \| 20 \| 295 \| 12 \| 394 \| Baseplate J/gp47 family protein \| Baseplate J/gp47 family protein; afdb-uniprot50 \| AF-A0A154LLK7-F1-MODEL\_V4 \| 1.0 \| 1.638e-12 \| 314 \| 0.159 \| 364 \| 209 \| 13 \| 20 \| 293 \| 8 \| 364 \| Baseplate\_J domain-containing protein \| Baseplate\_J domain-containing protein; afdb-uniprot50 \| AF-A0A3D1IYK7-F1-MODEL\_V4 \| 1.0 \| 1.408e-11 \| 314 \| 0.142 \| 392 \| 207 \| 18 \| 14 \| 295 \| 1 \| 373 \| Baseplate\_J domain-containing protein \| Baseplate\_J domain-containing protein; afdb-uniprot50 \| AF-A0A1L6I8I3-F1-MODEL\_V4 \| 1.0 \| 4.401e-13 \| 314 \| 0.144 \| 402 \| 205 \| 15 \| 20 \| 293 \| 4 \| 394 \| Baseplate\_J domain-containing protein \| Baseplate\_J domain-containing protein; afdb-uniprot50 \| AF-A0A1Y1QPN6-F1-MODEL\_V4 \| 1.0 \| 5.933e-13 \| 312 \| 0.142 \| 357 \| 212 \| 12 \| 12 \| 292 \| 2 \| 340 \| Baseplate\_J domain-containing protein \| Baseplate\_J domain-containing protein; afdb-uniprot50 \| AF-R6PWP6-F1-MODEL\_V4 \| 1.0 \| 1.255e-13 \| 312 \| 0.169 \| 359 \| 200 \| 14 \| 23 \| 293 \| 5 \| 353 \| Baseplate J family protein \| Baseplate J family protein; afdb-uniprot50 \| AF-A0A437M1E1-F1-MODEL\_V4 \| 1.0 \| 3.356e-12 \| 312 \| 0.148 \| 390 \| 206 \| 17 \| 14 \| 293 \| 1 \| 374 \| Baseplate\_J domain-containing protein \| Baseplate\_J domain-containing protein; afdb-uniprot50 \| AF-A0A5A9X997-F1-MODEL\_V4 \| 1.0 \| 8.73e-12 \| 312 \| 0.153 \| 385 \| 206 \| 18 \| 20 \| 295 \| 4 \| 377 \| Baseplate\_J domain-containing protein \| Baseplate\_J domain-containing protein; afdb-uniprot50 \| AF-A0A3D5BKD5-F1-MODEL\_V4 \| 1.0 \| 7.78e-14 \| 312 \| 0.152 \| 399 \| 219 \| 15 \| 14 \| 299 \| 3 \| 395 \| Baseplate\_J domain-containing protein \| Baseplate\_J domain-containing protein; afdb-uniprot50 \| AF-A0A5C7QAD1-F1-MODEL\_V4 \| 1.0 \| 3.997e-10 \| 312 \| 0.148 \| 209 \| 150 \| 10 \| 112 \| 295 \| 310 \| 515 \| Uncharacterized protein \| Uncharacterized protein; afdb-uniprot50 \| AF-A0A7Z7J4X3-F1-MODEL\_V4 \| 1.0 \| 2.62e-08 \| 311 \| 0.451 \| 124 \| 59 \| 2 \| 21 \| 137 \| 1 \| 122 \| Uncharacterized protein \| Uncharacterized protein; afdb-uniprot50 \| AF-M5DNC9-F1-MODEL\_V4 \| 1.0 \| 7.998e-13 \| 311 \| 0.149 \| 361 \| 214 \| 14 \| 13 \| 293 \| 1 \| 348 \| Baseplate J-like protein \| Baseplate J-like protein; afdb-uniprot50 \| AF-A0A381M382-F1-MODEL\_V4 \| 1.0 \| 2.859e-07 \| 310 \| 0.319 \| 94 \| 61 \| 1 \| 209 \| 299 \| 40 \| 133 \| Phage-related baseplate assembly protein \| Phage-related baseplate assembly protein; afdb-uniprot50 \| AF-A0A2V7X4R7-F1-MODEL\_V4 \| 1.0 \| 1.04e-09 \| 309 \| 0.177 \| 214 \| 147 \| 10 \| 112 \| 299 \| 201 \| 411 \| Baseplate\_J domain-containing protein \| Baseplate\_J domain-containing protein; afdb-uniprot50 \| AF-A0A174T558-F1-MODEL\_V4 \| 1.0 \| 1.952e-10 \| 309 \| 0.14 \| 220 \| 159 \| 10 \| 96 \| 296 \| 525 \| 733 \| Uncharacterized homolog of phage Mu protein gp47 \| Uncharacterized homolog of phage Mu protein gp47; afdb-uniprot50 \| AF-A0A7U6AEJ8-F1-MODEL\_V4 \| 1.0 \| 3.134e-08 \| 308 \| 0.454 \| 110 \| 60 \| 0 \| 135 \| 244 \| 2 \| 111 \| Baseplate\_J domain-containing protein \| Baseplate\_J domain-containing protein; afdb-uniprot50 \| AF-A0A2H0IXG9-F1-MODEL\_V4 \| 1.0 \| 1.205e-08 \| 308 \| 0.203 \| 182 \| 119 \| 8 \| 133 \| 293 \| 3 \| 179 \| Uncharacterized protein \| Uncharacterized protein; afdb-uniprot50 \| AF-A0A0A8WTQ2-F1-MODEL\_V4 \| 1.0 \| 8.491e-13 \| 308 \| 0.163 \| 324 \| 201 \| 14 \| 13 \| 296 \| 1 \| 294 \| Baseplate\_J domain-containing protein \| Baseplate\_J domain-containing protein; afdb-uniprot50 \| AF-M3JG00-F1-MODEL\_V4 \| 1.0 \| 1.21e-10 \| 308 \| 0.13 \| 391 \| 213 \| 16 \| 13 \| 296 \| 2 \| 372 \| Baseplate\_J domain-containing protein \| Baseplate\_J domain-containing protein; afdb-uniprot50 \| AF-A0A534UQ31-F1-MODEL\_V4 \| 1.0 \| 1.364e-10 \| 308 \| 0.179 \| 217 \| 145 \| 10 \| 110 \| 296 \| 778 \| 991 \| Putative baseplate assembly protein \| Putative baseplate assembly protein; afdb-uniprot50 \| AF-A0A379TSV1-F1-MODEL\_V4 \| 1.0 \| 3.631e-07 \| 307 \| 0.322 \| 90 \| 59 \| 2 \| 210 \| 298 \| 1 \| 89 \| Putative bacteriophage baseplate assembly protein \| Putative bacteriophage baseplate assembly protein; afdb-uniprot50 \| AF-A0A1G3LLV8-F1-MODEL\_V4 \| 1.0 \| 3.148e-10 \| 307 \| 0.119 \| 495 \| 217 \| 19 \| 12 \| 295 \| 7 \| 493 \| Baseplate\_J domain-containing protein \| Baseplate\_J domain-containing protein; afdb-uniprot50 \| AF-A0A150TFN7-F1-MODEL\_V4 \| 1.0 \| 1.631e-10 \| 307 \| 0.147 \| 223 \| 153 \| 10 \| 110 \| 300 \| 454 \| 671 \| Baseplate\_J domain-containing protein \| Baseplate\_J domain-containing protein; afdb-uniprot50 \| AF-A0A2I0CWU0-F1-MODEL\_V4 \| 1.0 \| 4.959e-13 \| 306 \| 0.147 \| 366 \| 204 \| 13 \| 22 \| 294 \| 14 \| 364 \| Baseplate\_J domain-containing protein \| Baseplate\_J domain-containing protein; afdb-uniprot50 \| AF-A0A7C9QTR2-F1-MODEL\_V4 \| 1.0 \| 4.505e-10 \| 306 \| 0.175 \| 211 \| 148 \| 11 \| 110 \| 296 \| 791 \| 999 \| Uncharacterized protein \| Uncharacterized protein; afdb-uniprot50 \| AF-A0A6N8PP11-F1-MODEL\_V4 \| 1.0 \| 3.222e-07 \| 305 \| 0.554 \| 83 \| 37 \| 0 \| 80 \| 162 \| 1 \| 83 \| Baseplate assembly protein \| Baseplate assembly protein; afdb-uniprot50 \| AF-E6LJD1-F1-MODEL\_V4 \| 1.0 \| 6.874e-12 \| 305 \| 0.127 \| 385 \| 212 \| 18 \| 22 \| 295 \| 12 \| 383 \| Baseplate\_J domain-containing protein \| Baseplate\_J domain-containing protein; afdb-uniprot50 \| AF-E3R381-F1-MODEL\_V4 \| 1.0 \| 4.803e-12 \| 305 \| 0.123 \| 397 \| 207 \| 20 \| 22 \| 296 \| 15 \| 392 \| Baseplate\_J domain-containing protein \| Baseplate\_J domain-containing protein; afdb-uniprot50 \| AF-A0A355JCM0-F1-MODEL\_V4 \| 1.0 \| 2.884e-11 \| 305 \| 0.136 \| 417 \| 213 \| 17 \| 20 \| 300 \| 9 \| 414 \| Baseplate\_J domain-containing protein \| Baseplate\_J domain-containing protein; afdb-uniprot50 \| AF-A0A2E1AI28-F1-MODEL\_V4 \| 1.0 \| 6.844e-10 \| 305 \| 0.189 \| 216 \| 146 \| 10 \| 111 \| 300 \| 431 \| 643 \| Putative baseplate assembly protein \| Putative baseplate assembly protein; afdb-uniprot50 \| AF-A0A3S4LRQ5-F1-MODEL\_V4 \| 1.0 \| 2.468e-08 \| 304 \| 0.342 \| 111 \| 71 \| 2 \| 191 \| 300 \| 2 \| 111 \| Baseplate protein \| Baseplate protein; afdb-uniprot50 \| AF-A0A317HNT2-F1-MODEL\_V4 \| 1.0 \| 1.501e-13 \| 304 \| 0.169 \| 349 \| 203 \| 17 \| 1 \| 294 \| 4 \| 320 \| Baseplate\_J domain-containing protein \| Baseplate\_J domain-containing protein; afdb-uniprot50 \| AF-A0A0E3NKG6-F1-MODEL\_V4 \| 1.0 \| 7.502e-11 \| 304 \| 0.163 \| 233 \| 151 \| 11 \| 110 \| 300 \| 579 \| 809 \| Baseplate\_J domain-containing protein \| Baseplate\_J domain-containing protein; afdb-uniprot50 \| AF-J9ZB16-F1-MODEL\_V4 \| 1.0 \| 3.356e-12 \| 303 \| 0.152 \| 368 \| 213 \| 12 \| 20 \| 295 \| 2 \| 362 \| Baseplate J family protein \| Baseplate J family protein; afdb-uniprot50 \| AF-A0A3R7UR03-F1-MODEL\_V4 \| 1.0 \| 1.952e-10 \| 303 \| 0.148 \| 222 \| 147 \| 10 \| 113 \| 294 \| 380 \| 599 \| Baseplate\_J domain-containing protein \| Baseplate\_J domain-containing protein; afdb-uniprot50 \| AF-A0A1H4QY22-F1-MODEL\_V4 \| 1.0 \| 3.435e-09 \| 302 \| 0.187 \| 171 \| 124 \| 8 \| 132 \| 295 \| 4 \| 166 \| Baseplate J-like protein \| Baseplate J-like protein; afdb-uniprot50 \| AF-A0A3F3A315-F1-MODEL\_V4 \| 1.0 \| 1.29e-12 \| 302 \| 0.116 \| 360 \| 225 \| 13 \| 23 \| 295 \| 1 \| 354 \| Baseplate\_J domain-containing protein \| Baseplate\_J domain-containing protein; afdb-uniprot50 \| AF-A0A7T8FM21-F1-MODEL\_V4 \| 1.0 \| 1.594e-13 \| 302 \| 0.141 \| 374 \| 216 \| 15 \| 12 \| 292 \| 3 \| 364 \| Baseplate J/gp47 family protein \| Baseplate J/gp47 family protein; afdb-uniprot50 \| AF-A0A1S1HMN1-F1-MODEL\_V4 \| 1.0 \| 3.997e-10 \| 302 \| 0.095 \| 378 \| 225 \| 16 \| 20 \| 294 \| 10 \| 373 \| Baseplate\_J domain-containing protein \| Baseplate\_J domain-containing protein; afdb-uniprot50 \| AF-A0A7Y6W8F8-F1-MODEL\_V4 \| 1.0 \| 1.846e-12 \| 302 \| 0.17 \| 404 \| 194 \| 17 \| 21 \| 293 \| 6 \| 399 \| Baseplate J/gp47 family protein \| Baseplate J/gp47 family protein; afdb-uniprot50 \| AF-A0A1D2R572-F1-MODEL\_V4 \| 1.0 \| 2.793e-10 \| 302 \| 0.164 \| 194 \| 140 \| 9 \| 111 \| 296 \| 333 \| 512 \| Uncharacterized protein \| Uncharacterized protein; afdb-uniprot50 \| AF-A0A431IBF5-F1-MODEL\_V4 \| 1.0 \| 2.345e-12 \| 301 \| 0.136 \| 375 \| 214 \| 18 \| 13 \| 293 \| 1 \| 359 \| Baseplate J protein \| Baseplate J protein; afdb-uniprot50 \| AF-A0A433WZS4-F1-MODEL\_V4 \| 1.0 \| 4.524e-12 \| 301 \| 0.176 \| 379 \| 198 \| 17 \| 13 \| 296 \| 3 \| 362 \| Baseplate\_J domain-containing protein \| Baseplate\_J domain-containing protein; afdb-uniprot50 \| AF-A0A838CGI4-F1-MODEL\_V4 \| 1.0 \| 1.435e-06 \| 300 \| 0.4 \| 95 \| 57 \| 0 \| 8 \| 102 \| 4 \| 98 \| Baseplate J protein \| Baseplate J protein; afdb-uniprot50 \| AF-A0A257PB54-F1-MODEL\_V4 \| 1.0 \| 1.326e-11 \| 299 \| 0.131 \| 366 \| 217 \| 15 \| 20 \| 293 \| 4 \| 360 \| Baseplate\_J domain-containing protein \| Baseplate\_J domain-containing protein; afdb-uniprot50 \| AF-A0A1Y3QLM5-F1-MODEL\_V4 \| 1.0 \| 5.933e-13 \| 299 \| 0.168 \| 398 \| 197 \| 18 \| 20 \| 295 \| 7 \| 392 \| Baseplate\_J domain-containing protein \| Baseplate\_J domain-containing protein; afdb-uniprot50 \| AF-A0A0F9SDC1-F1-MODEL\_V4 \| 1.0 \| 1.326e-11 \| 299 \| 0.122 \| 472 \| 211 \| 18 \| 22 \| 294 \| 5 \| 472 \| Baseplate\_J domain-containing protein \| Baseplate\_J domain-containing protein; afdb-uniprot50 \| AF-A0A0D5NAR8-F1-MODEL\_V4 \| 1.0 \| 1.364e-10 \| 299 \| 0.166 \| 228 \| 151 \| 10 \| 110 \| 300 \| 447 \| 672 \| Baseplate\_J domain-containing protein \| Baseplate\_J domain-containing protein; afdb-uniprot50 \| AF-A0A0S4LB89-F1-MODEL\_V4 \| 1.0 \| 3.547e-10 \| 298 \| 0.202 \| 227 \| 143 \| 9 \| 110 \| 300 \| 434 \| 658 \| Baseplate\_J domain-containing protein \| Baseplate\_J domain-containing protein; afdb-uniprot50 \| AF-A0A655DHR5-F1-MODEL\_V4 \| 1.0 \| 6.244e-09 \| 297 \| 0.323 \| 133 \| 87 \| 3 \| 170 \| 300 \| 3 \| 134 \| Phage baseplate protein \| Phage baseplate protein; afdb-uniprot50 \| AF-A0A164ZAU4-F1-MODEL\_V4 \| 1.0 \| 2.261e-09 \| 296 \| 0.201 \| 194 \| 134 \| 8 \| 116 \| 294 \| 12 \| 199 \| Phage baseplate \| Phage baseplate; afdb-uniprot50 \| AF-A0A824RZ05-F1-MODEL\_V4 \| 1.0 \| 1.78e-09 \| 296 \| 0.245 \| 236 \| 100 \| 4 \| 10 \| 167 \| 17 \| 252 \| Baseplate assembly protein \| Baseplate assembly protein; afdb-uniprot50 \| AF-A0A7K4BK92-F1-MODEL\_V4 \| 1.0 \| 1.906e-13 \| 296 \| 0.145 \| 393 \| 196 \| 20 \| 20 \| 295 \| 11 \| 380 \| Baseplate\_J domain-containing protein \| Baseplate\_J domain-containing protein; afdb-uniprot50 \| AF-A0A2T4N3C8-F1-MODEL\_V4 \| 1.0 \| 2.421e-13 \| 294 \| 0.138 \| 398 \| 223 \| 14 \| 11 \| 294 \| 2 \| 393 \| Baseplate\_J domain-containing protein \| Baseplate\_J domain-containing protein; afdb-uniprot50 \| AF-L0L0C7-F1-MODEL\_V4 \| 1.0 \| 5.077e-10 \| 294 \| 0.164 \| 225 \| 150 \| 9 \| 111 \| 299 \| 997 \| 1219 \| Baseplate\_J domain-containing protein \| Baseplate\_J domain-containing protein; afdb-uniprot50 \| AF-A0A1Z4C4M5-F1-MODEL\_V4 \| 1.0 \| 2.559e-11 \| 293 \| 0.152 \| 380 \| 200 \| 17 \| 20 \| 293 \| 3 \| 366 \| Baseplate\_J domain-containing protein \| Baseplate\_J domain-containing protein; afdb-uniprot50 \| AF-A0A5C7Q1H5-F1-MODEL\_V4 \| 1.0 \| 6.271e-11 \| 293 \| 0.152 \| 381 \| 208 \| 17 \| 20 \| 295 \| 3 \| 373 \| Baseplate\_J domain-containing protein \| Baseplate\_J domain-containing protein; afdb-uniprot50 \| AF-A0A5F1RNH5-F1-MODEL\_V4 \| 1.0 \| 3.075e-13 \| 293 \| 0.15 \| 386 \| 210 \| 18 \| 20 \| 296 \| 8 \| 384 \| Baseplate\_J domain-containing protein \| Baseplate\_J domain-containing protein; afdb-uniprot50 \| AF-A0A059V4B9-F1-MODEL\_V4 \| 1.0 \| 2.859e-07 \| 292 \| 0.261 \| 111 \| 81 \| 1 \| 191 \| 300 \| 6 \| 116 \| Phage-related baseplate assembly protein \| Phage-related baseplate assembly protein; afdb-uniprot50 \| AF-A0A1Z4HHP6-F1-MODEL\_V4 \| 1.0 \| 1.016e-12 \| 292 \| 0.185 \| 345 \| 197 \| 22 \| 13 \| 299 \| 3 \| 321 \| Baseplate\_J domain-containing protein \| Baseplate\_J domain-containing protein; afdb-uniprot50 \| AF-A0A1R0WGM4-F1-MODEL\_V4 \| 1.0 \| 1.369e-12 \| 291 \| 0.157 \| 355 \| 202 \| 16 \| 28 \| 296 \| 2 \| 345 \| Baseplate\_J domain-containing protein \| Baseplate\_J domain-containing protein; afdb-uniprot50 \| AF-A0A316PC92-F1-MODEL\_V4 \| 1.0 \| 6.686e-13 \| 291 \| 0.121 \| 355 \| 222 \| 11 \| 22 \| 292 \| 1 \| 349 \| Baseplate\_J domain-containing protein \| Baseplate\_J domain-containing protein; afdb-uniprot50 \| AF-A0A1G8B3H6-F1-MODEL\_V4 \| 1.0 \| 7.502e-11 \| 291 \| 0.139 \| 381 \| 209 \| 19 \| 20 \| 300 \| 3 \| 364 \| Baseplate J-like protein \| Baseplate J-like protein; afdb-uniprot50 \| AF-A0A662L870-F1-MODEL\_V4 \| 1.0 \| 4.505e-10 \| 291 \| 0.166 \| 210 \| 148 \| 10 \| 110 \| 294 \| 422 \| 629 \| Putative baseplate assembly protein \| Putative baseplate assembly protein; afdb-uniprot50 \| AF-A0A7W7H569-F1-MODEL\_V4 \| 1.0 \| 1.321e-09 \| 291 \| 0.181 \| 215 \| 144 \| 10 \| 110 \| 300 \| 966 \| 1172 \| Baseplate\_J domain-containing protein \| Baseplate\_J domain-containing protein; afdb-uniprot50 \| AF-A0A2X3LXX5-F1-MODEL\_V4 \| 1.0 \| 1.352e-06 \| 290 \| 0.345 \| 84 \| 55 \| 0 \| 216 \| 299 \| 2 \| 85 \| Baseplate assembly protein J \| Baseplate assembly protein J; afdb-uniprot50 \| AF-D4DSI2-F1-MODEL\_V4 \| 1.0 \| 1.069e-08 \| 290 \| 0.233 \| 154 \| 103 \| 7 \| 151 \| 294 \| 2 \| 150 \| Baseplate\_J domain-containing protein \| Baseplate\_J domain-containing protein; afdb-uniprot50 \| AF-A0A398BBK4-F1-MODEL\_V4 \| 1.0 \| 2.978e-12 \| 290 \| 0.138 \| 383 \| 213 \| 17 \| 20 \| 295 \| 8 \| 380 \| Baseplate J/gp47 family protein \| Baseplate J/gp47 family protein; afdb-uniprot50 \| AF-A0A0F7FBD4-F1-MODEL\_V4 \| 1.0 \| 1.838e-10 \| 290 \| 0.127 \| 391 \| 214 \| 19 \| 20 \| 295 \| 4 \| 382 \| Baseplate\_J domain-containing protein \| Baseplate\_J domain-containing protein; afdb-uniprot50 \| AF-A0A1G9EXR2-F1-MODEL\_V4 \| 1.0 \| 9.227e-10 \| 290 \| 0.221 \| 212 \| 134 \| 13 \| 110 \| 296 \| 870 \| 1075 \| Putative baseplate assembly protein \| Putative baseplate assembly protein; afdb-uniprot50 \| AF-A0A7R7TB47-F1-MODEL\_V4 \| 1.0 \| 5.907e-11 \| 289 \| 0.132 \| 384 \| 210 \| 17 \| 14 \| 294 \| 1 \| 364 \| Uncharacterized protein \| Uncharacterized protein; afdb-uniprot50 \| AF-H6NDS9-F1-MODEL\_V4 \| 1.0 \| 7.535e-13 \| 288 \| 0.157 \| 394 \| 204 \| 20 \| 20 \| 295 \| 8 \| 391 \| Baseplate protein \| Baseplate protein; afdb-uniprot50 \| AF-A0A1I1TQL3-F1-MODEL\_V4 \| 1.0 \| 1.215e-12 \| 288 \| 0.146 \| 396 \| 203 \| 17 \| 20 \| 296 \| 14 \| 393 \| Uncharacterized phage protein gp47/JayE \| Uncharacterized phage protein gp47/JayE; afdb-uniprot50 \| AF-Q2L2U7-F1-MODEL\_V4 \| 1.0 \| 1.677e-09 \| 288 \| 0.12 \| 415 \| 214 \| 19 \| 13 \| 295 \| 1 \| 396 \| Phage protein \| Phage protein; afdb-uniprot50 \| AF-A0A810LC53-F1-MODEL\_V4 \| 1.0 \| 3.148e-10 \| 288 \| 0.214 \| 224 \| 137 \| 14 \| 110 \| 300 \| 405 \| 622 \| Putative baseplate assembly protein \| Putative baseplate assembly protein; afdb-uniprot50 \| AF-A0A321LRK0-F1-MODEL\_V4 \| 1.0 \| 8.692e-10 \| 288 \| 0.158 \| 214 \| 147 \| 11 \| 111 \| 295 \| 557 \| 766 \| Putative baseplate assembly protein \| Putative baseplate assembly protein; afdb-uniprot50 \| AF-V4IZ11-F1-MODEL\_V4 \| 1.0 \| 2.19e-08 \| 287 \| 0.271 \| 151 \| 85 \| 3 \| 168 \| 295 \| 2 \| 150 \| Baseplate\_J domain-containing protein \| Baseplate\_J domain-containing protein; afdb-uniprot50 \| AF-A0A353REU8-F1-MODEL\_V4 \| 1.0 \| 1.244e-09 \| 287 \| 0.201 \| 199 \| 135 \| 10 \| 111 \| 294 \| 39 \| 228 \| Baseplate\_J domain-containing protein \| Baseplate\_J domain-containing protein; afdb-uniprot50 \| AF-A0A2V1HBX8-F1-MODEL\_V4 \| 1.0 \| 1.78e-09 \| 287 \| 0.103 \| 377 \| 219 \| 20 \| 22 \| 294 \| 12 \| 373 \| Baseplate\_J domain-containing protein \| Baseplate\_J domain-containing protein; afdb-uniprot50 \| AF-A0A1F5C406-F1-MODEL\_V4 \| 1.0 \| 1.898e-11 \| 287 \| 0.119 \| 469 \| 210 \| 18 \| 20 \| 292 \| 4 \| 465 \| Baseplate\_J domain-containing protein \| Baseplate\_J domain-containing protein; afdb-uniprot50 \| AF-A0A7G9KUY0-F1-MODEL\_V4 \| 1.0 \| 2.39e-07 \| 285 \| 0.549 \| 102 \| 45 \| 1 \| 1 \| 102 \| 1 \| 101 \| Baseplate assembly protein J \| Baseplate assembly protein J; afdb-uniprot50 \| AF-A0A432ULA7-F1-MODEL\_V4 \| 1.0 \| 1.587e-11 \| 285 \| 0.14 \| 350 \| 211 \| 14 \| 20 \| 293 \| 2 \| 337 \| Baseplate\_J domain-containing protein \| Baseplate\_J domain-containing protein; afdb-uniprot50 \| AF-A0A3A0GCC9-F1-MODEL\_V4 \| 1.0 \| 8.692e-10 \| 285 \| 0.169 \| 218 \| 148 \| 11 \| 110 \| 296 \| 527 \| 742 \| Putative baseplate assembly protein \| Putative baseplate assembly protein; afdb-uniprot50 \| AF-A0A239C2Z7-F1-MODEL\_V4 \| 1.0 \| 7.502e-11 \| 285 \| 0.191 \| 245 \| 136 \| 13 \| 110 \| 296 \| 507 \| 747 \| Putative baseplate assembly protein \| Putative baseplate assembly protein; afdb-uniprot50 \| AF-A0A1Z4HHL8-F1-MODEL\_V4 \| 1.0 \| 1.104e-09 \| 284 \| 0.144 \| 235 \| 145 \| 11 \| 110 \| 294 \| 844 \| 1072 \| Baseplate\_J domain-containing protein \| Baseplate\_J domain-containing protein; afdb-uniprot50 \| AF-A0A251ZY30-F1-MODEL\_V4 \| 1.0 \| 3.562e-12 \| 283 \| 0.132 \| 376 \| 215 \| 15 \| 16 \| 292 \| 1 \| 364 \| Baseplate\_J domain-containing protein \| Baseplate\_J domain-containing protein; afdb-uniprot50 \| AF-A0A7T4UTX7-F1-MODEL\_V4 \| 1.0 \| 1.408e-11 \| 283 \| 0.124 \| 386 \| 218 \| 19 \| 21 \| 296 \| 23 \| 398 \| Uncharacterized protein \| Uncharacterized protein; afdb-uniprot50 \| AF-G0ABR9-F1-MODEL\_V4 \| 1.0 \| 3.435e-09 \| 283 \| 0.153 \| 209 \| 153 \| 10 \| 110 \| 296 \| 824 \| 1030 \| Baseplate\_J domain-containing protein \| Baseplate\_J domain-containing protein; afdb-uniprot50 \| AF-I5B7H1-F1-MODEL\_V4 \| 1.0 \| 7.266e-10 \| 283 \| 0.164 \| 237 \| 155 \| 12 \| 101 \| 296 \| 794 \| 1028 \| Baseplate J-like protein \| Baseplate J-like protein; afdb-uniprot50 \| AF-A0A1Q6YE39-F1-MODEL\_V4 \| 1.0 \| 6.073e-10 \| 282 \| 0.165 \| 218 \| 148 \| 10 \| 110 \| 295 \| 442 \| 657 \| Putative baseplate assembly protein \| Putative baseplate assembly protein; afdb-uniprot50 \| AF-A0A7C6P3L4-F1-MODEL\_V4 \| 1.0 \| 9.796e-10 \| 282 \| 0.15 \| 212 \| 153 \| 10 \| 110 \| 298 \| 496 \| 703 \| Baseplate\_J domain-containing protein \| Baseplate\_J domain-containing protein; afdb-uniprot50 \| AF-A0A1S0V4H6-F1-MODEL\_V4 \| 1.0 \| 2.693e-07 \| 281 \| 0.504 \| 105 \| 52 \| 0 \| 1 \| 105 \| 1 \| 105 \| Uncharacterized protein \| Uncharacterized protein; afdb-uniprot50 \| AF-A0A7W7HM91-F1-MODEL\_V4 \| 1.0 \| 1.402e-09 \| 281 \| 0.161 \| 217 \| 148 \| 9 \| 110 \| 294 \| 396 \| 610 \| Putative phage baseplate assembly protein \| Putative phage baseplate assembly protein; afdb-uniprot50 \| AF-A0A3D4UXE9-F1-MODEL\_V4 \| 1.0 \| 4.917e-09 \| 280 \| 0.147 \| 217 \| 147 \| 9 \| 114 \| 294 \| 21 \| 235 \| Uncharacterized protein \| Uncharacterized protein; afdb-uniprot50 \| AF-A0A1P8MVC1-F1-MODEL\_V4 \| 1.0 \| 8.188e-10 \| 280 \| 0.179 \| 234 \| 137 \| 12 \| 110 \| 294 \| 426 \| 653 \| Putative baseplate assembly protein \| Putative baseplate assembly protein; afdb-uniprot50 \| AF-A0A7X8K102-F1-MODEL\_V4 \| 1.0 \| 1.321e-09 \| 280 \| 0.118 \| 211 \| 154 \| 10 \| 115 \| 298 \| 494 \| 699 \| Baseplate\_J domain-containing protein \| Baseplate\_J domain-containing protein; afdb-uniprot50 \| AF-A0A4U2ZFI7-F1-MODEL\_V4 \| 1.0 \| 1.454e-12 \| 278 \| 0.138 \| 382 \| 224 \| 14 \| 8 \| 296 \| 2 \| 371 \| Baseplate J/gp47 family protein \| Baseplate J/gp47 family protein; afdb-uniprot50 \| AF-A0A3M4S7G9-F1-MODEL\_V4 \| 1.0 \| 8.382e-07 \| 277 \| 0.438 \| 98 \| 55 \| 0 \| 101 \| 198 \| 2 \| 99 \| Phage baseplate assembly protein J \| Phage baseplate assembly protein J; afdb-uniprot50 \| AF-A0A2E6KN58-F1-MODEL\_V4 \| 1.0 \| 6.657e-11 \| 277 \| 0.11 \| 362 \| 224 \| 13 \| 20 \| 294 \| 10 \| 360 \| Baseplate J protein \| Baseplate J protein; afdb-uniprot50 \| AF-A0A6N4HYA5-F1-MODEL\_V4 \| 1.0 \| 9.569e-13 \| 277 \| 0.147 \| 408 \| 209 \| 18 \| 13 \| 296 \| 1 \| 393 \| Baseplate\_J domain-containing protein \| Baseplate\_J domain-containing protein; afdb-uniprot50 \| AF-A0A550JHH0-F1-MODEL\_V4 \| 1.0 \| 5.099e-12 \| 276 \| 0.189 \| 328 \| 189 \| 18 \| 13 \| 294 \| 1 \| 297 \| Baseplate\_J domain-containing protein \| Baseplate\_J domain-containing protein; afdb-uniprot50 \| AF-A0A150HQF5-F1-MODEL\_V4 \| 1.0 \| 1.177e-11 \| 276 \| 0.134 \| 380 \| 208 \| 13 \| 20 \| 293 \| 4 \| 368 \| Baseplate J-like protein \| Baseplate J-like protein; afdb-uniprot50 \| AF-A0A2N0F3D7-F1-MODEL\_V4 \| 1.0 \| 1.177e-11 \| 276 \| 0.121 \| 395 \| 215 \| 17 \| 20 \| 294 \| 9 \| 391 \| Putative phage protein gp47/JayE \| Putative phage protein gp47/JayE; afdb-uniprot50 \| AF-A0A535END5-F1-MODEL\_V4 \| 1.0 \| 1.78e-09 \| 276 \| 0.193 \| 232 \| 138 \| 13 \| 111 \| 300 \| 190 \| 414 \| Putative baseplate assembly protein \| Putative baseplate assembly protein; afdb-uniprot50 \| AF-A0A7C4C5M9-F1-MODEL\_V4 \| 1.0 \| 2.139e-11 \| 276 \| 0.12 \| 439 \| 192 \| 12 \| 42 \| 300 \| 97 \| 521 \| Baseplate\_J domain-containing protein \| Baseplate\_J domain-containing protein; afdb-uniprot50 \| AF-A0A6P2EU18-F1-MODEL\_V4 \| 1.0 \| 2.4e-09 \| 276 \| 0.172 \| 214 \| 144 \| 10 \| 110 \| 294 \| 501 \| 710 \| Baseplate J-like protein \| Baseplate J-like protein; afdb-uniprot50 \| AF-A0A2K3JD84-F1-MODEL\_V4 \| 1.0 \| 2.717e-11 \| 275 \| 0.114 \| 446 \| 218 \| 15 \| 16 \| 292 \| 1 \| 438 \| Uncharacterized protein \| Uncharacterized protein; afdb-uniprot50 \| AF-A0A4T3EY99-F1-MODEL\_V4 \| 1.0 \| 8.692e-10 \| 275 \| 0.162 \| 228 \| 150 \| 11 \| 110 \| 300 \| 442 \| 665 \| Putative baseplate assembly protein \| Putative baseplate assembly protein; afdb-uniprot50 \| AF-A0A376ZP12-F1-MODEL\_V4 \| 1.0 \| 2.181e-06 \| 274 \| 0.298 \| 87 \| 59 \| 1 \| 210 \| 296 \| 1 \| 85 \| Baseplate assembly protein J \| Baseplate assembly protein J; afdb-uniprot50 \| AF-A0A661L8M5-F1-MODEL\_V4 \| 1.0 \| 4.244e-10 \| 274 \| 0.164 \| 207 \| 137 \| 10 \| 116 \| 295 \| 18 \| 215 \| Baseplate\_J domain-containing protein \| Baseplate\_J domain-containing protein; afdb-uniprot50 \| AF-A0A4Q3RYT5-F1-MODEL\_V4 \| 1.0 \| 6.844e-10 \| 274 \| 0.204 \| 225 \| 145 \| 11 \| 102 \| 294 \| 407 \| 629 \| Putative baseplate assembly protein \| Putative baseplate assembly protein; afdb-uniprot50 \| AF-A0A4U9DA24-F1-MODEL\_V4 \| 1.0 \| 4.11e-09 \| 273 \| 0.189 \| 206 \| 136 \| 11 \| 113 \| 293 \| 25 \| 224 \| Uncharacterized homolog of phage Mu protein gp47 \| Uncharacterized homolog of phage Mu protein gp47; afdb-uniprot50 \| AF-A0A2R7QCT8-F1-MODEL\_V4 \| 1.0 \| 8.188e-10 \| 273 \| 0.186 \| 204 \| 138 \| 11 \| 110 \| 294 \| 115 \| 309 \| Baseplate\_J domain-containing protein \| Baseplate\_J domain-containing protein; afdb-uniprot50 \| AF-A0A6H5RIG3-F1-MODEL\_V4 \| 1.0 \| 1.96e-12 \| 273 \| 0.142 \| 387 \| 210 \| 18 \| 20 \| 296 \| 8 \| 382 \| Baseplate J/gp47 family protein \| Baseplate J/gp47 family protein; afdb-uniprot50 \| AF-A0A5D8QH11-F1-MODEL\_V4 \| 1.0 \| 1.205e-08 \| 273 \| 0.087 \| 479 \| 224 \| 19 \| 22 \| 296 \| 14 \| 483 \| Uncharacterized protein \| Uncharacterized protein; afdb-uniprot50 \| AF-I3ZGQ5-F1-MODEL\_V4 \| 1.0 \| 5.721e-10 \| 273 \| 0.207 \| 227 \| 136 \| 11 \| 111 \| 298 \| 588 \| 809 \| Putative phage Mu protein gp47-like protein \| Putative phage Mu protein gp47-like protein; afdb-uniprot50 \| AF-A0A1Z9HXQ8-F1-MODEL\_V4 \| 1.0 \| 4.363e-09 \| 272 \| 0.147 \| 217 \| 147 \| 9 \| 114 \| 294 \| 386 \| 600 \| Uncharacterized protein \| Uncharacterized protein; afdb-uniprot50 \| AF-A0A7Y4U5C5-F1-MODEL\_V4 \| 1.0 \| 2.13e-09 \| 272 \| 0.186 \| 225 \| 146 \| 10 \| 110 \| 300 \| 438 \| 659 \| Putative baseplate assembly protein \| Putative baseplate assembly protein; afdb-uniprot50 \| AF-A0A2I0F8R2-F1-MODEL\_V4 \| 1.0 \| 1.369e-12 \| 271 \| 0.125 \| 398 \| 223 \| 15 \| 16 \| 294 \| 1 \| 392 \| Baseplate\_J domain-containing protein \| Baseplate\_J domain-containing protein; afdb-uniprot50 \| AF-A0A2V7W195-F1-MODEL\_V4 \| 1.0 \| 2.19e-08 \| 271 \| 0.189 \| 216 \| 144 \| 11 \| 111 \| 299 \| 429 \| 640 \| Putative baseplate assembly protein \| Putative baseplate assembly protein; afdb-uniprot50 \| AF-A0A660N0R2-F1-MODEL\_V4 \| 1.0 \| 2.859e-07 \| 270 \| 0.306 \| 111 \| 75 \| 2 \| 190 \| 299 \| 1 \| 110 \| Baseplate assembly protein \| Baseplate assembly protein; afdb-uniprot50 \| AF-A0A2K2TVT0-F1-MODEL\_V4 \| 1.0 \| 2.884e-11 \| 270 \| 0.137 \| 394 \| 212 \| 17 \| 18 \| 295 \| 4 \| 385 \| Baseplate\_J domain-containing protein \| Baseplate\_J domain-containing protein; afdb-uniprot50 \| AF-A0A6I3MDQ0-F1-MODEL\_V4 \| 1.0 \| 2.072e-10 \| 270 \| 0.138 \| 252 \| 150 \| 14 \| 110 \| 296 \| 415 \| 664 \| Uncharacterized protein \| Uncharacterized protein; afdb-uniprot50 \| AF-A0A0U1DD70-F1-MODEL\_V4 \| 1.0 \| 1.239e-07 \| 269 \| 0.26 \| 115 \| 81 \| 2 \| 180 \| 294 \| 3 \| 113 \| Baseplate assembly protein GpJ \| Baseplate assembly protein GpJ; afdb-uniprot50 \| AF-A0A5T0H790-F1-MODEL\_V4 \| 1.0 \| 1.069e-08 \| 269 \| 0.23 \| 243 \| 107 \| 6 \| 1 \| 163 \| 3 \| 245 \| Baseplate assembly protein \| Baseplate assembly protein; afdb-uniprot50 \| AF-A0A522CNZ0-F1-MODEL\_V4 \| 1.0 \| 2.489e-12 \| 269 \| 0.135 \| 390 \| 215 \| 17 \| 20 \| 300 \| 3 \| 379 \| Baseplate\_J domain-containing protein \| Baseplate\_J domain-containing protein; afdb-uniprot50 \| AF-A0A7J3V0A5-F1-MODEL\_V4 \| 1.0 \| 1.21e-10 \| 269 \| 0.111 \| 438 \| 196 \| 15 \| 42 \| 300 \| 101 \| 524 \| DUF4815 domain-containing protein \| DUF4815 domain-containing protein; afdb-uniprot50 \| AF-A0A849PJE9-F1-MODEL\_V4 \| 1.0 \| 2.548e-09 \| 269 \| 0.162 \| 227 \| 149 \| 13 \| 110 \| 299 \| 424 \| 646 \| Putative baseplate assembly protein \| Putative baseplate assembly protein; afdb-uniprot50 \| AF-A0A133PS66-F1-MODEL\_V4 \| 1.0 \| 6.874e-12 \| 268 \| 0.109 \| 394 \| 219 \| 16 \| 20 \| 296 \| 22 \| 400 \| Baseplate\_J domain-containing protein \| Baseplate\_J domain-containing protein; afdb-uniprot50 \| AF-A0A3N5LBL1-F1-MODEL\_V4 \| 1.0 \| 3.663e-11 \| 268 \| 0.157 \| 368 \| 198 \| 13 \| 20 \| 280 \| 2 \| 364 \| Baseplate\_J domain-containing protein \| Baseplate\_J domain-containing protein; afdb-uniprot50 \| AF-A0A0D6KGS5-F1-MODEL\_V4 \| 1.0 \| 1.244e-09 \| 268 \| 0.141 \| 248 \| 152 \| 10 \| 110 \| 300 \| 951 \| 1194 \| Baseplate\_J domain-containing protein \| Baseplate\_J domain-containing protein; afdb-uniprot50 \| AF-A0A2D6RI92-F1-MODEL\_V4 \| 1.0 \| 2.13e-09 \| 267 \| 0.154 \| 226 \| 148 \| 11 \| 110 \| 294 \| 371 \| 594 \| Uncharacterized protein \| Uncharacterized protein; afdb-uniprot50 \| AF-A0A7Y3CTP7-F1-MODEL\_V4 \| 1.0 \| 6.447e-10 \| 266 \| 0.144 \| 443 \| 184 \| 18 \| 42 \| 293 \| 6 \| 444 \| Baseplate\_J domain-containing protein \| Baseplate\_J domain-containing protein; afdb-uniprot50 \| AF-A0A854ZKD2-F1-MODEL\_V4 \| 1.0 \| 2.4e-09 \| 266 \| 0.159 \| 220 \| 148 \| 11 \| 111 \| 295 \| 422 \| 639 \| Putative baseplate assembly protein \| Putative baseplate assembly protein; afdb-uniprot50 \| AF-A0A1V4VN91-F1-MODEL\_V4 \| 1.0 \| 1.677e-09 \| 266 \| 0.131 \| 229 \| 158 \| 11 \| 111 \| 300 \| 522 \| 748 \| Baseplate J-like protein \| Baseplate J-like protein; afdb-uniprot50 \| AF-A0A3C0F0C1-F1-MODEL\_V4 \| 1.0 \| 1.104e-09 \| 265 \| 0.15 \| 226 \| 146 \| 12 \| 110 \| 294 \| 350 \| 570 \| Uncharacterized protein \| Uncharacterized protein; afdb-uniprot50 \| AF-A0A7X2DY09-F1-MODEL\_V4 \| 1.0 \| 6.6e-07 \| 264 \| 0.505 \| 87 \| 43 \| 0 \| 143 \| 229 \| 2 \| 88 \| Baseplate assembly protein \| Baseplate assembly protein; afdb-uniprot50 \| AF-A0A399W6F7-F1-MODEL\_V4 \| 1.0 \| 1.04e-09 \| 264 \| 0.146 \| 252 \| 146 \| 13 \| 110 \| 296 \| 439 \| 686 \| Putative baseplate assembly protein \| Putative baseplate assembly protein; afdb-uniprot50 \| AF-A0A645HBP4-F1-MODEL\_V4 \| 1.0 \| 1.67e-07 \| 263 \| 0.21 \| 138 \| 103 \| 5 \| 158 \| 294 \| 4 \| 136 \| Baseplate\_J domain-containing protein \| Baseplate\_J domain-containing protein; afdb-uniprot50 \| AF-A0A3D1R508-F1-MODEL\_V4 \| 1.0 \| 2.19e-08 \| 263 \| 0.185 \| 210 \| 127 \| 11 \| 127 \| 295 \| 85 \| 291 \| T4-like baseplate wedge \| T4-like baseplate wedge; afdb-uniprot50 \| AF-A0A7T8EB75-F1-MODEL\_V4 \| 1.0 \| 9.267e-12 \| 263 \| 0.115 \| 400 \| 230 \| 21 \| 13 \| 295 \| 1 \| 393 \| Baseplate\_J domain-containing protein \| Baseplate\_J domain-containing protein; afdb-uniprot50 \| AF-A0A4R8GX56-F1-MODEL\_V4 \| 1.0 \| 8.223e-12 \| 263 \| 0.119 \| 401 \| 212 \| 18 \| 20 \| 296 \| 14 \| 397 \| Putative phage protein gp47/JayE \| Putative phage protein gp47/JayE; afdb-uniprot50 \| AF-A0A2W2DLG7-F1-MODEL\_V4 \| 1.0 \| 1.205e-08 \| 263 \| 0.215 \| 213 \| 137 \| 11 \| 111 \| 299 \| 406 \| 612 \| Baseplate\_J domain-containing protein \| Baseplate\_J domain-containing protein; afdb-uniprot50 \| AF-A0A6P1MHD6-F1-MODEL\_V4 \| 1.0 \| 4.11e-09 \| 263 \| 0.153 \| 209 \| 143 \| 10 \| 116 \| 296 \| 483 \| 685 \| Baseplate\_J domain-containing protein \| Baseplate\_J domain-containing protein; afdb-uniprot50 \| AF-A0A1T3NNP8-F1-MODEL\_V4 \| 1.0 \| 8.692e-10 \| 263 \| 0.161 \| 248 \| 144 \| 14 \| 111 \| 296 \| 486 \| 731 \| Baseplate\_J domain-containing protein \| Baseplate\_J domain-containing protein; afdb-uniprot50 \| AF-A0A1Z4PZG8-F1-MODEL\_V4 \| 1.0 \| 4.11e-09 \| 263 \| 0.157 \| 228 \| 142 \| 12 \| 113 \| 296 \| 642 \| 863 \| Baseplate\_J domain-containing protein \| Baseplate\_J domain-containing protein; afdb-uniprot50 \| AF-A0A783X0V2-F1-MODEL\_V4 \| 1.0 \| 1.14e-10 \| 262 \| 0.14 \| 385 \| 212 \| 17 \| 20 \| 293 \| 4 \| 380 \| Baseplate J/gp47 family protein \| Baseplate J/gp47 family protein; afdb-uniprot50 \| AF-A0A336NB80-F1-MODEL\_V4 \| 1.0 \| 3.733e-06 \| 261 \| 0.236 \| 93 \| 68 \| 3 \| 207 \| 298 \| 18 \| 108 \| Uncharacterized protein \| Uncharacterized protein; afdb-uniprot50 \| AF-A0A0R2IBD9-F1-MODEL\_V4 \| 1.0 \| 7.297e-12 \| 261 \| 0.118 \| 396 \| 218 \| 20 \| 20 \| 296 \| 8 \| 391 \| Phage Mu gp47 related protein \| Phage Mu gp47 related protein; afdb-uniprot50 \| AF-A0A1C7W237-F1-MODEL\_V4 \| 1.0 \| 6.244e-09 \| 261 \| 0.107 \| 411 \| 222 \| 20 \| 13 \| 296 \| 1 \| 393 \| Baseplate\_J domain-containing protein \| Baseplate\_J domain-containing protein; afdb-uniprot50 \| AF-A0A7C7CAA3-F1-MODEL\_V4 \| 1.0 \| 7.47e-09 \| 261 \| 0.152 \| 216 \| 143 \| 11 \| 116 \| 294 \| 389 \| 601 \| Uncharacterized protein \| Uncharacterized protein; afdb-uniprot50 \| AF-A0A7X5W2W2-F1-MODEL\_V4 \| 1.0 \| 1.89e-09 \| 261 \| 0.162 \| 227 \| 146 \| 10 \| 110 \| 294 \| 452 \| 676 \| Putative baseplate assembly protein \| Putative baseplate assembly protein; afdb-uniprot50 \| AF-A0A0E3WU79-F1-MODEL\_V4 \| 1.0 \| 1.58e-09 \| 261 \| 0.181 \| 226 \| 146 \| 13 \| 111 \| 300 \| 990 \| 1212 \| Baseplate\_J domain-containing protein \| Baseplate\_J domain-containing protein; afdb-uniprot50 \| AF-A0A6M0BMD7-F1-MODEL\_V4 \| 1.0 \| 6.244e-09 \| 260 \| 0.184 \| 211 \| 127 \| 11 \| 125 \| 296 \| 21 \| 225 \| Baseplate protein J \| Baseplate protein J; afdb-uniprot50 \| AF-A0A843FGQ1-F1-MODEL\_V4 \| 1.0 \| 3.161e-12 \| 260 \| 0.155 \| 392 \| 183 \| 16 \| 20 \| 270 \| 16 \| 400 \| Baseplate J/gp47 family protein \| Baseplate J/gp47 family protein; afdb-uniprot50 \| AF-A0A4Y7S2C9-F1-MODEL\_V4 \| 1.0 \| 5.882e-09 \| 259 \| 0.177 \| 214 \| 145 \| 13 \| 110 \| 300 \| 417 \| 622 \| Baseplate J-like protein \| Baseplate J-like protein; afdb-uniprot50 \| AF-A0A844MB01-F1-MODEL\_V4 \| 1.0 \| 1.89e-09 \| 259 \| 0.194 \| 221 \| 136 \| 10 \| 110 \| 295 \| 745 \| 958 \| Putative baseplate assembly protein \| Putative baseplate assembly protein; afdb-uniprot50 \| AF-A0A518BPM4-F1-MODEL\_V4 \| 1.0 \| 8.418e-09 \| 259 \| 0.169 \| 207 \| 145 \| 9 \| 113 \| 294 \| 754 \| 958 \| Uncharacterized protein \| Uncharacterized protein; afdb-uniprot50 \| AF-A1ZLI5-F1-MODEL\_V4 \| 1.0 \| 3.341e-10 \| 258 \| 0.136 \| 316 \| 182 \| 13 \| 20 \| 300 \| 1 \| 260 \| Uncharacterized protein \| Uncharacterized protein; afdb-uniprot50 \| AF-A0A662VR35-F1-MODEL\_V4 \| 1.0 \| 5.541e-09 \| 257 \| 0.175 \| 217 \| 140 \| 14 \| 110 \| 296 \| 96 \| 303 \| Putative baseplate assembly protein \| Putative baseplate assembly protein; afdb-uniprot50 \| AF-A0A1Z4TUW1-F1-MODEL\_V4 \| 1.0 \| 2.559e-11 \| 257 \| 0.145 \| 351 \| 206 \| 16 \| 13 \| 296 \| 3 \| 326 \| Baseplate\_J domain-containing protein \| Baseplate\_J domain-containing protein; afdb-uniprot50 \| AF-A0A510PDS8-F1-MODEL\_V4 \| 1.0 \| 4.363e-09 \| 257 \| 0.177 \| 231 \| 147 \| 9 \| 110 \| 299 \| 940 \| 1168 \| Putative baseplate assembly protein \| Putative baseplate assembly protein; afdb-uniprot50 \| AF-A0A418VPI8-F1-MODEL\_V4 \| 1.0 \| 3.236e-09 \| 256 \| 0.199 \| 211 \| 136 \| 11 \| 111 \| 295 \| 192 \| 395 \| Putative baseplate assembly protein \| Putative baseplate assembly protein; afdb-uniprot50 \| AF-A0A0Q1A8V2-F1-MODEL\_V4 \| 1.0 \| 6.629e-09 \| 256 \| 0.132 \| 226 \| 155 \| 9 \| 110 \| 296 \| 805 \| 1028 \| Baseplate\_J domain-containing protein \| Baseplate\_J domain-containing protein; afdb-uniprot50 \| AF-A0A2V2EBS4-F1-MODEL\_V4 \| 1.0 \| 2.015e-11 \| 255 \| 0.129 \| 386 \| 214 \| 20 \| 20 \| 295 \| 12 \| 385 \| Baseplate\_J domain-containing protein \| Baseplate\_J domain-containing protein; afdb-uniprot50 \| AF-A0A6C2C7Q0-F1-MODEL\_V4 \| 1.0 \| 3.062e-11 \| 255 \| 0.137 \| 393 \| 209 \| 17 \| 20 \| 293 \| 8 \| 389 \| Baseplate J/gp47 family protein \| Baseplate J/gp47 family protein; afdb-uniprot50 \| AF-A0A559TCY2-F1-MODEL\_V4 \| 1.0 \| 1.135e-08 \| 255 \| 0.116 \| 405 \| 218 \| 18 \| 12 \| 294 \| 5 \| 391 \| Baseplate J-like protein \| Baseplate J-like protein; afdb-uniprot50 \| AF-A0A5Q0TCA5-F1-MODEL\_V4 \| 1.0 \| 9.838e-12 \| 255 \| 0.124 \| 403 \| 218 \| 16 \| 16 \| 294 \| 1 \| 392 \| Baseplate\_J domain-containing protein \| Baseplate\_J domain-containing protein; afdb-uniprot50 \| AF-A0A2N9MSM6-F1-MODEL\_V4 \| 1.0 \| 2.325e-08 \| 254 \| 0.138 \| 217 \| 149 \| 8 \| 112 \| 296 \| 69 \| 279 \| Baseplate\_J domain-containing protein \| Baseplate\_J domain-containing protein; afdb-uniprot50 \| AF-C4K4X2-F1-MODEL\_V4 \| 1.0 \| 8.937e-09 \| 254 \| 0.196 \| 209 \| 135 \| 12 \| 111 \| 293 \| 102 \| 303 \| Phage baseplate J-like protein \| Phage baseplate J-like protein; afdb-uniprot50 \| AF-D1YVC1-F1-MODEL\_V4 \| 1.0 \| 9.528e-11 \| 254 \| 0.118 \| 431 \| 182 \| 13 \| 42 \| 294 \| 100 \| 510 \| Baseplate\_J domain-containing protein \| Baseplate\_J domain-containing protein; afdb-uniprot50 \| AF-A0A315XDX8-F1-MODEL\_V4 \| 1.0 \| 2.325e-08 \| 254 \| 0.142 \| 211 \| 148 \| 9 \| 113 \| 294 \| 738 \| 944 \| Putative baseplate assembly protein \| Putative baseplate assembly protein; afdb-uniprot50 \| AF-A0A318XA71-F1-MODEL\_V4 \| 1.0 \| 2.62e-08 \| 253 \| 0.172 \| 237 \| 144 \| 12 \| 110 \| 299 \| 443 \| 674 \| Putative phage baseplate assembly protein \| Putative phage baseplate assembly protein; afdb-uniprot50 \| AF-A0A661IT72-F1-MODEL\_V4 \| 1.0 \| 7.037e-09 \| 252 \| 0.105 \| 218 \| 159 \| 11 \| 111 \| 298 \| 723 \| 934 \| Putative baseplate assembly protein \| Putative baseplate assembly protein; afdb-uniprot50 \| AF-A0A069EWX3-F1-MODEL\_V4 \| 1.0 \| 2.139e-11 \| 251 \| 0.115 \| 400 \| 211 \| 18 \| 20 \| 295 \| 14 \| 394 \| Baseplate\_J domain-containing protein \| Baseplate\_J domain-containing protein; afdb-uniprot50 \| AF-C2Z1K9-F1-MODEL\_V4 \| 1.0 \| 1.788e-11 \| 251 \| 0.124 \| 411 \| 217 \| 23 \| 13 \| 296 \| 1 \| 395 \| Baseplate\_J domain-containing protein \| Baseplate\_J domain-containing protein; afdb-uniprot50 \| AF-A0A3M1QRQ3-F1-MODEL\_V4 \| 1.0 \| 2.063e-08 \| 251 \| 0.142 \| 217 \| 151 \| 12 \| 111 \| 300 \| 909 \| 1117 \| Putative baseplate assembly protein \| Putative baseplate assembly protein; afdb-uniprot50 \| AF-A0A1H4G0X1-F1-MODEL\_V4 \| 1.0 \| 1.58e-09 \| 250 \| 0.159 \| 244 \| 144 \| 12 \| 110 \| 294 \| 489 \| 730 \| Putative baseplate assembly protein \| Putative baseplate assembly protein; afdb-uniprot50 \| AF-A0A5B9EFW7-F1-MODEL\_V4 \| 1.0 \| 2.62e-08 \| 250 \| 0.172 \| 220 \| 138 \| 11 \| 112 \| 296 \| 773 \| 983 \| Putative baseplate assembly protein \| Putative baseplate assembly protein; afdb-uniprot50 \| AF-A0A6M0AC30-F1-MODEL\_V4 \| 1.0 \| 1.89e-09 \| 249 \| 0.183 \| 251 \| 141 \| 13 \| 101 \| 296 \| 213 \| 454 \| Putative baseplate assembly protein \| Putative baseplate assembly protein; afdb-uniprot50 \| AF-A0A3M3Y5F7-F1-MODEL\_V4 \| 1.0 \| 4.11e-09 \| 248 \| 0.14 \| 228 \| 153 \| 11 \| 102 \| 294 \| 14 \| 233 \| Baseplate\_J domain-containing protein \| Baseplate\_J domain-containing protein; afdb-uniprot50 \| AF-A0A1I0C9X9-F1-MODEL\_V4 \| 1.0 \| 1.831e-08 \| 248 \| 0.16 \| 212 \| 140 \| 12 \| 115 \| 296 \| 495 \| 698 \| Putative baseplate assembly protein \| Putative baseplate assembly protein; afdb-uniprot50 \| AF-A0A5D0UQC1-F1-MODEL\_V4 \| 1.0 \| 3.048e-09 \| 248 \| 0.216 \| 226 \| 134 \| 14 \| 110 \| 298 \| 616 \| 835 \| Uncharacterized protein \| Uncharacterized protein; afdb-uniprot50 \| AF-A0A6P0LHK6-F1-MODEL\_V4 \| 1.0 \| 7.037e-09 \| 248 \| 0.168 \| 232 \| 139 \| 14 \| 110 \| 296 \| 791 \| 1013 \| Putative baseplate assembly protein \| Putative baseplate assembly protein; afdb-uniprot50 \| AF-A0A1W1VPN7-F1-MODEL\_V4 \| 1.0 \| 8.455e-11 \| 247 \| 0.116 \| 396 \| 213 \| 18 \| 20 \| 294 \| 12 \| 391 \| Uncharacterized phage protein gp47/JayE \| Uncharacterized phage protein gp47/JayE; afdb-uniprot50 \| AF-A0A521U5U0-F1-MODEL\_V4 \| 1.0 \| 3.663e-11 \| 246 \| 0.161 \| 391 \| 199 \| 19 \| 20 \| 295 \| 13 \| 389 \| Baseplate\_J domain-containing protein \| Baseplate\_J domain-containing protein; afdb-uniprot50 \| AF-A0A6L9Z4K9-F1-MODEL\_V4 \| 1.0 \| 7.47e-09 \| 246 \| 0.177 \| 237 \| 139 \| 13 \| 110 \| 296 \| 345 \| 575 \| Putative baseplate assembly protein \| Putative baseplate assembly protein; afdb-uniprot50 \| AF-A0A850I740-F1-MODEL\_V4 \| 1.0 \| 2.19e-08 \| 245 \| 0.122 \| 221 \| 161 \| 12 \| 110 \| 299 \| 414 \| 632 \| Uncharacterized protein \| Uncharacterized protein; afdb-uniprot50 \| AF-A0A7Y5GDB4-F1-MODEL\_V4 \| 1.0 \| 1.677e-09 \| 245 \| 0.16 \| 243 \| 142 \| 14 \| 110 \| 294 \| 528 \| 766 \| Baseplate J/gp47 family protein \| Baseplate J/gp47 family protein; afdb-uniprot50 \| AF-C3X351-F1-MODEL\_V4 \| 1.0 \| 2.13e-09 \| 244 \| 0.13 \| 382 \| 206 \| 19 \| 20 \| 294 \| 4 \| 366 \| Baseplate\_J domain-containing protein \| Baseplate\_J domain-containing protein; afdb-uniprot50 \| AF-E3IXC3-F1-MODEL\_V4 \| 1.0 \| 2.705e-09 \| 244 \| 0.191 \| 266 \| 135 \| 11 \| 110 \| 298 \| 964 \| 1226 \| Baseplate J family protein \| Baseplate J family protein; afdb-uniprot50 \| AF-A0A2D6E3S6-F1-MODEL\_V4 \| 1.0 \| 1.007e-08 \| 243 \| 0.151 \| 224 \| 148 \| 12 \| 110 \| 294 \| 379 \| 599 \| Uncharacterized protein \| Uncharacterized protein; afdb-uniprot50 \| AF-A0A1W9V6S3-F1-MODEL\_V4 \| 1.0 \| 8.692e-10 \| 242 \| 0.102 \| 480 \| 211 \| 18 \| 22 \| 294 \| 13 \| 479 \| Baseplate\_J domain-containing protein \| Baseplate\_J domain-containing protein; afdb-uniprot50 \| AF-A0A4Y6PV84-F1-MODEL\_V4 \| 1.0 \| 1.205e-08 \| 242 \| 0.157 \| 260 \| 136 \| 10 \| 115 \| 294 \| 457 \| 713 \| Putative baseplate assembly protein \| Putative baseplate assembly protein; afdb-uniprot50 \| AF-A0A6P0ZY62-F1-MODEL\_V4 \| 1.0 \| 6.244e-09 \| 242 \| 0.185 \| 237 \| 136 \| 14 \| 110 \| 296 \| 936 \| 1165 \| Putative baseplate assembly protein \| Putative baseplate assembly protein; afdb-uniprot50 \| AF-A0A527W6E1-F1-MODEL\_V4 \| 1.0 \| 1.099e-07 \| 241 \| 0.304 \| 128 \| 86 \| 2 \| 171 \| 298 \| 7 \| 131 \| Uncharacterized protein \| Uncharacterized protein; afdb-uniprot50 \| AF-A0A061NPP2-F1-MODEL\_V4 \| 1.0 \| 4.762e-08 \| 240 \| 0.21 \| 252 \| 113 \| 8 \| 8 \| 174 \| 1 \| 251 \| Phage-related baseplate assembly protein \| Phage-related baseplate assembly protein; afdb-uniprot50 \| AF-A0A0P8CHE4-F1-MODEL\_V4 \| 1.0 \| 4.486e-08 \| 240 \| 0.125 \| 224 \| 156 \| 9 \| 111 \| 296 \| 870 \| 1091 \| Baseplate J-like protein \| Baseplate J-like protein; afdb-uniprot50 \| AF-A0A0F9A493-F1-MODEL\_V4 \| 1.0 \| 2.609e-06 \| 239 \| 0.19 \| 110 \| 87 \| 2 \| 187 \| 295 \| 2 \| 110 \| Uncharacterized protein \| Uncharacterized protein; afdb-uniprot50 \| AF-A0A1V3IJ84-F1-MODEL\_V4 \| 1.0 \| 5.565e-11 \| 239 \| 0.135 \| 383 \| 210 \| 16 \| 20 \| 294 \| 10 \| 379 \| Baseplate\_J domain-containing protein \| Baseplate\_J domain-containing protein; afdb-uniprot50 \| AF-A0A823TIV2-F1-MODEL\_V4 \| 1.0 \| 1.162e-05 \| 238 \| 0.355 \| 107 \| 65 \| 3 \| 1 \| 104 \| 1 \| 106 \| Baseplate assembly protein \| Baseplate assembly protein; afdb-uniprot50 \| AF-A0A1I4DEF2-F1-MODEL\_V4 \| 1.0 \| 4.225e-08 \| 238 \| 0.129 \| 208 \| 155 \| 9 \| 110 \| 295 \| 421 \| 624 \| Uncharacterized protein \| Uncharacterized protein; afdb-uniprot50 \| AF-A0A7C3XHE7-F1-MODEL\_V4 \| 1.0 \| 4.896e-07 \| 237 \| 0.202 \| 138 \| 96 \| 6 \| 152 \| 278 \| 18 \| 152 \| Baseplate\_J domain-containing protein \| Baseplate\_J domain-containing protein; afdb-uniprot50 \| AF-A0A544QQX2-F1-MODEL\_V4 \| 1.0 \| 7.502e-11 \| 237 \| 0.137 \| 423 \| 209 \| 23 \| 13 \| 295 \| 1 \| 407 \| Baseplate J family protein \| Baseplate J family protein; afdb-uniprot50 \| AF-A0A3A9ATA2-F1-MODEL\_V4 \| 1.0 \| 1.67e-07 \| 237 \| 0.08 \| 483 \| 230 \| 16 \| 20 \| 296 \| 12 \| 486 \| Uncharacterized protein \| Uncharacterized protein; afdb-uniprot50 \| AF-A0A1V5BAG1-F1-MODEL\_V4 \| 1.0 \| 4.486e-08 \| 237 \| 0.168 \| 232 \| 147 \| 13 \| 110 \| 300 \| 904 \| 1130 \| Baseplate J-like protein \| Baseplate J-like protein; afdb-uniprot50 \| AF-A0A825LZJ7-F1-MODEL\_V4 \| 1.0 \| 6.815e-08 \| 235 \| 0.24 \| 225 \| 85 \| 7 \| 77 \| 222 \| 1 \| 218 \| Baseplate assembly protein \| Baseplate assembly protein; afdb-uniprot50 \| AF-A0A2S9SX29-F1-MODEL\_V4 \| 1.0 \| 2.063e-08 \| 235 \| 0.175 \| 200 \| 138 \| 10 \| 111 \| 293 \| 53 \| 242 \| Baseplate\_J domain-containing protein \| Baseplate\_J domain-containing protein; afdb-uniprot50 \| AF-A0A2V2DR45-F1-MODEL\_V4 \| 1.0 \| 2.4e-09 \| 235 \| 0.134 \| 327 \| 191 \| 13 \| 52 \| 293 \| 2 \| 321 \| Baseplate J protein \| Baseplate J protein; afdb-uniprot50 \| AF-A0A897N0F2-F1-MODEL\_V4 \| 1.0 \| 3.148e-10 \| 235 \| 0.122 \| 433 \| 210 \| 20 \| 13 \| 296 \| 1 \| 412 \| Putative phage protein gp47/JayE \| Putative phage protein gp47/JayE; afdb-uniprot50 \| AF-A0A2P8VTN4-F1-MODEL\_V4 \| 1.0 \| 2.781e-08 \| 235 \| 0.167 \| 227 \| 137 \| 16 \| 110 \| 296 \| 647 \| 861 \| Uncharacterized protein \| Uncharacterized protein; afdb-uniprot50 \| AF-A0A841VH02-F1-MODEL\_V4 \| 1.0 \| 1.624e-08 \| 234 \| 0.15 \| 259 \| 145 \| 11 \| 111 \| 296 \| 259 \| 515 \| Baseplate J/gp47 family protein \| Baseplate J/gp47 family protein; afdb-uniprot50 \| AF-W2CWP2-F1-MODEL\_V4 \| 1.0 \| 9.149e-06 \| 233 \| 0.465 \| 101 \| 54 \| 0 \| 166 \| 266 \| 2 \| 102 \| Baseplate\_J domain-containing protein \| Baseplate\_J domain-containing protein; afdb-uniprot50 \| AF-A0A086ML19-F1-MODEL\_V4 \| 1.0 \| 1.943e-08 \| 233 \| 0.211 \| 222 \| 128 \| 14 \| 110 \| 294 \| 701 \| 912 \| Baseplate\_J domain-containing protein \| Baseplate\_J domain-containing protein; afdb-uniprot50 \| AF-A0A508B8W8-F1-MODEL\_V4 \| 1.0 \| 2.468e-08 \| 233 \| 0.135 \| 243 \| 145 \| 13 \| 111 \| 298 \| 812 \| 1044 \| Putative baseplate assembly protein \| Putative baseplate assembly protein; afdb-uniprot50 \| AF-A0A1B6BCR6-F1-MODEL\_V4 \| 1.0 \| 8.937e-09 \| 232 \| 0.124 \| 209 \| 151 \| 10 \| 112 \| 294 \| 406 \| 608 \| Uncharacterized protein \| Uncharacterized protein; afdb-uniprot50 \| AF-A0A4R2KD98-F1-MODEL\_V4 \| 1.0 \| 8.655e-08 \| 232 \| 0.163 \| 239 \| 142 \| 14 \| 110 \| 300 \| 720 \| 948 \| Putative phage baseplate assembly protein \| Putative phage baseplate assembly protein; afdb-uniprot50 \| AF-A0A836SK26-F1-MODEL\_V4 \| 1.0 \| 7.713e-10 \| 231 \| 0.122 \| 367 \| 226 \| 19 \| 12 \| 294 \| 2 \| 356 \| Uncharacterized protein \| Uncharacterized protein; afdb-uniprot50 \| AF-A0A610PH99-F1-MODEL\_V4 \| 1.0 \| 5.494e-05 \| 230 \| 0.385 \| 96 \| 57 \| 2 \| 10 \| 103 \| 17 \| 112 \| Baseplate assembly protein \| Baseplate assembly protein; afdb-uniprot50 \| AF-A0A6M0AIH3-F1-MODEL\_V4 \| 1.0 \| 5.055e-08 \| 230 \| 0.18 \| 199 \| 113 \| 12 \| 142 \| 296 \| 2 \| 194 \| Baseplate\_J domain-containing protein \| Baseplate\_J domain-containing protein; afdb-uniprot50 \| AF-A0A7U1HJI5-F1-MODEL\_V4 \| 1.0 \| 5.697e-08 \| 230 \| 0.187 \| 240 \| 138 \| 14 \| 110 \| 300 \| 652 \| 883 \| Uncharacterized protein \| Uncharacterized protein; afdb-uniprot50 \| AF-A0A3M1B3R2-F1-MODEL\_V4 \| 1.0 \| 5.697e-08 \| 229 \| 0.159 \| 207 \| 141 \| 10 \| 110 \| 294 \| 425 \| 620 \| Uncharacterized protein \| Uncharacterized protein; afdb-uniprot50 \| AF-A0A846BNC7-F1-MODEL\_V4 \| 1.0 \| 9.754e-08 \| 229 \| 0.115 \| 242 \| 155 \| 11 \| 111 \| 296 \| 966 \| 1204 \| Putative baseplate assembly protein \| Putative baseplate assembly protein; afdb-uniprot50 \| AF-A0A846TRD8-F1-MODEL\_V4 \| 1.0 \| 6.42e-08 \| 228 \| 0.159 \| 226 \| 143 \| 11 \| 114 \| 300 \| 704 \| 921 \| Uncharacterized protein \| Uncharacterized protein; afdb-uniprot50 \| AF-A0A3C0GH36-F1-MODEL\_V4 \| 1.0 \| 2.39e-07 \| 227 \| 0.159 \| 176 \| 113 \| 8 \| 152 \| 294 \| 17 \| 190 \| Uncharacterized protein \| Uncharacterized protein; afdb-uniprot50 \| AF-A0A3N9V016-F1-MODEL\_V4 \| 1.0 \| 4.11e-09 \| 223 \| 0.106 \| 500 \| 227 \| 22 \| 13 \| 296 \| 1 \| 496 \| Putative baseplate assembly protein \| Putative baseplate assembly protein; afdb-uniprot50 \| AF-A0A540WCV7-F1-MODEL\_V4 \| 1.0 \| 1.724e-08 \| 223 \| 0.198 \| 247 \| 140 \| 14 \| 101 \| 296 \| 406 \| 645 \| Putative baseplate assembly protein \| Putative baseplate assembly protein; afdb-uniprot50 \| AF-A0A3G2IBU6-F1-MODEL\_V4 \| 1.0 \| 0.000161 \| 222 \| 0.432 \| 74 \| 42 \| 0 \| 8 \| 81 \| 6 \| 79 \| Uncharacterized protein \| Uncharacterized protein; afdb-uniprot50 \| AF-A0A6L9YWI8-F1-MODEL\_V4 \| 1.0 \| 1.035e-07 \| 222 \| 0.171 \| 233 \| 134 \| 12 \| 116 \| 296 \| 26 \| 251 \| Baseplate\_J domain-containing protein \| Baseplate\_J domain-containing protein; afdb-uniprot50 \| AF-A0A5S3XXB9-F1-MODEL\_V4 \| 1.0 \| 1.624e-08 \| 222 \| 0.127 \| 211 \| 144 \| 14 \| 112 \| 295 \| 270 \| 467 \| Uncharacterized protein \| Uncharacterized protein; afdb-uniprot50 \| AF-G2G240-F1-MODEL\_V4 \| 1.0 \| 1.396e-07 \| 222 \| 0.183 \| 212 \| 138 \| 10 \| 111 \| 294 \| 433 \| 637 \| Baseplate\_J domain-containing protein \| Baseplate\_J domain-containing protein; afdb-uniprot50 \| AF-A0A1F8S3X9-F1-MODEL\_V4 \| 1.0 \| 7.235e-08 \| 222 \| 0.126 \| 253 \| 144 \| 11 \| 115 \| 299 \| 719 \| 962 \| Uncharacterized protein \| Uncharacterized protein; afdb-uniprot50 \| AF-A0A660E4D8-F1-MODEL\_V4 \| 1.0 \| 3.997e-10 \| 221 \| 0.125 \| 397 \| 215 \| 16 \| 20 \| 294 \| 11 \| 397 \| Baseplate\_J domain-containing protein \| Baseplate\_J domain-containing protein; afdb-uniprot50 \| AF-A0A226RDN1-F1-MODEL\_V4 \| 1.0 \| 1.172e-09 \| 220 \| 0.108 \| 404 \| 220 \| 17 \| 20 \| 293 \| 8 \| 401 \| Baseplate\_J domain-containing protein \| Baseplate\_J domain-containing protein; afdb-uniprot50 \| AF-A0A352AD62-F1-MODEL\_V4 \| 1.0 \| 4.896e-07 \| 220 \| 0.151 \| 224 \| 138 \| 15 \| 114 \| 296 \| 627 \| 839 \| Uncharacterized protein \| Uncharacterized protein; afdb-uniprot50 \| AF-A0A1Z3HRW5-F1-MODEL\_V4 \| 1.0 \| 1.67e-07 \| 219 \| 0.203 \| 231 \| 136 \| 13 \| 107 \| 296 \| 548 \| 771 \| Uncharacterized protein \| Uncharacterized protein; afdb-uniprot50 \| AF-I3IJH1-F1-MODEL\_V4 \| 1.0 \| 6.42e-08 \| 219 \| 0.162 \| 240 \| 142 \| 14 \| 110 \| 300 \| 635 \| 864 \| Uncharacterized protein \| Uncharacterized protein; afdb-uniprot50 \| AF-A0A3B0YCG0-F1-MODEL\_V4 \| 1.0 \| 1.167e-07 \| 219 \| 0.152 \| 236 \| 139 \| 15 \| 115 \| 300 \| 665 \| 889 \| Uncharacterized protein \| Uncharacterized protein; afdb-uniprot50 \| AF-A0A1Z4LU77-F1-MODEL\_V4 \| 1.0 \| 6.815e-08 \| 219 \| 0.126 \| 253 \| 149 \| 11 \| 111 \| 294 \| 817 \| 1066 \| Baseplate\_J domain-containing protein \| Baseplate\_J domain-containing protein; afdb-uniprot50 \| AF-A0A7W7V276-F1-MODEL\_V4 \| 1.0 \| 6.42e-08 \| 219 \| 0.142 \| 260 \| 144 \| 11 \| 110 \| 296 \| 952 \| 1205 \| Putative phage baseplate assembly protein \| Putative phage baseplate assembly protein; afdb-uniprot50 \| AF-A0A8A5XIZ9-F1-MODEL\_V4 \| 1.0 \| 2.682e-05 \| 218 \| 0.456 \| 92 \| 50 \| 0 \| 1 \| 92 \| 4 \| 95 \| Uncharacterized protein \| Uncharacterized protein; afdb-uniprot50 \| AF-A0A5M7L0X9-F1-MODEL\_V4 \| 1.0 \| 3.997e-10 \| 218 \| 0.135 \| 391 \| 211 \| 16 \| 21 \| 300 \| 11 \| 385 \| Baseplate J/gp47 family protein \| Baseplate J/gp47 family protein; afdb-uniprot50 \| AF-A0A5Q2NZU8-F1-MODEL\_V4 \| 1.0 \| 2.199e-10 \| 218 \| 0.135 \| 412 \| 206 \| 17 \| 20 \| 295 \| 8 \| 405 \| Baseplate\_J domain-containing protein \| Baseplate\_J domain-containing protein; afdb-uniprot50 \| AF-A0A2V9KQL4-F1-MODEL\_V4 \| 1.0 \| 4.896e-07 \| 218 \| 0.174 \| 218 \| 140 \| 13 \| 115 \| 300 \| 655 \| 864 \| Uncharacterized protein \| Uncharacterized protein; afdb-uniprot50 \| AF-A0A1W9S1W4-F1-MODEL\_V4 \| 1.0 \| 1.239e-07 \| 217 \| 0.123 \| 194 \| 148 \| 5 \| 15 \| 206 \| 1 \| 174 \| Uncharacterized protein \| Uncharacterized protein; afdb-uniprot50 \| AF-A0A178LTM7-F1-MODEL\_V4 \| 1.0 \| 5.197e-07 \| 217 \| 0.207 \| 222 \| 127 \| 12 \| 115 \| 296 \| 659 \| 871 \| Baseplate\_J domain-containing protein \| Baseplate\_J domain-containing protein; afdb-uniprot50 \| AF-A0A0D0H1J7-F1-MODEL\_V4 \| 1.0 \| 5.672e-06 \| 216 \| 0.375 \| 104 \| 65 \| 0 \| 123 \| 226 \| 6 \| 109 \| KY49.ctg7180000000025\_quiver, whole genome shotgun sequence \| KY49.ctg7180000000025\_quiver, whole genome shotgun sequence; afdb-uniprot50 \| AF-A0A6P0RHI9-F1-MODEL\_V4 \| 1.0 \| 1.239e-07 \| 216 \| 0.143 \| 258 \| 143 \| 11 \| 111 \| 294 \| 899 \| 1152 \| Baseplate protein J \| Baseplate protein J; afdb-uniprot50 \| AF-A0A354WSY4-F1-MODEL\_V4 \| 1.0 \| 1.011e-10 \| 215 \| 0.113 \| 398 \| 213 \| 18 \| 20 \| 293 \| 14 \| 395 \| Baseplate\_J domain-containing protein \| Baseplate\_J domain-containing protein; afdb-uniprot50 \| AF-A0A832EBT2-F1-MODEL\_V4 \| 1.0 \| 1.274e-06 \| 213 \| 0.132 \| 234 \| 150 \| 14 \| 110 \| 295 \| 675 \| 903 \| Uncharacterized protein \| Uncharacterized protein; afdb-uniprot50 \| AF-A0A0F9MQV1-F1-MODEL\_V4 \| 1.0 \| 5.366e-08 \| 212 \| 0.138 \| 282 \| 134 \| 10 \| 119 \| 294 \| 2 \| 280 \| Uncharacterized protein \| Uncharacterized protein; afdb-uniprot50 \| AF-A0A352X2K1-F1-MODEL\_V4 \| 1.0 \| 1.035e-07 \| 211 \| 0.136 \| 257 \| 165 \| 10 \| 98 \| 300 \| 628 \| 881 \| Putative baseplate assembly protein \| Putative baseplate assembly protein; afdb-uniprot50 \| AF-A0A497BHK9-F1-MODEL\_V4 \| 1.0 \| 2.793e-10 \| 210 \| 0.136 \| 389 \| 209 \| 19 \| 22 \| 295 \| 13 \| 389 \| Baseplate\_J domain-containing protein \| Baseplate\_J domain-containing protein; afdb-uniprot50 \| AF-A0A661PHR5-F1-MODEL\_V4 \| 1.0 \| 4.782e-10 \| 209 \| 0.127 \| 399 \| 211 \| 19 \| 20 \| 295 \| 10 \| 394 \| Baseplate\_J domain-containing protein \| Baseplate\_J domain-containing protein; afdb-uniprot50 \| AF-A0A1F1L5J3-F1-MODEL\_V4 \| 1.0 \| 4.11e-09 \| 206 \| 0.106 \| 393 \| 218 \| 19 \| 20 \| 294 \| 14 \| 391 \| Baseplate\_J domain-containing protein \| Baseplate\_J domain-containing protein; afdb-uniprot50 \| AF-A0A1C5V4K5-F1-MODEL\_V4 \| 1.0 \| 8.346e-05 \| 204 \| 0.126 \| 95 \| 81 \| 1 \| 207 \| 299 \| 20 \| 114 \| Uncharacterized protein \| Uncharacterized protein; afdb-uniprot50 \| AF-A0A367QYY9-F1-MODEL\_V4 \| 1.0 \| 7.68e-08 \| 204 \| 0.159 \| 232 \| 146 \| 11 \| 113 \| 298 \| 620 \| 848 \| Putative baseplate assembly protein \| Putative baseplate assembly protein; afdb-uniprot50 \| AF-A0A7X5IBM0-F1-MODEL\_V4 \| 1.0 \| 2.4e-09 \| 202 \| 0.102 \| 392 \| 223 \| 15 \| 20 \| 294 \| 13 \| 392 \| Uncharacterized protein \| Uncharacterized protein; afdb-uniprot50 \| AF-A0A7G2RX52-F1-MODEL\_V4 \| 1.0 \| 3.871e-09 \| 200 \| 0.118 \| 397 \| 214 \| 19 \| 20 \| 296 \| 11 \| 391 \| Baseplate\_J domain-containing protein \| Baseplate\_J domain-containing protein; afdb-uniprot50 \| AF-A0A5E4HHS7-F1-MODEL\_V4 \| 1.0 \| 2.537e-07 \| 200 \| 0.173 \| 236 \| 134 \| 18 \| 110 \| 299 \| 850 \| 1070 \| PA14 domain protein \| PA14 domain protein; afdb-uniprot50 \| AF-A0A7X8EVJ9-F1-MODEL\_V4 \| 1.0 \| 8.655e-08 \| 199 \| 0.099 \| 221 \| 157 \| 12 \| 111 \| 295 \| 495 \| 709 \| Uncharacterized protein \| Uncharacterized protein; afdb-uniprot50 \| AF-A0A7X7LHT1-F1-MODEL\_V4 \| 1.0 \| 2.13e-09 \| 197 \| 0.121 \| 305 \| 194 \| 18 \| 20 \| 295 \| 4 \| 263 \| Baseplate\_J domain-containing protein \| Baseplate\_J domain-containing protein; afdb-uniprot50 \| AF-A0A2U3KMN2-F1-MODEL\_V4 \| 1.0 \| 2.054e-06 \| 197 \| 0.121 \| 238 \| 144 \| 14 \| 110 \| 296 \| 640 \| 863 \| Uncharacterized protein \| Uncharacterized protein; afdb-uniprot50 \| AF-A0A5Y6SBS8-F1-MODEL\_V4 \| 1.0 \| 0.000171 \| 196 \| 0.349 \| 106 \| 61 \| 2 \| 1 \| 98 \| 1 \| 106 \| Baseplate assembly protein \| Baseplate assembly protein; afdb-uniprot50 \| AF-A0A806KJW5-F1-MODEL\_V4 \| 1.0 \| 1.358e-08 \| 196 \| 0.11 \| 489 \| 207 \| 21 \| 21 \| 300 \| 12 \| 481 \| Uncharacterized protein \| Uncharacterized protein; afdb-uniprot50 \| AF-A1ZMF0-F1-MODEL\_V4 \| 1.0 \| 4.632e-09 \| 195 \| 0.132 \| 309 \| 194 \| 17 \| 20 \| 294 \| 1 \| 269 \| Baseplate\_J domain-containing protein \| Baseplate\_J domain-containing protein; afdb-uniprot50 \| AF-N1WF81-F1-MODEL\_V4 \| 1.0 \| 6.629e-09 \| 194 \| 0.121 \| 411 \| 208 \| 22 \| 21 \| 296 \| 11 \| 403 \| Baseplate J-like protein \| Baseplate J-like protein; afdb-uniprot50 \| AF-N1ZQP9-F1-MODEL\_V4 \| 1.0 \| 4.612e-07 \| 194 \| 0.106 \| 207 \| 157 \| 10 \| 116 \| 298 \| 500 \| 702 \| Baseplate\_J domain-containing protein \| Baseplate\_J domain-containing protein; afdb-uniprot50 \| AF-A0A836ZSV8-F1-MODEL\_V4 \| 1.0 \| 0.0004189 \| 193 \| 0.338 \| 71 \| 47 \| 0 \| 210 \| 280 \| 2 \| 72 \| Baseplate assembly protein \| Baseplate assembly protein; afdb-uniprot50 \| AF-A0A431I9K0-F1-MODEL\_V4 \| 1.0 \| 1.358e-08 \| 190 \| 0.09 \| 475 \| 220 \| 15 \| 21 \| 293 \| 12 \| 476 \| Baseplate\_J domain-containing protein \| Baseplate\_J domain-containing protein; afdb-uniprot50 \| AF-A0A841JK07-F1-MODEL\_V4 \| 1.0 \| 1.943e-08 \| 189 \| 0.134 \| 313 \| 192 \| 19 \| 20 \| 295 \| 1 \| 271 \| Uncharacterized protein \| Uncharacterized protein; afdb-uniprot50 \| AF-A0A6P0JNB2-F1-MODEL\_V4 \| 1.0 \| 6.022e-06 \| 189 \| 0.167 \| 155 \| 105 \| 8 \| 159 \| 295 \| 149 \| 297 \| Uncharacterized protein \| Uncharacterized protein; afdb-uniprot50 \| AF-A0A1W6B8F3-F1-MODEL\_V4 \| 1.0 \| 3.532e-08 \| 189 \| 0.097 \| 402 \| 216 \| 21 \| 20 \| 295 \| 11 \| 391 \| Baseplate\_J domain-containing protein \| Baseplate\_J domain-containing protein; afdb-uniprot50 \| AF-A0A7X5QI95-F1-MODEL\_V4 \| 1.0 \| 6.392e-06 \| 184 \| 0.177 \| 163 \| 103 \| 6 \| 155 \| 293 \| 2 \| 157 \| Phage baseplate protein \| Phage baseplate protein; afdb-uniprot50 \| AF-F9D775-F1-MODEL\_V4 \| 1.0 \| 6.815e-08 \| 179 \| 0.134 \| 313 \| 197 \| 17 \| 21 \| 296 \| 1 \| 276 \| Uncharacterized protein \| Uncharacterized protein; afdb-uniprot50 \| AF-A0A1I1E3U7-F1-MODEL\_V4 \| 1.0 \| 6.047e-08 \| 178 \| 0.171 \| 309 \| 180 \| 17 \| 20 \| 295 \| 1 \| 266 \| Uncharacterized protein \| Uncharacterized protein; afdb-uniprot50 \| AF-A0A1B8SLD9-F1-MODEL\_V4 \| 1.0 \| 3.517e-06 \| 178 \| 0.086 \| 591 \| 219 \| 20 \| 20 \| 296 \| 1 \| 584 \| Baseplate\_J domain-containing protein \| Baseplate\_J domain-containing protein; afdb-uniprot50 \| AF-A0A8B3TC69-F1-MODEL\_V4 \| 1.0 \| 2.325e-08 \| 177 \| 0.119 \| 394 \| 213 \| 19 \| 20 \| 294 \| 9 \| 387 \| Uncharacterized protein \| Uncharacterized protein; afdb-uniprot50 \| AF-A0A0Q0ZIP1-F1-MODEL\_V4 \| 1.0 \| 9.188e-08 \| 177 \| 0.13 \| 346 \| 201 \| 19 \| 1 \| 295 \| 1 \| 297 \| Uncharacterized protein \| Uncharacterized protein; afdb-uniprot50 \| AF-A0A520BF88-F1-MODEL\_V4 \| 1.0 \| 8.153e-08 \| 176 \| 0.146 \| 362 \| 188 \| 21 \| 13 \| 295 \| 4 \| 323 \| Uncharacterized protein \| Uncharacterized protein; afdb-uniprot50 \| AF-A0A7V1ZEF9-F1-MODEL\_V4 \| 1.0 \| 4.896e-07 \| 176 \| 0.096 \| 260 \| 150 \| 13 \| 110 \| 294 \| 291 \| 540 \| Uncharacterized protein \| Uncharacterized protein; afdb-uniprot50 \| AF-A0A661FKB4-F1-MODEL\_V4 \| 1.0 \| 1.065e-06 \| 175 \| 0.177 \| 186 \| 109 \| 9 \| 151 \| 295 \| 802 \| 984 \| Baseplate\_J domain-containing protein \| Baseplate\_J domain-containing protein; afdb-uniprot50 \| AF-A0A1U7I513-F1-MODEL\_V4 \| 1.0 \| 4.466e-06 \| 174 \| 0.149 \| 167 \| 108 \| 9 \| 158 \| 296 \| 980 \| 1140 \| Baseplate\_J domain-containing protein \| Baseplate\_J domain-containing protein; afdb-uniprot50 \| AF-A0A7C1Z9L5-F1-MODEL\_V4 \| 1.0 \| 7.204e-06 \| 170 \| 0.097 \| 453 \| 206 \| 19 \| 1 \| 273 \| 1 \| 430 \| Uncharacterized protein \| Uncharacterized protein; afdb-uniprot50 \| AF-A0A257P8Z7-F1-MODEL\_V4 \| 1.0 \| 0.002232 \| 169 \| 0.261 \| 84 \| 62 \| 0 \| 20 \| 103 \| 4 \| 87 \| Baseplate protein \| Baseplate protein; afdb-uniprot50 \| AF-A0A848JKX1-F1-MODEL\_V4 \| 1.0 \| 2.39e-07 \| 165 \| 0.156 \| 339 \| 187 \| 17 \| 23 \| 300 \| 9 \| 309 \| Uncharacterized protein \| Uncharacterized protein; afdb-uniprot50 \| AF-K9PW54-F1-MODEL\_V4 \| 1.0 \| 3.517e-06 \| 164 \| 0.156 \| 269 \| 139 \| 12 \| 111 \| 296 \| 949 \| 1212 \| Baseplate\_J domain-containing protein \| Baseplate\_J domain-containing protein; afdb-uniprot50 \| AF-A0A2E4BK49-F1-MODEL\_V4 \| 1.0 \| 1.476e-05 \| 160 \| 0.136 \| 183 \| 130 \| 9 \| 112 \| 270 \| 297 \| 475 \| Uncharacterized protein \| Uncharacterized protein; afdb-uniprot50 \| AF-G6F3I2-F1-MODEL\_V4 \| 1.0 \| 4.486e-08 \| 159 \| 0.121 \| 470 \| 206 \| 20 \| 20 \| 295 \| 12 \| 468 \| Baseplate\_J domain-containing protein \| Baseplate\_J domain-containing protein; afdb-uniprot50 \| AF-A0A7S9CLQ2-F1-MODEL\_V4 \| 1.0 \| 1.035e-07 \| 158 \| 0.121 \| 370 \| 192 \| 17 \| 20 \| 270 \| 17 \| 372 \| Baseplate J/gp47 family protein \| Baseplate J/gp47 family protein; afdb-uniprot50 \| AF-A0A0F8Y9S2-F1-MODEL\_V4 \| 1.0 \| 0.006164 \| 157 \| 0.228 \| 92 \| 65 \| 2 \| 13 \| 104 \| 1 \| 86 \| Baseplate\_J domain-containing protein \| Baseplate\_J domain-containing protein; afdb-uniprot50 \| AF-A0A6N8HUB6-F1-MODEL\_V4 \| 1.0 \| 1.566e-05 \| 155 \| 0.136 \| 220 \| 142 \| 13 \| 110 \| 295 \| 725 \| 930 \| Baseplate\_J domain-containing protein \| Baseplate\_J domain-containing protein; afdb-uniprot50 \| AF-A0A1L8I3J9-F1-MODEL\_V4 \| 1.0 \| 0.0001517 \| 151 \| 0.172 \| 214 \| 100 \| 5 \| 20 \| 158 \| 12 \| 223 \| Baseplate\_J domain-containing protein \| Baseplate\_J domain-containing protein; afdb-uniprot50 \| AF-A0A3P7PM03-F1-MODEL\_V4 \| 1.0 \| 1.352e-06 \| 144 \| 0.087 \| 332 \| 215 \| 15 \| 13 \| 298 \| 215 \| 504 \| Uncharacterized protein \| Uncharacterized protein; afdb-uniprot50 \| AF-A0A3L7ZAM7-F1-MODEL\_V4 \| 1.0 \| 1.765e-05 \| 121 \| 0.096 \| 343 \| 191 \| 21 \| 20 \| 294 \| 1 \| 292 \| Uncharacterized protein \| Uncharacterized protein; afdb-uniprot50 \| AF-A0A2W4SAT6-F1-MODEL\_V4 \| 1.0 \| 0.001981 \| 119 \| 0.146 \| 212 \| 104 \| 5 \| 22 \| 158 \| 13 \| 222 \| Baseplate\_J domain-containing protein \| Baseplate\_J domain-containing protein; afdb-uniprot50 \| AF-A0A2G8MLJ1-F1-MODEL\_V4 \| 1.0 \| 0.0001517 \| 118 \| 0.115 \| 295 \| 154 \| 10 \| 20 \| 227 \| 17 \| 291 \| Baseplate\_J domain-containing protein \| Baseplate\_J domain-containing protein; afdb-uniprot50 \| AF-A0A6A8LR27-F1-MODEL\_V4 \| 1.0 \| 0.007829 \| 108 \| 0.115 \| 234 \| 110 \| 5 \| 20 \| 158 \| 8 \| 239 \| Baseplate\_J domain-containing protein \| Baseplate\_J domain-containing protein; afdb-uniprot50 \| AF-A0A3S4F3P3-F1-MODEL\_V4 \| 1.0 \| 0.000967 \| 101 \| 0.116 \| 317 \| 155 \| 11 \| 20 \| 225 \| 22 \| 324 \| Bacteriophage protein \| Bacteriophage protein; afdb-uniprot50 \| AF-A0A5Y6EZV3-F1-MODEL\_V4 \| 0.999 \| 0.0002927 \| 99 \| 0.111 \| 304 \| 164 \| 14 \| 8 \| 225 \| 2 \| 285 \| Baseplate\_J domain-containing protein \| Baseplate\_J domain-containing protein; afdb-uniprot50 \| AF-A0A849R7X1-F1-MODEL\_V4 \| 0.995 \| 0.005807 \| 88 \| 0.094 \| 498 \| 201 \| 20 \| 22 \| 278 \| 14 \| 502 \| Uncharacterized protein \| Uncharacterized protein; afdb-uniprot50 \| AF-A0A828Z879-F1-MODEL\_V4 \| 0.994 \| 0.004057 \| 87 \| 0.111 \| 322 \| 170 \| 10 \| 22 \| 242 \| 12 \| 318 \| Baseplate J-like protein \| Baseplate J-like protein; afdb-uniprot50 \| AF-A0A426VQ90-F1-MODEL\_V4 \| 0.986 \| 0.003195 \| 81 \| 0.098 \| 384 \| 158 \| 11 \| 20 \| 226 \| 9 \| 381 \| Uncharacterized protein \| Uncharacterized protein |
| Top keywords  (threshold 1.00e-02 (evalue)) | **Baseplate, assembly, domain\_containing, Baseplate\_J, J, gp47, Phage, J\_like, Phage\_related, Putative** |
| Output files | ../../similar\_structures/13\_FANPEZAQ\_CDS\_0013\_afdb-proteome\_foldseek.tsv ../../similar\_structures/13\_FANPEZAQ\_CDS\_0013\_afdb-uniprot50\_foldseek.tsv ../../similar\_structures/13\_FANPEZAQ\_CDS\_0013\_merged.svg ../../similar\_structures/13\_FANPEZAQ\_CDS\_0013\_pdb\_foldseek.tsv |

  
  
  

Return to summary | Go to previous | Go to next

  


---

**Sequence/structure alignments coloring**  
Each object in the alignment figures is colored according to its E-value following this color coding:

1e-100
10
