## Supplementary material for "Completing the BASEL phage collection to unlock hidden diversity for systematic exploration of phage-host interactions": Data S2: 14.html

FANPEZAQ\_CDS\_0014


Return to summary | Go to previous | Go to next

|  |  |
| --- | --- |
| FANPEZAQ\_CDS\_0014  Page creation date: 02 Sep 2024, 12:00  Project folder: n/a  Input sequences file: Escherichia\_virus\_HeidiAbel.gb | tail phage i p2 bacteriophage formation p2\_related p2\_like putative fragment domain\_containing tail\_p2\_i gpi duf2313 baseplate fiber i\_like j prophage assembly phage\_related tail\_like |

### Sequence information

|  |  |
| --- | --- |
| Name | FANPEZAQ\_CDS\_0014  14\_FANPEZAQ\_CDS\_0014 (pipeline id) |
| Imported annotations | Escherichia\_virus\_HeidiAbel Bas97 |
| Protein sequence | MSDLLPPNATAQERALANTIARISDVPLAARKMWNPDTIPANLLPWMAWAFSVDDWNNNW TEQEKRNVIKNSLFVHKHKGTLAAIKSAVEPLGYIIRIVEWWEDTPQAEPYTFRLEVGLL DKGVDESIYDQFTRLIETNKNLRSHMKALTIKSEINGVAYFGAGMVSGITTTVYPYIAEN LESTGGVFTAAAEQSVDAVSVYPFT |
| Number of residues | 205 |
| Molecular weight (Da) | 22921.75 |
| Output files | ../../query\_sequences/14\_FANPEZAQ\_CDS\_0014.fasta |

### Putative domain architecture and protein family

#### Search results (HHblits)^1^

|  |  |
| --- | --- |
| Domain family databases searched | Pfam, Ncbi-cd, Cath, Phrogs |
| Results, scheme(s)  (Top layers only; threshold 1.00e-03 (evalue)) | xml version="1.0" encoding="utf-8" standalone="no"?       2024-09-02T21:08:14.957402 image/svg+xml   Matplotlib v3.7.2, https://matplotlib.org/ |
| Results, table  (E-value ≤ 1.00e-03 (evalue)) | db \| id \| prob \| evalue \| pvalue \| score \| cols \| query \| query\_len \| template \| template\_len \| name \| description; pfam \| PF09684 \| 99.5 \| 2e-18 \| 3.9e-22 \| 125.7 \| 130 \| (7, 141) \| 205 \| (1, 138) \| 138 \| Tail\_P2\_I \| Phage tail protein (Tail\_P2\_I); pfam \| PF10076 \| 99.2 \| 3e-16 \| 6e-20 \| 118.6 \| 138 \| (9, 148) \| 205 \| (18, 185) \| 188 \| Phage\_Mu\_Gp48 \| Bacteriophage Mu-like, Gp48; pfam \| PF11041 \| 99.2 \| 1e-15 \| 2e-19 \| 116.0 \| 129 \| (8, 147) \| 205 \| (10, 180) \| 181 \| DUF2612 \| Protein of unknown function (DUF2612); phrogs \| 48 \| 100.0 \| 4.3e-47 \| 5.7e-51 \| 309.0 \| 201 \| (1, 204) \| 205 \| (7, 209) \| 211 \| tail protein \| tail protein; Category: tail; p200917 VI\_08608; phrogs \| 5726 \| 99.9 \| 4e-32 \| 4.7e-36 \| 196.5 \| 114 \| (2, 117) \| 205 \| (3, 122) \| 128 \| tail protein \| tail protein; Category: tail; p214505 VI\_09983; phrogs \| 9915 \| 99.9 \| 6.9e-31 \| 8e-35 \| 236.4 \| 167 \| (2, 169) \| 205 \| (7, 175) \| 1008 \| NA \| NA; Category: unknown function; p204625 VI\_11278; phrogs \| 14404 \| 99.9 \| 1.4e-27 \| 1.6e-31 \| 196.4 \| 152 \| (2, 157) \| 205 \| (12, 170) \| 378 \| baseplate protein \| baseplate protein; Category: tail; p127197 VI\_01263; phrogs \| 29945 \| 99.7 \| 5.3e-22 \| 5.9e-26 \| 158.2 \| 131 \| (25, 156) \| 205 \| (119, 256) \| 369 \| NA \| NA; Category: unknown function; p52203 VI\_08944; phrogs \| 1228 \| 99.6 \| 8.2e-20 \| 9.7e-24 \| 143.7 \| 117 \| (30, 153) \| 205 \| (43, 160) \| 218 \| tail protein \| tail protein; Category: tail; p284533 VI\_03834; phrogs \| 19272 \| 99.5 \| 4.5e-19 \| 5.1e-23 \| 141.9 \| 148 \| (2, 156) \| 205 \| (1, 149) \| 301 \| NA \| NA; Category: unknown function; p52189 VI\_08944; phrogs \| 34934 \| 99.4 \| 6.5e-18 \| 7.3e-22 \| 137.4 \| 96 \| (1, 96) \| 205 \| (2, 98) \| 392 \| tail protein \| tail protein; Category: tail; p102007 VI\_04294; phrogs \| 17612 \| 99.4 \| 4.3e-17 \| 4.9e-21 \| 133.3 \| 140 \| (3, 153) \| 205 \| (1, 141) \| 386 \| tail protein \| tail protein; Category: tail; p217156 VI\_00068; phrogs \| 8207 \| 98.8 \| 3.8e-13 \| 4.4e-17 \| 109.1 \| 115 \| (37, 158) \| 205 \| (20, 134) \| 348 \| NA \| NA; Category: unknown function; p142933 VI\_00095; phrogs \| 2871 \| 95.9 \| 5e-05 \| 5.9e-09 \| 69.2 \| 88 \| (24, 147) \| 205 \| (21, 108) \| 1049 \| tail protein \| tail protein; Category: tail; p28620 VI\_01314; phrogs \| 32946 \| 94.9 \| 0.00042 \| 4.6e-08 \| 49.3 \| 71 \| (28, 98) \| 205 \| (54, 127) \| 191 \| tail protein \| tail protein; Category: tail; KU234533\_p33 |
| Top keywords  (threshold 1.00e-03 (evalue)) | **tail, VI\_08944, Phage, Tail\_P2\_I, Bacteriophage, Mu\_like, Gp48, DUF2612, p200917, VI\_08608** |
| Output files | ../../domain\_architecture/14\_FANPEZAQ\_CDS\_0014\_cath.hhr ../../domain\_architecture/14\_FANPEZAQ\_CDS\_0014\_merged.svg ../../domain\_architecture/14\_FANPEZAQ\_CDS\_0014\_ncbi-cd.hhr ../../domain\_architecture/14\_FANPEZAQ\_CDS\_0014\_pfam.hhr ../../domain\_architecture/14\_FANPEZAQ\_CDS\_0014\_phrogs.hhr |

### Identical protein sequences/structures

#### Search results

|  |  |
| --- | --- |
| Protein sequence databases searched | Pdb, Swissprot, Refseq |
| Identical proteins found | -- |
| Top keywords | -- |
| Output files | -- |

### Similar protein sequences/structures

#### Sequence similarity search results (HHblits)^1^

|  |  |
| --- | --- |
| Sequence databases searched | Uniclust, Pdb70 |
| Results, scheme(s)  (Top layers only, threshold 1.00e-03 (evalue)) | xml version="1.0" encoding="utf-8" standalone="no"?       2024-09-02T21:08:34.315745 image/svg+xml   Matplotlib v3.7.2, https://matplotlib.org/ |
| Results, table(s)  (threshold 1.00e-03 (evalue)) | db \| id \| prob \| evalue \| pvalue \| score \| cols \| query \| query\_len \| template \| template\_len \| name \| description; uniclust \| UniRef100\_A0A009QFG0 \| 100.0 \| 1.9e-54 \| 4.2e-60 \| 357.7 \| 202 \| (1, 204) \| 205 \| (47, 248) \| 282 \| Phage tail protein I \| Phage tail protein I; uniclust \| UniRef100\_A0A024E9W6 \| 100.0 \| 2.1e-54 \| 4.5e-60 \| 354.2 \| 204 \| (1, 204) \| 205 \| (50, 253) \| 294 \| Tail protein I \| Tail protein I; uniclust \| UniRef100\_A0A1G7SAK3 \| 100.0 \| 8.4e-52 \| 1.8e-57 \| 333.0 \| 201 \| (1, 204) \| 205 \| (4, 204) \| 264 \| Phage tail protein, P2 protein I family \| Phage tail protein, P2 protein I family; uniclust \| UniRef100\_A0A077N9G9 \| 100.0 \| 2e-50 \| 4e-56 \| 318.1 \| 201 \| (2, 204) \| 205 \| (39, 239) \| 267 \| Tail protein I (GpI) \| Tail protein I (GpI); uniclust \| UniRef100\_A0A103EG21 \| 100.0 \| 4.7e-49 \| 9.4e-55 \| 309.7 \| 176 \| (1, 176) \| 205 \| (4, 179) \| 249 \| Phage tail protein \| Phage tail protein; uniclust \| UniRef100\_A0A066T5L9 \| 100.0 \| 9.9e-49 \| 2.1e-54 \| 312.3 \| 204 \| (1, 204) \| 205 \| (30, 235) \| 241 \| Phage tail protein I \| Phage tail protein I; uniclust \| UniRef100\_A0A066RUL7 \| 100.0 \| 1.7e-48 \| 3.6e-54 \| 320.2 \| 187 \| (1, 188) \| 205 \| (14, 202) \| 278 \| Tail protein \| Tail protein; uniclust \| UniRef100\_A0A1C3EBR0 \| 100.0 \| 7.2e-47 \| 1.5e-52 \| 308.1 \| 199 \| (2, 203) \| 205 \| (14, 219) \| 276 \| Phage tail protein I \| Phage tail protein I; uniclust \| UniRef100\_A0A0D7V3M6 \| 100.0 \| 1e-44 \| 1.9e-50 \| 278.5 \| 202 \| (2, 203) \| 205 \| (4, 206) \| 238 \| Phage tail protein I \| Phage tail protein I; uniclust \| UniRef100\_A0A016XH76 \| 100.0 \| 1.3e-44 \| 2.7e-50 \| 291.4 \| 169 \| (1, 170) \| 205 \| (26, 197) \| 256 \| Tail protein \| Tail protein; uniclust \| UniRef100\_A0A2U9LD38 \| 100.0 \| 2.8e-43 \| 5.5e-49 \| 279.0 \| 175 \| (2, 178) \| 205 \| (14, 188) \| 283 \| Phage tail protein I \| Phage tail protein I; uniclust \| UniRef100\_A0A377XEC8 \| 100.0 \| 1.9e-42 \| 3.8e-48 \| 256.2 \| 158 \| (1, 160) \| 205 \| (3, 160) \| 165 \| Tail protein I \| Tail protein I; uniclust \| UniRef100\_A0A0S1B435 \| 100.0 \| 1.9e-42 \| 4.1e-48 \| 282.8 \| 159 \| (2, 163) \| 205 \| (20, 185) \| 280 \| Phage P2 baseplate assembly gpI-like protein \| Phage P2 baseplate assembly gpI-like protein; uniclust \| UniRef100\_A0A0J8DB16 \| 100.0 \| 2.1e-42 \| 4.5e-48 \| 278.8 \| 202 \| (2, 205) \| 205 \| (17, 224) \| 259 \| Phage tail protein, P2 protein I family \| Phage tail protein, P2 protein I family; uniclust \| UniRef100\_A0A0A1AHN2 \| 100.0 \| 6e-42 \| 1.2e-47 \| 274.1 \| 203 \| (1, 205) \| 205 \| (10, 219) \| 257 \| Phage tail protein I \| Phage tail protein I; uniclust \| UniRef100\_A0A0B6CYD1 \| 100.0 \| 1.9e-41 \| 4.2e-47 \| 283.8 \| 164 \| (1, 165) \| 205 \| (16, 187) \| 299 \| Phage tail protein I \| Phage tail protein I; uniclust \| UniRef100\_A0A074LXC9 \| 100.0 \| 3.8e-41 \| 7.8e-47 \| 265.7 \| 199 \| (2, 204) \| 205 \| (19, 223) \| 224 \| Tail protein \| Tail protein; uniclust \| UniRef100\_A0A1W9H3W1 \| 100.0 \| 6.4e-41 \| 1.3e-46 \| 264.3 \| 153 \| (2, 154) \| 205 \| (14, 169) \| 245 \| Phage tail protein I \| Phage tail protein I; uniclust \| UniRef100\_A0A087NAA8 \| 100.0 \| 2e-40 \| 4e-46 \| 266.3 \| 170 \| (1, 171) \| 205 \| (46, 218) \| 271 \| Phage tail protein I \| Phage tail protein I; uniclust \| UniRef100\_A0A1X7CGW5 \| 100.0 \| 5.3e-40 \| 1.1e-45 \| 258.1 \| 164 \| (1, 173) \| 205 \| (20, 184) \| 247 \| Phage tail protein, P2 protein I family \| Phage tail protein, P2 protein I family; uniclust \| UniRef100\_A0A031FSG0 \| 100.0 \| 5.8e-40 \| 1.2e-45 \| 263.0 \| 202 \| (2, 205) \| 205 \| (11, 213) \| 244 \| Phage tail protein I \| Phage tail protein I; uniclust \| UniRef100\_A0A017HBC1 \| 100.0 \| 7.9e-40 \| 1.7e-45 \| 263.2 \| 164 \| (1, 169) \| 205 \| (32, 211) \| 234 \| Phage tail fiber protein \| Phage tail fiber protein; uniclust \| UniRef100\_A0A1G7HXV1 \| 100.0 \| 4.5e-39 \| 8.2e-45 \| 249.9 \| 201 \| (2, 204) \| 205 \| (3, 204) \| 295 \| Phage tail protein, P2 protein I family (Fragment) \| Phage tail protein, P2 protein I family (Fragment); uniclust \| UniRef100\_A0A1A9R6A9 \| 100.0 \| 2.3e-38 \| 4.6e-44 \| 260.7 \| 155 \| (1, 156) \| 205 \| (18, 178) \| 333 \| Phage tail protein I \| Phage tail protein I; uniclust \| UniRef100\_A0A061NN14 \| 100.0 \| 2.9e-38 \| 6.7e-44 \| 259.9 \| 199 \| (2, 204) \| 205 \| (15, 219) \| 249 \| Putative phage tail protein \| Putative phage tail protein; uniclust \| UniRef100\_A0A2X3LRF3 \| 100.0 \| 7.6e-38 \| 1.4e-43 \| 251.7 \| 186 \| (2, 189) \| 205 \| (169, 355) \| 362 \| Tail protein I \| Tail protein I; uniclust \| UniRef100\_UPI0002D8351F \| 100.0 \| 9.8e-38 \| 1.8e-43 \| 243.5 \| 203 \| (2, 204) \| 205 \| (4, 207) \| 267 \| phage tail protein I \| phage tail protein I; uniclust \| UniRef100\_A0A0F9Z9G5 \| 100.0 \| 1.2e-37 \| 2.4e-43 \| 251.7 \| 203 \| (1, 205) \| 205 \| (41, 247) \| 262 \| Phage tail protein \| Phage tail protein; uniclust \| UniRef100\_A0A0N8R6G9 \| 100.0 \| 2.2e-37 \| 4e-43 \| 236.5 \| 200 \| (1, 202) \| 205 \| (56, 255) \| 256 \| Tail protein I \| Tail protein I; uniclust \| UniRef100\_A0A239C7G7 \| 100.0 \| 2.4e-37 \| 5.1e-43 \| 246.2 \| 201 \| (2, 205) \| 205 \| (18, 224) \| 229 \| Phage tail protein, P2 protein I family \| Phage tail protein, P2 protein I family; uniclust \| UniRef100\_A0A0A8H9H6 \| 100.0 \| 2.7e-37 \| 5.9e-43 \| 251.0 \| 185 \| (1, 191) \| 205 \| (8, 194) \| 242 \| Phage P2 family tail protein \| Phage P2 family tail protein; uniclust \| UniRef100\_A0A1X3I4G6 \| 100.0 \| 3.4e-37 \| 6.3e-43 \| 235.2 \| 181 \| (2, 184) \| 205 \| (3, 183) \| 254 \| Putative phage tail protein \| Putative phage tail protein; uniclust \| UniRef100\_A0A134C9I1 \| 100.0 \| 3.1e-37 \| 6.5e-43 \| 247.0 \| 200 \| (3, 204) \| 205 \| (11, 214) \| 249 \| Phage tail protein I-like protein \| Phage tail protein I-like protein; uniclust \| UniRef100\_A0A375ABY7 \| 100.0 \| 9e-37 \| 1.6e-42 \| 251.3 \| 197 \| (2, 200) \| 205 \| (304, 500) \| 504 \| Phage baseplate assembly protein J \| Phage baseplate assembly protein J; uniclust \| UniRef100\_A0A8T5Z8P8 \| 100.0 \| 1.9e-36 \| 3.8e-42 \| 220.9 \| 139 \| (2, 140) \| 205 \| (7, 146) \| 148 \| Phage tail protein I (Fragment) \| Phage tail protein I (Fragment); uniclust \| UniRef100\_A0A077KVE4 \| 100.0 \| 2e-36 \| 4.1e-42 \| 231.0 \| 172 \| (1, 174) \| 205 \| (16, 187) \| 193 \| Phage tail protein I \| Phage tail protein I; uniclust \| UniRef100\_A0A1B9NST5 \| 100.0 \| 6.9e-36 \| 1.4e-41 \| 211.0 \| 105 \| (1, 105) \| 205 \| (6, 110) \| 115 \| Phage tail protein I \| Phage tail protein I; uniclust \| UniRef100\_A0A1E7HY16 \| 100.0 \| 8.7e-36 \| 1.7e-41 \| 216.5 \| 146 \| (2, 147) \| 205 \| (3, 148) \| 150 \| Phage tail protein I (Fragment) \| Phage tail protein I (Fragment); uniclust \| UniRef100\_A0A7D6BR59 \| 100.0 \| 1.4e-35 \| 2.6e-41 \| 243.8 \| 173 \| (2, 175) \| 205 \| (6, 179) \| 376 \| Phage tail protein I \| Phage tail protein I; uniclust \| UniRef100\_UPI001080C16A \| 100.0 \| 1.6e-35 \| 3.1e-41 \| 257.2 \| 202 \| (1, 203) \| 205 \| (5, 208) \| 631 \| phage tail protein I \| phage tail protein I; uniclust \| UniRef100\_UPI0011A48CC0 \| 100.0 \| 1.7e-35 \| 3.1e-41 \| 230.9 \| 185 \| (17, 203) \| 205 \| (117, 301) \| 301 \| phage tail protein I \| phage tail protein I; uniclust \| UniRef100\_A0A1J5B6D6 \| 100.0 \| 2e-35 \| 4e-41 \| 244.4 \| 159 \| (1, 159) \| 205 \| (7, 182) \| 341 \| Phage tail protein \| Phage tail protein; uniclust \| UniRef100\_A0A0H3ZS74 \| 100.0 \| 4e-35 \| 7.5e-41 \| 226.5 \| 183 \| (2, 186) \| 205 \| (4, 187) \| 237 \| Tail protein I \| Tail protein I; uniclust \| UniRef100\_A0A085ASA6 \| 100.0 \| 6.6e-35 \| 1.3e-40 \| 269.1 \| 201 \| (2, 203) \| 205 \| (168, 370) \| 1175 \| Tail protein I \| Tail protein I; uniclust \| UniRef100\_A0A975LVB5 \| 100.0 \| 8.3e-35 \| 1.6e-40 \| 223.6 \| 158 \| (2, 160) \| 205 \| (5, 163) \| 246 \| Phage tail protein I \| Phage tail protein I; uniclust \| UniRef100\_A0A0P7CQM9 \| 100.0 \| 1.3e-34 \| 2.5e-40 \| 228.4 \| 162 \| (1, 165) \| 205 \| (9, 176) \| 250 \| Phage tail protein \| Phage tail protein; uniclust \| UniRef100\_A0A0C5VFB1 \| 100.0 \| 1.3e-34 \| 2.7e-40 \| 236.1 \| 164 \| (2, 166) \| 205 \| (121, 293) \| 317 \| Bacteriophage P2-related tail formation protein \| Bacteriophage P2-related tail formation protein; uniclust \| UniRef100\_A0A285VTA9 \| 100.0 \| 2.2e-34 \| 4.5e-40 \| 244.7 \| 146 \| (1, 147) \| 205 \| (3, 181) \| 399 \| Phage tail protein, P2 protein I family \| Phage tail protein, P2 protein I family; uniclust \| UniRef100\_A0A3N6RV94 \| 100.0 \| 3.1e-34 \| 5.7e-40 \| 234.1 \| 189 \| (13, 203) \| 205 \| (230, 418) \| 418 \| Phage tail protein I \| Phage tail protein I; uniclust \| UniRef100\_A0A925L380 \| 100.0 \| 3.7e-34 \| 7.1e-40 \| 226.8 \| 170 \| (1, 171) \| 205 \| (6, 179) \| 276 \| Phage tail protein I \| Phage tail protein I; uniclust \| UniRef100\_A0A5D8SJY0 \| 100.0 \| 1e-33 \| 2e-39 \| 208.0 \| 152 \| (23, 176) \| 205 \| (7, 158) \| 159 \| Phage tail protein I (Fragment) \| Phage tail protein I (Fragment); uniclust \| UniRef100\_A0A0E1S4D3 \| 100.0 \| 1.4e-33 \| 2.7e-39 \| 205.8 \| 119 \| (2, 120) \| 205 \| (7, 125) \| 148 \| Phage tail protein I \| Phage tail protein I; uniclust \| UniRef100\_A0A1B3E8I5 \| 100.0 \| 1.5e-33 \| 2.8e-39 \| 222.1 \| 160 \| (1, 165) \| 205 \| (4, 170) \| 281 \| Phage tail protein I \| Phage tail protein I; uniclust \| UniRef100\_A0A1W1UNV3 \| 100.0 \| 1.6e-33 \| 3.2e-39 \| 216.5 \| 163 \| (2, 169) \| 205 \| (8, 187) \| 193 \| Phage tail protein, P2 protein I family \| Phage tail protein, P2 protein I family; uniclust \| UniRef100\_A0A840WZA1 \| 100.0 \| 1.8e-33 \| 3.6e-39 \| 222.1 \| 169 \| (2, 172) \| 205 \| (13, 188) \| 222 \| Phage tail P2-like protein \| Phage tail P2-like protein; uniclust \| UniRef100\_A0A1S1MWK0 \| 100.0 \| 2.6e-33 \| 5.1e-39 \| 216.4 \| 174 \| (30, 204) \| 205 \| (36, 209) \| 227 \| Phage tail protein I \| Phage tail protein I; uniclust \| UniRef100\_A0A329B4G1 \| 100.0 \| 2.7e-33 \| 5.3e-39 \| 199.6 \| 117 \| (1, 117) \| 205 \| (5, 121) \| 124 \| Phage tail P2-like protein (Fragment) \| Phage tail P2-like protein (Fragment); uniclust \| UniRef100\_A0A0T7DV65 \| 100.0 \| 4.2e-33 \| 8.4e-39 \| 220.2 \| 197 \| (4, 202) \| 205 \| (35, 238) \| 244 \| Phage tail protein I \| Phage tail protein I; uniclust \| UniRef100\_A0A547PS50 \| 100.0 \| 5.3e-33 \| 1.1e-38 \| 218.9 \| 180 \| (2, 183) \| 205 \| (6, 190) \| 230 \| Phage tail protein I \| Phage tail protein I; uniclust \| UniRef100\_A0A2X4TXF0 \| 100.0 \| 7.5e-33 \| 1.4e-38 \| 216.5 \| 173 \| (2, 176) \| 205 \| (3, 175) \| 302 \| Bacteriophage P2-related tail formation protein \| Bacteriophage P2-related tail formation protein; uniclust \| UniRef100\_UPI000DFE0774 \| 100.0 \| 1.7e-32 \| 3.1e-38 \| 215.2 \| 196 \| (2, 199) \| 205 \| (3, 200) \| 309 \| phage tail protein I \| phage tail protein I; uniclust \| UniRef100\_A0A085AFN4 \| 100.0 \| 1.7e-32 \| 3.7e-38 \| 220.7 \| 190 \| (2, 194) \| 205 \| (13, 206) \| 224 \| Tail protein I \| Tail protein I; uniclust \| UniRef100\_A0A1Q6JPU8 \| 100.0 \| 1.9e-32 \| 3.9e-38 \| 227.3 \| 199 \| (3, 203) \| 205 \| (18, 222) \| 310 \| Phage tail protein I \| Phage tail protein I; uniclust \| UniRef100\_A0A1C3FFM0 \| 99.9 \| 2.4e-32 \| 5.1e-38 \| 221.0 \| 172 \| (2, 178) \| 205 \| (20, 197) \| 253 \| Putative phage tail fiber protein \| Putative phage tail fiber protein; uniclust \| UniRef100\_UPI000C7DCBEE \| 99.9 \| 5.9e-32 \| 1.1e-37 \| 222.0 \| 170 \| (2, 172) \| 205 \| (7, 178) \| 412 \| phage tail protein I \| phage tail protein I; uniclust \| UniRef100\_A0A0C1QG10 \| 99.9 \| 7.4e-32 \| 1.4e-37 \| 216.0 \| 172 \| (2, 174) \| 205 \| (5, 198) \| 305 \| Phage tail protein I \| Phage tail protein I; uniclust \| UniRef100\_A0A939NK51 \| 99.9 \| 9.3e-32 \| 1.8e-37 \| 191.4 \| 122 \| (2, 125) \| 205 \| (3, 124) \| 128 \| Phage tail protein I \| Phage tail protein I; uniclust \| UniRef100\_A0A1C3ELC5 \| 99.9 \| 1e-31 \| 1.9e-37 \| 203.9 \| 157 \| (2, 159) \| 205 \| (3, 174) \| 231 \| Phage tail protein I \| Phage tail protein I; uniclust \| UniRef100\_A0A7H4P5M4 \| 99.9 \| 1.9e-31 \| 3.4e-37 \| 204.4 \| 170 \| (2, 172) \| 205 \| (7, 178) \| 253 \| Putative prophage tail protein \| Putative prophage tail protein; uniclust \| UniRef100\_A0A1H2N8K0 \| 99.9 \| 2.5e-31 \| 5.2e-37 \| 223.6 \| 152 \| (1, 152) \| 205 \| (6, 191) \| 371 \| Phage tail protein I \| Phage tail protein I; uniclust \| UniRef100\_A0A145VQK9 \| 99.9 \| 1e-30 \| 1.9e-36 \| 198.2 \| 153 \| (2, 155) \| 205 \| (6, 159) \| 232 \| Phage tail protein I \| Phage tail protein I; uniclust \| UniRef100\_UPI0008FEF499 \| 99.9 \| 1.3e-30 \| 2.6e-36 \| 228.6 \| 170 \| (2, 172) \| 205 \| (7, 178) \| 713 \| phage tail protein \| phage tail protein; uniclust \| UniRef100\_A0A1C6DZY7 \| 99.9 \| 1.5e-30 \| 3.2e-36 \| 214.4 \| 173 \| (2, 176) \| 205 \| (37, 215) \| 261 \| Bacteriophage P2-related tail formation protein \| Bacteriophage P2-related tail formation protein; uniclust \| UniRef100\_A0A074TKV9 \| 99.9 \| 2.1e-30 \| 4.3e-36 \| 207.1 \| 172 \| (2, 177) \| 205 \| (10, 195) \| 257 \| Tail protein \| Tail protein; uniclust \| UniRef100\_UPI0009B30C98 \| 99.9 \| 3.2e-30 \| 5.8e-36 \| 199.4 \| 191 \| (4, 194) \| 205 \| (71, 262) \| 269 \| phage tail protein I \| phage tail protein I; uniclust \| UniRef100\_UPI0006B4121B \| 99.9 \| 3.8e-30 \| 6.9e-36 \| 203.3 \| 169 \| (1, 169) \| 205 \| (1, 170) \| 322 \| phage tail protein I \| phage tail protein I; uniclust \| UniRef100\_A0A1D2QS86 \| 99.9 \| 3.9e-30 \| 7.8e-36 \| 193.4 \| 139 \| (26, 165) \| 205 \| (4, 145) \| 163 \| Phage tail protein I \| Phage tail protein I; uniclust \| UniRef100\_A0A1X7MNV1 \| 99.9 \| 7.5e-30 \| 1.4e-35 \| 183.2 \| 118 \| (2, 121) \| 205 \| (6, 123) \| 140 \| Phage tail protein I (Fragment) \| Phage tail protein I (Fragment); uniclust \| UniRef100\_A0A376RLD7 \| 99.9 \| 8.2e-30 \| 1.5e-35 \| 192.0 \| 126 \| (2, 129) \| 205 \| (86, 211) \| 219 \| Putative tail protein I (Gpi) \| Putative tail protein I (Gpi); uniclust \| UniRef100\_UPI000641FAAA \| 99.9 \| 9.7e-30 \| 1.8e-35 \| 181.4 \| 138 \| (27, 164) \| 205 \| (2, 139) \| 142 \| phage tail protein I \| phage tail protein I; uniclust \| UniRef100\_A0A142BH84 \| 99.9 \| 1e-29 \| 2e-35 \| 207.5 \| 120 \| (1, 120) \| 205 \| (1, 120) \| 352 \| Phage tail protein \| Phage tail protein; uniclust \| UniRef100\_A0A011MIC4 \| 99.9 \| 1.3e-29 \| 2.6e-35 \| 201.5 \| 148 \| (1, 153) \| 205 \| (21, 184) \| 214 \| Tail fiber protein \| Tail fiber protein; uniclust \| UniRef100\_A0A162A844 \| 99.9 \| 1.5e-29 \| 2.8e-35 \| 204.4 \| 163 \| (2, 167) \| 205 \| (9, 178) \| 305 \| Tail protein \| Tail protein; uniclust \| UniRef100\_UPI00073535E0 \| 99.9 \| 2.7e-29 \| 5.2e-35 \| 181.8 \| 131 \| (2, 135) \| 205 \| (6, 138) \| 140 \| phage tail protein I \| phage tail protein I; uniclust \| UniRef100\_A0A2A2C599 \| 99.9 \| 2.9e-29 \| 5.8e-35 \| 198.5 \| 196 \| (2, 198) \| 205 \| (11, 215) \| 228 \| Phage tail protein I \| Phage tail protein I; uniclust \| UniRef100\_D1Y2C1 \| 99.9 \| 3.9e-29 \| 7.6e-35 \| 195.7 \| 196 \| (3, 204) \| 205 \| (12, 215) \| 240 \| Phage tail protein \| Phage tail protein; uniclust \| UniRef100\_A0A7G2U2N6 \| 99.9 \| 4.8e-29 \| 8.8e-35 \| 178.4 \| 143 \| (2, 146) \| 205 \| (5, 152) \| 152 \| Phage tail protein I \| Phage tail protein I; uniclust \| UniRef100\_A0A7W6WC48 \| 99.9 \| 7.4e-29 \| 1.4e-34 \| 202.6 \| 152 \| (3, 154) \| 205 \| (4, 166) \| 348 \| Phage tail P2-like protein \| Phage tail P2-like protein; uniclust \| UniRef100\_UPI00044AA162 \| 99.9 \| 8e-29 \| 1.5e-34 \| 183.3 \| 148 \| (44, 193) \| 205 \| (4, 154) \| 161 \| phage tail protein I \| phage tail protein I; uniclust \| UniRef100\_UPI000DF97472 \| 99.9 \| 8.4e-29 \| 1.5e-34 \| 215.8 \| 152 \| (1, 153) \| 205 \| (2, 154) \| 925 \| phage tail protein I \| phage tail protein I; uniclust \| UniRef100\_A0A1X7NDR7 \| 99.9 \| 1e-28 \| 2.1e-34 \| 196.3 \| 149 \| (1, 152) \| 205 \| (4, 196) \| 238 \| Phage tail protein, P2 protein I family \| Phage tail protein, P2 protein I family; uniclust \| UniRef100\_A0A2G6HAI3 \| 99.9 \| 2.3e-28 \| 4.2e-34 \| 197.4 \| 158 \| (3, 163) \| 205 \| (6, 168) \| 385 \| Phage tail protein I (Fragment) \| Phage tail protein I (Fragment); uniclust \| UniRef100\_A0A8I1NYE7 \| 99.9 \| 2.9e-28 \| 5.6e-34 \| 188.0 \| 203 \| (2, 204) \| 205 \| (13, 220) \| 221 \| Phage tail protein I \| Phage tail protein I; uniclust \| UniRef100\_A0A167GZS5 \| 99.9 \| 3.1e-28 \| 5.8e-34 \| 194.6 \| 163 \| (2, 167) \| 205 \| (3, 172) \| 328 \| Tail protein \| Tail protein; uniclust \| UniRef100\_A0A149PPI8 \| 99.9 \| 3.5e-28 \| 6.5e-34 \| 189.6 \| 151 \| (2, 154) \| 205 \| (86, 239) \| 263 \| Tail protein \| Tail protein; uniclust \| UniRef100\_A0A137SPR3 \| 99.9 \| 4.7e-28 \| 9.9e-34 \| 202.5 \| 167 \| (2, 171) \| 205 \| (20, 200) \| 309 \| Phage tail protein I-like protein \| Phage tail protein I-like protein; uniclust \| UniRef100\_A0A078MHK2 \| 99.9 \| 5e-28 \| 1.1e-33 \| 202.9 \| 154 \| (2, 159) \| 205 \| (14, 173) \| 293 \| Phage tail protein (Tail\_P2\_I) \| Phage tail protein (Tail\_P2\_I); uniclust \| UniRef100\_A0A3N2E0S8 \| 99.9 \| 6.1e-28 \| 1.2e-33 \| 186.3 \| 150 \| (2, 152) \| 205 \| (13, 195) \| 234 \| Phage tail P2-like protein \| Phage tail P2-like protein; uniclust \| UniRef100\_UPI0005DFBD90 \| 99.9 \| 6.8e-28 \| 1.2e-33 \| 196.4 \| 149 \| (29, 179) \| 205 \| (265, 413) \| 417 \| phage tail protein I \| phage tail protein I; uniclust \| UniRef100\_A0A0F6A4X0 \| 99.9 \| 7.1e-28 \| 1.4e-33 \| 195.0 \| 150 \| (5, 157) \| 205 \| (11, 167) \| 302 \| Tail protein \| Tail protein; uniclust \| UniRef100\_A0A316MKT1 \| 99.9 \| 7.6e-28 \| 1.5e-33 \| 193.8 \| 164 \| (37, 204) \| 205 \| (75, 241) \| 278 \| Phage tail protein I \| Phage tail protein I; uniclust \| UniRef100\_UPI00210E7D19 \| 99.9 \| 9.4e-28 \| 1.7e-33 \| 191.4 \| 196 \| (2, 204) \| 205 \| (3, 199) \| 342 \| phage tail protein I \| phage tail protein I; uniclust \| UniRef100\_A0A447N0G4 \| 99.9 \| 9.4e-28 \| 1.8e-33 \| 162.5 \| 89 \| (1, 89) \| 205 \| (3, 91) \| 91 \| Orf38 p2 I-like tail protein \| Orf38 p2 I-like tail protein; uniclust \| UniRef100\_A0A968H8P4 \| 99.9 \| 1.2e-27 \| 2.2e-33 \| 183.7 \| 199 \| (4, 204) \| 205 \| (6, 210) \| 217 \| Phage tail protein I \| Phage tail protein I; uniclust \| UniRef100\_A0A178HP38 \| 99.9 \| 1.1e-27 \| 2.3e-33 \| 201.6 \| 153 \| (3, 155) \| 205 \| (7, 180) \| 343 \| Phage tail protein I \| Phage tail protein I; uniclust \| UniRef100\_A0A7W7N2Q5 \| 99.9 \| 1.5e-27 \| 2.7e-33 \| 189.7 \| 163 \| (2, 165) \| 205 \| (10, 175) \| 332 \| Phage tail P2-like protein \| Phage tail P2-like protein; uniclust \| UniRef100\_A0A376J7K4 \| 99.9 \| 1.7e-27 \| 3.2e-33 \| 167.5 \| 115 \| (42, 158) \| 205 \| (2, 116) \| 124 \| Putative tail protein I (Gpi) \| Putative tail protein I (Gpi); uniclust \| UniRef100\_A0A1S1MZR1 \| 99.9 \| 1.8e-27 \| 3.4e-33 \| 192.2 \| 149 \| (2, 153) \| 205 \| (8, 163) \| 313 \| Phage tail protein I \| Phage tail protein I; uniclust \| UniRef100\_UPI00235456C8 \| 99.9 \| 1.9e-27 \| 3.6e-33 \| 192.1 \| 142 \| (2, 150) \| 205 \| (6, 148) \| 384 \| phage tail protein I \| phage tail protein I; uniclust \| UniRef100\_A0A0A8RA06 \| 99.9 \| 3.1e-27 \| 5.9e-33 \| 213.4 \| 150 \| (1, 150) \| 205 \| (1, 150) \| 970 \| Putative bacteriophage protein \| Putative bacteriophage protein; uniclust \| UniRef100\_A0A269PJC4 \| 99.9 \| 3.8e-27 \| 6.9e-33 \| 174.0 \| 149 \| (29, 177) \| 205 \| (35, 183) \| 186 \| Phage tail protein I \| Phage tail protein I; uniclust \| UniRef100\_A0A1D2QMU0 \| 99.9 \| 4e-27 \| 8e-33 \| 174.6 \| 101 \| (1, 101) \| 205 \| (3, 104) \| 151 \| Phage tail protein I \| Phage tail protein I; uniclust \| UniRef100\_A0A0L8A9U9 \| 99.9 \| 5.9e-27 \| 1.1e-32 \| 194.0 \| 151 \| (2, 154) \| 205 \| (256, 409) \| 433 \| Tail protein \| Tail protein; uniclust \| UniRef100\_UPI0020352A08 \| 99.9 \| 7.7e-27 \| 1.4e-32 \| 191.2 \| 151 \| (2, 153) \| 205 \| (7, 159) \| 434 \| phage tail protein I \| phage tail protein I; uniclust \| UniRef100\_A0A059UZC6 \| 99.9 \| 7.6e-27 \| 1.6e-32 \| 198.8 \| 149 \| (3, 152) \| 205 \| (6, 195) \| 385 \| Tail protein I \| Tail protein I; uniclust \| UniRef100\_UPI0022A77364 \| 99.9 \| 1.9e-26 \| 3.4e-32 \| 172.1 \| 179 \| (24, 203) \| 205 \| (4, 183) \| 199 \| phage tail protein I \| phage tail protein I; uniclust \| UniRef100\_A0A5C7PF23 \| 99.9 \| 2.4e-26 \| 5e-32 \| 181.9 \| 149 \| (2, 153) \| 205 \| (12, 185) \| 211 \| Phage tail protein I \| Phage tail protein I; uniclust \| UniRef100\_A0A1X7L2M2 \| 99.9 \| 2.8e-26 \| 5.5e-32 \| 196.5 \| 146 \| (1, 147) \| 205 \| (8, 189) \| 496 \| Phage tail protein, P2 protein I family \| Phage tail protein, P2 protein I family; uniclust \| UniRef100\_A0A6L7TCB7 \| 99.9 \| 3e-26 \| 5.5e-32 \| 178.2 \| 146 \| (2, 153) \| 205 \| (101, 246) \| 272 \| Phage tail protein I \| Phage tail protein I; uniclust \| UniRef100\_A0A239CJC5 \| 99.9 \| 3.2e-26 \| 5.9e-32 \| 182.3 \| 152 \| (2, 154) \| 205 \| (1, 154) \| 332 \| Phage tail protein, P2 protein I family \| Phage tail protein, P2 protein I family; uniclust \| UniRef100\_A0A415HAB7 \| 99.9 \| 3.7e-26 \| 7.5e-32 \| 186.0 \| 161 \| (1, 166) \| 205 \| (14, 180) \| 277 \| Phage tail protein I \| Phage tail protein I; uniclust \| UniRef100\_A0A090V3M0 \| 99.9 \| 3.7e-26 \| 7.6e-32 \| 214.4 \| 156 \| (2, 159) \| 205 \| (3, 158) \| 1137 \| Tail protein \| Tail protein; uniclust \| UniRef100\_UPI0020300C06 \| 99.9 \| 5.6e-26 \| 1e-31 \| 163.5 \| 119 \| (2, 120) \| 205 \| (4, 122) \| 154 \| phage tail protein I \| phage tail protein I; uniclust \| UniRef100\_A0A0Q0ZJE5 \| 99.9 \| 5.5e-26 \| 1.1e-31 \| 182.3 \| 150 \| (1, 154) \| 205 \| (11, 167) \| 225 \| Phage tail protein \| Phage tail protein; uniclust \| UniRef100\_A0A0B7J8H6 \| 99.9 \| 5.9e-26 \| 1.2e-31 \| 185.2 \| 163 \| (2, 165) \| 205 \| (18, 181) \| 258 \| Phage tail protein I \| Phage tail protein I; uniclust \| UniRef100\_UPI0009E5A335 \| 99.9 \| 6.8e-26 \| 1.2e-31 \| 173.2 \| 151 \| (2, 153) \| 205 \| (5, 162) \| 237 \| phage tail protein I \| phage tail protein I; uniclust \| UniRef100\_A0A0F4QK22 \| 99.9 \| 7.4e-26 \| 1.5e-31 \| 190.0 \| 124 \| (28, 154) \| 205 \| (126, 260) \| 376 \| Phage tail protein I \| Phage tail protein I; uniclust \| UniRef100\_UPI0012EB4EFA \| 99.9 \| 8.3e-26 \| 1.5e-31 \| 185.7 \| 158 \| (2, 159) \| 205 \| (4, 162) \| 444 \| phage tail protein I \| phage tail protein I; uniclust \| UniRef100\_A0A1Y3DKT1 \| 99.9 \| 9.6e-26 \| 1.8e-31 \| 188.0 \| 174 \| (1, 176) \| 205 \| (1, 174) \| 434 \| Phage tail protein I \| Phage tail protein I; uniclust \| UniRef100\_A0A8S0FTH7 \| 99.9 \| 1.3e-25 \| 2.4e-31 \| 175.7 \| 144 \| (2, 145) \| 205 \| (138, 282) \| 285 \| Baseplate protein J-like domain-containing protein \| Baseplate protein J-like domain-containing protein; uniclust \| UniRef100\_A0A934JMB0 \| 99.9 \| 1.6e-25 \| 3.1e-31 \| 173.8 \| 193 \| (2, 195) \| 205 \| (13, 206) \| 220 \| Phage tail protein I \| Phage tail protein I; uniclust \| UniRef100\_A0A0E4G100 \| 99.9 \| 1.6e-25 \| 3.6e-31 \| 199.1 \| 95 \| (1, 95) \| 205 \| (18, 112) \| 426 \| Phage tail protein I \| Phage tail protein I; uniclust \| UniRef100\_UPI001B39611C \| 99.9 \| 2.2e-25 \| 4e-31 \| 178.3 \| 163 \| (2, 165) \| 205 \| (5, 174) \| 342 \| phage tail protein I \| phage tail protein I; uniclust \| UniRef100\_A0A1G7SDN3 \| 99.9 \| 3.4e-25 \| 6.2e-31 \| 169.6 \| 149 \| (2, 153) \| 205 \| (54, 218) \| 238 \| Phage tail protein, P2 protein I family \| Phage tail protein, P2 protein I family; uniclust \| UniRef100\_A0A348HHI4 \| 99.9 \| 4.5e-25 \| 8.3e-31 \| 171.3 \| 201 \| (2, 204) \| 205 \| (4, 206) \| 266 \| Bacteriophage P2-related tail formation protein \| Bacteriophage P2-related tail formation protein; uniclust \| UniRef100\_UPI0013D775D4 \| 99.9 \| 4.5e-25 \| 8.3e-31 \| 167.9 \| 148 \| (2, 155) \| 205 \| (3, 151) \| 227 \| phage tail protein I \| phage tail protein I; uniclust \| UniRef100\_A0A9E6RJL8 \| 99.9 \| 4.5e-25 \| 8.4e-31 \| 167.1 \| 198 \| (5, 202) \| 205 \| (6, 213) \| 219 \| Phage tail protein I \| Phage tail protein I; uniclust \| UniRef100\_A0A8T7D0V5 \| 99.8 \| 6.4e-25 \| 1.2e-30 \| 175.1 \| 159 \| (2, 161) \| 205 \| (3, 174) \| 332 \| Phage tail protein I \| Phage tail protein I; uniclust \| UniRef100\_UPI000F04D2C2 \| 99.8 \| 7.1e-25 \| 1.3e-30 \| 149.5 \| 109 \| (33, 141) \| 205 \| (1, 109) \| 109 \| phage tail protein I \| phage tail protein I; uniclust \| UniRef100\_A0A022PNX6 \| 99.8 \| 6.9e-25 \| 1.4e-30 \| 175.0 \| 160 \| (4, 168) \| 205 \| (24, 203) \| 219 \| Phage tail protein, P2 protein I family \| Phage tail protein, P2 protein I family; uniclust \| UniRef100\_UPI002264A9C4 \| 99.8 \| 9.1e-25 \| 1.7e-30 \| 152.0 \| 114 \| (2, 115) \| 205 \| (4, 117) \| 123 \| phage tail protein I \| phage tail protein I; uniclust \| UniRef100\_UPI0013A68137 \| 99.8 \| 9.2e-25 \| 1.7e-30 \| 167.0 \| 165 \| (23, 188) \| 205 \| (23, 190) \| 207 \| phage tail protein I \| phage tail protein I; uniclust \| UniRef100\_I3TTD2 \| 99.8 \| 1.2e-24 \| 2.2e-30 \| 170.3 \| 164 \| (2, 166) \| 205 \| (6, 175) \| 282 \| Gp15 protein \| Gp15 protein; uniclust \| UniRef100\_A0A541BHK6 \| 99.8 \| 1.3e-24 \| 2.5e-30 \| 173.9 \| 163 \| (3, 166) \| 205 \| (23, 196) \| 297 \| Phage tail protein I \| Phage tail protein I; uniclust \| UniRef100\_A0A3S4GL93 \| 99.8 \| 1.6e-24 \| 3e-30 \| 156.6 \| 104 \| (2, 105) \| 205 \| (16, 119) \| 147 \| Bacteriophage P2-related tail formation protein \| Bacteriophage P2-related tail formation protein; uniclust \| UniRef100\_UPI00069850DB \| 99.8 \| 1.6e-24 \| 3e-30 \| 173.7 \| 151 \| (2, 153) \| 205 \| (16, 202) \| 314 \| phage tail protein I \| phage tail protein I; uniclust \| UniRef100\_UPI000D57C2B6 \| 99.8 \| 1.7e-24 \| 3e-30 \| 157.4 \| 158 \| (44, 203) \| 205 \| (2, 163) \| 163 \| phage tail protein I \| phage tail protein I; uniclust \| UniRef100\_A0A7U9H2I1 \| 99.8 \| 1.7e-24 \| 3.2e-30 \| 169.4 \| 191 \| (12, 204) \| 205 \| (21, 217) \| 281 \| Phage tail protein I \| Phage tail protein I; uniclust \| UniRef100\_A0A0W8JFZ8 \| 99.8 \| 1.6e-24 \| 3.2e-30 \| 175.5 \| 163 \| (2, 165) \| 205 \| (29, 193) \| 270 \| Phage tail protein I \| Phage tail protein I; uniclust \| UniRef100\_A0A266LLA0 \| 99.8 \| 1.8e-24 \| 3.4e-30 \| 147.7 \| 90 \| (2, 91) \| 205 \| (4, 93) \| 95 \| Phage tail protein I (Fragment) \| Phage tail protein I (Fragment); uniclust \| UniRef100\_A0A927WW89 \| 99.8 \| 2e-24 \| 4e-30 \| 187.9 \| 174 \| (3, 180) \| 205 \| (12, 195) \| 583 \| Phage tail protein I \| Phage tail protein I; uniclust \| UniRef100\_A0A174T2Y7 \| 99.8 \| 2.3e-24 \| 4.5e-30 \| 170.1 \| 170 \| (2, 175) \| 205 \| (12, 187) \| 226 \| Bacteriophage P2-related tail formation protein \| Bacteriophage P2-related tail formation protein; uniclust \| UniRef100\_A0A1N7LRG2 \| 99.8 \| 2.8e-24 \| 5.3e-30 \| 163.3 \| 153 \| (2, 154) \| 205 \| (6, 160) \| 209 \| Phage tail protein, P2 protein I family \| Phage tail protein, P2 protein I family; uniclust \| UniRef100\_A0A173SBX4 \| 99.8 \| 3.2e-24 \| 6.4e-30 \| 185.6 \| 149 \| (2, 153) \| 205 \| (25, 179) \| 457 \| Bacteriophage P2-related tail formation protein \| Bacteriophage P2-related tail formation protein; uniclust \| UniRef100\_A0A428VMX7 \| 99.8 \| 3.4e-24 \| 6.5e-30 \| 180.3 \| 161 \| (2, 169) \| 205 \| (288, 465) \| 471 \| Phage tail protein I \| Phage tail protein I; uniclust \| UniRef100\_A0A2S7JR38 \| 99.8 \| 4e-24 \| 7.8e-30 \| 169.7 \| 152 \| (2, 154) \| 205 \| (7, 172) \| 249 \| Phage tail protein I \| Phage tail protein I; uniclust \| UniRef100\_A0A2D1CSN0 \| 99.8 \| 4.1e-24 \| 7.8e-30 \| 188.2 \| 142 \| (3, 144) \| 205 \| (5, 146) \| 733 \| Putative tail fiber protein \| Putative tail fiber protein; uniclust \| UniRef100\_A0A1E3G6D5 \| 99.8 \| 4.3e-24 \| 7.9e-30 \| 160.5 \| 162 \| (1, 164) \| 205 \| (1, 162) \| 205 \| Phage tail protein I \| Phage tail protein I; uniclust \| UniRef100\_A0A318KN12 \| 99.8 \| 6.9e-24 \| 1.3e-29 \| 170.9 \| 147 \| (3, 150) \| 205 \| (5, 152) \| 358 \| Phage tail P2-like protein (Fragment) \| Phage tail P2-like protein (Fragment); uniclust \| UniRef100\_A0A067ZG50 \| 99.8 \| 9.5e-24 \| 1.8e-29 \| 159.8 \| 172 \| (2, 175) \| 205 \| (10, 191) \| 192 \| Tail protein I \| Tail protein I; uniclust \| UniRef100\_A0A064AHQ3 \| 99.8 \| 2.3e-23 \| 4.6e-29 \| 165.0 \| 158 \| (3, 164) \| 205 \| (13, 178) \| 218 \| Phage tail protein \| Phage tail protein; uniclust \| UniRef100\_UPI001F0C5BA4 \| 99.8 \| 2.6e-23 \| 4.8e-29 \| 147.8 \| 104 \| (2, 105) \| 205 \| (1, 104) \| 139 \| phage tail protein I \| phage tail protein I; uniclust \| UniRef100\_A0A150MJA7 \| 99.8 \| 2.5e-23 \| 4.8e-29 \| 169.4 \| 149 \| (2, 154) \| 205 \| (10, 161) \| 298 \| Phage tail protein I \| Phage tail protein I; uniclust \| UniRef100\_A0A2D3WQP3 \| 99.8 \| 2.7e-23 \| 5.5e-29 \| 162.5 \| 150 \| (1, 153) \| 205 \| (5, 177) \| 201 \| Phage tail protein I \| Phage tail protein I; uniclust \| UniRef100\_UPI001D0F882A \| 99.8 \| 3.2e-23 \| 5.8e-29 \| 176.1 \| 152 \| (2, 154) \| 205 \| (5, 159) \| 603 \| phage tail protein I \| phage tail protein I; uniclust \| UniRef100\_A0A1T4WXE7 \| 99.8 \| 3.4e-23 \| 6.2e-29 \| 160.3 \| 154 \| (2, 157) \| 205 \| (6, 168) \| 253 \| Phage tail protein, P2 protein I family \| Phage tail protein, P2 protein I family; uniclust \| UniRef100\_A0A2B2B8I7 \| 99.8 \| 3.5e-23 \| 6.4e-29 \| 136.9 \| 85 \| (34, 120) \| 205 \| (1, 85) \| 90 \| Phage tail protein I (Fragment) \| Phage tail protein I (Fragment); uniclust \| UniRef100\_A0A812QW15 \| 99.8 \| 4e-23 \| 7.4e-29 \| 183.2 \| 146 \| (13, 159) \| 205 \| (289, 442) \| 1062 \| J protein \| J protein; uniclust \| UniRef100\_UPI000AD33DF5 \| 99.8 \| 4.4e-23 \| 8.2e-29 \| 147.9 \| 127 \| (47, 174) \| 205 \| (1, 129) \| 135 \| phage tail protein \| phage tail protein; uniclust \| UniRef100\_A0A016QLD8 \| 99.8 \| 4.2e-23 \| 8.8e-29 \| 180.6 \| 118 \| (29, 155) \| 205 \| (251, 370) \| 429 \| FHA domain-containing protein \| FHA domain-containing protein; uniclust \| UniRef100\_A0A1W9USS5 \| 99.8 \| 6.2e-23 \| 1.3e-28 \| 178.1 \| 124 \| (30, 155) \| 205 \| (213, 362) \| 397 \| Phage tail protein I \| Phage tail protein I; uniclust \| UniRef100\_A0A5Q2UG14 \| 99.8 \| 7.9e-23 \| 1.5e-28 \| 171.7 \| 152 \| (2, 154) \| 205 \| (7, 164) \| 428 \| Phage tail protein I \| Phage tail protein I; uniclust \| UniRef100\_A0A3C1WTS9 \| 99.8 \| 8.8e-23 \| 1.7e-28 \| 160.2 \| 187 \| (3, 194) \| 205 \| (18, 208) \| 247 \| Phage tail protein I \| Phage tail protein I; uniclust \| UniRef100\_A0A0A8JLX9 \| 99.8 \| 8.7e-23 \| 1.7e-28 \| 167.9 \| 148 \| (3, 153) \| 205 \| (11, 164) \| 348 \| Phage tail protein \| Phage tail protein; uniclust \| UniRef100\_A0A1H8VRL6 \| 99.8 \| 1.2e-22 \| 2.3e-28 \| 163.9 \| 147 \| (2, 153) \| 205 \| (39, 191) \| 315 \| Phage tail protein, P2 protein I family \| Phage tail protein, P2 protein I family; uniclust \| UniRef100\_A0A418VVG6 \| 99.8 \| 1.2e-22 \| 2.3e-28 \| 170.1 \| 152 \| (1, 152) \| 205 \| (1, 169) \| 515 \| Phage tail protein I \| Phage tail protein I; uniclust \| UniRef100\_A0A812RFY7 \| 99.8 \| 1.3e-22 \| 2.4e-28 \| 176.0 \| 150 \| (1, 153) \| 205 \| (266, 431) \| 772 \| J protein \| J protein; uniclust \| UniRef100\_A0A069Q1X1 \| 99.8 \| 1.5e-22 \| 3.2e-28 \| 170.0 \| 122 \| (29, 152) \| 205 \| (101, 240) \| 300 \| Uncharacterized protein \| Uncharacterized protein; uniclust \| UniRef100\_UPI0018DE8610 \| 99.8 \| 1.8e-22 \| 3.3e-28 \| 142.3 \| 130 \| (43, 172) \| 205 \| (1, 130) \| 131 \| phage tail protein I \| phage tail protein I; uniclust \| UniRef100\_UPI0005BBF981 \| 99.8 \| 2e-22 \| 3.8e-28 \| 140.7 \| 88 \| (33, 120) \| 205 \| (2, 89) \| 111 \| phage tail protein I \| phage tail protein I; uniclust \| UniRef100\_A0A1E7J923 \| 99.8 \| 1.8e-22 \| 3.8e-28 \| 174.1 \| 124 \| (29, 154) \| 205 \| (145, 288) \| 392 \| Phage tail protein I \| Phage tail protein I; uniclust \| UniRef100\_A0A966HEM8 \| 99.8 \| 2e-22 \| 3.8e-28 \| 164.4 \| 147 \| (3, 153) \| 205 \| (12, 165) \| 336 \| Phage tail protein I \| Phage tail protein I; uniclust \| UniRef100\_A0A0L6JHZ7 \| 99.8 \| 1.8e-22 \| 3.9e-28 \| 178.6 \| 172 \| (12, 189) \| 205 \| (211, 401) \| 442 \| Phage tail protein \| Phage tail protein; uniclust \| UniRef100\_A0A1Y1QXH2 \| 99.8 \| 2.3e-22 \| 4.3e-28 \| 154.5 \| 149 \| (2, 152) \| 205 \| (5, 155) \| 237 \| Phage tail protein I \| Phage tail protein I; uniclust \| UniRef100\_A0A376MUF5 \| 99.8 \| 3e-22 \| 5.4e-28 \| 143.2 \| 94 \| (32, 127) \| 205 \| (43, 136) \| 143 \| Putative tail protein I (Gpi) \| Putative tail protein I (Gpi); uniclust \| UniRef100\_A0A2X2DI53 \| 99.8 \| 2.8e-22 \| 5.5e-28 \| 144.8 \| 118 \| (85, 204) \| 205 \| (1, 118) \| 124 \| Phage tail protein \| Phage tail protein; uniclust \| UniRef100\_A0A521CHZ6 \| 99.8 \| 3.8e-22 \| 7.3e-28 \| 160.1 \| 150 \| (2, 153) \| 205 \| (4, 164) \| 287 \| Phage tail protein, P2 protein I family \| Phage tail protein, P2 protein I family; uniclust \| UniRef100\_UPI002244F0CF \| 99.8 \| 5.9e-22 \| 1.1e-27 \| 132.0 \| 87 \| (35, 121) \| 205 \| (2, 88) \| 93 \| phage tail protein I \| phage tail protein I; uniclust \| UniRef100\_UPI001CF0BE9A \| 99.8 \| 6.8e-22 \| 1.2e-27 \| 159.4 \| 161 \| (2, 169) \| 205 \| (167, 344) \| 350 \| phage tail protein I \| phage tail protein I; uniclust \| UniRef100\_A0A845R5N3 \| 99.8 \| 9.5e-22 \| 1.8e-27 \| 157.0 \| 197 \| (2, 200) \| 205 \| (10, 213) \| 250 \| Phage tail protein I \| Phage tail protein I; uniclust \| UniRef100\_A0A1C5WN41 \| 99.8 \| 9.4e-22 \| 1.9e-27 \| 164.9 \| 171 \| (3, 176) \| 205 \| (47, 222) \| 345 \| Bacteriophage P2-related tail formation protein \| Bacteriophage P2-related tail formation protein; uniclust \| UniRef100\_UPI00210D76C9 \| 99.8 \| 1.2e-21 \| 2.2e-27 \| 163.0 \| 152 \| (2, 155) \| 205 \| (14, 184) \| 471 \| phage tail protein I \| phage tail protein I; uniclust \| UniRef100\_A0A074LMF0 \| 99.8 \| 1.1e-21 \| 2.4e-27 \| 183.6 \| 158 \| (12, 175) \| 205 \| (496, 672) \| 713 \| Phage tail protein \| Phage tail protein; uniclust \| UniRef100\_A0A2J4JSF3 \| 99.7 \| 2.1e-21 \| 4e-27 \| 155.4 \| 160 \| (3, 164) \| 205 \| (11, 178) \| 255 \| Phage tail protein I \| Phage tail protein I; uniclust \| UniRef100\_UPI000DFB77B5 \| 99.7 \| 2.3e-21 \| 4.3e-27 \| 140.6 \| 128 \| (44, 172) \| 205 \| (10, 139) \| 147 \| phage tail protein I \| phage tail protein I; uniclust \| UniRef100\_A0A0C1IR17 \| 99.7 \| 2.1e-21 \| 4.3e-27 \| 158.1 \| 187 \| (2, 189) \| 205 \| (11, 206) \| 234 \| Phage tail protein I \| Phage tail protein I; uniclust \| UniRef100\_A0A1W2DZK8 \| 99.7 \| 2.9e-21 \| 5.7e-27 \| 154.3 \| 201 \| (3, 204) \| 205 \| (29, 238) \| 243 \| Phage tail protein, P2 protein I family \| Phage tail protein, P2 protein I family; uniclust \| UniRef100\_A0A847XBJ9 \| 99.7 \| 3.1e-21 \| 5.9e-27 \| 144.5 \| 132 \| (39, 173) \| 205 \| (1, 132) \| 173 \| Phage tail protein I \| Phage tail protein I; uniclust \| UniRef100\_A0A126QLJ0 \| 99.7 \| 2.9e-21 \| 6e-27 \| 158.9 \| 191 \| (13, 205) \| 205 \| (33, 224) \| 257 \| Uncharacterized protein \| Uncharacterized protein; uniclust \| UniRef100\_UPI0007DC4380 \| 99.7 \| 3.4e-21 \| 6.2e-27 \| 141.4 \| 148 \| (3, 152) \| 205 \| (4, 157) \| 168 \| phage tail protein I \| phage tail protein I; uniclust \| UniRef100\_A0A1W9VL85 \| 99.7 \| 3.1e-21 \| 6.3e-27 \| 165.2 \| 124 \| (30, 155) \| 205 \| (202, 351) \| 384 \| Phage tail protein I \| Phage tail protein I; uniclust \| UniRef100\_A0A023PZC5 \| 99.7 \| 3.5e-21 \| 6.7e-27 \| 155.0 \| 125 \| (28, 153) \| 205 \| (124, 257) \| 307 \| Tail protein I \| Tail protein I; uniclust \| UniRef100\_A0A080M1T9 \| 99.7 \| 3.2e-21 \| 6.7e-27 \| 158.2 \| 123 \| (29, 153) \| 205 \| (67, 198) \| 249 \| Phage tail protein domain protein \| Phage tail protein domain protein; uniclust \| UniRef100\_A0A6G5QFK7 \| 99.7 \| 3.5e-21 \| 7.3e-27 \| 152.7 \| 154 \| (45, 202) \| 205 \| (42, 195) \| 204 \| Phage P2 family tail protein \| Phage P2 family tail protein; uniclust \| UniRef100\_A0A6L3X1R5 \| 99.7 \| 4.9e-21 \| 9.6e-27 \| 132.8 \| 85 \| (2, 86) \| 205 \| (6, 92) \| 96 \| Phage tail protein I (Fragment) \| Phage tail protein I (Fragment); uniclust \| UniRef100\_A0A315X9K7 \| 99.7 \| 5.1e-21 \| 1e-26 \| 150.9 \| 121 \| (29, 153) \| 205 \| (73, 197) \| 212 \| Phage tail protein \| Phage tail protein; uniclust \| UniRef100\_UPI000471CB6A \| 99.7 \| 5.5e-21 \| 1e-26 \| 163.7 \| 150 \| (2, 152) \| 205 \| (7, 162) \| 530 \| phage tail protein I \| phage tail protein I; uniclust \| UniRef100\_A0A2P8QYP1 \| 99.7 \| 6.1e-21 \| 1.1e-26 \| 150.2 \| 186 \| (1, 192) \| 205 \| (1, 188) \| 282 \| Phage tail protein I \| Phage tail protein I; uniclust \| UniRef100\_UPI0013D51FFC \| 99.7 \| 6.2e-21 \| 1.1e-26 \| 147.0 \| 143 \| (30, 176) \| 205 \| (46, 188) \| 238 \| phage tail protein I \| phage tail protein I; uniclust \| UniRef100\_A0A973ALZ7 \| 99.7 \| 6.9e-21 \| 1.3e-26 \| 155.3 \| 152 \| (2, 153) \| 205 \| (3, 172) \| 383 \| Phage tail protein I \| Phage tail protein I; uniclust \| UniRef100\_UPI001E610C52 \| 99.7 \| 7.1e-21 \| 1.3e-26 \| 154.2 \| 145 \| (3, 151) \| 205 \| (12, 162) \| 360 \| phage tail protein I \| phage tail protein I; uniclust \| UniRef100\_A0A838RLH5 \| 99.7 \| 9.3e-21 \| 1.8e-26 \| 144.1 \| 121 \| (30, 154) \| 205 \| (21, 158) \| 176 \| Phage tail protein I \| Phage tail protein I; uniclust \| UniRef100\_A0A089JZ90 \| 99.7 \| 9.1e-21 \| 1.8e-26 \| 156.5 \| 122 \| (29, 153) \| 205 \| (64, 207) \| 289 \| Tail protein \| Tail protein; uniclust \| UniRef100\_UPI001F490D58 \| 99.7 \| 1.1e-20 \| 2.1e-26 \| 133.6 \| 118 \| (2, 119) \| 205 \| (6, 124) \| 131 \| phage tail protein I \| phage tail protein I; uniclust \| UniRef100\_A0A1X3J0Q7 \| 99.7 \| 1.1e-20 \| 2.1e-26 \| 131.8 \| 104 \| (71, 176) \| 205 \| (2, 105) \| 108 \| Tail protein I (GpI) \| Tail protein I (GpI); uniclust \| UniRef100\_UPI000E3DDFBE \| 99.7 \| 1.9e-20 \| 3.4e-26 \| 123.7 \| 82 \| (30, 111) \| 205 \| (6, 87) \| 87 \| phage tail protein I \| phage tail protein I; uniclust \| UniRef100\_A0A0H3ZYQ7 \| 99.7 \| 1.9e-20 \| 3.5e-26 \| 145.6 \| 194 \| (2, 196) \| 205 \| (9, 211) \| 220 \| Tail protein I \| Tail protein I; uniclust \| UniRef100\_UPI000BA1955F \| 99.7 \| 2.1e-20 \| 3.9e-26 \| 129.3 \| 97 \| (9, 105) \| 205 \| (5, 101) \| 106 \| phage tail protein I \| phage tail protein I; uniclust \| UniRef100\_A0A6N9P8X1 \| 99.7 \| 2.5e-20 \| 4.7e-26 \| 153.8 \| 152 \| (4, 158) \| 205 \| (16, 174) \| 357 \| Phage tail protein I \| Phage tail protein I; uniclust \| UniRef100\_F4BFR7 \| 99.7 \| 4.3e-20 \| 8e-26 \| 140.8 \| 165 \| (1, 171) \| 205 \| (1, 167) \| 195 \| Phage tail protein \| Phage tail protein; uniclust \| UniRef100\_A0A968XWL3 \| 99.7 \| 5e-20 \| 9.2e-26 \| 125.3 \| 102 \| (2, 103) \| 205 \| (1, 103) \| 103 \| Phage tail protein I \| Phage tail protein I; uniclust \| UniRef100\_A0A3A6E7Q8 \| 99.7 \| 5.8e-20 \| 1.1e-25 \| 150.3 \| 148 \| (3, 153) \| 205 \| (14, 168) \| 339 \| Phage tail protein I \| Phage tail protein I; uniclust \| UniRef100\_A0A1B9NSL6 \| 99.7 \| 6.3e-20 \| 1.2e-25 \| 133.3 \| 128 \| (75, 204) \| 205 \| (1, 128) \| 155 \| Phage tail protein I \| Phage tail protein I; uniclust \| UniRef100\_A0A6N7JGU4 \| 99.7 \| 6.5e-20 \| 1.2e-25 \| 154.2 \| 155 \| (2, 156) \| 205 \| (3, 170) \| 507 \| Phage tail protein I \| Phage tail protein I; uniclust \| UniRef100\_I7LG06 \| 99.7 \| 1.5e-19 \| 2.7e-25 \| 143.5 \| 147 \| (3, 153) \| 205 \| (11, 164) \| 295 \| Phage tail protein I \| Phage tail protein I; uniclust \| UniRef100\_A0A2A5JYU3 \| 99.7 \| 1.5e-19 \| 2.9e-25 \| 130.0 \| 130 \| (42, 175) \| 205 \| (7, 136) \| 138 \| Phage tail protein I \| Phage tail protein I; uniclust \| UniRef100\_UPI0009718AD0 \| 99.7 \| 1.7e-19 \| 3.2e-25 \| 129.0 \| 103 \| (3, 105) \| 205 \| (6, 109) \| 139 \| phage tail protein I \| phage tail protein I; uniclust \| UniRef100\_UPI00197B9417 \| 99.7 \| 1.8e-19 \| 3.3e-25 \| 139.8 \| 151 \| (2, 153) \| 205 \| (7, 167) \| 222 \| phage tail protein I \| phage tail protein I; uniclust \| UniRef100\_A0A1C6IIK7 \| 99.7 \| 1.8e-19 \| 3.6e-25 \| 151.1 \| 142 \| (10, 153) \| 205 \| (26, 173) \| 331 \| Bacteriophage P2-related tail formation protein \| Bacteriophage P2-related tail formation protein; uniclust \| UniRef100\_A0A0E3SE24 \| 99.7 \| 2e-19 \| 4.1e-25 \| 164.1 \| 158 \| (10, 174) \| 205 \| (447, 648) \| 709 \| NHL repeat domain protein \| NHL repeat domain protein; uniclust \| UniRef100\_A0A2S6N2X4 \| 99.6 \| 2.3e-19 \| 4.5e-25 \| 139.5 \| 195 \| (2, 204) \| 205 \| (9, 206) \| 207 \| Uncharacterized protein \| Uncharacterized protein; uniclust \| UniRef100\_A0A3D5LSM9 \| 99.6 \| 2.4e-19 \| 4.7e-25 \| 148.0 \| 158 \| (2, 164) \| 205 \| (18, 181) \| 352 \| Phage tail protein I \| Phage tail protein I; uniclust \| UniRef100\_A0A0C1MY76 \| 99.6 \| 2.3e-19 \| 4.7e-25 \| 158.6 \| 136 \| (13, 154) \| 205 \| (185, 349) \| 520 \| Phage tail protein \| Phage tail protein; uniclust \| UniRef100\_A0A0X8JTG7 \| 99.6 \| 2.6e-19 \| 5e-25 \| 140.4 \| 130 \| (32, 165) \| 205 \| (45, 174) \| 214 \| Phage tail protein \| Phage tail protein; uniclust \| UniRef100\_A0A0S2ZDH3 \| 99.6 \| 2.6e-19 \| 5.3e-25 \| 144.0 \| 143 \| (30, 178) \| 205 \| (49, 191) \| 225 \| Phage tail protein \| Phage tail protein; uniclust \| UniRef100\_A0A1G3UAH8 \| 99.6 \| 2.8e-19 \| 5.3e-25 \| 140.2 \| 145 \| (3, 153) \| 205 \| (2, 150) \| 245 \| Phage tail protein I \| Phage tail protein I; uniclust \| UniRef100\_A0A7X1ZFY9 \| 99.6 \| 3.1e-19 \| 5.7e-25 \| 134.7 \| 131 \| (73, 205) \| 205 \| (40, 170) \| 199 \| Phage tail protein I \| Phage tail protein I; uniclust \| UniRef100\_A0A6I2UKH8 \| 99.6 \| 3.2e-19 \| 5.8e-25 \| 142.0 \| 158 \| (31, 192) \| 205 \| (44, 201) \| 300 \| Phage tail protein I \| Phage tail protein I; uniclust \| UniRef100\_UPI001C7642B4 \| 99.6 \| 3.7e-19 \| 6.8e-25 \| 137.0 \| 111 \| (35, 150) \| 205 \| (48, 159) \| 230 \| phage tail protein I \| phage tail protein I; uniclust \| UniRef100\_A0A6N9P2M1 \| 99.6 \| 4.1e-19 \| 7.7e-25 \| 146.2 \| 140 \| (30, 172) \| 205 \| (49, 188) \| 344 \| Phage tail protein I \| Phage tail protein I; uniclust \| UniRef100\_A0A7U3VT62 \| 99.6 \| 4.4e-19 \| 8.2e-25 \| 149.5 \| 150 \| (1, 157) \| 205 \| (1, 162) \| 513 \| Fused protein containing phage-related tail protein GpI \| Fused protein containing phage-related tail protein GpI; uniclust \| UniRef100\_A0A073CHG0 \| 99.6 \| 4.5e-19 \| 9.1e-25 \| 153.3 \| 136 \| (13, 154) \| 205 \| (218, 382) \| 409 \| Phage tail protein \| Phage tail protein; uniclust \| UniRef100\_A0A268EA24 \| 99.6 \| 4.8e-19 \| 9.3e-25 \| 150.3 \| 146 \| (2, 153) \| 205 \| (16, 166) \| 402 \| Phage tail protein I \| Phage tail protein I; uniclust \| UniRef100\_A0A2Z5STI6 \| 99.6 \| 4.9e-19 \| 1e-24 \| 162.1 \| 167 \| (12, 185) \| 205 \| (483, 703) \| 733 \| Phage tail protein \| Phage tail protein; uniclust \| UniRef100\_A0A839IX64 \| 99.6 \| 5.7e-19 \| 1.1e-24 \| 139.6 \| 158 \| (2, 165) \| 205 \| (4, 162) \| 284 \| Phage tail protein I \| Phage tail protein I; uniclust \| UniRef100\_A0A318KXU1 \| 99.6 \| 6.4e-19 \| 1.2e-24 \| 143.1 \| 139 \| (13, 154) \| 205 \| (15, 158) \| 356 \| Phage tail P2-like protein \| Phage tail P2-like protein; uniclust \| UniRef100\_A0A928YGD5 \| 99.6 \| 6.8e-19 \| 1.3e-24 \| 136.7 \| 157 \| (3, 164) \| 205 \| (13, 175) \| 217 \| Phage tail protein I \| Phage tail protein I; uniclust \| UniRef100\_A0A1U7IY23 \| 99.6 \| 8.3e-19 \| 1.7e-24 \| 148.8 \| 119 \| (29, 153) \| 205 \| (68, 194) \| 360 \| Phage tail protein I \| Phage tail protein I; uniclust \| UniRef100\_A0A143XK95 \| 99.6 \| 1e-18 \| 2e-24 \| 145.1 \| 166 \| (2, 170) \| 205 \| (10, 182) \| 331 \| Phage tail protein (Tail\_P2\_I) \| Phage tail protein (Tail\_P2\_I); uniclust \| UniRef100\_A0A1C6EQC2 \| 99.6 \| 9.7e-19 \| 2e-24 \| 149.0 \| 151 \| (2, 154) \| 205 \| (10, 166) \| 337 \| Bacteriophage P2-related tail formation protein \| Bacteriophage P2-related tail formation protein; uniclust \| UniRef100\_A0A022MS75 \| 99.6 \| 9.8e-19 \| 2.1e-24 \| 148.9 \| 118 \| (29, 152) \| 205 \| (124, 253) \| 321 \| Tail protein \| Tail protein; uniclust \| UniRef100\_UPI0010454A05 \| 99.6 \| 1.3e-18 \| 2.4e-24 \| 137.8 \| 148 \| (3, 153) \| 205 \| (10, 170) \| 286 \| phage tail protein I \| phage tail protein I; uniclust \| UniRef100\_UPI00209B7E43 \| 99.6 \| 1.4e-18 \| 2.6e-24 \| 146.1 \| 146 \| (4, 150) \| 205 \| (5, 190) \| 491 \| phage tail protein I \| phage tail protein I; uniclust \| UniRef100\_A0A143HC79 \| 99.6 \| 1.5e-18 \| 2.8e-24 \| 135.4 \| 147 \| (3, 153) \| 205 \| (11, 160) \| 219 \| Phage tail protein I \| Phage tail protein I; uniclust \| UniRef100\_A0A4S2HBU2 \| 99.6 \| 1.6e-18 \| 2.9e-24 \| 138.3 \| 178 \| (4, 183) \| 205 \| (12, 196) \| 304 \| Phage tail protein I \| Phage tail protein I; uniclust \| UniRef100\_A0A3C2C7T1 \| 99.6 \| 1.7e-18 \| 3.3e-24 \| 144.5 \| 156 \| (12, 173) \| 205 \| (106, 293) \| 333 \| Phage tail protein (Fragment) \| Phage tail protein (Fragment); uniclust \| UniRef100\_UPI001CB6F6EA \| 99.6 \| 1.9e-18 \| 3.4e-24 \| 122.9 \| 92 \| (29, 120) \| 205 \| (34, 125) \| 132 \| phage tail protein I \| phage tail protein I; uniclust \| UniRef100\_A0A1H2SW83 \| 99.6 \| 2e-18 \| 3.6e-24 \| 121.6 \| 86 \| (31, 118) \| 205 \| (29, 114) \| 124 \| Phage tail protein, P2 protein I family \| Phage tail protein, P2 protein I family; uniclust \| UniRef100\_UPI0020A62330 \| 99.6 \| 2.3e-18 \| 4.2e-24 \| 125.9 \| 153 \| (47, 201) \| 205 \| (1, 154) \| 158 \| phage tail protein I \| phage tail protein I; uniclust \| UniRef100\_A0A844Q9W2 \| 99.6 \| 2.9e-18 \| 5.3e-24 \| 133.3 \| 195 \| (3, 198) \| 205 \| (32, 237) \| 245 \| Phage tail protein I \| Phage tail protein I; uniclust \| UniRef100\_A0A1C3HHS1 \| 99.6 \| 2.9e-18 \| 5.5e-24 \| 133.7 \| 136 \| (14, 153) \| 205 \| (31, 180) \| 216 \| Phage tail protein (Tail\_P2\_I) \| Phage tail protein (Tail\_P2\_I); uniclust \| UniRef100\_A0A0D6KHP6 \| 99.6 \| 2.8e-18 \| 5.5e-24 \| 151.9 \| 119 \| (29, 153) \| 205 \| (68, 194) \| 535 \| Putative bacteriophage tail protein \| Putative bacteriophage tail protein; uniclust \| UniRef100\_A0A5A8F1G5 \| 99.6 \| 3.9e-18 \| 7.2e-24 \| 135.3 \| 141 \| (8, 153) \| 205 \| (15, 156) \| 287 \| Phage tail protein I \| Phage tail protein I; uniclust \| UniRef100\_A0A136KPW8 \| 99.6 \| 3.6e-18 \| 7.3e-24 \| 146.0 \| 123 \| (28, 154) \| 205 \| (236, 375) \| 381 \| Phage tail protein \| Phage tail protein; uniclust \| UniRef100\_UPI0018A3C604 \| 99.6 \| 3.9e-18 \| 7.3e-24 \| 133.1 \| 170 \| (3, 176) \| 205 \| (7, 179) \| 236 \| phage tail protein I \| phage tail protein I; uniclust \| UniRef100\_UPI000887A604 \| 99.6 \| 4e-18 \| 7.4e-24 \| 127.1 \| 90 \| (1, 92) \| 205 \| (1, 90) \| 180 \| phage tail protein I \| phage tail protein I; uniclust \| UniRef100\_A0A6G5Y431 \| 99.6 \| 4.2e-18 \| 7.7e-24 \| 128.1 \| 121 \| (31, 153) \| 205 \| (46, 169) \| 191 \| Phage tail protein I \| Phage tail protein I; uniclust \| UniRef100\_A0A3A9DQT7 \| 99.6 \| 4.4e-18 \| 8.2e-24 \| 125.6 \| 120 \| (30, 151) \| 205 \| (46, 165) \| 168 \| Phage tail protein I (Fragment) \| Phage tail protein I (Fragment); uniclust \| UniRef100\_A0A2X0WX70 \| 99.6 \| 6.7e-18 \| 1.2e-23 \| 133.3 \| 165 \| (1, 166) \| 205 \| (10, 182) \| 275 \| Bacteriophage P2-related tail formation protein \| Bacteriophage P2-related tail formation protein; uniclust \| UniRef100\_A0A174TLI5 \| 99.6 \| 9.8e-18 \| 1.8e-23 \| 138.1 \| 148 \| (14, 164) \| 205 \| (50, 201) \| 366 \| Bacteriophage P2-related tail formation protein \| Bacteriophage P2-related tail formation protein; uniclust \| UniRef100\_A0A072YA28 \| 99.6 \| 8.8e-18 \| 1.9e-23 \| 138.7 \| 159 \| (33, 205) \| 205 \| (63, 225) \| 236 \| Phage protein \| Phage protein; uniclust \| UniRef100\_A0A1V0Q0B5 \| 99.5 \| 1.3e-17 \| 2.6e-23 \| 153.1 \| 164 \| (12, 181) \| 205 \| (473, 657) \| 690 \| NHL repeat domain protein \| NHL repeat domain protein; uniclust \| UniRef100\_UPI001BB07F4E \| 99.5 \| 1.4e-17 \| 2.6e-23 \| 143.3 \| 148 \| (3, 152) \| 205 \| (7, 164) \| 622 \| phage tail protein I \| phage tail protein I; uniclust \| UniRef100\_A0A927WKT3 \| 99.5 \| 1.8e-17 \| 3.3e-23 \| 133.7 \| 120 \| (35, 157) \| 205 \| (47, 166) \| 327 \| Phage tail protein I \| Phage tail protein I; uniclust \| UniRef100\_UPI00200D63BF \| 99.5 \| 2e-17 \| 3.7e-23 \| 114.6 \| 108 \| (53, 161) \| 205 \| (2, 109) \| 111 \| phage tail protein I \| phage tail protein I; uniclust \| UniRef100\_A0A6A0HXA4 \| 99.5 \| 2e-17 \| 4e-23 \| 149.8 \| 164 \| (12, 181) \| 205 \| (448, 664) \| 699 \| Phage tail protein \| Phage tail protein; uniclust \| UniRef100\_UPI001C0048D9 \| 99.5 \| 2.3e-17 \| 4.2e-23 \| 130.7 \| 120 \| (30, 153) \| 205 \| (66, 185) \| 280 \| phage tail protein I \| phage tail protein I; uniclust \| UniRef100\_A0A345DE45 \| 99.5 \| 2.5e-17 \| 4.6e-23 \| 125.6 \| 145 \| (2, 153) \| 205 \| (16, 182) \| 208 \| Phage tail protein I \| Phage tail protein I; uniclust \| UniRef100\_A0A1D9FX43 \| 99.5 \| 2.6e-17 \| 5.1e-23 \| 133.8 \| 122 \| (29, 153) \| 205 \| (56, 214) \| 261 \| Phage tail protein \| Phage tail protein; uniclust \| UniRef100\_UPI001CC0D908 \| 99.5 \| 3.1e-17 \| 5.8e-23 \| 132.9 \| 149 \| (2, 152) \| 205 \| (8, 157) \| 338 \| phage tail protein I \| phage tail protein I; uniclust \| UniRef100\_A0A0A8IL57 \| 99.5 \| 4.9e-17 \| 9.9e-23 \| 132.9 \| 96 \| (2, 101) \| 205 \| (37, 133) \| 248 \| Tail protein \| Tail protein; uniclust \| UniRef100\_UPI002020F9DF \| 99.5 \| 6.2e-17 \| 1.1e-22 \| 106.7 \| 75 \| (2, 76) \| 205 \| (7, 83) \| 83 \| phage tail protein I \| phage tail protein I; uniclust \| UniRef100\_UPI0018AC4BC9 \| 99.5 \| 6.5e-17 \| 1.2e-22 \| 118.7 \| 125 \| (73, 199) \| 205 \| (1, 126) \| 159 \| phage tail protein \| phage tail protein; uniclust \| UniRef100\_A0A1C4SGP1 \| 99.5 \| 7.6e-17 \| 1.5e-22 \| 125.7 \| 121 \| (29, 153) \| 205 \| (55, 185) \| 192 \| Phage tail protein, P2 protein I family (Fragment) \| Phage tail protein, P2 protein I family (Fragment); uniclust \| UniRef100\_A0A173R5F4 \| 99.5 \| 8.3e-17 \| 1.6e-22 \| 131.1 \| 147 \| (2, 152) \| 205 \| (12, 165) \| 301 \| Bacteriophage P2-related tail formation protein \| Bacteriophage P2-related tail formation protein; uniclust \| UniRef100\_A0A5M4DA93 \| 99.5 \| 9.6e-17 \| 1.8e-22 \| 129.3 \| 121 \| (29, 153) \| 205 \| (131, 268) \| 274 \| Phage tail protein \| Phage tail protein; uniclust \| UniRef100\_A0A3N5LKH2 \| 99.5 \| 9.8e-17 \| 1.9e-22 \| 133.7 \| 161 \| (12, 179) \| 205 \| (59, 282) \| 318 \| Phage tail protein I (Fragment) \| Phage tail protein I (Fragment); uniclust \| UniRef100\_A0A1G5H9F7 \| 99.5 \| 9e-17 \| 2e-22 \| 141.0 \| 145 \| (29, 182) \| 205 \| (194, 351) \| 374 \| Phage tail protein domain-containing protein \| Phage tail protein domain-containing protein; uniclust \| UniRef100\_A0A7M1L784 \| 99.5 \| 1.3e-16 \| 2.4e-22 \| 116.2 \| 143 \| (59, 204) \| 205 \| (6, 150) \| 150 \| Phage tail protein \| Phage tail protein; uniclust \| UniRef100\_A0A3M2ARC6 \| 99.5 \| 1.3e-16 \| 2.4e-22 \| 125.3 \| 115 \| (30, 152) \| 205 \| (92, 207) \| 210 \| Phage tail protein \| Phage tail protein; uniclust \| UniRef100\_A0A1N6X8I0 \| 99.5 \| 1.2e-16 \| 2.4e-22 \| 133.8 \| 123 \| (29, 153) \| 205 \| (106, 246) \| 277 \| Phage tail protein domain-containing protein \| Phage tail protein domain-containing protein; uniclust \| UniRef100\_A0A4U0YVX5 \| 99.5 \| 1.5e-16 \| 3e-22 \| 101.9 \| 58 \| (1, 58) \| 205 \| (6, 63) \| 64 \| Phage tail protein I (Fragment) \| Phage tail protein I (Fragment); uniclust \| UniRef100\_A4J3V2 \| 99.5 \| 1.9e-16 \| 3.6e-22 \| 137.2 \| 146 \| (3, 152) \| 205 \| (12, 164) \| 649 \| Phage tail protein I \| Phage tail protein I; uniclust \| UniRef100\_A0A062V414 \| 99.5 \| 2e-16 \| 3.9e-22 \| 131.9 \| 122 \| (30, 153) \| 205 \| (144, 282) \| 309 \| Phage tail protein \| Phage tail protein; uniclust \| UniRef100\_A0A3M5GB96 \| 99.5 \| 2.2e-16 \| 4.1e-22 \| 112.5 \| 81 \| (13, 95) \| 205 \| (45, 125) \| 130 \| Tail protein I \| Tail protein I; uniclust \| UniRef100\_A0A0F0HY11 \| 99.4 \| 3.1e-16 \| 6.7e-22 \| 147.4 \| 156 \| (12, 175) \| 205 \| (490, 691) \| 736 \| Uncharacterized protein \| Uncharacterized protein; uniclust \| UniRef100\_A0A3A8YWF4 \| 99.4 \| 3.6e-16 \| 6.9e-22 \| 123.7 \| 192 \| (5, 201) \| 205 \| (14, 211) \| 219 \| Phage tail protein I \| Phage tail protein I; uniclust \| UniRef100\_A0A8S5Q6A9 \| 99.4 \| 3.9e-16 \| 7.3e-22 \| 120.4 \| 166 \| (3, 172) \| 205 \| (14, 185) \| 222 \| Tail protein \| Tail protein; uniclust \| UniRef100\_A0A0Q8XJJ0 \| 99.4 \| 3.3e-16 \| 7.3e-22 \| 147.9 \| 159 \| (12, 176) \| 205 \| (454, 696) \| 748 \| Phage tail protein \| Phage tail protein; uniclust \| UniRef100\_A0A1M6N7M0 \| 99.4 \| 4e-16 \| 7.3e-22 \| 130.3 \| 118 \| (31, 152) \| 205 \| (41, 158) \| 435 \| Phage tail protein, P2 protein I family \| Phage tail protein, P2 protein I family; uniclust \| UniRef100\_A0A0A8WVT7 \| 99.4 \| 4.2e-16 \| 8.3e-22 \| 130.1 \| 88 \| (13, 101) \| 205 \| (27, 116) \| 316 \| Phage tail protein \| Phage tail protein; uniclust \| UniRef100\_A0A2E0L127 \| 99.4 \| 4.4e-16 \| 8.8e-22 \| 135.2 \| 116 \| (30, 154) \| 205 \| (234, 350) \| 408 \| FHA domain-containing protein \| FHA domain-containing protein; uniclust \| UniRef100\_A0A285NFM9 \| 99.4 \| 4.8e-16 \| 9.1e-22 \| 121.6 \| 175 \| (28, 205) \| 205 \| (45, 220) \| 221 \| Phage tail protein (Tail\_P2\_I) \| Phage tail protein (Tail\_P2\_I); uniclust \| UniRef100\_A0A0N8IBW6 \| 99.4 \| 5.2e-16 \| 9.5e-22 \| 117.9 \| 146 \| (4, 152) \| 205 \| (7, 168) \| 198 \| Phage tail protein \| Phage tail protein; uniclust \| UniRef100\_A0A3A9CZ71 \| 99.4 \| 5.3e-16 \| 9.9e-22 \| 130.3 \| 145 \| (3, 151) \| 205 \| (13, 164) \| 410 \| Phage tail protein \| Phage tail protein; uniclust \| UniRef100\_A0A523SGB7 \| 99.4 \| 5.9e-16 \| 1.1e-21 \| 123.8 \| 122 \| (29, 154) \| 205 \| (146, 283) \| 295 \| OmpR/PhoB-type domain-containing protein \| OmpR/PhoB-type domain-containing protein; uniclust \| UniRef100\_A0A014MDU0 \| 99.4 \| 5.8e-16 \| 1.2e-21 \| 141.3 \| 172 \| (2, 182) \| 205 \| (376, 609) \| 649 \| Tail protein \| Tail protein; uniclust \| UniRef100\_A0A4Y7S1X8 \| 99.4 \| 6.2e-16 \| 1.2e-21 \| 136.9 \| 154 \| (12, 171) \| 205 \| (379, 550) \| 593 \| Phage tail protein \| Phage tail protein; uniclust \| UniRef100\_A0A1Q6KUF2 \| 99.4 \| 6.3e-16 \| 1.2e-21 \| 126.1 \| 159 \| (3, 166) \| 205 \| (12, 175) \| 275 \| Phage tail protein I \| Phage tail protein I; uniclust \| UniRef100\_A0A7X4ZQB7 \| 99.4 \| 7.2e-16 \| 1.4e-21 \| 123.1 \| 171 \| (2, 175) \| 205 \| (10, 187) \| 252 \| Phage tail protein I \| Phage tail protein I; uniclust \| UniRef100\_A0A0Q0T5D2 \| 99.4 \| 7.4e-16 \| 1.4e-21 \| 108.9 \| 110 \| (65, 174) \| 205 \| (2, 111) \| 117 \| Phage tail protein \| Phage tail protein; uniclust \| UniRef100\_A0A095WFB1 \| 99.4 \| 9e-16 \| 1.8e-21 \| 121.9 \| 130 \| (30, 165) \| 205 \| (45, 174) \| 212 \| Phage tail protein I \| Phage tail protein I; uniclust \| UniRef100\_A0A2P9HMM9 \| 99.4 \| 9.3e-16 \| 1.9e-21 \| 134.9 \| 92 \| (2, 93) \| 205 \| (18, 111) \| 440 \| Phage tail protein I \| Phage tail protein I; uniclust \| UniRef100\_A0A069AQR6 \| 99.4 \| 8.7e-16 \| 1.9e-21 \| 127.1 \| 189 \| (3, 205) \| 205 \| (14, 211) \| 228 \| DUF2313 domain-containing protein \| DUF2313 domain-containing protein; uniclust \| UniRef100\_A0A6B9SWZ5 \| 99.4 \| 1e-15 \| 2e-21 \| 122.6 \| 141 \| (2, 151) \| 205 \| (4, 148) \| 221 \| Phage tail protein \| Phage tail protein; uniclust \| UniRef100\_A0A1M7YBP4 \| 99.4 \| 1.2e-15 \| 2.2e-21 \| 126.9 \| 145 \| (3, 152) \| 205 \| (11, 162) \| 339 \| Phage tail protein (Tail\_P2\_I) \| Phage tail protein (Tail\_P2\_I); uniclust \| UniRef100\_UPI00189D0918 \| 99.4 \| 1.2e-15 \| 2.3e-21 \| 122.8 \| 151 \| (2, 153) \| 205 \| (5, 158) \| 310 \| phage tail protein I \| phage tail protein I; uniclust \| UniRef100\_UPI001B3C6DE8 \| 99.4 \| 1.3e-15 \| 2.4e-21 \| 117.7 \| 148 \| (3, 154) \| 205 \| (11, 164) \| 222 \| phage tail protein \| phage tail protein; uniclust \| UniRef100\_A0A1G9EY78 \| 99.4 \| 1.1e-15 \| 2.4e-21 \| 143.3 \| 157 \| (12, 174) \| 205 \| (465, 683) \| 740 \| Phage tail protein domain-containing protein \| Phage tail protein domain-containing protein; uniclust \| UniRef100\_A0A099PD40 \| 99.4 \| 1.3e-15 \| 2.7e-21 \| 136.6 \| 159 \| (12, 175) \| 205 \| (221, 431) \| 537 \| Phage tail protein \| Phage tail protein; uniclust \| UniRef100\_A0A2N6CT40 \| 99.4 \| 1.7e-15 \| 3.2e-21 \| 125.7 \| 159 \| (28, 194) \| 205 \| (110, 315) \| 342 \| Phage tail protein I \| Phage tail protein I; uniclust \| UniRef100\_A0A6D0ZU35 \| 99.4 \| 2.4e-15 \| 4.5e-21 \| 110.6 \| 121 \| (63, 185) \| 205 \| (7, 127) \| 152 \| Phage tail protein I (Fragment) \| Phage tail protein I (Fragment); uniclust \| UniRef100\_A0A1C6BNG9 \| 99.4 \| 2.4e-15 \| 4.6e-21 \| 119.9 \| 172 \| (2, 175) \| 205 \| (10, 188) \| 250 \| Bacteriophage P2-related tail formation protein \| Bacteriophage P2-related tail formation protein; uniclust \| UniRef100\_A0A3A9EQX9 \| 99.4 \| 2.9e-15 \| 5.4e-21 \| 122.8 \| 128 \| (35, 164) \| 205 \| (54, 182) \| 346 \| Phage tail protein I \| Phage tail protein I; uniclust \| UniRef100\_UPI0009BF8F77 \| 99.4 \| 3.1e-15 \| 5.7e-21 \| 131.9 \| 119 \| (32, 151) \| 205 \| (34, 153) \| 767 \| phage tail protein I \| phage tail protein I; uniclust \| UniRef100\_UPI00202CF454 \| 99.4 \| 3.1e-15 \| 5.7e-21 \| 99.7 \| 74 \| (47, 120) \| 205 \| (1, 74) \| 86 \| phage tail protein I \| phage tail protein I; uniclust \| UniRef100\_A0A218Q5U8 \| 99.4 \| 3e-15 \| 5.8e-21 \| 125.4 \| 119 \| (29, 153) \| 205 \| (64, 190) \| 365 \| Phage tail protein \| Phage tail protein; uniclust \| UniRef100\_A0A1Q6PWS5 \| 99.3 \| 3.2e-15 \| 6.1e-21 \| 126.0 \| 136 \| (12, 153) \| 205 \| (23, 159) \| 369 \| Phage tail protein I \| Phage tail protein I; uniclust \| UniRef100\_UPI001F4B27F2 \| 99.3 \| 3.5e-15 \| 6.4e-21 \| 99.0 \| 76 \| (51, 126) \| 205 \| (2, 77) \| 84 \| phage tail protein I \| phage tail protein I; uniclust \| UniRef100\_Q9KW54 \| 99.3 \| 3.7e-15 \| 6.9e-21 \| 111.9 \| 128 \| (14, 144) \| 205 \| (18, 163) \| 164 \| Tail protein \| Tail protein; uniclust \| UniRef100\_A0A2W6SVJ0 \| 99.3 \| 3.8e-15 \| 7.4e-21 \| 128.0 \| 94 \| (1, 94) \| 205 \| (2, 96) \| 415 \| Phage tail protein I \| Phage tail protein I; uniclust \| UniRef100\_A0A1T4W599 \| 99.3 \| 4.3e-15 \| 8.5e-21 \| 126.6 \| 141 \| (2, 150) \| 205 \| (12, 155) \| 358 \| Phage tail protein (Tail\_P2\_I) \| Phage tail protein (Tail\_P2\_I); uniclust \| UniRef100\_UPI001FEBD5CB \| 99.3 \| 4.7e-15 \| 8.7e-21 \| 107.4 \| 112 \| (92, 205) \| 205 \| (1, 112) \| 141 \| phage tail protein \| phage tail protein; uniclust \| UniRef100\_A0A7T7D8U4 \| 99.3 \| 5.5e-15 \| 1e-20 \| 127.7 \| 191 \| (3, 196) \| 205 \| (390, 587) \| 596 \| Phage tail protein I \| Phage tail protein I; uniclust \| UniRef100\_UPI001C997A2E \| 99.3 \| 6.1e-15 \| 1.1e-20 \| 99.7 \| 85 \| (3, 87) \| 205 \| (6, 92) \| 93 \| phage tail protein I \| phage tail protein I; uniclust \| UniRef100\_A0A969G7N1 \| 99.3 \| 6.4e-15 \| 1.2e-20 \| 115.6 \| 117 \| (29, 153) \| 205 \| (18, 144) \| 247 \| Phage tail protein I \| Phage tail protein I; uniclust \| UniRef100\_UPI0021795371 \| 99.3 \| 6.6e-15 \| 1.3e-20 \| 110.3 \| 119 \| (2, 120) \| 205 \| (10, 134) \| 155 \| phage tail protein I \| phage tail protein I; uniclust \| UniRef100\_A0A2D3VXF3 \| 99.3 \| 7.5e-15 \| 1.4e-20 \| 110.0 \| 146 \| (2, 150) \| 205 \| (1, 147) \| 176 \| Phage tail protein I \| Phage tail protein I; uniclust \| UniRef100\_A0A3N5MGM2 \| 99.3 \| 9.6e-15 \| 1.8e-20 \| 119.1 \| 164 \| (12, 182) \| 205 \| (91, 306) \| 338 \| Phage tail protein I (Fragment) \| Phage tail protein I (Fragment); uniclust \| UniRef100\_A0A158HNT1 \| 99.3 \| 9.6e-15 \| 2e-20 \| 135.4 \| 157 \| (12, 174) \| 205 \| (428, 667) \| 710 \| Serine/threonine-protein kinase PknD \| Serine/threonine-protein kinase PknD; uniclust \| UniRef100\_D0LX69 \| 99.3 \| 1.3e-14 \| 2.4e-20 \| 122.1 \| 123 \| (30, 154) \| 205 \| (118, 260) \| 448 \| Phage tail protein \| Phage tail protein; uniclust \| UniRef100\_A0A021X9K5 \| 99.3 \| 1.2e-14 \| 2.5e-20 \| 127.9 \| 95 \| (5, 101) \| 205 \| (11, 109) \| 413 \| Phage tail protein I \| Phage tail protein I; uniclust \| UniRef100\_UPI000798D5FE \| 99.3 \| 1.3e-14 \| 2.5e-20 \| 100.0 \| 95 \| (80, 176) \| 205 \| (1, 95) \| 96 \| phage tail protein I \| phage tail protein I; uniclust \| UniRef100\_UPI001126ED48 \| 99.3 \| 1.4e-14 \| 2.6e-20 \| 118.8 \| 90 \| (3, 92) \| 205 \| (15, 106) \| 353 \| phage tail protein I \| phage tail protein I; uniclust \| UniRef100\_A0A160TAS9 \| 99.3 \| 1.3e-14 \| 2.7e-20 \| 135.7 \| 158 \| (12, 175) \| 205 \| (473, 695) \| 748 \| Phage tail protein \| Phage tail protein; uniclust \| UniRef100\_A0A655CJ05 \| 99.3 \| 1.7e-14 \| 3.4e-20 \| 108.9 \| 98 \| (3, 100) \| 205 \| (11, 114) \| 139 \| Phage Tail protein I \| Phage Tail protein I; uniclust \| UniRef100\_UPI0015626642 \| 99.3 \| 1.8e-14 \| 3.4e-20 \| 105.8 \| 114 \| (2, 121) \| 205 \| (10, 124) \| 133 \| phage tail protein \| phage tail protein; uniclust \| UniRef100\_UPI001EEDE119 \| 99.3 \| 1.9e-14 \| 3.4e-20 \| 104.2 \| 98 \| (50, 147) \| 205 \| (1, 98) \| 139 \| phage tail protein I \| phage tail protein I; uniclust \| UniRef100\_UPI001EEE1D57 \| 99.3 \| 1.9e-14 \| 3.5e-20 \| 94.7 \| 67 \| (23, 89) \| 205 \| (10, 77) \| 79 \| phage tail protein I \| phage tail protein I; uniclust \| UniRef100\_A0A6I1I488 \| 99.3 \| 2e-14 \| 3.7e-20 \| 102.2 \| 72 \| (4, 75) \| 205 \| (7, 78) \| 124 \| Phage tail protein I \| Phage tail protein I; uniclust \| UniRef100\_A0A535XYK6 \| 99.3 \| 2.1e-14 \| 3.8e-20 \| 110.3 \| 110 \| (30, 147) \| 205 \| (93, 203) \| 208 \| Phage tail protein I \| Phage tail protein I; uniclust \| UniRef100\_A0A7Z7WAU5 \| 99.3 \| 2.1e-14 \| 3.8e-20 \| 98.5 \| 90 \| (75, 165) \| 205 \| (1, 96) \| 100 \| Bacteriophage P2-related tail formation protein \| Bacteriophage P2-related tail formation protein; uniclust \| UniRef100\_A0A7X1ZG08 \| 99.3 \| 2.2e-14 \| 4e-20 \| 103.6 \| 72 \| (2, 73) \| 205 \| (60, 133) \| 137 \| Phage tail protein I \| Phage tail protein I; uniclust \| UniRef100\_A0A2T4UGC8 \| 99.3 \| 2.1e-14 \| 4.1e-20 \| 117.2 \| 121 \| (29, 154) \| 205 \| (98, 227) \| 253 \| Phage tail protein \| Phage tail protein; uniclust \| UniRef100\_A0A024LRF3 \| 99.3 \| 2.2e-14 \| 4.6e-20 \| 129.2 \| 145 \| (2, 153) \| 205 \| (49, 196) \| 468 \| Phage protein \| Phage protein; uniclust \| UniRef100\_A0A2T4JG36 \| 99.2 \| 3.3e-14 \| 6e-20 \| 114.6 \| 95 \| (2, 96) \| 205 \| (4, 98) \| 300 \| Phage tail protein I \| Phage tail protein I; uniclust \| UniRef100\_A0A238WY20 \| 99.2 \| 3.3e-14 \| 6.4e-20 \| 131.6 \| 162 \| (6, 171) \| 205 \| (663, 851) \| 910 \| Phage tail protein domain-containing protein \| Phage tail protein domain-containing protein; uniclust \| UniRef100\_A0A084SKL4 \| 99.2 \| 3.8e-14 \| 7.8e-20 \| 120.5 \| 115 \| (28, 152) \| 205 \| (84, 208) \| 318 \| Tail protein \| Tail protein; uniclust \| UniRef100\_A0A2T1ENV4 \| 99.2 \| 4.6e-14 \| 8.6e-20 \| 112.6 \| 121 \| (29, 153) \| 205 \| (57, 188) \| 252 \| Phage tail protein I \| Phage tail protein I; uniclust \| UniRef100\_A0A011UNJ4 \| 99.2 \| 5.7e-14 \| 1.1e-19 \| 115.6 \| 142 \| (2, 151) \| 205 \| (8, 152) \| 272 \| Phage tail protein \| Phage tail protein; uniclust \| UniRef100\_A0A607GXP1 \| 99.2 \| 5.9e-14 \| 1.1e-19 \| 100.8 \| 95 \| (69, 164) \| 205 \| (2, 99) \| 114 \| Phage tail protein \| Phage tail protein; uniclust \| UniRef100\_A0A4R3LI10 \| 99.2 \| 6e-14 \| 1.2e-19 \| 125.9 \| 140 \| (2, 151) \| 205 \| (8, 151) \| 532 \| P2-related tail formation protein \| P2-related tail formation protein; uniclust \| UniRef100\_A0A0P6W260 \| 99.2 \| 6.9e-14 \| 1.3e-19 \| 118.4 \| 90 \| (3, 92) \| 205 \| (15, 106) \| 465 \| Phage tail protein \| Phage tail protein; uniclust \| UniRef100\_A0A4Q8MC19 \| 99.2 \| 7.7e-14 \| 1.5e-19 \| 119.2 \| 94 \| (2, 95) \| 205 \| (3, 97) \| 422 \| Phage tail protein I \| Phage tail protein I; uniclust \| UniRef100\_A0A9D5W5Y6 \| 99.2 \| 7.8e-14 \| 1.5e-19 \| 109.0 \| 148 \| (2, 152) \| 205 \| (4, 188) \| 206 \| Phage tail protein \| Phage tail protein; uniclust \| UniRef100\_UPI001E4BC782 \| 99.2 \| 8.2e-14 \| 1.5e-19 \| 97.1 \| 71 \| (3, 73) \| 205 \| (8, 79) \| 104 \| phage tail protein I \| phage tail protein I; uniclust \| UniRef100\_A0A5M6I8Y5 \| 99.2 \| 9e-14 \| 1.6e-19 \| 103.1 \| 117 \| (85, 204) \| 205 \| (40, 156) \| 160 \| Phage tail protein I \| Phage tail protein I; uniclust \| UniRef100\_UPI000DE39F06 \| 99.2 \| 1.2e-13 \| 2.1e-19 \| 93.5 \| 78 \| (2, 79) \| 205 \| (3, 82) \| 91 \| phage tail protein I \| phage tail protein I; uniclust \| UniRef100\_A0A7X3ZHW1 \| 99.2 \| 1.6e-13 \| 2.9e-19 \| 113.5 \| 144 \| (3, 149) \| 205 \| (191, 340) \| 366 \| Phage tail protein I \| Phage tail protein I; uniclust \| UniRef100\_A0A1G6NCN4 \| 99.2 \| 1.4e-13 \| 3.1e-19 \| 130.7 \| 157 \| (11, 175) \| 205 \| (496, 715) \| 753 \| Phage tail protein domain-containing protein \| Phage tail protein domain-containing protein; uniclust \| UniRef100\_A0A381EC89 \| 99.2 \| 1.6e-13 \| 3.2e-19 \| 114.4 \| 146 \| (1, 153) \| 205 \| (2, 161) \| 298 \| Bacteriophage P2-related tail formation protein \| Bacteriophage P2-related tail formation protein; uniclust \| UniRef100\_A0A1D9LHX5 \| 99.2 \| 1.7e-13 \| 3.2e-19 \| 118.5 \| 145 \| (1, 150) \| 205 \| (2, 150) \| 444 \| Phage tail protein \| Phage tail protein; uniclust \| UniRef100\_A0A6B8KI89 \| 99.2 \| 2e-13 \| 3.7e-19 \| 110.1 \| 190 \| (2, 196) \| 205 \| (86, 278) \| 295 \| Phage tail protein I \| Phage tail protein I; uniclust \| UniRef100\_A0A1H3ZWU0 \| 99.1 \| 1.9e-13 \| 4e-19 \| 121.7 \| 96 \| (2, 99) \| 205 \| (15, 111) \| 413 \| Phage tail protein, P2 protein I family \| Phage tail protein, P2 protein I family; uniclust \| UniRef100\_A0A645H901 \| 99.1 \| 2.2e-13 \| 4.1e-19 \| 97.5 \| 119 \| (52, 174) \| 205 \| (4, 123) \| 127 \| Phage tail protein I \| Phage tail protein I; uniclust \| UniRef100\_UPI0013771264 \| 99.1 \| 2.3e-13 \| 4.3e-19 \| 114.1 \| 90 \| (2, 93) \| 205 \| (6, 95) \| 418 \| phage tail protein I \| phage tail protein I; uniclust \| UniRef100\_A0A072TF05 \| 99.1 \| 2.6e-13 \| 4.8e-19 \| 127.3 \| 148 \| (4, 153) \| 205 \| (1006, 1174) \| 1809 \| Bacteriophage regulatory protein, putative (Fragment) \| Bacteriophage regulatory protein, putative (Fragment); uniclust \| UniRef100\_UPI000F016F72 \| 99.1 \| 2.9e-13 \| 5.3e-19 \| 92.9 \| 89 \| (39, 128) \| 205 \| (3, 91) \| 98 \| phage tail protein I \| phage tail protein I; uniclust \| UniRef100\_A0A7W6E8X0 \| 99.1 \| 2.8e-13 \| 5.3e-19 \| 107.2 \| 97 \| (3, 103) \| 205 \| (6, 102) \| 214 \| P2-related tail formation protein \| P2-related tail formation protein; uniclust \| UniRef100\_A0A9D2DDK1 \| 99.1 \| 3.5e-13 \| 6.6e-19 \| 112.6 \| 139 \| (11, 151) \| 205 \| (15, 166) \| 357 \| Phage tail protein \| Phage tail protein; uniclust \| UniRef100\_A0A448QV26 \| 99.1 \| 3.9e-13 \| 7.2e-19 \| 105.5 \| 134 \| (15, 153) \| 205 \| (67, 217) \| 237 \| Bacteriophage P2-related tail formation protein \| Bacteriophage P2-related tail formation protein; uniclust \| UniRef100\_A0A031HNM3 \| 99.1 \| 3.5e-13 \| 7.3e-19 \| 117.7 \| 135 \| (2, 153) \| 205 \| (40, 178) \| 350 \| Phage tail protein \| Phage tail protein; uniclust \| UniRef100\_A0A1T2XBJ5 \| 99.1 \| 4.3e-13 \| 7.9e-19 \| 99.4 \| 135 \| (59, 195) \| 205 \| (8, 142) \| 158 \| Phage P2 family tail protein \| Phage P2 family tail protein; uniclust \| UniRef100\_A0A1C0V8J7 \| 99.1 \| 4.5e-13 \| 8.3e-19 \| 105.6 \| 151 \| (2, 153) \| 205 \| (2, 163) \| 244 \| Phage tail protein I \| Phage tail protein I; uniclust \| UniRef100\_A0A064AK86 \| 99.1 \| 4.7e-13 \| 8.6e-19 \| 100.8 \| 117 \| (36, 155) \| 205 \| (6, 122) \| 175 \| Tail protein \| Tail protein; uniclust \| UniRef100\_A0A2E2NRK6 \| 99.1 \| 4.4e-13 \| 8.6e-19 \| 117.3 \| 122 \| (30, 153) \| 205 \| (206, 366) \| 421 \| Uncharacterized protein \| Uncharacterized protein; uniclust \| UniRef100\_A0A2W5WSP6 \| 99.1 \| 4.7e-13 \| 8.9e-19 \| 114.4 \| 98 \| (3, 100) \| 205 \| (5, 103) \| 407 \| Phage tail protein I \| Phage tail protein I; uniclust \| UniRef100\_A0A1W1Z3V2 \| 99.1 \| 5.3e-13 \| 1.1e-18 \| 116.8 \| 93 \| (2, 95) \| 205 \| (6, 99) \| 427 \| Phage tail protein, P2 protein I family \| Phage tail protein, P2 protein I family; uniclust \| UniRef100\_A0A3P6JJC9 \| 99.1 \| 5.9e-13 \| 1.1e-18 \| 90.7 \| 77 \| (75, 153) \| 205 \| (1, 77) \| 94 \| Phage tail protein \| Phage tail protein; uniclust \| UniRef100\_A0A6I2UWE9 \| 99.1 \| 6.1e-13 \| 1.1e-18 \| 95.3 \| 69 \| (33, 101) \| 205 \| (1, 70) \| 126 \| Phage tail protein I (Fragment) \| Phage tail protein I (Fragment); uniclust \| UniRef100\_A0A1M6B964 \| 99.1 \| 6.1e-13 \| 1.2e-18 \| 115.9 \| 142 \| (1, 151) \| 205 \| (4, 148) \| 437 \| Phage tail protein (Tail\_P2\_I) \| Phage tail protein (Tail\_P2\_I); uniclust \| UniRef100\_A0A3A8UCJ4 \| 99.1 \| 6.4e-13 \| 1.2e-18 \| 115.2 \| 143 \| (5, 152) \| 205 \| (18, 167) \| 518 \| Phage tail protein I \| Phage tail protein I; uniclust \| UniRef100\_UPI000B329C9F \| 99.1 \| 7.7e-13 \| 1.5e-18 \| 100.1 \| 95 \| (54, 151) \| 205 \| (11, 121) \| 151 \| phage tail protein \| phage tail protein; uniclust \| UniRef100\_A0A968NW80 \| 99.1 \| 7.9e-13 \| 1.5e-18 \| 114.3 \| 135 \| (34, 174) \| 205 \| (240, 422) \| 465 \| Phage tail protein I \| Phage tail protein I; uniclust \| UniRef100\_A0A1C5SWA5 \| 99.1 \| 7.2e-13 \| 1.5e-18 \| 116.4 \| 158 \| (12, 174) \| 205 \| (175, 352) \| 373 \| Phage tail protein domain \| Phage tail protein domain; uniclust \| UniRef100\_A0A8H2LSC2 \| 99.1 \| 8.7e-13 \| 1.6e-18 \| 106.9 \| 68 \| (2, 69) \| 205 \| (6, 73) \| 303 \| Methyltransferase \| Methyltransferase; uniclust \| UniRef100\_A0A957FSB1 \| 99.1 \| 1.1e-12 \| 2.1e-18 \| 111.5 \| 114 \| (30, 153) \| 205 \| (337, 451) \| 464 \| Phage tail protein I \| Phage tail protein I; uniclust \| UniRef100\_A0A955Q574 \| 99.0 \| 1.3e-12 \| 2.4e-18 \| 102.0 \| 117 \| (35, 153) \| 205 \| (61, 196) \| 225 \| Uncharacterized protein \| Uncharacterized protein; uniclust \| UniRef100\_A0A1G3M9I7 \| 99.0 \| 1.4e-12 \| 2.7e-18 \| 105.0 \| 148 \| (27, 177) \| 205 \| (41, 191) \| 220 \| Uncharacterized protein \| Uncharacterized protein; uniclust \| UniRef100\_UPI000B5A0976 \| 99.0 \| 1.6e-12 \| 2.9e-18 \| 108.1 \| 151 \| (2, 153) \| 205 \| (1, 170) \| 374 \| phage tail protein \| phage tail protein; uniclust \| UniRef100\_A0A7X6FP85 \| 99.0 \| 1.6e-12 \| 3e-18 \| 91.6 \| 88 \| (2, 90) \| 205 \| (5, 93) \| 113 \| Phage tail protein I \| Phage tail protein I; uniclust \| UniRef100\_A0A0D8FVA9 \| 99.0 \| 1.7e-12 \| 3.4e-18 \| 107.2 \| 120 \| (29, 153) \| 205 \| (64, 188) \| 244 \| Phage tail protein \| Phage tail protein; uniclust \| UniRef100\_I3D8X8 \| 99.0 \| 1.8e-12 \| 3.6e-18 \| 95.7 \| 99 \| (1, 101) \| 205 \| (1, 108) \| 123 \| Phage tail protein \| Phage tail protein; uniclust \| UniRef100\_A0A0C1MU06 \| 99.0 \| 1.9e-12 \| 3.9e-18 \| 111.4 \| 143 \| (3, 153) \| 205 \| (18, 162) \| 327 \| Phage tail protein \| Phage tail protein; uniclust \| UniRef100\_A0A508B349 \| 99.0 \| 2.1e-12 \| 3.9e-18 \| 107.8 \| 120 \| (30, 152) \| 205 \| (197, 320) \| 387 \| Uncharacterized protein \| Uncharacterized protein; uniclust \| UniRef100\_A0A2W6YVX7 \| 99.0 \| 2.3e-12 \| 4.4e-18 \| 103.3 \| 197 \| (3, 204) \| 205 \| (23, 236) \| 238 \| Phage tail protein \| Phage tail protein; uniclust \| UniRef100\_A0A352AM66 \| 99.0 \| 2.4e-12 \| 4.5e-18 \| 94.6 \| 60 \| (29, 89) \| 205 \| (70, 129) \| 129 \| Phage tail protein (Fragment) \| Phage tail protein (Fragment); uniclust \| UniRef100\_A0A0Q7UJR8 \| 99.0 \| 2.5e-12 \| 4.8e-18 \| 104.3 \| 122 \| (28, 153) \| 205 \| (43, 172) \| 259 \| Phage tail protein \| Phage tail protein; uniclust \| UniRef100\_A0A0Q8B1P1 \| 99.0 \| 2.6e-12 \| 4.8e-18 \| 108.3 \| 94 \| (2, 95) \| 205 \| (13, 106) \| 423 \| Phage tail protein \| Phage tail protein; uniclust \| UniRef100\_A0A1Q3SJJ0 \| 99.0 \| 2.8e-12 \| 5.2e-18 \| 105.0 \| 121 \| (28, 152) \| 205 \| (60, 199) \| 325 \| Phage tail protein I \| Phage tail protein I; uniclust \| UniRef100\_A0A1X7Q586 \| 99.0 \| 3e-12 \| 6e-18 \| 111.4 \| 142 \| (1, 151) \| 205 \| (2, 145) \| 401 \| p2-related tail formation protein \| p2-related tail formation protein; uniclust \| UniRef100\_A0A2N5DAD0 \| 99.0 \| 3.5e-12 \| 7.1e-18 \| 120.2 \| 94 \| (10, 104) \| 205 \| (561, 658) \| 848 \| Uncharacterized protein \| Uncharacterized protein; uniclust \| UniRef100\_UPI001EDCD62C \| 99.0 \| 4.1e-12 \| 7.6e-18 \| 93.9 \| 117 \| (13, 142) \| 205 \| (13, 131) \| 151 \| phage tail protein \| phage tail protein; uniclust \| UniRef100\_A0A1I1V086 \| 99.0 \| 4.8e-12 \| 9.3e-18 \| 109.3 \| 93 \| (2, 94) \| 205 \| (5, 98) \| 404 \| Phage tail protein, P2 protein I family \| Phage tail protein, P2 protein I family; uniclust \| UniRef100\_A0A3D1NS20 \| 98.9 \| 5.3e-12 \| 9.8e-18 \| 97.4 \| 139 \| (11, 151) \| 205 \| (15, 166) \| 203 \| Phage tail protein \| Phage tail protein; uniclust \| UniRef100\_A0A3A4NQI1 \| 98.9 \| 6e-12 \| 1.1e-17 \| 98.2 \| 119 \| (28, 148) \| 205 \| (45, 182) \| 206 \| Phage tail protein I \| Phage tail protein I; uniclust \| UniRef100\_A0A4Y6PWX7 \| 98.9 \| 7e-12 \| 1.3e-17 \| 110.7 \| 126 \| (20, 147) \| 205 \| (432, 572) \| 670 \| Phage tail protein I \| Phage tail protein I; uniclust \| UniRef100\_UPI0021D519AB \| 98.9 \| 7.7e-12 \| 1.4e-17 \| 91.2 \| 132 \| (73, 204) \| 205 \| (5, 137) \| 138 \| phage tail protein \| phage tail protein; uniclust \| UniRef100\_UPI001C49953D \| 98.9 \| 1e-11 \| 1.9e-17 \| 95.5 \| 142 \| (2, 147) \| 205 \| (5, 168) \| 172 \| phage tail protein \| phage tail protein; uniclust \| UniRef100\_A0A929K3N6 \| 98.9 \| 1.1e-11 \| 2e-17 \| 92.3 \| 92 \| (12, 104) \| 205 \| (24, 119) \| 157 \| Phage tail protein I (Fragment) \| Phage tail protein I (Fragment); uniclust \| UniRef100\_A0A7C4AJB7 \| 98.9 \| 1.1e-11 \| 2e-17 \| 106.5 \| 94 \| (2, 95) \| 205 \| (4, 98) \| 397 \| Phage tail protein I (Fragment) \| Phage tail protein I (Fragment); uniclust \| UniRef100\_A0A3A9B594 \| 98.9 \| 1.1e-11 \| 2e-17 \| 100.3 \| 148 \| (3, 153) \| 205 \| (16, 174) \| 288 \| Phage tail protein I (Fragment) \| Phage tail protein I (Fragment); uniclust \| UniRef100\_A0A7C3GTL1 \| 98.9 \| 1.2e-11 \| 2.2e-17 \| 109.8 \| 140 \| (12, 153) \| 205 \| (464, 637) \| 707 \| Phage tail protein I \| Phage tail protein I; uniclust \| UniRef100\_UPI0021C5C126 \| 98.9 \| 1.2e-11 \| 2.2e-17 \| 81.9 \| 64 \| (38, 101) \| 205 \| (4, 70) \| 78 \| phage tail protein I \| phage tail protein I; uniclust \| UniRef100\_A0A411CW10 \| 98.9 \| 1.2e-11 \| 2.4e-17 \| 99.6 \| 116 \| (36, 153) \| 205 \| (58, 175) \| 205 \| Tail protein \| Tail protein; uniclust \| UniRef100\_A0A3P8M2T9 \| 98.9 \| 1.4e-11 \| 2.6e-17 \| 85.7 \| 66 \| (2, 67) \| 205 \| (3, 68) \| 103 \| Bacteriophage P2-related tail formation protein \| Bacteriophage P2-related tail formation protein; uniclust \| UniRef100\_UPI000417C6A3 \| 98.9 \| 1.5e-11 \| 2.7e-17 \| 103.8 \| 121 \| (33, 153) \| 205 \| (41, 180) \| 414 \| phage tail protein \| phage tail protein; uniclust \| UniRef100\_UPI00201AC909 \| 98.9 \| 1.6e-11 \| 2.9e-17 \| 77.7 \| 46 \| (30, 75) \| 205 \| (10, 55) \| 61 \| phage tail protein I \| phage tail protein I; uniclust \| UniRef100\_A0A969IZC1 \| 98.9 \| 1.7e-11 \| 3.2e-17 \| 98.0 \| 116 \| (29, 151) \| 205 \| (52, 175) \| 261 \| Phage tail protein I \| Phage tail protein I; uniclust \| UniRef100\_A0A087KKF1 \| 98.9 \| 1.7e-11 \| 3.2e-17 \| 87.8 \| 81 \| (5, 85) \| 205 \| (7, 91) \| 108 \| Tail fiber protein (Fragment) \| Tail fiber protein (Fragment); uniclust \| UniRef100\_A0A496RFZ9 \| 98.9 \| 1.7e-11 \| 3.3e-17 \| 109.3 \| 159 \| (12, 176) \| 205 \| (381, 549) \| 587 \| Phage tail protein (Fragment) \| Phage tail protein (Fragment); uniclust \| UniRef100\_A0A3D2IPZ0 \| 98.9 \| 1.8e-11 \| 3.4e-17 \| 97.2 \| 133 \| (30, 168) \| 205 \| (49, 181) \| 247 \| Phage tail protein I \| Phage tail protein I; uniclust \| UniRef100\_UPI000C6CA109 \| 98.9 \| 1.9e-11 \| 3.4e-17 \| 104.4 \| 141 \| (29, 175) \| 205 \| (254, 418) \| 460 \| phage tail protein \| phage tail protein; uniclust \| UniRef100\_A0A929PG22 \| 98.9 \| 1.9e-11 \| 3.5e-17 \| 99.5 \| 75 \| (29, 104) \| 205 \| (31, 106) \| 299 \| Uncharacterized protein \| Uncharacterized protein; uniclust \| UniRef100\_UPI00209C94E4 \| 98.9 \| 1.9e-11 \| 3.6e-17 \| 96.7 \| 135 \| (30, 171) \| 205 \| (54, 204) \| 240 \| phage tail protein \| phage tail protein; uniclust \| UniRef100\_UPI000E352F0F \| 98.9 \| 1.9e-11 \| 3.7e-17 \| 91.1 \| 84 \| (91, 176) \| 205 \| (5, 88) \| 137 \| phage tail protein I \| phage tail protein I; uniclust \| UniRef100\_UPI0006865A37 \| 98.9 \| 2.1e-11 \| 3.9e-17 \| 105.2 \| 141 \| (3, 145) \| 205 \| (12, 193) \| 511 \| phage tail protein \| phage tail protein; uniclust \| UniRef100\_A0A090SU32 \| 98.8 \| 2.4e-11 \| 4.5e-17 \| 78.9 \| 65 \| (19, 83) \| 205 \| (4, 69) \| 70 \| Uncharacterized protein \| Uncharacterized protein; uniclust \| UniRef100\_A0A6S6YHP9 \| 98.8 \| 2.4e-11 \| 4.5e-17 \| 99.4 \| 140 \| (2, 151) \| 205 \| (4, 147) \| 291 \| Phage tail protein \| Phage tail protein; uniclust \| UniRef100\_A0A928VPI1 \| 98.8 \| 2.8e-11 \| 5.2e-17 \| 102.8 \| 121 \| (30, 153) \| 205 \| (253, 412) \| 435 \| Uncharacterized protein \| Uncharacterized protein; uniclust \| UniRef100\_A0A1J0LXM1 \| 98.8 \| 2.6e-11 \| 5.2e-17 \| 103.2 \| 116 \| (35, 151) \| 205 \| (53, 183) \| 308 \| Phage tail protein (Tail\_P2\_I) \| Phage tail protein (Tail\_P2\_I); uniclust \| UniRef100\_A0A0Q6MMI2 \| 98.8 \| 2.7e-11 \| 5.8e-17 \| 117.9 \| 161 \| (12, 178) \| 205 \| (635, 867) \| 968 \| Phage tail protein \| Phage tail protein; uniclust \| UniRef100\_A0A9E6AVE8 \| 98.8 \| 3.2e-11 \| 5.8e-17 \| 96.0 \| 165 \| (3, 168) \| 205 \| (14, 186) \| 249 \| Phage tail protein I \| Phage tail protein I; uniclust \| UniRef100\_A0A1S2E3G0 \| 98.8 \| 3.1e-11 \| 5.8e-17 \| 99.8 \| 139 \| (13, 152) \| 205 \| (19, 166) \| 286 \| Phage tail protein I \| Phage tail protein I; uniclust \| UniRef100\_UPI001CF59578 \| 98.8 \| 3.2e-11 \| 5.9e-17 \| 101.4 \| 114 \| (29, 147) \| 205 \| (202, 333) \| 396 \| phage tail protein \| phage tail protein; uniclust \| UniRef100\_A0A2V8RW14 \| 98.8 \| 3.2e-11 \| 6.3e-17 \| 105.2 \| 136 \| (30, 171) \| 205 \| (195, 383) \| 431 \| Phage tail protein \| Phage tail protein; uniclust \| UniRef100\_UPI0003FB0752 \| 98.8 \| 3.5e-11 \| 6.4e-17 \| 96.1 \| 171 \| (30, 203) \| 205 \| (68, 240) \| 256 \| phage tail protein \| phage tail protein; uniclust \| UniRef100\_U1KSX1 \| 98.8 \| 3.7e-11 \| 6.7e-17 \| 92.0 \| 98 \| (53, 153) \| 205 \| (1, 105) \| 187 \| Phage tail protein I \| Phage tail protein I; uniclust \| UniRef100\_A0A2P8W8C8 \| 98.8 \| 3.6e-11 \| 6.9e-17 \| 110.7 \| 134 \| (12, 150) \| 205 \| (467, 644) \| 709 \| Phage tail protein (Fragment) \| Phage tail protein (Fragment); uniclust \| UniRef100\_UPI0018E0E16F \| 98.8 \| 3.9e-11 \| 7.1e-17 \| 102.2 \| 111 \| (36, 147) \| 205 \| (50, 196) \| 446 \| phage tail protein I \| phage tail protein I; uniclust \| UniRef100\_UPI000DE5AB00 \| 98.8 \| 4e-11 \| 7.4e-17 \| 86.9 \| 110 \| (1, 110) \| 205 \| (1, 116) \| 130 \| phage tail protein I \| phage tail protein I; uniclust \| UniRef100\_UPI000B064F58 \| 98.8 \| 4.1e-11 \| 7.6e-17 \| 85.2 \| 97 \| (13, 115) \| 205 \| (13, 111) \| 112 \| phage tail protein \| phage tail protein; uniclust \| UniRef100\_A0A1H0YPQ4 \| 98.8 \| 4.7e-11 \| 8.8e-17 \| 96.0 \| 202 \| (3, 204) \| 205 \| (9, 248) \| 250 \| Phage tail protein, P2 protein I family \| Phage tail protein, P2 protein I family; uniclust \| UniRef100\_A0A3A8IIH2 \| 98.8 \| 5.1e-11 \| 1e-16 \| 100.7 \| 122 \| (42, 168) \| 205 \| (134, 281) \| 298 \| Phage tail protein \| Phage tail protein; uniclust \| UniRef100\_UPI0016870817 \| 98.8 \| 5.6e-11 \| 1e-16 \| 107.9 \| 133 \| (12, 147) \| 205 \| (646, 832) \| 898 \| phage tail protein \| phage tail protein; uniclust \| UniRef100\_A0A0M2RHZ0 \| 98.8 \| 5.6e-11 \| 1.1e-16 \| 97.8 \| 115 \| (29, 152) \| 205 \| (52, 175) \| 266 \| Phage tail protein \| Phage tail protein; uniclust \| UniRef100\_UPI0021F725AA \| 98.8 \| 5.8e-11 \| 1.1e-16 \| 76.4 \| 56 \| (30, 85) \| 205 \| (10, 65) \| 66 \| phage tail protein I \| phage tail protein I; uniclust \| UniRef100\_A0A238TE88 \| 98.8 \| 5.7e-11 \| 1.1e-16 \| 87.5 \| 86 \| (65, 153) \| 205 \| (2, 103) \| 123 \| Uncharacterized protein \| Uncharacterized protein; uniclust \| UniRef100\_UPI0021AC41D2 \| 98.8 \| 6.8e-11 \| 1.2e-16 \| 95.6 \| 147 \| (4, 153) \| 205 \| (5, 164) \| 278 \| phage tail protein \| phage tail protein; uniclust \| UniRef100\_A0A4V2ZG90 \| 98.8 \| 6.9e-11 \| 1.3e-16 \| 106.1 \| 142 \| (30, 177) \| 205 \| (509, 676) \| 704 \| Phage tail protein \| Phage tail protein; uniclust \| UniRef100\_A0A0P8A262 \| 98.8 \| 7e-11 \| 1.3e-16 \| 93.6 \| 120 \| (45, 171) \| 205 \| (2, 171) \| 214 \| Uncharacterized protein \| Uncharacterized protein; uniclust \| UniRef100\_A0A968LQ35 \| 98.7 \| 8.3e-11 \| 1.5e-16 \| 92.6 \| 116 \| (30, 153) \| 205 \| (100, 215) \| 226 \| Phage tail protein I \| Phage tail protein I; uniclust \| UniRef100\_A0A059PCS6 \| 98.7 \| 8.6e-11 \| 1.6e-16 \| 101.5 \| 137 \| (1, 149) \| 205 \| (15, 153) \| 417 \| Tail I \| Tail I; uniclust \| UniRef100\_A0A938E8C9 \| 98.7 \| 9.1e-11 \| 1.7e-16 \| 89.0 \| 70 \| (30, 100) \| 205 \| (67, 140) \| 173 \| Uncharacterized protein (Fragment) \| Uncharacterized protein (Fragment); uniclust \| UniRef100\_A0A0B5FHF2 \| 98.7 \| 8.5e-11 \| 1.7e-16 \| 106.0 \| 101 \| (3, 104) \| 205 \| (11, 120) \| 529 \| Tail protein \| Tail protein; uniclust \| UniRef100\_A0A450WSG3 \| 98.7 \| 9.9e-11 \| 1.8e-16 \| 96.1 \| 145 \| (1, 150) \| 205 \| (1, 149) \| 315 \| P2-related tail formation protein \| P2-related tail formation protein; uniclust \| UniRef100\_A0A533MJS2 \| 98.7 \| 1e-10 \| 1.9e-16 \| 102.5 \| 89 \| (12, 101) \| 205 \| (333, 425) \| 582 \| Phage tail protein I \| Phage tail protein I; uniclust \| UniRef100\_A0A9E8EP50 \| 98.7 \| 1e-10 \| 1.9e-16 \| 94.2 \| 109 \| (39, 153) \| 205 \| (38, 164) \| 270 \| Phage tail protein \| Phage tail protein; uniclust \| UniRef100\_A0A124IN49 \| 98.7 \| 9.6e-11 \| 2e-16 \| 103.4 \| 152 \| (30, 188) \| 205 \| (196, 363) \| 381 \| Phage tail protein \| Phage tail protein; uniclust \| UniRef100\_UPI001FC81059 \| 98.7 \| 1.1e-10 \| 2.1e-16 \| 96.0 \| 119 \| (29, 153) \| 205 \| (153, 283) \| 321 \| phage tail protein \| phage tail protein; uniclust \| UniRef100\_UPI001FDF4D5B \| 98.7 \| 1.2e-10 \| 2.2e-16 \| 86.9 \| 69 \| (30, 98) \| 205 \| (33, 101) \| 155 \| phage tail protein I \| phage tail protein I; uniclust \| UniRef100\_A0A969BUE2 \| 98.7 \| 1.3e-10 \| 2.3e-16 \| 99.8 \| 117 \| (30, 151) \| 205 \| (240, 397) \| 466 \| Uncharacterized protein \| Uncharacterized protein; uniclust \| UniRef100\_A0A8J7KWD7 \| 98.7 \| 1.2e-10 \| 2.3e-16 \| 88.5 \| 110 \| (30, 151) \| 205 \| (42, 156) \| 158 \| Phage tail-like protein \| Phage tail-like protein; uniclust \| UniRef100\_A0A1J7BCP7 \| 98.7 \| 1.4e-10 \| 2.6e-16 \| 94.1 \| 120 \| (30, 153) \| 205 \| (60, 189) \| 252 \| Phage tail protein \| Phage tail protein; uniclust \| UniRef100\_A0A2W5C641 \| 98.7 \| 1.6e-10 \| 3.1e-16 \| 95.6 \| 139 \| (2, 147) \| 205 \| (5, 146) \| 266 \| Phage tail protein (Fragment) \| Phage tail protein (Fragment); uniclust \| UniRef100\_A0A8S5UB99 \| 98.7 \| 1.9e-10 \| 3.4e-16 \| 93.9 \| 108 \| (38, 151) \| 205 \| (75, 182) \| 295 \| Tail protein \| Tail protein; uniclust \| UniRef100\_U2F716 \| 98.7 \| 1.9e-10 \| 3.6e-16 \| 87.0 \| 118 \| (43, 167) \| 205 \| (1, 133) \| 143 \| Tail fiber \| Tail fiber; uniclust \| UniRef100\_A0A7C6KQ05 \| 98.7 \| 2e-10 \| 3.8e-16 \| 87.7 \| 133 \| (14, 150) \| 205 \| (21, 158) \| 180 \| Phage tail protein \| Phage tail protein; uniclust \| UniRef100\_A0A7J9Y123 \| 98.7 \| 2.2e-10 \| 4e-16 \| 92.2 \| 117 \| (30, 153) \| 205 \| (64, 181) \| 262 \| Phage tail protein \| Phage tail protein; uniclust \| UniRef100\_A0A0K2RLR3 \| 98.7 \| 2.3e-10 \| 4.3e-16 \| 91.4 \| 113 \| (30, 153) \| 205 \| (69, 194) \| 209 \| Phage tail protein \| Phage tail protein; uniclust \| UniRef100\_A0A7C4ELX4 \| 98.7 \| 2.4e-10 \| 4.4e-16 \| 88.6 \| 94 \| (2, 95) \| 205 \| (5, 102) \| 199 \| Phage tail protein (Fragment) \| Phage tail protein (Fragment); uniclust \| UniRef100\_UPI000DEEDDBB \| 98.7 \| 2.4e-10 \| 4.4e-16 \| 72.9 \| 61 \| (1, 61) \| 205 \| (1, 61) \| 62 \| phage tail protein I \| phage tail protein I; uniclust \| UniRef100\_A0A938CLU1 \| 98.7 \| 2.4e-10 \| 4.5e-16 \| 71.6 \| 56 \| (27, 82) \| 205 \| (2, 57) \| 57 \| Phage tail protein I \| Phage tail protein I; uniclust \| UniRef100\_A0A059IUT0 \| 98.7 \| 2.5e-10 \| 4.7e-16 \| 93.7 \| 137 \| (1, 149) \| 205 \| (15, 153) \| 269 \| Phage tail protein \| Phage tail protein; uniclust \| UniRef100\_A0A327JSB1 \| 98.6 \| 2.9e-10 \| 5.4e-16 \| 82.7 \| 89 \| (2, 91) \| 205 \| (6, 98) \| 130 \| Phage tail protein I (Fragment) \| Phage tail protein I (Fragment); uniclust \| UniRef100\_A0A371J258 \| 98.6 \| 3e-10 \| 5.6e-16 \| 82.4 \| 108 \| (2, 111) \| 205 \| (10, 123) \| 123 \| Phage tail protein (Fragment) \| Phage tail protein (Fragment); uniclust \| UniRef100\_A0A239INL8 \| 98.6 \| 3.2e-10 \| 6.3e-16 \| 94.7 \| 119 \| (29, 152) \| 205 \| (64, 187) \| 273 \| Phage tail protein domain-containing protein \| Phage tail protein domain-containing protein; uniclust \| UniRef100\_A0A212KXG2 \| 98.6 \| 3.3e-10 \| 6.3e-16 \| 100.3 \| 105 \| (40, 150) \| 205 \| (77, 181) \| 469 \| Phage tail protein \| Phage tail protein; uniclust \| UniRef100\_A0A4R7I2Z4 \| 98.6 \| 3.6e-10 \| 6.7e-16 \| 92.3 \| 121 \| (30, 154) \| 205 \| (51, 181) \| 271 \| Phage tail-like protein \| Phage tail-like protein; uniclust \| UniRef100\_A0A4R2JPC5 \| 98.6 \| 4.2e-10 \| 7.8e-16 \| 85.3 \| 116 \| (28, 151) \| 205 \| (41, 157) \| 158 \| Phage tail-like protein \| Phage tail-like protein; uniclust \| UniRef100\_UPI0022473D3F \| 98.6 \| 4.4e-10 \| 8.2e-16 \| 86.5 \| 110 \| (37, 147) \| 205 \| (27, 161) \| 188 \| phage tail protein \| phage tail protein; uniclust \| UniRef100\_A0A925IPT4 \| 98.6 \| 5.1e-10 \| 9.4e-16 \| 96.2 \| 115 \| (34, 153) \| 205 \| (207, 355) \| 463 \| Phage tail protein \| Phage tail protein; uniclust \| UniRef100\_UPI002356C388 \| 98.6 \| 5.2e-10 \| 9.5e-16 \| 85.0 \| 136 \| (3, 147) \| 205 \| (5, 144) \| 171 \| phage tail protein \| phage tail protein; uniclust \| UniRef100\_UPI001916A594 \| 98.6 \| 5.2e-10 \| 9.6e-16 \| 84.3 \| 139 \| (3, 153) \| 205 \| (6, 145) \| 162 \| phage tail protein \| phage tail protein; uniclust \| UniRef100\_G0FVT8 \| 98.6 \| 5.1e-10 \| 9.6e-16 \| 103.2 \| 120 \| (30, 151) \| 205 \| (711, 844) \| 847 \| Baseplate protein J-like domain-containing protein \| Baseplate protein J-like domain-containing protein; uniclust \| UniRef100\_UPI001EF24CD4 \| 98.6 \| 5.5e-10 \| 1e-15 \| 94.3 \| 132 \| (30, 163) \| 205 \| (180, 312) \| 391 \| phage tail protein \| phage tail protein; uniclust \| UniRef100\_A0A7W0VDI4 \| 98.6 \| 5.6e-10 \| 1e-15 \| 86.4 \| 70 \| (31, 101) \| 205 \| (118, 188) \| 194 \| PASTA domain-containing protein (Fragment) \| PASTA domain-containing protein (Fragment); uniclust \| UniRef100\_A0A1G9PB45 \| 98.6 \| 5.2e-10 \| 1.1e-15 \| 103.7 \| 63 \| (30, 92) \| 205 \| (74, 140) \| 561 \| Phage tail protein domain-containing protein \| Phage tail protein domain-containing protein; uniclust \| UniRef100\_A0A962F9E9 \| 98.6 \| 5.9e-10 \| 1.1e-15 \| 87.2 \| 148 \| (2, 153) \| 205 \| (8, 158) \| 209 \| Phage tail protein I \| Phage tail protein I; uniclust \| UniRef100\_UPI001E6408FD \| 98.6 \| 6e-10 \| 1.1e-15 \| 82.3 \| 134 \| (59, 204) \| 205 \| (6, 142) \| 142 \| DUF2313 domain-containing protein \| DUF2313 domain-containing protein |
| Top keywords  (threshold 1.00e-03 (evalue)) | **tail, Phage, I, Fragment, P2, P2\_related, formation, Bacteriophage, domain\_containing, Putative** |
| Output files | ../../similar\_sequences/14\_FANPEZAQ\_CDS\_0014\_merged.svg ../../similar\_sequences/14\_FANPEZAQ\_CDS\_0014\_pdb70.a3m ../../similar\_sequences/14\_FANPEZAQ\_CDS\_0014\_pdb70.hhr ../../similar\_sequences/14\_FANPEZAQ\_CDS\_0014\_uniclust.a3m ../../similar\_sequences/14\_FANPEZAQ\_CDS\_0014\_uniclust.hhr |

#### Structure prediction (AlphaFold)^2^

|  |  |
| --- | --- |
| Stats | xml version="1.0" encoding="utf-8" standalone="no"?       2024-09-02T21:09:12.909538 image/svg+xml   Matplotlib v3.7.2, https://matplotlib.org/ |
| Predicted structure | **NGL Viewer Controls:**  - Center: *Left-Click* - Rotate: *Left-Click + Drag* - Translate: *Right-Click + Drag* - Zoom: *Shift + Left-Click + Drag* |
| Output files | ../../predicted\_structures/14\_FANPEZAQ\_CDS\_0014/features.pkl ../../predicted\_structures/14\_FANPEZAQ\_CDS\_0014/ranked\_0.pdb ../../predicted\_structures/14\_FANPEZAQ\_CDS\_0014/ranked\_0\_plots.svg ../../predicted\_structures/14\_FANPEZAQ\_CDS\_0014/result\_model\_1\_ptm\_pred\_0.pkl |

#### Structure similarity search results (Foldseek)^3^

|  |  |
| --- | --- |
| Structure databases searched | Pdb, Afdb-proteome, Afdb-uniprot50 |
| Results, scheme(s)  (Top layers only, threshold 1.00e-02 (evalue)) | xml version="1.0" encoding="utf-8" standalone="no"?       2024-09-02T21:10:29.376195 image/svg+xml   Matplotlib v3.7.2, https://matplotlib.org/ |
| Results, table  (threshold 1.00e-02 (evalue)) | db \| id \| prob \| evalue \| bits \| fident \| alnlen \| mismatch \| gapopen \| qstart \| qend \| tstart \| tend \| name \| description; pdb \| 6U5B\_0 \| 1.0 \| 1.456e-10 \| 303 \| 0.348 \| 152 \| 97 \| 1 \| 1 \| 152 \| 1 \| 150 \| Tri2 PA0619 \| Tri2 PA0619; afdb-proteome \| AF-Q8ZMU4-F1-MODEL\_V4 \| 1.0 \| 7.453e-23 \| 701 \| 0.418 \| 203 \| 116 \| 1 \| 1 \| 203 \| 1 \| 201 \| Fels-2 prophage protein \| Fels-2 prophage protein; afdb-proteome \| AF-A0A0H3GQR7-F1-MODEL\_V4 \| 1.0 \| 1.573e-20 \| 600 \| 0.37 \| 200 \| 121 \| 2 \| 1 \| 197 \| 2 \| 199 \| Tail protein I \| Tail protein I; afdb-proteome \| AF-A0A0H3GWZ9-F1-MODEL\_V4 \| 1.0 \| 6.461e-19 \| 513 \| 0.359 \| 181 \| 114 \| 1 \| 5 \| 185 \| 1 \| 179 \| Tail protein I \| Tail protein I; afdb-proteome \| AF-Q8ZKK4-F1-MODEL\_V4 \| 1.0 \| 6.181e-16 \| 454 \| 0.317 \| 211 \| 134 \| 6 \| 1 \| 203 \| 1 \| 209 \| Putative phage tail protein \| Putative phage tail protein; afdb-proteome \| AF-G3XD92-F1-MODEL\_V4 \| 1.0 \| 9.347e-17 \| 443 \| 0.324 \| 179 \| 118 \| 2 \| 1 \| 178 \| 1 \| 177 \| Probable bacteriophage protein \| Probable bacteriophage protein; afdb-proteome \| AF-Q6NXT4-F1-MODEL\_V4 \| 0.817 \| 0.008268 \| 62 \| 0.128 \| 234 \| 147 \| 13 \| 8 \| 203 \| 169 \| 383 \| Zinc transporter 6 \| Zinc transporter 6; afdb-uniprot50 \| AF-A0A2Z3I623-F1-MODEL\_V4 \| 1.0 \| 4.101e-28 \| 866 \| 0.645 \| 203 \| 72 \| 0 \| 2 \| 204 \| 20 \| 222 \| Phage tail protein I \| Phage tail protein I; afdb-uniprot50 \| AF-C3X1Y3-F1-MODEL\_V4 \| 1.0 \| 1.568e-24 \| 765 \| 0.453 \| 205 \| 110 \| 1 \| 1 \| 205 \| 2 \| 204 \| Phage tail protein I \| Phage tail protein I; afdb-uniprot50 \| AF-A0A3S8ZPY7-F1-MODEL\_V4 \| 1.0 \| 8.51e-22 \| 730 \| 0.405 \| 180 \| 107 \| 0 \| 1 \| 180 \| 2 \| 181 \| Phage tail protein I \| Phage tail protein I; afdb-uniprot50 \| AF-A0A5C9AIB0-F1-MODEL\_V4 \| 1.0 \| 6.211e-22 \| 713 \| 0.435 \| 186 \| 103 \| 1 \| 1 \| 186 \| 1 \| 184 \| Phage tail protein I \| Phage tail protein I; afdb-uniprot50 \| AF-E4PPS5-F1-MODEL\_V4 \| 1.0 \| 9.652e-22 \| 712 \| 0.392 \| 191 \| 116 \| 0 \| 1 \| 191 \| 1 \| 191 \| Phage tail protein I \| Phage tail protein I; afdb-uniprot50 \| AF-A0A1N6I1G1-F1-MODEL\_V4 \| 1.0 \| 7.99e-22 \| 708 \| 0.388 \| 193 \| 117 \| 1 \| 4 \| 196 \| 3 \| 194 \| Phage tail protein, P2 protein I family \| Phage tail protein, P2 protein I family; afdb-uniprot50 \| AF-A0A4Y5WA04-F1-MODEL\_V4 \| 1.0 \| 2.188e-21 \| 705 \| 0.451 \| 175 \| 96 \| 0 \| 1 \| 175 \| 1 \| 175 \| Phage tail protein I \| Phage tail protein I; afdb-uniprot50 \| AF-A0A846QRB9-F1-MODEL\_V4 \| 1.0 \| 2.666e-23 \| 703 \| 0.443 \| 203 \| 113 \| 0 \| 1 \| 203 \| 2 \| 204 \| Phage tail P2-like protein \| Phage tail P2-like protein; afdb-uniprot50 \| AF-A0A0D7V3M6-F1-MODEL\_V4 \| 1.0 \| 1.065e-22 \| 703 \| 0.386 \| 207 \| 124 \| 2 \| 1 \| 204 \| 1 \| 207 \| Uncharacterized protein \| Uncharacterized protein; afdb-uniprot50 \| AF-A0A430HF69-F1-MODEL\_V4 \| 1.0 \| 1.541e-20 \| 703 \| 0.415 \| 178 \| 102 \| 2 \| 1 \| 176 \| 1 \| 178 \| Phage tail protein I \| Phage tail protein I; afdb-uniprot50 \| AF-Q8ZMU4-F1-MODEL\_V4 \| 1.0 \| 2.74e-22 \| 701 \| 0.418 \| 203 \| 116 \| 1 \| 1 \| 203 \| 1 \| 201 \| Fels-2 prophage protein \| Fels-2 prophage protein; afdb-uniprot50 \| AF-A0A5Y2X4S5-F1-MODEL\_V4 \| 1.0 \| 1.929e-21 \| 695 \| 0.46 \| 176 \| 91 \| 2 \| 1 \| 174 \| 1 \| 174 \| Phage tail protein I \| Phage tail protein I; afdb-uniprot50 \| AF-A0A7W5P3V3-F1-MODEL\_V4 \| 1.0 \| 6.437e-23 \| 689 \| 0.438 \| 203 \| 112 \| 1 \| 1 \| 203 \| 2 \| 202 \| Phage tail P2-like protein \| Phage tail P2-like protein; afdb-uniprot50 \| AF-A0A250DSF2-F1-MODEL\_V4 \| 1.0 \| 5.43e-20 \| 689 \| 0.382 \| 178 \| 107 \| 1 \| 1 \| 175 \| 1 \| 178 \| Phage tail protein I \| Phage tail protein I; afdb-uniprot50 \| AF-A0A328X6D0-F1-MODEL\_V4 \| 1.0 \| 2.331e-21 \| 685 \| 0.418 \| 184 \| 103 \| 2 \| 1 \| 181 \| 1 \| 183 \| Phage tail P2-like protein \| Phage tail P2-like protein; afdb-uniprot50 \| AF-A0A5Y8G7Y1-F1-MODEL\_V4 \| 1.0 \| 9.063e-22 \| 684 \| 0.448 \| 185 \| 98 \| 2 \| 1 \| 183 \| 1 \| 183 \| Phage tail protein I \| Phage tail protein I; afdb-uniprot50 \| AF-A0A7U7FJK2-F1-MODEL\_V4 \| 1.0 \| 2.13e-22 \| 684 \| 0.428 \| 203 \| 114 \| 1 \| 1 \| 203 \| 29 \| 229 \| Putative bacteriophage protein \| Putative bacteriophage protein; afdb-uniprot50 \| AF-A0A1Y3P6W7-F1-MODEL\_V4 \| 1.0 \| 1.701e-21 \| 683 \| 0.41 \| 185 \| 106 \| 2 \| 1 \| 185 \| 1 \| 182 \| Phage tail protein \| Phage tail protein; afdb-uniprot50 \| AF-A0A1I5KZ34-F1-MODEL\_V4 \| 1.0 \| 4.828e-22 \| 680 \| 0.414 \| 205 \| 118 \| 1 \| 1 \| 203 \| 1 \| 205 \| Phage tail protein, P2 protein I family \| Phage tail protein, P2 protein I family; afdb-uniprot50 \| AF-A0A137YJL7-F1-MODEL\_V4 \| 1.0 \| 1.276e-20 \| 678 \| 0.375 \| 176 \| 109 \| 1 \| 1 \| 175 \| 1 \| 176 \| Phage tail protein \| Phage tail protein; afdb-uniprot50 \| AF-A0A144SRX6-F1-MODEL\_V4 \| 1.0 \| 1.396e-19 \| 677 \| 0.474 \| 156 \| 78 \| 2 \| 1 \| 154 \| 1 \| 154 \| Phage Tail protein I \| Phage Tail protein I; afdb-uniprot50 \| AF-A0A1G7HXV1-F1-MODEL\_V4 \| 1.0 \| 5.832e-22 \| 677 \| 0.429 \| 205 \| 114 \| 2 \| 2 \| 205 \| 3 \| 205 \| Phage tail protein, P2 protein I family \| Phage tail protein, P2 protein I family; afdb-uniprot50 \| AF-A0A061JKL1-F1-MODEL\_V4 \| 1.0 \| 1.641e-20 \| 676 \| 0.357 \| 182 \| 116 \| 1 \| 1 \| 182 \| 1 \| 181 \| Uncharacterized protein \| Uncharacterized protein; afdb-uniprot50 \| AF-A0A1N6Q2Z7-F1-MODEL\_V4 \| 1.0 \| 2.716e-20 \| 675 \| 0.396 \| 179 \| 107 \| 1 \| 1 \| 178 \| 1 \| 179 \| Phage tail protein, P2 protein I family \| Phage tail protein, P2 protein I family; afdb-uniprot50 \| AF-A0A0N8R6G9-F1-MODEL\_V4 \| 1.0 \| 3.107e-22 \| 673 \| 0.413 \| 203 \| 117 \| 1 \| 1 \| 203 \| 56 \| 256 \| Tail protein I \| Tail protein I; afdb-uniprot50 \| AF-A0A0Q0CU47-F1-MODEL\_V4 \| 1.0 \| 1.701e-21 \| 670 \| 0.41 \| 190 \| 110 \| 1 \| 1 \| 190 \| 1 \| 188 \| Tail protein I \| Tail protein I; afdb-uniprot50 \| AF-A0A4Y6UC71-F1-MODEL\_V4 \| 1.0 \| 8.438e-20 \| 670 \| 0.346 \| 173 \| 112 \| 1 \| 4 \| 175 \| 18 \| 190 \| Phage tail protein I \| Phage tail protein I; afdb-uniprot50 \| AF-A0A1H0NP46-F1-MODEL\_V4 \| 1.0 \| 2.112e-20 \| 670 \| 0.388 \| 188 \| 115 \| 0 \| 5 \| 192 \| 1 \| 188 \| Phage tail protein, P2 protein I family \| Phage tail protein, P2 protein I family; afdb-uniprot50 \| AF-A0A6I3XGM5-F1-MODEL\_V4 \| 1.0 \| 2.461e-19 \| 666 \| 0.41 \| 178 \| 103 \| 2 \| 1 \| 178 \| 1 \| 176 \| Phage tail protein I \| Phage tail protein I; afdb-uniprot50 \| AF-A0A254PRB8-F1-MODEL\_V4 \| 1.0 \| 1.584e-19 \| 661 \| 0.359 \| 178 \| 112 \| 1 \| 1 \| 178 \| 6 \| 181 \| Phage tail protein I \| Phage tail protein I; afdb-uniprot50 \| AF-A0A395R2T8-F1-MODEL\_V4 \| 1.0 \| 4.787e-20 \| 659 \| 0.386 \| 181 \| 109 \| 1 \| 1 \| 179 \| 1 \| 181 \| Uncharacterized protein \| Uncharacterized protein; afdb-uniprot50 \| AF-A0A0S1B435-F1-MODEL\_V4 \| 1.0 \| 2.188e-21 \| 658 \| 0.371 \| 202 \| 127 \| 0 \| 4 \| 205 \| 3 \| 204 \| Phage P2 baseplate assembly gpI-like protein \| Phage P2 baseplate assembly gpI-like protein; afdb-uniprot50 \| AF-A0A485AEW3-F1-MODEL\_V4 \| 1.0 \| 3.622e-21 \| 657 \| 0.448 \| 194 \| 103 \| 2 \| 1 \| 192 \| 105 \| 296 \| Bacteriophage P2-related tail formation protein \| Bacteriophage P2-related tail formation protein; afdb-uniprot50 \| AF-A0A2N7UDJ2-F1-MODEL\_V4 \| 1.0 \| 1.701e-21 \| 654 \| 0.423 \| 203 \| 114 \| 3 \| 2 \| 203 \| 6 \| 206 \| Phage tail protein I \| Phage tail protein I; afdb-uniprot50 \| AF-A0A6B8QMK9-F1-MODEL\_V4 \| 1.0 \| 2.188e-21 \| 653 \| 0.366 \| 202 \| 128 \| 0 \| 4 \| 205 \| 9 \| 210 \| Phage tail protein I \| Phage tail protein I; afdb-uniprot50 \| AF-A0A1F1QBB0-F1-MODEL\_V4 \| 1.0 \| 9.063e-22 \| 652 \| 0.381 \| 202 \| 123 \| 1 \| 1 \| 202 \| 1 \| 200 \| Phage tail protein \| Phage tail protein; afdb-uniprot50 \| AF-A0A0D0TMA8-F1-MODEL\_V4 \| 1.0 \| 9.918e-21 \| 649 \| 0.432 \| 185 \| 103 \| 1 \| 1 \| 185 \| 1 \| 183 \| Phage tail protein \| Phage tail protein; afdb-uniprot50 \| AF-A0A3N6RV94-F1-MODEL\_V4 \| 1.0 \| 3.193e-21 \| 648 \| 0.428 \| 203 \| 114 \| 1 \| 1 \| 203 \| 182 \| 382 \| Phage tail protein I \| Phage tail protein I; afdb-uniprot50 \| AF-A0A2T5J1G3-F1-MODEL\_V4 \| 1.0 \| 1.276e-20 \| 644 \| 0.429 \| 205 \| 114 \| 3 \| 1 \| 205 \| 1 \| 202 \| Phage tail P2-like protein \| Phage tail P2-like protein; afdb-uniprot50 \| AF-A0A1X3ISP6-F1-MODEL\_V4 \| 1.0 \| 1.474e-17 \| 643 \| 0.496 \| 133 \| 65 \| 1 \| 22 \| 154 \| 8 \| 138 \| Phage tail protein I \| Phage tail protein I; afdb-uniprot50 \| AF-A0A7W6RFJ4-F1-MODEL\_V4 \| 1.0 \| 1.701e-21 \| 643 \| 0.404 \| 205 \| 119 \| 2 \| 2 \| 205 \| 5 \| 207 \| Phage tail P2-like protein \| Phage tail P2-like protein; afdb-uniprot50 \| AF-C1DS15-F1-MODEL\_V4 \| 1.0 \| 2.893e-20 \| 643 \| 0.383 \| 206 \| 123 \| 2 \| 1 \| 203 \| 1 \| 205 \| Phage P2 baseplate assembly gpI-like protein \| Phage P2 baseplate assembly gpI-like protein; afdb-uniprot50 \| AF-A0A7Z3GTI5-F1-MODEL\_V4 \| 1.0 \| 2.249e-20 \| 638 \| 0.424 \| 179 \| 100 \| 2 \| 1 \| 178 \| 1 \| 177 \| Phage tail protein I \| Phage tail protein I; afdb-uniprot50 \| AF-A0A5E9SCQ9-F1-MODEL\_V4 \| 1.0 \| 6.383e-21 \| 638 \| 0.421 \| 190 \| 108 \| 1 \| 14 \| 203 \| 2 \| 189 \| Phage tail protein I \| Phage tail protein I; afdb-uniprot50 \| AF-A0A143DE04-F1-MODEL\_V4 \| 1.0 \| 5.627e-21 \| 638 \| 0.371 \| 207 \| 127 \| 3 \| 1 \| 205 \| 2 \| 207 \| Uncharacterized protein \| Uncharacterized protein; afdb-uniprot50 \| AF-A0A1Y2SIU0-F1-MODEL\_V4 \| 1.0 \| 9.063e-22 \| 637 \| 0.436 \| 204 \| 112 \| 2 \| 1 \| 203 \| 1 \| 202 \| Phage tail protein I \| Phage tail protein I; afdb-uniprot50 \| AF-A0A2X4TXF0-F1-MODEL\_V4 \| 1.0 \| 6.159e-20 \| 637 \| 0.431 \| 176 \| 97 \| 2 \| 1 \| 175 \| 1 \| 174 \| Bacteriophage P2-related tail formation protein \| Bacteriophage P2-related tail formation protein; afdb-uniprot50 \| AF-A0A7W2HYN2-F1-MODEL\_V4 \| 1.0 \| 4.962e-21 \| 636 \| 0.376 \| 207 \| 125 \| 3 \| 1 \| 205 \| 1 \| 205 \| Phage tail protein I \| Phage tail protein I; afdb-uniprot50 \| AF-A0A2X3LRF3-F1-MODEL\_V4 \| 1.0 \| 7.71e-21 \| 633 \| 0.394 \| 195 \| 115 \| 2 \| 1 \| 194 \| 168 \| 360 \| Tail protein I \| Tail protein I; afdb-uniprot50 \| AF-A0A7C8HXE3-F1-MODEL\_V4 \| 1.0 \| 1.231e-19 \| 632 \| 0.369 \| 184 \| 112 \| 3 \| 1 \| 180 \| 1 \| 184 \| Uncharacterized protein \| Uncharacterized protein; afdb-uniprot50 \| AF-A0A5N9VY60-F1-MODEL\_V4 \| 1.0 \| 5.384e-18 \| 631 \| 0.42 \| 150 \| 85 \| 1 \| 24 \| 173 \| 2 \| 149 \| Phage tail protein I \| Phage tail protein I; afdb-uniprot50 \| AF-A0A6L2ZSC6-F1-MODEL\_V4 \| 1.0 \| 8.745e-21 \| 630 \| 0.442 \| 208 \| 112 \| 3 \| 1 \| 205 \| 6 \| 212 \| Phage tail protein I \| Phage tail protein I; afdb-uniprot50 \| AF-A0A2E3KNC1-F1-MODEL\_V4 \| 1.0 \| 1.276e-20 \| 626 \| 0.385 \| 192 \| 108 \| 3 \| 1 \| 188 \| 1 \| 186 \| Phage tail protein I \| Phage tail protein I; afdb-uniprot50 \| AF-A0A0H3ZIR2-F1-MODEL\_V4 \| 1.0 \| 2.112e-20 \| 621 \| 0.359 \| 203 \| 130 \| 0 \| 1 \| 203 \| 3 \| 205 \| Phage tail fiber protein \| Phage tail fiber protein; afdb-uniprot50 \| AF-A0A4U8YMN4-F1-MODEL\_V4 \| 1.0 \| 1.796e-19 \| 620 \| 0.393 \| 188 \| 109 \| 5 \| 1 \| 184 \| 1 \| 187 \| Tail p2 i: phage tail protein i \| Tail p2 i: phage tail protein i; afdb-uniprot50 \| AF-A0A2E9SIS4-F1-MODEL\_V4 \| 1.0 \| 1.796e-19 \| 620 \| 0.352 \| 193 \| 125 \| 0 \| 1 \| 193 \| 26 \| 218 \| Phage tail protein I \| Phage tail protein I; afdb-uniprot50 \| AF-A0A5D3J470-F1-MODEL\_V4 \| 1.0 \| 5.099e-20 \| 618 \| 0.379 \| 187 \| 111 \| 2 \| 1 \| 187 \| 2 \| 183 \| Phage tail protein I \| Phage tail protein I; afdb-uniprot50 \| AF-A0A7X4KDS0-F1-MODEL\_V4 \| 1.0 \| 4.92e-19 \| 617 \| 0.393 \| 178 \| 107 \| 1 \| 1 \| 178 \| 2 \| 178 \| Phage tail protein I \| Phage tail protein I; afdb-uniprot50 \| AF-A0A447P481-F1-MODEL\_V4 \| 1.0 \| 3.93e-18 \| 611 \| 0.428 \| 168 \| 92 \| 2 \| 33 \| 200 \| 1 \| 164 \| Putative phage tail protein \| Putative phage tail protein; afdb-uniprot50 \| AF-R6J7K3-F1-MODEL\_V4 \| 1.0 \| 9.234e-19 \| 610 \| 0.366 \| 183 \| 112 \| 3 \| 1 \| 181 \| 5 \| 185 \| Phage tail protein I \| Phage tail protein I; afdb-uniprot50 \| AF-G9Y651-F1-MODEL\_V4 \| 1.0 \| 3.824e-19 \| 609 \| 0.36 \| 183 \| 113 \| 3 \| 1 \| 180 \| 2 \| 183 \| Phage tail protein I \| Phage tail protein I; afdb-uniprot50 \| AF-A0A8A9XHG1-F1-MODEL\_V4 \| 1.0 \| 1.447e-20 \| 609 \| 0.365 \| 208 \| 127 \| 3 \| 1 \| 205 \| 1 \| 206 \| Phage tail protein I \| Phage tail protein I; afdb-uniprot50 \| AF-A0A7S9AD83-F1-MODEL\_V4 \| 1.0 \| 2.037e-19 \| 605 \| 0.351 \| 188 \| 115 \| 1 \| 1 \| 188 \| 1 \| 181 \| Phage tail protein I \| Phage tail protein I; afdb-uniprot50 \| AF-A0A0Q8Y302-F1-MODEL\_V4 \| 1.0 \| 3.193e-21 \| 603 \| 0.398 \| 206 \| 121 \| 1 \| 1 \| 203 \| 1 \| 206 \| Phage tail protein \| Phage tail protein; afdb-uniprot50 \| AF-K4Y7C4-F1-MODEL\_V4 \| 1.0 \| 8.597e-17 \| 600 \| 0.481 \| 133 \| 67 \| 1 \| 43 \| 175 \| 2 \| 132 \| Phage tail protein \| Phage tail protein; afdb-uniprot50 \| AF-A0A828QYR1-F1-MODEL\_V4 \| 1.0 \| 4.92e-19 \| 598 \| 0.42 \| 176 \| 100 \| 2 \| 28 \| 203 \| 1 \| 174 \| Phage tail protein I \| Phage tail protein I; afdb-uniprot50 \| AF-A0A809HAG4-F1-MODEL\_V4 \| 1.0 \| 7.645e-19 \| 595 \| 0.372 \| 177 \| 109 \| 2 \| 1 \| 176 \| 5 \| 180 \| Phage tail protein I \| Phage tail protein I; afdb-uniprot50 \| AF-A0A1Y3DKT1-F1-MODEL\_V4 \| 1.0 \| 2.461e-19 \| 594 \| 0.381 \| 181 \| 108 \| 3 \| 1 \| 179 \| 1 \| 179 \| Phage tail protein I \| Phage tail protein I; afdb-uniprot50 \| AF-A0A378Q2Y3-F1-MODEL\_V4 \| 1.0 \| 7.376e-18 \| 593 \| 0.373 \| 174 \| 107 \| 2 \| 2 \| 175 \| 19 \| 190 \| Bacteriophage P2-related tail formation protein \| Bacteriophage P2-related tail formation protein; afdb-uniprot50 \| AF-A0A2N0CX35-F1-MODEL\_V4 \| 1.0 \| 2.868e-18 \| 593 \| 0.397 \| 176 \| 104 \| 2 \| 1 \| 175 \| 5 \| 179 \| Phage tail protein I \| Phage tail protein I; afdb-uniprot50 \| AF-A0A8B5FYA0-F1-MODEL\_V4 \| 1.0 \| 1.672e-17 \| 591 \| 0.47 \| 153 \| 77 \| 2 \| 1 \| 151 \| 1 \| 151 \| Phage tail protein I \| Phage tail protein I; afdb-uniprot50 \| AF-A0A1F0H4V7-F1-MODEL\_V4 \| 1.0 \| 4.787e-20 \| 591 \| 0.331 \| 208 \| 131 \| 3 \| 1 \| 205 \| 4 \| 206 \| Uncharacterized protein \| Uncharacterized protein; afdb-uniprot50 \| AF-A0A4R2Z4J2-F1-MODEL\_V4 \| 1.0 \| 8.671e-19 \| 588 \| 0.448 \| 176 \| 94 \| 2 \| 1 \| 175 \| 1 \| 174 \| Phage tail P2-like protein \| Phage tail P2-like protein; afdb-uniprot50 \| AF-A0A6G8D863-F1-MODEL\_V4 \| 1.0 \| 2.791e-19 \| 587 \| 0.269 \| 208 \| 149 \| 2 \| 1 \| 205 \| 1 \| 208 \| Phage tail protein I \| Phage tail protein I; afdb-uniprot50 \| AF-A0A4P9VQK1-F1-MODEL\_V4 \| 1.0 \| 2.868e-18 \| 585 \| 0.411 \| 180 \| 101 \| 4 \| 1 \| 179 \| 1 \| 176 \| Phage tail protein I \| Phage tail protein I; afdb-uniprot50 \| AF-A0A5M7L3Q9-F1-MODEL\_V4 \| 1.0 \| 4.619e-19 \| 583 \| 0.395 \| 172 \| 102 \| 1 \| 4 \| 175 \| 9 \| 178 \| Phage tail protein I \| Phage tail protein I; afdb-uniprot50 \| AF-A0A7D5XQF7-F1-MODEL\_V4 \| 1.0 \| 7.239e-21 \| 581 \| 0.359 \| 203 \| 128 \| 1 \| 1 \| 203 \| 1 \| 201 \| Phage tail protein I \| Phage tail protein I; afdb-uniprot50 \| AF-A0A376J222-F1-MODEL\_V4 \| 1.0 \| 1.672e-17 \| 577 \| 0.48 \| 150 \| 76 \| 1 \| 26 \| 175 \| 7 \| 154 \| Tail protein I (GpI) \| Tail protein I (GpI); afdb-uniprot50 \| AF-N8XJQ6-F1-MODEL\_V4 \| 1.0 \| 4.337e-19 \| 576 \| 0.296 \| 206 \| 140 \| 3 \| 1 \| 205 \| 1 \| 202 \| Phage tail protein I \| Phage tail protein I; afdb-uniprot50 \| AF-A0A2T5HH08-F1-MODEL\_V4 \| 1.0 \| 1.311e-19 \| 572 \| 0.336 \| 211 \| 127 \| 4 \| 1 \| 205 \| 2 \| 205 \| Phage tail P2-like protein \| Phage tail P2-like protein; afdb-uniprot50 \| AF-A0A103EG21-F1-MODEL\_V4 \| 1.0 \| 8.986e-20 \| 571 \| 0.376 \| 202 \| 125 \| 1 \| 1 \| 201 \| 1 \| 202 \| Uncharacterized protein \| Uncharacterized protein; afdb-uniprot50 \| AF-A0A085ASA6-F1-MODEL\_V4 \| 1.0 \| 7.58e-17 \| 570 \| 0.385 \| 174 \| 100 \| 4 \| 1 \| 168 \| 1 \| 173 \| Tail protein I \| Tail protein I; afdb-uniprot50 \| AF-A0A2Y0P998-F1-MODEL\_V4 \| 1.0 \| 2.354e-16 \| 567 \| 0.372 \| 153 \| 94 \| 1 \| 1 \| 153 \| 13 \| 163 \| Putative tail protein I (Gpi) \| Putative tail protein I (Gpi); afdb-uniprot50 \| AF-A0A2U9LD38-F1-MODEL\_V4 \| 1.0 \| 3.372e-19 \| 567 \| 0.365 \| 205 \| 128 \| 1 \| 1 \| 205 \| 3 \| 205 \| Phage tail protein I \| Phage tail protein I; afdb-uniprot50 \| AF-A0A853IMJ7-F1-MODEL\_V4 \| 1.0 \| 1.897e-17 \| 566 \| 0.321 \| 171 \| 113 \| 3 \| 1 \| 169 \| 4 \| 173 \| Phage tail protein I \| Phage tail protein I; afdb-uniprot50 \| AF-A0A1Q5W016-F1-MODEL\_V4 \| 1.0 \| 4.457e-18 \| 564 \| 0.431 \| 176 \| 97 \| 2 \| 1 \| 175 \| 1 \| 174 \| Phage tail protein I \| Phage tail protein I; afdb-uniprot50 \| AF-A0A853IED5-F1-MODEL\_V4 \| 1.0 \| 1.83e-16 \| 563 \| 0.335 \| 155 \| 101 \| 2 \| 26 \| 179 \| 3 \| 156 \| Phage tail protein I \| Phage tail protein I; afdb-uniprot50 \| AF-V5ZB74-F1-MODEL\_V4 \| 1.0 \| 1.913e-19 \| 562 \| 0.36 \| 186 \| 116 \| 2 \| 1 \| 185 \| 1 \| 184 \| Tail protein I GpI \| Tail protein I GpI; afdb-uniprot50 \| AF-A0A4T2AB33-F1-MODEL\_V4 \| 1.0 \| 3.93e-18 \| 562 \| 0.321 \| 205 \| 133 \| 4 \| 3 \| 205 \| 4 \| 204 \| Phage tail protein I \| Phage tail protein I; afdb-uniprot50 \| AF-A0A7U1PSN1-F1-MODEL\_V4 \| 1.0 \| 4.073e-19 \| 561 \| 0.328 \| 201 \| 130 \| 3 \| 1 \| 198 \| 1 \| 199 \| Phage tail protein I \| Phage tail protein I; afdb-uniprot50 \| AF-A0A0M4U7R5-F1-MODEL\_V4 \| 1.0 \| 1.178e-16 \| 560 \| 0.32 \| 175 \| 117 \| 2 \| 1 \| 175 \| 3 \| 175 \| Phage tail protein \| Phage tail protein; afdb-uniprot50 \| AF-A0A829DXU4-F1-MODEL\_V4 \| 1.0 \| 2.599e-17 \| 559 \| 0.318 \| 179 \| 121 \| 1 \| 1 \| 179 \| 1 \| 178 \| Phage tail protein I \| Phage tail protein I; afdb-uniprot50 \| AF-A0A743CBY9-F1-MODEL\_V4 \| 1.0 \| 2.037e-19 \| 559 \| 0.311 \| 202 \| 138 \| 1 \| 1 \| 201 \| 3 \| 204 \| Phage tail protein I \| Phage tail protein I; afdb-uniprot50 \| AF-A0A0D8TVW9-F1-MODEL\_V4 \| 1.0 \| 1.076e-17 \| 557 \| 0.328 \| 189 \| 122 \| 2 \| 1 \| 189 \| 2 \| 185 \| Tail protein \| Tail protein; afdb-uniprot50 \| AF-A0A5E6YIM8-F1-MODEL\_V4 \| 1.0 \| 1.897e-17 \| 557 \| 0.379 \| 179 \| 103 \| 4 \| 1 \| 175 \| 3 \| 177 \| Uncharacterized protein \| Uncharacterized protein; afdb-uniprot50 \| AF-A0A2U3F145-F1-MODEL\_V4 \| 1.0 \| 2.094e-18 \| 555 \| 0.405 \| 180 \| 105 \| 1 \| 24 \| 203 \| 5 \| 182 \| Phage tail protein I \| Phage tail protein I; afdb-uniprot50 \| AF-A0A1E7RBV3-F1-MODEL\_V4 \| 1.0 \| 4.073e-19 \| 552 \| 0.357 \| 190 \| 111 \| 2 \| 1 \| 188 \| 2 \| 182 \| Phage tail protein I \| Phage tail protein I; afdb-uniprot50 \| AF-A0A145VQK9-F1-MODEL\_V4 \| 1.0 \| 9.489e-18 \| 551 \| 0.331 \| 202 \| 132 \| 3 \| 1 \| 201 \| 5 \| 204 \| Uncharacterized protein \| Uncharacterized protein; afdb-uniprot50 \| AF-A0A362XRN1-F1-MODEL\_V4 \| 1.0 \| 2.791e-19 \| 547 \| 0.412 \| 177 \| 104 \| 0 \| 28 \| 204 \| 1 \| 177 \| Phage tail P2-like protein \| Phage tail P2-like protein; afdb-uniprot50 \| AF-A0A828I9B9-F1-MODEL\_V4 \| 1.0 \| 2.058e-14 \| 547 \| 0.359 \| 128 \| 80 \| 1 \| 26 \| 153 \| 1 \| 126 \| Phage tail protein I \| Phage tail protein I; afdb-uniprot50 \| AF-A0A246KFX4-F1-MODEL\_V4 \| 1.0 \| 3.435e-16 \| 545 \| 0.322 \| 183 \| 117 \| 4 \| 1 \| 178 \| 8 \| 188 \| Bacteriophage P2-related tail formation protein \| Bacteriophage P2-related tail formation protein; afdb-uniprot50 \| AF-A0A7C1LB13-F1-MODEL\_V4 \| 1.0 \| 6.448e-16 \| 544 \| 0.344 \| 151 \| 95 \| 1 \| 38 \| 188 \| 1 \| 147 \| Phage tail protein I \| Phage tail protein I; afdb-uniprot50 \| AF-A0A4U8YL85-F1-MODEL\_V4 \| 1.0 \| 4.58e-17 \| 544 \| 0.32 \| 206 \| 137 \| 3 \| 1 \| 205 \| 3 \| 206 \| Tail p2 i: phage tail protein i \| Tail p2 i: phage tail protein i; afdb-uniprot50 \| AF-W0ITV5-F1-MODEL\_V4 \| 1.0 \| 3.93e-18 \| 543 \| 0.383 \| 201 \| 118 \| 3 \| 4 \| 203 \| 11 \| 206 \| Tail protein \| Tail protein; afdb-uniprot50 \| AF-L3CBD8-F1-MODEL\_V4 \| 1.0 \| 8.141e-19 \| 543 \| 0.349 \| 206 \| 129 \| 3 \| 1 \| 203 \| 1 \| 204 \| Phage tail protein I \| Phage tail protein I; afdb-uniprot50 \| AF-A0A2I8XW61-F1-MODEL\_V4 \| 1.0 \| 3.435e-16 \| 543 \| 0.369 \| 173 \| 106 \| 3 \| 2 \| 172 \| 7 \| 178 \| Phage tail protein I \| Phage tail protein I; afdb-uniprot50 \| AF-A0A6N3DZD4-F1-MODEL\_V4 \| 1.0 \| 4.419e-16 \| 540 \| 0.374 \| 171 \| 104 \| 3 \| 4 \| 172 \| 9 \| 178 \| Phage tail protein (Tail\_P2\_I) \| Phage tail protein (Tail\_P2\_I); afdb-uniprot50 \| AF-A0A5D3JC96-F1-MODEL\_V4 \| 1.0 \| 1.188e-18 \| 539 \| 0.331 \| 187 \| 120 \| 2 \| 1 \| 187 \| 2 \| 183 \| Phage tail protein I \| Phage tail protein I; afdb-uniprot50 \| AF-A0A5E4XFG2-F1-MODEL\_V4 \| 1.0 \| 1.178e-16 \| 538 \| 0.326 \| 187 \| 117 \| 4 \| 1 \| 185 \| 1 \| 180 \| Phage tail protein I \| Phage tail protein I; afdb-uniprot50 \| AF-A0A7H4P5M4-F1-MODEL\_V4 \| 1.0 \| 7.58e-17 \| 538 \| 0.367 \| 177 \| 107 \| 4 \| 1 \| 175 \| 6 \| 179 \| Putative prophage tail protein \| Putative prophage tail protein; afdb-uniprot50 \| AF-A0A7V8DKV8-F1-MODEL\_V4 \| 1.0 \| 2.599e-17 \| 535 \| 0.351 \| 188 \| 114 \| 4 \| 1 \| 186 \| 1 \| 182 \| Phage tail protein P2 protein I family \| Phage tail protein P2 protein I family; afdb-uniprot50 \| AF-A0A7Y3N1Q0-F1-MODEL\_V4 \| 1.0 \| 8.366e-18 \| 535 \| 0.27 \| 207 \| 149 \| 2 \| 1 \| 205 \| 5 \| 211 \| Phage tail protein I \| Phage tail protein I; afdb-uniprot50 \| AF-A0A849VNG3-F1-MODEL\_V4 \| 1.0 \| 1.136e-15 \| 534 \| 0.323 \| 164 \| 101 \| 3 \| 1 \| 157 \| 2 \| 162 \| Phage tail protein I \| Phage tail protein I; afdb-uniprot50 \| AF-R9ARZ6-F1-MODEL\_V4 \| 1.0 \| 3.792e-17 \| 533 \| 0.318 \| 176 \| 120 \| 0 \| 1 \| 176 \| 1 \| 176 \| Phage tail protein I \| Phage tail protein I; afdb-uniprot50 \| AF-A0A5M8FWX9-F1-MODEL\_V4 \| 1.0 \| 3.343e-17 \| 533 \| 0.276 \| 210 \| 145 \| 4 \| 1 \| 205 \| 1 \| 208 \| Phage tail protein I \| Phage tail protein I; afdb-uniprot50 \| AF-A0A348HHI4-F1-MODEL\_V4 \| 1.0 \| 1.614e-16 \| 533 \| 0.296 \| 209 \| 141 \| 5 \| 1 \| 205 \| 1 \| 207 \| Bacteriophage P2-related tail formation protein \| Bacteriophage P2-related tail formation protein; afdb-uniprot50 \| AF-A0A066RUL7-F1-MODEL\_V4 \| 1.0 \| 3.792e-17 \| 532 \| 0.286 \| 192 \| 133 \| 2 \| 13 \| 203 \| 2 \| 190 \| Tail protein \| Tail protein; afdb-uniprot50 \| AF-A0A6D0PQW3-F1-MODEL\_V4 \| 1.0 \| 1.178e-16 \| 531 \| 0.355 \| 177 \| 106 \| 5 \| 1 \| 171 \| 1 \| 175 \| Phage tail protein I \| Phage tail protein I; afdb-uniprot50 \| AF-F4V3B6-F1-MODEL\_V4 \| 1.0 \| 2.44e-17 \| 530 \| 0.324 \| 197 \| 126 \| 4 \| 1 \| 191 \| 1 \| 196 \| Tail protein I (GpI) \| Tail protein I (GpI); afdb-uniprot50 \| AF-G4CPH7-F1-MODEL\_V4 \| 1.0 \| 1.584e-19 \| 528 \| 0.385 \| 205 \| 123 \| 2 \| 1 \| 205 \| 3 \| 204 \| Phage tail protein I \| Phage tail protein I; afdb-uniprot50 \| AF-A0A1I1QCJ1-F1-MODEL\_V4 \| 1.0 \| 9.489e-18 \| 528 \| 0.294 \| 214 \| 139 \| 5 \| 1 \| 205 \| 1 \| 211 \| Phage tail protein, P2 protein I family \| Phage tail protein, P2 protein I family; afdb-uniprot50 \| AF-A0A547EQ25-F1-MODEL\_V4 \| 1.0 \| 4.337e-19 \| 526 \| 0.334 \| 206 \| 136 \| 1 \| 1 \| 205 \| 4 \| 209 \| Phage tail protein I \| Phage tail protein I; afdb-uniprot50 \| AF-A0A827H7B9-F1-MODEL\_V4 \| 1.0 \| 8.597e-17 \| 525 \| 0.352 \| 176 \| 111 \| 2 \| 2 \| 175 \| 6 \| 180 \| Phage tail protein I \| Phage tail protein I; afdb-uniprot50 \| AF-A0A702LDY9-F1-MODEL\_V4 \| 1.0 \| 1.254e-16 \| 523 \| 0.369 \| 176 \| 103 \| 4 \| 1 \| 170 \| 1 \| 174 \| Phage tail protein I \| Phage tail protein I; afdb-uniprot50 \| AF-A0A853IAZ5-F1-MODEL\_V4 \| 1.0 \| 9.489e-18 \| 522 \| 0.378 \| 198 \| 112 \| 4 \| 1 \| 191 \| 2 \| 195 \| Phage tail protein I \| Phage tail protein I; afdb-uniprot50 \| AF-A0A7Y7D7P6-F1-MODEL\_V4 \| 1.0 \| 1.423e-16 \| 521 \| 0.296 \| 182 \| 122 \| 3 \| 1 \| 180 \| 1 \| 178 \| Phage tail protein I \| Phage tail protein I; afdb-uniprot50 \| AF-A0A7X5J840-F1-MODEL\_V4 \| 1.0 \| 5.532e-17 \| 521 \| 0.269 \| 208 \| 147 \| 3 \| 1 \| 205 \| 2 \| 207 \| Phage tail protein I \| Phage tail protein I; afdb-uniprot50 \| AF-A1VSH3-F1-MODEL\_V4 \| 1.0 \| 8.671e-19 \| 520 \| 0.348 \| 172 \| 112 \| 0 \| 2 \| 173 \| 5 \| 176 \| Phage tail protein I \| Phage tail protein I; afdb-uniprot50 \| AF-A0A7V7NR87-F1-MODEL\_V4 \| 1.0 \| 2.076e-16 \| 520 \| 0.301 \| 196 \| 134 \| 2 \| 1 \| 195 \| 3 \| 196 \| Phage tail protein I \| Phage tail protein I; afdb-uniprot50 \| AF-A0A1X3I4G6-F1-MODEL\_V4 \| 1.0 \| 1.047e-18 \| 520 \| 0.342 \| 207 \| 125 \| 5 \| 1 \| 200 \| 2 \| 204 \| Putative phage tail protein \| Putative phage tail protein; afdb-uniprot50 \| AF-A0A4Q3CYC6-F1-MODEL\_V4 \| 1.0 \| 1.336e-16 \| 519 \| 0.38 \| 155 \| 96 \| 0 \| 48 \| 202 \| 1 \| 155 \| Phage tail protein I \| Phage tail protein I; afdb-uniprot50 \| AF-L0R6A3-F1-MODEL\_V4 \| 1.0 \| 6.503e-18 \| 519 \| 0.301 \| 209 \| 137 \| 3 \| 1 \| 205 \| 2 \| 205 \| Uncharacterized protein \| Uncharacterized protein; afdb-uniprot50 \| AF-A0A7H9K2Z1-F1-MODEL\_V4 \| 1.0 \| 4.878e-17 \| 519 \| 0.366 \| 180 \| 106 \| 4 \| 1 \| 174 \| 1 \| 178 \| Phage tail protein I \| Phage tail protein I; afdb-uniprot50 \| AF-A0A5S3R576-F1-MODEL\_V4 \| 1.0 \| 1.781e-17 \| 518 \| 0.331 \| 211 \| 133 \| 4 \| 1 \| 205 \| 1 \| 209 \| Phage tail protein I \| Phage tail protein I; afdb-uniprot50 \| AF-A0A726YCV2-F1-MODEL\_V4 \| 1.0 \| 4.495e-20 \| 518 \| 0.366 \| 224 \| 121 \| 4 \| 1 \| 205 \| 2 \| 223 \| Phage tail protein I \| Phage tail protein I; afdb-uniprot50 \| AF-A0A1N7LRG2-F1-MODEL\_V4 \| 1.0 \| 1.614e-16 \| 517 \| 0.251 \| 207 \| 150 \| 4 \| 1 \| 205 \| 5 \| 208 \| Phage tail protein, P2 protein I family \| Phage tail protein, P2 protein I family; afdb-uniprot50 \| AF-A0A2J5PKJ6-F1-MODEL\_V4 \| 1.0 \| 1.766e-15 \| 516 \| 0.383 \| 159 \| 95 \| 2 \| 4 \| 160 \| 9 \| 166 \| Phage tail protein I \| Phage tail protein I; afdb-uniprot50 \| AF-A0A2A3WJP6-F1-MODEL\_V4 \| 1.0 \| 7.58e-17 \| 516 \| 0.388 \| 180 \| 100 \| 5 \| 1 \| 170 \| 1 \| 180 \| Phage tail protein I \| Phage tail protein I; afdb-uniprot50 \| AF-A0A3E0X4D1-F1-MODEL\_V4 \| 1.0 \| 3.055e-18 \| 515 \| 0.34 \| 179 \| 115 \| 2 \| 1 \| 178 \| 2 \| 178 \| Phage tail protein I \| Phage tail protein I; afdb-uniprot50 \| AF-A0A7X9X6E3-F1-MODEL\_V4 \| 1.0 \| 1.336e-16 \| 515 \| 0.331 \| 184 \| 114 \| 4 \| 1 \| 180 \| 1 \| 179 \| Phage tail protein I \| Phage tail protein I; afdb-uniprot50 \| AF-A0A1W1XKA3-F1-MODEL\_V4 \| 1.0 \| 6.106e-18 \| 514 \| 0.382 \| 178 \| 106 \| 4 \| 1 \| 176 \| 1 \| 176 \| Phage tail protein, P2 protein I family \| Phage tail protein, P2 protein I family; afdb-uniprot50 \| AF-A0A6L9MLM2-F1-MODEL\_V4 \| 1.0 \| 4.878e-17 \| 514 \| 0.291 \| 206 \| 141 \| 3 \| 4 \| 205 \| 17 \| 221 \| Phage tail protein I \| Phage tail protein I; afdb-uniprot50 \| AF-A0A7K0GMX4-F1-MODEL\_V4 \| 1.0 \| 4.004e-15 \| 514 \| 0.339 \| 165 \| 100 \| 3 \| 1 \| 157 \| 1 \| 164 \| Phage tail protein I \| Phage tail protein I; afdb-uniprot50 \| AF-A0A2N3KX23-F1-MODEL\_V4 \| 1.0 \| 4.301e-17 \| 513 \| 0.338 \| 201 \| 130 \| 3 \| 4 \| 202 \| 3 \| 202 \| Phage tail protein I \| Phage tail protein I; afdb-uniprot50 \| AF-A0A4V3E318-F1-MODEL\_V4 \| 1.0 \| 1.038e-16 \| 513 \| 0.297 \| 195 \| 124 \| 5 \| 1 \| 191 \| 1 \| 186 \| Phage tail P2-like protein \| Phage tail P2-like protein; afdb-uniprot50 \| AF-A0A3Q8FAV9-F1-MODEL\_V4 \| 1.0 \| 4.073e-19 \| 513 \| 0.385 \| 184 \| 108 \| 1 \| 1 \| 184 \| 1 \| 179 \| Phage tail protein I \| Phage tail protein I; afdb-uniprot50 \| AF-A0A495BJ44-F1-MODEL\_V4 \| 1.0 \| 2.768e-17 \| 511 \| 0.314 \| 207 \| 129 \| 3 \| 4 \| 205 \| 5 \| 203 \| Phage tail P2-like protein \| Phage tail P2-like protein; afdb-uniprot50 \| AF-A0A5S3XWD0-F1-MODEL\_V4 \| 1.0 \| 5.151e-15 \| 510 \| 0.27 \| 174 \| 117 \| 3 \| 1 \| 167 \| 8 \| 178 \| Phage tail protein I \| Phage tail protein I; afdb-uniprot50 \| AF-A0A377XEC8-F1-MODEL\_V4 \| 1.0 \| 1.83e-16 \| 509 \| 0.414 \| 157 \| 86 \| 3 \| 1 \| 156 \| 1 \| 152 \| Tail protein I \| Tail protein I; afdb-uniprot50 \| AF-A0A2S6H5I0-F1-MODEL\_V4 \| 1.0 \| 1.21e-15 \| 508 \| 0.395 \| 167 \| 83 \| 4 \| 1 \| 151 \| 1 \| 165 \| Phage tail P2-like protein \| Phage tail P2-like protein; afdb-uniprot50 \| AF-A0A853SGW9-F1-MODEL\_V4 \| 1.0 \| 3.166e-19 \| 508 \| 0.364 \| 206 \| 130 \| 1 \| 1 \| 205 \| 3 \| 208 \| Phage tail protein (Tail\_P2\_I) \| Phage tail protein (Tail\_P2\_I); afdb-uniprot50 \| AF-A0A1Y2K0Y0-F1-MODEL\_V4 \| 1.0 \| 7.117e-17 \| 508 \| 0.33 \| 218 \| 129 \| 4 \| 1 \| 205 \| 2 \| 215 \| Putative phage tail protein I \| Putative phage tail protein I; afdb-uniprot50 \| AF-A0A5P9F0G1-F1-MODEL\_V4 \| 1.0 \| 5.012e-16 \| 507 \| 0.306 \| 186 \| 119 \| 5 \| 1 \| 181 \| 1 \| 181 \| Phage tail protein \| Phage tail protein; afdb-uniprot50 \| AF-A0A4P0YQA9-F1-MODEL\_V4 \| 1.0 \| 1.474e-17 \| 506 \| 0.345 \| 200 \| 125 \| 5 \| 1 \| 197 \| 1 \| 197 \| Tail protein I (GpI) from prophage \| Tail protein I (GpI) from prophage; afdb-uniprot50 \| AF-A0A5J6LD05-F1-MODEL\_V4 \| 1.0 \| 6.683e-17 \| 506 \| 0.309 \| 207 \| 136 \| 4 \| 1 \| 205 \| 1 \| 202 \| Phage tail protein I \| Phage tail protein I; afdb-uniprot50 \| AF-A0A2G2L0X1-F1-MODEL\_V4 \| 1.0 \| 1.396e-19 \| 506 \| 0.368 \| 209 \| 123 \| 6 \| 1 \| 205 \| 2 \| 205 \| Phage tail protein I \| Phage tail protein I; afdb-uniprot50 \| AF-A0A166Z6T7-F1-MODEL\_V4 \| 1.0 \| 1.672e-17 \| 506 \| 0.31 \| 222 \| 135 \| 5 \| 1 \| 205 \| 4 \| 224 \| Tail protein \| Tail protein; afdb-uniprot50 \| AF-A0A0B6CYD1-F1-MODEL\_V4 \| 1.0 \| 1.038e-16 \| 505 \| 0.336 \| 205 \| 125 \| 4 \| 1 \| 205 \| 1 \| 194 \| Phage tail protein I \| Phage tail protein I; afdb-uniprot50 \| AF-A0A2A3MGU4-F1-MODEL\_V4 \| 1.0 \| 3.253e-18 \| 505 \| 0.375 \| 173 \| 106 \| 1 \| 4 \| 174 \| 42 \| 214 \| Phage tail protein I \| Phage tail protein I; afdb-uniprot50 \| AF-A0A4R0DV06-F1-MODEL\_V4 \| 1.0 \| 1.719e-16 \| 504 \| 0.352 \| 176 \| 113 \| 1 \| 1 \| 176 \| 1 \| 175 \| Phage tail protein I \| Phage tail protein I; afdb-uniprot50 \| AF-A0A376SCH0-F1-MODEL\_V4 \| 1.0 \| 2.507e-16 \| 504 \| 0.397 \| 176 \| 104 \| 1 \| 1 \| 176 \| 3 \| 176 \| Tail protein \| Tail protein; afdb-uniprot50 \| AF-A0A1M3PG28-F1-MODEL\_V4 \| 1.0 \| 4.419e-16 \| 503 \| 0.346 \| 182 \| 108 \| 5 \| 4 \| 181 \| 3 \| 177 \| Phage tail protein I \| Phage tail protein I; afdb-uniprot50 \| AF-A0A1M7YYQ0-F1-MODEL\_V4 \| 1.0 \| 3.76e-15 \| 503 \| 0.354 \| 161 \| 95 \| 3 \| 1 \| 152 \| 1 \| 161 \| Phage tail protein (Tail\_P2\_I) \| Phage tail protein (Tail\_P2\_I); afdb-uniprot50 \| AF-A0A291LZR2-F1-MODEL\_V4 \| 1.0 \| 7.856e-18 \| 501 \| 0.306 \| 183 \| 125 \| 1 \| 1 \| 183 \| 1 \| 181 \| Phage tail protein I \| Phage tail protein I; afdb-uniprot50 \| AF-A0A2A4XSX7-F1-MODEL\_V4 \| 1.0 \| 1.881e-15 \| 501 \| 0.337 \| 166 \| 101 \| 3 \| 1 \| 159 \| 1 \| 164 \| Phage tail protein I \| Phage tail protein I; afdb-uniprot50 \| AF-A0A1A9KGZ0-F1-MODEL\_V4 \| 1.0 \| 1.557e-15 \| 501 \| 0.378 \| 156 \| 97 \| 0 \| 48 \| 203 \| 2 \| 157 \| Phage tail protein I \| Phage tail protein I; afdb-uniprot50 \| AF-A0A380C5Q3-F1-MODEL\_V4 \| 1.0 \| 7.58e-17 \| 500 \| 0.31 \| 206 \| 136 \| 2 \| 1 \| 201 \| 2 \| 206 \| Bacteriophage P2-related tail formation protein \| Bacteriophage P2-related tail formation protein; afdb-uniprot50 \| AF-A0A2G4U387-F1-MODEL\_V4 \| 1.0 \| 9.57e-20 \| 500 \| 0.377 \| 204 \| 124 \| 2 \| 1 \| 203 \| 17 \| 218 \| Phage tail protein I \| Phage tail protein I; afdb-uniprot50 \| AF-A0A5Q2KBP8-F1-MODEL\_V4 \| 1.0 \| 6.926e-18 \| 499 \| 0.395 \| 172 \| 102 \| 1 \| 4 \| 175 \| 37 \| 206 \| Phage tail protein I \| Phage tail protein I; afdb-uniprot50 \| AF-A0A853HRN2-F1-MODEL\_V4 \| 1.0 \| 6.683e-17 \| 498 \| 0.328 \| 198 \| 109 \| 5 \| 2 \| 179 \| 4 \| 197 \| Phage tail protein I \| Phage tail protein I; afdb-uniprot50 \| AF-A0A857EA04-F1-MODEL\_V4 \| 1.0 \| 4.747e-18 \| 496 \| 0.364 \| 181 \| 112 \| 2 \| 1 \| 180 \| 3 \| 181 \| Phage tail protein I \| Phage tail protein I; afdb-uniprot50 \| AF-A0A7J0BYT0-F1-MODEL\_V4 \| 1.0 \| 1.515e-16 \| 496 \| 0.352 \| 207 \| 129 \| 3 \| 1 \| 205 \| 1 \| 204 \| Uncharacterized protein \| Uncharacterized protein; afdb-uniprot50 \| AF-A0A6S5X5R7-F1-MODEL\_V4 \| 1.0 \| 5.532e-17 \| 496 \| 0.292 \| 226 \| 138 \| 4 \| 1 \| 205 \| 2 \| 226 \| Uncharacterized protein \| Uncharacterized protein; afdb-uniprot50 \| AF-A0A1V3IL14-F1-MODEL\_V4 \| 1.0 \| 5.055e-18 \| 495 \| 0.392 \| 176 \| 107 \| 0 \| 1 \| 176 \| 3 \| 178 \| Phage tail protein I \| Phage tail protein I; afdb-uniprot50 \| AF-A0A4S3LTA7-F1-MODEL\_V4 \| 1.0 \| 1.3e-17 \| 494 \| 0.397 \| 176 \| 103 \| 2 \| 1 \| 175 \| 1 \| 174 \| Phage tail protein I \| Phage tail protein I; afdb-uniprot50 \| AF-A0A0D8DEP2-F1-MODEL\_V4 \| 1.0 \| 2.211e-16 \| 494 \| 0.297 \| 212 \| 141 \| 2 \| 2 \| 205 \| 7 \| 218 \| Tail protein \| Tail protein; afdb-uniprot50 \| AF-A0A125AY37-F1-MODEL\_V4 \| 1.0 \| 4.707e-16 \| 493 \| 0.304 \| 184 \| 120 \| 3 \| 1 \| 180 \| 1 \| 180 \| Phage tail protein \| Phage tail protein; afdb-uniprot50 \| AF-A0A212KBJ6-F1-MODEL\_V4 \| 1.0 \| 9.751e-17 \| 493 \| 0.296 \| 233 \| 132 \| 5 \| 4 \| 205 \| 3 \| 234 \| Uncharacterized protein \| Uncharacterized protein; afdb-uniprot50 \| AF-A0A0C5VFB1-F1-MODEL\_V4 \| 1.0 \| 1.178e-16 \| 492 \| 0.31 \| 203 \| 129 \| 7 \| 1 \| 194 \| 125 \| 325 \| Bacteriophage P2-related tail formation protein \| Bacteriophage P2-related tail formation protein; afdb-uniprot50 \| AF-A0A6N6MG39-F1-MODEL\_V4 \| 1.0 \| 6.867e-16 \| 491 \| 0.322 \| 208 \| 135 \| 5 \| 3 \| 205 \| 11 \| 217 \| Phage tail protein I \| Phage tail protein I; afdb-uniprot50 \| AF-A0A269PIH6-F1-MODEL\_V4 \| 1.0 \| 1.178e-16 \| 491 \| 0.318 \| 226 \| 131 \| 6 \| 1 \| 205 \| 5 \| 228 \| Phage tail protein I \| Phage tail protein I; afdb-uniprot50 \| AF-A0A0H3ZYQ7-F1-MODEL\_V4 \| 1.0 \| 9.751e-17 \| 489 \| 0.242 \| 202 \| 146 \| 4 \| 4 \| 201 \| 11 \| 209 \| Tail protein I \| Tail protein I; afdb-uniprot50 \| AF-A0A0U3HNX6-F1-MODEL\_V4 \| 1.0 \| 9.751e-17 \| 489 \| 0.281 \| 217 \| 138 \| 5 \| 2 \| 201 \| 5 \| 220 \| Uncharacterized protein \| Uncharacterized protein; afdb-uniprot50 \| AF-A0A826V4P5-F1-MODEL\_V4 \| 1.0 \| 5.151e-15 \| 488 \| 0.489 \| 143 \| 69 \| 2 \| 1 \| 141 \| 1 \| 141 \| Phage tail protein I \| Phage tail protein I; afdb-uniprot50 \| AF-A0A545T778-F1-MODEL\_V4 \| 1.0 \| 4.878e-17 \| 488 \| 0.313 \| 214 \| 130 \| 4 \| 1 \| 202 \| 2 \| 210 \| Phage tail protein I \| Phage tail protein I; afdb-uniprot50 \| AF-A0A2G2D6J3-F1-MODEL\_V4 \| 1.0 \| 2.211e-16 \| 488 \| 0.309 \| 207 \| 133 \| 6 \| 5 \| 205 \| 26 \| 228 \| Phage tail protein I \| Phage tail protein I; afdb-uniprot50 \| AF-A0A432WB46-F1-MODEL\_V4 \| 1.0 \| 3.792e-17 \| 488 \| 0.312 \| 221 \| 136 \| 5 \| 1 \| 205 \| 2 \| 222 \| Phage tail protein I \| Phage tail protein I; afdb-uniprot50 \| AF-A0A2N3KSK0-F1-MODEL\_V4 \| 1.0 \| 3.315e-15 \| 487 \| 0.314 \| 191 \| 118 \| 6 \| 1 \| 179 \| 1 \| 190 \| Phage tail protein I \| Phage tail protein I; afdb-uniprot50 \| AF-A0A0E3UDX0-F1-MODEL\_V4 \| 1.0 \| 5.293e-14 \| 486 \| 0.35 \| 154 \| 96 \| 2 \| 13 \| 163 \| 2 \| 154 \| Tail protein \| Tail protein; afdb-uniprot50 \| AF-A0A1Z3UCW5-F1-MODEL\_V4 \| 1.0 \| 3.435e-16 \| 486 \| 0.308 \| 191 \| 119 \| 5 \| 1 \| 187 \| 2 \| 183 \| Phage tail protein I \| Phage tail protein I; afdb-uniprot50 \| AF-A0A7W5G584-F1-MODEL\_V4 \| 1.0 \| 6.626e-15 \| 486 \| 0.377 \| 167 \| 86 \| 5 \| 1 \| 156 \| 1 \| 160 \| Phage tail P2-like protein \| Phage tail P2-like protein; afdb-uniprot50 \| AF-Q1GH43-F1-MODEL\_V4 \| 1.0 \| 4.301e-17 \| 485 \| 0.344 \| 177 \| 115 \| 1 \| 1 \| 176 \| 5 \| 181 \| Phage tail protein I \| Phage tail protein I; afdb-uniprot50 \| AF-A0A6M3J2X1-F1-MODEL\_V4 \| 1.0 \| 2.923e-15 \| 485 \| 0.221 \| 203 \| 153 \| 4 \| 1 \| 202 \| 3 \| 201 \| Putative tail protein \| Putative tail protein; afdb-uniprot50 \| AF-A0A5N8ADV9-F1-MODEL\_V4 \| 1.0 \| 5.012e-16 \| 485 \| 0.324 \| 197 \| 117 \| 4 \| 1 \| 192 \| 1 \| 186 \| Phage tail protein I \| Phage tail protein I; afdb-uniprot50 \| AF-G2HXA1-F1-MODEL\_V4 \| 1.0 \| 1.766e-15 \| 483 \| 0.251 \| 187 \| 129 \| 3 \| 1 \| 183 \| 3 \| 182 \| Phage tail protein \| Phage tail protein; afdb-uniprot50 \| AF-A0A290WU73-F1-MODEL\_V4 \| 1.0 \| 5.734e-18 \| 483 \| 0.337 \| 231 \| 125 \| 2 \| 3 \| 205 \| 12 \| 242 \| Phage tail protein I \| Phage tail protein I; afdb-uniprot50 \| AF-E6L5Z8-F1-MODEL\_V4 \| 1.0 \| 4.15e-16 \| 482 \| 0.241 \| 211 \| 152 \| 3 \| 1 \| 205 \| 1 \| 209 \| Phage tail protein \| Phage tail protein; afdb-uniprot50 \| AF-A0A1C3EBR0-F1-MODEL\_V4 \| 1.0 \| 4.419e-16 \| 481 \| 0.269 \| 197 \| 131 \| 5 \| 4 \| 187 \| 3 \| 199 \| Phage tail protein I \| Phage tail protein I; afdb-uniprot50 \| AF-A0A2S2E575-F1-MODEL\_V4 \| 1.0 \| 4.419e-16 \| 481 \| 0.33 \| 206 \| 122 \| 5 \| 1 \| 194 \| 1 \| 202 \| Baseplate protein \| Baseplate protein; afdb-uniprot50 \| AF-A0A2G6IRQ3-F1-MODEL\_V4 \| 1.0 \| 8.295e-16 \| 481 \| 0.292 \| 229 \| 133 \| 6 \| 4 \| 205 \| 10 \| 236 \| Phage tail protein I \| Phage tail protein I; afdb-uniprot50 \| AF-A0A7W7N2Q5-F1-MODEL\_V4 \| 1.0 \| 2.133e-15 \| 481 \| 0.31 \| 200 \| 133 \| 4 \| 2 \| 197 \| 10 \| 208 \| Phage tail P2-like protein \| Phage tail P2-like protein; afdb-uniprot50 \| AF-A0A7W0BQ66-F1-MODEL\_V4 \| 1.0 \| 7.789e-16 \| 480 \| 0.301 \| 186 \| 123 \| 3 \| 1 \| 184 \| 3 \| 183 \| Phage tail P2-like protein \| Phage tail P2-like protein; afdb-uniprot50 \| AF-A0A3A9HV48-F1-MODEL\_V4 \| 1.0 \| 1.83e-16 \| 480 \| 0.348 \| 204 \| 124 \| 6 \| 4 \| 205 \| 5 \| 201 \| Phage tail protein I \| Phage tail protein I; afdb-uniprot50 \| AF-A0A516SLX1-F1-MODEL\_V4 \| 1.0 \| 2.577e-15 \| 479 \| 0.342 \| 207 \| 124 \| 5 \| 4 \| 205 \| 17 \| 216 \| Phage tail protein I \| Phage tail protein I; afdb-uniprot50 \| AF-A0A1N6M5D6-F1-MODEL\_V4 \| 1.0 \| 4.15e-16 \| 479 \| 0.241 \| 224 \| 150 \| 4 \| 1 \| 205 \| 4 \| 226 \| Phage tail protein (Tail\_P2\_I) \| Phage tail protein (Tail\_P2\_I); afdb-uniprot50 \| AF-A0A4R5W2H2-F1-MODEL\_V4 \| 1.0 \| 3.112e-15 \| 478 \| 0.314 \| 178 \| 115 \| 4 \| 4 \| 176 \| 10 \| 185 \| Phage tail protein I \| Phage tail protein I; afdb-uniprot50 \| AF-A0A354XLH7-F1-MODEL\_V4 \| 1.0 \| 2.076e-16 \| 478 \| 0.352 \| 213 \| 126 \| 6 \| 1 \| 205 \| 1 \| 209 \| Phage tail protein I \| Phage tail protein I; afdb-uniprot50 \| AF-F4HB19-F1-MODEL\_V4 \| 1.0 \| 1.614e-16 \| 477 \| 0.331 \| 178 \| 115 \| 3 \| 1 \| 176 \| 4 \| 179 \| Phage tail protein \| Phage tail protein; afdb-uniprot50 \| AF-A0A828HQ92-F1-MODEL\_V4 \| 1.0 \| 1.146e-17 \| 477 \| 0.323 \| 207 \| 136 \| 2 \| 1 \| 205 \| 2 \| 206 \| Phage tail protein I \| Phage tail protein I; afdb-uniprot50 \| AF-A0A285Z1J9-F1-MODEL\_V4 \| 1.0 \| 3.69e-18 \| 477 \| 0.318 \| 207 \| 132 \| 2 \| 1 \| 198 \| 4 \| 210 \| Phage tail protein, P2 protein I family \| Phage tail protein, P2 protein I family; afdb-uniprot50 \| AF-A0A1E3G6D5-F1-MODEL\_V4 \| 1.0 \| 2.419e-15 \| 476 \| 0.263 \| 205 \| 149 \| 2 \| 1 \| 205 \| 1 \| 203 \| Phage tail protein I \| Phage tail protein I; afdb-uniprot50 \| AF-A0A080K8W0-F1-MODEL\_V4 \| 1.0 \| 8.366e-18 \| 476 \| 0.362 \| 204 \| 126 \| 2 \| 2 \| 205 \| 4 \| 203 \| Bacteriophage P2-related tail formation protein \| Bacteriophage P2-related tail formation protein; afdb-uniprot50 \| AF-A0A3N2E0S8-F1-MODEL\_V4 \| 1.0 \| 3.003e-14 \| 475 \| 0.339 \| 162 \| 101 \| 3 \| 2 \| 163 \| 10 \| 165 \| Phage tail P2-like protein \| Phage tail P2-like protein; afdb-uniprot50 \| AF-A0A166WJW3-F1-MODEL\_V4 \| 1.0 \| 9.408e-16 \| 475 \| 0.273 \| 227 \| 142 \| 5 \| 1 \| 205 \| 3 \| 228 \| Uncharacterized protein \| Uncharacterized protein; afdb-uniprot50 \| AF-A0A4R5HGG6-F1-MODEL\_V4 \| 1.0 \| 1.168e-14 \| 474 \| 0.201 \| 213 \| 161 \| 4 \| 1 \| 205 \| 24 \| 235 \| Phage tail protein I \| Phage tail protein I; afdb-uniprot50 \| AF-A0A7G8SH16-F1-MODEL\_V4 \| 1.0 \| 3.69e-18 \| 473 \| 0.386 \| 176 \| 106 \| 2 \| 1 \| 175 \| 4 \| 178 \| Phage tail protein I \| Phage tail protein I; afdb-uniprot50 \| AF-A0A0E3BUJ7-F1-MODEL\_V4 \| 1.0 \| 1.221e-17 \| 471 \| 0.347 \| 213 \| 119 \| 7 \| 1 \| 203 \| 1 \| 203 \| Tail protein \| Tail protein; afdb-uniprot50 \| AF-A0A1I1QA98-F1-MODEL\_V4 \| 1.0 \| 3.435e-16 \| 471 \| 0.259 \| 239 \| 140 \| 5 \| 2 \| 205 \| 20 \| 256 \| Phage tail protein, P2 protein I family \| Phage tail protein, P2 protein I family; afdb-uniprot50 \| AF-A0A367M848-F1-MODEL\_V4 \| 1.0 \| 1.168e-14 \| 470 \| 0.339 \| 153 \| 91 \| 3 \| 4 \| 152 \| 5 \| 151 \| Phage tail protein I \| Phage tail protein I; afdb-uniprot50 \| AF-A0A369VQL1-F1-MODEL\_V4 \| 1.0 \| 1.719e-16 \| 470 \| 0.267 \| 183 \| 129 \| 3 \| 1 \| 179 \| 4 \| 185 \| Phage tail protein I \| Phage tail protein I; afdb-uniprot50 \| AF-A0A0L8A9U9-F1-MODEL\_V4 \| 1.0 \| 5.338e-16 \| 469 \| 0.336 \| 184 \| 113 \| 4 \| 2 \| 182 \| 256 \| 433 \| Tail protein \| Tail protein; afdb-uniprot50 \| AF-A0A5S3XJT3-F1-MODEL\_V4 \| 1.0 \| 2.82e-14 \| 467 \| 0.345 \| 162 \| 92 \| 6 \| 1 \| 152 \| 7 \| 164 \| Phage tail protein I \| Phage tail protein I; afdb-uniprot50 \| AF-A0A7T8NTY3-F1-MODEL\_V4 \| 1.0 \| 2.923e-15 \| 465 \| 0.275 \| 185 \| 119 \| 5 \| 4 \| 178 \| 7 \| 186 \| Phage tail protein I \| Phage tail protein I; afdb-uniprot50 \| AF-A0A1E3G7K1-F1-MODEL\_V4 \| 1.0 \| 5.012e-16 \| 464 \| 0.267 \| 213 \| 144 \| 4 \| 1 \| 201 \| 1 \| 213 \| Phage tail protein I \| Phage tail protein I; afdb-uniprot50 \| AF-A0A2A4ZMU3-F1-MODEL\_V4 \| 1.0 \| 2.003e-15 \| 464 \| 0.32 \| 215 \| 133 \| 6 \| 1 \| 205 \| 1 \| 212 \| Phage tail protein I \| Phage tail protein I; afdb-uniprot50 \| AF-A0A5S9Q3G5-F1-MODEL\_V4 \| 1.0 \| 5.293e-14 \| 463 \| 0.371 \| 159 \| 85 \| 4 \| 1 \| 151 \| 1 \| 152 \| Uncharacterized protein \| Uncharacterized protein; afdb-uniprot50 \| AF-A0A1X7MJV0-F1-MODEL\_V4 \| 1.0 \| 1.002e-15 \| 462 \| 0.275 \| 229 \| 139 \| 8 \| 1 \| 205 \| 5 \| 230 \| Phage tail protein I \| Phage tail protein I; afdb-uniprot50 \| AF-A0A0F9Z9G5-F1-MODEL\_V4 \| 1.0 \| 2.192e-14 \| 460 \| 0.281 \| 181 \| 125 \| 2 \| 1 \| 178 \| 2 \| 180 \| Uncharacterized protein \| Uncharacterized protein; afdb-uniprot50 \| AF-A0A269PJC4-F1-MODEL\_V4 \| 1.0 \| 3.896e-16 \| 459 \| 0.272 \| 169 \| 123 \| 0 \| 12 \| 180 \| 18 \| 186 \| Phage tail protein I \| Phage tail protein I; afdb-uniprot50 \| AF-B5JY67-F1-MODEL\_V4 \| 1.0 \| 1.528e-18 \| 459 \| 0.323 \| 198 \| 127 \| 3 \| 1 \| 196 \| 5 \| 197 \| Phage tail protein I \| Phage tail protein I; afdb-uniprot50 \| AF-A0A849VB51-F1-MODEL\_V4 \| 1.0 \| 1.278e-13 \| 455 \| 0.31 \| 164 \| 101 \| 5 \| 3 \| 155 \| 5 \| 167 \| Phage tail protein I \| Phage tail protein I; afdb-uniprot50 \| AF-A0A2D3VXF3-F1-MODEL\_V4 \| 1.0 \| 1.933e-14 \| 454 \| 0.215 \| 181 \| 134 \| 5 \| 1 \| 178 \| 1 \| 176 \| Phage tail protein I \| Phage tail protein I; afdb-uniprot50 \| AF-Q8ZKK4-F1-MODEL\_V4 \| 1.0 \| 2.272e-15 \| 454 \| 0.317 \| 211 \| 134 \| 6 \| 1 \| 203 \| 1 \| 209 \| Putative phage tail protein \| Putative phage tail protein; afdb-uniprot50 \| AF-A0A6M1CS17-F1-MODEL\_V4 \| 1.0 \| 3.97e-13 \| 453 \| 0.374 \| 131 \| 77 \| 2 \| 30 \| 158 \| 6 \| 133 \| Phage tail protein I \| Phage tail protein I; afdb-uniprot50 \| AF-A0A2A7UXW9-F1-MODEL\_V4 \| 1.0 \| 1.002e-15 \| 452 \| 0.312 \| 208 \| 135 \| 5 \| 1 \| 205 \| 41 \| 243 \| Phage tail protein I \| Phage tail protein I; afdb-uniprot50 \| AF-A0A6C8GFU6-F1-MODEL\_V4 \| 1.0 \| 2.82e-14 \| 451 \| 0.308 \| 172 \| 113 \| 3 \| 36 \| 203 \| 6 \| 175 \| Phage tail protein \| Phage tail protein; afdb-uniprot50 \| AF-A0A806LU75-F1-MODEL\_V4 \| 1.0 \| 1.002e-15 \| 451 \| 0.273 \| 194 \| 132 \| 2 \| 12 \| 205 \| 29 \| 213 \| Tail protein \| Tail protein; afdb-uniprot50 \| AF-A0A6L3Y824-F1-MODEL\_V4 \| 1.0 \| 1.658e-15 \| 451 \| 0.287 \| 205 \| 139 \| 5 \| 5 \| 205 \| 18 \| 219 \| Phage tail protein I \| Phage tail protein I; afdb-uniprot50 \| AF-A0A6L8GK81-F1-MODEL\_V4 \| 1.0 \| 2.898e-13 \| 450 \| 0.288 \| 156 \| 103 \| 5 \| 1 \| 154 \| 1 \| 150 \| Phage tail protein I \| Phage tail protein I; afdb-uniprot50 \| AF-A0A482INU7-F1-MODEL\_V4 \| 1.0 \| 4.541e-15 \| 450 \| 0.275 \| 214 \| 144 \| 4 \| 1 \| 205 \| 1 \| 212 \| Phage tail protein I \| Phage tail protein I; afdb-uniprot50 \| AF-A0A326GL50-F1-MODEL\_V4 \| 1.0 \| 8.295e-16 \| 450 \| 0.295 \| 186 \| 126 \| 3 \| 1 \| 182 \| 4 \| 188 \| Phage tail protein I \| Phage tail protein I; afdb-uniprot50 \| AF-B7WUI3-F1-MODEL\_V4 \| 1.0 \| 1.2e-13 \| 449 \| 0.263 \| 163 \| 110 \| 4 \| 24 \| 182 \| 6 \| 162 \| Phage tail protein I \| Phage tail protein I; afdb-uniprot50 \| AF-A0A3R9Y2T4-F1-MODEL\_V4 \| 1.0 \| 3.93e-18 \| 449 \| 0.333 \| 204 \| 132 \| 2 \| 3 \| 205 \| 2 \| 202 \| Phage tail protein I \| Phage tail protein I; afdb-uniprot50 \| AF-A0A5S3USV2-F1-MODEL\_V4 \| 1.0 \| 9.078e-15 \| 449 \| 0.25 \| 240 \| 143 \| 7 \| 1 \| 205 \| 2 \| 239 \| Phage tail protein I \| Phage tail protein I; afdb-uniprot50 \| AF-A0A3Q8E075-F1-MODEL\_V4 \| 1.0 \| 1.313e-12 \| 448 \| 0.314 \| 143 \| 93 \| 3 \| 20 \| 161 \| 3 \| 141 \| Phage tail protein I \| Phage tail protein I; afdb-uniprot50 \| AF-A0A510U3R5-F1-MODEL\_V4 \| 1.0 \| 2.133e-15 \| 448 \| 0.265 \| 211 \| 142 \| 6 \| 4 \| 202 \| 9 \| 218 \| Uncharacterized protein \| Uncharacterized protein; afdb-uniprot50 \| AF-A0A1G5B4I5-F1-MODEL\_V4 \| 1.0 \| 4.419e-16 \| 447 \| 0.306 \| 212 \| 129 \| 6 \| 3 \| 203 \| 12 \| 216 \| Phage tail protein, P2 protein I family \| Phage tail protein, P2 protein I family; afdb-uniprot50 \| AF-A0A375ABY7-F1-MODEL\_V4 \| 1.0 \| 4.073e-19 \| 447 \| 0.41 \| 200 \| 116 \| 1 \| 4 \| 203 \| 306 \| 503 \| Phage baseplate assembly protein J \| Phage baseplate assembly protein J; afdb-uniprot50 \| AF-A0A326L3P6-F1-MODEL\_V4 \| 1.0 \| 3.029e-16 \| 446 \| 0.352 \| 187 \| 112 \| 4 \| 4 \| 186 \| 21 \| 202 \| Phage tail protein I \| Phage tail protein I; afdb-uniprot50 \| AF-A0A0C6FTI9-F1-MODEL\_V4 \| 1.0 \| 5.486e-15 \| 446 \| 0.296 \| 216 \| 135 \| 7 \| 1 \| 201 \| 7 \| 220 \| Phage tail protein I \| Phage tail protein I; afdb-uniprot50 \| AF-A0A1U9RGY6-F1-MODEL\_V4 \| 1.0 \| 3.659e-16 \| 446 \| 0.283 \| 233 \| 131 \| 7 \| 1 \| 203 \| 2 \| 228 \| Phage tail protein I \| Phage tail protein I; afdb-uniprot50 \| AF-A0A4P9VF49-F1-MODEL\_V4 \| 1.0 \| 1.106e-16 \| 446 \| 0.33 \| 209 \| 134 \| 5 \| 1 \| 205 \| 1 \| 207 \| Phage tail protein I \| Phage tail protein I; afdb-uniprot50 \| AF-A0A3S7DS03-F1-MODEL\_V4 \| 1.0 \| 7.723e-14 \| 445 \| 0.295 \| 169 \| 109 \| 7 \| 4 \| 164 \| 2 \| 168 \| Phage tail protein I \| Phage tail protein I; afdb-uniprot50 \| AF-A0A2N1APA2-F1-MODEL\_V4 \| 1.0 \| 8.225e-14 \| 445 \| 0.266 \| 195 \| 112 \| 3 \| 1 \| 164 \| 1 \| 195 \| Phage tail protein I \| Phage tail protein I; afdb-uniprot50 \| AF-A0A1W9H3W1-F1-MODEL\_V4 \| 1.0 \| 1.361e-13 \| 444 \| 0.306 \| 163 \| 105 \| 6 \| 1 \| 157 \| 1 \| 161 \| Uncharacterized protein \| Uncharacterized protein; afdb-uniprot50 \| AF-R5H8L2-F1-MODEL\_V4 \| 1.0 \| 9.329e-14 \| 444 \| 0.278 \| 176 \| 121 \| 3 \| 1 \| 173 \| 42 \| 214 \| Putative phage tail protein \| Putative phage tail protein; afdb-uniprot50 \| AF-E2CN32-F1-MODEL\_V4 \| 1.0 \| 5.486e-15 \| 444 \| 0.29 \| 210 \| 127 \| 5 \| 4 \| 193 \| 10 \| 217 \| Putative phage tail protein \| Putative phage tail protein; afdb-uniprot50 \| AF-A0A350DIF7-F1-MODEL\_V4 \| 1.0 \| 2.67e-16 \| 444 \| 0.315 \| 219 \| 130 \| 5 \| 1 \| 203 \| 2 \| 216 \| Phage tail protein I \| Phage tail protein I; afdb-uniprot50 \| AF-G3XD92-F1-MODEL\_V4 \| 1.0 \| 3.435e-16 \| 443 \| 0.324 \| 179 \| 118 \| 2 \| 1 \| 178 \| 1 \| 177 \| Probable bacteriophage protein \| Probable bacteriophage protein; afdb-uniprot50 \| AF-A0A2R3PXS6-F1-MODEL\_V4 \| 1.0 \| 1.881e-15 \| 443 \| 0.329 \| 185 \| 113 \| 5 \| 2 \| 182 \| 82 \| 259 \| Phage tail protein I \| Phage tail protein I; afdb-uniprot50 \| AF-A0A3S8YQ57-F1-MODEL\_V4 \| 1.0 \| 2.272e-15 \| 443 \| 0.35 \| 197 \| 113 \| 5 \| 3 \| 192 \| 4 \| 192 \| Phage tail protein I \| Phage tail protein I; afdb-uniprot50 \| AF-A0A516SJT0-F1-MODEL\_V4 \| 1.0 \| 8.524e-15 \| 442 \| 0.33 \| 209 \| 127 \| 6 \| 2 \| 205 \| 3 \| 203 \| Phage tail protein I \| Phage tail protein I; afdb-uniprot50 \| AF-A0A1T4WXE7-F1-MODEL\_V4 \| 1.0 \| 1.557e-15 \| 442 \| 0.262 \| 221 \| 145 \| 7 \| 1 \| 203 \| 1 \| 221 \| Phage tail protein, P2 protein I family \| Phage tail protein, P2 protein I family; afdb-uniprot50 \| AF-A0A853HRM2-F1-MODEL\_V4 \| 1.0 \| 1.115e-18 \| 441 \| 0.402 \| 206 \| 119 \| 3 \| 1 \| 203 \| 2 \| 206 \| Phage tail protein I \| Phage tail protein I; afdb-uniprot50 \| AF-A0A1M7U2A5-F1-MODEL\_V4 \| 1.0 \| 5.151e-15 \| 441 \| 0.283 \| 208 \| 139 \| 6 \| 3 \| 203 \| 11 \| 215 \| Phage tail protein, P2 protein I family \| Phage tail protein, P2 protein I family; afdb-uniprot50 \| AF-A0A2C9ZYU5-F1-MODEL\_V4 \| 1.0 \| 8.225e-14 \| 441 \| 0.342 \| 152 \| 94 \| 3 \| 1 \| 148 \| 81 \| 230 \| Phage tail protein I \| Phage tail protein I; afdb-uniprot50 \| AF-A0A2C9D5L0-F1-MODEL\_V4 \| 1.0 \| 1.933e-14 \| 440 \| 0.274 \| 211 \| 141 \| 6 \| 4 \| 203 \| 20 \| 229 \| Phage tail protein I \| Phage tail protein I; afdb-uniprot50 \| AF-A0A1G7SAK3-F1-MODEL\_V4 \| 1.0 \| 1.373e-15 \| 440 \| 0.336 \| 217 \| 127 \| 5 \| 1 \| 205 \| 1 \| 212 \| Phage tail protein, P2 protein I family \| Phage tail protein, P2 protein I family; afdb-uniprot50 \| AF-A0A541BB46-F1-MODEL\_V4 \| 1.0 \| 2.67e-16 \| 440 \| 0.326 \| 202 \| 123 \| 5 \| 1 \| 192 \| 9 \| 207 \| Phage tail protein I \| Phage tail protein I; afdb-uniprot50 \| AF-A0A1C3EE42-F1-MODEL\_V4 \| 1.0 \| 1.411e-14 \| 439 \| 0.292 \| 198 \| 114 \| 6 \| 4 \| 176 \| 5 \| 201 \| Phage tail protein I \| Phage tail protein I; afdb-uniprot50 \| AF-A0A8A6BIR6-F1-MODEL\_V4 \| 1.0 \| 6.222e-15 \| 439 \| 0.256 \| 222 \| 136 \| 8 \| 4 \| 203 \| 9 \| 223 \| Phage tail protein I \| Phage tail protein I; afdb-uniprot50 \| AF-A0A2T4JDB8-F1-MODEL\_V4 \| 1.0 \| 1.751e-13 \| 438 \| 0.321 \| 168 \| 95 \| 4 \| 1 \| 152 \| 1 \| 165 \| Phage tail protein I \| Phage tail protein I; afdb-uniprot50 \| AF-A0A7Y8ADW1-F1-MODEL\_V4 \| 1.0 \| 2.335e-14 \| 438 \| 0.225 \| 213 \| 156 \| 4 \| 1 \| 205 \| 5 \| 216 \| Phage tail protein I \| Phage tail protein I; afdb-uniprot50 \| AF-A0A350LX93-F1-MODEL\_V4 \| 1.0 \| 5.195e-17 \| 437 \| 0.363 \| 212 \| 119 \| 5 \| 1 \| 202 \| 1 \| 206 \| Phage tail protein I \| Phage tail protein I; afdb-uniprot50 \| AF-A0A1W0C8U7-F1-MODEL\_V4 \| 1.0 \| 1.6e-14 \| 437 \| 0.253 \| 217 \| 149 \| 6 \| 1 \| 205 \| 1 \| 216 \| Phage tail protein I \| Phage tail protein I; afdb-uniprot50 \| AF-A0A0M9GJD3-F1-MODEL\_V4 \| 1.0 \| 1.704e-14 \| 436 \| 0.229 \| 209 \| 150 \| 6 \| 4 \| 203 \| 10 \| 216 \| Phage tail protein, P2 protein I family \| Phage tail protein, P2 protein I family; afdb-uniprot50 \| AF-A0A1M3AMT4-F1-MODEL\_V4 \| 1.0 \| 1.002e-15 \| 436 \| 0.269 \| 208 \| 146 \| 6 \| 4 \| 205 \| 15 \| 222 \| Phage tail protein I \| Phage tail protein I; afdb-uniprot50 \| AF-A0A109RVS0-F1-MODEL\_V4 \| 1.0 \| 3.628e-14 \| 435 \| 0.283 \| 212 \| 141 \| 7 \| 1 \| 205 \| 1 \| 208 \| Uncharacterized protein \| Uncharacterized protein; afdb-uniprot50 \| AF-A0A6S5P7S3-F1-MODEL\_V4 \| 1.0 \| 1.03e-14 \| 435 \| 0.229 \| 244 \| 144 \| 8 \| 1 \| 200 \| 1 \| 244 \| Uncharacterized protein \| Uncharacterized protein; afdb-uniprot50 \| AF-A0A8A6KEH0-F1-MODEL\_V4 \| 1.0 \| 2.399e-13 \| 434 \| 0.321 \| 165 \| 93 \| 4 \| 1 \| 152 \| 2 \| 160 \| Phage tail protein I \| Phage tail protein I; afdb-uniprot50 \| AF-A0A191WAB0-F1-MODEL\_V4 \| 1.0 \| 1.69e-12 \| 434 \| 0.201 \| 174 \| 131 \| 3 \| 1 \| 173 \| 7 \| 173 \| Phage tail protein \| Phage tail protein; afdb-uniprot50 \| AF-A0A7X8PBJ4-F1-MODEL\_V4 \| 1.0 \| 5.338e-16 \| 434 \| 0.351 \| 185 \| 110 \| 5 \| 1 \| 179 \| 1 \| 181 \| Phage tail protein I \| Phage tail protein I; afdb-uniprot50 \| AF-A0A2S7V2R9-F1-MODEL\_V4 \| 1.0 \| 7.723e-14 \| 434 \| 0.153 \| 196 \| 163 \| 3 \| 1 \| 195 \| 8 \| 201 \| Phage tail protein I \| Phage tail protein I; afdb-uniprot50 \| AF-A0A844Q9W2-F1-MODEL\_V4 \| 1.0 \| 1.168e-14 \| 434 \| 0.254 \| 208 \| 142 \| 6 \| 5 \| 202 \| 37 \| 241 \| Phage tail protein I \| Phage tail protein I; afdb-uniprot50 \| AF-G8PUM6-F1-MODEL\_V4 \| 1.0 \| 7.252e-14 \| 434 \| 0.271 \| 210 \| 144 \| 5 \| 4 \| 205 \| 10 \| 218 \| Phage tail protein I \| Phage tail protein I; afdb-uniprot50 \| AF-A0A6P2IUC4-F1-MODEL\_V4 \| 1.0 \| 4.228e-13 \| 433 \| 0.295 \| 149 \| 99 \| 2 \| 35 \| 180 \| 2 \| 147 \| Phage tail protein I \| Phage tail protein I; afdb-uniprot50 \| AF-A0A7G2U2N6-F1-MODEL\_V4 \| 1.0 \| 6.169e-13 \| 433 \| 0.326 \| 153 \| 93 \| 3 \| 1 \| 145 \| 1 \| 151 \| Phage tail protein I \| Phage tail protein I; afdb-uniprot50 \| AF-A0A541BHK6-F1-MODEL\_V4 \| 1.0 \| 5.151e-15 \| 433 \| 0.276 \| 224 \| 136 \| 8 \| 4 \| 205 \| 6 \| 225 \| Phage tail protein I \| Phage tail protein I; afdb-uniprot50 \| AF-A0A6N7BYW6-F1-MODEL\_V4 \| 1.0 \| 7.515e-15 \| 432 \| 0.262 \| 179 \| 127 \| 3 \| 1 \| 178 \| 2 \| 176 \| Uncharacterized protein \| Uncharacterized protein; afdb-uniprot50 \| AF-A0A7X4JXH6-F1-MODEL\_V4 \| 1.0 \| 2.948e-17 \| 432 \| 0.37 \| 205 \| 120 \| 6 \| 1 \| 203 \| 4 \| 201 \| Phage tail protein I \| Phage tail protein I; afdb-uniprot50 \| AF-A0A5S3YYC2-F1-MODEL\_V4 \| 1.0 \| 9.078e-15 \| 432 \| 0.223 \| 233 \| 150 \| 7 \| 1 \| 205 \| 2 \| 231 \| Phage tail protein I \| Phage tail protein I; afdb-uniprot50 \| AF-A0A5P9HBR8-F1-MODEL\_V4 \| 1.0 \| 9.078e-15 \| 432 \| 0.26 \| 200 \| 125 \| 6 \| 2 \| 184 \| 5 \| 198 \| Phage tail protein \| Phage tail protein; afdb-uniprot50 \| AF-A0A2A5BNC1-F1-MODEL\_V4 \| 1.0 \| 1.127e-13 \| 431 \| 0.325 \| 163 \| 93 \| 6 \| 1 \| 151 \| 1 \| 158 \| Phage tail protein I \| Phage tail protein I; afdb-uniprot50 \| AF-A0A285NMK5-F1-MODEL\_V4 \| 1.0 \| 2.192e-14 \| 431 \| 0.265 \| 196 \| 135 \| 3 \| 12 \| 205 \| 23 \| 211 \| Phage tail protein, P2 protein I family \| Phage tail protein, P2 protein I family; afdb-uniprot50 \| AF-A0A510U7K2-F1-MODEL\_V4 \| 1.0 \| 4.836e-15 \| 431 \| 0.268 \| 227 \| 140 \| 7 \| 4 \| 205 \| 7 \| 232 \| Uncharacterized protein \| Uncharacterized protein; afdb-uniprot50 \| AF-C5A8Q4-F1-MODEL\_V4 \| 1.0 \| 1.544e-13 \| 430 \| 0.31 \| 161 \| 103 \| 4 \| 1 \| 157 \| 1 \| 157 \| Phage-related tail protein \| Phage-related tail protein; afdb-uniprot50 \| AF-A0A7W4PUW2-F1-MODEL\_V4 \| 1.0 \| 4.878e-17 \| 430 \| 0.339 \| 212 \| 131 \| 3 \| 1 \| 205 \| 1 \| 210 \| Phage tail protein I \| Phage tail protein I; afdb-uniprot50 \| AF-A0A4R2P790-F1-MODEL\_V4 \| 1.0 \| 5.637e-14 \| 430 \| 0.288 \| 208 \| 138 \| 7 \| 1 \| 205 \| 1 \| 201 \| Phage tail P2-like protein \| Phage tail P2-like protein; afdb-uniprot50 \| AF-B7RNM5-F1-MODEL\_V4 \| 1.0 \| 1.325e-14 \| 429 \| 0.253 \| 213 \| 148 \| 4 \| 1 \| 205 \| 6 \| 215 \| Putative phage tail protein \| Putative phage tail protein; afdb-uniprot50 \| AF-A0A8B1NMW4-F1-MODEL\_V4 \| 1.0 \| 3.863e-14 \| 428 \| 0.372 \| 153 \| 91 \| 2 \| 42 \| 194 \| 2 \| 149 \| Uncharacterized protein \| Uncharacterized protein; afdb-uniprot50 \| AF-A0A847KR13-F1-MODEL\_V4 \| 1.0 \| 3.287e-13 \| 428 \| 0.293 \| 167 \| 111 \| 5 \| 4 \| 165 \| 17 \| 181 \| Phage tail protein I \| Phage tail protein I; afdb-uniprot50 \| AF-A0A5Z0QYI5-F1-MODEL\_V4 \| 1.0 \| 1.897e-17 \| 428 \| 0.356 \| 202 \| 125 \| 4 \| 1 \| 200 \| 1 \| 199 \| Phage tail protein I \| Phage tail protein I; afdb-uniprot50 \| AF-A0A8B2TV07-F1-MODEL\_V4 \| 1.0 \| 5.486e-15 \| 428 \| 0.22 \| 209 \| 151 \| 4 \| 1 \| 205 \| 1 \| 201 \| Phage tail protein I \| Phage tail protein I; afdb-uniprot50 \| AF-A0A1M3HHZ6-F1-MODEL\_V4 \| 1.0 \| 2.419e-15 \| 428 \| 0.306 \| 222 \| 128 \| 7 \| 1 \| 204 \| 1 \| 214 \| Phage tail protein I \| Phage tail protein I; afdb-uniprot50 \| AF-A0A3A3D8P6-F1-MODEL\_V4 \| 1.0 \| 8.524e-15 \| 428 \| 0.27 \| 174 \| 117 \| 3 \| 1 \| 167 \| 8 \| 178 \| Phage tail protein I \| Phage tail protein I; afdb-uniprot50 \| AF-A0A233HGS3-F1-MODEL\_V4 \| 1.0 \| 8.225e-14 \| 427 \| 0.206 \| 203 \| 152 \| 4 \| 1 \| 202 \| 7 \| 201 \| Phage tail protein \| Phage tail protein; afdb-uniprot50 \| AF-A0A238JAQ9-F1-MODEL\_V4 \| 1.0 \| 1.127e-13 \| 425 \| 0.3 \| 180 \| 106 \| 5 \| 1 \| 165 \| 1 \| 175 \| Phage tail protein (Tail\_P2\_I) \| Phage tail protein (Tail\_P2\_I); afdb-uniprot50 \| AF-A0A3S4IC27-F1-MODEL\_V4 \| 1.0 \| 3.198e-14 \| 425 \| 0.274 \| 211 \| 145 \| 4 \| 1 \| 205 \| 1 \| 209 \| Bacteriophage P2-related tail formation protein \| Bacteriophage P2-related tail formation protein; afdb-uniprot50 \| AF-A0A7Y1X1Y1-F1-MODEL\_V4 \| 1.0 \| 1.57e-17 \| 425 \| 0.334 \| 209 \| 133 \| 3 \| 1 \| 205 \| 1 \| 207 \| Phage tail protein I \| Phage tail protein I; afdb-uniprot50 \| AF-A0A7X6WY32-F1-MODEL\_V4 \| 1.0 \| 1.127e-13 \| 425 \| 0.243 \| 201 \| 113 \| 4 \| 1 \| 162 \| 2 \| 202 \| Phage tail protein I \| Phage tail protein I; afdb-uniprot50 \| AF-D0IJ10-F1-MODEL\_V4 \| 1.0 \| 6.57e-13 \| 424 \| 0.27 \| 159 \| 111 \| 3 \| 19 \| 176 \| 2 \| 156 \| Tail protein I \| Tail protein I; afdb-uniprot50 \| AF-A0A1B9NZR7-F1-MODEL\_V4 \| 1.0 \| 1.799e-12 \| 424 \| 0.158 \| 177 \| 141 \| 3 \| 1 \| 176 \| 7 \| 176 \| Uncharacterized protein \| Uncharacterized protein; afdb-uniprot50 \| AF-A0A1W0CDK0-F1-MODEL\_V4 \| 1.0 \| 5.151e-15 \| 424 \| 0.269 \| 215 \| 133 \| 8 \| 4 \| 205 \| 5 \| 208 \| Phage tail protein I \| Phage tail protein I; afdb-uniprot50 \| AF-A0A2S5F905-F1-MODEL\_V4 \| 1.0 \| 2.399e-13 \| 423 \| 0.294 \| 170 \| 105 \| 7 \| 4 \| 163 \| 2 \| 166 \| Phage tail protein I \| Phage tail protein I; afdb-uniprot50 \| AF-A0A840WZA1-F1-MODEL\_V4 \| 1.0 \| 8.759e-14 \| 423 \| 0.28 \| 200 \| 130 \| 5 \| 2 \| 192 \| 7 \| 201 \| Phage tail P2-like protein \| Phage tail P2-like protein; afdb-uniprot50 \| AF-A0A1H1G374-F1-MODEL\_V4 \| 1.0 \| 2.335e-14 \| 423 \| 0.25 \| 212 \| 148 \| 4 \| 4 \| 205 \| 7 \| 217 \| Phage tail protein, P2 protein I family \| Phage tail protein, P2 protein I family; afdb-uniprot50 \| AF-A0A5S9QVU4-F1-MODEL\_V4 \| 1.0 \| 4.503e-13 \| 422 \| 0.303 \| 178 \| 111 \| 5 \| 1 \| 167 \| 3 \| 178 \| Uncharacterized protein \| Uncharacterized protein; afdb-uniprot50 \| AF-A0A6N4CKX4-F1-MODEL\_V4 \| 1.0 \| 2.507e-16 \| 422 \| 0.354 \| 192 \| 114 \| 4 \| 4 \| 194 \| 3 \| 185 \| Uncharacterized protein \| Uncharacterized protein; afdb-uniprot50 \| AF-A0A4U8UH95-F1-MODEL\_V4 \| 1.0 \| 2.335e-14 \| 422 \| 0.242 \| 202 \| 146 \| 5 \| 1 \| 200 \| 2 \| 198 \| Phage tail protein I \| Phage tail protein I; afdb-uniprot50 \| AF-A0A1M3I801-F1-MODEL\_V4 \| 1.0 \| 5.151e-15 \| 422 \| 0.313 \| 188 \| 113 \| 6 \| 2 \| 181 \| 5 \| 184 \| Phage tail protein I \| Phage tail protein I; afdb-uniprot50 \| AF-A0A840LX24-F1-MODEL\_V4 \| 1.0 \| 1.45e-13 \| 422 \| 0.223 \| 215 \| 155 \| 7 \| 1 \| 205 \| 7 \| 219 \| Phage tail P2-like protein \| Phage tail P2-like protein; afdb-uniprot50 \| AF-A0A1N6M6A1-F1-MODEL\_V4 \| 1.0 \| 1.127e-13 \| 422 \| 0.208 \| 201 \| 155 \| 4 \| 4 \| 202 \| 15 \| 213 \| Phage tail protein (Tail\_P2\_I) \| Phage tail protein (Tail\_P2\_I); afdb-uniprot50 \| AF-A0A423PQM5-F1-MODEL\_V4 \| 1.0 \| 9.329e-14 \| 421 \| 0.287 \| 188 \| 117 \| 7 \| 2 \| 183 \| 8 \| 184 \| Tail protein \| Tail protein; afdb-uniprot50 \| AF-K5Z1Z7-F1-MODEL\_V4 \| 1.0 \| 2.399e-13 \| 421 \| 0.333 \| 171 \| 95 \| 4 \| 1 \| 155 \| 5 \| 172 \| Phage tail protein I \| Phage tail protein I; afdb-uniprot50 \| AF-A0A5C5NW28-F1-MODEL\_V4 \| 1.0 \| 1.865e-13 \| 421 \| 0.255 \| 184 \| 122 \| 5 \| 1 \| 170 \| 1 \| 183 \| Phage tail protein I \| Phage tail protein I; afdb-uniprot50 \| AF-A0A0C1G372-F1-MODEL\_V4 \| 1.0 \| 3.086e-13 \| 420 \| 0.349 \| 169 \| 90 \| 5 \| 1 \| 152 \| 1 \| 166 \| Tail protein \| Tail protein; afdb-uniprot50 \| AF-A0A261E3S3-F1-MODEL\_V4 \| 1.0 \| 8.004e-15 \| 420 \| 0.256 \| 207 \| 142 \| 6 \| 4 \| 199 \| 2 \| 207 \| Phage tail protein I \| Phage tail protein I; afdb-uniprot50 \| AF-A0A2T5NQ88-F1-MODEL\_V4 \| 1.0 \| 9.329e-14 \| 420 \| 0.27 \| 200 \| 140 \| 3 \| 10 \| 205 \| 5 \| 202 \| Phage tail protein I \| Phage tail protein I; afdb-uniprot50 \| AF-A0A2T6LZT5-F1-MODEL\_V4 \| 1.0 \| 1.6e-14 \| 419 \| 0.327 \| 168 \| 99 \| 5 \| 1 \| 158 \| 3 \| 166 \| Phage tail P2-like protein \| Phage tail P2-like protein; afdb-uniprot50 \| AF-A0A0L9YAM1-F1-MODEL\_V4 \| 1.0 \| 8.225e-14 \| 418 \| 0.237 \| 198 \| 144 \| 6 \| 11 \| 205 \| 24 \| 217 \| Phage tail protein I \| Phage tail protein I; afdb-uniprot50 \| AF-A0A413FZE3-F1-MODEL\_V4 \| 1.0 \| 2.721e-13 \| 417 \| 0.239 \| 171 \| 123 \| 3 \| 11 \| 180 \| 29 \| 193 \| Phage tail protein I \| Phage tail protein I; afdb-uniprot50 \| AF-A0A6L9FPZ5-F1-MODEL\_V4 \| 1.0 \| 1.865e-13 \| 417 \| 0.275 \| 185 \| 123 \| 5 \| 4 \| 183 \| 17 \| 195 \| Phage tail protein I \| Phage tail protein I; afdb-uniprot50 \| AF-A0A1R1MK88-F1-MODEL\_V4 \| 1.0 \| 1.986e-13 \| 417 \| 0.254 \| 181 \| 121 \| 4 \| 4 \| 172 \| 7 \| 185 \| Phage tail protein I \| Phage tail protein I; afdb-uniprot50 \| AF-A0A806DA82-F1-MODEL\_V4 \| 1.0 \| 5.637e-14 \| 416 \| 0.276 \| 188 \| 124 \| 5 \| 1 \| 182 \| 2 \| 183 \| Phage tail protein I \| Phage tail protein I; afdb-uniprot50 \| AF-E6X1N3-F1-MODEL\_V4 \| 1.0 \| 1.058e-13 \| 416 \| 0.247 \| 194 \| 130 \| 4 \| 1 \| 180 \| 2 \| 193 \| Uncharacterized protein \| Uncharacterized protein; afdb-uniprot50 \| AF-A0A1I5MPV2-F1-MODEL\_V4 \| 1.0 \| 1.097e-14 \| 416 \| 0.247 \| 214 \| 146 \| 6 \| 3 \| 205 \| 11 \| 220 \| Phage tail protein, P2 protein I family \| Phage tail protein, P2 protein I family; afdb-uniprot50 \| AF-A0A5C1DKB3-F1-MODEL\_V4 \| 1.0 \| 1.2e-13 \| 416 \| 0.276 \| 195 \| 133 \| 4 \| 1 \| 189 \| 1 \| 193 \| Phage tail protein I \| Phage tail protein I; afdb-uniprot50 \| AF-A0A4Y8REX3-F1-MODEL\_V4 \| 1.0 \| 4.264e-15 \| 414 \| 0.317 \| 214 \| 128 \| 8 \| 4 \| 203 \| 20 \| 229 \| Phage tail protein I \| Phage tail protein I; afdb-uniprot50 \| AF-A0A377AZI4-F1-MODEL\_V4 \| 1.0 \| 2.898e-13 \| 413 \| 0.401 \| 137 \| 80 \| 1 \| 56 \| 192 \| 4 \| 138 \| Tail protein I \| Tail protein I; afdb-uniprot50 \| AF-F3YY39-F1-MODEL\_V4 \| 1.0 \| 3.003e-14 \| 412 \| 0.313 \| 214 \| 131 \| 6 \| 4 \| 205 \| 13 \| 222 \| Phage tail protein I \| Phage tail protein I; afdb-uniprot50 \| AF-U3AMK5-F1-MODEL\_V4 \| 1.0 \| 7.723e-14 \| 412 \| 0.251 \| 211 \| 144 \| 5 \| 4 \| 203 \| 20 \| 227 \| Putative phage tail protein \| Putative phage tail protein; afdb-uniprot50 \| AF-A0A2P8QYP1-F1-MODEL\_V4 \| 1.0 \| 3.863e-14 \| 412 \| 0.242 \| 206 \| 150 \| 4 \| 1 \| 205 \| 1 \| 201 \| Phage tail protein I \| Phage tail protein I; afdb-uniprot50 \| AF-A0A827ZHP7-F1-MODEL\_V4 \| 1.0 \| 5.439e-13 \| 411 \| 0.504 \| 125 \| 60 \| 1 \| 1 \| 125 \| 2 \| 124 \| Phage tail protein I \| Phage tail protein I; afdb-uniprot50 \| AF-A0A7Y3Z485-F1-MODEL\_V4 \| 1.0 \| 4.503e-13 \| 411 \| 0.237 \| 177 \| 124 \| 4 \| 2 \| 176 \| 8 \| 175 \| Uncharacterized protein \| Uncharacterized protein; afdb-uniprot50 \| AF-A0A016XH76-F1-MODEL\_V4 \| 1.0 \| 2.507e-16 \| 411 \| 0.327 \| 186 \| 121 \| 3 \| 4 \| 186 \| 14 \| 198 \| Tail protein \| Tail protein; afdb-uniprot50 \| AF-A0A7X4WDI7-F1-MODEL\_V4 \| 1.0 \| 2.898e-13 \| 411 \| 0.196 \| 209 \| 157 \| 4 \| 1 \| 205 \| 4 \| 205 \| Phage tail protein \| Phage tail protein; afdb-uniprot50 \| AF-R6P8B0-F1-MODEL\_V4 \| 1.0 \| 7.252e-14 \| 411 \| 0.276 \| 188 \| 128 \| 4 \| 1 \| 183 \| 1 \| 185 \| Putative phage tail protein \| Putative phage tail protein; afdb-uniprot50 \| AF-A0A5S3YYF1-F1-MODEL\_V4 \| 1.0 \| 5.293e-14 \| 411 \| 0.276 \| 217 \| 133 \| 6 \| 12 \| 205 \| 18 \| 233 \| Phage tail protein I \| Phage tail protein I; afdb-uniprot50 \| AF-A0A1I7EL73-F1-MODEL\_V4 \| 1.0 \| 2.076e-16 \| 409 \| 0.325 \| 203 \| 125 \| 4 \| 1 \| 200 \| 1 \| 194 \| Phage tail protein, P2 protein I family \| Phage tail protein, P2 protein I family; afdb-uniprot50 \| AF-A0A359KDK8-F1-MODEL\_V4 \| 1.0 \| 3.406e-14 \| 409 \| 0.226 \| 212 \| 146 \| 7 \| 1 \| 197 \| 5 \| 213 \| Phage tail protein I \| Phage tail protein I; afdb-uniprot50 \| AF-A0A418VVG6-F1-MODEL\_V4 \| 1.0 \| 1.097e-14 \| 409 \| 0.283 \| 243 \| 134 \| 5 \| 1 \| 205 \| 1 \| 241 \| Phage tail protein I \| Phage tail protein I; afdb-uniprot50 \| AF-A0A399MBD2-F1-MODEL\_V4 \| 1.0 \| 1.127e-13 \| 408 \| 0.264 \| 208 \| 141 \| 7 \| 1 \| 200 \| 1 \| 204 \| Phage tail protein I \| Phage tail protein I; afdb-uniprot50 \| AF-A0A2E3Q532-F1-MODEL\_V4 \| 1.0 \| 1.69e-12 \| 408 \| 0.26 \| 196 \| 112 \| 4 \| 1 \| 163 \| 1 \| 196 \| Phage tail protein I \| Phage tail protein I; afdb-uniprot50 \| AF-R8AR41-F1-MODEL\_V4 \| 1.0 \| 1.865e-13 \| 407 \| 0.237 \| 206 \| 147 \| 4 \| 1 \| 203 \| 1 \| 199 \| Phage tail protein I \| Phage tail protein I; afdb-uniprot50 \| AF-A0A812QW15-F1-MODEL\_V4 \| 1.0 \| 3.003e-14 \| 407 \| 0.331 \| 172 \| 97 \| 4 \| 26 \| 180 \| 303 \| 473 \| J protein \| J protein; afdb-uniprot50 \| AF-A0A660NKJ4-F1-MODEL\_V4 \| 1.0 \| 7.452e-13 \| 406 \| 0.319 \| 169 \| 95 \| 5 \| 1 \| 152 \| 1 \| 166 \| Phage tail protein I \| Phage tail protein I; afdb-uniprot50 \| AF-I7E1B5-F1-MODEL\_V4 \| 1.0 \| 2.507e-16 \| 406 \| 0.358 \| 209 \| 115 \| 10 \| 1 \| 200 \| 1 \| 199 \| Phage tail protein I \| Phage tail protein I; afdb-uniprot50 \| AF-A0A1T4UVP0-F1-MODEL\_V4 \| 1.0 \| 5.486e-15 \| 406 \| 0.279 \| 215 \| 134 \| 8 \| 4 \| 205 \| 1 \| 207 \| Phage tail protein, P2 protein I family \| Phage tail protein, P2 protein I family; afdb-uniprot50 \| AF-A0A2G3J6I4-F1-MODEL\_V4 \| 1.0 \| 3.728e-13 \| 406 \| 0.231 \| 212 \| 153 \| 5 \| 1 \| 205 \| 1 \| 209 \| Phage tail protein \| Phage tail protein; afdb-uniprot50 \| AF-A0A840Q0S0-F1-MODEL\_V4 \| 1.0 \| 4.382e-14 \| 406 \| 0.266 \| 195 \| 135 \| 5 \| 12 \| 205 \| 28 \| 215 \| Phage tail P2-like protein \| Phage tail P2-like protein; afdb-uniprot50 \| AF-A0A7I9KDL4-F1-MODEL\_V4 \| 1.0 \| 6.003e-14 \| 405 \| 0.23 \| 200 \| 146 \| 4 \| 1 \| 198 \| 1 \| 194 \| Phage tail protein I \| Phage tail protein I; afdb-uniprot50 \| AF-A0A7M3MBM5-F1-MODEL\_V4 \| 1.0 \| 2.486e-14 \| 405 \| 0.266 \| 221 \| 137 \| 7 \| 4 \| 205 \| 12 \| 226 \| Phage tail protein I \| Phage tail protein I; afdb-uniprot50 \| AF-F2K226-F1-MODEL\_V4 \| 1.0 \| 3.287e-13 \| 405 \| 0.218 \| 220 \| 152 \| 6 \| 4 \| 205 \| 15 \| 232 \| Phage tail protein I \| Phage tail protein I; afdb-uniprot50 \| AF-A0A512JPD4-F1-MODEL\_V4 \| 1.0 \| 1.815e-14 \| 404 \| 0.24 \| 250 \| 142 \| 9 \| 1 \| 205 \| 1 \| 247 \| Uncharacterized protein \| Uncharacterized protein; afdb-uniprot50 \| AF-A0A855ISH0-F1-MODEL\_V4 \| 1.0 \| 5.107e-13 \| 403 \| 0.224 \| 169 \| 121 \| 3 \| 2 \| 166 \| 10 \| 172 \| Phage tail protein I \| Phage tail protein I; afdb-uniprot50 \| AF-A0A547PS50-F1-MODEL\_V4 \| 1.0 \| 9.935e-14 \| 403 \| 0.279 \| 215 \| 130 \| 8 \| 4 \| 200 \| 8 \| 215 \| Phage tail protein I \| Phage tail protein I; afdb-uniprot50 \| AF-W4M090-F1-MODEL\_V4 \| 1.0 \| 8.524e-15 \| 403 \| 0.31 \| 229 \| 131 \| 8 \| 1 \| 205 \| 1 \| 226 \| Uncharacterized protein \| Uncharacterized protein; afdb-uniprot50 \| AF-A0A162A844-F1-MODEL\_V4 \| 1.0 \| 6.394e-14 \| 403 \| 0.271 \| 173 \| 115 \| 4 \| 1 \| 165 \| 1 \| 170 \| Tail protein \| Tail protein; afdb-uniprot50 \| AF-A0A285M2N0-F1-MODEL\_V4 \| 1.0 \| 9.586e-13 \| 402 \| 0.281 \| 167 \| 110 \| 4 \| 2 \| 158 \| 5 \| 171 \| Phage tail protein, P2 protein I family \| Phage tail protein, P2 protein I family; afdb-uniprot50 \| AF-A0A839IX64-F1-MODEL\_V4 \| 1.0 \| 2.097e-11 \| 402 \| 0.254 \| 157 \| 108 \| 5 \| 1 \| 154 \| 1 \| 151 \| Phage tail protein I \| Phage tail protein I; afdb-uniprot50 \| AF-A0A1H7YJ96-F1-MODEL\_V4 \| 1.0 \| 2.115e-13 \| 401 \| 0.251 \| 195 \| 132 \| 5 \| 12 \| 205 \| 33 \| 214 \| Phage tail protein, P2 protein I family \| Phage tail protein, P2 protein I family; afdb-uniprot50 \| AF-A0A3S2V1V0-F1-MODEL\_V4 \| 1.0 \| 2.82e-14 \| 401 \| 0.234 \| 264 \| 141 \| 6 \| 1 \| 205 \| 7 \| 268 \| Phage tail protein I \| Phage tail protein I; afdb-uniprot50 \| AF-A0A239CJC5-F1-MODEL\_V4 \| 1.0 \| 2.844e-16 \| 401 \| 0.31 \| 203 \| 126 \| 5 \| 1 \| 200 \| 1 \| 192 \| Phage tail protein, P2 protein I family \| Phage tail protein, P2 protein I family; afdb-uniprot50 \| AF-A0A4R6M881-F1-MODEL\_V4 \| 1.0 \| 1.021e-12 \| 400 \| 0.331 \| 184 \| 111 \| 6 \| 3 \| 178 \| 11 \| 190 \| Phage tail P2-like protein \| Phage tail P2-like protein; afdb-uniprot50 \| AF-A0A1G8HUV2-F1-MODEL\_V4 \| 1.0 \| 1.69e-12 \| 400 \| 0.181 \| 209 \| 165 \| 6 \| 1 \| 205 \| 12 \| 218 \| Phage tail protein, P2 protein I family \| Phage tail protein, P2 protein I family; afdb-uniprot50 \| AF-A0A7L5Y1L5-F1-MODEL\_V4 \| 1.0 \| 2.335e-14 \| 399 \| 0.282 \| 184 \| 127 \| 4 \| 2 \| 182 \| 6 \| 187 \| Phage tail protein I \| Phage tail protein I; afdb-uniprot50 \| AF-A0A345J3J1-F1-MODEL\_V4 \| 1.0 \| 1.644e-13 \| 399 \| 0.243 \| 205 \| 149 \| 4 \| 1 \| 203 \| 4 \| 204 \| Phage tail protein, P2 family \| Phage tail protein, P2 family; afdb-uniprot50 \| AF-A0A855N9J0-F1-MODEL\_V4 \| 1.0 \| 4.503e-13 \| 399 \| 0.233 \| 206 \| 152 \| 4 \| 1 \| 205 \| 1 \| 201 \| Phage tail protein I \| Phage tail protein I; afdb-uniprot50 \| AF-F8FW00-F1-MODEL\_V4 \| 1.0 \| 1.45e-13 \| 398 \| 0.283 \| 219 \| 139 \| 8 \| 1 \| 205 \| 1 \| 215 \| Pyocin R2\_PP, tail formation protein \| Pyocin R2\_PP, tail formation protein; afdb-uniprot50 \| AF-A0A1C3FFM0-F1-MODEL\_V4 \| 1.0 \| 3.377e-12 \| 397 \| 0.289 \| 159 \| 106 \| 4 \| 1 \| 157 \| 3 \| 156 \| Putative phage tail fiber protein \| Putative phage tail fiber protein; afdb-uniprot50 \| AF-A0A0W8JFZ8-F1-MODEL\_V4 \| 1.0 \| 5.439e-13 \| 396 \| 0.2 \| 200 \| 149 \| 3 \| 16 \| 205 \| 2 \| 200 \| Uncharacterized protein \| Uncharacterized protein; afdb-uniprot50 \| AF-A0A2M7MVB7-F1-MODEL\_V4 \| 1.0 \| 8.225e-14 \| 396 \| 0.257 \| 229 \| 121 \| 7 \| 1 \| 181 \| 1 \| 228 \| Phage tail protein I \| Phage tail protein I; afdb-uniprot50 \| AF-A0A7U2UZK0-F1-MODEL\_V4 \| 1.0 \| 1.751e-13 \| 396 \| 0.246 \| 215 \| 148 \| 8 \| 4 \| 205 \| 10 \| 223 \| Phage tail protein I \| Phage tail protein I; afdb-uniprot50 \| AF-R7JD91-F1-MODEL\_V4 \| 1.0 \| 1.986e-13 \| 396 \| 0.278 \| 183 \| 118 \| 7 \| 2 \| 176 \| 43 \| 219 \| Putative phage tail protein \| Putative phage tail protein; afdb-uniprot50 \| AF-A0A4S2HBU2-F1-MODEL\_V4 \| 1.0 \| 5.248e-12 \| 396 \| 0.248 \| 161 \| 111 \| 6 \| 5 \| 158 \| 13 \| 170 \| Phage tail protein I \| Phage tail protein I; afdb-uniprot50 \| AF-A0A7J0BHY1-F1-MODEL\_V4 \| 1.0 \| 1.158e-12 \| 395 \| 0.257 \| 194 \| 138 \| 5 \| 13 \| 204 \| 28 \| 217 \| Phage tail protein I \| Phage tail protein I; afdb-uniprot50 \| AF-A0A855ITT2-F1-MODEL\_V4 \| 1.0 \| 8.452e-13 \| 395 \| 0.151 \| 205 \| 167 \| 4 \| 1 \| 200 \| 8 \| 210 \| Phage tail protein I \| Phage tail protein I; afdb-uniprot50 \| AF-A0A7U9H2I1-F1-MODEL\_V4 \| 1.0 \| 2.721e-13 \| 395 \| 0.252 \| 198 \| 138 \| 5 \| 11 \| 203 \| 24 \| 216 \| Phage tail protein I \| Phage tail protein I; afdb-uniprot50 \| AF-A0A1X3J174-F1-MODEL\_V4 \| 1.0 \| 3.471e-11 \| 394 \| 0.438 \| 105 \| 57 \| 1 \| 71 \| 175 \| 2 \| 104 \| Tail protein I (GpI) \| Tail protein I (GpI); afdb-uniprot50 \| AF-I3TTD2-F1-MODEL\_V4 \| 1.0 \| 2.253e-13 \| 394 \| 0.285 \| 221 \| 141 \| 5 \| 1 \| 205 \| 1 \| 220 \| Gp15 protein \| Gp15 protein; afdb-uniprot50 \| AF-A0A7W6REA6-F1-MODEL\_V4 \| 1.0 \| 1.03e-14 \| 393 \| 0.264 \| 227 \| 145 \| 6 \| 1 \| 205 \| 9 \| 235 \| Phage tail P2-like protein \| Phage tail P2-like protein; afdb-uniprot50 \| AF-A0A290TRG8-F1-MODEL\_V4 \| 1.0 \| 1.69e-12 \| 393 \| 0.245 \| 183 \| 123 \| 6 \| 14 \| 186 \| 108 \| 285 \| Uncharacterized protein \| Uncharacterized protein; afdb-uniprot50 \| AF-A0A5S3WQZ3-F1-MODEL\_V4 \| 1.0 \| 1.69e-12 \| 393 \| 0.311 \| 170 \| 100 \| 5 \| 11 \| 165 \| 104 \| 271 \| Phage tail protein I \| Phage tail protein I; afdb-uniprot50 \| AF-F3LHZ7-F1-MODEL\_V4 \| 1.0 \| 1.158e-12 \| 392 \| 0.287 \| 174 \| 102 \| 6 \| 1 \| 156 \| 1 \| 170 \| Tail protein I \| Tail protein I; afdb-uniprot50 \| AF-A0A7X6GUL0-F1-MODEL\_V4 \| 1.0 \| 3.171e-12 \| 392 \| 0.263 \| 171 \| 110 \| 6 \| 4 \| 162 \| 8 \| 174 \| Phage tail protein I \| Phage tail protein I; afdb-uniprot50 \| AF-U1JMH5-F1-MODEL\_V4 \| 1.0 \| 1.704e-14 \| 392 \| 0.29 \| 210 \| 138 \| 5 \| 1 \| 202 \| 1 \| 207 \| Putative prophage tail protein \| Putative prophage tail protein; afdb-uniprot50 \| AF-R9KAQ1-F1-MODEL\_V4 \| 1.0 \| 8.759e-14 \| 392 \| 0.255 \| 219 \| 141 \| 6 \| 3 \| 205 \| 21 \| 233 \| Phage tail protein I \| Phage tail protein I; afdb-uniprot50 \| AF-A0A891ZWB9-F1-MODEL\_V4 \| 1.0 \| 1.916e-12 \| 391 \| 0.228 \| 175 \| 125 \| 5 \| 11 \| 181 \| 3 \| 171 \| Bacteriophage P2-related tail formation protein \| Bacteriophage P2-related tail formation protein; afdb-uniprot50 \| AF-A0A522IBS6-F1-MODEL\_V4 \| 1.0 \| 3.287e-13 \| 391 \| 0.271 \| 225 \| 140 \| 7 \| 1 \| 203 \| 1 \| 223 \| Phage tail protein I \| Phage tail protein I; afdb-uniprot50 \| AF-A0A239EKX6-F1-MODEL\_V4 \| 1.0 \| 1.373e-15 \| 391 \| 0.272 \| 213 \| 138 \| 4 \| 1 \| 196 \| 4 \| 216 \| Phage tail protein, P2 protein I family \| Phage tail protein, P2 protein I family; afdb-uniprot50 \| AF-A0A318KN12-F1-MODEL\_V4 \| 1.0 \| 1.19e-11 \| 391 \| 0.335 \| 161 \| 105 \| 2 \| 4 \| 163 \| 6 \| 165 \| Phage tail P2-like protein \| Phage tail P2-like protein; afdb-uniprot50 \| AF-A0A5T0QSQ9-F1-MODEL\_V4 \| 1.0 \| 1.544e-13 \| 390 \| 0.234 \| 196 \| 140 \| 6 \| 1 \| 193 \| 1 \| 189 \| Phage tail protein I \| Phage tail protein I; afdb-uniprot50 \| AF-A0A376YFL4-F1-MODEL\_V4 \| 1.0 \| 5.685e-16 \| 390 \| 0.372 \| 196 \| 119 \| 2 \| 1 \| 194 \| 1 \| 194 \| Tail protein \| Tail protein; afdb-uniprot50 \| AF-A0A432PXG2-F1-MODEL\_V4 \| 1.0 \| 1.058e-13 \| 390 \| 0.258 \| 213 \| 142 \| 7 \| 4 \| 205 \| 6 \| 213 \| Phage tail protein I \| Phage tail protein I; afdb-uniprot50 \| AF-A0A6N6XVM0-F1-MODEL\_V4 \| 1.0 \| 3.97e-13 \| 389 \| 0.238 \| 189 \| 128 \| 7 \| 3 \| 181 \| 11 \| 193 \| Phage tail protein I \| Phage tail protein I; afdb-uniprot50 \| AF-A0A090IT51-F1-MODEL\_V4 \| 1.0 \| 9.586e-13 \| 388 \| 0.194 \| 206 \| 158 \| 3 \| 1 \| 205 \| 8 \| 206 \| Phage tail protein I \| Phage tail protein I; afdb-uniprot50 \| AF-A0A8B2QT57-F1-MODEL\_V4 \| 1.0 \| 1.127e-13 \| 387 \| 0.293 \| 201 \| 131 \| 7 \| 4 \| 202 \| 3 \| 194 \| Phage tail protein I \| Phage tail protein I; afdb-uniprot50 \| AF-A0A1E7HY16-F1-MODEL\_V4 \| 1.0 \| 7.19e-12 \| 386 \| 0.366 \| 139 \| 83 \| 2 \| 1 \| 139 \| 2 \| 135 \| Phage tail protein I \| Phage tail protein I; afdb-uniprot50 \| AF-B8GRZ9-F1-MODEL\_V4 \| 1.0 \| 2.192e-14 \| 386 \| 0.337 \| 157 \| 98 \| 3 \| 1 \| 153 \| 1 \| 155 \| Bacteriophage P2-related tail formation protein-like protein \| Bacteriophage P2-related tail formation protein-like protein; afdb-uniprot50 \| AF-A0A3S5I5C6-F1-MODEL\_V4 \| 1.0 \| 9.851e-12 \| 386 \| 0.307 \| 169 \| 100 \| 6 \| 11 \| 164 \| 104 \| 270 \| Phage tail protein I \| Phage tail protein I; afdb-uniprot50 \| AF-B9NP80-F1-MODEL\_V4 \| 1.0 \| 3.97e-13 \| 385 \| 0.265 \| 211 \| 145 \| 5 \| 1 \| 203 \| 4 \| 212 \| Pyocin R2\_PP, tail formation protein GpI \| Pyocin R2\_PP, tail formation protein GpI; afdb-uniprot50 \| AF-A0A7X7NDE5-F1-MODEL\_V4 \| 1.0 \| 2.041e-12 \| 385 \| 0.236 \| 190 \| 130 \| 6 \| 1 \| 184 \| 1 \| 181 \| Phage tail protein I \| Phage tail protein I; afdb-uniprot50 \| AF-A0A1Y4W9W6-F1-MODEL\_V4 \| 1.0 \| 9.001e-13 \| 385 \| 0.264 \| 204 \| 138 \| 7 \| 4 \| 198 \| 20 \| 220 \| Phage tail protein I \| Phage tail protein I; afdb-uniprot50 \| AF-A0A259C7Y6-F1-MODEL\_V4 \| 1.0 \| 3.97e-13 \| 384 \| 0.287 \| 160 \| 105 \| 5 \| 26 \| 178 \| 5 \| 162 \| Phage tail protein I \| Phage tail protein I; afdb-uniprot50 \| AF-A0A8A7F3K5-F1-MODEL\_V4 \| 1.0 \| 1.6e-14 \| 384 \| 0.262 \| 229 \| 143 \| 8 \| 1 \| 205 \| 1 \| 227 \| Phage tail protein I \| Phage tail protein I; afdb-uniprot50 \| AF-L0RFK5-F1-MODEL\_V4 \| 1.0 \| 1.361e-13 \| 383 \| 0.269 \| 267 \| 130 \| 9 \| 1 \| 205 \| 1 \| 264 \| Uncharacterized protein \| Uncharacterized protein; afdb-uniprot50 \| AF-A0A134C9I1-F1-MODEL\_V4 \| 1.0 \| 1.799e-12 \| 382 \| 0.245 \| 204 \| 141 \| 9 \| 5 \| 202 \| 13 \| 209 \| Phage tail protein I-like protein \| Phage tail protein I-like protein; afdb-uniprot50 \| AF-A0A1B3E8I5-F1-MODEL\_V4 \| 1.0 \| 1.087e-12 \| 382 \| 0.267 \| 258 \| 131 \| 7 \| 1 \| 205 \| 1 \| 253 \| Uncharacterized protein \| Uncharacterized protein; afdb-uniprot50 \| AF-A0A849VC39-F1-MODEL\_V4 \| 1.0 \| 1.865e-13 \| 381 \| 0.299 \| 167 \| 107 \| 3 \| 3 \| 162 \| 9 \| 172 \| Phage tail protein I \| Phage tail protein I; afdb-uniprot50 \| AF-A0A4R0F5F4-F1-MODEL\_V4 \| 1.0 \| 2.721e-13 \| 380 \| 0.267 \| 176 \| 127 \| 2 \| 1 \| 176 \| 1 \| 174 \| Phage tail protein I \| Phage tail protein I; afdb-uniprot50 \| AF-A0A2A3MPV6-F1-MODEL\_V4 \| 1.0 \| 2.555e-13 \| 380 \| 0.35 \| 180 \| 106 \| 6 \| 1 \| 173 \| 46 \| 221 \| Phage tail protein I \| Phage tail protein I; afdb-uniprot50 \| AF-A0A6N8I028-F1-MODEL\_V4 \| 1.0 \| 8.759e-14 \| 379 \| 0.278 \| 194 \| 124 \| 5 \| 14 \| 203 \| 35 \| 216 \| Phage tail protein \| Phage tail protein; afdb-uniprot50 \| AF-A0A2G2D3E6-F1-MODEL\_V4 \| 1.0 \| 2.625e-12 \| 379 \| 0.216 \| 194 \| 140 \| 5 \| 4 \| 187 \| 14 \| 205 \| Phage tail protein I \| Phage tail protein I; afdb-uniprot50 \| AF-A0A0A8JLX9-F1-MODEL\_V4 \| 1.0 \| 2.533e-11 \| 379 \| 0.246 \| 158 \| 110 \| 5 \| 4 \| 155 \| 12 \| 166 \| Phage tail protein \| Phage tail protein; afdb-uniprot50 \| AF-A0A353G6I2-F1-MODEL\_V4 \| 1.0 \| 4.192e-11 \| 377 \| 0.233 \| 167 \| 112 \| 8 \| 1 \| 152 \| 1 \| 166 \| Phage tail protein I \| Phage tail protein I; afdb-uniprot50 \| AF-A0A0Q2UZB0-F1-MODEL\_V4 \| 1.0 \| 1.021e-12 \| 377 \| 0.261 \| 214 \| 142 \| 7 \| 1 \| 203 \| 14 \| 222 \| Uncharacterized protein \| Uncharacterized protein; afdb-uniprot50 \| AF-A0A165W2J4-F1-MODEL\_V4 \| 1.0 \| 2.465e-12 \| 377 \| 0.286 \| 206 \| 139 \| 5 \| 4 \| 205 \| 10 \| 211 \| Uncharacterized protein \| Uncharacterized protein; afdb-uniprot50 \| AF-A0A2G6HAI3-F1-MODEL\_V4 \| 1.0 \| 6.626e-15 \| 377 \| 0.324 \| 234 \| 125 \| 9 \| 1 \| 205 \| 1 \| 230 \| Phage tail protein I \| Phage tail protein I; afdb-uniprot50 \| AF-A0A1C6D4J3-F1-MODEL\_V4 \| 1.0 \| 5.589e-12 \| 376 \| 0.28 \| 178 \| 113 \| 8 \| 4 \| 171 \| 15 \| 187 \| Bacteriophage P2-related tail formation protein \| Bacteriophage P2-related tail formation protein; afdb-uniprot50 \| AF-D5EF95-F1-MODEL\_V4 \| 1.0 \| 2.253e-13 \| 376 \| 0.285 \| 217 \| 134 \| 8 \| 3 \| 204 \| 8 \| 218 \| Phage tail protein I \| Phage tail protein I; afdb-uniprot50 \| AF-A0A1G7SDN3-F1-MODEL\_V4 \| 1.0 \| 9.25e-12 \| 376 \| 0.301 \| 169 \| 99 \| 4 \| 4 \| 156 \| 56 \| 221 \| Phage tail protein, P2 protein I family \| Phage tail protein, P2 protein I family; afdb-uniprot50 \| AF-G9PUJ6-F1-MODEL\_V4 \| 1.0 \| 1.45e-13 \| 375 \| 0.233 \| 210 \| 141 \| 6 \| 5 \| 203 \| 14 \| 214 \| Phage tail protein I \| Phage tail protein I; afdb-uniprot50 \| AF-A0A7U3VT62-F1-MODEL\_V4 \| 1.0 \| 7.657e-12 \| 375 \| 0.337 \| 169 \| 93 \| 5 \| 1 \| 157 \| 1 \| 162 \| Fused protein containing phage-related tail protein GpI \| Fused protein containing phage-related tail protein GpI; afdb-uniprot50 \| AF-A0A1C3E9B0-F1-MODEL\_V4 \| 1.0 \| 4.928e-12 \| 374 \| 0.179 \| 201 \| 155 \| 7 \| 1 \| 196 \| 7 \| 202 \| Phage tail protein I \| Phage tail protein I; afdb-uniprot50 \| AF-A0A087NAA8-F1-MODEL\_V4 \| 1.0 \| 6.169e-13 \| 374 \| 0.325 \| 181 \| 111 \| 5 \| 1 \| 174 \| 3 \| 179 \| Uncharacterized protein \| Uncharacterized protein; afdb-uniprot50 \| AF-E5Y5X1-F1-MODEL\_V4 \| 1.0 \| 6.394e-14 \| 374 \| 0.257 \| 233 \| 141 \| 8 \| 4 \| 205 \| 14 \| 245 \| Phage tail protein I \| Phage tail protein I; afdb-uniprot50 \| AF-A0A545T5T7-F1-MODEL\_V4 \| 1.0 \| 3.97e-13 \| 373 \| 0.308 \| 224 \| 106 \| 9 \| 3 \| 186 \| 14 \| 228 \| Phage tail protein I \| Phage tail protein I; afdb-uniprot50 \| AF-A0A1Q6PTY4-F1-MODEL\_V4 \| 1.0 \| 6.57e-13 \| 373 \| 0.246 \| 219 \| 144 \| 7 \| 4 \| 205 \| 14 \| 228 \| Phage tail protein I \| Phage tail protein I; afdb-uniprot50 \| AF-A0A1W1UNV3-F1-MODEL\_V4 \| 1.0 \| 3.377e-12 \| 371 \| 0.296 \| 172 \| 101 \| 4 \| 1 \| 155 \| 1 \| 169 \| Phage tail protein, P2 protein I family \| Phage tail protein, P2 protein I family; afdb-uniprot50 \| AF-A0A1Q6JPU8-F1-MODEL\_V4 \| 1.0 \| 2.379e-11 \| 371 \| 0.279 \| 154 \| 103 \| 4 \| 11 \| 163 \| 30 \| 176 \| Phage tail protein I \| Phage tail protein I; afdb-uniprot50 \| AF-A0A7X5B8H9-F1-MODEL\_V4 \| 1.0 \| 4.08e-12 \| 370 \| 0.222 \| 207 \| 154 \| 5 \| 1 \| 201 \| 7 \| 212 \| Phage tail protein \| Phage tail protein; afdb-uniprot50 \| AF-A0A2X0WX70-F1-MODEL\_V4 \| 1.0 \| 9.586e-13 \| 370 \| 0.208 \| 211 \| 156 \| 5 \| 4 \| 205 \| 13 \| 221 \| Bacteriophage P2-related tail formation protein \| Bacteriophage P2-related tail formation protein; afdb-uniprot50 \| AF-A0A0F6A4X0-F1-MODEL\_V4 \| 1.0 \| 3.728e-13 \| 370 \| 0.266 \| 165 \| 110 \| 5 \| 1 \| 157 \| 1 \| 162 \| Tail protein \| Tail protein; afdb-uniprot50 \| AF-A0A497UKP7-F1-MODEL\_V4 \| 1.0 \| 3.597e-12 \| 369 \| 0.247 \| 194 \| 125 \| 7 \| 1 \| 188 \| 1 \| 179 \| P2-related tail formation protein \| P2-related tail formation protein; afdb-uniprot50 \| AF-F9S7B6-F1-MODEL\_V4 \| 1.0 \| 4.345e-12 \| 369 \| 0.227 \| 180 \| 130 \| 6 \| 4 \| 176 \| 28 \| 205 \| Bacteriophage tail protein I \| Bacteriophage tail protein I; afdb-uniprot50 \| AF-A0A0J9EGJ6-F1-MODEL\_V4 \| 1.0 \| 2.721e-13 \| 368 \| 0.244 \| 229 \| 128 \| 7 \| 14 \| 200 \| 2 \| 227 \| Tail protein I \| Tail protein I; afdb-uniprot50 \| AF-A0A0H5CXM9-F1-MODEL\_V4 \| 1.0 \| 2.379e-11 \| 367 \| 0.264 \| 178 \| 112 \| 6 \| 1 \| 165 \| 5 \| 176 \| Phage tail protein I \| Phage tail protein I; afdb-uniprot50 \| AF-A0A1A9G374-F1-MODEL\_V4 \| 1.0 \| 1.19e-11 \| 367 \| 0.152 \| 243 \| 166 \| 7 \| 1 \| 205 \| 3 \| 243 \| Uncharacterized protein \| Uncharacterized protein; afdb-uniprot50 \| AF-A0A5Q2UG14-F1-MODEL\_V4 \| 1.0 \| 7.452e-13 \| 367 \| 0.273 \| 183 \| 119 \| 7 \| 5 \| 177 \| 1 \| 179 \| Phage tail protein I \| Phage tail protein I; afdb-uniprot50 \| AF-A0A2K2G515-F1-MODEL\_V4 \| 1.0 \| 2.898e-13 \| 366 \| 0.264 \| 189 \| 124 \| 6 \| 1 \| 180 \| 3 \| 185 \| Phage tail protein I \| Phage tail protein I; afdb-uniprot50 \| AF-A0A1I5W407-F1-MODEL\_V4 \| 1.0 \| 1.097e-14 \| 364 \| 0.295 \| 220 \| 128 \| 8 \| 1 \| 200 \| 1 \| 213 \| Phage tail protein, P2 protein I family \| Phage tail protein, P2 protein I family; afdb-uniprot50 \| AF-A0A1Q6JQM0-F1-MODEL\_V4 \| 1.0 \| 5.793e-13 \| 364 \| 0.273 \| 212 \| 136 \| 6 \| 5 \| 205 \| 15 \| 219 \| Phage tail protein I \| Phage tail protein I; afdb-uniprot50 \| AF-A0A3G6WST2-F1-MODEL\_V4 \| 1.0 \| 1.544e-13 \| 364 \| 0.378 \| 169 \| 86 \| 4 \| 1 \| 153 \| 34 \| 199 \| Phage tail protein I \| Phage tail protein I; afdb-uniprot50 \| AF-A0A1X7MNV1-F1-MODEL\_V4 \| 1.0 \| 7.869e-11 \| 363 \| 0.446 \| 121 \| 61 \| 2 \| 1 \| 117 \| 1 \| 119 \| Phage tail protein I \| Phage tail protein I; afdb-uniprot50 \| AF-A0A1B1NTF0-F1-MODEL\_V4 \| 1.0 \| 4.345e-12 \| 363 \| 0.234 \| 183 \| 129 \| 7 \| 3 \| 176 \| 19 \| 199 \| Baseplate protein \| Baseplate protein; afdb-uniprot50 \| AF-D8IV10-F1-MODEL\_V4 \| 1.0 \| 1.799e-12 \| 363 \| 0.206 \| 232 \| 156 \| 7 \| 2 \| 205 \| 6 \| 237 \| Tail-related protein \| Tail-related protein; afdb-uniprot50 \| AF-A0A6L7TCB7-F1-MODEL\_V4 \| 1.0 \| 1.865e-13 \| 363 \| 0.345 \| 171 \| 100 \| 5 \| 1 \| 167 \| 100 \| 262 \| Phage tail protein I \| Phage tail protein I; afdb-uniprot50 \| AF-A0A6C9G6K6-F1-MODEL\_V4 \| 1.0 \| 7.869e-11 \| 362 \| 0.514 \| 103 \| 50 \| 0 \| 1 \| 103 \| 1 \| 103 \| Phage tail protein I \| Phage tail protein I; afdb-uniprot50 \| AF-A0A290TLP4-F1-MODEL\_V4 \| 1.0 \| 6.222e-15 \| 362 \| 0.298 \| 181 \| 122 \| 3 \| 2 \| 179 \| 5 \| 183 \| Uncharacterized protein \| Uncharacterized protein; afdb-uniprot50 \| AF-A0A142BH84-F1-MODEL\_V4 \| 1.0 \| 2.041e-12 \| 362 \| 0.5 \| 116 \| 58 \| 0 \| 1 \| 116 \| 1 \| 116 \| Phage tail protein \| Phage tail protein; afdb-uniprot50 \| AF-A0A645F841-F1-MODEL\_V4 \| 1.0 \| 3.287e-13 \| 361 \| 0.278 \| 208 \| 132 \| 10 \| 3 \| 200 \| 11 \| 210 \| Uncharacterized protein \| Uncharacterized protein; afdb-uniprot50 \| AF-F4QJC1-F1-MODEL\_V4 \| 1.0 \| 3.003e-14 \| 360 \| 0.312 \| 189 \| 119 \| 6 \| 1 \| 185 \| 1 \| 182 \| Phage tail protein I \| Phage tail protein I; afdb-uniprot50 \| AF-A0A655UT13-F1-MODEL\_V4 \| 1.0 \| 2.233e-11 \| 360 \| 0.251 \| 171 \| 107 \| 5 \| 1 \| 155 \| 2 \| 167 \| Bacteriophage P2-related tail formation protein \| Bacteriophage P2-related tail formation protein; afdb-uniprot50 \| AF-A0A3D4MXN2-F1-MODEL\_V4 \| 1.0 \| 6.339e-12 \| 360 \| 0.279 \| 186 \| 114 \| 8 \| 4 \| 174 \| 39 \| 219 \| Phage tail protein I \| Phage tail protein I; afdb-uniprot50 \| AF-A0A6L7FZ02-F1-MODEL\_V4 \| 1.0 \| 2.978e-12 \| 360 \| 0.228 \| 223 \| 150 \| 8 \| 1 \| 205 \| 1 \| 219 \| Phage tail protein I \| Phage tail protein I; afdb-uniprot50 \| AF-F3YY51-F1-MODEL\_V4 \| 1.0 \| 1.233e-12 \| 357 \| 0.266 \| 206 \| 140 \| 6 \| 4 \| 203 \| 13 \| 213 \| Phage tail protein I \| Phage tail protein I; afdb-uniprot50 \| AF-A0A3S2Y7C0-F1-MODEL\_V4 \| 1.0 \| 1.986e-13 \| 357 \| 0.288 \| 184 \| 117 \| 4 \| 4 \| 184 \| 16 \| 188 \| Phage tail protein I \| Phage tail protein I; afdb-uniprot50 \| AF-A0A6L6VKC1-F1-MODEL\_V4 \| 1.0 \| 7.19e-12 \| 357 \| 0.191 \| 225 \| 162 \| 8 \| 1 \| 205 \| 1 \| 225 \| Phage tail protein I \| Phage tail protein I; afdb-uniprot50 \| AF-A0A161XW60-F1-MODEL\_V4 \| 1.0 \| 7.869e-11 \| 356 \| 0.277 \| 166 \| 107 \| 5 \| 3 \| 157 \| 98 \| 261 \| Tail protein \| Tail protein; afdb-uniprot50 \| AF-A0A3B6Y3G2-F1-MODEL\_V4 \| 1.0 \| 1.03e-14 \| 354 \| 0.275 \| 185 \| 123 \| 3 \| 4 \| 179 \| 1 \| 183 \| Phage tail protein I \| Phage tail protein I; afdb-uniprot50 \| AF-A0A1Y4W8I7-F1-MODEL\_V4 \| 1.0 \| 1.916e-12 \| 354 \| 0.258 \| 213 \| 135 \| 9 \| 3 \| 199 \| 11 \| 216 \| Phage tail protein I \| Phage tail protein I; afdb-uniprot50 \| AF-A0A3A9ENS4-F1-MODEL\_V4 \| 1.0 \| 1.158e-12 \| 354 \| 0.241 \| 240 \| 137 \| 10 \| 4 \| 203 \| 15 \| 249 \| Phage tail protein I \| Phage tail protein I; afdb-uniprot50 \| AF-A0A1Y3WRA3-F1-MODEL\_V4 \| 1.0 \| 1.849e-11 \| 353 \| 0.219 \| 187 \| 135 \| 7 \| 5 \| 183 \| 13 \| 196 \| Phage tail protein I \| Phage tail protein I; afdb-uniprot50 \| AF-A0A1G3UAH8-F1-MODEL\_V4 \| 1.0 \| 9.586e-13 \| 353 \| 0.242 \| 227 \| 138 \| 10 \| 4 \| 205 \| 3 \| 220 \| Phage tail protein I \| Phage tail protein I; afdb-uniprot50 \| AF-A0A7C9LKP3-F1-MODEL\_V4 \| 1.0 \| 9.505e-11 \| 351 \| 0.189 \| 179 \| 135 \| 6 \| 4 \| 177 \| 18 \| 191 \| Phage tail protein I \| Phage tail protein I; afdb-uniprot50 \| AF-A0A1B8QCU6-F1-MODEL\_V4 \| 1.0 \| 2.335e-14 \| 351 \| 0.284 \| 179 \| 123 \| 3 \| 4 \| 182 \| 19 \| 192 \| Phage tail protein I \| Phage tail protein I; afdb-uniprot50 \| AF-F7L031-F1-MODEL\_V4 \| 1.0 \| 7.657e-12 \| 351 \| 0.248 \| 189 \| 122 \| 5 \| 3 \| 175 \| 11 \| 195 \| Putative phage tail protein \| Putative phage tail protein; afdb-uniprot50 \| AF-A0A327Q855-F1-MODEL\_V4 \| 1.0 \| 4.627e-12 \| 351 \| 0.254 \| 220 \| 142 \| 9 \| 1 \| 205 \| 1 \| 213 \| Phage tail P2-like protein \| Phage tail P2-like protein; afdb-uniprot50 \| AF-A0A239C7G7-F1-MODEL\_V4 \| 1.0 \| 4.928e-12 \| 351 \| 0.263 \| 205 \| 134 \| 5 \| 4 \| 197 \| 12 \| 210 \| Phage tail protein, P2 protein I family \| Phage tail protein, P2 protein I family; afdb-uniprot50 \| AF-A0A135IPT9-F1-MODEL\_V4 \| 1.0 \| 6.626e-15 \| 351 \| 0.343 \| 224 \| 122 \| 9 \| 1 \| 205 \| 5 \| 222 \| Uncharacterized protein \| Uncharacterized protein; afdb-uniprot50 \| AF-A0A085AFN4-F1-MODEL\_V4 \| 1.0 \| 8.155e-12 \| 350 \| 0.256 \| 191 \| 123 \| 6 \| 1 \| 182 \| 2 \| 182 \| Tail protein I \| Tail protein I; afdb-uniprot50 \| AF-A0A1M7R7F1-F1-MODEL\_V4 \| 1.0 \| 7.19e-12 \| 350 \| 0.223 \| 201 \| 134 \| 8 \| 1 \| 192 \| 1 \| 188 \| P2-related tail formation protein \| P2-related tail formation protein; afdb-uniprot50 \| AF-A0A0A2SKX6-F1-MODEL\_V4 \| 1.0 \| 2.796e-12 \| 349 \| 0.254 \| 185 \| 120 \| 8 \| 3 \| 177 \| 11 \| 187 \| Uncharacterized protein \| Uncharacterized protein; afdb-uniprot50 \| AF-B6IMH1-F1-MODEL\_V4 \| 1.0 \| 1.531e-11 \| 349 \| 0.236 \| 224 \| 152 \| 6 \| 1 \| 205 \| 1 \| 224 \| Uncharacterized protein \| Uncharacterized protein; afdb-uniprot50 \| AF-A0A0J7Y5C0-F1-MODEL\_V4 \| 1.0 \| 1.087e-12 \| 348 \| 0.266 \| 229 \| 138 \| 4 \| 1 \| 200 \| 3 \| 230 \| Tail protein \| Tail protein; afdb-uniprot50 \| AF-A0A5S3YZW9-F1-MODEL\_V4 \| 1.0 \| 3.471e-11 \| 348 \| 0.255 \| 192 \| 125 \| 7 \| 5 \| 180 \| 97 \| 286 \| Phage tail protein I \| Phage tail protein I; afdb-uniprot50 \| AF-A0A2S9GTF6-F1-MODEL\_V4 \| 1.0 \| 3.377e-12 \| 347 \| 0.346 \| 150 \| 90 \| 4 \| 1 \| 149 \| 3 \| 145 \| Phage tail protein I \| Phage tail protein I; afdb-uniprot50 \| AF-A0A7T1BKU5-F1-MODEL\_V4 \| 1.0 \| 8.685e-12 \| 347 \| 0.278 \| 183 \| 119 \| 7 \| 5 \| 177 \| 11 \| 190 \| Phage tail protein I \| Phage tail protein I; afdb-uniprot50 \| AF-A0A3A9EMV3-F1-MODEL\_V4 \| 1.0 \| 5.393e-11 \| 346 \| 0.258 \| 147 \| 97 \| 5 \| 11 \| 154 \| 31 \| 168 \| Phage tail protein I \| Phage tail protein I; afdb-uniprot50 \| AF-A0A4P6B4N0-F1-MODEL\_V4 \| 1.0 \| 1.969e-11 \| 346 \| 0.268 \| 186 \| 116 \| 8 \| 3 \| 177 \| 11 \| 187 \| Phage tail protein \| Phage tail protein; afdb-uniprot50 \| AF-A0A1H9HC51-F1-MODEL\_V4 \| 1.0 \| 1.644e-13 \| 346 \| 0.289 \| 214 \| 135 \| 6 \| 1 \| 203 \| 3 \| 210 \| Phage tail protein, P2 protein I family \| Phage tail protein, P2 protein I family; afdb-uniprot50 \| AF-A0A6N9P645-F1-MODEL\_V4 \| 1.0 \| 1.302e-10 \| 346 \| 0.251 \| 163 \| 110 \| 5 \| 6 \| 163 \| 21 \| 176 \| Phage tail protein I \| Phage tail protein I; afdb-uniprot50 \| AF-A0A249DWE7-F1-MODEL\_V4 \| 1.0 \| 2.173e-12 \| 345 \| 0.315 \| 165 \| 98 \| 4 \| 1 \| 156 \| 2 \| 160 \| Phage tail protein I \| Phage tail protein I; afdb-uniprot50 \| AF-A0A1H0YPQ4-F1-MODEL\_V4 \| 1.0 \| 3.471e-11 \| 345 \| 0.211 \| 222 \| 156 \| 9 \| 1 \| 205 \| 1 \| 220 \| Phage tail protein, P2 protein I family \| Phage tail protein, P2 protein I family; afdb-uniprot50 \| AF-B6WRR5-F1-MODEL\_V4 \| 1.0 \| 3.171e-12 \| 345 \| 0.228 \| 206 \| 144 \| 5 \| 1 \| 205 \| 54 \| 245 \| Phage tail protein I-like protein \| Phage tail protein I-like protein; afdb-uniprot50 \| AF-A0A5C7PF23-F1-MODEL\_V4 \| 1.0 \| 1.399e-12 \| 344 \| 0.272 \| 209 \| 115 \| 6 \| 1 \| 175 \| 2 \| 207 \| Phage tail protein I \| Phage tail protein I; afdb-uniprot50 \| AF-A0A1Y1QXH2-F1-MODEL\_V4 \| 1.0 \| 1.9e-10 \| 344 \| 0.264 \| 151 \| 106 \| 4 \| 1 \| 148 \| 4 \| 152 \| Phage tail protein I \| Phage tail protein I; afdb-uniprot50 \| AF-A0A023PZC5-F1-MODEL\_V4 \| 1.0 \| 7.869e-11 \| 343 \| 0.269 \| 178 \| 118 \| 6 \| 1 \| 166 \| 97 \| 274 \| Tail protein I \| Tail protein I; afdb-uniprot50 \| AF-A0A2P8EI45-F1-MODEL\_V4 \| 1.0 \| 5.064e-11 \| 341 \| 0.231 \| 177 \| 124 \| 7 \| 4 \| 172 \| 13 \| 185 \| Phage tail P2-like protein \| Phage tail P2-like protein; afdb-uniprot50 \| AF-A0A351B539-F1-MODEL\_V4 \| 1.0 \| 9.586e-13 \| 341 \| 0.235 \| 289 \| 133 \| 8 \| 1 \| 205 \| 8 \| 292 \| Phage tail protein I \| Phage tail protein I; afdb-uniprot50 \| AF-A0A2J4RC72-F1-MODEL\_V4 \| 1.0 \| 2.024e-10 \| 340 \| 0.441 \| 120 \| 63 \| 2 \| 1 \| 117 \| 6 \| 124 \| Phage tail protein I \| Phage tail protein I; afdb-uniprot50 \| AF-A0A1E4MW51-F1-MODEL\_V4 \| 1.0 \| 4.465e-11 \| 340 \| 0.283 \| 141 \| 93 \| 3 \| 22 \| 157 \| 3 \| 140 \| Phage tail protein I \| Phage tail protein I; afdb-uniprot50 \| AF-A0A3T0N1K0-F1-MODEL\_V4 \| 1.0 \| 3.406e-14 \| 340 \| 0.275 \| 243 \| 133 \| 7 \| 1 \| 205 \| 4 \| 241 \| Phage tail protein I \| Phage tail protein I; afdb-uniprot50 \| AF-A0A5U3DU51-F1-MODEL\_V4 \| 1.0 \| 1.387e-10 \| 339 \| 0.391 \| 120 \| 71 \| 1 \| 75 \| 194 \| 2 \| 119 \| Phage tail protein I \| Phage tail protein I; afdb-uniprot50 \| AF-A0A5C8XTJ9-F1-MODEL\_V4 \| 1.0 \| 5.952e-12 \| 339 \| 0.361 \| 155 \| 95 \| 3 \| 53 \| 205 \| 3 \| 155 \| Phage tail protein I \| Phage tail protein I; afdb-uniprot50 \| AF-A0A6I2UKH8-F1-MODEL\_V4 \| 1.0 \| 7.19e-12 \| 339 \| 0.19 \| 210 \| 155 \| 8 \| 4 \| 205 \| 12 \| 214 \| Phage tail protein I \| Phage tail protein I; afdb-uniprot50 \| AF-A0A285M2Z8-F1-MODEL\_V4 \| 1.0 \| 4.588e-10 \| 338 \| 0.213 \| 164 \| 118 \| 5 \| 4 \| 157 \| 14 \| 176 \| Phage tail protein, P2 protein I family \| Phage tail protein, P2 protein I family; afdb-uniprot50 \| AF-A0A124INN1-F1-MODEL\_V4 \| 1.0 \| 5.248e-12 \| 337 \| 0.258 \| 186 \| 118 \| 7 \| 4 \| 178 \| 12 \| 188 \| Uncharacterized protein \| Uncharacterized protein; afdb-uniprot50 \| AF-A0A081B6D0-F1-MODEL\_V4 \| 1.0 \| 3.06e-11 \| 337 \| 0.284 \| 165 \| 99 \| 4 \| 1 \| 149 \| 1 \| 162 \| Phage tail protein I \| Phage tail protein I; afdb-uniprot50 \| AF-A0A413ED79-F1-MODEL\_V4 \| 1.0 \| 6.751e-12 \| 337 \| 0.221 \| 208 \| 149 \| 7 \| 3 \| 201 \| 11 \| 214 \| Phage tail protein I \| Phage tail protein I; afdb-uniprot50 \| AF-R9M1L6-F1-MODEL\_V4 \| 1.0 \| 5.064e-11 \| 337 \| 0.274 \| 171 \| 104 \| 7 \| 5 \| 163 \| 15 \| 177 \| Phage tail protein I \| Phage tail protein I; afdb-uniprot50 \| AF-A0A4R7SS34-F1-MODEL\_V4 \| 1.0 \| 1.573e-10 \| 336 \| 0.234 \| 162 \| 109 \| 5 \| 1 \| 151 \| 3 \| 160 \| Phage tail P2-like protein \| Phage tail P2-like protein; afdb-uniprot50 \| AF-A0A101K5W9-F1-MODEL\_V4 \| 1.0 \| 7.657e-12 \| 336 \| 0.235 \| 212 \| 137 \| 8 \| 3 \| 198 \| 15 \| 217 \| Phage tail protein \| Phage tail protein; afdb-uniprot50 \| AF-D8IV17-F1-MODEL\_V4 \| 1.0 \| 3.259e-11 \| 336 \| 0.209 \| 220 \| 154 \| 6 \| 4 \| 205 \| 9 \| 226 \| Phage-related tail protein \| Phage-related tail protein; afdb-uniprot50 \| AF-A0A318KXU1-F1-MODEL\_V4 \| 1.0 \| 6.117e-11 \| 335 \| 0.301 \| 166 \| 98 \| 6 \| 1 \| 156 \| 1 \| 158 \| Phage tail P2-like protein \| Phage tail P2-like protein; afdb-uniprot50 \| AF-E0NXI4-F1-MODEL\_V4 \| 1.0 \| 1.63e-11 \| 331 \| 0.23 \| 213 \| 143 \| 8 \| 3 \| 203 \| 12 \| 215 \| Phage tail protein \| Phage tail protein; afdb-uniprot50 \| AF-A0A3T0L2N1-F1-MODEL\_V4 \| 1.0 \| 3.728e-13 \| 331 \| 0.235 \| 212 \| 144 \| 7 \| 1 \| 205 \| 3 \| 203 \| Uncharacterized protein \| Uncharacterized protein; afdb-uniprot50 \| AF-A0A259U4Q0-F1-MODEL\_V4 \| 1.0 \| 2.041e-12 \| 330 \| 0.231 \| 207 \| 146 \| 7 \| 3 \| 201 \| 12 \| 213 \| Phage tail protein \| Phage tail protein; afdb-uniprot50 \| AF-A0A1M6SJT3-F1-MODEL\_V4 \| 1.0 \| 1.9e-10 \| 329 \| 0.231 \| 199 \| 141 \| 8 \| 5 \| 197 \| 12 \| 204 \| Phage tail protein, P2 protein I family \| Phage tail protein, P2 protein I family; afdb-uniprot50 \| AF-A0A376VMZ8-F1-MODEL\_V4 \| 1.0 \| 4.045e-10 \| 328 \| 0.552 \| 105 \| 45 \| 1 \| 1 \| 103 \| 1 \| 105 \| Tail protein I (GpI) \| Tail protein I (GpI); afdb-uniprot50 \| AF-A0A2V2FTF7-F1-MODEL\_V4 \| 1.0 \| 2.233e-11 \| 328 \| 0.241 \| 170 \| 113 \| 4 \| 12 \| 179 \| 37 \| 192 \| Phage tail protein I \| Phage tail protein I; afdb-uniprot50 \| AF-E2CGH4-F1-MODEL\_V4 \| 1.0 \| 2.533e-11 \| 328 \| 0.211 \| 213 \| 144 \| 7 \| 4 \| 194 \| 15 \| 225 \| Phage tail protein I \| Phage tail protein I; afdb-uniprot50 \| AF-A0A2S0PE89-F1-MODEL\_V4 \| 1.0 \| 7.19e-12 \| 328 \| 0.215 \| 237 \| 130 \| 8 \| 1 \| 190 \| 1 \| 228 \| Phage tail protein I \| Phage tail protein I; afdb-uniprot50 \| AF-A0A1B1A2A5-F1-MODEL\_V4 \| 1.0 \| 5.637e-14 \| 327 \| 0.302 \| 205 \| 132 \| 5 \| 9 \| 205 \| 4 \| 205 \| Uncharacterized protein \| Uncharacterized protein; afdb-uniprot50 \| AF-A0A845R5N3-F1-MODEL\_V4 \| 1.0 \| 7.869e-11 \| 327 \| 0.175 \| 233 \| 157 \| 9 \| 4 \| 204 \| 1 \| 230 \| Phage tail protein I \| Phage tail protein I; afdb-uniprot50 \| AF-K1KKI1-F1-MODEL\_V4 \| 1.0 \| 4.08e-12 \| 326 \| 0.244 \| 213 \| 143 \| 9 \| 3 \| 200 \| 12 \| 221 \| Phage tail protein I \| Phage tail protein I; afdb-uniprot50 \| AF-A0A827WDG4-F1-MODEL\_V4 \| 1.0 \| 4.886e-10 \| 325 \| 0.472 \| 110 \| 56 \| 1 \| 1 \| 110 \| 2 \| 109 \| Phage tail protein I \| Phage tail protein I; afdb-uniprot50 \| AF-A0A0X8JTG7-F1-MODEL\_V4 \| 1.0 \| 3.06e-11 \| 324 \| 0.197 \| 192 \| 139 \| 8 \| 14 \| 197 \| 28 \| 212 \| Phage tail protein \| Phage tail protein; afdb-uniprot50 \| AF-D4M9N0-F1-MODEL\_V4 \| 1.0 \| 9.001e-13 \| 324 \| 0.282 \| 237 \| 130 \| 11 \| 1 \| 205 \| 1 \| 229 \| Phage tail protein, P2 protein I family \| Phage tail protein, P2 protein I family; afdb-uniprot50 \| AF-A0A433LI31-F1-MODEL\_V4 \| 1.0 \| 6.938e-11 \| 323 \| 0.221 \| 190 \| 132 \| 6 \| 1 \| 184 \| 1 \| 180 \| Phage tail protein I \| Phage tail protein I; afdb-uniprot50 \| AF-A0A5M8P5X2-F1-MODEL\_V4 \| 1.0 \| 1.531e-11 \| 323 \| 0.215 \| 209 \| 151 \| 7 \| 5 \| 202 \| 31 \| 237 \| Phage tail protein I \| Phage tail protein I; afdb-uniprot50 \| AF-A0A0J2JHP4-F1-MODEL\_V4 \| 1.0 \| 1.784e-10 \| 322 \| 0.318 \| 138 \| 89 \| 3 \| 64 \| 198 \| 3 \| 138 \| Phage tail protein I \| Phage tail protein I; afdb-uniprot50 \| AF-A0A511Z872-F1-MODEL\_V4 \| 1.0 \| 2.097e-11 \| 322 \| 0.218 \| 215 \| 143 \| 7 \| 4 \| 205 \| 12 \| 214 \| Uncharacterized protein \| Uncharacterized protein; afdb-uniprot50 \| AF-A0A7C8GR62-F1-MODEL\_V4 \| 1.0 \| 2.698e-11 \| 322 \| 0.231 \| 212 \| 140 \| 9 \| 4 \| 205 \| 12 \| 210 \| Phage tail protein I \| Phage tail protein I; afdb-uniprot50 \| AF-A0A2S0JR95-F1-MODEL\_V4 \| 1.0 \| 1.849e-11 \| 321 \| 0.2 \| 224 \| 149 \| 8 \| 4 \| 202 \| 10 \| 228 \| Phage tail protein I \| Phage tail protein I; afdb-uniprot50 \| AF-A0A4P9VH54-F1-MODEL\_V4 \| 1.0 \| 1.302e-10 \| 320 \| 0.325 \| 120 \| 81 \| 0 \| 26 \| 145 \| 1 \| 120 \| Phage tail protein I \| Phage tail protein I; afdb-uniprot50 \| AF-A0A379ZTT6-F1-MODEL\_V4 \| 1.0 \| 2.772e-10 \| 320 \| 0.37 \| 135 \| 83 \| 1 \| 69 \| 203 \| 7 \| 139 \| Putative phage tail protein \| Putative phage tail protein; afdb-uniprot50 \| AF-A0A376WQ79-F1-MODEL\_V4 \| 1.0 \| 5.695e-09 \| 320 \| 0.525 \| 97 \| 46 \| 0 \| 1 \| 97 \| 3 \| 99 \| Tail protein \| Tail protein; afdb-uniprot50 \| AF-V4JM45-F1-MODEL\_V4 \| 1.0 \| 1.049e-11 \| 320 \| 0.323 \| 195 \| 90 \| 9 \| 2 \| 159 \| 14 \| 203 \| Tail protein \| Tail protein; afdb-uniprot50 \| AF-A0A777HJF3-F1-MODEL\_V4 \| 1.0 \| 3.144e-10 \| 319 \| 0.392 \| 125 \| 74 \| 1 \| 79 \| 203 \| 1 \| 123 \| Phage tail protein I \| Phage tail protein I; afdb-uniprot50 \| AF-A0A7W4RI07-F1-MODEL\_V4 \| 1.0 \| 1.799e-12 \| 319 \| 0.309 \| 171 \| 112 \| 4 \| 38 \| 205 \| 1 \| 168 \| Phage tail P2-like protein \| Phage tail P2-like protein; afdb-uniprot50 \| AF-J4UR00-F1-MODEL\_V4 \| 1.0 \| 5.064e-11 \| 319 \| 0.225 \| 182 \| 136 \| 4 \| 23 \| 203 \| 4 \| 181 \| Phage tail protein I-like protein \| Phage tail protein I-like protein; afdb-uniprot50 \| AF-A0A0X1U7S8-F1-MODEL\_V4 \| 1.0 \| 2.024e-10 \| 319 \| 0.201 \| 174 \| 128 \| 6 \| 10 \| 178 \| 20 \| 187 \| Phage tail protein \| Phage tail protein; afdb-uniprot50 \| AF-A0A6N7W593-F1-MODEL\_V4 \| 1.0 \| 5.064e-11 \| 319 \| 0.219 \| 214 \| 144 \| 8 \| 4 \| 205 \| 13 \| 215 \| Phage tail protein I \| Phage tail protein I; afdb-uniprot50 \| AF-A0A6B8KI89-F1-MODEL\_V4 \| 1.0 \| 1.35e-11 \| 319 \| 0.228 \| 206 \| 151 \| 7 \| 1 \| 203 \| 85 \| 285 \| Phage tail protein I \| Phage tail protein I; afdb-uniprot50 \| AF-A0A6N7JGU4-F1-MODEL\_V4 \| 1.0 \| 1.45e-13 \| 319 \| 0.253 \| 249 \| 142 \| 5 \| 1 \| 205 \| 2 \| 250 \| Phage tail protein I \| Phage tail protein I; afdb-uniprot50 \| AF-A0A143HC79-F1-MODEL\_V4 \| 1.0 \| 1.616e-09 \| 318 \| 0.23 \| 156 \| 107 \| 7 \| 9 \| 159 \| 19 \| 166 \| Uncharacterized protein \| Uncharacterized protein; afdb-uniprot50 \| AF-E3H9A4-F1-MODEL\_V4 \| 1.0 \| 3.259e-11 \| 318 \| 0.179 \| 184 \| 134 \| 6 \| 4 \| 179 \| 12 \| 186 \| Phage tail protein \| Phage tail protein; afdb-uniprot50 \| AF-A0A7C3CG54-F1-MODEL\_V4 \| 1.0 \| 8.612e-10 \| 318 \| 0.262 \| 164 \| 103 \| 5 \| 4 \| 155 \| 10 \| 167 \| Phage tail protein I \| Phage tail protein I; afdb-uniprot50 \| AF-A0A3A6E7Q8-F1-MODEL\_V4 \| 1.0 \| 4.192e-11 \| 318 \| 0.242 \| 239 \| 143 \| 9 \| 1 \| 205 \| 10 \| 244 \| Phage tail protein I \| Phage tail protein I; afdb-uniprot50 \| AF-A0A0B0HIN7-F1-MODEL\_V4 \| 1.0 \| 2.603e-10 \| 318 \| 0.227 \| 167 \| 112 \| 8 \| 4 \| 160 \| 12 \| 171 \| Phage tail protein (Tail\_P2\_I) \| Phage tail protein (Tail\_P2\_I); afdb-uniprot50 \| AF-B6VLW9-F1-MODEL\_V4 \| 1.0 \| 5.021e-09 \| 317 \| 0.49 \| 100 \| 51 \| 0 \| 1 \| 100 \| 2 \| 101 \| Putative bacteriophage protein \| Putative bacteriophage protein; afdb-uniprot50 \| AF-A0A1M7LUF7-F1-MODEL\_V4 \| 1.0 \| 2.097e-11 \| 317 \| 0.223 \| 188 \| 126 \| 6 \| 1 \| 178 \| 9 \| 186 \| Phage tail protein, P2 protein I family \| Phage tail protein, P2 protein I family; afdb-uniprot50 \| AF-A0A1H0P8S4-F1-MODEL\_V4 \| 1.0 \| 6.117e-11 \| 317 \| 0.206 \| 208 \| 153 \| 9 \| 5 \| 205 \| 12 \| 214 \| Phage tail protein, P2 protein I family \| Phage tail protein, P2 protein I family; afdb-uniprot50 \| AF-A0A376RLD7-F1-MODEL\_V4 \| 1.0 \| 8.925e-11 \| 316 \| 0.515 \| 130 \| 60 \| 2 \| 1 \| 129 \| 84 \| 211 \| Putative tail protein I (Gpi) \| Putative tail protein I (Gpi); afdb-uniprot50 \| AF-A0A0P8W181-F1-MODEL\_V4 \| 1.0 \| 5.393e-11 \| 316 \| 0.219 \| 205 \| 150 \| 5 \| 6 \| 205 \| 19 \| 218 \| Phage tail protein \| Phage tail protein; afdb-uniprot50 \| AF-A0A1M6GFJ1-F1-MODEL\_V4 \| 1.0 \| 3.259e-11 \| 316 \| 0.251 \| 215 \| 144 \| 7 \| 3 \| 205 \| 12 \| 221 \| Phage tail protein, P2 protein I family \| Phage tail protein, P2 protein I family; afdb-uniprot50 \| AF-A0A0U9HK81-F1-MODEL\_V4 \| 1.0 \| 2.873e-11 \| 315 \| 0.239 \| 217 \| 139 \| 8 \| 3 \| 205 \| 11 \| 215 \| Phage tail protein, P2 protein I family \| Phage tail protein, P2 protein I family; afdb-uniprot50 \| AF-A0A5B9Y8K3-F1-MODEL\_V4 \| 1.0 \| 2.155e-10 \| 314 \| 0.219 \| 182 \| 128 \| 7 \| 3 \| 176 \| 11 \| 186 \| Phage tail protein I \| Phage tail protein I; afdb-uniprot50 \| AF-A0A2E2N191-F1-MODEL\_V4 \| 1.0 \| 5.107e-13 \| 314 \| 0.261 \| 199 \| 133 \| 5 \| 10 \| 202 \| 1 \| 191 \| Phage tail protein I \| Phage tail protein I; afdb-uniprot50 \| AF-H1PQH3-F1-MODEL\_V4 \| 1.0 \| 5.902e-10 \| 314 \| 0.235 \| 174 \| 118 \| 4 \| 12 \| 180 \| 29 \| 192 \| Phage tail protein I \| Phage tail protein I; afdb-uniprot50 \| AF-A0A7U9X3N6-F1-MODEL\_V4 \| 1.0 \| 3.798e-10 \| 313 \| 0.257 \| 167 \| 111 \| 5 \| 3 \| 162 \| 11 \| 171 \| Uncharacterized protein \| Uncharacterized protein; afdb-uniprot50 \| AF-A0A162QMS0-F1-MODEL\_V4 \| 1.0 \| 3.471e-11 \| 313 \| 0.229 \| 218 \| 141 \| 9 \| 3 \| 203 \| 11 \| 218 \| Phage tail protein \| Phage tail protein; afdb-uniprot50 \| AF-A0A1C6EQC2-F1-MODEL\_V4 \| 1.0 \| 5.204e-10 \| 313 \| 0.184 \| 179 \| 134 \| 7 \| 5 \| 175 \| 13 \| 187 \| Bacteriophage P2-related tail formation protein \| Bacteriophage P2-related tail formation protein; afdb-uniprot50 \| AF-G4KQ65-F1-MODEL\_V4 \| 1.0 \| 4.465e-11 \| 312 \| 0.239 \| 184 \| 116 \| 9 \| 4 \| 174 \| 11 \| 183 \| Uncharacterized protein \| Uncharacterized protein; afdb-uniprot50 \| AF-A0A369BHI8-F1-MODEL\_V4 \| 1.0 \| 2.295e-10 \| 312 \| 0.231 \| 177 \| 122 \| 8 \| 6 \| 177 \| 20 \| 187 \| Phage tail P2-like protein \| Phage tail P2-like protein; afdb-uniprot50 \| AF-A0A844HQG5-F1-MODEL\_V4 \| 1.0 \| 2.873e-11 \| 312 \| 0.281 \| 188 \| 114 \| 3 \| 1 \| 170 \| 1 \| 185 \| Phage tail protein I \| Phage tail protein I; afdb-uniprot50 \| AF-A0A7X5KSS8-F1-MODEL\_V4 \| 1.0 \| 4.465e-11 \| 312 \| 0.207 \| 202 \| 151 \| 5 \| 3 \| 198 \| 11 \| 209 \| Uncharacterized protein \| Uncharacterized protein; afdb-uniprot50 \| AF-A0A1M7RI70-F1-MODEL\_V4 \| 1.0 \| 1.387e-10 \| 311 \| 0.199 \| 206 \| 147 \| 7 \| 1 \| 197 \| 1 \| 197 \| Phage tail protein (Tail\_P2\_I) \| Phage tail protein (Tail\_P2\_I); afdb-uniprot50 \| AF-A0A3S5D9L0-F1-MODEL\_V4 \| 1.0 \| 1.661e-08 \| 310 \| 0.471 \| 89 \| 45 \| 1 \| 35 \| 123 \| 2 \| 88 \| Phage tail protein I \| Phage tail protein I; afdb-uniprot50 \| AF-A0A2X2UK87-F1-MODEL\_V4 \| 1.0 \| 2.533e-11 \| 310 \| 0.227 \| 211 \| 134 \| 7 \| 12 \| 202 \| 30 \| 231 \| Phage tail protein \| Phage tail protein; afdb-uniprot50 \| AF-A0A173SBX4-F1-MODEL\_V4 \| 1.0 \| 2.097e-11 \| 310 \| 0.211 \| 208 \| 141 \| 6 \| 12 \| 205 \| 31 \| 229 \| Bacteriophage P2-related tail formation protein \| Bacteriophage P2-related tail formation protein; afdb-uniprot50 \| AF-A0A3S4HEN1-F1-MODEL\_V4 \| 1.0 \| 1.387e-10 \| 309 \| 0.189 \| 232 \| 156 \| 10 \| 1 \| 203 \| 2 \| 230 \| Bacteriophage P2-related tail formation protein \| Bacteriophage P2-related tail formation protein; afdb-uniprot50 \| AF-A0A7X5ICE8-F1-MODEL\_V4 \| 1.0 \| 1.338e-09 \| 309 \| 0.191 \| 167 \| 123 \| 7 \| 5 \| 163 \| 1 \| 163 \| Uncharacterized protein \| Uncharacterized protein; afdb-uniprot50 \| AF-A0A0C1IR17-F1-MODEL\_V4 \| 1.0 \| 6.938e-11 \| 307 \| 0.189 \| 211 \| 149 \| 6 \| 1 \| 190 \| 1 \| 210 \| Uncharacterized protein \| Uncharacterized protein; afdb-uniprot50 \| AF-A0A7Z0N0J2-F1-MODEL\_V4 \| 1.0 \| 1.19e-11 \| 306 \| 0.3 \| 190 \| 95 \| 6 \| 1 \| 157 \| 1 \| 185 \| Phage tail protein I \| Phage tail protein I; afdb-uniprot50 \| AF-A0A385Q1K2-F1-MODEL\_V4 \| 1.0 \| 1.387e-10 \| 305 \| 0.211 \| 180 \| 128 \| 8 \| 3 \| 175 \| 16 \| 188 \| Phage tail protein I \| Phage tail protein I; afdb-uniprot50 \| AF-A0A8A8BWX0-F1-MODEL\_V4 \| 1.0 \| 2.097e-11 \| 305 \| 0.289 \| 166 \| 98 \| 4 \| 1 \| 149 \| 2 \| 164 \| Phage tail protein I \| Phage tail protein I; afdb-uniprot50 \| AF-A0A0C9LZZ8-F1-MODEL\_V4 \| 1.0 \| 1.19e-11 \| 305 \| 0.266 \| 184 \| 110 \| 6 \| 5 \| 179 \| 1 \| 168 \| DNA, contig: SP610 \| DNA, contig: SP610; afdb-uniprot50 \| AF-A0A3P6KXK0-F1-MODEL\_V4 \| 1.0 \| 7.869e-11 \| 305 \| 0.234 \| 213 \| 141 \| 9 \| 1 \| 202 \| 9 \| 210 \| Phage tail protein (Tail\_P2\_I) \| Phage tail protein (Tail\_P2\_I); afdb-uniprot50 \| AF-A0A5A8F1G5-F1-MODEL\_V4 \| 1.0 \| 6.459e-09 \| 305 \| 0.301 \| 146 \| 94 \| 4 \| 11 \| 156 \| 22 \| 159 \| Phage tail protein I \| Phage tail protein I; afdb-uniprot50 \| AF-R5GZX5-F1-MODEL\_V4 \| 1.0 \| 5.393e-11 \| 304 \| 0.238 \| 172 \| 110 \| 6 \| 11 \| 176 \| 32 \| 188 \| Uncharacterized protein \| Uncharacterized protein; afdb-uniprot50 \| AF-A0A2V2GDM7-F1-MODEL\_V4 \| 1.0 \| 7.869e-11 \| 304 \| 0.198 \| 186 \| 128 \| 8 \| 5 \| 178 \| 18 \| 194 \| Phage tail protein I \| Phage tail protein I; afdb-uniprot50 \| AF-A0A3A9G247-F1-MODEL\_V4 \| 1.0 \| 2.379e-11 \| 304 \| 0.224 \| 249 \| 135 \| 10 \| 5 \| 205 \| 14 \| 252 \| Phage tail protein I \| Phage tail protein I; afdb-uniprot50 \| AF-A0A2W5JKU7-F1-MODEL\_V4 \| 1.0 \| 1.18e-09 \| 303 \| 0.304 \| 138 \| 88 \| 3 \| 48 \| 182 \| 2 \| 134 \| Phage tail protein I \| Phage tail protein I; afdb-uniprot50 \| AF-A0A7X1ZFY9-F1-MODEL\_V4 \| 1.0 \| 5.393e-11 \| 303 \| 0.342 \| 149 \| 90 \| 2 \| 55 \| 203 \| 28 \| 168 \| Phage tail protein I \| Phage tail protein I; afdb-uniprot50 \| AF-A0A167C5L9-F1-MODEL\_V4 \| 1.0 \| 2.079e-09 \| 302 \| 0.245 \| 163 \| 117 \| 5 \| 12 \| 173 \| 25 \| 182 \| Phage tail protein \| Phage tail protein; afdb-uniprot50 \| AF-A0A2Z6GC43-F1-MODEL\_V4 \| 1.0 \| 2.295e-10 \| 302 \| 0.294 \| 173 \| 101 \| 5 \| 3 \| 159 \| 2 \| 169 \| Tail protein \| Tail protein; afdb-uniprot50 \| AF-A0A4Q2KVQ7-F1-MODEL\_V4 \| 1.0 \| 9.851e-12 \| 302 \| 0.234 \| 179 \| 125 \| 5 \| 5 \| 175 \| 13 \| 187 \| Phage tail protein I \| Phage tail protein I; afdb-uniprot50 \| AF-A0A7G9GXJ3-F1-MODEL\_V4 \| 1.0 \| 2.233e-11 \| 302 \| 0.208 \| 211 \| 146 \| 12 \| 3 \| 205 \| 2 \| 199 \| Phage tail protein I \| Phage tail protein I; afdb-uniprot50 \| AF-A0A174UHU3-F1-MODEL\_V4 \| 1.0 \| 1.148e-10 \| 302 \| 0.211 \| 227 \| 141 \| 9 \| 5 \| 202 \| 15 \| 232 \| Bacteriophage P2-related tail formation protein \| Bacteriophage P2-related tail formation protein; afdb-uniprot50 \| AF-A0A825BCX2-F1-MODEL\_V4 \| 1.0 \| 2.079e-09 \| 301 \| 0.243 \| 144 \| 103 \| 3 \| 52 \| 193 \| 2 \| 141 \| Phage tail protein I \| Phage tail protein I; afdb-uniprot50 \| AF-A0A212J3U9-F1-MODEL\_V4 \| 1.0 \| 1.675e-10 \| 301 \| 0.267 \| 168 \| 104 \| 5 \| 1 \| 152 \| 2 \| 166 \| Phage tail protein I \| Phage tail protein I; afdb-uniprot50 \| AF-A0A1W2CVH4-F1-MODEL\_V4 \| 1.0 \| 2.952e-10 \| 301 \| 0.137 \| 232 \| 171 \| 7 \| 2 \| 205 \| 6 \| 236 \| Uncharacterized protein \| Uncharacterized protein; afdb-uniprot50 \| AF-A0A2T4JNB1-F1-MODEL\_V4 \| 1.0 \| 1.19e-11 \| 300 \| 0.32 \| 184 \| 105 \| 5 \| 1 \| 167 \| 2 \| 182 \| Phage tail protein I \| Phage tail protein I; afdb-uniprot50 \| AF-A0A414Q2J0-F1-MODEL\_V4 \| 1.0 \| 5.542e-10 \| 300 \| 0.21 \| 166 \| 110 \| 7 \| 5 \| 158 \| 15 \| 171 \| Phage tail protein I \| Phage tail protein I; afdb-uniprot50 \| AF-A0A1M4WE79-F1-MODEL\_V4 \| 1.0 \| 5.952e-12 \| 299 \| 0.296 \| 172 \| 102 \| 5 \| 1 \| 156 \| 1 \| 169 \| Phage tail protein, P2 protein I family \| Phage tail protein, P2 protein I family; afdb-uniprot50 \| AF-A0A4Z0R3W7-F1-MODEL\_V4 \| 1.0 \| 3.349e-10 \| 299 \| 0.215 \| 172 \| 123 \| 5 \| 12 \| 176 \| 38 \| 204 \| Phage tail protein I \| Phage tail protein I; afdb-uniprot50 \| AF-A0A1X7NDR7-F1-MODEL\_V4 \| 1.0 \| 5.393e-11 \| 299 \| 0.246 \| 219 \| 108 \| 8 \| 1 \| 165 \| 4 \| 219 \| Phage tail protein, P2 protein I family \| Phage tail protein, P2 protein I family; afdb-uniprot50 \| AF-A0A3C1WTS9-F1-MODEL\_V4 \| 1.0 \| 2.444e-10 \| 299 \| 0.235 \| 187 \| 130 \| 6 \| 11 \| 194 \| 28 \| 204 \| Phage tail protein I \| Phage tail protein I; afdb-uniprot50 \| AF-A0A853IIL6-F1-MODEL\_V4 \| 1.0 \| 2.581e-08 \| 298 \| 0.519 \| 104 \| 49 \| 1 \| 1 \| 104 \| 1 \| 103 \| Phage tail protein I \| Phage tail protein I; afdb-uniprot50 \| AF-A0A510KR89-F1-MODEL\_V4 \| 1.0 \| 4.588e-10 \| 298 \| 0.238 \| 180 \| 117 \| 6 \| 12 \| 184 \| 33 \| 199 \| Uncharacterized protein \| Uncharacterized protein; afdb-uniprot50 \| AF-C0QSD8-F1-MODEL\_V4 \| 1.0 \| 2.155e-10 \| 298 \| 0.153 \| 196 \| 159 \| 6 \| 12 \| 205 \| 29 \| 219 \| Uncharacterized protein \| Uncharacterized protein; afdb-uniprot50 \| AF-A0A266LJW6-F1-MODEL\_V4 \| 1.0 \| 5.347e-09 \| 297 \| 0.366 \| 112 \| 69 \| 2 \| 65 \| 175 \| 2 \| 112 \| Phage tail protein I \| Phage tail protein I; afdb-uniprot50 \| AF-A0A0N8IBW6-F1-MODEL\_V4 \| 1.0 \| 6.065e-09 \| 297 \| 0.225 \| 164 \| 108 \| 4 \| 4 \| 151 \| 7 \| 167 \| Uncharacterized protein \| Uncharacterized protein; afdb-uniprot50 \| AF-A0A2G6EZC5-F1-MODEL\_V4 \| 1.0 \| 1.518e-09 \| 297 \| 0.194 \| 190 \| 138 \| 6 \| 3 \| 182 \| 15 \| 199 \| Phage tail protein \| Phage tail protein; afdb-uniprot50 \| AF-A0A2A2IJZ7-F1-MODEL\_V4 \| 1.0 \| 5.064e-11 \| 297 \| 0.234 \| 205 \| 146 \| 7 \| 5 \| 202 \| 13 \| 213 \| Phage tail protein I \| Phage tail protein I; afdb-uniprot50 \| AF-A0A139D6J5-F1-MODEL\_V4 \| 1.0 \| 1.477e-10 \| 297 \| 0.207 \| 212 \| 149 \| 6 \| 5 \| 205 \| 14 \| 217 \| Uncharacterized protein \| Uncharacterized protein; afdb-uniprot50 \| AF-L8MFE9-F1-MODEL\_V4 \| 1.0 \| 5.743e-11 \| 296 \| 0.289 \| 166 \| 94 \| 5 \| 1 \| 147 \| 1 \| 161 \| Putative TAIL-RELATED PROTEIN (TAIL FORMATION-LIKE PROTEIN) \| Putative TAIL-RELATED PROTEIN (TAIL FORMATION-LIKE PROTEIN); afdb-uniprot50 \| AF-D1ALM0-F1-MODEL\_V4 \| 1.0 \| 4.755e-11 \| 296 \| 0.241 \| 215 \| 135 \| 8 \| 3 \| 202 \| 12 \| 213 \| Uncharacterized protein \| Uncharacterized protein; afdb-uniprot50 \| AF-A0A4Q3YY71-F1-MODEL\_V4 \| 1.0 \| 1.969e-11 \| 295 \| 0.252 \| 198 \| 120 \| 8 \| 2 \| 178 \| 6 \| 196 \| Phage tail protein I \| Phage tail protein I; afdb-uniprot50 \| AF-A0A4U7J706-F1-MODEL\_V4 \| 1.0 \| 2.024e-10 \| 294 \| 0.22 \| 186 \| 121 \| 11 \| 5 \| 177 \| 13 \| 187 \| Phage tail protein I \| Phage tail protein I; afdb-uniprot50 \| AF-A0A2N2S1K8-F1-MODEL\_V4 \| 1.0 \| 2.379e-11 \| 294 \| 0.309 \| 165 \| 94 \| 6 \| 2 \| 149 \| 4 \| 165 \| Phage tail protein I \| Phage tail protein I; afdb-uniprot50 \| AF-A0A447IP21-F1-MODEL\_V4 \| 1.0 \| 1.078e-10 \| 294 \| 0.241 \| 232 \| 112 \| 6 \| 1 \| 171 \| 1 \| 229 \| Phage tail protein (Tail\_P2\_I) \| Phage tail protein (Tail\_P2\_I); afdb-uniprot50 \| AF-A0A1Y4NK51-F1-MODEL\_V4 \| 1.0 \| 1.675e-10 \| 294 \| 0.208 \| 216 \| 148 \| 7 \| 3 \| 205 \| 26 \| 231 \| Phage tail protein I \| Phage tail protein I; afdb-uniprot50 \| AF-A0A6L3Y496-F1-MODEL\_V4 \| 1.0 \| 2.927e-08 \| 293 \| 0.33 \| 100 \| 64 \| 2 \| 78 \| 175 \| 1 \| 99 \| Phage tail protein I \| Phage tail protein I; afdb-uniprot50 \| AF-A0A064AK86-F1-MODEL\_V4 \| 1.0 \| 1.518e-09 \| 293 \| 0.177 \| 175 \| 133 \| 5 \| 35 \| 202 \| 5 \| 175 \| Tail protein \| Tail protein; afdb-uniprot50 \| AF-A0A2S0JKT1-F1-MODEL\_V4 \| 1.0 \| 4.886e-10 \| 293 \| 0.191 \| 225 \| 150 \| 9 \| 5 \| 202 \| 13 \| 232 \| Phage tail protein I \| Phage tail protein I; afdb-uniprot50 \| AF-A0A165PIY6-F1-MODEL\_V4 \| 1.0 \| 7.593e-10 \| 293 \| 0.199 \| 226 \| 159 \| 12 \| 1 \| 205 \| 1 \| 225 \| Phage tail protein \| Phage tail protein; afdb-uniprot50 \| AF-U2PI12-F1-MODEL\_V4 \| 1.0 \| 1.833e-09 \| 292 \| 0.196 \| 168 \| 115 \| 4 \| 12 \| 176 \| 37 \| 187 \| Phage tail protein \| Phage tail protein; afdb-uniprot50 \| AF-D9R641-F1-MODEL\_V4 \| 1.0 \| 1.425e-09 \| 290 \| 0.165 \| 175 \| 125 \| 7 \| 1 \| 163 \| 9 \| 174 \| Uncharacterized protein \| Uncharacterized protein; afdb-uniprot50 \| AF-A0A1T4QDN6-F1-MODEL\_V4 \| 1.0 \| 7.389e-11 \| 289 \| 0.217 \| 184 \| 127 \| 8 \| 4 \| 178 \| 9 \| 184 \| Phage tail protein, P2 protein I family \| Phage tail protein, P2 protein I family; afdb-uniprot50 \| AF-A0A2D3WQP3-F1-MODEL\_V4 \| 1.0 \| 7.129e-10 \| 289 \| 0.194 \| 190 \| 112 \| 5 \| 1 \| 152 \| 5 \| 191 \| Phage tail protein I \| Phage tail protein I; afdb-uniprot50 \| AF-A0A7T5EN83-F1-MODEL\_V4 \| 1.0 \| 1.148e-10 \| 288 \| 0.231 \| 186 \| 123 \| 7 \| 3 \| 177 \| 11 \| 187 \| Phage tail protein I \| Phage tail protein I; afdb-uniprot50 \| AF-A0A6C1BSQ9-F1-MODEL\_V4 \| 1.0 \| 1.387e-10 \| 288 \| 0.235 \| 217 \| 139 \| 8 \| 4 \| 205 \| 5 \| 209 \| Phage tail protein I \| Phage tail protein I; afdb-uniprot50 \| AF-A0A329TXY5-F1-MODEL\_V4 \| 1.0 \| 5.021e-09 \| 288 \| 0.217 \| 161 \| 114 \| 7 \| 5 \| 157 \| 13 \| 169 \| Phage tail protein I \| Phage tail protein I; afdb-uniprot50 \| AF-A0A316Q9A4-F1-MODEL\_V4 \| 1.0 \| 4.755e-11 \| 287 \| 0.235 \| 212 \| 142 \| 9 \| 5 \| 205 \| 12 \| 214 \| Phage tail protein I \| Phage tail protein I; afdb-uniprot50 \| AF-A0A7C5ZID1-F1-MODEL\_V4 \| 1.0 \| 3.665e-09 \| 287 \| 0.203 \| 157 \| 113 \| 7 \| 12 \| 164 \| 175 \| 323 \| Phage tail protein I \| Phage tail protein I; afdb-uniprot50 \| AF-A0A318EJ45-F1-MODEL\_V4 \| 1.0 \| 2.952e-10 \| 286 \| 0.191 \| 188 \| 134 \| 8 \| 3 \| 180 \| 11 \| 190 \| Tail protein P2 I \| Tail protein P2 I; afdb-uniprot50 \| AF-A0A2S6N2X4-F1-MODEL\_V4 \| 1.0 \| 6.938e-11 \| 286 \| 0.247 \| 210 \| 144 \| 10 \| 1 \| 205 \| 7 \| 207 \| Uncharacterized protein \| Uncharacterized protein; afdb-uniprot50 \| AF-A0A416B7R3-F1-MODEL\_V4 \| 1.0 \| 5.204e-10 \| 286 \| 0.215 \| 204 \| 147 \| 6 \| 5 \| 201 \| 13 \| 210 \| Uncharacterized protein \| Uncharacterized protein; afdb-uniprot50 \| AF-A0A7M1L784-F1-MODEL\_V4 \| 1.0 \| 2.512e-09 \| 285 \| 0.272 \| 154 \| 101 \| 6 \| 56 \| 203 \| 1 \| 149 \| Uncharacterized protein \| Uncharacterized protein; afdb-uniprot50 \| AF-A0A7U9RV01-F1-MODEL\_V4 \| 1.0 \| 1.784e-10 \| 285 \| 0.193 \| 207 \| 146 \| 9 \| 4 \| 199 \| 13 \| 209 \| Uncharacterized protein \| Uncharacterized protein; afdb-uniprot50 \| AF-A8SB35-F1-MODEL\_V4 \| 1.0 \| 1.338e-09 \| 284 \| 0.201 \| 208 \| 145 \| 9 \| 3 \| 197 \| 16 \| 215 \| Uncharacterized protein \| Uncharacterized protein; afdb-uniprot50 \| AF-A0A7X4ZQB7-F1-MODEL\_V4 \| 1.0 \| 4.886e-10 \| 284 \| 0.206 \| 218 \| 152 \| 7 \| 3 \| 202 \| 11 \| 225 \| Uncharacterized protein \| Uncharacterized protein; afdb-uniprot50 \| AF-A0A1M6LS85-F1-MODEL\_V4 \| 1.0 \| 1.148e-10 \| 284 \| 0.213 \| 229 \| 141 \| 9 \| 5 \| 203 \| 14 \| 233 \| Phage tail protein, P2 protein I family \| Phage tail protein, P2 protein I family; afdb-uniprot50 \| AF-A0A836NZX7-F1-MODEL\_V4 \| 1.0 \| 1.936e-07 \| 283 \| 0.465 \| 101 \| 54 \| 0 \| 1 \| 101 \| 2 \| 102 \| Tail protein \| Tail protein; afdb-uniprot50 \| AF-A0A1V4GEU5-F1-MODEL\_V4 \| 1.0 \| 4.045e-10 \| 283 \| 0.218 \| 183 \| 125 \| 8 \| 4 \| 175 \| 13 \| 188 \| Uncharacterized protein \| Uncharacterized protein; afdb-uniprot50 \| AF-A0A0Q2XP81-F1-MODEL\_V4 \| 1.0 \| 2.512e-09 \| 283 \| 0.196 \| 168 \| 113 \| 6 \| 4 \| 157 \| 13 \| 172 \| Uncharacterized protein \| Uncharacterized protein; afdb-uniprot50 \| AF-A0A373LAF9-F1-MODEL\_V4 \| 1.0 \| 9.767e-10 \| 283 \| 0.226 \| 212 \| 142 \| 8 \| 4 \| 202 \| 14 \| 216 \| Phage tail protein I \| Phage tail protein I; afdb-uniprot50 \| AF-A0A316MKT1-F1-MODEL\_V4 \| 1.0 \| 1.256e-09 \| 282 \| 0.207 \| 231 \| 152 \| 6 \| 4 \| 205 \| 13 \| 241 \| Phage tail protein I \| Phage tail protein I; afdb-uniprot50 \| AF-A0A3A6JUC8-F1-MODEL\_V4 \| 1.0 \| 1.108e-09 \| 282 \| 0.158 \| 196 \| 159 \| 3 \| 11 \| 205 \| 28 \| 218 \| Phage tail protein I \| Phage tail protein I; afdb-uniprot50 \| AF-A0A7X3ZHW1-F1-MODEL\_V4 \| 1.0 \| 3.903e-09 \| 281 \| 0.267 \| 161 \| 105 \| 5 \| 2 \| 155 \| 190 \| 344 \| Phage tail protein I \| Phage tail protein I; afdb-uniprot50 \| AF-A0A329MGA2-F1-MODEL\_V4 \| 1.0 \| 1.108e-09 \| 280 \| 0.215 \| 246 \| 144 \| 10 \| 3 \| 205 \| 40 \| 279 \| Uncharacterized protein \| Uncharacterized protein; afdb-uniprot50 \| AF-A0A1Y4BCC7-F1-MODEL\_V4 \| 1.0 \| 2.772e-10 \| 279 \| 0.239 \| 192 \| 116 \| 11 \| 4 \| 180 \| 11 \| 187 \| Phage tail protein I \| Phage tail protein I; afdb-uniprot50 \| AF-A0A377AJR9-F1-MODEL\_V4 \| 1.0 \| 2.137e-08 \| 278 \| 0.495 \| 103 \| 52 \| 0 \| 1 \| 103 \| 2 \| 104 \| Putative tail protein I (Gpi) \| Putative tail protein I (Gpi); afdb-uniprot50 \| AF-A0A3D2IPZ0-F1-MODEL\_V4 \| 1.0 \| 2.849e-09 \| 278 \| 0.199 \| 201 \| 141 \| 8 \| 5 \| 198 \| 18 \| 205 \| Uncharacterized protein \| Uncharacterized protein; afdb-uniprot50 \| AF-A0A7U5GZ11-F1-MODEL\_V4 \| 1.0 \| 1.223e-10 \| 278 \| 0.236 \| 211 \| 147 \| 7 \| 4 \| 205 \| 12 \| 217 \| Phage tail protein I \| Phage tail protein I; afdb-uniprot50 \| AF-A0A842J0C5-F1-MODEL\_V4 \| 1.0 \| 3.903e-09 \| 276 \| 0.245 \| 159 \| 106 \| 7 \| 12 \| 167 \| 28 \| 175 \| Uncharacterized protein \| Uncharacterized protein; afdb-uniprot50 \| AF-C3WBC4-F1-MODEL\_V4 \| 1.0 \| 7.593e-10 \| 276 \| 0.267 \| 157 \| 102 \| 5 \| 5 \| 153 \| 15 \| 166 \| Phage tail protein \| Phage tail protein; afdb-uniprot50 \| AF-A0A8B3RDD9-F1-MODEL\_V4 \| 1.0 \| 2.215e-09 \| 276 \| 0.223 \| 179 \| 108 \| 6 \| 4 \| 157 \| 6 \| 178 \| Phage tail protein I \| Phage tail protein I; afdb-uniprot50 \| AF-A0A3A9CSI0-F1-MODEL\_V4 \| 1.0 \| 6.065e-09 \| 276 \| 0.182 \| 175 \| 124 \| 6 \| 3 \| 167 \| 18 \| 183 \| Uncharacterized protein \| Uncharacterized protein; afdb-uniprot50 \| AF-B0PEZ8-F1-MODEL\_V4 \| 1.0 \| 9.767e-10 \| 276 \| 0.204 \| 205 \| 151 \| 6 \| 4 \| 201 \| 11 \| 210 \| Phage tail protein \| Phage tail protein; afdb-uniprot50 \| AF-A0A7U9MZ97-F1-MODEL\_V4 \| 1.0 \| 1.18e-09 \| 276 \| 0.213 \| 211 \| 150 \| 8 \| 4 \| 205 \| 11 \| 214 \| Uncharacterized protein \| Uncharacterized protein; afdb-uniprot50 \| AF-A0A348AJ17-F1-MODEL\_V4 \| 1.0 \| 7.869e-11 \| 276 \| 0.274 \| 226 \| 132 \| 10 \| 3 \| 205 \| 12 \| 228 \| Phage tail protein \| Phage tail protein; afdb-uniprot50 \| AF-A0A3S4GL93-F1-MODEL\_V4 \| 1.0 \| 2.424e-08 \| 275 \| 0.504 \| 101 \| 50 \| 0 \| 4 \| 104 \| 18 \| 118 \| Bacteriophage P2-related tail formation protein \| Bacteriophage P2-related tail formation protein; afdb-uniprot50 \| AF-A0A3A8WB53-F1-MODEL\_V4 \| 1.0 \| 7.593e-10 \| 275 \| 0.21 \| 219 \| 148 \| 10 \| 5 \| 205 \| 13 \| 224 \| Phage tail protein I \| Phage tail protein I; afdb-uniprot50 \| AF-A0A285NFM9-F1-MODEL\_V4 \| 1.0 \| 2.772e-10 \| 274 \| 0.227 \| 211 \| 146 \| 8 \| 6 \| 205 \| 16 \| 220 \| Phage tail protein (Tail\_P2\_I) \| Phage tail protein (Tail\_P2\_I); afdb-uniprot50 \| AF-A0A1Q6KUF2-F1-MODEL\_V4 \| 1.0 \| 1.338e-09 \| 274 \| 0.183 \| 223 \| 152 \| 11 \| 4 \| 205 \| 13 \| 226 \| Uncharacterized protein \| Uncharacterized protein; afdb-uniprot50 \| AF-A0A5S4VLN5-F1-MODEL\_V4 \| 1.0 \| 1.338e-09 \| 273 \| 0.216 \| 217 \| 151 \| 8 \| 3 \| 205 \| 11 \| 222 \| Uncharacterized protein \| Uncharacterized protein; afdb-uniprot50 \| AF-R5BTZ8-F1-MODEL\_V4 \| 1.0 \| 1.138e-08 \| 272 \| 0.245 \| 171 \| 117 \| 8 \| 3 \| 165 \| 11 \| 177 \| Uncharacterized protein \| Uncharacterized protein; afdb-uniprot50 \| AF-A0A078MHK2-F1-MODEL\_V4 \| 1.0 \| 2.079e-09 \| 272 \| 0.176 \| 232 \| 150 \| 8 \| 3 \| 198 \| 11 \| 237 \| Phage tail protein (Tail\_P2\_I) \| Phage tail protein (Tail\_P2\_I); afdb-uniprot50 \| AF-A0A3E2TEK9-F1-MODEL\_V4 \| 1.0 \| 2.952e-10 \| 272 \| 0.196 \| 204 \| 139 \| 6 \| 12 \| 205 \| 36 \| 224 \| Uncharacterized protein \| Uncharacterized protein; afdb-uniprot50 \| AF-A0A3D5LSM9-F1-MODEL\_V4 \| 1.0 \| 7.593e-10 \| 271 \| 0.206 \| 233 \| 140 \| 8 \| 3 \| 199 \| 11 \| 234 \| Uncharacterized protein \| Uncharacterized protein; afdb-uniprot50 \| AF-A0A416UB11-F1-MODEL\_V4 \| 1.0 \| 1.721e-09 \| 271 \| 0.153 \| 247 \| 157 \| 10 \| 1 \| 202 \| 9 \| 248 \| Phage tail protein \| Phage tail protein; afdb-uniprot50 \| AF-A0A7T7D8U4-F1-MODEL\_V4 \| 1.0 \| 2.849e-09 \| 271 \| 0.257 \| 214 \| 136 \| 10 \| 3 \| 200 \| 390 \| 596 \| Phage tail protein I \| Phage tail protein I; afdb-uniprot50 \| AF-A0A346QYG0-F1-MODEL\_V4 \| 1.0 \| 4.845e-08 \| 270 \| 0.45 \| 100 \| 54 \| 1 \| 2 \| 100 \| 19 \| 118 \| Phage tail protein I \| Phage tail protein I; afdb-uniprot50 \| AF-A0A6L9HFG1-F1-MODEL\_V4 \| 1.0 \| 8.612e-10 \| 269 \| 0.175 \| 234 \| 153 \| 10 \| 3 \| 205 \| 21 \| 245 \| Phage tail protein I \| Phage tail protein I; afdb-uniprot50 \| AF-A0A7V8HY86-F1-MODEL\_V4 \| 1.0 \| 3.231e-09 \| 268 \| 0.434 \| 115 \| 65 \| 0 \| 1 \| 115 \| 5 \| 119 \| Phage tail protein I \| Phage tail protein I; afdb-uniprot50 \| AF-D4K5G8-F1-MODEL\_V4 \| 1.0 \| 5.204e-10 \| 268 \| 0.22 \| 181 \| 122 \| 8 \| 3 \| 171 \| 11 \| 184 \| Phage tail protein (Tail\_P2\_I) \| Phage tail protein (Tail\_P2\_I); afdb-uniprot50 \| AF-A0A1G5JKM4-F1-MODEL\_V4 \| 1.0 \| 3.441e-09 \| 266 \| 0.164 \| 164 \| 127 \| 6 \| 12 \| 173 \| 30 \| 185 \| Phage tail protein (Tail\_P2\_I) \| Phage tail protein (Tail\_P2\_I); afdb-uniprot50 \| AF-A0A1H8VRL6-F1-MODEL\_V4 \| 1.0 \| 1.04e-09 \| 266 \| 0.18 \| 238 \| 149 \| 11 \| 3 \| 205 \| 11 \| 237 \| Phage tail protein, P2 protein I family \| Phage tail protein, P2 protein I family; afdb-uniprot50 \| AF-A0A7X4Z3Q6-F1-MODEL\_V4 \| 1.0 \| 7.326e-09 \| 266 \| 0.224 \| 187 \| 118 \| 10 \| 5 \| 169 \| 18 \| 199 \| Uncharacterized protein \| Uncharacterized protein; afdb-uniprot50 \| AF-A0A812RFY7-F1-MODEL\_V4 \| 1.0 \| 1.477e-10 \| 266 \| 0.317 \| 164 \| 97 \| 4 \| 1 \| 150 \| 266 \| 428 \| J protein \| J protein; afdb-uniprot50 \| AF-A0A8B4XHD1-F1-MODEL\_V4 \| 1.0 \| 6.404e-07 \| 265 \| 0.44 \| 93 \| 52 \| 0 \| 1 \| 93 \| 1 \| 93 \| Phage tail P2-like protein \| Phage tail P2-like protein; afdb-uniprot50 \| AF-A0A6N3FBW1-F1-MODEL\_V4 \| 1.0 \| 3.441e-09 \| 265 \| 0.197 \| 192 \| 128 \| 10 \| 4 \| 184 \| 14 \| 190 \| Phage tail protein (Tail\_P2\_I) \| Phage tail protein (Tail\_P2\_I); afdb-uniprot50 \| AF-A0A6N9P2M1-F1-MODEL\_V4 \| 1.0 \| 2.358e-09 \| 264 \| 0.201 \| 238 \| 151 \| 9 \| 1 \| 205 \| 18 \| 249 \| Uncharacterized protein \| Uncharacterized protein; afdb-uniprot50 \| AF-A0A1C6IIK7-F1-MODEL\_V4 \| 1.0 \| 2.952e-10 \| 264 \| 0.191 \| 225 \| 150 \| 10 \| 5 \| 205 \| 14 \| 230 \| Bacteriophage P2-related tail formation protein \| Bacteriophage P2-related tail formation protein; afdb-uniprot50 \| AF-A0A2E4F7R5-F1-MODEL\_V4 \| 1.0 \| 1.518e-09 \| 264 \| 0.152 \| 249 \| 163 \| 10 \| 3 \| 205 \| 79 \| 325 \| PASTA domain-containing protein \| PASTA domain-containing protein; afdb-uniprot50 \| AF-A0A7S7QIG1-F1-MODEL\_V4 \| 1.0 \| 6.232e-08 \| 264 \| 0.376 \| 101 \| 62 \| 1 \| 1 \| 100 \| 1 \| 101 \| Phage tail protein I \| Phage tail protein I; afdb-uniprot50 \| AF-A0A1Y3S126-F1-MODEL\_V4 \| 1.0 \| 4.588e-10 \| 263 \| 0.161 \| 254 \| 152 \| 10 \| 3 \| 205 \| 17 \| 260 \| Phage tail protein I \| Phage tail protein I; afdb-uniprot50 \| AF-A0A238JEV2-F1-MODEL\_V4 \| 1.0 \| 6.637e-08 \| 262 \| 0.242 \| 128 \| 92 \| 4 \| 1 \| 124 \| 7 \| 133 \| Phage tail protein (Tail\_P2\_I) \| Phage tail protein (Tail\_P2\_I); afdb-uniprot50 \| AF-A0A3A8UCJ4-F1-MODEL\_V4 \| 1.0 \| 3.231e-09 \| 261 \| 0.221 \| 212 \| 144 \| 9 \| 5 \| 200 \| 13 \| 219 \| Uncharacterized protein \| Uncharacterized protein; afdb-uniprot50 \| AF-A0A2N8HQL6-F1-MODEL\_V4 \| 1.0 \| 5.695e-09 \| 260 \| 0.178 \| 213 \| 156 \| 7 \| 4 \| 205 \| 11 \| 215 \| Phage tail protein I \| Phage tail protein I; afdb-uniprot50 \| AF-A0A7C8LUT2-F1-MODEL\_V4 \| 1.0 \| 3.231e-09 \| 259 \| 0.155 \| 199 \| 163 \| 4 \| 4 \| 200 \| 19 \| 214 \| Uncharacterized protein \| Uncharacterized protein; afdb-uniprot50 \| AF-A0A238WY20-F1-MODEL\_V4 \| 1.0 \| 4.427e-09 \| 259 \| 0.155 \| 244 \| 162 \| 7 \| 2 \| 205 \| 659 \| 898 \| Phage tail protein domain-containing protein \| Phage tail protein domain-containing protein; afdb-uniprot50 \| AF-A0A4P9VKB1-F1-MODEL\_V4 \| 1.0 \| 1.327e-07 \| 258 \| 0.408 \| 93 \| 55 \| 0 \| 1 \| 93 \| 2 \| 94 \| Phage tail protein I \| Phage tail protein I; afdb-uniprot50 \| AF-A0A1W9UFI2-F1-MODEL\_V4 \| 1.0 \| 1.108e-09 \| 258 \| 0.195 \| 230 \| 143 \| 10 \| 12 \| 205 \| 28 \| 251 \| Uncharacterized protein \| Uncharacterized protein; afdb-uniprot50 \| AF-A0A1H0BY95-F1-MODEL\_V4 \| 1.0 \| 1.464e-08 \| 258 \| 0.181 \| 243 \| 145 \| 7 \| 1 \| 197 \| 1 \| 235 \| Phage tail protein (Tail\_P2\_I) \| Phage tail protein (Tail\_P2\_I); afdb-uniprot50 \| AF-A0A1S1TLC3-F1-MODEL\_V4 \| 1.0 \| 1.603e-07 \| 258 \| 0.333 \| 102 \| 66 \| 2 \| 1 \| 100 \| 4 \| 105 \| Phage tail protein I \| Phage tail protein I; afdb-uniprot50 \| AF-A8GLQ8-F1-MODEL\_V4 \| 1.0 \| 8.612e-10 \| 255 \| 0.28 \| 171 \| 103 \| 5 \| 16 \| 182 \| 4 \| 158 \| Phage tail protein I \| Phage tail protein I; afdb-uniprot50 \| AF-A0A833LKH9-F1-MODEL\_V4 \| 1.0 \| 2.652e-07 \| 254 \| 0.32 \| 100 \| 67 \| 1 \| 1 \| 99 \| 2 \| 101 \| Phage tail protein I \| Phage tail protein I; afdb-uniprot50 \| AF-A0A3L7AJI5-F1-MODEL\_V4 \| 1.0 \| 4.389e-07 \| 254 \| 0.303 \| 99 \| 67 \| 2 \| 4 \| 100 \| 6 \| 104 \| Phage tail protein I \| Phage tail protein I; afdb-uniprot50 \| AF-A0A826R5C0-F1-MODEL\_V4 \| 1.0 \| 1.098e-07 \| 252 \| 0.417 \| 103 \| 58 \| 1 \| 1 \| 101 \| 1 \| 103 \| Phage tail protein I \| Phage tail protein I; afdb-uniprot50 \| AF-A0A7X2NLM7-F1-MODEL\_V4 \| 1.0 \| 1.769e-08 \| 252 \| 0.153 \| 248 \| 161 \| 12 \| 1 \| 202 \| 9 \| 253 \| Phage tail protein \| Phage tail protein; afdb-uniprot50 \| AF-A0A4S5JBM7-F1-MODEL\_V4 \| 1.0 \| 1.413e-07 \| 252 \| 0.346 \| 101 \| 65 \| 1 \| 1 \| 100 \| 3 \| 103 \| Phage tail protein I \| Phage tail protein I; afdb-uniprot50 \| AF-A0A417IXT6-F1-MODEL\_V4 \| 1.0 \| 3.766e-08 \| 251 \| 0.202 \| 158 \| 110 \| 4 \| 4 \| 152 \| 13 \| 163 \| Uncharacterized protein \| Uncharacterized protein; afdb-uniprot50 \| AF-A0A2G6EHZ3-F1-MODEL\_V4 \| 1.0 \| 2.952e-10 \| 251 \| 0.228 \| 171 \| 109 \| 4 \| 4 \| 155 \| 28 \| 194 \| Phage tail protein I \| Phage tail protein I; afdb-uniprot50 \| AF-A0A6N2Z557-F1-MODEL\_V4 \| 1.0 \| 6.694e-10 \| 251 \| 0.251 \| 207 \| 137 \| 9 \| 3 \| 200 \| 11 \| 208 \| Phage tail protein (Tail\_P2\_I) \| Phage tail protein (Tail\_P2\_I); afdb-uniprot50 \| AF-A7MW91-F1-MODEL\_V4 \| 1.0 \| 8.539e-08 \| 250 \| 0.193 \| 160 \| 120 \| 7 \| 53 \| 205 \| 2 \| 159 \| Uncharacterized protein \| Uncharacterized protein; afdb-uniprot50 \| AF-A0A6D0ZU35-F1-MODEL\_V4 \| 1.0 \| 3.118e-08 \| 249 \| 0.253 \| 146 \| 95 \| 5 \| 66 \| 201 \| 3 \| 144 \| Phage tail protein I \| Phage tail protein I; afdb-uniprot50 \| AF-A0A2P7V3T2-F1-MODEL\_V4 \| 1.0 \| 3.903e-09 \| 249 \| 0.243 \| 218 \| 129 \| 13 \| 3 \| 199 \| 11 \| 213 \| Phage tail protein I \| Phage tail protein I; afdb-uniprot50 \| AF-A0A7G9WG79-F1-MODEL\_V4 \| 1.0 \| 1.56e-08 \| 249 \| 0.183 \| 202 \| 151 \| 6 \| 4 \| 196 \| 14 \| 210 \| Phage tail protein I \| Phage tail protein I; afdb-uniprot50 \| AF-A0A367FXP5-F1-MODEL\_V4 \| 1.0 \| 1.212e-08 \| 249 \| 0.188 \| 260 \| 149 \| 9 \| 3 \| 205 \| 11 \| 265 \| Uncharacterized protein \| Uncharacterized protein; afdb-uniprot50 \| AF-I0GRI5-F1-MODEL\_V4 \| 1.0 \| 2.603e-10 \| 248 \| 0.22 \| 213 \| 145 \| 7 \| 3 \| 203 \| 11 \| 214 \| Uncharacterized protein \| Uncharacterized protein; afdb-uniprot50 \| AF-A0A1Y4BD08-F1-MODEL\_V4 \| 1.0 \| 3.665e-09 \| 248 \| 0.2 \| 229 \| 145 \| 10 \| 4 \| 203 \| 14 \| 233 \| Phage tail protein I \| Phage tail protein I; afdb-uniprot50 \| AF-A0A2U9T741-F1-MODEL\_V4 \| 1.0 \| 2.079e-09 \| 247 \| 0.19 \| 210 \| 136 \| 11 \| 16 \| 205 \| 61 \| 256 \| Phage tail protein \| Phage tail protein; afdb-uniprot50 \| AF-A0A4Z1QZ30-F1-MODEL\_V4 \| 1.0 \| 7.736e-07 \| 246 \| 0.306 \| 101 \| 68 \| 2 \| 1 \| 99 \| 17 \| 117 \| Phage tail protein I \| Phage tail protein I; afdb-uniprot50 \| AF-A0A5C8BFD6-F1-MODEL\_V4 \| 1.0 \| 1.769e-08 \| 244 \| 0.184 \| 211 \| 155 \| 7 \| 4 \| 202 \| 12 \| 217 \| Uncharacterized protein \| Uncharacterized protein; afdb-uniprot50 \| AF-R9KX12-F1-MODEL\_V4 \| 1.0 \| 3.903e-09 \| 244 \| 0.158 \| 221 \| 144 \| 9 \| 11 \| 204 \| 34 \| 239 \| Phage tail protein I \| Phage tail protein I; afdb-uniprot50 \| AF-A0A1C6DZY7-F1-MODEL\_V4 \| 1.0 \| 7.528e-08 \| 241 \| 0.191 \| 178 \| 124 \| 7 \| 5 \| 168 \| 14 \| 185 \| Bacteriophage P2-related tail formation protein \| Bacteriophage P2-related tail formation protein; afdb-uniprot50 \| AF-A0A1G6EJP1-F1-MODEL\_V4 \| 1.0 \| 1.505e-07 \| 241 \| 0.366 \| 112 \| 65 \| 2 \| 1 \| 111 \| 3 \| 109 \| Phage tail protein, P2 protein I family \| Phage tail protein, P2 protein I family; afdb-uniprot50 \| AF-Q72D30-F1-MODEL\_V4 \| 1.0 \| 3.118e-08 \| 238 \| 0.15 \| 206 \| 163 \| 7 \| 5 \| 205 \| 19 \| 217 \| Uncharacterized protein \| Uncharacterized protein; afdb-uniprot50 \| AF-A0A8B2NSS2-F1-MODEL\_V4 \| 1.0 \| 6.404e-07 \| 237 \| 0.373 \| 99 \| 61 \| 1 \| 3 \| 100 \| 2 \| 100 \| Phage tail protein I \| Phage tail protein I; afdb-uniprot50 \| AF-A0A7C3GTL1-F1-MODEL\_V4 \| 1.0 \| 1.661e-08 \| 237 \| 0.15 \| 233 \| 153 \| 9 \| 12 \| 205 \| 471 \| 697 \| Uncharacterized protein \| Uncharacterized protein; afdb-uniprot50 \| AF-G4KPI3-F1-MODEL\_V4 \| 1.0 \| 3.903e-09 \| 236 \| 0.206 \| 184 \| 134 \| 8 \| 3 \| 179 \| 16 \| 194 \| Uncharacterized protein \| Uncharacterized protein; afdb-uniprot50 \| AF-A0A2P1S8D0-F1-MODEL\_V4 \| 1.0 \| 6.694e-10 \| 236 \| 0.227 \| 211 \| 139 \| 9 \| 3 \| 198 \| 11 \| 212 \| Phage tail protein I \| Phage tail protein I; afdb-uniprot50 \| AF-A0A1Q6PXA8-F1-MODEL\_V4 \| 1.0 \| 2.358e-09 \| 236 \| 0.184 \| 211 \| 155 \| 8 \| 5 \| 202 \| 13 \| 219 \| Phage tail protein I \| Phage tail protein I; afdb-uniprot50 \| AF-A0A0N0N4N5-F1-MODEL\_V4 \| 1.0 \| 1.661e-08 \| 236 \| 0.175 \| 200 \| 152 \| 7 \| 15 \| 205 \| 34 \| 229 \| Phage tail protein \| Phage tail protein; afdb-uniprot50 \| AF-R9LUW9-F1-MODEL\_V4 \| 1.0 \| 1.291e-08 \| 236 \| 0.202 \| 212 \| 141 \| 9 \| 12 \| 205 \| 29 \| 230 \| Uncharacterized protein \| Uncharacterized protein; afdb-uniprot50 \| AF-A0A5C8S6W3-F1-MODEL\_V4 \| 1.0 \| 4.121e-07 \| 235 \| 0.362 \| 102 \| 63 \| 2 \| 1 \| 100 \| 8 \| 109 \| Phage tail protein I \| Phage tail protein I; afdb-uniprot50 \| AF-A0A2K8MAJ7-F1-MODEL\_V4 \| 1.0 \| 9.685e-08 \| 234 \| 0.154 \| 240 \| 165 \| 8 \| 2 \| 205 \| 658 \| 895 \| Uncharacterized protein \| Uncharacterized protein; afdb-uniprot50 \| AF-R6TSD6-F1-MODEL\_V4 \| 1.0 \| 2.581e-08 \| 233 \| 0.181 \| 231 \| 150 \| 11 \| 4 \| 205 \| 14 \| 234 \| Uncharacterized protein \| Uncharacterized protein; afdb-uniprot50 \| AF-H1CIQ0-F1-MODEL\_V4 \| 1.0 \| 9.424e-09 \| 232 \| 0.179 \| 229 \| 149 \| 9 \| 3 \| 200 \| 12 \| 232 \| Phage tail protein I \| Phage tail protein I; afdb-uniprot50 \| AF-A0A2T4JG36-F1-MODEL\_V4 \| 1.0 \| 1.989e-06 \| 232 \| 0.43 \| 93 \| 53 \| 0 \| 1 \| 93 \| 3 \| 95 \| Phage tail protein I \| Phage tail protein I; afdb-uniprot50 \| AF-A0A6F9X0W8-F1-MODEL\_V4 \| 1.0 \| 3.118e-08 \| 226 \| 0.177 \| 192 \| 143 \| 9 \| 5 \| 190 \| 19 \| 201 \| Uncharacterized protein \| Uncharacterized protein; afdb-uniprot50 \| AF-A0A502CNF8-F1-MODEL\_V4 \| 1.0 \| 1.546e-06 \| 226 \| 0.353 \| 99 \| 62 \| 2 \| 4 \| 100 \| 3 \| 101 \| Phage tail protein I \| Phage tail protein I; afdb-uniprot50 \| AF-A0A497FKG3-F1-MODEL\_V4 \| 1.0 \| 3.87e-07 \| 221 \| 0.193 \| 150 \| 108 \| 4 \| 44 \| 182 \| 3 \| 150 \| Phage tail protein I \| Phage tail protein I; afdb-uniprot50 \| AF-Q7NKR6-F1-MODEL\_V4 \| 1.0 \| 4.011e-08 \| 221 \| 0.157 \| 228 \| 147 \| 11 \| 12 \| 205 \| 183 \| 399 \| Gll1411 protein \| Gll1411 protein; afdb-uniprot50 \| AF-A0A7J9Y123-F1-MODEL\_V4 \| 1.0 \| 1.17e-07 \| 220 \| 0.137 \| 218 \| 152 \| 8 \| 11 \| 203 \| 49 \| 255 \| Phage tail protein \| Phage tail protein; afdb-uniprot50 \| AF-A0A854ZEC9-F1-MODEL\_V4 \| 1.0 \| 1.818e-07 \| 220 \| 0.172 \| 185 \| 138 \| 8 \| 12 \| 189 \| 493 \| 669 \| Uncharacterized protein \| Uncharacterized protein; afdb-uniprot50 \| AF-A0A1D2QMU0-F1-MODEL\_V4 \| 1.0 \| 1.246e-07 \| 219 \| 0.284 \| 151 \| 85 \| 6 \| 4 \| 136 \| 6 \| 151 \| Phage tail protein I \| Phage tail protein I; afdb-uniprot50 \| AF-A0A4U8S340-F1-MODEL\_V4 \| 1.0 \| 4.549e-08 \| 218 \| 0.175 \| 205 \| 151 \| 8 \| 4 \| 205 \| 3 \| 192 \| Uncharacterized protein \| Uncharacterized protein; afdb-uniprot50 \| AF-A0A6G5QFK7-F1-MODEL\_V4 \| 1.0 \| 4.157e-09 \| 218 \| 0.192 \| 208 \| 155 \| 7 \| 1 \| 205 \| 1 \| 198 \| Phage P2 family tail protein \| Phage P2 family tail protein; afdb-uniprot50 \| AF-A0A5W6Y441-F1-MODEL\_V4 \| 1.0 \| 2.825e-07 \| 215 \| 0.277 \| 126 \| 86 \| 3 \| 80 \| 203 \| 1 \| 123 \| Phage tail protein I \| Phage tail protein I; afdb-uniprot50 \| AF-S5MV89-F1-MODEL\_V4 \| 1.0 \| 7.528e-08 \| 215 \| 0.319 \| 122 \| 81 \| 1 \| 55 \| 176 \| 17 \| 136 \| Phage tail fiber \| Phage tail fiber; afdb-uniprot50 \| AF-A0A4D7B3A9-F1-MODEL\_V4 \| 1.0 \| 5.852e-08 \| 215 \| 0.371 \| 132 \| 69 \| 3 \| 4 \| 134 \| 11 \| 129 \| Phage tail protein I \| Phage tail protein I; afdb-uniprot50 \| AF-A0A6P0Y575-F1-MODEL\_V4 \| 1.0 \| 2.581e-08 \| 214 \| 0.152 \| 197 \| 137 \| 9 \| 1 \| 184 \| 52 \| 231 \| Phage tail protein \| Phage tail protein; afdb-uniprot50 \| AF-A0A6N8BB33-F1-MODEL\_V4 \| 1.0 \| 4.675e-07 \| 214 \| 0.127 \| 212 \| 155 \| 4 \| 12 \| 194 \| 29 \| 239 \| Uncharacterized protein \| Uncharacterized protein; afdb-uniprot50 \| AF-A0A4Q6CU04-F1-MODEL\_V4 \| 1.0 \| 3.008e-07 \| 213 \| 0.285 \| 133 \| 94 \| 1 \| 1 \| 133 \| 6 \| 137 \| Phage tail protein I \| Phage tail protein I; afdb-uniprot50 \| AF-A0A1I1V086-F1-MODEL\_V4 \| 1.0 \| 4.804e-06 \| 213 \| 0.297 \| 101 \| 69 \| 2 \| 2 \| 100 \| 4 \| 104 \| Phage tail protein, P2 protein I family \| Phage tail protein, P2 protein I family; afdb-uniprot50 \| AF-A0A2W5WSP6-F1-MODEL\_V4 \| 1.0 \| 5.116e-06 \| 211 \| 0.326 \| 104 \| 66 \| 3 \| 1 \| 100 \| 1 \| 104 \| Phage tail protein I \| Phage tail protein I; afdb-uniprot50 \| AF-A0A4R7I2Z4-F1-MODEL\_V4 \| 1.0 \| 1.413e-07 \| 207 \| 0.173 \| 219 \| 157 \| 10 \| 5 \| 205 \| 20 \| 232 \| Phage tail-like protein \| Phage tail-like protein; afdb-uniprot50 \| AF-A0A1X3J0Q7-F1-MODEL\_V4 \| 1.0 \| 1.754e-06 \| 206 \| 0.454 \| 88 \| 46 \| 1 \| 88 \| 175 \| 4 \| 89 \| Tail protein I (GpI) \| Tail protein I (GpI); afdb-uniprot50 \| AF-A0A1T2XBJ5-F1-MODEL\_V4 \| 1.0 \| 6.014e-07 \| 206 \| 0.217 \| 147 \| 111 \| 3 \| 54 \| 199 \| 3 \| 146 \| Phage P2 family tail protein \| Phage P2 family tail protein; afdb-uniprot50 \| AF-A0A827B048-F1-MODEL\_V4 \| 1.0 \| 1.28e-06 \| 205 \| 0.454 \| 88 \| 46 \| 1 \| 88 \| 175 \| 2 \| 87 \| Phage tail protein I \| Phage tail protein I; afdb-uniprot50 \| AF-U5QMV0-F1-MODEL\_V4 \| 1.0 \| 1.413e-07 \| 205 \| 0.156 \| 230 \| 151 \| 9 \| 12 \| 205 \| 180 \| 402 \| Phage tail protein \| Phage tail protein; afdb-uniprot50 \| AF-A0A1W1Z3V2-F1-MODEL\_V4 \| 1.0 \| 4.235e-06 \| 203 \| 0.318 \| 110 \| 73 \| 2 \| 4 \| 111 \| 8 \| 117 \| Phage tail protein, P2 protein I family \| Phage tail protein, P2 protein I family; afdb-uniprot50 \| AF-A0A1M4V0G4-F1-MODEL\_V4 \| 1.0 \| 1.603e-07 \| 201 \| 0.168 \| 214 \| 153 \| 10 \| 3 \| 205 \| 7 \| 206 \| Uncharacterized protein \| Uncharacterized protein; afdb-uniprot50 \| AF-A0A3A9FB62-F1-MODEL\_V4 \| 1.0 \| 1.868e-06 \| 199 \| 0.187 \| 165 \| 107 \| 9 \| 4 \| 149 \| 12 \| 168 \| Uncharacterized protein \| Uncharacterized protein; afdb-uniprot50 \| AF-A0A1C5HA19-F1-MODEL\_V4 \| 1.0 \| 6.232e-08 \| 198 \| 0.15 \| 233 \| 156 \| 11 \| 4 \| 205 \| 9 \| 230 \| Phage tail protein domain-containing protein \| Phage tail protein domain-containing protein; afdb-uniprot50 \| AF-A0A3L7JDQ4-F1-MODEL\_V4 \| 1.0 \| 7.009e-06 \| 198 \| 0.298 \| 104 \| 68 \| 2 \| 1 \| 99 \| 2 \| 105 \| Phage tail protein I \| Phage tail protein I; afdb-uniprot50 \| AF-A0A239INL8-F1-MODEL\_V4 \| 1.0 \| 9.685e-08 \| 196 \| 0.143 \| 202 \| 156 \| 6 \| 12 \| 202 \| 37 \| 232 \| Phage tail protein domain-containing protein \| Phage tail protein domain-containing protein; afdb-uniprot50 \| AF-A0A0E4G100-F1-MODEL\_V4 \| 1.0 \| 3.634e-07 \| 194 \| 0.264 \| 136 \| 82 \| 4 \| 4 \| 137 \| 9 \| 128 \| Uncharacterized protein \| Uncharacterized protein; afdb-uniprot50 \| AF-A0A0Q8B1P1-F1-MODEL\_V4 \| 1.0 \| 1.28e-06 \| 194 \| 0.294 \| 129 \| 77 \| 3 \| 4 \| 131 \| 15 \| 130 \| Uncharacterized protein \| Uncharacterized protein; afdb-uniprot50 \| AF-A0A2G2MJ30-F1-MODEL\_V4 \| 1.0 \| 8.239e-07 \| 192 \| 0.123 \| 227 \| 168 \| 12 \| 4 \| 205 \| 12 \| 232 \| Uncharacterized protein \| Uncharacterized protein; afdb-uniprot50 \| AF-A0A5F1S592-F1-MODEL\_V4 \| 1.0 \| 1.17e-07 \| 191 \| 0.251 \| 175 \| 106 \| 7 \| 2 \| 168 \| 6 \| 163 \| Phage tail protein I \| Phage tail protein I; afdb-uniprot50 \| AF-A0A1H4FZQ0-F1-MODEL\_V4 \| 1.0 \| 9.344e-07 \| 190 \| 0.103 \| 261 \| 156 \| 8 \| 14 \| 198 \| 466 \| 724 \| Phage tail protein domain-containing protein \| Phage tail protein domain-containing protein; afdb-uniprot50 \| AF-A0A439EP98-F1-MODEL\_V4 \| 1.0 \| 1.129e-06 \| 189 \| 0.265 \| 113 \| 73 \| 4 \| 75 \| 183 \| 2 \| 108 \| Phage tail protein I \| Phage tail protein I; afdb-uniprot50 \| AF-A0A4Q8MC19-F1-MODEL\_V4 \| 1.0 \| 0.000144 \| 189 \| 0.255 \| 90 \| 65 \| 2 \| 13 \| 100 \| 2 \| 91 \| Phage tail protein I \| Phage tail protein I; afdb-uniprot50 \| AF-A0A5M6I8Y5-F1-MODEL\_V4 \| 1.0 \| 2.903e-06 \| 187 \| 0.365 \| 115 \| 70 \| 1 \| 89 \| 203 \| 44 \| 155 \| Phage tail protein I \| Phage tail protein I; afdb-uniprot50 \| AF-A0A221SYK6-F1-MODEL\_V4 \| 1.0 \| 1.28e-06 \| 187 \| 0.157 \| 216 \| 154 \| 10 \| 5 \| 201 \| 150 \| 356 \| Phage tail protein \| Phage tail protein; afdb-uniprot50 \| AF-A0A1A6FKQ9-F1-MODEL\_V4 \| 1.0 \| 9.602e-06 \| 187 \| 0.298 \| 104 \| 67 \| 3 \| 1 \| 100 \| 1 \| 102 \| Phage tail protein I \| Phage tail protein I; afdb-uniprot50 \| AF-A0A0F4R541-F1-MODEL\_V4 \| 1.0 \| 3.634e-07 \| 186 \| 0.134 \| 215 \| 157 \| 11 \| 5 \| 202 \| 7 \| 209 \| Uncharacterized protein \| Uncharacterized protein; afdb-uniprot50 \| AF-A0A4P6YWR5-F1-MODEL\_V4 \| 1.0 \| 1.06e-06 \| 186 \| 0.116 \| 214 \| 170 \| 9 \| 1 \| 200 \| 9 \| 217 \| Uncharacterized protein \| Uncharacterized protein; afdb-uniprot50 \| AF-W1HTC4-F1-MODEL\_V4 \| 1.0 \| 4.51e-06 \| 184 \| 0.363 \| 99 \| 59 \| 2 \| 93 \| 189 \| 2 \| 98 \| Tail protein I \| Tail protein I; afdb-uniprot50 \| AF-A0A542PWC5-F1-MODEL\_V4 \| 1.0 \| 5.802e-06 \| 180 \| 0.113 \| 283 \| 158 \| 9 \| 5 \| 200 \| 18 \| 294 \| Uncharacterized protein \| Uncharacterized protein; afdb-uniprot50 \| AF-A0A5Z1E0Y7-F1-MODEL\_V4 \| 1.0 \| 1.316e-05 \| 179 \| 0.223 \| 121 \| 88 \| 3 \| 84 \| 202 \| 1 \| 117 \| Phage tail protein I \| Phage tail protein I; afdb-uniprot50 \| AF-A0A0F5VGY1-F1-MODEL\_V4 \| 1.0 \| 1.202e-06 \| 179 \| 0.164 \| 195 \| 144 \| 7 \| 12 \| 201 \| 27 \| 207 \| Phage protein \| Phage protein; afdb-uniprot50 \| AF-A0A021X9K5-F1-MODEL\_V4 \| 1.0 \| 1.754e-06 \| 176 \| 0.255 \| 141 \| 81 \| 5 \| 1 \| 134 \| 2 \| 125 \| Phage tail protein I \| Phage tail protein I; afdb-uniprot50 \| AF-A0A7T7VP87-F1-MODEL\_V4 \| 1.0 \| 4.549e-08 \| 175 \| 0.244 \| 213 \| 121 \| 7 \| 3 \| 200 \| 10 \| 197 \| Phage tail protein I \| Phage tail protein I; afdb-uniprot50 \| AF-A0A2U2DFB4-F1-MODEL\_V4 \| 1.0 \| 5.116e-06 \| 173 \| 0.276 \| 134 \| 80 \| 5 \| 1 \| 130 \| 5 \| 125 \| Phage tail protein I \| Phage tail protein I; afdb-uniprot50 \| AF-A0A3M0SWH5-F1-MODEL\_V4 \| 1.0 \| 9.344e-07 \| 170 \| 0.157 \| 203 \| 143 \| 13 \| 12 \| 205 \| 24 \| 207 \| DUF2313 domain-containing protein \| DUF2313 domain-containing protein; afdb-uniprot50 \| AF-A0A849PJG0-F1-MODEL\_V4 \| 1.0 \| 2.403e-06 \| 170 \| 0.14 \| 264 \| 151 \| 10 \| 5 \| 197 \| 140 \| 398 \| Uncharacterized protein \| Uncharacterized protein; afdb-uniprot50 \| AF-A0A2W6SVJ0-F1-MODEL\_V4 \| 1.0 \| 1.692e-05 \| 170 \| 0.251 \| 139 \| 86 \| 5 \| 1 \| 137 \| 1 \| 123 \| Phage tail protein I \| Phage tail protein I; afdb-uniprot50 \| AF-A0A1B9NSL6-F1-MODEL\_V4 \| 1.0 \| 7.464e-06 \| 169 \| 0.306 \| 111 \| 75 \| 1 \| 95 \| 205 \| 1 \| 109 \| Phage tail protein I \| Phage tail protein I; afdb-uniprot50 \| AF-A0A744CNX5-F1-MODEL\_V4 \| 1.0 \| 7.009e-06 \| 168 \| 0.268 \| 138 \| 91 \| 5 \| 3 \| 132 \| 11 \| 146 \| Phage tail protein I \| Phage tail protein I; afdb-uniprot50 \| AF-A0A6V6Y3Y7-F1-MODEL\_V4 \| 1.0 \| 5.802e-06 \| 168 \| 0.139 \| 208 \| 150 \| 10 \| 4 \| 197 \| 17 \| 209 \| Uncharacterized protein \| Uncharacterized protein; afdb-uniprot50 \| AF-A0A1S8T002-F1-MODEL\_V4 \| 1.0 \| 2.338e-07 \| 163 \| 0.19 \| 199 \| 139 \| 12 \| 11 \| 205 \| 48 \| 228 \| Uncharacterized protein \| Uncharacterized protein; afdb-uniprot50 \| AF-A0A450WI71-F1-MODEL\_V4 \| 1.0 \| 1.492e-05 \| 160 \| 0.139 \| 215 \| 167 \| 9 \| 1 \| 205 \| 3 \| 209 \| Concanavalin A-like lectin/glucanases superfamily protein \| Concanavalin A-like lectin/glucanases superfamily protein; afdb-uniprot50 \| AF-A0A2M9VEP7-F1-MODEL\_V4 \| 1.0 \| 1.246e-07 \| 159 \| 0.246 \| 203 \| 97 \| 2 \| 1 \| 203 \| 1 \| 147 \| Phage tail protein I \| Phage tail protein I; afdb-uniprot50 \| AF-U5MT65-F1-MODEL\_V4 \| 1.0 \| 1.098e-07 \| 159 \| 0.227 \| 211 \| 138 \| 14 \| 5 \| 205 \| 6 \| 201 \| Uncharacterized protein \| Uncharacterized protein; afdb-uniprot50 \| AF-A5ZQ01-F1-MODEL\_V4 \| 1.0 \| 3.292e-06 \| 158 \| 0.156 \| 198 \| 143 \| 10 \| 12 \| 204 \| 35 \| 213 \| Uncharacterized protein \| Uncharacterized protein; afdb-uniprot50 \| AF-A0A541BDT5-F1-MODEL\_V4 \| 1.0 \| 2.177e-05 \| 155 \| 0.138 \| 224 \| 160 \| 10 \| 1 \| 205 \| 3 \| 212 \| Uncharacterized protein \| Uncharacterized protein; afdb-uniprot50 \| AF-G7MCE0-F1-MODEL\_V4 \| 1.0 \| 2.49e-07 \| 151 \| 0.215 \| 200 \| 134 \| 12 \| 11 \| 205 \| 27 \| 208 \| Uncharacterized protein \| Uncharacterized protein; afdb-uniprot50 \| AF-A0A2P8EL31-F1-MODEL\_V4 \| 1.0 \| 5.116e-06 \| 149 \| 0.14 \| 206 \| 160 \| 7 \| 11 \| 205 \| 31 \| 230 \| Uncharacterized protein DUF2313 \| Uncharacterized protein DUF2313; afdb-uniprot50 \| AF-A0A4R3NVK9-F1-MODEL\_V4 \| 1.0 \| 3.506e-06 \| 144 \| 0.206 \| 232 \| 139 \| 9 \| 1 \| 205 \| 2 \| 215 \| Phage tail P2-like protein \| Phage tail P2-like protein; afdb-uniprot50 \| AF-A0A7Z0TG82-F1-MODEL\_V4 \| 1.0 \| 7.95e-06 \| 143 \| 0.186 \| 215 \| 142 \| 15 \| 3 \| 205 \| 9 \| 202 \| YmfQ family protein \| YmfQ family protein; afdb-uniprot50 \| AF-W4SMV0-F1-MODEL\_V4 \| 1.0 \| 0.0001192 \| 142 \| 0.243 \| 119 \| 78 \| 4 \| 69 \| 182 \| 3 \| 114 \| Phage-related tail protein \| Phage-related tail protein; afdb-uniprot50 \| AF-A0A7X5DCQ4-F1-MODEL\_V4 \| 1.0 \| 3.177e-05 \| 142 \| 0.123 \| 210 \| 162 \| 10 \| 5 \| 203 \| 14 \| 212 \| DUF2313 domain-containing protein \| DUF2313 domain-containing protein; afdb-uniprot50 \| AF-A0A1W7LUH0-F1-MODEL\_V4 \| 1.0 \| 6.581e-06 \| 140 \| 0.181 \| 198 \| 139 \| 13 \| 12 \| 205 \| 24 \| 202 \| Putative phage protein XkdU \| Putative phage protein XkdU; afdb-uniprot50 \| AF-A0A412NHA8-F1-MODEL\_V4 \| 1.0 \| 6.763e-05 \| 140 \| 0.136 \| 198 \| 150 \| 9 \| 14 \| 203 \| 34 \| 218 \| DUF2313 domain-containing protein \| DUF2313 domain-containing protein; afdb-uniprot50 \| AF-R9KLV1-F1-MODEL\_V4 \| 1.0 \| 9.602e-06 \| 140 \| 0.12 \| 208 \| 153 \| 12 \| 12 \| 205 \| 42 \| 233 \| Uncharacterized protein \| Uncharacterized protein; afdb-uniprot50 \| AF-A0A2I9CLJ2-F1-MODEL\_V4 \| 1.0 \| 3.837e-05 \| 137 \| 0.172 \| 215 \| 153 \| 9 \| 2 \| 202 \| 5 \| 208 \| Uncharacterized protein \| Uncharacterized protein; afdb-uniprot50 \| AF-A0A828HRI0-F1-MODEL\_V4 \| 1.0 \| 5.257e-05 \| 135 \| 0.253 \| 126 \| 79 \| 3 \| 93 \| 205 \| 6 \| 129 \| Phage tail protein I \| Phage tail protein I; afdb-uniprot50 \| AF-A0A704VV40-F1-MODEL\_V4 \| 1.0 \| 0.0001192 \| 132 \| 0.347 \| 95 \| 62 \| 0 \| 109 \| 203 \| 4 \| 98 \| Phage tail protein I \| Phage tail protein I; afdb-uniprot50 \| AF-A0A2A6Z7V1-F1-MODEL\_V4 \| 1.0 \| 0.0001192 \| 131 \| 0.113 \| 229 \| 169 \| 11 \| 4 \| 205 \| 19 \| 240 \| Uncharacterized protein \| Uncharacterized protein; afdb-uniprot50 \| AF-A0A494X1U6-F1-MODEL\_V4 \| 1.0 \| 1.692e-05 \| 131 \| 0.129 \| 232 \| 150 \| 12 \| 12 \| 202 \| 125 \| 345 \| Uncharacterized protein \| Uncharacterized protein; afdb-uniprot50 \| AF-A0A265Q495-F1-MODEL\_V4 \| 1.0 \| 6.18e-06 \| 128 \| 0.2 \| 220 \| 143 \| 12 \| 1 \| 205 \| 1 \| 202 \| Uncharacterized protein \| Uncharacterized protein; afdb-uniprot50 \| AF-A0A417ZEM8-F1-MODEL\_V4 \| 1.0 \| 0.000144 \| 125 \| 0.09 \| 210 \| 174 \| 6 \| 3 \| 205 \| 5 \| 204 \| Uncharacterized protein \| Uncharacterized protein; afdb-uniprot50 \| AF-A0A7U1GLI6-F1-MODEL\_V4 \| 1.0 \| 2.319e-05 \| 124 \| 0.153 \| 215 \| 157 \| 11 \| 5 \| 205 \| 8 \| 211 \| DUF2313 domain-containing protein \| DUF2313 domain-containing protein; afdb-uniprot50 \| AF-A0A0C1PS25-F1-MODEL\_V4 \| 1.0 \| 8.169e-05 \| 123 \| 0.141 \| 219 \| 156 \| 9 \| 3 \| 202 \| 19 \| 224 \| Uncharacterized protein \| Uncharacterized protein; afdb-uniprot50 \| AF-A0A7X5CZ78-F1-MODEL\_V4 \| 1.0 \| 0.0003702 \| 119 \| 0.127 \| 212 \| 153 \| 9 \| 14 \| 205 \| 34 \| 233 \| DUF2313 domain-containing protein \| DUF2313 domain-containing protein; afdb-uniprot50 \| AF-A0A7X5L3R8-F1-MODEL\_V4 \| 1.0 \| 6.35e-05 \| 114 \| 0.156 \| 211 \| 154 \| 11 \| 4 \| 203 \| 10 \| 207 \| DUF2313 domain-containing protein \| DUF2313 domain-containing protein; afdb-uniprot50 \| AF-A0A1V5NBE0-F1-MODEL\_V4 \| 1.0 \| 0.0002101 \| 112 \| 0.135 \| 199 \| 154 \| 9 \| 10 \| 205 \| 1 \| 184 \| Uncharacterized protein \| Uncharacterized protein; afdb-uniprot50 \| AF-A0A415TAE8-F1-MODEL\_V4 \| 1.0 \| 0.0002878 \| 102 \| 0.086 \| 209 \| 161 \| 11 \| 5 \| 197 \| 11 \| 205 \| DUF2313 domain-containing protein \| DUF2313 domain-containing protein; afdb-uniprot50 \| AF-A0A0P7H682-F1-MODEL\_V4 \| 1.0 \| 0.0001533 \| 100 \| 0.21 \| 219 \| 145 \| 7 \| 1 \| 205 \| 1 \| 205 \| Phage tail protein I \| Phage tail protein I; afdb-uniprot50 \| AF-A0A2G6AV33-F1-MODEL\_V4 \| 0.998 \| 0.0003065 \| 95 \| 0.172 \| 209 \| 146 \| 12 \| 6 \| 205 \| 27 \| 217 \| Uncharacterized protein \| Uncharacterized protein; afdb-uniprot50 \| AF-A0A6N9PSR9-F1-MODEL\_V4 \| 0.998 \| 0.000144 \| 93 \| 0.149 \| 208 \| 152 \| 9 \| 5 \| 201 \| 9 \| 202 \| DUF2313 domain-containing protein \| DUF2313 domain-containing protein; afdb-uniprot50 \| AF-A0A7X6N9Z3-F1-MODEL\_V4 \| 0.986 \| 0.003805 \| 81 \| 0.141 \| 212 \| 129 \| 11 \| 6 \| 203 \| 203 \| 375 \| Flagellar hook-length control protein FliK \| Flagellar hook-length control protein FliK; afdb-uniprot50 \| AF-A0A1I6IPJ1-F1-MODEL\_V4 \| 0.912 \| 0.005212 \| 68 \| 0.097 \| 194 \| 155 \| 7 \| 12 \| 200 \| 303 \| 481 \| TIGR00341 family protein \| TIGR00341 family protein; afdb-uniprot50 \| AF-A0A2E0A3F5-F1-MODEL\_V4 \| 0.663 \| 0.009784 \| 56 \| 0.125 \| 223 \| 148 \| 11 \| 1 \| 205 \| 267 \| 460 \| Flg\_hook domain-containing protein \| Flg\_hook domain-containing protein |
| Top keywords  (threshold 1.00e-02 (evalue)) | **tail, Phage, I, P2, P2\_like, Bacteriophage, formation, Putative, P2\_related, Tail\_P2\_I** |
| Output files | ../../similar\_structures/14\_FANPEZAQ\_CDS\_0014\_afdb-proteome\_foldseek.tsv ../../similar\_structures/14\_FANPEZAQ\_CDS\_0014\_afdb-uniprot50\_foldseek.tsv ../../similar\_structures/14\_FANPEZAQ\_CDS\_0014\_merged.svg ../../similar\_structures/14\_FANPEZAQ\_CDS\_0014\_pdb\_foldseek.tsv |

  
  
  

Return to summary | Go to previous | Go to next

  


---

**Sequence/structure alignments coloring**  
Each object in the alignment figures is colored according to its E-value following this color coding:

1e-100
10
