## Supplementary material for "Completing the BASEL phage collection to unlock hidden diversity for systematic exploration of phage-host interactions": Data S2: 15.html

FANPEZAQ\_CDS\_0015


Return to summary | Go to previous | Go to next

|  |  |
| --- | --- |
| FANPEZAQ\_CDS\_0015  Page creation date: 02 Sep 2024, 12:00  Project folder: n/a  Input sequences file: Escherichia\_virus\_HeidiAbel.gb | tail phage fiber fibre tail\_collar domain\_containing collar putative phage\_related repeat h fragment protein\_like variable bacteriophage gph repeat\_containing probable product fiber\_like duf3751 prophage dna inversion peptidase s74 duf2793 side adhesin pyocin short\_chain fatty acid transporter |

### Sequence information

|  |  |
| --- | --- |
| Name | FANPEZAQ\_CDS\_0015  15\_FANPEZAQ\_CDS\_0015 (pipeline id) |
| Imported annotations | Escherichia\_virus\_HeidiAbel Bas97 |
| Protein sequence | MATYYTLLTKIGQARIANAVALNQMVNLTYMAVGDGNGNPTTPNENQTALVREKYRATIN QLTVDPENPNYLVAEMIVPTTVGGWSIYEVGVFDDQNQLIAVANFPATYKPELVEGSGRD LIVRIIIQVSNTSVVTLKIDPAITLASQAWVAANYVKKITMAGGTTGQVLAKKSNANEDF QWVDPTAAVNVIVDSQTEEQTLASGQTIINLAKLTTQATAIYVEGVRLYPVTHYTINSDT RLTLKTPYPNGSKILAVQNDPTSTISLASTTQKGFTILSSATNSDDETKAATPKAVNDAV AALASTLGSASGRTVGNGANQIPDMNFFTSGTNWFKQPNGFIVQRGVSVAGAIGSPTFIT FPVPFTDYANVQIVVSYDNAEPYVTSIPTFAATPVNTGSGFSLMGSQVPGSNTTYARWIA IGK |
| Number of residues | 423 |
| Molecular weight (Da) | 45003.17 |
| Output files | ../../query\_sequences/15\_FANPEZAQ\_CDS\_0015.fasta |

### Putative domain architecture and protein family

#### Search results (HHblits)^1^

|  |  |
| --- | --- |
| Domain family databases searched | Pfam, Ncbi-cd, Cath, Phrogs |
| Results, scheme(s)  (Top layers only; threshold 1.00e-03 (evalue)) | xml version="1.0" encoding="utf-8" standalone="no"?       2024-09-02T21:08:15.225722 image/svg+xml   Matplotlib v3.7.2, https://matplotlib.org/ |
| Results, table  (E-value ≤ 1.00e-03 (evalue)) | db \| id \| prob \| evalue \| pvalue \| score \| cols \| query \| query\_len \| template \| template\_len \| name \| description; phrogs \| 6162 \| 100.0 \| 7.4e-53 \| 9.1e-57 \| 406.7 \| 313 \| (2, 423) \| 423 \| (3, 326) \| 326 \| tail fiber protein \| tail fiber protein; Category: tail; p300453 VI\_07590; phrogs \| 381 \| 100.0 \| 2.4e-51 \| 2.9e-55 \| 425.5 \| 203 \| (1, 304) \| 423 \| (2, 205) \| 748 \| tail fiber protein \| tail fiber protein; Category: tail; p250465 VI\_02803; phrogs \| 6202 \| 100.0 \| 9.1e-50 \| 1.1e-53 \| 394.5 \| 202 \| (2, 304) \| 423 \| (6, 208) \| 588 \| tail fiber protein \| tail fiber protein; Category: tail; p390163 VI\_08186; phrogs \| 327 \| 100.0 \| 2.2e-44 \| 2.6e-48 \| 361.4 \| 156 \| (1, 156) \| 423 \| (2, 157) \| 544 \| tail protein \| tail protein; Category: tail; p384764 VI\_09705; phrogs \| 988 \| 100.0 \| 8.5e-43 \| 1e-46 \| 350.9 \| 203 \| (1, 304) \| 423 \| (5, 208) \| 520 \| tail protein \| tail protein; Category: tail; p13596 VI\_07483; phrogs \| 4618 \| 100.0 \| 9.5e-37 \| 1.1e-40 \| 289.3 \| 204 \| (2, 308) \| 423 \| (1, 206) \| 365 \| tail collar fiber protein \| tail collar fiber protein; Category: tail; p204627 VI\_11278; phrogs \| 1780 \| 99.9 \| 4.7e-34 \| 5.9e-38 \| 259.7 \| 157 \| (1, 160) \| 423 \| (3, 159) \| 204 \| tail collar fiber protein \| tail collar fiber protein; Category: tail; p306437 VI\_06250; phrogs \| 11518 \| 99.9 \| 8.5e-32 \| 9.9e-36 \| 269.0 \| 153 \| (1, 153) \| 423 \| (1, 154) \| 648 \| tail fiber protein \| tail fiber protein; Category: tail; p203938 VI\_11941; phrogs \| 5181 \| 99.9 \| 6.7e-31 \| 7.9e-35 \| 263.6 \| 155 \| (1, 155) \| 423 \| (1, 156) \| 707 \| NA \| NA; Category: unknown function; p394343 VI\_08938; phrogs \| 2137 \| 99.9 \| 1e-30 \| 1.2e-34 \| 237.6 \| 154 \| (2, 155) \| 423 \| (4, 157) \| 281 \| tail fiber protein \| tail fiber protein; Category: tail; p58188 VI\_06099; phrogs \| 1162 \| 99.9 \| 3.1e-29 \| 3.7e-33 \| 236.4 \| 151 \| (2, 155) \| 423 \| (4, 154) \| 339 \| tail fiber protein \| tail fiber protein; Category: tail; p177110 VI\_06260; phrogs \| 14570 \| 99.9 \| 1.6e-27 \| 1.8e-31 \| 223.6 \| 154 \| (2, 155) \| 423 \| (6, 159) \| 540 \| NA \| NA; Category: unknown function; p191826 VI\_07392; phrogs \| 2557 \| 99.8 \| 3.2e-24 \| 4.2e-28 \| 182.9 \| 115 \| (304, 423) \| 423 \| (12, 129) \| 129 \| virion structural protein \| virion structural protein; Category: head and packaging; p173779 VI\_03738; phrogs \| 10153 \| 99.7 \| 2.1e-22 \| 2.3e-26 \| 180.1 \| 154 \| (2, 155) \| 423 \| (3, 156) \| 327 \| tail fiber protein \| tail fiber protein; Category: tail; p319252 VI\_06087; phrogs \| 27782 \| 99.6 \| 1.2e-20 \| 1.3e-24 \| 182.7 \| 155 \| (2, 156) \| 423 \| (3, 158) \| 641 \| NA \| NA; Category: unknown function; p342043 VI\_01070; phrogs \| 3737 \| 99.6 \| 1.3e-20 \| 1.6e-24 \| 167.4 \| 137 \| (1, 143) \| 423 \| (3, 139) \| 184 \| tail fiber protein \| tail fiber protein; Category: tail; p259608 VI\_12019; phrogs \| 30680 \| 99.6 \| 1.9e-20 \| 2.1e-24 \| 168.4 \| 154 \| (2, 155) \| 423 \| (1, 154) \| 338 \| NA \| NA; Category: unknown function; p169858 VI\_01362; phrogs \| 295 \| 99.6 \| 3.5e-20 \| 4.1e-24 \| 187.4 \| 194 \| (2, 304) \| 423 \| (171, 392) \| 813 \| tail fiber protein \| tail fiber protein; Category: tail; p109395 VI\_03810; phrogs \| 4890 \| 99.6 \| 6.5e-20 \| 7.3e-24 \| 149.2 \| 119 \| (304, 423) \| 423 \| (29, 153) \| 153 \| NA \| NA; Category: unknown function; p375303 VI\_02828; phrogs \| 14531 \| 99.5 \| 2.2e-19 \| 2.5e-23 \| 162.9 \| 173 \| (3, 183) \| 423 \| (5, 177) \| 359 \| NA \| NA; Category: unknown function; p181208 VI\_06278; phrogs \| 1396 \| 99.5 \| 2.3e-19 \| 3e-23 \| 168.5 \| 122 \| (294, 423) \| 423 \| (119, 261) \| 261 \| tail fiber protein \| tail fiber protein; Category: tail; p37946 VI\_07185; phrogs \| 9532 \| 99.4 \| 2.3e-17 \| 2.6e-21 \| 158.7 \| 154 \| (2, 155) \| 423 \| (3, 156) \| 759 \| tail fiber protein \| tail fiber protein; Category: tail; p329901 VI\_11396; phrogs \| 5075 \| 99.3 \| 2.2e-16 \| 2.5e-20 \| 144.8 \| 149 \| (2, 155) \| 423 \| (4, 153) \| 394 \| NA \| NA; Category: unknown function; p127048 VI\_01582; phrogs \| 7037 \| 99.2 \| 1.3e-15 \| 1.6e-19 \| 135.2 \| 146 \| (3, 152) \| 423 \| (4, 157) \| 185 \| tail fiber protein \| tail fiber protein; Category: tail; p172398 VI\_05814; phrogs \| 17585 \| 99.2 \| 1.9e-15 \| 2.2e-19 \| 145.5 \| 159 \| (4, 163) \| 423 \| (5, 163) \| 650 \| NA \| NA; Category: unknown function; p387047 VI\_09109; phrogs \| 23632 \| 99.0 \| 7.9e-14 \| 9.1e-18 \| 125.7 \| 152 \| (1, 156) \| 423 \| (1, 160) \| 290 \| NA \| NA; Category: unknown function; p130591 VI\_06329; phrogs \| 7489 \| 99.0 \| 8.2e-14 \| 9.5e-18 \| 137.0 \| 140 \| (4, 145) \| 423 \| (1, 147) \| 666 \| tail fiber protein \| tail fiber protein; Category: tail; p80663 VI\_09756; phrogs \| 25305 \| 98.9 \| 1.2e-13 \| 1.3e-17 \| 136.8 \| 159 \| (4, 163) \| 423 \| (5, 163) \| 870 \| tail fiber protein \| tail fiber protein; Category: tail; p176118 VI\_11367; phrogs \| 4209 \| 98.9 \| 1.9e-13 \| 2.3e-17 \| 130.1 \| 88 \| (329, 423) \| 423 \| (253, 341) \| 341 \| tail fiber protein \| tail fiber protein; Category: tail; p184403 VI\_05508; phrogs \| 8239 \| 98.8 \| 5.4e-13 \| 6.2e-17 \| 129.4 \| 140 \| (4, 145) \| 423 \| (2, 148) \| 654 \| tail fiber protein \| tail fiber protein; Category: tail; p255681 VI\_08125; phrogs \| 13714 \| 98.8 \| 8.6e-13 \| 1e-16 \| 113.6 \| 147 \| (3, 154) \| 423 \| (4, 156) \| 191 \| tail fiber protein \| tail fiber protein; Category: tail; p143339 VI\_09468; phrogs \| 4545 \| 98.7 \| 1.3e-12 \| 1.6e-16 \| 126.2 \| 138 \| (3, 145) \| 423 \| (2, 144) \| 512 \| NA \| NA; Category: unknown function; p341124 VI\_09299; phrogs \| 13388 \| 98.7 \| 1.6e-12 \| 1.9e-16 \| 105.2 \| 112 \| (4, 117) \| 423 \| (2, 118) \| 127 \| tail collar fiber protein \| tail collar fiber protein; Category: tail; p255876 VI\_08161; phrogs \| 15547 \| 98.1 \| 6.1e-10 \| 7e-14 \| 100.9 \| 103 \| (1, 111) \| 423 \| (1, 104) \| 323 \| NA \| NA; Category: unknown function; p391937 VI\_02991; phrogs \| 38337 \| 98.1 \| 9.4e-10 \| 1.1e-13 \| 103.7 \| 145 \| (4, 152) \| 423 \| (9, 160) \| 569 \| NA \| NA; Category: unknown function; p170432 VI\_01289; phrogs \| 19961 \| 98.1 \| 1e-09 \| 1.1e-13 \| 101.9 \| 150 \| (1, 155) \| 423 \| (1, 159) \| 439 \| tail fiber protein \| tail fiber protein; Category: tail; p321354 VI\_06335; phrogs \| 19921 \| 98.0 \| 2.1e-09 \| 2.4e-13 \| 104.6 \| 99 \| (7, 105) \| 423 \| (24, 133) \| 946 \| tail spike protein \| tail spike protein; Category: tail; NC\_027381\_p75; phrogs \| 8443 \| 97.6 \| 3e-08 \| 3.5e-12 \| 95.8 \| 106 \| (1, 114) \| 423 \| (1, 107) \| 530 \| tail fiber protein \| tail fiber protein; Category: tail; p142958 VI\_03022; phrogs \| 6683 \| 97.5 \| 5e-08 \| 5.8e-12 \| 88.9 \| 99 \| (317, 423) \| 423 \| (257, 359) \| 359 \| tail fiber protein \| tail fiber protein; Category: tail; NC\_010463\_p10; phrogs \| 4849 \| 97.5 \| 5.7e-08 \| 6.5e-12 \| 95.1 \| 105 \| (1, 111) \| 423 \| (1, 106) \| 790 \| tail protein \| tail protein; Category: tail; JX681814\_p23; phrogs \| 33377 \| 97.5 \| 6.1e-08 \| 6.8e-12 \| 88.2 \| 50 \| (330, 380) \| 423 \| (349, 409) \| 456 \| NA \| NA; Category: unknown function; p217392 VI\_05721; phrogs \| 25563 \| 97.5 \| 7e-08 \| 7.9e-12 \| 94.6 \| 102 \| (2, 109) \| 423 \| (1, 103) \| 829 \| NA \| NA; Category: unknown function; p252637 VI\_04196; phrogs \| 30004 \| 97.2 \| 3.6e-07 \| 4.1e-11 \| 81.2 \| 147 \| (5, 152) \| 423 \| (3, 155) \| 328 \| NA \| NA; Category: unknown function; p288306 VI\_01174; phrogs \| 10372 \| 97.2 \| 3.8e-07 \| 4.3e-11 \| 76.7 \| 43 \| (262, 304) \| 423 \| (62, 104) \| 200 \| NA \| NA; Category: unknown function; p28886 VI\_10606; phrogs \| 2631 \| 97.1 \| 6e-07 \| 6.9e-11 \| 84.6 \| 41 \| (264, 304) \| 423 \| (2, 42) \| 482 \| tail fiber protein \| tail fiber protein; Category: tail; p130317 VI\_07372; phrogs \| 6043 \| 97.1 \| 7e-07 \| 7.7e-11 \| 84.9 \| 139 \| (5, 145) \| 423 \| (2, 147) \| 814 \| NA \| NA; Category: unknown function; p336028 VI\_09513; phrogs \| 17472 \| 97.0 \| 7.5e-07 \| 8.7e-11 \| 88.2 \| 126 \| (4, 138) \| 423 \| (11, 138) \| 868 \| tail protein \| tail protein; Category: tail; KY290947\_p27; phrogs \| 36614 \| 96.8 \| 2.1e-06 \| 2.3e-10 \| 66.3 \| 40 \| (265, 304) \| 423 \| (3, 42) \| 131 \| NA \| NA; Category: unknown function; p237616 VI\_06905; phrogs \| 24573 \| 96.6 \| 4.3e-06 \| 4.8e-10 \| 79.0 \| 137 \| (6, 144) \| 423 \| (3, 146) \| 780 \| NA \| NA; Category: unknown function; p244508 VI\_06663; phrogs \| 8713 \| 94.8 \| 0.00048 \| 5.4e-08 \| 63.6 \| 40 \| (264, 303) \| 423 \| (352, 391) \| 541 \| tail fiber protein \| tail fiber protein; Category: tail; p399266 VI\_10066; phrogs \| 17623 \| 94.3 \| 0.00091 \| 1e-07 \| 65.8 \| 96 \| (5, 106) \| 423 \| (7, 103) \| 1121 \| tail collar fiber protein \| tail collar fiber protein; Category: tail; NC\_021343\_p35 |
| Top keywords  (threshold 1.00e-03 (evalue)) | **tail, fiber, collar, p300453, VI\_07590, p250465, VI\_02803, p390163, VI\_08186, p384764** |
| Output files | ../../domain\_architecture/15\_FANPEZAQ\_CDS\_0015\_cath.hhr ../../domain\_architecture/15\_FANPEZAQ\_CDS\_0015\_merged.svg ../../domain\_architecture/15\_FANPEZAQ\_CDS\_0015\_ncbi-cd.hhr ../../domain\_architecture/15\_FANPEZAQ\_CDS\_0015\_pfam.hhr ../../domain\_architecture/15\_FANPEZAQ\_CDS\_0015\_phrogs.hhr |

### Identical protein sequences/structures

#### Search results

|  |  |
| --- | --- |
| Protein sequence databases searched | Pdb, Swissprot, Refseq |
| Identical proteins found | -- |
| Top keywords | -- |
| Output files | -- |

### Similar protein sequences/structures

#### Sequence similarity search results (HHblits)^1^

|  |  |
| --- | --- |
| Sequence databases searched | Uniclust, Pdb70 |
| Results, scheme(s)  (Top layers only, threshold 1.00e-03 (evalue)) | xml version="1.0" encoding="utf-8" standalone="no"?       2024-09-02T21:08:34.936126 image/svg+xml   Matplotlib v3.7.2, https://matplotlib.org/ |
| Results, table(s)  (threshold 1.00e-03 (evalue)) | db \| id \| prob \| evalue \| pvalue \| score \| cols \| query \| query\_len \| template \| template\_len \| name \| description; uniclust \| UniRef100\_A0A381GJP7 \| 100.0 \| 1.2e-52 \| 2.3e-58 \| 364.7 \| 284 \| (1, 375) \| 423 \| (4, 351) \| 397 \| Phage tail fiber protein \| Phage tail fiber protein; uniclust \| UniRef100\_A0A1G7VLT1 \| 100.0 \| 7.7e-52 \| 1.5e-57 \| 363.3 \| 160 \| (1, 160) \| 423 \| (1, 160) \| 369 \| Phage tail-collar fibre protein \| Phage tail-collar fibre protein; uniclust \| UniRef100\_A0A0G2PMU5 \| 100.0 \| 4.6e-51 \| 8.7e-57 \| 374.3 \| 165 \| (1, 165) \| 423 \| (2, 166) \| 528 \| Tail fiber protein \| Tail fiber protein; uniclust \| UniRef100\_A0A261E4L1 \| 100.0 \| 1.3e-49 \| 2.5e-55 \| 346.0 \| 162 \| (1, 162) \| 423 \| (2, 163) \| 361 \| DNA inversion product \| DNA inversion product; uniclust \| UniRef100\_A0A088FQW7 \| 100.0 \| 5.2e-49 \| 1e-54 \| 364.8 \| 280 \| (1, 281) \| 423 \| (15, 299) \| 515 \| Tail protein \| Tail protein; uniclust \| UniRef100\_A0A080I6W6 \| 100.0 \| 3.8e-48 \| 7.1e-54 \| 335.9 \| 341 \| (2, 375) \| 423 \| (13, 358) \| 405 \| Phage tail-collar fiber family protein \| Phage tail-collar fiber family protein; uniclust \| UniRef100\_A0A061JKN2 \| 100.0 \| 9.7e-48 \| 1.8e-53 \| 352.0 \| 276 \| (1, 277) \| 423 \| (26, 304) \| 521 \| Tail protein \| Tail protein; uniclust \| UniRef100\_A0A024HA82 \| 100.0 \| 2.4e-47 \| 4.7e-53 \| 333.8 \| 155 \| (2, 156) \| 423 \| (6, 160) \| 358 \| Uncharacterized protein \| Uncharacterized protein; uniclust \| UniRef100\_A0A0E1LY00 \| 100.0 \| 5.6e-47 \| 1e-52 \| 343.6 \| 160 \| (1, 160) \| 423 \| (5, 165) \| 585 \| Putative phage tail fiber protein \| Putative phage tail fiber protein; uniclust \| UniRef100\_A0A080KKZ3 \| 100.0 \| 2.1e-46 \| 4e-52 \| 331.9 \| 158 \| (2, 159) \| 423 \| (3, 160) \| 343 \| Tail fiber protein \| Tail fiber protein; uniclust \| UniRef100\_A0A0J8YRQ6 \| 100.0 \| 3e-46 \| 5.7e-52 \| 329.5 \| 171 \| (2, 172) \| 423 \| (3, 173) \| 409 \| Phage tail protein \| Phage tail protein; uniclust \| UniRef100\_A0A827WUC6 \| 100.0 \| 3.2e-46 \| 5.8e-52 \| 304.5 \| 276 \| (2, 369) \| 423 \| (3, 278) \| 290 \| Phage tail protein (Fragment) \| Phage tail protein (Fragment); uniclust \| UniRef100\_A0A1B8TDC2 \| 100.0 \| 4.1e-46 \| 7.7e-52 \| 341.0 \| 154 \| (3, 156) \| 423 \| (4, 157) \| 566 \| Phage tail protein \| Phage tail protein; uniclust \| UniRef100\_A0A2N7UDN2 \| 100.0 \| 5.2e-46 \| 1e-51 \| 348.6 \| 160 \| (1, 160) \| 423 \| (15, 174) \| 523 \| Phage tail protein \| Phage tail protein; uniclust \| UniRef100\_A0A0M2WUJ2 \| 100.0 \| 1.1e-45 \| 2.1e-51 \| 345.0 \| 160 \| (1, 160) \| 423 \| (5, 164) \| 528 \| Uncharacterized protein \| Uncharacterized protein; uniclust \| UniRef100\_A0A022PCI5 \| 100.0 \| 1.4e-45 \| 2.6e-51 \| 328.1 \| 205 \| (2, 297) \| 423 \| (3, 207) \| 438 \| Phage-related tail fiber protein \| Phage-related tail fiber protein; uniclust \| UniRef100\_A0A1C3HKM5 \| 100.0 \| 2.9e-45 \| 5.5e-51 \| 357.2 \| 229 \| (1, 330) \| 423 \| (16, 262) \| 912 \| Phage tail fiber repeat protein \| Phage tail fiber repeat protein; uniclust \| UniRef100\_UPI0005E23890 \| 100.0 \| 3.8e-45 \| 7e-51 \| 311.9 \| 283 \| (2, 371) \| 423 \| (3, 337) \| 384 \| phage tail protein \| phage tail protein; uniclust \| UniRef100\_A0A144P4D2 \| 100.0 \| 9.8e-45 \| 1.9e-50 \| 353.5 \| 158 \| (2, 159) \| 423 \| (5, 163) \| 707 \| Tail Fiber protein \| Tail Fiber protein; uniclust \| UniRef100\_A0A1B8YGC4 \| 100.0 \| 3.9e-44 \| 7.6e-50 \| 340.4 \| 159 \| (2, 160) \| 423 \| (3, 161) \| 616 \| Phage tail fiber repeat protein \| Phage tail fiber repeat protein; uniclust \| UniRef100\_A0A7D7ENG5 \| 100.0 \| 5.7e-44 \| 1.1e-49 \| 321.1 \| 280 \| (2, 281) \| 423 \| (3, 282) \| 551 \| Tail fiber \| Tail fiber; uniclust \| UniRef100\_A0A0C2EE47 \| 100.0 \| 1.1e-43 \| 2.1e-49 \| 335.7 \| 157 \| (2, 158) \| 423 \| (10, 166) \| 603 \| Phage tail fiber protein \| Phage tail fiber protein; uniclust \| UniRef100\_A0A1G5RGJ6 \| 100.0 \| 1.2e-43 \| 2.3e-49 \| 304.5 \| 158 \| (2, 159) \| 423 \| (3, 160) \| 348 \| Phage tail fibre repeat-containing protein (Fragment) \| Phage tail fibre repeat-containing protein (Fragment); uniclust \| UniRef100\_UPI0005C6F866 \| 100.0 \| 3.6e-43 \| 6.8e-49 \| 308.4 \| 158 \| (3, 160) \| 423 \| (4, 161) \| 427 \| phage tail protein \| phage tail protein; uniclust \| UniRef100\_A0A077P0S8 \| 100.0 \| 4.4e-43 \| 8.3e-49 \| 318.9 \| 202 \| (1, 303) \| 423 \| (2, 203) \| 500 \| Uncharacterized protein \| Uncharacterized protein; uniclust \| UniRef100\_A0A1Q5TUM5 \| 100.0 \| 1.2e-42 \| 2.2e-48 \| 322.0 \| 159 \| (2, 160) \| 423 \| (3, 161) \| 573 \| Phage tail protein \| Phage tail protein; uniclust \| UniRef100\_A0A3L0W1T3 \| 100.0 \| 1.6e-42 \| 3e-48 \| 302.5 \| 160 \| (1, 160) \| 423 \| (2, 161) \| 368 \| Phage tail protein \| Phage tail protein; uniclust \| UniRef100\_A0A0K0I9Z1 \| 100.0 \| 2.1e-42 \| 4.1e-48 \| 337.0 \| 203 \| (1, 304) \| 423 \| (2, 205) \| 748 \| Phage tail protein \| Phage tail protein; uniclust \| UniRef100\_C6D9F9 \| 100.0 \| 4.1e-42 \| 7.8e-48 \| 317.8 \| 158 \| (2, 159) \| 423 \| (3, 160) \| 597 \| Tail fiber repeat 2 protein \| Tail fiber repeat 2 protein; uniclust \| UniRef100\_A0A2X7EVU3 \| 100.0 \| 6.3e-42 \| 1.2e-47 \| 310.3 \| 159 \| (2, 160) \| 423 \| (3, 161) \| 519 \| Phage tail collar domain-containing protein \| Phage tail collar domain-containing protein; uniclust \| UniRef100\_A0A0U4W4I1 \| 100.0 \| 9.8e-42 \| 1.8e-47 \| 308.5 \| 279 \| (3, 281) \| 423 \| (5, 290) \| 577 \| Ig-like domain-containing protein \| Ig-like domain-containing protein; uniclust \| UniRef100\_A0A0B6X5X6 \| 100.0 \| 2.2e-41 \| 4.1e-47 \| 292.7 \| 158 \| (2, 159) \| 423 \| (3, 160) \| 383 \| E14 prophage putative tail fiber protein (Modular protein) \| E14 prophage putative tail fiber protein (Modular protein); uniclust \| UniRef100\_A0A081RRN4 \| 100.0 \| 2.9e-41 \| 5.5e-47 \| 310.1 \| 159 \| (2, 160) \| 423 \| (3, 161) \| 684 \| Phage-related tail fiber protein \| Phage-related tail fiber protein; uniclust \| UniRef100\_A0A077NGB1 \| 100.0 \| 3.1e-41 \| 5.9e-47 \| 314.1 \| 159 \| (2, 160) \| 423 \| (3, 161) \| 765 \| Uncharacterized protein \| Uncharacterized protein; uniclust \| UniRef100\_A0A0M0TBG4 \| 100.0 \| 3.7e-41 \| 6.9e-47 \| 281.2 \| 204 \| (2, 301) \| 423 \| (3, 206) \| 274 \| Tail protein (Fragment) \| Tail protein (Fragment); uniclust \| UniRef100\_A0A0D5Y3B7 \| 100.0 \| 3.6e-41 \| 7e-47 \| 321.4 \| 157 \| (2, 158) \| 423 \| (22, 178) \| 498 \| Phage tail protein \| Phage tail protein; uniclust \| UniRef100\_A0A080KJ35 \| 100.0 \| 3.9e-41 \| 7.4e-47 \| 310.4 \| 203 \| (3, 302) \| 423 \| (16, 218) \| 530 \| Phage-related tail fiber protein \| Phage-related tail fiber protein; uniclust \| UniRef100\_A0A2G1CTX9 \| 100.0 \| 4.3e-41 \| 8.1e-47 \| 299.6 \| 173 \| (1, 173) \| 423 \| (5, 177) \| 391 \| Phage tail protein \| Phage tail protein; uniclust \| UniRef100\_A0A068QVC8 \| 100.0 \| 6.1e-41 \| 1.2e-46 \| 316.5 \| 158 \| (2, 159) \| 423 \| (3, 160) \| 696 \| Uncharacterized protein \| Uncharacterized protein; uniclust \| UniRef100\_A0A3D1C6R7 \| 100.0 \| 7.1e-41 \| 1.3e-46 \| 304.0 \| 171 \| (2, 172) \| 423 \| (10, 180) \| 511 \| Phage tail protein \| Phage tail protein; uniclust \| UniRef100\_A0A0A1ABU4 \| 100.0 \| 9.6e-41 \| 1.8e-46 \| 304.1 \| 200 \| (3, 303) \| 423 \| (5, 205) \| 539 \| Phage tail protein \| Phage tail protein; uniclust \| UniRef100\_A0A2X5BQE6 \| 100.0 \| 1.1e-40 \| 2.2e-46 \| 308.8 \| 160 \| (1, 160) \| 423 \| (4, 163) \| 540 \| Phage tail fibre repeat \| Phage tail fibre repeat; uniclust \| UniRef100\_A0A1D7Z5A8 \| 100.0 \| 2e-40 \| 3.8e-46 \| 299.2 \| 159 \| (2, 160) \| 423 \| (3, 161) \| 555 \| Phage tail protein \| Phage tail protein; uniclust \| UniRef100\_A0A0P7LYC5 \| 100.0 \| 3.9e-40 \| 7.4e-46 \| 303.1 \| 225 \| (2, 232) \| 423 \| (10, 236) \| 609 \| Phage tail protein \| Phage tail protein; uniclust \| UniRef100\_UPI000A487BB5 \| 100.0 \| 4.1e-40 \| 7.6e-46 \| 298.0 \| 214 \| (2, 305) \| 423 \| (3, 216) \| 578 \| phage tail protein \| phage tail protein; uniclust \| UniRef100\_UPI0004A05D1E \| 100.0 \| 6.7e-40 \| 1.2e-45 \| 295.7 \| 209 \| (2, 216) \| 423 \| (3, 211) \| 589 \| phage tail protein \| phage tail protein; uniclust \| UniRef100\_A0A1B1KLD7 \| 100.0 \| 1e-39 \| 1.9e-45 \| 305.5 \| 158 \| (2, 159) \| 423 \| (3, 160) \| 679 \| Uncharacterized protein \| Uncharacterized protein; uniclust \| UniRef100\_A0A376D432 \| 100.0 \| 1.1e-39 \| 2.1e-45 \| 282.2 \| 170 \| (2, 171) \| 423 \| (3, 172) \| 464 \| Putative variable tail fiber protein \| Putative variable tail fiber protein; uniclust \| UniRef100\_A0A0K9TAC2 \| 100.0 \| 1.2e-39 \| 2.2e-45 \| 279.0 \| 157 \| (3, 159) \| 423 \| (7, 164) \| 341 \| Phage tail fiber \| Phage tail fiber; uniclust \| UniRef100\_A0A068R2V7 \| 100.0 \| 1.3e-39 \| 2.4e-45 \| 283.6 \| 247 \| (2, 349) \| 423 \| (3, 278) \| 430 \| Uncharacterized protein \| Uncharacterized protein; uniclust \| UniRef100\_A0A1H5DGF8 \| 100.0 \| 1.4e-39 \| 2.7e-45 \| 311.9 \| 156 \| (2, 157) \| 423 \| (6, 161) \| 545 \| Phage tail-collar fibre protein \| Phage tail-collar fibre protein; uniclust \| UniRef100\_A0A5M7L9F8 \| 100.0 \| 2.1e-39 \| 3.9e-45 \| 281.1 \| 273 \| (3, 374) \| 423 \| (13, 346) \| 388 \| Phage tail protein \| Phage tail protein; uniclust \| UniRef100\_A0A1C4G344 \| 100.0 \| 2.3e-39 \| 4.3e-45 \| 305.5 \| 159 \| (2, 160) \| 423 \| (3, 162) \| 733 \| Phage tail fibre repeat-containing protein \| Phage tail fibre repeat-containing protein; uniclust \| UniRef100\_A0A078LPZ8 \| 100.0 \| 2.9e-39 \| 5.7e-45 \| 305.2 \| 169 \| (2, 170) \| 423 \| (10, 178) \| 566 \| Phage tail fiber protein \| Phage tail fiber protein; uniclust \| UniRef100\_UPI00042486BE \| 100.0 \| 3.2e-39 \| 5.9e-45 \| 296.7 \| 292 \| (2, 303) \| 423 \| (3, 315) \| 695 \| tail fiber protein \| tail fiber protein; uniclust \| UniRef100\_A0A0R9N0A5 \| 100.0 \| 3.2e-39 \| 6e-45 \| 295.1 \| 158 \| (3, 160) \| 423 \| (6, 163) \| 584 \| Phage tail protein \| Phage tail protein; uniclust \| UniRef100\_A0A2A2F373 \| 100.0 \| 7.4e-39 \| 1.4e-44 \| 266.5 \| 170 \| (1, 170) \| 423 \| (4, 173) \| 250 \| Phage tail protein \| Phage tail protein; uniclust \| UniRef100\_A0A014MD74 \| 100.0 \| 1.1e-38 \| 2.1e-44 \| 307.2 \| 178 \| (2, 180) \| 423 \| (95, 272) \| 647 \| Variable tail fiber protein \| Variable tail fiber protein; uniclust \| UniRef100\_A0A097R5S7 \| 100.0 \| 1.1e-38 \| 2.2e-44 \| 291.9 \| 171 \| (2, 172) \| 423 \| (17, 187) \| 428 \| Phage tail protein \| Phage tail protein; uniclust \| UniRef100\_D1NYY3 \| 100.0 \| 1.6e-38 \| 3.1e-44 \| 296.7 \| 155 \| (2, 156) \| 423 \| (1, 155) \| 501 \| Phage tail fiber repeat protein \| Phage tail fiber repeat protein; uniclust \| UniRef100\_A0A2S4QEE2 \| 100.0 \| 1.9e-38 \| 3.6e-44 \| 306.1 \| 299 \| (1, 304) \| 423 \| (2, 340) \| 691 \| Phage tail protein \| Phage tail protein; uniclust \| UniRef100\_A0A2I5TL70 \| 100.0 \| 2.6e-38 \| 5e-44 \| 287.2 \| 160 \| (2, 161) \| 423 \| (25, 184) \| 445 \| Phage tail protein \| Phage tail protein; uniclust \| UniRef100\_A0A0D8RJM6 \| 100.0 \| 2.7e-38 \| 5.1e-44 \| 257.9 \| 201 \| (2, 303) \| 423 \| (6, 207) \| 240 \| Phage tail protein \| Phage tail protein; uniclust \| UniRef100\_A0A0X4EUS0 \| 100.0 \| 3.5e-38 \| 6.8e-44 \| 310.2 \| 159 \| (2, 160) \| 423 \| (98, 257) \| 761 \| Phage tail protein \| Phage tail protein; uniclust \| UniRef100\_A0A1I6E807 \| 100.0 \| 3.9e-38 \| 7.5e-44 \| 306.7 \| 299 \| (4, 307) \| 423 \| (33, 370) \| 825 \| Phage tail fibre repeat-containing protein \| Phage tail fibre repeat-containing protein; uniclust \| UniRef100\_A0A080I1Q9 \| 100.0 \| 4.3e-38 \| 8.4e-44 \| 299.3 \| 172 \| (2, 173) \| 423 \| (6, 177) \| 599 \| Phage tail-collar fiber family protein \| Phage tail-collar fiber family protein; uniclust \| UniRef100\_A0A376PRJ2 \| 100.0 \| 4.9e-38 \| 9e-44 \| 253.0 \| 235 \| (2, 341) \| 423 \| (3, 237) \| 241 \| Phage tail collar protein \| Phage tail collar protein; uniclust \| UniRef100\_A0A6H2GG96 \| 100.0 \| 6.7e-38 \| 1.2e-43 \| 285.0 \| 158 \| (2, 159) \| 423 \| (3, 160) \| 694 \| Phage tail protein \| Phage tail protein; uniclust \| UniRef100\_A0A2D0ITF1 \| 100.0 \| 7e-38 \| 1.3e-43 \| 293.9 \| 158 \| (3, 160) \| 423 \| (4, 161) \| 857 \| Phage tail fiber protein \| Phage tail fiber protein; uniclust \| UniRef100\_A0A077PGX7 \| 100.0 \| 7.2e-38 \| 1.3e-43 \| 277.9 \| 154 \| (3, 156) \| 423 \| (4, 157) \| 512 \| Uncharacterized protein \| Uncharacterized protein; uniclust \| UniRef100\_A0A379VLX8 \| 100.0 \| 1.5e-37 \| 2.8e-43 \| 259.9 \| 171 \| (2, 172) \| 423 \| (3, 173) \| 335 \| Phage tail fiber protein \| Phage tail fiber protein; uniclust \| UniRef100\_A0A080HIT0 \| 100.0 \| 1.7e-37 \| 3.1e-43 \| 279.4 \| 170 \| (2, 171) \| 423 \| (3, 172) \| 499 \| Phage tail-collar fiber family protein \| Phage tail-collar fiber family protein; uniclust \| UniRef100\_A0A2R8CQR2 \| 100.0 \| 1.8e-37 \| 3.3e-43 \| 263.6 \| 160 \| (1, 160) \| 423 \| (1, 160) \| 364 \| Uncharacterized protein \| Uncharacterized protein; uniclust \| UniRef100\_A0A0T9KM71 \| 100.0 \| 2e-37 \| 3.7e-43 \| 270.8 \| 152 \| (3, 154) \| 423 \| (39, 190) \| 413 \| Variable tail fiber protein \| Variable tail fiber protein; uniclust \| UniRef100\_A0A1E7PZY5 \| 100.0 \| 2.4e-37 \| 4.7e-43 \| 262.0 \| 160 \| (1, 160) \| 423 \| (2, 161) \| 271 \| Phage tail protein \| Phage tail protein; uniclust \| UniRef100\_A0A806X791 \| 100.0 \| 3e-37 \| 5.7e-43 \| 272.2 \| 210 \| (1, 300) \| 423 \| (5, 214) \| 390 \| Phage tail collar domain-containing protein \| Phage tail collar domain-containing protein; uniclust \| UniRef100\_A0A2H4JF09 \| 100.0 \| 7.3e-37 \| 1.4e-42 \| 267.8 \| 157 \| (2, 158) \| 423 \| (6, 162) \| 416 \| Putative tail collar domain protein \| Putative tail collar domain protein; uniclust \| UniRef100\_A0A7V8UDY7 \| 100.0 \| 1.1e-36 \| 2.1e-42 \| 276.8 \| 155 \| (2, 156) \| 423 \| (6, 160) \| 572 \| Phage tail protein \| Phage tail protein; uniclust \| UniRef100\_A0A0A0GX64 \| 100.0 \| 1.3e-36 \| 2.4e-42 \| 299.1 \| 203 \| (1, 304) \| 423 \| (5, 208) \| 791 \| Putative tail fiber protein \| Putative tail fiber protein; uniclust \| UniRef100\_A0A1G9EQI8 \| 100.0 \| 1.9e-36 \| 3.5e-42 \| 272.8 \| 158 \| (1, 158) \| 423 \| (5, 162) \| 420 \| Phage-related tail fibre protein \| Phage-related tail fibre protein; uniclust \| UniRef100\_UPI001F4EC7D7 \| 100.0 \| 2e-36 \| 3.6e-42 \| 261.3 \| 157 \| (2, 158) \| 423 \| (3, 159) \| 428 \| phage tail protein \| phage tail protein; uniclust \| UniRef100\_A0A3S4G527 \| 100.0 \| 2.2e-36 \| 4.1e-42 \| 265.2 \| 244 \| (2, 302) \| 423 \| (3, 247) \| 426 \| Phage variable tail-fiber protein \| Phage variable tail-fiber protein; uniclust \| UniRef100\_A0A085GA02 \| 100.0 \| 2.3e-36 \| 4.4e-42 \| 285.9 \| 160 \| (1, 160) \| 423 \| (5, 165) \| 622 \| Putative tail fiber protein \| Putative tail fiber protein; uniclust \| UniRef100\_A0A085JCX9 \| 100.0 \| 3e-36 \| 5.9e-42 \| 296.7 \| 213 \| (1, 303) \| 423 \| (96, 308) \| 688 \| Tail fiber protein \| Tail fiber protein; uniclust \| UniRef100\_A0A3M2VPF6 \| 100.0 \| 3.7e-36 \| 7e-42 \| 262.9 \| 151 \| (2, 152) \| 423 \| (6, 156) \| 316 \| Putative Tail fiber protein H \| Putative Tail fiber protein H; uniclust \| UniRef100\_A0A0Q0EQE1 \| 100.0 \| 3.8e-36 \| 7.2e-42 \| 270.2 \| 156 \| (1, 156) \| 423 \| (5, 160) \| 434 \| Phage tail fiber protein h \| Phage tail fiber protein h; uniclust \| UniRef100\_A0A2K8QMP4 \| 100.0 \| 4.9e-36 \| 9.1e-42 \| 257.8 \| 170 \| (2, 171) \| 423 \| (3, 172) \| 383 \| Phage tail protein \| Phage tail protein; uniclust \| UniRef100\_UPI0010D6282B \| 100.0 \| 5.7e-36 \| 1.1e-41 \| 228.6 \| 153 \| (1, 153) \| 423 \| (2, 154) \| 169 \| phage tail protein \| phage tail protein; uniclust \| UniRef100\_A0A264VRT5 \| 100.0 \| 5.9e-36 \| 1.1e-41 \| 284.9 \| 156 \| (2, 157) \| 423 \| (3, 158) \| 568 \| Phage tail collar domain-containing protein \| Phage tail collar domain-containing protein; uniclust \| UniRef100\_A0A0D6BKK2 \| 100.0 \| 6e-36 \| 1.2e-41 \| 286.7 \| 157 \| (1, 157) \| 423 \| (5, 161) \| 605 \| Phage-related tail fibre protein-like protein \| Phage-related tail fibre protein-like protein; uniclust \| UniRef100\_A0A1H0SWR0 \| 100.0 \| 6.4e-36 \| 1.2e-41 \| 269.6 \| 154 \| (4, 157) \| 423 \| (5, 158) \| 549 \| Phage tail-collar fibre protein \| Phage tail-collar fibre protein; uniclust \| UniRef100\_A0A0E1ST60 \| 100.0 \| 6.5e-36 \| 1.3e-41 \| 295.6 \| 292 \| (3, 301) \| 423 \| (6, 336) \| 808 \| GpH \| GpH; uniclust \| UniRef100\_A0A068Z6W7 \| 100.0 \| 9.4e-36 \| 1.8e-41 \| 257.4 \| 169 \| (2, 170) \| 423 \| (3, 171) \| 410 \| Phage tail protein \| Phage tail protein; uniclust \| UniRef100\_UPI0009E43696 \| 100.0 \| 9.6e-36 \| 1.8e-41 \| 269.7 \| 169 \| (2, 171) \| 423 \| (3, 171) \| 583 \| phage tail protein \| phage tail protein; uniclust \| UniRef100\_A0A0P9YJT9 \| 100.0 \| 9.2e-36 \| 1.8e-41 \| 293.7 \| 159 \| (1, 159) \| 423 \| (5, 163) \| 710 \| Putative Tail fiber protein H \| Putative Tail fiber protein H; uniclust \| UniRef100\_A0A2K8VZI6 \| 100.0 \| 1.1e-35 \| 2e-41 \| 268.0 \| 153 \| (3, 155) \| 423 \| (30, 182) \| 542 \| Phage tail fiber protein \| Phage tail fiber protein; uniclust \| UniRef100\_V5YST5 \| 100.0 \| 1.1e-35 \| 2.1e-41 \| 259.9 \| 279 \| (1, 280) \| 423 \| (1, 279) \| 550 \| Putative tail-collar fibre protein \| Putative tail-collar fibre protein; uniclust \| UniRef100\_A0A1X7LWW5 \| 100.0 \| 1.2e-35 \| 2.3e-41 \| 269.1 \| 159 \| (3, 161) \| 423 \| (4, 162) \| 512 \| Phage tail fibre repeat-containing protein \| Phage tail fibre repeat-containing protein; uniclust \| UniRef100\_A0A024EJX5 \| 100.0 \| 1.2e-35 \| 2.4e-41 \| 293.9 \| 157 \| (1, 157) \| 423 \| (5, 161) \| 754 \| Tail fiber protein H, putative \| Tail fiber protein H, putative; uniclust \| UniRef100\_A0A3B7L361 \| 100.0 \| 1.3e-35 \| 2.4e-41 \| 273.2 \| 153 \| (1, 153) \| 423 \| (3, 155) \| 491 \| Phage tail protein \| Phage tail protein; uniclust \| UniRef100\_A0A747R6J3 \| 100.0 \| 1.4e-35 \| 2.7e-41 \| 255.4 \| 157 \| (2, 158) \| 423 \| (3, 159) \| 359 \| Phage tail protein \| Phage tail protein; uniclust \| UniRef100\_A0A0N9MHE6 \| 100.0 \| 1.5e-35 \| 3e-41 \| 290.9 \| 156 \| (2, 157) \| 423 \| (6, 161) \| 695 \| Phage tail fiber protein \| Phage tail fiber protein; uniclust \| UniRef100\_A0A379QSS5 \| 100.0 \| 2.2e-35 \| 4.1e-41 \| 262.4 \| 295 \| (3, 306) \| 423 \| (4, 313) \| 540 \| Tail protein \| Tail protein; uniclust \| UniRef100\_A0A1B9KN89 \| 100.0 \| 2.5e-35 \| 4.7e-41 \| 269.5 \| 156 \| (1, 156) \| 423 \| (2, 157) \| 619 \| Phage tail protein \| Phage tail protein; uniclust \| UniRef100\_A0A0Q8Y205 \| 100.0 \| 4.2e-35 \| 7.9e-41 \| 267.8 \| 158 \| (1, 158) \| 423 \| (2, 159) \| 505 \| Phage tail protein \| Phage tail protein; uniclust \| UniRef100\_A0A0H3F5Q0 \| 100.0 \| 6.9e-35 \| 1.3e-40 \| 277.6 \| 160 \| (1, 160) \| 423 \| (1, 160) \| 960 \| Tail fiber repeat 2 protein \| Tail fiber repeat 2 protein; uniclust \| UniRef100\_UPI00210164A2 \| 100.0 \| 9.3e-35 \| 1.7e-40 \| 239.9 \| 252 \| (2, 253) \| 423 \| (3, 261) \| 333 \| phage tail protein \| phage tail protein; uniclust \| UniRef100\_A0A248LIJ6 \| 100.0 \| 9.2e-35 \| 1.7e-40 \| 263.3 \| 154 \| (2, 155) \| 423 \| (3, 156) \| 521 \| Tail fiber protein 2 \| Tail fiber protein 2; uniclust \| UniRef100\_A0A5J2VSE0 \| 100.0 \| 9.5e-35 \| 1.8e-40 \| 266.5 \| 286 \| (2, 304) \| 423 \| (3, 293) \| 581 \| Phage tail protein \| Phage tail protein; uniclust \| UniRef100\_A0A0T9TRH4 \| 100.0 \| 9.2e-35 \| 1.8e-40 \| 278.7 \| 163 \| (2, 164) \| 423 \| (57, 219) \| 551 \| Tail fiber protein \| Tail fiber protein; uniclust \| UniRef100\_A0A1A0FJZ1 \| 100.0 \| 9.5e-35 \| 1.8e-40 \| 267.6 \| 158 \| (3, 160) \| 423 \| (7, 164) \| 418 \| Phage tail protein \| Phage tail protein; uniclust \| UniRef100\_A0A1G3JRW0 \| 100.0 \| 1.2e-34 \| 2.2e-40 \| 274.6 \| 240 \| (3, 246) \| 423 \| (6, 263) \| 629 \| Phage tail protein \| Phage tail protein; uniclust \| UniRef100\_A0A0W0MDX6 \| 100.0 \| 1.8e-34 \| 3.4e-40 \| 242.6 \| 160 \| (1, 160) \| 423 \| (1, 161) \| 312 \| Phage tail protein \| Phage tail protein; uniclust \| UniRef100\_A0A0H3NZ57 \| 100.0 \| 1.9e-34 \| 3.5e-40 \| 254.3 \| 157 \| (2, 158) \| 423 \| (3, 159) \| 514 \| Phage tail fiber protein \| Phage tail fiber protein; uniclust \| UniRef100\_A0A0T9KX86 \| 100.0 \| 2.4e-34 \| 4.4e-40 \| 268.1 \| 299 \| (2, 304) \| 423 \| (1, 338) \| 658 \| Tail fiber repeat 2-containing protein \| Tail fiber repeat 2-containing protein; uniclust \| UniRef100\_A0A1M4Y377 \| 100.0 \| 2.4e-34 \| 4.6e-40 \| 261.3 \| 160 \| (1, 160) \| 423 \| (1, 160) \| 542 \| Phage tail-collar fibre protein \| Phage tail-collar fibre protein; uniclust \| UniRef100\_A0A1Y2SIM3 \| 100.0 \| 3.1e-34 \| 5.6e-40 \| 258.9 \| 158 \| (3, 160) \| 423 \| (4, 161) \| 675 \| Tail protein \| Tail protein; uniclust \| UniRef100\_A0A0D1M6C4 \| 100.0 \| 4.5e-34 \| 8.6e-40 \| 272.2 \| 155 \| (2, 156) \| 423 \| (6, 160) \| 756 \| Phage tail collar domain-containing protein \| Phage tail collar domain-containing protein; uniclust \| UniRef100\_UPI000AC960B9 \| 100.0 \| 1.1e-33 \| 2.1e-39 \| 224.9 \| 156 \| (2, 157) \| 423 \| (3, 158) \| 244 \| phage tail protein \| phage tail protein; uniclust \| UniRef100\_A0A1C3H901 \| 100.0 \| 1.3e-33 \| 2.4e-39 \| 256.4 \| 161 \| (2, 162) \| 423 \| (3, 163) \| 655 \| Phage tail fiber repeat protein \| Phage tail fiber repeat protein; uniclust \| UniRef100\_A0A2C9P0H8 \| 100.0 \| 2.1e-33 \| 3.9e-39 \| 253.8 \| 157 \| (2, 158) \| 423 \| (5, 162) \| 581 \| Phage tail protein \| Phage tail protein; uniclust \| UniRef100\_A0A376DLB6 \| 100.0 \| 2.4e-33 \| 4.5e-39 \| 249.0 \| 156 \| (2, 157) \| 423 \| (3, 161) \| 430 \| Phage tail fibre repeat \| Phage tail fibre repeat; uniclust \| UniRef100\_A0A0H3NQY6 \| 100.0 \| 2.5e-33 \| 4.7e-39 \| 210.0 \| 131 \| (3, 133) \| 423 \| (7, 137) \| 139 \| Phage tail fiber protein \| Phage tail fiber protein; uniclust \| UniRef100\_A0A1I3XGI0 \| 100.0 \| 2.7e-33 \| 4.9e-39 \| 234.7 \| 152 \| (3, 154) \| 423 \| (4, 155) \| 372 \| Phage tail fibre repeat-containing protein (Fragment) \| Phage tail fibre repeat-containing protein (Fragment); uniclust \| UniRef100\_A0A2K9QFQ7 \| 100.0 \| 2.7e-33 \| 5.1e-39 \| 247.6 \| 168 \| (3, 170) \| 423 \| (4, 171) \| 474 \| Phage tail protein \| Phage tail protein; uniclust \| UniRef100\_A0A4P7L2V6 \| 100.0 \| 3.1e-33 \| 5.8e-39 \| 249.9 \| 154 \| (2, 155) \| 423 \| (1, 154) \| 508 \| Phage tail-collar fiber protein \| Phage tail-collar fiber protein; uniclust \| UniRef100\_A0A4R0FQS3 \| 100.0 \| 3.1e-33 \| 5.8e-39 \| 244.3 \| 157 \| (2, 160) \| 423 \| (3, 159) \| 399 \| Phage tail protein \| Phage tail protein; uniclust \| UniRef100\_A0A1H0YPL5 \| 100.0 \| 3.3e-33 \| 6.3e-39 \| 248.1 \| 165 \| (1, 165) \| 423 \| (3, 167) \| 405 \| Phage tail-collar fibre protein \| Phage tail-collar fibre protein; uniclust \| UniRef100\_A0A0A3AYU0 \| 100.0 \| 3.7e-33 \| 6.9e-39 \| 247.2 \| 150 \| (3, 152) \| 423 \| (4, 153) \| 466 \| Phage tail fiber protein \| Phage tail fiber protein; uniclust \| UniRef100\_A0A0A1HYZ6 \| 100.0 \| 3.9e-33 \| 7.2e-39 \| 251.2 \| 155 \| (1, 155) \| 423 \| (1, 156) \| 513 \| Prophage long tail fiber protein \| Prophage long tail fiber protein; uniclust \| UniRef100\_A0A143DGR5 \| 100.0 \| 3.8e-33 \| 7.4e-39 \| 257.5 \| 154 \| (1, 154) \| 423 \| (18, 171) \| 425 \| Phage tail collar domain-containing protein \| Phage tail collar domain-containing protein; uniclust \| UniRef100\_A0A0F7Y0J9 \| 100.0 \| 4.6e-33 \| 8.7e-39 \| 269.9 \| 155 \| (2, 156) \| 423 \| (6, 160) \| 1016 \| Phage-related tail fiber protein-like protein \| Phage-related tail fiber protein-like protein; uniclust \| UniRef100\_A0A6M0RCD0 \| 100.0 \| 4.9e-33 \| 9.4e-39 \| 234.5 \| 152 \| (2, 153) \| 423 \| (5, 156) \| 261 \| Phage tail protein (Fragment) \| Phage tail protein (Fragment); uniclust \| UniRef100\_A0A6D0UUI3 \| 100.0 \| 5.2e-33 \| 9.6e-39 \| 245.7 \| 157 \| (2, 158) \| 423 \| (3, 159) \| 576 \| Phage tail protein \| Phage tail protein; uniclust \| UniRef100\_UPI0002FB3713 \| 100.0 \| 6.3e-33 \| 1.2e-38 \| 224.9 \| 156 \| (2, 157) \| 423 \| (6, 161) \| 243 \| phage tail protein \| phage tail protein; uniclust \| UniRef100\_A0A376TTQ8 \| 100.0 \| 6.9e-33 \| 1.3e-38 \| 209.4 \| 155 \| (2, 156) \| 423 \| (3, 157) \| 168 \| Variable tail fiber protein \| Variable tail fiber protein; uniclust \| UniRef100\_A0A077NSD4 \| 100.0 \| 7.5e-33 \| 1.4e-38 \| 257.0 \| 158 \| (3, 160) \| 423 \| (4, 161) \| 922 \| Phage tail collar domain-containing protein \| Phage tail collar domain-containing protein; uniclust \| UniRef100\_A0A8T6A9G5 \| 100.0 \| 7.6e-33 \| 1.4e-38 \| 241.7 \| 158 \| (3, 160) \| 423 \| (7, 165) \| 478 \| Phage tail protein (Fragment) \| Phage tail protein (Fragment); uniclust \| UniRef100\_A0A077K818 \| 100.0 \| 7.6e-33 \| 1.4e-38 \| 255.3 \| 155 \| (3, 157) \| 423 \| (5, 159) \| 555 \| Phage tail collar domain-containing protein \| Phage tail collar domain-containing protein; uniclust \| UniRef100\_A0A1D3JVK3 \| 100.0 \| 8.6e-33 \| 1.6e-38 \| 251.0 \| 156 \| (2, 157) \| 423 \| (6, 161) \| 474 \| Phage tail protein \| Phage tail protein; uniclust \| UniRef100\_A0A2D0JYZ8 \| 100.0 \| 1e-32 \| 1.8e-38 \| 231.7 \| 204 \| (1, 301) \| 423 \| (1, 204) \| 376 \| Phage tail fiber protein \| Phage tail fiber protein; uniclust \| UniRef100\_A0A1N6MS83 \| 100.0 \| 1.2e-32 \| 2.3e-38 \| 241.3 \| 155 \| (1, 155) \| 423 \| (1, 155) \| 550 \| Uncharacterized protein \| Uncharacterized protein; uniclust \| UniRef100\_A0A4Y9K5F2 \| 100.0 \| 1.4e-32 \| 2.6e-38 \| 264.2 \| 152 \| (1, 152) \| 423 \| (3, 154) \| 667 \| Phage tail protein \| Phage tail protein; uniclust \| UniRef100\_UPI000A46B6C7 \| 100.0 \| 1.8e-32 \| 3.3e-38 \| 249.4 \| 159 \| (2, 160) \| 423 \| (6, 165) \| 653 \| phage tail protein \| phage tail protein; uniclust \| UniRef100\_A0A7S4LVA4 \| 100.0 \| 2.1e-32 \| 3.9e-38 \| 245.1 \| 160 \| (1, 160) \| 423 \| (1, 160) \| 587 \| Tail fiber protein \| Tail fiber protein; uniclust \| UniRef100\_A0A1B9JGB5 \| 100.0 \| 2.3e-32 \| 4.3e-38 \| 247.9 \| 254 \| (2, 300) \| 423 \| (3, 262) \| 665 \| Phage tail collar domain-containing protein \| Phage tail collar domain-containing protein; uniclust \| UniRef100\_UPI000445DE18 \| 100.0 \| 2.4e-32 \| 4.5e-38 \| 244.4 \| 159 \| (3, 161) \| 423 \| (7, 166) \| 482 \| tail fiber protein \| tail fiber protein; uniclust \| UniRef100\_A0A0A1AI95 \| 100.0 \| 2.5e-32 \| 4.7e-38 \| 257.0 \| 159 \| (1, 159) \| 423 \| (2, 161) \| 889 \| Phage tail protein \| Phage tail protein; uniclust \| UniRef100\_A0A0A7NV63 \| 99.9 \| 2.7e-32 \| 5.1e-38 \| 268.4 \| 159 \| (2, 160) \| 423 \| (295, 454) \| 1033 \| Tail fiber protein \| Tail fiber protein; uniclust \| UniRef100\_A0A101LTG1 \| 99.9 \| 3.3e-32 \| 6.2e-38 \| 250.0 \| 157 \| (3, 159) \| 423 \| (5, 162) \| 633 \| Phage tail protein \| Phage tail protein; uniclust \| UniRef100\_A0A0F3TBN3 \| 99.9 \| 3.3e-32 \| 6.2e-38 \| 264.0 \| 296 \| (4, 304) \| 423 \| (2, 336) \| 726 \| Phage tail protein \| Phage tail protein; uniclust \| UniRef100\_A0A0N9MHW1 \| 99.9 \| 4.1e-32 \| 7.7e-38 \| 271.0 \| 156 \| (2, 157) \| 423 \| (6, 161) \| 1312 \| Phage tail fiber protein \| Phage tail fiber protein; uniclust \| UniRef100\_A0A776QF13 \| 99.9 \| 5e-32 \| 9.3e-38 \| 231.5 \| 198 \| (2, 300) \| 423 \| (3, 201) \| 389 \| Phage tail protein (Fragment) \| Phage tail protein (Fragment); uniclust \| UniRef100\_A0A1N6I148 \| 99.9 \| 5.1e-32 \| 9.6e-38 \| 249.9 \| 164 \| (1, 164) \| 423 \| (2, 165) \| 519 \| Phage tail-collar fibre protein \| Phage tail-collar fibre protein; uniclust \| UniRef100\_A0A2U8I2Y4 \| 99.9 \| 5.5e-32 \| 1e-37 \| 235.4 \| 208 \| (3, 300) \| 423 \| (4, 211) \| 509 \| Phage tail protein \| Phage tail protein; uniclust \| UniRef100\_A0A0M1LJ12 \| 99.9 \| 5.5e-32 \| 1e-37 \| 229.7 \| 152 \| (3, 154) \| 423 \| (8, 159) \| 299 \| Phage tail protein \| Phage tail protein; uniclust \| UniRef100\_A0A858NPD6 \| 99.9 \| 6.1e-32 \| 1.1e-37 \| 248.3 \| 262 \| (1, 263) \| 423 \| (1, 262) \| 775 \| Tail fiber protein \| Tail fiber protein; uniclust \| UniRef100\_A0A398UJ84 \| 99.9 \| 6.5e-32 \| 1.2e-37 \| 242.4 \| 161 \| (1, 161) \| 423 \| (5, 166) \| 575 \| Phage tail protein (Fragment) \| Phage tail protein (Fragment); uniclust \| UniRef100\_UPI001BC85BAF \| 99.9 \| 8.4e-32 \| 1.6e-37 \| 246.4 \| 160 \| (1, 160) \| 423 \| (1, 160) \| 715 \| phage tail protein \| phage tail protein; uniclust \| UniRef100\_A0A3F3IWG0 \| 99.9 \| 1.1e-31 \| 2.1e-37 \| 245.0 \| 159 \| (2, 160) \| 423 \| (5, 164) \| 575 \| Bacteriophage T7 Gp17 C-terminal domain-containing protein \| Bacteriophage T7 Gp17 C-terminal domain-containing protein; uniclust \| UniRef100\_UPI001140E494 \| 99.9 \| 1.7e-31 \| 3.2e-37 \| 235.7 \| 160 \| (1, 160) \| 423 \| (1, 160) \| 550 \| phage tail protein \| phage tail protein; uniclust \| UniRef100\_G9Z5T4 \| 99.9 \| 1.9e-31 \| 3.6e-37 \| 259.1 \| 295 \| (4, 303) \| 423 \| (2, 335) \| 753 \| Phage tail collar domain protein \| Phage tail collar domain protein; uniclust \| UniRef100\_A0A0C1RAC0 \| 99.9 \| 2e-31 \| 3.9e-37 \| 247.4 \| 153 \| (2, 154) \| 423 \| (7, 159) \| 477 \| Phage tail-collar fiber family protein \| Phage tail-collar fiber family protein; uniclust \| UniRef100\_A0A1W1XL57 \| 99.9 \| 2.2e-31 \| 4e-37 \| 216.1 \| 157 \| (1, 157) \| 423 \| (2, 158) \| 234 \| Phage tail-collar fibre protein \| Phage tail-collar fibre protein; uniclust \| UniRef100\_A0A0H3FAF6 \| 99.9 \| 2.5e-31 \| 4.6e-37 \| 249.8 \| 160 \| (1, 160) \| 423 \| (1, 160) \| 795 \| Tail fiber repeat 2 protein \| Tail fiber repeat 2 protein; uniclust \| UniRef100\_A0A430HF82 \| 99.9 \| 2.5e-31 \| 4.6e-37 \| 214.7 \| 258 \| (2, 263) \| 423 \| (1, 258) \| 276 \| Phage tail protein \| Phage tail protein; uniclust \| UniRef100\_A0A068QZ58 \| 99.9 \| 2.8e-31 \| 5.3e-37 \| 256.5 \| 301 \| (2, 307) \| 423 \| (17, 356) \| 709 \| Bacteriophage T7 Gp17 C-terminal domain-containing protein \| Bacteriophage T7 Gp17 C-terminal domain-containing protein; uniclust \| UniRef100\_A0A5B8HYX9 \| 99.9 \| 2.9e-31 \| 5.3e-37 \| 236.3 \| 158 \| (3, 160) \| 423 \| (24, 181) \| 488 \| Phage tail protein \| Phage tail protein; uniclust \| UniRef100\_A0A077NM26 \| 99.9 \| 3e-31 \| 5.6e-37 \| 236.4 \| 154 \| (3, 156) \| 423 \| (4, 157) \| 641 \| Uncharacterized protein \| Uncharacterized protein; uniclust \| UniRef100\_UPI00098C1720 \| 99.9 \| 3.2e-31 \| 5.8e-37 \| 251.7 \| 159 \| (2, 160) \| 423 \| (3, 161) \| 1423 \| tail fiber protein \| tail fiber protein; uniclust \| UniRef100\_A0A1H5GVL6 \| 99.9 \| 3.1e-31 \| 5.8e-37 \| 238.0 \| 155 \| (2, 156) \| 423 \| (6, 160) \| 476 \| Phage-related tail fibre protein \| Phage-related tail fibre protein; uniclust \| UniRef100\_A0A2X1N770 \| 99.9 \| 3.3e-31 \| 6.1e-37 \| 245.6 \| 160 \| (1, 160) \| 423 \| (5, 165) \| 710 \| Tail fiber protein \| Tail fiber protein; uniclust \| UniRef100\_A0A0D4ZXS4 \| 99.9 \| 3.7e-31 \| 7e-37 \| 236.1 \| 163 \| (1, 163) \| 423 \| (17, 179) \| 374 \| Tail fiber protein \| Tail fiber protein; uniclust \| UniRef100\_UPI000AE7EBAD \| 99.9 \| 3.8e-31 \| 7.1e-37 \| 233.3 \| 155 \| (2, 156) \| 423 \| (6, 160) \| 434 \| phage tail protein \| phage tail protein; uniclust \| UniRef100\_A0A1H4Y3H3 \| 99.9 \| 3.8e-31 \| 7.1e-37 \| 253.1 \| 243 \| (1, 247) \| 423 \| (5, 264) \| 787 \| Phage Tail Collar Domain \| Phage Tail Collar Domain; uniclust \| UniRef100\_A0A433ZYK3 \| 99.9 \| 3.9e-31 \| 7.2e-37 \| 213.2 \| 156 \| (2, 157) \| 423 \| (3, 158) \| 249 \| Phage tail protein (Fragment) \| Phage tail protein (Fragment); uniclust \| UniRef100\_A0A1S6KZZ8 \| 99.9 \| 3.8e-31 \| 7.3e-37 \| 253.0 \| 169 \| (2, 170) \| 423 \| (51, 219) \| 738 \| Tail fiber protein \| Tail fiber protein; uniclust \| UniRef100\_UPI0005E045F7 \| 99.9 \| 4.3e-31 \| 7.8e-37 \| 222.5 \| 162 \| (2, 163) \| 423 \| (3, 164) \| 364 \| phage tail protein \| phage tail protein; uniclust \| UniRef100\_UPI001FC9489E \| 99.9 \| 5.4e-31 \| 9.9e-37 \| 212.1 \| 203 \| (1, 304) \| 423 \| (1, 203) \| 270 \| phage tail protein \| phage tail protein; uniclust \| UniRef100\_C5BEP6 \| 99.9 \| 5.8e-31 \| 1.1e-36 \| 224.7 \| 158 \| (2, 159) \| 423 \| (3, 160) \| 426 \| Phage tail fiber domain protein \| Phage tail fiber domain protein; uniclust \| UniRef100\_A0A024E8J4 \| 99.9 \| 6.1e-31 \| 1.2e-36 \| 245.4 \| 156 \| (2, 157) \| 423 \| (6, 161) \| 640 \| Tail fiber protein H, putative \| Tail fiber protein H, putative; uniclust \| UniRef100\_UPI000F950B8D \| 99.9 \| 6.5e-31 \| 1.2e-36 \| 228.0 \| 159 \| (1, 159) \| 423 \| (2, 160) \| 423 \| phage tail protein \| phage tail protein; uniclust \| UniRef100\_A0A484ZG34 \| 99.9 \| 1e-30 \| 1.8e-36 \| 215.3 \| 219 \| (2, 310) \| 423 \| (3, 221) \| 317 \| Phage tail fibre repeat \| Phage tail fibre repeat; uniclust \| UniRef100\_UPI000B30225B \| 99.9 \| 1e-30 \| 1.9e-36 \| 230.7 \| 158 \| (3, 160) \| 423 \| (4, 162) \| 575 \| tail fiber protein \| tail fiber protein; uniclust \| UniRef100\_A0A0Q8DAL0 \| 99.9 \| 1.1e-30 \| 2.1e-36 \| 239.3 \| 256 \| (2, 262) \| 423 \| (7, 279) \| 629 \| Phage tail protein \| Phage tail protein; uniclust \| UniRef100\_A0A836YZF5 \| 99.9 \| 1.2e-30 \| 2.2e-36 \| 229.1 \| 150 \| (3, 152) \| 423 \| (4, 153) \| 394 \| Tail fiber protein (Fragment) \| Tail fiber protein (Fragment); uniclust \| UniRef100\_A0A447PAU2 \| 99.9 \| 1.4e-30 \| 2.5e-36 \| 226.1 \| 170 \| (2, 171) \| 423 \| (3, 172) \| 446 \| Phage tail-like protein \| Phage tail-like protein; uniclust \| UniRef100\_A0A495BJ48 \| 99.9 \| 1.4e-30 \| 2.6e-36 \| 223.2 \| 155 \| (1, 155) \| 423 \| (2, 156) \| 380 \| Tail-collar fiber protein \| Tail-collar fiber protein; uniclust \| UniRef100\_A0A2X2BLJ3 \| 99.9 \| 1.6e-30 \| 3e-36 \| 212.0 \| 155 \| (2, 156) \| 423 \| (3, 157) \| 277 \| Phage variable tail fiber protein \| Phage variable tail fiber protein; uniclust \| UniRef100\_UPI001CE1390D \| 99.9 \| 2e-30 \| 3.6e-36 \| 241.8 \| 170 \| (2, 171) \| 423 \| (3, 172) \| 852 \| phage tail protein \| phage tail protein; uniclust \| UniRef100\_A0A3Z6GBS8 \| 99.9 \| 1.9e-30 \| 3.6e-36 \| 233.7 \| 159 \| (2, 160) \| 423 \| (5, 164) \| 563 \| Phage tail protein \| Phage tail protein; uniclust \| UniRef100\_UPI001FFE7AE7 \| 99.9 \| 2.3e-30 \| 4.2e-36 \| 216.7 \| 156 \| (3, 158) \| 423 \| (4, 159) \| 359 \| phage tail protein \| phage tail protein; uniclust \| UniRef100\_A0A1B9JGD4 \| 99.9 \| 2.4e-30 \| 4.5e-36 \| 221.5 \| 156 \| (3, 158) \| 423 \| (4, 159) \| 381 \| Phage tail collar domain-containing protein \| Phage tail collar domain-containing protein; uniclust \| UniRef100\_UPI0018A29913 \| 99.9 \| 2.8e-30 \| 5.2e-36 \| 223.8 \| 172 \| (2, 173) \| 423 \| (3, 174) \| 430 \| phage tail protein \| phage tail protein; uniclust \| UniRef100\_A0A072Y2M2 \| 99.9 \| 3e-30 \| 5.7e-36 \| 233.2 \| 153 \| (1, 153) \| 423 \| (3, 155) \| 441 \| Putative phage tail fiber protein \| Putative phage tail fiber protein; uniclust \| UniRef100\_A0A0D6BCT9 \| 99.9 \| 3.9e-30 \| 7.3e-36 \| 248.3 \| 243 \| (1, 247) \| 423 \| (5, 265) \| 1040 \| Phage-related tail fibre protein-like protein \| Phage-related tail fibre protein-like protein; uniclust \| UniRef100\_A0A2V1HJ68 \| 99.9 \| 3.9e-30 \| 7.3e-36 \| 229.5 \| 368 \| (4, 375) \| 423 \| (2, 484) \| 526 \| Phage tail protein \| Phage tail protein; uniclust \| UniRef100\_UPI00224B8CEA \| 99.9 \| 4.3e-30 \| 7.8e-36 \| 219.1 \| 160 \| (1, 160) \| 423 \| (1, 160) \| 419 \| phage tail protein \| phage tail protein; uniclust \| UniRef100\_A0A827QGW3 \| 99.9 \| 5.7e-30 \| 1.1e-35 \| 223.8 \| 158 \| (3, 160) \| 423 \| (4, 161) \| 364 \| Phage tail protein (Fragment) \| Phage tail protein (Fragment); uniclust \| UniRef100\_A0A0F4TBD1 \| 99.9 \| 6.1e-30 \| 1.1e-35 \| 235.5 \| 156 \| (2, 157) \| 423 \| (6, 161) \| 585 \| Phage tail protein \| Phage tail protein; uniclust \| UniRef100\_A0A8E1QKA4 \| 99.9 \| 6.6e-30 \| 1.2e-35 \| 210.9 \| 152 \| (2, 153) \| 423 \| (3, 154) \| 255 \| Tail fiber protein (Fragment) \| Tail fiber protein (Fragment); uniclust \| UniRef100\_A0A853HVL0 \| 99.9 \| 7.1e-30 \| 1.3e-35 \| 214.8 \| 163 \| (3, 165) \| 423 \| (4, 169) \| 332 \| Phage tail protein \| Phage tail protein; uniclust \| UniRef100\_UPI001AD852BE \| 99.9 \| 7.6e-30 \| 1.4e-35 \| 220.7 \| 160 \| (1, 160) \| 423 \| (1, 160) \| 474 \| phage tail protein \| phage tail protein; uniclust \| UniRef100\_A0A5U3F244 \| 99.9 \| 7.8e-30 \| 1.5e-35 \| 227.4 \| 319 \| (4, 333) \| 423 \| (2, 359) \| 431 \| Phage tail protein (Fragment) \| Phage tail protein (Fragment); uniclust \| UniRef100\_A0A5T5IEY5 \| 99.9 \| 8.3e-30 \| 1.6e-35 \| 223.5 \| 157 \| (2, 158) \| 423 \| (5, 162) \| 401 \| Phage tail protein (Fragment) \| Phage tail protein (Fragment); uniclust \| UniRef100\_A0A0S4I2P4 \| 99.9 \| 9e-30 \| 1.7e-35 \| 207.2 \| 157 \| (2, 158) \| 423 \| (6, 162) \| 255 \| Uncharacterized protein \| Uncharacterized protein; uniclust \| UniRef100\_UPI00041B59CC \| 99.9 \| 1e-29 \| 1.9e-35 \| 222.7 \| 170 \| (2, 171) \| 423 \| (3, 172) \| 474 \| phage tail protein \| phage tail protein; uniclust \| UniRef100\_A0A7X1Y572 \| 99.9 \| 1e-29 \| 1.9e-35 \| 231.6 \| 156 \| (2, 157) \| 423 \| (6, 161) \| 592 \| Phage tail protein \| Phage tail protein; uniclust \| UniRef100\_A0A2M8SLT6 \| 99.9 \| 1.3e-29 \| 2.3e-35 \| 220.3 \| 152 \| (2, 153) \| 423 \| (6, 157) \| 409 \| Phage tail protein \| Phage tail protein; uniclust \| UniRef100\_A0A0F4VR40 \| 99.9 \| 1.3e-29 \| 2.5e-35 \| 236.2 \| 221 \| (2, 230) \| 423 \| (7, 231) \| 497 \| Phage tail fiber protein \| Phage tail fiber protein; uniclust \| UniRef100\_A0A3S5ENC7 \| 99.9 \| 1.5e-29 \| 2.7e-35 \| 216.1 \| 158 \| (3, 160) \| 423 \| (4, 161) \| 383 \| Putative bacteriophage tail fiber protein \| Putative bacteriophage tail fiber protein; uniclust \| UniRef100\_UPI001E2832F3 \| 99.9 \| 1.6e-29 \| 2.9e-35 \| 216.0 \| 155 \| (2, 156) \| 423 \| (1, 155) \| 423 \| phage tail protein \| phage tail protein; uniclust \| UniRef100\_A0A1C6YVI6 \| 99.9 \| 1.9e-29 \| 3.5e-35 \| 236.7 \| 156 \| (1, 160) \| 423 \| (32, 190) \| 595 \| Phage-related tail fibre protein \| Phage-related tail fibre protein; uniclust \| UniRef100\_A0A084ZPT2 \| 99.9 \| 2.1e-29 \| 3.8e-35 \| 214.6 \| 155 \| (1, 155) \| 423 \| (2, 156) \| 330 \| Phage tail fiber protein \| Phage tail fiber protein; uniclust \| UniRef100\_UPI001EE069FF \| 99.9 \| 2.4e-29 \| 4.4e-35 \| 215.9 \| 158 \| (3, 160) \| 423 \| (4, 161) \| 441 \| phage tail protein \| phage tail protein; uniclust \| UniRef100\_UPI001D141119 \| 99.9 \| 2.8e-29 \| 5.2e-35 \| 211.6 \| 158 \| (2, 159) \| 423 \| (3, 160) \| 330 \| phage tail protein \| phage tail protein; uniclust \| UniRef100\_A0A4Q7BEF2 \| 99.9 \| 3.1e-29 \| 5.8e-35 \| 195.5 \| 158 \| (1, 158) \| 423 \| (2, 159) \| 186 \| Phage tail protein (Fragment) \| Phage tail protein (Fragment); uniclust \| UniRef100\_A0A1G4ZBJ6 \| 99.9 \| 3.1e-29 \| 6e-35 \| 244.6 \| 250 \| (1, 304) \| 423 \| (2, 253) \| 728 \| Phage T4 tail fibre \| Phage T4 tail fibre; uniclust \| UniRef100\_UPI001C266F20 \| 99.9 \| 3.4e-29 \| 6.2e-35 \| 216.5 \| 160 \| (1, 160) \| 423 \| (1, 160) \| 467 \| phage tail protein \| phage tail protein; uniclust \| UniRef100\_A0A1H0NP67 \| 99.9 \| 3.7e-29 \| 6.9e-35 \| 216.6 \| 154 \| (4, 157) \| 423 \| (5, 158) \| 388 \| Phage tail-collar fibre protein \| Phage tail-collar fibre protein; uniclust \| UniRef100\_UPI0009B80218 \| 99.9 \| 4.1e-29 \| 7.5e-35 \| 211.0 \| 167 \| (2, 168) \| 423 \| (3, 169) \| 382 \| phage tail protein \| phage tail protein; uniclust \| UniRef100\_A0A7V8F402 \| 99.9 \| 4.2e-29 \| 7.8e-35 \| 219.1 \| 155 \| (3, 157) \| 423 \| (4, 158) \| 479 \| Phage tail protein \| Phage tail protein; uniclust \| UniRef100\_A0A069A3R7 \| 99.9 \| 4.6e-29 \| 8.7e-35 \| 212.6 \| 152 \| (3, 154) \| 423 \| (8, 159) \| 301 \| Putative phage tail fiber protein \| Putative phage tail fiber protein; uniclust \| UniRef100\_UPI001476555E \| 99.9 \| 5e-29 \| 9.3e-35 \| 208.5 \| 157 \| (2, 158) \| 423 \| (6, 162) \| 297 \| phage tail protein \| phage tail protein; uniclust \| UniRef100\_A0A381C7I1 \| 99.9 \| 5.2e-29 \| 1e-34 \| 233.8 \| 149 \| (1, 149) \| 423 \| (2, 150) \| 504 \| Phage tail fibre repeat \| Phage tail fibre repeat; uniclust \| UniRef100\_A0A3M3BF19 \| 99.9 \| 5.5e-29 \| 1e-34 \| 181.2 \| 116 \| (2, 117) \| 423 \| (6, 121) \| 122 \| Tail fiber protein H (Fragment) \| Tail fiber protein H (Fragment); uniclust \| UniRef100\_UPI00201F34D1 \| 99.9 \| 5.8e-29 \| 1.1e-34 \| 221.7 \| 159 \| (2, 160) \| 423 \| (3, 162) \| 624 \| phage tail protein \| phage tail protein; uniclust \| UniRef100\_UPI000A9C88D5 \| 99.9 \| 6e-29 \| 1.1e-34 \| 223.6 \| 159 \| (2, 160) \| 423 \| (3, 161) \| 629 \| phage tail protein \| phage tail protein; uniclust \| UniRef100\_A0A1H7YIZ1 \| 99.9 \| 6.4e-29 \| 1.2e-34 \| 222.4 \| 157 \| (1, 157) \| 423 \| (2, 158) \| 404 \| Phage tail-collar fibre protein \| Phage tail-collar fibre protein; uniclust \| UniRef100\_A0A381I8S9 \| 99.9 \| 7.5e-29 \| 1.4e-34 \| 205.9 \| 154 \| (3, 156) \| 423 \| (8, 161) \| 266 \| Phage tail fiber protein \| Phage tail fiber protein; uniclust \| UniRef100\_UPI0009F41D80 \| 99.9 \| 7.5e-29 \| 1.4e-34 \| 231.1 \| 156 \| (2, 157) \| 423 \| (6, 161) \| 607 \| phage tail protein \| phage tail protein; uniclust \| UniRef100\_UPI000516E280 \| 99.9 \| 8.9e-29 \| 1.6e-34 \| 205.0 \| 155 \| (2, 156) \| 423 \| (6, 160) \| 308 \| phage tail protein \| phage tail protein; uniclust \| UniRef100\_A0A1I1UBJ2 \| 99.9 \| 9.3e-29 \| 1.7e-34 \| 212.8 \| 154 \| (1, 154) \| 423 \| (1, 154) \| 353 \| Phage tail-collar fibre protein \| Phage tail-collar fibre protein; uniclust \| UniRef100\_A0A0B5QHJ2 \| 99.9 \| 1e-28 \| 1.9e-34 \| 205.3 \| 150 \| (3, 152) \| 423 \| (14, 163) \| 280 \| Phage tail protein \| Phage tail protein; uniclust \| UniRef100\_UPI000DFE3857 \| 99.9 \| 1.2e-28 \| 2.1e-34 \| 183.7 \| 129 \| (2, 130) \| 423 \| (3, 131) \| 153 \| phage tail protein \| phage tail protein; uniclust \| UniRef100\_A0A2M8YKL8 \| 99.9 \| 1.2e-28 \| 2.3e-34 \| 201.6 \| 154 \| (1, 154) \| 423 \| (1, 154) \| 298 \| Tail-collar fiber protein \| Tail-collar fiber protein; uniclust \| UniRef100\_A0A080K6T3 \| 99.9 \| 1.3e-28 \| 2.3e-34 \| 180.6 \| 123 \| (2, 124) \| 423 \| (3, 125) \| 127 \| Tail fiber protein (Fragment) \| Tail fiber protein (Fragment); uniclust \| UniRef100\_UPI0009B7EA04 \| 99.9 \| 1.3e-28 \| 2.4e-34 \| 227.4 \| 163 \| (2, 164) \| 423 \| (3, 165) \| 813 \| phage tail protein \| phage tail protein; uniclust \| UniRef100\_UPI000B7CBA84 \| 99.9 \| 1.7e-28 \| 3e-34 \| 221.3 \| 157 \| (3, 159) \| 423 \| (7, 164) \| 700 \| tail fiber protein \| tail fiber protein; uniclust \| UniRef100\_UPI0012EBC142 \| 99.9 \| 1.9e-28 \| 3.4e-34 \| 213.8 \| 259 \| (4, 263) \| 423 \| (5, 263) \| 503 \| phage tail protein \| phage tail protein; uniclust \| UniRef100\_A0A0N0E1T2 \| 99.9 \| 1.8e-28 \| 3.4e-34 \| 241.3 \| 155 \| (1, 155) \| 423 \| (9, 163) \| 809 \| Phage-related tail fiber protein \| Phage-related tail fiber protein; uniclust \| UniRef100\_A0A0L8IPP7 \| 99.9 \| 2.6e-28 \| 4.8e-34 \| 223.9 \| 157 \| (1, 157) \| 423 \| (5, 161) \| 688 \| Tail fiber protein H \| Tail fiber protein H; uniclust \| UniRef100\_UPI0002E0FE29 \| 99.9 \| 2.8e-28 \| 5.1e-34 \| 207.1 \| 157 \| (2, 158) \| 423 \| (6, 162) \| 353 \| phage tail protein \| phage tail protein; uniclust \| UniRef100\_UPI0019693137 \| 99.9 \| 3e-28 \| 5.5e-34 \| 213.9 \| 165 \| (3, 167) \| 423 \| (4, 168) \| 535 \| phage tail protein \| phage tail protein; uniclust \| UniRef100\_UPI0005DFBD90 \| 99.9 \| 3.3e-28 \| 6.1e-34 \| 207.8 \| 154 \| (3, 156) \| 423 \| (4, 157) \| 417 \| phage tail protein I \| phage tail protein I; uniclust \| UniRef100\_UPI0002725035 \| 99.9 \| 3.3e-28 \| 6.1e-34 \| 206.7 \| 157 \| (2, 158) \| 423 \| (6, 162) \| 365 \| phage tail protein \| phage tail protein; uniclust \| UniRef100\_UPI001EFE032C \| 99.9 \| 3.3e-28 \| 6.2e-34 \| 216.5 \| 155 \| (2, 156) \| 423 \| (3, 157) \| 520 \| phage tail protein \| phage tail protein; uniclust \| UniRef100\_UPI001EFD6104 \| 99.9 \| 3.4e-28 \| 6.2e-34 \| 197.3 \| 155 \| (2, 156) \| 423 \| (3, 157) \| 278 \| phage tail protein \| phage tail protein; uniclust \| UniRef100\_UPI000C9DD6CC \| 99.9 \| 3.4e-28 \| 6.3e-34 \| 215.4 \| 184 \| (3, 186) \| 423 \| (4, 187) \| 582 \| phage tail protein \| phage tail protein; uniclust \| UniRef100\_A0A077QIY4 \| 99.9 \| 3.8e-28 \| 7.3e-34 \| 229.8 \| 295 \| (3, 303) \| 423 \| (1, 333) \| 574 \| Tail fiber protein \| Tail fiber protein; uniclust \| UniRef100\_A0A7Y8UX39 \| 99.9 \| 4.2e-28 \| 7.9e-34 \| 222.0 \| 155 \| (2, 156) \| 423 \| (6, 160) \| 619 \| Phage tail protein \| Phage tail protein; uniclust \| UniRef100\_A0A516S9W3 \| 99.9 \| 4.6e-28 \| 8.5e-34 \| 189.7 \| 183 \| (3, 185) \| 423 \| (6, 189) \| 204 \| Phage tail protein \| Phage tail protein; uniclust \| UniRef100\_UPI0022643D07 \| 99.9 \| 4.7e-28 \| 8.6e-34 \| 199.2 \| 205 \| (2, 307) \| 423 \| (5, 210) \| 308 \| phage tail protein \| phage tail protein; uniclust \| UniRef100\_A0A1N6Q2W5 \| 99.9 \| 4.6e-28 \| 8.6e-34 \| 190.8 \| 184 \| (2, 185) \| 423 \| (1, 185) \| 205 \| Phage tail-collar fibre protein \| Phage tail-collar fibre protein; uniclust \| UniRef100\_A0A3M5GCU5 \| 99.9 \| 5.7e-28 \| 1.1e-33 \| 188.2 \| 155 \| (2, 156) \| 423 \| (6, 160) \| 196 \| Phage-related tail fiber protein-like protein \| Phage-related tail fiber protein-like protein; uniclust \| UniRef100\_C3X1Y2 \| 99.9 \| 6.2e-28 \| 1.1e-33 \| 209.5 \| 183 \| (3, 185) \| 423 \| (4, 186) \| 480 \| Phage tail collar domain-containing protein \| Phage tail collar domain-containing protein; uniclust \| UniRef100\_A0A7X2MJW4 \| 99.9 \| 6.4e-28 \| 1.2e-33 \| 220.6 \| 299 \| (4, 307) \| 423 \| (2, 339) \| 635 \| Phage tail protein (Fragment) \| Phage tail protein (Fragment); uniclust \| UniRef100\_C6DCN9 \| 99.9 \| 6.5e-28 \| 1.2e-33 \| 217.5 \| 164 \| (2, 165) \| 423 \| (3, 166) \| 697 \| Phage-related tail fibre protein-like protein \| Phage-related tail fibre protein-like protein; uniclust \| UniRef100\_UPI001FF61CC4 \| 99.9 \| 7.7e-28 \| 1.4e-33 \| 211.2 \| 160 \| (1, 160) \| 423 \| (1, 160) \| 530 \| phage tail protein \| phage tail protein; uniclust \| UniRef100\_UPI0016768C5C \| 99.9 \| 7.8e-28 \| 1.4e-33 \| 212.1 \| 161 \| (1, 161) \| 423 \| (2, 164) \| 465 \| phage tail protein \| phage tail protein; uniclust \| UniRef100\_A0A1M7U2L6 \| 99.9 \| 8.5e-28 \| 1.6e-33 \| 205.3 \| 157 \| (3, 159) \| 423 \| (4, 160) \| 360 \| Phage tail-collar fibre protein \| Phage tail-collar fibre protein; uniclust \| UniRef100\_UPI0009BDDDCE \| 99.9 \| 1e-27 \| 1.9e-33 \| 215.6 \| 156 \| (2, 157) \| 423 \| (6, 161) \| 522 \| phage tail protein \| phage tail protein; uniclust \| UniRef100\_A0A077NG27 \| 99.9 \| 1e-27 \| 2e-33 \| 222.1 \| 140 \| (1, 145) \| 423 \| (1, 140) \| 457 \| Putative phage tail fiber protein \| Putative phage tail fiber protein; uniclust \| UniRef100\_UPI002017ADC9 \| 99.9 \| 1.2e-27 \| 2.2e-33 \| 219.4 \| 170 \| (2, 171) \| 423 \| (3, 172) \| 792 \| phage tail protein \| phage tail protein; uniclust \| UniRef100\_A0A0B6XER1 \| 99.9 \| 1.2e-27 \| 2.3e-33 \| 223.5 \| 153 \| (3, 160) \| 423 \| (4, 156) \| 738 \| Putative phage variable tail fiber protein \| Putative phage variable tail fiber protein; uniclust \| UniRef100\_A0A8S5N9Y2 \| 99.9 \| 1.3e-27 \| 2.3e-33 \| 226.3 \| 202 \| (1, 303) \| 423 \| (2, 209) \| 990 \| Baseplate wedge protein \| Baseplate wedge protein; uniclust \| UniRef100\_A0A4D6XN95 \| 99.9 \| 1.6e-27 \| 2.9e-33 \| 204.4 \| 154 \| (2, 155) \| 423 \| (6, 165) \| 428 \| Phage tail protein \| Phage tail protein; uniclust \| UniRef100\_A0A137SPV5 \| 99.9 \| 1.7e-27 \| 3.1e-33 \| 205.0 \| 164 \| (2, 169) \| 423 \| (7, 170) \| 294 \| Tail fiber protein \| Tail fiber protein; uniclust \| UniRef100\_A0A0N0FU17 \| 99.9 \| 1.8e-27 \| 3.3e-33 \| 211.8 \| 154 \| (2, 155) \| 423 \| (6, 159) \| 540 \| Tail fiber protein H \| Tail fiber protein H; uniclust \| UniRef100\_UPI000942066E \| 99.9 \| 1.8e-27 \| 3.3e-33 \| 209.8 \| 160 \| (2, 161) \| 423 \| (3, 162) \| 551 \| phage tail protein \| phage tail protein; uniclust \| UniRef100\_UPI0003EF618D \| 99.9 \| 1.8e-27 \| 3.5e-33 \| 205.0 \| 112 \| (60, 171) \| 423 \| (22, 133) \| 284 \| phage tail protein \| phage tail protein; uniclust \| UniRef100\_A0A077N2I9 \| 99.9 \| 1.9e-27 \| 3.6e-33 \| 234.0 \| 153 \| (3, 160) \| 423 \| (4, 156) \| 1303 \| Tail protein \| Tail protein; uniclust \| UniRef100\_UPI0003474E6B \| 99.9 \| 2.1e-27 \| 3.9e-33 \| 204.4 \| 158 \| (1, 158) \| 423 \| (1, 158) \| 441 \| phage tail protein \| phage tail protein; uniclust \| UniRef100\_A0A0A7RTP0 \| 99.9 \| 2.1e-27 \| 3.9e-33 \| 217.5 \| 152 \| (2, 153) \| 423 \| (3, 154) \| 603 \| Tail fiber protein \| Tail fiber protein; uniclust \| UniRef100\_A0A133PS36 \| 99.9 \| 2.2e-27 \| 4e-33 \| 204.5 \| 156 \| (1, 156) \| 423 \| (2, 157) \| 330 \| Uncharacterized protein \| Uncharacterized protein; uniclust \| UniRef100\_M1MEX9 \| 99.9 \| 2.2e-27 \| 4.2e-33 \| 216.4 \| 215 \| (2, 230) \| 423 \| (3, 217) \| 536 \| Putative phage tail fiber protein \| Putative phage tail fiber protein; uniclust \| UniRef100\_A0A1I0EJH2 \| 99.9 \| 2.4e-27 \| 4.4e-33 \| 190.3 \| 158 \| (2, 159) \| 423 \| (4, 161) \| 214 \| Phage tail-collar fibre protein \| Phage tail-collar fibre protein; uniclust \| UniRef100\_G7LSU4 \| 99.9 \| 2.5e-27 \| 4.5e-33 \| 213.2 \| 157 \| (1, 157) \| 423 \| (1, 157) \| 673 \| Uncharacterized protein \| Uncharacterized protein; uniclust \| UniRef100\_UPI001F5BB0E8 \| 99.9 \| 2.7e-27 \| 5e-33 \| 193.5 \| 155 \| (3, 157) \| 423 \| (4, 158) \| 292 \| phage tail protein \| phage tail protein; uniclust \| UniRef100\_A0A066RQJ3 \| 99.9 \| 2.7e-27 \| 5.1e-33 \| 218.3 \| 159 \| (1, 159) \| 423 \| (8, 166) \| 592 \| Tail fiber protein \| Tail fiber protein; uniclust \| UniRef100\_A0A351NNZ8 \| 99.9 \| 3.1e-27 \| 5.6e-33 \| 215.4 \| 169 \| (2, 170) \| 423 \| (3, 171) \| 775 \| Phage tail protein \| Phage tail protein; uniclust \| UniRef100\_UPI000F0649BD \| 99.9 \| 3.4e-27 \| 6.3e-33 \| 208.3 \| 157 \| (2, 158) \| 423 \| (6, 162) \| 481 \| phage tail protein \| phage tail protein; uniclust \| UniRef100\_A0A941I8B6 \| 99.9 \| 4e-27 \| 7.4e-33 \| 222.1 \| 153 \| (3, 155) \| 423 \| (4, 156) \| 914 \| Phage tail protein \| Phage tail protein; uniclust \| UniRef100\_A0A485CDL0 \| 99.9 \| 4.2e-27 \| 7.8e-33 \| 198.1 \| 157 \| (2, 158) \| 423 \| (4, 160) \| 282 \| Phage tail fibre repeat \| Phage tail fibre repeat; uniclust \| UniRef100\_UPI002280C8C9 \| 99.9 \| 4.5e-27 \| 8.2e-33 \| 204.1 \| 155 \| (3, 157) \| 423 \| (4, 158) \| 475 \| phage tail protein \| phage tail protein; uniclust \| UniRef100\_A0A1A9KIH8 \| 99.9 \| 4.5e-27 \| 8.3e-33 \| 217.4 \| 154 \| (1, 154) \| 423 \| (1, 154) \| 804 \| SGNH hydrolase-type esterase domain-containing protein \| SGNH hydrolase-type esterase domain-containing protein; uniclust \| UniRef100\_UPI000A85364E \| 99.9 \| 6.2e-27 \| 1.1e-32 \| 212.2 \| 260 \| (3, 263) \| 423 \| (4, 263) \| 725 \| phage tail protein \| phage tail protein; uniclust \| UniRef100\_A0A2T5HH07 \| 99.9 \| 6.6e-27 \| 1.2e-32 \| 194.5 \| 159 \| (2, 160) \| 423 \| (3, 163) \| 253 \| Tail-collar fiber protein \| Tail-collar fiber protein; uniclust \| UniRef100\_A0A1I6E7W8 \| 99.9 \| 6.7e-27 \| 1.2e-32 \| 202.8 \| 157 \| (2, 158) \| 423 \| (3, 159) \| 469 \| Phage-related tail fibre protein \| Phage-related tail fibre protein; uniclust \| UniRef100\_UPI000D348A11 \| 99.9 \| 6.7e-27 \| 1.2e-32 \| 201.2 \| 156 \| (1, 156) \| 423 \| (1, 156) \| 346 \| phage tail protein \| phage tail protein; uniclust \| UniRef100\_UPI000BE5AFC7 \| 99.9 \| 6.5e-27 \| 1.2e-32 \| 207.5 \| 247 \| (92, 374) \| 423 \| (17, 264) \| 310 \| phage tail protein \| phage tail protein; uniclust \| UniRef100\_A0A3M4NA57 \| 99.9 \| 8.2e-27 \| 1.5e-32 \| 203.0 \| 157 \| (2, 158) \| 423 \| (6, 162) \| 433 \| Tail fiber protein H (Fragment) \| Tail fiber protein H (Fragment); uniclust \| UniRef100\_UPI002162F542 \| 99.9 \| 8.6e-27 \| 1.6e-32 \| 211.9 \| 156 \| (1, 156) \| 423 \| (1, 156) \| 748 \| phage tail protein \| phage tail protein; uniclust \| UniRef100\_A0A0S4SND2 \| 99.9 \| 8.5e-27 \| 1.6e-32 \| 202.1 \| 154 \| (1, 157) \| 423 \| (2, 155) \| 309 \| Tail fiber protein H \| Tail fiber protein H; uniclust \| UniRef100\_A0A1S7SGS6 \| 99.9 \| 9.3e-27 \| 1.8e-32 \| 224.0 \| 153 \| (3, 155) \| 423 \| (19, 171) \| 585 \| DUF2793 domain-containing protein \| DUF2793 domain-containing protein; uniclust \| UniRef100\_UPI000DF33EE5 \| 99.9 \| 1.3e-26 \| 2.3e-32 \| 204.2 \| 160 \| (1, 160) \| 423 \| (2, 164) \| 539 \| phage tail protein \| phage tail protein; uniclust \| UniRef100\_A0A0C2RHK4 \| 99.9 \| 1.3e-26 \| 2.5e-32 \| 225.5 \| 157 \| (1, 157) \| 423 \| (2, 159) \| 926 \| Tail protein \| Tail protein; uniclust \| UniRef100\_UPI00168161AD \| 99.9 \| 1.5e-26 \| 2.7e-32 \| 183.3 \| 154 \| (3, 156) \| 423 \| (4, 157) \| 222 \| phage tail protein \| phage tail protein; uniclust \| UniRef100\_A0A0S1B3R8 \| 99.9 \| 1.5e-26 \| 2.9e-32 \| 177.9 \| 157 \| (1, 157) \| 423 \| (1, 158) \| 180 \| Side tail phage protein \| Side tail phage protein; uniclust \| UniRef100\_UPI0013B467C1 \| 99.9 \| 1.6e-26 \| 2.9e-32 \| 200.0 \| 155 \| (2, 156) \| 423 \| (3, 158) \| 434 \| phage tail protein \| phage tail protein; uniclust \| UniRef100\_A0A0E1PUY8 \| 99.9 \| 1.8e-26 \| 3.4e-32 \| 233.2 \| 159 \| (1, 161) \| 423 \| (2, 160) \| 966 \| Phage tail fiber protein \| Phage tail fiber protein; uniclust \| UniRef100\_A0A1H9AFH7 \| 99.9 \| 1.9e-26 \| 3.4e-32 \| 206.6 \| 156 \| (2, 157) \| 423 \| (6, 161) \| 544 \| Phage-related tail fibre protein \| Phage-related tail fibre protein; uniclust \| UniRef100\_UPI0009AC2AD7 \| 99.9 \| 1.9e-26 \| 3.5e-32 \| 215.1 \| 144 \| (1, 148) \| 423 \| (2, 145) \| 583 \| phage tail protein \| phage tail protein; uniclust \| UniRef100\_A0A318T0X1 \| 99.9 \| 1.9e-26 \| 3.7e-32 \| 220.5 \| 154 \| (2, 155) \| 423 \| (7, 160) \| 635 \| Uncharacterized protein DUF2793 \| Uncharacterized protein DUF2793; uniclust \| UniRef100\_A0A2T5NR62 \| 99.9 \| 2e-26 \| 3.8e-32 \| 208.5 \| 156 \| (3, 158) \| 423 \| (4, 160) \| 515 \| Phage tail protein \| Phage tail protein; uniclust \| UniRef100\_UPI00025CA96A \| 99.9 \| 2.1e-26 \| 4e-32 \| 214.8 \| 157 \| (3, 159) \| 423 \| (7, 164) \| 713 \| tail fiber protein \| tail fiber protein; uniclust \| UniRef100\_A0A285Z1M8 \| 99.9 \| 2.5e-26 \| 4.6e-32 \| 202.4 \| 156 \| (1, 156) \| 423 \| (1, 156) \| 417 \| Phage-related tail fibre protein \| Phage-related tail fibre protein; uniclust \| UniRef100\_A0A348HI80 \| 99.9 \| 2.5e-26 \| 4.6e-32 \| 194.8 \| 155 \| (1, 155) \| 423 \| (1, 155) \| 385 \| Phage-related tail fibre \| Phage-related tail fibre; uniclust \| UniRef100\_UPI001CC4633C \| 99.9 \| 2.6e-26 \| 4.8e-32 \| 210.4 \| 157 \| (3, 159) \| 423 \| (4, 160) \| 814 \| phage tail protein \| phage tail protein; uniclust \| UniRef100\_M8EFI1 \| 99.9 \| 2.6e-26 \| 4.8e-32 \| 202.5 \| 152 \| (3, 154) \| 423 \| (4, 155) \| 454 \| Phage-related tail fiber protein-like protein \| Phage-related tail fiber protein-like protein; uniclust \| UniRef100\_UPI00068A8B5B \| 99.9 \| 2.6e-26 \| 4.8e-32 \| 188.6 \| 152 \| (3, 154) \| 423 \| (4, 155) \| 266 \| phage tail protein \| phage tail protein; uniclust \| UniRef100\_A0A411WIN2 \| 99.9 \| 2.6e-26 \| 4.9e-32 \| 201.8 \| 145 \| (3, 153) \| 423 \| (4, 152) \| 396 \| Phage tail protein \| Phage tail protein; uniclust \| UniRef100\_A0A7V7KJT5 \| 99.9 \| 3.6e-26 \| 6.5e-32 \| 202.0 \| 154 \| (1, 154) \| 423 \| (1, 154) \| 553 \| Sialidase domain-containing protein \| Sialidase domain-containing protein; uniclust \| UniRef100\_UPI0005F91E19 \| 99.9 \| 3.6e-26 \| 6.6e-32 \| 201.5 \| 142 \| (2, 147) \| 423 \| (3, 144) \| 431 \| phage tail protein \| phage tail protein; uniclust \| UniRef100\_UPI001864AFE8 \| 99.9 \| 3.7e-26 \| 6.9e-32 \| 208.3 \| 152 \| (1, 152) \| 423 \| (2, 153) \| 564 \| phage tail protein \| phage tail protein; uniclust \| UniRef100\_A0A076LIZ9 \| 99.9 \| 3.9e-26 \| 7.3e-32 \| 214.7 \| 291 \| (3, 299) \| 423 \| (1, 328) \| 582 \| Phage Tail Collar domain protein \| Phage Tail Collar domain protein; uniclust \| UniRef100\_A0A1I9KF43 \| 99.9 \| 4.3e-26 \| 7.9e-32 \| 209.6 \| 154 \| (3, 156) \| 423 \| (4, 157) \| 839 \| Tail-collar fiber protein \| Tail-collar fiber protein; uniclust \| UniRef100\_D5EF94 \| 99.9 \| 4.4e-26 \| 8.1e-32 \| 201.7 \| 158 \| (3, 160) \| 423 \| (4, 161) \| 561 \| Phage-related tail fibre protein-like protein \| Phage-related tail fibre protein-like protein; uniclust \| UniRef100\_UPI001F14809F \| 99.9 \| 4.4e-26 \| 8.2e-32 \| 189.3 \| 159 \| (1, 159) \| 423 \| (1, 159) \| 325 \| phage tail protein \| phage tail protein; uniclust \| UniRef100\_A0A845BZU2 \| 99.9 \| 4.9e-26 \| 9e-32 \| 205.9 \| 156 \| (3, 158) \| 423 \| (4, 159) \| 701 \| Phage tail protein \| Phage tail protein; uniclust \| UniRef100\_A0A1S6TPA5 \| 99.9 \| 4.8e-26 \| 9.1e-32 \| 187.1 \| 151 \| (1, 154) \| 423 \| (3, 153) \| 227 \| Phage tail-collar fiber protein (DUF3751 domain) \| Phage tail-collar fiber protein (DUF3751 domain); uniclust \| UniRef100\_A0A2C9EHA1 \| 99.9 \| 4.9e-26 \| 9.1e-32 \| 195.7 \| 156 \| (2, 157) \| 423 \| (6, 161) \| 408 \| Putative tail fiber protein \| Putative tail fiber protein; uniclust \| UniRef100\_A0A2Z6AZ11 \| 99.9 \| 6.2e-26 \| 1.2e-31 \| 197.1 \| 148 \| (1, 148) \| 423 \| (2, 149) \| 348 \| Tail Collar domain protein \| Tail Collar domain protein; uniclust \| UniRef100\_UPI00068C6394 \| 99.9 \| 7.3e-26 \| 1.3e-31 \| 193.4 \| 154 \| (3, 156) \| 423 \| (7, 160) \| 407 \| phage tail protein \| phage tail protein; uniclust \| UniRef100\_UPI001F2CA588 \| 99.9 \| 8.6e-26 \| 1.6e-31 \| 189.7 \| 163 \| (2, 164) \| 423 \| (6, 168) \| 354 \| phage tail protein \| phage tail protein; uniclust \| UniRef100\_A0A0T9UVJ2 \| 99.9 \| 9e-26 \| 1.6e-31 \| 169.3 \| 129 \| (3, 131) \| 423 \| (4, 132) \| 163 \| Variable tail fiber protein \| Variable tail fiber protein; uniclust \| UniRef100\_A0A4U3FAV2 \| 99.9 \| 8.9e-26 \| 1.7e-31 \| 197.4 \| 156 \| (2, 157) \| 423 \| (3, 158) \| 354 \| Phage tail protein \| Phage tail protein; uniclust \| UniRef100\_UPI0017462346 \| 99.9 \| 9.1e-26 \| 1.7e-31 \| 184.4 \| 161 \| (1, 161) \| 423 \| (2, 166) \| 237 \| phage tail protein \| phage tail protein; uniclust \| UniRef100\_UPI001295318C \| 99.9 \| 9.3e-26 \| 1.7e-31 \| 208.8 \| 242 \| (2, 247) \| 423 \| (6, 264) \| 723 \| phage tail protein \| phage tail protein; uniclust \| UniRef100\_UPI0022E93E45 \| 99.9 \| 9.8e-26 \| 1.8e-31 \| 205.7 \| 153 \| (3, 155) \| 423 \| (4, 156) \| 767 \| phage tail protein \| phage tail protein; uniclust \| UniRef100\_A0A2W5JZT7 \| 99.9 \| 9.7e-26 \| 1.8e-31 \| 198.9 \| 156 \| (1, 159) \| 423 \| (3, 158) \| 370 \| Phage tail protein \| Phage tail protein; uniclust \| UniRef100\_A0A315WVX7 \| 99.9 \| 1e-25 \| 1.9e-31 \| 202.7 \| 155 \| (3, 157) \| 423 \| (7, 166) \| 561 \| Phage tail protein \| Phage tail protein; uniclust \| UniRef100\_UPI0013F605E0 \| 99.9 \| 1.1e-25 \| 2e-31 \| 196.4 \| 156 \| (3, 159) \| 423 \| (4, 160) \| 488 \| phage tail protein \| phage tail protein; uniclust \| UniRef100\_A0A5Q2KDB2 \| 99.9 \| 1.1e-25 \| 2e-31 \| 199.8 \| 200 \| (3, 303) \| 423 \| (4, 203) \| 575 \| Phage tail protein \| Phage tail protein; uniclust \| UniRef100\_A0A0G9KCC4 \| 99.9 \| 1.3e-25 \| 2.5e-31 \| 201.7 \| 155 \| (1, 158) \| 423 \| (1, 155) \| 405 \| Phage tail protein \| Phage tail protein; uniclust \| UniRef100\_A0A2N6J7T9 \| 99.9 \| 1.6e-25 \| 2.9e-31 \| 202.0 \| 156 \| (1, 156) \| 423 \| (1, 156) \| 628 \| Phage-related tail fiber protein-like protein \| Phage-related tail fiber protein-like protein; uniclust \| UniRef100\_UPI0022CCF947 \| 99.9 \| 1.8e-25 \| 3.3e-31 \| 206.9 \| 203 \| (1, 304) \| 423 \| (2, 207) \| 902 \| phage tail protein \| phage tail protein; uniclust \| UniRef100\_A0A0N9MHY1 \| 99.9 \| 1.9e-25 \| 3.7e-31 \| 217.4 \| 155 \| (2, 156) \| 423 \| (6, 160) \| 843 \| Phage tail collar domain-containing protein \| Phage tail collar domain-containing protein; uniclust \| UniRef100\_A0A0H3B1B9 \| 99.9 \| 1.9e-25 \| 3.7e-31 \| 213.1 \| 293 \| (3, 301) \| 423 \| (1, 334) \| 558 \| Tail fiber repeat 2 protein \| Tail fiber repeat 2 protein; uniclust \| UniRef100\_UPI001E3AD0D6 \| 99.9 \| 2e-25 \| 3.7e-31 \| 180.7 \| 155 \| (3, 157) \| 423 \| (4, 158) \| 268 \| phage tail protein \| phage tail protein; uniclust \| UniRef100\_A0A853I101 \| 99.9 \| 2.1e-25 \| 4e-31 \| 198.6 \| 155 \| (3, 157) \| 423 \| (4, 161) \| 435 \| Phage tail protein \| Phage tail protein; uniclust \| UniRef100\_UPI001AE086F0 \| 99.9 \| 2.3e-25 \| 4.2e-31 \| 204.3 \| 159 \| (1, 159) \| 423 \| (1, 159) \| 808 \| phage tail protein \| phage tail protein; uniclust \| UniRef100\_A0A0J8DFT2 \| 99.9 \| 2.8e-25 \| 5.2e-31 \| 185.3 \| 151 \| (3, 153) \| 423 \| (4, 154) \| 299 \| Phage tail-collar fiber protein \| Phage tail-collar fiber protein; uniclust \| UniRef100\_A0A377IMK2 \| 99.9 \| 4.2e-25 \| 7.7e-31 \| 206.7 \| 149 \| (3, 151) \| 423 \| (4, 152) \| 833 \| Putative phage-like tail fiber protein \| Putative phage-like tail fiber protein; uniclust \| UniRef100\_A0A265Q5Q8 \| 99.9 \| 4.3e-25 \| 7.9e-31 \| 197.9 \| 151 \| (2, 152) \| 423 \| (3, 153) \| 524 \| Phage tail protein \| Phage tail protein; uniclust \| UniRef100\_A0A380C811 \| 99.9 \| 4.5e-25 \| 8.4e-31 \| 184.9 \| 156 \| (1, 156) \| 423 \| (3, 164) \| 257 \| Phage tail protein \| Phage tail protein; uniclust \| UniRef100\_UPI000A4D95A8 \| 99.9 \| 4.6e-25 \| 8.4e-31 \| 202.1 \| 255 \| (3, 314) \| 423 \| (4, 262) \| 743 \| phage tail protein \| phage tail protein; uniclust \| UniRef100\_A0A921AXE4 \| 99.9 \| 4.7e-25 \| 8.7e-31 \| 181.2 \| 153 \| (3, 156) \| 423 \| (5, 157) \| 256 \| Phage tail protein \| Phage tail protein; uniclust \| UniRef100\_A0A1Q6KPQ8 \| 99.8 \| 5.7e-25 \| 1.1e-30 \| 185.8 \| 159 \| (3, 161) \| 423 \| (39, 197) \| 278 \| Phage tail protein \| Phage tail protein; uniclust \| UniRef100\_A0A6D2GE62 \| 99.8 \| 6.2e-25 \| 1.1e-30 \| 162.3 \| 129 \| (31, 159) \| 423 \| (1, 129) \| 148 \| Tail protein \| Tail protein; uniclust \| UniRef100\_A0A069B2A9 \| 99.8 \| 6.6e-25 \| 1.2e-30 \| 191.0 \| 154 \| (3, 156) \| 423 \| (25, 178) \| 371 \| Putative tail fiber protein \| Putative tail fiber protein; uniclust \| UniRef100\_A0A7Z7YC39 \| 99.8 \| 7e-25 \| 1.3e-30 \| 188.6 \| 153 \| (2, 154) \| 423 \| (3, 155) \| 426 \| Phage tail protein \| Phage tail protein; uniclust \| UniRef100\_A0A1K1QNR1 \| 99.8 \| 6.9e-25 \| 1.3e-30 \| 207.1 \| 153 \| (1, 153) \| 423 \| (1, 154) \| 682 \| Phage tail-collar fibre protein \| Phage tail-collar fibre protein; uniclust \| UniRef100\_UPI001FAEEA20 \| 99.8 \| 8e-25 \| 1.5e-30 \| 193.4 \| 155 \| (2, 156) \| 423 \| (6, 160) \| 544 \| phage tail protein \| phage tail protein; uniclust \| UniRef100\_A0A3M5NCX5 \| 99.8 \| 9.1e-25 \| 1.7e-30 \| 181.2 \| 156 \| (2, 157) \| 423 \| (6, 161) \| 318 \| Phage tail protein \| Phage tail protein; uniclust \| UniRef100\_A0A376FFE3 \| 99.8 \| 9.7e-25 \| 1.8e-30 \| 155.7 \| 112 \| (31, 142) \| 423 \| (1, 112) \| 121 \| Phage tail collar domain-containing protein \| Phage tail collar domain-containing protein; uniclust \| UniRef100\_A0A828ZHJ0 \| 99.8 \| 9.9e-25 \| 1.8e-30 \| 200.7 \| 159 \| (1, 159) \| 423 \| (1, 159) \| 592 \| Tail fiber protein (GpH) \| Tail fiber protein (GpH); uniclust \| UniRef100\_UPI00209EEE42 \| 99.8 \| 1e-24 \| 1.9e-30 \| 182.3 \| 160 \| (4, 163) \| 423 \| (5, 164) \| 338 \| phage tail protein \| phage tail protein; uniclust \| UniRef100\_A0A6I1HR03 \| 99.8 \| 1e-24 \| 1.9e-30 \| 201.0 \| 153 \| (3, 155) \| 423 \| (7, 159) \| 753 \| SGNH hydrolase-type esterase domain-containing protein \| SGNH hydrolase-type esterase domain-containing protein; uniclust \| UniRef100\_UPI001C450459 \| 99.8 \| 1.1e-24 \| 1.9e-30 \| 193.2 \| 158 \| (3, 160) \| 423 \| (25, 182) \| 558 \| phage tail protein \| phage tail protein; uniclust \| UniRef100\_A0A611ES10 \| 99.8 \| 1.2e-24 \| 2.2e-30 \| 166.6 \| 149 \| (59, 308) \| 423 \| (4, 152) \| 169 \| Phage tail protein (Fragment) \| Phage tail protein (Fragment); uniclust \| UniRef100\_UPI001FCEFC3F \| 99.8 \| 1.2e-24 \| 2.2e-30 \| 188.9 \| 193 \| (8, 301) \| 423 \| (1, 193) \| 462 \| phage tail protein \| phage tail protein; uniclust \| UniRef100\_B8DLJ2 \| 99.8 \| 1.4e-24 \| 2.7e-30 \| 197.4 \| 149 \| (4, 153) \| 423 \| (5, 154) \| 511 \| Tail fiber protein, putative \| Tail fiber protein, putative; uniclust \| UniRef100\_UPI00215BC1F7 \| 99.8 \| 1.5e-24 \| 2.7e-30 \| 195.8 \| 155 \| (2, 156) \| 423 \| (3, 157) \| 669 \| phage tail protein \| phage tail protein; uniclust \| UniRef100\_A0A4P2WWZ3 \| 99.8 \| 1.5e-24 \| 2.8e-30 \| 174.7 \| 157 \| (3, 160) \| 423 \| (5, 161) \| 225 \| Uncharacterized protein \| Uncharacterized protein; uniclust \| UniRef100\_A0A0A7RTQ0 \| 99.8 \| 1.5e-24 \| 2.8e-30 \| 191.9 \| 153 \| (1, 153) \| 423 \| (1, 153) \| 436 \| Tail-collar fiber family protein \| Tail-collar fiber family protein; uniclust \| UniRef100\_A0A8S5MN89 \| 99.8 \| 1.6e-24 \| 2.9e-30 \| 168.4 \| 155 \| (3, 157) \| 423 \| (4, 158) \| 201 \| Tail collar fiber protein \| Tail collar fiber protein; uniclust \| UniRef100\_A0A495AKT5 \| 99.8 \| 1.6e-24 \| 2.9e-30 \| 205.0 \| 282 \| (3, 304) \| 423 \| (129, 418) \| 907 \| Phage tail protein \| Phage tail protein; uniclust \| UniRef100\_A0A974X0L5 \| 99.8 \| 1.7e-24 \| 3.1e-30 \| 188.1 \| 153 \| (2, 154) \| 423 \| (1, 153) \| 463 \| Phage tail protein \| Phage tail protein; uniclust \| UniRef100\_UPI00158FB9FD \| 99.8 \| 1.7e-24 \| 3.1e-30 \| 159.5 \| 129 \| (3, 131) \| 423 \| (4, 132) \| 133 \| phage tail protein \| phage tail protein; uniclust \| UniRef100\_A0A078LT99 \| 99.8 \| 1.8e-24 \| 3.4e-30 \| 183.2 \| 163 \| (3, 165) \| 423 \| (7, 169) \| 339 \| Tail fiber protein H \| Tail fiber protein H; uniclust \| UniRef100\_UPI0018E08366 \| 99.8 \| 1.9e-24 \| 3.6e-30 \| 184.0 \| 157 \| (3, 159) \| 423 \| (5, 162) \| 391 \| phage tail protein \| phage tail protein; uniclust \| UniRef100\_A0A2H4J194 \| 99.8 \| 2e-24 \| 3.6e-30 \| 178.1 \| 157 \| (1, 157) \| 423 \| (1, 157) \| 303 \| Putative tail-collar fiber protein \| Putative tail-collar fiber protein; uniclust \| UniRef100\_A0A0H0GM37 \| 99.8 \| 2.3e-24 \| 4.3e-30 \| 205.5 \| 153 \| (3, 156) \| 423 \| (7, 160) \| 687 \| Phage tail protein \| Phage tail protein; uniclust \| UniRef100\_A0A2X1IXU3 \| 99.8 \| 2.4e-24 \| 4.5e-30 \| 163.2 \| 124 \| (41, 164) \| 423 \| (7, 130) \| 166 \| Variable tail fiber protein \| Variable tail fiber protein; uniclust \| UniRef100\_UPI001F1039DB \| 99.8 \| 2.4e-24 \| 4.5e-30 \| 198.7 \| 154 \| (3, 156) \| 423 \| (4, 157) \| 853 \| phage tail protein \| phage tail protein; uniclust \| UniRef100\_UPI00065A80D1 \| 99.8 \| 2.8e-24 \| 5.1e-30 \| 182.8 \| 155 \| (2, 156) \| 423 \| (86, 240) \| 384 \| phage tail protein I \| phage tail protein I; uniclust \| UniRef100\_A0A9E0KF93 \| 99.8 \| 2.9e-24 \| 5.3e-30 \| 182.2 \| 163 \| (1, 164) \| 423 \| (1, 163) \| 377 \| Phage tail protein \| Phage tail protein; uniclust \| UniRef100\_A0A1E3GML3 \| 99.8 \| 2.9e-24 \| 5.3e-30 \| 203.7 \| 156 \| (2, 157) \| 423 \| (4, 160) \| 768 \| Phage tail protein \| Phage tail protein; uniclust \| UniRef100\_A0A0D8DFZ0 \| 99.8 \| 3e-24 \| 5.5e-30 \| 173.1 \| 153 \| (2, 159) \| 423 \| (27, 179) \| 204 \| Tail fiber protein \| Tail fiber protein; uniclust \| UniRef100\_UPI0018A82FE3 \| 99.8 \| 3.1e-24 \| 5.8e-30 \| 154.9 \| 118 \| (1, 118) \| 423 \| (2, 119) \| 120 \| phage tail protein \| phage tail protein; uniclust \| UniRef100\_UPI001C5D3A71 \| 99.8 \| 3.5e-24 \| 6.5e-30 \| 170.4 \| 114 \| (47, 160) \| 423 \| (1, 114) \| 224 \| tail fiber protein \| tail fiber protein; uniclust \| UniRef100\_UPI001C474985 \| 99.8 \| 3.6e-24 \| 6.7e-30 \| 193.0 \| 158 \| (3, 160) \| 423 \| (25, 182) \| 655 \| phage tail protein \| phage tail protein; uniclust \| UniRef100\_A0A367LYA0 \| 99.8 \| 3.8e-24 \| 7e-30 \| 169.0 \| 155 \| (3, 157) \| 423 \| (7, 166) \| 198 \| Phage tail protein (Fragment) \| Phage tail protein (Fragment); uniclust \| UniRef100\_UPI001C48C03A \| 99.8 \| 4.2e-24 \| 7.7e-30 \| 155.0 \| 131 \| (14, 144) \| 423 \| (2, 132) \| 132 \| phage tail protein \| phage tail protein; uniclust \| UniRef100\_A0A2D1IR12 \| 99.8 \| 4.7e-24 \| 8.8e-30 \| 200.7 \| 157 \| (1, 159) \| 423 \| (2, 158) \| 604 \| Phage tail protein \| Phage tail protein; uniclust \| UniRef100\_A0A1Y5MYB7 \| 99.8 \| 4.7e-24 \| 9e-30 \| 187.8 \| 153 \| (2, 157) \| 423 \| (4, 157) \| 316 \| Phage tail protein \| Phage tail protein; uniclust \| UniRef100\_A0A3E1QH42 \| 99.8 \| 5.1e-24 \| 9.5e-30 \| 195.1 \| 150 \| (3, 152) \| 423 \| (4, 153) \| 636 \| Phage tail protein (Fragment) \| Phage tail protein (Fragment); uniclust \| UniRef100\_UPI001CA43773 \| 99.8 \| 5.5e-24 \| 1e-29 \| 192.6 \| 155 \| (2, 156) \| 423 \| (1, 156) \| 684 \| phage tail protein \| phage tail protein; uniclust \| UniRef100\_N8R6Q7 \| 99.8 \| 5.6e-24 \| 1e-29 \| 165.1 \| 153 \| (2, 154) \| 423 \| (1, 154) \| 198 \| Uncharacterized protein \| Uncharacterized protein; uniclust \| UniRef100\_A0A1B1KPJ7 \| 99.8 \| 5.6e-24 \| 1e-29 \| 185.2 \| 159 \| (3, 161) \| 423 \| (5, 163) \| 358 \| Phage tail protein \| Phage tail protein; uniclust \| UniRef100\_UPI000F76A94D \| 99.8 \| 5.8e-24 \| 1.1e-29 \| 170.7 \| 155 \| (1, 156) \| 423 \| (1, 159) \| 248 \| phage tail protein \| phage tail protein; uniclust \| UniRef100\_A0A3F3A8F4 \| 99.8 \| 5.7e-24 \| 1.1e-29 \| 191.7 \| 153 \| (1, 153) \| 423 \| (2, 154) \| 556 \| Putative phage tail fiber protein \| Putative phage tail fiber protein; uniclust \| UniRef100\_L1M5X1 \| 99.8 \| 7.2e-24 \| 1.3e-29 \| 193.5 \| 155 \| (2, 156) \| 423 \| (6, 160) \| 748 \| Tail fiber protein H \| Tail fiber protein H; uniclust \| UniRef100\_UPI001F2A79DD \| 99.8 \| 7.3e-24 \| 1.3e-29 \| 191.6 \| 153 \| (2, 154) \| 423 \| (6, 158) \| 673 \| phage tail protein \| phage tail protein; uniclust \| UniRef100\_A0A077NHN3 \| 99.8 \| 7.5e-24 \| 1.4e-29 \| 207.2 \| 152 \| (3, 159) \| 423 \| (4, 155) \| 1195 \| Tail protein \| Tail protein; uniclust \| UniRef100\_A0A2N2KCW4 \| 99.8 \| 8.5e-24 \| 1.6e-29 \| 168.0 \| 153 \| (1, 153) \| 423 \| (1, 153) \| 231 \| Phage tail protein (Fragment) \| Phage tail protein (Fragment); uniclust \| UniRef100\_UPI00190502AF \| 99.8 \| 8.6e-24 \| 1.6e-29 \| 192.6 \| 155 \| (1, 155) \| 423 \| (1, 159) \| 729 \| phage tail protein \| phage tail protein; uniclust \| UniRef100\_A0A290WU51 \| 99.8 \| 9e-24 \| 1.6e-29 \| 191.4 \| 150 \| (3, 152) \| 423 \| (4, 153) \| 686 \| SGNH hydrolase-type esterase domain-containing protein \| SGNH hydrolase-type esterase domain-containing protein; uniclust \| UniRef100\_UPI0018D16FB5 \| 99.8 \| 9.6e-24 \| 1.8e-29 \| 197.1 \| 153 \| (3, 155) \| 423 \| (5, 157) \| 668 \| phage tail protein \| phage tail protein; uniclust \| UniRef100\_UPI000240E034 \| 99.8 \| 1e-23 \| 1.9e-29 \| 178.8 \| 164 \| (2, 165) \| 423 \| (1, 164) \| 341 \| phage tail protein \| phage tail protein; uniclust \| UniRef100\_A0A1T4W1B3 \| 99.8 \| 1.1e-23 \| 2.1e-29 \| 180.9 \| 152 \| (3, 154) \| 423 \| (4, 155) \| 416 \| Phage tail-collar fibre protein \| Phage tail-collar fibre protein; uniclust \| UniRef100\_A0A5T3EMD8 \| 99.8 \| 1.1e-23 \| 2.1e-29 \| 151.6 \| 98 \| (2, 99) \| 423 \| (3, 101) \| 103 \| Phage tail protein (Fragment) \| Phage tail protein (Fragment); uniclust \| UniRef100\_A0A6G9RP95 \| 99.8 \| 1.1e-23 \| 2.1e-29 \| 196.5 \| 140 \| (1, 144) \| 423 \| (3, 142) \| 466 \| Phage tail protein \| Phage tail protein; uniclust \| UniRef100\_UPI001F07DAE0 \| 99.8 \| 1.1e-23 \| 2.1e-29 \| 178.8 \| 148 \| (2, 153) \| 423 \| (6, 153) \| 328 \| phage tail protein \| phage tail protein; uniclust \| UniRef100\_UPI001FEFA6B3 \| 99.8 \| 1.2e-23 \| 2.3e-29 \| 171.2 \| 154 \| (2, 155) \| 423 \| (3, 156) \| 274 \| phage tail protein \| phage tail protein; uniclust \| UniRef100\_A0A1X3I4D9 \| 99.8 \| 1.3e-23 \| 2.5e-29 \| 195.3 \| 147 \| (1, 154) \| 423 \| (8, 154) \| 442 \| Putative tail fiber protein (GpH) \| Putative tail fiber protein (GpH); uniclust \| UniRef100\_UPI001191EDC2 \| 99.8 \| 1.4e-23 \| 2.6e-29 \| 151.1 \| 116 \| (22, 137) \| 423 \| (10, 125) \| 126 \| phage tail protein \| phage tail protein; uniclust \| UniRef100\_A0A483Z142 \| 99.8 \| 1.4e-23 \| 2.6e-29 \| 169.5 \| 159 \| (1, 160) \| 423 \| (2, 160) \| 220 \| Phage tail protein (Fragment) \| Phage tail protein (Fragment); uniclust \| UniRef100\_A0A077N5A2 \| 99.8 \| 1.5e-23 \| 2.8e-29 \| 195.6 \| 275 \| (3, 281) \| 423 \| (1, 314) \| 657 \| Bacteriophage T7 Gp17 C-terminal domain-containing protein \| Bacteriophage T7 Gp17 C-terminal domain-containing protein; uniclust \| UniRef100\_UPI0015E27F6A \| 99.8 \| 1.6e-23 \| 3e-29 \| 185.6 \| 154 \| (3, 156) \| 423 \| (4, 157) \| 551 \| phage tail protein \| phage tail protein; uniclust \| UniRef100\_UPI000D6467B0 \| 99.8 \| 1.7e-23 \| 3.1e-29 \| 173.7 \| 157 \| (1, 157) \| 423 \| (1, 161) \| 278 \| phage tail protein \| phage tail protein; uniclust \| UniRef100\_A0A0J1NB15 \| 99.8 \| 1.7e-23 \| 3.1e-29 \| 196.2 \| 153 \| (2, 154) \| 423 \| (3, 155) \| 589 \| Phage tail protein \| Phage tail protein; uniclust \| UniRef100\_A0A0A8H9Y5 \| 99.8 \| 1.7e-23 \| 3.2e-29 \| 184.4 \| 154 \| (1, 157) \| 423 \| (3, 156) \| 348 \| Phage tail-collar fiber protein (DUF3751 domain) \| Phage tail-collar fiber protein (DUF3751 domain); uniclust \| UniRef100\_A0A239C905 \| 99.8 \| 1.9e-23 \| 3.5e-29 \| 185.6 \| 153 \| (2, 154) \| 423 \| (1, 153) \| 563 \| Phage tail-collar fibre protein \| Phage tail-collar fibre protein; uniclust \| UniRef100\_A0A8I1WMJ8 \| 99.8 \| 2e-23 \| 3.8e-29 \| 172.0 \| 160 \| (1, 160) \| 423 \| (1, 164) \| 298 \| Phage tail protein \| Phage tail protein; uniclust \| UniRef100\_UPI001F29E04B \| 99.8 \| 2.5e-23 \| 4.6e-29 \| 182.6 \| 149 \| (3, 151) \| 423 \| (4, 152) \| 500 \| phage tail protein \| phage tail protein; uniclust \| UniRef100\_A0A7Z2A8B6 \| 99.8 \| 2.5e-23 \| 4.8e-29 \| 187.0 \| 158 \| (1, 160) \| 423 \| (1, 158) \| 420 \| Phage tail protein \| Phage tail protein; uniclust \| UniRef100\_UPI001C106791 \| 99.8 \| 3e-23 \| 5.5e-29 \| 187.7 \| 151 \| (2, 152) \| 423 \| (3, 153) \| 669 \| phage tail protein \| phage tail protein; uniclust \| UniRef100\_A0A7Y8JX08 \| 99.8 \| 3.1e-23 \| 5.7e-29 \| 167.2 \| 156 \| (1, 156) \| 423 \| (1, 157) \| 244 \| Phage tail protein (Fragment) \| Phage tail protein (Fragment); uniclust \| UniRef100\_UPI0020A40C1E \| 99.8 \| 3.2e-23 \| 5.8e-29 \| 179.1 \| 157 \| (2, 160) \| 423 \| (3, 159) \| 434 \| phage tail protein \| phage tail protein; uniclust \| UniRef100\_UPI001C998269 \| 99.8 \| 3.2e-23 \| 5.9e-29 \| 189.9 \| 159 \| (1, 159) \| 423 \| (1, 163) \| 769 \| phage tail protein \| phage tail protein; uniclust \| UniRef100\_A0A077NV23 \| 99.8 \| 3.2e-23 \| 6.3e-29 \| 205.9 \| 244 \| (3, 251) \| 423 \| (1, 282) \| 689 \| Phage tail collar domain-containing protein \| Phage tail collar domain-containing protein; uniclust \| UniRef100\_A0A345J3J2 \| 99.8 \| 3.4e-23 \| 6.3e-29 \| 179.7 \| 156 \| (2, 157) \| 423 \| (1, 156) \| 450 \| Phage-related tail fiber protein \| Phage-related tail fiber protein; uniclust \| UniRef100\_A0A9D1QZY4 \| 99.8 \| 3.5e-23 \| 6.4e-29 \| 179.6 \| 161 \| (3, 163) \| 423 \| (4, 164) \| 417 \| Phage tail protein (Fragment) \| Phage tail protein (Fragment); uniclust \| UniRef100\_UPI00206B758F \| 99.8 \| 3.8e-23 \| 7e-29 \| 171.3 \| 152 \| (2, 153) \| 423 \| (4, 155) \| 310 \| phage tail protein \| phage tail protein; uniclust \| UniRef100\_A0A0J1LWH4 \| 99.8 \| 4.1e-23 \| 7.7e-29 \| 197.5 \| 212 \| (1, 308) \| 423 \| (96, 307) \| 716 \| Phage tail collar domain-containing protein \| Phage tail collar domain-containing protein; uniclust \| UniRef100\_A0A943L988 \| 99.8 \| 5e-23 \| 9.2e-29 \| 151.3 \| 136 \| (3, 138) \| 423 \| (5, 140) \| 141 \| Phage tail protein (Fragment) \| Phage tail protein (Fragment); uniclust \| UniRef100\_A0A846TIU6 \| 99.8 \| 5.4e-23 \| 1e-28 \| 167.9 \| 153 \| (3, 155) \| 423 \| (4, 156) \| 243 \| DUF2793 domain-containing protein \| DUF2793 domain-containing protein; uniclust \| UniRef100\_A0A022PLF6 \| 99.8 \| 5.4e-23 \| 1e-28 \| 199.6 \| 244 \| (3, 251) \| 423 \| (16, 297) \| 629 \| Phage tail-collar fiber protein \| Phage tail-collar fiber protein; uniclust \| UniRef100\_J6HCZ5 \| 99.8 \| 6e-23 \| 1.1e-28 \| 174.9 \| 152 \| (3, 154) \| 423 \| (4, 155) \| 384 \| Tail fiber protein \| Tail fiber protein; uniclust \| UniRef100\_UPI000B4A1C81 \| 99.8 \| 6.1e-23 \| 1.1e-28 \| 185.1 \| 152 \| (3, 154) \| 423 \| (4, 155) \| 647 \| phage tail protein \| phage tail protein; uniclust \| UniRef100\_A0A2D1CSN0 \| 99.8 \| 6.1e-23 \| 1.1e-28 \| 192.4 \| 154 \| (3, 156) \| 423 \| (135, 288) \| 733 \| Putative tail fiber protein \| Putative tail fiber protein; uniclust \| UniRef100\_UPI000AFA6AC9 \| 99.8 \| 6.5e-23 \| 1.2e-28 \| 190.2 \| 149 \| (3, 151) \| 423 \| (4, 152) \| 885 \| phage tail protein \| phage tail protein; uniclust \| UniRef100\_A0A0F9Z9H8 \| 99.8 \| 6.5e-23 \| 1.2e-28 \| 167.5 \| 152 \| (1, 159) \| 423 \| (7, 158) \| 234 \| Tail fiber-like protein \| Tail fiber-like protein; uniclust \| UniRef100\_A0A3G9G2L6 \| 99.8 \| 7.4e-23 \| 1.4e-28 \| 194.9 \| 153 \| (3, 156) \| 423 \| (5, 157) \| 735 \| Phage tail fiber protein \| Phage tail fiber protein; uniclust \| UniRef100\_UPI0021CF5D59 \| 99.8 \| 7.6e-23 \| 1.4e-28 \| 165.6 \| 123 \| (36, 158) \| 423 \| (2, 124) \| 260 \| phage tail protein \| phage tail protein; uniclust \| UniRef100\_A0A1G8FGR6 \| 99.8 \| 7.5e-23 \| 1.4e-28 \| 190.5 \| 158 \| (1, 158) \| 423 \| (2, 159) \| 657 \| Phage Tail Collar Domain \| Phage Tail Collar Domain; uniclust \| UniRef100\_A0A6B8QPL4 \| 99.8 \| 8.6e-23 \| 1.6e-28 \| 168.0 \| 153 \| (3, 155) \| 423 \| (46, 198) \| 292 \| Phage tail protein \| Phage tail protein; uniclust \| UniRef100\_UPI0006945938 \| 99.8 \| 8.7e-23 \| 1.6e-28 \| 178.7 \| 156 \| (1, 156) \| 423 \| (2, 159) \| 407 \| phage tail protein \| phage tail protein; uniclust \| UniRef100\_A0A1I5MQ45 \| 99.8 \| 9.3e-23 \| 1.7e-28 \| 183.1 \| 154 \| (1, 154) \| 423 \| (1, 154) \| 617 \| Chaperone of endosialidase \| Chaperone of endosialidase; uniclust \| UniRef100\_UPI000690FF63 \| 99.8 \| 9.3e-23 \| 1.7e-28 \| 177.2 \| 154 \| (3, 156) \| 423 \| (4, 157) \| 406 \| phage tail protein \| phage tail protein; uniclust \| UniRef100\_UPI00208DA219 \| 99.8 \| 9.4e-23 \| 1.7e-28 \| 159.5 \| 93 \| (71, 163) \| 423 \| (2, 94) \| 206 \| phage tail protein \| phage tail protein; uniclust \| UniRef100\_UPI0004CFD2CA \| 99.8 \| 9.7e-23 \| 1.8e-28 \| 171.2 \| 154 \| (2, 155) \| 423 \| (3, 156) \| 343 \| phage tail protein \| phage tail protein; uniclust \| UniRef100\_UPI001356777F \| 99.8 \| 1.1e-22 \| 2.1e-28 \| 180.2 \| 271 \| (4, 277) \| 423 \| (5, 276) \| 542 \| phage tail protein \| phage tail protein; uniclust \| UniRef100\_A0A2D3VW29 \| 99.8 \| 1.2e-22 \| 2.2e-28 \| 159.5 \| 153 \| (2, 154) \| 423 \| (1, 153) \| 211 \| Phage tail protein \| Phage tail protein; uniclust \| UniRef100\_UPI000693D28B \| 99.8 \| 1.3e-22 \| 2.4e-28 \| 173.0 \| 156 \| (1, 156) \| 423 \| (1, 156) \| 386 \| phage tail protein \| phage tail protein; uniclust \| UniRef100\_UPI000DFAA696 \| 99.8 \| 1.4e-22 \| 2.6e-28 \| 182.3 \| 154 \| (3, 156) \| 423 \| (7, 165) \| 510 \| phage tail protein \| phage tail protein; uniclust \| UniRef100\_G4CPH8 \| 99.8 \| 1.4e-22 \| 2.6e-28 \| 186.0 \| 154 \| (2, 155) \| 423 \| (3, 156) \| 779 \| Uncharacterized protein \| Uncharacterized protein; uniclust \| UniRef100\_A0A965NEJ7 \| 99.8 \| 1.5e-22 \| 2.7e-28 \| 147.8 \| 131 \| (2, 133) \| 423 \| (3, 133) \| 135 \| Uncharacterized protein (Fragment) \| Uncharacterized protein (Fragment); uniclust \| UniRef100\_A0A2G2D6M1 \| 99.8 \| 1.6e-22 \| 2.9e-28 \| 157.7 \| 154 \| (3, 157) \| 423 \| (22, 176) \| 187 \| Phage tail protein \| Phage tail protein; uniclust \| UniRef100\_A0A7C8HT98 \| 99.8 \| 1.6e-22 \| 2.9e-28 \| 184.3 \| 155 \| (1, 155) \| 423 \| (2, 156) \| 476 \| Phage tail collar domain-containing protein \| Phage tail collar domain-containing protein; uniclust \| UniRef100\_A0A1N6M5X0 \| 99.8 \| 1.5e-22 \| 3e-28 \| 186.4 \| 161 \| (2, 163) \| 423 \| (8, 168) \| 363 \| Phage Tail Collar Domain protein \| Phage Tail Collar Domain protein; uniclust \| UniRef100\_UPI001E52CB3C \| 99.8 \| 1.7e-22 \| 3.2e-28 \| 166.5 \| 204 \| (72, 277) \| 423 \| (5, 210) \| 296 \| phage tail protein \| phage tail protein; uniclust \| UniRef100\_UPI001967A92B \| 99.8 \| 2.3e-22 \| 4.2e-28 \| 175.4 \| 150 \| (3, 152) \| 423 \| (4, 153) \| 466 \| phage tail protein \| phage tail protein; uniclust \| UniRef100\_UPI0015D48CC2 \| 99.8 \| 2.4e-22 \| 4.4e-28 \| 139.5 \| 91 \| (23, 113) \| 423 \| (2, 92) \| 93 \| phage tail protein \| phage tail protein; uniclust \| UniRef100\_UPI000986ADFB \| 99.8 \| 2.8e-22 \| 5.1e-28 \| 169.2 \| 151 \| (2, 152) \| 423 \| (3, 153) \| 353 \| phage tail protein \| phage tail protein; uniclust \| UniRef100\_UPI0023545723 \| 99.8 \| 2.9e-22 \| 5.4e-28 \| 181.4 \| 154 \| (1, 154) \| 423 \| (1, 154) \| 667 \| phage tail protein \| phage tail protein; uniclust \| UniRef100\_A0A0A8JLH1 \| 99.8 \| 3.1e-22 \| 5.7e-28 \| 163.8 \| 157 \| (1, 157) \| 423 \| (1, 157) \| 280 \| Tail fiber protein \| Tail fiber protein; uniclust \| UniRef100\_UPI000E267FDC \| 99.8 \| 3.2e-22 \| 6e-28 \| 169.5 \| 136 \| (2, 144) \| 423 \| (1, 137) \| 294 \| phage tail protein \| phage tail protein; uniclust \| UniRef100\_UPI00069E8323 \| 99.8 \| 3.2e-22 \| 6e-28 \| 190.3 \| 151 \| (3, 154) \| 423 \| (5, 155) \| 779 \| phage tail protein \| phage tail protein; uniclust \| UniRef100\_UPI00068D786B \| 99.8 \| 3.4e-22 \| 6.2e-28 \| 148.9 \| 127 \| (2, 133) \| 423 \| (3, 129) \| 151 \| phage tail protein \| phage tail protein; uniclust \| UniRef100\_UPI0015AF74D6 \| 99.8 \| 3.4e-22 \| 6.2e-28 \| 158.5 \| 154 \| (1, 154) \| 423 \| (1, 154) \| 224 \| phage tail protein \| phage tail protein; uniclust \| UniRef100\_A0A4R4E5X6 \| 99.8 \| 3.8e-22 \| 6.9e-28 \| 181.0 \| 157 \| (1, 157) \| 423 \| (1, 157) \| 568 \| Trimeric autotransporter adhesin YadA-like head domain-containing protein \| Trimeric autotransporter adhesin YadA-like head domain-containing protein; uniclust \| UniRef100\_UPI00038C8FDF \| 99.8 \| 3.8e-22 \| 7e-28 \| 164.2 \| 151 \| (3, 153) \| 423 \| (5, 155) \| 268 \| phage tail protein \| phage tail protein; uniclust \| UniRef100\_A0A4Y9GWV8 \| 99.8 \| 3.9e-22 \| 7.3e-28 \| 183.5 \| 152 \| (1, 152) \| 423 \| (2, 153) \| 558 \| Phage tail protein (Fragment) \| Phage tail protein (Fragment); uniclust \| UniRef100\_A0A0A8TLD1 \| 99.8 \| 4.3e-22 \| 8.1e-28 \| 196.3 \| 157 \| (1, 159) \| 423 \| (2, 158) \| 998 \| Phage tail fiber protein \| Phage tail fiber protein; uniclust \| UniRef100\_UPI0014839BDF \| 99.8 \| 4.5e-22 \| 8.3e-28 \| 181.0 \| 155 \| (2, 156) \| 423 \| (6, 160) \| 698 \| phage tail protein \| phage tail protein; uniclust \| UniRef100\_A0A376RL24 \| 99.8 \| 4.7e-22 \| 8.8e-28 \| 157.9 \| 131 \| (73, 304) \| 423 \| (2, 132) \| 182 \| Bcv gene product \| Bcv gene product; uniclust \| UniRef100\_C6E6P3 \| 99.8 \| 4.9e-22 \| 9e-28 \| 160.3 \| 156 \| (2, 157) \| 423 \| (1, 157) \| 251 \| Phage-related tail fibre protein-like protein \| Phage-related tail fibre protein-like protein; uniclust \| UniRef100\_A0A2N2GS05 \| 99.8 \| 5.2e-22 \| 9.6e-28 \| 174.5 \| 153 \| (1, 153) \| 423 \| (1, 153) \| 497 \| Phage tail protein \| Phage tail protein; uniclust \| UniRef100\_UPI0008C2DC0F \| 99.8 \| 5.4e-22 \| 1e-27 \| 177.8 \| 157 \| (1, 157) \| 423 \| (1, 158) \| 537 \| phage tail protein \| phage tail protein; uniclust \| UniRef100\_A0A4P9VIJ6 \| 99.8 \| 5.5e-22 \| 1e-27 \| 156.2 \| 156 \| (3, 158) \| 423 \| (4, 162) \| 213 \| Phage tail protein \| Phage tail protein; uniclust \| UniRef100\_A0A7X2D3Y0 \| 99.8 \| 5.7e-22 \| 1e-27 \| 172.6 \| 152 \| (4, 155) \| 423 \| (5, 156) \| 454 \| Phage tail protein \| Phage tail protein; uniclust \| UniRef100\_A0A6M0SQX7 \| 99.8 \| 5.7e-22 \| 1.1e-27 \| 180.1 \| 153 \| (1, 154) \| 423 \| (2, 154) \| 574 \| Phage tail protein \| Phage tail protein; uniclust \| UniRef100\_A0A829RXG4 \| 99.8 \| 5.7e-22 \| 1.1e-27 \| 194.7 \| 156 \| (4, 160) \| 423 \| (6, 161) \| 802 \| Phage tail-collar fiber family protein \| Phage tail-collar fiber family protein; uniclust \| UniRef100\_A0A853IIN2 \| 99.8 \| 6.4e-22 \| 1.2e-27 \| 169.2 \| 161 \| (3, 163) \| 423 \| (6, 169) \| 349 \| Phage tail protein \| Phage tail protein; uniclust \| UniRef100\_A0A0M5MEM2 \| 99.8 \| 6.3e-22 \| 1.2e-27 \| 184.1 \| 153 \| (2, 157) \| 423 \| (1, 154) \| 480 \| Phage tail-collar fiber protein (DUF3751 domain) \| Phage tail-collar fiber protein (DUF3751 domain); uniclust \| UniRef100\_A0A2X2TYC3 \| 99.8 \| 6.7e-22 \| 1.2e-27 \| 170.7 \| 138 \| (1, 142) \| 423 \| (1, 138) \| 356 \| Phage tail fiber protein \| Phage tail fiber protein; uniclust \| UniRef100\_A0A7X7NDN0 \| 99.8 \| 7.4e-22 \| 1.4e-27 \| 147.8 \| 131 \| (2, 133) \| 423 \| (4, 134) \| 138 \| Phage tail protein \| Phage tail protein; uniclust \| UniRef100\_A0A1Y3CBA8 \| 99.8 \| 7.3e-22 \| 1.4e-27 \| 190.8 \| 159 \| (1, 161) \| 423 \| (2, 160) \| 838 \| Phage tail collar domain-containing protein \| Phage tail collar domain-containing protein; uniclust \| UniRef100\_UPI0004685D26 \| 99.8 \| 7.7e-22 \| 1.4e-27 \| 175.3 \| 156 \| (3, 158) \| 423 \| (5, 160) \| 517 \| phage tail protein \| phage tail protein; uniclust \| UniRef100\_A0A4P9VV53 \| 99.8 \| 7.6e-22 \| 1.4e-27 \| 175.8 \| 158 \| (3, 160) \| 423 \| (4, 164) \| 431 \| Phage tail protein \| Phage tail protein; uniclust \| UniRef100\_A0A949CHK2 \| 99.8 \| 7.8e-22 \| 1.5e-27 \| 182.1 \| 135 \| (3, 137) \| 423 \| (4, 139) \| 563 \| Phage tail protein \| Phage tail protein; uniclust \| UniRef100\_UPI0015E62003 \| 99.8 \| 8.3e-22 \| 1.5e-27 \| 132.8 \| 84 \| (74, 157) \| 423 \| (3, 86) \| 87 \| phage tail protein \| phage tail protein; uniclust \| UniRef100\_UPI0003D2F6E8 \| 99.8 \| 9.8e-22 \| 1.8e-27 \| 158.3 \| 139 \| (67, 306) \| 423 \| (3, 141) \| 228 \| phage tail protein \| phage tail protein; uniclust \| UniRef100\_A0A482IPK9 \| 99.8 \| 1.1e-21 \| 2e-27 \| 158.7 \| 156 \| (3, 158) \| 423 \| (4, 159) \| 255 \| Phage tail protein \| Phage tail protein; uniclust \| UniRef100\_A0A5M6I3L7 \| 99.8 \| 1.1e-21 \| 2.1e-27 \| 167.9 \| 156 \| (1, 156) \| 423 \| (3, 158) \| 312 \| Phage tail protein \| Phage tail protein; uniclust \| UniRef100\_A0A109RTW5 \| 99.8 \| 1.2e-21 \| 2.1e-27 \| 180.0 \| 158 \| (3, 161) \| 423 \| (4, 161) \| 685 \| Phage tail collar domain-containing protein \| Phage tail collar domain-containing protein; uniclust \| UniRef100\_E5Y5X2 \| 99.8 \| 1.2e-21 \| 2.1e-27 \| 182.3 \| 159 \| (3, 161) \| 423 \| (4, 162) \| 682 \| Phage tail collar domain-containing protein \| Phage tail collar domain-containing protein; uniclust \| UniRef100\_UPI001CF60DE5 \| 99.8 \| 1.2e-21 \| 2.2e-27 \| 182.3 \| 151 \| (3, 153) \| 423 \| (4, 154) \| 823 \| phage tail protein \| phage tail protein; uniclust \| UniRef100\_A0A8S0FS69 \| 99.8 \| 1.2e-21 \| 2.2e-27 \| 153.8 \| 137 \| (2, 142) \| 423 \| (4, 140) \| 165 \| Phage tail protein \| Phage tail protein; uniclust \| UniRef100\_A0A3D9UVK1 \| 99.8 \| 1.3e-21 \| 2.5e-27 \| 196.1 \| 153 \| (2, 159) \| 423 \| (3, 155) \| 2092 \| Tail fiber-like repeat protein \| Tail fiber-like repeat protein; uniclust \| UniRef100\_UPI001CA5DA19 \| 99.7 \| 1.5e-21 \| 2.7e-27 \| 180.8 \| 296 \| (5, 305) \| 423 \| (3, 337) \| 837 \| phage tail protein \| phage tail protein; uniclust \| UniRef100\_A0A4V1INX2 \| 99.7 \| 1.4e-21 \| 2.7e-27 \| 171.3 \| 155 \| (3, 157) \| 423 \| (4, 163) \| 391 \| Phage tail collar domain-containing protein \| Phage tail collar domain-containing protein; uniclust \| UniRef100\_UPI001B39AF5A \| 99.7 \| 1.5e-21 \| 2.7e-27 \| 168.2 \| 152 \| (3, 154) \| 423 \| (5, 156) \| 414 \| phage tail protein \| phage tail protein; uniclust \| UniRef100\_A0A380AJ63 \| 99.7 \| 1.6e-21 \| 3e-27 \| 186.5 \| 145 \| (1, 145) \| 423 \| (1, 145) \| 670 \| Caudovirales tail fibre assembly protein \| Caudovirales tail fibre assembly protein; uniclust \| UniRef100\_UPI001F48AD60 \| 99.7 \| 1.7e-21 \| 3.2e-27 \| 150.8 \| 154 \| (3, 156) \| 423 \| (4, 161) \| 190 \| phage tail protein \| phage tail protein; uniclust \| UniRef100\_UPI0021505079 \| 99.7 \| 1.9e-21 \| 3.4e-27 \| 180.3 \| 154 \| (3, 156) \| 423 \| (4, 157) \| 849 \| phage tail protein \| phage tail protein; uniclust \| UniRef100\_UPI0002CDE47E \| 99.7 \| 1.9e-21 \| 3.5e-27 \| 178.8 \| 153 \| (3, 156) \| 423 \| (5, 157) \| 606 \| phage tail protein \| phage tail protein; pdb70 \| 6CU2\_A \| 98.8 \| 1e-12 \| 1.1e-16 \| 120.4 \| 105 \| (309, 423) \| 423 \| (169, 273) \| 273 \| Probable bacteriophage protein \| 6CU2\_A Probable bacteriophage protein adhesin, tail fiber, fiber, phage; pdb70 \| 6CT8\_A \| 98.8 \| 1e-12 \| 1.1e-16 \| 120.4 \| 105 \| (309, 423) \| 423 \| (169, 273) \| 273 \| Probable bacteriophage protein \| 6CT8\_A Probable bacteriophage protein adhesin, tail fiber, fiber, phage HET: MSE; pdb70 \| 6CL6\_F \| 98.7 \| 3e-12 \| 3e-16 \| 123.9 \| 105 \| (309, 423) \| 423 \| (268, 372) \| 372 \| Tail fiber protein \| 6CL6\_F Tail fiber protein R-type pyocin, Pseudomonas aeruginosa, contractile HET: EDO; pdb70 \| 6CXB\_A \| 98.6 \| 9.4e-12 \| 1e-15 \| 105.5 \| 100 \| (322, 423) \| 423 \| (41, 146) \| 146 \| Probable bacteriophage protein \| 6CXB\_A Probable bacteriophage protein adhesin, tail fiber, fiber, phage; pdb70 \| 4MTM\_A \| 98.4 \| 6.1e-11 \| 6.7e-15 \| 101.7 \| 97 \| (321, 423) \| 423 \| (61, 164) \| 164 \| Putative tail fiber protein \| 4MTM\_A Putative tail fiber protein Lectin fold, Tail fiber, receptor HET: EDO, BR, GOL; pdb70 \| 6CL5\_D \| 97.8 \| 8.2e-09 \| 8.5e-13 \| 99.5 \| 99 \| (323, 423) \| 423 \| (280, 384) \| 384 \| Phage tail protein \| 6CL5\_D Phage tail protein R-type pyocin, Pseudomonas aeruginosa, contractile |
| Top keywords  (threshold 1.00e-03 (evalue)) | **tail, Phage, fiber, fibre, tail\_collar, Fragment, domain\_containing, collar, Putative, Phage\_related** |
| Output files | ../../similar\_sequences/15\_FANPEZAQ\_CDS\_0015\_merged.svg ../../similar\_sequences/15\_FANPEZAQ\_CDS\_0015\_pdb70.a3m ../../similar\_sequences/15\_FANPEZAQ\_CDS\_0015\_pdb70.hhr ../../similar\_sequences/15\_FANPEZAQ\_CDS\_0015\_uniclust.a3m ../../similar\_sequences/15\_FANPEZAQ\_CDS\_0015\_uniclust.hhr |

#### Structure prediction (AlphaFold)^2^

|  |  |
| --- | --- |
| Stats | xml version="1.0" encoding="utf-8" standalone="no"?       2024-09-02T21:09:14.212967 image/svg+xml   Matplotlib v3.7.2, https://matplotlib.org/ |
| Predicted structure | **NGL Viewer Controls:**  - Center: *Left-Click* - Rotate: *Left-Click + Drag* - Translate: *Right-Click + Drag* - Zoom: *Shift + Left-Click + Drag* |
| Output files | ../../predicted\_structures/15\_FANPEZAQ\_CDS\_0015/features.pkl ../../predicted\_structures/15\_FANPEZAQ\_CDS\_0015/ranked\_0.pdb ../../predicted\_structures/15\_FANPEZAQ\_CDS\_0015/ranked\_0\_plots.svg ../../predicted\_structures/15\_FANPEZAQ\_CDS\_0015/result\_model\_1\_ptm\_pred\_0.pkl |

#### Structure similarity search results (Foldseek)^3^

|  |  |
| --- | --- |
| Structure databases searched | Pdb, Afdb-proteome, Afdb-uniprot50 |
| Results, scheme(s)  (Top layers only, threshold 1.00e-02 (evalue)) | xml version="1.0" encoding="utf-8" standalone="no"?       2024-09-02T21:10:33.642177 image/svg+xml   Matplotlib v3.7.2, https://matplotlib.org/ |
| Results, table  (threshold 1.00e-02 (evalue)) | db \| id \| prob \| evalue \| bits \| fident \| alnlen \| mismatch \| gapopen \| qstart \| qend \| tstart \| tend \| name \| description; pdb \| 4MTM\_A \| 1.0 \| 5.97e-05 \| 171 \| 0.185 \| 167 \| 98 \| 11 \| 265 \| 423 \| 1 \| 137 \| Putative tail fiber protein \| Putative tail fiber protein; pdb \| 6CU2\_A \| 1.0 \| 0.0007118 \| 145 \| 0.191 \| 94 \| 62 \| 8 \| 334 \| 423 \| 166 \| 249 \| R2-type pyocin \| R2-type pyocin; pdb \| 6CL5\_F \| 1.0 \| 0.0006391 \| 136 \| 0.27 \| 100 \| 58 \| 9 \| 334 \| 423 \| 280 \| 374 \| Tail fiber protein \| Tail fiber protein; pdb \| 7RFV\_A \| 1.0 \| 0.003584 \| 128 \| 0.187 \| 80 \| 59 \| 4 \| 179 \| 258 \| 166 \| 239 \| Tailspike protein \| Tailspike protein; pdb \| 6CXB\_A \| 1.0 \| 0.002458 \| 121 \| 0.261 \| 107 \| 63 \| 10 \| 328 \| 423 \| 21 \| 122 \| R1-type pyocin tail fiber protein \| R1-type pyocin tail fiber protein; afdb-proteome \| AF-Q8ZKK5-F1-MODEL\_V4 \| 1.0 \| 2.829e-22 \| 488 \| 0.217 \| 551 \| 249 \| 24 \| 1 \| 412 \| 1 \| 508 \| Putative phage tail fiber protein H \| Putative phage tail fiber protein H; afdb-proteome \| AF-A0A0H3H1R2-F1-MODEL\_V4 \| 1.0 \| 1.45e-16 \| 414 \| 0.227 \| 334 \| 169 \| 14 \| 1 \| 258 \| 1 \| 321 \| Uncharacterized protein \| Uncharacterized protein; afdb-proteome \| AF-G3XD71-F1-MODEL\_V4 \| 1.0 \| 1.12e-19 \| 397 \| 0.151 \| 708 \| 295 \| 32 \| 1 \| 423 \| 5 \| 691 \| Probable bacteriophage protein \| Probable bacteriophage protein; afdb-proteome \| AF-A0A0H3GKQ5-F1-MODEL\_V4 \| 1.0 \| 3.083e-16 \| 393 \| 0.243 \| 341 \| 157 \| 14 \| 1 \| 258 \| 1 \| 323 \| Uncharacterized protein \| Uncharacterized protein; afdb-proteome \| AF-Q8ZMU5-F1-MODEL\_V4 \| 1.0 \| 5.949e-19 \| 378 \| 0.207 \| 583 \| 224 \| 23 \| 1 \| 421 \| 2 \| 508 \| Fels-2 prophage protein \| Fels-2 prophage protein; afdb-proteome \| AF-P76508-F1-MODEL\_V4 \| 1.0 \| 1.03e-06 \| 245 \| 0.287 \| 132 \| 82 \| 6 \| 296 \| 423 \| 49 \| 172 \| Putative uncharacterized protein YfdL \| Putative uncharacterized protein YfdL; afdb-proteome \| AF-P45581-F1-MODEL\_V4 \| 1.0 \| 8.547e-05 \| 206 \| 0.295 \| 88 \| 48 \| 6 \| 341 \| 423 \| 131 \| 209 \| Uncharacterized protein StfP \| Uncharacterized protein StfP; afdb-proteome \| AF-Q57793-F1-MODEL\_V4 \| 1.0 \| 0.0005634 \| 122 \| 0.156 \| 134 \| 88 \| 8 \| 307 \| 423 \| 26 \| 151 \| Uncharacterized protein MJ0347 \| Uncharacterized protein MJ0347; afdb-proteome \| AF-Q9I5S0-F1-MODEL\_V4 \| 0.923 \| 0.004607 \| 69 \| 0.114 \| 368 \| 169 \| 21 \| 193 \| 423 \| 1 \| 348 \| Uncharacterized protein \| Uncharacterized protein; afdb-proteome \| AF-Q9I5S3-F1-MODEL\_V4 \| 0.772 \| 0.006032 \| 60 \| 0.125 \| 389 \| 148 \| 24 \| 198 \| 423 \| 2 \| 361 \| Uncharacterized protein \| Uncharacterized protein; afdb-uniprot50 \| AF-A0A430HF82-F1-MODEL\_V4 \| 1.0 \| 1.815e-31 \| 1105 \| 0.473 \| 262 \| 126 \| 4 \| 4 \| 261 \| 3 \| 256 \| Uncharacterized protein \| Uncharacterized protein; afdb-uniprot50 \| AF-A0A2A3MGV4-F1-MODEL\_V4 \| 1.0 \| 7.366e-31 \| 1016 \| 0.456 \| 267 \| 138 \| 3 \| 1 \| 261 \| 1 \| 266 \| Phage tail protein \| Phage tail protein; afdb-uniprot50 \| AF-A0A250DS87-F1-MODEL\_V4 \| 1.0 \| 4.298e-31 \| 990 \| 0.43 \| 265 \| 146 \| 4 \| 1 \| 263 \| 1 \| 262 \| Phage tail protein \| Phage tail protein; afdb-uniprot50 \| AF-A0A0F9VFY7-F1-MODEL\_V4 \| 1.0 \| 1.505e-29 \| 962 \| 0.415 \| 265 \| 151 \| 3 \| 1 \| 263 \| 1 \| 263 \| Collar domain-containing protein \| Collar domain-containing protein; afdb-uniprot50 \| AF-A0A5E7KHM1-F1-MODEL\_V4 \| 1.0 \| 7.473e-30 \| 929 \| 0.436 \| 266 \| 143 \| 4 \| 1 \| 261 \| 1 \| 264 \| Collar domain-containing protein \| Collar domain-containing protein; afdb-uniprot50 \| AF-A0A432RGY3-F1-MODEL\_V4 \| 1.0 \| 2.674e-19 \| 857 \| 0.531 \| 158 \| 71 \| 1 \| 1 \| 155 \| 1 \| 158 \| Uncharacterized protein \| Uncharacterized protein; afdb-uniprot50 \| AF-A0A7V7KJT5-F1-MODEL\_V4 \| 1.0 \| 6.683e-19 \| 853 \| 0.503 \| 153 \| 76 \| 0 \| 1 \| 153 \| 1 \| 153 \| Sialidase domain-containing protein \| Sialidase domain-containing protein; afdb-uniprot50 \| AF-G4CPH8-F1-MODEL\_V4 \| 1.0 \| 1.5e-18 \| 848 \| 0.525 \| 154 \| 72 \| 1 \| 1 \| 153 \| 1 \| 154 \| Uncharacterized protein \| Uncharacterized protein; afdb-uniprot50 \| AF-A0A3N9U476-F1-MODEL\_V4 \| 1.0 \| 3.27e-20 \| 844 \| 0.465 \| 176 \| 90 \| 1 \| 1 \| 176 \| 1 \| 172 \| Collar domain-containing protein \| Collar domain-containing protein; afdb-uniprot50 \| AF-A5N994-F1-MODEL\_V4 \| 1.0 \| 3.899e-19 \| 843 \| 0.47 \| 153 \| 81 \| 0 \| 1 \| 153 \| 2 \| 154 \| Uncharacterized protein \| Uncharacterized protein; afdb-uniprot50 \| AF-A0A724JF68-F1-MODEL\_V4 \| 1.0 \| 7.855e-19 \| 842 \| 0.503 \| 157 \| 78 \| 0 \| 2 \| 158 \| 3 \| 159 \| Phage tail protein \| Phage tail protein; afdb-uniprot50 \| AF-A0A396RYI5-F1-MODEL\_V4 \| 1.0 \| 2.57e-18 \| 834 \| 0.458 \| 155 \| 84 \| 0 \| 2 \| 156 \| 6 \| 160 \| Uncharacterized protein \| Uncharacterized protein; afdb-uniprot50 \| AF-A0A4R1K4B3-F1-MODEL\_V4 \| 1.0 \| 9.233e-19 \| 832 \| 0.475 \| 162 \| 85 \| 0 \| 1 \| 162 \| 1 \| 162 \| Phage-related tail fiber protein \| Phage-related tail fiber protein; afdb-uniprot50 \| AF-A0A3Z6GCN4-F1-MODEL\_V4 \| 1.0 \| 6e-19 \| 830 \| 0.493 \| 158 \| 80 \| 0 \| 1 \| 158 \| 2 \| 159 \| Phage tail protein \| Phage tail protein; afdb-uniprot50 \| AF-A0A345LUV5-F1-MODEL\_V4 \| 1.0 \| 2.072e-18 \| 830 \| 0.519 \| 152 \| 73 \| 0 \| 4 \| 155 \| 3 \| 154 \| Collar domain-containing protein \| Collar domain-containing protein; afdb-uniprot50 \| AF-A0A5T7A7I4-F1-MODEL\_V4 \| 1.0 \| 3.748e-18 \| 828 \| 0.481 \| 158 \| 82 \| 0 \| 1 \| 158 \| 2 \| 159 \| Phage tail protein \| Phage tail protein; afdb-uniprot50 \| AF-A0A2P8VLU5-F1-MODEL\_V4 \| 1.0 \| 3.748e-18 \| 827 \| 0.487 \| 158 \| 81 \| 0 \| 1 \| 158 \| 2 \| 159 \| Phage tail protein \| Phage tail protein; afdb-uniprot50 \| AF-A0A6M0RCD0-F1-MODEL\_V4 \| 1.0 \| 1.258e-19 \| 825 \| 0.403 \| 166 \| 98 \| 1 \| 1 \| 166 \| 1 \| 165 \| Phage tail protein \| Phage tail protein; afdb-uniprot50 \| AF-A0A1I1WBP0-F1-MODEL\_V4 \| 1.0 \| 1.085e-18 \| 824 \| 0.478 \| 163 \| 83 \| 1 \| 1 \| 163 \| 1 \| 161 \| Phage tail-collar fibre protein \| Phage tail-collar fibre protein; afdb-uniprot50 \| AF-A0A080IG54-F1-MODEL\_V4 \| 1.0 \| 5.767e-18 \| 823 \| 0.481 \| 158 \| 82 \| 0 \| 1 \| 158 \| 2 \| 159 \| Phage tail-collar fiber family protein \| Phage tail-collar fiber family protein; afdb-uniprot50 \| AF-A0A5I1KPW3-F1-MODEL\_V4 \| 1.0 \| 1.763e-18 \| 820 \| 0.496 \| 157 \| 79 \| 0 \| 2 \| 158 \| 3 \| 159 \| Phage tail protein \| Phage tail protein; afdb-uniprot50 \| AF-A0A4R1TM91-F1-MODEL\_V4 \| 1.0 \| 1.145e-18 \| 819 \| 0.452 \| 157 \| 82 \| 2 \| 1 \| 153 \| 1 \| 157 \| Tail-collar fiber protein \| Tail-collar fiber protein; afdb-uniprot50 \| AF-G7LSU4-F1-MODEL\_V4 \| 1.0 \| 2.187e-18 \| 819 \| 0.512 \| 164 \| 79 \| 1 \| 1 \| 163 \| 1 \| 164 \| Uncharacterized protein \| Uncharacterized protein; afdb-uniprot50 \| AF-X6QCI0-F1-MODEL\_V4 \| 1.0 \| 7.155e-18 \| 816 \| 0.513 \| 150 \| 73 \| 0 \| 4 \| 153 \| 3 \| 152 \| Tail fiber protein \| Tail fiber protein; afdb-uniprot50 \| AF-D6IGH8-F1-MODEL\_V4 \| 1.0 \| 7.155e-18 \| 816 \| 0.481 \| 158 \| 82 \| 0 \| 1 \| 158 \| 2 \| 159 \| Phage tail fiber protein \| Phage tail fiber protein; afdb-uniprot50 \| AF-A0A7U3LRL7-F1-MODEL\_V4 \| 1.0 \| 8.749e-19 \| 814 \| 0.506 \| 160 \| 79 \| 0 \| 1 \| 160 \| 2 \| 161 \| Uncharacterized protein \| Uncharacterized protein; afdb-uniprot50 \| AF-A0A7T3MYV0-F1-MODEL\_V4 \| 1.0 \| 1.276e-18 \| 813 \| 0.437 \| 153 \| 86 \| 0 \| 1 \| 153 \| 2 \| 154 \| Phage tail protein \| Phage tail protein; afdb-uniprot50 \| AF-A0A5W6HQH5-F1-MODEL\_V4 \| 1.0 \| 9.885e-18 \| 813 \| 0.487 \| 158 \| 81 \| 0 \| 1 \| 158 \| 2 \| 159 \| Phage tail protein \| Phage tail protein; afdb-uniprot50 \| AF-A0A604P221-F1-MODEL\_V4 \| 1.0 \| 4.405e-18 \| 813 \| 0.496 \| 157 \| 79 \| 0 \| 2 \| 158 \| 3 \| 159 \| Phage tail protein \| Phage tail protein; afdb-uniprot50 \| AF-A0A6B3YWL7-F1-MODEL\_V4 \| 1.0 \| 1.963e-18 \| 811 \| 0.424 \| 153 \| 88 \| 0 \| 1 \| 153 \| 2 \| 154 \| Phage tail protein \| Phage tail protein; afdb-uniprot50 \| AF-A0A2S8QLP1-F1-MODEL\_V4 \| 1.0 \| 3.188e-18 \| 806 \| 0.453 \| 161 \| 88 \| 0 \| 1 \| 161 \| 2 \| 162 \| Uncharacterized protein \| Uncharacterized protein; afdb-uniprot50 \| AF-A0A5U9RX76-F1-MODEL\_V4 \| 1.0 \| 2.072e-18 \| 806 \| 0.493 \| 158 \| 80 \| 0 \| 1 \| 158 \| 2 \| 159 \| Phage tail protein \| Phage tail protein; afdb-uniprot50 \| AF-A0A292AIP5-F1-MODEL\_V4 \| 1.0 \| 1.043e-17 \| 806 \| 0.496 \| 155 \| 76 \| 1 \| 1 \| 155 \| 2 \| 154 \| Uncharacterized protein \| Uncharacterized protein; afdb-uniprot50 \| AF-A0A0U3LJE0-F1-MODEL\_V4 \| 1.0 \| 4.837e-19 \| 805 \| 0.448 \| 176 \| 97 \| 0 \| 1 \| 176 \| 1 \| 176 \| Bacteriophage variable tail fiber protein \| Bacteriophage variable tail fiber protein; afdb-uniprot50 \| AF-F3YY38-F1-MODEL\_V4 \| 1.0 \| 3.551e-18 \| 804 \| 0.519 \| 154 \| 73 \| 1 \| 1 \| 153 \| 2 \| 155 \| Uncharacterized protein \| Uncharacterized protein; afdb-uniprot50 \| AF-A0A5T6DFY3-F1-MODEL\_V4 \| 1.0 \| 1.67e-18 \| 803 \| 0.496 \| 157 \| 79 \| 0 \| 2 \| 158 \| 3 \| 159 \| Phage tail protein \| Phage tail protein; afdb-uniprot50 \| AF-A0A376D432-F1-MODEL\_V4 \| 1.0 \| 1.145e-18 \| 803 \| 0.481 \| 158 \| 82 \| 0 \| 1 \| 158 \| 2 \| 159 \| Putative variable tail fiber protein \| Putative variable tail fiber protein; afdb-uniprot50 \| AF-A0A447PAU2-F1-MODEL\_V4 \| 1.0 \| 1.5e-18 \| 803 \| 0.487 \| 158 \| 81 \| 0 \| 1 \| 158 \| 2 \| 159 \| Phage tail-like protein \| Phage tail-like protein; afdb-uniprot50 \| AF-A0A3N9UIV0-F1-MODEL\_V4 \| 1.0 \| 1.763e-18 \| 801 \| 0.458 \| 155 \| 84 \| 0 \| 1 \| 155 \| 1 \| 155 \| Uncharacterized protein \| Uncharacterized protein; afdb-uniprot50 \| AF-A0A348HHI2-F1-MODEL\_V4 \| 1.0 \| 2.218e-17 \| 798 \| 0.419 \| 155 \| 90 \| 0 \| 1 \| 155 \| 1 \| 155 \| Phage-related tail fibre protein \| Phage-related tail fibre protein; afdb-uniprot50 \| AF-A0A7L9R9Q1-F1-MODEL\_V4 \| 1.0 \| 2.072e-18 \| 798 \| 0.487 \| 162 \| 83 \| 0 \| 1 \| 162 \| 2 \| 163 \| Phage tail protein \| Phage tail protein; afdb-uniprot50 \| AF-A0A7D6C2F1-F1-MODEL\_V4 \| 1.0 \| 1.366e-17 \| 798 \| 0.432 \| 155 \| 88 \| 0 \| 1 \| 155 \| 2 \| 156 \| Phage tail protein \| Phage tail protein; afdb-uniprot50 \| AF-A0A5T2YBP8-F1-MODEL\_V4 \| 1.0 \| 2.471e-17 \| 797 \| 0.481 \| 158 \| 82 \| 0 \| 1 \| 158 \| 2 \| 159 \| Phage tail protein \| Phage tail protein; afdb-uniprot50 \| AF-A0A2Z4WEE0-F1-MODEL\_V4 \| 1.0 \| 3.188e-18 \| 796 \| 0.4 \| 155 \| 93 \| 0 \| 1 \| 155 \| 1 \| 155 \| Uncharacterized protein \| Uncharacterized protein; afdb-uniprot50 \| AF-A0A6L5EC52-F1-MODEL\_V4 \| 1.0 \| 1.86e-18 \| 795 \| 0.515 \| 159 \| 76 \| 1 \| 1 \| 158 \| 1 \| 159 \| Phage tail protein \| Phage tail protein; afdb-uniprot50 \| AF-A0A2G8C8T9-F1-MODEL\_V4 \| 1.0 \| 1.421e-18 \| 790 \| 0.496 \| 159 \| 79 \| 1 \| 1 \| 158 \| 1 \| 159 \| Phage tail protein \| Phage tail protein; afdb-uniprot50 \| AF-A0A0P9SFR0-F1-MODEL\_V4 \| 1.0 \| 2.102e-17 \| 790 \| 0.441 \| 154 \| 86 \| 0 \| 2 \| 155 \| 6 \| 159 \| Phage-related tail fiber protein-like protein \| Phage-related tail fiber protein-like protein; afdb-uniprot50 \| AF-A0A084ZPT2-F1-MODEL\_V4 \| 1.0 \| 7.155e-18 \| 788 \| 0.506 \| 156 \| 76 \| 1 \| 1 \| 155 \| 1 \| 156 \| Phage tail fiber protein \| Phage tail fiber protein; afdb-uniprot50 \| AF-A0A2R8CQR2-F1-MODEL\_V4 \| 1.0 \| 1.411e-26 \| 786 \| 0.235 \| 476 \| 203 \| 16 \| 1 \| 423 \| 1 \| 368 \| Uncharacterized protein \| Uncharacterized protein; afdb-uniprot50 \| AF-J6HCZ5-F1-MODEL\_V4 \| 1.0 \| 7.551e-18 \| 785 \| 0.434 \| 161 \| 91 \| 0 \| 1 \| 161 \| 2 \| 162 \| Tail fiber protein \| Tail fiber protein; afdb-uniprot50 \| AF-A0A5Y2E3R8-F1-MODEL\_V4 \| 1.0 \| 5.685e-19 \| 784 \| 0.429 \| 177 \| 96 \| 2 \| 1 \| 172 \| 1 \| 177 \| Phage tail protein \| Phage tail protein; afdb-uniprot50 \| AF-A0A447P489-F1-MODEL\_V4 \| 1.0 \| 5.851e-17 \| 783 \| 0.507 \| 136 \| 67 \| 0 \| 1 \| 136 \| 2 \| 137 \| Phage tail fiber protein \| Phage tail fiber protein; afdb-uniprot50 \| AF-A0A7U6FCV4-F1-MODEL\_V4 \| 1.0 \| 3.414e-17 \| 783 \| 0.424 \| 153 \| 88 \| 0 \| 1 \| 153 \| 5 \| 157 \| Uncharacterized protein \| Uncharacterized protein; afdb-uniprot50 \| AF-A0A714EVS6-F1-MODEL\_V4 \| 1.0 \| 4.649e-18 \| 783 \| 0.503 \| 157 \| 78 \| 0 \| 2 \| 158 \| 3 \| 159 \| Phage tail protein \| Phage tail protein; afdb-uniprot50 \| AF-A0A377Q9P4-F1-MODEL\_V4 \| 1.0 \| 1.366e-17 \| 783 \| 0.487 \| 154 \| 76 \| 1 \| 3 \| 153 \| 2 \| 155 \| Uncharacterized protein \| Uncharacterized protein; afdb-uniprot50 \| AF-C2BG12-F1-MODEL\_V4 \| 1.0 \| 4.235e-17 \| 782 \| 0.346 \| 153 \| 100 \| 0 \| 1 \| 153 \| 1 \| 153 \| Uncharacterized protein \| Uncharacterized protein; afdb-uniprot50 \| AF-A0A827L2X9-F1-MODEL\_V4 \| 1.0 \| 2.435e-18 \| 781 \| 0.49 \| 159 \| 80 \| 1 \| 1 \| 158 \| 1 \| 159 \| Phage tail protein \| Phage tail protein; afdb-uniprot50 \| AF-A0A3M6E5R0-F1-MODEL\_V4 \| 1.0 \| 3.234e-17 \| 780 \| 0.432 \| 155 \| 88 \| 0 \| 1 \| 155 \| 5 \| 159 \| Tail fiber protein H \| Tail fiber protein H; afdb-uniprot50 \| AF-A0A433ZYK3-F1-MODEL\_V4 \| 1.0 \| 4.174e-18 \| 779 \| 0.465 \| 161 \| 86 \| 0 \| 1 \| 161 \| 2 \| 162 \| Uncharacterized protein \| Uncharacterized protein; afdb-uniprot50 \| AF-D1P881-F1-MODEL\_V4 \| 1.0 \| 2.713e-18 \| 778 \| 0.453 \| 163 \| 87 \| 1 \| 4 \| 164 \| 3 \| 165 \| Uncharacterized protein \| Uncharacterized protein; afdb-uniprot50 \| AF-A0A506QSU0-F1-MODEL\_V4 \| 1.0 \| 5.253e-17 \| 777 \| 0.474 \| 158 \| 83 \| 0 \| 1 \| 158 \| 2 \| 159 \| Phage tail protein \| Phage tail protein; afdb-uniprot50 \| AF-A0A4P9VIJ6-F1-MODEL\_V4 \| 1.0 \| 4.649e-18 \| 777 \| 0.397 \| 166 \| 95 \| 3 \| 1 \| 161 \| 1 \| 166 \| Uncharacterized protein \| Uncharacterized protein; afdb-uniprot50 \| AF-A0A6G2EFZ9-F1-MODEL\_V4 \| 1.0 \| 1.86e-18 \| 775 \| 0.445 \| 166 \| 88 \| 1 \| 1 \| 162 \| 1 \| 166 \| Phage tail protein \| Phage tail protein; afdb-uniprot50 \| AF-A0A659NRL8-F1-MODEL\_V4 \| 1.0 \| 4.39e-22 \| 774 \| 0.321 \| 330 \| 111 \| 5 \| 2 \| 331 \| 3 \| 219 \| Phage tail protein \| Phage tail protein; afdb-uniprot50 \| AF-A0A5W0Y2X8-F1-MODEL\_V4 \| 1.0 \| 5.465e-18 \| 772 \| 0.481 \| 158 \| 82 \| 0 \| 1 \| 158 \| 2 \| 159 \| Phage tail protein \| Phage tail protein; afdb-uniprot50 \| AF-A0A080HXC3-F1-MODEL\_V4 \| 1.0 \| 1.192e-19 \| 771 \| 0.419 \| 193 \| 101 \| 2 \| 1 \| 184 \| 2 \| 192 \| Phage tail-collar fiber family protein \| Phage tail-collar fiber family protein; afdb-uniprot50 \| AF-F4VH24-F1-MODEL\_V4 \| 1.0 \| 1.294e-17 \| 771 \| 0.481 \| 158 \| 82 \| 0 \| 1 \| 158 \| 2 \| 159 \| Putative bacteriophage variable tail fiber protein H \| Putative bacteriophage variable tail fiber protein H; afdb-uniprot50 \| AF-A0A3N0FYV1-F1-MODEL\_V4 \| 1.0 \| 5.178e-18 \| 771 \| 0.484 \| 163 \| 83 \| 1 \| 1 \| 162 \| 1 \| 163 \| Phage tail protein \| Phage tail protein; afdb-uniprot50 \| AF-A0A0T9KM71-F1-MODEL\_V4 \| 1.0 \| 2.471e-17 \| 771 \| 0.465 \| 161 \| 86 \| 0 \| 1 \| 161 \| 2 \| 162 \| Variable tail fiber protein \| Variable tail fiber protein; afdb-uniprot50 \| AF-D4GNL8-F1-MODEL\_V4 \| 1.0 \| 8.531e-17 \| 771 \| 0.481 \| 158 \| 82 \| 0 \| 1 \| 158 \| 2 \| 159 \| H \| H; afdb-uniprot50 \| AF-E0SFL4-F1-MODEL\_V4 \| 1.0 \| 2.072e-18 \| 769 \| 0.5 \| 164 \| 79 \| 2 \| 1 \| 163 \| 93 \| 254 \| Tail fiber protein \| Tail fiber protein; afdb-uniprot50 \| AF-A0A5F1ELN5-F1-MODEL\_V4 \| 1.0 \| 9.233e-19 \| 768 \| 0.484 \| 159 \| 81 \| 1 \| 1 \| 158 \| 1 \| 159 \| Phage tail protein \| Phage tail protein; afdb-uniprot50 \| AF-A0A3M5N4H6-F1-MODEL\_V4 \| 1.0 \| 9.003e-17 \| 768 \| 0.441 \| 154 \| 86 \| 0 \| 2 \| 155 \| 6 \| 159 \| Phage-related tail fiber protein-like protein \| Phage-related tail fiber protein-like protein; afdb-uniprot50 \| AF-A0A0T9TRH4-F1-MODEL\_V4 \| 1.0 \| 2.874e-29 \| 766 \| 0.299 \| 468 \| 184 \| 15 \| 1 \| 423 \| 2 \| 370 \| Tail fiber protein \| Tail fiber protein; afdb-uniprot50 \| AF-A0A1M7FCH9-F1-MODEL\_V4 \| 1.0 \| 2.471e-17 \| 766 \| 0.477 \| 155 \| 81 \| 0 \| 1 \| 155 \| 2 \| 156 \| Phage tail-collar fibre protein \| Phage tail-collar fibre protein; afdb-uniprot50 \| AF-A0A385XY40-F1-MODEL\_V4 \| 1.0 \| 8.41e-18 \| 764 \| 0.49 \| 159 \| 80 \| 1 \| 1 \| 158 \| 1 \| 159 \| Collar domain-containing protein \| Collar domain-containing protein; afdb-uniprot50 \| AF-A0A741RNT9-F1-MODEL\_V4 \| 1.0 \| 1.605e-17 \| 763 \| 0.481 \| 158 \| 82 \| 0 \| 1 \| 158 \| 2 \| 159 \| Phage tail protein \| Phage tail protein; afdb-uniprot50 \| AF-A0A2T5HH07-F1-MODEL\_V4 \| 1.0 \| 3.414e-17 \| 761 \| 0.439 \| 157 \| 84 \| 2 \| 1 \| 153 \| 1 \| 157 \| Tail-collar fiber protein \| Tail-collar fiber protein; afdb-uniprot50 \| AF-A0A482IPK9-F1-MODEL\_V4 \| 1.0 \| 2.218e-17 \| 760 \| 0.359 \| 153 \| 98 \| 0 \| 1 \| 153 \| 2 \| 154 \| Uncharacterized protein \| Uncharacterized protein; afdb-uniprot50 \| AF-A0A2P7NQL6-F1-MODEL\_V4 \| 1.0 \| 6.516e-17 \| 758 \| 0.393 \| 155 \| 94 \| 0 \| 1 \| 155 \| 3 \| 157 \| Uncharacterized protein \| Uncharacterized protein; afdb-uniprot50 \| AF-W2C9W5-F1-MODEL\_V4 \| 1.0 \| 1.441e-17 \| 757 \| 0.509 \| 151 \| 74 \| 0 \| 1 \| 151 \| 2 \| 152 \| Phage tail protein \| Phage tail protein; afdb-uniprot50 \| AF-A0A741LDP6-F1-MODEL\_V4 \| 1.0 \| 8.084e-17 \| 757 \| 0.503 \| 151 \| 75 \| 0 \| 1 \| 151 \| 2 \| 152 \| Phage tail protein \| Phage tail protein; afdb-uniprot50 \| AF-A0A1S6TPA5-F1-MODEL\_V4 \| 1.0 \| 2.752e-17 \| 757 \| 0.352 \| 156 \| 98 \| 1 \| 3 \| 158 \| 2 \| 154 \| Phage tail-collar fiber protein (DUF3751 domain) \| Phage tail-collar fiber protein (DUF3751 domain); afdb-uniprot50 \| AF-A0A377LB56-F1-MODEL\_V4 \| 1.0 \| 1.162e-17 \| 757 \| 0.469 \| 162 \| 86 \| 0 \| 1 \| 162 \| 2 \| 163 \| Phage tail collar protein \| Phage tail collar protein; afdb-uniprot50 \| AF-A0A329VCK4-F1-MODEL\_V4 \| 1.0 \| 4.453e-21 \| 755 \| 0.343 \| 306 \| 100 \| 3 \| 4 \| 309 \| 3 \| 207 \| Phage tail protein \| Phage tail protein; afdb-uniprot50 \| AF-A0A2D5S2E6-F1-MODEL\_V4 \| 1.0 \| 6.877e-17 \| 753 \| 0.337 \| 151 \| 100 \| 0 \| 3 \| 153 \| 2 \| 152 \| Phage tail protein \| Phage tail protein; afdb-uniprot50 \| AF-A0A0S8ZMG3-F1-MODEL\_V4 \| 1.0 \| 1.179e-16 \| 751 \| 0.424 \| 153 \| 88 \| 0 \| 1 \| 153 \| 5 \| 157 \| Collar domain-containing protein \| Collar domain-containing protein; afdb-uniprot50 \| AF-V0XNW2-F1-MODEL\_V4 \| 1.0 \| 4.235e-17 \| 750 \| 0.448 \| 154 \| 84 \| 1 \| 1 \| 153 \| 2 \| 155 \| Uncharacterized protein \| Uncharacterized protein; afdb-uniprot50 \| AF-A0A1Y0FPI9-F1-MODEL\_V4 \| 1.0 \| 1.521e-17 \| 750 \| 0.468 \| 160 \| 83 \| 2 \| 1 \| 158 \| 1 \| 160 \| 6-phosphogluconolactonase \| 6-phosphogluconolactonase; afdb-uniprot50 \| AF-A0A5X2Z7L4-F1-MODEL\_V4 \| 1.0 \| 7.337e-20 \| 749 \| 0.322 \| 307 \| 107 \| 3 \| 1 \| 307 \| 2 \| 207 \| Phage tail protein \| Phage tail protein; afdb-uniprot50 \| AF-A0A839C2U0-F1-MODEL\_V4 \| 1.0 \| 4.716e-17 \| 749 \| 0.493 \| 158 \| 80 \| 0 \| 1 \| 158 \| 1 \| 158 \| Phage tail protein \| Phage tail protein; afdb-uniprot50 \| AF-A0A7W3F3N4-F1-MODEL\_V4 \| 1.0 \| 2.471e-17 \| 747 \| 0.487 \| 158 \| 81 \| 0 \| 1 \| 158 \| 2 \| 159 \| Phage tail protein \| Phage tail protein; afdb-uniprot50 \| AF-A0A522IBP7-F1-MODEL\_V4 \| 1.0 \| 2.471e-17 \| 746 \| 0.435 \| 163 \| 89 \| 2 \| 1 \| 162 \| 2 \| 162 \| Uncharacterized protein \| Uncharacterized protein; afdb-uniprot50 \| AF-A0A5S9NC05-F1-MODEL\_V4 \| 1.0 \| 7.258e-17 \| 746 \| 0.376 \| 154 \| 95 \| 1 \| 1 \| 153 \| 1 \| 154 \| Uncharacterized protein \| Uncharacterized protein; afdb-uniprot50 \| AF-A0A1I0EJH2-F1-MODEL\_V4 \| 1.0 \| 1.719e-16 \| 745 \| 0.418 \| 153 \| 88 \| 1 \| 1 \| 153 \| 1 \| 152 \| Phage tail-collar fibre protein \| Phage tail-collar fibre protein; afdb-uniprot50 \| AF-D7XVP6-F1-MODEL\_V4 \| 1.0 \| 1.516e-21 \| 745 \| 0.327 \| 324 \| 113 \| 5 \| 1 \| 324 \| 2 \| 220 \| Phage tail fiber repeat protein \| Phage tail fiber repeat protein; afdb-uniprot50 \| AF-A0A1G5RGJ6-F1-MODEL\_V4 \| 1.0 \| 2.742e-21 \| 744 \| 0.349 \| 323 \| 103 \| 5 \| 4 \| 326 \| 3 \| 218 \| Phage tail fibre repeat-containing protein \| Phage tail fibre repeat-containing protein; afdb-uniprot50 \| AF-A0A351NNZ8-F1-MODEL\_V4 \| 1.0 \| 2.102e-17 \| 744 \| 0.484 \| 161 \| 83 \| 0 \| 2 \| 162 \| 3 \| 163 \| Uncharacterized protein \| Uncharacterized protein; afdb-uniprot50 \| AF-A0A7A6RGS2-F1-MODEL\_V4 \| 1.0 \| 3.414e-17 \| 743 \| 0.453 \| 172 \| 94 \| 0 \| 1 \| 172 \| 2 \| 173 \| Phage tail protein \| Phage tail protein; afdb-uniprot50 \| AF-A0A417HJB2-F1-MODEL\_V4 \| 1.0 \| 2.904e-17 \| 743 \| 0.358 \| 156 \| 100 \| 0 \| 3 \| 158 \| 5 \| 160 \| Uncharacterized protein \| Uncharacterized protein; afdb-uniprot50 \| AF-A0A5U6FPA7-F1-MODEL\_V4 \| 1.0 \| 1.462e-16 \| 742 \| 0.503 \| 151 \| 75 \| 0 \| 1 \| 151 \| 2 \| 152 \| Phage tail protein \| Phage tail protein; afdb-uniprot50 \| AF-A0A1M5TXK3-F1-MODEL\_V4 \| 1.0 \| 1.653e-30 \| 742 \| 0.298 \| 469 \| 205 \| 18 \| 1 \| 423 \| 2 \| 392 \| Phage tail-collar fibre protein \| Phage tail-collar fibre protein; afdb-uniprot50 \| AF-D8J073-F1-MODEL\_V4 \| 1.0 \| 1.196e-30 \| 741 \| 0.28 \| 449 \| 249 \| 14 \| 1 \| 418 \| 3 \| 408 \| Bacteriophage variable tail fibre protein \| Bacteriophage variable tail fibre protein; afdb-uniprot50 \| AF-A0A1H5GVL6-F1-MODEL\_V4 \| 1.0 \| 6.516e-17 \| 741 \| 0.442 \| 156 \| 86 \| 1 \| 1 \| 155 \| 1 \| 156 \| Phage-related tail fibre protein \| Phage-related tail fibre protein; afdb-uniprot50 \| AF-A0A7X5IDT3-F1-MODEL\_V4 \| 1.0 \| 5.544e-17 \| 740 \| 0.31 \| 158 \| 109 \| 0 \| 1 \| 158 \| 1 \| 158 \| Peptidase S74 domain-containing protein \| Peptidase S74 domain-containing protein; afdb-uniprot50 \| AF-A0A6B4D2K2-F1-MODEL\_V4 \| 1.0 \| 6e-19 \| 738 \| 0.4 \| 195 \| 110 \| 2 \| 1 \| 188 \| 2 \| 196 \| Phage tail protein \| Phage tail protein; afdb-uniprot50 \| AF-A0A6G4CRK8-F1-MODEL\_V4 \| 1.0 \| 8.875e-18 \| 738 \| 0.465 \| 161 \| 86 \| 0 \| 1 \| 161 \| 2 \| 162 \| Phage tail protein \| Phage tail protein; afdb-uniprot50 \| AF-A0A2W5TDM3-F1-MODEL\_V4 \| 1.0 \| 9.003e-17 \| 738 \| 0.445 \| 157 \| 86 \| 1 \| 3 \| 158 \| 2 \| 158 \| Peptidase S74 domain-containing protein \| Peptidase S74 domain-containing protein; afdb-uniprot50 \| AF-A0A6A1RN19-F1-MODEL\_V4 \| 1.0 \| 1.058e-16 \| 738 \| 0.381 \| 160 \| 98 \| 1 \| 3 \| 162 \| 5 \| 163 \| Uncharacterized protein \| Uncharacterized protein; afdb-uniprot50 \| AF-A0A1H9EZ18-F1-MODEL\_V4 \| 1.0 \| 1.162e-17 \| 737 \| 0.493 \| 164 \| 81 \| 1 \| 1 \| 162 \| 2 \| 165 \| Phage Tail Collar Domain \| Phage Tail Collar Domain; afdb-uniprot50 \| AF-A0A8B5VHZ4-F1-MODEL\_V4 \| 1.0 \| 4.469e-17 \| 736 \| 0.444 \| 153 \| 85 \| 0 \| 1 \| 153 \| 2 \| 154 \| Phage tail protein \| Phage tail protein; afdb-uniprot50 \| AF-A0A806LIS8-F1-MODEL\_V4 \| 1.0 \| 2.471e-17 \| 736 \| 0.434 \| 161 \| 90 \| 1 \| 1 \| 161 \| 2 \| 161 \| Uncharacterized protein \| Uncharacterized protein; afdb-uniprot50 \| AF-A0A1W1XL57-F1-MODEL\_V4 \| 1.0 \| 3.802e-17 \| 734 \| 0.37 \| 162 \| 101 \| 1 \| 1 \| 161 \| 1 \| 162 \| Phage tail-collar fibre protein \| Phage tail-collar fibre protein; afdb-uniprot50 \| AF-N1ZHX9-F1-MODEL\_V4 \| 1.0 \| 1.629e-16 \| 734 \| 0.322 \| 158 \| 107 \| 0 \| 1 \| 158 \| 3 \| 160 \| Peptidase S74 domain-containing protein \| Peptidase S74 domain-containing protein; afdb-uniprot50 \| AF-A0A7W6RGC7-F1-MODEL\_V4 \| 1.0 \| 7.633e-21 \| 731 \| 0.423 \| 229 \| 110 \| 6 \| 1 \| 219 \| 1 \| 217 \| Phage-related tail fiber protein \| Phage-related tail fiber protein; afdb-uniprot50 \| AF-A0A418H7E0-F1-MODEL\_V4 \| 1.0 \| 1.244e-16 \| 728 \| 0.427 \| 159 \| 90 \| 1 \| 1 \| 158 \| 2 \| 160 \| Phage tail protein \| Phage tail protein; afdb-uniprot50 \| AF-A0A3D9EDX5-F1-MODEL\_V4 \| 1.0 \| 7.77e-16 \| 728 \| 0.5 \| 154 \| 76 \| 1 \| 1 \| 153 \| 1 \| 154 \| Tail-collar fiber protein \| Tail-collar fiber protein; afdb-uniprot50 \| AF-A0A1H4Y3H3-F1-MODEL\_V4 \| 1.0 \| 8.084e-17 \| 728 \| 0.44 \| 150 \| 84 \| 0 \| 4 \| 153 \| 8 \| 157 \| Phage Tail Collar Domain \| Phage Tail Collar Domain; afdb-uniprot50 \| AF-A0A5C9ABQ6-F1-MODEL\_V4 \| 1.0 \| 5.049e-16 \| 726 \| 0.507 \| 130 \| 64 \| 0 \| 1 \| 130 \| 2 \| 131 \| Phage tail protein \| Phage tail protein; afdb-uniprot50 \| AF-A0A0N9MHW1-F1-MODEL\_V4 \| 1.0 \| 1.386e-16 \| 726 \| 0.428 \| 154 \| 88 \| 0 \| 2 \| 155 \| 6 \| 159 \| Phage tail fiber protein \| Phage tail fiber protein; afdb-uniprot50 \| AF-A0A3M2VPF6-F1-MODEL\_V4 \| 1.0 \| 4.012e-17 \| 725 \| 0.403 \| 166 \| 99 \| 0 \| 1 \| 166 \| 5 \| 170 \| Putative Tail fiber protein H \| Putative Tail fiber protein H; afdb-uniprot50 \| AF-A0A3D4FGS6-F1-MODEL\_V4 \| 1.0 \| 2.072e-18 \| 724 \| 0.443 \| 178 \| 99 \| 0 \| 1 \| 178 \| 2 \| 179 \| Uncharacterized protein \| Uncharacterized protein; afdb-uniprot50 \| AF-A0A1I7J4R3-F1-MODEL\_V4 \| 1.0 \| 1.411e-26 \| 723 \| 0.252 \| 432 \| 221 \| 17 \| 1 \| 423 \| 1 \| 339 \| Phage tail-collar fibre protein \| Phage tail-collar fibre protein; afdb-uniprot50 \| AF-A0A3S6F0P7-F1-MODEL\_V4 \| 1.0 \| 1.105e-28 \| 723 \| 0.279 \| 473 \| 198 \| 11 \| 1 \| 423 \| 2 \| 381 \| Uncharacterized protein \| Uncharacterized protein; afdb-uniprot50 \| AF-A0A290WU51-F1-MODEL\_V4 \| 1.0 \| 1.101e-17 \| 722 \| 0.375 \| 181 \| 110 \| 1 \| 1 \| 178 \| 2 \| 182 \| Uncharacterized protein \| Uncharacterized protein; afdb-uniprot50 \| AF-A0A248XR00-F1-MODEL\_V4 \| 1.0 \| 2.645e-16 \| 722 \| 0.444 \| 153 \| 85 \| 0 \| 1 \| 153 \| 2 \| 154 \| Phage tail protein \| Phage tail protein; afdb-uniprot50 \| AF-D1NYY3-F1-MODEL\_V4 \| 1.0 \| 8.875e-18 \| 721 \| 0.472 \| 163 \| 84 \| 1 \| 4 \| 164 \| 3 \| 165 \| Uncharacterized protein \| Uncharacterized protein; afdb-uniprot50 \| AF-A0A7I0L7S4-F1-MODEL\_V4 \| 1.0 \| 8.2e-16 \| 721 \| 0.437 \| 151 \| 84 \| 1 \| 4 \| 153 \| 3 \| 153 \| Peptidase S74 domain-containing protein \| Peptidase S74 domain-containing protein; afdb-uniprot50 \| AF-A0A6N6MLW5-F1-MODEL\_V4 \| 1.0 \| 4.296e-16 \| 720 \| 0.364 \| 151 \| 96 \| 0 \| 1 \| 151 \| 1 \| 151 \| DUF2793 domain-containing protein \| DUF2793 domain-containing protein; afdb-uniprot50 \| AF-A0A6B8QPL4-F1-MODEL\_V4 \| 1.0 \| 2.218e-17 \| 720 \| 0.453 \| 161 \| 87 \| 1 \| 2 \| 162 \| 45 \| 204 \| Uncharacterized protein \| Uncharacterized protein; afdb-uniprot50 \| AF-A0A1I7C9J8-F1-MODEL\_V4 \| 1.0 \| 8.689e-27 \| 719 \| 0.263 \| 440 \| 203 \| 21 \| 1 \| 423 \| 1 \| 336 \| Phage tail-collar fibre protein \| Phage tail-collar fibre protein; afdb-uniprot50 \| AF-A0A7T4ZYC8-F1-MODEL\_V4 \| 1.0 \| 4.617e-26 \| 718 \| 0.275 \| 439 \| 186 \| 11 \| 1 \| 423 \| 2 \| 324 \| Phage tail protein \| Phage tail protein; afdb-uniprot50 \| AF-A0A839BSH3-F1-MODEL\_V4 \| 1.0 \| 7.004e-27 \| 718 \| 0.293 \| 447 \| 177 \| 13 \| 1 \| 423 \| 1 \| 332 \| Phage tail protein \| Phage tail protein; afdb-uniprot50 \| AF-A0A829HHD3-F1-MODEL\_V4 \| 1.0 \| 2.946e-16 \| 717 \| 0.345 \| 162 \| 105 \| 1 \| 1 \| 162 \| 2 \| 162 \| Uncharacterized protein \| Uncharacterized protein; afdb-uniprot50 \| AF-A0A418H6B6-F1-MODEL\_V4 \| 1.0 \| 6.264e-16 \| 715 \| 0.507 \| 132 \| 65 \| 0 \| 1 \| 132 \| 2 \| 133 \| Phage tail protein \| Phage tail protein; afdb-uniprot50 \| AF-A0A3S4I322-F1-MODEL\_V4 \| 1.0 \| 6.516e-17 \| 715 \| 0.456 \| 160 \| 85 \| 2 \| 1 \| 158 \| 1 \| 160 \| Uncharacterized protein \| Uncharacterized protein; afdb-uniprot50 \| AF-A0A411WMD6-F1-MODEL\_V4 \| 1.0 \| 2.516e-27 \| 715 \| 0.262 \| 530 \| 182 \| 14 \| 1 \| 423 \| 1 \| 428 \| Phage tail protein \| Phage tail protein; afdb-uniprot50 \| AF-A0A2N2GS05-F1-MODEL\_V4 \| 1.0 \| 3.109e-16 \| 715 \| 0.407 \| 162 \| 96 \| 0 \| 1 \| 162 \| 1 \| 162 \| Uncharacterized protein \| Uncharacterized protein; afdb-uniprot50 \| AF-A0A3S0LIM0-F1-MODEL\_V4 \| 1.0 \| 3.857e-16 \| 715 \| 0.327 \| 162 \| 108 \| 1 \| 1 \| 162 \| 2 \| 162 \| Alkaline phosphatase \| Alkaline phosphatase; afdb-uniprot50 \| AF-A0A776X733-F1-MODEL\_V4 \| 1.0 \| 2.187e-18 \| 713 \| 0.291 \| 298 \| 109 \| 4 \| 1 \| 297 \| 2 \| 198 \| Phage tail protein \| Phage tail protein; afdb-uniprot50 \| AF-A0A0N1HPL4-F1-MODEL\_V4 \| 1.0 \| 7.053e-19 \| 713 \| 0.405 \| 185 \| 103 \| 2 \| 1 \| 179 \| 2 \| 185 \| Tail protein \| Tail protein; afdb-uniprot50 \| AF-A0A375A5G5-F1-MODEL\_V4 \| 1.0 \| 4.649e-18 \| 712 \| 0.437 \| 185 \| 93 \| 2 \| 1 \| 174 \| 2 \| 186 \| Phage tail fiber protein \| Phage tail fiber protein; afdb-uniprot50 \| AF-A0A2N2KCW4-F1-MODEL\_V4 \| 1.0 \| 7.258e-17 \| 712 \| 0.398 \| 163 \| 98 \| 0 \| 1 \| 163 \| 1 \| 163 \| Uncharacterized protein \| Uncharacterized protein; afdb-uniprot50 \| AF-A0A560QZS3-F1-MODEL\_V4 \| 1.0 \| 1.914e-16 \| 712 \| 0.409 \| 154 \| 90 \| 1 \| 1 \| 153 \| 1 \| 154 \| Tail collar domain \| Tail collar domain; afdb-uniprot50 \| AF-A0A8B5S4N3-F1-MODEL\_V4 \| 1.0 \| 1.196e-15 \| 711 \| 0.346 \| 153 \| 99 \| 1 \| 1 \| 153 \| 2 \| 153 \| Uncharacterized protein \| Uncharacterized protein; afdb-uniprot50 \| AF-A0A506K1B3-F1-MODEL\_V4 \| 1.0 \| 5.935e-16 \| 711 \| 0.356 \| 160 \| 100 \| 2 \| 1 \| 158 \| 2 \| 160 \| Peptidase S74 domain-containing protein \| Peptidase S74 domain-containing protein; afdb-uniprot50 \| AF-A0A398PZR3-F1-MODEL\_V4 \| 1.0 \| 1.055e-20 \| 709 \| 0.301 \| 318 \| 142 \| 9 \| 1 \| 306 \| 2 \| 251 \| Phage tail protein \| Phage tail protein; afdb-uniprot50 \| AF-A0A7D6WRG1-F1-MODEL\_V4 \| 1.0 \| 1.505e-29 \| 709 \| 0.26 \| 507 \| 184 \| 13 \| 1 \| 423 \| 2 \| 401 \| Phage tail protein \| Phage tail protein; afdb-uniprot50 \| AF-A0A258E4J6-F1-MODEL\_V4 \| 1.0 \| 1.719e-16 \| 709 \| 0.402 \| 154 \| 91 \| 1 \| 1 \| 153 \| 1 \| 154 \| Uncharacterized protein \| Uncharacterized protein; afdb-uniprot50 \| AF-A0A774MST8-F1-MODEL\_V4 \| 1.0 \| 3.365e-18 \| 708 \| 0.28 \| 300 \| 112 \| 3 \| 1 \| 298 \| 2 \| 199 \| Phage tail protein \| Phage tail protein; afdb-uniprot50 \| AF-A0A0S4I2P4-F1-MODEL\_V4 \| 1.0 \| 8.41e-18 \| 708 \| 0.39 \| 182 \| 111 \| 0 \| 2 \| 183 \| 6 \| 187 \| Uncharacterized protein \| Uncharacterized protein; afdb-uniprot50 \| AF-A0A375G4X8-F1-MODEL\_V4 \| 1.0 \| 1.226e-17 \| 708 \| 0.365 \| 186 \| 114 \| 4 \| 1 \| 182 \| 1 \| 186 \| Peptidase S74 domain-containing protein \| Peptidase S74 domain-containing protein; afdb-uniprot50 \| AF-A0A6G6NFC5-F1-MODEL\_V4 \| 1.0 \| 3.234e-17 \| 707 \| 0.473 \| 173 \| 87 \| 2 \| 1 \| 173 \| 1 \| 169 \| Phage tail protein \| Phage tail protein; afdb-uniprot50 \| AF-A0A2D9EKL2-F1-MODEL\_V4 \| 1.0 \| 5.178e-18 \| 705 \| 0.407 \| 184 \| 105 \| 3 \| 1 \| 180 \| 1 \| 184 \| Uncharacterized protein \| Uncharacterized protein; afdb-uniprot50 \| AF-A0A7X1KJH1-F1-MODEL\_V4 \| 1.0 \| 5.465e-18 \| 704 \| 0.406 \| 187 \| 103 \| 3 \| 2 \| 180 \| 6 \| 192 \| Phage tail protein \| Phage tail protein; afdb-uniprot50 \| AF-A0A265Q5Q8-F1-MODEL\_V4 \| 1.0 \| 3.802e-17 \| 704 \| 0.45 \| 160 \| 86 \| 1 \| 1 \| 158 \| 2 \| 161 \| Uncharacterized protein \| Uncharacterized protein; afdb-uniprot50 \| AF-A0A2K4ISX8-F1-MODEL\_V4 \| 1.0 \| 2.471e-17 \| 704 \| 0.377 \| 172 \| 106 \| 1 \| 2 \| 172 \| 6 \| 177 \| Phage tail protein \| Phage tail protein; afdb-uniprot50 \| AF-A0A376ZKW5-F1-MODEL\_V4 \| 1.0 \| 1.629e-16 \| 703 \| 0.467 \| 154 \| 76 \| 1 \| 1 \| 148 \| 2 \| 155 \| Phage tail collar protein \| Phage tail collar protein; afdb-uniprot50 \| AF-A0A2G8D691-F1-MODEL\_V4 \| 1.0 \| 3.843e-20 \| 703 \| 0.27 \| 329 \| 134 \| 3 \| 1 \| 324 \| 2 \| 229 \| Phage tail protein \| Phage tail protein; afdb-uniprot50 \| AF-F3G6R6-F1-MODEL\_V4 \| 1.0 \| 2.904e-17 \| 702 \| 0.385 \| 187 \| 105 \| 3 \| 2 \| 181 \| 6 \| 189 \| Phage-related tail fiber protein-like protein \| Phage-related tail fiber protein-like protein; afdb-uniprot50 \| AF-A0A4S3LPH8-F1-MODEL\_V4 \| 1.0 \| 8.352e-26 \| 702 \| 0.315 \| 428 \| 180 \| 12 \| 1 \| 423 \| 2 \| 321 \| Phage tail protein \| Phage tail protein; afdb-uniprot50 \| AF-A0A7Y1JSW6-F1-MODEL\_V4 \| 1.0 \| 6.61e-16 \| 702 \| 0.393 \| 155 \| 94 \| 0 \| 1 \| 155 \| 5 \| 159 \| Phage tail protein \| Phage tail protein; afdb-uniprot50 \| AF-A0A1B9LPY6-F1-MODEL\_V4 \| 1.0 \| 2.733e-25 \| 700 \| 0.281 \| 434 \| 183 \| 16 \| 1 \| 423 \| 2 \| 317 \| Uncharacterized protein \| Uncharacterized protein; afdb-uniprot50 \| AF-A0A376J6K3-F1-MODEL\_V4 \| 1.0 \| 1.84e-15 \| 699 \| 0.412 \| 148 \| 86 \| 1 \| 1 \| 147 \| 5 \| 152 \| Phage protein gpH \| Phage protein gpH; afdb-uniprot50 \| AF-A0A4R7GU78-F1-MODEL\_V4 \| 1.0 \| 7.208e-25 \| 698 \| 0.279 \| 429 \| 179 \| 10 \| 1 \| 423 \| 2 \| 306 \| Tail-collar fiber protein \| Tail-collar fiber protein; afdb-uniprot50 \| AF-A0A377K993-F1-MODEL\_V4 \| 1.0 \| 7.053e-19 \| 698 \| 0.416 \| 209 \| 100 \| 3 \| 1 \| 187 \| 2 \| 210 \| Phage-related tail fiber protein-like protein \| Phage-related tail fiber protein-like protein; afdb-uniprot50 \| AF-A0A4R0HWF8-F1-MODEL\_V4 \| 1.0 \| 1.6e-21 \| 697 \| 0.312 \| 346 \| 115 \| 5 \| 1 \| 324 \| 1 \| 245 \| Phage tail protein \| Phage tail protein; afdb-uniprot50 \| AF-A0A1C7WI46-F1-MODEL\_V4 \| 1.0 \| 2.132e-16 \| 696 \| 0.372 \| 161 \| 98 \| 2 \| 1 \| 158 \| 1 \| 161 \| Uncharacterized protein \| Uncharacterized protein; afdb-uniprot50 \| AF-A0A2U1TNF9-F1-MODEL\_V4 \| 1.0 \| 2.57e-18 \| 695 \| 0.435 \| 193 \| 92 \| 1 \| 1 \| 176 \| 2 \| 194 \| Phage tail protein \| Phage tail protein; afdb-uniprot50 \| AF-F9MRF3-F1-MODEL\_V4 \| 1.0 \| 3.857e-16 \| 694 \| 0.366 \| 153 \| 97 \| 0 \| 1 \| 153 \| 2 \| 154 \| Uncharacterized protein \| Uncharacterized protein; afdb-uniprot50 \| AF-A0A4Y5WA69-F1-MODEL\_V4 \| 1.0 \| 6.045e-26 \| 694 \| 0.268 \| 439 \| 197 \| 19 \| 1 \| 423 \| 2 \| 332 \| Uncharacterized protein \| Uncharacterized protein; afdb-uniprot50 \| AF-A0A1A9KIH8-F1-MODEL\_V4 \| 1.0 \| 4.07e-16 \| 694 \| 0.421 \| 166 \| 96 \| 0 \| 1 \| 166 \| 1 \| 166 \| SGNH\_hydro domain-containing protein \| SGNH\_hydro domain-containing protein; afdb-uniprot50 \| AF-A0A3R8V706-F1-MODEL\_V4 \| 1.0 \| 1.929e-23 \| 693 \| 0.226 \| 442 \| 181 \| 13 \| 2 \| 423 \| 6 \| 306 \| Uncharacterized protein \| Uncharacterized protein; afdb-uniprot50 \| AF-A0A2D0LDE2-F1-MODEL\_V4 \| 1.0 \| 1.047e-28 \| 693 \| 0.252 \| 523 \| 187 \| 13 \| 1 \| 423 \| 2 \| 420 \| Phage tail protein \| Phage tail protein; afdb-uniprot50 \| AF-A0A161X4B1-F1-MODEL\_V4 \| 1.0 \| 4.533e-16 \| 693 \| 0.496 \| 153 \| 77 \| 0 \| 1 \| 153 \| 2 \| 154 \| SGNH\_hydro domain-containing protein \| SGNH\_hydro domain-containing protein; afdb-uniprot50 \| AF-A0A069A3R7-F1-MODEL\_V4 \| 1.0 \| 1.003e-16 \| 691 \| 0.397 \| 161 \| 97 \| 0 \| 1 \| 161 \| 6 \| 166 \| Putative phage tail fiber protein \| Putative phage tail fiber protein; afdb-uniprot50 \| AF-A0A3E0X1N8-F1-MODEL\_V4 \| 1.0 \| 2.409e-15 \| 691 \| 0.359 \| 153 \| 98 \| 0 \| 1 \| 153 \| 10 \| 162 \| Collar domain-containing protein \| Collar domain-containing protein; afdb-uniprot50 \| AF-A0A5E6S0H5-F1-MODEL\_V4 \| 1.0 \| 1.694e-17 \| 690 \| 0.373 \| 174 \| 106 \| 1 \| 1 \| 171 \| 5 \| 178 \| Uncharacterized protein \| Uncharacterized protein; afdb-uniprot50 \| AF-A0A3S4HKY0-F1-MODEL\_V4 \| 1.0 \| 8.381e-22 \| 689 \| 0.322 \| 350 \| 128 \| 6 \| 1 \| 331 \| 1 \| 260 \| Phage tail fibre repeat \| Phage tail fibre repeat; afdb-uniprot50 \| AF-A0A4U8UED2-F1-MODEL\_V4 \| 1.0 \| 1.483e-15 \| 689 \| 0.36 \| 158 \| 98 \| 1 \| 4 \| 161 \| 3 \| 157 \| Phage tail protein \| Phage tail protein; afdb-uniprot50 \| AF-A0A771BD76-F1-MODEL\_V4 \| 1.0 \| 4.358e-15 \| 689 \| 0.437 \| 151 \| 84 \| 1 \| 4 \| 153 \| 3 \| 153 \| Uncharacterized protein \| Uncharacterized protein; afdb-uniprot50 \| AF-A0A2N0DUE2-F1-MODEL\_V4 \| 1.0 \| 4.296e-16 \| 687 \| 0.402 \| 159 \| 94 \| 1 \| 1 \| 158 \| 1 \| 159 \| Tail collar domain \| Tail collar domain; afdb-uniprot50 \| AF-A0A625JTE0-F1-MODEL\_V4 \| 1.0 \| 5.329e-16 \| 686 \| 0.36 \| 158 \| 98 \| 1 \| 1 \| 158 \| 3 \| 157 \| Phage tail protein \| Phage tail protein; afdb-uniprot50 \| AF-A0A7X4KF01-F1-MODEL\_V4 \| 1.0 \| 5.049e-16 \| 684 \| 0.339 \| 165 \| 104 \| 3 \| 1 \| 163 \| 2 \| 163 \| Uncharacterized protein \| Uncharacterized protein; afdb-uniprot50 \| AF-J2PVQ7-F1-MODEL\_V4 \| 1.0 \| 5.049e-16 \| 684 \| 0.402 \| 159 \| 94 \| 1 \| 1 \| 158 \| 1 \| 159 \| Phage-related tail fiber protein \| Phage-related tail fiber protein; afdb-uniprot50 \| AF-A0A1G5MGW4-F1-MODEL\_V4 \| 1.0 \| 5.049e-16 \| 684 \| 0.396 \| 159 \| 95 \| 1 \| 1 \| 158 \| 1 \| 159 \| Phage-related tail fibre protein \| Phage-related tail fibre protein; afdb-uniprot50 \| AF-A0A2A2F373-F1-MODEL\_V4 \| 1.0 \| 1.565e-15 \| 683 \| 0.408 \| 159 \| 88 \| 2 \| 1 \| 153 \| 4 \| 162 \| Uncharacterized protein \| Uncharacterized protein; afdb-uniprot50 \| AF-A0A6L7E7U0-F1-MODEL\_V4 \| 1.0 \| 6.683e-19 \| 682 \| 0.401 \| 204 \| 96 \| 2 \| 1 \| 178 \| 2 \| 205 \| Phage tail protein \| Phage tail protein; afdb-uniprot50 \| AF-A0A3A6RIP6-F1-MODEL\_V4 \| 1.0 \| 3.602e-17 \| 681 \| 0.393 \| 188 \| 104 \| 2 \| 1 \| 178 \| 2 \| 189 \| Phage tail protein \| Phage tail protein; afdb-uniprot50 \| AF-A0A7W4RHL5-F1-MODEL\_V4 \| 1.0 \| 2.283e-15 \| 681 \| 0.41 \| 156 \| 91 \| 1 \| 1 \| 155 \| 2 \| 157 \| Uncharacterized protein \| Uncharacterized protein; afdb-uniprot50 \| AF-A0A516SJM3-F1-MODEL\_V4 \| 1.0 \| 1.346e-18 \| 680 \| 0.415 \| 195 \| 110 \| 2 \| 1 \| 191 \| 1 \| 195 \| Uncharacterized protein \| Uncharacterized protein; afdb-uniprot50 \| AF-A0A1Q6KPQ8-F1-MODEL\_V4 \| 1.0 \| 3.463e-16 \| 680 \| 0.339 \| 165 \| 106 \| 2 \| 1 \| 164 \| 1 \| 163 \| Uncharacterized protein \| Uncharacterized protein; afdb-uniprot50 \| AF-A0A2U2UFI2-F1-MODEL\_V4 \| 1.0 \| 3.899e-19 \| 679 \| 0.284 \| 330 \| 124 \| 5 \| 1 \| 329 \| 3 \| 221 \| Phage tail protein \| Phage tail protein; afdb-uniprot50 \| AF-A0A419AS73-F1-MODEL\_V4 \| 1.0 \| 2.498e-20 \| 679 \| 0.313 \| 325 \| 119 \| 5 \| 1 \| 324 \| 2 \| 223 \| Phage tail protein \| Phage tail protein; afdb-uniprot50 \| AF-C6E6P3-F1-MODEL\_V4 \| 1.0 \| 1.117e-16 \| 678 \| 0.39 \| 169 \| 97 \| 2 \| 4 \| 166 \| 3 \| 171 \| Phage-related tail fibre protein-like protein \| Phage-related tail fibre protein-like protein; afdb-uniprot50 \| AF-A0A7Z2QM66-F1-MODEL\_V4 \| 1.0 \| 8.2e-16 \| 678 \| 0.396 \| 159 \| 95 \| 1 \| 1 \| 158 \| 1 \| 159 \| Phage tail protein \| Phage tail protein; afdb-uniprot50 \| AF-A0A845RKE3-F1-MODEL\_V4 \| 1.0 \| 6.264e-16 \| 676 \| 0.326 \| 156 \| 105 \| 0 \| 1 \| 156 \| 29 \| 184 \| Uncharacterized protein \| Uncharacterized protein; afdb-uniprot50 \| AF-A0A1M7U2L6-F1-MODEL\_V4 \| 1.0 \| 1.043e-17 \| 676 \| 0.439 \| 189 \| 92 \| 2 \| 1 \| 175 \| 1 \| 189 \| Phage tail-collar fibre protein \| Phage tail-collar fibre protein; afdb-uniprot50 \| AF-A0A1Z1SPG3-F1-MODEL\_V4 \| 1.0 \| 7.66e-17 \| 676 \| 0.433 \| 173 \| 91 \| 1 \| 1 \| 166 \| 2 \| 174 \| Phage tail protein \| Phage tail protein; afdb-uniprot50 \| AF-A0A774SHB9-F1-MODEL\_V4 \| 1.0 \| 3.955e-18 \| 675 \| 0.298 \| 305 \| 110 \| 4 \| 1 \| 302 \| 1 \| 204 \| Phage tail protein \| Phage tail protein; afdb-uniprot50 \| AF-A0A1X3RYJ6-F1-MODEL\_V4 \| 1.0 \| 3.39e-25 \| 675 \| 0.266 \| 428 \| 200 \| 15 \| 1 \| 423 \| 1 \| 319 \| Phage tail protein \| Phage tail protein; afdb-uniprot50 \| AF-A0A828EEN1-F1-MODEL\_V4 \| 1.0 \| 2.674e-19 \| 675 \| 0.411 \| 209 \| 101 \| 3 \| 1 \| 187 \| 2 \| 210 \| Phage tail protein \| Phage tail protein; afdb-uniprot50 \| AF-A0A4Y8GSR2-F1-MODEL\_V4 \| 1.0 \| 4.567e-23 \| 675 \| 0.335 \| 331 \| 147 \| 8 \| 1 \| 323 \| 2 \| 267 \| Short-chain fatty acid transporter \| Short-chain fatty acid transporter; afdb-uniprot50 \| AF-C6DCN9-F1-MODEL\_V4 \| 1.0 \| 4.649e-18 \| 674 \| 0.441 \| 204 \| 88 \| 3 \| 1 \| 178 \| 2 \| 205 \| Phage-related tail fibre protein-like protein \| Phage-related tail fibre protein-like protein; afdb-uniprot50 \| AF-A0A542AK78-F1-MODEL\_V4 \| 1.0 \| 2.409e-15 \| 673 \| 0.353 \| 150 \| 96 \| 1 \| 4 \| 153 \| 3 \| 151 \| Tail-collar fiber protein \| Tail-collar fiber protein; afdb-uniprot50 \| AF-A0A824XYH8-F1-MODEL\_V4 \| 1.0 \| 8.2e-16 \| 673 \| 0.379 \| 158 \| 95 \| 1 \| 1 \| 158 \| 3 \| 157 \| Phage tail protein \| Phage tail protein; afdb-uniprot50 \| AF-A0A2C9EHA1-F1-MODEL\_V4 \| 1.0 \| 1.441e-17 \| 673 \| 0.363 \| 187 \| 111 \| 3 \| 4 \| 186 \| 8 \| 190 \| Putative tail fiber protein \| Putative tail fiber protein; afdb-uniprot50 \| AF-A0A072Y7K9-F1-MODEL\_V4 \| 1.0 \| 5.767e-18 \| 673 \| 0.338 \| 195 \| 120 \| 2 \| 1 \| 187 \| 2 \| 195 \| Tail fiber protein H \| Tail fiber protein H; afdb-uniprot50 \| AF-N8RAV5-F1-MODEL\_V4 \| 1.0 \| 2.132e-16 \| 673 \| 0.394 \| 175 \| 101 \| 2 \| 3 \| 172 \| 2 \| 176 \| Uncharacterized protein \| Uncharacterized protein; afdb-uniprot50 \| AF-A0A745FXY9-F1-MODEL\_V4 \| 1.0 \| 1.719e-16 \| 672 \| 0.368 \| 187 \| 105 \| 3 \| 1 \| 178 \| 2 \| 184 \| Phage tail protein \| Phage tail protein; afdb-uniprot50 \| AF-A0A380C811-F1-MODEL\_V4 \| 1.0 \| 1.992e-17 \| 672 \| 0.353 \| 181 \| 107 \| 3 \| 1 \| 176 \| 1 \| 176 \| Uncharacterized protein \| Uncharacterized protein; afdb-uniprot50 \| AF-A0A261E4L1-F1-MODEL\_V4 \| 1.0 \| 2.203e-25 \| 672 \| 0.278 \| 434 \| 207 \| 17 \| 1 \| 423 \| 2 \| 340 \| DNA inversion product \| DNA inversion product; afdb-uniprot50 \| AF-A0A556RUG3-F1-MODEL\_V4 \| 1.0 \| 3.054e-21 \| 672 \| 0.33 \| 306 \| 108 \| 1 \| 1 \| 306 \| 2 \| 210 \| Phage tail protein \| Phage tail protein; afdb-uniprot50 \| AF-A0A1E7RCE5-F1-MODEL\_V4 \| 1.0 \| 1.231e-28 \| 672 \| 0.282 \| 496 \| 230 \| 19 \| 4 \| 417 \| 3 \| 454 \| Phage tail protein \| Phage tail protein; afdb-uniprot50 \| AF-L1M5X1-F1-MODEL\_V4 \| 1.0 \| 4.235e-17 \| 672 \| 0.381 \| 186 \| 113 \| 2 \| 2 \| 186 \| 6 \| 190 \| Tail fiber protein H \| Tail fiber protein H; afdb-uniprot50 \| AF-A0A6H7NTP2-F1-MODEL\_V4 \| 1.0 \| 8.2e-16 \| 671 \| 0.392 \| 158 \| 93 \| 1 \| 1 \| 158 \| 3 \| 157 \| Phage tail protein \| Phage tail protein; afdb-uniprot50 \| AF-A0A624F4N8-F1-MODEL\_V4 \| 1.0 \| 1.133e-15 \| 671 \| 0.386 \| 158 \| 94 \| 1 \| 1 \| 158 \| 3 \| 157 \| Phage tail protein \| Phage tail protein; afdb-uniprot50 \| AF-A0A4Q0UTE8-F1-MODEL\_V4 \| 1.0 \| 2.791e-16 \| 671 \| 0.367 \| 177 \| 107 \| 3 \| 1 \| 176 \| 3 \| 175 \| Uncharacterized protein \| Uncharacterized protein; afdb-uniprot50 \| AF-A0A024HA82-F1-MODEL\_V4 \| 1.0 \| 5.142e-26 \| 671 \| 0.271 \| 445 \| 237 \| 20 \| 1 \| 423 \| 2 \| 381 \| Uncharacterized protein \| Uncharacterized protein; afdb-uniprot50 \| AF-A0A2Z6AZ11-F1-MODEL\_V4 \| 1.0 \| 3.451e-20 \| 670 \| 0.351 \| 236 \| 112 \| 6 \| 1 \| 219 \| 1 \| 212 \| Tail Collar domain protein \| Tail Collar domain protein; afdb-uniprot50 \| AF-A0A690UYS0-F1-MODEL\_V4 \| 1.0 \| 9.639e-16 \| 669 \| 0.386 \| 158 \| 94 \| 1 \| 1 \| 158 \| 3 \| 157 \| Phage tail protein \| Phage tail protein; afdb-uniprot50 \| AF-A0A0M0TBG4-F1-MODEL\_V4 \| 1.0 \| 2.242e-20 \| 668 \| 0.319 \| 307 \| 111 \| 4 \| 1 \| 306 \| 1 \| 210 \| Tail protein \| Tail protein; afdb-uniprot50 \| AF-A0A826S3K6-F1-MODEL\_V4 \| 1.0 \| 1.235e-24 \| 668 \| 0.276 \| 438 \| 184 \| 11 \| 1 \| 423 \| 2 \| 321 \| Phage tail protein \| Phage tail protein; afdb-uniprot50 \| AF-A0A2N0DX16-F1-MODEL\_V4 \| 1.0 \| 1.84e-15 \| 668 \| 0.409 \| 154 \| 91 \| 0 \| 2 \| 155 \| 6 \| 159 \| Phage tail protein \| Phage tail protein; afdb-uniprot50 \| AF-A0A1I3I6J5-F1-MODEL\_V4 \| 1.0 \| 6.132e-25 \| 667 \| 0.26 \| 438 \| 187 \| 17 \| 1 \| 423 \| 1 \| 316 \| Phage tail-collar fibre protein \| Phage tail-collar fibre protein; afdb-uniprot50 \| AF-A0A1B9JGD4-F1-MODEL\_V4 \| 1.0 \| 4.649e-18 \| 667 \| 0.394 \| 195 \| 95 \| 2 \| 1 \| 172 \| 2 \| 196 \| Collar domain-containing protein \| Collar domain-containing protein; afdb-uniprot50 \| AF-A0A4T2A2H5-F1-MODEL\_V4 \| 1.0 \| 3.439e-24 \| 667 \| 0.212 \| 451 \| 199 \| 15 \| 1 \| 423 \| 2 \| 324 \| DNA inversion product \| DNA inversion product; afdb-uniprot50 \| AF-A0A5C1A538-F1-MODEL\_V4 \| 1.0 \| 1.093e-25 \| 667 \| 0.225 \| 497 \| 210 \| 16 \| 1 \| 423 \| 1 \| 396 \| DNA inversion product \| DNA inversion product; afdb-uniprot50 \| AF-A0A0F4TBD1-F1-MODEL\_V4 \| 1.0 \| 2.542e-15 \| 667 \| 0.396 \| 154 \| 93 \| 0 \| 2 \| 155 \| 6 \| 159 \| Phage tail protein \| Phage tail protein; afdb-uniprot50 \| AF-A0A379UXV3-F1-MODEL\_V4 \| 1.0 \| 5.104e-19 \| 666 \| 0.411 \| 209 \| 101 \| 3 \| 1 \| 187 \| 2 \| 210 \| Phage tail fiber protein \| Phage tail fiber protein; afdb-uniprot50 \| AF-A0A447XE38-F1-MODEL\_V4 \| 1.0 \| 1.07e-19 \| 665 \| 0.308 \| 324 \| 123 \| 2 \| 1 \| 324 \| 2 \| 224 \| Tail fiber protein \| Tail fiber protein; afdb-uniprot50 \| AF-A0A721ZMC9-F1-MODEL\_V4 \| 1.0 \| 2.179e-22 \| 665 \| 0.304 \| 335 \| 162 \| 7 \| 2 \| 324 \| 3 \| 278 \| Phage tail protein \| Phage tail protein; afdb-uniprot50 \| AF-A0A221ZMW8-F1-MODEL\_V4 \| 1.0 \| 4.115e-19 \| 664 \| 0.414 \| 210 \| 100 \| 4 \| 1 \| 187 \| 1 \| 210 \| Phage tail fiber protein \| Phage tail fiber protein; afdb-uniprot50 \| AF-A0A7G5NY25-F1-MODEL\_V4 \| 1.0 \| 2.163e-15 \| 664 \| 0.375 \| 157 \| 98 \| 0 \| 2 \| 158 \| 7 \| 163 \| Phage tail protein \| Phage tail protein; afdb-uniprot50 \| AF-A0A3Q8FR82-F1-MODEL\_V4 \| 1.0 \| 2.48e-28 \| 664 \| 0.277 \| 429 \| 217 \| 20 \| 1 \| 407 \| 1 \| 358 \| Phage tail protein \| Phage tail protein; afdb-uniprot50 \| AF-A0A5P3ME88-F1-MODEL\_V4 \| 1.0 \| 4.716e-17 \| 664 \| 0.357 \| 190 \| 103 \| 4 \| 1 \| 172 \| 2 \| 190 \| Phage tail-collar fiber family protein \| Phage tail-collar fiber family protein; afdb-uniprot50 \| AF-A0A2N1CWE2-F1-MODEL\_V4 \| 1.0 \| 3.629e-24 \| 662 \| 0.242 \| 442 \| 180 \| 13 \| 2 \| 423 \| 6 \| 312 \| DNA inversion product \| DNA inversion product; afdb-uniprot50 \| AF-A0A6G5QMQ8-F1-MODEL\_V4 \| 1.0 \| 4.296e-16 \| 660 \| 0.365 \| 164 \| 98 \| 2 \| 3 \| 163 \| 2 \| 162 \| Phage tail-collar fiber protein (DUF3751 domain) \| Phage tail-collar fiber protein (DUF3751 domain); afdb-uniprot50 \| AF-A0A717EDS8-F1-MODEL\_V4 \| 1.0 \| 3.281e-16 \| 659 \| 0.347 \| 187 \| 109 \| 3 \| 1 \| 178 \| 5 \| 187 \| Phage tail protein \| Phage tail protein; afdb-uniprot50 \| AF-A0A1E7PZY5-F1-MODEL\_V4 \| 1.0 \| 2.155e-19 \| 659 \| 0.297 \| 329 \| 132 \| 8 \| 1 \| 324 \| 1 \| 235 \| Uncharacterized protein \| Uncharacterized protein; afdb-uniprot50 \| AF-A0A1B9LET1-F1-MODEL\_V4 \| 1.0 \| 8.352e-26 \| 659 \| 0.238 \| 473 \| 205 \| 13 \| 1 \| 423 \| 1 \| 368 \| Uncharacterized protein \| Uncharacterized protein; afdb-uniprot50 \| AF-A0A5B8HDY3-F1-MODEL\_V4 \| 1.0 \| 3.669e-27 \| 659 \| 0.265 \| 485 \| 201 \| 15 \| 1 \| 423 \| 1 \| 392 \| Phage tail protein \| Phage tail protein; afdb-uniprot50 \| AF-J2UA87-F1-MODEL\_V4 \| 1.0 \| 2.988e-15 \| 659 \| 0.389 \| 159 \| 96 \| 1 \| 1 \| 158 \| 1 \| 159 \| Phage-related tail fiber protein \| Phage-related tail fiber protein; afdb-uniprot50 \| AF-A0A4R7YKL0-F1-MODEL\_V4 \| 1.0 \| 1.782e-21 \| 658 \| 0.296 \| 320 \| 146 \| 11 \| 1 \| 306 \| 1 \| 255 \| Tail-collar fiber protein \| Tail-collar fiber protein; afdb-uniprot50 \| AF-A0A4R0GNU2-F1-MODEL\_V4 \| 1.0 \| 1.145e-18 \| 657 \| 0.423 \| 210 \| 98 \| 4 \| 1 \| 187 \| 1 \| 210 \| Phage tail protein \| Phage tail protein; afdb-uniprot50 \| AF-A0A6N8PLL2-F1-MODEL\_V4 \| 1.0 \| 7.443e-19 \| 655 \| 0.312 \| 333 \| 120 \| 7 \| 1 \| 330 \| 2 \| 228 \| Phage tail protein \| Phage tail protein; afdb-uniprot50 \| AF-A0A8B6JH22-F1-MODEL\_V4 \| 1.0 \| 5.935e-16 \| 655 \| 0.365 \| 164 \| 101 \| 1 \| 1 \| 161 \| 1 \| 164 \| Tail fiber protein \| Tail fiber protein; afdb-uniprot50 \| AF-A0A8A8PF66-F1-MODEL\_V4 \| 1.0 \| 5.31e-20 \| 655 \| 0.364 \| 236 \| 130 \| 5 \| 1 \| 223 \| 1 \| 229 \| Phage tail protein \| Phage tail protein; afdb-uniprot50 \| AF-A0A1I7HNR6-F1-MODEL\_V4 \| 1.0 \| 3.342e-26 \| 654 \| 0.239 \| 484 \| 199 \| 17 \| 1 \| 423 \| 2 \| 377 \| Phage tail fibre repeat-containing protein \| Phage tail fibre repeat-containing protein; afdb-uniprot50 \| AF-A0A1E3GML3-F1-MODEL\_V4 \| 1.0 \| 1.003e-16 \| 654 \| 0.39 \| 182 \| 99 \| 3 \| 3 \| 172 \| 2 \| 183 \| Uncharacterized protein \| Uncharacterized protein; afdb-uniprot50 \| AF-A0A5E8H5R3-F1-MODEL\_V4 \| 1.0 \| 4.854e-15 \| 653 \| 0.409 \| 154 \| 89 \| 2 \| 5 \| 157 \| 2 \| 154 \| Phage-related tail fiber protein \| Phage-related tail fiber protein; afdb-uniprot50 \| AF-A0A743JKX6-F1-MODEL\_V4 \| 1.0 \| 1.854e-22 \| 653 \| 0.312 \| 336 \| 154 \| 8 \| 1 \| 324 \| 1 \| 271 \| Phage tail protein \| Phage tail protein; afdb-uniprot50 \| AF-A0A7Z3MBJ3-F1-MODEL\_V4 \| 1.0 \| 1.222e-21 \| 653 \| 0.316 \| 325 \| 120 \| 3 \| 1 \| 324 \| 1 \| 224 \| Phage tail protein \| Phage tail protein; afdb-uniprot50 \| AF-A0A6V8GZ93-F1-MODEL\_V4 \| 1.0 \| 3.329e-15 \| 652 \| 0.373 \| 158 \| 96 \| 1 \| 1 \| 158 \| 3 \| 157 \| Phage tail protein \| Phage tail protein; afdb-uniprot50 \| AF-D6JBE2-F1-MODEL\_V4 \| 1.0 \| 1.834e-19 \| 652 \| 0.289 \| 307 \| 138 \| 8 \| 2 \| 296 \| 6 \| 244 \| GpH \| GpH; afdb-uniprot50 \| AF-A0A843NZ40-F1-MODEL\_V4 \| 1.0 \| 4.296e-16 \| 651 \| 0.402 \| 164 \| 92 \| 2 \| 1 \| 163 \| 2 \| 160 \| Phage tail protein \| Phage tail protein; afdb-uniprot50 \| AF-U6ZRT4-F1-MODEL\_V4 \| 1.0 \| 1.117e-16 \| 651 \| 0.412 \| 172 \| 98 \| 1 \| 4 \| 172 \| 8 \| 179 \| Collar domain-containing protein \| Collar domain-containing protein; afdb-uniprot50 \| AF-A0A7U5TL09-F1-MODEL\_V4 \| 1.0 \| 6.379e-26 \| 650 \| 0.283 \| 483 \| 188 \| 15 \| 1 \| 423 \| 2 \| 386 \| Phage tail protein \| Phage tail protein; afdb-uniprot50 \| AF-A0A5C7E1E3-F1-MODEL\_V4 \| 1.0 \| 5.122e-15 \| 649 \| 0.354 \| 158 \| 99 \| 1 \| 1 \| 158 \| 3 \| 157 \| Phage tail protein \| Phage tail protein; afdb-uniprot50 \| AF-A0A1Y4J5U1-F1-MODEL\_V4 \| 1.0 \| 7.882e-15 \| 649 \| 0.299 \| 157 \| 110 \| 0 \| 4 \| 160 \| 40 \| 196 \| Uncharacterized protein \| Uncharacterized protein; afdb-uniprot50 \| AF-A0A5C4YDH8-F1-MODEL\_V4 \| 1.0 \| 2.049e-15 \| 648 \| 0.392 \| 158 \| 93 \| 1 \| 1 \| 158 \| 3 \| 157 \| Phage tail protein \| Phage tail protein; afdb-uniprot50 \| AF-A0A7Y7YIA9-F1-MODEL\_V4 \| 1.0 \| 4.599e-15 \| 648 \| 0.422 \| 154 \| 89 \| 0 \| 2 \| 155 \| 6 \| 159 \| Phage tail protein \| Phage tail protein; afdb-uniprot50 \| AF-A0A381GJP7-F1-MODEL\_V4 \| 1.0 \| 4.343e-19 \| 647 \| 0.305 \| 314 \| 116 \| 4 \| 1 \| 308 \| 2 \| 219 \| Phage tail fiber protein \| Phage tail fiber protein; afdb-uniprot50 \| AF-A0A3Z0G4I0-F1-MODEL\_V4 \| 1.0 \| 4.803e-27 \| 647 \| 0.267 \| 493 \| 196 \| 14 \| 1 \| 423 \| 2 \| 399 \| Phage tail protein \| Phage tail protein; afdb-uniprot50 \| AF-A0A2D3VW29-F1-MODEL\_V4 \| 1.0 \| 3.513e-15 \| 646 \| 0.377 \| 151 \| 94 \| 0 \| 3 \| 153 \| 2 \| 152 \| Uncharacterized protein \| Uncharacterized protein; afdb-uniprot50 \| AF-A0A2N7P6T1-F1-MODEL\_V4 \| 1.0 \| 1.901e-24 \| 646 \| 0.214 \| 438 \| 226 \| 18 \| 3 \| 423 \| 2 \| 338 \| Uncharacterized protein \| Uncharacterized protein; afdb-uniprot50 \| AF-C1DBJ9-F1-MODEL\_V4 \| 1.0 \| 3.439e-24 \| 646 \| 0.252 \| 456 \| 197 \| 18 \| 2 \| 422 \| 3 \| 349 \| Probable phage tail fiber protein \| Probable phage tail fiber protein; afdb-uniprot50 \| AF-A0A542V4K6-F1-MODEL\_V4 \| 1.0 \| 2.516e-27 \| 646 \| 0.245 \| 505 \| 260 \| 24 \| 1 \| 423 \| 2 \| 467 \| Tail-collar fiber protein \| Tail-collar fiber protein; afdb-uniprot50 \| AF-A0A1S1HUY3-F1-MODEL\_V4 \| 1.0 \| 9.778e-15 \| 646 \| 0.36 \| 161 \| 101 \| 2 \| 1 \| 159 \| 1 \| 161 \| Uncharacterized protein \| Uncharacterized protein; afdb-uniprot50 \| AF-A0A0B5QHJ2-F1-MODEL\_V4 \| 1.0 \| 2.02e-16 \| 645 \| 0.342 \| 187 \| 115 \| 2 \| 1 \| 179 \| 1 \| 187 \| Phage tail protein \| Phage tail protein; afdb-uniprot50 \| AF-A0A376DLB6-F1-MODEL\_V4 \| 1.0 \| 2.978e-19 \| 644 \| 0.341 \| 249 \| 107 \| 5 \| 1 \| 221 \| 2 \| 221 \| Uncharacterized protein \| Uncharacterized protein; afdb-uniprot50 \| AF-A0A379QPW4-F1-MODEL\_V4 \| 1.0 \| 6.904e-28 \| 644 \| 0.256 \| 506 \| 244 \| 21 \| 1 \| 423 \| 1 \| 457 \| Tail protein \| Tail protein; afdb-uniprot50 \| AF-A0A318T0X1-F1-MODEL\_V4 \| 1.0 \| 3.109e-16 \| 644 \| 0.345 \| 162 \| 106 \| 0 \| 1 \| 162 \| 2 \| 163 \| Uncharacterized protein DUF2793 \| Uncharacterized protein DUF2793; afdb-uniprot50 \| AF-A0A1N6Q2W5-F1-MODEL\_V4 \| 1.0 \| 4.174e-18 \| 642 \| 0.36 \| 200 \| 113 \| 5 \| 3 \| 194 \| 2 \| 194 \| Phage tail-collar fibre protein \| Phage tail-collar fibre protein; afdb-uniprot50 \| AF-A0A5M0YJG9-F1-MODEL\_V4 \| 1.0 \| 3.513e-15 \| 641 \| 0.375 \| 152 \| 92 \| 1 \| 2 \| 153 \| 3 \| 151 \| Uncharacterized protein \| Uncharacterized protein; afdb-uniprot50 \| AF-A0A6C8Y4A3-F1-MODEL\_V4 \| 1.0 \| 4.518e-20 \| 641 \| 0.326 \| 315 \| 108 \| 4 \| 1 \| 308 \| 2 \| 219 \| Phage tail protein \| Phage tail protein; afdb-uniprot50 \| AF-A0A081RRN4-F1-MODEL\_V4 \| 1.0 \| 8.845e-22 \| 641 \| 0.331 \| 314 \| 132 \| 6 \| 4 \| 306 \| 3 \| 249 \| Phage-related tail fiber protein \| Phage-related tail fiber protein; afdb-uniprot50 \| AF-A0A0N1IDW0-F1-MODEL\_V4 \| 1.0 \| 9.639e-16 \| 640 \| 0.353 \| 164 \| 103 \| 1 \| 1 \| 161 \| 1 \| 164 \| Tail protein \| Tail protein; afdb-uniprot50 \| AF-A0A2X3H8W1-F1-MODEL\_V4 \| 1.0 \| 3.329e-15 \| 640 \| 0.401 \| 167 \| 92 \| 2 \| 1 \| 159 \| 1 \| 167 \| Tail fiber protein \| Tail fiber protein; afdb-uniprot50 \| AF-A0A1T4WZ85-F1-MODEL\_V4 \| 1.0 \| 2.636e-20 \| 640 \| 0.336 \| 244 \| 125 \| 10 \| 1 \| 223 \| 1 \| 228 \| Phage tail-collar fibre protein \| Phage tail-collar fibre protein; afdb-uniprot50 \| AF-A0A0S1B3R8-F1-MODEL\_V4 \| 1.0 \| 6.354e-15 \| 639 \| 0.378 \| 156 \| 96 \| 1 \| 1 \| 155 \| 1 \| 156 \| Side tail phage protein \| Side tail phage protein; afdb-uniprot50 \| AF-A0A4P5QDV9-F1-MODEL\_V4 \| 1.0 \| 1.213e-14 \| 639 \| 0.285 \| 161 \| 115 \| 0 \| 1 \| 161 \| 1 \| 161 \| Phage tail protein \| Phage tail protein; afdb-uniprot50 \| AF-A0A3D9DW60-F1-MODEL\_V4 \| 1.0 \| 1.431e-25 \| 639 \| 0.232 \| 490 \| 215 \| 19 \| 1 \| 423 \| 1 \| 396 \| Tail-collar fiber protein \| Tail-collar fiber protein; afdb-uniprot50 \| AF-A0A379X3P6-F1-MODEL\_V4 \| 1.0 \| 4.16e-22 \| 638 \| 0.298 \| 342 \| 155 \| 8 \| 1 \| 324 \| 1 \| 275 \| Gp19 \| Gp19; afdb-uniprot50 \| AF-A0A4P2WWZ3-F1-MODEL\_V4 \| 1.0 \| 3.414e-17 \| 637 \| 0.31 \| 203 \| 119 \| 5 \| 1 \| 183 \| 2 \| 203 \| Uncharacterized protein \| Uncharacterized protein; afdb-uniprot50 \| AF-A0A485CDL0-F1-MODEL\_V4 \| 1.0 \| 2.542e-15 \| 637 \| 0.38 \| 168 \| 98 \| 1 \| 1 \| 162 \| 2 \| 169 \| Uncharacterized protein \| Uncharacterized protein; afdb-uniprot50 \| AF-A0A2S0JB19-F1-MODEL\_V4 \| 1.0 \| 6.706e-15 \| 637 \| 0.358 \| 162 \| 98 \| 3 \| 1 \| 162 \| 1 \| 156 \| Uncharacterized protein \| Uncharacterized protein; afdb-uniprot50 \| AF-V4NM72-F1-MODEL\_V4 \| 1.0 \| 6.588e-20 \| 637 \| 0.287 \| 344 \| 126 \| 3 \| 1 \| 326 \| 1 \| 243 \| Uncharacterized protein \| Uncharacterized protein; afdb-uniprot50 \| AF-A0A1V0LL73-F1-MODEL\_V4 \| 1.0 \| 2.227e-28 \| 637 \| 0.271 \| 515 \| 228 \| 17 \| 1 \| 423 \| 2 \| 461 \| Phage tail protein \| Phage tail protein; afdb-uniprot50 \| AF-A0A2S4SSM1-F1-MODEL\_V4 \| 1.0 \| 6.087e-18 \| 637 \| 0.393 \| 211 \| 99 \| 3 \| 1 \| 182 \| 2 \| 212 \| Phage tail protein \| Phage tail protein; afdb-uniprot50 \| AF-A0A7U6ITD4-F1-MODEL\_V4 \| 1.0 \| 3.513e-15 \| 637 \| 0.343 \| 157 \| 98 \| 2 \| 1 \| 156 \| 2 \| 154 \| Uncharacterized protein \| Uncharacterized protein; afdb-uniprot50 \| AF-A0A380TT23-F1-MODEL\_V4 \| 1.0 \| 7.77e-16 \| 637 \| 0.409 \| 166 \| 93 \| 2 \| 3 \| 163 \| 2 \| 167 \| Plasmin and fibronectin-binding protein A \| Plasmin and fibronectin-binding protein A; afdb-uniprot50 \| AF-A0A369YED8-F1-MODEL\_V4 \| 1.0 \| 9.134e-16 \| 636 \| 0.375 \| 168 \| 100 \| 1 \| 1 \| 163 \| 1 \| 168 \| Uncharacterized protein \| Uncharacterized protein; afdb-uniprot50 \| AF-A5MYJ9-F1-MODEL\_V4 \| 1.0 \| 4.977e-17 \| 636 \| 0.369 \| 195 \| 114 \| 2 \| 1 \| 187 \| 2 \| 195 \| Predicted phage tail fibre protein \| Predicted phage tail fibre protein; afdb-uniprot50 \| AF-A0A0S4SND2-F1-MODEL\_V4 \| 1.0 \| 8.531e-17 \| 635 \| 0.397 \| 186 \| 85 \| 2 \| 1 \| 162 \| 1 \| 183 \| Tail fiber protein H \| Tail fiber protein H; afdb-uniprot50 \| AF-A0A6S4UU02-F1-MODEL\_V4 \| 1.0 \| 9.134e-16 \| 635 \| 0.31 \| 190 \| 112 \| 3 \| 1 \| 172 \| 2 \| 190 \| Mannuronan 5-epimerase \| Mannuronan 5-epimerase; afdb-uniprot50 \| AF-A0A6D0LVZ5-F1-MODEL\_V4 \| 1.0 \| 3.489e-23 \| 634 \| 0.29 \| 420 \| 127 \| 6 \| 2 \| 346 \| 3 \| 326 \| Phage tail protein \| Phage tail protein; afdb-uniprot50 \| AF-A0A1Q6AW37-F1-MODEL\_V4 \| 1.0 \| 2.172e-26 \| 634 \| 0.255 \| 505 \| 193 \| 19 \| 1 \| 423 \| 2 \| 405 \| Phage tail protein \| Phage tail protein; afdb-uniprot50 \| AF-C6D9F9-F1-MODEL\_V4 \| 1.0 \| 4.518e-20 \| 634 \| 0.332 \| 307 \| 103 \| 5 \| 1 \| 306 \| 1 \| 206 \| Tail fiber repeat 2 protein \| Tail fiber repeat 2 protein; afdb-uniprot50 \| AF-A0A2P8QYP4-F1-MODEL\_V4 \| 1.0 \| 3.065e-17 \| 633 \| 0.366 \| 199 \| 118 \| 2 \| 1 \| 194 \| 1 \| 196 \| Uncharacterized protein \| Uncharacterized protein; afdb-uniprot50 \| AF-A0A777TPN4-F1-MODEL\_V4 \| 1.0 \| 2.782e-20 \| 632 \| 0.277 \| 321 \| 159 \| 7 \| 1 \| 307 \| 2 \| 263 \| Short-chain fatty acid transporter \| Short-chain fatty acid transporter; afdb-uniprot50 \| AF-A0A7U4KUX0-F1-MODEL\_V4 \| 1.0 \| 3.212e-25 \| 632 \| 0.276 \| 485 \| 198 \| 14 \| 1 \| 423 \| 2 \| 395 \| Tail protein \| Tail protein; afdb-uniprot50 \| AF-A0A5T2TRA4-F1-MODEL\_V4 \| 1.0 \| 3.402e-21 \| 632 \| 0.326 \| 334 \| 144 \| 9 \| 1 \| 323 \| 2 \| 265 \| Uncharacterized protein \| Uncharacterized protein; afdb-uniprot50 \| AF-A0A8A5HZ74-F1-MODEL\_V4 \| 1.0 \| 1.549e-27 \| 630 \| 0.278 \| 532 \| 210 \| 20 \| 1 \| 423 \| 1 \| 467 \| Phage tail protein \| Phage tail protein; afdb-uniprot50 \| AF-A0A2R3MZM8-F1-MODEL\_V4 \| 1.0 \| 4.07e-16 \| 629 \| 0.312 \| 179 \| 115 \| 4 \| 1 \| 179 \| 2 \| 172 \| Uncharacterized protein \| Uncharacterized protein; afdb-uniprot50 \| AF-A0A823Q4U5-F1-MODEL\_V4 \| 1.0 \| 1.149e-14 \| 628 \| 0.375 \| 144 \| 85 \| 3 \| 1 \| 143 \| 3 \| 142 \| Phage tail protein \| Phage tail protein; afdb-uniprot50 \| AF-A0A3U9HGD2-F1-MODEL\_V4 \| 1.0 \| 6.379e-26 \| 628 \| 0.264 \| 496 \| 189 \| 18 \| 1 \| 423 \| 2 \| 394 \| Phage tail protein \| Phage tail protein; afdb-uniprot50 \| AF-A0A3E0X2Y9-F1-MODEL\_V4 \| 1.0 \| 8.2e-16 \| 627 \| 0.376 \| 178 \| 108 \| 2 \| 1 \| 177 \| 1 \| 176 \| Uncharacterized protein \| Uncharacterized protein; afdb-uniprot50 \| AF-A0A080HQP8-F1-MODEL\_V4 \| 1.0 \| 5.604e-20 \| 627 \| 0.322 \| 257 \| 130 \| 7 \| 1 \| 223 \| 2 \| 248 \| Phage tail-collar fiber family protein \| Phage tail-collar fiber family protein; afdb-uniprot50 \| AF-A0A447XW04-F1-MODEL\_V4 \| 1.0 \| 7.233e-21 \| 627 \| 0.326 \| 334 \| 144 \| 8 \| 1 \| 323 \| 2 \| 265 \| Tail fiber protein \| Tail fiber protein; afdb-uniprot50 \| AF-A0A6L6ITL6-F1-MODEL\_V4 \| 1.0 \| 1.538e-20 \| 626 \| 0.29 \| 351 \| 126 \| 6 \| 1 \| 324 \| 2 \| 256 \| Phage tail protein \| Phage tail protein; afdb-uniprot50 \| AF-A0A1S6U821-F1-MODEL\_V4 \| 1.0 \| 1.814e-16 \| 626 \| 0.326 \| 184 \| 116 \| 2 \| 3 \| 181 \| 1 \| 181 \| Phage tail collar fiber protein \| Phage tail collar fiber protein; afdb-uniprot50 \| AF-A0A7H9S4N1-F1-MODEL\_V4 \| 1.0 \| 1.361e-21 \| 625 \| 0.335 \| 334 \| 141 \| 9 \| 1 \| 323 \| 2 \| 265 \| Phage tail protein \| Phage tail protein; afdb-uniprot50 \| AF-A0A2A3VZP6-F1-MODEL\_V4 \| 1.0 \| 6.242e-20 \| 624 \| 0.29 \| 330 \| 150 \| 9 \| 1 \| 314 \| 1 \| 262 \| Phage tail protein \| Phage tail protein; afdb-uniprot50 \| AF-A0A2G2D6M1-F1-MODEL\_V4 \| 1.0 \| 3.329e-15 \| 623 \| 0.375 \| 165 \| 101 \| 2 \| 2 \| 165 \| 21 \| 184 \| Uncharacterized protein \| Uncharacterized protein; afdb-uniprot50 \| AF-A0A426VL03-F1-MODEL\_V4 \| 1.0 \| 1.017e-15 \| 623 \| 0.303 \| 181 \| 113 \| 4 \| 1 \| 178 \| 26 \| 196 \| Uncharacterized protein \| Uncharacterized protein; afdb-uniprot50 \| AF-W7PLX3-F1-MODEL\_V4 \| 1.0 \| 3.212e-25 \| 623 \| 0.202 \| 666 \| 178 \| 16 \| 1 \| 423 \| 1 \| 556 \| Side tail fiber protein \| Side tail fiber protein; afdb-uniprot50 \| AF-A0A2T9VBX3-F1-MODEL\_V4 \| 1.0 \| 3e-26 \| 622 \| 0.252 \| 495 \| 197 \| 18 \| 1 \| 423 \| 2 \| 395 \| Phage tail protein \| Phage tail protein; afdb-uniprot50 \| AF-A0A1T2X6C6-F1-MODEL\_V4 \| 1.0 \| 5.329e-16 \| 621 \| 0.329 \| 182 \| 106 \| 3 \| 6 \| 178 \| 4 \| 178 \| Phage tail-collar fiber protein (DUF3751 domain) \| Phage tail-collar fiber protein (DUF3751 domain); afdb-uniprot50 \| AF-A0A827GY52-F1-MODEL\_V4 \| 1.0 \| 4.205e-25 \| 621 \| 0.263 \| 501 \| 188 \| 21 \| 1 \| 423 \| 2 \| 399 \| Phage tail protein \| Phage tail protein; afdb-uniprot50 \| AF-A0A1S8PIF5-F1-MODEL\_V4 \| 1.0 \| 6.706e-15 \| 621 \| 0.414 \| 164 \| 93 \| 2 \| 1 \| 161 \| 1 \| 164 \| Glycosyl hydrolases family 43 \| Glycosyl hydrolases family 43; afdb-uniprot50 \| AF-A0A1H0YPL5-F1-MODEL\_V4 \| 1.0 \| 6.516e-17 \| 620 \| 0.32 \| 200 \| 118 \| 3 \| 3 \| 194 \| 5 \| 194 \| Phage tail-collar fibre protein \| Phage tail-collar fibre protein; afdb-uniprot50 \| AF-A0A4P8YDH1-F1-MODEL\_V4 \| 1.0 \| 1.646e-19 \| 619 \| 0.321 \| 252 \| 116 \| 5 \| 1 \| 221 \| 2 \| 229 \| Uncharacterized protein \| Uncharacterized protein; afdb-uniprot50 \| AF-A0A7X5J7V1-F1-MODEL\_V4 \| 1.0 \| 1.426e-14 \| 618 \| 0.373 \| 150 \| 92 \| 2 \| 5 \| 153 \| 2 \| 150 \| Uncharacterized protein \| Uncharacterized protein; afdb-uniprot50 \| AF-A0A6B2FQ21-F1-MODEL\_V4 \| 1.0 \| 9.993e-21 \| 618 \| 0.236 \| 432 \| 164 \| 15 \| 1 \| 423 \| 1 \| 275 \| Phage tail protein \| Phage tail protein; afdb-uniprot50 \| AF-A0A0R9N0A5-F1-MODEL\_V4 \| 1.0 \| 1.04e-21 \| 618 \| 0.316 \| 332 \| 154 \| 9 \| 1 \| 322 \| 1 \| 269 \| Phage tail protein \| Phage tail protein; afdb-uniprot50 \| AF-A0A4S2HC06-F1-MODEL\_V4 \| 1.0 \| 2.079e-14 \| 617 \| 0.32 \| 153 \| 104 \| 0 \| 1 \| 153 \| 2 \| 154 \| Uncharacterized protein \| Uncharacterized protein; afdb-uniprot50 \| AF-A0A421HAJ6-F1-MODEL\_V4 \| 1.0 \| 3.513e-15 \| 617 \| 0.395 \| 167 \| 93 \| 2 \| 1 \| 159 \| 1 \| 167 \| Phage tail protein \| Phage tail protein; afdb-uniprot50 \| AF-A0A0F6RH94-F1-MODEL\_V4 \| 1.0 \| 5.427e-26 \| 617 \| 0.239 \| 496 \| 207 \| 14 \| 1 \| 423 \| 2 \| 400 \| Tail protein \| Tail protein; afdb-uniprot50 \| AF-A0A4Q6YW23-F1-MODEL\_V4 \| 1.0 \| 7.363e-16 \| 615 \| 0.309 \| 181 \| 112 \| 4 \| 1 \| 178 \| 3 \| 173 \| Uncharacterized protein \| Uncharacterized protein; afdb-uniprot50 \| AF-A0A0A8FI71-F1-MODEL\_V4 \| 1.0 \| 6.401e-22 \| 615 \| 0.24 \| 433 \| 189 \| 17 \| 1 \| 422 \| 2 \| 305 \| Tail protein \| Tail protein; afdb-uniprot50 \| AF-A0A6L9JTE5-F1-MODEL\_V4 \| 1.0 \| 2.095e-21 \| 615 \| 0.35 \| 331 \| 126 \| 7 \| 4 \| 322 \| 3 \| 256 \| Phage tail protein \| Phage tail protein; afdb-uniprot50 \| AF-A0A653EAR8-F1-MODEL\_V4 \| 1.0 \| 2.358e-24 \| 615 \| 0.249 \| 453 \| 196 \| 19 \| 1 \| 423 \| 2 \| 340 \| Phage tail-collar fibre protein \| Phage tail-collar fibre protein; afdb-uniprot50 \| AF-A0A0U1KJN9-F1-MODEL\_V4 \| 1.0 \| 2.884e-25 \| 614 \| 0.249 \| 489 \| 207 \| 16 \| 1 \| 423 \| 2 \| 396 \| Tail fiber protein \| Tail fiber protein; afdb-uniprot50 \| AF-A0A849JYK8-F1-MODEL\_V4 \| 1.0 \| 4.145e-26 \| 614 \| 0.249 \| 497 \| 215 \| 16 \| 1 \| 423 \| 2 \| 414 \| Phage tail protein \| Phage tail protein; afdb-uniprot50 \| AF-C3X1Y2-F1-MODEL\_V4 \| 1.0 \| 1.121e-27 \| 614 \| 0.313 \| 501 \| 222 \| 19 \| 1 \| 412 \| 2 \| 469 \| Collar domain-containing protein \| Collar domain-containing protein; afdb-uniprot50 \| AF-A0A828AFE6-F1-MODEL\_V4 \| 1.0 \| 1.782e-21 \| 614 \| 0.273 \| 336 \| 167 \| 9 \| 1 \| 322 \| 2 \| 274 \| Phage tail protein \| Phage tail protein; afdb-uniprot50 \| AF-A0A4R5W1D0-F1-MODEL\_V4 \| 1.0 \| 1.101e-17 \| 614 \| 0.388 \| 216 \| 109 \| 6 \| 1 \| 194 \| 1 \| 215 \| Uncharacterized protein \| Uncharacterized protein; afdb-uniprot50 \| AF-A0A1R7Q9M1-F1-MODEL\_V4 \| 1.0 \| 1.426e-14 \| 614 \| 0.341 \| 158 \| 102 \| 2 \| 1 \| 158 \| 1 \| 156 \| Uncharacterized protein \| Uncharacterized protein; afdb-uniprot50 \| AF-A0A853IMF0-F1-MODEL\_V4 \| 1.0 \| 4.421e-14 \| 613 \| 0.368 \| 133 \| 81 \| 1 \| 1 \| 130 \| 1 \| 133 \| Phage tail protein \| Phage tail protein; afdb-uniprot50 \| AF-A0A826DDX5-F1-MODEL\_V4 \| 1.0 \| 1.213e-14 \| 613 \| 0.373 \| 158 \| 96 \| 1 \| 1 \| 158 \| 3 \| 157 \| Phage tail protein \| Phage tail protein; afdb-uniprot50 \| AF-H3R957-F1-MODEL\_V4 \| 1.0 \| 8.261e-23 \| 613 \| 0.259 \| 447 \| 186 \| 16 \| 1 \| 423 \| 1 \| 326 \| Uncharacterized protein \| Uncharacterized protein; afdb-uniprot50 \| AF-A0A6I3JS90-F1-MODEL\_V4 \| 1.0 \| 1.327e-19 \| 613 \| 0.323 \| 303 \| 107 \| 3 \| 4 \| 306 \| 3 \| 207 \| Collar domain-containing protein \| Collar domain-containing protein; afdb-uniprot50 \| AF-A0A811XII5-F1-MODEL\_V4 \| 1.0 \| 2.203e-25 \| 612 \| 0.223 \| 488 \| 213 \| 14 \| 1 \| 422 \| 2 \| 389 \| Phage tail fiber protein \| Phage tail fiber protein; afdb-uniprot50 \| AF-A0A423D1H4-F1-MODEL\_V4 \| 1.0 \| 8.814e-26 \| 612 \| 0.222 \| 490 \| 234 \| 19 \| 1 \| 423 \| 2 \| 411 \| Uncharacterized protein \| Uncharacterized protein; afdb-uniprot50 \| AF-A0A747Y1U2-F1-MODEL\_V4 \| 1.0 \| 5.31e-20 \| 611 \| 0.25 \| 351 \| 137 \| 7 \| 1 \| 326 \| 4 \| 253 \| Phage tail protein \| Phage tail protein; afdb-uniprot50 \| AF-A0A726YJW6-F1-MODEL\_V4 \| 1.0 \| 1.171e-24 \| 610 \| 0.238 \| 483 \| 197 \| 12 \| 4 \| 423 \| 14 \| 388 \| Uncharacterized protein \| Uncharacterized protein; afdb-uniprot50 \| AF-A0A2G0Q045-F1-MODEL\_V4 \| 1.0 \| 2.674e-19 \| 610 \| 0.316 \| 313 \| 113 \| 3 \| 1 \| 313 \| 2 \| 213 \| Tail protein \| Tail protein; afdb-uniprot50 \| AF-A0A447N0H8-F1-MODEL\_V4 \| 1.0 \| 2.367e-20 \| 609 \| 0.292 \| 338 \| 157 \| 10 \| 1 \| 323 \| 1 \| 271 \| Gp19 \| Gp19; afdb-uniprot50 \| AF-A0A7I9NTD9-F1-MODEL\_V4 \| 1.0 \| 1.676e-14 \| 609 \| 0.36 \| 158 \| 98 \| 1 \| 1 \| 158 \| 3 \| 157 \| Phage tail protein \| Phage tail protein; afdb-uniprot50 \| AF-A0A4P0YQ56-F1-MODEL\_V4 \| 1.0 \| 4.281e-20 \| 609 \| 0.261 \| 318 \| 156 \| 10 \| 1 \| 306 \| 1 \| 251 \| Putative phage tail fiber protein \| Putative phage tail fiber protein; afdb-uniprot50 \| AF-A0A857F214-F1-MODEL\_V4 \| 1.0 \| 3.451e-20 \| 609 \| 0.33 \| 306 \| 115 \| 4 \| 1 \| 306 \| 2 \| 217 \| Phage tail protein \| Phage tail protein; afdb-uniprot50 \| AF-A0A5X4GC92-F1-MODEL\_V4 \| 1.0 \| 6.401e-22 \| 609 \| 0.303 \| 349 \| 163 \| 9 \| 1 \| 328 \| 1 \| 290 \| Short-chain fatty acid transporter \| Short-chain fatty acid transporter; afdb-uniprot50 \| AF-A0A5Z0R0R1-F1-MODEL\_V4 \| 1.0 \| 8.319e-15 \| 609 \| 0.375 \| 157 \| 96 \| 2 \| 3 \| 159 \| 12 \| 166 \| Collar domain-containing protein \| Collar domain-containing protein; afdb-uniprot50 \| AF-A0A830Q9A3-F1-MODEL\_V4 \| 1.0 \| 6.221e-24 \| 608 \| 0.264 \| 457 \| 191 \| 14 \| 1 \| 423 \| 2 \| 347 \| Putative phage tail collar domain-containing protein \| Putative phage tail collar domain-containing protein; afdb-uniprot50 \| AF-C5BEP6-F1-MODEL\_V4 \| 1.0 \| 2.772e-24 \| 608 \| 0.223 \| 475 \| 206 \| 15 \| 1 \| 422 \| 2 \| 366 \| Phage tail fiber domain protein \| Phage tail fiber domain protein; afdb-uniprot50 \| AF-A0A6I2IHE3-F1-MODEL\_V4 \| 1.0 \| 3.476e-27 \| 608 \| 0.257 \| 509 \| 221 \| 20 \| 1 \| 423 \| 2 \| 439 \| Phage tail protein \| Phage tail protein; afdb-uniprot50 \| AF-A0A5X9L6P6-F1-MODEL\_V4 \| 1.0 \| 3.59e-21 \| 608 \| 0.314 \| 331 \| 153 \| 9 \| 2 \| 322 \| 3 \| 269 \| Phage tail protein \| Phage tail protein; afdb-uniprot50 \| AF-A0A5D3JF06-F1-MODEL\_V4 \| 1.0 \| 3.761e-14 \| 606 \| 0.417 \| 151 \| 87 \| 1 \| 1 \| 151 \| 2 \| 151 \| Phage tail protein \| Phage tail protein; afdb-uniprot50 \| AF-A0A826YSE8-F1-MODEL\_V4 \| 1.0 \| 4.22e-21 \| 606 \| 0.326 \| 334 \| 144 \| 8 \| 1 \| 323 \| 2 \| 265 \| Phage tail protein \| Phage tail protein; afdb-uniprot50 \| AF-A0A2X1PYX0-F1-MODEL\_V4 \| 1.0 \| 5.032e-20 \| 606 \| 0.293 \| 327 \| 150 \| 10 \| 1 \| 314 \| 1 \| 259 \| Bacteriophage tail protein \| Bacteriophage tail protein; afdb-uniprot50 \| AF-A0A024E8J4-F1-MODEL\_V4 \| 1.0 \| 2.579e-14 \| 606 \| 0.42 \| 145 \| 84 \| 0 \| 14 \| 158 \| 2 \| 146 \| Tail fiber protein H, putative \| Tail fiber protein H, putative; afdb-uniprot50 \| AF-A0A5H6KLG3-F1-MODEL\_V4 \| 1.0 \| 1.209e-18 \| 605 \| 0.267 \| 307 \| 123 \| 5 \| 1 \| 306 \| 4 \| 209 \| Phage tail protein \| Phage tail protein; afdb-uniprot50 \| AF-A0A2Y0PD72-F1-MODEL\_V4 \| 1.0 \| 8.595e-24 \| 605 \| 0.219 \| 483 \| 203 \| 12 \| 1 \| 423 \| 2 \| 370 \| Tail fiber domain-containing protein \| Tail fiber domain-containing protein; afdb-uniprot50 \| AF-A0A7W0B3B2-F1-MODEL\_V4 \| 1.0 \| 1.093e-25 \| 605 \| 0.236 \| 512 \| 248 \| 25 \| 1 \| 423 \| 1 \| 458 \| Phage-related tail fiber protein \| Phage-related tail fiber protein; afdb-uniprot50 \| AF-N1NKC6-F1-MODEL\_V4 \| 1.0 \| 1.381e-20 \| 605 \| 0.306 \| 330 \| 122 \| 5 \| 1 \| 324 \| 2 \| 230 \| Uncharacterized protein \| Uncharacterized protein; afdb-uniprot50 \| AF-A0A5U3G5W3-F1-MODEL\_V4 \| 1.0 \| 1.028e-18 \| 604 \| 0.244 \| 347 \| 138 \| 6 \| 1 \| 324 \| 1 \| 246 \| Uncharacterized protein \| Uncharacterized protein; afdb-uniprot50 \| AF-A0A4D7AWU2-F1-MODEL\_V4 \| 1.0 \| 6.108e-14 \| 604 \| 0.324 \| 154 \| 101 \| 3 \| 1 \| 153 \| 1 \| 152 \| DUF2793 domain-containing protein \| DUF2793 domain-containing protein; afdb-uniprot50 \| AF-A0A0U1HL65-F1-MODEL\_V4 \| 1.0 \| 1.141e-22 \| 604 \| 0.252 \| 491 \| 190 \| 20 \| 1 \| 423 \| 2 \| 383 \| Tail fiber protein \| Tail fiber protein; afdb-uniprot50 \| AF-D2BZE6-F1-MODEL\_V4 \| 1.0 \| 4.343e-19 \| 604 \| 0.31 \| 264 \| 138 \| 7 \| 1 \| 223 \| 2 \| 262 \| Phage-related tail fibre protein-like protein \| Phage-related tail fibre protein-like protein; afdb-uniprot50 \| AF-A0A433NXC7-F1-MODEL\_V4 \| 1.0 \| 9.469e-21 \| 602 \| 0.335 \| 331 \| 113 \| 6 \| 2 \| 332 \| 3 \| 226 \| Tail fiber protein \| Tail fiber protein; afdb-uniprot50 \| AF-N9QTJ7-F1-MODEL\_V4 \| 1.0 \| 6.264e-16 \| 602 \| 0.333 \| 207 \| 118 \| 5 \| 3 \| 190 \| 5 \| 210 \| Uncharacterized protein \| Uncharacterized protein; afdb-uniprot50 \| AF-A0A7Y1SG14-F1-MODEL\_V4 \| 1.0 \| 7.77e-16 \| 601 \| 0.351 \| 202 \| 111 \| 6 \| 1 \| 183 \| 1 \| 201 \| Phage tail protein \| Phage tail protein; afdb-uniprot50 \| AF-A0A7H4NYM5-F1-MODEL\_V4 \| 1.0 \| 7.855e-19 \| 601 \| 0.266 \| 334 \| 138 \| 7 \| 1 \| 324 \| 2 \| 238 \| Putative prophage tail fiber protein \| Putative prophage tail fiber protein; afdb-uniprot50 \| AF-A0A5M6I3L7-F1-MODEL\_V4 \| 1.0 \| 8.749e-19 \| 601 \| 0.287 \| 316 \| 135 \| 9 \| 1 \| 306 \| 1 \| 236 \| Uncharacterized protein \| Uncharacterized protein; afdb-uniprot50 \| AF-A0A827RXE4-F1-MODEL\_V4 \| 1.0 \| 1.516e-21 \| 600 \| 0.344 \| 334 \| 138 \| 10 \| 1 \| 324 \| 2 \| 264 \| Phage tail protein \| Phage tail protein; afdb-uniprot50 \| AF-A0A1G5QGK4-F1-MODEL\_V4 \| 1.0 \| 1.436e-21 \| 600 \| 0.338 \| 340 \| 118 \| 9 \| 4 \| 322 \| 3 \| 256 \| Phage tail fibre repeat-containing protein \| Phage tail fibre repeat-containing protein; afdb-uniprot50 \| AF-A0A077P0S8-F1-MODEL\_V4 \| 1.0 \| 2.926e-24 \| 600 \| 0.215 \| 622 \| 187 \| 15 \| 1 \| 423 \| 2 \| 521 \| Uncharacterized protein \| Uncharacterized protein; afdb-uniprot50 \| AF-A0A853HVL0-F1-MODEL\_V4 \| 1.0 \| 2.525e-23 \| 599 \| 0.217 \| 450 \| 207 \| 21 \| 1 \| 423 \| 1 \| 332 \| Phage tail protein \| Phage tail protein; afdb-uniprot50 \| AF-I6DBN4-F1-MODEL\_V4 \| 1.0 \| 8.502e-21 \| 598 \| 0.286 \| 342 \| 165 \| 12 \| 1 \| 308 \| 1 \| 297 \| Phage tail fiber repeat family protein \| Phage tail fiber repeat family protein; afdb-uniprot50 \| AF-A0A2H9EM17-F1-MODEL\_V4 \| 1.0 \| 3.342e-26 \| 598 \| 0.265 \| 535 \| 211 \| 22 \| 1 \| 423 \| 2 \| 466 \| Phage tail protein \| Phage tail protein; afdb-uniprot50 \| AF-J2V596-F1-MODEL\_V4 \| 1.0 \| 3.463e-16 \| 597 \| 0.356 \| 199 \| 119 \| 5 \| 2 \| 193 \| 6 \| 202 \| Uncharacterized protein \| Uncharacterized protein; afdb-uniprot50 \| AF-A0A7A4NFG1-F1-MODEL\_V4 \| 1.0 \| 2.367e-20 \| 596 \| 0.267 \| 344 \| 170 \| 9 \| 1 \| 332 \| 5 \| 278 \| Phage tail protein \| Phage tail protein; afdb-uniprot50 \| AF-F7SVK4-F1-MODEL\_V4 \| 1.0 \| 1.478e-19 \| 596 \| 0.337 \| 264 \| 134 \| 9 \| 1 \| 244 \| 1 \| 243 \| Uncharacterized protein \| Uncharacterized protein; afdb-uniprot50 \| AF-A0A1I3S1C5-F1-MODEL\_V4 \| 1.0 \| 6.588e-20 \| 595 \| 0.303 \| 330 \| 123 \| 6 \| 1 \| 324 \| 2 \| 230 \| Phage tail fibre repeat-containing protein \| Phage tail fibre repeat-containing protein; afdb-uniprot50 \| AF-A0A853I7M0-F1-MODEL\_V4 \| 1.0 \| 4.438e-25 \| 595 \| 0.211 \| 467 \| 248 \| 23 \| 1 \| 423 \| 1 \| 391 \| Phage tail protein \| Phage tail protein; afdb-uniprot50 \| AF-A0A066RYY3-F1-MODEL\_V4 \| 1.0 \| 2.636e-20 \| 595 \| 0.295 \| 305 \| 144 \| 8 \| 1 \| 259 \| 1 \| 280 \| Uncharacterized protein \| Uncharacterized protein; afdb-uniprot50 \| AF-A0A1C6Z3Q6-F1-MODEL\_V4 \| 1.0 \| 3.789e-21 \| 594 \| 0.313 \| 332 \| 155 \| 7 \| 1 \| 324 \| 2 \| 268 \| Phage tail fibre repeat-containing protein \| Phage tail fibre repeat-containing protein; afdb-uniprot50 \| AF-A0A1B9JGB9-F1-MODEL\_V4 \| 1.0 \| 3.999e-21 \| 593 \| 0.276 \| 355 \| 130 \| 3 \| 1 \| 343 \| 2 \| 241 \| Collar domain-containing protein \| Collar domain-containing protein; afdb-uniprot50 \| AF-A0A0T9UVJ2-F1-MODEL\_V4 \| 1.0 \| 6.803e-14 \| 592 \| 0.47 \| 134 \| 70 \| 1 \| 1 \| 133 \| 1 \| 134 \| Variable tail fiber protein \| Variable tail fiber protein; afdb-uniprot50 \| AF-A0A0B6FER7-F1-MODEL\_V4 \| 1.0 \| 8.261e-23 \| 592 \| 0.254 \| 436 \| 211 \| 21 \| 1 \| 423 \| 2 \| 336 \| Phage tail-collar fiber family protein \| Phage tail-collar fiber family protein; afdb-uniprot50 \| AF-A0A3S5DM66-F1-MODEL\_V4 \| 1.0 \| 4.101e-23 \| 592 \| 0.258 \| 449 \| 197 \| 17 \| 1 \| 408 \| 1 \| 354 \| Phage tail fiber protein H \| Phage tail fiber protein H; afdb-uniprot50 \| AF-A0A077PK87-F1-MODEL\_V4 \| 1.0 \| 4.056e-20 \| 592 \| 0.308 \| 331 \| 121 \| 6 \| 1 \| 324 \| 1 \| 230 \| E14 prophage putative tail fiber protein (Modular protein) \| E14 prophage putative tail fiber protein (Modular protein); afdb-uniprot50 \| AF-A0A833UC36-F1-MODEL\_V4 \| 1.0 \| 9.134e-16 \| 592 \| 0.368 \| 179 \| 103 \| 4 \| 1 \| 178 \| 1 \| 170 \| Uncharacterized protein \| Uncharacterized protein; afdb-uniprot50 \| AF-A0A831GBG9-F1-MODEL\_V4 \| 1.0 \| 7.882e-15 \| 591 \| 0.396 \| 169 \| 88 \| 5 \| 1 \| 159 \| 1 \| 165 \| Phage tail protein \| Phage tail protein; afdb-uniprot50 \| AF-A0A4T5FQZ6-F1-MODEL\_V4 \| 1.0 \| 1.113e-20 \| 591 \| 0.278 \| 359 \| 180 \| 12 \| 1 \| 324 \| 5 \| 319 \| Phage tail protein \| Phage tail protein; afdb-uniprot50 \| AF-A0A2X5PQJ0-F1-MODEL\_V4 \| 1.0 \| 3.928e-26 \| 591 \| 0.271 \| 537 \| 207 \| 24 \| 1 \| 423 \| 1 \| 467 \| Phage tail collar protein \| Phage tail collar protein; afdb-uniprot50 \| AF-A0A5M6IAF3-F1-MODEL\_V4 \| 1.0 \| 1.814e-16 \| 590 \| 0.418 \| 184 \| 104 \| 2 \| 1 \| 183 \| 1 \| 182 \| Uncharacterized protein \| Uncharacterized protein; afdb-uniprot50 \| AF-A0A1V2UTE0-F1-MODEL\_V4 \| 1.0 \| 4.784e-16 \| 590 \| 0.341 \| 208 \| 114 \| 5 \| 1 \| 206 \| 1 \| 187 \| Uncharacterized protein \| Uncharacterized protein; afdb-uniprot50 \| AF-A0A1A0FJZ1-F1-MODEL\_V4 \| 1.0 \| 3.83e-24 \| 589 \| 0.218 \| 481 \| 208 \| 17 \| 2 \| 423 \| 6 \| 377 \| Uncharacterized protein \| Uncharacterized protein; afdb-uniprot50 \| AF-R8WJW4-F1-MODEL\_V4 \| 1.0 \| 1.629e-16 \| 589 \| 0.425 \| 209 \| 98 \| 2 \| 1 \| 187 \| 2 \| 210 \| Uncharacterized protein \| Uncharacterized protein; afdb-uniprot50 \| AF-A0A7M3FHK4-F1-MODEL\_V4 \| 1.0 \| 2.683e-15 \| 589 \| 0.319 \| 197 \| 115 \| 4 \| 1 \| 179 \| 2 \| 197 \| Phage tail protein \| Phage tail protein; afdb-uniprot50 \| AF-A0A1G9YF31-F1-MODEL\_V4 \| 1.0 \| 5.728e-26 \| 588 \| 0.226 \| 512 \| 260 \| 30 \| 2 \| 423 \| 6 \| 471 \| Phage tail-collar fibre protein \| Phage tail-collar fibre protein; afdb-uniprot50 \| AF-A0A2X2BLJ3-F1-MODEL\_V4 \| 1.0 \| 2.752e-17 \| 587 \| 0.363 \| 223 \| 113 \| 3 \| 1 \| 194 \| 2 \| 224 \| Phage variable tail fiber protein \| Phage variable tail fiber protein; afdb-uniprot50 \| AF-A0A7Y8JX08-F1-MODEL\_V4 \| 1.0 \| 2.946e-16 \| 587 \| 0.341 \| 214 \| 112 \| 5 \| 1 \| 198 \| 1 \| 201 \| Phage tail protein \| Phage tail protein; afdb-uniprot50 \| AF-A0A3N6UMU5-F1-MODEL\_V4 \| 1.0 \| 7.233e-21 \| 587 \| 0.231 \| 453 \| 166 \| 15 \| 1 \| 423 \| 1 \| 301 \| Phage tail protein \| Phage tail protein; afdb-uniprot50 \| AF-A0A2V4E3U5-F1-MODEL\_V4 \| 1.0 \| 1.641e-23 \| 587 \| 0.271 \| 457 \| 188 \| 17 \| 1 \| 423 \| 2 \| 347 \| DNA inversion product \| DNA inversion product; afdb-uniprot50 \| AF-A0A6Y2VPA8-F1-MODEL\_V4 \| 1.0 \| 3.098e-20 \| 587 \| 0.27 \| 344 \| 169 \| 9 \| 1 \| 332 \| 5 \| 278 \| Phage tail protein \| Phage tail protein; afdb-uniprot50 \| AF-A0A6H8PL82-F1-MODEL\_V4 \| 1.0 \| 3.564e-14 \| 586 \| 0.381 \| 160 \| 94 \| 2 \| 1 \| 158 \| 1 \| 157 \| Phage tail protein \| Phage tail protein; afdb-uniprot50 \| AF-A0A5Y7WJU3-F1-MODEL\_V4 \| 1.0 \| 3.088e-24 \| 586 \| 0.209 \| 620 \| 187 \| 17 \| 1 \| 423 \| 2 \| 515 \| Phage tail protein \| Phage tail protein; afdb-uniprot50 \| AF-A0A1C4DJ79-F1-MODEL\_V4 \| 1.0 \| 3.223e-21 \| 585 \| 0.247 \| 437 \| 184 \| 14 \| 1 \| 423 \| 2 \| 307 \| Phage tail-collar fibre protein \| Phage tail-collar fibre protein; afdb-uniprot50 \| AF-A0A3X7LX61-F1-MODEL\_V4 \| 1.0 \| 1.908e-20 \| 585 \| 0.251 \| 346 \| 136 \| 5 \| 1 \| 324 \| 4 \| 248 \| Phage tail protein \| Phage tail protein; afdb-uniprot50 \| AF-A0A8A7FP84-F1-MODEL\_V4 \| 1.0 \| 6.588e-20 \| 585 \| 0.303 \| 330 \| 123 \| 6 \| 1 \| 324 \| 2 \| 230 \| Phage tail protein \| Phage tail protein; afdb-uniprot50 \| AF-A0A367LYA0-F1-MODEL\_V4 \| 1.0 \| 7.577e-14 \| 584 \| 0.371 \| 148 \| 88 \| 1 \| 6 \| 148 \| 3 \| 150 \| Phage tail protein \| Phage tail protein; afdb-uniprot50 \| AF-A0A827KPR0-F1-MODEL\_V4 \| 1.0 \| 1.694e-17 \| 584 \| 0.237 \| 312 \| 133 \| 6 \| 1 \| 308 \| 1 \| 211 \| Phage tail protein \| Phage tail protein; afdb-uniprot50 \| AF-A0A1H0FNR9-F1-MODEL\_V4 \| 1.0 \| 1.361e-21 \| 584 \| 0.22 \| 440 \| 198 \| 15 \| 1 \| 423 \| 1 \| 312 \| Phage tail-collar fibre protein \| Phage tail-collar fibre protein; afdb-uniprot50 \| AF-A0A443WD25-F1-MODEL\_V4 \| 1.0 \| 5.014e-24 \| 584 \| 0.277 \| 440 \| 200 \| 21 \| 2 \| 423 \| 3 \| 342 \| DNA inversion product \| DNA inversion product; afdb-uniprot50 \| AF-A0A140NE20-F1-MODEL\_V4 \| 1.0 \| 7.105e-26 \| 584 \| 0.243 \| 580 \| 241 \| 16 \| 1 \| 423 \| 2 \| 540 \| Phage-related tail fibre protein-like protein \| Phage-related tail fibre protein-like protein; afdb-uniprot50 \| AF-A0A2G2KYA8-F1-MODEL\_V4 \| 1.0 \| 3.899e-19 \| 584 \| 0.303 \| 264 \| 134 \| 9 \| 1 \| 257 \| 2 \| 222 \| Uncharacterized protein \| Uncharacterized protein; afdb-uniprot50 \| AF-A0A068QVC8-F1-MODEL\_V4 \| 1.0 \| 1.07e-19 \| 583 \| 0.306 \| 330 \| 122 \| 6 \| 1 \| 324 \| 2 \| 230 \| Uncharacterized protein \| Uncharacterized protein; afdb-uniprot50 \| AF-A0A7W4PXC5-F1-MODEL\_V4 \| 1.0 \| 4.174e-18 \| 583 \| 0.308 \| 253 \| 132 \| 9 \| 1 \| 247 \| 1 \| 216 \| Phage tail protein \| Phage tail protein; afdb-uniprot50 \| AF-A0A1B9JGB5-F1-MODEL\_V4 \| 1.0 \| 1.29e-21 \| 582 \| 0.295 \| 359 \| 154 \| 11 \| 1 \| 324 \| 1 \| 295 \| Collar domain-containing protein \| Collar domain-containing protein; afdb-uniprot50 \| AF-A0A1Q5TLV3-F1-MODEL\_V4 \| 1.0 \| 4.056e-20 \| 582 \| 0.298 \| 335 \| 125 \| 8 \| 1 \| 326 \| 1 \| 234 \| Tail protein \| Tail protein; afdb-uniprot50 \| AF-A0A2Z2JIM9-F1-MODEL\_V4 \| 1.0 \| 1.834e-19 \| 582 \| 0.271 \| 335 \| 162 \| 8 \| 1 \| 323 \| 5 \| 269 \| Phage tail fiber repeat protein \| Phage tail fiber repeat protein; afdb-uniprot50 \| AF-A0A7W3G1H3-F1-MODEL\_V4 \| 1.0 \| 1.105e-13 \| 581 \| 0.351 \| 148 \| 92 \| 1 \| 1 \| 148 \| 2 \| 145 \| Phage tail protein \| Phage tail protein; afdb-uniprot50 \| AF-A0A143DG88-F1-MODEL\_V4 \| 1.0 \| 1.07e-19 \| 581 \| 0.281 \| 316 \| 143 \| 9 \| 1 \| 299 \| 3 \| 251 \| Collar domain-containing protein \| Collar domain-containing protein; afdb-uniprot50 \| AF-A0A6I6IHZ7-F1-MODEL\_V4 \| 1.0 \| 2.627e-24 \| 581 \| 0.256 \| 491 \| 209 \| 15 \| 1 \| 423 \| 2 \| 404 \| Phage tail protein \| Phage tail protein; afdb-uniprot50 \| AF-A0A7A2HGT8-F1-MODEL\_V4 \| 1.0 \| 3.642e-20 \| 580 \| 0.274 \| 339 \| 160 \| 9 \| 1 \| 323 \| 1 \| 269 \| Phage tail protein \| Phage tail protein; afdb-uniprot50 \| AF-A0A3L2NND0-F1-MODEL\_V4 \| 1.0 \| 2.782e-20 \| 580 \| 0.299 \| 341 \| 159 \| 12 \| 1 \| 324 \| 1 \| 278 \| Short-chain fatty acid transporter \| Short-chain fatty acid transporter; afdb-uniprot50 \| AF-A0A1B9JGC2-F1-MODEL\_V4 \| 1.0 \| 1.985e-21 \| 580 \| 0.277 \| 367 \| 157 \| 10 \| 1 \| 322 \| 1 \| 304 \| Uncharacterized protein \| Uncharacterized protein; afdb-uniprot50 \| AF-A0A3E3I3G7-F1-MODEL\_V4 \| 1.0 \| 7.179e-14 \| 579 \| 0.341 \| 158 \| 96 \| 3 \| 1 \| 153 \| 1 \| 155 \| Uncharacterized protein \| Uncharacterized protein; afdb-uniprot50 \| AF-A0A2K1QCR8-F1-MODEL\_V4 \| 1.0 \| 2.333e-21 \| 579 \| 0.241 \| 439 \| 189 \| 13 \| 1 \| 422 \| 2 \| 313 \| Phage tail protein \| Phage tail protein; afdb-uniprot50 \| AF-A0A3N6UVH7-F1-MODEL\_V4 \| 1.0 \| 2.853e-22 \| 578 \| 0.26 \| 450 \| 196 \| 15 \| 1 \| 423 \| 2 \| 341 \| Phage tail protein \| Phage tail protein; afdb-uniprot50 \| AF-A0A7U7FJJ3-F1-MODEL\_V4 \| 1.0 \| 6.132e-25 \| 578 \| 0.238 \| 546 \| 197 \| 16 \| 1 \| 423 \| 2 \| 451 \| Hypothetical phage tail fiber protein \| Hypothetical phage tail fiber protein; afdb-uniprot50 \| AF-A0A377H523-F1-MODEL\_V4 \| 1.0 \| 6.83e-25 \| 578 \| 0.201 \| 572 \| 205 \| 16 \| 1 \| 423 \| 2 \| 470 \| Phage tail fibre repeat \| Phage tail fibre repeat; afdb-uniprot50 \| AF-A0A5C8XUF6-F1-MODEL\_V4 \| 1.0 \| 7.337e-20 \| 577 \| 0.254 \| 318 \| 158 \| 10 \| 1 \| 306 \| 1 \| 251 \| Uncharacterized protein \| Uncharacterized protein; afdb-uniprot50 \| AF-A0A7W5P3C6-F1-MODEL\_V4 \| 1.0 \| 1.533e-24 \| 577 \| 0.262 \| 506 \| 202 \| 16 \| 1 \| 423 \| 2 \| 419 \| Phage-related tail fiber protein \| Phage-related tail fiber protein; afdb-uniprot50 \| AF-A0A3G2IM13-F1-MODEL\_V4 \| 1.0 \| 1.56e-19 \| 577 \| 0.267 \| 352 \| 129 \| 2 \| 1 \| 324 \| 2 \| 252 \| Uncharacterized protein \| Uncharacterized protein; afdb-uniprot50 \| AF-A0A846QFW6-F1-MODEL\_V4 \| 1.0 \| 6.588e-20 \| 576 \| 0.332 \| 286 \| 142 \| 9 \| 1 \| 249 \| 1 \| 274 \| Phage-related tail fiber protein \| Phage-related tail fiber protein; afdb-uniprot50 \| AF-A0A1B8YGC4-F1-MODEL\_V4 \| 1.0 \| 2.095e-21 \| 576 \| 0.34 \| 344 \| 120 \| 6 \| 4 \| 326 \| 3 \| 260 \| Phage tail fiber repeat protein \| Phage tail fiber repeat protein; afdb-uniprot50 \| AF-A0A5T2CS27-F1-MODEL\_V4 \| 1.0 \| 2.616e-13 \| 575 \| 0.385 \| 127 \| 75 \| 1 \| 1 \| 127 \| 3 \| 126 \| Phage tail protein \| Phage tail protein; afdb-uniprot50 \| AF-A0A5W3T5U2-F1-MODEL\_V4 \| 1.0 \| 1.308e-20 \| 575 \| 0.288 \| 343 \| 156 \| 12 \| 1 \| 323 \| 1 \| 275 \| Phage tail protein \| Phage tail protein; afdb-uniprot50 \| AF-A0A827K3J7-F1-MODEL\_V4 \| 1.0 \| 7.053e-19 \| 575 \| 0.294 \| 309 \| 116 \| 4 \| 1 \| 308 \| 2 \| 209 \| Phage tail protein \| Phage tail protein; afdb-uniprot50 \| AF-A0A4R2GW55-F1-MODEL\_V4 \| 1.0 \| 5.871e-13 \| 575 \| 0.353 \| 147 \| 93 \| 2 \| 1 \| 146 \| 1 \| 146 \| Phage-related tail fiber protein \| Phage-related tail fiber protein; afdb-uniprot50 \| AF-A0A1Q8DQY0-F1-MODEL\_V4 \| 1.0 \| 7.996e-14 \| 574 \| 0.325 \| 160 \| 103 \| 3 \| 1 \| 158 \| 1 \| 157 \| Uncharacterized protein \| Uncharacterized protein; afdb-uniprot50 \| AF-A0A2V4EMJ8-F1-MODEL\_V4 \| 1.0 \| 2.401e-19 \| 574 \| 0.33 \| 284 \| 130 \| 9 \| 1 \| 237 \| 2 \| 272 \| Uncharacterized protein \| Uncharacterized protein; afdb-uniprot50 \| AF-A0A731FN07-F1-MODEL\_V4 \| 1.0 \| 5.747e-22 \| 574 \| 0.295 \| 366 \| 183 \| 9 \| 1 \| 321 \| 1 \| 336 \| Phage tail protein \| Phage tail protein; afdb-uniprot50 \| AF-A0A077QF37-F1-MODEL\_V4 \| 1.0 \| 2.674e-19 \| 574 \| 0.306 \| 330 \| 122 \| 5 \| 1 \| 324 \| 2 \| 230 \| Uncharacterized protein \| Uncharacterized protein; afdb-uniprot50 \| AF-A0A4R1NAG8-F1-MODEL\_V4 \| 1.0 \| 8.624e-20 \| 573 \| 0.304 \| 315 \| 113 \| 5 \| 1 \| 305 \| 1 \| 219 \| Tail fiber-like repeat protein \| Tail fiber-like repeat protein; afdb-uniprot50 \| AF-D4F114-F1-MODEL\_V4 \| 1.0 \| 8.624e-20 \| 573 \| 0.324 \| 330 \| 131 \| 5 \| 2 \| 331 \| 3 \| 240 \| Phage tail fiber repeat protein \| Phage tail fiber repeat protein; afdb-uniprot50 \| AF-A0A5W1P1H8-F1-MODEL\_V4 \| 1.0 \| 7.743e-20 \| 573 \| 0.282 \| 343 \| 158 \| 10 \| 1 \| 323 \| 1 \| 275 \| Phage tail protein \| Phage tail protein; afdb-uniprot50 \| AF-A0A3L5HA13-F1-MODEL\_V4 \| 1.0 \| 2.936e-20 \| 572 \| 0.283 \| 342 \| 158 \| 9 \| 1 \| 323 \| 2 \| 275 \| Phage tail protein \| Phage tail protein; afdb-uniprot50 \| AF-A0A3E2BGR8-F1-MODEL\_V4 \| 1.0 \| 7.443e-19 \| 572 \| 0.299 \| 307 \| 113 \| 5 \| 4 \| 306 \| 3 \| 211 \| Phage tail protein \| Phage tail protein; afdb-uniprot50 \| AF-A0A5U6Z5U0-F1-MODEL\_V4 \| 1.0 \| 2.087e-25 \| 572 \| 0.231 \| 562 \| 224 \| 17 \| 1 \| 423 \| 2 \| 494 \| Phage tail protein \| Phage tail protein; afdb-uniprot50 \| AF-A0A3G9G2L6-F1-MODEL\_V4 \| 1.0 \| 3.707e-15 \| 572 \| 0.32 \| 200 \| 117 \| 4 \| 1 \| 182 \| 3 \| 201 \| Phage tail fiber protein \| Phage tail fiber protein; afdb-uniprot50 \| AF-D3V0E0-F1-MODEL\_V4 \| 1.0 \| 2.534e-19 \| 572 \| 0.303 \| 330 \| 123 \| 5 \| 1 \| 324 \| 2 \| 230 \| E14 prophage putative tail fiber protein (Modular protein) \| E14 prophage putative tail fiber protein (Modular protein); afdb-uniprot50 \| AF-A0A5E6YEG9-F1-MODEL\_V4 \| 1.0 \| 2.392e-23 \| 571 \| 0.209 \| 454 \| 234 \| 20 \| 4 \| 423 \| 8 \| 370 \| Uncharacterized protein \| Uncharacterized protein; afdb-uniprot50 \| AF-F5MFA9-F1-MODEL\_V4 \| 1.0 \| 2.013e-20 \| 571 \| 0.28 \| 342 \| 159 \| 10 \| 1 \| 323 \| 2 \| 275 \| Phage tail fibre repeat family protein \| Phage tail fibre repeat family protein; afdb-uniprot50 \| AF-A0A8B2FET2-F1-MODEL\_V4 \| 1.0 \| 5.104e-19 \| 571 \| 0.306 \| 303 \| 112 \| 4 \| 4 \| 306 \| 3 \| 207 \| Phage tail protein \| Phage tail protein; afdb-uniprot50 \| AF-A0A2D0JSC8-F1-MODEL\_V4 \| 1.0 \| 1.646e-19 \| 571 \| 0.31 \| 335 \| 121 \| 8 \| 1 \| 326 \| 1 \| 234 \| Tail protein \| Tail protein; afdb-uniprot50 \| AF-A0A242PF12-F1-MODEL\_V4 \| 1.0 \| 6.153e-21 \| 571 \| 0.315 \| 345 \| 130 \| 5 \| 1 \| 344 \| 1 \| 240 \| Collar domain-containing protein \| Collar domain-containing protein; afdb-uniprot50 \| AF-R9B4A5-F1-MODEL\_V4 \| 1.0 \| 1.047e-13 \| 570 \| 0.254 \| 165 \| 113 \| 5 \| 1 \| 161 \| 2 \| 160 \| Collar domain-containing protein \| Collar domain-containing protein; afdb-uniprot50 \| AF-A0A376MUF9-F1-MODEL\_V4 \| 1.0 \| 1.381e-20 \| 570 \| 0.293 \| 341 \| 158 \| 9 \| 1 \| 322 \| 1 \| 277 \| Putative tail fiber protein (GpH) \| Putative tail fiber protein (GpH); afdb-uniprot50 \| AF-D3QUP0-F1-MODEL\_V4 \| 1.0 \| 9.851e-22 \| 570 \| 0.317 \| 343 \| 153 \| 7 \| 1 \| 332 \| 2 \| 274 \| DNA inversion product \| DNA inversion product; afdb-uniprot50 \| AF-A0A379F8Y3-F1-MODEL\_V4 \| 1.0 \| 8.531e-17 \| 570 \| 0.364 \| 222 \| 116 \| 4 \| 2 \| 207 \| 3 \| 215 \| Phage variable tail fiber protein \| Phage variable tail fiber protein; afdb-uniprot50 \| AF-A0A1C3K388-F1-MODEL\_V4 \| 1.0 \| 4.042e-24 \| 569 \| 0.252 \| 451 \| 229 \| 25 \| 3 \| 423 \| 2 \| 374 \| Phage tail fiber protein \| Phage tail fiber protein; afdb-uniprot50 \| AF-A0A389MCH5-F1-MODEL\_V4 \| 1.0 \| 6.683e-19 \| 569 \| 0.286 \| 304 \| 116 \| 3 \| 1 \| 304 \| 1 \| 203 \| Uncharacterized protein \| Uncharacterized protein; afdb-uniprot50 \| AF-A0A2N7UDN2-F1-MODEL\_V4 \| 1.0 \| 6.732e-26 \| 569 \| 0.215 \| 595 \| 268 \| 27 \| 1 \| 423 \| 1 \| 568 \| Uncharacterized protein \| Uncharacterized protein; afdb-uniprot50 \| AF-A0A3G2IDY2-F1-MODEL\_V4 \| 1.0 \| 1.56e-19 \| 569 \| 0.269 \| 352 \| 128 \| 4 \| 1 \| 324 \| 2 \| 252 \| Uncharacterized protein \| Uncharacterized protein; afdb-uniprot50 \| AF-A0A3M8C902-F1-MODEL\_V4 \| 1.0 \| 4.837e-19 \| 568 \| 0.333 \| 285 \| 133 \| 12 \| 1 \| 252 \| 2 \| 262 \| Phage tail protein \| Phage tail protein; afdb-uniprot50 \| AF-A0A2D0JYZ8-F1-MODEL\_V4 \| 1.0 \| 1.688e-21 \| 568 \| 0.288 \| 367 \| 141 \| 8 \| 1 \| 344 \| 1 \| 270 \| Phage tail fiber protein \| Phage tail fiber protein; afdb-uniprot50 \| AF-A0A5U3Y0T7-F1-MODEL\_V4 \| 1.0 \| 2.936e-20 \| 568 \| 0.251 \| 346 \| 136 \| 5 \| 1 \| 324 \| 4 \| 248 \| Phage tail protein \| Phage tail protein; afdb-uniprot50 \| AF-A0A7U0C064-F1-MODEL\_V4 \| 1.0 \| 3.642e-20 \| 568 \| 0.305 \| 344 \| 149 \| 9 \| 1 \| 324 \| 2 \| 275 \| Tail fiber protein \| Tail fiber protein; afdb-uniprot50 \| AF-A0A2D0IW09-F1-MODEL\_V4 \| 1.0 \| 5.032e-20 \| 568 \| 0.31 \| 329 \| 120 \| 5 \| 1 \| 324 \| 2 \| 228 \| Tail protein \| Tail protein; afdb-uniprot50 \| AF-A0A3D9ULH6-F1-MODEL\_V4 \| 1.0 \| 5.685e-19 \| 568 \| 0.309 \| 323 \| 121 \| 5 \| 1 \| 322 \| 2 \| 223 \| Tail fiber-like repeat protein \| Tail fiber-like repeat protein; afdb-uniprot50 \| AF-A0A7X7NDN0-F1-MODEL\_V4 \| 1.0 \| 5.871e-13 \| 567 \| 0.375 \| 128 \| 79 \| 1 \| 4 \| 131 \| 3 \| 129 \| Phage tail protein \| Phage tail protein; afdb-uniprot50 \| AF-A0A702QMM0-F1-MODEL\_V4 \| 1.0 \| 1.366e-17 \| 566 \| 0.23 \| 312 \| 134 \| 5 \| 1 \| 307 \| 1 \| 211 \| Phage tail protein \| Phage tail protein; afdb-uniprot50 \| AF-A0A2A7UY14-F1-MODEL\_V4 \| 1.0 \| 2.275e-19 \| 566 \| 0.216 \| 434 \| 165 \| 12 \| 1 \| 405 \| 1 \| 288 \| Uncharacterized protein \| Uncharacterized protein; afdb-uniprot50 \| AF-A0A391N963-F1-MODEL\_V4 \| 1.0 \| 1.125e-23 \| 566 \| 0.232 \| 465 \| 206 \| 21 \| 1 \| 423 \| 1 \| 356 \| Uncharacterized protein \| Uncharacterized protein; afdb-uniprot50 \| AF-A0A348HHI3-F1-MODEL\_V4 \| 1.0 \| 8.875e-18 \| 566 \| 0.301 \| 249 \| 129 \| 7 \| 1 \| 219 \| 1 \| 234 \| Phage-related tail fibre protein \| Phage-related tail fibre protein; afdb-uniprot50 \| AF-A6UZC4-F1-MODEL\_V4 \| 1.0 \| 2.392e-23 \| 566 \| 0.23 \| 473 \| 239 \| 19 \| 1 \| 423 \| 2 \| 399 \| GpH \| GpH; afdb-uniprot50 \| AF-L3C905-F1-MODEL\_V4 \| 1.0 \| 5.604e-20 \| 566 \| 0.264 \| 318 \| 155 \| 11 \| 1 \| 306 \| 1 \| 251 \| Uncharacterized protein \| Uncharacterized protein; afdb-uniprot50 \| AF-A0A2D0KDJ9-F1-MODEL\_V4 \| 1.0 \| 4.056e-20 \| 566 \| 0.299 \| 334 \| 125 \| 7 \| 1 \| 326 \| 2 \| 234 \| Tail protein \| Tail protein; afdb-uniprot50 \| AF-A0A0F7J9Z5-F1-MODEL\_V4 \| 1.0 \| 1.623e-20 \| 566 \| 0.338 \| 334 \| 140 \| 10 \| 1 \| 323 \| 2 \| 265 \| Variable tail fiber protein \| Variable tail fiber protein; afdb-uniprot50 \| AF-A0A829RXG4-F1-MODEL\_V4 \| 1.0 \| 6.61e-16 \| 565 \| 0.325 \| 212 \| 130 \| 4 \| 1 \| 212 \| 3 \| 201 \| Phage tail-collar fiber family protein \| Phage tail-collar fiber family protein; afdb-uniprot50 \| AF-A0A1V9IMJ2-F1-MODEL\_V4 \| 1.0 \| 1.258e-19 \| 564 \| 0.272 \| 305 \| 149 \| 12 \| 1 \| 258 \| 2 \| 280 \| Phage tail fiber protein \| Phage tail fiber protein; afdb-uniprot50 \| AF-A0A484ZG34-F1-MODEL\_V4 \| 1.0 \| 1.381e-20 \| 564 \| 0.253 \| 430 \| 188 \| 12 \| 1 \| 411 \| 2 \| 317 \| Phage tail fibre repeat \| Phage tail fibre repeat; afdb-uniprot50 \| AF-A0A078LT99-F1-MODEL\_V4 \| 1.0 \| 4.89e-22 \| 564 \| 0.213 \| 450 \| 206 \| 17 \| 4 \| 423 \| 8 \| 339 \| Tail fiber protein H \| Tail fiber protein H; afdb-uniprot50 \| AF-A0A2I8Q7M1-F1-MODEL\_V4 \| 1.0 \| 1.494e-22 \| 564 \| 0.241 \| 476 \| 216 \| 20 \| 1 \| 423 \| 2 \| 385 \| Phage tail protein \| Phage tail protein; afdb-uniprot50 \| AF-A0A7W4RI03-F1-MODEL\_V4 \| 1.0 \| 8.472e-25 \| 564 \| 0.245 \| 537 \| 259 \| 27 \| 1 \| 423 \| 1 \| 505 \| Uncharacterized protein \| Uncharacterized protein; afdb-uniprot50 \| AF-A0A840PS60-F1-MODEL\_V4 \| 1.0 \| 1.935e-19 \| 563 \| 0.271 \| 280 \| 160 \| 9 \| 1 \| 248 \| 2 \| 269 \| Phage-related tail fiber protein \| Phage-related tail fiber protein; afdb-uniprot50 \| AF-A0A5V1PKK2-F1-MODEL\_V4 \| 1.0 \| 6e-19 \| 563 \| 0.282 \| 308 \| 119 \| 4 \| 1 \| 307 \| 4 \| 210 \| Uncharacterized protein \| Uncharacterized protein; afdb-uniprot50 \| AF-A0A2X3E9B4-F1-MODEL\_V4 \| 1.0 \| 2.187e-18 \| 563 \| 0.333 \| 249 \| 126 \| 5 \| 1 \| 223 \| 1 \| 235 \| Uncharacterized protein \| Uncharacterized protein; afdb-uniprot50 \| AF-A0A2R8CKT4-F1-MODEL\_V4 \| 1.0 \| 1.109e-24 \| 563 \| 0.2 \| 545 \| 270 \| 22 \| 1 \| 423 \| 1 \| 501 \| Uncharacterized protein \| Uncharacterized protein; afdb-uniprot50 \| AF-A0A3D8XED5-F1-MODEL\_V4 \| 1.0 \| 4.281e-20 \| 562 \| 0.277 \| 342 \| 160 \| 9 \| 1 \| 323 \| 5 \| 278 \| Phage tail protein \| Phage tail protein; afdb-uniprot50 \| AF-A0A827CPD5-F1-MODEL\_V4 \| 1.0 \| 8.172e-20 \| 561 \| 0.277 \| 335 \| 160 \| 10 \| 1 \| 323 \| 5 \| 269 \| Phage tail protein \| Phage tail protein; afdb-uniprot50 \| AF-A0A502FV80-F1-MODEL\_V4 \| 1.0 \| 1.763e-18 \| 559 \| 0.338 \| 278 \| 109 \| 7 \| 1 \| 221 \| 2 \| 261 \| Phage tail protein \| Phage tail protein; afdb-uniprot50 \| AF-A0A7W9XSU5-F1-MODEL\_V4 \| 1.0 \| 7.208e-25 \| 559 \| 0.232 \| 537 \| 261 \| 21 \| 1 \| 423 \| 3 \| 502 \| Uncharacterized protein \| Uncharacterized protein; afdb-uniprot50 \| AF-A0A4R0E5Y1-F1-MODEL\_V4 \| 1.0 \| 9.399e-14 \| 558 \| 0.324 \| 157 \| 100 \| 3 \| 1 \| 155 \| 2 \| 154 \| Uncharacterized protein \| Uncharacterized protein; afdb-uniprot50 \| AF-U1ZYL0-F1-MODEL\_V4 \| 1.0 \| 1.158e-21 \| 558 \| 0.229 \| 448 \| 201 \| 17 \| 1 \| 423 \| 1 \| 329 \| Uncharacterized protein \| Uncharacterized protein; afdb-uniprot50 \| AF-W7PY95-F1-MODEL\_V4 \| 1.0 \| 3.054e-21 \| 558 \| 0.271 \| 364 \| 149 \| 16 \| 1 \| 259 \| 1 \| 353 \| Phage tail protein \| Phage tail protein; afdb-uniprot50 \| AF-W1J986-F1-MODEL\_V4 \| 1.0 \| 1.401e-19 \| 558 \| 0.306 \| 330 \| 122 \| 6 \| 1 \| 324 \| 2 \| 230 \| Collar domain-containing protein \| Collar domain-containing protein; afdb-uniprot50 \| AF-A0A7C8HVY4-F1-MODEL\_V4 \| 1.0 \| 1.346e-18 \| 557 \| 0.223 \| 371 \| 147 \| 6 \| 1 \| 342 \| 1 \| 259 \| Uncharacterized protein \| Uncharacterized protein; afdb-uniprot50 \| AF-A0A2I5TL70-F1-MODEL\_V4 \| 1.0 \| 7.969e-18 \| 556 \| 0.342 \| 248 \| 126 \| 4 \| 1 \| 221 \| 1 \| 238 \| Phage tail protein \| Phage tail protein; afdb-uniprot50 \| AF-A0A3G7TJE0-F1-MODEL\_V4 \| 1.0 \| 1.356e-25 \| 556 \| 0.23 \| 508 \| 281 \| 25 \| 2 \| 423 \| 6 \| 489 \| Phage tail fiber protein \| Phage tail fiber protein; afdb-uniprot50 \| AF-A0A8B4SMY3-F1-MODEL\_V4 \| 1.0 \| 3.776e-25 \| 556 \| 0.233 \| 562 \| 223 \| 18 \| 1 \| 423 \| 2 \| 494 \| Tail fiber protein \| Tail fiber protein; afdb-uniprot50 \| AF-A0A828ZHJ0-F1-MODEL\_V4 \| 1.0 \| 9.102e-20 \| 556 \| 0.243 \| 320 \| 153 \| 9 \| 1 \| 257 \| 1 \| 294 \| Tail fiber protein (GpH) \| Tail fiber protein (GpH); afdb-uniprot50 \| AF-A0A4Q7BEF2-F1-MODEL\_V4 \| 1.0 \| 1.867e-14 \| 555 \| 0.327 \| 171 \| 107 \| 4 \| 1 \| 166 \| 1 \| 168 \| Uncharacterized protein \| Uncharacterized protein; afdb-uniprot50 \| AF-C1DS14-F1-MODEL\_V4 \| 1.0 \| 1.032e-14 \| 555 \| 0.314 \| 197 \| 116 \| 5 \| 1 \| 178 \| 1 \| 197 \| Phage P2 tail fiber gpH-like protein \| Phage P2 tail fiber gpH-like protein; afdb-uniprot50 \| AF-A0A7T0RCA3-F1-MODEL\_V4 \| 1.0 \| 1.226e-17 \| 555 \| 0.322 \| 242 \| 132 \| 7 \| 2 \| 223 \| 6 \| 235 \| Phage tail protein \| Phage tail protein; afdb-uniprot50 \| AF-A0A7R7ZA19-F1-MODEL\_V4 \| 1.0 \| 3.776e-25 \| 555 \| 0.251 \| 489 \| 238 \| 22 \| 2 \| 423 \| 3 \| 430 \| Uncharacterized protein \| Uncharacterized protein; afdb-uniprot50 \| AF-A0A5U4CIE5-F1-MODEL\_V4 \| 1.0 \| 5.31e-20 \| 555 \| 0.251 \| 346 \| 136 \| 5 \| 1 \| 324 \| 4 \| 248 \| Phage tail protein \| Phage tail protein; afdb-uniprot50 \| AF-A5I4A1-F1-MODEL\_V4 \| 1.0 \| 5.851e-17 \| 555 \| 0.323 \| 229 \| 133 \| 4 \| 1 \| 216 \| 2 \| 221 \| Putative tail fiber protein \| Putative tail fiber protein; afdb-uniprot50 \| AF-A0A0A8JLH1-F1-MODEL\_V4 \| 1.0 \| 1.738e-19 \| 554 \| 0.253 \| 300 \| 161 \| 11 \| 1 \| 260 \| 1 \| 277 \| Tail fiber protein \| Tail fiber protein; afdb-uniprot50 \| AF-M8EFI1-F1-MODEL\_V4 \| 1.0 \| 3.629e-24 \| 554 \| 0.245 \| 460 \| 232 \| 21 \| 1 \| 407 \| 2 \| 399 \| Phage-related tail fiber protein-like protein \| Phage-related tail fiber protein-like protein; afdb-uniprot50 \| AF-A0A348D9H5-F1-MODEL\_V4 \| 1.0 \| 9.367e-18 \| 553 \| 0.27 \| 310 \| 123 \| 6 \| 1 \| 308 \| 1 \| 209 \| Uncharacterized protein \| Uncharacterized protein; afdb-uniprot50 \| AF-V4NRK3-F1-MODEL\_V4 \| 1.0 \| 1.634e-12 \| 552 \| 0.487 \| 119 \| 61 \| 0 \| 1 \| 119 \| 1 \| 119 \| Uncharacterized protein \| Uncharacterized protein; afdb-uniprot50 \| AF-A0A827S3F2-F1-MODEL\_V4 \| 1.0 \| 2.155e-19 \| 552 \| 0.248 \| 410 \| 139 \| 10 \| 1 \| 345 \| 2 \| 307 \| Phage tail protein \| Phage tail protein; afdb-uniprot50 \| AF-A0A810U842-F1-MODEL\_V4 \| 1.0 \| 1.431e-25 \| 552 \| 0.225 \| 598 \| 265 \| 27 \| 1 \| 423 \| 2 \| 576 \| Uncharacterized protein \| Uncharacterized protein; afdb-uniprot50 \| AF-A0A4S3LTE6-F1-MODEL\_V4 \| 1.0 \| 1.763e-18 \| 551 \| 0.28 \| 307 \| 120 \| 3 \| 1 \| 307 \| 2 \| 207 \| Uncharacterized protein \| Uncharacterized protein; afdb-uniprot50 \| AF-A0A7B5Z917-F1-MODEL\_V4 \| 1.0 \| 4.837e-19 \| 551 \| 0.274 \| 339 \| 133 \| 7 \| 1 \| 327 \| 1 \| 238 \| Phage tail protein \| Phage tail protein; afdb-uniprot50 \| AF-A0A7X1Y572-F1-MODEL\_V4 \| 1.0 \| 2.863e-18 \| 550 \| 0.276 \| 318 \| 149 \| 10 \| 2 \| 306 \| 6 \| 255 \| Uncharacterized protein \| Uncharacterized protein; afdb-uniprot50 \| AF-A0A077NGB1-F1-MODEL\_V4 \| 1.0 \| 1.085e-18 \| 550 \| 0.309 \| 330 \| 121 \| 6 \| 1 \| 324 \| 2 \| 230 \| Uncharacterized protein \| Uncharacterized protein; afdb-uniprot50 \| AF-A0A376TTQ8-F1-MODEL\_V4 \| 1.0 \| 3.075e-13 \| 549 \| 0.521 \| 119 \| 57 \| 0 \| 1 \| 119 \| 2 \| 120 \| Variable tail fiber protein \| Variable tail fiber protein; afdb-uniprot50 \| AF-A0A5C8XYZ1-F1-MODEL\_V4 \| 1.0 \| 2.179e-22 \| 549 \| 0.255 \| 474 \| 206 \| 22 \| 1 \| 423 \| 1 \| 378 \| Phage tail protein \| Phage tail protein; afdb-uniprot50 \| AF-A0A8A7FKW4-F1-MODEL\_V4 \| 1.0 \| 3.132e-23 \| 549 \| 0.203 \| 581 \| 197 \| 12 \| 1 \| 423 \| 1 \| 473 \| Phage tail protein \| Phage tail protein; afdb-uniprot50 \| AF-A0A4R6E2J7-F1-MODEL\_V4 \| 1.0 \| 1.617e-24 \| 549 \| 0.24 \| 557 \| 240 \| 21 \| 1 \| 423 \| 1 \| 508 \| Tail fiber-like repeat protein \| Tail fiber-like repeat protein; afdb-uniprot50 \| AF-A0A427Q396-F1-MODEL\_V4 \| 1.0 \| 2.489e-24 \| 549 \| 0.188 \| 573 \| 271 \| 25 \| 2 \| 423 \| 6 \| 535 \| Uncharacterized protein \| Uncharacterized protein; afdb-uniprot50 \| AF-A0A854BC76-F1-MODEL\_V4 \| 1.0 \| 2.203e-25 \| 548 \| 0.229 \| 496 \| 284 \| 22 \| 4 \| 423 \| 8 \| 481 \| Phage tail protein \| Phage tail protein; afdb-uniprot50 \| AF-A0A747EHS2-F1-MODEL\_V4 \| 1.0 \| 8.719e-23 \| 548 \| 0.177 \| 751 \| 175 \| 15 \| 1 \| 423 \| 1 \| 636 \| Phage tail protein \| Phage tail protein; afdb-uniprot50 \| AF-A0A6C8RYR2-F1-MODEL\_V4 \| 1.0 \| 2.57e-18 \| 547 \| 0.274 \| 393 \| 127 \| 10 \| 1 \| 329 \| 2 \| 300 \| Phage tail protein \| Phage tail protein; afdb-uniprot50 \| AF-A0A2D0LF95-F1-MODEL\_V4 \| 1.0 \| 1.956e-22 \| 547 \| 0.197 \| 604 \| 196 \| 13 \| 1 \| 423 \| 2 \| 497 \| Tail protein \| Tail protein; afdb-uniprot50 \| AF-A0A1V2GLB1-F1-MODEL\_V4 \| 1.0 \| 1.86e-18 \| 547 \| 0.276 \| 325 \| 130 \| 6 \| 1 \| 323 \| 1 \| 222 \| Phage tail protein \| Phage tail protein; afdb-uniprot50 \| AF-A0A2J9EBQ6-F1-MODEL\_V4 \| 1.0 \| 5.292e-24 \| 546 \| 0.243 \| 509 \| 222 \| 15 \| 1 \| 421 \| 2 \| 435 \| Phage tail protein \| Phage tail protein; afdb-uniprot50 \| AF-A0A752D550-F1-MODEL\_V4 \| 1.0 \| 2.148e-23 \| 546 \| 0.225 \| 595 \| 240 \| 24 \| 1 \| 423 \| 1 \| 546 \| Phage tail protein \| Phage tail protein; afdb-uniprot50 \| AF-A0A2X3JIY7-F1-MODEL\_V4 \| 1.0 \| 1.921e-12 \| 545 \| 0.376 \| 130 \| 76 \| 2 \| 3 \| 131 \| 7 \| 132 \| Phage protein gpH \| Phage protein gpH; afdb-uniprot50 \| AF-A0A377AV74-F1-MODEL\_V4 \| 1.0 \| 4.502e-24 \| 545 \| 0.251 \| 525 \| 192 \| 22 \| 1 \| 423 \| 1 \| 426 \| Phage tail collar domain-containing protein \| Phage tail collar domain-containing protein; afdb-uniprot50 \| AF-A0A659PJA5-F1-MODEL\_V4 \| 1.0 \| 4.82e-23 \| 545 \| 0.215 \| 543 \| 215 \| 22 \| 1 \| 423 \| 1 \| 452 \| Phage tail protein \| Phage tail protein; afdb-uniprot50 \| AF-A0A6A4RC06-F1-MODEL\_V4 \| 1.0 \| 2.627e-24 \| 545 \| 0.198 \| 573 \| 269 \| 27 \| 1 \| 423 \| 1 \| 532 \| Uncharacterized protein \| Uncharacterized protein; afdb-uniprot50 \| AF-A0A2V1Y908-F1-MODEL\_V4 \| 1.0 \| 6.31e-23 \| 543 \| 0.22 \| 477 \| 229 \| 17 \| 4 \| 423 \| 8 \| 398 \| Phage tail-collar fibre protein \| Phage tail-collar fibre protein; afdb-uniprot50 \| AF-A0A7W5DMQ9-F1-MODEL\_V4 \| 1.0 \| 1.956e-22 \| 543 \| 0.235 \| 502 \| 211 \| 25 \| 1 \| 423 \| 1 \| 408 \| Uncharacterized protein \| Uncharacterized protein; afdb-uniprot50 \| AF-A0A4U8YH51-F1-MODEL\_V4 \| 1.0 \| 2.926e-24 \| 542 \| 0.207 \| 468 \| 246 \| 17 \| 1 \| 423 \| 1 \| 388 \| Phage tail fibre protein \| Phage tail fibre protein; afdb-uniprot50 \| AF-A0A6N3HNW3-F1-MODEL\_V4 \| 1.0 \| 3.054e-21 \| 542 \| 0.173 \| 761 \| 213 \| 22 \| 1 \| 423 \| 1 \| 683 \| Phage tail fibre repeat protein \| Phage tail fibre repeat protein; afdb-uniprot50 \| AF-A0A3T2UQY7-F1-MODEL\_V4 \| 1.0 \| 1.623e-20 \| 542 \| 0.28 \| 364 \| 179 \| 16 \| 1 \| 322 \| 1 \| 323 \| Phage tail protein \| Phage tail protein; afdb-uniprot50 \| AF-A0A328TYP4-F1-MODEL\_V4 \| 1.0 \| 1.664e-22 \| 541 \| 0.235 \| 472 \| 220 \| 17 \| 1 \| 421 \| 2 \| 383 \| Phage tail-collar fiber family protein \| Phage tail-collar fiber family protein; afdb-uniprot50 \| AF-A0A4R0FQS3-F1-MODEL\_V4 \| 1.0 \| 4.042e-24 \| 541 \| 0.22 \| 485 \| 259 \| 25 \| 1 \| 423 \| 2 \| 429 \| Uncharacterized protein \| Uncharacterized protein; afdb-uniprot50 \| AF-A0A1I4ZMZ1-F1-MODEL\_V4 \| 1.0 \| 4.684e-25 \| 541 \| 0.231 \| 510 \| 246 \| 22 \| 4 \| 423 \| 8 \| 461 \| Phage tail-collar fibre protein \| Phage tail-collar fibre protein; afdb-uniprot50 \| AF-A0A0H3MMX0-F1-MODEL\_V4 \| 1.0 \| 4.96e-21 \| 541 \| 0.288 \| 357 \| 200 \| 12 \| 1 \| 324 \| 1 \| 336 \| Putative side tail phage protein \| Putative side tail phage protein; afdb-uniprot50 \| AF-A0A8B3UK72-F1-MODEL\_V4 \| 1.0 \| 2.155e-19 \| 541 \| 0.286 \| 335 \| 162 \| 8 \| 1 \| 323 \| 2 \| 271 \| Tail Fiber protein \| Tail Fiber protein; afdb-uniprot50 \| AF-B1LMZ0-F1-MODEL\_V4 \| 1.0 \| 2.435e-18 \| 541 \| 0.276 \| 325 \| 130 \| 6 \| 1 \| 323 \| 1 \| 222 \| Putative phage tail fiber protein \| Putative phage tail fiber protein; afdb-uniprot50 \| AF-A0A7Z8BNV6-F1-MODEL\_V4 \| 1.0 \| 3.551e-18 \| 541 \| 0.274 \| 324 \| 131 \| 5 \| 1 \| 323 \| 2 \| 222 \| Phage tail protein \| Phage tail protein; afdb-uniprot50 \| AF-A0A5K1MSE4-F1-MODEL\_V4 \| 1.0 \| 3.615e-13 \| 541 \| 0.352 \| 156 \| 98 \| 1 \| 1 \| 156 \| 2 \| 154 \| Uncharacterized protein \| Uncharacterized protein; afdb-uniprot50 \| AF-A0A5P2SMC4-F1-MODEL\_V4 \| 1.0 \| 4.751e-24 \| 540 \| 0.278 \| 478 \| 251 \| 24 \| 1 \| 423 \| 1 \| 439 \| Phage tail protein \| Phage tail protein; afdb-uniprot50 \| AF-A0A742CYR3-F1-MODEL\_V4 \| 1.0 \| 6.565e-24 \| 540 \| 0.239 \| 509 \| 227 \| 24 \| 1 \| 419 \| 1 \| 439 \| Phage tail protein \| Phage tail protein; afdb-uniprot50 \| AF-A0A5D4YFX8-F1-MODEL\_V4 \| 1.0 \| 5.368e-23 \| 540 \| 0.2 \| 629 \| 240 \| 22 \| 1 \| 423 \| 1 \| 572 \| Phage tail protein \| Phage tail protein; afdb-uniprot50 \| AF-A0A0L6FWL7-F1-MODEL\_V4 \| 1.0 \| 2.627e-24 \| 540 \| 0.181 \| 715 \| 280 \| 27 \| 1 \| 423 \| 5 \| 705 \| Collar domain-containing protein \| Collar domain-containing protein; afdb-uniprot50 \| AF-A0A398UJ84-F1-MODEL\_V4 \| 1.0 \| 2.713e-18 \| 539 \| 0.262 \| 324 \| 135 \| 5 \| 1 \| 323 \| 5 \| 225 \| Phage tail protein \| Phage tail protein; afdb-uniprot50 \| AF-A0A5W0I1C1-F1-MODEL\_V4 \| 1.0 \| 1.881e-21 \| 539 \| 0.281 \| 387 \| 183 \| 12 \| 1 \| 333 \| 1 \| 346 \| Phage tail protein \| Phage tail protein; afdb-uniprot50 \| AF-A0A6F9WXH3-F1-MODEL\_V4 \| 1.0 \| 2.479e-13 \| 539 \| 0.326 \| 156 \| 105 \| 0 \| 1 \| 156 \| 2 \| 157 \| Uncharacterized protein \| Uncharacterized protein; afdb-uniprot50 \| AF-A0A399Q0B5-F1-MODEL\_V4 \| 1.0 \| 3.489e-23 \| 539 \| 0.178 \| 548 \| 278 \| 29 \| 4 \| 423 \| 8 \| 511 \| Collar domain-containing protein \| Collar domain-containing protein; afdb-uniprot50 \| AF-A0A2S4QLD3-F1-MODEL\_V4 \| 1.0 \| 1.827e-23 \| 539 \| 0.233 \| 607 \| 258 \| 32 \| 1 \| 423 \| 1 \| 583 \| Uncharacterized protein \| Uncharacterized protein; afdb-uniprot50 \| AF-A0A080K6T3-F1-MODEL\_V4 \| 1.0 \| 2.654e-12 \| 538 \| 0.491 \| 116 \| 59 \| 0 \| 1 \| 116 \| 2 \| 117 \| Tail fiber protein \| Tail fiber protein; afdb-uniprot50 \| AF-A0A7R7HR43-F1-MODEL\_V4 \| 1.0 \| 2.148e-23 \| 538 \| 0.216 \| 577 \| 273 \| 24 \| 2 \| 423 \| 6 \| 558 \| Uncharacterized protein \| Uncharacterized protein; afdb-uniprot50 \| AF-D5EF94-F1-MODEL\_V4 \| 1.0 \| 6.242e-20 \| 538 \| 0.297 \| 306 \| 166 \| 10 \| 1 \| 258 \| 2 \| 306 \| Phage-related tail fibre protein-like protein \| Phage-related tail fibre protein-like protein; afdb-uniprot50 \| AF-A0A1E7WJE2-F1-MODEL\_V4 \| 1.0 \| 7.882e-15 \| 538 \| 0.301 \| 212 \| 126 \| 2 \| 1 \| 190 \| 4 \| 215 \| SGNH\_hydro domain-containing protein \| SGNH\_hydro domain-containing protein; afdb-uniprot50 \| AF-A0A376VKB4-F1-MODEL\_V4 \| 1.0 \| 5.032e-20 \| 537 \| 0.286 \| 359 \| 176 \| 13 \| 1 \| 324 \| 1 \| 314 \| Putative tail fiber protein (GpH) \| Putative tail fiber protein (GpH); afdb-uniprot50 \| AF-A0A837JCH1-F1-MODEL\_V4 \| 1.0 \| 5.329e-16 \| 537 \| 0.267 \| 232 \| 151 \| 5 \| 1 \| 223 \| 1 \| 222 \| Uncharacterized protein \| Uncharacterized protein; afdb-uniprot50 \| AF-A0A701UXD7-F1-MODEL\_V4 \| 1.0 \| 2.561e-22 \| 537 \| 0.254 \| 463 \| 223 \| 21 \| 1 \| 423 \| 1 \| 381 \| Phage tail protein \| Phage tail protein; afdb-uniprot50 \| AF-A0A6M0YIJ7-F1-MODEL\_V4 \| 1.0 \| 1.6e-21 \| 537 \| 0.209 \| 491 \| 211 \| 16 \| 2 \| 422 \| 3 \| 386 \| Phage tail protein \| Phage tail protein; afdb-uniprot50 \| AF-A0A2V4LES5-F1-MODEL\_V4 \| 1.0 \| 1.732e-23 \| 537 \| 0.226 \| 494 \| 259 \| 28 \| 1 \| 423 \| 3 \| 444 \| Uncharacterized protein \| Uncharacterized protein; afdb-uniprot50 \| AF-A0A8A2FEP4-F1-MODEL\_V4 \| 1.0 \| 3.59e-21 \| 537 \| 0.244 \| 413 \| 190 \| 17 \| 1 \| 361 \| 1 \| 343 \| Phage tail protein \| Phage tail protein; afdb-uniprot50 \| AF-A0A827VE52-F1-MODEL\_V4 \| 1.0 \| 2.782e-20 \| 537 \| 0.27 \| 366 \| 183 \| 13 \| 1 \| 333 \| 5 \| 319 \| Phage tail protein \| Phage tail protein; afdb-uniprot50 \| AF-A0A806Z174-F1-MODEL\_V4 \| 1.0 \| 3.629e-24 \| 537 \| 0.195 \| 695 \| 241 \| 23 \| 1 \| 423 \| 1 \| 649 \| Uncharacterized protein \| Uncharacterized protein; afdb-uniprot50 \| AF-R9K2R2-F1-MODEL\_V4 \| 1.0 \| 2.283e-15 \| 536 \| 0.242 \| 219 \| 129 \| 5 \| 1 \| 186 \| 1 \| 215 \| Uncharacterized protein \| Uncharacterized protein; afdb-uniprot50 \| AF-A0A5Y3PXB4-F1-MODEL\_V4 \| 1.0 \| 2.978e-19 \| 536 \| 0.25 \| 340 \| 134 \| 7 \| 1 \| 334 \| 4 \| 228 \| Phage tail protein \| Phage tail protein; afdb-uniprot50 \| AF-A0A7B6C8V7-F1-MODEL\_V4 \| 1.0 \| 4.056e-20 \| 536 \| 0.271 \| 364 \| 187 \| 10 \| 1 \| 332 \| 5 \| 322 \| Phage tail protein \| Phage tail protein; afdb-uniprot50 \| AF-A0A6C8Y9W3-F1-MODEL\_V4 \| 1.0 \| 1.025e-22 \| 535 \| 0.245 \| 469 \| 211 \| 18 \| 1 \| 423 \| 1 \| 372 \| Uncharacterized protein \| Uncharacterized protein; afdb-uniprot50 \| AF-A0A1E3XL37-F1-MODEL\_V4 \| 1.0 \| 5.604e-20 \| 535 \| 0.253 \| 332 \| 146 \| 7 \| 1 \| 322 \| 1 \| 240 \| Uncharacterized protein \| Uncharacterized protein; afdb-uniprot50 \| AF-A0A349XXC0-F1-MODEL\_V4 \| 1.0 \| 1.565e-15 \| 534 \| 0.292 \| 219 \| 116 \| 4 \| 1 \| 183 \| 1 \| 216 \| Phage tail protein \| Phage tail protein; afdb-uniprot50 \| AF-G9Z0C3-F1-MODEL\_V4 \| 1.0 \| 3.212e-25 \| 534 \| 0.255 \| 563 \| 257 \| 24 \| 1 \| 423 \| 2 \| 542 \| Uncharacterized protein \| Uncharacterized protein; afdb-uniprot50 \| AF-A0A375ABE7-F1-MODEL\_V4 \| 1.0 \| 3.317e-19 \| 533 \| 0.281 \| 316 \| 145 \| 9 \| 1 \| 251 \| 2 \| 300 \| Phage tail fiber protein \| Phage tail fiber protein; afdb-uniprot50 \| AF-A0A444RCG1-F1-MODEL\_V4 \| 1.0 \| 3.748e-18 \| 532 \| 0.266 \| 394 \| 131 \| 9 \| 1 \| 330 \| 2 \| 301 \| Phage tail protein \| Phage tail protein; afdb-uniprot50 \| AF-A0A3S4G527-F1-MODEL\_V4 \| 1.0 \| 2.636e-20 \| 532 \| 0.292 \| 359 \| 172 \| 14 \| 1 \| 324 \| 1 \| 312 \| Phage variable tail-fiber protein \| Phage variable tail-fiber protein; afdb-uniprot50 \| AF-A0A5U3EXM5-F1-MODEL\_V4 \| 1.0 \| 9.573e-24 \| 531 \| 0.232 \| 494 \| 231 \| 23 \| 1 \| 423 \| 2 \| 418 \| Uncharacterized protein \| Uncharacterized protein; afdb-uniprot50 \| AF-A0A2C6DJ56-F1-MODEL\_V4 \| 1.0 \| 1.538e-20 \| 531 \| 0.266 \| 394 \| 155 \| 12 \| 1 \| 377 \| 2 \| 278 \| Uncharacterized protein \| Uncharacterized protein; afdb-uniprot50 \| AF-A0A370S8A4-F1-MODEL\_V4 \| 1.0 \| 1.361e-21 \| 531 \| 0.245 \| 464 \| 213 \| 18 \| 1 \| 423 \| 1 \| 368 \| Tail fiber-like repeat protein \| Tail fiber-like repeat protein; afdb-uniprot50 \| AF-W2CTP2-F1-MODEL\_V4 \| 1.0 \| 6.539e-13 \| 530 \| 0.52 \| 125 \| 60 \| 0 \| 27 \| 151 \| 2 \| 126 \| Phage tail protein \| Phage tail protein; afdb-uniprot50 \| AF-A0A064DL20-F1-MODEL\_V4 \| 1.0 \| 6.565e-24 \| 530 \| 0.269 \| 501 \| 218 \| 22 \| 1 \| 423 \| 2 \| 432 \| Uncharacterized protein \| Uncharacterized protein; afdb-uniprot50 \| AF-A0A5S4U4U6-F1-MODEL\_V4 \| 1.0 \| 2.926e-24 \| 529 \| 0.258 \| 495 \| 218 \| 24 \| 1 \| 423 \| 2 \| 419 \| Phage tail protein \| Phage tail protein; afdb-uniprot50 \| AF-A0A1V2G5C9-F1-MODEL\_V4 \| 1.0 \| 5.292e-24 \| 529 \| 0.244 \| 528 \| 218 \| 24 \| 1 \| 423 \| 1 \| 452 \| Phage tail protein \| Phage tail protein; afdb-uniprot50 \| AF-A0A7Y9B7N8-F1-MODEL\_V4 \| 1.0 \| 1.376e-24 \| 529 \| 0.245 \| 600 \| 239 \| 25 \| 1 \| 423 \| 2 \| 564 \| Phage tail protein \| Phage tail protein; afdb-uniprot50 \| AF-A0A626YJZ8-F1-MODEL\_V4 \| 1.0 \| 2.401e-19 \| 529 \| 0.245 \| 346 \| 138 \| 7 \| 1 \| 324 \| 4 \| 248 \| Phage tail protein \| Phage tail protein; afdb-uniprot50 \| AF-A0A7U1Z2W3-F1-MODEL\_V4 \| 1.0 \| 2.187e-18 \| 529 \| 0.265 \| 339 \| 131 \| 6 \| 1 \| 326 \| 1 \| 234 \| Phage tail protein \| Phage tail protein; afdb-uniprot50 \| AF-I3DCB6-F1-MODEL\_V4 \| 1.0 \| 1.623e-20 \| 528 \| 0.245 \| 359 \| 172 \| 12 \| 1 \| 322 \| 2 \| 298 \| Phage tail fiber repeat protein \| Phage tail fiber repeat protein; afdb-uniprot50 \| AF-A0A0G9KCC4-F1-MODEL\_V4 \| 1.0 \| 8.531e-17 \| 528 \| 0.3 \| 236 \| 140 \| 5 \| 1 \| 223 \| 1 \| 224 \| Uncharacterized protein \| Uncharacterized protein; afdb-uniprot50 \| AF-A0A5Y2KP10-F1-MODEL\_V4 \| 1.0 \| 9.711e-23 \| 528 \| 0.201 \| 581 \| 233 \| 21 \| 1 \| 423 \| 1 \| 508 \| Phage tail protein \| Phage tail protein; afdb-uniprot50 \| AF-A0A1Q9QWT4-F1-MODEL\_V4 \| 1.0 \| 1.641e-23 \| 528 \| 0.184 \| 574 \| 278 \| 22 \| 4 \| 423 \| 8 \| 545 \| Uncharacterized protein \| Uncharacterized protein; afdb-uniprot50 \| AF-A0A4D6XN95-F1-MODEL\_V4 \| 1.0 \| 4.82e-23 \| 527 \| 0.216 \| 495 \| 245 \| 23 \| 1 \| 423 \| 5 \| 428 \| Uncharacterized protein \| Uncharacterized protein; afdb-uniprot50 \| AF-A0A4Y9GYI2-F1-MODEL\_V4 \| 1.0 \| 5.104e-19 \| 527 \| 0.232 \| 331 \| 153 \| 7 \| 1 \| 322 \| 2 \| 240 \| Uncharacterized protein \| Uncharacterized protein; afdb-uniprot50 \| AF-A0A2G1CTX9-F1-MODEL\_V4 \| 1.0 \| 9.573e-24 \| 526 \| 0.237 \| 479 \| 229 \| 20 \| 4 \| 423 \| 3 \| 404 \| Uncharacterized protein \| Uncharacterized protein; afdb-uniprot50 \| AF-A0A6G4BMA1-F1-MODEL\_V4 \| 1.0 \| 4.82e-23 \| 526 \| 0.232 \| 517 \| 229 \| 22 \| 1 \| 423 \| 2 \| 444 \| Phage tail protein \| Phage tail protein; afdb-uniprot50 \| AF-A0A062L6U3-F1-MODEL\_V4 \| 1.0 \| 5.271e-13 \| 526 \| 0.354 \| 161 \| 103 \| 1 \| 2 \| 162 \| 3 \| 162 \| Phage tail-collar fiber family protein \| Phage tail-collar fiber family protein; afdb-uniprot50 \| AF-D1Y2C2-F1-MODEL\_V4 \| 1.0 \| 2.148e-23 \| 525 \| 0.207 \| 521 \| 265 \| 18 \| 1 \| 423 \| 1 \| 471 \| Uncharacterized protein \| Uncharacterized protein; afdb-uniprot50 \| AF-A0A827W3S7-F1-MODEL\_V4 \| 1.0 \| 4.281e-20 \| 524 \| 0.281 \| 370 \| 175 \| 12 \| 1 \| 335 \| 5 \| 318 \| Uncharacterized protein \| Uncharacterized protein; afdb-uniprot50 \| AF-G5QJV8-F1-MODEL\_V4 \| 1.0 \| 4.16e-22 \| 524 \| 0.214 \| 481 \| 212 \| 15 \| 1 \| 423 \| 4 \| 376 \| Phage tail fiber protein \| Phage tail fiber protein; afdb-uniprot50 \| AF-A0A7H9FY19-F1-MODEL\_V4 \| 1.0 \| 3.489e-23 \| 524 \| 0.263 \| 485 \| 232 \| 25 \| 1 \| 422 \| 1 \| 423 \| Phage tail protein \| Phage tail protein; afdb-uniprot50 \| AF-A0A1I4WBK0-F1-MODEL\_V4 \| 1.0 \| 3.885e-23 \| 524 \| 0.198 \| 646 \| 256 \| 23 \| 2 \| 423 \| 6 \| 613 \| Phage tail-collar fibre protein \| Phage tail-collar fibre protein; afdb-uniprot50 \| AF-A0A807Z4S1-F1-MODEL\_V4 \| 1.0 \| 1.317e-12 \| 523 \| 0.314 \| 159 \| 101 \| 3 \| 1 \| 152 \| 3 \| 160 \| Uncharacterized protein \| Uncharacterized protein; afdb-uniprot50 \| AF-R7ZJ78-F1-MODEL\_V4 \| 1.0 \| 6.332e-19 \| 523 \| 0.276 \| 322 \| 148 \| 12 \| 1 \| 258 \| 1 \| 301 \| Phage-related tail fiber protein-like protein \| Phage-related tail fiber protein-like protein; afdb-uniprot50 \| AF-A0A3M5NCX5-F1-MODEL\_V4 \| 1.0 \| 1.346e-18 \| 523 \| 0.275 \| 312 \| 155 \| 11 \| 2 \| 247 \| 6 \| 312 \| Uncharacterized protein \| Uncharacterized protein; afdb-uniprot50 \| AF-A0A5E1A2X0-F1-MODEL\_V4 \| 1.0 \| 1.003e-16 \| 523 \| 0.316 \| 250 \| 130 \| 12 \| 1 \| 223 \| 2 \| 237 \| Phage tail collar domain-containing protein \| Phage tail collar domain-containing protein; afdb-uniprot50 \| AF-A0A4Z0B2H6-F1-MODEL\_V4 \| 1.0 \| 4.82e-23 \| 523 \| 0.213 \| 530 \| 261 \| 24 \| 2 \| 423 \| 6 \| 487 \| Uncharacterized protein \| Uncharacterized protein; afdb-uniprot50 \| AF-A4P0H3-F1-MODEL\_V4 \| 1.0 \| 3.143e-19 \| 523 \| 0.234 \| 332 \| 152 \| 5 \| 1 \| 322 \| 2 \| 241 \| Probable tail fiber protein \| Probable tail fiber protein; afdb-uniprot50 \| AF-A0A1B7IQU1-F1-MODEL\_V4 \| 1.0 \| 1.294e-17 \| 523 \| 0.281 \| 320 \| 124 \| 7 \| 1 \| 316 \| 1 \| 218 \| Phage tail fiber protein \| Phage tail fiber protein; afdb-uniprot50 \| AF-A0A1G5B425-F1-MODEL\_V4 \| 1.0 \| 2.102e-17 \| 522 \| 0.281 \| 270 \| 144 \| 12 \| 1 \| 244 \| 1 \| 246 \| Phage Tail Collar Domain \| Phage Tail Collar Domain; afdb-uniprot50 \| AF-A0A7S4LVA4-F1-MODEL\_V4 \| 1.0 \| 1.583e-18 \| 522 \| 0.267 \| 344 \| 133 \| 3 \| 1 \| 326 \| 1 \| 243 \| Uncharacterized protein \| Uncharacterized protein; afdb-uniprot50 \| AF-G5S7G0-F1-MODEL\_V4 \| 1.0 \| 7.053e-19 \| 522 \| 0.236 \| 346 \| 141 \| 7 \| 1 \| 324 \| 4 \| 248 \| Phage tail fiber protein \| Phage tail fiber protein; afdb-uniprot50 \| AF-A0A1B8TDC2-F1-MODEL\_V4 \| 1.0 \| 1.025e-22 \| 521 \| 0.241 \| 564 \| 239 \| 29 \| 1 \| 423 \| 1 \| 516 \| Phage tail protein \| Phage tail protein; afdb-uniprot50 \| AF-A0A7I9ALZ6-F1-MODEL\_V4 \| 1.0 \| 2.148e-23 \| 520 \| 0.246 \| 523 \| 243 \| 23 \| 1 \| 421 \| 1 \| 474 \| Putative tail fiber protein (GpH),Phage tail fibre repeat \| Putative tail fiber protein (GpH),Phage tail fibre repeat; afdb-uniprot50 \| AF-A0A7H8UZA3-F1-MODEL\_V4 \| 1.0 \| 2.042e-19 \| 520 \| 0.239 \| 334 \| 147 \| 9 \| 1 \| 322 \| 3 \| 241 \| Phage tail protein \| Phage tail protein; afdb-uniprot50 \| AF-A0A837CRU8-F1-MODEL\_V4 \| 1.0 \| 7.388e-12 \| 519 \| 0.504 \| 113 \| 55 \| 1 \| 1 \| 112 \| 1 \| 113 \| Phage tail fiber protein \| Phage tail fiber protein; afdb-uniprot50 \| AF-A4N472-F1-MODEL\_V4 \| 1.0 \| 2.978e-19 \| 519 \| 0.24 \| 332 \| 150 \| 8 \| 1 \| 322 \| 2 \| 241 \| Probable tail fiber protein \| Probable tail fiber protein; afdb-uniprot50 \| AF-A0A6X8RGM9-F1-MODEL\_V4 \| 1.0 \| 2.333e-21 \| 519 \| 0.179 \| 715 \| 214 \| 20 \| 1 \| 423 \| 1 \| 634 \| Phage tail protein \| Phage tail protein; afdb-uniprot50 \| AF-A0A5S3V1C2-F1-MODEL\_V4 \| 1.0 \| 5.705e-15 \| 519 \| 0.259 \| 212 \| 139 \| 5 \| 2 \| 203 \| 4 \| 207 \| Uncharacterized protein \| Uncharacterized protein; afdb-uniprot50 \| AF-A0A2G4UEQ4-F1-MODEL\_V4 \| 1.0 \| 6.683e-19 \| 518 \| 0.228 \| 438 \| 184 \| 12 \| 1 \| 413 \| 2 \| 310 \| Phage tail protein \| Phage tail protein; afdb-uniprot50 \| AF-A0A4S3Y564-F1-MODEL\_V4 \| 1.0 \| 1.01e-23 \| 518 \| 0.212 \| 570 \| 260 \| 29 \| 2 \| 423 \| 6 \| 534 \| Phage tail protein \| Phage tail protein; afdb-uniprot50 \| AF-A0A3D1C6R7-F1-MODEL\_V4 \| 1.0 \| 2.358e-24 \| 518 \| 0.246 \| 572 \| 248 \| 23 \| 1 \| 423 \| 2 \| 539 \| Phage tail protein \| Phage tail protein; afdb-uniprot50 \| AF-A0A1I5L005-F1-MODEL\_V4 \| 1.0 \| 1.226e-17 \| 517 \| 0.285 \| 291 \| 145 \| 10 \| 1 \| 246 \| 1 \| 273 \| Phage tail-collar fibre protein \| Phage tail-collar fibre protein; afdb-uniprot50 \| AF-A0A3R9RZV6-F1-MODEL\_V4 \| 1.0 \| 1.406e-15 \| 517 \| 0.271 \| 236 \| 128 \| 6 \| 1 \| 194 \| 1 \| 234 \| Uncharacterized protein \| Uncharacterized protein; afdb-uniprot50 \| AF-A0A5S4U6Y9-F1-MODEL\_V4 \| 1.0 \| 4.837e-19 \| 516 \| 0.246 \| 328 \| 167 \| 9 \| 1 \| 255 \| 2 \| 322 \| Phage tail protein \| Phage tail protein; afdb-uniprot50 \| AF-A0A0J8YRQ6-F1-MODEL\_V4 \| 1.0 \| 1.082e-22 \| 516 \| 0.248 \| 500 \| 200 \| 22 \| 1 \| 423 \| 2 \| 402 \| Uncharacterized protein \| Uncharacterized protein; afdb-uniprot50 \| AF-A0A5E4SRD1-F1-MODEL\_V4 \| 1.0 \| 6.087e-18 \| 516 \| 0.245 \| 318 \| 155 \| 10 \| 1 \| 253 \| 2 \| 299 \| Uncharacterized protein \| Uncharacterized protein; afdb-uniprot50 \| AF-A0A5Z2TND1-F1-MODEL\_V4 \| 1.0 \| 1.025e-22 \| 516 \| 0.232 \| 534 \| 247 \| 25 \| 1 \| 422 \| 1 \| 483 \| Phage tail protein \| Phage tail protein; afdb-uniprot50 \| AF-A0A7Y8F9X6-F1-MODEL\_V4 \| 1.0 \| 2.148e-23 \| 516 \| 0.173 \| 653 \| 293 \| 27 \| 2 \| 423 \| 6 \| 642 \| Phage tail protein \| Phage tail protein; afdb-uniprot50 \| AF-A0A4S2QLK6-F1-MODEL\_V4 \| 1.0 \| 7.053e-19 \| 516 \| 0.265 \| 331 \| 142 \| 7 \| 1 \| 322 \| 2 \| 240 \| Uncharacterized protein \| Uncharacterized protein; afdb-uniprot50 \| AF-A0A4R8JQJ0-F1-MODEL\_V4 \| 1.0 \| 2.782e-20 \| 515 \| 0.246 \| 438 \| 217 \| 14 \| 1 \| 417 \| 2 \| 347 \| Tail fiber-like repeat protein \| Tail fiber-like repeat protein; afdb-uniprot50 \| AF-A0A6N4NZG6-F1-MODEL\_V4 \| 1.0 \| 2.607e-17 \| 515 \| 0.276 \| 325 \| 127 \| 8 \| 1 \| 323 \| 1 \| 219 \| Phage tail protein \| Phage tail protein; afdb-uniprot50 \| AF-A0A7Y1CCL3-F1-MODEL\_V4 \| 1.0 \| 3.365e-18 \| 514 \| 0.263 \| 296 \| 147 \| 8 \| 2 \| 235 \| 6 \| 292 \| Phage tail protein \| Phage tail protein; afdb-uniprot50 \| AF-A0A5M9IW52-F1-MODEL\_V4 \| 1.0 \| 4.82e-23 \| 514 \| 0.187 \| 539 \| 279 \| 25 \| 1 \| 423 \| 1 \| 496 \| Collar domain-containing protein \| Collar domain-containing protein; afdb-uniprot50 \| AF-A0A336NWA0-F1-MODEL\_V4 \| 1.0 \| 7.053e-19 \| 514 \| 0.29 \| 351 \| 126 \| 5 \| 1 \| 324 \| 2 \| 256 \| Phage tail fiber protein \| Phage tail fiber protein; afdb-uniprot50 \| AF-A0A403NCR6-F1-MODEL\_V4 \| 1.0 \| 1.162e-17 \| 514 \| 0.242 \| 346 \| 139 \| 5 \| 1 \| 324 \| 2 \| 246 \| Phage tail protein \| Phage tail protein; afdb-uniprot50 \| AF-A0A625UNB3-F1-MODEL\_V4 \| 1.0 \| 1.24e-20 \| 513 \| 0.241 \| 455 \| 221 \| 16 \| 1 \| 411 \| 2 \| 376 \| Phage tail protein \| Phage tail protein; afdb-uniprot50 \| AF-A0A2V4KQ48-F1-MODEL\_V4 \| 1.0 \| 6.242e-20 \| 513 \| 0.238 \| 377 \| 174 \| 11 \| 1 \| 272 \| 5 \| 373 \| Uncharacterized protein \| Uncharacterized protein; afdb-uniprot50 \| AF-A0A701ZGL9-F1-MODEL\_V4 \| 1.0 \| 1.158e-21 \| 512 \| 0.24 \| 475 \| 208 \| 19 \| 1 \| 413 \| 1 \| 384 \| Phage tail protein \| Phage tail protein; afdb-uniprot50 \| AF-A0A1X3I4D9-F1-MODEL\_V4 \| 1.0 \| 1.985e-21 \| 511 \| 0.254 \| 500 \| 181 \| 18 \| 1 \| 417 \| 1 \| 391 \| Putative tail fiber protein (GpH) \| Putative tail fiber protein (GpH); afdb-uniprot50 \| AF-A0A2U0W405-F1-MODEL\_V4 \| 1.0 \| 2.148e-23 \| 511 \| 0.218 \| 513 \| 270 \| 24 \| 1 \| 423 \| 1 \| 472 \| Tail collar domain \| Tail collar domain; afdb-uniprot50 \| AF-F7U608-F1-MODEL\_V4 \| 1.0 \| 2.832e-15 \| 511 \| 0.289 \| 221 \| 118 \| 4 \| 1 \| 185 \| 1 \| 218 \| Phage-related tail fibre protein-like protein \| Phage-related tail fibre protein-like protein; afdb-uniprot50 \| AF-A0A829QY27-F1-MODEL\_V4 \| 1.0 \| 7.417e-23 \| 511 \| 0.211 \| 620 \| 254 \| 29 \| 4 \| 423 \| 3 \| 587 \| Tail fiber protein \| Tail fiber protein; afdb-uniprot50 \| AF-A0A4Q2B3R1-F1-MODEL\_V4 \| 1.0 \| 3.377e-14 \| 510 \| 0.314 \| 197 \| 116 \| 5 \| 2 \| 194 \| 11 \| 192 \| Uncharacterized protein \| Uncharacterized protein; afdb-uniprot50 \| AF-A0A826UY05-F1-MODEL\_V4 \| 1.0 \| 2.065e-22 \| 510 \| 0.224 \| 494 \| 245 \| 24 \| 1 \| 421 \| 2 \| 430 \| Phage tail protein \| Phage tail protein; afdb-uniprot50 \| AF-A0A1N6MS83-F1-MODEL\_V4 \| 1.0 \| 7.941e-22 \| 510 \| 0.247 \| 452 \| 179 \| 15 \| 1 \| 412 \| 1 \| 331 \| Uncharacterized protein \| Uncharacterized protein; afdb-uniprot50 \| AF-A0A1G4ZBJ6-F1-MODEL\_V4 \| 1.0 \| 7.312e-24 \| 510 \| 0.189 \| 627 \| 256 \| 29 \| 1 \| 423 \| 2 \| 580 \| Phage T4 tail fibre \| Phage T4 tail fibre; afdb-uniprot50 \| AF-A0A722H4T9-F1-MODEL\_V4 \| 1.0 \| 1.014e-19 \| 510 \| 0.255 \| 418 \| 183 \| 17 \| 1 \| 362 \| 2 \| 347 \| Phage tail protein \| Phage tail protein; afdb-uniprot50 \| AF-B3Y0S1-F1-MODEL\_V4 \| 1.0 \| 6.424e-18 \| 509 \| 0.271 \| 324 \| 132 \| 7 \| 1 \| 323 \| 5 \| 225 \| Putative phage tail fiber protein \| Putative phage tail fiber protein; afdb-uniprot50 \| AF-A0A5C7CK81-F1-MODEL\_V4 \| 1.0 \| 4.16e-22 \| 509 \| 0.265 \| 449 \| 200 \| 19 \| 1 \| 405 \| 2 \| 364 \| Uncharacterized protein \| Uncharacterized protein; afdb-uniprot50 \| AF-A0A1I3XGI0-F1-MODEL\_V4 \| 1.0 \| 3.27e-20 \| 508 \| 0.234 \| 473 \| 168 \| 14 \| 1 \| 417 \| 2 \| 336 \| Phage tail fibre repeat-containing protein \| Phage tail fibre repeat-containing protein; afdb-uniprot50 \| AF-A0A285Z1M8-F1-MODEL\_V4 \| 1.0 \| 4.328e-23 \| 508 \| 0.217 \| 478 \| 239 \| 19 \| 1 \| 423 \| 1 \| 398 \| Phage-related tail fibre protein \| Phage-related tail fibre protein; afdb-uniprot50 \| AF-A0A1J0EGB1-F1-MODEL\_V4 \| 1.0 \| 1.416e-22 \| 508 \| 0.221 \| 560 \| 254 \| 27 \| 1 \| 423 \| 3 \| 517 \| Phage tail protein \| Phage tail protein; afdb-uniprot50 \| AF-A0A778CR53-F1-MODEL\_V4 \| 1.0 \| 9.851e-22 \| 508 \| 0.237 \| 485 \| 219 \| 20 \| 1 \| 417 \| 2 \| 403 \| Phage tail protein \| Phage tail protein; afdb-uniprot50 \| AF-A0A2X2TYC3-F1-MODEL\_V4 \| 1.0 \| 1.605e-17 \| 507 \| 0.241 \| 307 \| 137 \| 5 \| 1 \| 307 \| 1 \| 211 \| Phage tail fiber protein \| Phage tail fiber protein; afdb-uniprot50 \| AF-A0A6G8F290-F1-MODEL\_V4 \| 1.0 \| 3.188e-18 \| 507 \| 0.237 \| 291 \| 153 \| 8 \| 1 \| 246 \| 7 \| 273 \| Tail protein \| Tail protein; afdb-uniprot50 \| AF-A0A7T9YBY6-F1-MODEL\_V4 \| 1.0 \| 3.353e-22 \| 507 \| 0.27 \| 458 \| 170 \| 14 \| 1 \| 417 \| 1 \| 335 \| Phage tail protein \| Phage tail protein; afdb-uniprot50 \| AF-W1J3T6-F1-MODEL\_V4 \| 1.0 \| 1.07e-19 \| 506 \| 0.282 \| 390 \| 154 \| 8 \| 1 \| 377 \| 2 \| 278 \| Uncharacterized protein \| Uncharacterized protein; afdb-uniprot50 \| AF-A0A2V5BHI2-F1-MODEL\_V4 \| 1.0 \| 1.757e-22 \| 506 \| 0.222 \| 557 \| 256 \| 24 \| 2 \| 423 \| 6 \| 520 \| Phage-related tail fiber protein \| Phage-related tail fiber protein; afdb-uniprot50 \| AF-A0A316MM73-F1-MODEL\_V4 \| 1.0 \| 6.779e-18 \| 505 \| 0.219 \| 342 \| 198 \| 12 \| 1 \| 323 \| 4 \| 295 \| Uncharacterized protein \| Uncharacterized protein; afdb-uniprot50 \| AF-A0A836YZF5-F1-MODEL\_V4 \| 1.0 \| 1.421e-18 \| 505 \| 0.258 \| 337 \| 145 \| 7 \| 1 \| 324 \| 2 \| 246 \| Tail fiber protein \| Tail fiber protein; afdb-uniprot50 \| AF-A0A376LG26-F1-MODEL\_V4 \| 1.0 \| 1.516e-21 \| 505 \| 0.23 \| 460 \| 243 \| 19 \| 1 \| 408 \| 2 \| 402 \| Putative side tail phage protein \| Putative side tail phage protein; afdb-uniprot50 \| AF-A0A5T5I9M4-F1-MODEL\_V4 \| 1.0 \| 1.222e-21 \| 505 \| 0.199 \| 512 \| 237 \| 17 \| 1 \| 421 \| 2 \| 431 \| Phage tail protein \| Phage tail protein; afdb-uniprot50 \| AF-A0A2M8YKL8-F1-MODEL\_V4 \| 1.0 \| 1.583e-18 \| 504 \| 0.318 \| 298 \| 143 \| 8 \| 1 \| 246 \| 1 \| 290 \| Tail-collar fiber protein \| Tail-collar fiber protein; afdb-uniprot50 \| AF-A0A853I4D4-F1-MODEL\_V4 \| 1.0 \| 1.5e-18 \| 504 \| 0.268 \| 317 \| 155 \| 12 \| 1 \| 258 \| 3 \| 301 \| Phage tail protein \| Phage tail protein; afdb-uniprot50 \| AF-A0A753WFH7-F1-MODEL\_V4 \| 1.0 \| 5.747e-22 \| 504 \| 0.207 \| 579 \| 216 \| 23 \| 1 \| 421 \| 1 \| 494 \| Phage tail protein \| Phage tail protein; afdb-uniprot50 \| AF-A0A1M4Y377-F1-MODEL\_V4 \| 1.0 \| 3.353e-22 \| 504 \| 0.193 \| 574 \| 263 \| 33 \| 1 \| 421 \| 1 \| 527 \| Phage tail-collar fibre protein \| Phage tail-collar fibre protein; afdb-uniprot50 \| AF-A0A7X5D8F7-F1-MODEL\_V4 \| 1.0 \| 6.087e-18 \| 502 \| 0.206 \| 348 \| 188 \| 13 \| 1 \| 318 \| 4 \| 293 \| Uncharacterized protein \| Uncharacterized protein; afdb-uniprot50 \| AF-A0A1B9JGD0-F1-MODEL\_V4 \| 1.0 \| 6.242e-20 \| 502 \| 0.274 \| 372 \| 139 \| 7 \| 1 \| 349 \| 1 \| 264 \| Collar domain-containing protein \| Collar domain-containing protein; afdb-uniprot50 \| AF-A0A172YYW4-F1-MODEL\_V4 \| 1.0 \| 1.757e-22 \| 502 \| 0.221 \| 546 \| 263 \| 26 \| 2 \| 423 \| 6 \| 513 \| Putative phage tail fiber protein \| Putative phage tail fiber protein; afdb-uniprot50 \| AF-A0A776LTW8-F1-MODEL\_V4 \| 1.0 \| 3.885e-23 \| 502 \| 0.273 \| 439 \| 173 \| 17 \| 1 \| 377 \| 1 \| 355 \| Phage tail protein \| Phage tail protein; afdb-uniprot50 \| AF-A0A0W0MX65-F1-MODEL\_V4 \| 1.0 \| 3.177e-22 \| 502 \| 0.224 \| 569 \| 245 \| 29 \| 2 \| 422 \| 6 \| 525 \| Uncharacterized protein \| Uncharacterized protein; afdb-uniprot50 \| AF-A0A243TCZ2-F1-MODEL\_V4 \| 1.0 \| 8.875e-18 \| 502 \| 0.253 \| 308 \| 134 \| 5 \| 1 \| 308 \| 1 \| 212 \| Phage tail protein \| Phage tail protein; afdb-uniprot50 \| AF-A0A1W7LPS7-F1-MODEL\_V4 \| 1.0 \| 6.175e-17 \| 502 \| 0.274 \| 266 \| 146 \| 9 \| 1 \| 255 \| 1 \| 230 \| Uncharacterized protein \| Uncharacterized protein; afdb-uniprot50 \| AF-A0A1D3JVK3-F1-MODEL\_V4 \| 1.0 \| 3.011e-22 \| 501 \| 0.179 \| 518 \| 275 \| 21 \| 2 \| 423 \| 6 \| 469 \| Uncharacterized protein \| Uncharacterized protein; afdb-uniprot50 \| AF-A0A1B7HT63-F1-MODEL\_V4 \| 1.0 \| 1.29e-21 \| 501 \| 0.248 \| 462 \| 224 \| 17 \| 1 \| 403 \| 2 \| 399 \| Phage tail fiber protein \| Phage tail fiber protein; afdb-uniprot50 \| AF-A0A4P9VLW7-F1-MODEL\_V4 \| 1.0 \| 4.837e-19 \| 500 \| 0.227 \| 427 \| 180 \| 15 \| 2 \| 419 \| 3 \| 288 \| Uncharacterized protein \| Uncharacterized protein; afdb-uniprot50 \| AF-M1RN38-F1-MODEL\_V4 \| 1.0 \| 9.102e-20 \| 500 \| 0.259 \| 359 \| 161 \| 11 \| 1 \| 261 \| 2 \| 353 \| Uncharacterized protein \| Uncharacterized protein; afdb-uniprot50 \| AF-A0A3F3I8L4-F1-MODEL\_V4 \| 1.0 \| 2.095e-21 \| 500 \| 0.192 \| 567 \| 203 \| 16 \| 1 \| 423 \| 4 \| 459 \| Uncharacterized protein \| Uncharacterized protein; afdb-uniprot50 \| AF-A0A1H0SWR0-F1-MODEL\_V4 \| 1.0 \| 9.851e-22 \| 500 \| 0.188 \| 615 \| 279 \| 18 \| 1 \| 423 \| 1 \| 587 \| Phage tail-collar fibre protein \| Phage tail-collar fibre protein; afdb-uniprot50 \| AF-A0A1Q5VBN3-F1-MODEL\_V4 \| 1.0 \| 1.881e-21 \| 500 \| 0.262 \| 438 \| 179 \| 10 \| 1 \| 414 \| 1 \| 318 \| Uncharacterized protein \| Uncharacterized protein; afdb-uniprot50 \| AF-A0A0M7E7A2-F1-MODEL\_V4 \| 1.0 \| 6.31e-23 \| 499 \| 0.241 \| 522 \| 214 \| 20 \| 1 \| 423 \| 2 \| 440 \| Variable tail fiber protein \| Variable tail fiber protein; afdb-uniprot50 \| AF-A0A7T9UBZ9-F1-MODEL\_V4 \| 1.0 \| 1.117e-16 \| 499 \| 0.267 \| 258 \| 168 \| 8 \| 1 \| 238 \| 1 \| 257 \| Phage tail protein \| Phage tail protein; afdb-uniprot50 \| AF-N9L859-F1-MODEL\_V4 \| 1.0 \| 5.049e-16 \| 499 \| 0.3 \| 256 \| 134 \| 8 \| 3 \| 246 \| 2 \| 224 \| Uncharacterized protein \| Uncharacterized protein; afdb-uniprot50 \| AF-A0A1I1UBJ2-F1-MODEL\_V4 \| 1.0 \| 6.952e-20 \| 498 \| 0.261 \| 382 \| 187 \| 19 \| 1 \| 322 \| 1 \| 347 \| Phage tail-collar fibre protein \| Phage tail-collar fibre protein; afdb-uniprot50 \| AF-A0A285M2J9-F1-MODEL\_V4 \| 1.0 \| 3.27e-20 \| 498 \| 0.21 \| 455 \| 237 \| 22 \| 1 \| 423 \| 1 \| 365 \| Phage tail-collar fibre protein \| Phage tail-collar fibre protein; afdb-uniprot50 \| AF-A0A556S900-F1-MODEL\_V4 \| 1.0 \| 5.747e-22 \| 498 \| 0.26 \| 488 \| 171 \| 13 \| 1 \| 409 \| 2 \| 378 \| Phage tail protein \| Phage tail protein; afdb-uniprot50 \| AF-A0A2S5C7R9-F1-MODEL\_V4 \| 1.0 \| 2.936e-20 \| 498 \| 0.175 \| 719 \| 227 \| 21 \| 1 \| 423 \| 1 \| 649 \| Phage tail protein \| Phage tail protein; afdb-uniprot50 \| AF-A0A5T8G1R8-F1-MODEL\_V4 \| 1.0 \| 7.525e-22 \| 498 \| 0.205 \| 491 \| 204 \| 15 \| 1 \| 423 \| 4 \| 376 \| Phage tail protein \| Phage tail protein; afdb-uniprot50 \| AF-A0A4U3FAV2-F1-MODEL\_V4 \| 1.0 \| 4.056e-20 \| 497 \| 0.25 \| 440 \| 212 \| 17 \| 1 \| 413 \| 2 \| 350 \| Uncharacterized protein \| Uncharacterized protein; afdb-uniprot50 \| AF-A0A7D6BYB5-F1-MODEL\_V4 \| 1.0 \| 2.853e-22 \| 497 \| 0.217 \| 528 \| 198 \| 14 \| 1 \| 406 \| 2 \| 436 \| Phage tail protein \| Phage tail protein; afdb-uniprot50 \| AF-A0A829CQA7-F1-MODEL\_V4 \| 1.0 \| 8.845e-22 \| 497 \| 0.167 \| 680 \| 236 \| 21 \| 1 \| 423 \| 1 \| 607 \| Phage tail fiber repeat family protein \| Phage tail fiber repeat family protein; afdb-uniprot50 \| AF-A0A1V0BI47-F1-MODEL\_V4 \| 1.0 \| 1.141e-22 \| 497 \| 0.161 \| 779 \| 268 \| 30 \| 1 \| 423 \| 2 \| 750 \| Uncharacterized protein \| Uncharacterized protein; afdb-uniprot50 \| AF-A0A1C3HKM5-F1-MODEL\_V4 \| 1.0 \| 1.24e-20 \| 497 \| 0.248 \| 451 \| 174 \| 13 \| 1 \| 417 \| 1 \| 320 \| Phage tail fiber repeat protein \| Phage tail fiber repeat protein; afdb-uniprot50 \| AF-A0A1Y2S899-F1-MODEL\_V4 \| 1.0 \| 3.098e-20 \| 496 \| 0.239 \| 501 \| 172 \| 17 \| 1 \| 416 \| 2 \| 378 \| Tail protein \| Tail protein; afdb-uniprot50 \| AF-A0A3A9I3B0-F1-MODEL\_V4 \| 1.0 \| 5.329e-16 \| 496 \| 0.265 \| 252 \| 152 \| 6 \| 1 \| 221 \| 1 \| 250 \| Collar domain-containing protein \| Collar domain-containing protein; afdb-uniprot50 \| AF-A0A855M4U4-F1-MODEL\_V4 \| 1.0 \| 1.04e-21 \| 496 \| 0.242 \| 541 \| 179 \| 16 \| 1 \| 423 \| 1 \| 428 \| Phage tail protein \| Phage tail protein; afdb-uniprot50 \| AF-A0A2K9M2M1-F1-MODEL\_V4 \| 1.0 \| 1.097e-21 \| 496 \| 0.179 \| 662 \| 281 \| 26 \| 2 \| 423 \| 6 \| 644 \| Phage tail protein \| Phage tail protein; afdb-uniprot50 \| AF-A0A7U9G0P5-F1-MODEL\_V4 \| 1.0 \| 1.077e-11 \| 495 \| 0.496 \| 127 \| 64 \| 0 \| 32 \| 158 \| 2 \| 128 \| Putative tail fiber protein (GpH) \| Putative tail fiber protein (GpH); afdb-uniprot50 \| AF-A0A483Z189-F1-MODEL\_V4 \| 1.0 \| 1.82e-12 \| 494 \| 0.416 \| 137 \| 77 \| 3 \| 23 \| 158 \| 4 \| 138 \| Phage tail protein \| Phage tail protein; afdb-uniprot50 \| AF-A0A2S8JIV1-F1-MODEL\_V4 \| 1.0 \| 7.577e-14 \| 494 \| 0.308 \| 198 \| 111 \| 5 \| 1 \| 176 \| 1 \| 194 \| Phage tail protein \| Phage tail protein; afdb-uniprot50 \| AF-A0A5U6G5Z8-F1-MODEL\_V4 \| 1.0 \| 7.633e-21 \| 494 \| 0.23 \| 481 \| 206 \| 18 \| 1 \| 405 \| 1 \| 393 \| Collar domain-containing protein \| Collar domain-containing protein; afdb-uniprot50 \| AF-A0A2U3F153-F1-MODEL\_V4 \| 1.0 \| 2.211e-21 \| 494 \| 0.252 \| 467 \| 164 \| 13 \| 1 \| 409 \| 2 \| 341 \| Uncharacterized protein \| Uncharacterized protein; afdb-uniprot50 \| AF-L0MEM1-F1-MODEL\_V4 \| 1.0 \| 1.688e-21 \| 493 \| 0.171 \| 695 \| 192 \| 15 \| 1 \| 423 \| 1 \| 583 \| Phage-related tail fiber protein \| Phage-related tail fiber protein; afdb-uniprot50 \| AF-A0A1W5DTY4-F1-MODEL\_V4 \| 1.0 \| 1.175e-20 \| 492 \| 0.267 \| 452 \| 167 \| 10 \| 1 \| 417 \| 2 \| 324 \| Phage tail fiber repeat protein \| Phage tail fiber repeat protein; afdb-uniprot50 \| AF-A0A6I1H834-F1-MODEL\_V4 \| 1.0 \| 8.719e-23 \| 492 \| 0.227 \| 563 \| 240 \| 28 \| 1 \| 423 \| 1 \| 508 \| Phage tail protein \| Phage tail protein; afdb-uniprot50 \| AF-A0A5P2SLL5-F1-MODEL\_V4 \| 1.0 \| 5.387e-19 \| 491 \| 0.297 \| 373 \| 156 \| 17 \| 1 \| 298 \| 2 \| 343 \| Phage tail protein \| Phage tail protein; afdb-uniprot50 \| AF-A0A1G3JRW0-F1-MODEL\_V4 \| 1.0 \| 5.16e-22 \| 491 \| 0.228 \| 543 \| 259 \| 22 \| 1 \| 423 \| 4 \| 506 \| Uncharacterized protein \| Uncharacterized protein; afdb-uniprot50 \| AF-A0A2U0SUN8-F1-MODEL\_V4 \| 1.0 \| 1.577e-22 \| 491 \| 0.206 \| 590 \| 272 \| 24 \| 1 \| 423 \| 1 \| 561 \| Tail-collar fiber protein \| Tail-collar fiber protein; afdb-uniprot50 \| AF-A0A5U6SMK2-F1-MODEL\_V4 \| 1.0 \| 1.782e-21 \| 491 \| 0.205 \| 481 \| 215 \| 16 \| 1 \| 423 \| 4 \| 375 \| Phage tail protein \| Phage tail protein; afdb-uniprot50 \| AF-A0A2U8I2Y4-F1-MODEL\_V4 \| 1.0 \| 2.462e-21 \| 490 \| 0.24 \| 529 \| 192 \| 18 \| 1 \| 421 \| 1 \| 427 \| Uncharacterized protein \| Uncharacterized protein; afdb-uniprot50 \| AF-A0A7L7BSE0-F1-MODEL\_V4 \| 1.0 \| 1.401e-19 \| 490 \| 0.239 \| 451 \| 209 \| 17 \| 1 \| 405 \| 2 \| 364 \| Phage tail protein \| Phage tail protein; afdb-uniprot50 \| AF-A0A1X3K047-F1-MODEL\_V4 \| 1.0 \| 3.735e-22 \| 489 \| 0.205 \| 556 \| 243 \| 25 \| 1 \| 423 \| 2 \| 491 \| Putative phage tail fiber protein H \| Putative phage tail fiber protein H; afdb-uniprot50 \| AF-A0A854J8Q1-F1-MODEL\_V4 \| 1.0 \| 3.098e-20 \| 489 \| 0.247 \| 453 \| 171 \| 13 \| 1 \| 417 \| 1 \| 319 \| Uncharacterized protein \| Uncharacterized protein; afdb-uniprot50 \| AF-D2MWT7-F1-MODEL\_V4 \| 1.0 \| 1.914e-16 \| 488 \| 0.244 \| 298 \| 131 \| 8 \| 1 \| 248 \| 3 \| 256 \| Tail fiber protein H, putative \| Tail fiber protein H, putative; afdb-uniprot50 \| AF-A0A125V1R9-F1-MODEL\_V4 \| 1.0 \| 3.501e-19 \| 488 \| 0.22 \| 359 \| 190 \| 8 \| 1 \| 306 \| 1 \| 322 \| Phage tail fiber protein \| Phage tail fiber protein; afdb-uniprot50 \| AF-A0A447QPL2-F1-MODEL\_V4 \| 1.0 \| 1.24e-20 \| 488 \| 0.236 \| 456 \| 174 \| 13 \| 1 \| 409 \| 1 \| 329 \| Phage tail fibre repeat \| Phage tail fibre repeat; afdb-uniprot50 \| AF-Q8ZKK5-F1-MODEL\_V4 \| 1.0 \| 1.04e-21 \| 488 \| 0.217 \| 551 \| 249 \| 24 \| 1 \| 412 \| 1 \| 508 \| Putative phage tail fiber protein H \| Putative phage tail fiber protein H; afdb-uniprot50 \| AF-A0A8B3U7S3-F1-MODEL\_V4 \| 1.0 \| 1.175e-20 \| 488 \| 0.184 \| 725 \| 236 \| 25 \| 1 \| 423 \| 1 \| 671 \| Tail Fiber protein \| Tail Fiber protein; afdb-uniprot50 \| AF-A0A4U9D1H6-F1-MODEL\_V4 \| 1.0 \| 2.914e-13 \| 487 \| 0.349 \| 186 \| 109 \| 3 \| 1 \| 176 \| 1 \| 184 \| Uncharacterized protein \| Uncharacterized protein; afdb-uniprot50 \| AF-G1UYT0-F1-MODEL\_V4 \| 1.0 \| 5.387e-19 \| 487 \| 0.277 \| 335 \| 179 \| 13 \| 1 \| 296 \| 1 \| 311 \| Uncharacterized protein \| Uncharacterized protein; afdb-uniprot50 \| AF-A0A0T9T666-F1-MODEL\_V4 \| 1.0 \| 5.767e-18 \| 487 \| 0.29 \| 331 \| 145 \| 9 \| 1 \| 309 \| 2 \| 264 \| Variable tail fiber protein \| Variable tail fiber protein; afdb-uniprot50 \| AF-A0A031FSS2-F1-MODEL\_V4 \| 1.0 \| 7.525e-22 \| 487 \| 0.216 \| 555 \| 260 \| 25 \| 1 \| 423 \| 1 \| 512 \| Prophage long tail fiber protein H \| Prophage long tail fiber protein H; afdb-uniprot50 \| AF-A0A831HRR1-F1-MODEL\_V4 \| 1.0 \| 1.436e-21 \| 487 \| 0.222 \| 517 \| 264 \| 25 \| 1 \| 421 \| 1 \| 475 \| Phage tail protein \| Phage tail protein; afdb-uniprot50 \| AF-A0A2N7RX48-F1-MODEL\_V4 \| 1.0 \| 4.16e-22 \| 486 \| 0.221 \| 491 \| 240 \| 23 \| 1 \| 423 \| 1 \| 417 \| Uncharacterized protein \| Uncharacterized protein; afdb-uniprot50 \| AF-A0A454TSP7-F1-MODEL\_V4 \| 1.0 \| 7.258e-17 \| 486 \| 0.293 \| 296 \| 152 \| 12 \| 1 \| 247 \| 1 \| 288 \| Phage tail protein \| Phage tail protein; afdb-uniprot50 \| AF-A0A380Q8N6-F1-MODEL\_V4 \| 1.0 \| 3.353e-22 \| 486 \| 0.239 \| 510 \| 269 \| 21 \| 1 \| 423 \| 2 \| 479 \| Tail fiber protein \| Tail fiber protein; afdb-uniprot50 \| AF-A0A376U5P4-F1-MODEL\_V4 \| 1.0 \| 1.058e-16 \| 485 \| 0.299 \| 344 \| 151 \| 16 \| 1 \| 324 \| 2 \| 275 \| Tail fiber protein \| Tail fiber protein; afdb-uniprot50 \| AF-H8NR92-F1-MODEL\_V4 \| 1.0 \| 2.863e-18 \| 485 \| 0.287 \| 320 \| 156 \| 8 \| 1 \| 254 \| 2 \| 315 \| Tail Collar domain-containing protein \| Tail Collar domain-containing protein; afdb-uniprot50 \| AF-A0A5Y2LQP0-F1-MODEL\_V4 \| 1.0 \| 5.235e-21 \| 485 \| 0.252 \| 464 \| 183 \| 13 \| 1 \| 405 \| 1 \| 359 \| Collar domain-containing protein \| Collar domain-containing protein; afdb-uniprot50 \| AF-A0A6N8PUZ8-F1-MODEL\_V4 \| 1.0 \| 8.056e-21 \| 485 \| 0.244 \| 474 \| 221 \| 20 \| 1 \| 413 \| 1 \| 398 \| Phage tail protein \| Phage tail protein; afdb-uniprot50 \| AF-A0A7Z0UXS0-F1-MODEL\_V4 \| 1.0 \| 1.421e-18 \| 485 \| 0.249 \| 349 \| 188 \| 12 \| 1 \| 324 \| 1 \| 300 \| Phage tail fiber protein \| Phage tail fiber protein; afdb-uniprot50 \| AF-A0A7V8F402-F1-MODEL\_V4 \| 1.0 \| 2.713e-18 \| 484 \| 0.302 \| 350 \| 170 \| 15 \| 1 \| 309 \| 1 \| 317 \| Uncharacterized protein \| Uncharacterized protein; afdb-uniprot50 \| AF-A0A144L7H0-F1-MODEL\_V4 \| 1.0 \| 8.845e-22 \| 484 \| 0.219 \| 607 \| 186 \| 23 \| 1 \| 423 \| 1 \| 503 \| Phage tail fiber protein \| Phage tail fiber protein; afdb-uniprot50 \| AF-A0A1Y3L4U4-F1-MODEL\_V4 \| 1.0 \| 3.539e-22 \| 484 \| 0.188 \| 674 \| 254 \| 34 \| 2 \| 423 \| 6 \| 638 \| Phage tail protein \| Phage tail protein; afdb-uniprot50 \| AF-A0A7U5ZFC7-F1-MODEL\_V4 \| 1.0 \| 2.211e-21 \| 484 \| 0.258 \| 460 \| 229 \| 18 \| 1 \| 413 \| 2 \| 396 \| Phage tail protein \| Phage tail protein; afdb-uniprot50 \| AF-A0A844Q9H3-F1-MODEL\_V4 \| 1.0 \| 1.308e-20 \| 483 \| 0.215 \| 505 \| 214 \| 24 \| 1 \| 422 \| 3 \| 408 \| Uncharacterized protein \| Uncharacterized protein; afdb-uniprot50 \| AF-A0A7M2QP04-F1-MODEL\_V4 \| 1.0 \| 1.555e-23 \| 483 \| 0.181 \| 591 \| 261 \| 24 \| 4 \| 423 \| 3 \| 541 \| Tail fiber protein \| Tail fiber protein; afdb-uniprot50 \| AF-A0A2J9H2E6-F1-MODEL\_V4 \| 1.0 \| 5.915e-20 \| 483 \| 0.221 \| 487 \| 183 \| 12 \| 1 \| 409 \| 1 \| 369 \| Phage tail protein \| Phage tail protein; afdb-uniprot50 \| AF-A0A143HDH4-F1-MODEL\_V4 \| 1.0 \| 5.196e-14 \| 482 \| 0.31 \| 193 \| 109 \| 5 \| 1 \| 177 \| 1 \| 185 \| Uncharacterized protein \| Uncharacterized protein; afdb-uniprot50 \| AF-A0A1N6I148-F1-MODEL\_V4 \| 1.0 \| 2.179e-22 \| 482 \| 0.198 \| 548 \| 273 \| 27 \| 1 \| 423 \| 2 \| 508 \| Phage tail-collar fibre protein \| Phage tail-collar fibre protein; afdb-uniprot50 \| AF-A0A1Q5TR71-F1-MODEL\_V4 \| 1.0 \| 1.538e-20 \| 482 \| 0.261 \| 448 \| 173 \| 16 \| 1 \| 408 \| 2 \| 331 \| Tail protein \| Tail protein; afdb-uniprot50 \| AF-A0A822NDL8-F1-MODEL\_V4 \| 1.0 \| 8.972e-21 \| 482 \| 0.258 \| 434 \| 185 \| 9 \| 1 \| 400 \| 2 \| 332 \| Uncharacterized protein \| Uncharacterized protein; afdb-uniprot50 \| AF-A0A2C9NUX8-F1-MODEL\_V4 \| 1.0 \| 4.012e-17 \| 482 \| 0.244 \| 307 \| 136 \| 5 \| 1 \| 307 \| 1 \| 211 \| Phage tail protein \| Phage tail protein; afdb-uniprot50 \| AF-K7ZZ34-F1-MODEL\_V4 \| 1.0 \| 7.797e-12 \| 481 \| 0.504 \| 121 \| 58 \| 1 \| 1 \| 119 \| 2 \| 122 \| Phage tail fiber protein \| Phage tail fiber protein; afdb-uniprot50 \| AF-N8R6Q7-F1-MODEL\_V4 \| 1.0 \| 5.705e-15 \| 481 \| 0.365 \| 197 \| 121 \| 3 \| 4 \| 198 \| 3 \| 197 \| Uncharacterized protein \| Uncharacterized protein; afdb-uniprot50 \| AF-A0A1Y5MYB7-F1-MODEL\_V4 \| 1.0 \| 2.607e-17 \| 481 \| 0.206 \| 315 \| 181 \| 15 \| 3 \| 259 \| 2 \| 305 \| Uncharacterized protein \| Uncharacterized protein; afdb-uniprot50 \| AF-A0A6B3ZPL8-F1-MODEL\_V4 \| 1.0 \| 1.129e-19 \| 481 \| 0.205 \| 369 \| 181 \| 12 \| 1 \| 257 \| 2 \| 370 \| Phage tail protein \| Phage tail protein; afdb-uniprot50 \| AF-A0A7U5SYK9-F1-MODEL\_V4 \| 1.0 \| 5.425e-11 \| 481 \| 0.248 \| 153 \| 108 \| 3 \| 6 \| 155 \| 10 \| 158 \| Phage tail protein \| Phage tail protein; afdb-uniprot50 \| AF-A0A4Y6UDH3-F1-MODEL\_V4 \| 1.0 \| 3.353e-22 \| 481 \| 0.206 \| 518 \| 271 \| 28 \| 1 \| 423 \| 18 \| 490 \| Uncharacterized protein \| Uncharacterized protein; afdb-uniprot50 \| AF-A0A377LGA7-F1-MODEL\_V4 \| 1.0 \| 6.066e-22 \| 481 \| 0.254 \| 492 \| 266 \| 26 \| 1 \| 423 \| 5 \| 464 \| Tail fiber protein \| Tail fiber protein; afdb-uniprot50 \| AF-A0A825RJE7-F1-MODEL\_V4 \| 1.0 \| 5.465e-18 \| 480 \| 0.237 \| 320 \| 169 \| 11 \| 1 \| 259 \| 3 \| 308 \| Phage tail protein \| Phage tail protein; afdb-uniprot50 \| AF-A0A4Y9K5F2-F1-MODEL\_V4 \| 1.0 \| 4.907e-18 \| 480 \| 0.259 \| 332 \| 144 \| 8 \| 1 \| 322 \| 2 \| 241 \| Uncharacterized protein \| Uncharacterized protein; afdb-uniprot50 \| AF-A0A6H9S1K4-F1-MODEL\_V4 \| 1.0 \| 4.87e-11 \| 479 \| 0.428 \| 112 \| 64 \| 0 \| 1 \| 112 \| 5 \| 116 \| Phage tail protein \| Phage tail protein; afdb-uniprot50 \| AF-A0A1G7D1W6-F1-MODEL\_V4 \| 1.0 \| 1.014e-19 \| 479 \| 0.253 \| 410 \| 145 \| 11 \| 1 \| 405 \| 2 \| 255 \| Phage tail-collar fibre protein \| Phage tail-collar fibre protein; afdb-uniprot50 \| AF-A0A379CXC0-F1-MODEL\_V4 \| 1.0 \| 1.29e-21 \| 479 \| 0.21 \| 599 \| 260 \| 32 \| 1 \| 423 \| 2 \| 563 \| Phage tail fibre repeat \| Phage tail fibre repeat; afdb-uniprot50 \| AF-A0A2D1CSN0-F1-MODEL\_V4 \| 1.0 \| 5.525e-21 \| 479 \| 0.186 \| 629 \| 276 \| 28 \| 2 \| 423 \| 6 \| 605 \| Putative tail fiber protein \| Putative tail fiber protein; afdb-uniprot50 \| AF-A0A1Y2SIM3-F1-MODEL\_V4 \| 1.0 \| 1.24e-20 \| 479 \| 0.254 \| 452 \| 178 \| 14 \| 1 \| 412 \| 1 \| 333 \| Tail protein \| Tail protein; afdb-uniprot50 \| AF-A0A4U6GR68-F1-MODEL\_V4 \| 1.0 \| 3.021e-18 \| 478 \| 0.276 \| 405 \| 147 \| 14 \| 1 \| 331 \| 2 \| 334 \| Phage tail protein \| Phage tail protein; afdb-uniprot50 \| AF-A0A5N7JNT7-F1-MODEL\_V4 \| 1.0 \| 6.494e-21 \| 478 \| 0.205 \| 530 \| 257 \| 24 \| 1 \| 423 \| 3 \| 475 \| Phage tail protein \| Phage tail protein; afdb-uniprot50 \| AF-A0A766JGR3-F1-MODEL\_V4 \| 1.0 \| 1.478e-19 \| 478 \| 0.211 \| 486 \| 210 \| 17 \| 1 \| 417 \| 5 \| 386 \| Phage tail protein \| Phage tail protein; afdb-uniprot50 \| AF-A0A077NM26-F1-MODEL\_V4 \| 1.0 \| 4.768e-20 \| 478 \| 0.257 \| 431 \| 177 \| 12 \| 1 \| 412 \| 1 \| 307 \| Uncharacterized protein \| Uncharacterized protein; afdb-uniprot50 \| AF-A0A754E741-F1-MODEL\_V4 \| 1.0 \| 2.936e-20 \| 477 \| 0.152 \| 713 \| 201 \| 17 \| 1 \| 423 \| 4 \| 602 \| Uncharacterized protein \| Uncharacterized protein; afdb-uniprot50 \| AF-A0A2G8N573-F1-MODEL\_V4 \| 1.0 \| 6.066e-22 \| 477 \| 0.207 \| 666 \| 260 \| 24 \| 2 \| 423 \| 6 \| 647 \| Phage tail protein \| Phage tail protein; afdb-uniprot50 \| AF-A4WEL3-F1-MODEL\_V4 \| 1.0 \| 3.365e-18 \| 476 \| 0.27 \| 340 \| 158 \| 12 \| 1 \| 255 \| 2 \| 336 \| Phage Tail Collar domain protein \| Phage Tail Collar domain protein; afdb-uniprot50 \| AF-A0A1H6YID8-F1-MODEL\_V4 \| 1.0 \| 1.24e-20 \| 476 \| 0.208 \| 479 \| 243 \| 21 \| 2 \| 423 \| 6 \| 405 \| Phage tail-collar fibre protein \| Phage tail-collar fibre protein; afdb-uniprot50 \| AF-A0A1B9KN89-F1-MODEL\_V4 \| 1.0 \| 1.757e-22 \| 476 \| 0.213 \| 619 \| 250 \| 27 \| 1 \| 423 \| 1 \| 578 \| Uncharacterized protein \| Uncharacterized protein; afdb-uniprot50 \| AF-A0A1G7S9W8-F1-MODEL\_V4 \| 1.0 \| 1.175e-20 \| 476 \| 0.205 \| 448 \| 233 \| 20 \| 1 \| 423 \| 1 \| 350 \| Uncharacterized protein \| Uncharacterized protein; afdb-uniprot50 \| AF-A0A4U9HA58-F1-MODEL\_V4 \| 1.0 \| 3.27e-20 \| 475 \| 0.253 \| 415 \| 157 \| 12 \| 1 \| 363 \| 1 \| 314 \| Phage tail fibre repeat \| Phage tail fibre repeat; afdb-uniprot50 \| AF-A0A0T9UMM8-F1-MODEL\_V4 \| 1.0 \| 1.381e-20 \| 475 \| 0.244 \| 470 \| 175 \| 14 \| 1 \| 405 \| 1 \| 355 \| Variable tail fiber protein \| Variable tail fiber protein; afdb-uniprot50 \| AF-A0A1N6MV94-F1-MODEL\_V4 \| 1.0 \| 1.129e-19 \| 475 \| 0.232 \| 474 \| 176 \| 11 \| 1 \| 417 \| 2 \| 344 \| Uncharacterized protein \| Uncharacterized protein; afdb-uniprot50 \| AF-Q1I687-F1-MODEL\_V4 \| 1.0 \| 2.333e-21 \| 475 \| 0.144 \| 908 \| 278 \| 29 \| 1 \| 423 \| 5 \| 898 \| Putative phage variable tail fibre protein \| Putative phage variable tail fibre protein; afdb-uniprot50 \| AF-A0A5T5IEY5-F1-MODEL\_V4 \| 1.0 \| 5.915e-20 \| 474 \| 0.214 \| 509 \| 187 \| 18 \| 1 \| 419 \| 4 \| 389 \| Phage tail protein \| Phage tail protein; afdb-uniprot50 \| AF-A0A068R2V7-F1-MODEL\_V4 \| 1.0 \| 1.457e-20 \| 474 \| 0.209 \| 540 \| 192 \| 16 \| 1 \| 420 \| 2 \| 426 \| Uncharacterized protein \| Uncharacterized protein; afdb-uniprot50 \| AF-A0A7D5LM12-F1-MODEL\_V4 \| 1.0 \| 1.436e-21 \| 474 \| 0.225 \| 537 \| 238 \| 26 \| 4 \| 423 \| 8 \| 483 \| Phage tail protein \| Phage tail protein; afdb-uniprot50 \| AF-A0A6D0PJP5-F1-MODEL\_V4 \| 1.0 \| 4.7e-21 \| 474 \| 0.19 \| 600 \| 243 \| 26 \| 1 \| 421 \| 2 \| 537 \| Phage tail protein \| Phage tail protein; afdb-uniprot50 \| AF-A0A236MFK9-F1-MODEL\_V4 \| 1.0 \| 1.07e-19 \| 474 \| 0.225 \| 444 \| 179 \| 15 \| 1 \| 392 \| 5 \| 335 \| Phage tail protein \| Phage tail protein; afdb-uniprot50 \| AF-A0A1B2FA13-F1-MODEL\_V4 \| 1.0 \| 1.985e-21 \| 474 \| 0.206 \| 615 \| 243 \| 31 \| 2 \| 423 \| 6 \| 568 \| Uncharacterized protein \| Uncharacterized protein; afdb-uniprot50 \| AF-A0A7I7E0P7-F1-MODEL\_V4 \| 1.0 \| 2.013e-20 \| 474 \| 0.261 \| 401 \| 209 \| 14 \| 1 \| 361 \| 2 \| 355 \| Uncharacterized protein \| Uncharacterized protein; afdb-uniprot50 \| AF-A0A7Y0N7P7-F1-MODEL\_V4 \| 1.0 \| 8.624e-20 \| 473 \| 0.226 \| 371 \| 206 \| 12 \| 1 \| 324 \| 1 \| 337 \| Phage tail protein \| Phage tail protein; afdb-uniprot50 \| AF-W1J3V9-F1-MODEL\_V4 \| 1.0 \| 1.713e-20 \| 473 \| 0.199 \| 586 \| 200 \| 16 \| 1 \| 422 \| 1 \| 481 \| Uncharacterized protein \| Uncharacterized protein; afdb-uniprot50 \| AF-A0A1H8TEQ6-F1-MODEL\_V4 \| 1.0 \| 3.789e-21 \| 473 \| 0.219 \| 566 \| 249 \| 29 \| 1 \| 423 \| 1 \| 516 \| Phage tail-collar fibre protein \| Phage tail-collar fibre protein; afdb-uniprot50 \| AF-A0A2D1IR12-F1-MODEL\_V4 \| 1.0 \| 4.633e-22 \| 473 \| 0.194 \| 577 \| 263 \| 25 \| 1 \| 423 \| 1 \| 529 \| Uncharacterized protein \| Uncharacterized protein; afdb-uniprot50 \| AF-A0A5A9YWL3-F1-MODEL\_V4 \| 1.0 \| 9.535e-13 \| 472 \| 0.295 \| 169 \| 105 \| 3 \| 24 \| 183 \| 6 \| 169 \| Phage tail protein \| Phage tail protein; afdb-uniprot50 \| AF-A0A495BJ48-F1-MODEL\_V4 \| 1.0 \| 3.843e-20 \| 472 \| 0.22 \| 498 \| 202 \| 20 \| 1 \| 423 \| 1 \| 387 \| Tail-collar fiber protein \| Tail-collar fiber protein; afdb-uniprot50 \| AF-A0A482IM57-F1-MODEL\_V4 \| 1.0 \| 8.749e-19 \| 472 \| 0.214 \| 368 \| 174 \| 15 \| 1 \| 260 \| 1 \| 361 \| Uncharacterized protein \| Uncharacterized protein; afdb-uniprot50 \| AF-A0A4R7GFV2-F1-MODEL\_V4 \| 1.0 \| 3.843e-20 \| 472 \| 0.225 \| 511 \| 223 \| 21 \| 1 \| 421 \| 2 \| 429 \| Tail-collar fiber protein \| Tail-collar fiber protein; afdb-uniprot50 \| AF-A0A3F3P2W1-F1-MODEL\_V4 \| 1.0 \| 1.516e-21 \| 472 \| 0.222 \| 518 \| 269 \| 22 \| 1 \| 421 \| 1 \| 481 \| Phage tail protein \| Phage tail protein; afdb-uniprot50 \| AF-A0A198GSC6-F1-MODEL\_V4 \| 1.0 \| 6.31e-23 \| 472 \| 0.216 \| 573 \| 263 \| 24 \| 1 \| 423 \| 2 \| 538 \| Phage tail fiber protein \| Phage tail fiber protein; afdb-uniprot50 \| AF-A0A376M6X6-F1-MODEL\_V4 \| 1.0 \| 9.993e-21 \| 472 \| 0.234 \| 504 \| 235 \| 25 \| 1 \| 392 \| 5 \| 469 \| Tail fiber protein (Gph) \| Tail fiber protein (Gph); afdb-uniprot50 \| AF-A0A7A6VQX8-F1-MODEL\_V4 \| 1.0 \| 1.623e-20 \| 472 \| 0.245 \| 480 \| 215 \| 23 \| 1 \| 392 \| 2 \| 422 \| Phage tail protein \| Phage tail protein; afdb-uniprot50 \| AF-N9CZ59-F1-MODEL\_V4 \| 1.0 \| 1.313e-16 \| 472 \| 0.215 \| 329 \| 173 \| 11 \| 1 \| 278 \| 2 \| 296 \| Uncharacterized protein \| Uncharacterized protein; afdb-uniprot50 \| AF-A0A853IIN2-F1-MODEL\_V4 \| 1.0 \| 4.405e-18 \| 471 \| 0.265 \| 339 \| 149 \| 19 \| 1 \| 258 \| 1 \| 320 \| Phage tail protein \| Phage tail protein; afdb-uniprot50 \| AF-A0A315WVX7-F1-MODEL\_V4 \| 1.0 \| 3.999e-21 \| 471 \| 0.218 \| 567 \| 264 \| 28 \| 1 \| 423 \| 1 \| 532 \| Phage tail protein \| Phage tail protein; afdb-uniprot50 \| AF-A0A4P6GED1-F1-MODEL\_V4 \| 1.0 \| 5.747e-22 \| 471 \| 0.178 \| 644 \| 283 \| 34 \| 1 \| 423 \| 1 \| 619 \| Phage tail protein \| Phage tail protein; afdb-uniprot50 \| AF-A0A7V8EWA4-F1-MODEL\_V4 \| 1.0 \| 2.211e-21 \| 471 \| 0.155 \| 655 \| 285 \| 26 \| 2 \| 423 \| 6 \| 625 \| Phage tail protein \| Phage tail protein; afdb-uniprot50 \| AF-A0A1Y3CBA8-F1-MODEL\_V4 \| 1.0 \| 1.588e-14 \| 471 \| 0.256 \| 238 \| 153 \| 6 \| 1 \| 223 \| 1 \| 229 \| Collar domain-containing protein \| Collar domain-containing protein; afdb-uniprot50 \| AF-A0A380PQB1-F1-MODEL\_V4 \| 1.0 \| 2.401e-19 \| 470 \| 0.254 \| 464 \| 182 \| 15 \| 1 \| 417 \| 2 \| 348 \| Tail fiber protein \| Tail fiber protein; afdb-uniprot50 \| AF-A0A5T8J817-F1-MODEL\_V4 \| 1.0 \| 1.24e-20 \| 470 \| 0.211 \| 483 \| 211 \| 16 \| 1 \| 423 \| 4 \| 376 \| Phage tail protein \| Phage tail protein; afdb-uniprot50 \| AF-A0A7I7ESP2-F1-MODEL\_V4 \| 1.0 \| 3.843e-20 \| 469 \| 0.266 \| 461 \| 220 \| 21 \| 1 \| 418 \| 2 \| 387 \| gp37\_C domain-containing protein \| gp37\_C domain-containing protein; afdb-uniprot50 \| AF-R5ITN8-F1-MODEL\_V4 \| 1.0 \| 2.194e-14 \| 468 \| 0.247 \| 234 \| 137 \| 7 \| 2 \| 198 \| 6 \| 237 \| Possible tail fiber protein \| Possible tail fiber protein; afdb-uniprot50 \| AF-A0A0M9GBQ6-F1-MODEL\_V4 \| 1.0 \| 3.054e-21 \| 468 \| 0.235 \| 514 \| 254 \| 31 \| 2 \| 423 \| 6 \| 472 \| Phage-related tail fiber protein \| Phage-related tail fiber protein; afdb-uniprot50 \| AF-A0A3S6EZT4-F1-MODEL\_V4 \| 1.0 \| 1.808e-20 \| 467 \| 0.216 \| 508 \| 207 \| 21 \| 1 \| 421 \| 2 \| 405 \| Uncharacterized protein \| Uncharacterized protein; afdb-uniprot50 \| AF-B8DLJ2-F1-MODEL\_V4 \| 1.0 \| 5.446e-22 \| 467 \| 0.196 \| 533 \| 290 \| 20 \| 1 \| 423 \| 2 \| 505 \| Tail fiber protein, putative \| Tail fiber protein, putative; afdb-uniprot50 \| AF-A0A6I1L6W4-F1-MODEL\_V4 \| 1.0 \| 1.782e-21 \| 467 \| 0.197 \| 576 \| 275 \| 27 \| 1 \| 423 \| 1 \| 541 \| Uncharacterized protein \| Uncharacterized protein; afdb-uniprot50 \| AF-A0A4U3LF57-F1-MODEL\_V4 \| 1.0 \| 2.498e-20 \| 467 \| 0.23 \| 446 \| 209 \| 17 \| 1 \| 405 \| 2 \| 354 \| Phage tail protein \| Phage tail protein; afdb-uniprot50 \| AF-H8NWL9-F1-MODEL\_V4 \| 1.0 \| 8.624e-20 \| 467 \| 0.249 \| 441 \| 188 \| 12 \| 1 \| 412 \| 1 \| 327 \| Tail fiber repeat 2 protein \| Tail fiber repeat 2 protein; afdb-uniprot50 \| AF-A0A825XI72-F1-MODEL\_V4 \| 1.0 \| 4.907e-18 \| 466 \| 0.24 \| 328 \| 167 \| 12 \| 1 \| 258 \| 1 \| 316 \| Phage tail protein \| Phage tail protein; afdb-uniprot50 \| AF-A0A2X4TXE4-F1-MODEL\_V4 \| 1.0 \| 4.056e-20 \| 466 \| 0.231 \| 453 \| 179 \| 13 \| 1 \| 417 \| 2 \| 321 \| Phage tail fibre repeat \| Phage tail fibre repeat; afdb-uniprot50 \| AF-A0A737EIK7-F1-MODEL\_V4 \| 1.0 \| 4.22e-21 \| 465 \| 0.228 \| 495 \| 228 \| 21 \| 1 \| 423 \| 1 \| 413 \| Phage tail protein \| Phage tail protein; afdb-uniprot50 \| AF-A0A362XHI5-F1-MODEL\_V4 \| 1.0 \| 5.685e-19 \| 465 \| 0.225 \| 425 \| 171 \| 12 \| 1 \| 380 \| 1 \| 312 \| Tail-collar fiber protein \| Tail-collar fiber protein; afdb-uniprot50 \| AF-A0A756ZEH1-F1-MODEL\_V4 \| 1.0 \| 9.334e-22 \| 465 \| 0.227 \| 584 \| 242 \| 32 \| 2 \| 423 \| 3 \| 539 \| Phage tail protein \| Phage tail protein; afdb-uniprot50 \| AF-A0A7X1BUI6-F1-MODEL\_V4 \| 1.0 \| 8.502e-21 \| 465 \| 0.225 \| 505 \| 219 \| 16 \| 1 \| 411 \| 2 \| 428 \| Phage tail protein \| Phage tail protein; afdb-uniprot50 \| AF-A0A383RYZ5-F1-MODEL\_V4 \| 1.0 \| 5.235e-21 \| 465 \| 0.191 \| 759 \| 252 \| 28 \| 2 \| 423 \| 6 \| 739 \| Putative tail fiber protein \| Putative tail fiber protein; afdb-uniprot50 \| AF-A0A7W3F2A9-F1-MODEL\_V4 \| 1.0 \| 1.457e-20 \| 464 \| 0.243 \| 477 \| 222 \| 22 \| 1 \| 417 \| 2 \| 399 \| Phage tail protein \| Phage tail protein; afdb-uniprot50 \| AF-A0A263JA61-F1-MODEL\_V4 \| 1.0 \| 1.113e-20 \| 464 \| 0.251 \| 421 \| 196 \| 14 \| 1 \| 409 \| 1 \| 314 \| Uncharacterized protein \| Uncharacterized protein; afdb-uniprot50 \| AF-A0A3A9I672-F1-MODEL\_V4 \| 1.0 \| 4.469e-17 \| 463 \| 0.181 \| 292 \| 173 \| 10 \| 3 \| 258 \| 16 \| 277 \| Uncharacterized protein \| Uncharacterized protein; afdb-uniprot50 \| AF-A0A6X8FNY2-F1-MODEL\_V4 \| 1.0 \| 3.143e-19 \| 463 \| 0.239 \| 468 \| 202 \| 19 \| 1 \| 412 \| 1 \| 370 \| Phage tail protein \| Phage tail protein; afdb-uniprot50 \| AF-A0A6G5QFT1-F1-MODEL\_V4 \| 1.0 \| 1.04e-21 \| 463 \| 0.193 \| 542 \| 268 \| 22 \| 4 \| 423 \| 3 \| 497 \| Phage tail-collar fiber protein (DUF3751 domain) \| Phage tail-collar fiber protein (DUF3751 domain); afdb-uniprot50 \| AF-A0A7Y8XK01-F1-MODEL\_V4 \| 1.0 \| 1.623e-20 \| 463 \| 0.252 \| 491 \| 183 \| 16 \| 1 \| 418 \| 2 \| 381 \| Phage tail protein \| Phage tail protein; afdb-uniprot50 \| AF-A0A656VM29-F1-MODEL\_V4 \| 1.0 \| 8.624e-20 \| 463 \| 0.254 \| 448 \| 175 \| 13 \| 1 \| 417 \| 1 \| 320 \| Uncharacterized protein \| Uncharacterized protein; afdb-uniprot50 \| AF-A0A3S4IZL9-F1-MODEL\_V4 \| 1.0 \| 7.337e-20 \| 461 \| 0.258 \| 468 \| 175 \| 18 \| 1 \| 417 \| 2 \| 348 \| Tail fiber protein \| Tail fiber protein; afdb-uniprot50 \| AF-A0A7Z7L7Z7-F1-MODEL\_V4 \| 1.0 \| 2.242e-20 \| 461 \| 0.218 \| 494 \| 252 \| 21 \| 1 \| 421 \| 5 \| 437 \| Putative tail fiber protein \| Putative tail fiber protein; afdb-uniprot50 \| AF-A0A364GR27-F1-MODEL\_V4 \| 1.0 \| 2.013e-20 \| 460 \| 0.208 \| 642 \| 221 \| 23 \| 1 \| 423 \| 1 \| 574 \| Tail-collar fiber protein \| Tail-collar fiber protein; afdb-uniprot50 \| AF-A0A8B5NF75-F1-MODEL\_V4 \| 1.0 \| 7.743e-20 \| 460 \| 0.211 \| 530 \| 202 \| 17 \| 1 \| 423 \| 1 \| 421 \| Phage tail protein \| Phage tail protein; afdb-uniprot50 \| AF-A0A828HMS4-F1-MODEL\_V4 \| 1.0 \| 2.742e-21 \| 459 \| 0.252 \| 491 \| 234 \| 25 \| 2 \| 408 \| 4 \| 445 \| Uncharacterized protein \| Uncharacterized protein; afdb-uniprot50 \| AF-A0A855N7K5-F1-MODEL\_V4 \| 1.0 \| 5.329e-16 \| 459 \| 0.22 \| 268 \| 178 \| 9 \| 3 \| 247 \| 2 \| 261 \| Uncharacterized protein \| Uncharacterized protein; afdb-uniprot50 \| AF-A0A5R8ZE78-F1-MODEL\_V4 \| 1.0 \| 3.955e-18 \| 459 \| 0.229 \| 375 \| 180 \| 11 \| 1 \| 272 \| 5 \| 373 \| Collar domain-containing protein \| Collar domain-containing protein; afdb-uniprot50 \| AF-A0A7A3AF76-F1-MODEL\_V4 \| 1.0 \| 3.899e-19 \| 459 \| 0.238 \| 457 \| 170 \| 17 \| 1 \| 408 \| 1 \| 328 \| Phage tail protein \| Phage tail protein; afdb-uniprot50 \| AF-A0A852Q0N8-F1-MODEL\_V4 \| 1.0 \| 1.192e-19 \| 459 \| 0.25 \| 443 \| 177 \| 12 \| 1 \| 418 \| 1 \| 313 \| Phage tail protein \| Phage tail protein; afdb-uniprot50 \| AF-A0A747DAY4-F1-MODEL\_V4 \| 1.0 \| 9.502e-17 \| 458 \| 0.217 \| 304 \| 164 \| 8 \| 1 \| 258 \| 32 \| 307 \| Collar domain-containing protein \| Collar domain-containing protein; afdb-uniprot50 \| AF-A0A2X5DKB2-F1-MODEL\_V4 \| 1.0 \| 7.053e-19 \| 458 \| 0.227 \| 435 \| 180 \| 13 \| 1 \| 405 \| 2 \| 310 \| Side tail phage protein \| Side tail phage protein; afdb-uniprot50 \| AF-A0A7X5IEB0-F1-MODEL\_V4 \| 1.0 \| 5.705e-15 \| 458 \| 0.218 \| 247 \| 177 \| 3 \| 1 \| 240 \| 3 \| 240 \| Peptidase S74 domain-containing protein \| Peptidase S74 domain-containing protein; afdb-uniprot50 \| AF-A0A4R7GCL9-F1-MODEL\_V4 \| 1.0 \| 1.258e-19 \| 458 \| 0.211 \| 516 \| 194 \| 13 \| 1 \| 414 \| 1 \| 405 \| Tail fiber protein gp37 \| Tail fiber protein gp37; afdb-uniprot50 \| AF-A0A2J9QNN2-F1-MODEL\_V4 \| 1.0 \| 2.125e-20 \| 458 \| 0.216 \| 438 \| 188 \| 15 \| 1 \| 392 \| 1 \| 329 \| Phage tail protein \| Phage tail protein; afdb-uniprot50 \| AF-A0A370T1V6-F1-MODEL\_V4 \| 1.0 \| 1.381e-20 \| 458 \| 0.253 \| 450 \| 175 \| 14 \| 1 \| 417 \| 1 \| 322 \| Phage-related tail fiber protein \| Phage-related tail fiber protein; afdb-uniprot50 \| AF-A0A827ZET3-F1-MODEL\_V4 \| 1.0 \| 5.685e-19 \| 457 \| 0.231 \| 458 \| 174 \| 17 \| 1 \| 409 \| 1 \| 329 \| Phage tail protein \| Phage tail protein; afdb-uniprot50 \| AF-A0A1I7H4F9-F1-MODEL\_V4 \| 1.0 \| 4.518e-20 \| 457 \| 0.145 \| 699 \| 282 \| 30 \| 1 \| 423 \| 1 \| 659 \| Phage Tail Collar Domain \| Phage Tail Collar Domain; afdb-uniprot50 \| AF-A0A1B8PH65-F1-MODEL\_V4 \| 1.0 \| 2.367e-20 \| 457 \| 0.153 \| 718 \| 256 \| 24 \| 1 \| 423 \| 2 \| 662 \| Collar domain-containing protein \| Collar domain-containing protein; afdb-uniprot50 \| AF-A0A0C1UK48-F1-MODEL\_V4 \| 1.0 \| 8.348e-11 \| 456 \| 0.435 \| 108 \| 61 \| 0 \| 1 \| 108 \| 2 \| 109 \| Phage tail-collar fiber family protein \| Phage tail-collar fiber family protein; afdb-uniprot50 \| AF-A0A564M9T5-F1-MODEL\_V4 \| 1.0 \| 1.327e-19 \| 456 \| 0.22 \| 468 \| 213 \| 17 \| 1 \| 422 \| 2 \| 363 \| Uncharacterized protein \| Uncharacterized protein; afdb-uniprot50 \| AF-A0A7W7KQY3-F1-MODEL\_V4 \| 1.0 \| 1.538e-20 \| 456 \| 0.186 \| 704 \| 264 \| 28 \| 1 \| 421 \| 1 \| 678 \| Uncharacterized protein \| Uncharacterized protein; afdb-uniprot50 \| AF-A0A558FD57-F1-MODEL\_V4 \| 1.0 \| 1.262e-15 \| 455 \| 0.237 \| 290 \| 162 \| 10 \| 1 \| 248 \| 1 \| 273 \| Uncharacterized protein \| Uncharacterized protein; afdb-uniprot50 \| AF-A0A4R2YQ67-F1-MODEL\_V4 \| 1.0 \| 1.834e-19 \| 455 \| 0.235 \| 437 \| 193 \| 13 \| 1 \| 415 \| 1 \| 318 \| Tail fiber-like repeat protein \| Tail fiber-like repeat protein; afdb-uniprot50 \| AF-A0A2C8EXU4-F1-MODEL\_V4 \| 1.0 \| 4.7e-21 \| 454 \| 0.173 \| 657 \| 255 \| 20 \| 4 \| 423 \| 8 \| 613 \| Putative Phage-related tail fiber protein \| Putative Phage-related tail fiber protein; afdb-uniprot50 \| AF-A0A1I7E346-F1-MODEL\_V4 \| 1.0 \| 4.281e-20 \| 454 \| 0.218 \| 499 \| 254 \| 24 \| 1 \| 420 \| 1 \| 442 \| Phage tail fibre repeat-containing protein \| Phage tail fibre repeat-containing protein; afdb-uniprot50 \| AF-A0A7Z7YC39-F1-MODEL\_V4 \| 1.0 \| 4.583e-19 \| 453 \| 0.227 \| 395 \| 158 \| 15 \| 1 \| 255 \| 2 \| 389 \| Uncharacterized protein \| Uncharacterized protein; afdb-uniprot50 \| AF-A0A377PB01-F1-MODEL\_V4 \| 1.0 \| 1.158e-21 \| 453 \| 0.211 \| 590 \| 252 \| 24 \| 2 \| 423 \| 5 \| 548 \| Phage tail fibre repeat \| Phage tail fibre repeat; afdb-uniprot50 \| AF-N1ZEN3-F1-MODEL\_V4 \| 1.0 \| 1.196e-15 \| 453 \| 0.22 \| 281 \| 172 \| 9 \| 1 \| 250 \| 1 \| 265 \| Peptidase S74 domain-containing protein \| Peptidase S74 domain-containing protein; afdb-uniprot50 \| AF-A0A031GY26-F1-MODEL\_V4 \| 1.0 \| 9.993e-21 \| 453 \| 0.178 \| 711 \| 281 \| 30 \| 1 \| 423 \| 1 \| 696 \| Phage-related tail fiber protein-like protein \| Phage-related tail fiber protein-like protein; afdb-uniprot50 \| AF-A0A560Z585-F1-MODEL\_V4 \| 1.0 \| 2.013e-20 \| 452 \| 0.204 \| 533 \| 246 \| 26 \| 2 \| 422 \| 6 \| 472 \| Tail-collar fiber protein \| Tail-collar fiber protein; afdb-uniprot50 \| AF-A0A8B2ZR20-F1-MODEL\_V4 \| 1.0 \| 1.244e-16 \| 452 \| 0.266 \| 319 \| 153 \| 11 \| 1 \| 306 \| 1 \| 251 \| Uncharacterized protein \| Uncharacterized protein; afdb-uniprot50 \| AF-A0A7Y0ZFL0-F1-MODEL\_V4 \| 1.0 \| 2.367e-20 \| 452 \| 0.184 \| 651 \| 287 \| 28 \| 2 \| 423 \| 6 \| 641 \| Uncharacterized protein \| Uncharacterized protein; afdb-uniprot50 \| AF-A0A1M4V0T2-F1-MODEL\_V4 \| 1.0 \| 4.281e-20 \| 451 \| 0.225 \| 483 \| 224 \| 23 \| 1 \| 423 \| 2 \| 394 \| Phage tail-collar fibre protein \| Phage tail-collar fibre protein; afdb-uniprot50 \| AF-A0A2A2PHW8-F1-MODEL\_V4 \| 1.0 \| 2.242e-20 \| 451 \| 0.191 \| 679 \| 265 \| 27 \| 2 \| 423 \| 6 \| 657 \| Uncharacterized protein \| Uncharacterized protein; afdb-uniprot50 \| AF-A0A845KEW0-F1-MODEL\_V4 \| 1.0 \| 1.014e-19 \| 451 \| 0.26 \| 449 \| 195 \| 18 \| 1 \| 406 \| 1 \| 355 \| Phage tail protein \| Phage tail protein; afdb-uniprot50 \| AF-A0A4S1DI76-F1-MODEL\_V4 \| 1.0 \| 2.57e-18 \| 451 \| 0.209 \| 396 \| 174 \| 16 \| 1 \| 259 \| 1 \| 394 \| Uncharacterized protein \| Uncharacterized protein; afdb-uniprot50 \| AF-A0A0H3ZQZ8-F1-MODEL\_V4 \| 1.0 \| 3.211e-10 \| 450 \| 0.211 \| 151 \| 112 \| 3 \| 6 \| 153 \| 10 \| 156 \| Phage tail fiber protein \| Phage tail fiber protein; afdb-uniprot50 \| AF-A0A7X2D3Y0-F1-MODEL\_V4 \| 1.0 \| 4.281e-20 \| 450 \| 0.201 \| 506 \| 256 \| 27 \| 1 \| 421 \| 1 \| 443 \| Uncharacterized protein \| Uncharacterized protein; afdb-uniprot50 \| AF-H2IRJ0-F1-MODEL\_V4 \| 1.0 \| 1.538e-20 \| 450 \| 0.188 \| 620 \| 264 \| 27 \| 1 \| 423 \| 1 \| 578 \| Phage-related tail fiber protein \| Phage-related tail fiber protein; afdb-uniprot50 \| AF-A0A822X4Z2-F1-MODEL\_V4 \| 1.0 \| 5.465e-18 \| 450 \| 0.25 \| 355 \| 164 \| 12 \| 1 \| 326 \| 2 \| 283 \| Tail Fiber protein \| Tail Fiber protein; afdb-uniprot50 \| AF-A0A1S7SGS6-F1-MODEL\_V4 \| 1.0 \| 3.955e-18 \| 449 \| 0.215 \| 389 \| 167 \| 14 \| 1 \| 258 \| 1 \| 382 \| Uncharacterized protein \| Uncharacterized protein; afdb-uniprot50 \| AF-A0A853I101-F1-MODEL\_V4 \| 1.0 \| 5.387e-19 \| 449 \| 0.204 \| 513 \| 244 \| 26 \| 1 \| 422 \| 1 \| 440 \| Phage tail protein \| Phage tail protein; afdb-uniprot50 \| AF-A0A7U7FC33-F1-MODEL\_V4 \| 1.0 \| 7.337e-20 \| 449 \| 0.244 \| 507 \| 177 \| 16 \| 1 \| 413 \| 2 \| 396 \| Putative phage-related protein \| Putative phage-related protein; afdb-uniprot50 \| AF-A0A850F4G9-F1-MODEL\_V4 \| 1.0 \| 5.83e-21 \| 449 \| 0.182 \| 549 \| 271 \| 21 \| 1 \| 423 \| 2 \| 498 \| Phage tail protein \| Phage tail protein; afdb-uniprot50 \| AF-A0A826V725-F1-MODEL\_V4 \| 1.0 \| 1.175e-20 \| 449 \| 0.211 \| 653 \| 248 \| 31 \| 1 \| 421 \| 1 \| 618 \| Phage tail protein \| Phage tail protein; afdb-uniprot50 \| AF-A0A2V5B0I4-F1-MODEL\_V4 \| 1.0 \| 7.633e-21 \| 449 \| 0.185 \| 669 \| 287 \| 33 \| 1 \| 423 \| 1 \| 657 \| Tail-collar fiber protein \| Tail-collar fiber protein; afdb-uniprot50 \| AF-A0A7M1NUJ9-F1-MODEL\_V4 \| 1.0 \| 9.606e-20 \| 448 \| 0.176 \| 647 \| 227 \| 28 \| 1 \| 422 \| 2 \| 567 \| Phage tail protein \| Phage tail protein; afdb-uniprot50 \| AF-A0A5R9QTW1-F1-MODEL\_V4 \| 1.0 \| 2.598e-21 \| 448 \| 0.227 \| 598 \| 265 \| 24 \| 1 \| 423 \| 1 \| 576 \| Phage tail protein \| Phage tail protein; afdb-uniprot50 \| AF-A0A826TQX2-F1-MODEL\_V4 \| 1.0 \| 1.738e-19 \| 448 \| 0.242 \| 487 \| 192 \| 17 \| 1 \| 417 \| 2 \| 381 \| Phage tail protein \| Phage tail protein; afdb-uniprot50 \| AF-A0A1T0AVV7-F1-MODEL\_V4 \| 1.0 \| 5.31e-20 \| 448 \| 0.235 \| 459 \| 195 \| 16 \| 1 \| 418 \| 1 \| 344 \| Collar domain-containing protein \| Collar domain-containing protein; afdb-uniprot50 \| AF-A0A0C1RAC0-F1-MODEL\_V4 \| 1.0 \| 2.978e-19 \| 447 \| 0.222 \| 413 \| 164 \| 13 \| 1 \| 258 \| 1 \| 411 \| Phage tail-collar fiber family protein \| Phage tail-collar fiber family protein; afdb-uniprot50 \| AF-A0A3G6ZV45-F1-MODEL\_V4 \| 1.0 \| 2.636e-20 \| 447 \| 0.179 \| 630 \| 285 \| 29 \| 2 \| 423 \| 6 \| 611 \| Phage tail fiber protein \| Phage tail fiber protein; afdb-uniprot50 \| AF-G9PUJ7-F1-MODEL\_V4 \| 1.0 \| 2.02e-16 \| 446 \| 0.248 \| 322 \| 155 \| 13 \| 5 \| 260 \| 3 \| 303 \| Uncharacterized protein \| Uncharacterized protein; afdb-uniprot50 \| AF-A0A379Z2I6-F1-MODEL\_V4 \| 1.0 \| 3.843e-20 \| 446 \| 0.232 \| 456 \| 175 \| 14 \| 1 \| 417 \| 2 \| 321 \| Phage tail fibre repeat \| Phage tail fibre repeat; afdb-uniprot50 \| AF-A0A4R2YZB4-F1-MODEL\_V4 \| 1.0 \| 3.098e-20 \| 446 \| 0.192 \| 625 \| 267 \| 26 \| 1 \| 420 \| 1 \| 592 \| Tail fiber-like repeat protein \| Tail fiber-like repeat protein; afdb-uniprot50 \| AF-A0A5U3EWV3-F1-MODEL\_V4 \| 1.0 \| 3.843e-20 \| 445 \| 0.227 \| 480 \| 212 \| 14 \| 1 \| 412 \| 2 \| 390 \| Phage tail protein \| Phage tail protein; afdb-uniprot50 \| AF-A0A0E3CGK1-F1-MODEL\_V4 \| 1.0 \| 1.129e-19 \| 445 \| 0.221 \| 491 \| 215 \| 24 \| 1 \| 377 \| 1 \| 438 \| Uncharacterized protein \| Uncharacterized protein; afdb-uniprot50 \| AF-A0A0B6XER1-F1-MODEL\_V4 \| 1.0 \| 2.02e-16 \| 445 \| 0.247 \| 311 \| 117 \| 8 \| 1 \| 306 \| 2 \| 200 \| Putative phage variable tail fiber protein \| Putative phage variable tail fiber protein; afdb-uniprot50 \| AF-A0A2D0KQU5-F1-MODEL\_V4 \| 1.0 \| 2.367e-20 \| 445 \| 0.208 \| 676 \| 262 \| 32 \| 1 \| 423 \| 2 \| 657 \| Phage tail fiber protein \| Phage tail fiber protein; afdb-uniprot50 \| AF-A0A152BV35-F1-MODEL\_V4 \| 1.0 \| 1.028e-18 \| 445 \| 0.207 \| 486 \| 212 \| 16 \| 1 \| 417 \| 5 \| 386 \| Phage tail protein \| Phage tail protein; afdb-uniprot50 \| AF-A7GFS8-F1-MODEL\_V4 \| 1.0 \| 2.308e-18 \| 444 \| 0.227 \| 395 \| 165 \| 10 \| 1 \| 258 \| 2 \| 393 \| Tail fiber protein H \| Tail fiber protein H; afdb-uniprot50 \| AF-A0A5K4Y4X7-F1-MODEL\_V4 \| 1.0 \| 5.915e-20 \| 444 \| 0.216 \| 540 \| 211 \| 27 \| 1 \| 421 \| 1 \| 447 \| Uncharacterized protein \| Uncharacterized protein; afdb-uniprot50 \| AF-B1JB17-F1-MODEL\_V4 \| 1.0 \| 7.743e-20 \| 444 \| 0.176 \| 737 \| 266 \| 26 \| 1 \| 423 \| 5 \| 714 \| Phage-related tail fibre protein-like protein \| Phage-related tail fibre protein-like protein; afdb-uniprot50 \| AF-A0A0A1HV98-F1-MODEL\_V4 \| 1.0 \| 2.636e-20 \| 444 \| 0.16 \| 743 \| 274 \| 25 \| 2 \| 423 \| 6 \| 719 \| Phage tail fiber protein \| Phage tail fiber protein; afdb-uniprot50 \| AF-A0A7Y8TQ40-F1-MODEL\_V4 \| 1.0 \| 4.977e-17 \| 443 \| 0.224 \| 321 \| 171 \| 14 \| 1 \| 259 \| 1 \| 305 \| Phage tail protein \| Phage tail protein; afdb-uniprot50 \| AF-A0A2W0FKV2-F1-MODEL\_V4 \| 1.0 \| 4.453e-21 \| 443 \| 0.216 \| 535 \| 252 \| 27 \| 2 \| 423 \| 6 \| 486 \| Phage tail protein \| Phage tail protein; afdb-uniprot50 \| AF-A0A1B1KLD7-F1-MODEL\_V4 \| 1.0 \| 4.056e-20 \| 443 \| 0.236 \| 448 \| 183 \| 15 \| 1 \| 417 \| 2 \| 321 \| Uncharacterized protein \| Uncharacterized protein; afdb-uniprot50 \| AF-A0A3I8FWP0-F1-MODEL\_V4 \| 1.0 \| 3.451e-20 \| 442 \| 0.185 \| 562 \| 244 \| 27 \| 1 \| 420 \| 4 \| 493 \| Uncharacterized protein \| Uncharacterized protein; afdb-uniprot50 \| AF-A0A3G7JDV4-F1-MODEL\_V4 \| 1.0 \| 1.56e-19 \| 442 \| 0.203 \| 526 \| 258 \| 21 \| 2 \| 410 \| 6 \| 487 \| Phage tail fiber protein \| Phage tail fiber protein; afdb-uniprot50 \| AF-A0A7G8SK23-F1-MODEL\_V4 \| 1.0 \| 1.276e-18 \| 442 \| 0.227 \| 449 \| 193 \| 11 \| 1 \| 412 \| 1 \| 332 \| Phage tail protein \| Phage tail protein; afdb-uniprot50 \| AF-A0A3V2YUH3-F1-MODEL\_V4 \| 1.0 \| 2.367e-20 \| 442 \| 0.201 \| 644 \| 252 \| 31 \| 1 \| 423 \| 1 \| 602 \| Uncharacterized protein \| Uncharacterized protein; afdb-uniprot50 \| AF-A0A1H8UEW5-F1-MODEL\_V4 \| 1.0 \| 1.209e-18 \| 442 \| 0.25 \| 455 \| 224 \| 15 \| 1 \| 405 \| 2 \| 389 \| Phage-related tail fibre protein \| Phage-related tail fibre protein; afdb-uniprot50 \| AF-A0A822N451-F1-MODEL\_V4 \| 1.0 \| 2.072e-18 \| 442 \| 0.233 \| 480 \| 168 \| 14 \| 1 \| 417 \| 2 \| 344 \| Tail fiber protein XnpH1 of the Xenorhabdicin, a R-type bacteriocin \| Tail fiber protein XnpH1 of the Xenorhabdicin, a R-type bacteriocin; afdb-uniprot50 \| AF-A0A2L0TU26-F1-MODEL\_V4 \| 1.0 \| 2.367e-20 \| 441 \| 0.233 \| 506 \| 241 \| 18 \| 2 \| 416 \| 3 \| 452 \| Phage tail protein \| Phage tail protein; afdb-uniprot50 \| AF-H8NT04-F1-MODEL\_V4 \| 1.0 \| 5.387e-19 \| 441 \| 0.229 \| 445 \| 192 \| 12 \| 1 \| 412 \| 1 \| 327 \| Tail fiber repeat 2 protein \| Tail fiber repeat 2 protein; afdb-uniprot50 \| AF-A0A5S3XN15-F1-MODEL\_V4 \| 1.0 \| 2.873e-14 \| 441 \| 0.218 \| 266 \| 174 \| 9 \| 1 \| 249 \| 3 \| 251 \| Uncharacterized protein \| Uncharacterized protein; afdb-uniprot50 \| AF-A0A840UDD6-F1-MODEL\_V4 \| 1.0 \| 1.478e-19 \| 440 \| 0.22 \| 480 \| 222 \| 23 \| 2 \| 417 \| 3 \| 394 \| Phage-related tail fiber protein \| Phage-related tail fiber protein; afdb-uniprot50 \| AF-A0A2A2MBN2-F1-MODEL\_V4 \| 1.0 \| 8.502e-21 \| 439 \| 0.254 \| 492 \| 204 \| 19 \| 1 \| 409 \| 1 \| 412 \| Uncharacterized protein \| Uncharacterized protein; afdb-uniprot50 \| AF-J2Y2U3-F1-MODEL\_V4 \| 1.0 \| 2.042e-19 \| 439 \| 0.167 \| 667 \| 284 \| 30 \| 1 \| 423 \| 1 \| 639 \| Putative phage tail protein \| Putative phage tail protein; afdb-uniprot50 \| AF-A0A7B5VCM3-F1-MODEL\_V4 \| 1.0 \| 2.791e-16 \| 438 \| 0.241 \| 298 \| 159 \| 15 \| 1 \| 256 \| 1 \| 273 \| Tail fiber \| Tail fiber; afdb-uniprot50 \| AF-A0A6G4HNB6-F1-MODEL\_V4 \| 1.0 \| 8.875e-18 \| 438 \| 0.221 \| 357 \| 180 \| 10 \| 1 \| 260 \| 1 \| 356 \| Phage tail protein \| Phage tail protein; afdb-uniprot50 \| AF-A0A1I1KYG8-F1-MODEL\_V4 \| 1.0 \| 3.021e-18 \| 438 \| 0.209 \| 488 \| 233 \| 21 \| 1 \| 423 \| 1 \| 400 \| Phage tail-collar fibre protein \| Phage tail-collar fibre protein; afdb-uniprot50 \| AF-A0A2W7IK50-F1-MODEL\_V4 \| 1.0 \| 5.604e-20 \| 438 \| 0.216 \| 512 \| 220 \| 31 \| 1 \| 423 \| 1 \| 420 \| Tail-collar fiber protein \| Tail-collar fiber protein; afdb-uniprot50 \| AF-A0A5B0C2P7-F1-MODEL\_V4 \| 1.0 \| 1.175e-20 \| 438 \| 0.18 \| 709 \| 281 \| 30 \| 2 \| 423 \| 6 \| 701 \| Phage tail protein \| Phage tail protein; afdb-uniprot50 \| AF-A0A827LBC4-F1-MODEL\_V4 \| 1.0 \| 2.822e-19 \| 438 \| 0.219 \| 470 \| 204 \| 19 \| 1 \| 423 \| 5 \| 358 \| Phage tail protein \| Phage tail protein; afdb-uniprot50 \| AF-A0A5V0QDA0-F1-MODEL\_V4 \| 1.0 \| 2.791e-16 \| 437 \| 0.228 \| 306 \| 164 \| 12 \| 1 \| 260 \| 1 \| 280 \| Collar domain-containing protein \| Collar domain-containing protein; afdb-uniprot50 \| AF-A0A7G9WG78-F1-MODEL\_V4 \| 1.0 \| 5.14e-11 \| 436 \| 0.404 \| 121 \| 72 \| 0 \| 28 \| 148 \| 1 \| 121 \| Phage tail protein \| Phage tail protein; afdb-uniprot50 \| AF-A0A2I4NEU5-F1-MODEL\_V4 \| 1.0 \| 1.808e-20 \| 436 \| 0.21 \| 500 \| 263 \| 25 \| 1 \| 422 \| 2 \| 447 \| Phage tail-collar fiber family protein \| Phage tail-collar fiber family protein; afdb-uniprot50 \| AF-A0A429K136-F1-MODEL\_V4 \| 1.0 \| 1.196e-15 \| 436 \| 0.2 \| 339 \| 177 \| 12 \| 1 \| 259 \| 1 \| 325 \| Uncharacterized protein \| Uncharacterized protein; afdb-uniprot50 \| AF-A0A6D0S3Z5-F1-MODEL\_V4 \| 1.0 \| 1.963e-18 \| 436 \| 0.213 \| 534 \| 195 \| 18 \| 1 \| 423 \| 2 \| 421 \| Phage tail protein \| Phage tail protein; afdb-uniprot50 \| AF-A0A8A6J8G1-F1-MODEL\_V4 \| 1.0 \| 2.435e-18 \| 436 \| 0.236 \| 453 \| 177 \| 10 \| 1 \| 417 \| 2 \| 321 \| Phage tail protein \| Phage tail protein; afdb-uniprot50 \| AF-R6P5W1-F1-MODEL\_V4 \| 1.0 \| 6e-19 \| 435 \| 0.168 \| 451 \| 222 \| 18 \| 4 \| 423 \| 3 \| 331 \| Putative tail fiber-related protein \| Putative tail fiber-related protein; afdb-uniprot50 \| AF-A0A3V5LF01-F1-MODEL\_V4 \| 1.0 \| 4.583e-19 \| 435 \| 0.224 \| 494 \| 225 \| 19 \| 1 \| 403 \| 1 \| 427 \| Phage tail protein \| Phage tail protein; afdb-uniprot50 \| AF-A0A6B4FV55-F1-MODEL\_V4 \| 1.0 \| 2.978e-19 \| 434 \| 0.201 \| 507 \| 222 \| 24 \| 1 \| 420 \| 2 \| 412 \| Phage tail protein \| Phage tail protein; afdb-uniprot50 \| AF-A0A4V1INX2-F1-MODEL\_V4 \| 1.0 \| 8.172e-20 \| 434 \| 0.209 \| 510 \| 214 \| 22 \| 1 \| 416 \| 1 \| 415 \| Uncharacterized protein \| Uncharacterized protein; afdb-uniprot50 \| AF-A0A7T7R8V5-F1-MODEL\_V4 \| 1.0 \| 6.779e-18 \| 433 \| 0.197 \| 451 \| 230 \| 21 \| 1 \| 423 \| 1 \| 347 \| Phage tail protein \| Phage tail protein; afdb-uniprot50 \| AF-A0A1X7LWW5-F1-MODEL\_V4 \| 1.0 \| 7.053e-19 \| 433 \| 0.192 \| 567 \| 199 \| 20 \| 1 \| 419 \| 1 \| 456 \| Phage tail fibre repeat-containing protein \| Phage tail fibre repeat-containing protein; afdb-uniprot50 \| AF-A0A2K4JEE0-F1-MODEL\_V4 \| 1.0 \| 3.098e-20 \| 433 \| 0.19 \| 572 \| 252 \| 26 \| 2 \| 421 \| 6 \| 518 \| Phage tail protein \| Phage tail protein; afdb-uniprot50 \| AF-A0A2N6J7T9-F1-MODEL\_V4 \| 1.0 \| 2.978e-19 \| 433 \| 0.224 \| 521 \| 221 \| 25 \| 1 \| 422 \| 1 \| 437 \| Phage-related tail fiber protein-like protein \| Phage-related tail fiber protein-like protein; afdb-uniprot50 \| AF-A0A857ER02-F1-MODEL\_V4 \| 1.0 \| 1.457e-20 \| 433 \| 0.167 \| 663 \| 286 \| 24 \| 1 \| 423 \| 1 \| 637 \| Uncharacterized protein \| Uncharacterized protein; afdb-uniprot50 \| AF-A0A8B6AAK0-F1-MODEL\_V4 \| 1.0 \| 1.478e-19 \| 433 \| 0.174 \| 746 \| 274 \| 27 \| 1 \| 423 \| 2 \| 728 \| Phage-related tail fiber protein-like protein \| Phage-related tail fiber protein-like protein; afdb-uniprot50 \| AF-A0A4U3C4N5-F1-MODEL\_V4 \| 1.0 \| 1.209e-18 \| 433 \| 0.189 \| 438 \| 235 \| 17 \| 1 \| 421 \| 91 \| 425 \| DUF1566 domain-containing protein \| DUF1566 domain-containing protein; afdb-uniprot50 \| AF-A0A269PIY6-F1-MODEL\_V4 \| 1.0 \| 2.02e-16 \| 433 \| 0.174 \| 338 \| 179 \| 13 \| 4 \| 258 \| 3 \| 323 \| Uncharacterized protein \| Uncharacterized protein; afdb-uniprot50 \| AF-A0A261REI1-F1-MODEL\_V4 \| 1.0 \| 2.341e-17 \| 432 \| 0.219 \| 456 \| 209 \| 21 \| 1 \| 412 \| 1 \| 353 \| Uncharacterized protein \| Uncharacterized protein; afdb-uniprot50 \| AF-A0A1I6E7W8-F1-MODEL\_V4 \| 1.0 \| 1.145e-18 \| 432 \| 0.195 \| 569 \| 210 \| 26 \| 1 \| 423 \| 2 \| 468 \| Phage-related tail fibre protein \| Phage-related tail fibre protein; afdb-uniprot50 \| AF-A0A1D7Z5A8-F1-MODEL\_V4 \| 1.0 \| 6e-19 \| 432 \| 0.211 \| 642 \| 177 \| 20 \| 1 \| 423 \| 2 \| 532 \| Phage tail protein \| Phage tail protein; afdb-uniprot50 \| AF-A0A853I581-F1-MODEL\_V4 \| 1.0 \| 6.952e-20 \| 431 \| 0.177 \| 512 \| 237 \| 25 \| 2 \| 423 \| 3 \| 420 \| Phage tail protein \| Phage tail protein; afdb-uniprot50 \| AF-A0A7J0BJF8-F1-MODEL\_V4 \| 1.0 \| 9.993e-21 \| 431 \| 0.189 \| 558 \| 264 \| 27 \| 1 \| 423 \| 1 \| 504 \| Tail fiber protein \| Tail fiber protein; afdb-uniprot50 \| AF-A0A2T5NWN9-F1-MODEL\_V4 \| 1.0 \| 3.098e-20 \| 430 \| 0.19 \| 561 \| 266 \| 31 \| 1 \| 420 \| 1 \| 514 \| Uncharacterized protein \| Uncharacterized protein; afdb-uniprot50 \| AF-A0A7M1L6P4-F1-MODEL\_V4 \| 1.0 \| 8.172e-20 \| 429 \| 0.194 \| 560 \| 266 \| 25 \| 1 \| 418 \| 1 \| 517 \| Phage tail protein \| Phage tail protein; afdb-uniprot50 \| AF-A0A329CFM1-F1-MODEL\_V4 \| 1.0 \| 2.341e-17 \| 428 \| 0.263 \| 372 \| 183 \| 16 \| 1 \| 323 \| 5 \| 334 \| Tail-collar fiber protein \| Tail-collar fiber protein; afdb-uniprot50 \| AF-A0A376LF19-F1-MODEL\_V4 \| 1.0 \| 3.694e-19 \| 428 \| 0.203 \| 535 \| 200 \| 18 \| 1 \| 423 \| 1 \| 421 \| Phage tail fiber protein H (GpH) \| Phage tail fiber protein H (GpH); afdb-uniprot50 \| AF-A0A4R6PI57-F1-MODEL\_V4 \| 1.0 \| 2.978e-19 \| 426 \| 0.188 \| 663 \| 280 \| 31 \| 1 \| 423 \| 1 \| 645 \| Phage-related tail fiber protein \| Phage-related tail fiber protein; afdb-uniprot50 \| AF-A0A3B0C125-F1-MODEL\_V4 \| 1.0 \| 2.349e-13 \| 426 \| 0.258 \| 213 \| 133 \| 7 \| 1 \| 194 \| 1 \| 207 \| Uncharacterized protein \| Uncharacterized protein; afdb-uniprot50 \| AF-A0A376FFE3-F1-MODEL\_V4 \| 1.0 \| 2.086e-10 \| 425 \| 0.517 \| 116 \| 55 \| 1 \| 32 \| 147 \| 2 \| 116 \| Phage tail collar domain-containing protein \| Phage tail collar domain-containing protein; afdb-uniprot50 \| AF-A0A285M654-F1-MODEL\_V4 \| 1.0 \| 2.308e-18 \| 425 \| 0.191 \| 418 \| 175 \| 13 \| 1 \| 258 \| 1 \| 415 \| Phage tail-collar fibre protein \| Phage tail-collar fibre protein; afdb-uniprot50 \| AF-A0A0N9MHE6-F1-MODEL\_V4 \| 1.0 \| 2.102e-17 \| 425 \| 0.286 \| 363 \| 183 \| 19 \| 2 \| 324 \| 6 \| 332 \| Phage tail fiber protein \| Phage tail fiber protein; afdb-uniprot50 \| AF-A0A1X1A8V7-F1-MODEL\_V4 \| 1.0 \| 2.752e-17 \| 425 \| 0.267 \| 396 \| 178 \| 16 \| 1 \| 311 \| 5 \| 373 \| Uncharacterized protein \| Uncharacterized protein; afdb-uniprot50 \| AF-A0A1W7LRA1-F1-MODEL\_V4 \| 1.0 \| 9.606e-20 \| 424 \| 0.177 \| 513 \| 276 \| 27 \| 1 \| 421 \| 1 \| 459 \| Putative phage tail fiber protein \| Putative phage tail fiber protein; afdb-uniprot50 \| AF-A0A1H2M4C6-F1-MODEL\_V4 \| 1.0 \| 3.694e-19 \| 424 \| 0.258 \| 461 \| 224 \| 20 \| 2 \| 405 \| 6 \| 405 \| Phage tail-collar fibre protein \| Phage tail-collar fibre protein; afdb-uniprot50 \| AF-A0A5E1AHB0-F1-MODEL\_V4 \| 1.0 \| 7.443e-19 \| 424 \| 0.231 \| 475 \| 252 \| 17 \| 1 \| 405 \| 2 \| 433 \| Tail Fiber protein \| Tail Fiber protein; afdb-uniprot50 \| AF-A0A5N9IEZ9-F1-MODEL\_V4 \| 1.0 \| 9.233e-19 \| 424 \| 0.154 \| 813 \| 269 \| 29 \| 2 \| 423 \| 6 \| 790 \| Phage tail protein \| Phage tail protein; afdb-uniprot50 \| AF-A0A2X5NM48-F1-MODEL\_V4 \| 1.0 \| 5.032e-20 \| 423 \| 0.159 \| 519 \| 261 \| 22 \| 3 \| 423 \| 2 \| 443 \| Uncharacterized protein \| Uncharacterized protein; afdb-uniprot50 \| AF-A0A315T3S6-F1-MODEL\_V4 \| 1.0 \| 1.014e-19 \| 423 \| 0.157 \| 817 \| 278 \| 35 \| 1 \| 423 \| 5 \| 804 \| Tail-collar fiber protein \| Tail-collar fiber protein; afdb-uniprot50 \| AF-A0A377IMK2-F1-MODEL\_V4 \| 1.0 \| 8.624e-20 \| 423 \| 0.222 \| 494 \| 242 \| 27 \| 1 \| 422 \| 1 \| 424 \| Putative phage-like tail fiber protein \| Putative phage-like tail fiber protein; afdb-uniprot50 \| AF-N9JQ34-F1-MODEL\_V4 \| 1.0 \| 6.087e-18 \| 422 \| 0.207 \| 434 \| 245 \| 16 \| 1 \| 423 \| 1 \| 346 \| Uncharacterized protein \| Uncharacterized protein; afdb-uniprot50 \| AF-A0A348HI80-F1-MODEL\_V4 \| 1.0 \| 1.583e-18 \| 422 \| 0.226 \| 495 \| 190 \| 21 \| 1 \| 417 \| 1 \| 380 \| Phage-related tail fibre \| Phage-related tail fibre; afdb-uniprot50 \| AF-A0A3E0FEW6-F1-MODEL\_V4 \| 1.0 \| 3.899e-19 \| 421 \| 0.212 \| 531 \| 249 \| 22 \| 4 \| 411 \| 8 \| 492 \| Tail-collar fiber protein \| Tail-collar fiber protein; afdb-uniprot50 \| AF-A0A1X0VZ24-F1-MODEL\_V4 \| 1.0 \| 1.629e-16 \| 420 \| 0.232 \| 322 \| 173 \| 13 \| 1 \| 252 \| 1 \| 318 \| Phage tail protein \| Phage tail protein; afdb-uniprot50 \| AF-A0A1B5ER81-F1-MODEL\_V4 \| 1.0 \| 6.242e-20 \| 420 \| 0.175 \| 702 \| 280 \| 33 \| 2 \| 423 \| 6 \| 688 \| Uncharacterized protein \| Uncharacterized protein; afdb-uniprot50 \| AF-A0A446AZN6-F1-MODEL\_V4 \| 1.0 \| 5.387e-19 \| 420 \| 0.16 \| 775 \| 286 \| 31 \| 2 \| 423 \| 6 \| 768 \| Uncharacterized protein \| Uncharacterized protein; afdb-uniprot50 \| AF-A0A1Q6JPI1-F1-MODEL\_V4 \| 1.0 \| 4.296e-16 \| 419 \| 0.197 \| 349 \| 192 \| 13 \| 4 \| 313 \| 15 \| 314 \| Uncharacterized protein \| Uncharacterized protein; afdb-uniprot50 \| AF-A0A2G8BXY9-F1-MODEL\_V4 \| 1.0 \| 3.551e-18 \| 419 \| 0.213 \| 472 \| 208 \| 21 \| 1 \| 423 \| 1 \| 358 \| Phage tail protein \| Phage tail protein; afdb-uniprot50 \| AF-A0A0U4W4I1-F1-MODEL\_V4 \| 1.0 \| 5.104e-19 \| 419 \| 0.218 \| 572 \| 252 \| 29 \| 1 \| 423 \| 2 \| 527 \| Ig-like domain-containing protein \| Ig-like domain-containing protein; afdb-uniprot50 \| AF-A0A246GVB8-F1-MODEL\_V4 \| 1.0 \| 4.649e-18 \| 419 \| 0.229 \| 419 \| 180 \| 15 \| 1 \| 311 \| 5 \| 388 \| Collar domain-containing protein \| Collar domain-containing protein; afdb-uniprot50 \| AF-A0A3S5ENC7-F1-MODEL\_V4 \| 1.0 \| 1.788e-17 \| 418 \| 0.24 \| 486 \| 187 \| 17 \| 2 \| 412 \| 3 \| 381 \| Putative bacteriophage tail fiber protein \| Putative bacteriophage tail fiber protein; afdb-uniprot50 \| AF-A0A1H9AFH7-F1-MODEL\_V4 \| 1.0 \| 1.313e-16 \| 418 \| 0.274 \| 361 \| 179 \| 16 \| 2 \| 311 \| 6 \| 334 \| Phage-related tail fibre protein \| Phage-related tail fibre protein; afdb-uniprot50 \| AF-A0A077P9Q6-F1-MODEL\_V4 \| 1.0 \| 4.07e-16 \| 418 \| 0.24 \| 333 \| 134 \| 9 \| 1 \| 326 \| 2 \| 222 \| Uncharacterized protein \| Uncharacterized protein; afdb-uniprot50 \| AF-J2MLE3-F1-MODEL\_V4 \| 1.0 \| 7.855e-19 \| 417 \| 0.253 \| 450 \| 236 \| 13 \| 2 \| 417 \| 6 \| 389 \| Putative phage protein \| Putative phage protein; afdb-uniprot50 \| AF-V8RA43-F1-MODEL\_V4 \| 1.0 \| 8.749e-19 \| 417 \| 0.14 \| 842 \| 267 \| 35 \| 1 \| 423 \| 1 \| 804 \| Tail protein \| Tail protein; afdb-uniprot50 \| AF-A0A0E1PUY8-F1-MODEL\_V4 \| 1.0 \| 6.588e-20 \| 417 \| 0.193 \| 485 \| 267 \| 23 \| 1 \| 423 \| 1 \| 423 \| Uncharacterized protein \| Uncharacterized protein; afdb-uniprot50 \| AF-A0A5C8BE21-F1-MODEL\_V4 \| 1.0 \| 6.446e-14 \| 416 \| 0.245 \| 248 \| 140 \| 5 \| 3 \| 217 \| 2 \| 235 \| Uncharacterized protein \| Uncharacterized protein; afdb-uniprot50 \| AF-A0A806X791-F1-MODEL\_V4 \| 1.0 \| 7.969e-18 \| 416 \| 0.213 \| 436 \| 242 \| 10 \| 1 \| 416 \| 2 \| 356 \| Uncharacterized protein \| Uncharacterized protein; afdb-uniprot50 \| AF-A0A828NYP4-F1-MODEL\_V4 \| 1.0 \| 4.649e-18 \| 416 \| 0.189 \| 464 \| 201 \| 16 \| 1 \| 409 \| 4 \| 347 \| Phage tail protein \| Phage tail protein; afdb-uniprot50 \| AF-A0A1K2AJW9-F1-MODEL\_V4 \| 1.0 \| 1.9e-09 \| 416 \| 0.401 \| 127 \| 76 \| 0 \| 32 \| 158 \| 2 \| 128 \| Phage tail-collar fibre protein \| Phage tail-collar fibre protein; afdb-uniprot50 \| AF-A0A0N9MHY1-F1-MODEL\_V4 \| 1.0 \| 2.904e-17 \| 416 \| 0.269 \| 378 \| 181 \| 16 \| 2 \| 306 \| 6 \| 361 \| Collar domain-containing protein \| Collar domain-containing protein; afdb-uniprot50 \| AF-A0A853I1G7-F1-MODEL\_V4 \| 1.0 \| 9.233e-19 \| 415 \| 0.174 \| 544 \| 213 \| 23 \| 2 \| 422 \| 3 \| 433 \| Phage tail protein \| Phage tail protein; afdb-uniprot50 \| AF-A0A1S8TWQ6-F1-MODEL\_V4 \| 1.0 \| 1.56e-19 \| 415 \| 0.201 \| 531 \| 260 \| 30 \| 1 \| 423 \| 2 \| 476 \| Uncharacterized protein \| Uncharacterized protein; afdb-uniprot50 \| AF-A0A8B5G8Z2-F1-MODEL\_V4 \| 1.0 \| 2.234e-09 \| 414 \| 0.48 \| 102 \| 53 \| 0 \| 1 \| 102 \| 2 \| 103 \| Phage tail protein \| Phage tail protein; afdb-uniprot50 \| AF-A0A2T5NR62-F1-MODEL\_V4 \| 1.0 \| 1.56e-19 \| 414 \| 0.193 \| 544 \| 217 \| 28 \| 1 \| 422 \| 1 \| 444 \| Uncharacterized protein \| Uncharacterized protein; afdb-uniprot50 \| AF-A0A3M2YA40-F1-MODEL\_V4 \| 1.0 \| 1.788e-17 \| 414 \| 0.243 \| 399 \| 199 \| 19 \| 1 \| 350 \| 5 \| 349 \| Uncharacterized protein \| Uncharacterized protein; afdb-uniprot50 \| AF-A0A3G6Z2B6-F1-MODEL\_V4 \| 1.0 \| 2.636e-20 \| 413 \| 0.176 \| 572 \| 255 \| 25 \| 1 \| 423 \| 1 \| 505 \| Uncharacterized protein \| Uncharacterized protein; afdb-uniprot50 \| AF-A0A1I5MQ45-F1-MODEL\_V4 \| 1.0 \| 9.745e-19 \| 413 \| 0.202 \| 513 \| 244 \| 22 \| 1 \| 412 \| 1 \| 449 \| Chaperone of endosialidase \| Chaperone of endosialidase; afdb-uniprot50 \| AF-A0A4V3CGG4-F1-MODEL\_V4 \| 1.0 \| 2.674e-19 \| 412 \| 0.189 \| 534 \| 246 \| 29 \| 1 \| 405 \| 1 \| 476 \| Tail-collar fiber protein \| Tail-collar fiber protein; afdb-uniprot50 \| AF-A0A285M2V3-F1-MODEL\_V4 \| 1.0 \| 7.443e-19 \| 410 \| 0.202 \| 500 \| 204 \| 19 \| 2 \| 422 \| 4 \| 387 \| Phage tail-collar fibre protein \| Phage tail-collar fibre protein; afdb-uniprot50 \| AF-A0A484QR79-F1-MODEL\_V4 \| 1.0 \| 4.837e-19 \| 410 \| 0.202 \| 548 \| 237 \| 30 \| 3 \| 423 \| 2 \| 476 \| Phage tail fiber protein \| Phage tail fiber protein; afdb-uniprot50 \| AF-A0A3E1QH42-F1-MODEL\_V4 \| 1.0 \| 7.443e-19 \| 410 \| 0.213 \| 441 \| 216 \| 20 \| 1 \| 415 \| 2 \| 337 \| Phage tail protein \| Phage tail protein; afdb-uniprot50 \| AF-A0A1W9UG34-F1-MODEL\_V4 \| 1.0 \| 1.346e-18 \| 409 \| 0.191 \| 495 \| 212 \| 23 \| 1 \| 423 \| 1 \| 379 \| Uncharacterized protein \| Uncharacterized protein; afdb-uniprot50 \| AF-A0A1H8JBA9-F1-MODEL\_V4 \| 1.0 \| 9.035e-13 \| 409 \| 0.318 \| 207 \| 121 \| 6 \| 18 \| 211 \| 2 \| 201 \| Phage tail-collar fibre protein \| Phage tail-collar fibre protein; afdb-uniprot50 \| AF-S4YR67-F1-MODEL\_V4 \| 1.0 \| 4.405e-18 \| 408 \| 0.228 \| 420 \| 197 \| 10 \| 1 \| 412 \| 2 \| 302 \| Uncharacterized protein \| Uncharacterized protein; afdb-uniprot50 \| AF-A0A5Q2KDB2-F1-MODEL\_V4 \| 1.0 \| 7.443e-19 \| 408 \| 0.229 \| 461 \| 201 \| 18 \| 1 \| 413 \| 1 \| 355 \| Uncharacterized protein \| Uncharacterized protein; afdb-uniprot50 \| AF-A0A5U3F095-F1-MODEL\_V4 \| 1.0 \| 1.192e-19 \| 408 \| 0.169 \| 719 \| 264 \| 33 \| 1 \| 423 \| 4 \| 685 \| Phage tail protein \| Phage tail protein; afdb-uniprot50 \| AF-A0A3Q8TZG5-F1-MODEL\_V4 \| 1.0 \| 1.213e-14 \| 408 \| 0.265 \| 290 \| 165 \| 11 \| 1 \| 255 \| 5 \| 281 \| Collar domain-containing protein \| Collar domain-containing protein; afdb-uniprot50 \| AF-A0A380T4V8-F1-MODEL\_V4 \| 1.0 \| 7.443e-19 \| 408 \| 0.174 \| 786 \| 276 \| 32 \| 2 \| 423 \| 6 \| 782 \| Putative tail fiber protein \| Putative tail fiber protein; afdb-uniprot50 \| AF-A0A5E4UH46-F1-MODEL\_V4 \| 1.0 \| 3.021e-18 \| 407 \| 0.228 \| 434 \| 218 \| 21 \| 1 \| 392 \| 2 \| 360 \| Uncharacterized protein \| Uncharacterized protein; afdb-uniprot50 \| AF-A0A1N6I1A6-F1-MODEL\_V4 \| 1.0 \| 8.2e-16 \| 406 \| 0.239 \| 351 \| 154 \| 12 \| 1 \| 255 \| 1 \| 334 \| Phage tail-collar fibre protein \| Phage tail-collar fibre protein; afdb-uniprot50 \| AF-A0A4U8YRJ4-F1-MODEL\_V4 \| 1.0 \| 3.857e-16 \| 406 \| 0.236 \| 342 \| 170 \| 11 \| 1 \| 257 \| 1 \| 336 \| Phage tail fibre protein \| Phage tail fibre protein; afdb-uniprot50 \| AF-Q1QXR9-F1-MODEL\_V4 \| 1.0 \| 1.763e-18 \| 406 \| 0.215 \| 551 \| 255 \| 26 \| 1 \| 422 \| 1 \| 502 \| Phage-related tail fibre protein-like protein \| Phage-related tail fibre protein-like protein; afdb-uniprot50 \| AF-E1W2D8-F1-MODEL\_V4 \| 1.0 \| 2.275e-19 \| 406 \| 0.206 \| 509 \| 209 \| 16 \| 1 \| 413 \| 2 \| 411 \| Probable tail fiber protein \| Probable tail fiber protein; afdb-uniprot50 \| AF-A0A1H1IHT2-F1-MODEL\_V4 \| 1.0 \| 1.145e-18 \| 406 \| 0.165 \| 696 \| 288 \| 27 \| 2 \| 423 \| 6 \| 682 \| Phage tail-collar fibre protein \| Phage tail-collar fibre protein; afdb-uniprot50 \| AF-A0A616NKZ7-F1-MODEL\_V4 \| 1.0 \| 1.085e-18 \| 406 \| 0.244 \| 494 \| 212 \| 23 \| 2 \| 423 \| 3 \| 407 \| Phage tail protein \| Phage tail protein; afdb-uniprot50 \| AF-A0A6I1HR03-F1-MODEL\_V4 \| 1.0 \| 6.424e-18 \| 406 \| 0.142 \| 771 \| 285 \| 27 \| 1 \| 422 \| 5 \| 747 \| Uncharacterized protein \| Uncharacterized protein; afdb-uniprot50 \| AF-A0A2D0LG90-F1-MODEL\_V4 \| 1.0 \| 9.265e-15 \| 406 \| 0.227 \| 316 \| 117 \| 8 \| 1 \| 306 \| 2 \| 200 \| Tail protein \| Tail protein; afdb-uniprot50 \| AF-A0A5Y3B2B3-F1-MODEL\_V4 \| 1.0 \| 3.414e-17 \| 405 \| 0.187 \| 453 \| 247 \| 18 \| 1 \| 420 \| 1 \| 365 \| Uncharacterized protein \| Uncharacterized protein; afdb-uniprot50 \| AF-A0A3M0SWC3-F1-MODEL\_V4 \| 1.0 \| 3.143e-19 \| 405 \| 0.187 \| 591 \| 249 \| 30 \| 1 \| 422 \| 1 \| 529 \| Uncharacterized protein \| Uncharacterized protein; afdb-uniprot50 \| AF-A0A2W5T9N3-F1-MODEL\_V4 \| 1.0 \| 7.363e-16 \| 404 \| 0.238 \| 310 \| 170 \| 13 \| 1 \| 258 \| 2 \| 297 \| Uncharacterized protein \| Uncharacterized protein; afdb-uniprot50 \| AF-A0A2X4TX84-F1-MODEL\_V4 \| 1.0 \| 2.187e-18 \| 404 \| 0.226 \| 500 \| 197 \| 21 \| 4 \| 419 \| 3 \| 396 \| Uncharacterized protein \| Uncharacterized protein; afdb-uniprot50 \| AF-N2IMU0-F1-MODEL\_V4 \| 1.0 \| 8.749e-19 \| 404 \| 0.23 \| 600 \| 226 \| 34 \| 1 \| 422 \| 1 \| 542 \| Uncharacterized protein \| Uncharacterized protein; afdb-uniprot50 \| AF-E5Y5X2-F1-MODEL\_V4 \| 1.0 \| 1.276e-18 \| 404 \| 0.209 \| 478 \| 246 \| 21 \| 1 \| 415 \| 2 \| 410 \| Uncharacterized protein \| Uncharacterized protein; afdb-uniprot50 \| AF-A0A7Z2TBJ7-F1-MODEL\_V4 \| 1.0 \| 2.863e-18 \| 404 \| 0.156 \| 781 \| 283 \| 37 \| 1 \| 423 \| 1 \| 763 \| Phage tail protein \| Phage tail protein; afdb-uniprot50 \| AF-A0A8A7WRM1-F1-MODEL\_V4 \| 1.0 \| 1.014e-19 \| 403 \| 0.251 \| 466 \| 230 \| 24 \| 1 \| 417 \| 1 \| 396 \| Phage tail protein \| Phage tail protein; afdb-uniprot50 \| AF-A0A3M4NA57-F1-MODEL\_V4 \| 1.0 \| 3.802e-17 \| 401 \| 0.244 \| 405 \| 201 \| 18 \| 1 \| 355 \| 5 \| 354 \| Tail fiber protein H \| Tail fiber protein H; afdb-uniprot50 \| AF-A0A1G9EQI8-F1-MODEL\_V4 \| 1.0 \| 4.343e-19 \| 401 \| 0.197 \| 581 \| 267 \| 29 \| 2 \| 416 \| 6 \| 552 \| Phage-related tail fibre protein \| Phage-related tail fibre protein; afdb-uniprot50 \| AF-A0A221FN03-F1-MODEL\_V4 \| 1.0 \| 2.506e-16 \| 400 \| 0.197 \| 364 \| 181 \| 12 \| 1 \| 258 \| 1 \| 359 \| Phage tail protein \| Phage tail protein; afdb-uniprot50 \| AF-A0A2D0ITF1-F1-MODEL\_V4 \| 1.0 \| 9.885e-18 \| 400 \| 0.234 \| 473 \| 175 \| 10 \| 1 \| 417 \| 2 \| 343 \| Phage tail fiber protein \| Phage tail fiber protein; afdb-uniprot50 \| AF-A0A7T1Y6B1-F1-MODEL\_V4 \| 1.0 \| 6.087e-18 \| 399 \| 0.228 \| 477 \| 214 \| 23 \| 2 \| 416 \| 6 \| 390 \| Phage tail protein \| Phage tail protein; afdb-uniprot50 \| AF-A0A855K828-F1-MODEL\_V4 \| 1.0 \| 1.763e-18 \| 399 \| 0.196 \| 515 \| 243 \| 21 \| 1 \| 421 \| 1 \| 438 \| Uncharacterized protein \| Uncharacterized protein; afdb-uniprot50 \| AF-A0A371YQG3-F1-MODEL\_V4 \| 1.0 \| 3.281e-16 \| 399 \| 0.194 \| 390 \| 181 \| 15 \| 1 \| 260 \| 2 \| 388 \| Uncharacterized protein \| Uncharacterized protein; afdb-uniprot50 \| AF-A0A3L0W1T3-F1-MODEL\_V4 \| 1.0 \| 3.551e-18 \| 398 \| 0.231 \| 466 \| 219 \| 20 \| 1 \| 416 \| 2 \| 378 \| Phage tail protein \| Phage tail protein; afdb-uniprot50 \| AF-A0A2J9GZ53-F1-MODEL\_V4 \| 1.0 \| 2.042e-19 \| 398 \| 0.221 \| 505 \| 261 \| 17 \| 1 \| 414 \| 4 \| 467 \| gp37\_C domain-containing protein \| gp37\_C domain-containing protein; afdb-uniprot50 \| AF-A0A0E3BR31-F1-MODEL\_V4 \| 1.0 \| 1.028e-18 \| 398 \| 0.236 \| 477 \| 232 \| 23 \| 1 \| 412 \| 1 \| 410 \| Tail protein \| Tail protein; afdb-uniprot50 \| AF-N9SPH7-F1-MODEL\_V4 \| 1.0 \| 6.087e-18 \| 398 \| 0.169 \| 577 \| 236 \| 25 \| 2 \| 421 \| 4 \| 494 \| Uncharacterized protein \| Uncharacterized protein; afdb-uniprot50 \| AF-A0A2K9QFQ7-F1-MODEL\_V4 \| 1.0 \| 9.233e-19 \| 397 \| 0.186 \| 567 \| 217 \| 23 \| 1 \| 421 \| 2 \| 469 \| Phage tail protein \| Phage tail protein; afdb-uniprot50 \| AF-A0A5C6YJN9-F1-MODEL\_V4 \| 1.0 \| 1.117e-16 \| 397 \| 0.237 \| 391 \| 206 \| 18 \| 1 \| 349 \| 10 \| 350 \| Uncharacterized protein \| Uncharacterized protein; afdb-uniprot50 \| AF-G3XD71-F1-MODEL\_V4 \| 1.0 \| 4.115e-19 \| 397 \| 0.151 \| 708 \| 295 \| 32 \| 1 \| 423 \| 5 \| 691 \| Probable bacteriophage protein \| Probable bacteriophage protein; afdb-uniprot50 \| AF-A0A3A8ECP1-F1-MODEL\_V4 \| 1.0 \| 1.196e-15 \| 396 \| 0.16 \| 437 \| 215 \| 20 \| 1 \| 421 \| 2 \| 302 \| Uncharacterized protein \| Uncharacterized protein; afdb-uniprot50 \| AF-A0A0M5MEM2-F1-MODEL\_V4 \| 1.0 \| 7.443e-19 \| 396 \| 0.18 \| 544 \| 265 \| 30 \| 3 \| 417 \| 2 \| 493 \| Phage tail-collar fiber protein (DUF3751 domain) \| Phage tail-collar fiber protein (DUF3751 domain); afdb-uniprot50 \| AF-A0A4P8SH52-F1-MODEL\_V4 \| 1.0 \| 6.683e-19 \| 396 \| 0.205 \| 593 \| 241 \| 27 \| 1 \| 421 \| 2 \| 536 \| Uncharacterized protein \| Uncharacterized protein; afdb-uniprot50 \| AF-A0A7L7MRD4-F1-MODEL\_V4 \| 1.0 \| 2.116e-09 \| 395 \| 0.376 \| 109 \| 66 \| 2 \| 1 \| 108 \| 1 \| 108 \| Uncharacterized protein \| Uncharacterized protein; afdb-uniprot50 \| AF-A0A2I1RIW9-F1-MODEL\_V4 \| 1.0 \| 1.047e-13 \| 395 \| 0.234 \| 269 \| 174 \| 10 \| 1 \| 261 \| 1 \| 245 \| Uncharacterized protein \| Uncharacterized protein; afdb-uniprot50 \| AF-A0A7Z2A8B6-F1-MODEL\_V4 \| 1.0 \| 1.67e-18 \| 395 \| 0.195 \| 471 \| 245 \| 24 \| 1 \| 422 \| 1 \| 386 \| Uncharacterized protein \| Uncharacterized protein; afdb-uniprot50 \| AF-A0A2K8VZI6-F1-MODEL\_V4 \| 1.0 \| 1.276e-18 \| 395 \| 0.2 \| 524 \| 213 \| 30 \| 1 \| 422 \| 2 \| 421 \| Phage tail fiber protein \| Phage tail fiber protein; afdb-uniprot50 \| AF-A0A345J3J2-F1-MODEL\_V4 \| 1.0 \| 2.042e-19 \| 395 \| 0.195 \| 502 \| 238 \| 26 \| 4 \| 412 \| 3 \| 431 \| Phage-related tail fiber protein \| Phage-related tail fiber protein; afdb-uniprot50 \| AF-A0A3P6JSJ9-F1-MODEL\_V4 \| 1.0 \| 9.233e-19 \| 395 \| 0.205 \| 541 \| 237 \| 25 \| 1 \| 421 \| 2 \| 469 \| Putative phage tail fiber protein \| Putative phage tail fiber protein; afdb-uniprot50 \| AF-A0A3F3A8F4-F1-MODEL\_V4 \| 1.0 \| 3.501e-19 \| 395 \| 0.219 \| 492 \| 252 \| 25 \| 1 \| 413 \| 1 \| 439 \| Putative phage tail fiber protein \| Putative phage tail fiber protein; afdb-uniprot50 \| AF-A0A5B8I5F0-F1-MODEL\_V4 \| 1.0 \| 3.021e-18 \| 395 \| 0.227 \| 496 \| 218 \| 19 \| 1 \| 385 \| 1 \| 442 \| Phage tail protein \| Phage tail protein; afdb-uniprot50 \| AF-A0A840MLL2-F1-MODEL\_V4 \| 1.0 \| 1.5e-18 \| 394 \| 0.209 \| 520 \| 252 \| 26 \| 1 \| 423 \| 1 \| 458 \| Phage-related tail fiber protein \| Phage-related tail fiber protein; afdb-uniprot50 \| AF-A0A0M4TW35-F1-MODEL\_V4 \| 1.0 \| 1.676e-14 \| 393 \| 0.238 \| 285 \| 181 \| 12 \| 3 \| 275 \| 2 \| 262 \| Uncharacterized protein \| Uncharacterized protein; afdb-uniprot50 \| AF-A0A0H3GKQ5-F1-MODEL\_V4 \| 1.0 \| 1.133e-15 \| 393 \| 0.243 \| 341 \| 157 \| 14 \| 1 \| 258 \| 1 \| 323 \| Uncharacterized protein \| Uncharacterized protein; afdb-uniprot50 \| AF-A0A072Y2M2-F1-MODEL\_V4 \| 1.0 \| 2.187e-18 \| 393 \| 0.204 \| 522 \| 235 \| 26 \| 1 \| 423 \| 2 \| 441 \| Putative phage tail fiber protein \| Putative phage tail fiber protein; afdb-uniprot50 \| AF-A0A248LIJ6-F1-MODEL\_V4 \| 1.0 \| 2.155e-19 \| 393 \| 0.204 \| 612 \| 226 \| 30 \| 2 \| 420 \| 3 \| 546 \| Tail fiber protein 2 \| Tail fiber protein 2; afdb-uniprot50 \| AF-A0A826U4X0-F1-MODEL\_V4 \| 1.0 \| 1.101e-17 \| 393 \| 0.216 \| 484 \| 231 \| 21 \| 1 \| 422 \| 1 \| 398 \| Phage tail protein \| Phage tail protein; afdb-uniprot50 \| AF-A0A6H3LMB7-F1-MODEL\_V4 \| 1.0 \| 2.771e-09 \| 392 \| 0.372 \| 129 \| 81 \| 0 \| 33 \| 161 \| 1 \| 129 \| Phage tail protein \| Phage tail protein; afdb-uniprot50 \| AF-A0A6S7CIQ6-F1-MODEL\_V4 \| 1.0 \| 5.178e-18 \| 392 \| 0.231 \| 462 \| 234 \| 25 \| 1 \| 404 \| 1 \| 399 \| Uncharacterized protein \| Uncharacterized protein; afdb-uniprot50 \| AF-A0A2K9CH40-F1-MODEL\_V4 \| 1.0 \| 4.405e-18 \| 392 \| 0.148 \| 849 \| 257 \| 32 \| 1 \| 423 \| 5 \| 813 \| Uncharacterized protein \| Uncharacterized protein; afdb-uniprot50 \| AF-A0A1B2QZE5-F1-MODEL\_V4 \| 1.0 \| 3.188e-18 \| 391 \| 0.229 \| 506 \| 223 \| 29 \| 1 \| 413 \| 1 \| 432 \| Uncharacterized protein \| Uncharacterized protein; afdb-uniprot50 \| AF-A0A7Y8ZMJ1-F1-MODEL\_V4 \| 1.0 \| 7.053e-19 \| 391 \| 0.199 \| 511 \| 272 \| 27 \| 1 \| 422 \| 1 \| 463 \| Phage-related tail fiber protein \| Phage-related tail fiber protein; afdb-uniprot50 \| AF-A0A1S8T0S9-F1-MODEL\_V4 \| 1.0 \| 2.401e-19 \| 391 \| 0.193 \| 549 \| 267 \| 28 \| 1 \| 423 \| 2 \| 500 \| Uncharacterized protein \| Uncharacterized protein; afdb-uniprot50 \| AF-A0A0D6BCT9-F1-MODEL\_V4 \| 1.0 \| 1.583e-18 \| 391 \| 0.151 \| 814 \| 283 \| 36 \| 2 \| 423 \| 6 \| 803 \| Phage-related tail fibre protein-like protein \| Phage-related tail fibre protein-like protein; afdb-uniprot50 \| AF-A0A0A8FH85-F1-MODEL\_V4 \| 1.0 \| 4.405e-18 \| 390 \| 0.21 \| 518 \| 257 \| 20 \| 1 \| 416 \| 1 \| 468 \| Collar domain-containing protein \| Collar domain-containing protein; afdb-uniprot50 \| AF-A0A240TRE0-F1-MODEL\_V4 \| 1.0 \| 1.763e-18 \| 389 \| 0.222 \| 499 \| 214 \| 24 \| 1 \| 416 \| 1 \| 408 \| Uncharacterized protein \| Uncharacterized protein; afdb-uniprot50 \| AF-A0A1N6MZM9-F1-MODEL\_V4 \| 1.0 \| 6.706e-15 \| 389 \| 0.233 \| 334 \| 121 \| 11 \| 1 \| 315 \| 2 \| 219 \| Uncharacterized protein \| Uncharacterized protein; afdb-uniprot50 \| AF-B2TMU2-F1-MODEL\_V4 \| 1.0 \| 7.155e-18 \| 388 \| 0.197 \| 521 \| 261 \| 27 \| 1 \| 423 \| 2 \| 463 \| Putative tail fiber protein \| Putative tail fiber protein; afdb-uniprot50 \| AF-A0A239C905-F1-MODEL\_V4 \| 1.0 \| 6.683e-19 \| 388 \| 0.171 \| 612 \| 250 \| 28 \| 4 \| 423 \| 3 \| 549 \| Phage tail-collar fibre protein \| Phage tail-collar fibre protein; afdb-uniprot50 \| AF-A0A7Y3ILG7-F1-MODEL\_V4 \| 1.0 \| 1.846e-11 \| 387 \| 0.212 \| 221 \| 134 \| 9 \| 6 \| 204 \| 10 \| 212 \| Uncharacterized protein \| Uncharacterized protein; afdb-uniprot50 \| AF-A7ZNA3-F1-MODEL\_V4 \| 1.0 \| 6.706e-15 \| 387 \| 0.217 \| 322 \| 168 \| 15 \| 1 \| 258 \| 3 \| 304 \| Uncharacterized protein \| Uncharacterized protein; afdb-uniprot50 \| AF-A0A3M5P7X5-F1-MODEL\_V4 \| 1.0 \| 2.187e-18 \| 387 \| 0.182 \| 538 \| 213 \| 18 \| 2 \| 423 \| 6 \| 432 \| Phage tail fiber protein h \| Phage tail fiber protein h; afdb-uniprot50 \| AF-A0A157QPJ3-F1-MODEL\_V4 \| 1.0 \| 1.441e-17 \| 387 \| 0.232 \| 404 \| 211 \| 15 \| 1 \| 351 \| 1 \| 358 \| Tail fiber protein \| Tail fiber protein; afdb-uniprot50 \| AF-A0A0E3M8H3-F1-MODEL\_V4 \| 1.0 \| 3.748e-18 \| 386 \| 0.207 \| 454 \| 262 \| 16 \| 1 \| 406 \| 2 \| 405 \| Putative phage tail fiber protein \| Putative phage tail fiber protein; afdb-uniprot50 \| AF-M1MEX9-F1-MODEL\_V4 \| 1.0 \| 4.174e-18 \| 386 \| 0.182 \| 538 \| 290 \| 28 \| 1 \| 423 \| 1 \| 503 \| Putative phage tail fiber protein \| Putative phage tail fiber protein; afdb-uniprot50 \| AF-Q4QK27-F1-MODEL\_V4 \| 1.0 \| 1.043e-17 \| 386 \| 0.214 \| 439 \| 218 \| 21 \| 1 \| 415 \| 2 \| 337 \| Probable tail fiber protein \| Probable tail fiber protein; afdb-uniprot50 \| AF-N9KFA7-F1-MODEL\_V4 \| 1.0 \| 8.779e-15 \| 385 \| 0.183 \| 337 \| 168 \| 16 \| 2 \| 258 \| 3 \| 312 \| Uncharacterized protein \| Uncharacterized protein; afdb-uniprot50 \| AF-A0A0P9N6Y3-F1-MODEL\_V4 \| 1.0 \| 3.602e-17 \| 385 \| 0.214 \| 550 \| 255 \| 26 \| 2 \| 423 \| 6 \| 506 \| Tail fiber protein H \| Tail fiber protein H; afdb-uniprot50 \| AF-A0A1C4DMT0-F1-MODEL\_V4 \| 1.0 \| 7.855e-19 \| 384 \| 0.203 \| 579 \| 216 \| 28 \| 1 \| 421 \| 2 \| 493 \| Phage tail-collar fibre protein \| Phage tail-collar fibre protein; afdb-uniprot50 \| AF-A0A509JDS2-F1-MODEL\_V4 \| 1.0 \| 3.365e-18 \| 384 \| 0.163 \| 717 \| 282 \| 31 \| 1 \| 423 \| 1 \| 693 \| Phage tail protein \| Phage tail protein; afdb-uniprot50 \| AF-A0A0H3ZXX7-F1-MODEL\_V4 \| 1.0 \| 3.365e-18 \| 384 \| 0.217 \| 520 \| 242 \| 28 \| 1 \| 417 \| 1 \| 458 \| Tail fiber protein \| Tail fiber protein; afdb-uniprot50 \| AF-A0A0P9YJT9-F1-MODEL\_V4 \| 1.0 \| 4.907e-18 \| 383 \| 0.208 \| 570 \| 252 \| 28 \| 1 \| 407 \| 5 \| 538 \| Putative Tail fiber protein H \| Putative Tail fiber protein H; afdb-uniprot50 \| AF-A0A725RYD2-F1-MODEL\_V4 \| 1.0 \| 1.028e-18 \| 383 \| 0.181 \| 705 \| 261 \| 33 \| 1 \| 421 \| 1 \| 673 \| Phage tail protein \| Phage tail protein; afdb-uniprot50 \| AF-A0A2N8B960-F1-MODEL\_V4 \| 1.0 \| 1.887e-17 \| 382 \| 0.191 \| 537 \| 226 \| 27 \| 2 \| 421 \| 6 \| 451 \| Phage tail protein \| Phage tail protein; afdb-uniprot50 \| AF-A0A2K4GCZ3-F1-MODEL\_V4 \| 1.0 \| 2.102e-17 \| 382 \| 0.207 \| 487 \| 261 \| 20 \| 2 \| 411 \| 6 \| 444 \| Uncharacterized protein \| Uncharacterized protein; afdb-uniprot50 \| AF-A0A6D2GE62-F1-MODEL\_V4 \| 1.0 \| 1.153e-10 \| 381 \| 0.434 \| 145 \| 77 \| 1 \| 32 \| 176 \| 2 \| 141 \| Tail protein \| Tail protein; afdb-uniprot50 \| AF-A0A127MM56-F1-MODEL\_V4 \| 1.0 \| 1.366e-17 \| 380 \| 0.259 \| 490 \| 217 \| 28 \| 1 \| 417 \| 1 \| 417 \| Uncharacterized protein \| Uncharacterized protein; afdb-uniprot50 \| AF-A0A423IG29-F1-MODEL\_V4 \| 1.0 \| 2.471e-17 \| 380 \| 0.202 \| 450 \| 247 \| 20 \| 1 \| 410 \| 5 \| 382 \| Uncharacterized protein \| Uncharacterized protein; afdb-uniprot50 \| AF-B5JYG6-F1-MODEL\_V4 \| 1.0 \| 5.122e-15 \| 379 \| 0.436 \| 188 \| 101 \| 4 \| 83 \| 268 \| 2 \| 186 \| Phage Tail Collar Domain family \| Phage Tail Collar Domain family; afdb-uniprot50 \| AF-A0A6S5JPB2-F1-MODEL\_V4 \| 1.0 \| 4.649e-18 \| 379 \| 0.228 \| 529 \| 210 \| 25 \| 1 \| 412 \| 2 \| 449 \| Uncharacterized protein \| Uncharacterized protein; afdb-uniprot50 \| AF-A0A1G6JDL2-F1-MODEL\_V4 \| 1.0 \| 1.244e-16 \| 379 \| 0.147 \| 575 \| 230 \| 22 \| 1 \| 421 \| 3 \| 471 \| Phage tail-collar fibre protein \| Phage tail-collar fibre protein; afdb-uniprot50 \| AF-A0A855HWY9-F1-MODEL\_V4 \| 1.0 \| 6.779e-18 \| 379 \| 0.191 \| 518 \| 261 \| 18 \| 1 \| 412 \| 2 \| 467 \| Uncharacterized protein \| Uncharacterized protein; afdb-uniprot50 \| AF-A0A8B6KUE2-F1-MODEL\_V4 \| 1.0 \| 6.087e-18 \| 378 \| 0.218 \| 508 \| 255 \| 22 \| 1 \| 418 \| 1 \| 456 \| Variable tail fiber protein \| Variable tail fiber protein; afdb-uniprot50 \| AF-Q8ZMU5-F1-MODEL\_V4 \| 1.0 \| 2.187e-18 \| 378 \| 0.207 \| 583 \| 224 \| 23 \| 1 \| 421 \| 2 \| 508 \| Fels-2 prophage protein \| Fels-2 prophage protein; afdb-uniprot50 \| AF-A0A5F0JKE5-F1-MODEL\_V4 \| 1.0 \| 2.341e-17 \| 378 \| 0.159 \| 683 \| 252 \| 35 \| 2 \| 423 \| 6 \| 627 \| Uncharacterized protein \| Uncharacterized protein; afdb-uniprot50 \| AF-A0A7Y8Z4M8-F1-MODEL\_V4 \| 1.0 \| 9.885e-18 \| 378 \| 0.163 \| 569 \| 251 \| 30 \| 1 \| 422 \| 1 \| 491 \| Uncharacterized protein \| Uncharacterized protein; afdb-uniprot50 \| AF-A0A1I5D5R7-F1-MODEL\_V4 \| 1.0 \| 6.354e-15 \| 378 \| 0.23 \| 325 \| 138 \| 9 \| 1 \| 323 \| 2 \| 216 \| Phage tail fibre repeat-containing protein \| Phage tail fibre repeat-containing protein; afdb-uniprot50 \| AF-A0A7X1KID4-F1-MODEL\_V4 \| 1.0 \| 3.021e-18 \| 377 \| 0.19 \| 551 \| 234 \| 24 \| 4 \| 423 \| 8 \| 477 \| Phage tail protein \| Phage tail protein; afdb-uniprot50 \| AF-A0A6I5RT98-F1-MODEL\_V4 \| 1.0 \| 1.652e-15 \| 377 \| 0.259 \| 378 \| 193 \| 13 \| 2 \| 310 \| 6 \| 365 \| Uncharacterized protein \| Uncharacterized protein; afdb-uniprot50 \| AF-A0A353G7W2-F1-MODEL\_V4 \| 1.0 \| 4.129e-15 \| 376 \| 0.224 \| 347 \| 165 \| 11 \| 1 \| 306 \| 2 \| 285 \| Uncharacterized protein \| Uncharacterized protein; afdb-uniprot50 \| AF-A0A1H8SZT3-F1-MODEL\_V4 \| 1.0 \| 4.649e-18 \| 375 \| 0.197 \| 568 \| 274 \| 27 \| 1 \| 421 \| 1 \| 533 \| Phage-related tail fibre protein \| Phage-related tail fibre protein; afdb-uniprot50 \| AF-A0A3G2IKC5-F1-MODEL\_V4 \| 1.0 \| 1.887e-17 \| 375 \| 0.171 \| 566 \| 245 \| 25 \| 1 \| 412 \| 1 \| 496 \| Uncharacterized protein \| Uncharacterized protein; afdb-uniprot50 \| AF-A0A379YEC2-F1-MODEL\_V4 \| 1.0 \| 2.506e-16 \| 373 \| 0.154 \| 485 \| 247 \| 25 \| 1 \| 420 \| 1 \| 387 \| Uncharacterized protein \| Uncharacterized protein; afdb-uniprot50 \| AF-A0A4P7L2V6-F1-MODEL\_V4 \| 1.0 \| 2.57e-18 \| 373 \| 0.231 \| 513 \| 237 \| 23 \| 4 \| 416 \| 3 \| 458 \| Phage tail-collar fiber protein \| Phage tail-collar fiber protein; afdb-uniprot50 \| AF-A0A2T5NRB6-F1-MODEL\_V4 \| 1.0 \| 6.877e-17 \| 373 \| 0.218 \| 498 \| 239 \| 26 \| 1 \| 423 \| 2 \| 424 \| Uncharacterized protein \| Uncharacterized protein; afdb-uniprot50 \| AF-A0A7Y1MB50-F1-MODEL\_V4 \| 1.0 \| 1.605e-17 \| 373 \| 0.187 \| 544 \| 268 \| 25 \| 2 \| 412 \| 6 \| 508 \| Phage tail protein \| Phage tail protein; afdb-uniprot50 \| AF-F9GSW5-F1-MODEL\_V4 \| 1.0 \| 7.443e-19 \| 373 \| 0.126 \| 746 \| 289 \| 27 \| 1 \| 423 \| 2 \| 707 \| Phage tail collar domain-containing protein \| Phage tail collar domain-containing protein; afdb-uniprot50 \| AF-A0A855ISI7-F1-MODEL\_V4 \| 1.0 \| 4.682e-10 \| 372 \| 0.185 \| 199 \| 126 \| 5 \| 6 \| 172 \| 10 \| 204 \| Uncharacterized protein \| Uncharacterized protein; afdb-uniprot50 \| AF-A0A0A8TLD1-F1-MODEL\_V4 \| 1.0 \| 9.885e-18 \| 372 \| 0.187 \| 559 \| 246 \| 31 \| 1 \| 412 \| 1 \| 498 \| Phage tail fiber protein \| Phage tail fiber protein; afdb-uniprot50 \| AF-A0A6G9RP95-F1-MODEL\_V4 \| 1.0 \| 3.109e-16 \| 371 \| 0.166 \| 480 \| 215 \| 20 \| 1 \| 421 \| 3 \| 356 \| Phage tail protein \| Phage tail protein; afdb-uniprot50 \| AF-A0A6M0YIR0-F1-MODEL\_V4 \| 1.0 \| 1.313e-16 \| 371 \| 0.179 \| 452 \| 246 \| 20 \| 1 \| 416 \| 2 \| 364 \| Uncharacterized protein \| Uncharacterized protein; afdb-uniprot50 \| AF-A0A7C8HXB6-F1-MODEL\_V4 \| 1.0 \| 1.605e-17 \| 371 \| 0.217 \| 533 \| 231 \| 23 \| 1 \| 414 \| 1 \| 466 \| Uncharacterized protein \| Uncharacterized protein; afdb-uniprot50 \| AF-A0A708DGG9-F1-MODEL\_V4 \| 1.0 \| 4.405e-18 \| 371 \| 0.196 \| 576 \| 222 \| 28 \| 1 \| 421 \| 1 \| 490 \| Uncharacterized protein \| Uncharacterized protein; afdb-uniprot50 \| AF-A0A1N7LRE7-F1-MODEL\_V4 \| 1.0 \| 1.39e-12 \| 371 \| 0.19 \| 315 \| 147 \| 8 \| 1 \| 310 \| 1 \| 212 \| Phage tail-collar fibre protein \| Phage tail-collar fibre protein; afdb-uniprot50 \| AF-A0A2H5E1S9-F1-MODEL\_V4 \| 1.0 \| 5.253e-17 \| 371 \| 0.212 \| 457 \| 235 \| 20 \| 2 \| 418 \| 3 \| 374 \| Uncharacterized protein \| Uncharacterized protein; afdb-uniprot50 \| AF-A0A379CAC0-F1-MODEL\_V4 \| 1.0 \| 2.375e-16 \| 371 \| 0.178 \| 414 \| 177 \| 10 \| 1 \| 412 \| 2 \| 254 \| Phage tail fibre repeat \| Phage tail fibre repeat; afdb-uniprot50 \| AF-M1MI28-F1-MODEL\_V4 \| 1.0 \| 6.087e-18 \| 371 \| 0.159 \| 582 \| 247 \| 25 \| 1 \| 422 \| 2 \| 501 \| Putative tail fiber protein \| Putative tail fiber protein; afdb-uniprot50 \| AF-A0A7W4RSH8-F1-MODEL\_V4 \| 1.0 \| 3.463e-16 \| 370 \| 0.202 \| 459 \| 231 \| 16 \| 1 \| 416 \| 1 \| 367 \| Phage-related tail fiber protein \| Phage-related tail fiber protein; afdb-uniprot50 \| AF-B1B8U4-F1-MODEL\_V4 \| 1.0 \| 1.226e-17 \| 370 \| 0.178 \| 559 \| 241 \| 21 \| 1 \| 421 \| 1 \| 479 \| Putative phage tail fiber protein \| Putative phage tail fiber protein; afdb-uniprot50 \| AF-A0A4P7LY25-F1-MODEL\_V4 \| 1.0 \| 7.155e-18 \| 370 \| 0.184 \| 705 \| 259 \| 37 \| 1 \| 421 \| 1 \| 673 \| Phage tail protein \| Phage tail protein; afdb-uniprot50 \| AF-A0A368TXZ3-F1-MODEL\_V4 \| 1.0 \| 7.66e-17 \| 369 \| 0.167 \| 536 \| 218 \| 19 \| 2 \| 416 \| 6 \| 434 \| Collar domain-containing protein \| Collar domain-containing protein; afdb-uniprot50 \| AF-A0A840M5X8-F1-MODEL\_V4 \| 1.0 \| 3.748e-18 \| 369 \| 0.161 \| 601 \| 277 \| 28 \| 1 \| 421 \| 2 \| 555 \| Uncharacterized protein \| Uncharacterized protein; afdb-uniprot50 \| AF-A0A488A766-F1-MODEL\_V4 \| 1.0 \| 7.308e-09 \| 368 \| 0.385 \| 114 \| 63 \| 2 \| 1 \| 111 \| 1 \| 110 \| Tail fiber protein \| Tail fiber protein; afdb-uniprot50 \| AF-A0A379FY21-F1-MODEL\_V4 \| 1.0 \| 1.226e-17 \| 367 \| 0.194 \| 612 \| 248 \| 31 \| 4 \| 421 \| 3 \| 563 \| Uncharacterized protein \| Uncharacterized protein; afdb-uniprot50 \| AF-A0A8B4MIL4-F1-MODEL\_V4 \| 1.0 \| 6.087e-18 \| 367 \| 0.192 \| 536 \| 272 \| 24 \| 1 \| 412 \| 1 \| 499 \| Bifunctional phage-like tail fiber protein/heme utilization protein \| Bifunctional phage-like tail fiber protein/heme utilization protein; afdb-uniprot50 \| AF-B0P727-F1-MODEL\_V4 \| 1.0 \| 2.791e-16 \| 366 \| 0.146 \| 497 \| 242 \| 23 \| 3 \| 420 \| 2 \| 395 \| Uncharacterized protein \| Uncharacterized protein; afdb-uniprot50 \| AF-A0A5T0CRZ1-F1-MODEL\_V4 \| 1.0 \| 6.264e-16 \| 366 \| 0.186 \| 402 \| 175 \| 14 \| 2 \| 258 \| 4 \| 398 \| Phage tail protein \| Phage tail protein; afdb-uniprot50 \| AF-A0A7Z1GL46-F1-MODEL\_V4 \| 1.0 \| 3.065e-17 \| 365 \| 0.196 \| 581 \| 267 \| 32 \| 2 \| 422 \| 6 \| 546 \| Tail collar domain \| Tail collar domain; afdb-uniprot50 \| AF-A0A6P2SH75-F1-MODEL\_V4 \| 1.0 \| 7.258e-17 \| 364 \| 0.216 \| 555 \| 257 \| 26 \| 1 \| 420 \| 2 \| 513 \| Uncharacterized protein \| Uncharacterized protein; afdb-uniprot50 \| AF-A0A380AJ63-F1-MODEL\_V4 \| 1.0 \| 2.791e-16 \| 364 \| 0.16 \| 516 \| 259 \| 24 \| 1 \| 414 \| 1 \| 444 \| Caudovirales tail fibre assembly protein \| Caudovirales tail fibre assembly protein; afdb-uniprot50 \| AF-A0A845BZU2-F1-MODEL\_V4 \| 1.0 \| 1.366e-17 \| 364 \| 0.204 \| 573 \| 264 \| 29 \| 1 \| 421 \| 1 \| 533 \| Uncharacterized protein \| Uncharacterized protein; afdb-uniprot50 \| AF-A0A3M3YCB8-F1-MODEL\_V4 \| 1.0 \| 7.258e-17 \| 364 \| 0.214 \| 521 \| 263 \| 27 \| 1 \| 412 \| 1 \| 483 \| Uncharacterized protein \| Uncharacterized protein; afdb-uniprot50 \| AF-N8XQE4-F1-MODEL\_V4 \| 1.0 \| 2.713e-18 \| 362 \| 0.171 \| 612 \| 267 \| 28 \| 1 \| 421 \| 4 \| 566 \| Pectate\_lyase\_3 domain-containing protein \| Pectate\_lyase\_3 domain-containing protein; afdb-uniprot50 \| AF-A0A378NBM0-F1-MODEL\_V4 \| 1.0 \| 1.992e-17 \| 362 \| 0.19 \| 578 \| 223 \| 27 \| 1 \| 422 \| 2 \| 490 \| Catalytic beta propeller domain of bacteriophage endosialidase \| Catalytic beta propeller domain of bacteriophage endosialidase; afdb-uniprot50 \| AF-E6WD86-F1-MODEL\_V4 \| 1.0 \| 1.605e-17 \| 361 \| 0.174 \| 597 \| 216 \| 25 \| 1 \| 421 \| 1 \| 496 \| Phage-related tail fibre protein-like protein \| Phage-related tail fibre protein-like protein; afdb-uniprot50 \| AF-A0A2S9X9N0-F1-MODEL\_V4 \| 1.0 \| 1.462e-16 \| 361 \| 0.153 \| 708 \| 249 \| 31 \| 1 \| 423 \| 1 \| 642 \| Uncharacterized protein \| Uncharacterized protein; afdb-uniprot50 \| AF-A0A6C2DDF4-F1-MODEL\_V4 \| 1.0 \| 3.365e-18 \| 361 \| 0.165 \| 683 \| 290 \| 36 \| 1 \| 423 \| 2 \| 664 \| gp37\_C domain-containing protein \| gp37\_C domain-containing protein; afdb-uniprot50 \| AF-A0A2T5KPN3-F1-MODEL\_V4 \| 1.0 \| 1.543e-16 \| 361 \| 0.21 \| 494 \| 253 \| 24 \| 1 \| 411 \| 5 \| 444 \| Tail collar domain \| Tail collar domain; afdb-uniprot50 \| AF-A0A530M9E5-F1-MODEL\_V4 \| 1.0 \| 1.162e-17 \| 360 \| 0.199 \| 506 \| 230 \| 23 \| 1 \| 413 \| 2 \| 425 \| Phage tail protein \| Phage tail protein; afdb-uniprot50 \| AF-A0A7Y8FLP5-F1-MODEL\_V4 \| 1.0 \| 6.516e-17 \| 360 \| 0.205 \| 512 \| 268 \| 26 \| 1 \| 412 \| 1 \| 473 \| Phage tail protein \| Phage tail protein; afdb-uniprot50 \| AF-A0A7U9D8U8-F1-MODEL\_V4 \| 1.0 \| 2.102e-17 \| 358 \| 0.183 \| 517 \| 266 \| 29 \| 1 \| 418 \| 2 \| 461 \| Uncharacterized protein \| Uncharacterized protein; afdb-uniprot50 \| AF-A0A483W7T5-F1-MODEL\_V4 \| 1.0 \| 1.992e-17 \| 358 \| 0.203 \| 635 \| 228 \| 31 \| 1 \| 422 \| 1 \| 570 \| Phage tail protein \| Phage tail protein; afdb-uniprot50 \| AF-A0A383RR92-F1-MODEL\_V4 \| 1.0 \| 6.877e-17 \| 358 \| 0.218 \| 567 \| 239 \| 29 \| 2 \| 409 \| 6 \| 527 \| Putative tail fiber protein \| Putative tail fiber protein; afdb-uniprot50 \| AF-A0A5E4YCM3-F1-MODEL\_V4 \| 1.0 \| 4.469e-17 \| 357 \| 0.171 \| 611 \| 241 \| 30 \| 1 \| 422 \| 2 \| 535 \| Uncharacterized protein \| Uncharacterized protein; afdb-uniprot50 \| AF-W7S9K3-F1-MODEL\_V4 \| 1.0 \| 1.366e-17 \| 356 \| 0.207 \| 458 \| 251 \| 22 \| 1 \| 405 \| 2 \| 400 \| Tail fiber protein \| Tail fiber protein; afdb-uniprot50 \| AF-A0A857E7Z2-F1-MODEL\_V4 \| 1.0 \| 1.543e-16 \| 356 \| 0.169 \| 454 \| 245 \| 17 \| 1 \| 421 \| 1 \| 355 \| Uncharacterized protein \| Uncharacterized protein; afdb-uniprot50 \| AF-A0A378YNC7-F1-MODEL\_V4 \| 1.0 \| 1.117e-16 \| 355 \| 0.205 \| 517 \| 206 \| 26 \| 1 \| 417 \| 2 \| 413 \| Uncharacterized protein \| Uncharacterized protein; afdb-uniprot50 \| AF-A0A5E5AEQ9-F1-MODEL\_V4 \| 1.0 \| 6.516e-17 \| 354 \| 0.2 \| 550 \| 222 \| 27 \| 1 \| 422 \| 2 \| 461 \| Tail fiber protein \| Tail fiber protein; afdb-uniprot50 \| AF-A0A5U0PXD3-F1-MODEL\_V4 \| 1.0 \| 2.752e-17 \| 354 \| 0.173 \| 537 \| 240 \| 24 \| 2 \| 421 \| 9 \| 458 \| Uncharacterized protein \| Uncharacterized protein; afdb-uniprot50 \| AF-A0A4V1IMV1-F1-MODEL\_V4 \| 1.0 \| 1.992e-17 \| 353 \| 0.181 \| 555 \| 290 \| 22 \| 1 \| 420 \| 1 \| 525 \| Uncharacterized protein \| Uncharacterized protein; afdb-uniprot50 \| AF-A0A239SWC2-F1-MODEL\_V4 \| 1.0 \| 2.946e-16 \| 353 \| 0.214 \| 457 \| 233 \| 26 \| 1 \| 392 \| 2 \| 397 \| Uncharacterized protein \| Uncharacterized protein; afdb-uniprot50 \| AF-A0A1C7C2V4-F1-MODEL\_V4 \| 1.0 \| 4.469e-17 \| 352 \| 0.147 \| 798 \| 286 \| 34 \| 1 \| 423 \| 1 \| 778 \| Uncharacterized protein \| Uncharacterized protein; afdb-uniprot50 \| AF-A0A5E7VM25-F1-MODEL\_V4 \| 1.0 \| 2.02e-16 \| 351 \| 0.2 \| 529 \| 282 \| 21 \| 2 \| 422 \| 6 \| 501 \| Uncharacterized protein \| Uncharacterized protein; afdb-uniprot50 \| AF-A0A6G8D7U0-F1-MODEL\_V4 \| 1.0 \| 2.163e-15 \| 349 \| 0.185 \| 480 \| 253 \| 20 \| 1 \| 423 \| 2 \| 400 \| Phage tail protein \| Phage tail protein; afdb-uniprot50 \| AF-A0A1Y3CH38-F1-MODEL\_V4 \| 1.0 \| 1.179e-16 \| 348 \| 0.18 \| 487 \| 277 \| 25 \| 1 \| 413 \| 1 \| 439 \| Collar domain-containing protein \| Collar domain-containing protein; afdb-uniprot50 \| AF-A0A6N7BXN9-F1-MODEL\_V4 \| 1.0 \| 1.231e-13 \| 347 \| 0.203 \| 325 \| 185 \| 11 \| 1 \| 256 \| 1 \| 320 \| Uncharacterized protein \| Uncharacterized protein; afdb-uniprot50 \| AF-A0A5E4XZ01-F1-MODEL\_V4 \| 1.0 \| 5.329e-16 \| 343 \| 0.196 \| 499 \| 245 \| 28 \| 1 \| 413 \| 2 \| 430 \| Uncharacterized protein \| Uncharacterized protein; afdb-uniprot50 \| AF-A0A177RPE9-F1-MODEL\_V4 \| 1.0 \| 6.175e-17 \| 342 \| 0.207 \| 486 \| 252 \| 23 \| 1 \| 412 \| 1 \| 427 \| Tail protein \| Tail protein; afdb-uniprot50 \| AF-A0A241WFG7-F1-MODEL\_V4 \| 1.0 \| 6.108e-14 \| 341 \| 0.209 \| 334 \| 169 \| 16 \| 2 \| 255 \| 3 \| 321 \| Uncharacterized protein \| Uncharacterized protein; afdb-uniprot50 \| AF-A0A1W2DYX6-F1-MODEL\_V4 \| 1.0 \| 1.914e-16 \| 340 \| 0.185 \| 554 \| 251 \| 30 \| 1 \| 422 \| 1 \| 485 \| Uncharacterized protein \| Uncharacterized protein; afdb-uniprot50 \| AF-X6QAX6-F1-MODEL\_V4 \| 1.0 \| 1.942e-15 \| 338 \| 0.186 \| 462 \| 232 \| 23 \| 1 \| 419 \| 1 \| 361 \| Tail fiber protein \| Tail fiber protein; afdb-uniprot50 \| AF-A0A1H5GLX5-F1-MODEL\_V4 \| 1.0 \| 4.296e-16 \| 338 \| 0.19 \| 536 \| 244 \| 27 \| 1 \| 423 \| 1 \| 459 \| Phage Tail Collar Domain \| Phage Tail Collar Domain; afdb-uniprot50 \| AF-A0A6M0CVM5-F1-MODEL\_V4 \| 1.0 \| 3.463e-16 \| 338 \| 0.208 \| 480 \| 227 \| 26 \| 2 \| 378 \| 6 \| 435 \| Phage tail protein \| Phage tail protein; afdb-uniprot50 \| AF-C3KCU2-F1-MODEL\_V4 \| 1.0 \| 6.976e-16 \| 337 \| 0.157 \| 678 \| 288 \| 31 \| 1 \| 421 \| 1 \| 651 \| Putative phage tail fiber-related protein \| Putative phage tail fiber-related protein; afdb-uniprot50 \| AF-A0A367MED4-F1-MODEL\_V4 \| 1.0 \| 2.163e-15 \| 336 \| 0.194 \| 478 \| 232 \| 25 \| 1 \| 415 \| 1 \| 388 \| Uncharacterized protein \| Uncharacterized protein; afdb-uniprot50 \| AF-A0A2T5J1H4-F1-MODEL\_V4 \| 1.0 \| 9.134e-16 \| 331 \| 0.178 \| 465 \| 261 \| 20 \| 1 \| 421 \| 1 \| 388 \| Tail-collar fiber protein \| Tail-collar fiber protein; afdb-uniprot50 \| AF-V8RCP1-F1-MODEL\_V4 \| 1.0 \| 1.942e-15 \| 331 \| 0.192 \| 494 \| 252 \| 24 \| 2 \| 423 \| 6 \| 424 \| Phage tail protein \| Phage tail protein; afdb-uniprot50 \| AF-A0A1B8QCU5-F1-MODEL\_V4 \| 1.0 \| 4.599e-15 \| 331 \| 0.187 \| 431 \| 208 \| 20 \| 1 \| 407 \| 1 \| 313 \| Uncharacterized protein \| Uncharacterized protein; afdb-uniprot50 \| AF-A0A2T0YWF7-F1-MODEL\_V4 \| 1.0 \| 1.133e-15 \| 330 \| 0.213 \| 446 \| 221 \| 23 \| 32 \| 423 \| 2 \| 371 \| Tail-collar fiber protein \| Tail-collar fiber protein; afdb-uniprot50 \| AF-A0A6B3J8L4-F1-MODEL\_V4 \| 1.0 \| 9.639e-16 \| 330 \| 0.169 \| 525 \| 258 \| 25 \| 1 \| 423 \| 1 \| 449 \| Uncharacterized protein \| Uncharacterized protein; afdb-uniprot50 \| AF-A0A1N6UIC0-F1-MODEL\_V4 \| 1.0 \| 5.049e-16 \| 329 \| 0.189 \| 527 \| 233 \| 25 \| 2 \| 421 \| 6 \| 445 \| Phage tail-collar fibre protein \| Phage tail-collar fibre protein; afdb-uniprot50 \| AF-A0A376P7J4-F1-MODEL\_V4 \| 1.0 \| 6.633e-12 \| 329 \| 0.256 \| 292 \| 114 \| 4 \| 32 \| 323 \| 2 \| 190 \| Putative phage tail fiber protein \| Putative phage tail fiber protein; afdb-uniprot50 \| AF-A0A330EWK9-F1-MODEL\_V4 \| 1.0 \| 1.074e-15 \| 327 \| 0.162 \| 510 \| 252 \| 20 \| 1 \| 421 \| 2 \| 425 \| Phage tail protein \| Phage tail protein; afdb-uniprot50 \| AF-A0A6M0SQX7-F1-MODEL\_V4 \| 1.0 \| 6.976e-16 \| 327 \| 0.162 \| 626 \| 267 \| 26 \| 1 \| 423 \| 1 \| 571 \| Uncharacterized protein \| Uncharacterized protein; afdb-uniprot50 \| AF-A0A1G5ACS0-F1-MODEL\_V4 \| 1.0 \| 5.544e-17 \| 323 \| 0.167 \| 555 \| 294 \| 28 \| 1 \| 423 \| 1 \| 519 \| Phage tail-collar fibre protein \| Phage tail-collar fibre protein; afdb-uniprot50 \| AF-A0A1C3CVN3-F1-MODEL\_V4 \| 1.0 \| 1.744e-15 \| 320 \| 0.169 \| 556 \| 280 \| 31 \| 1 \| 421 \| 1 \| 509 \| Uncharacterized protein \| Uncharacterized protein; afdb-uniprot50 \| AF-A0A368V9P8-F1-MODEL\_V4 \| 1.0 \| 1.386e-16 \| 315 \| 0.186 \| 516 \| 256 \| 24 \| 17 \| 421 \| 2 \| 464 \| Tail-collar fiber protein \| Tail-collar fiber protein; afdb-uniprot50 \| AF-A0A2W6YCW4-F1-MODEL\_V4 \| 1.0 \| 9.778e-15 \| 314 \| 0.165 \| 495 \| 269 \| 21 \| 1 \| 421 \| 1 \| 425 \| Uncharacterized protein \| Uncharacterized protein; afdb-uniprot50 \| AF-A0A150HM63-F1-MODEL\_V4 \| 1.0 \| 1.406e-15 \| 314 \| 0.17 \| 552 \| 268 \| 27 \| 1 \| 402 \| 1 \| 512 \| Uncharacterized protein \| Uncharacterized protein; afdb-uniprot50 \| AF-U1JBI7-F1-MODEL\_V4 \| 1.0 \| 1.089e-14 \| 312 \| 0.179 \| 467 \| 273 \| 27 \| 1 \| 414 \| 3 \| 412 \| Putative side tail phage protein \| Putative side tail phage protein; afdb-uniprot50 \| AF-A0A7Y3T9N6-F1-MODEL\_V4 \| 1.0 \| 3.154e-15 \| 311 \| 0.153 \| 535 \| 271 \| 23 \| 1 \| 423 \| 1 \| 465 \| DUF2793 domain-containing protein \| DUF2793 domain-containing protein; afdb-uniprot50 \| AF-U5MTV4-F1-MODEL\_V4 \| 1.0 \| 3.377e-14 \| 310 \| 0.125 \| 783 \| 261 \| 24 \| 1 \| 422 \| 2 \| 721 \| H: tail fiber protein \| H: tail fiber protein; afdb-uniprot50 \| AF-A0A6P1LU34-F1-MODEL\_V4 \| 1.0 \| 2.914e-13 \| 309 \| 0.148 \| 459 \| 225 \| 21 \| 1 \| 421 \| 2 \| 332 \| Uncharacterized protein \| Uncharacterized protein; afdb-uniprot50 \| AF-A0A2U8DMY3-F1-MODEL\_V4 \| 1.0 \| 6.706e-15 \| 306 \| 0.193 \| 512 \| 265 \| 23 \| 1 \| 418 \| 2 \| 459 \| Uncharacterized protein \| Uncharacterized protein; afdb-uniprot50 \| AF-A0A241V3C4-F1-MODEL\_V4 \| 1.0 \| 1.942e-15 \| 306 \| 0.2 \| 518 \| 261 \| 20 \| 1 \| 412 \| 2 \| 472 \| Uncharacterized protein \| Uncharacterized protein; afdb-uniprot50 \| AF-A0A1C3CUP3-F1-MODEL\_V4 \| 1.0 \| 6.021e-15 \| 305 \| 0.187 \| 475 \| 265 \| 25 \| 1 \| 423 \| 1 \| 406 \| Uncharacterized protein \| Uncharacterized protein; afdb-uniprot50 \| AF-A0A7Z2B2X5-F1-MODEL\_V4 \| 1.0 \| 3.513e-15 \| 303 \| 0.16 \| 581 \| 247 \| 25 \| 1 \| 421 \| 2 \| 501 \| Uncharacterized protein \| Uncharacterized protein; afdb-uniprot50 \| AF-A0A376H5A1-F1-MODEL\_V4 \| 1.0 \| 1.426e-14 \| 301 \| 0.161 \| 527 \| 227 \| 25 \| 1 \| 406 \| 2 \| 434 \| Phage tail protein \| Phage tail protein; afdb-uniprot50 \| AF-A0A5M8P6A1-F1-MODEL\_V4 \| 1.0 \| 1.28e-14 \| 288 \| 0.214 \| 452 \| 257 \| 25 \| 2 \| 413 \| 5 \| 398 \| Uncharacterized protein \| Uncharacterized protein; afdb-uniprot50 \| AF-S7JLI8-F1-MODEL\_V4 \| 1.0 \| 5.067e-12 \| 283 \| 0.161 \| 365 \| 205 \| 14 \| 6 \| 348 \| 10 \| 295 \| Tail fiber protein \| Tail fiber protein; afdb-uniprot50 \| AF-S4YWB1-F1-MODEL\_V4 \| 1.0 \| 1.105e-13 \| 279 \| 0.174 \| 447 \| 240 \| 19 \| 3 \| 422 \| 2 \| 346 \| Uncharacterized protein \| Uncharacterized protein; afdb-uniprot50 \| AF-G2HXA2-F1-MODEL\_V4 \| 1.0 \| 4.421e-14 \| 277 \| 0.149 \| 574 \| 271 \| 32 \| 1 \| 421 \| 1 \| 510 \| Phage tail fiber protein \| Phage tail fiber protein; afdb-uniprot50 \| AF-A0A3B0BXV7-F1-MODEL\_V4 \| 1.0 \| 1.371e-13 \| 271 \| 0.202 \| 410 \| 234 \| 20 \| 5 \| 404 \| 3 \| 329 \| Uncharacterized protein \| Uncharacterized protein; afdb-uniprot50 \| AF-A0A329MQ29-F1-MODEL\_V4 \| 1.0 \| 2.663e-08 \| 266 \| 0.216 \| 162 \| 119 \| 4 \| 6 \| 166 \| 5 \| 159 \| Uncharacterized protein \| Uncharacterized protein; afdb-uniprot50 \| AF-A0A849B068-F1-MODEL\_V4 \| 1.0 \| 2.722e-14 \| 255 \| 0.171 \| 495 \| 272 \| 24 \| 1 \| 420 \| 1 \| 432 \| Uncharacterized protein \| Uncharacterized protein; afdb-uniprot50 \| AF-A0A1K1T5P9-F1-MODEL\_V4 \| 1.0 \| 2.551e-11 \| 220 \| 0.235 \| 438 \| 212 \| 21 \| 32 \| 411 \| 2 \| 374 \| Phage tail-collar fibre protein \| Phage tail-collar fibre protein; afdb-uniprot50 \| AF-A0A378E622-F1-MODEL\_V4 \| 1.0 \| 8.14e-09 \| 217 \| 0.211 \| 303 \| 138 \| 12 \| 32 \| 258 \| 2 \| 279 \| Tail fiber protein \| Tail fiber protein |
| Top keywords  (threshold 1.00e-02 (evalue)) | **tail, Phage, fiber, fibre, tail\_collar, domain\_containing, Collar, Putative, Phage\_related, repeat** |
| Output files | ../../similar\_structures/15\_FANPEZAQ\_CDS\_0015\_afdb-proteome\_foldseek.tsv ../../similar\_structures/15\_FANPEZAQ\_CDS\_0015\_afdb-uniprot50\_foldseek.tsv ../../similar\_structures/15\_FANPEZAQ\_CDS\_0015\_merged.svg ../../similar\_structures/15\_FANPEZAQ\_CDS\_0015\_pdb\_foldseek.tsv |

  
  
  

Return to summary | Go to previous | Go to next

  


---

**Sequence/structure alignments coloring**  
Each object in the alignment figures is colored according to its E-value following this color coding:

1e-100
10
