## Supplementary material for "Completing the BASEL phage collection to unlock hidden diversity for systematic exploration of phage-host interactions": Data S2: 16.html

FANPEZAQ\_CDS\_0016


Return to summary | Go to previous | Go to next

|  |  |
| --- | --- |
| FANPEZAQ\_CDS\_0016  Page creation date: 02 Sep 2024, 12:00  Project folder: n/a  Input sequences file: Escherichia\_virus\_HeidiAbel.gb | tail assembly fiber phage chaperone caudovirales putative fibre lambda gpk prophage virus gp38 domain\_containing fragment hypothetical duf4376 bacteriophage cps\_53 kple1 g homolog from lambdoid |

### Sequence information

|  |  |
| --- | --- |
| Name | FANPEZAQ\_CDS\_0016  16\_FANPEZAQ\_CDS\_0016 (pipeline id) |
| Imported annotations | Escherichia\_virus\_HeidiAbel Bas97 |
| Protein sequence | MSKYMFSPSTNAFYPVALQSIYEEANMWPNDGVGCEEDVFSEFTGVPPEGKRRGSDKDGN PAWVDVPPLTKEQQIANAKSMQQYLIDQANDFMNSKQWPGKAAIGRLKDAELNQYKIWLD YLDALYAVDTSTAPEITWPEQPQ |
| Number of residues | 143 |
| Molecular weight (Da) | 16173.84 |
| Output files | ../../query\_sequences/16\_FANPEZAQ\_CDS\_0016.fasta |

### Putative domain architecture and protein family

#### Search results (HHblits)^1^

|  |  |
| --- | --- |
| Domain family databases searched | Pfam, Ncbi-cd, Cath, Phrogs |
| Results, scheme(s)  (Top layers only; threshold 1.00e-03 (evalue)) | xml version="1.0" encoding="utf-8" standalone="no"?       2024-09-02T21:08:15.476921 image/svg+xml   Matplotlib v3.7.2, https://matplotlib.org/ |
| Results, table  (E-value ≤ 1.00e-03 (evalue)) | db \| id \| prob \| evalue \| pvalue \| score \| cols \| query \| query\_len \| template \| template\_len \| name \| description; pfam \| PF02413 \| 98.9 \| 1.1e-13 \| 2.3e-17 \| 88.7 \| 94 \| (44, 143) \| 143 \| (27, 120) \| 120 \| Caudo\_TAP \| Caudovirales tail fibre assembly protein, lambda gpK; pfam \| PF16778 \| 97.4 \| 4.9e-08 \| 9.9e-12 \| 53.6 \| 56 \| (77, 143) \| 143 \| (3, 60) \| 60 \| Phage\_tail\_APC \| Phage tail assembly chaperone protein; cath \| 2kz6A01 \| 99.1 \| 4.5e-15 \| 7.7e-19 \| 95.0 \| 73 \| (28, 126) \| 143 \| (11, 83) \| 83 \| Uncharacterized protein \| CATHCODE: 6.10.140.1310 NAME: Uncharacterized protein. Chain: a. Engineered: yes SOURCE: Chromobacterium violaceum. Organism\_taxid: 536. Gene: cv\_0426. Expressed in: escherichia coli. Expression\_system\_taxid: 562. Expression\_system\_vector: p15tv lic CLASS: Special, ARCH: Helix non-globular, TOPOL: Helix Hairpins, HOMOL: Helix Hairpins; phrogs \| 188 \| 100.0 \| 6.4e-39 \| 8.4e-43 \| 225.6 \| 136 \| (1, 143) \| 143 \| (4, 140) \| 140 \| tail fiber assembly protein \| tail fiber assembly protein; Category: tail; p268800 VI\_09964; phrogs \| 5572 \| 99.7 \| 8.3e-22 \| 9.7e-26 \| 123.0 \| 75 \| (67, 143) \| 143 \| (4, 78) \| 80 \| NA \| NA; Category: unknown function; p62117 VI\_01572; phrogs \| 105 \| 99.7 \| 2.5e-21 \| 3.1e-25 \| 141.8 \| 80 \| (62, 143) \| 143 \| (124, 203) \| 203 \| tail fiber assembly \| tail fiber assembly; Category: tail; p319838 VI\_03642; phrogs \| 3792 \| 99.6 \| 5.7e-21 \| 6.6e-25 \| 120.8 \| 80 \| (56, 143) \| 143 \| (9, 88) \| 88 \| tail assembly chaperone \| tail assembly chaperone; Category: tail; p125307 VI\_08164; phrogs \| 136 \| 99.5 \| 1e-18 \| 1.3e-22 \| 125.3 \| 81 \| (55, 143) \| 143 \| (95, 175) \| 175 \| tail fiber assembly protein \| tail fiber assembly protein; Category: tail; p240703 VI\_01691; phrogs \| 5844 \| 99.1 \| 1.2e-14 \| 1.3e-18 \| 101.5 \| 111 \| (30, 142) \| 143 \| (81, 193) \| 199 \| NA \| NA; Category: unknown function; p304883 VI\_09788; phrogs \| 19350 \| 98.3 \| 1.1e-10 \| 1.3e-14 \| 67.4 \| 65 \| (1, 65) \| 143 \| (1, 66) \| 69 \| NA \| NA; Category: unknown function; p37945 VI\_07185; phrogs \| 5400 \| 98.3 \| 1.2e-10 \| 1.4e-14 \| 69.5 \| 59 \| (3, 68) \| 143 \| (1, 59) \| 74 \| NA \| NA; Category: unknown function; p80665 VI\_09756; phrogs \| 1795 \| 98.1 \| 1.2e-09 \| 1.4e-13 \| 73.9 \| 61 \| (78, 143) \| 143 \| (70, 133) \| 136 \| tail fiber protein \| tail fiber protein; Category: tail; KX660669\_p44; phrogs \| 8746 \| 98.0 \| 1.3e-09 \| 1.6e-13 \| 74.1 \| 61 \| (73, 143) \| 143 \| (92, 154) \| 154 \| tail fiber assembly protein \| tail fiber assembly protein; Category: tail; KP343639\_p7; phrogs \| 4628 \| 98.0 \| 2.5e-09 \| 2.8e-13 \| 70.0 \| 64 \| (70, 143) \| 143 \| (58, 123) \| 124 \| tail fiber assembly \| tail fiber assembly; Category: tail; NC\_026587\_p82; phrogs \| 24907 \| 97.6 \| 3.4e-08 \| 3.9e-12 \| 68.1 \| 66 \| (76, 143) \| 143 \| (130, 197) \| 206 \| NA \| NA; Category: unknown function; p115387 VI\_09903; phrogs \| 8716 \| 97.5 \| 5.7e-08 \| 6.7e-12 \| 63.2 \| 58 \| (71, 143) \| 143 \| (56, 115) \| 115 \| tail fiber protein \| tail fiber protein; Category: tail; NC\_020838\_p5; phrogs \| 37801 \| 97.4 \| 1.2e-07 \| 1.3e-11 \| 54.4 \| 36 \| (2, 37) \| 143 \| (1, 36) \| 73 \| NA \| NA; Category: unknown function; p165538 VI\_11050 |
| Top keywords  (threshold 1.00e-03 (evalue)) | **tail, assembly, fiber, Helix, chaperone, Hairpins, Caudovirales, fibre, lambda, gpK** |
| Output files | ../../domain\_architecture/16\_FANPEZAQ\_CDS\_0016\_cath.hhr ../../domain\_architecture/16\_FANPEZAQ\_CDS\_0016\_merged.svg ../../domain\_architecture/16\_FANPEZAQ\_CDS\_0016\_ncbi-cd.hhr ../../domain\_architecture/16\_FANPEZAQ\_CDS\_0016\_pfam.hhr ../../domain\_architecture/16\_FANPEZAQ\_CDS\_0016\_phrogs.hhr |

### Identical protein sequences/structures

#### Search results

|  |  |
| --- | --- |
| Protein sequence databases searched | Pdb, Swissprot, Refseq |
| Identical proteins found | -- |
| Top keywords | -- |
| Output files | -- |

### Similar protein sequences/structures

#### Sequence similarity search results (HHblits)^1^

|  |  |
| --- | --- |
| Sequence databases searched | Uniclust, Pdb70 |
| Results, scheme(s)  (Top layers only, threshold 1.00e-03 (evalue)) | xml version="1.0" encoding="utf-8" standalone="no"?       2024-09-02T21:08:35.691326 image/svg+xml   Matplotlib v3.7.2, https://matplotlib.org/ |
| Results, table(s)  (threshold 1.00e-03 (evalue)) | db \| id \| prob \| evalue \| pvalue \| score \| cols \| query \| query\_len \| template \| template\_len \| name \| description; uniclust \| UniRef100\_A0A071M0C5 \| 100.0 \| 1.3e-50 \| 2.5e-56 \| 261.9 \| 141 \| (1, 143) \| 143 \| (16, 156) \| 172 \| Tail fiber assembly protein \| Tail fiber assembly protein; uniclust \| UniRef100\_A0A345CNR7 \| 100.0 \| 3.3e-47 \| 6.2e-53 \| 234.6 \| 135 \| (1, 143) \| 143 \| (3, 137) \| 143 \| Tail fiber assembly protein \| Tail fiber assembly protein; uniclust \| UniRef100\_A0A022PLD3 \| 100.0 \| 1e-46 \| 1.9e-52 \| 251.1 \| 141 \| (1, 143) \| 143 \| (51, 191) \| 214 \| Caudovirales tail fiber assembly protein \| Caudovirales tail fiber assembly protein; uniclust \| UniRef100\_A0A0M7GTK9 \| 100.0 \| 5.2e-45 \| 9.9e-51 \| 229.7 \| 136 \| (1, 143) \| 143 \| (5, 142) \| 151 \| Bacteriophage tail assembly protein \| Bacteriophage tail assembly protein; uniclust \| UniRef100\_A0A077Q6L9 \| 100.0 \| 6e-45 \| 1.1e-50 \| 234.1 \| 143 \| (1, 143) \| 143 \| (17, 159) \| 177 \| Putative tail fiber assembly protein \| Putative tail fiber assembly protein; uniclust \| UniRef100\_A0A080IB21 \| 100.0 \| 2.7e-43 \| 5.3e-49 \| 235.4 \| 141 \| (1, 143) \| 143 \| (26, 166) \| 193 \| Caudovirales tail fiber assembly family protein \| Caudovirales tail fiber assembly family protein; uniclust \| UniRef100\_A0A022PIU1 \| 100.0 \| 9.4e-43 \| 1.8e-48 \| 237.7 \| 141 \| (1, 143) \| 143 \| (55, 195) \| 224 \| Caudovirales tail fiber assembly protein \| Caudovirales tail fiber assembly protein; uniclust \| UniRef100\_A0A2X1RV26 \| 100.0 \| 1.4e-42 \| 2.6e-48 \| 216.5 \| 141 \| (1, 143) \| 143 \| (1, 142) \| 146 \| Caudovirales tail fibre assembly protein \| Caudovirales tail fibre assembly protein; uniclust \| UniRef100\_A0A376YI45 \| 100.0 \| 3.6e-42 \| 6.7e-48 \| 214.1 \| 143 \| (1, 143) \| 143 \| (11, 153) \| 153 \| CPS-53 (KpLE1) prophage protein \| CPS-53 (KpLE1) prophage protein; uniclust \| UniRef100\_A0A0E8XEB7 \| 100.0 \| 9.8e-41 \| 1.9e-46 \| 217.2 \| 141 \| (1, 143) \| 143 \| (16, 157) \| 172 \| Tail assembly chaperone gp38 \| Tail assembly chaperone gp38; uniclust \| UniRef100\_A0A0H2VSV7 \| 100.0 \| 6.5e-40 \| 1.2e-45 \| 206.1 \| 136 \| (1, 143) \| 143 \| (37, 172) \| 188 \| Putative phage tail fibre protein \| Putative phage tail fibre protein; uniclust \| UniRef100\_A0A0V9JY68 \| 100.0 \| 7e-40 \| 1.3e-45 \| 207.7 \| 140 \| (1, 143) \| 143 \| (19, 158) \| 159 \| Phage tail protein \| Phage tail protein; uniclust \| UniRef100\_A0A071M4A9 \| 100.0 \| 9.5e-40 \| 1.8e-45 \| 212.7 \| 135 \| (1, 143) \| 143 \| (24, 158) \| 194 \| Tail fiber assembly protein \| Tail fiber assembly protein; uniclust \| UniRef100\_A0A071LRY7 \| 100.0 \| 4.7e-39 \| 8.9e-45 \| 199.8 \| 136 \| (1, 143) \| 143 \| (1, 136) \| 140 \| Tail fiber assembly protein \| Tail fiber assembly protein; uniclust \| UniRef100\_A0A8R2D774 \| 100.0 \| 4.5e-38 \| 8.5e-44 \| 195.6 \| 140 \| (2, 143) \| 143 \| (1, 143) \| 144 \| Tail fiber assembly protein \| Tail fiber assembly protein; uniclust \| UniRef100\_A0A3F3IA78 \| 100.0 \| 5.7e-35 \| 1.1e-40 \| 183.7 \| 134 \| (1, 143) \| 143 \| (4, 137) \| 155 \| Tail fiber assembly protein \| Tail fiber assembly protein; uniclust \| UniRef100\_A0A0D0PTQ5 \| 100.0 \| 1.1e-33 \| 2.2e-39 \| 177.9 \| 133 \| (1, 143) \| 143 \| (1, 135) \| 140 \| Tail fiber assembly protein \| Tail fiber assembly protein; uniclust \| UniRef100\_A0A4D7IX74 \| 100.0 \| 3.7e-33 \| 6.8e-39 \| 176.6 \| 138 \| (3, 142) \| 143 \| (59, 197) \| 197 \| Phage tail protein \| Phage tail protein; uniclust \| UniRef100\_A0A080KGQ0 \| 100.0 \| 9.8e-33 \| 1.9e-38 \| 176.0 \| 133 \| (1, 143) \| 143 \| (16, 148) \| 153 \| Caudovirales tail fiber assembly protein \| Caudovirales tail fiber assembly protein; uniclust \| UniRef100\_A0A031FRK0 \| 100.0 \| 1.4e-32 \| 2.7e-38 \| 176.8 \| 135 \| (1, 143) \| 143 \| (10, 146) \| 164 \| Prophage tail fimber assembly protein \| Prophage tail fimber assembly protein; uniclust \| UniRef100\_A0A080HNE4 \| 100.0 \| 2.2e-32 \| 4.3e-38 \| 181.2 \| 134 \| (1, 143) \| 143 \| (23, 159) \| 172 \| Caudovirales tail fiber assembly family protein \| Caudovirales tail fiber assembly family protein; uniclust \| UniRef100\_A0A127QUQ2 \| 99.9 \| 3.5e-32 \| 6.7e-38 \| 173.1 \| 137 \| (1, 143) \| 143 \| (4, 149) \| 154 \| Caudovirales tail fiber assembly family protein \| Caudovirales tail fiber assembly family protein; uniclust \| UniRef100\_A0A024EI51 \| 99.9 \| 3.6e-31 \| 7e-37 \| 179.1 \| 135 \| (1, 143) \| 143 \| (34, 170) \| 203 \| Phage tail protein \| Phage tail protein; uniclust \| UniRef100\_A0A068QUZ6 \| 99.9 \| 3.6e-31 \| 7e-37 \| 169.8 \| 135 \| (1, 143) \| 143 \| (2, 136) \| 148 \| Putative tail fiber assembly protein \| Putative tail fiber assembly protein; uniclust \| UniRef100\_A0A0A0YSY3 \| 99.9 \| 4.1e-31 \| 7.7e-37 \| 171.4 \| 135 \| (3, 143) \| 143 \| (68, 202) \| 220 \| Tail fiber assembly protein \| Tail fiber assembly protein; uniclust \| UniRef100\_A0A1Q6AW63 \| 99.9 \| 6.1e-31 \| 1.1e-36 \| 160.6 \| 137 \| (4, 143) \| 143 \| (6, 143) \| 147 \| Phage tail protein \| Phage tail protein; uniclust \| UniRef100\_UPI0002D7ACE8 \| 99.9 \| 1.3e-30 \| 2.5e-36 \| 162.6 \| 136 \| (1, 143) \| 143 \| (2, 139) \| 142 \| tail assembly chaperone \| tail assembly chaperone; uniclust \| UniRef100\_A0A0A6DH01 \| 99.9 \| 1.8e-30 \| 3.4e-36 \| 165.6 \| 134 \| (1, 142) \| 143 \| (5, 140) \| 156 \| Phage tail protein \| Phage tail protein; uniclust \| UniRef100\_A0A2L1UQL2 \| 99.9 \| 2e-30 \| 3.7e-36 \| 159.7 \| 132 \| (2, 143) \| 143 \| (1, 132) \| 134 \| Phage tail protein \| Phage tail protein; uniclust \| UniRef100\_A0A2D0IKG5 \| 99.9 \| 5.2e-30 \| 9.5e-36 \| 155.2 \| 135 \| (3, 143) \| 143 \| (15, 149) \| 149 \| Tail assembly chaperone \| Tail assembly chaperone; uniclust \| UniRef100\_A0A0N0XL62 \| 99.9 \| 2e-29 \| 3.9e-35 \| 162.3 \| 128 \| (1, 143) \| 143 \| (5, 134) \| 146 \| Caudovirales tail fiber assembly protein \| Caudovirales tail fiber assembly protein; uniclust \| UniRef100\_A0A1W0D5R5 \| 99.9 \| 2.2e-29 \| 4e-35 \| 153.8 \| 135 \| (3, 143) \| 143 \| (1, 137) \| 143 \| Phage tail protein \| Phage tail protein; uniclust \| UniRef100\_A0A1M3PGE6 \| 99.9 \| 2.9e-29 \| 5.5e-35 \| 155.3 \| 126 \| (1, 143) \| 143 \| (1, 128) \| 133 \| Phage tail protein \| Phage tail protein; uniclust \| UniRef100\_A0A0N9WEK9 \| 99.9 \| 6.4e-29 \| 1.2e-34 \| 156.0 \| 135 \| (1, 143) \| 143 \| (1, 137) \| 151 \| Phage tail protein \| Phage tail protein; uniclust \| UniRef100\_A0A2S5GQX3 \| 99.9 \| 8.9e-29 \| 1.7e-34 \| 159.4 \| 135 \| (3, 143) \| 143 \| (62, 198) \| 201 \| Tail fiber assembly protein \| Tail fiber assembly protein; uniclust \| UniRef100\_A0A031IRP6 \| 99.9 \| 1.5e-28 \| 2.8e-34 \| 165.1 \| 141 \| (1, 143) \| 143 \| (3, 198) \| 210 \| Prophage PSPPH06 tail fiber protein \| Prophage PSPPH06 tail fiber protein; uniclust \| UniRef100\_A0A2S7HKK0 \| 99.9 \| 3e-28 \| 5.6e-34 \| 146.4 \| 138 \| (3, 143) \| 143 \| (1, 138) \| 142 \| Phage tail protein \| Phage tail protein; uniclust \| UniRef100\_A0A1L4BM55 \| 99.9 \| 1.2e-27 \| 2.3e-33 \| 144.9 \| 121 \| (22, 143) \| 143 \| (3, 123) \| 127 \| Caudovirales tail fiber assembly protein \| Caudovirales tail fiber assembly protein; uniclust \| UniRef100\_A0A1Q5TJA7 \| 99.9 \| 1.8e-27 \| 3.3e-33 \| 152.7 \| 76 \| (66, 143) \| 143 \| (64, 139) \| 162 \| Tail fiber assembly protein \| Tail fiber assembly protein; uniclust \| UniRef100\_A0A501S8D9 \| 99.9 \| 3.8e-27 \| 7.2e-33 \| 146.8 \| 140 \| (1, 143) \| 143 \| (1, 140) \| 142 \| Tail fiber assembly protein \| Tail fiber assembly protein; uniclust \| UniRef100\_UPI0015857BD7 \| 99.9 \| 4.1e-27 \| 7.5e-33 \| 149.7 \| 142 \| (2, 143) \| 143 \| (70, 211) \| 211 \| tail fiber assembly protein \| tail fiber assembly protein; uniclust \| UniRef100\_A0A104MR71 \| 99.9 \| 4.5e-27 \| 8.4e-33 \| 147.8 \| 133 \| (1, 143) \| 143 \| (1, 139) \| 149 \| Tail fiber assembly protein \| Tail fiber assembly protein; uniclust \| UniRef100\_A0A0K8QR29 \| 99.9 \| 4.8e-27 \| 9.1e-33 \| 152.2 \| 127 \| (1, 143) \| 143 \| (16, 144) \| 160 \| Tail fiber assembly protein \| Tail fiber assembly protein; uniclust \| UniRef100\_A0A077P958 \| 99.9 \| 6.9e-27 \| 1.3e-32 \| 140.4 \| 91 \| (50, 143) \| 143 \| (17, 107) \| 114 \| Putative tail fiber chaperone Rac prophage \| Putative tail fiber chaperone Rac prophage; uniclust \| UniRef100\_A0A1G9YD25 \| 99.9 \| 1.2e-26 \| 2.3e-32 \| 145.2 \| 112 \| (30, 143) \| 143 \| (20, 132) \| 142 \| Virus tail fibre assembly protein, lambda gpK \| Virus tail fibre assembly protein, lambda gpK; uniclust \| UniRef100\_A0A0B1YWB2 \| 99.9 \| 1.2e-26 \| 2.3e-32 \| 151.4 \| 136 \| (1, 143) \| 143 \| (7, 144) \| 171 \| Phage tail protein \| Phage tail protein; uniclust \| UniRef100\_A0A022PH83 \| 99.9 \| 2e-26 \| 3.8e-32 \| 137.6 \| 95 \| (46, 143) \| 143 \| (4, 98) \| 110 \| Caudovirales tail fiber assembly protein \| Caudovirales tail fiber assembly protein; uniclust \| UniRef100\_A0A0E1SIM2 \| 99.9 \| 5.7e-26 \| 1.1e-31 \| 150.2 \| 114 \| (28, 143) \| 143 \| (45, 161) \| 184 \| Tail assembly chaperone \| Tail assembly chaperone; uniclust \| UniRef100\_A0A2D0KV06 \| 99.9 \| 6.6e-26 \| 1.2e-31 \| 161.5 \| 138 \| (2, 143) \| 143 \| (421, 561) \| 561 \| Tail assembly chaperone \| Tail assembly chaperone; uniclust \| UniRef100\_A0A3A3ZKX8 \| 99.9 \| 1.6e-25 \| 3.1e-31 \| 133.0 \| 91 \| (1, 91) \| 143 \| (2, 93) \| 98 \| Tail fiber assembly protein \| Tail fiber assembly protein; uniclust \| UniRef100\_A0A014N9D4 \| 99.9 \| 2.4e-25 \| 4.4e-31 \| 140.8 \| 139 \| (1, 143) \| 143 \| (35, 175) \| 176 \| Tail fiber assembly protein \| Tail fiber assembly protein; uniclust \| UniRef100\_A0A2K8QS96 \| 99.9 \| 2.4e-25 \| 4.5e-31 \| 133.8 \| 123 \| (2, 143) \| 143 \| (1, 126) \| 128 \| Phage tail protein \| Phage tail protein; uniclust \| UniRef100\_A0A158S497 \| 99.9 \| 4.2e-25 \| 8.1e-31 \| 139.5 \| 126 \| (2, 143) \| 143 \| (1, 128) \| 129 \| Tail fiber assembly protein \| Tail fiber assembly protein; uniclust \| UniRef100\_A0A369F960 \| 99.9 \| 5.1e-25 \| 9.7e-31 \| 140.5 \| 131 \| (1, 143) \| 143 \| (6, 137) \| 151 \| Tail fiber assembly protein \| Tail fiber assembly protein; uniclust \| UniRef100\_A0A208ZVY5 \| 99.8 \| 6.1e-25 \| 1.1e-30 \| 128.9 \| 76 \| (66, 143) \| 143 \| (15, 90) \| 94 \| Phage tail protein \| Phage tail protein; uniclust \| UniRef100\_A0A1H1NDT3 \| 99.8 \| 7.5e-25 \| 1.4e-30 \| 139.7 \| 126 \| (4, 143) \| 143 \| (1, 130) \| 143 \| Virus tail fibre assembly protein, lambda gpK \| Virus tail fibre assembly protein, lambda gpK; uniclust \| UniRef100\_A0A3V9FI80 \| 99.8 \| 9.1e-25 \| 1.7e-30 \| 137.4 \| 136 \| (2, 143) \| 143 \| (10, 145) \| 149 \| Phage tail protein \| Phage tail protein; uniclust \| UniRef100\_A0A0T9UQ83 \| 99.8 \| 9.5e-25 \| 1.8e-30 \| 130.7 \| 94 \| (48, 143) \| 143 \| (13, 106) \| 110 \| Tail assembly chaperone gp38 \| Tail assembly chaperone gp38; uniclust \| UniRef100\_A0A1G5QUE3 \| 99.8 \| 1.2e-24 \| 2.2e-30 \| 130.6 \| 100 \| (1, 100) \| 143 \| (6, 105) \| 109 \| Virus tail fibre assembly protein, lambda gpK \| Virus tail fibre assembly protein, lambda gpK; uniclust \| UniRef100\_UPI000792F1CB \| 99.8 \| 1.2e-24 \| 2.2e-30 \| 131.9 \| 140 \| (2, 143) \| 143 \| (1, 144) \| 144 \| tail fiber assembly protein \| tail fiber assembly protein; uniclust \| UniRef100\_A0A2S1BAI7 \| 99.8 \| 1.3e-24 \| 2.5e-30 \| 136.0 \| 104 \| (37, 143) \| 143 \| (36, 140) \| 144 \| Phage tail protein \| Phage tail protein; uniclust \| UniRef100\_A0A0U3TL14 \| 99.8 \| 1.4e-24 \| 2.7e-30 \| 138.1 \| 131 \| (1, 143) \| 143 \| (19, 149) \| 157 \| Tail fiber assembly protein \| Tail fiber assembly protein; uniclust \| UniRef100\_A0A0W1G7X1 \| 99.8 \| 1.5e-24 \| 2.9e-30 \| 140.4 \| 130 \| (1, 143) \| 143 \| (3, 154) \| 163 \| Phage tail protein \| Phage tail protein; uniclust \| UniRef100\_A0A2X4UV66 \| 99.8 \| 1.7e-24 \| 3.3e-30 \| 132.5 \| 94 \| (1, 95) \| 143 \| (3, 96) \| 116 \| Uncharacterized protein \| Uncharacterized protein; uniclust \| UniRef100\_A0A2J4V2S3 \| 99.8 \| 2.7e-24 \| 5.1e-30 \| 133.7 \| 81 \| (61, 143) \| 143 \| (46, 126) \| 132 \| Tail fiber assembly protein (Fragment) \| Tail fiber assembly protein (Fragment); uniclust \| UniRef100\_A0A0H5HS46 \| 99.8 \| 3.1e-24 \| 5.9e-30 \| 142.2 \| 137 \| (1, 143) \| 143 \| (31, 170) \| 212 \| Putative tail fiber assembly protein \| Putative tail fiber assembly protein; uniclust \| UniRef100\_A0A4R4JRM7 \| 99.8 \| 8.3e-24 \| 1.6e-29 \| 127.4 \| 74 \| (68, 143) \| 143 \| (37, 110) \| 117 \| Tail fiber assembly protein \| Tail fiber assembly protein; uniclust \| UniRef100\_A0A1S2TBU1 \| 99.8 \| 8.5e-24 \| 1.6e-29 \| 134.6 \| 137 \| (3, 142) \| 143 \| (27, 166) \| 174 \| Phage tail protein \| Phage tail protein; uniclust \| UniRef100\_A0A411WM04 \| 99.8 \| 1.9e-23 \| 3.7e-29 \| 128.3 \| 110 \| (27, 143) \| 143 \| (11, 121) \| 124 \| Tail fiber assembly protein \| Tail fiber assembly protein; uniclust \| UniRef100\_A0A2D0J0R2 \| 99.8 \| 2.3e-23 \| 4.3e-29 \| 141.1 \| 76 \| (66, 143) \| 143 \| (154, 229) \| 230 \| Phage tail fiber assembly \| Phage tail fiber assembly; uniclust \| UniRef100\_A0A1X0VZT8 \| 99.8 \| 2.8e-23 \| 5.3e-29 \| 132.8 \| 134 \| (4, 143) \| 143 \| (1, 154) \| 155 \| Phage tail protein \| Phage tail protein; uniclust \| UniRef100\_A0A7U5JH60 \| 99.8 \| 3.3e-23 \| 6.2e-29 \| 134.7 \| 135 \| (1, 140) \| 143 \| (21, 157) \| 161 \| Tail fiber assembly protein \| Tail fiber assembly protein; uniclust \| UniRef100\_A0A2L1I8I3 \| 99.8 \| 4e-23 \| 7.6e-29 \| 131.1 \| 77 \| (65, 143) \| 143 \| (56, 134) \| 141 \| Phage tail protein \| Phage tail protein; uniclust \| UniRef100\_A0A229A5L0 \| 99.8 \| 5.8e-23 \| 1.1e-28 \| 126.1 \| 142 \| (2, 143) \| 143 \| (7, 148) \| 152 \| Tail fiber assembly protein \| Tail fiber assembly protein; uniclust \| UniRef100\_A0A2U8I8G2 \| 99.8 \| 7.4e-23 \| 1.4e-28 \| 120.1 \| 84 \| (58, 143) \| 143 \| (5, 89) \| 94 \| Phage tail protein \| Phage tail protein; uniclust \| UniRef100\_A0A2X1MUF5 \| 99.8 \| 1.7e-22 \| 3.2e-28 \| 128.8 \| 85 \| (56, 143) \| 143 \| (63, 147) \| 157 \| Tail fiber assembly protein (Gpg) \| Tail fiber assembly protein (Gpg); uniclust \| UniRef100\_A0A7D6TWW6 \| 99.8 \| 2.3e-22 \| 4.3e-28 \| 126.3 \| 120 \| (1, 123) \| 143 \| (1, 120) \| 184 \| Tail fiber assembly protein \| Tail fiber assembly protein; uniclust \| UniRef100\_A0A336QKS4 \| 99.8 \| 2.3e-22 \| 4.4e-28 \| 122.8 \| 74 \| (67, 143) \| 143 \| (41, 114) \| 116 \| Phage tail fiber assembly protein \| Phage tail fiber assembly protein; uniclust \| UniRef100\_A0A068Z7G0 \| 99.8 \| 2.6e-22 \| 5e-28 \| 130.7 \| 72 \| (70, 143) \| 143 \| (80, 151) \| 157 \| Tail fiber assembly protein \| Tail fiber assembly protein; uniclust \| UniRef100\_A0A076LFJ0 \| 99.8 \| 3.2e-22 \| 6e-28 \| 117.9 \| 86 \| (58, 143) \| 143 \| (2, 87) \| 96 \| Phage tail fiber assembly protein \| Phage tail fiber assembly protein; uniclust \| UniRef100\_UPI000983374A \| 99.8 \| 3.6e-22 \| 6.8e-28 \| 120.2 \| 105 \| (34, 143) \| 143 \| (11, 116) \| 117 \| tail fiber assembly protein \| tail fiber assembly protein; uniclust \| UniRef100\_A0A0K1QN38 \| 99.8 \| 4.1e-22 \| 7.7e-28 \| 126.4 \| 76 \| (66, 143) \| 143 \| (49, 126) \| 143 \| Phage tail protein \| Phage tail protein; uniclust \| UniRef100\_A0A423XTX7 \| 99.8 \| 9.9e-22 \| 1.8e-27 \| 120.5 \| 137 \| (1, 143) \| 143 \| (1, 143) \| 146 \| Phage tail protein \| Phage tail protein; uniclust \| UniRef100\_A0A0H3AZX4 \| 99.8 \| 9.7e-22 \| 1.8e-27 \| 130.4 \| 73 \| (69, 143) \| 143 \| (111, 183) \| 197 \| Tail assembly chaperone gp38 \| Tail assembly chaperone gp38; uniclust \| UniRef100\_A0A2C5TSB9 \| 99.8 \| 1.1e-21 \| 2e-27 \| 113.5 \| 94 \| (48, 143) \| 143 \| (3, 96) \| 96 \| Phage tail protein \| Phage tail protein; uniclust \| UniRef100\_UPI000A9D9905 \| 99.7 \| 1.7e-21 \| 3.2e-27 \| 117.8 \| 121 \| (2, 143) \| 143 \| (1, 123) \| 123 \| phage tail assembly chaperone \| phage tail assembly chaperone; uniclust \| UniRef100\_A0A548QSH7 \| 99.7 \| 2.2e-21 \| 4e-27 \| 117.7 \| 131 \| (1, 141) \| 143 \| (3, 133) \| 137 \| Tail fiber assembly protein (Fragment) \| Tail fiber assembly protein (Fragment); uniclust \| UniRef100\_A0A090V754 \| 99.7 \| 3.6e-21 \| 6.7e-27 \| 113.3 \| 83 \| (59, 143) \| 143 \| (6, 90) \| 94 \| Predicted tail fiber assembly protein \| Predicted tail fiber assembly protein; uniclust \| UniRef100\_A0A5C5DE35 \| 99.7 \| 3.7e-21 \| 6.9e-27 \| 120.5 \| 138 \| (1, 143) \| 143 \| (1, 141) \| 145 \| Tail fiber assembly protein \| Tail fiber assembly protein; uniclust \| UniRef100\_A0A2T8WWY8 \| 99.7 \| 4.1e-21 \| 7.7e-27 \| 126.8 \| 68 \| (74, 143) \| 143 \| (112, 179) \| 194 \| Tail fiber assembly protein (Fragment) \| Tail fiber assembly protein (Fragment); uniclust \| UniRef100\_A0A0N9ZQ35 \| 99.7 \| 4.4e-21 \| 8.2e-27 \| 121.5 \| 134 \| (2, 143) \| 143 \| (10, 145) \| 156 \| Phage tail protein \| Phage tail protein; uniclust \| UniRef100\_E9CQ84 \| 99.7 \| 6.3e-21 \| 1.2e-26 \| 110.4 \| 80 \| (62, 143) \| 143 \| (5, 84) \| 87 \| Putative phage tail assembly protein (Fragment) \| Putative phage tail assembly protein (Fragment); uniclust \| UniRef100\_A0A0J1NAY6 \| 99.7 \| 6.3e-21 \| 1.2e-26 \| 128.8 \| 70 \| (72, 143) \| 143 \| (113, 182) \| 190 \| Tail fiber assembly protein \| Tail fiber assembly protein; uniclust \| UniRef100\_A0A1C7W7E1 \| 99.7 \| 6.8e-21 \| 1.3e-26 \| 110.4 \| 72 \| (1, 72) \| 143 \| (1, 72) \| 80 \| Phage tail protein \| Phage tail protein; uniclust \| UniRef100\_A0A077NP04 \| 99.7 \| 8.5e-21 \| 1.6e-26 \| 126.9 \| 80 \| (62, 143) \| 143 \| (118, 197) \| 201 \| Tail fiber assembly protein \| Tail fiber assembly protein; uniclust \| UniRef100\_A0A2T8TCL9 \| 99.7 \| 1.2e-20 \| 2.3e-26 \| 122.4 \| 132 \| (2, 143) \| 143 \| (1, 169) \| 180 \| Tail assembly chaperone \| Tail assembly chaperone; uniclust \| UniRef100\_UPI00094363AD \| 99.7 \| 1.7e-20 \| 3.2e-26 \| 116.7 \| 106 \| (2, 112) \| 143 \| (12, 117) \| 121 \| tail fiber assembly protein \| tail fiber assembly protein; uniclust \| UniRef100\_A0A3N2RWI5 \| 99.7 \| 2.1e-20 \| 3.9e-26 \| 121.0 \| 131 \| (3, 143) \| 143 \| (33, 184) \| 188 \| Tail fiber assembly protein \| Tail fiber assembly protein; uniclust \| UniRef100\_A0A2E3R781 \| 99.7 \| 2.7e-20 \| 4.9e-26 \| 114.3 \| 126 \| (1, 143) \| 143 \| (7, 135) \| 138 \| Phage tail protein \| Phage tail protein; uniclust \| UniRef100\_A0A1D8UTE2 \| 99.7 \| 2.8e-20 \| 5.3e-26 \| 109.5 \| 70 \| (1, 70) \| 143 \| (6, 75) \| 83 \| Uncharacterized protein \| Uncharacterized protein; uniclust \| UniRef100\_UPI001D07D678 \| 99.7 \| 3.3e-20 \| 6.2e-26 \| 115.0 \| 139 \| (2, 142) \| 143 \| (1, 159) \| 159 \| tail fiber assembly protein \| tail fiber assembly protein; uniclust \| UniRef100\_A0A2T6LTZ2 \| 99.7 \| 3.5e-20 \| 6.5e-26 \| 115.2 \| 127 \| (3, 143) \| 143 \| (1, 127) \| 134 \| Virus tail fiber assembly protein lambda gpK \| Virus tail fiber assembly protein lambda gpK; uniclust \| UniRef100\_A0A0T9T931 \| 99.7 \| 3.8e-20 \| 7.2e-26 \| 118.7 \| 110 \| (27, 143) \| 143 \| (44, 157) \| 160 \| Tail fiber assembly protein G \| Tail fiber assembly protein G; uniclust \| UniRef100\_UPI001322C4D5 \| 99.7 \| 4.2e-20 \| 7.8e-26 \| 106.3 \| 74 \| (68, 143) \| 143 \| (13, 86) \| 89 \| tail fiber assembly protein \| tail fiber assembly protein; uniclust \| UniRef100\_A0A0T9T1K4 \| 99.7 \| 4.5e-20 \| 8.5e-26 \| 123.2 \| 85 \| (57, 143) \| 143 \| (124, 208) \| 225 \| Tail fiber assembly protein G \| Tail fiber assembly protein G; uniclust \| UniRef100\_A0A022PI45 \| 99.7 \| 5e-20 \| 9.8e-26 \| 128.7 \| 80 \| (62, 143) \| 143 \| (134, 213) \| 230 \| Caudovirales tail fiber assembly protein \| Caudovirales tail fiber assembly protein; uniclust \| UniRef100\_A0A1Z1STM3 \| 99.7 \| 6e-20 \| 1.1e-25 \| 113.7 \| 133 \| (3, 143) \| 143 \| (1, 134) \| 134 \| Tail fiber assembly protein \| Tail fiber assembly protein; uniclust \| UniRef100\_A0A022PBR1 \| 99.7 \| 6e-20 \| 1.2e-25 \| 128.5 \| 80 \| (62, 143) \| 143 \| (155, 234) \| 245 \| Bacteriophage tail assembly protein \| Bacteriophage tail assembly protein; uniclust \| UniRef100\_A0A0T9R7L9 \| 99.7 \| 7.9e-20 \| 1.5e-25 \| 123.0 \| 128 \| (1, 143) \| 143 \| (35, 174) \| 202 \| Tail fiber assembly protein G \| Tail fiber assembly protein G; uniclust \| UniRef100\_A0A1Q6B6N7 \| 99.7 \| 8.5e-20 \| 1.6e-25 \| 107.6 \| 84 \| (58, 143) \| 143 \| (29, 112) \| 116 \| Phage tail protein \| Phage tail protein; uniclust \| UniRef100\_A0A127Q904 \| 99.7 \| 8.8e-20 \| 1.7e-25 \| 123.8 \| 104 \| (38, 143) \| 143 \| (86, 196) \| 213 \| Caudovirales tail fiber assembly family protein \| Caudovirales tail fiber assembly family protein; uniclust \| UniRef100\_A0A0A1IVW2 \| 99.7 \| 1e-19 \| 1.9e-25 \| 119.7 \| 75 \| (67, 143) \| 143 \| (76, 150) \| 161 \| Tail fiber component \| Tail fiber component; uniclust \| UniRef100\_A0A068R0K4 \| 99.7 \| 1.2e-19 \| 2.3e-25 \| 121.8 \| 72 \| (67, 143) \| 143 \| (110, 181) \| 205 \| Tail fiber assembly protein \| Tail fiber assembly protein; uniclust \| UniRef100\_A0A0A0CIE4 \| 99.7 \| 1.2e-19 \| 2.3e-25 \| 115.5 \| 86 \| (56, 143) \| 143 \| (46, 131) \| 143 \| Tail assembly protein (Fragment) \| Tail assembly protein (Fragment); uniclust \| UniRef100\_A0A024L1Y3 \| 99.7 \| 1.4e-19 \| 2.8e-25 \| 129.0 \| 81 \| (61, 143) \| 143 \| (149, 229) \| 266 \| Tail fiber assembly protein \| Tail fiber assembly protein; uniclust \| UniRef100\_UPI00131B923D \| 99.7 \| 1.8e-19 \| 3.3e-25 \| 109.5 \| 126 \| (4, 142) \| 143 \| (1, 127) \| 130 \| tail fiber assembly protein \| tail fiber assembly protein; uniclust \| UniRef100\_UPI001FC7C713 \| 99.7 \| 1.9e-19 \| 3.5e-25 \| 110.1 \| 114 \| (26, 143) \| 143 \| (34, 148) \| 149 \| tail fiber assembly protein \| tail fiber assembly protein; uniclust \| UniRef100\_UPI000664EF55 \| 99.6 \| 2.4e-19 \| 4.4e-25 \| 101.0 \| 80 \| (4, 88) \| 143 \| (1, 80) \| 82 \| tail fiber assembly protein \| tail fiber assembly protein; uniclust \| UniRef100\_A0A0H3I5Y1 \| 99.6 \| 2.5e-19 \| 4.7e-25 \| 113.9 \| 81 \| (61, 143) \| 143 \| (46, 128) \| 145 \| Tail fiber chaperone, Qin prophage \| Tail fiber chaperone, Qin prophage; uniclust \| UniRef100\_A0A0T9RSX1 \| 99.6 \| 2.8e-19 \| 5.2e-25 \| 101.8 \| 75 \| (67, 143) \| 143 \| (6, 80) \| 84 \| Tail fiber assembly protein p37 \| Tail fiber assembly protein p37; uniclust \| UniRef100\_A0A2S8PUH6 \| 99.6 \| 3.1e-19 \| 5.7e-25 \| 96.6 \| 59 \| (82, 142) \| 143 \| (2, 60) \| 64 \| Phage tail protein (Fragment) \| Phage tail protein (Fragment); uniclust \| UniRef100\_A0A0D1LLW8 \| 99.6 \| 3.3e-19 \| 6.3e-25 \| 109.5 \| 72 \| (70, 143) \| 143 \| (33, 106) \| 117 \| Tail fiber protein \| Tail fiber protein; uniclust \| UniRef100\_A0A1X8WD02 \| 99.6 \| 4.1e-19 \| 7.7e-25 \| 102.6 \| 79 \| (60, 143) \| 143 \| (11, 89) \| 91 \| Caudovirales tail fiber assembly protein \| Caudovirales tail fiber assembly protein; uniclust \| UniRef100\_A0A2R7PZL5 \| 99.6 \| 4.2e-19 \| 7.9e-25 \| 109.4 \| 66 \| (76, 143) \| 143 \| (52, 119) \| 121 \| Phage tail protein (Fragment) \| Phage tail protein (Fragment); uniclust \| UniRef100\_A0A851FVU6 \| 99.6 \| 5.2e-19 \| 9.7e-25 \| 109.8 \| 134 \| (2, 143) \| 143 \| (1, 135) \| 142 \| Tail fiber assembly protein \| Tail fiber assembly protein; uniclust \| UniRef100\_UPI000C786B2F \| 99.6 \| 6.9e-19 \| 1.3e-24 \| 105.0 \| 113 \| (29, 143) \| 143 \| (6, 122) \| 123 \| tail fiber assembly protein \| tail fiber assembly protein; uniclust \| UniRef100\_A0A418GGH7 \| 99.6 \| 7.2e-19 \| 1.3e-24 \| 109.5 \| 80 \| (61, 142) \| 143 \| (63, 142) \| 144 \| Phage tail protein (Fragment) \| Phage tail protein (Fragment); uniclust \| UniRef100\_A0A2E0R6L6 \| 99.6 \| 7e-19 \| 1.3e-24 \| 113.7 \| 122 \| (2, 143) \| 143 \| (4, 127) \| 148 \| Phage tail protein \| Phage tail protein; uniclust \| UniRef100\_A0A0P9WFS5 \| 99.6 \| 7.1e-19 \| 1.4e-24 \| 113.9 \| 81 \| (61, 143) \| 143 \| (68, 150) \| 157 \| Tail fiber assembly protein \| Tail fiber assembly protein; uniclust \| UniRef100\_UPI000560D39E \| 99.6 \| 7.4e-19 \| 1.4e-24 \| 111.1 \| 101 \| (30, 143) \| 143 \| (45, 147) \| 147 \| tail fiber assembly protein \| tail fiber assembly protein; uniclust \| UniRef100\_A0A0N1JDF2 \| 99.6 \| 7.6e-19 \| 1.4e-24 \| 112.3 \| 111 \| (30, 142) \| 143 \| (29, 141) \| 154 \| Prophage PSPPH06 \| Prophage PSPPH06; uniclust \| UniRef100\_A0A060AGS0 \| 99.6 \| 8.4e-19 \| 1.7e-24 \| 123.5 \| 74 \| (68, 143) \| 143 \| (127, 200) \| 232 \| Putative tail fiber assembly protein \| Putative tail fiber assembly protein; uniclust \| UniRef100\_A0A485I3W6 \| 99.6 \| 1.2e-18 \| 2.2e-24 \| 109.6 \| 134 \| (2, 142) \| 143 \| (6, 143) \| 167 \| Tail fiber assembly protein \| Tail fiber assembly protein; uniclust \| UniRef100\_A0A0H3KZW0 \| 99.6 \| 1.4e-18 \| 2.7e-24 \| 117.3 \| 82 \| (60, 143) \| 143 \| (123, 204) \| 208 \| Tail fiber assembly protein \| Tail fiber assembly protein; uniclust \| UniRef100\_A0A158FMJ5 \| 99.6 \| 1.5e-18 \| 2.7e-24 \| 111.0 \| 74 \| (67, 143) \| 143 \| (70, 143) \| 156 \| Caudovirales tail fiber assembly protein \| Caudovirales tail fiber assembly protein; uniclust \| UniRef100\_UPI000695DC3B \| 99.6 \| 1.8e-18 \| 3.3e-24 \| 103.1 \| 75 \| (67, 143) \| 143 \| (46, 120) \| 120 \| tail fiber assembly protein \| tail fiber assembly protein; uniclust \| UniRef100\_A0A376EWH3 \| 99.6 \| 2.3e-18 \| 4.3e-24 \| 103.5 \| 86 \| (56, 143) \| 143 \| (12, 103) \| 105 \| Phage tail assembly chaperone gp38 \| Phage tail assembly chaperone gp38; uniclust \| UniRef100\_UPI00068E9C67 \| 99.6 \| 2.7e-18 \| 4.9e-24 \| 102.7 \| 80 \| (62, 143) \| 143 \| (44, 123) \| 124 \| tail fiber assembly protein \| tail fiber assembly protein; uniclust \| UniRef100\_A0A2A9KIG4 \| 99.6 \| 3.1e-18 \| 5.9e-24 \| 110.2 \| 84 \| (58, 143) \| 143 \| (59, 148) \| 153 \| Virus tail fiber assembly protein lambda gpK \| Virus tail fiber assembly protein lambda gpK; uniclust \| UniRef100\_UPI00210832D5 \| 99.6 \| 3.8e-18 \| 7e-24 \| 108.7 \| 76 \| (66, 143) \| 143 \| (118, 193) \| 193 \| tail fiber assembly protein \| tail fiber assembly protein; uniclust \| UniRef100\_A0A1G7SHF9 \| 99.6 \| 4.3e-18 \| 8.4e-24 \| 117.0 \| 90 \| (1, 90) \| 143 \| (9, 99) \| 207 \| Uncharacterized protein (Fragment) \| Uncharacterized protein (Fragment); uniclust \| UniRef100\_A0A336P4J4 \| 99.6 \| 5.1e-18 \| 9.6e-24 \| 100.2 \| 91 \| (49, 143) \| 143 \| (7, 98) \| 102 \| Phage tail fiber assembly protein \| Phage tail fiber assembly protein; uniclust \| UniRef100\_A0A2J9ED04 \| 99.6 \| 5.1e-18 \| 9.8e-24 \| 110.7 \| 101 \| (30, 143) \| 143 \| (51, 153) \| 158 \| Tail fiber assembly protein \| Tail fiber assembly protein; uniclust \| UniRef100\_UPI00200582ED \| 99.6 \| 5.8e-18 \| 1.1e-23 \| 105.5 \| 93 \| (2, 94) \| 143 \| (1, 93) \| 165 \| tail fiber assembly protein \| tail fiber assembly protein; uniclust \| UniRef100\_A0A1F4JLP3 \| 99.6 \| 6.7e-18 \| 1.3e-23 \| 107.1 \| 124 \| (2, 142) \| 143 \| (15, 140) \| 146 \| Phage tail protein \| Phage tail protein; uniclust \| UniRef100\_UPI0004B0C64D \| 99.5 \| 1.3e-17 \| 2.3e-23 \| 97.3 \| 92 \| (46, 143) \| 143 \| (2, 93) \| 96 \| tail fiber assembly protein \| tail fiber assembly protein; uniclust \| UniRef100\_A0A0T9RKV0 \| 99.5 \| 1.4e-17 \| 2.6e-23 \| 106.2 \| 87 \| (50, 143) \| 143 \| (53, 140) \| 151 \| Putative phage tail fiber assembly protein \| Putative phage tail fiber assembly protein; uniclust \| UniRef100\_UPI0007359171 \| 99.5 \| 1.6e-17 \| 3e-23 \| 103.1 \| 139 \| (2, 143) \| 143 \| (1, 141) \| 141 \| hypothetical protein \| hypothetical protein; uniclust \| UniRef100\_A0A1C7WD91 \| 99.5 \| 1.6e-17 \| 3e-23 \| 116.8 \| 83 \| (58, 143) \| 143 \| (213, 295) \| 296 \| Phage tail protein \| Phage tail protein; uniclust \| UniRef100\_A0A5T9PXD7 \| 99.5 \| 1.6e-17 \| 3.1e-23 \| 105.2 \| 84 \| (57, 143) \| 143 \| (46, 129) \| 134 \| Tail fiber assembly protein \| Tail fiber assembly protein; uniclust \| UniRef100\_A0A2A2PEQ9 \| 99.5 \| 2.2e-17 \| 4.1e-23 \| 92.6 \| 60 \| (82, 143) \| 143 \| (9, 68) \| 71 \| Phage tail protein \| Phage tail protein; uniclust \| UniRef100\_A0A022PAN9 \| 99.5 \| 2.3e-17 \| 4.3e-23 \| 115.8 \| 72 \| (70, 143) \| 143 \| (138, 209) \| 252 \| Caudovirales tail fiber assembly protein \| Caudovirales tail fiber assembly protein; uniclust \| UniRef100\_A0A081RRP7 \| 99.5 \| 2.4e-17 \| 4.5e-23 \| 106.5 \| 74 \| (68, 143) \| 143 \| (97, 170) \| 171 \| Caudovirales tail fiber assembly protein \| Caudovirales tail fiber assembly protein; uniclust \| UniRef100\_A0A4R4K582 \| 99.5 \| 2.5e-17 \| 4.6e-23 \| 97.2 \| 73 \| (69, 143) \| 143 \| (14, 86) \| 95 \| Phage tail protein \| Phage tail protein; uniclust \| UniRef100\_A0A081RR89 \| 99.5 \| 4.9e-17 \| 9.1e-23 \| 99.5 \| 63 \| (78, 143) \| 143 \| (54, 116) \| 124 \| Caudovirales tail fiber assembly protein \| Caudovirales tail fiber assembly protein; uniclust \| UniRef100\_A0A1H8RBN5 \| 99.5 \| 4.7e-17 \| 9.1e-23 \| 108.5 \| 113 \| (2, 121) \| 143 \| (5, 124) \| 168 \| Phage tail assembly chaperone protein \| Phage tail assembly chaperone protein; uniclust \| UniRef100\_A0A5K4YG87 \| 99.5 \| 6.4e-17 \| 1.2e-22 \| 91.7 \| 75 \| (1, 75) \| 143 \| (1, 76) \| 79 \| Uncharacterized protein \| Uncharacterized protein; uniclust \| UniRef100\_A0A2Z3I402 \| 99.5 \| 6.5e-17 \| 1.2e-22 \| 105.1 \| 81 \| (61, 143) \| 143 \| (98, 180) \| 181 \| Phage tail protein \| Phage tail protein; uniclust \| UniRef100\_UPI001D06A5C9 \| 99.5 \| 7.5e-17 \| 1.4e-22 \| 88.9 \| 71 \| (60, 132) \| 143 \| (4, 74) \| 78 \| tail fiber assembly protein \| tail fiber assembly protein; uniclust \| UniRef100\_A0A0F0FPS3 \| 99.5 \| 8.9e-17 \| 1.7e-22 \| 97.6 \| 81 \| (61, 143) \| 143 \| (43, 125) \| 125 \| Tail fiber assembly protein \| Tail fiber assembly protein; uniclust \| UniRef100\_A0A0F7Y2W8 \| 99.5 \| 9.6e-17 \| 1.8e-22 \| 105.8 \| 68 \| (74, 143) \| 143 \| (116, 185) \| 186 \| Phage tail protein \| Phage tail protein; uniclust \| UniRef100\_A0A0A1FEP8 \| 99.5 \| 1e-16 \| 2e-22 \| 113.8 \| 71 \| (71, 143) \| 143 \| (173, 245) \| 261 \| Phage tail fiber assembly protein \| Phage tail fiber assembly protein; uniclust \| UniRef100\_UPI0021BD8A01 \| 99.5 \| 1.5e-16 \| 2.7e-22 \| 94.4 \| 67 \| (75, 143) \| 143 \| (42, 108) \| 108 \| tail fiber assembly protein \| tail fiber assembly protein; uniclust \| UniRef100\_A0A5N3CRX4 \| 99.5 \| 1.5e-16 \| 2.9e-22 \| 93.7 \| 70 \| (1, 70) \| 143 \| (9, 79) \| 86 \| Phage tail protein \| Phage tail protein; uniclust \| UniRef100\_A0A0C5DSP9 \| 99.5 \| 1.7e-16 \| 3.3e-22 \| 108.2 \| 72 \| (70, 143) \| 143 \| (116, 189) \| 208 \| Tail fiber assembly protein \| Tail fiber assembly protein; uniclust \| UniRef100\_A0A2X2RKD0 \| 99.5 \| 1.9e-16 \| 3.4e-22 \| 94.9 \| 85 \| (50, 143) \| 143 \| (26, 110) \| 114 \| Phage tail fiber assembly protein \| Phage tail fiber assembly protein; uniclust \| UniRef100\_A0A0K4WW72 \| 99.4 \| 2.5e-16 \| 4.7e-22 \| 92.5 \| 85 \| (1, 85) \| 143 \| (1, 85) \| 111 \| Tail fiber assembly protein \| Tail fiber assembly protein; uniclust \| UniRef100\_A0A2N1H596 \| 99.4 \| 3.3e-16 \| 6.2e-22 \| 100.1 \| 133 \| (2, 143) \| 143 \| (1, 137) \| 147 \| Phage tail protein \| Phage tail protein; uniclust \| UniRef100\_A0A743TIT6 \| 99.4 \| 3.4e-16 \| 6.2e-22 \| 91.5 \| 92 \| (46, 143) \| 143 \| (6, 97) \| 99 \| Tail fiber assembly protein (Fragment) \| Tail fiber assembly protein (Fragment); uniclust \| UniRef100\_A0A5V7Y4W7 \| 99.4 \| 3.5e-16 \| 6.4e-22 \| 91.8 \| 81 \| (58, 143) \| 143 \| (18, 98) \| 101 \| Tail fiber assembly protein \| Tail fiber assembly protein; uniclust \| UniRef100\_UPI001E56960D \| 99.4 \| 3.7e-16 \| 6.7e-22 \| 92.3 \| 97 \| (44, 143) \| 143 \| (3, 99) \| 110 \| tail fiber assembly protein \| tail fiber assembly protein; uniclust \| UniRef100\_A0A2I8Q8G5 \| 99.4 \| 3.9e-16 \| 7.1e-22 \| 93.8 \| 69 \| (73, 143) \| 143 \| (57, 125) \| 127 \| Phage tail protein \| Phage tail protein; uniclust \| UniRef100\_UPI00224466F1 \| 99.4 \| 4e-16 \| 7.4e-22 \| 94.7 \| 128 \| (2, 142) \| 143 \| (1, 130) \| 135 \| phage tail assembly chaperone \| phage tail assembly chaperone; uniclust \| UniRef100\_C4K6I6 \| 99.4 \| 4.2e-16 \| 7.8e-22 \| 82.2 \| 55 \| (87, 143) \| 143 \| (2, 56) \| 57 \| Putative tail fiber assembly protein \| Putative tail fiber assembly protein; uniclust \| UniRef100\_A0A225SMD4 \| 99.4 \| 4.3e-16 \| 8e-22 \| 102.4 \| 72 \| (70, 143) \| 143 \| (86, 159) \| 184 \| Phage tail protein \| Phage tail protein; uniclust \| UniRef100\_A0A380AA26 \| 99.4 \| 5.1e-16 \| 9.6e-22 \| 89.0 \| 68 \| (1, 68) \| 143 \| (1, 69) \| 80 \| Tail fiber assembly protein \| Tail fiber assembly protein; uniclust \| UniRef100\_A0A1Q8SPJ2 \| 99.4 \| 5.9e-16 \| 1.1e-21 \| 103.0 \| 88 \| (2, 89) \| 143 \| (8, 96) \| 213 \| DUF4376 domain-containing protein \| DUF4376 domain-containing protein; uniclust \| UniRef100\_UPI0003FF973E \| 99.4 \| 6.2e-16 \| 1.2e-21 \| 88.1 \| 67 \| (2, 68) \| 143 \| (3, 69) \| 79 \| hypothetical protein \| hypothetical protein; uniclust \| UniRef100\_UPI001F296025 \| 99.4 \| 6.9e-16 \| 1.3e-21 \| 94.2 \| 79 \| (65, 143) \| 143 \| (49, 127) \| 129 \| tail fiber assembly protein \| tail fiber assembly protein; uniclust \| UniRef100\_UPI00067AAE6F \| 99.4 \| 7.5e-16 \| 1.4e-21 \| 91.6 \| 100 \| (3, 114) \| 143 \| (1, 101) \| 112 \| tail fiber assembly protein \| tail fiber assembly protein; uniclust \| UniRef100\_A0A1G7PVA6 \| 99.4 \| 7.5e-16 \| 1.5e-21 \| 102.2 \| 64 \| (2, 70) \| 143 \| (1, 64) \| 160 \| Uncharacterized protein \| Uncharacterized protein; uniclust \| UniRef100\_A0A5T6NIG6 \| 99.4 \| 8e-16 \| 1.5e-21 \| 86.8 \| 70 \| (69, 143) \| 143 \| (6, 75) \| 78 \| Tail fiber assembly protein \| Tail fiber assembly protein; uniclust \| UniRef100\_A0A6I3WIV8 \| 99.4 \| 8.2e-16 \| 1.5e-21 \| 98.7 \| 67 \| (75, 143) \| 143 \| (85, 153) \| 157 \| Tail fiber assembly protein \| Tail fiber assembly protein; uniclust \| UniRef100\_UPI001883862A \| 99.4 \| 8.5e-16 \| 1.6e-21 \| 97.8 \| 137 \| (3, 142) \| 143 \| (35, 179) \| 183 \| tail fiber assembly protein \| tail fiber assembly protein; uniclust \| UniRef100\_A0A0W0MTR4 \| 99.4 \| 8.9e-16 \| 1.7e-21 \| 96.0 \| 70 \| (72, 143) \| 143 \| (69, 140) \| 140 \| Phage tail protein (Fragment) \| Phage tail protein (Fragment); uniclust \| UniRef100\_A0A080KAQ9 \| 99.4 \| 9e-16 \| 1.7e-21 \| 94.7 \| 65 \| (77, 143) \| 143 \| (50, 114) \| 124 \| Caudovirales tail fiber assembly protein (Fragment) \| Caudovirales tail fiber assembly protein (Fragment); uniclust \| UniRef100\_A0A432UNX0 \| 99.4 \| 1.2e-15 \| 2.2e-21 \| 100.2 \| 73 \| (2, 74) \| 143 \| (1, 74) \| 194 \| Tail fiber assembly protein \| Tail fiber assembly protein; uniclust \| UniRef100\_A0A0N9WN70 \| 99.4 \| 1.4e-15 \| 2.5e-21 \| 100.5 \| 136 \| (4, 143) \| 143 \| (1, 236) \| 239 \| Tail fiber assembly protein \| Tail fiber assembly protein; uniclust \| UniRef100\_A0A2R8CKR9 \| 99.4 \| 1.4e-15 \| 2.5e-21 \| 101.3 \| 67 \| (2, 68) \| 143 \| (3, 69) \| 202 \| Uncharacterized protein \| Uncharacterized protein; uniclust \| UniRef100\_A0A0A0CJA3 \| 99.4 \| 1.4e-15 \| 2.5e-21 \| 80.6 \| 44 \| (100, 143) \| 143 \| (9, 52) \| 55 \| Tail assembly protein (Fragment) \| Tail assembly protein (Fragment); uniclust \| UniRef100\_A0A1H3DNJ2 \| 99.4 \| 1.5e-15 \| 2.8e-21 \| 90.5 \| 66 \| (75, 142) \| 143 \| (38, 105) \| 107 \| Virus tail fibre assembly protein, lambda gpK \| Virus tail fibre assembly protein, lambda gpK; uniclust \| UniRef100\_A0A1D9LI34 \| 99.4 \| 1.6e-15 \| 3.1e-21 \| 96.9 \| 68 \| (2, 74) \| 143 \| (1, 68) \| 149 \| Tail fiber assembly protein \| Tail fiber assembly protein; uniclust \| UniRef100\_UPI0009EA3590 \| 99.4 \| 1.7e-15 \| 3.2e-21 \| 97.2 \| 68 \| (74, 143) \| 143 \| (95, 164) \| 164 \| tail fiber assembly protein \| tail fiber assembly protein; uniclust \| UniRef100\_UPI0002198843 \| 99.4 \| 1.9e-15 \| 3.6e-21 \| 99.7 \| 63 \| (2, 69) \| 143 \| (1, 63) \| 183 \| DUF4376 domain-containing protein \| DUF4376 domain-containing protein; uniclust \| UniRef100\_UPI000DDB98E6 \| 99.4 \| 3e-15 \| 5.5e-21 \| 89.3 \| 99 \| (1, 102) \| 143 \| (3, 103) \| 107 \| tail fiber assembly protein \| tail fiber assembly protein; uniclust \| UniRef100\_A0A1C4G4F0 \| 99.3 \| 3.3e-15 \| 6.1e-21 \| 100.2 \| 66 \| (73, 143) \| 143 \| (138, 203) \| 216 \| Virus tail fibre assembly protein, lambda gpK \| Virus tail fibre assembly protein, lambda gpK; uniclust \| UniRef100\_A0A0A7S0J4 \| 99.3 \| 3.6e-15 \| 6.8e-21 \| 92.2 \| 74 \| (67, 143) \| 143 \| (42, 115) \| 117 \| Caudovirales tail fiber assembly protein \| Caudovirales tail fiber assembly protein; uniclust \| UniRef100\_UPI00208F983D \| 99.3 \| 3.9e-15 \| 7.1e-21 \| 91.4 \| 81 \| (60, 142) \| 143 \| (62, 142) \| 143 \| tail fiber assembly protein \| tail fiber assembly protein; uniclust \| UniRef100\_D4I2C0 \| 99.3 \| 3.9e-15 \| 7.2e-21 \| 85.7 \| 56 \| (86, 143) \| 143 \| (2, 57) \| 97 \| Tail fiber assembly protein homolog from lambdoid prophage DLP12 \| Tail fiber assembly protein homolog from lambdoid prophage DLP12; uniclust \| UniRef100\_A0A0F4T3H0 \| 99.3 \| 4e-15 \| 7.6e-21 \| 97.7 \| 74 \| (66, 143) \| 143 \| (70, 144) \| 150 \| Tail fiber assembly protein \| Tail fiber assembly protein; uniclust \| UniRef100\_A0A087NAA6 \| 99.3 \| 4e-15 \| 7.6e-21 \| 99.0 \| 71 \| (1, 77) \| 143 \| (10, 80) \| 170 \| Uncharacterized protein \| Uncharacterized protein; uniclust \| UniRef100\_UPI001EE2F813 \| 99.3 \| 4.1e-15 \| 7.6e-21 \| 104.2 \| 73 \| (69, 143) \| 143 \| (288, 360) \| 360 \| tail fiber assembly protein \| tail fiber assembly protein; uniclust \| UniRef100\_A0A2N8AP07 \| 99.3 \| 4.8e-15 \| 8.8e-21 \| 98.1 \| 134 \| (2, 142) \| 143 \| (96, 231) \| 240 \| Phage tail protein \| Phage tail protein; uniclust \| UniRef100\_A0A8S5V3M4 \| 99.3 \| 4.8e-15 \| 8.9e-21 \| 94.5 \| 112 \| (10, 143) \| 143 \| (23, 137) \| 156 \| Tail assembly chaperone protein \| Tail assembly chaperone protein; uniclust \| UniRef100\_A0A225SMJ4 \| 99.3 \| 5.1e-15 \| 9.4e-21 \| 90.8 \| 66 \| (2, 72) \| 143 \| (1, 66) \| 129 \| Phage tail protein \| Phage tail protein; uniclust \| UniRef100\_A0A031FRM8 \| 99.3 \| 5.4e-15 \| 1e-20 \| 99.0 \| 79 \| (62, 143) \| 143 \| (86, 168) \| 171 \| Caudovirales tail fiber assembly protein \| Caudovirales tail fiber assembly protein; uniclust \| UniRef100\_A0A077F610 \| 99.3 \| 6.5e-15 \| 1.2e-20 \| 102.3 \| 70 \| (72, 143) \| 143 \| (158, 229) \| 246 \| Phage tail fiber assembly protein \| Phage tail fiber assembly protein; uniclust \| UniRef100\_G9E597 \| 99.3 \| 7e-15 \| 1.3e-20 \| 92.0 \| 127 \| (1, 143) \| 143 \| (19, 147) \| 151 \| Tail fiber assembly protein \| Tail fiber assembly protein; uniclust \| UniRef100\_A0A5C7CFK8 \| 99.3 \| 7.1e-15 \| 1.3e-20 \| 94.3 \| 76 \| (66, 143) \| 143 \| (100, 175) \| 191 \| Tail fiber assembly protein \| Tail fiber assembly protein; uniclust \| UniRef100\_UPI000680C4D0 \| 99.3 \| 7.6e-15 \| 1.4e-20 \| 98.3 \| 84 \| (58, 143) \| 143 \| (177, 260) \| 260 \| tail fiber assembly protein \| tail fiber assembly protein; uniclust \| UniRef100\_A0A024E8T8 \| 99.3 \| 8.7e-15 \| 1.6e-20 \| 97.9 \| 70 \| (72, 143) \| 143 \| (116, 187) \| 200 \| Tail fiber assembly protein \| Tail fiber assembly protein; uniclust \| UniRef100\_UPI00068D9C5E \| 99.3 \| 9.1e-15 \| 1.7e-20 \| 91.8 \| 65 \| (76, 143) \| 143 \| (66, 130) \| 131 \| tail fiber assembly protein \| tail fiber assembly protein; uniclust \| UniRef100\_A0A022PFM9 \| 99.3 \| 9.8e-15 \| 1.8e-20 \| 108.2 \| 84 \| (58, 143) \| 143 \| (450, 533) \| 534 \| Phage-related tail fiber protein \| Phage-related tail fiber protein; uniclust \| UniRef100\_A0A192A522 \| 99.3 \| 2e-14 \| 3.8e-20 \| 100.7 \| 76 \| (2, 77) \| 143 \| (4, 105) \| 240 \| DUF4376 domain-containing protein \| DUF4376 domain-containing protein; uniclust \| UniRef100\_A0A7Y0Z1A9 \| 99.3 \| 2.1e-14 \| 3.9e-20 \| 85.6 \| 67 \| (75, 143) \| 143 \| (37, 105) \| 105 \| Tail fiber assembly protein \| Tail fiber assembly protein; uniclust \| UniRef100\_UPI0002EF02CE \| 99.3 \| 2.2e-14 \| 4e-20 \| 81.4 \| 80 \| (2, 82) \| 143 \| (1, 80) \| 81 \| hypothetical protein \| hypothetical protein; uniclust \| UniRef100\_A0A246GTR2 \| 99.3 \| 2.4e-14 \| 4.6e-20 \| 93.9 \| 128 \| (2, 143) \| 143 \| (1, 133) \| 145 \| Phage tail protein \| Phage tail protein; uniclust \| UniRef100\_A0A0S4UUF1 \| 99.3 \| 2.7e-14 \| 5.1e-20 \| 98.4 \| 69 \| (72, 143) \| 143 \| (152, 220) \| 228 \| Integrase/recombinase \| Integrase/recombinase; uniclust \| UniRef100\_A0A0H3PBT9 \| 99.2 \| 2.8e-14 \| 5.4e-20 \| 102.2 \| 79 \| (63, 143) \| 143 \| (129, 207) \| 256 \| Uncharacterized protein (Fragment) \| Uncharacterized protein (Fragment); uniclust \| UniRef100\_UPI00077BD77D \| 99.2 \| 3.2e-14 \| 5.8e-20 \| 91.0 \| 94 \| (2, 95) \| 143 \| (1, 94) \| 184 \| tail fiber assembly protein \| tail fiber assembly protein; uniclust \| UniRef100\_A0A077NN52 \| 99.2 \| 3.3e-14 \| 6.1e-20 \| 88.8 \| 81 \| (61, 143) \| 143 \| (72, 152) \| 152 \| Tail fiber assembly protein \| Tail fiber assembly protein; uniclust \| UniRef100\_A0A2A5RDH4 \| 99.2 \| 3.3e-14 \| 6.3e-20 \| 92.5 \| 74 \| (68, 143) \| 143 \| (69, 146) \| 146 \| Phage tail protein \| Phage tail protein; uniclust \| UniRef100\_UPI00190065B7 \| 99.2 \| 3.5e-14 \| 6.5e-20 \| 87.6 \| 130 \| (1, 130) \| 143 \| (1, 132) \| 145 \| hypothetical protein \| hypothetical protein; uniclust \| UniRef100\_A0A1A9VZ06 \| 99.2 \| 3.6e-14 \| 6.7e-20 \| 102.2 \| 73 \| (69, 143) \| 143 \| (403, 475) \| 479 \| Bacteriophage P22 tailspike N-terminal domain-containing protein \| Bacteriophage P22 tailspike N-terminal domain-containing protein; uniclust \| UniRef100\_A0A344SLS6 \| 99.2 \| 3.7e-14 \| 6.9e-20 \| 81.0 \| 70 \| (69, 143) \| 143 \| (5, 74) \| 80 \| Tail fiber assembly protein \| Tail fiber assembly protein; uniclust \| UniRef100\_A0A6L7C6M0 \| 99.2 \| 4.3e-14 \| 8e-20 \| 86.4 \| 64 \| (78, 143) \| 143 \| (68, 131) \| 131 \| Tail fiber assembly protein \| Tail fiber assembly protein; uniclust \| UniRef100\_UPI001883B016 \| 99.2 \| 4.4e-14 \| 8e-20 \| 101.8 \| 137 \| (3, 142) \| 143 \| (333, 477) \| 481 \| tail fiber assembly protein \| tail fiber assembly protein; uniclust \| UniRef100\_A0A7L6CMI7 \| 99.2 \| 4.5e-14 \| 8.4e-20 \| 86.3 \| 64 \| (76, 143) \| 143 \| (48, 112) \| 113 \| Tail fiber assembly protein \| Tail fiber assembly protein; uniclust \| UniRef100\_A0A023WVR1 \| 99.2 \| 4.5e-14 \| 8.6e-20 \| 91.6 \| 61 \| (2, 68) \| 143 \| (1, 61) \| 137 \| Phage tail protein \| Phage tail protein; uniclust \| UniRef100\_A0A0Q8NI86 \| 99.2 \| 4.9e-14 \| 9.4e-20 \| 91.8 \| 107 \| (11, 143) \| 143 \| (26, 134) \| 150 \| Phage tail protein \| Phage tail protein; uniclust \| UniRef100\_A0A1C1W012 \| 99.2 \| 6.1e-14 \| 1.1e-19 \| 92.9 \| 110 \| (31, 142) \| 143 \| (84, 195) \| 203 \| Phage tail protein \| Phage tail protein; uniclust \| UniRef100\_A0A2L0M0L4 \| 99.2 \| 6.2e-14 \| 1.1e-19 \| 82.3 \| 73 \| (69, 143) \| 143 \| (26, 100) \| 106 \| Phage tail protein \| Phage tail protein; uniclust \| UniRef100\_A0A2V1HJ93 \| 99.2 \| 6.9e-14 \| 1.3e-19 \| 85.8 \| 74 \| (67, 142) \| 143 \| (46, 121) \| 122 \| Phage tail protein \| Phage tail protein; uniclust \| UniRef100\_A0A0L1KFJ7 \| 99.2 \| 6.9e-14 \| 1.3e-19 \| 91.8 \| 59 \| (1, 64) \| 143 \| (5, 64) \| 156 \| Putative tail fiber assembly protein \| Putative tail fiber assembly protein; uniclust \| UniRef100\_A0A0K1QML5 \| 99.2 \| 7e-14 \| 1.3e-19 \| 97.7 \| 79 \| (63, 143) \| 143 \| (150, 230) \| 255 \| Tail fiber assembly protein \| Tail fiber assembly protein; uniclust \| UniRef100\_A0A077NTU9 \| 99.2 \| 7.2e-14 \| 1.3e-19 \| 75.5 \| 47 \| (92, 143) \| 143 \| (7, 53) \| 59 \| Tail fiber assembly protein \| Tail fiber assembly protein; uniclust \| UniRef100\_A0A261SNT8 \| 99.2 \| 1e-13 \| 1.9e-19 \| 91.5 \| 133 \| (2, 143) \| 143 \| (4, 151) \| 154 \| Phage tail protein \| Phage tail protein; uniclust \| UniRef100\_UPI000190DFD3 \| 99.2 \| 1.1e-13 \| 2e-19 \| 79.4 \| 72 \| (70, 143) \| 143 \| (7, 79) \| 83 \| tail fiber assembly protein \| tail fiber assembly protein; uniclust \| UniRef100\_UPI0022FDBD1A \| 99.2 \| 1.1e-13 \| 2.1e-19 \| 81.4 \| 72 \| (70, 143) \| 143 \| (35, 107) \| 107 \| tail fiber assembly protein \| tail fiber assembly protein; uniclust \| UniRef100\_UPI001101862B \| 99.2 \| 1.3e-13 \| 2.4e-19 \| 76.1 \| 62 \| (77, 143) \| 143 \| (3, 64) \| 65 \| tail fiber assembly protein \| tail fiber assembly protein; uniclust \| UniRef100\_UPI001124FB43 \| 99.2 \| 1.3e-13 \| 2.5e-19 \| 79.3 \| 84 \| (58, 143) \| 143 \| (3, 87) \| 89 \| tail fiber assembly protein \| tail fiber assembly protein; uniclust \| UniRef100\_A0A0A8VHP2 \| 99.2 \| 1.3e-13 \| 2.5e-19 \| 90.3 \| 133 \| (1, 142) \| 143 \| (1, 139) \| 169 \| Putative tail fiber assembly protein \| Putative tail fiber assembly protein; uniclust \| UniRef100\_UPI001256250B \| 99.2 \| 1.5e-13 \| 2.7e-19 \| 82.1 \| 82 \| (1, 90) \| 143 \| (1, 82) \| 109 \| tail fiber assembly protein \| tail fiber assembly protein; uniclust \| UniRef100\_A0A0J7J5L4 \| 99.2 \| 1.4e-13 \| 2.7e-19 \| 95.2 \| 63 \| (1, 68) \| 143 \| (6, 68) \| 202 \| Uncharacterized protein \| Uncharacterized protein; uniclust \| UniRef100\_A0A1Z9Q5B5 \| 99.2 \| 1.5e-13 \| 2.9e-19 \| 88.2 \| 118 \| (4, 141) \| 143 \| (1, 122) \| 137 \| Uncharacterized protein \| Uncharacterized protein; uniclust \| UniRef100\_A0A1B9KI83 \| 99.2 \| 1.6e-13 \| 3e-19 \| 79.8 \| 72 \| (69, 143) \| 143 \| (12, 83) \| 85 \| Tail fiber assembly protein \| Tail fiber assembly protein; uniclust \| UniRef100\_UPI001CCF0462 \| 99.2 \| 1.8e-13 \| 3.3e-19 \| 73.6 \| 59 \| (83, 143) \| 143 \| (3, 63) \| 63 \| tail fiber assembly protein \| tail fiber assembly protein; uniclust \| UniRef100\_A0A031GY32 \| 99.2 \| 1.9e-13 \| 3.7e-19 \| 92.1 \| 87 \| (1, 88) \| 143 \| (8, 98) \| 183 \| Uncharacterized protein \| Uncharacterized protein; uniclust \| UniRef100\_A0A1C7WAJ7 \| 99.1 \| 2.2e-13 \| 4.1e-19 \| 77.0 \| 64 \| (78, 143) \| 143 \| (5, 68) \| 75 \| Phage tail protein \| Phage tail protein; uniclust \| UniRef100\_A0A2N8B7K5 \| 99.1 \| 2.5e-13 \| 4.7e-19 \| 85.4 \| 69 \| (72, 142) \| 143 \| (71, 141) \| 147 \| Phage tail protein \| Phage tail protein; uniclust \| UniRef100\_UPI00164D51E1 \| 99.1 \| 2.9e-13 \| 5.2e-19 \| 85.1 \| 110 \| (30, 141) \| 143 \| (47, 158) \| 160 \| phage tail assembly chaperone \| phage tail assembly chaperone; uniclust \| UniRef100\_A0A3S0YDP2 \| 99.1 \| 2.8e-13 \| 5.3e-19 \| 92.9 \| 88 \| (1, 89) \| 143 \| (1, 133) \| 240 \| DUF4376 domain-containing protein \| DUF4376 domain-containing protein; uniclust \| UniRef100\_A0A7W4JZ22 \| 99.1 \| 3.1e-13 \| 5.8e-19 \| 85.7 \| 125 \| (3, 142) \| 143 \| (7, 133) \| 139 \| Tail fiber assembly protein \| Tail fiber assembly protein; uniclust \| UniRef100\_UPI00080DBA8F \| 99.1 \| 3.2e-13 \| 5.8e-19 \| 87.3 \| 128 \| (4, 142) \| 143 \| (58, 185) \| 193 \| tail fiber assembly protein \| tail fiber assembly protein; uniclust \| UniRef100\_A0A6N9S5D4 \| 99.1 \| 3.2e-13 \| 6e-19 \| 88.2 \| 67 \| (75, 143) \| 143 \| (141, 207) \| 209 \| Chromosome partitioning protein ParB \| Chromosome partitioning protein ParB; uniclust \| UniRef100\_UPI0023423F92 \| 99.1 \| 3.9e-13 \| 7.1e-19 \| 84.8 \| 56 \| (85, 142) \| 143 \| (105, 160) \| 163 \| tail fiber assembly protein \| tail fiber assembly protein; uniclust \| UniRef100\_A0A024E960 \| 99.1 \| 3.9e-13 \| 7.5e-19 \| 91.2 \| 72 \| (70, 143) \| 143 \| (79, 152) \| 171 \| Phage tail protein \| Phage tail protein; uniclust \| UniRef100\_A0A7J0BMD4 \| 99.1 \| 4.3e-13 \| 8.1e-19 \| 82.9 \| 60 \| (2, 68) \| 143 \| (1, 60) \| 129 \| Uncharacterized protein \| Uncharacterized protein; uniclust \| UniRef100\_A0A1X3J0U4 \| 99.1 \| 4.8e-13 \| 8.9e-19 \| 64.4 \| 33 \| (111, 143) \| 143 \| (2, 34) \| 35 \| Tail fiber assembly protein-like protein \| Tail fiber assembly protein-like protein; uniclust \| UniRef100\_UPI001C47AEF2 \| 99.1 \| 4.9e-13 \| 9.1e-19 \| 81.9 \| 124 \| (2, 143) \| 143 \| (1, 130) \| 135 \| phage tail assembly chaperone \| phage tail assembly chaperone; uniclust \| UniRef100\_UPI001FD41782 \| 99.1 \| 5.3e-13 \| 9.7e-19 \| 75.7 \| 58 \| (83, 143) \| 143 \| (29, 86) \| 86 \| tail fiber assembly protein \| tail fiber assembly protein; uniclust \| UniRef100\_UPI0011B1F365 \| 99.1 \| 5.7e-13 \| 1.1e-18 \| 80.8 \| 73 \| (69, 143) \| 143 \| (24, 101) \| 112 \| tail fiber assembly protein \| tail fiber assembly protein; uniclust \| UniRef100\_A0A5V6PU83 \| 99.1 \| 5.8e-13 \| 1.1e-18 \| 85.6 \| 110 \| (1, 118) \| 143 \| (1, 111) \| 184 \| Tail fiber assembly protein \| Tail fiber assembly protein; uniclust \| UniRef100\_A0A376D3X0 \| 99.1 \| 5.9e-13 \| 1.1e-18 \| 77.8 \| 93 \| (2, 101) \| 143 \| (1, 93) \| 101 \| Tail assembly chaperone gp38 \| Tail assembly chaperone gp38; uniclust \| UniRef100\_A0A2J7VZP8 \| 99.1 \| 6.2e-13 \| 1.1e-18 \| 85.2 \| 65 \| (77, 143) \| 143 \| (114, 180) \| 180 \| Uncharacterized protein \| Uncharacterized protein; uniclust \| UniRef100\_A0A192CNA1 \| 99.1 \| 6.1e-13 \| 1.1e-18 \| 84.1 \| 70 \| (69, 143) \| 143 \| (69, 138) \| 140 \| Tail fiber assembly protein \| Tail fiber assembly protein; uniclust \| UniRef100\_UPI001F210AA1 \| 99.1 \| 6.3e-13 \| 1.2e-18 \| 82.0 \| 67 \| (74, 142) \| 143 \| (72, 140) \| 141 \| tail fiber assembly protein \| tail fiber assembly protein; uniclust \| UniRef100\_A0A351RD40 \| 99.1 \| 7.4e-13 \| 1.4e-18 \| 83.2 \| 83 \| (58, 142) \| 143 \| (47, 130) \| 130 \| Tail fiber assembly protein \| Tail fiber assembly protein; uniclust \| UniRef100\_A0A1C4BVV1 \| 99.1 \| 7.8e-13 \| 1.4e-18 \| 71.7 \| 56 \| (2, 63) \| 143 \| (1, 56) \| 60 \| Phage protein \| Phage protein; uniclust \| UniRef100\_A0A0H8H6L6 \| 99.1 \| 8.2e-13 \| 1.5e-18 \| 87.0 \| 69 \| (73, 143) \| 143 \| (104, 178) \| 183 \| Tail fiber assembly protein \| Tail fiber assembly protein; uniclust \| UniRef100\_A0A379CM87 \| 99.1 \| 8.7e-13 \| 1.6e-18 \| 90.0 \| 80 \| (62, 143) \| 143 \| (127, 206) \| 210 \| Caudovirales tail fibre assembly protein \| Caudovirales tail fibre assembly protein; uniclust \| UniRef100\_A0A1H1V7V6 \| 99.1 \| 8.7e-13 \| 1.7e-18 \| 83.3 \| 81 \| (60, 143) \| 143 \| (30, 112) \| 122 \| Virus tail fibre assembly protein, lambda gpK \| Virus tail fibre assembly protein, lambda gpK; uniclust \| UniRef100\_A0A1A9RM66 \| 99.1 \| 8.9e-13 \| 1.7e-18 \| 90.5 \| 67 \| (2, 74) \| 143 \| (18, 84) \| 228 \| Uncharacterized protein (Fragment) \| Uncharacterized protein (Fragment); uniclust \| UniRef100\_A0A0T9RR95 \| 99.1 \| 1.1e-12 \| 2e-18 \| 67.7 \| 44 \| (100, 143) \| 143 \| (4, 47) \| 48 \| Tail assembly chaperone gp38 \| Tail assembly chaperone gp38; uniclust \| UniRef100\_A0A0K4CN17 \| 99.0 \| 1.1e-12 \| 2.1e-18 \| 87.1 \| 84 \| (58, 143) \| 143 \| (115, 199) \| 203 \| Tail fiber assembly protein \| Tail fiber assembly protein; uniclust \| UniRef100\_UPI001F1FB5E9 \| 99.0 \| 1.2e-12 \| 2.2e-18 \| 81.6 \| 131 \| (2, 143) \| 143 \| (1, 149) \| 149 \| phage tail assembly chaperone \| phage tail assembly chaperone; uniclust \| UniRef100\_A0A1Q5TR83 \| 99.0 \| 1.3e-12 \| 2.4e-18 \| 88.8 \| 77 \| (65, 143) \| 143 \| (106, 182) \| 196 \| Tail assembly chaperone \| Tail assembly chaperone; uniclust \| UniRef100\_A0A3J2DEH3 \| 99.0 \| 1.3e-12 \| 2.4e-18 \| 77.4 \| 65 \| (77, 143) \| 143 \| (8, 78) \| 89 \| Phage tail protein (Fragment) \| Phage tail protein (Fragment); uniclust \| UniRef100\_A0A2X1PXX8 \| 99.0 \| 1.3e-12 \| 2.5e-18 \| 80.1 \| 81 \| (61, 143) \| 143 \| (27, 108) \| 116 \| Tail fiber assembly protein from lambdoid prophage \| Tail fiber assembly protein from lambdoid prophage; uniclust \| UniRef100\_E0M480 \| 99.0 \| 1.4e-12 \| 2.6e-18 \| 86.7 \| 76 \| (66, 143) \| 143 \| (119, 194) \| 196 \| Tail assembly chaperone gp38 \| Tail assembly chaperone gp38; uniclust \| UniRef100\_A0A5D8M825 \| 99.0 \| 1.4e-12 \| 2.6e-18 \| 64.3 \| 34 \| (110, 143) \| 143 \| (2, 35) \| 38 \| Tail fiber assembly protein \| Tail fiber assembly protein; uniclust \| UniRef100\_UPI000B7D225B \| 99.0 \| 1.5e-12 \| 2.7e-18 \| 80.7 \| 124 \| (9, 141) \| 143 \| (9, 132) \| 133 \| tail fiber assembly protein \| tail fiber assembly protein; uniclust \| UniRef100\_UPI00138F4819 \| 99.0 \| 1.6e-12 \| 2.9e-18 \| 76.7 \| 69 \| (70, 143) \| 143 \| (6, 74) \| 106 \| tail fiber assembly protein \| tail fiber assembly protein; uniclust \| UniRef100\_UPI001B8D2E1C \| 99.0 \| 1.7e-12 \| 3.1e-18 \| 83.3 \| 105 \| (36, 143) \| 143 \| (68, 180) \| 180 \| tail fiber assembly protein \| tail fiber assembly protein; uniclust \| UniRef100\_A0A090NC98 \| 99.0 \| 1.7e-12 \| 3.2e-18 \| 64.7 \| 31 \| (113, 143) \| 143 \| (8, 38) \| 40 \| Terminase large subunit \| Terminase large subunit; uniclust \| UniRef100\_UPI001E48B2CF \| 99.0 \| 2e-12 \| 3.6e-18 \| 75.3 \| 70 \| (72, 143) \| 143 \| (27, 96) \| 98 \| tail fiber assembly protein \| tail fiber assembly protein; uniclust \| UniRef100\_UPI0018F0CC46 \| 99.0 \| 2.2e-12 \| 4e-18 \| 78.8 \| 67 \| (74, 143) \| 143 \| (65, 131) \| 131 \| tail fiber assembly protein \| tail fiber assembly protein; uniclust \| UniRef100\_A0A8G2C8I7 \| 99.0 \| 2.6e-12 \| 4.8e-18 \| 80.9 \| 117 \| (2, 127) \| 143 \| (1, 121) \| 159 \| Caudovirales tail fibre assembly protein \| Caudovirales tail fibre assembly protein; uniclust \| UniRef100\_A0A315BKG8 \| 99.0 \| 3e-12 \| 5.6e-18 \| 76.9 \| 75 \| (3, 82) \| 143 \| (1, 75) \| 118 \| Phage tail protein \| Phage tail protein; uniclust \| UniRef100\_UPI002097513D \| 99.0 \| 3.4e-12 \| 6.2e-18 \| 80.1 \| 77 \| (64, 143) \| 143 \| (63, 139) \| 150 \| tail fiber assembly protein \| tail fiber assembly protein; uniclust \| UniRef100\_A0A0F4VEV3 \| 99.0 \| 3.4e-12 \| 6.4e-18 \| 82.1 \| 123 \| (2, 142) \| 143 \| (4, 127) \| 131 \| Phage tail protein \| Phage tail protein; uniclust \| UniRef100\_A0A066SRH2 \| 99.0 \| 4e-12 \| 7.3e-18 \| 75.4 \| 56 \| (85, 142) \| 143 \| (31, 86) \| 101 \| Phage tail protein \| Phage tail protein; uniclust \| UniRef100\_A0A376YM78 \| 99.0 \| 4.8e-12 \| 8.9e-18 \| 66.4 \| 46 \| (93, 143) \| 143 \| (2, 47) \| 51 \| Tail fiber assembly protein \| Tail fiber assembly protein; uniclust \| UniRef100\_UPI0013C49323 \| 99.0 \| 5e-12 \| 9.2e-18 \| 75.2 \| 75 \| (67, 143) \| 143 \| (36, 110) \| 110 \| tail fiber assembly protein \| tail fiber assembly protein; uniclust \| UniRef100\_A0A0T9UB25 \| 99.0 \| 5.1e-12 \| 9.5e-18 \| 73.2 \| 66 \| (76, 143) \| 143 \| (3, 73) \| 85 \| Putative phage tail fiber assembly protein \| Putative phage tail fiber assembly protein; uniclust \| UniRef100\_A0A221FYR6 \| 99.0 \| 5.2e-12 \| 9.5e-18 \| 67.2 \| 58 \| (82, 143) \| 143 \| (2, 59) \| 60 \| Tail fiber assembly protein \| Tail fiber assembly protein; uniclust \| UniRef100\_J8UT63 \| 99.0 \| 5.2e-12 \| 9.6e-18 \| 71.8 \| 63 \| (79, 143) \| 143 \| (15, 79) \| 85 \| Tail fiber assembly protein \| Tail fiber assembly protein; uniclust \| UniRef100\_A0A242NTF1 \| 98.9 \| 5.3e-12 \| 9.8e-18 \| 76.1 \| 104 \| (35, 142) \| 143 \| (3, 109) \| 110 \| Tail fiber assembly protein \| Tail fiber assembly protein; uniclust \| UniRef100\_UPI001C7D3E88 \| 98.9 \| 5.4e-12 \| 1e-17 \| 78.6 \| 135 \| (1, 142) \| 143 \| (1, 147) \| 147 \| tail fiber assembly protein \| tail fiber assembly protein; uniclust \| UniRef100\_UPI001FD7E4A7 \| 98.9 \| 5.6e-12 \| 1e-17 \| 72.1 \| 73 \| (67, 141) \| 143 \| (12, 84) \| 88 \| tail fiber assembly protein \| tail fiber assembly protein; uniclust \| UniRef100\_A0A1H4Y2W8 \| 98.9 \| 5.9e-12 \| 1.1e-17 \| 85.5 \| 70 \| (1, 75) \| 143 \| (3, 74) \| 201 \| Phage tail protein \| Phage tail protein; uniclust \| UniRef100\_A0A370S0C2 \| 98.9 \| 6e-12 \| 1.1e-17 \| 79.2 \| 66 \| (76, 143) \| 143 \| (56, 123) \| 132 \| Virus tail fiber assembly protein lambda gpK \| Virus tail fiber assembly protein lambda gpK; uniclust \| UniRef100\_A0A076G4D7 \| 98.9 \| 6.1e-12 \| 1.2e-17 \| 88.0 \| 66 \| (76, 143) \| 143 \| (113, 187) \| 213 \| Uncharacterized protein \| Uncharacterized protein; uniclust \| UniRef100\_E0SM49 \| 98.9 \| 6.6e-12 \| 1.2e-17 \| 74.6 \| 59 \| (73, 133) \| 143 \| (25, 83) \| 105 \| Tail fiber assembly protein \| Tail fiber assembly protein; uniclust \| UniRef100\_A0A1A9WW69 \| 98.9 \| 7.1e-12 \| 1.3e-17 \| 92.0 \| 72 \| (69, 142) \| 143 \| (98, 171) \| 529 \| Tail fiber assembly protein \| Tail fiber assembly protein; uniclust \| UniRef100\_UPI0015624E47 \| 98.9 \| 7.4e-12 \| 1.4e-17 \| 76.6 \| 64 \| (78, 143) \| 143 \| (67, 130) \| 130 \| tail fiber assembly protein \| tail fiber assembly protein; uniclust \| UniRef100\_A0A7X5ZI37 \| 98.9 \| 7.7e-12 \| 1.4e-17 \| 75.9 \| 74 \| (67, 143) \| 143 \| (47, 120) \| 124 \| Tail fiber assembly protein \| Tail fiber assembly protein; uniclust \| UniRef100\_A0A8I0PZM9 \| 98.9 \| 8.9e-12 \| 1.7e-17 \| 69.6 \| 62 \| (80, 143) \| 143 \| (5, 66) \| 70 \| Phage tail protein \| Phage tail protein; uniclust \| UniRef100\_UPI001BA53F6E \| 98.9 \| 9.1e-12 \| 1.7e-17 \| 68.9 \| 67 \| (73, 141) \| 143 \| (6, 72) \| 73 \| tail fiber assembly protein \| tail fiber assembly protein; uniclust \| UniRef100\_A0A3L2VMZ7 \| 98.9 \| 9.8e-12 \| 1.8e-17 \| 63.9 \| 39 \| (105, 143) \| 143 \| (2, 40) \| 50 \| Tail fiber assembly protein \| Tail fiber assembly protein; uniclust \| UniRef100\_A0A478GUA7 \| 98.9 \| 9.8e-12 \| 1.8e-17 \| 75.3 \| 69 \| (73, 143) \| 143 \| (19, 93) \| 100 \| Phage tail fiber assembly protein \| Phage tail fiber assembly protein; uniclust \| UniRef100\_UPI00039E0AEB \| 98.9 \| 1e-11 \| 1.9e-17 \| 83.5 \| 66 \| (2, 68) \| 143 \| (3, 68) \| 206 \| hypothetical protein \| hypothetical protein; uniclust \| UniRef100\_UPI000BCBA4DC \| 98.9 \| 1.1e-11 \| 1.9e-17 \| 70.0 \| 61 \| (77, 139) \| 143 \| (21, 81) \| 81 \| tail fiber assembly protein \| tail fiber assembly protein; uniclust \| UniRef100\_UPI001CDACA06 \| 98.9 \| 1.2e-11 \| 2.3e-17 \| 71.1 \| 80 \| (31, 110) \| 143 \| (8, 88) \| 90 \| tail fiber assembly protein \| tail fiber assembly protein; uniclust \| UniRef100\_A0A235HBB2 \| 98.9 \| 1.4e-11 \| 2.5e-17 \| 83.3 \| 83 \| (2, 89) \| 143 \| (1, 87) \| 193 \| DUF4376 domain-containing protein \| DUF4376 domain-containing protein; uniclust \| UniRef100\_A0A2V4F8Z2 \| 98.9 \| 1.5e-11 \| 2.7e-17 \| 70.2 \| 60 \| (2, 66) \| 143 \| (3, 63) \| 70 \| Uncharacterized protein \| Uncharacterized protein; uniclust \| UniRef100\_A0A176NIZ0 \| 98.9 \| 1.6e-11 \| 3e-17 \| 81.3 \| 80 \| (62, 143) \| 143 \| (58, 139) \| 147 \| Phage tail assembly chaperone protein \| Phage tail assembly chaperone protein; uniclust \| UniRef100\_A0A066T5Y8 \| 98.9 \| 1.6e-11 \| 3e-17 \| 81.1 \| 66 \| (75, 142) \| 143 \| (129, 194) \| 197 \| Phage tail protein \| Phage tail protein; uniclust \| UniRef100\_UPI0021E99FC3 \| 98.9 \| 1.7e-11 \| 3.2e-17 \| 75.4 \| 65 \| (77, 143) \| 143 \| (68, 134) \| 134 \| tail fiber assembly protein \| tail fiber assembly protein; uniclust \| UniRef100\_A0A953LSW1 \| 98.9 \| 1.8e-11 \| 3.2e-17 \| 76.6 \| 70 \| (4, 78) \| 143 \| (1, 70) \| 148 \| Uncharacterized protein (Fragment) \| Uncharacterized protein (Fragment); uniclust \| UniRef100\_A0A141RRG9 \| 98.9 \| 1.8e-11 \| 3.3e-17 \| 68.5 \| 65 \| (70, 136) \| 143 \| (12, 76) \| 77 \| Caudovirales tail fibre assembly protein \| Caudovirales tail fibre assembly protein; uniclust \| UniRef100\_A0A366F728 \| 98.9 \| 1.9e-11 \| 3.5e-17 \| 76.0 \| 99 \| (32, 143) \| 143 \| (17, 117) \| 120 \| Phage tail assembly chaperone \| Phage tail assembly chaperone; uniclust \| UniRef100\_A0A388SKY9 \| 98.8 \| 2.4e-11 \| 4.5e-17 \| 77.9 \| 68 \| (2, 75) \| 143 \| (1, 68) \| 152 \| Putative tail fiber assembly protein p37 \| Putative tail fiber assembly protein p37; uniclust \| UniRef100\_UPI0020A40FBA \| 98.8 \| 2.5e-11 \| 4.5e-17 \| 69.4 \| 64 \| (76, 142) \| 143 \| (23, 86) \| 86 \| tail fiber assembly protein \| tail fiber assembly protein; uniclust \| UniRef100\_A0A2G0Y6Y2 \| 98.8 \| 2.5e-11 \| 4.6e-17 \| 81.5 \| 64 \| (77, 142) \| 143 \| (155, 220) \| 228 \| Phage tail protein \| Phage tail protein; uniclust \| UniRef100\_A0A078LUM4 \| 98.8 \| 2.5e-11 \| 4.7e-17 \| 84.7 \| 63 \| (1, 69) \| 143 \| (2, 64) \| 203 \| Putative tail fiber assembly-like protein \| Putative tail fiber assembly-like protein; uniclust \| UniRef100\_Q9B026 \| 98.8 \| 2.7e-11 \| 4.9e-17 \| 75.8 \| 72 \| (70, 143) \| 143 \| (73, 144) \| 147 \| Probable tail fiber assembly protein \| Probable tail fiber assembly protein; uniclust \| UniRef100\_UPI001C256683 \| 98.8 \| 3.1e-11 \| 5.7e-17 \| 74.4 \| 103 \| (30, 143) \| 143 \| (23, 133) \| 134 \| tail fiber assembly protein \| tail fiber assembly protein; uniclust \| UniRef100\_UPI002157772D \| 98.8 \| 3.5e-11 \| 6.4e-17 \| 63.6 \| 51 \| (47, 97) \| 143 \| (5, 55) \| 57 \| tail fiber assembly protein \| tail fiber assembly protein; uniclust \| UniRef100\_A0A3R0JSW1 \| 98.8 \| 3.6e-11 \| 6.6e-17 \| 66.6 \| 64 \| (1, 64) \| 143 \| (4, 67) \| 70 \| GST N-terminal domain-containing protein \| GST N-terminal domain-containing protein; uniclust \| UniRef100\_A0A1G9YTU7 \| 98.8 \| 3.6e-11 \| 6.6e-17 \| 76.3 \| 98 \| (30, 143) \| 143 \| (46, 144) \| 144 \| Phage tail assembly chaperone protein \| Phage tail assembly chaperone protein; uniclust \| UniRef100\_A0A5D4YEH1 \| 98.8 \| 3.9e-11 \| 7.2e-17 \| 73.2 \| 73 \| (66, 143) \| 143 \| (38, 110) \| 114 \| Tail fiber assembly protein \| Tail fiber assembly protein; uniclust \| UniRef100\_A0A0D9API2 \| 98.8 \| 4.4e-11 \| 8.2e-17 \| 81.5 \| 69 \| (1, 75) \| 143 \| (11, 81) \| 213 \| Phage tail protein \| Phage tail protein; uniclust \| UniRef100\_A0A2E3R7C7 \| 98.8 \| 5.5e-11 \| 1e-16 \| 71.4 \| 89 \| (50, 143) \| 143 \| (7, 100) \| 102 \| Tail fiber assembly protein \| Tail fiber assembly protein; uniclust \| UniRef100\_A0A0F6B2I6 \| 98.8 \| 6.3e-11 \| 1.2e-16 \| 84.3 \| 77 \| (62, 143) \| 143 \| (299, 375) \| 378 \| Phage side tail fiber assembly protein \| Phage side tail fiber assembly protein; uniclust \| UniRef100\_UPI0012642E8B \| 98.8 \| 6.3e-11 \| 1.2e-16 \| 67.2 \| 55 \| (87, 143) \| 143 \| (25, 80) \| 82 \| tail fiber assembly protein \| tail fiber assembly protein; uniclust \| UniRef100\_A0A2D5R456 \| 98.8 \| 6.7e-11 \| 1.3e-16 \| 75.8 \| 98 \| (27, 143) \| 143 \| (36, 135) \| 136 \| Phage tail protein \| Phage tail protein; uniclust \| UniRef100\_A0A7X0SZA7 \| 98.7 \| 7.9e-11 \| 1.5e-16 \| 65.4 \| 44 \| (100, 143) \| 143 \| (7, 50) \| 69 \| Tail fiber assembly protein \| Tail fiber assembly protein; uniclust \| UniRef100\_A0A0W8JAN2 \| 98.7 \| 7.9e-11 \| 1.5e-16 \| 76.3 \| 122 \| (1, 131) \| 143 \| (2, 126) \| 147 \| Phage tail protein \| Phage tail protein; uniclust \| UniRef100\_UPI000CF05EB2 \| 98.7 \| 8.2e-11 \| 1.5e-16 \| 79.3 \| 60 \| (80, 141) \| 143 \| (184, 243) \| 244 \| IS5 family transposase \| IS5 family transposase; uniclust \| UniRef100\_A0A3A6R813 \| 98.7 \| 9e-11 \| 1.7e-16 \| 76.1 \| 115 \| (29, 143) \| 143 \| (38, 171) \| 185 \| Tail fiber assembly protein \| Tail fiber assembly protein; uniclust \| UniRef100\_A0A399PZJ5 \| 98.7 \| 9.4e-11 \| 1.7e-16 \| 74.3 \| 60 \| (3, 70) \| 143 \| (1, 60) \| 133 \| Phage tail protein \| Phage tail protein; uniclust \| UniRef100\_A0A0P1FIZ6 \| 98.7 \| 9.3e-11 \| 1.8e-16 \| 78.2 \| 65 \| (68, 143) \| 143 \| (76, 142) \| 148 \| Caudovirales tail fibre assembly protein \| Caudovirales tail fibre assembly protein; uniclust \| UniRef100\_A0A1H4V2G1 \| 98.7 \| 9.8e-11 \| 1.8e-16 \| 74.9 \| 117 \| (2, 128) \| 143 \| (1, 123) \| 133 \| Uncharacterized protein \| Uncharacterized protein; uniclust \| UniRef100\_A0A8I1G1G7 \| 98.7 \| 1e-10 \| 1.9e-16 \| 63.4 \| 55 \| (3, 63) \| 143 \| (1, 56) \| 61 \| Uncharacterized protein (Fragment) \| Uncharacterized protein (Fragment); uniclust \| UniRef100\_A0A0H3NQF5 \| 98.7 \| 1.2e-10 \| 2.2e-16 \| 67.9 \| 68 \| (73, 143) \| 143 \| (22, 89) \| 89 \| Tail fiber assembly protein \| Tail fiber assembly protein; uniclust \| UniRef100\_B4RM01 \| 98.7 \| 1.2e-10 \| 2.3e-16 \| 75.5 \| 58 \| (2, 68) \| 143 \| (1, 58) \| 157 \| Phage associated protein \| Phage associated protein; uniclust \| UniRef100\_UPI001FCA6AA2 \| 98.7 \| 1.3e-10 \| 2.3e-16 \| 75.2 \| 50 \| (91, 142) \| 143 \| (131, 180) \| 180 \| DUF1983 domain-containing protein \| DUF1983 domain-containing protein; uniclust \| UniRef100\_UPI000272327A \| 98.7 \| 1.5e-10 \| 2.8e-16 \| 67.7 \| 56 \| (4, 65) \| 143 \| (1, 56) \| 97 \| hypothetical protein \| hypothetical protein; uniclust \| UniRef100\_UPI0020C96B49 \| 98.7 \| 1.6e-10 \| 2.9e-16 \| 66.4 \| 67 \| (72, 142) \| 143 \| (21, 87) \| 87 \| tail fiber assembly protein \| tail fiber assembly protein; uniclust \| UniRef100\_A0A2S8YX24 \| 98.7 \| 1.6e-10 \| 3e-16 \| 73.2 \| 60 \| (82, 143) \| 143 \| (32, 93) \| 157 \| Phage tail protein \| Phage tail protein; uniclust \| UniRef100\_UPI0006877F13 \| 98.7 \| 1.7e-10 \| 3.1e-16 \| 64.7 \| 64 \| (1, 69) \| 143 \| (1, 64) \| 76 \| hypothetical protein \| hypothetical protein; uniclust \| UniRef100\_A0A2D0YIG7 \| 98.7 \| 1.7e-10 \| 3.2e-16 \| 75.2 \| 115 \| (1, 130) \| 143 \| (15, 131) \| 150 \| Tail fibers protein \| Tail fibers protein; uniclust \| UniRef100\_UPI001F153A13 \| 98.7 \| 1.8e-10 \| 3.4e-16 \| 70.7 \| 60 \| (81, 143) \| 143 \| (68, 127) \| 128 \| tail fiber assembly protein \| tail fiber assembly protein; uniclust \| UniRef100\_A0A078LGE4 \| 98.7 \| 2e-10 \| 3.7e-16 \| 68.6 \| 56 \| (74, 143) \| 143 \| (20, 77) \| 86 \| Uncharacterized protein \| Uncharacterized protein; uniclust \| UniRef100\_A0A0J6KHP8 \| 98.7 \| 2e-10 \| 3.8e-16 \| 73.8 \| 55 \| (78, 143) \| 143 \| (60, 116) \| 123 \| Phage tail protein \| Phage tail protein; uniclust \| UniRef100\_UPI001E61745F \| 98.7 \| 2.1e-10 \| 3.9e-16 \| 73.4 \| 72 \| (1, 77) \| 143 \| (1, 72) \| 167 \| hypothetical protein \| hypothetical protein; uniclust \| UniRef100\_A0A080IVY2 \| 98.7 \| 2.2e-10 \| 4.1e-16 \| 79.2 \| 69 \| (73, 143) \| 143 \| (116, 190) \| 216 \| Caudovirales tail fiber assembly family protein \| Caudovirales tail fiber assembly family protein; uniclust \| UniRef100\_UPI00227A8565 \| 98.7 \| 2.5e-10 \| 4.5e-16 \| 71.4 \| 68 \| (70, 142) \| 143 \| (76, 143) \| 143 \| tail fiber assembly protein \| tail fiber assembly protein; uniclust \| UniRef100\_A0A447L1E5 \| 98.6 \| 2.8e-10 \| 5.1e-16 \| 76.3 \| 73 \| (69, 143) \| 143 \| (156, 228) \| 229 \| Caudovirales tail fibre assembly protein \| Caudovirales tail fibre assembly protein; uniclust \| UniRef100\_A0A0T9KYK9 \| 98.6 \| 2.9e-10 \| 5.3e-16 \| 73.9 \| 71 \| (71, 143) \| 143 \| (88, 158) \| 165 \| Tail fiber assembly protein G \| Tail fiber assembly protein G; uniclust \| UniRef100\_UPI0009BA29CF \| 98.6 \| 3e-10 \| 5.5e-16 \| 80.8 \| 64 \| (78, 143) \| 143 \| (304, 369) \| 369 \| tail fiber assembly protein \| tail fiber assembly protein; uniclust \| UniRef100\_A0A2C6DTU1 \| 98.6 \| 3e-10 \| 5.6e-16 \| 55.4 \| 29 \| (115, 143) \| 143 \| (2, 30) \| 36 \| Tail fiber assembly protein \| Tail fiber assembly protein; uniclust \| UniRef100\_A0A0T9P2Y6 \| 98.6 \| 3.1e-10 \| 5.7e-16 \| 73.7 \| 71 \| (71, 143) \| 143 \| (92, 162) \| 164 \| Tail fiber assembly protein G \| Tail fiber assembly protein G; uniclust \| UniRef100\_UPI001061FCA3 \| 98.6 \| 3.3e-10 \| 6e-16 \| 71.0 \| 55 \| (86, 142) \| 143 \| (88, 142) \| 144 \| tail fiber assembly protein \| tail fiber assembly protein; uniclust \| UniRef100\_UPI00046AFBFA \| 98.6 \| 3.3e-10 \| 6.1e-16 \| 72.8 \| 62 \| (80, 143) \| 143 \| (99, 162) \| 162 \| tail fiber assembly protein \| tail fiber assembly protein; uniclust \| UniRef100\_A0A3D1IY78 \| 98.6 \| 3.5e-10 \| 6.4e-16 \| 66.1 \| 87 \| (2, 91) \| 143 \| (1, 94) \| 95 \| Uncharacterized protein (Fragment) \| Uncharacterized protein (Fragment); uniclust \| UniRef100\_Q126B3 \| 98.6 \| 3.7e-10 \| 6.8e-16 \| 75.4 \| 82 \| (4, 85) \| 143 \| (1, 112) \| 223 \| Phage tail protein \| Phage tail protein; uniclust \| UniRef100\_A0A0J6LG96 \| 98.6 \| 3.7e-10 \| 7e-16 \| 69.3 \| 73 \| (69, 143) \| 143 \| (19, 93) \| 102 \| Phage-like protein \| Phage-like protein; uniclust \| UniRef100\_UPI001BD46F23 \| 98.6 \| 3.9e-10 \| 7.2e-16 \| 74.1 \| 71 \| (70, 142) \| 143 \| (106, 178) \| 180 \| tail fiber assembly protein \| tail fiber assembly protein; uniclust \| UniRef100\_J3CFF5 \| 98.6 \| 4.4e-10 \| 8.1e-16 \| 64.0 \| 82 \| (60, 143) \| 143 \| (1, 82) \| 82 \| Caudovirales tail fiber assembly protein \| Caudovirales tail fiber assembly protein; uniclust \| UniRef100\_UPI00192B3CC5 \| 98.6 \| 4.5e-10 \| 8.3e-16 \| 86.9 \| 69 \| (72, 142) \| 143 \| (796, 864) \| 867 \| phage tail protein \| phage tail protein; uniclust \| UniRef100\_A0A2A2T4I5 \| 98.6 \| 4.3e-10 \| 8.4e-16 \| 75.8 \| 66 \| (67, 143) \| 143 \| (78, 144) \| 156 \| Phage tail protein \| Phage tail protein; uniclust \| UniRef100\_UPI001BE0504F \| 98.6 \| 5e-10 \| 9.2e-16 \| 64.9 \| 57 \| (85, 143) \| 143 \| (30, 86) \| 90 \| tail fiber assembly protein \| tail fiber assembly protein; uniclust \| UniRef100\_A0A5J6WBQ9 \| 98.6 \| 5.3e-10 \| 9.8e-16 \| 71.5 \| 85 \| (57, 143) \| 143 \| (55, 145) \| 146 \| Tail fiber assembly protein \| Tail fiber assembly protein; uniclust \| UniRef100\_UPI001F5BCB89 \| 98.6 \| 5.8e-10 \| 1.1e-15 \| 63.7 \| 68 \| (63, 132) \| 143 \| (4, 71) \| 83 \| tail fiber assembly protein \| tail fiber assembly protein; uniclust \| UniRef100\_A0A732CWE2 \| 98.6 \| 6.6e-10 \| 1.2e-15 \| 55.9 \| 38 \| (100, 137) \| 143 \| (4, 41) \| 41 \| Tail fiber assembly protein \| Tail fiber assembly protein; uniclust \| UniRef100\_A0A1W0CCP9 \| 98.6 \| 6.7e-10 \| 1.3e-15 \| 63.8 \| 63 \| (79, 143) \| 143 \| (2, 66) \| 72 \| Phage tail protein \| Phage tail protein; uniclust \| UniRef100\_UPI001F44B6A4 \| 98.6 \| 7e-10 \| 1.3e-15 \| 73.2 \| 60 \| (82, 143) \| 143 \| (143, 202) \| 202 \| tail fiber assembly protein \| tail fiber assembly protein; uniclust \| UniRef100\_UPI00117AE413 \| 98.6 \| 7.1e-10 \| 1.3e-15 \| 65.1 \| 69 \| (73, 143) \| 143 \| (27, 95) \| 97 \| tail fiber assembly protein \| tail fiber assembly protein; uniclust \| UniRef100\_A0A8S5R0T0 \| 98.6 \| 7.1e-10 \| 1.3e-15 \| 76.9 \| 68 \| (65, 143) \| 143 \| (122, 195) \| 203 \| Tail assembly chaperone protein \| Tail assembly chaperone protein; uniclust \| UniRef100\_UPI0020A688E0 \| 98.6 \| 7.6e-10 \| 1.4e-15 \| 58.2 \| 52 \| (88, 142) \| 143 \| (3, 54) \| 54 \| tail fiber assembly protein \| tail fiber assembly protein; uniclust \| UniRef100\_A0A077LR29 \| 98.5 \| 7.6e-10 \| 1.4e-15 \| 76.0 \| 67 \| (75, 143) \| 143 \| (115, 183) \| 199 \| Tail fiber assembly protein \| Tail fiber assembly protein; uniclust \| UniRef100\_UPI000DEF41CE \| 98.5 \| 8.2e-10 \| 1.5e-15 \| 72.2 \| 112 \| (24, 143) \| 143 \| (54, 170) \| 170 \| phage tail assembly chaperone \| phage tail assembly chaperone; uniclust \| UniRef100\_A0A0H5AKI0 \| 98.5 \| 8e-10 \| 1.5e-15 \| 66.1 \| 70 \| (72, 143) \| 143 \| (4, 75) \| 85 \| Phage-like protein \| Phage-like protein; uniclust \| UniRef100\_A0A3W0FGG3 \| 98.5 \| 8.2e-10 \| 1.5e-15 \| 57.6 \| 41 \| (100, 143) \| 143 \| (4, 44) \| 47 \| Tail fiber assembly protein \| Tail fiber assembly protein; uniclust \| UniRef100\_A0A377HWU3 \| 98.5 \| 9.4e-10 \| 1.7e-15 \| 66.3 \| 58 \| (2, 68) \| 143 \| (1, 58) \| 104 \| Putative tail fiber assembly protein \| Putative tail fiber assembly protein; uniclust \| UniRef100\_A0A381C907 \| 98.5 \| 1.1e-09 \| 2e-15 \| 63.7 \| 62 \| (24, 86) \| 143 \| (30, 91) \| 91 \| Caudovirales tail fibre assembly protein \| Caudovirales tail fibre assembly protein; uniclust \| UniRef100\_A0A1H2HWK3 \| 98.5 \| 1.1e-09 \| 2.1e-15 \| 72.5 \| 74 \| (67, 142) \| 143 \| (69, 144) \| 158 \| Phage tail assembly chaperone protein \| Phage tail assembly chaperone protein; uniclust \| UniRef100\_A0A090K647 \| 98.5 \| 1.1e-09 \| 2.1e-15 \| 66.9 \| 58 \| (74, 143) \| 143 \| (23, 82) \| 89 \| Uncharacterized phage protein \| Uncharacterized phage protein; uniclust \| UniRef100\_A0A2A2GZS3 \| 98.5 \| 1.2e-09 \| 2.2e-15 \| 61.6 \| 61 \| (1, 66) \| 143 \| (5, 65) \| 70 \| Uncharacterized protein \| Uncharacterized protein; uniclust \| UniRef100\_A0A977GFE5 \| 98.5 \| 1.2e-09 \| 2.2e-15 \| 61.7 \| 56 \| (86, 143) \| 143 \| (14, 78) \| 78 \| Tail fiber assembly protein \| Tail fiber assembly protein; uniclust \| UniRef100\_UPI00191DD07F \| 98.5 \| 1.5e-09 \| 2.7e-15 \| 62.4 \| 69 \| (73, 143) \| 143 \| (13, 82) \| 85 \| tail fiber assembly protein \| tail fiber assembly protein; uniclust \| UniRef100\_UPI001E5F398A \| 98.5 \| 1.5e-09 \| 2.7e-15 \| 68.2 \| 80 \| (62, 143) \| 143 \| (64, 143) \| 143 \| tail fiber assembly protein \| tail fiber assembly protein; uniclust \| UniRef100\_UPI000FAB35A7 \| 98.5 \| 1.6e-09 \| 2.9e-15 \| 59.1 \| 54 \| (69, 124) \| 143 \| (9, 62) \| 64 \| tail fiber assembly protein \| tail fiber assembly protein; uniclust \| UniRef100\_A0A2S9RN83 \| 98.5 \| 1.6e-09 \| 3e-15 \| 65.5 \| 58 \| (2, 68) \| 143 \| (4, 61) \| 106 \| Phage tail protein \| Phage tail protein; uniclust \| UniRef100\_A0A0A1I628 \| 98.5 \| 1.6e-09 \| 3.1e-15 \| 74.3 \| 58 \| (76, 143) \| 143 \| (87, 146) \| 166 \| Tail fiber assembly protein \| Tail fiber assembly protein; uniclust \| UniRef100\_A0A382DWT1 \| 98.5 \| 1.7e-09 \| 3.1e-15 \| 68.7 \| 58 \| (75, 143) \| 143 \| (64, 123) \| 126 \| Phage tail protein (Fragment) \| Phage tail protein (Fragment); uniclust \| UniRef100\_UPI00111B018F \| 98.5 \| 1.7e-09 \| 3.1e-15 \| 64.0 \| 97 \| (1, 98) \| 143 \| (1, 98) \| 100 \| tail fiber assembly protein \| tail fiber assembly protein; uniclust \| UniRef100\_A0A0B7JBQ8 \| 98.5 \| 1.6e-09 \| 3.1e-15 \| 70.1 \| 61 \| (72, 143) \| 143 \| (49, 111) \| 118 \| Phage tail protein \| Phage tail protein; uniclust \| UniRef100\_A0A5J6B771 \| 98.5 \| 1.7e-09 \| 3.1e-15 \| 68.8 \| 60 \| (4, 68) \| 143 \| (1, 60) \| 156 \| Tail fibers protein \| Tail fibers protein; uniclust \| UniRef100\_A0A3P3QNW6 \| 98.5 \| 1.7e-09 \| 3.2e-15 \| 64.5 \| 70 \| (69, 143) \| 143 \| (16, 87) \| 88 \| Phage tail protein \| Phage tail protein; uniclust \| UniRef100\_UPI0009416878 \| 98.5 \| 2e-09 \| 3.6e-15 \| 65.1 \| 62 \| (77, 143) \| 143 \| (39, 100) \| 105 \| tail fiber assembly protein \| tail fiber assembly protein; uniclust \| UniRef100\_UPI0022FED9A9 \| 98.5 \| 2e-09 \| 3.7e-15 \| 55.9 \| 49 \| (92, 143) \| 143 \| (2, 50) \| 50 \| tail fiber assembly protein \| tail fiber assembly protein; uniclust \| UniRef100\_A0A2D4ZXJ1 \| 98.5 \| 2e-09 \| 3.9e-15 \| 73.1 \| 60 \| (73, 143) \| 143 \| (96, 157) \| 158 \| Tail fiber assembly protein \| Tail fiber assembly protein; uniclust \| UniRef100\_A0A023Q0D1 \| 98.5 \| 2.1e-09 \| 4e-15 \| 65.9 \| 58 \| (75, 143) \| 143 \| (24, 83) \| 90 \| Phage tail protein \| Phage tail protein; uniclust \| UniRef100\_A0A553ZT71 \| 98.5 \| 2.2e-09 \| 4e-15 \| 53.0 \| 34 \| (110, 143) \| 143 \| (1, 34) \| 38 \| Tail fiber assembly protein (Fragment) \| Tail fiber assembly protein (Fragment); uniclust \| UniRef100\_A0A6J5MT50 \| 98.4 \| 2.2e-09 \| 4.1e-15 \| 70.6 \| 57 \| (74, 143) \| 143 \| (81, 139) \| 141 \| Phage tail assembly chaperone protein \| Phage tail assembly chaperone protein; uniclust \| UniRef100\_A0A5E7AJE0 \| 98.4 \| 2.4e-09 \| 4.5e-15 \| 71.2 \| 62 \| (1, 68) \| 143 \| (1, 62) \| 187 \| DUF4376 domain-containing protein \| DUF4376 domain-containing protein; uniclust \| UniRef100\_U1ZF67 \| 98.4 \| 2.6e-09 \| 4.7e-15 \| 60.8 \| 69 \| (2, 76) \| 143 \| (1, 71) \| 80 \| Uncharacterized protein (Fragment) \| Uncharacterized protein (Fragment); uniclust \| UniRef100\_A0A1Z8T124 \| 98.4 \| 2.5e-09 \| 4.7e-15 \| 65.9 \| 58 \| (74, 143) \| 143 \| (39, 97) \| 99 \| Phage tail protein \| Phage tail protein; uniclust \| UniRef100\_A0A826QMU1 \| 98.4 \| 2.6e-09 \| 4.8e-15 \| 61.1 \| 64 \| (78, 143) \| 143 \| (3, 72) \| 77 \| Tail fiber assembly protein \| Tail fiber assembly protein; uniclust \| UniRef100\_A0A0H0K3J8 \| 98.4 \| 3e-09 \| 5.5e-15 \| 60.8 \| 66 \| (1, 66) \| 143 \| (1, 67) \| 74 \| CPS-53 (KpLE1) prophage protein \| CPS-53 (KpLE1) prophage protein; uniclust \| UniRef100\_A0A096GRC5 \| 98.4 \| 2.9e-09 \| 5.5e-15 \| 70.7 \| 58 \| (75, 143) \| 143 \| (91, 149) \| 153 \| Tail fiber assembly protein \| Tail fiber assembly protein; uniclust \| UniRef100\_A0A418X1B6 \| 98.4 \| 3.1e-09 \| 5.7e-15 \| 72.0 \| 75 \| (3, 77) \| 143 \| (1, 126) \| 214 \| Uncharacterized protein \| Uncharacterized protein; uniclust \| UniRef100\_A0A380X5D8 \| 98.4 \| 3.1e-09 \| 5.7e-15 \| 63.1 \| 58 \| (3, 67) \| 143 \| (1, 58) \| 101 \| Tail fiber assembly protein \| Tail fiber assembly protein; uniclust \| UniRef100\_A0A2G8MZ68 \| 98.4 \| 3.1e-09 \| 5.9e-15 \| 74.3 \| 60 \| (2, 68) \| 143 \| (1, 60) \| 203 \| Phage tail protein \| Phage tail protein; uniclust \| UniRef100\_A0A7Y8YCH4 \| 98.4 \| 3.4e-09 \| 6.3e-15 \| 55.7 \| 45 \| (95, 141) \| 143 \| (6, 50) \| 53 \| Tail fiber assembly protein \| Tail fiber assembly protein; uniclust \| UniRef100\_UPI000288F1B1 \| 98.4 \| 3.4e-09 \| 6.4e-15 \| 67.2 \| 109 \| (27, 143) \| 143 \| (15, 128) \| 128 \| hypothetical protein \| hypothetical protein; uniclust \| UniRef100\_A0A286QHW2 \| 98.4 \| 3.4e-09 \| 6.4e-15 \| 70.5 \| 68 \| (74, 143) \| 143 \| (113, 180) \| 180 \| Putative tail fiber assembly protein \| Putative tail fiber assembly protein; uniclust \| UniRef100\_A0A1R0GHS9 \| 98.4 \| 3.5e-09 \| 6.6e-15 \| 73.6 \| 79 \| (63, 143) \| 143 \| (112, 192) \| 200 \| Phage tail protein \| Phage tail protein; uniclust \| UniRef100\_A0A855K755 \| 98.4 \| 3.8e-09 \| 7.1e-15 \| 65.9 \| 75 \| (66, 143) \| 143 \| (50, 124) \| 125 \| Tail fiber assembly protein \| Tail fiber assembly protein; uniclust \| UniRef100\_A0A763STN1 \| 98.4 \| 3.9e-09 \| 7.1e-15 \| 54.0 \| 41 \| (93, 135) \| 143 \| (1, 41) \| 46 \| Tail fiber assembly protein \| Tail fiber assembly protein; uniclust \| UniRef100\_UPI002023B97A \| 98.4 \| 4.2e-09 \| 7.7e-15 \| 58.1 \| 49 \| (77, 127) \| 143 \| (15, 63) \| 66 \| tail fiber assembly protein \| tail fiber assembly protein; uniclust \| UniRef100\_A0A167MRB5 \| 98.4 \| 4e-09 \| 7.7e-15 \| 68.6 \| 62 \| (71, 143) \| 143 \| (59, 122) \| 125 \| Tail fiber assembly protein \| Tail fiber assembly protein; uniclust \| UniRef100\_A0A6M4XJ55 \| 98.4 \| 4.2e-09 \| 7.7e-15 \| 58.4 \| 60 \| (3, 67) \| 143 \| (1, 60) \| 64 \| Uncharacterized protein \| Uncharacterized protein; uniclust \| UniRef100\_UPI001F0AB459 \| 98.4 \| 4.3e-09 \| 7.9e-15 \| 57.5 \| 63 \| (63, 125) \| 143 \| (2, 64) \| 64 \| tail fiber assembly protein \| tail fiber assembly protein; uniclust \| UniRef100\_UPI001CC0D473 \| 98.4 \| 4.5e-09 \| 8.3e-15 \| 57.4 \| 57 \| (2, 64) \| 143 \| (1, 57) \| 64 \| hypothetical protein \| hypothetical protein; uniclust \| UniRef100\_UPI001BD5EA33 \| 98.4 \| 4.5e-09 \| 8.3e-15 \| 58.4 \| 65 \| (77, 143) \| 143 \| (6, 70) \| 70 \| tail fiber assembly protein \| tail fiber assembly protein; uniclust \| UniRef100\_A0A2S9X7U2 \| 98.4 \| 4.5e-09 \| 8.3e-15 \| 75.9 \| 69 \| (64, 143) \| 143 \| (219, 289) \| 295 \| Phage tail protein \| Phage tail protein; uniclust \| UniRef100\_A0A0C5EHZ3 \| 98.4 \| 4.8e-09 \| 9e-15 \| 64.3 \| 61 \| (72, 143) \| 143 \| (29, 91) \| 95 \| Phage tail assembly chaperone protein \| Phage tail assembly chaperone protein; uniclust \| UniRef100\_A0A0H4VYX9 \| 98.4 \| 4.9e-09 \| 9e-15 \| 69.4 \| 88 \| (3, 95) \| 143 \| (42, 131) \| 201 \| Tail assembly chaperone \| Tail assembly chaperone; uniclust \| UniRef100\_UPI001F55841F \| 98.4 \| 4.9e-09 \| 9.1e-15 \| 57.9 \| 56 \| (77, 143) \| 143 \| (4, 61) \| 63 \| tail fiber assembly protein \| tail fiber assembly protein; uniclust \| UniRef100\_A0A6S6RXB7 \| 98.4 \| 5e-09 \| 9.3e-15 \| 56.5 \| 54 \| (57, 112) \| 143 \| (6, 59) \| 60 \| Uncharacterized protein \| Uncharacterized protein; uniclust \| UniRef100\_A0A2D4QWA0 \| 98.4 \| 4.9e-09 \| 9.3e-15 \| 64.5 \| 53 \| (77, 143) \| 143 \| (37, 91) \| 98 \| Phage tail protein \| Phage tail protein; uniclust \| UniRef100\_UPI002301F898 \| 98.4 \| 5.3e-09 \| 9.7e-15 \| 65.6 \| 66 \| (1, 66) \| 143 \| (1, 66) \| 140 \| hypothetical protein \| hypothetical protein; uniclust \| UniRef100\_A0A411GJZ7 \| 98.4 \| 5.6e-09 \| 1e-14 \| 64.4 \| 79 \| (60, 143) \| 143 \| (40, 118) \| 122 \| Uncharacterized protein \| Uncharacterized protein; uniclust \| UniRef100\_A0A126QKP2 \| 98.3 \| 6e-09 \| 1.1e-14 \| 70.5 \| 62 \| (71, 143) \| 143 \| (88, 151) \| 163 \| Phage tail protein \| Phage tail protein; uniclust \| UniRef100\_A0A1B7JW24 \| 98.3 \| 6.4e-09 \| 1.2e-14 \| 84.7 \| 80 \| (62, 143) \| 143 \| (672, 753) \| 754 \| Phage minor tail protein \| Phage minor tail protein; uniclust \| UniRef100\_B0VK51 \| 98.3 \| 6.7e-09 \| 1.2e-14 \| 70.1 \| 58 \| (4, 66) \| 143 \| (1, 58) \| 228 \| Uncharacterized protein 51 \| Uncharacterized protein 51; uniclust \| UniRef100\_Q7N1H8 \| 98.3 \| 6.8e-09 \| 1.2e-14 \| 71.2 \| 58 \| (75, 134) \| 143 \| (140, 197) \| 258 \| Photorhabdus luminescens subsp. laumondii TTO1 complete genome; segment 12/17 \| Photorhabdus luminescens subsp. laumondii TTO1 complete genome; segment 12/17; uniclust \| UniRef100\_A0A022PK72 \| 98.3 \| 6.8e-09 \| 1.3e-14 \| 79.1 \| 71 \| (71, 143) \| 143 \| (394, 464) \| 569 \| Phage tail protein \| Phage tail protein; uniclust \| UniRef100\_UPI0009339C22 \| 98.3 \| 6.9e-09 \| 1.3e-14 \| 60.9 \| 75 \| (10, 84) \| 143 \| (8, 82) \| 90 \| tail fiber assembly protein \| tail fiber assembly protein; uniclust \| UniRef100\_A0A0J6K2V7 \| 98.3 \| 8.2e-09 \| 1.6e-14 \| 63.7 \| 61 \| (72, 143) \| 143 \| (29, 91) \| 94 \| Phage tail protein \| Phage tail protein; uniclust \| UniRef100\_UPI000984B09A \| 98.3 \| 8.7e-09 \| 1.6e-14 \| 66.4 \| 68 \| (74, 143) \| 143 \| (64, 133) \| 134 \| phage tail assembly chaperone \| phage tail assembly chaperone; uniclust \| UniRef100\_A0A1A8TEZ4 \| 98.3 \| 8.8e-09 \| 1.6e-14 \| 69.0 \| 63 \| (69, 143) \| 143 \| (99, 163) \| 168 \| Caudovirales tail fibre assembly protein \| Caudovirales tail fibre assembly protein; uniclust \| UniRef100\_A0A2S3X1I2 \| 98.3 \| 9.2e-09 \| 1.7e-14 \| 80.1 \| 63 \| (79, 143) \| 143 \| (783, 847) \| 847 \| Phage tail protein \| Phage tail protein; uniclust \| UniRef100\_UPI0018F81469 \| 98.3 \| 9.4e-09 \| 1.7e-14 \| 56.7 \| 63 \| (2, 69) \| 143 \| (1, 63) \| 67 \| hypothetical protein \| hypothetical protein; uniclust \| UniRef100\_A0A1L5C071 \| 98.3 \| 9.1e-09 \| 1.7e-14 \| 69.9 \| 65 \| (70, 143) \| 143 \| (74, 141) \| 161 \| Putative tail fiber assembly protein \| Putative tail fiber assembly protein; uniclust \| UniRef100\_A0A6N6K3R1 \| 98.3 \| 9.5e-09 \| 1.7e-14 \| 59.3 \| 84 \| (4, 93) \| 143 \| (1, 84) \| 85 \| Phage tail protein \| Phage tail protein; uniclust \| UniRef100\_A0A7G2IV99 \| 98.3 \| 9.4e-09 \| 1.8e-14 \| 56.9 \| 47 \| (75, 123) \| 143 \| (12, 58) \| 60 \| Tail fiber assembly protein \| Tail fiber assembly protein; uniclust \| UniRef100\_A0A3N6V447 \| 98.3 \| 9.6e-09 \| 1.8e-14 \| 61.8 \| 68 \| (74, 143) \| 143 \| (33, 106) \| 108 \| Tail fiber assembly protein \| Tail fiber assembly protein; uniclust \| UniRef100\_A0A1N6I101 \| 98.3 \| 9.6e-09 \| 1.8e-14 \| 65.7 \| 117 \| (2, 127) \| 143 \| (1, 126) \| 140 \| Uncharacterized protein \| Uncharacterized protein; uniclust \| UniRef100\_A0A2S3X8M0 \| 98.3 \| 1e-08 \| 1.9e-14 \| 78.2 \| 63 \| (79, 143) \| 143 \| (603, 667) \| 667 \| Phage tail protein \| Phage tail protein; uniclust \| UniRef100\_A0A0U5EW02 \| 98.3 \| 1.1e-08 \| 2.1e-14 \| 69.7 \| 53 \| (2, 61) \| 143 \| (20, 72) \| 184 \| Uncharacterized protein \| Uncharacterized protein; uniclust \| UniRef100\_UPI000484336E \| 98.3 \| 1.2e-08 \| 2.2e-14 \| 62.7 \| 59 \| (83, 143) \| 143 \| (61, 121) \| 121 \| tail fiber assembly protein \| tail fiber assembly protein; uniclust \| UniRef100\_UPI0003353EC4 \| 98.3 \| 1.2e-08 \| 2.2e-14 \| 64.6 \| 110 \| (32, 142) \| 143 \| (14, 131) \| 132 \| tail fiber assembly protein \| tail fiber assembly protein; uniclust \| UniRef100\_A0A1A9VZ50 \| 98.3 \| 1.2e-08 \| 2.2e-14 \| 73.1 \| 68 \| (74, 143) \| 143 \| (293, 360) \| 363 \| Tail fiber assembly protein \| Tail fiber assembly protein; uniclust \| UniRef100\_UPI001C97D201 \| 98.3 \| 1.2e-08 \| 2.2e-14 \| 57.6 \| 44 \| (3, 46) \| 143 \| (1, 44) \| 75 \| hypothetical protein \| hypothetical protein; uniclust \| UniRef100\_UPI0015A437FB \| 98.3 \| 1.2e-08 \| 2.2e-14 \| 56.2 \| 60 \| (2, 67) \| 143 \| (1, 60) \| 66 \| hypothetical protein \| hypothetical protein; uniclust \| UniRef100\_UPI001F4303E0 \| 98.3 \| 1.2e-08 \| 2.2e-14 \| 54.6 \| 53 \| (89, 143) \| 143 \| (4, 56) \| 57 \| tail fiber assembly protein \| tail fiber assembly protein; uniclust \| UniRef100\_A0A443IQF6 \| 98.3 \| 1.2e-08 \| 2.2e-14 \| 68.4 \| 61 \| (72, 143) \| 143 \| (84, 146) \| 153 \| Tail fiber assembly protein \| Tail fiber assembly protein; uniclust \| UniRef100\_A0A1I1B6J1 \| 98.3 \| 1.3e-08 \| 2.4e-14 \| 68.5 \| 62 \| (71, 143) \| 143 \| (84, 146) \| 156 \| Phage tail assembly chaperone protein \| Phage tail assembly chaperone protein; uniclust \| UniRef100\_A0A345CT33 \| 98.3 \| 1.3e-08 \| 2.4e-14 \| 56.7 \| 59 \| (2, 67) \| 143 \| (1, 59) \| 66 \| Uncharacterized protein \| Uncharacterized protein; uniclust \| UniRef100\_UPI001F45C5CD \| 98.3 \| 1.4e-08 \| 2.5e-14 \| 58.3 \| 65 \| (66, 132) \| 143 \| (9, 74) \| 82 \| tail fiber assembly protein \| tail fiber assembly protein; uniclust \| UniRef100\_UPI0003061D81 \| 98.3 \| 1.4e-08 \| 2.6e-14 \| 53.1 \| 33 \| (110, 142) \| 143 \| (19, 51) \| 51 \| tail fiber assembly protein \| tail fiber assembly protein; uniclust \| UniRef100\_A0A0U1HXL6 \| 98.2 \| 1.5e-08 \| 2.7e-14 \| 63.5 \| 71 \| (71, 143) \| 143 \| (26, 101) \| 122 \| Putative phage tail fiber assembly protein \| Putative phage tail fiber assembly protein; uniclust \| UniRef100\_A0A0U1ZGY3 \| 98.2 \| 1.4e-08 \| 2.7e-14 \| 69.0 \| 59 \| (77, 143) \| 143 \| (92, 152) \| 167 \| Uncharacterized protein \| Uncharacterized protein; uniclust \| UniRef100\_A0A076GCL9 \| 98.2 \| 1.5e-08 \| 2.9e-14 \| 70.0 \| 69 \| (73, 143) \| 143 \| (112, 180) \| 186 \| Putative tail fiber assembly protein \| Putative tail fiber assembly protein; uniclust \| UniRef100\_A0A0H0G3F6 \| 98.2 \| 1.6e-08 \| 2.9e-14 \| 58.7 \| 46 \| (85, 132) \| 143 \| (12, 57) \| 82 \| Tail fiber assembly protein \| Tail fiber assembly protein; uniclust \| UniRef100\_UPI00143FE79E \| 98.2 \| 1.6e-08 \| 2.9e-14 \| 70.5 \| 66 \| (1, 67) \| 143 \| (84, 149) \| 289 \| tail fiber assembly protein \| tail fiber assembly protein; uniclust \| UniRef100\_A0A0T9V566 \| 98.2 \| 1.7e-08 \| 3.2e-14 \| 67.1 \| 132 \| (1, 142) \| 143 \| (2, 137) \| 143 \| Tail fiber assembly protein \| Tail fiber assembly protein; uniclust \| UniRef100\_A0A0A0HKV0 \| 98.2 \| 1.8e-08 \| 3.3e-14 \| 59.9 \| 62 \| (71, 143) \| 143 \| (14, 77) \| 80 \| Caudovirales tail fiber assembly protein \| Caudovirales tail fiber assembly protein; uniclust \| UniRef100\_A0A2X2RPY1 \| 98.2 \| 1.8e-08 \| 3.3e-14 \| 65.9 \| 64 \| (76, 141) \| 143 \| (114, 177) \| 180 \| Caudovirales tail fibre assembly protein \| Caudovirales tail fibre assembly protein; uniclust \| UniRef100\_A0A376J3U5 \| 98.2 \| 1.9e-08 \| 3.6e-14 \| 64.8 \| 72 \| (69, 143) \| 143 \| (73, 144) \| 144 \| Tail fiber assembly protein (Gpg) \| Tail fiber assembly protein (Gpg); uniclust \| UniRef100\_A0A7L6U9J7 \| 98.2 \| 2e-08 \| 3.8e-14 \| 54.9 \| 49 \| (92, 142) \| 143 \| (14, 62) \| 63 \| Tail fiber assembly protein \| Tail fiber assembly protein; uniclust \| UniRef100\_UPI00214F3669 \| 98.2 \| 2.1e-08 \| 3.8e-14 \| 58.4 \| 52 \| (87, 143) \| 143 \| (5, 56) \| 88 \| tail fiber assembly protein \| tail fiber assembly protein; uniclust \| UniRef100\_A0A8T6AMS3 \| 98.2 \| 2.1e-08 \| 3.9e-14 \| 54.6 \| 30 \| (113, 142) \| 143 \| (33, 62) \| 62 \| Tail fiber assembly protein \| Tail fiber assembly protein; uniclust \| UniRef100\_A0A444RIF9 \| 98.2 \| 2.1e-08 \| 3.9e-14 \| 62.9 \| 73 \| (69, 143) \| 143 \| (47, 126) \| 126 \| Tail fiber assembly protein \| Tail fiber assembly protein; uniclust \| UniRef100\_A0A0D5Y4U0 \| 98.2 \| 2.3e-08 \| 4.2e-14 \| 59.5 \| 57 \| (85, 143) \| 143 \| (32, 97) \| 98 \| Phage tail protein \| Phage tail protein; uniclust \| UniRef100\_UPI001ED8DAE3 \| 98.2 \| 2.3e-08 \| 4.3e-14 \| 77.3 \| 68 \| (74, 143) \| 143 \| (593, 660) \| 660 \| tail fiber assembly protein \| tail fiber assembly protein; uniclust \| UniRef100\_A0A2D6P997 \| 98.2 \| 2.3e-08 \| 4.4e-14 \| 63.7 \| 58 \| (75, 143) \| 143 \| (63, 121) \| 122 \| Tail fiber assembly protein \| Tail fiber assembly protein; uniclust \| UniRef100\_A0A2N7P764 \| 98.2 \| 2.6e-08 \| 4.9e-14 \| 63.2 \| 93 \| (4, 96) \| 143 \| (1, 94) \| 147 \| Uncharacterized protein \| Uncharacterized protein; uniclust \| UniRef100\_A0A2X1PJL4 \| 98.2 \| 2.7e-08 \| 5e-14 \| 60.0 \| 65 \| (27, 91) \| 143 \| (41, 105) \| 108 \| Putative phage tail assembly chaperone \| Putative phage tail assembly chaperone; uniclust \| UniRef100\_UPI001B82413B \| 98.2 \| 2.8e-08 \| 5.1e-14 \| 56.9 \| 56 \| (3, 64) \| 143 \| (1, 56) \| 80 \| hypothetical protein \| hypothetical protein; uniclust \| UniRef100\_UPI0004661D35 \| 98.2 \| 2.8e-08 \| 5.1e-14 \| 59.8 \| 70 \| (72, 143) \| 143 \| (16, 86) \| 93 \| phage tail assembly chaperone \| phage tail assembly chaperone; uniclust \| UniRef100\_UPI0021155A40 \| 98.2 \| 2.9e-08 \| 5.4e-14 \| 56.2 \| 64 \| (77, 142) \| 143 \| (4, 71) \| 72 \| tail fiber assembly protein \| tail fiber assembly protein; uniclust \| UniRef100\_C1D954 \| 98.2 \| 3.1e-08 \| 5.7e-14 \| 69.0 \| 65 \| (4, 74) \| 143 \| (1, 65) \| 283 \| HsdM \| HsdM; uniclust \| UniRef100\_A0A4R4J6I0 \| 98.2 \| 3.2e-08 \| 5.9e-14 \| 48.5 \| 28 \| (115, 142) \| 143 \| (9, 36) \| 37 \| Tail fiber assembly protein \| Tail fiber assembly protein; uniclust \| UniRef100\_A0A1A9WSI7 \| 98.2 \| 3.4e-08 \| 6.2e-14 \| 74.9 \| 78 \| (5, 82) \| 143 \| (509, 587) \| 604 \| Uncharacterized protein \| Uncharacterized protein; uniclust \| UniRef100\_A0A0F7Y2W3 \| 98.2 \| 3.3e-08 \| 6.3e-14 \| 77.3 \| 62 \| (80, 143) \| 143 \| (430, 493) \| 493 \| Tail fiber protein \| Tail fiber protein; uniclust \| UniRef100\_A0A484QPN5 \| 98.1 \| 3.4e-08 \| 6.4e-14 \| 63.2 \| 79 \| (63, 143) \| 143 \| (51, 131) \| 139 \| Tail fiber assembly protein \| Tail fiber assembly protein; uniclust \| UniRef100\_A0A2M7HGM4 \| 98.1 \| 3.5e-08 \| 6.4e-14 \| 57.5 \| 81 \| (47, 143) \| 143 \| (4, 84) \| 88 \| Phage tail protein \| Phage tail protein; uniclust \| UniRef100\_UPI001F25180A \| 98.1 \| 3.6e-08 \| 6.6e-14 \| 57.2 \| 72 \| (70, 143) \| 143 \| (8, 86) \| 86 \| tail fiber assembly protein \| tail fiber assembly protein; uniclust \| UniRef100\_A0A8T5ZT04 \| 98.1 \| 3.8e-08 \| 7e-14 \| 47.1 \| 30 \| (114, 143) \| 143 \| (3, 32) \| 33 \| Tail fiber assembly protein \| Tail fiber assembly protein; uniclust \| UniRef100\_UPI000734FD4D \| 98.1 \| 4.1e-08 \| 7.6e-14 \| 70.6 \| 81 \| (60, 142) \| 143 \| (134, 216) \| 294 \| tail fiber assembly protein \| tail fiber assembly protein; uniclust \| UniRef100\_S2KT03 \| 98.1 \| 4.3e-08 \| 7.8e-14 \| 64.1 \| 65 \| (76, 142) \| 143 \| (104, 176) \| 176 \| Phage tail fiber assembly protein \| Phage tail fiber assembly protein; uniclust \| UniRef100\_A0A5A9DBI3 \| 98.1 \| 4.3e-08 \| 8e-14 \| 48.9 \| 33 \| (109, 141) \| 143 \| (5, 37) \| 40 \| Tail fiber assembly protein \| Tail fiber assembly protein; uniclust \| UniRef100\_UPI0021CFB339 \| 98.1 \| 4.3e-08 \| 8e-14 \| 58.6 \| 64 \| (75, 143) \| 143 \| (38, 101) \| 102 \| tail fiber assembly protein \| tail fiber assembly protein; uniclust \| UniRef100\_A0A411H8C5 \| 98.1 \| 4.4e-08 \| 8.2e-14 \| 61.8 \| 64 \| (77, 143) \| 143 \| (69, 132) \| 134 \| Tail fiber assembly protein \| Tail fiber assembly protein; pdb70 \| 5YVQ\_B \| 99.2 \| 2.3e-15 \| 2.3e-19 \| 112.6 \| 75 \| (64, 143) \| 143 \| (101, 175) \| 175 \| Gene product J \| 5YVQ\_B Gene product J bacteriophage, VIRAL PROTEIN HET: GOL; pdb70 \| 2KZ6\_A \| 98.5 \| 2.6e-11 \| 2.5e-15 \| 84.2 \| 63 \| (70, 143) \| 143 \| (38, 102) \| 102 \| Uncharacterized protein \| 2KZ6\_A Uncharacterized protein protein of unknown function, Structural |
| Top keywords  (threshold 1.00e-03 (evalue)) | **tail, assembly, fiber, Phage, chaperone, Caudovirales, Fragment, Putative, fibre, hypothetical** |
| Output files | ../../similar\_sequences/16\_FANPEZAQ\_CDS\_0016\_merged.svg ../../similar\_sequences/16\_FANPEZAQ\_CDS\_0016\_pdb70.a3m ../../similar\_sequences/16\_FANPEZAQ\_CDS\_0016\_pdb70.hhr ../../similar\_sequences/16\_FANPEZAQ\_CDS\_0016\_uniclust.a3m ../../similar\_sequences/16\_FANPEZAQ\_CDS\_0016\_uniclust.hhr |

#### Structure prediction (AlphaFold)^2^

|  |  |
| --- | --- |
| Stats | xml version="1.0" encoding="utf-8" standalone="no"?       2024-09-02T21:09:15.217395 image/svg+xml   Matplotlib v3.7.2, https://matplotlib.org/ |
| Predicted structure | **NGL Viewer Controls:**  - Center: *Left-Click* - Rotate: *Left-Click + Drag* - Translate: *Right-Click + Drag* - Zoom: *Shift + Left-Click + Drag* |
| Output files | ../../predicted\_structures/16\_FANPEZAQ\_CDS\_0016/features.pkl ../../predicted\_structures/16\_FANPEZAQ\_CDS\_0016/ranked\_0.pdb ../../predicted\_structures/16\_FANPEZAQ\_CDS\_0016/ranked\_0\_plots.svg ../../predicted\_structures/16\_FANPEZAQ\_CDS\_0016/result\_model\_1\_ptm\_pred\_0.pkl |

#### Structure similarity search results (Foldseek)^3^

|  |  |
| --- | --- |
| Structure databases searched | Pdb, Afdb-proteome, Afdb-uniprot50 |
| Results, scheme(s)  (Top layers only, threshold 1.00e-02 (evalue)) | xml version="1.0" encoding="utf-8" standalone="no"?       2024-09-02T21:10:34.592546 image/svg+xml   Matplotlib v3.7.2, https://matplotlib.org/ |
| Results, table  (threshold 1.00e-02 (evalue)) | db \| id \| prob \| evalue \| bits \| fident \| alnlen \| mismatch \| gapopen \| qstart \| qend \| tstart \| tend \| name \| description; afdb-proteome \| AF-P77656-F1-MODEL\_V4 \| 1.0 \| 9.056e-18 \| 683 \| 0.564 \| 140 \| 61 \| 0 \| 4 \| 143 \| 3 \| 142 \| Uncharacterized protein YfdK \| Uncharacterized protein YfdK; afdb-proteome \| AF-Q8ZNJ2-F1-MODEL\_V4 \| 1.0 \| 6.388e-13 \| 448 \| 0.429 \| 142 \| 79 \| 2 \| 1 \| 141 \| 1 \| 141 \| Putative cytoplasmic protein \| Putative cytoplasmic protein; afdb-proteome \| AF-P09154-F1-MODEL\_V4 \| 1.0 \| 1.137e-09 \| 313 \| 0.368 \| 141 \| 78 \| 4 \| 3 \| 142 \| 2 \| 132 \| Uncharacterized protein YmfS \| Uncharacterized protein YmfS; afdb-proteome \| AF-G3XD08-F1-MODEL\_V4 \| 1.0 \| 8.406e-08 \| 260 \| 0.276 \| 123 \| 87 \| 2 \| 4 \| 126 \| 5 \| 125 \| Uncharacterized protein \| Uncharacterized protein; afdb-proteome \| AF-Q8ZMU7-F1-MODEL\_V4 \| 1.0 \| 4.528e-07 \| 202 \| 0.212 \| 146 \| 97 \| 6 \| 1 \| 142 \| 1 \| 132 \| Fels-2 prophage protein \| Fels-2 prophage protein; afdb-proteome \| AF-P09153-F1-MODEL\_V4 \| 1.0 \| 4.276e-06 \| 176 \| 0.256 \| 148 \| 77 \| 8 \| 2 \| 143 \| 79 \| 199 \| Prophage tail fiber assembly protein homolog TfaE \| Prophage tail fiber assembly protein homolog TfaE; afdb-proteome \| AF-P77326-F1-MODEL\_V4 \| 1.0 \| 6.248e-05 \| 171 \| 0.34 \| 100 \| 57 \| 5 \| 46 \| 143 \| 21 \| 113 \| Putative protein TfaS \| Putative protein TfaS; afdb-proteome \| AF-P77699-F1-MODEL\_V4 \| 1.0 \| 3.794e-05 \| 161 \| 0.208 \| 144 \| 84 \| 5 \| 3 \| 142 \| 67 \| 184 \| Protein TfaD \| Protein TfaD; afdb-proteome \| AF-P76155-F1-MODEL\_V4 \| 1.0 \| 0.0001406 \| 140 \| 0.205 \| 146 \| 82 \| 5 \| 3 \| 142 \| 74 \| 191 \| Prophage tail fiber assembly protein homolog TfaQ \| Prophage tail fiber assembly protein homolog TfaQ; afdb-proteome \| AF-Q8ZQ80-F1-MODEL\_V4 \| 1.0 \| 0.0001804 \| 140 \| 0.164 \| 146 \| 91 \| 6 \| 2 \| 143 \| 73 \| 191 \| Gifsy-2 prophage tail fiber assembly like-protein \| Gifsy-2 prophage tail fiber assembly like-protein; afdb-proteome \| AF-P40784-F1-MODEL\_V4 \| 1.0 \| 0.000192 \| 138 \| 0.178 \| 146 \| 89 \| 7 \| 2 \| 143 \| 73 \| 191 \| Tail fiber assembly protein homolog from lambdoid prophage Fels-1 \| Tail fiber assembly protein homolog from lambdoid prophage Fels-1; afdb-proteome \| AF-P77163-F1-MODEL\_V4 \| 1.0 \| 0.0002315 \| 136 \| 0.205 \| 146 \| 82 \| 5 \| 3 \| 142 \| 74 \| 191 \| Prophage tail fiber assembly protein homolog TfaR \| Prophage tail fiber assembly protein homolog TfaR; afdb-proteome \| AF-Q8ZN15-F1-MODEL\_V4 \| 1.0 \| 0.0006279 \| 127 \| 0.157 \| 146 \| 90 \| 6 \| 2 \| 143 \| 73 \| 189 \| Gifsy-1 prophage protein \| Gifsy-1 prophage protein; afdb-proteome \| AF-Q8ZNJ1-F1-MODEL\_V4 \| 1.0 \| 0.0002971 \| 122 \| 0.155 \| 161 \| 100 \| 7 \| 5 \| 143 \| 28 \| 174 \| Putative tail fiber assembly protein \| Putative tail fiber assembly protein; afdb-proteome \| AF-H9L459-F1-MODEL\_V4 \| 1.0 \| 0.001703 \| 112 \| 0.136 \| 147 \| 92 \| 7 \| 3 \| 143 \| 173 \| 290 \| Phage-tail assembly protein-like protein \| Phage-tail assembly protein-like protein; afdb-uniprot50 \| AF-A0A2Y9TWM4-F1-MODEL\_V4 \| 1.0 \| 4.014e-17 \| 676 \| 0.46 \| 141 \| 76 \| 0 \| 3 \| 143 \| 2 \| 142 \| Tail fiber assembly protein \| Tail fiber assembly protein; afdb-uniprot50 \| AF-A0A5Z5REI7-F1-MODEL\_V4 \| 1.0 \| 3.752e-18 \| 672 \| 0.566 \| 143 \| 61 \| 1 \| 1 \| 143 \| 1 \| 142 \| Tail assembly chaperone \| Tail assembly chaperone; afdb-uniprot50 \| AF-A0A3P5HII9-F1-MODEL\_V4 \| 1.0 \| 7.93e-18 \| 664 \| 0.566 \| 143 \| 61 \| 1 \| 1 \| 143 \| 1 \| 142 \| Prophage tail fiber assembly protein TfaE \| Prophage tail fiber assembly protein TfaE; afdb-uniprot50 \| AF-A0A1X3I4W0-F1-MODEL\_V4 \| 1.0 \| 1.306e-17 \| 661 \| 0.573 \| 143 \| 60 \| 1 \| 1 \| 143 \| 24 \| 165 \| Caudovirales tail fiber assembly protein \| Caudovirales tail fiber assembly protein; afdb-uniprot50 \| AF-A0A639BZQ5-F1-MODEL\_V4 \| 1.0 \| 6.61e-17 \| 658 \| 0.552 \| 143 \| 63 \| 1 \| 1 \| 143 \| 1 \| 142 \| Tail fiber assembly protein \| Tail fiber assembly protein; afdb-uniprot50 \| AF-A0A828GQS5-F1-MODEL\_V4 \| 1.0 \| 5.151e-17 \| 653 \| 0.51 \| 143 \| 70 \| 0 \| 1 \| 143 \| 1 \| 143 \| Tail fiber assembly protein \| Tail fiber assembly protein; afdb-uniprot50 \| AF-A0A3A6RJK8-F1-MODEL\_V4 \| 1.0 \| 1.023e-16 \| 651 \| 0.51 \| 143 \| 70 \| 0 \| 1 \| 143 \| 1 \| 143 \| Tail fiber assembly protein \| Tail fiber assembly protein; afdb-uniprot50 \| AF-A0A6I6DJ31-F1-MODEL\_V4 \| 1.0 \| 6.211e-17 \| 645 \| 0.564 \| 140 \| 60 \| 1 \| 4 \| 143 \| 4 \| 142 \| Tail fiber assembly protein \| Tail fiber assembly protein; afdb-uniprot50 \| AF-A0A8A5IWD8-F1-MODEL\_V4 \| 1.0 \| 1.583e-16 \| 637 \| 0.517 \| 143 \| 69 \| 0 \| 1 \| 143 \| 1 \| 143 \| Tail fiber assembly protein \| Tail fiber assembly protein; afdb-uniprot50 \| AF-A0A0Q4MYZ8-F1-MODEL\_V4 \| 1.0 \| 1.028e-15 \| 620 \| 0.442 \| 140 \| 76 \| 1 \| 4 \| 143 \| 2 \| 139 \| Uncharacterized protein \| Uncharacterized protein; afdb-uniprot50 \| AF-A0A411WHE0-F1-MODEL\_V4 \| 1.0 \| 1.028e-15 \| 618 \| 0.439 \| 141 \| 79 \| 0 \| 3 \| 143 \| 11 \| 151 \| Tail fiber assembly protein \| Tail fiber assembly protein; afdb-uniprot50 \| AF-A0A0H2V5L4-F1-MODEL\_V4 \| 1.0 \| 3.579e-15 \| 610 \| 0.423 \| 144 \| 82 \| 1 \| 1 \| 143 \| 11 \| 154 \| Uncharacterized protein \| Uncharacterized protein; afdb-uniprot50 \| AF-A0A7U2JL75-F1-MODEL\_V4 \| 1.0 \| 1.404e-15 \| 607 \| 0.454 \| 143 \| 78 \| 0 \| 1 \| 143 \| 1 \| 143 \| Tail fiber assembly protein \| Tail fiber assembly protein; afdb-uniprot50 \| AF-A0A433ZQC5-F1-MODEL\_V4 \| 1.0 \| 9.075e-16 \| 604 \| 0.489 \| 141 \| 69 \| 2 \| 3 \| 143 \| 2 \| 139 \| Uncharacterized protein \| Uncharacterized protein; afdb-uniprot50 \| AF-A0A0Q4MZH0-F1-MODEL\_V4 \| 1.0 \| 1.033e-14 \| 584 \| 0.417 \| 139 \| 79 \| 1 \| 4 \| 142 \| 2 \| 138 \| Uncharacterized protein \| Uncharacterized protein; afdb-uniprot50 \| AF-A0A827CVF2-F1-MODEL\_V4 \| 1.0 \| 2.313e-15 \| 569 \| 0.556 \| 142 \| 59 \| 3 \| 4 \| 143 \| 3 \| 142 \| Tail fiber assembly protein \| Tail fiber assembly protein; afdb-uniprot50 \| AF-A0A826JA78-F1-MODEL\_V4 \| 1.0 \| 9.12e-15 \| 568 \| 0.482 \| 141 \| 72 \| 1 \| 4 \| 143 \| 3 \| 143 \| Tail assembly chaperone \| Tail assembly chaperone; afdb-uniprot50 \| AF-A0A6N8NTV2-F1-MODEL\_V4 \| 1.0 \| 6.242e-16 \| 564 \| 0.526 \| 133 \| 63 \| 0 \| 1 \| 133 \| 1 \| 133 \| Tail fiber assembly protein \| Tail fiber assembly protein; afdb-uniprot50 \| AF-A0A6N4ED12-F1-MODEL\_V4 \| 1.0 \| 5.894e-15 \| 564 \| 0.363 \| 143 \| 89 \| 1 \| 1 \| 143 \| 11 \| 151 \| Phage tail protein \| Phage tail protein; afdb-uniprot50 \| AF-A0A377ALH1-F1-MODEL\_V4 \| 1.0 \| 6.677e-15 \| 561 \| 0.592 \| 125 \| 51 \| 0 \| 19 \| 143 \| 2 \| 126 \| CPS-53 (KpLE1) prophage protein \| CPS-53 (KpLE1) prophage protein; afdb-uniprot50 \| AF-A0A6G6JM87-F1-MODEL\_V4 \| 1.0 \| 1.1e-14 \| 560 \| 0.442 \| 140 \| 75 \| 2 \| 4 \| 143 \| 2 \| 138 \| Tail fiber assembly protein \| Tail fiber assembly protein; afdb-uniprot50 \| AF-A0A6D0JBZ2-F1-MODEL\_V4 \| 1.0 \| 7.107e-15 \| 558 \| 0.531 \| 141 \| 59 \| 2 \| 4 \| 143 \| 5 \| 139 \| Tail fiber assembly protein \| Tail fiber assembly protein; afdb-uniprot50 \| AF-A0A2U3F206-F1-MODEL\_V4 \| 1.0 \| 2.324e-14 \| 556 \| 0.394 \| 142 \| 84 \| 1 \| 1 \| 142 \| 1 \| 140 \| Tail fiber assembly protein \| Tail fiber assembly protein; afdb-uniprot50 \| AF-A0A6I2IQI1-F1-MODEL\_V4 \| 1.0 \| 2.803e-14 \| 552 \| 0.364 \| 140 \| 89 \| 0 \| 4 \| 143 \| 3 \| 142 \| Tail fiber assembly protein \| Tail fiber assembly protein; afdb-uniprot50 \| AF-A0A6B2FQM4-F1-MODEL\_V4 \| 1.0 \| 1.591e-15 \| 550 \| 0.398 \| 143 \| 84 \| 1 \| 1 \| 143 \| 7 \| 147 \| Tail fiber assembly protein \| Tail fiber assembly protein; afdb-uniprot50 \| AF-A0A732IA80-F1-MODEL\_V4 \| 1.0 \| 4.913e-14 \| 549 \| 0.446 \| 141 \| 76 \| 2 \| 4 \| 143 \| 5 \| 144 \| Tail fiber assembly protein \| Tail fiber assembly protein; afdb-uniprot50 \| AF-A0A411WM32-F1-MODEL\_V4 \| 1.0 \| 1.033e-14 \| 547 \| 0.321 \| 143 \| 95 \| 1 \| 1 \| 143 \| 14 \| 154 \| Tail fiber assembly protein \| Tail fiber assembly protein; afdb-uniprot50 \| AF-A0A4R4JDX7-F1-MODEL\_V4 \| 1.0 \| 1.502e-14 \| 546 \| 0.394 \| 142 \| 84 \| 1 \| 1 \| 142 \| 3 \| 142 \| Phage tail protein \| Phage tail protein; afdb-uniprot50 \| AF-A0A7Z3MDV7-F1-MODEL\_V4 \| 1.0 \| 1.502e-14 \| 545 \| 0.365 \| 145 \| 88 \| 2 \| 1 \| 143 \| 1 \| 143 \| Phage tail assembly chaperone gp38 \| Phage tail assembly chaperone gp38; afdb-uniprot50 \| AF-A0A064DHX9-F1-MODEL\_V4 \| 1.0 \| 2.474e-14 \| 542 \| 0.391 \| 143 \| 83 \| 3 \| 2 \| 143 \| 22 \| 161 \| Uncharacterized protein \| Uncharacterized protein; afdb-uniprot50 \| AF-A0A411WII7-F1-MODEL\_V4 \| 1.0 \| 7.143e-14 \| 534 \| 0.375 \| 144 \| 87 \| 2 \| 1 \| 143 \| 1 \| 142 \| Tail fiber assembly protein \| Tail fiber assembly protein; afdb-uniprot50 \| AF-A0A7Y7YPN7-F1-MODEL\_V4 \| 1.0 \| 4.913e-14 \| 532 \| 0.378 \| 140 \| 83 \| 2 \| 4 \| 143 \| 2 \| 137 \| Tail fiber assembly protein \| Tail fiber assembly protein; afdb-uniprot50 \| AF-A0A4V2X8Y2-F1-MODEL\_V4 \| 1.0 \| 6.305e-14 \| 532 \| 0.382 \| 141 \| 84 \| 2 \| 4 \| 143 \| 2 \| 140 \| Phage tail protein \| Phage tail protein; afdb-uniprot50 \| AF-A0A077QBZ2-F1-MODEL\_V4 \| 1.0 \| 7.143e-14 \| 530 \| 0.378 \| 140 \| 84 \| 2 \| 4 \| 143 \| 2 \| 138 \| Uncharacterized protein \| Uncharacterized protein; afdb-uniprot50 \| AF-A0A077NV63-F1-MODEL\_V4 \| 1.0 \| 9.756e-14 \| 524 \| 0.371 \| 140 \| 85 \| 2 \| 4 \| 143 \| 2 \| 138 \| Uncharacterized protein \| Uncharacterized protein; afdb-uniprot50 \| AF-A0A3D1C8C5-F1-MODEL\_V4 \| 1.0 \| 1.71e-13 \| 522 \| 0.386 \| 145 \| 85 \| 3 \| 1 \| 143 \| 1 \| 143 \| Tail fiber assembly protein \| Tail fiber assembly protein; afdb-uniprot50 \| AF-A0A7L6U240-F1-MODEL\_V4 \| 1.0 \| 3.175e-14 \| 521 \| 0.425 \| 141 \| 78 \| 2 \| 4 \| 143 \| 2 \| 140 \| Tail assembly chaperone \| Tail assembly chaperone; afdb-uniprot50 \| AF-A0A0H0RZ18-F1-MODEL\_V4 \| 1.0 \| 3.597e-14 \| 519 \| 0.397 \| 141 \| 84 \| 1 \| 4 \| 143 \| 3 \| 143 \| Phage tail protein \| Phage tail protein; afdb-uniprot50 \| AF-A0A376ST16-F1-MODEL\_V4 \| 1.0 \| 2.062e-13 \| 519 \| 0.507 \| 126 \| 62 \| 0 \| 1 \| 126 \| 1 \| 126 \| Tail fiber assembly protein \| Tail fiber assembly protein; afdb-uniprot50 \| AF-A0A1I3WQ77-F1-MODEL\_V4 \| 1.0 \| 3.175e-14 \| 518 \| 0.335 \| 143 \| 93 \| 1 \| 1 \| 143 \| 3 \| 143 \| Virus tail fibre assembly protein, lambda gpK \| Virus tail fibre assembly protein, lambda gpK; afdb-uniprot50 \| AF-A0A1V2BDV7-F1-MODEL\_V4 \| 1.0 \| 2.336e-13 \| 514 \| 0.378 \| 140 \| 85 \| 1 \| 4 \| 143 \| 2 \| 139 \| Uncharacterized protein \| Uncharacterized protein; afdb-uniprot50 \| AF-A0A2X1RV26-F1-MODEL\_V4 \| 1.0 \| 6.745e-13 \| 511 \| 0.342 \| 143 \| 92 \| 1 \| 1 \| 143 \| 1 \| 141 \| Caudovirales tail fibre assembly protein \| Caudovirales tail fibre assembly protein; afdb-uniprot50 \| AF-A0A7X5FY77-F1-MODEL\_V4 \| 1.0 \| 1.333e-13 \| 510 \| 0.393 \| 145 \| 84 \| 3 \| 1 \| 143 \| 1 \| 143 \| Tail fiber assembly protein \| Tail fiber assembly protein; afdb-uniprot50 \| AF-A0A0P0QDS8-F1-MODEL\_V4 \| 1.0 \| 2.998e-13 \| 507 \| 0.335 \| 140 \| 91 \| 1 \| 4 \| 143 \| 3 \| 140 \| Tail fiber assembly protein \| Tail fiber assembly protein; afdb-uniprot50 \| AF-A0A2X4V2H3-F1-MODEL\_V4 \| 1.0 \| 9.756e-14 \| 505 \| 0.356 \| 143 \| 89 \| 2 \| 1 \| 143 \| 1 \| 140 \| Caudovirales tail fibre assembly protein \| Caudovirales tail fibre assembly protein; afdb-uniprot50 \| AF-A0A351NIR9-F1-MODEL\_V4 \| 1.0 \| 3.396e-13 \| 504 \| 0.35 \| 140 \| 87 \| 3 \| 4 \| 143 \| 3 \| 138 \| Phage tail protein \| Phage tail protein; afdb-uniprot50 \| AF-A0A377AI92-F1-MODEL\_V4 \| 1.0 \| 7.143e-14 \| 504 \| 0.57 \| 128 \| 54 \| 1 \| 16 \| 143 \| 6 \| 132 \| CPS-53 (KpLE1) prophage protein \| CPS-53 (KpLE1) prophage protein; afdb-uniprot50 \| AF-A0A254PQ93-F1-MODEL\_V4 \| 1.0 \| 1.607e-13 \| 503 \| 0.349 \| 143 \| 89 \| 3 \| 1 \| 142 \| 2 \| 141 \| Phage tail protein \| Phage tail protein; afdb-uniprot50 \| AF-A0A3D1C5Q5-F1-MODEL\_V4 \| 1.0 \| 3.847e-13 \| 499 \| 0.333 \| 144 \| 93 \| 2 \| 1 \| 143 \| 1 \| 142 \| Phage tail protein \| Phage tail protein; afdb-uniprot50 \| AF-A0A2S8PTN0-F1-MODEL\_V4 \| 1.0 \| 5.954e-13 \| 497 \| 0.335 \| 140 \| 90 \| 2 \| 4 \| 142 \| 2 \| 139 \| Phage tail protein \| Phage tail protein; afdb-uniprot50 \| AF-A0A482PI23-F1-MODEL\_V4 \| 1.0 \| 5.566e-14 \| 497 \| 0.423 \| 144 \| 75 \| 2 \| 1 \| 143 \| 1 \| 137 \| Tail fiber assembly protein \| Tail fiber assembly protein; afdb-uniprot50 \| AF-A0A794RD75-F1-MODEL\_V4 \| 1.0 \| 2.195e-13 \| 496 \| 0.576 \| 125 \| 51 \| 2 \| 21 \| 143 \| 2 \| 126 \| Tail fiber assembly protein \| Tail fiber assembly protein; afdb-uniprot50 \| AF-A0A0H3FAT1-F1-MODEL\_V4 \| 1.0 \| 2.998e-13 \| 491 \| 0.354 \| 141 \| 88 \| 2 \| 4 \| 143 \| 3 \| 141 \| Tail assembly chaperone gp38 \| Tail assembly chaperone gp38; afdb-uniprot50 \| AF-A0A370S9Q0-F1-MODEL\_V4 \| 1.0 \| 6.745e-13 \| 490 \| 0.288 \| 142 \| 99 \| 1 \| 2 \| 143 \| 20 \| 159 \| Virus tail fiber assembly protein lambda gpK \| Virus tail fiber assembly protein lambda gpK; afdb-uniprot50 \| AF-A0A1C4G0U3-F1-MODEL\_V4 \| 1.0 \| 7.564e-15 \| 487 \| 0.45 \| 140 \| 74 \| 1 \| 4 \| 143 \| 2 \| 138 \| Virus tail fibre assembly protein, lambda gpK \| Virus tail fibre assembly protein, lambda gpK; afdb-uniprot50 \| AF-A0A857DVJ6-F1-MODEL\_V4 \| 1.0 \| 7.641e-13 \| 487 \| 0.392 \| 140 \| 82 \| 2 \| 4 \| 143 \| 2 \| 138 \| Tail fiber assembly protein \| Tail fiber assembly protein; afdb-uniprot50 \| AF-A0A7U5UFF6-F1-MODEL\_V4 \| 1.0 \| 1.111e-12 \| 483 \| 0.349 \| 143 \| 90 \| 2 \| 2 \| 143 \| 17 \| 157 \| Tail assembly chaperone \| Tail assembly chaperone; afdb-uniprot50 \| AF-A4W7Q4-F1-MODEL\_V4 \| 1.0 \| 4.963e-12 \| 481 \| 0.335 \| 140 \| 91 \| 1 \| 4 \| 143 \| 3 \| 140 \| Phage tail assembly chaperone gp38 \| Phage tail assembly chaperone gp38; afdb-uniprot50 \| AF-A0A831I510-F1-MODEL\_V4 \| 1.0 \| 1.044e-12 \| 478 \| 0.433 \| 143 \| 73 \| 1 \| 1 \| 143 \| 1 \| 135 \| Tail fiber assembly protein \| Tail fiber assembly protein; afdb-uniprot50 \| AF-W0HMV2-F1-MODEL\_V4 \| 1.0 \| 3.396e-13 \| 478 \| 0.321 \| 143 \| 95 \| 1 \| 1 \| 143 \| 1 \| 141 \| Uncharacterized protein \| Uncharacterized protein; afdb-uniprot50 \| AF-A0A4D7IX74-F1-MODEL\_V4 \| 1.0 \| 1.044e-12 \| 478 \| 0.412 \| 143 \| 80 \| 3 \| 2 \| 142 \| 57 \| 197 \| Phage tail protein \| Phage tail protein; afdb-uniprot50 \| AF-A0A087KXL6-F1-MODEL\_V4 \| 1.0 \| 6.337e-13 \| 477 \| 0.34 \| 144 \| 89 \| 4 \| 1 \| 143 \| 1 \| 139 \| Uncharacterized protein \| Uncharacterized protein; afdb-uniprot50 \| AF-A0A7U5TH15-F1-MODEL\_V4 \| 1.0 \| 9.213e-13 \| 476 \| 0.358 \| 145 \| 88 \| 4 \| 1 \| 143 \| 1 \| 142 \| Uncharacterized protein \| Uncharacterized protein; afdb-uniprot50 \| AF-A0A2A2HLN3-F1-MODEL\_V4 \| 1.0 \| 1.719e-12 \| 476 \| 0.34 \| 141 \| 90 \| 2 \| 4 \| 143 \| 2 \| 140 \| Phage tail protein \| Phage tail protein; afdb-uniprot50 \| AF-A0A6L9JIU7-F1-MODEL\_V4 \| 1.0 \| 1.719e-12 \| 474 \| 0.326 \| 141 \| 92 \| 2 \| 4 \| 143 \| 2 \| 140 \| Tail fiber assembly protein \| Tail fiber assembly protein; afdb-uniprot50 \| AF-A0A4R6PIE0-F1-MODEL\_V4 \| 1.0 \| 1.339e-12 \| 474 \| 0.364 \| 140 \| 87 \| 1 \| 4 \| 143 \| 3 \| 140 \| Virus tail fiber assembly protein lambda gpK \| Virus tail fiber assembly protein lambda gpK; afdb-uniprot50 \| AF-A0A5U1N744-F1-MODEL\_V4 \| 1.0 \| 6.337e-13 \| 474 \| 0.453 \| 141 \| 74 \| 3 \| 4 \| 142 \| 3 \| 142 \| Tail fiber assembly protein \| Tail fiber assembly protein; afdb-uniprot50 \| AF-A0A0A8VIE2-F1-MODEL\_V4 \| 1.0 \| 8.132e-13 \| 473 \| 0.335 \| 143 \| 92 \| 2 \| 1 \| 143 \| 1 \| 140 \| Tail fiber assembly protein p37 \| Tail fiber assembly protein p37; afdb-uniprot50 \| AF-A0A2V2BI76-F1-MODEL\_V4 \| 1.0 \| 1.24e-15 \| 472 \| 0.461 \| 143 \| 77 \| 0 \| 1 \| 143 \| 1 \| 143 \| Virus tail fiber assembly protein lambda gpK \| Virus tail fiber assembly protein lambda gpK; afdb-uniprot50 \| AF-U3U6G6-F1-MODEL\_V4 \| 1.0 \| 1.719e-12 \| 472 \| 0.355 \| 138 \| 86 \| 2 \| 4 \| 140 \| 2 \| 137 \| Uncharacterized protein \| Uncharacterized protein; afdb-uniprot50 \| AF-A0A826NIC5-F1-MODEL\_V4 \| 1.0 \| 2.195e-13 \| 471 \| 0.374 \| 139 \| 85 \| 1 \| 2 \| 140 \| 3 \| 139 \| Tail fiber assembly protein \| Tail fiber assembly protein; afdb-uniprot50 \| AF-A0A3S5G566-F1-MODEL\_V4 \| 1.0 \| 2.66e-12 \| 469 \| 0.345 \| 142 \| 89 \| 3 \| 1 \| 140 \| 1 \| 140 \| Tail fiber assembly protein \| Tail fiber assembly protein; afdb-uniprot50 \| AF-A0A811D365-F1-MODEL\_V4 \| 1.0 \| 3.013e-12 \| 465 \| 0.35 \| 140 \| 86 \| 2 \| 4 \| 143 \| 2 \| 136 \| Uncharacterized protein \| Uncharacterized protein; afdb-uniprot50 \| AF-A0A765X9T6-F1-MODEL\_V4 \| 1.0 \| 5.594e-13 \| 465 \| 0.46 \| 141 \| 69 \| 1 \| 3 \| 143 \| 2 \| 135 \| Tail fiber assembly protein \| Tail fiber assembly protein; afdb-uniprot50 \| AF-A0A7U5G183-F1-MODEL\_V4 \| 1.0 \| 7.215e-12 \| 461 \| 0.333 \| 144 \| 90 \| 4 \| 1 \| 143 \| 1 \| 139 \| Uncharacterized protein \| Uncharacterized protein; afdb-uniprot50 \| AF-H2IRJ1-F1-MODEL\_V4 \| 1.0 \| 3.867e-12 \| 461 \| 0.358 \| 145 \| 89 \| 3 \| 1 \| 143 \| 1 \| 143 \| Caudovirales tail fiber assembly protein \| Caudovirales tail fiber assembly protein; afdb-uniprot50 \| AF-A0A5B8I5R9-F1-MODEL\_V4 \| 1.0 \| 5.594e-13 \| 459 \| 0.356 \| 146 \| 88 \| 4 \| 1 \| 143 \| 1 \| 143 \| Tail assembly chaperone \| Tail assembly chaperone; afdb-uniprot50 \| AF-A0A6B0TPS5-F1-MODEL\_V4 \| 1.0 \| 6.369e-12 \| 457 \| 0.335 \| 143 \| 93 \| 1 \| 1 \| 143 \| 9 \| 149 \| Tail fiber assembly protein \| Tail fiber assembly protein; afdb-uniprot50 \| AF-A0A3N6SH38-F1-MODEL\_V4 \| 1.0 \| 7.679e-12 \| 456 \| 0.324 \| 145 \| 92 \| 5 \| 1 \| 143 \| 1 \| 141 \| Phage tail protein \| Phage tail protein; afdb-uniprot50 \| AF-A0A6I6IFF1-F1-MODEL\_V4 \| 1.0 \| 7.179e-13 \| 455 \| 0.375 \| 149 \| 83 \| 3 \| 3 \| 143 \| 2 \| 148 \| Tail fiber assembly protein \| Tail fiber assembly protein; afdb-uniprot50 \| AF-A0A3Q8DJE8-F1-MODEL\_V4 \| 1.0 \| 9.855e-12 \| 452 \| 0.392 \| 140 \| 79 \| 3 \| 4 \| 143 \| 4 \| 137 \| Phage tail protein \| Phage tail protein; afdb-uniprot50 \| AF-A0A2I8Q7J9-F1-MODEL\_V4 \| 1.0 \| 1.829e-12 \| 452 \| 0.349 \| 143 \| 89 \| 3 \| 1 \| 143 \| 11 \| 149 \| Phage tail protein \| Phage tail protein; afdb-uniprot50 \| AF-A0A7I6TII6-F1-MODEL\_V4 \| 1.0 \| 3.413e-12 \| 451 \| 0.298 \| 144 \| 98 \| 2 \| 1 \| 143 \| 1 \| 142 \| Tail assembly chaperone \| Tail assembly chaperone; afdb-uniprot50 \| AF-R8ANC1-F1-MODEL\_V4 \| 1.0 \| 2.817e-13 \| 451 \| 0.384 \| 143 \| 86 \| 2 \| 1 \| 143 \| 2 \| 142 \| Putative tail fiber assembly protein \| Putative tail fiber assembly protein; afdb-uniprot50 \| AF-A0A857EZE8-F1-MODEL\_V4 \| 1.0 \| 1.615e-12 \| 449 \| 0.357 \| 140 \| 87 \| 2 \| 4 \| 143 \| 3 \| 139 \| Tail fiber assembly protein \| Tail fiber assembly protein; afdb-uniprot50 \| AF-A0A3G2I9A2-F1-MODEL\_V4 \| 1.0 \| 8.173e-12 \| 448 \| 0.342 \| 140 \| 87 \| 2 \| 4 \| 143 \| 2 \| 136 \| Tail fiber assembly protein \| Tail fiber assembly protein; afdb-uniprot50 \| AF-Q8ZNJ2-F1-MODEL\_V4 \| 1.0 \| 2.348e-12 \| 448 \| 0.429 \| 142 \| 79 \| 2 \| 1 \| 141 \| 1 \| 141 \| Putative cytoplasmic protein \| Putative cytoplasmic protein; afdb-uniprot50 \| AF-A0A4P7KSD4-F1-MODEL\_V4 \| 1.0 \| 2.499e-12 \| 442 \| 0.312 \| 141 \| 94 \| 2 \| 4 \| 143 \| 9 \| 147 \| Caudovirales tail fiber assembly protein, lambda gpK \| Caudovirales tail fiber assembly protein, lambda gpK; afdb-uniprot50 \| AF-A0A7X1F2W4-F1-MODEL\_V4 \| 1.0 \| 2.083e-11 \| 439 \| 0.328 \| 140 \| 91 \| 2 \| 4 \| 143 \| 2 \| 138 \| Tail fiber assembly protein \| Tail fiber assembly protein; afdb-uniprot50 \| AF-A0A859TBI1-F1-MODEL\_V4 \| 1.0 \| 6.779e-12 \| 435 \| 0.333 \| 141 \| 91 \| 2 \| 3 \| 143 \| 2 \| 139 \| Tail fiber assembly protein \| Tail fiber assembly protein; afdb-uniprot50 \| AF-A0A0A3ZBR4-F1-MODEL\_V4 \| 1.0 \| 8.699e-12 \| 433 \| 0.342 \| 143 \| 85 \| 3 \| 4 \| 143 \| 3 \| 139 \| Uncharacterized protein \| Uncharacterized protein; afdb-uniprot50 \| AF-A0A564UCV1-F1-MODEL\_V4 \| 1.0 \| 6.014e-11 \| 431 \| 0.546 \| 108 \| 48 \| 1 \| 37 \| 143 \| 3 \| 110 \| Caudovirales tail fibre assembly protein \| Caudovirales tail fibre assembly protein; afdb-uniprot50 \| AF-A0A7D7ACY9-F1-MODEL\_V4 \| 1.0 \| 3.223e-11 \| 429 \| 0.347 \| 141 \| 88 \| 3 \| 4 \| 143 \| 3 \| 140 \| Tail fiber assembly protein \| Tail fiber assembly protein; afdb-uniprot50 \| AF-A0A212I9D1-F1-MODEL\_V4 \| 1.0 \| 1.525e-11 \| 428 \| 0.338 \| 142 \| 89 \| 3 \| 4 \| 143 \| 3 \| 141 \| Uncharacterized protein \| Uncharacterized protein; afdb-uniprot50 \| AF-A0A7I6VNU6-F1-MODEL\_V4 \| 1.0 \| 4.663e-12 \| 423 \| 0.335 \| 143 \| 91 \| 3 \| 1 \| 143 \| 1 \| 139 \| Tail assembly chaperone \| Tail assembly chaperone; afdb-uniprot50 \| AF-A0A7W5JQR9-F1-MODEL\_V4 \| 1.0 \| 2.217e-11 \| 420 \| 0.278 \| 140 \| 99 \| 1 \| 4 \| 143 \| 3 \| 140 \| Uncharacterized protein \| Uncharacterized protein; afdb-uniprot50 \| AF-A0A344S5G9-F1-MODEL\_V4 \| 1.0 \| 3.431e-11 \| 420 \| 0.345 \| 139 \| 87 \| 3 \| 3 \| 140 \| 2 \| 137 \| Phage tail protein \| Phage tail protein; afdb-uniprot50 \| AF-A0A6M5D0K4-F1-MODEL\_V4 \| 1.0 \| 2.36e-11 \| 419 \| 0.304 \| 148 \| 91 \| 3 \| 4 \| 143 \| 3 \| 146 \| Phage tail protein \| Phage tail protein; afdb-uniprot50 \| AF-A0A826S2B1-F1-MODEL\_V4 \| 1.0 \| 1.957e-11 \| 417 \| 0.34 \| 141 \| 84 \| 3 \| 4 \| 143 \| 22 \| 154 \| Tail fiber assembly protein \| Tail fiber assembly protein; afdb-uniprot50 \| AF-A0A7A3AEP4-F1-MODEL\_V4 \| 1.0 \| 6.369e-12 \| 416 \| 0.404 \| 141 \| 77 \| 2 \| 4 \| 143 \| 5 \| 139 \| Tail fiber assembly protein \| Tail fiber assembly protein; afdb-uniprot50 \| AF-Q2NQF0-F1-MODEL\_V4 \| 1.0 \| 6.014e-11 \| 415 \| 0.378 \| 132 \| 79 \| 2 \| 4 \| 135 \| 12 \| 140 \| Caudovirales tail fibre assembly protein \| Caudovirales tail fibre assembly protein; afdb-uniprot50 \| AF-A0A0J1N7J0-F1-MODEL\_V4 \| 1.0 \| 8.743e-11 \| 414 \| 0.29 \| 141 \| 98 \| 1 \| 3 \| 143 \| 2 \| 140 \| Uncharacterized protein \| Uncharacterized protein; afdb-uniprot50 \| AF-A0A810TM02-F1-MODEL\_V4 \| 1.0 \| 4.403e-11 \| 414 \| 0.305 \| 144 \| 94 \| 3 \| 1 \| 143 \| 1 \| 139 \| Phage tail fiber assembly protein \| Phage tail fiber assembly protein; afdb-uniprot50 \| AF-A0A1R0Q9D4-F1-MODEL\_V4 \| 1.0 \| 4.136e-11 \| 412 \| 0.34 \| 138 \| 84 \| 3 \| 4 \| 141 \| 2 \| 132 \| Phage tail protein \| Phage tail protein; afdb-uniprot50 \| AF-A0A5D4YFI5-F1-MODEL\_V4 \| 1.0 \| 5.309e-11 \| 410 \| 0.307 \| 140 \| 95 \| 1 \| 4 \| 143 \| 3 \| 140 \| Tail fiber assembly protein \| Tail fiber assembly protein; afdb-uniprot50 \| AF-A0A377F879-F1-MODEL\_V4 \| 1.0 \| 1.353e-10 \| 409 \| 0.491 \| 118 \| 60 \| 0 \| 26 \| 143 \| 23 \| 140 \| Putative phage tail assembly chaperone \| Putative phage tail assembly chaperone; afdb-uniprot50 \| AF-D2TYN1-F1-MODEL\_V4 \| 1.0 \| 4.136e-11 \| 406 \| 0.297 \| 141 \| 95 \| 3 \| 4 \| 143 \| 2 \| 139 \| Caudovirales tail fiber assembly protein, lambda gpK \| Caudovirales tail fiber assembly protein, lambda gpK; afdb-uniprot50 \| AF-A0A2D0LEZ5-F1-MODEL\_V4 \| 1.0 \| 2.983e-14 \| 405 \| 0.342 \| 146 \| 91 \| 2 \| 1 \| 143 \| 2 \| 145 \| Phage tail fiber assembly \| Phage tail fiber assembly; afdb-uniprot50 \| AF-A0A3N0FQC3-F1-MODEL\_V4 \| 1.0 \| 1.44e-10 \| 401 \| 0.335 \| 140 \| 85 \| 2 \| 4 \| 143 \| 5 \| 136 \| Phage tail protein \| Phage tail protein; afdb-uniprot50 \| AF-A0A3M4RIN5-F1-MODEL\_V4 \| 1.0 \| 2.083e-11 \| 400 \| 0.286 \| 143 \| 98 \| 3 \| 1 \| 142 \| 1 \| 140 \| Phage tail fiber assembly \| Phage tail fiber assembly; afdb-uniprot50 \| AF-A0A830Q7F2-F1-MODEL\_V4 \| 1.0 \| 2.512e-11 \| 400 \| 0.351 \| 145 \| 89 \| 4 \| 1 \| 143 \| 1 \| 142 \| Putative phage tail fiber assembly protein \| Putative phage tail fiber assembly protein; afdb-uniprot50 \| AF-A0A1H8UDQ4-F1-MODEL\_V4 \| 1.0 \| 2.093e-10 \| 398 \| 0.298 \| 144 \| 91 \| 3 \| 4 \| 143 \| 3 \| 140 \| Virus tail fibre assembly protein, lambda gpK \| Virus tail fibre assembly protein, lambda gpK; afdb-uniprot50 \| AF-A0A1Q6B0D0-F1-MODEL\_V4 \| 1.0 \| 7.251e-11 \| 398 \| 0.359 \| 142 \| 87 \| 3 \| 3 \| 143 \| 5 \| 143 \| Phage tail protein \| Phage tail protein; afdb-uniprot50 \| AF-A0A7L6YDG5-F1-MODEL\_V4 \| 1.0 \| 6.401e-11 \| 397 \| 0.292 \| 147 \| 93 \| 4 \| 1 \| 143 \| 4 \| 143 \| Tail fiber assembly protein \| Tail fiber assembly protein; afdb-uniprot50 \| AF-A0A7L6CIZ9-F1-MODEL\_V4 \| 1.0 \| 1.054e-10 \| 395 \| 0.342 \| 143 \| 88 \| 5 \| 4 \| 143 \| 4 \| 143 \| Tail fiber assembly protein \| Tail fiber assembly protein; afdb-uniprot50 \| AF-A0A4V3F6N1-F1-MODEL\_V4 \| 1.0 \| 1.533e-10 \| 394 \| 0.32 \| 125 \| 82 \| 2 \| 4 \| 127 \| 10 \| 132 \| Virus tail fiber assembly protein lambda gpK \| Virus tail fiber assembly protein lambda gpK; afdb-uniprot50 \| AF-A0A5F1T5X9-F1-MODEL\_V4 \| 1.0 \| 7.288e-10 \| 394 \| 0.33 \| 142 \| 89 \| 3 \| 2 \| 143 \| 23 \| 158 \| Tail fiber assembly protein \| Tail fiber assembly protein; afdb-uniprot50 \| AF-A0A2D0KV06-F1-MODEL\_V4 \| 1.0 \| 2.831e-12 \| 393 \| 0.356 \| 143 \| 85 \| 3 \| 4 \| 143 \| 423 \| 561 \| Tail assembly chaperone \| Tail assembly chaperone; afdb-uniprot50 \| AF-A0A6N8NR58-F1-MODEL\_V4 \| 1.0 \| 3.256e-09 \| 389 \| 0.623 \| 93 \| 34 \| 1 \| 51 \| 143 \| 1 \| 92 \| Tail fiber assembly protein \| Tail fiber assembly protein; afdb-uniprot50 \| AF-T0QGQ9-F1-MODEL\_V4 \| 1.0 \| 3.044e-10 \| 386 \| 0.366 \| 120 \| 74 \| 1 \| 1 \| 120 \| 3 \| 120 \| Putative tail fiber protein \| Putative tail fiber protein; afdb-uniprot50 \| AF-A0A1Q5V7C8-F1-MODEL\_V4 \| 1.0 \| 4.686e-11 \| 385 \| 0.323 \| 142 \| 90 \| 4 \| 4 \| 143 \| 2 \| 139 \| Uncharacterized protein \| Uncharacterized protein; afdb-uniprot50 \| AF-A0A4T4IVD4-F1-MODEL\_V4 \| 1.0 \| 4.136e-11 \| 385 \| 0.291 \| 144 \| 97 \| 3 \| 1 \| 143 \| 1 \| 140 \| Tail assembly chaperone \| Tail assembly chaperone; afdb-uniprot50 \| AF-A0A209ALQ9-F1-MODEL\_V4 \| 1.0 \| 2.073e-12 \| 385 \| 0.388 \| 144 \| 81 \| 4 \| 4 \| 143 \| 13 \| 153 \| Tail fiber assembly protein \| Tail fiber assembly protein; afdb-uniprot50 \| AF-A0A744A7C6-F1-MODEL\_V4 \| 1.0 \| 2.384e-09 \| 383 \| 0.541 \| 96 \| 43 \| 1 \| 47 \| 142 \| 1 \| 95 \| Tail fiber assembly protein \| Tail fiber assembly protein; afdb-uniprot50 \| AF-A0A654DRC8-F1-MODEL\_V4 \| 1.0 \| 2.998e-13 \| 378 \| 0.335 \| 140 \| 88 \| 3 \| 4 \| 143 \| 2 \| 136 \| Uncharacterized protein \| Uncharacterized protein; afdb-uniprot50 \| AF-A0A3S6EWT1-F1-MODEL\_V4 \| 1.0 \| 3.906e-10 \| 377 \| 0.302 \| 142 \| 94 \| 4 \| 4 \| 143 \| 2 \| 140 \| Uncharacterized protein \| Uncharacterized protein; afdb-uniprot50 \| AF-A0A1M5U130-F1-MODEL\_V4 \| 1.0 \| 5.594e-13 \| 377 \| 0.371 \| 140 \| 85 \| 2 \| 4 \| 143 \| 3 \| 139 \| Virus tail fibre assembly protein, lambda gpK \| Virus tail fibre assembly protein, lambda gpK; afdb-uniprot50 \| AF-A0A085JCX8-F1-MODEL\_V4 \| 1.0 \| 6.847e-10 \| 376 \| 0.292 \| 140 \| 96 \| 2 \| 4 \| 143 \| 2 \| 138 \| Tail fiber assembly protein \| Tail fiber assembly protein; afdb-uniprot50 \| AF-A0A068Z8F6-F1-MODEL\_V4 \| 1.0 \| 2.817e-13 \| 374 \| 0.349 \| 143 \| 88 \| 2 \| 4 \| 143 \| 3 \| 143 \| Tail fiber assembly protein \| Tail fiber assembly protein; afdb-uniprot50 \| AF-A0A5U3DTV8-F1-MODEL\_V4 \| 1.0 \| 1.122e-10 \| 373 \| 0.321 \| 140 \| 90 \| 2 \| 4 \| 143 \| 3 \| 137 \| Tail fiber assembly protein \| Tail fiber assembly protein; afdb-uniprot50 \| AF-A0A495AL45-F1-MODEL\_V4 \| 1.0 \| 1.194e-10 \| 372 \| 0.316 \| 142 \| 91 \| 3 \| 4 \| 143 \| 2 \| 139 \| Tail fiber assembly protein \| Tail fiber assembly protein; afdb-uniprot50 \| AF-A0A7U4I2C5-F1-MODEL\_V4 \| 1.0 \| 8.743e-11 \| 372 \| 0.3 \| 140 \| 96 \| 1 \| 4 \| 143 \| 3 \| 140 \| Uncharacterized protein \| Uncharacterized protein; afdb-uniprot50 \| AF-A0A806YUH0-F1-MODEL\_V4 \| 1.0 \| 2.073e-12 \| 372 \| 0.349 \| 143 \| 88 \| 3 \| 4 \| 143 \| 3 \| 143 \| Uncharacterized protein \| Uncharacterized protein; afdb-uniprot50 \| AF-A0A5A9D4L2-F1-MODEL\_V4 \| 1.0 \| 2.687e-10 \| 372 \| 0.351 \| 145 \| 83 \| 5 \| 1 \| 140 \| 41 \| 179 \| Tail fiber assembly protein \| Tail fiber assembly protein; afdb-uniprot50 \| AF-A0A5B0K5E4-F1-MODEL\_V4 \| 1.0 \| 2.348e-12 \| 370 \| 0.286 \| 143 \| 97 \| 3 \| 4 \| 143 \| 9 \| 149 \| Tail fiber assembly protein \| Tail fiber assembly protein; afdb-uniprot50 \| AF-A0A3D3HLN8-F1-MODEL\_V4 \| 1.0 \| 1.977e-09 \| 368 \| 0.441 \| 111 \| 59 \| 2 \| 33 \| 143 \| 5 \| 112 \| Phage tail protein \| Phage tail protein; afdb-uniprot50 \| AF-A0A5V3MS23-F1-MODEL\_V4 \| 1.0 \| 1.36e-09 \| 368 \| 0.264 \| 140 \| 101 \| 1 \| 4 \| 143 \| 3 \| 140 \| Tail fiber assembly protein \| Tail fiber assembly protein; afdb-uniprot50 \| AF-A0A8B3UIH1-F1-MODEL\_V4 \| 1.0 \| 2.524e-10 \| 368 \| 0.316 \| 142 \| 86 \| 4 \| 4 \| 140 \| 3 \| 138 \| Tail assembly chaperone gp38 \| Tail assembly chaperone gp38; afdb-uniprot50 \| AF-A0A7L6DU13-F1-MODEL\_V4 \| 1.0 \| 6.044e-10 \| 368 \| 0.321 \| 146 \| 93 \| 5 \| 1 \| 143 \| 1 \| 143 \| Tail fiber assembly protein \| Tail fiber assembly protein; afdb-uniprot50 \| AF-A0A376U5W9-F1-MODEL\_V4 \| 1.0 \| 6.498e-08 \| 367 \| 0.612 \| 80 \| 31 \| 0 \| 64 \| 143 \| 2 \| 81 \| Putative phage tail assembly chaperone \| Putative phage tail assembly chaperone; afdb-uniprot50 \| AF-A0A701UP36-F1-MODEL\_V4 \| 1.0 \| 1.54e-09 \| 366 \| 0.312 \| 144 \| 92 \| 5 \| 4 \| 143 \| 3 \| 143 \| Tail assembly chaperone \| Tail assembly chaperone; afdb-uniprot50 \| AF-A0A0P8I676-F1-MODEL\_V4 \| 1.0 \| 1.36e-09 \| 365 \| 0.364 \| 118 \| 72 \| 2 \| 27 \| 143 \| 1 \| 116 \| Phage tail protein \| Phage tail protein; afdb-uniprot50 \| AF-A0A2N5DU56-F1-MODEL\_V4 \| 1.0 \| 3.413e-12 \| 362 \| 0.354 \| 144 \| 90 \| 2 \| 1 \| 143 \| 9 \| 150 \| Phage tail protein \| Phage tail protein; afdb-uniprot50 \| AF-A0A069J7E8-F1-MODEL\_V4 \| 1.0 \| 1.54e-09 \| 361 \| 0.51 \| 98 \| 48 \| 0 \| 4 \| 101 \| 3 \| 100 \| Uncharacterized protein \| Uncharacterized protein; afdb-uniprot50 \| AF-A0A0T9UK09-F1-MODEL\_V4 \| 1.0 \| 2.831e-12 \| 361 \| 0.378 \| 148 \| 79 \| 4 \| 3 \| 143 \| 2 \| 143 \| Tail assembly chaperone gp38 \| Tail assembly chaperone gp38; afdb-uniprot50 \| AF-A0A501S8D9-F1-MODEL\_V4 \| 1.0 \| 8.743e-11 \| 360 \| 0.3 \| 143 \| 91 \| 3 \| 4 \| 143 \| 3 \| 139 \| Tail fiber assembly protein \| Tail fiber assembly protein; afdb-uniprot50 \| AF-A0A376WPX8-F1-MODEL\_V4 \| 1.0 \| 1.006e-07 \| 359 \| 0.637 \| 80 \| 29 \| 0 \| 64 \| 143 \| 7 \| 86 \| CPS-53 (KpLE1) prophage protein \| CPS-53 (KpLE1) prophage protein; afdb-uniprot50 \| AF-A0A0A2V4M0-F1-MODEL\_V4 \| 1.0 \| 2.36e-11 \| 359 \| 0.285 \| 147 \| 96 \| 4 \| 1 \| 143 \| 1 \| 142 \| Uncharacterized protein yfdK \| Uncharacterized protein yfdK; afdb-uniprot50 \| AF-A0A849JYU2-F1-MODEL\_V4 \| 1.0 \| 1.271e-10 \| 358 \| 0.307 \| 143 \| 97 \| 1 \| 1 \| 143 \| 2 \| 142 \| Tail fiber assembly protein \| Tail fiber assembly protein; afdb-uniprot50 \| AF-A0A2V1HEP7-F1-MODEL\_V4 \| 1.0 \| 1.271e-10 \| 354 \| 0.316 \| 142 \| 91 \| 3 \| 3 \| 143 \| 5 \| 141 \| Phage tail protein \| Phage tail protein; afdb-uniprot50 \| AF-A0A3W4K3C3-F1-MODEL\_V4 \| 1.0 \| 3.707e-08 \| 353 \| 0.584 \| 89 \| 37 \| 0 \| 55 \| 143 \| 1 \| 89 \| Phage tail protein \| Phage tail protein; afdb-uniprot50 \| AF-A0A827UX27-F1-MODEL\_V4 \| 1.0 \| 6.847e-10 \| 353 \| 0.257 \| 140 \| 102 \| 1 \| 4 \| 143 \| 3 \| 140 \| Tail fiber assembly protein \| Tail fiber assembly protein; afdb-uniprot50 \| AF-A0A7Y6NBC3-F1-MODEL\_V4 \| 1.0 \| 4.447e-09 \| 352 \| 0.302 \| 139 \| 90 \| 2 \| 4 \| 142 \| 3 \| 134 \| Tail fiber assembly protein \| Tail fiber assembly protein; afdb-uniprot50 \| AF-A0A6N3HR67-F1-MODEL\_V4 \| 1.0 \| 3.926e-09 \| 351 \| 0.291 \| 144 \| 86 \| 4 \| 4 \| 143 \| 3 \| 134 \| Caudovirales tail fibre assembly protein \| Caudovirales tail fibre assembly protein; afdb-uniprot50 \| AF-A0A424WAM4-F1-MODEL\_V4 \| 1.0 \| 1.745e-09 \| 348 \| 0.286 \| 143 \| 97 \| 3 \| 4 \| 143 \| 29 \| 169 \| Tail fiber assembly protein \| Tail fiber assembly protein; afdb-uniprot50 \| AF-A0A7D7HTP8-F1-MODEL\_V4 \| 1.0 \| 3.906e-10 \| 346 \| 0.244 \| 143 \| 103 \| 2 \| 1 \| 143 \| 4 \| 141 \| Tail assembly chaperone \| Tail assembly chaperone; afdb-uniprot50 \| AF-A0A837LIM0-F1-MODEL\_V4 \| 1.0 \| 3.028e-11 \| 346 \| 0.364 \| 159 \| 71 \| 5 \| 4 \| 143 \| 3 \| 150 \| Uncharacterized protein \| Uncharacterized protein; afdb-uniprot50 \| AF-A0A5C5DE35-F1-MODEL\_V4 \| 1.0 \| 1.06e-09 \| 344 \| 0.234 \| 145 \| 105 \| 4 \| 1 \| 142 \| 1 \| 142 \| Tail fiber assembly protein \| Tail fiber assembly protein; afdb-uniprot50 \| AF-A0A8A6CV85-F1-MODEL\_V4 \| 1.0 \| 5.736e-08 \| 343 \| 0.511 \| 88 \| 43 \| 0 \| 56 \| 143 \| 5 \| 92 \| Tail fiber assembly protein \| Tail fiber assembly protein; afdb-uniprot50 \| AF-A0A410XDP5-F1-MODEL\_V4 \| 1.0 \| 2.36e-11 \| 343 \| 0.276 \| 141 \| 99 \| 2 \| 4 \| 143 \| 2 \| 140 \| Phage tail protein \| Phage tail protein; afdb-uniprot50 \| AF-A0A376FB91-F1-MODEL\_V4 \| 1.0 \| 7.361e-08 \| 342 \| 0.515 \| 66 \| 32 \| 0 \| 3 \| 68 \| 2 \| 67 \| Phage tail assembly chaperone gp38 \| Phage tail assembly chaperone gp38; afdb-uniprot50 \| AF-A0A5P0JJ43-F1-MODEL\_V4 \| 1.0 \| 4.178e-09 \| 340 \| 0.344 \| 119 \| 74 \| 2 \| 4 \| 122 \| 2 \| 116 \| Tail fiber assembly protein \| Tail fiber assembly protein; afdb-uniprot50 \| AF-Q2NWF3-F1-MODEL\_V4 \| 1.0 \| 1.548e-08 \| 339 \| 0.333 \| 114 \| 74 \| 1 \| 1 \| 114 \| 19 \| 130 \| Uncharacterized protein \| Uncharacterized protein; afdb-uniprot50 \| AF-A0A7U5JH60-F1-MODEL\_V4 \| 1.0 \| 9.954e-10 \| 339 \| 0.314 \| 124 \| 82 \| 2 \| 4 \| 127 \| 3 \| 123 \| Uncharacterized protein \| Uncharacterized protein; afdb-uniprot50 \| AF-W1J7J0-F1-MODEL\_V4 \| 1.0 \| 5.013e-10 \| 338 \| 0.301 \| 146 \| 91 \| 4 \| 1 \| 143 \| 3 \| 140 \| Uncharacterized protein \| Uncharacterized protein; afdb-uniprot50 \| AF-A0A2Y0RH22-F1-MODEL\_V4 \| 1.0 \| 1.128e-09 \| 338 \| 0.298 \| 144 \| 86 \| 3 \| 4 \| 143 \| 3 \| 135 \| Tail assembly chaperone gp38 \| Tail assembly chaperone gp38; afdb-uniprot50 \| AF-A0A1I6E7B8-F1-MODEL\_V4 \| 1.0 \| 9.353e-10 \| 337 \| 0.28 \| 139 \| 96 \| 3 \| 4 \| 140 \| 3 \| 139 \| Virus tail fibre assembly protein, lambda gpK \| Virus tail fibre assembly protein, lambda gpK; afdb-uniprot50 \| AF-A0A5X3P5F8-F1-MODEL\_V4 \| 1.0 \| 4.71e-10 \| 336 \| 0.5 \| 138 \| 50 \| 3 \| 3 \| 140 \| 2 \| 120 \| Tail fiber assembly protein \| Tail fiber assembly protein; afdb-uniprot50 \| AF-D2U4A6-F1-MODEL\_V4 \| 1.0 \| 4.447e-09 \| 336 \| 0.283 \| 127 \| 88 \| 2 \| 4 \| 129 \| 12 \| 136 \| Phage tail assembly protein \| Phage tail assembly protein; afdb-uniprot50 \| AF-A0A0U1HYV0-F1-MODEL\_V4 \| 1.0 \| 4.447e-09 \| 336 \| 0.308 \| 146 \| 89 \| 5 \| 3 \| 143 \| 2 \| 140 \| Phage tail protein \| Phage tail protein; afdb-uniprot50 \| AF-A0A7X5FY66-F1-MODEL\_V4 \| 1.0 \| 1.2e-09 \| 335 \| 0.263 \| 144 \| 91 \| 4 \| 4 \| 143 \| 3 \| 135 \| Tail fiber assembly protein \| Tail fiber assembly protein; afdb-uniprot50 \| AF-A0A2D0IKG5-F1-MODEL\_V4 \| 1.0 \| 2.384e-09 \| 331 \| 0.282 \| 145 \| 90 \| 4 \| 3 \| 143 \| 15 \| 149 \| Tail assembly chaperone \| Tail assembly chaperone; afdb-uniprot50 \| AF-A0A777TWY1-F1-MODEL\_V4 \| 1.0 \| 7.398e-07 \| 329 \| 0.657 \| 70 \| 24 \| 0 \| 74 \| 143 \| 2 \| 71 \| Tail fiber assembly protein \| Tail fiber assembly protein; afdb-uniprot50 \| AF-A0A5D4YD83-F1-MODEL\_V4 \| 1.0 \| 6.847e-10 \| 329 \| 0.321 \| 143 \| 86 \| 5 \| 4 \| 143 \| 2 \| 136 \| Tail fiber assembly protein \| Tail fiber assembly protein; afdb-uniprot50 \| AF-W0BPG2-F1-MODEL\_V4 \| 1.0 \| 1.857e-09 \| 329 \| 0.279 \| 143 \| 98 \| 4 \| 3 \| 143 \| 2 \| 141 \| Uncharacterized protein \| Uncharacterized protein; afdb-uniprot50 \| AF-A0A376YI45-F1-MODEL\_V4 \| 1.0 \| 5.038e-09 \| 327 \| 0.44 \| 100 \| 56 \| 0 \| 4 \| 103 \| 3 \| 102 \| CPS-53 (KpLE1) prophage protein \| CPS-53 (KpLE1) prophage protein; afdb-uniprot50 \| AF-A0A2D0LAF6-F1-MODEL\_V4 \| 1.0 \| 6.075e-09 \| 327 \| 0.279 \| 143 \| 91 \| 4 \| 1 \| 143 \| 1 \| 131 \| Tail assembly chaperone \| Tail assembly chaperone; afdb-uniprot50 \| AF-A0A5R8TTA1-F1-MODEL\_V4 \| 1.0 \| 1.145e-06 \| 325 \| 0.657 \| 70 \| 24 \| 0 \| 74 \| 143 \| 2 \| 71 \| Tail fiber assembly protein \| Tail fiber assembly protein; afdb-uniprot50 \| AF-A0A7Y8YCI4-F1-MODEL\_V4 \| 1.0 \| 1.284e-08 \| 324 \| 0.271 \| 140 \| 94 \| 2 \| 4 \| 143 \| 2 \| 133 \| Tail fiber assembly protein \| Tail fiber assembly protein; afdb-uniprot50 \| AF-A0A411WM65-F1-MODEL\_V4 \| 1.0 \| 1.867e-08 \| 324 \| 0.292 \| 140 \| 90 \| 3 \| 4 \| 143 \| 3 \| 133 \| Tail fiber assembly protein \| Tail fiber assembly protein; afdb-uniprot50 \| AF-A0A7M3J414-F1-MODEL\_V4 \| 1.0 \| 2.384e-09 \| 323 \| 0.555 \| 108 \| 47 \| 1 \| 1 \| 108 \| 1 \| 107 \| Tail fiber assembly protein \| Tail fiber assembly protein; afdb-uniprot50 \| AF-A0A6N9SA34-F1-MODEL\_V4 \| 1.0 \| 2.7e-09 \| 323 \| 0.451 \| 124 \| 61 \| 1 \| 3 \| 126 \| 2 \| 118 \| Tail fiber assembly protein \| Tail fiber assembly protein; afdb-uniprot50 \| AF-A0A857F028-F1-MODEL\_V4 \| 1.0 \| 3.688e-09 \| 323 \| 0.346 \| 124 \| 78 \| 2 \| 20 \| 143 \| 2 \| 122 \| Tail fiber assembly protein \| Tail fiber assembly protein; afdb-uniprot50 \| AF-A0A847KVC2-F1-MODEL\_V4 \| 1.0 \| 3.965e-07 \| 319 \| 0.447 \| 67 \| 36 \| 1 \| 1 \| 66 \| 5 \| 71 \| Uncharacterized protein \| Uncharacterized protein; afdb-uniprot50 \| AF-A0A7U4HR21-F1-MODEL\_V4 \| 1.0 \| 2.251e-08 \| 319 \| 0.248 \| 145 \| 93 \| 4 \| 3 \| 143 \| 4 \| 136 \| Uncharacterized protein \| Uncharacterized protein; afdb-uniprot50 \| AF-A0A4C6ZPY9-F1-MODEL\_V4 \| 1.0 \| 9.4e-09 \| 319 \| 0.273 \| 139 \| 94 \| 2 \| 4 \| 142 \| 19 \| 150 \| Phage tail fiber assembly protein \| Phage tail fiber assembly protein; afdb-uniprot50 \| AF-A0A807LDM0-F1-MODEL\_V4 \| 1.0 \| 6.075e-09 \| 317 \| 0.266 \| 150 \| 90 \| 6 \| 4 \| 143 \| 3 \| 142 \| Uncharacterized protein \| Uncharacterized protein; afdb-uniprot50 \| AF-A0A7D6TWW6-F1-MODEL\_V4 \| 1.0 \| 6.044e-10 \| 317 \| 0.377 \| 127 \| 76 \| 2 \| 1 \| 127 \| 1 \| 124 \| Tail fiber assembly protein \| Tail fiber assembly protein; afdb-uniprot50 \| AF-A0A7L6EL48-F1-MODEL\_V4 \| 1.0 \| 1.987e-08 \| 314 \| 0.255 \| 145 \| 94 \| 5 \| 4 \| 143 \| 3 \| 138 \| Tail fiber assembly protein \| Tail fiber assembly protein; afdb-uniprot50 \| AF-A0A7Z2EIW0-F1-MODEL\_V4 \| 1.0 \| 6.136e-07 \| 313 \| 0.512 \| 82 \| 38 \| 1 \| 64 \| 143 \| 1 \| 82 \| Tail fiber assembly protein \| Tail fiber assembly protein; afdb-uniprot50 \| AF-A0A5E8P1V0-F1-MODEL\_V4 \| 1.0 \| 1.754e-08 \| 313 \| 0.322 \| 124 \| 81 \| 2 \| 20 \| 143 \| 5 \| 125 \| Phage tail protein \| Phage tail protein; afdb-uniprot50 \| AF-P09154-F1-MODEL\_V4 \| 1.0 \| 4.178e-09 \| 313 \| 0.368 \| 141 \| 78 \| 4 \| 3 \| 142 \| 2 \| 132 \| Uncharacterized protein YmfS \| Uncharacterized protein YmfS; afdb-uniprot50 \| AF-A0A829ITR5-F1-MODEL\_V4 \| 1.0 \| 2.575e-06 \| 312 \| 0.589 \| 73 \| 30 \| 0 \| 71 \| 143 \| 16 \| 88 \| Uncharacterized protein \| Uncharacterized protein; afdb-uniprot50 \| AF-A0A7U6IRQ5-F1-MODEL\_V4 \| 1.0 \| 5.65e-11 \| 312 \| 0.393 \| 145 \| 81 \| 5 \| 3 \| 143 \| 2 \| 143 \| Tail assembly chaperone \| Tail assembly chaperone; afdb-uniprot50 \| AF-A0A7W3BQ09-F1-MODEL\_V4 \| 1.0 \| 6.531e-07 \| 309 \| 0.409 \| 66 \| 39 \| 0 \| 1 \| 66 \| 1 \| 66 \| Uncharacterized protein \| Uncharacterized protein; afdb-uniprot50 \| AF-A0A243UU47-F1-MODEL\_V4 \| 1.0 \| 1.997e-07 \| 308 \| 0.435 \| 78 \| 44 \| 0 \| 4 \| 81 \| 3 \| 80 \| Phage tail fiber assembly protein \| Phage tail fiber assembly protein; afdb-uniprot50 \| AF-A0A5B9ATE9-F1-MODEL\_V4 \| 1.0 \| 4.757e-08 \| 308 \| 0.266 \| 139 \| 95 \| 2 \| 5 \| 143 \| 2 \| 133 \| Tail fiber assembly protein \| Tail fiber assembly protein; afdb-uniprot50 \| AF-A0A1Q5VEU1-F1-MODEL\_V4 \| 1.0 \| 8.787e-10 \| 308 \| 0.267 \| 146 \| 95 \| 6 \| 1 \| 142 \| 3 \| 140 \| Uncharacterized protein \| Uncharacterized protein; afdb-uniprot50 \| AF-A0A847KQA9-F1-MODEL\_V4 \| 1.0 \| 5.089e-07 \| 307 \| 0.458 \| 72 \| 37 \| 1 \| 1 \| 70 \| 6 \| 77 \| Tail fiber assembly protein \| Tail fiber assembly protein; afdb-uniprot50 \| AF-A0A1Z1STM3-F1-MODEL\_V4 \| 1.0 \| 5.389e-08 \| 307 \| 0.256 \| 117 \| 83 \| 3 \| 28 \| 143 \| 4 \| 117 \| Uncharacterized protein \| Uncharacterized protein; afdb-uniprot50 \| AF-A0A336NXC7-F1-MODEL\_V4 \| 1.0 \| 1.278e-09 \| 306 \| 0.358 \| 145 \| 66 \| 3 \| 1 \| 143 \| 3 \| 122 \| Phage tail fiber assembly protein \| Phage tail fiber assembly protein; afdb-uniprot50 \| AF-A0A5K6DP22-F1-MODEL\_V4 \| 1.0 \| 5.707e-09 \| 306 \| 0.241 \| 141 \| 104 \| 2 \| 3 \| 143 \| 9 \| 146 \| Uncharacterized protein \| Uncharacterized protein; afdb-uniprot50 \| AF-A0A5A9CAD4-F1-MODEL\_V4 \| 1.0 \| 1.284e-08 \| 305 \| 0.415 \| 113 \| 52 \| 1 \| 45 \| 143 \| 1 \| 113 \| Tail fiber assembly protein \| Tail fiber assembly protein; afdb-uniprot50 \| AF-A0A837F9A4-F1-MODEL\_V4 \| 1.0 \| 7.796e-09 \| 305 \| 0.326 \| 141 \| 84 \| 4 \| 4 \| 143 \| 3 \| 133 \| Uncharacterized protein \| Uncharacterized protein; afdb-uniprot50 \| AF-A0A376WQ73-F1-MODEL\_V4 \| 1.0 \| 6.136e-07 \| 304 \| 0.417 \| 67 \| 39 \| 0 \| 4 \| 70 \| 3 \| 69 \| CPS-53 (KpLE1) prophage protein \| CPS-53 (KpLE1) prophage protein; afdb-uniprot50 \| AF-A0A1N6MWN0-F1-MODEL\_V4 \| 1.0 \| 6.105e-08 \| 304 \| 0.312 \| 112 \| 74 \| 2 \| 3 \| 114 \| 2 \| 110 \| Uncharacterized protein \| Uncharacterized protein; afdb-uniprot50 \| AF-A0A741M1S9-F1-MODEL\_V4 \| 1.0 \| 8.831e-09 \| 301 \| 0.253 \| 138 \| 97 \| 2 \| 4 \| 141 \| 2 \| 133 \| Tail fiber assembly protein \| Tail fiber assembly protein; afdb-uniprot50 \| AF-A0A5M7L6C3-F1-MODEL\_V4 \| 1.0 \| 1.367e-08 \| 301 \| 0.276 \| 152 \| 92 \| 7 \| 1 \| 142 \| 1 \| 144 \| Tail fiber assembly protein \| Tail fiber assembly protein; afdb-uniprot50 \| AF-A0A7W3H786-F1-MODEL\_V4 \| 1.0 \| 1.011e-06 \| 300 \| 0.393 \| 66 \| 40 \| 0 \| 1 \| 66 \| 1 \| 66 \| Uncharacterized protein \| Uncharacterized protein; afdb-uniprot50 \| AF-A0A6C6ZQ08-F1-MODEL\_V4 \| 1.0 \| 2.115e-08 \| 300 \| 0.482 \| 114 \| 57 \| 2 \| 27 \| 139 \| 1 \| 113 \| Tail fiber assembly protein \| Tail fiber assembly protein; afdb-uniprot50 \| AF-A0A1V8NRE6-F1-MODEL\_V4 \| 1.0 \| 4.157e-10 \| 300 \| 0.361 \| 163 \| 74 \| 6 \| 4 \| 143 \| 2 \| 157 \| Uncharacterized protein \| Uncharacterized protein; afdb-uniprot50 \| AF-A0A2L0TKF4-F1-MODEL\_V4 \| 1.0 \| 1.867e-08 \| 300 \| 0.244 \| 143 \| 102 \| 3 \| 1 \| 143 \| 27 \| 163 \| Phage tail protein \| Phage tail protein; afdb-uniprot50 \| AF-A0A1C7W7E1-F1-MODEL\_V4 \| 1.0 \| 6.951e-07 \| 297 \| 0.465 \| 73 \| 38 \| 1 \| 1 \| 73 \| 1 \| 72 \| Uncharacterized protein \| Uncharacterized protein; afdb-uniprot50 \| AF-W1IUZ4-F1-MODEL\_V4 \| 1.0 \| 1.373e-07 \| 295 \| 0.368 \| 103 \| 65 \| 0 \| 41 \| 143 \| 2 \| 104 \| Putative tail fiber assembly protein \| Putative tail fiber assembly protein; afdb-uniprot50 \| AF-A0A5H5WYN2-F1-MODEL\_V4 \| 1.0 \| 4.47e-08 \| 295 \| 0.49 \| 104 \| 52 \| 1 \| 37 \| 140 \| 3 \| 105 \| Tail fiber assembly protein \| Tail fiber assembly protein; afdb-uniprot50 \| AF-A0A763SVG3-F1-MODEL\_V4 \| 1.0 \| 6.564e-06 \| 294 \| 0.671 \| 67 \| 22 \| 0 \| 77 \| 143 \| 2 \| 68 \| Tail fiber assembly protein \| Tail fiber assembly protein; afdb-uniprot50 \| AF-A0A4U6GPM8-F1-MODEL\_V4 \| 1.0 \| 1.2e-09 \| 293 \| 0.316 \| 142 \| 74 \| 3 \| 1 \| 141 \| 1 \| 120 \| Tail fiber assembly protein \| Tail fiber assembly protein; afdb-uniprot50 \| AF-A0A7M2VBL5-F1-MODEL\_V4 \| 1.0 \| 8.256e-10 \| 291 \| 0.304 \| 141 \| 87 \| 4 \| 4 \| 143 \| 3 \| 133 \| Tail fiber assembly protein \| Tail fiber assembly protein; afdb-uniprot50 \| AF-A0A0M0T7T7-F1-MODEL\_V4 \| 1.0 \| 3.67e-10 \| 291 \| 0.319 \| 147 \| 88 \| 4 \| 4 \| 142 \| 7 \| 149 \| Uncharacterized protein \| Uncharacterized protein; afdb-uniprot50 \| AF-A0A402TSN0-F1-MODEL\_V4 \| 1.0 \| 2.007e-06 \| 290 \| 0.402 \| 67 \| 39 \| 1 \| 4 \| 69 \| 3 \| 69 \| Tail fiber assembly protein \| Tail fiber assembly protein; afdb-uniprot50 \| AF-A0A7I6QQY2-F1-MODEL\_V4 \| 1.0 \| 2.42e-06 \| 290 \| 0.402 \| 67 \| 40 \| 0 \| 4 \| 70 \| 3 \| 69 \| Uncharacterized protein \| Uncharacterized protein; afdb-uniprot50 \| AF-A0A7I8Z101-F1-MODEL\_V4 \| 1.0 \| 1.664e-06 \| 289 \| 0.347 \| 69 \| 44 \| 1 \| 1 \| 69 \| 1 \| 68 \| Hypothetical protein \| Hypothetical protein; afdb-uniprot50 \| AF-A0A3S6EV28-F1-MODEL\_V4 \| 1.0 \| 3.272e-08 \| 289 \| 0.27 \| 144 \| 100 \| 4 \| 1 \| 143 \| 1 \| 140 \| Uncharacterized protein \| Uncharacterized protein; afdb-uniprot50 \| AF-A0A7U6IVZ9-F1-MODEL\_V4 \| 1.0 \| 3.483e-08 \| 287 \| 0.441 \| 120 \| 58 \| 2 \| 24 \| 143 \| 14 \| 124 \| Phage tail protein \| Phage tail protein; afdb-uniprot50 \| AF-A0A6G3I5P6-F1-MODEL\_V4 \| 1.0 \| 4.492e-07 \| 283 \| 0.435 \| 78 \| 44 \| 0 \| 4 \| 81 \| 3 \| 80 \| Tail fiber assembly protein \| Tail fiber assembly protein; afdb-uniprot50 \| AF-A0A0V9JY68-F1-MODEL\_V4 \| 1.0 \| 9.447e-08 \| 282 \| 0.235 \| 140 \| 101 \| 3 \| 4 \| 142 \| 3 \| 137 \| Uncharacterized protein \| Uncharacterized protein; afdb-uniprot50 \| AF-A0A5Q2VF37-F1-MODEL\_V4 \| 1.0 \| 2.575e-06 \| 281 \| 0.402 \| 67 \| 40 \| 0 \| 4 \| 70 \| 3 \| 69 \| Uncharacterized protein \| Uncharacterized protein; afdb-uniprot50 \| AF-A0A5N9VV86-F1-MODEL\_V4 \| 1.0 \| 2.903e-07 \| 281 \| 0.414 \| 99 \| 56 \| 1 \| 4 \| 102 \| 2 \| 98 \| Tail fiber assembly protein \| Tail fiber assembly protein; afdb-uniprot50 \| AF-A0A1C0U180-F1-MODEL\_V4 \| 1.0 \| 8.381e-07 \| 280 \| 0.377 \| 98 \| 59 \| 1 \| 46 \| 143 \| 1 \| 96 \| Caudovirales tail fiber assembly protein \| Caudovirales tail fiber assembly protein; afdb-uniprot50 \| AF-A0A198GQI9-F1-MODEL\_V4 \| 1.0 \| 1.07e-07 \| 280 \| 0.244 \| 139 \| 98 \| 3 \| 5 \| 143 \| 2 \| 133 \| Tail fiber assembly protein \| Tail fiber assembly protein; afdb-uniprot50 \| AF-A0A4P9D4P7-F1-MODEL\_V4 \| 1.0 \| 3.707e-08 \| 280 \| 0.504 \| 119 \| 47 \| 4 \| 31 \| 143 \| 19 \| 131 \| Phage tail protein \| Phage tail protein; afdb-uniprot50 \| AF-A0A2N0VZQ8-F1-MODEL\_V4 \| 1.0 \| 3.726e-07 \| 279 \| 0.251 \| 139 \| 94 \| 3 \| 5 \| 143 \| 2 \| 130 \| Phage tail protein \| Phage tail protein; afdb-uniprot50 \| AF-A0A0H5HS46-F1-MODEL\_V4 \| 1.0 \| 1.54e-09 \| 279 \| 0.299 \| 147 \| 86 \| 7 \| 4 \| 143 \| 2 \| 138 \| Putative tail fiber assembly protein \| Putative tail fiber assembly protein; afdb-uniprot50 \| AF-A0A827R0P6-F1-MODEL\_V4 \| 1.0 \| 3.09e-07 \| 278 \| 0.489 \| 92 \| 46 \| 1 \| 1 \| 92 \| 1 \| 91 \| Tail fiber assembly protein \| Tail fiber assembly protein; afdb-uniprot50 \| AF-A0A5M7KZH6-F1-MODEL\_V4 \| 1.0 \| 5.736e-08 \| 276 \| 0.258 \| 147 \| 96 \| 5 \| 4 \| 143 \| 5 \| 145 \| Tail fiber assembly protein \| Tail fiber assembly protein; afdb-uniprot50 \| AF-A0A1D8UTE2-F1-MODEL\_V4 \| 1.0 \| 6.986e-06 \| 273 \| 0.38 \| 71 \| 43 \| 1 \| 1 \| 70 \| 2 \| 72 \| Uncharacterized protein \| Uncharacterized protein; afdb-uniprot50 \| AF-J2U9X8-F1-MODEL\_V4 \| 1.0 \| 8.297e-09 \| 271 \| 0.268 \| 145 \| 89 \| 6 \| 3 \| 143 \| 2 \| 133 \| Caudovirales tail fiber assembly protein \| Caudovirales tail fiber assembly protein; afdb-uniprot50 \| AF-A0A5V6PU83-F1-MODEL\_V4 \| 1.0 \| 9.954e-10 \| 270 \| 0.3 \| 143 \| 87 \| 5 \| 1 \| 142 \| 1 \| 131 \| Uncharacterized protein \| Uncharacterized protein; afdb-uniprot50 \| AF-A0A7I6ZLB1-F1-MODEL\_V4 \| 1.0 \| 8.423e-06 \| 268 \| 0.363 \| 66 \| 42 \| 0 \| 3 \| 68 \| 2 \| 67 \| Uncharacterized protein \| Uncharacterized protein; afdb-uniprot50 \| AF-A0A2S9ASE0-F1-MODEL\_V4 \| 1.0 \| 2.408e-07 \| 268 \| 0.238 \| 147 \| 93 \| 5 \| 4 \| 143 \| 39 \| 173 \| Phage tail protein \| Phage tail protein; afdb-uniprot50 \| AF-A0A6M5DB00-F1-MODEL\_V4 \| 1.0 \| 1.133e-08 \| 268 \| 0.315 \| 146 \| 72 \| 6 \| 2 \| 143 \| 77 \| 198 \| Phage tail protein \| Phage tail protein; afdb-uniprot50 \| AF-A0A1G5QUE3-F1-MODEL\_V4 \| 1.0 \| 1.218e-06 \| 267 \| 0.433 \| 83 \| 46 \| 1 \| 4 \| 85 \| 2 \| 84 \| Virus tail fibre assembly protein, lambda gpK \| Virus tail fibre assembly protein, lambda gpK; afdb-uniprot50 \| AF-E0LV13-F1-MODEL\_V4 \| 1.0 \| 3.926e-09 \| 267 \| 0.312 \| 141 \| 89 \| 5 \| 4 \| 143 \| 3 \| 136 \| Tail assembly chaperone gp38 \| Tail assembly chaperone gp38; afdb-uniprot50 \| AF-A0A7W4P415-F1-MODEL\_V4 \| 1.0 \| 2.874e-09 \| 266 \| 0.297 \| 148 \| 81 \| 8 \| 1 \| 143 \| 2 \| 131 \| Uncharacterized protein \| Uncharacterized protein; afdb-uniprot50 \| AF-A0A7V7JCF9-F1-MODEL\_V4 \| 1.0 \| 9.954e-10 \| 265 \| 0.344 \| 148 \| 80 \| 7 \| 3 \| 143 \| 2 \| 139 \| Phage tail protein \| Phage tail protein; afdb-uniprot50 \| AF-A0A7D7KF73-F1-MODEL\_V4 \| 1.0 \| 1.139e-07 \| 264 \| 0.286 \| 143 \| 83 \| 5 \| 5 \| 143 \| 2 \| 129 \| Tail fiber assembly protein \| Tail fiber assembly protein; afdb-uniprot50 \| AF-A0A2C5TSB9-F1-MODEL\_V4 \| 1.0 \| 1.477e-05 \| 263 \| 0.412 \| 80 \| 45 \| 1 \| 64 \| 143 \| 9 \| 86 \| Phage tail protein \| Phage tail protein; afdb-uniprot50 \| AF-A0A2I0FUR0-F1-MODEL\_V4 \| 1.0 \| 1.133e-08 \| 263 \| 0.255 \| 141 \| 102 \| 2 \| 3 \| 142 \| 6 \| 144 \| Phage tail protein \| Phage tail protein; afdb-uniprot50 \| AF-A0A5F1BTK6-F1-MODEL\_V4 \| 1.0 \| 2.903e-07 \| 262 \| 0.284 \| 123 \| 86 \| 2 \| 4 \| 126 \| 5 \| 125 \| Phage tail protein \| Phage tail protein; afdb-uniprot50 \| AF-A0A6B1U976-F1-MODEL\_V4 \| 1.0 \| 6.951e-07 \| 260 \| 0.242 \| 140 \| 95 \| 3 \| 4 \| 143 \| 2 \| 130 \| Tail fiber assembly protein \| Tail fiber assembly protein; afdb-uniprot50 \| AF-G3XD08-F1-MODEL\_V4 \| 1.0 \| 3.09e-07 \| 260 \| 0.276 \| 123 \| 87 \| 2 \| 4 \| 126 \| 5 \| 125 \| Uncharacterized protein \| Uncharacterized protein; afdb-uniprot50 \| AF-A0A379QMG1-F1-MODEL\_V4 \| 1.0 \| 2.115e-08 \| 259 \| 0.289 \| 128 \| 74 \| 4 \| 1 \| 123 \| 1 \| 116 \| Caudovirales tail fibre assembly protein \| Caudovirales tail fibre assembly protein; afdb-uniprot50 \| AF-A0A2T4Y153-F1-MODEL\_V4 \| 1.0 \| 1.867e-08 \| 259 \| 0.3 \| 130 \| 82 \| 4 \| 1 \| 126 \| 1 \| 125 \| Uncharacterized protein \| Uncharacterized protein; afdb-uniprot50 \| AF-A0A7X3M9Z9-F1-MODEL\_V4 \| 1.0 \| 6.465e-09 \| 259 \| 0.288 \| 142 \| 91 \| 6 \| 3 \| 143 \| 21 \| 153 \| Tail fiber assembly protein \| Tail fiber assembly protein; afdb-uniprot50 \| AF-A0A5N3CRX4-F1-MODEL\_V4 \| 1.0 \| 1.387e-05 \| 258 \| 0.414 \| 70 \| 40 \| 1 \| 1 \| 69 \| 1 \| 70 \| Phage tail protein \| Phage tail protein; afdb-uniprot50 \| AF-A0A6B1U9P0-F1-MODEL\_V4 \| 1.0 \| 5.794e-06 \| 258 \| 0.394 \| 71 \| 42 \| 1 \| 1 \| 70 \| 1 \| 71 \| Phage tail protein \| Phage tail protein; afdb-uniprot50 \| AF-A0A345CNV0-F1-MODEL\_V4 \| 1.0 \| 1.997e-07 \| 258 \| 0.221 \| 149 \| 99 \| 6 \| 4 \| 143 \| 3 \| 143 \| Phage tail protein \| Phage tail protein; afdb-uniprot50 \| AF-A0A380SAU4-F1-MODEL\_V4 \| 1.0 \| 1.225e-05 \| 256 \| 0.376 \| 69 \| 42 \| 1 \| 1 \| 69 \| 1 \| 68 \| Uncharacterized protein \| Uncharacterized protein; afdb-uniprot50 \| AF-A0A4R7NR00-F1-MODEL\_V4 \| 1.0 \| 1.772e-06 \| 255 \| 0.302 \| 86 \| 59 \| 1 \| 5 \| 89 \| 2 \| 87 \| Uncharacterized protein DUF4376 \| Uncharacterized protein DUF4376; afdb-uniprot50 \| AF-A0A6S5KC07-F1-MODEL\_V4 \| 1.0 \| 1.876e-07 \| 254 \| 0.289 \| 145 \| 87 \| 6 \| 1 \| 143 \| 1 \| 131 \| Uncharacterized protein \| Uncharacterized protein; afdb-uniprot50 \| AF-A0A2K4PA33-F1-MODEL\_V4 \| 1.0 \| 1.07e-07 \| 253 \| 0.35 \| 120 \| 75 \| 2 \| 1 \| 120 \| 4 \| 120 \| Phage tail fiber assembly protein \| Phage tail fiber assembly protein; afdb-uniprot50 \| AF-A0A3V3S8C9-F1-MODEL\_V4 \| 1.0 \| 2.741e-06 \| 253 \| 0.285 \| 119 \| 79 \| 4 \| 28 \| 143 \| 10 \| 125 \| Tail assembly chaperone \| Tail assembly chaperone; afdb-uniprot50 \| AF-A0A6H1FWY0-F1-MODEL\_V4 \| 1.0 \| 1.455e-08 \| 253 \| 0.273 \| 128 \| 85 \| 3 \| 20 \| 143 \| 6 \| 129 \| Tail fiber assembly protein \| Tail fiber assembly protein; afdb-uniprot50 \| AF-A0A8B3T6D4-F1-MODEL\_V4 \| 1.0 \| 4.199e-08 \| 252 \| 0.279 \| 118 \| 80 \| 3 \| 4 \| 120 \| 3 \| 116 \| Uncharacterized protein \| Uncharacterized protein; afdb-uniprot50 \| AF-A0A855M959-F1-MODEL\_V4 \| 1.0 \| 4.47e-08 \| 251 \| 0.27 \| 144 \| 89 \| 4 \| 4 \| 143 \| 2 \| 133 \| Uncharacterized protein \| Uncharacterized protein; afdb-uniprot50 \| AF-A0A4Q1B5G0-F1-MODEL\_V4 \| 1.0 \| 6.564e-06 \| 249 \| 0.388 \| 72 \| 43 \| 1 \| 4 \| 74 \| 9 \| 80 \| Uncharacterized protein \| Uncharacterized protein; afdb-uniprot50 \| AF-A0A1U9Q763-F1-MODEL\_V4 \| 1.0 \| 6.531e-07 \| 249 \| 0.255 \| 141 \| 96 \| 6 \| 5 \| 139 \| 2 \| 139 \| Uncharacterized protein \| Uncharacterized protein; afdb-uniprot50 \| AF-A0A5B0C0W6-F1-MODEL\_V4 \| 1.0 \| 1.011e-06 \| 249 \| 0.241 \| 124 \| 90 \| 4 \| 4 \| 126 \| 28 \| 148 \| Tail fiber assembly protein \| Tail fiber assembly protein; afdb-uniprot50 \| AF-A0A352MQ92-F1-MODEL\_V4 \| 1.0 \| 6.951e-07 \| 247 \| 0.343 \| 102 \| 63 \| 3 \| 4 \| 105 \| 3 \| 100 \| Phage tail protein \| Phage tail protein; afdb-uniprot50 \| AF-U2MIZ1-F1-MODEL\_V4 \| 1.0 \| 6.167e-06 \| 246 \| 0.318 \| 91 \| 60 \| 1 \| 53 \| 143 \| 2 \| 90 \| Tail fiber assembly like-protein \| Tail fiber assembly like-protein; afdb-uniprot50 \| AF-A0A5Y3W528-F1-MODEL\_V4 \| 1.0 \| 8.921e-07 \| 246 \| 0.369 \| 92 \| 53 \| 2 \| 4 \| 94 \| 2 \| 89 \| Phage tail protein \| Phage tail protein; afdb-uniprot50 \| AF-A0A731EUV7-F1-MODEL\_V4 \| 1.0 \| 4.781e-07 \| 246 \| 0.376 \| 109 \| 66 \| 2 \| 1 \| 108 \| 1 \| 108 \| Tail fiber assembly protein \| Tail fiber assembly protein; afdb-uniprot50 \| AF-A0A2K9QKP4-F1-MODEL\_V4 \| 1.0 \| 2.888e-08 \| 245 \| 0.262 \| 145 \| 95 \| 4 \| 4 \| 143 \| 2 \| 139 \| Uncharacterized protein \| Uncharacterized protein; afdb-uniprot50 \| AF-A0A4D0B479-F1-MODEL\_V4 \| 1.0 \| 8.381e-07 \| 244 \| 0.282 \| 117 \| 81 \| 2 \| 28 \| 143 \| 11 \| 125 \| Phage tail fiber assembly protein \| Phage tail fiber assembly protein; afdb-uniprot50 \| AF-A0A7Y9CEQ5-F1-MODEL\_V4 \| 1.0 \| 1.455e-08 \| 244 \| 0.285 \| 140 \| 97 \| 2 \| 4 \| 143 \| 3 \| 139 \| Tail fiber assembly protein \| Tail fiber assembly protein; afdb-uniprot50 \| AF-A0A5X9FP93-F1-MODEL\_V4 \| 1.0 \| 9.011e-05 \| 243 \| 0.716 \| 60 \| 17 \| 0 \| 84 \| 143 \| 2 \| 61 \| Tail fiber assembly protein \| Tail fiber assembly protein; afdb-uniprot50 \| AF-A0A0H3AZJ6-F1-MODEL\_V4 \| 1.0 \| 7.796e-09 \| 243 \| 0.295 \| 149 \| 90 \| 7 \| 4 \| 143 \| 3 \| 145 \| Uncharacterized protein \| Uncharacterized protein; afdb-uniprot50 \| AF-A0A743U4J6-F1-MODEL\_V4 \| 1.0 \| 3.289e-07 \| 242 \| 0.244 \| 127 \| 92 \| 2 \| 1 \| 127 \| 1 \| 123 \| Tail assembly chaperone \| Tail assembly chaperone; afdb-uniprot50 \| AF-A0A6N3R610-F1-MODEL\_V4 \| 1.0 \| 7.473e-05 \| 240 \| 0.701 \| 57 \| 17 \| 0 \| 86 \| 142 \| 2 \| 58 \| Uncharacterized protein \| Uncharacterized protein; afdb-uniprot50 \| AF-W1F7R7-F1-MODEL\_V4 \| 1.0 \| 1.477e-05 \| 240 \| 0.323 \| 99 \| 64 \| 2 \| 45 \| 143 \| 1 \| 96 \| Phage tail fiber assembly protein \| Phage tail fiber assembly protein; afdb-uniprot50 \| AF-A0A7T5A242-F1-MODEL\_V4 \| 1.0 \| 1.469e-06 \| 238 \| 0.29 \| 117 \| 76 \| 3 \| 28 \| 143 \| 1 \| 111 \| Tail fiber assembly protein \| Tail fiber assembly protein; afdb-uniprot50 \| AF-A0A411WM04-F1-MODEL\_V4 \| 1.0 \| 7.361e-08 \| 238 \| 0.28 \| 125 \| 84 \| 3 \| 19 \| 143 \| 3 \| 121 \| Tail fiber assembly protein \| Tail fiber assembly protein; afdb-uniprot50 \| AF-A0A2U3F338-F1-MODEL\_V4 \| 1.0 \| 1.772e-06 \| 237 \| 0.212 \| 141 \| 100 \| 4 \| 4 \| 143 \| 5 \| 135 \| Uncharacterized protein \| Uncharacterized protein; afdb-uniprot50 \| AF-A0A433ZYI6-F1-MODEL\_V4 \| 1.0 \| 0.0001394 \| 236 \| 0.44 \| 75 \| 40 \| 1 \| 69 \| 143 \| 1 \| 73 \| Uncharacterized protein \| Uncharacterized protein; afdb-uniprot50 \| AF-A0A428L7S5-F1-MODEL\_V4 \| 1.0 \| 1.145e-06 \| 236 \| 0.241 \| 149 \| 97 \| 5 \| 1 \| 143 \| 1 \| 139 \| Tail assembly chaperone \| Tail assembly chaperone; afdb-uniprot50 \| AF-A0A6S7DGX0-F1-MODEL\_V4 \| 1.0 \| 3.105e-06 \| 236 \| 0.214 \| 149 \| 101 \| 5 \| 1 \| 143 \| 60 \| 198 \| Uncharacterized protein \| Uncharacterized protein; afdb-uniprot50 \| AF-A0A7Y0ZWM6-F1-MODEL\_V4 \| 1.0 \| 5.765e-07 \| 235 \| 0.27 \| 144 \| 95 \| 5 \| 4 \| 142 \| 2 \| 140 \| Tail fiber assembly protein \| Tail fiber assembly protein; afdb-uniprot50 \| AF-A0A828KRF7-F1-MODEL\_V4 \| 1.0 \| 3.105e-06 \| 234 \| 0.466 \| 90 \| 44 \| 3 \| 4 \| 91 \| 3 \| 90 \| Tail fiber assembly protein \| Tail fiber assembly protein; afdb-uniprot50 \| AF-A0A0A8FI31-F1-MODEL\_V4 \| 1.0 \| 1.876e-07 \| 233 \| 0.28 \| 139 \| 87 \| 2 \| 5 \| 143 \| 2 \| 127 \| Tail fiber protein \| Tail fiber protein; afdb-uniprot50 \| AF-A0A793IKX6-F1-MODEL\_V4 \| 1.0 \| 5.471e-05 \| 232 \| 0.369 \| 65 \| 41 \| 0 \| 1 \| 65 \| 1 \| 65 \| Tail fiber assembly protein \| Tail fiber assembly protein; afdb-uniprot50 \| AF-A0A2H5DVM4-F1-MODEL\_V4 \| 1.0 \| 3.985e-06 \| 231 \| 0.217 \| 147 \| 100 \| 4 \| 1 \| 143 \| 3 \| 138 \| Uncharacterized protein \| Uncharacterized protein; afdb-uniprot50 \| AF-A0A7Z2J1J3-F1-MODEL\_V4 \| 1.0 \| 1.139e-07 \| 231 \| 0.275 \| 145 \| 98 \| 4 \| 4 \| 143 \| 3 \| 145 \| Tail fiber assembly protein \| Tail fiber assembly protein; afdb-uniprot50 \| AF-A0A0F1B154-F1-MODEL\_V4 \| 1.0 \| 5.794e-06 \| 230 \| 0.214 \| 140 \| 97 \| 3 \| 4 \| 143 \| 2 \| 128 \| Uncharacterized protein \| Uncharacterized protein; afdb-uniprot50 \| AF-A0A1G7SVK8-F1-MODEL\_V4 \| 1.0 \| 2.125e-07 \| 229 \| 0.235 \| 157 \| 86 \| 5 \| 4 \| 126 \| 3 \| 159 \| Uncharacterized protein \| Uncharacterized protein; afdb-uniprot50 \| AF-A0A763CKN9-F1-MODEL\_V4 \| 1.0 \| 1.656e-07 \| 229 \| 0.181 \| 193 \| 100 \| 5 \| 4 \| 141 \| 2 \| 191 \| DUF4376 domain-containing protein \| DUF4376 domain-containing protein; afdb-uniprot50 \| AF-A0A6M5HRU9-F1-MODEL\_V4 \| 1.0 \| 0.0001394 \| 228 \| 0.375 \| 64 \| 39 \| 1 \| 3 \| 66 \| 2 \| 64 \| Uncharacterized protein \| Uncharacterized protein; afdb-uniprot50 \| AF-A0A0M7GTK9-F1-MODEL\_V4 \| 1.0 \| 1.469e-06 \| 228 \| 0.243 \| 152 \| 92 \| 6 \| 1 \| 142 \| 1 \| 139 \| Bacteriophage tail assembly protein \| Bacteriophage tail assembly protein; afdb-uniprot50 \| AF-A0A2N7P764-F1-MODEL\_V4 \| 1.0 \| 8.381e-07 \| 228 \| 0.231 \| 147 \| 98 \| 5 \| 5 \| 143 \| 2 \| 141 \| Uncharacterized protein \| Uncharacterized protein; afdb-uniprot50 \| AF-A0A0N0XL62-F1-MODEL\_V4 \| 1.0 \| 6.564e-06 \| 227 \| 0.213 \| 145 \| 100 \| 5 \| 1 \| 142 \| 1 \| 134 \| Caudovirales tail fiber assembly protein \| Caudovirales tail fiber assembly protein; afdb-uniprot50 \| AF-A0A1H0BDJ1-F1-MODEL\_V4 \| 1.0 \| 9.543e-06 \| 226 \| 0.182 \| 148 \| 106 \| 6 \| 1 \| 143 \| 1 \| 138 \| Virus tail fibre assembly protein, lambda gpK \| Virus tail fibre assembly protein, lambda gpK; afdb-uniprot50 \| AF-A0A5E8P460-F1-MODEL\_V4 \| 1.0 \| 7.914e-06 \| 225 \| 0.333 \| 90 \| 60 \| 0 \| 1 \| 90 \| 1 \| 90 \| Phage tail protein \| Phage tail protein; afdb-uniprot50 \| AF-A0A844U1B6-F1-MODEL\_V4 \| 1.0 \| 3.744e-06 \| 225 \| 0.231 \| 147 \| 101 \| 4 \| 1 \| 143 \| 1 \| 139 \| Phage tail protein \| Phage tail protein; afdb-uniprot50 \| AF-E3G2V8-F1-MODEL\_V4 \| 1.0 \| 1.772e-06 \| 224 \| 0.25 \| 148 \| 86 \| 8 \| 1 \| 143 \| 1 \| 128 \| Tail assembly chaperone gp38 \| Tail assembly chaperone gp38; afdb-uniprot50 \| AF-A0A077Q6L9-F1-MODEL\_V4 \| 1.0 \| 3.985e-06 \| 223 \| 0.371 \| 97 \| 56 \| 2 \| 50 \| 143 \| 2 \| 96 \| Putative tail fiber assembly protein \| Putative tail fiber assembly protein; afdb-uniprot50 \| AF-A0A077P958-F1-MODEL\_V4 \| 1.0 \| 0.0001681 \| 222 \| 0.333 \| 78 \| 50 \| 1 \| 66 \| 143 \| 2 \| 77 \| Putative tail fiber chaperone Rac prophage \| Putative tail fiber chaperone Rac prophage; afdb-uniprot50 \| AF-A0A8B2Y1X9-F1-MODEL\_V4 \| 1.0 \| 5.089e-07 \| 222 \| 0.246 \| 146 \| 101 \| 6 \| 1 \| 143 \| 1 \| 140 \| Phage tail protein \| Phage tail protein; afdb-uniprot50 \| AF-A0A643FMC7-F1-MODEL\_V4 \| 1.0 \| 7.874e-07 \| 221 \| 0.258 \| 143 \| 91 \| 4 \| 3 \| 143 \| 2 \| 131 \| Phage tail assembly chaperone \| Phage tail assembly chaperone; afdb-uniprot50 \| AF-W1EMA3-F1-MODEL\_V4 \| 1.0 \| 7.914e-06 \| 221 \| 0.229 \| 144 \| 93 \| 7 \| 4 \| 143 \| 3 \| 132 \| Uncharacterized protein \| Uncharacterized protein; afdb-uniprot50 \| AF-A0A2N8KM43-F1-MODEL\_V4 \| 1.0 \| 2.918e-06 \| 221 \| 0.256 \| 144 \| 88 \| 6 \| 4 \| 143 \| 3 \| 131 \| Tail assembly chaperone \| Tail assembly chaperone; afdb-uniprot50 \| AF-A0A377LDC8-F1-MODEL\_V4 \| 1.0 \| 1.218e-06 \| 220 \| 0.541 \| 85 \| 39 \| 0 \| 20 \| 104 \| 3 \| 87 \| CPS-53 (KpLE1) prophage protein \| CPS-53 (KpLE1) prophage protein; afdb-uniprot50 \| AF-A0A1I7J4J1-F1-MODEL\_V4 \| 1.0 \| 3.483e-08 \| 220 \| 0.238 \| 176 \| 80 \| 7 \| 5 \| 126 \| 2 \| 177 \| DUF4376 domain-containing protein \| DUF4376 domain-containing protein; afdb-uniprot50 \| AF-A0A2Z6EW35-F1-MODEL\_V4 \| 1.0 \| 2.741e-06 \| 218 \| 0.239 \| 146 \| 94 \| 6 \| 4 \| 143 \| 3 \| 137 \| Uncharacterized protein \| Uncharacterized protein; afdb-uniprot50 \| AF-A0A1H9W380-F1-MODEL\_V4 \| 1.0 \| 5.736e-08 \| 218 \| 0.238 \| 193 \| 87 \| 7 \| 4 \| 141 \| 3 \| 190 \| DUF4376 domain-containing protein \| DUF4376 domain-containing protein; afdb-uniprot50 \| AF-A0A2W5F2S4-F1-MODEL\_V4 \| 1.0 \| 5.794e-06 \| 217 \| 0.246 \| 142 \| 94 \| 5 \| 5 \| 142 \| 2 \| 134 \| Phage tail protein \| Phage tail protein; afdb-uniprot50 \| AF-A0A076LFJ0-F1-MODEL\_V4 \| 1.0 \| 0.0004285 \| 216 \| 0.373 \| 67 \| 40 \| 1 \| 77 \| 143 \| 2 \| 66 \| Phage tail fiber assembly protein \| Phage tail fiber assembly protein; afdb-uniprot50 \| AF-H3RFA6-F1-MODEL\_V4 \| 1.0 \| 0.0005853 \| 216 \| 0.342 \| 73 \| 46 \| 1 \| 71 \| 143 \| 8 \| 78 \| Uncharacterized protein \| Uncharacterized protein; afdb-uniprot50 \| AF-A0A849JQA2-F1-MODEL\_V4 \| 1.0 \| 1.076e-06 \| 216 \| 0.273 \| 146 \| 78 \| 7 \| 4 \| 143 \| 3 \| 126 \| Tail assembly chaperone \| Tail assembly chaperone; afdb-uniprot50 \| AF-A0A0D8L6B2-F1-MODEL\_V4 \| 1.0 \| 1.297e-06 \| 216 \| 0.218 \| 165 \| 95 \| 8 \| 3 \| 143 \| 21 \| 175 \| Uncharacterized protein \| Uncharacterized protein; afdb-uniprot50 \| AF-A0A447UEL9-F1-MODEL\_V4 \| 1.0 \| 6.167e-06 \| 215 \| 0.192 \| 140 \| 102 \| 3 \| 4 \| 143 \| 2 \| 130 \| Phage tail fiber assembly protein \| Phage tail fiber assembly protein; afdb-uniprot50 \| AF-A0A2I8DIP5-F1-MODEL\_V4 \| 1.0 \| 6.564e-06 \| 215 \| 0.244 \| 147 \| 95 \| 6 \| 4 \| 143 \| 24 \| 161 \| Phage tail protein \| Phage tail protein; afdb-uniprot50 \| AF-X1XFW3-F1-MODEL\_V4 \| 1.0 \| 2.136e-06 \| 215 \| 0.248 \| 141 \| 85 \| 5 \| 17 \| 142 \| 577 \| 711 \| DNA\_circ\_N domain-containing protein \| DNA\_circ\_N domain-containing protein; afdb-uniprot50 \| AF-A0A7X2R448-F1-MODEL\_V4 \| 1.0 \| 0.0002027 \| 213 \| 0.376 \| 77 \| 46 \| 1 \| 66 \| 142 \| 11 \| 85 \| Tail fiber assembly protein \| Tail fiber assembly protein; afdb-uniprot50 \| AF-A0A6G2GGE2-F1-MODEL\_V4 \| 1.0 \| 2.932e-05 \| 212 \| 0.317 \| 85 \| 57 \| 1 \| 4 \| 88 \| 2 \| 85 \| Tail fiber assembly protein \| Tail fiber assembly protein; afdb-uniprot50 \| AF-A0A4D8QSV8-F1-MODEL\_V4 \| 1.0 \| 1.016e-05 \| 210 \| 0.215 \| 144 \| 95 \| 5 \| 4 \| 142 \| 3 \| 133 \| Phage tail protein \| Phage tail protein; afdb-uniprot50 \| AF-A0A0D0PTQ5-F1-MODEL\_V4 \| 1.0 \| 2.285e-05 \| 210 \| 0.204 \| 142 \| 100 \| 4 \| 4 \| 143 \| 3 \| 133 \| Contig\_41, whole genome shotgun sequence \| Contig\_41, whole genome shotgun sequence; afdb-uniprot50 \| AF-A0A1J0EK84-F1-MODEL\_V4 \| 1.0 \| 8.381e-07 \| 210 \| 0.251 \| 147 \| 96 \| 4 \| 3 \| 143 \| 2 \| 140 \| Uncharacterized protein \| Uncharacterized protein; afdb-uniprot50 \| AF-A0A0M7JWH2-F1-MODEL\_V4 \| 1.0 \| 4.515e-06 \| 208 \| 0.282 \| 131 \| 83 \| 5 \| 1 \| 126 \| 1 \| 125 \| Caudovirales tail fibre assembly protein \| Caudovirales tail fibre assembly protein; afdb-uniprot50 \| AF-A0A0H2VSV7-F1-MODEL\_V4 \| 1.0 \| 7.914e-06 \| 208 \| 0.233 \| 124 \| 88 \| 2 \| 4 \| 127 \| 35 \| 151 \| Putative phage tail fibre protein \| Putative phage tail fibre protein; afdb-uniprot50 \| AF-A0A6H1S397-F1-MODEL\_V4 \| 1.0 \| 1.212e-07 \| 208 \| 0.214 \| 168 \| 93 \| 7 \| 1 \| 133 \| 84 \| 247 \| Uncharacterized protein \| Uncharacterized protein; afdb-uniprot50 \| AF-A0A7H8I663-F1-MODEL\_V4 \| 1.0 \| 0.0004854 \| 206 \| 0.3 \| 80 \| 52 \| 2 \| 66 \| 143 \| 1 \| 78 \| Tail fiber assembly protein \| Tail fiber assembly protein; afdb-uniprot50 \| AF-A0A775TK41-F1-MODEL\_V4 \| 1.0 \| 0.0003553 \| 205 \| 0.345 \| 81 \| 50 \| 2 \| 64 \| 143 \| 9 \| 87 \| Phage tail protein \| Phage tail protein; afdb-uniprot50 \| AF-A0A1I4ZNN3-F1-MODEL\_V4 \| 1.0 \| 3.744e-06 \| 205 \| 0.238 \| 151 \| 94 \| 6 \| 4 \| 143 \| 2 \| 142 \| Uncharacterized protein \| Uncharacterized protein; afdb-uniprot50 \| AF-A0A854BW95-F1-MODEL\_V4 \| 1.0 \| 1.297e-06 \| 204 \| 0.237 \| 143 \| 96 \| 5 \| 4 \| 142 \| 2 \| 135 \| Phage tail protein \| Phage tail protein; afdb-uniprot50 \| AF-A0A0T9RNX7-F1-MODEL\_V4 \| 1.0 \| 2.262e-07 \| 204 \| 0.258 \| 151 \| 95 \| 6 \| 1 \| 143 \| 2 \| 143 \| Putative tail fiber assembly protein \| Putative tail fiber assembly protein; afdb-uniprot50 \| AF-A0A6M5CYI8-F1-MODEL\_V4 \| 1.0 \| 0.001588 \| 203 \| 0.342 \| 70 \| 46 \| 0 \| 74 \| 143 \| 2 \| 71 \| Uncharacterized protein \| Uncharacterized protein; afdb-uniprot50 \| AF-A0A2C5TPI7-F1-MODEL\_V4 \| 1.0 \| 8.423e-06 \| 203 \| 0.26 \| 138 \| 77 \| 3 \| 4 \| 141 \| 2 \| 114 \| Phage tail protein \| Phage tail protein; afdb-uniprot50 \| AF-W1J6H1-F1-MODEL\_V4 \| 1.0 \| 5.444e-06 \| 203 \| 0.223 \| 143 \| 80 \| 4 \| 4 \| 143 \| 2 \| 116 \| Putaive tail fiber assembly protein \| Putaive tail fiber assembly protein; afdb-uniprot50 \| AF-A0A1S8CP93-F1-MODEL\_V4 \| 1.0 \| 1.772e-06 \| 203 \| 0.205 \| 141 \| 102 \| 5 \| 4 \| 143 \| 3 \| 134 \| Uncharacterized protein \| Uncharacterized protein; afdb-uniprot50 \| AF-G5QY84-F1-MODEL\_V4 \| 1.0 \| 0.0005499 \| 202 \| 0.343 \| 64 \| 42 \| 0 \| 4 \| 67 \| 2 \| 65 \| Uncharacterized protein \| Uncharacterized protein; afdb-uniprot50 \| AF-C5AKX8-F1-MODEL\_V4 \| 1.0 \| 2.432e-05 \| 202 \| 0.201 \| 144 \| 99 \| 5 \| 4 \| 143 \| 6 \| 137 \| Uncharacterized protein \| Uncharacterized protein; afdb-uniprot50 \| AF-A0A1B8HT06-F1-MODEL\_V4 \| 1.0 \| 2.273e-06 \| 202 \| 0.275 \| 145 \| 75 \| 5 \| 2 \| 142 \| 61 \| 179 \| Uncharacterized protein \| Uncharacterized protein; afdb-uniprot50 \| AF-A0A855M0Y7-F1-MODEL\_V4 \| 1.0 \| 4.242e-06 \| 202 \| 0.236 \| 148 \| 79 \| 7 \| 2 \| 143 \| 77 \| 196 \| Tail assembly chaperone \| Tail assembly chaperone; afdb-uniprot50 \| AF-A0A0M0T628-F1-MODEL\_V4 \| 1.0 \| 3.5e-07 \| 201 \| 0.279 \| 129 \| 87 \| 3 \| 1 \| 126 \| 1 \| 126 \| Uncharacterized protein \| Uncharacterized protein; afdb-uniprot50 \| AF-R8WJ06-F1-MODEL\_V4 \| 1.0 \| 1.664e-06 \| 201 \| 0.263 \| 144 \| 89 \| 4 \| 3 \| 142 \| 2 \| 132 \| Uncharacterized protein \| Uncharacterized protein; afdb-uniprot50 \| AF-A0A379QQG2-F1-MODEL\_V4 \| 1.0 \| 2.563e-07 \| 201 \| 0.223 \| 161 \| 89 \| 4 \| 1 \| 127 \| 1 \| 159 \| Phage tail fiber assembly \| Phage tail fiber assembly; afdb-uniprot50 \| AF-A0A826XDH3-F1-MODEL\_V4 \| 1.0 \| 0.000158 \| 200 \| 0.276 \| 94 \| 61 \| 4 \| 52 \| 143 \| 45 \| 133 \| Tail fiber assembly protein \| Tail fiber assembly protein; afdb-uniprot50 \| AF-A0A1B8HTT9-F1-MODEL\_V4 \| 1.0 \| 5.444e-06 \| 199 \| 0.277 \| 148 \| 66 \| 5 \| 4 \| 143 \| 2 \| 116 \| Uncharacterized protein \| Uncharacterized protein; afdb-uniprot50 \| AF-A0A432UNX0-F1-MODEL\_V4 \| 1.0 \| 2.563e-07 \| 199 \| 0.218 \| 192 \| 95 \| 8 \| 4 \| 140 \| 3 \| 194 \| Uncharacterized protein \| Uncharacterized protein; afdb-uniprot50 \| AF-A0A8A5H0Q5-F1-MODEL\_V4 \| 1.0 \| 1.673e-05 \| 198 \| 0.264 \| 117 \| 73 \| 4 \| 28 \| 143 \| 24 \| 128 \| Tail fiber assembly protein \| Tail fiber assembly protein; afdb-uniprot50 \| AF-A0A1A9WSI7-F1-MODEL\_V4 \| 1.0 \| 0.0001394 \| 198 \| 0.282 \| 85 \| 60 \| 1 \| 2 \| 85 \| 506 \| 590 \| Uncharacterized protein \| Uncharacterized protein; afdb-uniprot50 \| AF-A0A763SYU9-F1-MODEL\_V4 \| 1.0 \| 0.0004026 \| 197 \| 0.369 \| 65 \| 40 \| 1 \| 4 \| 68 \| 3 \| 66 \| Tail assembly chaperone \| Tail assembly chaperone; afdb-uniprot50 \| AF-A0A177SRB6-F1-MODEL\_V4 \| 1.0 \| 9.543e-06 \| 197 \| 0.229 \| 144 \| 97 \| 6 \| 3 \| 142 \| 2 \| 135 \| Phage tail protein \| Phage tail protein; afdb-uniprot50 \| AF-E0LW01-F1-MODEL\_V4 \| 1.0 \| 9.543e-06 \| 196 \| 0.183 \| 147 \| 88 \| 7 \| 2 \| 143 \| 12 \| 131 \| Tail assembly chaperone gp38 \| Tail assembly chaperone gp38; afdb-uniprot50 \| AF-A0A7W7ING5-F1-MODEL\_V4 \| 1.0 \| 1.673e-05 \| 196 \| 0.236 \| 152 \| 85 \| 8 \| 4 \| 142 \| 3 \| 136 \| Uncharacterized protein \| Uncharacterized protein; afdb-uniprot50 \| AF-A0A1Q8EUF2-F1-MODEL\_V4 \| 1.0 \| 4.537e-05 \| 195 \| 0.227 \| 123 \| 87 \| 2 \| 4 \| 126 \| 3 \| 117 \| Uncharacterized protein \| Uncharacterized protein; afdb-uniprot50 \| AF-D8MKJ6-F1-MODEL\_V4 \| 1.0 \| 6.986e-06 \| 195 \| 0.205 \| 146 \| 86 \| 6 \| 2 \| 143 \| 77 \| 196 \| Phage tail assembly chaperone gp38 \| Phage tail assembly chaperone gp38; afdb-uniprot50 \| AF-A0A1J7C8B7-F1-MODEL\_V4 \| 1.0 \| 4.242e-06 \| 195 \| 0.219 \| 146 \| 82 \| 6 \| 2 \| 142 \| 73 \| 191 \| Phage tail protein \| Phage tail protein; afdb-uniprot50 \| AF-A0A6L9LZN9-F1-MODEL\_V4 \| 1.0 \| 1.78e-05 \| 193 \| 0.248 \| 145 \| 90 \| 7 \| 4 \| 142 \| 3 \| 134 \| Phage tail protein \| Phage tail protein; afdb-uniprot50 \| AF-A0A2V1Y818-F1-MODEL\_V4 \| 1.0 \| 5.14e-05 \| 193 \| 0.171 \| 146 \| 102 \| 5 \| 4 \| 143 \| 2 \| 134 \| Virus tail fiber assembly protein lambda gpK \| Virus tail fiber assembly protein lambda gpK; afdb-uniprot50 \| AF-A0A1G9YD25-F1-MODEL\_V4 \| 1.0 \| 0.0001086 \| 191 \| 0.242 \| 140 \| 93 \| 3 \| 4 \| 143 \| 3 \| 129 \| Virus tail fibre assembly protein, lambda gpK \| Virus tail fibre assembly protein, lambda gpK; afdb-uniprot50 \| AF-A0A4Q9ESD9-F1-MODEL\_V4 \| 1.0 \| 2.42e-06 \| 191 \| 0.231 \| 164 \| 87 \| 6 \| 4 \| 142 \| 2 \| 151 \| Tail fiber assembly protein \| Tail fiber assembly protein; afdb-uniprot50 \| AF-A0A2U3F166-F1-MODEL\_V4 \| 1.0 \| 3.518e-06 \| 191 \| 0.226 \| 146 \| 91 \| 8 \| 4 \| 142 \| 5 \| 135 \| Phage tail protein \| Phage tail protein; afdb-uniprot50 \| AF-A0A209A5U3-F1-MODEL\_V4 \| 1.0 \| 2.147e-05 \| 190 \| 0.232 \| 146 \| 91 \| 7 \| 1 \| 142 \| 1 \| 129 \| Uncharacterized protein \| Uncharacterized protein; afdb-uniprot50 \| AF-A0A379FUE8-F1-MODEL\_V4 \| 1.0 \| 0.001026 \| 189 \| 0.383 \| 73 \| 43 \| 1 \| 70 \| 142 \| 4 \| 74 \| Bacteriophage tail assembly protein \| Bacteriophage tail assembly protein; afdb-uniprot50 \| AF-A0A3D1C7H0-F1-MODEL\_V4 \| 1.0 \| 0.0004026 \| 189 \| 0.28 \| 89 \| 62 \| 1 \| 55 \| 143 \| 4 \| 90 \| Phage tail protein \| Phage tail protein; afdb-uniprot50 \| AF-A0A2J9HPR0-F1-MODEL\_V4 \| 1.0 \| 0.0001156 \| 189 \| 0.226 \| 115 \| 78 \| 4 \| 33 \| 143 \| 4 \| 111 \| Tail assembly chaperone \| Tail assembly chaperone; afdb-uniprot50 \| AF-A0A7W8U3D6-F1-MODEL\_V4 \| 1.0 \| 3.518e-06 \| 189 \| 0.254 \| 157 \| 77 \| 7 \| 4 \| 126 \| 2 \| 152 \| Peptidoglycan/xylan/chitin deacetylase (PgdA/CDA1 family) \| Peptidoglycan/xylan/chitin deacetylase (PgdA/CDA1 family); afdb-uniprot50 \| AF-A0A3L1P348-F1-MODEL\_V4 \| 1.0 \| 0.000456 \| 188 \| 0.344 \| 90 \| 57 \| 1 \| 54 \| 143 \| 1 \| 88 \| Tail fiber assembly protein \| Tail fiber assembly protein; afdb-uniprot50 \| AF-A0A411SXP7-F1-MODEL\_V4 \| 1.0 \| 1.477e-05 \| 188 \| 0.208 \| 144 \| 96 \| 6 \| 2 \| 143 \| 6 \| 133 \| Uncharacterized protein \| Uncharacterized protein; afdb-uniprot50 \| AF-A0A7H9BG12-F1-MODEL\_V4 \| 1.0 \| 2.432e-05 \| 188 \| 0.221 \| 131 \| 86 \| 2 \| 5 \| 126 \| 2 \| 125 \| Uncharacterized protein \| Uncharacterized protein; afdb-uniprot50 \| AF-A0A7X5TMC5-F1-MODEL\_V4 \| 1.0 \| 0.0002158 \| 187 \| 0.282 \| 99 \| 69 \| 1 \| 45 \| 143 \| 8 \| 104 \| Phage tail protein \| Phage tail protein; afdb-uniprot50 \| AF-D0ZBI3-F1-MODEL\_V4 \| 1.0 \| 1.572e-05 \| 187 \| 0.204 \| 147 \| 85 \| 7 \| 2 \| 143 \| 73 \| 192 \| Phage tail assembly chaperone gp38 \| Phage tail assembly chaperone gp38; afdb-uniprot50 \| AF-W1ITK8-F1-MODEL\_V4 \| 1.0 \| 0.001588 \| 186 \| 0.315 \| 76 \| 50 \| 1 \| 68 \| 143 \| 13 \| 86 \| Putaive tail fiber assembly protein \| Putaive tail fiber assembly protein; afdb-uniprot50 \| AF-A0A0J7J7Q7-F1-MODEL\_V4 \| 1.0 \| 7.398e-07 \| 186 \| 0.217 \| 156 \| 88 \| 4 \| 5 \| 126 \| 2 \| 157 \| Uncharacterized protein \| Uncharacterized protein; afdb-uniprot50 \| AF-A0A618PKJ7-F1-MODEL\_V4 \| 1.0 \| 3.322e-05 \| 185 \| 0.238 \| 105 \| 80 \| 0 \| 1 \| 105 \| 2 \| 106 \| Tail fiber assembly protein \| Tail fiber assembly protein; afdb-uniprot50 \| AF-A0A376DH18-F1-MODEL\_V4 \| 1.0 \| 0.0001484 \| 185 \| 0.278 \| 104 \| 66 \| 4 \| 42 \| 143 \| 21 \| 117 \| Caudovirales tail fibre assembly protein \| Caudovirales tail fibre assembly protein; afdb-uniprot50 \| AF-A0A0U1HLA4-F1-MODEL\_V4 \| 1.0 \| 0.001914 \| 184 \| 0.276 \| 76 \| 53 \| 1 \| 68 \| 143 \| 6 \| 79 \| Tail assembly chaperone gp38 \| Tail assembly chaperone gp38; afdb-uniprot50 \| AF-A0A3A3ZKX8-F1-MODEL\_V4 \| 1.0 \| 0.0002444 \| 184 \| 0.32 \| 78 \| 53 \| 0 \| 4 \| 81 \| 3 \| 80 \| Uncharacterized protein \| Uncharacterized protein; afdb-uniprot50 \| AF-A0A838YFR4-F1-MODEL\_V4 \| 1.0 \| 9.591e-05 \| 184 \| 0.229 \| 144 \| 91 \| 6 \| 3 \| 143 \| 2 \| 128 \| Uncharacterized protein \| Uncharacterized protein; afdb-uniprot50 \| AF-A0A1M3PGE6-F1-MODEL\_V4 \| 1.0 \| 2.147e-05 \| 184 \| 0.248 \| 149 \| 86 \| 6 \| 1 \| 143 \| 1 \| 129 \| Uncharacterized protein \| Uncharacterized protein; afdb-uniprot50 \| AF-A0A5A9DGY0-F1-MODEL\_V4 \| 1.0 \| 0.0001394 \| 183 \| 0.352 \| 102 \| 59 \| 6 \| 45 \| 143 \| 52 \| 149 \| Tail fiber assembly protein \| Tail fiber assembly protein; afdb-uniprot50 \| AF-A0A420WVJ7-F1-MODEL\_V4 \| 1.0 \| 7.874e-07 \| 183 \| 0.229 \| 161 \| 85 \| 6 \| 5 \| 126 \| 2 \| 162 \| Uncharacterized protein \| Uncharacterized protein; afdb-uniprot50 \| AF-A0A7I6QR37-F1-MODEL\_V4 \| 1.0 \| 0.0004026 \| 182 \| 0.232 \| 99 \| 68 \| 4 \| 46 \| 143 \| 16 \| 107 \| Uncharacterized protein \| Uncharacterized protein; afdb-uniprot50 \| AF-A0A5Y9TWT7-F1-MODEL\_V4 \| 1.0 \| 4.829e-05 \| 182 \| 0.231 \| 147 \| 83 \| 6 \| 2 \| 143 \| 79 \| 200 \| Tail fiber assembly protein \| Tail fiber assembly protein; afdb-uniprot50 \| AF-A0A855SIT2-F1-MODEL\_V4 \| 1.0 \| 0.001492 \| 181 \| 0.392 \| 56 \| 34 \| 0 \| 1 \| 56 \| 1 \| 56 \| Tail fiber assembly protein \| Tail fiber assembly protein; afdb-uniprot50 \| AF-A0A080I832-F1-MODEL\_V4 \| 1.0 \| 0.0001789 \| 181 \| 0.295 \| 88 \| 59 \| 2 \| 1 \| 88 \| 1 \| 85 \| Tail assembly chaperone gp38 domain protein \| Tail assembly chaperone gp38 domain protein; afdb-uniprot50 \| AF-A0A8A5IKF3-F1-MODEL\_V4 \| 1.0 \| 0.0002947 \| 181 \| 0.245 \| 102 \| 69 \| 4 \| 45 \| 143 \| 6 \| 102 \| Tail fiber assembly protein \| Tail fiber assembly protein; afdb-uniprot50 \| AF-A0A0T9KM84-F1-MODEL\_V4 \| 1.0 \| 0.0007057 \| 179 \| 0.329 \| 82 \| 52 \| 2 \| 62 \| 143 \| 11 \| 89 \| Putative phage tail fiber assembly protein \| Putative phage tail fiber assembly protein; afdb-uniprot50 \| AF-J2SUJ2-F1-MODEL\_V4 \| 1.0 \| 2.147e-05 \| 179 \| 0.21 \| 147 \| 96 \| 6 \| 3 \| 142 \| 2 \| 135 \| Caudovirales tail fiber assembly protein \| Caudovirales tail fiber assembly protein; afdb-uniprot50 \| AF-A0A0H3I5Y1-F1-MODEL\_V4 \| 1.0 \| 5.471e-05 \| 179 \| 0.251 \| 147 \| 86 \| 8 \| 1 \| 143 \| 1 \| 127 \| Tail fiber chaperone, Qin prophage \| Tail fiber chaperone, Qin prophage; afdb-uniprot50 \| AF-A0A2W7L5R6-F1-MODEL\_V4 \| 1.0 \| 0.0001789 \| 179 \| 0.233 \| 154 \| 99 \| 7 \| 4 \| 142 \| 3 \| 152 \| Phage tail assembly chaperone \| Phage tail assembly chaperone; afdb-uniprot50 \| AF-A0A4U2U962-F1-MODEL\_V4 \| 1.0 \| 2.588e-05 \| 179 \| 0.183 \| 147 \| 88 \| 7 \| 2 \| 143 \| 60 \| 179 \| Tail fiber assembly protein \| Tail fiber assembly protein; afdb-uniprot50 \| AF-A0A2R8CKR9-F1-MODEL\_V4 \| 1.0 \| 6.136e-07 \| 179 \| 0.302 \| 139 \| 80 \| 5 \| 3 \| 140 \| 2 \| 124 \| Uncharacterized protein \| Uncharacterized protein; afdb-uniprot50 \| AF-A0A2K9MAQ1-F1-MODEL\_V4 \| 1.0 \| 0.000158 \| 179 \| 0.182 \| 148 \| 102 \| 5 \| 2 \| 143 \| 72 \| 206 \| Uncharacterized protein \| Uncharacterized protein; afdb-uniprot50 \| AF-A0A7Y9WJF3-F1-MODEL\_V4 \| 1.0 \| 4.537e-05 \| 178 \| 0.226 \| 146 \| 93 \| 8 \| 5 \| 143 \| 2 \| 134 \| Uncharacterized protein \| Uncharacterized protein; afdb-uniprot50 \| AF-U3TVP9-F1-MODEL\_V4 \| 1.0 \| 0.0007057 \| 178 \| 0.242 \| 99 \| 67 \| 4 \| 45 \| 143 \| 86 \| 176 \| Uncharacterized protein \| Uncharacterized protein; afdb-uniprot50 \| AF-A0A4P7L7D3-F1-MODEL\_V4 \| 1.0 \| 2.432e-05 \| 178 \| 0.231 \| 147 \| 81 \| 7 \| 2 \| 143 \| 87 \| 206 \| Uncharacterized protein \| Uncharacterized protein; afdb-uniprot50 \| AF-D2U2G7-F1-MODEL\_V4 \| 1.0 \| 2.932e-05 \| 177 \| 0.238 \| 147 \| 80 \| 7 \| 2 \| 143 \| 58 \| 177 \| Phage tail fiber assembly protein \| Phage tail fiber assembly protein; afdb-uniprot50 \| AF-A0A5C4RHG7-F1-MODEL\_V4 \| 1.0 \| 2.588e-05 \| 177 \| 0.226 \| 146 \| 83 \| 7 \| 2 \| 143 \| 75 \| 194 \| Tail fiber assembly protein \| Tail fiber assembly protein; afdb-uniprot50 \| AF-A0A484QG78-F1-MODEL\_V4 \| 1.0 \| 0.000131 \| 176 \| 0.195 \| 148 \| 99 \| 7 \| 4 \| 143 \| 3 \| 138 \| Tail fiber assembly protein \| Tail fiber assembly protein; afdb-uniprot50 \| AF-A0A7X5QHH8-F1-MODEL\_V4 \| 1.0 \| 3.322e-05 \| 176 \| 0.226 \| 146 \| 83 \| 6 \| 2 \| 143 \| 753 \| 872 \| Phage-tail\_3 domain-containing protein \| Phage-tail\_3 domain-containing protein; afdb-uniprot50 \| AF-R9VMU8-F1-MODEL\_V4 \| 1.0 \| 0.0005853 \| 175 \| 0.273 \| 95 \| 63 \| 2 \| 53 \| 143 \| 50 \| 142 \| Uncharacterized protein \| Uncharacterized protein; afdb-uniprot50 \| AF-A0A5C7CFK8-F1-MODEL\_V4 \| 1.0 \| 0.002962 \| 175 \| 0.278 \| 79 \| 54 \| 2 \| 66 \| 143 \| 99 \| 175 \| Tail fiber assembly protein \| Tail fiber assembly protein; afdb-uniprot50 \| AF-A0A7H8I670-F1-MODEL\_V4 \| 1.0 \| 6.198e-05 \| 175 \| 0.191 \| 146 \| 88 \| 7 \| 2 \| 143 \| 79 \| 198 \| Tail fiber assembly protein \| Tail fiber assembly protein; afdb-uniprot50 \| AF-A0A1G5QW41-F1-MODEL\_V4 \| 1.0 \| 2.932e-05 \| 175 \| 0.238 \| 147 \| 80 \| 6 \| 2 \| 143 \| 410 \| 529 \| Phage Tail Collar Domain \| Phage Tail Collar Domain; afdb-uniprot50 \| AF-A0A833NNS7-F1-MODEL\_V4 \| 1.0 \| 0.0003553 \| 174 \| 0.265 \| 98 \| 67 \| 2 \| 46 \| 143 \| 22 \| 114 \| Tail fiber assembly protein \| Tail fiber assembly protein; afdb-uniprot50 \| AF-A0A223W0F0-F1-MODEL\_V4 \| 1.0 \| 0.0004026 \| 174 \| 0.266 \| 109 \| 73 \| 3 \| 37 \| 143 \| 26 \| 129 \| Uncharacterized protein \| Uncharacterized protein; afdb-uniprot50 \| AF-A0A7H8UFS0-F1-MODEL\_V4 \| 1.0 \| 7.954e-05 \| 174 \| 0.314 \| 108 \| 57 \| 5 \| 30 \| 133 \| 26 \| 120 \| Tail fiber assembly protein \| Tail fiber assembly protein; afdb-uniprot50 \| AF-A0A2G0WBJ8-F1-MODEL\_V4 \| 1.0 \| 0.0001021 \| 173 \| 0.253 \| 146 \| 89 \| 7 \| 4 \| 142 \| 2 \| 134 \| Uncharacterized protein \| Uncharacterized protein; afdb-uniprot50 \| AF-E9TCL4-F1-MODEL\_V4 \| 1.0 \| 0.0004026 \| 172 \| 0.285 \| 98 \| 65 \| 3 \| 46 \| 143 \| 24 \| 116 \| Caudovirales tail fiber assembly protein \| Caudovirales tail fiber assembly protein; afdb-uniprot50 \| AF-A0A857BPZ5-F1-MODEL\_V4 \| 1.0 \| 7.021e-05 \| 172 \| 0.217 \| 147 \| 83 \| 6 \| 2 \| 143 \| 79 \| 198 \| Tail fiber assembly protein \| Tail fiber assembly protein; afdb-uniprot50 \| AF-T0QEZ4-F1-MODEL\_V4 \| 1.0 \| 0.001092 \| 171 \| 0.242 \| 99 \| 68 \| 4 \| 45 \| 143 \| 1 \| 92 \| Uncharacterized protein \| Uncharacterized protein; afdb-uniprot50 \| AF-A0A5N7JNQ3-F1-MODEL\_V4 \| 1.0 \| 9.591e-05 \| 171 \| 0.237 \| 143 \| 95 \| 5 \| 4 \| 142 \| 37 \| 169 \| Phage tail protein \| Phage tail protein; afdb-uniprot50 \| AF-A0A6I4LQ79-F1-MODEL\_V4 \| 1.0 \| 0.000663 \| 170 \| 0.265 \| 98 \| 67 \| 2 \| 46 \| 143 \| 1 \| 93 \| Tail fiber assembly protein \| Tail fiber assembly protein; afdb-uniprot50 \| AF-A0A403LVI5-F1-MODEL\_V4 \| 1.0 \| 0.000456 \| 170 \| 0.285 \| 98 \| 65 \| 3 \| 46 \| 143 \| 9 \| 101 \| Tail fiber assembly protein \| Tail fiber assembly protein; afdb-uniprot50 \| AF-A0A2I8Q8G5-F1-MODEL\_V4 \| 1.0 \| 0.0009639 \| 170 \| 0.305 \| 85 \| 55 \| 3 \| 60 \| 143 \| 44 \| 125 \| Uncharacterized protein \| Uncharacterized protein; afdb-uniprot50 \| AF-A0A7X5TM28-F1-MODEL\_V4 \| 1.0 \| 5.14e-05 \| 170 \| 0.217 \| 147 \| 83 \| 7 \| 2 \| 143 \| 89 \| 208 \| Phage tail protein \| Phage tail protein; afdb-uniprot50 \| AF-A0A6N8PP82-F1-MODEL\_V4 \| 1.0 \| 0.006663 \| 169 \| 0.454 \| 44 \| 24 \| 0 \| 1 \| 44 \| 1 \| 44 \| Tail fiber assembly protein \| Tail fiber assembly protein; afdb-uniprot50 \| AF-A0A208ZVY5-F1-MODEL\_V4 \| 1.0 \| 0.005526 \| 169 \| 0.26 \| 73 \| 52 \| 1 \| 71 \| 143 \| 4 \| 74 \| Phage tail protein \| Phage tail protein; afdb-uniprot50 \| AF-A0A411H8C5-F1-MODEL\_V4 \| 1.0 \| 0.0006229 \| 169 \| 0.265 \| 98 \| 58 \| 3 \| 46 \| 143 \| 26 \| 109 \| Uncharacterized protein \| Uncharacterized protein; afdb-uniprot50 \| AF-A0A1G5QVS6-F1-MODEL\_V4 \| 1.0 \| 0.0001789 \| 169 \| 0.226 \| 146 \| 83 \| 6 \| 2 \| 143 \| 6 \| 125 \| Virus tail fibre assembly protein, lambda gpK \| Virus tail fibre assembly protein, lambda gpK; afdb-uniprot50 \| AF-A0A3L2NWY4-F1-MODEL\_V4 \| 1.0 \| 0.0005166 \| 169 \| 0.265 \| 98 \| 67 \| 2 \| 46 \| 143 \| 35 \| 127 \| Tail fiber assembly protein \| Tail fiber assembly protein; afdb-uniprot50 \| AF-A0A345EKD5-F1-MODEL\_V4 \| 1.0 \| 9.591e-05 \| 169 \| 0.252 \| 119 \| 76 \| 6 \| 28 \| 143 \| 27 \| 135 \| Phage tail protein \| Phage tail protein; afdb-uniprot50 \| AF-F3YY47-F1-MODEL\_V4 \| 1.0 \| 0.0002947 \| 169 \| 0.195 \| 148 \| 102 \| 4 \| 4 \| 143 \| 3 \| 141 \| Prophage PSPPH06, putative tail fiber protein \| Prophage PSPPH06, putative tail fiber protein; afdb-uniprot50 \| AF-A0A855V8H1-F1-MODEL\_V4 \| 1.0 \| 0.0001156 \| 169 \| 0.208 \| 144 \| 105 \| 4 \| 4 \| 142 \| 3 \| 142 \| Phage tail protein \| Phage tail protein; afdb-uniprot50 \| AF-A0A736RL65-F1-MODEL\_V4 \| 1.0 \| 4.537e-05 \| 169 \| 0.214 \| 154 \| 88 \| 9 \| 1 \| 143 \| 44 \| 175 \| Tail fiber assembly protein \| Tail fiber assembly protein; afdb-uniprot50 \| AF-A0A849JE86-F1-MODEL\_V4 \| 1.0 \| 4.005e-05 \| 169 \| 0.2 \| 145 \| 86 \| 7 \| 2 \| 142 \| 82 \| 200 \| Phage tail protein \| Phage tail protein; afdb-uniprot50 \| AF-A0A1Q6B6N7-F1-MODEL\_V4 \| 1.0 \| 0.0009639 \| 168 \| 0.271 \| 92 \| 60 \| 3 \| 53 \| 142 \| 3 \| 89 \| Phage tail protein \| Phage tail protein; afdb-uniprot50 \| AF-A0A6F8VBJ7-F1-MODEL\_V4 \| 1.0 \| 0.0002296 \| 168 \| 0.222 \| 144 \| 88 \| 5 \| 5 \| 143 \| 2 \| 126 \| Uncharacterized protein \| Uncharacterized protein; afdb-uniprot50 \| AF-A0A7Z7PAS9-F1-MODEL\_V4 \| 1.0 \| 1.895e-05 \| 168 \| 0.22 \| 154 \| 90 \| 9 \| 1 \| 143 \| 41 \| 175 \| Bacteriophage tail assembly protein \| Bacteriophage tail assembly protein; afdb-uniprot50 \| AF-A0A2D0IJS5-F1-MODEL\_V4 \| 1.0 \| 4.005e-05 \| 168 \| 0.239 \| 146 \| 81 \| 7 \| 2 \| 143 \| 78 \| 197 \| Tail assembly protein \| Tail assembly protein; afdb-uniprot50 \| AF-A0A5C4RJQ8-F1-MODEL\_V4 \| 1.0 \| 6.597e-05 \| 168 \| 0.227 \| 145 \| 84 \| 6 \| 2 \| 143 \| 79 \| 198 \| Tail fiber assembly protein \| Tail fiber assembly protein; afdb-uniprot50 \| AF-A0A7X0T437-F1-MODEL\_V4 \| 1.0 \| 9.591e-05 \| 168 \| 0.191 \| 146 \| 88 \| 7 \| 2 \| 143 \| 81 \| 200 \| Tail fiber assembly protein \| Tail fiber assembly protein; afdb-uniprot50 \| AF-E7CGJ6-F1-MODEL\_V4 \| 1.0 \| 9.011e-05 \| 167 \| 0.201 \| 144 \| 85 \| 5 \| 3 \| 142 \| 114 \| 231 \| Phage-related tail fiber assembly protein G \| Phage-related tail fiber assembly protein G; afdb-uniprot50 \| AF-A0A250BJZ7-F1-MODEL\_V4 \| 1.0 \| 4.005e-05 \| 166 \| 0.227 \| 145 \| 81 \| 6 \| 2 \| 142 \| 176 \| 293 \| Phage tail protein \| Phage tail protein; afdb-uniprot50 \| AF-A0A6B1U8Y3-F1-MODEL\_V4 \| 1.0 \| 4.005e-05 \| 165 \| 0.208 \| 139 \| 93 \| 4 \| 4 \| 142 \| 3 \| 124 \| Tail fiber assembly protein \| Tail fiber assembly protein; afdb-uniprot50 \| AF-A0A4P8SN34-F1-MODEL\_V4 \| 1.0 \| 4.263e-05 \| 165 \| 0.206 \| 155 \| 87 \| 10 \| 1 \| 141 \| 61 \| 193 \| Tail assembly chaperone \| Tail assembly chaperone; afdb-uniprot50 \| AF-A0A443ZGX7-F1-MODEL\_V4 \| 1.0 \| 0.0002769 \| 164 \| 0.202 \| 143 \| 101 \| 6 \| 4 \| 142 \| 3 \| 136 \| Phage tail protein \| Phage tail protein; afdb-uniprot50 \| AF-A0A430BB89-F1-MODEL\_V4 \| 1.0 \| 3.105e-06 \| 164 \| 0.217 \| 161 \| 96 \| 4 \| 4 \| 141 \| 3 \| 156 \| Uncharacterized protein \| Uncharacterized protein; afdb-uniprot50 \| AF-A0A097QXF3-F1-MODEL\_V4 \| 1.0 \| 6.198e-05 \| 164 \| 0.207 \| 154 \| 89 \| 9 \| 1 \| 143 \| 44 \| 175 \| Uncharacterized protein \| Uncharacterized protein; afdb-uniprot50 \| AF-A0A422ZK35-F1-MODEL\_V4 \| 1.0 \| 0.0001394 \| 164 \| 0.213 \| 145 \| 86 \| 6 \| 2 \| 143 \| 73 \| 192 \| Tail assembly chaperone \| Tail assembly chaperone; afdb-uniprot50 \| AF-Q7N2W3-F1-MODEL\_V4 \| 1.0 \| 0.0009639 \| 164 \| 0.27 \| 100 \| 60 \| 3 \| 55 \| 143 \| 374 \| 471 \| Uncharacterized protein \| Uncharacterized protein; afdb-uniprot50 \| AF-A0A376EYU4-F1-MODEL\_V4 \| 1.0 \| 0.007092 \| 163 \| 0.416 \| 48 \| 28 \| 0 \| 4 \| 51 \| 3 \| 50 \| Phage tail assembly chaperone gp38 \| Phage tail assembly chaperone gp38; afdb-uniprot50 \| AF-A0A2U3TGW6-F1-MODEL\_V4 \| 1.0 \| 0.0002027 \| 163 \| 0.202 \| 148 \| 92 \| 8 \| 1 \| 142 \| 1 \| 128 \| Uncharacterized protein \| Uncharacterized protein; afdb-uniprot50 \| AF-A0A851GEG3-F1-MODEL\_V4 \| 1.0 \| 4.829e-05 \| 163 \| 0.208 \| 149 \| 88 \| 8 \| 4 \| 143 \| 50 \| 177 \| Tail fiber assembly protein \| Tail fiber assembly protein; afdb-uniprot50 \| AF-A0A4R7P8A5-F1-MODEL\_V4 \| 1.0 \| 5.823e-05 \| 163 \| 0.219 \| 146 \| 82 \| 7 \| 2 \| 142 \| 88 \| 206 \| Virus tail fiber assembly protein lambda gpK \| Virus tail fiber assembly protein lambda gpK; afdb-uniprot50 \| AF-A0A2A2GZI5-F1-MODEL\_V4 \| 1.0 \| 0.0001231 \| 163 \| 0.204 \| 147 \| 85 \| 7 \| 2 \| 143 \| 87 \| 206 \| Uncharacterized protein \| Uncharacterized protein; afdb-uniprot50 \| AF-A0A7U4N670-F1-MODEL\_V4 \| 1.0 \| 6.597e-05 \| 162 \| 0.206 \| 155 \| 90 \| 9 \| 1 \| 143 \| 40 \| 173 \| Uncharacterized protein \| Uncharacterized protein; afdb-uniprot50 \| AF-D3V998-F1-MODEL\_V4 \| 1.0 \| 0.000456 \| 161 \| 0.267 \| 101 \| 69 \| 3 \| 43 \| 143 \| 5 \| 100 \| Uncharacterized protein \| Uncharacterized protein; afdb-uniprot50 \| AF-A0A0T9UVT2-F1-MODEL\_V4 \| 1.0 \| 9.591e-05 \| 161 \| 0.294 \| 95 \| 62 \| 4 \| 4 \| 95 \| 2 \| 94 \| Uncharacterized protein \| Uncharacterized protein; afdb-uniprot50 \| AF-A0A077P8M4-F1-MODEL\_V4 \| 1.0 \| 0.0001021 \| 161 \| 0.179 \| 156 \| 91 \| 10 \| 1 \| 143 \| 45 \| 176 \| Putative tail fiber assembly protein \| Putative tail fiber assembly protein; afdb-uniprot50 \| AF-A0A3D9UGT7-F1-MODEL\_V4 \| 1.0 \| 9.011e-05 \| 161 \| 0.236 \| 144 \| 85 \| 6 \| 1 \| 143 \| 58 \| 177 \| Virus tail fiber assembly protein lambda gpK \| Virus tail fiber assembly protein lambda gpK; afdb-uniprot50 \| AF-P77699-F1-MODEL\_V4 \| 1.0 \| 0.0001394 \| 161 \| 0.208 \| 144 \| 84 \| 5 \| 3 \| 142 \| 67 \| 184 \| Protein TfaD \| Protein TfaD; afdb-uniprot50 \| AF-A0A2G8BY00-F1-MODEL\_V4 \| 1.0 \| 0.000158 \| 161 \| 0.171 \| 146 \| 91 \| 5 \| 2 \| 143 \| 76 \| 195 \| Uncharacterized protein \| Uncharacterized protein; afdb-uniprot50 \| AF-W7NUC6-F1-MODEL\_V4 \| 1.0 \| 0.0001681 \| 161 \| 0.172 \| 145 \| 92 \| 7 \| 2 \| 143 \| 79 \| 198 \| Tail fiber assembly protein \| Tail fiber assembly protein; afdb-uniprot50 \| AF-A0A7Z3MKM2-F1-MODEL\_V4 \| 1.0 \| 9.591e-05 \| 160 \| 0.185 \| 156 \| 92 \| 10 \| 1 \| 143 \| 2 \| 135 \| Tail fiber assembly protein \| Tail fiber assembly protein; afdb-uniprot50 \| AF-A0A1X1A8Y4-F1-MODEL\_V4 \| 1.0 \| 0.0007511 \| 160 \| 0.178 \| 123 \| 93 \| 2 \| 4 \| 126 \| 5 \| 119 \| Phage tail protein \| Phage tail protein; afdb-uniprot50 \| AF-A0A0M0T4V1-F1-MODEL\_V4 \| 1.0 \| 0.0001086 \| 160 \| 0.227 \| 145 \| 84 \| 7 \| 2 \| 143 \| 78 \| 197 \| Tail assembly protein \| Tail assembly protein; afdb-uniprot50 \| AF-A0A7D6NVZ1-F1-MODEL\_V4 \| 1.0 \| 0.0001789 \| 159 \| 0.259 \| 127 \| 67 \| 3 \| 17 \| 143 \| 54 \| 153 \| Tail fiber assembly protein \| Tail fiber assembly protein; afdb-uniprot50 \| AF-A0A6N8QLY1-F1-MODEL\_V4 \| 1.0 \| 0.000131 \| 159 \| 0.188 \| 143 \| 86 \| 5 \| 3 \| 141 \| 69 \| 185 \| Tail fiber assembly protein \| Tail fiber assembly protein; afdb-uniprot50 \| AF-A0A455W0B7-F1-MODEL\_V4 \| 1.0 \| 0.0005166 \| 158 \| 0.225 \| 120 \| 77 \| 5 \| 25 \| 143 \| 22 \| 126 \| Uncharacterized protein \| Uncharacterized protein; afdb-uniprot50 \| AF-A0A4P6GKN4-F1-MODEL\_V4 \| 1.0 \| 0.0003782 \| 158 \| 0.182 \| 137 \| 94 \| 6 \| 3 \| 135 \| 2 \| 124 \| Phage tail protein \| Phage tail protein; afdb-uniprot50 \| AF-A0A455VVP5-F1-MODEL\_V4 \| 1.0 \| 0.0001789 \| 158 \| 0.179 \| 145 \| 91 \| 7 \| 2 \| 143 \| 33 \| 152 \| Prophage tail fiber assembly protein homolog TfaE \| Prophage tail fiber assembly protein homolog TfaE; afdb-uniprot50 \| AF-A0A2K4IBZ4-F1-MODEL\_V4 \| 1.0 \| 5.471e-05 \| 158 \| 0.206 \| 145 \| 100 \| 6 \| 3 \| 143 \| 2 \| 135 \| Phage tail protein \| Phage tail protein; afdb-uniprot50 \| AF-A0A1B9L6I4-F1-MODEL\_V4 \| 1.0 \| 0.0001484 \| 158 \| 0.217 \| 147 \| 83 \| 6 \| 2 \| 143 \| 82 \| 201 \| Uncharacterized protein \| Uncharacterized protein; afdb-uniprot50 \| AF-A0A7X2R1K4-F1-MODEL\_V4 \| 1.0 \| 0.0002027 \| 157 \| 0.261 \| 126 \| 66 \| 3 \| 17 \| 142 \| 37 \| 135 \| Tail fiber assembly protein \| Tail fiber assembly protein; afdb-uniprot50 \| AF-A0A0L1C3B6-F1-MODEL\_V4 \| 1.0 \| 0.0004854 \| 156 \| 0.285 \| 119 \| 72 \| 7 \| 28 \| 143 \| 27 \| 135 \| Tail fiber assembly protein \| Tail fiber assembly protein; afdb-uniprot50 \| AF-A0A4S3LMC6-F1-MODEL\_V4 \| 1.0 \| 8.466e-05 \| 156 \| 0.206 \| 155 \| 90 \| 9 \| 1 \| 143 \| 38 \| 171 \| Tail fiber assembly protein \| Tail fiber assembly protein; afdb-uniprot50 \| AF-A0A1Y0KR81-F1-MODEL\_V4 \| 1.0 \| 1.225e-05 \| 156 \| 0.181 \| 154 \| 93 \| 5 \| 4 \| 126 \| 2 \| 153 \| Uncharacterized protein \| Uncharacterized protein; afdb-uniprot50 \| AF-A0A2D0J0R2-F1-MODEL\_V4 \| 1.0 \| 0.0002601 \| 156 \| 0.232 \| 129 \| 72 \| 3 \| 15 \| 143 \| 112 \| 213 \| Phage tail fiber assembly \| Phage tail fiber assembly; afdb-uniprot50 \| AF-A0A4P7L774-F1-MODEL\_V4 \| 1.0 \| 0.0001394 \| 156 \| 0.21 \| 147 \| 84 \| 7 \| 2 \| 143 \| 97 \| 216 \| Uncharacterized protein \| Uncharacterized protein; afdb-uniprot50 \| AF-L4J2A0-F1-MODEL\_V4 \| 1.0 \| 0.0001905 \| 156 \| 0.229 \| 148 \| 81 \| 8 \| 2 \| 143 \| 79 \| 199 \| Phage tail fiber assembly protein \| Phage tail fiber assembly protein; afdb-uniprot50 \| AF-A0A2Z4RQA3-F1-MODEL\_V4 \| 1.0 \| 0.0002947 \| 155 \| 0.254 \| 122 \| 84 \| 4 \| 5 \| 126 \| 2 \| 116 \| Phage tail protein \| Phage tail protein; afdb-uniprot50 \| AF-A0A3B0MF92-F1-MODEL\_V4 \| 1.0 \| 0.0002296 \| 155 \| 0.197 \| 152 \| 92 \| 9 \| 1 \| 143 \| 39 \| 169 \| Uncharacterized protein \| Uncharacterized protein; afdb-uniprot50 \| AF-A7MLN9-F1-MODEL\_V4 \| 1.0 \| 0.0002601 \| 155 \| 0.166 \| 144 \| 92 \| 6 \| 2 \| 142 \| 75 \| 193 \| Uncharacterized protein \| Uncharacterized protein; afdb-uniprot50 \| AF-A0A2D0IXB9-F1-MODEL\_V4 \| 1.0 \| 0.0002027 \| 155 \| 0.19 \| 147 \| 87 \| 7 \| 2 \| 143 \| 87 \| 206 \| Tail fiber protein of a prophage \| Tail fiber protein of a prophage; afdb-uniprot50 \| AF-A0A1H9EZ77-F1-MODEL\_V4 \| 1.0 \| 7.954e-05 \| 155 \| 0.16 \| 199 \| 89 \| 7 \| 4 \| 143 \| 26 \| 205 \| Virus tail fibre assembly protein, lambda gpK \| Virus tail fibre assembly protein, lambda gpK; afdb-uniprot50 \| AF-A0A2P5IM29-F1-MODEL\_V4 \| 1.0 \| 0.0001681 \| 154 \| 0.278 \| 115 \| 69 \| 6 \| 28 \| 140 \| 27 \| 129 \| Phage tail protein \| Phage tail protein; afdb-uniprot50 \| AF-A0A5U6ST04-F1-MODEL\_V4 \| 1.0 \| 0.0001086 \| 154 \| 0.19 \| 147 \| 95 \| 7 \| 1 \| 143 \| 11 \| 137 \| Tail fiber assembly protein \| Tail fiber assembly protein; afdb-uniprot50 \| AF-F2K8I0-F1-MODEL\_V4 \| 1.0 \| 6.597e-05 \| 154 \| 0.203 \| 123 \| 91 \| 4 \| 4 \| 126 \| 3 \| 118 \| Conserved hypothetical phage protein Putative tail fiber assembly protein \| Conserved hypothetical phage protein Putative tail fiber assembly protein; afdb-uniprot50 \| AF-A0A2S8QNI7-F1-MODEL\_V4 \| 1.0 \| 0.0002947 \| 154 \| 0.209 \| 148 \| 83 \| 7 \| 2 \| 143 \| 25 \| 144 \| Phage tail protein \| Phage tail protein; afdb-uniprot50 \| AF-A0A3N8BA59-F1-MODEL\_V4 \| 1.0 \| 0.0002296 \| 154 \| 0.221 \| 158 \| 89 \| 11 \| 1 \| 142 \| 1 \| 140 \| Phage tail protein \| Phage tail protein; afdb-uniprot50 \| AF-A0A080KEX6-F1-MODEL\_V4 \| 1.0 \| 0.0002027 \| 154 \| 0.225 \| 142 \| 88 \| 5 \| 2 \| 143 \| 31 \| 150 \| Caudovirales tail fiber assembly protein \| Caudovirales tail fiber assembly protein; afdb-uniprot50 \| AF-A0A6S7EC07-F1-MODEL\_V4 \| 1.0 \| 8.466e-05 \| 154 \| 0.251 \| 139 \| 87 \| 6 \| 1 \| 127 \| 1 \| 134 \| Uncharacterized protein \| Uncharacterized protein; afdb-uniprot50 \| AF-A0A5F1I731-F1-MODEL\_V4 \| 1.0 \| 0.0001484 \| 154 \| 0.211 \| 151 \| 91 \| 8 \| 1 \| 143 \| 39 \| 169 \| Tail assembly chaperone \| Tail assembly chaperone; afdb-uniprot50 \| AF-A0A379FLW3-F1-MODEL\_V4 \| 1.0 \| 0.0003137 \| 154 \| 0.236 \| 127 \| 74 \| 2 \| 17 \| 143 \| 119 \| 222 \| Bacteriophage tail assembly protein \| Bacteriophage tail assembly protein; afdb-uniprot50 \| AF-A0A346B7G2-F1-MODEL\_V4 \| 1.0 \| 0.0003339 \| 153 \| 0.194 \| 149 \| 104 \| 7 \| 1 \| 142 \| 1 \| 140 \| Phage tail protein \| Phage tail protein; afdb-uniprot50 \| AF-A0A6P1Q4F9-F1-MODEL\_V4 \| 1.0 \| 0.000131 \| 153 \| 0.197 \| 162 \| 96 \| 8 \| 5 \| 143 \| 25 \| 175 \| Uncharacterized protein \| Uncharacterized protein; afdb-uniprot50 \| AF-A0A2L2B7F2-F1-MODEL\_V4 \| 1.0 \| 0.0002947 \| 153 \| 0.247 \| 89 \| 58 \| 3 \| 4 \| 89 \| 3 \| 85 \| DUF4376 domain-containing protein \| DUF4376 domain-containing protein; afdb-uniprot50 \| AF-A0A7X5TFD8-F1-MODEL\_V4 \| 1.0 \| 0.0002027 \| 153 \| 0.21 \| 147 \| 84 \| 7 \| 2 \| 143 \| 78 \| 197 \| Phage tail protein \| Phage tail protein; afdb-uniprot50 \| AF-A0A2T4HXL0-F1-MODEL\_V4 \| 1.0 \| 0.0002027 \| 153 \| 0.205 \| 146 \| 84 \| 7 \| 2 \| 142 \| 87 \| 205 \| Phage tail protein \| Phage tail protein; afdb-uniprot50 \| AF-A0A3A6RRJ9-F1-MODEL\_V4 \| 1.0 \| 0.007092 \| 152 \| 0.293 \| 75 \| 51 \| 1 \| 66 \| 140 \| 9 \| 81 \| Uncharacterized protein \| Uncharacterized protein; afdb-uniprot50 \| AF-A0A1U6INB7-F1-MODEL\_V4 \| 1.0 \| 0.001317 \| 152 \| 0.179 \| 145 \| 94 \| 7 \| 4 \| 143 \| 3 \| 127 \| Phage tail assembly chaperone protein \| Phage tail assembly chaperone protein; afdb-uniprot50 \| AF-A0A837WAY8-F1-MODEL\_V4 \| 1.0 \| 0.0007057 \| 152 \| 0.197 \| 142 \| 100 \| 6 \| 4 \| 143 \| 3 \| 132 \| Phage tail protein \| Phage tail protein; afdb-uniprot50 \| AF-A0A2D0IW15-F1-MODEL\_V4 \| 1.0 \| 0.0002444 \| 152 \| 0.2 \| 155 \| 91 \| 10 \| 1 \| 143 \| 38 \| 171 \| Tail assembly chaperone \| Tail assembly chaperone; afdb-uniprot50 \| AF-A0A2S8Q9I9-F1-MODEL\_V4 \| 1.0 \| 0.0002769 \| 152 \| 0.219 \| 146 \| 84 \| 7 \| 2 \| 143 \| 61 \| 180 \| Phage tail protein \| Phage tail protein; afdb-uniprot50 \| AF-A0A2S5K611-F1-MODEL\_V4 \| 1.0 \| 0.0003553 \| 152 \| 0.142 \| 147 \| 94 \| 7 \| 2 \| 143 \| 78 \| 197 \| Phage tail protein \| Phage tail protein; afdb-uniprot50 \| AF-A0A447L1E5-F1-MODEL\_V4 \| 1.0 \| 0.0005499 \| 152 \| 0.169 \| 124 \| 90 \| 6 \| 25 \| 143 \| 113 \| 228 \| Caudovirales tail fibre assembly protein \| Caudovirales tail fibre assembly protein; afdb-uniprot50 \| AF-A0A2P1VLZ0-F1-MODEL\_V4 \| 1.0 \| 0.0002296 \| 151 \| 0.205 \| 146 \| 86 \| 5 \| 2 \| 143 \| 80 \| 199 \| Phage tail protein \| Phage tail protein; afdb-uniprot50 \| AF-A0A375AA84-F1-MODEL\_V4 \| 1.0 \| 0.0003553 \| 151 \| 0.162 \| 148 \| 90 \| 7 \| 2 \| 143 \| 83 \| 202 \| Tail fiber assembly protein \| Tail fiber assembly protein; afdb-uniprot50 \| AF-V0AZV2-F1-MODEL\_V4 \| 1.0 \| 0.0002296 \| 151 \| 0.222 \| 144 \| 80 \| 7 \| 2 \| 140 \| 82 \| 198 \| Caudovirales tail fiber assembly protein \| Caudovirales tail fiber assembly protein; afdb-uniprot50 \| AF-A0A336QKS4-F1-MODEL\_V4 \| 1.0 \| 0.000663 \| 150 \| 0.209 \| 143 \| 81 \| 5 \| 4 \| 143 \| 3 \| 116 \| Phage tail fiber assembly protein \| Phage tail fiber assembly protein; afdb-uniprot50 \| AF-A0A119AU52-F1-MODEL\_V4 \| 1.0 \| 0.000456 \| 150 \| 0.18 \| 144 \| 102 \| 6 \| 4 \| 140 \| 5 \| 139 \| Uncharacterized protein \| Uncharacterized protein; afdb-uniprot50 \| AF-A0A4Z0TBQ0-F1-MODEL\_V4 \| 1.0 \| 0.0005166 \| 150 \| 0.194 \| 144 \| 86 \| 5 \| 3 \| 142 \| 27 \| 144 \| Phage tail protein \| Phage tail protein; afdb-uniprot50 \| AF-A0A7V8IHT1-F1-MODEL\_V4 \| 1.0 \| 0.0002444 \| 150 \| 0.191 \| 146 \| 86 \| 6 \| 2 \| 142 \| 86 \| 204 \| Tail assembly protein \| Tail assembly protein; afdb-uniprot50 \| AF-A0A376D3X0-F1-MODEL\_V4 \| 1.0 \| 0.0002027 \| 149 \| 0.363 \| 99 \| 56 \| 1 \| 3 \| 101 \| 2 \| 93 \| Tail assembly chaperone gp38 \| Tail assembly chaperone gp38; afdb-uniprot50 \| AF-A0A2P9EJZ1-F1-MODEL\_V4 \| 1.0 \| 0.002783 \| 149 \| 0.302 \| 86 \| 57 \| 2 \| 55 \| 140 \| 29 \| 111 \| Tail fiber assembly protein homolog from lambdoid prophage DLP12 \| Tail fiber assembly protein homolog from lambdoid prophage DLP12; afdb-uniprot50 \| AF-A0A330EV64-F1-MODEL\_V4 \| 1.0 \| 0.0006229 \| 149 \| 0.21 \| 157 \| 85 \| 8 \| 1 \| 143 \| 1 \| 132 \| Uncharacterized protein \| Uncharacterized protein; afdb-uniprot50 \| AF-A0A4R5L703-F1-MODEL\_V4 \| 1.0 \| 0.001401 \| 149 \| 0.21 \| 152 \| 98 \| 7 \| 1 \| 142 \| 1 \| 140 \| Phage tail protein \| Phage tail protein; afdb-uniprot50 \| AF-A0A315WUN1-F1-MODEL\_V4 \| 1.0 \| 0.001401 \| 149 \| 0.186 \| 123 \| 92 \| 3 \| 4 \| 126 \| 7 \| 121 \| Phage tail protein \| Phage tail protein; afdb-uniprot50 \| AF-A0A2A7U735-F1-MODEL\_V4 \| 1.0 \| 0.0003782 \| 149 \| 0.207 \| 154 \| 89 \| 8 \| 1 \| 143 \| 39 \| 170 \| Tail assembly chaperone \| Tail assembly chaperone; afdb-uniprot50 \| AF-A0A379GMK5-F1-MODEL\_V4 \| 1.0 \| 0.0002947 \| 149 \| 0.198 \| 146 \| 87 \| 7 \| 1 \| 142 \| 86 \| 205 \| Bacteriophage tail assembly protein \| Bacteriophage tail assembly protein; afdb-uniprot50 \| AF-Q7N5D1-F1-MODEL\_V4 \| 1.0 \| 0.0003339 \| 149 \| 0.19 \| 147 \| 87 \| 7 \| 2 \| 143 \| 374 \| 493 \| Collar domain-containing protein \| Collar domain-containing protein; afdb-uniprot50 \| AF-A0A7D6IG48-F1-MODEL\_V4 \| 1.0 \| 0.0005499 \| 148 \| 0.209 \| 143 \| 99 \| 4 \| 5 \| 143 \| 2 \| 134 \| Tail fiber assembly protein \| Tail fiber assembly protein; afdb-uniprot50 \| AF-A0A261SNT8-F1-MODEL\_V4 \| 1.0 \| 0.0004285 \| 148 \| 0.218 \| 119 \| 81 \| 5 \| 29 \| 143 \| 33 \| 143 \| Uncharacterized protein \| Uncharacterized protein; afdb-uniprot50 \| AF-A0A077NP04-F1-MODEL\_V4 \| 1.0 \| 0.0004026 \| 148 \| 0.163 \| 147 \| 91 \| 7 \| 2 \| 143 \| 62 \| 181 \| Uncharacterized protein \| Uncharacterized protein; afdb-uniprot50 \| AF-A0A379DRV6-F1-MODEL\_V4 \| 1.0 \| 0.000456 \| 148 \| 0.189 \| 148 \| 86 \| 7 \| 2 \| 143 \| 77 \| 196 \| Bacteriophage tail assembly protein \| Bacteriophage tail assembly protein; afdb-uniprot50 \| AF-A0A1B9MVS0-F1-MODEL\_V4 \| 1.0 \| 0.0003782 \| 148 \| 0.197 \| 147 \| 86 \| 7 \| 2 \| 143 \| 83 \| 202 \| Uncharacterized protein \| Uncharacterized protein; afdb-uniprot50 \| AF-A0A077NJ56-F1-MODEL\_V4 \| 1.0 \| 0.0003137 \| 147 \| 0.243 \| 148 \| 75 \| 10 \| 4 \| 143 \| 5 \| 123 \| Uncharacterized protein \| Uncharacterized protein; afdb-uniprot50 \| AF-W1IQA1-F1-MODEL\_V4 \| 1.0 \| 0.001401 \| 147 \| 0.195 \| 123 \| 82 \| 5 \| 25 \| 143 \| 28 \| 137 \| Putative tail fiber assembly protein \| Putative tail fiber assembly protein; afdb-uniprot50 \| AF-A0A068Z1I4-F1-MODEL\_V4 \| 1.0 \| 0.0002947 \| 147 \| 0.212 \| 141 \| 87 \| 6 \| 2 \| 141 \| 70 \| 187 \| Phage tail assembly protein \| Phage tail assembly protein; afdb-uniprot50 \| AF-A0A7X1BIN5-F1-MODEL\_V4 \| 1.0 \| 0.0004285 \| 147 \| 0.182 \| 148 \| 87 \| 6 \| 2 \| 143 \| 77 \| 196 \| Tail fiber assembly protein \| Tail fiber assembly protein; afdb-uniprot50 \| AF-A0A1A9VZ50-F1-MODEL\_V4 \| 1.0 \| 0.000456 \| 147 \| 0.178 \| 146 \| 90 \| 6 \| 2 \| 143 \| 241 \| 360 \| Uncharacterized protein \| Uncharacterized protein; afdb-uniprot50 \| AF-A0A7Y0VF10-F1-MODEL\_V4 \| 1.0 \| 0.001317 \| 146 \| 0.204 \| 142 \| 99 \| 4 \| 3 \| 142 \| 2 \| 131 \| Phage tail protein \| Phage tail protein; afdb-uniprot50 \| AF-A0A7M2QM89-F1-MODEL\_V4 \| 1.0 \| 0.0004285 \| 146 \| 0.187 \| 144 \| 85 \| 7 \| 2 \| 140 \| 82 \| 198 \| Uncharacterized protein \| Uncharacterized protein; afdb-uniprot50 \| AF-A1JND9-F1-MODEL\_V4 \| 1.0 \| 0.0003137 \| 146 \| 0.19 \| 147 \| 86 \| 7 \| 2 \| 143 \| 86 \| 204 \| Putative phage tail fiber assembly protein \| Putative phage tail fiber assembly protein; afdb-uniprot50 \| AF-A0A3R8W675-F1-MODEL\_V4 \| 1.0 \| 0.0004026 \| 146 \| 0.191 \| 146 \| 88 \| 7 \| 2 \| 143 \| 87 \| 206 \| Tail fiber assembly protein \| Tail fiber assembly protein; afdb-uniprot50 \| AF-A0A2N4VAI3-F1-MODEL\_V4 \| 1.0 \| 1.303e-05 \| 145 \| 0.221 \| 140 \| 89 \| 5 \| 1 \| 127 \| 1 \| 133 \| Uncharacterized protein \| Uncharacterized protein; afdb-uniprot50 \| AF-A0A410UEB7-F1-MODEL\_V4 \| 1.0 \| 0.003153 \| 145 \| 0.197 \| 86 \| 65 \| 2 \| 4 \| 89 \| 3 \| 84 \| Uncharacterized protein \| Uncharacterized protein; afdb-uniprot50 \| AF-A0A6L6FGL7-F1-MODEL\_V4 \| 1.0 \| 0.0002947 \| 145 \| 0.206 \| 145 \| 85 \| 7 \| 2 \| 142 \| 65 \| 183 \| Tail fiber assembly protein \| Tail fiber assembly protein; afdb-uniprot50 \| AF-A0A2N5EJ48-F1-MODEL\_V4 \| 1.0 \| 0.0004026 \| 145 \| 0.21 \| 147 \| 84 \| 7 \| 2 \| 143 \| 96 \| 215 \| Phage tail protein \| Phage tail protein; afdb-uniprot50 \| AF-A0A658A1F9-F1-MODEL\_V4 \| 1.0 \| 0.005192 \| 144 \| 0.257 \| 66 \| 47 \| 2 \| 2 \| 66 \| 19 \| 83 \| Uncharacterized protein \| Uncharacterized protein; afdb-uniprot50 \| AF-A0A423NVP6-F1-MODEL\_V4 \| 1.0 \| 0.0002947 \| 144 \| 0.174 \| 143 \| 103 \| 6 \| 4 \| 142 \| 3 \| 134 \| Uncharacterized protein \| Uncharacterized protein; afdb-uniprot50 \| AF-A0A554PRY0-F1-MODEL\_V4 \| 1.0 \| 0.001026 \| 144 \| 0.167 \| 143 \| 105 \| 4 \| 4 \| 142 \| 2 \| 134 \| Phage tail protein \| Phage tail protein; afdb-uniprot50 \| AF-A0A344UII6-F1-MODEL\_V4 \| 1.0 \| 1.78e-05 \| 144 \| 0.248 \| 149 \| 92 \| 7 \| 4 \| 137 \| 3 \| 146 \| Uncharacterized protein \| Uncharacterized protein; afdb-uniprot50 \| AF-B8DLJ3-F1-MODEL\_V4 \| 1.0 \| 0.0006229 \| 144 \| 0.169 \| 159 \| 101 \| 5 \| 4 \| 140 \| 3 \| 152 \| Uncharacterized protein \| Uncharacterized protein; afdb-uniprot50 \| AF-A0A379Z175-F1-MODEL\_V4 \| 1.0 \| 0.0002769 \| 144 \| 0.162 \| 154 \| 96 \| 9 \| 1 \| 142 \| 40 \| 172 \| Caudovirales tail fibre assembly protein \| Caudovirales tail fibre assembly protein; afdb-uniprot50 \| AF-A0A1C0U9B5-F1-MODEL\_V4 \| 1.0 \| 0.0007994 \| 144 \| 0.193 \| 155 \| 94 \| 9 \| 1 \| 143 \| 42 \| 177 \| Caudovirales tail fiber assembly protein \| Caudovirales tail fiber assembly protein; afdb-uniprot50 \| AF-A0A2A7T5Q6-F1-MODEL\_V4 \| 1.0 \| 0.0004026 \| 144 \| 0.198 \| 146 \| 85 \| 7 \| 3 \| 143 \| 74 \| 192 \| Tail assembly chaperone \| Tail assembly chaperone; afdb-uniprot50 \| AF-U7QV10-F1-MODEL\_V4 \| 1.0 \| 0.0007057 \| 143 \| 0.169 \| 153 \| 88 \| 8 \| 2 \| 143 \| 59 \| 183 \| Uncharacterized protein \| Uncharacterized protein; afdb-uniprot50 \| AF-A0A4U9HJ43-F1-MODEL\_V4 \| 1.0 \| 0.0004854 \| 141 \| 0.208 \| 149 \| 81 \| 8 \| 2 \| 143 \| 71 \| 189 \| Caudovirales tail fibre assembly protein \| Caudovirales tail fibre assembly protein; afdb-uniprot50 \| AF-A0A556RUH4-F1-MODEL\_V4 \| 1.0 \| 0.007092 \| 140 \| 0.204 \| 98 \| 71 \| 3 \| 46 \| 143 \| 1 \| 91 \| Tail fiber assembly protein \| Tail fiber assembly protein; afdb-uniprot50 \| AF-A0A1B3Z882-F1-MODEL\_V4 \| 1.0 \| 0.000131 \| 139 \| 0.19 \| 131 \| 94 \| 4 \| 4 \| 127 \| 3 \| 128 \| Uncharacterized protein \| Uncharacterized protein; afdb-uniprot50 \| AF-A0A7V8TLT1-F1-MODEL\_V4 \| 1.0 \| 0.0005166 \| 139 \| 0.219 \| 155 \| 86 \| 10 \| 1 \| 143 \| 38 \| 169 \| Tail fiber assembly protein \| Tail fiber assembly protein; afdb-uniprot50 \| AF-A0A4V2W777-F1-MODEL\_V4 \| 1.0 \| 0.001237 \| 139 \| 0.156 \| 147 \| 92 \| 7 \| 1 \| 142 \| 175 \| 294 \| Virus tail fiber assembly protein lambda gpK \| Virus tail fiber assembly protein lambda gpK; afdb-uniprot50 \| AF-A0A315BKG8-F1-MODEL\_V4 \| 1.0 \| 0.0005853 \| 138 \| 0.24 \| 137 \| 84 \| 6 \| 4 \| 140 \| 2 \| 118 \| Uncharacterized protein \| Uncharacterized protein; afdb-uniprot50 \| AF-A0A3G5LCQ7-F1-MODEL\_V4 \| 1.0 \| 0.0005499 \| 138 \| 0.216 \| 125 \| 67 \| 6 \| 45 \| 143 \| 3 \| 122 \| Phage tail protein \| Phage tail protein; afdb-uniprot50 \| AF-A0A2N8AP07-F1-MODEL\_V4 \| 1.0 \| 0.002615 \| 138 \| 0.146 \| 143 \| 109 \| 6 \| 4 \| 142 \| 98 \| 231 \| Uncharacterized protein \| Uncharacterized protein; afdb-uniprot50 \| AF-A0A485A861-F1-MODEL\_V4 \| 1.0 \| 0.0004026 \| 138 \| 0.193 \| 145 \| 92 \| 9 \| 4 \| 143 \| 618 \| 742 \| Caudovirales tail fibre assembly protein \| Caudovirales tail fibre assembly protein; afdb-uniprot50 \| AF-U1TQF0-F1-MODEL\_V4 \| 1.0 \| 0.0008509 \| 137 \| 0.179 \| 139 \| 105 \| 4 \| 5 \| 143 \| 2 \| 131 \| Uncharacterized protein \| Uncharacterized protein; afdb-uniprot50 \| AF-A0A3D9YP94-F1-MODEL\_V4 \| 1.0 \| 0.001237 \| 137 \| 0.214 \| 149 \| 88 \| 8 \| 5 \| 142 \| 2 \| 132 \| Uncharacterized protein \| Uncharacterized protein; afdb-uniprot50 \| AF-A0A1S8CIT8-F1-MODEL\_V4 \| 1.0 \| 0.001588 \| 137 \| 0.171 \| 146 \| 91 \| 7 \| 2 \| 143 \| 77 \| 196 \| Uncharacterized protein \| Uncharacterized protein; afdb-uniprot50 \| AF-S1IG16-F1-MODEL\_V4 \| 1.0 \| 0.0003137 \| 136 \| 0.206 \| 145 \| 88 \| 9 \| 4 \| 140 \| 6 \| 131 \| Uncharacterized protein \| Uncharacterized protein; afdb-uniprot50 \| AF-A0A1N6MS03-F1-MODEL\_V4 \| 1.0 \| 0.0009639 \| 136 \| 0.165 \| 163 \| 97 \| 8 \| 5 \| 143 \| 22 \| 169 \| Uncharacterized protein \| Uncharacterized protein; afdb-uniprot50 \| AF-A0A7T9HE50-F1-MODEL\_V4 \| 1.0 \| 0.0006229 \| 135 \| 0.209 \| 124 \| 80 \| 4 \| 4 \| 126 \| 3 \| 109 \| Tail fiber assembly protein \| Tail fiber assembly protein; afdb-uniprot50 \| AF-A0A3S9K111-F1-MODEL\_V4 \| 1.0 \| 0.0008509 \| 135 \| 0.185 \| 140 \| 97 \| 5 \| 4 \| 141 \| 3 \| 127 \| Phage tail protein \| Phage tail protein; afdb-uniprot50 \| AF-A0A2V4KJM6-F1-MODEL\_V4 \| 1.0 \| 0.001026 \| 135 \| 0.167 \| 137 \| 102 \| 6 \| 4 \| 140 \| 3 \| 127 \| Uncharacterized protein \| Uncharacterized protein; afdb-uniprot50 \| AF-D4BTZ7-F1-MODEL\_V4 \| 1.0 \| 0.0007057 \| 135 \| 0.18 \| 150 \| 97 \| 8 \| 1 \| 143 \| 39 \| 169 \| Caudovirales tail fiber assembly protein \| Caudovirales tail fiber assembly protein; afdb-uniprot50 \| AF-A0A2J9H2E1-F1-MODEL\_V4 \| 1.0 \| 0.001092 \| 135 \| 0.171 \| 146 \| 91 \| 7 \| 2 \| 143 \| 74 \| 193 \| Phage tail protein \| Phage tail protein; afdb-uniprot50 \| AF-U1ZF67-F1-MODEL\_V4 \| 1.0 \| 0.002783 \| 133 \| 0.262 \| 80 \| 51 \| 2 \| 4 \| 81 \| 3 \| 76 \| Uncharacterized protein \| Uncharacterized protein; afdb-uniprot50 \| AF-A0A1F8VMN7-F1-MODEL\_V4 \| 1.0 \| 0.001026 \| 132 \| 0.184 \| 130 \| 91 \| 5 \| 5 \| 130 \| 2 \| 120 \| Uncharacterized protein \| Uncharacterized protein; afdb-uniprot50 \| AF-A0A085H8Z3-F1-MODEL\_V4 \| 1.0 \| 0.001492 \| 132 \| 0.193 \| 145 \| 87 \| 6 \| 3 \| 143 \| 78 \| 196 \| Tail fiber assembly protein \| Tail fiber assembly protein; afdb-uniprot50 \| AF-A0A225SMJ4-F1-MODEL\_V4 \| 1.0 \| 0.0006229 \| 131 \| 0.217 \| 124 \| 78 \| 5 \| 5 \| 127 \| 2 \| 107 \| Uncharacterized protein \| Uncharacterized protein; afdb-uniprot50 \| AF-A0A3S0J0M3-F1-MODEL\_V4 \| 1.0 \| 0.003571 \| 131 \| 0.136 \| 146 \| 104 \| 7 \| 4 \| 142 \| 3 \| 133 \| Phage tail protein \| Phage tail protein; afdb-uniprot50 \| AF-A0A345CRH0-F1-MODEL\_V4 \| 1.0 \| 2.432e-05 \| 131 \| 0.219 \| 132 \| 87 \| 7 \| 4 \| 127 \| 3 \| 126 \| Uncharacterized protein \| Uncharacterized protein; afdb-uniprot50 \| AF-A0A1G6QQ09-F1-MODEL\_V4 \| 1.0 \| 0.003801 \| 131 \| 0.167 \| 155 \| 88 \| 5 \| 4 \| 126 \| 3 \| 148 \| Uncharacterized protein \| Uncharacterized protein; afdb-uniprot50 \| AF-A0A2A2PTQ0-F1-MODEL\_V4 \| 1.0 \| 0.003801 \| 130 \| 0.165 \| 109 \| 86 \| 3 \| 28 \| 135 \| 1 \| 105 \| Uncharacterized protein \| Uncharacterized protein; afdb-uniprot50 \| AF-A0A7D6E693-F1-MODEL\_V4 \| 1.0 \| 0.001162 \| 130 \| 0.186 \| 123 \| 93 \| 3 \| 4 \| 126 \| 2 \| 117 \| Phage tail protein \| Phage tail protein; afdb-uniprot50 \| AF-A0A1B4FRS3-F1-MODEL\_V4 \| 1.0 \| 0.001798 \| 130 \| 0.192 \| 135 \| 94 \| 6 \| 12 \| 142 \| 16 \| 139 \| Phage tail protein \| Phage tail protein; afdb-uniprot50 \| AF-V0ATP8-F1-MODEL\_V4 \| 1.0 \| 0.0003137 \| 130 \| 0.218 \| 151 \| 87 \| 9 \| 1 \| 143 \| 40 \| 167 \| Caudovirales tail fiber assembly protein \| Caudovirales tail fiber assembly protein; afdb-uniprot50 \| AF-A0A2C8EYE8-F1-MODEL\_V4 \| 1.0 \| 0.004583 \| 130 \| 0.186 \| 123 \| 90 \| 5 \| 27 \| 143 \| 78 \| 196 \| Uncharacterized protein \| Uncharacterized protein; afdb-uniprot50 \| AF-A0A7W3CXQ1-F1-MODEL\_V4 \| 1.0 \| 0.0007057 \| 129 \| 0.186 \| 134 \| 91 \| 4 \| 4 \| 133 \| 3 \| 122 \| Uncharacterized protein \| Uncharacterized protein; afdb-uniprot50 \| AF-A0A2K4MJ83-F1-MODEL\_V4 \| 1.0 \| 0.0001156 \| 129 \| 0.197 \| 147 \| 95 \| 5 \| 4 \| 142 \| 2 \| 133 \| Uncharacterized protein \| Uncharacterized protein; afdb-uniprot50 \| AF-A0A291MWI8-F1-MODEL\_V4 \| 1.0 \| 3.536e-05 \| 129 \| 0.2 \| 165 \| 94 \| 6 \| 4 \| 141 \| 3 \| 156 \| Uncharacterized protein \| Uncharacterized protein; afdb-uniprot50 \| AF-A0A0U1KG08-F1-MODEL\_V4 \| 1.0 \| 0.001237 \| 128 \| 0.227 \| 145 \| 90 \| 8 \| 4 \| 143 \| 3 \| 130 \| Putative phage tail fiber assembly protein \| Putative phage tail fiber assembly protein; afdb-uniprot50 \| AF-A0A5P1DHX9-F1-MODEL\_V4 \| 1.0 \| 0.005526 \| 128 \| 0.171 \| 99 \| 80 \| 2 \| 28 \| 126 \| 49 \| 145 \| Phage tail protein \| Phage tail protein; afdb-uniprot50 \| AF-A0A6L6H6J7-F1-MODEL\_V4 \| 1.0 \| 0.001092 \| 128 \| 0.155 \| 167 \| 98 \| 6 \| 14 \| 143 \| 13 \| 173 \| Tail fiber assembly protein \| Tail fiber assembly protein; afdb-uniprot50 \| AF-A0A3M4M4E6-F1-MODEL\_V4 \| 1.0 \| 0.0002601 \| 128 \| 0.2 \| 110 \| 69 \| 3 \| 4 \| 99 \| 3 \| 107 \| Uncharacterized protein \| Uncharacterized protein; afdb-uniprot50 \| AF-A0A5N7JQ80-F1-MODEL\_V4 \| 1.0 \| 0.003801 \| 127 \| 0.191 \| 120 \| 88 \| 3 \| 3 \| 122 \| 2 \| 112 \| Phage tail protein \| Phage tail protein; afdb-uniprot50 \| AF-A0A7W2PYW0-F1-MODEL\_V4 \| 1.0 \| 0.006261 \| 127 \| 0.144 \| 125 \| 95 \| 3 \| 4 \| 126 \| 5 \| 119 \| Phage tail protein \| Phage tail protein; afdb-uniprot50 \| AF-A0A0C1W5C6-F1-MODEL\_V4 \| 1.0 \| 0.003355 \| 126 \| 0.237 \| 118 \| 82 \| 4 \| 28 \| 140 \| 15 \| 129 \| Uncharacterized protein \| Uncharacterized protein; afdb-uniprot50 \| AF-A0A7D4DZS1-F1-MODEL\_V4 \| 1.0 \| 0.005882 \| 125 \| 0.133 \| 142 \| 94 \| 6 \| 4 \| 143 \| 3 \| 117 \| Uncharacterized protein \| Uncharacterized protein; afdb-uniprot50 \| AF-A0A1S1V1A2-F1-MODEL\_V4 \| 1.0 \| 0.005526 \| 125 \| 0.222 \| 108 \| 80 \| 3 \| 28 \| 135 \| 1 \| 104 \| Uncharacterized protein \| Uncharacterized protein; afdb-uniprot50 \| AF-A0A2U8TMA5-F1-MODEL\_V4 \| 1.0 \| 0.005526 \| 125 \| 0.165 \| 127 \| 92 \| 4 \| 3 \| 126 \| 2 \| 117 \| Phage tail protein \| Phage tail protein; afdb-uniprot50 \| AF-F3YW31-F1-MODEL\_V4 \| 1.0 \| 0.002168 \| 125 \| 0.226 \| 128 \| 80 \| 3 \| 1 \| 127 \| 1 \| 110 \| Uncharacterized protein \| Uncharacterized protein; afdb-uniprot50 \| AF-A0A6N0JI93-F1-MODEL\_V4 \| 1.0 \| 0.001401 \| 124 \| 0.164 \| 152 \| 88 \| 6 \| 4 \| 142 \| 3 \| 128 \| Uncharacterized protein \| Uncharacterized protein; afdb-uniprot50 \| AF-A0A109L8T1-F1-MODEL\_V4 \| 1.0 \| 0.00169 \| 124 \| 0.164 \| 146 \| 101 \| 9 \| 4 \| 142 \| 3 \| 134 \| Uncharacterized protein \| Uncharacterized protein; afdb-uniprot50 \| AF-A0A418WP23-F1-MODEL\_V4 \| 1.0 \| 0.0002769 \| 124 \| 0.172 \| 151 \| 106 \| 4 \| 4 \| 143 \| 3 \| 145 \| Uncharacterized protein \| Uncharacterized protein; afdb-uniprot50 \| AF-A0A1D9BC70-F1-MODEL\_V4 \| 1.0 \| 0.0003553 \| 124 \| 0.142 \| 183 \| 102 \| 5 \| 4 \| 140 \| 2 \| 175 \| DUF4376 domain-containing protein \| DUF4376 domain-containing protein; afdb-uniprot50 \| AF-A0A5E7Q4J6-F1-MODEL\_V4 \| 1.0 \| 0.0005166 \| 123 \| 0.208 \| 139 \| 93 \| 6 \| 5 \| 135 \| 2 \| 131 \| Uncharacterized protein \| Uncharacterized protein; afdb-uniprot50 \| AF-A0A5S9HN81-F1-MODEL\_V4 \| 1.0 \| 0.0007511 \| 123 \| 0.198 \| 141 \| 92 \| 5 \| 4 \| 129 \| 3 \| 137 \| Uncharacterized protein \| Uncharacterized protein; afdb-uniprot50 \| AF-A0A377N9X0-F1-MODEL\_V4 \| 1.0 \| 0.002615 \| 123 \| 0.174 \| 149 \| 87 \| 7 \| 2 \| 143 \| 70 \| 189 \| Caudovirales tail fibre assembly protein \| Caudovirales tail fibre assembly protein; afdb-uniprot50 \| AF-A0A4R6YXJ0-F1-MODEL\_V4 \| 1.0 \| 0.0007994 \| 123 \| 0.183 \| 158 \| 97 \| 5 \| 1 \| 126 \| 1 \| 158 \| Uncharacterized protein \| Uncharacterized protein; afdb-uniprot50 \| AF-A0A250KYB2-F1-MODEL\_V4 \| 1.0 \| 0.001492 \| 121 \| 0.16 \| 193 \| 94 \| 5 \| 4 \| 138 \| 2 \| 184 \| DUF4376 domain-containing protein \| DUF4376 domain-containing protein; afdb-uniprot50 \| AF-Q1I688-F1-MODEL\_V4 \| 1.0 \| 0.002037 \| 120 \| 0.181 \| 143 \| 102 \| 6 \| 4 \| 142 \| 2 \| 133 \| Putative phage protein \| Putative phage protein; afdb-uniprot50 \| AF-A0A2A7T6M2-F1-MODEL\_V4 \| 1.0 \| 0.004878 \| 120 \| 0.177 \| 118 \| 77 \| 6 \| 46 \| 143 \| 29 \| 146 \| Phage tail protein \| Phage tail protein; afdb-uniprot50 \| AF-A0A806X527-F1-MODEL\_V4 \| 1.0 \| 0.002308 \| 120 \| 0.193 \| 145 \| 69 \| 4 \| 45 \| 143 \| 53 \| 195 \| Phage tail protein \| Phage tail protein; afdb-uniprot50 \| AF-A0A7V8RB58-F1-MODEL\_V4 \| 1.0 \| 0.0002947 \| 119 \| 0.223 \| 152 \| 98 \| 6 \| 4 \| 143 \| 3 \| 146 \| Uncharacterized protein \| Uncharacterized protein; afdb-uniprot50 \| AF-A0A0T9T410-F1-MODEL\_V4 \| 1.0 \| 0.004583 \| 119 \| 0.164 \| 146 \| 103 \| 7 \| 4 \| 143 \| 32 \| 164 \| Tail fiber assembly protein G \| Tail fiber assembly protein G; afdb-uniprot50 \| AF-A0A2N1H596-F1-MODEL\_V4 \| 1.0 \| 0.003571 \| 118 \| 0.217 \| 124 \| 80 \| 6 \| 28 \| 143 \| 1 \| 115 \| Phage tail protein \| Phage tail protein; afdb-uniprot50 \| AF-A0A261SHH5-F1-MODEL\_V4 \| 1.0 \| 0.0004285 \| 118 \| 0.18 \| 144 \| 94 \| 6 \| 4 \| 143 \| 3 \| 126 \| Uncharacterized protein \| Uncharacterized protein; afdb-uniprot50 \| AF-A0A537MFJ5-F1-MODEL\_V4 \| 1.0 \| 0.0005166 \| 118 \| 0.195 \| 138 \| 89 \| 7 \| 1 \| 126 \| 1 \| 128 \| Uncharacterized protein \| Uncharacterized protein; afdb-uniprot50 \| AF-A0A2V3Y7F0-F1-MODEL\_V4 \| 1.0 \| 0.002037 \| 118 \| 0.2 \| 140 \| 87 \| 8 \| 4 \| 135 \| 3 \| 125 \| Phage tail assembly chaperone \| Phage tail assembly chaperone; afdb-uniprot50 \| AF-A0A5E6WPB4-F1-MODEL\_V4 \| 1.0 \| 0.004583 \| 118 \| 0.255 \| 86 \| 58 \| 1 \| 4 \| 89 \| 2 \| 81 \| Uncharacterized protein \| Uncharacterized protein; afdb-uniprot50 \| AF-A0A5P0J6I2-F1-MODEL\_V4 \| 1.0 \| 0.0001484 \| 116 \| 0.234 \| 132 \| 80 \| 5 \| 3 \| 127 \| 2 \| 119 \| Uncharacterized protein \| Uncharacterized protein; afdb-uniprot50 \| AF-A0A5C7Q2A1-F1-MODEL\_V4 \| 1.0 \| 0.002168 \| 116 \| 0.195 \| 169 \| 95 \| 8 \| 1 \| 140 \| 1 \| 157 \| Uncharacterized protein \| Uncharacterized protein; afdb-uniprot50 \| AF-A0A1C7WD91-F1-MODEL\_V4 \| 1.0 \| 0.001401 \| 116 \| 0.154 \| 201 \| 92 \| 7 \| 2 \| 142 \| 71 \| 253 \| Uncharacterized protein \| Uncharacterized protein; afdb-uniprot50 \| AF-A0A083ZV50-F1-MODEL\_V4 \| 1.0 \| 0.004046 \| 115 \| 0.159 \| 163 \| 80 \| 4 \| 28 \| 142 \| 48 \| 201 \| Caudovirales tail fiber assembly protein \| Caudovirales tail fiber assembly protein; afdb-uniprot50 \| AF-A0A0E8XEB7-F1-MODEL\_V4 \| 1.0 \| 0.0004854 \| 114 \| 0.242 \| 161 \| 82 \| 8 \| 2 \| 143 \| 18 \| 157 \| Tail assembly chaperone gp38 \| Tail assembly chaperone gp38; afdb-uniprot50 \| AF-A0A1V5QIZ0-F1-MODEL\_V4 \| 1.0 \| 0.002457 \| 112 \| 0.16 \| 187 \| 95 \| 6 \| 5 \| 138 \| 2 \| 179 \| DUF4376 domain-containing protein \| DUF4376 domain-containing protein; afdb-uniprot50 \| AF-H9BMI8-F1-MODEL\_V4 \| 1.0 \| 0.006663 \| 111 \| 0.194 \| 159 \| 93 \| 10 \| 4 \| 143 \| 6 \| 148 \| Uncharacterized protein \| Uncharacterized protein; afdb-uniprot50 \| AF-A0A1A9VZ06-F1-MODEL\_V4 \| 1.0 \| 0.007092 \| 110 \| 0.188 \| 159 \| 87 \| 9 \| 5 \| 142 \| 337 \| 474 \| Head\_binding domain-containing protein \| Head\_binding domain-containing protein; afdb-uniprot50 \| AF-A0A4R3J9P7-F1-MODEL\_V4 \| 1.0 \| 0.006261 \| 108 \| 0.146 \| 198 \| 101 \| 6 \| 4 \| 139 \| 3 \| 194 \| Uncharacterized protein \| Uncharacterized protein; afdb-uniprot50 \| AF-A0A6G6ZV78-F1-MODEL\_V4 \| 1.0 \| 0.004306 \| 105 \| 0.142 \| 169 \| 103 \| 7 \| 4 \| 143 \| 3 \| 158 \| Phage tail protein \| Phage tail protein; afdb-uniprot50 \| AF-A0A2T6LN68-F1-MODEL\_V4 \| 1.0 \| 0.0002158 \| 105 \| 0.197 \| 142 \| 89 \| 5 \| 1 \| 127 \| 1 \| 132 \| Uncharacterized protein \| Uncharacterized protein; afdb-uniprot50 \| AF-A0A2V2AD22-F1-MODEL\_V4 \| 1.0 \| 0.008034 \| 105 \| 0.245 \| 102 \| 70 \| 4 \| 27 \| 126 \| 81 \| 177 \| Uncharacterized protein \| Uncharacterized protein; afdb-uniprot50 \| AF-A0A1I2QMV9-F1-MODEL\_V4 \| 1.0 \| 0.0006229 \| 104 \| 0.181 \| 138 \| 91 \| 5 \| 4 \| 134 \| 2 \| 124 \| Uncharacterized protein \| Uncharacterized protein; afdb-uniprot50 \| AF-A0A377QEM3-F1-MODEL\_V4 \| 1.0 \| 0.002783 \| 104 \| 0.171 \| 134 \| 89 \| 5 \| 1 \| 126 \| 1 \| 120 \| Uncharacterized protein \| Uncharacterized protein; afdb-uniprot50 \| AF-A0A427DLX8-F1-MODEL\_V4 \| 1.0 \| 0.0002947 \| 102 \| 0.209 \| 148 \| 92 \| 8 \| 4 \| 142 \| 2 \| 133 \| Uncharacterized protein \| Uncharacterized protein; afdb-uniprot50 \| AF-A0A7T7KK36-F1-MODEL\_V4 \| 1.0 \| 0.005526 \| 101 \| 0.159 \| 188 \| 102 \| 8 \| 4 \| 140 \| 2 \| 184 \| Uncharacterized protein \| Uncharacterized protein; afdb-uniprot50 \| AF-A0A354UB04-F1-MODEL\_V4 \| 1.0 \| 0.009688 \| 100 \| 0.205 \| 146 \| 96 \| 5 \| 3 \| 138 \| 2 \| 137 \| Uncharacterized protein \| Uncharacterized protein; afdb-uniprot50 \| AF-A0A2S9AZ00-F1-MODEL\_V4 \| 1.0 \| 0.003801 \| 100 \| 0.201 \| 124 \| 74 \| 4 \| 3 \| 109 \| 2 \| 117 \| Uncharacterized protein \| Uncharacterized protein; afdb-uniprot50 \| AF-A0A7Z8DSQ9-F1-MODEL\_V4 \| 1.0 \| 0.0008509 \| 100 \| 0.163 \| 196 \| 90 \| 6 \| 1 \| 141 \| 10 \| 186 \| DUF4376 domain-containing protein \| DUF4376 domain-containing protein; afdb-uniprot50 \| AF-A0A845BIX0-F1-MODEL\_V4 \| 0.999 \| 0.005526 \| 99 \| 0.125 \| 262 \| 105 \| 8 \| 2 \| 141 \| 21 \| 280 \| DUF4376 domain-containing protein \| DUF4376 domain-containing protein; afdb-uniprot50 \| AF-A0A557QX70-F1-MODEL\_V4 \| 0.999 \| 0.006261 \| 98 \| 0.163 \| 184 \| 96 \| 6 \| 4 \| 139 \| 3 \| 176 \| Uncharacterized protein \| Uncharacterized protein; afdb-uniprot50 \| AF-A0A142BH82-F1-MODEL\_V4 \| 0.999 \| 0.0009056 \| 97 \| 0.17 \| 176 \| 88 \| 8 \| 4 \| 130 \| 3 \| 169 \| Uncharacterized protein \| Uncharacterized protein; afdb-uniprot50 \| AF-A0A4R2MGV0-F1-MODEL\_V4 \| 0.999 \| 0.002962 \| 96 \| 0.205 \| 136 \| 80 \| 6 \| 4 \| 127 \| 3 \| 122 \| Uncharacterized protein \| Uncharacterized protein; afdb-uniprot50 \| AF-A0A7I8EBT6-F1-MODEL\_V4 \| 0.999 \| 0.002308 \| 96 \| 0.16 \| 187 \| 98 \| 6 \| 12 \| 141 \| 2 \| 186 \| Uncharacterized protein \| Uncharacterized protein; afdb-uniprot50 \| AF-W9BTD0-F1-MODEL\_V4 \| 0.997 \| 0.001588 \| 92 \| 0.175 \| 165 \| 101 \| 10 \| 2 \| 143 \| 11 \| 163 \| Uncharacterized protein \| Uncharacterized protein; afdb-uniprot50 \| AF-A0A1Y0N4F4-F1-MODEL\_V4 \| 0.996 \| 0.007092 \| 90 \| 0.183 \| 131 \| 80 \| 6 \| 5 \| 127 \| 2 \| 113 \| Uncharacterized protein \| Uncharacterized protein; afdb-uniprot50 \| AF-A0A4D8RIN6-F1-MODEL\_V4 \| 0.996 \| 0.008034 \| 90 \| 0.161 \| 198 \| 98 \| 8 \| 2 \| 140 \| 329 \| 517 \| DUF4376 domain-containing protein \| DUF4376 domain-containing protein; afdb-uniprot50 \| AF-A0A2W0FDX5-F1-MODEL\_V4 \| 0.961 \| 0.003153 \| 74 \| 0.232 \| 116 \| 66 \| 6 \| 28 \| 134 \| 5 \| 106 \| Phage tail protein \| Phage tail protein |
| Top keywords  (threshold 1.00e-02 (evalue)) | **tail, assembly, fiber, Phage, chaperone, Caudovirales, fibre, Putative, prophage, lambda** |
| Output files | ../../similar\_structures/16\_FANPEZAQ\_CDS\_0016\_afdb-proteome\_foldseek.tsv ../../similar\_structures/16\_FANPEZAQ\_CDS\_0016\_afdb-uniprot50\_foldseek.tsv ../../similar\_structures/16\_FANPEZAQ\_CDS\_0016\_merged.svg ../../similar\_structures/16\_FANPEZAQ\_CDS\_0016\_pdb\_foldseek.tsv |

  
  
  

Return to summary | Go to previous | Go to next

  


---

**Sequence/structure alignments coloring**  
Each object in the alignment figures is colored according to its E-value following this color coding:

1e-100
10
