## Supplementary material for "Completing the BASEL phage collection to unlock hidden diversity for systematic exploration of phage-host interactions": Data S2: 17.html

FANPEZAQ\_CDS\_0017


Return to summary | Go to previous | Go to next

|  |  |
| --- | --- |
| FANPEZAQ\_CDS\_0017  Page creation date: 02 Sep 2024, 12:00  Project folder: n/a  Input sequences file: Escherichia\_virus\_HeidiAbel.gb | tail collar domain\_containing phage fiber microcystin\_dependent hypothetical baseplate t4 wedge gp6 complex tube putative baseplate\_tail pre\_attachment fragment phage\_related side microcystin dependent prophage fibre short protein\_like repeat acid transporter structural fatty short\_chain containing n\_terminal bacteriophage long |

### Sequence information

|  |  |
| --- | --- |
| Name | FANPEZAQ\_CDS\_0017  17\_FANPEZAQ\_CDS\_0017 (pipeline id) |
| Imported annotations | Escherichia\_virus\_HeidiAbel Bas97 |
| Protein sequence | MSQILEFWKGLSGAVLPFAGANAPTGWLMCDGSAVSRTTFAELFASIGTTYGTGDGSTTF NLPDLRGRVAAGKDNMGGTPANRLTAAGAGITGTALGATGGSQTHTMTTGQMPAHSHGVN DPTHAHSVYDPTHTHAVYDPGHVHSYGRVTTGNGQGSDIGTANNHVTANTGAATTGISLY ASATGIGIYGAATGISIQNQGGGAAHNITQPTIVLNHIIKI |
| Number of residues | 221 |
| Molecular weight (Da) | 22212.26 |
| Output files | ../../query\_sequences/17\_FANPEZAQ\_CDS\_0017.fasta |

### Putative domain architecture and protein family

#### Search results (HHblits)^1^

|  |  |
| --- | --- |
| Domain family databases searched | Pfam, Ncbi-cd, Cath, Phrogs |
| Results, scheme(s)  (Top layers only; threshold 1.00e-03 (evalue)) | xml version="1.0" encoding="utf-8" standalone="no"?       2024-09-02T21:08:15.827230 image/svg+xml   Matplotlib v3.7.2, https://matplotlib.org/ |
| Results, table  (E-value ≤ 1.00e-03 (evalue)) | db \| id \| prob \| evalue \| pvalue \| score \| cols \| query \| query\_len \| template \| template\_len \| name \| description; phrogs \| 8763 \| 99.9 \| 4.4e-34 \| 5.4e-38 \| 233.1 \| 170 \| (28, 221) \| 221 \| (1, 170) \| 170 \| tail protein \| tail protein; Category: tail; NC\_023006\_p17; phrogs \| 8469 \| 99.9 \| 9.4e-32 \| 1.1e-35 \| 230.8 \| 168 \| (9, 221) \| 221 \| (114, 290) \| 292 \| tail fiber protein \| tail fiber protein; Category: tail; NC\_019917\_p61; phrogs \| 523 \| 99.9 \| 1.3e-28 \| 1.6e-32 \| 204.1 \| 89 \| (6, 123) \| 221 \| (40, 128) \| 183 \| tail collar fiber protein \| tail collar fiber protein; Category: tail; p284593 VI\_03272; phrogs \| 1800 \| 99.8 \| 1.3e-26 \| 1.6e-30 \| 193.7 \| 155 \| (8, 221) \| 221 \| (70, 224) \| 224 \| tail collar fiber protein \| tail collar fiber protein; Category: tail; KJ019100\_p265; phrogs \| 1858 \| 99.8 \| 9.8e-24 \| 1.3e-27 \| 182.0 \| 86 \| (8, 122) \| 221 \| (103, 188) \| 256 \| tail fiber protein \| tail fiber protein; Category: tail; p362065 VI\_01943; phrogs \| 9116 \| 99.7 \| 4.7e-23 \| 5.4e-27 \| 182.6 \| 100 \| (9, 121) \| 221 \| (58, 157) \| 443 \| tail protein \| tail protein; Category: tail; KU160641\_p20; phrogs \| 2097 \| 99.7 \| 7.3e-23 \| 8.8e-27 \| 185.7 \| 117 \| (8, 145) \| 221 \| (255, 371) \| 454 \| tail fiber protein \| tail fiber protein; Category: tail; p203462 VI\_04227; phrogs \| 31611 \| 99.7 \| 3.4e-22 \| 3.8e-26 \| 171.9 \| 198 \| (7, 221) \| 221 \| (156, 384) \| 384 \| tail collar protein \| tail collar protein; Category: tail; KU234532\_p33; phrogs \| 14307 \| 99.7 \| 6.7e-22 \| 7.7e-26 \| 141.2 \| 81 \| (6, 115) \| 221 \| (3, 83) \| 83 \| NA \| NA; Category: unknown function; p28392 VI\_07752; phrogs \| 20472 \| 99.5 \| 1.7e-18 \| 1.9e-22 \| 147.4 \| 86 \| (7, 121) \| 221 \| (186, 271) \| 331 \| NA \| NA; Category: unknown function; p276897 VI\_03881; phrogs \| 25747 \| 99.4 \| 7.2e-18 \| 8e-22 \| 153.9 \| 95 \| (11, 124) \| 221 \| (679, 773) \| 901 \| tail fiber protein \| tail fiber protein; Category: tail; KR052480\_p88; phrogs \| 295 \| 99.4 \| 1.6e-17 \| 1.9e-21 \| 156.2 \| 108 \| (5, 146) \| 221 \| (624, 746) \| 813 \| tail fiber protein \| tail fiber protein; Category: tail; p109395 VI\_03810; phrogs \| 4849 \| 99.4 \| 2.5e-17 \| 2.9e-21 \| 152.3 \| 96 \| (7, 121) \| 221 \| (550, 662) \| 790 \| tail protein \| tail protein; Category: tail; JX681814\_p23; phrogs \| 3100 \| 99.3 \| 9.5e-17 \| 1.1e-20 \| 143.0 \| 100 \| (7, 140) \| 221 \| (249, 363) \| 436 \| tail fiber protein \| tail fiber protein; Category: tail; p172841 VI\_09914; phrogs \| 5795 \| 99.3 \| 9.4e-17 \| 1.2e-20 \| 134.3 \| 89 \| (7, 122) \| 221 \| (62, 152) \| 215 \| tail protein \| tail protein; Category: tail; JN254801\_p12; phrogs \| 327 \| 99.3 \| 1.7e-16 \| 2.1e-20 \| 146.2 \| 87 \| (5, 120) \| 221 \| (381, 467) \| 544 \| tail protein \| tail protein; Category: tail; p384764 VI\_09705; phrogs \| 6256 \| 99.1 \| 6.9e-15 \| 8.2e-19 \| 134.5 \| 108 \| (4, 147) \| 221 \| (251, 379) \| 579 \| tail fiber protein \| tail fiber protein; Category: tail; KY883648\_p41; phrogs \| 6459 \| 99.1 \| 1.1e-14 \| 1.2e-18 \| 110.4 \| 66 \| (27, 121) \| 221 \| (3, 68) \| 121 \| tail collar protein \| tail collar protein; Category: tail; p30245 VI\_11136; phrogs \| 19618 \| 99.0 \| 6.4e-14 \| 7.2e-18 \| 121.0 \| 90 \| (7, 120) \| 221 \| (133, 222) \| 387 \| tail collar protein \| tail collar protein; Category: tail; p51395 VI\_03026; phrogs \| 12846 \| 98.9 \| 2.7e-13 \| 3e-17 \| 119.8 \| 91 \| (5, 121) \| 221 \| (150, 240) \| 544 \| tail fiber protein; Ig-domain containing \| tail fiber protein; Ig-domain containing; Category: tail; MF979563\_p29; phrogs \| 4711 \| 98.7 \| 3.9e-12 \| 4.5e-16 \| 111.4 \| 101 \| (9, 121) \| 221 \| (223, 400) \| 461 \| NA \| NA; Category: unknown function; p26028 VI\_12353; phrogs \| 7765 \| 98.6 \| 1.1e-11 \| 1.3e-15 \| 110.7 \| 102 \| (8, 121) \| 221 \| (340, 517) \| 578 \| NA \| NA; Category: unknown function; p66770 VI\_12458; phrogs \| 38759 \| 98.1 \| 9.8e-10 \| 1.1e-13 \| 100.0 \| 78 \| (37, 139) \| 221 \| (648, 727) \| 787 \| NA \| NA; Category: unknown function; p217390 VI\_05721; phrogs \| 9370 \| 98.0 \| 1.4e-09 \| 1.5e-13 \| 94.2 \| 100 \| (10, 121) \| 221 \| (226, 401) \| 458 \| tail fiber protein \| tail fiber protein; Category: tail; p122631 VI\_12395; phrogs \| 26622 \| 97.9 \| 4.8e-09 \| 5.4e-13 \| 94.5 \| 84 \| (9, 119) \| 221 \| (174, 257) \| 691 \| tail fiber protein \| tail fiber protein; Category: tail; p171543 VI\_11794; phrogs \| 2631 \| 97.6 \| 3.9e-08 \| 4.6e-12 \| 86.8 \| 85 \| (7, 125) \| 221 \| (337, 421) \| 482 \| tail fiber protein \| tail fiber protein; Category: tail; p130317 VI\_07372; phrogs \| 8239 \| 97.5 \| 5.3e-08 \| 6.1e-12 \| 87.8 \| 84 \| (6, 124) \| 221 \| (442, 525) \| 654 \| tail fiber protein \| tail fiber protein; Category: tail; p255681 VI\_08125; phrogs \| 20242 \| 97.4 \| 8e-08 \| 9.2e-12 \| 89.5 \| 75 \| (14, 118) \| 221 \| (913, 987) \| 1001 \| tail fiber protein \| tail fiber protein; Category: tail; NC\_014660\_p244; phrogs \| 6921 \| 97.3 \| 2.2e-07 \| 2.6e-11 \| 86.7 \| 67 \| (8, 74) \| 221 \| (234, 326) \| 767 \| tail fiber protein \| tail fiber protein; Category: tail; KM979355\_p107; phrogs \| 4591 \| 97.2 \| 3.7e-07 \| 4.3e-11 \| 81.5 \| 53 \| (2, 54) \| 221 \| (5, 59) \| 567 \| short tail fiber protein \| short tail fiber protein; Category: tail; KX349292\_p35; phrogs \| 7489 \| 97.0 \| 9.6e-07 \| 1.1e-10 \| 80.2 \| 42 \| (8, 49) \| 221 \| (497, 539) \| 666 \| tail fiber protein \| tail fiber protein; Category: tail; p80663 VI\_09756; phrogs \| 24267 \| 96.7 \| 3.1e-06 \| 3.5e-10 \| 68.7 \| 29 \| (93, 121) \| 221 \| (185, 213) \| 266 \| NA \| NA; Category: unknown function; p146200 VI\_04950; phrogs \| 4924 \| 96.6 \| 4.7e-06 \| 5.6e-10 \| 78.4 \| 67 \| (6, 74) \| 221 \| (537, 617) \| 981 \| short tail fiber protein \| short tail fiber protein; Category: tail; NC\_031900\_p33; phrogs \| 38064 \| 96.6 \| 5.5e-06 \| 6.2e-10 \| 67.4 \| 62 \| (3, 64) \| 221 \| (213, 276) \| 295 \| tail collar protein \| tail collar protein; Category: tail; NC\_029002\_p19; phrogs \| 31602 \| 96.2 \| 1.9e-05 \| 2.2e-09 \| 67.7 \| 67 \| (7, 73) \| 221 \| (11, 97) \| 428 \| virion structural protein \| virion structural protein; Category: head and packaging; NC\_013085\_p212; phrogs \| 2631 \| 95.7 \| 8.8e-05 \| 1e-08 \| 64.9 \| 33 \| (4, 36) \| 221 \| (179, 225) \| 482 \| tail fiber protein \| tail fiber protein; Category: tail; p130317 VI\_07372; phrogs \| 13173 \| 94.7 \| 0.0005 \| 5.8e-08 \| 63.3 \| 95 \| (12, 122) \| 221 \| (445, 557) \| 920 \| tail fiber protein \| tail fiber protein; Category: tail; MG201401\_p40 |
| Top keywords  (threshold 1.00e-03 (evalue)) | **tail, fiber, collar, p130317, VI\_07372, short, NC\_023006\_p17, NC\_019917\_p61, p284593, VI\_03272** |
| Output files | ../../domain\_architecture/17\_FANPEZAQ\_CDS\_0017\_cath.hhr ../../domain\_architecture/17\_FANPEZAQ\_CDS\_0017\_merged.svg ../../domain\_architecture/17\_FANPEZAQ\_CDS\_0017\_ncbi-cd.hhr ../../domain\_architecture/17\_FANPEZAQ\_CDS\_0017\_pfam.hhr ../../domain\_architecture/17\_FANPEZAQ\_CDS\_0017\_phrogs.hhr |

### Identical protein sequences/structures

#### Search results

|  |  |
| --- | --- |
| Protein sequence databases searched | Pdb, Swissprot, Refseq |
| Identical proteins found | -- |
| Top keywords | -- |
| Output files | -- |

### Similar protein sequences/structures

#### Sequence similarity search results (HHblits)^1^

|  |  |
| --- | --- |
| Sequence databases searched | Uniclust, Pdb70 |
| Results, scheme(s)  (Top layers only, threshold 1.00e-03 (evalue)) | xml version="1.0" encoding="utf-8" standalone="no"?       2024-09-02T21:08:36.325454 image/svg+xml   Matplotlib v3.7.2, https://matplotlib.org/ |
| Results, table(s)  (threshold 1.00e-03 (evalue)) | db \| id \| prob \| evalue \| pvalue \| score \| cols \| query \| query\_len \| template \| template\_len \| name \| description; uniclust \| UniRef100\_A0A0P0EA18 \| 100.0 \| 8e-41 \| 1.5e-46 \| 242.0 \| 180 \| (6, 221) \| 221 \| (63, 242) \| 247 \| Phage tail collar domain-containing protein \| Phage tail collar domain-containing protein; uniclust \| UniRef100\_A0A0M4T0P4 \| 100.0 \| 8.7e-38 \| 1.6e-43 \| 241.3 \| 200 \| (8, 221) \| 221 \| (157, 367) \| 368 \| Phage tail collar domain-containing protein \| Phage tail collar domain-containing protein; uniclust \| UniRef100\_A0A0M5IIJ7 \| 100.0 \| 1.6e-37 \| 2.9e-43 \| 234.2 \| 171 \| (9, 221) \| 221 \| (127, 297) \| 302 \| Microcystin-dependent protein-like protein \| Microcystin-dependent protein-like protein; uniclust \| UniRef100\_A0A813X408 \| 100.0 \| 1.4e-36 \| 2.7e-42 \| 222.5 \| 195 \| (8, 221) \| 221 \| (59, 256) \| 270 \| Phage tail collar domain-containing protein \| Phage tail collar domain-containing protein; uniclust \| UniRef100\_A0A1C2DSX6 \| 100.0 \| 6.8e-36 \| 1.3e-41 \| 214.0 \| 113 \| (6, 118) \| 221 \| (30, 142) \| 218 \| Phage tail collar domain-containing protein \| Phage tail collar domain-containing protein; uniclust \| UniRef100\_A0A0B3RIK5 \| 100.0 \| 3.2e-35 \| 6e-41 \| 217.5 \| 186 \| (7, 221) \| 221 \| (78, 263) \| 265 \| Phage Tail Collar \| Phage Tail Collar; uniclust \| UniRef100\_A0A1V1P4C0 \| 100.0 \| 8.1e-34 \| 1.5e-39 \| 219.8 \| 175 \| (10, 221) \| 221 \| (208, 386) \| 386 \| Microcystin-dependent protein-like protein \| Microcystin-dependent protein-like protein; uniclust \| UniRef100\_A0A1F3ABU4 \| 100.0 \| 8e-34 \| 1.5e-39 \| 199.8 \| 100 \| (8, 119) \| 221 \| (19, 118) \| 176 \| Phage tail collar domain-containing protein \| Phage tail collar domain-containing protein; uniclust \| UniRef100\_A0A813P3R8 \| 100.0 \| 9.5e-34 \| 1.8e-39 \| 199.8 \| 174 \| (9, 221) \| 221 \| (38, 211) \| 220 \| Phage tail collar domain-containing protein \| Phage tail collar domain-containing protein; uniclust \| UniRef100\_A0A1V4VVI6 \| 100.0 \| 3.4e-33 \| 6.4e-39 \| 213.1 \| 178 \| (7, 221) \| 221 \| (205, 382) \| 386 \| Phage Tail Collar Domain protein \| Phage Tail Collar Domain protein; uniclust \| UniRef100\_A0A3D0M2I4 \| 100.0 \| 4.4e-33 \| 8.2e-39 \| 199.0 \| 99 \| (8, 119) \| 221 \| (71, 169) \| 230 \| Phage tail collar domain-containing protein \| Phage tail collar domain-containing protein; uniclust \| UniRef100\_A0A6J5M5U3 \| 100.0 \| 1.6e-32 \| 3e-38 \| 197.3 \| 94 \| (8, 119) \| 221 \| (72, 165) \| 238 \| MdpB Microcystin-dependent protein \| MdpB Microcystin-dependent protein; uniclust \| UniRef100\_A0A074J4T7 \| 100.0 \| 2.6e-32 \| 4.8e-38 \| 188.4 \| 100 \| (8, 119) \| 221 \| (48, 147) \| 194 \| Phage tail collar domain-containing protein \| Phage tail collar domain-containing protein; uniclust \| UniRef100\_A0A2E7V2E5 \| 99.9 \| 3.8e-32 \| 7e-38 \| 203.8 \| 108 \| (7, 117) \| 221 \| (152, 259) \| 325 \| Phage tail collar domain-containing protein \| Phage tail collar domain-containing protein; uniclust \| UniRef100\_A0A0H4IKC1 \| 99.9 \| 4.2e-32 \| 7.8e-38 \| 198.2 \| 181 \| (8, 221) \| 221 \| (82, 262) \| 264 \| Putative tail collar protein \| Putative tail collar protein; uniclust \| UniRef100\_A0A0D7Q2R9 \| 99.9 \| 7.7e-32 \| 1.4e-37 \| 207.0 \| 116 \| (9, 126) \| 221 \| (133, 248) \| 335 \| Phage tail collar domain-containing protein \| Phage tail collar domain-containing protein; uniclust \| UniRef100\_A0A813VP22 \| 99.9 \| 1.2e-31 \| 2.1e-37 \| 200.4 \| 196 \| (7, 221) \| 221 \| (136, 339) \| 346 \| Phage tail collar domain-containing protein \| Phage tail collar domain-containing protein; uniclust \| UniRef100\_A0A177Q1Y6 \| 99.9 \| 5.3e-31 \| 9.9e-37 \| 195.1 \| 108 \| (8, 119) \| 221 \| (132, 239) \| 301 \| Phage tail collar domain-containing protein \| Phage tail collar domain-containing protein; uniclust \| UniRef100\_A0A814BNA3 \| 99.9 \| 7.4e-31 \| 1.4e-36 \| 190.3 \| 176 \| (7, 221) \| 221 \| (104, 279) \| 280 \| Phage tail collar domain-containing protein \| Phage tail collar domain-containing protein; uniclust \| UniRef100\_A0A814NW00 \| 99.9 \| 8.6e-31 \| 1.6e-36 \| 216.5 \| 198 \| (10, 221) \| 221 \| (585, 787) \| 792 \| Phage tail collar domain-containing protein \| Phage tail collar domain-containing protein; uniclust \| UniRef100\_A0A6N4T611 \| 99.9 \| 1.1e-30 \| 2e-36 \| 178.0 \| 90 \| (8, 119) \| 221 \| (21, 110) \| 168 \| Phage tail collar domain-containing protein \| Phage tail collar domain-containing protein; uniclust \| UniRef100\_A0A2E4T1Y0 \| 99.9 \| 1.4e-30 \| 2.6e-36 \| 189.3 \| 178 \| (10, 221) \| 221 \| (108, 286) \| 286 \| Phage tail collar domain-containing protein \| Phage tail collar domain-containing protein; uniclust \| UniRef100\_A0A6C0EEV3 \| 99.9 \| 2e-30 \| 3.7e-36 \| 191.4 \| 173 \| (8, 221) \| 221 \| (176, 348) \| 349 \| Phage tail collar domain-containing protein \| Phage tail collar domain-containing protein; uniclust \| UniRef100\_A0A060DEV3 \| 99.9 \| 4.2e-30 \| 7.9e-36 \| 177.4 \| 158 \| (7, 221) \| 221 \| (28, 186) \| 186 \| Phage tail collar domain-containing protein \| Phage tail collar domain-containing protein; uniclust \| UniRef100\_A0A1G0C5V8 \| 99.9 \| 8e-30 \| 1.5e-35 \| 193.7 \| 157 \| (10, 221) \| 221 \| (116, 276) \| 280 \| Phage tail collar domain-containing protein \| Phage tail collar domain-containing protein; uniclust \| UniRef100\_A0A0Q6Z9B8 \| 99.9 \| 9.4e-30 \| 1.7e-35 \| 196.1 \| 111 \| (9, 120) \| 221 \| (202, 313) \| 387 \| Phage tail collar domain-containing protein \| Phage tail collar domain-containing protein; uniclust \| UniRef100\_A0A074J2N7 \| 99.9 \| 1.3e-29 \| 2.4e-35 \| 178.8 \| 97 \| (10, 120) \| 221 \| (6, 102) \| 188 \| Phage tail collar domain-containing protein \| Phage tail collar domain-containing protein; uniclust \| UniRef100\_A0A814FG22 \| 99.9 \| 1.6e-29 \| 3e-35 \| 183.5 \| 197 \| (8, 221) \| 221 \| (41, 245) \| 255 \| Phage tail collar domain-containing protein \| Phage tail collar domain-containing protein; uniclust \| UniRef100\_A0A1F3PPS7 \| 99.9 \| 1.6e-29 \| 3.1e-35 \| 196.5 \| 159 \| (9, 221) \| 221 \| (156, 317) \| 319 \| Phage tail collar domain-containing protein \| Phage tail collar domain-containing protein; uniclust \| UniRef100\_A0A813X8J9 \| 99.9 \| 2e-29 \| 3.8e-35 \| 198.7 \| 196 \| (9, 221) \| 221 \| (262, 465) \| 466 \| Phage tail collar domain-containing protein \| Phage tail collar domain-containing protein; uniclust \| UniRef100\_A0A0B4EK47 \| 99.9 \| 2.9e-29 \| 5.4e-35 \| 184.8 \| 99 \| (9, 120) \| 221 \| (59, 157) \| 237 \| Phage tail collar domain-containing protein \| Phage tail collar domain-containing protein; uniclust \| UniRef100\_A0A0L0ENS5 \| 99.9 \| 5.2e-29 \| 9.8e-35 \| 187.1 \| 98 \| (9, 119) \| 221 \| (50, 147) \| 249 \| Phage tail collar domain-containing protein \| Phage tail collar domain-containing protein; uniclust \| UniRef100\_A0A063KYT2 \| 99.9 \| 5.5e-29 \| 1e-34 \| 184.2 \| 99 \| (9, 120) \| 221 \| (49, 147) \| 233 \| Phage tail protein \| Phage tail protein; uniclust \| UniRef100\_A0A0Q0XB69 \| 99.9 \| 7.2e-29 \| 1.3e-34 \| 178.6 \| 97 \| (10, 119) \| 221 \| (45, 141) \| 219 \| Phage tail collar domain-containing protein \| Phage tail collar domain-containing protein; uniclust \| UniRef100\_A0A010YCA6 \| 99.9 \| 8e-29 \| 1.5e-34 \| 183.5 \| 99 \| (9, 120) \| 221 \| (30, 128) \| 228 \| Microcystin-dependent protein \| Microcystin-dependent protein; uniclust \| UniRef100\_A0A0C3ENX2 \| 99.9 \| 1.1e-28 \| 2.1e-34 \| 180.4 \| 97 \| (9, 118) \| 221 \| (32, 128) \| 213 \| Tail protein \| Tail protein; uniclust \| UniRef100\_A0A077L9W7 \| 99.9 \| 1.4e-28 \| 2.7e-34 \| 180.8 \| 100 \| (8, 120) \| 221 \| (26, 125) \| 249 \| Tail Collar domain protein \| Tail Collar domain protein; uniclust \| UniRef100\_A0A0D6AS89 \| 99.9 \| 1.5e-28 \| 2.8e-34 \| 180.6 \| 92 \| (7, 120) \| 221 \| (52, 145) \| 216 \| Microcystin dependent protein \| Microcystin dependent protein; uniclust \| UniRef100\_A0A0U4JQ24 \| 99.9 \| 4.3e-28 \| 7.9e-34 \| 186.0 \| 97 \| (10, 119) \| 221 \| (59, 155) \| 455 \| Tail protein \| Tail protein; uniclust \| UniRef100\_A0A0A1DL90 \| 99.9 \| 4.5e-28 \| 8.3e-34 \| 166.9 \| 99 \| (9, 120) \| 221 \| (12, 110) \| 180 \| Microcystin dependent protein \| Microcystin dependent protein; uniclust \| UniRef100\_A0A5C7MBK6 \| 99.9 \| 6.6e-28 \| 1.2e-33 \| 182.3 \| 182 \| (9, 221) \| 221 \| (204, 385) \| 387 \| Tail fiber protein \| Tail fiber protein; uniclust \| UniRef100\_A0A157ZM93 \| 99.9 \| 8.8e-28 \| 1.6e-33 \| 175.4 \| 170 \| (9, 221) \| 221 \| (83, 266) \| 269 \| Trimeric autotransporter adhesin YadA-like stalk domain-containing protein \| Trimeric autotransporter adhesin YadA-like stalk domain-containing protein; uniclust \| UniRef100\_A0A023Y4H2 \| 99.9 \| 9e-28 \| 1.7e-33 \| 179.6 \| 98 \| (9, 119) \| 221 \| (51, 148) \| 242 \| Phage tail collar domain-containing protein \| Phage tail collar domain-containing protein; uniclust \| UniRef100\_A0A160TGR6 \| 99.9 \| 9.5e-28 \| 1.8e-33 \| 178.0 \| 99 \| (9, 120) \| 221 \| (34, 132) \| 243 \| Microcystin dependent protein \| Microcystin dependent protein; uniclust \| UniRef100\_A0A814Z5L0 \| 99.9 \| 1e-27 \| 1.9e-33 \| 177.1 \| 189 \| (8, 221) \| 221 \| (117, 320) \| 322 \| Phage tail collar domain-containing protein \| Phage tail collar domain-containing protein; uniclust \| UniRef100\_A0A0Q6S010 \| 99.9 \| 1.1e-27 \| 2e-33 \| 179.1 \| 97 \| (10, 119) \| 221 \| (41, 137) \| 247 \| Phage tail collar domain-containing protein \| Phage tail collar domain-containing protein; uniclust \| UniRef100\_A0A073AGH6 \| 99.9 \| 1.1e-27 \| 2e-33 \| 176.6 \| 94 \| (9, 120) \| 221 \| (48, 141) \| 227 \| Phage tail protein \| Phage tail protein; uniclust \| UniRef100\_UPI000693AA4D \| 99.9 \| 1.1e-27 \| 2.1e-33 \| 184.8 \| 117 \| (9, 138) \| 221 \| (377, 501) \| 614 \| tail fiber protein \| tail fiber protein; uniclust \| UniRef100\_A0A814K3I7 \| 99.9 \| 1.4e-27 \| 2.5e-33 \| 186.3 \| 189 \| (11, 221) \| 221 \| (286, 484) \| 489 \| Phage tail collar domain-containing protein \| Phage tail collar domain-containing protein; uniclust \| UniRef100\_A0A072T796 \| 99.9 \| 1.5e-27 \| 2.7e-33 \| 180.1 \| 98 \| (9, 119) \| 221 \| (56, 153) \| 267 \| Tail collar protein \| Tail collar protein; uniclust \| UniRef100\_A0A010ZGK4 \| 99.9 \| 1.5e-27 \| 2.9e-33 \| 180.6 \| 98 \| (9, 119) \| 221 \| (70, 167) \| 255 \| Microcystin-dependent protein \| Microcystin-dependent protein; uniclust \| UniRef100\_A0A0A8WR50 \| 99.9 \| 1.8e-27 \| 3.5e-33 \| 184.4 \| 145 \| (9, 221) \| 221 \| (143, 290) \| 291 \| Phage Tail Collar Domain protein \| Phage Tail Collar Domain protein; uniclust \| UniRef100\_A0A8S5MVZ6 \| 99.9 \| 2.1e-27 \| 3.9e-33 \| 170.3 \| 92 \| (8, 120) \| 221 \| (88, 179) \| 242 \| Baseplate structural protein \| Baseplate structural protein; uniclust \| UniRef100\_A0A011A738 \| 99.9 \| 2.1e-27 \| 3.9e-33 \| 182.1 \| 98 \| (9, 119) \| 221 \| (67, 164) \| 292 \| Microcystin-dependent protein \| Microcystin-dependent protein; uniclust \| UniRef100\_A0A813NFR6 \| 99.9 \| 2.3e-27 \| 4.2e-33 \| 195.9 \| 195 \| (9, 221) \| 221 \| (499, 702) \| 707 \| Phage tail collar domain-containing protein \| Phage tail collar domain-containing protein; uniclust \| UniRef100\_A0A1E4LGS2 \| 99.9 \| 2.5e-27 \| 4.6e-33 \| 186.8 \| 124 \| (9, 139) \| 221 \| (224, 347) \| 432 \| Phage tail collar domain-containing protein \| Phage tail collar domain-containing protein; uniclust \| UniRef100\_UPI0018C60E32 \| 99.9 \| 2.7e-27 \| 5e-33 \| 174.5 \| 175 \| (7, 221) \| 221 \| (204, 378) \| 378 \| tail fiber protein \| tail fiber protein; uniclust \| UniRef100\_A0A1G8XFZ8 \| 99.9 \| 3.1e-27 \| 5.7e-33 \| 179.8 \| 172 \| (7, 221) \| 221 \| (148, 321) \| 323 \| Microcystin-dependent protein \| Microcystin-dependent protein; uniclust \| UniRef100\_A0A1G6EJM0 \| 99.9 \| 3.5e-27 \| 6.4e-33 \| 170.4 \| 192 \| (7, 220) \| 221 \| (117, 313) \| 316 \| Microcystin-dependent protein \| Microcystin-dependent protein; uniclust \| UniRef100\_A0A2I7SF00 \| 99.9 \| 4.3e-27 \| 8.1e-33 \| 178.3 \| 144 \| (9, 221) \| 221 \| (146, 291) \| 293 \| Phage tail collar domain-containing protein \| Phage tail collar domain-containing protein; uniclust \| UniRef100\_A0A0F9MAR7 \| 99.9 \| 5e-27 \| 9.4e-33 \| 167.6 \| 70 \| (7, 119) \| 221 \| (27, 98) \| 178 \| Phage tail collar domain-containing protein \| Phage tail collar domain-containing protein; uniclust \| UniRef100\_A0A150TEA7 \| 99.9 \| 5.1e-27 \| 9.4e-33 \| 190.2 \| 164 \| (8, 221) \| 221 \| (499, 664) \| 664 \| Phage tail collar domain-containing protein \| Phage tail collar domain-containing protein; uniclust \| UniRef100\_A0A0P9DVX3 \| 99.9 \| 7.3e-27 \| 1.4e-32 \| 183.5 \| 97 \| (10, 119) \| 221 \| (215, 311) \| 389 \| Phage tail collar domain-containing protein \| Phage tail collar domain-containing protein; uniclust \| UniRef100\_A0A163AZI2 \| 99.9 \| 9.7e-27 \| 1.8e-32 \| 180.0 \| 153 \| (9, 221) \| 221 \| (232, 384) \| 387 \| Phage tail collar domain-containing protein \| Phage tail collar domain-containing protein; uniclust \| UniRef100\_A0A2D6U0S1 \| 99.9 \| 1.9e-26 \| 3.5e-32 \| 172.4 \| 109 \| (11, 119) \| 221 \| (77, 191) \| 271 \| Phage tail collar domain-containing protein \| Phage tail collar domain-containing protein; uniclust \| UniRef100\_A0A9E8ELC9 \| 99.9 \| 2e-26 \| 3.7e-32 \| 162.4 \| 97 \| (10, 119) \| 221 \| (108, 204) \| 273 \| Tail fiber protein \| Tail fiber protein; uniclust \| UniRef100\_A0A522ETL0 \| 99.9 \| 2.8e-26 \| 5.2e-32 \| 165.1 \| 106 \| (9, 119) \| 221 \| (83, 188) \| 243 \| Tail fiber protein \| Tail fiber protein; uniclust \| UniRef100\_A0A1H3XG57 \| 99.9 \| 2.9e-26 \| 5.5e-32 \| 161.2 \| 97 \| (11, 120) \| 221 \| (10, 106) \| 182 \| Microcystin-dependent protein \| Microcystin-dependent protein; uniclust \| UniRef100\_A0A815GMY2 \| 99.9 \| 3.3e-26 \| 6.2e-32 \| 149.1 \| 121 \| (7, 144) \| 221 \| (14, 134) \| 135 \| Phage tail collar domain-containing protein \| Phage tail collar domain-containing protein; uniclust \| UniRef100\_A0A0R3DQ44 \| 99.9 \| 3.3e-26 \| 6.2e-32 \| 179.2 \| 112 \| (8, 119) \| 221 \| (196, 308) \| 383 \| Phage tail collar domain-containing protein \| Phage tail collar domain-containing protein; uniclust \| UniRef100\_A0A7C5Z090 \| 99.9 \| 5.8e-26 \| 1.1e-31 \| 163.2 \| 102 \| (6, 120) \| 221 \| (165, 266) \| 315 \| Phage tail collar domain-containing protein \| Phage tail collar domain-containing protein; uniclust \| UniRef100\_A0A173LYL4 \| 99.9 \| 5.9e-26 \| 1.1e-31 \| 182.7 \| 104 \| (9, 127) \| 221 \| (542, 645) \| 704 \| Collagen triple helix repeat protein \| Collagen triple helix repeat protein; uniclust \| UniRef100\_A0A1Z8ZQ15 \| 99.9 \| 5.8e-26 \| 1.1e-31 \| 175.1 \| 139 \| (8, 221) \| 221 \| (212, 352) \| 353 \| Phage tail collar domain-containing protein \| Phage tail collar domain-containing protein; uniclust \| UniRef100\_A0A2W6Y1Z1 \| 99.9 \| 6.1e-26 \| 1.1e-31 \| 172.7 \| 168 \| (11, 221) \| 221 \| (124, 302) \| 312 \| Phage tail protein \| Phage tail protein; uniclust \| UniRef100\_A0A813ML39 \| 99.9 \| 1.1e-25 \| 2e-31 \| 181.2 \| 194 \| (10, 220) \| 221 \| (548, 751) \| 753 \| Phage tail collar domain-containing protein \| Phage tail collar domain-containing protein; uniclust \| UniRef100\_A0A940VND6 \| 99.9 \| 1.1e-25 \| 2.1e-31 \| 168.9 \| 95 \| (9, 119) \| 221 \| (270, 364) \| 416 \| Phage tail collar domain-containing protein \| Phage tail collar domain-containing protein; uniclust \| UniRef100\_A0A0Q5Z2P4 \| 99.9 \| 1.4e-25 \| 2.7e-31 \| 175.5 \| 104 \| (10, 119) \| 221 \| (148, 251) \| 421 \| Phage tail collar domain-containing protein \| Phage tail collar domain-containing protein; uniclust \| UniRef100\_A0A0C2D114 \| 99.9 \| 1.5e-25 \| 2.8e-31 \| 164.2 \| 143 \| (8, 221) \| 221 \| (70, 214) \| 216 \| Phage tail fiber protein \| Phage tail fiber protein; uniclust \| UniRef100\_A0A0P9DVX3 \| 99.9 \| 1.6e-25 \| 3.1e-31 \| 176.1 \| 92 \| (10, 119) \| 221 \| (47, 138) \| 389 \| Phage tail collar domain-containing protein \| Phage tail collar domain-containing protein; uniclust \| UniRef100\_A0A8S5RTE1 \| 99.9 \| 1.7e-25 \| 3.1e-31 \| 159.9 \| 98 \| (7, 121) \| 221 \| (86, 183) \| 249 \| Tail collar domain \| Tail collar domain; uniclust \| UniRef100\_A0A369RCN8 \| 99.9 \| 2.6e-25 \| 4.9e-31 \| 158.9 \| 98 \| (9, 119) \| 221 \| (115, 212) \| 275 \| Phage tail collar domain-containing protein \| Phage tail collar domain-containing protein; uniclust \| UniRef100\_A0A031GQD8 \| 99.9 \| 3.3e-25 \| 6.1e-31 \| 154.6 \| 102 \| (9, 119) \| 221 \| (3, 104) \| 177 \| Microcystin dependent protein \| Microcystin dependent protein; uniclust \| UniRef100\_A0A060DHI4 \| 99.9 \| 4.1e-25 \| 7.6e-31 \| 166.4 \| 99 \| (10, 121) \| 221 \| (47, 145) \| 267 \| Phage tail collar domain-containing protein \| Phage tail collar domain-containing protein; uniclust \| UniRef100\_A0A8T3YWH3 \| 99.9 \| 4.3e-25 \| 7.9e-31 \| 158.5 \| 94 \| (9, 126) \| 221 \| (88, 181) \| 241 \| Tail fiber protein \| Tail fiber protein; uniclust \| UniRef100\_A0A0A1FCT1 \| 99.9 \| 4.2e-25 \| 7.9e-31 \| 155.3 \| 98 \| (9, 119) \| 221 \| (8, 105) \| 177 \| Microcystin dependent protein \| Microcystin dependent protein; uniclust \| UniRef100\_UPI0020119B83 \| 99.8 \| 5.8e-25 \| 1.1e-30 \| 157.1 \| 99 \| (9, 119) \| 221 \| (155, 253) \| 299 \| tail fiber protein \| tail fiber protein; uniclust \| UniRef100\_A0A813PLM5 \| 99.8 \| 5.9e-25 \| 1.1e-30 \| 154.4 \| 193 \| (12, 221) \| 221 \| (19, 219) \| 220 \| Uncharacterized protein \| Uncharacterized protein; uniclust \| UniRef100\_A0A0Q5ZY68 \| 99.8 \| 6.1e-25 \| 1.1e-30 \| 167.9 \| 112 \| (6, 118) \| 221 \| (136, 247) \| 316 \| Phage tail collar domain-containing protein \| Phage tail collar domain-containing protein; uniclust \| UniRef100\_A0A0A1PS49 \| 99.8 \| 1.3e-24 \| 2.5e-30 \| 163.9 \| 185 \| (7, 221) \| 221 \| (194, 379) \| 379 \| Phage Tail Collar Domain protein \| Phage Tail Collar Domain protein; uniclust \| UniRef100\_A0A4Y6CGU0 \| 99.8 \| 1.4e-24 \| 2.6e-30 \| 180.5 \| 151 \| (8, 221) \| 221 \| (546, 698) \| 718 \| Phage tail collar domain-containing protein \| Phage tail collar domain-containing protein; uniclust \| UniRef100\_UPI0018C7C22C \| 99.8 \| 1.4e-24 \| 2.6e-30 \| 158.7 \| 177 \| (8, 221) \| 221 \| (181, 357) \| 357 \| tail fiber protein \| tail fiber protein; uniclust \| UniRef100\_A0A6L5FV64 \| 99.8 \| 1.5e-24 \| 2.9e-30 \| 155.0 \| 98 \| (9, 120) \| 221 \| (111, 208) \| 268 \| Phage tail collar domain-containing protein \| Phage tail collar domain-containing protein; uniclust \| UniRef100\_A0A481Z0P9 \| 99.8 \| 1.6e-24 \| 3.1e-30 \| 158.2 \| 74 \| (9, 120) \| 221 \| (96, 169) \| 231 \| Tail collar domain protein \| Tail collar domain protein; uniclust \| UniRef100\_A0A3N6N9Q9 \| 99.8 \| 1.8e-24 \| 3.3e-30 \| 163.8 \| 97 \| (10, 119) \| 221 \| (139, 235) \| 316 \| Phage tail collar domain-containing protein \| Phage tail collar domain-containing protein; uniclust \| UniRef100\_A0A059FRJ9 \| 99.8 \| 1.9e-24 \| 3.5e-30 \| 166.6 \| 98 \| (9, 119) \| 221 \| (60, 157) \| 272 \| Phage tail collar domain-containing protein \| Phage tail collar domain-containing protein; uniclust \| UniRef100\_A0A943I3K4 \| 99.8 \| 3e-24 \| 5.5e-30 \| 143.2 \| 88 \| (11, 115) \| 221 \| (4, 91) \| 165 \| Tail fiber protein \| Tail fiber protein; uniclust \| UniRef100\_A0A3M1ZWN2 \| 99.8 \| 4e-24 \| 7.3e-30 \| 153.6 \| 97 \| (10, 119) \| 221 \| (34, 131) \| 269 \| Phage tail collar domain-containing protein \| Phage tail collar domain-containing protein; uniclust \| UniRef100\_A0A1X7I1H1 \| 99.8 \| 4.3e-24 \| 8e-30 \| 166.0 \| 93 \| (9, 119) \| 221 \| (181, 273) \| 346 \| Microcystin-dependent protein \| Microcystin-dependent protein; uniclust \| UniRef100\_A0A5K0UBR7 \| 99.8 \| 4.6e-24 \| 8.6e-30 \| 158.2 \| 72 \| (8, 119) \| 221 \| (94, 165) \| 264 \| Uncharacterized protein \| Uncharacterized protein; uniclust \| UniRef100\_A0A0F8YBQ7 \| 99.8 \| 5.8e-24 \| 1.1e-29 \| 154.4 \| 93 \| (9, 120) \| 221 \| (74, 166) \| 227 \| Phage tail collar domain-containing protein (Fragment) \| Phage tail collar domain-containing protein (Fragment); uniclust \| UniRef100\_A0A1Y4GY33 \| 99.8 \| 6.1e-24 \| 1.1e-29 \| 157.7 \| 97 \| (8, 121) \| 221 \| (184, 280) \| 339 \| Phage tail collar domain-containing protein \| Phage tail collar domain-containing protein; uniclust \| UniRef100\_A0A7V2Y2J7 \| 99.8 \| 6.6e-24 \| 1.2e-29 \| 160.9 \| 112 \| (7, 120) \| 221 \| (150, 261) \| 330 \| Phage tail collar domain-containing protein \| Phage tail collar domain-containing protein; uniclust \| UniRef100\_A0A516KT37 \| 99.8 \| 6.7e-24 \| 1.2e-29 \| 160.8 \| 99 \| (9, 120) \| 221 \| (263, 363) \| 430 \| Minor tail protein \| Minor tail protein; uniclust \| UniRef100\_A0A6C0HZC1 \| 99.8 \| 6.9e-24 \| 1.3e-29 \| 151.5 \| 191 \| (8, 221) \| 221 \| (42, 256) \| 256 \| Phage tail collar domain-containing protein \| Phage tail collar domain-containing protein; uniclust \| UniRef100\_A0A1X7I1H1 \| 99.8 \| 6.9e-24 \| 1.3e-29 \| 164.9 \| 98 \| (9, 119) \| 221 \| (7, 104) \| 346 \| Microcystin-dependent protein \| Microcystin-dependent protein; uniclust \| UniRef100\_A0A1F3IL82 \| 99.8 \| 7.1e-24 \| 1.3e-29 \| 160.8 \| 86 \| (9, 118) \| 221 \| (141, 228) \| 319 \| Phage tail collar domain-containing protein \| Phage tail collar domain-containing protein; uniclust \| UniRef100\_A0A937XD69 \| 99.8 \| 7.5e-24 \| 1.4e-29 \| 144.5 \| 146 \| (7, 221) \| 221 \| (65, 210) \| 211 \| Tail fiber protein \| Tail fiber protein; uniclust \| UniRef100\_UPI000A039BE3 \| 99.8 \| 8.3e-24 \| 1.5e-29 \| 157.5 \| 102 \| (9, 122) \| 221 \| (239, 340) \| 412 \| tail fiber protein \| tail fiber protein; uniclust \| UniRef100\_A0A7G9A427 \| 99.8 \| 9.3e-24 \| 1.7e-29 \| 157.7 \| 197 \| (8, 221) \| 221 \| (222, 423) \| 424 \| Phage tail collar domain-containing protein \| Phage tail collar domain-containing protein; uniclust \| UniRef100\_A0A7K0RNX1 \| 99.8 \| 1.1e-23 \| 2e-29 \| 162.1 \| 105 \| (10, 120) \| 221 \| (344, 448) \| 562 \| Phage tail collar domain-containing protein \| Phage tail collar domain-containing protein; uniclust \| UniRef100\_A0A386HRX7 \| 99.8 \| 1.2e-23 \| 2.2e-29 \| 158.7 \| 69 \| (9, 120) \| 221 \| (158, 228) \| 295 \| Tail fiber protein \| Tail fiber protein; uniclust \| UniRef100\_A0A2M7SZQ3 \| 99.8 \| 1.3e-23 \| 2.5e-29 \| 165.3 \| 106 \| (8, 119) \| 221 \| (249, 357) \| 438 \| Phage tail collar domain-containing protein \| Phage tail collar domain-containing protein; uniclust \| UniRef100\_A0A2S5DRN7 \| 99.8 \| 1.8e-23 \| 3.3e-29 \| 150.1 \| 170 \| (9, 221) \| 221 \| (104, 281) \| 282 \| Phage tail protein \| Phage tail protein; uniclust \| UniRef100\_A0A150WBY6 \| 99.8 \| 1.8e-23 \| 3.5e-29 \| 157.1 \| 110 \| (6, 120) \| 221 \| (86, 195) \| 257 \| Phage tail collar domain-containing protein \| Phage tail collar domain-containing protein; uniclust \| UniRef100\_A0A3A8TU84 \| 99.8 \| 2.1e-23 \| 4e-29 \| 142.0 \| 97 \| (10, 119) \| 221 \| (4, 100) \| 144 \| Phage tail protein (Fragment) \| Phage tail protein (Fragment); uniclust \| UniRef100\_A0A1A6CD73 \| 99.8 \| 2.4e-23 \| 4.7e-29 \| 164.1 \| 142 \| (11, 221) \| 221 \| (144, 287) \| 291 \| Phage tail collar domain-containing protein \| Phage tail collar domain-containing protein; uniclust \| UniRef100\_A0A8J7Q9N7 \| 99.8 \| 2.6e-23 \| 4.7e-29 \| 151.2 \| 104 \| (8, 124) \| 221 \| (120, 224) \| 289 \| Tail fiber protein \| Tail fiber protein; uniclust \| UniRef100\_A0A974Z143 \| 99.8 \| 2.6e-23 \| 4.8e-29 \| 160.3 \| 99 \| (8, 119) \| 221 \| (387, 485) \| 538 \| Tail fiber protein \| Tail fiber protein; uniclust \| UniRef100\_A0A2L1HSI4 \| 99.8 \| 2.6e-23 \| 4.9e-29 \| 157.0 \| 172 \| (8, 221) \| 221 \| (178, 349) \| 349 \| Phage tail collar domain-containing protein \| Phage tail collar domain-containing protein; uniclust \| UniRef100\_A0A969SMF4 \| 99.8 \| 3.9e-23 \| 7.1e-29 \| 155.7 \| 195 \| (10, 220) \| 221 \| (246, 450) \| 453 \| Phage tail collar domain-containing protein \| Phage tail collar domain-containing protein; uniclust \| UniRef100\_A0A3D1PJI0 \| 99.8 \| 4.1e-23 \| 7.6e-29 \| 150.2 \| 99 \| (8, 119) \| 221 \| (44, 142) \| 337 \| Phage tail collar domain-containing protein \| Phage tail collar domain-containing protein; uniclust \| UniRef100\_A0A661AU03 \| 99.8 \| 4.1e-23 \| 7.6e-29 \| 162.9 \| 171 \| (9, 221) \| 221 \| (552, 722) \| 722 \| Phage tail collar domain-containing protein (Fragment) \| Phage tail collar domain-containing protein (Fragment); uniclust \| UniRef100\_A0A1D9LMZ6 \| 99.8 \| 5.4e-23 \| 1e-28 \| 152.3 \| 94 \| (8, 120) \| 221 \| (99, 192) \| 252 \| Phage tail collar domain-containing protein \| Phage tail collar domain-containing protein; uniclust \| UniRef100\_E3T4R8 \| 99.8 \| 6.5e-23 \| 1.2e-28 \| 142.9 \| 99 \| (9, 119) \| 221 \| (77, 175) \| 242 \| Phage tail collar domain-containing protein \| Phage tail collar domain-containing protein; uniclust \| UniRef100\_A0A1Z9FAS1 \| 99.8 \| 6.8e-23 \| 1.3e-28 \| 169.4 \| 107 \| (9, 119) \| 221 \| (493, 601) \| 679 \| Phage tail collar domain-containing protein \| Phage tail collar domain-containing protein; uniclust \| UniRef100\_A0A1Z7ZRI5 \| 99.8 \| 6.9e-23 \| 1.3e-28 \| 159.7 \| 107 \| (8, 120) \| 221 \| (125, 231) \| 382 \| Phage tail collar domain-containing protein \| Phage tail collar domain-containing protein; uniclust \| UniRef100\_A0A098S053 \| 99.8 \| 1e-22 \| 1.9e-28 \| 147.3 \| 92 \| (9, 119) \| 221 \| (65, 156) \| 219 \| Phage tail collar domain-containing protein \| Phage tail collar domain-containing protein; uniclust \| UniRef100\_D3F201 \| 99.8 \| 1.2e-22 \| 2.1e-28 \| 147.4 \| 184 \| (18, 221) \| 221 \| (73, 260) \| 262 \| Tail Collar domain protein \| Tail Collar domain protein; uniclust \| UniRef100\_A0A7Y7P107 \| 99.8 \| 1.2e-22 \| 2.3e-28 \| 161.4 \| 108 \| (8, 120) \| 221 \| (213, 321) \| 380 \| Tail fiber protein \| Tail fiber protein; uniclust \| UniRef100\_A0A2E1UKG0 \| 99.8 \| 1.3e-22 \| 2.3e-28 \| 159.8 \| 169 \| (8, 221) \| 221 \| (305, 490) \| 491 \| Major tropism determinant N-terminal domain-containing protein (Fragment) \| Major tropism determinant N-terminal domain-containing protein (Fragment); uniclust \| UniRef100\_A0A844YWZ8 \| 99.8 \| 1.3e-22 \| 2.3e-28 \| 146.4 \| 93 \| (8, 119) \| 221 \| (115, 207) \| 254 \| Phage tail protein \| Phage tail protein; uniclust \| UniRef100\_A0A6P1ZJ46 \| 99.8 \| 1.3e-22 \| 2.4e-28 \| 161.9 \| 185 \| (5, 221) \| 221 \| (248, 438) \| 439 \| Phage tail collar domain-containing protein \| Phage tail collar domain-containing protein; uniclust \| UniRef100\_A0A1V5QDU0 \| 99.8 \| 1.4e-22 \| 2.6e-28 \| 171.9 \| 109 \| (8, 120) \| 221 \| (1889, 1997) \| 2045 \| Phage Tail Collar Domain protein \| Phage Tail Collar Domain protein; uniclust \| UniRef100\_A0A1Q8TCI9 \| 99.8 \| 1.4e-22 \| 2.7e-28 \| 152.6 \| 88 \| (13, 119) \| 221 \| (108, 195) \| 290 \| Phage tail collar domain-containing protein \| Phage tail collar domain-containing protein; uniclust \| UniRef100\_A0A2A3DSC8 \| 99.8 \| 1.7e-22 \| 3.1e-28 \| 153.3 \| 90 \| (10, 118) \| 221 \| (131, 220) \| 282 \| Phage tail collar domain-containing protein \| Phage tail collar domain-containing protein; uniclust \| UniRef100\_A0A143DGR5 \| 99.8 \| 1.7e-22 \| 3.1e-28 \| 162.2 \| 93 \| (7, 118) \| 221 \| (266, 358) \| 425 \| Phage tail collar domain-containing protein \| Phage tail collar domain-containing protein; uniclust \| UniRef100\_A0A074JN26 \| 99.8 \| 2.1e-22 \| 3.9e-28 \| 146.9 \| 98 \| (10, 120) \| 221 \| (54, 151) \| 231 \| Phage tail collar domain-containing protein \| Phage tail collar domain-containing protein; uniclust \| UniRef100\_A0A158D1B4 \| 99.8 \| 2.2e-22 \| 4e-28 \| 151.3 \| 193 \| (7, 221) \| 221 \| (172, 373) \| 373 \| Phage tail collar domain-containing protein \| Phage tail collar domain-containing protein; uniclust \| UniRef100\_D0LMW0 \| 99.8 \| 3e-22 \| 5.5e-28 \| 141.4 \| 95 \| (13, 119) \| 221 \| (113, 207) \| 264 \| Tail Collar domain protein \| Tail Collar domain protein; uniclust \| UniRef100\_A0A820P0S5 \| 99.8 \| 3.1e-22 \| 5.7e-28 \| 153.4 \| 194 \| (13, 220) \| 221 \| (250, 447) \| 449 \| Phage tail collar domain-containing protein \| Phage tail collar domain-containing protein; uniclust \| UniRef100\_A0A8S5PNQ0 \| 99.8 \| 3.8e-22 \| 6.9e-28 \| 140.7 \| 105 \| (7, 126) \| 221 \| (63, 167) \| 231 \| Baseplate wedge protein \| Baseplate wedge protein; uniclust \| UniRef100\_UPI000CE3D78C \| 99.8 \| 3.8e-22 \| 7e-28 \| 149.2 \| 100 \| (8, 120) \| 221 \| (269, 368) \| 419 \| tail fiber protein \| tail fiber protein; uniclust \| UniRef100\_J0MTG7 \| 99.8 \| 4.4e-22 \| 8.2e-28 \| 145.0 \| 87 \| (9, 119) \| 221 \| (160, 246) \| 310 \| Phage tail collar domain protein \| Phage tail collar domain protein; uniclust \| UniRef100\_A0A945NP07 \| 99.8 \| 5.6e-22 \| 1e-27 \| 151.9 \| 99 \| (9, 122) \| 221 \| (309, 407) \| 520 \| Phage tail collar domain-containing protein \| Phage tail collar domain-containing protein; uniclust \| UniRef100\_A0A358QWR4 \| 99.8 \| 5.9e-22 \| 1.1e-27 \| 140.5 \| 120 \| (8, 221) \| 221 \| (123, 244) \| 244 \| Phage tail collar domain-containing protein \| Phage tail collar domain-containing protein; uniclust \| UniRef100\_A0A813U148 \| 99.8 \| 6.4e-22 \| 1.2e-27 \| 154.6 \| 194 \| (9, 221) \| 221 \| (432, 633) \| 633 \| Phage tail collar domain-containing protein (Fragment) \| Phage tail collar domain-containing protein (Fragment); uniclust \| UniRef100\_A0A3B0MDZ4 \| 99.8 \| 7e-22 \| 1.3e-27 \| 147.2 \| 77 \| (9, 85) \| 221 \| (149, 225) \| 315 \| Phage tail collar domain-containing protein \| Phage tail collar domain-containing protein; uniclust \| UniRef100\_A0A2W4QXT5 \| 99.8 \| 7.1e-22 \| 1.3e-27 \| 138.3 \| 149 \| (8, 221) \| 221 \| (99, 247) \| 247 \| Uncharacterized protein \| Uncharacterized protein; uniclust \| UniRef100\_A0A024LSG2 \| 99.8 \| 8.4e-22 \| 1.6e-27 \| 153.1 \| 92 \| (9, 119) \| 221 \| (178, 269) \| 345 \| Putative phage tail fiber protein \| Putative phage tail fiber protein; uniclust \| UniRef100\_A0A0S4U5F8 \| 99.8 \| 8.6e-22 \| 1.6e-27 \| 145.2 \| 91 \| (10, 119) \| 221 \| (73, 163) \| 236 \| Putative phage tail collar protein \| Putative phage tail collar protein; uniclust \| UniRef100\_A0A1G7NQ56 \| 99.8 \| 9.2e-22 \| 1.7e-27 \| 146.0 \| 67 \| (8, 76) \| 221 \| (77, 144) \| 250 \| Phage Tail Collar Domain \| Phage Tail Collar Domain; uniclust \| UniRef100\_A0A0G1QR78 \| 99.8 \| 1e-21 \| 1.9e-27 \| 139.6 \| 99 \| (9, 119) \| 221 \| (131, 229) \| 276 \| Tail Collar domain protein \| Tail Collar domain protein; uniclust \| UniRef100\_A0A1G7VYT4 \| 99.8 \| 1.1e-21 \| 2e-27 \| 156.7 \| 92 \| (10, 119) \| 221 \| (4, 95) \| 518 \| Microcystin-dependent protein \| Microcystin-dependent protein; uniclust \| UniRef100\_A0A291LA10 \| 99.8 \| 1.2e-21 \| 2.3e-27 \| 132.5 \| 92 \| (9, 119) \| 221 \| (35, 126) \| 176 \| Putative tail-collar fiber protein \| Putative tail-collar fiber protein; uniclust \| UniRef100\_A0A516M2K7 \| 99.8 \| 1.3e-21 \| 2.5e-27 \| 150.6 \| 167 \| (6, 221) \| 221 \| (283, 449) \| 449 \| Putative tail fiber protein \| Putative tail fiber protein; uniclust \| UniRef100\_A0A0A0DZA9 \| 99.7 \| 1.8e-21 \| 3.3e-27 \| 149.7 \| 97 \| (11, 120) \| 221 \| (59, 156) \| 297 \| Phage tail collar domain-containing protein \| Phage tail collar domain-containing protein; uniclust \| UniRef100\_A0A6G6K382 \| 99.7 \| 1.8e-21 \| 3.3e-27 \| 150.1 \| 111 \| (9, 119) \| 221 \| (252, 365) \| 419 \| Phage tail collar domain-containing protein \| Phage tail collar domain-containing protein; uniclust \| UniRef100\_A0A351KZF5 \| 99.7 \| 1.9e-21 \| 3.4e-27 \| 153.0 \| 164 \| (8, 221) \| 221 \| (510, 677) \| 677 \| Phage tail collar domain-containing protein \| Phage tail collar domain-containing protein; uniclust \| UniRef100\_A0A1G6T0X4 \| 99.7 \| 1.9e-21 \| 3.5e-27 \| 140.9 \| 97 \| (10, 119) \| 221 \| (21, 117) \| 224 \| Microcystin-dependent protein \| Microcystin-dependent protein; uniclust \| UniRef100\_UPI0010939542 \| 99.7 \| 2.1e-21 \| 3.8e-27 \| 138.7 \| 96 \| (8, 103) \| 221 \| (106, 201) \| 284 \| tail fiber protein \| tail fiber protein; uniclust \| UniRef100\_A0A5J4Y931 \| 99.7 \| 2.5e-21 \| 4.5e-27 \| 140.6 \| 93 \| (21, 126) \| 221 \| (162, 254) \| 323 \| Phage tail collar domain-containing protein \| Phage tail collar domain-containing protein; uniclust \| UniRef100\_A0A1F7BR55 \| 99.7 \| 2.6e-21 \| 4.8e-27 \| 148.5 \| 110 \| (10, 119) \| 221 \| (365, 474) \| 523 \| Phage tail collar domain-containing protein \| Phage tail collar domain-containing protein; uniclust \| UniRef100\_E4T5W3 \| 99.7 \| 2.7e-21 \| 5e-27 \| 155.9 \| 96 \| (11, 119) \| 221 \| (739, 838) \| 898 \| Tail Collar domain protein \| Tail Collar domain protein; uniclust \| UniRef100\_A0A936WGZ5 \| 99.7 \| 2.8e-21 \| 5.2e-27 \| 148.0 \| 98 \| (9, 119) \| 221 \| (288, 388) \| 449 \| Tail fiber protein \| Tail fiber protein; uniclust \| UniRef100\_A0A1G3BQZ8 \| 99.7 \| 2.8e-21 \| 5.2e-27 \| 143.7 \| 107 \| (8, 119) \| 221 \| (39, 151) \| 227 \| Phage tail collar domain-containing protein \| Phage tail collar domain-containing protein; uniclust \| UniRef100\_A0A9E0UQ04 \| 99.7 \| 2.9e-21 \| 5.4e-27 \| 137.2 \| 97 \| (11, 120) \| 221 \| (75, 171) \| 242 \| Phage tail protein \| Phage tail protein; uniclust \| UniRef100\_A0A1G7VYT4 \| 99.7 \| 3e-21 \| 5.5e-27 \| 154.4 \| 96 \| (10, 119) \| 221 \| (173, 268) \| 518 \| Microcystin-dependent protein \| Microcystin-dependent protein; uniclust \| UniRef100\_A0A958X9M4 \| 99.7 \| 3.2e-21 \| 5.8e-27 \| 155.6 \| 157 \| (8, 221) \| 221 \| (581, 737) \| 737 \| Uncharacterized protein \| Uncharacterized protein; uniclust \| UniRef100\_A0A0N9MHY1 \| 99.7 \| 3.2e-21 \| 5.9e-27 \| 162.5 \| 193 \| (9, 221) \| 221 \| (651, 843) \| 843 \| Phage tail collar domain-containing protein \| Phage tail collar domain-containing protein; uniclust \| UniRef100\_A0A1G0CK00 \| 99.7 \| 3.4e-21 \| 6.4e-27 \| 133.5 \| 98 \| (9, 119) \| 221 \| (6, 103) \| 185 \| Phage tail protein \| Phage tail protein; uniclust \| UniRef100\_A0A5C7LV16 \| 99.7 \| 3.5e-21 \| 6.4e-27 \| 154.7 \| 174 \| (8, 221) \| 221 \| (97, 270) \| 856 \| Phage tail collar domain-containing protein \| Phage tail collar domain-containing protein; uniclust \| UniRef100\_A0A0D0N756 \| 99.7 \| 3.4e-21 \| 6.4e-27 \| 139.2 \| 102 \| (9, 119) \| 221 \| (29, 138) \| 202 \| Phage tail collar domain-containing protein \| Phage tail collar domain-containing protein; uniclust \| UniRef100\_A0A815YNV7 \| 99.7 \| 4e-21 \| 7.3e-27 \| 134.7 \| 124 \| (7, 147) \| 221 \| (118, 241) \| 248 \| Phage tail collar domain-containing protein \| Phage tail collar domain-containing protein; uniclust \| UniRef100\_A0A2W7PA10 \| 99.7 \| 4e-21 \| 7.6e-27 \| 141.1 \| 92 \| (9, 119) \| 221 \| (77, 168) \| 224 \| Tail collar domain \| Tail collar domain; uniclust \| UniRef100\_A0A2K9MAN8 \| 99.7 \| 4.7e-21 \| 8.6e-27 \| 132.2 \| 103 \| (10, 119) \| 221 \| (13, 115) \| 221 \| Phage tail collar domain-containing protein \| Phage tail collar domain-containing protein; uniclust \| UniRef100\_A0A845XUZ4 \| 99.7 \| 4.7e-21 \| 8.7e-27 \| 146.6 \| 103 \| (8, 127) \| 221 \| (339, 441) \| 507 \| Phage tail collar domain-containing protein \| Phage tail collar domain-containing protein; uniclust \| UniRef100\_A0A1Q4DT89 \| 99.7 \| 4.7e-21 \| 8.7e-27 \| 138.3 \| 185 \| (8, 221) \| 221 \| (33, 227) \| 227 \| Phage tail collar domain-containing protein \| Phage tail collar domain-containing protein; uniclust \| UniRef100\_A0A966D4Q4 \| 99.7 \| 5.2e-21 \| 9.7e-27 \| 139.7 \| 77 \| (9, 85) \| 221 \| (40, 116) \| 262 \| Phage tail collar domain-containing protein \| Phage tail collar domain-containing protein; uniclust \| UniRef100\_A0A0Q6ZRG1 \| 99.7 \| 5.3e-21 \| 1e-26 \| 147.9 \| 94 \| (10, 122) \| 221 \| (166, 259) \| 316 \| Phage tail collar domain-containing protein \| Phage tail collar domain-containing protein; uniclust \| UniRef100\_A0A835XN54 \| 99.7 \| 5.7e-21 \| 1.1e-26 \| 152.1 \| 99 \| (9, 119) \| 221 \| (447, 545) \| 631 \| Phage tail collar domain-containing protein \| Phage tail collar domain-containing protein; uniclust \| UniRef100\_A0A0P7BJN4 \| 99.7 \| 6e-21 \| 1.1e-26 \| 148.6 \| 91 \| (9, 119) \| 221 \| (458, 548) \| 599 \| Phage tail collar domain-containing protein \| Phage tail collar domain-containing protein; uniclust \| UniRef100\_A0A815F2F0 \| 99.7 \| 6.2e-21 \| 1.2e-26 \| 144.6 \| 128 \| (8, 148) \| 221 \| (172, 300) \| 355 \| Phage tail collar domain-containing protein \| Phage tail collar domain-containing protein; uniclust \| UniRef100\_A0A3M9M907 \| 99.7 \| 6.6e-21 \| 1.2e-26 \| 140.5 \| 177 \| (7, 221) \| 221 \| (137, 317) \| 317 \| Phage tail collar domain-containing protein \| Phage tail collar domain-containing protein; uniclust \| UniRef100\_A0A6B3UTZ7 \| 99.7 \| 6.6e-21 \| 1.2e-26 \| 142.6 \| 79 \| (7, 85) \| 221 \| (159, 237) \| 312 \| Phage tail protein \| Phage tail protein; uniclust \| UniRef100\_UPI001F3D999F \| 99.7 \| 8.5e-21 \| 1.6e-26 \| 139.8 \| 141 \| (8, 221) \| 221 \| (218, 360) \| 360 \| hypothetical protein \| hypothetical protein; uniclust \| UniRef100\_A0A1X7F9Q3 \| 99.7 \| 9.3e-21 \| 1.7e-26 \| 139.0 \| 75 \| (9, 83) \| 221 \| (77, 151) \| 219 \| Phage Tail Collar Domain \| Phage Tail Collar Domain; uniclust \| UniRef100\_A0A899ISM6 \| 99.7 \| 9.7e-21 \| 1.8e-26 \| 134.7 \| 76 \| (8, 83) \| 221 \| (29, 104) \| 207 \| Tail protein \| Tail protein; uniclust \| UniRef100\_A0A0B9H1Q2 \| 99.7 \| 1e-20 \| 1.9e-26 \| 139.9 \| 98 \| (9, 119) \| 221 \| (51, 148) \| 252 \| Phage tail collar domain-containing protein \| Phage tail collar domain-containing protein; uniclust \| UniRef100\_A0A927Y330 \| 99.7 \| 1e-20 \| 1.9e-26 \| 152.8 \| 108 \| (8, 128) \| 221 \| (648, 755) \| 901 \| Fibronectin type-III domain-containing protein \| Fibronectin type-III domain-containing protein; uniclust \| UniRef100\_A0A0M2NH56 \| 99.7 \| 1e-20 \| 1.9e-26 \| 133.3 \| 99 \| (6, 119) \| 221 \| (107, 207) \| 256 \| Microcystin dependent protein \| Microcystin dependent protein; uniclust \| UniRef100\_A0A120FHB0 \| 99.7 \| 1e-20 \| 1.9e-26 \| 143.9 \| 176 \| (8, 221) \| 221 \| (209, 384) \| 384 \| Phage tail collar domain-containing protein \| Phage tail collar domain-containing protein; uniclust \| UniRef100\_A0A814HPR8 \| 99.7 \| 1e-20 \| 1.9e-26 \| 134.8 \| 161 \| (8, 221) \| 221 \| (41, 215) \| 219 \| Phage tail collar domain-containing protein \| Phage tail collar domain-containing protein; uniclust \| UniRef100\_A0A225DUJ5 \| 99.7 \| 1.2e-20 \| 2.1e-26 \| 140.0 \| 125 \| (7, 145) \| 221 \| (105, 242) \| 326 \| Phage tail fiber \| Phage tail fiber; uniclust \| UniRef100\_UPI0009F51E47 \| 99.7 \| 1.2e-20 \| 2.2e-26 \| 142.4 \| 95 \| (10, 119) \| 221 \| (296, 390) \| 444 \| tail fiber protein \| tail fiber protein; uniclust \| UniRef100\_A0A350MM43 \| 99.7 \| 1.2e-20 \| 2.3e-26 \| 134.2 \| 69 \| (8, 119) \| 221 \| (93, 163) \| 215 \| Tail fiber protein (Fragment) \| Tail fiber protein (Fragment); uniclust \| UniRef100\_A0A815DS73 \| 99.7 \| 1.3e-20 \| 2.4e-26 \| 120.8 \| 108 \| (7, 126) \| 221 \| (25, 132) \| 137 \| Phage tail collar domain-containing protein \| Phage tail collar domain-containing protein; uniclust \| UniRef100\_A0A815JLE8 \| 99.7 \| 1.3e-20 \| 2.4e-26 \| 145.5 \| 194 \| (10, 221) \| 221 \| (271, 473) \| 474 \| Phage tail collar domain-containing protein \| Phage tail collar domain-containing protein; uniclust \| UniRef100\_A0A162G8C3 \| 99.7 \| 1.4e-20 \| 2.7e-26 \| 138.4 \| 69 \| (8, 77) \| 221 \| (54, 122) \| 224 \| Phage tail collar domain-containing protein \| Phage tail collar domain-containing protein; uniclust \| UniRef100\_A0A0Q2UK27 \| 99.7 \| 1.6e-20 \| 2.9e-26 \| 129.7 \| 94 \| (7, 119) \| 221 \| (23, 116) \| 166 \| Phage tail collar domain-containing protein \| Phage tail collar domain-containing protein; uniclust \| UniRef100\_A0A0F9B578 \| 99.7 \| 1.6e-20 \| 3e-26 \| 136.7 \| 98 \| (11, 119) \| 221 \| (110, 210) \| 262 \| Phage tail collar domain-containing protein \| Phage tail collar domain-containing protein; uniclust \| UniRef100\_A0A1H0FQQ1 \| 99.7 \| 1.6e-20 \| 3e-26 \| 127.8 \| 97 \| (10, 119) \| 221 \| (4, 100) \| 176 \| Microcystin-dependent protein \| Microcystin-dependent protein; uniclust \| UniRef100\_A0A0H2MCN6 \| 99.7 \| 1.7e-20 \| 3.2e-26 \| 139.4 \| 197 \| (2, 221) \| 221 \| (63, 280) \| 280 \| Minor tail protein \| Minor tail protein; uniclust \| UniRef100\_A0A816HQP8 \| 99.7 \| 1.8e-20 \| 3.3e-26 \| 118.9 \| 112 \| (23, 147) \| 221 \| (3, 114) \| 129 \| Phage tail collar domain-containing protein (Fragment) \| Phage tail collar domain-containing protein (Fragment); uniclust \| UniRef100\_A0A354T924 \| 99.7 \| 1.8e-20 \| 3.4e-26 \| 139.0 \| 110 \| (7, 121) \| 221 \| (118, 228) \| 279 \| Phage tail collar domain-containing protein \| Phage tail collar domain-containing protein; uniclust \| UniRef100\_A0A814J2Q1 \| 99.7 \| 2.1e-20 \| 3.8e-26 \| 156.9 \| 190 \| (9, 221) \| 221 \| (1103, 1304) \| 1304 \| Elongation of very long chain fatty acids protein \| Elongation of very long chain fatty acids protein; uniclust \| UniRef100\_A0A7W8G0N9 \| 99.7 \| 2.2e-20 \| 4.1e-26 \| 125.7 \| 90 \| (12, 119) \| 221 \| (6, 95) \| 168 \| Microcystin-dependent protein \| Microcystin-dependent protein; uniclust \| UniRef100\_A0A819QFI6 \| 99.7 \| 2.3e-20 \| 4.2e-26 \| 121.1 \| 120 \| (11, 149) \| 221 \| (2, 126) \| 147 \| Phage tail collar domain-containing protein (Fragment) \| Phage tail collar domain-containing protein (Fragment); uniclust \| UniRef100\_A0A964YWP9 \| 99.7 \| 2.5e-20 \| 4.6e-26 \| 137.7 \| 184 \| (10, 215) \| 221 \| (173, 358) \| 367 \| Phage tail collar domain-containing protein \| Phage tail collar domain-containing protein; uniclust \| UniRef100\_A0A1M4TNW1 \| 99.7 \| 2.5e-20 \| 4.7e-26 \| 151.7 \| 84 \| (9, 119) \| 221 \| (566, 649) \| 728 \| Microcystin-dependent protein \| Microcystin-dependent protein; uniclust \| UniRef100\_A0A9E9KT91 \| 99.7 \| 2.6e-20 \| 4.8e-26 \| 139.5 \| 76 \| (10, 85) \| 221 \| (138, 213) \| 352 \| Phage tail protein \| Phage tail protein; uniclust \| UniRef100\_A0A968JEA1 \| 99.7 \| 2.9e-20 \| 5.4e-26 \| 134.0 \| 65 \| (9, 73) \| 221 \| (72, 136) \| 300 \| Tail fiber protein \| Tail fiber protein; uniclust \| UniRef100\_A0A9E3L5V0 \| 99.7 \| 3.3e-20 \| 6e-26 \| 136.4 \| 107 \| (9, 119) \| 221 \| (147, 255) \| 329 \| Tail fiber protein \| Tail fiber protein; uniclust \| UniRef100\_A0A0J5GKU1 \| 99.7 \| 3.3e-20 \| 6.2e-26 \| 129.2 \| 93 \| (8, 119) \| 221 \| (22, 114) \| 172 \| Phage tail collar domain-containing protein \| Phage tail collar domain-containing protein; uniclust \| UniRef100\_A0A1I3Z6T6 \| 99.7 \| 3.4e-20 \| 6.3e-26 \| 122.3 \| 98 \| (9, 119) \| 221 \| (17, 114) \| 135 \| Phage Tail Collar Domain (Fragment) \| Phage Tail Collar Domain (Fragment); uniclust \| UniRef100\_A0A090N7J5 \| 99.7 \| 3.4e-20 \| 6.4e-26 \| 148.9 \| 189 \| (7, 221) \| 221 \| (205, 417) \| 456 \| Microcystin-dependent protein \| Microcystin-dependent protein; uniclust \| UniRef100\_A0A1G0LDM4 \| 99.7 \| 3.4e-20 \| 6.4e-26 \| 138.1 \| 74 \| (8, 120) \| 221 \| (159, 234) \| 306 \| Phage tail collar domain-containing protein \| Phage tail collar domain-containing protein; uniclust \| UniRef100\_A0A480A6W0 \| 99.7 \| 3.5e-20 \| 6.5e-26 \| 148.6 \| 145 \| (11, 221) \| 221 \| (548, 695) \| 696 \| Phage tail collar domain-containing protein \| Phage tail collar domain-containing protein; uniclust \| UniRef100\_A0A1V5X7T8 \| 99.7 \| 3.6e-20 \| 6.6e-26 \| 129.3 \| 151 \| (8, 221) \| 221 \| (84, 237) \| 237 \| Phage Tail Collar Domain protein \| Phage Tail Collar Domain protein; uniclust \| UniRef100\_A0A7C8AIK9 \| 99.7 \| 3.6e-20 \| 6.6e-26 \| 139.5 \| 107 \| (9, 120) \| 221 \| (176, 284) \| 338 \| Tail fiber protein \| Tail fiber protein; uniclust \| UniRef100\_A0A1F3PPY1 \| 99.7 \| 4e-20 \| 7.4e-26 \| 140.5 \| 89 \| (8, 120) \| 221 \| (212, 300) \| 378 \| Uncharacterized protein \| Uncharacterized protein; uniclust \| UniRef100\_A0A947MTH0 \| 99.7 \| 4.4e-20 \| 8.1e-26 \| 141.2 \| 101 \| (7, 119) \| 221 \| (229, 329) \| 497 \| Tail fiber protein \| Tail fiber protein; uniclust \| UniRef100\_UPI00158EA18A \| 99.7 \| 4.8e-20 \| 8.7e-26 \| 122.1 \| 151 \| (24, 221) \| 221 \| (10, 165) \| 166 \| hypothetical protein \| hypothetical protein; uniclust \| UniRef100\_A0A1G4PCJ5 \| 99.7 \| 5.1e-20 \| 9.5e-26 \| 137.2 \| 70 \| (8, 120) \| 221 \| (126, 197) \| 250 \| Microcystin-dependent protein \| Microcystin-dependent protein; uniclust \| UniRef100\_A0A2E4EDR2 \| 99.7 \| 5.5e-20 \| 1e-25 \| 119.1 \| 98 \| (9, 119) \| 221 \| (24, 121) \| 124 \| Phage tail collar domain-containing protein \| Phage tail collar domain-containing protein; uniclust \| UniRef100\_UPI001D053F05 \| 99.7 \| 5.6e-20 \| 1e-25 \| 144.5 \| 99 \| (9, 120) \| 221 \| (471, 569) \| 651 \| uncharacterized protein \| uncharacterized protein; uniclust \| UniRef100\_A0A846FV65 \| 99.7 \| 5.8e-20 \| 1.1e-25 \| 136.5 \| 92 \| (9, 119) \| 221 \| (225, 316) \| 381 \| Phage tail collar domain-containing protein \| Phage tail collar domain-containing protein; uniclust \| UniRef100\_A0A0Q4T3S1 \| 99.7 \| 6.2e-20 \| 1.1e-25 \| 140.2 \| 107 \| (13, 119) \| 221 \| (148, 255) \| 423 \| Phage tail collar domain-containing protein \| Phage tail collar domain-containing protein; uniclust \| UniRef100\_A0A1H8IGX2 \| 99.7 \| 6.3e-20 \| 1.2e-25 \| 150.2 \| 75 \| (10, 84) \| 221 \| (285, 359) \| 462 \| Phage Tail Collar Domain \| Phage Tail Collar Domain; uniclust \| UniRef100\_A0A941A575 \| 99.7 \| 7.2e-20 \| 1.3e-25 \| 128.3 \| 94 \| (10, 119) \| 221 \| (48, 156) \| 215 \| Phage tail collar domain-containing protein \| Phage tail collar domain-containing protein; uniclust \| UniRef100\_A0A212J771 \| 99.7 \| 8.2e-20 \| 1.5e-25 \| 135.3 \| 96 \| (9, 119) \| 221 \| (132, 229) \| 301 \| Phage tail collar domain-containing protein \| Phage tail collar domain-containing protein; uniclust \| UniRef100\_A9AVC5 \| 99.7 \| 8.7e-20 \| 1.6e-25 \| 151.1 \| 152 \| (7, 221) \| 221 \| (782, 935) \| 935 \| Tail Collar domain protein \| Tail Collar domain protein; uniclust \| UniRef100\_A0A820I8T0 \| 99.7 \| 8.7e-20 \| 1.6e-25 \| 122.4 \| 125 \| (10, 147) \| 221 \| (11, 136) \| 180 \| Phage tail collar domain-containing protein (Fragment) \| Phage tail collar domain-containing protein (Fragment); uniclust \| UniRef100\_A0A822BDI4 \| 99.7 \| 9.2e-20 \| 1.7e-25 \| 121.7 \| 120 \| (9, 146) \| 221 \| (26, 146) \| 174 \| Phage tail collar domain-containing protein (Fragment) \| Phage tail collar domain-containing protein (Fragment); uniclust \| UniRef100\_A0A323UBG8 \| 99.7 \| 9.8e-20 \| 1.8e-25 \| 132.6 \| 97 \| (4, 119) \| 221 \| (103, 201) \| 322 \| Phage tail collar domain-containing protein \| Phage tail collar domain-containing protein; uniclust \| UniRef100\_A0A0C1EKM6 \| 99.7 \| 9.9e-20 \| 1.8e-25 \| 140.6 \| 92 \| (9, 119) \| 221 \| (204, 295) \| 358 \| Phage tail collar domain-containing protein \| Phage tail collar domain-containing protein; uniclust \| UniRef100\_A0A163Z5D0 \| 99.7 \| 1e-19 \| 1.9e-25 \| 139.9 \| 65 \| (8, 74) \| 221 \| (114, 178) \| 308 \| Phage tail collar domain-containing protein \| Phage tail collar domain-containing protein; uniclust \| UniRef100\_A0A819MQL5 \| 99.7 \| 1.1e-19 \| 2e-25 \| 129.6 \| 126 \| (11, 149) \| 221 \| (49, 175) \| 228 \| Phage tail collar domain-containing protein \| Phage tail collar domain-containing protein; uniclust \| UniRef100\_A0A1D7Y2A0 \| 99.7 \| 1.2e-19 \| 2.3e-25 \| 139.0 \| 159 \| (9, 221) \| 221 \| (132, 303) \| 309 \| Phage tail collar domain-containing protein \| Phage tail collar domain-containing protein; uniclust \| UniRef100\_A0A2N2CG59 \| 99.7 \| 1.3e-19 \| 2.4e-25 \| 124.1 \| 144 \| (8, 221) \| 221 \| (61, 206) \| 206 \| Phage tail collar domain-containing protein \| Phage tail collar domain-containing protein; uniclust \| UniRef100\_A0A4Y5NZ21 \| 99.7 \| 1.4e-19 \| 2.6e-25 \| 127.7 \| 96 \| (10, 119) \| 221 \| (74, 170) \| 225 \| Tail fiber \| Tail fiber; uniclust \| UniRef100\_A0A3S9QK74 \| 99.7 \| 1.4e-19 \| 2.6e-25 \| 131.4 \| 98 \| (8, 118) \| 221 \| (133, 232) \| 303 \| Phage tail collar domain-containing protein \| Phage tail collar domain-containing protein; uniclust \| UniRef100\_A0A349QFL3 \| 99.7 \| 1.4e-19 \| 2.6e-25 \| 127.0 \| 93 \| (8, 119) \| 221 \| (25, 135) \| 186 \| Phage tail collar domain-containing protein \| Phage tail collar domain-containing protein; uniclust \| UniRef100\_UPI001FB54CC0 \| 99.7 \| 1.5e-19 \| 2.8e-25 \| 133.3 \| 100 \| (7, 120) \| 221 \| (195, 294) \| 356 \| tail fiber protein \| tail fiber protein; uniclust \| UniRef100\_A0A4V0I998 \| 99.7 \| 1.5e-19 \| 2.8e-25 \| 147.6 \| 100 \| (8, 119) \| 221 \| (505, 604) \| 1012 \| Phage tail collar domain-containing protein \| Phage tail collar domain-containing protein; uniclust \| UniRef100\_A0A1V5WWW4 \| 99.7 \| 1.9e-19 \| 3.4e-25 \| 130.6 \| 98 \| (9, 119) \| 221 \| (158, 256) \| 311 \| Phage Tail Collar Domain protein \| Phage Tail Collar Domain protein; uniclust \| UniRef100\_UPI001EF32E70 \| 99.7 \| 2e-19 \| 3.7e-25 \| 123.7 \| 166 \| (12, 221) \| 221 \| (47, 212) \| 212 \| tail fiber protein \| tail fiber protein; uniclust \| UniRef100\_A0A077PNS6 \| 99.7 \| 2e-19 \| 3.7e-25 \| 131.9 \| 90 \| (4, 119) \| 221 \| (10, 107) \| 190 \| Phage tail collar domain-containing protein \| Phage tail collar domain-containing protein; uniclust \| UniRef100\_A0A0G4FSR3 \| 99.7 \| 2.1e-19 \| 3.9e-25 \| 140.9 \| 98 \| (10, 119) \| 221 \| (479, 576) \| 625 \| Phage tail collar domain-containing protein \| Phage tail collar domain-containing protein; uniclust \| UniRef100\_A0A2A2TNN2 \| 99.7 \| 2.1e-19 \| 4e-25 \| 152.8 \| 158 \| (8, 221) \| 221 \| (762, 924) \| 924 \| LamG-like jellyroll fold domain-containing protein \| LamG-like jellyroll fold domain-containing protein; uniclust \| UniRef100\_I0KE34 \| 99.7 \| 2.3e-19 \| 4.3e-25 \| 124.4 \| 97 \| (11, 120) \| 221 \| (53, 149) \| 224 \| Tail collar domain protein \| Tail collar domain protein; uniclust \| UniRef100\_A0A813P951 \| 99.6 \| 2.4e-19 \| 4.5e-25 \| 140.1 \| 108 \| (9, 128) \| 221 \| (397, 504) \| 579 \| LamG-like jellyroll fold domain-containing protein \| LamG-like jellyroll fold domain-containing protein; uniclust \| UniRef100\_A0A1H8SWG1 \| 99.6 \| 2.5e-19 \| 4.5e-25 \| 127.3 \| 85 \| (7, 119) \| 221 \| (125, 211) \| 265 \| Microcystin-dependent protein \| Microcystin-dependent protein; uniclust \| UniRef100\_A0A923UU29 \| 99.6 \| 2.7e-19 \| 5e-25 \| 128.4 \| 80 \| (6, 85) \| 221 \| (39, 118) \| 231 \| Tail fiber protein \| Tail fiber protein; uniclust \| UniRef100\_A0A937CVC5 \| 99.6 \| 3.3e-19 \| 6.1e-25 \| 131.8 \| 119 \| (8, 137) \| 221 \| (175, 293) \| 363 \| Tail fiber protein \| Tail fiber protein; uniclust \| UniRef100\_A0A388TAF9 \| 99.6 \| 3.3e-19 \| 6.2e-25 \| 125.7 \| 145 \| (9, 221) \| 221 \| (25, 176) \| 177 \| Phage tail fiber protein (Fragment) \| Phage tail fiber protein (Fragment); uniclust \| UniRef100\_A0A959MGB6 \| 99.6 \| 3.4e-19 \| 6.3e-25 \| 130.7 \| 100 \| (11, 120) \| 221 \| (177, 276) \| 340 \| Tail fiber protein \| Tail fiber protein; uniclust \| UniRef100\_A0A8J7LAJ2 \| 99.6 \| 3.5e-19 \| 6.5e-25 \| 128.3 \| 97 \| (8, 119) \| 221 \| (140, 236) \| 294 \| Tail fiber protein \| Tail fiber protein; uniclust \| UniRef100\_A0A062GTY4 \| 99.6 \| 3.5e-19 \| 6.5e-25 \| 129.1 \| 77 \| (7, 84) \| 221 \| (70, 146) \| 218 \| Phage Tail Collar domain protein \| Phage Tail Collar domain protein; uniclust \| UniRef100\_A0A923NCB6 \| 99.6 \| 3.7e-19 \| 6.9e-25 \| 142.7 \| 99 \| (9, 120) \| 221 \| (617, 716) \| 796 \| Tail fiber protein \| Tail fiber protein; uniclust \| UniRef100\_A0A235AB52 \| 99.6 \| 3.8e-19 \| 7.1e-25 \| 150.0 \| 97 \| (8, 117) \| 221 \| (479, 575) \| 677 \| Phage tail collar domain-containing protein \| Phage tail collar domain-containing protein; uniclust \| UniRef100\_A0A074JA67 \| 99.6 \| 4.1e-19 \| 7.6e-25 \| 126.0 \| 91 \| (10, 119) \| 221 \| (52, 142) \| 207 \| Phage tail collar domain-containing protein \| Phage tail collar domain-containing protein; uniclust \| UniRef100\_A0A1V4BWF5 \| 99.6 \| 4.3e-19 \| 7.9e-25 \| 127.5 \| 70 \| (8, 120) \| 221 \| (89, 160) \| 258 \| Uncharacterized protein \| Uncharacterized protein; uniclust \| UniRef100\_A0A1G1PZ25 \| 99.6 \| 5.1e-19 \| 9.5e-25 \| 131.1 \| 80 \| (7, 119) \| 221 \| (145, 226) \| 278 \| Prepilin-type N-terminal cleavage/methylation domain-containing protein \| Prepilin-type N-terminal cleavage/methylation domain-containing protein; uniclust \| UniRef100\_A0A1W9JSL6 \| 99.6 \| 5.2e-19 \| 9.6e-25 \| 131.5 \| 74 \| (12, 85) \| 221 \| (136, 209) \| 312 \| Phage tail collar domain-containing protein \| Phage tail collar domain-containing protein; uniclust \| UniRef100\_A0A0A1FH46 \| 99.6 \| 5.8e-19 \| 1.1e-24 \| 133.4 \| 80 \| (6, 85) \| 221 \| (150, 229) \| 320 \| Phage tail fiber protein \| Phage tail fiber protein; uniclust \| UniRef100\_A0A832HQN6 \| 99.6 \| 5.9e-19 \| 1.1e-24 \| 122.0 \| 76 \| (8, 122) \| 221 \| (51, 126) \| 191 \| Phage tail collar domain-containing protein \| Phage tail collar domain-containing protein; uniclust \| UniRef100\_A0A120IHK0 \| 99.6 \| 6.2e-19 \| 1.1e-24 \| 125.1 \| 90 \| (11, 119) \| 221 \| (38, 127) \| 187 \| Phage tail collar domain-containing protein \| Phage tail collar domain-containing protein; uniclust \| UniRef100\_UPI000C7DD9BD \| 99.6 \| 6.7e-19 \| 1.2e-24 \| 133.5 \| 95 \| (8, 119) \| 221 \| (298, 392) \| 447 \| tail fiber protein \| tail fiber protein; uniclust \| UniRef100\_A0A1Q6UCI5 \| 99.6 \| 6.7e-19 \| 1.3e-24 \| 126.7 \| 77 \| (8, 84) \| 221 \| (36, 112) \| 195 \| Phage tail collar domain-containing protein \| Phage tail collar domain-containing protein; uniclust \| UniRef100\_A0A2E5T0N1 \| 99.6 \| 7.7e-19 \| 1.4e-24 \| 137.4 \| 92 \| (9, 122) \| 221 \| (291, 382) \| 426 \| Phage tail collar domain-containing protein \| Phage tail collar domain-containing protein; uniclust \| UniRef100\_A0A3M0CFX7 \| 99.6 \| 8.2e-19 \| 1.5e-24 \| 124.5 \| 99 \| (10, 120) \| 221 \| (7, 105) \| 209 \| Microcystin-dependent protein \| Microcystin-dependent protein; uniclust \| UniRef100\_A0A1H9HCA5 \| 99.6 \| 8.3e-19 \| 1.5e-24 \| 133.1 \| 94 \| (7, 119) \| 221 \| (199, 292) \| 345 \| Microcystin-dependent protein \| Microcystin-dependent protein; uniclust \| UniRef100\_A0A433WUA5 \| 99.6 \| 8.6e-19 \| 1.6e-24 \| 131.3 \| 53 \| (7, 75) \| 221 \| (60, 114) \| 261 \| Phage tail collar domain-containing protein \| Phage tail collar domain-containing protein; uniclust \| UniRef100\_A0A095S9W8 \| 99.6 \| 8.8e-19 \| 1.6e-24 \| 125.0 \| 66 \| (9, 75) \| 221 \| (29, 94) \| 170 \| Phage tail Collar \| Phage tail Collar; uniclust \| UniRef100\_A0A6C0DW18 \| 99.6 \| 9.8e-19 \| 1.8e-24 \| 118.4 \| 94 \| (13, 119) \| 221 \| (37, 130) \| 188 \| Phage tail collar domain-containing protein \| Phage tail collar domain-containing protein; uniclust \| UniRef100\_A0A2N2SSE1 \| 99.6 \| 9.8e-19 \| 1.8e-24 \| 124.7 \| 95 \| (8, 108) \| 221 \| (75, 169) \| 271 \| Phage tail protein \| Phage tail protein; uniclust \| UniRef100\_A0A083UIH7 \| 99.6 \| 1e-18 \| 1.9e-24 \| 146.2 \| 76 \| (10, 85) \| 221 \| (353, 428) \| 636 \| Tail fiber protein \| Tail fiber protein; uniclust \| UniRef100\_A0A6M5Z7F0 \| 99.6 \| 1.1e-18 \| 2e-24 \| 135.2 \| 102 \| (7, 119) \| 221 \| (391, 492) \| 544 \| Phage tail collar domain-containing protein \| Phage tail collar domain-containing protein; uniclust \| UniRef100\_A0A2N2HV87 \| 99.6 \| 1.1e-18 \| 2.1e-24 \| 128.8 \| 109 \| (11, 119) \| 221 \| (174, 287) \| 354 \| Phage tail collar domain-containing protein \| Phage tail collar domain-containing protein; uniclust \| UniRef100\_UPI000E5A2982 \| 99.6 \| 1.2e-18 \| 2.2e-24 \| 127.9 \| 94 \| (22, 119) \| 221 \| (174, 267) \| 337 \| tail fiber protein \| tail fiber protein; uniclust \| UniRef100\_A0A1Y1Q2G4 \| 99.6 \| 1.2e-18 \| 2.3e-24 \| 132.3 \| 91 \| (10, 119) \| 221 \| (71, 161) \| 326 \| Phage tail collar domain-containing protein \| Phage tail collar domain-containing protein; uniclust \| UniRef100\_A0A6C0JGK4 \| 99.6 \| 1.2e-18 \| 2.3e-24 \| 121.6 \| 97 \| (9, 119) \| 221 \| (76, 172) \| 232 \| Phage tail collar domain-containing protein \| Phage tail collar domain-containing protein; uniclust \| UniRef100\_A0A926SHI8 \| 99.6 \| 1.3e-18 \| 2.3e-24 \| 124.3 \| 73 \| (10, 119) \| 221 \| (140, 214) \| 272 \| Tail fiber protein \| Tail fiber protein; uniclust \| UniRef100\_A0A969MIB8 \| 99.6 \| 1.3e-18 \| 2.4e-24 \| 139.3 \| 98 \| (10, 119) \| 221 \| (514, 611) \| 767 \| Uncharacterized protein \| Uncharacterized protein; uniclust \| UniRef100\_A0A0Q0Y1I2 \| 99.6 \| 1.3e-18 \| 2.4e-24 \| 133.9 \| 102 \| (8, 119) \| 221 \| (54, 174) \| 268 \| Putative rhizosphere induced protein RhiB \| Putative rhizosphere induced protein RhiB; uniclust \| UniRef100\_A0A088FQW7 \| 99.6 \| 1.3e-18 \| 2.4e-24 \| 143.0 \| 93 \| (8, 119) \| 221 \| (316, 408) \| 515 \| Tail protein \| Tail protein; uniclust \| UniRef100\_UPI0015862BE5 \| 99.6 \| 1.4e-18 \| 2.5e-24 \| 121.2 \| 71 \| (8, 78) \| 221 \| (11, 81) \| 182 \| tail fiber protein \| tail fiber protein; uniclust \| UniRef100\_A0A821B254 \| 99.6 \| 1.4e-18 \| 2.5e-24 \| 126.6 \| 121 \| (11, 147) \| 221 \| (106, 228) \| 318 \| Phage tail collar domain-containing protein \| Phage tail collar domain-containing protein; uniclust \| UniRef100\_A0A075X7J1 \| 99.6 \| 1.6e-18 \| 2.9e-24 \| 128.5 \| 149 \| (7, 221) \| 221 \| (160, 313) \| 313 \| Phage tail collar domain-containing protein \| Phage tail collar domain-containing protein; uniclust \| UniRef100\_A0A2S9GTJ9 \| 99.6 \| 1.6e-18 \| 3e-24 \| 123.2 \| 90 \| (8, 119) \| 221 \| (17, 108) \| 205 \| Phage Tail Collar Domain \| Phage Tail Collar Domain; uniclust \| UniRef100\_A0A930BHL5 \| 99.6 \| 1.6e-18 \| 3e-24 \| 113.3 \| 56 \| (21, 119) \| 221 \| (7, 62) \| 128 \| Uncharacterized protein \| Uncharacterized protein; uniclust \| UniRef100\_A0A369KVY1 \| 99.6 \| 1.6e-18 \| 3e-24 \| 131.8 \| 104 \| (8, 119) \| 221 \| (80, 186) \| 314 \| Phage tail collar domain-containing protein \| Phage tail collar domain-containing protein; uniclust \| UniRef100\_A0A7M2CD16 \| 99.6 \| 1.6e-18 \| 3e-24 \| 124.0 \| 67 \| (9, 119) \| 221 \| (54, 120) \| 188 \| Phage tail collar domain-containing protein \| Phage tail collar domain-containing protein; uniclust \| UniRef100\_UPI001591FAB7 \| 99.6 \| 1.7e-18 \| 3.1e-24 \| 125.2 \| 192 \| (8, 221) \| 221 \| (89, 284) \| 284 \| tail fiber protein \| tail fiber protein; uniclust \| UniRef100\_A0A942VA29 \| 99.6 \| 1.7e-18 \| 3.1e-24 \| 127.1 \| 99 \| (9, 119) \| 221 \| (104, 202) \| 293 \| Tail fiber protein \| Tail fiber protein; uniclust \| UniRef100\_A0A3M1QYC2 \| 99.6 \| 1.7e-18 \| 3.2e-24 \| 127.3 \| 105 \| (9, 119) \| 221 \| (132, 238) \| 306 \| Phage tail collar domain-containing protein (Fragment) \| Phage tail collar domain-containing protein (Fragment); uniclust \| UniRef100\_A0A136Q4S0 \| 99.6 \| 1.9e-18 \| 3.5e-24 \| 136.7 \| 95 \| (10, 119) \| 221 \| (144, 240) \| 567 \| Phage Tail Collar domain protein \| Phage Tail Collar domain protein; uniclust \| UniRef100\_X0Z504 \| 99.6 \| 2e-18 \| 3.7e-24 \| 120.4 \| 98 \| (9, 119) \| 221 \| (76, 173) \| 229 \| Phage tail collar domain-containing protein (Fragment) \| Phage tail collar domain-containing protein (Fragment); uniclust \| UniRef100\_A0A212BGA2 \| 99.6 \| 2e-18 \| 3.7e-24 \| 121.8 \| 74 \| (12, 85) \| 221 \| (67, 140) \| 249 \| Phage tail collar domain-containing protein \| Phage tail collar domain-containing protein; uniclust \| UniRef100\_A0A085FNZ1 \| 99.6 \| 2.1e-18 \| 4e-24 \| 141.8 \| 95 \| (20, 114) \| 221 \| (233, 393) \| 486 \| Uncharacterized protein \| Uncharacterized protein; uniclust \| UniRef100\_A0A4P6FA25 \| 99.6 \| 2.4e-18 \| 4.5e-24 \| 116.7 \| 86 \| (22, 120) \| 221 \| (7, 92) \| 190 \| Phage tail collar domain-containing protein \| Phage tail collar domain-containing protein; uniclust \| UniRef100\_A0A0J5P3M7 \| 99.6 \| 2.4e-18 \| 4.6e-24 \| 122.4 \| 65 \| (7, 84) \| 221 \| (16, 80) \| 160 \| Phage tail collar domain-containing protein \| Phage tail collar domain-containing protein; uniclust \| UniRef100\_A0A1W6NZ62 \| 99.6 \| 2.5e-18 \| 4.6e-24 \| 130.8 \| 101 \| (9, 119) \| 221 \| (278, 378) \| 456 \| Phage tail collar domain-containing protein \| Phage tail collar domain-containing protein; uniclust \| UniRef100\_A0A934ZLY9 \| 99.6 \| 2.6e-18 \| 4.8e-24 \| 125.6 \| 114 \| (8, 126) \| 221 \| (145, 258) \| 315 \| Tail fiber protein \| Tail fiber protein; uniclust \| UniRef100\_UPI00168913C5 \| 99.6 \| 2.7e-18 \| 4.9e-24 \| 137.3 \| 97 \| (9, 119) \| 221 \| (556, 659) \| 748 \| tail fiber protein \| tail fiber protein; uniclust \| UniRef100\_A0A926VM20 \| 99.6 \| 2.7e-18 \| 5e-24 \| 128.5 \| 85 \| (8, 119) \| 221 \| (252, 336) \| 394 \| Tail fiber protein \| Tail fiber protein; uniclust \| UniRef100\_A0A072T5I3 \| 99.6 \| 2.7e-18 \| 5e-24 \| 115.8 \| 94 \| (10, 119) \| 221 \| (3, 96) \| 175 \| Phage tail protein \| Phage tail protein; uniclust \| UniRef100\_A0A820RLX9 \| 99.6 \| 2.9e-18 \| 5.3e-24 \| 111.0 \| 98 \| (8, 119) \| 221 \| (25, 122) \| 141 \| Phage tail collar domain-containing protein \| Phage tail collar domain-containing protein; uniclust \| UniRef100\_A0A0E9MQR3 \| 99.6 \| 3.1e-18 \| 5.8e-24 \| 143.8 \| 94 \| (11, 119) \| 221 \| (359, 452) \| 693 \| PEP-CTERM protein-sorting domain-containing protein \| PEP-CTERM protein-sorting domain-containing protein; uniclust \| UniRef100\_A0A977UYB6 \| 99.6 \| 3.6e-18 \| 6.7e-24 \| 140.1 \| 76 \| (8, 83) \| 221 \| (829, 904) \| 1018 \| Phage tail protein \| Phage tail protein; uniclust \| UniRef100\_A0A6L4A8E9 \| 99.6 \| 3.7e-18 \| 6.8e-24 \| 122.5 \| 100 \| (8, 119) \| 221 \| (47, 146) \| 281 \| Phage tail collar domain-containing protein \| Phage tail collar domain-containing protein; uniclust \| UniRef100\_A0A1G7VYT4 \| 99.6 \| 4.7e-18 \| 8.8e-24 \| 136.9 \| 98 \| (10, 119) \| 221 \| (345, 442) \| 518 \| Microcystin-dependent protein \| Microcystin-dependent protein; uniclust \| UniRef100\_A0A0F1B0R1 \| 99.6 \| 4.9e-18 \| 9.2e-24 \| 123.7 \| 72 \| (4, 85) \| 221 \| (38, 109) \| 194 \| Phage tail collar domain-containing protein \| Phage tail collar domain-containing protein; uniclust \| UniRef100\_A0A2D9HKR9 \| 99.6 \| 5.4e-18 \| 1e-23 \| 116.5 \| 94 \| (7, 103) \| 221 \| (74, 167) \| 206 \| Phage tail collar domain-containing protein \| Phage tail collar domain-containing protein; uniclust \| UniRef100\_A0A2A5VXT6 \| 99.6 \| 5.4e-18 \| 1e-23 \| 135.5 \| 103 \| (8, 118) \| 221 \| (268, 376) \| 435 \| Phage tail collar domain-containing protein \| Phage tail collar domain-containing protein; uniclust \| UniRef100\_A0A6C0JIE1 \| 99.6 \| 6.3e-18 \| 1.2e-23 \| 118.9 \| 95 \| (9, 119) \| 221 \| (12, 118) \| 196 \| Phage tail collar domain-containing protein \| Phage tail collar domain-containing protein; uniclust \| UniRef100\_A0A7C7C9X1 \| 99.6 \| 6.5e-18 \| 1.2e-23 \| 118.8 \| 94 \| (13, 127) \| 221 \| (87, 182) \| 242 \| Tail fiber protein \| Tail fiber protein; uniclust \| UniRef100\_A0A257UMQ9 \| 99.6 \| 6.6e-18 \| 1.2e-23 \| 118.5 \| 96 \| (12, 119) \| 221 \| (69, 165) \| 238 \| Phage tail collar domain-containing protein \| Phage tail collar domain-containing protein; uniclust \| UniRef100\_A0A1V5WG08 \| 99.6 \| 7.1e-18 \| 1.3e-23 \| 118.2 \| 79 \| (8, 121) \| 221 \| (90, 168) \| 235 \| Phage tail collar domain-containing protein \| Phage tail collar domain-containing protein; uniclust \| UniRef100\_A0A2D5S2C0 \| 99.6 \| 7e-18 \| 1.3e-23 \| 119.6 \| 78 \| (8, 85) \| 221 \| (16, 93) \| 177 \| Phage tail protein (Fragment) \| Phage tail protein (Fragment); uniclust \| UniRef100\_A0A845WN63 \| 99.6 \| 7.2e-18 \| 1.3e-23 \| 121.2 \| 88 \| (9, 119) \| 221 \| (136, 224) \| 282 \| Phage tail collar domain-containing protein \| Phage tail collar domain-containing protein; uniclust \| UniRef100\_A0A553JK32 \| 99.6 \| 7.2e-18 \| 1.3e-23 \| 131.7 \| 75 \| (9, 126) \| 221 \| (324, 398) \| 483 \| Phage tail collar domain-containing protein \| Phage tail collar domain-containing protein; uniclust \| UniRef100\_K7XWX8 \| 99.6 \| 7.5e-18 \| 1.4e-23 \| 125.3 \| 77 \| (9, 85) \| 221 \| (199, 275) \| 370 \| Phage tail fiber protein \| Phage tail fiber protein; uniclust \| UniRef100\_A0A800M671 \| 99.6 \| 7.6e-18 \| 1.4e-23 \| 120.9 \| 96 \| (11, 119) \| 221 \| (114, 209) \| 279 \| Phage tail protein \| Phage tail protein; uniclust \| UniRef100\_D1BW55 \| 99.6 \| 7.7e-18 \| 1.4e-23 \| 127.9 \| 85 \| (22, 119) \| 221 \| (236, 320) \| 443 \| Tail Collar domain protein \| Tail Collar domain protein; uniclust \| UniRef100\_A0A967KK39 \| 99.6 \| 7.9e-18 \| 1.4e-23 \| 126.3 \| 96 \| (11, 119) \| 221 \| (6, 101) \| 399 \| Phage tail protein \| Phage tail protein; uniclust \| UniRef100\_A0A0E9MQR3 \| 99.6 \| 7.8e-18 \| 1.5e-23 \| 141.6 \| 96 \| (11, 120) \| 221 \| (501, 596) \| 693 \| PEP-CTERM protein-sorting domain-containing protein \| PEP-CTERM protein-sorting domain-containing protein; uniclust \| UniRef100\_A0A2V9WHC2 \| 99.6 \| 8.1e-18 \| 1.5e-23 \| 122.6 \| 99 \| (7, 126) \| 221 \| (130, 230) \| 315 \| Phage tail collar domain-containing protein \| Phage tail collar domain-containing protein; uniclust \| UniRef100\_A0A969GKG4 \| 99.6 \| 8.8e-18 \| 1.6e-23 \| 123.9 \| 84 \| (9, 119) \| 221 \| (205, 288) \| 346 \| Phage tail collar domain-containing protein \| Phage tail collar domain-containing protein; uniclust \| UniRef100\_A0A8T6N729 \| 99.6 \| 9e-18 \| 1.6e-23 \| 109.7 \| 67 \| (9, 75) \| 221 \| (27, 93) \| 148 \| Phage tail collar domain-containing protein (Fragment) \| Phage tail collar domain-containing protein (Fragment); uniclust \| UniRef100\_A0A3B8Q1P4 \| 99.6 \| 9.9e-18 \| 1.8e-23 \| 124.7 \| 68 \| (9, 119) \| 221 \| (252, 319) \| 371 \| Uncharacterized protein \| Uncharacterized protein; uniclust \| UniRef100\_A0A8S0HYQ4 \| 99.6 \| 1.1e-17 \| 1.9e-23 \| 118.6 \| 127 \| (8, 221) \| 221 \| (115, 243) \| 243 \| Phage tail collar domain-containing protein \| Phage tail collar domain-containing protein; uniclust \| UniRef100\_A0A5B8XJ48 \| 99.6 \| 1.1e-17 \| 2e-23 \| 130.8 \| 96 \| (12, 119) \| 221 \| (434, 529) \| 580 \| Phage tail collar domain protein \| Phage tail collar domain protein; uniclust \| UniRef100\_A0A965JC12 \| 99.6 \| 1.1e-17 \| 2e-23 \| 122.2 \| 76 \| (7, 82) \| 221 \| (89, 164) \| 281 \| Phage tail collar domain-containing protein \| Phage tail collar domain-containing protein; uniclust \| UniRef100\_A0A959IFW3 \| 99.6 \| 1.1e-17 \| 2e-23 \| 133.1 \| 84 \| (8, 119) \| 221 \| (561, 644) \| 701 \| Tail fiber protein \| Tail fiber protein; uniclust \| UniRef100\_A0A7C5IR70 \| 99.6 \| 1.1e-17 \| 2.1e-23 \| 130.0 \| 146 \| (11, 221) \| 221 \| (405, 550) \| 550 \| Phage tail collar domain-containing protein \| Phage tail collar domain-containing protein; uniclust \| UniRef100\_UPI0018DC42F4 \| 99.5 \| 1.2e-17 \| 2.1e-23 \| 129.1 \| 93 \| (22, 128) \| 221 \| (361, 453) \| 519 \| tail fiber protein \| tail fiber protein; uniclust \| UniRef100\_A0A1F9IXH9 \| 99.5 \| 1.2e-17 \| 2.2e-23 \| 115.8 \| 105 \| (10, 118) \| 221 \| (65, 170) \| 217 \| Phage tail collar domain-containing protein \| Phage tail collar domain-containing protein; uniclust \| UniRef100\_A0A955C5E6 \| 99.5 \| 1.2e-17 \| 2.2e-23 \| 126.5 \| 101 \| (11, 119) \| 221 \| (265, 368) \| 431 \| Tail fiber protein \| Tail fiber protein; uniclust \| UniRef100\_A0A022PFL8 \| 99.5 \| 1.2e-17 \| 2.2e-23 \| 119.1 \| 84 \| (7, 119) \| 221 \| (50, 133) \| 206 \| Phage-related tail fiber protein \| Phage-related tail fiber protein; uniclust \| UniRef100\_A0A2E1T177 \| 99.5 \| 1.3e-17 \| 2.4e-23 \| 128.2 \| 89 \| (8, 139) \| 221 \| (108, 198) \| 278 \| Uncharacterized protein \| Uncharacterized protein; uniclust \| UniRef100\_A0A3S0C1T6 \| 99.5 \| 1.3e-17 \| 2.4e-23 \| 125.8 \| 67 \| (9, 75) \| 221 \| (83, 149) \| 354 \| Tail fiber protein \| Tail fiber protein; uniclust \| UniRef100\_A0A9E0PA08 \| 99.5 \| 1.4e-17 \| 2.6e-23 \| 122.8 \| 111 \| (8, 119) \| 221 \| (182, 295) \| 344 \| Tail fiber protein \| Tail fiber protein; uniclust \| UniRef100\_A0A0D6GNQ8 \| 99.5 \| 1.4e-17 \| 2.6e-23 \| 126.1 \| 72 \| (4, 85) \| 221 \| (115, 186) \| 281 \| Tail fiber protein \| Tail fiber protein; uniclust \| UniRef100\_A0A2X1ULZ5 \| 99.5 \| 1.5e-17 \| 2.7e-23 \| 123.8 \| 98 \| (9, 145) \| 221 \| (75, 174) \| 258 \| Phage tail fibre repeat \| Phage tail fibre repeat; uniclust \| UniRef100\_A0A6C0JYK5 \| 99.5 \| 1.5e-17 \| 2.8e-23 \| 140.5 \| 149 \| (8, 221) \| 221 \| (1367, 1519) \| 1521 \| Trimeric autotransporter adhesin YadA-like head domain-containing protein \| Trimeric autotransporter adhesin YadA-like head domain-containing protein; uniclust \| UniRef100\_UPI001E32DA79 \| 99.5 \| 1.5e-17 \| 2.8e-23 \| 115.5 \| 98 \| (9, 119) \| 221 \| (121, 218) \| 221 \| tail fiber protein \| tail fiber protein; uniclust \| UniRef100\_UPI001FC84116 \| 99.5 \| 1.6e-17 \| 2.9e-23 \| 131.5 \| 92 \| (8, 117) \| 221 \| (462, 553) \| 624 \| tail fiber protein \| tail fiber protein; uniclust \| UniRef100\_A0A0E3P079 \| 99.5 \| 1.6e-17 \| 3e-23 \| 130.8 \| 98 \| (9, 119) \| 221 \| (331, 428) \| 431 \| Microcystin dependent protein \| Microcystin dependent protein; uniclust \| UniRef100\_UPI0004242599 \| 99.5 \| 1.7e-17 \| 3.1e-23 \| 119.8 \| 77 \| (8, 84) \| 221 \| (111, 187) \| 291 \| tail fiber protein \| tail fiber protein; uniclust \| UniRef100\_A0A2E4L524 \| 99.5 \| 1.8e-17 \| 3.3e-23 \| 135.2 \| 87 \| (9, 118) \| 221 \| (789, 875) \| 927 \| Phage tail collar domain-containing protein \| Phage tail collar domain-containing protein; uniclust \| UniRef100\_A0A938GHZ7 \| 99.5 \| 1.9e-17 \| 3.5e-23 \| 131.6 \| 72 \| (8, 120) \| 221 \| (372, 445) \| 514 \| Uncharacterized protein \| Uncharacterized protein; uniclust \| UniRef100\_A0A1H1FGW5 \| 99.5 \| 1.9e-17 \| 3.5e-23 \| 126.0 \| 112 \| (8, 119) \| 221 \| (199, 375) \| 449 \| Phage Tail Collar Domain \| Phage Tail Collar Domain; uniclust \| UniRef100\_A0A1K0IP40 \| 99.5 \| 2.1e-17 \| 3.9e-23 \| 136.7 \| 91 \| (10, 119) \| 221 \| (406, 496) \| 548 \| Phage tail collar domain-containing protein \| Phage tail collar domain-containing protein; uniclust \| UniRef100\_A0A7C3YVS4 \| 99.5 \| 2.3e-17 \| 4.2e-23 \| 112.4 \| 90 \| (11, 119) \| 221 \| (46, 135) \| 192 \| Phage tail collar domain-containing protein \| Phage tail collar domain-containing protein; uniclust \| UniRef100\_A0A835XXH8 \| 99.5 \| 2.3e-17 \| 4.3e-23 \| 128.3 \| 109 \| (7, 127) \| 221 \| (356, 464) \| 550 \| Phage tail collar domain-containing protein \| Phage tail collar domain-containing protein; uniclust \| UniRef100\_A0A375EED5 \| 99.5 \| 2.4e-17 \| 4.4e-23 \| 125.2 \| 94 \| (11, 121) \| 221 \| (4, 97) \| 440 \| Phage tail collar domain-containing protein \| Phage tail collar domain-containing protein; uniclust \| UniRef100\_A0A418RLT6 \| 99.5 \| 2.4e-17 \| 4.5e-23 \| 122.0 \| 64 \| (8, 119) \| 221 \| (105, 170) \| 247 \| Tail fiber protein \| Tail fiber protein; uniclust \| UniRef100\_A0A1X3I976 \| 99.5 \| 2.6e-17 \| 4.8e-23 \| 113.5 \| 72 \| (4, 85) \| 221 \| (16, 87) \| 166 \| Side tail fiber protein-like protein (Fragment) \| Side tail fiber protein-like protein (Fragment); uniclust \| UniRef100\_A0A024L5N7 \| 99.5 \| 2.6e-17 \| 4.9e-23 \| 126.8 \| 87 \| (4, 119) \| 221 \| (112, 198) \| 289 \| Phage tail protein \| Phage tail protein; uniclust \| UniRef100\_A0A2G1DGN7 \| 99.5 \| 2.7e-17 \| 5.1e-23 \| 122.9 \| 77 \| (8, 84) \| 221 \| (165, 241) \| 317 \| Phage tail collar domain-containing protein \| Phage tail collar domain-containing protein; uniclust \| UniRef100\_A0A1G7NGA5 \| 99.5 \| 2.7e-17 \| 5.1e-23 \| 125.5 \| 76 \| (9, 84) \| 221 \| (113, 202) \| 333 \| Phage Tail Collar Domain \| Phage Tail Collar Domain; uniclust \| UniRef100\_A0A1I5LHR7 \| 99.5 \| 2.9e-17 \| 5.3e-23 \| 109.5 \| 67 \| (10, 119) \| 221 \| (40, 108) \| 167 \| Microcystin-dependent protein \| Microcystin-dependent protein; uniclust \| UniRef100\_A0A816UR79 \| 99.5 \| 2.9e-17 \| 5.3e-23 \| 109.6 \| 145 \| (24, 221) \| 221 \| (21, 166) \| 168 \| Phage tail collar domain-containing protein \| Phage tail collar domain-containing protein; uniclust \| UniRef100\_A0A370CFE4 \| 99.5 \| 2.9e-17 \| 5.4e-23 \| 126.6 \| 100 \| (9, 119) \| 221 \| (231, 331) \| 382 \| Phage tail collar domain-containing protein \| Phage tail collar domain-containing protein; uniclust \| UniRef100\_K8GNE3 \| 99.5 \| 3e-17 \| 5.4e-23 \| 130.9 \| 196 \| (8, 221) \| 221 \| (480, 703) \| 703 \| Phage tail collar family protein \| Phage tail collar family protein; uniclust \| UniRef100\_A0A7R9WFI0 \| 99.5 \| 3e-17 \| 5.5e-23 \| 118.5 \| 97 \| (11, 120) \| 221 \| (48, 144) \| 257 \| Phage tail collar domain-containing protein \| Phage tail collar domain-containing protein; uniclust \| UniRef100\_A0A346HW44 \| 99.5 \| 3.2e-17 \| 6e-23 \| 133.8 \| 110 \| (9, 137) \| 221 \| (275, 384) \| 574 \| Putative tail-fiber protein \| Putative tail-fiber protein; uniclust \| UniRef100\_A0A9D1L441 \| 99.5 \| 3.3e-17 \| 6.1e-23 \| 127.0 \| 98 \| (9, 119) \| 221 \| (239, 336) \| 527 \| Tail fiber protein \| Tail fiber protein; uniclust \| UniRef100\_A0A819C810 \| 99.5 \| 3.5e-17 \| 6.3e-23 \| 137.1 \| 198 \| (10, 220) \| 221 \| (352, 562) \| 1314 \| RNA helicase \| RNA helicase; uniclust \| UniRef100\_A0A6C0HLH9 \| 99.5 \| 3.5e-17 \| 6.4e-23 \| 119.4 \| 88 \| (8, 119) \| 221 \| (145, 233) \| 312 \| Phage tail collar domain-containing protein \| Phage tail collar domain-containing protein; uniclust \| UniRef100\_A0A022PDV2 \| 99.5 \| 3.6e-17 \| 6.8e-23 \| 128.9 \| 87 \| (6, 118) \| 221 \| (173, 267) \| 353 \| Phage tail collar family protein \| Phage tail collar family protein; uniclust \| UniRef100\_A0A7X5TNA6 \| 99.5 \| 3.8e-17 \| 7.1e-23 \| 115.2 \| 83 \| (8, 119) \| 221 \| (52, 134) \| 188 \| Phage tail collar domain-containing protein \| Phage tail collar domain-containing protein; uniclust \| UniRef100\_A0A1J5P6P6 \| 99.5 \| 3.9e-17 \| 7.1e-23 \| 117.9 \| 100 \| (16, 119) \| 221 \| (111, 210) \| 288 \| Phage terminase large subunit \| Phage terminase large subunit; uniclust \| UniRef100\_A0A6C0KWH9 \| 99.5 \| 4.1e-17 \| 7.5e-23 \| 132.0 \| 164 \| (10, 221) \| 221 \| (667, 831) \| 831 \| Phage tail collar domain-containing protein \| Phage tail collar domain-containing protein; uniclust \| UniRef100\_UPI000E5A64BF \| 99.5 \| 4.3e-17 \| 7.9e-23 \| 122.4 \| 105 \| (9, 119) \| 221 \| (236, 341) \| 394 \| tail fiber protein \| tail fiber protein; uniclust \| UniRef100\_A0A098UFE1 \| 99.5 \| 5.1e-17 \| 9.6e-23 \| 127.5 \| 98 \| (9, 119) \| 221 \| (74, 171) \| 305 \| Phage tail collar domain-containing protein \| Phage tail collar domain-containing protein; uniclust \| UniRef100\_A0A024FDZ3 \| 99.5 \| 5.3e-17 \| 1e-22 \| 128.1 \| 99 \| (6, 136) \| 221 \| (161, 260) \| 331 \| Phage tail fibers \| Phage tail fibers; uniclust \| UniRef100\_UPI001AFA07E9 \| 99.5 \| 5.5e-17 \| 1e-22 \| 126.6 \| 162 \| (8, 221) \| 221 \| (395, 561) \| 561 \| hypothetical protein \| hypothetical protein; uniclust \| UniRef100\_A0A972W8A9 \| 99.5 \| 5.9e-17 \| 1.1e-22 \| 120.7 \| 107 \| (8, 119) \| 221 \| (211, 321) \| 368 \| Tail fiber protein \| Tail fiber protein; uniclust \| UniRef100\_A0A1Y5K9L9 \| 99.5 \| 6.1e-17 \| 1.1e-22 \| 116.9 \| 101 \| (10, 123) \| 221 \| (5, 105) \| 233 \| Phage tail collar domain-containing protein \| Phage tail collar domain-containing protein; uniclust \| UniRef100\_F2Y385 \| 99.5 \| 6.2e-17 \| 1.1e-22 \| 117.7 \| 150 \| (8, 221) \| 221 \| (154, 303) \| 303 \| Putative phage tail collar domain family protein \| Putative phage tail collar domain family protein; uniclust \| UniRef100\_A0A5A9GP38 \| 99.5 \| 6.3e-17 \| 1.2e-22 \| 118.8 \| 109 \| (8, 124) \| 221 \| (89, 204) \| 256 \| Tail fiber protein \| Tail fiber protein; uniclust \| UniRef100\_A0A5C5PXD1 \| 99.5 \| 6.4e-17 \| 1.2e-22 \| 122.9 \| 90 \| (10, 118) \| 221 \| (185, 274) \| 335 \| Phage tail collar domain-containing protein \| Phage tail collar domain-containing protein; uniclust \| UniRef100\_A0A0F9I8V3 \| 99.5 \| 6.4e-17 \| 1.2e-22 \| 115.4 \| 69 \| (7, 118) \| 221 \| (26, 100) \| 184 \| Phage tail collar domain-containing protein \| Phage tail collar domain-containing protein; uniclust \| UniRef100\_A0A937FTQ8 \| 99.5 \| 6.6e-17 \| 1.2e-22 \| 122.0 \| 178 \| (10, 221) \| 221 \| (206, 409) \| 410 \| Uncharacterized protein \| Uncharacterized protein; uniclust \| UniRef100\_A0A835XV31 \| 99.5 \| 6.6e-17 \| 1.2e-22 \| 137.9 \| 108 \| (8, 127) \| 221 \| (1556, 1663) \| 1750 \| EF-hand domain-containing protein \| EF-hand domain-containing protein; uniclust \| UniRef100\_A0A815B3Y9 \| 99.5 \| 6.6e-17 \| 1.2e-22 \| 129.3 \| 196 \| (9, 221) \| 221 \| (380, 583) \| 585 \| Phage tail collar domain-containing protein \| Phage tail collar domain-containing protein; uniclust \| UniRef100\_A0A2Z6B2C1 \| 99.5 \| 6.6e-17 \| 1.2e-22 \| 117.3 \| 74 \| (8, 83) \| 221 \| (119, 192) \| 257 \| Putative phage tail fiber-related protein \| Putative phage tail fiber-related protein; uniclust \| UniRef100\_A0A077PK15 \| 99.5 \| 7.6e-17 \| 1.4e-22 \| 124.4 \| 69 \| (7, 85) \| 221 \| (160, 228) \| 318 \| Putative E14 prophage tail fiber protein (Modular protein) \| Putative E14 prophage tail fiber protein (Modular protein); uniclust \| UniRef100\_UPI0020139BF7 \| 99.5 \| 7.7e-17 \| 1.4e-22 \| 122.6 \| 131 \| (9, 146) \| 221 \| (204, 334) \| 440 \| tail fiber protein \| tail fiber protein; uniclust \| UniRef100\_A0A6J5LZ45 \| 99.5 \| 7.8e-17 \| 1.4e-22 \| 116.2 \| 108 \| (12, 119) \| 221 \| (124, 232) \| 285 \| MdpB Microcystin-dependent protein \| MdpB Microcystin-dependent protein; uniclust \| UniRef100\_A0A0N9MHW1 \| 99.5 \| 7.8e-17 \| 1.5e-22 \| 142.1 \| 169 \| (10, 221) \| 221 \| (1144, 1312) \| 1312 \| Phage tail fiber protein \| Phage tail fiber protein; uniclust \| UniRef100\_U3A9G3 \| 99.5 \| 8.1e-17 \| 1.5e-22 \| 128.2 \| 157 \| (9, 221) \| 221 \| (531, 687) \| 687 \| Phage tail collar domain-containing protein \| Phage tail collar domain-containing protein; uniclust \| UniRef100\_Q55EP2 \| 99.5 \| 8.1e-17 \| 1.5e-22 \| 107.4 \| 90 \| (10, 124) \| 221 \| (25, 114) \| 166 \| Phage tail collar domain-containing protein \| Phage tail collar domain-containing protein; uniclust \| UniRef100\_A0A5Q2W2K9 \| 99.5 \| 8.2e-17 \| 1.5e-22 \| 127.2 \| 146 \| (8, 221) \| 221 \| (405, 634) \| 634 \| Tail collar \| Tail collar; uniclust \| UniRef100\_A0A845XBF3 \| 99.5 \| 8.4e-17 \| 1.6e-22 \| 132.6 \| 76 \| (8, 121) \| 221 \| (456, 541) \| 610 \| Tail fiber protein \| Tail fiber protein; uniclust \| UniRef100\_A0A6J5S6B7 \| 99.5 \| 8.6e-17 \| 1.6e-22 \| 123.9 \| 112 \| (8, 119) \| 221 \| (299, 415) \| 494 \| Phage tail collar domain containing protein \| Phage tail collar domain containing protein; uniclust \| UniRef100\_A0A1Y1IPX0 \| 99.5 \| 9.8e-17 \| 1.8e-22 \| 111.9 \| 65 \| (38, 120) \| 221 \| (29, 93) \| 164 \| Phage tail collar domain-containing protein \| Phage tail collar domain-containing protein; uniclust \| UniRef100\_A0A1E4J0W6 \| 99.5 \| 1e-16 \| 1.9e-22 \| 119.4 \| 116 \| (11, 145) \| 221 \| (121, 236) \| 337 \| Phage tail collar domain-containing protein \| Phage tail collar domain-containing protein; uniclust \| UniRef100\_A0A0H5Q3U7 \| 99.5 \| 1e-16 \| 1.9e-22 \| 116.3 \| 89 \| (11, 120) \| 221 \| (71, 159) \| 213 \| Phage tail collar domain-containing protein \| Phage tail collar domain-containing protein; uniclust \| UniRef100\_A0A1G3JKE1 \| 99.5 \| 1e-16 \| 1.9e-22 \| 121.9 \| 78 \| (8, 85) \| 221 \| (137, 214) \| 286 \| Peptidase S74 domain-containing protein \| Peptidase S74 domain-containing protein; uniclust \| UniRef100\_A0A814Z794 \| 99.5 \| 1e-16 \| 1.9e-22 \| 116.6 \| 122 \| (9, 145) \| 221 \| (119, 241) \| 252 \| Phage tail collar domain-containing protein \| Phage tail collar domain-containing protein; uniclust \| UniRef100\_A0A835XQ12 \| 99.5 \| 1.1e-16 \| 2e-22 \| 120.8 \| 98 \| (10, 119) \| 221 \| (235, 335) \| 409 \| Phage tail collar domain-containing protein \| Phage tail collar domain-containing protein; uniclust \| UniRef100\_A0A2I7SAV6 \| 99.5 \| 1.1e-16 \| 2e-22 \| 118.2 \| 87 \| (8, 119) \| 221 \| (121, 207) \| 268 \| Short tail fiber protein \| Short tail fiber protein; uniclust \| UniRef100\_A0A3M1AQT8 \| 99.5 \| 1.1e-16 \| 2e-22 \| 132.3 \| 195 \| (9, 221) \| 221 \| (853, 1066) \| 1066 \| Phage tail collar domain-containing protein \| Phage tail collar domain-containing protein; uniclust \| UniRef100\_A0A814VVY1 \| 99.5 \| 1.1e-16 \| 2e-22 \| 124.1 \| 92 \| (10, 119) \| 221 \| (330, 421) \| 423 \| Phage tail collar domain-containing protein \| Phage tail collar domain-containing protein; uniclust \| UniRef100\_A0A1M6Z489 \| 99.5 \| 1.1e-16 \| 2.1e-22 \| 114.9 \| 104 \| (9, 118) \| 221 \| (114, 217) \| 264 \| Phage Tail Collar Domain \| Phage Tail Collar Domain; uniclust \| UniRef100\_A0A2S4LWD3 \| 99.5 \| 1.1e-16 \| 2.1e-22 \| 141.0 \| 190 \| (10, 221) \| 221 \| (844, 1066) \| 1067 \| Microcystin-dependent protein \| Microcystin-dependent protein; uniclust \| UniRef100\_A0A522BAB4 \| 99.5 \| 1.2e-16 \| 2.1e-22 \| 113.3 \| 97 \| (11, 120) \| 221 \| (98, 194) \| 249 \| Phage tail collar domain-containing protein \| Phage tail collar domain-containing protein; uniclust \| UniRef100\_A0A133XLD8 \| 99.5 \| 1.2e-16 \| 2.2e-22 \| 116.7 \| 78 \| (8, 85) \| 221 \| (124, 201) \| 268 \| Phage tail collar domain-containing protein \| Phage tail collar domain-containing protein; uniclust \| UniRef100\_A0A376BTN0 \| 99.5 \| 1.2e-16 \| 2.3e-22 \| 132.3 \| 94 \| (7, 119) \| 221 \| (507, 600) \| 669 \| Phage Tail Collar Domain \| Phage Tail Collar Domain; uniclust \| UniRef100\_A0A6J5QPP2 \| 99.5 \| 1.3e-16 \| 2.4e-22 \| 118.2 \| 93 \| (10, 121) \| 221 \| (137, 229) \| 348 \| Phage tail collar domain containing protein \| Phage tail collar domain containing protein; uniclust \| UniRef100\_A0A150PN60 \| 99.5 \| 1.4e-16 \| 2.5e-22 \| 120.1 \| 69 \| (8, 119) \| 221 \| (273, 343) \| 404 \| Phage tail collar domain-containing protein \| Phage tail collar domain-containing protein; uniclust \| UniRef100\_A0A966QWS9 \| 99.5 \| 1.4e-16 \| 2.5e-22 \| 118.8 \| 69 \| (8, 76) \| 221 \| (208, 277) \| 368 \| Phage tail collar domain-containing protein \| Phage tail collar domain-containing protein; uniclust \| UniRef100\_A0A2E6GQF0 \| 99.5 \| 1.4e-16 \| 2.6e-22 \| 118.0 \| 98 \| (10, 119) \| 221 \| (199, 297) \| 351 \| Phage tail collar domain-containing protein \| Phage tail collar domain-containing protein; uniclust \| UniRef100\_A0A821TQI6 \| 99.5 \| 1.4e-16 \| 2.6e-22 \| 115.8 \| 106 \| (25, 149) \| 221 \| (14, 119) \| 279 \| Tesmin/TSO1-like CXC domain-containing protein (Fragment) \| Tesmin/TSO1-like CXC domain-containing protein (Fragment); uniclust \| UniRef100\_A0A1G3HMJ8 \| 99.5 \| 1.4e-16 \| 2.7e-22 \| 125.7 \| 75 \| (10, 84) \| 221 \| (174, 248) \| 324 \| Phage tail collar domain-containing protein \| Phage tail collar domain-containing protein; uniclust \| UniRef100\_A0A158DQ00 \| 99.5 \| 1.4e-16 \| 2.7e-22 \| 123.6 \| 72 \| (9, 119) \| 221 \| (79, 152) \| 278 \| Phage-related tail fiber protein \| Phage-related tail fiber protein; uniclust \| UniRef100\_A0A822B9W3 \| 99.5 \| 1.5e-16 \| 2.8e-22 \| 103.2 \| 96 \| (8, 119) \| 221 \| (32, 132) \| 140 \| Phage tail collar domain-containing protein \| Phage tail collar domain-containing protein; uniclust \| UniRef100\_A0A084EG14 \| 99.5 \| 1.6e-16 \| 2.9e-22 \| 110.7 \| 97 \| (10, 119) \| 221 \| (14, 111) \| 195 \| Tail collar domain-containing protein \| Tail collar domain-containing protein; uniclust \| UniRef100\_UPI001C0EAFD0 \| 99.5 \| 1.6e-16 \| 3e-22 \| 125.9 \| 151 \| (8, 221) \| 221 \| (495, 647) \| 647 \| hypothetical protein \| hypothetical protein; uniclust \| UniRef100\_A0A2R4H0T7 \| 99.5 \| 1.6e-16 \| 3e-22 \| 122.7 \| 97 \| (10, 119) \| 221 \| (265, 445) \| 501 \| Phage tail collar domain-containing protein \| Phage tail collar domain-containing protein; uniclust \| UniRef100\_A0A1Y4GXK4 \| 99.5 \| 1.7e-16 \| 3e-22 \| 113.0 \| 94 \| (9, 119) \| 221 \| (94, 187) \| 245 \| Phage tail collar domain-containing protein \| Phage tail collar domain-containing protein; uniclust \| UniRef100\_UPI001D104AA5 \| 99.5 \| 1.7e-16 \| 3.1e-22 \| 113.5 \| 100 \| (9, 119) \| 221 \| (77, 176) \| 265 \| tail fiber protein \| tail fiber protein; uniclust \| UniRef100\_A0A1C3YQ05 \| 99.5 \| 1.7e-16 \| 3.1e-22 \| 121.9 \| 78 \| (8, 85) \| 221 \| (151, 228) \| 314 \| Phage tail fibre repeat-containing protein \| Phage tail fibre repeat-containing protein; uniclust \| UniRef100\_A0A0G3CP38 \| 99.5 \| 1.8e-16 \| 3.4e-22 \| 111.7 \| 73 \| (4, 86) \| 221 \| (15, 87) \| 179 \| Phage tail collar domain-containing protein \| Phage tail collar domain-containing protein; uniclust \| UniRef100\_A0A074VFM7 \| 99.5 \| 2.1e-16 \| 3.9e-22 \| 127.9 \| 78 \| (8, 85) \| 221 \| (226, 303) \| 411 \| Phage Tail Collar Domain \| Phage Tail Collar Domain; uniclust \| UniRef100\_UPI000532B154 \| 99.5 \| 2.2e-16 \| 4e-22 \| 112.9 \| 83 \| (8, 119) \| 221 \| (72, 154) \| 218 \| tail fiber protein \| tail fiber protein; uniclust \| UniRef100\_A0A7W4IK36 \| 99.5 \| 2.2e-16 \| 4e-22 \| 125.4 \| 107 \| (20, 133) \| 221 \| (3, 109) \| 659 \| Phage tail collar domain-containing protein \| Phage tail collar domain-containing protein; uniclust \| UniRef100\_A0A4R1FMB3 \| 99.5 \| 2.2e-16 \| 4.1e-22 \| 130.3 \| 92 \| (9, 119) \| 221 \| (668, 759) \| 817 \| Tail collar domain \| Tail collar domain; uniclust \| UniRef100\_A0A817Z300 \| 99.4 \| 2.5e-16 \| 4.6e-22 \| 120.2 \| 198 \| (10, 221) \| 221 \| (243, 448) \| 449 \| Phage tail collar domain-containing protein \| Phage tail collar domain-containing protein; uniclust \| UniRef100\_A0A8S2DDE4 \| 99.4 \| 2.6e-16 \| 4.7e-22 \| 126.2 \| 177 \| (15, 220) \| 221 \| (2, 189) \| 728 \| Amino acid permease/ SLC12A domain-containing protein \| Amino acid permease/ SLC12A domain-containing protein; uniclust \| UniRef100\_A0A2D8EVB5 \| 99.4 \| 2.6e-16 \| 4.8e-22 \| 124.9 \| 75 \| (9, 126) \| 221 \| (508, 586) \| 652 \| Major tropism determinant N-terminal domain-containing protein \| Major tropism determinant N-terminal domain-containing protein; uniclust \| UniRef100\_A0A1X0TJA2 \| 99.4 \| 2.6e-16 \| 4.8e-22 \| 114.2 \| 76 \| (10, 85) \| 221 \| (66, 141) \| 224 \| Phage tail collar domain-containing protein \| Phage tail collar domain-containing protein; uniclust \| UniRef100\_A0A7L4ZU76 \| 99.4 \| 2.7e-16 \| 4.9e-22 \| 116.9 \| 95 \| (11, 119) \| 221 \| (4, 98) \| 356 \| Phage tail collar domain-containing protein \| Phage tail collar domain-containing protein; uniclust \| UniRef100\_A0A8S5VPB3 \| 99.4 \| 2.9e-16 \| 5.4e-22 \| 116.1 \| 40 \| (9, 74) \| 221 \| (34, 75) \| 262 \| Tail fiber protein \| Tail fiber protein; uniclust \| UniRef100\_A0A6J5PAS1 \| 99.4 \| 3e-16 \| 5.4e-22 \| 113.2 \| 93 \| (7, 119) \| 221 \| (140, 232) \| 281 \| Phage tail collar domain containing protein \| Phage tail collar domain containing protein; uniclust \| UniRef100\_A0A534A2U7 \| 99.4 \| 3.1e-16 \| 5.6e-22 \| 105.1 \| 86 \| (23, 119) \| 221 \| (36, 121) \| 170 \| Phage tail collar domain-containing protein \| Phage tail collar domain-containing protein; uniclust \| UniRef100\_A0A3M2BJC7 \| 99.4 \| 3.1e-16 \| 5.8e-22 \| 106.3 \| 76 \| (9, 84) \| 221 \| (92, 167) \| 183 \| Tail fiber protein (Fragment) \| Tail fiber protein (Fragment); uniclust \| UniRef100\_A0A957SMD6 \| 99.4 \| 3.2e-16 \| 5.9e-22 \| 125.9 \| 67 \| (10, 119) \| 221 \| (617, 685) \| 742 \| Uncharacterized protein \| Uncharacterized protein; uniclust \| UniRef100\_A0A212LXT2 \| 99.4 \| 3.3e-16 \| 6e-22 \| 114.8 \| 74 \| (12, 119) \| 221 \| (140, 218) \| 273 \| Phage Tail Collar domain protein \| Phage Tail Collar domain protein; uniclust \| UniRef100\_A0A2Z6AZ11 \| 99.4 \| 3.6e-16 \| 6.7e-22 \| 119.8 \| 74 \| (10, 83) \| 221 \| (187, 260) \| 348 \| Tail Collar domain protein \| Tail Collar domain protein; uniclust \| UniRef100\_A0A3D4BUX3 \| 99.4 \| 3.7e-16 \| 6.9e-22 \| 92.9 \| 64 \| (10, 74) \| 221 \| (5, 68) \| 72 \| Phage tail protein (Fragment) \| Phage tail protein (Fragment); uniclust \| UniRef100\_A0A963Z1H4 \| 99.4 \| 3.9e-16 \| 7.1e-22 \| 111.0 \| 60 \| (17, 76) \| 221 \| (27, 86) \| 252 \| Tail fiber protein \| Tail fiber protein; uniclust \| UniRef100\_A0A1B8YGC4 \| 99.4 \| 4.2e-16 \| 7.8e-22 \| 131.8 \| 68 \| (8, 85) \| 221 \| (470, 537) \| 616 \| Phage tail fiber repeat protein \| Phage tail fiber repeat protein; uniclust \| UniRef100\_UPI001FFB714D \| 99.4 \| 4.3e-16 \| 7.8e-22 \| 112.2 \| 91 \| (10, 119) \| 221 \| (108, 198) \| 277 \| phage tail protein \| phage tail protein; uniclust \| UniRef100\_A0A0K4CUX8 \| 99.4 \| 4.2e-16 \| 7.9e-22 \| 107.1 \| 64 \| (12, 85) \| 221 \| (8, 71) \| 148 \| Alternative C-terminus of phage tail fiber protein, fused to tail fiber protein by inversion \| Alternative C-terminus of phage tail fiber protein, fused to tail fiber protein by inversion; uniclust \| UniRef100\_A0A0T9PN75 \| 99.4 \| 4.5e-16 \| 8.4e-22 \| 127.5 \| 87 \| (7, 119) \| 221 \| (270, 364) \| 447 \| Phage Tail Collar Domain \| Phage Tail Collar Domain; uniclust \| UniRef100\_A0A0A2S0D5 \| 99.4 \| 4.6e-16 \| 8.5e-22 \| 110.2 \| 86 \| (4, 118) \| 221 \| (34, 119) \| 186 \| Phage tail protein \| Phage tail protein; uniclust \| UniRef100\_A0A6J7VL93 \| 99.4 \| 4.6e-16 \| 8.5e-22 \| 116.0 \| 98 \| (22, 119) \| 221 \| (207, 307) \| 363 \| Phage tail collar domain containing protein \| Phage tail collar domain containing protein; uniclust \| UniRef100\_A0A5E7XPA5 \| 99.4 \| 4.7e-16 \| 8.6e-22 \| 111.9 \| 80 \| (10, 128) \| 221 \| (110, 191) \| 275 \| Phage tail collar domain-containing protein \| Phage tail collar domain-containing protein; uniclust \| UniRef100\_A0A071MNQ9 \| 99.4 \| 4.8e-16 \| 9e-22 \| 129.9 \| 191 \| (9, 221) \| 221 \| (430, 652) \| 653 \| Tail protein \| Tail protein; uniclust \| UniRef100\_UPI0006A999EC \| 99.4 \| 5e-16 \| 9.2e-22 \| 115.3 \| 73 \| (8, 119) \| 221 \| (191, 265) \| 352 \| tail fiber protein \| tail fiber protein; uniclust \| UniRef100\_A0A0F8WDG3 \| 99.4 \| 5.1e-16 \| 9.7e-22 \| 107.2 \| 55 \| (9, 106) \| 221 \| (19, 75) \| 135 \| Phage tail collar domain-containing protein \| Phage tail collar domain-containing protein; uniclust \| UniRef100\_UPI001FC8B715 \| 99.4 \| 5.3e-16 \| 9.7e-22 \| 99.3 \| 67 \| (9, 75) \| 221 \| (47, 113) \| 128 \| phage tail protein \| phage tail protein; uniclust \| UniRef100\_C6ABW9 \| 99.4 \| 5.3e-16 \| 9.7e-22 \| 117.5 \| 75 \| (9, 83) \| 221 \| (220, 294) \| 370 \| Phage tail collar protein \| Phage tail collar protein; uniclust \| UniRef100\_UPI000736F1C9 \| 99.4 \| 5.6e-16 \| 1e-21 \| 119.7 \| 73 \| (13, 85) \| 221 \| (335, 407) \| 496 \| tail fiber protein \| tail fiber protein; uniclust \| UniRef100\_A0A821Z358 \| 99.4 \| 5.7e-16 \| 1e-21 \| 102.3 \| 93 \| (11, 119) \| 221 \| (2, 99) \| 154 \| Phage tail collar domain-containing protein \| Phage tail collar domain-containing protein; uniclust \| UniRef100\_A0A954EH12 \| 99.4 \| 6e-16 \| 1.1e-21 \| 116.6 \| 100 \| (10, 121) \| 221 \| (114, 213) \| 395 \| Tail fiber protein (Fragment) \| Tail fiber protein (Fragment); uniclust \| UniRef100\_A0A061JKN2 \| 99.4 \| 6e-16 \| 1.1e-21 \| 126.9 \| 76 \| (10, 85) \| 221 \| (327, 402) \| 521 \| Tail protein \| Tail protein; uniclust \| UniRef100\_A0A0K8JHT0 \| 99.4 \| 6.3e-16 \| 1.2e-21 \| 118.7 \| 92 \| (9, 119) \| 221 \| (193, 285) \| 469 \| Phage tail collar domain-containing protein \| Phage tail collar domain-containing protein; uniclust \| UniRef100\_A0A022PG07 \| 99.4 \| 6.3e-16 \| 1.2e-21 \| 124.6 \| 88 \| (6, 119) \| 221 \| (256, 351) \| 434 \| Phage tail collar family protein \| Phage tail collar family protein; uniclust \| UniRef100\_A0A6M0S0R8 \| 99.4 \| 6.5e-16 \| 1.2e-21 \| 99.9 \| 62 \| (15, 119) \| 221 \| (2, 65) \| 125 \| Tail fiber protein \| Tail fiber protein; uniclust \| UniRef100\_A0A5E4KZA2 \| 99.4 \| 6.7e-16 \| 1.2e-21 \| 111.2 \| 67 \| (10, 119) \| 221 \| (117, 185) \| 277 \| Phage Tail Collar Domain protein \| Phage Tail Collar Domain protein; uniclust \| UniRef100\_A0A081RQS4 \| 99.4 \| 6.9e-16 \| 1.3e-21 \| 123.6 \| 83 \| (8, 119) \| 221 \| (331, 413) \| 474 \| Phage-related tail fiber protein (Fragment) \| Phage-related tail fiber protein (Fragment); uniclust \| UniRef100\_A0A6J5QSS9 \| 99.4 \| 7.2e-16 \| 1.3e-21 \| 113.0 \| 102 \| (7, 119) \| 221 \| (109, 210) \| 317 \| MdpB Microcystin-dependent protein \| MdpB Microcystin-dependent protein; uniclust \| UniRef100\_A0A1S8LFT6 \| 99.4 \| 7.1e-16 \| 1.3e-21 \| 112.1 \| 66 \| (10, 75) \| 221 \| (23, 88) \| 178 \| Phage tail collar domain protein \| Phage tail collar domain protein; uniclust \| UniRef100\_A0A7T8IWR9 \| 99.4 \| 7.4e-16 \| 1.4e-21 \| 120.8 \| 82 \| (9, 119) \| 221 \| (423, 506) \| 568 \| Tail fiber protein \| Tail fiber protein; uniclust \| UniRef100\_A0A2E0ZP65 \| 99.4 \| 7.8e-16 \| 1.4e-21 \| 100.8 \| 75 \| (9, 83) \| 221 \| (20, 94) \| 146 \| Phage tail collar domain-containing protein \| Phage tail collar domain-containing protein; uniclust \| UniRef100\_A0A4U6CQW2 \| 99.4 \| 7.8e-16 \| 1.4e-21 \| 117.5 \| 98 \| (9, 119) \| 221 \| (238, 335) \| 342 \| Phage tail collar domain-containing protein \| Phage tail collar domain-containing protein; uniclust \| UniRef100\_A0A6P2HBE9 \| 99.4 \| 7.8e-16 \| 1.5e-21 \| 112.9 \| 171 \| (9, 218) \| 221 \| (54, 228) \| 233 \| Tail fiber protein \| Tail fiber protein; uniclust \| UniRef100\_A0A957NRE5 \| 99.4 \| 8.1e-16 \| 1.5e-21 \| 115.6 \| 67 \| (10, 119) \| 221 \| (265, 333) \| 388 \| Uncharacterized protein (Fragment) \| Uncharacterized protein (Fragment); uniclust \| UniRef100\_A0A228IJ11 \| 99.4 \| 8.3e-16 \| 1.5e-21 \| 122.6 \| 174 \| (5, 221) \| 221 \| (305, 527) \| 531 \| Tail fiber protein \| Tail fiber protein; uniclust \| UniRef100\_A0A973FDY3 \| 99.4 \| 8.9e-16 \| 1.6e-21 \| 109.1 \| 182 \| (10, 221) \| 221 \| (70, 251) \| 251 \| Tail fiber protein \| Tail fiber protein; uniclust \| UniRef100\_A0A1X3DKU1 \| 99.4 \| 8.9e-16 \| 1.7e-21 \| 112.8 \| 76 \| (8, 83) \| 221 \| (80, 155) \| 256 \| Phage tail collar domain-containing protein \| Phage tail collar domain-containing protein; uniclust \| UniRef100\_A0A835ZHE3 \| 99.4 \| 9.7e-16 \| 1.8e-21 \| 111.3 \| 79 \| (24, 119) \| 221 \| (16, 94) \| 294 \| Phage tail collar domain-containing protein \| Phage tail collar domain-containing protein; uniclust \| UniRef100\_A0A1G5MZE4 \| 99.4 \| 1.2e-15 \| 2.2e-21 \| 126.1 \| 74 \| (12, 85) \| 221 \| (662, 735) \| 825 \| Phage Tail Collar Domain \| Phage Tail Collar Domain; uniclust \| UniRef100\_A0A1V5WU45 \| 99.4 \| 1.2e-15 \| 2.2e-21 \| 108.1 \| 125 \| (11, 143) \| 221 \| (8, 138) \| 210 \| Phage Tail Collar Domain protein \| Phage Tail Collar Domain protein; uniclust \| UniRef100\_A0A2N9Y859 \| 99.4 \| 1.2e-15 \| 2.2e-21 \| 120.4 \| 78 \| (8, 85) \| 221 \| (157, 234) \| 390 \| Phage tail protein \| Phage tail protein; uniclust \| UniRef100\_A0A090ZDZ3 \| 99.4 \| 1.3e-15 \| 2.4e-21 \| 102.9 \| 91 \| (10, 119) \| 221 \| (38, 128) \| 130 \| Phage Tail Collar domain protein \| Phage Tail Collar domain protein; uniclust \| UniRef100\_A0A2Z2P0U1 \| 99.4 \| 1.3e-15 \| 2.5e-21 \| 112.1 \| 67 \| (9, 117) \| 221 \| (67, 133) \| 193 \| Phage tail collar domain-containing protein \| Phage tail collar domain-containing protein; uniclust \| UniRef100\_A0A849R972 \| 99.4 \| 1.4e-15 \| 2.6e-21 \| 109.8 \| 67 \| (8, 119) \| 221 \| (164, 232) \| 280 \| Tail fiber protein \| Tail fiber protein; uniclust \| UniRef100\_A0A1I2LDU5 \| 99.4 \| 1.4e-15 \| 2.6e-21 \| 119.0 \| 94 \| (7, 119) \| 221 \| (305, 398) \| 450 \| Microcystin-dependent protein \| Microcystin-dependent protein; uniclust \| UniRef100\_UPI001F3862FB \| 99.4 \| 1.4e-15 \| 2.6e-21 \| 112.5 \| 92 \| (9, 119) \| 221 \| (175, 266) \| 338 \| phage tail protein \| phage tail protein; uniclust \| UniRef100\_A0A815V480 \| 99.4 \| 1.5e-15 \| 2.7e-21 \| 123.1 \| 195 \| (10, 221) \| 221 \| (500, 702) \| 803 \| Phage tail collar domain-containing protein \| Phage tail collar domain-containing protein; uniclust \| UniRef100\_A0A069PMJ9 \| 99.4 \| 1.6e-15 \| 3e-21 \| 124.4 \| 76 \| (10, 85) \| 221 \| (294, 385) \| 521 \| Tail protein \| Tail protein; uniclust \| UniRef100\_A0A956Q788 \| 99.4 \| 1.7e-15 \| 3.1e-21 \| 108.9 \| 112 \| (8, 119) \| 221 \| (83, 221) \| 270 \| Uncharacterized protein \| Uncharacterized protein; uniclust \| UniRef100\_A0A2I2L5J8 \| 99.4 \| 1.7e-15 \| 3.1e-21 \| 125.6 \| 168 \| (8, 221) \| 221 \| (890, 1059) \| 1059 \| Collagen-like protein with Phage Tail Collar Domain \| Collagen-like protein with Phage Tail Collar Domain; uniclust \| UniRef100\_A0A0A0CRJ5 \| 99.4 \| 1.8e-15 \| 3.3e-21 \| 108.0 \| 67 \| (7, 83) \| 221 \| (52, 118) \| 195 \| Phage tail collar domain-containing protein \| Phage tail collar domain-containing protein; uniclust \| UniRef100\_A0A0H3GJI2 \| 99.4 \| 1.8e-15 \| 3.3e-21 \| 117.8 \| 75 \| (11, 85) \| 221 \| (233, 307) \| 385 \| Phage tail collar domain-containing protein \| Phage tail collar domain-containing protein; uniclust \| UniRef100\_UPI00210D7C8A \| 99.4 \| 1.8e-15 \| 3.4e-21 \| 91.8 \| 83 \| (24, 119) \| 221 \| (1, 92) \| 94 \| tail fiber protein \| tail fiber protein; uniclust \| UniRef100\_A0A928V6C5 \| 99.4 \| 2.2e-15 \| 3.9e-21 \| 117.8 \| 70 \| (10, 119) \| 221 \| (401, 472) \| 544 \| FHA domain-containing protein \| FHA domain-containing protein; uniclust \| UniRef100\_A0A516LLH4 \| 99.4 \| 2.3e-15 \| 4.2e-21 \| 97.2 \| 80 \| (8, 87) \| 221 \| (15, 94) \| 133 \| Putative tail fiber protein \| Putative tail fiber protein; uniclust \| UniRef100\_UPI0004113DCB \| 99.4 \| 2.3e-15 \| 4.3e-21 \| 119.3 \| 147 \| (9, 221) \| 221 \| (476, 627) \| 627 \| hypothetical protein \| hypothetical protein; uniclust \| UniRef100\_A0A1G8CBW8 \| 99.4 \| 2.4e-15 \| 4.4e-21 \| 104.9 \| 77 \| (9, 85) \| 221 \| (23, 99) \| 181 \| Phage Tail Collar Domain \| Phage Tail Collar Domain; uniclust \| UniRef100\_A0A6V8M1A9 \| 99.4 \| 2.4e-15 \| 4.5e-21 \| 110.4 \| 101 \| (7, 119) \| 221 \| (76, 178) \| 243 \| Phage tail collar domain-containing protein \| Phage tail collar domain-containing protein; uniclust \| UniRef100\_A0A1I3YTE4 \| 99.4 \| 2.5e-15 \| 4.6e-21 \| 116.6 \| 83 \| (8, 119) \| 221 \| (227, 309) \| 358 \| Phage Tail Collar Domain (Fragment) \| Phage Tail Collar Domain (Fragment); uniclust \| UniRef100\_UPI001F13FAFE \| 99.4 \| 2.5e-15 \| 4.6e-21 \| 104.6 \| 79 \| (7, 85) \| 221 \| (72, 150) \| 215 \| phage tail protein \| phage tail protein; uniclust \| UniRef100\_A0A1H0KC46 \| 99.4 \| 2.6e-15 \| 4.8e-21 \| 111.5 \| 105 \| (7, 119) \| 221 \| (36, 151) \| 215 \| Microcystin-dependent protein \| Microcystin-dependent protein; pdb70 \| 1PDI\_F \| 99.4 \| 3.5e-17 \| 3.3e-21 \| 142.9 \| 112 \| (8, 123) \| 221 \| (90, 204) \| 278 \| Short tail fiber protein \| 1PDI\_F Short tail fiber protein STRUCTURAL PROTEIN; pdb70 \| 5IV5\_AI \| 99.3 \| 4.4e-16 \| 4.1e-20 \| 146.1 \| 112 \| (8, 123) \| 221 \| (339, 453) \| 527 \| Baseplate wedge protein gp6, Baseplate \| 5IV5\_AI Baseplate wedge protein gp6, Baseplate T4, baseplate-tail tube complex, pre-attachment; pdb70 \| 5IV5\_AJ \| 99.3 \| 4.4e-16 \| 4.1e-20 \| 146.1 \| 112 \| (8, 123) \| 221 \| (339, 453) \| 527 \| Baseplate wedge protein gp6, Baseplate \| 5IV5\_AJ Baseplate wedge protein gp6, Baseplate T4, baseplate-tail tube complex, pre-attachment; pdb70 \| 5IV5\_BA \| 99.3 \| 4.4e-16 \| 4.1e-20 \| 146.1 \| 112 \| (8, 123) \| 221 \| (339, 453) \| 527 \| Baseplate wedge protein gp6, Baseplate \| 5IV5\_BA Baseplate wedge protein gp6, Baseplate T4, baseplate-tail tube complex, pre-attachment; pdb70 \| 5IV5\_DB \| 99.3 \| 4.4e-16 \| 4.1e-20 \| 146.1 \| 112 \| (8, 123) \| 221 \| (339, 453) \| 527 \| Baseplate wedge protein gp6, Baseplate \| 5IV5\_DB Baseplate wedge protein gp6, Baseplate T4, baseplate-tail tube complex, pre-attachment; pdb70 \| 5IV5\_DC \| 99.3 \| 4.4e-16 \| 4.1e-20 \| 146.1 \| 112 \| (8, 123) \| 221 \| (339, 453) \| 527 \| Baseplate wedge protein gp6, Baseplate \| 5IV5\_DC Baseplate wedge protein gp6, Baseplate T4, baseplate-tail tube complex, pre-attachment; pdb70 \| 5IV5\_DD \| 99.3 \| 4.4e-16 \| 4.1e-20 \| 146.1 \| 112 \| (8, 123) \| 221 \| (339, 453) \| 527 \| Baseplate wedge protein gp6, Baseplate \| 5IV5\_DD Baseplate wedge protein gp6, Baseplate T4, baseplate-tail tube complex, pre-attachment; pdb70 \| 5IV5\_FE \| 99.3 \| 4.4e-16 \| 4.1e-20 \| 146.1 \| 112 \| (8, 123) \| 221 \| (339, 453) \| 527 \| Baseplate wedge protein gp6, Baseplate \| 5IV5\_FE Baseplate wedge protein gp6, Baseplate T4, baseplate-tail tube complex, pre-attachment; pdb70 \| 5IV5\_FF \| 99.3 \| 4.4e-16 \| 4.1e-20 \| 146.1 \| 112 \| (8, 123) \| 221 \| (339, 453) \| 527 \| Baseplate wedge protein gp6, Baseplate \| 5IV5\_FF Baseplate wedge protein gp6, Baseplate T4, baseplate-tail tube complex, pre-attachment; pdb70 \| 5IV5\_FG \| 99.3 \| 4.4e-16 \| 4.1e-20 \| 146.1 \| 112 \| (8, 123) \| 221 \| (339, 453) \| 527 \| Baseplate wedge protein gp6, Baseplate \| 5IV5\_FG Baseplate wedge protein gp6, Baseplate T4, baseplate-tail tube complex, pre-attachment; pdb70 \| 5IV5\_HH \| 99.3 \| 4.4e-16 \| 4.1e-20 \| 146.1 \| 112 \| (8, 123) \| 221 \| (339, 453) \| 527 \| Baseplate wedge protein gp6, Baseplate \| 5IV5\_HH Baseplate wedge protein gp6, Baseplate T4, baseplate-tail tube complex, pre-attachment; pdb70 \| 5IV5\_HJ \| 99.3 \| 4.4e-16 \| 4.1e-20 \| 146.1 \| 112 \| (8, 123) \| 221 \| (339, 453) \| 527 \| Baseplate wedge protein gp6, Baseplate \| 5IV5\_HJ Baseplate wedge protein gp6, Baseplate T4, baseplate-tail tube complex, pre-attachment; pdb70 \| 5IV5\_O \| 99.3 \| 4.4e-16 \| 4.1e-20 \| 146.1 \| 112 \| (8, 123) \| 221 \| (339, 453) \| 527 \| Baseplate wedge protein gp6, Baseplate \| 5IV5\_O Baseplate wedge protein gp6, Baseplate T4, baseplate-tail tube complex, pre-attachment; pdb70 \| 5IV5\_P \| 99.3 \| 4.4e-16 \| 4.1e-20 \| 146.1 \| 112 \| (8, 123) \| 221 \| (339, 453) \| 527 \| Baseplate wedge protein gp6, Baseplate \| 5IV5\_P Baseplate wedge protein gp6, Baseplate T4, baseplate-tail tube complex, pre-attachment; pdb70 \| 5IV5\_Q \| 99.3 \| 4.4e-16 \| 4.1e-20 \| 146.1 \| 112 \| (8, 123) \| 221 \| (339, 453) \| 527 \| Baseplate wedge protein gp6, Baseplate \| 5IV5\_Q Baseplate wedge protein gp6, Baseplate T4, baseplate-tail tube complex, pre-attachment; pdb70 \| 5IV5\_n \| 99.3 \| 4.4e-16 \| 4.1e-20 \| 146.1 \| 112 \| (8, 123) \| 221 \| (339, 453) \| 527 \| Baseplate wedge protein gp6, Baseplate \| 5IV5\_n Baseplate wedge protein gp6, Baseplate T4, baseplate-tail tube complex, pre-attachment; pdb70 \| 1OCY\_A \| 99.2 \| 6.1e-16 \| 6.1e-20 \| 127.3 \| 111 \| (7, 121) \| 221 \| (9, 122) \| 198 \| BACTERIOPHAGE T4 SHORT TAIL FIBRE \| 1OCY\_A BACTERIOPHAGE T4 SHORT TAIL FIBRE STRUCTURAL PROTEIN, FIBROUS PROTEIN, LIPO-POLYSACCHARIDE HET: CIT, SO4; pdb70 \| 2XGF\_A \| 99.1 \| 1.7e-14 \| 1.6e-18 \| 122.8 \| 80 \| (7, 121) \| 221 \| (26, 105) \| 242 \| LONG TAIL FIBER PROTEIN P37 \| 2XGF\_A LONG TAIL FIBER PROTEIN P37 VIRAL PROTEIN, FIBER PROTEIN; pdb70 \| 5LYE\_A \| 98.4 \| 5.4e-11 \| 5e-15 \| 104.9 \| 65 \| (7, 73) \| 221 \| (254, 319) \| 322 \| Gp12 \| 5LYE\_A Gp12 STRUCTURAL PROTEIN, GENE PRODUCT 12; pdb70 \| 2FKK\_A \| 98.0 \| 2.8e-09 \| 2.7e-13 \| 88.1 \| 55 \| (66, 121) \| 221 \| (87, 145) \| 206 \| Baseplate structural protein Gp10 \| 2FKK\_A Baseplate structural protein Gp10 Bacteriophage T4 Baseplate HET: EDO; pdb70 \| 5IV5\_HI \| 97.3 \| 1.7e-07 \| 1.6e-11 \| 86.7 \| 112 \| (8, 123) \| 221 \| (339, 453) \| 527 \| Baseplate wedge protein gp6, Baseplate \| 5IV5\_HI Baseplate wedge protein gp6, Baseplate T4, baseplate-tail tube complex, pre-attachment; pdb70 \| 5HX2\_H \| 97.0 \| 1.2e-06 \| 1e-10 \| 82.2 \| 55 \| (66, 121) \| 221 \| (483, 541) \| 602 \| Baseplate wedge protein gp7, Baseplate \| 5HX2\_H Baseplate wedge protein gp7, Baseplate T4, baseplate, complex, VIRAL PROTEIN; pdb70 \| 5IV5\_AC \| 97.0 \| 1.2e-06 \| 1e-10 \| 82.2 \| 55 \| (66, 121) \| 221 \| (483, 541) \| 602 \| Baseplate wedge protein gp6, Baseplate \| 5IV5\_AC Baseplate wedge protein gp6, Baseplate T4, baseplate-tail tube complex, pre-attachment; pdb70 \| 5IV5\_AD \| 97.0 \| 1.2e-06 \| 1e-10 \| 82.2 \| 55 \| (66, 121) \| 221 \| (483, 541) \| 602 \| Baseplate wedge protein gp6, Baseplate \| 5IV5\_AD Baseplate wedge protein gp6, Baseplate T4, baseplate-tail tube complex, pre-attachment; pdb70 \| 5IV5\_AE \| 97.0 \| 1.2e-06 \| 1e-10 \| 82.2 \| 55 \| (66, 121) \| 221 \| (483, 541) \| 602 \| Baseplate wedge protein gp6, Baseplate \| 5IV5\_AE Baseplate wedge protein gp6, Baseplate T4, baseplate-tail tube complex, pre-attachment; pdb70 \| 5IV5\_CF \| 97.0 \| 1.2e-06 \| 1e-10 \| 82.2 \| 55 \| (66, 121) \| 221 \| (483, 541) \| 602 \| Baseplate wedge protein gp6, Baseplate \| 5IV5\_CF Baseplate wedge protein gp6, Baseplate T4, baseplate-tail tube complex, pre-attachment; pdb70 \| 5IV5\_CG \| 97.0 \| 1.2e-06 \| 1e-10 \| 82.2 \| 55 \| (66, 121) \| 221 \| (483, 541) \| 602 \| Baseplate wedge protein gp6, Baseplate \| 5IV5\_CG Baseplate wedge protein gp6, Baseplate T4, baseplate-tail tube complex, pre-attachment; pdb70 \| 5IV5\_CH \| 97.0 \| 1.2e-06 \| 1e-10 \| 82.2 \| 55 \| (66, 121) \| 221 \| (483, 541) \| 602 \| Baseplate wedge protein gp6, Baseplate \| 5IV5\_CH Baseplate wedge protein gp6, Baseplate T4, baseplate-tail tube complex, pre-attachment; pdb70 \| 5IV5\_EI \| 97.0 \| 1.2e-06 \| 1e-10 \| 82.2 \| 55 \| (66, 121) \| 221 \| (483, 541) \| 602 \| Baseplate wedge protein gp6, Baseplate \| 5IV5\_EI Baseplate wedge protein gp6, Baseplate T4, baseplate-tail tube complex, pre-attachment; pdb70 \| 5IV5\_EJ \| 97.0 \| 1.2e-06 \| 1e-10 \| 82.2 \| 55 \| (66, 121) \| 221 \| (483, 541) \| 602 \| Baseplate wedge protein gp6, Baseplate \| 5IV5\_EJ Baseplate wedge protein gp6, Baseplate T4, baseplate-tail tube complex, pre-attachment; pdb70 \| 5IV5\_FA \| 97.0 \| 1.2e-06 \| 1e-10 \| 82.2 \| 55 \| (66, 121) \| 221 \| (483, 541) \| 602 \| Baseplate wedge protein gp6, Baseplate \| 5IV5\_FA Baseplate wedge protein gp6, Baseplate T4, baseplate-tail tube complex, pre-attachment; pdb70 \| 5IV5\_HB \| 97.0 \| 1.2e-06 \| 1e-10 \| 82.2 \| 55 \| (66, 121) \| 221 \| (483, 541) \| 602 \| Baseplate wedge protein gp6, Baseplate \| 5IV5\_HB Baseplate wedge protein gp6, Baseplate T4, baseplate-tail tube complex, pre-attachment; pdb70 \| 5IV5\_HC \| 97.0 \| 1.2e-06 \| 1e-10 \| 82.2 \| 55 \| (66, 121) \| 221 \| (483, 541) \| 602 \| Baseplate wedge protein gp6, Baseplate \| 5IV5\_HC Baseplate wedge protein gp6, Baseplate T4, baseplate-tail tube complex, pre-attachment; pdb70 \| 5IV5\_HD \| 97.0 \| 1.2e-06 \| 1e-10 \| 82.2 \| 55 \| (66, 121) \| 221 \| (483, 541) \| 602 \| Baseplate wedge protein gp6, Baseplate \| 5IV5\_HD Baseplate wedge protein gp6, Baseplate T4, baseplate-tail tube complex, pre-attachment; pdb70 \| 5IV5\_I \| 97.0 \| 1.2e-06 \| 1e-10 \| 82.2 \| 55 \| (66, 121) \| 221 \| (483, 541) \| 602 \| Baseplate wedge protein gp6, Baseplate \| 5IV5\_I Baseplate wedge protein gp6, Baseplate T4, baseplate-tail tube complex, pre-attachment; pdb70 \| 5IV5\_J \| 97.0 \| 1.2e-06 \| 1e-10 \| 82.2 \| 55 \| (66, 121) \| 221 \| (483, 541) \| 602 \| Baseplate wedge protein gp6, Baseplate \| 5IV5\_J Baseplate wedge protein gp6, Baseplate T4, baseplate-tail tube complex, pre-attachment; pdb70 \| 5IV5\_K \| 97.0 \| 1.2e-06 \| 1e-10 \| 82.2 \| 55 \| (66, 121) \| 221 \| (483, 541) \| 602 \| Baseplate wedge protein gp6, Baseplate \| 5IV5\_K Baseplate wedge protein gp6, Baseplate T4, baseplate-tail tube complex, pre-attachment; pdb70 \| 5IV7\_FC \| 97.0 \| 1.2e-06 \| 1e-10 \| 82.2 \| 55 \| (66, 121) \| 221 \| (483, 541) \| 602 \| Baseplate wedge protein gp6, Baseplate \| 5IV7\_FC Baseplate wedge protein gp6, Baseplate T4, baseplate, post-attachment, bacteriophage, bacterial |
| Top keywords  (threshold 1.00e-03 (evalue)) | **tail, Phage, collar, domain\_containing, fiber, Baseplate, T4, wedge, gp6, complex** |
| Output files | ../../similar\_sequences/17\_FANPEZAQ\_CDS\_0017\_merged.svg ../../similar\_sequences/17\_FANPEZAQ\_CDS\_0017\_pdb70.a3m ../../similar\_sequences/17\_FANPEZAQ\_CDS\_0017\_pdb70.hhr ../../similar\_sequences/17\_FANPEZAQ\_CDS\_0017\_uniclust.a3m ../../similar\_sequences/17\_FANPEZAQ\_CDS\_0017\_uniclust.hhr |

#### Structure prediction (AlphaFold)^2^

|  |  |
| --- | --- |
| Stats | xml version="1.0" encoding="utf-8" standalone="no"?       2024-09-02T21:09:16.168346 image/svg+xml   Matplotlib v3.7.2, https://matplotlib.org/ |
| Predicted structure | **NGL Viewer Controls:**  - Center: *Left-Click* - Rotate: *Left-Click + Drag* - Translate: *Right-Click + Drag* - Zoom: *Shift + Left-Click + Drag* |
| Output files | ../../predicted\_structures/17\_FANPEZAQ\_CDS\_0017/features.pkl ../../predicted\_structures/17\_FANPEZAQ\_CDS\_0017/ranked\_0.pdb ../../predicted\_structures/17\_FANPEZAQ\_CDS\_0017/ranked\_0\_plots.svg ../../predicted\_structures/17\_FANPEZAQ\_CDS\_0017/result\_model\_1\_ptm\_pred\_0.pkl |

#### Structure similarity search results (Foldseek)^3^

|  |  |
| --- | --- |
| Structure databases searched | Pdb, Afdb-proteome, Afdb-uniprot50 |
| Results, scheme(s)  (Top layers only, threshold 1.00e-02 (evalue)) | xml version="1.0" encoding="utf-8" standalone="no"?       2024-09-02T21:10:37.577455 image/svg+xml   Matplotlib v3.7.2, https://matplotlib.org/ |
| Results, table  (threshold 1.00e-02 (evalue)) | db \| id \| prob \| evalue \| bits \| fident \| alnlen \| mismatch \| gapopen \| qstart \| qend \| tstart \| tend \| name \| description; pdb \| 1OCY\_A \| 1.0 \| 0.0001505 \| 113 \| 0.18 \| 238 \| 110 \| 10 \| 12 \| 220 \| 14 \| 195 \| BACTERIOPHAGE T4 SHORT TAIL FIBRE \| BACTERIOPHAGE T4 SHORT TAIL FIBRE; pdb \| 1PDI\_A \| 1.0 \| 0.0001695 \| 111 \| 0.236 \| 237 \| 120 \| 16 \| 1 \| 220 \| 83 \| 275 \| Short tail fiber protein \| Short tail fiber protein; pdb \| 5IV5\_O \| 1.0 \| 0.0001337 \| 110 \| 0.207 \| 231 \| 127 \| 12 \| 5 \| 220 \| 334 \| 523 \| Tail tube protein gp19 \| Tail tube protein gp19; pdb \| 2XGF\_B \| 1.0 \| 8.824e-05 \| 106 \| 0.191 \| 246 \| 128 \| 11 \| 11 \| 220 \| 4 \| 214 \| LONG TAIL FIBER PROTEIN P37 \| LONG TAIL FIBER PROTEIN P37; pdb \| 6OV6\_C \| 0.986 \| 0.001717 \| 81 \| 0.135 \| 237 \| 112 \| 16 \| 1 \| 221 \| 84 \| 243 \| C24 PROTEIN \| C24 PROTEIN; pdb \| 6OV6\_A \| 0.967 \| 0.003108 \| 75 \| 0.14 \| 235 \| 113 \| 15 \| 1 \| 221 \| 95 \| 254 \| C24 PROTEIN \| C24 PROTEIN; pdb \| 6OV6\_B \| 0.956 \| 0.004438 \| 73 \| 0.16 \| 231 \| 113 \| 16 \| 1 \| 221 \| 80 \| 239 \| C24 PROTEIN \| C24 PROTEIN; afdb-proteome \| AF-A0A0H3GJI2-F1-MODEL\_V4 \| 1.0 \| 1.608e-08 \| 198 \| 0.251 \| 223 \| 77 \| 9 \| 10 \| 221 \| 231 \| 374 \| Collar domain-containing protein \| Collar domain-containing protein; afdb-proteome \| AF-Q8ZQF5-F1-MODEL\_V4 \| 1.0 \| 7.814e-05 \| 115 \| 0.174 \| 212 \| 142 \| 8 \| 11 \| 220 \| 608 \| 788 \| Putative Fels-1 prophage minor tail protein \| Putative Fels-1 prophage minor tail protein; afdb-proteome \| AF-Q8ZQ81-F1-MODEL\_V4 \| 1.0 \| 0.0002562 \| 115 \| 0.155 \| 212 \| 146 \| 9 \| 11 \| 220 \| 630 \| 810 \| Gifsy-2 prophage probable tail fiber protein \| Gifsy-2 prophage probable tail fiber protein; afdb-proteome \| AF-P77515-F1-MODEL\_V4 \| 0.988 \| 0.0005881 \| 82 \| 0.202 \| 242 \| 144 \| 10 \| 12 \| 220 \| 93 \| 318 \| Prophage side tail fiber protein homolog StfQ \| Prophage side tail fiber protein homolog StfQ; afdb-uniprot50 \| AF-A0A7W4VIA3-F1-MODEL\_V4 \| 1.0 \| 3.91e-22 \| 772 \| 0.489 \| 231 \| 107 \| 5 \| 1 \| 221 \| 1 \| 230 \| Microcystin-dependent protein \| Microcystin-dependent protein; afdb-uniprot50 \| AF-A0A512NCR2-F1-MODEL\_V4 \| 1.0 \| 1.571e-19 \| 622 \| 0.479 \| 223 \| 75 \| 9 \| 3 \| 221 \| 22 \| 207 \| Tail protein \| Tail protein; afdb-uniprot50 \| AF-A0A5C7Q4X8-F1-MODEL\_V4 \| 1.0 \| 4.111e-18 \| 585 \| 0.433 \| 212 \| 80 \| 6 \| 12 \| 221 \| 20 \| 193 \| Phage tail protein \| Phage tail protein; afdb-uniprot50 \| AF-A0A522VPY7-F1-MODEL\_V4 \| 1.0 \| 3.441e-18 \| 583 \| 0.459 \| 211 \| 63 \| 3 \| 11 \| 221 \| 61 \| 220 \| Tail fiber protein \| Tail fiber protein; afdb-uniprot50 \| AF-A0A6A7M412-F1-MODEL\_V4 \| 1.0 \| 4.852e-19 \| 556 \| 0.466 \| 223 \| 75 \| 10 \| 2 \| 221 \| 65 \| 246 \| Collar domain-containing protein \| Collar domain-containing protein; afdb-uniprot50 \| AF-A0A5C7MBC1-F1-MODEL\_V4 \| 1.0 \| 6.23e-18 \| 553 \| 0.459 \| 211 \| 59 \| 5 \| 11 \| 221 \| 108 \| 263 \| Phage tail protein \| Phage tail protein; afdb-uniprot50 \| AF-A0A523UNM2-F1-MODEL\_V4 \| 1.0 \| 5.871e-18 \| 552 \| 0.364 \| 211 \| 104 \| 7 \| 11 \| 221 \| 35 \| 215 \| Collar domain-containing protein \| Collar domain-containing protein; afdb-uniprot50 \| AF-A0A7L7MLE0-F1-MODEL\_V4 \| 1.0 \| 1.61e-17 \| 546 \| 0.377 \| 220 \| 111 \| 6 \| 1 \| 220 \| 54 \| 247 \| Tail fiber protein \| Tail fiber protein; afdb-uniprot50 \| AF-A0A5C7LV16-F1-MODEL\_V4 \| 1.0 \| 2.271e-18 \| 543 \| 0.466 \| 210 \| 72 \| 4 \| 11 \| 220 \| 100 \| 269 \| Collar domain-containing protein \| Collar domain-containing protein; afdb-uniprot50 \| AF-A0A516H786-F1-MODEL\_V4 \| 1.0 \| 3.484e-17 \| 536 \| 0.438 \| 212 \| 67 \| 5 \| 11 \| 221 \| 66 \| 226 \| Collar domain-containing protein \| Collar domain-containing protein; afdb-uniprot50 \| AF-A0A844QL60-F1-MODEL\_V4 \| 1.0 \| 2.3e-17 \| 534 \| 0.416 \| 221 \| 90 \| 7 \| 2 \| 220 \| 81 \| 264 \| Uncharacterized protein \| Uncharacterized protein; afdb-uniprot50 \| AF-A0A5C7PH71-F1-MODEL\_V4 \| 1.0 \| 2.194e-16 \| 526 \| 0.382 \| 212 \| 109 \| 4 \| 11 \| 221 \| 40 \| 230 \| Tail fiber protein \| Tail fiber protein; afdb-uniprot50 \| AF-H0HNH5-F1-MODEL\_V4 \| 1.0 \| 1.076e-16 \| 520 \| 0.403 \| 213 \| 93 \| 5 \| 11 \| 221 \| 206 \| 386 \| Microcystin-dependent protein-like protein \| Microcystin-dependent protein-like protein; afdb-uniprot50 \| AF-A0A6G6K382-F1-MODEL\_V4 \| 1.0 \| 1.076e-16 \| 520 \| 0.381 \| 223 \| 83 \| 4 \| 3 \| 221 \| 251 \| 422 \| Collar domain-containing protein \| Collar domain-containing protein; afdb-uniprot50 \| AF-A0A1Y2J6J7-F1-MODEL\_V4 \| 1.0 \| 2.622e-16 \| 517 \| 0.402 \| 216 \| 88 \| 8 \| 12 \| 221 \| 25 \| 205 \| Collar domain-containing protein \| Collar domain-containing protein; afdb-uniprot50 \| AF-A0A060DEV3-F1-MODEL\_V4 \| 1.0 \| 2.358e-15 \| 514 \| 0.355 \| 211 \| 79 \| 8 \| 12 \| 221 \| 13 \| 167 \| Collar domain-containing protein \| Collar domain-containing protein; afdb-uniprot50 \| AF-D5BQR2-F1-MODEL\_V4 \| 1.0 \| 7.103e-17 \| 511 \| 0.386 \| 212 \| 87 \| 6 \| 11 \| 221 \| 119 \| 288 \| Microcystin-dependent protein-like protein \| Microcystin-dependent protein-like protein; afdb-uniprot50 \| AF-A0A1M3FXB5-F1-MODEL\_V4 \| 1.0 \| 5.945e-17 \| 501 \| 0.413 \| 225 \| 106 \| 7 \| 11 \| 221 \| 93 \| 305 \| Collar domain-containing protein \| Collar domain-containing protein; afdb-uniprot50 \| AF-A0A1F7BR55-F1-MODEL\_V4 \| 1.0 \| 6.02e-16 \| 501 \| 0.386 \| 220 \| 81 \| 3 \| 2 \| 221 \| 358 \| 523 \| Collar domain-containing protein \| Collar domain-containing protein; afdb-uniprot50 \| AF-A0A516IWE1-F1-MODEL\_V4 \| 1.0 \| 2.471e-16 \| 499 \| 0.398 \| 216 \| 89 \| 8 \| 12 \| 221 \| 149 \| 329 \| Phage tail protein \| Phage tail protein; afdb-uniprot50 \| AF-B5ZFN5-F1-MODEL\_V4 \| 1.0 \| 1.002e-17 \| 496 \| 0.454 \| 211 \| 94 \| 6 \| 12 \| 220 \| 86 \| 277 \| Tail Collar domain protein \| Tail Collar domain protein; afdb-uniprot50 \| AF-A0A6H1ZGL2-F1-MODEL\_V4 \| 1.0 \| 1.731e-16 \| 493 \| 0.426 \| 211 \| 96 \| 5 \| 11 \| 221 \| 152 \| 337 \| Putative tail collar protein \| Putative tail collar protein; afdb-uniprot50 \| AF-A0A1G6EJM0-F1-MODEL\_V4 \| 1.0 \| 3.973e-16 \| 486 \| 0.409 \| 210 \| 105 \| 4 \| 12 \| 220 \| 122 \| 313 \| Microcystin-dependent protein \| Microcystin-dependent protein; afdb-uniprot50 \| AF-A0A840HY65-F1-MODEL\_V4 \| 1.0 \| 4.474e-16 \| 484 \| 0.444 \| 209 \| 80 \| 5 \| 12 \| 220 \| 53 \| 225 \| Microcystin-dependent protein \| Microcystin-dependent protein; afdb-uniprot50 \| AF-A0A1X3FVC4-F1-MODEL\_V4 \| 1.0 \| 4.474e-16 \| 476 \| 0.443 \| 212 \| 81 \| 6 \| 12 \| 220 \| 150 \| 327 \| Phage tail protein \| Phage tail protein; afdb-uniprot50 \| AF-A0A4Q7TER5-F1-MODEL\_V4 \| 1.0 \| 4.474e-16 \| 475 \| 0.4 \| 215 \| 95 \| 8 \| 10 \| 220 \| 200 \| 384 \| Microcystin-dependent protein \| Microcystin-dependent protein; afdb-uniprot50 \| AF-A0A836CG45-F1-MODEL\_V4 \| 1.0 \| 4.377e-13 \| 464 \| 0.386 \| 212 \| 70 \| 4 \| 11 \| 221 \| 2 \| 154 \| Uncharacterized protein \| Uncharacterized protein; afdb-uniprot50 \| AF-A0A225DUJ5-F1-MODEL\_V4 \| 1.0 \| 3.744e-16 \| 463 \| 0.421 \| 211 \| 92 \| 5 \| 11 \| 221 \| 130 \| 310 \| Phage tail fiber \| Phage tail fiber; afdb-uniprot50 \| AF-A0A1E4LGS2-F1-MODEL\_V4 \| 1.0 \| 5.346e-16 \| 458 \| 0.409 \| 210 \| 101 \| 4 \| 11 \| 220 \| 178 \| 364 \| Collar domain-containing protein \| Collar domain-containing protein; afdb-uniprot50 \| AF-A0A7J4F737-F1-MODEL\_V4 \| 1.0 \| 4.023e-15 \| 457 \| 0.386 \| 212 \| 75 \| 6 \| 11 \| 221 \| 100 \| 257 \| Collar domain-containing protein \| Collar domain-containing protein; afdb-uniprot50 \| AF-A0A2E7V2E5-F1-MODEL\_V4 \| 1.0 \| 7.729e-15 \| 450 \| 0.345 \| 211 \| 100 \| 7 \| 11 \| 221 \| 135 \| 307 \| Collar domain-containing protein \| Collar domain-containing protein; afdb-uniprot50 \| AF-A0A7C1IMI0-F1-MODEL\_V4 \| 1.0 \| 7.284e-15 \| 447 \| 0.374 \| 211 \| 76 \| 5 \| 11 \| 221 \| 137 \| 291 \| Collar domain-containing protein \| Collar domain-containing protein; afdb-uniprot50 \| AF-A0A4Q1VB42-F1-MODEL\_V4 \| 1.0 \| 1.399e-14 \| 446 \| 0.331 \| 214 \| 109 \| 6 \| 12 \| 221 \| 152 \| 335 \| Phage tail protein \| Phage tail protein; afdb-uniprot50 \| AF-A0A814Y0G9-F1-MODEL\_V4 \| 1.0 \| 1.319e-14 \| 445 \| 0.361 \| 213 \| 93 \| 5 \| 11 \| 221 \| 108 \| 279 \| Hypothetical protein \| Hypothetical protein; afdb-uniprot50 \| AF-A0A3D0M2I4-F1-MODEL\_V4 \| 1.0 \| 1.774e-14 \| 442 \| 0.336 \| 220 \| 86 \| 4 \| 2 \| 221 \| 57 \| 216 \| Uncharacterized protein \| Uncharacterized protein; afdb-uniprot50 \| AF-A0A815SI59-F1-MODEL\_V4 \| 1.0 \| 1.672e-14 \| 440 \| 0.376 \| 210 \| 92 \| 4 \| 11 \| 220 \| 399 \| 569 \| Hypothetical protein \| Hypothetical protein; afdb-uniprot50 \| AF-A0A815N1T3-F1-MODEL\_V4 \| 1.0 \| 2.418e-13 \| 439 \| 0.35 \| 211 \| 94 \| 4 \| 11 \| 221 \| 2 \| 169 \| Hypothetical protein \| Hypothetical protein; afdb-uniprot50 \| AF-A0A7C1UDS7-F1-MODEL\_V4 \| 1.0 \| 2.388e-14 \| 437 \| 0.35 \| 214 \| 81 \| 5 \| 8 \| 221 \| 134 \| 289 \| Collar domain-containing protein \| Collar domain-containing protein; afdb-uniprot50 \| AF-A0A819R841-F1-MODEL\_V4 \| 1.0 \| 7.284e-15 \| 436 \| 0.333 \| 219 \| 120 \| 8 \| 11 \| 220 \| 308 \| 509 \| Hypothetical protein \| Hypothetical protein; afdb-uniprot50 \| AF-G9RT16-F1-MODEL\_V4 \| 1.0 \| 1.693e-13 \| 433 \| 0.369 \| 211 \| 67 \| 6 \| 11 \| 221 \| 11 \| 155 \| Collar domain-containing protein \| Collar domain-containing protein; afdb-uniprot50 \| AF-W9GZ80-F1-MODEL\_V4 \| 1.0 \| 1.883e-14 \| 433 \| 0.369 \| 219 \| 87 \| 8 \| 4 \| 220 \| 95 \| 264 \| Collar domain-containing protein \| Collar domain-containing protein; afdb-uniprot50 \| AF-A0A817QHN6-F1-MODEL\_V4 \| 1.0 \| 2.094e-15 \| 433 \| 0.36 \| 219 \| 114 \| 8 \| 11 \| 220 \| 373 \| 574 \| Hypothetical protein \| Hypothetical protein; afdb-uniprot50 \| AF-A0A821J8J7-F1-MODEL\_V4 \| 1.0 \| 9.801e-15 \| 432 \| 0.334 \| 218 \| 120 \| 7 \| 11 \| 220 \| 191 \| 391 \| Hypothetical protein \| Hypothetical protein; afdb-uniprot50 \| AF-A0A7T7YU12-F1-MODEL\_V4 \| 1.0 \| 6.55e-14 \| 430 \| 0.393 \| 216 \| 88 \| 5 \| 12 \| 220 \| 109 \| 288 \| Tail fiber protein \| Tail fiber protein; afdb-uniprot50 \| AF-A0A820NDG9-F1-MODEL\_V4 \| 1.0 \| 2.818e-15 \| 429 \| 0.361 \| 221 \| 111 \| 11 \| 11 \| 220 \| 104 \| 305 \| Hypothetical protein \| Hypothetical protein; afdb-uniprot50 \| AF-A0A1V1P4C0-F1-MODEL\_V4 \| 1.0 \| 1.774e-14 \| 429 \| 0.369 \| 211 \| 97 \| 6 \| 11 \| 221 \| 189 \| 363 \| Microcystin-dependent protein-like protein \| Microcystin-dependent protein-like protein; afdb-uniprot50 \| AF-A0A0D6AS89-F1-MODEL\_V4 \| 1.0 \| 6.779e-16 \| 428 \| 0.341 \| 240 \| 105 \| 8 \| 3 \| 221 \| 37 \| 244 \| Microcystin dependent protein \| Microcystin dependent protein; afdb-uniprot50 \| AF-A0A177Q1Y6-F1-MODEL\_V4 \| 1.0 \| 2.534e-14 \| 428 \| 0.342 \| 228 \| 88 \| 8 \| 1 \| 221 \| 81 \| 253 \| Collar domain-containing protein \| Collar domain-containing protein; afdb-uniprot50 \| AF-A0A846V389-F1-MODEL\_V4 \| 1.0 \| 2.25e-14 \| 428 \| 0.303 \| 224 \| 120 \| 8 \| 1 \| 220 \| 140 \| 331 \| Microcystin-dependent protein \| Microcystin-dependent protein; afdb-uniprot50 \| AF-A0A6B2LL29-F1-MODEL\_V4 \| 1.0 \| 1.352e-12 \| 427 \| 0.317 \| 211 \| 83 \| 6 \| 11 \| 221 \| 3 \| 152 \| Collar domain-containing protein \| Collar domain-containing protein; afdb-uniprot50 \| AF-A0A815B3Y9-F1-MODEL\_V4 \| 1.0 \| 4.53e-15 \| 427 \| 0.343 \| 233 \| 119 \| 13 \| 3 \| 220 \| 369 \| 582 \| Hypothetical protein \| Hypothetical protein; afdb-uniprot50 \| AF-A0A815K0V8-F1-MODEL\_V4 \| 1.0 \| 6.468e-15 \| 423 \| 0.32 \| 225 \| 120 \| 11 \| 11 \| 220 \| 184 \| 390 \| Hypothetical protein \| Hypothetical protein; afdb-uniprot50 \| AF-A0A813P3R8-F1-MODEL\_V4 \| 1.0 \| 1.053e-13 \| 422 \| 0.368 \| 225 \| 93 \| 7 \| 3 \| 220 \| 17 \| 199 \| Hypothetical protein \| Hypothetical protein; afdb-uniprot50 \| AF-A0A816E9L0-F1-MODEL\_V4 \| 1.0 \| 3.791e-15 \| 422 \| 0.336 \| 220 \| 117 \| 9 \| 11 \| 220 \| 481 \| 681 \| Hypothetical protein \| Hypothetical protein; afdb-uniprot50 \| AF-A0A6C0ALD8-F1-MODEL\_V4 \| 1.0 \| 9.353e-14 \| 420 \| 0.345 \| 211 \| 107 \| 7 \| 12 \| 221 \| 49 \| 229 \| Collar domain-containing protein \| Collar domain-containing protein; afdb-uniprot50 \| AF-A0A1I7KTY4-F1-MODEL\_V4 \| 1.0 \| 1.485e-14 \| 420 \| 0.448 \| 212 \| 76 \| 9 \| 11 \| 220 \| 199 \| 371 \| Microcystin-dependent protein \| Microcystin-dependent protein; afdb-uniprot50 \| AF-A0A818BYP4-F1-MODEL\_V4 \| 1.0 \| 6.864e-15 \| 420 \| 0.327 \| 220 \| 119 \| 9 \| 11 \| 220 \| 407 \| 607 \| Hypothetical protein \| Hypothetical protein; afdb-uniprot50 \| AF-A0A7W4IKD8-F1-MODEL\_V4 \| 1.0 \| 5.101e-15 \| 418 \| 0.409 \| 220 \| 102 \| 7 \| 8 \| 220 \| 140 \| 338 \| Tail fiber protein \| Tail fiber protein; afdb-uniprot50 \| AF-A0A816GY78-F1-MODEL\_V4 \| 1.0 \| 5.744e-15 \| 417 \| 0.337 \| 222 \| 113 \| 9 \| 11 \| 220 \| 57 \| 256 \| Hypothetical protein \| Hypothetical protein; afdb-uniprot50 \| AF-A0A2M7SZQ3-F1-MODEL\_V4 \| 1.0 \| 1.973e-15 \| 416 \| 0.386 \| 220 \| 91 \| 9 \| 3 \| 221 \| 234 \| 410 \| Collar domain-containing protein \| Collar domain-containing protein; afdb-uniprot50 \| AF-A0A4Y6CGU0-F1-MODEL\_V4 \| 1.0 \| 1.417e-13 \| 413 \| 0.355 \| 211 \| 87 \| 4 \| 11 \| 221 \| 522 \| 683 \| Collar domain-containing protein \| Collar domain-containing protein; afdb-uniprot50 \| AF-A0A820E4P5-F1-MODEL\_V4 \| 1.0 \| 2.723e-13 \| 411 \| 0.364 \| 211 \| 95 \| 7 \| 11 \| 221 \| 25 \| 196 \| Hypothetical protein \| Hypothetical protein; afdb-uniprot50 \| AF-A0A1J5P6P6-F1-MODEL\_V4 \| 1.0 \| 9.924e-14 \| 411 \| 0.427 \| 213 \| 88 \| 7 \| 12 \| 221 \| 106 \| 287 \| Phage terminase large subunit \| Phage terminase large subunit; afdb-uniprot50 \| AF-A0A822CIC7-F1-MODEL\_V4 \| 1.0 \| 9.236e-15 \| 411 \| 0.337 \| 219 \| 119 \| 8 \| 11 \| 220 \| 132 \| 333 \| Hypothetical protein \| Hypothetical protein; afdb-uniprot50 \| AF-A0A821HM98-F1-MODEL\_V4 \| 1.0 \| 8.704e-15 \| 411 \| 0.319 \| 222 \| 118 \| 9 \| 11 \| 219 \| 231 \| 432 \| Hypothetical protein \| Hypothetical protein; afdb-uniprot50 \| AF-A0A0P0EA18-F1-MODEL\_V4 \| 1.0 \| 1.693e-13 \| 407 \| 0.317 \| 230 \| 91 \| 6 \| 1 \| 221 \| 18 \| 190 \| Collar domain-containing protein \| Collar domain-containing protein; afdb-uniprot50 \| AF-A0A5J4Y931-F1-MODEL\_V4 \| 1.0 \| 1.907e-13 \| 404 \| 0.334 \| 212 \| 98 \| 5 \| 12 \| 221 \| 153 \| 323 \| Collar domain-containing protein \| Collar domain-containing protein; afdb-uniprot50 \| AF-A0A2E3QAG3-F1-MODEL\_V4 \| 1.0 \| 2.049e-12 \| 402 \| 0.291 \| 223 \| 99 \| 7 \| 1 \| 221 \| 10 \| 175 \| Phage tail protein \| Phage tail protein; afdb-uniprot50 \| AF-A0A3S0TER0-F1-MODEL\_V4 \| 1.0 \| 7.827e-14 \| 400 \| 0.377 \| 209 \| 83 \| 6 \| 12 \| 220 \| 488 \| 649 \| Collar domain-containing protein \| Collar domain-containing protein; afdb-uniprot50 \| AF-A0A4Q3WNL6-F1-MODEL\_V4 \| 1.0 \| 3.618e-14 \| 399 \| 0.33 \| 227 \| 95 \| 6 \| 3 \| 221 \| 239 \| 416 \| Tail fiber protein \| Tail fiber protein; afdb-uniprot50 \| AF-A0A820D1C7-F1-MODEL\_V4 \| 1.0 \| 2.689e-14 \| 397 \| 0.328 \| 222 \| 116 \| 11 \| 12 \| 220 \| 171 \| 372 \| Hypothetical protein \| Hypothetical protein; afdb-uniprot50 \| AF-A0A817IDX0-F1-MODEL\_V4 \| 1.0 \| 8.813e-14 \| 397 \| 0.348 \| 218 \| 117 \| 8 \| 11 \| 220 \| 399 \| 599 \| Hypothetical protein \| Hypothetical protein; afdb-uniprot50 \| AF-A0A821YM97-F1-MODEL\_V4 \| 1.0 \| 7.376e-14 \| 396 \| 0.283 \| 219 \| 131 \| 7 \| 11 \| 220 \| 31 \| 232 \| Hypothetical protein \| Hypothetical protein; afdb-uniprot50 \| AF-A0A6C0HZC1-F1-MODEL\_V4 \| 1.0 \| 4.868e-14 \| 396 \| 0.302 \| 228 \| 124 \| 6 \| 12 \| 221 \| 46 \| 256 \| Collar domain-containing protein \| Collar domain-containing protein; afdb-uniprot50 \| AF-A0A816FNH1-F1-MODEL\_V4 \| 1.0 \| 5.482e-14 \| 395 \| 0.321 \| 224 \| 115 \| 10 \| 11 \| 219 \| 431 \| 632 \| Hypothetical protein \| Hypothetical protein; afdb-uniprot50 \| AF-A0A7V3QID4-F1-MODEL\_V4 \| 1.0 \| 2.023e-13 \| 392 \| 0.341 \| 211 \| 78 \| 5 \| 11 \| 221 \| 207 \| 356 \| Collar domain-containing protein \| Collar domain-containing protein; afdb-uniprot50 \| AF-A0A821GMF9-F1-MODEL\_V4 \| 1.0 \| 7.827e-14 \| 391 \| 0.324 \| 219 \| 113 \| 9 \| 11 \| 220 \| 15 \| 207 \| Hypothetical protein \| Hypothetical protein; afdb-uniprot50 \| AF-A0A818P2B2-F1-MODEL\_V4 \| 1.0 \| 6.173e-14 \| 390 \| 0.339 \| 233 \| 103 \| 7 \| 1 \| 221 \| 46 \| 239 \| Hypothetical protein \| Hypothetical protein; afdb-uniprot50 \| AF-A0A814FG22-F1-MODEL\_V4 \| 1.0 \| 1.319e-14 \| 390 \| 0.336 \| 220 \| 116 \| 10 \| 11 \| 220 \| 49 \| 248 \| Hypothetical protein \| Hypothetical protein; afdb-uniprot50 \| AF-A0A6N6JSU8-F1-MODEL\_V4 \| 1.0 \| 4.645e-13 \| 388 \| 0.354 \| 217 \| 101 \| 9 \| 11 \| 221 \| 25 \| 208 \| Collar domain-containing protein \| Collar domain-containing protein; afdb-uniprot50 \| AF-A0A815V480-F1-MODEL\_V4 \| 1.0 \| 2.023e-13 \| 388 \| 0.322 \| 217 \| 122 \| 9 \| 12 \| 220 \| 502 \| 701 \| Hypothetical protein \| Hypothetical protein; afdb-uniprot50 \| AF-A0A4Q8ARL3-F1-MODEL\_V4 \| 1.0 \| 1.274e-12 \| 387 \| 0.31 \| 229 \| 89 \| 6 \| 3 \| 221 \| 26 \| 195 \| Microcystin-dependent protein \| Microcystin-dependent protein; afdb-uniprot50 \| AF-A0A3M1NVN9-F1-MODEL\_V4 \| 1.0 \| 2.723e-13 \| 387 \| 0.364 \| 269 \| 112 \| 11 \| 2 \| 220 \| 59 \| 318 \| Collar domain-containing protein \| Collar domain-containing protein; afdb-uniprot50 \| AF-A0A158D1B4-F1-MODEL\_V4 \| 1.0 \| 1.09e-15 \| 385 \| 0.414 \| 229 \| 104 \| 6 \| 11 \| 221 \| 176 \| 392 \| Phage tail collar domain-containing protein \| Phage tail collar domain-containing protein; afdb-uniprot50 \| AF-A0A4Q0AIU5-F1-MODEL\_V4 \| 1.0 \| 2.336e-11 \| 384 \| 0.293 \| 211 \| 88 \| 8 \| 11 \| 221 \| 2 \| 151 \| Collar domain-containing protein \| Collar domain-containing protein; afdb-uniprot50 \| AF-A0A822DWL8-F1-MODEL\_V4 \| 1.0 \| 1.907e-13 \| 384 \| 0.322 \| 220 \| 120 \| 8 \| 11 \| 220 \| 12 \| 212 \| Hypothetical protein \| Hypothetical protein; afdb-uniprot50 \| AF-A0A818VB53-F1-MODEL\_V4 \| 1.0 \| 1.053e-13 \| 384 \| 0.309 \| 233 \| 126 \| 9 \| 2 \| 220 \| 11 \| 222 \| Hypothetical protein \| Hypothetical protein; afdb-uniprot50 \| AF-A0A818SKZ2-F1-MODEL\_V4 \| 1.0 \| 1.053e-13 \| 384 \| 0.327 \| 220 \| 112 \| 9 \| 12 \| 220 \| 33 \| 227 \| Hypothetical protein \| Hypothetical protein; afdb-uniprot50 \| AF-A0A815JLE8-F1-MODEL\_V4 \| 1.0 \| 8.813e-14 \| 384 \| 0.353 \| 218 \| 115 \| 9 \| 11 \| 220 \| 272 \| 471 \| Hypothetical protein \| Hypothetical protein; afdb-uniprot50 \| AF-A0A1Y4GYF8-F1-MODEL\_V4 \| 1.0 \| 2.336e-11 \| 381 \| 0.308 \| 211 \| 85 \| 4 \| 11 \| 221 \| 9 \| 158 \| Collar domain-containing protein \| Collar domain-containing protein; afdb-uniprot50 \| AF-A0A819LSZ1-F1-MODEL\_V4 \| 1.0 \| 7.469e-13 \| 379 \| 0.343 \| 230 \| 95 \| 8 \| 3 \| 221 \| 30 \| 214 \| Hypothetical protein \| Hypothetical protein; afdb-uniprot50 \| AF-A0A369RCN8-F1-MODEL\_V4 \| 1.0 \| 1.066e-12 \| 379 \| 0.319 \| 210 \| 81 \| 4 \| 12 \| 221 \| 73 \| 220 \| Collar domain-containing protein \| Collar domain-containing protein; afdb-uniprot50 \| AF-A0A1C3XJM3-F1-MODEL\_V4 \| 1.0 \| 6.193e-09 \| 379 \| 0.579 \| 88 \| 36 \| 1 \| 16 \| 102 \| 128 \| 215 \| Phage Tail Collar Domain \| Phage Tail Collar Domain; afdb-uniprot50 \| AF-A0A559TE11-F1-MODEL\_V4 \| 1.0 \| 9.924e-14 \| 378 \| 0.384 \| 229 \| 109 \| 9 \| 2 \| 221 \| 183 \| 388 \| Microcystin-dependent protein \| Microcystin-dependent protein; afdb-uniprot50 \| AF-A0A815U1H6-F1-MODEL\_V4 \| 1.0 \| 3.452e-13 \| 378 \| 0.31 \| 219 \| 125 \| 8 \| 11 \| 220 \| 514 \| 715 \| Hypothetical protein \| Hypothetical protein; afdb-uniprot50 \| AF-A0A2N2HV87-F1-MODEL\_V4 \| 1.0 \| 5.482e-14 \| 375 \| 0.346 \| 222 \| 93 \| 8 \| 11 \| 221 \| 174 \| 354 \| Uncharacterized protein \| Uncharacterized protein; afdb-uniprot50 \| AF-A0A7C5Z090-F1-MODEL\_V4 \| 1.0 \| 4.125e-13 \| 374 \| 0.371 \| 221 \| 74 \| 7 \| 1 \| 221 \| 160 \| 315 \| Collar domain-containing protein \| Collar domain-containing protein; afdb-uniprot50 \| AF-A0A840BX98-F1-MODEL\_V4 \| 1.0 \| 7.926e-13 \| 374 \| 0.333 \| 225 \| 113 \| 9 \| 2 \| 220 \| 296 \| 489 \| Microcystin-dependent protein \| Microcystin-dependent protein; afdb-uniprot50 \| AF-A0A816MB52-F1-MODEL\_V4 \| 1.0 \| 2.023e-13 \| 374 \| 0.323 \| 229 \| 115 \| 12 \| 11 \| 220 \| 505 \| 712 \| Hypothetical protein \| Hypothetical protein; afdb-uniprot50 \| AF-A0A816Z751-F1-MODEL\_V4 \| 1.0 \| 1.186e-13 \| 373 \| 0.314 \| 229 \| 110 \| 10 \| 11 \| 221 \| 2 \| 201 \| Hypothetical protein \| Hypothetical protein; afdb-uniprot50 \| AF-A0A835XXH8-F1-MODEL\_V4 \| 1.0 \| 1.132e-12 \| 373 \| 0.313 \| 230 \| 115 \| 8 \| 2 \| 220 \| 350 \| 547 \| Uncharacterized protein \| Uncharacterized protein; afdb-uniprot50 \| AF-A0A521K5D5-F1-MODEL\_V4 \| 1.0 \| 1.335e-13 \| 372 \| 0.361 \| 216 \| 83 \| 6 \| 12 \| 221 \| 49 \| 215 \| Collar domain-containing protein \| Collar domain-containing protein; afdb-uniprot50 \| AF-A0A820QPV6-F1-MODEL\_V4 \| 1.0 \| 4.929e-13 \| 372 \| 0.313 \| 217 \| 127 \| 8 \| 11 \| 220 \| 92 \| 293 \| Hypothetical protein \| Hypothetical protein; afdb-uniprot50 \| AF-A0A814LGM5-F1-MODEL\_V4 \| 1.0 \| 6.633e-13 \| 371 \| 0.35 \| 211 \| 96 \| 8 \| 12 \| 221 \| 48 \| 218 \| Hypothetical protein \| Hypothetical protein; afdb-uniprot50 \| AF-A0A1F3B1K2-F1-MODEL\_V4 \| 1.0 \| 1.931e-12 \| 370 \| 0.316 \| 212 \| 79 \| 5 \| 11 \| 221 \| 374 \| 520 \| Collar domain-containing protein \| Collar domain-containing protein; afdb-uniprot50 \| AF-A0A815QNH7-F1-MODEL\_V4 \| 1.0 \| 3.663e-13 \| 369 \| 0.327 \| 217 \| 122 \| 7 \| 11 \| 220 \| 297 \| 496 \| Hypothetical protein \| Hypothetical protein; afdb-uniprot50 \| AF-A0A814Z5Y2-F1-MODEL\_V4 \| 1.0 \| 7.376e-14 \| 368 \| 0.313 \| 220 \| 121 \| 11 \| 11 \| 220 \| 124 \| 323 \| Hypothetical protein \| Hypothetical protein; afdb-uniprot50 \| AF-A0A4D7B3R7-F1-MODEL\_V4 \| 1.0 \| 1.931e-12 \| 366 \| 0.35 \| 214 \| 114 \| 9 \| 11 \| 221 \| 125 \| 316 \| Tail fiber protein \| Tail fiber protein; afdb-uniprot50 \| AF-A0A3N7JCP5-F1-MODEL\_V4 \| 1.0 \| 3.496e-12 \| 364 \| 0.271 \| 214 \| 128 \| 6 \| 10 \| 220 \| 3 \| 191 \| Phage tail protein \| Phage tail protein; afdb-uniprot50 \| AF-A0A2D9HKR9-F1-MODEL\_V4 \| 1.0 \| 2.049e-12 \| 364 \| 0.355 \| 211 \| 54 \| 4 \| 11 \| 221 \| 78 \| 206 \| Collar domain-containing protein \| Collar domain-containing protein; afdb-uniprot50 \| AF-A0A3D1PJI0-F1-MODEL\_V4 \| 1.0 \| 2.174e-12 \| 364 \| 0.325 \| 209 \| 78 \| 5 \| 12 \| 220 \| 48 \| 193 \| Collar domain-containing protein \| Collar domain-containing protein; afdb-uniprot50 \| AF-A0A813ML39-F1-MODEL\_V4 \| 1.0 \| 1.258e-13 \| 363 \| 0.319 \| 219 \| 124 \| 9 \| 11 \| 220 \| 515 \| 717 \| Hypothetical protein \| Hypothetical protein; afdb-uniprot50 \| AF-A0A821Z962-F1-MODEL\_V4 \| 1.0 \| 2.889e-13 \| 362 \| 0.297 \| 225 \| 122 \| 9 \| 11 \| 220 \| 220 \| 423 \| Hypothetical protein \| Hypothetical protein; afdb-uniprot50 \| AF-A0A819Y0C9-F1-MODEL\_V4 \| 1.0 \| 1.258e-13 \| 360 \| 0.299 \| 227 \| 115 \| 9 \| 11 \| 221 \| 50 \| 248 \| Hypothetical protein \| Hypothetical protein; afdb-uniprot50 \| AF-A0A817VC28-F1-MODEL\_V4 \| 1.0 \| 1.005e-12 \| 359 \| 0.319 \| 216 \| 119 \| 8 \| 12 \| 220 \| 29 \| 223 \| Hypothetical protein \| Hypothetical protein; afdb-uniprot50 \| AF-A0A1Y4GY33-F1-MODEL\_V4 \| 1.0 \| 8.41e-13 \| 359 \| 0.321 \| 224 \| 88 \| 6 \| 2 \| 221 \| 174 \| 337 \| Collar domain-containing protein \| Collar domain-containing protein; afdb-uniprot50 \| AF-A0A6A7MGA2-F1-MODEL\_V4 \| 1.0 \| 7.127e-12 \| 359 \| 0.312 \| 211 \| 107 \| 4 \| 11 \| 221 \| 171 \| 343 \| Collar domain-containing protein \| Collar domain-containing protein; afdb-uniprot50 \| AF-A0A7W2NTN7-F1-MODEL\_V4 \| 1.0 \| 8.516e-12 \| 357 \| 0.281 \| 217 \| 103 \| 8 \| 10 \| 220 \| 3 \| 172 \| Phage tail protein \| Phage tail protein; afdb-uniprot50 \| AF-A0A5S3SAM7-F1-MODEL\_V4 \| 1.0 \| 8.026e-12 \| 355 \| 0.41 \| 200 \| 80 \| 9 \| 24 \| 220 \| 1 \| 165 \| Phage tail protein \| Phage tail protein; afdb-uniprot50 \| AF-A0A817QZ18-F1-MODEL\_V4 \| 1.0 \| 9.151e-11 \| 355 \| 0.311 \| 209 \| 103 \| 5 \| 14 \| 221 \| 4 \| 172 \| Hypothetical protein \| Hypothetical protein; afdb-uniprot50 \| AF-A0A6N3F5J0-F1-MODEL\_V4 \| 1.0 \| 1.274e-12 \| 355 \| 0.323 \| 213 \| 74 \| 9 \| 11 \| 221 \| 49 \| 193 \| Phage Tail Collar Domain protein \| Phage Tail Collar Domain protein; afdb-uniprot50 \| AF-A0A1V5DL01-F1-MODEL\_V4 \| 1.0 \| 2.418e-13 \| 355 \| 0.374 \| 219 \| 87 \| 10 \| 11 \| 221 \| 160 \| 336 \| Phage Tail Collar Domain protein \| Phage Tail Collar Domain protein; afdb-uniprot50 \| AF-A0A7V8IVS3-F1-MODEL\_V4 \| 1.0 \| 8.925e-13 \| 355 \| 0.331 \| 211 \| 80 \| 8 \| 12 \| 221 \| 256 \| 406 \| Collar domain-containing protein \| Collar domain-containing protein; afdb-uniprot50 \| AF-E4T5W3-F1-MODEL\_V4 \| 1.0 \| 2.448e-12 \| 354 \| 0.362 \| 215 \| 78 \| 7 \| 11 \| 221 \| 739 \| 898 \| Tail Collar domain protein \| Tail Collar domain protein; afdb-uniprot50 \| AF-A0A512NCS0-F1-MODEL\_V4 \| 1.0 \| 3.452e-13 \| 353 \| 0.338 \| 213 \| 97 \| 7 \| 12 \| 221 \| 145 \| 316 \| Collar domain-containing protein \| Collar domain-containing protein; afdb-uniprot50 \| AF-A0A1F3ABU4-F1-MODEL\_V4 \| 1.0 \| 2.366e-10 \| 352 \| 0.314 \| 207 \| 79 \| 6 \| 16 \| 221 \| 1 \| 145 \| Collar domain-containing protein \| Collar domain-containing protein; afdb-uniprot50 \| AF-A0A2I0N9H0-F1-MODEL\_V4 \| 1.0 \| 2.598e-12 \| 352 \| 0.285 \| 224 \| 111 \| 11 \| 1 \| 221 \| 168 \| 345 \| Collar domain-containing protein \| Collar domain-containing protein; afdb-uniprot50 \| AF-A0A6N4RF73-F1-MODEL\_V4 \| 1.0 \| 1.258e-13 \| 351 \| 0.344 \| 215 \| 109 \| 7 \| 11 \| 221 \| 39 \| 225 \| Tail fiber protein \| Tail fiber protein; afdb-uniprot50 \| AF-A0A257UMQ9-F1-MODEL\_V4 \| 1.0 \| 2.174e-12 \| 351 \| 0.378 \| 214 \| 87 \| 8 \| 11 \| 221 \| 68 \| 238 \| Collar domain-containing protein \| Collar domain-containing protein; afdb-uniprot50 \| AF-A0A2A6FNG7-F1-MODEL\_V4 \| 1.0 \| 1.453e-11 \| 350 \| 0.311 \| 225 \| 101 \| 6 \| 3 \| 221 \| 11 \| 187 \| Collar domain-containing protein \| Collar domain-containing protein; afdb-uniprot50 \| AF-A0A817WT08-F1-MODEL\_V4 \| 1.0 \| 5.231e-13 \| 348 \| 0.308 \| 227 \| 113 \| 7 \| 12 \| 221 \| 55 \| 254 \| Hypothetical protein \| Hypothetical protein; afdb-uniprot50 \| AF-A0A7V2Y2J7-F1-MODEL\_V4 \| 1.0 \| 1.435e-12 \| 348 \| 0.297 \| 215 \| 86 \| 10 \| 11 \| 221 \| 161 \| 314 \| Collar domain-containing protein \| Collar domain-containing protein; afdb-uniprot50 \| AF-A0A0G1QRC3-F1-MODEL\_V4 \| 1.0 \| 8.516e-12 \| 347 \| 0.331 \| 211 \| 74 \| 7 \| 11 \| 221 \| 133 \| 276 \| Tail Collar domain protein \| Tail Collar domain protein; afdb-uniprot50 \| AF-A0A1V4VVI6-F1-MODEL\_V4 \| 1.0 \| 3.936e-12 \| 346 \| 0.304 \| 223 \| 92 \| 10 \| 2 \| 221 \| 219 \| 381 \| Phage Tail Collar Domain protein \| Phage Tail Collar Domain protein; afdb-uniprot50 \| AF-A0A661AU03-F1-MODEL\_V4 \| 1.0 \| 6.716e-12 \| 345 \| 0.341 \| 211 \| 97 \| 7 \| 11 \| 221 \| 554 \| 722 \| Collar domain-containing protein \| Collar domain-containing protein; afdb-uniprot50 \| AF-A0A816AZ21-F1-MODEL\_V4 \| 1.0 \| 7.469e-13 \| 344 \| 0.286 \| 220 \| 125 \| 8 \| 11 \| 220 \| 262 \| 459 \| Hypothetical protein \| Hypothetical protein; afdb-uniprot50 \| AF-A0A813U148-F1-MODEL\_V4 \| 1.0 \| 1.186e-13 \| 343 \| 0.298 \| 241 \| 124 \| 10 \| 2 \| 221 \| 300 \| 516 \| Hypothetical protein \| Hypothetical protein; afdb-uniprot50 \| AF-A0A818QJV6-F1-MODEL\_V4 \| 1.0 \| 1.005e-12 \| 342 \| 0.266 \| 221 \| 128 \| 9 \| 11 \| 220 \| 450 \| 647 \| Hypothetical protein \| Hypothetical protein; afdb-uniprot50 \| AF-A0A816H4T3-F1-MODEL\_V4 \| 1.0 \| 3.104e-12 \| 341 \| 0.297 \| 222 \| 124 \| 8 \| 11 \| 220 \| 352 \| 553 \| Hypothetical protein \| Hypothetical protein; afdb-uniprot50 \| AF-A0A0Q4T3S1-F1-MODEL\_V4 \| 1.0 \| 3.71e-12 \| 340 \| 0.334 \| 284 \| 108 \| 11 \| 12 \| 221 \| 125 \| 401 \| Collar domain-containing protein \| Collar domain-containing protein; afdb-uniprot50 \| AF-A0A431IBK6-F1-MODEL\_V4 \| 1.0 \| 1.955e-11 \| 339 \| 0.299 \| 224 \| 82 \| 7 \| 2 \| 221 \| 29 \| 181 \| Collar domain-containing protein \| Collar domain-containing protein; afdb-uniprot50 \| AF-A0A086D2K0-F1-MODEL\_V4 \| 1.0 \| 3.756e-11 \| 339 \| 0.276 \| 224 \| 105 \| 8 \| 11 \| 220 \| 6 \| 186 \| Tail protein \| Tail protein; afdb-uniprot50 \| AF-A0A6M5Z7F0-F1-MODEL\_V4 \| 1.0 \| 1.216e-11 \| 339 \| 0.364 \| 211 \| 73 \| 8 \| 11 \| 221 \| 395 \| 544 \| Collar domain-containing protein \| Collar domain-containing protein; afdb-uniprot50 \| AF-A0A228K8W8-F1-MODEL\_V4 \| 1.0 \| 1.736e-11 \| 338 \| 0.269 \| 234 \| 115 \| 8 \| 1 \| 221 \| 94 \| 284 \| Uncharacterized protein \| Uncharacterized protein; afdb-uniprot50 \| AF-A0A656QC36-F1-MODEL\_V4 \| 1.0 \| 1.08e-11 \| 337 \| 0.359 \| 220 \| 93 \| 7 \| 9 \| 220 \| 104 \| 283 \| Uncharacterized protein \| Uncharacterized protein; afdb-uniprot50 \| AF-A0A7S6S8F8-F1-MODEL\_V4 \| 1.0 \| 3.144e-11 \| 336 \| 0.294 \| 221 \| 123 \| 8 \| 11 \| 220 \| 4 \| 202 \| Phage tail protein \| Phage tail protein; afdb-uniprot50 \| AF-A0A813XM12-F1-MODEL\_V4 \| 1.0 \| 4.587e-14 \| 336 \| 0.333 \| 240 \| 117 \| 12 \| 2 \| 221 \| 73 \| 289 \| Hypothetical protein \| Hypothetical protein; afdb-uniprot50 \| AF-A0A815S9R3-F1-MODEL\_V4 \| 1.0 \| 1.931e-12 \| 335 \| 0.283 \| 236 \| 127 \| 9 \| 11 \| 220 \| 337 \| 556 \| Hypothetical protein \| Hypothetical protein; afdb-uniprot50 \| AF-A0A2V9WHC2-F1-MODEL\_V4 \| 1.0 \| 1.82e-12 \| 334 \| 0.372 \| 215 \| 93 \| 7 \| 12 \| 220 \| 135 \| 313 \| Collar domain-containing protein \| Collar domain-containing protein; afdb-uniprot50 \| AF-D0LMW0-F1-MODEL\_V4 \| 1.0 \| 1.842e-11 \| 333 \| 0.291 \| 223 \| 96 \| 7 \| 2 \| 220 \| 98 \| 262 \| Tail Collar domain protein \| Tail Collar domain protein; afdb-uniprot50 \| AF-A0A0F0DN26-F1-MODEL\_V4 \| 1.0 \| 9.151e-11 \| 332 \| 0.278 \| 223 \| 106 \| 7 \| 11 \| 220 \| 6 \| 186 \| Tail protein \| Tail protein; afdb-uniprot50 \| AF-A0A819AK34-F1-MODEL\_V4 \| 1.0 \| 1.132e-12 \| 332 \| 0.313 \| 230 \| 127 \| 10 \| 2 \| 221 \| 98 \| 306 \| Hypothetical protein \| Hypothetical protein; afdb-uniprot50 \| AF-A0A2L2NDQ4-F1-MODEL\_V4 \| 1.0 \| 2.889e-13 \| 332 \| 0.328 \| 256 \| 112 \| 7 \| 11 \| 221 \| 160 \| 400 \| Phage tail collar domain protein \| Phage tail collar domain protein; afdb-uniprot50 \| AF-A0A367Q7H7-F1-MODEL\_V4 \| 1.0 \| 3.54e-11 \| 331 \| 0.305 \| 219 \| 86 \| 6 \| 4 \| 220 \| 312 \| 466 \| Collar domain-containing protein \| Collar domain-containing protein; afdb-uniprot50 \| AF-A0A561GTC3-F1-MODEL\_V4 \| 1.0 \| 3.54e-11 \| 330 \| 0.294 \| 221 \| 116 \| 11 \| 10 \| 220 \| 3 \| 193 \| Microcystin-dependent protein \| Microcystin-dependent protein; afdb-uniprot50 \| AF-A0A1Y2LD10-F1-MODEL\_V4 \| 1.0 \| 8.127e-11 \| 328 \| 0.298 \| 211 \| 77 \| 5 \| 12 \| 221 \| 52 \| 192 \| Collar domain-containing protein \| Collar domain-containing protein; afdb-uniprot50 \| AF-D9SMM5-F1-MODEL\_V4 \| 1.0 \| 1.842e-11 \| 328 \| 0.275 \| 225 \| 110 \| 7 \| 11 \| 220 \| 4 \| 190 \| Tail Collar domain protein \| Tail Collar domain protein; afdb-uniprot50 \| AF-A0A199XMG2-F1-MODEL\_V4 \| 1.0 \| 6.409e-11 \| 327 \| 0.304 \| 217 \| 93 \| 9 \| 12 \| 220 \| 5 \| 171 \| Phage tail collar domain protein \| Phage tail collar domain protein; afdb-uniprot50 \| AF-A0A818C432-F1-MODEL\_V4 \| 1.0 \| 4.992e-12 \| 327 \| 0.295 \| 237 \| 121 \| 8 \| 3 \| 220 \| 14 \| 223 \| Hypothetical protein \| Hypothetical protein; afdb-uniprot50 \| AF-A0A6L4A8E9-F1-MODEL\_V4 \| 1.0 \| 7.563e-12 \| 327 \| 0.319 \| 210 \| 89 \| 7 \| 12 \| 220 \| 51 \| 207 \| Collar domain-containing protein \| Collar domain-containing protein; afdb-uniprot50 \| AF-A0A3B9AMF4-F1-MODEL\_V4 \| 1.0 \| 2.827e-10 \| 325 \| 0.291 \| 216 \| 99 \| 6 \| 10 \| 220 \| 3 \| 169 \| Phage tail protein \| Phage tail protein; afdb-uniprot50 \| AF-X0Z504-F1-MODEL\_V4 \| 1.0 \| 2.962e-11 \| 325 \| 0.317 \| 211 \| 85 \| 6 \| 11 \| 221 \| 78 \| 229 \| Collar domain-containing protein \| Collar domain-containing protein; afdb-uniprot50 \| AF-A0A4Y8ADI2-F1-MODEL\_V4 \| 1.0 \| 1.955e-11 \| 323 \| 0.295 \| 223 \| 104 \| 9 \| 11 \| 220 \| 5 \| 187 \| Phage tail protein \| Phage tail protein; afdb-uniprot50 \| AF-A0A6J5DWF4-F1-MODEL\_V4 \| 1.0 \| 1.657e-10 \| 322 \| 0.282 \| 230 \| 111 \| 8 \| 3 \| 221 \| 79 \| 265 \| Uncharacterized protein \| Uncharacterized protein; afdb-uniprot50 \| AF-A0A820P0S5-F1-MODEL\_V4 \| 1.0 \| 2.598e-12 \| 322 \| 0.355 \| 211 \| 107 \| 8 \| 20 \| 220 \| 224 \| 415 \| Hypothetical protein \| Hypothetical protein; afdb-uniprot50 \| AF-A0A845XUZ4-F1-MODEL\_V4 \| 1.0 \| 1.542e-11 \| 322 \| 0.342 \| 210 \| 93 \| 6 \| 12 \| 221 \| 343 \| 507 \| Uncharacterized protein \| Uncharacterized protein; afdb-uniprot50 \| AF-A0A832E9G7-F1-MODEL\_V4 \| 1.0 \| 5.118e-10 \| 321 \| 0.278 \| 212 \| 96 \| 7 \| 10 \| 220 \| 3 \| 158 \| Phage tail protein \| Phage tail protein; afdb-uniprot50 \| AF-A0A2U8WK05-F1-MODEL\_V4 \| 1.0 \| 5.692e-11 \| 321 \| 0.227 \| 237 \| 131 \| 9 \| 11 \| 220 \| 4 \| 215 \| Collar domain-containing protein \| Collar domain-containing protein; afdb-uniprot50 \| AF-A0A816ETV2-F1-MODEL\_V4 \| 1.0 \| 5.551e-13 \| 321 \| 0.314 \| 232 \| 124 \| 10 \| 1 \| 221 \| 66 \| 273 \| Hypothetical protein \| Hypothetical protein; afdb-uniprot50 \| AF-A0A1J5PBH2-F1-MODEL\_V4 \| 1.0 \| 6.04e-11 \| 321 \| 0.288 \| 215 \| 118 \| 10 \| 12 \| 221 \| 138 \| 322 \| Phage tail collar domain protein \| Phage tail collar domain protein; afdb-uniprot50 \| AF-A0A1Q3W4R7-F1-MODEL\_V4 \| 1.0 \| 2.229e-10 \| 320 \| 0.202 \| 276 \| 129 \| 6 \| 11 \| 220 \| 218 \| 468 \| Uncharacterized protein \| Uncharacterized protein; afdb-uniprot50 \| AF-A0A1W2GA44-F1-MODEL\_V4 \| 1.0 \| 1.387e-10 \| 319 \| 0.227 \| 224 \| 135 \| 8 \| 11 \| 220 \| 4 \| 203 \| Microcystin-dependent protein \| Microcystin-dependent protein; afdb-uniprot50 \| AF-A0A815G9G8-F1-MODEL\_V4 \| 1.0 \| 7.038e-13 \| 319 \| 0.311 \| 225 \| 114 \| 10 \| 12 \| 221 \| 100 \| 298 \| Hypothetical protein \| Hypothetical protein; afdb-uniprot50 \| AF-D1BW55-F1-MODEL\_V4 \| 1.0 \| 2.075e-11 \| 319 \| 0.302 \| 245 \| 124 \| 9 \| 1 \| 220 \| 215 \| 437 \| Tail Collar domain protein \| Tail Collar domain protein; afdb-uniprot50 \| AF-A0A3A4PGH4-F1-MODEL\_V4 \| 1.0 \| 8.733e-10 \| 318 \| 0.266 \| 210 \| 103 \| 5 \| 11 \| 220 \| 5 \| 163 \| Phage tail protein \| Phage tail protein; afdb-uniprot50 \| AF-A9H7F1-F1-MODEL\_V4 \| 1.0 \| 5.231e-13 \| 318 \| 0.468 \| 177 \| 72 \| 5 \| 28 \| 198 \| 1 \| 161 \| Putative membrane protein \| Putative membrane protein; afdb-uniprot50 \| AF-A0A3E0DWJ8-F1-MODEL\_V4 \| 1.0 \| 3.183e-10 \| 318 \| 0.276 \| 213 \| 111 \| 7 \| 10 \| 220 \| 3 \| 174 \| Microcystin-dependent protein \| Microcystin-dependent protein; afdb-uniprot50 \| AF-A0A1X7NXK8-F1-MODEL\_V4 \| 1.0 \| 5.054e-11 \| 317 \| 0.216 \| 236 \| 137 \| 9 \| 11 \| 220 \| 4 \| 217 \| Microcystin-dependent protein \| Microcystin-dependent protein; afdb-uniprot50 \| AF-A0A0D7Q2R9-F1-MODEL\_V4 \| 1.0 \| 4.489e-11 \| 317 \| 0.251 \| 219 \| 116 \| 9 \| 12 \| 220 \| 120 \| 300 \| Collar domain-containing protein \| Collar domain-containing protein; afdb-uniprot50 \| AF-A0A0G4FSR3-F1-MODEL\_V4 \| 1.0 \| 1.636e-11 \| 317 \| 0.261 \| 226 \| 95 \| 11 \| 2 \| 221 \| 466 \| 625 \| Collar domain-containing protein \| Collar domain-containing protein; afdb-uniprot50 \| AF-A0A4R2H649-F1-MODEL\_V4 \| 1.0 \| 1.758e-10 \| 316 \| 0.303 \| 221 \| 89 \| 9 \| 11 \| 220 \| 4 \| 170 \| Microcystin-dependent protein \| Microcystin-dependent protein; afdb-uniprot50 \| AF-A0A4V0I5Q5-F1-MODEL\_V4 \| 1.0 \| 5.364e-11 \| 314 \| 0.336 \| 211 \| 75 \| 7 \| 11 \| 220 \| 287 \| 433 \| Uncharacterized protein \| Uncharacterized protein; afdb-uniprot50 \| AF-A0A2D8BVW2-F1-MODEL\_V4 \| 1.0 \| 1.842e-11 \| 313 \| 0.273 \| 223 \| 89 \| 7 \| 12 \| 221 \| 24 \| 186 \| Collar domain-containing protein \| Collar domain-containing protein; afdb-uniprot50 \| AF-A0A316AC07-F1-MODEL\_V4 \| 1.0 \| 5.692e-11 \| 313 \| 0.313 \| 223 \| 101 \| 8 \| 2 \| 220 \| 21 \| 195 \| Microcystin-dependent protein \| Microcystin-dependent protein; afdb-uniprot50 \| AF-A0A2N2SSE1-F1-MODEL\_V4 \| 1.0 \| 3.496e-12 \| 313 \| 0.378 \| 219 \| 103 \| 8 \| 11 \| 221 \| 78 \| 271 \| Phage tail protein \| Phage tail protein; afdb-uniprot50 \| AF-A0A1W6NZ62-F1-MODEL\_V4 \| 1.0 \| 2.336e-11 \| 313 \| 0.293 \| 225 \| 116 \| 7 \| 4 \| 220 \| 265 \| 454 \| Collar domain-containing protein \| Collar domain-containing protein; afdb-uniprot50 \| AF-A0A154VNW1-F1-MODEL\_V4 \| 1.0 \| 9.711e-11 \| 312 \| 0.208 \| 235 \| 149 \| 6 \| 11 \| 220 \| 38 \| 260 \| Collar domain-containing protein \| Collar domain-containing protein; afdb-uniprot50 \| AF-A0A820N1U4-F1-MODEL\_V4 \| 1.0 \| 4.704e-12 \| 312 \| 0.283 \| 236 \| 126 \| 11 \| 2 \| 220 \| 79 \| 288 \| Hypothetical protein \| Hypothetical protein; afdb-uniprot50 \| AF-A0A4R6F7Q8-F1-MODEL\_V4 \| 1.0 \| 7.658e-11 \| 312 \| 0.308 \| 246 \| 101 \| 8 \| 1 \| 220 \| 102 \| 304 \| Uncharacterized protein \| Uncharacterized protein; afdb-uniprot50 \| AF-A0A835XN54-F1-MODEL\_V4 \| 1.0 \| 1.842e-11 \| 312 \| 0.316 \| 224 \| 110 \| 8 \| 3 \| 221 \| 436 \| 621 \| Uncharacterized protein \| Uncharacterized protein; afdb-uniprot50 \| AF-A0A819RFW5-F1-MODEL\_V4 \| 1.0 \| 6.716e-12 \| 311 \| 0.301 \| 222 \| 114 \| 7 \| 12 \| 220 \| 37 \| 230 \| Hypothetical protein \| Hypothetical protein; afdb-uniprot50 \| AF-A0A821GTZ3-F1-MODEL\_V4 \| 1.0 \| 1.523e-12 \| 311 \| 0.28 \| 225 \| 120 \| 10 \| 11 \| 220 \| 165 \| 362 \| Hypothetical protein \| Hypothetical protein; afdb-uniprot50 \| AF-A0A1J5HLH8-F1-MODEL\_V4 \| 1.0 \| 1.018e-11 \| 311 \| 0.339 \| 212 \| 104 \| 8 \| 12 \| 221 \| 439 \| 616 \| Collar domain-containing protein \| Collar domain-containing protein; afdb-uniprot50 \| AF-A0A328LF96-F1-MODEL\_V4 \| 1.0 \| 5.118e-10 \| 310 \| 0.246 \| 211 \| 110 \| 7 \| 11 \| 220 \| 5 \| 167 \| Microcystin-dependent protein \| Microcystin-dependent protein; afdb-uniprot50 \| AF-A0A199XM71-F1-MODEL\_V4 \| 1.0 \| 2.664e-10 \| 309 \| 0.295 \| 213 \| 105 \| 6 \| 10 \| 220 \| 3 \| 172 \| Phage tail collar domain protein \| Phage tail collar domain protein; afdb-uniprot50 \| AF-A0A814GRD0-F1-MODEL\_V4 \| 1.0 \| 3.336e-11 \| 309 \| 0.313 \| 217 \| 111 \| 8 \| 14 \| 220 \| 1 \| 189 \| Hypothetical protein \| Hypothetical protein; afdb-uniprot50 \| AF-A0A503X4M5-F1-MODEL\_V4 \| 1.0 \| 1.758e-10 \| 309 \| 0.243 \| 234 \| 123 \| 9 \| 12 \| 221 \| 112 \| 315 \| Uncharacterized protein \| Uncharacterized protein; afdb-uniprot50 \| AF-A0A7Z0J594-F1-MODEL\_V4 \| 1.0 \| 7.658e-11 \| 309 \| 0.309 \| 210 \| 83 \| 7 \| 12 \| 221 \| 409 \| 556 \| Microcystin-dependent protein \| Microcystin-dependent protein; afdb-uniprot50 \| AF-A0A173LYL4-F1-MODEL\_V4 \| 1.0 \| 3.104e-12 \| 309 \| 0.378 \| 219 \| 106 \| 11 \| 12 \| 221 \| 578 \| 775 \| Collagen triple helix repeat protein \| Collagen triple helix repeat protein; afdb-uniprot50 \| AF-A0A0Q4QY73-F1-MODEL\_V4 \| 1.0 \| 1.107e-09 \| 308 \| 0.289 \| 211 \| 99 \| 6 \| 11 \| 220 \| 4 \| 164 \| Collar domain-containing protein \| Collar domain-containing protein; afdb-uniprot50 \| AF-A0A1E3L3M0-F1-MODEL\_V4 \| 1.0 \| 1.175e-09 \| 308 \| 0.248 \| 213 \| 113 \| 6 \| 11 \| 220 \| 5 \| 173 \| Collar domain-containing protein \| Collar domain-containing protein; afdb-uniprot50 \| AF-A0A4S8NSY4-F1-MODEL\_V4 \| 1.0 \| 4.763e-11 \| 308 \| 0.232 \| 237 \| 139 \| 9 \| 11 \| 220 \| 47 \| 267 \| Collar domain-containing protein \| Collar domain-containing protein; afdb-uniprot50 \| AF-D3F201-F1-MODEL\_V4 \| 1.0 \| 2.075e-11 \| 307 \| 0.359 \| 220 \| 112 \| 10 \| 12 \| 221 \| 61 \| 261 \| Tail Collar domain protein \| Tail Collar domain protein; afdb-uniprot50 \| AF-A0A816HBL7-F1-MODEL\_V4 \| 1.0 \| 2.926e-12 \| 307 \| 0.32 \| 225 \| 100 \| 10 \| 11 \| 220 \| 194 \| 380 \| Hypothetical protein \| Hypothetical protein; afdb-uniprot50 \| AF-A0A4V2W2R3-F1-MODEL\_V4 \| 1.0 \| 1.107e-09 \| 306 \| 0.275 \| 214 \| 92 \| 7 \| 11 \| 220 \| 4 \| 158 \| Microcystin-dependent protein \| Microcystin-dependent protein; afdb-uniprot50 \| AF-A0A1Y6CS75-F1-MODEL\_V4 \| 1.0 \| 4.545e-10 \| 306 \| 0.226 \| 225 \| 114 \| 9 \| 11 \| 220 \| 6 \| 185 \| Microcystin-dependent protein \| Microcystin-dependent protein; afdb-uniprot50 \| AF-A0A3D6BMR4-F1-MODEL\_V4 \| 1.0 \| 3.54e-11 \| 306 \| 0.29 \| 231 \| 136 \| 7 \| 1 \| 221 \| 4 \| 216 \| Collar domain-containing protein \| Collar domain-containing protein; afdb-uniprot50 \| AF-A0A1Y4GXK4-F1-MODEL\_V4 \| 1.0 \| 7.658e-11 \| 306 \| 0.289 \| 211 \| 89 \| 6 \| 11 \| 221 \| 83 \| 232 \| Collar domain-containing protein \| Collar domain-containing protein; afdb-uniprot50 \| AF-A0A5C1ACN0-F1-MODEL\_V4 \| 1.0 \| 8.229e-10 \| 306 \| 0.261 \| 252 \| 102 \| 6 \| 2 \| 221 \| 337 \| 536 \| Uncharacterized protein \| Uncharacterized protein; afdb-uniprot50 \| AF-A0A371WXZ3-F1-MODEL\_V4 \| 1.0 \| 4.489e-11 \| 305 \| 0.276 \| 213 \| 124 \| 8 \| 11 \| 220 \| 4 \| 189 \| Phage tail protein \| Phage tail protein; afdb-uniprot50 \| AF-A0A816TS03-F1-MODEL\_V4 \| 1.0 \| 1.369e-11 \| 305 \| 0.289 \| 228 \| 118 \| 10 \| 11 \| 220 \| 427 \| 628 \| Hypothetical protein \| Hypothetical protein; afdb-uniprot50 \| AF-A0A818AVD1-F1-MODEL\_V4 \| 1.0 \| 1.715e-12 \| 305 \| 0.277 \| 220 \| 127 \| 9 \| 11 \| 220 \| 494 \| 691 \| Hypothetical protein \| Hypothetical protein; afdb-uniprot50 \| AF-A0A3S0R516-F1-MODEL\_V4 \| 1.0 \| 1.657e-10 \| 304 \| 0.269 \| 223 \| 127 \| 7 \| 11 \| 220 \| 5 \| 204 \| Phage tail protein \| Phage tail protein; afdb-uniprot50 \| AF-A0A835XPQ2-F1-MODEL\_V4 \| 1.0 \| 2.336e-11 \| 304 \| 0.318 \| 220 \| 116 \| 7 \| 3 \| 220 \| 440 \| 627 \| Uncharacterized protein \| Uncharacterized protein; afdb-uniprot50 \| AF-A0A397M176-F1-MODEL\_V4 \| 1.0 \| 4.545e-10 \| 303 \| 0.265 \| 211 \| 111 \| 6 \| 11 \| 220 \| 5 \| 172 \| Microcystin-dependent protein \| Microcystin-dependent protein; afdb-uniprot50 \| AF-A0A7W0BUS8-F1-MODEL\_V4 \| 1.0 \| 3.038e-09 \| 302 \| 0.247 \| 210 \| 109 \| 5 \| 11 \| 220 \| 5 \| 165 \| Microcystin-dependent protein \| Microcystin-dependent protein; afdb-uniprot50 \| AF-G8THK5-F1-MODEL\_V4 \| 1.0 \| 8.733e-10 \| 302 \| 0.281 \| 213 \| 101 \| 7 \| 12 \| 220 \| 5 \| 169 \| Tail Collar domain protein \| Tail Collar domain protein; afdb-uniprot50 \| AF-A0A5N1IZX8-F1-MODEL\_V4 \| 1.0 \| 1.404e-09 \| 301 \| 0.278 \| 212 \| 108 \| 5 \| 11 \| 220 \| 4 \| 172 \| Phage tail protein \| Phage tail protein; afdb-uniprot50 \| AF-A0A4Q3L744-F1-MODEL\_V4 \| 1.0 \| 1.323e-09 \| 301 \| 0.294 \| 214 \| 115 \| 9 \| 11 \| 220 \| 4 \| 185 \| Phage tail protein \| Phage tail protein; afdb-uniprot50 \| AF-A0A3B8I8V5-F1-MODEL\_V4 \| 1.0 \| 7.308e-10 \| 301 \| 0.27 \| 218 \| 121 \| 8 \| 10 \| 220 \| 3 \| 189 \| Phage tail protein \| Phage tail protein; afdb-uniprot50 \| AF-A0A7K1Y201-F1-MODEL\_V4 \| 1.0 \| 6.116e-10 \| 301 \| 0.262 \| 217 \| 125 \| 9 \| 11 \| 220 \| 4 \| 192 \| Phage tail protein \| Phage tail protein; afdb-uniprot50 \| AF-G3ISC7-F1-MODEL\_V4 \| 1.0 \| 9.711e-11 \| 301 \| 0.296 \| 229 \| 123 \| 9 \| 11 \| 220 \| 5 \| 214 \| Tail Collar domain protein \| Tail Collar domain protein; afdb-uniprot50 \| AF-A0A6A8A4D1-F1-MODEL\_V4 \| 1.0 \| 2.101e-10 \| 301 \| 0.244 \| 237 \| 136 \| 11 \| 11 \| 220 \| 70 \| 290 \| Collar domain-containing protein \| Collar domain-containing protein; afdb-uniprot50 \| AF-A0A2E0J1D8-F1-MODEL\_V4 \| 1.0 \| 1.758e-10 \| 301 \| 0.253 \| 221 \| 87 \| 5 \| 11 \| 221 \| 215 \| 367 \| Collar domain-containing protein \| Collar domain-containing protein; afdb-uniprot50 \| AF-A0A4R6FF43-F1-MODEL\_V4 \| 1.0 \| 2.127e-09 \| 300 \| 0.275 \| 214 \| 100 \| 7 \| 10 \| 220 \| 3 \| 164 \| Microcystin-dependent protein \| Microcystin-dependent protein; afdb-uniprot50 \| AF-A0A6L5E183-F1-MODEL\_V4 \| 1.0 \| 1.247e-09 \| 300 \| 0.274 \| 215 \| 130 \| 6 \| 12 \| 221 \| 5 \| 198 \| Phage tail protein \| Phage tail protein; afdb-uniprot50 \| AF-A0A2D8SKJ2-F1-MODEL\_V4 \| 1.0 \| 1.561e-10 \| 300 \| 0.281 \| 227 \| 116 \| 9 \| 12 \| 220 \| 5 \| 202 \| Phage tail protein \| Phage tail protein; afdb-uniprot50 \| AF-A0A7Z3TTQ5-F1-MODEL\_V4 \| 1.0 \| 1.561e-10 \| 300 \| 0.202 \| 237 \| 148 \| 8 \| 11 \| 220 \| 29 \| 251 \| Tail fiber protein \| Tail fiber protein; afdb-uniprot50 \| AF-A0A2P8G260-F1-MODEL\_V4 \| 1.0 \| 1.49e-09 \| 299 \| 0.271 \| 210 \| 100 \| 5 \| 11 \| 220 \| 4 \| 160 \| Microcystin-dependent protein \| Microcystin-dependent protein; afdb-uniprot50 \| AF-A0A3B1BHP0-F1-MODEL\_V4 \| 1.0 \| 2.229e-10 \| 299 \| 0.298 \| 211 \| 117 \| 8 \| 11 \| 220 \| 5 \| 185 \| Microcystin dependent protein \| Microcystin dependent protein; afdb-uniprot50 \| AF-A0A0U3HN04-F1-MODEL\_V4 \| 1.0 \| 5.763e-10 \| 299 \| 0.226 \| 212 \| 142 \| 6 \| 11 \| 221 \| 6 \| 196 \| Collar domain-containing protein \| Collar domain-containing protein; afdb-uniprot50 \| AF-A0A522BAB4-F1-MODEL\_V4 \| 1.0 \| 8.229e-10 \| 299 \| 0.363 \| 212 \| 72 \| 7 \| 12 \| 221 \| 99 \| 249 \| Collar domain-containing protein \| Collar domain-containing protein; afdb-uniprot50 \| AF-A0A157ZM93-F1-MODEL\_V4 \| 1.0 \| 6.409e-11 \| 299 \| 0.282 \| 237 \| 110 \| 8 \| 1 \| 221 \| 76 \| 268 \| YadA\_stalk domain-containing protein \| YadA\_stalk domain-containing protein; afdb-uniprot50 \| AF-A0A817VYB8-F1-MODEL\_V4 \| 1.0 \| 1.471e-10 \| 299 \| 0.266 \| 221 \| 124 \| 9 \| 1 \| 220 \| 252 \| 435 \| Hypothetical protein \| Hypothetical protein; afdb-uniprot50 \| AF-A0A815HLR0-F1-MODEL\_V4 \| 1.0 \| 1.616e-12 \| 299 \| 0.303 \| 237 \| 128 \| 9 \| 2 \| 221 \| 417 \| 633 \| Hypothetical protein \| Hypothetical protein; afdb-uniprot50 \| AF-A0A286G5D7-F1-MODEL\_V4 \| 1.0 \| 6.49e-10 \| 298 \| 0.266 \| 210 \| 108 \| 7 \| 11 \| 220 \| 4 \| 167 \| Microcystin-dependent protein \| Microcystin-dependent protein; afdb-uniprot50 \| AF-A0A6N8HFD9-F1-MODEL\_V4 \| 1.0 \| 1.581e-09 \| 298 \| 0.254 \| 212 \| 113 \| 7 \| 10 \| 220 \| 4 \| 171 \| Phage tail protein \| Phage tail protein; afdb-uniprot50 \| AF-A0A2P8D2J1-F1-MODEL\_V4 \| 1.0 \| 1.49e-09 \| 298 \| 0.292 \| 212 \| 108 \| 5 \| 10 \| 220 \| 3 \| 173 \| Microcystin-dependent protein \| Microcystin-dependent protein; afdb-uniprot50 \| AF-A0A173M9U9-F1-MODEL\_V4 \| 1.0 \| 1.231e-10 \| 298 \| 0.278 \| 219 \| 117 \| 10 \| 12 \| 220 \| 5 \| 192 \| Microcystin-dependent protein \| Microcystin-dependent protein; afdb-uniprot50 \| AF-A0A1B9VN74-F1-MODEL\_V4 \| 1.0 \| 9.151e-11 \| 298 \| 0.229 \| 235 \| 137 \| 9 \| 11 \| 220 \| 33 \| 248 \| Tail Collar domain protein \| Tail Collar domain protein; afdb-uniprot50 \| AF-A0A2E0M2L0-F1-MODEL\_V4 \| 1.0 \| 1.758e-10 \| 298 \| 0.282 \| 251 \| 124 \| 8 \| 9 \| 220 \| 18 \| 251 \| Collar domain-containing protein \| Collar domain-containing protein; afdb-uniprot50 \| AF-A0A1V9FKC3-F1-MODEL\_V4 \| 1.0 \| 6.887e-10 \| 297 \| 0.251 \| 211 \| 113 \| 7 \| 12 \| 220 \| 6 \| 173 \| Collar domain-containing protein \| Collar domain-containing protein; afdb-uniprot50 \| AF-A0A4Z0NDY8-F1-MODEL\_V4 \| 1.0 \| 2.229e-10 \| 297 \| 0.283 \| 289 \| 116 \| 9 \| 11 \| 221 \| 143 \| 418 \| Collar domain-containing protein \| Collar domain-containing protein; afdb-uniprot50 \| AF-A0A7W4IK36-F1-MODEL\_V4 \| 1.0 \| 1.453e-11 \| 296 \| 0.424 \| 205 \| 98 \| 6 \| 18 \| 220 \| 1 \| 187 \| Collar domain-containing protein \| Collar domain-containing protein; afdb-uniprot50 \| AF-A0A2P8D2H1-F1-MODEL\_V4 \| 1.0 \| 2.127e-09 \| 295 \| 0.267 \| 217 \| 105 \| 7 \| 10 \| 220 \| 3 \| 171 \| Microcystin-dependent protein \| Microcystin-dependent protein; afdb-uniprot50 \| AF-A0A3N7CVV8-F1-MODEL\_V4 \| 1.0 \| 1.581e-09 \| 295 \| 0.255 \| 219 \| 110 \| 8 \| 11 \| 220 \| 5 \| 179 \| Phage tail protein \| Phage tail protein; afdb-uniprot50 \| AF-A0A1M6T5E4-F1-MODEL\_V4 \| 1.0 \| 7.658e-11 \| 295 \| 0.288 \| 225 \| 130 \| 6 \| 11 \| 220 \| 4 \| 213 \| Microcystin-dependent protein \| Microcystin-dependent protein; afdb-uniprot50 \| AF-A0A6L5FV64-F1-MODEL\_V4 \| 1.0 \| 1.842e-11 \| 295 \| 0.327 \| 235 \| 113 \| 11 \| 3 \| 221 \| 45 \| 250 \| Collar domain-containing protein \| Collar domain-containing protein; afdb-uniprot50 \| AF-A0A6L8KW90-F1-MODEL\_V4 \| 1.0 \| 1.107e-09 \| 295 \| 0.234 \| 247 \| 138 \| 10 \| 11 \| 220 \| 34 \| 266 \| Phage tail protein \| Phage tail protein; afdb-uniprot50 \| AF-A0A835XQ12-F1-MODEL\_V4 \| 1.0 \| 2.962e-11 \| 295 \| 0.273 \| 227 \| 117 \| 11 \| 3 \| 221 \| 223 \| 409 \| Uncharacterized protein \| Uncharacterized protein; afdb-uniprot50 \| AF-A0A5R8ZI20-F1-MODEL\_V4 \| 1.0 \| 2.698e-09 \| 294 \| 0.261 \| 214 \| 102 \| 7 \| 11 \| 220 \| 4 \| 165 \| Phage tail protein \| Phage tail protein; afdb-uniprot50 \| AF-A0A4Q0RK75-F1-MODEL\_V4 \| 1.0 \| 1.889e-09 \| 294 \| 0.256 \| 218 \| 102 \| 9 \| 11 \| 220 \| 4 \| 169 \| Phage tail protein \| Phage tail protein; afdb-uniprot50 \| AF-A0A1I1NQ92-F1-MODEL\_V4 \| 1.0 \| 3.804e-10 \| 294 \| 0.282 \| 216 \| 117 \| 10 \| 11 \| 220 \| 4 \| 187 \| Microcystin-dependent protein \| Microcystin-dependent protein; afdb-uniprot50 \| AF-A0A521AUE6-F1-MODEL\_V4 \| 1.0 \| 1.107e-09 \| 294 \| 0.273 \| 216 \| 136 \| 8 \| 10 \| 220 \| 3 \| 202 \| Microcystin-dependent protein \| Microcystin-dependent protein; afdb-uniprot50 \| AF-A0A1G8MR82-F1-MODEL\_V4 \| 1.0 \| 8.624e-11 \| 294 \| 0.282 \| 258 \| 129 \| 9 \| 3 \| 221 \| 28 \| 268 \| Phage Tail Collar Domain \| Phage Tail Collar Domain; afdb-uniprot50 \| AF-E5APE0-F1-MODEL\_V4 \| 1.0 \| 6.49e-10 \| 294 \| 0.318 \| 226 \| 102 \| 7 \| 2 \| 220 \| 161 \| 341 \| PHAGE TAIL PROTEIN \| PHAGE TAIL PROTEIN; afdb-uniprot50 \| AF-A0A1I7LHC6-F1-MODEL\_V4 \| 1.0 \| 2.51e-10 \| 294 \| 0.293 \| 279 \| 121 \| 12 \| 11 \| 221 \| 143 \| 413 \| Microcystin-dependent protein \| Microcystin-dependent protein; afdb-uniprot50 \| AF-A0A819YKS0-F1-MODEL\_V4 \| 1.0 \| 2.792e-11 \| 293 \| 0.253 \| 225 \| 126 \| 9 \| 11 \| 220 \| 6 \| 203 \| Hypothetical protein \| Hypothetical protein; afdb-uniprot50 \| AF-A0A2E0ZP65-F1-MODEL\_V4 \| 1.0 \| 2.732e-08 \| 292 \| 0.566 \| 90 \| 33 \| 1 \| 2 \| 85 \| 7 \| 96 \| Collar domain-containing protein \| Collar domain-containing protein; afdb-uniprot50 \| AF-A0A7W7Z7K7-F1-MODEL\_V4 \| 1.0 \| 1.889e-09 \| 292 \| 0.251 \| 211 \| 108 \| 7 \| 11 \| 220 \| 5 \| 166 \| Microcystin-dependent protein \| Microcystin-dependent protein; afdb-uniprot50 \| AF-A0A1V9FKH6-F1-MODEL\_V4 \| 1.0 \| 5.183e-09 \| 292 \| 0.243 \| 218 \| 110 \| 8 \| 10 \| 221 \| 3 \| 171 \| Collar domain-containing protein \| Collar domain-containing protein; afdb-uniprot50 \| AF-A0A6C0DW18-F1-MODEL\_V4 \| 1.0 \| 5.763e-10 \| 292 \| 0.28 \| 214 \| 88 \| 7 \| 11 \| 220 \| 35 \| 186 \| Collar domain-containing protein \| Collar domain-containing protein; afdb-uniprot50 \| AF-A0A7C3YVS4-F1-MODEL\_V4 \| 1.0 \| 1.231e-10 \| 292 \| 0.276 \| 210 \| 88 \| 9 \| 12 \| 221 \| 47 \| 192 \| Collar domain-containing protein \| Collar domain-containing protein; afdb-uniprot50 \| AF-A0A7H9DPU9-F1-MODEL\_V4 \| 1.0 \| 1.49e-09 \| 292 \| 0.278 \| 219 \| 111 \| 7 \| 11 \| 220 \| 5 \| 185 \| Phage tail protein \| Phage tail protein; afdb-uniprot50 \| AF-A0A1Q3GJM7-F1-MODEL\_V4 \| 1.0 \| 6.49e-10 \| 292 \| 0.274 \| 226 \| 110 \| 10 \| 10 \| 220 \| 3 \| 189 \| Collar domain-containing protein \| Collar domain-containing protein; afdb-uniprot50 \| AF-A0A3T0W1B7-F1-MODEL\_V4 \| 1.0 \| 6.887e-10 \| 292 \| 0.235 \| 217 \| 133 \| 8 \| 11 \| 220 \| 6 \| 196 \| Phage tail protein \| Phage tail protein; afdb-uniprot50 \| AF-A0A521AUD9-F1-MODEL\_V4 \| 1.0 \| 4.036e-10 \| 292 \| 0.281 \| 224 \| 127 \| 9 \| 10 \| 220 \| 4 \| 206 \| Microcystin-dependent protein \| Microcystin-dependent protein; afdb-uniprot50 \| AF-A0A850KVP1-F1-MODEL\_V4 \| 1.0 \| 2.366e-10 \| 292 \| 0.246 \| 215 \| 117 \| 9 \| 11 \| 220 \| 38 \| 212 \| Tail fiber protein \| Tail fiber protein; afdb-uniprot50 \| AF-A0A821ML04-F1-MODEL\_V4 \| 1.0 \| 3.804e-10 \| 292 \| 0.254 \| 212 \| 110 \| 10 \| 11 \| 220 \| 195 \| 360 \| Hypothetical protein \| Hypothetical protein; afdb-uniprot50 \| AF-A0A1V2H5U5-F1-MODEL\_V4 \| 1.0 \| 1.561e-10 \| 292 \| 0.319 \| 213 \| 118 \| 9 \| 12 \| 220 \| 213 \| 402 \| Collar domain-containing protein \| Collar domain-containing protein; afdb-uniprot50 \| AF-A0A1E4J0W6-F1-MODEL\_V4 \| 1.0 \| 1.352e-12 \| 291 \| 0.344 \| 247 \| 121 \| 9 \| 1 \| 221 \| 106 \| 337 \| Collar domain-containing protein \| Collar domain-containing protein; afdb-uniprot50 \| AF-A0A2E5YUW9-F1-MODEL\_V4 \| 1.0 \| 1.369e-11 \| 291 \| 0.265 \| 241 \| 124 \| 5 \| 11 \| 221 \| 148 \| 365 \| Collar domain-containing protein \| Collar domain-containing protein; afdb-uniprot50 \| AF-A0A258K539-F1-MODEL\_V4 \| 1.0 \| 2.396e-09 \| 290 \| 0.25 \| 220 \| 103 \| 7 \| 11 \| 221 \| 4 \| 170 \| Collar domain-containing protein \| Collar domain-containing protein; afdb-uniprot50 \| AF-A0A1T4L5J9-F1-MODEL\_V4 \| 1.0 \| 2.258e-09 \| 290 \| 0.279 \| 215 \| 102 \| 7 \| 11 \| 220 \| 5 \| 171 \| Microcystin-dependent protein \| Microcystin-dependent protein; afdb-uniprot50 \| AF-A0A6L9IQ28-F1-MODEL\_V4 \| 1.0 \| 2.366e-10 \| 290 \| 0.276 \| 213 \| 117 \| 9 \| 11 \| 220 \| 21 \| 199 \| Phage tail protein \| Phage tail protein; afdb-uniprot50 \| AF-A0A3N6N9Q9-F1-MODEL\_V4 \| 1.0 \| 3e-10 \| 290 \| 0.259 \| 216 \| 105 \| 7 \| 9 \| 220 \| 133 \| 297 \| Collar domain-containing protein \| Collar domain-containing protein; afdb-uniprot50 \| AF-A6EAC0-F1-MODEL\_V4 \| 1.0 \| 1.247e-09 \| 289 \| 0.252 \| 214 \| 115 \| 6 \| 11 \| 220 \| 4 \| 176 \| Microcystin-dependent protein \| Microcystin-dependent protein; afdb-uniprot50 \| AF-A0A7U0PKD0-F1-MODEL\_V4 \| 1.0 \| 6.801e-11 \| 289 \| 0.272 \| 224 \| 127 \| 11 \| 11 \| 220 \| 5 \| 206 \| Phage tail protein \| Phage tail protein; afdb-uniprot50 \| AF-A0A5C7MBK6-F1-MODEL\_V4 \| 1.0 \| 2.51e-10 \| 289 \| 0.279 \| 236 \| 99 \| 9 \| 12 \| 221 \| 276 \| 466 \| Uncharacterized protein \| Uncharacterized protein; afdb-uniprot50 \| AF-A0A1H7BRU9-F1-MODEL\_V4 \| 1.0 \| 1.043e-09 \| 288 \| 0.271 \| 217 \| 102 \| 9 \| 11 \| 220 \| 4 \| 171 \| Microcystin-dependent protein \| Microcystin-dependent protein; afdb-uniprot50 \| AF-A0A1G4QLN4-F1-MODEL\_V4 \| 1.0 \| 1.107e-09 \| 288 \| 0.276 \| 210 \| 116 \| 7 \| 11 \| 220 \| 4 \| 177 \| Microcystin-dependent protein \| Microcystin-dependent protein; afdb-uniprot50 \| AF-A0A7W9ZEX5-F1-MODEL\_V4 \| 1.0 \| 1.49e-09 \| 288 \| 0.269 \| 215 \| 111 \| 10 \| 11 \| 220 \| 4 \| 177 \| Microcystin-dependent protein \| Microcystin-dependent protein; afdb-uniprot50 \| AF-F0SE37-F1-MODEL\_V4 \| 1.0 \| 1.107e-09 \| 288 \| 0.263 \| 224 \| 118 \| 8 \| 11 \| 221 \| 4 \| 193 \| Tail Collar domain protein \| Tail Collar domain protein; afdb-uniprot50 \| AF-A0A1G6Y0D3-F1-MODEL\_V4 \| 1.0 \| 1.758e-10 \| 288 \| 0.207 \| 212 \| 130 \| 7 \| 11 \| 220 \| 40 \| 215 \| Microcystin-dependent protein \| Microcystin-dependent protein; afdb-uniprot50 \| AF-A0A1Y4GLN6-F1-MODEL\_V4 \| 1.0 \| 1.458e-06 \| 287 \| 0.538 \| 65 \| 30 \| 0 \| 12 \| 76 \| 25 \| 89 \| Collar domain-containing protein \| Collar domain-containing protein; afdb-uniprot50 \| AF-D8GPZ1-F1-MODEL\_V4 \| 1.0 \| 7.755e-10 \| 287 \| 0.274 \| 222 \| 137 \| 6 \| 11 \| 220 \| 5 \| 214 \| Collar domain-containing protein \| Collar domain-containing protein; afdb-uniprot50 \| AF-K8GNE3-F1-MODEL\_V4 \| 1.0 \| 8.516e-12 \| 287 \| 0.334 \| 254 \| 118 \| 8 \| 1 \| 221 \| 468 \| 703 \| Phage tail collar family protein \| Phage tail collar family protein; afdb-uniprot50 \| AF-A0A2C6C321-F1-MODEL\_V4 \| 1.0 \| 1.175e-09 \| 286 \| 0.24 \| 212 \| 121 \| 7 \| 10 \| 220 \| 3 \| 175 \| Phage tail protein \| Phage tail protein; afdb-uniprot50 \| AF-A0A444WG53-F1-MODEL\_V4 \| 1.0 \| 4.087e-09 \| 286 \| 0.291 \| 213 \| 107 \| 5 \| 11 \| 220 \| 5 \| 176 \| Tail Collar domain protein \| Tail Collar domain protein; afdb-uniprot50 \| AF-A0A7T8SK15-F1-MODEL\_V4 \| 1.0 \| 8.229e-10 \| 286 \| 0.273 \| 216 \| 107 \| 9 \| 12 \| 220 \| 9 \| 181 \| Tail fiber protein \| Tail fiber protein; afdb-uniprot50 \| AF-A0A7Y4RAK2-F1-MODEL\_V4 \| 1.0 \| 9.711e-11 \| 286 \| 0.292 \| 239 \| 141 \| 8 \| 2 \| 221 \| 64 \| 293 \| Tail fiber protein \| Tail fiber protein; afdb-uniprot50 \| AF-A0A222EY63-F1-MODEL\_V4 \| 1.0 \| 1.889e-09 \| 285 \| 0.289 \| 211 \| 79 \| 6 \| 11 \| 221 \| 21 \| 160 \| Collar domain-containing protein \| Collar domain-containing protein; afdb-uniprot50 \| AF-A0A817V4J7-F1-MODEL\_V4 \| 1.0 \| 1.107e-09 \| 285 \| 0.305 \| 219 \| 104 \| 6 \| 11 \| 221 \| 17 \| 195 \| Hypothetical protein \| Hypothetical protein; afdb-uniprot50 \| AF-A0A7L4ZNP4-F1-MODEL\_V4 \| 1.0 \| 5.118e-10 \| 285 \| 0.254 \| 224 \| 112 \| 9 \| 2 \| 220 \| 21 \| 194 \| Collar domain-containing protein \| Collar domain-containing protein; afdb-uniprot50 \| AF-A0A1F9IXH9-F1-MODEL\_V4 \| 1.0 \| 5.054e-11 \| 285 \| 0.33 \| 230 \| 84 \| 6 \| 2 \| 221 \| 48 \| 217 \| Collar domain-containing protein \| Collar domain-containing protein; afdb-uniprot50 \| AF-A0A222FDY7-F1-MODEL\_V4 \| 1.0 \| 2.542e-09 \| 285 \| 0.223 \| 224 \| 136 \| 8 \| 11 \| 220 \| 29 \| 228 \| Phage tail protein \| Phage tail protein; afdb-uniprot50 \| AF-A0A833JCH6-F1-MODEL\_V4 \| 1.0 \| 2.51e-10 \| 285 \| 0.235 \| 259 \| 146 \| 12 \| 2 \| 221 \| 68 \| 313 \| Uncharacterized protein \| Uncharacterized protein; afdb-uniprot50 \| AF-A0A2U2BCA2-F1-MODEL\_V4 \| 1.0 \| 3.585e-10 \| 284 \| 0.278 \| 233 \| 118 \| 7 \| 2 \| 220 \| 24 \| 220 \| Phage tail protein \| Phage tail protein; afdb-uniprot50 \| AF-A0A1H8W0U3-F1-MODEL\_V4 \| 1.0 \| 6.193e-09 \| 283 \| 0.235 \| 212 \| 111 \| 6 \| 10 \| 220 \| 3 \| 164 \| Microcystin-dependent protein \| Microcystin-dependent protein; afdb-uniprot50 \| AF-A0A519S7T0-F1-MODEL\_V4 \| 1.0 \| 7.4e-09 \| 283 \| 0.257 \| 214 \| 105 \| 10 \| 11 \| 220 \| 4 \| 167 \| Phage tail protein \| Phage tail protein; afdb-uniprot50 \| AF-A0A3S1AVP9-F1-MODEL\_V4 \| 1.0 \| 2.127e-09 \| 283 \| 0.265 \| 222 \| 115 \| 6 \| 11 \| 220 \| 5 \| 190 \| Phage tail protein \| Phage tail protein; afdb-uniprot50 \| AF-A0A3S9QK74-F1-MODEL\_V4 \| 1.0 \| 1.03e-10 \| 283 \| 0.252 \| 214 \| 111 \| 10 \| 11 \| 221 \| 136 \| 303 \| Collar domain-containing protein \| Collar domain-containing protein; afdb-uniprot50 \| AF-A0A1I5JPD8-F1-MODEL\_V4 \| 1.0 \| 2.258e-09 \| 283 \| 0.28 \| 214 \| 123 \| 7 \| 12 \| 220 \| 171 \| 358 \| Uncharacterized protein \| Uncharacterized protein; afdb-uniprot50 \| AF-A0A2N9X676-F1-MODEL\_V4 \| 1.0 \| 1.657e-10 \| 283 \| 0.281 \| 213 \| 119 \| 8 \| 11 \| 221 \| 245 \| 425 \| Collar domain-containing protein \| Collar domain-containing protein; afdb-uniprot50 \| AF-A0A848FSQ8-F1-MODEL\_V4 \| 1.0 \| 5.5e-09 \| 282 \| 0.24 \| 212 \| 114 \| 5 \| 12 \| 220 \| 5 \| 172 \| Phage tail protein \| Phage tail protein; afdb-uniprot50 \| AF-A0A4Q7N378-F1-MODEL\_V4 \| 1.0 \| 3.63e-09 \| 282 \| 0.267 \| 217 \| 106 \| 7 \| 10 \| 220 \| 3 \| 172 \| Microcystin-dependent protein \| Microcystin-dependent protein; afdb-uniprot50 \| AF-A0A2E3FMJ4-F1-MODEL\_V4 \| 1.0 \| 3.378e-10 \| 282 \| 0.328 \| 210 \| 102 \| 8 \| 11 \| 220 \| 504 \| 674 \| Collar domain-containing protein \| Collar domain-containing protein; afdb-uniprot50 \| AF-A0A7S7U734-F1-MODEL\_V4 \| 1.0 \| 8.843e-09 \| 281 \| 0.255 \| 215 \| 102 \| 9 \| 11 \| 220 \| 5 \| 166 \| Phage tail protein \| Phage tail protein; afdb-uniprot50 \| AF-A0A521AU55-F1-MODEL\_V4 \| 1.0 \| 3.183e-10 \| 281 \| 0.252 \| 222 \| 131 \| 10 \| 11 \| 220 \| 13 \| 211 \| Microcystin-dependent protein \| Microcystin-dependent protein; afdb-uniprot50 \| AF-A0A348AH31-F1-MODEL\_V4 \| 1.0 \| 7.4e-09 \| 280 \| 0.284 \| 211 \| 103 \| 7 \| 11 \| 220 \| 4 \| 167 \| Phage Tail Collar Domain protein \| Phage Tail Collar Domain protein; afdb-uniprot50 \| AF-A0A1C2GP00-F1-MODEL\_V4 \| 1.0 \| 9.834e-10 \| 280 \| 0.248 \| 213 \| 120 \| 9 \| 11 \| 220 \| 5 \| 180 \| Collar domain-containing protein \| Collar domain-containing protein; afdb-uniprot50 \| AF-F0SE36-F1-MODEL\_V4 \| 1.0 \| 6.887e-10 \| 280 \| 0.297 \| 222 \| 110 \| 9 \| 11 \| 220 \| 5 \| 192 \| Tail Collar domain protein \| Tail Collar domain protein; afdb-uniprot50 \| AF-F8GEA0-F1-MODEL\_V4 \| 1.0 \| 4.087e-09 \| 280 \| 0.227 \| 229 \| 130 \| 7 \| 11 \| 220 \| 69 \| 269 \| Tail Collar domain protein \| Tail Collar domain protein; afdb-uniprot50 \| AF-A0A2T9K1M7-F1-MODEL\_V4 \| 1.0 \| 1.107e-09 \| 279 \| 0.281 \| 224 \| 116 \| 8 \| 11 \| 220 \| 4 \| 196 \| Phage tail protein \| Phage tail protein; afdb-uniprot50 \| AF-A0A3G8WSK6-F1-MODEL\_V4 \| 1.0 \| 3.421e-09 \| 279 \| 0.266 \| 225 \| 120 \| 10 \| 10 \| 220 \| 3 \| 196 \| Phage tail protein \| Phage tail protein; afdb-uniprot50 \| AF-A0A2E4T1Y0-F1-MODEL\_V4 \| 1.0 \| 8.516e-12 \| 279 \| 0.295 \| 230 \| 115 \| 9 \| 11 \| 221 \| 118 \| 319 \| Collar domain-containing protein \| Collar domain-containing protein; afdb-uniprot50 \| AF-A0A818V6Z8-F1-MODEL\_V4 \| 1.0 \| 4.823e-10 \| 279 \| 0.269 \| 223 \| 128 \| 10 \| 11 \| 220 \| 389 \| 589 \| Hypothetical protein \| Hypothetical protein; afdb-uniprot50 \| AF-A0A1J0WLQ5-F1-MODEL\_V4 \| 1.0 \| 2.698e-09 \| 278 \| 0.248 \| 217 \| 108 \| 9 \| 12 \| 220 \| 14 \| 183 \| Collar domain-containing protein \| Collar domain-containing protein; afdb-uniprot50 \| AF-A0A818YA82-F1-MODEL\_V4 \| 1.0 \| 5.118e-10 \| 277 \| 0.235 \| 225 \| 135 \| 9 \| 11 \| 220 \| 98 \| 300 \| Hypothetical protein \| Hypothetical protein; afdb-uniprot50 \| AF-A0A1F3K302-F1-MODEL\_V4 \| 1.0 \| 5.763e-10 \| 276 \| 0.279 \| 222 \| 124 \| 8 \| 10 \| 220 \| 3 \| 199 \| Collar domain-containing protein \| Collar domain-containing protein; afdb-uniprot50 \| AF-W6RSW9-F1-MODEL\_V4 \| 1.0 \| 3.144e-11 \| 276 \| 0.282 \| 244 \| 130 \| 11 \| 5 \| 221 \| 36 \| 261 \| Putative tail fiber protein \| Putative tail fiber protein; afdb-uniprot50 \| AF-A0A7H1JCB2-F1-MODEL\_V4 \| 1.0 \| 1.98e-10 \| 276 \| 0.252 \| 242 \| 124 \| 9 \| 11 \| 220 \| 41 \| 257 \| Tail fiber protein \| Tail fiber protein; afdb-uniprot50 \| AF-A0A553JK32-F1-MODEL\_V4 \| 1.0 \| 1.247e-09 \| 276 \| 0.316 \| 224 \| 108 \| 6 \| 2 \| 220 \| 297 \| 480 \| Collar domain-containing protein \| Collar domain-containing protein; afdb-uniprot50 \| AF-A0A7C8AIS3-F1-MODEL\_V4 \| 1.0 \| 1.404e-09 \| 275 \| 0.242 \| 231 \| 113 \| 8 \| 1 \| 221 \| 164 \| 342 \| Collar domain-containing protein \| Collar domain-containing protein; afdb-uniprot50 \| AF-A0A849EVQ5-F1-MODEL\_V4 \| 1.0 \| 3.421e-09 \| 274 \| 0.271 \| 221 \| 111 \| 9 \| 11 \| 220 \| 6 \| 187 \| Phage tail protein \| Phage tail protein; afdb-uniprot50 \| AF-A0A7W6LK96-F1-MODEL\_V4 \| 1.0 \| 6.887e-10 \| 274 \| 0.2 \| 235 \| 148 \| 8 \| 11 \| 220 \| 41 \| 260 \| Microcystin-dependent protein \| Microcystin-dependent protein; afdb-uniprot50 \| AF-A0A423L7N4-F1-MODEL\_V4 \| 1.0 \| 9.037e-12 \| 274 \| 0.318 \| 226 \| 119 \| 9 \| 12 \| 221 \| 176 \| 382 \| Collar domain-containing protein \| Collar domain-containing protein; afdb-uniprot50 \| AF-A0A5B8XJ48-F1-MODEL\_V4 \| 1.0 \| 1.889e-09 \| 274 \| 0.301 \| 212 \| 79 \| 5 \| 11 \| 220 \| 433 \| 577 \| Phage tail collar domain protein \| Phage tail collar domain protein; afdb-uniprot50 \| AF-A0A395JR72-F1-MODEL\_V4 \| 1.0 \| 2.863e-09 \| 273 \| 0.282 \| 223 \| 108 \| 9 \| 11 \| 221 \| 5 \| 187 \| Phage tail protein \| Phage tail protein; afdb-uniprot50 \| AF-A0A3B7MPY4-F1-MODEL\_V4 \| 1.0 \| 4.884e-09 \| 272 \| 0.287 \| 216 \| 99 \| 7 \| 12 \| 220 \| 5 \| 172 \| Phage tail protein \| Phage tail protein; afdb-uniprot50 \| AF-A0A2S5DKY5-F1-MODEL\_V4 \| 1.0 \| 2.315e-07 \| 272 \| 0.45 \| 100 \| 40 \| 3 \| 9 \| 106 \| 23 \| 109 \| Phage tail protein \| Phage tail protein; afdb-uniprot50 \| AF-A0A359L8X4-F1-MODEL\_V4 \| 1.0 \| 2.698e-09 \| 272 \| 0.259 \| 231 \| 115 \| 11 \| 11 \| 220 \| 4 \| 199 \| Phage tail protein \| Phage tail protein; afdb-uniprot50 \| AF-A0A0Q6ZRG1-F1-MODEL\_V4 \| 1.0 \| 4.823e-10 \| 272 \| 0.273 \| 223 \| 112 \| 10 \| 1 \| 221 \| 156 \| 330 \| Collar domain-containing protein \| Collar domain-containing protein; afdb-uniprot50 \| AF-A0A6N1X655-F1-MODEL\_V4 \| 1.0 \| 1.657e-10 \| 272 \| 0.293 \| 215 \| 120 \| 9 \| 11 \| 221 \| 201 \| 387 \| Tail fiber protein \| Tail fiber protein; afdb-uniprot50 \| AF-A0A2W5K0J5-F1-MODEL\_V4 \| 1.0 \| 1.453e-11 \| 272 \| 0.325 \| 227 \| 116 \| 6 \| 9 \| 221 \| 228 \| 431 \| Collar domain-containing protein \| Collar domain-containing protein; afdb-uniprot50 \| AF-A0A152A3A6-F1-MODEL\_V4 \| 1.0 \| 3.852e-09 \| 272 \| 0.247 \| 214 \| 120 \| 6 \| 10 \| 221 \| 442 \| 616 \| Collar domain-containing protein \| Collar domain-containing protein; afdb-uniprot50 \| AF-A0A1H2GA33-F1-MODEL\_V4 \| 1.0 \| 4.036e-10 \| 271 \| 0.261 \| 237 \| 110 \| 9 \| 11 \| 220 \| 34 \| 232 \| Microcystin-dependent protein \| Microcystin-dependent protein; afdb-uniprot50 \| AF-A0A1W2F866-F1-MODEL\_V4 \| 1.0 \| 2.664e-10 \| 271 \| 0.339 \| 224 \| 87 \| 8 \| 1 \| 221 \| 123 \| 288 \| Phage Tail Collar Domain \| Phage Tail Collar Domain; afdb-uniprot50 \| AF-L0H2G4-F1-MODEL\_V4 \| 1.0 \| 8.843e-09 \| 270 \| 0.245 \| 216 \| 111 \| 8 \| 11 \| 220 \| 6 \| 175 \| Microcystin-dependent protein \| Microcystin-dependent protein; afdb-uniprot50 \| AF-A0A2D3RDN9-F1-MODEL\_V4 \| 1.0 \| 3.038e-09 \| 270 \| 0.284 \| 211 \| 102 \| 8 \| 11 \| 220 \| 32 \| 194 \| Collar domain-containing protein \| Collar domain-containing protein; afdb-uniprot50 \| AF-A0A0M9DM99-F1-MODEL\_V4 \| 1.0 \| 7.755e-10 \| 270 \| 0.261 \| 241 \| 124 \| 9 \| 11 \| 220 \| 4 \| 221 \| Phage tail collar protein \| Phage tail collar protein; afdb-uniprot50 \| AF-A0A178MM09-F1-MODEL\_V4 \| 1.0 \| 5.763e-10 \| 270 \| 0.224 \| 218 \| 136 \| 7 \| 11 \| 220 \| 31 \| 223 \| Collar domain-containing protein \| Collar domain-containing protein; afdb-uniprot50 \| AF-A0A839IKX9-F1-MODEL\_V4 \| 1.0 \| 7.4e-09 \| 269 \| 0.213 \| 220 \| 125 \| 5 \| 11 \| 220 \| 4 \| 185 \| Tail fiber protein \| Tail fiber protein; afdb-uniprot50 \| AF-A0A1K1QMP3-F1-MODEL\_V4 \| 1.0 \| 1.34e-08 \| 269 \| 0.276 \| 224 \| 106 \| 10 \| 11 \| 220 \| 4 \| 185 \| Microcystin-dependent protein \| Microcystin-dependent protein; afdb-uniprot50 \| AF-A0A4Z1BHZ3-F1-MODEL\_V4 \| 1.0 \| 6.572e-09 \| 269 \| 0.235 \| 234 \| 104 \| 7 \| 11 \| 220 \| 4 \| 186 \| Phage tail protein \| Phage tail protein; afdb-uniprot50 \| AF-A0A2N3KBQ2-F1-MODEL\_V4 \| 1.0 \| 1.889e-09 \| 269 \| 0.261 \| 218 \| 128 \| 7 \| 10 \| 220 \| 3 \| 194 \| Collar domain-containing protein \| Collar domain-containing protein; afdb-uniprot50 \| AF-A0A1H8GZ50-F1-MODEL\_V4 \| 1.0 \| 2.479e-11 \| 269 \| 0.268 \| 235 \| 123 \| 11 \| 11 \| 220 \| 4 \| 214 \| Microcystin-dependent protein \| Microcystin-dependent protein; afdb-uniprot50 \| AF-A0A4R7BEN1-F1-MODEL\_V4 \| 1.0 \| 1.98e-10 \| 269 \| 0.327 \| 214 \| 102 \| 9 \| 12 \| 221 \| 241 \| 416 \| Microcystin-dependent protein \| Microcystin-dependent protein; afdb-uniprot50 \| AF-A0A2N3T4D1-F1-MODEL\_V4 \| 1.0 \| 6.193e-09 \| 268 \| 0.262 \| 213 \| 117 \| 8 \| 11 \| 220 \| 4 \| 179 \| Microcystin-dependent protein \| Microcystin-dependent protein; afdb-uniprot50 \| AF-A0A162JQP9-F1-MODEL\_V4 \| 1.0 \| 1.404e-09 \| 268 \| 0.24 \| 225 \| 115 \| 9 \| 11 \| 220 \| 5 \| 188 \| Collar domain-containing protein \| Collar domain-containing protein; afdb-uniprot50 \| AF-A0A4Q5WXF2-F1-MODEL\_V4 \| 1.0 \| 4.545e-10 \| 267 \| 0.282 \| 230 \| 119 \| 9 \| 11 \| 220 \| 5 \| 208 \| Phage tail protein \| Phage tail protein; afdb-uniprot50 \| AF-A0A212BGA2-F1-MODEL\_V4 \| 1.0 \| 2.366e-10 \| 267 \| 0.336 \| 217 \| 107 \| 11 \| 10 \| 221 \| 65 \| 249 \| Collar domain-containing protein \| Collar domain-containing protein; afdb-uniprot50 \| AF-A0A4D8R397-F1-MODEL\_V4 \| 1.0 \| 1.175e-09 \| 267 \| 0.242 \| 239 \| 131 \| 11 \| 11 \| 220 \| 31 \| 248 \| Collar domain-containing protein \| Collar domain-containing protein; afdb-uniprot50 \| AF-A0A800M671-F1-MODEL\_V4 \| 1.0 \| 1.78e-09 \| 267 \| 0.279 \| 211 \| 97 \| 8 \| 11 \| 220 \| 114 \| 270 \| Phage tail protein \| Phage tail protein; afdb-uniprot50 \| AF-A0NQ95-F1-MODEL\_V4 \| 1.0 \| 1.387e-10 \| 267 \| 0.27 \| 229 \| 128 \| 9 \| 2 \| 221 \| 131 \| 329 \| Putative tail fiber-related protein \| Putative tail fiber-related protein; afdb-uniprot50 \| AF-A0A844A272-F1-MODEL\_V4 \| 1.0 \| 1.889e-09 \| 267 \| 0.3 \| 233 \| 128 \| 8 \| 1 \| 221 \| 245 \| 454 \| Tail fiber protein \| Tail fiber protein; afdb-uniprot50 \| AF-A0A4S2GXX0-F1-MODEL\_V4 \| 1.0 \| 5.836e-09 \| 266 \| 0.251 \| 211 \| 108 \| 7 \| 11 \| 220 \| 29 \| 190 \| Phage tail protein \| Phage tail protein; afdb-uniprot50 \| AF-A0A177MPX3-F1-MODEL\_V4 \| 1.0 \| 5.836e-09 \| 266 \| 0.251 \| 215 \| 121 \| 6 \| 11 \| 220 \| 19 \| 198 \| Collar domain-containing protein \| Collar domain-containing protein; afdb-uniprot50 \| AF-A0A2H0IRE9-F1-MODEL\_V4 \| 1.0 \| 4.087e-09 \| 266 \| 0.256 \| 230 \| 118 \| 7 \| 11 \| 220 \| 4 \| 200 \| Phage tail protein \| Phage tail protein; afdb-uniprot50 \| AF-A0A845L736-F1-MODEL\_V4 \| 1.0 \| 6.887e-10 \| 266 \| 0.305 \| 229 \| 113 \| 11 \| 11 \| 220 \| 4 \| 205 \| Uncharacterized protein \| Uncharacterized protein; afdb-uniprot50 \| AF-F7P1I1-F1-MODEL\_V4 \| 1.0 \| 1.509e-08 \| 265 \| 0.216 \| 217 \| 116 \| 6 \| 11 \| 220 \| 4 \| 173 \| Microcystin-dependent protein \| Microcystin-dependent protein; afdb-uniprot50 \| AF-A0A2E4PG81-F1-MODEL\_V4 \| 1.0 \| 9.958e-09 \| 265 \| 0.229 \| 218 \| 112 \| 8 \| 11 \| 220 \| 6 \| 175 \| Phage tail protein \| Phage tail protein; afdb-uniprot50 \| AF-A0A653P8Z9-F1-MODEL\_V4 \| 1.0 \| 9.834e-10 \| 265 \| 0.242 \| 231 \| 124 \| 10 \| 11 \| 220 \| 33 \| 233 \| Collar domain-containing protein \| Collar domain-containing protein; afdb-uniprot50 \| AF-A0A2S1SFX4-F1-MODEL\_V4 \| 1.0 \| 7.853e-09 \| 264 \| 0.257 \| 214 \| 110 \| 9 \| 11 \| 220 \| 6 \| 174 \| Phage tail protein \| Phage tail protein; afdb-uniprot50 \| AF-A0A4Q6AGP9-F1-MODEL\_V4 \| 1.0 \| 7.4e-09 \| 264 \| 0.255 \| 215 \| 112 \| 6 \| 10 \| 220 \| 3 \| 173 \| Phage tail protein \| Phage tail protein; afdb-uniprot50 \| AF-A0A7H9DR54-F1-MODEL\_V4 \| 1.0 \| 2.005e-09 \| 264 \| 0.278 \| 219 \| 113 \| 11 \| 11 \| 220 \| 4 \| 186 \| Phage tail protein \| Phage tail protein; afdb-uniprot50 \| AF-A0A1Y4GJI2-F1-MODEL\_V4 \| 1.0 \| 1.164e-05 \| 263 \| 0.516 \| 60 \| 29 \| 0 \| 12 \| 71 \| 23 \| 82 \| Collar domain-containing protein \| Collar domain-containing protein; afdb-uniprot50 \| AF-A0A383RWZ2-F1-MODEL\_V4 \| 1.0 \| 3.223e-09 \| 263 \| 0.263 \| 216 \| 110 \| 7 \| 11 \| 220 \| 4 \| 176 \| Collar domain-containing protein \| Collar domain-containing protein; afdb-uniprot50 \| AF-A0A291QWR7-F1-MODEL\_V4 \| 1.0 \| 4.603e-09 \| 263 \| 0.263 \| 216 \| 122 \| 8 \| 11 \| 220 \| 4 \| 188 \| Phage tail protein \| Phage tail protein; afdb-uniprot50 \| AF-A0A0G4ICD0-F1-MODEL\_V4 \| 1.0 \| 2.396e-09 \| 263 \| 0.27 \| 211 \| 103 \| 7 \| 11 \| 220 \| 42 \| 202 \| Collar domain-containing protein \| Collar domain-containing protein; afdb-uniprot50 \| AF-A0A7Y7EEW4-F1-MODEL\_V4 \| 1.0 \| 4.603e-09 \| 263 \| 0.217 \| 234 \| 140 \| 13 \| 11 \| 220 \| 7 \| 221 \| Tail fiber protein \| Tail fiber protein; afdb-uniprot50 \| AF-A0A2Z4PXV1-F1-MODEL\_V4 \| 1.0 \| 1.78e-09 \| 263 \| 0.222 \| 238 \| 138 \| 10 \| 11 \| 220 \| 13 \| 231 \| Collar domain-containing protein \| Collar domain-containing protein; afdb-uniprot50 \| AF-U1ID74-F1-MODEL\_V4 \| 1.0 \| 6.116e-10 \| 263 \| 0.287 \| 216 \| 126 \| 7 \| 12 \| 220 \| 98 \| 292 \| NitT/TauT family transport system permease \| NitT/TauT family transport system permease; afdb-uniprot50 \| AF-A0A7W6ICU7-F1-MODEL\_V4 \| 1.0 \| 1.758e-10 \| 263 \| 0.266 \| 233 \| 118 \| 9 \| 11 \| 220 \| 358 \| 560 \| Microcystin-dependent protein \| Microcystin-dependent protein; afdb-uniprot50 \| AF-A0A2K9MAN8-F1-MODEL\_V4 \| 1.0 \| 1.955e-11 \| 262 \| 0.321 \| 233 \| 131 \| 7 \| 1 \| 221 \| 4 \| 221 \| Collar domain-containing protein \| Collar domain-containing protein; afdb-uniprot50 \| AF-A0A3D9L8W2-F1-MODEL\_V4 \| 1.0 \| 2.542e-09 \| 262 \| 0.246 \| 227 \| 135 \| 11 \| 11 \| 220 \| 5 \| 212 \| Microcystin-dependent protein \| Microcystin-dependent protein; afdb-uniprot50 \| AF-A0A2L1HSI4-F1-MODEL\_V4 \| 1.0 \| 1.678e-09 \| 262 \| 0.256 \| 234 \| 125 \| 11 \| 1 \| 221 \| 151 \| 348 \| Collar domain-containing protein \| Collar domain-containing protein; afdb-uniprot50 \| AF-A0A5B8UCR2-F1-MODEL\_V4 \| 1.0 \| 1.175e-09 \| 261 \| 0.311 \| 231 \| 89 \| 11 \| 3 \| 220 \| 35 \| 208 \| Tail fiber protein \| Tail fiber protein; afdb-uniprot50 \| AF-A0A7V1GDM0-F1-MODEL\_V4 \| 1.0 \| 3.038e-09 \| 261 \| 0.267 \| 232 \| 134 \| 10 \| 11 \| 220 \| 6 \| 223 \| Phage tail protein \| Phage tail protein; afdb-uniprot50 \| AF-A0A433BEE5-F1-MODEL\_V4 \| 1.0 \| 3.63e-09 \| 260 \| 0.233 \| 214 \| 109 \| 8 \| 11 \| 220 \| 13 \| 175 \| Collar domain-containing protein \| Collar domain-containing protein; afdb-uniprot50 \| AF-A0A7T4YH49-F1-MODEL\_V4 \| 1.0 \| 1.175e-09 \| 260 \| 0.29 \| 217 \| 91 \| 11 \| 11 \| 221 \| 42 \| 201 \| Tail fiber protein \| Tail fiber protein; afdb-uniprot50 \| AF-A0A0F9W1V5-F1-MODEL\_V4 \| 1.0 \| 2.396e-09 \| 259 \| 0.29 \| 217 \| 80 \| 7 \| 12 \| 220 \| 22 \| 172 \| Collar domain-containing protein \| Collar domain-containing protein; afdb-uniprot50 \| AF-A0A6J4FWR0-F1-MODEL\_V4 \| 1.0 \| 4.337e-09 \| 259 \| 0.283 \| 215 \| 114 \| 9 \| 11 \| 220 \| 5 \| 184 \| Putative Microcystin dependent protein \| Putative Microcystin dependent protein; afdb-uniprot50 \| AF-A0A1V4X328-F1-MODEL\_V4 \| 1.0 \| 2.51e-10 \| 259 \| 0.342 \| 213 \| 98 \| 8 \| 12 \| 221 \| 228 \| 401 \| Phage Tail Collar Domain protein \| Phage Tail Collar Domain protein; afdb-uniprot50 \| AF-A0A6N1WZL5-F1-MODEL\_V4 \| 1.0 \| 4.823e-10 \| 259 \| 0.297 \| 215 \| 119 \| 6 \| 11 \| 221 \| 824 \| 1010 \| Tail fiber protein \| Tail fiber protein; afdb-uniprot50 \| AF-A1AK23-F1-MODEL\_V4 \| 1.0 \| 7.4e-09 \| 258 \| 0.25 \| 224 \| 127 \| 8 \| 11 \| 220 \| 4 \| 200 \| Phage Tail Collar domain protein \| Phage Tail Collar domain protein; afdb-uniprot50 \| AF-A0A848FK93-F1-MODEL\_V4 \| 1.0 \| 3.038e-09 \| 258 \| 0.235 \| 212 \| 123 \| 7 \| 11 \| 220 \| 29 \| 203 \| Phage tail protein \| Phage tail protein; afdb-uniprot50 \| AF-A0A251WMF7-F1-MODEL\_V4 \| 1.0 \| 3.464e-08 \| 257 \| 0.242 \| 214 \| 88 \| 8 \| 11 \| 220 \| 5 \| 148 \| Collar domain-containing protein \| Collar domain-containing protein; afdb-uniprot50 \| AF-L0H3J2-F1-MODEL\_V4 \| 1.0 \| 2.574e-08 \| 257 \| 0.267 \| 213 \| 108 \| 12 \| 11 \| 220 \| 5 \| 172 \| Microcystin-dependent protein \| Microcystin-dependent protein; afdb-uniprot50 \| AF-A0A1V9FC14-F1-MODEL\_V4 \| 1.0 \| 6.193e-09 \| 257 \| 0.23 \| 217 \| 116 \| 8 \| 10 \| 220 \| 4 \| 175 \| Collar domain-containing protein \| Collar domain-containing protein; afdb-uniprot50 \| AF-A0A351HL10-F1-MODEL\_V4 \| 1.0 \| 4.603e-09 \| 257 \| 0.266 \| 218 \| 133 \| 8 \| 10 \| 220 \| 3 \| 200 \| Phage tail protein \| Phage tail protein; afdb-uniprot50 \| AF-A0A2D8AUD5-F1-MODEL\_V4 \| 1.0 \| 1.231e-10 \| 257 \| 0.263 \| 224 \| 121 \| 8 \| 12 \| 221 \| 24 \| 217 \| Collar domain-containing protein \| Collar domain-containing protein; afdb-uniprot50 \| AF-A0A2E3CYM6-F1-MODEL\_V4 \| 1.0 \| 5.5e-09 \| 257 \| 0.265 \| 211 \| 129 \| 6 \| 11 \| 220 \| 34 \| 219 \| Phage tail protein \| Phage tail protein; afdb-uniprot50 \| AF-A0A538RHD6-F1-MODEL\_V4 \| 1.0 \| 1.323e-09 \| 257 \| 0.296 \| 219 \| 114 \| 10 \| 11 \| 221 \| 56 \| 242 \| Collar domain-containing protein \| Collar domain-containing protein; afdb-uniprot50 \| AF-A0A3E0WQ94-F1-MODEL\_V4 \| 1.0 \| 3.378e-10 \| 257 \| 0.274 \| 237 \| 116 \| 12 \| 1 \| 221 \| 279 \| 475 \| Collar domain-containing protein \| Collar domain-containing protein; afdb-uniprot50 \| AF-A0A3M0CW52-F1-MODEL\_V4 \| 1.0 \| 1.34e-08 \| 256 \| 0.231 \| 220 \| 123 \| 8 \| 11 \| 221 \| 7 \| 189 \| Microcystin-dependent protein \| Microcystin-dependent protein; afdb-uniprot50 \| AF-A0A090VAI2-F1-MODEL\_V4 \| 1.0 \| 1.121e-08 \| 256 \| 0.278 \| 223 \| 106 \| 8 \| 11 \| 220 \| 28 \| 208 \| Microcystin dependent protein \| Microcystin dependent protein; afdb-uniprot50 \| AF-A0A1M3LS74-F1-MODEL\_V4 \| 1.0 \| 1.803e-08 \| 255 \| 0.26 \| 211 \| 115 \| 9 \| 11 \| 220 \| 5 \| 175 \| Collar domain-containing protein \| Collar domain-containing protein; afdb-uniprot50 \| AF-A8HTM6-F1-MODEL\_V4 \| 1.0 \| 2.574e-08 \| 255 \| 0.225 \| 235 \| 125 \| 10 \| 12 \| 221 \| 148 \| 350 \| Microcystin-dependent protein \| Microcystin-dependent protein; afdb-uniprot50 \| AF-A0A7Z6VNA8-F1-MODEL\_V4 \| 1.0 \| 5.183e-09 \| 255 \| 0.283 \| 226 \| 129 \| 10 \| 2 \| 221 \| 456 \| 654 \| Phage tail protein \| Phage tail protein; afdb-uniprot50 \| AF-A0A2E1HKS8-F1-MODEL\_V4 \| 1.0 \| 3.464e-08 \| 254 \| 0.24 \| 212 \| 115 \| 6 \| 11 \| 220 \| 6 \| 173 \| Phage tail protein \| Phage tail protein; afdb-uniprot50 \| AF-A0A7Z0I2P9-F1-MODEL\_V4 \| 1.0 \| 8.333e-09 \| 254 \| 0.234 \| 213 \| 122 \| 6 \| 11 \| 220 \| 40 \| 214 \| Tail fiber protein \| Tail fiber protein; afdb-uniprot50 \| AF-A0A1J0EHY7-F1-MODEL\_V4 \| 1.0 \| 2.792e-11 \| 254 \| 0.33 \| 227 \| 115 \| 9 \| 12 \| 221 \| 131 \| 337 \| Collar domain-containing protein \| Collar domain-containing protein; afdb-uniprot50 \| AF-A0A354WRB2-F1-MODEL\_V4 \| 1.0 \| 1.263e-08 \| 253 \| 0.285 \| 210 \| 87 \| 6 \| 12 \| 221 \| 22 \| 168 \| Collar domain-containing protein \| Collar domain-containing protein; afdb-uniprot50 \| AF-A0A1P8X333-F1-MODEL\_V4 \| 1.0 \| 3.076e-08 \| 253 \| 0.248 \| 217 \| 108 \| 8 \| 11 \| 220 \| 6 \| 174 \| Collar domain-containing protein \| Collar domain-containing protein; afdb-uniprot50 \| AF-A0A3N5E5J5-F1-MODEL\_V4 \| 1.0 \| 2.154e-08 \| 253 \| 0.254 \| 216 \| 117 \| 9 \| 11 \| 220 \| 5 \| 182 \| Phage tail protein \| Phage tail protein; afdb-uniprot50 \| AF-A0A233RHW0-F1-MODEL\_V4 \| 1.0 \| 1.121e-08 \| 253 \| 0.229 \| 222 \| 128 \| 10 \| 11 \| 220 \| 5 \| 195 \| Collar domain-containing protein \| Collar domain-containing protein; afdb-uniprot50 \| AF-A0A0U3F2K0-F1-MODEL\_V4 \| 1.0 \| 5.836e-09 \| 253 \| 0.269 \| 215 \| 98 \| 9 \| 11 \| 220 \| 43 \| 203 \| Collar domain-containing protein \| Collar domain-containing protein; afdb-uniprot50 \| AF-A0A6N6VYM1-F1-MODEL\_V4 \| 1.0 \| 5.118e-10 \| 253 \| 0.277 \| 245 \| 144 \| 8 \| 2 \| 221 \| 45 \| 281 \| Collar domain-containing protein \| Collar domain-containing protein; afdb-uniprot50 \| AF-B5JYG6-F1-MODEL\_V4 \| 1.0 \| 3.421e-09 \| 253 \| 0.26 \| 234 \| 124 \| 10 \| 1 \| 220 \| 200 \| 398 \| Phage Tail Collar Domain family \| Phage Tail Collar Domain family; afdb-uniprot50 \| AF-A0A366FYJ6-F1-MODEL\_V4 \| 1.0 \| 1.247e-09 \| 253 \| 0.323 \| 213 \| 89 \| 8 \| 11 \| 221 \| 646 \| 805 \| Tail collar domain \| Tail collar domain; afdb-uniprot50 \| AF-A0A1Y0FCL5-F1-MODEL\_V4 \| 1.0 \| 6.974e-09 \| 252 \| 0.251 \| 223 \| 110 \| 7 \| 3 \| 221 \| 24 \| 193 \| Collar domain-containing protein \| Collar domain-containing protein; afdb-uniprot50 \| AF-I3TXM0-F1-MODEL\_V4 \| 1.0 \| 2.396e-09 \| 252 \| 0.256 \| 222 \| 115 \| 10 \| 11 \| 220 \| 39 \| 222 \| Tail Collar domain protein \| Tail Collar domain protein; afdb-uniprot50 \| AF-A0A7C1MNT2-F1-MODEL\_V4 \| 1.0 \| 4.036e-10 \| 252 \| 0.302 \| 225 \| 119 \| 11 \| 11 \| 220 \| 68 \| 269 \| Tail fiber protein \| Tail fiber protein; afdb-uniprot50 \| AF-A0A2H5E1S5-F1-MODEL\_V4 \| 1.0 \| 3.378e-10 \| 252 \| 0.301 \| 219 \| 126 \| 6 \| 5 \| 221 \| 218 \| 411 \| Collar domain-containing protein \| Collar domain-containing protein; afdb-uniprot50 \| AF-A0A382GIC3-F1-MODEL\_V4 \| 1.0 \| 1.323e-09 \| 251 \| 0.278 \| 223 \| 93 \| 9 \| 8 \| 221 \| 29 \| 192 \| Collar domain-containing protein \| Collar domain-containing protein; afdb-uniprot50 \| AF-A0A4R1A3G5-F1-MODEL\_V4 \| 1.0 \| 2.542e-09 \| 251 \| 0.244 \| 237 \| 121 \| 8 \| 2 \| 220 \| 19 \| 215 \| Phage tail protein \| Phage tail protein; afdb-uniprot50 \| AF-N9TWJ4-F1-MODEL\_V4 \| 1.0 \| 3.223e-09 \| 251 \| 0.269 \| 223 \| 124 \| 10 \| 11 \| 220 \| 29 \| 225 \| Tail collar domain-containing protein \| Tail collar domain-containing protein; afdb-uniprot50 \| AF-A0A419H9E0-F1-MODEL\_V4 \| 1.0 \| 1.404e-09 \| 251 \| 0.229 \| 253 \| 132 \| 10 \| 11 \| 220 \| 24 \| 256 \| Collar domain-containing protein \| Collar domain-containing protein; afdb-uniprot50 \| AF-A0A0Q5F3R8-F1-MODEL\_V4 \| 1.0 \| 2.863e-09 \| 251 \| 0.201 \| 248 \| 135 \| 12 \| 11 \| 220 \| 40 \| 262 \| Collar domain-containing protein \| Collar domain-containing protein; afdb-uniprot50 \| AF-A0A1I4FJV1-F1-MODEL\_V4 \| 1.0 \| 5.183e-09 \| 250 \| 0.277 \| 220 \| 106 \| 8 \| 11 \| 220 \| 29 \| 205 \| Microcystin-dependent protein \| Microcystin-dependent protein; afdb-uniprot50 \| AF-A0A1S8LIP9-F1-MODEL\_V4 \| 1.0 \| 6.974e-09 \| 250 \| 0.227 \| 237 \| 131 \| 10 \| 11 \| 220 \| 4 \| 215 \| Phage tail collar domain protein \| Phage tail collar domain protein; afdb-uniprot50 \| AF-A0A819NXH0-F1-MODEL\_V4 \| 1.0 \| 1.094e-10 \| 250 \| 0.262 \| 248 \| 123 \| 14 \| 2 \| 220 \| 13 \| 229 \| Hypothetical protein \| Hypothetical protein; afdb-uniprot50 \| AF-A0A2S1XFQ2-F1-MODEL\_V4 \| 1.0 \| 7.755e-10 \| 250 \| 0.215 \| 237 \| 140 \| 8 \| 11 \| 220 \| 31 \| 248 \| Collar domain-containing protein \| Collar domain-containing protein; afdb-uniprot50 \| AF-F4KQ07-F1-MODEL\_V4 \| 1.0 \| 2.127e-09 \| 250 \| 0.243 \| 234 \| 116 \| 13 \| 2 \| 221 \| 129 \| 315 \| Tail Collar domain protein \| Tail Collar domain protein; afdb-uniprot50 \| AF-A0A257ULB7-F1-MODEL\_V4 \| 1.0 \| 1.561e-10 \| 250 \| 0.368 \| 244 \| 116 \| 10 \| 12 \| 220 \| 133 \| 373 \| Collar domain-containing protein \| Collar domain-containing protein; afdb-uniprot50 \| AF-A0A1G5E937-F1-MODEL\_V4 \| 1.0 \| 1.699e-08 \| 249 \| 0.231 \| 229 \| 115 \| 8 \| 11 \| 220 \| 4 \| 190 \| Microcystin-dependent protein \| Microcystin-dependent protein; afdb-uniprot50 \| AF-A0A7U3BZ46-F1-MODEL\_V4 \| 1.0 \| 1.581e-09 \| 249 \| 0.303 \| 224 \| 93 \| 8 \| 1 \| 220 \| 142 \| 306 \| Collar domain-containing protein \| Collar domain-containing protein; afdb-uniprot50 \| AF-A0A814Z5L0-F1-MODEL\_V4 \| 1.0 \| 3.183e-10 \| 249 \| 0.291 \| 230 \| 116 \| 10 \| 12 \| 220 \| 111 \| 314 \| Hypothetical protein \| Hypothetical protein; afdb-uniprot50 \| AF-A0A2T4JN94-F1-MODEL\_V4 \| 1.0 \| 1.404e-09 \| 249 \| 0.255 \| 219 \| 100 \| 8 \| 6 \| 221 \| 209 \| 367 \| Collar domain-containing protein \| Collar domain-containing protein; afdb-uniprot50 \| AF-A0A2K8QRG9-F1-MODEL\_V4 \| 1.0 \| 3.076e-08 \| 248 \| 0.209 \| 267 \| 138 \| 10 \| 11 \| 220 \| 37 \| 287 \| Collar domain-containing protein \| Collar domain-containing protein; afdb-uniprot50 \| AF-A0A8A7KLE0-F1-MODEL\_V4 \| 1.0 \| 9.958e-09 \| 247 \| 0.278 \| 223 \| 93 \| 8 \| 3 \| 221 \| 38 \| 196 \| Phage tail protein \| Phage tail protein; afdb-uniprot50 \| AF-A0A835ZKI7-F1-MODEL\_V4 \| 1.0 \| 2.698e-09 \| 247 \| 0.293 \| 225 \| 120 \| 10 \| 12 \| 220 \| 25 \| 226 \| Uncharacterized protein \| Uncharacterized protein; afdb-uniprot50 \| AF-B8FJJ3-F1-MODEL\_V4 \| 1.0 \| 4.884e-09 \| 247 \| 0.259 \| 216 \| 86 \| 8 \| 12 \| 221 \| 117 \| 264 \| Tail Collar domain protein \| Tail Collar domain protein; afdb-uniprot50 \| AF-A0A1X0T8R2-F1-MODEL\_V4 \| 1.0 \| 4.392e-08 \| 246 \| 0.251 \| 215 \| 109 \| 8 \| 11 \| 220 \| 19 \| 186 \| Phage tail collar domain-containing protein \| Phage tail collar domain-containing protein; afdb-uniprot50 \| AF-A0A1G0MZ06-F1-MODEL\_V4 \| 1.0 \| 6.655e-08 \| 246 \| 0.236 \| 224 \| 130 \| 9 \| 11 \| 220 \| 4 \| 200 \| Collar domain-containing protein \| Collar domain-containing protein; afdb-uniprot50 \| AF-A0A0B3RIK5-F1-MODEL\_V4 \| 1.0 \| 3.421e-09 \| 246 \| 0.293 \| 276 \| 121 \| 9 \| 3 \| 221 \| 27 \| 285 \| Phage Tail Collar \| Phage Tail Collar; afdb-uniprot50 \| AF-A0A7Y7P107-F1-MODEL\_V4 \| 1.0 \| 2.005e-09 \| 246 \| 0.215 \| 237 \| 107 \| 10 \| 1 \| 221 \| 122 \| 295 \| Tail fiber protein \| Tail fiber protein; afdb-uniprot50 \| AF-A0A7C1Q6C6-F1-MODEL\_V4 \| 1.0 \| 1.49e-09 \| 245 \| 0.276 \| 235 \| 101 \| 13 \| 1 \| 221 \| 50 \| 229 \| Collar domain-containing protein \| Collar domain-containing protein; afdb-uniprot50 \| AF-A0A7Z2VX81-F1-MODEL\_V4 \| 1.0 \| 2.827e-10 \| 245 \| 0.218 \| 252 \| 135 \| 12 \| 11 \| 220 \| 33 \| 264 \| Phage tail protein \| Phage tail protein; afdb-uniprot50 \| AF-A0A1R3EDN7-F1-MODEL\_V4 \| 1.0 \| 1.509e-08 \| 244 \| 0.238 \| 222 \| 137 \| 10 \| 13 \| 220 \| 2 \| 205 \| Collar domain-containing protein \| Collar domain-containing protein; afdb-uniprot50 \| AF-A0A120IHK0-F1-MODEL\_V4 \| 1.0 \| 3.676e-08 \| 243 \| 0.23 \| 213 \| 101 \| 8 \| 11 \| 221 \| 24 \| 175 \| Collar domain-containing protein \| Collar domain-containing protein; afdb-uniprot50 \| AF-A0A1J4ST39-F1-MODEL\_V4 \| 1.0 \| 1.422e-08 \| 243 \| 0.254 \| 216 \| 110 \| 9 \| 11 \| 220 \| 5 \| 175 \| Collar domain-containing protein \| Collar domain-containing protein; afdb-uniprot50 \| AF-C6DJW4-F1-MODEL\_V4 \| 1.0 \| 2.03e-08 \| 243 \| 0.192 \| 229 \| 139 \| 6 \| 11 \| 220 \| 35 \| 236 \| Tail Collar domain protein \| Tail Collar domain protein; afdb-uniprot50 \| AF-A0A1B9VN93-F1-MODEL\_V4 \| 1.0 \| 8.333e-09 \| 243 \| 0.173 \| 242 \| 143 \| 11 \| 11 \| 220 \| 35 \| 251 \| Tail Collar domain protein \| Tail Collar domain protein; afdb-uniprot50 \| AF-A0A661ZRY0-F1-MODEL\_V4 \| 1.0 \| 2.005e-09 \| 243 \| 0.272 \| 220 \| 133 \| 6 \| 11 \| 221 \| 94 \| 295 \| Collar domain-containing protein \| Collar domain-containing protein; afdb-uniprot50 \| AF-A0A3S9H6F8-F1-MODEL\_V4 \| 1.0 \| 7.853e-09 \| 243 \| 0.264 \| 219 \| 95 \| 5 \| 3 \| 221 \| 143 \| 295 \| Collar domain-containing protein \| Collar domain-containing protein; afdb-uniprot50 \| AF-A0A3S9SJT2-F1-MODEL\_V4 \| 1.0 \| 2.542e-09 \| 243 \| 0.319 \| 219 \| 72 \| 9 \| 5 \| 221 \| 183 \| 326 \| Phage tail protein \| Phage tail protein; afdb-uniprot50 \| AF-A0A3B0MDZ4-F1-MODEL\_V4 \| 1.0 \| 7.217e-11 \| 243 \| 0.283 \| 236 \| 128 \| 8 \| 3 \| 221 \| 155 \| 366 \| Collar domain-containing protein \| Collar domain-containing protein; afdb-uniprot50 \| AF-A0A7W3S4S1-F1-MODEL\_V4 \| 1.0 \| 1.19e-08 \| 243 \| 0.302 \| 281 \| 120 \| 11 \| 11 \| 221 \| 145 \| 419 \| Microcystin-dependent protein \| Microcystin-dependent protein; afdb-uniprot50 \| AF-A0A411MQE7-F1-MODEL\_V4 \| 1.0 \| 1.121e-08 \| 243 \| 0.267 \| 217 \| 127 \| 7 \| 11 \| 221 \| 256 \| 446 \| Collar domain-containing protein \| Collar domain-containing protein; afdb-uniprot50 \| AF-L8MKX1-F1-MODEL\_V4 \| 1.0 \| 2.664e-10 \| 243 \| 0.317 \| 217 \| 111 \| 9 \| 10 \| 221 \| 342 \| 526 \| Phage tail fiber protein \| Phage tail fiber protein; afdb-uniprot50 \| AF-A0A819H8M6-F1-MODEL\_V4 \| 1.0 \| 5.431e-10 \| 243 \| 0.338 \| 201 \| 104 \| 9 \| 11 \| 198 \| 481 \| 665 \| Hypothetical protein \| Hypothetical protein; afdb-uniprot50 \| AF-A0A7V8INF4-F1-MODEL\_V4 \| 1.0 \| 8.954e-08 \| 242 \| 0.252 \| 210 \| 99 \| 6 \| 11 \| 220 \| 1 \| 152 \| Collar domain-containing protein \| Collar domain-containing protein; afdb-uniprot50 \| AF-A0A8A6KKJ5-F1-MODEL\_V4 \| 1.0 \| 4.504e-06 \| 242 \| 0.437 \| 80 \| 43 \| 1 \| 2 \| 79 \| 72 \| 151 \| Tail fiber protein \| Tail fiber protein; afdb-uniprot50 \| AF-A0A4D8PL44-F1-MODEL\_V4 \| 1.0 \| 2.005e-09 \| 242 \| 0.235 \| 242 \| 129 \| 11 \| 11 \| 220 \| 31 \| 248 \| Collar domain-containing protein \| Collar domain-containing protein; afdb-uniprot50 \| AF-A0A0G9JNT3-F1-MODEL\_V4 \| 1.0 \| 2.698e-09 \| 242 \| 0.284 \| 225 \| 104 \| 12 \| 3 \| 221 \| 158 \| 331 \| Collar domain-containing protein \| Collar domain-containing protein; afdb-uniprot50 \| AF-A0A125Q5U9-F1-MODEL\_V4 \| 1.0 \| 1.803e-08 \| 242 \| 0.253 \| 233 \| 131 \| 11 \| 7 \| 221 \| 139 \| 346 \| Collar domain-containing protein \| Collar domain-containing protein; afdb-uniprot50 \| AF-A0A7V9P271-F1-MODEL\_V4 \| 1.0 \| 2.005e-09 \| 242 \| 0.274 \| 211 \| 107 \| 5 \| 11 \| 221 \| 256 \| 420 \| Tail fiber protein \| Tail fiber protein; afdb-uniprot50 \| AF-A0A6L3AU25-F1-MODEL\_V4 \| 1.0 \| 9.384e-09 \| 241 \| 0.176 \| 238 \| 148 \| 11 \| 11 \| 220 \| 31 \| 248 \| Phage tail protein \| Phage tail protein; afdb-uniprot50 \| AF-A0A3N5NQW9-F1-MODEL\_V4 \| 1.0 \| 8.333e-09 \| 241 \| 0.268 \| 212 \| 65 \| 6 \| 12 \| 221 \| 266 \| 389 \| Tail fiber protein \| Tail fiber protein; afdb-uniprot50 \| AF-A0A150J402-F1-MODEL\_V4 \| 1.0 \| 1.057e-08 \| 241 \| 0.215 \| 218 \| 95 \| 5 \| 12 \| 221 \| 267 \| 416 \| Phage Tail Collar Domain protein \| Phage Tail Collar Domain protein; afdb-uniprot50 \| AF-A0A1S8LYQ0-F1-MODEL\_V4 \| 1.0 \| 2.732e-08 \| 240 \| 0.259 \| 212 \| 93 \| 6 \| 9 \| 220 \| 30 \| 177 \| Phage tail collar domain protein \| Phage tail collar domain protein; afdb-uniprot50 \| AF-A0A3S0V0B8-F1-MODEL\_V4 \| 1.0 \| 1.057e-08 \| 240 \| 0.227 \| 246 \| 131 \| 10 \| 18 \| 220 \| 2 \| 231 \| Collar domain-containing protein \| Collar domain-containing protein; afdb-uniprot50 \| AF-A0A1M6BZM6-F1-MODEL\_V4 \| 1.0 \| 9.502e-08 \| 239 \| 0.22 \| 213 \| 118 \| 6 \| 11 \| 220 \| 4 \| 171 \| Phage Tail Collar Domain protein \| Phage Tail Collar Domain protein; afdb-uniprot50 \| AF-A0A7W9FLI3-F1-MODEL\_V4 \| 1.0 \| 6.572e-09 \| 239 \| 0.261 \| 218 \| 124 \| 10 \| 11 \| 220 \| 8 \| 196 \| Microcystin-dependent protein \| Microcystin-dependent protein; afdb-uniprot50 \| AF-A0A2U2HDJ5-F1-MODEL\_V4 \| 1.0 \| 6.193e-09 \| 239 \| 0.263 \| 239 \| 110 \| 12 \| 11 \| 220 \| 4 \| 205 \| Phage tail protein \| Phage tail protein; afdb-uniprot50 \| AF-A0A4Y7XFR7-F1-MODEL\_V4 \| 1.0 \| 1.581e-09 \| 239 \| 0.268 \| 227 \| 116 \| 9 \| 2 \| 221 \| 49 \| 232 \| Collar domain-containing protein \| Collar domain-containing protein; afdb-uniprot50 \| AF-A0A2E5YV74-F1-MODEL\_V4 \| 1.0 \| 9.267e-10 \| 239 \| 0.237 \| 269 \| 123 \| 9 \| 11 \| 221 \| 147 \| 391 \| Collar domain-containing protein \| Collar domain-containing protein; afdb-uniprot50 \| AF-A0A833JB53-F1-MODEL\_V4 \| 1.0 \| 2.863e-09 \| 238 \| 0.216 \| 249 \| 151 \| 11 \| 3 \| 220 \| 38 \| 273 \| Uncharacterized protein \| Uncharacterized protein; afdb-uniprot50 \| AF-A0A446AXF9-F1-MODEL\_V4 \| 1.0 \| 4.087e-09 \| 238 \| 0.269 \| 219 \| 126 \| 8 \| 11 \| 221 \| 116 \| 308 \| Collar domain-containing protein \| Collar domain-containing protein; afdb-uniprot50 \| AF-A0A366D7H9-F1-MODEL\_V4 \| 1.0 \| 1.121e-08 \| 238 \| 0.253 \| 221 \| 88 \| 7 \| 2 \| 221 \| 356 \| 500 \| Microcystin-dependent protein \| Microcystin-dependent protein; afdb-uniprot50 \| AF-A0A1H2FLZ8-F1-MODEL\_V4 \| 1.0 \| 3.223e-09 \| 238 \| 0.276 \| 210 \| 94 \| 6 \| 11 \| 220 \| 268 \| 419 \| Microcystin-dependent protein \| Microcystin-dependent protein; afdb-uniprot50 \| AF-A0A820I8T0-F1-MODEL\_V4 \| 1.0 \| 2.542e-09 \| 237 \| 0.391 \| 184 \| 85 \| 8 \| 11 \| 182 \| 12 \| 180 \| Hypothetical protein \| Hypothetical protein; afdb-uniprot50 \| AF-A0A495JVM4-F1-MODEL\_V4 \| 1.0 \| 1.422e-08 \| 237 \| 0.236 \| 228 \| 115 \| 10 \| 9 \| 220 \| 11 \| 195 \| Tail collar domain \| Tail collar domain; afdb-uniprot50 \| AF-A0A560CBU6-F1-MODEL\_V4 \| 1.0 \| 3.852e-09 \| 237 \| 0.217 \| 235 \| 142 \| 9 \| 11 \| 220 \| 31 \| 248 \| Microcystin-dependent protein \| Microcystin-dependent protein; afdb-uniprot50 \| AF-A0A2A5VXT6-F1-MODEL\_V4 \| 1.0 \| 1.509e-08 \| 237 \| 0.235 \| 229 \| 112 \| 7 \| 1 \| 221 \| 246 \| 419 \| Collar domain-containing protein \| Collar domain-containing protein; afdb-uniprot50 \| AF-A0A098S053-F1-MODEL\_V4 \| 1.0 \| 5.248e-08 \| 236 \| 0.244 \| 225 \| 97 \| 10 \| 11 \| 221 \| 14 \| 179 \| Collar domain-containing protein \| Collar domain-containing protein; afdb-uniprot50 \| AF-A0A5C7LB37-F1-MODEL\_V4 \| 1.0 \| 5.183e-09 \| 236 \| 0.207 \| 222 \| 125 \| 12 \| 3 \| 221 \| 51 \| 224 \| Tail fiber protein \| Tail fiber protein; afdb-uniprot50 \| AF-A0A2R7UAN8-F1-MODEL\_V4 \| 1.0 \| 2.127e-09 \| 236 \| 0.3 \| 220 \| 113 \| 9 \| 12 \| 221 \| 46 \| 234 \| Collar domain-containing protein \| Collar domain-containing protein; afdb-uniprot50 \| AF-A0A1X3DKU1-F1-MODEL\_V4 \| 1.0 \| 3.038e-09 \| 236 \| 0.3 \| 220 \| 116 \| 8 \| 3 \| 221 \| 61 \| 243 \| Collar domain-containing protein \| Collar domain-containing protein; afdb-uniprot50 \| AF-A0A448VHU0-F1-MODEL\_V4 \| 1.0 \| 4.337e-09 \| 236 \| 0.246 \| 231 \| 111 \| 11 \| 1 \| 221 \| 255 \| 432 \| Putative phage tail fiber protein \| Putative phage tail fiber protein; afdb-uniprot50 \| AF-A0A480ATV8-F1-MODEL\_V4 \| 1.0 \| 7.4e-09 \| 235 \| 0.286 \| 220 \| 100 \| 8 \| 4 \| 221 \| 64 \| 228 \| Tail protein \| Tail protein; afdb-uniprot50 \| AF-A0A533TV88-F1-MODEL\_V4 \| 1.0 \| 9.958e-09 \| 235 \| 0.225 \| 262 \| 131 \| 13 \| 11 \| 220 \| 39 \| 280 \| Tail fiber protein \| Tail fiber protein; afdb-uniprot50 \| AF-A0A4R7MXL7-F1-MODEL\_V4 \| 1.0 \| 8.843e-09 \| 235 \| 0.291 \| 223 \| 108 \| 10 \| 1 \| 221 \| 593 \| 767 \| Tail collar domain \| Tail collar domain; afdb-uniprot50 \| AF-A0A1X0ZXQ4-F1-MODEL\_V4 \| 1.0 \| 7.4e-09 \| 235 \| 0.274 \| 222 \| 110 \| 10 \| 1 \| 221 \| 701 \| 872 \| Collar domain-containing protein \| Collar domain-containing protein; afdb-uniprot50 \| AF-A0A367V8J5-F1-MODEL\_V4 \| 1.0 \| 2.574e-08 \| 234 \| 0.25 \| 220 \| 105 \| 9 \| 1 \| 220 \| 14 \| 173 \| Collar domain-containing protein \| Collar domain-containing protein; afdb-uniprot50 \| AF-A0A821SA43-F1-MODEL\_V4 \| 1.0 \| 5.836e-09 \| 234 \| 0.286 \| 206 \| 111 \| 9 \| 23 \| 219 \| 14 \| 192 \| Hypothetical protein \| Hypothetical protein; afdb-uniprot50 \| AF-A0A7X4H185-F1-MODEL\_V4 \| 1.0 \| 7.308e-10 \| 234 \| 0.23 \| 243 \| 130 \| 12 \| 11 \| 220 \| 4 \| 222 \| Collar domain-containing protein \| Collar domain-containing protein; afdb-uniprot50 \| AF-A0A2R6VDR8-F1-MODEL\_V4 \| 1.0 \| 9.958e-09 \| 234 \| 0.224 \| 227 \| 138 \| 9 \| 11 \| 220 \| 34 \| 239 \| Collar domain-containing protein \| Collar domain-containing protein; afdb-uniprot50 \| AF-A0A323UBG8-F1-MODEL\_V4 \| 1.0 \| 8.333e-09 \| 234 \| 0.257 \| 245 \| 130 \| 10 \| 3 \| 220 \| 101 \| 320 \| Collar domain-containing protein \| Collar domain-containing protein; afdb-uniprot50 \| AF-A0A8A3LGF1-F1-MODEL\_V4 \| 1.0 \| 1.175e-09 \| 234 \| 0.316 \| 212 \| 94 \| 10 \| 11 \| 221 \| 181 \| 342 \| Tail fiber protein \| Tail fiber protein; afdb-uniprot50 \| AF-A0A378UDT9-F1-MODEL\_V4 \| 1.0 \| 1.057e-08 \| 234 \| 0.265 \| 211 \| 95 \| 8 \| 11 \| 221 \| 429 \| 579 \| Putative phage tail fiber protein \| Putative phage tail fiber protein; afdb-uniprot50 \| AF-A0A1E4UZN2-F1-MODEL\_V4 \| 1.0 \| 1.34e-08 \| 233 \| 0.25 \| 216 \| 121 \| 10 \| 11 \| 220 \| 34 \| 214 \| Collar domain-containing protein \| Collar domain-containing protein; afdb-uniprot50 \| AF-A0A222FGG6-F1-MODEL\_V4 \| 1.0 \| 4.946e-08 \| 233 \| 0.233 \| 227 \| 125 \| 9 \| 11 \| 220 \| 54 \| 248 \| Phage tail protein \| Phage tail protein; afdb-uniprot50 \| AF-I8TWE2-F1-MODEL\_V4 \| 1.0 \| 2.154e-08 \| 233 \| 0.244 \| 229 \| 103 \| 6 \| 3 \| 221 \| 114 \| 282 \| Tail Collar domain protein \| Tail Collar domain protein; afdb-uniprot50 \| AF-A0A5P3MW99-F1-MODEL\_V4 \| 1.0 \| 1.78e-09 \| 233 \| 0.28 \| 232 \| 92 \| 9 \| 2 \| 221 \| 147 \| 315 \| Tail fiber protein \| Tail fiber protein; afdb-uniprot50 \| AF-A0A031IW68-F1-MODEL\_V4 \| 1.0 \| 9.384e-09 \| 233 \| 0.297 \| 222 \| 110 \| 12 \| 1 \| 221 \| 600 \| 776 \| Tail fiber protein H \| Tail fiber protein H; afdb-uniprot50 \| AF-A0A7H9BG73-F1-MODEL\_V4 \| 1.0 \| 1.121e-08 \| 232 \| 0.285 \| 214 \| 91 \| 11 \| 11 \| 221 \| 25 \| 179 \| Tail fiber protein \| Tail fiber protein; afdb-uniprot50 \| AF-A6FCS9-F1-MODEL\_V4 \| 1.0 \| 2.899e-08 \| 232 \| 0.217 \| 239 \| 120 \| 11 \| 11 \| 220 \| 4 \| 204 \| Collar domain-containing protein \| Collar domain-containing protein; afdb-uniprot50 \| AF-A0A5C6V133-F1-MODEL\_V4 \| 1.0 \| 1.121e-08 \| 232 \| 0.288 \| 215 \| 108 \| 9 \| 11 \| 221 \| 128 \| 301 \| Tail fiber protein \| Tail fiber protein; afdb-uniprot50 \| AF-A0A498REH0-F1-MODEL\_V4 \| 1.0 \| 1.34e-08 \| 232 \| 0.279 \| 211 \| 91 \| 6 \| 11 \| 221 \| 253 \| 402 \| Collar domain-containing protein \| Collar domain-containing protein; afdb-uniprot50 \| AF-A0A359D4J8-F1-MODEL\_V4 \| 1.0 \| 2.426e-08 \| 232 \| 0.229 \| 218 \| 93 \| 7 \| 12 \| 221 \| 379 \| 529 \| Collar domain-containing protein \| Collar domain-containing protein; afdb-uniprot50 \| AF-A0A1M7JT24-F1-MODEL\_V4 \| 1.0 \| 6.655e-08 \| 231 \| 0.233 \| 227 \| 116 \| 11 \| 11 \| 220 \| 4 \| 189 \| Microcystin-dependent protein \| Microcystin-dependent protein; afdb-uniprot50 \| AF-A0A821FDS7-F1-MODEL\_V4 \| 1.0 \| 1.422e-08 \| 231 \| 0.41 \| 173 \| 81 \| 8 \| 11 \| 172 \| 49 \| 211 \| Hypothetical protein \| Hypothetical protein; afdb-uniprot50 \| AF-A0A0Q2UK27-F1-MODEL\_V4 \| 1.0 \| 2.456e-07 \| 230 \| 0.222 \| 220 \| 96 \| 9 \| 3 \| 220 \| 17 \| 163 \| Collar domain-containing protein \| Collar domain-containing protein; afdb-uniprot50 \| AF-A0A2U1XUJ9-F1-MODEL\_V4 \| 1.0 \| 1.205e-07 \| 230 \| 0.251 \| 231 \| 89 \| 7 \| 2 \| 221 \| 18 \| 175 \| Collar domain-containing protein \| Collar domain-containing protein; afdb-uniprot50 \| AF-A0A1D9LMZ6-F1-MODEL\_V4 \| 1.0 \| 8.333e-09 \| 230 \| 0.283 \| 229 \| 84 \| 10 \| 2 \| 221 \| 39 \| 196 \| Collar domain-containing protein \| Collar domain-containing protein; afdb-uniprot50 \| AF-A0A661CQ93-F1-MODEL\_V4 \| 1.0 \| 2.005e-09 \| 230 \| 0.235 \| 221 \| 119 \| 7 \| 11 \| 220 \| 13 \| 194 \| Collar domain-containing protein \| Collar domain-containing protein; afdb-uniprot50 \| AF-A0A1X7F9Q3-F1-MODEL\_V4 \| 1.0 \| 9.384e-09 \| 230 \| 0.283 \| 222 \| 83 \| 8 \| 4 \| 221 \| 73 \| 222 \| Phage Tail Collar Domain \| Phage Tail Collar Domain; afdb-uniprot50 \| AF-A0A7W3FCC0-F1-MODEL\_V4 \| 1.0 \| 1.601e-08 \| 230 \| 0.272 \| 224 \| 110 \| 13 \| 11 \| 220 \| 30 \| 214 \| Tail fiber protein \| Tail fiber protein; afdb-uniprot50 \| AF-A0A2R4VPM6-F1-MODEL\_V4 \| 1.0 \| 3.676e-08 \| 230 \| 0.241 \| 215 \| 129 \| 9 \| 11 \| 220 \| 35 \| 220 \| Collar domain-containing protein \| Collar domain-containing protein; afdb-uniprot50 \| AF-A0A318V634-F1-MODEL\_V4 \| 1.0 \| 5.91e-08 \| 230 \| 0.213 \| 230 \| 146 \| 8 \| 11 \| 220 \| 38 \| 252 \| Microcystin-dependent protein \| Microcystin-dependent protein; afdb-uniprot50 \| AF-A0A3C2AA29-F1-MODEL\_V4 \| 1.0 \| 8.333e-09 \| 230 \| 0.251 \| 219 \| 94 \| 6 \| 3 \| 221 \| 136 \| 284 \| Collar domain-containing protein \| Collar domain-containing protein; afdb-uniprot50 \| AF-A0A651FYR1-F1-MODEL\_V4 \| 1.0 \| 1.008e-07 \| 230 \| 0.217 \| 234 \| 124 \| 9 \| 12 \| 221 \| 114 \| 312 \| Uncharacterized protein \| Uncharacterized protein; afdb-uniprot50 \| AF-A0A7X4KFU6-F1-MODEL\_V4 \| 1.0 \| 8.733e-10 \| 229 \| 0.247 \| 238 \| 133 \| 11 \| 11 \| 220 \| 4 \| 223 \| Collar domain-containing protein \| Collar domain-containing protein; afdb-uniprot50 \| AF-A0A522ETL0-F1-MODEL\_V4 \| 1.0 \| 9.384e-09 \| 229 \| 0.252 \| 214 \| 86 \| 6 \| 12 \| 221 \| 110 \| 253 \| Tail fiber protein \| Tail fiber protein; afdb-uniprot50 \| AF-A0A1U9MIU8-F1-MODEL\_V4 \| 1.0 \| 8.333e-09 \| 229 \| 0.241 \| 211 \| 108 \| 9 \| 12 \| 221 \| 154 \| 313 \| Phage Tail Collar Domain \| Phage Tail Collar Domain; afdb-uniprot50 \| AF-A0A0M0H0I1-F1-MODEL\_V4 \| 1.0 \| 7.952e-08 \| 228 \| 0.245 \| 212 \| 89 \| 7 \| 11 \| 221 \| 14 \| 155 \| Phage Tail Collar Domain \| Phage Tail Collar Domain; afdb-uniprot50 \| AF-A0A0J5P3M7-F1-MODEL\_V4 \| 1.0 \| 2.574e-08 \| 228 \| 0.279 \| 215 \| 99 \| 10 \| 11 \| 221 \| 14 \| 176 \| Collar domain-containing protein \| Collar domain-containing protein; afdb-uniprot50 \| AF-A0A2Z6AZX8-F1-MODEL\_V4 \| 1.0 \| 1.913e-08 \| 228 \| 0.244 \| 225 \| 112 \| 11 \| 1 \| 221 \| 37 \| 207 \| Phage Tail Collar Domain protein \| Phage Tail Collar Domain protein; afdb-uniprot50 \| AF-A0A239KLT7-F1-MODEL\_V4 \| 1.0 \| 1.678e-09 \| 228 \| 0.232 \| 254 \| 138 \| 12 \| 11 \| 220 \| 4 \| 244 \| Microcystin-dependent protein \| Microcystin-dependent protein; afdb-uniprot50 \| AF-A0A2S5KL32-F1-MODEL\_V4 \| 1.0 \| 7.952e-08 \| 228 \| 0.23 \| 243 \| 132 \| 11 \| 11 \| 220 \| 51 \| 271 \| Phage tail protein \| Phage tail protein; afdb-uniprot50 \| AF-A0A1V2R607-F1-MODEL\_V4 \| 1.0 \| 3.676e-08 \| 227 \| 0.189 \| 248 \| 139 \| 11 \| 11 \| 220 \| 35 \| 258 \| Phage tail protein \| Phage tail protein; afdb-uniprot50 \| AF-A0A2A6HIY1-F1-MODEL\_V4 \| 1.0 \| 6.193e-09 \| 227 \| 0.303 \| 227 \| 99 \| 10 \| 1 \| 221 \| 183 \| 356 \| Collar domain-containing protein \| Collar domain-containing protein; afdb-uniprot50 \| AF-A0A7G8ZSD1-F1-MODEL\_V4 \| 1.0 \| 1.19e-08 \| 227 \| 0.265 \| 222 \| 99 \| 10 \| 12 \| 221 \| 246 \| 415 \| Tail fiber protein \| Tail fiber protein; afdb-uniprot50 \| AF-A0A833JEF0-F1-MODEL\_V4 \| 1.0 \| 7.952e-08 \| 226 \| 0.22 \| 222 \| 104 \| 10 \| 2 \| 221 \| 47 \| 201 \| Uncharacterized protein \| Uncharacterized protein; afdb-uniprot50 \| AF-A0A560GUX4-F1-MODEL\_V4 \| 1.0 \| 1.889e-09 \| 226 \| 0.223 \| 269 \| 134 \| 12 \| 11 \| 220 \| 5 \| 257 \| Microcystin-dependent protein \| Microcystin-dependent protein; afdb-uniprot50 \| AF-A0A0U1KXP1-F1-MODEL\_V4 \| 1.0 \| 9.958e-09 \| 226 \| 0.295 \| 210 \| 90 \| 7 \| 12 \| 221 \| 172 \| 323 \| Phage tail fibers \| Phage tail fibers; afdb-uniprot50 \| AF-A0A846QFW6-F1-MODEL\_V4 \| 1.0 \| 3.676e-08 \| 226 \| 0.27 \| 211 \| 80 \| 6 \| 11 \| 221 \| 317 \| 453 \| Phage-related tail fiber protein \| Phage-related tail fiber protein; afdb-uniprot50 \| AF-A0A662BD73-F1-MODEL\_V4 \| 1.0 \| 6.271e-08 \| 225 \| 0.276 \| 210 \| 75 \| 7 \| 12 \| 221 \| 39 \| 171 \| Phage tail protein \| Phage tail protein; afdb-uniprot50 \| AF-A0A2N3PZP5-F1-MODEL\_V4 \| 1.0 \| 7.4e-09 \| 225 \| 0.242 \| 256 \| 129 \| 10 \| 11 \| 220 \| 4 \| 240 \| Collar domain-containing protein \| Collar domain-containing protein; afdb-uniprot50 \| AF-A0A7S7WJF1-F1-MODEL\_V4 \| 1.0 \| 2.286e-08 \| 225 \| 0.266 \| 214 \| 104 \| 7 \| 9 \| 221 \| 143 \| 304 \| Collar domain-containing protein \| Collar domain-containing protein; afdb-uniprot50 \| AF-A0A7Y8AZA7-F1-MODEL\_V4 \| 1.0 \| 1.422e-08 \| 225 \| 0.274 \| 211 \| 89 \| 7 \| 12 \| 221 \| 190 \| 337 \| Tail fiber protein \| Tail fiber protein; afdb-uniprot50 \| AF-A0A7Z0F930-F1-MODEL\_V4 \| 1.0 \| 4.337e-09 \| 225 \| 0.247 \| 218 \| 131 \| 8 \| 9 \| 221 \| 161 \| 350 \| Phage-related tail fiber protein \| Phage-related tail fiber protein; afdb-uniprot50 \| AF-C8PDQ5-F1-MODEL\_V4 \| 1.0 \| 1.34e-08 \| 225 \| 0.237 \| 227 \| 108 \| 11 \| 1 \| 221 \| 224 \| 391 \| Phage Tail Collar Domain protein \| Phage Tail Collar Domain protein; afdb-uniprot50 \| AF-A0A0F9S6G2-F1-MODEL\_V4 \| 1.0 \| 2.732e-08 \| 225 \| 0.28 \| 214 \| 84 \| 6 \| 12 \| 220 \| 313 \| 461 \| Collar domain-containing protein \| Collar domain-containing protein; afdb-uniprot50 \| AF-A0A378TUW3-F1-MODEL\_V4 \| 1.0 \| 2.426e-08 \| 225 \| 0.285 \| 221 \| 105 \| 11 \| 3 \| 221 \| 343 \| 512 \| Putative phage tail fiber protein \| Putative phage tail fiber protein; afdb-uniprot50 \| AF-A0A2E9QLA8-F1-MODEL\_V4 \| 1.0 \| 3.076e-08 \| 224 \| 0.236 \| 241 \| 128 \| 13 \| 3 \| 221 \| 106 \| 312 \| Collar domain-containing protein \| Collar domain-containing protein; afdb-uniprot50 \| AF-A0A250DS87-F1-MODEL\_V4 \| 1.0 \| 7.853e-09 \| 224 \| 0.271 \| 232 \| 124 \| 11 \| 1 \| 220 \| 290 \| 488 \| Phage tail protein \| Phage tail protein; afdb-uniprot50 \| AF-A0A1H6MNN9-F1-MODEL\_V4 \| 1.0 \| 1.263e-08 \| 224 \| 0.294 \| 221 \| 105 \| 9 \| 2 \| 221 \| 455 \| 625 \| Microcystin-dependent protein \| Microcystin-dependent protein; afdb-uniprot50 \| AF-A0A0L6FYH4-F1-MODEL\_V4 \| 1.0 \| 9.267e-10 \| 224 \| 0.285 \| 210 \| 130 \| 6 \| 12 \| 221 \| 493 \| 682 \| Phage tail protein \| Phage tail protein; afdb-uniprot50 \| AF-A0A1G3BQZ8-F1-MODEL\_V4 \| 1.0 \| 6.193e-09 \| 223 \| 0.285 \| 221 \| 107 \| 7 \| 3 \| 221 \| 32 \| 203 \| Collar domain-containing protein \| Collar domain-containing protein; afdb-uniprot50 \| AF-A0A559ZVU9-F1-MODEL\_V4 \| 1.0 \| 2.286e-08 \| 223 \| 0.255 \| 223 \| 108 \| 11 \| 11 \| 220 \| 19 \| 196 \| Tail collar domain \| Tail collar domain; afdb-uniprot50 \| AF-A0A5S9PGT0-F1-MODEL\_V4 \| 1.0 \| 1.34e-08 \| 223 \| 0.235 \| 225 \| 121 \| 10 \| 11 \| 220 \| 37 \| 225 \| Collar domain-containing protein \| Collar domain-containing protein; afdb-uniprot50 \| AF-E2PCZ5-F1-MODEL\_V4 \| 1.0 \| 2.574e-08 \| 223 \| 0.295 \| 220 \| 101 \| 9 \| 5 \| 221 \| 141 \| 309 \| Phage Tail Collar Domain protein \| Phage Tail Collar Domain protein; afdb-uniprot50 \| AF-A0A075KIT9-F1-MODEL\_V4 \| 1.0 \| 6.193e-09 \| 223 \| 0.287 \| 219 \| 99 \| 6 \| 3 \| 221 \| 156 \| 317 \| Tail Collar domain protein \| Tail Collar domain protein; afdb-uniprot50 \| AF-A0A0Q4WR10-F1-MODEL\_V4 \| 1.0 \| 4.884e-09 \| 223 \| 0.381 \| 215 \| 118 \| 6 \| 12 \| 216 \| 108 \| 317 \| Collar domain-containing protein \| Collar domain-containing protein; afdb-uniprot50 \| AF-A0A4R6DCB1-F1-MODEL\_V4 \| 1.0 \| 2.426e-08 \| 223 \| 0.245 \| 220 \| 117 \| 9 \| 2 \| 221 \| 235 \| 405 \| Phage tail fiber protein \| Phage tail fiber protein; afdb-uniprot50 \| AF-A0A6L5BI96-F1-MODEL\_V4 \| 1.0 \| 2.863e-09 \| 223 \| 0.28 \| 214 \| 86 \| 7 \| 10 \| 221 \| 295 \| 442 \| Collar domain-containing protein \| Collar domain-containing protein; afdb-uniprot50 \| AF-A0A2N8HQI7-F1-MODEL\_V4 \| 1.0 \| 1.121e-08 \| 223 \| 0.274 \| 215 \| 95 \| 8 \| 12 \| 221 \| 321 \| 479 \| Collar domain-containing protein \| Collar domain-containing protein; afdb-uniprot50 \| AF-A0A1H5GLX5-F1-MODEL\_V4 \| 1.0 \| 3.264e-08 \| 223 \| 0.262 \| 221 \| 114 \| 10 \| 2 \| 221 \| 315 \| 487 \| Phage Tail Collar Domain \| Phage Tail Collar Domain; afdb-uniprot50 \| AF-A0A7W2L2C3-F1-MODEL\_V4 \| 1.0 \| 7.494e-08 \| 223 \| 0.245 \| 220 \| 122 \| 9 \| 3 \| 221 \| 384 \| 560 \| Tail fiber protein \| Tail fiber protein; afdb-uniprot50 \| AF-A0A498DEB1-F1-MODEL\_V4 \| 1.0 \| 1.07e-07 \| 222 \| 0.247 \| 222 \| 97 \| 8 \| 12 \| 220 \| 23 \| 187 \| Collar domain-containing protein \| Collar domain-containing protein; afdb-uniprot50 \| AF-A0A509KW42-F1-MODEL\_V4 \| 1.0 \| 2.574e-08 \| 222 \| 0.231 \| 220 \| 95 \| 5 \| 1 \| 220 \| 51 \| 196 \| Collar domain-containing protein \| Collar domain-containing protein; afdb-uniprot50 \| AF-A0A381TMQ4-F1-MODEL\_V4 \| 1.0 \| 8.843e-09 \| 222 \| 0.252 \| 214 \| 82 \| 9 \| 11 \| 221 \| 69 \| 207 \| Uncharacterized protein \| Uncharacterized protein; afdb-uniprot50 \| AF-A0A344UNR3-F1-MODEL\_V4 \| 1.0 \| 5.248e-08 \| 222 \| 0.264 \| 223 \| 102 \| 7 \| 1 \| 221 \| 162 \| 324 \| Collar domain-containing protein \| Collar domain-containing protein; afdb-uniprot50 \| AF-A0A3S0C1T6-F1-MODEL\_V4 \| 1.0 \| 9.384e-09 \| 222 \| 0.276 \| 300 \| 117 \| 12 \| 3 \| 221 \| 73 \| 353 \| Tail fiber protein \| Tail fiber protein; afdb-uniprot50 \| AF-A0A1K0IP40-F1-MODEL\_V4 \| 1.0 \| 2.732e-08 \| 222 \| 0.27 \| 218 \| 90 \| 9 \| 6 \| 221 \| 320 \| 470 \| Collar domain-containing protein \| Collar domain-containing protein; afdb-uniprot50 \| AF-A0A154VGC7-F1-MODEL\_V4 \| 1.0 \| 6.271e-08 \| 221 \| 0.25 \| 220 \| 107 \| 10 \| 1 \| 220 \| 14 \| 175 \| Collar domain-containing protein \| Collar domain-containing protein; afdb-uniprot50 \| AF-A0A7Z0FDF9-F1-MODEL\_V4 \| 1.0 \| 4.139e-08 \| 221 \| 0.236 \| 224 \| 128 \| 10 \| 11 \| 220 \| 6 \| 200 \| Microcystin-dependent protein \| Microcystin-dependent protein; afdb-uniprot50 \| AF-A0A844YWZ8-F1-MODEL\_V4 \| 1.0 \| 1.057e-08 \| 221 \| 0.279 \| 211 \| 76 \| 6 \| 11 \| 221 \| 96 \| 230 \| Phage tail protein \| Phage tail protein; afdb-uniprot50 \| AF-E2CN30-F1-MODEL\_V4 \| 1.0 \| 1.34e-08 \| 221 \| 0.257 \| 225 \| 86 \| 9 \| 3 \| 221 \| 116 \| 265 \| Tail fiber protein gpH \| Tail fiber protein gpH; afdb-uniprot50 \| AF-A0A1Z7ZRI5-F1-MODEL\_V4 \| 1.0 \| 1.107e-09 \| 221 \| 0.293 \| 259 \| 131 \| 10 \| 11 \| 221 \| 95 \| 349 \| Collar domain-containing protein \| Collar domain-containing protein; afdb-uniprot50 \| AF-A0A348HHI3-F1-MODEL\_V4 \| 1.0 \| 7.853e-09 \| 221 \| 0.275 \| 225 \| 103 \| 8 \| 2 \| 221 \| 204 \| 373 \| Phage-related tail fibre protein \| Phage-related tail fibre protein; afdb-uniprot50 \| AF-A0A376BTN0-F1-MODEL\_V4 \| 1.0 \| 2.574e-08 \| 221 \| 0.312 \| 221 \| 82 \| 9 \| 1 \| 221 \| 479 \| 629 \| Phage Tail Collar Domain \| Phage Tail Collar Domain; afdb-uniprot50 \| AF-A0A845XCJ3-F1-MODEL\_V4 \| 1.0 \| 5.248e-08 \| 220 \| 0.254 \| 212 \| 103 \| 8 \| 11 \| 221 \| 25 \| 182 \| Uncharacterized protein \| Uncharacterized protein; afdb-uniprot50 \| AF-A0A7H1CRS4-F1-MODEL\_V4 \| 1.0 \| 1.357e-07 \| 220 \| 0.257 \| 210 \| 124 \| 5 \| 11 \| 220 \| 5 \| 182 \| Collar domain-containing protein \| Collar domain-containing protein; afdb-uniprot50 \| AF-N9NNT1-F1-MODEL\_V4 \| 1.0 \| 3.676e-08 \| 220 \| 0.285 \| 221 \| 81 \| 8 \| 2 \| 221 \| 61 \| 205 \| Collar domain-containing protein \| Collar domain-containing protein; afdb-uniprot50 \| AF-A0A6I1ND50-F1-MODEL\_V4 \| 1.0 \| 8.843e-09 \| 220 \| 0.246 \| 211 \| 85 \| 6 \| 11 \| 221 \| 212 \| 348 \| Collar domain-containing protein \| Collar domain-containing protein; afdb-uniprot50 \| AF-A0A509LEA5-F1-MODEL\_V4 \| 1.0 \| 3.852e-09 \| 220 \| 0.288 \| 222 \| 86 \| 11 \| 12 \| 220 \| 195 \| 357 \| Collar domain-containing protein \| Collar domain-containing protein; afdb-uniprot50 \| AF-A0A0F8Z7I6-F1-MODEL\_V4 \| 1.0 \| 3.95e-07 \| 219 \| 0.212 \| 212 \| 90 \| 6 \| 12 \| 221 \| 21 \| 157 \| Collar domain-containing protein \| Collar domain-containing protein; afdb-uniprot50 \| AF-A0A5J6QYJ5-F1-MODEL\_V4 \| 1.0 \| 3.223e-09 \| 219 \| 0.311 \| 231 \| 131 \| 8 \| 1 \| 221 \| 37 \| 249 \| Tail fiber protein \| Tail fiber protein; afdb-uniprot50 \| AF-A0A430HC49-F1-MODEL\_V4 \| 1.0 \| 1.34e-08 \| 219 \| 0.227 \| 251 \| 119 \| 14 \| 11 \| 220 \| 55 \| 271 \| Phage tail protein \| Phage tail protein; afdb-uniprot50 \| AF-A0A263JBT7-F1-MODEL\_V4 \| 1.0 \| 1.509e-08 \| 219 \| 0.269 \| 219 \| 95 \| 10 \| 12 \| 221 \| 131 \| 293 \| Collar domain-containing protein \| Collar domain-containing protein; afdb-uniprot50 \| AF-C3X8Y3-F1-MODEL\_V4 \| 1.0 \| 2.574e-08 \| 218 \| 0.257 \| 225 \| 93 \| 6 \| 1 \| 221 \| 116 \| 270 \| Phage Tail Collar Domain protein \| Phage Tail Collar Domain protein; afdb-uniprot50 \| AF-A0A2E2N138-F1-MODEL\_V4 \| 1.0 \| 5.984e-07 \| 217 \| 0.241 \| 211 \| 87 \| 8 \| 11 \| 221 \| 1 \| 138 \| Collar domain-containing protein \| Collar domain-containing protein; afdb-uniprot50 \| AF-G9PUJ8-F1-MODEL\_V4 \| 1.0 \| 1.528e-07 \| 217 \| 0.211 \| 217 \| 100 \| 9 \| 11 \| 221 \| 16 \| 167 \| Collar domain-containing protein \| Collar domain-containing protein; afdb-uniprot50 \| AF-A0A498RAD1-F1-MODEL\_V4 \| 1.0 \| 5.91e-08 \| 217 \| 0.25 \| 212 \| 88 \| 6 \| 11 \| 221 \| 46 \| 187 \| Collar domain-containing protein \| Collar domain-containing protein; afdb-uniprot50 \| AF-A0A4D8QIM4-F1-MODEL\_V4 \| 1.0 \| 2.426e-08 \| 217 \| 0.25 \| 251 \| 112 \| 12 \| 11 \| 220 \| 4 \| 219 \| Phage tail protein \| Phage tail protein; afdb-uniprot50 \| AF-A0A351LDF1-F1-MODEL\_V4 \| 1.0 \| 2.03e-08 \| 217 \| 0.267 \| 247 \| 137 \| 10 \| 3 \| 221 \| 61 \| 291 \| Collar domain-containing protein \| Collar domain-containing protein; afdb-uniprot50 \| AF-A0A258ZJM9-F1-MODEL\_V4 \| 1.0 \| 3.076e-08 \| 217 \| 0.287 \| 212 \| 82 \| 8 \| 11 \| 221 \| 228 \| 371 \| Collar domain-containing protein \| Collar domain-containing protein; afdb-uniprot50 \| AF-A0A7U2B2P4-F1-MODEL\_V4 \| 1.0 \| 1.357e-07 \| 216 \| 0.24 \| 237 \| 88 \| 11 \| 1 \| 221 \| 24 \| 184 \| Tail fiber protein \| Tail fiber protein; afdb-uniprot50 \| AF-C4GFX3-F1-MODEL\_V4 \| 1.0 \| 1.803e-08 \| 216 \| 0.242 \| 227 \| 110 \| 9 \| 2 \| 221 \| 139 \| 310 \| Phage Tail Collar Domain protein \| Phage Tail Collar Domain protein; afdb-uniprot50 \| AF-A0A7X3H8V8-F1-MODEL\_V4 \| 1.0 \| 4.946e-08 \| 216 \| 0.261 \| 222 \| 97 \| 8 \| 1 \| 221 \| 175 \| 330 \| Collar domain-containing protein \| Collar domain-containing protein; afdb-uniprot50 \| AF-A0A4R1FMB3-F1-MODEL\_V4 \| 1.0 \| 3.464e-08 \| 216 \| 0.239 \| 221 \| 95 \| 6 \| 1 \| 221 \| 585 \| 732 \| Tail collar domain \| Tail collar domain; afdb-uniprot50 \| AF-I3CVE0-F1-MODEL\_V4 \| 1.0 \| 5.569e-08 \| 215 \| 0.26 \| 223 \| 100 \| 9 \| 2 \| 221 \| 28 \| 188 \| Collar domain-containing protein \| Collar domain-containing protein; afdb-uniprot50 \| AF-A0A1E4LLZ9-F1-MODEL\_V4 \| 1.0 \| 1.057e-08 \| 215 \| 0.228 \| 219 \| 135 \| 10 \| 11 \| 220 \| 11 \| 204 \| Collar domain-containing protein \| Collar domain-containing protein; afdb-uniprot50 \| AF-A0A2D5I712-F1-MODEL\_V4 \| 1.0 \| 2.366e-10 \| 215 \| 0.311 \| 225 \| 127 \| 7 \| 12 \| 221 \| 17 \| 228 \| Collar domain-containing protein \| Collar domain-containing protein; afdb-uniprot50 \| AF-A0A1H9M0F5-F1-MODEL\_V4 \| 1.0 \| 2.181e-07 \| 215 \| 0.208 \| 230 \| 123 \| 8 \| 10 \| 220 \| 38 \| 227 \| Microcystin-dependent protein \| Microcystin-dependent protein; afdb-uniprot50 \| AF-A0A2V2BG48-F1-MODEL\_V4 \| 1.0 \| 4.392e-08 \| 215 \| 0.19 \| 262 \| 142 \| 13 \| 11 \| 220 \| 35 \| 278 \| Microcystin-dependent protein \| Microcystin-dependent protein; afdb-uniprot50 \| AF-A0A0B6CXA9-F1-MODEL\_V4 \| 1.0 \| 4.946e-08 \| 215 \| 0.227 \| 220 \| 89 \| 7 \| 3 \| 221 \| 186 \| 325 \| Phage Tail Collar domain protein \| Phage Tail Collar domain protein; afdb-uniprot50 \| AF-A0A2N9X5V1-F1-MODEL\_V4 \| 1.0 \| 2.574e-08 \| 215 \| 0.261 \| 218 \| 90 \| 10 \| 11 \| 221 \| 182 \| 335 \| Collar domain-containing protein \| Collar domain-containing protein; afdb-uniprot50 \| AF-A0A1V5JDC5-F1-MODEL\_V4 \| 1.0 \| 5.248e-08 \| 215 \| 0.29 \| 217 \| 80 \| 9 \| 11 \| 220 \| 228 \| 377 \| Phage Tail Collar Domain protein \| Phage Tail Collar Domain protein; afdb-uniprot50 \| AF-A0A7Z1M4M9-F1-MODEL\_V4 \| 1.0 \| 1.34e-08 \| 215 \| 0.298 \| 221 \| 106 \| 7 \| 1 \| 221 \| 217 \| 388 \| Short tail fiber-like protein \| Short tail fiber-like protein; afdb-uniprot50 \| AF-A0A6M1M2X3-F1-MODEL\_V4 \| 1.0 \| 7.588e-07 \| 214 \| 0.195 \| 220 \| 102 \| 9 \| 12 \| 221 \| 4 \| 158 \| Tail fiber protein \| Tail fiber protein; afdb-uniprot50 \| AF-A0A661CMF7-F1-MODEL\_V4 \| 1.0 \| 4.661e-08 \| 214 \| 0.224 \| 245 \| 120 \| 11 \| 1 \| 221 \| 1 \| 199 \| Collar domain-containing protein \| Collar domain-containing protein; afdb-uniprot50 \| AF-Q8XYR6-F1-MODEL\_V4 \| 1.0 \| 8.843e-09 \| 214 \| 0.272 \| 220 \| 109 \| 8 \| 3 \| 221 \| 56 \| 225 \| Probable phage tail collar protein \| Probable phage tail collar protein; afdb-uniprot50 \| AF-A0A841R3L5-F1-MODEL\_V4 \| 1.0 \| 9.958e-09 \| 214 \| 0.237 \| 215 \| 134 \| 7 \| 12 \| 221 \| 53 \| 242 \| Microcystin-dependent protein \| Microcystin-dependent protein; afdb-uniprot50 \| AF-A0A4Q0ZFJ9-F1-MODEL\_V4 \| 1.0 \| 5.91e-08 \| 214 \| 0.224 \| 214 \| 97 \| 5 \| 12 \| 221 \| 169 \| 317 \| Collar domain-containing protein \| Collar domain-containing protein; afdb-uniprot50 \| AF-A0A348HHI2-F1-MODEL\_V4 \| 1.0 \| 3.264e-08 \| 213 \| 0.28 \| 210 \| 82 \| 9 \| 12 \| 221 \| 277 \| 417 \| Phage-related tail fibre protein \| Phage-related tail fibre protein; afdb-uniprot50 \| AF-A0A3E0X1N8-F1-MODEL\_V4 \| 1.0 \| 2.396e-09 \| 213 \| 0.257 \| 225 \| 110 \| 8 \| 9 \| 221 \| 250 \| 429 \| Collar domain-containing protein \| Collar domain-containing protein; afdb-uniprot50 \| AF-A0A840RBH8-F1-MODEL\_V4 \| 1.0 \| 8.052e-07 \| 212 \| 0.186 \| 214 \| 105 \| 6 \| 11 \| 221 \| 26 \| 173 \| Microcystin-dependent protein \| Microcystin-dependent protein; afdb-uniprot50 \| AF-A4TXJ3-F1-MODEL\_V4 \| 1.0 \| 2.766e-07 \| 212 \| 0.2 \| 229 \| 129 \| 9 \| 11 \| 220 \| 34 \| 227 \| Phage Tail Collar \| Phage Tail Collar; afdb-uniprot50 \| AF-A0A0A1FH46-F1-MODEL\_V4 \| 1.0 \| 5.248e-08 \| 212 \| 0.253 \| 221 \| 95 \| 10 \| 2 \| 221 \| 98 \| 249 \| Phage tail fiber protein \| Phage tail fiber protein; afdb-uniprot50 \| AF-A0A7D7SIN8-F1-MODEL\_V4 \| 1.0 \| 3.9e-08 \| 212 \| 0.275 \| 225 \| 102 \| 12 \| 5 \| 221 \| 210 \| 381 \| Tail fiber protein \| Tail fiber protein; afdb-uniprot50 \| AF-A0A522WRR5-F1-MODEL\_V4 \| 1.0 \| 6.271e-08 \| 211 \| 0.257 \| 214 \| 128 \| 9 \| 11 \| 220 \| 5 \| 191 \| Collar domain-containing protein \| Collar domain-containing protein; afdb-uniprot50 \| AF-A0A1B6BGB3-F1-MODEL\_V4 \| 1.0 \| 7.062e-08 \| 211 \| 0.222 \| 225 \| 132 \| 11 \| 11 \| 220 \| 4 \| 200 \| Microcystin dependent protein \| Microcystin dependent protein; afdb-uniprot50 \| AF-A0A2K1FR86-F1-MODEL\_V4 \| 1.0 \| 3.676e-08 \| 211 \| 0.26 \| 223 \| 91 \| 8 \| 2 \| 221 \| 63 \| 214 \| Collar domain-containing protein \| Collar domain-containing protein; afdb-uniprot50 \| AF-A0A424HEY5-F1-MODEL\_V4 \| 1.0 \| 6.193e-09 \| 211 \| 0.247 \| 222 \| 113 \| 7 \| 7 \| 221 \| 102 \| 276 \| Tail fiber protein \| Tail fiber protein; afdb-uniprot50 \| AF-A0A5C7LV11-F1-MODEL\_V4 \| 1.0 \| 2.426e-08 \| 211 \| 0.273 \| 219 \| 93 \| 6 \| 3 \| 221 \| 153 \| 305 \| Collar domain-containing protein \| Collar domain-containing protein; afdb-uniprot50 \| AF-X2H9P4-F1-MODEL\_V4 \| 1.0 \| 3.264e-08 \| 211 \| 0.283 \| 212 \| 91 \| 9 \| 12 \| 221 \| 204 \| 356 \| Phage tail fiber protein \| Phage tail fiber protein; afdb-uniprot50 \| AF-A0A251WMU1-F1-MODEL\_V4 \| 1.0 \| 1.008e-07 \| 210 \| 0.27 \| 211 \| 100 \| 8 \| 11 \| 220 \| 5 \| 162 \| Collar domain-containing protein \| Collar domain-containing protein; afdb-uniprot50 \| AF-A0A4Q0HQ98-F1-MODEL\_V4 \| 1.0 \| 2.154e-08 \| 210 \| 0.252 \| 234 \| 142 \| 9 \| 3 \| 221 \| 2 \| 217 \| Collar domain-containing protein \| Collar domain-containing protein; afdb-uniprot50 \| AF-A5ENH3-F1-MODEL\_V4 \| 1.0 \| 2.426e-08 \| 210 \| 0.404 \| 205 \| 86 \| 8 \| 16 \| 213 \| 116 \| 291 \| Putative phage tail Collar domain protein \| Putative phage tail Collar domain protein; afdb-uniprot50 \| AF-A0A816MWF3-F1-MODEL\_V4 \| 1.0 \| 8.545e-07 \| 210 \| 0.395 \| 149 \| 66 \| 5 \| 11 \| 152 \| 186 \| 317 \| Hypothetical protein \| Hypothetical protein; afdb-uniprot50 \| AF-F3YY59-F1-MODEL\_V4 \| 1.0 \| 1.803e-08 \| 210 \| 0.252 \| 218 \| 91 \| 9 \| 5 \| 221 \| 281 \| 427 \| Tail Collar domain protein \| Tail Collar domain protein; afdb-uniprot50 \| AF-A0A1H9AFH7-F1-MODEL\_V4 \| 1.0 \| 2.732e-08 \| 210 \| 0.256 \| 226 \| 115 \| 9 \| 1 \| 221 \| 367 \| 544 \| Phage-related tail fibre protein \| Phage-related tail fibre protein; afdb-uniprot50 \| AF-A0A4R1SCQ5-F1-MODEL\_V4 \| 1.0 \| 1.601e-08 \| 210 \| 0.308 \| 211 \| 78 \| 8 \| 11 \| 221 \| 428 \| 570 \| Phage-related tail fiber protein \| Phage-related tail fiber protein; afdb-uniprot50 \| AF-A0A4R1G1E3-F1-MODEL\_V4 \| 1.0 \| 4.946e-08 \| 210 \| 0.262 \| 221 \| 90 \| 7 \| 1 \| 221 \| 554 \| 701 \| Tail collar domain \| Tail collar domain; afdb-uniprot50 \| AF-A0A4P7BDA5-F1-MODEL\_V4 \| 1.0 \| 2.154e-08 \| 209 \| 0.225 \| 239 \| 122 \| 9 \| 11 \| 220 \| 4 \| 208 \| Phage tail protein \| Phage tail protein; afdb-uniprot50 \| AF-A0A6C0BHW3-F1-MODEL\_V4 \| 1.0 \| 3.076e-08 \| 209 \| 0.236 \| 241 \| 142 \| 10 \| 1 \| 220 \| 8 \| 227 \| Collar domain-containing protein \| Collar domain-containing protein; afdb-uniprot50 \| AF-A0A1W0CQA8-F1-MODEL\_V4 \| 1.0 \| 1.19e-08 \| 209 \| 0.339 \| 221 \| 79 \| 9 \| 1 \| 221 \| 171 \| 324 \| Collar domain-containing protein \| Collar domain-containing protein; afdb-uniprot50 \| AF-A0A0N9MHW1-F1-MODEL\_V4 \| 1.0 \| 2.286e-08 \| 209 \| 0.25 \| 220 \| 117 \| 11 \| 3 \| 221 \| 978 \| 1150 \| Phage tail fiber protein \| Phage tail fiber protein; afdb-uniprot50 \| AF-A0A133XLD8-F1-MODEL\_V4 \| 1.0 \| 1.528e-07 \| 208 \| 0.242 \| 210 \| 94 \| 6 \| 11 \| 220 \| 77 \| 221 \| Collar domain-containing protein \| Collar domain-containing protein; afdb-uniprot50 \| AF-A0A7Y2P2Q3-F1-MODEL\_V4 \| 1.0 \| 4.087e-09 \| 208 \| 0.223 \| 251 \| 115 \| 13 \| 11 \| 220 \| 4 \| 215 \| Collar domain-containing protein \| Collar domain-containing protein; afdb-uniprot50 \| AF-A0A3G9GF41-F1-MODEL\_V4 \| 1.0 \| 2.732e-08 \| 208 \| 0.246 \| 211 \| 88 \| 7 \| 11 \| 221 \| 187 \| 326 \| Phage tail fiber protein \| Phage tail fiber protein; afdb-uniprot50 \| AF-A0A4P7LA05-F1-MODEL\_V4 \| 1.0 \| 1.621e-07 \| 208 \| 0.233 \| 227 \| 116 \| 9 \| 1 \| 221 \| 192 \| 366 \| Phage Tail Collar Domain protein \| Phage Tail Collar Domain protein; afdb-uniprot50 \| AF-A0A1S1CJX7-F1-MODEL\_V4 \| 1.0 \| 2.181e-07 \| 208 \| 0.265 \| 211 \| 80 \| 5 \| 11 \| 221 \| 373 \| 508 \| Collar domain-containing protein \| Collar domain-containing protein; afdb-uniprot50 \| AF-A0A2N2HSX5-F1-MODEL\_V4 \| 1.0 \| 5.91e-08 \| 207 \| 0.273 \| 212 \| 81 \| 6 \| 11 \| 221 \| 58 \| 197 \| Collar domain-containing protein \| Collar domain-containing protein; afdb-uniprot50 \| AF-A0A2W7PA10-F1-MODEL\_V4 \| 1.0 \| 3.464e-08 \| 207 \| 0.274 \| 211 \| 80 \| 6 \| 11 \| 221 \| 79 \| 216 \| Tail collar domain \| Tail collar domain; afdb-uniprot50 \| AF-C3X8U2-F1-MODEL\_V4 \| 1.0 \| 5.248e-08 \| 207 \| 0.224 \| 223 \| 117 \| 10 \| 3 \| 221 \| 96 \| 266 \| Phage Tail Collar Domain protein \| Phage Tail Collar Domain protein; afdb-uniprot50 \| AF-A0A1C3YPF7-F1-MODEL\_V4 \| 1.0 \| 7.494e-08 \| 207 \| 0.245 \| 212 \| 101 \| 8 \| 11 \| 221 \| 132 \| 285 \| Phage Tail Collar Domain \| Phage Tail Collar Domain; afdb-uniprot50 \| AF-A0A1X3GF36-F1-MODEL\_V4 \| 1.0 \| 8.052e-07 \| 207 \| 0.213 \| 243 \| 129 \| 9 \| 12 \| 221 \| 104 \| 317 \| Uncharacterized protein \| Uncharacterized protein; afdb-uniprot50 \| AF-A0A4Y9GKA5-F1-MODEL\_V4 \| 1.0 \| 8.954e-08 \| 207 \| 0.254 \| 232 \| 100 \| 11 \| 2 \| 221 \| 177 \| 347 \| Tail fiber protein \| Tail fiber protein; afdb-uniprot50 \| AF-A0A1Q6U9D9-F1-MODEL\_V4 \| 1.0 \| 2.154e-08 \| 207 \| 0.233 \| 227 \| 84 \| 11 \| 12 \| 221 \| 208 \| 361 \| Collar domain-containing protein \| Collar domain-containing protein; afdb-uniprot50 \| AF-A0A1E7RCE5-F1-MODEL\_V4 \| 1.0 \| 7.952e-08 \| 207 \| 0.232 \| 215 \| 100 \| 9 \| 11 \| 221 \| 303 \| 456 \| Phage tail protein \| Phage tail protein; afdb-uniprot50 \| AF-U0ZCR8-F1-MODEL\_V4 \| 1.0 \| 4.946e-08 \| 207 \| 0.265 \| 222 \| 99 \| 7 \| 1 \| 221 \| 307 \| 465 \| Collar domain-containing protein \| Collar domain-containing protein; afdb-uniprot50 \| AF-A0A3A9HSP0-F1-MODEL\_V4 \| 1.0 \| 3.305e-07 \| 206 \| 0.263 \| 209 \| 95 \| 7 \| 12 \| 220 \| 24 \| 173 \| Tail fiber protein \| Tail fiber protein; afdb-uniprot50 \| AF-A0A367VXU5-F1-MODEL\_V4 \| 1.0 \| 4.72e-07 \| 206 \| 0.244 \| 213 \| 94 \| 9 \| 12 \| 220 \| 25 \| 174 \| Collar domain-containing protein \| Collar domain-containing protein; afdb-uniprot50 \| AF-A0A822CRA9-F1-MODEL\_V4 \| 1.0 \| 5.984e-07 \| 206 \| 0.365 \| 164 \| 82 \| 5 \| 11 \| 166 \| 222 \| 371 \| Hypothetical protein \| Hypothetical protein; afdb-uniprot50 \| AF-A0A7Y8AQ99-F1-MODEL\_V4 \| 1.0 \| 1.44e-07 \| 206 \| 0.257 \| 210 \| 95 \| 8 \| 12 \| 221 \| 300 \| 448 \| Tail fiber protein \| Tail fiber protein; afdb-uniprot50 \| AF-A0A2T5UPX7-F1-MODEL\_V4 \| 1.0 \| 1.205e-07 \| 206 \| 0.251 \| 215 \| 91 \| 9 \| 11 \| 220 \| 184 \| 333 \| Microcystin-dependent protein \| Microcystin-dependent protein; afdb-uniprot50 \| AF-H8FV07-F1-MODEL\_V4 \| 1.0 \| 2.732e-08 \| 206 \| 0.285 \| 210 \| 73 \| 7 \| 12 \| 220 \| 484 \| 617 \| Collar domain-containing protein \| Collar domain-containing protein; afdb-uniprot50 \| AF-A0A7X7EMF1-F1-MODEL\_V4 \| 1.0 \| 1.205e-07 \| 205 \| 0.253 \| 221 \| 92 \| 9 \| 3 \| 221 \| 90 \| 239 \| Tail fiber protein \| Tail fiber protein; afdb-uniprot50 \| AF-A0A4R4G488-F1-MODEL\_V4 \| 1.0 \| 6.655e-08 \| 205 \| 0.244 \| 221 \| 92 \| 9 \| 2 \| 221 \| 150 \| 296 \| Collar domain-containing protein \| Collar domain-containing protein; afdb-uniprot50 \| AF-A0A4R7WSX9-F1-MODEL\_V4 \| 1.0 \| 6.655e-08 \| 205 \| 0.245 \| 212 \| 111 \| 7 \| 11 \| 221 \| 153 \| 316 \| Tail collar domain \| Tail collar domain; afdb-uniprot50 \| AF-A0A2K2G606-F1-MODEL\_V4 \| 1.0 \| 2.181e-07 \| 205 \| 0.245 \| 220 \| 107 \| 9 \| 3 \| 221 \| 194 \| 355 \| Collar domain-containing protein \| Collar domain-containing protein; afdb-uniprot50 \| AF-C6ABW9-F1-MODEL\_V4 \| 1.0 \| 9.502e-08 \| 205 \| 0.207 \| 226 \| 106 \| 9 \| 2 \| 221 \| 212 \| 370 \| Phage tail collar protein \| Phage tail collar protein; afdb-uniprot50 \| AF-A0A818RWQ3-F1-MODEL\_V4 \| 1.0 \| 2.154e-08 \| 205 \| 0.281 \| 206 \| 130 \| 7 \| 11 \| 214 \| 31 \| 220 \| Hypothetical protein \| Hypothetical protein; afdb-uniprot50 \| AF-A0A291LYY5-F1-MODEL\_V4 \| 1.0 \| 5.569e-08 \| 205 \| 0.247 \| 222 \| 93 \| 7 \| 1 \| 221 \| 296 \| 444 \| Collar domain-containing protein \| Collar domain-containing protein; afdb-uniprot50 \| AF-A0A2N6FYL6-F1-MODEL\_V4 \| 1.0 \| 2.315e-07 \| 204 \| 0.208 \| 221 \| 118 \| 7 \| 2 \| 221 \| 47 \| 211 \| Collar domain-containing protein \| Collar domain-containing protein; afdb-uniprot50 \| AF-C4GGG9-F1-MODEL\_V4 \| 1.0 \| 7.062e-08 \| 204 \| 0.242 \| 227 \| 110 \| 10 \| 2 \| 221 \| 98 \| 269 \| Phage Tail Collar Domain protein \| Phage Tail Collar Domain protein; afdb-uniprot50 \| AF-A0A286B0I3-F1-MODEL\_V4 \| 1.0 \| 5.91e-08 \| 204 \| 0.208 \| 216 \| 126 \| 7 \| 11 \| 221 \| 301 \| 476 \| Phage Tail Collar Domain \| Phage Tail Collar Domain; afdb-uniprot50 \| AF-U1BY63-F1-MODEL\_V4 \| 1.0 \| 5.248e-08 \| 204 \| 0.258 \| 213 \| 91 \| 6 \| 9 \| 221 \| 424 \| 569 \| Collar domain-containing protein \| Collar domain-containing protein; afdb-uniprot50 \| AF-A0A446AZ89-F1-MODEL\_V4 \| 1.0 \| 2.732e-08 \| 204 \| 0.307 \| 221 \| 115 \| 8 \| 11 \| 221 \| 562 \| 754 \| Collar domain-containing protein \| Collar domain-containing protein; afdb-uniprot50 \| AF-A0A3M5GHQ5-F1-MODEL\_V4 \| 1.0 \| 2.181e-07 \| 203 \| 0.222 \| 225 \| 127 \| 9 \| 12 \| 221 \| 5 \| 196 \| Phage tail Collar \| Phage tail Collar; afdb-uniprot50 \| AF-A0A1V5LAC3-F1-MODEL\_V4 \| 1.0 \| 1.72e-07 \| 203 \| 0.2 \| 224 \| 110 \| 10 \| 3 \| 221 \| 50 \| 209 \| Phage Tail Collar Domain protein \| Phage Tail Collar Domain protein; afdb-uniprot50 \| AF-A0A7X8LQL7-F1-MODEL\_V4 \| 1.0 \| 3.464e-08 \| 203 \| 0.253 \| 217 \| 88 \| 6 \| 11 \| 221 \| 70 \| 218 \| Phage tail protein \| Phage tail protein; afdb-uniprot50 \| AF-A0A5C5PXD1-F1-MODEL\_V4 \| 1.0 \| 4.139e-08 \| 203 \| 0.24 \| 216 \| 93 \| 7 \| 11 \| 221 \| 92 \| 241 \| Collar domain-containing protein \| Collar domain-containing protein; afdb-uniprot50 \| AF-A0A5E6WIQ1-F1-MODEL\_V4 \| 1.0 \| 3.464e-08 \| 203 \| 0.27 \| 229 \| 106 \| 11 \| 1 \| 221 \| 107 \| 282 \| Collar domain-containing protein \| Collar domain-containing protein; afdb-uniprot50 \| AF-A0A7H1M8U3-F1-MODEL\_V4 \| 1.0 \| 4.661e-08 \| 203 \| 0.28 \| 228 \| 112 \| 9 \| 2 \| 221 \| 103 \| 286 \| Phage Tail Collar domain protein \| Phage Tail Collar domain protein; afdb-uniprot50 \| AF-A0A2N9XQU6-F1-MODEL\_V4 \| 1.0 \| 1.601e-08 \| 203 \| 0.251 \| 219 \| 96 \| 12 \| 11 \| 221 \| 140 \| 298 \| Collar domain-containing protein \| Collar domain-containing protein; afdb-uniprot50 \| AF-A0A3M9M907-F1-MODEL\_V4 \| 1.0 \| 1.205e-07 \| 203 \| 0.273 \| 238 \| 97 \| 13 \| 2 \| 221 \| 131 \| 310 \| Collar domain-containing protein \| Collar domain-containing protein; afdb-uniprot50 \| AF-A0A162GDK8-F1-MODEL\_V4 \| 1.0 \| 1.135e-07 \| 203 \| 0.273 \| 212 \| 93 \| 7 \| 12 \| 220 \| 461 \| 614 \| Collar domain-containing protein \| Collar domain-containing protein; afdb-uniprot50 \| AF-A0A5A7M8N1-F1-MODEL\_V4 \| 1.0 \| 5.91e-08 \| 203 \| 0.224 \| 223 \| 112 \| 10 \| 4 \| 220 \| 1057 \| 1224 \| Collar domain-containing protein \| Collar domain-containing protein; afdb-uniprot50 \| AF-A0A367WHK4-F1-MODEL\_V4 \| 1.0 \| 7.062e-08 \| 202 \| 0.263 \| 209 \| 96 \| 7 \| 12 \| 220 \| 51 \| 201 \| Collar domain-containing protein \| Collar domain-containing protein; afdb-uniprot50 \| AF-A0A2L2BGH7-F1-MODEL\_V4 \| 1.0 \| 5.569e-08 \| 202 \| 0.267 \| 213 \| 89 \| 7 \| 9 \| 221 \| 109 \| 254 \| Collar domain-containing protein \| Collar domain-containing protein; afdb-uniprot50 \| AF-A0A5M6CCN1-F1-MODEL\_V4 \| 1.0 \| 8.333e-09 \| 202 \| 0.252 \| 250 \| 133 \| 11 \| 3 \| 220 \| 34 \| 261 \| Collar domain-containing protein \| Collar domain-containing protein; afdb-uniprot50 \| AF-A0A1V5RYX1-F1-MODEL\_V4 \| 1.0 \| 1.803e-08 \| 202 \| 0.3 \| 210 \| 111 \| 9 \| 11 \| 220 \| 6 \| 179 \| Phage Tail Collar Domain protein \| Phage Tail Collar Domain protein; afdb-uniprot50 \| AF-A0A1W9JSL6-F1-MODEL\_V4 \| 1.0 \| 2.766e-07 \| 202 \| 0.267 \| 217 \| 120 \| 10 \| 9 \| 221 \| 133 \| 314 \| Collar domain-containing protein \| Collar domain-containing protein; afdb-uniprot50 \| AF-A0A817X4U4-F1-MODEL\_V4 \| 1.0 \| 2.766e-07 \| 202 \| 0.237 \| 223 \| 94 \| 8 \| 2 \| 221 \| 191 \| 340 \| Hypothetical protein \| Hypothetical protein; afdb-uniprot50 \| AF-A0A7U3LRL7-F1-MODEL\_V4 \| 1.0 \| 2.315e-07 \| 202 \| 0.233 \| 227 \| 98 \| 11 \| 4 \| 221 \| 200 \| 359 \| Uncharacterized protein \| Uncharacterized protein; afdb-uniprot50 \| AF-A0A6I6QS22-F1-MODEL\_V4 \| 1.0 \| 5.569e-08 \| 202 \| 0.271 \| 210 \| 83 \| 7 \| 12 \| 221 \| 418 \| 557 \| Collar domain-containing protein \| Collar domain-containing protein; afdb-uniprot50 \| AF-A0A0N9MHY1-F1-MODEL\_V4 \| 1.0 \| 2.03e-08 \| 202 \| 0.283 \| 219 \| 124 \| 9 \| 11 \| 221 \| 555 \| 748 \| Collar domain-containing protein \| Collar domain-containing protein; afdb-uniprot50 \| AF-A0A512NQJ7-F1-MODEL\_V4 \| 1.0 \| 2.972e-06 \| 201 \| 0.386 \| 132 \| 70 \| 3 \| 28 \| 159 \| 1 \| 121 \| Collar domain-containing protein \| Collar domain-containing protein; afdb-uniprot50 \| AF-A0A7W6MM04-F1-MODEL\_V4 \| 1.0 \| 3.347e-06 \| 201 \| 0.195 \| 220 \| 100 \| 10 \| 12 \| 220 \| 4 \| 157 \| Collar domain-containing protein \| Collar domain-containing protein; afdb-uniprot50 \| AF-A0A1Y4GG17-F1-MODEL\_V4 \| 1.0 \| 2.181e-07 \| 201 \| 0.227 \| 211 \| 114 \| 9 \| 11 \| 220 \| 25 \| 187 \| Collar domain-containing protein \| Collar domain-containing protein; afdb-uniprot50 \| AF-A0A077L9W7-F1-MODEL\_V4 \| 1.0 \| 1.621e-07 \| 201 \| 0.199 \| 221 \| 153 \| 5 \| 11 \| 220 \| 5 \| 212 \| Tail Collar domain protein \| Tail Collar domain protein; afdb-uniprot50 \| AF-A0A486SU05-F1-MODEL\_V4 \| 1.0 \| 5.569e-08 \| 201 \| 0.234 \| 230 \| 98 \| 10 \| 1 \| 221 \| 119 \| 279 \| Phage Tail Collar Domain \| Phage Tail Collar Domain; afdb-uniprot50 \| AF-A0A1H9HCA5-F1-MODEL\_V4 \| 1.0 \| 2.607e-07 \| 201 \| 0.224 \| 218 \| 96 \| 10 \| 3 \| 220 \| 167 \| 311 \| Microcystin-dependent protein \| Microcystin-dependent protein; afdb-uniprot50 \| AF-A0A7D5QQI0-F1-MODEL\_V4 \| 1.0 \| 1.07e-07 \| 201 \| 0.312 \| 211 \| 70 \| 7 \| 11 \| 221 \| 323 \| 458 \| Tail fiber protein \| Tail fiber protein; afdb-uniprot50 \| AF-A0A202B4T0-F1-MODEL\_V4 \| 1.0 \| 7.062e-08 \| 200 \| 0.234 \| 230 \| 95 \| 7 \| 9 \| 221 \| 66 \| 231 \| Collar domain-containing protein \| Collar domain-containing protein; afdb-uniprot50 \| AF-A0A1W9LKS6-F1-MODEL\_V4 \| 1.0 \| 1.826e-07 \| 200 \| 0.269 \| 226 \| 127 \| 7 \| 9 \| 221 \| 122 \| 322 \| Collar domain-containing protein \| Collar domain-containing protein; afdb-uniprot50 \| AF-A0A7W6RGC7-F1-MODEL\_V4 \| 1.0 \| 3.076e-08 \| 200 \| 0.253 \| 229 \| 96 \| 9 \| 3 \| 221 \| 192 \| 355 \| Phage-related tail fiber protein \| Phage-related tail fiber protein; afdb-uniprot50 \| AF-A0A7Y1MB50-F1-MODEL\_V4 \| 1.0 \| 1.279e-07 \| 200 \| 0.255 \| 211 \| 84 \| 5 \| 11 \| 221 \| 515 \| 652 \| Phage tail protein \| Phage tail protein; afdb-uniprot50 \| AF-A0A8A9YBB3-F1-MODEL\_V4 \| 1.0 \| 4.661e-08 \| 200 \| 0.245 \| 220 \| 123 \| 9 \| 12 \| 221 \| 646 \| 832 \| Phage tail protein \| Phage tail protein; afdb-uniprot50 \| AF-A0A6N4RFI0-F1-MODEL\_V4 \| 1.0 \| 1.528e-07 \| 199 \| 0.341 \| 208 \| 110 \| 6 \| 16 \| 214 \| 3 \| 192 \| Tail fiber protein \| Tail fiber protein; afdb-uniprot50 \| AF-A0A6C0KMM6-F1-MODEL\_V4 \| 1.0 \| 6.271e-08 \| 199 \| 0.24 \| 212 \| 116 \| 10 \| 11 \| 220 \| 46 \| 214 \| Collar domain-containing protein \| Collar domain-containing protein; afdb-uniprot50 \| AF-A0A5S9HP15-F1-MODEL\_V4 \| 1.0 \| 2.607e-07 \| 199 \| 0.254 \| 224 \| 95 \| 8 \| 5 \| 221 \| 121 \| 279 \| Collar domain-containing protein \| Collar domain-containing protein; afdb-uniprot50 \| AF-A0A353PD69-F1-MODEL\_V4 \| 1.0 \| 1.135e-07 \| 199 \| 0.228 \| 232 \| 127 \| 11 \| 11 \| 221 \| 89 \| 289 \| Collar domain-containing protein \| Collar domain-containing protein; afdb-uniprot50 \| AF-A0A515EKF3-F1-MODEL\_V4 \| 1.0 \| 3.9e-08 \| 199 \| 0.298 \| 211 \| 84 \| 8 \| 11 \| 221 \| 179 \| 325 \| Collar domain-containing protein \| Collar domain-containing protein; afdb-uniprot50 \| AF-A0A454TSP7-F1-MODEL\_V4 \| 1.0 \| 3.9e-08 \| 199 \| 0.283 \| 212 \| 95 \| 8 \| 11 \| 221 \| 291 \| 446 \| Phage tail protein \| Phage tail protein; afdb-uniprot50 \| AF-A0A0F9VFY7-F1-MODEL\_V4 \| 1.0 \| 2.315e-07 \| 199 \| 0.256 \| 218 \| 98 \| 10 \| 5 \| 221 \| 294 \| 448 \| Collar domain-containing protein \| Collar domain-containing protein; afdb-uniprot50 \| AF-A0A076PYE4-F1-MODEL\_V4 \| 1.0 \| 3.9e-08 \| 199 \| 0.269 \| 226 \| 107 \| 8 \| 1 \| 220 \| 908 \| 1081 \| Collar domain-containing protein \| Collar domain-containing protein; afdb-uniprot50 \| AF-A0A7C5BRR0-F1-MODEL\_V4 \| 1.0 \| 1.44e-07 \| 198 \| 0.242 \| 210 \| 90 \| 6 \| 12 \| 221 \| 54 \| 194 \| Uncharacterized protein \| Uncharacterized protein; afdb-uniprot50 \| AF-C3X3R4-F1-MODEL\_V4 \| 1.0 \| 7.062e-08 \| 198 \| 0.228 \| 236 \| 89 \| 11 \| 2 \| 220 \| 70 \| 229 \| Collar domain-containing protein \| Collar domain-containing protein; afdb-uniprot50 \| AF-A0A1Q6U9C4-F1-MODEL\_V4 \| 1.0 \| 5.639e-07 \| 198 \| 0.243 \| 222 \| 98 \| 8 \| 2 \| 220 \| 91 \| 245 \| Collar domain-containing protein \| Collar domain-containing protein; afdb-uniprot50 \| AF-A0A2E3Q9T6-F1-MODEL\_V4 \| 1.0 \| 5.91e-08 \| 198 \| 0.268 \| 212 \| 90 \| 9 \| 11 \| 221 \| 105 \| 252 \| Collar domain-containing protein \| Collar domain-containing protein; afdb-uniprot50 \| AF-A0A0H3GJI2-F1-MODEL\_V4 \| 1.0 \| 5.91e-08 \| 198 \| 0.251 \| 223 \| 77 \| 9 \| 10 \| 221 \| 231 \| 374 \| Collar domain-containing protein \| Collar domain-containing protein; afdb-uniprot50 \| AF-A0A212J3U4-F1-MODEL\_V4 \| 1.0 \| 1.07e-07 \| 198 \| 0.246 \| 215 \| 91 \| 8 \| 11 \| 221 \| 267 \| 414 \| Collar domain-containing protein \| Collar domain-containing protein; afdb-uniprot50 \| AF-A0A840MKP4-F1-MODEL\_V4 \| 1.0 \| 6.271e-08 \| 198 \| 0.274 \| 211 \| 94 \| 6 \| 11 \| 221 \| 285 \| 436 \| Phage-related tail fiber protein \| Phage-related tail fiber protein; afdb-uniprot50 \| AF-A0A2S8IPM3-F1-MODEL\_V4 \| 1.0 \| 6.655e-08 \| 198 \| 0.242 \| 256 \| 136 \| 13 \| 1 \| 220 \| 421 \| 654 \| Uncharacterized protein \| Uncharacterized protein; afdb-uniprot50 \| AF-A0A7W8S7J7-F1-MODEL\_V4 \| 1.0 \| 3.507e-07 \| 198 \| 0.198 \| 257 \| 147 \| 12 \| 1 \| 220 \| 487 \| 721 \| Uncharacterized protein \| Uncharacterized protein; afdb-uniprot50 \| AF-A0A077LFW5-F1-MODEL\_V4 \| 1.0 \| 7.952e-08 \| 197 \| 0.242 \| 235 \| 118 \| 9 \| 11 \| 220 \| 5 \| 204 \| Tail collar domain-containing protein \| Tail collar domain-containing protein; afdb-uniprot50 \| AF-A0A143DC03-F1-MODEL\_V4 \| 1.0 \| 2.056e-07 \| 197 \| 0.248 \| 221 \| 97 \| 9 \| 1 \| 220 \| 116 \| 268 \| Collar domain-containing protein \| Collar domain-containing protein; afdb-uniprot50 \| AF-A0A2I1XBM6-F1-MODEL\_V4 \| 1.0 \| 6.655e-08 \| 197 \| 0.259 \| 216 \| 94 \| 10 \| 12 \| 221 \| 186 \| 341 \| Phage tail protein \| Phage tail protein; afdb-uniprot50 \| AF-A0A0S9N328-F1-MODEL\_V4 \| 1.0 \| 6.271e-08 \| 197 \| 0.257 \| 237 \| 142 \| 12 \| 1 \| 221 \| 226 \| 444 \| Collar domain-containing protein \| Collar domain-containing protein; afdb-uniprot50 \| AF-A0A381F1Q4-F1-MODEL\_V4 \| 1.0 \| 3.464e-08 \| 197 \| 0.272 \| 213 \| 116 \| 8 \| 11 \| 221 \| 271 \| 446 \| Phage Tail Collar Domain \| Phage Tail Collar Domain; afdb-uniprot50 \| AF-A0A1H8IGX2-F1-MODEL\_V4 \| 1.0 \| 1.528e-07 \| 197 \| 0.231 \| 220 \| 97 \| 6 \| 2 \| 221 \| 304 \| 451 \| Phage Tail Collar Domain \| Phage Tail Collar Domain; afdb-uniprot50 \| AF-A0A7W6RFU8-F1-MODEL\_V4 \| 1.0 \| 7.588e-07 \| 196 \| 0.222 \| 211 \| 121 \| 9 \| 11 \| 220 \| 41 \| 209 \| Microcystin-dependent protein \| Microcystin-dependent protein; afdb-uniprot50 \| AF-A0A6A7Y806-F1-MODEL\_V4 \| 1.0 \| 7.494e-08 \| 196 \| 0.228 \| 214 \| 140 \| 8 \| 10 \| 220 \| 20 \| 211 \| Collar domain-containing protein \| Collar domain-containing protein; afdb-uniprot50 \| AF-A0A177NMW9-F1-MODEL\_V4 \| 1.0 \| 6.655e-08 \| 196 \| 0.259 \| 239 \| 119 \| 11 \| 11 \| 220 \| 40 \| 249 \| Collar domain-containing protein \| Collar domain-containing protein; afdb-uniprot50 \| AF-A0A3Q9BAD5-F1-MODEL\_V4 \| 1.0 \| 2.181e-07 \| 196 \| 0.216 \| 226 \| 110 \| 8 \| 3 \| 221 \| 163 \| 328 \| Collar domain-containing protein \| Collar domain-containing protein; afdb-uniprot50 \| AF-A0A496KQQ3-F1-MODEL\_V4 \| 1.0 \| 3.95e-07 \| 196 \| 0.274 \| 211 \| 78 \| 9 \| 11 \| 221 \| 226 \| 361 \| Collar domain-containing protein \| Collar domain-containing protein; afdb-uniprot50 \| AF-A0A530YYK5-F1-MODEL\_V4 \| 1.0 \| 5.569e-08 \| 195 \| 0.251 \| 227 \| 113 \| 6 \| 3 \| 221 \| 70 \| 247 \| Collar domain-containing protein \| Collar domain-containing protein; afdb-uniprot50 \| AF-A0A6L5HA01-F1-MODEL\_V4 \| 1.0 \| 6.739e-07 \| 195 \| 0.247 \| 222 \| 93 \| 6 \| 1 \| 221 \| 110 \| 258 \| Collar domain-containing protein \| Collar domain-containing protein; afdb-uniprot50 \| AF-A0A509LD13-F1-MODEL\_V4 \| 1.0 \| 5.008e-07 \| 195 \| 0.252 \| 214 \| 85 \| 6 \| 12 \| 220 \| 46 \| 189 \| Collar domain-containing protein \| Collar domain-containing protein; afdb-uniprot50 \| AF-A0A2A3MGV4-F1-MODEL\_V4 \| 1.0 \| 1.621e-07 \| 195 \| 0.233 \| 210 \| 88 \| 6 \| 12 \| 221 \| 307 \| 443 \| Phage tail protein \| Phage tail protein; afdb-uniprot50 \| AF-F4KS41-F1-MODEL\_V4 \| 1.0 \| 5.984e-07 \| 195 \| 0.243 \| 218 \| 104 \| 11 \| 11 \| 221 \| 312 \| 475 \| Tail Collar domain protein \| Tail Collar domain protein; afdb-uniprot50 \| AF-A0A840MLL2-F1-MODEL\_V4 \| 1.0 \| 1.44e-07 \| 195 \| 0.225 \| 235 \| 124 \| 7 \| 1 \| 221 \| 469 \| 659 \| Phage-related tail fiber protein \| Phage-related tail fiber protein; afdb-uniprot50 \| AF-A0A378UFL6-F1-MODEL\_V4 \| 1.0 \| 6.655e-08 \| 195 \| 0.262 \| 221 \| 112 \| 8 \| 2 \| 221 \| 532 \| 702 \| Putative phage tail fiber protein \| Putative phage tail fiber protein; afdb-uniprot50 \| AF-A0A496KG49-F1-MODEL\_V4 \| 1.0 \| 2.154e-08 \| 194 \| 0.279 \| 222 \| 101 \| 10 \| 11 \| 221 \| 106 \| 279 \| Collar domain-containing protein \| Collar domain-containing protein; afdb-uniprot50 \| AF-A0A7Z7RIG7-F1-MODEL\_V4 \| 1.0 \| 1.44e-07 \| 194 \| 0.216 \| 222 \| 112 \| 9 \| 1 \| 221 \| 144 \| 304 \| Phage Tail Collar Domain \| Phage Tail Collar Domain; afdb-uniprot50 \| AF-A0A6P1ZJ46-F1-MODEL\_V4 \| 1.0 \| 1.44e-07 \| 194 \| 0.287 \| 216 \| 112 \| 9 \| 12 \| 221 \| 204 \| 383 \| Collar domain-containing protein \| Collar domain-containing protein; afdb-uniprot50 \| AF-A0A378UGB4-F1-MODEL\_V4 \| 1.0 \| 1.135e-07 \| 194 \| 0.225 \| 222 \| 110 \| 8 \| 11 \| 220 \| 471 \| 642 \| Putative phage tail fiber protein \| Putative phage tail fiber protein; afdb-uniprot50 \| AF-A0A7Y1CQT0-F1-MODEL\_V4 \| 1.0 \| 1.528e-07 \| 194 \| 0.256 \| 218 \| 87 \| 6 \| 11 \| 221 \| 503 \| 652 \| Collar domain-containing protein \| Collar domain-containing protein; afdb-uniprot50 \| AF-R7I616-F1-MODEL\_V4 \| 1.0 \| 6.655e-08 \| 193 \| 0.217 \| 230 \| 97 \| 9 \| 2 \| 221 \| 94 \| 250 \| Collar domain-containing protein \| Collar domain-containing protein; afdb-uniprot50 \| AF-A0A2H0ZA63-F1-MODEL\_V4 \| 1.0 \| 5.314e-07 \| 193 \| 0.235 \| 212 \| 92 \| 8 \| 11 \| 221 \| 129 \| 271 \| Collar domain-containing protein \| Collar domain-containing protein; afdb-uniprot50 \| AF-A0A817YYG9-F1-MODEL\_V4 \| 1.0 \| 1.19e-08 \| 193 \| 0.32 \| 206 \| 97 \| 6 \| 20 \| 216 \| 395 \| 566 \| Hypothetical protein \| Hypothetical protein; afdb-uniprot50 \| AF-G9PUK2-F1-MODEL\_V4 \| 1.0 \| 2.082e-06 \| 192 \| 0.238 \| 214 \| 96 \| 10 \| 12 \| 220 \| 24 \| 175 \| Collar domain-containing protein \| Collar domain-containing protein; afdb-uniprot50 \| AF-A0A285UDY9-F1-MODEL\_V4 \| 1.0 \| 3.464e-08 \| 192 \| 0.303 \| 231 \| 121 \| 11 \| 11 \| 221 \| 156 \| 366 \| Tail collar domain \| Tail collar domain; afdb-uniprot50 \| AF-C3X912-F1-MODEL\_V4 \| 1.0 \| 2.456e-07 \| 192 \| 0.224 \| 218 \| 95 \| 10 \| 11 \| 221 \| 286 \| 436 \| Phage Tail Collar Domain protein \| Phage Tail Collar Domain protein; afdb-uniprot50 \| AF-C5B185-F1-MODEL\_V4 \| 1.0 \| 3.305e-07 \| 192 \| 0.235 \| 212 \| 91 \| 6 \| 11 \| 221 \| 308 \| 449 \| Collar domain-containing protein \| Collar domain-containing protein; afdb-uniprot50 \| AF-A0A5E7KHM1-F1-MODEL\_V4 \| 1.0 \| 4.946e-08 \| 192 \| 0.302 \| 251 \| 120 \| 9 \| 12 \| 221 \| 305 \| 541 \| Collar domain-containing protein \| Collar domain-containing protein; afdb-uniprot50 \| AF-A0A1G9EQI8-F1-MODEL\_V4 \| 1.0 \| 3.305e-07 \| 192 \| 0.2 \| 215 \| 108 \| 10 \| 11 \| 221 \| 401 \| 555 \| Phage-related tail fibre protein \| Phage-related tail fibre protein; afdb-uniprot50 \| AF-A0A7T9LJD7-F1-MODEL\_V4 \| 1.0 \| 2.181e-07 \| 191 \| 0.251 \| 223 \| 98 \| 11 \| 3 \| 221 \| 13 \| 170 \| Tail fiber protein \| Tail fiber protein; afdb-uniprot50 \| AF-A0A815S8I3-F1-MODEL\_V4 \| 1.0 \| 1.295e-06 \| 191 \| 0.355 \| 169 \| 91 \| 5 \| 11 \| 171 \| 16 \| 174 \| Hypothetical protein \| Hypothetical protein; afdb-uniprot50 \| AF-A0A0P9CHF4-F1-MODEL\_V4 \| 1.0 \| 8.545e-07 \| 191 \| 0.218 \| 220 \| 115 \| 9 \| 4 \| 221 \| 70 \| 234 \| Collar domain-containing protein \| Collar domain-containing protein; afdb-uniprot50 \| AF-A0A6B3UTZ7-F1-MODEL\_V4 \| 1.0 \| 1.135e-07 \| 191 \| 0.283 \| 212 \| 79 \| 7 \| 11 \| 221 \| 106 \| 245 \| Phage tail protein \| Phage tail protein; afdb-uniprot50 \| AF-A0A662DPJ2-F1-MODEL\_V4 \| 1.0 \| 1.44e-07 \| 191 \| 0.264 \| 246 \| 130 \| 13 \| 2 \| 221 \| 50 \| 270 \| Collar domain-containing protein \| Collar domain-containing protein; afdb-uniprot50 \| AF-A0A2S9XA95-F1-MODEL\_V4 \| 1.0 \| 2.456e-07 \| 191 \| 0.246 \| 223 \| 104 \| 9 \| 1 \| 221 \| 158 \| 318 \| Collar domain-containing protein \| Collar domain-containing protein; afdb-uniprot50 \| AF-A0A2K9CDC4-F1-MODEL\_V4 \| 1.0 \| 2.315e-07 \| 191 \| 0.241 \| 232 \| 108 \| 8 \| 1 \| 221 \| 145 \| 319 \| Collar domain-containing protein \| Collar domain-containing protein; afdb-uniprot50 \| AF-A0A2N9Y0U7-F1-MODEL\_V4 \| 1.0 \| 3.115e-07 \| 191 \| 0.243 \| 222 \| 93 \| 9 \| 11 \| 221 \| 172 \| 329 \| Collar domain-containing protein \| Collar domain-containing protein; afdb-uniprot50 \| AF-A0A2W0GCJ0-F1-MODEL\_V4 \| 1.0 \| 8.954e-08 \| 191 \| 0.284 \| 211 \| 89 \| 7 \| 11 \| 221 \| 233 \| 381 \| Phage tail protein \| Phage tail protein; afdb-uniprot50 \| AF-A0A3B8Q380-F1-MODEL\_V4 \| 1.0 \| 3.95e-07 \| 191 \| 0.215 \| 232 \| 119 \| 10 \| 1 \| 221 \| 220 \| 399 \| Collar domain-containing protein \| Collar domain-containing protein; afdb-uniprot50 \| AF-V9H7W8-F1-MODEL\_V4 \| 1.0 \| 3.95e-07 \| 191 \| 0.25 \| 228 \| 108 \| 11 \| 2 \| 221 \| 249 \| 421 \| Collar domain-containing protein \| Collar domain-containing protein; afdb-uniprot50 \| AF-A0A2K4ISX8-F1-MODEL\_V4 \| 1.0 \| 5.314e-07 \| 191 \| 0.284 \| 211 \| 90 \| 9 \| 11 \| 221 \| 508 \| 657 \| Phage tail protein \| Phage tail protein; afdb-uniprot50 \| AF-A0A3S0SEE9-F1-MODEL\_V4 \| 1.0 \| 8.954e-08 \| 190 \| 0.277 \| 227 \| 101 \| 9 \| 4 \| 221 \| 91 \| 263 \| Collar domain-containing protein \| Collar domain-containing protein; afdb-uniprot50 \| AF-A0A835ZHE3-F1-MODEL\_V4 \| 1.0 \| 5.984e-07 \| 190 \| 0.284 \| 225 \| 120 \| 10 \| 12 \| 220 \| 4 \| 203 \| Uncharacterized protein \| Uncharacterized protein; afdb-uniprot50 \| AF-A0A3C1RWZ2-F1-MODEL\_V4 \| 1.0 \| 7.062e-08 \| 190 \| 0.261 \| 268 \| 147 \| 9 \| 3 \| 221 \| 49 \| 314 \| Collar domain-containing protein \| Collar domain-containing protein; afdb-uniprot50 \| AF-A0A5M8P654-F1-MODEL\_V4 \| 1.0 \| 5.569e-08 \| 190 \| 0.254 \| 216 \| 110 \| 11 \| 11 \| 221 \| 171 \| 340 \| Phage tail protein \| Phage tail protein; afdb-uniprot50 \| AF-A0A3C0UIR2-F1-MODEL\_V4 \| 1.0 \| 9.502e-08 \| 190 \| 0.271 \| 210 \| 87 \| 9 \| 12 \| 220 \| 168 \| 312 \| Collar domain-containing protein \| Collar domain-containing protein; afdb-uniprot50 \| AF-A0A143DG88-F1-MODEL\_V4 \| 1.0 \| 6.739e-07 \| 190 \| 0.291 \| 158 \| 83 \| 7 \| 11 \| 162 \| 255 \| 389 \| Collar domain-containing protein \| Collar domain-containing protein; afdb-uniprot50 \| AF-A0A822BJ94-F1-MODEL\_V4 \| 1.0 \| 2.209e-06 \| 190 \| 0.368 \| 163 \| 83 \| 5 \| 11 \| 166 \| 407 \| 556 \| Hypothetical protein \| Hypothetical protein; afdb-uniprot50 \| AF-A0A1S8LA53-F1-MODEL\_V4 \| 1.0 \| 1.937e-07 \| 190 \| 0.247 \| 210 \| 91 \| 7 \| 12 \| 221 \| 669 \| 811 \| Phage tail collar domain protein \| Phage tail collar domain protein; afdb-uniprot50 \| AF-A0A1Y4GGY2-F1-MODEL\_V4 \| 1.0 \| 1.07e-07 \| 189 \| 0.256 \| 222 \| 104 \| 10 \| 12 \| 220 \| 24 \| 197 \| Collar domain-containing protein \| Collar domain-containing protein; afdb-uniprot50 \| AF-A0A292RR50-F1-MODEL\_V4 \| 1.0 \| 2.181e-07 \| 188 \| 0.168 \| 237 \| 99 \| 11 \| 3 \| 221 \| 49 \| 205 \| Collar domain-containing protein \| Collar domain-containing protein; afdb-uniprot50 \| AF-A0A222FFN0-F1-MODEL\_V4 \| 1.0 \| 4.448e-07 \| 188 \| 0.19 \| 226 \| 140 \| 10 \| 11 \| 220 \| 30 \| 228 \| Collar domain-containing protein \| Collar domain-containing protein; afdb-uniprot50 \| AF-C3X3V5-F1-MODEL\_V4 \| 1.0 \| 3.676e-08 \| 188 \| 0.226 \| 225 \| 114 \| 8 \| 3 \| 220 \| 87 \| 258 \| Collar domain-containing protein \| Collar domain-containing protein; afdb-uniprot50 \| AF-K9F0F3-F1-MODEL\_V4 \| 1.0 \| 1.357e-07 \| 188 \| 0.27 \| 248 \| 139 \| 11 \| 3 \| 220 \| 31 \| 266 \| Microcystin-dependent protein \| Microcystin-dependent protein; afdb-uniprot50 \| AF-A0A1Y6D0G3-F1-MODEL\_V4 \| 1.0 \| 3.722e-07 \| 188 \| 0.246 \| 211 \| 98 \| 7 \| 12 \| 221 \| 166 \| 316 \| Phage Tail Collar Domain \| Phage Tail Collar Domain; afdb-uniprot50 \| AF-A0A5B7ZM63-F1-MODEL\_V4 \| 1.0 \| 9.502e-08 \| 188 \| 0.232 \| 237 \| 89 \| 11 \| 3 \| 221 \| 179 \| 340 \| Tail fiber protein \| Tail fiber protein; afdb-uniprot50 \| AF-A0A7U2FZZ3-F1-MODEL\_V4 \| 1.0 \| 2.181e-07 \| 188 \| 0.226 \| 230 \| 101 \| 10 \| 1 \| 221 \| 254 \| 415 \| Tail fiber protein \| Tail fiber protein; afdb-uniprot50 \| AF-A0A7S7WIR9-F1-MODEL\_V4 \| 1.0 \| 1.135e-07 \| 187 \| 0.272 \| 224 \| 91 \| 11 \| 10 \| 221 \| 80 \| 243 \| Collar domain-containing protein \| Collar domain-containing protein; afdb-uniprot50 \| AF-A0A135WED4-F1-MODEL\_V4 \| 1.0 \| 1.205e-07 \| 187 \| 0.254 \| 232 \| 92 \| 13 \| 11 \| 221 \| 147 \| 318 \| Collar domain-containing protein \| Collar domain-containing protein; afdb-uniprot50 \| AF-U1AK99-F1-MODEL\_V4 \| 1.0 \| 3.305e-07 \| 187 \| 0.263 \| 228 \| 91 \| 10 \| 2 \| 221 \| 263 \| 421 \| Collar domain-containing protein \| Collar domain-containing protein; afdb-uniprot50 \| AF-A0A0K8JHT0-F1-MODEL\_V4 \| 1.0 \| 1.07e-07 \| 187 \| 0.264 \| 212 \| 90 \| 9 \| 12 \| 220 \| 196 \| 344 \| Collar domain-containing protein \| Collar domain-containing protein; afdb-uniprot50 \| AF-A7INV5-F1-MODEL\_V4 \| 1.0 \| 4.448e-07 \| 187 \| 0.203 \| 211 \| 94 \| 7 \| 11 \| 221 \| 356 \| 492 \| Tail Collar domain protein \| Tail Collar domain protein; afdb-uniprot50 \| AF-A0A0D9LFX9-F1-MODEL\_V4 \| 1.0 \| 1.72e-07 \| 186 \| 0.303 \| 211 \| 70 \| 9 \| 12 \| 221 \| 206 \| 340 \| Phage Tail Collar domain protein \| Phage Tail Collar domain protein; afdb-uniprot50 \| AF-A0A381EVQ5-F1-MODEL\_V4 \| 1.0 \| 6.271e-08 \| 186 \| 0.265 \| 230 \| 96 \| 9 \| 11 \| 221 \| 190 \| 365 \| Phage Tail Collar Domain \| Phage Tail Collar Domain; afdb-uniprot50 \| AF-C3X3K6-F1-MODEL\_V4 \| 1.0 \| 3.507e-07 \| 186 \| 0.227 \| 237 \| 104 \| 11 \| 2 \| 220 \| 198 \| 373 \| Collar domain-containing protein \| Collar domain-containing protein; afdb-uniprot50 \| AF-A0A1Q3ZE37-F1-MODEL\_V4 \| 1.0 \| 1.458e-06 \| 186 \| 0.202 \| 217 \| 102 \| 8 \| 12 \| 220 \| 456 \| 609 \| Collar domain-containing protein \| Collar domain-containing protein; afdb-uniprot50 \| AF-A0A817T034-F1-MODEL\_V4 \| 1.0 \| 2.574e-08 \| 186 \| 0.312 \| 211 \| 106 \| 10 \| 16 \| 216 \| 420 \| 601 \| Hypothetical protein \| Hypothetical protein; afdb-uniprot50 \| AF-A0A353WK11-F1-MODEL\_V4 \| 1.0 \| 2.766e-07 \| 185 \| 0.258 \| 217 \| 85 \| 7 \| 6 \| 221 \| 85 \| 226 \| Collar domain-containing protein \| Collar domain-containing protein; afdb-uniprot50 \| AF-C3X3V2-F1-MODEL\_V4 \| 1.0 \| 1.72e-07 \| 185 \| 0.227 \| 233 \| 92 \| 10 \| 3 \| 220 \| 129 \| 288 \| Collar domain-containing protein \| Collar domain-containing protein; afdb-uniprot50 \| AF-C3X8V2-F1-MODEL\_V4 \| 1.0 \| 2.607e-07 \| 185 \| 0.243 \| 226 \| 96 \| 9 \| 3 \| 220 \| 70 \| 228 \| Phage Tail Collar Domain protein \| Phage Tail Collar Domain protein; afdb-uniprot50 \| AF-A0A352DBM0-F1-MODEL\_V4 \| 1.0 \| 3.722e-07 \| 185 \| 0.219 \| 214 \| 103 \| 7 \| 9 \| 221 \| 351 \| 501 \| Collar domain-containing protein \| Collar domain-containing protein; afdb-uniprot50 \| AF-A0A3M2DZ18-F1-MODEL\_V4 \| 1.0 \| 1.07e-07 \| 184 \| 0.276 \| 235 \| 109 \| 8 \| 1 \| 220 \| 223 \| 411 \| Tail fiber protein \| Tail fiber protein; afdb-uniprot50 \| AF-Q7P176-F1-MODEL\_V4 \| 1.0 \| 5.008e-07 \| 184 \| 0.206 \| 223 \| 115 \| 10 \| 1 \| 221 \| 273 \| 435 \| Probable bacteriophge tail fiber protein \| Probable bacteriophge tail fiber protein; afdb-uniprot50 \| AF-A0A7R7UYC3-F1-MODEL\_V4 \| 1.0 \| 4.448e-07 \| 184 \| 0.279 \| 211 \| 84 \| 7 \| 11 \| 221 \| 453 \| 595 \| Collar domain-containing protein \| Collar domain-containing protein; afdb-uniprot50 \| AF-A0A2N3LS20-F1-MODEL\_V4 \| 1.0 \| 1.083e-06 \| 183 \| 0.203 \| 211 \| 91 \| 8 \| 13 \| 221 \| 176 \| 311 \| Collar domain-containing protein \| Collar domain-containing protein; afdb-uniprot50 \| AF-A0A822BDI4-F1-MODEL\_V4 \| 1.0 \| 9.067e-07 \| 182 \| 0.349 \| 166 \| 71 \| 7 \| 12 \| 162 \| 29 \| 172 \| Hypothetical protein \| Hypothetical protein; afdb-uniprot50 \| AF-A0A821UTN8-F1-MODEL\_V4 \| 1.0 \| 1.295e-06 \| 182 \| 0.261 \| 172 \| 78 \| 6 \| 11 \| 179 \| 348 \| 473 \| Hypothetical protein \| Hypothetical protein; afdb-uniprot50 \| AF-A0A3N9U476-F1-MODEL\_V4 \| 1.0 \| 1.742e-06 \| 182 \| 0.218 \| 211 \| 101 \| 8 \| 11 \| 221 \| 362 \| 508 \| Collar domain-containing protein \| Collar domain-containing protein; afdb-uniprot50 \| AF-A0A7W9ZHL4-F1-MODEL\_V4 \| 1.0 \| 1.295e-06 \| 181 \| 0.24 \| 212 \| 122 \| 7 \| 11 \| 220 \| 5 \| 179 \| Microcystin-dependent protein \| Microcystin-dependent protein; afdb-uniprot50 \| AF-A0A661CNJ0-F1-MODEL\_V4 \| 1.0 \| 3.115e-07 \| 181 \| 0.229 \| 222 \| 122 \| 8 \| 12 \| 220 \| 13 \| 198 \| Collar domain-containing protein \| Collar domain-containing protein; afdb-uniprot50 \| AF-A0A6C0KKB8-F1-MODEL\_V4 \| 1.0 \| 2.456e-07 \| 181 \| 0.266 \| 214 \| 113 \| 7 \| 11 \| 221 \| 31 \| 203 \| Collar domain-containing protein \| Collar domain-containing protein; afdb-uniprot50 \| AF-A0A7W9FLH6-F1-MODEL\_V4 \| 1.0 \| 1.15e-06 \| 181 \| 0.204 \| 220 \| 139 \| 8 \| 10 \| 220 \| 14 \| 206 \| Microcystin-dependent protein \| Microcystin-dependent protein; afdb-uniprot50 \| AF-C3X192-F1-MODEL\_V4 \| 1.0 \| 3.95e-07 \| 181 \| 0.245 \| 228 \| 100 \| 9 \| 1 \| 220 \| 67 \| 230 \| Collar domain-containing protein \| Collar domain-containing protein; afdb-uniprot50 \| AF-A0A2S5DKP9-F1-MODEL\_V4 \| 1.0 \| 2.607e-07 \| 181 \| 0.231 \| 225 \| 107 \| 9 \| 2 \| 221 \| 218 \| 381 \| Collar domain-containing protein \| Collar domain-containing protein; afdb-uniprot50 \| AF-A0A2M9R257-F1-MODEL\_V4 \| 1.0 \| 8.843e-09 \| 181 \| 0.286 \| 269 \| 125 \| 14 \| 11 \| 221 \| 149 \| 408 \| Collar domain-containing protein \| Collar domain-containing protein; afdb-uniprot50 \| AF-C3X7Z8-F1-MODEL\_V4 \| 1.0 \| 4.448e-07 \| 181 \| 0.244 \| 209 \| 94 \| 7 \| 12 \| 220 \| 201 \| 345 \| Phage Tail Collar Domain protein \| Phage Tail Collar Domain protein; afdb-uniprot50 \| AF-A0A7Y6S8C6-F1-MODEL\_V4 \| 1.0 \| 2.209e-06 \| 180 \| 0.29 \| 200 \| 90 \| 8 \| 22 \| 221 \| 2 \| 149 \| Tail fiber protein \| Tail fiber protein; afdb-uniprot50 \| AF-A0A177M7V7-F1-MODEL\_V4 \| 1.0 \| 5.984e-07 \| 180 \| 0.221 \| 239 \| 128 \| 13 \| 11 \| 220 \| 5 \| 214 \| Collar domain-containing protein \| Collar domain-containing protein; afdb-uniprot50 \| AF-A0A816UWW5-F1-MODEL\_V4 \| 1.0 \| 4.779e-06 \| 180 \| 0.228 \| 171 \| 89 \| 4 \| 11 \| 179 \| 180 \| 309 \| Hypothetical protein \| Hypothetical protein; afdb-uniprot50 \| AF-C3X8I7-F1-MODEL\_V4 \| 1.0 \| 6.35e-07 \| 180 \| 0.23 \| 226 \| 97 \| 9 \| 2 \| 220 \| 96 \| 251 \| Phage Tail Collar Domain protein \| Phage Tail Collar Domain protein; afdb-uniprot50 \| AF-A0A143DGR5-F1-MODEL\_V4 \| 1.0 \| 4.191e-07 \| 180 \| 0.218 \| 224 \| 92 \| 8 \| 11 \| 220 \| 329 \| 483 \| Collar domain-containing protein \| Collar domain-containing protein; afdb-uniprot50 \| AF-A0A820VF90-F1-MODEL\_V4 \| 1.0 \| 1.458e-06 \| 179 \| 0.256 \| 222 \| 115 \| 8 \| 11 \| 220 \| 1 \| 184 \| Hypothetical protein \| Hypothetical protein; afdb-uniprot50 \| AF-A0A074JSZ8-F1-MODEL\_V4 \| 1.0 \| 2.766e-07 \| 179 \| 0.235 \| 221 \| 91 \| 7 \| 12 \| 220 \| 57 \| 211 \| Collar domain-containing protein \| Collar domain-containing protein; afdb-uniprot50 \| AF-A0A1G8DPL3-F1-MODEL\_V4 \| 1.0 \| 2.286e-08 \| 179 \| 0.245 \| 253 \| 130 \| 15 \| 11 \| 220 \| 4 \| 238 \| Microcystin-dependent protein \| Microcystin-dependent protein; afdb-uniprot50 \| AF-A0A356FYY9-F1-MODEL\_V4 \| 1.0 \| 1.826e-07 \| 178 \| 0.198 \| 237 \| 108 \| 11 \| 2 \| 221 \| 87 \| 258 \| Collar domain-containing protein \| Collar domain-containing protein; afdb-uniprot50 \| AF-A0A6P0WRL1-F1-MODEL\_V4 \| 1.0 \| 3.115e-07 \| 178 \| 0.24 \| 254 \| 150 \| 5 \| 6 \| 220 \| 67 \| 316 \| Collar domain-containing protein \| Collar domain-containing protein; afdb-uniprot50 \| AF-A0A336NEY6-F1-MODEL\_V4 \| 1.0 \| 5.008e-07 \| 178 \| 0.211 \| 218 \| 120 \| 7 \| 12 \| 221 \| 160 \| 333 \| Phage Tail Collar Domain \| Phage Tail Collar Domain; afdb-uniprot50 \| AF-C3X3R8-F1-MODEL\_V4 \| 1.0 \| 6.35e-07 \| 178 \| 0.218 \| 233 \| 95 \| 9 \| 3 \| 220 \| 78 \| 238 \| Collar domain-containing protein \| Collar domain-containing protein; afdb-uniprot50 \| AF-A0A8B2QXZ8-F1-MODEL\_V4 \| 1.0 \| 2.315e-07 \| 178 \| 0.231 \| 212 \| 95 \| 8 \| 11 \| 221 \| 571 \| 715 \| Tail collar domain \| Tail collar domain; afdb-uniprot50 \| AF-C3XAA4-F1-MODEL\_V4 \| 1.0 \| 1.083e-06 \| 177 \| 0.207 \| 217 \| 81 \| 8 \| 11 \| 220 \| 15 \| 147 \| Phage Tail Collar Domain protein \| Phage Tail Collar Domain protein; afdb-uniprot50 \| AF-A0A7Z7RHC2-F1-MODEL\_V4 \| 1.0 \| 1.021e-06 \| 177 \| 0.224 \| 218 \| 103 \| 8 \| 11 \| 221 \| 586 \| 744 \| Phage Tail Collar Domain \| Phage Tail Collar Domain; afdb-uniprot50 \| AF-A0A521ZHZ2-F1-MODEL\_V4 \| 1.0 \| 1.642e-06 \| 176 \| 0.242 \| 227 \| 99 \| 13 \| 2 \| 220 \| 211 \| 372 \| Tail fiber protein \| Tail fiber protein; afdb-uniprot50 \| AF-C3X8V7-F1-MODEL\_V4 \| 1.0 \| 1.22e-06 \| 175 \| 0.236 \| 224 \| 99 \| 8 \| 3 \| 220 \| 181 \| 338 \| Phage Tail Collar Domain protein \| Phage Tail Collar Domain protein; afdb-uniprot50 \| AF-A0A7Z0JL73-F1-MODEL\_V4 \| 1.0 \| 1.021e-06 \| 175 \| 0.208 \| 254 \| 142 \| 15 \| 3 \| 220 \| 693 \| 923 \| Microcystin-dependent protein \| Microcystin-dependent protein; afdb-uniprot50 \| AF-A0A6P3BUP6-F1-MODEL\_V4 \| 1.0 \| 9.622e-07 \| 174 \| 0.237 \| 240 \| 150 \| 8 \| 3 \| 220 \| 93 \| 321 \| Putative phage tail protein \| Putative phage tail protein; afdb-uniprot50 \| AF-A0A3R8V3H5-F1-MODEL\_V4 \| 1.0 \| 7.952e-08 \| 174 \| 0.284 \| 271 \| 122 \| 11 \| 11 \| 221 \| 68 \| 326 \| Tail fiber protein \| Tail fiber protein; afdb-uniprot50 \| AF-J1JT46-F1-MODEL\_V4 \| 1.0 \| 3.722e-07 \| 174 \| 0.195 \| 256 \| 141 \| 9 \| 1 \| 221 \| 149 \| 374 \| Collar domain-containing protein \| Collar domain-containing protein; afdb-uniprot50 \| AF-C3X1Y2-F1-MODEL\_V4 \| 1.0 \| 1.374e-06 \| 174 \| 0.255 \| 211 \| 92 \| 7 \| 12 \| 220 \| 191 \| 338 \| Collar domain-containing protein \| Collar domain-containing protein; afdb-uniprot50 \| AF-A0A2W5JYE1-F1-MODEL\_V4 \| 1.0 \| 5.314e-07 \| 173 \| 0.236 \| 220 \| 94 \| 7 \| 1 \| 220 \| 154 \| 299 \| Collar domain-containing protein \| Collar domain-containing protein; afdb-uniprot50 \| AF-A0A661P6R3-F1-MODEL\_V4 \| 1.0 \| 1.008e-07 \| 173 \| 0.303 \| 231 \| 113 \| 10 \| 3 \| 214 \| 63 \| 264 \| Collar domain-containing protein \| Collar domain-containing protein; afdb-uniprot50 \| AF-A0A0C4Y828-F1-MODEL\_V4 \| 1.0 \| 4.448e-07 \| 173 \| 0.195 \| 246 \| 162 \| 8 \| 1 \| 220 \| 286 \| 521 \| Phage-related tail fiber protein \| Phage-related tail fiber protein; afdb-uniprot50 \| AF-A0A251X838-F1-MODEL\_V4 \| 1.0 \| 2.801e-06 \| 172 \| 0.184 \| 244 \| 136 \| 14 \| 3 \| 221 \| 149 \| 354 \| Collar domain-containing protein \| Collar domain-containing protein; afdb-uniprot50 \| AF-A0A7W8WHR4-F1-MODEL\_V4 \| 1.0 \| 5.984e-07 \| 172 \| 0.189 \| 259 \| 149 \| 11 \| 2 \| 220 \| 457 \| 694 \| Microcystin-dependent protein \| Microcystin-dependent protein; afdb-uniprot50 \| AF-A0A1H9M0F1-F1-MODEL\_V4 \| 1.0 \| 3.154e-06 \| 171 \| 0.193 \| 227 \| 129 \| 10 \| 8 \| 220 \| 31 \| 217 \| Microcystin-dependent protein \| Microcystin-dependent protein; afdb-uniprot50 \| AF-A0A3B9Q8G5-F1-MODEL\_V4 \| 1.0 \| 7.151e-07 \| 171 \| 0.247 \| 226 \| 80 \| 8 \| 11 \| 221 \| 152 \| 302 \| Collar domain-containing protein \| Collar domain-containing protein; afdb-uniprot50 \| AF-A0A819J9H2-F1-MODEL\_V4 \| 1.0 \| 4.191e-07 \| 171 \| 0.217 \| 221 \| 91 \| 9 \| 11 \| 220 \| 183 \| 332 \| Hypothetical protein \| Hypothetical protein; afdb-uniprot50 \| AF-A0A7X3HHB3-F1-MODEL\_V4 \| 1.0 \| 3.722e-07 \| 170 \| 0.204 \| 250 \| 155 \| 13 \| 1 \| 220 \| 207 \| 442 \| Uncharacterized protein \| Uncharacterized protein; afdb-uniprot50 \| AF-A0A660N0Q8-F1-MODEL\_V4 \| 1.0 \| 5.382e-06 \| 169 \| 0.254 \| 220 \| 101 \| 8 \| 16 \| 221 \| 2 \| 172 \| Collar domain-containing protein \| Collar domain-containing protein; afdb-uniprot50 \| AF-A0A3D5BA67-F1-MODEL\_V4 \| 1.0 \| 8.052e-07 \| 169 \| 0.217 \| 221 \| 97 \| 10 \| 3 \| 220 \| 92 \| 239 \| Collar domain-containing protein \| Collar domain-containing protein; afdb-uniprot50 \| AF-A0A6P0WKN5-F1-MODEL\_V4 \| 1.0 \| 5.314e-07 \| 169 \| 0.252 \| 222 \| 121 \| 7 \| 3 \| 214 \| 51 \| 237 \| Tail fiber protein \| Tail fiber protein; afdb-uniprot50 \| AF-A0A2E1KV02-F1-MODEL\_V4 \| 1.0 \| 5.248e-08 \| 169 \| 0.336 \| 229 \| 110 \| 8 \| 11 \| 214 \| 49 \| 260 \| Uncharacterized protein \| Uncharacterized protein; afdb-uniprot50 \| AF-A0A2N8HS50-F1-MODEL\_V4 \| 1.0 \| 1.458e-06 \| 168 \| 0.21 \| 214 \| 104 \| 8 \| 11 \| 221 \| 95 \| 246 \| Collar domain-containing protein \| Collar domain-containing protein; afdb-uniprot50 \| AF-A0A820D142-F1-MODEL\_V4 \| 1.0 \| 5.984e-07 \| 168 \| 0.317 \| 192 \| 106 \| 6 \| 19 \| 210 \| 28 \| 194 \| Hypothetical protein \| Hypothetical protein; afdb-uniprot50 \| AF-C3X910-F1-MODEL\_V4 \| 1.0 \| 3.722e-07 \| 168 \| 0.264 \| 212 \| 86 \| 9 \| 12 \| 220 \| 265 \| 409 \| Phage Tail Collar Domain protein \| Phage Tail Collar Domain protein; afdb-uniprot50 \| AF-A0A381EST0-F1-MODEL\_V4 \| 1.0 \| 3.95e-07 \| 167 \| 0.233 \| 240 \| 91 \| 10 \| 11 \| 221 \| 235 \| 410 \| Phage Tail Collar Domain \| Phage Tail Collar Domain; afdb-uniprot50 \| AF-A0A2S9X7Q4-F1-MODEL\_V4 \| 1.0 \| 1.44e-07 \| 167 \| 0.284 \| 211 \| 103 \| 11 \| 11 \| 221 \| 392 \| 554 \| Collar domain-containing protein \| Collar domain-containing protein; afdb-uniprot50 \| AF-A0A5S9PHC1-F1-MODEL\_V4 \| 1.0 \| 7.241e-06 \| 166 \| 0.192 \| 223 \| 132 \| 10 \| 9 \| 220 \| 27 \| 212 \| Collar domain-containing protein \| Collar domain-containing protein; afdb-uniprot50 \| AF-A0A7Y8KXK1-F1-MODEL\_V4 \| 1.0 \| 6.739e-07 \| 166 \| 0.216 \| 226 \| 120 \| 12 \| 2 \| 220 \| 129 \| 304 \| Tail fiber protein \| Tail fiber protein; afdb-uniprot50 \| AF-A0A1X0TJA2-F1-MODEL\_V4 \| 1.0 \| 1.567e-05 \| 165 \| 0.364 \| 118 \| 60 \| 4 \| 20 \| 136 \| 1 \| 104 \| Collar domain-containing protein \| Collar domain-containing protein; afdb-uniprot50 \| AF-A0A6L5JUV3-F1-MODEL\_V4 \| 1.0 \| 1.235e-05 \| 165 \| 0.228 \| 206 \| 83 \| 6 \| 15 \| 220 \| 35 \| 164 \| Collar domain-containing protein \| Collar domain-containing protein; afdb-uniprot50 \| AF-A0A1Y1Q392-F1-MODEL\_V4 \| 1.0 \| 2.519e-05 \| 165 \| 0.325 \| 166 \| 87 \| 6 \| 2 \| 162 \| 54 \| 199 \| Collar domain-containing protein \| Collar domain-containing protein; afdb-uniprot50 \| AF-A0A7V8JNI0-F1-MODEL\_V4 \| 1.0 \| 3.676e-08 \| 165 \| 0.266 \| 270 \| 138 \| 8 \| 1 \| 221 \| 172 \| 430 \| Collar domain-containing protein \| Collar domain-containing protein; afdb-uniprot50 \| AF-A0A816C073-F1-MODEL\_V4 \| 1.0 \| 1.295e-06 \| 165 \| 0.307 \| 195 \| 110 \| 6 \| 20 \| 214 \| 42 \| 211 \| Hypothetical protein \| Hypothetical protein; afdb-uniprot50 \| AF-A0A1Y4GG09-F1-MODEL\_V4 \| 1.0 \| 1.22e-06 \| 164 \| 0.205 \| 238 \| 93 \| 10 \| 12 \| 220 \| 24 \| 194 \| Collar domain-containing protein \| Collar domain-containing protein; afdb-uniprot50 \| AF-A0A522VZY9-F1-MODEL\_V4 \| 1.0 \| 1.15e-06 \| 164 \| 0.219 \| 232 \| 138 \| 9 \| 12 \| 220 \| 2 \| 213 \| Tail fiber protein \| Tail fiber protein; afdb-uniprot50 \| AF-A0A2D5I6Z8-F1-MODEL\_V4 \| 1.0 \| 1.826e-07 \| 164 \| 0.268 \| 227 \| 137 \| 9 \| 11 \| 220 \| 25 \| 239 \| Collar domain-containing protein \| Collar domain-containing protein; afdb-uniprot50 \| AF-A0A7Y5MLB7-F1-MODEL\_V4 \| 1.0 \| 9.622e-07 \| 163 \| 0.259 \| 227 \| 117 \| 12 \| 6 \| 221 \| 175 \| 361 \| Tail fiber protein \| Tail fiber protein; afdb-uniprot50 \| AF-A0A3A9HNI7-F1-MODEL\_V4 \| 1.0 \| 2.64e-06 \| 162 \| 0.252 \| 218 \| 92 \| 9 \| 11 \| 220 \| 20 \| 174 \| Tail fiber protein \| Tail fiber protein; afdb-uniprot50 \| AF-A0A7H2RJ25-F1-MODEL\_V4 \| 1.0 \| 5.382e-06 \| 162 \| 0.165 \| 218 \| 116 \| 7 \| 3 \| 220 \| 64 \| 215 \| Tail fiber protein \| Tail fiber protein; afdb-uniprot50 \| AF-A0A6I2KGM5-F1-MODEL\_V4 \| 1.0 \| 2.607e-07 \| 162 \| 0.214 \| 270 \| 132 \| 12 \| 11 \| 220 \| 79 \| 328 \| Tail fiber protein \| Tail fiber protein; afdb-uniprot50 \| AF-A0A7T1Y6B1-F1-MODEL\_V4 \| 1.0 \| 4.779e-06 \| 162 \| 0.203 \| 221 \| 106 \| 9 \| 11 \| 221 \| 233 \| 393 \| Phage tail protein \| Phage tail protein; afdb-uniprot50 \| AF-A0A448V8J3-F1-MODEL\_V4 \| 1.0 \| 6.739e-07 \| 162 \| 0.208 \| 292 \| 135 \| 11 \| 12 \| 221 \| 159 \| 436 \| Phage Tail Collar Domain \| Phage Tail Collar Domain; afdb-uniprot50 \| AF-A0A1Z7Z823-F1-MODEL\_V4 \| 1.0 \| 6.35e-07 \| 162 \| 0.281 \| 213 \| 81 \| 6 \| 9 \| 221 \| 418 \| 558 \| Collar domain-containing protein \| Collar domain-containing protein; afdb-uniprot50 \| AF-A0A077LCE0-F1-MODEL\_V4 \| 1.0 \| 7.588e-07 \| 161 \| 0.238 \| 235 \| 119 \| 11 \| 11 \| 220 \| 5 \| 204 \| Phage Tail Collar domain protein \| Phage Tail Collar domain protein; afdb-uniprot50 \| AF-A0A6L6YMH3-F1-MODEL\_V4 \| 1.0 \| 2.64e-06 \| 160 \| 0.198 \| 227 \| 106 \| 8 \| 3 \| 221 \| 114 \| 272 \| Collar domain-containing protein \| Collar domain-containing protein; afdb-uniprot50 \| AF-A9ITY1-F1-MODEL\_V4 \| 1.0 \| 1.872e-05 \| 160 \| 0.168 \| 244 \| 119 \| 5 \| 2 \| 221 \| 116 \| 299 \| Phage related protein \| Phage related protein; afdb-uniprot50 \| AF-Q0BEK5-F1-MODEL\_V4 \| 1.0 \| 2.209e-06 \| 159 \| 0.243 \| 246 \| 142 \| 9 \| 1 \| 220 \| 506 \| 733 \| Phage Tail Collar domain protein \| Phage Tail Collar domain protein; afdb-uniprot50 \| AF-A0A7C4AHI4-F1-MODEL\_V4 \| 1.0 \| 2.972e-06 \| 155 \| 0.221 \| 212 \| 93 \| 8 \| 11 \| 220 \| 79 \| 220 \| Collar domain-containing protein \| Collar domain-containing protein; afdb-uniprot50 \| AF-A0A246KV99-F1-MODEL\_V4 \| 1.0 \| 3.507e-07 \| 155 \| 0.29 \| 248 \| 135 \| 10 \| 11 \| 221 \| 246 \| 489 \| Collar domain-containing protein \| Collar domain-containing protein; afdb-uniprot50 \| AF-A0A1Y4GRC1-F1-MODEL\_V4 \| 1.0 \| 1.235e-05 \| 154 \| 0.22 \| 222 \| 94 \| 11 \| 11 \| 221 \| 23 \| 176 \| Collar domain-containing protein \| Collar domain-containing protein; afdb-uniprot50 \| AF-A0A368TXZ3-F1-MODEL\_V4 \| 1.0 \| 4.244e-06 \| 154 \| 0.216 \| 217 \| 112 \| 9 \| 11 \| 221 \| 273 \| 437 \| Collar domain-containing protein \| Collar domain-containing protein; afdb-uniprot50 \| AF-A0A1Y5MHL2-F1-MODEL\_V4 \| 1.0 \| 1.742e-06 \| 154 \| 0.21 \| 223 \| 97 \| 10 \| 11 \| 221 \| 507 \| 662 \| Collar domain-containing protein \| Collar domain-containing protein; afdb-uniprot50 \| AF-A0A817QMQ8-F1-MODEL\_V4 \| 1.0 \| 5.639e-07 \| 152 \| 0.293 \| 218 \| 126 \| 10 \| 11 \| 221 \| 20 \| 216 \| Hypothetical protein \| Hypothetical protein; afdb-uniprot50 \| AF-C3X9T8-F1-MODEL\_V4 \| 1.0 \| 7.241e-06 \| 151 \| 0.23 \| 217 \| 83 \| 11 \| 12 \| 220 \| 196 \| 336 \| Phage Tail Collar Domain protein \| Phage Tail Collar Domain protein; afdb-uniprot50 \| AF-A0A1G9MSY8-F1-MODEL\_V4 \| 1.0 \| 1.476e-05 \| 150 \| 0.187 \| 315 \| 134 \| 16 \| 2 \| 221 \| 103 \| 390 \| Uncharacterized protein \| Uncharacterized protein; afdb-uniprot50 \| AF-A0A1A9RLT3-F1-MODEL\_V4 \| 1.0 \| 1.097e-05 \| 150 \| 0.218 \| 211 \| 87 \| 8 \| 12 \| 220 \| 155 \| 289 \| Uncharacterized protein \| Uncharacterized protein; afdb-uniprot50 \| AF-C1D6M7-F1-MODEL\_V4 \| 1.0 \| 3.154e-06 \| 149 \| 0.239 \| 209 \| 101 \| 10 \| 12 \| 220 \| 105 \| 255 \| Phage-related protein \| Phage-related protein; afdb-uniprot50 \| AF-A0A662BH77-F1-MODEL\_V4 \| 1.0 \| 1.391e-05 \| 149 \| 0.211 \| 227 \| 130 \| 10 \| 3 \| 220 \| 354 \| 540 \| Uncharacterized protein \| Uncharacterized protein; afdb-uniprot50 \| AF-A0A437M1B9-F1-MODEL\_V4 \| 1.0 \| 2.108e-05 \| 149 \| 0.229 \| 309 \| 135 \| 14 \| 1 \| 221 \| 337 \| 630 \| Uncharacterized protein \| Uncharacterized protein; afdb-uniprot50 \| AF-A0A560C055-F1-MODEL\_V4 \| 1.0 \| 6.824e-06 \| 147 \| 0.209 \| 229 \| 77 \| 5 \| 3 \| 221 \| 728 \| 862 \| Tail collar domain \| Tail collar domain; afdb-uniprot50 \| AF-I4FLZ9-F1-MODEL\_V4 \| 1.0 \| 4.05e-05 \| 147 \| 0.244 \| 213 \| 113 \| 6 \| 11 \| 221 \| 782 \| 948 \| Uncharacterized protein \| Uncharacterized protein; afdb-uniprot50 \| AF-A0A349P466-F1-MODEL\_V4 \| 1.0 \| 1.567e-05 \| 146 \| 0.206 \| 218 \| 131 \| 9 \| 12 \| 220 \| 94 \| 278 \| Uncharacterized protein \| Uncharacterized protein; afdb-uniprot50 \| AF-A0A3S4IBW8-F1-MODEL\_V4 \| 1.0 \| 2.972e-06 \| 146 \| 0.272 \| 224 \| 120 \| 9 \| 9 \| 214 \| 323 \| 521 \| Phage Tail Collar Domain \| Phage Tail Collar Domain; afdb-uniprot50 \| AF-A0A4Q3MBS5-F1-MODEL\_V4 \| 1.0 \| 8.545e-07 \| 145 \| 0.225 \| 239 \| 129 \| 9 \| 12 \| 220 \| 5 \| 217 \| Collar domain-containing protein \| Collar domain-containing protein; afdb-uniprot50 \| AF-A0A818P2E2-F1-MODEL\_V4 \| 1.0 \| 3.194e-05 \| 142 \| 0.267 \| 183 \| 94 \| 7 \| 2 \| 161 \| 25 \| 190 \| Hypothetical protein \| Hypothetical protein; afdb-uniprot50 \| AF-A0A813X408-F1-MODEL\_V4 \| 1.0 \| 3.552e-06 \| 142 \| 0.37 \| 170 \| 76 \| 5 \| 16 \| 174 \| 67 \| 216 \| Hypothetical protein \| Hypothetical protein; afdb-uniprot50 \| AF-A0A1H1GAM2-F1-MODEL\_V4 \| 1.0 \| 0.0001409 \| 142 \| 0.2 \| 335 \| 141 \| 14 \| 12 \| 221 \| 107 \| 439 \| Uncharacterized protein \| Uncharacterized protein; afdb-uniprot50 \| AF-A0A292S0E4-F1-MODEL\_V4 \| 1.0 \| 2.519e-05 \| 141 \| 0.203 \| 226 \| 98 \| 8 \| 3 \| 220 \| 78 \| 229 \| Collar domain-containing protein \| Collar domain-containing protein; afdb-uniprot50 \| AF-A0A3C0GBI9-F1-MODEL\_V4 \| 1.0 \| 1.135e-07 \| 141 \| 0.323 \| 229 \| 123 \| 10 \| 14 \| 214 \| 1 \| 225 \| Collar domain-containing protein \| Collar domain-containing protein; afdb-uniprot50 \| AF-A0A2S9X536-F1-MODEL\_V4 \| 1.0 \| 8.154e-06 \| 141 \| 0.234 \| 235 \| 127 \| 12 \| 3 \| 220 \| 283 \| 481 \| Collar domain-containing protein \| Collar domain-containing protein; afdb-uniprot50 \| AF-A0A821TQI6-F1-MODEL\_V4 \| 1.0 \| 1.642e-06 \| 139 \| 0.34 \| 191 \| 95 \| 8 \| 25 \| 214 \| 13 \| 173 \| Hypothetical protein \| Hypothetical protein; afdb-uniprot50 \| AF-A0A1I1D6G6-F1-MODEL\_V4 \| 1.0 \| 2.082e-06 \| 139 \| 0.221 \| 221 \| 125 \| 11 \| 11 \| 221 \| 93 \| 276 \| Phage Tail Collar Domain \| Phage Tail Collar Domain; afdb-uniprot50 \| AF-U1BY59-F1-MODEL\_V4 \| 1.0 \| 4.779e-06 \| 136 \| 0.275 \| 189 \| 96 \| 6 \| 11 \| 188 \| 389 \| 547 \| Collar domain-containing protein \| Collar domain-containing protein; afdb-uniprot50 \| AF-A0A5V1PKQ6-F1-MODEL\_V4 \| 1.0 \| 9.866e-05 \| 135 \| 0.176 \| 227 \| 147 \| 10 \| 3 \| 220 \| 86 \| 281 \| Tail fiber protein \| Tail fiber protein; afdb-uniprot50 \| AF-A0A2D6P7B0-F1-MODEL\_V4 \| 1.0 \| 1.391e-05 \| 133 \| 0.309 \| 213 \| 110 \| 8 \| 12 \| 214 \| 116 \| 301 \| Collar domain-containing protein \| Collar domain-containing protein; afdb-uniprot50 \| AF-A0A806X791-F1-MODEL\_V4 \| 1.0 \| 5.783e-05 \| 133 \| 0.2 \| 234 \| 143 \| 9 \| 2 \| 220 \| 154 \| 358 \| Uncharacterized protein \| Uncharacterized protein; afdb-uniprot50 \| AF-A0A5V0QBY5-F1-MODEL\_V4 \| 1.0 \| 9.866e-05 \| 131 \| 0.171 \| 228 \| 147 \| 11 \| 3 \| 220 \| 115 \| 310 \| Phage tail protein \| Phage tail protein; afdb-uniprot50 \| AF-A0A6C8YBT0-F1-MODEL\_V4 \| 1.0 \| 0.0001111 \| 130 \| 0.18 \| 211 \| 142 \| 7 \| 11 \| 220 \| 166 \| 346 \| Tail fiber protein \| Tail fiber protein; afdb-uniprot50 \| AF-A0A060HJ12-F1-MODEL\_V4 \| 1.0 \| 1.849e-06 \| 129 \| 0.273 \| 205 \| 129 \| 5 \| 11 \| 214 \| 137 \| 322 \| Tail protein \| Tail protein; afdb-uniprot50 \| AF-A0A5V0QBB4-F1-MODEL\_V4 \| 1.0 \| 0.0002551 \| 127 \| 0.171 \| 222 \| 150 \| 9 \| 3 \| 220 \| 115 \| 306 \| Phage tail protein \| Phage tail protein; afdb-uniprot50 \| AF-A0A5U0QPU6-F1-MODEL\_V4 \| 1.0 \| 9.298e-05 \| 127 \| 0.183 \| 224 \| 138 \| 10 \| 5 \| 220 \| 197 \| 383 \| Tail fiber protein \| Tail fiber protein; afdb-uniprot50 \| AF-A0A410UF18-F1-MODEL\_V4 \| 1.0 \| 1.458e-06 \| 127 \| 0.295 \| 227 \| 122 \| 8 \| 11 \| 214 \| 183 \| 394 \| Collar domain-containing protein \| Collar domain-containing protein; afdb-uniprot50 \| AF-A0A1E7YUN4-F1-MODEL\_V4 \| 1.0 \| 7.333e-05 \| 126 \| 0.2 \| 239 \| 136 \| 11 \| 2 \| 220 \| 9 \| 212 \| Long tail fiber protein p37 \| Long tail fiber protein p37; afdb-uniprot50 \| AF-A0A821RQV6-F1-MODEL\_V4 \| 1.0 \| 2.519e-05 \| 126 \| 0.275 \| 200 \| 107 \| 5 \| 19 \| 212 \| 51 \| 218 \| Hypothetical protein \| Hypothetical protein; afdb-uniprot50 \| AF-A0A610C2R2-F1-MODEL\_V4 \| 1.0 \| 0.0001586 \| 126 \| 0.185 \| 221 \| 147 \| 8 \| 3 \| 220 \| 290 \| 480 \| Tail fiber protein \| Tail fiber protein; afdb-uniprot50 \| AF-A0A1C7W6K6-F1-MODEL\_V4 \| 1.0 \| 0.0002707 \| 125 \| 0.219 \| 214 \| 130 \| 7 \| 12 \| 220 \| 85 \| 266 \| Collar domain-containing protein \| Collar domain-containing protein; afdb-uniprot50 \| AF-A0A6L4BC09-F1-MODEL\_V4 \| 1.0 \| 9.298e-05 \| 125 \| 0.316 \| 155 \| 93 \| 5 \| 16 \| 167 \| 75 \| 219 \| Tail fiber protein \| Tail fiber protein; afdb-uniprot50 \| AF-A0A821JPY2-F1-MODEL\_V4 \| 1.0 \| 3.01e-05 \| 124 \| 0.266 \| 206 \| 109 \| 8 \| 16 \| 214 \| 139 \| 309 \| Hypothetical protein \| Hypothetical protein; afdb-uniprot50 \| AF-A0A376IY78-F1-MODEL\_V4 \| 1.0 \| 0.0001896 \| 124 \| 0.169 \| 230 \| 141 \| 10 \| 3 \| 220 \| 217 \| 408 \| Phage tail fibre repeat \| Phage tail fibre repeat; afdb-uniprot50 \| AF-A0A719G1Y9-F1-MODEL\_V4 \| 1.0 \| 0.0006214 \| 123 \| 0.146 \| 211 \| 149 \| 8 \| 11 \| 220 \| 27 \| 207 \| Tail fiber protein \| Tail fiber protein; afdb-uniprot50 \| AF-A0A639YPN2-F1-MODEL\_V4 \| 1.0 \| 0.0001586 \| 122 \| 0.201 \| 228 \| 138 \| 10 \| 2 \| 220 \| 141 \| 333 \| Tail fiber protein \| Tail fiber protein; afdb-uniprot50 \| AF-A0A1J7C8B9-F1-MODEL\_V4 \| 1.0 \| 0.0002135 \| 121 \| 0.192 \| 228 \| 140 \| 12 \| 2 \| 220 \| 77 \| 269 \| Collar domain-containing protein \| Collar domain-containing protein; afdb-uniprot50 \| AF-H3R9T3-F1-MODEL\_V4 \| 1.0 \| 8.762e-05 \| 120 \| 0.211 \| 227 \| 133 \| 11 \| 1 \| 220 \| 155 \| 342 \| Phage tail collar protein \| Phage tail collar protein; afdb-uniprot50 \| AF-A0A379QX05-F1-MODEL\_V4 \| 1.0 \| 9.298e-05 \| 120 \| 0.213 \| 220 \| 129 \| 11 \| 11 \| 220 \| 234 \| 419 \| DNA recombinase-like protein \| DNA recombinase-like protein; afdb-uniprot50 \| AF-A0A741N024-F1-MODEL\_V4 \| 1.0 \| 0.0003865 \| 120 \| 0.18 \| 211 \| 142 \| 7 \| 11 \| 220 \| 518 \| 698 \| Collar domain-containing protein \| Collar domain-containing protein; afdb-uniprot50 \| AF-A0A833J123-F1-MODEL\_V4 \| 1.0 \| 0.0002404 \| 119 \| 0.164 \| 231 \| 146 \| 11 \| 2 \| 220 \| 155 \| 350 \| Phage tail protein \| Phage tail protein; afdb-uniprot50 \| AF-A0A5U0QU90-F1-MODEL\_V4 \| 1.0 \| 0.0008872 \| 119 \| 0.162 \| 221 \| 152 \| 9 \| 3 \| 220 \| 177 \| 367 \| Tail fiber protein \| Tail fiber protein; afdb-uniprot50 \| AF-A0A707P6F2-F1-MODEL\_V4 \| 1.0 \| 0.0008872 \| 118 \| 0.151 \| 211 \| 148 \| 7 \| 11 \| 220 \| 14 \| 194 \| Tail fiber protein \| Tail fiber protein; afdb-uniprot50 \| AF-V7IMA5-F1-MODEL\_V4 \| 1.0 \| 0.0002135 \| 118 \| 0.2 \| 215 \| 133 \| 8 \| 11 \| 220 \| 290 \| 470 \| Phage Tail Collar domain protein \| Phage Tail Collar domain protein; afdb-uniprot50 \| AF-A0A439EPD8-F1-MODEL\_V4 \| 1.0 \| 6.512e-05 \| 117 \| 0.28 \| 214 \| 102 \| 10 \| 11 \| 207 \| 17 \| 195 \| Phage tail protein \| Phage tail protein; afdb-uniprot50 \| AF-A0A759ZPD9-F1-MODEL\_V4 \| 1.0 \| 0.0009415 \| 117 \| 0.155 \| 212 \| 146 \| 9 \| 11 \| 220 \| 179 \| 359 \| Collar domain-containing protein \| Collar domain-containing protein; afdb-uniprot50 \| AF-A0A602CNX1-F1-MODEL\_V4 \| 1.0 \| 0.0004101 \| 117 \| 0.169 \| 212 \| 143 \| 8 \| 11 \| 220 \| 463 \| 643 \| Phage tail protein \| Phage tail protein; afdb-uniprot50 \| AF-A0A3U5FGR7-F1-MODEL\_V4 \| 1.0 \| 0.0007425 \| 117 \| 0.169 \| 213 \| 142 \| 10 \| 11 \| 220 \| 630 \| 810 \| Phage tail protein \| Phage tail protein; afdb-uniprot50 \| AF-A0A2E1UKG0-F1-MODEL\_V4 \| 1.0 \| 0.0003048 \| 116 \| 0.247 \| 226 \| 112 \| 10 \| 11 \| 221 \| 315 \| 497 \| Mtd\_N domain-containing protein \| Mtd\_N domain-containing protein; afdb-uniprot50 \| AF-A0A706NAW6-F1-MODEL\_V4 \| 1.0 \| 0.0005518 \| 116 \| 0.169 \| 212 \| 143 \| 8 \| 11 \| 220 \| 360 \| 540 \| Phage tail protein \| Phage tail protein; afdb-uniprot50 \| AF-A0A762B2E8-F1-MODEL\_V4 \| 1.0 \| 0.00106 \| 116 \| 0.16 \| 212 \| 145 \| 9 \| 11 \| 220 \| 510 \| 690 \| Phage tail protein \| Phage tail protein; afdb-uniprot50 \| AF-A0A085HGE9-F1-MODEL\_V4 \| 1.0 \| 0.0002551 \| 115 \| 0.212 \| 226 \| 138 \| 9 \| 3 \| 220 \| 92 \| 285 \| Phage tail fiber protein \| Phage tail fiber protein; afdb-uniprot50 \| AF-A0A778CR53-F1-MODEL\_V4 \| 1.0 \| 6.136e-05 \| 115 \| 0.194 \| 236 \| 138 \| 12 \| 11 \| 220 \| 416 \| 625 \| Phage tail protein \| Phage tail protein; afdb-uniprot50 \| AF-Q8ZQ81-F1-MODEL\_V4 \| 1.0 \| 0.0009415 \| 115 \| 0.155 \| 212 \| 146 \| 9 \| 11 \| 220 \| 630 \| 810 \| Gifsy-2 prophage probable tail fiber protein \| Gifsy-2 prophage probable tail fiber protein; afdb-uniprot50 \| AF-H3RF68-F1-MODEL\_V4 \| 1.0 \| 0.0002404 \| 114 \| 0.187 \| 224 \| 142 \| 10 \| 2 \| 220 \| 106 \| 294 \| Putative tail fiber protein \| Putative tail fiber protein; afdb-uniprot50 \| AF-A0A701J2Z3-F1-MODEL\_V4 \| 1.0 \| 0.001125 \| 114 \| 0.156 \| 211 \| 147 \| 7 \| 11 \| 220 \| 160 \| 340 \| Phage tail protein \| Phage tail protein; afdb-uniprot50 \| AF-A0A619AEI5-F1-MODEL\_V4 \| 1.0 \| 0.0002265 \| 114 \| 0.195 \| 215 \| 134 \| 8 \| 11 \| 220 \| 237 \| 417 \| Tail fiber protein \| Tail fiber protein; afdb-uniprot50 \| AF-A0A6H2Y0W8-F1-MODEL\_V4 \| 1.0 \| 0.0005518 \| 114 \| 0.204 \| 215 \| 131 \| 9 \| 11 \| 220 \| 267 \| 446 \| Shikimate transporter \| Shikimate transporter; afdb-uniprot50 \| AF-A0A757ZPK5-F1-MODEL\_V4 \| 1.0 \| 0.0002707 \| 114 \| 0.204 \| 215 \| 132 \| 8 \| 11 \| 220 \| 509 \| 689 \| Phage tail protein \| Phage tail protein; afdb-uniprot50 \| AF-A0A738DCD2-F1-MODEL\_V4 \| 1.0 \| 0.0007879 \| 113 \| 0.165 \| 212 \| 144 \| 8 \| 11 \| 220 \| 165 \| 345 \| Tail fiber protein \| Tail fiber protein; afdb-uniprot50 \| AF-A0A757D915-F1-MODEL\_V4 \| 1.0 \| 0.0005518 \| 113 \| 0.165 \| 212 \| 144 \| 8 \| 11 \| 220 \| 267 \| 447 \| Phage tail protein \| Phage tail protein; afdb-uniprot50 \| AF-A0A5U6SMK2-F1-MODEL\_V4 \| 1.0 \| 0.0005856 \| 113 \| 0.174 \| 212 \| 140 \| 8 \| 12 \| 220 \| 429 \| 608 \| Phage tail protein \| Phage tail protein; afdb-uniprot50 \| AF-A0A3V2NRU9-F1-MODEL\_V4 \| 1.0 \| 0.001514 \| 112 \| 0.161 \| 211 \| 146 \| 8 \| 11 \| 220 \| 220 \| 400 \| Shikimate transporter \| Shikimate transporter; afdb-uniprot50 \| AF-A0A707VIZ3-F1-MODEL\_V4 \| 1.0 \| 0.0008872 \| 112 \| 0.165 \| 212 \| 144 \| 8 \| 11 \| 220 \| 339 \| 519 \| Phage tail protein \| Phage tail protein; afdb-uniprot50 \| AF-A0A604P221-F1-MODEL\_V4 \| 1.0 \| 0.0002265 \| 112 \| 0.195 \| 220 \| 128 \| 10 \| 11 \| 220 \| 419 \| 599 \| Phage tail protein \| Phage tail protein; afdb-uniprot50 \| AF-A0A2X4T8S9-F1-MODEL\_V4 \| 1.0 \| 0.0003642 \| 111 \| 0.19 \| 221 \| 133 \| 12 \| 11 \| 220 \| 102 \| 287 \| DNA recombinase-like protein \| DNA recombinase-like protein; afdb-uniprot50 \| AF-A0A820G8G8-F1-MODEL\_V4 \| 1.0 \| 0.0001786 \| 111 \| 0.232 \| 181 \| 95 \| 6 \| 11 \| 162 \| 179 \| 344 \| Hypothetical protein \| Hypothetical protein; afdb-uniprot50 \| AF-A0A7Z0Y4I4-F1-MODEL\_V4 \| 1.0 \| 0.0003642 \| 111 \| 0.195 \| 215 \| 134 \| 8 \| 11 \| 220 \| 198 \| 378 \| Phage tail protein \| Phage tail protein; afdb-uniprot50 \| AF-A0A5W2GRF4-F1-MODEL\_V4 \| 1.0 \| 0.0009415 \| 110 \| 0.156 \| 211 \| 147 \| 7 \| 11 \| 220 \| 284 \| 464 \| Phage tail protein \| Phage tail protein; afdb-uniprot50 \| AF-A0A5Z0R0R1-F1-MODEL\_V4 \| 1.0 \| 0.0002872 \| 110 \| 0.213 \| 220 \| 129 \| 12 \| 11 \| 220 \| 318 \| 503 \| Collar domain-containing protein \| Collar domain-containing protein; afdb-uniprot50 \| AF-A0A731WFF8-F1-MODEL\_V4 \| 1.0 \| 0.0001251 \| 110 \| 0.213 \| 215 \| 130 \| 11 \| 11 \| 220 \| 341 \| 521 \| Collar domain-containing protein \| Collar domain-containing protein; afdb-uniprot50 \| AF-G5RWL7-F1-MODEL\_V4 \| 1.0 \| 0.0002265 \| 109 \| 0.185 \| 221 \| 129 \| 9 \| 11 \| 220 \| 83 \| 263 \| Tail fiber assembly like-protein \| Tail fiber assembly like-protein; afdb-uniprot50 \| AF-A0A600RD36-F1-MODEL\_V4 \| 1.0 \| 0.0003865 \| 109 \| 0.209 \| 220 \| 125 \| 11 \| 11 \| 220 \| 95 \| 275 \| Tail fiber protein \| Tail fiber protein; afdb-uniprot50 \| AF-A0A5Y2ZE76-F1-MODEL\_V4 \| 1.0 \| 0.001194 \| 109 \| 0.16 \| 212 \| 145 \| 8 \| 11 \| 220 \| 163 \| 343 \| Tail fiber protein \| Tail fiber protein; afdb-uniprot50 \| AF-A0A708RGM0-F1-MODEL\_V4 \| 1.0 \| 0.00106 \| 109 \| 0.165 \| 212 \| 144 \| 8 \| 11 \| 220 \| 339 \| 519 \| Phage tail protein \| Phage tail protein; afdb-uniprot50 \| AF-A0A6C0BYV0-F1-MODEL\_V4 \| 1.0 \| 0.001514 \| 108 \| 0.216 \| 240 \| 120 \| 10 \| 12 \| 221 \| 85 \| 286 \| Uncharacterized protein \| Uncharacterized protein; afdb-uniprot50 \| AF-A0A7Y5M3U8-F1-MODEL\_V4 \| 1.0 \| 0.001194 \| 108 \| 0.158 \| 214 \| 144 \| 8 \| 11 \| 220 \| 206 \| 387 \| Tail fiber protein \| Tail fiber protein; afdb-uniprot50 \| AF-A0A7H8WZS4-F1-MODEL\_V4 \| 1.0 \| 0.0006214 \| 108 \| 0.177 \| 225 \| 126 \| 11 \| 11 \| 220 \| 512 \| 692 \| Phage tail protein \| Phage tail protein; afdb-uniprot50 \| AF-A0A827KJI0-F1-MODEL\_V4 \| 1.0 \| 0.001344 \| 107 \| 0.176 \| 221 \| 143 \| 9 \| 7 \| 220 \| 6 \| 194 \| Tail fiber protein \| Tail fiber protein; afdb-uniprot50 \| AF-A0A855SIZ9-F1-MODEL\_V4 \| 1.0 \| 0.0007425 \| 107 \| 0.181 \| 232 \| 136 \| 11 \| 11 \| 220 \| 17 \| 216 \| Phage tail protein \| Phage tail protein; afdb-uniprot50 \| AF-A0A736VG58-F1-MODEL\_V4 \| 1.0 \| 0.0004101 \| 107 \| 0.198 \| 217 \| 128 \| 11 \| 12 \| 220 \| 297 \| 475 \| Tail fiber protein \| Tail fiber protein; afdb-uniprot50 \| AF-A0A5Y2LQP0-F1-MODEL\_V4 \| 1.0 \| 0.0004101 \| 107 \| 0.209 \| 215 \| 131 \| 9 \| 11 \| 220 \| 364 \| 544 \| Collar domain-containing protein \| Collar domain-containing protein; afdb-uniprot50 \| AF-A0A2E9W8F9-F1-MODEL\_V4 \| 1.0 \| 0.0005518 \| 106 \| 0.22 \| 204 \| 139 \| 6 \| 13 \| 214 \| 8 \| 193 \| Collar domain-containing protein \| Collar domain-containing protein; afdb-uniprot50 \| AF-A0A5T3NHC4-F1-MODEL\_V4 \| 1.0 \| 0.003475 \| 106 \| 0.137 \| 211 \| 151 \| 7 \| 11 \| 220 \| 525 \| 705 \| Collar domain-containing protein \| Collar domain-containing protein; afdb-uniprot50 \| AF-A0A0W7WFQ0-F1-MODEL\_V4 \| 1.0 \| 0.0004352 \| 105 \| 0.191 \| 235 \| 143 \| 7 \| 11 \| 214 \| 30 \| 248 \| Collar domain-containing protein \| Collar domain-containing protein; afdb-uniprot50 \| AF-A0A5A9C9Z6-F1-MODEL\_V4 \| 1.0 \| 0.0008361 \| 105 \| 0.188 \| 233 \| 136 \| 10 \| 11 \| 220 \| 671 \| 873 \| Tail fiber protein \| Tail fiber protein; afdb-uniprot50 \| AF-A0A714KQT2-F1-MODEL\_V4 \| 1.0 \| 0.0009415 \| 104 \| 0.171 \| 221 \| 132 \| 10 \| 11 \| 220 \| 27 \| 207 \| Tail fiber protein \| Tail fiber protein; afdb-uniprot50 \| AF-A0A707NFK7-F1-MODEL\_V4 \| 1.0 \| 0.0008361 \| 104 \| 0.175 \| 216 \| 137 \| 11 \| 11 \| 220 \| 280 \| 460 \| Phage tail protein \| Phage tail protein; afdb-uniprot50 \| AF-A0A6J7EVX5-F1-MODEL\_V4 \| 1.0 \| 4.504e-06 \| 104 \| 0.405 \| 227 \| 106 \| 4 \| 13 \| 221 \| 347 \| 562 \| Unannotated protein \| Unannotated protein; afdb-uniprot50 \| AF-A0A5E8KNR2-F1-MODEL\_V4 \| 1.0 \| 0.0001683 \| 103 \| 0.263 \| 201 \| 95 \| 7 \| 16 \| 216 \| 2 \| 149 \| Collar domain-containing protein \| Collar domain-containing protein; afdb-uniprot50 \| AF-L3Q3I4-F1-MODEL\_V4 \| 1.0 \| 0.001427 \| 103 \| 0.197 \| 228 \| 130 \| 13 \| 8 \| 220 \| 243 \| 432 \| Collar domain-containing protein \| Collar domain-containing protein; afdb-uniprot50 \| AF-A0A2I5TL70-F1-MODEL\_V4 \| 1.0 \| 0.0009415 \| 102 \| 0.177 \| 236 \| 155 \| 9 \| 1 \| 220 \| 208 \| 420 \| Phage tail protein \| Phage tail protein; afdb-uniprot50 \| AF-A0A4Y8GSR2-F1-MODEL\_V4 \| 1.0 \| 0.0006594 \| 101 \| 0.188 \| 234 \| 152 \| 8 \| 3 \| 220 \| 290 \| 501 \| Short-chain fatty acid transporter \| Short-chain fatty acid transporter; afdb-uniprot50 \| AF-A0A717LCK5-F1-MODEL\_V4 \| 1.0 \| 0.00106 \| 100 \| 0.183 \| 224 \| 126 \| 12 \| 11 \| 220 \| 29 \| 209 \| Tail fiber protein \| Tail fiber protein; afdb-uniprot50 \| AF-A0A479JBS8-F1-MODEL\_V4 \| 1.0 \| 0.00052 \| 100 \| 0.222 \| 247 \| 148 \| 11 \| 1 \| 220 \| 134 \| 363 \| Phage tail fiber protein \| Phage tail fiber protein; afdb-uniprot50 \| AF-A0A815QGL9-F1-MODEL\_V4 \| 1.0 \| 0.0001683 \| 100 \| 0.183 \| 223 \| 136 \| 7 \| 2 \| 214 \| 646 \| 832 \| Hypothetical protein \| Hypothetical protein; afdb-uniprot50 \| AF-A0A485DQI3-F1-MODEL\_V4 \| 0.999 \| 0.0004101 \| 99 \| 0.205 \| 214 \| 136 \| 8 \| 11 \| 220 \| 260 \| 443 \| Phage lambda-related protein \| Phage lambda-related protein; afdb-uniprot50 \| AF-I6CNP4-F1-MODEL\_V4 \| 0.999 \| 0.001809 \| 98 \| 0.188 \| 239 \| 136 \| 11 \| 11 \| 220 \| 42 \| 251 \| Long tail fiber protein p37 \| Long tail fiber protein p37; afdb-uniprot50 \| AF-A0A328TN35-F1-MODEL\_V4 \| 0.999 \| 0.001809 \| 98 \| 0.207 \| 255 \| 143 \| 13 \| 2 \| 221 \| 34 \| 264 \| Phage tail fiber repeat family protein \| Phage tail fiber repeat family protein; afdb-uniprot50 \| AF-D2TUQ3-F1-MODEL\_V4 \| 0.999 \| 0.0002404 \| 98 \| 0.166 \| 246 \| 143 \| 14 \| 3 \| 220 \| 69 \| 280 \| Putative phage tail fibre protein \| Putative phage tail fibre protein; afdb-uniprot50 \| AF-A0A8A7UB48-F1-MODEL\_V4 \| 0.999 \| 0.001606 \| 98 \| 0.184 \| 239 \| 137 \| 10 \| 11 \| 220 \| 96 \| 305 \| Tail fiber protein \| Tail fiber protein; afdb-uniprot50 \| AF-A0A828NYP4-F1-MODEL\_V4 \| 0.999 \| 0.003086 \| 96 \| 0.189 \| 222 \| 137 \| 12 \| 8 \| 220 \| 362 \| 549 \| Phage tail protein \| Phage tail protein; afdb-uniprot50 \| AF-A0A0T9T368-F1-MODEL\_V4 \| 0.998 \| 0.00106 \| 95 \| 0.199 \| 236 \| 136 \| 11 \| 1 \| 220 \| 86 \| 284 \| Phage tail protein \| Phage tail protein; afdb-uniprot50 \| AF-A0A8B4KTQ3-F1-MODEL\_V4 \| 0.998 \| 0.00593 \| 95 \| 0.171 \| 256 \| 135 \| 10 \| 12 \| 220 \| 452 \| 677 \| PPE-repeat protein \| PPE-repeat protein; afdb-uniprot50 \| AF-A0A830SMV6-F1-MODEL\_V4 \| 0.998 \| 0.002583 \| 94 \| 0.184 \| 239 \| 137 \| 10 \| 11 \| 220 \| 205 \| 414 \| Short-chain fatty acid transporter \| Short-chain fatty acid transporter; afdb-uniprot50 \| AF-A0A243TCZ2-F1-MODEL\_V4 \| 0.998 \| 0.0009415 \| 94 \| 0.193 \| 238 \| 129 \| 12 \| 11 \| 220 \| 386 \| 588 \| Phage tail protein \| Phage tail protein; afdb-uniprot50 \| AF-A0A1F3CD68-F1-MODEL\_V4 \| 0.997 \| 0.002908 \| 92 \| 0.22 \| 240 \| 114 \| 11 \| 12 \| 221 \| 388 \| 584 \| Uncharacterized protein \| Uncharacterized protein; afdb-uniprot50 \| AF-A0A8B4K950-F1-MODEL\_V4 \| 0.997 \| 0.0004618 \| 92 \| 0.239 \| 242 \| 136 \| 11 \| 11 \| 220 \| 924 \| 1149 \| Putative tail fiber protein \| Putative tail fiber protein; afdb-uniprot50 \| AF-A0A828H8Q0-F1-MODEL\_V4 \| 0.997 \| 0.0003234 \| 92 \| 0.237 \| 240 \| 139 \| 11 \| 11 \| 220 \| 949 \| 1174 \| Short-chain fatty acid transporter \| Short-chain fatty acid transporter; afdb-uniprot50 \| AF-W0BPN1-F1-MODEL\_V4 \| 0.997 \| 0.0007425 \| 91 \| 0.2 \| 220 \| 143 \| 8 \| 12 \| 220 \| 176 \| 373 \| Collar domain-containing protein \| Collar domain-containing protein; afdb-uniprot50 \| AF-A0A4U3LF57-F1-MODEL\_V4 \| 0.997 \| 0.001194 \| 91 \| 0.232 \| 219 \| 141 \| 10 \| 11 \| 220 \| 359 \| 559 \| Phage tail protein \| Phage tail protein; afdb-uniprot50 \| AF-A0A7A3AF76-F1-MODEL\_V4 \| 0.997 \| 0.001919 \| 91 \| 0.2 \| 220 \| 146 \| 8 \| 11 \| 220 \| 668 \| 867 \| Phage tail protein \| Phage tail protein; afdb-uniprot50 \| AF-A0A827K9D2-F1-MODEL\_V4 \| 0.996 \| 0.001344 \| 90 \| 0.175 \| 228 \| 143 \| 8 \| 11 \| 220 \| 87 \| 287 \| Tail fiber protein \| Tail fiber protein; afdb-uniprot50 \| AF-A0A1E7Z390-F1-MODEL\_V4 \| 0.996 \| 0.002161 \| 90 \| 0.208 \| 249 \| 150 \| 11 \| 2 \| 221 \| 76 \| 306 \| Phage tail protein \| Phage tail protein; afdb-uniprot50 \| AF-A0A7Z3MBJ3-F1-MODEL\_V4 \| 0.996 \| 0.001514 \| 90 \| 0.215 \| 246 \| 151 \| 9 \| 1 \| 220 \| 200 \| 429 \| Phage tail protein \| Phage tail protein; afdb-uniprot50 \| AF-A0A2J9H2E6-F1-MODEL\_V4 \| 0.996 \| 0.002294 \| 90 \| 0.194 \| 231 \| 136 \| 10 \| 11 \| 220 \| 468 \| 669 \| Phage tail protein \| Phage tail protein; afdb-uniprot50 \| AF-A0A7T7XQ37-F1-MODEL\_V4 \| 0.996 \| 0.003086 \| 90 \| 0.214 \| 257 \| 116 \| 13 \| 1 \| 220 \| 540 \| 747 \| Uncharacterized protein \| Uncharacterized protein; afdb-uniprot50 \| AF-A0A2J0PYI7-F1-MODEL\_V4 \| 0.996 \| 0.001267 \| 90 \| 0.196 \| 254 \| 141 \| 10 \| 11 \| 220 \| 572 \| 806 \| Collar domain-containing protein \| Collar domain-containing protein; afdb-uniprot50 \| AF-K4VSX7-F1-MODEL\_V4 \| 0.996 \| 0.001427 \| 89 \| 0.2 \| 224 \| 141 \| 7 \| 11 \| 220 \| 101 \| 300 \| Major tail fiber protein \| Major tail fiber protein; afdb-uniprot50 \| AF-A0A090R0D4-F1-MODEL\_V4 \| 0.996 \| 0.001919 \| 89 \| 0.157 \| 242 \| 134 \| 10 \| 1 \| 220 \| 4 \| 197 \| Phage tail fibers \| Phage tail fibers; afdb-uniprot50 \| AF-A0A5F0PX95-F1-MODEL\_V4 \| 0.996 \| 0.002037 \| 89 \| 0.176 \| 226 \| 145 \| 8 \| 11 \| 220 \| 189 \| 389 \| Tail fiber protein \| Tail fiber protein; afdb-uniprot50 \| AF-A0A4C8VIP7-F1-MODEL\_V4 \| 0.996 \| 0.002583 \| 89 \| 0.175 \| 211 \| 150 \| 6 \| 11 \| 220 \| 650 \| 837 \| Phage side tail fiber protein \| Phage side tail fiber protein; afdb-uniprot50 \| AF-A0A517N136-F1-MODEL\_V4 \| 0.995 \| 0.003475 \| 88 \| 0.262 \| 217 \| 120 \| 7 \| 17 \| 214 \| 958 \| 1153 \| Phage Tail Collar Domain protein \| Phage Tail Collar Domain protein; afdb-uniprot50 \| AF-A0A2P5GM50-F1-MODEL\_V4 \| 0.994 \| 0.00106 \| 87 \| 0.206 \| 228 \| 136 \| 9 \| 12 \| 220 \| 225 \| 426 \| Collar domain-containing protein \| Collar domain-containing protein; afdb-uniprot50 \| AF-A0A236MFK9-F1-MODEL\_V4 \| 0.994 \| 0.003688 \| 87 \| 0.187 \| 262 \| 129 \| 12 \| 11 \| 220 \| 317 \| 546 \| Phage tail protein \| Phage tail protein; afdb-uniprot50 \| AF-A0A7W4KMJ0-F1-MODEL\_V4 \| 0.994 \| 0.001514 \| 87 \| 0.182 \| 225 \| 145 \| 7 \| 11 \| 220 \| 453 \| 653 \| Tail fiber protein \| Tail fiber protein; afdb-uniprot50 \| AF-A0A1Q4PEJ7-F1-MODEL\_V4 \| 0.994 \| 0.001705 \| 87 \| 0.21 \| 214 \| 137 \| 10 \| 11 \| 220 \| 501 \| 686 \| Phage tail protein \| Phage tail protein; afdb-uniprot50 \| AF-A0A8B5NF75-F1-MODEL\_V4 \| 0.994 \| 0.002161 \| 87 \| 0.174 \| 223 \| 149 \| 7 \| 11 \| 220 \| 670 \| 870 \| Phage tail protein \| Phage tail protein; afdb-uniprot50 \| AF-A0A827FX06-F1-MODEL\_V4 \| 0.994 \| 0.002583 \| 87 \| 0.195 \| 220 \| 147 \| 7 \| 11 \| 220 \| 942 \| 1141 \| Phage tail protein \| Phage tail protein; afdb-uniprot50 \| AF-A0A5P0PY25-F1-MODEL\_V4 \| 0.993 \| 0.0007879 \| 86 \| 0.212 \| 235 \| 133 \| 10 \| 12 \| 220 \| 2 \| 210 \| Tail fiber protein \| Tail fiber protein; afdb-uniprot50 \| AF-A0A7A6N7A4-F1-MODEL\_V4 \| 0.993 \| 0.002741 \| 86 \| 0.179 \| 228 \| 142 \| 8 \| 11 \| 220 \| 251 \| 451 \| Tail fiber protein \| Tail fiber protein; afdb-uniprot50 \| AF-A0A2X5DKB2-F1-MODEL\_V4 \| 0.993 \| 0.002037 \| 86 \| 0.232 \| 219 \| 141 \| 10 \| 11 \| 220 \| 315 \| 515 \| Side tail phage protein \| Side tail phage protein; afdb-uniprot50 \| AF-D6I5M1-F1-MODEL\_V4 \| 0.993 \| 0.0005856 \| 86 \| 0.229 \| 231 \| 135 \| 10 \| 12 \| 220 \| 342 \| 551 \| Collar domain-containing protein \| Collar domain-containing protein; afdb-uniprot50 \| AF-A0A1X3JXT7-F1-MODEL\_V4 \| 0.993 \| 0.002908 \| 86 \| 0.214 \| 242 \| 140 \| 11 \| 12 \| 220 \| 419 \| 643 \| Side tail fiber protein-like protein \| Side tail fiber protein-like protein; afdb-uniprot50 \| AF-A0A478KQU1-F1-MODEL\_V4 \| 0.993 \| 0.0007879 \| 86 \| 0.232 \| 241 \| 137 \| 10 \| 12 \| 220 \| 611 \| 835 \| Phage side tail fiber protein \| Phage side tail fiber protein; afdb-uniprot50 \| AF-A0A2Z2JLN6-F1-MODEL\_V4 \| 0.993 \| 0.002294 \| 86 \| 0.2 \| 219 \| 147 \| 6 \| 11 \| 220 \| 748 \| 947 \| Putative membrane protein \| Putative membrane protein; afdb-uniprot50 \| AF-A0A827JSA7-F1-MODEL\_V4 \| 0.992 \| 0.001344 \| 85 \| 0.185 \| 248 \| 145 \| 10 \| 11 \| 220 \| 23 \| 251 \| Tail fiber protein \| Tail fiber protein; afdb-uniprot50 \| AF-A0A7D6J6I4-F1-MODEL\_V4 \| 0.992 \| 0.004407 \| 85 \| 0.217 \| 239 \| 143 \| 10 \| 12 \| 220 \| 539 \| 763 \| Tail fiber protein \| Tail fiber protein; afdb-uniprot50 \| AF-A0A827LBC4-F1-MODEL\_V4 \| 0.992 \| 0.007519 \| 85 \| 0.204 \| 235 \| 127 \| 13 \| 11 \| 220 \| 616 \| 815 \| Phage tail protein \| Phage tail protein; afdb-uniprot50 \| AF-A0A1R0G2E0-F1-MODEL\_V4 \| 0.992 \| 0.00106 \| 85 \| 0.181 \| 243 \| 139 \| 10 \| 11 \| 220 \| 698 \| 913 \| Prophage tail fiber N-terminal domain-containing protein \| Prophage tail fiber N-terminal domain-containing protein; afdb-uniprot50 \| AF-A0A3P5DNK9-F1-MODEL\_V4 \| 0.992 \| 0.001705 \| 85 \| 0.208 \| 249 \| 146 \| 11 \| 5 \| 220 \| 722 \| 952 \| Uncharacterized protein \| Uncharacterized protein; afdb-uniprot50 \| AF-A0A2X8FBW3-F1-MODEL\_V4 \| 0.992 \| 0.003913 \| 85 \| 0.231 \| 238 \| 141 \| 10 \| 12 \| 220 \| 760 \| 984 \| Putative tail fiber protein \| Putative tail fiber protein; afdb-uniprot50 \| AF-A0A0H3MMX0-F1-MODEL\_V4 \| 0.991 \| 0.004407 \| 84 \| 0.16 \| 237 \| 158 \| 9 \| 1 \| 220 \| 312 \| 524 \| Putative side tail phage protein \| Putative side tail phage protein; afdb-uniprot50 \| AF-A0A6N6XUS5-F1-MODEL\_V4 \| 0.991 \| 0.001514 \| 84 \| 0.215 \| 241 \| 143 \| 11 \| 11 \| 220 \| 585 \| 810 \| Short-chain fatty acid transporter \| Short-chain fatty acid transporter; afdb-uniprot50 \| AF-A0A377NAA6-F1-MODEL\_V4 \| 0.991 \| 0.003086 \| 84 \| 0.17 \| 229 \| 149 \| 9 \| 11 \| 220 \| 650 \| 856 \| Tail fiber protein \| Tail fiber protein; afdb-uniprot50 \| AF-A0A828UHZ0-F1-MODEL\_V4 \| 0.99 \| 0.003475 \| 83 \| 0.187 \| 224 \| 144 \| 7 \| 11 \| 220 \| 403 \| 602 \| Phage tail collar domain protein \| Phage tail collar domain protein; afdb-uniprot50 \| AF-A0A8B4JKQ6-F1-MODEL\_V4 \| 0.99 \| 0.001344 \| 83 \| 0.198 \| 252 \| 139 \| 12 \| 11 \| 220 \| 384 \| 614 \| Tail fiber protein \| Tail fiber protein; afdb-uniprot50 \| AF-A0A379R3M2-F1-MODEL\_V4 \| 0.99 \| 0.002294 \| 83 \| 0.204 \| 220 \| 146 \| 8 \| 11 \| 220 \| 524 \| 724 \| Gifsy-2 prophage tail fiber protein \| Gifsy-2 prophage tail fiber protein; afdb-uniprot50 \| AF-A0A6C8XTX3-F1-MODEL\_V4 \| 0.99 \| 0.003688 \| 83 \| 0.163 \| 232 \| 142 \| 8 \| 11 \| 220 \| 608 \| 809 \| Phage tail protein \| Phage tail protein; afdb-uniprot50 \| AF-W0BR44-F1-MODEL\_V4 \| 0.988 \| 0.001919 \| 82 \| 0.208 \| 221 \| 141 \| 8 \| 12 \| 220 \| 364 \| 562 \| Collar domain-containing protein \| Collar domain-containing protein; afdb-uniprot50 \| AF-A0A793GNT0-F1-MODEL\_V4 \| 0.986 \| 0.001606 \| 81 \| 0.235 \| 234 \| 130 \| 12 \| 12 \| 220 \| 33 \| 242 \| Tail fiber protein \| Tail fiber protein; afdb-uniprot50 \| AF-F4V1R4-F1-MODEL\_V4 \| 0.986 \| 0.001606 \| 81 \| 0.226 \| 234 \| 132 \| 11 \| 12 \| 220 \| 143 \| 352 \| Side tail fiber protein lambdoid prophage Qin-like protein \| Side tail fiber protein lambdoid prophage Qin-like protein; afdb-uniprot50 \| AF-A0A8B4L7W3-F1-MODEL\_V4 \| 0.986 \| 0.001705 \| 81 \| 0.229 \| 231 \| 135 \| 10 \| 12 \| 220 \| 183 \| 392 \| Phage tail domain-containing protein \| Phage tail domain-containing protein; afdb-uniprot50 \| AF-A0A1X3M1P4-F1-MODEL\_V4 \| 0.986 \| 0.004962 \| 81 \| 0.207 \| 227 \| 134 \| 10 \| 12 \| 220 \| 250 \| 448 \| Side tail fiber protein-like protein \| Side tail fiber protein-like protein; afdb-uniprot50 \| AF-A0A7U0GZW4-F1-MODEL\_V4 \| 0.986 \| 0.003475 \| 81 \| 0.205 \| 239 \| 147 \| 9 \| 12 \| 220 \| 241 \| 466 \| Phage tail tape measure protein \| Phage tail tape measure protein; afdb-uniprot50 \| AF-A0A826TQX2-F1-MODEL\_V4 \| 0.986 \| 0.002583 \| 81 \| 0.194 \| 221 \| 144 \| 9 \| 12 \| 220 \| 505 \| 703 \| Phage tail protein \| Phage tail protein; afdb-uniprot50 \| AF-A0A7B5PWV2-F1-MODEL\_V4 \| 0.986 \| 0.004407 \| 81 \| 0.207 \| 222 \| 140 \| 10 \| 12 \| 220 \| 544 \| 742 \| Collar domain-containing protein \| Collar domain-containing protein; afdb-uniprot50 \| AF-B2TVZ0-F1-MODEL\_V4 \| 0.984 \| 0.002294 \| 80 \| 0.198 \| 252 \| 139 \| 11 \| 11 \| 220 \| 209 \| 439 \| Side tail fiber protein \| Side tail fiber protein; afdb-uniprot50 \| AF-A0A376VIM7-F1-MODEL\_V4 \| 0.984 \| 0.002037 \| 80 \| 0.194 \| 252 \| 140 \| 12 \| 11 \| 220 \| 273 \| 503 \| Side tail fiber protein from bacteriophage origin \| Side tail fiber protein from bacteriophage origin; afdb-uniprot50 \| AF-A0A6L9FK66-F1-MODEL\_V4 \| 0.984 \| 0.006677 \| 80 \| 0.188 \| 234 \| 152 \| 9 \| 1 \| 220 \| 309 \| 518 \| Uncharacterized protein \| Uncharacterized protein; afdb-uniprot50 \| AF-A0A646JEL0-F1-MODEL\_V4 \| 0.984 \| 0.001809 \| 80 \| 0.183 \| 245 \| 145 \| 10 \| 12 \| 220 \| 626 \| 851 \| Phage tail protein \| Phage tail protein; afdb-uniprot50 \| AF-A0A4S4IN19-F1-MODEL\_V4 \| 0.984 \| 0.006677 \| 80 \| 0.227 \| 246 \| 137 \| 12 \| 12 \| 220 \| 804 \| 1033 \| Qin prophage, putative side tail fiber assembly protein \| Qin prophage, putative side tail fiber assembly protein; afdb-uniprot50 \| AF-A0A376SP24-F1-MODEL\_V4 \| 0.981 \| 0.002294 \| 79 \| 0.235 \| 234 \| 130 \| 12 \| 12 \| 220 \| 108 \| 317 \| Prophage tail fiber domain-containing protein \| Prophage tail fiber domain-containing protein; afdb-uniprot50 \| AF-A0A2X6JPY3-F1-MODEL\_V4 \| 0.981 \| 0.002294 \| 79 \| 0.225 \| 226 \| 142 \| 9 \| 12 \| 220 \| 461 \| 670 \| Prophage tail fiber N-terminal domain-containing protein \| Prophage tail fiber N-terminal domain-containing protein; afdb-uniprot50 \| AF-A0A2X2HTT9-F1-MODEL\_V4 \| 0.981 \| 0.006677 \| 79 \| 0.219 \| 228 \| 142 \| 9 \| 11 \| 220 \| 712 \| 921 \| Side tail fiber protein \| Side tail fiber protein; afdb-uniprot50 \| AF-A0A775EW34-F1-MODEL\_V4 \| 0.978 \| 0.002908 \| 78 \| 0.234 \| 235 \| 131 \| 12 \| 11 \| 220 \| 54 \| 264 \| Tail fiber protein \| Tail fiber protein; afdb-uniprot50 \| AF-A0A6C0LBP0-F1-MODEL\_V4 \| 0.978 \| 0.001606 \| 78 \| 0.197 \| 269 \| 141 \| 13 \| 12 \| 221 \| 15 \| 267 \| Collar domain-containing protein \| Collar domain-containing protein; afdb-uniprot50 \| AF-A0A5X4GC92-F1-MODEL\_V4 \| 0.978 \| 0.007519 \| 78 \| 0.221 \| 248 \| 147 \| 10 \| 3 \| 220 \| 253 \| 484 \| Short-chain fatty acid transporter \| Short-chain fatty acid transporter; afdb-uniprot50 \| AF-A0A810TS17-F1-MODEL\_V4 \| 0.978 \| 0.002741 \| 78 \| 0.221 \| 235 \| 134 \| 11 \| 11 \| 220 \| 297 \| 507 \| Short-chain dehydrogenase \| Short-chain dehydrogenase; afdb-uniprot50 \| AF-A0A4B9BPF7-F1-MODEL\_V4 \| 0.978 \| 0.002434 \| 78 \| 0.222 \| 229 \| 139 \| 9 \| 12 \| 220 \| 305 \| 514 \| Phage tail protein \| Phage tail protein; afdb-uniprot50 \| AF-A0A6D0EXL5-F1-MODEL\_V4 \| 0.978 \| 0.002161 \| 78 \| 0.216 \| 236 \| 133 \| 11 \| 11 \| 220 \| 449 \| 658 \| Tail fiber protein \| Tail fiber protein; afdb-uniprot50 \| AF-A0A152BV35-F1-MODEL\_V4 \| 0.978 \| 0.004676 \| 78 \| 0.195 \| 225 \| 139 \| 8 \| 12 \| 220 \| 510 \| 708 \| Phage tail protein \| Phage tail protein; afdb-uniprot50 \| AF-A0A836ZDU5-F1-MODEL\_V4 \| 0.975 \| 0.003475 \| 77 \| 0.227 \| 237 \| 129 \| 12 \| 11 \| 220 \| 319 \| 528 \| Phage tail fiber repeat family protein \| Phage tail fiber repeat family protein; afdb-uniprot50 \| AF-A0A222QUV6-F1-MODEL\_V4 \| 0.971 \| 0.007979 \| 76 \| 0.209 \| 229 \| 131 \| 11 \| 12 \| 220 \| 338 \| 536 \| Putative tail fiber protein \| Putative tail fiber protein; afdb-uniprot50 \| AF-A0A377F1I4-F1-MODEL\_V4 \| 0.971 \| 0.005266 \| 76 \| 0.209 \| 248 \| 139 \| 11 \| 12 \| 220 \| 349 \| 578 \| Phage tail protein \| Phage tail protein; afdb-uniprot50 \| AF-Q7BTW5-F1-MODEL\_V4 \| 0.971 \| 0.004962 \| 76 \| 0.217 \| 258 \| 135 \| 10 \| 11 \| 220 \| 620 \| 858 \| Tail fiber protein homolog \| Tail fiber protein homolog; afdb-uniprot50 \| AF-A0A6N8NGX5-F1-MODEL\_V4 \| 0.967 \| 0.003913 \| 75 \| 0.235 \| 234 \| 130 \| 12 \| 12 \| 220 \| 365 \| 574 \| Collar domain-containing protein \| Collar domain-containing protein; afdb-uniprot50 \| AF-A0A854RAR2-F1-MODEL\_V4 \| 0.961 \| 0.007979 \| 74 \| 0.198 \| 252 \| 139 \| 13 \| 11 \| 220 \| 36 \| 266 \| Phage tail protein \| Phage tail protein; afdb-uniprot50 \| AF-A0A7U8W607-F1-MODEL\_V4 \| 0.956 \| 0.007979 \| 73 \| 0.183 \| 251 \| 144 \| 10 \| 11 \| 220 \| 213 \| 443 \| Tail protein 2 \| Tail protein 2; afdb-uniprot50 \| AF-A0A7B3JM82-F1-MODEL\_V4 \| 0.956 \| 0.006292 \| 73 \| 0.19 \| 252 \| 141 \| 9 \| 11 \| 220 \| 678 \| 908 \| Uncharacterized protein \| Uncharacterized protein |
| Top keywords  (threshold 1.00e-02 (evalue)) | **Collar, tail, domain\_containing, Phage, fiber, Hypothetical, Microcystin\_dependent, Putative, Side, Phage\_related** |
| Output files | ../../similar\_structures/17\_FANPEZAQ\_CDS\_0017\_afdb-proteome\_foldseek.tsv ../../similar\_structures/17\_FANPEZAQ\_CDS\_0017\_afdb-uniprot50\_foldseek.tsv ../../similar\_structures/17\_FANPEZAQ\_CDS\_0017\_merged.svg ../../similar\_structures/17\_FANPEZAQ\_CDS\_0017\_pdb\_foldseek.tsv |

  
  
  

Return to summary | Go to previous | Go to next

  


---

**Sequence/structure alignments coloring**  
Each object in the alignment figures is colored according to its E-value following this color coding:

1e-100
10
