## Supplementary material for "Completing the BASEL phage collection to unlock hidden diversity for systematic exploration of phage-host interactions": Data S2: 18.html

FANPEZAQ\_CDS\_0018


Return to summary | Go to previous | Go to next

|  |  |
| --- | --- |
| FANPEZAQ\_CDS\_0018  Page creation date: 02 Sep 2024, 12:00  Project folder: n/a  Input sequences file: Escherichia\_virus\_HeidiAbel.gb | tail phage sheath domain\_containing fragment subtilisin\_like fi c\_terminal phage\_sheath\_1c major phage\_sheath\_1 putative prophage monomer outer membrane secretion hypothetical contractile ragb susd nutrient uptake structural system bacteriophage afp3 type vi afp2 phage\_related afp1 anti\_feeding afp |

### Sequence information

|  |  |
| --- | --- |
| Name | FANPEZAQ\_CDS\_0018  18\_FANPEZAQ\_CDS\_0018 (pipeline id) |
| Imported annotations | Escherichia\_virus\_HeidiAbel Bas97 |
| Protein sequence | MAGTDFLHGVEVLEIDSGSRVIQTVKSSVIGIVGTAPNADPDAFPLNTPTLIAGSQREAA KLDMLGTGEGTLPKALDAILDQIGAVVIVVRVEEGESDTATLANIMGGVNASTGKYEGVH ALLGAESVVGYSPRILCVPGFTHTRVQDGVTAISIGNGGSGYTEAPTITLAGGGGGSGAT AQATVVGGVITKITVTNAGMGYTEAPTVTITGGNGTGATATASVGIAANAVVSELLGIAN RLRAVIIADGPNTNDADAIAYRKDFGSKRVYVVDPKVLVLDEDGTTITDWASPRVAGLIA RIDNERGFWWSPSNQEIYGIQGTARTIDFTLGDTSSRANLLNENEVTTIIRQDGFRLWGN RTTSSDPKWAFLCVVRTADIINDSLLRAHLWAVDRGITKQYVNDVIEGVNAYLRNLVSLG AILGGTCWADPDLNTADQIADGKVYFDFDFTPVYPAEHIIFRSQLTNDYISEVFE |
| Number of residues | 475 |
| Molecular weight (Da) | 49868.35 |
| Output files | ../../query\_sequences/18\_FANPEZAQ\_CDS\_0018.fasta |

### Putative domain architecture and protein family

#### Search results (HHblits)^1^

|  |  |
| --- | --- |
| Domain family databases searched | Pfam, Ncbi-cd, Cath, Phrogs |
| Results, scheme(s)  (Top layers only; threshold 1.00e-03 (evalue)) | xml version="1.0" encoding="utf-8" standalone="no"?       2024-09-02T21:08:16.093537 image/svg+xml   Matplotlib v3.7.2, https://matplotlib.org/ |
| Results, table  (E-value ≤ 1.00e-03 (evalue)) | db \| id \| prob \| evalue \| pvalue \| score \| cols \| query \| query\_len \| template \| template\_len \| name \| description; pfam \| PF17541 \| 99.8 \| 3.7e-26 \| 8.1e-30 \| 231.7 \| 231 \| (229, 472) \| 475 \| (173, 422) \| 430 \| TssC \| Type VI secretion system, TssC, VipB; pfam \| PF10758 \| 99.5 \| 8.8e-19 \| 1.8e-22 \| 178.3 \| 175 \| (287, 469) \| 475 \| (244, 435) \| 436 \| DUF2586 \| Protein of unknown function (DUF2586); pfam \| PF17482 \| 98.0 \| 9.7e-10 \| 2.2e-13 \| 83.6 \| 97 \| (366, 465) \| 475 \| (3, 99) \| 100 \| Phage\_sheath\_1C \| Phage tail sheath C-terminal domain; pfam \| PF05943 \| 97.8 \| 3.7e-09 \| 7.6e-13 \| 100.2 \| 130 \| (229, 363) \| 475 \| (107, 301) \| 302 \| VipB \| EvpB/VC\_A0108, tail sheath N-terminal domain; pfam \| PF18945 \| 96.6 \| 2.5e-06 \| 5.2e-10 \| 68.2 \| 85 \| (371, 467) \| 475 \| (15, 110) \| 112 \| VipB\_2 \| EvpB/VC\_A0108, tail sheath gpW/gp25-like domain; cath \| 3foaB01 \| 99.9 \| 2e-27 \| 3.9e-31 \| 207.7 \| 199 \| (26, 327) \| 475 \| (2, 230) \| 231 \| Tail sheath protein gp18 \| CATHCODE: 3.40.50.11780 NAME: Tail sheath protein gp18. Chain: a, b, c, d. Fragment: deletion mutantgp18m: unp residues 1-510. Engineered: yes. Mutation: yes SOURCE: Enterobacteria phage t4. Bacteriophage t4. Organism\_taxid: 10665. Gene: 18, gb aaa32541. Expressed in: escherichia coli. Expression\_system\_taxid: 562. CLASS: Alpha Beta, ARCH: 3-Layer(aba) Sandwich, TOPOL: Rossmann fold, HOMOL: Rossmann fold; phrogs \| 23 \| 100.0 \| 2.6e-57 \| 3.2e-61 \| 471.2 \| 439 \| (3, 474) \| 475 \| (5, 523) \| 530 \| tail sheath \| tail sheath; Category: tail; NC\_017972\_p61; phrogs \| 279 \| 100.0 \| 9.5e-45 \| 1.2e-48 \| 368.3 \| 417 \| (4, 465) \| 475 \| (10, 472) \| 473 \| tail sheath \| tail sheath; Category: tail; p47709 VI\_05997; phrogs \| 13288 \| 99.9 \| 7.5e-29 \| 8.5e-33 \| 207.2 \| 195 \| (280, 474) \| 475 \| (4, 204) \| 208 \| NA \| NA; Category: unknown function; p370787 VI\_12002; phrogs \| 1036 \| 99.9 \| 2.6e-27 \| 3.1e-31 \| 231.4 \| 421 \| (9, 471) \| 475 \| (15, 518) \| 529 \| tail sheath \| tail sheath; Category: tail; p43549 VI\_12111; phrogs \| 17610 \| 99.8 \| 2.4e-24 \| 2.7e-28 \| 185.3 \| 227 \| (245, 472) \| 475 \| (2, 250) \| 257 \| tail sheath \| tail sheath; Category: tail; p306105 VI\_06232; phrogs \| 10581 \| 99.7 \| 3e-22 \| 3.4e-26 \| 196.7 \| 204 \| (265, 468) \| 475 \| (673, 887) \| 888 \| tail sheath \| tail sheath; Category: tail; NC\_019526\_p41; phrogs \| 907 \| 99.6 \| 6.2e-21 \| 7.3e-25 \| 181.1 \| 184 \| (287, 472) \| 475 \| (171, 368) \| 370 \| tail sheath \| tail sheath; Category: tail; NC\_009542\_p28; phrogs \| 10244 \| 99.6 \| 4.6e-20 \| 5.3e-24 \| 166.9 \| 323 \| (9, 467) \| 475 \| (11, 346) \| 348 \| tail sheath \| tail sheath; Category: tail; p54311 VI\_12213; phrogs \| 15814 \| 99.1 \| 3.8e-15 \| 4.2e-19 \| 138.1 \| 294 \| (162, 471) \| 475 \| (411, 730) \| 738 \| NA \| NA; Category: unknown function; p401786 VI\_06225; phrogs \| 16736 \| 99.1 \| 6.5e-15 \| 7.3e-19 \| 108.5 \| 100 \| (374, 473) \| 475 \| (2, 101) \| 108 \| NA \| NA; Category: unknown function; p404698 VI\_10663; phrogs \| 30392 \| 99.0 \| 2.8e-14 \| 3.1e-18 \| 131.6 \| 279 \| (162, 469) \| 475 \| (478, 769) \| 774 \| tail sheath \| tail sheath; Category: tail; NC\_029002\_p92; phrogs \| 16991 \| 98.9 \| 1.2e-13 \| 1.4e-17 \| 132.6 \| 168 \| (285, 465) \| 475 \| (545, 715) \| 717 \| tail sheath \| tail sheath; Category: tail; NC\_009760\_p40; phrogs \| 11242 \| 98.9 \| 2.8e-13 \| 3.2e-17 \| 101.6 \| 94 \| (2, 100) \| 475 \| (1, 95) \| 112 \| NA \| NA; Category: unknown function; p435652 VI\_11847; phrogs \| 1480 \| 98.8 \| 9e-13 \| 1.1e-16 \| 122.7 \| 313 \| (9, 464) \| 475 \| (5, 354) \| 355 \| tail sheath \| tail sheath; Category: tail; NC\_029116\_p9; phrogs \| 20240 \| 98.7 \| 1.8e-12 \| 2e-16 \| 108.3 \| 228 \| (2, 263) \| 475 \| (1, 249) \| 272 \| NA \| NA; Category: unknown function; p370788 VI\_12002; phrogs \| 34521 \| 98.7 \| 3.3e-12 \| 3.7e-16 \| 122.0 \| 204 \| (268, 473) \| 475 \| (734, 940) \| 944 \| tail sheath \| tail sheath; Category: tail; LC168164\_p135; phrogs \| 19666 \| 98.6 \| 7e-12 \| 7.8e-16 \| 120.1 \| 233 \| (227, 465) \| 475 \| (743, 985) \| 987 \| tail sheath \| tail sheath; Category: tail; MF360957\_p13; phrogs \| 165 \| 98.0 \| 1.3e-09 \| 1.6e-13 \| 106.5 \| 364 \| (9, 427) \| 475 \| (19, 433) \| 473 \| tail sheath \| tail sheath; Category: tail; p220370 VI\_01324; phrogs \| 12477 \| 97.0 \| 1e-06 \| 1.2e-10 \| 54.0 \| 48 \| (3, 50) \| 475 \| (1, 48) \| 48 \| NA \| NA; Category: unknown function; p311722 VI\_10874; phrogs \| 38392 \| 96.7 \| 3.8e-06 \| 4.3e-10 \| 70.7 \| 92 \| (2, 101) \| 475 \| (1, 94) \| 329 \| tail sheath \| tail sheath; Category: tail; MF001355\_p127; phrogs \| 20365 \| 95.6 \| 0.0001 \| 1.1e-08 \| 45.7 \| 50 \| (424, 473) \| 475 \| (2, 51) \| 58 \| NA \| NA; Category: unknown function; p352521 VI\_10849 |
| Top keywords  (threshold 1.00e-03 (evalue)) | **tail, sheath, Phage, EvpB, VC\_A0108, yes, t4, Rossmann, fold, VI\_12002** |
| Output files | ../../domain\_architecture/18\_FANPEZAQ\_CDS\_0018\_cath.hhr ../../domain\_architecture/18\_FANPEZAQ\_CDS\_0018\_merged.svg ../../domain\_architecture/18\_FANPEZAQ\_CDS\_0018\_ncbi-cd.hhr ../../domain\_architecture/18\_FANPEZAQ\_CDS\_0018\_pfam.hhr ../../domain\_architecture/18\_FANPEZAQ\_CDS\_0018\_phrogs.hhr |

### Identical protein sequences/structures

#### Search results

|  |  |
| --- | --- |
| Protein sequence databases searched | Pdb, Swissprot, Refseq |
| Identical proteins found | -- |
| Top keywords | -- |
| Output files | -- |

### Similar protein sequences/structures

#### Sequence similarity search results (HHblits)^1^

|  |  |
| --- | --- |
| Sequence databases searched | Uniclust, Pdb70 |
| Results, scheme(s)  (Top layers only, threshold 1.00e-03 (evalue)) | xml version="1.0" encoding="utf-8" standalone="no"?       2024-09-02T21:08:37.015043 image/svg+xml   Matplotlib v3.7.2, https://matplotlib.org/ |
| Results, table(s)  (threshold 1.00e-03 (evalue)) | db \| id \| prob \| evalue \| pvalue \| score \| cols \| query \| query\_len \| template \| template\_len \| name \| description; uniclust \| UniRef100\_A0A088FVH5 \| 100.0 \| 3.3e-79 \| 6.5e-85 \| 563.7 \| 392 \| (1, 475) \| 475 \| (21, 412) \| 424 \| Tail protein \| Tail protein; uniclust \| UniRef100\_A0A009Q2W1 \| 100.0 \| 2.3e-74 \| 4.5e-80 \| 555.2 \| 385 \| (1, 475) \| 475 \| (82, 468) \| 525 \| Phage tail sheath family protein \| Phage tail sheath family protein; uniclust \| UniRef100\_A0A095ULV3 \| 100.0 \| 1.4e-72 \| 2.7e-78 \| 527.7 \| 364 \| (1, 468) \| 475 \| (52, 437) \| 446 \| Phage tail sheath protein \| Phage tail sheath protein; uniclust \| UniRef100\_A0A2D7W1J0 \| 100.0 \| 6.9e-72 \| 1.3e-77 \| 520.6 \| 441 \| (1, 468) \| 475 \| (1, 507) \| 513 \| Phage tail protein \| Phage tail protein; uniclust \| UniRef100\_A0A024E974 \| 100.0 \| 8.4e-71 \| 1.7e-76 \| 523.6 \| 385 \| (1, 475) \| 475 \| (72, 456) \| 475 \| Major tail sheath protein \| Major tail sheath protein; uniclust \| UniRef100\_A0A068TBL7 \| 100.0 \| 1.9e-70 \| 3.7e-76 \| 519.2 \| 383 \| (6, 475) \| 475 \| (29, 415) \| 440 \| Phage tail sheath protein \| Phage tail sheath protein; uniclust \| UniRef100\_A0A094QDP2 \| 100.0 \| 2e-70 \| 3.9e-76 \| 517.9 \| 233 \| (229, 467) \| 475 \| (282, 530) \| 544 \| Putative phage tail sheath protein \| Putative phage tail sheath protein; uniclust \| UniRef100\_A0A061YI42 \| 100.0 \| 2.1e-70 \| 4.1e-76 \| 522.2 \| 381 \| (1, 475) \| 475 \| (1, 495) \| 507 \| Phage tail protein \| Phage tail protein; uniclust \| UniRef100\_A0A062UZH4 \| 100.0 \| 2.7e-69 \| 5.3e-75 \| 525.6 \| 356 \| (1, 469) \| 475 \| (125, 501) \| 536 \| Phage tail sheath protein FI \| Phage tail sheath protein FI; uniclust \| UniRef100\_A0A011NBA1 \| 100.0 \| 3.7e-67 \| 7.2e-73 \| 482.8 \| 384 \| (1, 474) \| 475 \| (2, 387) \| 400 \| Tail sheath protein \| Tail sheath protein; uniclust \| UniRef100\_A0A0P6X2M2 \| 100.0 \| 3.9e-67 \| 7.4e-73 \| 489.6 \| 235 \| (229, 469) \| 475 \| (290, 538) \| 552 \| Tail protein \| Tail protein; uniclust \| UniRef100\_A0A016QTR2 \| 100.0 \| 1.1e-66 \| 2.2e-72 \| 523.5 \| 447 \| (7, 470) \| 475 \| (74, 587) \| 629 \| Phage tail sheath protein \| Phage tail sheath protein; uniclust \| UniRef100\_A0A2X2RMN3 \| 100.0 \| 3.5e-66 \| 6.7e-72 \| 460.8 \| 375 \| (2, 474) \| 475 \| (1, 375) \| 383 \| Phage tail sheath protein \| Phage tail sheath protein; uniclust \| UniRef100\_A0A0K1RXU5 \| 100.0 \| 2.4e-65 \| 4.6e-71 \| 470.3 \| 361 \| (2, 467) \| 475 \| (1, 418) \| 432 \| Phage tail sheath protein FI \| Phage tail sheath protein FI; uniclust \| UniRef100\_A0A0B2YHC3 \| 100.0 \| 4.4e-65 \| 8.5e-71 \| 476.9 \| 360 \| (7, 467) \| 475 \| (31, 418) \| 458 \| Tail protein \| Tail protein; uniclust \| UniRef100\_A0A011NH28 \| 100.0 \| 2.6e-64 \| 5.2e-70 \| 501.9 \| 233 \| (230, 468) \| 475 \| (380, 623) \| 643 \| Major tail sheath protein \| Major tail sheath protein; uniclust \| UniRef100\_A0A059VPR9 \| 100.0 \| 3e-64 \| 5.8e-70 \| 459.2 \| 364 \| (7, 467) \| 475 \| (21, 422) \| 432 \| Putative phage tail sheath protein \| Putative phage tail sheath protein; uniclust \| UniRef100\_A0A011MIB3 \| 100.0 \| 5e-64 \| 9.7e-70 \| 482.3 \| 380 \| (1, 474) \| 475 \| (53, 525) \| 538 \| Phage tail protein \| Phage tail protein; uniclust \| UniRef100\_A0A1G6NJX4 \| 100.0 \| 8.8e-64 \| 1.7e-69 \| 479.3 \| 357 \| (5, 469) \| 475 \| (112, 486) \| 502 \| Phage tail sheath protein FI \| Phage tail sheath protein FI; uniclust \| UniRef100\_A0A0D0HFB9 \| 100.0 \| 1.3e-62 \| 2.5e-68 \| 443.6 \| 384 \| (1, 474) \| 475 \| (1, 386) \| 393 \| Phage tail sheath family protein \| Phage tail sheath family protein; uniclust \| UniRef100\_A0A1Q7VC09 \| 100.0 \| 4.2e-62 \| 8.2e-68 \| 477.1 \| 238 \| (225, 468) \| 475 \| (422, 673) \| 680 \| Phage tail protein \| Phage tail protein; uniclust \| UniRef100\_A0A0R3MVK4 \| 100.0 \| 5.3e-62 \| 9.8e-68 \| 433.5 \| 468 \| (1, 475) \| 475 \| (1, 469) \| 469 \| Phage tail protein \| Phage tail protein; uniclust \| UniRef100\_A0A1J5DEX6 \| 100.0 \| 7.4e-62 \| 1.4e-67 \| 462.9 \| 233 \| (229, 468) \| 475 \| (337, 579) \| 591 \| Tail sheath protein C-terminal domain-containing protein \| Tail sheath protein C-terminal domain-containing protein; uniclust \| UniRef100\_A0A099PD65 \| 100.0 \| 1.3e-60 \| 2.7e-66 \| 482.7 \| 237 \| (229, 471) \| 475 \| (461, 712) \| 745 \| Tail sheath protein C-terminal domain-containing protein \| Tail sheath protein C-terminal domain-containing protein; uniclust \| UniRef100\_A0A1E7Q659 \| 100.0 \| 4.2e-59 \| 7.9e-65 \| 412.6 \| 358 \| (2, 467) \| 475 \| (1, 375) \| 377 \| Phage tail protein \| Phage tail protein; uniclust \| UniRef100\_A0A090AMB4 \| 100.0 \| 9e-59 \| 1.8e-64 \| 467.2 \| 237 \| (227, 469) \| 475 \| (417, 673) \| 689 \| Putative phage tail sheath protein \| Putative phage tail sheath protein; uniclust \| UniRef100\_A0A014LUF2 \| 100.0 \| 5.2e-58 \| 1e-63 \| 457.6 \| 445 \| (2, 468) \| 475 \| (155, 677) \| 748 \| Tail protein \| Tail protein; uniclust \| UniRef100\_A0A0C1QXU3 \| 100.0 \| 1.1e-57 \| 2.3e-63 \| 466.5 \| 235 \| (229, 469) \| 475 \| (558, 804) \| 810 \| Tail protein \| Tail protein; uniclust \| UniRef100\_A0A059WKP9 \| 100.0 \| 1.2e-57 \| 2.5e-63 \| 445.8 \| 429 \| (1, 475) \| 475 \| (3, 488) \| 501 \| Tail sheath protein \| Tail sheath protein; uniclust \| UniRef100\_A0A060H1U5 \| 100.0 \| 3.1e-56 \| 5.6e-62 \| 384.5 \| 392 \| (2, 474) \| 475 \| (5, 396) \| 396 \| Tail protein \| Tail protein; uniclust \| UniRef100\_A0A011MDR6 \| 100.0 \| 3.7e-56 \| 7.5e-62 \| 453.7 \| 232 \| (231, 468) \| 475 \| (486, 730) \| 737 \| Major tail sheath protein \| Major tail sheath protein; uniclust \| UniRef100\_A0A022PIB5 \| 100.0 \| 7e-56 \| 1.3e-61 \| 394.9 \| 366 \| (19, 474) \| 475 \| (18, 388) \| 394 \| Phage tail sheath protein FI \| Phage tail sheath protein FI; uniclust \| UniRef100\_A0A1A7PR96 \| 100.0 \| 7.2e-56 \| 1.4e-61 \| 406.5 \| 429 \| (2, 473) \| 475 \| (1, 473) \| 478 \| Phage tail protein (Fragment) \| Phage tail protein (Fragment); uniclust \| UniRef100\_A0A0F0HJ39 \| 100.0 \| 9.2e-56 \| 1.7e-61 \| 403.6 \| 369 \| (1, 468) \| 475 \| (1, 495) \| 501 \| Phage tail protein (Fragment) \| Phage tail protein (Fragment); uniclust \| UniRef100\_A0A069PUW5 \| 100.0 \| 1.5e-55 \| 2.9e-61 \| 430.5 \| 234 \| (229, 468) \| 475 \| (427, 689) \| 701 \| Tail protein \| Tail protein; uniclust \| UniRef100\_A0A812WA44 \| 100.0 \| 2.9e-55 \| 5.2e-61 \| 404.1 \| 442 \| (2, 467) \| 475 \| (1, 513) \| 780 \| 18 protein \| 18 protein; uniclust \| UniRef100\_A0A085ART8 \| 100.0 \| 1e-54 \| 2e-60 \| 404.6 \| 368 \| (4, 471) \| 475 \| (51, 532) \| 546 \| Phage tail sheath monomer \| Phage tail sheath monomer; uniclust \| UniRef100\_A0A022PCH2 \| 100.0 \| 1.1e-54 \| 2.1e-60 \| 415.3 \| 348 \| (1, 469) \| 475 \| (60, 425) \| 449 \| Phage tail sheath protein FI \| Phage tail sheath protein FI; uniclust \| UniRef100\_A0A011PSV2 \| 100.0 \| 3.1e-54 \| 6.1e-60 \| 437.5 \| 181 \| (287, 468) \| 475 \| (412, 602) \| 768 \| Tail sheath protein \| Tail sheath protein; uniclust \| UniRef100\_A0A5A9EN65 \| 100.0 \| 6.6e-54 \| 1.2e-59 \| 394.8 \| 462 \| (2, 475) \| 475 \| (1, 617) \| 617 \| Phage tail protein \| Phage tail protein; uniclust \| UniRef100\_A0A1A9VKH7 \| 100.0 \| 9.6e-54 \| 1.8e-59 \| 380.4 \| 375 \| (4, 462) \| 475 \| (2, 383) \| 535 \| Phage tail protein \| Phage tail protein; uniclust \| UniRef100\_A0A1G3FX38 \| 100.0 \| 9.9e-54 \| 1.9e-59 \| 404.7 \| 358 \| (7, 468) \| 475 \| (77, 459) \| 472 \| Phage tail protein (Fragment) \| Phage tail protein (Fragment); uniclust \| UniRef100\_A0A1D2QMV4 \| 100.0 \| 1.7e-53 \| 3.2e-59 \| 386.2 \| 379 \| (1, 475) \| 475 \| (15, 395) \| 399 \| Phage tail protein \| Phage tail protein; uniclust \| UniRef100\_A0A2M9PCU5 \| 100.0 \| 3.1e-53 \| 5.8e-59 \| 371.2 \| 246 \| (230, 475) \| 475 \| (55, 301) \| 315 \| Phage tail protein (Fragment) \| Phage tail protein (Fragment); uniclust \| UniRef100\_A0A0P9N092 \| 100.0 \| 4.4e-53 \| 8.2e-59 \| 373.0 \| 385 \| (1, 474) \| 475 \| (29, 418) \| 421 \| Major tail sheath protein \| Major tail sheath protein; uniclust \| UniRef100\_A0A080LVP7 \| 100.0 \| 7.4e-53 \| 1.4e-58 \| 380.9 \| 233 \| (231, 470) \| 475 \| (114, 359) \| 372 \| Tail sheath protein \| Tail sheath protein; uniclust \| UniRef100\_A0A0A8WRE5 \| 100.0 \| 9.7e-53 \| 1.8e-58 \| 387.6 \| 353 \| (7, 470) \| 475 \| (11, 381) \| 521 \| Phage tail sheath protein \| Phage tail sheath protein; uniclust \| UniRef100\_A0A031JIL8 \| 100.0 \| 1.1e-52 \| 2.3e-58 \| 433.0 \| 235 \| (230, 470) \| 475 \| (520, 775) \| 782 \| Phage tail sheath protein \| Phage tail sheath protein; uniclust \| UniRef100\_A0A812QV43 \| 100.0 \| 1.7e-52 \| 3.1e-58 \| 396.7 \| 386 \| (2, 474) \| 475 \| (1, 390) \| 1104 \| GpFI protein \| GpFI protein; uniclust \| UniRef100\_A0A1F3S987 \| 100.0 \| 2e-52 \| 4e-58 \| 395.4 \| 355 \| (2, 467) \| 475 \| (1, 443) \| 447 \| Tail sheath protein C-terminal domain-containing protein \| Tail sheath protein C-terminal domain-containing protein; uniclust \| UniRef100\_A0A7R9HD16 \| 100.0 \| 2.4e-52 \| 4.4e-58 \| 367.9 \| 378 \| (2, 466) \| 475 \| (100, 484) \| 487 \| Phage tail sheath protein \| Phage tail sheath protein; uniclust \| UniRef100\_A0A011TTH8 \| 100.0 \| 9.1e-52 \| 1.8e-57 \| 395.0 \| 458 \| (2, 475) \| 475 \| (1, 476) \| 492 \| Tail sheath protein \| Tail sheath protein; uniclust \| UniRef100\_A0A2U3LUJ3 \| 100.0 \| 1.4e-51 \| 2.6e-57 \| 380.6 \| 234 \| (229, 468) \| 475 \| (206, 451) \| 477 \| Phage tail sheath protein \| Phage tail sheath protein; uniclust \| UniRef100\_A0A022PDU0 \| 100.0 \| 1.4e-51 \| 2.8e-57 \| 401.1 \| 177 \| (286, 468) \| 475 \| (272, 458) \| 520 \| Phage tail sheath protein FI \| Phage tail sheath protein FI; uniclust \| UniRef100\_A0A356K0G5 \| 100.0 \| 2.3e-51 \| 4.3e-57 \| 360.3 \| 296 \| (1, 396) \| 475 \| (1, 325) \| 325 \| Phage tail protein (Fragment) \| Phage tail protein (Fragment); uniclust \| UniRef100\_A0A1H3QMK3 \| 100.0 \| 3.3e-51 \| 6.2e-57 \| 392.6 \| 232 \| (230, 467) \| 475 \| (550, 805) \| 809 \| Uncharacterized protein \| Uncharacterized protein; uniclust \| UniRef100\_A0A0Q8LR77 \| 100.0 \| 3.4e-51 \| 6.5e-57 \| 401.5 \| 232 \| (230, 467) \| 475 \| (621, 867) \| 870 \| Phage tail protein \| Phage tail protein; uniclust \| UniRef100\_A0A1Q3SJK3 \| 100.0 \| 3.6e-51 \| 6.7e-57 \| 374.8 \| 233 \| (229, 467) \| 475 \| (364, 611) \| 615 \| Phage tail protein \| Phage tail protein; uniclust \| UniRef100\_A0A7R9HA11 \| 100.0 \| 6.3e-51 \| 1.2e-56 \| 392.6 \| 364 \| (15, 467) \| 475 \| (627, 991) \| 1133 \| Uncharacterized protein \| Uncharacterized protein; uniclust \| UniRef100\_A0A0F3IT60 \| 100.0 \| 2e-50 \| 3.9e-56 \| 380.9 \| 362 \| (5, 467) \| 475 \| (5, 500) \| 504 \| Phage tail protein (Fragment) \| Phage tail protein (Fragment); uniclust \| UniRef100\_A0A2V9GXL0 \| 100.0 \| 2.4e-50 \| 4.6e-56 \| 388.0 \| 234 \| (229, 468) \| 475 \| (530, 782) \| 786 \| Phage tail protein \| Phage tail protein; uniclust \| UniRef100\_A0A1A1YQW6 \| 100.0 \| 1.7e-49 \| 3.3e-55 \| 402.4 \| 234 \| (229, 468) \| 475 \| (859, 1114) \| 1121 \| Phage tail protein \| Phage tail protein; uniclust \| UniRef100\_A0A5Q4D9T1 \| 100.0 \| 1.8e-49 \| 3.4e-55 \| 360.3 \| 359 \| (7, 468) \| 475 \| (14, 401) \| 436 \| Phage tail sheath family protein \| Phage tail sheath family protein; uniclust \| UniRef100\_A0A061JGM3 \| 100.0 \| 3.9e-49 \| 7.3e-55 \| 341.2 \| 304 \| (2, 397) \| 475 \| (1, 305) \| 306 \| Major tail sheath protein \| Major tail sheath protein; uniclust \| UniRef100\_A0A2P8VUG9 \| 100.0 \| 7.1e-49 \| 1.3e-54 \| 364.5 \| 233 \| (229, 467) \| 475 \| (293, 536) \| 541 \| Phage tail protein \| Phage tail protein; uniclust \| UniRef100\_A0A519FU64 \| 100.0 \| 1.8e-48 \| 3.4e-54 \| 343.4 \| 305 \| (86, 475) \| 475 \| (13, 318) \| 324 \| Phage tail sheath family protein (Fragment) \| Phage tail sheath family protein (Fragment); uniclust \| UniRef100\_A0A0B5FRP8 \| 100.0 \| 2.9e-48 \| 5.4e-54 \| 362.1 \| 423 \| (2, 468) \| 475 \| (1, 501) \| 513 \| Tail protein \| Tail protein; uniclust \| UniRef100\_A0A1Q7MAI6 \| 100.0 \| 3.9e-48 \| 7.2e-54 \| 343.8 \| 369 \| (3, 464) \| 475 \| (2, 381) \| 515 \| Phage tail protein \| Phage tail protein; uniclust \| UniRef100\_A0A4Q5SQU1 \| 100.0 \| 3.8e-48 \| 7.4e-54 \| 344.1 \| 234 \| (230, 469) \| 475 \| (47, 304) \| 307 \| Phage tail sheath family protein (Fragment) \| Phage tail sheath family protein (Fragment); uniclust \| UniRef100\_A0A6J5N355 \| 100.0 \| 3.9e-48 \| 7.5e-54 \| 367.6 \| 433 \| (2, 469) \| 475 \| (1, 503) \| 511 \| Tail sheath protein \| Tail sheath protein; uniclust \| UniRef100\_A0A7J6YK69 \| 100.0 \| 5.1e-48 \| 9.4e-54 \| 383.6 \| 384 \| (2, 468) \| 475 \| (1278, 1668) \| 2523 \| Uncharacterized protein \| Uncharacterized protein; uniclust \| UniRef100\_A0A2W5Z8T4 \| 100.0 \| 1.4e-47 \| 2.5e-53 \| 343.7 \| 347 \| (6, 468) \| 475 \| (6, 376) \| 565 \| AAA+ ATPase domain-containing protein \| AAA+ ATPase domain-containing protein; uniclust \| UniRef100\_A0A367ZMD7 \| 100.0 \| 1.4e-47 \| 2.7e-53 \| 358.7 \| 235 \| (228, 468) \| 475 \| (356, 607) \| 621 \| Phage tail sheath protein FI \| Phage tail sheath protein FI; uniclust \| UniRef100\_A0A2X3JUE5 \| 100.0 \| 2.2e-47 \| 4.3e-53 \| 355.1 \| 343 \| (2, 435) \| 475 \| (1, 349) \| 394 \| Major tail sheath protein FI \| Major tail sheath protein FI; uniclust \| UniRef100\_A0A0D6L3S9 \| 100.0 \| 6.8e-47 \| 1.3e-52 \| 351.5 \| 368 \| (2, 466) \| 475 \| (3, 466) \| 470 \| Uncharacterized protein (Fragment) \| Uncharacterized protein (Fragment); uniclust \| UniRef100\_A0A0W8I564 \| 100.0 \| 1.3e-46 \| 2.5e-52 \| 354.8 \| 334 \| (18, 469) \| 475 \| (25, 377) \| 508 \| Tail sheath protein subtilisin-like domain-containing protein \| Tail sheath protein subtilisin-like domain-containing protein; uniclust \| UniRef100\_A0A2E2NTG5 \| 100.0 \| 1.6e-46 \| 2.9e-52 \| 352.6 \| 229 \| (230, 468) \| 475 \| (329, 593) \| 600 \| Phage tail protein \| Phage tail protein; uniclust \| UniRef100\_A0A956NBW1 \| 100.0 \| 3.3e-46 \| 6.1e-52 \| 340.0 \| 336 \| (18, 468) \| 475 \| (53, 405) \| 521 \| Phage tail sheath subtilisin-like domain-containing protein \| Phage tail sheath subtilisin-like domain-containing protein; uniclust \| UniRef100\_A0A074M4V8 \| 100.0 \| 7e-46 \| 1.4e-51 \| 356.3 \| 409 \| (7, 468) \| 475 \| (12, 470) \| 481 \| Phage tail protein \| Phage tail protein; uniclust \| UniRef100\_A0A1D8TT95 \| 100.0 \| 1.1e-45 \| 2e-51 \| 352.3 \| 232 \| (229, 466) \| 475 \| (536, 788) \| 792 \| Phage tail protein \| Phage tail protein; uniclust \| UniRef100\_A0A0J0Y2J6 \| 100.0 \| 1.3e-45 \| 2.5e-51 \| 358.4 \| 179 \| (287, 466) \| 475 \| (516, 704) \| 708 \| Phage tail sheath protein \| Phage tail sheath protein; uniclust \| UniRef100\_A0A0Q8XJ68 \| 100.0 \| 3.2e-45 \| 6.2e-51 \| 360.8 \| 234 \| (229, 471) \| 475 \| (515, 767) \| 783 \| Tail sheath protein C-terminal domain-containing protein \| Tail sheath protein C-terminal domain-containing protein; uniclust \| UniRef100\_A0A0P0QJ01 \| 100.0 \| 5.3e-45 \| 1e-50 \| 342.5 \| 376 \| (7, 469) \| 475 \| (16, 416) \| 433 \| Phage tail sheath family protein \| Phage tail sheath family protein; uniclust \| UniRef100\_A0A350QJ90 \| 100.0 \| 1.3e-44 \| 2.5e-50 \| 327.4 \| 233 \| (230, 468) \| 475 \| (119, 369) \| 409 \| Phage tail sheath family protein (Fragment) \| Phage tail sheath family protein (Fragment); uniclust \| UniRef100\_A0A014LI00 \| 100.0 \| 2.3e-44 \| 4.5e-50 \| 342.4 \| 420 \| (7, 472) \| 475 \| (31, 524) \| 528 \| Tail protein \| Tail protein; uniclust \| UniRef100\_A0A1U7N9U3 \| 100.0 \| 5.5e-44 \| 1e-49 \| 326.7 \| 378 \| (1, 466) \| 475 \| (1, 476) \| 480 \| Phage tail protein \| Phage tail protein; uniclust \| UniRef100\_A0A410G7I7 \| 100.0 \| 5.7e-44 \| 1.1e-49 \| 345.3 \| 231 \| (231, 468) \| 475 \| (533, 785) \| 789 \| Phage tail sheath family protein \| Phage tail sheath family protein; uniclust \| UniRef100\_A0A2V9K9Q7 \| 100.0 \| 8.7e-44 \| 1.7e-49 \| 335.5 \| 235 \| (229, 470) \| 475 \| (242, 498) \| 500 \| Phage tail protein \| Phage tail protein; uniclust \| UniRef100\_A0A0T9RBT0 \| 100.0 \| 2.5e-43 \| 4.7e-49 \| 301.3 \| 236 \| (240, 475) \| 475 \| (3, 240) \| 247 \| Phage tail sheath monomer \| Phage tail sheath monomer; uniclust \| UniRef100\_A0A0N0JDZ8 \| 100.0 \| 1e-42 \| 2e-48 \| 349.9 \| 236 \| (229, 470) \| 475 \| (420, 667) \| 729 \| Tail sheath protein C-terminal domain-containing protein \| Tail sheath protein C-terminal domain-containing protein; uniclust \| UniRef100\_A0A2N5D506 \| 100.0 \| 1.4e-42 \| 2.6e-48 \| 309.1 \| 346 \| (1, 466) \| 475 \| (1, 367) \| 371 \| Phage tail sheath family protein \| Phage tail sheath family protein; uniclust \| UniRef100\_A0A1A9VZ70 \| 100.0 \| 2.3e-42 \| 4.3e-48 \| 310.2 \| 289 \| (101, 474) \| 475 \| (52, 354) \| 359 \| Tail sheath protein subtilisin-like domain-containing protein \| Tail sheath protein subtilisin-like domain-containing protein; uniclust \| UniRef100\_A0A0L9YBE2 \| 100.0 \| 2.3e-42 \| 4.5e-48 \| 333.7 \| 377 \| (1, 475) \| 475 \| (1, 490) \| 497 \| Phage tail sheath protein \| Phage tail sheath protein; uniclust \| UniRef100\_A0A952H001 \| 100.0 \| 3.3e-42 \| 6.1e-48 \| 307.8 \| 233 \| (229, 467) \| 475 \| (218, 461) \| 466 \| Phage tail sheath family protein \| Phage tail sheath family protein; uniclust \| UniRef100\_A0A031FQW7 \| 100.0 \| 4.4e-42 \| 8.5e-48 \| 322.4 \| 366 \| (7, 475) \| 475 \| (14, 390) \| 414 \| Phage tail sheath monomer \| Phage tail sheath monomer; uniclust \| UniRef100\_A0A0S7Z2I8 \| 100.0 \| 7.2e-42 \| 1.4e-47 \| 340.4 \| 437 \| (6, 468) \| 475 \| (80, 582) \| 655 \| Tail sheath protein subtilisin-like domain-containing protein \| Tail sheath protein subtilisin-like domain-containing protein; uniclust \| UniRef100\_A0A5B7V907 \| 100.0 \| 7.8e-42 \| 1.4e-47 \| 302.7 \| 232 \| (231, 468) \| 475 \| (231, 483) \| 488 \| Phage tail sheath protein \| Phage tail sheath protein; uniclust \| UniRef100\_A0A0Q6AB86 \| 100.0 \| 1.5e-41 \| 2.9e-47 \| 328.5 \| 180 \| (286, 466) \| 475 \| (451, 640) \| 644 \| VWFA domain-containing protein \| VWFA domain-containing protein; uniclust \| UniRef100\_A0A2E2NTG3 \| 100.0 \| 1.9e-41 \| 3.6e-47 \| 311.5 \| 228 \| (230, 468) \| 475 \| (149, 395) \| 401 \| Phage tail sheath family protein \| Phage tail sheath family protein; uniclust \| UniRef100\_A0A143G8E1 \| 100.0 \| 3.7e-41 \| 7.1e-47 \| 294.1 \| 274 \| (1, 364) \| 475 \| (3, 283) \| 285 \| Phage tail protein \| Phage tail protein; uniclust \| UniRef100\_A0A1B7IQW9 \| 100.0 \| 6.3e-41 \| 1.2e-46 \| 308.6 \| 248 \| (227, 474) \| 475 \| (395, 644) \| 649 \| Phage tail sheath monomer \| Phage tail sheath monomer; uniclust \| UniRef100\_A0A1B1NT66 \| 100.0 \| 2.2e-40 \| 4.1e-46 \| 300.9 \| 376 \| (4, 474) \| 475 \| (1, 426) \| 434 \| Putative prophage major tail sheath protein \| Putative prophage major tail sheath protein; uniclust \| UniRef100\_A0A090GUB3 \| 100.0 \| 4.6e-40 \| 8.8e-46 \| 301.7 \| 395 \| (2, 474) \| 475 \| (1, 420) \| 435 \| Phage tail protein \| Phage tail protein; uniclust \| UniRef100\_A0A010SSQ8 \| 100.0 \| 8e-40 \| 1.6e-45 \| 319.5 \| 430 \| (7, 466) \| 475 \| (71, 553) \| 609 \| Phage tail protein \| Phage tail protein; uniclust \| UniRef100\_A0A4V1UFD5 \| 100.0 \| 1.1e-39 \| 2.1e-45 \| 298.7 \| 236 \| (229, 470) \| 475 \| (204, 461) \| 464 \| Phage tail sheath family protein \| Phage tail sheath family protein; uniclust \| UniRef100\_A0A1C4YCD9 \| 100.0 \| 1.3e-39 \| 2.4e-45 \| 284.2 \| 198 \| (268, 471) \| 475 \| (16, 233) \| 270 \| Phage tail sheath protein \| Phage tail sheath protein; uniclust \| UniRef100\_A0A3M2D088 \| 100.0 \| 1.4e-39 \| 2.7e-45 \| 266.8 \| 176 \| (289, 470) \| 475 \| (1, 184) \| 193 \| Phage tail sheath family protein (Fragment) \| Phage tail sheath family protein (Fragment); uniclust \| UniRef100\_A0A2A7RYU0 \| 100.0 \| 1.5e-39 \| 2.8e-45 \| 281.3 \| 310 \| (3, 402) \| 475 \| (1, 320) \| 339 \| Tail sheath protein subtilisin-like domain-containing protein \| Tail sheath protein subtilisin-like domain-containing protein; uniclust \| UniRef100\_A0A077KVI5 \| 100.0 \| 2.3e-39 \| 4.3e-45 \| 278.3 \| 239 \| (230, 473) \| 475 \| (4, 244) \| 262 \| Tail sheath protein subtilisin-like domain-containing protein \| Tail sheath protein subtilisin-like domain-containing protein; uniclust \| UniRef100\_A0A1C3NFI6 \| 100.0 \| 2.4e-39 \| 4.8e-45 \| 312.4 \| 238 \| (225, 468) \| 475 \| (265, 515) \| 530 \| Phage tail sheath protein FI \| Phage tail sheath protein FI; uniclust \| UniRef100\_A0A0Q8AN78 \| 100.0 \| 2.6e-39 \| 4.9e-45 \| 289.8 \| 388 \| (1, 474) \| 475 \| (1, 400) \| 413 \| Phage tail protein \| Phage tail protein; uniclust \| UniRef100\_A0A1A7R3U7 \| 100.0 \| 3.4e-39 \| 6.4e-45 \| 303.9 \| 372 \| (2, 468) \| 475 \| (1, 463) \| 727 \| Phage tail sheath protein FI \| Phage tail sheath protein FI; uniclust \| UniRef100\_A0A1V5CVP8 \| 100.0 \| 6.5e-39 \| 1.2e-44 \| 292.7 \| 234 \| (228, 467) \| 475 \| (249, 502) \| 507 \| Phage tail sheath protein \| Phage tail sheath protein; uniclust \| UniRef100\_A0A963BNV3 \| 100.0 \| 7e-39 \| 1.3e-44 \| 282.7 \| 233 \| (229, 467) \| 475 \| (218, 461) \| 464 \| Phage tail sheath family protein \| Phage tail sheath family protein; uniclust \| UniRef100\_A0A0A8H9Y0 \| 100.0 \| 7.6e-39 \| 1.5e-44 \| 294.6 \| 366 \| (5, 474) \| 475 \| (3, 377) \| 385 \| Major tail sheath protein \| Major tail sheath protein; uniclust \| UniRef100\_A0A1Q7BHJ3 \| 100.0 \| 8.2e-39 \| 1.5e-44 \| 285.6 \| 334 \| (30, 464) \| 475 \| (2, 353) \| 516 \| Phage tail protein \| Phage tail protein; uniclust \| UniRef100\_A0A136QAK0 \| 100.0 \| 1.5e-38 \| 2.8e-44 \| 262.3 \| 243 \| (233, 475) \| 475 \| (3, 246) \| 249 \| Tail protein (Fragment) \| Tail protein (Fragment); uniclust \| UniRef100\_A0A2D5XZB8 \| 100.0 \| 1.6e-38 \| 3.1e-44 \| 304.8 \| 229 \| (235, 469) \| 475 \| (416, 670) \| 677 \| Tail sheath protein C-terminal domain-containing protein \| Tail sheath protein C-terminal domain-containing protein; uniclust \| UniRef100\_A0A150S1N9 \| 100.0 \| 1.7e-38 \| 3.2e-44 \| 292.8 \| 399 \| (7, 471) \| 475 \| (11, 452) \| 462 \| Tail sheath protein subtilisin-like domain-containing protein \| Tail sheath protein subtilisin-like domain-containing protein; uniclust \| UniRef100\_A0A8S5LXM1 \| 100.0 \| 1.8e-38 \| 3.4e-44 \| 289.1 \| 431 \| (2, 467) \| 475 \| (1, 496) \| 500 \| Tail sheath protein \| Tail sheath protein; uniclust \| UniRef100\_A0A7V9BVW5 \| 100.0 \| 1.9e-38 \| 3.6e-44 \| 291.3 \| 227 \| (232, 468) \| 475 \| (326, 587) \| 593 \| Phage tail sheath family protein \| Phage tail sheath family protein; uniclust \| UniRef100\_A0A0M3V6G8 \| 100.0 \| 1.9e-38 \| 3.7e-44 \| 299.2 \| 414 \| (7, 466) \| 475 \| (17, 476) \| 509 \| Tail sheath protein subtilisin-like domain-containing protein \| Tail sheath protein subtilisin-like domain-containing protein; uniclust \| UniRef100\_A0A433VYQ0 \| 100.0 \| 2.9e-38 \| 5.3e-44 \| 303.0 \| 231 \| (231, 467) \| 475 \| (893, 1149) \| 1173 \| Inter-alpha-trypsin inhibitor domain-containing protein \| Inter-alpha-trypsin inhibitor domain-containing protein; uniclust \| UniRef100\_A0A134CAP7 \| 100.0 \| 5.6e-38 \| 1.1e-43 \| 295.0 \| 281 \| (101, 475) \| 475 \| (126, 425) \| 430 \| Phage tail protein (Fragment) \| Phage tail protein (Fragment); uniclust \| UniRef100\_A0A1X7M8N6 \| 100.0 \| 7.8e-38 \| 1.5e-43 \| 268.6 \| 227 \| (1, 315) \| 475 \| (1, 228) \| 231 \| Phage major tail sheath protein \| Phage major tail sheath protein; uniclust \| UniRef100\_A0A7C3H8R7 \| 100.0 \| 9.4e-38 \| 1.7e-43 \| 281.0 \| 233 \| (229, 467) \| 475 \| (247, 499) \| 504 \| Phage tail sheath family protein (Fragment) \| Phage tail sheath family protein (Fragment); uniclust \| UniRef100\_A0A0M2S0L1 \| 100.0 \| 9.6e-38 \| 1.9e-43 \| 311.0 \| 237 \| (228, 469) \| 475 \| (523, 779) \| 786 \| Tail sheath protein subtilisin-like domain-containing protein \| Tail sheath protein subtilisin-like domain-containing protein; uniclust \| UniRef100\_A0A0Q4LKW2 \| 100.0 \| 1.7e-37 \| 3.2e-43 \| 260.7 \| 182 \| (286, 468) \| 475 \| (15, 206) \| 213 \| Tail sheath protein C-terminal domain-containing protein \| Tail sheath protein C-terminal domain-containing protein; uniclust \| UniRef100\_A0A5D0URY7 \| 100.0 \| 4.3e-37 \| 8e-43 \| 290.0 \| 225 \| (235, 467) \| 475 \| (556, 800) \| 804 \| Phage tail sheath family protein \| Phage tail sheath family protein; uniclust \| UniRef100\_A0A7Y6Z423 \| 100.0 \| 5e-37 \| 9.3e-43 \| 283.1 \| 233 \| (229, 467) \| 475 \| (295, 553) \| 557 \| Phage tail sheath family protein \| Phage tail sheath family protein; uniclust \| UniRef100\_A0A3N5KUP2 \| 100.0 \| 7.6e-37 \| 1.4e-42 \| 253.9 \| 190 \| (269, 465) \| 475 \| (4, 201) \| 214 \| Phage tail protein (Fragment) \| Phage tail protein (Fragment); uniclust \| UniRef100\_A0A285NJC0 \| 100.0 \| 9.1e-37 \| 1.7e-42 \| 268.1 \| 374 \| (3, 474) \| 475 \| (1, 391) \| 399 \| Tail sheath protein subtilisin-like domain-containing protein \| Tail sheath protein subtilisin-like domain-containing protein; uniclust \| UniRef100\_A0A0A7HAU9 \| 100.0 \| 9.5e-37 \| 1.9e-42 \| 293.1 \| 426 \| (1, 466) \| 475 \| (1, 482) \| 490 \| Tail sheath protein \| Tail sheath protein; uniclust \| UniRef100\_A0A2S9WYU3 \| 100.0 \| 1e-36 \| 2e-42 \| 250.1 \| 157 \| (306, 468) \| 475 \| (1, 165) \| 184 \| Phage tail protein (Fragment) \| Phage tail protein (Fragment); uniclust \| UniRef100\_A0A6N7C2F0 \| 100.0 \| 2e-36 \| 3.7e-42 \| 263.4 \| 233 \| (231, 469) \| 475 \| (25, 269) \| 286 \| Uncharacterized protein \| Uncharacterized protein; uniclust \| UniRef100\_A0A2S7K9H5 \| 100.0 \| 3.6e-36 \| 6.7e-42 \| 266.5 \| 345 \| (7, 465) \| 475 \| (106, 469) \| 473 \| Phage tail protein \| Phage tail protein; uniclust \| UniRef100\_A0A2G6CJB9 \| 100.0 \| 4.4e-36 \| 8.3e-42 \| 277.4 \| 244 \| (226, 473) \| 475 \| (203, 474) \| 481 \| Phage tail protein \| Phage tail protein; uniclust \| UniRef100\_A0A6J7X5P4 \| 100.0 \| 4.9e-36 \| 9e-42 \| 272.9 \| 232 \| (231, 468) \| 475 \| (345, 604) \| 610 \| Phage tail sheath C-terminal domain containing protein \| Phage tail sheath C-terminal domain containing protein; uniclust \| UniRef100\_A0A0F9ST52 \| 100.0 \| 1.1e-35 \| 2.1e-41 \| 295.7 \| 242 \| (229, 474) \| 475 \| (517, 786) \| 796 \| Tail sheath protein C-terminal domain-containing protein \| Tail sheath protein C-terminal domain-containing protein; uniclust \| UniRef100\_A0A1S6HVK6 \| 100.0 \| 1.3e-35 \| 2.5e-41 \| 261.9 \| 180 \| (286, 467) \| 475 \| (77, 266) \| 279 \| Phage tail sheath protein \| Phage tail sheath protein; uniclust \| UniRef100\_A0A6G9ZDK1 \| 100.0 \| 1.5e-35 \| 2.8e-41 \| 273.1 \| 230 \| (231, 466) \| 475 \| (447, 691) \| 695 \| Phage tail protein \| Phage tail protein; uniclust \| UniRef100\_A0A1C3FCE3 \| 100.0 \| 1.5e-35 \| 2.8e-41 \| 250.7 \| 209 \| (266, 474) \| 475 \| (11, 225) \| 232 \| Phage tail sheath monomer \| Phage tail sheath monomer; uniclust \| UniRef100\_A0A6V8KE03 \| 100.0 \| 2e-35 \| 3.8e-41 \| 260.4 \| 270 \| (2, 388) \| 475 \| (9, 343) \| 345 \| Tail sheath protein subtilisin-like domain-containing protein \| Tail sheath protein subtilisin-like domain-containing protein; uniclust \| UniRef100\_A0A3A0B079 \| 100.0 \| 2e-35 \| 3.8e-41 \| 270.2 \| 236 \| (229, 470) \| 475 \| (257, 510) \| 523 \| Phage tail sheath family protein (Fragment) \| Phage tail sheath family protein (Fragment); uniclust \| UniRef100\_UPI001E51F530 \| 100.0 \| 3.6e-35 \| 6.6e-41 \| 271.0 \| 343 \| (16, 469) \| 475 \| (238, 609) \| 646 \| phage tail sheath subtilisin-like domain-containing protein \| phage tail sheath subtilisin-like domain-containing protein; uniclust \| UniRef100\_A0A1Z4RAC9 \| 100.0 \| 5.1e-35 \| 9.8e-41 \| 268.8 \| 360 \| (3, 466) \| 475 \| (8, 384) \| 398 \| Phage tail sheath protein fi-like protein \| Phage tail sheath protein fi-like protein; uniclust \| UniRef100\_A0A1Q7YH55 \| 100.0 \| 5.3e-35 \| 9.9e-41 \| 266.9 \| 433 \| (7, 464) \| 475 \| (10, 494) \| 508 \| Tail sheath protein subtilisin-like domain-containing protein \| Tail sheath protein subtilisin-like domain-containing protein; uniclust \| UniRef100\_A0A2K8ZCA4 \| 100.0 \| 5.9e-35 \| 1.2e-40 \| 281.9 \| 197 \| (270, 467) \| 475 \| (301, 509) \| 514 \| Phage tail protein \| Phage tail protein; uniclust \| UniRef100\_A0A518BPN9 \| 100.0 \| 8.9e-35 \| 1.6e-40 \| 270.0 \| 232 \| (229, 466) \| 475 \| (489, 744) \| 750 \| Phage tail sheath protein \| Phage tail sheath protein; uniclust \| UniRef100\_A0A0A0YVK8 \| 100.0 \| 1.3e-34 \| 2.5e-40 \| 279.9 \| 235 \| (229, 469) \| 475 \| (307, 559) \| 575 \| Tail sheath protein \| Tail sheath protein; uniclust \| UniRef100\_A0A7Z2GDE3 \| 100.0 \| 1.4e-34 \| 2.7e-40 \| 282.0 \| 232 \| (230, 467) \| 475 \| (717, 979) \| 983 \| BIG2 domain-containing protein \| BIG2 domain-containing protein; uniclust \| UniRef100\_A0A8X6MUK6 \| 100.0 \| 2.6e-34 \| 4.7e-40 \| 256.5 \| 212 \| (230, 445) \| 475 \| (45, 256) \| 498 \| Putative prophage major tail sheath protein \| Putative prophage major tail sheath protein; uniclust \| UniRef100\_A0A0T7DUW1 \| 100.0 \| 3.1e-34 \| 5.8e-40 \| 272.8 \| 232 \| (242, 474) \| 475 \| (352, 604) \| 612 \| Putative bacteriophage tail sheath protein \| Putative bacteriophage tail sheath protein; uniclust \| UniRef100\_A0A2E7CJ71 \| 100.0 \| 3.8e-34 \| 7.1e-40 \| 271.9 \| 195 \| (267, 467) \| 475 \| (426, 635) \| 643 \| Phage tail sheath family protein \| Phage tail sheath family protein; uniclust \| UniRef100\_A0A1W0CRX8 \| 100.0 \| 6.1e-34 \| 1.2e-39 \| 267.1 \| 176 \| (287, 468) \| 475 \| (375, 560) \| 562 \| Phage tail protein \| Phage tail protein; uniclust \| UniRef100\_A0A5N8UQV4 \| 100.0 \| 8.5e-34 \| 1.6e-39 \| 247.6 \| 230 \| (231, 468) \| 475 \| (110, 366) \| 367 \| Phage tail sheath family protein \| Phage tail sheath family protein; uniclust \| UniRef100\_A0A2S2DPT2 \| 100.0 \| 8.6e-34 \| 1.6e-39 \| 265.7 \| 362 \| (7, 466) \| 475 \| (296, 819) \| 823 \| Tail sheath protein C-terminal domain-containing protein \| Tail sheath protein C-terminal domain-containing protein; uniclust \| UniRef100\_A0A2A8B6F9 \| 100.0 \| 1.6e-33 \| 3.1e-39 \| 226.4 \| 168 \| (295, 468) \| 475 \| (1, 177) \| 181 \| Phage tail sheath protein \| Phage tail sheath protein; uniclust \| UniRef100\_UPI00035D90AA \| 100.0 \| 3.5e-33 \| 6.6e-39 \| 271.1 \| 231 \| (231, 467) \| 475 \| (157, 427) \| 837 \| phage tail sheath C-terminal domain-containing protein \| phage tail sheath C-terminal domain-containing protein; uniclust \| UniRef100\_A0A126V0R7 \| 100.0 \| 4.8e-33 \| 9.1e-39 \| 257.6 \| 225 \| (231, 465) \| 475 \| (264, 505) \| 511 \| Phage tail protein \| Phage tail protein; uniclust \| UniRef100\_A0A3S4DC63 \| 100.0 \| 8.4e-33 \| 1.6e-38 \| 228.1 \| 196 \| (268, 468) \| 475 \| (7, 213) \| 228 \| Phage tail sheath protein \| Phage tail sheath protein; uniclust \| UniRef100\_A0A838FVS3 \| 100.0 \| 1e-32 \| 1.9e-38 \| 253.3 \| 333 \| (26, 466) \| 475 \| (32, 382) \| 465 \| Uncharacterized protein \| Uncharacterized protein; uniclust \| UniRef100\_A0A1M3D377 \| 100.0 \| 1.2e-32 \| 2.3e-38 \| 246.9 \| 354 \| (5, 461) \| 475 \| (30, 428) \| 430 \| Tail sheath protein C-terminal domain-containing protein \| Tail sheath protein C-terminal domain-containing protein; uniclust \| UniRef100\_A0A6J5S5A9 \| 100.0 \| 1.3e-32 \| 2.5e-38 \| 256.9 \| 231 \| (230, 466) \| 475 \| (532, 780) \| 783 \| Tail sheath protein \| Tail sheath protein; uniclust \| UniRef100\_A0A3N5DYH3 \| 100.0 \| 1.4e-32 \| 2.6e-38 \| 258.8 \| 180 \| (286, 466) \| 475 \| (325, 514) \| 518 \| Phage tail sheath family protein \| Phage tail sheath family protein; uniclust \| UniRef100\_A0A4Q6CEC2 \| 100.0 \| 1.6e-32 \| 3.1e-38 \| 233.4 \| 237 \| (230, 474) \| 475 \| (7, 245) \| 249 \| Phage tail protein (Fragment) \| Phage tail protein (Fragment); uniclust \| UniRef100\_A0A385ISI4 \| 100.0 \| 1.9e-32 \| 3.7e-38 \| 270.9 \| 236 \| (229, 470) \| 475 \| (359, 610) \| 622 \| Putative tail sheath protein \| Putative tail sheath protein; uniclust \| UniRef100\_A0A7K2KKT6 \| 99.9 \| 3.3e-32 \| 6e-38 \| 229.6 \| 167 \| (231, 403) \| 475 \| (131, 305) \| 306 \| Phage tail sheath family protein (Fragment) \| Phage tail sheath family protein (Fragment); uniclust \| UniRef100\_A0A2D6SUT0 \| 99.9 \| 7.2e-32 \| 1.3e-37 \| 247.8 \| 224 \| (230, 459) \| 475 \| (226, 470) \| 553 \| Phage tail protein \| Phage tail protein; uniclust \| UniRef100\_A0A353GA80 \| 99.9 \| 8.2e-32 \| 1.5e-37 \| 233.0 \| 292 \| (2, 381) \| 475 \| (1, 323) \| 325 \| Phage tail protein (Fragment) \| Phage tail protein (Fragment); uniclust \| UniRef100\_A0A925M9A4 \| 99.9 \| 9.8e-32 \| 1.8e-37 \| 229.7 \| 183 \| (286, 469) \| 475 \| (72, 264) \| 273 \| Phage tail sheath family protein \| Phage tail sheath family protein; uniclust \| UniRef100\_A0A0N7C9H8 \| 99.9 \| 1.2e-31 \| 2.3e-37 \| 251.4 \| 233 \| (231, 469) \| 475 \| (161, 411) \| 427 \| Tail sheath protein \| Tail sheath protein; uniclust \| UniRef100\_A0A2X2BJR7 \| 99.9 \| 2.5e-31 \| 4.8e-37 \| 223.4 \| 221 \| (115, 421) \| 475 \| (18, 240) \| 241 \| Major tail sheath protein \| Major tail sheath protein; uniclust \| UniRef100\_A0A954RU13 \| 99.9 \| 2.6e-31 \| 4.8e-37 \| 243.4 \| 235 \| (232, 467) \| 475 \| (360, 624) \| 627 \| Phage tail sheath family protein \| Phage tail sheath family protein; uniclust \| UniRef100\_A0A662Q4X3 \| 99.9 \| 6.4e-31 \| 1.2e-36 \| 236.7 \| 190 \| (267, 468) \| 475 \| (312, 511) \| 526 \| Uncharacterized protein \| Uncharacterized protein; uniclust \| UniRef100\_A0A960XNF8 \| 99.9 \| 7.4e-31 \| 1.4e-36 \| 213.3 \| 194 \| (268, 467) \| 475 \| (14, 229) \| 232 \| Phage tail sheath family protein \| Phage tail sheath family protein; uniclust \| UniRef100\_A0A2W4QEG8 \| 99.9 \| 2.2e-30 \| 4.2e-36 \| 243.7 \| 175 \| (286, 466) \| 475 \| (356, 545) \| 549 \| Tail sheath protein C-terminal domain-containing protein \| Tail sheath protein C-terminal domain-containing protein; uniclust \| UniRef100\_A0A958D3A0 \| 99.9 \| 2.6e-30 \| 4.7e-36 \| 208.6 \| 186 \| (231, 422) \| 475 \| (19, 220) \| 220 \| Phage tail sheath family protein (Fragment) \| Phage tail sheath family protein (Fragment); uniclust \| UniRef100\_A0A953FW81 \| 99.9 \| 2.8e-30 \| 5.2e-36 \| 232.7 \| 345 \| (7, 466) \| 475 \| (155, 523) \| 526 \| Tail sheath protein C-terminal domain-containing protein \| Tail sheath protein C-terminal domain-containing protein; uniclust \| UniRef100\_A0A7V9U445 \| 99.9 \| 3.3e-30 \| 6.2e-36 \| 220.8 \| 200 \| (264, 469) \| 475 \| (42, 251) \| 259 \| Phage tail sheath family protein \| Phage tail sheath family protein; uniclust \| UniRef100\_A0A955QN58 \| 99.9 \| 3.7e-30 \| 6.8e-36 \| 212.2 \| 225 \| (237, 467) \| 475 \| (7, 251) \| 255 \| Phage tail sheath family protein \| Phage tail sheath family protein; uniclust \| UniRef100\_A0A951KAP4 \| 99.9 \| 4.3e-30 \| 7.9e-36 \| 240.1 \| 235 \| (229, 469) \| 475 \| (499, 759) \| 765 \| Phage tail sheath family protein \| Phage tail sheath family protein; uniclust \| UniRef100\_A0A957BWX3 \| 99.9 \| 4.4e-30 \| 8e-36 \| 233.4 \| 366 \| (2, 465) \| 475 \| (1, 409) \| 569 \| TIR domain-containing protein \| TIR domain-containing protein; uniclust \| UniRef100\_A0A0M7QCF0 \| 99.9 \| 6.8e-30 \| 1.3e-35 \| 230.1 \| 233 \| (231, 469) \| 475 \| (33, 283) \| 295 \| Putative tail sheath monomer [Shigella phage Shfl2] \| Putative tail sheath monomer [Shigella phage Shfl2]; uniclust \| UniRef100\_UPI001ABB0727 \| 99.9 \| 1.4e-29 \| 2.6e-35 \| 237.8 \| 174 \| (287, 466) \| 475 \| (498, 685) \| 689 \| phage tail sheath C-terminal domain-containing protein \| phage tail sheath C-terminal domain-containing protein; uniclust \| UniRef100\_A0A7X8L470 \| 99.9 \| 1.5e-29 \| 2.8e-35 \| 229.6 \| 227 \| (229, 464) \| 475 \| (322, 557) \| 559 \| Phage tail sheath family protein \| Phage tail sheath family protein; uniclust \| UniRef100\_A0A353BET0 \| 99.9 \| 1.7e-29 \| 3e-35 \| 235.4 \| 232 \| (229, 466) \| 475 \| (478, 729) \| 734 \| Tail sheath protein C-terminal domain-containing protein \| Tail sheath protein C-terminal domain-containing protein; uniclust \| UniRef100\_A0A7X5ZX35 \| 99.9 \| 3.4e-29 \| 6.3e-35 \| 226.4 \| 227 \| (235, 467) \| 475 \| (282, 531) \| 536 \| Phage tail sheath family protein \| Phage tail sheath family protein; uniclust \| UniRef100\_A0A2W1BAD9 \| 99.9 \| 4.3e-29 \| 8e-35 \| 228.0 \| 425 \| (3, 466) \| 475 \| (1, 450) \| 460 \| Tail sheath protein C-terminal domain-containing protein \| Tail sheath protein C-terminal domain-containing protein; uniclust \| UniRef100\_A0A2D7XCW1 \| 99.9 \| 4.3e-29 \| 8.4e-35 \| 234.4 \| 235 \| (229, 469) \| 475 \| (155, 421) \| 426 \| Phage tail protein \| Phage tail protein; uniclust \| UniRef100\_A0A023W640 \| 99.9 \| 4.4e-29 \| 8.8e-35 \| 257.0 \| 233 \| (231, 469) \| 475 \| (410, 661) \| 699 \| Tail sheath protein \| Tail sheath protein; uniclust \| UniRef100\_A0A1I4LMX5 \| 99.9 \| 5.1e-29 \| 1e-34 \| 235.6 \| 293 \| (1, 352) \| 475 \| (1, 383) \| 400 \| Phage tail protein \| Phage tail protein; uniclust \| UniRef100\_UPI000A44B44F \| 99.9 \| 7.1e-29 \| 1.3e-34 \| 203.4 \| 181 \| (231, 421) \| 475 \| (33, 243) \| 243 \| phage tail sheath subtilisin-like domain-containing protein \| phage tail sheath subtilisin-like domain-containing protein; uniclust \| UniRef100\_A0A011NTQ5 \| 99.9 \| 1.3e-28 \| 2.5e-34 \| 262.3 \| 236 \| (229, 470) \| 475 \| (671, 936) \| 946 \| Phage tail sheath protein \| Phage tail sheath protein; uniclust \| UniRef100\_A0A849TMA6 \| 99.9 \| 1.4e-28 \| 2.7e-34 \| 234.8 \| 432 \| (7, 467) \| 475 \| (16, 501) \| 635 \| Phage tail protein \| Phage tail protein; uniclust \| UniRef100\_A0A9E1IB57 \| 99.9 \| 1.5e-28 \| 2.7e-34 \| 218.2 \| 194 \| (268, 467) \| 475 \| (235, 443) \| 451 \| Phage tail sheath family protein \| Phage tail sheath family protein; uniclust \| UniRef100\_A0A2D5F6D1 \| 99.9 \| 2.2e-28 \| 4.1e-34 \| 233.2 \| 238 \| (227, 468) \| 475 \| (425, 679) \| 686 \| Tail sheath protein C-terminal domain-containing protein \| Tail sheath protein C-terminal domain-containing protein; uniclust \| UniRef100\_A0A011NTT3 \| 99.9 \| 3.2e-28 \| 6.1e-34 \| 238.9 \| 219 \| (229, 464) \| 475 \| (330, 558) \| 560 \| Phage tail sheath protein \| Phage tail sheath protein; uniclust \| UniRef100\_A0A0Q8V0M7 \| 99.9 \| 3.7e-28 \| 7.2e-34 \| 242.1 \| 239 \| (229, 473) \| 475 \| (449, 709) \| 713 \| Tail sheath protein C-terminal domain-containing protein \| Tail sheath protein C-terminal domain-containing protein; uniclust \| UniRef100\_A0A429I985 \| 99.9 \| 4e-28 \| 7.5e-34 \| 223.7 \| 427 \| (2, 468) \| 475 \| (18, 506) \| 514 \| Uncharacterized protein (Fragment) \| Uncharacterized protein (Fragment); uniclust \| UniRef100\_UPI0020BD5A97 \| 99.9 \| 4.5e-28 \| 8.3e-34 \| 197.4 \| 188 \| (106, 377) \| 475 \| (43, 231) \| 231 \| phage tail sheath subtilisin-like domain-containing protein \| phage tail sheath subtilisin-like domain-containing protein; uniclust \| UniRef100\_A0A1Y3AZE0 \| 99.9 \| 4.6e-28 \| 8.4e-34 \| 246.9 \| 322 \| (6, 454) \| 475 \| (933, 1270) \| 3230 \| Chitinase-like protein \| Chitinase-like protein; uniclust \| UniRef100\_UPI0022869F9D \| 99.9 \| 5.2e-28 \| 9.6e-34 \| 206.0 \| 177 \| (286, 466) \| 475 \| (127, 315) \| 319 \| phage tail sheath subtilisin-like domain-containing protein \| phage tail sheath subtilisin-like domain-containing protein; uniclust \| UniRef100\_A0A0B0SCR7 \| 99.9 \| 5.6e-28 \| 1.1e-33 \| 236.9 \| 428 \| (2, 472) \| 475 \| (1, 470) \| 487 \| DUF2586 family protein \| DUF2586 family protein; uniclust \| UniRef100\_UPI0009DD7DD7 \| 99.9 \| 6.9e-28 \| 1.3e-33 \| 219.6 \| 222 \| (234, 466) \| 475 \| (332, 568) \| 572 \| phage tail sheath C-terminal domain-containing protein \| phage tail sheath C-terminal domain-containing protein; uniclust \| UniRef100\_A0A952SK04 \| 99.9 \| 8.5e-28 \| 1.6e-33 \| 219.9 \| 235 \| (231, 469) \| 475 \| (325, 588) \| 593 \| Phage tail sheath family protein \| Phage tail sheath family protein; uniclust \| UniRef100\_A0A1Z4PN29 \| 99.9 \| 9.1e-28 \| 1.7e-33 \| 216.0 \| 355 \| (6, 466) \| 475 \| (3, 369) \| 376 \| Putative prophage major tail sheath protein \| Putative prophage major tail sheath protein; uniclust \| UniRef100\_A0A447T5K7 \| 99.9 \| 9.5e-28 \| 1.8e-33 \| 211.8 \| 320 \| (2, 363) \| 475 \| (1, 364) \| 377 \| Phage tail sheath protein \| Phage tail sheath protein; uniclust \| UniRef100\_A0A2D5FCB7 \| 99.9 \| 1.1e-27 \| 2e-33 \| 212.4 \| 233 \| (231, 469) \| 475 \| (86, 337) \| 341 \| Phage tail protein (Fragment) \| Phage tail protein (Fragment); uniclust \| UniRef100\_A0A0Q6MQW8 \| 99.9 \| 1.1e-27 \| 2.1e-33 \| 239.1 \| 202 \| (266, 468) \| 475 \| (351, 563) \| 580 \| Phage tail protein \| Phage tail protein; uniclust \| UniRef100\_A0A800F6T4 \| 99.9 \| 1.4e-27 \| 2.7e-33 \| 203.9 \| 177 \| (287, 466) \| 475 \| (96, 283) \| 291 \| Phage tail sheath family protein \| Phage tail sheath family protein; uniclust \| UniRef100\_UPI0013159F4D \| 99.9 \| 1.5e-27 \| 2.7e-33 \| 196.0 \| 191 \| (266, 467) \| 475 \| (38, 241) \| 244 \| phage tail sheath C-terminal domain-containing protein \| phage tail sheath C-terminal domain-containing protein; uniclust \| UniRef100\_E2CFK5 \| 99.9 \| 1.7e-27 \| 3.1e-33 \| 211.8 \| 378 \| (5, 474) \| 475 \| (6, 387) \| 398 \| Putative tail sheath protein \| Putative tail sheath protein; uniclust \| UniRef100\_A0A081C205 \| 99.9 \| 1.9e-27 \| 3.6e-33 \| 216.5 \| 339 \| (5, 465) \| 475 \| (10, 360) \| 361 \| Phage tail sheath protein \| Phage tail sheath protein; uniclust \| UniRef100\_A0A1F7S379 \| 99.9 \| 2.2e-27 \| 4.1e-33 \| 211.6 \| 240 \| (229, 468) \| 475 \| (185, 448) \| 464 \| Phage tail protein (Fragment) \| Phage tail protein (Fragment); uniclust \| UniRef100\_A0A064A4E1 \| 99.9 \| 2.6e-27 \| 4.9e-33 \| 200.9 \| 222 \| (243, 475) \| 475 \| (12, 250) \| 251 \| Phage tail protein (Fragment) \| Phage tail protein (Fragment); uniclust \| UniRef100\_UPI0005EF3930 \| 99.9 \| 3e-27 \| 5.5e-33 \| 195.4 \| 247 \| (42, 388) \| 475 \| (6, 253) \| 254 \| phage tail sheath subtilisin-like domain-containing protein \| phage tail sheath subtilisin-like domain-containing protein; uniclust \| UniRef100\_A0A403T4T3 \| 99.9 \| 3e-27 \| 5.6e-33 \| 221.2 \| 284 \| (1, 374) \| 475 \| (1, 290) \| 751 \| Tail sheath protein subtilisin-like domain-containing protein \| Tail sheath protein subtilisin-like domain-containing protein; uniclust \| UniRef100\_A0A6P0I039 \| 99.9 \| 5.6e-27 \| 1e-32 \| 205.3 \| 175 \| (286, 466) \| 475 \| (148, 336) \| 340 \| Phage tail sheath family protein \| Phage tail sheath family protein; uniclust \| UniRef100\_A0A0Q5TAS8 \| 99.9 \| 6.9e-27 \| 1.3e-32 \| 207.7 \| 179 \| (287, 466) \| 475 \| (183, 371) \| 375 \| Tail sheath protein C-terminal domain-containing protein \| Tail sheath protein C-terminal domain-containing protein; uniclust \| UniRef100\_UPI00211EA2BE \| 99.9 \| 8.6e-27 \| 1.6e-32 \| 180.3 \| 136 \| (296, 437) \| 475 \| (3, 148) \| 164 \| phage tail sheath subtilisin-like domain-containing protein \| phage tail sheath subtilisin-like domain-containing protein; uniclust \| UniRef100\_A0A2T4JE69 \| 99.9 \| 8.6e-27 \| 1.6e-32 \| 193.4 \| 206 \| (114, 397) \| 475 \| (14, 220) \| 226 \| Phage tail protein (Fragment) \| Phage tail protein (Fragment); uniclust \| UniRef100\_A0A849EW92 \| 99.9 \| 9.2e-27 \| 1.7e-32 \| 218.3 \| 230 \| (231, 467) \| 475 \| (507, 758) \| 761 \| Phage tail sheath family protein \| Phage tail sheath family protein; uniclust \| UniRef100\_A0A813CV22 \| 99.9 \| 9.4e-27 \| 1.7e-32 \| 242.1 \| 226 \| (233, 464) \| 475 \| (292, 546) \| 7673 \| FtsH protein \| FtsH protein; uniclust \| UniRef100\_A0A955ZJD4 \| 99.9 \| 9.4e-27 \| 1.8e-32 \| 217.2 \| 234 \| (229, 468) \| 475 \| (324, 586) \| 597 \| Phage tail sheath subtilisin-like domain-containing protein \| Phage tail sheath subtilisin-like domain-containing protein; uniclust \| UniRef100\_A0A959ZZ97 \| 99.9 \| 9.9e-27 \| 1.9e-32 \| 218.7 \| 429 \| (2, 467) \| 475 \| (1, 474) \| 479 \| Uncharacterized protein \| Uncharacterized protein; uniclust \| UniRef100\_A0A813BIE1 \| 99.9 \| 1.1e-26 \| 2.1e-32 \| 232.3 \| 220 \| (232, 457) \| 475 \| (291, 539) \| 1968 \| FtsH protein \| FtsH protein; uniclust \| UniRef100\_A0A1Q7Z022 \| 99.9 \| 1.4e-26 \| 2.6e-32 \| 203.1 \| 231 \| (232, 467) \| 475 \| (146, 391) \| 399 \| Uncharacterized protein \| Uncharacterized protein; uniclust \| UniRef100\_UPI000D34BEAD \| 99.9 \| 2.3e-26 \| 4.3e-32 \| 197.4 \| 204 \| (257, 467) \| 475 \| (112, 324) \| 334 \| phage tail sheath C-terminal domain-containing protein \| phage tail sheath C-terminal domain-containing protein; uniclust \| UniRef100\_A0A1Y3AZE0 \| 99.9 \| 2.7e-26 \| 4.9e-32 \| 234.6 \| 161 \| (286, 452) \| 475 \| (1543, 1712) \| 3230 \| Chitinase-like protein \| Chitinase-like protein; uniclust \| UniRef100\_A0A3M1ZRW3 \| 99.9 \| 2.8e-26 \| 5.2e-32 \| 213.6 \| 235 \| (231, 469) \| 475 \| (445, 699) \| 702 \| Phage tail sheath family protein (Fragment) \| Phage tail sheath family protein (Fragment); uniclust \| UniRef100\_A0A6J5KV31 \| 99.9 \| 4.2e-26 \| 8.1e-32 \| 228.4 \| 235 \| (229, 469) \| 475 \| (544, 788) \| 794 \| Tail sheath protein \| Tail sheath protein; uniclust \| UniRef100\_A0A0A8J9Q3 \| 99.9 \| 4.1e-26 \| 8.1e-32 \| 240.0 \| 237 \| (229, 469) \| 475 \| (752, 1020) \| 1043 \| Tail sheath protein \| Tail sheath protein; uniclust \| UniRef100\_A0A376TUN6 \| 99.9 \| 4.7e-26 \| 9.2e-32 \| 194.5 \| 172 \| (229, 400) \| 475 \| (17, 190) \| 202 \| Tail sheath protein \| Tail sheath protein; uniclust \| UniRef100\_A0A1Y3TJF4 \| 99.9 \| 6.1e-26 \| 1.1e-31 \| 205.0 \| 379 \| (4, 474) \| 475 \| (1, 484) \| 490 \| Phage tail protein \| Phage tail protein; uniclust \| UniRef100\_A0A1G6LSA3 \| 99.9 \| 6e-26 \| 1.1e-31 \| 208.4 \| 351 \| (4, 466) \| 475 \| (1, 366) \| 373 \| Phage tail sheath protein \| Phage tail sheath protein; uniclust \| UniRef100\_A0A0B0HE64 \| 99.9 \| 8.3e-26 \| 1.6e-31 \| 204.7 \| 274 \| (4, 374) \| 475 \| (1, 371) \| 383 \| Phage tail sheath protein \| Phage tail sheath protein; uniclust \| UniRef100\_A0A956GJ63 \| 99.9 \| 9.1e-26 \| 1.7e-31 \| 195.3 \| 232 \| (233, 470) \| 475 \| (43, 305) \| 311 \| Phage tail sheath family protein \| Phage tail sheath family protein; uniclust \| UniRef100\_A0A0F0HL05 \| 99.9 \| 9.7e-26 \| 1.8e-31 \| 193.1 \| 234 \| (231, 470) \| 475 \| (14, 278) \| 289 \| Tail protein (Fragment) \| Tail protein (Fragment); uniclust \| UniRef100\_A0A257MWQ1 \| 99.9 \| 1.4e-25 \| 2.7e-31 \| 179.8 \| 126 \| (343, 468) \| 475 \| (13, 148) \| 152 \| Putative phage tail sheath protein FI (Fragment) \| Putative phage tail sheath protein FI (Fragment); uniclust \| UniRef100\_UPI001F56A12B \| 99.9 \| 1.4e-25 \| 2.7e-31 \| 210.9 \| 243 \| (230, 472) \| 475 \| (24, 267) \| 772 \| phage tail tape measure protein \| phage tail tape measure protein; uniclust \| UniRef100\_A0A0S8A5K3 \| 99.9 \| 1.5e-25 \| 2.8e-31 \| 168.2 \| 122 \| (273, 400) \| 475 \| (1, 125) \| 126 \| Tail sheath protein subtilisin-like domain-containing protein \| Tail sheath protein subtilisin-like domain-containing protein; uniclust \| UniRef100\_A0A090ZAL6 \| 99.9 \| 1.6e-25 \| 3.1e-31 \| 219.7 \| 408 \| (7, 467) \| 475 \| (15, 465) \| 475 \| Phage tail sheath family protein \| Phage tail sheath family protein; uniclust \| UniRef100\_A0A8X6HJI7 \| 99.9 \| 2e-25 \| 3.7e-31 \| 197.8 \| 217 \| (22, 325) \| 475 \| (175, 392) \| 428 \| Phage tail protein \| Phage tail protein; uniclust \| UniRef100\_A0A1Z9H610 \| 99.9 \| 2e-25 \| 3.9e-31 \| 236.3 \| 236 \| (228, 469) \| 475 \| (709, 977) \| 982 \| Tail sheath protein C-terminal domain-containing protein \| Tail sheath protein C-terminal domain-containing protein; uniclust \| UniRef100\_UPI000FEEF31E \| 99.9 \| 2.1e-25 \| 4e-31 \| 186.3 \| 221 \| (33, 342) \| 475 \| (2, 224) \| 224 \| phage tail sheath subtilisin-like domain-containing protein \| phage tail sheath subtilisin-like domain-containing protein; uniclust \| UniRef100\_A0A523DTF3 \| 99.9 \| 2.2e-25 \| 4.1e-31 \| 169.3 \| 131 \| (288, 424) \| 475 \| (2, 136) \| 146 \| Phage tail sheath family protein \| Phage tail sheath family protein; uniclust \| UniRef100\_UPI00138F8D04 \| 99.9 \| 2.3e-25 \| 4.1e-31 \| 198.9 \| 233 \| (230, 468) \| 475 \| (199, 439) \| 456 \| phage tail sheath subtilisin-like domain-containing protein \| phage tail sheath subtilisin-like domain-containing protein; uniclust \| UniRef100\_A0A941J0D5 \| 99.9 \| 2.4e-25 \| 4.5e-31 \| 185.4 \| 229 \| (229, 466) \| 475 \| (13, 258) \| 264 \| Phage tail sheath family protein \| Phage tail sheath family protein; uniclust \| UniRef100\_A0A370LRA8 \| 99.9 \| 3.1e-25 \| 5.7e-31 \| 214.2 \| 237 \| (229, 469) \| 475 \| (647, 902) \| 911 \| Tail sheath protein C-terminal domain-containing protein \| Tail sheath protein C-terminal domain-containing protein; uniclust \| UniRef100\_A0A4Q3AFM2 \| 99.9 \| 3.2e-25 \| 6.2e-31 \| 211.6 \| 260 \| (5, 363) \| 475 \| (9, 405) \| 453 \| Phage tail protein (Fragment) \| Phage tail protein (Fragment); uniclust \| UniRef100\_A0A0F9FWT7 \| 99.9 \| 3.7e-25 \| 7.2e-31 \| 219.8 \| 232 \| (229, 466) \| 475 \| (400, 641) \| 655 \| Tail sheath protein subtilisin-like domain-containing protein (Fragment) \| Tail sheath protein subtilisin-like domain-containing protein (Fragment); uniclust \| UniRef100\_A0A2Z4FJ67 \| 99.9 \| 4.2e-25 \| 7.7e-31 \| 199.0 \| 333 \| (19, 467) \| 475 \| (126, 487) \| 495 \| Tail sheath protein subtilisin-like domain-containing protein \| Tail sheath protein subtilisin-like domain-containing protein; uniclust \| UniRef100\_A0A7C4ZXL6 \| 99.9 \| 4.2e-25 \| 7.8e-31 \| 197.7 \| 338 \| (1, 464) \| 475 \| (1, 358) \| 360 \| Phage tail protein \| Phage tail protein; uniclust \| UniRef100\_UPI000AE5212C \| 99.9 \| 4.8e-25 \| 8.9e-31 \| 181.2 \| 199 \| (87, 371) \| 475 \| (5, 211) \| 213 \| phage tail sheath subtilisin-like domain-containing protein \| phage tail sheath subtilisin-like domain-containing protein; uniclust \| UniRef100\_A0A950FF97 \| 99.8 \| 6e-25 \| 1.1e-30 \| 205.7 \| 244 \| (230, 473) \| 475 \| (441, 716) \| 725 \| Phage tail sheath subtilisin-like domain-containing protein \| Phage tail sheath subtilisin-like domain-containing protein; uniclust \| UniRef100\_A0A3C1Q7E4 \| 99.8 \| 6.1e-25 \| 1.1e-30 \| 188.8 \| 205 \| (261, 475) \| 475 \| (64, 279) \| 280 \| Phage tail protein \| Phage tail protein; uniclust \| UniRef100\_A0A2W6P9S6 \| 99.8 \| 8.1e-25 \| 1.5e-30 \| 176.1 \| 184 \| (290, 474) \| 475 \| (2, 185) \| 191 \| Phage tail sheath family protein (Fragment) \| Phage tail sheath family protein (Fragment); uniclust \| UniRef100\_A0A448JE63 \| 99.8 \| 8.6e-25 \| 1.6e-30 \| 195.3 \| 238 \| (231, 474) \| 475 \| (66, 305) \| 317 \| Major tail sheath protein \| Major tail sheath protein; uniclust \| UniRef100\_A0A1F3RZR2 \| 99.8 \| 1.1e-24 \| 1.9e-30 \| 193.6 \| 180 \| (286, 466) \| 475 \| (243, 431) \| 434 \| Uncharacterized protein \| Uncharacterized protein; uniclust \| UniRef100\_A0A6M0F244 \| 99.8 \| 1.3e-24 \| 2.4e-30 \| 187.8 \| 180 \| (286, 466) \| 475 \| (151, 340) \| 344 \| Tail sheath protein C-terminal domain-containing protein \| Tail sheath protein C-terminal domain-containing protein; uniclust \| UniRef100\_A0A965TRY7 \| 99.8 \| 1.3e-24 \| 2.4e-30 \| 199.1 \| 205 \| (258, 466) \| 475 \| (359, 573) \| 576 \| Phage tail sheath family protein \| Phage tail sheath family protein; uniclust \| UniRef100\_A0A929F6P8 \| 99.8 \| 1.5e-24 \| 2.7e-30 \| 184.5 \| 163 \| (231, 399) \| 475 \| (126, 305) \| 305 \| Phage tail sheath family protein (Fragment) \| Phage tail sheath family protein (Fragment); uniclust \| UniRef100\_A0A080LR57 \| 99.8 \| 1.6e-24 \| 2.9e-30 \| 168.2 \| 117 \| (353, 469) \| 475 \| (7, 133) \| 136 \| Tail sheath protein \| Tail sheath protein; uniclust \| UniRef100\_A0A327J8X6 \| 99.8 \| 2e-24 \| 3.6e-30 \| 193.9 \| 427 \| (6, 469) \| 475 \| (1, 468) \| 475 \| Tail sheath protein subtilisin-like domain-containing protein \| Tail sheath protein subtilisin-like domain-containing protein; uniclust \| UniRef100\_A0A350EYQ4 \| 99.8 \| 2e-24 \| 3.6e-30 \| 172.6 \| 145 \| (317, 467) \| 475 \| (6, 161) \| 197 \| Tail sheath protein C-terminal domain-containing protein (Fragment) \| Tail sheath protein C-terminal domain-containing protein (Fragment); uniclust \| UniRef100\_A0A370E172 \| 99.8 \| 2e-24 \| 3.6e-30 \| 189.3 \| 180 \| (286, 466) \| 475 \| (193, 382) \| 386 \| Phage tail protein \| Phage tail protein; uniclust \| UniRef100\_A0A1Y3AZE0 \| 99.8 \| 2e-24 \| 3.6e-30 \| 221.4 \| 371 \| (3, 466) \| 475 \| (1726, 2118) \| 3230 \| Chitinase-like protein \| Chitinase-like protein; uniclust \| UniRef100\_UPI001EDA5694 \| 99.8 \| 2.2e-24 \| 4e-30 \| 186.3 \| 326 \| (8, 468) \| 475 \| (2, 340) \| 343 \| hypothetical protein \| hypothetical protein; uniclust \| UniRef100\_A0A0F3RPG1 \| 99.8 \| 2.2e-24 \| 4e-30 \| 173.7 \| 191 \| (2, 275) \| 475 \| (1, 192) \| 192 \| Phage tail sheath family protein \| Phage tail sheath family protein; uniclust \| UniRef100\_A0A085K2M7 \| 99.8 \| 2.7e-24 \| 5.2e-30 \| 183.6 \| 188 \| (1, 278) \| 475 \| (11, 199) \| 201 \| Phage tail protein \| Phage tail protein; uniclust \| UniRef100\_A0A1J6QRE2 \| 99.8 \| 3.3e-24 \| 6.2e-30 \| 190.1 \| 372 \| (7, 474) \| 475 \| (4, 390) \| 397 \| Phage tail protein \| Phage tail protein; uniclust \| UniRef100\_A0A8S0FXS2 \| 99.8 \| 3.7e-24 \| 7.2e-30 \| 189.4 \| 161 \| (231, 391) \| 475 \| (57, 231) \| 232 \| Tail sheath protein subtilisin-like domain-containing protein \| Tail sheath protein subtilisin-like domain-containing protein; uniclust \| UniRef100\_A0A2E6EJU7 \| 99.8 \| 4.1e-24 \| 7.6e-30 \| 192.5 \| 193 \| (266, 468) \| 475 \| (281, 484) \| 487 \| Phage tail protein \| Phage tail protein; uniclust \| UniRef100\_A0A1Q8A397 \| 99.8 \| 4.4e-24 \| 8.3e-30 \| 164.5 \| 105 \| (364, 468) \| 475 \| (2, 112) \| 133 \| Tail sheath protein C-terminal domain-containing protein \| Tail sheath protein C-terminal domain-containing protein; uniclust \| UniRef100\_A0A925P9Z3 \| 99.8 \| 4.8e-24 \| 9.2e-30 \| 177.6 \| 159 \| (306, 470) \| 475 \| (10, 175) \| 182 \| Phage tail sheath protein \| Phage tail sheath protein; uniclust \| UniRef100\_A0A355DNH3 \| 99.8 \| 5.3e-24 \| 9.9e-30 \| 194.1 \| 128 \| (230, 363) \| 475 \| (301, 455) \| 464 \| Phage tail protein (Fragment) \| Phage tail protein (Fragment); uniclust \| UniRef100\_A0A833AQN6 \| 99.8 \| 5.6e-24 \| 1.1e-29 \| 217.7 \| 236 \| (229, 470) \| 475 \| (691, 967) \| 974 \| Phage tail sheath family protein \| Phage tail sheath family protein; uniclust \| UniRef100\_UPI0015C33B2A \| 99.8 \| 6.9e-24 \| 1.3e-29 \| 161.0 \| 129 \| (285, 419) \| 475 \| (9, 141) \| 143 \| phage tail sheath subtilisin-like domain-containing protein \| phage tail sheath subtilisin-like domain-containing protein; uniclust \| UniRef100\_A0A376YN89 \| 99.8 \| 7.9e-24 \| 1.4e-29 \| 169.7 \| 171 \| (260, 430) \| 475 \| (3, 174) \| 200 \| Major tail sheath protein FI \| Major tail sheath protein FI; uniclust \| UniRef100\_A0A1C0V8T7 \| 99.8 \| 1.1e-23 \| 2e-29 \| 196.7 \| 244 \| (229, 473) \| 475 \| (428, 686) \| 688 \| Tail sheath protein subtilisin-like domain-containing protein \| Tail sheath protein subtilisin-like domain-containing protein; uniclust \| UniRef100\_UPI00116C6DAE \| 99.8 \| 1.1e-23 \| 2.1e-29 \| 164.5 \| 147 \| (291, 437) \| 475 \| (8, 154) \| 158 \| phage tail sheath subtilisin-like domain-containing protein \| phage tail sheath subtilisin-like domain-containing protein; uniclust \| UniRef100\_A0A7W1Q988 \| 99.8 \| 1.2e-23 \| 2.3e-29 \| 180.9 \| 239 \| (3, 363) \| 475 \| (2, 249) \| 329 \| Phage tail sheath family protein \| Phage tail sheath family protein; uniclust \| UniRef100\_A0A292GIW4 \| 99.8 \| 1.3e-23 \| 2.6e-29 \| 204.2 \| 241 \| (229, 469) \| 475 \| (374, 636) \| 649 \| Tail sheath protein \| Tail sheath protein; uniclust \| UniRef100\_A0A2D6PBJ4 \| 99.8 \| 1.6e-23 \| 3e-29 \| 197.6 \| 237 \| (229, 469) \| 475 \| (394, 647) \| 652 \| Tail sheath protein C-terminal domain-containing protein \| Tail sheath protein C-terminal domain-containing protein; uniclust \| UniRef100\_UPI0022347FD7 \| 99.8 \| 1.6e-23 \| 3e-29 \| 163.6 \| 141 \| (231, 371) \| 475 \| (28, 169) \| 170 \| phage tail sheath subtilisin-like domain-containing protein \| phage tail sheath subtilisin-like domain-containing protein; uniclust \| UniRef100\_A0A266LLW5 \| 99.8 \| 1.8e-23 \| 3.3e-29 \| 161.5 \| 143 \| (231, 373) \| 475 \| (4, 147) \| 149 \| Phage tail protein (Fragment) \| Phage tail protein (Fragment); uniclust \| UniRef100\_A0A6S6PL19 \| 99.8 \| 1.9e-23 \| 3.5e-29 \| 192.8 \| 429 \| (7, 466) \| 475 \| (17, 487) \| 497 \| Phage tail protein \| Phage tail protein; uniclust \| UniRef100\_A0A412P9T8 \| 99.8 \| 1.9e-23 \| 3.6e-29 \| 187.2 \| 114 \| (287, 403) \| 475 \| (265, 382) \| 384 \| Phage tail sheath family protein \| Phage tail sheath family protein; uniclust \| UniRef100\_A0A3A8PQG2 \| 99.8 \| 1.9e-23 \| 3.6e-29 \| 194.5 \| 232 \| (230, 467) \| 475 \| (202, 472) \| 481 \| Phage tail sheath family protein (Fragment) \| Phage tail sheath family protein (Fragment); uniclust \| UniRef100\_A0A813AAM4 \| 99.8 \| 2e-23 \| 3.7e-29 \| 204.7 \| 234 \| (229, 468) \| 475 \| (1012, 1266) \| 1276 \| 18 protein \| 18 protein; uniclust \| UniRef100\_A0A0C5ADZ8 \| 99.8 \| 2e-23 \| 3.8e-29 \| 208.8 \| 232 \| (232, 469) \| 475 \| (494, 744) \| 763 \| Tail sheath monomer \| Tail sheath monomer; uniclust \| UniRef100\_A0A3A8JLS6 \| 99.8 \| 2.1e-23 \| 3.9e-29 \| 182.1 \| 241 \| (230, 474) \| 475 \| (74, 342) \| 347 \| Phage tail sheath family protein \| Phage tail sheath family protein; uniclust \| UniRef100\_A0A382AQG4 \| 99.8 \| 3.2e-23 \| 5.9e-29 \| 192.6 \| 234 \| (229, 468) \| 475 \| (251, 494) \| 500 \| Tail sheath protein C-terminal domain-containing protein (Fragment) \| Tail sheath protein C-terminal domain-containing protein (Fragment); uniclust \| UniRef100\_A0A268THL2 \| 99.8 \| 4e-23 \| 7.6e-29 \| 189.0 \| 357 \| (5, 474) \| 475 \| (3, 362) \| 374 \| Phage tail protein \| Phage tail protein; uniclust \| UniRef100\_A0A5F1HTD7 \| 99.8 \| 5.6e-23 \| 1e-28 \| 179.1 \| 174 \| (287, 466) \| 475 \| (168, 354) \| 361 \| Uncharacterized protein \| Uncharacterized protein; uniclust \| UniRef100\_A0A1H9D327 \| 99.8 \| 5.4e-23 \| 1e-28 \| 204.7 \| 236 \| (229, 470) \| 475 \| (621, 887) \| 894 \| IPT/TIG domain-containing protein \| IPT/TIG domain-containing protein; uniclust \| UniRef100\_A0A1Z8RYF4 \| 99.8 \| 6e-23 \| 1.2e-28 \| 218.7 \| 234 \| (231, 470) \| 475 \| (746, 1004) \| 1008 \| Tail sheath protein C-terminal domain-containing protein \| Tail sheath protein C-terminal domain-containing protein; uniclust \| UniRef100\_A0A059WTY0 \| 99.8 \| 6.9e-23 \| 1.3e-28 \| 163.4 \| 136 \| (231, 372) \| 475 \| (51, 192) \| 192 \| Phage tail sheath protein (Fragment) \| Phage tail sheath protein (Fragment); uniclust \| UniRef100\_A0A3E1Y3Z4 \| 99.8 \| 7.1e-23 \| 1.3e-28 \| 188.4 \| 179 \| (287, 466) \| 475 \| (394, 582) \| 586 \| Phage tail sheath family protein \| Phage tail sheath family protein; uniclust \| UniRef100\_A0A7C6A7S9 \| 99.8 \| 7.4e-23 \| 1.4e-28 \| 184.7 \| 431 \| (2, 469) \| 475 \| (1, 479) \| 487 \| Tail sheath protein C-terminal domain-containing protein \| Tail sheath protein C-terminal domain-containing protein; uniclust \| UniRef100\_A0A0N8TDW0 \| 99.8 \| 9.2e-23 \| 1.7e-28 \| 165.1 \| 128 \| (340, 467) \| 475 \| (24, 161) \| 165 \| Phage tail sheath protein FI \| Phage tail sheath protein FI; uniclust \| UniRef100\_UPI0009B437D5 \| 99.8 \| 1.2e-22 \| 2.1e-28 \| 190.4 \| 233 \| (229, 468) \| 475 \| (425, 685) \| 702 \| phage tail sheath family protein \| phage tail sheath family protein; uniclust \| UniRef100\_A0A5Z1S5A9 \| 99.8 \| 1.2e-22 \| 2.3e-28 \| 171.8 \| 215 \| (257, 474) \| 475 \| (10, 226) \| 233 \| Phage tail sheath family protein (Fragment) \| Phage tail sheath family protein (Fragment); uniclust \| UniRef100\_A0A0G9K7V9 \| 99.8 \| 1.3e-22 \| 2.4e-28 \| 187.1 \| 237 \| (231, 472) \| 475 \| (140, 379) \| 388 \| Tail protein \| Tail protein; uniclust \| UniRef100\_A0A5C7JAB2 \| 99.8 \| 1.4e-22 \| 2.6e-28 \| 192.6 \| 237 \| (229, 469) \| 475 \| (310, 575) \| 580 \| Tail sheath protein C-terminal domain-containing protein \| Tail sheath protein C-terminal domain-containing protein; uniclust \| UniRef100\_A0A3N5G9W8 \| 99.8 \| 1.4e-22 \| 2.7e-28 \| 197.4 \| 218 \| (229, 464) \| 475 \| (323, 551) \| 553 \| Phage tail sheath protein (Fragment) \| Phage tail sheath protein (Fragment); uniclust \| UniRef100\_UPI00143D32B4 \| 99.8 \| 1.8e-22 \| 3.3e-28 \| 174.8 \| 180 \| (287, 468) \| 475 \| (151, 340) \| 342 \| phage tail sheath family protein \| phage tail sheath family protein; uniclust \| UniRef100\_A0A2V8N5I3 \| 99.8 \| 2.6e-22 \| 4.7e-28 \| 170.3 \| 197 \| (266, 468) \| 475 \| (78, 283) \| 291 \| Tail sheath protein C-terminal domain-containing protein \| Tail sheath protein C-terminal domain-containing protein; uniclust \| UniRef100\_A0A075E197 \| 99.8 \| 2.6e-22 \| 5e-28 \| 198.3 \| 233 \| (232, 467) \| 475 \| (395, 642) \| 660 \| Tail sheath family protein \| Tail sheath family protein; uniclust \| UniRef100\_A0A085EKS0 \| 99.8 \| 2.6e-22 \| 5.2e-28 \| 193.9 \| 363 \| (2, 469) \| 475 \| (14, 411) \| 419 \| Tail sheath protein \| Tail sheath protein; uniclust \| UniRef100\_A0A4R3Z7M7 \| 99.8 \| 3.1e-22 \| 5.6e-28 \| 154.7 \| 140 \| (231, 378) \| 475 \| (12, 156) \| 157 \| Tail sheath protein \| Tail sheath protein; uniclust \| UniRef100\_A0A7I8MZ51 \| 99.8 \| 3.1e-22 \| 5.8e-28 \| 182.1 \| 230 \| (233, 468) \| 475 \| (271, 511) \| 520 \| Tail sheath protein C-terminal domain-containing protein \| Tail sheath protein C-terminal domain-containing protein; uniclust \| UniRef100\_A0A0H4WYB2 \| 99.8 \| 4e-22 \| 7.8e-28 \| 194.1 \| 400 \| (7, 465) \| 475 \| (36, 490) \| 493 \| Phage tail sheath protein \| Phage tail sheath protein; uniclust \| UniRef100\_A0A081NYA3 \| 99.8 \| 6.6e-22 \| 1.3e-27 \| 196.2 \| 427 \| (5, 470) \| 475 \| (3, 468) \| 481 \| Tail sheath protein subtilisin-like domain-containing protein \| Tail sheath protein subtilisin-like domain-containing protein; uniclust \| UniRef100\_A0A6A1TYH7 \| 99.8 \| 8.7e-22 \| 1.6e-27 \| 169.9 \| 243 \| (225, 474) \| 475 \| (69, 313) \| 329 \| Phage tail protein \| Phage tail protein; uniclust \| UniRef100\_UPI0006204962 \| 99.8 \| 9.9e-22 \| 1.8e-27 \| 168.8 \| 258 \| (5, 362) \| 475 \| (3, 271) \| 271 \| phage tail protein \| phage tail protein; uniclust \| UniRef100\_A0A015U7W7 \| 99.8 \| 1.1e-21 \| 2.3e-27 \| 195.7 \| 368 \| (1, 471) \| 475 \| (20, 418) \| 427 \| DUF2586 family protein \| DUF2586 family protein; uniclust \| UniRef100\_A0A061JGT4 \| 99.7 \| 1.4e-21 \| 2.6e-27 \| 153.9 \| 115 \| (360, 474) \| 475 \| (8, 122) \| 138 \| Major tail sheath protein \| Major tail sheath protein; uniclust \| UniRef100\_J2WTG1 \| 99.7 \| 1.6e-21 \| 2.9e-27 \| 179.4 \| 228 \| (231, 466) \| 475 \| (323, 559) \| 570 \| Phage tail sheath protein FI (Fragment) \| Phage tail sheath protein FI (Fragment); uniclust \| UniRef100\_A0A4Q3UNL2 \| 99.7 \| 1.6e-21 \| 2.9e-27 \| 171.4 \| 149 \| (227, 381) \| 475 \| (227, 380) \| 380 \| Phage tail sheath family protein (Fragment) \| Phage tail sheath family protein (Fragment); uniclust \| UniRef100\_A0A842WMT0 \| 99.7 \| 2e-21 \| 3.6e-27 \| 187.3 \| 233 \| (230, 465) \| 475 \| (680, 929) \| 949 \| Tail sheath protein subtilisin-like domain-containing protein \| Tail sheath protein subtilisin-like domain-containing protein; uniclust \| UniRef100\_A0A2E3R271 \| 99.7 \| 2.1e-21 \| 3.9e-27 \| 172.9 \| 377 \| (7, 474) \| 475 \| (3, 423) \| 424 \| Tail sheath protein subtilisin-like domain-containing protein \| Tail sheath protein subtilisin-like domain-containing protein; uniclust \| UniRef100\_A0A2D6AJR1 \| 99.7 \| 2.2e-21 \| 4.2e-27 \| 185.1 \| 176 \| (287, 468) \| 475 \| (231, 412) \| 452 \| Phage tail protein \| Phage tail protein; uniclust \| UniRef100\_A0A956ILR2 \| 99.7 \| 3.1e-21 \| 5.7e-27 \| 169.2 \| 231 \| (231, 467) \| 475 \| (100, 359) \| 371 \| Phage tail sheath family protein \| Phage tail sheath family protein; uniclust \| UniRef100\_UPI001B8D85E7 \| 99.7 \| 3.2e-21 \| 5.8e-27 \| 184.7 \| 131 \| (287, 423) \| 475 \| (678, 812) \| 876 \| phage tail sheath family protein \| phage tail sheath family protein; uniclust \| UniRef100\_A0A6A4R6J7 \| 99.7 \| 3.5e-21 \| 6.5e-27 \| 170.2 \| 286 \| (7, 394) \| 475 \| (12, 307) \| 311 \| Phage tail protein (Fragment) \| Phage tail protein (Fragment); uniclust \| UniRef100\_A0A6M0FV63 \| 99.7 \| 3.6e-21 \| 6.8e-27 \| 173.2 \| 344 \| (7, 466) \| 475 \| (4, 362) \| 367 \| Tail sheath protein subtilisin-like domain-containing protein \| Tail sheath protein subtilisin-like domain-containing protein; uniclust \| UniRef100\_A0A976JLS4 \| 99.7 \| 3.7e-21 \| 6.8e-27 \| 171.0 \| 233 \| (230, 465) \| 475 \| (152, 408) \| 415 \| Phage tail sheath family protein \| Phage tail sheath family protein; uniclust \| UniRef100\_A0A4S2S3L3 \| 99.7 \| 3.9e-21 \| 7.2e-27 \| 184.7 \| 233 \| (232, 470) \| 475 \| (668, 912) \| 916 \| Phage tail sheath family protein \| Phage tail sheath family protein; uniclust \| UniRef100\_A0A7K1RD86 \| 99.7 \| 4.7e-21 \| 8.7e-27 \| 181.2 \| 342 \| (43, 470) \| 475 \| (384, 743) \| 756 \| Phage tail protein \| Phage tail protein; uniclust \| UniRef100\_A0A2W6WG49 \| 99.7 \| 4.8e-21 \| 8.9e-27 \| 161.6 \| 243 \| (3, 474) \| 475 \| (1, 243) \| 249 \| Tail sheath protein C-terminal domain-containing protein \| Tail sheath protein C-terminal domain-containing protein; uniclust \| UniRef100\_A0A0F8XJ22 \| 99.7 \| 5e-21 \| 9.5e-27 \| 157.4 \| 155 \| (309, 469) \| 475 \| (3, 163) \| 176 \| Tail sheath protein C-terminal domain-containing protein (Fragment) \| Tail sheath protein C-terminal domain-containing protein (Fragment); uniclust \| UniRef100\_A0A957QB84 \| 99.7 \| 5.4e-21 \| 9.8e-27 \| 172.3 \| 235 \| (229, 469) \| 475 \| (179, 458) \| 466 \| Phage tail sheath family protein \| Phage tail sheath family protein; uniclust \| UniRef100\_A0A292SAA6 \| 99.7 \| 1.2e-20 \| 2.2e-26 \| 173.0 \| 201 \| (266, 469) \| 475 \| (239, 443) \| 451 \| Tail sheath protein C-terminal domain-containing protein \| Tail sheath protein C-terminal domain-containing protein; uniclust \| UniRef100\_A0A924Q0Y1 \| 99.7 \| 1.5e-20 \| 2.8e-26 \| 170.1 \| 208 \| (265, 473) \| 475 \| (107, 327) \| 340 \| Phage tail protein \| Phage tail protein; uniclust \| UniRef100\_E3BEQ6 \| 99.7 \| 1.8e-20 \| 3.2e-26 \| 163.8 \| 215 \| (240, 466) \| 475 \| (127, 353) \| 356 \| Uncharacterized protein \| Uncharacterized protein; uniclust \| UniRef100\_A0A096CJ44 \| 99.7 \| 1.8e-20 \| 3.5e-26 \| 180.1 \| 322 \| (7, 466) \| 475 \| (56, 394) \| 395 \| Tail sheath protein C-terminal domain-containing protein \| Tail sheath protein C-terminal domain-containing protein; uniclust \| UniRef100\_A0A3N7HIZ4 \| 99.7 \| 2.1e-20 \| 4.2e-26 \| 188.0 \| 198 \| (265, 467) \| 475 \| (406, 611) \| 622 \| Phage tail sheath protein \| Phage tail sheath protein; uniclust \| UniRef100\_A0A2S5F8Z6 \| 99.7 \| 2.4e-20 \| 4.4e-26 \| 168.4 \| 240 \| (234, 474) \| 475 \| (204, 462) \| 469 \| Phage tail protein \| Phage tail protein; uniclust \| UniRef100\_A0A011PBZ6 \| 99.7 \| 2.5e-20 \| 4.7e-26 \| 177.0 \| 204 \| (266, 471) \| 475 \| (298, 509) \| 519 \| Phage tail sheath protein \| Phage tail sheath protein; uniclust \| UniRef100\_A0A482MLF8 \| 99.7 \| 3.6e-20 \| 6.8e-26 \| 179.2 \| 234 \| (229, 468) \| 475 \| (397, 642) \| 657 \| Tail sheath protein \| Tail sheath protein; uniclust \| UniRef100\_A0A257GTL1 \| 99.7 \| 3.5e-20 \| 6.8e-26 \| 171.0 \| 188 \| (1, 280) \| 475 \| (1, 222) \| 303 \| Phage tail protein (Fragment) \| Phage tail protein (Fragment); uniclust \| UniRef100\_A0A923X4B7 \| 99.7 \| 3.7e-20 \| 6.9e-26 \| 139.2 \| 102 \| (366, 467) \| 475 \| (4, 111) \| 120 \| Phage tail sheath family protein \| Phage tail sheath family protein; uniclust \| UniRef100\_A0A1X7MDT6 \| 99.7 \| 3.9e-20 \| 7.2e-26 \| 140.4 \| 123 \| (300, 422) \| 475 \| (1, 123) \| 138 \| Phage tail sheath protein (Fragment) \| Phage tail sheath protein (Fragment); uniclust \| UniRef100\_A0A7J4G8F5 \| 99.7 \| 4e-20 \| 7.5e-26 \| 171.5 \| 233 \| (231, 469) \| 475 \| (241, 493) \| 502 \| Tail sheath protein subtilisin-like domain-containing protein (Fragment) \| Tail sheath protein subtilisin-like domain-containing protein (Fragment); uniclust \| UniRef100\_A0A1Q7C2B6 \| 99.7 \| 4.1e-20 \| 7.8e-26 \| 144.8 \| 115 \| (353, 467) \| 475 \| (3, 123) \| 129 \| Uncharacterized protein \| Uncharacterized protein; uniclust \| UniRef100\_A0A972PJ70 \| 99.7 \| 5e-20 \| 9.2e-26 \| 144.1 \| 140 \| (336, 475) \| 475 \| (20, 164) \| 165 \| Phage tail sheath family protein \| Phage tail sheath family protein; uniclust \| UniRef100\_A0A2E6W8A2 \| 99.7 \| 5e-20 \| 9.4e-26 \| 179.2 \| 239 \| (229, 469) \| 475 \| (392, 660) \| 668 \| Tail sheath protein C-terminal domain-containing protein (Fragment) \| Tail sheath protein C-terminal domain-containing protein (Fragment); uniclust \| UniRef100\_UPI000470A430 \| 99.7 \| 5.1e-20 \| 9.5e-26 \| 155.9 \| 221 \| (2, 311) \| 475 \| (1, 259) \| 259 \| phage tail sheath family protein \| phage tail sheath family protein; uniclust \| UniRef100\_A0A193QGV7 \| 99.7 \| 5.5e-20 \| 1e-25 \| 148.9 \| 198 \| (260, 473) \| 475 \| (3, 201) \| 205 \| Phage tail sheath protein \| Phage tail sheath protein; uniclust \| UniRef100\_A0A3C1KVB2 \| 99.7 \| 5.9e-20 \| 1.1e-25 \| 169.4 \| 119 \| (229, 354) \| 475 \| (340, 460) \| 461 \| Tail sheath protein subtilisin-like domain-containing protein (Fragment) \| Tail sheath protein subtilisin-like domain-containing protein (Fragment); uniclust \| UniRef100\_UPI0020B107FC \| 99.7 \| 8.1e-20 \| 1.5e-25 \| 174.4 \| 428 \| (7, 465) \| 475 \| (15, 497) \| 818 \| hypothetical protein \| hypothetical protein; uniclust \| UniRef100\_A0A1I5H9Z7 \| 99.7 \| 1e-19 \| 1.9e-25 \| 168.5 \| 367 \| (16, 471) \| 475 \| (15, 409) \| 424 \| Phage tail sheath protein FI \| Phage tail sheath protein FI; uniclust \| UniRef100\_A0A6P0LXV0 \| 99.7 \| 1e-19 \| 1.9e-25 \| 145.9 \| 130 \| (1, 144) \| 475 \| (1, 136) \| 171 \| Phage tail sheath family protein (Fragment) \| Phage tail sheath family protein (Fragment); uniclust \| UniRef100\_A0A350EYQ5 \| 99.7 \| 1e-19 \| 1.9e-25 \| 148.6 \| 201 \| (4, 316) \| 475 \| (3, 213) \| 216 \| Phage tail sheath family protein (Fragment) \| Phage tail sheath family protein (Fragment); uniclust \| UniRef100\_A0A1G7 \| 99.7 \| 1e-19 \| 1.9e-25 \| 163.4 \| 232 \| (231, 468) \| 475 \| (79, 341) \| 353 \| Tail tube glycoprotein (Fragment) \| Tail tube glycoprotein (Fragment); uniclust \| UniRef100\_A0A1V5KBQ7 \| 99.7 \| 1.6e-19 \| 2.8e-25 \| 173.9 \| 234 \| (229, 466) \| 475 \| (625, 883) \| 894 \| Phage tail sheath protein \| Phage tail sheath protein; uniclust \| UniRef100\_A0A926CBS6 \| 99.7 \| 1.6e-19 \| 2.9e-25 \| 148.8 \| 195 \| (267, 468) \| 475 \| (19, 220) \| 228 \| Phage tail sheath family protein \| Phage tail sheath family protein; uniclust \| UniRef100\_UPI000A73B8BF \| 99.7 \| 1.6e-19 \| 3e-25 \| 147.4 \| 193 \| (2, 282) \| 475 \| (1, 215) \| 215 \| phage tail sheath subtilisin-like domain-containing protein \| phage tail sheath subtilisin-like domain-containing protein; uniclust \| UniRef100\_A0A2A4T3R4 \| 99.7 \| 1.6e-19 \| 3.1e-25 \| 187.7 \| 232 \| (231, 469) \| 475 \| (784, 1035) \| 1041 \| Tail sheath protein C-terminal domain-containing protein \| Tail sheath protein C-terminal domain-containing protein; uniclust \| UniRef100\_A0A950BPE5 \| 99.7 \| 2e-19 \| 3.8e-25 \| 158.2 \| 311 \| (2, 328) \| 475 \| (1, 363) \| 370 \| Phage tail sheath family protein (Fragment) \| Phage tail sheath family protein (Fragment); uniclust \| UniRef100\_A0A497CTY6 \| 99.6 \| 2.5e-19 \| 4.7e-25 \| 164.5 \| 127 \| (229, 361) \| 475 \| (357, 499) \| 499 \| Peptidase C30 domain-containing protein (Fragment) \| Peptidase C30 domain-containing protein (Fragment); uniclust \| UniRef100\_A0A2D6E608 \| 99.6 \| 2.7e-19 \| 5.1e-25 \| 176.5 \| 240 \| (229, 474) \| 475 \| (529, 800) \| 801 \| Tail sheath protein C-terminal domain-containing protein \| Tail sheath protein C-terminal domain-containing protein; uniclust \| UniRef100\_A0A1L6I8K2 \| 99.6 \| 3.2e-19 \| 5.9e-25 \| 164.7 \| 280 \| (168, 466) \| 475 \| (97, 391) \| 405 \| Phage tail protein \| Phage tail protein; uniclust \| UniRef100\_A0A661H7Y5 \| 99.6 \| 5.7e-19 \| 1.1e-24 \| 141.5 \| 129 \| (337, 467) \| 475 \| (20, 158) \| 164 \| Tail sheath protein C-terminal domain-containing protein \| Tail sheath protein C-terminal domain-containing protein; uniclust \| UniRef100\_A0A261KJC2 \| 99.6 \| 5.8e-19 \| 1.1e-24 \| 165.9 \| 91 \| (1, 94) \| 475 \| (1, 94) \| 327 \| Tail protein (Fragment) \| Tail protein (Fragment); uniclust \| UniRef100\_A0A150TG83 \| 99.6 \| 6.2e-19 \| 1.2e-24 \| 169.0 \| 197 \| (268, 470) \| 475 \| (477, 685) \| 699 \| Tail sheath protein C-terminal domain-containing protein \| Tail sheath protein C-terminal domain-containing protein; uniclust \| UniRef100\_A0A0H2YXP9 \| 99.6 \| 6.6e-19 \| 1.2e-24 \| 144.6 \| 137 \| (3, 144) \| 475 \| (1, 143) \| 185 \| GpF1 \| GpF1; uniclust \| UniRef100\_A0A068ZW53 \| 99.6 \| 6.4e-19 \| 1.3e-24 \| 173.8 \| 414 \| (6, 463) \| 475 \| (34, 502) \| 508 \| Phage tail sheath protein \| Phage tail sheath protein; uniclust \| UniRef100\_A0A7X6FQF0 \| 99.6 \| 7e-19 \| 1.3e-24 \| 148.4 \| 238 \| (2, 251) \| 475 \| (1, 247) \| 266 \| Phage tail protein \| Phage tail protein; uniclust \| UniRef100\_A0A951GZA7 \| 99.6 \| 7.3e-19 \| 1.3e-24 \| 157.6 \| 163 \| (231, 399) \| 475 \| (232, 427) \| 428 \| Phage tail sheath family protein (Fragment) \| Phage tail sheath family protein (Fragment); uniclust \| UniRef100\_J2VU54 \| 99.6 \| 7.2e-19 \| 1.4e-24 \| 157.2 \| 73 \| (287, 365) \| 475 \| (239, 313) \| 318 \| Uncharacterized protein (Fragment) \| Uncharacterized protein (Fragment); uniclust \| UniRef100\_A0A8J7Q6J0 \| 99.6 \| 8.9e-19 \| 1.6e-24 \| 148.1 \| 116 \| (353, 468) \| 475 \| (8, 129) \| 270 \| Phage tail sheath family protein \| Phage tail sheath family protein; uniclust \| UniRef100\_A0A5D8RP33 \| 99.6 \| 9e-19 \| 1.7e-24 \| 135.2 \| 117 \| (357, 473) \| 475 \| (1, 120) \| 129 \| Phage tail sheath family protein (Fragment) \| Phage tail sheath family protein (Fragment); uniclust \| UniRef100\_A0A6V8KHY3 \| 99.6 \| 8.9e-19 \| 1.7e-24 \| 133.3 \| 95 \| (374, 468) \| 475 \| (1, 101) \| 110 \| Tail sheath protein C-terminal domain-containing protein \| Tail sheath protein C-terminal domain-containing protein; uniclust \| UniRef100\_A0A6P0JV64 \| 99.6 \| 9.9e-19 \| 1.9e-24 \| 155.3 \| 220 \| (2, 323) \| 475 \| (1, 304) \| 310 \| Phage tail sheath family protein (Fragment) \| Phage tail sheath family protein (Fragment); uniclust \| UniRef100\_UPI000615A201 \| 99.6 \| 1.1e-18 \| 2e-24 \| 162.3 \| 202 \| (1, 303) \| 475 \| (1, 316) \| 338 \| phage tail sheath C-terminal domain-containing protein \| phage tail sheath C-terminal domain-containing protein; uniclust \| UniRef100\_UPI00156E4C90 \| 99.6 \| 1.3e-18 \| 2.4e-24 \| 134.0 \| 126 \| (269, 394) \| 475 \| (3, 129) \| 130 \| phage tail sheath subtilisin-like domain-containing protein \| phage tail sheath subtilisin-like domain-containing protein; uniclust \| UniRef100\_A0A4P9VIU9 \| 99.6 \| 1.3e-18 \| 2.5e-24 \| 137.7 \| 130 \| (2, 137) \| 475 \| (1, 130) \| 141 \| Phage tail protein \| Phage tail protein; uniclust \| UniRef100\_A0A7L5Y128 \| 99.6 \| 1.8e-18 \| 3.2e-24 \| 163.7 \| 337 \| (43, 466) \| 475 \| (331, 687) \| 713 \| Right-handed parallel beta-helix repeat-containing protein \| Right-handed parallel beta-helix repeat-containing protein; uniclust \| UniRef100\_A0A4U9D4B3 \| 99.6 \| 1.8e-18 \| 3.3e-24 \| 147.4 \| 194 \| (32, 315) \| 475 \| (1, 199) \| 285 \| Phage tail sheath protein \| Phage tail sheath protein; uniclust \| UniRef100\_A0A3B8PUP4 \| 99.6 \| 2.1e-18 \| 3.9e-24 \| 156.9 \| 140 \| (229, 374) \| 475 \| (333, 485) \| 485 \| Phage tail protein (Fragment) \| Phage tail protein (Fragment); uniclust \| UniRef100\_A0A955R619 \| 99.6 \| 3.3e-18 \| 6e-24 \| 145.6 \| 232 \| (229, 467) \| 475 \| (12, 272) \| 281 \| Phage tail sheath family protein \| Phage tail sheath family protein; uniclust \| UniRef100\_UPI000D149F10 \| 99.6 \| 3.3e-18 \| 6.1e-24 \| 140.3 \| 150 \| (267, 426) \| 475 \| (58, 210) \| 217 \| phage tail sheath C-terminal domain-containing protein \| phage tail sheath C-terminal domain-containing protein; uniclust \| UniRef100\_A0A099P8J3 \| 99.6 \| 3.2e-18 \| 6.1e-24 \| 130.7 \| 87 \| (1, 95) \| 475 \| (1, 90) \| 106 \| Uncharacterized protein (Fragment) \| Uncharacterized protein (Fragment); uniclust \| UniRef100\_A0A379QW15 \| 99.6 \| 3.6e-18 \| 6.8e-24 \| 133.0 \| 128 \| (346, 474) \| 475 \| (2, 134) \| 140 \| Tail protein \| Tail protein; uniclust \| UniRef100\_A0A7S5T3D9 \| 99.6 \| 3.7e-18 \| 6.9e-24 \| 165.7 \| 229 \| (229, 463) \| 475 \| (572, 817) \| 833 \| Tail sheath protein subtilisin-like domain-containing protein \| Tail sheath protein subtilisin-like domain-containing protein; uniclust \| UniRef100\_A0A660SQZ6 \| 99.6 \| 4e-18 \| 7.4e-24 \| 149.2 \| 234 \| (231, 468) \| 475 \| (91, 341) \| 347 \| Tail sheath protein C-terminal domain-containing protein (Fragment) \| Tail sheath protein C-terminal domain-containing protein (Fragment); uniclust \| UniRef100\_UPI000A41458E \| 99.6 \| 4.9e-18 \| 9e-24 \| 133.6 \| 135 \| (266, 400) \| 475 \| (21, 161) \| 163 \| phage tail sheath subtilisin-like domain-containing protein \| phage tail sheath subtilisin-like domain-containing protein; uniclust \| UniRef100\_A0A9C8Q3C5 \| 99.6 \| 5.4e-18 \| 9.9e-24 \| 134.7 \| 137 \| (337, 473) \| 475 \| (31, 168) \| 177 \| Phage tail sheath family protein \| Phage tail sheath family protein; uniclust \| UniRef100\_UPI00158EB90D \| 99.6 \| 5.9e-18 \| 1.1e-23 \| 150.2 \| 225 \| (231, 464) \| 475 \| (120, 365) \| 387 \| phage tail sheath family protein \| phage tail sheath family protein; uniclust \| UniRef100\_A0A963ER11 \| 99.6 \| 7.9e-18 \| 1.5e-23 \| 144.6 \| 224 \| (237, 470) \| 475 \| (7, 243) \| 259 \| Uncharacterized protein \| Uncharacterized protein; uniclust \| UniRef100\_A0A0A0BPC6 \| 99.6 \| 1e-17 \| 1.9e-23 \| 147.8 \| 208 \| (259, 475) \| 475 \| (43, 265) \| 269 \| Tail protein \| Tail protein; uniclust \| UniRef100\_A0A0F8VRZ6 \| 99.6 \| 1.1e-17 \| 2e-23 \| 131.1 \| 127 \| (343, 469) \| 475 \| (3, 132) \| 139 \| Tail sheath protein C-terminal domain-containing protein (Fragment) \| Tail sheath protein C-terminal domain-containing protein (Fragment); uniclust \| UniRef100\_A0A1Z4K7L4 \| 99.5 \| 1.1e-17 \| 2.1e-23 \| 150.5 \| 224 \| (230, 463) \| 475 \| (127, 367) \| 394 \| Phage tail sheath protein fi-like protein \| Phage tail sheath protein fi-like protein; uniclust \| UniRef100\_A0A6G5QYL8 \| 99.5 \| 1.4e-17 \| 2.6e-23 \| 156.6 \| 240 \| (231, 474) \| 475 \| (411, 653) \| 658 \| Phage tail sheath protein FI (Integrase domain) \| Phage tail sheath protein FI (Integrase domain); uniclust \| UniRef100\_UPI00223C8DA3 \| 99.5 \| 2.1e-17 \| 3.8e-23 \| 145.1 \| 200 \| (266, 470) \| 475 \| (137, 344) \| 350 \| hypothetical protein \| hypothetical protein; uniclust \| UniRef100\_A0A8S5MEQ3 \| 99.5 \| 2.1e-17 \| 3.8e-23 \| 157.8 \| 235 \| (229, 469) \| 475 \| (514, 764) \| 768 \| Tail sheath protein \| Tail sheath protein; uniclust \| UniRef100\_UPI001C5FF8DD \| 99.5 \| 2.1e-17 \| 3.9e-23 \| 115.9 \| 85 \| (289, 373) \| 475 \| (3, 87) \| 87 \| phage tail sheath subtilisin-like domain-containing protein \| phage tail sheath subtilisin-like domain-containing protein; uniclust \| UniRef100\_A0A0F9PP94 \| 99.5 \| 2.1e-17 \| 4e-23 \| 166.9 \| 231 \| (229, 465) \| 475 \| (504, 752) \| 761 \| Tail sheath protein C-terminal domain-containing protein \| Tail sheath protein C-terminal domain-containing protein; uniclust \| UniRef100\_A0A7Y0GV13 \| 99.5 \| 3.3e-17 \| 6.2e-23 \| 152.8 \| 90 \| (229, 318) \| 475 \| (388, 483) \| 487 \| Phage tail sheath family protein (Fragment) \| Phage tail sheath family protein (Fragment); uniclust \| UniRef100\_A0A0C2VDI1 \| 99.5 \| 3.6e-17 \| 7.1e-23 \| 166.0 \| 229 \| (228, 467) \| 475 \| (322, 569) \| 591 \| Tail sheath protein subtilisin-like domain-containing protein \| Tail sheath protein subtilisin-like domain-containing protein; uniclust \| UniRef100\_A0A2H0PCS0 \| 99.5 \| 4e-17 \| 7.3e-23 \| 155.2 \| 244 \| (229, 474) \| 475 \| (453, 724) \| 728 \| Tail sheath protein C-terminal domain-containing protein \| Tail sheath protein C-terminal domain-containing protein; uniclust \| UniRef100\_A0A0F9KEX1 \| 99.5 \| 4e-17 \| 7.4e-23 \| 158.4 \| 226 \| (243, 468) \| 475 \| (681, 922) \| 926 \| Tail sheath protein C-terminal domain-containing protein (Fragment) \| Tail sheath protein C-terminal domain-containing protein (Fragment); uniclust \| UniRef100\_A0A6V8LKN3 \| 99.5 \| 4.4e-17 \| 8.2e-23 \| 125.4 \| 117 \| (353, 469) \| 475 \| (15, 138) \| 144 \| Tail sheath protein C-terminal domain-containing protein \| Tail sheath protein C-terminal domain-containing protein; uniclust \| UniRef100\_A0A4Q5HCU8 \| 99.5 \| 4.8e-17 \| 8.7e-23 \| 122.9 \| 113 \| (287, 402) \| 475 \| (10, 126) \| 129 \| Phage tail sheath family protein \| Phage tail sheath family protein; uniclust \| UniRef100\_A0A1F3SEE9 \| 99.5 \| 6e-17 \| 1.1e-22 \| 143.3 \| 322 \| (22, 463) \| 475 \| (23, 365) \| 370 \| Tail sheath protein C-terminal domain-containing protein \| Tail sheath protein C-terminal domain-containing protein; uniclust \| UniRef100\_A0A434RRN7 \| 99.5 \| 6.7e-17 \| 1.2e-22 \| 139.6 \| 241 \| (230, 471) \| 475 \| (40, 293) \| 307 \| Phage tail protein (Fragment) \| Phage tail protein (Fragment); uniclust \| UniRef100\_A0A7W0HPR7 \| 99.5 \| 8.2e-17 \| 1.5e-22 \| 134.3 \| 202 \| (29, 360) \| 475 \| (4, 211) \| 238 \| Tail sheath protein subtilisin-like domain-containing protein \| Tail sheath protein subtilisin-like domain-containing protein; uniclust \| UniRef100\_A0A1H2UW00 \| 99.5 \| 8.1e-17 \| 1.5e-22 \| 123.1 \| 112 \| (363, 474) \| 475 \| (2, 114) \| 122 \| Phage tail sheath protein \| Phage tail sheath protein; uniclust \| UniRef100\_A0A2D7XGG3 \| 99.5 \| 9.7e-17 \| 1.8e-22 \| 155.1 \| 196 \| (266, 467) \| 475 \| (582, 784) \| 793 \| Tail sheath protein C-terminal domain-containing protein \| Tail sheath protein C-terminal domain-containing protein; uniclust \| UniRef100\_UPI0013CE83EC \| 99.5 \| 1e-16 \| 1.8e-22 \| 117.8 \| 104 \| (259, 362) \| 475 \| (2, 106) \| 106 \| phage tail sheath subtilisin-like domain-containing protein \| phage tail sheath subtilisin-like domain-containing protein; uniclust \| UniRef100\_A0A024AZG0 \| 99.5 \| 1.2e-16 \| 2.4e-22 \| 163.5 \| 227 \| (229, 471) \| 475 \| (337, 581) \| 615 \| Tail sheath protein \| Tail sheath protein; uniclust \| UniRef100\_UPI001D12A7C3 \| 99.5 \| 1.3e-16 \| 2.4e-22 \| 143.0 \| 305 \| (7, 430) \| 475 \| (44, 368) \| 410 \| phage tail sheath C-terminal domain-containing protein \| phage tail sheath C-terminal domain-containing protein; uniclust \| UniRef100\_A0A1Z8RYI1 \| 99.5 \| 1.4e-16 \| 2.7e-22 \| 156.9 \| 200 \| (265, 470) \| 475 \| (514, 720) \| 723 \| Tail sheath protein subtilisin-like domain-containing protein (Fragment) \| Tail sheath protein subtilisin-like domain-containing protein (Fragment); uniclust \| UniRef100\_A0A0F6W355 \| 99.5 \| 1.6e-16 \| 3e-22 \| 156.4 \| 422 \| (5, 472) \| 475 \| (1, 481) \| 489 \| Tail sheath protein subtilisin-like domain-containing protein \| Tail sheath protein subtilisin-like domain-containing protein; uniclust \| UniRef100\_A0A0Q0D1V1 \| 99.5 \| 1.7e-16 \| 3.1e-22 \| 121.1 \| 118 \| (357, 474) \| 475 \| (1, 118) \| 122 \| Major tail sheath protein \| Major tail sheath protein; uniclust \| UniRef100\_A0A7S5RC19 \| 99.5 \| 2.1e-16 \| 3.8e-22 \| 151.4 \| 233 \| (231, 469) \| 475 \| (516, 761) \| 773 \| Tail sheath protein \| Tail sheath protein; uniclust \| UniRef100\_A0A2Z3EDS5 \| 99.4 \| 2.5e-16 \| 4.6e-22 \| 150.0 \| 237 \| (229, 468) \| 475 \| (459, 712) \| 727 \| Tail sheath protein \| Tail sheath protein; uniclust \| UniRef100\_A0A4V6JGZ6 \| 99.4 \| 2.8e-16 \| 5.1e-22 \| 138.8 \| 259 \| (41, 390) \| 475 \| (75, 351) \| 359 \| Phage tail sheath protein \| Phage tail sheath protein; uniclust \| UniRef100\_UPI00096A297B \| 99.4 \| 3e-16 \| 5.5e-22 \| 132.8 \| 191 \| (18, 321) \| 475 \| (9, 206) \| 261 \| phage tail sheath family protein \| phage tail sheath family protein; uniclust \| UniRef100\_UPI001922CB15 \| 99.4 \| 3.7e-16 \| 6.7e-22 \| 127.6 \| 184 \| (41, 324) \| 475 \| (13, 197) \| 204 \| phage tail sheath subtilisin-like domain-containing protein \| phage tail sheath subtilisin-like domain-containing protein; uniclust \| UniRef100\_UPI00197F3314 \| 99.4 \| 3.9e-16 \| 7.2e-22 \| 116.7 \| 116 \| (359, 474) \| 475 \| (2, 118) \| 120 \| phage tail sheath C-terminal domain-containing protein \| phage tail sheath C-terminal domain-containing protein; uniclust \| UniRef100\_A0A1M5WCL1 \| 99.4 \| 4.1e-16 \| 7.8e-22 \| 124.8 \| 86 \| (2, 95) \| 475 \| (1, 92) \| 136 \| Phage tail sheath protein \| Phage tail sheath protein; uniclust \| UniRef100\_A0A0T9RBS8 \| 99.4 \| 4.9e-16 \| 9.2e-22 \| 119.0 \| 90 \| (2, 95) \| 475 \| (1, 90) \| 118 \| Phage tail sheath monomer \| Phage tail sheath monomer; uniclust \| UniRef100\_UPI00234B027B \| 99.4 \| 6.6e-16 \| 1.2e-21 \| 133.5 \| 139 \| (2, 145) \| 475 \| (1, 139) \| 302 \| ATP-binding protein \| ATP-binding protein; uniclust \| UniRef100\_A0A257LFR8 \| 99.4 \| 6.3e-16 \| 1.2e-21 \| 142.4 \| 85 \| (5, 97) \| 475 \| (8, 94) \| 272 \| Phage tail protein (Fragment) \| Phage tail protein (Fragment); uniclust \| UniRef100\_A0A3D0UP20 \| 99.4 \| 6.7e-16 \| 1.2e-21 \| 115.3 \| 88 \| (310, 403) \| 475 \| (1, 90) \| 119 \| Tail sheath protein subtilisin-like domain-containing protein (Fragment) \| Tail sheath protein subtilisin-like domain-containing protein (Fragment); uniclust \| UniRef100\_A0A1C4JDU7 \| 99.4 \| 6.7e-16 \| 1.3e-21 \| 135.9 \| 87 \| (2, 96) \| 475 \| (1, 89) \| 233 \| Phage tail sheath protein \| Phage tail sheath protein; uniclust \| UniRef100\_UPI0018EC0130 \| 99.4 \| 7.4e-16 \| 1.4e-21 \| 110.4 \| 92 \| (289, 381) \| 475 \| (1, 95) \| 95 \| phage tail sheath subtilisin-like domain-containing protein \| phage tail sheath subtilisin-like domain-containing protein; uniclust \| UniRef100\_A0A3B9JCS2 \| 99.4 \| 8.1e-16 \| 1.5e-21 \| 121.1 \| 91 \| (1, 94) \| 475 \| (1, 103) \| 125 \| Phage tail sheath family protein (Fragment) \| Phage tail sheath family protein (Fragment); uniclust \| UniRef100\_A0A1M6F1L4 \| 99.4 \| 8.6e-16 \| 1.6e-21 \| 136.5 \| 122 \| (2, 142) \| 475 \| (1, 159) \| 268 \| Phage tail sheath protein \| Phage tail sheath protein; uniclust \| UniRef100\_A0A0B0HIM0 \| 99.4 \| 1e-15 \| 1.9e-21 \| 117.3 \| 109 \| (367, 475) \| 475 \| (4, 112) \| 114 \| Phage tail sheath protein \| Phage tail sheath protein; uniclust \| UniRef100\_A0A2I7QP51 \| 99.4 \| 1.3e-15 \| 2.3e-21 \| 155.2 \| 231 \| (229, 464) \| 475 \| (1030, 1276) \| 1283 \| Tail sheath protein \| Tail sheath protein; uniclust \| UniRef100\_A0A484YNN0 \| 99.4 \| 1.4e-15 \| 2.5e-21 \| 122.0 \| 134 \| (233, 366) \| 475 \| (3, 138) \| 180 \| Tail Sheath protein \| Tail Sheath protein; uniclust \| UniRef100\_A0A2E2FJ22 \| 99.4 \| 1.9e-15 \| 3.5e-21 \| 149.0 \| 239 \| (229, 469) \| 475 \| (360, 637) \| 649 \| Tail sheath protein subtilisin-like domain-containing protein \| Tail sheath protein subtilisin-like domain-containing protein; uniclust \| UniRef100\_A0A0P7JRX0 \| 99.4 \| 2.1e-15 \| 3.9e-21 \| 139.1 \| 404 \| (2, 466) \| 475 \| (1, 428) \| 434 \| Tail sheath protein subtilisin-like domain-containing protein \| Tail sheath protein subtilisin-like domain-containing protein; uniclust \| UniRef100\_A0A2X3JRA6 \| 99.4 \| 2.2e-15 \| 4e-21 \| 117.7 \| 141 \| (238, 379) \| 475 \| (8, 150) \| 153 \| Major tail sheath protein FI \| Major tail sheath protein FI; uniclust \| UniRef100\_A0A1H5ZKW0 \| 99.4 \| 2.2e-15 \| 4.1e-21 \| 144.5 \| 384 \| (7, 466) \| 475 \| (10, 426) \| 440 \| Phage tail sheath protein \| Phage tail sheath protein; uniclust \| UniRef100\_A0A1H8RZB9 \| 99.4 \| 2.3e-15 \| 4.4e-21 \| 146.1 \| 411 \| (1, 468) \| 475 \| (1, 441) \| 451 \| Phage tail sheath protein \| Phage tail sheath protein; uniclust \| UniRef100\_X1DII4 \| 99.4 \| 2.8e-15 \| 5.2e-21 \| 113.6 \| 85 \| (343, 427) \| 475 \| (17, 104) \| 128 \| Tail sheath protein C-terminal domain-containing protein (Fragment) \| Tail sheath protein C-terminal domain-containing protein (Fragment); uniclust \| UniRef100\_A0A7V9TCY7 \| 99.4 \| 3.1e-15 \| 5.7e-21 \| 113.1 \| 83 \| (4, 94) \| 475 \| (4, 88) \| 117 \| Phage tail sheath family protein (Fragment) \| Phage tail sheath family protein (Fragment); uniclust \| UniRef100\_UPI0015CF41A0 \| 99.3 \| 3.8e-15 \| 7.1e-21 \| 108.5 \| 83 \| (287, 376) \| 475 \| (9, 94) \| 94 \| phage tail sheath subtilisin-like domain-containing protein \| phage tail sheath subtilisin-like domain-containing protein; uniclust \| UniRef100\_A0A076YKJ9 \| 99.3 \| 4.6e-15 \| 8.6e-21 \| 149.0 \| 236 \| (232, 470) \| 475 \| (582, 831) \| 839 \| Putative tail sheath protein \| Putative tail sheath protein; uniclust \| UniRef100\_A0A6B3GW63 \| 99.3 \| 5.3e-15 \| 9.8e-21 \| 119.5 \| 98 \| (231, 328) \| 475 \| (80, 183) \| 187 \| Phage tail sheath family protein (Fragment) \| Phage tail sheath family protein (Fragment); uniclust \| UniRef100\_UPI0006CC31C6 \| 99.3 \| 5.6e-15 \| 1e-20 \| 112.1 \| 110 \| (337, 446) \| 475 \| (12, 126) \| 128 \| hypothetical protein \| hypothetical protein; uniclust \| UniRef100\_A0A946FFJ9 \| 99.3 \| 5.8e-15 \| 1.1e-20 \| 135.0 \| 87 \| (2, 96) \| 475 \| (1, 89) \| 370 \| Uncharacterized protein (Fragment) \| Uncharacterized protein (Fragment); uniclust \| UniRef100\_A0A7C3UJZ4 \| 99.3 \| 5.9e-15 \| 1.1e-20 \| 147.5 \| 201 \| (263, 470) \| 475 \| (566, 770) \| 776 \| Tail sheath protein C-terminal domain-containing protein \| Tail sheath protein C-terminal domain-containing protein; uniclust \| UniRef100\_UPI001D0F62D8 \| 99.3 \| 6.2e-15 \| 1.1e-20 \| 126.2 \| 180 \| (231, 416) \| 475 \| (61, 244) \| 275 \| phage tail sheath subtilisin-like domain-containing protein \| phage tail sheath subtilisin-like domain-containing protein; uniclust \| UniRef100\_A0A5C7M3L5 \| 99.3 \| 7.6e-15 \| 1.4e-20 \| 132.7 \| 132 \| (229, 363) \| 475 \| (273, 418) \| 420 \| Tail sheath protein subtilisin-like domain-containing protein (Fragment) \| Tail sheath protein subtilisin-like domain-containing protein (Fragment); uniclust \| UniRef100\_A0A644XLY2 \| 99.3 \| 8.2e-15 \| 1.5e-20 \| 113.4 \| 123 \| (353, 475) \| 475 \| (3, 129) \| 132 \| Prophage major tail sheath protein \| Prophage major tail sheath protein; uniclust \| UniRef100\_W4MDK3 \| 99.3 \| 8.9e-15 \| 1.6e-20 \| 121.1 \| 186 \| (2, 285) \| 475 \| (1, 193) \| 218 \| Tail sheath protein subtilisin-like domain-containing protein \| Tail sheath protein subtilisin-like domain-containing protein; uniclust \| UniRef100\_A0A0U2YXC1 \| 99.3 \| 9.8e-15 \| 1.9e-20 \| 143.6 \| 90 \| (1, 98) \| 475 \| (1, 92) \| 476 \| Phage tail sheath protein (Fragment) \| Phage tail sheath protein (Fragment); uniclust \| UniRef100\_A0A1Z8WSY0 \| 99.3 \| 1e-14 \| 1.9e-20 \| 152.1 \| 199 \| (265, 469) \| 475 \| (1466, 1672) \| 1675 \| Tail sheath protein C-terminal domain-containing protein \| Tail sheath protein C-terminal domain-containing protein; uniclust \| UniRef100\_A0A8S0FU88 \| 99.3 \| 1.2e-14 \| 2.3e-20 \| 109.2 \| 105 \| (286, 390) \| 475 \| (16, 120) \| 121 \| Tail sheath protein subtilisin-like domain-containing protein \| Tail sheath protein subtilisin-like domain-containing protein; uniclust \| UniRef100\_A0A1Y3CXG9 \| 99.3 \| 1.3e-14 \| 2.4e-20 \| 132.4 \| 187 \| (3, 279) \| 475 \| (1, 194) \| 343 \| FHA domain-containing protein \| FHA domain-containing protein; uniclust \| UniRef100\_A0A967K953 \| 99.3 \| 1.4e-14 \| 2.5e-20 \| 96.0 \| 63 \| (355, 417) \| 475 \| (1, 63) \| 63 \| Phage tail sheath family protein (Fragment) \| Phage tail sheath family protein (Fragment); uniclust \| UniRef100\_A0A327KJ47 \| 99.3 \| 1.5e-14 \| 2.8e-20 \| 126.7 \| 314 \| (2, 326) \| 475 \| (2, 325) \| 326 \| Phage tail protein (Fragment) \| Phage tail protein (Fragment); uniclust \| UniRef100\_A0A3C2AP21 \| 99.3 \| 1.6e-14 \| 3e-20 \| 145.2 \| 203 \| (265, 468) \| 475 \| (851, 1064) \| 1069 \| Uncharacterized protein (Fragment) \| Uncharacterized protein (Fragment); uniclust \| UniRef100\_A0A3B8IT26 \| 99.3 \| 2e-14 \| 3.7e-20 \| 133.5 \| 121 \| (287, 408) \| 475 \| (404, 527) \| 528 \| Tail sheath protein subtilisin-like domain-containing protein (Fragment) \| Tail sheath protein subtilisin-like domain-containing protein (Fragment); uniclust \| UniRef100\_H6WFT7 \| 99.3 \| 2.7e-14 \| 4.9e-20 \| 142.1 \| 191 \| (268, 468) \| 475 \| (909, 1111) \| 1137 \| Tail sheath protein \| Tail sheath protein; uniclust \| UniRef100\_A0A068NXW1 \| 99.3 \| 2.7e-14 \| 5.3e-20 \| 138.9 \| 331 \| (7, 467) \| 475 \| (23, 394) \| 398 \| Tail sheath protein subtilisin-like domain-containing protein \| Tail sheath protein subtilisin-like domain-containing protein; uniclust \| UniRef100\_A0A524QBZ3 \| 99.2 \| 2.9e-14 \| 5.4e-20 \| 135.4 \| 87 \| (2, 95) \| 475 \| (44, 137) \| 550 \| Phage tail sheath family protein (Fragment) \| Phage tail sheath family protein (Fragment); uniclust \| UniRef100\_A0A926AT07 \| 99.2 \| 2.9e-14 \| 5.5e-20 \| 132.3 \| 85 \| (4, 96) \| 475 \| (4, 90) \| 308 \| Phage tail sheath family protein (Fragment) \| Phage tail sheath family protein (Fragment); uniclust \| UniRef100\_A0A327JRX8 \| 99.2 \| 3e-14 \| 5.5e-20 \| 114.5 \| 156 \| (307, 464) \| 475 \| (5, 160) \| 178 \| Tail sheath protein C-terminal domain-containing protein (Fragment) \| Tail sheath protein C-terminal domain-containing protein (Fragment); uniclust \| UniRef100\_A0A356X1N2 \| 99.2 \| 3e-14 \| 5.6e-20 \| 117.0 \| 173 \| (295, 469) \| 475 \| (2, 182) \| 187 \| Phage tail protein (Fragment) \| Phage tail protein (Fragment); uniclust \| UniRef100\_A0A0C3J1K3 \| 99.2 \| 3.1e-14 \| 6e-20 \| 142.8 \| 186 \| (267, 470) \| 475 \| (301, 507) \| 523 \| Phage tail sheath family protein \| Phage tail sheath family protein; uniclust \| UniRef100\_A0A6V6ZU17 \| 99.2 \| 3.3e-14 \| 6.1e-20 \| 134.9 \| 317 \| (1, 330) \| 475 \| (1, 332) \| 649 \| Uncharacterized protein \| Uncharacterized protein; uniclust \| UniRef100\_A0A075BS62 \| 99.2 \| 3.5e-14 \| 6.5e-20 \| 140.1 \| 232 \| (229, 468) \| 475 \| (522, 768) \| 776 \| Putative tail sheath protein \| Putative tail sheath protein; uniclust \| UniRef100\_A0A239K6F1 \| 99.2 \| 3.9e-14 \| 7.4e-20 \| 118.6 \| 88 \| (1, 96) \| 475 \| (1, 90) \| 187 \| Phage tail sheath protein \| Phage tail sheath protein; uniclust \| UniRef100\_A0A370E0M7 \| 99.2 \| 3.9e-14 \| 7.4e-20 \| 118.1 \| 123 \| (2, 144) \| 475 \| (3, 149) \| 160 \| Uncharacterized protein \| Uncharacterized protein; uniclust \| UniRef100\_A0A931KX23 \| 99.2 \| 4.2e-14 \| 7.8e-20 \| 132.4 \| 396 \| (19, 467) \| 475 \| (15, 457) \| 462 \| Uncharacterized protein \| Uncharacterized protein; uniclust \| UniRef100\_A0A956AK97 \| 99.2 \| 4.8e-14 \| 8.9e-20 \| 115.9 \| 176 \| (245, 424) \| 475 \| (2, 196) \| 205 \| Phage tail protein (Fragment) \| Phage tail protein (Fragment); uniclust \| UniRef100\_A0A4Y8ZK08 \| 99.2 \| 4.7e-14 \| 8.9e-20 \| 116.8 \| 83 \| (6, 96) \| 475 \| (6, 90) \| 175 \| Phage tail sheath family protein (Fragment) \| Phage tail sheath family protein (Fragment); uniclust \| UniRef100\_A0A7K0P9W3 \| 99.2 \| 4.9e-14 \| 9.1e-20 \| 133.1 \| 232 \| (230, 466) \| 475 \| (227, 480) \| 486 \| Tail sheath protein subtilisin-like domain-containing protein \| Tail sheath protein subtilisin-like domain-containing protein; uniclust \| UniRef100\_A0A1I1S0H2 \| 99.2 \| 4.9e-14 \| 9.2e-20 \| 105.6 \| 87 \| (381, 467) \| 475 \| (2, 95) \| 99 \| Phage tail sheath protein \| Phage tail sheath protein; uniclust \| UniRef100\_A0A6J5RUV9 \| 99.2 \| 5.1e-14 \| 9.3e-20 \| 125.8 \| 336 \| (8, 464) \| 475 \| (3, 362) \| 373 \| Uncharacterized protein \| Uncharacterized protein; uniclust \| UniRef100\_A0A368A7G8 \| 99.2 \| 5.1e-14 \| 9.4e-20 \| 115.4 \| 74 \| (260, 334) \| 475 \| (110, 183) \| 201 \| Phage tail sheath protein FI \| Phage tail sheath protein FI; uniclust \| UniRef100\_A0A839HMA6 \| 99.2 \| 5.2e-14 \| 9.5e-20 \| 128.8 \| 180 \| (287, 467) \| 475 \| (259, 454) \| 457 \| Phage tail sheath family protein \| Phage tail sheath family protein; uniclust \| UniRef100\_A0A935W4P1 \| 99.2 \| 5.4e-14 \| 1e-19 \| 110.7 \| 105 \| (364, 468) \| 475 \| (23, 133) \| 140 \| Uncharacterized protein \| Uncharacterized protein; uniclust \| UniRef100\_A0A7C5GVL1 \| 99.2 \| 6e-14 \| 1.1e-19 \| 127.8 \| 126 \| (1, 144) \| 475 \| (1, 159) \| 325 \| Phage tail protein (Fragment) \| Phage tail protein (Fragment); uniclust \| UniRef100\_A0A2E7WB84 \| 99.2 \| 7e-14 \| 1.3e-19 \| 109.8 \| 112 \| (358, 469) \| 475 \| (2, 114) \| 124 \| Tail sheath protein C-terminal domain-containing protein (Fragment) \| Tail sheath protein C-terminal domain-containing protein (Fragment); uniclust \| UniRef100\_A0A450ZA88 \| 99.2 \| 7.7e-14 \| 1.4e-19 \| 117.3 \| 122 \| (230, 351) \| 475 \| (27, 150) \| 208 \| Tail sheath protein subtilisin-like domain-containing protein \| Tail sheath protein subtilisin-like domain-containing protein; uniclust \| UniRef100\_A0A3A8EJ92 \| 99.2 \| 7.7e-14 \| 1.5e-19 \| 113.8 \| 136 \| (2, 142) \| 475 \| (1, 139) \| 150 \| Phage tail protein (Fragment) \| Phage tail protein (Fragment); uniclust \| UniRef100\_A0A0G3M576 \| 99.2 \| 8.3e-14 \| 1.7e-19 \| 138.8 \| 368 \| (4, 468) \| 475 \| (2, 412) \| 415 \| DUF2586 family protein \| DUF2586 family protein; uniclust \| UniRef100\_UPI001872E667 \| 99.2 \| 9.2e-14 \| 1.7e-19 \| 115.8 \| 122 \| (353, 474) \| 475 \| (97, 218) \| 223 \| P2 family phage major capsid protein \| P2 family phage major capsid protein; uniclust \| UniRef100\_A0A0F9KVW3 \| 99.2 \| 9.2e-14 \| 1.7e-19 \| 138.0 \| 223 \| (244, 467) \| 475 \| (829, 1073) \| 1079 \| Tail sheath protein C-terminal domain-containing protein (Fragment) \| Tail sheath protein C-terminal domain-containing protein (Fragment); uniclust \| UniRef100\_UPI00193ADC5D \| 99.2 \| 9.6e-14 \| 1.8e-19 \| 110.3 \| 145 \| (326, 474) \| 475 \| (8, 159) \| 164 \| hypothetical protein \| hypothetical protein; uniclust \| UniRef100\_UPI00047DF1B9 \| 99.2 \| 1.3e-13 \| 2.4e-19 \| 127.1 \| 233 \| (229, 467) \| 475 \| (220, 474) \| 482 \| hypothetical protein \| hypothetical protein; uniclust \| UniRef100\_UPI001FEE2E79 \| 99.2 \| 1.7e-13 \| 3.1e-19 \| 113.2 \| 133 \| (340, 473) \| 475 \| (58, 193) \| 208 \| hypothetical protein \| hypothetical protein; uniclust \| UniRef100\_A0A523V8M4 \| 99.2 \| 1.8e-13 \| 3.3e-19 \| 125.8 \| 236 \| (225, 466) \| 475 \| (206, 460) \| 467 \| Tail sheath protein subtilisin-like domain-containing protein (Fragment) \| Tail sheath protein subtilisin-like domain-containing protein (Fragment); uniclust \| UniRef100\_UPI001F1A55FA \| 99.2 \| 1.9e-13 \| 3.5e-19 \| 108.3 \| 102 \| (230, 331) \| 475 \| (10, 112) \| 160 \| phage tail sheath subtilisin-like domain-containing protein \| phage tail sheath subtilisin-like domain-containing protein; uniclust \| UniRef100\_UPI001CC11497 \| 99.1 \| 2.2e-13 \| 4.1e-19 \| 103.6 \| 94 \| (372, 465) \| 475 \| (19, 118) \| 126 \| phage tail sheath C-terminal domain-containing protein \| phage tail sheath C-terminal domain-containing protein; uniclust \| UniRef100\_UPI001EEA4F30 \| 99.1 \| 2.3e-13 \| 4.2e-19 \| 98.4 \| 81 \| (308, 388) \| 475 \| (1, 81) \| 96 \| phage tail sheath subtilisin-like domain-containing protein \| phage tail sheath subtilisin-like domain-containing protein; uniclust \| UniRef100\_K8GN70 \| 99.1 \| 2.4e-13 \| 4.4e-19 \| 133.8 \| 214 \| (244, 467) \| 475 \| (704, 927) \| 939 \| Phage tail sheath protein FI \| Phage tail sheath protein FI; uniclust \| UniRef100\_A0A950PI68 \| 99.1 \| 2.8e-13 \| 5.2e-19 \| 107.8 \| 100 \| (365, 464) \| 475 \| (57, 161) \| 164 \| Phage tail sheath family protein \| Phage tail sheath family protein; uniclust \| UniRef100\_A0A3Q9LGI5 \| 99.1 \| 2.8e-13 \| 5.2e-19 \| 112.3 \| 167 \| (62, 315) \| 475 \| (6, 180) \| 192 \| Tail sheath protein subtilisin-like domain-containing protein \| Tail sheath protein subtilisin-like domain-containing protein; uniclust \| UniRef100\_A0A3B0XDA8 \| 99.1 \| 3.3e-13 \| 6.2e-19 \| 113.8 \| 87 \| (1, 95) \| 475 \| (1, 89) \| 196 \| Uncharacterized protein (Fragment) \| Uncharacterized protein (Fragment); uniclust \| UniRef100\_UPI001FCE0DAA \| 99.1 \| 3.9e-13 \| 7.2e-19 \| 111.9 \| 122 \| (225, 346) \| 475 \| (94, 217) \| 218 \| phage tail sheath subtilisin-like domain-containing protein \| phage tail sheath subtilisin-like domain-containing protein; pdb70 \| 6RBN\_D \| 100.0 \| 1.6e-43 \| 1.2e-47 \| 348.8 \| 351 \| (1, 468) \| 475 \| (1, 450) \| 451 \| Afp1, Afp2, Afp3, Afp16 \| 6RBN\_D Afp1, Afp2, Afp3, Afp16 Anti-feeding prophage, secretion system, AFP; pdb70 \| 6RAO\_D \| 100.0 \| 2.1e-43 \| 1.6e-47 \| 347.9 \| 351 \| (1, 468) \| 475 \| (1, 450) \| 451 \| Afp1, Afp2, Afp3, Afp5, Afp9 \| 6RAO\_D Afp1, Afp2, Afp3, Afp5, Afp9 Anti-feeding prophage, secretion system, AFP; pdb70 \| 3J9Q\_B \| 100.0 \| 2.9e-42 \| 2.2e-46 \| 336.4 \| 382 \| (5, 475) \| 475 \| (2, 384) \| 386 \| sheath, tube \| 3J9Q\_B sheath, tube pyocin, bacteriocin, sheath, tube, STRUCTURAL; pdb70 \| 6J0B\_J \| 100.0 \| 9.1e-40 \| 7e-44 \| 314.9 \| 337 \| (1, 468) \| 475 \| (1, 353) \| 355 \| Pvc2, Conserved hypothetical phage tail \| 6J0B\_J Pvc2, Conserved hypothetical phage tail assembly, Photorhabdus asymbiotica, PVC, contractile; pdb70 \| 6J0C\_c \| 100.0 \| 9.1e-40 \| 7e-44 \| 314.9 \| 337 \| (1, 468) \| 475 \| (1, 353) \| 355 \| Pvc2 \| 6J0C\_c Pvc2 assembly, Photorhabdus asymbiotica, PVC, contractile; pdb70 \| 6J0N\_u \| 100.0 \| 2.7e-39 \| 2e-43 \| 318.8 \| 351 \| (1, 468) \| 475 \| (1, 439) \| 440 \| Pvc1, Pvc9, Pvc11, Pvc12, Pvc4 \| 6J0N\_u Pvc1, Pvc9, Pvc11, Pvc12, Pvc4 assembly, Photorhabdus asymbiotica, PVC, contractile; pdb70 \| 6RAO\_C \| 100.0 \| 5.2e-39 \| 4e-43 \| 309.4 \| 336 \| (1, 468) \| 475 \| (1, 353) \| 354 \| Afp1, Afp2, Afp3, Afp5, Afp9 \| 6RAO\_C Afp1, Afp2, Afp3, Afp5, Afp9 Anti-feeding prophage, secretion system, AFP; pdb70 \| 6RBN\_C \| 100.0 \| 5.2e-39 \| 4e-43 \| 309.4 \| 336 \| (1, 468) \| 475 \| (1, 353) \| 354 \| Afp1, Afp2, Afp3, Afp16 \| 6RBN\_C Afp1, Afp2, Afp3, Afp16 Anti-feeding prophage, secretion system, AFP; pdb70 \| 6RAO\_E \| 100.0 \| 1.9e-37 \| 1.5e-41 \| 305.2 \| 376 \| (4, 468) \| 475 \| (2, 405) \| 417 \| Afp1, Afp2, Afp3, Afp5, Afp9 \| 6RAO\_E Afp1, Afp2, Afp3, Afp5, Afp9 Anti-feeding prophage, secretion system, AFP; pdb70 \| 6J0N\_V \| 100.0 \| 3.4e-36 \| 2.6e-40 \| 295.5 \| 369 \| (4, 468) \| 475 \| (2, 405) \| 410 \| Pvc1, Pvc9, Pvc11, Pvc12, Pvc4 \| 6J0N\_V Pvc1, Pvc9, Pvc11, Pvc12, Pvc4 assembly, Photorhabdus asymbiotica, PVC, contractile; pdb70 \| 3HXL\_A \| 99.9 \| 1e-33 \| 7.8e-38 \| 280.9 \| 394 \| (4, 469) \| 475 \| (10, 441) \| 446 \| uncharacterized protein DSY3957 \| 3HXL\_A uncharacterized protein DSY3957 alpha-beta three-domained protein., Structural Genomics; pdb70 \| 3LML\_A \| 99.9 \| 6.9e-33 \| 5.3e-37 \| 275.8 \| 396 \| (3, 469) \| 475 \| (11, 455) \| 460 \| Lin1278 protein \| 3LML\_A Lin1278 protein Structural Genomics, PSI-2, Protein Structure; pdb70 \| 3LML\_B \| 99.9 \| 6.9e-33 \| 5.3e-37 \| 275.8 \| 396 \| (3, 469) \| 475 \| (11, 455) \| 460 \| Lin1278 protein \| 3LML\_B Lin1278 protein Structural Genomics, PSI-2, Protein Structure HET: MSE; pdb70 \| 3J2M\_V \| 99.9 \| 8e-33 \| 6e-37 \| 281.0 \| 239 \| (230, 474) \| 475 \| (388, 655) \| 659 \| Tail connector protein Gp15, Tail \| 3J2M\_V Tail connector protein Gp15, Tail bacteriophage T4, phage tail terminator; pdb70 \| 5LI4\_B \| 99.9 \| 3.9e-27 \| 3e-31 \| 240.3 \| 248 \| (201, 470) \| 475 \| (319, 585) \| 587 \| Uncharacterized protein \| 5LI4\_B Uncharacterized protein polyvalent staphylococcal bactoriophage, Myoviridae, tail; pdb70 \| 6GKW\_A \| 99.9 \| 6.5e-27 \| 5e-31 \| 224.8 \| 316 \| (3, 464) \| 475 \| (3, 355) \| 356 \| Putative phage XkdK-like protein \| 6GKW\_A Putative phage XkdK-like protein Diffocin Sheath, structural protein HET: MSE; pdb70 \| 3FOA\_C \| 98.8 \| 6.7e-13 \| 5e-17 \| 129.6 \| 84 \| (3, 95) \| 475 \| (1, 86) \| 510 \| Tail sheath protein Gp18 \| 3FOA\_C Tail sheath protein Gp18 alpha-beta, viral structural protein, bacteriophage; pdb70 \| 3FOI\_B \| 98.8 \| 6.7e-13 \| 5e-17 \| 129.6 \| 84 \| (3, 95) \| 475 \| (1, 86) \| 510 \| Tail sheath protein Gp18 \| 3FOI\_B Tail sheath protein Gp18 alpha-beta, viral structural protein, bacteriophage |
| Top keywords  (threshold 1.00e-03 (evalue)) | **tail, sheath, Phage, domain\_containing, Fragment, subtilisin\_like, C\_terminal, Major, FI, Putative** |
| Output files | ../../similar\_sequences/18\_FANPEZAQ\_CDS\_0018\_merged.svg ../../similar\_sequences/18\_FANPEZAQ\_CDS\_0018\_pdb70.a3m ../../similar\_sequences/18\_FANPEZAQ\_CDS\_0018\_pdb70.hhr ../../similar\_sequences/18\_FANPEZAQ\_CDS\_0018\_uniclust.a3m ../../similar\_sequences/18\_FANPEZAQ\_CDS\_0018\_uniclust.hhr |

#### Structure prediction (AlphaFold)^2^

|  |  |
| --- | --- |
| Stats | xml version="1.0" encoding="utf-8" standalone="no"?       2024-09-02T21:09:17.817824 image/svg+xml   Matplotlib v3.7.2, https://matplotlib.org/ |
| Predicted structure | **NGL Viewer Controls:**  - Center: *Left-Click* - Rotate: *Left-Click + Drag* - Translate: *Right-Click + Drag* - Zoom: *Shift + Left-Click + Drag* |
| Output files | ../../predicted\_structures/18\_FANPEZAQ\_CDS\_0018/features.pkl ../../predicted\_structures/18\_FANPEZAQ\_CDS\_0018/ranked\_0.pdb ../../predicted\_structures/18\_FANPEZAQ\_CDS\_0018/ranked\_0\_plots.svg ../../predicted\_structures/18\_FANPEZAQ\_CDS\_0018/result\_model\_1\_ptm\_pred\_0.pkl |

#### Structure similarity search results (Foldseek)^3^

|  |  |
| --- | --- |
| Structure databases searched | Pdb, Afdb-proteome, Afdb-uniprot50 |
| Results, scheme(s)  (Top layers only, threshold 1.00e-02 (evalue)) | xml version="1.0" encoding="utf-8" standalone="no"?       2024-09-02T21:10:41.882959 image/svg+xml   Matplotlib v3.7.2, https://matplotlib.org/ |
| Results, table  (threshold 1.00e-02 (evalue)) | db \| id \| prob \| evalue \| bits \| fident \| alnlen \| mismatch \| gapopen \| qstart \| qend \| tstart \| tend \| name \| description; pdb \| 6U5B\_N \| 1.0 \| 2.288e-35 \| 1224 \| 0.372 \| 473 \| 201 \| 8 \| 5 \| 473 \| 1 \| 381 \| Sheath PA0622 \| Sheath PA0622; pdb \| 8BKY\_A \| 1.0 \| 9.482e-19 \| 574 \| 0.228 \| 490 \| 229 \| 19 \| 3 \| 466 \| 1 \| 367 \| Phage tail sheath protein \| Phage tail sheath protein; pdb \| 6RAO\_D \| 1.0 \| 7.019e-18 \| 536 \| 0.161 \| 484 \| 256 \| 17 \| 5 \| 466 \| 2 \| 357 \| Afp3 \| Afp3; pdb \| 6RC8\_B \| 1.0 \| 4.835e-16 \| 477 \| 0.163 \| 477 \| 263 \| 20 \| 5 \| 468 \| 2 \| 355 \| Afp3 \| Afp3; pdb \| 6J0N\_H \| 1.0 \| 1.908e-15 \| 452 \| 0.137 \| 493 \| 256 \| 21 \| 3 \| 468 \| 2 \| 352 \| Pvc2 \| Pvc2; pdb \| 6J0N\_V \| 1.0 \| 8.089e-16 \| 449 \| 0.134 \| 514 \| 269 \| 22 \| 1 \| 468 \| 5 \| 388 \| Pvc3 \| Pvc3; pdb \| 8BL4\_D \| 1.0 \| 5.341e-15 \| 437 \| 0.199 \| 457 \| 246 \| 23 \| 12 \| 448 \| 8 \| 364 \| Phage tail protein \| Phage tail protein; pdb \| 6RGL\_A \| 1.0 \| 2.166e-12 \| 384 \| 0.141 \| 481 \| 256 \| 20 \| 7 \| 466 \| 4 \| 348 \| Afp2 \| Afp2; pdb \| 6J0N\_T \| 1.0 \| 2.363e-14 \| 374 \| 0.142 \| 526 \| 254 \| 20 \| 9 \| 468 \| 5 \| 399 \| Pvc3 \| Pvc3; pdb \| 7AEK\_3A \| 1.0 \| 3.373e-11 \| 365 \| 0.229 \| 192 \| 136 \| 5 \| 285 \| 466 \| 465 \| 654 \| Putative phage tail sheath protein FI \| Putative phage tail sheath protein FI; pdb \| 7AE0\_3A \| 1.0 \| 2.649e-14 \| 363 \| 0.142 \| 611 \| 274 \| 25 \| 4 \| 466 \| 1 \| 509 \| Phage tail protein \| Phage tail protein; pdb \| 7B5H\_AO \| 1.0 \| 5.572e-14 \| 360 \| 0.144 \| 581 \| 270 \| 25 \| 3 \| 466 \| 1 \| 471 \| All3325 protein \| All3325 protein; pdb \| 7B5H\_AP \| 1.0 \| 1.295e-12 \| 317 \| 0.147 \| 585 \| 271 \| 29 \| 3 \| 466 \| 1 \| 478 \| All3325 protein \| All3325 protein; pdb \| 6RGL\_D \| 1.0 \| 3.945e-10 \| 283 \| 0.156 \| 505 \| 264 \| 25 \| 1 \| 468 \| 4 \| 383 \| Afp4 \| Afp4; pdb \| 8JAN\_N \| 1.0 \| 6.8e-12 \| 251 \| 0.131 \| 578 \| 322 \| 38 \| 7 \| 466 \| 12 \| 527 \| Gp22 \| Gp22; pdb \| 6GKW\_A \| 1.0 \| 2.492e-06 \| 177 \| 0.117 \| 468 \| 220 \| 27 \| 28 \| 453 \| 23 \| 339 \| Putative phage XkdK-like protein \| Putative phage XkdK-like protein; pdb \| 8AU1\_A \| 1.0 \| 3.718e-06 \| 157 \| 0.106 \| 282 \| 193 \| 14 \| 231 \| 472 \| 425 \| 687 \| Putative tail sheath protein \| Putative tail sheath protein; pdb \| 5N8N\_B \| 1.0 \| 4.674e-06 \| 150 \| 0.11 \| 327 \| 187 \| 22 \| 231 \| 467 \| 141 \| 453 \| EvpB family type VI secretion protein \| EvpB family type VI secretion protein; pdb \| 5URX\_1B \| 1.0 \| 6.222e-06 \| 140 \| 0.116 \| 334 \| 180 \| 21 \| 231 \| 468 \| 114 \| 428 \| TssC \| TssC; pdb \| 7KH1\_A6 \| 1.0 \| 2.099e-06 \| 130 \| 0.107 \| 603 \| 294 \| 44 \| 1 \| 465 \| 1 \| 497 \| tail sheath protein, gp6 \| tail sheath protein, gp6; pdb \| 3J9O\_D \| 1.0 \| 0.0001026 \| 128 \| 0.07 \| 340 \| 201 \| 18 \| 231 \| 474 \| 103 \| 423 \| Intracellular growth locus protein B \| Intracellular growth locus protein B; pdb \| 3J9G\_B \| 1.0 \| 4.605e-05 \| 127 \| 0.083 \| 334 \| 191 \| 20 \| 231 \| 468 \| 114 \| 428 \| VipB \| VipB; pdb \| 8HDR\_M \| 1.0 \| 9.148e-05 \| 125 \| 0.111 \| 509 \| 268 \| 31 \| 8 \| 466 \| 8 \| 381 \| Pam3 sheath protein \| Pam3 sheath protein; pdb \| 5MXN\_A \| 1.0 \| 0.0001924 \| 113 \| 0.088 \| 328 \| 196 \| 18 \| 231 \| 468 \| 158 \| 472 \| Type VI secretion protein \| Type VI secretion protein; pdb \| 5OJQ\_D \| 1.0 \| 0.0001924 \| 113 \| 0.088 \| 328 \| 196 \| 18 \| 231 \| 468 \| 158 \| 472 \| Type VI secretion protein \| Type VI secretion protein; pdb \| 8FVG\_A \| 1.0 \| 4.949e-06 \| 108 \| 0.087 \| 586 \| 321 \| 36 \| 1 \| 466 \| 1 \| 492 \| Sheath protein gp31 \| Sheath protein gp31; pdb \| 3J2M\_U \| 0.999 \| 0.0009014 \| 98 \| 0.098 \| 649 \| 248 \| 30 \| 26 \| 422 \| 1 \| 564 \| Tail sheath protein Gp18 \| Tail sheath protein Gp18; pdb \| 8ENV\_E \| 0.997 \| 2.914e-05 \| 91 \| 0.096 \| 593 \| 304 \| 39 \| 6 \| 467 \| 10 \| 501 \| Structural protein gp45 \| Structural protein gp45; afdb-proteome \| AF-G3XD39-F1-MODEL\_V4 \| 1.0 \| 1.989e-44 \| 1677 \| 0.372 \| 473 \| 201 \| 8 \| 5 \| 473 \| 2 \| 382 \| Probable bacteriophage protein \| Probable bacteriophage protein; afdb-proteome \| AF-A0A0H3GRZ6-F1-MODEL\_V4 \| 1.0 \| 2.361e-44 \| 1663 \| 0.339 \| 477 \| 220 \| 8 \| 3 \| 474 \| 1 \| 387 \| Putative prophage tail sheath \| Putative prophage tail sheath; afdb-proteome \| AF-A0A0H3GQS2-F1-MODEL\_V4 \| 1.0 \| 8.541e-42 \| 1591 \| 0.323 \| 476 \| 229 \| 6 \| 1 \| 474 \| 1 \| 385 \| Phage tail sheath protein \| Phage tail sheath protein; afdb-proteome \| AF-Q8ZMV0-F1-MODEL\_V4 \| 1.0 \| 2.133e-41 \| 1578 \| 0.327 \| 474 \| 227 \| 5 \| 3 \| 474 \| 2 \| 385 \| Fels-2 prophage protein \| Fels-2 prophage protein; afdb-proteome \| AF-Q8ZKJ2-F1-MODEL\_V4 \| 1.0 \| 5.16e-35 \| 1011 \| 0.252 \| 547 \| 261 \| 21 \| 1 \| 474 \| 1 \| 472 \| Putative phage tail sheath protein \| Putative phage tail sheath protein; afdb-proteome \| AF-P44233-F1-MODEL\_V4 \| 1.0 \| 3.131e-09 \| 194 \| 0.103 \| 579 \| 302 \| 35 \| 2 \| 466 \| 11 \| 486 \| Mu-like prophage FluMu tail sheath protein \| Mu-like prophage FluMu tail sheath protein; afdb-proteome \| AF-A0A0H3GRW7-F1-MODEL\_V4 \| 1.0 \| 2.382e-06 \| 181 \| 0.104 \| 517 \| 270 \| 27 \| 6 \| 468 \| 1 \| 378 \| Putative bacteriophage protein \| Putative bacteriophage protein; afdb-proteome \| AF-Q9I748-F1-MODEL\_V4 \| 1.0 \| 5.086e-07 \| 168 \| 0.104 \| 335 \| 196 \| 21 \| 231 \| 475 \| 178 \| 498 \| Type VI secretion system sheath protein TssC1 \| Type VI secretion system sheath protein TssC1; afdb-proteome \| AF-Q8ZRL7-F1-MODEL\_V4 \| 1.0 \| 6.77e-07 \| 166 \| 0.119 \| 343 \| 185 \| 23 \| 231 \| 475 \| 179 \| 502 \| Putative cytoplasmic protein \| Putative cytoplasmic protein; afdb-proteome \| AF-Q9I1B3-F1-MODEL\_V4 \| 1.0 \| 1.01e-06 \| 156 \| 0.119 \| 334 \| 177 \| 19 \| 231 \| 468 \| 175 \| 487 \| Uricase PuuD \| Uricase PuuD; afdb-proteome \| AF-A0A0H3GSE9-F1-MODEL\_V4 \| 1.0 \| 5.306e-06 \| 152 \| 0.091 \| 328 \| 194 \| 21 \| 230 \| 468 \| 196 \| 508 \| EvpB/family type VI secretion protein \| EvpB/family type VI secretion protein; afdb-proteome \| AF-Q9I367-F1-MODEL\_V4 \| 1.0 \| 5.618e-06 \| 142 \| 0.093 \| 331 \| 191 \| 19 \| 231 \| 468 \| 173 \| 487 \| Uncharacterized protein \| Uncharacterized protein; afdb-uniprot50 \| AF-A0A1T3BL82-F1-MODEL\_V4 \| 1.0 \| 1.03e-70 \| 2600 \| 0.585 \| 473 \| 185 \| 7 \| 1 \| 473 \| 1 \| 462 \| Uncharacterized protein \| Uncharacterized protein; afdb-uniprot50 \| AF-A0A2G6HAE9-F1-MODEL\_V4 \| 1.0 \| 1.716e-62 \| 2511 \| 0.573 \| 471 \| 122 \| 3 \| 5 \| 475 \| 2 \| 393 \| Phage tail protein \| Phage tail protein; afdb-uniprot50 \| AF-V7HPP7-F1-MODEL\_V4 \| 1.0 \| 1.745e-67 \| 2446 \| 0.606 \| 478 \| 168 \| 10 \| 1 \| 473 \| 1 \| 463 \| Tail sheath protein \| Tail sheath protein; afdb-uniprot50 \| AF-A0A522WEK9-F1-MODEL\_V4 \| 1.0 \| 1.982e-64 \| 2352 \| 0.522 \| 475 \| 212 \| 9 \| 3 \| 474 \| 2 \| 464 \| Uncharacterized protein \| Uncharacterized protein; afdb-uniprot50 \| AF-A0A7S8HDT1-F1-MODEL\_V4 \| 1.0 \| 1.982e-64 \| 2330 \| 0.488 \| 473 \| 232 \| 5 \| 3 \| 474 \| 2 \| 465 \| Phage tail sheath subtilisin-like domain-containing protein \| Phage tail sheath subtilisin-like domain-containing protein; afdb-uniprot50 \| AF-A0A212KBR8-F1-MODEL\_V4 \| 1.0 \| 5.548e-64 \| 2330 \| 0.511 \| 475 \| 217 \| 9 \| 3 \| 474 \| 2 \| 464 \| Putative phage tail protein \| Putative phage tail protein; afdb-uniprot50 \| AF-A0A3R8TA92-F1-MODEL\_V4 \| 1.0 \| 1.287e-58 \| 2302 \| 0.539 \| 471 \| 132 \| 7 \| 3 \| 473 \| 1 \| 386 \| Phage tail protein \| Phage tail protein; afdb-uniprot50 \| AF-A0A524RVX4-F1-MODEL\_V4 \| 1.0 \| 9.543e-62 \| 2282 \| 0.466 \| 476 \| 238 \| 7 \| 5 \| 473 \| 4 \| 470 \| Phage tail protein \| Phage tail protein; afdb-uniprot50 \| AF-A0A5M6IF21-F1-MODEL\_V4 \| 1.0 \| 2.523e-61 \| 2272 \| 0.564 \| 473 \| 127 \| 4 \| 1 \| 473 \| 52 \| 445 \| Phage tail protein \| Phage tail protein; afdb-uniprot50 \| AF-A0A2V3UCK1-F1-MODEL\_V4 \| 1.0 \| 1.978e-60 \| 2261 \| 0.465 \| 481 \| 238 \| 9 \| 3 \| 474 \| 2 \| 472 \| Uncharacterized protein \| Uncharacterized protein; afdb-uniprot50 \| AF-A0A5A9EN65-F1-MODEL\_V4 \| 1.0 \| 2.125e-61 \| 2242 \| 0.433 \| 625 \| 193 \| 10 \| 3 \| 475 \| 2 \| 617 \| Phage tail protein \| Phage tail protein; afdb-uniprot50 \| AF-K5XWL1-F1-MODEL\_V4 \| 1.0 \| 9.956e-61 \| 2238 \| 0.508 \| 482 \| 217 \| 7 \| 3 \| 474 \| 2 \| 473 \| Phage\_sheath\_1 domain-containing protein \| Phage\_sheath\_1 domain-containing protein; afdb-uniprot50 \| AF-A0A812QV43-F1-MODEL\_V4 \| 1.0 \| 7.37e-60 \| 2168 \| 0.554 \| 474 \| 125 \| 3 \| 1 \| 473 \| 1 \| 389 \| GpFI protein \| GpFI protein; afdb-uniprot50 \| AF-Q31HT2-F1-MODEL\_V4 \| 1.0 \| 8.352e-53 \| 2082 \| 0.431 \| 473 \| 179 \| 4 \| 3 \| 473 \| 2 \| 386 \| Phage tail sheath protein \| Phage tail sheath protein; afdb-uniprot50 \| AF-A0A1M5PX14-F1-MODEL\_V4 \| 1.0 \| 1.362e-58 \| 2076 \| 0.429 \| 477 \| 259 \| 8 \| 1 \| 474 \| 1 \| 467 \| Uncharacterized protein \| Uncharacterized protein; afdb-uniprot50 \| AF-A0A7W6RFE6-F1-MODEL\_V4 \| 1.0 \| 7.788e-56 \| 2065 \| 0.425 \| 482 \| 252 \| 12 \| 5 \| 474 \| 3 \| 471 \| Phage\_sheath\_1 domain-containing protein \| Phage\_sheath\_1 domain-containing protein; afdb-uniprot50 \| AF-A0A1A9VKH7-F1-MODEL\_V4 \| 1.0 \| 1.249e-56 \| 2030 \| 0.505 \| 465 \| 141 \| 5 \| 5 \| 463 \| 3 \| 384 \| Uncharacterized protein \| Uncharacterized protein; afdb-uniprot50 \| AF-E2CJT2-F1-MODEL\_V4 \| 1.0 \| 1.731e-51 \| 2025 \| 0.405 \| 471 \| 190 \| 5 \| 3 \| 473 \| 2 \| 382 \| Phage tail sheath protein \| Phage tail sheath protein; afdb-uniprot50 \| AF-A0A7V7KJN4-F1-MODEL\_V4 \| 1.0 \| 2.09e-56 \| 2014 \| 0.448 \| 587 \| 128 \| 5 \| 3 \| 475 \| 1 \| 505 \| Phage tail protein \| Phage tail protein; afdb-uniprot50 \| AF-A0A8A8PHF1-F1-MODEL\_V4 \| 1.0 \| 4.151e-56 \| 2007 \| 0.453 \| 587 \| 125 \| 5 \| 3 \| 475 \| 1 \| 505 \| Phage tail protein \| Phage tail protein; afdb-uniprot50 \| AF-A0A149SWC0-F1-MODEL\_V4 \| 1.0 \| 5.296e-57 \| 1990 \| 0.441 \| 491 \| 247 \| 13 \| 3 \| 474 \| 2 \| 484 \| Uncharacterized protein \| Uncharacterized protein; afdb-uniprot50 \| AF-A0A2W5KVX7-F1-MODEL\_V4 \| 1.0 \| 1.734e-55 \| 1983 \| 0.429 \| 584 \| 136 \| 5 \| 5 \| 474 \| 2 \| 502 \| Phage tail protein \| Phage tail protein; afdb-uniprot50 \| AF-F4BFQ2-F1-MODEL\_V4 \| 1.0 \| 2.695e-50 \| 1977 \| 0.421 \| 474 \| 181 \| 8 \| 5 \| 475 \| 2 \| 385 \| Uncharacterized protein \| Uncharacterized protein; afdb-uniprot50 \| AF-A0A7R9HD16-F1-MODEL\_V4 \| 1.0 \| 2.741e-55 \| 1974 \| 0.504 \| 472 \| 142 \| 5 \| 2 \| 467 \| 100 \| 485 \| Hypothetical protein \| Hypothetical protein; afdb-uniprot50 \| AF-A0A167FZT2-F1-MODEL\_V4 \| 1.0 \| 2.241e-53 \| 1955 \| 0.476 \| 510 \| 147 \| 10 \| 3 \| 473 \| 1 \| 429 \| Tail protein \| Tail protein; afdb-uniprot50 \| AF-A0A6N7JIY2-F1-MODEL\_V4 \| 1.0 \| 1.262e-49 \| 1932 \| 0.416 \| 471 \| 188 \| 6 \| 5 \| 474 \| 2 \| 386 \| Phage tail protein \| Phage tail protein; afdb-uniprot50 \| AF-A0A7R9HA11-F1-MODEL\_V4 \| 1.0 \| 6.104e-55 \| 1920 \| 0.507 \| 467 \| 136 \| 7 \| 13 \| 474 \| 422 \| 799 \| Hypothetical protein \| Hypothetical protein; afdb-uniprot50 \| AF-A0A432QUT0-F1-MODEL\_V4 \| 1.0 \| 5.444e-55 \| 1918 \| 0.4 \| 475 \| 271 \| 8 \| 1 \| 474 \| 1 \| 462 \| Uncharacterized protein \| Uncharacterized protein; afdb-uniprot50 \| AF-A0A345DE55-F1-MODEL\_V4 \| 1.0 \| 1.109e-48 \| 1915 \| 0.388 \| 479 \| 201 \| 5 \| 1 \| 475 \| 1 \| 391 \| Prophage major tail sheath protein \| Prophage major tail sheath protein; afdb-uniprot50 \| AF-A0A239EII1-F1-MODEL\_V4 \| 1.0 \| 9.915e-53 \| 1902 \| 0.412 \| 587 \| 146 \| 4 \| 3 \| 474 \| 2 \| 504 \| Uncharacterized protein \| Uncharacterized protein; afdb-uniprot50 \| AF-A0A410UFB0-F1-MODEL\_V4 \| 1.0 \| 1.705e-50 \| 1900 \| 0.449 \| 478 \| 180 \| 7 \| 1 \| 473 \| 1 \| 400 \| Phage tail protein \| Phage tail protein; afdb-uniprot50 \| AF-A0A423PRQ5-F1-MODEL\_V4 \| 1.0 \| 1.779e-49 \| 1898 \| 0.401 \| 478 \| 207 \| 8 \| 1 \| 474 \| 1 \| 403 \| Tail protein \| Tail protein; afdb-uniprot50 \| AF-A0A6M6IDP4-F1-MODEL\_V4 \| 1.0 \| 3.534e-49 \| 1882 \| 0.536 \| 416 \| 119 \| 6 \| 3 \| 418 \| 2 \| 343 \| Phage tail protein \| Phage tail protein; afdb-uniprot50 \| AF-A0A0B6D4V4-F1-MODEL\_V4 \| 1.0 \| 1.455e-47 \| 1852 \| 0.365 \| 473 \| 209 \| 5 \| 3 \| 474 \| 1 \| 383 \| Phage tail sheath family protein \| Phage tail sheath family protein; afdb-uniprot50 \| AF-A0A2N1ANM3-F1-MODEL\_V4 \| 1.0 \| 1.521e-50 \| 1849 \| 0.507 \| 475 \| 140 \| 5 \| 1 \| 475 \| 1 \| 381 \| Phage tail protein \| Phage tail protein; afdb-uniprot50 \| AF-A0A3N2E0U0-F1-MODEL\_V4 \| 1.0 \| 1.607e-46 \| 1825 \| 0.372 \| 470 \| 206 \| 4 \| 5 \| 474 \| 2 \| 382 \| Uncharacterized protein \| Uncharacterized protein; afdb-uniprot50 \| AF-A0A1G7PXY9-F1-MODEL\_V4 \| 1.0 \| 5.121e-47 \| 1797 \| 0.398 \| 474 \| 191 \| 7 \| 5 \| 474 \| 2 \| 385 \| Uncharacterized protein \| Uncharacterized protein; afdb-uniprot50 \| AF-A0A5P9F317-F1-MODEL\_V4 \| 1.0 \| 3.798e-50 \| 1787 \| 0.354 \| 589 \| 179 \| 6 \| 1 \| 475 \| 1 \| 502 \| Phage tail sheath protein \| Phage tail sheath protein; afdb-uniprot50 \| AF-A0A6L3Y4M8-F1-MODEL\_V4 \| 1.0 \| 4.918e-52 \| 1782 \| 0.412 \| 480 \| 258 \| 14 \| 1 \| 474 \| 1 \| 462 \| Phage tail protein \| Phage tail protein; afdb-uniprot50 \| AF-A0A1Y1QXK5-F1-MODEL\_V4 \| 1.0 \| 1.496e-45 \| 1777 \| 0.348 \| 471 \| 214 \| 6 \| 5 \| 474 \| 2 \| 380 \| Phage\_sheath\_1 domain-containing protein \| Phage\_sheath\_1 domain-containing protein; afdb-uniprot50 \| AF-A0A2E3N1I9-F1-MODEL\_V4 \| 1.0 \| 2.977e-49 \| 1774 \| 0.435 \| 482 \| 178 \| 11 \| 1 \| 475 \| 1 \| 395 \| Phage tail protein \| Phage tail protein; afdb-uniprot50 \| AF-A0A258L5K8-F1-MODEL\_V4 \| 1.0 \| 1.377e-51 \| 1774 \| 0.379 \| 496 \| 269 \| 14 \| 1 \| 474 \| 1 \| 479 \| Uncharacterized protein \| Uncharacterized protein; afdb-uniprot50 \| AF-A0A8B2NUE5-F1-MODEL\_V4 \| 1.0 \| 3.247e-51 \| 1760 \| 0.345 \| 492 \| 291 \| 12 \| 1 \| 474 \| 1 \| 479 \| Phage tail protein \| Phage tail protein; afdb-uniprot50 \| AF-A0A3G4V9H2-F1-MODEL\_V4 \| 1.0 \| 2.363e-45 \| 1754 \| 0.336 \| 475 \| 225 \| 5 \| 3 \| 475 \| 2 \| 388 \| Phage tail protein \| Phage tail protein; afdb-uniprot50 \| AF-A0A2N3KSI6-F1-MODEL\_V4 \| 1.0 \| 8.807e-45 \| 1742 \| 0.334 \| 478 \| 227 \| 4 \| 1 \| 475 \| 3 \| 392 \| Phage tail protein \| Phage tail protein; afdb-uniprot50 \| AF-A0A847KPL0-F1-MODEL\_V4 \| 1.0 \| 7.232e-51 \| 1737 \| 0.374 \| 475 \| 271 \| 14 \| 5 \| 474 \| 2 \| 455 \| Phage tail protein \| Phage tail protein; afdb-uniprot50 \| AF-A0A2S6N2V9-F1-MODEL\_V4 \| 1.0 \| 6.249e-45 \| 1689 \| 0.343 \| 477 \| 224 \| 9 \| 4 \| 474 \| 3 \| 396 \| Uncharacterized protein \| Uncharacterized protein; afdb-uniprot50 \| AF-A0A258CZQ8-F1-MODEL\_V4 \| 1.0 \| 3.199e-50 \| 1686 \| 0.321 \| 489 \| 300 \| 14 \| 1 \| 474 \| 1 \| 472 \| Phage tail protein \| Phage tail protein; afdb-uniprot50 \| AF-A0A1D2QS78-F1-MODEL\_V4 \| 1.0 \| 4.134e-48 \| 1672 \| 0.403 \| 552 \| 156 \| 7 \| 5 \| 475 \| 2 \| 460 \| Uncharacterized protein \| Uncharacterized protein; afdb-uniprot50 \| AF-A0A2W5K1C0-F1-MODEL\_V4 \| 1.0 \| 5.186e-44 \| 1670 \| 0.523 \| 403 \| 117 \| 4 \| 72 \| 474 \| 3 \| 330 \| Phage tail protein \| Phage tail protein; afdb-uniprot50 \| AF-A0A7T8NTW5-F1-MODEL\_V4 \| 1.0 \| 3.68e-44 \| 1669 \| 0.337 \| 480 \| 228 \| 8 \| 1 \| 474 \| 1 \| 396 \| Phage tail sheath subtilisin-like domain-containing protein \| Phage tail sheath subtilisin-like domain-containing protein; afdb-uniprot50 \| AF-A0A6L7FX52-F1-MODEL\_V4 \| 1.0 \| 5.492e-44 \| 1644 \| 0.352 \| 476 \| 215 \| 9 \| 1 \| 474 \| 1 \| 385 \| Phage tail protein \| Phage tail protein; afdb-uniprot50 \| AF-A0A774N4X1-F1-MODEL\_V4 \| 1.0 \| 2.394e-42 \| 1634 \| 0.32 \| 474 \| 230 \| 5 \| 3 \| 474 \| 2 \| 385 \| Phage tail protein \| Phage tail protein; afdb-uniprot50 \| AF-A0A4R1K4A7-F1-MODEL\_V4 \| 1.0 \| 8.568e-47 \| 1617 \| 0.359 \| 587 \| 163 \| 8 \| 3 \| 475 \| 1 \| 488 \| Uncharacterized protein \| Uncharacterized protein; afdb-uniprot50 \| AF-A0A285ZDJ9-F1-MODEL\_V4 \| 1.0 \| 6.329e-42 \| 1601 \| 0.348 \| 477 \| 214 \| 7 \| 3 \| 475 \| 1 \| 384 \| Uncharacterized protein \| Uncharacterized protein; afdb-uniprot50 \| AF-A0A482NKP0-F1-MODEL\_V4 \| 1.0 \| 2.104e-41 \| 1596 \| 0.316 \| 474 \| 232 \| 5 \| 3 \| 474 \| 1 \| 384 \| Phage tail sheath protein \| Phage tail sheath protein; afdb-uniprot50 \| AF-A0A7G7RNU6-F1-MODEL\_V4 \| 1.0 \| 3.783e-42 \| 1583 \| 0.321 \| 479 \| 227 \| 7 \| 3 \| 473 \| 2 \| 390 \| Phage tail sheath protein \| Phage tail sheath protein; afdb-uniprot50 \| AF-A0A367WTL7-F1-MODEL\_V4 \| 1.0 \| 3.52e-41 \| 1581 \| 0.312 \| 474 \| 233 \| 6 \| 3 \| 474 \| 2 \| 384 \| Tail sheath protein \| Tail sheath protein; afdb-uniprot50 \| AF-G8LQ42-F1-MODEL\_V4 \| 1.0 \| 1.933e-43 \| 1580 \| 0.329 \| 480 \| 222 \| 6 \| 5 \| 474 \| 39 \| 428 \| Tail Sheath Protein \| Tail Sheath Protein; afdb-uniprot50 \| AF-Q8ZMV0-F1-MODEL\_V4 \| 1.0 \| 7.839e-41 \| 1578 \| 0.327 \| 474 \| 227 \| 5 \| 3 \| 474 \| 2 \| 385 \| Fels-2 prophage protein \| Fels-2 prophage protein; afdb-uniprot50 \| AF-A0A2W6YVE1-F1-MODEL\_V4 \| 1.0 \| 1.077e-46 \| 1578 \| 0.349 \| 498 \| 286 \| 16 \| 3 \| 474 \| 1 \| 486 \| Phage tail protein \| Phage tail protein; afdb-uniprot50 \| AF-Q5D5G3-F1-MODEL\_V4 \| 1.0 \| 3.093e-40 \| 1570 \| 0.464 \| 377 \| 115 \| 5 \| 19 \| 395 \| 2 \| 291 \| Phage tail sheath protein \| Phage tail sheath protein; afdb-uniprot50 \| AF-A0A1N7LRF5-F1-MODEL\_V4 \| 1.0 \| 9.854e-41 \| 1562 \| 0.314 \| 477 \| 233 \| 8 \| 1 \| 474 \| 3 \| 388 \| Uncharacterized protein \| Uncharacterized protein; afdb-uniprot50 \| AF-A0A350LX90-F1-MODEL\_V4 \| 1.0 \| 1.059e-41 \| 1555 \| 0.305 \| 472 \| 240 \| 8 \| 4 \| 474 \| 3 \| 387 \| Phage tail protein \| Phage tail protein; afdb-uniprot50 \| AF-A0A836RQY2-F1-MODEL\_V4 \| 1.0 \| 2.606e-40 \| 1538 \| 0.397 \| 428 \| 168 \| 5 \| 48 \| 473 \| 1 \| 340 \| Phage tail protein \| Phage tail protein; afdb-uniprot50 \| AF-A0A3R1AVG7-F1-MODEL\_V4 \| 1.0 \| 2.073e-40 \| 1536 \| 0.314 \| 477 \| 234 \| 5 \| 1 \| 474 \| 1 \| 387 \| Phage tail sheath protein \| Phage tail sheath protein; afdb-uniprot50 \| AF-A0A806D7P7-F1-MODEL\_V4 \| 1.0 \| 6.144e-40 \| 1534 \| 0.284 \| 471 \| 244 \| 5 \| 5 \| 474 \| 2 \| 380 \| Tail sheath protein \| Tail sheath protein; afdb-uniprot50 \| AF-A0A7W4X2I7-F1-MODEL\_V4 \| 1.0 \| 2.135e-42 \| 1528 \| 0.508 \| 362 \| 101 \| 5 \| 3 \| 364 \| 2 \| 286 \| Phage tail sheath protein FI \| Phage tail sheath protein FI; afdb-uniprot50 \| AF-A0A3M4LWQ9-F1-MODEL\_V4 \| 1.0 \| 1.933e-43 \| 1519 \| 0.366 \| 477 \| 206 \| 7 \| 2 \| 474 \| 73 \| 457 \| Major tail sheath protein \| Major tail sheath protein; afdb-uniprot50 \| AF-A0A1B2R8T9-F1-MODEL\_V4 \| 1.0 \| 7.295e-40 \| 1501 \| 0.292 \| 476 \| 243 \| 7 \| 3 \| 474 \| 2 \| 387 \| Phage tail protein \| Phage tail protein; afdb-uniprot50 \| AF-A0A376QG66-F1-MODEL\_V4 \| 1.0 \| 2.016e-42 \| 1500 \| 0.333 \| 480 \| 222 \| 8 \| 3 \| 474 \| 1 \| 390 \| Major tail sheath protein FI \| Major tail sheath protein FI; afdb-uniprot50 \| AF-A0A369Z1S6-F1-MODEL\_V4 \| 1.0 \| 1.449e-39 \| 1498 \| 0.28 \| 475 \| 249 \| 6 \| 3 \| 474 \| 2 \| 386 \| Phage tail protein \| Phage tail protein; afdb-uniprot50 \| AF-A0A2G3K3C7-F1-MODEL\_V4 \| 1.0 \| 2.227e-41 \| 1496 \| 0.294 \| 510 \| 232 \| 8 \| 3 \| 474 \| 2 \| 421 \| Phage tail protein \| Phage tail protein; afdb-uniprot50 \| AF-A0A0C2IBA1-F1-MODEL\_V4 \| 1.0 \| 1.698e-42 \| 1493 \| 0.327 \| 473 \| 226 \| 5 \| 4 \| 474 \| 84 \| 466 \| Phage tail sheath monomer \| Phage tail sheath monomer; afdb-uniprot50 \| AF-A0A0K1NER8-F1-MODEL\_V4 \| 1.0 \| 1.849e-40 \| 1492 \| 0.291 \| 481 \| 241 \| 7 \| 3 \| 474 \| 1 \| 390 \| Uncharacterized protein \| Uncharacterized protein; afdb-uniprot50 \| AF-A0A0Q8XV39-F1-MODEL\_V4 \| 1.0 \| 2.718e-39 \| 1491 \| 0.3 \| 479 \| 239 \| 8 \| 1 \| 474 \| 1 \| 388 \| Phage tail protein \| Phage tail protein; afdb-uniprot50 \| AF-A0A348HI83-F1-MODEL\_V4 \| 1.0 \| 1.428e-38 \| 1483 \| 0.3 \| 476 \| 240 \| 6 \| 1 \| 474 \| 1 \| 385 \| Phage tail sheath protein FI \| Phage tail sheath protein FI; afdb-uniprot50 \| AF-A0A4P9VGN2-F1-MODEL\_V4 \| 1.0 \| 4.816e-39 \| 1478 \| 0.305 \| 478 \| 230 \| 9 \| 3 \| 474 \| 2 \| 383 \| Phage tail protein \| Phage tail protein; afdb-uniprot50 \| AF-A0A2M8RTB4-F1-MODEL\_V4 \| 1.0 \| 2.718e-39 \| 1476 \| 0.271 \| 475 \| 253 \| 7 \| 3 \| 474 \| 2 \| 386 \| Phage tail protein \| Phage tail protein; afdb-uniprot50 \| AF-A0A6B8KJH9-F1-MODEL\_V4 \| 1.0 \| 1.043e-40 \| 1475 \| 0.319 \| 485 \| 237 \| 12 \| 1 \| 474 \| 1 \| 403 \| Phage tail protein \| Phage tail protein; afdb-uniprot50 \| AF-A0A1A9RDH2-F1-MODEL\_V4 \| 1.0 \| 1.449e-39 \| 1472 \| 0.297 \| 474 \| 240 \| 5 \| 3 \| 474 \| 2 \| 384 \| Phage tail protein \| Phage tail protein; afdb-uniprot50 \| AF-B1H6B6-F1-MODEL\_V4 \| 1.0 \| 2.389e-38 \| 1470 \| 0.305 \| 451 \| 221 \| 5 \| 25 \| 473 \| 1 \| 361 \| Phage tail sheath protein \| Phage tail sheath protein; afdb-uniprot50 \| AF-A0A7W7Z2X1-F1-MODEL\_V4 \| 1.0 \| 3.186e-42 \| 1468 \| 0.307 \| 485 \| 310 \| 16 \| 1 \| 474 \| 1 \| 470 \| Uncharacterized protein \| Uncharacterized protein; afdb-uniprot50 \| AF-A0A2D9ER23-F1-MODEL\_V4 \| 1.0 \| 2.131e-38 \| 1458 \| 0.298 \| 476 \| 240 \| 7 \| 3 \| 474 \| 1 \| 386 \| Phage tail protein \| Phage tail protein; afdb-uniprot50 \| AF-A0A377H6F5-F1-MODEL\_V4 \| 1.0 \| 6.054e-39 \| 1457 \| 0.306 \| 483 \| 234 \| 8 \| 3 \| 474 \| 2 \| 394 \| Phage tail sheath protein \| Phage tail sheath protein; afdb-uniprot50 \| AF-A0A1T4W573-F1-MODEL\_V4 \| 1.0 \| 1.601e-38 \| 1456 \| 0.292 \| 478 \| 239 \| 7 \| 3 \| 474 \| 2 \| 386 \| Uncharacterized protein \| Uncharacterized protein; afdb-uniprot50 \| AF-A0A379C9Z4-F1-MODEL\_V4 \| 1.0 \| 1.136e-38 \| 1453 \| 0.272 \| 473 \| 251 \| 5 \| 3 \| 474 \| 2 \| 382 \| Phage tail sheath protein \| Phage tail sheath protein; afdb-uniprot50 \| AF-A0A017HDD9-F1-MODEL\_V4 \| 1.0 \| 8.678e-44 \| 1451 \| 0.334 \| 479 \| 290 \| 15 \| 3 \| 474 \| 2 \| 458 \| Phage\_sheath\_1 domain-containing protein \| Phage\_sheath\_1 domain-containing protein; afdb-uniprot50 \| AF-A0A545T5U0-F1-MODEL\_V4 \| 1.0 \| 7.096e-42 \| 1449 \| 0.344 \| 540 \| 182 \| 9 \| 5 \| 475 \| 3 \| 439 \| Phage tail protein \| Phage tail protein; afdb-uniprot50 \| AF-A0A7W2BMI1-F1-MODEL\_V4 \| 1.0 \| 1.625e-39 \| 1441 \| 0.287 \| 487 \| 245 \| 10 \| 3 \| 474 \| 2 \| 401 \| Phage tail sheath subtilisin-like domain-containing protein \| Phage tail sheath subtilisin-like domain-containing protein; afdb-uniprot50 \| AF-A0A7W3DV09-F1-MODEL\_V4 \| 1.0 \| 1.013e-38 \| 1439 \| 0.283 \| 484 \| 243 \| 9 \| 3 \| 474 \| 2 \| 393 \| Phage tail sheath subtilisin-like domain-containing protein \| Phage tail sheath subtilisin-like domain-containing protein; afdb-uniprot50 \| AF-A0A8B3UBL2-F1-MODEL\_V4 \| 1.0 \| 2.162e-39 \| 1435 \| 0.3 \| 482 \| 234 \| 10 \| 1 \| 474 \| 16 \| 402 \| Major tail sheath protein FI \| Major tail sheath protein FI; afdb-uniprot50 \| AF-A0A0Q8AN78-F1-MODEL\_V4 \| 1.0 \| 6.054e-39 \| 1425 \| 0.267 \| 486 \| 258 \| 8 \| 1 \| 474 \| 1 \| 400 \| Uncharacterized protein \| Uncharacterized protein; afdb-uniprot50 \| AF-A0A3P8KCN2-F1-MODEL\_V4 \| 1.0 \| 6.316e-38 \| 1408 \| 0.301 \| 478 \| 236 \| 9 \| 3 \| 474 \| 2 \| 387 \| Phage tail sheath protein \| Phage tail sheath protein; afdb-uniprot50 \| AF-A0A779QIX5-F1-MODEL\_V4 \| 1.0 \| 6.688e-38 \| 1404 \| 0.303 \| 464 \| 222 \| 10 \| 19 \| 474 \| 2 \| 372 \| Phage tail protein \| Phage tail protein; afdb-uniprot50 \| AF-A0A2W5GCK7-F1-MODEL\_V4 \| 1.0 \| 2.016e-42 \| 1404 \| 0.375 \| 421 \| 245 \| 8 \| 55 \| 474 \| 4 \| 407 \| Phage tail protein \| Phage tail protein; afdb-uniprot50 \| AF-A0A286BYN5-F1-MODEL\_V4 \| 1.0 \| 1.105e-40 \| 1404 \| 0.324 \| 472 \| 227 \| 6 \| 3 \| 472 \| 1 \| 382 \| Phage tail tape measure protein, TP901 family, core region \| Phage tail tape measure protein, TP901 family, core region; afdb-uniprot50 \| AF-A0A7X3ZJQ5-F1-MODEL\_V4 \| 1.0 \| 2.016e-42 \| 1398 \| 0.304 \| 598 \| 205 \| 14 \| 6 \| 475 \| 1 \| 515 \| Phage tail sheath family protein \| Phage tail sheath family protein; afdb-uniprot50 \| AF-A0A084ZPR3-F1-MODEL\_V4 \| 1.0 \| 6.054e-39 \| 1394 \| 0.311 \| 481 \| 231 \| 12 \| 1 \| 474 \| 17 \| 404 \| Phage tail sheath monomer \| Phage tail sheath monomer; afdb-uniprot50 \| AF-A0A0P6WF34-F1-MODEL\_V4 \| 1.0 \| 2.227e-41 \| 1383 \| 0.268 \| 502 \| 328 \| 12 \| 1 \| 474 \| 1 \| 491 \| Uncharacterized protein \| Uncharacterized protein; afdb-uniprot50 \| AF-A0A8B5G355-F1-MODEL\_V4 \| 1.0 \| 4.675e-37 \| 1376 \| 0.332 \| 445 \| 199 \| 8 \| 3 \| 439 \| 1 \| 355 \| Phage tail protein \| Phage tail protein; afdb-uniprot50 \| AF-A0A4R7GG91-F1-MODEL\_V4 \| 1.0 \| 3.317e-37 \| 1376 \| 0.275 \| 483 \| 248 \| 10 \| 3 \| 474 \| 2 \| 393 \| Phage\_sheath\_1 domain-containing protein \| Phage\_sheath\_1 domain-containing protein; afdb-uniprot50 \| AF-A0A5T6QRP6-F1-MODEL\_V4 \| 1.0 \| 2.456e-36 \| 1375 \| 0.328 \| 439 \| 203 \| 6 \| 3 \| 439 \| 2 \| 350 \| Phage tail protein \| Phage tail protein; afdb-uniprot50 \| AF-A0A4P9VGL6-F1-MODEL\_V4 \| 1.0 \| 1.218e-35 \| 1365 \| 0.614 \| 244 \| 93 \| 1 \| 231 \| 473 \| 12 \| 255 \| Uncharacterized protein \| Uncharacterized protein; afdb-uniprot50 \| AF-A0A2X5SGI3-F1-MODEL\_V4 \| 1.0 \| 1.292e-39 \| 1364 \| 0.275 \| 573 \| 212 \| 8 \| 5 \| 474 \| 2 \| 474 \| Phage tail sheath protein \| Phage tail sheath protein; afdb-uniprot50 \| AF-V4PWM7-F1-MODEL\_V4 \| 1.0 \| 5.469e-36 \| 1363 \| 0.28 \| 470 \| 231 \| 7 \| 6 \| 474 \| 1 \| 364 \| Tail sheath protein \| Tail sheath protein; afdb-uniprot50 \| AF-A0A794DQD8-F1-MODEL\_V4 \| 1.0 \| 3.275e-40 \| 1363 \| 0.282 \| 592 \| 207 \| 10 \| 1 \| 474 \| 1 \| 492 \| Phage tail protein \| Phage tail protein; afdb-uniprot50 \| AF-A0A2A4XUJ8-F1-MODEL\_V4 \| 1.0 \| 4.675e-37 \| 1362 \| 0.238 \| 477 \| 273 \| 9 \| 3 \| 474 \| 4 \| 395 \| Uncharacterized protein \| Uncharacterized protein; afdb-uniprot50 \| AF-A0A7W6F1M4-F1-MODEL\_V4 \| 1.0 \| 6.59e-37 \| 1361 \| 0.243 \| 476 \| 265 \| 7 \| 3 \| 474 \| 1 \| 385 \| Phage\_sheath\_1 domain-containing protein \| Phage\_sheath\_1 domain-containing protein; afdb-uniprot50 \| AF-A0A6N7F0I1-F1-MODEL\_V4 \| 1.0 \| 1.15e-35 \| 1350 \| 0.25 \| 471 \| 254 \| 5 \| 3 \| 472 \| 2 \| 374 \| Phage tail protein \| Phage tail protein; afdb-uniprot50 \| AF-A0A375AAB9-F1-MODEL\_V4 \| 1.0 \| 1.203e-38 \| 1349 \| 0.307 \| 478 \| 235 \| 9 \| 1 \| 474 \| 47 \| 432 \| Phage tail sheath monomer \| Phage tail sheath monomer; afdb-uniprot50 \| AF-A0A767WHR7-F1-MODEL\_V4 \| 1.0 \| 5.165e-36 \| 1348 \| 0.331 \| 432 \| 191 \| 8 \| 20 \| 443 \| 1 \| 342 \| Phage tail sheath protein \| Phage tail sheath protein; afdb-uniprot50 \| AF-A0A0S2G9I8-F1-MODEL\_V4 \| 1.0 \| 2.038e-35 \| 1348 \| 0.315 \| 444 \| 211 \| 5 \| 33 \| 474 \| 2 \| 354 \| Major tail sheath protein \| Major tail sheath protein; afdb-uniprot50 \| AF-A0A4R7SNF9-F1-MODEL\_V4 \| 1.0 \| 1.119e-37 \| 1347 \| 0.286 \| 481 \| 246 \| 11 \| 1 \| 474 \| 1 \| 391 \| Phage\_sheath\_1 domain-containing protein \| Phage\_sheath\_1 domain-containing protein; afdb-uniprot50 \| AF-A0A1I6JLA7-F1-MODEL\_V4 \| 1.0 \| 1.645e-36 \| 1346 \| 0.288 \| 472 \| 241 \| 7 \| 5 \| 474 \| 2 \| 380 \| Phage\_sheath\_1 domain-containing protein \| Phage\_sheath\_1 domain-containing protein; afdb-uniprot50 \| AF-A0A717IAE2-F1-MODEL\_V4 \| 1.0 \| 5.165e-36 \| 1345 \| 0.317 \| 438 \| 207 \| 5 \| 39 \| 474 \| 2 \| 349 \| Phage tail sheath protein \| Phage tail sheath protein; afdb-uniprot50 \| AF-A0A1Y3NT02-F1-MODEL\_V4 \| 1.0 \| 2.753e-36 \| 1339 \| 0.352 \| 454 \| 186 \| 11 \| 1 \| 443 \| 1 \| 357 \| Phage tail protein \| Phage tail protein; afdb-uniprot50 \| AF-A0A6N8Q1J1-F1-MODEL\_V4 \| 1.0 \| 4.287e-35 \| 1337 \| 0.32 \| 424 \| 196 \| 5 \| 3 \| 424 \| 2 \| 335 \| Phage tail protein \| Phage tail protein; afdb-uniprot50 \| AF-A0A8A7KVG0-F1-MODEL\_V4 \| 1.0 \| 1.218e-35 \| 1335 \| 0.275 \| 475 \| 248 \| 7 \| 4 \| 474 \| 5 \| 387 \| Phage tail protein \| Phage tail protein; afdb-uniprot50 \| AF-A0A4R3EE36-F1-MODEL\_V4 \| 1.0 \| 2.19e-36 \| 1324 \| 0.301 \| 485 \| 234 \| 16 \| 1 \| 474 \| 1 \| 391 \| Uncharacterized protein \| Uncharacterized protein; afdb-uniprot50 \| AF-A0A6I2IBC8-F1-MODEL\_V4 \| 1.0 \| 7.187e-39 \| 1321 \| 0.277 \| 590 \| 208 \| 10 \| 5 \| 474 \| 1 \| 491 \| Phage tail protein \| Phage tail protein; afdb-uniprot50 \| AF-A0A6L8HKK1-F1-MODEL\_V4 \| 1.0 \| 6.519e-44 \| 1309 \| 0.361 \| 534 \| 206 \| 15 \| 3 \| 475 \| 1 \| 460 \| Phage tail protein \| Phage tail protein; afdb-uniprot50 \| AF-A0A7V8DAB9-F1-MODEL\_V4 \| 1.0 \| 1.925e-35 \| 1308 \| 0.274 \| 474 \| 240 \| 8 \| 5 \| 474 \| 2 \| 375 \| Phage tail sheath monomer \| Phage tail sheath monomer; afdb-uniprot50 \| AF-A0A528LBW2-F1-MODEL\_V4 \| 1.0 \| 4.548e-39 \| 1304 \| 0.281 \| 501 \| 317 \| 20 \| 1 \| 474 \| 1 \| 485 \| Phage tail protein \| Phage tail protein; afdb-uniprot50 \| AF-A0A212S7J7-F1-MODEL\_V4 \| 1.0 \| 3.18e-38 \| 1303 \| 0.29 \| 493 \| 310 \| 18 \| 3 \| 474 \| 1 \| 474 \| Uncharacterized protein \| Uncharacterized protein; afdb-uniprot50 \| AF-A0A2S8RS47-F1-MODEL\_V4 \| 1.0 \| 2.959e-37 \| 1302 \| 0.297 \| 470 \| 234 \| 8 \| 1 \| 468 \| 1 \| 376 \| Uncharacterized protein \| Uncharacterized protein; afdb-uniprot50 \| AF-A0A1I7J4F6-F1-MODEL\_V4 \| 1.0 \| 5.32e-38 \| 1301 \| 0.277 \| 588 \| 220 \| 9 \| 1 \| 474 \| 1 \| 497 \| Uncharacterized protein \| Uncharacterized protein; afdb-uniprot50 \| AF-A0A704CVA7-F1-MODEL\_V4 \| 1.0 \| 1.2e-34 \| 1297 \| 0.315 \| 437 \| 207 \| 5 \| 40 \| 474 \| 1 \| 347 \| Phage tail sheath protein \| Phage tail sheath protein; afdb-uniprot50 \| AF-A0A256CAX3-F1-MODEL\_V4 \| 1.0 \| 1.133e-34 \| 1292 \| 0.247 \| 477 \| 264 \| 9 \| 3 \| 474 \| 2 \| 388 \| Phage\_sheath\_1 domain-containing protein \| Phage\_sheath\_1 domain-containing protein; afdb-uniprot50 \| AF-U4T476-F1-MODEL\_V4 \| 1.0 \| 1.531e-35 \| 1283 \| 0.253 \| 489 \| 253 \| 13 \| 3 \| 473 \| 1 \| 395 \| Major tail sheath protein \| Major tail sheath protein; afdb-uniprot50 \| AF-A0A1D2QMV4-F1-MODEL\_V4 \| 1.0 \| 1.791e-34 \| 1281 \| 0.273 \| 476 \| 247 \| 10 \| 1 \| 474 \| 1 \| 379 \| Uncharacterized protein \| Uncharacterized protein; afdb-uniprot50 \| AF-A0A2T5J1F6-F1-MODEL\_V4 \| 1.0 \| 8.902e-38 \| 1278 \| 0.256 \| 600 \| 228 \| 8 \| 3 \| 474 \| 2 \| 511 \| Uncharacterized protein DUF3383 \| Uncharacterized protein DUF3383; afdb-uniprot50 \| AF-A0A7Y1CWN9-F1-MODEL\_V4 \| 1.0 \| 3.989e-34 \| 1268 \| 0.384 \| 385 \| 151 \| 3 \| 92 \| 475 \| 7 \| 306 \| Phage tail sheath family protein \| Phage tail sheath family protein; afdb-uniprot50 \| AF-A0A7C2J5J0-F1-MODEL\_V4 \| 1.0 \| 1.49e-37 \| 1265 \| 0.275 \| 559 \| 226 \| 16 \| 1 \| 475 \| 1 \| 464 \| Phage tail sheath family protein \| Phage tail sheath family protein; afdb-uniprot50 \| AF-A0A5M7LJZ8-F1-MODEL\_V4 \| 1.0 \| 3.719e-37 \| 1248 \| 0.29 \| 486 \| 240 \| 11 \| 1 \| 474 \| 82 \| 474 \| Phage tail protein \| Phage tail protein; afdb-uniprot50 \| AF-E2CFK5-F1-MODEL\_V4 \| 1.0 \| 7.067e-34 \| 1246 \| 0.253 \| 478 \| 260 \| 13 \| 1 \| 474 \| 1 \| 385 \| Putative tail sheath protein \| Putative tail sheath protein; afdb-uniprot50 \| AF-A0A7V8JNR8-F1-MODEL\_V4 \| 1.0 \| 3.461e-36 \| 1245 \| 0.628 \| 296 \| 99 \| 1 \| 190 \| 474 \| 1 \| 296 \| Putative prophage major tail sheath protein \| Putative prophage major tail sheath protein; afdb-uniprot50 \| AF-A0A246KG12-F1-MODEL\_V4 \| 1.0 \| 3.36e-34 \| 1237 \| 0.261 \| 482 \| 243 \| 11 \| 1 \| 474 \| 1 \| 377 \| Phage tail sheath protein \| Phage tail sheath protein; afdb-uniprot50 \| AF-A0A2W6XD46-F1-MODEL\_V4 \| 1.0 \| 9.96e-34 \| 1228 \| 0.252 \| 476 \| 258 \| 7 \| 1 \| 474 \| 1 \| 380 \| Phage tail protein \| Phage tail protein; afdb-uniprot50 \| AF-A0A2W5VPE1-F1-MODEL\_V4 \| 1.0 \| 1.117e-33 \| 1227 \| 0.244 \| 483 \| 245 \| 11 \| 3 \| 474 \| 2 \| 375 \| Phage tail protein \| Phage tail protein; afdb-uniprot50 \| AF-A0A2E4CSI8-F1-MODEL\_V4 \| 1.0 \| 4.607e-36 \| 1221 \| 0.244 \| 585 \| 232 \| 7 \| 5 \| 474 \| 2 \| 491 \| Phage tail protein \| Phage tail protein; afdb-uniprot50 \| AF-A0A4D7B771-F1-MODEL\_V4 \| 1.0 \| 3.311e-33 \| 1217 \| 0.232 \| 486 \| 274 \| 10 \| 1 \| 474 \| 1 \| 399 \| Uncharacterized protein \| Uncharacterized protein; afdb-uniprot50 \| AF-A0A6C1BRA6-F1-MODEL\_V4 \| 1.0 \| 3.41e-35 \| 1216 \| 0.274 \| 507 \| 245 \| 16 \| 3 \| 474 \| 1 \| 419 \| Phage tail sheath family protein \| Phage tail sheath family protein; afdb-uniprot50 \| AF-A0A6L5IG60-F1-MODEL\_V4 \| 1.0 \| 1.618e-31 \| 1205 \| 0.5 \| 246 \| 122 \| 1 \| 230 \| 474 \| 24 \| 269 \| Phage tail sheath family protein \| Phage tail sheath family protein; afdb-uniprot50 \| AF-A0A4D7DC88-F1-MODEL\_V4 \| 1.0 \| 1.979e-33 \| 1204 \| 0.273 \| 482 \| 229 \| 10 \| 6 \| 474 \| 1 \| 374 \| Phage tail protein \| Phage tail protein; afdb-uniprot50 \| AF-A0A2X5PXW7-F1-MODEL\_V4 \| 1.0 \| 1.642e-32 \| 1186 \| 0.311 \| 433 \| 200 \| 8 \| 50 \| 474 \| 1 \| 343 \| Major tail sheath protein FI \| Major tail sheath protein FI; afdb-uniprot50 \| AF-A0A7B6D7K3-F1-MODEL\_V4 \| 1.0 \| 7.708e-36 \| 1185 \| 0.265 \| 564 \| 229 \| 15 \| 1 \| 472 \| 1 \| 470 \| Phage tail sheath family protein \| Phage tail sheath family protein; afdb-uniprot50 \| AF-A0A4U0YPU6-F1-MODEL\_V4 \| 1.0 \| 1.893e-30 \| 1184 \| 0.502 \| 245 \| 121 \| 1 \| 231 \| 474 \| 2 \| 246 \| Phage tail protein \| Phage tail protein; afdb-uniprot50 \| AF-A0A3D9UGV7-F1-MODEL\_V4 \| 1.0 \| 1.618e-31 \| 1180 \| 0.312 \| 400 \| 187 \| 4 \| 77 \| 474 \| 2 \| 315 \| Uncharacterized protein \| Uncharacterized protein; afdb-uniprot50 \| AF-A0A2X2RMN3-F1-MODEL\_V4 \| 1.0 \| 1.921e-31 \| 1176 \| 0.309 \| 426 \| 193 \| 7 \| 50 \| 474 \| 1 \| 326 \| Phage tail sheath protein \| Phage tail sheath protein; afdb-uniprot50 \| AF-A0A336QN02-F1-MODEL\_V4 \| 1.0 \| 6.303e-34 \| 1174 \| 0.32 \| 434 \| 194 \| 6 \| 42 \| 474 \| 67 \| 400 \| Phage tail sheath protein \| Phage tail sheath protein; afdb-uniprot50 \| AF-A0A4V3BMQ0-F1-MODEL\_V4 \| 1.0 \| 6.889e-40 \| 1172 \| 0.25 \| 660 \| 290 \| 21 \| 3 \| 474 \| 1 \| 643 \| Phage tail sheath protein FI \| Phage tail sheath protein FI; afdb-uniprot50 \| AF-A0A6J4N9R5-F1-MODEL\_V4 \| 1.0 \| 2.872e-35 \| 1168 \| 0.254 \| 565 \| 239 \| 15 \| 1 \| 471 \| 1 \| 477 \| Phage tail sheath monomer \| Phage tail sheath monomer; afdb-uniprot50 \| AF-A0A376ZHD7-F1-MODEL\_V4 \| 1.0 \| 1.252e-33 \| 1164 \| 0.296 \| 479 \| 229 \| 14 \| 1 \| 468 \| 1 \| 382 \| Major tail sheath protein FI \| Major tail sheath protein FI; afdb-uniprot50 \| AF-A0A7Z7Y9X6-F1-MODEL\_V4 \| 1.0 \| 1.509e-34 \| 1160 \| 0.26 \| 591 \| 225 \| 12 \| 1 \| 475 \| 1 \| 495 \| Phage tail protein \| Phage tail protein; afdb-uniprot50 \| AF-A0A6M1HTF4-F1-MODEL\_V4 \| 1.0 \| 6.384e-31 \| 1150 \| 0.333 \| 393 \| 167 \| 8 \| 3 \| 390 \| 1 \| 303 \| Phage tail sheath protein \| Phage tail sheath protein; afdb-uniprot50 \| AF-A0A371WU78-F1-MODEL\_V4 \| 1.0 \| 7.067e-34 \| 1144 \| 0.208 \| 608 \| 329 \| 20 \| 3 \| 474 \| 1 \| 591 \| Uncharacterized protein \| Uncharacterized protein; afdb-uniprot50 \| AF-A0A1M5AJX8-F1-MODEL\_V4 \| 1.0 \| 3.454e-32 \| 1141 \| 0.245 \| 480 \| 253 \| 12 \| 4 \| 474 \| 5 \| 384 \| Uncharacterized protein \| Uncharacterized protein; afdb-uniprot50 \| AF-A0A444K860-F1-MODEL\_V4 \| 1.0 \| 1.2e-34 \| 1141 \| 0.257 \| 498 \| 298 \| 14 \| 3 \| 474 \| 1 \| 452 \| Phage tail protein \| Phage tail protein; afdb-uniprot50 \| AF-A0A3M2ZQ51-F1-MODEL\_V4 \| 1.0 \| 3.603e-31 \| 1139 \| 0.345 \| 400 \| 168 \| 7 \| 5 \| 399 \| 2 \| 312 \| Phage\_sheath\_1 domain-containing protein \| Phage\_sheath\_1 domain-containing protein; afdb-uniprot50 \| AF-A0A136QAK0-F1-MODEL\_V4 \| 1.0 \| 1.114e-29 \| 1135 \| 0.51 \| 243 \| 118 \| 1 \| 233 \| 474 \| 3 \| 245 \| Uncharacterized protein \| Uncharacterized protein; afdb-uniprot50 \| AF-A0A2G2D3E0-F1-MODEL\_V4 \| 1.0 \| 3.872e-32 \| 1133 \| 0.258 \| 500 \| 251 \| 15 \| 3 \| 474 \| 2 \| 409 \| Phage tail protein \| Phage tail protein; afdb-uniprot50 \| AF-A0A7Y7RNU4-F1-MODEL\_V4 \| 1.0 \| 6.861e-32 \| 1130 \| 0.236 \| 483 \| 261 \| 11 \| 1 \| 474 \| 1 \| 384 \| Phage tail protein \| Phage tail protein; afdb-uniprot50 \| AF-A0A485D547-F1-MODEL\_V4 \| 1.0 \| 6.76e-31 \| 1125 \| 0.297 \| 393 \| 183 \| 6 \| 89 \| 474 \| 2 \| 308 \| Phage tail sheath protein \| Phage tail sheath protein; afdb-uniprot50 \| AF-A0A061JGM3-F1-MODEL\_V4 \| 1.0 \| 1.921e-31 \| 1124 \| 0.498 \| 331 \| 82 \| 4 \| 3 \| 333 \| 2 \| 248 \| Major tail sheath protein \| Major tail sheath protein; afdb-uniprot50 \| AF-A0A2A5BNT9-F1-MODEL\_V4 \| 1.0 \| 2.866e-31 \| 1117 \| 0.25 \| 476 \| 246 \| 13 \| 5 \| 475 \| 2 \| 371 \| Phage tail protein \| Phage tail protein; afdb-uniprot50 \| AF-A0A2X3JUE5-F1-MODEL\_V4 \| 1.0 \| 3.214e-31 \| 1116 \| 0.309 \| 423 \| 194 \| 11 \| 1 \| 417 \| 1 \| 331 \| Major tail sheath protein FI \| Major tail sheath protein FI; afdb-uniprot50 \| AF-B3QTJ6-F1-MODEL\_V4 \| 1.0 \| 3.712e-33 \| 1108 \| 0.249 \| 521 \| 236 \| 17 \| 1 \| 475 \| 1 \| 412 \| Tail sheath protein \| Tail sheath protein; afdb-uniprot50 \| AF-A0A285NJC0-F1-MODEL\_V4 \| 1.0 \| 2.034e-31 \| 1104 \| 0.221 \| 496 \| 257 \| 12 \| 3 \| 474 \| 1 \| 391 \| Phage\_sheath\_1 domain-containing protein \| Phage\_sheath\_1 domain-containing protein; afdb-uniprot50 \| AF-A0A1J5HQE5-F1-MODEL\_V4 \| 1.0 \| 2.122e-30 \| 1102 \| 0.244 \| 491 \| 257 \| 13 \| 1 \| 474 \| 1 \| 394 \| Phage\_sheath\_1C domain-containing protein \| Phage\_sheath\_1C domain-containing protein; afdb-uniprot50 \| AF-A0A1T4UYU3-F1-MODEL\_V4 \| 1.0 \| 6.384e-31 \| 1101 \| 0.215 \| 483 \| 268 \| 14 \| 1 \| 474 \| 5 \| 385 \| Phage tail sheath protein \| Phage tail sheath protein; afdb-uniprot50 \| AF-A0A7K1PIA1-F1-MODEL\_V4 \| 1.0 \| 2.28e-31 \| 1098 \| 0.253 \| 498 \| 259 \| 14 \| 3 \| 474 \| 1 \| 411 \| Phage tail protein \| Phage tail protein; afdb-uniprot50 \| AF-A0A3D1NSE5-F1-MODEL\_V4 \| 1.0 \| 9.527e-31 \| 1096 \| 0.253 \| 494 \| 248 \| 14 \| 3 \| 471 \| 1 \| 398 \| Phage\_sheath\_1 domain-containing protein \| Phage\_sheath\_1 domain-containing protein; afdb-uniprot50 \| AF-A0A839IZ43-F1-MODEL\_V4 \| 1.0 \| 1.528e-31 \| 1093 \| 0.234 \| 482 \| 255 \| 12 \| 1 \| 474 \| 9 \| 384 \| Phage tail protein \| Phage tail protein; afdb-uniprot50 \| AF-A0A5M8P666-F1-MODEL\_V4 \| 1.0 \| 1.443e-31 \| 1086 \| 0.232 \| 507 \| 268 \| 15 \| 3 \| 474 \| 1 \| 421 \| Phage tail protein \| Phage tail protein; afdb-uniprot50 \| AF-A0A355TSY3-F1-MODEL\_V4 \| 1.0 \| 7.373e-33 \| 1086 \| 0.238 \| 557 \| 228 \| 13 \| 20 \| 471 \| 2 \| 467 \| Phage tail protein \| Phage tail protein; afdb-uniprot50 \| AF-A0A0Q3EW34-F1-MODEL\_V4 \| 1.0 \| 8.498e-31 \| 1078 \| 0.242 \| 483 \| 254 \| 19 \| 3 \| 474 \| 2 \| 383 \| Phage\_sheath\_1 domain-containing protein \| Phage\_sheath\_1 domain-containing protein; afdb-uniprot50 \| AF-A0A846VLL6-F1-MODEL\_V4 \| 1.0 \| 7.907e-30 \| 1075 \| 0.218 \| 495 \| 279 \| 14 \| 1 \| 474 \| 1 \| 408 \| Uncharacterized protein \| Uncharacterized protein; afdb-uniprot50 \| AF-A0A3L7AKN3-F1-MODEL\_V4 \| 1.0 \| 3.815e-31 \| 1064 \| 0.259 \| 498 \| 257 \| 15 \| 3 \| 474 \| 1 \| 412 \| Phage tail protein \| Phage tail protein; afdb-uniprot50 \| AF-A0A2D2CYV4-F1-MODEL\_V4 \| 1.0 \| 1.896e-34 \| 1053 \| 0.231 \| 610 \| 325 \| 21 \| 3 \| 474 \| 1 \| 604 \| Uncharacterized protein \| Uncharacterized protein; afdb-uniprot50 \| AF-A0A5S9NEJ8-F1-MODEL\_V4 \| 1.0 \| 3.55e-30 \| 1052 \| 0.239 \| 505 \| 264 \| 14 \| 3 \| 474 \| 2 \| 419 \| Prophage major tail sheath protein \| Prophage major tail sheath protein; afdb-uniprot50 \| AF-A0A318TT75-F1-MODEL\_V4 \| 1.0 \| 1.323e-29 \| 1049 \| 0.22 \| 494 \| 279 \| 13 \| 3 \| 474 \| 1 \| 410 \| Uncharacterized protein \| Uncharacterized protein; afdb-uniprot50 \| AF-A0A022PIB5-F1-MODEL\_V4 \| 1.0 \| 1.36e-27 \| 1044 \| 0.373 \| 246 \| 152 \| 2 \| 231 \| 474 \| 28 \| 273 \| Phage tail sheath protein FI \| Phage tail sheath protein FI; afdb-uniprot50 \| AF-A0A3S4JT82-F1-MODEL\_V4 \| 1.0 \| 1.735e-28 \| 1043 \| 0.309 \| 378 \| 169 \| 5 \| 3 \| 378 \| 1 \| 288 \| Phage tail sheath protein \| Phage tail sheath protein; afdb-uniprot50 \| AF-A0A0Q2Y3L7-F1-MODEL\_V4 \| 1.0 \| 7.053e-30 \| 1034 \| 0.215 \| 505 \| 275 \| 16 \| 1 \| 473 \| 1 \| 416 \| Uncharacterized protein \| Uncharacterized protein; afdb-uniprot50 \| AF-A0A3A6S7R0-F1-MODEL\_V4 \| 1.0 \| 7.791e-29 \| 1030 \| 0.31 \| 400 \| 178 \| 11 \| 1 \| 394 \| 1 \| 308 \| Phage tail sheath family protein \| Phage tail sheath family protein; afdb-uniprot50 \| AF-A0A1T4WVG7-F1-MODEL\_V4 \| 1.0 \| 1.098e-28 \| 1028 \| 0.224 \| 489 \| 253 \| 13 \| 3 \| 475 \| 2 \| 380 \| Phage\_sheath\_1 domain-containing protein \| Phage\_sheath\_1 domain-containing protein; afdb-uniprot50 \| AF-A0A4R6E0C1-F1-MODEL\_V4 \| 1.0 \| 1.917e-27 \| 1026 \| 0.298 \| 355 \| 163 \| 3 \| 119 \| 472 \| 8 \| 277 \| Tail sheath protein \| Tail sheath protein; afdb-uniprot50 \| AF-A0A0Q1F5U2-F1-MODEL\_V4 \| 1.0 \| 5.378e-31 \| 1024 \| 0.234 \| 562 \| 243 \| 18 \| 5 \| 473 \| 2 \| 469 \| Phage\_sheath\_1 domain-containing protein \| Phage\_sheath\_1 domain-containing protein; afdb-uniprot50 \| AF-A0A5P0JHK8-F1-MODEL\_V4 \| 1.0 \| 1.066e-26 \| 1020 \| 0.371 \| 242 \| 150 \| 2 \| 235 \| 474 \| 3 \| 244 \| Phage tail protein \| Phage tail protein; afdb-uniprot50 \| AF-A0A2R8CQQ9-F1-MODEL\_V4 \| 1.0 \| 2.31e-28 \| 1018 \| 0.243 \| 477 \| 191 \| 7 \| 1 \| 474 \| 1 \| 310 \| Prophage major tail sheath protein \| Prophage major tail sheath protein; afdb-uniprot50 \| AF-A0A2Y0G935-F1-MODEL\_V4 \| 1.0 \| 1.945e-28 \| 1017 \| 0.28 \| 396 \| 186 \| 9 \| 88 \| 474 \| 2 \| 307 \| Major tail sheath protein FI \| Major tail sheath protein FI; afdb-uniprot50 \| AF-A0A0T7FB95-F1-MODEL\_V4 \| 1.0 \| 4.93e-29 \| 1011 \| 0.233 \| 489 \| 268 \| 18 \| 1 \| 473 \| 1 \| 398 \| Phage tail sheath protein \| Phage tail sheath protein; afdb-uniprot50 \| AF-A0A6A1TYH7-F1-MODEL\_V4 \| 1.0 \| 2.154e-31 \| 1007 \| 0.371 \| 320 \| 190 \| 7 \| 157 \| 474 \| 3 \| 313 \| Uncharacterized protein \| Uncharacterized protein; afdb-uniprot50 \| AF-A0A2S5J5L4-F1-MODEL\_V4 \| 1.0 \| 1.231e-28 \| 1000 \| 0.21 \| 493 \| 259 \| 15 \| 1 \| 474 \| 1 \| 382 \| Phage tail protein \| Phage tail protein; afdb-uniprot50 \| AF-S6SL27-F1-MODEL\_V4 \| 1.0 \| 2e-26 \| 999 \| 0.531 \| 224 \| 104 \| 1 \| 253 \| 475 \| 1 \| 224 \| Major tail sheath protein \| Major tail sheath protein; afdb-uniprot50 \| AF-A0A437NT20-F1-MODEL\_V4 \| 1.0 \| 9.249e-29 \| 999 \| 0.191 \| 497 \| 292 \| 14 \| 3 \| 474 \| 1 \| 412 \| Uncharacterized protein \| Uncharacterized protein; afdb-uniprot50 \| AF-A0A327QFK8-F1-MODEL\_V4 \| 1.0 \| 1.787e-30 \| 996 \| 0.22 \| 566 \| 252 \| 16 \| 3 \| 472 \| 2 \| 474 \| Phage\_sheath\_1 domain-containing protein \| Phage\_sheath\_1 domain-containing protein; afdb-uniprot50 \| AF-A0A7K1RD86-F1-MODEL\_V4 \| 1.0 \| 4.796e-31 \| 996 \| 0.245 \| 506 \| 264 \| 15 \| 1 \| 474 \| 328 \| 747 \| Uncharacterized protein \| Uncharacterized protein; afdb-uniprot50 \| AF-A0A6D0KAC9-F1-MODEL\_V4 \| 1.0 \| 8.498e-31 \| 985 \| 0.257 \| 525 \| 174 \| 8 \| 5 \| 412 \| 2 \| 427 \| Phage tail protein \| Phage tail protein; afdb-uniprot50 \| AF-A0A4U8UCG0-F1-MODEL\_V4 \| 1.0 \| 6.198e-29 \| 979 \| 0.25 \| 496 \| 245 \| 22 \| 1 \| 474 \| 1 \| 391 \| Phage tail sheath family protein \| Phage tail sheath family protein; afdb-uniprot50 \| AF-A0A2X3F9K0-F1-MODEL\_V4 \| 1.0 \| 1.398e-25 \| 978 \| 0.357 \| 238 \| 151 \| 2 \| 239 \| 474 \| 2 \| 239 \| Phage tail sheath protein \| Phage tail sheath protein; afdb-uniprot50 \| AF-A0A4R2GWM4-F1-MODEL\_V4 \| 1.0 \| 4.092e-28 \| 970 \| 0.199 \| 511 \| 285 \| 18 \| 1 \| 474 \| 1 \| 424 \| Phage\_sheath\_1 domain-containing protein \| Phage\_sheath\_1 domain-containing protein; afdb-uniprot50 \| AF-A0A2C9D5D6-F1-MODEL\_V4 \| 1.0 \| 1.591e-26 \| 965 \| 0.202 \| 490 \| 279 \| 15 \| 3 \| 474 \| 1 \| 396 \| Major tail sheath protein \| Major tail sheath protein; afdb-uniprot50 \| AF-A0A418G9S2-F1-MODEL\_V4 \| 1.0 \| 6.746e-27 \| 964 \| 0.277 \| 393 \| 183 \| 8 \| 95 \| 472 \| 1 \| 307 \| Phage tail sheath family protein \| Phage tail sheath family protein; afdb-uniprot50 \| AF-A0A7W4PVA6-F1-MODEL\_V4 \| 1.0 \| 3.914e-25 \| 948 \| 0.34 \| 247 \| 160 \| 3 \| 230 \| 474 \| 11 \| 256 \| Phage tail sheath family protein \| Phage tail sheath family protein; afdb-uniprot50 \| AF-A0A1F1HV50-F1-MODEL\_V4 \| 1.0 \| 5.221e-29 \| 944 \| 0.209 \| 562 \| 256 \| 18 \| 1 \| 474 \| 1 \| 461 \| Phage tail protein \| Phage tail protein; afdb-uniprot50 \| AF-A0A379CWG5-F1-MODEL\_V4 \| 1.0 \| 2.118e-26 \| 943 \| 0.315 \| 365 \| 152 \| 7 \| 3 \| 359 \| 1 \| 275 \| Phage tail sheath protein \| Phage tail sheath protein; afdb-uniprot50 \| AF-A0A4D9Y687-F1-MODEL\_V4 \| 1.0 \| 4.716e-26 \| 942 \| 0.324 \| 373 \| 154 \| 8 \| 3 \| 367 \| 1 \| 283 \| Phage tail protein \| Phage tail protein; afdb-uniprot50 \| AF-A0A1M7RIE6-F1-MODEL\_V4 \| 1.0 \| 7.358e-29 \| 941 \| 0.215 \| 566 \| 257 \| 19 \| 1 \| 473 \| 1 \| 472 \| Phage\_sheath\_1 domain-containing protein \| Phage\_sheath\_1 domain-containing protein; afdb-uniprot50 \| AF-A0A611ESW8-F1-MODEL\_V4 \| 1.0 \| 4.269e-27 \| 939 \| 0.306 \| 382 \| 167 \| 8 \| 35 \| 408 \| 2 \| 293 \| Phage tail protein \| Phage tail protein; afdb-uniprot50 \| AF-A0A369RI45-F1-MODEL\_V4 \| 1.0 \| 2.897e-24 \| 938 \| 0.723 \| 177 \| 49 \| 0 \| 291 \| 467 \| 12 \| 188 \| Phage tail sheath protein \| Phage tail sheath protein; afdb-uniprot50 \| AF-A0A4Q6D3G5-F1-MODEL\_V4 \| 1.0 \| 3.491e-25 \| 931 \| 0.386 \| 251 \| 144 \| 4 \| 227 \| 475 \| 4 \| 246 \| Phage\_sheath\_1 domain-containing protein \| Phage\_sheath\_1 domain-containing protein; afdb-uniprot50 \| AF-A0A327J8X6-F1-MODEL\_V4 \| 1.0 \| 1.462e-28 \| 928 \| 0.172 \| 561 \| 285 \| 16 \| 6 \| 474 \| 1 \| 473 \| Phage\_sheath\_1 domain-containing protein \| Phage\_sheath\_1 domain-containing protein; afdb-uniprot50 \| AF-A0A531KHM3-F1-MODEL\_V4 \| 1.0 \| 1.437e-23 \| 921 \| 0.685 \| 175 \| 55 \| 0 \| 301 \| 475 \| 2 \| 176 \| Phage tail protein \| Phage tail protein; afdb-uniprot50 \| AF-A0A833LH34-F1-MODEL\_V4 \| 1.0 \| 1.459e-24 \| 913 \| 0.328 \| 262 \| 161 \| 4 \| 227 \| 474 \| 27 \| 287 \| Phage tail protein \| Phage tail protein; afdb-uniprot50 \| AF-F4QJB7-F1-MODEL\_V4 \| 1.0 \| 3.44e-24 \| 911 \| 0.333 \| 237 \| 150 \| 2 \| 239 \| 474 \| 4 \| 233 \| Major tail sheath protein \| Major tail sheath protein; afdb-uniprot50 \| AF-A0A0T9LR91-F1-MODEL\_V4 \| 1.0 \| 1.096e-24 \| 901 \| 0.248 \| 474 \| 166 \| 5 \| 1 \| 474 \| 1 \| 284 \| Major tail sheath protein \| Major tail sheath protein; afdb-uniprot50 \| AF-A0A2D3VMA0-F1-MODEL\_V4 \| 1.0 \| 4.144e-25 \| 896 \| 0.207 \| 477 \| 265 \| 14 \| 5 \| 474 \| 2 \| 372 \| Phage tail protein \| Phage tail protein; afdb-uniprot50 \| AF-A0A7L5XUF0-F1-MODEL\_V4 \| 1.0 \| 5.682e-27 \| 896 \| 0.196 \| 519 \| 281 \| 18 \| 3 \| 474 \| 1 \| 430 \| Phage tail protein \| Phage tail protein; afdb-uniprot50 \| AF-A0A836P084-F1-MODEL\_V4 \| 1.0 \| 9.095e-24 \| 892 \| 0.339 \| 224 \| 147 \| 1 \| 231 \| 453 \| 8 \| 231 \| Tail sheath protein \| Tail sheath protein; afdb-uniprot50 \| AF-A0A496L9D9-F1-MODEL\_V4 \| 1.0 \| 1.889e-26 \| 885 \| 0.257 \| 443 \| 196 \| 7 \| 7 \| 415 \| 1 \| 344 \| Phage tail protein \| Phage tail protein; afdb-uniprot50 \| AF-A0A266LLW9-F1-MODEL\_V4 \| 1.0 \| 4.388e-25 \| 880 \| 0.292 \| 366 \| 162 \| 7 \| 1 \| 362 \| 1 \| 273 \| Phage tail protein \| Phage tail protein; afdb-uniprot50 \| AF-A0A1H0YQB3-F1-MODEL\_V4 \| 1.0 \| 1.112e-25 \| 877 \| 0.172 \| 515 \| 295 \| 14 \| 3 \| 474 \| 1 \| 427 \| Uncharacterized protein \| Uncharacterized protein; afdb-uniprot50 \| AF-A0A3M4S0Q3-F1-MODEL\_V4 \| 1.0 \| 2.897e-24 \| 871 \| 0.304 \| 364 \| 144 \| 8 \| 10 \| 372 \| 1 \| 256 \| Major tail sheath protein \| Major tail sheath protein; afdb-uniprot50 \| AF-A0A0H3ZR03-F1-MODEL\_V4 \| 1.0 \| 9.23e-25 \| 864 \| 0.185 \| 479 \| 278 \| 15 \| 1 \| 474 \| 1 \| 372 \| Phage tail sheath monomer \| Phage tail sheath monomer; afdb-uniprot50 \| AF-A0A7C4DFG7-F1-MODEL\_V4 \| 1.0 \| 5.599e-26 \| 864 \| 0.222 \| 503 \| 262 \| 18 \| 5 \| 475 \| 2 \| 407 \| Phage tail sheath protein \| Phage tail sheath protein; afdb-uniprot50 \| AF-A0A0D6ARN2-F1-MODEL\_V4 \| 1.0 \| 5.694e-31 \| 864 \| 0.235 \| 590 \| 299 \| 26 \| 5 \| 474 \| 2 \| 559 \| Phage tail sheath monomer \| Phage tail sheath monomer; afdb-uniprot50 \| AF-A0A6L5VN99-F1-MODEL\_V4 \| 1.0 \| 1.502e-26 \| 862 \| 0.283 \| 385 \| 175 \| 8 \| 103 \| 472 \| 89 \| 387 \| Phage tail sheath family protein \| Phage tail sheath family protein; afdb-uniprot50 \| AF-A0A2T4JE69-F1-MODEL\_V4 \| 1.0 \| 1.357e-23 \| 860 \| 0.361 \| 293 \| 104 \| 5 \| 100 \| 391 \| 2 \| 212 \| Phage tail protein \| Phage tail protein; afdb-uniprot50 \| AF-A0A1E7Q659-F1-MODEL\_V4 \| 1.0 \| 2.941e-25 \| 860 \| 0.19 \| 498 \| 267 \| 16 \| 3 \| 467 \| 2 \| 396 \| Uncharacterized protein \| Uncharacterized protein; afdb-uniprot50 \| AF-A0A2X4TXD9-F1-MODEL\_V4 \| 1.0 \| 1.377e-24 \| 857 \| 0.288 \| 357 \| 162 \| 5 \| 5 \| 359 \| 3 \| 269 \| Phage tail sheath protein \| Phage tail sheath protein; afdb-uniprot50 \| AF-A0A6H7P0U8-F1-MODEL\_V4 \| 1.0 \| 7.891e-26 \| 856 \| 0.198 \| 509 \| 276 \| 12 \| 5 \| 474 \| 2 \| 417 \| Phage tail sheath family protein \| Phage tail sheath family protein; afdb-uniprot50 \| AF-V0YRL2-F1-MODEL\_V4 \| 1.0 \| 1.78e-22 \| 855 \| 0.405 \| 202 \| 119 \| 1 \| 274 \| 474 \| 2 \| 203 \| Phage tail sheath protein \| Phage tail sheath protein; afdb-uniprot50 \| AF-A0A6L8LZU4-F1-MODEL\_V4 \| 1.0 \| 2.941e-25 \| 854 \| 0.185 \| 481 \| 278 \| 16 \| 3 \| 474 \| 4 \| 379 \| Uncharacterized protein \| Uncharacterized protein; afdb-uniprot50 \| AF-A0A8B2TR80-F1-MODEL\_V4 \| 1.0 \| 1.066e-26 \| 854 \| 0.185 \| 550 \| 246 \| 14 \| 28 \| 474 \| 7 \| 457 \| Uncharacterized protein \| Uncharacterized protein; afdb-uniprot50 \| AF-A0A1V4ZC03-F1-MODEL\_V4 \| 1.0 \| 3.857e-24 \| 848 \| 0.161 \| 478 \| 271 \| 13 \| 3 \| 468 \| 2 \| 361 \| Phage tail sheath protein \| Phage tail sheath protein; afdb-uniprot50 \| AF-D1P897-F1-MODEL\_V4 \| 1.0 \| 4.445e-22 \| 847 \| 0.368 \| 209 \| 131 \| 1 \| 267 \| 474 \| 1 \| 209 \| Phage tail sheath protein \| Phage tail sheath protein; afdb-uniprot50 \| AF-A0A7X6FQ87-F1-MODEL\_V4 \| 1.0 \| 3.339e-22 \| 844 \| 0.386 \| 215 \| 129 \| 2 \| 262 \| 474 \| 4 \| 217 \| Phage\_sheath\_1 domain-containing protein \| Phage\_sheath\_1 domain-containing protein; afdb-uniprot50 \| AF-A0A5R8Y7Q1-F1-MODEL\_V4 \| 1.0 \| 1.096e-24 \| 840 \| 0.198 \| 499 \| 286 \| 20 \| 3 \| 474 \| 1 \| 412 \| Uncharacterized protein \| Uncharacterized protein; afdb-uniprot50 \| AF-A0A715MBH1-F1-MODEL\_V4 \| 1.0 \| 2.509e-22 \| 835 \| 0.297 \| 339 \| 146 \| 6 \| 39 \| 375 \| 1 \| 249 \| Phage tail protein \| Phage tail protein; afdb-uniprot50 \| AF-A0A690AYC2-F1-MODEL\_V4 \| 1.0 \| 9.368e-26 \| 833 \| 0.18 \| 515 \| 282 \| 15 \| 5 \| 474 \| 2 \| 421 \| Phage tail sheath family protein \| Phage tail sheath family protein; afdb-uniprot50 \| AF-A0A2A5M6V2-F1-MODEL\_V4 \| 1.0 \| 5.755e-24 \| 828 \| 0.207 \| 468 \| 238 \| 12 \| 33 \| 474 \| 2 \| 362 \| Phage tail protein \| Phage tail protein; afdb-uniprot50 \| AF-A0A7I9TNA5-F1-MODEL\_V4 \| 1.0 \| 5.21e-25 \| 821 \| 0.202 \| 519 \| 268 \| 16 \| 1 \| 474 \| 1 \| 418 \| Phage tail sheath family protein \| Phage tail sheath family protein; afdb-uniprot50 \| AF-A0A824RG65-F1-MODEL\_V4 \| 1.0 \| 1.282e-23 \| 817 \| 0.226 \| 407 \| 215 \| 7 \| 71 \| 474 \| 30 \| 339 \| Phage tail sheath family protein \| Phage tail sheath family protein; afdb-uniprot50 \| AF-A0A7K3ZN16-F1-MODEL\_V4 \| 1.0 \| 1.143e-23 \| 817 \| 0.19 \| 478 \| 251 \| 14 \| 2 \| 467 \| 3 \| 356 \| Phage tail sheath family protein \| Phage tail sheath family protein; afdb-uniprot50 \| AF-A0A446CQT9-F1-MODEL\_V4 \| 1.0 \| 8.589e-24 \| 815 \| 0.188 \| 483 \| 257 \| 14 \| 1 \| 466 \| 1 \| 365 \| Prophage major tail sheath protein \| Prophage major tail sheath protein; afdb-uniprot50 \| AF-A0A6G5QYL8-F1-MODEL\_V4 \| 1.0 \| 3.154e-22 \| 814 \| 0.307 \| 247 \| 164 \| 5 \| 231 \| 474 \| 411 \| 653 \| Phage tail sheath protein FI (Integrase domain) \| Phage tail sheath protein FI (Integrase domain); afdb-uniprot50 \| AF-A0A534UC15-F1-MODEL\_V4 \| 1.0 \| 1.611e-23 \| 812 \| 0.175 \| 473 \| 281 \| 16 \| 1 \| 460 \| 1 \| 377 \| Phage tail sheath family protein \| Phage tail sheath family protein; afdb-uniprot50 \| AF-A0A6V6ZU17-F1-MODEL\_V4 \| 1.0 \| 3.815e-31 \| 810 \| 0.294 \| 434 \| 248 \| 14 \| 1 \| 419 \| 1 \| 391 \| Uncharacterized protein \| Uncharacterized protein; afdb-uniprot50 \| AF-A0A4Q6CEC2-F1-MODEL\_V4 \| 1.0 \| 1.318e-21 \| 805 \| 0.366 \| 224 \| 135 \| 3 \| 254 \| 475 \| 11 \| 229 \| Phage tail protein \| Phage tail protein; afdb-uniprot50 \| AF-A0A1Z4PN29-F1-MODEL\_V4 \| 1.0 \| 1.282e-23 \| 805 \| 0.179 \| 478 \| 269 \| 14 \| 5 \| 466 \| 2 \| 372 \| Putative prophage major tail sheath protein \| Putative prophage major tail sheath protein; afdb-uniprot50 \| AF-A0A521PE06-F1-MODEL\_V4 \| 1.0 \| 8.112e-24 \| 804 \| 0.185 \| 485 \| 263 \| 17 \| 3 \| 468 \| 2 \| 373 \| Phage tail sheath family protein \| Phage tail sheath family protein; afdb-uniprot50 \| AF-A0A349XNZ8-F1-MODEL\_V4 \| 1.0 \| 1.522e-23 \| 803 \| 0.255 \| 399 \| 193 \| 12 \| 93 \| 474 \| 14 \| 325 \| Phage tail protein \| Phage tail protein; afdb-uniprot50 \| AF-A0A1J5D0C9-F1-MODEL\_V4 \| 1.0 \| 1.941e-24 \| 801 \| 0.207 \| 492 \| 255 \| 19 \| 1 \| 466 \| 1 \| 383 \| Phage tail protein \| Phage tail protein; afdb-uniprot50 \| AF-A0A6I7PVR7-F1-MODEL\_V4 \| 1.0 \| 5.355e-23 \| 800 \| 0.175 \| 483 \| 274 \| 17 \| 3 \| 468 \| 2 \| 377 \| Phage tail sheath family protein \| Phage tail sheath family protein; afdb-uniprot50 \| AF-A0A6G3I5H9-F1-MODEL\_V4 \| 1.0 \| 4.379e-21 \| 798 \| 0.378 \| 214 \| 127 \| 3 \| 265 \| 472 \| 15 \| 228 \| Phage tail sheath family protein \| Phage tail sheath family protein; afdb-uniprot50 \| AF-A0A4Q0ZK21-F1-MODEL\_V4 \| 1.0 \| 1.885e-22 \| 795 \| 0.182 \| 478 \| 272 \| 14 \| 2 \| 474 \| 5 \| 368 \| Phage tail protein \| Phage tail protein; afdb-uniprot50 \| AF-A0A6M0L0M0-F1-MODEL\_V4 \| 1.0 \| 5.058e-23 \| 794 \| 0.181 \| 491 \| 271 \| 20 \| 3 \| 468 \| 2 \| 386 \| Phage tail sheath family protein \| Phage tail sheath family protein; afdb-uniprot50 \| AF-A0A6A4R6J7-F1-MODEL\_V4 \| 1.0 \| 8.961e-23 \| 792 \| 0.218 \| 407 \| 211 \| 10 \| 1 \| 398 \| 1 \| 309 \| Phage tail protein \| Phage tail protein; afdb-uniprot50 \| AF-A0A1I6LKB0-F1-MODEL\_V4 \| 1.0 \| 2.696e-23 \| 788 \| 0.187 \| 480 \| 252 \| 11 \| 5 \| 467 \| 3 \| 361 \| Phage tail sheath protein \| Phage tail sheath protein; afdb-uniprot50 \| AF-A0A1H2U7X5-F1-MODEL\_V4 \| 1.0 \| 1.706e-23 \| 788 \| 0.183 \| 486 \| 259 \| 18 \| 4 \| 468 \| 5 \| 373 \| Uncharacterized protein \| Uncharacterized protein; afdb-uniprot50 \| AF-A0A823XP83-F1-MODEL\_V4 \| 1.0 \| 1.048e-21 \| 786 \| 0.217 \| 377 \| 196 \| 6 \| 100 \| 474 \| 3 \| 282 \| Phage tail sheath family protein \| Phage tail sheath family protein; afdb-uniprot50 \| AF-A0A1E3GNZ3-F1-MODEL\_V4 \| 1.0 \| 2.271e-23 \| 786 \| 0.173 \| 485 \| 268 \| 14 \| 3 \| 466 \| 2 \| 374 \| Phage tail sheath protein \| Phage tail sheath protein; afdb-uniprot50 \| AF-A0A285MD48-F1-MODEL\_V4 \| 1.0 \| 1.211e-23 \| 785 \| 0.165 \| 503 \| 295 \| 20 \| 3 \| 474 \| 2 \| 410 \| Uncharacterized protein \| Uncharacterized protein; afdb-uniprot50 \| AF-A0A5T0J8E0-F1-MODEL\_V4 \| 1.0 \| 4.379e-21 \| 777 \| 0.28 \| 246 \| 172 \| 3 \| 231 \| 474 \| 26 \| 268 \| Phage tail sheath family protein \| Phage tail sheath family protein; afdb-uniprot50 \| AF-A0A1I5RNY6-F1-MODEL\_V4 \| 1.0 \| 2.546e-23 \| 773 \| 0.17 \| 493 \| 278 \| 16 \| 5 \| 474 \| 6 \| 390 \| Uncharacterized protein \| Uncharacterized protein; afdb-uniprot50 \| AF-W9GDP8-F1-MODEL\_V4 \| 1.0 \| 5.133e-24 \| 772 \| 0.188 \| 472 \| 253 \| 16 \| 19 \| 466 \| 3 \| 368 \| Phage tail sheath protein \| Phage tail sheath protein; afdb-uniprot50 \| AF-A0A5Y3QFL3-F1-MODEL\_V4 \| 1.0 \| 2.472e-21 \| 768 \| 0.308 \| 327 \| 132 \| 7 \| 76 \| 394 \| 2 \| 242 \| Phage tail protein \| Phage tail protein; afdb-uniprot50 \| AF-A0A0F3RPG1-F1-MODEL\_V4 \| 1.0 \| 2.771e-21 \| 765 \| 0.439 \| 273 \| 71 \| 4 \| 3 \| 275 \| 2 \| 192 \| Phage tail sheath family protein \| Phage tail sheath family protein; afdb-uniprot50 \| AF-A0A1G8HX37-F1-MODEL\_V4 \| 1.0 \| 2.205e-21 \| 763 \| 0.205 \| 408 \| 219 \| 8 \| 75 \| 474 \| 2 \| 312 \| Uncharacterized protein \| Uncharacterized protein; afdb-uniprot50 \| AF-A0A436QUU1-F1-MODEL\_V4 \| 1.0 \| 1.632e-20 \| 762 \| 0.288 \| 260 \| 171 \| 4 \| 228 \| 474 \| 23 \| 281 \| Phage tail protein \| Phage tail protein; afdb-uniprot50 \| AF-A0A6L3F9E2-F1-MODEL\_V4 \| 1.0 \| 2.271e-23 \| 762 \| 0.209 \| 502 \| 253 \| 20 \| 1 \| 468 \| 7 \| 398 \| Phage tail sheath family protein \| Phage tail sheath family protein; afdb-uniprot50 \| AF-A0A7T8BA55-F1-MODEL\_V4 \| 1.0 \| 1.913e-23 \| 759 \| 0.173 \| 513 \| 269 \| 20 \| 3 \| 468 \| 2 \| 406 \| Phage tail sheath family protein \| Phage tail sheath family protein; afdb-uniprot50 \| AF-A0A2U2MY54-F1-MODEL\_V4 \| 1.0 \| 1.611e-23 \| 757 \| 0.182 \| 500 \| 273 \| 16 \| 3 \| 467 \| 2 \| 400 \| Uncharacterized protein \| Uncharacterized protein; afdb-uniprot50 \| AF-E6X1M8-F1-MODEL\_V4 \| 1.0 \| 3.29e-21 \| 756 \| 0.191 \| 485 \| 261 \| 17 \| 5 \| 474 \| 2 \| 370 \| Major tail sheath protein \| Major tail sheath protein; afdb-uniprot50 \| AF-A0A530RAK9-F1-MODEL\_V4 \| 1.0 \| 1.519e-19 \| 754 \| 0.688 \| 154 \| 48 \| 0 \| 320 \| 473 \| 1 \| 154 \| Phage tail protein \| Phage tail protein; afdb-uniprot50 \| AF-A0A7C3RPS7-F1-MODEL\_V4 \| 1.0 \| 5.755e-24 \| 754 \| 0.235 \| 441 \| 205 \| 11 \| 81 \| 475 \| 9 \| 363 \| Phage tail sheath family protein \| Phage tail sheath family protein; afdb-uniprot50 \| AF-A0A7V8JNF3-F1-MODEL\_V4 \| 1.0 \| 1.861e-25 \| 753 \| 0.57 \| 228 \| 92 \| 3 \| 3 \| 226 \| 2 \| 227 \| Putative prophage major tail sheath protein \| Putative prophage major tail sheath protein; afdb-uniprot50 \| AF-A0A1M6GFB7-F1-MODEL\_V4 \| 1.0 \| 3.589e-23 \| 753 \| 0.198 \| 524 \| 245 \| 18 \| 3 \| 467 \| 2 \| 409 \| Uncharacterized protein \| Uncharacterized protein; afdb-uniprot50 \| AF-A0A537ZZS7-F1-MODEL\_V4 \| 1.0 \| 1.754e-21 \| 750 \| 0.196 \| 463 \| 234 \| 14 \| 3 \| 456 \| 2 \| 335 \| Phage tail sheath family protein \| Phage tail sheath family protein; afdb-uniprot50 \| AF-A0A6G5QNE9-F1-MODEL\_V4 \| 1.0 \| 7.024e-22 \| 750 \| 0.169 \| 478 \| 276 \| 15 \| 5 \| 474 \| 2 \| 366 \| Phage tail sheath protein FI \| Phage tail sheath protein FI; afdb-uniprot50 \| AF-A0A3S7DS92-F1-MODEL\_V4 \| 1.0 \| 2.276e-27 \| 750 \| 0.191 \| 621 \| 304 \| 28 \| 3 \| 474 \| 1 \| 572 \| Phage tail protein \| Phage tail protein; afdb-uniprot50 \| AF-A0A6N8U467-F1-MODEL\_V4 \| 1.0 \| 8.463e-23 \| 748 \| 0.186 \| 483 \| 259 \| 16 \| 7 \| 467 \| 1 \| 371 \| Uncharacterized protein \| Uncharacterized protein; afdb-uniprot50 \| AF-A0A7M3MAW8-F1-MODEL\_V4 \| 1.0 \| 2.025e-23 \| 748 \| 0.205 \| 512 \| 269 \| 19 \| 3 \| 475 \| 1 \| 413 \| Phage tail sheath family protein \| Phage tail sheath family protein; afdb-uniprot50 \| AF-A0A6L9FQ12-F1-MODEL\_V4 \| 1.0 \| 7.676e-28 \| 748 \| 0.184 \| 623 \| 326 \| 30 \| 3 \| 474 \| 1 \| 592 \| Phage tail protein \| Phage tail protein; afdb-uniprot50 \| AF-A0A7Y4DJI9-F1-MODEL\_V4 \| 1.0 \| 1.996e-22 \| 746 \| 0.14 \| 478 \| 295 \| 15 \| 1 \| 474 \| 3 \| 368 \| Uncharacterized protein \| Uncharacterized protein; afdb-uniprot50 \| AF-A0A524IP71-F1-MODEL\_V4 \| 1.0 \| 1.996e-22 \| 744 \| 0.177 \| 489 \| 268 \| 16 \| 1 \| 466 \| 1 \| 378 \| Phage tail sheath family protein \| Phage tail sheath family protein; afdb-uniprot50 \| AF-A0A1C0V8T7-F1-MODEL\_V4 \| 1.0 \| 4.52e-27 \| 744 \| 0.187 \| 730 \| 287 \| 30 \| 1 \| 471 \| 2 \| 684 \| Phage\_sheath\_1 domain-containing protein \| Phage\_sheath\_1 domain-containing protein; afdb-uniprot50 \| AF-A0A1Q7X247-F1-MODEL\_V4 \| 1.0 \| 6.732e-23 \| 741 \| 0.188 \| 473 \| 243 \| 16 \| 3 \| 464 \| 24 \| 366 \| Uncharacterized protein \| Uncharacterized protein; afdb-uniprot50 \| AF-A0A432UGQ1-F1-MODEL\_V4 \| 1.0 \| 2.405e-23 \| 737 \| 0.204 \| 571 \| 257 \| 22 \| 5 \| 475 \| 2 \| 474 \| Phage tail sheath family protein \| Phage tail sheath family protein; afdb-uniprot50 \| AF-A0A1Q7MAI6-F1-MODEL\_V4 \| 1.0 \| 1.02e-23 \| 735 \| 0.171 \| 497 \| 293 \| 18 \| 1 \| 474 \| 1 \| 401 \| Uncharacterized protein \| Uncharacterized protein; afdb-uniprot50 \| AF-A0A377I6M5-F1-MODEL\_V4 \| 1.0 \| 2.978e-22 \| 732 \| 0.238 \| 398 \| 193 \| 12 \| 93 \| 473 \| 31 \| 335 \| Tail sheath-like protein \| Tail sheath-like protein; afdb-uniprot50 \| AF-A0A6P0MAY6-F1-MODEL\_V4 \| 1.0 \| 1.78e-22 \| 730 \| 0.18 \| 481 \| 262 \| 17 \| 5 \| 466 \| 2 \| 369 \| Uncharacterized protein \| Uncharacterized protein; afdb-uniprot50 \| AF-A0A1J4YMP9-F1-MODEL\_V4 \| 1.0 \| 1.564e-21 \| 727 \| 0.171 \| 485 \| 270 \| 21 \| 5 \| 468 \| 3 \| 376 \| Uncharacterized protein \| Uncharacterized protein; afdb-uniprot50 \| AF-A0A376YN89-F1-MODEL\_V4 \| 1.0 \| 1.541e-20 \| 724 \| 0.372 \| 196 \| 122 \| 1 \| 259 \| 453 \| 2 \| 197 \| Major tail sheath protein FI \| Major tail sheath protein FI; afdb-uniprot50 \| AF-A0A660N068-F1-MODEL\_V4 \| 1.0 \| 3.8e-23 \| 723 \| 0.207 \| 507 \| 217 \| 16 \| 1 \| 419 \| 1 \| 410 \| Phage tail protein \| Phage tail protein; afdb-uniprot50 \| AF-A0A4R9F3E7-F1-MODEL\_V4 \| 1.0 \| 2.771e-21 \| 720 \| 0.187 \| 469 \| 250 \| 19 \| 15 \| 468 \| 2 \| 354 \| Phage tail sheath family protein \| Phage tail sheath family protein; afdb-uniprot50 \| AF-A0A0F0GLD7-F1-MODEL\_V4 \| 1.0 \| 3.543e-26 \| 720 \| 0.197 \| 548 \| 304 \| 18 \| 3 \| 467 \| 2 \| 496 \| Tail protein \| Tail protein; afdb-uniprot50 \| AF-A0A353GA80-F1-MODEL\_V4 \| 1.0 \| 6.633e-22 \| 719 \| 0.251 \| 414 \| 188 \| 15 \| 1 \| 382 \| 1 \| 324 \| Phage tail protein \| Phage tail protein; afdb-uniprot50 \| AF-A0A403T4T3-F1-MODEL\_V4 \| 1.0 \| 4.454e-26 \| 719 \| 0.256 \| 449 \| 208 \| 9 \| 1 \| 419 \| 1 \| 353 \| Phage\_sheath\_1 domain-containing protein \| Phage\_sheath\_1 domain-containing protein; afdb-uniprot50 \| AF-A0A6B9YRM7-F1-MODEL\_V4 \| 1.0 \| 5.277e-22 \| 715 \| 0.158 \| 506 \| 278 \| 22 \| 1 \| 468 \| 8 \| 403 \| Phage tail sheath family protein \| Phage tail sheath family protein; afdb-uniprot50 \| AF-B6WRQ3-F1-MODEL\_V4 \| 1.0 \| 1.416e-22 \| 715 \| 0.171 \| 590 \| 279 \| 20 \| 1 \| 475 \| 1 \| 495 \| Uncharacterized protein \| Uncharacterized protein; afdb-uniprot50 \| AF-A0A0D0X3W4-F1-MODEL\_V4 \| 1.0 \| 1.784e-26 \| 715 \| 0.201 \| 556 \| 307 \| 25 \| 3 \| 468 \| 2 \| 510 \| Tail protein \| Tail protein; afdb-uniprot50 \| AF-A0A523TY78-F1-MODEL\_V4 \| 1.0 \| 9.919e-26 \| 712 \| 0.173 \| 547 \| 314 \| 23 \| 1 \| 468 \| 1 \| 488 \| Phage tail sheath family protein \| Phage tail sheath family protein; afdb-uniprot50 \| AF-A0A7Y5XG39-F1-MODEL\_V4 \| 1.0 \| 3.062e-20 \| 711 \| 0.277 \| 252 \| 162 \| 8 \| 230 \| 467 \| 73 \| 318 \| Phage tail sheath family protein \| Phage tail sheath family protein; afdb-uniprot50 \| AF-A0A369BHJ8-F1-MODEL\_V4 \| 1.0 \| 1.126e-22 \| 710 \| 0.197 \| 506 \| 270 \| 25 \| 5 \| 474 \| 2 \| 407 \| Phage tail sheath protein FI \| Phage tail sheath protein FI; afdb-uniprot50 \| AF-A0A535YX28-F1-MODEL\_V4 \| 1.0 \| 1.112e-25 \| 706 \| 0.2 \| 565 \| 286 \| 24 \| 3 \| 467 \| 2 \| 500 \| Phage tail sheath family protein \| Phage tail sheath family protein; afdb-uniprot50 \| AF-A0A2X2BJR7-F1-MODEL\_V4 \| 1.0 \| 1.279e-19 \| 704 \| 0.383 \| 193 \| 117 \| 2 \| 230 \| 420 \| 27 \| 219 \| Major tail sheath protein \| Major tail sheath protein; afdb-uniprot50 \| AF-A0A1Q9Q7N6-F1-MODEL\_V4 \| 1.0 \| 4.637e-21 \| 703 \| 0.184 \| 505 \| 254 \| 13 \| 3 \| 468 \| 2 \| 387 \| Uncharacterized protein \| Uncharacterized protein; afdb-uniprot50 \| AF-A0A7Y1Y9C5-F1-MODEL\_V4 \| 1.0 \| 1.754e-21 \| 700 \| 0.165 \| 490 \| 277 \| 20 \| 1 \| 466 \| 3 \| 384 \| Phage tail sheath family protein \| Phage tail sheath family protein; afdb-uniprot50 \| AF-A0A3P1YCU6-F1-MODEL\_V4 \| 1.0 \| 2.109e-18 \| 699 \| 0.336 \| 202 \| 132 \| 2 \| 274 \| 474 \| 1 \| 201 \| Phage tail sheath family protein \| Phage tail sheath family protein; afdb-uniprot50 \| AF-A0A774NAH9-F1-MODEL\_V4 \| 1.0 \| 5.424e-20 \| 699 \| 0.269 \| 341 \| 154 \| 7 \| 88 \| 420 \| 2 \| 255 \| Phage tail protein \| Phage tail protein; afdb-uniprot50 \| AF-A0A7H8K818-F1-MODEL\_V4 \| 1.0 \| 1.05e-25 \| 698 \| 0.199 \| 556 \| 309 \| 21 \| 3 \| 468 \| 2 \| 511 \| Phage tail sheath family protein \| Phage tail sheath family protein; afdb-uniprot50 \| AF-A0A1X7M8N6-F1-MODEL\_V4 \| 1.0 \| 1.992e-18 \| 697 \| 0.397 \| 254 \| 65 \| 2 \| 30 \| 283 \| 2 \| 167 \| Phage major tail sheath protein \| Phage major tail sheath protein; afdb-uniprot50 \| AF-A0A177NQ44-F1-MODEL\_V4 \| 1.0 \| 3.107e-21 \| 697 \| 0.197 \| 502 \| 251 \| 18 \| 3 \| 467 \| 2 \| 388 \| Phage tail protein \| Phage tail protein; afdb-uniprot50 \| AF-A0A7K1H3K3-F1-MODEL\_V4 \| 1.0 \| 8.355e-26 \| 697 \| 0.184 \| 557 \| 314 \| 24 \| 3 \| 468 \| 2 \| 508 \| Phage tail sheath family protein \| Phage tail sheath family protein; afdb-uniprot50 \| AF-A0A849Q2Q2-F1-MODEL\_V4 \| 1.0 \| 3.536e-22 \| 693 \| 0.17 \| 504 \| 262 \| 22 \| 1 \| 467 \| 1 \| 385 \| Uncharacterized protein \| Uncharacterized protein; afdb-uniprot50 \| AF-A0A0G9K7V9-F1-MODEL\_V4 \| 1.0 \| 8.095e-20 \| 693 \| 0.152 \| 491 \| 285 \| 17 \| 5 \| 474 \| 2 \| 382 \| Tail protein \| Tail protein; afdb-uniprot50 \| AF-A0A833EFL3-F1-MODEL\_V4 \| 1.0 \| 1.158e-20 \| 693 \| 0.195 \| 502 \| 269 \| 20 \| 3 \| 475 \| 1 \| 396 \| Phage tail sheath protein \| Phage tail sheath protein; afdb-uniprot50 \| AF-A0A4Q6GVN0-F1-MODEL\_V4 \| 1.0 \| 4.379e-21 \| 692 \| 0.154 \| 499 \| 277 \| 19 \| 1 \| 468 \| 2 \| 386 \| Phage tail sheath family protein \| Phage tail sheath family protein; afdb-uniprot50 \| AF-A0A6G7Z242-F1-MODEL\_V4 \| 1.0 \| 6.185e-25 \| 692 \| 0.193 \| 573 \| 300 \| 23 \| 3 \| 468 \| 2 \| 519 \| Phage tail sheath family protein \| Phage tail sheath family protein; afdb-uniprot50 \| AF-A0A1E4GSC1-F1-MODEL\_V4 \| 1.0 \| 3.906e-21 \| 691 \| 0.184 \| 487 \| 266 \| 18 \| 3 \| 468 \| 2 \| 378 \| Uncharacterized protein \| Uncharacterized protein; afdb-uniprot50 \| AF-A0A538DVN2-F1-MODEL\_V4 \| 1.0 \| 2.935e-21 \| 689 \| 0.195 \| 492 \| 260 \| 20 \| 3 \| 468 \| 2 \| 383 \| Phage tail sheath family protein \| Phage tail sheath family protein; afdb-uniprot50 \| AF-L1P591-F1-MODEL\_V4 \| 1.0 \| 7.532e-19 \| 687 \| 0.215 \| 376 \| 188 \| 10 \| 113 \| 474 \| 2 \| 284 \| Phage\_sheath\_1 domain-containing protein \| Phage\_sheath\_1 domain-containing protein; afdb-uniprot50 \| AF-A0A2E3R271-F1-MODEL\_V4 \| 1.0 \| 4.706e-22 \| 687 \| 0.176 \| 521 \| 283 \| 24 \| 3 \| 475 \| 2 \| 424 \| Phage\_sheath\_1 domain-containing protein \| Phage\_sheath\_1 domain-containing protein; afdb-uniprot50 \| AF-A0A812WA44-F1-MODEL\_V4 \| 1.0 \| 1.247e-25 \| 687 \| 0.207 \| 569 \| 290 \| 28 \| 3 \| 467 \| 2 \| 513 \| 18 protein \| 18 protein; afdb-uniprot50 \| AF-A0A7X5APW2-F1-MODEL\_V4 \| 1.0 \| 3.29e-21 \| 685 \| 0.171 \| 496 \| 264 \| 21 \| 3 \| 466 \| 2 \| 382 \| Phage\_sheath\_1 domain-containing protein \| Phage\_sheath\_1 domain-containing protein; afdb-uniprot50 \| AF-A0A5Q4D9T1-F1-MODEL\_V4 \| 1.0 \| 7.875e-22 \| 685 \| 0.182 \| 499 \| 269 \| 23 \| 1 \| 466 \| 7 \| 399 \| Phage tail sheath family protein \| Phage tail sheath family protein; afdb-uniprot50 \| AF-A0A859FDD1-F1-MODEL\_V4 \| 1.0 \| 5.671e-23 \| 685 \| 0.178 \| 576 \| 263 \| 21 \| 5 \| 474 \| 2 \| 473 \| Phage tail sheath family protein \| Phage tail sheath family protein; afdb-uniprot50 \| AF-A0A3N5RBQ4-F1-MODEL\_V4 \| 1.0 \| 1.97e-25 \| 685 \| 0.188 \| 561 \| 305 \| 26 \| 1 \| 467 \| 1 \| 504 \| Phage tail sheath family protein \| Phage tail sheath family protein; afdb-uniprot50 \| AF-A0A2W5Z8T4-F1-MODEL\_V4 \| 1.0 \| 1.588e-22 \| 685 \| 0.186 \| 494 \| 257 \| 17 \| 1 \| 467 \| 1 \| 376 \| AAA domain-containing protein \| AAA domain-containing protein; afdb-uniprot50 \| AF-A0A150R6N1-F1-MODEL\_V4 \| 1.0 \| 7.22e-20 \| 685 \| 0.281 \| 252 \| 160 \| 9 \| 231 \| 468 \| 690 \| 934 \| Uncharacterized protein \| Uncharacterized protein; afdb-uniprot50 \| AF-A0A1X4I4Z3-F1-MODEL\_V4 \| 1.0 \| 1.279e-19 \| 684 \| 0.3 \| 253 \| 156 \| 9 \| 230 \| 467 \| 65 \| 311 \| Phage tail protein \| Phage tail protein; afdb-uniprot50 \| AF-A0A3B9JCN2-F1-MODEL\_V4 \| 1.0 \| 4.252e-19 \| 682 \| 0.277 \| 252 \| 162 \| 8 \| 230 \| 467 \| 104 \| 349 \| Phage tail protein \| Phage tail protein; afdb-uniprot50 \| AF-A0A3S0CXJ3-F1-MODEL\_V4 \| 1.0 \| 2.021e-19 \| 682 \| 0.266 \| 251 \| 165 \| 8 \| 231 \| 468 \| 111 \| 355 \| Phage tail sheath family protein \| Phage tail sheath family protein; afdb-uniprot50 \| AF-D4K2N0-F1-MODEL\_V4 \| 1.0 \| 2.3e-20 \| 682 \| 0.165 \| 494 \| 272 \| 17 \| 5 \| 472 \| 1 \| 380 \| Phage tail sheath protein FI \| Phage tail sheath protein FI; afdb-uniprot50 \| AF-A0A4R5TTN3-F1-MODEL\_V4 \| 1.0 \| 3.689e-21 \| 681 \| 0.188 \| 503 \| 257 \| 18 \| 3 \| 468 \| 2 \| 390 \| Phage tail sheath family protein \| Phage tail sheath family protein; afdb-uniprot50 \| AF-A0A2X0WI40-F1-MODEL\_V4 \| 1.0 \| 4.983e-22 \| 680 \| 0.164 \| 594 \| 275 \| 26 \| 3 \| 475 \| 1 \| 493 \| Phage tail sheath protein \| Phage tail sheath protein; afdb-uniprot50 \| AF-A0A1G5B506-F1-MODEL\_V4 \| 1.0 \| 3.536e-22 \| 679 \| 0.174 \| 583 \| 264 \| 21 \| 5 \| 475 \| 2 \| 478 \| Uncharacterized protein \| Uncharacterized protein; afdb-uniprot50 \| AF-A0A0S8BQ23-F1-MODEL\_V4 \| 1.0 \| 2.477e-25 \| 678 \| 0.202 \| 563 \| 302 \| 27 \| 1 \| 467 \| 1 \| 512 \| Uncharacterized protein \| Uncharacterized protein; afdb-uniprot50 \| AF-V2YG99-F1-MODEL\_V4 \| 1.0 \| 5.047e-19 \| 676 \| 0.258 \| 251 \| 167 \| 7 \| 231 \| 468 \| 88 \| 332 \| Uncharacterized protein \| Uncharacterized protein; afdb-uniprot50 \| AF-V6U633-F1-MODEL\_V4 \| 1.0 \| 3.581e-19 \| 676 \| 0.28 \| 253 \| 162 \| 8 \| 230 \| 468 \| 135 \| 381 \| Putative phage tail sheath protein \| Putative phage tail sheath protein; afdb-uniprot50 \| AF-A0A497BTG9-F1-MODEL\_V4 \| 1.0 \| 1.459e-24 \| 676 \| 0.175 \| 580 \| 297 \| 21 \| 1 \| 468 \| 1 \| 511 \| Phage tail sheath family protein \| Phage tail sheath family protein; afdb-uniprot50 \| AF-A0A7C5IAH6-F1-MODEL\_V4 \| 1.0 \| 7.976e-19 \| 675 \| 0.235 \| 251 \| 174 \| 7 \| 230 \| 469 \| 168 \| 411 \| Phage tail sheath family protein \| Phage tail sheath family protein; afdb-uniprot50 \| AF-A0A0Q7MZB4-F1-MODEL\_V4 \| 1.0 \| 8.233e-25 \| 675 \| 0.186 \| 575 \| 299 \| 25 \| 1 \| 468 \| 1 \| 513 \| Phage tail protein \| Phage tail protein; afdb-uniprot50 \| AF-A0A6G2PQT4-F1-MODEL\_V4 \| 1.0 \| 3.017e-19 \| 674 \| 0.273 \| 252 \| 161 \| 7 \| 230 \| 468 \| 48 \| 290 \| Phage tail sheath family protein \| Phage tail sheath family protein; afdb-uniprot50 \| AF-A0A848Y6F5-F1-MODEL\_V4 \| 1.0 \| 3.581e-19 \| 674 \| 0.259 \| 258 \| 166 \| 7 \| 230 \| 468 \| 66 \| 317 \| Phage tail sheath family protein \| Phage tail sheath family protein; afdb-uniprot50 \| AF-M0BGK5-F1-MODEL\_V4 \| 1.0 \| 2.14e-19 \| 673 \| 0.256 \| 250 \| 165 \| 7 \| 231 \| 468 \| 268 \| 508 \| Phage tail sheath protein FI \| Phage tail sheath protein FI; afdb-uniprot50 \| AF-A0A651E436-F1-MODEL\_V4 \| 1.0 \| 5.659e-19 \| 673 \| 0.274 \| 251 \| 164 \| 7 \| 230 \| 468 \| 288 \| 532 \| Phage tail sheath family protein \| Phage tail sheath family protein; afdb-uniprot50 \| AF-A0A5J6QAB9-F1-MODEL\_V4 \| 1.0 \| 3.696e-25 \| 673 \| 0.169 \| 633 \| 314 \| 29 \| 1 \| 468 \| 1 \| 586 \| Uncharacterized protein \| Uncharacterized protein; afdb-uniprot50 \| AF-A0A6G9ZDV3-F1-MODEL\_V4 \| 1.0 \| 1.499e-22 \| 672 \| 0.177 \| 534 \| 265 \| 18 \| 5 \| 468 \| 1 \| 430 \| Uncharacterized protein \| Uncharacterized protein; afdb-uniprot50 \| AF-A0A4U7JHW2-F1-MODEL\_V4 \| 1.0 \| 5.992e-19 \| 671 \| 0.261 \| 252 \| 167 \| 8 \| 230 \| 468 \| 311 \| 556 \| Phage tail sheath family protein \| Phage tail sheath family protein; afdb-uniprot50 \| AF-A0A7V3SDC0-F1-MODEL\_V4 \| 1.0 \| 6.833e-24 \| 669 \| 0.168 \| 576 \| 303 \| 23 \| 1 \| 468 \| 1 \| 508 \| Phage tail sheath family protein \| Phage tail sheath family protein; afdb-uniprot50 \| AF-A0A447JQB1-F1-MODEL\_V4 \| 1.0 \| 2.731e-20 \| 667 \| 0.268 \| 358 \| 161 \| 8 \| 90 \| 431 \| 19 \| 291 \| Tail protein \| Tail protein; afdb-uniprot50 \| AF-A0A1V3NDB0-F1-MODEL\_V4 \| 1.0 \| 3.242e-20 \| 666 \| 0.175 \| 473 \| 256 \| 14 \| 9 \| 468 \| 31 \| 382 \| Phage\_sheath\_1C domain-containing protein \| Phage\_sheath\_1C domain-containing protein; afdb-uniprot50 \| AF-A0A7C7E3G3-F1-MODEL\_V4 \| 1.0 \| 4.015e-19 \| 665 \| 0.254 \| 255 \| 168 \| 9 \| 230 \| 468 \| 158 \| 406 \| Phage tail sheath family protein \| Phage tail sheath family protein; afdb-uniprot50 \| AF-A0A1V4VN97-F1-MODEL\_V4 \| 1.0 \| 8.233e-25 \| 665 \| 0.179 \| 567 \| 294 \| 24 \| 5 \| 467 \| 3 \| 501 \| Phage tail sheath protein \| Phage tail sheath protein; afdb-uniprot50 \| AF-A0A2V7W469-F1-MODEL\_V4 \| 1.0 \| 9.63e-24 \| 663 \| 0.152 \| 598 \| 306 \| 19 \| 3 \| 468 \| 2 \| 530 \| Uncharacterized protein \| Uncharacterized protein; afdb-uniprot50 \| AF-A0A656TYE7-F1-MODEL\_V4 \| 1.0 \| 7.976e-19 \| 662 \| 0.257 \| 256 \| 165 \| 7 \| 232 \| 468 \| 12 \| 261 \| Uncharacterized protein \| Uncharacterized protein; afdb-uniprot50 \| AF-A0A1Q7S8Q3-F1-MODEL\_V4 \| 1.0 \| 1.062e-18 \| 662 \| 0.268 \| 253 \| 164 \| 8 \| 230 \| 467 \| 162 \| 408 \| Phage tail protein \| Phage tail protein; afdb-uniprot50 \| AF-A0A1S2NW23-F1-MODEL\_V4 \| 1.0 \| 2.082e-21 \| 662 \| 0.185 \| 495 \| 268 \| 20 \| 20 \| 466 \| 2 \| 409 \| Uncharacterized protein \| Uncharacterized protein; afdb-uniprot50 \| AF-A0A2S5F8Z6-F1-MODEL\_V4 \| 1.0 \| 6.264e-22 \| 661 \| 0.215 \| 408 \| 201 \| 9 \| 86 \| 474 \| 155 \| 462 \| Phage tail protein \| Phage tail protein; afdb-uniprot50 \| AF-A0A1Z8SQC7-F1-MODEL\_V4 \| 1.0 \| 3.642e-24 \| 660 \| 0.165 \| 669 \| 328 \| 25 \| 3 \| 473 \| 2 \| 637 \| Uncharacterized protein \| Uncharacterized protein; afdb-uniprot50 \| AF-A0A6V8M398-F1-MODEL\_V4 \| 1.0 \| 6.345e-19 \| 659 \| 0.269 \| 252 \| 166 \| 8 \| 229 \| 468 \| 73 \| 318 \| Uncharacterized protein \| Uncharacterized protein; afdb-uniprot50 \| AF-B3PD96-F1-MODEL\_V4 \| 1.0 \| 2.177e-24 \| 658 \| 0.166 \| 589 \| 304 \| 26 \| 3 \| 468 \| 2 \| 526 \| Phage tail sheath protein \| Phage tail sheath protein; afdb-uniprot50 \| AF-A0A7U6QNP6-F1-MODEL\_V4 \| 1.0 \| 2.33e-17 \| 657 \| 0.312 \| 205 \| 133 \| 4 \| 276 \| 473 \| 5 \| 208 \| Phage\_sheath\_1 domain-containing protein \| Phage\_sheath\_1 domain-containing protein; afdb-uniprot50 \| AF-A0A2P1PP13-F1-MODEL\_V4 \| 1.0 \| 1.703e-19 \| 657 \| 0.154 \| 484 \| 275 \| 21 \| 11 \| 468 \| 4 \| 379 \| Uncharacterized protein \| Uncharacterized protein; afdb-uniprot50 \| AF-A0A1Y3X9S1-F1-MODEL\_V4 \| 1.0 \| 1.459e-24 \| 657 \| 0.161 \| 562 \| 306 \| 20 \| 11 \| 468 \| 1 \| 500 \| Uncharacterized protein \| Uncharacterized protein; afdb-uniprot50 \| AF-A0A3G2R5A9-F1-MODEL\_V4 \| 1.0 \| 4.084e-24 \| 657 \| 0.169 \| 550 \| 324 \| 26 \| 5 \| 466 \| 3 \| 507 \| Phage tail sheath family protein \| Phage tail sheath family protein; afdb-uniprot50 \| AF-A0A7X0FS83-F1-MODEL\_V4 \| 1.0 \| 2.266e-19 \| 656 \| 0.171 \| 456 \| 238 \| 18 \| 28 \| 466 \| 4 \| 336 \| Uncharacterized protein \| Uncharacterized protein; afdb-uniprot50 \| AF-E1QHA9-F1-MODEL\_V4 \| 1.0 \| 1.11e-21 \| 655 \| 0.188 \| 583 \| 260 \| 24 \| 3 \| 474 \| 1 \| 481 \| Uncharacterized protein \| Uncharacterized protein; afdb-uniprot50 \| AF-A0A1H8VSD3-F1-MODEL\_V4 \| 1.0 \| 1.094e-20 \| 654 \| 0.192 \| 578 \| 262 \| 22 \| 5 \| 475 \| 2 \| 481 \| Uncharacterized protein \| Uncharacterized protein; afdb-uniprot50 \| AF-A5UTU6-F1-MODEL\_V4 \| 1.0 \| 4.848e-24 \| 653 \| 0.185 \| 581 \| 290 \| 28 \| 5 \| 467 \| 3 \| 517 \| Phage tail sheath protein \| Phage tail sheath protein; afdb-uniprot50 \| AF-A0A351G7S8-F1-MODEL\_V4 \| 1.0 \| 1.301e-24 \| 653 \| 0.178 \| 600 \| 294 \| 29 \| 5 \| 467 \| 3 \| 540 \| Phage tail protein \| Phage tail protein; afdb-uniprot50 \| AF-A0A4Q3UGF8-F1-MODEL\_V4 \| 1.0 \| 1.062e-18 \| 652 \| 0.226 \| 260 \| 171 \| 8 \| 231 \| 467 \| 118 \| 370 \| Phage tail sheath family protein \| Phage tail sheath family protein; afdb-uniprot50 \| AF-A0A7C3BTZ0-F1-MODEL\_V4 \| 1.0 \| 1.992e-18 \| 652 \| 0.265 \| 252 \| 165 \| 8 \| 230 \| 467 \| 207 \| 452 \| Phage tail sheath family protein \| Phage tail sheath family protein; afdb-uniprot50 \| AF-A0A0A2WM66-F1-MODEL\_V4 \| 1.0 \| 4.848e-24 \| 651 \| 0.177 \| 559 \| 306 \| 20 \| 1 \| 468 \| 4 \| 499 \| Phage tail sheath protein FI \| Phage tail sheath protein FI; afdb-uniprot50 \| AF-A0A2W4URD9-F1-MODEL\_V4 \| 1.0 \| 4.569e-20 \| 650 \| 0.191 \| 486 \| 249 \| 16 \| 3 \| 467 \| 2 \| 364 \| Phage tail sheath family protein \| Phage tail sheath family protein; afdb-uniprot50 \| AF-A0A5E4K8W2-F1-MODEL\_V4 \| 1.0 \| 3.147e-18 \| 649 \| 0.242 \| 252 \| 172 \| 7 \| 230 \| 468 \| 59 \| 304 \| Phage tail sheath protein subtilisin-like domain protein \| Phage tail sheath protein subtilisin-like domain protein; afdb-uniprot50 \| AF-A0A2V8QNT6-F1-MODEL\_V4 \| 1.0 \| 1.26e-18 \| 649 \| 0.238 \| 260 \| 168 \| 8 \| 231 \| 467 \| 58 \| 310 \| Uncharacterized protein \| Uncharacterized protein; afdb-uniprot50 \| AF-A0A811B248-F1-MODEL\_V4 \| 1.0 \| 9.095e-24 \| 649 \| 0.171 \| 576 \| 301 \| 21 \| 5 \| 468 \| 3 \| 513 \| Tail protein \| Tail protein; afdb-uniprot50 \| AF-A0A150MJ89-F1-MODEL\_V4 \| 1.0 \| 2.3e-20 \| 648 \| 0.156 \| 580 \| 282 \| 19 \| 5 \| 475 \| 2 \| 482 \| Uncharacterized protein \| Uncharacterized protein; afdb-uniprot50 \| AF-U3TVR5-F1-MODEL\_V4 \| 1.0 \| 3.681e-17 \| 647 \| 0.364 \| 203 \| 121 \| 4 \| 278 \| 473 \| 2 \| 203 \| Tail sheath protein \| Tail sheath protein; afdb-uniprot50 \| AF-G9PUI4-F1-MODEL\_V4 \| 1.0 \| 2.172e-20 \| 646 \| 0.173 \| 588 \| 271 \| 23 \| 5 \| 475 \| 2 \| 491 \| Uncharacterized protein \| Uncharacterized protein; afdb-uniprot50 \| AF-A0A261QLY6-F1-MODEL\_V4 \| 1.0 \| 1.244e-21 \| 645 \| 0.262 \| 454 \| 150 \| 17 \| 3 \| 365 \| 2 \| 361 \| Phage tail protein \| Phage tail protein; afdb-uniprot50 \| AF-A0A0M2VSK3-F1-MODEL\_V4 \| 1.0 \| 4.983e-22 \| 645 \| 0.186 \| 578 \| 260 \| 23 \| 5 \| 472 \| 2 \| 479 \| Phage tail protein \| Phage tail protein; afdb-uniprot50 \| AF-A0A1B6ASG7-F1-MODEL\_V4 \| 1.0 \| 3.068e-24 \| 645 \| 0.19 \| 625 \| 300 \| 27 \| 3 \| 467 \| 2 \| 580 \| Putative phage tail sheath protein \| Putative phage tail sheath protein; afdb-uniprot50 \| AF-A0A4R1QQL4-F1-MODEL\_V4 \| 1.0 \| 2.855e-23 \| 644 \| 0.155 \| 565 \| 317 \| 27 \| 1 \| 468 \| 10 \| 511 \| Uncharacterized protein \| Uncharacterized protein; afdb-uniprot50 \| AF-A0A7X5DZ34-F1-MODEL\_V4 \| 1.0 \| 9.095e-24 \| 643 \| 0.162 \| 629 \| 304 \| 26 \| 3 \| 468 \| 1 \| 569 \| Phage tail sheath family protein \| Phage tail sheath family protein; afdb-uniprot50 \| AF-A0A3M1GT53-F1-MODEL\_V4 \| 1.0 \| 5.516e-25 \| 643 \| 0.144 \| 621 \| 333 \| 27 \| 3 \| 467 \| 2 \| 579 \| Phage tail sheath family protein \| Phage tail sheath family protein; afdb-uniprot50 \| AF-A0A523UW76-F1-MODEL\_V4 \| 1.0 \| 9.63e-24 \| 642 \| 0.189 \| 554 \| 296 \| 23 \| 5 \| 468 \| 3 \| 493 \| Phage tail sheath family protein \| Phage tail sheath family protein; afdb-uniprot50 \| AF-A0A1G6NJX4-F1-MODEL\_V4 \| 1.0 \| 1.124e-18 \| 641 \| 0.261 \| 249 \| 165 \| 8 \| 231 \| 468 \| 165 \| 405 \| Uncharacterized protein \| Uncharacterized protein; afdb-uniprot50 \| AF-A0A1C0AIV2-F1-MODEL\_V4 \| 1.0 \| 6.44e-20 \| 640 \| 0.214 \| 467 \| 229 \| 21 \| 10 \| 457 \| 1 \| 348 \| Phage\_sheath\_1 domain-containing protein \| Phage\_sheath\_1 domain-containing protein; afdb-uniprot50 \| AF-A0A7H8TM34-F1-MODEL\_V4 \| 1.0 \| 5.266e-18 \| 640 \| 0.259 \| 254 \| 167 \| 8 \| 230 \| 468 \| 289 \| 536 \| Phage tail sheath family protein \| Phage tail sheath family protein; afdb-uniprot50 \| AF-A0A1P8MVD1-F1-MODEL\_V4 \| 1.0 \| 1.678e-18 \| 640 \| 0.258 \| 263 \| 164 \| 9 \| 230 \| 468 \| 871 \| 1126 \| Uncharacterized protein \| Uncharacterized protein; afdb-uniprot50 \| AF-A0A0B2YW47-F1-MODEL\_V4 \| 1.0 \| 3.062e-20 \| 639 \| 0.187 \| 490 \| 260 \| 23 \| 1 \| 461 \| 1 \| 381 \| Uncharacterized protein \| Uncharacterized protein; afdb-uniprot50 \| AF-A0A417HJA2-F1-MODEL\_V4 \| 1.0 \| 1.632e-20 \| 639 \| 0.175 \| 574 \| 270 \| 22 \| 3 \| 473 \| 1 \| 473 \| Phage tail protein \| Phage tail protein; afdb-uniprot50 \| AF-A0A2E8RZ20-F1-MODEL\_V4 \| 1.0 \| 1.282e-23 \| 638 \| 0.159 \| 632 \| 316 \| 29 \| 1 \| 468 \| 2 \| 581 \| Uncharacterized protein \| Uncharacterized protein; afdb-uniprot50 \| AF-A0A448QV01-F1-MODEL\_V4 \| 1.0 \| 1.474e-17 \| 637 \| 0.273 \| 249 \| 161 \| 8 \| 239 \| 474 \| 2 \| 243 \| Phage tail sheath protein \| Phage tail sheath protein; afdb-uniprot50 \| AF-A0A844UR49-F1-MODEL\_V4 \| 1.0 \| 7.743e-17 \| 636 \| 0.328 \| 198 \| 131 \| 2 \| 278 \| 474 \| 2 \| 198 \| Uncharacterized protein \| Uncharacterized protein; afdb-uniprot50 \| AF-A0A3M1TA51-F1-MODEL\_V4 \| 1.0 \| 7.235e-24 \| 636 \| 0.17 \| 634 \| 310 \| 31 \| 1 \| 471 \| 66 \| 646 \| Phage tail sheath family protein \| Phage tail sheath family protein; afdb-uniprot50 \| AF-A0A4Q7FYR0-F1-MODEL\_V4 \| 1.0 \| 1.357e-23 \| 635 \| 0.237 \| 433 \| 218 \| 18 \| 5 \| 337 \| 3 \| 423 \| Uncharacterized protein \| Uncharacterized protein; afdb-uniprot50 \| AF-A0A3N5NWC7-F1-MODEL\_V4 \| 1.0 \| 7.422e-18 \| 634 \| 0.288 \| 253 \| 160 \| 8 \| 230 \| 468 \| 21 \| 267 \| Phage tail sheath family protein \| Phage tail sheath family protein; afdb-uniprot50 \| AF-A0A349GR39-F1-MODEL\_V4 \| 1.0 \| 3.906e-21 \| 634 \| 0.19 \| 576 \| 259 \| 26 \| 5 \| 474 \| 1 \| 474 \| Phage tail protein \| Phage tail protein; afdb-uniprot50 \| AF-A0A4P8XM20-F1-MODEL\_V4 \| 1.0 \| 5.829e-21 \| 633 \| 0.17 \| 582 \| 272 \| 22 \| 1 \| 475 \| 1 \| 478 \| Uncharacterized protein \| Uncharacterized protein; afdb-uniprot50 \| AF-F2NYF2-F1-MODEL\_V4 \| 1.0 \| 4.023e-23 \| 633 \| 0.163 \| 556 \| 303 \| 23 \| 5 \| 468 \| 1 \| 486 \| Tail sheath protein \| Tail sheath protein; afdb-uniprot50 \| AF-I5B7I5-F1-MODEL\_V4 \| 1.0 \| 5.435e-24 \| 633 \| 0.179 \| 625 \| 326 \| 32 \| 5 \| 468 \| 3 \| 601 \| Phage tail sheath protein FI \| Phage tail sheath protein FI; afdb-uniprot50 \| AF-A0A1T5DNU0-F1-MODEL\_V4 \| 1.0 \| 1.437e-23 \| 632 \| 0.166 \| 638 \| 314 \| 26 \| 1 \| 467 \| 9 \| 599 \| Uncharacterized protein \| Uncharacterized protein; afdb-uniprot50 \| AF-A0A1B8VUJ7-F1-MODEL\_V4 \| 1.0 \| 9.754e-21 \| 631 \| 0.173 \| 582 \| 271 \| 23 \| 6 \| 475 \| 1 \| 484 \| Phage tail protein \| Phage tail protein; afdb-uniprot50 \| AF-A0A6L3F7Z3-F1-MODEL\_V4 \| 1.0 \| 1.653e-17 \| 630 \| 0.26 \| 250 \| 167 \| 7 \| 231 \| 468 \| 288 \| 531 \| Uncharacterized protein \| Uncharacterized protein; afdb-uniprot50 \| AF-A0A124U4Q1-F1-MODEL\_V4 \| 1.0 \| 4.777e-23 \| 629 \| 0.158 \| 586 \| 313 \| 28 \| 1 \| 466 \| 1 \| 526 \| Phage tail protein \| Phage tail protein; afdb-uniprot50 \| AF-A0A524Q8Z7-F1-MODEL\_V4 \| 1.0 \| 5.576e-18 \| 628 \| 0.248 \| 250 \| 165 \| 8 \| 231 \| 467 \| 87 \| 326 \| Phage tail sheath family protein \| Phage tail sheath family protein; afdb-uniprot50 \| AF-A0A2E4F959-F1-MODEL\_V4 \| 1.0 \| 5.755e-24 \| 628 \| 0.189 \| 622 \| 307 \| 26 \| 3 \| 470 \| 2 \| 580 \| Uncharacterized protein \| Uncharacterized protein; afdb-uniprot50 \| AF-A0A1V4IW92-F1-MODEL\_V4 \| 1.0 \| 1.094e-20 \| 626 \| 0.156 \| 583 \| 279 \| 20 \| 5 \| 474 \| 2 \| 484 \| Uncharacterized protein \| Uncharacterized protein; afdb-uniprot50 \| AF-A0A6B8RL73-F1-MODEL\_V4 \| 1.0 \| 1.033e-20 \| 623 \| 0.166 \| 571 \| 278 \| 21 \| 5 \| 474 \| 2 \| 475 \| Phage tail sheath protein \| Phage tail sheath protein; afdb-uniprot50 \| AF-A0A318XBR6-F1-MODEL\_V4 \| 1.0 \| 7.548e-23 \| 623 \| 0.168 \| 593 \| 305 \| 24 \| 1 \| 468 \| 22 \| 551 \| Uncharacterized protein \| Uncharacterized protein; afdb-uniprot50 \| AF-A0A1G9NRA0-F1-MODEL\_V4 \| 1.0 \| 6.732e-23 \| 623 \| 0.165 \| 573 \| 308 \| 28 \| 1 \| 467 \| 64 \| 572 \| Uncharacterized protein \| Uncharacterized protein; afdb-uniprot50 \| AF-S4XSN5-F1-MODEL\_V4 \| 1.0 \| 2.271e-23 \| 623 \| 0.166 \| 765 \| 288 \| 32 \| 3 \| 468 \| 2 \| 715 \| Uncharacterized protein \| Uncharacterized protein; afdb-uniprot50 \| AF-A0A1I6IEN1-F1-MODEL\_V4 \| 1.0 \| 1.474e-17 \| 623 \| 0.235 \| 251 \| 171 \| 7 \| 229 \| 467 \| 734 \| 975 \| Phage tail sheath protein FI \| Phage tail sheath protein FI; afdb-uniprot50 \| AF-A0A810LEP1-F1-MODEL\_V4 \| 1.0 \| 4.569e-20 \| 622 \| 0.202 \| 480 \| 272 \| 23 \| 4 \| 468 \| 1 \| 384 \| Uncharacterized protein \| Uncharacterized protein; afdb-uniprot50 \| AF-A0A413ED89-F1-MODEL\_V4 \| 1.0 \| 5.743e-20 \| 622 \| 0.169 \| 585 \| 273 \| 25 \| 1 \| 475 \| 1 \| 482 \| Phage tail sheath family protein \| Phage tail sheath family protein; afdb-uniprot50 \| AF-A0A0Q7BE77-F1-MODEL\_V4 \| 1.0 \| 9.63e-24 \| 622 \| 0.169 \| 648 \| 292 \| 28 \| 5 \| 468 \| 3 \| 588 \| Uncharacterized protein \| Uncharacterized protein; afdb-uniprot50 \| AF-R5BL49-F1-MODEL\_V4 \| 1.0 \| 1.519e-19 \| 621 \| 0.184 \| 580 \| 264 \| 21 \| 5 \| 474 \| 2 \| 482 \| Uncharacterized protein \| Uncharacterized protein; afdb-uniprot50 \| AF-A0A7C5ZLD0-F1-MODEL\_V4 \| 1.0 \| 5.355e-23 \| 620 \| 0.17 \| 594 \| 310 \| 22 \| 1 \| 468 \| 1 \| 537 \| Phage tail sheath family protein \| Phage tail sheath family protein; afdb-uniprot50 \| AF-A0A6N7AQF3-F1-MODEL\_V4 \| 1.0 \| 4.127e-17 \| 619 \| 0.611 \| 152 \| 59 \| 0 \| 324 \| 475 \| 3 \| 154 \| Major tail sheath protein \| Major tail sheath protein; afdb-uniprot50 \| AF-A0A414Q2K8-F1-MODEL\_V4 \| 1.0 \| 1.728e-20 \| 618 \| 0.166 \| 571 \| 273 \| 26 \| 5 \| 474 \| 2 \| 470 \| Phage tail sheath family protein \| Phage tail sheath family protein; afdb-uniprot50 \| AF-A0A851IXL8-F1-MODEL\_V4 \| 1.0 \| 3.635e-20 \| 618 \| 0.145 \| 569 \| 293 \| 19 \| 5 \| 474 \| 2 \| 476 \| Phage tail sheath protein \| Phage tail sheath protein; afdb-uniprot50 \| AF-A0A4U9UQL5-F1-MODEL\_V4 \| 1.0 \| 2.435e-20 \| 617 \| 0.165 \| 567 \| 259 \| 20 \| 23 \| 475 \| 1 \| 467 \| Phage-related contractile tail sheath protein \| Phage-related contractile tail sheath protein; afdb-uniprot50 \| AF-A0A7Z9GAM2-F1-MODEL\_V4 \| 1.0 \| 1.437e-23 \| 617 \| 0.152 \| 634 \| 316 \| 30 \| 1 \| 467 \| 1 \| 579 \| Phage tail sheath family protein \| Phage tail sheath family protein; afdb-uniprot50 \| AF-A0A4Q7AXR1-F1-MODEL\_V4 \| 1.0 \| 1.277e-15 \| 615 \| 0.32 \| 181 \| 123 \| 0 \| 294 \| 474 \| 15 \| 195 \| Phage tail protein \| Phage tail protein; afdb-uniprot50 \| AF-A0A853IM30-F1-MODEL\_V4 \| 1.0 \| 3.898e-17 \| 615 \| 0.28 \| 300 \| 124 \| 6 \| 102 \| 399 \| 2 \| 211 \| Phage tail sheath subtilisin-like domain-containing protein \| Phage tail sheath subtilisin-like domain-containing protein; afdb-uniprot50 \| AF-A0A3N5AEN8-F1-MODEL\_V4 \| 1.0 \| 7.114e-19 \| 615 \| 0.182 \| 494 \| 249 \| 22 \| 1 \| 468 \| 1 \| 365 \| Uncharacterized protein \| Uncharacterized protein; afdb-uniprot50 \| AF-A0A454VXY5-F1-MODEL\_V4 \| 1.0 \| 2.467e-17 \| 614 \| 0.275 \| 247 \| 159 \| 8 \| 230 \| 462 \| 19 \| 259 \| Phage tail sheath family protein \| Phage tail sheath family protein; afdb-uniprot50 \| AF-F8KPL7-F1-MODEL\_V4 \| 1.0 \| 3.101e-17 \| 614 \| 0.218 \| 252 \| 189 \| 5 \| 225 \| 474 \| 27 \| 272 \| Major tail sheath protein \| Major tail sheath protein; afdb-uniprot50 \| AF-A0A1C6BNB5-F1-MODEL\_V4 \| 1.0 \| 2.3e-20 \| 614 \| 0.166 \| 566 \| 279 \| 20 \| 5 \| 474 \| 2 \| 470 \| Uncharacterized protein \| Uncharacterized protein; afdb-uniprot50 \| AF-A0A3C0WX80-F1-MODEL\_V4 \| 1.0 \| 3.642e-24 \| 614 \| 0.167 \| 555 \| 302 \| 27 \| 5 \| 468 \| 3 \| 488 \| Phage tail sheath family protein \| Phage tail sheath family protein; afdb-uniprot50 \| AF-A0A1N6X8E9-F1-MODEL\_V4 \| 1.0 \| 5.671e-23 \| 614 \| 0.166 \| 614 \| 330 \| 26 \| 1 \| 468 \| 3 \| 580 \| Uncharacterized protein \| Uncharacterized protein; afdb-uniprot50 \| AF-A0A7X3MK04-F1-MODEL\_V4 \| 1.0 \| 8.463e-23 \| 612 \| 0.168 \| 634 \| 300 \| 27 \| 3 \| 468 \| 2 \| 576 \| Phage tail sheath family protein \| Phage tail sheath family protein; afdb-uniprot50 \| AF-A0A2W6ZWR9-F1-MODEL\_V4 \| 1.0 \| 3.635e-20 \| 612 \| 0.21 \| 385 \| 191 \| 12 \| 102 \| 468 \| 562 \| 851 \| Phage tail protein \| Phage tail protein; afdb-uniprot50 \| AF-A0A1A1YQW6-F1-MODEL\_V4 \| 1.0 \| 1.392e-17 \| 612 \| 0.241 \| 261 \| 168 \| 8 \| 231 \| 468 \| 857 \| 1110 \| Uncharacterized protein \| Uncharacterized protein; afdb-uniprot50 \| AF-A0A2E1AI36-F1-MODEL\_V4 \| 1.0 \| 3.589e-23 \| 611 \| 0.182 \| 575 \| 287 \| 29 \| 5 \| 468 \| 3 \| 505 \| Phage tail protein \| Phage tail protein; afdb-uniprot50 \| AF-A0A1Y4HZF3-F1-MODEL\_V4 \| 1.0 \| 3.201e-23 \| 611 \| 0.164 \| 609 \| 314 \| 29 \| 3 \| 467 \| 2 \| 559 \| Phage tail protein \| Phage tail protein; afdb-uniprot50 \| AF-A0A2N2DHN9-F1-MODEL\_V4 \| 1.0 \| 4.023e-23 \| 610 \| 0.168 \| 619 \| 320 \| 31 \| 5 \| 467 \| 3 \| 582 \| Phage tail protein \| Phage tail protein; afdb-uniprot50 \| AF-A0A7V9U445-F1-MODEL\_V4 \| 1.0 \| 3.898e-17 \| 609 \| 0.254 \| 236 \| 158 \| 8 \| 245 \| 468 \| 12 \| 241 \| Phage tail sheath family protein \| Phage tail sheath family protein; afdb-uniprot50 \| AF-X0PXM3-F1-MODEL\_V4 \| 1.0 \| 2.2e-17 \| 609 \| 0.222 \| 261 \| 173 \| 8 \| 231 \| 468 \| 870 \| 1123 \| Uncharacterized protein \| Uncharacterized protein; afdb-uniprot50 \| AF-A0A3A6RI96-F1-MODEL\_V4 \| 1.0 \| 9.192e-17 \| 608 \| 0.276 \| 297 \| 121 \| 7 \| 77 \| 366 \| 2 \| 211 \| Phage tail sheath family protein \| Phage tail sheath family protein; afdb-uniprot50 \| AF-A0A416S5I6-F1-MODEL\_V4 \| 1.0 \| 1.416e-22 \| 608 \| 0.165 \| 557 \| 312 \| 27 \| 3 \| 468 \| 2 \| 496 \| Phage tail sheath family protein \| Phage tail sheath family protein; afdb-uniprot50 \| AF-A0A5C5ZX69-F1-MODEL\_V4 \| 1.0 \| 3.589e-23 \| 608 \| 0.167 \| 789 \| 320 \| 34 \| 3 \| 468 \| 2 \| 776 \| Phage tail sheath protein \| Phage tail sheath protein; afdb-uniprot50 \| AF-A0A836ZRP0-F1-MODEL\_V4 \| 1.0 \| 1.224e-16 \| 607 \| 0.32 \| 175 \| 118 \| 1 \| 231 \| 404 \| 8 \| 182 \| Tail sheath protein \| Tail sheath protein; afdb-uniprot50 \| AF-A0A133NQP6-F1-MODEL\_V4 \| 1.0 \| 8.095e-20 \| 606 \| 0.165 \| 579 \| 277 \| 21 \| 3 \| 474 \| 1 \| 480 \| Uncharacterized protein \| Uncharacterized protein; afdb-uniprot50 \| AF-A0A7C1G8D3-F1-MODEL\_V4 \| 1.0 \| 1.885e-22 \| 606 \| 0.187 \| 596 \| 295 \| 24 \| 1 \| 468 \| 1 \| 534 \| Phage tail sheath family protein \| Phage tail sheath family protein; afdb-uniprot50 \| AF-A0A0J6J0B1-F1-MODEL\_V4 \| 1.0 \| 4.777e-23 \| 606 \| 0.169 \| 591 \| 306 \| 23 \| 1 \| 468 \| 14 \| 542 \| Uncharacterized protein \| Uncharacterized protein; afdb-uniprot50 \| AF-A0A6L9A8T7-F1-MODEL\_V4 \| 1.0 \| 3.433e-20 \| 605 \| 0.174 \| 556 \| 253 \| 17 \| 30 \| 475 \| 2 \| 461 \| Phage tail sheath family protein \| Phage tail sheath family protein; afdb-uniprot50 \| AF-A0A3A0B079-F1-MODEL\_V4 \| 1.0 \| 4.628e-17 \| 605 \| 0.226 \| 256 \| 174 \| 6 \| 231 \| 468 \| 242 \| 491 \| Uncharacterized protein \| Uncharacterized protein; afdb-uniprot50 \| AF-A0A535I9P1-F1-MODEL\_V4 \| 1.0 \| 2.369e-22 \| 604 \| 0.193 \| 588 \| 281 \| 24 \| 1 \| 468 \| 1 \| 514 \| Phage tail sheath family protein \| Phage tail sheath family protein; afdb-uniprot50 \| AF-A0A2N0KBH5-F1-MODEL\_V4 \| 1.0 \| 1.996e-22 \| 603 \| 0.17 \| 635 \| 299 \| 30 \| 3 \| 468 \| 1 \| 576 \| Phage tail sheath family protein \| Phage tail sheath family protein; afdb-uniprot50 \| AF-A0A2E1AI38-F1-MODEL\_V4 \| 1.0 \| 6.633e-22 \| 602 \| 0.135 \| 603 \| 319 \| 24 \| 3 \| 467 \| 2 \| 539 \| Uncharacterized protein \| Uncharacterized protein; afdb-uniprot50 \| AF-A0A7I8MYS9-F1-MODEL\_V4 \| 1.0 \| 1.499e-22 \| 601 \| 0.158 \| 656 \| 320 \| 33 \| 3 \| 468 \| 2 \| 615 \| Phage tail sheath protein FI \| Phage tail sheath protein FI; afdb-uniprot50 \| AF-A0A2V9PGB7-F1-MODEL\_V4 \| 1.0 \| 1.156e-16 \| 600 \| 0.233 \| 231 \| 161 \| 5 \| 245 \| 466 \| 12 \| 235 \| Uncharacterized protein \| Uncharacterized protein; afdb-uniprot50 \| AF-A0A660S5R2-F1-MODEL\_V4 \| 1.0 \| 1.108e-17 \| 600 \| 0.137 \| 481 \| 259 \| 19 \| 1 \| 466 \| 1 \| 340 \| Phage tail sheath protein \| Phage tail sheath protein; afdb-uniprot50 \| AF-A0A7W7XTS9-F1-MODEL\_V4 \| 1.0 \| 1.909e-19 \| 600 \| 0.163 \| 477 \| 266 \| 15 \| 3 \| 455 \| 2 \| 369 \| Uncharacterized protein \| Uncharacterized protein; afdb-uniprot50 \| AF-Q2N8P2-F1-MODEL\_V4 \| 1.0 \| 1.588e-22 \| 600 \| 0.168 \| 658 \| 315 \| 33 \| 3 \| 468 \| 2 \| 619 \| Phage tail sheath protein, putative \| Phage tail sheath protein, putative; afdb-uniprot50 \| AF-A0A1Y6BIP2-F1-MODEL\_V4 \| 1.0 \| 8.463e-23 \| 600 \| 0.172 \| 679 \| 304 \| 28 \| 1 \| 468 \| 1 \| 632 \| Phage\_sheath\_1C domain-containing protein \| Phage\_sheath\_1C domain-containing protein; afdb-uniprot50 \| AF-A0A1M5CL98-F1-MODEL\_V4 \| 1.0 \| 1.126e-22 \| 599 \| 0.175 \| 594 \| 292 \| 26 \| 3 \| 468 \| 2 \| 525 \| Uncharacterized protein \| Uncharacterized protein; afdb-uniprot50 \| AF-A0A497VVB2-F1-MODEL\_V4 \| 1.0 \| 1.416e-22 \| 599 \| 0.148 \| 658 \| 309 \| 30 \| 5 \| 468 \| 3 \| 602 \| Uncharacterized protein \| Uncharacterized protein; afdb-uniprot50 \| AF-A0A1H8WFQ1-F1-MODEL\_V4 \| 1.0 \| 9.33e-18 \| 599 \| 0.241 \| 253 \| 166 \| 9 \| 231 \| 468 \| 736 \| 977 \| Phage tail sheath protein FI \| Phage tail sheath protein FI; afdb-uniprot50 \| AF-A0A351BHJ8-F1-MODEL\_V4 \| 1.0 \| 4.37e-17 \| 598 \| 0.266 \| 259 \| 164 \| 8 \| 230 \| 468 \| 31 \| 283 \| Phage tail sheath family protein \| Phage tail sheath family protein; afdb-uniprot50 \| AF-A0A143PL89-F1-MODEL\_V4 \| 1.0 \| 5.916e-22 \| 598 \| 0.153 \| 579 \| 312 \| 22 \| 5 \| 468 \| 3 \| 518 \| Major tail sheath protein \| Major tail sheath protein; afdb-uniprot50 \| AF-A0A117NJJ3-F1-MODEL\_V4 \| 1.0 \| 1.193e-22 \| 598 \| 0.164 \| 577 \| 322 \| 27 \| 3 \| 470 \| 1 \| 526 \| Uncharacterized protein \| Uncharacterized protein; afdb-uniprot50 \| AF-A0A2E5HMA5-F1-MODEL\_V4 \| 1.0 \| 1.499e-22 \| 598 \| 0.149 \| 647 \| 312 \| 29 \| 3 \| 468 \| 2 \| 590 \| Uncharacterized protein \| Uncharacterized protein; afdb-uniprot50 \| AF-A0A1H6TSH5-F1-MODEL\_V4 \| 1.0 \| 1.885e-22 \| 598 \| 0.169 \| 671 \| 301 \| 31 \| 3 \| 468 \| 2 \| 620 \| Uncharacterized protein \| Uncharacterized protein; afdb-uniprot50 \| AF-A0A3A8B9H2-F1-MODEL\_V4 \| 1.0 \| 1.263e-22 \| 598 \| 0.152 \| 710 \| 327 \| 30 \| 3 \| 468 \| 2 \| 680 \| Uncharacterized protein \| Uncharacterized protein; afdb-uniprot50 \| AF-A0A7Z7I241-F1-MODEL\_V4 \| 1.0 \| 2.33e-17 \| 598 \| 0.255 \| 262 \| 164 \| 10 \| 231 \| 468 \| 895 \| 1149 \| Phage tail sheath protein FI \| Phage tail sheath protein FI; afdb-uniprot50 \| AF-A0A7S9UQU7-F1-MODEL\_V4 \| 1.0 \| 7.313e-17 \| 597 \| 0.234 \| 260 \| 169 \| 8 \| 231 \| 467 \| 266 \| 518 \| Phage tail sheath family protein \| Phage tail sheath family protein; afdb-uniprot50 \| AF-A0A1Q3SJK3-F1-MODEL\_V4 \| 1.0 \| 5.355e-23 \| 597 \| 0.161 \| 637 \| 330 \| 25 \| 10 \| 468 \| 1 \| 611 \| Uncharacterized protein \| Uncharacterized protein; afdb-uniprot50 \| AF-W4RWM2-F1-MODEL\_V4 \| 1.0 \| 3.574e-15 \| 596 \| 0.331 \| 166 \| 111 \| 0 \| 309 \| 474 \| 2 \| 167 \| Tail sheath protein \| Tail sheath protein; afdb-uniprot50 \| AF-A0A3S9PC88-F1-MODEL\_V4 \| 1.0 \| 3.898e-17 \| 595 \| 0.25 \| 256 \| 169 \| 9 \| 230 \| 468 \| 144 \| 393 \| Phage tail sheath family protein \| Phage tail sheath family protein; afdb-uniprot50 \| AF-A0A2K4ZHW7-F1-MODEL\_V4 \| 1.0 \| 2.172e-20 \| 594 \| 0.175 \| 586 \| 261 \| 28 \| 5 \| 473 \| 2 \| 482 \| Uncharacterized protein \| Uncharacterized protein; afdb-uniprot50 \| AF-A0A7V6E9A4-F1-MODEL\_V4 \| 1.0 \| 1.996e-22 \| 594 \| 0.133 \| 624 \| 343 \| 27 \| 1 \| 467 \| 6 \| 588 \| Phage tail sheath family protein \| Phage tail sheath family protein; afdb-uniprot50 \| AF-C7RKK7-F1-MODEL\_V4 \| 1.0 \| 2.978e-22 \| 594 \| 0.171 \| 654 \| 316 \| 34 \| 3 \| 468 \| 2 \| 617 \| Tail sheath protein \| Tail sheath protein; afdb-uniprot50 \| AF-A0A1Q7BLH0-F1-MODEL\_V4 \| 1.0 \| 1.126e-22 \| 594 \| 0.139 \| 746 \| 319 \| 29 \| 5 \| 468 \| 3 \| 707 \| Uncharacterized protein \| Uncharacterized protein; afdb-uniprot50 \| AF-A0A1L6L570-F1-MODEL\_V4 \| 1.0 \| 4.983e-22 \| 593 \| 0.156 \| 689 \| 311 \| 31 \| 1 \| 468 \| 145 \| 783 \| Phage tail sheath protein FI \| Phage tail sheath protein FI; afdb-uniprot50 \| AF-A0A011MDR6-F1-MODEL\_V4 \| 1.0 \| 1.608e-19 \| 593 \| 0.206 \| 387 \| 196 \| 12 \| 102 \| 468 \| 706 \| 1001 \| Major tail sheath protein \| Major tail sheath protein; afdb-uniprot50 \| AF-A0A317HHF4-F1-MODEL\_V4 \| 1.0 \| 3.792e-19 \| 592 \| 0.148 \| 491 \| 280 \| 19 \| 3 \| 467 \| 15 \| 393 \| Phage tail sheath family protein \| Phage tail sheath family protein; afdb-uniprot50 \| AF-A0A158AZ89-F1-MODEL\_V4 \| 1.0 \| 2.14e-19 \| 592 \| 0.165 \| 509 \| 275 \| 17 \| 1 \| 468 \| 15 \| 414 \| Tail sheath protein \| Tail sheath protein; afdb-uniprot50 \| AF-A0A3N5PYF2-F1-MODEL\_V4 \| 1.0 \| 3.681e-17 \| 592 \| 0.242 \| 260 \| 168 \| 9 \| 231 \| 468 \| 360 \| 612 \| Phage tail sheath family protein \| Phage tail sheath family protein; afdb-uniprot50 \| AF-A0A1V0Q0C4-F1-MODEL\_V4 \| 1.0 \| 3.107e-21 \| 592 \| 0.139 \| 730 \| 335 \| 25 \| 5 \| 467 \| 1 \| 703 \| Phage tail sheath protein FI \| Phage tail sheath protein FI; afdb-uniprot50 \| AF-A0A7W0WXT9-F1-MODEL\_V4 \| 1.0 \| 1.413e-18 \| 591 \| 0.2 \| 429 \| 205 \| 16 \| 1 \| 399 \| 1 \| 321 \| Phage tail sheath family protein \| Phage tail sheath family protein; afdb-uniprot50 \| AF-A0A1X4H0I2-F1-MODEL\_V4 \| 1.0 \| 4.9e-17 \| 591 \| 0.213 \| 258 \| 177 \| 8 \| 230 \| 467 \| 99 \| 350 \| Uncharacterized protein \| Uncharacterized protein; afdb-uniprot50 \| AF-A0A1W9H7R4-F1-MODEL\_V4 \| 1.0 \| 1.046e-17 \| 590 \| 0.192 \| 446 \| 233 \| 20 \| 46 \| 466 \| 1 \| 344 \| Phage\_sheath\_1C domain-containing protein \| Phage\_sheath\_1C domain-containing protein; afdb-uniprot50 \| AF-A0A525C5G4-F1-MODEL\_V4 \| 1.0 \| 1.78e-22 \| 590 \| 0.158 \| 581 \| 307 \| 26 \| 5 \| 466 \| 3 \| 520 \| Phage tail sheath family protein \| Phage tail sheath family protein; afdb-uniprot50 \| AF-D0LXB9-F1-MODEL\_V4 \| 1.0 \| 6.264e-22 \| 590 \| 0.158 \| 624 \| 326 \| 30 \| 1 \| 467 \| 2 \| 583 \| Tail sheath protein \| Tail sheath protein; afdb-uniprot50 \| AF-A0A845WCQ0-F1-MODEL\_V4 \| 1.0 \| 5.344e-19 \| 589 \| 0.162 \| 485 \| 268 \| 21 \| 1 \| 466 \| 1 \| 366 \| Phage tail sheath protein \| Phage tail sheath protein; afdb-uniprot50 \| AF-A0A6G4XFE5-F1-MODEL\_V4 \| 1.0 \| 1.885e-22 \| 589 \| 0.155 \| 567 \| 303 \| 18 \| 1 \| 467 \| 7 \| 497 \| Phage tail sheath family protein \| Phage tail sheath family protein; afdb-uniprot50 \| AF-A0A1H2U4F8-F1-MODEL\_V4 \| 1.0 \| 1.588e-22 \| 589 \| 0.178 \| 661 \| 310 \| 32 \| 3 \| 468 \| 1 \| 623 \| Uncharacterized protein \| Uncharacterized protein; afdb-uniprot50 \| AF-X5R2Z1-F1-MODEL\_V4 \| 1.0 \| 2.813e-22 \| 587 \| 0.176 \| 579 \| 294 \| 23 \| 3 \| 468 \| 2 \| 510 \| Tail protein \| Tail protein; afdb-uniprot50 \| AF-A0A4D9Y1N5-F1-MODEL\_V4 \| 1.0 \| 6.332e-15 \| 585 \| 0.416 \| 149 \| 87 \| 0 \| 326 \| 474 \| 1 \| 149 \| Phage tail protein \| Phage tail protein; afdb-uniprot50 \| AF-A0A7T9VKM0-F1-MODEL\_V4 \| 1.0 \| 9.59e-16 \| 585 \| 0.327 \| 186 \| 120 \| 2 \| 294 \| 474 \| 7 \| 192 \| Phage tail protein \| Phage tail protein; afdb-uniprot50 \| AF-A0A3M1BL67-F1-MODEL\_V4 \| 1.0 \| 2.233e-18 \| 585 \| 0.247 \| 380 \| 167 \| 15 \| 3 \| 351 \| 1 \| 292 \| Phage tail sheath family protein \| Phage tail sheath family protein; afdb-uniprot50 \| AF-A0A562GH79-F1-MODEL\_V4 \| 1.0 \| 1.244e-21 \| 585 \| 0.181 \| 534 \| 305 \| 23 \| 1 \| 468 \| 1 \| 468 \| Phage\_sheath\_1 domain-containing protein \| Phage\_sheath\_1 domain-containing protein; afdb-uniprot50 \| AF-A0A1V5CVP8-F1-MODEL\_V4 \| 1.0 \| 5.188e-17 \| 584 \| 0.273 \| 256 \| 160 \| 11 \| 231 \| 466 \| 177 \| 426 \| Phage tail sheath protein \| Phage tail sheath protein; afdb-uniprot50 \| AF-A0A5Q0GXI4-F1-MODEL\_V4 \| 1.0 \| 8.338e-22 \| 584 \| 0.16 \| 581 \| 314 \| 20 \| 3 \| 467 \| 15 \| 537 \| Phage tail sheath family protein \| Phage tail sheath family protein; afdb-uniprot50 \| AF-A0A4Q9KMC0-F1-MODEL\_V4 \| 1.0 \| 1.395e-21 \| 584 \| 0.147 \| 707 \| 336 \| 30 \| 3 \| 468 \| 2 \| 682 \| Phage tail protein \| Phage tail protein; afdb-uniprot50 \| AF-I5B7I6-F1-MODEL\_V4 \| 1.0 \| 2.509e-22 \| 584 \| 0.163 \| 732 \| 311 \| 33 \| 3 \| 468 \| 2 \| 697 \| Phage tail sheath protein FI \| Phage tail sheath protein FI; afdb-uniprot50 \| AF-A0A1M6SAY6-F1-MODEL\_V4 \| 1.0 \| 8.942e-19 \| 583 \| 0.155 \| 586 \| 276 \| 25 \| 4 \| 475 \| 1 \| 481 \| Uncharacterized protein \| Uncharacterized protein; afdb-uniprot50 \| AF-R7Z895-F1-MODEL\_V4 \| 1.0 \| 5.123e-20 \| 582 \| 0.173 \| 581 \| 266 \| 27 \| 5 \| 475 \| 1 \| 477 \| Uncharacterized protein \| Uncharacterized protein; afdb-uniprot50 \| AF-A0A1V4GEV1-F1-MODEL\_V4 \| 1.0 \| 6.44e-20 \| 582 \| 0.168 \| 589 \| 264 \| 25 \| 5 \| 474 \| 2 \| 483 \| Uncharacterized protein \| Uncharacterized protein; afdb-uniprot50 \| AF-A0A4Q3UKU1-F1-MODEL\_V4 \| 1.0 \| 8.338e-22 \| 582 \| 0.151 \| 653 \| 317 \| 29 \| 3 \| 467 \| 2 \| 605 \| Phage\_sheath\_1C domain-containing protein \| Phage\_sheath\_1C domain-containing protein; afdb-uniprot50 \| AF-A0A2V2S183-F1-MODEL\_V4 \| 1.0 \| 2.509e-22 \| 582 \| 0.157 \| 712 \| 323 \| 32 \| 5 \| 467 \| 3 \| 686 \| Uncharacterized protein \| Uncharacterized protein; afdb-uniprot50 \| AF-A0A2S0JKR3-F1-MODEL\_V4 \| 1.0 \| 9.076e-20 \| 581 \| 0.175 \| 581 \| 267 \| 26 \| 1 \| 475 \| 1 \| 475 \| Phage tail protein \| Phage tail protein; afdb-uniprot50 \| AF-A0A2R6JS05-F1-MODEL\_V4 \| 1.0 \| 8.338e-22 \| 580 \| 0.153 \| 625 \| 307 \| 29 \| 5 \| 468 \| 3 \| 566 \| Phage tail protein \| Phage tail protein; afdb-uniprot50 \| AF-A0A4P5W0E4-F1-MODEL\_V4 \| 1.0 \| 3.792e-19 \| 579 \| 0.171 \| 496 \| 275 \| 21 \| 6 \| 468 \| 1 \| 393 \| Uncharacterized protein \| Uncharacterized protein; afdb-uniprot50 \| AF-A0A524Q0G6-F1-MODEL\_V4 \| 1.0 \| 1.933e-16 \| 578 \| 0.25 \| 244 \| 164 \| 7 \| 235 \| 467 \| 3 \| 238 \| Phage tail sheath family protein \| Phage tail sheath family protein; afdb-uniprot50 \| AF-A0A6S7AUX4-F1-MODEL\_V4 \| 1.0 \| 4.706e-22 \| 578 \| 0.153 \| 769 \| 323 \| 32 \| 3 \| 468 \| 2 \| 745 \| Uncharacterized protein \| Uncharacterized protein; afdb-uniprot50 \| AF-A0A496NWC5-F1-MODEL\_V4 \| 1.0 \| 3.194e-19 \| 577 \| 0.153 \| 580 \| 283 \| 20 \| 3 \| 474 \| 1 \| 480 \| Phage tail sheath family protein \| Phage tail sheath family protein; afdb-uniprot50 \| AF-A0A7W0PUB1-F1-MODEL\_V4 \| 1.0 \| 6.805e-16 \| 577 \| 0.23 \| 234 \| 165 \| 5 \| 231 \| 455 \| 272 \| 499 \| Phage tail sheath family protein \| Phage tail sheath family protein; afdb-uniprot50 \| AF-A0A827A5S8-F1-MODEL\_V4 \| 1.0 \| 2.69e-19 \| 576 \| 0.23 \| 456 \| 135 \| 8 \| 5 \| 343 \| 2 \| 358 \| Phage tail protein \| Phage tail protein; afdb-uniprot50 \| AF-A0A5A8F1E7-F1-MODEL\_V4 \| 1.0 \| 3.147e-18 \| 576 \| 0.174 \| 499 \| 260 \| 22 \| 5 \| 474 \| 2 \| 377 \| Uncharacterized protein \| Uncharacterized protein; afdb-uniprot50 \| AF-A0A849EW92-F1-MODEL\_V4 \| 1.0 \| 5.188e-17 \| 576 \| 0.254 \| 259 \| 166 \| 10 \| 231 \| 468 \| 507 \| 759 \| Phage tail sheath family protein \| Phage tail sheath family protein; afdb-uniprot50 \| AF-A0A4Q9TLA8-F1-MODEL\_V4 \| 1.0 \| 5.344e-19 \| 575 \| 0.157 \| 576 \| 281 \| 25 \| 1 \| 475 \| 1 \| 472 \| Phage tail sheath family protein \| Phage tail sheath family protein; afdb-uniprot50 \| AF-J2WTG1-F1-MODEL\_V4 \| 1.0 \| 1.091e-16 \| 575 \| 0.217 \| 248 \| 177 \| 7 \| 230 \| 468 \| 322 \| 561 \| Phage tail sheath protein FI \| Phage tail sheath protein FI; afdb-uniprot50 \| AF-S7TBH7-F1-MODEL\_V4 \| 1.0 \| 3.906e-21 \| 574 \| 0.159 \| 557 \| 332 \| 26 \| 3 \| 473 \| 2 \| 508 \| Phage tail sheath protein \| Phage tail sheath protein; afdb-uniprot50 \| AF-A0A7I8MZ51-F1-MODEL\_V4 \| 1.0 \| 3.898e-17 \| 574 \| 0.247 \| 246 \| 168 \| 8 \| 234 \| 468 \| 272 \| 511 \| Phage\_sheath\_1C domain-containing protein \| Phage\_sheath\_1C domain-containing protein; afdb-uniprot50 \| AF-A0A1N6L5N5-F1-MODEL\_V4 \| 1.0 \| 7.875e-22 \| 573 \| 0.166 \| 560 \| 313 \| 22 \| 1 \| 466 \| 1 \| 500 \| Uncharacterized protein \| Uncharacterized protein; afdb-uniprot50 \| AF-A0A4Q7LV90-F1-MODEL\_V4 \| 1.0 \| 3.906e-21 \| 573 \| 0.185 \| 603 \| 304 \| 28 \| 1 \| 467 \| 11 \| 562 \| Uncharacterized protein \| Uncharacterized protein; afdb-uniprot50 \| AF-A0A7T9J4P3-F1-MODEL\_V4 \| 1.0 \| 3.29e-21 \| 572 \| 0.148 \| 560 \| 326 \| 27 \| 1 \| 472 \| 3 \| 499 \| Phage tail sheath family protein \| Phage tail sheath family protein; afdb-uniprot50 \| AF-A0A661R710-F1-MODEL\_V4 \| 1.0 \| 3.536e-22 \| 572 \| 0.16 \| 637 \| 333 \| 33 \| 3 \| 468 \| 2 \| 607 \| Phage tail sheath family protein \| Phage tail sheath family protein; afdb-uniprot50 \| AF-A0A7Y5WMY0-F1-MODEL\_V4 \| 1.0 \| 6.633e-22 \| 572 \| 0.157 \| 717 \| 333 \| 32 \| 3 \| 467 \| 1 \| 698 \| Uncharacterized protein \| Uncharacterized protein; afdb-uniprot50 \| AF-I4IAB3-F1-MODEL\_V4 \| 1.0 \| 3.477e-17 \| 572 \| 0.225 \| 257 \| 172 \| 8 \| 231 \| 466 \| 687 \| 937 \| Putative phage tail sheath protein (Modular protein) \| Putative phage tail sheath protein (Modular protein); afdb-uniprot50 \| AF-A0A5T1QWS3-F1-MODEL\_V4 \| 1.0 \| 2.2e-17 \| 571 \| 0.242 \| 330 \| 152 \| 6 \| 71 \| 397 \| 51 \| 285 \| Phage tail sheath family protein \| Phage tail sheath family protein; afdb-uniprot50 \| AF-A0A2E7CK62-F1-MODEL\_V4 \| 1.0 \| 2.109e-18 \| 571 \| 0.153 \| 489 \| 294 \| 21 \| 1 \| 467 \| 2 \| 392 \| Uncharacterized protein \| Uncharacterized protein; afdb-uniprot50 \| AF-A0A5M9U7D7-F1-MODEL\_V4 \| 1.0 \| 1.26e-18 \| 571 \| 0.154 \| 583 \| 284 \| 19 \| 3 \| 475 \| 1 \| 484 \| Phage tail sheath family protein \| Phage tail sheath family protein; afdb-uniprot50 \| AF-A0A7V6Q5J0-F1-MODEL\_V4 \| 1.0 \| 1.141e-19 \| 570 \| 0.18 \| 586 \| 259 \| 28 \| 1 \| 474 \| 1 \| 476 \| Uncharacterized protein \| Uncharacterized protein; afdb-uniprot50 \| AF-A0A2E0KUM3-F1-MODEL\_V4 \| 1.0 \| 2.205e-21 \| 570 \| 0.153 \| 698 \| 318 \| 31 \| 3 \| 468 \| 1 \| 657 \| Uncharacterized protein \| Uncharacterized protein; afdb-uniprot50 \| AF-A0A1C3FCE3-F1-MODEL\_V4 \| 1.0 \| 7.099e-15 \| 569 \| 0.375 \| 181 \| 106 \| 3 \| 299 \| 473 \| 2 \| 181 \| Phage tail sheath monomer \| Phage tail sheath monomer; afdb-uniprot50 \| AF-A0A3R8ZYP4-F1-MODEL\_V4 \| 1.0 \| 1.538e-16 \| 569 \| 0.333 \| 186 \| 120 \| 3 \| 254 \| 438 \| 44 \| 226 \| Phage\_sheath\_1 domain-containing protein \| Phage\_sheath\_1 domain-containing protein; afdb-uniprot50 \| AF-A0A7W4H212-F1-MODEL\_V4 \| 1.0 \| 4.767e-19 \| 569 \| 0.198 \| 463 \| 237 \| 19 \| 11 \| 456 \| 3 \| 348 \| Phage tail sheath subtilisin-like domain-containing protein \| Phage tail sheath subtilisin-like domain-containing protein; afdb-uniprot50 \| AF-A0A7T7XS11-F1-MODEL\_V4 \| 1.0 \| 4.828e-16 \| 569 \| 0.264 \| 242 \| 158 \| 7 \| 230 \| 457 \| 228 \| 463 \| Phage tail sheath family protein \| Phage tail sheath family protein; afdb-uniprot50 \| AF-A0A2N1QG40-F1-MODEL\_V4 \| 1.0 \| 5.047e-19 \| 569 \| 0.164 \| 584 \| 274 \| 22 \| 5 \| 474 \| 2 \| 485 \| Phage tail protein \| Phage tail protein; afdb-uniprot50 \| AF-A0A494XZK3-F1-MODEL\_V4 \| 1.0 \| 3.681e-17 \| 568 \| 0.15 \| 492 \| 250 \| 20 \| 1 \| 468 \| 1 \| 348 \| Phage tail sheath family protein \| Phage tail sheath family protein; afdb-uniprot50 \| AF-A0A4R1HQ95-F1-MODEL\_V4 \| 1.0 \| 6.264e-22 \| 568 \| 0.146 \| 671 \| 327 \| 36 \| 1 \| 467 \| 1 \| 629 \| Uncharacterized protein \| Uncharacterized protein; afdb-uniprot50 \| AF-A0A5X8YUE9-F1-MODEL\_V4 \| 1.0 \| 1.001e-14 \| 567 \| 0.329 \| 173 \| 115 \| 1 \| 302 \| 474 \| 2 \| 173 \| Phage tail sheath family protein \| Phage tail sheath family protein; afdb-uniprot50 \| AF-A0A3M1CYL5-F1-MODEL\_V4 \| 1.0 \| 4.252e-19 \| 566 \| 0.148 \| 498 \| 283 \| 22 \| 7 \| 468 \| 1 \| 393 \| Phage tail sheath family protein \| Phage tail sheath family protein; afdb-uniprot50 \| AF-A0A832W4U3-F1-MODEL\_V4 \| 1.0 \| 2.364e-18 \| 566 \| 0.192 \| 384 \| 209 \| 7 \| 95 \| 467 \| 239 \| 532 \| Phage tail sheath family protein \| Phage tail sheath family protein; afdb-uniprot50 \| AF-A0A6G9ZD31-F1-MODEL\_V4 \| 1.0 \| 4.306e-16 \| 565 \| 0.241 \| 253 \| 171 \| 8 \| 231 \| 468 \| 157 \| 403 \| Uncharacterized protein \| Uncharacterized protein; afdb-uniprot50 \| AF-A0A7V1ZW97-F1-MODEL\_V4 \| 1.0 \| 8.829e-22 \| 565 \| 0.147 \| 625 \| 326 \| 28 \| 3 \| 468 \| 2 \| 578 \| Phage tail sheath family protein \| Phage tail sheath family protein; afdb-uniprot50 \| AF-W5WF51-F1-MODEL\_V4 \| 1.0 \| 1.966e-21 \| 565 \| 0.148 \| 652 \| 329 \| 28 \| 1 \| 468 \| 2 \| 610 \| Uncharacterized protein \| Uncharacterized protein; afdb-uniprot50 \| AF-A0A3A0GJR6-F1-MODEL\_V4 \| 1.0 \| 2.771e-21 \| 565 \| 0.153 \| 744 \| 302 \| 37 \| 1 \| 466 \| 1 \| 694 \| Phage tail sheath family protein \| Phage tail sheath family protein; afdb-uniprot50 \| AF-A0A6P0TJE0-F1-MODEL\_V4 \| 1.0 \| 1.724e-16 \| 564 \| 0.203 \| 275 \| 176 \| 9 \| 231 \| 468 \| 240 \| 508 \| Phage tail sheath family protein \| Phage tail sheath family protein; afdb-uniprot50 \| AF-A0A7V5V361-F1-MODEL\_V4 \| 1.0 \| 1.477e-21 \| 564 \| 0.177 \| 661 \| 318 \| 34 \| 5 \| 466 \| 3 \| 636 \| Phage tail sheath family protein \| Phage tail sheath family protein; afdb-uniprot50 \| AF-A0A2K8MIB7-F1-MODEL\_V4 \| 1.0 \| 1.678e-18 \| 564 \| 0.212 \| 381 \| 188 \| 12 \| 101 \| 467 \| 647 \| 929 \| Uncharacterized protein \| Uncharacterized protein; afdb-uniprot50 \| AF-A0A0Q6V5H7-F1-MODEL\_V4 \| 1.0 \| 1.11e-21 \| 563 \| 0.144 \| 673 \| 343 \| 31 \| 1 \| 467 \| 1 \| 646 \| Uncharacterized protein \| Uncharacterized protein; afdb-uniprot50 \| AF-A0A1Q6YDS8-F1-MODEL\_V4 \| 1.0 \| 1.656e-21 \| 563 \| 0.158 \| 731 \| 321 \| 32 \| 3 \| 468 \| 2 \| 702 \| Uncharacterized protein \| Uncharacterized protein; afdb-uniprot50 \| AF-A0A826YSY8-F1-MODEL\_V4 \| 1.0 \| 1.206e-15 \| 562 \| 0.424 \| 158 \| 90 \| 1 \| 272 \| 428 \| 5 \| 162 \| Phage tail protein \| Phage tail protein; afdb-uniprot50 \| AF-A0A511MZM7-F1-MODEL\_V4 \| 1.0 \| 1.564e-21 \| 562 \| 0.145 \| 733 \| 322 \| 35 \| 3 \| 468 \| 2 \| 696 \| Uncharacterized protein \| Uncharacterized protein; afdb-uniprot50 \| AF-A0A4Y8ND74-F1-MODEL\_V4 \| 1.0 \| 2.364e-18 \| 561 \| 0.182 \| 389 \| 208 \| 11 \| 93 \| 468 \| 226 \| 517 \| Phage tail sheath family protein \| Phage tail sheath family protein; afdb-uniprot50 \| AF-A0A7U3YQ73-F1-MODEL\_V4 \| 1.0 \| 7.875e-22 \| 561 \| 0.17 \| 661 \| 316 \| 40 \| 3 \| 466 \| 1 \| 625 \| Phage tail sheath protein \| Phage tail sheath protein; afdb-uniprot50 \| AF-A0A6J4YL04-F1-MODEL\_V4 \| 1.0 \| 6.906e-17 \| 561 \| 0.249 \| 269 \| 163 \| 10 \| 231 \| 466 \| 739 \| 1001 \| Phage tail sheath protein FI \| Phage tail sheath protein FI; afdb-uniprot50 \| AF-A0A2U3LUJ3-F1-MODEL\_V4 \| 1.0 \| 1.724e-16 \| 560 \| 0.241 \| 265 \| 172 \| 9 \| 227 \| 468 \| 96 \| 354 \| Phage tail sheath protein \| Phage tail sheath protein; afdb-uniprot50 \| AF-A0A7V8YCS0-F1-MODEL\_V4 \| 1.0 \| 1.992e-18 \| 560 \| 0.131 \| 503 \| 291 \| 26 \| 1 \| 467 \| 2 \| 394 \| Phage tail sheath family protein \| Phage tail sheath family protein; afdb-uniprot50 \| AF-A0A1Q4ZX14-F1-MODEL\_V4 \| 1.0 \| 1.372e-16 \| 560 \| 0.251 \| 262 \| 165 \| 9 \| 230 \| 466 \| 727 \| 982 \| Uncharacterized protein \| Uncharacterized protein; afdb-uniprot50 \| AF-A0A1G9EWK1-F1-MODEL\_V4 \| 1.0 \| 1.564e-21 \| 559 \| 0.145 \| 696 \| 313 \| 35 \| 1 \| 467 \| 1 \| 643 \| Uncharacterized protein \| Uncharacterized protein; afdb-uniprot50 \| AF-A0A1Q6YE31-F1-MODEL\_V4 \| 1.0 \| 7.437e-22 \| 558 \| 0.156 \| 709 \| 328 \| 35 \| 3 \| 467 \| 2 \| 684 \| Uncharacterized protein \| Uncharacterized protein; afdb-uniprot50 \| AF-A0A399ZLA4-F1-MODEL\_V4 \| 1.0 \| 3.689e-21 \| 558 \| 0.141 \| 867 \| 315 \| 34 \| 5 \| 468 \| 3 \| 842 \| Uncharacterized protein \| Uncharacterized protein; afdb-uniprot50 \| AF-A0A1Y3SWA1-F1-MODEL\_V4 \| 1.0 \| 1.26e-18 \| 556 \| 0.172 \| 575 \| 271 \| 25 \| 6 \| 474 \| 1 \| 476 \| Phage tail protein \| Phage tail protein; afdb-uniprot50 \| AF-A0A6N8HUV9-F1-MODEL\_V4 \| 1.0 \| 2.771e-21 \| 556 \| 0.184 \| 557 \| 284 \| 29 \| 12 \| 463 \| 1 \| 491 \| Phage tail sheath protein subtilisin-like domain protein \| Phage tail sheath protein subtilisin-like domain protein; afdb-uniprot50 \| AF-A0A7C6A7S9-F1-MODEL\_V4 \| 1.0 \| 4.637e-21 \| 555 \| 0.157 \| 551 \| 308 \| 25 \| 1 \| 468 \| 1 \| 478 \| Phage\_sheath\_1C domain-containing protein \| Phage\_sheath\_1C domain-containing protein; afdb-uniprot50 \| AF-C4ZMC0-F1-MODEL\_V4 \| 1.0 \| 9.754e-21 \| 555 \| 0.149 \| 770 \| 303 \| 37 \| 3 \| 468 \| 2 \| 723 \| Tail sheath protein \| Tail sheath protein; afdb-uniprot50 \| AF-A0A3B0QYN2-F1-MODEL\_V4 \| 1.0 \| 1.937e-20 \| 554 \| 0.148 \| 572 \| 314 \| 24 \| 3 \| 467 \| 1 \| 506 \| Phage tail sheath protein FI \| Phage tail sheath protein FI; afdb-uniprot50 \| AF-A0A1I3I3B9-F1-MODEL\_V4 \| 1.0 \| 2.205e-21 \| 554 \| 0.165 \| 628 \| 304 \| 32 \| 3 \| 468 \| 2 \| 571 \| Uncharacterized protein \| Uncharacterized protein; afdb-uniprot50 \| AF-A0A0S4LDL9-F1-MODEL\_V4 \| 1.0 \| 3.107e-21 \| 554 \| 0.149 \| 696 \| 324 \| 34 \| 1 \| 467 \| 1 \| 657 \| Putative Phage tail sheath protein \| Putative Phage tail sheath protein; afdb-uniprot50 \| AF-A0A6C9SSS4-F1-MODEL\_V4 \| 1.0 \| 4.18e-14 \| 553 \| 0.421 \| 147 \| 85 \| 0 \| 328 \| 474 \| 1 \| 147 \| Phage tail protein \| Phage tail protein; afdb-uniprot50 \| AF-A0A5F1HSS1-F1-MODEL\_V4 \| 1.0 \| 2.078e-17 \| 553 \| 0.156 \| 493 \| 255 \| 20 \| 1 \| 468 \| 11 \| 367 \| Phage tail protein \| Phage tail protein; afdb-uniprot50 \| AF-A0A5C8AYR4-F1-MODEL\_V4 \| 1.0 \| 7.976e-19 \| 553 \| 0.156 \| 580 \| 279 \| 25 \| 3 \| 474 \| 1 \| 477 \| Uncharacterized protein \| Uncharacterized protein; afdb-uniprot50 \| AF-A0A401Z4M6-F1-MODEL\_V4 \| 1.0 \| 1.315e-17 \| 553 \| 0.231 \| 281 \| 166 \| 9 \| 230 \| 466 \| 462 \| 736 \| Tail protein \| Tail protein; afdb-uniprot50 \| AF-A0A3A9D0K2-F1-MODEL\_V4 \| 1.0 \| 1.335e-18 \| 552 \| 0.153 \| 591 \| 280 \| 27 \| 3 \| 474 \| 2 \| 490 \| Phage tail sheath family protein \| Phage tail sheath family protein; afdb-uniprot50 \| AF-A0A3D4UZL2-F1-MODEL\_V4 \| 1.0 \| 2.017e-15 \| 552 \| 0.184 \| 249 \| 187 \| 8 \| 230 \| 468 \| 248 \| 490 \| Uncharacterized protein \| Uncharacterized protein; afdb-uniprot50 \| AF-A0A329LW71-F1-MODEL\_V4 \| 1.0 \| 2.771e-21 \| 552 \| 0.145 \| 832 \| 325 \| 33 \| 5 \| 468 \| 3 \| 816 \| Phage tail protein \| Phage tail protein; afdb-uniprot50 \| AF-A0A4Q3MWA4-F1-MODEL\_V4 \| 1.0 \| 3.484e-21 \| 551 \| 0.16 \| 616 \| 296 \| 31 \| 9 \| 468 \| 8 \| 558 \| Phage tail sheath family protein \| Phage tail sheath family protein; afdb-uniprot50 \| AF-A0A7S9UGE8-F1-MODEL\_V4 \| 1.0 \| 1.728e-20 \| 551 \| 0.136 \| 702 \| 322 \| 24 \| 9 \| 468 \| 8 \| 666 \| Phage tail sheath family protein \| Phage tail sheath family protein; afdb-uniprot50 \| AF-A0A3N5L8K0-F1-MODEL\_V4 \| 1.0 \| 8.216e-21 \| 551 \| 0.157 \| 755 \| 308 \| 34 \| 5 \| 459 \| 3 \| 729 \| Phage tail protein \| Phage tail protein; afdb-uniprot50 \| AF-A0A1Y4I4D9-F1-MODEL\_V4 \| 1.0 \| 5.829e-21 \| 550 \| 0.185 \| 540 \| 302 \| 28 \| 5 \| 475 \| 2 \| 472 \| Uncharacterized protein \| Uncharacterized protein; afdb-uniprot50 \| AF-A0A2S7K9H5-F1-MODEL\_V4 \| 1.0 \| 1.909e-19 \| 550 \| 0.195 \| 482 \| 247 \| 22 \| 9 \| 467 \| 108 \| 471 \| Uncharacterized protein \| Uncharacterized protein; afdb-uniprot50 \| AF-A0A1P8MV71-F1-MODEL\_V4 \| 1.0 \| 1.226e-20 \| 550 \| 0.131 \| 708 \| 314 \| 24 \| 3 \| 468 \| 2 \| 650 \| Uncharacterized protein \| Uncharacterized protein; afdb-uniprot50 \| AF-A0A4Q7YEW2-F1-MODEL\_V4 \| 1.0 \| 2.771e-21 \| 549 \| 0.155 \| 559 \| 320 \| 22 \| 1 \| 468 \| 6 \| 503 \| Phage\_sheath\_1 domain-containing protein \| Phage\_sheath\_1 domain-containing protein; afdb-uniprot50 \| AF-A0A5B7V9X8-F1-MODEL\_V4 \| 1.0 \| 3.792e-19 \| 549 \| 0.193 \| 429 \| 215 \| 14 \| 56 \| 468 \| 230 \| 543 \| Phage tail sheath protein \| Phage tail sheath protein; afdb-uniprot50 \| AF-A0A0D8CYY8-F1-MODEL\_V4 \| 1.0 \| 8.338e-22 \| 549 \| 0.17 \| 647 \| 325 \| 37 \| 3 \| 466 \| 1 \| 618 \| Uncharacterized protein \| Uncharacterized protein; afdb-uniprot50 \| AF-E3HBL6-F1-MODEL\_V4 \| 1.0 \| 5.904e-18 \| 548 \| 0.15 \| 590 \| 274 \| 24 \| 5 \| 475 \| 1 \| 482 \| Uncharacterized protein \| Uncharacterized protein; afdb-uniprot50 \| AF-A0A7Z6LAT1-F1-MODEL\_V4 \| 1.0 \| 3.689e-21 \| 547 \| 0.266 \| 364 \| 198 \| 14 \| 162 \| 474 \| 91 \| 436 \| Phage tail protein \| Phage tail protein; afdb-uniprot50 \| AF-A0A4V6JGZ6-F1-MODEL\_V4 \| 1.0 \| 1.19e-18 \| 546 \| 0.24 \| 429 \| 179 \| 12 \| 2 \| 398 \| 46 \| 359 \| Phage tail sheath protein \| Phage tail sheath protein; afdb-uniprot50 \| AF-A0A1L6M114-F1-MODEL\_V4 \| 1.0 \| 2.766e-17 \| 545 \| 0.164 \| 500 \| 267 \| 17 \| 2 \| 468 \| 7 \| 388 \| Phage tail sheath protein FI \| Phage tail sheath protein FI; afdb-uniprot50 \| AF-A0A0Q8V3E8-F1-MODEL\_V4 \| 1.0 \| 7.759e-21 \| 545 \| 0.152 \| 675 \| 317 \| 30 \| 3 \| 466 \| 2 \| 632 \| Uncharacterized protein \| Uncharacterized protein; afdb-uniprot50 \| AF-A0A5C6M4S1-F1-MODEL\_V4 \| 1.0 \| 9.754e-21 \| 544 \| 0.134 \| 715 \| 327 \| 37 \| 1 \| 467 \| 1 \| 671 \| Tail protein \| Tail protein; afdb-uniprot50 \| AF-A0A1L6L1T8-F1-MODEL\_V4 \| 1.0 \| 2.082e-21 \| 544 \| 0.141 \| 784 \| 337 \| 33 \| 3 \| 467 \| 1 \| 767 \| Phage tail sheath protein FI \| Phage tail sheath protein FI; afdb-uniprot50 \| AF-A0A3N5XL57-F1-MODEL\_V4 \| 1.0 \| 5.505e-21 \| 544 \| 0.128 \| 795 \| 337 \| 35 \| 5 \| 468 \| 3 \| 772 \| Uncharacterized protein \| Uncharacterized protein; afdb-uniprot50 \| AF-U2DD43-F1-MODEL\_V4 \| 1.0 \| 1.992e-18 \| 543 \| 0.149 \| 575 \| 288 \| 20 \| 5 \| 474 \| 1 \| 479 \| Uncharacterized protein \| Uncharacterized protein; afdb-uniprot50 \| AF-A0A4P7HLX1-F1-MODEL\_V4 \| 1.0 \| 6.264e-22 \| 543 \| 0.181 \| 595 \| 302 \| 26 \| 3 \| 468 \| 2 \| 540 \| Phage tail sheath family protein \| Phage tail sheath family protein; afdb-uniprot50 \| AF-A0A2V6EZ80-F1-MODEL\_V4 \| 1.0 \| 3.29e-21 \| 543 \| 0.149 \| 797 \| 327 \| 37 \| 5 \| 467 \| 3 \| 782 \| Uncharacterized protein \| Uncharacterized protein; afdb-uniprot50 \| AF-A0A428MM72-F1-MODEL\_V4 \| 1.0 \| 5.199e-21 \| 542 \| 0.149 \| 717 \| 324 \| 33 \| 3 \| 468 \| 2 \| 683 \| Uncharacterized protein \| Uncharacterized protein; afdb-uniprot50 \| AF-A0A7W1N6A9-F1-MODEL\_V4 \| 1.0 \| 6.252e-18 \| 541 \| 0.143 \| 502 \| 280 \| 22 \| 6 \| 468 \| 1 \| 391 \| Phage tail sheath family protein \| Phage tail sheath family protein; afdb-uniprot50 \| AF-A0A1U7IY13-F1-MODEL\_V4 \| 1.0 \| 7.976e-19 \| 541 \| 0.169 \| 519 \| 259 \| 25 \| 1 \| 462 \| 1 \| 404 \| Phage\_sheath\_1 domain-containing protein \| Phage\_sheath\_1 domain-containing protein; afdb-uniprot50 \| AF-A0A4R1J9T0-F1-MODEL\_V4 \| 1.0 \| 2.435e-20 \| 541 \| 0.144 \| 604 \| 320 \| 33 \| 1 \| 466 \| 1 \| 545 \| Uncharacterized protein \| Uncharacterized protein; afdb-uniprot50 \| AF-A0A7Y4NFZ4-F1-MODEL\_V4 \| 1.0 \| 5.199e-21 \| 541 \| 0.144 \| 788 \| 333 \| 34 \| 3 \| 467 \| 1 \| 769 \| Uncharacterized protein \| Uncharacterized protein; afdb-uniprot50 \| AF-A0A656TEJ8-F1-MODEL\_V4 \| 1.0 \| 3.574e-15 \| 540 \| 0.292 \| 212 \| 134 \| 6 \| 267 \| 468 \| 1 \| 206 \| Tail protein \| Tail protein; afdb-uniprot50 \| AF-A0A2T9JXK8-F1-MODEL\_V4 \| 1.0 \| 4.445e-22 \| 540 \| 0.15 \| 718 \| 310 \| 30 \| 1 \| 468 \| 18 \| 685 \| Uncharacterized protein \| Uncharacterized protein; afdb-uniprot50 \| AF-A0A3G7BLM4-F1-MODEL\_V4 \| 1.0 \| 2.651e-18 \| 539 \| 0.153 \| 554 \| 265 \| 24 \| 3 \| 468 \| 2 \| 439 \| Phage tail sheath protein FI \| Phage tail sheath protein FI; afdb-uniprot50 \| AF-A0A1M7R6C0-F1-MODEL\_V4 \| 1.0 \| 1.564e-21 \| 539 \| 0.146 \| 798 \| 314 \| 38 \| 1 \| 466 \| 1 \| 763 \| Uncharacterized protein \| Uncharacterized protein; afdb-uniprot50 \| AF-A0A5S4GTI6-F1-MODEL\_V4 \| 1.0 \| 3.062e-20 \| 538 \| 0.153 \| 643 \| 325 \| 28 \| 1 \| 468 \| 44 \| 641 \| Phage tail sheath family protein \| Phage tail sheath family protein; afdb-uniprot50 \| AF-M2QID3-F1-MODEL\_V4 \| 1.0 \| 8.095e-20 \| 538 \| 0.131 \| 798 \| 309 \| 28 \| 5 \| 468 \| 3 \| 750 \| Phage tail sheath protein FI \| Phage tail sheath protein FI; afdb-uniprot50 \| AF-A0A7W7LTD8-F1-MODEL\_V4 \| 1.0 \| 3.529e-18 \| 537 \| 0.192 \| 421 \| 222 \| 14 \| 68 \| 468 \| 158 \| 480 \| Uncharacterized protein \| Uncharacterized protein; afdb-uniprot50 \| AF-A0A538QBQ6-F1-MODEL\_V4 \| 1.0 \| 6.819e-20 \| 537 \| 0.167 \| 579 \| 324 \| 31 \| 1 \| 474 \| 1 \| 526 \| Phage\_sheath\_1 domain-containing protein \| Phage\_sheath\_1 domain-containing protein; afdb-uniprot50 \| AF-A0A6I8LL75-F1-MODEL\_V4 \| 1.0 \| 5.199e-21 \| 537 \| 0.178 \| 660 \| 322 \| 31 \| 3 \| 463 \| 2 \| 639 \| Phage tail sheath protein FI \| Phage tail sheath protein FI; afdb-uniprot50 \| AF-A0A839MYJ4-F1-MODEL\_V4 \| 1.0 \| 3.849e-20 \| 537 \| 0.16 \| 686 \| 319 \| 27 \| 2 \| 468 \| 4 \| 651 \| Uncharacterized protein \| Uncharacterized protein; afdb-uniprot50 \| AF-D1JAL9-F1-MODEL\_V4 \| 1.0 \| 5.829e-21 \| 537 \| 0.127 \| 899 \| 329 \| 35 \| 3 \| 468 \| 2 \| 877 \| Uncharacterized protein \| Uncharacterized protein; afdb-uniprot50 \| AF-A0A6G9ZDL1-F1-MODEL\_V4 \| 1.0 \| 6.62e-18 \| 536 \| 0.182 \| 421 \| 227 \| 18 \| 72 \| 467 \| 247 \| 575 \| Uncharacterized protein \| Uncharacterized protein; afdb-uniprot50 \| AF-J2GKS0-F1-MODEL\_V4 \| 1.0 \| 3.242e-20 \| 536 \| 0.137 \| 771 \| 323 \| 31 \| 3 \| 468 \| 2 \| 735 \| Phage tail sheath protein FI \| Phage tail sheath protein FI; afdb-uniprot50 \| AF-A0A7Y9CZI0-F1-MODEL\_V4 \| 1.0 \| 3.956e-18 \| 534 \| 0.166 \| 576 \| 271 \| 27 \| 5 \| 474 \| 2 \| 474 \| Phage tail sheath protein FI \| Phage tail sheath protein FI; afdb-uniprot50 \| AF-A0A661ZAW2-F1-MODEL\_V4 \| 1.0 \| 3.235e-16 \| 534 \| 0.219 \| 260 \| 181 \| 8 \| 231 \| 468 \| 332 \| 591 \| Uncharacterized protein \| Uncharacterized protein; afdb-uniprot50 \| AF-A0A4Z0M8Z8-F1-MODEL\_V4 \| 1.0 \| 1.937e-20 \| 534 \| 0.159 \| 697 \| 316 \| 29 \| 3 \| 468 \| 1 \| 658 \| Uncharacterized protein \| Uncharacterized protein; afdb-uniprot50 \| AF-A0A1L6LX19-F1-MODEL\_V4 \| 1.0 \| 8.321e-18 \| 533 \| 0.147 \| 521 \| 275 \| 20 \| 5 \| 468 \| 1 \| 409 \| Phage tail sheath protein FI \| Phage tail sheath protein FI; afdb-uniprot50 \| AF-A0A1C6D4Q3-F1-MODEL\_V4 \| 1.0 \| 8.216e-21 \| 532 \| 0.183 \| 546 \| 292 \| 30 \| 6 \| 473 \| 1 \| 470 \| Phage tail sheath protein \| Phage tail sheath protein; afdb-uniprot50 \| AF-E3IXB2-F1-MODEL\_V4 \| 1.0 \| 5.576e-18 \| 531 \| 0.148 \| 506 \| 273 \| 19 \| 6 \| 467 \| 1 \| 392 \| Phage tail sheath protein fi-like protein \| Phage tail sheath protein fi-like protein; afdb-uniprot50 \| AF-A0A399IIQ1-F1-MODEL\_V4 \| 1.0 \| 2.651e-18 \| 531 \| 0.146 \| 566 \| 277 \| 22 \| 17 \| 474 \| 2 \| 469 \| Phage tail sheath protein \| Phage tail sheath protein; afdb-uniprot50 \| AF-A0A3D8JWB0-F1-MODEL\_V4 \| 1.0 \| 3.062e-20 \| 531 \| 0.144 \| 628 \| 332 \| 31 \| 3 \| 466 \| 2 \| 587 \| Phage tail sheath family protein \| Phage tail sheath family protein; afdb-uniprot50 \| AF-A0A2V6FAV4-F1-MODEL\_V4 \| 1.0 \| 4.569e-20 \| 531 \| 0.127 \| 811 \| 337 \| 30 \| 3 \| 468 \| 2 \| 786 \| Uncharacterized protein \| Uncharacterized protein; afdb-uniprot50 \| AF-A0A4R5H846-F1-MODEL\_V4 \| 1.0 \| 2.731e-20 \| 531 \| 0.133 \| 846 \| 315 \| 36 \| 3 \| 468 \| 2 \| 809 \| Phage tail sheath protein \| Phage tail sheath protein; afdb-uniprot50 \| AF-A0A2T1ENU3-F1-MODEL\_V4 \| 1.0 \| 1.031e-16 \| 530 \| 0.158 \| 474 \| 262 \| 19 \| 1 \| 455 \| 2 \| 357 \| Phage tail sheath family protein \| Phage tail sheath family protein; afdb-uniprot50 \| AF-A0A1M6SKF9-F1-MODEL\_V4 \| 1.0 \| 8.095e-20 \| 530 \| 0.181 \| 550 \| 299 \| 31 \| 5 \| 475 \| 2 \| 479 \| Uncharacterized protein \| Uncharacterized protein; afdb-uniprot50 \| AF-A0A2L2XDC6-F1-MODEL\_V4 \| 1.0 \| 7.645e-20 \| 528 \| 0.178 \| 510 \| 281 \| 23 \| 25 \| 466 \| 1 \| 440 \| Phage tail sheath protein FI \| Phage tail sheath protein FI; afdb-uniprot50 \| AF-A0A630BY26-F1-MODEL\_V4 \| 1.0 \| 3.326e-14 \| 527 \| 0.273 \| 201 \| 141 \| 3 \| 276 \| 474 \| 4 \| 201 \| Phage tail sheath family protein \| Phage tail sheath family protein; afdb-uniprot50 \| AF-A0A828D4K5-F1-MODEL\_V4 \| 1.0 \| 1.001e-14 \| 527 \| 0.225 \| 301 \| 135 \| 6 \| 75 \| 372 \| 16 \| 221 \| Phage tail sheath family protein \| Phage tail sheath family protein; afdb-uniprot50 \| AF-A0A6H9V561-F1-MODEL\_V4 \| 1.0 \| 2.731e-20 \| 526 \| 0.13 \| 895 \| 320 \| 34 \| 1 \| 468 \| 1 \| 863 \| Uncharacterized protein \| Uncharacterized protein; afdb-uniprot50 \| AF-A0A1L6M381-F1-MODEL\_V4 \| 1.0 \| 2.2e-17 \| 525 \| 0.152 \| 513 \| 266 \| 23 \| 1 \| 467 \| 2 \| 391 \| Phage tail sheath protein FI \| Phage tail sheath protein FI; afdb-uniprot50 \| AF-R7HVY5-F1-MODEL\_V4 \| 1.0 \| 2.467e-17 \| 525 \| 0.148 \| 572 \| 286 \| 22 \| 5 \| 474 \| 1 \| 473 \| Uncharacterized protein \| Uncharacterized protein; afdb-uniprot50 \| AF-A0A5B7V907-F1-MODEL\_V4 \| 1.0 \| 5.732e-16 \| 525 \| 0.223 \| 260 \| 171 \| 9 \| 230 \| 466 \| 230 \| 481 \| Phage tail sheath protein \| Phage tail sheath protein; afdb-uniprot50 \| AF-A0A150TH68-F1-MODEL\_V4 \| 1.0 \| 1.298e-20 \| 525 \| 0.187 \| 596 \| 293 \| 26 \| 6 \| 468 \| 1 \| 537 \| Uncharacterized protein \| Uncharacterized protein; afdb-uniprot50 \| AF-A0A2D6MDR9-F1-MODEL\_V4 \| 1.0 \| 1.909e-19 \| 525 \| 0.145 \| 727 \| 309 \| 28 \| 3 \| 474 \| 2 \| 670 \| Uncharacterized protein \| Uncharacterized protein; afdb-uniprot50 \| AF-A0A1H6TUH7-F1-MODEL\_V4 \| 1.0 \| 1.909e-19 \| 524 \| 0.148 \| 699 \| 312 \| 26 \| 1 \| 468 \| 6 \| 651 \| Uncharacterized protein \| Uncharacterized protein; afdb-uniprot50 \| AF-A0A832EAA6-F1-MODEL\_V4 \| 1.0 \| 1.298e-20 \| 524 \| 0.135 \| 776 \| 333 \| 33 \| 1 \| 466 \| 1 \| 748 \| Uncharacterized protein \| Uncharacterized protein; afdb-uniprot50 \| AF-Q1D3W0-F1-MODEL\_V4 \| 1.0 \| 3.849e-20 \| 524 \| 0.152 \| 822 \| 309 \| 38 \| 1 \| 468 \| 4 \| 791 \| Putative Phage tail sheath protein \| Putative Phage tail sheath protein; afdb-uniprot50 \| AF-A0A376J7I4-F1-MODEL\_V4 \| 1.0 \| 8.304e-14 \| 523 \| 0.363 \| 146 \| 90 \| 3 \| 238 \| 381 \| 2 \| 146 \| Major tail sheath protein FI \| Major tail sheath protein FI; afdb-uniprot50 \| AF-A0A3D4L0R8-F1-MODEL\_V4 \| 1.0 \| 8.095e-20 \| 523 \| 0.16 \| 550 \| 305 \| 29 \| 5 \| 474 \| 2 \| 474 \| Phage tail protein \| Phage tail protein; afdb-uniprot50 \| AF-A0A6G9ZDK1-F1-MODEL\_V4 \| 1.0 \| 4.973e-18 \| 523 \| 0.176 \| 419 \| 228 \| 16 \| 70 \| 468 \| 372 \| 693 \| Uncharacterized protein \| Uncharacterized protein; afdb-uniprot50 \| AF-A4X0M3-F1-MODEL\_V4 \| 1.0 \| 6.44e-20 \| 523 \| 0.153 \| 842 \| 315 \| 35 \| 1 \| 468 \| 1 \| 818 \| Uncharacterized protein \| Uncharacterized protein; afdb-uniprot50 \| AF-A0A0F9KEX1-F1-MODEL\_V4 \| 1.0 \| 2.017e-15 \| 523 \| 0.207 \| 260 \| 180 \| 9 \| 231 \| 468 \| 667 \| 922 \| Phage\_sheath\_1C domain-containing protein \| Phage\_sheath\_1C domain-containing protein; afdb-uniprot50 \| AF-A0A690ZQB1-F1-MODEL\_V4 \| 1.0 \| 7.406e-14 \| 522 \| 0.35 \| 154 \| 97 \| 2 \| 230 \| 381 \| 38 \| 190 \| Phage tail sheath family protein \| Phage tail sheath family protein; afdb-uniprot50 \| AF-A0A1H0JRF2-F1-MODEL\_V4 \| 1.0 \| 2.891e-20 \| 522 \| 0.154 \| 621 \| 305 \| 34 \| 9 \| 466 \| 8 \| 571 \| Uncharacterized protein \| Uncharacterized protein; afdb-uniprot50 \| AF-A0A266LKS8-F1-MODEL\_V4 \| 1.0 \| 7.842e-14 \| 521 \| 0.507 \| 136 \| 67 \| 0 \| 339 \| 474 \| 2 \| 137 \| Phage tail protein \| Phage tail protein; afdb-uniprot50 \| AF-A0A378N170-F1-MODEL\_V4 \| 1.0 \| 5.255e-14 \| 521 \| 0.317 \| 173 \| 114 \| 3 \| 304 \| 474 \| 10 \| 180 \| Phage tail sheath protein \| Phage tail sheath protein; afdb-uniprot50 \| AF-A0A6N7C2F0-F1-MODEL\_V4 \| 1.0 \| 2.645e-14 \| 521 \| 0.239 \| 234 \| 161 \| 5 \| 245 \| 467 \| 6 \| 233 \| Phage\_sheath\_1C domain-containing protein \| Phage\_sheath\_1C domain-containing protein; afdb-uniprot50 \| AF-A0A1I3W1P1-F1-MODEL\_V4 \| 1.0 \| 5.424e-20 \| 521 \| 0.146 \| 794 \| 334 \| 26 \| 3 \| 468 \| 2 \| 779 \| Uncharacterized protein \| Uncharacterized protein; afdb-uniprot50 \| AF-A0A365VQG7-F1-MODEL\_V4 \| 1.0 \| 7.842e-14 \| 520 \| 0.335 \| 143 \| 93 \| 2 \| 231 \| 371 \| 51 \| 193 \| Phage tail protein \| Phage tail protein; afdb-uniprot50 \| AF-A0A160T8X3-F1-MODEL\_V4 \| 1.0 \| 1.608e-19 \| 520 \| 0.156 \| 583 \| 297 \| 25 \| 3 \| 455 \| 2 \| 519 \| Putative Tail sheath protein \| Putative Tail sheath protein; afdb-uniprot50 \| AF-A0A840BSA8-F1-MODEL\_V4 \| 1.0 \| 1.018e-19 \| 520 \| 0.138 \| 778 \| 339 \| 35 \| 3 \| 468 \| 2 \| 759 \| Uncharacterized protein \| Uncharacterized protein; afdb-uniprot50 \| AF-A0A5X8YWD7-F1-MODEL\_V4 \| 1.0 \| 2.196e-13 \| 519 \| 0.333 \| 165 \| 109 \| 1 \| 310 \| 474 \| 3 \| 166 \| Phage tail sheath family protein \| Phage tail sheath family protein; afdb-uniprot50 \| AF-A0A1G5B734-F1-MODEL\_V4 \| 1.0 \| 1.046e-17 \| 519 \| 0.142 \| 532 \| 268 \| 19 \| 1 \| 457 \| 8 \| 426 \| Phage\_sheath\_1 domain-containing protein \| Phage\_sheath\_1 domain-containing protein; afdb-uniprot50 \| AF-A0A661RQ90-F1-MODEL\_V4 \| 1.0 \| 2.651e-18 \| 519 \| 0.144 \| 519 \| 281 \| 23 \| 1 \| 467 \| 10 \| 417 \| LTD domain-containing protein \| LTD domain-containing protein; afdb-uniprot50 \| AF-A9CK92-F1-MODEL\_V4 \| 1.0 \| 3.107e-21 \| 519 \| 0.284 \| 355 \| 207 \| 12 \| 77 \| 419 \| 2 \| 321 \| Uncharacterized protein \| Uncharacterized protein; afdb-uniprot50 \| AF-A0A2T5JUP8-F1-MODEL\_V4 \| 1.0 \| 2.504e-18 \| 519 \| 0.13 \| 757 \| 336 \| 28 \| 1 \| 468 \| 1 \| 723 \| Tail sheath protein \| Tail sheath protein; afdb-uniprot50 \| AF-A0A5J6PCS3-F1-MODEL\_V4 \| 1.0 \| 1.018e-19 \| 519 \| 0.135 \| 784 \| 322 \| 35 \| 3 \| 468 \| 1 \| 746 \| Phage tail protein \| Phage tail protein; afdb-uniprot50 \| AF-A0A662Q4X3-F1-MODEL\_V4 \| 1.0 \| 3.635e-20 \| 518 \| 0.169 \| 579 \| 302 \| 31 \| 1 \| 468 \| 1 \| 511 \| Phage\_sheath\_1 domain-containing protein \| Phage\_sheath\_1 domain-containing protein; afdb-uniprot50 \| AF-A0A417XSR8-F1-MODEL\_V4 \| 1.0 \| 2.541e-19 \| 518 \| 0.132 \| 901 \| 303 \| 34 \| 3 \| 468 \| 1 \| 857 \| Uncharacterized protein \| Uncharacterized protein; afdb-uniprot50 \| AF-A0A351U536-F1-MODEL\_V4 \| 1.0 \| 1.001e-14 \| 516 \| 0.28 \| 221 \| 138 \| 8 \| 260 \| 465 \| 6 \| 220 \| Phage tail protein \| Phage tail protein; afdb-uniprot50 \| AF-A0A074LXD5-F1-MODEL\_V4 \| 1.0 \| 4.315e-20 \| 516 \| 0.189 \| 538 \| 297 \| 28 \| 5 \| 474 \| 1 \| 467 \| Uncharacterized protein \| Uncharacterized protein; afdb-uniprot50 \| AF-A0A261GTB9-F1-MODEL\_V4 \| 1.0 \| 5.344e-19 \| 516 \| 0.151 \| 593 \| 313 \| 32 \| 1 \| 467 \| 1 \| 529 \| Uncharacterized protein \| Uncharacterized protein; afdb-uniprot50 \| AF-A0A6L9G0N3-F1-MODEL\_V4 \| 1.0 \| 7.976e-19 \| 516 \| 0.131 \| 769 \| 335 \| 28 \| 1 \| 468 \| 1 \| 737 \| Phage tail sheath protein \| Phage tail sheath protein; afdb-uniprot50 \| AF-A0A3A8JXS8-F1-MODEL\_V4 \| 1.0 \| 2.541e-19 \| 515 \| 0.158 \| 601 \| 304 \| 31 \| 5 \| 467 \| 2 \| 538 \| Phage tail sheath family protein \| Phage tail sheath family protein; afdb-uniprot50 \| AF-A0A7G9YSQ3-F1-MODEL\_V4 \| 1.0 \| 1.803e-19 \| 515 \| 0.132 \| 818 \| 326 \| 32 \| 3 \| 467 \| 2 \| 788 \| Uncharacterized protein \| Uncharacterized protein; afdb-uniprot50 \| AF-A0A1F7S379-F1-MODEL\_V4 \| 1.0 \| 5.647e-15 \| 514 \| 0.186 \| 263 \| 190 \| 9 \| 230 \| 468 \| 186 \| 448 \| Uncharacterized protein \| Uncharacterized protein; afdb-uniprot50 \| AF-A0A2W5QE72-F1-MODEL\_V4 \| 1.0 \| 7.22e-20 \| 514 \| 0.157 \| 624 \| 320 \| 31 \| 1 \| 467 \| 32 \| 606 \| Phage tail sheath family protein \| Phage tail sheath family protein; afdb-uniprot50 \| AF-A0A7Y6QLZ2-F1-MODEL\_V4 \| 1.0 \| 2.467e-17 \| 514 \| 0.187 \| 411 \| 210 \| 12 \| 84 \| 467 \| 733 \| 1046 \| Phage tail sheath subtilisin-like domain-containing protein \| Phage tail sheath subtilisin-like domain-containing protein; afdb-uniprot50 \| AF-U2KP13-F1-MODEL\_V4 \| 1.0 \| 9.211e-21 \| 512 \| 0.161 \| 612 \| 296 \| 31 \| 5 \| 455 \| 3 \| 558 \| Phage tail sheath protein \| Phage tail sheath protein; afdb-uniprot50 \| AF-A0A090AMB4-F1-MODEL\_V4 \| 1.0 \| 1.018e-19 \| 512 \| 0.158 \| 613 \| 336 \| 32 \| 3 \| 467 \| 2 \| 582 \| Putative phage tail sheath protein \| Putative phage tail sheath protein; afdb-uniprot50 \| AF-A0A4Y6PV73-F1-MODEL\_V4 \| 1.0 \| 1.803e-19 \| 512 \| 0.143 \| 821 \| 326 \| 35 \| 5 \| 468 \| 3 \| 802 \| Uncharacterized protein \| Uncharacterized protein; afdb-uniprot50 \| AF-A0A7G9WG91-F1-MODEL\_V4 \| 1.0 \| 1.75e-17 \| 511 \| 0.138 \| 583 \| 296 \| 26 \| 1 \| 475 \| 1 \| 484 \| Phage tail sheath family protein \| Phage tail sheath family protein; afdb-uniprot50 \| AF-A0A2T5RE85-F1-MODEL\_V4 \| 1.0 \| 2.4e-19 \| 511 \| 0.144 \| 669 \| 321 \| 31 \| 3 \| 474 \| 2 \| 615 \| Tail sheath protein \| Tail sheath protein; afdb-uniprot50 \| AF-A0A2K8U9B6-F1-MODEL\_V4 \| 1.0 \| 2.541e-19 \| 510 \| 0.141 \| 712 \| 327 \| 29 \| 1 \| 468 \| 2 \| 672 \| Uncharacterized protein \| Uncharacterized protein; afdb-uniprot50 \| AF-A0A2E4ZEU9-F1-MODEL\_V4 \| 1.0 \| 1.608e-19 \| 509 \| 0.156 \| 556 \| 307 \| 24 \| 3 \| 468 \| 1 \| 484 \| Uncharacterized protein \| Uncharacterized protein; afdb-uniprot50 \| AF-J9Z8U4-F1-MODEL\_V4 \| 1.0 \| 1.124e-18 \| 509 \| 0.145 \| 551 \| 328 \| 22 \| 1 \| 468 \| 4 \| 494 \| Uncharacterized protein \| Uncharacterized protein; afdb-uniprot50 \| AF-A0A1D2QTZ4-F1-MODEL\_V4 \| 1.0 \| 1.354e-19 \| 508 \| 0.141 \| 709 \| 333 \| 30 \| 1 \| 468 \| 1 \| 674 \| Phage\_sheath\_1C domain-containing protein \| Phage\_sheath\_1C domain-containing protein; afdb-uniprot50 \| AF-A0A2X4UYA7-F1-MODEL\_V4 \| 1.0 \| 8.554e-16 \| 505 \| 0.154 \| 484 \| 258 \| 22 \| 1 \| 466 \| 1 \| 351 \| Phage tail sheath protein \| Phage tail sheath protein; afdb-uniprot50 \| AF-A0A1C6IJ19-F1-MODEL\_V4 \| 1.0 \| 1.208e-19 \| 505 \| 0.172 \| 555 \| 302 \| 31 \| 5 \| 475 \| 2 \| 482 \| Phage tail sheath protein \| Phage tail sheath protein; afdb-uniprot50 \| AF-A0A1V5QKC2-F1-MODEL\_V4 \| 1.0 \| 1.674e-14 \| 505 \| 0.182 \| 268 \| 194 \| 10 \| 226 \| 474 \| 225 \| 486 \| Phage tail sheath protein \| Phage tail sheath protein; afdb-uniprot50 \| AF-A0A1S6FHM7-F1-MODEL\_V4 \| 1.0 \| 2.972e-18 \| 505 \| 0.121 \| 740 \| 356 \| 33 \| 1 \| 467 \| 1 \| 719 \| Phage\_sheath\_1C domain-containing protein \| Phage\_sheath\_1C domain-containing protein; afdb-uniprot50 \| AF-A0A2R2Q2H2-F1-MODEL\_V4 \| 1.0 \| 4.697e-18 \| 504 \| 0.159 \| 572 \| 255 \| 22 \| 1 \| 468 \| 1 \| 450 \| AfpX3 \| AfpX3; afdb-uniprot50 \| AF-A0A845SZJ5-F1-MODEL\_V4 \| 1.0 \| 8.942e-19 \| 504 \| 0.167 \| 556 \| 300 \| 31 \| 1 \| 475 \| 1 \| 474 \| Phage tail protein \| Phage tail protein; afdb-uniprot50 \| AF-A0A8A6HCG5-F1-MODEL\_V4 \| 1.0 \| 5.047e-19 \| 504 \| 0.148 \| 689 \| 332 \| 30 \| 2 \| 468 \| 4 \| 659 \| Uncharacterized protein \| Uncharacterized protein; afdb-uniprot50 \| AF-A0A4R8CM40-F1-MODEL\_V4 \| 1.0 \| 5.047e-19 \| 504 \| 0.138 \| 700 \| 326 \| 31 \| 2 \| 468 \| 9 \| 664 \| Uncharacterized protein \| Uncharacterized protein; afdb-uniprot50 \| AF-A0A523INP5-F1-MODEL\_V4 \| 1.0 \| 1.703e-19 \| 504 \| 0.155 \| 668 \| 324 \| 32 \| 10 \| 464 \| 1 \| 641 \| Phage tail sheath family protein \| Phage tail sheath family protein; afdb-uniprot50 \| AF-A0A522C7H8-F1-MODEL\_V4 \| 1.0 \| 3.898e-17 \| 503 \| 0.143 \| 510 \| 278 \| 24 \| 1 \| 468 \| 7 \| 399 \| Phage tail sheath family protein \| Phage tail sheath family protein; afdb-uniprot50 \| AF-A0A382G038-F1-MODEL\_V4 \| 1.0 \| 9.449e-15 \| 503 \| 0.178 \| 247 \| 189 \| 7 \| 231 \| 468 \| 186 \| 427 \| Uncharacterized protein \| Uncharacterized protein; afdb-uniprot50 \| AF-A0A266LLW5-F1-MODEL\_V4 \| 1.0 \| 1.221e-12 \| 502 \| 0.504 \| 121 \| 59 \| 1 \| 243 \| 362 \| 2 \| 122 \| Phage tail protein \| Phage tail protein; afdb-uniprot50 \| AF-V0XQG6-F1-MODEL\_V4 \| 1.0 \| 2.196e-13 \| 502 \| 0.272 \| 257 \| 100 \| 4 \| 113 \| 367 \| 2 \| 173 \| Phage\_sheath\_1 domain-containing protein \| Phage\_sheath\_1 domain-containing protein; afdb-uniprot50 \| AF-A0A1W0CS69-F1-MODEL\_V4 \| 1.0 \| 4.007e-15 \| 501 \| 0.158 \| 487 \| 250 \| 19 \| 1 \| 468 \| 1 \| 346 \| Phage tail protein \| Phage tail protein; afdb-uniprot50 \| AF-U2PM90-F1-MODEL\_V4 \| 1.0 \| 8.554e-16 \| 500 \| 0.169 \| 406 \| 226 \| 12 \| 90 \| 474 \| 19 \| 334 \| Uncharacterized protein \| Uncharacterized protein; afdb-uniprot50 \| AF-A0A1J5DEX6-F1-MODEL\_V4 \| 1.0 \| 7.645e-20 \| 499 \| 0.148 \| 614 \| 314 \| 30 \| 3 \| 457 \| 2 \| 565 \| Uncharacterized protein \| Uncharacterized protein; afdb-uniprot50 \| AF-A0A850NPQ6-F1-MODEL\_V4 \| 1.0 \| 7.532e-19 \| 498 \| 0.149 \| 550 \| 325 \| 28 \| 2 \| 468 \| 6 \| 495 \| Phage tail protein \| Phage tail protein; afdb-uniprot50 \| AF-D5CT51-F1-MODEL\_V4 \| 1.0 \| 2.233e-18 \| 498 \| 0.144 \| 554 \| 328 \| 23 \| 2 \| 468 \| 10 \| 504 \| Uncharacterized protein \| Uncharacterized protein; afdb-uniprot50 \| AF-A0A4Y6PVA2-F1-MODEL\_V4 \| 1.0 \| 2.364e-18 \| 498 \| 0.137 \| 686 \| 333 \| 30 \| 1 \| 455 \| 1 \| 658 \| Phage tail sheath family protein \| Phage tail sheath family protein; afdb-uniprot50 \| AF-A0A662L329-F1-MODEL\_V4 \| 1.0 \| 6.819e-20 \| 497 \| 0.187 \| 495 \| 267 \| 20 \| 3 \| 421 \| 2 \| 437 \| Phage tail protein \| Phage tail protein; afdb-uniprot50 \| AF-A0A3T1B9R3-F1-MODEL\_V4 \| 1.0 \| 4.315e-20 \| 497 \| 0.143 \| 598 \| 337 \| 23 \| 1 \| 466 \| 4 \| 558 \| Uncharacterized protein \| Uncharacterized protein; afdb-uniprot50 \| AF-A0A2S3UN30-F1-MODEL\_V4 \| 1.0 \| 2.4e-19 \| 497 \| 0.143 \| 642 \| 328 \| 28 \| 5 \| 466 \| 2 \| 601 \| Tail sheath protein \| Tail sheath protein; afdb-uniprot50 \| AF-A0A7Z7I242-F1-MODEL\_V4 \| 1.0 \| 9.469e-19 \| 497 \| 0.141 \| 728 \| 315 \| 28 \| 6 \| 468 \| 1 \| 683 \| Phage tail sheath protein \| Phage tail sheath protein; afdb-uniprot50 \| AF-A0A1Y4T0D0-F1-MODEL\_V4 \| 1.0 \| 6.069e-16 \| 496 \| 0.186 \| 413 \| 212 \| 15 \| 90 \| 474 \| 44 \| 360 \| Uncharacterized protein \| Uncharacterized protein; afdb-uniprot50 \| AF-A0A4C2EPL5-F1-MODEL\_V4 \| 1.0 \| 3.581e-19 \| 496 \| 0.13 \| 888 \| 335 \| 34 \| 3 \| 468 \| 2 \| 873 \| Uncharacterized protein \| Uncharacterized protein; afdb-uniprot50 \| AF-A0A430VBI2-F1-MODEL\_V4 \| 1.0 \| 2.228e-14 \| 496 \| 0.141 \| 255 \| 194 \| 13 \| 231 \| 468 \| 705 \| 951 \| Uncharacterized protein \| Uncharacterized protein; afdb-uniprot50 \| AF-A0A1C5SXM5-F1-MODEL\_V4 \| 1.0 \| 9.61e-20 \| 495 \| 0.162 \| 608 \| 279 \| 25 \| 23 \| 463 \| 2 \| 545 \| Phage tail sheath protein \| Phage tail sheath protein; afdb-uniprot50 \| AF-A0A2T6N8W8-F1-MODEL\_V4 \| 1.0 \| 9.076e-20 \| 495 \| 0.144 \| 759 \| 317 \| 31 \| 3 \| 468 \| 13 \| 731 \| Phage\_sheath\_1C domain-containing protein \| Phage\_sheath\_1C domain-containing protein; afdb-uniprot50 \| AF-A0A1V5BE78-F1-MODEL\_V4 \| 1.0 \| 9.61e-20 \| 495 \| 0.131 \| 824 \| 333 \| 37 \| 1 \| 466 \| 1 \| 799 \| Phage tail sheath protein \| Phage tail sheath protein; afdb-uniprot50 \| AF-A0A4Q5RRV7-F1-MODEL\_V4 \| 1.0 \| 3.235e-16 \| 494 \| 0.135 \| 634 \| 274 \| 24 \| 1 \| 466 \| 1 \| 528 \| Phage tail sheath family protein \| Phage tail sheath family protein; afdb-uniprot50 \| AF-A0A6N8I017-F1-MODEL\_V4 \| 1.0 \| 5.743e-20 \| 493 \| 0.162 \| 559 \| 303 \| 31 \| 1 \| 474 \| 1 \| 479 \| Uncharacterized protein \| Uncharacterized protein; afdb-uniprot50 \| AF-A0A3C1WS05-F1-MODEL\_V4 \| 1.0 \| 1.629e-16 \| 493 \| 0.151 \| 588 \| 284 \| 22 \| 1 \| 475 \| 1 \| 486 \| Phage tail protein \| Phage tail protein; afdb-uniprot50 \| AF-A0A840MS80-F1-MODEL\_V4 \| 1.0 \| 1.992e-18 \| 493 \| 0.135 \| 687 \| 341 \| 27 \| 1 \| 468 \| 5 \| 657 \| Uncharacterized protein \| Uncharacterized protein; afdb-uniprot50 \| AF-A0A7J4LL39-F1-MODEL\_V4 \| 1.0 \| 5.047e-19 \| 493 \| 0.15 \| 710 \| 333 \| 39 \| 5 \| 471 \| 3 \| 685 \| Phage tail sheath family protein \| Phage tail sheath family protein; afdb-uniprot50 \| AF-A0A447PAZ2-F1-MODEL\_V4 \| 1.0 \| 5.178e-13 \| 492 \| 0.375 \| 144 \| 88 \| 2 \| 230 \| 371 \| 8 \| 151 \| Major tail sheath protein \| Major tail sheath protein; afdb-uniprot50 \| AF-A0A382DIG5-F1-MODEL\_V4 \| 1.0 \| 2.645e-14 \| 492 \| 0.176 \| 266 \| 192 \| 11 \| 230 \| 474 \| 60 \| 319 \| Uncharacterized protein \| Uncharacterized protein; afdb-uniprot50 \| AF-A0A2G2EDZ2-F1-MODEL\_V4 \| 1.0 \| 1.538e-16 \| 492 \| 0.145 \| 577 \| 282 \| 27 \| 1 \| 468 \| 1 \| 475 \| Uncharacterized protein \| Uncharacterized protein; afdb-uniprot50 \| AF-A0A2D7CYR9-F1-MODEL\_V4 \| 1.0 \| 1.26e-18 \| 492 \| 0.151 \| 642 \| 318 \| 34 \| 3 \| 475 \| 2 \| 585 \| Uncharacterized protein \| Uncharacterized protein; afdb-uniprot50 \| AF-A0A518BPN9-F1-MODEL\_V4 \| 1.0 \| 2.266e-19 \| 492 \| 0.137 \| 757 \| 349 \| 31 \| 3 \| 467 \| 2 \| 746 \| Phage tail sheath protein \| Phage tail sheath protein; afdb-uniprot50 \| AF-A0A525HH54-F1-MODEL\_V4 \| 1.0 \| 1.258e-14 \| 492 \| 0.215 \| 264 \| 172 \| 8 \| 230 \| 465 \| 606 \| 862 \| Uncharacterized protein \| Uncharacterized protein; afdb-uniprot50 \| AF-A0A064A4E1-F1-MODEL\_V4 \| 1.0 \| 1.988e-14 \| 491 \| 0.282 \| 237 \| 141 \| 11 \| 254 \| 474 \| 26 \| 249 \| Phage tail protein \| Phage tail protein; afdb-uniprot50 \| AF-A0A2D5FCB7-F1-MODEL\_V4 \| 1.0 \| 2.105e-14 \| 491 \| 0.202 \| 262 \| 184 \| 10 \| 226 \| 468 \| 34 \| 289 \| Uncharacterized protein \| Uncharacterized protein; afdb-uniprot50 \| AF-A0A3M0WVE2-F1-MODEL\_V4 \| 1.0 \| 4.315e-20 \| 491 \| 0.178 \| 511 \| 260 \| 22 \| 1 \| 415 \| 1 \| 447 \| Phage tail sheath family protein \| Phage tail sheath family protein; afdb-uniprot50 \| AF-A0A4U3M6X8-F1-MODEL\_V4 \| 1.0 \| 6.345e-19 \| 491 \| 0.132 \| 567 \| 329 \| 24 \| 3 \| 468 \| 2 \| 506 \| Phage tail sheath family protein \| Phage tail sheath family protein; afdb-uniprot50 \| AF-A0A2D7XGG3-F1-MODEL\_V4 \| 1.0 \| 6.332e-15 \| 491 \| 0.166 \| 277 \| 193 \| 9 \| 229 \| 473 \| 522 \| 792 \| Uncharacterized protein \| Uncharacterized protein; afdb-uniprot50 \| AF-A0A2E0KX46-F1-MODEL\_V4 \| 1.0 \| 1.332e-14 \| 491 \| 0.176 \| 261 \| 186 \| 9 \| 229 \| 466 \| 571 \| 825 \| Uncharacterized protein \| Uncharacterized protein; afdb-uniprot50 \| AF-A0A2D6ANC4-F1-MODEL\_V4 \| 1.0 \| 1.493e-14 \| 491 \| 0.13 \| 275 \| 207 \| 8 \| 225 \| 473 \| 853 \| 1121 \| Uncharacterized protein \| Uncharacterized protein; afdb-uniprot50 \| AF-A0A368A7G8-F1-MODEL\_V4 \| 1.0 \| 2.536e-15 \| 490 \| 0.288 \| 353 \| 98 \| 7 \| 1 \| 352 \| 1 \| 201 \| Phage tail sheath protein FI \| Phage tail sheath protein FI; afdb-uniprot50 \| AF-A0A2E4W7V1-F1-MODEL\_V4 \| 1.0 \| 1.853e-17 \| 490 \| 0.11 \| 727 \| 326 \| 29 \| 1 \| 475 \| 1 \| 658 \| Phage tail protein \| Phage tail protein; afdb-uniprot50 \| AF-A0A2W5F6C6-F1-MODEL\_V4 \| 1.0 \| 4.252e-19 \| 490 \| 0.151 \| 705 \| 310 \| 30 \| 1 \| 467 \| 1 \| 654 \| Uncharacterized protein \| Uncharacterized protein; afdb-uniprot50 \| AF-A0A7Y5LPM5-F1-MODEL\_V4 \| 1.0 \| 2.4e-19 \| 490 \| 0.141 \| 684 \| 332 \| 30 \| 1 \| 459 \| 1 \| 653 \| Phage tail sheath family protein \| Phage tail sheath family protein; afdb-uniprot50 \| AF-A0A399YVR8-F1-MODEL\_V4 \| 1.0 \| 2.849e-19 \| 490 \| 0.142 \| 838 \| 325 \| 40 \| 1 \| 466 \| 1 \| 816 \| Uncharacterized protein \| Uncharacterized protein; afdb-uniprot50 \| AF-A0A329WRQ2-F1-MODEL\_V4 \| 1.0 \| 8.681e-17 \| 489 \| 0.145 \| 579 \| 263 \| 23 \| 1 \| 467 \| 1 \| 459 \| Phage tail protein \| Phage tail protein; afdb-uniprot50 \| AF-A0A0F9KVW3-F1-MODEL\_V4 \| 1.0 \| 4.493e-15 \| 489 \| 0.191 \| 261 \| 188 \| 8 \| 231 \| 468 \| 814 \| 1074 \| Phage\_sheath\_1C domain-containing protein \| Phage\_sheath\_1C domain-containing protein; afdb-uniprot50 \| AF-A8T9K8-F1-MODEL\_V4 \| 1.0 \| 1.206e-15 \| 488 \| 0.158 \| 497 \| 246 \| 22 \| 1 \| 468 \| 1 \| 354 \| Phage\_sheath\_1C domain-containing protein \| Phage\_sheath\_1C domain-containing protein; afdb-uniprot50 \| AF-R7MS92-F1-MODEL\_V4 \| 1.0 \| 1.206e-15 \| 488 \| 0.179 \| 413 \| 216 \| 14 \| 85 \| 475 \| 68 \| 379 \| Uncharacterized protein \| Uncharacterized protein; afdb-uniprot50 \| AF-A0A4P8QGP8-F1-MODEL\_V4 \| 1.0 \| 6.345e-19 \| 488 \| 0.142 \| 568 \| 319 \| 28 \| 1 \| 468 \| 4 \| 503 \| Phage tail protein \| Phage tail protein; afdb-uniprot50 \| AF-A0A0F5K3T8-F1-MODEL\_V4 \| 1.0 \| 6.805e-16 \| 486 \| 0.147 \| 516 \| 261 \| 23 \| 3 \| 468 \| 2 \| 388 \| Uncharacterized protein \| Uncharacterized protein; afdb-uniprot50 \| AF-A0A842J0D4-F1-MODEL\_V4 \| 1.0 \| 6.718e-19 \| 486 \| 0.178 \| 561 \| 290 \| 36 \| 5 \| 475 \| 2 \| 481 \| Phage tail protein \| Phage tail protein; afdb-uniprot50 \| AF-A0A143HDI3-F1-MODEL\_V4 \| 1.0 \| 2.725e-16 \| 486 \| 0.165 \| 586 \| 273 \| 25 \| 1 \| 474 \| 1 \| 482 \| Uncharacterized protein \| Uncharacterized protein; afdb-uniprot50 \| AF-A0A257PQZ1-F1-MODEL\_V4 \| 1.0 \| 8.942e-19 \| 485 \| 0.137 \| 540 \| 308 \| 28 \| 19 \| 468 \| 28 \| 499 \| Uncharacterized protein \| Uncharacterized protein; afdb-uniprot50 \| AF-A0A3N5PP70-F1-MODEL\_V4 \| 1.0 \| 4.015e-19 \| 484 \| 0.15 \| 679 \| 282 \| 27 \| 3 \| 442 \| 2 \| 624 \| Phage tail sheath family protein \| Phage tail sheath family protein; afdb-uniprot50 \| AF-A0A2D6E403-F1-MODEL\_V4 \| 1.0 \| 1.877e-14 \| 484 \| 0.134 \| 268 \| 201 \| 9 \| 225 \| 467 \| 773 \| 1034 \| Uncharacterized protein \| Uncharacterized protein; afdb-uniprot50 \| AF-A0A7J4GTR8-F1-MODEL\_V4 \| 1.0 \| 1.312e-13 \| 483 \| 0.16 \| 255 \| 188 \| 9 \| 241 \| 475 \| 150 \| 398 \| Uncharacterized protein \| Uncharacterized protein; afdb-uniprot50 \| AF-K6Y8P6-F1-MODEL\_V4 \| 1.0 \| 1.19e-18 \| 483 \| 0.135 \| 709 \| 346 \| 33 \| 1 \| 468 \| 1 \| 683 \| Uncharacterized protein \| Uncharacterized protein; afdb-uniprot50 \| AF-A0A2D9CBG6-F1-MODEL\_V4 \| 1.0 \| 3.095e-13 \| 482 \| 0.157 \| 247 \| 192 \| 7 \| 231 \| 467 \| 133 \| 373 \| Phage\_sheath\_1 domain-containing protein \| Phage\_sheath\_1 domain-containing protein; afdb-uniprot50 \| AF-A0A7Y4SSP2-F1-MODEL\_V4 \| 1.0 \| 7.22e-20 \| 482 \| 0.172 \| 667 \| 317 \| 29 \| 5 \| 465 \| 3 \| 640 \| Phage tail protein \| Phage tail protein; afdb-uniprot50 \| AF-A0A3N5STT9-F1-MODEL\_V4 \| 1.0 \| 1.41e-14 \| 481 \| 0.186 \| 241 \| 175 \| 8 \| 231 \| 466 \| 208 \| 432 \| Phage tail protein \| Phage tail protein; afdb-uniprot50 \| AF-A0A4U7J714-F1-MODEL\_V4 \| 1.0 \| 7.532e-19 \| 481 \| 0.169 \| 568 \| 298 \| 32 \| 1 \| 475 \| 1 \| 487 \| Uncharacterized protein \| Uncharacterized protein; afdb-uniprot50 \| AF-A0A432LJ81-F1-MODEL\_V4 \| 1.0 \| 2.295e-16 \| 481 \| 0.132 \| 610 \| 281 \| 22 \| 1 \| 468 \| 1 \| 504 \| Phage tail sheath family protein \| Phage tail sheath family protein; afdb-uniprot50 \| AF-A0A7X8L470-F1-MODEL\_V4 \| 1.0 \| 6.345e-19 \| 481 \| 0.134 \| 579 \| 315 \| 24 \| 1 \| 466 \| 54 \| 559 \| Phage tail sheath family protein \| Phage tail sheath family protein; afdb-uniprot50 \| AF-A0A4Q7WMN5-F1-MODEL\_V4 \| 1.0 \| 2.972e-18 \| 481 \| 0.155 \| 688 \| 323 \| 33 \| 1 \| 467 \| 3 \| 653 \| Phage\_sheath\_1 domain-containing protein \| Phage\_sheath\_1 domain-containing protein; afdb-uniprot50 \| AF-A0A2D9C4T6-F1-MODEL\_V4 \| 1.0 \| 2.364e-18 \| 481 \| 0.143 \| 737 \| 341 \| 34 \| 1 \| 468 \| 1 \| 715 \| Phage\_sheath\_1C domain-containing protein \| Phage\_sheath\_1C domain-containing protein; afdb-uniprot50 \| AF-A0A2E7WA37-F1-MODEL\_V4 \| 1.0 \| 2.645e-14 \| 481 \| 0.149 \| 268 \| 198 \| 8 \| 225 \| 468 \| 703 \| 964 \| Uncharacterized protein \| Uncharacterized protein; afdb-uniprot50 \| AF-A0A832XY63-F1-MODEL\_V4 \| 1.0 \| 1.988e-14 \| 480 \| 0.169 \| 266 \| 194 \| 9 \| 230 \| 474 \| 82 \| 341 \| Uncharacterized protein \| Uncharacterized protein; afdb-uniprot50 \| AF-A0A2V9GSQ5-F1-MODEL\_V4 \| 1.0 \| 4.189e-18 \| 480 \| 0.14 \| 535 \| 328 \| 29 \| 1 \| 468 \| 1 \| 470 \| Phage tail sheath protein \| Phage tail sheath protein; afdb-uniprot50 \| AF-A0A7G7M251-F1-MODEL\_V4 \| 1.0 \| 3.141e-14 \| 479 \| 0.167 \| 461 \| 155 \| 4 \| 5 \| 465 \| 2 \| 233 \| Phage\_sheath\_1C domain-containing protein \| Phage\_sheath\_1C domain-containing protein; afdb-uniprot50 \| AF-A0A5F1HY87-F1-MODEL\_V4 \| 1.0 \| 1.372e-16 \| 479 \| 0.155 \| 509 \| 264 \| 24 \| 1 \| 466 \| 66 \| 451 \| Phage tail protein \| Phage tail protein; afdb-uniprot50 \| AF-A0A4Y9M3D3-F1-MODEL\_V4 \| 1.0 \| 4.973e-18 \| 479 \| 0.136 \| 694 \| 329 \| 29 \| 1 \| 468 \| 1 \| 649 \| Phage tail sheath family protein \| Phage tail sheath family protein; afdb-uniprot50 \| AF-A0A1W1X8C9-F1-MODEL\_V4 \| 1.0 \| 5.992e-19 \| 479 \| 0.134 \| 828 \| 323 \| 36 \| 1 \| 466 \| 1 \| 796 \| Uncharacterized protein \| Uncharacterized protein; afdb-uniprot50 \| AF-A0A2E1QDI5-F1-MODEL\_V4 \| 1.0 \| 7.099e-15 \| 479 \| 0.172 \| 273 \| 194 \| 8 \| 231 \| 473 \| 811 \| 1081 \| Phage\_sheath\_1C domain-containing protein \| Phage\_sheath\_1C domain-containing protein; afdb-uniprot50 \| AF-A0A2E6H1W9-F1-MODEL\_V4 \| 1.0 \| 3.729e-14 \| 479 \| 0.144 \| 270 \| 199 \| 8 \| 225 \| 468 \| 915 \| 1178 \| Phage\_sheath\_1 domain-containing protein \| Phage\_sheath\_1 domain-containing protein; afdb-uniprot50 \| AF-A0A7Y6QLZ9-F1-MODEL\_V4 \| 1.0 \| 1.584e-18 \| 478 \| 0.153 \| 677 \| 311 \| 28 \| 5 \| 455 \| 4 \| 644 \| Phage tail sheath family protein \| Phage tail sheath family protein; afdb-uniprot50 \| AF-M0DAR5-F1-MODEL\_V4 \| 1.0 \| 2.504e-18 \| 478 \| 0.137 \| 785 \| 327 \| 34 \| 3 \| 468 \| 2 \| 755 \| Phage tail sheath protein FI \| Phage tail sheath protein FI; afdb-uniprot50 \| AF-A0A3D3QRZ8-F1-MODEL\_V4 \| 1.0 \| 4.973e-18 \| 476 \| 0.218 \| 366 \| 201 \| 20 \| 166 \| 468 \| 26 \| 369 \| Uncharacterized protein \| Uncharacterized protein; afdb-uniprot50 \| AF-A0A7K0CGL4-F1-MODEL\_V4 \| 1.0 \| 3.332e-18 \| 475 \| 0.154 \| 701 \| 305 \| 26 \| 1 \| 468 \| 1 \| 646 \| Phage\_sheath\_1 domain-containing protein \| Phage\_sheath\_1 domain-containing protein; afdb-uniprot50 \| AF-A0A651HNK2-F1-MODEL\_V4 \| 1.0 \| 2.504e-18 \| 475 \| 0.149 \| 754 \| 317 \| 31 \| 3 \| 468 \| 1 \| 717 \| Phage tail sheath protein \| Phage tail sheath protein; afdb-uniprot50 \| AF-A0A7W4X2D6-F1-MODEL\_V4 \| 1.0 \| 4.55e-12 \| 474 \| 0.789 \| 95 \| 20 \| 0 \| 381 \| 475 \| 2 \| 96 \| Phage tail sheath protein FI \| Phage tail sheath protein FI; afdb-uniprot50 \| AF-A0A3P6KDJ0-F1-MODEL\_V4 \| 1.0 \| 5.732e-16 \| 474 \| 0.185 \| 427 \| 216 \| 15 \| 84 \| 474 \| 32 \| 362 \| Phage tail sheath protein FI \| Phage tail sheath protein FI; afdb-uniprot50 \| AF-K4ZEC4-F1-MODEL\_V4 \| 1.0 \| 3.332e-18 \| 474 \| 0.215 \| 394 \| 217 \| 19 \| 148 \| 475 \| 64 \| 431 \| Phage tail sheath protein FI \| Phage tail sheath protein FI; afdb-uniprot50 \| AF-A0A0Q9UDN4-F1-MODEL\_V4 \| 1.0 \| 1.124e-18 \| 474 \| 0.138 \| 723 \| 324 \| 32 \| 1 \| 468 \| 3 \| 681 \| Uncharacterized protein \| Uncharacterized protein; afdb-uniprot50 \| AF-A0A2D7BTI7-F1-MODEL\_V4 \| 1.0 \| 1.471e-13 \| 473 \| 0.17 \| 276 \| 194 \| 10 \| 231 \| 475 \| 558 \| 829 \| Phage\_sheath\_1C domain-containing protein \| Phage\_sheath\_1C domain-containing protein; afdb-uniprot50 \| AF-A0A7C3F0E2-F1-MODEL\_V4 \| 1.0 \| 1.105e-13 \| 473 \| 0.145 \| 261 \| 194 \| 8 \| 231 \| 468 \| 642 \| 896 \| Phage\_sheath\_1C domain-containing protein \| Phage\_sheath\_1C domain-containing protein; afdb-uniprot50 \| AF-A0A3C1Q7E4-F1-MODEL\_V4 \| 1.0 \| 1.17e-13 \| 471 \| 0.202 \| 247 \| 168 \| 11 \| 245 \| 472 \| 15 \| 251 \| Uncharacterized protein \| Uncharacterized protein; afdb-uniprot50 \| AF-A0A7Z9UQ35-F1-MODEL\_V4 \| 1.0 \| 1.105e-13 \| 471 \| 0.18 \| 260 \| 185 \| 10 \| 230 \| 468 \| 136 \| 388 \| Uncharacterized protein \| Uncharacterized protein; afdb-uniprot50 \| AF-A0A382CH64-F1-MODEL\_V4 \| 1.0 \| 1.558e-13 \| 471 \| 0.201 \| 238 \| 176 \| 7 \| 229 \| 457 \| 417 \| 649 \| Uncharacterized protein \| Uncharacterized protein; afdb-uniprot50 \| AF-A0A545UE66-F1-MODEL\_V4 \| 1.0 \| 5.047e-19 \| 471 \| 0.14 \| 834 \| 332 \| 36 \| 1 \| 467 \| 1 \| 816 \| Uncharacterized protein \| Uncharacterized protein; afdb-uniprot50 \| AF-A0A484YNN0-F1-MODEL\_V4 \| 1.0 \| 2.074e-13 \| 470 \| 0.3 \| 170 \| 117 \| 2 \| 232 \| 399 \| 2 \| 171 \| Tail Sheath protein \| Tail Sheath protein; afdb-uniprot50 \| AF-M1YXK8-F1-MODEL\_V4 \| 1.0 \| 5.266e-18 \| 470 \| 0.151 \| 819 \| 310 \| 40 \| 5 \| 466 \| 3 \| 793 \| Uncharacterized protein \| Uncharacterized protein; afdb-uniprot50 \| AF-A0A2A4T3R4-F1-MODEL\_V4 \| 1.0 \| 1.17e-13 \| 470 \| 0.169 \| 253 \| 189 \| 11 \| 231 \| 468 \| 712 \| 958 \| Uncharacterized protein \| Uncharacterized protein; afdb-uniprot50 \| AF-A0A7C7C6D0-F1-MODEL\_V4 \| 1.0 \| 8.304e-14 \| 470 \| 0.137 \| 270 \| 195 \| 8 \| 231 \| 468 \| 697 \| 960 \| Uncharacterized protein \| Uncharacterized protein; afdb-uniprot50 \| AF-A0A2D6E3Y1-F1-MODEL\_V4 \| 1.0 \| 8.304e-14 \| 470 \| 0.141 \| 269 \| 200 \| 9 \| 225 \| 468 \| 787 \| 1049 \| Phage\_sheath\_1 domain-containing protein \| Phage\_sheath\_1 domain-containing protein; afdb-uniprot50 \| AF-A0A1C5DEJ3-F1-MODEL\_V4 \| 1.0 \| 4.767e-19 \| 469 \| 0.194 \| 489 \| 263 \| 22 \| 71 \| 467 \| 3 \| 452 \| Uncharacterized protein \| Uncharacterized protein; afdb-uniprot50 \| AF-A0A401XA82-F1-MODEL\_V4 \| 1.0 \| 2.2e-17 \| 469 \| 0.144 \| 545 \| 310 \| 26 \| 7 \| 468 \| 15 \| 485 \| Phage tail sheath protein \| Phage tail sheath protein; afdb-uniprot50 \| AF-A0A382AQG4-F1-MODEL\_V4 \| 1.0 \| 4.687e-14 \| 469 \| 0.205 \| 248 \| 184 \| 9 \| 229 \| 466 \| 269 \| 513 \| Uncharacterized protein \| Uncharacterized protein; afdb-uniprot50 \| AF-A0A1Z8SSC5-F1-MODEL\_V4 \| 1.0 \| 2.462e-13 \| 469 \| 0.162 \| 258 \| 191 \| 8 \| 231 \| 467 \| 634 \| 887 \| Phage\_sheath\_1C domain-containing protein \| Phage\_sheath\_1C domain-containing protein; afdb-uniprot50 \| AF-A0A2V9GXL0-F1-MODEL\_V4 \| 1.0 \| 5.266e-18 \| 468 \| 0.143 \| 797 \| 312 \| 36 \| 3 \| 468 \| 1 \| 757 \| Phage tail protein \| Phage tail protein; afdb-uniprot50 \| AF-A0A2E8NWC6-F1-MODEL\_V4 \| 1.0 \| 1.258e-14 \| 468 \| 0.208 \| 268 \| 181 \| 8 \| 231 \| 468 \| 794 \| 1060 \| Phage\_sheath\_1C domain-containing protein \| Phage\_sheath\_1C domain-containing protein; afdb-uniprot50 \| AF-X0TSA3-F1-MODEL\_V4 \| 1.0 \| 1.044e-13 \| 467 \| 0.176 \| 272 \| 192 \| 12 \| 228 \| 474 \| 107 \| 371 \| Uncharacterized protein \| Uncharacterized protein; afdb-uniprot50 \| AF-A0A7C5MPJ1-F1-MODEL\_V4 \| 1.0 \| 9.469e-19 \| 467 \| 0.154 \| 616 \| 312 \| 26 \| 3 \| 466 \| 2 \| 560 \| Phage tail sheath protein \| Phage tail sheath protein; afdb-uniprot50 \| AF-A0A0F9MLP2-F1-MODEL\_V4 \| 1.0 \| 1.471e-13 \| 467 \| 0.149 \| 247 \| 196 \| 7 \| 231 \| 468 \| 343 \| 584 \| Phage\_sheath\_1 domain-containing protein \| Phage\_sheath\_1 domain-containing protein; afdb-uniprot50 \| AF-A0A7K4MRN1-F1-MODEL\_V4 \| 1.0 \| 3.47e-13 \| 466 \| 0.159 \| 269 \| 198 \| 12 \| 227 \| 474 \| 241 \| 502 \| Phage tail sheath subtilisin-like domain-containing protein \| Phage tail sheath subtilisin-like domain-containing protein; afdb-uniprot50 \| AF-A0A1Z9M029-F1-MODEL\_V4 \| 1.0 \| 3.521e-14 \| 466 \| 0.187 \| 278 \| 192 \| 10 \| 231 \| 475 \| 868 \| 1144 \| Phage\_sheath\_1C domain-containing protein \| Phage\_sheath\_1C domain-containing protein; afdb-uniprot50 \| AF-A0A497PR65-F1-MODEL\_V4 \| 1.0 \| 1.221e-12 \| 465 \| 0.194 \| 237 \| 169 \| 7 \| 247 \| 467 \| 13 \| 243 \| Phage\_sheath\_1 domain-containing protein \| Phage\_sheath\_1 domain-containing protein; afdb-uniprot50 \| AF-A0A2D4QWX6-F1-MODEL\_V4 \| 1.0 \| 6.252e-18 \| 465 \| 0.16 \| 609 \| 342 \| 33 \| 1 \| 474 \| 4 \| 577 \| Uncharacterized protein \| Uncharacterized protein; afdb-uniprot50 \| AF-A0A538QL19-F1-MODEL\_V4 \| 1.0 \| 1.496e-18 \| 465 \| 0.131 \| 847 \| 325 \| 41 \| 1 \| 468 \| 1 \| 815 \| Phage tail sheath family protein \| Phage tail sheath family protein; afdb-uniprot50 \| AF-A0A3S4W452-F1-MODEL\_V4 \| 1.0 \| 4.243e-15 \| 463 \| 0.198 \| 352 \| 159 \| 7 \| 71 \| 391 \| 15 \| 274 \| Major tail sheath protein \| Major tail sheath protein; afdb-uniprot50 \| AF-A0A0N7JW40-F1-MODEL\_V4 \| 1.0 \| 1.799e-15 \| 463 \| 0.145 \| 515 \| 282 \| 24 \| 1 \| 457 \| 6 \| 420 \| Phage tail sheath protein FI \| Phage tail sheath protein FI; afdb-uniprot50 \| AF-A0A831UJ70-F1-MODEL\_V4 \| 1.0 \| 1.678e-18 \| 463 \| 0.163 \| 662 \| 322 \| 39 \| 5 \| 465 \| 3 \| 633 \| Phage tail sheath family protein \| Phage tail sheath family protein; afdb-uniprot50 \| AF-A0A2D7XGW4-F1-MODEL\_V4 \| 1.0 \| 6.995e-14 \| 463 \| 0.166 \| 277 \| 193 \| 10 \| 229 \| 473 \| 381 \| 651 \| Uncharacterized protein \| Uncharacterized protein; afdb-uniprot50 \| AF-A0A5C7XHQ9-F1-MODEL\_V4 \| 1.0 \| 1.392e-17 \| 463 \| 0.129 \| 885 \| 336 \| 39 \| 3 \| 468 \| 1 \| 869 \| Uncharacterized protein \| Uncharacterized protein; afdb-uniprot50 \| AF-A0A1X7MDT6-F1-MODEL\_V4 \| 1.0 \| 1.389e-13 \| 462 \| 0.708 \| 120 \| 35 \| 0 \| 301 \| 420 \| 2 \| 121 \| Phage tail sheath protein \| Phage tail sheath protein; afdb-uniprot50 \| AF-A0A7Y2ZSL3-F1-MODEL\_V4 \| 1.0 \| 1.584e-18 \| 462 \| 0.143 \| 669 \| 334 \| 37 \| 1 \| 473 \| 2 \| 627 \| Uncharacterized protein \| Uncharacterized protein; afdb-uniprot50 \| AF-A0A2D7M0S1-F1-MODEL\_V4 \| 1.0 \| 9.879e-18 \| 462 \| 0.135 \| 714 \| 318 \| 25 \| 1 \| 467 \| 1 \| 661 \| Uncharacterized protein \| Uncharacterized protein; afdb-uniprot50 \| AF-J0UZD7-F1-MODEL\_V4 \| 1.0 \| 2.074e-13 \| 462 \| 0.15 \| 253 \| 191 \| 10 \| 233 \| 468 \| 424 \| 669 \| Phage tail sheath protein FI \| Phage tail sheath protein FI; afdb-uniprot50 \| AF-A0A848UQ52-F1-MODEL\_V4 \| 1.0 \| 2.233e-18 \| 461 \| 0.14 \| 614 \| 330 \| 32 \| 1 \| 467 \| 1 \| 563 \| Uncharacterized protein \| Uncharacterized protein; afdb-uniprot50 \| AF-A0A356TQ48-F1-MODEL\_V4 \| 1.0 \| 6.906e-17 \| 461 \| 0.132 \| 769 \| 323 \| 32 \| 5 \| 474 \| 2 \| 725 \| Phage tail protein \| Phage tail protein; afdb-uniprot50 \| AF-A0A524BCG6-F1-MODEL\_V4 \| 1.0 \| 3.89e-13 \| 460 \| 0.169 \| 254 \| 186 \| 8 \| 230 \| 468 \| 282 \| 525 \| Uncharacterized protein \| Uncharacterized protein; afdb-uniprot50 \| AF-A0A356KTV5-F1-MODEL\_V4 \| 1.0 \| 2.541e-19 \| 460 \| 0.147 \| 588 \| 279 \| 25 \| 1 \| 424 \| 1 \| 529 \| Uncharacterized protein \| Uncharacterized protein; afdb-uniprot50 \| AF-A0A5J4SH86-F1-MODEL\_V4 \| 1.0 \| 9.858e-14 \| 460 \| 0.142 \| 260 \| 192 \| 10 \| 231 \| 468 \| 783 \| 1033 \| Uncharacterized protein \| Uncharacterized protein; afdb-uniprot50 \| AF-A0A497CSG5-F1-MODEL\_V4 \| 1.0 \| 6.509e-13 \| 459 \| 0.205 \| 219 \| 159 \| 7 \| 265 \| 474 \| 3 \| 215 \| Phage tail protein \| Phage tail protein; afdb-uniprot50 \| AF-A0A2P5K775-F1-MODEL\_V4 \| 1.0 \| 6.805e-16 \| 459 \| 0.146 \| 512 \| 277 \| 28 \| 3 \| 468 \| 2 \| 399 \| Tail sheath protein \| Tail sheath protein; afdb-uniprot50 \| AF-A0A3B8M4T7-F1-MODEL\_V4 \| 1.0 \| 7.532e-19 \| 458 \| 0.183 \| 435 \| 232 \| 20 \| 144 \| 467 \| 12 \| 434 \| Phage tail sheath family protein \| Phage tail sheath family protein; afdb-uniprot50 \| AF-A0A6M3K7K1-F1-MODEL\_V4 \| 1.0 \| 4.127e-17 \| 458 \| 0.12 \| 615 \| 355 \| 35 \| 3 \| 474 \| 2 \| 573 \| Putative tail sheath protein \| Putative tail sheath protein; afdb-uniprot50 \| AF-A0A660SQZ6-F1-MODEL\_V4 \| 1.0 \| 2.074e-13 \| 457 \| 0.146 \| 260 \| 197 \| 10 \| 228 \| 468 \| 88 \| 341 \| Uncharacterized protein \| Uncharacterized protein; afdb-uniprot50 \| AF-Q1H2H8-F1-MODEL\_V4 \| 1.0 \| 1.206e-15 \| 457 \| 0.131 \| 563 \| 280 \| 25 \| 3 \| 468 \| 2 \| 452 \| Phage\_sheath\_1C domain-containing protein \| Phage\_sheath\_1C domain-containing protein; afdb-uniprot50 \| AF-A0A1Z8S9S0-F1-MODEL\_V4 \| 1.0 \| 2.462e-13 \| 456 \| 0.174 \| 270 \| 187 \| 9 \| 231 \| 468 \| 735 \| 1000 \| Phage\_sheath\_1C domain-containing protein \| Phage\_sheath\_1C domain-containing protein; afdb-uniprot50 \| AF-A0A8B2R094-F1-MODEL\_V4 \| 1.0 \| 2.257e-11 \| 455 \| 0.409 \| 127 \| 75 \| 0 \| 348 \| 474 \| 1 \| 127 \| Phage tail protein \| Phage tail protein; afdb-uniprot50 \| AF-A0A1I3LPK1-F1-MODEL\_V4 \| 1.0 \| 1.015e-15 \| 455 \| 0.133 \| 583 \| 258 \| 23 \| 1 \| 468 \| 1 \| 451 \| Uncharacterized protein \| Uncharacterized protein; afdb-uniprot50 \| AF-A0A176RSK4-F1-MODEL\_V4 \| 1.0 \| 1.105e-13 \| 454 \| 0.188 \| 244 \| 180 \| 9 \| 231 \| 466 \| 4 \| 237 \| Phage tail sheath protein \| Phage tail sheath protein; afdb-uniprot50 \| AF-A0A4R2C040-F1-MODEL\_V4 \| 1.0 \| 1.962e-17 \| 454 \| 0.14 \| 684 \| 304 \| 29 \| 2 \| 455 \| 3 \| 632 \| Uncharacterized protein \| Uncharacterized protein; afdb-uniprot50 \| AF-A0A5P3VE82-F1-MODEL\_V4 \| 1.0 \| 8.321e-18 \| 454 \| 0.129 \| 728 \| 322 \| 32 \| 6 \| 468 \| 1 \| 681 \| Phage tail sheath family protein \| Phage tail sheath family protein; afdb-uniprot50 \| AF-A0A2E7N1B0-F1-MODEL\_V4 \| 1.0 \| 7.406e-14 \| 453 \| 0.168 \| 273 \| 199 \| 7 \| 231 \| 475 \| 227 \| 499 \| Phage\_sheath\_1C domain-containing protein \| Phage\_sheath\_1C domain-containing protein; afdb-uniprot50 \| AF-S4XY80-F1-MODEL\_V4 \| 1.0 \| 4.189e-18 \| 453 \| 0.19 \| 702 \| 313 \| 29 \| 5 \| 468 \| 3 \| 686 \| Uncharacterized protein \| Uncharacterized protein; afdb-uniprot50 \| AF-A0A2D7XGK1-F1-MODEL\_V4 \| 1.0 \| 1.849e-13 \| 453 \| 0.151 \| 278 \| 196 \| 9 \| 225 \| 468 \| 833 \| 1104 \| Uncharacterized protein \| Uncharacterized protein; afdb-uniprot50 \| AF-A0A3B8QSR2-F1-MODEL\_V4 \| 1.0 \| 1.65e-13 \| 452 \| 0.143 \| 279 \| 198 \| 9 \| 225 \| 468 \| 108 \| 380 \| Uncharacterized protein \| Uncharacterized protein; afdb-uniprot50 \| AF-A0A6N0ZC76-F1-MODEL\_V4 \| 1.0 \| 1.933e-16 \| 452 \| 0.142 \| 561 \| 325 \| 31 \| 7 \| 467 \| 1 \| 505 \| Phage tail protein \| Phage tail protein; afdb-uniprot50 \| AF-A0A4Q9KLK9-F1-MODEL\_V4 \| 1.0 \| 2.504e-18 \| 452 \| 0.155 \| 680 \| 328 \| 28 \| 1 \| 456 \| 1 \| 657 \| Phage tail sheath family protein \| Phage tail sheath family protein; afdb-uniprot50 \| AF-A0A7J4ULP8-F1-MODEL\_V4 \| 1.0 \| 5.178e-13 \| 451 \| 0.174 \| 264 \| 186 \| 8 \| 231 \| 468 \| 124 \| 381 \| Phage tail protein \| Phage tail protein; afdb-uniprot50 \| AF-A0A842PA51-F1-MODEL\_V4 \| 1.0 \| 2.607e-13 \| 451 \| 0.167 \| 275 \| 192 \| 11 \| 230 \| 473 \| 280 \| 548 \| Phage tail sheath subtilisin-like domain-containing protein \| Phage tail sheath subtilisin-like domain-containing protein; afdb-uniprot50 \| AF-A0A524MJ07-F1-MODEL\_V4 \| 1.0 \| 1.992e-18 \| 451 \| 0.152 \| 623 \| 323 \| 31 \| 1 \| 468 \| 1 \| 573 \| Phage\_sheath\_1C domain-containing protein \| Phage\_sheath\_1C domain-containing protein; afdb-uniprot50 \| AF-A0A1Z9EFV8-F1-MODEL\_V4 \| 1.0 \| 5.805e-13 \| 451 \| 0.168 \| 267 \| 195 \| 11 \| 230 \| 475 \| 663 \| 923 \| Uncharacterized protein \| Uncharacterized protein; afdb-uniprot50 \| AF-A0A2D8CBU3-F1-MODEL\_V4 \| 1.0 \| 3.674e-13 \| 451 \| 0.155 \| 276 \| 195 \| 9 \| 225 \| 468 \| 861 \| 1130 \| Phage\_sheath\_1C domain-containing protein \| Phage\_sheath\_1C domain-containing protein; afdb-uniprot50 \| AF-A0A660RI94-F1-MODEL\_V4 \| 1.0 \| 4.127e-17 \| 450 \| 0.16 \| 510 \| 304 \| 21 \| 3 \| 456 \| 1 \| 442 \| Phage\_sheath\_1C domain-containing protein \| Phage\_sheath\_1C domain-containing protein; afdb-uniprot50 \| AF-A0A2D8ENN3-F1-MODEL\_V4 \| 1.0 \| 3.89e-13 \| 450 \| 0.145 \| 289 \| 196 \| 10 \| 231 \| 474 \| 308 \| 590 \| Uncharacterized protein \| Uncharacterized protein; afdb-uniprot50 \| AF-A0A7Z2GQZ2-F1-MODEL\_V4 \| 1.0 \| 2.078e-17 \| 449 \| 0.193 \| 423 \| 263 \| 27 \| 68 \| 461 \| 135 \| 508 \| Uncharacterized protein \| Uncharacterized protein; afdb-uniprot50 \| AF-A0A2E9U1Y6-F1-MODEL\_V4 \| 1.0 \| 1.962e-17 \| 449 \| 0.125 \| 748 \| 355 \| 34 \| 1 \| 475 \| 1 \| 722 \| Phage tail protein \| Phage tail protein; afdb-uniprot50 \| AF-A0A376ZSE8-F1-MODEL\_V4 \| 1.0 \| 2.291e-12 \| 448 \| 0.344 \| 148 \| 92 \| 3 \| 252 \| 398 \| 7 \| 150 \| Major tail sheath protein FI \| Major tail sheath protein FI; afdb-uniprot50 \| AF-A0A849TMA6-F1-MODEL\_V4 \| 1.0 \| 9.733e-17 \| 448 \| 0.148 \| 558 \| 307 \| 34 \| 7 \| 468 \| 14 \| 499 \| Phage tail protein \| Phage tail protein; afdb-uniprot50 \| AF-A0A353BET0-F1-MODEL\_V4 \| 1.0 \| 1.075e-15 \| 448 \| 0.17 \| 423 \| 210 \| 16 \| 73 \| 468 \| 424 \| 732 \| Uncharacterized protein \| Uncharacterized protein; afdb-uniprot50 \| AF-A0A511H440-F1-MODEL\_V4 \| 1.0 \| 7.422e-18 \| 448 \| 0.158 \| 794 \| 290 \| 36 \| 25 \| 468 \| 1 \| 765 \| Uncharacterized protein \| Uncharacterized protein; afdb-uniprot50 \| AF-A0A6C9Q6A8-F1-MODEL\_V4 \| 1.0 \| 8.906e-11 \| 447 \| 0.41 \| 117 \| 69 \| 0 \| 358 \| 474 \| 3 \| 119 \| Phage tail protein \| Phage tail protein; afdb-uniprot50 \| AF-A0A3N5HQ21-F1-MODEL\_V4 \| 1.0 \| 4.015e-19 \| 447 \| 0.177 \| 534 \| 270 \| 26 \| 3 \| 420 \| 1 \| 481 \| Uncharacterized protein \| Uncharacterized protein; afdb-uniprot50 \| AF-A0A519CSN0-F1-MODEL\_V4 \| 1.0 \| 1.881e-18 \| 447 \| 0.127 \| 907 \| 342 \| 35 \| 3 \| 468 \| 2 \| 899 \| Phage\_sheath\_1C domain-containing protein \| Phage\_sheath\_1C domain-containing protein; afdb-uniprot50 \| AF-A0A3D1A0D5-F1-MODEL\_V4 \| 1.0 \| 2.196e-13 \| 447 \| 0.154 \| 278 \| 195 \| 9 \| 225 \| 468 \| 751 \| 1022 \| Uncharacterized protein \| Uncharacterized protein; afdb-uniprot50 \| AF-A0A2E6A8L5-F1-MODEL\_V4 \| 1.0 \| 2.462e-13 \| 447 \| 0.156 \| 275 \| 195 \| 9 \| 225 \| 468 \| 766 \| 1034 \| Uncharacterized protein \| Uncharacterized protein; afdb-uniprot50 \| AF-A0A3C2A712-F1-MODEL\_V4 \| 1.0 \| 7.406e-14 \| 446 \| 0.178 \| 241 \| 177 \| 10 \| 230 \| 466 \| 84 \| 307 \| Phage tail protein \| Phage tail protein; afdb-uniprot50 \| AF-L9PQK2-F1-MODEL\_V4 \| 1.0 \| 2.928e-17 \| 446 \| 0.143 \| 647 \| 293 \| 28 \| 19 \| 468 \| 28 \| 610 \| Uncharacterized protein \| Uncharacterized protein; afdb-uniprot50 \| AF-A0A2D5MP23-F1-MODEL\_V4 \| 1.0 \| 1.653e-17 \| 446 \| 0.129 \| 679 \| 347 \| 38 \| 3 \| 474 \| 2 \| 643 \| Uncharacterized protein \| Uncharacterized protein; afdb-uniprot50 \| AF-A0A2T1DKD7-F1-MODEL\_V4 \| 1.0 \| 6.62e-18 \| 445 \| 0.142 \| 610 \| 293 \| 27 \| 3 \| 452 \| 2 \| 541 \| Phage tail sheath protein \| Phage tail sheath protein; afdb-uniprot50 \| AF-A0A5C5ZXM5-F1-MODEL\_V4 \| 1.0 \| 3.529e-18 \| 445 \| 0.144 \| 597 \| 321 \| 25 \| 1 \| 455 \| 1 \| 549 \| Phage tail sheath protein \| Phage tail sheath protein; afdb-uniprot50 \| AF-A0A2D6FIM5-F1-MODEL\_V4 \| 1.0 \| 2.76e-13 \| 444 \| 0.145 \| 274 \| 196 \| 9 \| 227 \| 468 \| 700 \| 967 \| Phage\_sheath\_1 domain-containing protein \| Phage\_sheath\_1 domain-containing protein; afdb-uniprot50 \| AF-A0A5D0S8K9-F1-MODEL\_V4 \| 1.0 \| 1.453e-16 \| 443 \| 0.202 \| 385 \| 226 \| 16 \| 114 \| 468 \| 423 \| 756 \| Phage tail sheath family protein \| Phage tail sheath family protein; afdb-uniprot50 \| AF-A0A1X7MDW3-F1-MODEL\_V4 \| 1.0 \| 1.428e-11 \| 442 \| 0.705 \| 95 \| 28 \| 0 \| 379 \| 473 \| 2 \| 96 \| Phage-related contractile tail sheath protein \| Phage-related contractile tail sheath protein; afdb-uniprot50 \| AF-A0A1M4WNH5-F1-MODEL\_V4 \| 1.0 \| 7.743e-17 \| 442 \| 0.14 \| 570 \| 320 \| 27 \| 1 \| 468 \| 1 \| 502 \| Phage tail sheath protein \| Phage tail sheath protein; afdb-uniprot50 \| AF-A0A2D5XZB8-F1-MODEL\_V4 \| 1.0 \| 2.364e-18 \| 442 \| 0.153 \| 643 \| 351 \| 38 \| 3 \| 468 \| 2 \| 627 \| Phage\_sheath\_1C domain-containing protein \| Phage\_sheath\_1C domain-containing protein; afdb-uniprot50 \| AF-A0A2D5YHB3-F1-MODEL\_V4 \| 1.0 \| 1.826e-16 \| 442 \| 0.11 \| 766 \| 322 \| 32 \| 5 \| 474 \| 2 \| 703 \| Uncharacterized protein \| Uncharacterized protein; afdb-uniprot50 \| AF-A0A5C7LNH5-F1-MODEL\_V4 \| 1.0 \| 6.509e-13 \| 442 \| 0.13 \| 269 \| 203 \| 9 \| 225 \| 468 \| 730 \| 992 \| Phage\_sheath\_1 domain-containing protein \| Phage\_sheath\_1 domain-containing protein; afdb-uniprot50 \| AF-A0A376X2C2-F1-MODEL\_V4 \| 1.0 \| 2.257e-11 \| 441 \| 0.351 \| 148 \| 91 \| 2 \| 330 \| 472 \| 1 \| 148 \| Phage-related contractile tail sheath protein \| Phage-related contractile tail sheath protein; afdb-uniprot50 \| AF-A0A1V5Z4B1-F1-MODEL\_V4 \| 1.0 \| 1.046e-17 \| 441 \| 0.149 \| 496 \| 289 \| 20 \| 46 \| 453 \| 1 \| 451 \| Phage tail sheath protein \| Phage tail sheath protein; afdb-uniprot50 \| AF-G4KQ75-F1-MODEL\_V4 \| 1.0 \| 3.283e-17 \| 441 \| 0.172 \| 556 \| 297 \| 31 \| 5 \| 474 \| 2 \| 480 \| Putative tail sheath protein \| Putative tail sheath protein; afdb-uniprot50 \| AF-A0A7Z1V658-F1-MODEL\_V4 \| 1.0 \| 2.928e-17 \| 441 \| 0.136 \| 535 \| 307 \| 31 \| 19 \| 468 \| 28 \| 492 \| Phage tail protein \| Phage tail protein; afdb-uniprot50 \| AF-A0A2A4ZDU8-F1-MODEL\_V4 \| 1.0 \| 8.199e-17 \| 441 \| 0.133 \| 673 \| 347 \| 35 \| 3 \| 475 \| 2 \| 638 \| Uncharacterized protein \| Uncharacterized protein; afdb-uniprot50 \| AF-A0A2E6PTS5-F1-MODEL\_V4 \| 1.0 \| 4.361e-13 \| 441 \| 0.141 \| 275 \| 196 \| 10 \| 228 \| 468 \| 708 \| 976 \| Uncharacterized protein \| Uncharacterized protein; afdb-uniprot50 \| AF-A0A2D6X5Z5-F1-MODEL\_V4 \| 1.0 \| 2.922e-13 \| 441 \| 0.161 \| 278 \| 192 \| 8 \| 225 \| 467 \| 726 \| 997 \| Uncharacterized protein \| Uncharacterized protein; afdb-uniprot50 \| AF-A0A5C7LM99-F1-MODEL\_V4 \| 1.0 \| 5.483e-13 \| 441 \| 0.133 \| 270 \| 202 \| 11 \| 225 \| 468 \| 748 \| 1011 \| Uncharacterized protein \| Uncharacterized protein; afdb-uniprot50 \| AF-A0A6C7KTB5-F1-MODEL\_V4 \| 1.0 \| 1.428e-11 \| 438 \| 0.279 \| 186 \| 129 \| 3 \| 291 \| 474 \| 8 \| 190 \| Phage tail sheath family protein \| Phage tail sheath family protein; afdb-uniprot50 \| AF-A0A382KT14-F1-MODEL\_V4 \| 1.0 \| 1.153e-12 \| 438 \| 0.18 \| 260 \| 186 \| 10 \| 237 \| 475 \| 3 \| 256 \| Uncharacterized protein \| Uncharacterized protein; afdb-uniprot50 \| AF-A0A011RAT7-F1-MODEL\_V4 \| 1.0 \| 5.413e-16 \| 438 \| 0.138 \| 570 \| 321 \| 32 \| 7 \| 467 \| 1 \| 509 \| Phage tail sheath protein \| Phage tail sheath protein; afdb-uniprot50 \| AF-A0A099PD65-F1-MODEL\_V4 \| 1.0 \| 1.933e-16 \| 438 \| 0.153 \| 417 \| 264 \| 19 \| 119 \| 468 \| 234 \| 628 \| Phage\_sheath\_1C domain-containing protein \| Phage\_sheath\_1C domain-containing protein; afdb-uniprot50 \| AF-A0A1Z9F154-F1-MODEL\_V4 \| 1.0 \| 6.16e-17 \| 438 \| 0.127 \| 829 \| 347 \| 34 \| 3 \| 474 \| 2 \| 810 \| Uncharacterized protein \| Uncharacterized protein; afdb-uniprot50 \| AF-A0A563CW44-F1-MODEL\_V4 \| 1.0 \| 2.607e-13 \| 438 \| 0.165 \| 260 \| 186 \| 12 \| 231 \| 468 \| 910 \| 1160 \| Phage\_sheath\_1 domain-containing protein \| Phage\_sheath\_1 domain-containing protein; afdb-uniprot50 \| AF-A0A0Q9M0T6-F1-MODEL\_V4 \| 1.0 \| 2.2e-17 \| 437 \| 0.144 \| 685 \| 320 \| 31 \| 1 \| 455 \| 1 \| 649 \| Uncharacterized protein \| Uncharacterized protein; afdb-uniprot50 \| AF-A0A810LBR4-F1-MODEL\_V4 \| 1.0 \| 1.156e-16 \| 437 \| 0.126 \| 932 \| 316 \| 37 \| 3 \| 467 \| 1 \| 900 \| Uncharacterized protein \| Uncharacterized protein; afdb-uniprot50 \| AF-A0A5N9VYH6-F1-MODEL\_V4 \| 1.0 \| 6.057e-12 \| 436 \| 0.429 \| 135 \| 77 \| 0 \| 303 \| 437 \| 1 \| 135 \| Phage tail protein \| Phage tail protein; afdb-uniprot50 \| AF-A0A5C7LEF6-F1-MODEL\_V4 \| 1.0 \| 1.153e-12 \| 436 \| 0.13 \| 269 \| 203 \| 8 \| 225 \| 468 \| 740 \| 1002 \| Phage\_sheath\_1 domain-containing protein \| Phage\_sheath\_1 domain-containing protein; afdb-uniprot50 \| AF-A0A800EES6-F1-MODEL\_V4 \| 1.0 \| 5.483e-13 \| 435 \| 0.133 \| 278 \| 201 \| 10 \| 225 \| 468 \| 79 \| 350 \| Phage\_sheath\_1C domain-containing protein \| Phage\_sheath\_1C domain-containing protein; afdb-uniprot50 \| AF-A0A842WMT0-F1-MODEL\_V4 \| 1.0 \| 1.674e-14 \| 434 \| 0.138 \| 398 \| 221 \| 15 \| 95 \| 468 \| 633 \| 932 \| Uncharacterized protein \| Uncharacterized protein; afdb-uniprot50 \| AF-A0A2D6E5M5-F1-MODEL\_V4 \| 1.0 \| 1.65e-13 \| 434 \| 0.153 \| 281 \| 196 \| 11 \| 225 \| 468 \| 750 \| 1025 \| Uncharacterized protein \| Uncharacterized protein; afdb-uniprot50 \| AF-A0A521CKY5-F1-MODEL\_V4 \| 1.0 \| 1.224e-16 \| 433 \| 0.136 \| 556 \| 309 \| 26 \| 9 \| 468 \| 2 \| 482 \| Uncharacterized protein \| Uncharacterized protein; afdb-uniprot50 \| AF-A0A2D5F6D1-F1-MODEL\_V4 \| 1.0 \| 1.372e-16 \| 433 \| 0.11 \| 752 \| 321 \| 33 \| 1 \| 468 \| 2 \| 689 \| Uncharacterized protein \| Uncharacterized protein; afdb-uniprot50 \| AF-A0A3D5RSJ9-F1-MODEL\_V4 \| 1.0 \| 3.277e-13 \| 433 \| 0.127 \| 275 \| 200 \| 10 \| 228 \| 468 \| 599 \| 867 \| Uncharacterized protein \| Uncharacterized protein; afdb-uniprot50 \| AF-A0A3N5GCM3-F1-MODEL\_V4 \| 1.0 \| 7.298e-13 \| 432 \| 0.194 \| 221 \| 151 \| 8 \| 259 \| 466 \| 11 \| 217 \| Phage tail sheath protein \| Phage tail sheath protein; afdb-uniprot50 \| AF-A0A6I5RGG6-F1-MODEL\_V4 \| 1.0 \| 4.56e-16 \| 432 \| 0.15 \| 586 \| 316 \| 30 \| 1 \| 468 \| 1 \| 522 \| Uncharacterized protein \| Uncharacterized protein; afdb-uniprot50 \| AF-A0A7J4UP87-F1-MODEL\_V4 \| 1.0 \| 5.817e-17 \| 432 \| 0.141 \| 698 \| 348 \| 33 \| 3 \| 474 \| 2 \| 673 \| Uncharacterized protein \| Uncharacterized protein; afdb-uniprot50 \| AF-A0A2K3J0B0-F1-MODEL\_V4 \| 1.0 \| 8.681e-17 \| 432 \| 0.134 \| 704 \| 348 \| 36 \| 3 \| 473 \| 4 \| 678 \| Uncharacterized protein \| Uncharacterized protein; afdb-uniprot50 \| AF-A0A3D4UU50-F1-MODEL\_V4 \| 1.0 \| 5.494e-17 \| 431 \| 0.128 \| 750 \| 335 \| 38 \| 1 \| 468 \| 1 \| 713 \| Uncharacterized protein \| Uncharacterized protein; afdb-uniprot50 \| AF-A0A1V5KBQ7-F1-MODEL\_V4 \| 1.0 \| 1.877e-14 \| 431 \| 0.159 \| 401 \| 209 \| 14 \| 94 \| 466 \| 583 \| 883 \| Phage tail sheath protein \| Phage tail sheath protein; afdb-uniprot50 \| AF-A0A0Q8V0M7-F1-MODEL\_V4 \| 1.0 \| 8.681e-17 \| 429 \| 0.121 \| 747 \| 337 \| 32 \| 1 \| 468 \| 1 \| 706 \| Phage\_sheath\_1C domain-containing protein \| Phage\_sheath\_1C domain-containing protein; afdb-uniprot50 \| AF-A0A101JEY1-F1-MODEL\_V4 \| 1.0 \| 1.826e-16 \| 429 \| 0.127 \| 745 \| 344 \| 33 \| 5 \| 468 \| 2 \| 721 \| Phage\_sheath\_1 domain-containing protein \| Phage\_sheath\_1 domain-containing protein; afdb-uniprot50 \| AF-A0A696JDI9-F1-MODEL\_V4 \| 1.0 \| 4.234e-11 \| 428 \| 0.255 \| 168 \| 122 \| 2 \| 307 \| 474 \| 2 \| 166 \| Phage tail sheath family protein \| Phage tail sheath family protein; afdb-uniprot50 \| AF-A0A0F0HL05-F1-MODEL\_V4 \| 1.0 \| 9.174e-13 \| 428 \| 0.195 \| 266 \| 177 \| 9 \| 233 \| 467 \| 17 \| 276 \| Uncharacterized protein \| Uncharacterized protein; afdb-uniprot50 \| AF-A0A7X7WC07-F1-MODEL\_V4 \| 1.0 \| 1.296e-16 \| 428 \| 0.165 \| 526 \| 303 \| 25 \| 1 \| 468 \| 3 \| 450 \| Phage\_sheath\_1C domain-containing protein \| Phage\_sheath\_1C domain-containing protein; afdb-uniprot50 \| AF-K8GN70-F1-MODEL\_V4 \| 1.0 \| 2.568e-12 \| 428 \| 0.213 \| 253 \| 174 \| 9 \| 231 \| 468 \| 686 \| 928 \| Phage tail sheath protein FI \| Phage tail sheath protein FI; afdb-uniprot50 \| AF-A0A2D6WY17-F1-MODEL\_V4 \| 1.0 \| 1.352e-15 \| 427 \| 0.114 \| 828 \| 325 \| 30 \| 3 \| 475 \| 2 \| 775 \| Uncharacterized protein \| Uncharacterized protein; afdb-uniprot50 \| AF-A0A2D6MI23-F1-MODEL\_V4 \| 1.0 \| 8.182e-13 \| 427 \| 0.147 \| 278 \| 197 \| 10 \| 225 \| 468 \| 773 \| 1044 \| Uncharacterized protein \| Uncharacterized protein; afdb-uniprot50 \| AF-A0A6P2CYQ9-F1-MODEL\_V4 \| 1.0 \| 2.078e-17 \| 426 \| 0.14 \| 610 \| 329 \| 31 \| 3 \| 459 \| 2 \| 568 \| Phage\_sheath\_1C domain-containing protein \| Phage\_sheath\_1C domain-containing protein; afdb-uniprot50 \| AF-A0A1A9F3T7-F1-MODEL\_V4 \| 1.0 \| 1.296e-16 \| 425 \| 0.146 \| 790 \| 314 \| 32 \| 3 \| 463 \| 1 \| 758 \| Uncharacterized protein \| Uncharacterized protein; afdb-uniprot50 \| AF-A0A833DC75-F1-MODEL\_V4 \| 1.0 \| 6.906e-17 \| 424 \| 0.128 \| 683 \| 352 \| 34 \| 1 \| 474 \| 4 \| 651 \| Uncharacterized protein \| Uncharacterized protein; afdb-uniprot50 \| AF-A0A800EC41-F1-MODEL\_V4 \| 1.0 \| 2.291e-12 \| 423 \| 0.14 \| 277 \| 199 \| 9 \| 225 \| 468 \| 132 \| 402 \| Phage\_sheath\_1C domain-containing protein \| Phage\_sheath\_1C domain-containing protein; afdb-uniprot50 \| AF-A0A6W2FV74-F1-MODEL\_V4 \| 1.0 \| 8.536e-12 \| 422 \| 0.225 \| 270 \| 112 \| 5 \| 92 \| 359 \| 9 \| 183 \| Phage tail sheath family protein \| Phage tail sheath family protein; afdb-uniprot50 \| AF-D4EA08-F1-MODEL\_V4 \| 1.0 \| 1.699e-15 \| 422 \| 0.133 \| 639 \| 273 \| 24 \| 1 \| 468 \| 1 \| 529 \| Phage\_sheath\_1C domain-containing protein \| Phage\_sheath\_1C domain-containing protein; afdb-uniprot50 \| AF-A0A2D6E5Y4-F1-MODEL\_V4 \| 1.0 \| 5.178e-13 \| 422 \| 0.164 \| 274 \| 187 \| 10 \| 231 \| 468 \| 762 \| 1029 \| Phage\_sheath\_1 domain-containing protein \| Phage\_sheath\_1 domain-containing protein; afdb-uniprot50 \| AF-A0A661IMA0-F1-MODEL\_V4 \| 1.0 \| 7.406e-14 \| 421 \| 0.117 \| 475 \| 283 \| 23 \| 6 \| 468 \| 1 \| 351 \| Uncharacterized protein \| Uncharacterized protein; afdb-uniprot50 \| AF-A0A2D6E608-F1-MODEL\_V4 \| 1.0 \| 4.89e-13 \| 421 \| 0.147 \| 284 \| 198 \| 13 \| 230 \| 475 \| 616 \| 893 \| Phage\_sheath\_1C domain-containing protein \| Phage\_sheath\_1C domain-containing protein; afdb-uniprot50 \| AF-A0A3C0GEV4-F1-MODEL\_V4 \| 1.0 \| 1.822e-12 \| 421 \| 0.145 \| 274 \| 192 \| 10 \| 225 \| 462 \| 640 \| 907 \| Phage\_sheath\_1C domain-containing protein \| Phage\_sheath\_1C domain-containing protein; afdb-uniprot50 \| AF-A0A1N7K568-F1-MODEL\_V4 \| 1.0 \| 7.959e-15 \| 420 \| 0.133 \| 563 \| 280 \| 24 \| 1 \| 466 \| 16 \| 467 \| Uncharacterized protein \| Uncharacterized protein; afdb-uniprot50 \| AF-C7BIE9-F1-MODEL\_V4 \| 1.0 \| 8.554e-16 \| 420 \| 0.127 \| 614 \| 266 \| 23 \| 1 \| 468 \| 1 \| 490 \| Uncharacterized protein \| Uncharacterized protein; afdb-uniprot50 \| AF-A0A7C1VR73-F1-MODEL\_V4 \| 1.0 \| 4.56e-16 \| 420 \| 0.129 \| 631 \| 336 \| 37 \| 1 \| 468 \| 1 \| 580 \| Phage\_sheath\_1 domain-containing protein \| Phage\_sheath\_1 domain-containing protein; afdb-uniprot50 \| AF-A0A2H3KZU4-F1-MODEL\_V4 \| 1.0 \| 1.724e-16 \| 420 \| 0.153 \| 503 \| 292 \| 24 \| 46 \| 467 \| 167 \| 616 \| Uncharacterized protein \| Uncharacterized protein; afdb-uniprot50 \| AF-A0A5C7JJ84-F1-MODEL\_V4 \| 1.0 \| 2.43e-16 \| 420 \| 0.129 \| 712 \| 320 \| 31 \| 1 \| 468 \| 5 \| 660 \| Phage tail sheath family protein \| Phage tail sheath family protein; afdb-uniprot50 \| AF-A0A3M2DCM6-F1-MODEL\_V4 \| 1.0 \| 1.075e-15 \| 419 \| 0.127 \| 541 \| 331 \| 25 \| 7 \| 460 \| 1 \| 487 \| Phage tail sheath protein \| Phage tail sheath protein; afdb-uniprot50 \| AF-A0A5C7LE80-F1-MODEL\_V4 \| 1.0 \| 2.568e-12 \| 419 \| 0.129 \| 270 \| 203 \| 9 \| 225 \| 468 \| 456 \| 719 \| Uncharacterized protein \| Uncharacterized protein; afdb-uniprot50 \| AF-A0A3M2VUF5-F1-MODEL\_V4 \| 1.0 \| 2.531e-11 \| 418 \| 0.284 \| 165 \| 111 \| 3 \| 312 \| 474 \| 1 \| 160 \| Putative Phage tail sheath protein \| Putative Phage tail sheath protein; afdb-uniprot50 \| AF-A0A350YB92-F1-MODEL\_V4 \| 1.0 \| 9.733e-17 \| 418 \| 0.148 \| 525 \| 301 \| 22 \| 3 \| 452 \| 2 \| 455 \| Phage tail protein \| Phage tail protein; afdb-uniprot50 \| AF-A0A5R8TFK6-F1-MODEL\_V4 \| 1.0 \| 6.704e-15 \| 418 \| 0.137 \| 587 \| 267 \| 26 \| 5 \| 468 \| 2 \| 471 \| Phage tail sheath family protein \| Phage tail sheath family protein; afdb-uniprot50 \| AF-A0A381RCS0-F1-MODEL\_V4 \| 1.0 \| 5.732e-16 \| 418 \| 0.121 \| 776 \| 357 \| 42 \| 1 \| 475 \| 1 \| 752 \| Uncharacterized protein \| Uncharacterized protein; afdb-uniprot50 \| AF-A0A094J1V3-F1-MODEL\_V4 \| 1.0 \| 8.554e-16 \| 417 \| 0.12 \| 800 \| 323 \| 31 \| 3 \| 474 \| 2 \| 748 \| Phage tail sheath protein \| Phage tail sheath protein; afdb-uniprot50 \| AF-A0A6N8K8V9-F1-MODEL\_V4 \| 1.0 \| 1.105e-13 \| 416 \| 0.268 \| 287 \| 108 \| 6 \| 43 \| 326 \| 127 \| 314 \| Phage tail protein \| Phage tail protein; afdb-uniprot50 \| AF-A0A524QSS2-F1-MODEL\_V4 \| 1.0 \| 1.089e-12 \| 416 \| 0.181 \| 265 \| 179 \| 12 \| 232 \| 468 \| 208 \| 462 \| Uncharacterized protein \| Uncharacterized protein; afdb-uniprot50 \| AF-A0A2D6F2P0-F1-MODEL\_V4 \| 1.0 \| 1.153e-12 \| 416 \| 0.16 \| 268 \| 195 \| 9 \| 225 \| 468 \| 337 \| 598 \| Phage\_sheath\_1C domain-containing protein \| Phage\_sheath\_1C domain-containing protein; afdb-uniprot50 \| AF-A0A7C3HFY0-F1-MODEL\_V4 \| 1.0 \| 1.139e-15 \| 415 \| 0.144 \| 588 \| 328 \| 27 \| 2 \| 467 \| 51 \| 585 \| Uncharacterized protein \| Uncharacterized protein; afdb-uniprot50 \| AF-A0A2A5GVC5-F1-MODEL\_V4 \| 1.0 \| 2.43e-16 \| 415 \| 0.125 \| 692 \| 352 \| 36 \| 3 \| 473 \| 4 \| 663 \| Uncharacterized protein \| Uncharacterized protein; afdb-uniprot50 \| AF-A0A1L6L565-F1-MODEL\_V4 \| 1.0 \| 1.853e-17 \| 415 \| 0.149 \| 462 \| 277 \| 25 \| 100 \| 468 \| 493 \| 931 \| Phage tail sheath protein FI \| Phage tail sheath protein FI; afdb-uniprot50 \| AF-R6PS34-F1-MODEL\_V4 \| 1.0 \| 1.45e-12 \| 414 \| 0.21 \| 252 \| 173 \| 9 \| 245 \| 475 \| 15 \| 261 \| Uncharacterized protein \| Uncharacterized protein; afdb-uniprot50 \| AF-A0A2E6W8A2-F1-MODEL\_V4 \| 1.0 \| 4.058e-12 \| 413 \| 0.133 \| 278 \| 201 \| 8 \| 225 \| 468 \| 363 \| 634 \| Uncharacterized protein \| Uncharacterized protein; afdb-uniprot50 \| AF-A0A1Z8RYI1-F1-MODEL\_V4 \| 1.0 \| 3.228e-12 \| 411 \| 0.144 \| 270 \| 199 \| 11 \| 225 \| 468 \| 400 \| 663 \| Phage\_sheath\_1 domain-containing protein \| Phage\_sheath\_1 domain-containing protein; afdb-uniprot50 \| AF-A0A1Q7BLH3-F1-MODEL\_V4 \| 1.0 \| 5.494e-17 \| 410 \| 0.172 \| 550 \| 282 \| 32 \| 3 \| 455 \| 2 \| 475 \| Phage\_sheath\_1 domain-containing protein \| Phage\_sheath\_1 domain-containing protein; afdb-uniprot50 \| AF-A0A8A6KD18-F1-MODEL\_V4 \| 1.0 \| 4.56e-16 \| 410 \| 0.108 \| 781 \| 344 \| 33 \| 5 \| 468 \| 2 \| 746 \| Phage tail sheath family protein \| Phage tail sheath family protein; afdb-uniprot50 \| AF-A0A849QUP3-F1-MODEL\_V4 \| 1.0 \| 4.9e-17 \| 409 \| 0.163 \| 490 \| 259 \| 25 \| 3 \| 398 \| 2 \| 434 \| Phage tail sheath family protein \| Phage tail sheath family protein; afdb-uniprot50 \| AF-A0A2D8FVW0-F1-MODEL\_V4 \| 1.0 \| 3.419e-12 \| 409 \| 0.136 \| 279 \| 200 \| 8 \| 225 \| 468 \| 579 \| 851 \| Phage\_sheath\_1C domain-containing protein \| Phage\_sheath\_1C domain-containing protein; afdb-uniprot50 \| AF-A0A5F1HTD7-F1-MODEL\_V4 \| 1.0 \| 4.618e-13 \| 408 \| 0.19 \| 315 \| 162 \| 12 \| 231 \| 468 \| 61 \| 359 \| Phage\_sheath\_1C domain-containing protein \| Phage\_sheath\_1C domain-containing protein; afdb-uniprot50 \| AF-A0A3M2BQ74-F1-MODEL\_V4 \| 1.0 \| 5.72e-12 \| 408 \| 0.141 \| 276 \| 194 \| 9 \| 231 \| 467 \| 508 \| 779 \| Uncharacterized protein \| Uncharacterized protein; afdb-uniprot50 \| AF-A0A382PEL7-F1-MODEL\_V4 \| 1.0 \| 4.483e-11 \| 407 \| 0.169 \| 230 \| 175 \| 7 \| 255 \| 474 \| 8 \| 231 \| Uncharacterized protein \| Uncharacterized protein; afdb-uniprot50 \| AF-A0A853I305-F1-MODEL\_V4 \| 1.0 \| 4.426e-14 \| 407 \| 0.121 \| 594 \| 295 \| 25 \| 3 \| 468 \| 2 \| 496 \| Phage tail sheath family protein \| Phage tail sheath family protein; afdb-uniprot50 \| AF-A0A1Z8THJ1-F1-MODEL\_V4 \| 1.0 \| 5.72e-12 \| 407 \| 0.168 \| 273 \| 192 \| 11 \| 231 \| 474 \| 289 \| 555 \| Uncharacterized protein \| Uncharacterized protein; afdb-uniprot50 \| AF-A0A2E7TM91-F1-MODEL\_V4 \| 1.0 \| 5.402e-12 \| 407 \| 0.184 \| 265 \| 191 \| 10 \| 230 \| 474 \| 410 \| 669 \| Uncharacterized protein \| Uncharacterized protein; afdb-uniprot50 \| AF-A0A2E3KCZ5-F1-MODEL\_V4 \| 1.0 \| 1.629e-16 \| 407 \| 0.124 \| 770 \| 351 \| 31 \| 1 \| 473 \| 1 \| 743 \| Uncharacterized protein \| Uncharacterized protein; afdb-uniprot50 \| AF-A0A3S4VXL9-F1-MODEL\_V4 \| 1.0 \| 8.182e-13 \| 406 \| 0.195 \| 251 \| 115 \| 3 \| 119 \| 367 \| 64 \| 229 \| Major tail sheath protein \| Major tail sheath protein; afdb-uniprot50 \| AF-A0A2D6AJR1-F1-MODEL\_V4 \| 1.0 \| 4.687e-14 \| 406 \| 0.133 \| 523 \| 274 \| 25 \| 5 \| 468 \| 1 \| 403 \| Uncharacterized protein \| Uncharacterized protein; afdb-uniprot50 \| AF-A0A7V8IUR4-F1-MODEL\_V4 \| 1.0 \| 2.612e-17 \| 406 \| 0.155 \| 534 \| 266 \| 25 \| 1 \| 413 \| 1 \| 470 \| Phage tail sheath protein \| Phage tail sheath protein; afdb-uniprot50 \| AF-A0A1Z9GEN2-F1-MODEL\_V4 \| 1.0 \| 1.156e-16 \| 406 \| 0.112 \| 687 \| 361 \| 41 \| 1 \| 474 \| 1 \| 651 \| Uncharacterized protein \| Uncharacterized protein; afdb-uniprot50 \| AF-A0A2N6N302-F1-MODEL\_V4 \| 1.0 \| 3.62e-12 \| 406 \| 0.247 \| 202 \| 129 \| 9 \| 283 \| 467 \| 477 \| 672 \| Phage tail protein \| Phage tail protein; afdb-uniprot50 \| AF-A0A1R3VFK8-F1-MODEL\_V4 \| 1.0 \| 2.725e-16 \| 406 \| 0.139 \| 783 \| 317 \| 32 \| 5 \| 457 \| 4 \| 759 \| Putative Phage tail sheath protein \| Putative Phage tail sheath protein; afdb-uniprot50 \| AF-A0A0R0ABK8-F1-MODEL\_V4 \| 1.0 \| 3.088e-09 \| 405 \| 0.54 \| 87 \| 38 \| 2 \| 140 \| 226 \| 103 \| 187 \| Uncharacterized protein \| Uncharacterized protein; afdb-uniprot50 \| AF-A0A7Y6NZ27-F1-MODEL\_V4 \| 1.0 \| 6.069e-16 \| 404 \| 0.106 \| 674 \| 350 \| 38 \| 1 \| 474 \| 1 \| 621 \| Phage tail sheath subtilisin-like domain-containing protein \| Phage tail sheath subtilisin-like domain-containing protein; afdb-uniprot50 \| AF-A0A4P2R373-F1-MODEL\_V4 \| 1.0 \| 1.799e-15 \| 402 \| 0.131 \| 561 \| 312 \| 26 \| 10 \| 468 \| 65 \| 551 \| Uncharacterized protein \| Uncharacterized protein; afdb-uniprot50 \| AF-A0A2D8EUZ2-F1-MODEL\_V4 \| 1.0 \| 6.426e-16 \| 401 \| 0.115 \| 703 \| 366 \| 37 \| 3 \| 474 \| 2 \| 679 \| Uncharacterized protein \| Uncharacterized protein; afdb-uniprot50 \| AF-A0A1H9D327-F1-MODEL\_V4 \| 1.0 \| 1.156e-16 \| 401 \| 0.15 \| 477 \| 272 \| 26 \| 99 \| 468 \| 462 \| 912 \| IPT/TIG domain-containing protein \| IPT/TIG domain-containing protein; afdb-uniprot50 \| AF-A0A2V2GPW4-F1-MODEL\_V4 \| 1.0 \| 6.16e-17 \| 400 \| 0.171 \| 560 \| 295 \| 29 \| 5 \| 474 \| 2 \| 482 \| Phage tail protein \| Phage tail protein; afdb-uniprot50 \| AF-A0A7V9L2C0-F1-MODEL\_V4 \| 1.0 \| 2.33e-17 \| 400 \| 0.158 \| 548 \| 272 \| 28 \| 1 \| 401 \| 2 \| 507 \| Phage tail sheath family protein \| Phage tail sheath family protein; afdb-uniprot50 \| AF-A0A0F9QR86-F1-MODEL\_V4 \| 1.0 \| 1.255e-10 \| 399 \| 0.375 \| 152 \| 93 \| 2 \| 324 \| 474 \| 19 \| 169 \| Phage\_sheath\_1C domain-containing protein \| Phage\_sheath\_1C domain-containing protein; afdb-uniprot50 \| AF-A0A1E7IPR4-F1-MODEL\_V4 \| 1.0 \| 1.091e-16 \| 399 \| 0.137 \| 711 \| 341 \| 37 \| 5 \| 466 \| 3 \| 689 \| Phage\_sheath\_1C domain-containing protein \| Phage\_sheath\_1C domain-containing protein; afdb-uniprot50 \| AF-A0A370LRA8-F1-MODEL\_V4 \| 1.0 \| 4.828e-16 \| 399 \| 0.125 \| 824 \| 341 \| 39 \| 3 \| 474 \| 10 \| 805 \| Uncharacterized protein \| Uncharacterized protein; afdb-uniprot50 \| AF-A0A3M1ZRW3-F1-MODEL\_V4 \| 1.0 \| 5.494e-17 \| 398 \| 0.174 \| 470 \| 240 \| 19 \| 141 \| 471 \| 241 \| 701 \| Phage tail sheath family protein \| Phage tail sheath family protein; afdb-uniprot50 \| AF-A0A7V7ZI79-F1-MODEL\_V4 \| 1.0 \| 1.001e-14 \| 398 \| 0.144 \| 366 \| 237 \| 16 \| 151 \| 468 \| 570 \| 907 \| Phage tail sheath family protein \| Phage tail sheath family protein; afdb-uniprot50 \| AF-A0A327JRX8-F1-MODEL\_V4 \| 1.0 \| 4.677e-10 \| 396 \| 0.255 \| 168 \| 123 \| 1 \| 307 \| 474 \| 5 \| 170 \| Phage\_sheath\_1C domain-containing protein \| Phage\_sheath\_1C domain-containing protein; afdb-uniprot50 \| AF-A0A7C7GUK5-F1-MODEL\_V4 \| 1.0 \| 1.139e-15 \| 394 \| 0.132 \| 740 \| 349 \| 38 \| 3 \| 473 \| 2 \| 717 \| Uncharacterized protein \| Uncharacterized protein; afdb-uniprot50 \| AF-A0A843FVB4-F1-MODEL\_V4 \| 1.0 \| 1.601e-11 \| 394 \| 0.14 \| 263 \| 199 \| 10 \| 227 \| 468 \| 619 \| 875 \| Uncharacterized protein \| Uncharacterized protein; afdb-uniprot50 \| AF-A0A0Q8FLN8-F1-MODEL\_V4 \| 1.0 \| 1.826e-16 \| 394 \| 0.15 \| 498 \| 276 \| 25 \| 96 \| 468 \| 436 \| 911 \| Phage\_sheath\_1C domain-containing protein \| Phage\_sheath\_1C domain-containing protein; afdb-uniprot50 \| AF-A0A4Q9KAR5-F1-MODEL\_V4 \| 1.0 \| 2.725e-16 \| 393 \| 0.118 \| 752 \| 326 \| 35 \| 1 \| 468 \| 6 \| 704 \| Phage tail sheath protein \| Phage tail sheath protein; afdb-uniprot50 \| AF-J2ZV19-F1-MODEL\_V4 \| 1.0 \| 6.069e-16 \| 392 \| 0.141 \| 475 \| 285 \| 20 \| 100 \| 468 \| 465 \| 922 \| Phage tail sheath protein FI \| Phage tail sheath protein FI; afdb-uniprot50 \| AF-A0A813AAM4-F1-MODEL\_V4 \| 1.0 \| 6.426e-16 \| 392 \| 0.133 \| 796 \| 334 \| 34 \| 1 \| 468 \| 499 \| 1266 \| 18 protein \| 18 protein; afdb-uniprot50 \| AF-A0A2D6PBJ4-F1-MODEL\_V4 \| 1.0 \| 6.805e-16 \| 391 \| 0.127 \| 598 \| 313 \| 29 \| 1 \| 453 \| 1 \| 534 \| Uncharacterized protein \| Uncharacterized protein; afdb-uniprot50 \| AF-A0A2W4LBW3-F1-MODEL\_V4 \| 1.0 \| 7.959e-15 \| 391 \| 0.16 \| 506 \| 290 \| 25 \| 46 \| 468 \| 101 \| 554 \| Phage tail sheath protein \| Phage tail sheath protein; afdb-uniprot50 \| AF-A0A2D9Z9K2-F1-MODEL\_V4 \| 1.0 \| 5.98e-15 \| 391 \| 0.102 \| 781 \| 364 \| 34 \| 1 \| 475 \| 1 \| 750 \| Uncharacterized protein \| Uncharacterized protein; afdb-uniprot50 \| AF-A0A1I3BRN0-F1-MODEL\_V4 \| 1.0 \| 1.224e-16 \| 390 \| 0.132 \| 765 \| 331 \| 39 \| 3 \| 467 \| 2 \| 733 \| Phage\_sheath\_1 domain-containing protein \| Phage\_sheath\_1 domain-containing protein; afdb-uniprot50 \| AF-A0A2E3G893-F1-MODEL\_V4 \| 1.0 \| 3.188e-15 \| 389 \| 0.13 \| 619 \| 350 \| 32 \| 3 \| 474 \| 2 \| 578 \| Uncharacterized protein \| Uncharacterized protein; afdb-uniprot50 \| AF-A0A662ABX1-F1-MODEL\_V4 \| 1.0 \| 4.243e-15 \| 388 \| 0.118 \| 710 \| 320 \| 28 \| 1 \| 468 \| 1 \| 646 \| Uncharacterized protein \| Uncharacterized protein; afdb-uniprot50 \| AF-A0A7J4UPE8-F1-MODEL\_V4 \| 1.0 \| 2.136e-15 \| 388 \| 0.133 \| 698 \| 338 \| 36 \| 3 \| 475 \| 2 \| 657 \| Uncharacterized protein \| Uncharacterized protein; afdb-uniprot50 \| AF-A0A356X586-F1-MODEL\_V4 \| 1.0 \| 1.153e-12 \| 388 \| 0.141 \| 304 \| 195 \| 14 \| 225 \| 468 \| 400 \| 697 \| Phage\_sheath\_1C domain-containing protein \| Phage\_sheath\_1C domain-containing protein; afdb-uniprot50 \| AF-A0A849BHN9-F1-MODEL\_V4 \| 1.0 \| 4.05e-08 \| 388 \| 0.551 \| 78 \| 34 \| 1 \| 147 \| 224 \| 595 \| 671 \| Uncharacterized protein \| Uncharacterized protein; afdb-uniprot50 \| AF-A0A6G2FUR4-F1-MODEL\_V4 \| 1.0 \| 5.112e-16 \| 386 \| 0.181 \| 419 \| 242 \| 18 \| 138 \| 463 \| 14 \| 424 \| Uncharacterized protein \| Uncharacterized protein; afdb-uniprot50 \| AF-A0A2V8SH17-F1-MODEL\_V4 \| 1.0 \| 2.168e-16 \| 386 \| 0.168 \| 468 \| 245 \| 19 \| 132 \| 467 \| 28 \| 483 \| Phage tail sheath family protein \| Phage tail sheath family protein; afdb-uniprot50 \| AF-A0A292SAA6-F1-MODEL\_V4 \| 1.0 \| 6.057e-12 \| 385 \| 0.17 \| 287 \| 185 \| 9 \| 230 \| 468 \| 161 \| 442 \| Phage\_sheath\_1C domain-containing protein \| Phage\_sheath\_1C domain-containing protein; afdb-uniprot50 \| AF-A0A2A7RYU0-F1-MODEL\_V4 \| 1.0 \| 4.747e-11 \| 384 \| 0.199 \| 316 \| 161 \| 9 \| 107 \| 418 \| 2 \| 229 \| Phage\_sheath\_1 domain-containing protein \| Phage\_sheath\_1 domain-containing protein; afdb-uniprot50 \| AF-A0A0F0HLZ5-F1-MODEL\_V4 \| 1.0 \| 3.776e-11 \| 383 \| 0.282 \| 195 \| 125 \| 4 \| 283 \| 468 \| 3 \| 191 \| Tail protein \| Tail protein; afdb-uniprot50 \| AF-A0A7I8DID5-F1-MODEL\_V4 \| 1.0 \| 2.136e-15 \| 383 \| 0.144 \| 559 \| 311 \| 30 \| 6 \| 475 \| 1 \| 480 \| Uncharacterized protein \| Uncharacterized protein; afdb-uniprot50 \| AF-A0A1X7MP25-F1-MODEL\_V4 \| 1.0 \| 1.31e-09 \| 381 \| 0.678 \| 84 \| 27 \| 0 \| 390 \| 473 \| 1 \| 84 \| Phage-related contractile tail sheath protein \| Phage-related contractile tail sheath protein; afdb-uniprot50 \| AF-A0A800EG04-F1-MODEL\_V4 \| 1.0 \| 2.531e-11 \| 381 \| 0.129 \| 248 \| 186 \| 10 \| 245 \| 468 \| 5 \| 246 \| Phage\_sheath\_1 domain-containing protein \| Phage\_sheath\_1 domain-containing protein; afdb-uniprot50 \| AF-A0A5C7JAB2-F1-MODEL\_V4 \| 1.0 \| 2.685e-15 \| 381 \| 0.128 \| 636 \| 334 \| 36 \| 3 \| 468 \| 2 \| 586 \| Phage\_sheath\_1C domain-containing protein \| Phage\_sheath\_1C domain-containing protein; afdb-uniprot50 \| AF-A0A736IA12-F1-MODEL\_V4 \| 1.0 \| 8.775e-10 \| 378 \| 0.351 \| 131 \| 82 \| 2 \| 345 \| 474 \| 2 \| 130 \| Phage tail sheath family protein \| Phage tail sheath family protein; afdb-uniprot50 \| AF-A0A1Q7YAH8-F1-MODEL\_V4 \| 1.0 \| 1.258e-14 \| 378 \| 0.136 \| 511 \| 296 \| 24 \| 46 \| 468 \| 197 \| 649 \| Uncharacterized protein \| Uncharacterized protein; afdb-uniprot50 \| AF-A0A730JWC2-F1-MODEL\_V4 \| 1.0 \| 4.953e-10 \| 375 \| 0.472 \| 110 \| 58 \| 0 \| 340 \| 449 \| 2 \| 111 \| Phage tail protein \| Phage tail protein; afdb-uniprot50 \| AF-A0A6V8EZA9-F1-MODEL\_V4 \| 1.0 \| 7.099e-15 \| 375 \| 0.1 \| 624 \| 349 \| 34 \| 15 \| 466 \| 2 \| 584 \| Uncharacterized protein \| Uncharacterized protein; afdb-uniprot50 \| AF-A0A6N6X9E8-F1-MODEL\_V4 \| 1.0 \| 8.646e-09 \| 374 \| 0.432 \| 111 \| 63 \| 0 \| 364 \| 474 \| 1 \| 111 \| Phage tail protein \| Phage tail protein; afdb-uniprot50 \| AF-A0A2E6EJU7-F1-MODEL\_V4 \| 1.0 \| 2.136e-15 \| 372 \| 0.137 \| 590 \| 274 \| 23 \| 1 \| 468 \| 8 \| 484 \| Uncharacterized protein \| Uncharacterized protein; afdb-uniprot50 \| AF-A0A559QHL7-F1-MODEL\_V4 \| 1.0 \| 8.793e-14 \| 372 \| 0.134 \| 578 \| 325 \| 30 \| 1 \| 468 \| 1 \| 512 \| Phage tail sheath protein \| Phage tail sheath protein; afdb-uniprot50 \| AF-A0A651EMP1-F1-MODEL\_V4 \| 1.0 \| 1.849e-13 \| 372 \| 0.159 \| 383 \| 259 \| 14 \| 99 \| 455 \| 460 \| 805 \| Uncharacterized protein \| Uncharacterized protein; afdb-uniprot50 \| AF-I4HNE8-F1-MODEL\_V4 \| 1.0 \| 4.007e-15 \| 371 \| 0.147 \| 399 \| 237 \| 19 \| 142 \| 466 \| 6 \| 375 \| Uncharacterized protein \| Uncharacterized protein; afdb-uniprot50 \| AF-A0A193QGV7-F1-MODEL\_V4 \| 1.0 \| 3.141e-14 \| 370 \| 0.285 \| 217 \| 138 \| 2 \| 259 \| 474 \| 2 \| 202 \| Phage tail sheath protein \| Phage tail sheath protein; afdb-uniprot50 \| AF-A0A3N8BHF0-F1-MODEL\_V4 \| 1.0 \| 4.067e-16 \| 370 \| 0.152 \| 557 \| 299 \| 31 \| 5 \| 468 \| 2 \| 478 \| Phage tail sheath family protein \| Phage tail sheath family protein; afdb-uniprot50 \| AF-Q2N8P0-F1-MODEL\_V4 \| 1.0 \| 3.784e-15 \| 370 \| 0.159 \| 470 \| 255 \| 23 \| 113 \| 468 \| 482 \| 925 \| Phage\_sheath\_1C domain-containing protein \| Phage\_sheath\_1C domain-containing protein; afdb-uniprot50 \| AF-A0A2D6WY00-F1-MODEL\_V4 \| 1.0 \| 7.406e-14 \| 368 \| 0.128 \| 568 \| 279 \| 23 \| 5 \| 468 \| 1 \| 456 \| Uncharacterized protein \| Uncharacterized protein; afdb-uniprot50 \| AF-A0A450ZA88-F1-MODEL\_V4 \| 1.0 \| 1.49e-10 \| 367 \| 0.214 \| 256 \| 114 \| 3 \| 121 \| 375 \| 3 \| 172 \| Phage\_sheath\_1 domain-containing protein \| Phage\_sheath\_1 domain-containing protein; afdb-uniprot50 \| AF-A0A3D4UXI7-F1-MODEL\_V4 \| 1.0 \| 5.805e-13 \| 366 \| 0.121 \| 534 \| 267 \| 24 \| 5 \| 468 \| 1 \| 402 \| Uncharacterized protein \| Uncharacterized protein; afdb-uniprot50 \| AF-A0A5J6PAG1-F1-MODEL\_V4 \| 1.0 \| 7.175e-08 \| 365 \| 0.44 \| 84 \| 45 \| 2 \| 144 \| 226 \| 120 \| 202 \| Phage tail protein \| Phage tail protein; afdb-uniprot50 \| AF-A0A823Z769-F1-MODEL\_V4 \| 1.0 \| 1.073e-11 \| 365 \| 0.192 \| 358 \| 164 \| 12 \| 1 \| 336 \| 1 \| 255 \| Phage tail sheath family protein \| Phage tail sheath family protein; afdb-uniprot50 \| AF-A0A011NTT3-F1-MODEL\_V4 \| 1.0 \| 4.243e-15 \| 365 \| 0.117 \| 683 \| 323 \| 30 \| 3 \| 452 \| 2 \| 637 \| Phage tail sheath protein \| Phage tail sheath protein; afdb-uniprot50 \| AF-A0A0F9G5K0-F1-MODEL\_V4 \| 1.0 \| 6.319e-11 \| 364 \| 0.141 \| 269 \| 194 \| 13 \| 233 \| 473 \| 207 \| 466 \| Uncharacterized protein \| Uncharacterized protein; afdb-uniprot50 \| AF-A0A3C1WGA5-F1-MODEL\_V4 \| 1.0 \| 7.598e-08 \| 364 \| 0.542 \| 83 \| 37 \| 1 \| 142 \| 224 \| 114 \| 195 \| Uncharacterized protein \| Uncharacterized protein; afdb-uniprot50 \| AF-T5KPS4-F1-MODEL\_V4 \| 1.0 \| 2.563e-08 \| 363 \| 0.494 \| 87 \| 38 \| 3 \| 139 \| 224 \| 13 \| 94 \| Uncharacterized protein \| Uncharacterized protein; afdb-uniprot50 \| AF-A0A2V8S9Y8-F1-MODEL\_V4 \| 1.0 \| 8.906e-11 \| 363 \| 0.17 \| 247 \| 176 \| 10 \| 231 \| 456 \| 228 \| 466 \| Uncharacterized protein \| Uncharacterized protein; afdb-uniprot50 \| AF-A0A2E3GP57-F1-MODEL\_V4 \| 1.0 \| 3.521e-14 \| 363 \| 0.132 \| 725 \| 327 \| 33 \| 3 \| 473 \| 4 \| 680 \| Uncharacterized protein \| Uncharacterized protein; afdb-uniprot50 \| AF-A0A4Q3AL90-F1-MODEL\_V4 \| 1.0 \| 3.134e-10 \| 362 \| 0.173 \| 219 \| 161 \| 7 \| 265 \| 468 \| 3 \| 216 \| Phage\_sheath\_1C domain-containing protein \| Phage\_sheath\_1C domain-containing protein; afdb-uniprot50 \| AF-A0A843FPA9-F1-MODEL\_V4 \| 1.0 \| 2.493e-10 \| 362 \| 0.144 \| 250 \| 187 \| 10 \| 234 \| 468 \| 801 \| 1038 \| Uncharacterized protein \| Uncharacterized protein; afdb-uniprot50 \| AF-A0A2E7URM1-F1-MODEL\_V4 \| 1.0 \| 2.498e-14 \| 361 \| 0.158 \| 447 \| 227 \| 21 \| 72 \| 466 \| 277 \| 626 \| Uncharacterized protein \| Uncharacterized protein; afdb-uniprot50 \| AF-A0A1Z9STS7-F1-MODEL\_V4 \| 1.0 \| 3.729e-14 \| 360 \| 0.118 \| 774 \| 361 \| 36 \| 3 \| 474 \| 2 \| 755 \| Uncharacterized protein \| Uncharacterized protein; afdb-uniprot50 \| AF-E3BL79-F1-MODEL\_V4 \| 1.0 \| 8.062e-12 \| 359 \| 0.101 \| 510 \| 279 \| 23 \| 1 \| 466 \| 3 \| 377 \| Uncharacterized protein \| Uncharacterized protein; afdb-uniprot50 \| AF-A0A2D8ERL6-F1-MODEL\_V4 \| 1.0 \| 1.17e-13 \| 359 \| 0.124 \| 531 \| 268 \| 26 \| 15 \| 466 \| 11 \| 423 \| Phage\_sheath\_1C domain-containing protein \| Phage\_sheath\_1C domain-containing protein; afdb-uniprot50 \| AF-A0A2S3UNF6-F1-MODEL\_V4 \| 1.0 \| 4.18e-14 \| 359 \| 0.11 \| 802 \| 351 \| 36 \| 5 \| 466 \| 1 \| 779 \| Phage tail sheath protein FI \| Phage tail sheath protein FI; afdb-uniprot50 \| AF-A0A496KHE0-F1-MODEL\_V4 \| 1.0 \| 1.622e-08 \| 358 \| 0.436 \| 94 \| 51 \| 2 \| 136 \| 227 \| 663 \| 756 \| Uncharacterized protein \| Uncharacterized protein; afdb-uniprot50 \| AF-A0A7Y5K224-F1-MODEL\_V4 \| 1.0 \| 5.636e-11 \| 357 \| 0.195 \| 235 \| 153 \| 9 \| 253 \| 467 \| 20 \| 238 \| Phage tail sheath family protein \| Phage tail sheath family protein; afdb-uniprot50 \| AF-I5B7I1-F1-MODEL\_V4 \| 1.0 \| 2.536e-15 \| 357 \| 0.152 \| 624 \| 291 \| 39 \| 5 \| 437 \| 1 \| 577 \| Phage tail sheath protein FI \| Phage tail sheath protein FI; afdb-uniprot50 \| AF-A0A1A7PRB7-F1-MODEL\_V4 \| 1.0 \| 2.457e-09 \| 355 \| 0.318 \| 154 \| 95 \| 4 \| 327 \| 474 \| 1 \| 150 \| Phage tail protein \| Phage tail protein; afdb-uniprot50 \| AF-D3AR67-F1-MODEL\_V4 \| 1.0 \| 4.56e-16 \| 355 \| 0.16 \| 493 \| 253 \| 23 \| 121 \| 463 \| 14 \| 495 \| Phage tail sheath protein \| Phage tail sheath protein; afdb-uniprot50 \| AF-A0A2E6FCL3-F1-MODEL\_V4 \| 1.0 \| 3.948e-14 \| 355 \| 0.133 \| 622 \| 354 \| 36 \| 3 \| 473 \| 2 \| 589 \| Uncharacterized protein \| Uncharacterized protein; afdb-uniprot50 \| AF-A0A382CP73-F1-MODEL\_V4 \| 1.0 \| 3.326e-14 \| 355 \| 0.111 \| 655 \| 343 \| 35 \| 3 \| 445 \| 2 \| 629 \| Uncharacterized protein \| Uncharacterized protein; afdb-uniprot50 \| AF-A0A2E0HZL7-F1-MODEL\_V4 \| 1.0 \| 1.17e-13 \| 355 \| 0.123 \| 832 \| 340 \| 41 \| 1 \| 474 \| 1 \| 800 \| Uncharacterized protein \| Uncharacterized protein; afdb-uniprot50 \| AF-A0A6P1B3L2-F1-MODEL\_V4 \| 1.0 \| 1.717e-08 \| 354 \| 0.802 \| 86 \| 17 \| 0 \| 362 \| 447 \| 1 \| 86 \| Phage tail sheath family protein \| Phage tail sheath family protein; afdb-uniprot50 \| AF-A0A1G8EJS8-F1-MODEL\_V4 \| 1.0 \| 1.332e-14 \| 354 \| 0.12 \| 549 \| 327 \| 22 \| 1 \| 468 \| 13 \| 486 \| Phage tail sheath protein \| Phage tail sheath protein; afdb-uniprot50 \| AF-A0A328FDX7-F1-MODEL\_V4 \| 1.0 \| 1.747e-13 \| 353 \| 0.125 \| 572 \| 337 \| 28 \| 7 \| 468 \| 1 \| 519 \| Uncharacterized protein \| Uncharacterized protein; afdb-uniprot50 \| AF-A0A2D0N1H2-F1-MODEL\_V4 \| 1.0 \| 1.203e-11 \| 353 \| 0.097 \| 798 \| 287 \| 24 \| 1 \| 466 \| 3 \| 698 \| Uncharacterized protein \| Uncharacterized protein; afdb-uniprot50 \| AF-A0A7K4MQW1-F1-MODEL\_V4 \| 1.0 \| 9.31e-14 \| 353 \| 0.1 \| 768 \| 376 \| 39 \| 3 \| 474 \| 2 \| 750 \| Phage tail sheath subtilisin-like domain-containing protein \| Phage tail sheath subtilisin-like domain-containing protein; afdb-uniprot50 \| AF-A0A2N2MEV8-F1-MODEL\_V4 \| 1.0 \| 9.449e-15 \| 351 \| 0.14 \| 492 \| 261 \| 21 \| 102 \| 455 \| 500 \| 967 \| Uncharacterized protein \| Uncharacterized protein; afdb-uniprot50 \| AF-A0A3A8PQG2-F1-MODEL\_V4 \| 1.0 \| 6.147e-13 \| 350 \| 0.153 \| 392 \| 233 \| 18 \| 159 \| 468 \| 23 \| 397 \| Uncharacterized protein \| Uncharacterized protein; afdb-uniprot50 \| AF-A0A418HAB9-F1-MODEL\_V4 \| 1.0 \| 1.468e-09 \| 349 \| 0.436 \| 119 \| 66 \| 1 \| 278 \| 395 \| 1 \| 119 \| Phage tail protein \| Phage tail protein; afdb-uniprot50 \| AF-A0A0Q0D1V1-F1-MODEL\_V4 \| 1.0 \| 1.026e-08 \| 347 \| 0.484 \| 99 \| 51 \| 0 \| 376 \| 474 \| 2 \| 100 \| Major tail sheath protein \| Major tail sheath protein; afdb-uniprot50 \| AF-A0A3M1BNH9-F1-MODEL\_V4 \| 1.0 \| 3.27e-09 \| 347 \| 0.333 \| 135 \| 82 \| 4 \| 346 \| 474 \| 2 \| 134 \| Phage tail sheath family protein \| Phage tail sheath family protein; afdb-uniprot50 \| AF-A0A3M1QC03-F1-MODEL\_V4 \| 1.0 \| 2.191e-09 \| 346 \| 0.238 \| 180 \| 129 \| 2 \| 291 \| 468 \| 9 \| 182 \| Phage tail sheath family protein \| Phage tail sheath family protein; afdb-uniprot50 \| AF-A0A6M1Q203-F1-MODEL\_V4 \| 1.0 \| 2.998e-07 \| 345 \| 0.415 \| 77 \| 43 \| 1 \| 148 \| 224 \| 2 \| 76 \| RagB/SusD family nutrient uptake outer membrane protein \| RagB/SusD family nutrient uptake outer membrane protein; afdb-uniprot50 \| AF-A0A383D6D3-F1-MODEL\_V4 \| 1.0 \| 1.255e-10 \| 345 \| 0.176 \| 204 \| 152 \| 7 \| 229 \| 424 \| 39 \| 234 \| Phage\_sheath\_1 domain-containing protein \| Phage\_sheath\_1 domain-containing protein; afdb-uniprot50 \| AF-A0A2D0AK03-F1-MODEL\_V4 \| 1.0 \| 5.091e-08 \| 345 \| 0.5 \| 86 \| 41 \| 2 \| 140 \| 224 \| 99 \| 183 \| Phage tail protein \| Phage tail protein; afdb-uniprot50 \| AF-A0A5E4M1E6-F1-MODEL\_V4 \| 1.0 \| 3.095e-13 \| 345 \| 0.136 \| 448 \| 228 \| 16 \| 84 \| 468 \| 699 \| 1050 \| Phage\_sheath\_1C domain-containing protein \| Phage\_sheath\_1C domain-containing protein; afdb-uniprot50 \| AF-A0A845AZ49-F1-MODEL\_V4 \| 1.0 \| 2.607e-13 \| 343 \| 0.122 \| 603 \| 327 \| 33 \| 7 \| 467 \| 3 \| 545 \| Uncharacterized protein \| Uncharacterized protein; afdb-uniprot50 \| AF-A0A7V2ED64-F1-MODEL\_V4 \| 1.0 \| 5.027e-11 \| 343 \| 0.148 \| 296 \| 198 \| 12 \| 227 \| 473 \| 859 \| 1149 \| Phage\_sheath\_1C domain-containing protein \| Phage\_sheath\_1C domain-containing protein; afdb-uniprot50 \| AF-A0A1S6HM92-F1-MODEL\_V4 \| 1.0 \| 5.255e-14 \| 341 \| 0.178 \| 542 \| 283 \| 27 \| 1 \| 468 \| 1 \| 453 \| Phage tail sheath protein \| Phage tail sheath protein; afdb-uniprot50 \| AF-R9L6C8-F1-MODEL\_V4 \| 1.0 \| 3.088e-09 \| 339 \| 0.235 \| 208 \| 137 \| 9 \| 278 \| 472 \| 2 \| 200 \| Uncharacterized protein \| Uncharacterized protein; afdb-uniprot50 \| AF-A0A7C5N1C4-F1-MODEL\_V4 \| 1.0 \| 1.103e-09 \| 338 \| 0.21 \| 195 \| 128 \| 8 \| 288 \| 468 \| 461 \| 643 \| Phage tail sheath family protein \| Phage tail sheath family protein; afdb-uniprot50 \| AF-A0A854M7T7-F1-MODEL\_V4 \| 1.0 \| 2.966e-14 \| 336 \| 0.123 \| 687 \| 332 \| 34 \| 3 \| 474 \| 4 \| 635 \| Uncharacterized protein \| Uncharacterized protein; afdb-uniprot50 \| AF-A0A7X7X247-F1-MODEL\_V4 \| 1.0 \| 5.255e-14 \| 335 \| 0.14 \| 528 \| 280 \| 26 \| 1 \| 424 \| 44 \| 501 \| Uncharacterized protein \| Uncharacterized protein; afdb-uniprot50 \| AF-A0A1F1BZS2-F1-MODEL\_V4 \| 1.0 \| 1.071e-07 \| 335 \| 0.369 \| 92 \| 56 \| 2 \| 137 \| 226 \| 361 \| 452 \| Uncharacterized protein \| Uncharacterized protein; afdb-uniprot50 \| AF-N6YII7-F1-MODEL\_V4 \| 1.0 \| 4.172e-10 \| 334 \| 0.134 \| 261 \| 185 \| 9 \| 231 \| 452 \| 131 \| 389 \| Uncharacterized protein \| Uncharacterized protein; afdb-uniprot50 \| AF-A0A1H8JA80-F1-MODEL\_V4 \| 1.0 \| 7.614e-12 \| 334 \| 0.14 \| 519 \| 222 \| 17 \| 68 \| 468 \| 251 \| 663 \| Phage\_sheath\_1C domain-containing protein \| Phage\_sheath\_1C domain-containing protein; afdb-uniprot50 \| AF-A0A496KYN9-F1-MODEL\_V4 \| 1.0 \| 4.541e-08 \| 333 \| 0.385 \| 96 \| 57 \| 2 \| 133 \| 226 \| 712 \| 807 \| Uncharacterized protein \| Uncharacterized protein; afdb-uniprot50 \| AF-A0A2E0S344-F1-MODEL\_V4 \| 1.0 \| 6.509e-13 \| 332 \| 0.116 \| 809 \| 342 \| 41 \| 1 \| 474 \| 1 \| 771 \| Uncharacterized protein \| Uncharacterized protein; afdb-uniprot50 \| AF-A0A1U7CXB8-F1-MODEL\_V4 \| 1.0 \| 1.134e-07 \| 331 \| 0.505 \| 89 \| 39 \| 2 \| 140 \| 224 \| 79 \| 166 \| Uncharacterized protein \| Uncharacterized protein; afdb-uniprot50 \| AF-A0A3N5W9U7-F1-MODEL\_V4 \| 1.0 \| 7.927e-07 \| 331 \| 0.53 \| 83 \| 38 \| 1 \| 144 \| 226 \| 212 \| 293 \| C-type lectin domain-containing protein \| C-type lectin domain-containing protein; afdb-uniprot50 \| AF-A0A4P5QN36-F1-MODEL\_V4 \| 1.0 \| 3.361e-07 \| 330 \| 0.463 \| 82 \| 42 \| 2 \| 142 \| 223 \| 123 \| 202 \| Uncharacterized protein \| Uncharacterized protein; afdb-uniprot50 \| AF-A0A3N5MEB4-F1-MODEL\_V4 \| 1.0 \| 1.326e-06 \| 330 \| 0.576 \| 78 \| 32 \| 1 \| 147 \| 224 \| 323 \| 399 \| Fibronectin type-III domain-containing protein \| Fibronectin type-III domain-containing protein; afdb-uniprot50 \| AF-A0A1F1HV94-F1-MODEL\_V4 \| 1.0 \| 4.808e-08 \| 330 \| 0.387 \| 98 \| 56 \| 3 \| 134 \| 227 \| 631 \| 728 \| Uncharacterized protein \| Uncharacterized protein; afdb-uniprot50 \| AF-A0A419GQ10-F1-MODEL\_V4 \| 1.0 \| 1.792e-07 \| 327 \| 0.522 \| 90 \| 40 \| 2 \| 139 \| 225 \| 96 \| 185 \| Uncharacterized protein \| Uncharacterized protein; afdb-uniprot50 \| AF-A0A6L6S241-F1-MODEL\_V4 \| 1.0 \| 3.833e-12 \| 327 \| 0.114 \| 444 \| 235 \| 16 \| 83 \| 467 \| 541 \| 885 \| Phage\_sheath\_1C domain-containing protein \| Phage\_sheath\_1C domain-containing protein; afdb-uniprot50 \| AF-A0A843GVB5-F1-MODEL\_V4 \| 1.0 \| 2.1e-10 \| 326 \| 0.158 \| 253 \| 174 \| 10 \| 245 \| 468 \| 12 \| 254 \| Uncharacterized protein \| Uncharacterized protein; afdb-uniprot50 \| AF-A0A2D8EN41-F1-MODEL\_V4 \| 1.0 \| 1.329e-10 \| 323 \| 0.099 \| 522 \| 282 \| 18 \| 2 \| 468 \| 7 \| 395 \| Uncharacterized protein \| Uncharacterized protein; afdb-uniprot50 \| AF-A0A450Z311-F1-MODEL\_V4 \| 1.0 \| 6.4e-08 \| 322 \| 0.376 \| 93 \| 58 \| 0 \| 381 \| 473 \| 2 \| 94 \| Phage\_sheath\_1C domain-containing protein \| Phage\_sheath\_1C domain-containing protein; afdb-uniprot50 \| AF-X0YSG3-F1-MODEL\_V4 \| 1.0 \| 4.11e-09 \| 322 \| 0.185 \| 200 \| 146 \| 9 \| 231 \| 419 \| 11 \| 204 \| Phage\_sheath\_1 domain-containing protein \| Phage\_sheath\_1 domain-containing protein; afdb-uniprot50 \| AF-A0A2W7DZU1-F1-MODEL\_V4 \| 1.0 \| 7.943e-11 \| 321 \| 0.197 \| 309 \| 150 \| 12 \| 1 \| 300 \| 1 \| 220 \| Uncharacterized protein \| Uncharacterized protein; afdb-uniprot50 \| AF-A0A3S0E3K7-F1-MODEL\_V4 \| 1.0 \| 2.219e-06 \| 321 \| 0.448 \| 78 \| 42 \| 1 \| 147 \| 224 \| 119 \| 195 \| Fibronectin type-III domain-containing protein \| Fibronectin type-III domain-containing protein; afdb-uniprot50 \| AF-A0A1G3MC68-F1-MODEL\_V4 \| 1.0 \| 6.509e-13 \| 319 \| 0.121 \| 427 \| 264 \| 18 \| 140 \| 474 \| 376 \| 783 \| Uncharacterized protein \| Uncharacterized protein; afdb-uniprot50 \| AF-A0A1F7WZM2-F1-MODEL\_V4 \| 1.0 \| 9.713e-13 \| 318 \| 0.114 \| 532 \| 326 \| 24 \| 1 \| 455 \| 1 \| 464 \| Uncharacterized protein \| Uncharacterized protein; afdb-uniprot50 \| AF-A0A661IMM3-F1-MODEL\_V4 \| 1.0 \| 3.134e-10 \| 318 \| 0.131 \| 266 \| 186 \| 12 \| 231 \| 466 \| 463 \| 713 \| Uncharacterized protein \| Uncharacterized protein; afdb-uniprot50 \| AF-A0A1Y6CY82-F1-MODEL\_V4 \| 1.0 \| 6.044e-08 \| 317 \| 0.373 \| 99 \| 61 \| 1 \| 376 \| 473 \| 2 \| 100 \| Phage tail sheath protein \| Phage tail sheath protein; afdb-uniprot50 \| AF-A0A7V8NSJ0-F1-MODEL\_V4 \| 1.0 \| 3.277e-13 \| 317 \| 0.142 \| 542 \| 284 \| 29 \| 48 \| 456 \| 2 \| 495 \| Phage tail sheath subtilisin-like domain-containing protein \| Phage tail sheath subtilisin-like domain-containing protein; afdb-uniprot50 \| AF-A0A0G1W3T5-F1-MODEL\_V4 \| 1.0 \| 3.411e-08 \| 316 \| 0.398 \| 103 \| 57 \| 2 \| 145 \| 243 \| 182 \| 283 \| Autotransporter-associated beta strand repeat protein \| Autotransporter-associated beta strand repeat protein; afdb-uniprot50 \| AF-A0A2V3RDB6-F1-MODEL\_V4 \| 1.0 \| 1.897e-07 \| 315 \| 0.305 \| 118 \| 77 \| 2 \| 358 \| 471 \| 2 \| 118 \| Tail sheath protein \| Tail sheath protein; afdb-uniprot50 \| AF-A0A225DFW4-F1-MODEL\_V4 \| 1.0 \| 1.598e-07 \| 315 \| 0.425 \| 94 \| 46 \| 3 \| 140 \| 226 \| 191 \| 283 \| Putative major tail protein \| Putative major tail protein; afdb-uniprot50 \| AF-A0A7X1EF97-F1-MODEL\_V4 \| 1.0 \| 4.541e-08 \| 314 \| 0.346 \| 101 \| 65 \| 1 \| 260 \| 359 \| 2 \| 102 \| Phage tail sheath subtilisin-like domain-containing protein \| Phage tail sheath subtilisin-like domain-containing protein; afdb-uniprot50 \| AF-A0A1I1EU22-F1-MODEL\_V4 \| 1.0 \| 7.842e-14 \| 313 \| 0.112 \| 533 \| 311 \| 30 \| 1 \| 466 \| 4 \| 441 \| Phage tail sheath protein \| Phage tail sheath protein; afdb-uniprot50 \| AF-A0A1H2UW00-F1-MODEL\_V4 \| 1.0 \| 3.174e-07 \| 312 \| 0.363 \| 99 \| 63 \| 0 \| 376 \| 474 \| 7 \| 105 \| Phage tail sheath protein \| Phage tail sheath protein; afdb-uniprot50 \| AF-A0A2D6XA84-F1-MODEL\_V4 \| 1.0 \| 8.393e-07 \| 310 \| 0.487 \| 80 \| 39 \| 2 \| 147 \| 224 \| 21 \| 100 \| Uncharacterized protein \| Uncharacterized protein; afdb-uniprot50 \| AF-A0A4T3EYF2-F1-MODEL\_V4 \| 1.0 \| 1.293e-12 \| 310 \| 0.172 \| 429 \| 243 \| 21 \| 79 \| 457 \| 386 \| 752 \| Phage\_sheath\_1 domain-containing protein \| Phage\_sheath\_1 domain-containing protein; afdb-uniprot50 \| AF-A0A3N7HIZ4-F1-MODEL\_V4 \| 1.0 \| 4.297e-12 \| 309 \| 0.134 \| 497 \| 274 \| 23 \| 72 \| 467 \| 201 \| 642 \| Uncharacterized protein \| Uncharacterized protein; afdb-uniprot50 \| AF-A0A2T6GH30-F1-MODEL\_V4 \| 1.0 \| 2.563e-08 \| 306 \| 0.423 \| 118 \| 45 \| 5 \| 109 \| 226 \| 99 \| 193 \| Phage tail protein \| Phage tail protein; afdb-uniprot50 \| AF-A0A1Z9B2F7-F1-MODEL\_V4 \| 1.0 \| 3.128e-06 \| 306 \| 0.382 \| 81 \| 48 \| 1 \| 146 \| 226 \| 613 \| 691 \| Uncharacterized protein \| Uncharacterized protein; afdb-uniprot50 \| AF-A0A381W6D4-F1-MODEL\_V4 \| 1.0 \| 1.487e-06 \| 303 \| 0.38 \| 84 \| 51 \| 1 \| 142 \| 224 \| 134 \| 217 \| DUF4815 domain-containing protein \| DUF4815 domain-containing protein; afdb-uniprot50 \| AF-A0A7Y7P2S0-F1-MODEL\_V4 \| 1.0 \| 1.535e-12 \| 303 \| 0.131 \| 823 \| 324 \| 37 \| 1 \| 468 \| 1 \| 787 \| Phage tail sheath subtilisin-like domain-containing protein \| Phage tail sheath subtilisin-like domain-containing protein; afdb-uniprot50 \| AF-A0A540WH21-F1-MODEL\_V4 \| 1.0 \| 4.474e-07 \| 302 \| 0.306 \| 98 \| 68 \| 0 \| 377 \| 474 \| 4 \| 101 \| Phage\_sheath\_1C domain-containing protein \| Phage\_sheath\_1C domain-containing protein; afdb-uniprot50 \| AF-A1B8I0-F1-MODEL\_V4 \| 1.0 \| 1.692e-07 \| 299 \| 0.447 \| 96 \| 50 \| 2 \| 132 \| 225 \| 384 \| 478 \| Uncharacterized protein \| Uncharacterized protein; afdb-uniprot50 \| AF-A0A7W0VBV0-F1-MODEL\_V4 \| 1.0 \| 8.287e-10 \| 298 \| 0.169 \| 366 \| 176 \| 13 \| 1 \| 342 \| 1 \| 262 \| Phage tail sheath family protein \| Phage tail sheath family protein; afdb-uniprot50 \| AF-A0A2V7WD79-F1-MODEL\_V4 \| 1.0 \| 1.509e-07 \| 298 \| 0.444 \| 99 \| 44 \| 2 \| 139 \| 227 \| 294 \| 391 \| Uncharacterized protein \| Uncharacterized protein; afdb-uniprot50 \| AF-A0A3P6JQ12-F1-MODEL\_V4 \| 1.0 \| 2.039e-08 \| 297 \| 0.208 \| 192 \| 131 \| 6 \| 290 \| 467 \| 7 \| 191 \| Phage tail sheath protein FI \| Phage tail sheath protein FI; afdb-uniprot50 \| AF-A0A3N5G9W8-F1-MODEL\_V4 \| 1.0 \| 1.312e-13 \| 296 \| 0.152 \| 426 \| 228 \| 18 \| 145 \| 452 \| 36 \| 446 \| Uncharacterized protein \| Uncharacterized protein; afdb-uniprot50 \| AF-A0A1F5C1Z1-F1-MODEL\_V4 \| 1.0 \| 4.11e-09 \| 295 \| 0.134 \| 253 \| 187 \| 11 \| 232 \| 466 \| 6 \| 244 \| Uncharacterized protein \| Uncharacterized protein; afdb-uniprot50 \| AF-G4KPJ2-F1-MODEL\_V4 \| 1.0 \| 3.612e-08 \| 293 \| 0.235 \| 178 \| 118 \| 4 \| 308 \| 474 \| 2 \| 172 \| Putative tail sheath protein \| Putative tail sheath protein; afdb-uniprot50 \| AF-A0A843GHW6-F1-MODEL\_V4 \| 1.0 \| 4.808e-08 \| 290 \| 0.135 \| 236 \| 173 \| 9 \| 254 \| 467 \| 2 \| 228 \| Uncharacterized protein \| Uncharacterized protein; afdb-uniprot50 \| AF-A0A4Q3NY20-F1-MODEL\_V4 \| 1.0 \| 1.822e-12 \| 290 \| 0.147 \| 624 \| 351 \| 37 \| 1 \| 474 \| 1 \| 593 \| Uncharacterized protein \| Uncharacterized protein; afdb-uniprot50 \| AF-I0QL77-F1-MODEL\_V4 \| 1.0 \| 1.201e-07 \| 289 \| 0.471 \| 87 \| 45 \| 1 \| 291 \| 376 \| 8 \| 94 \| Major tail sheath protein \| Major tail sheath protein; afdb-uniprot50 \| AF-A0A6V8GYL1-F1-MODEL\_V4 \| 1.0 \| 2.831e-07 \| 288 \| 0.232 \| 129 \| 97 \| 1 \| 346 \| 474 \| 5 \| 131 \| Phage tail sheath family protein \| Phage tail sheath family protein; afdb-uniprot50 \| AF-A0A3S0LFZ3-F1-MODEL\_V4 \| 1.0 \| 6.057e-12 \| 288 \| 0.136 \| 446 \| 260 \| 21 \| 116 \| 455 \| 346 \| 772 \| Uncharacterized protein \| Uncharacterized protein; afdb-uniprot50 \| AF-A0A2D6U3C2-F1-MODEL\_V4 \| 1.0 \| 3.932e-06 \| 286 \| 0.385 \| 83 \| 49 \| 1 \| 145 \| 225 \| 55 \| 137 \| Phage tail protein \| Phage tail protein; afdb-uniprot50 \| AF-A0A842NAQ9-F1-MODEL\_V4 \| 1.0 \| 5.402e-12 \| 286 \| 0.112 \| 731 \| 318 \| 37 \| 3 \| 423 \| 2 \| 711 \| Phage tail sheath subtilisin-like domain-containing protein \| Phage tail sheath subtilisin-like domain-containing protein; afdb-uniprot50 \| AF-A0A2E4BMY0-F1-MODEL\_V4 \| 1.0 \| 2.043e-12 \| 286 \| 0.121 \| 445 \| 265 \| 21 \| 145 \| 473 \| 543 \| 977 \| Uncharacterized protein \| Uncharacterized protein; afdb-uniprot50 \| AF-A0A0Q4JAE9-F1-MODEL\_V4 \| 1.0 \| 2.096e-06 \| 284 \| 0.206 \| 97 \| 77 \| 0 \| 377 \| 473 \| 13 \| 109 \| Uncharacterized protein \| Uncharacterized protein; afdb-uniprot50 \| AF-A0A825L246-F1-MODEL\_V4 \| 1.0 \| 1.349e-11 \| 284 \| 0.137 \| 408 \| 242 \| 25 \| 138 \| 457 \| 174 \| 559 \| Uncharacterized protein \| Uncharacterized protein; afdb-uniprot50 \| AF-A0A1Z8Q4K5-F1-MODEL\_V4 \| 1.0 \| 4.163e-06 \| 282 \| 0.481 \| 81 \| 40 \| 1 \| 146 \| 224 \| 1010 \| 1090 \| Uncharacterized protein \| Uncharacterized protein; afdb-uniprot50 \| AF-A0A2E7T9B3-F1-MODEL\_V4 \| 1.0 \| 9.411e-07 \| 281 \| 0.413 \| 92 \| 49 \| 2 \| 136 \| 225 \| 128 \| 216 \| Uncharacterized protein \| Uncharacterized protein; afdb-uniprot50 \| AF-A0A661GUA0-F1-MODEL\_V4 \| 1.0 \| 2.291e-12 \| 280 \| 0.13 \| 498 \| 300 \| 30 \| 48 \| 465 \| 9 \| 453 \| Uncharacterized protein \| Uncharacterized protein; afdb-uniprot50 \| AF-A0A2V8RQP8-F1-MODEL\_V4 \| 1.0 \| 4.474e-07 \| 280 \| 0.423 \| 104 \| 54 \| 4 \| 126 \| 226 \| 167 \| 267 \| Uncharacterized protein \| Uncharacterized protein; afdb-uniprot50 \| AF-A0A3B9Z7Q0-F1-MODEL\_V4 \| 1.0 \| 7.614e-12 \| 279 \| 0.104 \| 844 \| 354 \| 39 \| 2 \| 468 \| 10 \| 828 \| Phage\_sheath\_1C domain-containing protein \| Phage\_sheath\_1C domain-containing protein; afdb-uniprot50 \| AF-G5LWT8-F1-MODEL\_V4 \| 1.0 \| 3.312e-06 \| 278 \| 0.293 \| 92 \| 65 \| 0 \| 382 \| 473 \| 8 \| 99 \| Major tail sheath protein \| Major tail sheath protein; afdb-uniprot50 \| AF-A0A5U2F9H2-F1-MODEL\_V4 \| 1.0 \| 3.312e-06 \| 277 \| 0.323 \| 99 \| 67 \| 0 \| 376 \| 474 \| 12 \| 110 \| Phage tail sheath family protein \| Phage tail sheath family protein; afdb-uniprot50 \| AF-A0A1V5B852-F1-MODEL\_V4 \| 1.0 \| 1.822e-12 \| 276 \| 0.126 \| 666 \| 317 \| 38 \| 20 \| 468 \| 28 \| 645 \| Phage tail sheath protein \| Phage tail sheath protein; afdb-uniprot50 \| AF-A0A7Z1HP85-F1-MODEL\_V4 \| 1.0 \| 3.507e-06 \| 274 \| 0.383 \| 86 \| 53 \| 0 \| 389 \| 474 \| 2 \| 87 \| Phage tail protein \| Phage tail protein; afdb-uniprot50 \| AF-A0A3S4J167-F1-MODEL\_V4 \| 1.0 \| 1.487e-06 \| 272 \| 0.428 \| 98 \| 56 \| 0 \| 350 \| 447 \| 2 \| 99 \| Major tail sheath protein \| Major tail sheath protein; afdb-uniprot50 \| AF-R7JPV7-F1-MODEL\_V4 \| 1.0 \| 1.667e-06 \| 272 \| 0.286 \| 129 \| 86 \| 3 \| 350 \| 473 \| 1 \| 128 \| Phage-related contractile tail sheath protein \| Phage-related contractile tail sheath protein; afdb-uniprot50 \| AF-A0A0B0HIM0-F1-MODEL\_V4 \| 1.0 \| 8.27e-06 \| 270 \| 0.305 \| 95 \| 66 \| 0 \| 381 \| 475 \| 5 \| 99 \| Uncharacterized protein \| Uncharacterized protein; afdb-uniprot50 \| AF-A0A447T5K7-F1-MODEL\_V4 \| 1.0 \| 6.226e-10 \| 270 \| 0.225 \| 270 \| 108 \| 7 \| 85 \| 344 \| 82 \| 260 \| Phage tail sheath protein \| Phage tail sheath protein; afdb-uniprot50 \| AF-A0A7Z2GRK3-F1-MODEL\_V4 \| 1.0 \| 1.49e-10 \| 270 \| 0.192 \| 359 \| 166 \| 15 \| 1 \| 275 \| 1 \| 319 \| Uncharacterized protein \| Uncharacterized protein; afdb-uniprot50 \| AF-A0A376YIM8-F1-MODEL\_V4 \| 1.0 \| 2.596e-05 \| 269 \| 0.354 \| 79 \| 51 \| 0 \| 376 \| 454 \| 9 \| 87 \| Phage-related contractile tail sheath protein \| Phage-related contractile tail sheath protein; afdb-uniprot50 \| AF-A0A0C2Z8E2-F1-MODEL\_V4 \| 1.0 \| 8.887e-07 \| 269 \| 0.258 \| 124 \| 90 \| 1 \| 351 \| 474 \| 1 \| 122 \| Uncharacterized protein \| Uncharacterized protein; afdb-uniprot50 \| AF-A0A2N7QTB6-F1-MODEL\_V4 \| 1.0 \| 4.474e-07 \| 266 \| 0.369 \| 111 \| 51 \| 5 \| 129 \| 225 \| 516 \| 621 \| Beta\_helix domain-containing protein \| Beta\_helix domain-containing protein; afdb-uniprot50 \| AF-A0A5U2F6F1-F1-MODEL\_V4 \| 1.0 \| 1.667e-06 \| 265 \| 0.336 \| 98 \| 64 \| 1 \| 260 \| 356 \| 2 \| 99 \| Phage tail sheath family protein \| Phage tail sheath family protein; afdb-uniprot50 \| AF-A0A2V8JS33-F1-MODEL\_V4 \| 1.0 \| 2.596e-05 \| 263 \| 0.433 \| 83 \| 44 \| 3 \| 145 \| 226 \| 225 \| 305 \| Uncharacterized protein \| Uncharacterized protein; afdb-uniprot50 \| AF-A0A4Q3L8D9-F1-MODEL\_V4 \| 1.0 \| 6.592e-10 \| 262 \| 0.138 \| 333 \| 220 \| 15 \| 186 \| 474 \| 462 \| 771 \| Uncharacterized protein \| Uncharacterized protein; afdb-uniprot50 \| AF-A0A644XLY2-F1-MODEL\_V4 \| 1.0 \| 1.04e-05 \| 258 \| 0.214 \| 121 \| 91 \| 1 \| 358 \| 474 \| 8 \| 128 \| Uncharacterized protein \| Uncharacterized protein; afdb-uniprot50 \| AF-A0A2S9JTK1-F1-MODEL\_V4 \| 1.0 \| 1.425e-07 \| 258 \| 0.303 \| 132 \| 59 \| 5 \| 116 \| 225 \| 394 \| 514 \| RagB/SusD family nutrient uptake outer membrane protein \| RagB/SusD family nutrient uptake outer membrane protein; afdb-uniprot50 \| AF-A0A7K4MSZ8-F1-MODEL\_V4 \| 1.0 \| 6.677e-07 \| 257 \| 0.316 \| 120 \| 56 \| 4 \| 109 \| 224 \| 117 \| 214 \| DUF4815 domain-containing protein \| DUF4815 domain-containing protein; afdb-uniprot50 \| AF-A0A5D4H7Y3-F1-MODEL\_V4 \| 1.0 \| 2.009e-07 \| 256 \| 0.278 \| 133 \| 61 \| 5 \| 116 \| 225 \| 358 \| 478 \| RagB/SusD family nutrient uptake outer membrane protein \| RagB/SusD family nutrient uptake outer membrane protein; afdb-uniprot50 \| AF-A0A3S4F1N2-F1-MODEL\_V4 \| 1.0 \| 7.376e-06 \| 255 \| 0.38 \| 84 \| 52 \| 0 \| 390 \| 473 \| 1 \| 84 \| Major tail sheath protein \| Major tail sheath protein; afdb-uniprot50 \| AF-A0A382MVY2-F1-MODEL\_V4 \| 1.0 \| 6.213e-06 \| 255 \| 0.42 \| 88 \| 46 \| 1 \| 142 \| 224 \| 39 \| 126 \| Uncharacterized protein \| Uncharacterized protein; afdb-uniprot50 \| AF-A0A5E9SB03-F1-MODEL\_V4 \| 1.0 \| 4.943e-06 \| 253 \| 0.3 \| 90 \| 63 \| 0 \| 385 \| 474 \| 2 \| 91 \| Phage tail sheath family protein \| Phage tail sheath family protein; afdb-uniprot50 \| AF-A0A0L1JK15-F1-MODEL\_V4 \| 1.0 \| 2.187e-05 \| 252 \| 0.411 \| 85 \| 47 \| 3 \| 140 \| 222 \| 374 \| 457 \| Uncharacterized protein \| Uncharacterized protein; afdb-uniprot50 \| AF-A0A2G4KB82-F1-MODEL\_V4 \| 1.0 \| 3.874e-05 \| 249 \| 0.437 \| 80 \| 45 \| 0 \| 144 \| 223 \| 269 \| 348 \| Uncharacterized protein \| Uncharacterized protein; afdb-uniprot50 \| AF-A0A1W9F593-F1-MODEL\_V4 \| 1.0 \| 9.964e-07 \| 247 \| 0.448 \| 98 \| 54 \| 0 \| 321 \| 418 \| 2 \| 99 \| Phage\_sheath\_1C domain-containing protein \| Phage\_sheath\_1C domain-containing protein; afdb-uniprot50 \| AF-A0A4Q7FYN1-F1-MODEL\_V4 \| 1.0 \| 3.312e-06 \| 245 \| 0.26 \| 150 \| 97 \| 2 \| 326 \| 474 \| 1 \| 137 \| Uncharacterized protein \| Uncharacterized protein; afdb-uniprot50 \| AF-B5JF22-F1-MODEL\_V4 \| 1.0 \| 3.082e-05 \| 244 \| 0.494 \| 91 \| 38 \| 3 \| 140 \| 224 \| 641 \| 729 \| Uncharacterized protein \| Uncharacterized protein; afdb-uniprot50 \| AF-A0A495J6Q7-F1-MODEL\_V4 \| 1.0 \| 2.488e-06 \| 243 \| 0.348 \| 109 \| 56 \| 2 \| 119 \| 226 \| 412 \| 506 \| Putative outer membrane starch-binding protein \| Putative outer membrane starch-binding protein; afdb-uniprot50 \| AF-A0A0F8VRZ6-F1-MODEL\_V4 \| 1.0 \| 2.91e-05 \| 241 \| 0.241 \| 112 \| 84 \| 1 \| 358 \| 468 \| 4 \| 115 \| Phage\_sheath\_1C domain-containing protein \| Phage\_sheath\_1C domain-containing protein; afdb-uniprot50 \| AF-A0A2D6AVV5-F1-MODEL\_V4 \| 1.0 \| 2.096e-06 \| 241 \| 0.192 \| 166 \| 124 \| 4 \| 308 \| 468 \| 2 \| 162 \| Uncharacterized protein \| Uncharacterized protein; afdb-uniprot50 \| AF-A0A4R4G448-F1-MODEL\_V4 \| 1.0 \| 6.966e-06 \| 239 \| 0.467 \| 77 \| 41 \| 0 \| 398 \| 474 \| 1 \| 77 \| Phage tail sheath family protein \| Phage tail sheath family protein; afdb-uniprot50 \| AF-A0A6M1E7R6-F1-MODEL\_V4 \| 1.0 \| 1.447e-08 \| 239 \| 0.284 \| 158 \| 76 \| 6 \| 116 \| 242 \| 379 \| 530 \| RagB/SusD family nutrient uptake outer membrane protein \| RagB/SusD family nutrient uptake outer membrane protein; afdb-uniprot50 \| AF-G5R8L3-F1-MODEL\_V4 \| 1.0 \| 2.452e-05 \| 237 \| 0.32 \| 81 \| 55 \| 0 \| 393 \| 473 \| 1 \| 81 \| Major tail sheath protein \| Major tail sheath protein; afdb-uniprot50 \| AF-A0A377U4U1-F1-MODEL\_V4 \| 1.0 \| 2.315e-05 \| 237 \| 0.35 \| 80 \| 52 \| 0 \| 395 \| 474 \| 1 \| 80 \| Phage tail sheath protein \| Phage tail sheath protein; afdb-uniprot50 \| AF-A0A356X1N2-F1-MODEL\_V4 \| 1.0 \| 6.579e-06 \| 235 \| 0.179 \| 184 \| 137 \| 5 \| 295 \| 468 \| 2 \| 181 \| Phage tail protein \| Phage tail protein; afdb-uniprot50 \| AF-A0A827DWH9-F1-MODEL\_V4 \| 1.0 \| 1.384e-05 \| 234 \| 0.402 \| 72 \| 43 \| 0 \| 403 \| 474 \| 4 \| 75 \| Phage tail protein \| Phage tail protein; afdb-uniprot50 \| AF-A0A7K4MS80-F1-MODEL\_V4 \| 1.0 \| 4.541e-08 \| 231 \| 0.286 \| 164 \| 63 \| 8 \| 93 \| 231 \| 84 \| 218 \| DUF4815 domain-containing protein \| DUF4815 domain-containing protein; afdb-uniprot50 \| AF-A0A7X5N4I8-F1-MODEL\_V4 \| 1.0 \| 5.542e-06 \| 228 \| 0.34 \| 97 \| 64 \| 0 \| 327 \| 423 \| 1 \| 97 \| Phage tail protein \| Phage tail protein; afdb-uniprot50 \| AF-A0A7C3WVH1-F1-MODEL\_V4 \| 1.0 \| 6.226e-10 \| 228 \| 0.118 \| 488 \| 312 \| 32 \| 49 \| 466 \| 519 \| 958 \| Uncharacterized protein \| Uncharacterized protein; afdb-uniprot50 \| AF-A0A317II01-F1-MODEL\_V4 \| 1.0 \| 8.757e-06 \| 225 \| 0.317 \| 104 \| 61 \| 3 \| 121 \| 223 \| 57 \| 151 \| Uncharacterized protein \| Uncharacterized protein; afdb-uniprot50 \| AF-A0A350DHD4-F1-MODEL\_V4 \| 1.0 \| 3.263e-05 \| 224 \| 0.388 \| 72 \| 44 \| 0 \| 403 \| 474 \| 5 \| 76 \| Phage tail protein \| Phage tail protein; afdb-uniprot50 \| AF-A0A3A9EM69-F1-MODEL\_V4 \| 1.0 \| 0.0002154 \| 218 \| 0.208 \| 96 \| 76 \| 0 \| 378 \| 473 \| 2 \| 97 \| Uncharacterized protein \| Uncharacterized protein; afdb-uniprot50 \| AF-R6I7Y6-F1-MODEL\_V4 \| 1.0 \| 0.0001714 \| 215 \| 0.236 \| 93 \| 71 \| 0 \| 382 \| 474 \| 6 \| 98 \| Uncharacterized protein \| Uncharacterized protein; afdb-uniprot50 \| AF-A0A653AK93-F1-MODEL\_V4 \| 1.0 \| 5.781e-05 \| 214 \| 0.206 \| 131 \| 99 \| 3 \| 349 \| 474 \| 2 \| 132 \| Tail sheath protein \| Tail sheath protein; afdb-uniprot50 \| AF-A0A4U9HES6-F1-MODEL\_V4 \| 1.0 \| 2.749e-05 \| 211 \| 0.523 \| 65 \| 31 \| 0 \| 410 \| 474 \| 2 \| 66 \| Phage tail sheath protein \| Phage tail sheath protein; afdb-uniprot50 \| AF-U2KYU4-F1-MODEL\_V4 \| 1.0 \| 3.514e-10 \| 210 \| 0.101 \| 643 \| 327 \| 31 \| 30 \| 473 \| 3 \| 593 \| Uncharacterized protein \| Uncharacterized protein; afdb-uniprot50 \| AF-A0A6C7S0W9-F1-MODEL\_V4 \| 1.0 \| 0.0002558 \| 209 \| 0.209 \| 110 \| 85 \| 1 \| 361 \| 470 \| 2 \| 109 \| Phage tail sheath family protein \| Phage tail sheath family protein; afdb-uniprot50 \| AF-A0A1W1Z5G4-F1-MODEL\_V4 \| 1.0 \| 5.016e-07 \| 209 \| 0.264 \| 155 \| 75 \| 6 \| 116 \| 243 \| 385 \| 527 \| Starch-binding associating with outer membrane \| Starch-binding associating with outer membrane; afdb-uniprot50 \| AF-A0A382Q741-F1-MODEL\_V4 \| 1.0 \| 0.0001714 \| 207 \| 0.182 \| 126 \| 101 \| 2 \| 350 \| 473 \| 1 \| 126 \| Phage\_sheath\_1C domain-containing protein \| Phage\_sheath\_1C domain-containing protein; afdb-uniprot50 \| AF-A0A0F8XJ22-F1-MODEL\_V4 \| 1.0 \| 0.0001529 \| 206 \| 0.145 \| 131 \| 109 \| 2 \| 341 \| 468 \| 2 \| 132 \| Uncharacterized protein \| Uncharacterized protein; afdb-uniprot50 \| AF-A0A519WUA4-F1-MODEL\_V4 \| 1.0 \| 2.525e-07 \| 205 \| 0.26 \| 161 \| 74 \| 9 \| 116 \| 243 \| 388 \| 536 \| RagB/SusD family nutrient uptake outer membrane protein \| RagB/SusD family nutrient uptake outer membrane protein; afdb-uniprot50 \| AF-A0A376KHV7-F1-MODEL\_V4 \| 1.0 \| 1.166e-05 \| 204 \| 0.275 \| 120 \| 62 \| 2 \| 273 \| 367 \| 3 \| 122 \| Major tail sheath protein FI \| Major tail sheath protein FI; afdb-uniprot50 \| AF-X0XYY1-F1-MODEL\_V4 \| 1.0 \| 5.46e-05 \| 204 \| 0.155 \| 148 \| 122 \| 2 \| 324 \| 468 \| 1 \| 148 \| Phage\_sheath\_1C domain-containing protein \| Phage\_sheath\_1C domain-containing protein; afdb-uniprot50 \| AF-A0A6M1N703-F1-MODEL\_V4 \| 1.0 \| 1.219e-08 \| 197 \| 0.241 \| 207 \| 98 \| 12 \| 61 \| 243 \| 377 \| 548 \| RagB/SusD family nutrient uptake outer membrane protein \| RagB/SusD family nutrient uptake outer membrane protein; afdb-uniprot50 \| AF-A0A353G6H1-F1-MODEL\_V4 \| 1.0 \| 0.0002154 \| 196 \| 0.365 \| 82 \| 52 \| 0 \| 393 \| 474 \| 4 \| 85 \| Phage tail protein \| Phage tail protein; afdb-uniprot50 \| AF-A0A1V6IIL8-F1-MODEL\_V4 \| 1.0 \| 9.674e-05 \| 196 \| 0.134 \| 194 \| 158 \| 6 \| 282 \| 470 \| 246 \| 434 \| Uncharacterized protein \| Uncharacterized protein; afdb-uniprot50 \| AF-A0A495J3X6-F1-MODEL\_V4 \| 1.0 \| 1.692e-07 \| 196 \| 0.3 \| 180 \| 81 \| 7 \| 83 \| 243 \| 356 \| 509 \| Putative outer membrane starch-binding protein \| Putative outer membrane starch-binding protein; afdb-uniprot50 \| AF-U2QTT3-F1-MODEL\_V4 \| 1.0 \| 1.575e-06 \| 192 \| 0.109 \| 501 \| 282 \| 28 \| 4 \| 466 \| 7 \| 381 \| Uncharacterized protein \| Uncharacterized protein; afdb-uniprot50 \| AF-A8GDF6-F1-MODEL\_V4 \| 1.0 \| 4.737e-07 \| 190 \| 0.21 \| 247 \| 97 \| 3 \| 94 \| 339 \| 1 \| 150 \| Phage tail sheath protein FI-like protein \| Phage tail sheath protein FI-like protein; afdb-uniprot50 \| AF-A0A5Z1TET5-F1-MODEL\_V4 \| 1.0 \| 0.0009531 \| 189 \| 0.177 \| 96 \| 77 \| 1 \| 379 \| 474 \| 2 \| 95 \| Phage tail sheath family protein \| Phage tail sheath family protein; afdb-uniprot50 \| AF-A0A7C2S225-F1-MODEL\_V4 \| 1.0 \| 0.0009002 \| 186 \| 0.193 \| 119 \| 93 \| 2 \| 358 \| 473 \| 3 \| 121 \| Phage tail protein \| Phage tail protein; afdb-uniprot50 \| AF-A0A350PJK4-F1-MODEL\_V4 \| 1.0 \| 0.0008501 \| 185 \| 0.169 \| 112 \| 92 \| 1 \| 358 \| 468 \| 2 \| 113 \| Phage\_sheath\_1C domain-containing protein \| Phage\_sheath\_1C domain-containing protein; afdb-uniprot50 \| AF-A0A356WZX7-F1-MODEL\_V4 \| 1.0 \| 0.0003036 \| 185 \| 0.154 \| 136 \| 107 \| 3 \| 341 \| 468 \| 3 \| 138 \| Uncharacterized protein \| Uncharacterized protein; afdb-uniprot50 \| AF-A0A519XZM0-F1-MODEL\_V4 \| 1.0 \| 8.887e-07 \| 183 \| 0.256 \| 179 \| 81 \| 7 \| 61 \| 226 \| 139 \| 278 \| RagB/SusD family nutrient uptake outer membrane protein \| RagB/SusD family nutrient uptake outer membrane protein; afdb-uniprot50 \| AF-A0A519SBQ2-F1-MODEL\_V4 \| 1.0 \| 9.964e-07 \| 183 \| 0.245 \| 179 \| 83 \| 7 \| 61 \| 226 \| 361 \| 500 \| DUF5017 domain-containing protein \| DUF5017 domain-containing protein; afdb-uniprot50 \| AF-A0A519MRJ7-F1-MODEL\_V4 \| 1.0 \| 1.055e-06 \| 179 \| 0.245 \| 179 \| 83 \| 8 \| 61 \| 226 \| 280 \| 419 \| RagB/SusD family nutrient uptake outer membrane protein \| RagB/SusD family nutrient uptake outer membrane protein; afdb-uniprot50 \| AF-A0A7W4KR57-F1-MODEL\_V4 \| 1.0 \| 0.0002558 \| 178 \| 0.325 \| 80 \| 54 \| 0 \| 382 \| 461 \| 2 \| 81 \| Phage tail sheath family protein \| Phage tail sheath family protein; afdb-uniprot50 \| AF-A0A248LIR3-F1-MODEL\_V4 \| 1.0 \| 0.0001529 \| 177 \| 0.426 \| 61 \| 35 \| 0 \| 413 \| 473 \| 28 \| 88 \| Phage tail sheath protein \| Phage tail sheath protein; afdb-uniprot50 \| AF-A0A350PHK9-F1-MODEL\_V4 \| 1.0 \| 0.001689 \| 173 \| 0.178 \| 95 \| 78 \| 0 \| 374 \| 468 \| 5 \| 99 \| Phage tail protein \| Phage tail protein; afdb-uniprot50 \| AF-A0A4R0MPD7-F1-MODEL\_V4 \| 1.0 \| 3.043e-08 \| 170 \| 0.225 \| 231 \| 104 \| 11 \| 113 \| 288 \| 399 \| 609 \| RagB/SusD family nutrient uptake outer membrane protein \| RagB/SusD family nutrient uptake outer membrane protein; afdb-uniprot50 \| AF-A0A0F9LR86-F1-MODEL\_V4 \| 1.0 \| 0.003761 \| 168 \| 0.156 \| 115 \| 96 \| 1 \| 353 \| 466 \| 6 \| 120 \| Uncharacterized protein \| Uncharacterized protein; afdb-uniprot50 \| AF-A0A496KP74-F1-MODEL\_V4 \| 1.0 \| 0.001343 \| 166 \| 0.277 \| 72 \| 52 \| 0 \| 403 \| 474 \| 4 \| 75 \| Phage tail protein \| Phage tail protein |
| Top keywords  (threshold 1.00e-02 (evalue)) | **tail, Phage, sheath, domain\_containing, Phage\_sheath\_1C, Phage\_sheath\_1, FI, Major, Putative, subtilisin\_like** |
| Output files | ../../similar\_structures/18\_FANPEZAQ\_CDS\_0018\_afdb-proteome\_foldseek.tsv ../../similar\_structures/18\_FANPEZAQ\_CDS\_0018\_afdb-uniprot50\_foldseek.tsv ../../similar\_structures/18\_FANPEZAQ\_CDS\_0018\_merged.svg ../../similar\_structures/18\_FANPEZAQ\_CDS\_0018\_pdb\_foldseek.tsv |

  
  
  

Return to summary | Go to previous | Go to next

  


---

**Sequence/structure alignments coloring**  
Each object in the alignment figures is colored according to its E-value following this color coding:

1e-100
10
