## Supplementary material for "Completing the BASEL phage collection to unlock hidden diversity for systematic exploration of phage-host interactions": Data S2: 19.html

FANPEZAQ\_CDS\_0019


Return to summary | Go to previous | Go to next

|  |  |
| --- | --- |
| FANPEZAQ\_CDS\_0019  Page creation date: 02 Sep 2024, 12:00  Project folder: n/a  Input sequences file: Escherichia\_virus\_HeidiAbel.gb | tail phage tube major fii putative fragment contractile prophage p2 sheath bacteriophage hypothetical core head closure domain\_containing from structural tp901\_1 connector mumc02 phage\_like probable protoporphyrinogen oxidase pyocin bacteriocin tail\_like gpfi fii\_like afp1 baseplate hcp gp53 fels\_2 p2w3 conserved region |

### Sequence information

|  |  |
| --- | --- |
| Name | FANPEZAQ\_CDS\_0019  19\_FANPEZAQ\_CDS\_0019 (pipeline id) |
| Imported annotations | Escherichia\_virus\_HeidiAbel Bas97 |
| Protein sequence | MAARDVLKNINLFVDGRGYAGQVESVTPPTLTLQTEEFRAGGMDAPIDVTLGMEKLEASF ALRAYDSDVLALFGLSEGASVPLTFRGALESYDGSVKAVVMNMRGKITSMEPGTWTPGEL SSLNVTVSLNYFKQQHNGKTIHEIDVENMVRIINGVDTLAAIRSALAM |
| Number of residues | 168 |
| Molecular weight (Da) | 18113.49 |
| Output files | ../../query\_sequences/19\_FANPEZAQ\_CDS\_0019.fasta |

### Putative domain architecture and protein family

#### Search results (HHblits)^1^

|  |  |
| --- | --- |
| Domain family databases searched | Pfam, Ncbi-cd, Cath, Phrogs |
| Results, scheme(s)  (Top layers only; threshold 1.00e-03 (evalue)) | xml version="1.0" encoding="utf-8" standalone="no"?       2024-09-02T21:08:16.313371 image/svg+xml   Matplotlib v3.7.2, https://matplotlib.org/ |
| Results, table  (E-value ≤ 1.00e-03 (evalue)) | db \| id \| prob \| evalue \| pvalue \| score \| cols \| query \| query\_len \| template \| template\_len \| name \| description; pfam \| PF04985 \| 100.0 \| 1.3e-42 \| 2.5e-46 \| 248.0 \| 159 \| (5, 167) \| 168 \| (2, 161) \| 161 \| Phage\_tube \| Phage tail tube protein FII; phrogs \| 55 \| 100.0 \| 3e-52 \| 3.9e-56 \| 320.5 \| 166 \| (2, 168) \| 168 \| (1, 171) \| 172 \| head closure \| head closure; Category: connector; p363376 VI\_00073; phrogs \| 13676 \| 99.9 \| 2.7e-27 \| 3e-31 \| 157.0 \| 89 \| (80, 168) \| 168 \| (3, 92) \| 92 \| head closure \| head closure; Category: connector; p268213 VI\_02784; phrogs \| 15831 \| 99.2 \| 6.8e-16 \| 7.8e-20 \| 89.6 \| 45 \| (1, 45) \| 168 \| (1, 45) \| 46 \| head closure \| head closure; Category: connector; p388731 VI\_08178 |
| Top keywords  (threshold 1.00e-03 (evalue)) | **head, closure, connector, Phage, tail, tube, FII, p363376, VI\_00073, p268213** |
| Output files | ../../domain\_architecture/19\_FANPEZAQ\_CDS\_0019\_cath.hhr ../../domain\_architecture/19\_FANPEZAQ\_CDS\_0019\_merged.svg ../../domain\_architecture/19\_FANPEZAQ\_CDS\_0019\_ncbi-cd.hhr ../../domain\_architecture/19\_FANPEZAQ\_CDS\_0019\_pfam.hhr ../../domain\_architecture/19\_FANPEZAQ\_CDS\_0019\_phrogs.hhr |

### Identical protein sequences/structures

#### Search results

|  |  |
| --- | --- |
| Protein sequence databases searched | Pdb, Swissprot, Refseq |
| Identical proteins found | Refseq  - YP\_009168894.1: Head closure (Escherichia phage vB\_EcoM\_ECO1230-10) - YP\_009600711.1: Head closure (Escherichia phage vB\_EcoM\_ECOO78) - ADE87917.1: Putative phage tail protein (Escherichia phage vB\_EcoM\_ECO1230-10) - ARM70443.1: Putative tail protein (Escherichia phage vB\_EcoM\_ECOO78) - QIG65610.1: Major tail tube protein (Salmonella phage PT1) |
| Top keywords | **tail, Head, closure, Putative, phage, Major, tube** |
| Output files | ../../identical\_sequences/19\_FANPEZAQ\_CDS\_0019\_refseq.fasta |

### Similar protein sequences/structures

#### Sequence similarity search results (HHblits)^1^

|  |  |
| --- | --- |
| Sequence databases searched | Uniclust, Pdb70 |
| Results, scheme(s)  (Top layers only, threshold 1.00e-03 (evalue)) | xml version="1.0" encoding="utf-8" standalone="no"?       2024-09-02T21:08:37.339295 image/svg+xml   Matplotlib v3.7.2, https://matplotlib.org/ |
| Results, table(s)  (threshold 1.00e-03 (evalue)) | db \| id \| prob \| evalue \| pvalue \| score \| cols \| query \| query\_len \| template \| template\_len \| name \| description; uniclust \| UniRef100\_A0A0B4BLQ2 \| 100.0 \| 3.6e-56 \| 7.9e-62 \| 333.3 \| 168 \| (1, 168) \| 168 \| (8, 175) \| 189 \| Major tail tube protein \| Major tail tube protein; uniclust \| UniRef100\_A0A024E873 \| 100.0 \| 6.1e-54 \| 1.4e-59 \| 328.6 \| 168 \| (1, 168) \| 168 \| (22, 189) \| 201 \| Major tail tube protein \| Major tail tube protein; uniclust \| UniRef100\_A0A009PKY6 \| 100.0 \| 2.6e-52 \| 5.5e-58 \| 321.7 \| 168 \| (1, 168) \| 168 \| (44, 214) \| 239 \| Phage major tail tube protein \| Phage major tail tube protein; uniclust \| UniRef100\_A0A017HBX3 \| 100.0 \| 2.8e-47 \| 6.1e-53 \| 285.6 \| 165 \| (2, 168) \| 168 \| (26, 190) \| 192 \| Phage major tail tube protein \| Phage major tail tube protein; uniclust \| UniRef100\_A0A075WT57 \| 100.0 \| 7.7e-46 \| 1.6e-51 \| 273.1 \| 167 \| (1, 168) \| 168 \| (3, 173) \| 174 \| Tail protein \| Tail protein; uniclust \| UniRef100\_A0A0U1DAP0 \| 100.0 \| 5.5e-45 \| 1e-50 \| 261.6 \| 167 \| (2, 168) \| 168 \| (16, 183) \| 183 \| Phage tail tube protein \| Phage tail tube protein; uniclust \| UniRef100\_A0A011NCB3 \| 100.0 \| 8.1e-45 \| 1.7e-50 \| 273.2 \| 164 \| (4, 168) \| 168 \| (24, 191) \| 201 \| Phage major tail tube protein \| Phage major tail tube protein; uniclust \| UniRef100\_A0A061NPQ1 \| 100.0 \| 3.3e-43 \| 7.4e-49 \| 267.6 \| 164 \| (1, 168) \| 168 \| (20, 190) \| 192 \| Phage major tail tube protein \| Phage major tail tube protein; uniclust \| UniRef100\_A0A0X1U7S9 \| 100.0 \| 1.2e-41 \| 2.6e-47 \| 254.4 \| 166 \| (1, 168) \| 168 \| (11, 183) \| 183 \| Phage tail tube protein FII \| Phage tail tube protein FII; uniclust \| UniRef100\_A0A0E3Y6F4 \| 100.0 \| 1.6e-41 \| 3.5e-47 \| 252.4 \| 164 \| (1, 168) \| 168 \| (7, 177) \| 178 \| Major tail tube protein \| Major tail tube protein; uniclust \| UniRef100\_A0A0P7HQ39 \| 100.0 \| 1.2e-40 \| 2.6e-46 \| 234.2 \| 129 \| (40, 168) \| 168 \| (1, 130) \| 131 \| Major tail tube protein \| Major tail tube protein; uniclust \| UniRef100\_A0A8X6GSQ5 \| 100.0 \| 1.9e-40 \| 3.5e-46 \| 244.0 \| 162 \| (2, 163) \| 168 \| (1, 163) \| 234 \| Phage tail tube protein FII \| Phage tail tube protein FII; uniclust \| UniRef100\_A0A165XH78 \| 100.0 \| 3.2e-40 \| 6.7e-46 \| 244.8 \| 166 \| (3, 168) \| 168 \| (10, 178) \| 180 \| Phage tail tube protein FII \| Phage tail tube protein FII; uniclust \| UniRef100\_A0A0D7WZ75 \| 100.0 \| 3.8e-40 \| 8.6e-46 \| 248.3 \| 164 \| (1, 168) \| 168 \| (9, 179) \| 181 \| Phage tail protein \| Phage tail protein; uniclust \| UniRef100\_A0A066RR28 \| 100.0 \| 9e-39 \| 1.8e-44 \| 231.1 \| 168 \| (1, 168) \| 168 \| (2, 171) \| 172 \| Major tail tube protein \| Major tail tube protein; uniclust \| UniRef100\_A0A081MYF2 \| 100.0 \| 8e-37 \| 1.6e-42 \| 220.8 \| 166 \| (2, 168) \| 168 \| (4, 170) \| 170 \| Phage major tail tube protein \| Phage major tail tube protein; uniclust \| UniRef100\_A0A2M7MWC6 \| 100.0 \| 1.2e-36 \| 2.1e-42 \| 223.1 \| 166 \| (2, 167) \| 168 \| (1, 166) \| 235 \| Phage major tail tube protein (Fragment) \| Phage major tail tube protein (Fragment); uniclust \| UniRef100\_A0A2X2BQ70 \| 100.0 \| 2e-36 \| 4.1e-42 \| 215.5 \| 139 \| (1, 140) \| 168 \| (1, 142) \| 145 \| Major tail tube protein \| Major tail tube protein; uniclust \| UniRef100\_A0A450Z351 \| 100.0 \| 4.7e-36 \| 9.4e-42 \| 208.8 \| 125 \| (1, 125) \| 168 \| (3, 127) \| 129 \| Phage tail tube protein FII \| Phage tail tube protein FII; uniclust \| UniRef100\_A0A8X6MUK6 \| 100.0 \| 6.6e-36 \| 1.2e-41 \| 238.1 \| 165 \| (3, 167) \| 168 \| (264, 429) \| 498 \| Putative prophage major tail sheath protein \| Putative prophage major tail sheath protein; uniclust \| UniRef100\_A0A3G2IAH9 \| 100.0 \| 1.7e-35 \| 3.3e-41 \| 215.9 \| 162 \| (4, 167) \| 168 \| (9, 173) \| 176 \| Phage tail protein \| Phage tail protein; uniclust \| UniRef100\_A0A061JHN2 \| 100.0 \| 2.2e-34 \| 4e-40 \| 207.1 \| 146 \| (2, 148) \| 168 \| (1, 146) \| 201 \| Major tail tube protein \| Major tail tube protein; uniclust \| UniRef100\_A0A1Y6CXU4 \| 100.0 \| 2.4e-34 \| 4.7e-40 \| 207.7 \| 168 \| (1, 168) \| 168 \| (3, 172) \| 173 \| Phage major tail tube protein \| Phage major tail tube protein; uniclust \| UniRef100\_A0A087M4E0 \| 100.0 \| 4.8e-34 \| 1e-39 \| 214.5 \| 165 \| (4, 168) \| 168 \| (8, 179) \| 185 \| Tail protein \| Tail protein; uniclust \| UniRef100\_A0A061K3S3 \| 100.0 \| 8.2e-34 \| 1.7e-39 \| 209.2 \| 166 \| (1, 168) \| 168 \| (6, 173) \| 174 \| Phage tail protein \| Phage tail protein; uniclust \| UniRef100\_A0A1V3VT23 \| 99.9 \| 2.8e-32 \| 5.4e-38 \| 196.8 \| 166 \| (1, 168) \| 168 \| (1, 169) \| 171 \| Phage tail protein (Fragment) \| Phage tail protein (Fragment); uniclust \| UniRef100\_A0A376W1N8 \| 99.9 \| 5.3e-32 \| 1e-37 \| 192.1 \| 148 \| (21, 168) \| 168 \| (2, 153) \| 153 \| Major tail sheath protein FII from prophage \| Major tail sheath protein FII from prophage; uniclust \| UniRef100\_A0A2W6PC71 \| 99.9 \| 2.6e-31 \| 5.1e-37 \| 186.4 \| 129 \| (38, 168) \| 168 \| (1, 132) \| 134 \| Phage tail protein (Fragment) \| Phage tail protein (Fragment); uniclust \| UniRef100\_A0A8X6WS87 \| 99.9 \| 1.4e-30 \| 2.5e-36 \| 191.1 \| 144 \| (25, 168) \| 168 \| (84, 228) \| 228 \| Tail tube protein \| Tail tube protein; uniclust \| UniRef100\_A0A261QLN0 \| 99.9 \| 1.8e-30 \| 3.6e-36 \| 184.9 \| 139 \| (28, 167) \| 168 \| (1, 143) \| 148 \| Phage major tail tube protein (Fragment) \| Phage major tail tube protein (Fragment); uniclust \| UniRef100\_A0A1A9VKH7 \| 99.9 \| 2.2e-30 \| 4e-36 \| 208.7 \| 152 \| (17, 168) \| 168 \| (383, 535) \| 535 \| Phage tail protein \| Phage tail protein; uniclust \| UniRef100\_A0A0F4NK74 \| 99.9 \| 6.8e-30 \| 1.4e-35 \| 187.7 \| 163 \| (1, 167) \| 168 \| (7, 172) \| 173 \| Phage major tail tube protein \| Phage major tail tube protein; uniclust \| UniRef100\_A0A0C2UPV3 \| 99.9 \| 9.3e-30 \| 1.9e-35 \| 186.4 \| 156 \| (1, 166) \| 168 \| (4, 160) \| 161 \| Major tail tube protein \| Major tail tube protein; uniclust \| UniRef100\_A0A085AFM9 \| 99.9 \| 1.1e-29 \| 2.4e-35 \| 188.6 \| 167 \| (1, 168) \| 168 \| (1, 167) \| 168 \| Phage major tail tube protein \| Phage major tail tube protein; uniclust \| UniRef100\_A0A0A8H9N3 \| 99.9 \| 8.3e-29 \| 1.6e-34 \| 180.1 \| 164 \| (2, 168) \| 168 \| (8, 174) \| 174 \| Phage contractile tail tube protein, P2 family \| Phage contractile tail tube protein, P2 family; uniclust \| UniRef100\_A0A031FRM0 \| 99.9 \| 1.7e-28 \| 3.4e-34 \| 181.0 \| 166 \| (1, 167) \| 168 \| (2, 167) \| 170 \| Putative phage tail-like protein \| Putative phage tail-like protein; uniclust \| UniRef100\_A0A095WG55 \| 99.9 \| 2e-28 \| 4e-34 \| 178.4 \| 162 \| (2, 168) \| 168 \| (6, 175) \| 176 \| Phage major tail tube protein \| Phage major tail tube protein; uniclust \| UniRef100\_A0A0Q6RZW5 \| 99.9 \| 5.3e-28 \| 1.1e-33 \| 180.4 \| 166 \| (2, 167) \| 168 \| (4, 171) \| 184 \| Phage tail protein \| Phage tail protein; uniclust \| UniRef100\_UPI00031064D1 \| 99.9 \| 9.2e-28 \| 1.7e-33 \| 163.4 \| 123 \| (2, 126) \| 168 \| (1, 123) \| 125 \| phage major tail tube protein \| phage major tail tube protein; uniclust \| UniRef100\_A0A0S2ZDF0 \| 99.9 \| 2.2e-27 \| 4.5e-33 \| 174.0 \| 161 \| (1, 167) \| 168 \| (1, 169) \| 169 \| Phage tail protein \| Phage tail protein; uniclust \| UniRef100\_E1SFY3 \| 99.9 \| 2.6e-27 \| 4.8e-33 \| 182.6 \| 153 \| (1, 156) \| 168 \| (1, 156) \| 335 \| Major tail tube protein (Protein FII) \| Major tail tube protein (Protein FII); uniclust \| UniRef100\_A0A345DE54 \| 99.9 \| 1.1e-26 \| 1.9e-32 \| 164.5 \| 167 \| (2, 168) \| 168 \| (1, 169) \| 170 \| Phage tail tube protein FII \| Phage tail tube protein FII; uniclust \| UniRef100\_A0A1S6TP66 \| 99.9 \| 1.4e-26 \| 2.7e-32 \| 171.3 \| 161 \| (3, 167) \| 168 \| (7, 168) \| 175 \| Phage major tail tube protein \| Phage major tail tube protein; uniclust \| UniRef100\_A0A450Z316 \| 99.9 \| 2.1e-26 \| 4.2e-32 \| 151.2 \| 82 \| (87, 168) \| 168 \| (2, 84) \| 85 \| Phage tail tube protein FII \| Phage tail tube protein FII; uniclust \| UniRef100\_A0A0C1J3Q2 \| 99.9 \| 2.7e-26 \| 5.8e-32 \| 171.7 \| 156 \| (6, 162) \| 168 \| (9, 173) \| 174 \| Uncharacterized protein \| Uncharacterized protein; uniclust \| UniRef100\_A0A059IUF5 \| 99.9 \| 3.2e-26 \| 6.1e-32 \| 153.4 \| 106 \| (2, 107) \| 168 \| (1, 107) \| 110 \| Contractile tail tube protein \| Contractile tail tube protein; uniclust \| UniRef100\_A0A0A8ILB8 \| 99.9 \| 3.1e-26 \| 6.5e-32 \| 171.0 \| 149 \| (16, 168) \| 168 \| (21, 172) \| 178 \| Tail tube \| Tail tube; uniclust \| UniRef100\_A0A173R5V1 \| 99.9 \| 5e-26 \| 1e-31 \| 168.0 \| 155 \| (5, 166) \| 168 \| (10, 170) \| 170 \| Phage major tail tube protein \| Phage major tail tube protein; uniclust \| UniRef100\_A0A0S4XLP2 \| 99.9 \| 6.7e-26 \| 1.4e-31 \| 167.2 \| 159 \| (5, 166) \| 168 \| (10, 169) \| 172 \| Phage major tail tube protein \| Phage major tail tube protein; uniclust \| UniRef100\_A0A248LI16 \| 99.9 \| 9.6e-26 \| 1.8e-31 \| 148.1 \| 104 \| (2, 168) \| 168 \| (1, 104) \| 104 \| Phage tail protein \| Phage tail protein; uniclust \| UniRef100\_Q2NU86 \| 99.9 \| 1.2e-25 \| 2.5e-31 \| 154.9 \| 112 \| (1, 114) \| 168 \| (1, 114) \| 115 \| Hypothetical phage protein \| Hypothetical phage protein; uniclust \| UniRef100\_J0PYV7 \| 99.8 \| 5.5e-25 \| 1e-30 \| 150.8 \| 130 \| (36, 168) \| 168 \| (2, 131) \| 131 \| Phage major tail tube protein \| Phage major tail tube protein; uniclust \| UniRef100\_A0A379YDV4 \| 99.8 \| 2.1e-24 \| 4.3e-30 \| 161.8 \| 156 \| (3, 160) \| 168 \| (1, 159) \| 191 \| Phage major tail tube protein \| Phage major tail tube protein; uniclust \| UniRef100\_A0A8D8JHV8 \| 99.8 \| 3.8e-24 \| 7.3e-30 \| 143.7 \| 105 \| (64, 168) \| 168 \| (3, 107) \| 107 \| Tail tube protein \| Tail tube protein; uniclust \| UniRef100\_A0A1H2FLU6 \| 99.8 \| 7.1e-24 \| 1.4e-29 \| 156.9 \| 167 \| (1, 167) \| 168 \| (3, 174) \| 188 \| Phage tail tube protein FII \| Phage tail tube protein FII; uniclust \| UniRef100\_UPI00082ED17C \| 99.8 \| 1e-23 \| 2e-29 \| 151.5 \| 163 \| (3, 168) \| 168 \| (2, 167) \| 168 \| phage major tail tube protein \| phage major tail tube protein; uniclust \| UniRef100\_A0A1E7NW44 \| 99.8 \| 1.6e-23 \| 3.2e-29 \| 153.5 \| 162 \| (1, 168) \| 168 \| (3, 166) \| 166 \| Phage tail protein \| Phage tail protein; uniclust \| UniRef100\_A0A645DSK8 \| 99.8 \| 3.8e-23 \| 7.6e-29 \| 141.1 \| 90 \| (79, 168) \| 168 \| (13, 107) \| 107 \| Phage tail tube protein FII \| Phage tail tube protein FII; uniclust \| UniRef100\_UPI001FCC480A \| 99.8 \| 4.6e-23 \| 8.4e-29 \| 154.3 \| 143 \| (23, 167) \| 168 \| (106, 248) \| 249 \| phage major tail tube protein \| phage major tail tube protein; uniclust \| UniRef100\_UPI000B6EE99C \| 99.8 \| 5.5e-23 \| 1.1e-28 \| 141.0 \| 113 \| (1, 113) \| 168 \| (1, 116) \| 120 \| phage major tail tube protein \| phage major tail tube protein; uniclust \| UniRef100\_A0A173SC52 \| 99.8 \| 2.8e-22 \| 5.1e-28 \| 150.4 \| 157 \| (7, 167) \| 168 \| (65, 227) \| 251 \| Phage major tail tube protein \| Phage major tail tube protein; uniclust \| UniRef100\_A0A2A4XT18 \| 99.8 \| 5.2e-22 \| 9.8e-28 \| 143.0 \| 166 \| (3, 168) \| 168 \| (2, 167) \| 167 \| Phage tail protein \| Phage tail protein; uniclust \| UniRef100\_A0A1N7LR57 \| 99.8 \| 1.2e-21 \| 2.3e-27 \| 140.4 \| 160 \| (6, 168) \| 168 \| (12, 172) \| 173 \| Phage major tail tube protein \| Phage major tail tube protein; uniclust \| UniRef100\_A0A378PU11 \| 99.7 \| 1.9e-21 \| 3.4e-27 \| 136.4 \| 115 \| (51, 166) \| 168 \| (12, 127) \| 152 \| Phage major tail tube protein \| Phage major tail tube protein; uniclust \| UniRef100\_D1AFA7 \| 99.7 \| 3.6e-21 \| 6.7e-27 \| 137.6 \| 161 \| (3, 166) \| 168 \| (5, 172) \| 173 \| Major tail tube protein \| Major tail tube protein; uniclust \| UniRef100\_A0A1E3G652 \| 99.7 \| 3.6e-21 \| 6.7e-27 \| 136.9 \| 166 \| (1, 168) \| 168 \| (1, 166) \| 167 \| Phage major tail tube protein \| Phage major tail tube protein; uniclust \| UniRef100\_UPI001F575EED \| 99.7 \| 4e-21 \| 7.6e-27 \| 138.7 \| 166 \| (1, 168) \| 168 \| (3, 170) \| 171 \| phage major tail tube protein \| phage major tail tube protein; uniclust \| UniRef100\_A0A7I8DLE1 \| 99.7 \| 6.2e-21 \| 1.2e-26 \| 138.6 \| 143 \| (22, 167) \| 168 \| (26, 174) \| 175 \| Bacteriophage protein \| Bacteriophage protein; uniclust \| UniRef100\_A0A9E6RJX5 \| 99.7 \| 1.1e-20 \| 2e-26 \| 134.9 \| 162 \| (4, 167) \| 168 \| (3, 168) \| 169 \| Phage major tail tube protein \| Phage major tail tube protein; uniclust \| UniRef100\_A0A1D2QMU1 \| 99.7 \| 1.6e-20 \| 3e-26 \| 136.2 \| 164 \| (2, 167) \| 168 \| (4, 168) \| 172 \| Phage major tail tube protein \| Phage major tail tube protein; uniclust \| UniRef100\_A0A1Y1QXH8 \| 99.7 \| 1.9e-20 \| 3.4e-26 \| 133.9 \| 162 \| (2, 167) \| 168 \| (8, 170) \| 171 \| Phage tail protein \| Phage tail protein; uniclust \| UniRef100\_A0A2T5U148 \| 99.7 \| 2.6e-20 \| 5.2e-26 \| 120.3 \| 73 \| (96, 168) \| 168 \| (3, 75) \| 76 \| P2 family phage contractile tail tube protein \| P2 family phage contractile tail tube protein; uniclust \| UniRef100\_A0A0B6D149 \| 99.7 \| 6.6e-20 \| 1.2e-25 \| 131.2 \| 159 \| (5, 168) \| 168 \| (11, 171) \| 171 \| Phage tail tube FII family protein \| Phage tail tube FII family protein; uniclust \| UniRef100\_A0A6P0MC46 \| 99.7 \| 9.9e-20 \| 2e-25 \| 135.8 \| 163 \| (5, 168) \| 168 \| (8, 176) \| 178 \| Phage tail protein \| Phage tail protein; uniclust \| UniRef100\_A0A1X7MCI7 \| 99.7 \| 1.1e-19 \| 2.2e-25 \| 119.2 \| 67 \| (2, 69) \| 168 \| (15, 81) \| 85 \| Phage major tail tube protein (Fragment) \| Phage major tail tube protein (Fragment); uniclust \| UniRef100\_A0A090KJ19 \| 99.7 \| 1.3e-19 \| 2.8e-25 \| 136.0 \| 158 \| (1, 162) \| 168 \| (3, 160) \| 165 \| Putative phage tail tube protein \| Putative phage tail tube protein; uniclust \| UniRef100\_A0A328TPU4 \| 99.7 \| 1.7e-19 \| 3.3e-25 \| 127.1 \| 119 \| (4, 123) \| 168 \| (7, 129) \| 133 \| Phage major tail tube protein \| Phage major tail tube protein; uniclust \| UniRef100\_A0A2E3R286 \| 99.7 \| 2.1e-19 \| 3.9e-25 \| 128.8 \| 162 \| (3, 166) \| 168 \| (4, 172) \| 172 \| Phage tail protein \| Phage tail protein; uniclust \| UniRef100\_A0A812QV43 \| 99.6 \| 5.4e-19 \| 1e-24 \| 153.1 \| 143 \| (18, 160) \| 168 \| (388, 531) \| 1104 \| GpFI protein \| GpFI protein; uniclust \| UniRef100\_A0A2A5BNC9 \| 99.6 \| 6.7e-19 \| 1.2e-24 \| 126.4 \| 166 \| (3, 168) \| 168 \| (3, 171) \| 173 \| Phage tail protein \| Phage tail protein; uniclust \| UniRef100\_A0A376FP60 \| 99.6 \| 1.1e-18 \| 2.1e-24 \| 108.9 \| 66 \| (103, 168) \| 168 \| (1, 66) \| 66 \| Phage major tail tube protein \| Phage major tail tube protein; uniclust \| UniRef100\_A0A2X2J512 \| 99.6 \| 1.9e-18 \| 3.6e-24 \| 114.1 \| 73 \| (96, 168) \| 168 \| (11, 84) \| 87 \| Tail tube protein \| Tail tube protein; uniclust \| UniRef100\_A0A0T7E1W1 \| 99.6 \| 1.9e-18 \| 3.7e-24 \| 125.4 \| 158 \| (9, 168) \| 168 \| (8, 165) \| 166 \| Phage tail protein \| Phage tail protein; uniclust \| UniRef100\_UPI00056FD72B \| 99.6 \| 2.3e-18 \| 4.3e-24 \| 123.3 \| 140 \| (1, 141) \| 168 \| (1, 144) \| 150 \| phage major tail tube protein \| phage major tail tube protein; uniclust \| UniRef100\_A0A7R9HA11 \| 99.6 \| 2.5e-18 \| 4.7e-24 \| 151.4 \| 166 \| (2, 168) \| 168 \| (953, 1133) \| 1133 \| Uncharacterized protein \| Uncharacterized protein; uniclust \| UniRef100\_A0A369RL94 \| 99.6 \| 6.5e-18 \| 1.2e-23 \| 120.3 \| 128 \| (40, 168) \| 168 \| (22, 151) \| 151 \| Phage major tail tube protein \| Phage major tail tube protein; uniclust \| UniRef100\_A0A078LLW6 \| 99.6 \| 7.2e-18 \| 1.3e-23 \| 122.9 \| 104 \| (1, 104) \| 168 \| (1, 107) \| 189 \| Phage P2 FII-like protein \| Phage P2 FII-like protein; uniclust \| UniRef100\_A0A0F9Z9G8 \| 99.6 \| 8e-18 \| 1.5e-23 \| 118.6 \| 111 \| (4, 115) \| 168 \| (6, 120) \| 137 \| Prophage MuMc02, major tail tube protein \| Prophage MuMc02, major tail tube protein; uniclust \| UniRef100\_UPI001FFFF203 \| 99.5 \| 2.3e-17 \| 4.3e-23 \| 110.3 \| 95 \| (3, 97) \| 168 \| (5, 99) \| 107 \| phage major tail tube protein \| phage major tail tube protein; uniclust \| UniRef100\_UPI0021E519AF \| 99.5 \| 3e-17 \| 5.4e-23 \| 107.1 \| 77 \| (92, 168) \| 168 \| (16, 92) \| 92 \| phage major tail tube protein \| phage major tail tube protein; uniclust \| UniRef100\_A0A450SXT5 \| 99.5 \| 3.1e-17 \| 5.7e-23 \| 110.6 \| 85 \| (43, 127) \| 168 \| (1, 85) \| 112 \| Uncharacterized protein \| Uncharacterized protein; uniclust \| UniRef100\_A0A2I7R3K6 \| 99.5 \| 4.5e-17 \| 8.3e-23 \| 117.1 \| 166 \| (2, 168) \| 168 \| (3, 170) \| 170 \| Tail tube protein \| Tail tube protein; uniclust \| UniRef100\_A0A485CDJ1 \| 99.5 \| 8e-17 \| 1.5e-22 \| 120.4 \| 105 \| (61, 165) \| 168 \| (2, 107) \| 225 \| Phage major tail tube protein \| Phage major tail tube protein; uniclust \| UniRef100\_A0A327JAQ8 \| 99.5 \| 9.4e-17 \| 1.7e-22 \| 115.5 \| 164 \| (3, 168) \| 168 \| (1, 169) \| 170 \| Phage tail protein \| Phage tail protein; uniclust \| UniRef100\_A0A022PER4 \| 99.5 \| 9.3e-17 \| 1.9e-22 \| 106.7 \| 65 \| (1, 65) \| 168 \| (10, 76) \| 83 \| Phage tail tube protein FII \| Phage tail tube protein FII; uniclust \| UniRef100\_A0A0N1CCE3 \| 99.5 \| 1.1e-16 \| 2.1e-22 \| 119.1 \| 157 \| (5, 161) \| 168 \| (3, 163) \| 177 \| Phage tail protein \| Phage tail protein; uniclust \| UniRef100\_A0A645CLD3 \| 99.5 \| 2.1e-16 \| 3.8e-22 \| 107.4 \| 110 \| (56, 168) \| 168 \| (1, 115) \| 116 \| Phage tail tube protein FII \| Phage tail tube protein FII; uniclust \| UniRef100\_UPI001F3D1B67 \| 99.5 \| 2e-16 \| 3.8e-22 \| 115.1 \| 165 \| (1, 166) \| 168 \| (1, 165) \| 166 \| phage major tail tube protein \| phage major tail tube protein; uniclust \| UniRef100\_A0A1I1EZL7 \| 99.4 \| 2.3e-16 \| 4.6e-22 \| 118.1 \| 160 \| (6, 168) \| 168 \| (8, 179) \| 181 \| Phage tail tube protein FII \| Phage tail tube protein FII; uniclust \| UniRef100\_A0A285IVC6 \| 99.4 \| 2.9e-16 \| 5.7e-22 \| 96.5 \| 54 \| (1, 54) \| 168 \| (2, 55) \| 58 \| Phage contractile tail tube protein, P2 family (Fragment) \| Phage contractile tail tube protein, P2 family (Fragment); uniclust \| UniRef100\_A0A1Q6PWR7 \| 99.4 \| 3e-16 \| 5.9e-22 \| 116.0 \| 152 \| (9, 166) \| 168 \| (11, 170) \| 171 \| Phage tail protein \| Phage tail protein; uniclust \| UniRef100\_A0A355TQV7 \| 99.4 \| 4.9e-16 \| 9e-22 \| 115.7 \| 162 \| (5, 167) \| 168 \| (48, 214) \| 214 \| Phage tail protein \| Phage tail protein; uniclust \| UniRef100\_A0A0T7DUY0 \| 99.4 \| 4.9e-16 \| 1e-21 \| 117.8 \| 147 \| (6, 155) \| 168 \| (15, 167) \| 180 \| Phage tail protein \| Phage tail protein; uniclust \| UniRef100\_UPI0021B36975 \| 99.4 \| 7.3e-16 \| 1.3e-21 \| 105.6 \| 94 \| (23, 116) \| 168 \| (19, 113) \| 121 \| phage major tail tube protein \| phage major tail tube protein; uniclust \| UniRef100\_A0A1I5H9X3 \| 99.4 \| 7.8e-16 \| 1.5e-21 \| 114.0 \| 147 \| (17, 168) \| 168 \| (22, 172) \| 173 \| Phage tail tube protein FII \| Phage tail tube protein FII; uniclust \| UniRef100\_UPI00201CE701 \| 99.4 \| 1.3e-15 \| 2.4e-21 \| 106.2 \| 130 \| (37, 167) \| 168 \| (2, 134) \| 135 \| phage major tail tube protein \| phage major tail tube protein; uniclust \| UniRef100\_E8KG78 \| 99.4 \| 2.5e-15 \| 4.7e-21 \| 102.7 \| 89 \| (4, 93) \| 168 \| (6, 94) \| 109 \| Putative phage major tail tube protein \| Putative phage major tail tube protein; uniclust \| UniRef100\_A4P152 \| 99.3 \| 3.4e-15 \| 6.3e-21 \| 100.9 \| 91 \| (1, 92) \| 168 \| (1, 91) \| 110 \| Uncharacterized protein \| Uncharacterized protein; uniclust \| UniRef100\_UPI0015930F13 \| 99.3 \| 4.3e-15 \| 8.2e-21 \| 97.2 \| 80 \| (10, 90) \| 168 \| (1, 80) \| 82 \| phage major tail tube protein \| phage major tail tube protein; uniclust \| UniRef100\_A0A930QI81 \| 99.3 \| 8.3e-15 \| 1.5e-20 \| 91.6 \| 70 \| (99, 168) \| 168 \| (1, 70) \| 70 \| Phage major tail tube protein \| Phage major tail tube protein; uniclust \| UniRef100\_A0A1Y0FNC6 \| 99.3 \| 8.1e-15 \| 1.6e-20 \| 95.3 \| 72 \| (96, 167) \| 168 \| (5, 76) \| 78 \| FAD-dependent pyridine nucleotide-disulfide oxidoreductase \| FAD-dependent pyridine nucleotide-disulfide oxidoreductase; uniclust \| UniRef100\_A0A7J6YK69 \| 99.3 \| 1e-14 \| 1.9e-20 \| 133.5 \| 86 \| (7, 92) \| 168 \| (1664, 1749) \| 2523 \| Uncharacterized protein \| Uncharacterized protein; uniclust \| UniRef100\_UPI001904BA8E \| 99.3 \| 1.1e-14 \| 2e-20 \| 92.9 \| 69 \| (4, 72) \| 168 \| (7, 75) \| 78 \| phage major tail tube protein \| phage major tail tube protein; uniclust \| UniRef100\_UPI00036492D7 \| 99.3 \| 1.5e-14 \| 2.7e-20 \| 93.9 \| 62 \| (1, 62) \| 168 \| (1, 62) \| 86 \| phage major tail tube protein \| phage major tail tube protein; uniclust \| UniRef100\_A0A962FNC8 \| 99.3 \| 1.6e-14 \| 3e-20 \| 104.6 \| 163 \| (5, 168) \| 168 \| (3, 168) \| 170 \| Phage major tail tube protein \| Phage major tail tube protein; uniclust \| UniRef100\_A0A377U0N6 \| 99.3 \| 2.2e-14 \| 4.1e-20 \| 91.5 \| 65 \| (1, 65) \| 168 \| (1, 67) \| 70 \| Phage major tail tube protein \| Phage major tail tube protein; uniclust \| UniRef100\_A0A1T4WVE5 \| 99.3 \| 2.9e-14 \| 5.3e-20 \| 102.7 \| 145 \| (2, 147) \| 168 \| (4, 151) \| 163 \| Phage tail tube protein FII \| Phage tail tube protein FII; uniclust \| UniRef100\_UPI000971A697 \| 99.2 \| 4e-14 \| 7.3e-20 \| 102.6 \| 167 \| (2, 168) \| 168 \| (1, 169) \| 170 \| phage major tail tube protein \| phage major tail tube protein; uniclust \| UniRef100\_A0A8B2NVA0 \| 99.2 \| 7.3e-14 \| 1.3e-19 \| 101.5 \| 156 \| (4, 162) \| 168 \| (8, 166) \| 172 \| Phage major tail tube protein \| Phage major tail tube protein; uniclust \| UniRef100\_A0A2T4JJM7 \| 99.2 \| 7.6e-14 \| 1.4e-19 \| 90.6 \| 74 \| (95, 168) \| 168 \| (12, 85) \| 85 \| Phage major tail tube protein (Fragment) \| Phage major tail tube protein (Fragment); uniclust \| UniRef100\_A0A9E6DQQ8 \| 99.2 \| 1.2e-13 \| 2.2e-19 \| 99.8 \| 156 \| (3, 160) \| 168 \| (4, 163) \| 165 \| Phage major tail tube protein \| Phage major tail tube protein; uniclust \| UniRef100\_A0A4Q6CYM3 \| 99.1 \| 2e-13 \| 3.7e-19 \| 93.2 \| 108 \| (61, 168) \| 168 \| (6, 114) \| 114 \| Phage major tail tube protein \| Phage major tail tube protein; uniclust \| UniRef100\_UPI00226CFAB6 \| 99.1 \| 2.4e-13 \| 4.4e-19 \| 97.7 \| 156 \| (10, 168) \| 168 \| (1, 158) \| 158 \| phage major tail tube protein \| phage major tail tube protein; uniclust \| UniRef100\_A0A836RL47 \| 99.1 \| 2.9e-13 \| 5.3e-19 \| 90.5 \| 89 \| (1, 93) \| 168 \| (1, 91) \| 100 \| Phage tail protein (Fragment) \| Phage tail protein (Fragment); uniclust \| UniRef100\_UPI001BB6B4AF \| 99.1 \| 3.1e-13 \| 5.7e-19 \| 94.9 \| 129 \| (35, 168) \| 168 \| (1, 135) \| 135 \| phage major tail tube protein \| phage major tail tube protein; uniclust \| UniRef100\_A0A418HAQ3 \| 99.1 \| 5.2e-13 \| 9.8e-19 \| 80.6 \| 53 \| (116, 168) \| 168 \| (1, 53) \| 53 \| Phage tail protein (Fragment) \| Phage tail protein (Fragment); uniclust \| UniRef100\_UPI001E503DE8 \| 99.1 \| 7.7e-13 \| 1.4e-18 \| 87.3 \| 87 \| (82, 168) \| 168 \| (2, 92) \| 92 \| phage major tail tube protein \| phage major tail tube protein; uniclust \| UniRef100\_UPI002284FB3C \| 99.1 \| 8.2e-13 \| 1.5e-18 \| 102.4 \| 128 \| (7, 138) \| 168 \| (138, 273) \| 273 \| phage major tail tube protein \| phage major tail tube protein; uniclust \| UniRef100\_UPI0021F6DC2B \| 99.1 \| 9e-13 \| 1.6e-18 \| 86.2 \| 69 \| (100, 168) \| 168 \| (19, 87) \| 87 \| phage major tail tube protein \| phage major tail tube protein; uniclust \| UniRef100\_A0A377U351 \| 99.0 \| 1.7e-12 \| 3.1e-18 \| 81.9 \| 68 \| (82, 149) \| 168 \| (2, 70) \| 71 \| Phage major tail tube protein \| Phage major tail tube protein; uniclust \| UniRef100\_A0A6L3XLT6 \| 99.0 \| 1.9e-12 \| 3.8e-18 \| 92.8 \| 110 \| (3, 114) \| 168 \| (5, 114) \| 128 \| Phage tail protein (Fragment) \| Phage tail protein (Fragment); uniclust \| UniRef100\_A0A2G6CDE2 \| 99.0 \| 3.7e-12 \| 6.8e-18 \| 86.8 \| 103 \| (43, 149) \| 168 \| (1, 104) \| 109 \| Uncharacterized protein \| Uncharacterized protein; uniclust \| UniRef100\_A0A376X2D4 \| 99.0 \| 4.1e-12 \| 7.7e-18 \| 78.9 \| 54 \| (5, 58) \| 168 \| (8, 61) \| 62 \| Phage major tail tube protein \| Phage major tail tube protein; uniclust \| UniRef100\_A0A840FUU9 \| 99.0 \| 4.2e-12 \| 7.8e-18 \| 79.4 \| 61 \| (108, 168) \| 168 \| (2, 62) \| 67 \| P2 family phage contractile tail tube protein \| P2 family phage contractile tail tube protein; uniclust \| UniRef100\_Q31HT1 \| 99.0 \| 5.2e-12 \| 9.5e-18 \| 92.0 \| 159 \| (7, 168) \| 168 \| (9, 167) \| 167 \| Phage tail tube protein \| Phage tail tube protein; uniclust \| UniRef100\_UPI0022E25BA4 \| 98.9 \| 5.6e-12 \| 1e-17 \| 80.1 \| 71 \| (98, 168) \| 168 \| (2, 72) \| 73 \| phage major tail tube protein \| phage major tail tube protein; uniclust \| UniRef100\_A0A4Q6CFC6 \| 98.9 \| 9.1e-12 \| 1.7e-17 \| 83.9 \| 75 \| (1, 75) \| 168 \| (1, 75) \| 101 \| Uncharacterized protein (Fragment) \| Uncharacterized protein (Fragment); uniclust \| UniRef100\_UPI00037521CA \| 98.9 \| 9.9e-12 \| 1.8e-17 \| 86.9 \| 116 \| (3, 118) \| 168 \| (2, 120) \| 121 \| phage major tail tube protein \| phage major tail tube protein; uniclust \| UniRef100\_A0A5E6QXM5 \| 98.9 \| 1e-11 \| 1.9e-17 \| 84.7 \| 92 \| (77, 168) \| 168 \| (4, 99) \| 100 \| Uncharacterized protein \| Uncharacterized protein; uniclust \| UniRef100\_A0A4D7B2K3 \| 98.9 \| 1.1e-11 \| 2e-17 \| 90.7 \| 157 \| (7, 164) \| 168 \| (7, 166) \| 171 \| Phage tail protein \| Phage tail protein; uniclust \| UniRef100\_A0A317H401 \| 98.9 \| 1.1e-11 \| 2.1e-17 \| 88.7 \| 121 \| (46, 166) \| 168 \| (4, 126) \| 148 \| Uncharacterized protein (Fragment) \| Uncharacterized protein (Fragment); uniclust \| UniRef100\_UPI0021B4D849 \| 98.9 \| 1.2e-11 \| 2.1e-17 \| 90.2 \| 155 \| (8, 168) \| 168 \| (10, 165) \| 166 \| phage major tail tube protein \| phage major tail tube protein; uniclust \| UniRef100\_A0A656GGP8 \| 98.9 \| 1.9e-11 \| 3.5e-17 \| 72.4 \| 41 \| (128, 168) \| 168 \| (10, 50) \| 50 \| Major tail tube protein (Fragment) \| Major tail tube protein (Fragment); uniclust \| UniRef100\_A0A0H3PCB3 \| 98.8 \| 3.6e-11 \| 6.9e-17 \| 70.8 \| 42 \| (126, 167) \| 168 \| (2, 43) \| 44 \| Uncharacterized protein \| Uncharacterized protein; uniclust \| UniRef100\_A0A376J2P6 \| 98.8 \| 3.8e-11 \| 7.2e-17 \| 71.9 \| 43 \| (1, 43) \| 168 \| (1, 43) \| 49 \| Phage major tail tube protein FII \| Phage major tail tube protein FII; uniclust \| UniRef100\_A0A6M0FV91 \| 98.8 \| 4.8e-11 \| 9e-17 \| 89.0 \| 142 \| (18, 160) \| 168 \| (26, 174) \| 176 \| DUF1326 domain-containing protein \| DUF1326 domain-containing protein; uniclust \| UniRef100\_A0A0H3G198 \| 98.8 \| 4.8e-11 \| 9.2e-17 \| 87.2 \| 139 \| (1, 168) \| 168 \| (1, 140) \| 148 \| Major tail tube protein \| Major tail tube protein; uniclust \| UniRef100\_UPI000E0283AC \| 98.7 \| 1e-10 \| 1.9e-16 \| 78.6 \| 69 \| (2, 71) \| 168 \| (7, 77) \| 90 \| phage major tail tube protein \| phage major tail tube protein; uniclust \| UniRef100\_A0A285NFT0 \| 98.7 \| 1.4e-10 \| 2.6e-16 \| 85.6 \| 140 \| (10, 151) \| 168 \| (12, 158) \| 164 \| Phage tail tube protein FII \| Phage tail tube protein FII; uniclust \| UniRef100\_A0A451A1E0 \| 98.7 \| 1.8e-10 \| 3.3e-16 \| 67.4 \| 42 \| (127, 168) \| 168 \| (5, 46) \| 46 \| Phage tail tube protein FII \| Phage tail tube protein FII; uniclust \| UniRef100\_A0A376P9I8 \| 98.7 \| 2.3e-10 \| 4.3e-16 \| 73.0 \| 50 \| (1, 50) \| 168 \| (1, 50) \| 72 \| Major tail tube protein FII \| Major tail tube protein FII; uniclust \| UniRef100\_UPI00204C4744 \| 98.7 \| 2.4e-10 \| 4.4e-16 \| 74.7 \| 61 \| (1, 61) \| 168 \| (1, 61) \| 82 \| phage major tail tube protein \| phage major tail tube protein; uniclust \| UniRef100\_A0A419A4I5 \| 98.7 \| 2.5e-10 \| 4.6e-16 \| 81.2 \| 106 \| (10, 132) \| 168 \| (3, 108) \| 136 \| Phage tail protein \| Phage tail protein; uniclust \| UniRef100\_UPI000D9096AA \| 98.6 \| 2.8e-10 \| 5.2e-16 \| 84.8 \| 134 \| (2, 137) \| 168 \| (2, 137) \| 186 \| phage major tail tube protein \| phage major tail tube protein; uniclust \| UniRef100\_UPI00159035F8 \| 98.6 \| 3e-10 \| 5.5e-16 \| 77.8 \| 103 \| (65, 167) \| 168 \| (2, 106) \| 108 \| phage major tail tube protein \| phage major tail tube protein; uniclust \| UniRef100\_UPI001E291166 \| 98.6 \| 3.6e-10 \| 6.7e-16 \| 79.0 \| 83 \| (2, 88) \| 168 \| (38, 122) \| 122 \| phage major tail tube protein \| phage major tail tube protein; uniclust \| UniRef100\_UPI001C69230E \| 98.6 \| 5e-10 \| 9.2e-16 \| 75.6 \| 93 \| (76, 168) \| 168 \| (4, 98) \| 98 \| phage major tail tube protein \| phage major tail tube protein; uniclust \| UniRef100\_A0A6L5WKW9 \| 98.6 \| 5.5e-10 \| 1e-15 \| 79.1 \| 115 \| (49, 163) \| 168 \| (12, 129) \| 131 \| Phage tail protein (Fragment) \| Phage tail protein (Fragment); uniclust \| UniRef100\_UPI000CDE1566 \| 98.5 \| 8.7e-10 \| 1.6e-15 \| 71.1 \| 61 \| (107, 167) \| 168 \| (10, 70) \| 71 \| phage major tail tube protein \| phage major tail tube protein; uniclust \| UniRef100\_UPI001BA0B277 \| 98.5 \| 9.6e-10 \| 1.8e-15 \| 68.0 \| 59 \| (109, 167) \| 168 \| (1, 59) \| 60 \| phage major tail tube protein \| phage major tail tube protein; uniclust \| UniRef100\_A0A3A9EEY6 \| 98.5 \| 1.3e-09 \| 2.4e-15 \| 69.9 \| 67 \| (101, 167) \| 168 \| (6, 72) \| 73 \| Phage tail protein (Fragment) \| Phage tail protein (Fragment); uniclust \| UniRef100\_A0A954BDX2 \| 98.5 \| 1.5e-09 \| 2.8e-15 \| 77.4 \| 121 \| (47, 167) \| 168 \| (4, 128) \| 135 \| Phage major tail tube protein \| Phage major tail tube protein; uniclust \| UniRef100\_A0A4Q0U6A3 \| 98.4 \| 2.3e-09 \| 4.3e-15 \| 76.2 \| 111 \| (53, 166) \| 168 \| (17, 132) \| 132 \| Phage tail protein \| Phage tail protein; uniclust \| UniRef100\_A0A376DH64 \| 98.4 \| 4.8e-09 \| 8.9e-15 \| 63.1 \| 46 \| (123, 168) \| 168 \| (2, 48) \| 49 \| Major tail sheath protein FII from prophage \| Major tail sheath protein FII from prophage; uniclust \| UniRef100\_UPI00048CC744 \| 98.4 \| 4.9e-09 \| 9.1e-15 \| 70.3 \| 86 \| (3, 88) \| 168 \| (1, 86) \| 92 \| phage major tail tube protein \| phage major tail tube protein; uniclust \| UniRef100\_UPI002357F360 \| 98.3 \| 7.5e-09 \| 1.4e-14 \| 69.1 \| 78 \| (38, 116) \| 168 \| (8, 89) \| 89 \| phage major tail tube protein \| phage major tail tube protein; uniclust \| UniRef100\_E2CFK4 \| 98.3 \| 7.4e-09 \| 1.4e-14 \| 77.6 \| 159 \| (6, 167) \| 168 \| (6, 167) \| 169 \| Uncharacterized protein \| Uncharacterized protein; uniclust \| UniRef100\_A0A090SU12 \| 98.3 \| 8.2e-09 \| 1.5e-14 \| 72.9 \| 75 \| (3, 77) \| 168 \| (1, 75) \| 124 \| Uncharacterized protein \| Uncharacterized protein; uniclust \| UniRef100\_A0A6D1AG81 \| 98.3 \| 9.4e-09 \| 1.7e-14 \| 63.7 \| 41 \| (120, 160) \| 168 \| (2, 42) \| 59 \| Phage tail protein (Fragment) \| Phage tail protein (Fragment); uniclust \| UniRef100\_A0A2W5HIC4 \| 98.3 \| 1.1e-08 \| 2e-14 \| 61.9 \| 41 \| (1, 41) \| 168 \| (1, 41) \| 52 \| Phage tail protein (Fragment) \| Phage tail protein (Fragment); uniclust \| UniRef100\_A0A7Y5Z3W4 \| 98.2 \| 1.5e-08 \| 2.8e-14 \| 55.7 \| 33 \| (136, 168) \| 168 \| (1, 33) \| 33 \| Phage tail protein \| Phage tail protein; uniclust \| UniRef100\_UPI00190410E6 \| 98.2 \| 2e-08 \| 3.7e-14 \| 65.5 \| 41 \| (128, 168) \| 168 \| (37, 77) \| 77 \| phage major tail tube protein \| phage major tail tube protein; uniclust \| UniRef100\_A0A376J1C5 \| 98.2 \| 2.8e-08 \| 5.2e-14 \| 68.5 \| 81 \| (44, 125) \| 168 \| (2, 86) \| 99 \| Phage major tail tube protein FII \| Phage major tail tube protein FII; uniclust \| UniRef100\_UPI000421B9C0 \| 98.2 \| 3.2e-08 \| 6e-14 \| 65.2 \| 65 \| (28, 93) \| 168 \| (1, 65) \| 76 \| phage major tail tube protein \| phage major tail tube protein; uniclust \| UniRef100\_UPI0021195812 \| 98.1 \| 4.1e-08 \| 7.6e-14 \| 67.0 \| 94 \| (29, 123) \| 168 \| (1, 98) \| 99 \| phage major tail tube protein \| phage major tail tube protein; uniclust \| UniRef100\_A0A0T6ZCB2 \| 98.1 \| 4.9e-08 \| 8.9e-14 \| 69.4 \| 116 \| (46, 166) \| 168 \| (3, 126) \| 126 \| Uncharacterized protein \| Uncharacterized protein; uniclust \| UniRef100\_UPI00201D4D24 \| 98.1 \| 5e-08 \| 9.2e-14 \| 60.9 \| 52 \| (24, 75) \| 168 \| (2, 55) \| 60 \| phage major tail tube protein \| phage major tail tube protein; uniclust \| UniRef100\_UPI001A9CAF53 \| 98.1 \| 5.4e-08 \| 9.9e-14 \| 64.8 \| 75 \| (94, 168) \| 168 \| (11, 85) \| 85 \| phage major tail tube protein \| phage major tail tube protein; uniclust \| UniRef100\_UPI0012C62FDD \| 98.1 \| 6.3e-08 \| 1.2e-13 \| 64.1 \| 70 \| (97, 166) \| 168 \| (7, 78) \| 80 \| phage major tail tube protein \| phage major tail tube protein; uniclust \| UniRef100\_A0A3P6JSU6 \| 98.0 \| 8.2e-08 \| 1.5e-13 \| 65.1 \| 72 \| (4, 75) \| 168 \| (2, 78) \| 94 \| Putative phage major tail tube protein \| Putative phage major tail tube protein; uniclust \| UniRef100\_A0A448QUZ8 \| 98.0 \| 1e-07 \| 1.9e-13 \| 59.6 \| 46 \| (4, 49) \| 168 \| (5, 50) \| 60 \| Phage major tail tube protein \| Phage major tail tube protein; uniclust \| UniRef100\_UPI0019699564 \| 98.0 \| 1.2e-07 \| 2.2e-13 \| 62.4 \| 65 \| (103, 167) \| 168 \| (4, 70) \| 72 \| phage major tail tube protein \| phage major tail tube protein; uniclust \| UniRef100\_U2EIY6 \| 98.0 \| 1.4e-07 \| 2.5e-13 \| 57.9 \| 53 \| (115, 167) \| 168 \| (1, 53) \| 54 \| Uncharacterized protein \| Uncharacterized protein; uniclust \| UniRef100\_UPI001E51626C \| 97.9 \| 1.7e-07 \| 3.2e-13 \| 60.7 \| 56 \| (19, 74) \| 168 \| (9, 64) \| 72 \| phage major tail tube protein \| phage major tail tube protein; uniclust \| UniRef100\_UPI0013ED546A \| 97.9 \| 2.4e-07 \| 4.4e-13 \| 70.6 \| 112 \| (24, 137) \| 168 \| (4, 122) \| 195 \| phage major tail tube protein \| phage major tail tube protein; uniclust \| UniRef100\_UPI0013D30FFB \| 97.9 \| 3.1e-07 \| 5.6e-13 \| 55.8 \| 48 \| (121, 168) \| 168 \| (4, 51) \| 51 \| phage major tail tube protein \| phage major tail tube protein; uniclust \| UniRef100\_A0A2X3LT05 \| 97.8 \| 3.7e-07 \| 6.8e-13 \| 53.3 \| 40 \| (103, 142) \| 168 \| (1, 40) \| 42 \| Major tail tube protein FII \| Major tail tube protein FII; uniclust \| UniRef100\_UPI000F6D0689 \| 97.8 \| 4.3e-07 \| 7.9e-13 \| 55.6 \| 45 \| (124, 168) \| 168 \| (8, 52) \| 53 \| phage major tail tube protein \| phage major tail tube protein; uniclust \| UniRef100\_UPI000360283B \| 97.8 \| 5.6e-07 \| 1e-12 \| 56.7 \| 40 \| (1, 40) \| 168 \| (1, 40) \| 61 \| phage major tail tube protein \| phage major tail tube protein; uniclust \| UniRef100\_UPI000F0131DD \| 97.8 \| 5.8e-07 \| 1.1e-12 \| 63.2 \| 108 \| (57, 164) \| 168 \| (1, 108) \| 113 \| phage major tail tube protein \| phage major tail tube protein; uniclust \| UniRef100\_A0A840E036 \| 97.8 \| 7e-07 \| 1.3e-12 \| 62.7 \| 105 \| (64, 168) \| 168 \| (4, 111) \| 111 \| Uncharacterized protein \| Uncharacterized protein; uniclust \| UniRef100\_A0A109RVA8 \| 97.8 \| 7.1e-07 \| 1.3e-12 \| 57.6 \| 43 \| (125, 167) \| 168 \| (26, 68) \| 69 \| Uncharacterized protein \| Uncharacterized protein; uniclust \| UniRef100\_A0A376YLN8 \| 97.7 \| 8.6e-07 \| 1.6e-12 \| 48.3 \| 30 \| (1, 30) \| 168 \| (1, 30) \| 30 \| Phage major tail tube protein FII \| Phage major tail tube protein FII; uniclust \| UniRef100\_UPI00210EA4CD \| 97.7 \| 8.6e-07 \| 1.6e-12 \| 61.2 \| 79 \| (34, 113) \| 168 \| (4, 83) \| 100 \| phage major tail tube protein \| phage major tail tube protein; uniclust \| UniRef100\_UPI001C8BCB0F \| 97.7 \| 8.8e-07 \| 1.6e-12 \| 57.7 \| 64 \| (7, 73) \| 168 \| (5, 69) \| 72 \| phage major tail tube protein \| phage major tail tube protein; uniclust \| UniRef100\_UPI0009B7D3B4 \| 97.7 \| 1.1e-06 \| 1.9e-12 \| 56.1 \| 39 \| (24, 62) \| 168 \| (2, 40) \| 64 \| phage major tail tube protein \| phage major tail tube protein; uniclust \| UniRef100\_A0A853IEB6 \| 97.7 \| 1.1e-06 \| 2.1e-12 \| 51.2 \| 36 \| (133, 168) \| 168 \| (6, 41) \| 41 \| Phage major tail tube protein \| Phage major tail tube protein; uniclust \| UniRef100\_UPI0021C9E4C1 \| 97.7 \| 1.3e-06 \| 2.4e-12 \| 58.1 \| 72 \| (23, 94) \| 168 \| (3, 74) \| 81 \| phage major tail tube protein \| phage major tail tube protein; uniclust \| UniRef100\_A0A7H4Z562 \| 97.6 \| 1.5e-06 \| 2.8e-12 \| 59.4 \| 82 \| (4, 88) \| 168 \| (6, 90) \| 94 \| Phage tail protein (Fragment) \| Phage tail protein (Fragment); uniclust \| UniRef100\_A0A4Y8VEF2 \| 97.6 \| 1.7e-06 \| 3e-12 \| 59.6 \| 55 \| (113, 167) \| 168 \| (27, 81) \| 97 \| DUF2790 domain-containing protein \| DUF2790 domain-containing protein; uniclust \| UniRef100\_A0A7U9RYY6 \| 97.6 \| 2e-06 \| 3.6e-12 \| 62.5 \| 119 \| (45, 166) \| 168 \| (9, 134) \| 134 \| Uncharacterized protein \| Uncharacterized protein; uniclust \| UniRef100\_UPI00215029C9 \| 97.5 \| 2.9e-06 \| 5.4e-12 \| 56.1 \| 67 \| (61, 127) \| 168 \| (4, 71) \| 77 \| phage major tail tube protein \| phage major tail tube protein; uniclust \| UniRef100\_A0A143DGN7 \| 97.5 \| 3e-06 \| 5.5e-12 \| 56.9 \| 41 \| (1, 41) \| 168 \| (1, 41) \| 84 \| Uncharacterized protein \| Uncharacterized protein; uniclust \| UniRef100\_A0A378TY62 \| 97.5 \| 3.3e-06 \| 6.1e-12 \| 48.0 \| 33 \| (1, 33) \| 168 \| (1, 33) \| 36 \| Phage-like protein \| Phage-like protein; uniclust \| UniRef100\_A0A527W089 \| 97.5 \| 4.3e-06 \| 7.8e-12 \| 56.3 \| 75 \| (94, 168) \| 168 \| (8, 83) \| 84 \| Uncharacterized protein \| Uncharacterized protein; uniclust \| UniRef100\_A0A2G2DI71 \| 97.5 \| 4.5e-06 \| 8.3e-12 \| 56.9 \| 74 \| (93, 166) \| 168 \| (14, 88) \| 90 \| Uncharacterized protein (Fragment) \| Uncharacterized protein (Fragment); uniclust \| UniRef100\_UPI000B1E361C \| 97.4 \| 5.1e-06 \| 9.4e-12 \| 53.7 \| 55 \| (4, 66) \| 168 \| (6, 60) \| 67 \| phage major tail tube protein \| phage major tail tube protein; uniclust \| UniRef100\_UPI000AF19DF4 \| 97.4 \| 5.8e-06 \| 1.1e-11 \| 60.0 \| 67 \| (24, 90) \| 168 \| (51, 118) \| 131 \| phage major tail tube protein \| phage major tail tube protein; uniclust \| UniRef100\_UPI001E2BAB95 \| 97.4 \| 6.4e-06 \| 1.2e-11 \| 53.3 \| 58 \| (45, 103) \| 168 \| (2, 63) \| 66 \| phage major tail tube protein \| phage major tail tube protein; uniclust \| UniRef100\_A0A1T3BR98 \| 97.4 \| 8.5e-06 \| 1.6e-11 \| 52.0 \| 47 \| (3, 49) \| 168 \| (6, 52) \| 62 \| Uncharacterized protein \| Uncharacterized protein; uniclust \| UniRef100\_I0QL77 \| 97.3 \| 9.6e-06 \| 1.8e-11 \| 59.6 \| 44 \| (123, 166) \| 168 \| (95, 138) \| 140 \| Major tail sheath protein \| Major tail sheath protein; uniclust \| UniRef100\_A0A850F557 \| 97.3 \| 1.1e-05 \| 2e-11 \| 57.0 \| 86 \| (79, 165) \| 168 \| (11, 98) \| 101 \| Phage major tail tube protein \| Phage major tail tube protein; uniclust \| UniRef100\_UPI001F4B2576 \| 97.3 \| 1.2e-05 \| 2.2e-11 \| 54.1 \| 72 \| (95, 166) \| 168 \| (11, 82) \| 82 \| phage major tail tube protein \| phage major tail tube protein; uniclust \| UniRef100\_A0A377HW70 \| 97.3 \| 1.4e-05 \| 2.5e-11 \| 51.6 \| 47 \| (1, 47) \| 168 \| (1, 47) \| 65 \| Phage major tail tube protein \| Phage major tail tube protein; uniclust \| UniRef100\_UPI0022470942 \| 97.2 \| 2.3e-05 \| 4.2e-11 \| 50.3 \| 36 \| (1, 36) \| 168 \| (1, 36) \| 63 \| phage major tail tube protein \| phage major tail tube protein; uniclust \| UniRef100\_A0A3A8EJ92 \| 97.1 \| 2.6e-05 \| 5e-11 \| 60.1 \| 86 \| (83, 168) \| 168 \| (65, 150) \| 150 \| Phage tail protein (Fragment) \| Phage tail protein (Fragment); uniclust \| UniRef100\_A0A448QV03 \| 97.1 \| 3.9e-05 \| 7.2e-11 \| 53.5 \| 92 \| (50, 142) \| 168 \| (2, 97) \| 98 \| Phage major tail tube protein \| Phage major tail tube protein; uniclust \| UniRef100\_A0A8S0FSJ8 \| 97.1 \| 4e-05 \| 7.3e-11 \| 49.6 \| 62 \| (1, 63) \| 168 \| (1, 62) \| 65 \| Uncharacterized protein \| Uncharacterized protein; uniclust \| UniRef100\_UPI00116EFC7B \| 97.0 \| 4.5e-05 \| 8.3e-11 \| 53.3 \| 45 \| (124, 168) \| 168 \| (50, 94) \| 99 \| phage major tail tube protein \| phage major tail tube protein; uniclust \| UniRef100\_A0A836W881 \| 97.0 \| 4.9e-05 \| 9e-11 \| 48.5 \| 57 \| (3, 60) \| 168 \| (4, 60) \| 60 \| Phage tail protein (Fragment) \| Phage tail protein (Fragment); uniclust \| UniRef100\_A0A6L5WIA9 \| 97.0 \| 5e-05 \| 9.2e-11 \| 46.3 \| 39 \| (3, 41) \| 168 \| (5, 43) \| 48 \| Phage tail protein \| Phage tail protein; uniclust \| UniRef100\_UPI001F0E249E \| 97.0 \| 6.4e-05 \| 1.2e-10 \| 46.7 \| 35 \| (133, 167) \| 168 \| (2, 36) \| 52 \| phage major tail tube protein \| phage major tail tube protein; uniclust \| UniRef100\_A0A853IE67 \| 97.0 \| 6.5e-05 \| 1.2e-10 \| 42.4 \| 33 \| (136, 168) \| 168 \| (1, 33) \| 33 \| Phage major tail tube protein \| Phage major tail tube protein; uniclust \| UniRef100\_R9J2X8 \| 96.9 \| 9.1e-05 \| 1.7e-10 \| 51.2 \| 72 \| (79, 150) \| 168 \| (13, 89) \| 91 \| Uncharacterized protein \| Uncharacterized protein; uniclust \| UniRef100\_UPI0005CAA7C2 \| 96.8 \| 0.00011 \| 2.1e-10 \| 54.1 \| 88 \| (5, 94) \| 168 \| (8, 96) \| 134 \| phage major tail tube protein \| phage major tail tube protein; uniclust \| UniRef100\_R7BVV5 \| 96.8 \| 0.00015 \| 2.7e-10 \| 47.1 \| 38 \| (131, 168) \| 168 \| (24, 61) \| 62 \| Major tail tube protein \| Major tail tube protein; uniclust \| UniRef100\_A0A4Q3YXT5 \| 96.8 \| 0.00015 \| 2.7e-10 \| 41.6 \| 33 \| (136, 168) \| 168 \| (3, 35) \| 35 \| Uncharacterized protein \| Uncharacterized protein; uniclust \| UniRef100\_UPI0018E48B0B \| 96.7 \| 0.00019 \| 3.4e-10 \| 47.6 \| 60 \| (101, 160) \| 168 \| (8, 67) \| 71 \| phage major tail tube protein \| phage major tail tube protein; uniclust \| UniRef100\_A0A6I4YK80 \| 96.7 \| 0.00022 \| 3.9e-10 \| 53.0 \| 47 \| (122, 168) \| 168 \| (92, 138) \| 138 \| GIY-YIG nuclease family protein \| GIY-YIG nuclease family protein; uniclust \| UniRef100\_A0A0H3ZYX4 \| 96.6 \| 0.00035 \| 6.5e-10 \| 50.3 \| 106 \| (48, 156) \| 168 \| (4, 109) \| 113 \| Uncharacterized protein \| Uncharacterized protein; uniclust \| UniRef100\_UPI000870CDAC \| 96.6 \| 0.00036 \| 6.7e-10 \| 53.2 \| 64 \| (97, 160) \| 168 \| (96, 161) \| 162 \| phage major tail tube protein \| phage major tail tube protein; uniclust \| UniRef100\_J9FNQ2 \| 96.6 \| 0.00038 \| 6.9e-10 \| 47.3 \| 70 \| (97, 166) \| 168 \| (10, 79) \| 80 \| Phage major tail tube protein (Fragment) \| Phage major tail tube protein (Fragment); uniclust \| UniRef100\_UPI0014783C6E \| 96.5 \| 0.00041 \| 7.5e-10 \| 49.9 \| 58 \| (111, 168) \| 168 \| (53, 111) \| 111 \| phage major tail tube protein \| phage major tail tube protein; uniclust \| UniRef100\_A0A376J940 \| 96.5 \| 0.00049 \| 9e-10 \| 46.6 \| 67 \| (60, 128) \| 168 \| (4, 70) \| 78 \| Major tail tube protein FII \| Major tail tube protein FII; uniclust \| UniRef100\_UPI0021BD40D7 \| 96.5 \| 0.00055 \| 1e-09 \| 50.7 \| 58 \| (36, 93) \| 168 \| (60, 119) \| 131 \| phage major tail tube protein \| phage major tail tube protein; pdb70 \| 3J9Q\_l \| 100.0 \| 3.3e-49 \| 2.8e-53 \| 292.7 \| 168 \| (1, 168) \| 168 \| (1, 168) \| 168 \| sheath, tube \| 3J9Q\_l sheath, tube pyocin, bacteriocin, sheath, tube, STRUCTURAL; pdb70 \| 5W5E\_c \| 100.0 \| 3.3e-49 \| 2.8e-53 \| 292.7 \| 168 \| (1, 168) \| 168 \| (1, 168) \| 168 \| FIIR2 protein \| 5W5E\_c FIIR2 protein pyocin, bacteriocin, STRUCTURAL PROTEIN |
| Top keywords  (threshold 1.00e-03 (evalue)) | **tail, Phage, tube, major, FII, Fragment, sheath, Putative, contractile, P2** |
| Output files | ../../similar\_sequences/19\_FANPEZAQ\_CDS\_0019\_merged.svg ../../similar\_sequences/19\_FANPEZAQ\_CDS\_0019\_pdb70.a3m ../../similar\_sequences/19\_FANPEZAQ\_CDS\_0019\_pdb70.hhr ../../similar\_sequences/19\_FANPEZAQ\_CDS\_0019\_uniclust.a3m ../../similar\_sequences/19\_FANPEZAQ\_CDS\_0019\_uniclust.hhr |

#### Structure prediction (AlphaFold)^2^

|  |  |
| --- | --- |
| Stats | xml version="1.0" encoding="utf-8" standalone="no"?       2024-09-02T21:09:18.882232 image/svg+xml   Matplotlib v3.7.2, https://matplotlib.org/ |
| Predicted structure | **NGL Viewer Controls:**  - Center: *Left-Click* - Rotate: *Left-Click + Drag* - Translate: *Right-Click + Drag* - Zoom: *Shift + Left-Click + Drag* |
| Output files | ../../predicted\_structures/19\_FANPEZAQ\_CDS\_0019/features.pkl ../../predicted\_structures/19\_FANPEZAQ\_CDS\_0019/ranked\_0.pdb ../../predicted\_structures/19\_FANPEZAQ\_CDS\_0019/ranked\_0\_plots.svg ../../predicted\_structures/19\_FANPEZAQ\_CDS\_0019/result\_model\_1\_ptm\_pred\_0.pkl |

#### Structure similarity search results (Foldseek)^3^

|  |  |
| --- | --- |
| Structure databases searched | Pdb, Afdb-proteome, Afdb-uniprot50 |
| Results, scheme(s)  (Top layers only, threshold 1.00e-02 (evalue)) | xml version="1.0" encoding="utf-8" standalone="no"?       2024-09-02T21:10:44.149803 image/svg+xml   Matplotlib v3.7.2, https://matplotlib.org/ |
| Results, table  (threshold 1.00e-02 (evalue)) | db \| id \| prob \| evalue \| bits \| fident \| alnlen \| mismatch \| gapopen \| qstart \| qend \| tstart \| tend \| name \| description; pdb \| 6U5F\_M \| 1.0 \| 7.011e-16 \| 693 \| 0.393 \| 165 \| 96 \| 2 \| 6 \| 168 \| 4 \| 166 \| Tube PA0623 \| Tube PA0623; pdb \| 7B5H\_AN \| 1.0 \| 0.0001664 \| 174 \| 0.123 \| 138 \| 104 \| 10 \| 9 \| 135 \| 9 \| 140 \| All3324 protein \| All3324 protein; pdb \| 6RAO\_A \| 1.0 \| 0.0002933 \| 163 \| 0.117 \| 136 \| 105 \| 9 \| 9 \| 135 \| 17 \| 146 \| Afp1 \| Afp1; pdb \| 7AEB\_Y \| 1.0 \| 0.0003424 \| 151 \| 0.166 \| 144 \| 99 \| 9 \| 7 \| 135 \| 7 \| 144 \| Phospholipid/glycerol acyltransferase \| Phospholipid/glycerol acyltransferase; pdb \| 5IV5\_W \| 1.0 \| 0.0001501 \| 145 \| 0.161 \| 136 \| 86 \| 8 \| 22 \| 136 \| 62 \| 190 \| Baseplate wedge protein gp7 \| Baseplate wedge protein gp7; pdb \| 7ADZ\_1A \| 1.0 \| 0.002081 \| 133 \| 0.087 \| 148 \| 111 \| 10 \| 1 \| 135 \| 3 \| 139 \| Phage tail protein \| Phage tail protein; pdb \| 6J0N\_N \| 1.0 \| 0.000912 \| 131 \| 0.102 \| 137 \| 109 \| 8 \| 10 \| 136 \| 11 \| 143 \| Pvc5 \| Pvc5; pdb \| 7KJK\_A7 \| 1.0 \| 0.003485 \| 123 \| 0.138 \| 144 \| 104 \| 10 \| 6 \| 135 \| 4 \| 141 \| Tail tube protein \| Tail tube protein; pdb \| 8GRA\_C \| 1.0 \| 0.001065 \| 122 \| 0.136 \| 132 \| 93 \| 10 \| 9 \| 135 \| 1 \| 116 \| Bacterodales T6SS protein TssD (Hcp) \| Bacterodales T6SS protein TssD (Hcp); pdb \| 6BDC\_A \| 1.0 \| 0.002835 \| 121 \| 0.156 \| 134 \| 94 \| 10 \| 10 \| 140 \| 3 \| 120 \| Hcp1 \| Hcp1; pdb \| 6V8I\_EA \| 1.0 \| 0.005 \| 111 \| 0.094 \| 148 \| 105 \| 11 \| 5 \| 136 \| 6 \| 140 \| Major Tail Protein, gp53 \| Major Tail Protein, gp53; pdb \| 6RAO\_F \| 1.0 \| 0.001011 \| 109 \| 0.109 \| 164 \| 109 \| 11 \| 1 \| 147 \| 5 \| 148 \| Afp5 \| Afp5; pdb \| 2X8K\_B \| 1.0 \| 0.006812 \| 106 \| 0.114 \| 148 \| 91 \| 8 \| 8 \| 135 \| 1 \| 128 \| HYPOTHETICAL PROTEIN 19.1 \| HYPOTHETICAL PROTEIN 19.1; pdb \| 2X8K\_C \| 1.0 \| 0.007552 \| 105 \| 0.126 \| 142 \| 97 \| 8 \| 8 \| 135 \| 1 \| 129 \| HYPOTHETICAL PROTEIN 19.1 \| HYPOTHETICAL PROTEIN 19.1; pdb \| 6V8I\_EB \| 1.0 \| 0.003863 \| 104 \| 0.108 \| 147 \| 112 \| 11 \| 1 \| 136 \| 2 \| 140 \| Major Tail Protein, gp53 \| Major Tail Protein, gp53; pdb \| 5IV5\_R \| 1.0 \| 0.002429 \| 103 \| 0.138 \| 159 \| 99 \| 13 \| 1 \| 132 \| 14 \| 161 \| Tail tube protein gp19 \| Tail tube protein gp19; pdb \| 8JAN\_L \| 0.998 \| 0.007552 \| 95 \| 0.13 \| 146 \| 107 \| 9 \| 1 \| 129 \| 13 \| 155 \| BplB \| BplB; pdb \| 6J0N\_B \| 0.998 \| 0.003143 \| 95 \| 0.094 \| 159 \| 105 \| 11 \| 1 \| 129 \| 1 \| 150 \| Pvc7 \| Pvc7; pdb \| 7FCF\_B \| 0.996 \| 0.008816 \| 90 \| 0.122 \| 163 \| 104 \| 12 \| 9 \| 152 \| 1 \| 143 \| Fimbrial protein \| Fimbrial protein; pdb \| 7AEB\_M \| 0.981 \| 0.009773 \| 79 \| 0.088 \| 159 \| 97 \| 11 \| 7 \| 129 \| 5 \| 151 \| Phage tail protein \| Phage tail protein; pdb \| 7ZQP\_D \| 0.887 \| 0.006145 \| 66 \| 0.177 \| 180 \| 91 \| 15 \| 4 \| 160 \| 46 \| 191 \| Probable baseplate hub protein \| Probable baseplate hub protein; pdb \| 8BFK\_L \| 0.855 \| 0.005543 \| 64 \| 0.089 \| 191 \| 112 \| 12 \| 3 \| 136 \| 45 \| 230 \| Putative virion structural protein \| Putative virion structural protein; afdb-proteome \| AF-Q9I5S9-F1-MODEL\_V4 \| 1.0 \| 1.257e-17 \| 817 \| 0.392 \| 168 \| 101 \| 1 \| 1 \| 168 \| 1 \| 167 \| Probable bacteriophage protein \| Probable bacteriophage protein; afdb-proteome \| AF-Q4AAY4-F1-MODEL\_V4 \| 1.0 \| 1.493e-16 \| 751 \| 0.356 \| 171 \| 107 \| 2 \| 1 \| 168 \| 1 \| 171 \| Fels-2 prophage protein \| Fels-2 prophage protein; afdb-proteome \| AF-A0A0H3GX02-F1-MODEL\_V4 \| 1.0 \| 3.778e-16 \| 737 \| 0.343 \| 172 \| 108 \| 4 \| 1 \| 168 \| 1 \| 171 \| Putative prophage tail tube protein \| Putative prophage tail tube protein; afdb-proteome \| AF-A0A0H3GQH1-F1-MODEL\_V4 \| 1.0 \| 5.706e-16 \| 733 \| 0.309 \| 171 \| 115 \| 2 \| 1 \| 168 \| 1 \| 171 \| Phage major tail tube protein \| Phage major tail tube protein; afdb-proteome \| AF-Q8ZKJ3-F1-MODEL\_V4 \| 1.0 \| 6.661e-16 \| 672 \| 0.24 \| 175 \| 125 \| 4 \| 1 \| 168 \| 1 \| 174 \| Putative phage tail core protein \| Putative phage tail core protein; afdb-proteome \| AF-G3XD38-F1-MODEL\_V4 \| 1.0 \| 3.543e-05 \| 182 \| 0.146 \| 157 \| 103 \| 10 \| 8 \| 135 \| 6 \| 160 \| Uncharacterized protein \| Uncharacterized protein; afdb-proteome \| AF-A0A3P7DZT8-F1-MODEL\_V4 \| 1.0 \| 0.0003425 \| 144 \| 0.12 \| 158 \| 106 \| 12 \| 1 \| 135 \| 13 \| 160 \| Uncharacterized protein \| Uncharacterized protein; afdb-proteome \| AF-A0A133CKK6-F1-MODEL\_V4 \| 1.0 \| 0.0004914 \| 125 \| 0.147 \| 142 \| 92 \| 8 \| 10 \| 135 \| 5 \| 133 \| Phage tail protein \| Phage tail protein; afdb-proteome \| AF-P71389-F1-MODEL\_V4 \| 1.0 \| 0.007175 \| 106 \| 0.102 \| 137 \| 95 \| 8 \| 1 \| 129 \| 1 \| 117 \| Mu-like prophage FluMu DNA circularization protein \| Mu-like prophage FluMu DNA circularization protein; afdb-proteome \| AF-Q2FX62-F1-MODEL\_V4 \| 1.0 \| 0.008818 \| 102 \| 0.127 \| 149 \| 99 \| 13 \| 5 \| 136 \| 9 \| 143 \| Phage structural protein, putative \| Phage structural protein, putative; afdb-proteome \| AF-Q2FYC9-F1-MODEL\_V4 \| 0.996 \| 0.008818 \| 89 \| 0.11 \| 145 \| 92 \| 10 \| 10 \| 133 \| 5 \| 133 \| Holin-like protein \| Holin-like protein; afdb-uniprot50 \| AF-A0A0M1I4B3-F1-MODEL\_V4 \| 1.0 \| 2.239e-23 \| 1126 \| 0.566 \| 166 \| 72 \| 0 \| 3 \| 168 \| 5 \| 170 \| Major tail tube protein \| Major tail tube protein; afdb-uniprot50 \| AF-A0A1H2ZHT2-F1-MODEL\_V4 \| 1.0 \| 3.442e-22 \| 1057 \| 0.505 \| 168 \| 82 \| 1 \| 1 \| 168 \| 1 \| 167 \| Uncharacterized protein \| Uncharacterized protein; afdb-uniprot50 \| AF-A0A6A4RBN6-F1-MODEL\_V4 \| 1.0 \| 5.475e-22 \| 1056 \| 0.523 \| 168 \| 80 \| 0 \| 1 \| 168 \| 1 \| 168 \| Phage major tail tube protein \| Phage major tail tube protein; afdb-uniprot50 \| AF-A0A1T4WZ67-F1-MODEL\_V4 \| 1.0 \| 1.315e-21 \| 1022 \| 0.517 \| 168 \| 81 \| 0 \| 1 \| 168 \| 1 \| 168 \| Uncharacterized protein \| Uncharacterized protein; afdb-uniprot50 \| AF-A0A1H9YDK4-F1-MODEL\_V4 \| 1.0 \| 3.327e-21 \| 1022 \| 0.452 \| 168 \| 92 \| 0 \| 1 \| 168 \| 1 \| 168 \| Uncharacterized protein \| Uncharacterized protein; afdb-uniprot50 \| AF-A0A4T2A7H2-F1-MODEL\_V4 \| 1.0 \| 1.563e-20 \| 995 \| 0.449 \| 169 \| 92 \| 1 \| 1 \| 168 \| 1 \| 169 \| Phage major tail tube protein \| Phage major tail tube protein; afdb-uniprot50 \| AF-A0A7C8LY52-F1-MODEL\_V4 \| 1.0 \| 1.824e-20 \| 978 \| 0.476 \| 168 \| 87 \| 1 \| 1 \| 167 \| 1 \| 168 \| Uncharacterized protein \| Uncharacterized protein; afdb-uniprot50 \| AF-A0A7S8C720-F1-MODEL\_V4 \| 1.0 \| 2.242e-20 \| 976 \| 0.464 \| 168 \| 89 \| 1 \| 1 \| 168 \| 1 \| 167 \| Phage major tail tube protein \| Phage major tail tube protein; afdb-uniprot50 \| AF-Q73HI5-F1-MODEL\_V4 \| 1.0 \| 1.089e-20 \| 974 \| 0.47 \| 168 \| 86 \| 2 \| 1 \| 168 \| 1 \| 165 \| Prophage P2W3, contractile tail tube protein \| Prophage P2W3, contractile tail tube protein; afdb-uniprot50 \| AF-A0A7Y6Z4U2-F1-MODEL\_V4 \| 1.0 \| 1.271e-20 \| 968 \| 0.485 \| 169 \| 86 \| 1 \| 1 \| 168 \| 1 \| 169 \| Phage major tail tube protein \| Phage major tail tube protein; afdb-uniprot50 \| AF-A0A7Z0MZH7-F1-MODEL\_V4 \| 1.0 \| 4.614e-20 \| 967 \| 0.415 \| 166 \| 96 \| 1 \| 4 \| 168 \| 3 \| 168 \| Phage major tail tube protein \| Phage major tail tube protein; afdb-uniprot50 \| AF-A0A149SW17-F1-MODEL\_V4 \| 1.0 \| 3.953e-20 \| 963 \| 0.386 \| 168 \| 103 \| 0 \| 1 \| 168 \| 2 \| 169 \| Uncharacterized protein \| Uncharacterized protein; afdb-uniprot50 \| AF-A0A4U8YKH6-F1-MODEL\_V4 \| 1.0 \| 3.754e-20 \| 958 \| 0.45 \| 162 \| 89 \| 0 \| 7 \| 168 \| 8 \| 169 \| Tail tube: phage major tail tube protein \| Tail tube: phage major tail tube protein; afdb-uniprot50 \| AF-A0A2D8Q7A4-F1-MODEL\_V4 \| 1.0 \| 3.386e-20 \| 948 \| 0.416 \| 168 \| 96 \| 2 \| 1 \| 168 \| 2 \| 167 \| Phage major tail tube protein \| Phage major tail tube protein; afdb-uniprot50 \| AF-A0A178GTC8-F1-MODEL\_V4 \| 1.0 \| 2.901e-20 \| 945 \| 0.456 \| 171 \| 89 \| 3 \| 1 \| 168 \| 1 \| 170 \| Phage tail protein \| Phage tail protein; afdb-uniprot50 \| AF-A0A081N7T9-F1-MODEL\_V4 \| 1.0 \| 7.727e-20 \| 936 \| 0.402 \| 169 \| 99 \| 2 \| 1 \| 168 \| 1 \| 168 \| Uncharacterized protein \| Uncharacterized protein; afdb-uniprot50 \| AF-A0A6I2KEU0-F1-MODEL\_V4 \| 1.0 \| 1.053e-19 \| 933 \| 0.534 \| 144 \| 67 \| 0 \| 2 \| 145 \| 3 \| 146 \| Phage major tail tube protein \| Phage major tail tube protein; afdb-uniprot50 \| AF-A0A6M5E4Y8-F1-MODEL\_V4 \| 1.0 \| 6.97e-20 \| 932 \| 0.476 \| 170 \| 85 \| 3 \| 1 \| 168 \| 1 \| 168 \| Phage major tail tube protein \| Phage major tail tube protein; afdb-uniprot50 \| AF-A0A0H3K123-F1-MODEL\_V4 \| 1.0 \| 1.59e-19 \| 930 \| 0.463 \| 166 \| 89 \| 0 \| 3 \| 168 \| 2 \| 167 \| Phage tail tube protein FII \| Phage tail tube protein FII; afdb-uniprot50 \| AF-B8GS08-F1-MODEL\_V4 \| 1.0 \| 1.763e-19 \| 923 \| 0.446 \| 168 \| 89 \| 2 \| 1 \| 168 \| 1 \| 164 \| Contractile tail tube protein \| Contractile tail tube protein; afdb-uniprot50 \| AF-A0A853ILE9-F1-MODEL\_V4 \| 1.0 \| 1.59e-19 \| 923 \| 0.392 \| 168 \| 101 \| 1 \| 1 \| 168 \| 1 \| 167 \| Phage major tail tube protein \| Phage major tail tube protein; afdb-uniprot50 \| AF-A0A3N4UQP0-F1-MODEL\_V4 \| 1.0 \| 2.529e-19 \| 917 \| 0.437 \| 169 \| 93 \| 2 \| 1 \| 168 \| 1 \| 168 \| Uncharacterized protein \| Uncharacterized protein; afdb-uniprot50 \| AF-A0A4R3LLQ7-F1-MODEL\_V4 \| 1.0 \| 5.772e-19 \| 900 \| 0.464 \| 168 \| 90 \| 0 \| 1 \| 168 \| 2 \| 169 \| Uncharacterized protein \| Uncharacterized protein; afdb-uniprot50 \| AF-A0A6L2ZQ79-F1-MODEL\_V4 \| 1.0 \| 4.945e-19 \| 896 \| 0.423 \| 170 \| 95 \| 2 \| 1 \| 168 \| 2 \| 170 \| Phage major tail tube protein \| Phage major tail tube protein; afdb-uniprot50 \| AF-A0A450W6Q5-F1-MODEL\_V4 \| 1.0 \| 4.023e-19 \| 894 \| 0.423 \| 170 \| 95 \| 2 \| 1 \| 168 \| 1 \| 169 \| Uncharacterized protein \| Uncharacterized protein; afdb-uniprot50 \| AF-A0A4Q4L4P4-F1-MODEL\_V4 \| 1.0 \| 1.128e-18 \| 893 \| 0.41 \| 168 \| 98 \| 1 \| 1 \| 168 \| 1 \| 167 \| Phage major tail tube protein \| Phage major tail tube protein; afdb-uniprot50 \| AF-A0A7X2D4H3-F1-MODEL\_V4 \| 1.0 \| 1.794e-18 \| 889 \| 0.44 \| 161 \| 89 \| 1 \| 7 \| 167 \| 14 \| 173 \| Phage major tail tube protein \| Phage major tail tube protein; afdb-uniprot50 \| AF-A0A7X3TYV3-F1-MODEL\_V4 \| 1.0 \| 6.077e-19 \| 888 \| 0.422 \| 168 \| 95 \| 2 \| 2 \| 168 \| 3 \| 169 \| Phage major tail tube protein \| Phage major tail tube protein; afdb-uniprot50 \| AF-A0A3N8AXB8-F1-MODEL\_V4 \| 1.0 \| 1.188e-18 \| 887 \| 0.446 \| 168 \| 92 \| 1 \| 1 \| 168 \| 1 \| 167 \| Phage major tail tube protein \| Phage major tail tube protein; afdb-uniprot50 \| AF-A0A3N2E2A2-F1-MODEL\_V4 \| 1.0 \| 1.317e-18 \| 880 \| 0.416 \| 168 \| 93 \| 2 \| 1 \| 168 \| 2 \| 164 \| Uncharacterized protein \| Uncharacterized protein; afdb-uniprot50 \| AF-A0A423MS82-F1-MODEL\_V4 \| 1.0 \| 1.704e-18 \| 880 \| 0.404 \| 168 \| 99 \| 1 \| 1 \| 168 \| 1 \| 167 \| Phage major tail tube protein \| Phage major tail tube protein; afdb-uniprot50 \| AF-A0A836CEB5-F1-MODEL\_V4 \| 1.0 \| 1.794e-18 \| 877 \| 0.301 \| 166 \| 114 \| 1 \| 2 \| 167 \| 5 \| 168 \| Phage major tail tube protein \| Phage major tail tube protein; afdb-uniprot50 \| AF-A0A257ZWP2-F1-MODEL\_V4 \| 1.0 \| 1.072e-18 \| 874 \| 0.335 \| 167 \| 109 \| 1 \| 2 \| 168 \| 4 \| 168 \| Phage tail protein \| Phage tail protein; afdb-uniprot50 \| AF-A0A081B6C6-F1-MODEL\_V4 \| 1.0 \| 1.704e-18 \| 873 \| 0.384 \| 169 \| 99 \| 3 \| 1 \| 168 \| 1 \| 165 \| Phage major tail tube protein \| Phage major tail tube protein; afdb-uniprot50 \| AF-A0A5A9EMS2-F1-MODEL\_V4 \| 1.0 \| 1.704e-18 \| 871 \| 0.41 \| 168 \| 98 \| 1 \| 1 \| 168 \| 1 \| 167 \| Phage major tail tube protein \| Phage major tail tube protein; afdb-uniprot50 \| AF-A0A1E3G6Z6-F1-MODEL\_V4 \| 1.0 \| 1.989e-18 \| 870 \| 0.386 \| 168 \| 99 \| 2 \| 1 \| 168 \| 3 \| 166 \| Phage major tail tube protein \| Phage major tail tube protein; afdb-uniprot50 \| AF-A0A2V3UAX1-F1-MODEL\_V4 \| 1.0 \| 2.574e-18 \| 869 \| 0.373 \| 166 \| 104 \| 0 \| 3 \| 168 \| 1 \| 166 \| P2 family phage contractile tail tube protein \| P2 family phage contractile tail tube protein; afdb-uniprot50 \| AF-A0A1M7YZ05-F1-MODEL\_V4 \| 1.0 \| 2.445e-18 \| 856 \| 0.438 \| 171 \| 90 \| 3 \| 1 \| 167 \| 1 \| 169 \| Phage tail tube protein FII \| Phage tail tube protein FII; afdb-uniprot50 \| AF-A0A0D8CN41-F1-MODEL\_V4 \| 1.0 \| 1.46e-18 \| 853 \| 0.366 \| 169 \| 104 \| 3 \| 1 \| 168 \| 1 \| 167 \| Tail protein \| Tail protein; afdb-uniprot50 \| AF-A0A348FYG6-F1-MODEL\_V4 \| 1.0 \| 9.343e-18 \| 853 \| 0.353 \| 167 \| 108 \| 0 \| 2 \| 168 \| 3 \| 169 \| Tail protein \| Tail protein; afdb-uniprot50 \| AF-A0A5P9F0T7-F1-MODEL\_V4 \| 1.0 \| 3.331e-18 \| 851 \| 0.388 \| 170 \| 101 \| 3 \| 1 \| 168 \| 1 \| 169 \| Phage tail tube protein FII \| Phage tail tube protein FII; afdb-uniprot50 \| AF-A0A0J1K2H5-F1-MODEL\_V4 \| 1.0 \| 4.311e-18 \| 849 \| 0.372 \| 169 \| 104 \| 1 \| 1 \| 167 \| 1 \| 169 \| Major tail tube protein \| Major tail tube protein; afdb-uniprot50 \| AF-A0A2G6CS51-F1-MODEL\_V4 \| 1.0 \| 5.874e-18 \| 848 \| 0.366 \| 169 \| 104 \| 3 \| 1 \| 168 \| 1 \| 167 \| Phage major tail tube protein \| Phage major tail tube protein; afdb-uniprot50 \| AF-A0A7Y8ENP0-F1-MODEL\_V4 \| 1.0 \| 1.209e-17 \| 842 \| 0.392 \| 168 \| 101 \| 1 \| 1 \| 168 \| 1 \| 167 \| Phage major tail tube protein \| Phage major tail tube protein; afdb-uniprot50 \| AF-A0A4P9VQM1-F1-MODEL\_V4 \| 1.0 \| 5.874e-18 \| 841 \| 0.385 \| 171 \| 102 \| 2 \| 1 \| 168 \| 2 \| 172 \| Phage major tail tube protein \| Phage major tail tube protein; afdb-uniprot50 \| AF-A0A2W5DEX1-F1-MODEL\_V4 \| 1.0 \| 1.148e-17 \| 840 \| 0.416 \| 168 \| 98 \| 0 \| 1 \| 168 \| 1 \| 168 \| Phage major tail tube protein \| Phage major tail tube protein; afdb-uniprot50 \| AF-A0A7Y8ED06-F1-MODEL\_V4 \| 1.0 \| 1.411e-17 \| 839 \| 0.41 \| 168 \| 98 \| 1 \| 1 \| 168 \| 1 \| 167 \| Phage major tail tube protein \| Phage major tail tube protein; afdb-uniprot50 \| AF-A0A2A7UY84-F1-MODEL\_V4 \| 1.0 \| 8.004e-18 \| 839 \| 0.382 \| 170 \| 102 \| 3 \| 1 \| 168 \| 1 \| 169 \| Phage major tail tube protein \| Phage major tail tube protein; afdb-uniprot50 \| AF-A0A3S4BVE8-F1-MODEL\_V4 \| 1.0 \| 8.004e-18 \| 838 \| 0.446 \| 168 \| 90 \| 1 \| 1 \| 168 \| 3 \| 167 \| Phage major tail tube protein \| Phage major tail tube protein; afdb-uniprot50 \| AF-A0A4R2P5B2-F1-MODEL\_V4 \| 1.0 \| 1.036e-17 \| 837 \| 0.388 \| 170 \| 99 \| 3 \| 1 \| 168 \| 1 \| 167 \| Uncharacterized protein \| Uncharacterized protein; afdb-uniprot50 \| AF-A0A522WEK0-F1-MODEL\_V4 \| 1.0 \| 1.273e-17 \| 833 \| 0.375 \| 168 \| 101 \| 2 \| 2 \| 168 \| 3 \| 167 \| Phage major tail tube protein \| Phage major tail tube protein; afdb-uniprot50 \| AF-A0A2E9SIR0-F1-MODEL\_V4 \| 1.0 \| 1.209e-17 \| 831 \| 0.415 \| 171 \| 96 \| 3 \| 1 \| 168 \| 1 \| 170 \| Phage major tail tube protein \| Phage major tail tube protein; afdb-uniprot50 \| AF-A0A2G6IRW7-F1-MODEL\_V4 \| 1.0 \| 1.036e-17 \| 830 \| 0.284 \| 165 \| 115 \| 3 \| 6 \| 168 \| 4 \| 167 \| Uncharacterized protein \| Uncharacterized protein; afdb-uniprot50 \| AF-A0A2S6N2V7-F1-MODEL\_V4 \| 1.0 \| 1.647e-17 \| 827 \| 0.325 \| 163 \| 110 \| 0 \| 3 \| 165 \| 5 \| 167 \| Uncharacterized protein \| Uncharacterized protein; afdb-uniprot50 \| AF-A0A0J9EDK5-F1-MODEL\_V4 \| 1.0 \| 2.62e-17 \| 824 \| 0.351 \| 168 \| 107 \| 2 \| 3 \| 168 \| 1 \| 168 \| Phage major tail tube protein \| Phage major tail tube protein; afdb-uniprot50 \| AF-A0A7X3ZJY9-F1-MODEL\_V4 \| 1.0 \| 1.036e-17 \| 824 \| 0.417 \| 170 \| 96 \| 3 \| 1 \| 168 \| 1 \| 169 \| Phage major tail tube protein \| Phage major tail tube protein; afdb-uniprot50 \| AF-S6I331-F1-MODEL\_V4 \| 1.0 \| 2.905e-17 \| 816 \| 0.408 \| 169 \| 98 \| 2 \| 1 \| 168 \| 1 \| 168 \| Phage major tail tube protein \| Phage major tail tube protein; afdb-uniprot50 \| AF-A0A7T8NU05-F1-MODEL\_V4 \| 1.0 \| 3.759e-17 \| 814 \| 0.343 \| 169 \| 107 \| 2 \| 1 \| 168 \| 3 \| 168 \| Phage major tail tube protein \| Phage major tail tube protein; afdb-uniprot50 \| AF-A0A7X9ZX79-F1-MODEL\_V4 \| 1.0 \| 5.979e-17 \| 813 \| 0.343 \| 169 \| 110 \| 1 \| 1 \| 168 \| 1 \| 169 \| Phage major tail tube protein \| Phage major tail tube protein; afdb-uniprot50 \| AF-A0A432QTQ3-F1-MODEL\_V4 \| 1.0 \| 3.958e-17 \| 813 \| 0.293 \| 167 \| 118 \| 0 \| 1 \| 167 \| 2 \| 168 \| Uncharacterized protein \| Uncharacterized protein; afdb-uniprot50 \| AF-A0A1I5DR52-F1-MODEL\_V4 \| 1.0 \| 4.388e-17 \| 807 \| 0.37 \| 170 \| 104 \| 3 \| 1 \| 168 \| 1 \| 169 \| Uncharacterized protein \| Uncharacterized protein; afdb-uniprot50 \| AF-A0A4V1IPB1-F1-MODEL\_V4 \| 1.0 \| 3.22e-17 \| 803 \| 0.396 \| 164 \| 96 \| 3 \| 6 \| 168 \| 4 \| 165 \| Phage major tail tube protein \| Phage major tail tube protein; afdb-uniprot50 \| AF-A0A379C9M6-F1-MODEL\_V4 \| 1.0 \| 1.169e-16 \| 802 \| 0.319 \| 169 \| 114 \| 1 \| 1 \| 168 \| 1 \| 169 \| Phage major tail tube protein \| Phage major tail tube protein; afdb-uniprot50 \| AF-A0A3D9EDI9-F1-MODEL\_V4 \| 1.0 \| 6.628e-17 \| 801 \| 0.368 \| 171 \| 105 \| 2 \| 1 \| 168 \| 1 \| 171 \| Uncharacterized protein \| Uncharacterized protein; afdb-uniprot50 \| AF-A0A1Y6CXV9-F1-MODEL\_V4 \| 1.0 \| 2.363e-17 \| 800 \| 0.329 \| 170 \| 111 \| 2 \| 1 \| 168 \| 1 \| 169 \| Uncharacterized protein \| Uncharacterized protein; afdb-uniprot50 \| AF-A0A524RVY1-F1-MODEL\_V4 \| 1.0 \| 5.678e-17 \| 798 \| 0.396 \| 169 \| 96 \| 4 \| 1 \| 168 \| 4 \| 167 \| Phage major tail tube protein \| Phage major tail tube protein; afdb-uniprot50 \| AF-A0A2J8GXU0-F1-MODEL\_V4 \| 1.0 \| 1.486e-17 \| 797 \| 0.41 \| 168 \| 93 \| 4 \| 3 \| 168 \| 5 \| 168 \| Phage major tail tube protein \| Phage major tail tube protein; afdb-uniprot50 \| AF-C6ABW7-F1-MODEL\_V4 \| 1.0 \| 6.979e-17 \| 796 \| 0.363 \| 168 \| 104 \| 1 \| 1 \| 168 \| 4 \| 168 \| Phage tail tube protein FII \| Phage tail tube protein FII; afdb-uniprot50 \| AF-A0A6B8KGJ9-F1-MODEL\_V4 \| 1.0 \| 1.512e-16 \| 792 \| 0.278 \| 169 \| 121 \| 1 \| 1 \| 168 \| 5 \| 173 \| Uncharacterized protein \| Uncharacterized protein; afdb-uniprot50 \| AF-A0A061JHN2-F1-MODEL\_V4 \| 1.0 \| 5.979e-17 \| 784 \| 0.442 \| 147 \| 80 \| 2 \| 1 \| 147 \| 1 \| 145 \| Major tail tube protein \| Major tail tube protein; afdb-uniprot50 \| AF-A0A2M7MWC6-F1-MODEL\_V4 \| 1.0 \| 1.765e-16 \| 783 \| 0.391 \| 166 \| 100 \| 1 \| 1 \| 166 \| 1 \| 165 \| Phage major tail tube protein \| Phage major tail tube protein; afdb-uniprot50 \| AF-A0A3D1C6N3-F1-MODEL\_V4 \| 1.0 \| 2.285e-16 \| 779 \| 0.333 \| 171 \| 111 \| 2 \| 1 \| 168 \| 1 \| 171 \| Phage major tail tube protein \| Phage major tail tube protein; afdb-uniprot50 \| AF-A0A0C5VGC6-F1-MODEL\_V4 \| 1.0 \| 1.923e-17 \| 779 \| 0.381 \| 173 \| 91 \| 3 \| 1 \| 158 \| 1 \| 172 \| Phage tail tube protein FII \| Phage tail tube protein FII; afdb-uniprot50 \| AF-A0A6H9GVB3-F1-MODEL\_V4 \| 1.0 \| 1.364e-16 \| 778 \| 0.35 \| 171 \| 106 \| 3 \| 1 \| 168 \| 1 \| 169 \| Major tail tube protein \| Major tail tube protein; afdb-uniprot50 \| AF-A0A6F8NL50-F1-MODEL\_V4 \| 1.0 \| 1.957e-16 \| 777 \| 0.329 \| 164 \| 109 \| 1 \| 6 \| 168 \| 11 \| 174 \| Bacteriophage P2 major tail sheath protein GPFII \| Bacteriophage P2 major tail sheath protein GPFII; afdb-uniprot50 \| AF-A0A4U0YZA9-F1-MODEL\_V4 \| 1.0 \| 2.17e-16 \| 775 \| 0.325 \| 169 \| 109 \| 2 \| 1 \| 168 \| 1 \| 165 \| Phage major tail tube protein \| Phage major tail tube protein; afdb-uniprot50 \| AF-A0A1E3G652-F1-MODEL\_V4 \| 1.0 \| 1.436e-16 \| 773 \| 0.281 \| 167 \| 117 \| 3 \| 1 \| 167 \| 2 \| 165 \| Uncharacterized protein \| Uncharacterized protein; afdb-uniprot50 \| AF-A0A7K0GNR9-F1-MODEL\_V4 \| 1.0 \| 7.348e-17 \| 772 \| 0.459 \| 148 \| 79 \| 1 \| 21 \| 168 \| 1 \| 147 \| Phage major tail tube protein \| Phage major tail tube protein; afdb-uniprot50 \| AF-A0A1A9VKH7-F1-MODEL\_V4 \| 1.0 \| 4.388e-17 \| 772 \| 0.45 \| 153 \| 83 \| 1 \| 17 \| 168 \| 383 \| 535 \| Uncharacterized protein \| Uncharacterized protein; afdb-uniprot50 \| AF-A0A1M3PG22-F1-MODEL\_V4 \| 1.0 \| 2.17e-16 \| 771 \| 0.333 \| 171 \| 111 \| 2 \| 1 \| 168 \| 1 \| 171 \| Phage major tail tube protein \| Phage major tail tube protein; afdb-uniprot50 \| AF-A0A537MFH6-F1-MODEL\_V4 \| 1.0 \| 3.826e-16 \| 769 \| 0.335 \| 167 \| 109 \| 2 \| 1 \| 166 \| 1 \| 166 \| Phage major tail tube protein \| Phage major tail tube protein; afdb-uniprot50 \| AF-A0A380TT15-F1-MODEL\_V4 \| 1.0 \| 4.951e-16 \| 768 \| 0.331 \| 169 \| 111 \| 2 \| 1 \| 168 \| 1 \| 168 \| Phage major tail tube protein \| Phage major tail tube protein; afdb-uniprot50 \| AF-A0A1A9RES3-F1-MODEL\_V4 \| 1.0 \| 1.512e-16 \| 768 \| 0.319 \| 169 \| 112 \| 3 \| 1 \| 168 \| 2 \| 168 \| Phage major tail tube protein \| Phage major tail tube protein; afdb-uniprot50 \| AF-A0A840G0D8-F1-MODEL\_V4 \| 1.0 \| 3.826e-16 \| 767 \| 0.347 \| 170 \| 108 \| 3 \| 1 \| 168 \| 1 \| 169 \| Uncharacterized protein \| Uncharacterized protein; afdb-uniprot50 \| AF-A0A212KMZ0-F1-MODEL\_V4 \| 1.0 \| 9.509e-17 \| 763 \| 0.361 \| 163 \| 97 \| 4 \| 7 \| 168 \| 10 \| 166 \| Prophage P2W3, contractile tail tube protein \| Prophage P2W3, contractile tail tube protein; afdb-uniprot50 \| AF-A0A5S9Q1W1-F1-MODEL\_V4 \| 1.0 \| 5.489e-16 \| 763 \| 0.335 \| 170 \| 111 \| 1 \| 1 \| 168 \| 2 \| 171 \| Uncharacterized protein \| Uncharacterized protein; afdb-uniprot50 \| AF-A0A5M6I344-F1-MODEL\_V4 \| 1.0 \| 3.278e-16 \| 760 \| 0.345 \| 168 \| 107 \| 2 \| 1 \| 166 \| 23 \| 189 \| Phage major tail tube protein \| Phage major tail tube protein; afdb-uniprot50 \| AF-C8N757-F1-MODEL\_V4 \| 1.0 \| 6.746e-16 \| 757 \| 0.359 \| 167 \| 105 \| 2 \| 1 \| 166 \| 1 \| 166 \| Phage major tail tube protein \| Phage major tail tube protein; afdb-uniprot50 \| AF-A0A2A5BNC9-F1-MODEL\_V4 \| 1.0 \| 4.702e-16 \| 757 \| 0.272 \| 169 \| 120 \| 3 \| 1 \| 166 \| 1 \| 169 \| Uncharacterized protein \| Uncharacterized protein; afdb-uniprot50 \| AF-A0A3N2E0L8-F1-MODEL\_V4 \| 1.0 \| 1.11e-16 \| 756 \| 0.361 \| 163 \| 98 \| 3 \| 1 \| 158 \| 1 \| 162 \| Uncharacterized protein \| Uncharacterized protein; afdb-uniprot50 \| AF-A0A318L0T4-F1-MODEL\_V4 \| 1.0 \| 6.407e-16 \| 755 \| 0.284 \| 169 \| 120 \| 1 \| 1 \| 168 \| 1 \| 169 \| Uncharacterized protein \| Uncharacterized protein; afdb-uniprot50 \| AF-A0A1Z3N265-F1-MODEL\_V4 \| 1.0 \| 8.73e-16 \| 755 \| 0.321 \| 171 \| 113 \| 2 \| 1 \| 168 \| 1 \| 171 \| Phage major tail tube protein \| Phage major tail tube protein; afdb-uniprot50 \| AF-A0A161YC88-F1-MODEL\_V4 \| 1.0 \| 4.028e-16 \| 755 \| 0.313 \| 172 \| 112 \| 4 \| 1 \| 168 \| 2 \| 171 \| Major tail tube protein \| Major tail tube protein; afdb-uniprot50 \| AF-A0A2E5BU07-F1-MODEL\_V4 \| 1.0 \| 7.874e-16 \| 754 \| 0.305 \| 170 \| 115 \| 2 \| 1 \| 168 \| 1 \| 169 \| Phage major tail tube protein \| Phage major tail tube protein; afdb-uniprot50 \| AF-A0A0K1NFI5-F1-MODEL\_V4 \| 1.0 \| 8.73e-16 \| 753 \| 0.313 \| 169 \| 114 \| 2 \| 1 \| 168 \| 1 \| 168 \| Phage tail protein \| Phage tail protein; afdb-uniprot50 \| AF-A0A857E651-F1-MODEL\_V4 \| 1.0 \| 3.451e-16 \| 753 \| 0.323 \| 170 \| 110 \| 4 \| 1 \| 167 \| 1 \| 168 \| Phage tail protein \| Phage tail protein; afdb-uniprot50 \| AF-A0A4Q3U3Y4-F1-MODEL\_V4 \| 1.0 \| 7.479e-16 \| 751 \| 0.309 \| 168 \| 113 \| 3 \| 1 \| 166 \| 1 \| 167 \| Phage major tail tube protein \| Phage major tail tube protein; afdb-uniprot50 \| AF-Q4AAY4-F1-MODEL\_V4 \| 1.0 \| 5.489e-16 \| 751 \| 0.356 \| 171 \| 107 \| 2 \| 1 \| 168 \| 1 \| 171 \| Fels-2 prophage protein \| Fels-2 prophage protein; afdb-uniprot50 \| AF-W9BWM4-F1-MODEL\_V4 \| 1.0 \| 1.073e-15 \| 747 \| 0.339 \| 171 \| 108 \| 3 \| 1 \| 168 \| 1 \| 169 \| Major tail tube protein \| Major tail tube protein; afdb-uniprot50 \| AF-A0A837E5C9-F1-MODEL\_V4 \| 1.0 \| 6.407e-16 \| 746 \| 0.301 \| 169 \| 113 \| 2 \| 1 \| 168 \| 2 \| 166 \| Uncharacterized protein \| Uncharacterized protein; afdb-uniprot50 \| AF-A0A6N7JGS1-F1-MODEL\_V4 \| 1.0 \| 9.192e-16 \| 746 \| 0.301 \| 169 \| 113 \| 3 \| 1 \| 168 \| 3 \| 167 \| Uncharacterized protein \| Uncharacterized protein; afdb-uniprot50 \| AF-A0A326GGT4-F1-MODEL\_V4 \| 1.0 \| 1.189e-15 \| 745 \| 0.297 \| 168 \| 116 \| 2 \| 1 \| 167 \| 1 \| 167 \| Phage major tail tube protein \| Phage major tail tube protein; afdb-uniprot50 \| AF-A0A659Q6F3-F1-MODEL\_V4 \| 1.0 \| 7.874e-16 \| 743 \| 0.323 \| 173 \| 110 \| 4 \| 1 \| 168 \| 1 \| 171 \| Phage major tail tube protein \| Phage major tail tube protein; afdb-uniprot50 \| AF-A0A3R8V2Y2-F1-MODEL\_V4 \| 1.0 \| 1.539e-15 \| 743 \| 0.315 \| 171 \| 114 \| 2 \| 1 \| 168 \| 1 \| 171 \| Phage major tail tube protein \| Phage major tail tube protein; afdb-uniprot50 \| AF-A0A2E4CSP6-F1-MODEL\_V4 \| 1.0 \| 1.621e-15 \| 742 \| 0.325 \| 169 \| 112 \| 2 \| 1 \| 168 \| 1 \| 168 \| Phage major tail tube protein \| Phage major tail tube protein; afdb-uniprot50 \| AF-H6SQI7-F1-MODEL\_V4 \| 1.0 \| 1.319e-15 \| 742 \| 0.352 \| 170 \| 105 \| 4 \| 1 \| 167 \| 1 \| 168 \| Phage major tail tube protein (Probable bacteriophage tail tube protein FII) \| Phage major tail tube protein (Probable bacteriophage tail tube protein FII); afdb-uniprot50 \| AF-A0A3V8HSW3-F1-MODEL\_V4 \| 1.0 \| 1.621e-15 \| 742 \| 0.343 \| 172 \| 109 \| 3 \| 1 \| 168 \| 1 \| 172 \| Phage major tail tube protein \| Phage major tail tube protein; afdb-uniprot50 \| AF-A0A2R7NI64-F1-MODEL\_V4 \| 1.0 \| 1.621e-15 \| 740 \| 0.313 \| 172 \| 113 \| 4 \| 1 \| 168 \| 2 \| 172 \| Phage major tail tube protein \| Phage major tail tube protein; afdb-uniprot50 \| AF-A0A5S9HN48-F1-MODEL\_V4 \| 1.0 \| 1.252e-15 \| 739 \| 0.295 \| 169 \| 118 \| 1 \| 1 \| 168 \| 4 \| 172 \| Major tail tube protein \| Major tail tube protein; afdb-uniprot50 \| AF-A0A1D2QSB5-F1-MODEL\_V4 \| 1.0 \| 1.436e-16 \| 739 \| 0.347 \| 187 \| 97 \| 4 \| 1 \| 163 \| 1 \| 186 \| Uncharacterized protein \| Uncharacterized protein; afdb-uniprot50 \| AF-A0A345DE54-F1-MODEL\_V4 \| 1.0 \| 5.489e-16 \| 738 \| 0.266 \| 169 \| 121 \| 3 \| 1 \| 167 \| 1 \| 168 \| Uncharacterized protein \| Uncharacterized protein; afdb-uniprot50 \| AF-A0A326L1L7-F1-MODEL\_V4 \| 1.0 \| 3.009e-15 \| 738 \| 0.305 \| 170 \| 116 \| 2 \| 1 \| 168 \| 1 \| 170 \| Phage major tail tube protein \| Phage major tail tube protein; afdb-uniprot50 \| AF-A0A7X4FXW3-F1-MODEL\_V4 \| 1.0 \| 5.779e-16 \| 737 \| 0.313 \| 172 \| 111 \| 5 \| 1 \| 168 \| 1 \| 169 \| Phage major tail tube protein \| Phage major tail tube protein; afdb-uniprot50 \| AF-A0A0H3GX02-F1-MODEL\_V4 \| 1.0 \| 1.388e-15 \| 737 \| 0.343 \| 172 \| 108 \| 4 \| 1 \| 168 \| 1 \| 171 \| Putative prophage tail tube protein \| Putative prophage tail tube protein; afdb-uniprot50 \| AF-A0A3B7Q0S0-F1-MODEL\_V4 \| 1.0 \| 2.097e-15 \| 732 \| 0.323 \| 170 \| 111 \| 4 \| 1 \| 168 \| 1 \| 168 \| Phage major tail tube protein \| Phage major tail tube protein; afdb-uniprot50 \| AF-A0A5E4XFQ3-F1-MODEL\_V4 \| 1.0 \| 1.436e-16 \| 732 \| 0.343 \| 169 \| 108 \| 2 \| 2 \| 168 \| 4 \| 171 \| Phage major tail tube protein \| Phage major tail tube protein; afdb-uniprot50 \| AF-A0A0Q2YZZ0-F1-MODEL\_V4 \| 1.0 \| 1.892e-15 \| 729 \| 0.269 \| 167 \| 118 \| 2 \| 2 \| 168 \| 4 \| 166 \| Uncharacterized protein \| Uncharacterized protein; afdb-uniprot50 \| AF-E2CJT1-F1-MODEL\_V4 \| 1.0 \| 2.448e-15 \| 728 \| 0.267 \| 168 \| 122 \| 1 \| 1 \| 168 \| 1 \| 167 \| Phage tail tube protein FII \| Phage tail tube protein FII; afdb-uniprot50 \| AF-A0A379YDU2-F1-MODEL\_V4 \| 1.0 \| 1.073e-15 \| 727 \| 0.329 \| 170 \| 109 \| 4 \| 1 \| 167 \| 1 \| 168 \| Phage major tail tube protein \| Phage major tail tube protein; afdb-uniprot50 \| AF-A0A2A2B589-F1-MODEL\_V4 \| 1.0 \| 1.189e-15 \| 723 \| 0.304 \| 171 \| 116 \| 2 \| 1 \| 168 \| 1 \| 171 \| Phage major tail tube protein \| Phage major tail tube protein; afdb-uniprot50 \| AF-A0A2I1RIY9-F1-MODEL\_V4 \| 1.0 \| 2.325e-15 \| 720 \| 0.345 \| 171 \| 107 \| 4 \| 1 \| 167 \| 1 \| 170 \| Phage major tail tube protein \| Phage major tail tube protein; afdb-uniprot50 \| AF-A0A743CC81-F1-MODEL\_V4 \| 1.0 \| 1.13e-15 \| 719 \| 0.349 \| 169 \| 107 \| 3 \| 1 \| 167 \| 1 \| 168 \| Phage tail protein \| Phage tail protein; afdb-uniprot50 \| AF-F4BFQ1-F1-MODEL\_V4 \| 1.0 \| 1.992e-15 \| 717 \| 0.291 \| 161 \| 109 \| 3 \| 3 \| 162 \| 2 \| 158 \| Phage major tail tube protein \| Phage major tail tube protein; afdb-uniprot50 \| AF-A0A246KGG0-F1-MODEL\_V4 \| 1.0 \| 8.438e-15 \| 714 \| 0.275 \| 167 \| 120 \| 1 \| 1 \| 166 \| 1 \| 167 \| Phage major tail tube protein \| Phage major tail tube protein; afdb-uniprot50 \| AF-A0A2G6EWZ7-F1-MODEL\_V4 \| 1.0 \| 6.193e-15 \| 714 \| 0.272 \| 169 \| 122 \| 1 \| 1 \| 168 \| 1 \| 169 \| Phage major tail tube protein \| Phage major tail tube protein; afdb-uniprot50 \| AF-A0A0U1DAP0-F1-MODEL\_V4 \| 1.0 \| 1.992e-15 \| 712 \| 0.446 \| 141 \| 77 \| 1 \| 29 \| 168 \| 2 \| 142 \| Phage tail tube protein \| Phage tail tube protein; afdb-uniprot50 \| AF-A0A359KE96-F1-MODEL\_V4 \| 1.0 \| 6.193e-15 \| 712 \| 0.278 \| 169 \| 117 \| 3 \| 1 \| 166 \| 1 \| 167 \| Phage major tail tube protein \| Phage major tail tube protein; afdb-uniprot50 \| AF-A0A7Y3N2Q7-F1-MODEL\_V4 \| 1.0 \| 3.009e-15 \| 710 \| 0.272 \| 169 \| 121 \| 2 \| 1 \| 168 \| 3 \| 170 \| Phage major tail tube protein \| Phage major tail tube protein; afdb-uniprot50 \| AF-A0A1X3I4U3-F1-MODEL\_V4 \| 1.0 \| 6.866e-15 \| 709 \| 0.304 \| 171 \| 116 \| 2 \| 1 \| 168 \| 3 \| 173 \| Phage major tail tube protein \| Phage major tail tube protein; afdb-uniprot50 \| AF-A0A7C2APA8-F1-MODEL\_V4 \| 1.0 \| 7.479e-16 \| 702 \| 0.293 \| 174 \| 114 \| 5 \| 1 \| 168 \| 1 \| 171 \| Phage major tail tube protein \| Phage major tail tube protein; afdb-uniprot50 \| AF-A0A258L6Q4-F1-MODEL\_V4 \| 1.0 \| 1.13e-15 \| 701 \| 0.301 \| 169 \| 112 \| 3 \| 2 \| 168 \| 5 \| 169 \| Phage tail protein \| Phage tail protein; afdb-uniprot50 \| AF-R7JNY0-F1-MODEL\_V4 \| 1.0 \| 2.448e-15 \| 700 \| 0.283 \| 166 \| 114 \| 3 \| 7 \| 168 \| 6 \| 170 \| Phage tail core protein \| Phage tail core protein; afdb-uniprot50 \| AF-A0A7G7RNU7-F1-MODEL\_V4 \| 1.0 \| 2.448e-15 \| 700 \| 0.335 \| 173 \| 108 \| 3 \| 1 \| 168 \| 1 \| 171 \| Phage major tail tube protein \| Phage major tail tube protein; afdb-uniprot50 \| AF-A0A6M8T223-F1-MODEL\_V4 \| 1.0 \| 2.325e-15 \| 699 \| 0.225 \| 173 \| 127 \| 4 \| 1 \| 168 \| 1 \| 171 \| Phage major tail tube protein \| Phage major tail tube protein; afdb-uniprot50 \| AF-A0A0F9QWB3-F1-MODEL\_V4 \| 1.0 \| 8.438e-15 \| 698 \| 0.253 \| 166 \| 122 \| 1 \| 2 \| 167 \| 4 \| 167 \| Uncharacterized protein \| Uncharacterized protein; afdb-uniprot50 \| AF-A0A089PTU9-F1-MODEL\_V4 \| 1.0 \| 4.317e-15 \| 696 \| 0.343 \| 172 \| 106 \| 6 \| 1 \| 168 \| 1 \| 169 \| Tail protein \| Tail protein; afdb-uniprot50 \| AF-A0A6C1BNV4-F1-MODEL\_V4 \| 1.0 \| 3.336e-15 \| 695 \| 0.242 \| 165 \| 121 \| 3 \| 7 \| 168 \| 7 \| 170 \| Phage major tail tube protein \| Phage major tail tube protein; afdb-uniprot50 \| AF-A0A827QKM1-F1-MODEL\_V4 \| 1.0 \| 4.317e-15 \| 694 \| 0.305 \| 170 \| 113 \| 4 \| 1 \| 167 \| 1 \| 168 \| Phage major tail tube protein \| Phage major tail tube protein; afdb-uniprot50 \| AF-A0A2D5S2E1-F1-MODEL\_V4 \| 1.0 \| 7.612e-15 \| 693 \| 0.282 \| 170 \| 117 \| 5 \| 2 \| 168 \| 3 \| 170 \| Uncharacterized protein \| Uncharacterized protein; afdb-uniprot50 \| AF-A0A2E6BPJ5-F1-MODEL\_V4 \| 1.0 \| 1.275e-14 \| 690 \| 0.295 \| 169 \| 117 \| 2 \| 1 \| 168 \| 1 \| 168 \| Phage major tail tube protein \| Phage major tail tube protein; afdb-uniprot50 \| AF-A0A806CP35-F1-MODEL\_V4 \| 1.0 \| 6.866e-15 \| 688 \| 0.3 \| 170 \| 114 \| 4 \| 1 \| 167 \| 1 \| 168 \| Phage major tail tube protein \| Phage major tail tube protein; afdb-uniprot50 \| AF-A0A450YW94-F1-MODEL\_V4 \| 1.0 \| 1.621e-15 \| 684 \| 0.24 \| 166 \| 121 \| 3 \| 7 \| 168 \| 10 \| 174 \| Uncharacterized protein \| Uncharacterized protein; afdb-uniprot50 \| AF-A0A0Q8B4N4-F1-MODEL\_V4 \| 1.0 \| 6.521e-15 \| 679 \| 0.255 \| 168 \| 121 \| 4 \| 3 \| 168 \| 1 \| 166 \| Uncharacterized protein \| Uncharacterized protein; afdb-uniprot50 \| AF-A0A809HDP3-F1-MODEL\_V4 \| 1.0 \| 8.438e-15 \| 677 \| 0.27 \| 170 \| 120 \| 3 \| 1 \| 168 \| 1 \| 168 \| Phage tail protein \| Phage tail protein; afdb-uniprot50 \| AF-A0A1N7LR57-F1-MODEL\_V4 \| 1.0 \| 1.829e-14 \| 676 \| 0.251 \| 167 \| 120 \| 3 \| 2 \| 167 \| 9 \| 171 \| Uncharacterized protein \| Uncharacterized protein; afdb-uniprot50 \| AF-A0A7W6RG44-F1-MODEL\_V4 \| 1.0 \| 2.624e-14 \| 673 \| 0.254 \| 169 \| 123 \| 3 \| 3 \| 168 \| 1 \| 169 \| Uncharacterized protein \| Uncharacterized protein; afdb-uniprot50 \| AF-A0A4R6E272-F1-MODEL\_V4 \| 1.0 \| 1.925e-14 \| 672 \| 0.295 \| 169 \| 114 \| 4 \| 1 \| 167 \| 1 \| 166 \| Uncharacterized protein \| Uncharacterized protein; afdb-uniprot50 \| AF-R7MC56-F1-MODEL\_V4 \| 1.0 \| 6.866e-15 \| 672 \| 0.229 \| 174 \| 127 \| 4 \| 1 \| 168 \| 1 \| 173 \| Prophage MuMc02 major tail tube protein \| Prophage MuMc02 major tail tube protein; afdb-uniprot50 \| AF-Q8ZKJ3-F1-MODEL\_V4 \| 1.0 \| 2.448e-15 \| 672 \| 0.24 \| 175 \| 125 \| 4 \| 1 \| 168 \| 1 \| 174 \| Putative phage tail core protein \| Putative phage tail core protein; afdb-uniprot50 \| AF-A0A085AFM9-F1-MODEL\_V4 \| 1.0 \| 2.909e-14 \| 670 \| 0.245 \| 167 \| 124 \| 2 \| 1 \| 167 \| 2 \| 166 \| Phage major tail tube protein \| Phage major tail tube protein; afdb-uniprot50 \| AF-A0A4Q6D1V8-F1-MODEL\_V4 \| 1.0 \| 1.275e-14 \| 666 \| 0.29 \| 155 \| 108 \| 1 \| 1 \| 153 \| 1 \| 155 \| Uncharacterized protein \| Uncharacterized protein; afdb-uniprot50 \| AF-Q31HT1-F1-MODEL\_V4 \| 1.0 \| 2.366e-14 \| 666 \| 0.265 \| 162 \| 116 \| 3 \| 7 \| 168 \| 9 \| 167 \| Phage tail tube protein \| Phage tail tube protein; afdb-uniprot50 \| AF-A0A1C0V8D9-F1-MODEL\_V4 \| 1.0 \| 9.85e-15 \| 666 \| 0.224 \| 165 \| 123 \| 3 \| 7 \| 167 \| 7 \| 170 \| Uncharacterized protein \| Uncharacterized protein; afdb-uniprot50 \| AF-A0A482IPK0-F1-MODEL\_V4 \| 1.0 \| 1.488e-14 \| 664 \| 0.226 \| 172 \| 127 \| 4 \| 1 \| 168 \| 4 \| 173 \| Phage major tail tube protein \| Phage major tail tube protein; afdb-uniprot50 \| AF-A0A4Q0GQ17-F1-MODEL\_V4 \| 1.0 \| 2.909e-14 \| 664 \| 0.374 \| 155 \| 94 \| 2 \| 17 \| 168 \| 33 \| 187 \| Phage major tail tube protein \| Phage major tail tube protein; afdb-uniprot50 \| AF-F2K1Q9-F1-MODEL\_V4 \| 1.0 \| 1.342e-14 \| 663 \| 0.236 \| 169 \| 122 \| 6 \| 1 \| 168 \| 3 \| 165 \| Major tail tube protein \| Major tail tube protein; afdb-uniprot50 \| AF-U0ZR36-F1-MODEL\_V4 \| 1.0 \| 5.039e-15 \| 663 \| 0.236 \| 173 \| 125 \| 4 \| 1 \| 168 \| 1 \| 171 \| Tail protein \| Tail protein; afdb-uniprot50 \| AF-A0A256CAU6-F1-MODEL\_V4 \| 1.0 \| 1.567e-14 \| 662 \| 0.307 \| 169 \| 114 \| 3 \| 1 \| 167 \| 3 \| 170 \| Uncharacterized protein \| Uncharacterized protein; afdb-uniprot50 \| AF-E2CN26-F1-MODEL\_V4 \| 1.0 \| 1.037e-14 \| 660 \| 0.264 \| 170 \| 120 \| 5 \| 1 \| 167 \| 1 \| 168 \| Phage tail tube protein FII \| Phage tail tube protein FII; afdb-uniprot50 \| AF-A0A1J5HZS6-F1-MODEL\_V4 \| 1.0 \| 1.737e-14 \| 660 \| 0.224 \| 174 \| 127 \| 5 \| 1 \| 167 \| 1 \| 173 \| Uncharacterized protein \| Uncharacterized protein; afdb-uniprot50 \| AF-A0A1B9NSN8-F1-MODEL\_V4 \| 1.0 \| 9.043e-14 \| 659 \| 0.343 \| 172 \| 107 \| 4 \| 1 \| 168 \| 1 \| 170 \| Phage major tail tube protein \| Phage major tail tube protein; afdb-uniprot50 \| AF-A0A1H2FLU6-F1-MODEL\_V4 \| 1.0 \| 2.492e-14 \| 659 \| 0.232 \| 168 \| 124 \| 5 \| 6 \| 168 \| 4 \| 171 \| Uncharacterized protein \| Uncharacterized protein; afdb-uniprot50 \| AF-A0A1E4AKG0-F1-MODEL\_V4 \| 1.0 \| 7.229e-15 \| 659 \| 0.274 \| 175 \| 119 \| 4 \| 1 \| 168 \| 1 \| 174 \| Phage major tail tube protein \| Phage major tail tube protein; afdb-uniprot50 \| AF-J5KT56-F1-MODEL\_V4 \| 1.0 \| 3.575e-14 \| 658 \| 0.209 \| 167 \| 126 \| 4 \| 1 \| 166 \| 2 \| 163 \| Phage tail tube protein FII \| Phage tail tube protein FII; afdb-uniprot50 \| AF-A0A3S5C3B8-F1-MODEL\_V4 \| 1.0 \| 2.624e-14 \| 655 \| 0.197 \| 167 \| 129 \| 3 \| 6 \| 168 \| 7 \| 172 \| Phage-like protein \| Phage-like protein; afdb-uniprot50 \| AF-W6W214-F1-MODEL\_V4 \| 1.0 \| 1.092e-14 \| 652 \| 0.298 \| 171 \| 113 \| 5 \| 3 \| 168 \| 1 \| 169 \| Major tail tube protein \| Major tail tube protein; afdb-uniprot50 \| AF-A0A1C3EBQ2-F1-MODEL\_V4 \| 1.0 \| 5.128e-14 \| 650 \| 0.263 \| 167 \| 117 \| 6 \| 6 \| 168 \| 8 \| 172 \| Uncharacterized protein \| Uncharacterized protein; afdb-uniprot50 \| AF-A0A327Q9I2-F1-MODEL\_V4 \| 1.0 \| 2.762e-14 \| 649 \| 0.227 \| 167 \| 123 \| 4 \| 7 \| 168 \| 8 \| 173 \| Uncharacterized protein \| Uncharacterized protein; afdb-uniprot50 \| AF-E1SFY3-F1-MODEL\_V4 \| 1.0 \| 6.637e-14 \| 649 \| 0.35 \| 157 \| 96 \| 3 \| 1 \| 154 \| 1 \| 154 \| Major tail tube protein (Protein FII) \| Major tail tube protein (Protein FII); afdb-uniprot50 \| AF-B6JEF9-F1-MODEL\_V4 \| 1.0 \| 1.003e-13 \| 645 \| 0.289 \| 169 \| 113 \| 4 \| 6 \| 167 \| 4 \| 172 \| Putative phage tail tube protein FII \| Putative phage tail tube protein FII; afdb-uniprot50 \| AF-A0A163V2U5-F1-MODEL\_V4 \| 1.0 \| 1.366e-13 \| 643 \| 0.212 \| 165 \| 128 \| 2 \| 6 \| 168 \| 4 \| 168 \| Uncharacterized protein \| Uncharacterized protein; afdb-uniprot50 \| AF-A0A377JJD6-F1-MODEL\_V4 \| 1.0 \| 5.128e-14 \| 643 \| 0.184 \| 173 \| 135 \| 4 \| 1 \| 168 \| 1 \| 172 \| Tail core-like protein \| Tail core-like protein; afdb-uniprot50 \| AF-B6IMG1-F1-MODEL\_V4 \| 1.0 \| 2.624e-14 \| 643 \| 0.218 \| 174 \| 129 \| 5 \| 1 \| 168 \| 1 \| 173 \| Phage major tail tube protein \| Phage major tail tube protein; afdb-uniprot50 \| AF-A0A1T2YJS6-F1-MODEL\_V4 \| 1.0 \| 9.043e-14 \| 642 \| 0.232 \| 168 \| 127 \| 2 \| 1 \| 168 \| 2 \| 167 \| Phage tail protein \| Phage tail protein; afdb-uniprot50 \| AF-A0A4V6YSQ4-F1-MODEL\_V4 \| 1.0 \| 5.686e-14 \| 642 \| 0.209 \| 167 \| 127 \| 4 \| 1 \| 166 \| 7 \| 169 \| Uncharacterized protein \| Uncharacterized protein; afdb-uniprot50 \| AF-A0A1N6M6E1-F1-MODEL\_V4 \| 1.0 \| 6.637e-14 \| 641 \| 0.216 \| 166 \| 128 \| 2 \| 2 \| 167 \| 3 \| 166 \| Major tail tube protein \| Major tail tube protein; afdb-uniprot50 \| AF-A0A7C4AK38-F1-MODEL\_V4 \| 1.0 \| 1.567e-14 \| 640 \| 0.228 \| 175 \| 126 \| 6 \| 1 \| 168 \| 2 \| 174 \| Uncharacterized protein \| Uncharacterized protein; afdb-uniprot50 \| AF-A0A7L5Y1E9-F1-MODEL\_V4 \| 1.0 \| 1.514e-13 \| 640 \| 0.207 \| 169 \| 130 \| 3 \| 3 \| 167 \| 1 \| 169 \| Phage major tail tube protein \| Phage major tail tube protein; afdb-uniprot50 \| AF-A0A1R4LKF5-F1-MODEL\_V4 \| 1.0 \| 1.003e-13 \| 639 \| 0.221 \| 167 \| 128 \| 2 \| 2 \| 168 \| 3 \| 167 \| Phage tail tube protein FII \| Phage tail tube protein FII; afdb-uniprot50 \| AF-A0A379AQE4-F1-MODEL\_V4 \| 1.0 \| 5.128e-14 \| 639 \| 0.18 \| 166 \| 131 \| 3 \| 7 \| 168 \| 9 \| 173 \| Phage major tail tube protein \| Phage major tail tube protein; afdb-uniprot50 \| AF-A0A348HHH9-F1-MODEL\_V4 \| 1.0 \| 1.056e-13 \| 638 \| 0.233 \| 163 \| 124 \| 1 \| 6 \| 168 \| 7 \| 168 \| Phage tail tube protein FII \| Phage tail tube protein FII; afdb-uniprot50 \| AF-A0A3G2IKK8-F1-MODEL\_V4 \| 1.0 \| 7.358e-14 \| 637 \| 0.253 \| 158 \| 114 \| 3 \| 6 \| 160 \| 3 \| 159 \| Uncharacterized protein \| Uncharacterized protein; afdb-uniprot50 \| AF-A0A7C3RQ40-F1-MODEL\_V4 \| 1.0 \| 1.413e-14 \| 633 \| 0.238 \| 176 \| 122 \| 6 \| 1 \| 168 \| 1 \| 172 \| Phage major tail tube protein \| Phage major tail tube protein; afdb-uniprot50 \| AF-A0A1Y1QXH8-F1-MODEL\_V4 \| 1.0 \| 2.288e-13 \| 631 \| 0.251 \| 163 \| 117 \| 2 \| 7 \| 168 \| 13 \| 171 \| Uncharacterized protein \| Uncharacterized protein; afdb-uniprot50 \| AF-A0A7W2BMA7-F1-MODEL\_V4 \| 1.0 \| 9.521e-14 \| 631 \| 0.258 \| 174 \| 121 \| 7 \| 1 \| 168 \| 1 \| 172 \| Phage major tail tube protein \| Phage major tail tube protein; afdb-uniprot50 \| AF-G2HXA5-F1-MODEL\_V4 \| 1.0 \| 3.395e-14 \| 630 \| 0.285 \| 161 \| 108 \| 5 \| 7 \| 166 \| 10 \| 164 \| Phage tail protein \| Phage tail protein; afdb-uniprot50 \| AF-A0A3G2IAV8-F1-MODEL\_V4 \| 1.0 \| 7.747e-14 \| 630 \| 0.269 \| 163 \| 115 \| 3 \| 7 \| 167 \| 5 \| 165 \| Uncharacterized protein \| Uncharacterized protein; afdb-uniprot50 \| AF-A0A1G3UAL9-F1-MODEL\_V4 \| 1.0 \| 1.96e-13 \| 628 \| 0.191 \| 162 \| 127 \| 3 \| 6 \| 166 \| 9 \| 167 \| Uncharacterized protein \| Uncharacterized protein; afdb-uniprot50 \| AF-A0A3T0L2U1-F1-MODEL\_V4 \| 1.0 \| 7.358e-14 \| 625 \| 0.21 \| 166 \| 125 \| 4 \| 1 \| 166 \| 1 \| 160 \| Uncharacterized protein \| Uncharacterized protein; afdb-uniprot50 \| AF-A0A6I0CJR1-F1-MODEL\_V4 \| 1.0 \| 6.303e-14 \| 625 \| 0.216 \| 171 \| 128 \| 4 \| 1 \| 167 \| 4 \| 172 \| Phage major tail tube protein \| Phage major tail tube protein; afdb-uniprot50 \| AF-A0A1R1MK86-F1-MODEL\_V4 \| 1.0 \| 5.128e-14 \| 624 \| 0.268 \| 175 \| 119 \| 6 \| 1 \| 168 \| 2 \| 174 \| Uncharacterized protein \| Uncharacterized protein; afdb-uniprot50 \| AF-A0A2N3KSL5-F1-MODEL\_V4 \| 1.0 \| 1.111e-13 \| 622 \| 0.269 \| 171 \| 116 \| 6 \| 1 \| 168 \| 1 \| 165 \| Uncharacterized protein \| Uncharacterized protein; afdb-uniprot50 \| AF-A0A327JM09-F1-MODEL\_V4 \| 1.0 \| 1.297e-13 \| 622 \| 0.204 \| 171 \| 130 \| 6 \| 1 \| 166 \| 1 \| 170 \| Phage tail protein \| Phage tail protein; afdb-uniprot50 \| AF-A0A828HN09-F1-MODEL\_V4 \| 1.0 \| 4.033e-13 \| 621 \| 0.26 \| 173 \| 119 \| 5 \| 1 \| 168 \| 1 \| 169 \| Phage tail protein \| Phage tail protein; afdb-uniprot50 \| AF-A0A7U7YUD0-F1-MODEL\_V4 \| 1.0 \| 1.17e-13 \| 621 \| 0.244 \| 172 \| 120 \| 6 \| 2 \| 168 \| 5 \| 171 \| Phage tail protein \| Phage tail protein; afdb-uniprot50 \| AF-A0A075KC25-F1-MODEL\_V4 \| 1.0 \| 4.173e-14 \| 620 \| 0.231 \| 173 \| 122 \| 6 \| 2 \| 167 \| 6 \| 174 \| Major tail tube protein \| Major tail tube protein; afdb-uniprot50 \| AF-A0A1T4W4F8-F1-MODEL\_V4 \| 1.0 \| 3.117e-13 \| 619 \| 0.227 \| 167 \| 125 \| 3 \| 3 \| 168 \| 2 \| 165 \| Uncharacterized protein \| Uncharacterized protein; afdb-uniprot50 \| AF-A0A6A4R9K0-F1-MODEL\_V4 \| 1.0 \| 1.232e-13 \| 619 \| 0.232 \| 168 \| 127 \| 2 \| 1 \| 168 \| 2 \| 167 \| Phage tail protein \| Phage tail protein; afdb-uniprot50 \| AF-A0A2W5VKB0-F1-MODEL\_V4 \| 1.0 \| 3.638e-13 \| 619 \| 0.264 \| 170 \| 117 \| 5 \| 1 \| 166 \| 2 \| 167 \| Phage tail protein \| Phage tail protein; afdb-uniprot50 \| AF-G4CJF7-F1-MODEL\_V4 \| 1.0 \| 9.521e-14 \| 618 \| 0.208 \| 173 \| 131 \| 4 \| 1 \| 168 \| 1 \| 172 \| Phage tail core protein \| Phage tail core protein; afdb-uniprot50 \| AF-E0NXJ4-F1-MODEL\_V4 \| 1.0 \| 6.637e-14 \| 618 \| 0.291 \| 175 \| 111 \| 6 \| 2 \| 168 \| 18 \| 187 \| Phage tail tube protein FII \| Phage tail tube protein FII; afdb-uniprot50 \| AF-A0A6G5QXU3-F1-MODEL\_V4 \| 1.0 \| 2.063e-13 \| 617 \| 0.247 \| 170 \| 121 \| 5 \| 2 \| 168 \| 6 \| 171 \| Phage tail tube protein FII \| Phage tail tube protein FII; afdb-uniprot50 \| AF-A0A1T4PVU5-F1-MODEL\_V4 \| 1.0 \| 5.686e-14 \| 617 \| 0.255 \| 176 \| 118 \| 5 \| 1 \| 168 \| 1 \| 171 \| Uncharacterized protein \| Uncharacterized protein; afdb-uniprot50 \| AF-C9MV95-F1-MODEL\_V4 \| 1.0 \| 1.65e-14 \| 617 \| 0.209 \| 172 \| 125 \| 5 \| 4 \| 168 \| 7 \| 174 \| Putative phage major tail tube protein \| Putative phage major tail tube protein; afdb-uniprot50 \| AF-A0A5S9QVA0-F1-MODEL\_V4 \| 1.0 \| 2.67e-13 \| 615 \| 0.291 \| 168 \| 115 \| 3 \| 1 \| 167 \| 1 \| 165 \| Uncharacterized protein \| Uncharacterized protein; afdb-uniprot50 \| AF-R5BQ89-F1-MODEL\_V4 \| 1.0 \| 3.764e-14 \| 615 \| 0.248 \| 177 \| 120 \| 7 \| 1 \| 168 \| 1 \| 173 \| Major tail tube protein \| Major tail tube protein; afdb-uniprot50 \| AF-A0A377AJ06-F1-MODEL\_V4 \| 1.0 \| 3.575e-14 \| 615 \| 0.324 \| 179 \| 107 \| 6 \| 1 \| 167 \| 1 \| 177 \| Phage major tail tube protein \| Phage major tail tube protein; afdb-uniprot50 \| AF-A0A316Q9G3-F1-MODEL\_V4 \| 1.0 \| 3.575e-14 \| 612 \| 0.211 \| 175 \| 128 \| 4 \| 1 \| 168 \| 4 \| 175 \| Phage tail protein \| Phage tail protein; afdb-uniprot50 \| AF-A0A414Q2K2-F1-MODEL\_V4 \| 1.0 \| 5.986e-14 \| 611 \| 0.24 \| 175 \| 123 \| 5 \| 1 \| 168 \| 1 \| 172 \| Phage tail protein \| Phage tail protein; afdb-uniprot50 \| AF-A0A1Q8Q269-F1-MODEL\_V4 \| 1.0 \| 8.588e-14 \| 611 \| 0.227 \| 176 \| 122 \| 7 \| 1 \| 168 \| 6 \| 175 \| Uncharacterized protein \| Uncharacterized protein; afdb-uniprot50 \| AF-A0A0T9RBM0-F1-MODEL\_V4 \| 1.0 \| 1.514e-13 \| 610 \| 0.339 \| 174 \| 109 \| 3 \| 1 \| 168 \| 1 \| 174 \| Major tail tube protein \| Major tail tube protein; afdb-uniprot50 \| AF-A0A376JS17-F1-MODEL\_V4 \| 1.0 \| 3.282e-13 \| 608 \| 0.333 \| 153 \| 96 \| 5 \| 1 \| 148 \| 1 \| 152 \| Phage major tail tube protein FII \| Phage major tail tube protein FII; afdb-uniprot50 \| AF-A0A839IVJ8-F1-MODEL\_V4 \| 1.0 \| 4.472e-13 \| 608 \| 0.239 \| 167 \| 124 \| 3 \| 2 \| 168 \| 3 \| 166 \| Phage major tail tube protein \| Phage major tail tube protein; afdb-uniprot50 \| AF-A0A7X5EZ69-F1-MODEL\_V4 \| 1.0 \| 1.96e-13 \| 608 \| 0.171 \| 181 \| 135 \| 4 \| 3 \| 168 \| 1 \| 181 \| Uncharacterized protein \| Uncharacterized protein; afdb-uniprot50 \| AF-A0A2P8EI37-F1-MODEL\_V4 \| 1.0 \| 7.747e-14 \| 607 \| 0.222 \| 175 \| 123 \| 7 \| 1 \| 168 \| 1 \| 169 \| Uncharacterized protein \| Uncharacterized protein; afdb-uniprot50 \| AF-A0A417ISQ5-F1-MODEL\_V4 \| 1.0 \| 1.438e-13 \| 606 \| 0.245 \| 175 \| 121 \| 5 \| 1 \| 168 \| 1 \| 171 \| Phage tail protein \| Phage tail protein; afdb-uniprot50 \| AF-A0A7U7ZB02-F1-MODEL\_V4 \| 1.0 \| 2.96e-13 \| 606 \| 0.244 \| 172 \| 120 \| 6 \| 1 \| 168 \| 4 \| 169 \| Phage tail protein \| Phage tail protein; afdb-uniprot50 \| AF-A0A6N7VZ75-F1-MODEL\_V4 \| 1.0 \| 1.438e-13 \| 605 \| 0.264 \| 174 \| 115 \| 5 \| 1 \| 167 \| 2 \| 169 \| Phage tail protein \| Phage tail protein; afdb-uniprot50 \| AF-A0A7K1RPH2-F1-MODEL\_V4 \| 1.0 \| 4.472e-13 \| 604 \| 0.2 \| 175 \| 132 \| 5 \| 1 \| 167 \| 1 \| 175 \| Phage tail protein \| Phage tail protein; afdb-uniprot50 \| AF-A0A4P7L0L1-F1-MODEL\_V4 \| 1.0 \| 4.957e-13 \| 603 \| 0.348 \| 149 \| 94 \| 2 \| 23 \| 168 \| 2 \| 150 \| Phage tail tube protein FII \| Phage tail tube protein FII; afdb-uniprot50 \| AF-A0A4U8S350-F1-MODEL\_V4 \| 1.0 \| 1.514e-13 \| 603 \| 0.218 \| 169 \| 124 \| 5 \| 1 \| 166 \| 3 \| 166 \| Phage tail protein \| Phage tail protein; afdb-uniprot50 \| AF-A0A7C8KSM3-F1-MODEL\_V4 \| 1.0 \| 1.679e-13 \| 603 \| 0.217 \| 175 \| 127 \| 4 \| 1 \| 168 \| 4 \| 175 \| Phage tail protein \| Phage tail protein; afdb-uniprot50 \| AF-A0A4V2ZH72-F1-MODEL\_V4 \| 1.0 \| 4.033e-13 \| 602 \| 0.222 \| 166 \| 127 \| 2 \| 1 \| 166 \| 2 \| 165 \| Phage tail protein \| Phage tail protein; afdb-uniprot50 \| AF-A0A0B6D149-F1-MODEL\_V4 \| 1.0 \| 1.438e-13 \| 600 \| 0.264 \| 170 \| 117 \| 6 \| 1 \| 168 \| 8 \| 171 \| Phage tail tube FII family protein \| Phage tail tube FII family protein; afdb-uniprot50 \| AF-A0A4U8YRD6-F1-MODEL\_V4 \| 1.0 \| 1.679e-13 \| 600 \| 0.232 \| 176 \| 123 \| 7 \| 1 \| 168 \| 1 \| 172 \| Tail tube protein \| Tail tube protein; afdb-uniprot50 \| AF-A0A413EDA9-F1-MODEL\_V4 \| 1.0 \| 8.157e-14 \| 600 \| 0.262 \| 175 \| 118 \| 5 \| 1 \| 168 \| 2 \| 172 \| Phage tail protein \| Phage tail protein; afdb-uniprot50 \| AF-A0A662Z5W3-F1-MODEL\_V4 \| 1.0 \| 1.594e-13 \| 599 \| 0.232 \| 176 \| 122 \| 6 \| 1 \| 168 \| 5 \| 175 \| Uncharacterized protein \| Uncharacterized protein; afdb-uniprot50 \| AF-A0A840PWU5-F1-MODEL\_V4 \| 1.0 \| 3.282e-13 \| 598 \| 0.211 \| 175 \| 126 \| 5 \| 1 \| 168 \| 6 \| 175 \| Uncharacterized protein \| Uncharacterized protein; afdb-uniprot50 \| AF-A0A7W6IIC9-F1-MODEL\_V4 \| 1.0 \| 6.415e-13 \| 597 \| 0.231 \| 177 \| 125 \| 7 \| 1 \| 168 \| 1 \| 175 \| Uncharacterized protein \| Uncharacterized protein; afdb-uniprot50 \| AF-A0A212LD32-F1-MODEL\_V4 \| 1.0 \| 7.488e-13 \| 596 \| 0.192 \| 166 \| 131 \| 3 \| 6 \| 168 \| 8 \| 173 \| Uncharacterized protein \| Uncharacterized protein; afdb-uniprot50 \| AF-R5BKU1-F1-MODEL\_V4 \| 1.0 \| 2.67e-13 \| 596 \| 0.255 \| 176 \| 117 \| 6 \| 2 \| 168 \| 5 \| 175 \| Uncharacterized protein \| Uncharacterized protein; afdb-uniprot50 \| AF-A0A7X2TC80-F1-MODEL\_V4 \| 1.0 \| 6.637e-14 \| 594 \| 0.255 \| 172 \| 119 \| 5 \| 1 \| 165 \| 1 \| 170 \| Phage tail protein \| Phage tail protein; afdb-uniprot50 \| AF-A0A0D6ARX0-F1-MODEL\_V4 \| 1.0 \| 9.043e-14 \| 593 \| 0.23 \| 156 \| 114 \| 4 \| 7 \| 157 \| 5 \| 159 \| Putative phage tail core protein \| Putative phage tail core protein; afdb-uniprot50 \| AF-A0A268TWR0-F1-MODEL\_V4 \| 1.0 \| 8.302e-13 \| 593 \| 0.17 \| 170 \| 132 \| 5 \| 1 \| 166 \| 6 \| 170 \| Uncharacterized protein \| Uncharacterized protein; afdb-uniprot50 \| AF-A0A1E4M047-F1-MODEL\_V4 \| 1.0 \| 8.741e-13 \| 592 \| 0.213 \| 169 \| 126 \| 4 \| 6 \| 167 \| 6 \| 174 \| Uncharacterized protein \| Uncharacterized protein; afdb-uniprot50 \| AF-A0A376RTK4-F1-MODEL\_V4 \| 1.0 \| 1.02e-12 \| 591 \| 0.324 \| 151 \| 95 \| 4 \| 1 \| 146 \| 1 \| 149 \| Phage major tail tube protein \| Phage major tail tube protein; afdb-uniprot50 \| AF-A0A2A4XT18-F1-MODEL\_V4 \| 1.0 \| 4.957e-13 \| 591 \| 0.282 \| 170 \| 116 \| 4 \| 1 \| 168 \| 1 \| 166 \| Uncharacterized protein \| Uncharacterized protein; afdb-uniprot50 \| AF-A0A413FZE2-F1-MODEL\_V4 \| 1.0 \| 2.536e-13 \| 591 \| 0.226 \| 172 \| 122 \| 5 \| 1 \| 166 \| 4 \| 170 \| Uncharacterized protein \| Uncharacterized protein; afdb-uniprot50 \| AF-A0A1M7RI12-F1-MODEL\_V4 \| 1.0 \| 1.366e-13 \| 591 \| 0.205 \| 170 \| 123 \| 5 \| 7 \| 168 \| 9 \| 174 \| Uncharacterized protein \| Uncharacterized protein; afdb-uniprot50 \| AF-B0UK47-F1-MODEL\_V4 \| 1.0 \| 8.741e-13 \| 588 \| 0.228 \| 175 \| 128 \| 5 \| 1 \| 168 \| 1 \| 175 \| Major tail tube protein \| Major tail tube protein; afdb-uniprot50 \| AF-A0A2G1CVI0-F1-MODEL\_V4 \| 1.0 \| 1.438e-13 \| 587 \| 0.233 \| 171 \| 120 \| 6 \| 1 \| 166 \| 7 \| 171 \| Uncharacterized protein \| Uncharacterized protein; afdb-uniprot50 \| AF-U4T7P1-F1-MODEL\_V4 \| 1.0 \| 2.173e-13 \| 587 \| 0.248 \| 165 \| 117 \| 4 \| 7 \| 166 \| 11 \| 173 \| Putative bacteriophage tail core protein \| Putative bacteriophage tail core protein; afdb-uniprot50 \| AF-A0A355TQV7-F1-MODEL\_V4 \| 1.0 \| 1.861e-13 \| 586 \| 0.242 \| 165 \| 119 \| 4 \| 7 \| 166 \| 50 \| 213 \| Uncharacterized protein \| Uncharacterized protein; afdb-uniprot50 \| AF-A0A1A6FJS5-F1-MODEL\_V4 \| 1.0 \| 1.464e-12 \| 584 \| 0.215 \| 176 \| 130 \| 5 \| 1 \| 168 \| 1 \| 176 \| Uncharacterized protein \| Uncharacterized protein; afdb-uniprot50 \| AF-A0A385Q1F7-F1-MODEL\_V4 \| 1.0 \| 1.111e-13 \| 583 \| 0.255 \| 172 \| 119 \| 4 \| 1 \| 165 \| 1 \| 170 \| Phage tail protein \| Phage tail protein; afdb-uniprot50 \| AF-A0A1Q5XTX7-F1-MODEL\_V4 \| 1.0 \| 3.117e-13 \| 582 \| 0.238 \| 176 \| 121 \| 6 \| 1 \| 168 \| 1 \| 171 \| Phage tail protein \| Phage tail protein; afdb-uniprot50 \| AF-A0A1L8CRR8-F1-MODEL\_V4 \| 1.0 \| 3.455e-13 \| 582 \| 0.219 \| 173 \| 124 \| 5 \| 3 \| 168 \| 4 \| 172 \| Phage tail protein \| Phage tail protein; afdb-uniprot50 \| AF-A0A4Q2SZY2-F1-MODEL\_V4 \| 1.0 \| 1.995e-12 \| 582 \| 0.228 \| 175 \| 125 \| 6 \| 1 \| 167 \| 1 \| 173 \| Uncharacterized protein \| Uncharacterized protein; afdb-uniprot50 \| AF-A0A098MDI9-F1-MODEL\_V4 \| 1.0 \| 2.96e-13 \| 581 \| 0.227 \| 176 \| 121 \| 7 \| 1 \| 168 \| 4 \| 172 \| Tail protein \| Tail protein; afdb-uniprot50 \| AF-A0A1S0V4H2-F1-MODEL\_V4 \| 1.0 \| 1.894e-12 \| 580 \| 0.474 \| 118 \| 62 \| 0 \| 51 \| 168 \| 1 \| 118 \| Uncharacterized protein \| Uncharacterized protein; afdb-uniprot50 \| AF-A0A7M3MBM7-F1-MODEL\_V4 \| 1.0 \| 3.282e-13 \| 580 \| 0.222 \| 175 \| 123 \| 7 \| 1 \| 167 \| 2 \| 171 \| Phage tail protein \| Phage tail protein; afdb-uniprot50 \| AF-A0A3L7AKR1-F1-MODEL\_V4 \| 1.0 \| 3.703e-12 \| 579 \| 0.21 \| 176 \| 130 \| 5 \| 1 \| 168 \| 4 \| 178 \| Phage tail protein \| Phage tail protein; afdb-uniprot50 \| AF-A0A2D3WHP5-F1-MODEL\_V4 \| 1.0 \| 9.204e-13 \| 577 \| 0.213 \| 169 \| 126 \| 5 \| 1 \| 166 \| 5 \| 169 \| Uncharacterized protein \| Uncharacterized protein; afdb-uniprot50 \| AF-A0A2N0CX46-F1-MODEL\_V4 \| 1.0 \| 6.093e-13 \| 576 \| 0.295 \| 169 \| 108 \| 5 \| 1 \| 167 \| 1 \| 160 \| Phage tail protein \| Phage tail protein; afdb-uniprot50 \| AF-A0A2E3R286-F1-MODEL\_V4 \| 1.0 \| 1.056e-13 \| 574 \| 0.192 \| 171 \| 131 \| 4 \| 1 \| 165 \| 2 \| 171 \| Uncharacterized protein \| Uncharacterized protein; afdb-uniprot50 \| AF-B3QTJ5-F1-MODEL\_V4 \| 1.0 \| 3.638e-13 \| 573 \| 0.237 \| 164 \| 119 \| 4 \| 1 \| 160 \| 4 \| 165 \| Phage major tail tube protein \| Phage major tail tube protein; afdb-uniprot50 \| AF-A0A1D2QMU1-F1-MODEL\_V4 \| 1.0 \| 2.1e-12 \| 573 \| 0.222 \| 171 \| 126 \| 4 \| 1 \| 168 \| 3 \| 169 \| Uncharacterized protein \| Uncharacterized protein; afdb-uniprot50 \| AF-A0A064AK82-F1-MODEL\_V4 \| 1.0 \| 2.96e-13 \| 573 \| 0.218 \| 174 \| 124 \| 5 \| 1 \| 166 \| 2 \| 171 \| Major tail tube protein \| Major tail tube protein; afdb-uniprot50 \| AF-D1AFA7-F1-MODEL\_V4 \| 1.0 \| 2.811e-13 \| 573 \| 0.208 \| 173 \| 127 \| 4 \| 1 \| 166 \| 3 \| 172 \| Major tail tube protein \| Major tail tube protein; afdb-uniprot50 \| AF-A0A1Q6JPJ0-F1-MODEL\_V4 \| 1.0 \| 1.366e-13 \| 573 \| 0.263 \| 171 \| 116 \| 5 \| 1 \| 165 \| 8 \| 174 \| Phage tail protein \| Phage tail protein; afdb-uniprot50 \| AF-G1UXP6-F1-MODEL\_V4 \| 1.0 \| 8.741e-13 \| 572 \| 0.217 \| 179 \| 124 \| 7 \| 1 \| 168 \| 5 \| 178 \| Uncharacterized protein \| Uncharacterized protein; afdb-uniprot50 \| AF-A0A496JWV3-F1-MODEL\_V4 \| 1.0 \| 1.623e-12 \| 571 \| 0.267 \| 146 \| 103 \| 3 \| 25 \| 168 \| 2 \| 145 \| Phage tail protein \| Phage tail protein; afdb-uniprot50 \| AF-A0A2P7V3M3-F1-MODEL\_V4 \| 1.0 \| 1.131e-12 \| 570 \| 0.228 \| 175 \| 122 \| 6 \| 2 \| 168 \| 5 \| 174 \| Phage tail protein \| Phage tail protein; afdb-uniprot50 \| AF-A0A7C3GHU1-F1-MODEL\_V4 \| 1.0 \| 3.117e-13 \| 570 \| 0.224 \| 174 \| 128 \| 5 \| 1 \| 168 \| 4 \| 176 \| Phage major tail tube protein \| Phage major tail tube protein; afdb-uniprot50 \| AF-A0A2E2N2N1-F1-MODEL\_V4 \| 1.0 \| 5.22e-13 \| 569 \| 0.209 \| 172 \| 125 \| 5 \| 1 \| 165 \| 2 \| 169 \| Phage tail protein \| Phage tail protein; afdb-uniprot50 \| AF-A0A0C2UPV3-F1-MODEL\_V4 \| 1.0 \| 2.536e-13 \| 568 \| 0.233 \| 163 \| 111 \| 7 \| 6 \| 166 \| 9 \| 159 \| Uncharacterized protein \| Uncharacterized protein; afdb-uniprot50 \| AF-A0A5N3PH51-F1-MODEL\_V4 \| 1.0 \| 2.581e-12 \| 567 \| 0.204 \| 176 \| 128 \| 6 \| 1 \| 167 \| 4 \| 176 \| Uncharacterized protein \| Uncharacterized protein; afdb-uniprot50 \| AF-A0A3G2R410-F1-MODEL\_V4 \| 1.0 \| 7.885e-13 \| 566 \| 0.208 \| 173 \| 126 \| 6 \| 1 \| 166 \| 1 \| 169 \| Phage tail protein \| Phage tail protein; afdb-uniprot50 \| AF-A0A4P8G1X0-F1-MODEL\_V4 \| 1.0 \| 2.328e-12 \| 565 \| 0.226 \| 159 \| 122 \| 1 \| 9 \| 167 \| 1 \| 158 \| Uncharacterized protein \| Uncharacterized protein; afdb-uniprot50 \| AF-A0A0X1U7S9-F1-MODEL\_V4 \| 1.0 \| 2.96e-13 \| 563 \| 0.267 \| 172 \| 116 \| 6 \| 1 \| 165 \| 3 \| 171 \| Phage tail tube protein FII \| Phage tail tube protein FII; afdb-uniprot50 \| AF-A0A8B2NVA0-F1-MODEL\_V4 \| 1.0 \| 2.211e-12 \| 563 \| 0.203 \| 172 \| 126 \| 5 \| 2 \| 168 \| 6 \| 171 \| Uncharacterized protein \| Uncharacterized protein; afdb-uniprot50 \| AF-A0A0D7X7K2-F1-MODEL\_V4 \| 1.0 \| 5.22e-13 \| 563 \| 0.232 \| 176 \| 120 \| 8 \| 1 \| 168 \| 6 \| 174 \| Uncharacterized protein \| Uncharacterized protein; afdb-uniprot50 \| AF-B6WRQ4-F1-MODEL\_V4 \| 1.0 \| 3.282e-13 \| 560 \| 0.145 \| 172 \| 139 \| 4 \| 1 \| 167 \| 22 \| 190 \| Putative phage major tail tube protein \| Putative phage major tail tube protein; afdb-uniprot50 \| AF-A0A316DHI7-F1-MODEL\_V4 \| 1.0 \| 9.69e-13 \| 558 \| 0.215 \| 176 \| 123 \| 8 \| 1 \| 168 \| 4 \| 172 \| Uncharacterized protein \| Uncharacterized protein; afdb-uniprot50 \| AF-A0A812QV43-F1-MODEL\_V4 \| 1.0 \| 3.517e-12 \| 558 \| 0.345 \| 136 \| 88 \| 1 \| 23 \| 157 \| 393 \| 528 \| GpFI protein \| GpFI protein; afdb-uniprot50 \| AF-A0A327JAQ8-F1-MODEL\_V4 \| 1.0 \| 1.995e-12 \| 557 \| 0.22 \| 168 \| 122 \| 6 \| 6 \| 167 \| 4 \| 168 \| Uncharacterized protein \| Uncharacterized protein; afdb-uniprot50 \| AF-A0A7J0BHY9-F1-MODEL\_V4 \| 1.0 \| 1.39e-12 \| 557 \| 0.193 \| 176 \| 130 \| 6 \| 1 \| 168 \| 1 \| 172 \| Uncharacterized protein \| Uncharacterized protein; afdb-uniprot50 \| AF-A0A5N3S1K2-F1-MODEL\_V4 \| 1.0 \| 1.541e-12 \| 555 \| 0.179 \| 162 \| 130 \| 2 \| 1 \| 162 \| 1 \| 159 \| Uncharacterized protein \| Uncharacterized protein; afdb-uniprot50 \| AF-A0A373CZ49-F1-MODEL\_V4 \| 1.0 \| 6.093e-13 \| 555 \| 0.273 \| 172 \| 116 \| 5 \| 1 \| 165 \| 4 \| 173 \| Phage tail protein \| Phage tail protein; afdb-uniprot50 \| AF-D6KLG9-F1-MODEL\_V4 \| 1.0 \| 1.799e-12 \| 554 \| 0.221 \| 176 \| 122 \| 6 \| 3 \| 168 \| 1 \| 171 \| Putative phage major tail tube protein \| Putative phage major tail tube protein; afdb-uniprot50 \| AF-A0A1V4GES7-F1-MODEL\_V4 \| 1.0 \| 8.302e-13 \| 554 \| 0.238 \| 176 \| 120 \| 6 \| 1 \| 167 \| 2 \| 172 \| Uncharacterized protein \| Uncharacterized protein; afdb-uniprot50 \| AF-A0A285MEY3-F1-MODEL\_V4 \| 1.0 \| 2.861e-12 \| 553 \| 0.226 \| 168 \| 121 \| 4 \| 6 \| 168 \| 8 \| 171 \| Phage tail tube protein FII \| Phage tail tube protein FII; afdb-uniprot50 \| AF-A0A268ELC0-F1-MODEL\_V4 \| 1.0 \| 1.074e-12 \| 553 \| 0.215 \| 176 \| 124 \| 7 \| 1 \| 167 \| 1 \| 171 \| Uncharacterized protein \| Uncharacterized protein; afdb-uniprot50 \| AF-A0A329UB98-F1-MODEL\_V4 \| 1.0 \| 8.741e-13 \| 553 \| 0.17 \| 170 \| 133 \| 4 \| 2 \| 165 \| 9 \| 176 \| Phage tail protein \| Phage tail protein; afdb-uniprot50 \| AF-A0A845STF4-F1-MODEL\_V4 \| 1.0 \| 2.581e-12 \| 550 \| 0.205 \| 175 \| 126 \| 5 \| 1 \| 168 \| 4 \| 172 \| Phage tail protein \| Phage tail protein; afdb-uniprot50 \| AF-A0A2N0CX99-F1-MODEL\_V4 \| 1.0 \| 2.328e-12 \| 548 \| 0.301 \| 166 \| 105 \| 5 \| 1 \| 164 \| 1 \| 157 \| Phage tail protein \| Phage tail protein; afdb-uniprot50 \| AF-A0A1B9NZR8-F1-MODEL\_V4 \| 1.0 \| 2.581e-12 \| 548 \| 0.175 \| 165 \| 133 \| 2 \| 1 \| 165 \| 1 \| 162 \| Uncharacterized protein \| Uncharacterized protein; afdb-uniprot50 \| AF-A0A0M7ALA7-F1-MODEL\_V4 \| 1.0 \| 7.621e-12 \| 548 \| 0.173 \| 173 \| 132 \| 6 \| 2 \| 168 \| 3 \| 170 \| Major tail tube protein \| Major tail tube protein; afdb-uniprot50 \| AF-A0A6L9HFW7-F1-MODEL\_V4 \| 1.0 \| 2.718e-12 \| 547 \| 0.194 \| 175 \| 130 \| 6 \| 1 \| 168 \| 2 \| 172 \| Phage tail protein \| Phage tail protein; afdb-uniprot50 \| AF-A0A2Y0TYM0-F1-MODEL\_V4 \| 1.0 \| 1.709e-12 \| 547 \| 0.278 \| 169 \| 109 \| 4 \| 1 \| 165 \| 1 \| 160 \| Major tail tube protein FII \| Major tail tube protein FII; afdb-uniprot50 \| AF-A0A1B8VX25-F1-MODEL\_V4 \| 1.0 \| 2.718e-12 \| 546 \| 0.181 \| 176 \| 132 \| 6 \| 1 \| 168 \| 1 \| 172 \| Uncharacterized protein \| Uncharacterized protein; afdb-uniprot50 \| AF-A0A5A8F086-F1-MODEL\_V4 \| 1.0 \| 1.32e-12 \| 545 \| 0.173 \| 173 \| 136 \| 5 \| 1 \| 168 \| 1 \| 171 \| Uncharacterized protein \| Uncharacterized protein; afdb-uniprot50 \| AF-G4KQ74-F1-MODEL\_V4 \| 1.0 \| 2.1e-12 \| 544 \| 0.205 \| 175 \| 124 \| 6 \| 1 \| 165 \| 2 \| 171 \| Putative tail tube protein \| Putative tail tube protein; afdb-uniprot50 \| AF-A0A1N7A8E0-F1-MODEL\_V4 \| 1.0 \| 1.995e-12 \| 544 \| 0.194 \| 175 \| 126 \| 7 \| 1 \| 167 \| 4 \| 171 \| Uncharacterized protein \| Uncharacterized protein; afdb-uniprot50 \| AF-A0A5S4VNC9-F1-MODEL\_V4 \| 1.0 \| 2.96e-13 \| 543 \| 0.229 \| 170 \| 124 \| 4 \| 1 \| 165 \| 3 \| 170 \| Phage tail protein \| Phage tail protein; afdb-uniprot50 \| AF-A0A163TB63-F1-MODEL\_V4 \| 1.0 \| 4.551e-12 \| 543 \| 0.217 \| 175 \| 126 \| 6 \| 1 \| 168 \| 1 \| 171 \| Bacteriophage protein \| Bacteriophage protein; afdb-uniprot50 \| AF-R5GLL1-F1-MODEL\_V4 \| 1.0 \| 3.34e-12 \| 543 \| 0.202 \| 173 \| 128 \| 5 \| 1 \| 167 \| 2 \| 170 \| Putative phage major tail tube protein \| Putative phage major tail tube protein; afdb-uniprot50 \| AF-A0A7X6JBC6-F1-MODEL\_V4 \| 1.0 \| 5.593e-12 \| 542 \| 0.383 \| 133 \| 79 \| 1 \| 36 \| 168 \| 2 \| 131 \| Phage major tail tube protein \| Phage major tail tube protein; afdb-uniprot50 \| AF-E6X1M7-F1-MODEL\_V4 \| 1.0 \| 2.718e-12 \| 542 \| 0.218 \| 165 \| 118 \| 5 \| 6 \| 166 \| 9 \| 166 \| Major tail tube protein \| Major tail tube protein; afdb-uniprot50 \| AF-A0A3D5LTH1-F1-MODEL\_V4 \| 1.0 \| 2.288e-13 \| 542 \| 0.217 \| 170 \| 126 \| 5 \| 2 \| 165 \| 5 \| 173 \| Phage tail protein \| Phage tail protein; afdb-uniprot50 \| AF-A0A3C1WQU9-F1-MODEL\_V4 \| 1.0 \| 1.131e-12 \| 542 \| 0.232 \| 172 \| 122 \| 5 \| 2 \| 165 \| 10 \| 179 \| Phage tail protein \| Phage tail protein; afdb-uniprot50 \| AF-A0A611ERL2-F1-MODEL\_V4 \| 1.0 \| 3.172e-12 \| 541 \| 0.34 \| 150 \| 91 \| 5 \| 1 \| 144 \| 1 \| 148 \| Phage major tail tube protein \| Phage major tail tube protein; afdb-uniprot50 \| AF-D6LHL5-F1-MODEL\_V4 \| 1.0 \| 4.957e-13 \| 541 \| 0.212 \| 174 \| 123 \| 8 \| 1 \| 166 \| 1 \| 168 \| Phage-related contractile tail tube protein \| Phage-related contractile tail tube protein; afdb-uniprot50 \| AF-A0A833A242-F1-MODEL\_V4 \| 1.0 \| 1.32e-12 \| 540 \| 0.195 \| 174 \| 131 \| 5 \| 1 \| 168 \| 2 \| 172 \| Phage tail protein \| Phage tail protein; afdb-uniprot50 \| AF-A0A844Q9G1-F1-MODEL\_V4 \| 1.0 \| 3.013e-12 \| 538 \| 0.259 \| 177 \| 116 \| 8 \| 1 \| 167 \| 1 \| 172 \| Phage tail protein \| Phage tail protein; afdb-uniprot50 \| AF-A0A1T4WVE5-F1-MODEL\_V4 \| 1.0 \| 9.367e-12 \| 537 \| 0.201 \| 149 \| 115 \| 4 \| 1 \| 146 \| 3 \| 150 \| Uncharacterized protein \| Uncharacterized protein; afdb-uniprot50 \| AF-A0A1I5MNT8-F1-MODEL\_V4 \| 1.0 \| 4.033e-13 \| 537 \| 0.235 \| 170 \| 124 \| 4 \| 1 \| 165 \| 1 \| 169 \| Uncharacterized protein \| Uncharacterized protein; afdb-uniprot50 \| AF-A0A174PJS3-F1-MODEL\_V4 \| 1.0 \| 1.541e-12 \| 536 \| 0.217 \| 175 \| 123 \| 8 \| 1 \| 165 \| 1 \| 171 \| Phage major tail tube protein \| Phage major tail tube protein; afdb-uniprot50 \| AF-A0A660NLN2-F1-MODEL\_V4 \| 1.0 \| 1.415e-11 \| 534 \| 0.663 \| 116 \| 39 \| 0 \| 1 \| 116 \| 1 \| 116 \| Phage major tail tube protein \| Phage major tail tube protein; afdb-uniprot50 \| AF-A0A2X1PJG7-F1-MODEL\_V4 \| 1.0 \| 3.703e-12 \| 534 \| 0.312 \| 157 \| 101 \| 5 \| 15 \| 167 \| 2 \| 155 \| Major tail tube protein FII \| Major tail tube protein FII; afdb-uniprot50 \| AF-A0A1A9AX50-F1-MODEL\_V4 \| 1.0 \| 2.766e-11 \| 532 \| 0.216 \| 162 \| 124 \| 3 \| 2 \| 162 \| 3 \| 162 \| Phage tail protein \| Phage tail protein; afdb-uniprot50 \| AF-A0A842IW21-F1-MODEL\_V4 \| 1.0 \| 8.302e-13 \| 532 \| 0.215 \| 172 \| 126 \| 5 \| 1 \| 168 \| 2 \| 168 \| Phage major tail tube protein \| Phage major tail tube protein; afdb-uniprot50 \| AF-A0A078MHI9-F1-MODEL\_V4 \| 1.0 \| 3.703e-12 \| 532 \| 0.184 \| 173 \| 129 \| 6 \| 3 \| 168 \| 1 \| 168 \| Phage tail tube protein FII \| Phage tail tube protein FII; afdb-uniprot50 \| AF-A0A2D3W2J9-F1-MODEL\_V4 \| 1.0 \| 3.703e-12 \| 528 \| 0.26 \| 161 \| 108 \| 6 \| 7 \| 166 \| 10 \| 160 \| Uncharacterized protein \| Uncharacterized protein; afdb-uniprot50 \| AF-A0A061NPQ1-F1-MODEL\_V4 \| 1.0 \| 2.451e-12 \| 528 \| 0.238 \| 172 \| 118 \| 6 \| 1 \| 165 \| 4 \| 169 \| Phage major tail tube protein \| Phage major tail tube protein; afdb-uniprot50 \| AF-A0A4V2KL57-F1-MODEL\_V4 \| 1.0 \| 1.212e-11 \| 528 \| 0.217 \| 175 \| 125 \| 6 \| 1 \| 168 \| 2 \| 171 \| Phage tail protein \| Phage tail protein; afdb-uniprot50 \| AF-G9PUI5-F1-MODEL\_V4 \| 1.0 \| 6.201e-12 \| 528 \| 0.195 \| 174 \| 129 \| 6 \| 1 \| 167 \| 2 \| 171 \| Phage major tail tube protein \| Phage major tail tube protein; afdb-uniprot50 \| AF-A0A5T0IXF5-F1-MODEL\_V4 \| 1.0 \| 1.831e-11 \| 526 \| 0.245 \| 155 \| 108 \| 5 \| 18 \| 168 \| 2 \| 151 \| Phage tail protein \| Phage tail protein; afdb-uniprot50 \| AF-A0A6L5TDP9-F1-MODEL\_V4 \| 1.0 \| 2.451e-12 \| 526 \| 0.235 \| 170 \| 121 \| 5 \| 2 \| 165 \| 5 \| 171 \| Phage tail protein \| Phage tail protein; afdb-uniprot50 \| AF-A0A6N8K4V4-F1-MODEL\_V4 \| 1.0 \| 1.928e-11 \| 525 \| 0.331 \| 145 \| 90 \| 4 \| 29 \| 168 \| 1 \| 143 \| Phage major tail tube protein \| Phage major tail tube protein; afdb-uniprot50 \| AF-R7HWW0-F1-MODEL\_V4 \| 1.0 \| 2.451e-12 \| 525 \| 0.201 \| 174 \| 128 \| 5 \| 1 \| 165 \| 1 \| 172 \| Phage tail tube protein FII \| Phage tail tube protein FII; afdb-uniprot50 \| AF-A0A2V2FVX9-F1-MODEL\_V4 \| 1.0 \| 1.276e-11 \| 525 \| 0.198 \| 176 \| 129 \| 6 \| 1 \| 168 \| 2 \| 173 \| Uncharacterized protein \| Uncharacterized protein; afdb-uniprot50 \| AF-A0A0D2GJP7-F1-MODEL\_V4 \| 1.0 \| 1.569e-11 \| 525 \| 0.22 \| 172 \| 123 \| 4 \| 4 \| 168 \| 7 \| 174 \| Tail protein \| Tail protein; afdb-uniprot50 \| AF-A0A1T2XA25-F1-MODEL\_V4 \| 1.0 \| 8.449e-12 \| 524 \| 0.193 \| 176 \| 128 \| 7 \| 1 \| 168 \| 6 \| 175 \| Phage tail protein \| Phage tail protein; afdb-uniprot50 \| AF-E3HBL5-F1-MODEL\_V4 \| 1.0 \| 7.621e-12 \| 521 \| 0.232 \| 172 \| 120 \| 6 \| 4 \| 167 \| 5 \| 172 \| Major tail tube protein \| Major tail tube protein; afdb-uniprot50 \| AF-A0A7I8DLE1-F1-MODEL\_V4 \| 1.0 \| 7.621e-12 \| 521 \| 0.203 \| 177 \| 129 \| 6 \| 1 \| 168 \| 2 \| 175 \| Bacteriophage protein \| Bacteriophage protein; afdb-uniprot50 \| AF-A0A8B2TUX0-F1-MODEL\_V4 \| 1.0 \| 5.593e-12 \| 520 \| 0.252 \| 170 \| 121 \| 3 \| 1 \| 168 \| 1 \| 166 \| Phage tail protein \| Phage tail protein; afdb-uniprot50 \| AF-E5VKJ4-F1-MODEL\_V4 \| 1.0 \| 1.464e-12 \| 520 \| 0.222 \| 171 \| 127 \| 4 \| 1 \| 165 \| 2 \| 172 \| Phage major tail tube protein \| Phage major tail tube protein; afdb-uniprot50 \| AF-A0A3D4L176-F1-MODEL\_V4 \| 1.0 \| 8.896e-12 \| 516 \| 0.204 \| 176 \| 127 \| 7 \| 2 \| 168 \| 4 \| 175 \| Phage tail protein \| Phage tail protein; afdb-uniprot50 \| AF-A0A6I0DW72-F1-MODEL\_V4 \| 1.0 \| 3.399e-11 \| 514 \| 0.207 \| 169 \| 124 \| 5 \| 6 \| 168 \| 6 \| 170 \| Uncharacterized protein \| Uncharacterized protein; afdb-uniprot50 \| AF-A0A087M4E0-F1-MODEL\_V4 \| 1.0 \| 5.994e-11 \| 514 \| 0.174 \| 178 \| 134 \| 7 \| 1 \| 167 \| 1 \| 176 \| Uncharacterized protein \| Uncharacterized protein; afdb-uniprot50 \| AF-A0A7T5JQ37-F1-MODEL\_V4 \| 1.0 \| 2.137e-11 \| 511 \| 0.237 \| 177 \| 121 \| 6 \| 1 \| 168 \| 2 \| 173 \| Phage major tail tube protein \| Phage major tail tube protein; afdb-uniprot50 \| AF-D1Y708-F1-MODEL\_V4 \| 1.0 \| 1.739e-11 \| 510 \| 0.241 \| 174 \| 119 \| 6 \| 1 \| 166 \| 2 \| 170 \| Putative phage major tail tube protein \| Putative phage major tail tube protein; afdb-uniprot50 \| AF-A0A7X4ZQ89-F1-MODEL\_V4 \| 1.0 \| 1.212e-11 \| 509 \| 0.223 \| 152 \| 108 \| 4 \| 22 \| 165 \| 4 \| 153 \| Phage tail protein \| Phage tail protein; afdb-uniprot50 \| AF-A0A5C8BE13-F1-MODEL\_V4 \| 1.0 \| 1.541e-12 \| 506 \| 0.191 \| 172 \| 130 \| 6 \| 1 \| 165 \| 1 \| 170 \| Uncharacterized protein \| Uncharacterized protein; afdb-uniprot50 \| AF-A0A6P0MHV0-F1-MODEL\_V4 \| 1.0 \| 4.632e-11 \| 506 \| 0.193 \| 176 \| 132 \| 5 \| 1 \| 168 \| 1 \| 174 \| Uncharacterized protein \| Uncharacterized protein; afdb-uniprot50 \| AF-E6L5Z4-F1-MODEL\_V4 \| 1.0 \| 3.579e-11 \| 505 \| 0.207 \| 169 \| 124 \| 7 \| 3 \| 168 \| 1 \| 162 \| Phage major tail tube protein \| Phage major tail tube protein; afdb-uniprot50 \| AF-A0A349HHP9-F1-MODEL\_V4 \| 1.0 \| 1.928e-11 \| 505 \| 0.168 \| 178 \| 133 \| 8 \| 1 \| 168 \| 1 \| 173 \| Phage tail protein \| Phage tail protein; afdb-uniprot50 \| AF-A0A3A9BH43-F1-MODEL\_V4 \| 1.0 \| 1.344e-11 \| 505 \| 0.175 \| 171 \| 129 \| 6 \| 4 \| 165 \| 9 \| 176 \| Uncharacterized protein \| Uncharacterized protein; afdb-uniprot50 \| AF-A0A4R2GWP5-F1-MODEL\_V4 \| 1.0 \| 6.875e-12 \| 504 \| 0.215 \| 167 \| 120 \| 6 \| 2 \| 158 \| 3 \| 168 \| Phage tail tube protein FII \| Phage tail tube protein FII; afdb-uniprot50 \| AF-A0A173SC52-F1-MODEL\_V4 \| 1.0 \| 1.831e-11 \| 504 \| 0.207 \| 169 \| 124 \| 5 \| 4 \| 166 \| 62 \| 226 \| Phage major tail tube protein \| Phage major tail tube protein; afdb-uniprot50 \| AF-A0A1W2EJX0-F1-MODEL\_V4 \| 1.0 \| 4.632e-11 \| 502 \| 0.182 \| 175 \| 130 \| 7 \| 1 \| 168 \| 4 \| 172 \| Uncharacterized protein \| Uncharacterized protein; afdb-uniprot50 \| AF-E2CJG9-F1-MODEL\_V4 \| 1.0 \| 2.495e-11 \| 501 \| 0.18 \| 166 \| 125 \| 6 \| 10 \| 168 \| 4 \| 165 \| Major tail tube protein \| Major tail tube protein; afdb-uniprot50 \| AF-A0A139D6K0-F1-MODEL\_V4 \| 1.0 \| 4.551e-12 \| 501 \| 0.188 \| 170 \| 132 \| 4 \| 1 \| 165 \| 3 \| 171 \| Major tail tube protein \| Major tail tube protein; afdb-uniprot50 \| AF-A0A5M9TYS6-F1-MODEL\_V4 \| 1.0 \| 2.627e-11 \| 501 \| 0.175 \| 177 \| 132 \| 7 \| 1 \| 168 \| 1 \| 172 \| Uncharacterized protein \| Uncharacterized protein; afdb-uniprot50 \| AF-A0A1V3VT23-F1-MODEL\_V4 \| 1.0 \| 4.877e-11 \| 500 \| 0.288 \| 135 \| 91 \| 3 \| 21 \| 152 \| 2 \| 134 \| Phage tail protein \| Phage tail protein; afdb-uniprot50 \| AF-S9TYX5-F1-MODEL\_V4 \| 1.0 \| 3.399e-11 \| 500 \| 0.165 \| 175 \| 132 \| 7 \| 3 \| 168 \| 5 \| 174 \| Major tail tube protein \| Major tail tube protein; afdb-uniprot50 \| AF-A0A1I5RNP2-F1-MODEL\_V4 \| 1.0 \| 4.399e-11 \| 499 \| 0.203 \| 167 \| 123 \| 6 \| 1 \| 166 \| 1 \| 158 \| Phage contractile tail tube protein, P2 family \| Phage contractile tail tube protein, P2 family; afdb-uniprot50 \| AF-A0A553SNF7-F1-MODEL\_V4 \| 1.0 \| 1.276e-11 \| 499 \| 0.209 \| 172 \| 126 \| 4 \| 1 \| 165 \| 1 \| 169 \| Phage tail protein \| Phage tail protein; afdb-uniprot50 \| AF-A0A6N9P8Y1-F1-MODEL\_V4 \| 1.0 \| 7.621e-12 \| 499 \| 0.196 \| 173 \| 132 \| 4 \| 1 \| 168 \| 1 \| 171 \| Uncharacterized protein \| Uncharacterized protein; afdb-uniprot50 \| AF-A0A7W6PS12-F1-MODEL\_V4 \| 1.0 \| 6.997e-11 \| 498 \| 0.184 \| 168 \| 126 \| 4 \| 6 \| 168 \| 8 \| 169 \| Uncharacterized protein \| Uncharacterized protein; afdb-uniprot50 \| AF-A0A316ML96-F1-MODEL\_V4 \| 1.0 \| 4.551e-12 \| 498 \| 0.156 \| 172 \| 139 \| 4 \| 1 \| 167 \| 1 \| 171 \| Phage tail protein \| Phage tail protein; afdb-uniprot50 \| AF-E2CFK4-F1-MODEL\_V4 \| 1.0 \| 2.137e-11 \| 495 \| 0.192 \| 171 \| 130 \| 5 \| 1 \| 167 \| 1 \| 167 \| Uncharacterized protein \| Uncharacterized protein; afdb-uniprot50 \| AF-A0A7G9GXI4-F1-MODEL\_V4 \| 1.0 \| 7.238e-12 \| 495 \| 0.189 \| 174 \| 128 \| 7 \| 1 \| 166 \| 1 \| 169 \| Phage major tail tube protein \| Phage major tail tube protein; afdb-uniprot50 \| AF-A0A369BH51-F1-MODEL\_V4 \| 1.0 \| 1.212e-11 \| 495 \| 0.18 \| 177 \| 131 \| 6 \| 1 \| 166 \| 1 \| 174 \| Uncharacterized protein \| Uncharacterized protein; afdb-uniprot50 \| AF-A0A7J5WEE2-F1-MODEL\_V4 \| 1.0 \| 7.757e-11 \| 495 \| 0.222 \| 175 \| 122 \| 8 \| 2 \| 167 \| 6 \| 175 \| Major tail tube \| Major tail tube; afdb-uniprot50 \| AF-A0A285M2K2-F1-MODEL\_V4 \| 1.0 \| 3.399e-11 \| 492 \| 0.238 \| 176 \| 119 \| 8 \| 1 \| 168 \| 1 \| 169 \| Phage contractile tail tube protein, P2 family \| Phage contractile tail tube protein, P2 family; afdb-uniprot50 \| AF-A0A3A9CS87-F1-MODEL\_V4 \| 1.0 \| 4.551e-12 \| 492 \| 0.2 \| 170 \| 125 \| 8 \| 3 \| 165 \| 7 \| 172 \| Phage tail protein \| Phage tail protein; afdb-uniprot50 \| AF-F5S5D1-F1-MODEL\_V4 \| 1.0 \| 1.299e-10 \| 491 \| 0.294 \| 136 \| 94 \| 2 \| 1 \| 135 \| 1 \| 135 \| Uncharacterized protein \| Uncharacterized protein; afdb-uniprot50 \| AF-A0A7C4DIZ6-F1-MODEL\_V4 \| 1.0 \| 3.579e-11 \| 491 \| 0.236 \| 148 \| 105 \| 4 \| 28 \| 168 \| 2 \| 148 \| Uncharacterized protein \| Uncharacterized protein; afdb-uniprot50 \| AF-A0A1Q6PWR7-F1-MODEL\_V4 \| 1.0 \| 1.739e-11 \| 491 \| 0.209 \| 172 \| 122 \| 6 \| 3 \| 166 \| 5 \| 170 \| Uncharacterized protein \| Uncharacterized protein; afdb-uniprot50 \| AF-A0A3M4BUI2-F1-MODEL\_V4 \| 1.0 \| 3.46e-10 \| 484 \| 0.383 \| 120 \| 74 \| 0 \| 49 \| 168 \| 2 \| 121 \| Major tail tube protein \| Major tail tube protein; afdb-uniprot50 \| AF-A0A4V6EN13-F1-MODEL\_V4 \| 1.0 \| 1.739e-11 \| 484 \| 0.19 \| 173 \| 131 \| 6 \| 1 \| 165 \| 1 \| 172 \| Phage major tail tube protein \| Phage major tail tube protein; afdb-uniprot50 \| AF-A0A1M3AIS0-F1-MODEL\_V4 \| 1.0 \| 3.46e-10 \| 483 \| 0.201 \| 154 \| 117 \| 3 \| 8 \| 155 \| 2 \| 155 \| Phage tail protein \| Phage tail protein; afdb-uniprot50 \| AF-A0A1H1D1J4-F1-MODEL\_V4 \| 1.0 \| 3.399e-11 \| 481 \| 0.132 \| 166 \| 133 \| 6 \| 2 \| 158 \| 6 \| 169 \| Uncharacterized protein \| Uncharacterized protein; afdb-uniprot50 \| AF-A0A143HCW7-F1-MODEL\_V4 \| 1.0 \| 2.451e-12 \| 480 \| 0.224 \| 169 \| 125 \| 4 \| 2 \| 165 \| 4 \| 171 \| Uncharacterized protein \| Uncharacterized protein; afdb-uniprot50 \| AF-A0A0S4XLP2-F1-MODEL\_V4 \| 1.0 \| 4.399e-11 \| 478 \| 0.177 \| 169 \| 127 \| 5 \| 3 \| 168 \| 5 \| 164 \| Uncharacterized protein \| Uncharacterized protein; afdb-uniprot50 \| AF-A0A4D7B2K3-F1-MODEL\_V4 \| 1.0 \| 4.399e-11 \| 478 \| 0.222 \| 171 \| 127 \| 6 \| 1 \| 167 \| 1 \| 169 \| Uncharacterized protein \| Uncharacterized protein; afdb-uniprot50 \| AF-A0A5M8P5M4-F1-MODEL\_V4 \| 1.0 \| 4.399e-11 \| 476 \| 0.175 \| 160 \| 123 \| 5 \| 7 \| 158 \| 10 \| 168 \| Uncharacterized protein \| Uncharacterized protein; afdb-uniprot50 \| AF-A0A1Q6KUB1-F1-MODEL\_V4 \| 1.0 \| 9.863e-12 \| 476 \| 0.173 \| 173 \| 133 \| 5 \| 2 \| 165 \| 5 \| 176 \| Uncharacterized protein \| Uncharacterized protein; afdb-uniprot50 \| AF-A0A5R8Y7D8-F1-MODEL\_V4 \| 1.0 \| 1.113e-10 \| 473 \| 0.18 \| 177 \| 128 \| 9 \| 1 \| 167 \| 1 \| 170 \| Uncharacterized protein \| Uncharacterized protein; afdb-uniprot50 \| AF-A0A285M8B1-F1-MODEL\_V4 \| 1.0 \| 2.29e-10 \| 469 \| 0.177 \| 175 \| 130 \| 8 \| 1 \| 166 \| 1 \| 170 \| Phage tail tube protein FII \| Phage tail tube protein FII; afdb-uniprot50 \| AF-A0A4R3EGB1-F1-MODEL\_V4 \| 1.0 \| 5.994e-11 \| 469 \| 0.205 \| 180 \| 126 \| 7 \| 1 \| 167 \| 1 \| 176 \| Uncharacterized protein \| Uncharacterized protein; afdb-uniprot50 \| AF-A0A0Q7A9A4-F1-MODEL\_V4 \| 1.0 \| 3.643e-10 \| 468 \| 0.173 \| 179 \| 132 \| 8 \| 1 \| 167 \| 1 \| 175 \| Uncharacterized protein \| Uncharacterized protein; afdb-uniprot50 \| AF-A0A448J7T4-F1-MODEL\_V4 \| 1.0 \| 5.794e-10 \| 459 \| 0.222 \| 144 \| 104 \| 5 \| 1 \| 140 \| 4 \| 143 \| Tail protein \| Tail protein; afdb-uniprot50 \| AF-A0A268TJI4-F1-MODEL\_V4 \| 1.0 \| 8.312e-10 \| 447 \| 0.161 \| 149 \| 117 \| 5 \| 1 \| 146 \| 5 \| 148 \| Uncharacterized protein \| Uncharacterized protein; afdb-uniprot50 \| AF-A0A1A9G1E2-F1-MODEL\_V4 \| 1.0 \| 1.057e-10 \| 446 \| 0.173 \| 167 \| 127 \| 6 \| 1 \| 158 \| 1 \| 165 \| Phage tail protein \| Phage tail protein; afdb-uniprot50 \| AF-A0A1C6BND7-F1-MODEL\_V4 \| 1.0 \| 2.815e-10 \| 442 \| 0.192 \| 171 \| 126 \| 7 \| 2 \| 165 \| 5 \| 170 \| Phage major tail tube protein \| Phage major tail tube protein; afdb-uniprot50 \| AF-A0A7Y9D078-F1-MODEL\_V4 \| 1.0 \| 3.769e-11 \| 440 \| 0.215 \| 172 \| 124 \| 6 \| 2 \| 165 \| 3 \| 171 \| P2 family phage contractile tail tube protein \| P2 family phage contractile tail tube protein; afdb-uniprot50 \| AF-A0A0Q2UUR3-F1-MODEL\_V4 \| 1.0 \| 1.516e-10 \| 439 \| 0.161 \| 167 \| 130 \| 5 \| 1 \| 158 \| 1 \| 166 \| Uncharacterized protein \| Uncharacterized protein; afdb-uniprot50 \| AF-A0A413PG50-F1-MODEL\_V4 \| 1.0 \| 2.29e-10 \| 438 \| 0.173 \| 173 \| 127 \| 8 \| 1 \| 165 \| 3 \| 167 \| Protoporphyrinogen oxidase \| Protoporphyrinogen oxidase; afdb-uniprot50 \| AF-E3H9B4-F1-MODEL\_V4 \| 1.0 \| 5.503e-10 \| 438 \| 0.16 \| 168 \| 128 \| 5 \| 1 \| 159 \| 3 \| 166 \| Major tail tube protein \| Major tail tube protein; afdb-uniprot50 \| AF-A0A2G6EZB0-F1-MODEL\_V4 \| 1.0 \| 1.597e-10 \| 436 \| 0.224 \| 178 \| 110 \| 9 \| 3 \| 166 \| 1 \| 164 \| Phage tail protein \| Phage tail protein; afdb-uniprot50 \| AF-A0A2K1DEW6-F1-MODEL\_V4 \| 1.0 \| 7.498e-10 \| 434 \| 0.212 \| 160 \| 115 \| 7 \| 1 \| 157 \| 1 \| 152 \| Uncharacterized protein \| Uncharacterized protein; afdb-uniprot50 \| AF-J0PYV7-F1-MODEL\_V4 \| 1.0 \| 3.344e-09 \| 431 \| 0.341 \| 126 \| 80 \| 1 \| 43 \| 168 \| 1 \| 123 \| Phage major tail tube protein \| Phage major tail tube protein; afdb-uniprot50 \| AF-G1WGC6-F1-MODEL\_V4 \| 1.0 \| 1.172e-10 \| 431 \| 0.192 \| 171 \| 125 \| 8 \| 1 \| 165 \| 4 \| 167 \| Uncharacterized protein \| Uncharacterized protein; afdb-uniprot50 \| AF-A0A345J3J5-F1-MODEL\_V4 \| 1.0 \| 2.455e-09 \| 430 \| 0.208 \| 168 \| 124 \| 7 \| 1 \| 166 \| 1 \| 161 \| Phage tail tube protein \| Phage tail tube protein; afdb-uniprot50 \| AF-A0A0C1J3Q2-F1-MODEL\_V4 \| 1.0 \| 1.322e-09 \| 428 \| 0.156 \| 160 \| 125 \| 5 \| 8 \| 158 \| 8 \| 166 \| Uncharacterized protein \| Uncharacterized protein; afdb-uniprot50 \| AF-A0A379X2H6-F1-MODEL\_V4 \| 1.0 \| 2.103e-09 \| 426 \| 0.242 \| 140 \| 101 \| 3 \| 33 \| 168 \| 1 \| 139 \| Tail protein \| Tail protein; afdb-uniprot50 \| AF-A0A377HSE9-F1-MODEL\_V4 \| 1.0 \| 3.904e-09 \| 425 \| 0.392 \| 125 \| 71 \| 3 \| 47 \| 168 \| 1 \| 123 \| Phage major tail tube protein \| Phage major tail tube protein; afdb-uniprot50 \| AF-A0A370DZ67-F1-MODEL\_V4 \| 1.0 \| 4.11e-09 \| 420 \| 0.46 \| 102 \| 55 \| 0 \| 67 \| 168 \| 7 \| 108 \| Uncharacterized protein \| Uncharacterized protein; afdb-uniprot50 \| AF-A0A6L8LZW8-F1-MODEL\_V4 \| 1.0 \| 1.466e-09 \| 420 \| 0.183 \| 158 \| 119 \| 8 \| 3 \| 157 \| 2 \| 152 \| Uncharacterized protein \| Uncharacterized protein; afdb-uniprot50 \| AF-A8TC06-F1-MODEL\_V4 \| 1.0 \| 4.328e-09 \| 418 \| 0.195 \| 138 \| 110 \| 1 \| 30 \| 167 \| 1 \| 137 \| Uncharacterized protein \| Uncharacterized protein; afdb-uniprot50 \| AF-A0A7Y3Z3M6-F1-MODEL\_V4 \| 1.0 \| 1.543e-09 \| 413 \| 0.178 \| 157 \| 123 \| 5 \| 1 \| 157 \| 1 \| 151 \| Uncharacterized protein \| Uncharacterized protein; afdb-uniprot50 \| AF-A0A2N8HQH0-F1-MODEL\_V4 \| 1.0 \| 4.039e-10 \| 413 \| 0.213 \| 173 \| 120 \| 10 \| 1 \| 165 \| 4 \| 168 \| Protoporphyrinogen oxidase \| Protoporphyrinogen oxidase; afdb-uniprot50 \| AF-A0A5S9NBV0-F1-MODEL\_V4 \| 1.0 \| 1.322e-09 \| 409 \| 0.193 \| 160 \| 119 \| 5 \| 8 \| 158 \| 10 \| 168 \| Uncharacterized protein \| Uncharacterized protein; afdb-uniprot50 \| AF-A0A285NFT0-F1-MODEL\_V4 \| 1.0 \| 9.215e-10 \| 405 \| 0.197 \| 162 \| 118 \| 7 \| 1 \| 153 \| 1 \| 159 \| Phage tail tube protein FII \| Phage tail tube protein FII; afdb-uniprot50 \| AF-A0A378URC9-F1-MODEL\_V4 \| 1.0 \| 3.708e-09 \| 404 \| 0.151 \| 139 \| 115 \| 2 \| 33 \| 168 \| 3 \| 141 \| Phage-like protein \| Phage-like protein; afdb-uniprot50 \| AF-A0A845Y1I5-F1-MODEL\_V4 \| 1.0 \| 3.836e-10 \| 404 \| 0.174 \| 172 \| 124 \| 7 \| 6 \| 160 \| 4 \| 174 \| Uncharacterized protein \| Uncharacterized protein; afdb-uniprot50 \| AF-A0A2W7RI33-F1-MODEL\_V4 \| 1.0 \| 2.14e-08 \| 401 \| 0.297 \| 121 \| 80 \| 2 \| 49 \| 168 \| 1 \| 117 \| Uncharacterized protein \| Uncharacterized protein; afdb-uniprot50 \| AF-F8KPL6-F1-MODEL\_V4 \| 1.0 \| 1.625e-09 \| 399 \| 0.181 \| 143 \| 112 \| 3 \| 27 \| 166 \| 2 \| 142 \| Uncharacterized protein \| Uncharacterized protein; afdb-uniprot50 \| AF-A0A7G9WG90-F1-MODEL\_V4 \| 1.0 \| 7.498e-10 \| 397 \| 0.142 \| 176 \| 128 \| 6 \| 1 \| 165 \| 6 \| 169 \| Phage major tail tube protein \| Phage major tail tube protein; afdb-uniprot50 \| AF-A0A2J8HSA6-F1-MODEL\_V4 \| 1.0 \| 2.214e-09 \| 393 \| 0.215 \| 158 \| 116 \| 5 \| 1 \| 157 \| 1 \| 151 \| Uncharacterized protein \| Uncharacterized protein; afdb-uniprot50 \| AF-A0A735IT95-F1-MODEL\_V4 \| 1.0 \| 2.63e-08 \| 390 \| 0.305 \| 118 \| 78 \| 4 \| 54 \| 168 \| 1 \| 117 \| Phage major tail tube protein \| Phage major tail tube protein; afdb-uniprot50 \| AF-A0A348SUU8-F1-MODEL\_V4 \| 1.0 \| 2.77e-08 \| 390 \| 0.272 \| 125 \| 87 \| 4 \| 47 \| 168 \| 1 \| 124 \| Phage tail protein \| Phage tail protein; afdb-uniprot50 \| AF-A0A358PQ20-F1-MODEL\_V4 \| 1.0 \| 1.193e-09 \| 390 \| 0.189 \| 174 \| 124 \| 8 \| 1 \| 165 \| 5 \| 170 \| Protoporphyrinogen oxidase \| Protoporphyrinogen oxidase; afdb-uniprot50 \| AF-A0A6L9ABJ3-F1-MODEL\_V4 \| 1.0 \| 1.214e-08 \| 387 \| 0.209 \| 129 \| 97 \| 3 \| 7 \| 131 \| 9 \| 136 \| Phage major tail tube protein \| Phage major tail tube protein; afdb-uniprot50 \| AF-A0A3A9EJ85-F1-MODEL\_V4 \| 1.0 \| 4.11e-09 \| 387 \| 0.175 \| 148 \| 116 \| 3 \| 25 \| 167 \| 1 \| 147 \| Phage tail protein \| Phage tail protein; afdb-uniprot50 \| AF-A0A1C6JGP5-F1-MODEL\_V4 \| 1.0 \| 1.322e-09 \| 387 \| 0.136 \| 169 \| 135 \| 7 \| 1 \| 164 \| 6 \| 168 \| Phage tail tube protein FII \| Phage tail tube protein FII; afdb-uniprot50 \| AF-A0A378PU11-F1-MODEL\_V4 \| 1.0 \| 6.884e-09 \| 378 \| 0.34 \| 132 \| 81 \| 4 \| 38 \| 167 \| 1 \| 128 \| Phage major tail tube protein \| Phage major tail tube protein; afdb-uniprot50 \| AF-A0A3E2BFZ8-F1-MODEL\_V4 \| 1.0 \| 4.405e-08 \| 376 \| 0.255 \| 133 \| 94 \| 4 \| 1 \| 129 \| 1 \| 132 \| Phage tail protein \| Phage tail protein; afdb-uniprot50 \| AF-A0A1Z4PN25-F1-MODEL\_V4 \| 1.0 \| 3.176e-09 \| 375 \| 0.174 \| 189 \| 123 \| 5 \| 1 \| 168 \| 1 \| 177 \| Uncharacterized protein \| Uncharacterized protein; afdb-uniprot50 \| AF-A0A541BHJ3-F1-MODEL\_V4 \| 1.0 \| 7.248e-09 \| 367 \| 0.2 \| 165 \| 115 \| 9 \| 1 \| 158 \| 1 \| 155 \| Uncharacterized protein \| Uncharacterized protein; afdb-uniprot50 \| AF-A0A2X2BQ70-F1-MODEL\_V4 \| 1.0 \| 3.584e-08 \| 365 \| 0.296 \| 118 \| 76 \| 4 \| 23 \| 135 \| 4 \| 119 \| Major tail tube protein \| Major tail tube protein; afdb-uniprot50 \| AF-A0A419A4I5-F1-MODEL\_V4 \| 1.0 \| 2.373e-08 \| 361 \| 0.403 \| 129 \| 60 \| 2 \| 8 \| 136 \| 1 \| 112 \| Phage tail protein \| Phage tail protein; afdb-uniprot50 \| AF-A0A5M7P9N9-F1-MODEL\_V4 \| 1.0 \| 9.546e-08 \| 354 \| 0.341 \| 129 \| 75 \| 4 \| 1 \| 124 \| 1 \| 124 \| Phage tail protein \| Phage tail protein; afdb-uniprot50 \| AF-A0A5Y7FW73-F1-MODEL\_V4 \| 1.0 \| 3.29e-07 \| 346 \| 0.227 \| 132 \| 92 \| 6 \| 1 \| 128 \| 4 \| 129 \| Phage tail protein \| Phage tail protein; afdb-uniprot50 \| AF-A0A317H401-F1-MODEL\_V4 \| 1.0 \| 7.767e-08 \| 341 \| 0.236 \| 127 \| 87 \| 4 \| 47 \| 167 \| 5 \| 127 \| Uncharacterized protein \| Uncharacterized protein; afdb-uniprot50 \| AF-A0A645CLD3-F1-MODEL\_V4 \| 1.0 \| 1.37e-07 \| 334 \| 0.284 \| 116 \| 75 \| 3 \| 57 \| 167 \| 2 \| 114 \| Uncharacterized protein \| Uncharacterized protein; afdb-uniprot50 \| AF-A0A059IUF5-F1-MODEL\_V4 \| 1.0 \| 4.97e-07 \| 331 \| 0.585 \| 82 \| 34 \| 0 \| 6 \| 87 \| 5 \| 86 \| Contractile tail tube protein \| Contractile tail tube protein; afdb-uniprot50 \| AF-A0A607GZE9-F1-MODEL\_V4 \| 1.0 \| 1.965e-07 \| 331 \| 0.242 \| 136 \| 91 \| 5 \| 39 \| 168 \| 1 \| 130 \| Phage tail protein \| Phage tail protein; afdb-uniprot50 \| AF-A0A6L5WKW9-F1-MODEL\_V4 \| 1.0 \| 1.37e-07 \| 331 \| 0.223 \| 130 \| 98 \| 3 \| 38 \| 164 \| 1 \| 130 \| Phage tail protein \| Phage tail protein; afdb-uniprot50 \| AF-A0A450SXT5-F1-MODEL\_V4 \| 1.0 \| 1.866e-07 \| 330 \| 0.505 \| 93 \| 46 \| 0 \| 43 \| 135 \| 1 \| 93 \| Uncharacterized protein \| Uncharacterized protein; afdb-uniprot50 \| AF-A0A376DET1-F1-MODEL\_V4 \| 1.0 \| 1.023e-06 \| 328 \| 0.327 \| 122 \| 74 \| 4 \| 1 \| 118 \| 1 \| 118 \| Major tail sheath protein FII from prophage \| Major tail sheath protein FII from prophage; afdb-uniprot50 \| AF-A0A8A8MDP2-F1-MODEL\_V4 \| 1.0 \| 1.023e-06 \| 321 \| 0.211 \| 137 \| 101 \| 5 \| 36 \| 168 \| 2 \| 135 \| Uncharacterized protein \| Uncharacterized protein; afdb-uniprot50 \| AF-A0A3A6RI32-F1-MODEL\_V4 \| 1.0 \| 1.627e-06 \| 318 \| 0.297 \| 121 \| 81 \| 3 \| 1 \| 119 \| 1 \| 119 \| Phage tail protein \| Phage tail protein; afdb-uniprot50 \| AF-A0A6S6NIH2-F1-MODEL\_V4 \| 1.0 \| 6.654e-08 \| 318 \| 0.159 \| 138 \| 108 \| 4 \| 29 \| 158 \| 2 \| 139 \| Uncharacterized protein \| Uncharacterized protein; afdb-uniprot50 \| AF-A0A5C9AFX2-F1-MODEL\_V4 \| 1.0 \| 8.764e-07 \| 316 \| 0.33 \| 106 \| 66 \| 3 \| 66 \| 168 \| 1 \| 104 \| Phage tail protein \| Phage tail protein; afdb-uniprot50 \| AF-A0A1C6YVF4-F1-MODEL\_V4 \| 1.0 \| 2.178e-07 \| 314 \| 0.321 \| 115 \| 75 \| 2 \| 53 \| 165 \| 1 \| 114 \| Uncharacterized protein \| Uncharacterized protein; afdb-uniprot50 \| AF-A0A379ZTI5-F1-MODEL\_V4 \| 1.0 \| 1.545e-06 \| 311 \| 0.322 \| 124 \| 77 \| 4 \| 1 \| 119 \| 1 \| 122 \| Phage major tail tube protein FII \| Phage major tail tube protein FII; afdb-uniprot50 \| AF-A0A369RL94-F1-MODEL\_V4 \| 1.0 \| 6.319e-08 \| 309 \| 0.367 \| 106 \| 62 \| 2 \| 64 \| 168 \| 46 \| 147 \| Phage major tail tube protein \| Phage major tail tube protein; afdb-uniprot50 \| AF-A0A4Q6CYM3-F1-MODEL\_V4 \| 1.0 \| 1.194e-06 \| 302 \| 0.208 \| 115 \| 88 \| 2 \| 55 \| 168 \| 2 \| 114 \| Uncharacterized protein \| Uncharacterized protein; afdb-uniprot50 \| AF-A0A7I6S727-F1-MODEL\_V4 \| 1.0 \| 6.217e-06 \| 301 \| 0.333 \| 117 \| 75 \| 2 \| 1 \| 114 \| 1 \| 117 \| Uncharacterized protein \| Uncharacterized protein; afdb-uniprot50 \| AF-A0A078LLW6-F1-MODEL\_V4 \| 1.0 \| 1.899e-06 \| 288 \| 0.382 \| 123 \| 65 \| 6 \| 1 \| 118 \| 1 \| 117 \| Phage P2 FII-like protein \| Phage P2 FII-like protein; afdb-uniprot50 \| AF-A0A5Y6SC09-F1-MODEL\_V4 \| 1.0 \| 4.334e-06 \| 287 \| 0.24 \| 100 \| 73 \| 2 \| 72 \| 168 \| 2 \| 101 \| Phage tail protein \| Phage tail protein; afdb-uniprot50 \| AF-A0A4Z0KQR6-F1-MODEL\_V4 \| 1.0 \| 1.419e-05 \| 284 \| 0.375 \| 72 \| 45 \| 0 \| 97 \| 168 \| 4 \| 75 \| Phage tail protein \| Phage tail protein; afdb-uniprot50 \| AF-A0A828B488-F1-MODEL\_V4 \| 1.0 \| 8.045e-06 \| 284 \| 0.313 \| 102 \| 66 \| 3 \| 1 \| 100 \| 1 \| 100 \| Phage tail protein \| Phage tail protein; afdb-uniprot50 \| AF-A0A2T4JJM7-F1-MODEL\_V4 \| 1.0 \| 1.096e-05 \| 282 \| 0.309 \| 84 \| 57 \| 1 \| 86 \| 168 \| 2 \| 85 \| Phage major tail tube protein \| Phage major tail tube protein; afdb-uniprot50 \| AF-A0A485CDJ1-F1-MODEL\_V4 \| 1.0 \| 9.227e-07 \| 281 \| 0.271 \| 114 \| 72 \| 6 \| 59 \| 167 \| 2 \| 109 \| Phage major tail tube protein \| Phage major tail tube protein; afdb-uniprot50 \| AF-A0A244C1R5-F1-MODEL\_V4 \| 1.0 \| 2.725e-06 \| 279 \| 0.438 \| 89 \| 50 \| 0 \| 3 \| 91 \| 2 \| 90 \| Phage tail protein \| Phage tail protein; afdb-uniprot50 \| AF-A0A7I9KEL5-F1-MODEL\_V4 \| 1.0 \| 1.836e-05 \| 272 \| 0.24 \| 108 \| 77 \| 3 \| 64 \| 168 \| 3 \| 108 \| Phage tail protein \| Phage tail protein; afdb-uniprot50 \| AF-A0A3P6KKF7-F1-MODEL\_V4 \| 1.0 \| 3.909e-06 \| 269 \| 0.311 \| 106 \| 72 \| 1 \| 64 \| 168 \| 6 \| 111 \| Major tail tube protein \| Major tail tube protein; afdb-uniprot50 \| AF-A0A0F9Z9G8-F1-MODEL\_V4 \| 1.0 \| 7.641e-06 \| 268 \| 0.241 \| 112 \| 79 \| 4 \| 7 \| 114 \| 9 \| 118 \| Prophage MuMc02, major tail tube protein \| Prophage MuMc02, major tail tube protein; afdb-uniprot50 \| AF-A0A4Q0U6A3-F1-MODEL\_V4 \| 1.0 \| 1.804e-06 \| 268 \| 0.19 \| 131 \| 102 \| 3 \| 38 \| 165 \| 2 \| 131 \| Uncharacterized protein \| Uncharacterized protein; afdb-uniprot50 \| AF-A4P152-F1-MODEL\_V4 \| 1.0 \| 1.494e-05 \| 262 \| 0.402 \| 92 \| 54 \| 1 \| 1 \| 92 \| 1 \| 91 \| Uncharacterized protein \| Uncharacterized protein; afdb-uniprot50 \| AF-A0A829E329-F1-MODEL\_V4 \| 1.0 \| 1.096e-05 \| 262 \| 0.181 \| 116 \| 90 \| 3 \| 33 \| 144 \| 1 \| 115 \| Phage major tail tube protein \| Phage major tail tube protein; afdb-uniprot50 \| AF-A0A7Z7RJ53-F1-MODEL\_V4 \| 1.0 \| 4.644e-05 \| 258 \| 0.292 \| 82 \| 57 \| 1 \| 87 \| 167 \| 2 \| 83 \| Phage major tail tube protein \| Phage major tail tube protein; afdb-uniprot50 \| AF-A0A2G6CDE2-F1-MODEL\_V4 \| 1.0 \| 2.256e-05 \| 253 \| 0.241 \| 112 \| 78 \| 3 \| 43 \| 153 \| 1 \| 106 \| Uncharacterized protein \| Uncharacterized protein; afdb-uniprot50 \| AF-A0A7U9RYY6-F1-MODEL\_V4 \| 1.0 \| 7.257e-06 \| 252 \| 0.181 \| 138 \| 102 \| 7 \| 34 \| 165 \| 1 \| 133 \| Uncharacterized protein \| Uncharacterized protein; afdb-uniprot50 \| AF-A0A607GXK8-F1-MODEL\_V4 \| 1.0 \| 3.978e-05 \| 251 \| 0.25 \| 112 \| 77 \| 5 \| 59 \| 168 \| 3 \| 109 \| Phage tail protein \| Phage tail protein; afdb-uniprot50 \| AF-A0A7W4KR61-F1-MODEL\_V4 \| 1.0 \| 5.708e-05 \| 247 \| 0.344 \| 93 \| 58 \| 2 \| 1 \| 91 \| 1 \| 92 \| Phage major tail tube protein \| Phage major tail tube protein; afdb-uniprot50 \| AF-A0A645DSK8-F1-MODEL\_V4 \| 1.0 \| 2.035e-05 \| 245 \| 0.276 \| 94 \| 64 \| 2 \| 76 \| 165 \| 3 \| 96 \| Uncharacterized protein \| Uncharacterized protein; afdb-uniprot50 \| AF-A0A826K908-F1-MODEL\_V4 \| 1.0 \| 0.0001116 \| 237 \| 0.333 \| 96 \| 61 \| 2 \| 1 \| 93 \| 1 \| 96 \| Phage tail protein \| Phage tail protein; afdb-uniprot50 \| AF-A0A2W6PC71-F1-MODEL\_V4 \| 1.0 \| 8.188e-05 \| 236 \| 0.264 \| 106 \| 73 \| 4 \| 64 \| 167 \| 4 \| 106 \| Phage tail protein \| Phage tail protein; afdb-uniprot50 \| AF-A0A0D1VL43-F1-MODEL\_V4 \| 1.0 \| 4.804e-06 \| 235 \| 0.151 \| 145 \| 107 \| 8 \| 1 \| 135 \| 4 \| 142 \| Phage tail tube protein \| Phage tail tube protein; afdb-uniprot50 \| AF-A0A7K0ZTX3-F1-MODEL\_V4 \| 1.0 \| 5.326e-06 \| 234 \| 0.147 \| 142 \| 103 \| 6 \| 1 \| 132 \| 1 \| 134 \| Uncharacterized protein \| Uncharacterized protein; afdb-uniprot50 \| AF-A0A163TDC7-F1-MODEL\_V4 \| 1.0 \| 0.0001685 \| 231 \| 0.238 \| 84 \| 62 \| 2 \| 87 \| 168 \| 2 \| 85 \| Bacteriophage protein \| Bacteriophage protein; afdb-uniprot50 \| AF-A0A0T6ZCB2-F1-MODEL\_V4 \| 1.0 \| 1.419e-05 \| 229 \| 0.2 \| 130 \| 95 \| 6 \| 40 \| 165 \| 1 \| 125 \| Uncharacterized protein \| Uncharacterized protein; afdb-uniprot50 \| AF-A0A1S7MQV2-F1-MODEL\_V4 \| 1.0 \| 7.015e-05 \| 228 \| 0.195 \| 123 \| 84 \| 4 \| 61 \| 168 \| 2 \| 124 \| Uncharacterized protein \| Uncharacterized protein; afdb-uniprot50 \| AF-A0A450Z316-F1-MODEL\_V4 \| 1.0 \| 0.0002972 \| 225 \| 0.467 \| 62 \| 33 \| 0 \| 107 \| 168 \| 2 \| 63 \| Uncharacterized protein \| Uncharacterized protein; afdb-uniprot50 \| AF-A0A799JWC1-F1-MODEL\_V4 \| 1.0 \| 0.0001175 \| 225 \| 0.174 \| 109 \| 85 \| 3 \| 64 \| 168 \| 2 \| 109 \| Phage major tail tube protein \| Phage major tail tube protein; afdb-uniprot50 \| AF-A0A645IWR8-F1-MODEL\_V4 \| 1.0 \| 5.421e-05 \| 221 \| 0.277 \| 90 \| 62 \| 2 \| 79 \| 165 \| 7 \| 96 \| Uncharacterized protein \| Uncharacterized protein; afdb-uniprot50 \| AF-A0A850F557-F1-MODEL\_V4 \| 1.0 \| 7.015e-05 \| 221 \| 0.229 \| 96 \| 71 \| 2 \| 75 \| 168 \| 7 \| 101 \| Phage major tail tube protein \| Phage major tail tube protein; afdb-uniprot50 \| AF-A0A0H3PB23-F1-MODEL\_V4 \| 1.0 \| 0.0001371 \| 219 \| 0.185 \| 124 \| 95 \| 4 \| 1 \| 119 \| 1 \| 123 \| Uncharacterized protein \| Uncharacterized protein; afdb-uniprot50 \| AF-Q2NU86-F1-MODEL\_V4 \| 1.0 \| 0.0004049 \| 216 \| 0.237 \| 101 \| 73 \| 3 \| 1 \| 98 \| 1 \| 100 \| Hypothetical phage protein \| Hypothetical phage protein; afdb-uniprot50 \| AF-A0A376ZNY1-F1-MODEL\_V4 \| 1.0 \| 0.000152 \| 213 \| 0.273 \| 95 \| 65 \| 3 \| 74 \| 167 \| 3 \| 94 \| Major tail tube protein FII \| Major tail tube protein FII; afdb-uniprot50 \| AF-A0A447QPL3-F1-MODEL\_V4 \| 1.0 \| 0.0008775 \| 212 \| 0.378 \| 66 \| 41 \| 0 \| 103 \| 168 \| 1 \| 66 \| Phage major tail tube protein \| Phage major tail tube protein; afdb-uniprot50 \| AF-A0A7C5N8L8-F1-MODEL\_V4 \| 1.0 \| 0.0001371 \| 212 \| 0.154 \| 142 \| 103 \| 7 \| 3 \| 135 \| 2 \| 135 \| Uncharacterized protein \| Uncharacterized protein; afdb-uniprot50 \| AF-A0A2G2DI71-F1-MODEL\_V4 \| 1.0 \| 0.0003652 \| 211 \| 0.229 \| 87 \| 65 \| 2 \| 84 \| 168 \| 4 \| 90 \| Uncharacterized protein \| Uncharacterized protein; afdb-uniprot50 \| AF-A0A2U2AQA0-F1-MODEL\_V4 \| 1.0 \| 1.419e-05 \| 210 \| 0.109 \| 155 \| 108 \| 9 \| 1 \| 129 \| 1 \| 151 \| Uncharacterized protein \| Uncharacterized protein; afdb-uniprot50 \| AF-A0A6L3XLT6-F1-MODEL\_V4 \| 1.0 \| 0.0002296 \| 209 \| 0.277 \| 90 \| 64 \| 1 \| 1 \| 90 \| 1 \| 89 \| Phage tail protein \| Phage tail protein; afdb-uniprot50 \| AF-A0A376JCW1-F1-MODEL\_V4 \| 1.0 \| 0.0002296 \| 209 \| 0.205 \| 117 \| 88 \| 3 \| 7 \| 119 \| 9 \| 124 \| Phage major tail tube protein \| Phage major tail tube protein; afdb-uniprot50 \| AF-A0A519ED62-F1-MODEL\_V4 \| 1.0 \| 4.89e-05 \| 209 \| 0.139 \| 151 \| 112 \| 7 \| 1 \| 136 \| 1 \| 148 \| Uncharacterized protein \| Uncharacterized protein; afdb-uniprot50 \| AF-A0A7W4VT46-F1-MODEL\_V4 \| 1.0 \| 3.408e-05 \| 204 \| 0.174 \| 143 \| 98 \| 9 \| 1 \| 132 \| 1 \| 134 \| Uncharacterized protein \| Uncharacterized protein; afdb-uniprot50 \| AF-A0A1E5G2B2-F1-MODEL\_V4 \| 1.0 \| 6.328e-05 \| 204 \| 0.109 \| 146 \| 109 \| 10 \| 1 \| 135 \| 8 \| 143 \| Uncharacterized protein \| Uncharacterized protein; afdb-uniprot50 \| AF-A0A376ZEC3-F1-MODEL\_V4 \| 1.0 \| 0.0007915 \| 203 \| 0.297 \| 74 \| 51 \| 1 \| 95 \| 167 \| 3 \| 76 \| Major tail tube protein FII \| Major tail tube protein FII; afdb-uniprot50 \| AF-A0A538F034-F1-MODEL\_V4 \| 1.0 \| 3.978e-05 \| 201 \| 0.124 \| 137 \| 105 \| 9 \| 9 \| 135 \| 13 \| 144 \| Phage tail protein \| Phage tail protein; afdb-uniprot50 \| AF-E8KG78-F1-MODEL\_V4 \| 1.0 \| 0.000152 \| 199 \| 0.188 \| 106 \| 82 \| 3 \| 1 \| 104 \| 4 \| 107 \| Putative phage major tail tube protein \| Putative phage major tail tube protein; afdb-uniprot50 \| AF-A0A1I4SH27-F1-MODEL\_V4 \| 1.0 \| 2.256e-05 \| 199 \| 0.164 \| 140 \| 88 \| 8 \| 9 \| 129 \| 7 \| 136 \| Uncharacterized protein \| Uncharacterized protein; afdb-uniprot50 \| AF-A0A561K6F5-F1-MODEL\_V4 \| 1.0 \| 0.0008334 \| 198 \| 0.362 \| 80 \| 50 \| 1 \| 1 \| 79 \| 1 \| 80 \| P2 family phage contractile tail tube protein \| P2 family phage contractile tail tube protein; afdb-uniprot50 \| AF-A0A4D9X464-F1-MODEL\_V4 \| 1.0 \| 0.0002296 \| 198 \| 0.244 \| 90 \| 65 \| 2 \| 7 \| 94 \| 9 \| 97 \| Phage major tail tube protein \| Phage major tail tube protein; afdb-uniprot50 \| AF-A0A1F2QQE5-F1-MODEL\_V4 \| 1.0 \| 0.0001868 \| 193 \| 0.16 \| 143 \| 100 \| 10 \| 1 \| 135 \| 1 \| 131 \| Uncharacterized protein \| Uncharacterized protein; afdb-uniprot50 \| AF-A0A448QV03-F1-MODEL\_V4 \| 1.0 \| 0.0008775 \| 192 \| 0.195 \| 97 \| 73 \| 3 \| 49 \| 141 \| 1 \| 96 \| Phage major tail tube protein \| Phage major tail tube protein; afdb-uniprot50 \| AF-A0A0F9F7Z1-F1-MODEL\_V4 \| 1.0 \| 0.0001371 \| 192 \| 0.136 \| 147 \| 102 \| 9 \| 1 \| 136 \| 1 \| 133 \| Uncharacterized protein \| Uncharacterized protein; afdb-uniprot50 \| AF-A0A3C1F3W3-F1-MODEL\_V4 \| 1.0 \| 0.0002296 \| 192 \| 0.165 \| 145 \| 99 \| 9 \| 1 \| 136 \| 1 \| 132 \| Uncharacterized protein \| Uncharacterized protein; afdb-uniprot50 \| AF-A0A2T5U148-F1-MODEL\_V4 \| 1.0 \| 0.003025 \| 191 \| 0.323 \| 65 \| 44 \| 0 \| 103 \| 167 \| 1 \| 65 \| P2 family phage contractile tail tube protein \| P2 family phage contractile tail tube protein; afdb-uniprot50 \| AF-A0A2W7QPU3-F1-MODEL\_V4 \| 1.0 \| 0.002002 \| 191 \| 0.384 \| 78 \| 48 \| 0 \| 1 \| 78 \| 1 \| 78 \| Uncharacterized protein \| Uncharacterized protein; afdb-uniprot50 \| AF-A0A4T0UJL6-F1-MODEL\_V4 \| 1.0 \| 0.0001175 \| 190 \| 0.132 \| 143 \| 103 \| 8 \| 1 \| 132 \| 1 \| 133 \| Uncharacterized protein \| Uncharacterized protein; afdb-uniprot50 \| AF-A0A0H3ZYX4-F1-MODEL\_V4 \| 1.0 \| 8.622e-05 \| 189 \| 0.173 \| 115 \| 90 \| 4 \| 43 \| 157 \| 1 \| 110 \| Uncharacterized protein \| Uncharacterized protein; afdb-uniprot50 \| AF-A0A1S2JZI5-F1-MODEL\_V4 \| 1.0 \| 0.0002546 \| 189 \| 0.158 \| 145 \| 104 \| 9 \| 1 \| 135 \| 1 \| 137 \| Uncharacterized protein \| Uncharacterized protein; afdb-uniprot50 \| AF-A0A176XNG2-F1-MODEL\_V4 \| 1.0 \| 9.078e-05 \| 188 \| 0.168 \| 148 \| 94 \| 8 \| 1 \| 129 \| 1 \| 138 \| Uncharacterized protein \| Uncharacterized protein; afdb-uniprot50 \| AF-A0A3A9WR79-F1-MODEL\_V4 \| 1.0 \| 0.001136 \| 186 \| 0.102 \| 137 \| 108 \| 9 \| 9 \| 137 \| 18 \| 147 \| Phage tail protein \| Phage tail protein; afdb-uniprot50 \| AF-A0A1G6LSJ2-F1-MODEL\_V4 \| 1.0 \| 0.0002071 \| 185 \| 0.131 \| 144 \| 103 \| 9 \| 1 \| 132 \| 1 \| 134 \| Uncharacterized protein \| Uncharacterized protein; afdb-uniprot50 \| AF-A0A1I0EEG8-F1-MODEL\_V4 \| 1.0 \| 0.0008334 \| 185 \| 0.103 \| 145 \| 109 \| 10 \| 1 \| 135 \| 1 \| 134 \| Uncharacterized protein \| Uncharacterized protein; afdb-uniprot50 \| AF-A0A1Z8QIQ5-F1-MODEL\_V4 \| 1.0 \| 0.001078 \| 185 \| 0.123 \| 138 \| 104 \| 9 \| 10 \| 135 \| 23 \| 155 \| Uncharacterized protein \| Uncharacterized protein; afdb-uniprot50 \| AF-R6J7P7-F1-MODEL\_V4 \| 1.0 \| 0.0002181 \| 184 \| 0.151 \| 145 \| 110 \| 8 \| 1 \| 136 \| 1 \| 141 \| Uncharacterized protein \| Uncharacterized protein; afdb-uniprot50 \| AF-A0A377U351-F1-MODEL\_V4 \| 1.0 \| 0.002461 \| 183 \| 0.347 \| 69 \| 44 \| 1 \| 82 \| 149 \| 2 \| 70 \| Phage major tail tube protein \| Phage major tail tube protein; afdb-uniprot50 \| AF-A0A527W089-F1-MODEL\_V4 \| 1.0 \| 0.003914 \| 183 \| 0.16 \| 81 \| 66 \| 2 \| 89 \| 167 \| 2 \| 82 \| Uncharacterized protein \| Uncharacterized protein; afdb-uniprot50 \| AF-A0A1U7GJ66-F1-MODEL\_V4 \| 1.0 \| 0.0002681 \| 183 \| 0.163 \| 147 \| 107 \| 7 \| 1 \| 136 \| 1 \| 142 \| Uncharacterized protein \| Uncharacterized protein; afdb-uniprot50 \| AF-A0A376W1N8-F1-MODEL\_V4 \| 1.0 \| 0.001902 \| 181 \| 0.216 \| 97 \| 70 \| 4 \| 43 \| 136 \| 1 \| 94 \| Major tail sheath protein FII from prophage \| Major tail sheath protein FII from prophage; afdb-uniprot50 \| AF-A1ZC06-F1-MODEL\_V4 \| 1.0 \| 0.0003846 \| 181 \| 0.17 \| 135 \| 98 \| 8 \| 9 \| 136 \| 18 \| 145 \| Afp1 \| Afp1; afdb-uniprot50 \| AF-A0A1S1NS96-F1-MODEL\_V4 \| 1.0 \| 6.662e-05 \| 181 \| 0.133 \| 180 \| 117 \| 11 \| 1 \| 160 \| 1 \| 161 \| Uncharacterized protein \| Uncharacterized protein; afdb-uniprot50 \| AF-A0A354NH67-F1-MODEL\_V4 \| 1.0 \| 0.0008775 \| 180 \| 0.139 \| 136 \| 103 \| 7 \| 9 \| 135 \| 8 \| 138 \| Uncharacterized protein \| Uncharacterized protein; afdb-uniprot50 \| AF-A0A3S9YSX8-F1-MODEL\_V4 \| 1.0 \| 0.0004489 \| 178 \| 0.13 \| 153 \| 107 \| 8 \| 1 \| 135 \| 1 \| 145 \| Uncharacterized protein \| Uncharacterized protein; afdb-uniprot50 \| AF-A0A416PS04-F1-MODEL\_V4 \| 1.0 \| 0.0001371 \| 178 \| 0.163 \| 141 \| 100 \| 10 \| 8 \| 139 \| 17 \| 148 \| Uncharacterized protein \| Uncharacterized protein; afdb-uniprot50 \| AF-A0A1Q7Q1G9-F1-MODEL\_V4 \| 1.0 \| 0.0002681 \| 178 \| 0.091 \| 142 \| 115 \| 9 \| 1 \| 136 \| 21 \| 154 \| Uncharacterized protein \| Uncharacterized protein; afdb-uniprot50 \| AF-A0A0F9HCI3-F1-MODEL\_V4 \| 1.0 \| 0.001629 \| 177 \| 0.122 \| 139 \| 104 \| 10 \| 10 \| 136 \| 15 \| 147 \| Uncharacterized protein \| Uncharacterized protein; afdb-uniprot50 \| AF-A0A2N2JDW8-F1-MODEL\_V4 \| 1.0 \| 0.001902 \| 177 \| 0.08 \| 136 \| 110 \| 8 \| 9 \| 135 \| 68 \| 197 \| Phage tail protein \| Phage tail protein; afdb-uniprot50 \| AF-A0A3A9EEY6-F1-MODEL\_V4 \| 1.0 \| 0.008931 \| 176 \| 0.25 \| 68 \| 51 \| 0 \| 101 \| 168 \| 6 \| 73 \| Phage tail protein \| Phage tail protein; afdb-uniprot50 \| AF-A0A261SH18-F1-MODEL\_V4 \| 1.0 \| 0.0001967 \| 176 \| 0.156 \| 147 \| 97 \| 8 \| 1 \| 129 \| 1 \| 138 \| Uncharacterized protein \| Uncharacterized protein; afdb-uniprot50 \| AF-A0A7Y0JM30-F1-MODEL\_V4 \| 1.0 \| 0.0008334 \| 175 \| 0.16 \| 143 \| 100 \| 9 \| 3 \| 135 \| 2 \| 134 \| Uncharacterized protein \| Uncharacterized protein; afdb-uniprot50 \| AF-A0A2L2BC49-F1-MODEL\_V4 \| 1.0 \| 0.0001175 \| 175 \| 0.143 \| 146 \| 93 \| 9 \| 3 \| 129 \| 5 \| 137 \| Uncharacterized protein \| Uncharacterized protein; afdb-uniprot50 \| AF-A0A3A9VWV0-F1-MODEL\_V4 \| 1.0 \| 0.001902 \| 175 \| 0.109 \| 137 \| 107 \| 9 \| 9 \| 137 \| 18 \| 147 \| Phage tail protein \| Phage tail protein; afdb-uniprot50 \| AF-A0A1S1HPU9-F1-MODEL\_V4 \| 1.0 \| 0.0007517 \| 175 \| 0.08 \| 161 \| 117 \| 8 \| 1 \| 141 \| 1 \| 150 \| Phage tail protein \| Phage tail protein; afdb-uniprot50 \| AF-A0A413W1T0-F1-MODEL\_V4 \| 1.0 \| 0.0003652 \| 174 \| 0.136 \| 146 \| 110 \| 9 \| 1 \| 135 \| 1 \| 141 \| Phage tail protein \| Phage tail protein; afdb-uniprot50 \| AF-A0A8B5RKL2-F1-MODEL\_V4 \| 1.0 \| 0.009403 \| 173 \| 0.285 \| 63 \| 45 \| 0 \| 106 \| 168 \| 1 \| 63 \| Phage tail protein \| Phage tail protein; afdb-uniprot50 \| AF-A0A5W2M6M4-F1-MODEL\_V4 \| 1.0 \| 0.007267 \| 173 \| 0.343 \| 67 \| 43 \| 1 \| 103 \| 168 \| 3 \| 69 \| Phage tail protein \| Phage tail protein; afdb-uniprot50 \| AF-A0A7Y2IRB2-F1-MODEL\_V4 \| 1.0 \| 0.0005809 \| 173 \| 0.155 \| 135 \| 94 \| 9 \| 10 \| 135 \| 9 \| 132 \| Uncharacterized protein \| Uncharacterized protein; afdb-uniprot50 \| AF-A0A6G9IB21-F1-MODEL\_V4 \| 1.0 \| 0.0005809 \| 173 \| 0.09 \| 155 \| 110 \| 10 \| 1 \| 136 \| 1 \| 143 \| Phage tail protein \| Phage tail protein; afdb-uniprot50 \| AF-A0A4S1E497-F1-MODEL\_V4 \| 1.0 \| 0.0004489 \| 173 \| 0.12 \| 149 \| 101 \| 6 \| 1 \| 129 \| 1 \| 139 \| Uncharacterized protein \| Uncharacterized protein; afdb-uniprot50 \| AF-X1I0E2-F1-MODEL\_V4 \| 1.0 \| 0.0004726 \| 172 \| 0.125 \| 144 \| 106 \| 7 \| 3 \| 135 \| 2 \| 136 \| Uncharacterized protein \| Uncharacterized protein; afdb-uniprot50 \| AF-A0A5V4ZED2-F1-MODEL\_V4 \| 1.0 \| 0.0007915 \| 171 \| 0.107 \| 139 \| 106 \| 7 \| 9 \| 137 \| 12 \| 142 \| Uncharacterized protein \| Uncharacterized protein; afdb-uniprot50 \| AF-A0A1L6M2V1-F1-MODEL\_V4 \| 1.0 \| 0.001396 \| 171 \| 0.101 \| 138 \| 107 \| 8 \| 9 \| 135 \| 22 \| 153 \| Uncharacterized protein \| Uncharacterized protein; afdb-uniprot50 \| AF-A0A327S9J4-F1-MODEL\_V4 \| 1.0 \| 0.001469 \| 170 \| 0.104 \| 144 \| 109 \| 12 \| 1 \| 135 \| 1 \| 133 \| TP901-1 family phage major tail protein \| TP901-1 family phage major tail protein; afdb-uniprot50 \| AF-A0A2E6EJY7-F1-MODEL\_V4 \| 1.0 \| 0.002728 \| 169 \| 0.071 \| 139 \| 114 \| 8 \| 9 \| 135 \| 15 \| 150 \| Phage tail protein \| Phage tail protein; afdb-uniprot50 \| AF-A0A6D1E7W5-F1-MODEL\_V4 \| 1.0 \| 0.001078 \| 168 \| 0.071 \| 154 \| 112 \| 9 \| 1 \| 136 \| 2 \| 142 \| Phage tail protein \| Phage tail protein; afdb-uniprot50 \| AF-A0A2W4S886-F1-MODEL\_V4 \| 1.0 \| 0.00222 \| 167 \| 0.094 \| 138 \| 108 \| 10 \| 9 \| 135 \| 11 \| 142 \| Phage tail protein \| Phage tail protein; afdb-uniprot50 \| AF-A0A853IAX0-F1-MODEL\_V4 \| 1.0 \| 0.0004049 \| 167 \| 0.109 \| 146 \| 111 \| 11 \| 1 \| 136 \| 17 \| 153 \| Phage tail protein \| Phage tail protein; afdb-uniprot50 \| AF-A0A0G3CNN1-F1-MODEL\_V4 \| 1.0 \| 0.0009239 \| 167 \| 0.108 \| 156 \| 107 \| 9 \| 1 \| 136 \| 1 \| 144 \| Uncharacterized protein \| Uncharacterized protein; afdb-uniprot50 \| AF-A0A4R5EZG6-F1-MODEL\_V4 \| 1.0 \| 0.0004489 \| 165 \| 0.176 \| 147 \| 95 \| 12 \| 1 \| 135 \| 1 \| 133 \| Uncharacterized protein \| Uncharacterized protein; afdb-uniprot50 \| AF-T0QQH1-F1-MODEL\_V4 \| 1.0 \| 0.001806 \| 165 \| 0.114 \| 131 \| 109 \| 5 \| 10 \| 136 \| 9 \| 136 \| Uncharacterized protein \| Uncharacterized protein; afdb-uniprot50 \| AF-A0A534PKH6-F1-MODEL\_V4 \| 1.0 \| 0.0007517 \| 165 \| 0.117 \| 145 \| 108 \| 9 \| 1 \| 135 \| 8 \| 142 \| Phage tail protein \| Phage tail protein; afdb-uniprot50 \| AF-A0A497Y9K5-F1-MODEL\_V4 \| 1.0 \| 0.003185 \| 165 \| 0.104 \| 134 \| 108 \| 8 \| 9 \| 136 \| 25 \| 152 \| Phage tail protein \| Phage tail protein; afdb-uniprot50 \| AF-A0A2E0EK74-F1-MODEL\_V4 \| 1.0 \| 0.001259 \| 165 \| 0.106 \| 159 \| 113 \| 9 \| 1 \| 135 \| 1 \| 154 \| Uncharacterized protein \| Uncharacterized protein; afdb-uniprot50 \| AF-A0A090SU12-F1-MODEL\_V4 \| 1.0 \| 0.008931 \| 162 \| 0.293 \| 75 \| 51 \| 1 \| 1 \| 75 \| 1 \| 73 \| Uncharacterized protein \| Uncharacterized protein; afdb-uniprot50 \| AF-A0A1C6AJU2-F1-MODEL\_V4 \| 1.0 \| 0.001469 \| 161 \| 0.158 \| 139 \| 107 \| 7 \| 5 \| 136 \| 11 \| 146 \| Conserved hypothetical phage tail region protein \| Conserved hypothetical phage tail region protein; afdb-uniprot50 \| AF-A0A7V6WL49-F1-MODEL\_V4 \| 1.0 \| 0.001547 \| 160 \| 0.16 \| 125 \| 88 \| 8 \| 22 \| 136 \| 30 \| 147 \| Uncharacterized protein \| Uncharacterized protein; afdb-uniprot50 \| AF-A0A419GEK8-F1-MODEL\_V4 \| 1.0 \| 0.001259 \| 160 \| 0.131 \| 183 \| 106 \| 9 \| 5 \| 136 \| 5 \| 185 \| Uncharacterized protein \| Uncharacterized protein; afdb-uniprot50 \| AF-A0A3G9IH27-F1-MODEL\_V4 \| 1.0 \| 0.001469 \| 159 \| 0.143 \| 139 \| 105 \| 7 \| 1 \| 132 \| 1 \| 132 \| Uncharacterized protein \| Uncharacterized protein; afdb-uniprot50 \| AF-A0A3R0Y9I4-F1-MODEL\_V4 \| 1.0 \| 0.002108 \| 159 \| 0.108 \| 138 \| 105 \| 7 \| 10 \| 137 \| 13 \| 142 \| Big-1 domain-containing protein \| Big-1 domain-containing protein; afdb-uniprot50 \| AF-A0A4P5Y9X1-F1-MODEL\_V4 \| 1.0 \| 0.003717 \| 158 \| 0.101 \| 138 \| 113 \| 6 \| 1 \| 135 \| 4 \| 133 \| Uncharacterized protein \| Uncharacterized protein; afdb-uniprot50 \| AF-A0A7X9N1M4-F1-MODEL\_V4 \| 1.0 \| 0.0003295 \| 157 \| 0.138 \| 173 \| 106 \| 12 \| 10 \| 167 \| 13 \| 157 \| Phage major tail protein, TP901-1 family \| Phage major tail protein, TP901-1 family; afdb-uniprot50 \| AF-A0A2E7URN5-F1-MODEL\_V4 \| 1.0 \| 0.002337 \| 156 \| 0.096 \| 155 \| 108 \| 10 \| 1 \| 135 \| 7 \| 149 \| Uncharacterized protein \| Uncharacterized protein; afdb-uniprot50 \| AF-E8X0R1-F1-MODEL\_V4 \| 1.0 \| 0.003185 \| 153 \| 0.17 \| 141 \| 102 \| 6 \| 9 \| 135 \| 16 \| 155 \| Uncharacterized protein \| Uncharacterized protein; afdb-uniprot50 \| AF-A0A6P2BDC7-F1-MODEL\_V4 \| 1.0 \| 0.002873 \| 152 \| 0.136 \| 132 \| 104 \| 6 \| 9 \| 135 \| 10 \| 136 \| Uncharacterized protein \| Uncharacterized protein; afdb-uniprot50 \| AF-A0A6M1N9D7-F1-MODEL\_V4 \| 1.0 \| 0.00353 \| 152 \| 0.095 \| 146 \| 108 \| 9 \| 10 \| 136 \| 9 \| 149 \| Uncharacterized protein \| Uncharacterized protein; afdb-uniprot50 \| AF-A0A1Y4GV23-F1-MODEL\_V4 \| 1.0 \| 0.0009239 \| 152 \| 0.121 \| 140 \| 101 \| 8 \| 1 \| 135 \| 1 \| 123 \| Uncharacterized protein \| Uncharacterized protein; afdb-uniprot50 \| AF-A0A0T2MKH8-F1-MODEL\_V4 \| 1.0 \| 0.00222 \| 151 \| 0.151 \| 132 \| 98 \| 9 \| 10 \| 136 \| 9 \| 131 \| Uncharacterized protein \| Uncharacterized protein; afdb-uniprot50 \| AF-A0A1V6HRI2-F1-MODEL\_V4 \| 1.0 \| 0.003025 \| 150 \| 0.165 \| 139 \| 93 \| 10 \| 3 \| 129 \| 2 \| 129 \| Uncharacterized protein \| Uncharacterized protein; afdb-uniprot50 \| AF-A0A521U4D5-F1-MODEL\_V4 \| 1.0 \| 0.002873 \| 149 \| 0.173 \| 150 \| 107 \| 10 \| 1 \| 141 \| 1 \| 142 \| Uncharacterized protein \| Uncharacterized protein; afdb-uniprot50 \| AF-A0A0F5Q2G1-F1-MODEL\_V4 \| 1.0 \| 0.0004726 \| 149 \| 0.147 \| 149 \| 102 \| 11 \| 1 \| 141 \| 31 \| 162 \| Uncharacterized protein \| Uncharacterized protein; afdb-uniprot50 \| AF-A0A2A7U7Q2-F1-MODEL\_V4 \| 1.0 \| 0.008482 \| 149 \| 0.063 \| 141 \| 110 \| 8 \| 5 \| 129 \| 8 \| 142 \| Uncharacterized protein \| Uncharacterized protein; afdb-uniprot50 \| AF-A0A840ENA2-F1-MODEL\_V4 \| 1.0 \| 0.003717 \| 147 \| 0.108 \| 138 \| 100 \| 10 \| 10 \| 140 \| 11 \| 132 \| Uncharacterized protein \| Uncharacterized protein; afdb-uniprot50 \| AF-A0A6B9HSX7-F1-MODEL\_V4 \| 1.0 \| 0.001902 \| 147 \| 0.131 \| 145 \| 94 \| 13 \| 9 \| 134 \| 12 \| 143 \| Phage major tail protein, TP901-1 family \| Phage major tail protein, TP901-1 family; afdb-uniprot50 \| AF-A0A2D7ZXH8-F1-MODEL\_V4 \| 1.0 \| 0.001715 \| 146 \| 0.14 \| 142 \| 100 \| 10 \| 1 \| 135 \| 1 \| 127 \| Uncharacterized protein \| Uncharacterized protein; afdb-uniprot50 \| AF-F6GGQ2-F1-MODEL\_V4 \| 1.0 \| 0.001547 \| 146 \| 0.069 \| 144 \| 112 \| 12 \| 7 \| 147 \| 4 \| 128 \| Uncharacterized protein \| Uncharacterized protein; afdb-uniprot50 \| AF-A0A7W7CHS3-F1-MODEL\_V4 \| 1.0 \| 0.0005517 \| 146 \| 0.13 \| 138 \| 103 \| 9 \| 1 \| 129 \| 9 \| 138 \| Phage tail-like protein \| Phage tail-like protein; afdb-uniprot50 \| AF-A0A1E3GXP7-F1-MODEL\_V4 \| 1.0 \| 0.008056 \| 146 \| 0.082 \| 146 \| 113 \| 5 \| 9 \| 136 \| 7 \| 149 \| Uncharacterized protein \| Uncharacterized protein; afdb-uniprot50 \| AF-D9WWW1-F1-MODEL\_V4 \| 1.0 \| 0.003353 \| 146 \| 0.125 \| 152 \| 103 \| 9 \| 10 \| 140 \| 117 \| 259 \| Uncharacterized protein \| Uncharacterized protein; afdb-uniprot50 \| AF-A0A7L5UII7-F1-MODEL\_V4 \| 1.0 \| 0.002002 \| 145 \| 0.104 \| 143 \| 106 \| 13 \| 5 \| 136 \| 12 \| 143 \| Phage tail protein \| Phage tail protein; afdb-uniprot50 \| AF-A0A376W7L3-F1-MODEL\_V4 \| 1.0 \| 0.00222 \| 144 \| 0.095 \| 157 \| 105 \| 10 \| 1 \| 136 \| 2 \| 142 \| Major tail subunit encoded within prophage CP-933V \| Major tail subunit encoded within prophage CP-933V; afdb-uniprot50 \| AF-A0A1F9ZG88-F1-MODEL\_V4 \| 1.0 \| 0.002461 \| 144 \| 0.096 \| 145 \| 111 \| 9 \| 1 \| 135 \| 15 \| 149 \| Uncharacterized protein \| Uncharacterized protein; afdb-uniprot50 \| AF-A0A1H7T315-F1-MODEL\_V4 \| 1.0 \| 0.003185 \| 144 \| 0.14 \| 128 \| 95 \| 9 \| 10 \| 135 \| 5 \| 119 \| Uncharacterized protein \| Uncharacterized protein; afdb-uniprot50 \| AF-A0A0K5ZHP5-F1-MODEL\_V4 \| 1.0 \| 0.004339 \| 144 \| 0.071 \| 154 \| 111 \| 11 \| 1 \| 136 \| 20 \| 159 \| Phage major tail subunit \| Phage major tail subunit; afdb-uniprot50 \| AF-A0A521X7C1-F1-MODEL\_V4 \| 1.0 \| 0.0002546 \| 143 \| 0.148 \| 141 \| 99 \| 9 \| 6 \| 140 \| 3 \| 128 \| Uncharacterized protein \| Uncharacterized protein; afdb-uniprot50 \| AF-A0A0S7YR27-F1-MODEL\_V4 \| 1.0 \| 0.001902 \| 143 \| 0.126 \| 142 \| 104 \| 7 \| 1 \| 129 \| 1 \| 135 \| Uncharacterized protein \| Uncharacterized protein; afdb-uniprot50 \| AF-A0A3M2CPG9-F1-MODEL\_V4 \| 1.0 \| 0.003717 \| 143 \| 0.162 \| 129 \| 95 \| 9 \| 8 \| 129 \| 1 \| 123 \| Uncharacterized protein \| Uncharacterized protein; afdb-uniprot50 \| AF-A0A1G2VKT4-F1-MODEL\_V4 \| 1.0 \| 0.005912 \| 143 \| 0.102 \| 136 \| 112 \| 8 \| 1 \| 129 \| 633 \| 765 \| Kazal-like domain-containing protein \| Kazal-like domain-containing protein; afdb-uniprot50 \| AF-A0A2N1DEF5-F1-MODEL\_V4 \| 1.0 \| 0.002108 \| 142 \| 0.104 \| 144 \| 111 \| 7 \| 1 \| 136 \| 9 \| 142 \| Phage tail protein \| Phage tail protein; afdb-uniprot50 \| AF-A0A396BX50-F1-MODEL\_V4 \| 1.0 \| 0.0007517 \| 141 \| 0.118 \| 135 \| 98 \| 11 \| 10 \| 140 \| 5 \| 122 \| Type VI secretion system needle protein Hcp \| Type VI secretion system needle protein Hcp; afdb-uniprot50 \| AF-A0A7Y1UT33-F1-MODEL\_V4 \| 1.0 \| 0.008482 \| 141 \| 0.105 \| 133 \| 110 \| 7 \| 9 \| 135 \| 18 \| 147 \| Phage tail protein \| Phage tail protein; afdb-uniprot50 \| AF-E6M101-F1-MODEL\_V4 \| 1.0 \| 0.002108 \| 141 \| 0.098 \| 153 \| 99 \| 8 \| 10 \| 137 \| 10 \| 148 \| Uncharacterized protein \| Uncharacterized protein; afdb-uniprot50 \| AF-A0A1G7J1E5-F1-MODEL\_V4 \| 1.0 \| 0.0008775 \| 141 \| 0.093 \| 149 \| 107 \| 11 \| 1 \| 144 \| 1 \| 126 \| Uncharacterized protein \| Uncharacterized protein; afdb-uniprot50 \| AF-A0A4U7JH93-F1-MODEL\_V4 \| 1.0 \| 0.004121 \| 140 \| 0.088 \| 135 \| 110 \| 7 \| 1 \| 129 \| 6 \| 133 \| Phage tail protein \| Phage tail protein; afdb-uniprot50 \| AF-A0A6J4RD76-F1-MODEL\_V4 \| 1.0 \| 0.002591 \| 140 \| 0.167 \| 131 \| 95 \| 8 \| 5 \| 129 \| 11 \| 133 \| Uncharacterized protein \| Uncharacterized protein; afdb-uniprot50 \| AF-A0A852QIZ0-F1-MODEL\_V4 \| 1.0 \| 0.005065 \| 139 \| 0.08 \| 124 \| 105 \| 4 \| 18 \| 135 \| 49 \| 169 \| Uncharacterized protein \| Uncharacterized protein; afdb-uniprot50 \| AF-A0A1E8CW82-F1-MODEL\_V4 \| 1.0 \| 0.0003652 \| 139 \| 0.178 \| 140 \| 93 \| 11 \| 6 \| 136 \| 1 \| 127 \| ATPase\_AAA\_core domain-containing protein \| ATPase\_AAA\_core domain-containing protein; afdb-uniprot50 \| AF-A0A2G6JA31-F1-MODEL\_V4 \| 1.0 \| 0.006901 \| 138 \| 0.076 \| 144 \| 115 \| 9 \| 1 \| 135 \| 10 \| 144 \| Phage tail protein \| Phage tail protein; afdb-uniprot50 \| AF-A0A5E6QXM5-F1-MODEL\_V4 \| 1.0 \| 0.007267 \| 137 \| 0.204 \| 83 \| 64 \| 1 \| 88 \| 168 \| 2 \| 84 \| Uncharacterized protein \| Uncharacterized protein; afdb-uniprot50 \| AF-A0A831QR42-F1-MODEL\_V4 \| 1.0 \| 0.002591 \| 137 \| 0.148 \| 141 \| 90 \| 13 \| 1 \| 134 \| 1 \| 118 \| Uncharacterized protein \| Uncharacterized protein; afdb-uniprot50 \| AF-A0A1N6FWD3-F1-MODEL\_V4 \| 1.0 \| 0.004121 \| 137 \| 0.138 \| 130 \| 103 \| 7 \| 10 \| 135 \| 9 \| 133 \| Conserved hypothetical phage tail region protein \| Conserved hypothetical phage tail region protein; afdb-uniprot50 \| AF-A0A5C5ZW62-F1-MODEL\_V4 \| 1.0 \| 0.004569 \| 137 \| 0.128 \| 132 \| 95 \| 8 \| 10 \| 130 \| 9 \| 131 \| Uncharacterized protein \| Uncharacterized protein; afdb-uniprot50 \| AF-A0A366ZZU0-F1-MODEL\_V4 \| 1.0 \| 0.007267 \| 137 \| 0.131 \| 122 \| 94 \| 5 \| 18 \| 135 \| 42 \| 155 \| Uncharacterized protein \| Uncharacterized protein; afdb-uniprot50 \| AF-A0A1H7UBE6-F1-MODEL\_V4 \| 1.0 \| 0.003914 \| 136 \| 0.104 \| 143 \| 104 \| 11 \| 10 \| 144 \| 30 \| 156 \| Uncharacterized protein \| Uncharacterized protein; afdb-uniprot50 \| AF-A0A6H1ZDK9-F1-MODEL\_V4 \| 1.0 \| 0.003914 \| 135 \| 0.14 \| 142 \| 102 \| 10 \| 3 \| 136 \| 5 \| 134 \| Uncharacterized protein \| Uncharacterized protein; afdb-uniprot50 \| AF-A0A1Y4DW26-F1-MODEL\_V4 \| 1.0 \| 0.004121 \| 135 \| 0.139 \| 151 \| 97 \| 11 \| 6 \| 138 \| 2 \| 137 \| Uncharacterized protein \| Uncharacterized protein; afdb-uniprot50 \| AF-A0A1G6Z760-F1-MODEL\_V4 \| 1.0 \| 0.001259 \| 135 \| 0.166 \| 144 \| 90 \| 16 \| 10 \| 146 \| 5 \| 125 \| Uncharacterized protein \| Uncharacterized protein; afdb-uniprot50 \| AF-A0A3N0E5J3-F1-MODEL\_V4 \| 1.0 \| 0.003185 \| 134 \| 0.116 \| 137 \| 100 \| 11 \| 7 \| 140 \| 4 \| 122 \| Uncharacterized protein \| Uncharacterized protein; afdb-uniprot50 \| AF-A0A2W7NBR6-F1-MODEL\_V4 \| 1.0 \| 0.004121 \| 132 \| 0.098 \| 183 \| 122 \| 12 \| 1 \| 168 \| 3 \| 157 \| TP901-1 family phage major tail protein \| TP901-1 family phage major tail protein; afdb-uniprot50 \| AF-A0A553DU56-F1-MODEL\_V4 \| 1.0 \| 0.003717 \| 131 \| 0.125 \| 144 \| 100 \| 14 \| 1 \| 140 \| 1 \| 122 \| Uncharacterized protein \| Uncharacterized protein; afdb-uniprot50 \| AF-A0A5C1AH35-F1-MODEL\_V4 \| 1.0 \| 0.003914 \| 130 \| 0.137 \| 124 \| 89 \| 6 \| 22 \| 135 \| 22 \| 137 \| Uncharacterized protein \| Uncharacterized protein; afdb-uniprot50 \| AF-A0A4P6U525-F1-MODEL\_V4 \| 1.0 \| 0.007267 \| 130 \| 0.141 \| 141 \| 93 \| 7 \| 10 \| 129 \| 15 \| 148 \| Uncharacterized protein \| Uncharacterized protein; afdb-uniprot50 \| AF-A0A6B8KDY4-F1-MODEL\_V4 \| 1.0 \| 0.008056 \| 129 \| 0.081 \| 159 \| 111 \| 11 \| 1 \| 135 \| 3 \| 150 \| Uncharacterized protein \| Uncharacterized protein; afdb-uniprot50 \| AF-A0A0F9A0R1-F1-MODEL\_V4 \| 1.0 \| 0.001469 \| 128 \| 0.153 \| 150 \| 96 \| 13 \| 1 \| 136 \| 1 \| 133 \| Uncharacterized protein \| Uncharacterized protein; afdb-uniprot50 \| AF-A0A6N7RZ20-F1-MODEL\_V4 \| 1.0 \| 0.0007517 \| 128 \| 0.121 \| 173 \| 127 \| 10 \| 1 \| 160 \| 1 \| 161 \| Uncharacterized protein \| Uncharacterized protein; afdb-uniprot50 \| AF-A0A4P7X061-F1-MODEL\_V4 \| 1.0 \| 0.004339 \| 128 \| 0.079 \| 139 \| 105 \| 9 \| 10 \| 143 \| 5 \| 125 \| Uncharacterized protein \| Uncharacterized protein; afdb-uniprot50 \| AF-A0A329N2I3-F1-MODEL\_V4 \| 1.0 \| 0.005333 \| 126 \| 0.138 \| 144 \| 98 \| 14 \| 1 \| 140 \| 1 \| 122 \| Uncharacterized protein \| Uncharacterized protein; afdb-uniprot50 \| AF-A0A4P6YRT8-F1-MODEL\_V4 \| 1.0 \| 0.006901 \| 125 \| 0.094 \| 137 \| 101 \| 8 \| 9 \| 135 \| 11 \| 134 \| Phage tail protein \| Phage tail protein; afdb-uniprot50 \| AF-A0A662BGP6-F1-MODEL\_V4 \| 1.0 \| 0.007267 \| 124 \| 0.117 \| 145 \| 108 \| 9 \| 1 \| 135 \| 42 \| 176 \| Uncharacterized protein \| Uncharacterized protein; afdb-uniprot50 \| AF-A0A7H8PN01-F1-MODEL\_V4 \| 1.0 \| 0.008931 \| 122 \| 0.14 \| 135 \| 94 \| 11 \| 10 \| 136 \| 5 \| 125 \| Phage tail protein \| Phage tail protein; afdb-uniprot50 \| AF-A0A7Y2EDK0-F1-MODEL\_V4 \| 1.0 \| 0.005333 \| 122 \| 0.127 \| 149 \| 94 \| 12 \| 1 \| 135 \| 11 \| 137 \| Phage tail protein \| Phage tail protein; afdb-uniprot50 \| AF-A0A1F1DKW7-F1-MODEL\_V4 \| 1.0 \| 0.007651 \| 122 \| 0.096 \| 156 \| 100 \| 10 \| 1 \| 129 \| 7 \| 148 \| Uncharacterized protein \| Uncharacterized protein; afdb-uniprot50 \| AF-A0A540QCB7-F1-MODEL\_V4 \| 1.0 \| 0.002728 \| 121 \| 0.122 \| 196 \| 121 \| 11 \| 1 \| 168 \| 4 \| 176 \| Uncharacterized protein \| Uncharacterized protein; afdb-uniprot50 \| AF-A0A1K1N5G7-F1-MODEL\_V4 \| 1.0 \| 0.005333 \| 120 \| 0.109 \| 137 \| 101 \| 10 \| 7 \| 140 \| 4 \| 122 \| Uncharacterized protein \| Uncharacterized protein; afdb-uniprot50 \| AF-A0A1G7HEH5-F1-MODEL\_V4 \| 1.0 \| 0.006555 \| 119 \| 0.123 \| 138 \| 98 \| 12 \| 7 \| 140 \| 4 \| 122 \| Uncharacterized protein \| Uncharacterized protein; afdb-uniprot50 \| AF-A0A2D6AJM6-F1-MODEL\_V4 \| 1.0 \| 0.007267 \| 119 \| 0.125 \| 152 \| 109 \| 12 \| 1 \| 140 \| 1 \| 140 \| Uncharacterized protein \| Uncharacterized protein; afdb-uniprot50 \| AF-A0A3A9VUY5-F1-MODEL\_V4 \| 1.0 \| 0.00222 \| 119 \| 0.157 \| 140 \| 95 \| 10 \| 5 \| 140 \| 30 \| 150 \| Uncharacterized protein \| Uncharacterized protein; afdb-uniprot50 \| AF-A0A3A6P734-F1-MODEL\_V4 \| 1.0 \| 0.008056 \| 118 \| 0.108 \| 138 \| 98 \| 11 \| 4 \| 135 \| 7 \| 125 \| Uncharacterized protein \| Uncharacterized protein; afdb-uniprot50 \| AF-A0A1J0LW85-F1-MODEL\_V4 \| 1.0 \| 0.008056 \| 117 \| 0.167 \| 137 \| 95 \| 11 \| 4 \| 132 \| 9 \| 134 \| Uncharacterized protein \| Uncharacterized protein; afdb-uniprot50 \| AF-A0A2D6X4M8-F1-MODEL\_V4 \| 1.0 \| 0.008931 \| 113 \| 0.138 \| 152 \| 94 \| 12 \| 1 \| 129 \| 4 \| 141 \| Uncharacterized protein \| Uncharacterized protein; afdb-uniprot50 \| AF-A0A1I0J0M0-F1-MODEL\_V4 \| 1.0 \| 0.006555 \| 113 \| 0.124 \| 153 \| 104 \| 11 \| 6 \| 135 \| 9 \| 154 \| Phage tail tube protein \| Phage tail tube protein; afdb-uniprot50 \| AF-A0A0K1EBJ0-F1-MODEL\_V4 \| 1.0 \| 0.00481 \| 113 \| 0.14 \| 142 \| 87 \| 7 \| 11 \| 136 \| 13 \| 135 \| Uncharacterized protein \| Uncharacterized protein; afdb-uniprot50 \| AF-A0A7U9NPC4-F1-MODEL\_V4 \| 1.0 \| 0.008931 \| 113 \| 0.167 \| 143 \| 91 \| 9 \| 7 \| 133 \| 4 \| 134 \| Uncharacterized protein \| Uncharacterized protein; afdb-uniprot50 \| AF-A0A0F9EL08-F1-MODEL\_V4 \| 1.0 \| 0.006225 \| 111 \| 0.138 \| 152 \| 110 \| 11 \| 1 \| 136 \| 79 \| 225 \| Uncharacterized protein \| Uncharacterized protein; afdb-uniprot50 \| AF-I0K9T2-F1-MODEL\_V4 \| 1.0 \| 0.002728 \| 110 \| 0.187 \| 144 \| 83 \| 12 \| 1 \| 132 \| 1 \| 122 \| PaaR repeat-containing protein \| PaaR repeat-containing protein; afdb-uniprot50 \| AF-A0A2W2GU04-F1-MODEL\_V4 \| 1.0 \| 0.009901 \| 109 \| 0.134 \| 156 \| 95 \| 9 \| 1 \| 129 \| 6 \| 148 \| Uncharacterized protein \| Uncharacterized protein; afdb-uniprot50 \| AF-A0A7L4Q6N2-F1-MODEL\_V4 \| 1.0 \| 0.008056 \| 108 \| 0.106 \| 132 \| 90 \| 8 \| 11 \| 129 \| 5 \| 121 \| Uncharacterized protein \| Uncharacterized protein; afdb-uniprot50 \| AF-A0A021VU20-F1-MODEL\_V4 \| 1.0 \| 0.009403 \| 108 \| 0.131 \| 152 \| 93 \| 10 \| 10 \| 137 \| 18 \| 154 \| Uncharacterized protein \| Uncharacterized protein; afdb-uniprot50 \| AF-A0A6M3IF35-F1-MODEL\_V4 \| 1.0 \| 0.00481 \| 107 \| 0.12 \| 150 \| 106 \| 11 \| 1 \| 135 \| 1 \| 139 \| Uncharacterized protein \| Uncharacterized protein; afdb-uniprot50 \| AF-A0A518ICN5-F1-MODEL\_V4 \| 0.999 \| 0.004569 \| 99 \| 0.131 \| 145 \| 87 \| 7 \| 1 \| 129 \| 1 \| 122 \| Uncharacterized protein \| Uncharacterized protein; afdb-uniprot50 \| AF-A0A512H4S7-F1-MODEL\_V4 \| 0.998 \| 0.008931 \| 94 \| 0.129 \| 154 \| 104 \| 11 \| 4 \| 135 \| 11 \| 156 \| Uncharacterized protein \| Uncharacterized protein |
| Top keywords  (threshold 1.00e-02 (evalue)) | **tail, Phage, tube, major, FII, Putative, prophage, contractile, P2, Bacteriophage** |
| Output files | ../../similar\_structures/19\_FANPEZAQ\_CDS\_0019\_afdb-proteome\_foldseek.tsv ../../similar\_structures/19\_FANPEZAQ\_CDS\_0019\_afdb-uniprot50\_foldseek.tsv ../../similar\_structures/19\_FANPEZAQ\_CDS\_0019\_merged.svg ../../similar\_structures/19\_FANPEZAQ\_CDS\_0019\_pdb\_foldseek.tsv |

  
  
  

Return to summary | Go to previous | Go to next

  


---

**Sequence/structure alignments coloring**  
Each object in the alignment figures is colored according to its E-value following this color coding:

1e-100
10
