## Supplementary material for "Completing the BASEL phage collection to unlock hidden diversity for systematic exploration of phage-host interactions": Data S2: 20.html

FANPEZAQ\_CDS\_0020


Return to summary | Go to previous | Go to next

|  |  |
| --- | --- |
| FANPEZAQ\_CDS\_0020  Page creation date: 02 Sep 2024, 12:00  Project folder: n/a  Input sequences file: Escherichia\_virus\_HeidiAbel.gb | tail phage assembly or e chaperone 14\_like gp41 flumu mu\_like prophage putative fragment bacteriophage hypothetical related phage\_related domain\_containing tube small phage\_like phage\_tac\_7 containing hypothethical major whole genome shotgun sequence arsr transcriptional regulator |

### Sequence information

|  |  |
| --- | --- |
| Name | FANPEZAQ\_CDS\_0020  20\_FANPEZAQ\_CDS\_0020 (pipeline id) |
| Imported annotations | Escherichia\_virus\_HeidiAbel Bas97 |
| Protein sequence | MAKLPDYLKFNHETGHCDITLSRPLKIAGVDVPVLRMREPTVADQEIASDMGGSDAAREI QTFANLCEQAPDDIRRLPLKDYKRLQAAYVNFID |
| Number of residues | 94 |
| Molecular weight (Da) | 10588.98 |
| Output files | ../../query\_sequences/20\_FANPEZAQ\_CDS\_0020.fasta |

### Putative domain architecture and protein family

#### Search results (HHblits)^1^

|  |  |
| --- | --- |
| Domain family databases searched | Pfam, Ncbi-cd, Cath, Phrogs |
| Results, scheme(s)  (Top layers only; threshold 1.00e-03 (evalue)) | xml version="1.0" encoding="utf-8" standalone="no"?       2024-09-02T21:08:16.522205 image/svg+xml   Matplotlib v3.7.2, https://matplotlib.org/ |
| Results, table  (E-value ≤ 1.00e-03 (evalue)) | db \| id \| prob \| evalue \| pvalue \| score \| cols \| query \| query\_len \| template \| template\_len \| name \| description; pfam \| PF10109 \| 99.5 \| 1.1e-19 \| 2.2e-23 \| 102.9 \| 77 \| (18, 94) \| 94 \| (1, 84) \| 84 \| Phage\_TAC\_7 \| Phage tail assembly chaperone proteins, E, or 41 or 14; phrogs \| 103 \| 99.7 \| 3.3e-23 \| 4.1e-27 \| 128.6 \| 84 \| (11, 94) \| 94 \| (10, 97) \| 108 \| tail protein \| tail protein; Category: tail; NC\_003278\_p26; phrogs \| 166 \| 99.6 \| 4.9e-21 \| 6.4e-25 \| 116.5 \| 79 \| (16, 94) \| 94 \| (2, 89) \| 98 \| tail protein \| tail protein; Category: tail; p131196 VI\_08606; phrogs \| 25559 \| 99.1 \| 1.3e-14 \| 1.5e-18 \| 89.6 \| 92 \| (1, 93) \| 94 \| (19, 121) \| 131 \| NA \| NA; Category: unknown function; p249034 VI\_04203; phrogs \| 3140 \| 99.0 \| 3.8e-14 \| 4.6e-18 \| 84.7 \| 77 \| (17, 93) \| 94 \| (2, 90) \| 92 \| tail assembly chaperone \| tail assembly chaperone; Category: tail; NC\_024369\_p21; phrogs \| 4648 \| 98.0 \| 2.2e-09 \| 2.6e-13 \| 67.4 \| 81 \| (13, 93) \| 94 \| (53, 139) \| 139 \| NA \| NA; Category: unknown function; KY421186\_p67; phrogs \| 17026 \| 97.9 \| 3.4e-09 \| 3.9e-13 \| 68.0 \| 77 \| (12, 94) \| 94 \| (94, 170) \| 180 \| NA \| NA; Category: unknown function; p161962 VI\_09828; phrogs \| 7608 \| 97.5 \| 5e-08 \| 5.7e-12 \| 57.4 \| 72 \| (17, 91) \| 94 \| (2, 81) \| 100 \| NA \| NA; Category: unknown function; p416168 VI\_03036; phrogs \| 9738 \| 96.4 \| 1e-05 \| 1.1e-09 \| 50.0 \| 82 \| (13, 94) \| 94 \| (6, 131) \| 155 \| NA \| NA; Category: unknown function; p338877 VI\_04185; phrogs \| 30387 \| 96.3 \| 1.3e-05 \| 1.5e-09 \| 49.1 \| 74 \| (7, 84) \| 94 \| (63, 136) \| 151 \| NA \| NA; Category: unknown function; p193301 VI\_04290; phrogs \| 24302 \| 96.1 \| 2.6e-05 \| 3e-09 \| 45.6 \| 78 \| (16, 93) \| 94 \| (3, 94) \| 106 \| NA \| NA; Category: unknown function; NC\_025427\_p15; phrogs \| 30394 \| 95.2 \| 0.00024 \| 2.7e-08 \| 36.2 \| 30 \| (64, 93) \| 94 \| (2, 31) \| 56 \| NA \| NA; Category: unknown function; p376042 VI\_09042 |
| Top keywords  (threshold 1.00e-03 (evalue)) | **tail, assembly, chaperone, or, Phage, proteins, E, NC\_003278\_p26, p131196, VI\_08606** |
| Output files | ../../domain\_architecture/20\_FANPEZAQ\_CDS\_0020\_cath.hhr ../../domain\_architecture/20\_FANPEZAQ\_CDS\_0020\_merged.svg ../../domain\_architecture/20\_FANPEZAQ\_CDS\_0020\_ncbi-cd.hhr ../../domain\_architecture/20\_FANPEZAQ\_CDS\_0020\_pfam.hhr ../../domain\_architecture/20\_FANPEZAQ\_CDS\_0020\_phrogs.hhr |

### Identical protein sequences/structures

#### Search results

|  |  |
| --- | --- |
| Protein sequence databases searched | Pdb, Swissprot, Refseq |
| Identical proteins found | -- |
| Top keywords | -- |
| Output files | -- |

### Similar protein sequences/structures

#### Sequence similarity search results (HHblits)^1^

|  |  |
| --- | --- |
| Sequence databases searched | Uniclust, Pdb70 |
| Results, scheme(s)  (Top layers only, threshold 1.00e-03 (evalue)) | xml version="1.0" encoding="utf-8" standalone="no"?       2024-09-02T21:08:37.930161 image/svg+xml   Matplotlib v3.7.2, https://matplotlib.org/ |
| Results, table(s)  (threshold 1.00e-03 (evalue)) | db \| id \| prob \| evalue \| pvalue \| score \| cols \| query \| query\_len \| template \| template\_len \| name \| description; uniclust \| UniRef100\_A0A023Q1E1 \| 99.9 \| 1.9e-26 \| 4.1e-32 \| 146.9 \| 91 \| (2, 94) \| 94 \| (16, 106) \| 137 \| Phage tail assembly protein \| Phage tail assembly protein; uniclust \| UniRef100\_A0A0K2IH21 \| 99.8 \| 8.1e-24 \| 1.7e-29 \| 133.4 \| 92 \| (2, 94) \| 94 \| (16, 107) \| 133 \| Phage tail assembly protein \| Phage tail assembly protein; uniclust \| UniRef100\_A0A069PKK6 \| 99.8 \| 1.3e-23 \| 2.6e-29 \| 131.8 \| 82 \| (13, 94) \| 94 \| (20, 102) \| 129 \| Phage tail assembly protein \| Phage tail assembly protein; uniclust \| UniRef100\_A0A024HBU0 \| 99.8 \| 6e-23 \| 1.2e-28 \| 128.1 \| 90 \| (2, 93) \| 94 \| (14, 103) \| 122 \| Phage tail assembly protein \| Phage tail assembly protein; uniclust \| UniRef100\_A0A060H358 \| 99.8 \| 8e-23 \| 1.6e-28 \| 126.7 \| 93 \| (1, 94) \| 94 \| (29, 121) \| 126 \| Phage tail assembly protein \| Phage tail assembly protein; uniclust \| UniRef100\_A0A3M1X2P8 \| 99.8 \| 7.9e-23 \| 1.6e-28 \| 122.4 \| 82 \| (13, 94) \| 94 \| (6, 87) \| 95 \| Phage tail assembly protein \| Phage tail assembly protein; uniclust \| UniRef100\_A0A0P8Y9Q0 \| 99.8 \| 1.2e-22 \| 2.5e-28 \| 121.6 \| 93 \| (1, 94) \| 94 \| (3, 96) \| 96 \| Phage tail assembly protein \| Phage tail assembly protein; uniclust \| UniRef100\_A0A1G7S922 \| 99.7 \| 7.5e-21 \| 1.5e-26 \| 115.8 \| 83 \| (12, 94) \| 94 \| (5, 87) \| 103 \| Phage tail assembly chaperone protein, E, or 41 or 14 \| Phage tail assembly chaperone protein, E, or 41 or 14; uniclust \| UniRef100\_A0A212KJV9 \| 99.7 \| 1.9e-20 \| 4.2e-26 \| 113.5 \| 82 \| (12, 94) \| 94 \| (2, 84) \| 96 \| Phage tail assembly protein \| Phage tail assembly protein; uniclust \| UniRef100\_A0A0C1H2X0 \| 99.7 \| 4.1e-20 \| 8.3e-26 \| 115.0 \| 79 \| (14, 93) \| 94 \| (28, 107) \| 122 \| Phage tail assembly protein \| Phage tail assembly protein; uniclust \| UniRef100\_A0A0M4L5V1 \| 99.7 \| 5.2e-20 \| 1.1e-25 \| 110.8 \| 80 \| (13, 94) \| 94 \| (7, 86) \| 99 \| Uncharacterized protein \| Uncharacterized protein; uniclust \| UniRef100\_A0A021XB90 \| 99.7 \| 1.2e-19 \| 2.6e-25 \| 115.8 \| 82 \| (12, 94) \| 94 \| (19, 100) \| 137 \| Mu-like prophage FluMu protein gp41 \| Mu-like prophage FluMu protein gp41; uniclust \| UniRef100\_A0A024E9W1 \| 99.6 \| 2.6e-19 \| 5.3e-25 \| 112.3 \| 88 \| (2, 91) \| 94 \| (18, 106) \| 125 \| Phage tail assembly protein \| Phage tail assembly protein; uniclust \| UniRef100\_A0A011LYC3 \| 99.6 \| 5.4e-19 \| 1.1e-24 \| 111.6 \| 79 \| (14, 92) \| 94 \| (18, 97) \| 129 \| Phage tail protein \| Phage tail protein; uniclust \| UniRef100\_A0A239LH38 \| 99.6 \| 7.3e-19 \| 1.5e-24 \| 105.6 \| 80 \| (15, 94) \| 94 \| (4, 84) \| 93 \| Phage tail assembly chaperone protein, E, or 41 or 14 \| Phage tail assembly chaperone protein, E, or 41 or 14; uniclust \| UniRef100\_A0A024E9U8 \| 99.6 \| 8.7e-19 \| 1.8e-24 \| 111.1 \| 82 \| (12, 93) \| 94 \| (8, 98) \| 128 \| Sigma-factor domain-containing protein \| Sigma-factor domain-containing protein; uniclust \| UniRef100\_A0A1X7MB04 \| 99.6 \| 1e-18 \| 1.9e-24 \| 100.5 \| 79 \| (16, 94) \| 94 \| (3, 81) \| 85 \| Uncharacterized protein (Fragment) \| Uncharacterized protein (Fragment); uniclust \| UniRef100\_A0A085ARU0 \| 99.6 \| 1e-18 \| 2.1e-24 \| 110.8 \| 82 \| (12, 93) \| 94 \| (17, 99) \| 136 \| Phage tail assembly protein \| Phage tail assembly protein; uniclust \| UniRef100\_A0A143DC33 \| 99.6 \| 1.1e-18 \| 2.2e-24 \| 106.7 \| 81 \| (14, 94) \| 94 \| (3, 90) \| 104 \| Phage tail protein \| Phage tail protein; uniclust \| UniRef100\_A0A023WUD5 \| 99.6 \| 1.1e-18 \| 2.3e-24 \| 112.4 \| 82 \| (13, 94) \| 94 \| (37, 126) \| 141 \| Phage tail assembly protein \| Phage tail assembly protein; uniclust \| UniRef100\_A0A0P1FH37 \| 99.6 \| 1.2e-18 \| 2.4e-24 \| 105.8 \| 83 \| (12, 94) \| 94 \| (3, 88) \| 106 \| Phage tail protein E \| Phage tail protein E; uniclust \| UniRef100\_A0A0K1JXV3 \| 99.6 \| 6.9e-18 \| 1.4e-23 \| 105.5 \| 88 \| (3, 92) \| 94 \| (6, 95) \| 123 \| Phage tail assembly protein \| Phage tail assembly protein; uniclust \| UniRef100\_A0A0Q4II11 \| 99.6 \| 7.1e-18 \| 1.5e-23 \| 104.1 \| 87 \| (4, 94) \| 94 \| (9, 95) \| 107 \| Phage tail protein \| Phage tail protein; uniclust \| UniRef100\_A0A239C819 \| 99.6 \| 8.3e-18 \| 1.6e-23 \| 101.9 \| 80 \| (15, 94) \| 94 \| (26, 105) \| 110 \| Phage tail assembly chaperone protein, E, or 41 or 14 \| Phage tail assembly chaperone protein, E, or 41 or 14; uniclust \| UniRef100\_A0A4U0QBM5 \| 99.5 \| 1.1e-17 \| 2.3e-23 \| 99.7 \| 82 \| (12, 94) \| 94 \| (4, 85) \| 93 \| Phage tail assembly protein \| Phage tail assembly protein; uniclust \| UniRef100\_A0A0C5VFB7 \| 99.5 \| 1.6e-17 \| 3e-23 \| 100.7 \| 82 \| (13, 94) \| 94 \| (18, 99) \| 106 \| Mu-like prophage FluMu protein gp41 \| Mu-like prophage FluMu protein gp41; uniclust \| UniRef100\_A0A7L8G6A2 \| 99.5 \| 1.6e-17 \| 3.1e-23 \| 104.2 \| 91 \| (2, 93) \| 94 \| (6, 96) \| 145 \| Tape measure chaperone \| Tape measure chaperone; uniclust \| UniRef100\_A0A0R3MUP2 \| 99.5 \| 1.6e-17 \| 3.1e-23 \| 101.8 \| 79 \| (15, 93) \| 94 \| (19, 97) \| 109 \| Phage tail assembly protein \| Phage tail assembly protein; uniclust \| UniRef100\_A0A022PMF2 \| 99.5 \| 1.5e-17 \| 3.1e-23 \| 104.6 \| 81 \| (13, 93) \| 94 \| (15, 96) \| 123 \| Mu-like prophage FluMu protein gp41 \| Mu-like prophage FluMu protein gp41; uniclust \| UniRef100\_A0A212KXH5 \| 99.5 \| 3.2e-17 \| 6.4e-23 \| 101.5 \| 81 \| (13, 93) \| 94 \| (3, 83) \| 119 \| Phage tail assembly protein \| Phage tail assembly protein; uniclust \| UniRef100\_A0A0N0XKQ2 \| 99.4 \| 3.8e-16 \| 7.4e-22 \| 94.2 \| 78 \| (16, 93) \| 94 \| (3, 89) \| 99 \| Phage tail protein E \| Phage tail protein E; uniclust \| UniRef100\_A0A011UPH2 \| 99.4 \| 4.1e-16 \| 8.3e-22 \| 96.5 \| 82 \| (12, 93) \| 94 \| (9, 90) \| 111 \| Phage tail assembly protein \| Phage tail assembly protein; uniclust \| UniRef100\_A0A017HBQ7 \| 99.4 \| 4.2e-16 \| 8.6e-22 \| 97.1 \| 81 \| (13, 94) \| 94 \| (8, 88) \| 115 \| Phage tail assembly protein \| Phage tail assembly protein; uniclust \| UniRef100\_A0A149SWH3 \| 99.4 \| 5e-16 \| 9.6e-22 \| 94.5 \| 92 \| (2, 93) \| 94 \| (14, 110) \| 111 \| Phage tail assembly protein \| Phage tail assembly protein; uniclust \| UniRef100\_A0A143DG95 \| 99.4 \| 8.3e-16 \| 1.7e-21 \| 95.6 \| 79 \| (16, 94) \| 94 \| (9, 91) \| 115 \| Phage tail protein \| Phage tail protein; uniclust \| UniRef100\_A0A1I3SW87 \| 99.4 \| 8.4e-16 \| 1.7e-21 \| 92.1 \| 80 \| (14, 93) \| 94 \| (3, 84) \| 93 \| Phage tail assembly chaperone protein, E, or 41 or 14 \| Phage tail assembly chaperone protein, E, or 41 or 14; uniclust \| UniRef100\_A0A1M5PX63 \| 99.4 \| 8.5e-16 \| 1.7e-21 \| 94.2 \| 83 \| (12, 94) \| 94 \| (11, 94) \| 103 \| Phage tail assembly chaperone protein, E, or 41 or 14 \| Phage tail assembly chaperone protein, E, or 41 or 14; uniclust \| UniRef100\_A0A4S5JA13 \| 99.4 \| 1.7e-15 \| 3.4e-21 \| 98.3 \| 80 \| (13, 94) \| 94 \| (37, 116) \| 148 \| Phage tail assembly protein \| Phage tail assembly protein; uniclust \| UniRef100\_A0A0U5I607 \| 99.4 \| 2e-15 \| 3.8e-21 \| 93.8 \| 83 \| (12, 94) \| 94 \| (6, 89) \| 119 \| Uncharacterized protein \| Uncharacterized protein; uniclust \| UniRef100\_A0A2W5N4W8 \| 99.4 \| 2.8e-15 \| 5.1e-21 \| 91.5 \| 89 \| (3, 93) \| 94 \| (35, 123) \| 124 \| Phage tail assembly protein \| Phage tail assembly protein; uniclust \| UniRef100\_A0A066ZM25 \| 99.4 \| 2.9e-15 \| 5.6e-21 \| 90.6 \| 81 \| (13, 93) \| 94 \| (14, 96) \| 99 \| Phage tail assembly protein \| Phage tail assembly protein; uniclust \| UniRef100\_A0A2J8GXT3 \| 99.3 \| 3.3e-15 \| 6.6e-21 \| 92.4 \| 78 \| (16, 93) \| 94 \| (10, 100) \| 113 \| Phage tail assembly protein \| Phage tail assembly protein; uniclust \| UniRef100\_A0A0H3WRU3 \| 99.3 \| 4.7e-15 \| 9.4e-21 \| 92.7 \| 80 \| (13, 93) \| 94 \| (18, 101) \| 117 \| Phage tail assembly protein \| Phage tail assembly protein; uniclust \| UniRef100\_A0A031IT93 \| 99.3 \| 5.3e-15 \| 1.1e-20 \| 99.5 \| 90 \| (2, 93) \| 94 \| (8, 98) \| 200 \| Phage tail assembly protein \| Phage tail assembly protein; uniclust \| UniRef100\_A0A0C4YA46 \| 99.3 \| 5.4e-15 \| 1.1e-20 \| 96.3 \| 81 \| (13, 93) \| 94 \| (38, 122) \| 144 \| Tail protein \| Tail protein; uniclust \| UniRef100\_A0A2G6CS65 \| 99.3 \| 7.9e-15 \| 1.6e-20 \| 85.9 \| 71 \| (15, 94) \| 94 \| (3, 73) \| 79 \| Phage tail assembly protein \| Phage tail assembly protein; uniclust \| UniRef100\_A0A6I4YRP3 \| 99.3 \| 1.1e-14 \| 2.1e-20 \| 90.4 \| 91 \| (2, 94) \| 94 \| (8, 108) \| 115 \| Phage tail assembly protein \| Phage tail assembly protein; uniclust \| UniRef100\_A0A031IT93 \| 99.3 \| 1.1e-14 \| 2.2e-20 \| 98.0 \| 88 \| (2, 91) \| 94 \| (105, 193) \| 200 \| Phage tail assembly protein \| Phage tail assembly protein; uniclust \| UniRef100\_A0A151FEY6 \| 99.3 \| 1.1e-14 \| 2.3e-20 \| 92.0 \| 81 \| (13, 93) \| 94 \| (19, 110) \| 126 \| Phage tail assembly protein \| Phage tail assembly protein; uniclust \| UniRef100\_A0A2V3VR35 \| 99.3 \| 1.2e-14 \| 2.4e-20 \| 89.0 \| 82 \| (13, 94) \| 94 \| (9, 94) \| 103 \| Tail assembly chaperone E/41/14-like protein \| Tail assembly chaperone E/41/14-like protein; uniclust \| UniRef100\_A0A1I3TBZ1 \| 99.3 \| 1.5e-14 \| 2.9e-20 \| 92.4 \| 81 \| (13, 93) \| 94 \| (28, 125) \| 138 \| Phage tail assembly chaperone protein, E, or 41 or 14 \| Phage tail assembly chaperone protein, E, or 41 or 14; uniclust \| UniRef100\_A0A061YFP8 \| 99.3 \| 1.6e-14 \| 3.1e-20 \| 86.5 \| 77 \| (16, 92) \| 94 \| (8, 84) \| 97 \| Phage tail assembly protein \| Phage tail assembly protein; uniclust \| UniRef100\_A0A014N9C7 \| 99.3 \| 1.5e-14 \| 3.2e-20 \| 96.7 \| 78 \| (16, 93) \| 94 \| (55, 136) \| 174 \| Mu-like prophage FluMu gp41 family protein \| Mu-like prophage FluMu gp41 family protein; uniclust \| UniRef100\_A0A1G3UAH2 \| 99.3 \| 1.8e-14 \| 3.5e-20 \| 83.8 \| 66 \| (25, 94) \| 94 \| (8, 73) \| 75 \| Phage tail protein \| Phage tail protein; uniclust \| UniRef100\_A0A066TS77 \| 99.3 \| 1.9e-14 \| 3.7e-20 \| 90.6 \| 75 \| (16, 90) \| 94 \| (39, 115) \| 125 \| Phage tail assembly protein \| Phage tail assembly protein; uniclust \| UniRef100\_A0A098TF13 \| 99.3 \| 2.2e-14 \| 4.6e-20 \| 92.2 \| 83 \| (12, 94) \| 94 \| (12, 107) \| 132 \| Uncharacterized protein \| Uncharacterized protein; uniclust \| UniRef100\_A0A0A1HVW3 \| 99.3 \| 2.4e-14 \| 4.9e-20 \| 92.9 \| 79 \| (15, 93) \| 94 \| (36, 118) \| 141 \| Putative phage tail protein \| Putative phage tail protein; uniclust \| UniRef100\_A0A1G5SVM9 \| 99.3 \| 2.5e-14 \| 5e-20 \| 89.7 \| 79 \| (16, 94) \| 94 \| (25, 103) \| 117 \| Phage tail assembly chaperone protein, E, or 41 or 14 \| Phage tail assembly chaperone protein, E, or 41 or 14; uniclust \| UniRef100\_A0A259MGY1 \| 99.2 \| 4.5e-14 \| 8.9e-20 \| 86.4 \| 78 \| (16, 93) \| 94 \| (2, 82) \| 101 \| Phage tail assembly protein \| Phage tail assembly protein; uniclust \| UniRef100\_A0A009QFF4 \| 99.2 \| 4.8e-14 \| 9.8e-20 \| 92.9 \| 82 \| (13, 94) \| 94 \| (41, 126) \| 157 \| Mu-like prophage FluMu gp41 family protein \| Mu-like prophage FluMu gp41 family protein; uniclust \| UniRef100\_A0A165RMJ9 \| 99.2 \| 5e-14 \| 9.9e-20 \| 86.6 \| 80 \| (13, 93) \| 94 \| (5, 84) \| 107 \| Phage tail assembly chaperone protein, E, or 41 or 14 \| Phage tail assembly chaperone protein, E, or 41 or 14; uniclust \| UniRef100\_A0A7J0BVD3 \| 99.2 \| 5.4e-14 \| 1e-19 \| 86.6 \| 82 \| (13, 94) \| 94 \| (25, 106) \| 129 \| Phage tail assembly protein \| Phage tail assembly protein; uniclust \| UniRef100\_A0A084EX45 \| 99.2 \| 5.5e-14 \| 1.1e-19 \| 91.1 \| 82 \| (13, 94) \| 94 \| (41, 126) \| 146 \| Phage tail protein E \| Phage tail protein E; uniclust \| UniRef100\_A0A2T1B5W6 \| 99.2 \| 6e-14 \| 1.2e-19 \| 86.0 \| 81 \| (14, 94) \| 94 \| (15, 101) \| 111 \| Tail assembly chaperone E/41/14-like protein \| Tail assembly chaperone E/41/14-like protein; uniclust \| UniRef100\_A0A087NA98 \| 99.2 \| 5.5e-14 \| 1.2e-19 \| 90.3 \| 79 \| (15, 93) \| 94 \| (26, 108) \| 130 \| Phage-related tail protein \| Phage-related tail protein; uniclust \| UniRef100\_A0A1F8WN21 \| 99.2 \| 6.9e-14 \| 1.3e-19 \| 83.9 \| 79 \| (14, 94) \| 94 \| (4, 82) \| 94 \| Phage tail assembly protein \| Phage tail assembly protein; uniclust \| UniRef100\_A0A248LIV2 \| 99.2 \| 8.8e-14 \| 1.6e-19 \| 82.2 \| 77 \| (15, 91) \| 94 \| (2, 78) \| 98 \| Phage\_TAC\_7 domain containing protein \| Phage\_TAC\_7 domain containing protein; uniclust \| UniRef100\_A0A1Y2K105 \| 99.2 \| 1e-13 \| 1.9e-19 \| 79.2 \| 78 \| (16, 94) \| 94 \| (3, 80) \| 80 \| Phage tail assembly protein \| Phage tail assembly protein; uniclust \| UniRef100\_A0A1A9R7D8 \| 99.2 \| 1.1e-13 \| 2.3e-19 \| 88.2 \| 80 \| (14, 93) \| 94 \| (23, 106) \| 129 \| Phage tail protein \| Phage tail protein; uniclust \| UniRef100\_A0A370DZC3 \| 99.2 \| 1.3e-13 \| 2.5e-19 \| 82.9 \| 77 \| (16, 93) \| 94 \| (24, 100) \| 100 \| Phage tail assembly protein \| Phage tail assembly protein; uniclust \| UniRef100\_A0A5E4XFR1 \| 99.2 \| 1.3e-13 \| 2.5e-19 \| 84.9 \| 80 \| (15, 94) \| 94 \| (27, 110) \| 119 \| Phage tail assembly protein \| Phage tail assembly protein; uniclust \| UniRef100\_A0A9D2KQ00 \| 99.2 \| 1.4e-13 \| 2.6e-19 \| 81.1 \| 79 \| (15, 94) \| 94 \| (2, 81) \| 90 \| Phage tail assembly protein \| Phage tail assembly protein; uniclust \| UniRef100\_A0A071MEA2 \| 99.2 \| 1.3e-13 \| 2.6e-19 \| 91.2 \| 79 \| (15, 93) \| 94 \| (67, 149) \| 165 \| Phage tail protein \| Phage tail protein; uniclust \| UniRef100\_UPI00192BA721 \| 99.2 \| 1.5e-13 \| 2.8e-19 \| 85.0 \| 82 \| (12, 93) \| 94 \| (3, 84) \| 123 \| phage tail assembly protein \| phage tail assembly protein; uniclust \| UniRef100\_A0A1B8QCY3 \| 99.2 \| 1.4e-13 \| 2.9e-19 \| 86.5 \| 83 \| (12, 94) \| 94 \| (15, 101) \| 117 \| Phage tail protein E \| Phage tail protein E; uniclust \| UniRef100\_A0A0M7MLL6 \| 99.1 \| 2.1e-13 \| 4.2e-19 \| 86.4 \| 80 \| (15, 94) \| 94 \| (15, 101) \| 124 \| Phage tail assembly protein \| Phage tail assembly protein; uniclust \| UniRef100\_A0A074LRJ5 \| 99.1 \| 2.1e-13 \| 4.5e-19 \| 88.3 \| 80 \| (14, 93) \| 94 \| (21, 112) \| 131 \| Phage tail assembly protein \| Phage tail assembly protein; uniclust \| UniRef100\_A0A1I7ND34 \| 99.1 \| 2.7e-13 \| 5e-19 \| 84.5 \| 78 \| (16, 94) \| 94 \| (4, 81) \| 137 \| Phage tail assembly chaperone protein, E, or 41 or 14 \| Phage tail assembly chaperone protein, E, or 41 or 14; uniclust \| UniRef100\_A0A068Z8E9 \| 99.1 \| 2.8e-13 \| 5.7e-19 \| 89.0 \| 82 \| (13, 94) \| 94 \| (48, 133) \| 154 \| Phage tail assembly protein \| Phage tail assembly protein; uniclust \| UniRef100\_A0A1M6QZE2 \| 99.1 \| 3.1e-13 \| 5.7e-19 \| 83.6 \| 78 \| (15, 92) \| 94 \| (30, 108) \| 127 \| Phage tail assembly chaperone protein, E, or 41 or 14 \| Phage tail assembly chaperone protein, E, or 41 or 14; uniclust \| UniRef100\_A0A1F3A562 \| 99.1 \| 4.2e-13 \| 8e-19 \| 87.3 \| 81 \| (14, 94) \| 94 \| (8, 101) \| 165 \| Phage tail assembly protein \| Phage tail assembly protein; uniclust \| UniRef100\_A0A450ZAK9 \| 99.1 \| 4.5e-13 \| 8.3e-19 \| 80.4 \| 82 \| (13, 94) \| 94 \| (14, 99) \| 107 \| Phage tail assembly chaperone protein, E, or 41 or 14 \| Phage tail assembly chaperone protein, E, or 41 or 14; uniclust \| UniRef100\_A0A0J6K5A0 \| 99.1 \| 5.7e-13 \| 1.1e-18 \| 79.6 \| 80 \| (15, 94) \| 94 \| (12, 93) \| 93 \| Phage tail assembly protein \| Phage tail assembly protein; uniclust \| UniRef100\_A0A067ZIM0 \| 99.1 \| 6.2e-13 \| 1.3e-18 \| 85.1 \| 78 \| (16, 93) \| 94 \| (4, 93) \| 123 \| Phage tail assembly protein \| Phage tail assembly protein; uniclust \| UniRef100\_A0A0N1C7F5 \| 99.1 \| 6.1e-13 \| 1.3e-18 \| 90.0 \| 80 \| (14, 94) \| 94 \| (65, 144) \| 177 \| Phage tail assembly protein \| Phage tail assembly protein; uniclust \| UniRef100\_A0A1H7TYI9 \| 99.1 \| 6.6e-13 \| 1.3e-18 \| 83.7 \| 81 \| (13, 93) \| 94 \| (12, 94) \| 124 \| Phage tail assembly chaperone protein, E, or 41 or 14 \| Phage tail assembly chaperone protein, E, or 41 or 14; uniclust \| UniRef100\_A0A068T8B4 \| 99.1 \| 6.4e-13 \| 1.3e-18 \| 90.3 \| 82 \| (13, 94) \| 94 \| (76, 157) \| 187 \| Phage tail assembly protein \| Phage tail assembly protein; uniclust \| UniRef100\_A0A1M3BPS8 \| 99.1 \| 7.1e-13 \| 1.5e-18 \| 84.1 \| 80 \| (15, 94) \| 94 \| (20, 104) \| 121 \| Phage tail assembly protein \| Phage tail assembly protein; uniclust \| UniRef100\_A0A524RVY5 \| 99.1 \| 8.6e-13 \| 1.6e-18 \| 80.3 \| 92 \| (3, 94) \| 94 \| (9, 104) \| 116 \| Phage tail assembly protein \| Phage tail assembly protein; uniclust \| UniRef100\_A0A016XHR3 \| 99.1 \| 9.8e-13 \| 1.9e-18 \| 84.0 \| 80 \| (14, 93) \| 94 \| (32, 115) \| 132 \| Phage tail assembly protein \| Phage tail assembly protein; uniclust \| UniRef100\_A0A0F3IN91 \| 99.1 \| 1e-12 \| 2e-18 \| 79.2 \| 80 \| (14, 94) \| 94 \| (2, 83) \| 95 \| Phage tail assembly protein (Fragment) \| Phage tail assembly protein (Fragment); uniclust \| UniRef100\_UPI000B7E693B \| 99.0 \| 1.2e-12 \| 2.3e-18 \| 76.3 \| 74 \| (20, 93) \| 94 \| (2, 83) \| 86 \| phage tail assembly protein \| phage tail assembly protein; uniclust \| UniRef100\_A0A0A3ASC6 \| 99.0 \| 1.2e-12 \| 2.5e-18 \| 81.7 \| 77 \| (16, 92) \| 94 \| (19, 100) \| 111 \| Phage tail protein \| Phage tail protein; uniclust \| UniRef100\_A0A3N7CHT4 \| 99.0 \| 1.3e-12 \| 2.5e-18 \| 79.4 \| 82 \| (13, 94) \| 94 \| (5, 93) \| 101 \| Phage tail assembly protein \| Phage tail assembly protein; uniclust \| UniRef100\_A0A0G9KCB9 \| 99.0 \| 1.4e-12 \| 2.8e-18 \| 80.3 \| 78 \| (16, 94) \| 94 \| (30, 107) \| 109 \| Phage tail assembly protein \| Phage tail assembly protein; uniclust \| UniRef100\_UPI0021AA529F \| 99.0 \| 1.7e-12 \| 3.1e-18 \| 83.0 \| 79 \| (12, 90) \| 94 \| (14, 92) \| 156 \| phage tail assembly protein \| phage tail assembly protein; uniclust \| UniRef100\_UPI00082D28E9 \| 99.0 \| 1.7e-12 \| 3.2e-18 \| 76.7 \| 81 \| (12, 93) \| 94 \| (3, 83) \| 96 \| phage tail assembly protein \| phage tail assembly protein; uniclust \| UniRef100\_A0A137SPX7 \| 99.0 \| 2.6e-12 \| 5.3e-18 \| 85.1 \| 79 \| (15, 93) \| 94 \| (52, 144) \| 157 \| Phage tail assembly protein \| Phage tail assembly protein; uniclust \| UniRef100\_A0A0N0MC10 \| 99.0 \| 3.4e-12 \| 6.6e-18 \| 83.3 \| 79 \| (16, 94) \| 94 \| (47, 125) \| 152 \| Uncharacterized protein \| Uncharacterized protein; uniclust \| UniRef100\_A0A502FV57 \| 99.0 \| 4e-12 \| 7.7e-18 \| 76.9 \| 76 \| (16, 93) \| 94 \| (11, 86) \| 95 \| Phage tail assembly protein \| Phage tail assembly protein; uniclust \| UniRef100\_A0A1V0DX85 \| 99.0 \| 4e-12 \| 8.1e-18 \| 85.0 \| 82 \| (13, 94) \| 94 \| (25, 110) \| 167 \| Tapemeasure chaperone \| Tapemeasure chaperone; uniclust \| UniRef100\_A0A650EN83 \| 99.0 \| 4.3e-12 \| 8.6e-18 \| 79.0 \| 77 \| (18, 94) \| 94 \| (3, 83) \| 111 \| Phage tail assembly protein \| Phage tail assembly protein; uniclust \| UniRef100\_A0A554XFV5 \| 99.0 \| 4.9e-12 \| 9e-18 \| 74.9 \| 87 \| (1, 91) \| 94 \| (1, 88) \| 96 \| Phage tail assembly chaperone protein, E, or 41 or 14 \| Phage tail assembly chaperone protein, E, or 41 or 14; uniclust \| UniRef100\_A0A0Q4I3S2 \| 99.0 \| 4.6e-12 \| 9.4e-18 \| 84.3 \| 78 \| (16, 93) \| 94 \| (55, 136) \| 163 \| Phage tail protein \| Phage tail protein; uniclust \| UniRef100\_UPI0009D9DDE6 \| 99.0 \| 5.2e-12 \| 9.6e-18 \| 75.9 \| 89 \| (3, 92) \| 94 \| (5, 99) \| 101 \| phage tail assembly protein \| phage tail assembly protein; uniclust \| UniRef100\_A0A066RUL2 \| 98.9 \| 5.3e-12 \| 1e-17 \| 81.4 \| 80 \| (14, 93) \| 94 \| (10, 98) \| 141 \| Phage tail assembly protein \| Phage tail assembly protein; uniclust \| UniRef100\_Q65WG3 \| 98.9 \| 7.7e-12 \| 1.4e-17 \| 77.4 \| 75 \| (17, 92) \| 94 \| (46, 121) \| 127 \| Phage tail assembly protein \| Phage tail assembly protein; uniclust \| UniRef100\_A0A070A7W1 \| 98.9 \| 6.8e-12 \| 1.4e-17 \| 81.4 \| 91 \| (2, 94) \| 94 \| (22, 124) \| 133 \| Phage tail assembly protein \| Phage tail assembly protein; uniclust \| UniRef100\_A0A1A9VKI4 \| 98.9 \| 8.6e-12 \| 1.6e-17 \| 82.2 \| 81 \| (14, 94) \| 94 \| (111, 191) \| 197 \| Phage tail assembly protein \| Phage tail assembly protein; uniclust \| UniRef100\_A0A366IL82 \| 98.9 \| 8.9e-12 \| 1.7e-17 \| 78.6 \| 79 \| (15, 93) \| 94 \| (17, 99) \| 125 \| Tail assembly chaperone E/41/14-like protein \| Tail assembly chaperone E/41/14-like protein; uniclust \| UniRef100\_A0A1H9Q8X8 \| 98.9 \| 1e-11 \| 1.9e-17 \| 76.1 \| 81 \| (13, 93) \| 94 \| (14, 105) \| 111 \| Phage tail assembly chaperone protein, E, or 41 or 14 \| Phage tail assembly chaperone protein, E, or 41 or 14; uniclust \| UniRef100\_A0A1A6FM26 \| 98.9 \| 1e-11 \| 2.2e-17 \| 82.5 \| 79 \| (12, 94) \| 94 \| (33, 112) \| 152 \| Uncharacterized protein \| Uncharacterized protein; uniclust \| UniRef100\_A0A089JI40 \| 98.9 \| 1.1e-11 \| 2.3e-17 \| 80.0 \| 78 \| (16, 93) \| 94 \| (24, 116) \| 131 \| Phage tail assembly protein \| Phage tail assembly protein; uniclust \| UniRef100\_A0A2P5MUZ4 \| 98.9 \| 1.1e-11 \| 2.3e-17 \| 77.9 \| 74 \| (17, 92) \| 94 \| (3, 77) \| 110 \| Phage tail assembly protein \| Phage tail assembly protein; uniclust \| UniRef100\_UPI000C6D6F4B \| 98.9 \| 1.3e-11 \| 2.4e-17 \| 74.8 \| 78 \| (17, 94) \| 94 \| (2, 80) \| 105 \| phage tail assembly protein \| phage tail assembly protein; uniclust \| UniRef100\_UPI001F58CA0F \| 98.9 \| 1.4e-11 \| 2.6e-17 \| 76.8 \| 81 \| (13, 93) \| 94 \| (11, 93) \| 120 \| phage tail assembly protein \| phage tail assembly protein; uniclust \| UniRef100\_A0A0B4B3T3 \| 98.9 \| 1.4e-11 \| 2.6e-17 \| 73.6 \| 89 \| (2, 90) \| 94 \| (3, 94) \| 98 \| Phage tail assembly protein \| Phage tail assembly protein; uniclust \| UniRef100\_A0A5S9R620 \| 98.9 \| 1.4e-11 \| 2.7e-17 \| 73.8 \| 77 \| (18, 94) \| 94 \| (4, 82) \| 94 \| Phage tail assembly protein \| Phage tail assembly protein; uniclust \| UniRef100\_A0A1N7LRA5 \| 98.9 \| 1.6e-11 \| 3.1e-17 \| 77.5 \| 85 \| (8, 93) \| 94 \| (29, 113) \| 129 \| Phage tail assembly chaperone protein, E, or 41 or 14 \| Phage tail assembly chaperone protein, E, or 41 or 14; uniclust \| UniRef100\_A0A077KTP9 \| 98.9 \| 1.7e-11 \| 3.4e-17 \| 78.2 \| 81 \| (13, 93) \| 94 \| (28, 112) \| 126 \| Phage tail assembly protein \| Phage tail assembly protein; uniclust \| UniRef100\_A0A212KBP9 \| 98.9 \| 1.9e-11 \| 3.6e-17 \| 75.2 \| 77 \| (17, 93) \| 94 \| (5, 84) \| 112 \| Uncharacterized protein \| Uncharacterized protein; uniclust \| UniRef100\_A0A8S5PW02 \| 98.9 \| 1.9e-11 \| 3.6e-17 \| 76.4 \| 78 \| (13, 90) \| 94 \| (3, 86) \| 115 \| Tail assembly chaperone protein \| Tail assembly chaperone protein; uniclust \| UniRef100\_UPI0013ED4072 \| 98.9 \| 2.1e-11 \| 3.8e-17 \| 72.8 \| 82 \| (12, 93) \| 94 \| (13, 94) \| 99 \| phage tail assembly protein \| phage tail assembly protein; uniclust \| UniRef100\_A0A087KRL7 \| 98.8 \| 2.1e-11 \| 4e-17 \| 75.1 \| 78 \| (17, 94) \| 94 \| (2, 87) \| 107 \| Phage small tail protein E \| Phage small tail protein E; uniclust \| UniRef100\_A0A174U989 \| 98.8 \| 2.1e-11 \| 4.1e-17 \| 80.6 \| 80 \| (14, 93) \| 94 \| (56, 147) \| 157 \| Phage tail assembly protein \| Phage tail assembly protein; uniclust \| UniRef100\_A0A0K6HHB3 \| 98.8 \| 2.2e-11 \| 4.8e-17 \| 81.1 \| 83 \| (12, 94) \| 94 \| (24, 120) \| 153 \| Uncharacterized protein \| Uncharacterized protein; uniclust \| UniRef100\_A0A085G9Z8 \| 98.8 \| 3e-11 \| 6e-17 \| 76.8 \| 80 \| (15, 94) \| 94 \| (22, 105) \| 124 \| Tail protein \| Tail protein; uniclust \| UniRef100\_A0A916UDS7 \| 98.8 \| 3.4e-11 \| 6.4e-17 \| 73.2 \| 79 \| (16, 94) \| 94 \| (19, 98) \| 102 \| Uncharacterized protein \| Uncharacterized protein; uniclust \| UniRef100\_A0A6M8HND0 \| 98.8 \| 3.6e-11 \| 6.8e-17 \| 74.1 \| 82 \| (13, 94) \| 94 \| (16, 103) \| 111 \| Phage tail assembly protein \| Phage tail assembly protein; uniclust \| UniRef100\_A0A5A8F0S7 \| 98.8 \| 3.9e-11 \| 7.4e-17 \| 70.7 \| 69 \| (20, 93) \| 94 \| (6, 77) \| 84 \| Phage tail assembly protein \| Phage tail assembly protein; uniclust \| UniRef100\_S7TEH7 \| 98.8 \| 4.1e-11 \| 7.5e-17 \| 70.2 \| 76 \| (18, 93) \| 94 \| (6, 84) \| 88 \| Mu-like prophage FluMu protein gp41 \| Mu-like prophage FluMu protein gp41; uniclust \| UniRef100\_A0A6N8BZ30 \| 98.8 \| 4.3e-11 \| 8.4e-17 \| 73.2 \| 75 \| (16, 94) \| 94 \| (16, 92) \| 100 \| Phage tail assembly protein \| Phage tail assembly protein; uniclust \| UniRef100\_UPI00138AC42E \| 98.8 \| 5.4e-11 \| 9.9e-17 \| 69.1 \| 76 \| (18, 94) \| 94 \| (4, 79) \| 83 \| phage tail assembly protein \| phage tail assembly protein; uniclust \| UniRef100\_A0A2A4R5T3 \| 98.8 \| 4.8e-11 \| 1e-16 \| 80.1 \| 89 \| (3, 94) \| 94 \| (50, 151) \| 163 \| Phage tail assembly protein \| Phage tail assembly protein; uniclust \| UniRef100\_A0A431JGG1 \| 98.8 \| 5.3e-11 \| 1e-16 \| 74.6 \| 79 \| (14, 94) \| 94 \| (41, 119) \| 119 \| Phage tail assembly protein \| Phage tail assembly protein; uniclust \| UniRef100\_A0A1Y6CZP6 \| 98.8 \| 5.3e-11 \| 1e-16 \| 75.4 \| 79 \| (16, 94) \| 94 \| (4, 91) \| 121 \| Phage tail assembly chaperone protein, E, or 41 or 14 \| Phage tail assembly chaperone protein, E, or 41 or 14; uniclust \| UniRef100\_UPI0006B454B7 \| 98.8 \| 6.7e-11 \| 1.2e-16 \| 70.1 \| 80 \| (13, 93) \| 94 \| (4, 83) \| 94 \| phage tail assembly protein \| phage tail assembly protein; uniclust \| UniRef100\_A0A8X6G6V4 \| 98.8 \| 6.8e-11 \| 1.3e-16 \| 76.1 \| 80 \| (15, 94) \| 94 \| (69, 148) \| 154 \| Phage tail assembly protein \| Phage tail assembly protein; uniclust \| UniRef100\_A0A1J5B7U6 \| 98.8 \| 6.7e-11 \| 1.3e-16 \| 70.6 \| 76 \| (16, 91) \| 94 \| (3, 81) \| 87 \| Phage tail assembly protein \| Phage tail assembly protein; uniclust \| UniRef100\_A0A0J7LYN2 \| 98.8 \| 6.4e-11 \| 1.3e-16 \| 77.2 \| 80 \| (14, 94) \| 94 \| (40, 126) \| 138 \| Mu-like prophage FluMu protein gp41 \| Mu-like prophage FluMu protein gp41; uniclust \| UniRef100\_A0A3M5CVP7 \| 98.8 \| 7.3e-11 \| 1.4e-16 \| 67.2 \| 64 \| (2, 67) \| 94 \| (5, 69) \| 70 \| Uncharacterized protein \| Uncharacterized protein; uniclust \| UniRef100\_A0A0D2HN99 \| 98.7 \| 8e-11 \| 1.5e-16 \| 68.1 \| 77 \| (17, 93) \| 94 \| (3, 80) \| 81 \| Phage tail assembly protein \| Phage tail assembly protein; uniclust \| UniRef100\_A0A0W1G7Y5 \| 98.7 \| 9.6e-11 \| 1.9e-16 \| 72.1 \| 77 \| (16, 93) \| 94 \| (6, 89) \| 99 \| Phage tail assembly protein \| Phage tail assembly protein; uniclust \| UniRef100\_A0A1N6NVV8 \| 98.7 \| 9.8e-11 \| 1.9e-16 \| 77.3 \| 90 \| (2, 94) \| 94 \| (46, 148) \| 157 \| Phage tail assembly chaperone protein, E, or 41 or 14 \| Phage tail assembly chaperone protein, E, or 41 or 14; uniclust \| UniRef100\_A0A0B0HZT8 \| 98.7 \| 1e-10 \| 2e-16 \| 74.5 \| 79 \| (16, 94) \| 94 \| (25, 120) \| 123 \| Phage tail assembly protein \| Phage tail assembly protein; uniclust \| UniRef100\_A0A0D8ID76 \| 98.7 \| 9.7e-11 \| 2e-16 \| 76.6 \| 81 \| (13, 93) \| 94 \| (34, 127) \| 134 \| Mu-like prophage FluMu protein gp41 \| Mu-like prophage FluMu protein gp41; uniclust \| UniRef100\_A0A317H3T5 \| 98.7 \| 1.1e-10 \| 2.2e-16 \| 76.8 \| 78 \| (16, 93) \| 94 \| (57, 144) \| 153 \| PqqD family protein \| PqqD family protein; uniclust \| UniRef100\_A0A059V670 \| 98.7 \| 1.2e-10 \| 2.3e-16 \| 76.2 \| 88 \| (3, 92) \| 94 \| (7, 95) \| 150 \| Phage tail assembly protein \| Phage tail assembly protein; uniclust \| UniRef100\_A0A068Z6X2 \| 98.7 \| 1.2e-10 \| 2.5e-16 \| 75.5 \| 81 \| (14, 94) \| 94 \| (23, 109) \| 130 \| Phage tail assembly protein \| Phage tail assembly protein; uniclust \| UniRef100\_A0A0D5LRC8 \| 98.7 \| 1.3e-10 \| 2.6e-16 \| 78.3 \| 90 \| (2, 94) \| 94 \| (58, 159) \| 172 \| Tail assembly chaperone E/41/14-like protein \| Tail assembly chaperone E/41/14-like protein; uniclust \| UniRef100\_UPI0013D24EDB \| 98.7 \| 1.6e-10 \| 3e-16 \| 71.5 \| 79 \| (13, 91) \| 94 \| (19, 97) \| 122 \| phage tail assembly protein \| phage tail assembly protein; uniclust \| UniRef100\_A0A2S6QN92 \| 98.7 \| 1.7e-10 \| 3e-16 \| 73.4 \| 71 \| (15, 85) \| 94 \| (3, 74) \| 147 \| Uncharacterized protein (Fragment) \| Uncharacterized protein (Fragment); uniclust \| UniRef100\_A0A349HHP8 \| 98.7 \| 1.7e-10 \| 3.1e-16 \| 71.9 \| 79 \| (12, 90) \| 94 \| (20, 100) \| 128 \| Phage tail assembly protein \| Phage tail assembly protein; uniclust \| UniRef100\_A0A378YUI0 \| 98.7 \| 1.7e-10 \| 3.2e-16 \| 76.3 \| 78 \| (16, 93) \| 94 \| (3, 90) \| 176 \| Phage tail assembly protein \| Phage tail assembly protein; uniclust \| UniRef100\_UPI001CCE5A3E \| 98.7 \| 1.8e-10 \| 3.2e-16 \| 73.2 \| 77 \| (16, 92) \| 94 \| (10, 86) \| 145 \| phage tail assembly protein \| phage tail assembly protein; uniclust \| UniRef100\_A0A151FHH4 \| 98.7 \| 1.7e-10 \| 3.5e-16 \| 73.4 \| 81 \| (12, 92) \| 94 \| (4, 99) \| 114 \| Phage tail assembly protein \| Phage tail assembly protein; uniclust \| UniRef100\_A0A336NB71 \| 98.7 \| 1.8e-10 \| 3.5e-16 \| 80.5 \| 73 \| (17, 91) \| 94 \| (7, 79) \| 248 \| Phage related protein \| Phage related protein; uniclust \| UniRef100\_UPI002247CADB \| 98.7 \| 2.1e-10 \| 3.9e-16 \| 71.7 \| 82 \| (13, 94) \| 94 \| (41, 122) \| 130 \| phage tail assembly protein \| phage tail assembly protein; uniclust \| UniRef100\_A0A0J8DFL1 \| 98.7 \| 2e-10 \| 4.1e-16 \| 75.0 \| 78 \| (16, 93) \| 94 \| (26, 115) \| 134 \| Uncharacterized protein \| Uncharacterized protein; uniclust \| UniRef100\_A0A139D6Z5 \| 98.7 \| 2e-10 \| 4.1e-16 \| 75.0 \| 79 \| (15, 93) \| 94 \| (41, 132) \| 137 \| Phage tail assembly protein \| Phage tail assembly protein; uniclust \| UniRef100\_A0A0S4VLK3 \| 98.7 \| 2.2e-10 \| 4.2e-16 \| 70.2 \| 78 \| (16, 93) \| 94 \| (6, 84) \| 109 \| Phage tail assembly protein \| Phage tail assembly protein; uniclust \| UniRef100\_A0A133ZYZ6 \| 98.7 \| 2e-10 \| 4.2e-16 \| 74.6 \| 78 \| (16, 93) \| 94 \| (33, 123) \| 130 \| Phage tail assembly protein \| Phage tail assembly protein; uniclust \| UniRef100\_A0A2E6KNH6 \| 98.7 \| 2.4e-10 \| 4.4e-16 \| 67.4 \| 79 \| (16, 94) \| 94 \| (7, 87) \| 90 \| Phage tail assembly protein \| Phage tail assembly protein; uniclust \| UniRef100\_UPI002101B5F4 \| 98.7 \| 2.4e-10 \| 4.5e-16 \| 75.2 \| 82 \| (12, 93) \| 94 \| (59, 141) \| 187 \| phage tail assembly protein \| phage tail assembly protein; uniclust \| UniRef100\_A0A158D0Y5 \| 98.7 \| 2.4e-10 \| 4.6e-16 \| 77.2 \| 78 \| (14, 92) \| 94 \| (2, 83) \| 187 \| Phage tail assembly protein \| Phage tail assembly protein; uniclust \| UniRef100\_A0A212KMT6 \| 98.6 \| 2.6e-10 \| 4.8e-16 \| 74.3 \| 78 \| (16, 93) \| 94 \| (84, 161) \| 174 \| Phage tail assembly protein \| Phage tail assembly protein; uniclust \| UniRef100\_UPI001FF44E79 \| 98.6 \| 2.6e-10 \| 4.9e-16 \| 67.5 \| 76 \| (15, 90) \| 94 \| (3, 79) \| 92 \| phage tail assembly protein \| phage tail assembly protein; uniclust \| UniRef100\_A0A0Q4XXX4 \| 98.6 \| 2.5e-10 \| 5.1e-16 \| 74.7 \| 82 \| (13, 94) \| 94 \| (10, 105) \| 134 \| Uncharacterized protein \| Uncharacterized protein; uniclust \| UniRef100\_A0A0Q7A331 \| 98.6 \| 2.9e-10 \| 5.7e-16 \| 73.4 \| 82 \| (12, 94) \| 94 \| (22, 105) \| 129 \| Phage tail assembly protein \| Phage tail assembly protein; uniclust \| UniRef100\_UPI0022384EAA \| 98.6 \| 3.2e-10 \| 5.8e-16 \| 71.3 \| 82 \| (12, 93) \| 94 \| (38, 120) \| 134 \| phage tail assembly protein \| phage tail assembly protein; uniclust \| UniRef100\_A0A212LR71 \| 98.6 \| 2.9e-10 \| 5.8e-16 \| 75.6 \| 81 \| (12, 94) \| 94 \| (43, 130) \| 156 \| Uncharacterized protein \| Uncharacterized protein; uniclust \| UniRef100\_A0A1E2RY06 \| 98.6 \| 3.2e-10 \| 6.2e-16 \| 72.0 \| 83 \| (12, 94) \| 94 \| (27, 121) \| 125 \| Phage tail protein E \| Phage tail protein E; uniclust \| UniRef100\_A0A102DFQ3 \| 98.6 \| 3.3e-10 \| 6.3e-16 \| 70.9 \| 79 \| (14, 94) \| 94 \| (42, 120) \| 120 \| Phage tail assembly protein \| Phage tail assembly protein; uniclust \| UniRef100\_A0A935YDJ1 \| 98.6 \| 4e-10 \| 7.3e-16 \| 67.9 \| 80 \| (15, 94) \| 94 \| (6, 93) \| 102 \| Phage tail assembly protein \| Phage tail assembly protein; uniclust \| UniRef100\_A0A1G7D511 \| 98.6 \| 3.7e-10 \| 7.3e-16 \| 70.8 \| 82 \| (13, 94) \| 94 \| (4, 90) \| 110 \| Phage tail assembly chaperone protein, E, or 41 or 14 \| Phage tail assembly chaperone protein, E, or 41 or 14; uniclust \| UniRef100\_UPI00197C731A \| 98.6 \| 4e-10 \| 7.4e-16 \| 70.6 \| 82 \| (12, 93) \| 94 \| (25, 109) \| 131 \| phage tail assembly protein \| phage tail assembly protein; uniclust \| UniRef100\_A0A1C7L9J9 \| 98.6 \| 4e-10 \| 8e-16 \| 72.7 \| 80 \| (14, 93) \| 94 \| (31, 114) \| 127 \| Tail E family protein \| Tail E family protein; uniclust \| UniRef100\_A0A7M3MAZ0 \| 98.6 \| 4.4e-10 \| 8.2e-16 \| 68.2 \| 76 \| (15, 90) \| 94 \| (6, 84) \| 102 \| Phage tail assembly protein \| Phage tail assembly protein; uniclust \| UniRef100\_A0A849MU39 \| 98.6 \| 4.4e-10 \| 8.2e-16 \| 68.3 \| 80 \| (15, 94) \| 94 \| (14, 99) \| 107 \| Phage tail assembly protein \| Phage tail assembly protein; uniclust \| UniRef100\_UPI0020CEC014 \| 98.6 \| 4.8e-10 \| 8.8e-16 \| 69.1 \| 86 \| (8, 93) \| 94 \| (21, 110) \| 117 \| phage tail assembly protein \| phage tail assembly protein; uniclust \| UniRef100\_A0A929HY00 \| 98.6 \| 4.8e-10 \| 8.8e-16 \| 66.1 \| 78 \| (13, 92) \| 94 \| (11, 88) \| 89 \| Phage tail assembly protein \| Phage tail assembly protein; uniclust \| UniRef100\_A0A0A2SKX1 \| 98.6 \| 4.5e-10 \| 9e-16 \| 72.9 \| 79 \| (15, 93) \| 94 \| (35, 124) \| 137 \| Phage tail assembly protein \| Phage tail assembly protein; uniclust \| UniRef100\_A0A094ZEZ2 \| 98.6 \| 4.4e-10 \| 9.1e-16 \| 79.1 \| 75 \| (18, 92) \| 94 \| (31, 111) \| 223 \| Phage protein \| Phage protein; uniclust \| UniRef100\_A0A2R8CKK2 \| 98.6 \| 5.1e-10 \| 9.5e-16 \| 73.3 \| 78 \| (16, 93) \| 94 \| (71, 152) \| 162 \| Phage tail protein E \| Phage tail protein E; uniclust \| UniRef100\_UPI00156FF9AB \| 98.6 \| 5.3e-10 \| 1e-15 \| 68.6 \| 82 \| (13, 94) \| 94 \| (11, 96) \| 105 \| phage tail assembly protein \| phage tail assembly protein; uniclust \| UniRef100\_A0A077PQ35 \| 98.6 \| 5.4e-10 \| 1e-15 \| 68.1 \| 80 \| (14, 93) \| 94 \| (3, 86) \| 96 \| Phage tail assembly protein \| Phage tail assembly protein; uniclust \| UniRef100\_A0A212S820 \| 98.6 \| 5.3e-10 \| 1e-15 \| 72.9 \| 82 \| (13, 94) \| 94 \| (30, 116) \| 141 \| Phage tail assembly chaperone protein, E, or 41 or 14 \| Phage tail assembly chaperone protein, E, or 41 or 14; uniclust \| UniRef100\_A0A031FSF6 \| 98.6 \| 5e-10 \| 1.1e-15 \| 78.4 \| 78 \| (16, 94) \| 94 \| (116, 193) \| 209 \| Phage-like protein \| Phage-like protein; uniclust \| UniRef100\_UPI001364C58B \| 98.6 \| 6.1e-10 \| 1.1e-15 \| 67.4 \| 78 \| (14, 92) \| 94 \| (7, 88) \| 95 \| phage tail assembly protein \| phage tail assembly protein; uniclust \| UniRef100\_A0A4Q8MDQ4 \| 98.6 \| 6e-10 \| 1.2e-15 \| 68.1 \| 77 \| (17, 93) \| 94 \| (2, 86) \| 95 \| Phage tail assembly protein \| Phage tail assembly protein; uniclust \| UniRef100\_A0A5C7NIW4 \| 98.6 \| 6.9e-10 \| 1.3e-15 \| 66.2 \| 77 \| (17, 94) \| 94 \| (2, 86) \| 95 \| Phage tail assembly protein \| Phage tail assembly protein; uniclust \| UniRef100\_A0A074TD79 \| 98.6 \| 7.1e-10 \| 1.4e-15 \| 68.7 \| 74 \| (16, 90) \| 94 \| (4, 77) \| 102 \| Phage tail assembly protein \| Phage tail assembly protein; uniclust \| UniRef100\_A0A0A8GWA2 \| 98.6 \| 7.2e-10 \| 1.4e-15 \| 66.0 \| 62 \| (33, 94) \| 94 \| (12, 73) \| 82 \| Phage tail assembly protein \| Phage tail assembly protein; uniclust \| UniRef100\_A0A1H6QNL8 \| 98.5 \| 8.1e-10 \| 1.5e-15 \| 73.8 \| 77 \| (15, 92) \| 94 \| (3, 83) \| 186 \| Phage tail assembly chaperone protein, E, or 41 or 14 \| Phage tail assembly chaperone protein, E, or 41 or 14; uniclust \| UniRef100\_A0A7W4ICA4 \| 98.5 \| 7.7e-10 \| 1.5e-15 \| 73.1 \| 90 \| (2, 93) \| 94 \| (38, 139) \| 147 \| Phage tail assembly protein \| Phage tail assembly protein; uniclust \| UniRef100\_A0A0C6FCI1 \| 98.5 \| 8.7e-10 \| 1.7e-15 \| 74.6 \| 83 \| (12, 94) \| 94 \| (70, 154) \| 182 \| Uncharacterized protein \| Uncharacterized protein; uniclust \| UniRef100\_A0A173SBH0 \| 98.5 \| 9e-10 \| 1.8e-15 \| 72.4 \| 79 \| (15, 93) \| 94 \| (54, 144) \| 149 \| Phage tail assembly protein \| Phage tail assembly protein; uniclust \| UniRef100\_A0A4Z0E5Q8 \| 98.5 \| 1.1e-09 \| 2e-15 \| 64.9 \| 76 \| (15, 90) \| 94 \| (3, 78) \| 83 \| Phage tail assembly protein \| Phage tail assembly protein; uniclust \| UniRef100\_A0A016XIN1 \| 98.5 \| 1.1e-09 \| 2.1e-15 \| 67.7 \| 79 \| (13, 93) \| 94 \| (8, 86) \| 119 \| Phage tail assembly protein \| Phage tail assembly protein; uniclust \| UniRef100\_A0A925L1X7 \| 98.5 \| 1.1e-09 \| 2.1e-15 \| 69.2 \| 86 \| (8, 94) \| 94 \| (14, 100) \| 128 \| Phage tail assembly protein \| Phage tail assembly protein; uniclust \| UniRef100\_A0A094ZEZ2 \| 98.5 \| 1.3e-09 \| 2.6e-15 \| 76.9 \| 78 \| (16, 93) \| 94 \| (128, 210) \| 223 \| Phage protein \| Phage protein; uniclust \| UniRef100\_A0NQ99 \| 98.5 \| 1.5e-09 \| 2.7e-15 \| 67.8 \| 79 \| (14, 93) \| 94 \| (5, 84) \| 121 \| Tail assembly chaperone \| Tail assembly chaperone; uniclust \| UniRef100\_A0A090G422 \| 98.5 \| 1.5e-09 \| 2.8e-15 \| 69.6 \| 72 \| (16, 91) \| 94 \| (49, 120) \| 134 \| Phage tail assembly protein \| Phage tail assembly protein; uniclust \| UniRef100\_UPI0013D729BF \| 98.5 \| 1.5e-09 \| 2.9e-15 \| 66.9 \| 78 \| (16, 93) \| 94 \| (10, 97) \| 106 \| phage tail assembly protein \| phage tail assembly protein; uniclust \| UniRef100\_A0A8J2Z5X2 \| 98.5 \| 1.6e-09 \| 3e-15 \| 65.9 \| 74 \| (17, 91) \| 94 \| (4, 78) \| 91 \| Tail assembly chaperone E/41/14-like protein \| Tail assembly chaperone E/41/14-like protein; uniclust \| UniRef100\_A0A7J5WEE9 \| 98.5 \| 1.7e-09 \| 3.1e-15 \| 65.5 \| 73 \| (18, 90) \| 94 \| (25, 98) \| 103 \| Phage tail assembly protein \| Phage tail assembly protein; uniclust \| UniRef100\_A0A423PRR2 \| 98.5 \| 1.7e-09 \| 3.3e-15 \| 68.2 \| 80 \| (15, 94) \| 94 \| (28, 109) \| 116 \| Phage tail assembly protein \| Phage tail assembly protein; uniclust \| UniRef100\_A0A0H5CY95 \| 98.5 \| 1.7e-09 \| 3.4e-15 \| 69.5 \| 79 \| (15, 93) \| 94 \| (3, 102) \| 122 \| Phage tail assembly protein \| Phage tail assembly protein; uniclust \| UniRef100\_A0A1G9AZL3 \| 98.5 \| 2e-09 \| 3.8e-15 \| 67.9 \| 88 \| (3, 91) \| 94 \| (10, 108) \| 122 \| Phage tail assembly chaperone protein, E, or 41 or 14 \| Phage tail assembly chaperone protein, E, or 41 or 14; uniclust \| UniRef100\_A0A5P9JUS1 \| 98.5 \| 2e-09 \| 3.8e-15 \| 64.6 \| 80 \| (14, 93) \| 94 \| (5, 86) \| 90 \| Uncharacterized protein \| Uncharacterized protein; uniclust \| UniRef100\_A0A061NN23 \| 98.5 \| 2e-09 \| 3.8e-15 \| 67.4 \| 79 \| (16, 94) \| 94 \| (25, 115) \| 116 \| Phage protein \| Phage protein; uniclust \| UniRef100\_A0A940MVV2 \| 98.5 \| 2.1e-09 \| 4e-15 \| 66.5 \| 70 \| (23, 93) \| 94 \| (26, 95) \| 105 \| Phage tail assembly protein \| Phage tail assembly protein; uniclust \| UniRef100\_A0A4D7BGG9 \| 98.5 \| 2.1e-09 \| 4e-15 \| 69.5 \| 81 \| (13, 93) \| 94 \| (7, 101) \| 133 \| Phage tail assembly protein \| Phage tail assembly protein; uniclust \| UniRef100\_UPI0004B2DC3D \| 98.4 \| 2.6e-09 \| 4.7e-15 \| 67.6 \| 76 \| (17, 92) \| 94 \| (3, 82) \| 136 \| phage tail assembly protein \| phage tail assembly protein; uniclust \| UniRef100\_UPI00143B4262 \| 98.4 \| 2.5e-09 \| 4.7e-15 \| 66.0 \| 76 \| (16, 91) \| 94 \| (3, 86) \| 110 \| phage tail assembly protein \| phage tail assembly protein; uniclust \| UniRef100\_A0A1I7NC51 \| 98.4 \| 2.7e-09 \| 5.3e-15 \| 69.1 \| 83 \| (12, 94) \| 94 \| (21, 105) \| 135 \| Phage tail assembly chaperone protein, E, or 41 or 14 \| Phage tail assembly chaperone protein, E, or 41 or 14; uniclust \| UniRef100\_A0A1B9VJR2 \| 98.4 \| 2.8e-09 \| 5.6e-15 \| 69.0 \| 90 \| (4, 94) \| 94 \| (16, 117) \| 130 \| Mu-like prophage FluMu protein gp41 \| Mu-like prophage FluMu protein gp41; uniclust \| UniRef100\_A0A1M6ZC09 \| 98.4 \| 3.1e-09 \| 6e-15 \| 63.2 \| 76 \| (18, 94) \| 94 \| (2, 79) \| 83 \| Phage tail assembly chaperone protein, E, or 41 or 14 \| Phage tail assembly chaperone protein, E, or 41 or 14; uniclust \| UniRef100\_A0A0U3UX96 \| 98.4 \| 3.4e-09 \| 6.4e-15 \| 66.1 \| 79 \| (16, 94) \| 94 \| (5, 91) \| 111 \| Phage small tail protein E \| Phage small tail protein E; uniclust \| UniRef100\_A0A2S4LWK7 \| 98.4 \| 4.6e-09 \| 8.5e-15 \| 69.1 \| 79 \| (15, 93) \| 94 \| (83, 165) \| 179 \| Tail assembly chaperone E/41/14-like protein \| Tail assembly chaperone E/41/14-like protein; uniclust \| UniRef100\_A0A098ARP9 \| 98.4 \| 4.9e-09 \| 9e-15 \| 58.2 \| 57 \| (18, 74) \| 94 \| (3, 59) \| 61 \| Putative phage related protein \| Putative phage related protein; uniclust \| UniRef100\_A0A2E2IRC7 \| 98.4 \| 5.1e-09 \| 1e-14 \| 65.9 \| 75 \| (16, 90) \| 94 \| (5, 93) \| 108 \| Phage tail assembly protein \| Phage tail assembly protein; uniclust \| UniRef100\_UPI001BCB7C8E \| 98.4 \| 5.6e-09 \| 1.1e-14 \| 64.4 \| 83 \| (12, 94) \| 94 \| (3, 95) \| 103 \| phage tail assembly protein \| phage tail assembly protein; uniclust \| UniRef100\_A0A1I2QQM4 \| 98.3 \| 5.8e-09 \| 1.1e-14 \| 64.3 \| 82 \| (13, 94) \| 94 \| (16, 98) \| 107 \| Phage tail assembly chaperone protein, E, or 41 or 14 \| Phage tail assembly chaperone protein, E, or 41 or 14; uniclust \| UniRef100\_A0A8J7LX76 \| 98.3 \| 6e-09 \| 1.1e-14 \| 67.1 \| 83 \| (12, 94) \| 94 \| (29, 115) \| 137 \| Phage tail assembly protein \| Phage tail assembly protein; uniclust \| UniRef100\_A0A031FSF6 \| 98.3 \| 5.6e-09 \| 1.2e-14 \| 73.5 \| 79 \| (15, 93) \| 94 \| (18, 98) \| 209 \| Phage-like protein \| Phage-like protein; uniclust \| UniRef100\_A0A349GR41 \| 98.3 \| 6.3e-09 \| 1.2e-14 \| 66.7 \| 81 \| (13, 93) \| 94 \| (30, 123) \| 125 \| Phage tail assembly protein \| Phage tail assembly protein; uniclust \| UniRef100\_A0A2S6HV68 \| 98.3 \| 6.2e-09 \| 1.3e-14 \| 67.1 \| 78 \| (16, 93) \| 94 \| (25, 116) \| 120 \| Tail assembly chaperone E/41/14-like protein \| Tail assembly chaperone E/41/14-like protein; uniclust \| UniRef100\_UPI0007377FCF \| 98.3 \| 6.6e-09 \| 1.3e-14 \| 63.5 \| 70 \| (13, 82) \| 94 \| (18, 91) \| 92 \| phage tail assembly protein \| phage tail assembly protein; uniclust \| UniRef100\_UPI001FB7CC94 \| 98.3 \| 7e-09 \| 1.3e-14 \| 63.6 \| 76 \| (17, 92) \| 94 \| (8, 86) \| 109 \| phage tail assembly protein \| phage tail assembly protein; uniclust \| UniRef100\_A0A256CAU0 \| 98.3 \| 7.1e-09 \| 1.3e-14 \| 63.7 \| 79 \| (15, 93) \| 94 \| (3, 85) \| 100 \| Phage tail protein \| Phage tail protein; uniclust \| UniRef100\_UPI0014125629 \| 98.3 \| 7.2e-09 \| 1.3e-14 \| 65.2 \| 82 \| (12, 94) \| 94 \| (23, 113) \| 121 \| hypothetical protein \| hypothetical protein; uniclust \| UniRef100\_A0A1V4VYZ4 \| 98.3 \| 7e-09 \| 1.4e-14 \| 65.6 \| 77 \| (17, 93) \| 94 \| (3, 91) \| 110 \| Phage tail protein E \| Phage tail protein E; uniclust \| UniRef100\_A0A1X3S1E1 \| 98.3 \| 7.6e-09 \| 1.5e-14 \| 66.5 \| 79 \| (16, 94) \| 94 \| (5, 91) \| 133 \| Phage tail protein \| Phage tail protein; uniclust \| UniRef100\_A0A7X4CN54 \| 98.3 \| 8.3e-09 \| 1.5e-14 \| 63.5 \| 78 \| (16, 93) \| 94 \| (33, 111) \| 112 \| Phage tail assembly protein \| Phage tail assembly protein; uniclust \| UniRef100\_E2CN25 \| 98.3 \| 8.2e-09 \| 1.6e-14 \| 65.3 \| 79 \| (16, 94) \| 94 \| (4, 87) \| 123 \| Phage-related protein \| Phage-related protein; uniclust \| UniRef100\_A0A1P8QQ42 \| 98.3 \| 9e-09 \| 1.7e-14 \| 63.4 \| 82 \| (12, 93) \| 94 \| (29, 110) \| 112 \| Uncharacterized protein \| Uncharacterized protein; uniclust \| UniRef100\_A0A1H2FLU3 \| 98.3 \| 8.8e-09 \| 1.7e-14 \| 67.4 \| 83 \| (12, 94) \| 94 \| (7, 129) \| 141 \| Phage tail assembly chaperone protein, E, or 41 or 14 \| Phage tail assembly chaperone protein, E, or 41 or 14; uniclust \| UniRef100\_A0A0C2VFH9 \| 98.3 \| 9.5e-09 \| 1.7e-14 \| 60.7 \| 77 \| (15, 94) \| 94 \| (4, 82) \| 86 \| Phage tail assembly protein \| Phage tail assembly protein; uniclust \| UniRef100\_A0A922XTU6 \| 98.3 \| 9.8e-09 \| 1.8e-14 \| 62.2 \| 82 \| (13, 94) \| 94 \| (10, 92) \| 101 \| Uncharacterized protein \| Uncharacterized protein; uniclust \| UniRef100\_A0A2S5N1P1 \| 98.3 \| 9.3e-09 \| 1.8e-14 \| 68.3 \| 82 \| (13, 94) \| 94 \| (54, 136) \| 157 \| Phage tail assembly protein \| Phage tail assembly protein; uniclust \| UniRef100\_A0A6L6YK12 \| 98.3 \| 1.1e-08 \| 2e-14 \| 62.3 \| 79 \| (16, 94) \| 94 \| (3, 86) \| 96 \| Phage tail assembly protein \| Phage tail assembly protein; uniclust \| UniRef100\_A0A2E2N165 \| 98.3 \| 1.1e-08 \| 2e-14 \| 65.7 \| 79 \| (15, 93) \| 94 \| (2, 93) \| 125 \| Phage tail assembly protein \| Phage tail assembly protein; uniclust \| UniRef100\_A0A1M3BNG8 \| 98.3 \| 1.1e-08 \| 2.1e-14 \| 66.3 \| 79 \| (12, 93) \| 94 \| (39, 117) \| 126 \| Phage tail assembly protein \| Phage tail assembly protein; uniclust \| UniRef100\_A0A2N6FYM4 \| 98.3 \| 1.2e-08 \| 2.1e-14 \| 60.1 \| 77 \| (16, 93) \| 94 \| (5, 83) \| 84 \| Phage tail assembly protein \| Phage tail assembly protein; uniclust \| UniRef100\_UPI0003802596 \| 98.3 \| 1.2e-08 \| 2.2e-14 \| 60.0 \| 78 \| (17, 94) \| 94 \| (2, 80) \| 84 \| phage tail assembly protein \| phage tail assembly protein; uniclust \| UniRef100\_A0A844Q6C1 \| 98.3 \| 1.2e-08 \| 2.3e-14 \| 63.6 \| 77 \| (17, 93) \| 94 \| (2, 92) \| 105 \| Phage tail assembly protein \| Phage tail assembly protein; uniclust \| UniRef100\_A0A1I5RNJ4 \| 98.3 \| 1.3e-08 \| 2.4e-14 \| 61.1 \| 80 \| (15, 94) \| 94 \| (2, 89) \| 89 \| Phage tail assembly chaperone protein, E, or 41 or 14 \| Phage tail assembly chaperone protein, E, or 41 or 14; uniclust \| UniRef100\_A0A0D8L4Q2 \| 98.3 \| 1.3e-08 \| 2.5e-14 \| 64.4 \| 88 \| (4, 93) \| 94 \| (17, 108) \| 120 \| Phage tail protein \| Phage tail protein; uniclust \| UniRef100\_A0A1Y1S1F4 \| 98.3 \| 1.3e-08 \| 2.5e-14 \| 63.6 \| 81 \| (12, 94) \| 94 \| (42, 123) \| 123 \| Phage tail protein \| Phage tail protein; uniclust \| UniRef100\_UPI0018AD3EFF \| 98.3 \| 1.4e-08 \| 2.6e-14 \| 55.9 \| 52 \| (38, 90) \| 94 \| (1, 52) \| 58 \| phage tail assembly protein \| phage tail assembly protein; uniclust \| UniRef100\_A0A0C3RQ40 \| 98.3 \| 1.2e-08 \| 2.6e-14 \| 66.7 \| 80 \| (12, 91) \| 94 \| (17, 106) \| 126 \| Phage protein \| Phage protein; uniclust \| UniRef100\_UPI00037665EA \| 98.2 \| 1.5e-08 \| 2.8e-14 \| 63.6 \| 74 \| (17, 90) \| 94 \| (4, 78) \| 113 \| phage tail assembly protein \| phage tail assembly protein; uniclust \| UniRef100\_A0A2D8T8N2 \| 98.2 \| 1.5e-08 \| 2.9e-14 \| 64.1 \| 85 \| (3, 93) \| 94 \| (11, 99) \| 109 \| Phage tail assembly protein \| Phage tail assembly protein; uniclust \| UniRef100\_UPI000D381B68 \| 98.2 \| 1.6e-08 \| 2.9e-14 \| 62.1 \| 62 \| (33, 94) \| 94 \| (43, 104) \| 109 \| phage tail assembly protein \| phage tail assembly protein; uniclust \| UniRef100\_UPI00094A100B \| 98.2 \| 1.6e-08 \| 3e-14 \| 66.2 \| 79 \| (15, 93) \| 94 \| (59, 139) \| 149 \| phage tail assembly protein \| phage tail assembly protein; uniclust \| UniRef100\_A0A4R5W1E0 \| 98.2 \| 1.6e-08 \| 3e-14 \| 62.9 \| 78 \| (13, 90) \| 94 \| (10, 88) \| 118 \| Phage tail assembly protein \| Phage tail assembly protein; uniclust \| UniRef100\_UPI001030D5AE \| 98.2 \| 1.6e-08 \| 3e-14 \| 62.1 \| 76 \| (15, 91) \| 94 \| (29, 104) \| 109 \| phage tail assembly protein \| phage tail assembly protein; uniclust \| UniRef100\_UPI001903CD40 \| 98.2 \| 1.8e-08 \| 3.3e-14 \| 62.7 \| 76 \| (17, 93) \| 94 \| (13, 88) \| 118 \| phage tail assembly protein \| phage tail assembly protein; uniclust \| UniRef100\_UPI0002A3D1A3 \| 98.2 \| 1.8e-08 \| 3.5e-14 \| 60.7 \| 62 \| (32, 93) \| 94 \| (14, 75) \| 86 \| phage tail assembly protein \| phage tail assembly protein; uniclust \| UniRef100\_A0A7J6YK69 \| 98.2 \| 2.1e-08 \| 3.9e-14 \| 83.0 \| 75 \| (16, 90) \| 94 \| (1760, 1834) \| 2523 \| Uncharacterized protein \| Uncharacterized protein; uniclust \| UniRef100\_A0A8G2CCC5 \| 98.2 \| 2.2e-08 \| 4.1e-14 \| 61.5 \| 76 \| (16, 91) \| 94 \| (6, 84) \| 109 \| Phage tail assembly chaperone protein, E, or 41 or 14 \| Phage tail assembly chaperone protein, E, or 41 or 14; uniclust \| UniRef100\_A0A1I9KFV2 \| 98.2 \| 2.2e-08 \| 4.4e-14 \| 70.7 \| 75 \| (18, 94) \| 94 \| (127, 203) \| 216 \| Uncharacterized protein \| Uncharacterized protein; uniclust \| UniRef100\_A0A066QH52 \| 98.2 \| 2.6e-08 \| 5.1e-14 \| 65.5 \| 79 \| (16, 94) \| 94 \| (27, 111) \| 136 \| Phage tail E family protein \| Phage tail E family protein; uniclust \| UniRef100\_A0A1X7I5S5 \| 98.2 \| 2.9e-08 \| 5.2e-14 \| 62.1 \| 74 \| (20, 93) \| 94 \| (13, 94) \| 121 \| Phage tail assembly chaperone protein, E, or 41 or 14 \| Phage tail assembly chaperone protein, E, or 41 or 14; uniclust \| UniRef100\_A0A4P9VF09 \| 98.2 \| 3e-08 \| 5.5e-14 \| 54.0 \| 40 \| (18, 57) \| 94 \| (4, 43) \| 54 \| Phage tail assembly protein \| Phage tail assembly protein; uniclust \| UniRef100\_UPI001FD41C6C \| 98.2 \| 3.2e-08 \| 5.8e-14 \| 58.8 \| 73 \| (16, 89) \| 94 \| (3, 75) \| 88 \| phage tail assembly protein \| phage tail assembly protein; uniclust \| UniRef100\_A0A258L5N6 \| 98.2 \| 3.1e-08 \| 6.1e-14 \| 69.4 \| 78 \| (16, 94) \| 94 \| (115, 192) \| 223 \| Phage tail assembly protein \| Phage tail assembly protein; uniclust \| UniRef100\_Q31HT0 \| 98.1 \| 3.5e-08 \| 6.4e-14 \| 56.1 \| 64 \| (30, 94) \| 94 \| (5, 68) \| 68 \| Phage tail assembly protein \| Phage tail assembly protein; uniclust \| UniRef100\_A0A2J0QXA4 \| 98.1 \| 3.5e-08 \| 6.6e-14 \| 61.0 \| 79 \| (16, 94) \| 94 \| (20, 102) \| 107 \| Phage tail assembly protein \| Phage tail assembly protein; uniclust \| UniRef100\_A0A0M7ALB4 \| 98.1 \| 3.9e-08 \| 7.2e-14 \| 59.9 \| 80 \| (14, 93) \| 94 \| (19, 99) \| 102 \| Tail assembly chaperone E/41/14-like protein \| Tail assembly chaperone E/41/14-like protein; uniclust \| UniRef100\_A0A9D1VF36 \| 98.1 \| 4.2e-08 \| 7.8e-14 \| 60.5 \| 79 \| (13, 93) \| 94 \| (21, 100) \| 110 \| Phage tail assembly protein \| Phage tail assembly protein; uniclust \| UniRef100\_A0A3A6N335 \| 98.1 \| 4.1e-08 \| 7.9e-14 \| 61.9 \| 82 \| (13, 94) \| 94 \| (6, 97) \| 108 \| Phage tail assembly protein \| Phage tail assembly protein; uniclust \| UniRef100\_A0A661TQP7 \| 98.1 \| 4.7e-08 \| 8.7e-14 \| 59.5 \| 81 \| (13, 93) \| 94 \| (3, 93) \| 101 \| Phage tail assembly protein \| Phage tail assembly protein; uniclust \| UniRef100\_A0A2T1B6V7 \| 98.1 \| 5e-08 \| 9.4e-14 \| 66.3 \| 79 \| (15, 93) \| 94 \| (95, 175) \| 205 \| Tail assembly chaperone E/41/14-like protein \| Tail assembly chaperone E/41/14-like protein; uniclust \| UniRef100\_A0A1X0SVA2 \| 98.1 \| 4.8e-08 \| 9.5e-14 \| 62.4 \| 80 \| (15, 94) \| 94 \| (2, 90) \| 115 \| Uncharacterized protein \| Uncharacterized protein; uniclust \| UniRef100\_A0A0Q8AP47 \| 98.1 \| 5.3e-08 \| 1e-13 \| 61.0 \| 75 \| (16, 90) \| 94 \| (3, 80) \| 108 \| Phage tail protein \| Phage tail protein; uniclust \| UniRef100\_A0A0G0BNF2 \| 98.1 \| 5.5e-08 \| 1.1e-13 \| 61.3 \| 62 \| (31, 92) \| 94 \| (15, 83) \| 106 \| Phage tail assembly protein \| Phage tail assembly protein; uniclust \| UniRef100\_A0A212KMT6 \| 98.1 \| 6.1e-08 \| 1.1e-13 \| 64.0 \| 75 \| (18, 92) \| 94 \| (3, 77) \| 174 \| Phage tail assembly protein \| Phage tail assembly protein; uniclust \| UniRef100\_A0A7X6FP61 \| 98.1 \| 6.2e-08 \| 1.1e-13 \| 54.4 \| 56 \| (15, 71) \| 94 \| (6, 61) \| 63 \| Phage tail assembly protein \| Phage tail assembly protein; uniclust \| UniRef100\_A0A3N2E0T6 \| 98.1 \| 6.4e-08 \| 1.2e-13 \| 59.1 \| 69 \| (15, 83) \| 94 \| (20, 89) \| 103 \| Tail assembly chaperone E/41/14-like protein \| Tail assembly chaperone E/41/14-like protein; uniclust \| UniRef100\_A0A961TLJ2 \| 98.1 \| 6.7e-08 \| 1.2e-13 \| 60.6 \| 78 \| (15, 93) \| 94 \| (27, 104) \| 121 \| Phage tail assembly protein \| Phage tail assembly protein; uniclust \| UniRef100\_A0A1B1KPJ6 \| 98.1 \| 7.1e-08 \| 1.3e-13 \| 61.3 \| 78 \| (16, 93) \| 94 \| (25, 106) \| 120 \| Tail protein \| Tail protein; uniclust \| UniRef100\_A0A1H1G2N2 \| 98.1 \| 7e-08 \| 1.3e-13 \| 61.3 \| 79 \| (16, 94) \| 94 \| (3, 86) \| 118 \| Phage tail assembly chaperone protein, E, or 41 or 14 \| Phage tail assembly chaperone protein, E, or 41 or 14; uniclust \| UniRef100\_A0A1V3RSD9 \| 98.1 \| 7.6e-08 \| 1.4e-13 \| 58.7 \| 76 \| (16, 91) \| 94 \| (16, 97) \| 102 \| Phage tail assembly protein \| Phage tail assembly protein; uniclust \| UniRef100\_A0A6B8KJI9 \| 98.0 \| 8e-08 \| 1.5e-13 \| 60.6 \| 75 \| (16, 94) \| 94 \| (52, 126) \| 126 \| Phage tail assembly protein \| Phage tail assembly protein; uniclust \| UniRef100\_A0A266LMF2 \| 98.0 \| 7.9e-08 \| 1.5e-13 \| 57.2 \| 64 \| (28, 91) \| 94 \| (3, 67) \| 79 \| Phage tail assembly protein (Fragment) \| Phage tail assembly protein (Fragment); uniclust \| UniRef100\_UPI0009EC53EA \| 98.0 \| 8.7e-08 \| 1.6e-13 \| 68.9 \| 70 \| (15, 84) \| 94 \| (21, 101) \| 344 \| phage tail assembly protein \| phage tail assembly protein; uniclust \| UniRef100\_UPI00056E07A5 \| 98.0 \| 8.7e-08 \| 1.6e-13 \| 55.3 \| 56 \| (16, 72) \| 94 \| (16, 72) \| 73 \| phage tail assembly protein \| phage tail assembly protein; uniclust \| UniRef100\_A0A2I7R3J1 \| 98.0 \| 8.8e-08 \| 1.6e-13 \| 56.5 \| 80 \| (15, 94) \| 94 \| (2, 83) \| 83 \| Putative tail protein \| Putative tail protein; uniclust \| UniRef100\_A0A1G7ZBM7 \| 98.0 \| 8.7e-08 \| 1.7e-13 \| 65.5 \| 79 \| (13, 92) \| 94 \| (13, 91) \| 196 \| Phage tail assembly chaperone protein, E, or 41 or 14 \| Phage tail assembly chaperone protein, E, or 41 or 14; uniclust \| UniRef100\_A0A7G6RKV0 \| 98.0 \| 9.1e-08 \| 1.8e-13 \| 60.1 \| 74 \| (16, 93) \| 94 \| (19, 97) \| 107 \| Phage tail assembly protein \| Phage tail assembly protein; uniclust \| UniRef100\_A0A0P7I4K8 \| 98.0 \| 9.9e-08 \| 1.8e-13 \| 63.2 \| 81 \| (14, 94) \| 94 \| (28, 108) \| 162 \| Tail assembly chaperone \| Tail assembly chaperone; uniclust \| UniRef100\_UPI00234F299C \| 98.0 \| 1.1e-07 \| 2e-13 \| 59.3 \| 79 \| (15, 93) \| 94 \| (3, 85) \| 117 \| phage tail assembly protein \| phage tail assembly protein; uniclust \| UniRef100\_A0A2D3VZ53 \| 98.0 \| 1.2e-07 \| 2.1e-13 \| 58.1 \| 59 \| (33, 92) \| 94 \| (38, 96) \| 103 \| Phage tail assembly protein \| Phage tail assembly protein; uniclust \| UniRef100\_A0A0A8IK84 \| 98.0 \| 1.1e-07 \| 2.2e-13 \| 64.3 \| 70 \| (16, 89) \| 94 \| (73, 142) \| 153 \| Uncharacterized protein \| Uncharacterized protein; uniclust \| UniRef100\_F3YY69 \| 98.0 \| 1.2e-07 \| 2.2e-13 \| 58.7 \| 75 \| (16, 90) \| 94 \| (4, 86) \| 111 \| Phage tail assembly protein \| Phage tail assembly protein; uniclust \| UniRef100\_A0A2N9Y8Z8 \| 98.0 \| 1.2e-07 \| 2.4e-13 \| 63.8 \| 81 \| (14, 94) \| 94 \| (8, 127) \| 151 \| Phage tail assembly protein \| Phage tail assembly protein; uniclust \| UniRef100\_A0A9E7YFR7 \| 98.0 \| 1.3e-07 \| 2.4e-13 \| 60.3 \| 76 \| (14, 93) \| 94 \| (16, 91) \| 135 \| Uncharacterized protein \| Uncharacterized protein; uniclust \| UniRef100\_A0A7J0BJG9 \| 98.0 \| 1.3e-07 \| 2.4e-13 \| 58.8 \| 76 \| (15, 90) \| 94 \| (4, 82) \| 112 \| Phage tail assembly protein \| Phage tail assembly protein; uniclust \| UniRef100\_UPI0012BCC767 \| 98.0 \| 1.3e-07 \| 2.4e-13 \| 56.9 \| 79 \| (16, 94) \| 94 \| (4, 82) \| 93 \| phage tail assembly protein \| phage tail assembly protein; uniclust \| UniRef100\_UPI002147F80F \| 98.0 \| 1.4e-07 \| 2.5e-13 \| 54.5 \| 61 \| (33, 94) \| 94 \| (4, 64) \| 73 \| phage tail assembly protein \| phage tail assembly protein; uniclust \| UniRef100\_A0A4R3J9N6 \| 98.0 \| 1.5e-07 \| 2.8e-13 \| 56.9 \| 78 \| (17, 94) \| 94 \| (5, 92) \| 96 \| Tail assembly chaperone E/41/14-like protein \| Tail assembly chaperone E/41/14-like protein; uniclust \| UniRef100\_A0A258L5N6 \| 98.0 \| 1.5e-07 \| 2.9e-13 \| 66.2 \| 77 \| (15, 91) \| 94 \| (21, 99) \| 223 \| Phage tail assembly protein \| Phage tail assembly protein; uniclust \| UniRef100\_UPI00174E383D \| 98.0 \| 1.7e-07 \| 3.1e-13 \| 59.0 \| 78 \| (17, 94) \| 94 \| (4, 94) \| 122 \| phage tail assembly protein \| phage tail assembly protein; uniclust \| UniRef100\_UPI002100D467 \| 97.9 \| 1.7e-07 \| 3.1e-13 \| 58.5 \| 81 \| (13, 94) \| 94 \| (16, 96) \| 108 \| phage tail assembly protein \| phage tail assembly protein; uniclust \| UniRef100\_A0A077L031 \| 97.9 \| 1.7e-07 \| 3.2e-13 \| 58.7 \| 79 \| (16, 94) \| 94 \| (17, 98) \| 104 \| Phage tail assembly protein \| Phage tail assembly protein; uniclust \| UniRef100\_A0A1S6U833 \| 97.9 \| 1.7e-07 \| 3.4e-13 \| 56.7 \| 61 \| (33, 94) \| 94 \| (13, 73) \| 82 \| Putative Mu-like phage protein \| Putative Mu-like phage protein; uniclust \| UniRef100\_A0A1G8JZK0 \| 97.9 \| 1.8e-07 \| 3.4e-13 \| 61.3 \| 78 \| (16, 93) \| 94 \| (53, 134) \| 147 \| Phage tail assembly chaperone protein, E, or 41 or 14 \| Phage tail assembly chaperone protein, E, or 41 or 14; uniclust \| UniRef100\_A0A0P0URT8 \| 97.9 \| 2e-07 \| 3.7e-13 \| 58.2 \| 74 \| (16, 89) \| 94 \| (3, 78) \| 117 \| Phage-related terminase \| Phage-related terminase; uniclust \| UniRef100\_A0A1X7MF74 \| 97.9 \| 2.1e-07 \| 3.9e-13 \| 50.2 \| 42 \| (53, 94) \| 94 \| (9, 50) \| 50 \| Phage tail assembly chaperone protein, E, or 41 or 14 \| Phage tail assembly chaperone protein, E, or 41 or 14; uniclust \| UniRef100\_A0A7W6WBB0 \| 97.9 \| 2.2e-07 \| 4.1e-13 \| 57.7 \| 76 \| (16, 91) \| 94 \| (13, 91) \| 112 \| Tail assembly chaperone E/41/14-like protein \| Tail assembly chaperone E/41/14-like protein; uniclust \| UniRef100\_A0A068CE07 \| 97.9 \| 2.1e-07 \| 4.2e-13 \| 61.9 \| 82 \| (13, 94) \| 94 \| (6, 101) \| 142 \| Uncharacterized protein \| Uncharacterized protein; uniclust \| UniRef100\_A0A095CPF9 \| 97.9 \| 2.4e-07 \| 4.4e-13 \| 58.0 \| 75 \| (14, 89) \| 94 \| (14, 88) \| 112 \| Uncharacterized protein \| Uncharacterized protein; uniclust \| UniRef100\_A0A0K1NGL7 \| 97.9 \| 2.4e-07 \| 4.6e-13 \| 55.6 \| 66 \| (28, 93) \| 94 \| (3, 72) \| 83 \| Phage tail protein \| Phage tail protein; uniclust \| UniRef100\_A0A1I1EZQ6 \| 97.9 \| 2.5e-07 \| 4.7e-13 \| 60.4 \| 78 \| (16, 93) \| 94 \| (49, 126) \| 145 \| Phage tail assembly protein \| Phage tail assembly protein; uniclust \| UniRef100\_UPI00131AA224 \| 97.9 \| 2.8e-07 \| 5.1e-13 \| 56.4 \| 78 \| (13, 90) \| 94 \| (7, 95) \| 102 \| phage tail assembly protein \| phage tail assembly protein; uniclust \| UniRef100\_A0A151FYQ4 \| 97.9 \| 3.1e-07 \| 5.9e-13 \| 60.1 \| 78 \| (16, 93) \| 94 \| (52, 139) \| 146 \| Phage tail assembly protein \| Phage tail assembly protein; uniclust \| UniRef100\_A0A380ALS8 \| 97.9 \| 3.3e-07 \| 6.1e-13 \| 50.4 \| 39 \| (13, 51) \| 94 \| (14, 52) \| 55 \| Uncharacterized protein \| Uncharacterized protein; uniclust \| UniRef100\_UPI00137719BA \| 97.8 \| 3.5e-07 \| 6.5e-13 \| 56.2 \| 74 \| (16, 90) \| 94 \| (6, 83) \| 104 \| phage tail assembly protein \| phage tail assembly protein; uniclust \| UniRef100\_A0A380ZEC4 \| 97.8 \| 3.5e-07 \| 6.6e-13 \| 51.2 \| 46 \| (21, 68) \| 94 \| (11, 57) \| 57 \| Uncharacterized protein \| Uncharacterized protein; uniclust \| UniRef100\_A0A1V5WR09 \| 97.8 \| 3.6e-07 \| 6.6e-13 \| 55.3 \| 78 \| (17, 94) \| 94 \| (7, 84) \| 95 \| Phage tail protein E \| Phage tail protein E; uniclust \| UniRef100\_A0A418VWW7 \| 97.8 \| 3.7e-07 \| 6.9e-13 \| 54.0 \| 62 \| (32, 93) \| 94 \| (2, 67) \| 77 \| Phage tail assembly protein \| Phage tail assembly protein; uniclust \| UniRef100\_UPI002243AD49 \| 97.8 \| 3.8e-07 \| 7e-13 \| 61.3 \| 79 \| (15, 93) \| 94 \| (96, 178) \| 190 \| phage tail assembly protein \| phage tail assembly protein; uniclust \| UniRef100\_A0A395D213 \| 97.8 \| 3.7e-07 \| 7.1e-13 \| 57.9 \| 78 \| (12, 92) \| 94 \| (32, 109) \| 116 \| Uncharacterized protein \| Uncharacterized protein; uniclust \| UniRef100\_A0A9E3GWE3 \| 97.8 \| 4e-07 \| 7.4e-13 \| 56.1 \| 76 \| (18, 93) \| 94 \| (3, 79) \| 105 \| Phage tail assembly protein \| Phage tail assembly protein; uniclust \| UniRef100\_UPI001CD5C949 \| 97.8 \| 4.2e-07 \| 7.7e-13 \| 59.2 \| 83 \| (12, 94) \| 94 \| (16, 98) \| 152 \| phage tail assembly protein \| phage tail assembly protein; uniclust \| UniRef100\_A0A5C7PFE2 \| 97.8 \| 4.4e-07 \| 8e-13 \| 56.2 \| 80 \| (15, 94) \| 94 \| (13, 95) \| 109 \| Phage tail assembly protein \| Phage tail assembly protein; uniclust \| UniRef100\_A0A1M6LDM2 \| 97.8 \| 4.6e-07 \| 8.9e-13 \| 57.6 \| 69 \| (24, 93) \| 94 \| (26, 94) \| 108 \| Phage tail assembly chaperone protein, E, or 41 or 14 \| Phage tail assembly chaperone protein, E, or 41 or 14; uniclust \| UniRef100\_UPI001FEFFE9E \| 97.8 \| 5.1e-07 \| 9.4e-13 \| 60.5 \| 78 \| (16, 93) \| 94 \| (42, 129) \| 186 \| phage tail assembly protein \| phage tail assembly protein; uniclust \| UniRef100\_A0A0D0GXK6 \| 97.8 \| 5e-07 \| 9.7e-13 \| 56.9 \| 80 \| (15, 94) \| 94 \| (13, 105) \| 105 \| Mu-like prophage FluMu gp41 family protein \| Mu-like prophage FluMu gp41 family protein; uniclust \| UniRef100\_UPI00226FF933 \| 97.8 \| 5.4e-07 \| 1e-12 \| 59.0 \| 80 \| (12, 92) \| 94 \| (68, 147) \| 157 \| phage tail assembly protein \| phage tail assembly protein; uniclust \| UniRef100\_A0A2S7JR59 \| 97.8 \| 5.6e-07 \| 1e-12 \| 56.3 \| 80 \| (14, 93) \| 94 \| (12, 93) \| 110 \| Phage tail assembly protein \| Phage tail assembly protein; uniclust \| UniRef100\_A0A011LWS3 \| 97.8 \| 5.5e-07 \| 1.1e-12 \| 57.1 \| 77 \| (16, 92) \| 94 \| (4, 84) \| 108 \| Tail protein \| Tail protein; uniclust \| UniRef100\_A0A6M3ZU73 \| 97.8 \| 5.8e-07 \| 1.1e-12 \| 60.0 \| 74 \| (20, 93) \| 94 \| (2, 84) \| 174 \| Phage tail assembly protein \| Phage tail assembly protein; uniclust \| UniRef100\_A0A1Q6KUF5 \| 97.8 \| 5.8e-07 \| 1.1e-12 \| 59.6 \| 80 \| (13, 92) \| 94 \| (38, 130) \| 136 \| Uncharacterized protein \| Uncharacterized protein; uniclust \| UniRef100\_UPI002094E750 \| 97.8 \| 6.3e-07 \| 1.2e-12 \| 54.1 \| 73 \| (18, 90) \| 94 \| (3, 84) \| 92 \| phage tail assembly protein \| phage tail assembly protein; uniclust \| UniRef100\_A0A1L4D148 \| 97.8 \| 6.4e-07 \| 1.2e-12 \| 55.6 \| 78 \| (16, 93) \| 94 \| (19, 96) \| 106 \| Phage tail assembly protein \| Phage tail assembly protein; uniclust \| UniRef100\_A6WZ02 \| 97.8 \| 6.6e-07 \| 1.2e-12 \| 56.0 \| 73 \| (17, 89) \| 94 \| (2, 92) \| 110 \| Phage tail assembly protein \| Phage tail assembly protein; uniclust \| UniRef100\_A0A1X7MFT2 \| 97.8 \| 7.1e-07 \| 1.3e-12 \| 46.2 \| 35 \| (16, 50) \| 94 \| (3, 37) \| 40 \| Phage tail assembly protein (Fragment) \| Phage tail assembly protein (Fragment); uniclust \| UniRef100\_UPI001CBE614B \| 97.7 \| 7.1e-07 \| 1.3e-12 \| 53.7 \| 61 \| (2, 64) \| 94 \| (5, 66) \| 90 \| phage tail assembly protein \| phage tail assembly protein; uniclust \| UniRef100\_UPI0021F431C5 \| 97.7 \| 7.3e-07 \| 1.3e-12 \| 57.9 \| 78 \| (13, 90) \| 94 \| (36, 114) \| 148 \| phage tail assembly protein \| phage tail assembly protein; uniclust \| UniRef100\_A0A944GS05 \| 97.7 \| 7.5e-07 \| 1.4e-12 \| 58.5 \| 80 \| (12, 92) \| 94 \| (57, 136) \| 158 \| Phage tail assembly protein \| Phage tail assembly protein; uniclust \| UniRef100\_A0A512JPF4 \| 97.7 \| 7.3e-07 \| 1.4e-12 \| 59.4 \| 83 \| (12, 94) \| 94 \| (52, 137) \| 159 \| Uncharacterized protein \| Uncharacterized protein; uniclust \| UniRef100\_UPI0022DE3CC5 \| 97.7 \| 7.8e-07 \| 1.4e-12 \| 49.2 \| 56 \| (37, 94) \| 94 \| (1, 56) \| 56 \| phage tail assembly protein \| phage tail assembly protein; uniclust \| UniRef100\_A0A257KLL7 \| 97.7 \| 7.5e-07 \| 1.4e-12 \| 55.4 \| 76 \| (13, 93) \| 94 \| (8, 83) \| 95 \| Phage tail assembly protein \| Phage tail assembly protein; uniclust \| UniRef100\_A0A431IB45 \| 97.7 \| 8e-07 \| 1.5e-12 \| 57.7 \| 76 \| (12, 90) \| 94 \| (66, 141) \| 147 \| Phage tail assembly protein \| Phage tail assembly protein; uniclust \| UniRef100\_A0A0U2ATQ0 \| 97.7 \| 7.9e-07 \| 1.5e-12 \| 55.1 \| 81 \| (14, 94) \| 94 \| (7, 91) \| 101 \| Phage tail assembly protein \| Phage tail assembly protein; uniclust \| UniRef100\_UPI0013C53023 \| 97.7 \| 8.4e-07 \| 1.5e-12 \| 50.5 \| 58 \| (16, 73) \| 94 \| (7, 64) \| 65 \| phage tail assembly protein \| phage tail assembly protein; uniclust \| UniRef100\_A0A1X9SVU8 \| 97.7 \| 8.1e-07 \| 1.6e-12 \| 55.7 \| 62 \| (33, 94) \| 94 \| (40, 102) \| 102 \| FluMu gp41 family protein \| FluMu gp41 family protein; uniclust \| UniRef100\_A0A094ZPX7 \| 97.7 \| 8.2e-07 \| 1.6e-12 \| 57.9 \| 82 \| (12, 94) \| 94 \| (22, 109) \| 125 \| Phage tail assembly protein \| Phage tail assembly protein; uniclust \| UniRef100\_A0A6N9P3E0 \| 97.7 \| 9e-07 \| 1.7e-12 \| 58.8 \| 80 \| (14, 93) \| 94 \| (76, 167) \| 172 \| Phage tail assembly protein \| Phage tail assembly protein; uniclust \| UniRef100\_A0A1D8UTE9 \| 97.7 \| 8.8e-07 \| 1.7e-12 \| 56.6 \| 81 \| (13, 93) \| 94 \| (13, 102) \| 115 \| Phage tail assembly protein \| Phage tail assembly protein; uniclust \| UniRef100\_E9L500 \| 97.7 \| 9.2e-07 \| 1.7e-12 \| 54.8 \| 61 \| (33, 93) \| 94 \| (37, 97) \| 104 \| Phage tail assembly protein \| Phage tail assembly protein; uniclust \| UniRef100\_A0A948AK24 \| 97.7 \| 9.3e-07 \| 1.7e-12 \| 56.9 \| 81 \| (13, 93) \| 94 \| (42, 126) \| 138 \| Phage tail assembly protein \| Phage tail assembly protein; uniclust \| UniRef100\_A0A176Z3V3 \| 97.7 \| 9.4e-07 \| 1.8e-12 \| 56.2 \| 76 \| (16, 91) \| 94 \| (3, 93) \| 110 \| Phage tail assembly protein \| Phage tail assembly protein; uniclust \| UniRef100\_K0JHK7 \| 97.7 \| 1e-06 \| 2e-12 \| 54.8 \| 81 \| (13, 93) \| 94 \| (4, 85) \| 101 \| Phage protein \| Phage protein; uniclust \| UniRef100\_A0A0J9EDU6 \| 97.7 \| 1.1e-06 \| 2e-12 \| 51.3 \| 59 \| (35, 94) \| 94 \| (1, 59) \| 75 \| Phage protein \| Phage protein; uniclust \| UniRef100\_A0A7L5Y296 \| 97.7 \| 1.1e-06 \| 2.1e-12 \| 60.7 \| 77 \| (16, 94) \| 94 \| (77, 153) \| 185 \| Uncharacterized protein \| Uncharacterized protein; uniclust \| UniRef100\_A0A1Q9A2K6 \| 97.7 \| 1.1e-06 \| 2.1e-12 \| 58.5 \| 75 \| (14, 91) \| 94 \| (73, 147) \| 157 \| Phage tail assembly protein \| Phage tail assembly protein; uniclust \| UniRef100\_UPI0003955618 \| 97.7 \| 1.2e-06 \| 2.2e-12 \| 55.7 \| 81 \| (13, 93) \| 94 \| (31, 115) \| 126 \| phage tail assembly protein \| phage tail assembly protein; uniclust \| UniRef100\_UPI001F249746 \| 97.7 \| 1.2e-06 \| 2.2e-12 \| 55.1 \| 62 \| (18, 80) \| 94 \| (2, 64) \| 107 \| phage tail assembly protein \| phage tail assembly protein; uniclust \| UniRef100\_A0A0S4V3E5 \| 97.7 \| 1.3e-06 \| 2.4e-12 \| 56.3 \| 74 \| (16, 90) \| 94 \| (6, 83) \| 131 \| Hypothethical protein \| Hypothethical protein; uniclust \| UniRef100\_R7I6D8 \| 97.6 \| 1.4e-06 \| 2.6e-12 \| 53.4 \| 79 \| (16, 94) \| 94 \| (3, 87) \| 95 \| Phage tail assembly protein \| Phage tail assembly protein; uniclust \| UniRef100\_UPI0021CEA9E1 \| 97.6 \| 1.4e-06 \| 2.6e-12 \| 52.7 \| 74 \| (18, 91) \| 94 \| (2, 75) \| 92 \| phage tail assembly protein \| phage tail assembly protein; uniclust \| UniRef100\_A0A968GD13 \| 97.6 \| 1.3e-06 \| 2.6e-12 \| 58.7 \| 85 \| (9, 93) \| 94 \| (62, 149) \| 151 \| Uncharacterized protein \| Uncharacterized protein; uniclust \| UniRef100\_A0A8I2FQ78 \| 97.6 \| 1.4e-06 \| 2.6e-12 \| 53.2 \| 81 \| (13, 94) \| 94 \| (17, 98) \| 98 \| Phage tail assembly protein \| Phage tail assembly protein; uniclust \| UniRef100\_A0A1M5RFG1 \| 97.6 \| 1.4e-06 \| 2.7e-12 \| 52.7 \| 76 \| (18, 94) \| 94 \| (2, 78) \| 87 \| Phage tail assembly chaperone protein, E, or 41 or 14 \| Phage tail assembly chaperone protein, E, or 41 or 14; uniclust \| UniRef100\_A0A7V8JT65 \| 97.6 \| 1.4e-06 \| 2.7e-12 \| 53.5 \| 78 \| (16, 94) \| 94 \| (6, 83) \| 95 \| Phage tail assembly protein \| Phage tail assembly protein; uniclust \| UniRef100\_A0A3A6P6S0 \| 97.6 \| 1.5e-06 \| 2.8e-12 \| 54.8 \| 83 \| (12, 94) \| 94 \| (14, 107) \| 120 \| Phage tail assembly protein \| Phage tail assembly protein; uniclust \| UniRef100\_A0A437N1X6 \| 97.6 \| 1.5e-06 \| 2.8e-12 \| 53.9 \| 77 \| (17, 93) \| 94 \| (12, 92) \| 108 \| Phage tail assembly protein \| Phage tail assembly protein; uniclust \| UniRef100\_A0A3A9B6H8 \| 97.6 \| 1.5e-06 \| 2.9e-12 \| 56.5 \| 82 \| (12, 93) \| 94 \| (24, 118) \| 122 \| Phage tail assembly protein \| Phage tail assembly protein; uniclust \| UniRef100\_A0A9D8STM0 \| 97.6 \| 1.6e-06 \| 3e-12 \| 52.8 \| 76 \| (18, 93) \| 94 \| (2, 78) \| 89 \| Uncharacterized protein \| Uncharacterized protein; uniclust \| UniRef100\_A0A7H1NUI4 \| 97.6 \| 1.8e-06 \| 3.4e-12 \| 55.5 \| 80 \| (14, 93) \| 94 \| (26, 110) \| 123 \| Phage tail assembly chaperone protein \| Phage tail assembly chaperone protein; uniclust \| UniRef100\_A0A225SNI8 \| 97.6 \| 1.9e-06 \| 3.6e-12 \| 54.0 \| 76 \| (18, 93) \| 94 \| (28, 105) \| 115 \| Phage tail assembly protein \| Phage tail assembly protein; uniclust \| UniRef100\_A0A1F4HNB5 \| 97.6 \| 2e-06 \| 3.7e-12 \| 54.0 \| 77 \| (18, 94) \| 94 \| (24, 107) \| 115 \| Phage tail protein \| Phage tail protein; uniclust \| UniRef100\_A0A8J7UJ23 \| 97.6 \| 2e-06 \| 3.7e-12 \| 51.4 \| 74 \| (18, 91) \| 94 \| (3, 78) \| 85 \| Phage tail assembly protein \| Phage tail assembly protein; uniclust \| UniRef100\_UPI001D10B551 \| 97.6 \| 2.1e-06 \| 3.8e-12 \| 57.4 \| 79 \| (16, 94) \| 94 \| (71, 153) \| 176 \| phage tail assembly protein \| phage tail assembly protein; uniclust \| UniRef100\_A0A1G3JSY5 \| 97.6 \| 1.9e-06 \| 3.9e-12 \| 53.8 \| 65 \| (30, 94) \| 94 \| (8, 76) \| 90 \| Phage tail protein \| Phage tail protein; uniclust \| UniRef100\_A0A433X0G4 \| 97.6 \| 2.1e-06 \| 3.9e-12 \| 55.9 \| 90 \| (2, 93) \| 94 \| (38, 139) \| 147 \| Phage tail protein \| Phage tail protein; uniclust \| UniRef100\_A0A076LV38 \| 97.6 \| 2.1e-06 \| 3.9e-12 \| 53.4 \| 81 \| (14, 94) \| 94 \| (5, 89) \| 98 \| Putative bacteriophage protein \| Putative bacteriophage protein; uniclust \| UniRef100\_A0A0D2JFD8 \| 97.6 \| 2.2e-06 \| 4e-12 \| 52.2 \| 79 \| (15, 93) \| 94 \| (4, 92) \| 95 \| Phage tail assembly protein \| Phage tail assembly protein; uniclust \| UniRef100\_A0A174TN41 \| 97.6 \| 2e-06 \| 4e-12 \| 57.5 \| 81 \| (13, 93) \| 94 \| (33, 134) \| 139 \| Uncharacterized protein \| Uncharacterized protein; uniclust \| UniRef100\_A0A5M9U1M0 \| 97.6 \| 2.1e-06 \| 4.1e-12 \| 55.5 \| 78 \| (16, 93) \| 94 \| (20, 114) \| 118 \| Phage tail assembly protein \| Phage tail assembly protein; uniclust \| UniRef100\_A0A165XHA6 \| 97.6 \| 2.3e-06 \| 4.2e-12 \| 56.7 \| 75 \| (13, 94) \| 94 \| (70, 149) \| 165 \| Phage tail assembly protein \| Phage tail assembly protein; uniclust \| UniRef100\_UPI0016750E23 \| 97.6 \| 2.6e-06 \| 4.7e-12 \| 54.0 \| 82 \| (12, 93) \| 94 \| (27, 112) \| 121 \| phage tail assembly protein \| phage tail assembly protein; uniclust \| UniRef100\_A0A5N8AC27 \| 97.6 \| 2.6e-06 \| 4.8e-12 \| 55.1 \| 78 \| (16, 93) \| 94 \| (43, 124) \| 139 \| Phage tail assembly protein \| Phage tail assembly protein; uniclust \| UniRef100\_A0A367Q8P7 \| 97.5 \| 2.8e-06 \| 5.3e-12 \| 59.1 \| 82 \| (2, 93) \| 94 \| (109, 197) \| 203 \| Uncharacterized protein \| Uncharacterized protein; uniclust \| UniRef100\_A0A0Q6BTN7 \| 97.5 \| 2.8e-06 \| 5.5e-12 \| 52.3 \| 66 \| (17, 86) \| 94 \| (11, 76) \| 89 \| Phage tail assembly protein \| Phage tail assembly protein; uniclust \| UniRef100\_A0A6L8HXB9 \| 97.5 \| 3.1e-06 \| 5.7e-12 \| 60.1 \| 77 \| (16, 93) \| 94 \| (22, 99) \| 280 \| SMP-30/Gluconolactonase/LRE-like region domain-containing protein \| SMP-30/Gluconolactonase/LRE-like region domain-containing protein; uniclust \| UniRef100\_UPI001378214B \| 97.5 \| 3.1e-06 \| 5.7e-12 \| 54.2 \| 80 \| (14, 93) \| 94 \| (26, 109) \| 129 \| phage tail assembly protein \| phage tail assembly protein; uniclust \| UniRef100\_A0A9E9B835 \| 97.5 \| 3.2e-06 \| 5.8e-12 \| 55.0 \| 74 \| (14, 91) \| 94 \| (62, 135) \| 143 \| Phage tail assembly protein \| Phage tail assembly protein; uniclust \| UniRef100\_S9S610 \| 97.5 \| 3.3e-06 \| 6.1e-12 \| 52.6 \| 77 \| (16, 92) \| 94 \| (14, 92) \| 109 \| Putative phage related protein \| Putative phage related protein; uniclust \| UniRef100\_A0A4S8Q2I0 \| 97.5 \| 3.3e-06 \| 6.3e-12 \| 52.1 \| 77 \| (17, 93) \| 94 \| (2, 78) \| 93 \| Uncharacterized protein \| Uncharacterized protein; uniclust \| UniRef100\_A0A564WJQ0 \| 97.5 \| 3.5e-06 \| 6.4e-12 \| 52.1 \| 79 \| (16, 94) \| 94 \| (2, 85) \| 103 \| Uncharacterized protein \| Uncharacterized protein; uniclust \| UniRef100\_A0A485CDJ1 \| 97.5 \| 3.6e-06 \| 6.7e-12 \| 58.2 \| 81 \| (13, 93) \| 94 \| (129, 213) \| 225 \| Phage major tail tube protein \| Phage major tail tube protein; uniclust \| UniRef100\_A0A0B1YRZ3 \| 97.5 \| 3.5e-06 \| 6.7e-12 \| 53.4 \| 81 \| (13, 93) \| 94 \| (3, 92) \| 108 \| Phage tail assembly protein \| Phage tail assembly protein; uniclust \| UniRef100\_A0A961Y6K2 \| 97.5 \| 3.9e-06 \| 7.2e-12 \| 54.1 \| 73 \| (18, 94) \| 94 \| (5, 81) \| 128 \| Phage tail assembly protein \| Phage tail assembly protein; uniclust \| UniRef100\_A0A176EWX2 \| 97.5 \| 4.3e-06 \| 7.9e-12 \| 50.4 \| 72 \| (18, 89) \| 94 \| (3, 75) \| 86 \| Phage tail protein \| Phage tail protein; uniclust \| UniRef100\_A0A1I4TJ36 \| 97.5 \| 4e-06 \| 8e-12 \| 55.3 \| 77 \| (16, 93) \| 94 \| (3, 96) \| 127 \| Uncharacterized protein \| Uncharacterized protein; uniclust \| UniRef100\_A0A7W6RF20 \| 97.5 \| 4.3e-06 \| 8.1e-12 \| 54.7 \| 83 \| (12, 94) \| 94 \| (15, 102) \| 132 \| Tail assembly chaperone E/41/14-like protein \| Tail assembly chaperone E/41/14-like protein; uniclust \| UniRef100\_A0A1X3I4M1 \| 97.4 \| 5e-06 \| 9.3e-12 \| 52.8 \| 80 \| (15, 94) \| 94 \| (27, 112) \| 113 \| Phage tail protein E \| Phage tail protein E; uniclust \| UniRef100\_A0A8B2QYK0 \| 97.4 \| 5.1e-06 \| 9.4e-12 \| 41.3 \| 32 \| (62, 93) \| 94 \| (1, 32) \| 32 \| Phage tail assembly protein \| Phage tail assembly protein; uniclust \| UniRef100\_A0A1E3G5Y7 \| 97.4 \| 5.8e-06 \| 1.1e-11 \| 48.9 \| 72 \| (17, 93) \| 94 \| (3, 74) \| 78 \| Phage tail assembly protein \| Phage tail assembly protein; uniclust \| UniRef100\_A0A4D7YHU8 \| 97.4 \| 5.8e-06 \| 1.1e-11 \| 48.2 \| 60 \| (33, 93) \| 94 \| (2, 61) \| 72 \| Phage tail assembly protein \| Phage tail assembly protein; uniclust \| UniRef100\_UPI0006863B40 \| 97.4 \| 6.2e-06 \| 1.1e-11 \| 56.4 \| 76 \| (16, 93) \| 94 \| (113, 188) \| 203 \| phage tail assembly protein \| phage tail assembly protein; uniclust \| UniRef100\_UPI0015D41224 \| 97.4 \| 6.3e-06 \| 1.2e-11 \| 52.5 \| 73 \| (18, 92) \| 94 \| (2, 75) \| 123 \| phage tail assembly protein \| phage tail assembly protein; uniclust \| UniRef100\_A0A1X0WB04 \| 97.4 \| 5.9e-06 \| 1.2e-11 \| 57.2 \| 83 \| (12, 94) \| 94 \| (83, 169) \| 183 \| Uncharacterized protein \| Uncharacterized protein; uniclust \| UniRef100\_A0A2S4MDR7 \| 97.4 \| 6.2e-06 \| 1.2e-11 \| 52.9 \| 79 \| (16, 94) \| 94 \| (23, 107) \| 118 \| Tail assembly chaperone E/41/14-like protein \| Tail assembly chaperone E/41/14-like protein; uniclust \| UniRef100\_A0A2A4XUJ7 \| 97.4 \| 6.3e-06 \| 1.2e-11 \| 52.0 \| 77 \| (17, 93) \| 94 \| (4, 89) \| 106 \| Phage tail assembly protein \| Phage tail assembly protein; uniclust \| UniRef100\_UPI001FB9F85B \| 97.4 \| 6.6e-06 \| 1.2e-11 \| 57.4 \| 82 \| (13, 94) \| 94 \| (113, 200) \| 226 \| hypothetical protein \| hypothetical protein; uniclust \| UniRef100\_A0A087M272 \| 97.4 \| 6.7e-06 \| 1.3e-11 \| 51.3 \| 78 \| (15, 93) \| 94 \| (13, 91) \| 100 \| Phage tail assembly protein \| Phage tail assembly protein; uniclust \| UniRef100\_A0A097P6T1 \| 97.4 \| 7.1e-06 \| 1.3e-11 \| 53.7 \| 77 \| (18, 94) \| 94 \| (3, 91) \| 147 \| Tail assembly chaperone \| Tail assembly chaperone; uniclust \| UniRef100\_A0A7C1VCE0 \| 97.4 \| 7.3e-06 \| 1.3e-11 \| 52.8 \| 79 \| (15, 93) \| 94 \| (53, 133) \| 133 \| Phage tail assembly protein \| Phage tail assembly protein; uniclust \| UniRef100\_A0A150HJQ6 \| 97.4 \| 7.2e-06 \| 1.4e-11 \| 53.1 \| 80 \| (15, 94) \| 94 \| (29, 114) \| 126 \| Phage tail protein E \| Phage tail protein E; uniclust \| UniRef100\_A0A1T4WVF8 \| 97.4 \| 8.1e-06 \| 1.5e-11 \| 55.4 \| 70 \| (17, 92) \| 94 \| (113, 183) \| 189 \| Phage tail assembly chaperone protein, E, or 41 or 14 \| Phage tail assembly chaperone protein, E, or 41 or 14; uniclust \| UniRef100\_A0A078L7Q4 \| 97.4 \| 8.1e-06 \| 1.5e-11 \| 55.5 \| 80 \| (14, 93) \| 94 \| (28, 115) \| 167 \| Phage tail assembly protein \| Phage tail assembly protein; uniclust \| UniRef100\_A0A0A3TNQ9 \| 97.4 \| 8.2e-06 \| 1.6e-11 \| 50.9 \| 77 \| (17, 93) \| 94 \| (2, 91) \| 97 \| Phage tail assembly protein \| Phage tail assembly protein; uniclust \| UniRef100\_A0A4R2P5C3 \| 97.4 \| 8.6e-06 \| 1.6e-11 \| 51.4 \| 81 \| (13, 93) \| 94 \| (22, 106) \| 115 \| Tail assembly chaperone E/41/14-like protein \| Tail assembly chaperone E/41/14-like protein; uniclust \| UniRef100\_D8JWC8 \| 97.3 \| 8.8e-06 \| 1.6e-11 \| 52.6 \| 74 \| (19, 92) \| 94 \| (34, 126) \| 134 \| Phage protein \| Phage protein; uniclust \| UniRef100\_UPI0013EA8D57 \| 97.3 \| 9e-06 \| 1.6e-11 \| 51.9 \| 79 \| (13, 92) \| 94 \| (35, 123) \| 123 \| phage tail assembly protein \| phage tail assembly protein; uniclust \| UniRef100\_A0A2S6N2U8 \| 97.3 \| 9.5e-06 \| 1.7e-11 \| 50.8 \| 74 \| (17, 92) \| 94 \| (33, 106) \| 109 \| Phage tail assembly protein \| Phage tail assembly protein; uniclust \| UniRef100\_A0A4Q2U0L3 \| 97.3 \| 9.5e-06 \| 1.8e-11 \| 51.7 \| 80 \| (14, 93) \| 94 \| (11, 102) \| 122 \| Phage tail assembly protein \| Phage tail assembly protein; uniclust \| UniRef100\_A0A0U1DB82 \| 97.3 \| 9.5e-06 \| 1.8e-11 \| 45.1 \| 45 \| (50, 94) \| 94 \| (5, 49) \| 55 \| Putative phage related protein \| Putative phage related protein; uniclust \| UniRef100\_S5Y192 \| 97.3 \| 9.6e-06 \| 1.8e-11 \| 51.0 \| 73 \| (18, 90) \| 94 \| (2, 80) \| 112 \| Uncharacterized protein \| Uncharacterized protein; uniclust \| UniRef100\_A0A560T0I5 \| 97.3 \| 1.1e-05 \| 2.1e-11 \| 50.6 \| 76 \| (16, 91) \| 94 \| (4, 80) \| 104 \| Tail assembly chaperone E/41/14-like protein \| Tail assembly chaperone E/41/14-like protein; uniclust \| UniRef100\_A0A7W6EB82 \| 97.3 \| 1.2e-05 \| 2.1e-11 \| 53.1 \| 68 \| (16, 87) \| 94 \| (75, 142) \| 154 \| Phage tail assembly protein \| Phage tail assembly protein; uniclust \| UniRef100\_UPI001EDF0F93 \| 97.3 \| 1.2e-05 \| 2.1e-11 \| 46.4 \| 56 \| (37, 92) \| 94 \| (1, 56) \| 67 \| phage tail assembly protein \| phage tail assembly protein; uniclust \| UniRef100\_UPI00051682BF \| 97.3 \| 1.2e-05 \| 2.1e-11 \| 46.2 \| 46 \| (3, 50) \| 94 \| (12, 57) \| 65 \| phage tail assembly protein \| phage tail assembly protein; uniclust \| UniRef100\_UPI002226FE03 \| 97.3 \| 1.2e-05 \| 2.1e-11 \| 51.9 \| 77 \| (16, 92) \| 94 \| (31, 118) \| 132 \| phage tail assembly protein \| phage tail assembly protein; uniclust \| UniRef100\_A0A2A5EJW3 \| 97.3 \| 1.2e-05 \| 2.3e-11 \| 54.4 \| 77 \| (15, 91) \| 94 \| (5, 96) \| 186 \| Phage tail assembly protein \| Phage tail assembly protein; uniclust \| UniRef100\_A0A7X6FRD4 \| 97.3 \| 1.3e-05 \| 2.3e-11 \| 47.3 \| 33 \| (18, 50) \| 94 \| (3, 35) \| 76 \| Phage tail assembly protein \| Phage tail assembly protein; uniclust \| UniRef100\_A0A0S4XLQ8 \| 97.3 \| 1.3e-05 \| 2.4e-11 \| 49.5 \| 61 \| (33, 94) \| 94 \| (21, 81) \| 90 \| Phage tail assembly protein \| Phage tail assembly protein; uniclust \| UniRef100\_A0A526RPY9 \| 97.3 \| 1.4e-05 \| 2.5e-11 \| 48.6 \| 70 \| (17, 90) \| 94 \| (6, 75) \| 90 \| Phage tail assembly protein (Fragment) \| Phage tail assembly protein (Fragment); uniclust \| UniRef100\_A0A2G1CU23 \| 97.3 \| 1.4e-05 \| 2.5e-11 \| 49.1 \| 77 \| (16, 93) \| 94 \| (13, 89) \| 96 \| Phage tail assembly protein \| Phage tail assembly protein; uniclust \| UniRef100\_UPI000277D392 \| 97.3 \| 1.4e-05 \| 2.5e-11 \| 51.1 \| 39 \| (15, 53) \| 94 \| (2, 40) \| 123 \| phage tail assembly protein \| phage tail assembly protein; uniclust \| UniRef100\_A0A317H578 \| 97.3 \| 1.4e-05 \| 2.6e-11 \| 53.4 \| 76 \| (13, 91) \| 94 \| (8, 83) \| 167 \| Phage tail assembly protein \| Phage tail assembly protein; uniclust \| UniRef100\_A0A2S3W4R7 \| 97.2 \| 1.5e-05 \| 2.8e-11 \| 55.5 \| 80 \| (14, 93) \| 94 \| (97, 182) \| 191 \| Uncharacterized protein \| Uncharacterized protein; uniclust \| UniRef100\_UPI0006528B58 \| 97.2 \| 1.6e-05 \| 2.9e-11 \| 46.2 \| 34 \| (18, 51) \| 94 \| (3, 36) \| 69 \| phage tail assembly protein \| phage tail assembly protein; uniclust \| UniRef100\_A0A1C6E0X6 \| 97.2 \| 1.5e-05 \| 3e-11 \| 57.5 \| 78 \| (17, 94) \| 94 \| (45, 137) \| 237 \| Phage tail assembly protein \| Phage tail assembly protein; uniclust \| UniRef100\_A0A8X6GSQ5 \| 97.2 \| 1.6e-05 \| 3e-11 \| 56.0 \| 65 \| (30, 94) \| 94 \| (164, 228) \| 234 \| Phage tail tube protein FII \| Phage tail tube protein FII; uniclust \| UniRef100\_UPI001C12BF21 \| 97.2 \| 1.8e-05 \| 3.3e-11 \| 47.9 \| 72 \| (21, 93) \| 94 \| (1, 73) \| 87 \| phage tail assembly protein \| phage tail assembly protein; uniclust \| UniRef100\_A0A0B7J8H1 \| 97.2 \| 1.7e-05 \| 3.3e-11 \| 55.8 \| 80 \| (14, 94) \| 94 \| (100, 180) \| 196 \| Phage tail assembly protein \| Phage tail assembly protein; uniclust \| UniRef100\_A0A859FB61 \| 97.2 \| 1.9e-05 \| 3.4e-11 \| 52.7 \| 81 \| (14, 94) \| 94 \| (71, 163) \| 164 \| Phage tail assembly protein \| Phage tail assembly protein; uniclust \| UniRef100\_A0A1G7PX14 \| 97.2 \| 1.9e-05 \| 3.4e-11 \| 51.5 \| 81 \| (12, 94) \| 94 \| (33, 114) \| 139 \| Phage tail assembly chaperone protein, E, or 41 or 14 \| Phage tail assembly chaperone protein, E, or 41 or 14; uniclust \| UniRef100\_UPI00186A80F2 \| 97.2 \| 1.9e-05 \| 3.4e-11 \| 48.1 \| 73 \| (17, 90) \| 94 \| (3, 75) \| 91 \| phage tail assembly protein \| phage tail assembly protein; uniclust \| UniRef100\_A0A085AFM8 \| 97.2 \| 1.9e-05 \| 3.6e-11 \| 55.6 \| 76 \| (16, 91) \| 94 \| (4, 87) \| 209 \| Phage protein \| Phage protein; uniclust \| UniRef100\_A0A812QV43 \| 97.2 \| 2.1e-05 \| 3.8e-11 \| 63.9 \| 75 \| (16, 91) \| 94 \| (1023, 1097) \| 1104 \| GpFI protein \| GpFI protein; uniclust \| UniRef100\_UPI00235E67D6 \| 97.2 \| 2.2e-05 \| 4e-11 \| 47.7 \| 74 \| (17, 91) \| 94 \| (2, 75) \| 89 \| phage tail assembly protein \| phage tail assembly protein; uniclust \| UniRef100\_A0A1L3SPW8 \| 97.2 \| 2.2e-05 \| 4.2e-11 \| 51.1 \| 79 \| (16, 94) \| 94 \| (2, 94) \| 119 \| Uncharacterized protein \| Uncharacterized protein; uniclust \| UniRef100\_A0A067W7B7 \| 97.1 \| 2.6e-05 \| 4.7e-11 \| 45.5 \| 56 \| (25, 82) \| 94 \| (15, 70) \| 70 \| Uncharacterized protein \| Uncharacterized protein; uniclust \| UniRef100\_A0A6I3KEY4 \| 97.1 \| 2.6e-05 \| 4.8e-11 \| 46.4 \| 63 \| (16, 88) \| 94 \| (3, 66) \| 79 \| Phage tail assembly chaperone protein, E, or 41 or 14 \| Phage tail assembly chaperone protein, E, or 41 or 14; uniclust \| UniRef100\_A0A261QN74 \| 97.1 \| 2.9e-05 \| 5.5e-11 \| 48.2 \| 39 \| (56, 94) \| 94 \| (25, 63) \| 89 \| Phage tail assembly protein \| Phage tail assembly protein; uniclust \| UniRef100\_A0A9D1NLP2 \| 97.1 \| 3e-05 \| 5.5e-11 \| 47.7 \| 74 \| (17, 90) \| 94 \| (3, 80) \| 95 \| Phage tail assembly protein \| Phage tail assembly protein; uniclust \| UniRef100\_A0A8S7UTI4 \| 97.1 \| 3.1e-05 \| 5.6e-11 \| 41.6 \| 35 \| (17, 51) \| 94 \| (2, 36) \| 45 \| Phage tail assembly protein (Fragment) \| Phage tail assembly protein (Fragment); uniclust \| UniRef100\_A0A1W0CCH5 \| 97.1 \| 3.4e-05 \| 6.3e-11 \| 47.4 \| 77 \| (16, 92) \| 94 \| (3, 89) \| 95 \| Phage tail assembly protein \| Phage tail assembly protein; uniclust \| UniRef100\_A0A2U1TJU4 \| 97.1 \| 3.4e-05 \| 6.4e-11 \| 48.9 \| 80 \| (15, 94) \| 94 \| (5, 90) \| 105 \| Phage tail protein \| Phage tail protein; uniclust \| UniRef100\_A0A094JIN5 \| 97.1 \| 3.5e-05 \| 6.5e-11 \| 45.6 \| 62 \| (18, 79) \| 94 \| (2, 73) \| 76 \| Uncharacterized protein \| Uncharacterized protein; uniclust \| UniRef100\_A0A6L3YW32 \| 97.1 \| 3.7e-05 \| 6.9e-11 \| 51.5 \| 33 \| (18, 50) \| 94 \| (3, 35) \| 155 \| Phage tail assembly protein \| Phage tail assembly protein; uniclust \| UniRef100\_A0A9E8T1G1 \| 97.1 \| 3.8e-05 \| 7e-11 \| 50.7 \| 74 \| (18, 91) \| 94 \| (41, 116) \| 149 \| Phage tail assembly protein \| Phage tail assembly protein; uniclust \| UniRef100\_A0A0H3ZVR6 \| 97.1 \| 3.5e-05 \| 7.1e-11 \| 54.9 \| 61 \| (33, 94) \| 94 \| (130, 190) \| 203 \| Phage tail assembly protein \| Phage tail assembly protein; uniclust \| UniRef100\_A0A255XY73 \| 97.1 \| 4e-05 \| 7.3e-11 \| 59.5 \| 81 \| (13, 94) \| 94 \| (505, 587) \| 595 \| Uncharacterized protein \| Uncharacterized protein; uniclust \| UniRef100\_UPI00082FD7B7 \| 97.1 \| 4e-05 \| 7.3e-11 \| 50.0 \| 73 \| (16, 88) \| 94 \| (5, 85) \| 137 \| phage tail assembly protein \| phage tail assembly protein; uniclust \| UniRef100\_A0A2H5DVZ8 \| 97.1 \| 4e-05 \| 7.3e-11 \| 51.0 \| 78 \| (13, 90) \| 94 \| (64, 151) \| 158 \| Phage tail assembly protein \| Phage tail assembly protein; uniclust \| UniRef100\_UPI001CC07461 \| 97.1 \| 4.1e-05 \| 7.6e-11 \| 49.9 \| 75 \| (16, 90) \| 94 \| (4, 79) \| 137 \| phage tail assembly protein \| phage tail assembly protein; uniclust \| UniRef100\_A0A252BR55 \| 97.1 \| 4e-05 \| 7.7e-11 \| 51.2 \| 88 \| (2, 94) \| 94 \| (33, 124) \| 140 \| Phage tail assembly protein \| Phage tail assembly protein; uniclust \| UniRef100\_A0A482IXJ4 \| 97.1 \| 4.3e-05 \| 7.8e-11 \| 50.5 \| 74 \| (17, 90) \| 94 \| (42, 116) \| 149 \| Phage tail assembly protein \| Phage tail assembly protein; uniclust \| UniRef100\_UPI0022AFC0B4 \| 97.0 \| 4.3e-05 \| 7.9e-11 \| 48.6 \| 80 \| (14, 94) \| 94 \| (28, 107) \| 116 \| hypothetical protein \| hypothetical protein; uniclust \| UniRef100\_A0A2R7ILE3 \| 97.0 \| 4.5e-05 \| 8.4e-11 \| 49.1 \| 76 \| (14, 90) \| 94 \| (2, 87) \| 114 \| Phage tail assembly protein \| Phage tail assembly protein; uniclust \| UniRef100\_UPI00226FC3D2 \| 97.0 \| 4.6e-05 \| 8.5e-11 \| 50.1 \| 79 \| (15, 93) \| 94 \| (3, 98) \| 145 \| hypothetical protein \| hypothetical protein; uniclust \| UniRef100\_A0A1G0GGY7 \| 97.0 \| 4.5e-05 \| 8.8e-11 \| 49.6 \| 78 \| (14, 93) \| 94 \| (4, 91) \| 113 \| Phage tail assembly protein \| Phage tail assembly protein; uniclust \| UniRef100\_UPI000A356193 \| 97.0 \| 4.9e-05 \| 9e-11 \| 43.3 \| 59 \| (36, 94) \| 94 \| (2, 61) \| 61 \| phage tail assembly protein \| phage tail assembly protein; uniclust \| UniRef100\_A0A072TPP1 \| 97.0 \| 5e-05 \| 9.1e-11 \| 60.0 \| 80 \| (14, 93) \| 94 \| (625, 710) \| 726 \| Bacteriophage mu tail sheath protein \| Bacteriophage mu tail sheath protein; uniclust \| UniRef100\_A0A0B1Q7C3 \| 97.0 \| 4.9e-05 \| 9.5e-11 \| 49.2 \| 78 \| (17, 94) \| 94 \| (3, 88) \| 113 \| Uncharacterized protein \| Uncharacterized protein; uniclust \| UniRef100\_A0A9D9A8Q3 \| 97.0 \| 5.5e-05 \| 1e-10 \| 50.0 \| 78 \| (16, 94) \| 94 \| (43, 121) \| 148 \| Phage tail assembly protein \| Phage tail assembly protein; uniclust \| UniRef100\_A0A444L4U4 \| 97.0 \| 5.6e-05 \| 1e-10 \| 47.4 \| 67 \| (20, 86) \| 94 \| (2, 84) \| 105 \| Phage tail assembly protein \| Phage tail assembly protein; uniclust \| UniRef100\_A0A858X005 \| 97.0 \| 6.1e-05 \| 1.2e-10 \| 49.2 \| 77 \| (17, 93) \| 94 \| (2, 101) \| 125 \| Phage tail assembly protein \| Phage tail assembly protein; uniclust \| UniRef100\_A0A1J5HSC3 \| 97.0 \| 6.4e-05 \| 1.2e-10 \| 48.2 \| 79 \| (16, 94) \| 94 \| (6, 102) \| 121 \| Phage tail protein \| Phage tail protein; uniclust \| UniRef100\_UPI000E25BC78 \| 97.0 \| 6.5e-05 \| 1.2e-10 \| 48.2 \| 73 \| (16, 89) \| 94 \| (4, 76) \| 122 \| hypothetical protein \| hypothetical protein; uniclust \| UniRef100\_UPI00082C10DB \| 97.0 \| 6.6e-05 \| 1.2e-10 \| 52.9 \| 77 \| (17, 93) \| 94 \| (45, 126) \| 239 \| phage tail assembly protein \| phage tail assembly protein; uniclust \| UniRef100\_A0A432QTZ1 \| 97.0 \| 6.7e-05 \| 1.2e-10 \| 51.0 \| 77 \| (16, 92) \| 94 \| (11, 88) \| 182 \| Phage tail assembly protein \| Phage tail assembly protein; uniclust \| UniRef100\_A0A1Z4RAB7 \| 97.0 \| 6.8e-05 \| 1.2e-10 \| 46.9 \| 60 \| (35, 94) \| 94 \| (33, 100) \| 103 \| Phage tail assembly protein \| Phage tail assembly protein; uniclust \| UniRef100\_A0A1E3WJ29 \| 97.0 \| 6.8e-05 \| 1.2e-10 \| 47.6 \| 77 \| (17, 93) \| 94 \| (27, 105) \| 114 \| Phage tail assembly protein \| Phage tail assembly protein; uniclust \| UniRef100\_UPI001BA5F6AD \| 97.0 \| 6.9e-05 \| 1.3e-10 \| 48.0 \| 73 \| (13, 86) \| 94 \| (34, 106) \| 121 \| hypothetical protein \| hypothetical protein; uniclust \| UniRef100\_UPI00234EDE79 \| 96.9 \| 7e-05 \| 1.3e-10 \| 50.6 \| 89 \| (3, 94) \| 94 \| (65, 166) \| 173 \| phage tail assembly protein \| phage tail assembly protein; uniclust \| UniRef100\_A0A4Q3U4W6 \| 96.9 \| 7.2e-05 \| 1.3e-10 \| 48.7 \| 77 \| (17, 93) \| 94 \| (40, 120) \| 133 \| Phage tail assembly protein \| Phage tail assembly protein; uniclust \| UniRef100\_UPI001A5E140C \| 96.9 \| 7.4e-05 \| 1.4e-10 \| 48.5 \| 78 \| (16, 93) \| 94 \| (27, 123) \| 130 \| phage tail assembly protein \| phage tail assembly protein; uniclust \| UniRef100\_A0A3G7TJ80 \| 96.9 \| 7.6e-05 \| 1.4e-10 \| 49.0 \| 76 \| (15, 90) \| 94 \| (3, 79) \| 136 \| Phage tail assembly protein \| Phage tail assembly protein; uniclust \| UniRef100\_A0A2W5AE53 \| 96.9 \| 7.9e-05 \| 1.4e-10 \| 46.2 \| 77 \| (17, 93) \| 94 \| (3, 85) \| 98 \| Phage tail assembly protein \| Phage tail assembly protein; uniclust \| UniRef100\_A0A5C7JB85 \| 96.9 \| 8.7e-05 \| 1.6e-10 \| 45.7 \| 74 \| (18, 92) \| 94 \| (5, 79) \| 94 \| Phage tail assembly protein \| Phage tail assembly protein; uniclust \| UniRef100\_V4NZH5 \| 96.9 \| 8.8e-05 \| 1.6e-10 \| 46.0 \| 74 \| (16, 90) \| 94 \| (13, 95) \| 97 \| Phage protein \| Phage protein; uniclust \| UniRef100\_UPI0016521F9B \| 96.9 \| 8.8e-05 \| 1.6e-10 \| 51.5 \| 79 \| (15, 93) \| 94 \| (77, 156) \| 210 \| phage tail assembly protein \| phage tail assembly protein; uniclust \| UniRef100\_A0A090IQA6 \| 96.9 \| 8.5e-05 \| 1.6e-10 \| 53.6 \| 80 \| (14, 94) \| 94 \| (143, 223) \| 236 \| Uncharacterized phage protein \| Uncharacterized phage protein; uniclust \| UniRef100\_A0A268TN79 \| 96.9 \| 8.5e-05 \| 1.6e-10 \| 46.1 \| 62 \| (33, 94) \| 94 \| (18, 81) \| 86 \| Phage tail assembly protein \| Phage tail assembly protein; uniclust \| UniRef100\_A0RRS4 \| 96.9 \| 9e-05 \| 1.7e-10 \| 47.2 \| 62 \| (33, 94) \| 94 \| (46, 108) \| 108 \| Phage tail assembly protein \| Phage tail assembly protein; uniclust \| UniRef100\_A0A2W6XD89 \| 96.9 \| 9.7e-05 \| 1.8e-10 \| 47.9 \| 78 \| (16, 93) \| 94 \| (30, 111) \| 124 \| Phage tail assembly protein \| Phage tail assembly protein; uniclust \| UniRef100\_X1CAN2 \| 96.9 \| 9.8e-05 \| 1.8e-10 \| 46.9 \| 80 \| (13, 93) \| 94 \| (32, 113) \| 113 \| Uncharacterized protein \| Uncharacterized protein; uniclust \| UniRef100\_A0A358C7S8 \| 96.9 \| 9.9e-05 \| 1.8e-10 \| 46.5 \| 79 \| (16, 94) \| 94 \| (5, 91) \| 107 \| Phage tail assembly protein \| Phage tail assembly protein; uniclust \| UniRef100\_UPI00082C10DB \| 96.9 \| 0.0001 \| 1.9e-10 \| 52.0 \| 79 \| (15, 93) \| 94 \| (137, 219) \| 239 \| phage tail assembly protein \| phage tail assembly protein; uniclust \| UniRef100\_UPI00200A87DE \| 96.9 \| 0.00011 \| 2e-10 \| 50.1 \| 77 \| (14, 92) \| 94 \| (94, 170) \| 180 \| phage tail assembly protein \| phage tail assembly protein |
| Top keywords  (threshold 1.00e-03 (evalue)) | **tail, Phage, assembly, or, E, chaperone, 14\_like, Mu\_like, FluMu, gp41** |
| Output files | ../../similar\_sequences/20\_FANPEZAQ\_CDS\_0020\_merged.svg ../../similar\_sequences/20\_FANPEZAQ\_CDS\_0020\_pdb70.a3m ../../similar\_sequences/20\_FANPEZAQ\_CDS\_0020\_pdb70.hhr ../../similar\_sequences/20\_FANPEZAQ\_CDS\_0020\_uniclust.a3m ../../similar\_sequences/20\_FANPEZAQ\_CDS\_0020\_uniclust.hhr |

#### Structure prediction (AlphaFold)^2^

|  |  |
| --- | --- |
| Stats | xml version="1.0" encoding="utf-8" standalone="no"?       2024-09-02T21:09:19.650353 image/svg+xml   Matplotlib v3.7.2, https://matplotlib.org/ |
| Predicted structure | **NGL Viewer Controls:**  - Center: *Left-Click* - Rotate: *Left-Click + Drag* - Translate: *Right-Click + Drag* - Zoom: *Shift + Left-Click + Drag* |
| Output files | ../../predicted\_structures/20\_FANPEZAQ\_CDS\_0020/features.pkl ../../predicted\_structures/20\_FANPEZAQ\_CDS\_0020/ranked\_0.pdb ../../predicted\_structures/20\_FANPEZAQ\_CDS\_0020/ranked\_0\_plots.svg ../../predicted\_structures/20\_FANPEZAQ\_CDS\_0020/result\_model\_1\_ptm\_pred\_0.pkl |

#### Structure similarity search results (Foldseek)^3^

|  |  |
| --- | --- |
| Structure databases searched | Pdb, Afdb-proteome, Afdb-uniprot50 |
| Results, scheme(s)  (Top layers only, threshold 1.00e-02 (evalue)) | xml version="1.0" encoding="utf-8" standalone="no"?       2024-09-02T21:10:44.797899 image/svg+xml   Matplotlib v3.7.2, https://matplotlib.org/ |
| Results, table  (threshold 1.00e-02 (evalue)) | db \| id \| prob \| evalue \| bits \| fident \| alnlen \| mismatch \| gapopen \| qstart \| qend \| tstart \| tend \| name \| description; afdb-proteome \| AF-G3XCX2-F1-MODEL\_V4 \| 1.0 \| 1.424e-08 \| 314 \| 0.387 \| 93 \| 54 \| 2 \| 2 \| 93 \| 5 \| 95 \| Uncharacterized protein \| Uncharacterized protein; afdb-proteome \| AF-Q8ZMV1-F1-MODEL\_V4 \| 1.0 \| 0.00306 \| 127 \| 0.13 \| 92 \| 70 \| 3 \| 10 \| 94 \| 1 \| 89 \| Fels-2 prophage protein \| Fels-2 prophage protein; afdb-proteome \| AF-A0A0H3GM31-F1-MODEL\_V4 \| 1.0 \| 0.003962 \| 125 \| 0.147 \| 88 \| 63 \| 3 \| 15 \| 94 \| 5 \| 88 \| Phage tail protein E \| Phage tail protein E; afdb-uniprot50 \| AF-A0A653KXL9-F1-MODEL\_V4 \| 1.0 \| 9.457e-13 \| 514 \| 0.663 \| 95 \| 30 \| 2 \| 1 \| 94 \| 1 \| 94 \| Phage tail assembly protein \| Phage tail assembly protein; afdb-uniprot50 \| AF-A0A7R8GJ05-F1-MODEL\_V4 \| 1.0 \| 2.337e-12 \| 506 \| 0.623 \| 93 \| 34 \| 1 \| 2 \| 94 \| 25 \| 116 \| Uncharacterized protein \| Uncharacterized protein; afdb-uniprot50 \| AF-K5YQ81-F1-MODEL\_V4 \| 1.0 \| 2.019e-10 \| 422 \| 0.494 \| 93 \| 45 \| 1 \| 2 \| 94 \| 16 \| 106 \| Uncharacterized protein \| Uncharacterized protein; afdb-uniprot50 \| AF-A0A2Z3I5X9-F1-MODEL\_V4 \| 1.0 \| 3.174e-10 \| 408 \| 0.489 \| 96 \| 45 \| 2 \| 1 \| 94 \| 1 \| 94 \| Phage tail assembly protein \| Phage tail assembly protein; afdb-uniprot50 \| AF-A0A0D0QRI8-F1-MODEL\_V4 \| 1.0 \| 7.352e-10 \| 399 \| 0.616 \| 86 \| 32 \| 1 \| 9 \| 94 \| 2 \| 86 \| Contig\_41, whole genome shotgun sequence \| Contig\_41, whole genome shotgun sequence; afdb-uniprot50 \| AF-A0A315BIP8-F1-MODEL\_V4 \| 1.0 \| 1.817e-09 \| 398 \| 0.56 \| 82 \| 36 \| 0 \| 13 \| 94 \| 10 \| 91 \| Uncharacterized protein \| Uncharacterized protein; afdb-uniprot50 \| AF-A0A7V8FKG1-F1-MODEL\_V4 \| 1.0 \| 1.083e-09 \| 393 \| 0.473 \| 95 \| 48 \| 2 \| 1 \| 94 \| 1 \| 94 \| Uncharacterized protein \| Uncharacterized protein; afdb-uniprot50 \| AF-A0A158E8B3-F1-MODEL\_V4 \| 1.0 \| 7.352e-10 \| 391 \| 0.482 \| 85 \| 42 \| 2 \| 6 \| 90 \| 14 \| 96 \| Uncharacterized protein \| Uncharacterized protein; afdb-uniprot50 \| AF-B8KZU3-F1-MODEL\_V4 \| 1.0 \| 1.938e-09 \| 386 \| 0.473 \| 93 \| 47 \| 1 \| 2 \| 94 \| 6 \| 96 \| Uncharacterized protein \| Uncharacterized protein; afdb-uniprot50 \| AF-A0A2W5N4W8-F1-MODEL\_V4 \| 1.0 \| 7.058e-09 \| 368 \| 0.478 \| 92 \| 46 \| 2 \| 3 \| 94 \| 35 \| 124 \| Phage tail assembly protein \| Phage tail assembly protein; afdb-uniprot50 \| AF-A0A143DHP2-F1-MODEL\_V4 \| 1.0 \| 1.04e-08 \| 359 \| 0.518 \| 79 \| 38 \| 0 \| 16 \| 94 \| 5 \| 83 \| Uncharacterized protein \| Uncharacterized protein; afdb-uniprot50 \| AF-A0A410UF81-F1-MODEL\_V4 \| 1.0 \| 4.31e-08 \| 334 \| 0.384 \| 91 \| 54 \| 1 \| 2 \| 92 \| 30 \| 118 \| Phage tail assembly protein \| Phage tail assembly protein; afdb-uniprot50 \| AF-A0A7G7TJ95-F1-MODEL\_V4 \| 1.0 \| 6.352e-08 \| 333 \| 0.354 \| 93 \| 58 \| 1 \| 2 \| 94 \| 5 \| 95 \| Phage tail assembly protein \| Phage tail assembly protein; afdb-uniprot50 \| AF-A0A2U1XZ20-F1-MODEL\_V4 \| 1.0 \| 1.674e-07 \| 329 \| 0.384 \| 78 \| 48 \| 0 \| 15 \| 92 \| 5 \| 82 \| Uncharacterized protein \| Uncharacterized protein; afdb-uniprot50 \| AF-A0A2K4GEU8-F1-MODEL\_V4 \| 1.0 \| 4.598e-08 \| 327 \| 0.378 \| 95 \| 56 \| 2 \| 1 \| 94 \| 8 \| 100 \| Phage tail assembly protein \| Phage tail assembly protein; afdb-uniprot50 \| AF-A0A5S9Q2R5-F1-MODEL\_V4 \| 1.0 \| 1.293e-07 \| 321 \| 0.412 \| 80 \| 47 \| 0 \| 15 \| 94 \| 4 \| 83 \| Uncharacterized protein \| Uncharacterized protein; afdb-uniprot50 \| AF-A0A7Y7J7Q5-F1-MODEL\_V4 \| 1.0 \| 6.352e-08 \| 321 \| 0.34 \| 94 \| 59 \| 2 \| 2 \| 94 \| 5 \| 96 \| Phage tail assembly protein \| Phage tail assembly protein; afdb-uniprot50 \| AF-A0A4Q3YYA7-F1-MODEL\_V4 \| 1.0 \| 5.232e-08 \| 319 \| 0.397 \| 93 \| 51 \| 3 \| 3 \| 94 \| 10 \| 98 \| Phage tail assembly protein \| Phage tail assembly protein; afdb-uniprot50 \| AF-A0A2E3N1U5-F1-MODEL\_V4 \| 1.0 \| 1.905e-07 \| 314 \| 0.373 \| 83 \| 52 \| 0 \| 12 \| 94 \| 4 \| 86 \| Uncharacterized protein \| Uncharacterized protein; afdb-uniprot50 \| AF-A0A0M1I5Y4-F1-MODEL\_V4 \| 1.0 \| 9.985e-08 \| 314 \| 0.442 \| 95 \| 49 \| 3 \| 2 \| 94 \| 4 \| 96 \| Uncharacterized protein \| Uncharacterized protein; afdb-uniprot50 \| AF-A0A2K9M0P8-F1-MODEL\_V4 \| 1.0 \| 9.36e-08 \| 313 \| 0.308 \| 94 \| 62 \| 2 \| 2 \| 94 \| 4 \| 95 \| Phage tail assembly protein \| Phage tail assembly protein; afdb-uniprot50 \| AF-A0A291LZ34-F1-MODEL\_V4 \| 1.0 \| 2.033e-07 \| 312 \| 0.519 \| 77 \| 37 \| 0 \| 16 \| 92 \| 4 \| 80 \| Uncharacterized protein \| Uncharacterized protein; afdb-uniprot50 \| AF-A0A149SWH3-F1-MODEL\_V4 \| 1.0 \| 1.379e-07 \| 310 \| 0.309 \| 97 \| 62 \| 2 \| 3 \| 94 \| 6 \| 102 \| Uncharacterized protein \| Uncharacterized protein; afdb-uniprot50 \| AF-A0A0P8Y9Q0-F1-MODEL\_V4 \| 1.0 \| 9.985e-08 \| 307 \| 0.41 \| 95 \| 53 \| 3 \| 1 \| 94 \| 1 \| 93 \| Uncharacterized protein \| Uncharacterized protein; afdb-uniprot50 \| AF-A0A6I2J072-F1-MODEL\_V4 \| 1.0 \| 2.808e-07 \| 306 \| 0.341 \| 85 \| 56 \| 0 \| 10 \| 94 \| 1 \| 85 \| Phage tail assembly protein \| Phage tail assembly protein; afdb-uniprot50 \| AF-A0A443ZHA6-F1-MODEL\_V4 \| 1.0 \| 1.57e-07 \| 306 \| 0.382 \| 94 \| 55 \| 2 \| 2 \| 94 \| 5 \| 96 \| Phage tail assembly protein \| Phage tail assembly protein; afdb-uniprot50 \| AF-A0A1G6JG81-F1-MODEL\_V4 \| 1.0 \| 1.905e-07 \| 305 \| 0.423 \| 92 \| 49 \| 2 \| 5 \| 94 \| 11 \| 100 \| Phage tail assembly chaperone protein, E, or 41 or 14 \| Phage tail assembly chaperone protein, E, or 41 or 14; afdb-uniprot50 \| AF-A0A3M4SLW6-F1-MODEL\_V4 \| 1.0 \| 1.786e-07 \| 304 \| 0.351 \| 94 \| 58 \| 2 \| 2 \| 94 \| 7 \| 98 \| Uncharacterized protein \| Uncharacterized protein; afdb-uniprot50 \| AF-A0A3M2V2F9-F1-MODEL\_V4 \| 1.0 \| 2.168e-07 \| 304 \| 0.329 \| 94 \| 60 \| 2 \| 2 \| 94 \| 11 \| 102 \| Uncharacterized protein \| Uncharacterized protein; afdb-uniprot50 \| AF-B7RNN7-F1-MODEL\_V4 \| 1.0 \| 3.636e-07 \| 301 \| 0.414 \| 94 \| 53 \| 2 \| 2 \| 94 \| 4 \| 96 \| Uncharacterized protein \| Uncharacterized protein; afdb-uniprot50 \| AF-A0A515BE98-F1-MODEL\_V4 \| 1.0 \| 2.995e-07 \| 300 \| 0.378 \| 95 \| 55 \| 3 \| 2 \| 94 \| 3 \| 95 \| Phage tail assembly protein \| Phage tail assembly protein; afdb-uniprot50 \| AF-A0A1T4W2E5-F1-MODEL\_V4 \| 1.0 \| 2.995e-07 \| 298 \| 0.379 \| 79 \| 49 \| 0 \| 16 \| 94 \| 2 \| 80 \| Phage tail assembly chaperone protein, E, or 41 or 14 \| Phage tail assembly chaperone protein, E, or 41 or 14; afdb-uniprot50 \| AF-A0A1N7LRA5-F1-MODEL\_V4 \| 1.0 \| 5.023e-07 \| 295 \| 0.318 \| 91 \| 61 \| 1 \| 3 \| 93 \| 5 \| 94 \| Phage tail assembly chaperone protein, E, or 41 or 14 \| Phage tail assembly chaperone protein, E, or 41 or 14; afdb-uniprot50 \| AF-A0A0C1KHK2-F1-MODEL\_V4 \| 1.0 \| 2.168e-07 \| 294 \| 0.268 \| 93 \| 65 \| 2 \| 2 \| 93 \| 7 \| 97 \| Uncharacterized protein \| Uncharacterized protein; afdb-uniprot50 \| AF-A0A554X0U8-F1-MODEL\_V4 \| 1.0 \| 8.985e-07 \| 291 \| 0.389 \| 77 \| 47 \| 0 \| 18 \| 94 \| 7 \| 83 \| Phage tail assembly chaperone protein E, or 41 or 14 \| Phage tail assembly chaperone protein E, or 41 or 14; afdb-uniprot50 \| AF-A0A6A4RGH5-F1-MODEL\_V4 \| 1.0 \| 7.896e-07 \| 291 \| 0.336 \| 95 \| 61 \| 2 \| 1 \| 94 \| 1 \| 94 \| Phage tail assembly protein \| Phage tail assembly protein; afdb-uniprot50 \| AF-A0A0Q8Z8Q7-F1-MODEL\_V4 \| 1.0 \| 5.023e-07 \| 289 \| 0.4 \| 90 \| 51 \| 2 \| 4 \| 92 \| 1 \| 88 \| Uncharacterized protein \| Uncharacterized protein; afdb-uniprot50 \| AF-A0A2S0PEE6-F1-MODEL\_V4 \| 1.0 \| 3.195e-07 \| 289 \| 0.447 \| 76 \| 42 \| 0 \| 18 \| 93 \| 4 \| 79 \| Phage tail assembly protein \| Phage tail assembly protein; afdb-uniprot50 \| AF-A0A266NDD3-F1-MODEL\_V4 \| 1.0 \| 4.709e-07 \| 289 \| 0.351 \| 94 \| 58 \| 2 \| 2 \| 94 \| 7 \| 98 \| Uncharacterized protein \| Uncharacterized protein; afdb-uniprot50 \| AF-A0A2N8BUS6-F1-MODEL\_V4 \| 1.0 \| 7.402e-07 \| 288 \| 0.358 \| 92 \| 56 \| 2 \| 2 \| 92 \| 7 \| 96 \| Phage tail assembly protein \| Phage tail assembly protein; afdb-uniprot50 \| AF-A0A0C5VFB7-F1-MODEL\_V4 \| 1.0 \| 1.023e-06 \| 287 \| 0.369 \| 84 \| 53 \| 0 \| 11 \| 94 \| 15 \| 98 \| Uncharacterized protein \| Uncharacterized protein; afdb-uniprot50 \| AF-A0A1Q6U7M3-F1-MODEL\_V4 \| 1.0 \| 7.896e-07 \| 286 \| 0.35 \| 80 \| 52 \| 0 \| 15 \| 94 \| 3 \| 82 \| Uncharacterized protein \| Uncharacterized protein; afdb-uniprot50 \| AF-A0A2R3H0M0-F1-MODEL\_V4 \| 1.0 \| 8.423e-07 \| 286 \| 0.36 \| 86 \| 54 \| 1 \| 10 \| 94 \| 1 \| 86 \| Uncharacterized protein \| Uncharacterized protein; afdb-uniprot50 \| AF-A0A4V6IKT7-F1-MODEL\_V4 \| 1.0 \| 5.023e-07 \| 284 \| 0.441 \| 77 \| 43 \| 0 \| 18 \| 94 \| 5 \| 81 \| Bacteriophage tail protein gp41 putative \| Bacteriophage tail protein gp41 putative; afdb-uniprot50 \| AF-A0A2V4FUR3-F1-MODEL\_V4 \| 1.0 \| 6.939e-07 \| 284 \| 0.312 \| 96 \| 62 \| 3 \| 1 \| 94 \| 1 \| 94 \| Phage tail assembly protein \| Phage tail assembly protein; afdb-uniprot50 \| AF-A0A7J0BX99-F1-MODEL\_V4 \| 1.0 \| 1.164e-06 \| 283 \| 0.454 \| 77 \| 42 \| 0 \| 18 \| 94 \| 4 \| 80 \| Uncharacterized protein \| Uncharacterized protein; afdb-uniprot50 \| AF-A0A248LIV2-F1-MODEL\_V4 \| 1.0 \| 1.413e-06 \| 280 \| 0.379 \| 79 \| 49 \| 0 \| 16 \| 94 \| 3 \| 81 \| Phage\_TAC\_7 domain containing protein \| Phage\_TAC\_7 domain containing protein; afdb-uniprot50 \| AF-A0A239C819-F1-MODEL\_V4 \| 1.0 \| 1.951e-06 \| 276 \| 0.341 \| 79 \| 52 \| 0 \| 16 \| 94 \| 2 \| 80 \| Phage tail assembly chaperone protein, E, or 41 or 14 \| Phage tail assembly chaperone protein, E, or 41 or 14; afdb-uniprot50 \| AF-H0SKG3-F1-MODEL\_V4 \| 1.0 \| 3.272e-06 \| 276 \| 0.346 \| 78 \| 51 \| 0 \| 15 \| 92 \| 11 \| 88 \| Uncharacterized protein \| Uncharacterized protein; afdb-uniprot50 \| AF-A0A5E7KKS2-F1-MODEL\_V4 \| 1.0 \| 2.369e-06 \| 276 \| 0.284 \| 95 \| 65 \| 2 \| 1 \| 94 \| 3 \| 95 \| Uncharacterized protein \| Uncharacterized protein; afdb-uniprot50 \| AF-A0A7W5B8R9-F1-MODEL\_V4 \| 1.0 \| 1.413e-06 \| 275 \| 0.347 \| 92 \| 57 \| 2 \| 3 \| 93 \| 2 \| 91 \| Uncharacterized protein \| Uncharacterized protein; afdb-uniprot50 \| AF-A0A4Y6UC53-F1-MODEL\_V4 \| 1.0 \| 1.507e-06 \| 275 \| 0.31 \| 100 \| 61 \| 4 \| 2 \| 94 \| 14 \| 112 \| Phage tail assembly protein \| Phage tail assembly protein; afdb-uniprot50 \| AF-A0A743CDA4-F1-MODEL\_V4 \| 1.0 \| 2.369e-06 \| 274 \| 0.27 \| 85 \| 62 \| 0 \| 10 \| 94 \| 1 \| 85 \| Phage tail assembly protein \| Phage tail assembly protein; afdb-uniprot50 \| AF-A0A4U1KTT9-F1-MODEL\_V4 \| 1.0 \| 1.091e-06 \| 272 \| 0.329 \| 91 \| 58 \| 2 \| 3 \| 92 \| 10 \| 98 \| Phage tail assembly protein \| Phage tail assembly protein; afdb-uniprot50 \| AF-C6BVX2-F1-MODEL\_V4 \| 1.0 \| 2.369e-06 \| 270 \| 0.415 \| 77 \| 45 \| 0 \| 18 \| 94 \| 3 \| 79 \| Uncharacterized protein \| Uncharacterized protein; afdb-uniprot50 \| AF-A0A849VJT3-F1-MODEL\_V4 \| 1.0 \| 4.52e-06 \| 267 \| 0.428 \| 77 \| 44 \| 0 \| 18 \| 94 \| 5 \| 81 \| Phage tail assembly protein \| Phage tail assembly protein; afdb-uniprot50 \| AF-A0A7C8LWZ0-F1-MODEL\_V4 \| 1.0 \| 1.829e-06 \| 266 \| 0.329 \| 79 \| 52 \| 1 \| 16 \| 94 \| 3 \| 80 \| Uncharacterized protein \| Uncharacterized protein; afdb-uniprot50 \| AF-A0A8A5W0U8-F1-MODEL\_V4 \| 1.0 \| 5.853e-06 \| 266 \| 0.337 \| 77 \| 51 \| 0 \| 18 \| 94 \| 3 \| 79 \| Phage tail assembly protein \| Phage tail assembly protein; afdb-uniprot50 \| AF-A0A1Y3NT57-F1-MODEL\_V4 \| 1.0 \| 5.144e-06 \| 261 \| 0.279 \| 93 \| 64 \| 2 \| 2 \| 93 \| 12 \| 102 \| Uncharacterized protein \| Uncharacterized protein; afdb-uniprot50 \| AF-A0A554XCC7-F1-MODEL\_V4 \| 1.0 \| 4.52e-06 \| 260 \| 0.32 \| 78 \| 52 \| 1 \| 16 \| 93 \| 2 \| 78 \| Phage tail assembly chaperone protein, E, or 41 or 14 \| Phage tail assembly chaperone protein, E, or 41 or 14; afdb-uniprot50 \| AF-A0A853I2L0-F1-MODEL\_V4 \| 1.0 \| 7.58e-06 \| 260 \| 0.289 \| 76 \| 54 \| 0 \| 18 \| 93 \| 4 \| 79 \| Phage tail assembly protein \| Phage tail assembly protein; afdb-uniprot50 \| AF-A0A516SAU7-F1-MODEL\_V4 \| 1.0 \| 2.527e-06 \| 260 \| 0.279 \| 93 \| 63 \| 2 \| 1 \| 93 \| 1 \| 89 \| Phage tail assembly protein \| Phage tail assembly protein; afdb-uniprot50 \| AF-A0A1H9YDQ2-F1-MODEL\_V4 \| 1.0 \| 4.52e-06 \| 258 \| 0.367 \| 79 \| 50 \| 0 \| 16 \| 94 \| 3 \| 81 \| Phage tail assembly chaperone protein, E, or 41 or 14 \| Phage tail assembly chaperone protein, E, or 41 or 14; afdb-uniprot50 \| AF-A0A3A9HX51-F1-MODEL\_V4 \| 1.0 \| 6.244e-06 \| 258 \| 0.381 \| 76 \| 47 \| 0 \| 18 \| 93 \| 6 \| 81 \| Phage tail assembly protein \| Phage tail assembly protein; afdb-uniprot50 \| AF-A0A161YCG8-F1-MODEL\_V4 \| 1.0 \| 8.626e-06 \| 258 \| 0.342 \| 76 \| 50 \| 0 \| 18 \| 93 \| 4 \| 79 \| Uncharacterized protein \| Uncharacterized protein; afdb-uniprot50 \| AF-A1W3J4-F1-MODEL\_V4 \| 1.0 \| 1.164e-06 \| 258 \| 0.326 \| 101 \| 61 \| 1 \| 1 \| 94 \| 1 \| 101 \| Uncharacterized protein \| Uncharacterized protein; afdb-uniprot50 \| AF-A0A3G2IAI1-F1-MODEL\_V4 \| 1.0 \| 7.106e-06 \| 258 \| 0.302 \| 86 \| 59 \| 1 \| 10 \| 94 \| 1 \| 86 \| Phage tail assembly protein \| Phage tail assembly protein; afdb-uniprot50 \| AF-A0A2N0CXC7-F1-MODEL\_V4 \| 1.0 \| 6.244e-06 \| 254 \| 0.302 \| 86 \| 59 \| 1 \| 10 \| 94 \| 1 \| 86 \| Phage tail assembly protein \| Phage tail assembly protein; afdb-uniprot50 \| AF-A0A5A9EPG0-F1-MODEL\_V4 \| 1.0 \| 5.144e-06 \| 252 \| 0.487 \| 80 \| 38 \| 1 \| 18 \| 94 \| 2 \| 81 \| Phage tail assembly protein \| Phage tail assembly protein; afdb-uniprot50 \| AF-A0A212KXH5-F1-MODEL\_V4 \| 1.0 \| 5.144e-06 \| 252 \| 0.32 \| 78 \| 53 \| 0 \| 15 \| 92 \| 5 \| 82 \| Uncharacterized protein \| Uncharacterized protein; afdb-uniprot50 \| AF-A0A1N6I0V2-F1-MODEL\_V4 \| 1.0 \| 5.144e-06 \| 251 \| 0.329 \| 79 \| 52 \| 1 \| 16 \| 94 \| 3 \| 80 \| Phage tail assembly chaperone protein, E, or 41 or 14 \| Phage tail assembly chaperone protein, E, or 41 or 14; afdb-uniprot50 \| AF-A0A6N7JJ06-F1-MODEL\_V4 \| 1.0 \| 8.086e-06 \| 249 \| 0.341 \| 79 \| 52 \| 0 \| 16 \| 94 \| 2 \| 80 \| Phage tail assembly protein \| Phage tail assembly protein; afdb-uniprot50 \| AF-A0A5Q0THE6-F1-MODEL\_V4 \| 1.0 \| 2.527e-06 \| 249 \| 0.255 \| 94 \| 70 \| 0 \| 1 \| 94 \| 5 \| 98 \| Uncharacterized protein \| Uncharacterized protein; afdb-uniprot50 \| AF-A0A2W4T4A0-F1-MODEL\_V4 \| 1.0 \| 9.202e-06 \| 247 \| 0.192 \| 83 \| 67 \| 0 \| 11 \| 93 \| 1 \| 83 \| Uncharacterized protein \| Uncharacterized protein; afdb-uniprot50 \| AF-G4CJF6-F1-MODEL\_V4 \| 1.0 \| 9.202e-06 \| 247 \| 0.313 \| 83 \| 55 \| 2 \| 13 \| 94 \| 13 \| 94 \| Uncharacterized protein \| Uncharacterized protein; afdb-uniprot50 \| AF-A0A1E3LH51-F1-MODEL\_V4 \| 1.0 \| 9.816e-06 \| 244 \| 0.395 \| 81 \| 48 \| 1 \| 13 \| 92 \| 6 \| 86 \| Uncharacterized protein \| Uncharacterized protein; afdb-uniprot50 \| AF-A0A125GZ96-F1-MODEL\_V4 \| 1.0 \| 1.271e-05 \| 243 \| 0.412 \| 80 \| 46 \| 1 \| 16 \| 94 \| 3 \| 82 \| Uncharacterized protein \| Uncharacterized protein; afdb-uniprot50 \| AF-A0A5E4XFR1-F1-MODEL\_V4 \| 1.0 \| 3.491e-06 \| 242 \| 0.27 \| 100 \| 67 \| 2 \| 1 \| 94 \| 10 \| 109 \| Uncharacterized protein \| Uncharacterized protein; afdb-uniprot50 \| AF-A0A7X3U083-F1-MODEL\_V4 \| 1.0 \| 6.661e-06 \| 241 \| 0.325 \| 83 \| 55 \| 1 \| 10 \| 92 \| 1 \| 82 \| Phage tail assembly protein \| Phage tail assembly protein; afdb-uniprot50 \| AF-A0A5V6NLU6-F1-MODEL\_V4 \| 1.0 \| 2.426e-05 \| 241 \| 0.275 \| 80 \| 57 \| 1 \| 16 \| 94 \| 9 \| 88 \| Phage tail assembly protein \| Phage tail assembly protein; afdb-uniprot50 \| AF-A0A7Y8GD21-F1-MODEL\_V4 \| 1.0 \| 1.271e-05 \| 241 \| 0.311 \| 93 \| 61 \| 3 \| 1 \| 92 \| 6 \| 96 \| Phage tail assembly protein \| Phage tail assembly protein; afdb-uniprot50 \| AF-A0A1C3ELA5-F1-MODEL\_V4 \| 1.0 \| 4.339e-05 \| 239 \| 0.223 \| 76 \| 59 \| 0 \| 18 \| 93 \| 3 \| 78 \| Uncharacterized protein \| Uncharacterized protein; afdb-uniprot50 \| AF-A0A103RA62-F1-MODEL\_V4 \| 1.0 \| 1.873e-05 \| 238 \| 0.412 \| 80 \| 46 \| 1 \| 16 \| 94 \| 3 \| 82 \| Uncharacterized protein \| Uncharacterized protein; afdb-uniprot50 \| AF-A0A345DE52-F1-MODEL\_V4 \| 1.0 \| 8.086e-06 \| 237 \| 0.255 \| 86 \| 64 \| 0 \| 9 \| 94 \| 1 \| 86 \| Uncharacterized protein \| Uncharacterized protein; afdb-uniprot50 \| AF-A0A495BJ37-F1-MODEL\_V4 \| 1.0 \| 5.487e-06 \| 237 \| 0.41 \| 78 \| 45 \| 1 \| 16 \| 92 \| 2 \| 79 \| Tail assembly chaperone E/41/14-like protein \| Tail assembly chaperone E/41/14-like protein; afdb-uniprot50 \| AF-V4PY80-F1-MODEL\_V4 \| 1.0 \| 1.446e-05 \| 236 \| 0.309 \| 84 \| 56 \| 2 \| 10 \| 91 \| 1 \| 84 \| Phage tail protein \| Phage tail protein; afdb-uniprot50 \| AF-A0A5P9F0G6-F1-MODEL\_V4 \| 1.0 \| 7.58e-06 \| 236 \| 0.279 \| 86 \| 53 \| 1 \| 18 \| 94 \| 5 \| 90 \| Uncharacterized protein \| Uncharacterized protein; afdb-uniprot50 \| AF-A0A2P5LZ45-F1-MODEL\_V4 \| 1.0 \| 5.853e-06 \| 235 \| 0.407 \| 81 \| 45 \| 2 \| 14 \| 94 \| 3 \| 80 \| Uncharacterized protein \| Uncharacterized protein; afdb-uniprot50 \| AF-A0A1X7L2K2-F1-MODEL\_V4 \| 1.0 \| 9.816e-06 \| 235 \| 0.289 \| 83 \| 55 \| 1 \| 16 \| 94 \| 2 \| 84 \| Phage tail assembly chaperone protein, E, or 41 or 14 \| Phage tail assembly chaperone protein, E, or 41 or 14; afdb-uniprot50 \| AF-G1USF7-F1-MODEL\_V4 \| 1.0 \| 2.274e-05 \| 234 \| 0.294 \| 78 \| 55 \| 0 \| 15 \| 92 \| 5 \| 82 \| Uncharacterized protein \| Uncharacterized protein; afdb-uniprot50 \| AF-A0A4T2ACC4-F1-MODEL\_V4 \| 1.0 \| 2.76e-05 \| 234 \| 0.32 \| 81 \| 55 \| 0 \| 14 \| 94 \| 4 \| 84 \| Phage tail assembly protein \| Phage tail assembly protein; afdb-uniprot50 \| AF-A0A0W0IQA4-F1-MODEL\_V4 \| 1.0 \| 1.543e-05 \| 233 \| 0.352 \| 85 \| 54 \| 1 \| 10 \| 93 \| 1 \| 85 \| Uncharacterized protein \| Uncharacterized protein; afdb-uniprot50 \| AF-A0A3G2IKE7-F1-MODEL\_V4 \| 1.0 \| 1.047e-05 \| 232 \| 0.313 \| 99 \| 63 \| 1 \| 1 \| 94 \| 1 \| 99 \| Phage tail assembly protein \| Phage tail assembly protein; afdb-uniprot50 \| AF-A0A2S2E568-F1-MODEL\_V4 \| 1.0 \| 5.268e-05 \| 231 \| 0.363 \| 77 \| 49 \| 0 \| 18 \| 94 \| 9 \| 85 \| Uncharacterized protein \| Uncharacterized protein; afdb-uniprot50 \| AF-F2BEW6-F1-MODEL\_V4 \| 1.0 \| 3.575e-05 \| 230 \| 0.277 \| 83 \| 58 \| 2 \| 13 \| 94 \| 28 \| 109 \| Uncharacterized protein \| Uncharacterized protein; afdb-uniprot50 \| AF-A0A853I978-F1-MODEL\_V4 \| 1.0 \| 6.821e-05 \| 229 \| 0.253 \| 79 \| 59 \| 0 \| 16 \| 94 \| 2 \| 80 \| Phage tail assembly protein \| Phage tail assembly protein; afdb-uniprot50 \| AF-A0A644XKN4-F1-MODEL\_V4 \| 1.0 \| 2.945e-05 \| 229 \| 0.256 \| 78 \| 58 \| 0 \| 15 \| 92 \| 5 \| 82 \| Uncharacterized protein \| Uncharacterized protein; afdb-uniprot50 \| AF-A0A554XFV5-F1-MODEL\_V4 \| 1.0 \| 8.086e-06 \| 229 \| 0.354 \| 93 \| 55 \| 2 \| 1 \| 92 \| 1 \| 89 \| Phage tail assembly chaperone protein, E, or 41 or 14 \| Phage tail assembly chaperone protein, E, or 41 or 14; afdb-uniprot50 \| AF-A0A1B6VVH9-F1-MODEL\_V4 \| 1.0 \| 2.76e-05 \| 228 \| 0.337 \| 83 \| 53 \| 2 \| 13 \| 94 \| 13 \| 94 \| Uncharacterized protein \| Uncharacterized protein; afdb-uniprot50 \| AF-A8U2I1-F1-MODEL\_V4 \| 1.0 \| 4.938e-05 \| 227 \| 0.237 \| 80 \| 61 \| 0 \| 14 \| 93 \| 9 \| 88 \| Uncharacterized protein \| Uncharacterized protein; afdb-uniprot50 \| AF-A0A1A9VKI4-F1-MODEL\_V4 \| 1.0 \| 7.762e-05 \| 227 \| 0.298 \| 77 \| 54 \| 0 \| 16 \| 92 \| 113 \| 189 \| Uncharacterized protein \| Uncharacterized protein; afdb-uniprot50 \| AF-A0A1H8TDI4-F1-MODEL\_V4 \| 1.0 \| 9.202e-06 \| 226 \| 0.315 \| 95 \| 62 \| 2 \| 1 \| 94 \| 1 \| 93 \| Phage tail assembly chaperone protein, E, or 41 or 14 \| Phage tail assembly chaperone protein, E, or 41 or 14; afdb-uniprot50 \| AF-A0A198XEB0-F1-MODEL\_V4 \| 1.0 \| 6.394e-05 \| 224 \| 0.303 \| 79 \| 53 \| 2 \| 15 \| 92 \| 18 \| 95 \| Uncharacterized protein \| Uncharacterized protein; afdb-uniprot50 \| AF-G3IRG0-F1-MODEL\_V4 \| 1.0 \| 6.661e-06 \| 224 \| 0.322 \| 93 \| 52 \| 3 \| 2 \| 94 \| 6 \| 87 \| Uncharacterized protein \| Uncharacterized protein; afdb-uniprot50 \| AF-A0A0J1HAC3-F1-MODEL\_V4 \| 1.0 \| 2.76e-05 \| 222 \| 0.297 \| 84 \| 59 \| 0 \| 9 \| 92 \| 1 \| 84 \| Uncharacterized protein \| Uncharacterized protein; afdb-uniprot50 \| AF-A0A4Y9WRE6-F1-MODEL\_V4 \| 1.0 \| 0.0001144 \| 221 \| 0.262 \| 80 \| 58 \| 1 \| 16 \| 94 \| 11 \| 90 \| Phage tail assembly protein \| Phage tail assembly protein; afdb-uniprot50 \| AF-A0A316MPU8-F1-MODEL\_V4 \| 1.0 \| 2.76e-05 \| 221 \| 0.341 \| 85 \| 50 \| 2 \| 16 \| 94 \| 4 \| 88 \| Uncharacterized protein \| Uncharacterized protein; afdb-uniprot50 \| AF-A0A1F1HUP3-F1-MODEL\_V4 \| 1.0 \| 5.619e-05 \| 220 \| 0.294 \| 78 \| 53 \| 2 \| 18 \| 94 \| 16 \| 92 \| Uncharacterized protein \| Uncharacterized protein; afdb-uniprot50 \| AF-A0A7J0BVD3-F1-MODEL\_V4 \| 1.0 \| 4.629e-05 \| 220 \| 0.337 \| 77 \| 51 \| 0 \| 18 \| 94 \| 30 \| 106 \| Uncharacterized protein \| Uncharacterized protein; afdb-uniprot50 \| AF-A0A1W0CDM4-F1-MODEL\_V4 \| 1.0 \| 4.339e-05 \| 219 \| 0.407 \| 76 \| 44 \| 1 \| 18 \| 92 \| 8 \| 83 \| Uncharacterized protein \| Uncharacterized protein; afdb-uniprot50 \| AF-A0A258KUC8-F1-MODEL\_V4 \| 1.0 \| 0.0001389 \| 219 \| 0.215 \| 79 \| 60 \| 1 \| 16 \| 92 \| 3 \| 81 \| Uncharacterized protein \| Uncharacterized protein; afdb-uniprot50 \| AF-A0A7S6MHP3-F1-MODEL\_V4 \| 1.0 \| 4.629e-05 \| 219 \| 0.317 \| 82 \| 55 \| 1 \| 12 \| 92 \| 12 \| 93 \| Phage tail assembly protein \| Phage tail assembly protein; afdb-uniprot50 \| AF-A0A2G6CS65-F1-MODEL\_V4 \| 1.0 \| 4.938e-05 \| 218 \| 0.35 \| 77 \| 40 \| 2 \| 18 \| 94 \| 4 \| 70 \| Uncharacterized protein \| Uncharacterized protein; afdb-uniprot50 \| AF-A0A0D2HN99-F1-MODEL\_V4 \| 1.0 \| 8.833e-05 \| 218 \| 0.307 \| 78 \| 53 \| 1 \| 18 \| 94 \| 4 \| 81 \| Uncharacterized protein \| Uncharacterized protein; afdb-uniprot50 \| AF-A0A399Q037-F1-MODEL\_V4 \| 1.0 \| 6.821e-05 \| 218 \| 0.345 \| 81 \| 52 \| 1 \| 14 \| 93 \| 5 \| 85 \| Phage tail assembly protein \| Phage tail assembly protein; afdb-uniprot50 \| AF-A0A0F0DXF6-F1-MODEL\_V4 \| 1.0 \| 4.629e-05 \| 218 \| 0.379 \| 79 \| 48 \| 1 \| 16 \| 93 \| 3 \| 81 \| Uncharacterized protein \| Uncharacterized protein; afdb-uniprot50 \| AF-A0A228R688-F1-MODEL\_V4 \| 1.0 \| 7.277e-05 \| 216 \| 0.333 \| 78 \| 51 \| 1 \| 16 \| 92 \| 3 \| 80 \| Uncharacterized protein \| Uncharacterized protein; afdb-uniprot50 \| AF-A0A2V3R3G9-F1-MODEL\_V4 \| 1.0 \| 1.356e-05 \| 216 \| 0.263 \| 95 \| 69 \| 1 \| 1 \| 94 \| 6 \| 100 \| Tail assembly chaperone E/41/14-like protein \| Tail assembly chaperone E/41/14-like protein; afdb-uniprot50 \| AF-A0A6L6JDE6-F1-MODEL\_V4 \| 1.0 \| 2.76e-05 \| 215 \| 0.361 \| 94 \| 48 \| 4 \| 1 \| 94 \| 7 \| 88 \| Uncharacterized protein \| Uncharacterized protein; afdb-uniprot50 \| AF-A0A1N7DI63-F1-MODEL\_V4 \| 1.0 \| 0.0001144 \| 215 \| 0.32 \| 78 \| 51 \| 2 \| 16 \| 92 \| 22 \| 98 \| Phage tail assembly chaperone protein, E, or 41 or 14 \| Phage tail assembly chaperone protein, E, or 41 or 14; afdb-uniprot50 \| AF-A0A081N7U0-F1-MODEL\_V4 \| 1.0 \| 7.277e-05 \| 214 \| 0.207 \| 77 \| 60 \| 1 \| 18 \| 94 \| 6 \| 81 \| Uncharacterized protein \| Uncharacterized protein; afdb-uniprot50 \| AF-A0A258KTP8-F1-MODEL\_V4 \| 1.0 \| 0.0002183 \| 214 \| 0.227 \| 79 \| 59 \| 1 \| 16 \| 92 \| 4 \| 82 \| Uncharacterized protein \| Uncharacterized protein; afdb-uniprot50 \| AF-A0A450W6P6-F1-MODEL\_V4 \| 1.0 \| 2.76e-05 \| 214 \| 0.25 \| 92 \| 61 \| 1 \| 11 \| 94 \| 22 \| 113 \| Phage tail assembly chaperone protein, E, or 41 or 14 \| Phage tail assembly chaperone protein, E, or 41 or 14; afdb-uniprot50 \| AF-A0A4P5VJP3-F1-MODEL\_V4 \| 1.0 \| 2.274e-05 \| 213 \| 0.25 \| 92 \| 64 \| 2 \| 1 \| 92 \| 2 \| 88 \| Uncharacterized protein \| Uncharacterized protein; afdb-uniprot50 \| AF-A0A1B4S7U1-F1-MODEL\_V4 \| 1.0 \| 3.813e-05 \| 213 \| 0.349 \| 83 \| 47 \| 2 \| 16 \| 94 \| 3 \| 82 \| Uncharacterized protein \| Uncharacterized protein; afdb-uniprot50 \| AF-A0A1N7JL23-F1-MODEL\_V4 \| 1.0 \| 9.202e-06 \| 213 \| 0.297 \| 94 \| 58 \| 1 \| 1 \| 94 \| 9 \| 94 \| Phage tail assembly chaperone protein, E, or 41 or 14 \| Phage tail assembly chaperone protein, E, or 41 or 14; afdb-uniprot50 \| AF-A0A349HHP8-F1-MODEL\_V4 \| 1.0 \| 4.068e-05 \| 213 \| 0.294 \| 78 \| 53 \| 1 \| 15 \| 90 \| 23 \| 100 \| Uncharacterized protein \| Uncharacterized protein; afdb-uniprot50 \| AF-A0A2N1WD23-F1-MODEL\_V4 \| 1.0 \| 4.339e-05 \| 212 \| 0.292 \| 89 \| 58 \| 2 \| 11 \| 94 \| 3 \| 91 \| Phage tail assembly protein \| Phage tail assembly protein; afdb-uniprot50 \| AF-A0A6L8HHU5-F1-MODEL\_V4 \| 1.0 \| 7.277e-05 \| 212 \| 0.333 \| 75 \| 49 \| 1 \| 18 \| 92 \| 25 \| 98 \| Phage tail assembly protein \| Phage tail assembly protein; afdb-uniprot50 \| AF-A0A6B3LE99-F1-MODEL\_V4 \| 1.0 \| 1.356e-05 \| 211 \| 0.293 \| 92 \| 57 \| 1 \| 1 \| 92 \| 4 \| 87 \| Phage tail assembly protein \| Phage tail assembly protein; afdb-uniprot50 \| AF-A0A081B6C5-F1-MODEL\_V4 \| 1.0 \| 0.0001144 \| 209 \| 0.285 \| 77 \| 54 \| 1 \| 18 \| 94 \| 3 \| 78 \| Conserved protein \| Conserved protein; afdb-uniprot50 \| AF-A0A378WGB6-F1-MODEL\_V4 \| 1.0 \| 0.0001389 \| 209 \| 0.271 \| 81 \| 57 \| 2 \| 15 \| 94 \| 13 \| 92 \| Phage protein \| Phage protein; afdb-uniprot50 \| AF-A0A4R1K4S3-F1-MODEL\_V4 \| 1.0 \| 0.0001302 \| 209 \| 0.258 \| 85 \| 63 \| 0 \| 10 \| 94 \| 1 \| 85 \| Tail assembly chaperone E/41/14-like protein \| Tail assembly chaperone E/41/14-like protein; afdb-uniprot50 \| AF-A0A376BTM2-F1-MODEL\_V4 \| 1.0 \| 0.0001144 \| 209 \| 0.285 \| 84 \| 57 \| 3 \| 13 \| 94 \| 11 \| 93 \| Uncharacterized protein \| Uncharacterized protein; afdb-uniprot50 \| AF-W0IYF1-F1-MODEL\_V4 \| 1.0 \| 0.0001072 \| 208 \| 0.333 \| 78 \| 51 \| 1 \| 18 \| 94 \| 3 \| 80 \| Uncharacterized protein \| Uncharacterized protein; afdb-uniprot50 \| AF-A0A3S1CIX7-F1-MODEL\_V4 \| 1.0 \| 7.762e-05 \| 208 \| 0.209 \| 86 \| 66 \| 1 \| 10 \| 93 \| 1 \| 86 \| Phage tail assembly protein \| Phage tail assembly protein; afdb-uniprot50 \| AF-A0A522WEL1-F1-MODEL\_V4 \| 1.0 \| 0.0001302 \| 208 \| 0.337 \| 80 \| 52 \| 1 \| 16 \| 94 \| 2 \| 81 \| Phage tail assembly protein \| Phage tail assembly protein; afdb-uniprot50 \| AF-A0A1B4SJY4-F1-MODEL\_V4 \| 1.0 \| 0.0002329 \| 208 \| 0.311 \| 77 \| 52 \| 1 \| 18 \| 93 \| 2 \| 78 \| Uncharacterized protein \| Uncharacterized protein; afdb-uniprot50 \| AF-A0A420FTB7-F1-MODEL\_V4 \| 1.0 \| 1.543e-05 \| 208 \| 0.285 \| 98 \| 66 \| 2 \| 1 \| 94 \| 1 \| 98 \| Uncharacterized protein \| Uncharacterized protein; afdb-uniprot50 \| AF-Q65WG3-F1-MODEL\_V4 \| 1.0 \| 0.000158 \| 208 \| 0.337 \| 77 \| 49 \| 2 \| 18 \| 93 \| 47 \| 122 \| Uncharacterized protein \| Uncharacterized protein; afdb-uniprot50 \| AF-A0A3G4V6S2-F1-MODEL\_V4 \| 1.0 \| 7.762e-05 \| 207 \| 0.295 \| 88 \| 53 \| 1 \| 16 \| 94 \| 6 \| 93 \| Phage tail assembly protein \| Phage tail assembly protein; afdb-uniprot50 \| AF-A0A2W4TDV9-F1-MODEL\_V4 \| 1.0 \| 0.0002484 \| 206 \| 0.246 \| 77 \| 58 \| 0 \| 18 \| 94 \| 5 \| 81 \| Phage tail assembly protein \| Phage tail assembly protein; afdb-uniprot50 \| AF-A0A7M3MAZ0-F1-MODEL\_V4 \| 1.0 \| 4.629e-05 \| 206 \| 0.257 \| 97 \| 60 \| 2 \| 1 \| 94 \| 1 \| 88 \| Uncharacterized protein \| Uncharacterized protein; afdb-uniprot50 \| AF-A0A2R2IRA1-F1-MODEL\_V4 \| 1.0 \| 4.068e-05 \| 205 \| 0.255 \| 94 \| 69 \| 1 \| 1 \| 93 \| 1 \| 94 \| Uncharacterized protein \| Uncharacterized protein; afdb-uniprot50 \| AF-A0A1A9KIB7-F1-MODEL\_V4 \| 1.0 \| 6.821e-05 \| 205 \| 0.311 \| 93 \| 59 \| 3 \| 1 \| 93 \| 1 \| 88 \| Uncharacterized protein \| Uncharacterized protein; afdb-uniprot50 \| AF-A0A5A8F0S7-F1-MODEL\_V4 \| 1.0 \| 0.0002329 \| 204 \| 0.265 \| 79 \| 54 \| 1 \| 16 \| 94 \| 2 \| 76 \| Phage tail assembly protein \| Phage tail assembly protein; afdb-uniprot50 \| AF-A0A3D5IQS9-F1-MODEL\_V4 \| 1.0 \| 4.629e-05 \| 204 \| 0.223 \| 85 \| 65 \| 1 \| 9 \| 93 \| 5 \| 88 \| Phage tail assembly protein \| Phage tail assembly protein; afdb-uniprot50 \| AF-A0A2N3KSE8-F1-MODEL\_V4 \| 1.0 \| 5.268e-05 \| 203 \| 0.229 \| 96 \| 64 \| 3 \| 2 \| 93 \| 23 \| 112 \| Uncharacterized protein \| Uncharacterized protein; afdb-uniprot50 \| AF-E6L5Z3-F1-MODEL\_V4 \| 1.0 \| 3.141e-05 \| 202 \| 0.285 \| 84 \| 59 \| 1 \| 9 \| 92 \| 7 \| 89 \| Uncharacterized protein \| Uncharacterized protein; afdb-uniprot50 \| AF-A0A5C4PTD6-F1-MODEL\_V4 \| 1.0 \| 0.000265 \| 202 \| 0.324 \| 77 \| 50 \| 2 \| 18 \| 93 \| 13 \| 88 \| Phage tail assembly protein \| Phage tail assembly protein; afdb-uniprot50 \| AF-A0A0U2B5L0-F1-MODEL\_V4 \| 1.0 \| 4.629e-05 \| 202 \| 0.257 \| 101 \| 67 \| 3 \| 1 \| 94 \| 1 \| 100 \| Uncharacterized protein \| Uncharacterized protein; afdb-uniprot50 \| AF-A0A0A2YRD7-F1-MODEL\_V4 \| 1.0 \| 0.0001072 \| 201 \| 0.289 \| 83 \| 58 \| 1 \| 9 \| 90 \| 2 \| 84 \| Uncharacterized protein \| Uncharacterized protein; afdb-uniprot50 \| AF-A0A348FYG5-F1-MODEL\_V4 \| 1.0 \| 0.0002329 \| 201 \| 0.22 \| 77 \| 60 \| 0 \| 18 \| 94 \| 6 \| 82 \| Uncharacterized protein \| Uncharacterized protein; afdb-uniprot50 \| AF-A0A7J0BJG9-F1-MODEL\_V4 \| 1.0 \| 6.394e-05 \| 201 \| 0.41 \| 78 \| 41 \| 2 \| 16 \| 89 \| 5 \| 81 \| Uncharacterized protein \| Uncharacterized protein; afdb-uniprot50 \| AF-A0A524RVY5-F1-MODEL\_V4 \| 1.0 \| 5.994e-05 \| 201 \| 0.247 \| 105 \| 67 \| 5 \| 1 \| 94 \| 1 \| 104 \| Phage tail assembly protein \| Phage tail assembly protein; afdb-uniprot50 \| AF-A0A165NWZ7-F1-MODEL\_V4 \| 1.0 \| 0.0001686 \| 200 \| 0.269 \| 78 \| 57 \| 0 \| 15 \| 92 \| 2 \| 79 \| Uncharacterized protein \| Uncharacterized protein; afdb-uniprot50 \| AF-A0A0U5I607-F1-MODEL\_V4 \| 1.0 \| 2.76e-05 \| 200 \| 0.258 \| 93 \| 61 \| 2 \| 4 \| 94 \| 1 \| 87 \| Uncharacterized protein \| Uncharacterized protein; afdb-uniprot50 \| AF-A0A1Y1S1F4-F1-MODEL\_V4 \| 1.0 \| 0.0001918 \| 200 \| 0.179 \| 89 \| 70 \| 3 \| 7 \| 94 \| 37 \| 123 \| Uncharacterized protein \| Uncharacterized protein; afdb-uniprot50 \| AF-A0A1Y2K105-F1-MODEL\_V4 \| 1.0 \| 0.0001918 \| 199 \| 0.272 \| 77 \| 55 \| 1 \| 18 \| 94 \| 5 \| 80 \| Uncharacterized protein \| Uncharacterized protein; afdb-uniprot50 \| AF-A0A1M6YPI3-F1-MODEL\_V4 \| 1.0 \| 4.938e-05 \| 198 \| 0.257 \| 97 \| 69 \| 2 \| 1 \| 94 \| 2 \| 98 \| Phage tail assembly chaperone protein, E, or 41 or 14 \| Phage tail assembly chaperone protein, E, or 41 or 14; afdb-uniprot50 \| AF-A0A2A5DQ32-F1-MODEL\_V4 \| 1.0 \| 0.0005057 \| 197 \| 0.151 \| 79 \| 66 \| 1 \| 16 \| 94 \| 3 \| 80 \| Uncharacterized protein \| Uncharacterized protein; afdb-uniprot50 \| AF-A0A837E6Q2-F1-MODEL\_V4 \| 1.0 \| 0.0002329 \| 195 \| 0.346 \| 78 \| 50 \| 1 \| 18 \| 94 \| 2 \| 79 \| Uncharacterized protein \| Uncharacterized protein; afdb-uniprot50 \| AF-A0A1M3AJN0-F1-MODEL\_V4 \| 1.0 \| 0.000158 \| 195 \| 0.282 \| 78 \| 55 \| 1 \| 16 \| 93 \| 3 \| 79 \| Uncharacterized protein \| Uncharacterized protein; afdb-uniprot50 \| AF-E4ZB37-F1-MODEL\_V4 \| 1.0 \| 0.0002046 \| 194 \| 0.262 \| 80 \| 57 \| 2 \| 16 \| 94 \| 33 \| 111 \| Uncharacterized protein \| Uncharacterized protein; afdb-uniprot50 \| AF-A0A1I7ND34-F1-MODEL\_V4 \| 1.0 \| 0.0001798 \| 194 \| 0.298 \| 77 \| 53 \| 1 \| 16 \| 92 \| 4 \| 79 \| Phage tail assembly chaperone protein, E, or 41 or 14 \| Phage tail assembly chaperone protein, E, or 41 or 14; afdb-uniprot50 \| AF-A0A212KMT6-F1-MODEL\_V4 \| 1.0 \| 0.0004166 \| 194 \| 0.293 \| 75 \| 53 \| 0 \| 20 \| 94 \| 88 \| 162 \| Uncharacterized protein \| Uncharacterized protein; afdb-uniprot50 \| AF-A0A4P9VU53-F1-MODEL\_V4 \| 1.0 \| 0.0005754 \| 193 \| 0.219 \| 73 \| 57 \| 0 \| 16 \| 88 \| 2 \| 74 \| Phage tail assembly protein \| Phage tail assembly protein; afdb-uniprot50 \| AF-S7TEH7-F1-MODEL\_V4 \| 1.0 \| 0.000122 \| 193 \| 0.321 \| 87 \| 54 \| 3 \| 11 \| 94 \| 1 \| 85 \| Mu-like prophage FluMu protein gp41 \| Mu-like prophage FluMu protein gp41; afdb-uniprot50 \| AF-A0A6L5JVI4-F1-MODEL\_V4 \| 1.0 \| 0.0001072 \| 193 \| 0.226 \| 84 \| 64 \| 1 \| 10 \| 93 \| 1 \| 83 \| Uncharacterized protein \| Uncharacterized protein; afdb-uniprot50 \| AF-A0A6I3KFP2-F1-MODEL\_V4 \| 1.0 \| 0.0002329 \| 191 \| 0.237 \| 80 \| 60 \| 1 \| 14 \| 93 \| 4 \| 82 \| Uncharacterized protein \| Uncharacterized protein; afdb-uniprot50 \| AF-A0A2D2C1C9-F1-MODEL\_V4 \| 1.0 \| 0.0002183 \| 191 \| 0.217 \| 78 \| 60 \| 1 \| 15 \| 92 \| 4 \| 80 \| Uncharacterized protein \| Uncharacterized protein; afdb-uniprot50 \| AF-A0A212KBP9-F1-MODEL\_V4 \| 1.0 \| 0.000265 \| 191 \| 0.378 \| 82 \| 48 \| 3 \| 14 \| 92 \| 2 \| 83 \| Uncharacterized protein \| Uncharacterized protein; afdb-uniprot50 \| AF-J0QC11-F1-MODEL\_V4 \| 1.0 \| 0.0004166 \| 190 \| 0.189 \| 79 \| 62 \| 1 \| 15 \| 93 \| 5 \| 81 \| Uncharacterized protein \| Uncharacterized protein; afdb-uniprot50 \| AF-A0A450ZAK9-F1-MODEL\_V4 \| 1.0 \| 7.277e-05 \| 190 \| 0.232 \| 86 \| 58 \| 3 \| 15 \| 94 \| 16 \| 99 \| Phage tail assembly chaperone protein, E, or 41 or 14 \| Phage tail assembly chaperone protein, E, or 41 or 14; afdb-uniprot50 \| AF-L0R6A7-F1-MODEL\_V4 \| 1.0 \| 0.0004166 \| 189 \| 0.337 \| 80 \| 44 \| 3 \| 15 \| 94 \| 2 \| 72 \| Uncharacterized protein \| Uncharacterized protein; afdb-uniprot50 \| AF-A0A4R6U4B4-F1-MODEL\_V4 \| 1.0 \| 0.0002827 \| 189 \| 0.269 \| 78 \| 56 \| 1 \| 18 \| 94 \| 2 \| 79 \| Tail assembly chaperone E/41/14-like protein \| Tail assembly chaperone E/41/14-like protein; afdb-uniprot50 \| AF-A0A445MWH4-F1-MODEL\_V4 \| 1.0 \| 0.0002484 \| 188 \| 0.23 \| 78 \| 58 \| 2 \| 18 \| 94 \| 8 \| 84 \| Uncharacterized protein \| Uncharacterized protein; afdb-uniprot50 \| AF-A0A522IQG1-F1-MODEL\_V4 \| 1.0 \| 0.000848 \| 188 \| 0.294 \| 78 \| 54 \| 1 \| 18 \| 94 \| 2 \| 79 \| Phage tail assembly protein \| Phage tail assembly protein; afdb-uniprot50 \| AF-A0A809Z9Z9-F1-MODEL\_V4 \| 1.0 \| 0.0001798 \| 188 \| 0.211 \| 85 \| 66 \| 1 \| 10 \| 93 \| 1 \| 85 \| Uncharacterized protein \| Uncharacterized protein; afdb-uniprot50 \| AF-A0A837E323-F1-MODEL\_V4 \| 1.0 \| 0.0007452 \| 188 \| 0.197 \| 76 \| 60 \| 1 \| 18 \| 93 \| 6 \| 80 \| Uncharacterized protein \| Uncharacterized protein; afdb-uniprot50 \| AF-G1UYU1-F1-MODEL\_V4 \| 1.0 \| 0.0005754 \| 187 \| 0.253 \| 83 \| 56 \| 1 \| 16 \| 92 \| 3 \| 85 \| Uncharacterized protein \| Uncharacterized protein; afdb-uniprot50 \| AF-A0A3M4ZXY2-F1-MODEL\_V4 \| 1.0 \| 0.0003905 \| 186 \| 0.43 \| 72 \| 40 \| 1 \| 23 \| 93 \| 1 \| 72 \| Uncharacterized protein \| Uncharacterized protein; afdb-uniprot50 \| AF-Q602Z2-F1-MODEL\_V4 \| 1.0 \| 0.0004444 \| 186 \| 0.41 \| 78 \| 45 \| 1 \| 18 \| 94 \| 3 \| 80 \| Uncharacterized protein \| Uncharacterized protein; afdb-uniprot50 \| AF-A0A1W2F849-F1-MODEL\_V4 \| 1.0 \| 0.0003432 \| 185 \| 0.3 \| 70 \| 48 \| 1 \| 18 \| 87 \| 5 \| 73 \| Phage tail assembly chaperone protein, E, or 41 or 14 \| Phage tail assembly chaperone protein, E, or 41 or 14; afdb-uniprot50 \| AF-A0A0J6K5A0-F1-MODEL\_V4 \| 1.0 \| 0.0001072 \| 183 \| 0.234 \| 94 \| 62 \| 3 \| 2 \| 92 \| 4 \| 90 \| Uncharacterized protein \| Uncharacterized protein; afdb-uniprot50 \| AF-A0A1H1FGP4-F1-MODEL\_V4 \| 1.0 \| 0.000474 \| 183 \| 0.177 \| 79 \| 64 \| 1 \| 10 \| 88 \| 1 \| 78 \| Phage tail assembly chaperone protein, E, or 41 or 14 \| Phage tail assembly chaperone protein, E, or 41 or 14; afdb-uniprot50 \| AF-A0A066RUL2-F1-MODEL\_V4 \| 1.0 \| 0.0004444 \| 183 \| 0.275 \| 87 \| 53 \| 2 \| 18 \| 94 \| 9 \| 95 \| Uncharacterized protein \| Uncharacterized protein; afdb-uniprot50 \| AF-A0A317H578-F1-MODEL\_V4 \| 1.0 \| 0.0001798 \| 183 \| 0.2 \| 85 \| 62 \| 3 \| 8 \| 91 \| 4 \| 83 \| Uncharacterized protein \| Uncharacterized protein; afdb-uniprot50 \| AF-A0A7W8GZY1-F1-MODEL\_V4 \| 1.0 \| 0.0006986 \| 181 \| 0.24 \| 75 \| 56 \| 1 \| 18 \| 92 \| 8 \| 81 \| Uncharacterized protein \| Uncharacterized protein; afdb-uniprot50 \| AF-A0A2G1CU23-F1-MODEL\_V4 \| 1.0 \| 0.0002046 \| 181 \| 0.202 \| 89 \| 70 \| 1 \| 4 \| 92 \| 1 \| 88 \| Uncharacterized protein \| Uncharacterized protein; afdb-uniprot50 \| AF-A0A7J5WEE9-F1-MODEL\_V4 \| 1.0 \| 9.423e-05 \| 181 \| 0.244 \| 90 \| 67 \| 1 \| 1 \| 89 \| 8 \| 97 \| Uncharacterized protein \| Uncharacterized protein; afdb-uniprot50 \| AF-A0A1F3A562-F1-MODEL\_V4 \| 1.0 \| 0.000848 \| 181 \| 0.23 \| 78 \| 59 \| 1 \| 16 \| 92 \| 5 \| 82 \| Uncharacterized protein \| Uncharacterized protein; afdb-uniprot50 \| AF-A0A833IQP1-F1-MODEL\_V4 \| 1.0 \| 0.0001481 \| 180 \| 0.204 \| 93 \| 63 \| 3 \| 1 \| 93 \| 1 \| 82 \| Phage tail assembly protein \| Phage tail assembly protein; afdb-uniprot50 \| AF-A0A6N8BZ30-F1-MODEL\_V4 \| 1.0 \| 0.0005057 \| 180 \| 0.176 \| 85 \| 69 \| 1 \| 10 \| 93 \| 1 \| 85 \| Phage tail assembly protein \| Phage tail assembly protein; afdb-uniprot50 \| AF-A0A6P2SQ52-F1-MODEL\_V4 \| 1.0 \| 0.001333 \| 179 \| 0.269 \| 78 \| 56 \| 1 \| 18 \| 94 \| 2 \| 79 \| Uncharacterized protein \| Uncharacterized protein; afdb-uniprot50 \| AF-A0A5C8S8T5-F1-MODEL\_V4 \| 1.0 \| 0.0002484 \| 179 \| 0.139 \| 86 \| 73 \| 1 \| 9 \| 93 \| 1 \| 86 \| Phage tail assembly protein \| Phage tail assembly protein; afdb-uniprot50 \| AF-A0A6L9FIH6-F1-MODEL\_V4 \| 1.0 \| 0.000474 \| 179 \| 0.271 \| 92 \| 52 \| 2 \| 18 \| 94 \| 3 \| 94 \| Uncharacterized protein \| Uncharacterized protein; afdb-uniprot50 \| AF-A0A4V2RF86-F1-MODEL\_V4 \| 1.0 \| 0.0003905 \| 178 \| 0.246 \| 77 \| 57 \| 1 \| 16 \| 92 \| 2 \| 77 \| Tail assembly chaperone E/41/14-like protein \| Tail assembly chaperone E/41/14-like protein; afdb-uniprot50 \| AF-A0A4U8YLQ2-F1-MODEL\_V4 \| 1.0 \| 0.0005754 \| 178 \| 0.253 \| 79 \| 57 \| 2 \| 16 \| 93 \| 2 \| 79 \| Bacteriophage tail protein gp41 putative \| Bacteriophage tail protein gp41 putative; afdb-uniprot50 \| AF-A0A0E3BR39-F1-MODEL\_V4 \| 1.0 \| 0.001098 \| 177 \| 0.231 \| 82 \| 62 \| 1 \| 12 \| 92 \| 5 \| 86 \| Uncharacterized protein \| Uncharacterized protein; afdb-uniprot50 \| AF-Q1QI92-F1-MODEL\_V4 \| 1.0 \| 0.0005394 \| 177 \| 0.155 \| 77 \| 64 \| 1 \| 16 \| 92 \| 5 \| 80 \| Uncharacterized protein \| Uncharacterized protein; afdb-uniprot50 \| AF-A0A4R2GHJ3-F1-MODEL\_V4 \| 1.0 \| 0.0005394 \| 176 \| 0.25 \| 76 \| 56 \| 1 \| 18 \| 93 \| 3 \| 77 \| Tail assembly chaperone E/41/14-like protein \| Tail assembly chaperone E/41/14-like protein; afdb-uniprot50 \| AF-A0A1H9Q8X8-F1-MODEL\_V4 \| 1.0 \| 0.0002827 \| 176 \| 0.188 \| 106 \| 74 \| 2 \| 1 \| 94 \| 1 \| 106 \| Phage tail assembly chaperone protein, E, or 41 or 14 \| Phage tail assembly chaperone protein, E, or 41 or 14; afdb-uniprot50 \| AF-A0A6M7VG49-F1-MODEL\_V4 \| 1.0 \| 0.001333 \| 175 \| 0.162 \| 86 \| 71 \| 1 \| 9 \| 94 \| 1 \| 85 \| Phage tail assembly protein \| Phage tail assembly protein; afdb-uniprot50 \| AF-A0A1A9VZ72-F1-MODEL\_V4 \| 1.0 \| 0.001171 \| 175 \| 0.189 \| 79 \| 63 \| 1 \| 16 \| 93 \| 4 \| 82 \| Uncharacterized protein \| Uncharacterized protein; afdb-uniprot50 \| AF-A0A1M4WE05-F1-MODEL\_V4 \| 1.0 \| 0.0009046 \| 175 \| 0.189 \| 74 \| 59 \| 1 \| 15 \| 88 \| 7 \| 79 \| Phage tail assembly chaperone protein, E, or 41 or 14 \| Phage tail assembly chaperone protein, E, or 41 or 14; afdb-uniprot50 \| AF-A0A2N6FYM4-F1-MODEL\_V4 \| 1.0 \| 0.0002183 \| 174 \| 0.182 \| 93 \| 65 \| 3 \| 4 \| 94 \| 1 \| 84 \| Uncharacterized protein \| Uncharacterized protein; afdb-uniprot50 \| AF-A0A4Q0USL0-F1-MODEL\_V4 \| 1.0 \| 0.000122 \| 174 \| 0.191 \| 94 \| 75 \| 1 \| 1 \| 94 \| 5 \| 97 \| Uncharacterized protein \| Uncharacterized protein; afdb-uniprot50 \| AF-A0A2D3VZ53-F1-MODEL\_V4 \| 1.0 \| 0.0003432 \| 174 \| 0.28 \| 82 \| 50 \| 3 \| 13 \| 94 \| 26 \| 98 \| Uncharacterized protein \| Uncharacterized protein; afdb-uniprot50 \| AF-A0A837JCA3-F1-MODEL\_V4 \| 1.0 \| 0.000265 \| 174 \| 0.241 \| 91 \| 62 \| 2 \| 2 \| 92 \| 22 \| 105 \| Uncharacterized protein \| Uncharacterized protein; afdb-uniprot50 \| AF-F3YY69-F1-MODEL\_V4 \| 1.0 \| 0.0007452 \| 174 \| 0.267 \| 86 \| 55 \| 2 \| 14 \| 91 \| 2 \| 87 \| Uncharacterized protein \| Uncharacterized protein; afdb-uniprot50 \| AF-A0A847KS38-F1-MODEL\_V4 \| 1.0 \| 0.000848 \| 173 \| 0.227 \| 79 \| 60 \| 1 \| 16 \| 94 \| 3 \| 80 \| Phage tail assembly protein \| Phage tail assembly protein; afdb-uniprot50 \| AF-A0A212KJV9-F1-MODEL\_V4 \| 1.0 \| 0.0005754 \| 172 \| 0.262 \| 80 \| 57 \| 1 \| 16 \| 93 \| 4 \| 83 \| Uncharacterized protein \| Uncharacterized protein; afdb-uniprot50 \| AF-D5RTF6-F1-MODEL\_V4 \| 1.0 \| 0.0002827 \| 172 \| 0.197 \| 86 \| 68 \| 1 \| 9 \| 94 \| 1 \| 85 \| Uncharacterized protein \| Uncharacterized protein; afdb-uniprot50 \| AF-A0A7R7YEW5-F1-MODEL\_V4 \| 1.0 \| 0.0004166 \| 171 \| 0.233 \| 90 \| 57 \| 2 \| 16 \| 93 \| 6 \| 95 \| Uncharacterized protein \| Uncharacterized protein; afdb-uniprot50 \| AF-E5Y5W3-F1-MODEL\_V4 \| 1.0 \| 0.000965 \| 171 \| 0.265 \| 83 \| 55 \| 1 \| 16 \| 92 \| 4 \| 86 \| Uncharacterized protein \| Uncharacterized protein; afdb-uniprot50 \| AF-A0A2N3KJK8-F1-MODEL\_V4 \| 1.0 \| 0.0007452 \| 171 \| 0.186 \| 102 \| 70 \| 5 \| 3 \| 92 \| 11 \| 111 \| Uncharacterized protein \| Uncharacterized protein; afdb-uniprot50 \| AF-U6ZUR1-F1-MODEL\_V4 \| 1.0 \| 0.001841 \| 171 \| 0.194 \| 77 \| 61 \| 1 \| 18 \| 94 \| 102 \| 177 \| Uncharacterized protein \| Uncharacterized protein; afdb-uniprot50 \| AF-A0A2P7PD21-F1-MODEL\_V4 \| 1.0 \| 0.001422 \| 170 \| 0.194 \| 77 \| 60 \| 1 \| 16 \| 92 \| 5 \| 79 \| Phage tail assembly protein \| Phage tail assembly protein; afdb-uniprot50 \| AF-A0A2C5TPJ7-F1-MODEL\_V4 \| 1.0 \| 0.0006139 \| 169 \| 0.188 \| 85 \| 64 \| 2 \| 15 \| 94 \| 3 \| 87 \| Phage tail assembly protein \| Phage tail assembly protein; afdb-uniprot50 \| AF-A0A847G972-F1-MODEL\_V4 \| 1.0 \| 0.001171 \| 169 \| 0.272 \| 77 \| 53 \| 2 \| 16 \| 89 \| 18 \| 94 \| Phage tail assembly protein \| Phage tail assembly protein; afdb-uniprot50 \| AF-A0A165RMJ9-F1-MODEL\_V4 \| 1.0 \| 0.001029 \| 169 \| 0.171 \| 76 \| 62 \| 1 \| 10 \| 85 \| 1 \| 75 \| Uncharacterized protein \| Uncharacterized protein; afdb-uniprot50 \| AF-A0A827DRS4-F1-MODEL\_V4 \| 1.0 \| 0.0006548 \| 168 \| 0.244 \| 86 \| 56 \| 2 \| 1 \| 85 \| 4 \| 81 \| Phage tail assembly protein \| Phage tail assembly protein; afdb-uniprot50 \| AF-U3TX64-F1-MODEL\_V4 \| 1.0 \| 0.001517 \| 168 \| 0.215 \| 79 \| 61 \| 1 \| 16 \| 93 \| 2 \| 80 \| Uncharacterized protein \| Uncharacterized protein; afdb-uniprot50 \| AF-F7U3Q5-F1-MODEL\_V4 \| 1.0 \| 0.000965 \| 168 \| 0.181 \| 77 \| 62 \| 1 \| 16 \| 92 \| 2 \| 77 \| Uncharacterized protein \| Uncharacterized protein; afdb-uniprot50 \| AF-A0A827G1L9-F1-MODEL\_V4 \| 1.0 \| 0.0001686 \| 168 \| 0.197 \| 91 \| 65 \| 3 \| 9 \| 93 \| 2 \| 90 \| Phage tail assembly protein \| Phage tail assembly protein; afdb-uniprot50 \| AF-A0A087KTW1-F1-MODEL\_V4 \| 1.0 \| 0.001171 \| 168 \| 0.204 \| 83 \| 62 \| 2 \| 15 \| 93 \| 4 \| 86 \| Uncharacterized protein \| Uncharacterized protein; afdb-uniprot50 \| AF-A0A1D9BCU0-F1-MODEL\_V4 \| 1.0 \| 0.0006139 \| 168 \| 0.235 \| 85 \| 58 \| 2 \| 16 \| 93 \| 2 \| 86 \| Uncharacterized protein \| Uncharacterized protein; afdb-uniprot50 \| AF-A0A7W6WBB0-F1-MODEL\_V4 \| 1.0 \| 0.0002827 \| 168 \| 0.235 \| 89 \| 63 \| 2 \| 10 \| 93 \| 5 \| 93 \| Uncharacterized protein \| Uncharacterized protein; afdb-uniprot50 \| AF-A0A3M1X2P8-F1-MODEL\_V4 \| 1.0 \| 0.0006139 \| 167 \| 0.164 \| 91 \| 67 \| 2 \| 3 \| 93 \| 1 \| 82 \| Phage tail assembly protein \| Phage tail assembly protein; afdb-uniprot50 \| AF-A0A4D8R9U9-F1-MODEL\_V4 \| 1.0 \| 0.0002183 \| 167 \| 0.223 \| 94 \| 62 \| 2 \| 1 \| 94 \| 1 \| 83 \| Phage tail assembly protein \| Phage tail assembly protein; afdb-uniprot50 \| AF-A0A7U0N692-F1-MODEL\_V4 \| 1.0 \| 0.001618 \| 167 \| 0.172 \| 81 \| 65 \| 1 \| 15 \| 93 \| 6 \| 86 \| Phage tail assembly protein \| Phage tail assembly protein; afdb-uniprot50 \| AF-A0A017HBQ7-F1-MODEL\_V4 \| 1.0 \| 0.000848 \| 166 \| 0.157 \| 76 \| 63 \| 1 \| 10 \| 85 \| 1 \| 75 \| Uncharacterized protein \| Uncharacterized protein; afdb-uniprot50 \| AF-A0A6J5FMZ1-F1-MODEL\_V4 \| 1.0 \| 0.001098 \| 166 \| 0.212 \| 80 \| 61 \| 1 \| 15 \| 92 \| 3 \| 82 \| Uncharacterized protein \| Uncharacterized protein; afdb-uniprot50 \| AF-A0A7C1VCE0-F1-MODEL\_V4 \| 1.0 \| 0.001333 \| 166 \| 0.164 \| 85 \| 69 \| 2 \| 7 \| 89 \| 45 \| 129 \| Phage tail assembly protein \| Phage tail assembly protein; afdb-uniprot50 \| AF-A0A4U0QBM5-F1-MODEL\_V4 \| 1.0 \| 0.001098 \| 165 \| 0.266 \| 75 \| 54 \| 1 \| 18 \| 92 \| 10 \| 83 \| Phage tail assembly protein \| Phage tail assembly protein; afdb-uniprot50 \| AF-A0A840C379-F1-MODEL\_V4 \| 1.0 \| 0.002544 \| 165 \| 0.189 \| 79 \| 62 \| 1 \| 16 \| 92 \| 2 \| 80 \| Uncharacterized protein \| Uncharacterized protein; afdb-uniprot50 \| AF-A0A6F8URG2-F1-MODEL\_V4 \| 1.0 \| 0.001171 \| 165 \| 0.202 \| 79 \| 62 \| 1 \| 10 \| 88 \| 1 \| 78 \| Uncharacterized protein \| Uncharacterized protein; afdb-uniprot50 \| AF-A0A855R6X1-F1-MODEL\_V4 \| 1.0 \| 0.001098 \| 165 \| 0.243 \| 78 \| 57 \| 1 \| 15 \| 90 \| 3 \| 80 \| Phage tail assembly protein \| Phage tail assembly protein; afdb-uniprot50 \| AF-A0A484GB85-F1-MODEL\_V4 \| 1.0 \| 0.0006986 \| 165 \| 0.259 \| 81 \| 54 \| 2 \| 18 \| 93 \| 3 \| 82 \| Phage tail assembly protein \| Phage tail assembly protein; afdb-uniprot50 \| AF-A0A2S9IP70-F1-MODEL\_V4 \| 1.0 \| 0.002235 \| 165 \| 0.202 \| 69 \| 54 \| 1 \| 15 \| 83 \| 5 \| 72 \| Phage tail assembly protein \| Phage tail assembly protein; afdb-uniprot50 \| AF-A0A0Q2UGR9-F1-MODEL\_V4 \| 1.0 \| 0.0005754 \| 165 \| 0.191 \| 94 \| 70 \| 2 \| 1 \| 94 \| 1 \| 88 \| Uncharacterized protein \| Uncharacterized protein; afdb-uniprot50 \| AF-A0A4R2D341-F1-MODEL\_V4 \| 1.0 \| 0.001726 \| 164 \| 0.197 \| 76 \| 60 \| 1 \| 18 \| 93 \| 3 \| 77 \| Tail assembly chaperone E/41/14-like protein \| Tail assembly chaperone E/41/14-like protein; afdb-uniprot50 \| AF-A0A1F8WN21-F1-MODEL\_V4 \| 1.0 \| 0.001098 \| 164 \| 0.2 \| 80 \| 60 \| 2 \| 16 \| 94 \| 3 \| 79 \| Uncharacterized protein \| Uncharacterized protein; afdb-uniprot50 \| AF-A0A2U1TU93-F1-MODEL\_V4 \| 1.0 \| 0.001841 \| 164 \| 0.168 \| 83 \| 65 \| 2 \| 15 \| 93 \| 4 \| 86 \| Phage tail assembly protein \| Phage tail assembly protein; afdb-uniprot50 \| AF-H8FUZ8-F1-MODEL\_V4 \| 1.0 \| 0.0005754 \| 164 \| 0.176 \| 85 \| 63 \| 1 \| 16 \| 93 \| 5 \| 89 \| Putative bacteriophage protein \| Putative bacteriophage protein; afdb-uniprot50 \| AF-A0A4Z0W7G4-F1-MODEL\_V4 \| 1.0 \| 0.001964 \| 164 \| 0.197 \| 81 \| 63 \| 1 \| 15 \| 93 \| 27 \| 107 \| Phage tail assembly protein \| Phage tail assembly protein; afdb-uniprot50 \| AF-A0A4R5W1E0-F1-MODEL\_V4 \| 1.0 \| 0.004551 \| 164 \| 0.271 \| 81 \| 58 \| 1 \| 15 \| 94 \| 12 \| 92 \| Phage tail assembly protein \| Phage tail assembly protein; afdb-uniprot50 \| AF-A0A8B3NM35-F1-MODEL\_V4 \| 1.0 \| 0.0005754 \| 164 \| 0.213 \| 89 \| 65 \| 2 \| 6 \| 94 \| 24 \| 107 \| Phage tail assembly protein \| Phage tail assembly protein; afdb-uniprot50 \| AF-A0A854G7K2-F1-MODEL\_V4 \| 1.0 \| 0.005524 \| 163 \| 0.246 \| 77 \| 57 \| 1 \| 18 \| 93 \| 2 \| 78 \| Uncharacterized protein \| Uncharacterized protein; afdb-uniprot50 \| AF-A0A2T3GP98-F1-MODEL\_V4 \| 1.0 \| 0.001726 \| 162 \| 0.194 \| 77 \| 61 \| 1 \| 16 \| 92 \| 4 \| 79 \| Phage tail assembly protein \| Phage tail assembly protein; afdb-uniprot50 \| AF-A0A344UIJ8-F1-MODEL\_V4 \| 1.0 \| 0.003514 \| 162 \| 0.259 \| 77 \| 56 \| 1 \| 18 \| 93 \| 2 \| 78 \| Phage tail assembly protein \| Phage tail assembly protein; afdb-uniprot50 \| AF-A0A346QYG8-F1-MODEL\_V4 \| 1.0 \| 0.0004166 \| 162 \| 0.216 \| 83 \| 60 \| 2 \| 14 \| 92 \| 4 \| 85 \| Phage tail assembly protein \| Phage tail assembly protein; afdb-uniprot50 \| AF-A0A2K9P758-F1-MODEL\_V4 \| 1.0 \| 0.001517 \| 162 \| 0.136 \| 88 \| 72 \| 2 \| 10 \| 93 \| 1 \| 88 \| Uncharacterized protein \| Uncharacterized protein; afdb-uniprot50 \| AF-A0A7Z0QG19-F1-MODEL\_V4 \| 1.0 \| 0.0006548 \| 162 \| 0.225 \| 93 \| 61 \| 1 \| 11 \| 92 \| 7 \| 99 \| Phage tail assembly protein \| Phage tail assembly protein; afdb-uniprot50 \| AF-A0A1K0J3G7-F1-MODEL\_V4 \| 1.0 \| 0.002096 \| 162 \| 0.2 \| 80 \| 63 \| 1 \| 14 \| 92 \| 4 \| 83 \| Uncharacterized protein \| Uncharacterized protein; afdb-uniprot50 \| AF-W0BWL8-F1-MODEL\_V4 \| 1.0 \| 0.0005394 \| 162 \| 0.189 \| 95 \| 69 \| 3 \| 4 \| 93 \| 1 \| 92 \| Uncharacterized protein \| Uncharacterized protein; afdb-uniprot50 \| AF-A0A2S7JR59-F1-MODEL\_V4 \| 1.0 \| 0.0003661 \| 162 \| 0.229 \| 96 \| 70 \| 2 \| 1 \| 94 \| 2 \| 95 \| Phage tail assembly protein \| Phage tail assembly protein; afdb-uniprot50 \| AF-A0A349GR41-F1-MODEL\_V4 \| 1.0 \| 0.000474 \| 162 \| 0.198 \| 101 \| 66 \| 4 \| 6 \| 93 \| 25 \| 123 \| Uncharacterized protein \| Uncharacterized protein; afdb-uniprot50 \| AF-A0A5Q8CAB4-F1-MODEL\_V4 \| 1.0 \| 0.0004444 \| 162 \| 0.225 \| 93 \| 63 \| 3 \| 2 \| 94 \| 32 \| 115 \| Phage tail assembly protein \| Phage tail assembly protein; afdb-uniprot50 \| AF-A0A2V3UB04-F1-MODEL\_V4 \| 1.0 \| 0.003749 \| 161 \| 0.216 \| 74 \| 58 \| 0 \| 18 \| 91 \| 3 \| 76 \| Tail assembly chaperone E/41/14-like protein \| Tail assembly chaperone E/41/14-like protein; afdb-uniprot50 \| AF-A0A564G557-F1-MODEL\_V4 \| 1.0 \| 0.001029 \| 161 \| 0.197 \| 76 \| 60 \| 1 \| 18 \| 92 \| 14 \| 89 \| Uncharacterized protein \| Uncharacterized protein; afdb-uniprot50 \| AF-A0A6L9MMR9-F1-MODEL\_V4 \| 1.0 \| 0.002235 \| 161 \| 0.189 \| 79 \| 63 \| 1 \| 10 \| 88 \| 1 \| 78 \| Phage tail assembly protein \| Phage tail assembly protein; afdb-uniprot50 \| AF-A0A853T2H5-F1-MODEL\_V4 \| 1.0 \| 0.0009046 \| 161 \| 0.222 \| 72 \| 53 \| 2 \| 15 \| 85 \| 14 \| 83 \| Phage tail assembly chaperone protein, E, or 41 or 14 \| Phage tail assembly chaperone protein, E, or 41 or 14; afdb-uniprot50 \| AF-A0A2J8GXT3-F1-MODEL\_V4 \| 1.0 \| 0.002385 \| 161 \| 0.162 \| 86 \| 63 \| 1 \| 18 \| 94 \| 3 \| 88 \| Uncharacterized protein \| Uncharacterized protein; afdb-uniprot50 \| AF-A0A3N2E0T6-F1-MODEL\_V4 \| 1.0 \| 0.0009046 \| 161 \| 0.253 \| 79 \| 54 \| 2 \| 18 \| 91 \| 23 \| 101 \| Tail assembly chaperone E/41/14-like protein \| Tail assembly chaperone E/41/14-like protein; afdb-uniprot50 \| AF-A0A4Z1R4X7-F1-MODEL\_V4 \| 1.0 \| 0.0004166 \| 161 \| 0.224 \| 98 \| 69 \| 2 \| 2 \| 94 \| 4 \| 99 \| Phage tail assembly protein \| Phage tail assembly protein; afdb-uniprot50 \| AF-E7H537-F1-MODEL\_V4 \| 1.0 \| 0.00125 \| 160 \| 0.226 \| 84 \| 57 \| 1 \| 18 \| 93 \| 2 \| 85 \| Uncharacterized protein \| Uncharacterized protein; afdb-uniprot50 \| AF-A0A102DFQ3-F1-MODEL\_V4 \| 1.0 \| 0.0002329 \| 160 \| 0.21 \| 100 \| 69 \| 3 \| 1 \| 94 \| 4 \| 99 \| Uncharacterized protein \| Uncharacterized protein; afdb-uniprot50 \| AF-A0A7L9Q1H1-F1-MODEL\_V4 \| 1.0 \| 0.000965 \| 159 \| 0.252 \| 95 \| 66 \| 3 \| 1 \| 94 \| 1 \| 91 \| Phage tail assembly protein \| Phage tail assembly protein; afdb-uniprot50 \| AF-X0YLY6-F1-MODEL\_V4 \| 1.0 \| 0.0007949 \| 159 \| 0.151 \| 86 \| 71 \| 1 \| 9 \| 94 \| 2 \| 85 \| Uncharacterized protein \| Uncharacterized protein; afdb-uniprot50 \| AF-A0A1S7Q6W0-F1-MODEL\_V4 \| 1.0 \| 0.002235 \| 159 \| 0.223 \| 76 \| 59 \| 0 \| 18 \| 93 \| 3 \| 78 \| Uncharacterized protein \| Uncharacterized protein; afdb-uniprot50 \| AF-A0A143DC33-F1-MODEL\_V4 \| 1.0 \| 0.001098 \| 159 \| 0.252 \| 87 \| 58 \| 2 \| 14 \| 93 \| 3 \| 89 \| Uncharacterized protein \| Uncharacterized protein; afdb-uniprot50 \| AF-A0A502GP93-F1-MODEL\_V4 \| 1.0 \| 0.0007949 \| 159 \| 0.218 \| 96 \| 70 \| 3 \| 3 \| 93 \| 37 \| 132 \| Phage tail assembly protein \| Phage tail assembly protein; afdb-uniprot50 \| AF-A0A7V8JT65-F1-MODEL\_V4 \| 1.0 \| 0.003088 \| 158 \| 0.126 \| 79 \| 68 \| 1 \| 14 \| 92 \| 4 \| 81 \| Uncharacterized protein \| Uncharacterized protein; afdb-uniprot50 \| AF-A0A258KWE3-F1-MODEL\_V4 \| 1.0 \| 0.003749 \| 158 \| 0.16 \| 81 \| 66 \| 1 \| 16 \| 94 \| 3 \| 83 \| Uncharacterized protein \| Uncharacterized protein; afdb-uniprot50 \| AF-A0A323UKS3-F1-MODEL\_V4 \| 1.0 \| 0.000474 \| 158 \| 0.18 \| 94 \| 73 \| 2 \| 1 \| 94 \| 1 \| 90 \| Phage tail assembly protein \| Phage tail assembly protein; afdb-uniprot50 \| AF-A0A7W4ICA4-F1-MODEL\_V4 \| 1.0 \| 0.0006548 \| 158 \| 0.25 \| 104 \| 65 \| 5 \| 3 \| 94 \| 36 \| 138 \| Phage tail assembly protein \| Phage tail assembly protein; afdb-uniprot50 \| AF-A0A316GTU0-F1-MODEL\_V4 \| 1.0 \| 0.001841 \| 157 \| 0.207 \| 77 \| 60 \| 1 \| 18 \| 93 \| 3 \| 79 \| Tail assembly chaperone E/41/14-like protein \| Tail assembly chaperone E/41/14-like protein; afdb-uniprot50 \| AF-R5EZ88-F1-MODEL\_V4 \| 1.0 \| 0.001333 \| 157 \| 0.178 \| 84 \| 61 \| 2 \| 18 \| 93 \| 7 \| 90 \| Putative bacteriophage protein \| Putative bacteriophage protein; afdb-uniprot50 \| AF-A0A4Q1B5V9-F1-MODEL\_V4 \| 1.0 \| 0.004266 \| 157 \| 0.189 \| 79 \| 63 \| 1 \| 16 \| 93 \| 15 \| 93 \| Uncharacterized protein \| Uncharacterized protein; afdb-uniprot50 \| AF-A0A0M3AY55-F1-MODEL\_V4 \| 1.0 \| 0.003999 \| 156 \| 0.166 \| 72 \| 59 \| 1 \| 16 \| 87 \| 5 \| 75 \| Uncharacterized protein \| Uncharacterized protein; afdb-uniprot50 \| AF-A0A1X7F9Z0-F1-MODEL\_V4 \| 1.0 \| 0.001618 \| 156 \| 0.192 \| 78 \| 62 \| 1 \| 15 \| 92 \| 10 \| 86 \| Phage tail assembly chaperone protein, E, or 41 or 14 \| Phage tail assembly chaperone protein, E, or 41 or 14; afdb-uniprot50 \| AF-A0A629KGM0-F1-MODEL\_V4 \| 1.0 \| 0.003749 \| 156 \| 0.246 \| 77 \| 57 \| 1 \| 18 \| 93 \| 8 \| 84 \| Phage tail assembly protein \| Phage tail assembly protein; afdb-uniprot50 \| AF-A0A2A2H0C8-F1-MODEL\_V4 \| 1.0 \| 0.003749 \| 156 \| 0.197 \| 81 \| 64 \| 1 \| 14 \| 93 \| 3 \| 83 \| Uncharacterized protein \| Uncharacterized protein; afdb-uniprot50 \| AF-A0A847S1J7-F1-MODEL\_V4 \| 1.0 \| 0.004551 \| 156 \| 0.259 \| 77 \| 56 \| 1 \| 18 \| 93 \| 2 \| 78 \| Phage tail assembly protein \| Phage tail assembly protein; afdb-uniprot50 \| AF-A0A109JFT6-F1-MODEL\_V4 \| 1.0 \| 0.001029 \| 156 \| 0.232 \| 86 \| 55 \| 1 \| 18 \| 92 \| 14 \| 99 \| Uncharacterized protein \| Uncharacterized protein; afdb-uniprot50 \| AF-E3H9B2-F1-MODEL\_V4 \| 1.0 \| 0.0007949 \| 156 \| 0.15 \| 106 \| 75 \| 4 \| 1 \| 93 \| 7 \| 110 \| Uncharacterized protein \| Uncharacterized protein; afdb-uniprot50 \| AF-A0A1Y2JNK8-F1-MODEL\_V4 \| 1.0 \| 0.001726 \| 156 \| 0.168 \| 95 \| 68 \| 1 \| 11 \| 94 \| 21 \| 115 \| Uncharacterized protein \| Uncharacterized protein; afdb-uniprot50 \| AF-A0A0J7LYN2-F1-MODEL\_V4 \| 1.0 \| 0.001517 \| 156 \| 0.128 \| 101 \| 74 \| 5 \| 3 \| 92 \| 16 \| 113 \| Mu-like prophage FluMu protein gp41 \| Mu-like prophage FluMu protein gp41; afdb-uniprot50 \| AF-A0A5Y9K6W5-F1-MODEL\_V4 \| 1.0 \| 0.002544 \| 155 \| 0.302 \| 76 \| 45 \| 2 \| 18 \| 93 \| 5 \| 72 \| Phage tail assembly protein \| Phage tail assembly protein; afdb-uniprot50 \| AF-I3D8M1-F1-MODEL\_V4 \| 1.0 \| 0.002235 \| 155 \| 0.202 \| 79 \| 58 \| 3 \| 16 \| 92 \| 3 \| 78 \| Phage tail protein E \| Phage tail protein E; afdb-uniprot50 \| AF-A0A0D2JFD8-F1-MODEL\_V4 \| 1.0 \| 0.0007949 \| 155 \| 0.238 \| 88 \| 57 \| 3 \| 16 \| 93 \| 5 \| 92 \| Uncharacterized protein \| Uncharacterized protein; afdb-uniprot50 \| AF-A0A418W4D9-F1-MODEL\_V4 \| 1.0 \| 0.001098 \| 155 \| 0.202 \| 89 \| 67 \| 2 \| 9 \| 93 \| 2 \| 90 \| Phage tail assembly protein \| Phage tail assembly protein; afdb-uniprot50 \| AF-A0A258L5N6-F1-MODEL\_V4 \| 1.0 \| 0.005178 \| 155 \| 0.189 \| 79 \| 63 \| 1 \| 15 \| 93 \| 122 \| 199 \| Uncharacterized protein \| Uncharacterized protein; afdb-uniprot50 \| AF-A0A1G3UAH2-F1-MODEL\_V4 \| 1.0 \| 0.002544 \| 154 \| 0.285 \| 77 \| 45 \| 3 \| 18 \| 94 \| 3 \| 69 \| Uncharacterized protein \| Uncharacterized protein; afdb-uniprot50 \| AF-A0A3G2V508-F1-MODEL\_V4 \| 1.0 \| 0.0005394 \| 154 \| 0.223 \| 94 \| 66 \| 2 \| 4 \| 93 \| 1 \| 91 \| Phage tail assembly protein \| Phage tail assembly protein; afdb-uniprot50 \| AF-A0A850I2F0-F1-MODEL\_V4 \| 1.0 \| 0.000848 \| 154 \| 0.192 \| 104 \| 65 \| 2 \| 2 \| 94 \| 34 \| 129 \| Phage tail assembly protein \| Phage tail assembly protein; afdb-uniprot50 \| AF-A0A2W6YN90-F1-MODEL\_V4 \| 1.0 \| 0.0006986 \| 154 \| 0.193 \| 93 \| 73 \| 1 \| 2 \| 92 \| 38 \| 130 \| Uncharacterized protein \| Uncharacterized protein; afdb-uniprot50 \| AF-A0A0F4NK84-F1-MODEL\_V4 \| 1.0 \| 0.002544 \| 153 \| 0.277 \| 72 \| 48 \| 2 \| 23 \| 94 \| 2 \| 69 \| Uncharacterized protein \| Uncharacterized protein; afdb-uniprot50 \| AF-Q7NY03-F1-MODEL\_V4 \| 1.0 \| 0.009882 \| 153 \| 0.22 \| 77 \| 59 \| 1 \| 18 \| 93 \| 3 \| 79 \| Uncharacterized protein \| Uncharacterized protein; afdb-uniprot50 \| AF-A0A5P9JUS1-F1-MODEL\_V4 \| 1.0 \| 0.002385 \| 153 \| 0.188 \| 85 \| 65 \| 2 \| 10 \| 91 \| 1 \| 84 \| Uncharacterized protein \| Uncharacterized protein; afdb-uniprot50 \| AF-A0A430BCQ8-F1-MODEL\_V4 \| 1.0 \| 0.0006548 \| 153 \| 0.186 \| 91 \| 67 \| 2 \| 4 \| 94 \| 7 \| 90 \| Uncharacterized protein \| Uncharacterized protein; afdb-uniprot50 \| AF-A0A077PQ35-F1-MODEL\_V4 \| 1.0 \| 0.003999 \| 153 \| 0.18 \| 83 \| 64 \| 2 \| 15 \| 93 \| 4 \| 86 \| Uncharacterized protein \| Uncharacterized protein; afdb-uniprot50 \| AF-A0A6B2KN65-F1-MODEL\_V4 \| 1.0 \| 0.001422 \| 153 \| 0.147 \| 88 \| 65 \| 1 \| 16 \| 93 \| 3 \| 90 \| Phage tail assembly protein \| Phage tail assembly protein; afdb-uniprot50 \| AF-S9S610-F1-MODEL\_V4 \| 1.0 \| 0.0009046 \| 153 \| 0.266 \| 90 \| 59 \| 3 \| 8 \| 92 \| 5 \| 92 \| Putative phage related protein \| Putative phage related protein; afdb-uniprot50 \| AF-A0A423PRR2-F1-MODEL\_V4 \| 1.0 \| 0.004854 \| 153 \| 0.175 \| 80 \| 63 \| 2 \| 15 \| 92 \| 28 \| 106 \| Uncharacterized protein \| Uncharacterized protein; afdb-uniprot50 \| AF-A0A5M8ZBZ0-F1-MODEL\_V4 \| 1.0 \| 0.0005754 \| 153 \| 0.206 \| 97 \| 67 \| 6 \| 6 \| 94 \| 11 \| 105 \| Uncharacterized protein \| Uncharacterized protein; afdb-uniprot50 \| AF-A0A7W8PBC6-F1-MODEL\_V4 \| 1.0 \| 0.005178 \| 152 \| 0.185 \| 81 \| 60 \| 3 \| 16 \| 94 \| 2 \| 78 \| Uncharacterized protein \| Uncharacterized protein; afdb-uniprot50 \| AF-I9WY64-F1-MODEL\_V4 \| 1.0 \| 0.002895 \| 152 \| 0.246 \| 77 \| 56 \| 1 \| 18 \| 94 \| 6 \| 80 \| Uncharacterized protein \| Uncharacterized protein; afdb-uniprot50 \| AF-A0A1Y5TZY3-F1-MODEL\_V4 \| 1.0 \| 0.003294 \| 152 \| 0.246 \| 81 \| 54 \| 2 \| 16 \| 92 \| 5 \| 82 \| Phage tail protein E \| Phage tail protein E; afdb-uniprot50 \| AF-A0A2I8DJT9-F1-MODEL\_V4 \| 1.0 \| 0.004551 \| 152 \| 0.202 \| 79 \| 63 \| 0 \| 16 \| 94 \| 4 \| 82 \| Phage tail assembly protein \| Phage tail assembly protein; afdb-uniprot50 \| AF-A0A5E5PBU8-F1-MODEL\_V4 \| 1.0 \| 0.003514 \| 152 \| 0.214 \| 84 \| 62 \| 1 \| 15 \| 94 \| 5 \| 88 \| Phage tail assembly protein \| Phage tail assembly protein; afdb-uniprot50 \| AF-A0A645HXF0-F1-MODEL\_V4 \| 1.0 \| 0.003749 \| 152 \| 0.222 \| 81 \| 60 \| 2 \| 14 \| 92 \| 2 \| 81 \| Uncharacterized protein \| Uncharacterized protein; afdb-uniprot50 \| AF-A0A521D126-F1-MODEL\_V4 \| 1.0 \| 0.001618 \| 152 \| 0.193 \| 93 \| 64 \| 2 \| 3 \| 92 \| 1 \| 85 \| Phage tail assembly chaperone protein, E, or 41 or 14 \| Phage tail assembly chaperone protein, E, or 41 or 14; afdb-uniprot50 \| AF-A0A5B9CYR7-F1-MODEL\_V4 \| 1.0 \| 0.003749 \| 152 \| 0.219 \| 73 \| 55 \| 1 \| 16 \| 88 \| 6 \| 76 \| Phage tail assembly protein \| Phage tail assembly protein; afdb-uniprot50 \| AF-A0A5N7YGG3-F1-MODEL\_V4 \| 1.0 \| 0.006286 \| 151 \| 0.421 \| 57 \| 33 \| 0 \| 37 \| 93 \| 3 \| 59 \| Phage tail assembly protein \| Phage tail assembly protein; afdb-uniprot50 \| AF-A0A258D055-F1-MODEL\_V4 \| 1.0 \| 0.003749 \| 151 \| 0.202 \| 69 \| 53 \| 2 \| 16 \| 83 \| 4 \| 71 \| Uncharacterized protein \| Uncharacterized protein; afdb-uniprot50 \| AF-A0A1P8QQ42-F1-MODEL\_V4 \| 1.0 \| 0.001171 \| 151 \| 0.189 \| 95 \| 74 \| 1 \| 1 \| 92 \| 1 \| 95 \| Uncharacterized protein \| Uncharacterized protein; afdb-uniprot50 \| AF-A0A516W9Z3-F1-MODEL\_V4 \| 1.0 \| 0.002235 \| 151 \| 0.159 \| 88 \| 65 \| 1 \| 14 \| 92 \| 2 \| 89 \| Phage tail assembly protein \| Phage tail assembly protein; afdb-uniprot50 \| AF-A0A411WIN1-F1-MODEL\_V4 \| 1.0 \| 0.002895 \| 150 \| 0.301 \| 83 \| 51 \| 2 \| 1 \| 82 \| 1 \| 77 \| Phage tail assembly protein \| Phage tail assembly protein; afdb-uniprot50 \| AF-A0A2M8G048-F1-MODEL\_V4 \| 1.0 \| 0.002714 \| 150 \| 0.209 \| 81 \| 59 \| 2 \| 15 \| 93 \| 4 \| 81 \| Phage tail assembly protein \| Phage tail assembly protein; afdb-uniprot50 \| AF-A0A4Y9GJY5-F1-MODEL\_V4 \| 1.0 \| 0.007631 \| 150 \| 0.217 \| 78 \| 60 \| 1 \| 13 \| 89 \| 11 \| 88 \| Phage tail assembly protein \| Phage tail assembly protein; afdb-uniprot50 \| AF-A0A7Y3JAD0-F1-MODEL\_V4 \| 1.0 \| 0.0007949 \| 150 \| 0.206 \| 92 \| 64 \| 3 \| 2 \| 92 \| 4 \| 87 \| Phage tail assembly protein \| Phage tail assembly protein; afdb-uniprot50 \| AF-A0A849MU39-F1-MODEL\_V4 \| 1.0 \| 0.00125 \| 150 \| 0.19 \| 84 \| 62 \| 3 \| 16 \| 93 \| 15 \| 98 \| Phage tail assembly protein \| Phage tail assembly protein; afdb-uniprot50 \| AF-A0A2U8WC22-F1-MODEL\_V4 \| 1.0 \| 0.0006548 \| 150 \| 0.223 \| 103 \| 64 \| 3 \| 4 \| 92 \| 1 \| 101 \| Uncharacterized protein \| Uncharacterized protein; afdb-uniprot50 \| AF-A0A482IXJ4-F1-MODEL\_V4 \| 1.0 \| 0.003088 \| 150 \| 0.295 \| 71 \| 48 \| 2 \| 18 \| 86 \| 43 \| 113 \| Phage tail assembly protein \| Phage tail assembly protein; afdb-uniprot50 \| AF-A0A7W6EB82-F1-MODEL\_V4 \| 1.0 \| 0.0005394 \| 150 \| 0.202 \| 94 \| 66 \| 3 \| 1 \| 92 \| 61 \| 147 \| Uncharacterized protein \| Uncharacterized protein; afdb-uniprot50 \| AF-A0A4Q0ZNJ1-F1-MODEL\_V4 \| 1.0 \| 0.001964 \| 149 \| 0.267 \| 71 \| 48 \| 2 \| 24 \| 94 \| 6 \| 72 \| Uncharacterized protein \| Uncharacterized protein; afdb-uniprot50 \| AF-A0A2A7V0W4-F1-MODEL\_V4 \| 1.0 \| 0.002895 \| 149 \| 0.17 \| 88 \| 64 \| 1 \| 15 \| 93 \| 2 \| 89 \| Phage tail assembly protein \| Phage tail assembly protein; afdb-uniprot50 \| AF-A0A2N5C424-F1-MODEL\_V4 \| 1.0 \| 0.002385 \| 149 \| 0.202 \| 84 \| 63 \| 2 \| 15 \| 94 \| 11 \| 94 \| Phage tail assembly protein \| Phage tail assembly protein; afdb-uniprot50 \| AF-A0A4U5ZRD0-F1-MODEL\_V4 \| 1.0 \| 0.007154 \| 149 \| 0.171 \| 70 \| 57 \| 1 \| 18 \| 87 \| 56 \| 124 \| Phage tail assembly protein \| Phage tail assembly protein; afdb-uniprot50 \| AF-A0A6S5Y2Z1-F1-MODEL\_V4 \| 1.0 \| 0.001422 \| 148 \| 0.234 \| 94 \| 68 \| 2 \| 1 \| 92 \| 1 \| 92 \| Uncharacterized protein \| Uncharacterized protein; afdb-uniprot50 \| AF-A0A4R7WAS9-F1-MODEL\_V4 \| 1.0 \| 0.001726 \| 148 \| 0.2 \| 85 \| 59 \| 1 \| 18 \| 93 \| 6 \| 90 \| Tail assembly chaperone E/41/14-like protein \| Tail assembly chaperone E/41/14-like protein; afdb-uniprot50 \| AF-A0A1W1Z5N4-F1-MODEL\_V4 \| 1.0 \| 0.001726 \| 148 \| 0.119 \| 84 \| 66 \| 3 \| 4 \| 87 \| 1 \| 76 \| Phage tail assembly chaperone protein, E, or 41 or 14 \| Phage tail assembly chaperone protein, E, or 41 or 14; afdb-uniprot50 \| AF-A0A4S5JA13-F1-MODEL\_V4 \| 1.0 \| 0.001333 \| 148 \| 0.195 \| 92 \| 72 \| 1 \| 1 \| 92 \| 30 \| 119 \| Phage tail assembly protein \| Phage tail assembly protein; afdb-uniprot50 \| AF-A0A174ERE2-F1-MODEL\_V4 \| 1.0 \| 0.000265 \| 148 \| 0.231 \| 108 \| 68 \| 4 \| 1 \| 93 \| 36 \| 143 \| Uncharacterized protein \| Uncharacterized protein; afdb-uniprot50 \| AF-A0A154IH79-F1-MODEL\_V4 \| 1.0 \| 0.004854 \| 147 \| 0.197 \| 76 \| 60 \| 1 \| 18 \| 93 \| 3 \| 77 \| Uncharacterized protein \| Uncharacterized protein; afdb-uniprot50 \| AF-A0A6L5PED5-F1-MODEL\_V4 \| 1.0 \| 0.002895 \| 147 \| 0.148 \| 81 \| 65 \| 2 \| 15 \| 91 \| 4 \| 84 \| Uncharacterized protein \| Uncharacterized protein; afdb-uniprot50 \| AF-B6WS21-F1-MODEL\_V4 \| 1.0 \| 0.002096 \| 147 \| 0.23 \| 78 \| 56 \| 3 \| 18 \| 93 \| 5 \| 80 \| Phage tail protein E \| Phage tail protein E; afdb-uniprot50 \| AF-A0A6N9TJJ6-F1-MODEL\_V4 \| 1.0 \| 0.002714 \| 147 \| 0.228 \| 83 \| 57 \| 2 \| 15 \| 93 \| 2 \| 81 \| Phage tail assembly protein \| Phage tail assembly protein; afdb-uniprot50 \| AF-A0A2U0TF96-F1-MODEL\_V4 \| 1.0 \| 0.008141 \| 147 \| 0.2 \| 75 \| 59 \| 1 \| 18 \| 92 \| 6 \| 79 \| Tail assembly chaperone E/41/14-like protein \| Tail assembly chaperone E/41/14-like protein; afdb-uniprot50 \| AF-E9CK62-F1-MODEL\_V4 \| 1.0 \| 0.002544 \| 147 \| 0.209 \| 86 \| 60 \| 1 \| 16 \| 93 \| 2 \| 87 \| Uncharacterized protein \| Uncharacterized protein; afdb-uniprot50 \| AF-A0A1E2RY06-F1-MODEL\_V4 \| 1.0 \| 0.001333 \| 147 \| 0.209 \| 105 \| 71 \| 2 \| 1 \| 93 \| 2 \| 106 \| Uncharacterized protein \| Uncharacterized protein; afdb-uniprot50 \| AF-A0A4V2RX89-F1-MODEL\_V4 \| 1.0 \| 0.0009046 \| 147 \| 0.191 \| 94 \| 73 \| 1 \| 2 \| 92 \| 31 \| 124 \| Tail assembly chaperone E/41/14-like protein \| Tail assembly chaperone E/41/14-like protein; afdb-uniprot50 \| AF-A0A600X9S7-F1-MODEL\_V4 \| 1.0 \| 0.004266 \| 146 \| 0.233 \| 77 \| 58 \| 1 \| 18 \| 93 \| 4 \| 80 \| Phage tail assembly protein \| Phage tail assembly protein; afdb-uniprot50 \| AF-A0A5E4Z4W9-F1-MODEL\_V4 \| 1.0 \| 0.005893 \| 145 \| 0.214 \| 84 \| 62 \| 1 \| 15 \| 94 \| 5 \| 88 \| Phage tail assembly protein \| Phage tail assembly protein; afdb-uniprot50 \| AF-A0A2G0Q146-F1-MODEL\_V4 \| 1.0 \| 0.004854 \| 145 \| 0.129 \| 85 \| 68 \| 2 \| 15 \| 93 \| 3 \| 87 \| Uncharacterized protein \| Uncharacterized protein; afdb-uniprot50 \| AF-A0A2M9VHD7-F1-MODEL\_V4 \| 1.0 \| 0.002544 \| 145 \| 0.183 \| 87 \| 62 \| 1 \| 16 \| 93 \| 3 \| 89 \| Phage tail assembly protein \| Phage tail assembly protein; afdb-uniprot50 \| AF-A0A1X0TET7-F1-MODEL\_V4 \| 1.0 \| 0.003999 \| 145 \| 0.244 \| 90 \| 57 \| 3 \| 16 \| 94 \| 3 \| 92 \| Uncharacterized protein \| Uncharacterized protein; afdb-uniprot50 \| AF-A0A2A5EJW3-F1-MODEL\_V4 \| 1.0 \| 0.002714 \| 145 \| 0.189 \| 95 \| 62 \| 2 \| 11 \| 90 \| 1 \| 95 \| Uncharacterized protein \| Uncharacterized protein; afdb-uniprot50 \| AF-A0A1G7ZBM7-F1-MODEL\_V4 \| 1.0 \| 0.0009046 \| 145 \| 0.186 \| 91 \| 71 \| 3 \| 3 \| 92 \| 2 \| 90 \| Phage tail assembly chaperone protein, E, or 41 or 14 \| Phage tail assembly chaperone protein, E, or 41 or 14; afdb-uniprot50 \| AF-A0A098ARP9-F1-MODEL\_V4 \| 1.0 \| 0.009264 \| 144 \| 0.339 \| 56 \| 37 \| 0 \| 18 \| 73 \| 3 \| 58 \| Putative phage related protein \| Putative phage related protein; afdb-uniprot50 \| AF-A0A6G5QYA4-F1-MODEL\_V4 \| 1.0 \| 0.0006548 \| 144 \| 0.214 \| 98 \| 65 \| 4 \| 1 \| 94 \| 1 \| 90 \| Uncharacterized protein \| Uncharacterized protein; afdb-uniprot50 \| AF-A0A1C3WNV3-F1-MODEL\_V4 \| 1.0 \| 0.002714 \| 144 \| 0.191 \| 89 \| 61 \| 1 \| 15 \| 92 \| 7 \| 95 \| Phage tail assembly chaperone protein, E, or 41 or 14 \| Phage tail assembly chaperone protein, E, or 41 or 14; afdb-uniprot50 \| AF-A0A743BZ38-F1-MODEL\_V4 \| 1.0 \| 0.005178 \| 144 \| 0.233 \| 77 \| 58 \| 1 \| 18 \| 93 \| 4 \| 80 \| Phage tail assembly protein \| Phage tail assembly protein; afdb-uniprot50 \| AF-A0A0A8JLJ0-F1-MODEL\_V4 \| 1.0 \| 0.002235 \| 144 \| 0.221 \| 95 \| 61 \| 4 \| 12 \| 93 \| 25 \| 119 \| Uncharacterized protein \| Uncharacterized protein; afdb-uniprot50 \| AF-A0A4R3EF76-F1-MODEL\_V4 \| 1.0 \| 0.000848 \| 144 \| 0.206 \| 92 \| 71 \| 2 \| 2 \| 92 \| 31 \| 121 \| Tail assembly chaperone E/41/14-like protein \| Tail assembly chaperone E/41/14-like protein; afdb-uniprot50 \| AF-R9K258-F1-MODEL\_V4 \| 1.0 \| 0.0007452 \| 144 \| 0.2 \| 105 \| 71 \| 4 \| 1 \| 93 \| 39 \| 142 \| Uncharacterized protein \| Uncharacterized protein; afdb-uniprot50 \| AF-A0A259CE66-F1-MODEL\_V4 \| 1.0 \| 0.007631 \| 143 \| 0.213 \| 75 \| 58 \| 1 \| 18 \| 92 \| 3 \| 76 \| Uncharacterized protein \| Uncharacterized protein; afdb-uniprot50 \| AF-A0A2S7JEN3-F1-MODEL\_V4 \| 1.0 \| 0.002385 \| 143 \| 0.159 \| 88 \| 62 \| 2 \| 18 \| 93 \| 6 \| 93 \| Phage tail assembly protein \| Phage tail assembly protein; afdb-uniprot50 \| AF-A0A0D1LLV6-F1-MODEL\_V4 \| 1.0 \| 0.003088 \| 143 \| 0.195 \| 87 \| 61 \| 1 \| 16 \| 93 \| 3 \| 89 \| ArsR family transcriptional regulator \| ArsR family transcriptional regulator; afdb-uniprot50 \| AF-A0A509MUH0-F1-MODEL\_V4 \| 1.0 \| 0.006286 \| 143 \| 0.127 \| 94 \| 74 \| 2 \| 7 \| 94 \| 31 \| 122 \| Uncharacterized protein \| Uncharacterized protein; afdb-uniprot50 \| AF-A0A840C0Q8-F1-MODEL\_V4 \| 1.0 \| 0.003088 \| 143 \| 0.225 \| 93 \| 71 \| 1 \| 1 \| 92 \| 27 \| 119 \| Uncharacterized protein \| Uncharacterized protein; afdb-uniprot50 \| AF-A0A606FEL0-F1-MODEL\_V4 \| 1.0 \| 0.005893 \| 142 \| 0.236 \| 76 \| 57 \| 1 \| 18 \| 92 \| 4 \| 79 \| Phage tail assembly protein \| Phage tail assembly protein; afdb-uniprot50 \| AF-A0A547PS60-F1-MODEL\_V4 \| 1.0 \| 0.008684 \| 142 \| 0.185 \| 70 \| 56 \| 1 \| 18 \| 87 \| 6 \| 74 \| Uncharacterized protein \| Uncharacterized protein; afdb-uniprot50 \| AF-A0A502FV57-F1-MODEL\_V4 \| 1.0 \| 0.000965 \| 142 \| 0.206 \| 92 \| 69 \| 2 \| 4 \| 94 \| 1 \| 89 \| Phage tail assembly protein \| Phage tail assembly protein; afdb-uniprot50 \| AF-A0A2V3VR35-F1-MODEL\_V4 \| 1.0 \| 0.004266 \| 142 \| 0.193 \| 88 \| 62 \| 2 \| 14 \| 92 \| 2 \| 89 \| Tail assembly chaperone E/41/14-like protein \| Tail assembly chaperone E/41/14-like protein; afdb-uniprot50 \| AF-A0A249A392-F1-MODEL\_V4 \| 1.0 \| 0.007631 \| 142 \| 0.197 \| 86 \| 64 \| 3 \| 13 \| 93 \| 12 \| 97 \| Phage tail assembly protein \| Phage tail assembly protein; afdb-uniprot50 \| AF-A0A2C9D5V1-F1-MODEL\_V4 \| 1.0 \| 0.003088 \| 142 \| 0.176 \| 85 \| 63 \| 4 \| 15 \| 94 \| 5 \| 87 \| Uncharacterized protein \| Uncharacterized protein; afdb-uniprot50 \| AF-A0A7X5CQL9-F1-MODEL\_V4 \| 1.0 \| 0.0009046 \| 142 \| 0.237 \| 97 \| 62 \| 3 \| 9 \| 93 \| 34 \| 130 \| Phage tail assembly protein \| Phage tail assembly protein; afdb-uniprot50 \| AF-A0A174U989-F1-MODEL\_V4 \| 1.0 \| 0.0006139 \| 142 \| 0.144 \| 104 \| 77 \| 3 \| 2 \| 93 \| 28 \| 131 \| Uncharacterized protein \| Uncharacterized protein; afdb-uniprot50 \| AF-A0A4Y9VR20-F1-MODEL\_V4 \| 1.0 \| 0.003999 \| 141 \| 0.209 \| 86 \| 58 \| 2 \| 18 \| 93 \| 2 \| 87 \| Phage tail assembly protein \| Phage tail assembly protein; afdb-uniprot50 \| AF-A0A450UE46-F1-MODEL\_V4 \| 1.0 \| 0.005893 \| 141 \| 0.223 \| 76 \| 58 \| 1 \| 18 \| 93 \| 3 \| 77 \| Mu-like prophage FluMu protein gp41 \| Mu-like prophage FluMu protein gp41; afdb-uniprot50 \| AF-A0A5E8GXQ8-F1-MODEL\_V4 \| 1.0 \| 0.0009046 \| 141 \| 0.206 \| 97 \| 72 \| 3 \| 1 \| 92 \| 5 \| 101 \| Uncharacterized protein \| Uncharacterized protein; afdb-uniprot50 \| AF-A0A2S1BAT4-F1-MODEL\_V4 \| 1.0 \| 0.009264 \| 140 \| 0.166 \| 78 \| 60 \| 2 \| 16 \| 88 \| 4 \| 81 \| Phage tail assembly protein \| Phage tail assembly protein; afdb-uniprot50 \| AF-A0A1F9FW38-F1-MODEL\_V4 \| 1.0 \| 0.008684 \| 140 \| 0.219 \| 73 \| 53 \| 2 \| 18 \| 89 \| 7 \| 76 \| Uncharacterized protein \| Uncharacterized protein; afdb-uniprot50 \| AF-A0A837WJ78-F1-MODEL\_V4 \| 1.0 \| 0.005524 \| 140 \| 0.188 \| 85 \| 61 \| 2 \| 16 \| 92 \| 3 \| 87 \| Uncharacterized protein \| Uncharacterized protein; afdb-uniprot50 \| AF-A0A2A5DI31-F1-MODEL\_V4 \| 1.0 \| 0.004266 \| 140 \| 0.13 \| 84 \| 63 \| 2 \| 15 \| 93 \| 8 \| 86 \| Uncharacterized protein \| Uncharacterized protein; afdb-uniprot50 \| AF-A0A6L7G199-F1-MODEL\_V4 \| 1.0 \| 0.007154 \| 140 \| 0.171 \| 76 \| 60 \| 2 \| 18 \| 91 \| 6 \| 80 \| Uncharacterized protein \| Uncharacterized protein; afdb-uniprot50 \| AF-A0A4Y8RG68-F1-MODEL\_V4 \| 1.0 \| 0.001964 \| 140 \| 0.155 \| 90 \| 68 \| 3 \| 1 \| 90 \| 1 \| 82 \| Phage tail assembly protein \| Phage tail assembly protein; afdb-uniprot50 \| AF-A0A7H1NUI4-F1-MODEL\_V4 \| 1.0 \| 0.002385 \| 140 \| 0.163 \| 98 \| 71 \| 2 \| 1 \| 93 \| 15 \| 106 \| Phage tail assembly chaperone protein \| Phage tail assembly chaperone protein; afdb-uniprot50 \| AF-A8I7U6-F1-MODEL\_V4 \| 1.0 \| 0.0005394 \| 140 \| 0.178 \| 95 \| 71 \| 3 \| 2 \| 92 \| 48 \| 139 \| Uncharacterized protein \| Uncharacterized protein; afdb-uniprot50 \| AF-A0A4P2SNB1-F1-MODEL\_V4 \| 1.0 \| 0.004854 \| 139 \| 0.182 \| 93 \| 68 \| 1 \| 10 \| 94 \| 1 \| 93 \| Uncharacterized protein \| Uncharacterized protein; afdb-uniprot50 \| AF-A0A1H7ZZV8-F1-MODEL\_V4 \| 1.0 \| 0.001618 \| 139 \| 0.142 \| 105 \| 75 \| 4 \| 1 \| 93 \| 4 \| 105 \| Uncharacterized protein \| Uncharacterized protein; afdb-uniprot50 \| AF-A0A1X7I5S5-F1-MODEL\_V4 \| 1.0 \| 0.003088 \| 139 \| 0.176 \| 85 \| 61 \| 4 \| 18 \| 93 \| 10 \| 94 \| Phage tail assembly chaperone protein, E, or 41 or 14 \| Phage tail assembly chaperone protein, E, or 41 or 14; afdb-uniprot50 \| AF-A0A1M3E4Y4-F1-MODEL\_V4 \| 1.0 \| 0.003088 \| 139 \| 0.166 \| 90 \| 71 \| 4 \| 7 \| 94 \| 45 \| 132 \| Uncharacterized protein \| Uncharacterized protein; afdb-uniprot50 \| AF-A0A4R6EIC9-F1-MODEL\_V4 \| 1.0 \| 0.003514 \| 138 \| 0.139 \| 93 \| 68 \| 3 \| 10 \| 94 \| 1 \| 89 \| Tail assembly chaperone E/41/14-like protein \| Tail assembly chaperone E/41/14-like protein; afdb-uniprot50 \| AF-A0A4R5M9C3-F1-MODEL\_V4 \| 1.0 \| 0.008141 \| 138 \| 0.173 \| 104 \| 75 \| 4 \| 1 \| 93 \| 3 \| 106 \| Phage tail assembly protein \| Phage tail assembly protein; afdb-uniprot50 \| AF-X1XFW3-F1-MODEL\_V4 \| 1.0 \| 0.005893 \| 138 \| 0.211 \| 85 \| 59 \| 1 \| 18 \| 94 \| 101 \| 185 \| DNA\_circ\_N domain-containing protein \| DNA\_circ\_N domain-containing protein; afdb-uniprot50 \| AF-A0A806VH15-F1-MODEL\_V4 \| 1.0 \| 0.002096 \| 137 \| 0.204 \| 93 \| 65 \| 3 \| 1 \| 91 \| 8 \| 93 \| Uncharacterized protein \| Uncharacterized protein; afdb-uniprot50 \| AF-A0A0Q8AKH2-F1-MODEL\_V4 \| 1.0 \| 0.008141 \| 137 \| 0.188 \| 85 \| 60 \| 1 \| 18 \| 93 \| 4 \| 88 \| ArsR family transcriptional regulator \| ArsR family transcriptional regulator; afdb-uniprot50 \| AF-A0A509BPY4-F1-MODEL\_V4 \| 1.0 \| 0.003749 \| 137 \| 0.225 \| 93 \| 68 \| 2 \| 2 \| 93 \| 11 \| 100 \| Uncharacterized protein \| Uncharacterized protein; afdb-uniprot50 \| AF-A0A2D0IHJ0-F1-MODEL\_V4 \| 1.0 \| 0.003999 \| 137 \| 0.211 \| 90 \| 58 \| 2 \| 18 \| 94 \| 5 \| 94 \| Uncharacterized protein \| Uncharacterized protein; afdb-uniprot50 \| AF-A0A1H1JSE7-F1-MODEL\_V4 \| 1.0 \| 0.003088 \| 137 \| 0.149 \| 107 \| 77 \| 4 \| 1 \| 94 \| 1 \| 106 \| Phage tail assembly chaperone protein, E, or 41 or 14 \| Phage tail assembly chaperone protein, E, or 41 or 14; afdb-uniprot50 \| AF-A0A3A6P0B7-F1-MODEL\_V4 \| 1.0 \| 0.004854 \| 137 \| 0.178 \| 101 \| 70 \| 6 \| 3 \| 92 \| 1 \| 99 \| Uncharacterized protein \| Uncharacterized protein; afdb-uniprot50 \| AF-A0A1Z1SQQ7-F1-MODEL\_V4 \| 1.0 \| 0.008684 \| 137 \| 0.16 \| 100 \| 72 \| 4 \| 2 \| 94 \| 10 \| 104 \| Phage tail protein \| Phage tail protein; afdb-uniprot50 \| AF-A0A3A6P6S0-F1-MODEL\_V4 \| 1.0 \| 0.005178 \| 137 \| 0.166 \| 90 \| 64 \| 2 \| 15 \| 93 \| 17 \| 106 \| Phage tail assembly protein \| Phage tail assembly protein; afdb-uniprot50 \| AF-A0A5M8C2B4-F1-MODEL\_V4 \| 1.0 \| 0.001171 \| 137 \| 0.183 \| 98 \| 67 \| 4 \| 9 \| 93 \| 30 \| 127 \| Phage tail assembly protein \| Phage tail assembly protein; afdb-uniprot50 \| AF-A0A658Z224-F1-MODEL\_V4 \| 1.0 \| 0.005178 \| 137 \| 0.159 \| 88 \| 66 \| 1 \| 14 \| 93 \| 3 \| 90 \| Bacteriophage protein \| Bacteriophage protein; afdb-uniprot50 \| AF-A0A4U8UC82-F1-MODEL\_V4 \| 1.0 \| 0.005893 \| 136 \| 0.319 \| 72 \| 45 \| 2 \| 23 \| 94 \| 3 \| 70 \| Phage tail assembly protein \| Phage tail assembly protein; afdb-uniprot50 \| AF-A0A5U1J7E5-F1-MODEL\_V4 \| 1.0 \| 0.008684 \| 136 \| 0.232 \| 73 \| 55 \| 1 \| 18 \| 89 \| 6 \| 78 \| Phage tail assembly protein \| Phage tail assembly protein; afdb-uniprot50 \| AF-A0A5E7ADD5-F1-MODEL\_V4 \| 1.0 \| 0.006286 \| 136 \| 0.2 \| 85 \| 59 \| 1 \| 18 \| 93 \| 6 \| 90 \| Uncharacterized protein \| Uncharacterized protein; afdb-uniprot50 \| AF-E5F1U9-F1-MODEL\_V4 \| 1.0 \| 0.006286 \| 136 \| 0.291 \| 72 \| 48 \| 1 \| 22 \| 93 \| 29 \| 97 \| Uncharacterized protein \| Uncharacterized protein; afdb-uniprot50 \| AF-A0A7W9QVC4-F1-MODEL\_V4 \| 1.0 \| 0.006286 \| 136 \| 0.208 \| 96 \| 66 \| 5 \| 2 \| 93 \| 8 \| 97 \| Uncharacterized protein \| Uncharacterized protein; afdb-uniprot50 \| AF-A0A411T3J1-F1-MODEL\_V4 \| 1.0 \| 0.003749 \| 136 \| 0.279 \| 86 \| 52 \| 2 \| 18 \| 93 \| 9 \| 94 \| Phage tail assembly protein \| Phage tail assembly protein; afdb-uniprot50 \| AF-A0A0E1GP01-F1-MODEL\_V4 \| 1.0 \| 0.001726 \| 136 \| 0.175 \| 97 \| 69 \| 4 \| 1 \| 94 \| 20 \| 108 \| Uncharacterized protein \| Uncharacterized protein; afdb-uniprot50 \| AF-A0A431JGG1-F1-MODEL\_V4 \| 1.0 \| 0.003294 \| 136 \| 0.16 \| 87 \| 70 \| 2 \| 9 \| 94 \| 33 \| 117 \| Phage tail assembly protein \| Phage tail assembly protein; afdb-uniprot50 \| AF-A0A4Q2SXK7-F1-MODEL\_V4 \| 1.0 \| 0.001618 \| 136 \| 0.21 \| 95 \| 68 \| 3 \| 2 \| 94 \| 65 \| 154 \| Phage tail assembly protein \| Phage tail assembly protein; afdb-uniprot50 \| AF-A0A1S1D167-F1-MODEL\_V4 \| 1.0 \| 0.006706 \| 135 \| 0.195 \| 87 \| 59 \| 2 \| 15 \| 93 \| 2 \| 85 \| Uncharacterized protein \| Uncharacterized protein; afdb-uniprot50 \| AF-A0A650EN83-F1-MODEL\_V4 \| 1.0 \| 0.004551 \| 135 \| 0.172 \| 81 \| 63 \| 1 \| 18 \| 94 \| 7 \| 87 \| Uncharacterized protein \| Uncharacterized protein; afdb-uniprot50 \| AF-A0A7U4R2Y7-F1-MODEL\_V4 \| 1.0 \| 0.005178 \| 135 \| 0.165 \| 103 \| 77 \| 3 \| 1 \| 94 \| 1 \| 103 \| Uncharacterized protein \| Uncharacterized protein; afdb-uniprot50 \| AF-A0A239JSH3-F1-MODEL\_V4 \| 1.0 \| 0.001618 \| 135 \| 0.172 \| 110 \| 72 \| 6 \| 1 \| 92 \| 1 \| 109 \| Phage tail assembly chaperone protein, E, or 41 or 14 \| Phage tail assembly chaperone protein, E, or 41 or 14; afdb-uniprot50 \| AF-A0A504U8T9-F1-MODEL\_V4 \| 1.0 \| 0.005178 \| 135 \| 0.141 \| 78 \| 64 \| 1 \| 15 \| 92 \| 35 \| 109 \| Phage tail assembly protein \| Phage tail assembly protein; afdb-uniprot50 \| AF-H4FBM5-F1-MODEL\_V4 \| 1.0 \| 0.002544 \| 135 \| 0.184 \| 92 \| 67 \| 3 \| 1 \| 92 \| 28 \| 111 \| Uncharacterized protein \| Uncharacterized protein; afdb-uniprot50 \| AF-A0A0F6B300-F1-MODEL\_V4 \| 1.0 \| 0.001964 \| 135 \| 0.197 \| 86 \| 59 \| 4 \| 16 \| 92 \| 5 \| 89 \| Uncharacterized protein \| Uncharacterized protein; afdb-uniprot50 \| AF-A0A431MY95-F1-MODEL\_V4 \| 1.0 \| 0.001964 \| 134 \| 0.209 \| 86 \| 61 \| 3 \| 9 \| 94 \| 2 \| 80 \| Phage tail assembly protein \| Phage tail assembly protein; afdb-uniprot50 \| AF-R7I6D8-F1-MODEL\_V4 \| 1.0 \| 0.005178 \| 134 \| 0.21 \| 76 \| 54 \| 3 \| 16 \| 85 \| 3 \| 78 \| Uncharacterized protein \| Uncharacterized protein; afdb-uniprot50 \| AF-A0A345CNU2-F1-MODEL\_V4 \| 1.0 \| 0.003514 \| 134 \| 0.149 \| 87 \| 64 \| 3 \| 15 \| 94 \| 7 \| 90 \| Phage tail assembly protein \| Phage tail assembly protein; afdb-uniprot50 \| AF-K0JHK7-F1-MODEL\_V4 \| 1.0 \| 0.004266 \| 134 \| 0.159 \| 88 \| 69 \| 2 \| 10 \| 94 \| 1 \| 86 \| Uncharacterized protein \| Uncharacterized protein; afdb-uniprot50 \| AF-A0A418X1D6-F1-MODEL\_V4 \| 1.0 \| 0.005178 \| 134 \| 0.183 \| 87 \| 59 \| 2 \| 18 \| 92 \| 2 \| 88 \| Phage tail assembly protein \| Phage tail assembly protein; afdb-uniprot50 \| AF-A0A0S4V3E5-F1-MODEL\_V4 \| 1.0 \| 0.009264 \| 134 \| 0.172 \| 81 \| 60 \| 3 \| 16 \| 91 \| 6 \| 84 \| Hypothethical protein \| Hypothethical protein; afdb-uniprot50 \| AF-A0A1I1EZQ6-F1-MODEL\_V4 \| 1.0 \| 0.005178 \| 134 \| 0.141 \| 85 \| 73 \| 0 \| 3 \| 87 \| 36 \| 120 \| Uncharacterized protein \| Uncharacterized protein; afdb-uniprot50 \| AF-A0A0Q6C6X2-F1-MODEL\_V4 \| 1.0 \| 0.003514 \| 134 \| 0.172 \| 93 \| 65 \| 3 \| 1 \| 92 \| 67 \| 148 \| Uncharacterized protein \| Uncharacterized protein; afdb-uniprot50 \| AF-A0A7G6U2B7-F1-MODEL\_V4 \| 1.0 \| 0.006706 \| 133 \| 0.157 \| 76 \| 59 \| 2 \| 18 \| 93 \| 6 \| 76 \| Phage tail assembly protein \| Phage tail assembly protein; afdb-uniprot50 \| AF-A0A522K8G1-F1-MODEL\_V4 \| 1.0 \| 0.007154 \| 133 \| 0.193 \| 93 \| 71 \| 2 \| 3 \| 92 \| 2 \| 93 \| Phage tail assembly protein \| Phage tail assembly protein; afdb-uniprot50 \| AF-A0A5M8P9I7-F1-MODEL\_V4 \| 1.0 \| 0.005178 \| 133 \| 0.157 \| 121 \| 64 \| 4 \| 10 \| 92 \| 1 \| 121 \| Uncharacterized protein \| Uncharacterized protein; afdb-uniprot50 \| AF-A0A285T5R0-F1-MODEL\_V4 \| 1.0 \| 0.00125 \| 133 \| 0.202 \| 99 \| 69 \| 3 \| 1 \| 92 \| 40 \| 135 \| Uncharacterized protein \| Uncharacterized protein; afdb-uniprot50 \| AF-A0A7W7KJ99-F1-MODEL\_V4 \| 1.0 \| 0.007154 \| 132 \| 0.202 \| 84 \| 62 \| 3 \| 15 \| 94 \| 3 \| 85 \| Uncharacterized protein \| Uncharacterized protein; afdb-uniprot50 \| AF-A0A291PD75-F1-MODEL\_V4 \| 1.0 \| 0.003088 \| 132 \| 0.177 \| 96 \| 69 \| 5 \| 5 \| 93 \| 11 \| 103 \| Uncharacterized protein \| Uncharacterized protein; afdb-uniprot50 \| AF-A0A3R9CVH5-F1-MODEL\_V4 \| 1.0 \| 0.008141 \| 132 \| 0.17 \| 88 \| 61 \| 3 \| 15 \| 94 \| 18 \| 101 \| Phage tail assembly protein \| Phage tail assembly protein; afdb-uniprot50 \| AF-A0A016XIN1-F1-MODEL\_V4 \| 1.0 \| 0.009882 \| 132 \| 0.157 \| 89 \| 69 \| 3 \| 9 \| 94 \| 2 \| 87 \| Uncharacterized protein \| Uncharacterized protein; afdb-uniprot50 \| AF-A0A3R7HUR4-F1-MODEL\_V4 \| 1.0 \| 0.006706 \| 132 \| 0.243 \| 82 \| 58 \| 3 \| 15 \| 94 \| 5 \| 84 \| Tail assembly chaperone E/41/14-like protein \| Tail assembly chaperone E/41/14-like protein; afdb-uniprot50 \| AF-A0A7Y6Z4K4-F1-MODEL\_V4 \| 1.0 \| 0.003514 \| 131 \| 0.172 \| 93 \| 67 \| 4 \| 9 \| 94 \| 2 \| 91 \| Phage tail assembly protein \| Phage tail assembly protein; afdb-uniprot50 \| AF-A0A2W5S0Y9-F1-MODEL\_V4 \| 1.0 \| 0.009264 \| 131 \| 0.175 \| 91 \| 63 \| 3 \| 12 \| 94 \| 14 \| 100 \| Phage tail assembly protein \| Phage tail assembly protein; afdb-uniprot50 \| AF-A0A225SNI8-F1-MODEL\_V4 \| 1.0 \| 0.003294 \| 131 \| 0.2 \| 95 \| 70 \| 4 \| 2 \| 93 \| 14 \| 105 \| Uncharacterized protein \| Uncharacterized protein; afdb-uniprot50 \| AF-A0A7Y6T7N0-F1-MODEL\_V4 \| 1.0 \| 0.003749 \| 131 \| 0.205 \| 102 \| 67 \| 2 \| 2 \| 92 \| 12 \| 110 \| Phage tail assembly protein \| Phage tail assembly protein; afdb-uniprot50 \| AF-A0A3A9EEN6-F1-MODEL\_V4 \| 1.0 \| 0.001517 \| 131 \| 0.154 \| 97 \| 70 \| 4 \| 9 \| 93 \| 35 \| 131 \| Phage tail assembly protein \| Phage tail assembly protein; afdb-uniprot50 \| AF-A0A485CDJ1-F1-MODEL\_V4 \| 1.0 \| 0.004854 \| 131 \| 0.148 \| 101 \| 75 \| 4 \| 1 \| 94 \| 118 \| 214 \| Phage major tail tube protein \| Phage major tail tube protein; afdb-uniprot50 \| AF-A0A0D0GXK6-F1-MODEL\_V4 \| 1.0 \| 0.005524 \| 130 \| 0.175 \| 108 \| 73 \| 4 \| 1 \| 94 \| 1 \| 106 \| KY49.ctg7180000000025\_quiver, whole genome shotgun sequence \| KY49.ctg7180000000025\_quiver, whole genome shotgun sequence; afdb-uniprot50 \| AF-N8V954-F1-MODEL\_V4 \| 1.0 \| 0.009264 \| 130 \| 0.188 \| 90 \| 61 \| 3 \| 13 \| 94 \| 14 \| 99 \| Uncharacterized protein \| Uncharacterized protein; afdb-uniprot50 \| AF-E2CHK2-F1-MODEL\_V4 \| 1.0 \| 0.001333 \| 130 \| 0.195 \| 97 \| 69 \| 3 \| 2 \| 92 \| 25 \| 118 \| Uncharacterized protein \| Uncharacterized protein; afdb-uniprot50 \| AF-A0A5P8MUZ6-F1-MODEL\_V4 \| 1.0 \| 0.001841 \| 130 \| 0.178 \| 95 \| 72 \| 3 \| 1 \| 92 \| 48 \| 139 \| Uncharacterized protein \| Uncharacterized protein; afdb-uniprot50 \| AF-A0A6I5WQG3-F1-MODEL\_V4 \| 1.0 \| 0.008684 \| 129 \| 0.162 \| 86 \| 67 \| 3 \| 13 \| 93 \| 14 \| 99 \| Phage tail assembly protein \| Phage tail assembly protein; afdb-uniprot50 \| AF-A0A6N9P3E0-F1-MODEL\_V4 \| 1.0 \| 0.001964 \| 129 \| 0.2 \| 105 \| 69 \| 5 \| 1 \| 93 \| 66 \| 167 \| Phage tail assembly protein \| Phage tail assembly protein; afdb-uniprot50 \| AF-A0A1X9T1B4-F1-MODEL\_V4 \| 1.0 \| 0.006286 \| 128 \| 0.189 \| 95 \| 65 \| 4 \| 1 \| 94 \| 1 \| 84 \| FluMu gp41 family protein \| FluMu gp41 family protein; afdb-uniprot50 \| AF-A0A0Q4II11-F1-MODEL\_V4 \| 1.0 \| 0.003294 \| 128 \| 0.177 \| 96 \| 75 \| 2 \| 1 \| 94 \| 2 \| 95 \| Uncharacterized protein \| Uncharacterized protein; afdb-uniprot50 \| AF-A0A841S0F6-F1-MODEL\_V4 \| 1.0 \| 0.007154 \| 128 \| 0.147 \| 102 \| 75 \| 3 \| 1 \| 94 \| 8 \| 105 \| Uncharacterized protein \| Uncharacterized protein; afdb-uniprot50 \| AF-A0A6A1TS96-F1-MODEL\_V4 \| 1.0 \| 0.003749 \| 128 \| 0.186 \| 102 \| 70 \| 2 \| 1 \| 92 \| 45 \| 143 \| Phage tail assembly protein \| Phage tail assembly protein; afdb-uniprot50 \| AF-A0A2W6TML0-F1-MODEL\_V4 \| 1.0 \| 0.004551 \| 128 \| 0.184 \| 92 \| 74 \| 1 \| 2 \| 92 \| 67 \| 158 \| Uncharacterized protein \| Uncharacterized protein; afdb-uniprot50 \| AF-A0A736RG71-F1-MODEL\_V4 \| 1.0 \| 0.003999 \| 127 \| 0.195 \| 87 \| 60 \| 4 \| 16 \| 93 \| 2 \| 87 \| Phage tail assembly protein \| Phage tail assembly protein; afdb-uniprot50 \| AF-A0A1P8JYY7-F1-MODEL\_V4 \| 1.0 \| 0.005893 \| 127 \| 0.177 \| 96 \| 66 \| 3 \| 9 \| 93 \| 5 \| 98 \| Uncharacterized protein \| Uncharacterized protein; afdb-uniprot50 \| AF-A0A2U1SE14-F1-MODEL\_V4 \| 1.0 \| 0.008684 \| 127 \| 0.142 \| 98 \| 71 \| 3 \| 9 \| 93 \| 2 \| 99 \| Uncharacterized protein \| Uncharacterized protein; afdb-uniprot50 \| AF-A0A0D7PAE5-F1-MODEL\_V4 \| 1.0 \| 0.007154 \| 127 \| 0.166 \| 108 \| 76 \| 3 \| 1 \| 94 \| 1 \| 108 \| Uncharacterized protein \| Uncharacterized protein; afdb-uniprot50 \| AF-A0A7U5GYV5-F1-MODEL\_V4 \| 1.0 \| 0.001726 \| 127 \| 0.135 \| 111 \| 75 \| 5 \| 1 \| 93 \| 1 \| 108 \| Uncharacterized protein \| Uncharacterized protein; afdb-uniprot50 \| AF-L0NDQ1-F1-MODEL\_V4 \| 1.0 \| 0.005524 \| 127 \| 0.223 \| 94 \| 67 \| 3 \| 2 \| 92 \| 31 \| 121 \| Uncharacterized protein \| Uncharacterized protein; afdb-uniprot50 \| AF-A0A0P6VMZ8-F1-MODEL\_V4 \| 1.0 \| 0.003088 \| 127 \| 0.191 \| 94 \| 64 \| 4 \| 1 \| 88 \| 1 \| 88 \| Uncharacterized protein \| Uncharacterized protein; afdb-uniprot50 \| AF-A0A432UNV4-F1-MODEL\_V4 \| 1.0 \| 0.009264 \| 126 \| 0.155 \| 90 \| 68 \| 2 \| 10 \| 94 \| 1 \| 87 \| Phage tail assembly protein \| Phage tail assembly protein; afdb-uniprot50 \| AF-A0A6M8HND0-F1-MODEL\_V4 \| 1.0 \| 0.003999 \| 126 \| 0.175 \| 97 \| 66 \| 3 \| 4 \| 94 \| 8 \| 96 \| Phage tail assembly protein \| Phage tail assembly protein; afdb-uniprot50 \| AF-A0A1Q6JQ84-F1-MODEL\_V4 \| 1.0 \| 0.002235 \| 126 \| 0.196 \| 107 \| 73 \| 4 \| 1 \| 94 \| 27 \| 133 \| Uncharacterized protein \| Uncharacterized protein; afdb-uniprot50 \| AF-A0A829WCQ7-F1-MODEL\_V4 \| 1.0 \| 0.001964 \| 126 \| 0.194 \| 108 \| 71 \| 5 \| 2 \| 93 \| 30 \| 137 \| Uncharacterized protein \| Uncharacterized protein; afdb-uniprot50 \| AF-A0A2G6IRS6-F1-MODEL\_V4 \| 1.0 \| 0.003514 \| 126 \| 0.229 \| 96 \| 66 \| 3 \| 2 \| 92 \| 14 \| 106 \| Uncharacterized protein \| Uncharacterized protein; afdb-uniprot50 \| AF-A0A257KLL7-F1-MODEL\_V4 \| 1.0 \| 0.006706 \| 125 \| 0.207 \| 77 \| 56 \| 3 \| 18 \| 94 \| 2 \| 73 \| Uncharacterized protein \| Uncharacterized protein; afdb-uniprot50 \| AF-A0A378Q4F7-F1-MODEL\_V4 \| 1.0 \| 0.007631 \| 125 \| 0.2 \| 90 \| 58 \| 4 \| 14 \| 94 \| 7 \| 91 \| Phage tail protein E \| Phage tail protein E; afdb-uniprot50 \| AF-A0A2N3D8J3-F1-MODEL\_V4 \| 1.0 \| 0.009264 \| 125 \| 0.148 \| 101 \| 72 \| 4 \| 2 \| 94 \| 3 \| 97 \| Phage tail assembly protein \| Phage tail assembly protein; afdb-uniprot50 \| AF-A0A2S6QN92-F1-MODEL\_V4 \| 1.0 \| 0.004551 \| 125 \| 0.271 \| 70 \| 50 \| 1 \| 14 \| 82 \| 2 \| 71 \| Uncharacterized protein \| Uncharacterized protein; afdb-uniprot50 \| AF-A0A327K060-F1-MODEL\_V4 \| 1.0 \| 0.003294 \| 124 \| 0.244 \| 94 \| 64 \| 3 \| 3 \| 92 \| 2 \| 92 \| Uncharacterized protein \| Uncharacterized protein; afdb-uniprot50 \| AF-C1DBJ2-F1-MODEL\_V4 \| 1.0 \| 0.005893 \| 123 \| 0.172 \| 87 \| 62 \| 3 \| 15 \| 94 \| 2 \| 85 \| Uncharacterized protein \| Uncharacterized protein; afdb-uniprot50 \| AF-A0A547PW59-F1-MODEL\_V4 \| 1.0 \| 0.005524 \| 123 \| 0.207 \| 106 \| 66 \| 5 \| 1 \| 94 \| 3 \| 102 \| Phage tail assembly protein \| Phage tail assembly protein; afdb-uniprot50 \| AF-A0A7T3DHK5-F1-MODEL\_V4 \| 1.0 \| 0.008141 \| 123 \| 0.128 \| 101 \| 76 \| 5 \| 1 \| 94 \| 27 \| 122 \| Phage tail assembly protein \| Phage tail assembly protein; afdb-uniprot50 \| AF-A0A348HI85-F1-MODEL\_V4 \| 1.0 \| 0.008141 \| 122 \| 0.128 \| 101 \| 71 \| 4 \| 1 \| 93 \| 1 \| 92 \| Small-conductance mechanosensitive channel \| Small-conductance mechanosensitive channel; afdb-uniprot50 \| AF-A0A3L7JF17-F1-MODEL\_V4 \| 1.0 \| 0.004551 \| 122 \| 0.215 \| 93 \| 65 \| 4 \| 2 \| 92 \| 50 \| 136 \| Phage tail assembly protein \| Phage tail assembly protein; afdb-uniprot50 \| AF-A0A2C9D6L2-F1-MODEL\_V4 \| 1.0 \| 0.007631 \| 122 \| 0.18 \| 94 \| 70 \| 3 \| 3 \| 92 \| 61 \| 151 \| Uncharacterized protein \| Uncharacterized protein; afdb-uniprot50 \| AF-A0A2N3KX48-F1-MODEL\_V4 \| 1.0 \| 0.005178 \| 121 \| 0.197 \| 96 \| 57 \| 4 \| 10 \| 93 \| 1 \| 88 \| Phage tail protein \| Phage tail protein; afdb-uniprot50 \| AF-A0A806LZ96-F1-MODEL\_V4 \| 1.0 \| 0.007154 \| 121 \| 0.178 \| 95 \| 59 \| 4 \| 15 \| 93 \| 16 \| 107 \| Uncharacterized protein \| Uncharacterized protein; afdb-uniprot50 \| AF-A0NQ99-F1-MODEL\_V4 \| 1.0 \| 0.003749 \| 121 \| 0.185 \| 97 \| 65 \| 3 \| 10 \| 93 \| 1 \| 96 \| Uncharacterized protein \| Uncharacterized protein; afdb-uniprot50 \| AF-A0A2W5EHM5-F1-MODEL\_V4 \| 1.0 \| 0.008684 \| 121 \| 0.187 \| 96 \| 69 \| 3 \| 1 \| 92 \| 50 \| 140 \| Uncharacterized protein \| Uncharacterized protein; afdb-uniprot50 \| AF-A0A7W4JQX5-F1-MODEL\_V4 \| 1.0 \| 0.003514 \| 121 \| 0.25 \| 96 \| 64 \| 3 \| 2 \| 91 \| 14 \| 107 \| Phage tail assembly protein \| Phage tail assembly protein; afdb-uniprot50 \| AF-A0A410VCZ1-F1-MODEL\_V4 \| 1.0 \| 0.009264 \| 120 \| 0.228 \| 83 \| 59 \| 3 \| 12 \| 94 \| 7 \| 84 \| Uncharacterized protein \| Uncharacterized protein; afdb-uniprot50 \| AF-A0A653HUE2-F1-MODEL\_V4 \| 1.0 \| 0.009264 \| 120 \| 0.166 \| 90 \| 65 \| 3 \| 12 \| 94 \| 16 \| 102 \| Uncharacterized protein \| Uncharacterized protein; afdb-uniprot50 \| AF-A0A1M5NLQ3-F1-MODEL\_V4 \| 1.0 \| 0.004266 \| 120 \| 0.141 \| 106 \| 78 \| 4 \| 1 \| 93 \| 12 \| 117 \| Uncharacterized protein \| Uncharacterized protein; afdb-uniprot50 \| AF-A0A212LD12-F1-MODEL\_V4 \| 1.0 \| 0.005893 \| 120 \| 0.147 \| 95 \| 73 \| 3 \| 2 \| 92 \| 44 \| 134 \| Uncharacterized protein \| Uncharacterized protein; afdb-uniprot50 \| AF-A0A2E4CSJ3-F1-MODEL\_V4 \| 1.0 \| 0.008684 \| 119 \| 0.158 \| 101 \| 67 \| 4 \| 2 \| 94 \| 15 \| 105 \| Phage tail protein \| Phage tail protein; afdb-uniprot50 \| AF-A0A6B8KJI9-F1-MODEL\_V4 \| 1.0 \| 0.004266 \| 119 \| 0.19 \| 100 \| 69 \| 4 \| 1 \| 94 \| 33 \| 126 \| Uncharacterized protein \| Uncharacterized protein; afdb-uniprot50 \| AF-A0A7T9VKM3-F1-MODEL\_V4 \| 1.0 \| 0.008141 \| 119 \| 0.161 \| 93 \| 68 \| 3 \| 9 \| 94 \| 25 \| 114 \| Phage tail assembly protein \| Phage tail assembly protein; afdb-uniprot50 \| AF-A0A7L5Y296-F1-MODEL\_V4 \| 1.0 \| 0.008684 \| 119 \| 0.153 \| 98 \| 75 \| 3 \| 1 \| 93 \| 34 \| 128 \| Uncharacterized protein \| Uncharacterized protein; afdb-uniprot50 \| AF-A0A6N7EZF4-F1-MODEL\_V4 \| 1.0 \| 0.009264 \| 118 \| 0.186 \| 102 \| 65 \| 4 \| 1 \| 94 \| 12 \| 103 \| Phage tail assembly protein \| Phage tail assembly protein; afdb-uniprot50 \| AF-A0A2X0YJY8-F1-MODEL\_V4 \| 1.0 \| 0.005524 \| 118 \| 0.166 \| 108 \| 74 \| 4 \| 2 \| 93 \| 16 \| 123 \| Uncharacterized protein \| Uncharacterized protein; afdb-uniprot50 \| AF-A0A2M9WHJ3-F1-MODEL\_V4 \| 1.0 \| 0.009264 \| 118 \| 0.182 \| 93 \| 65 \| 4 \| 11 \| 92 \| 1 \| 93 \| Uncharacterized protein \| Uncharacterized protein; afdb-uniprot50 \| AF-B1HTN1-F1-MODEL\_V4 \| 1.0 \| 0.008141 \| 118 \| 0.144 \| 104 \| 78 \| 2 \| 1 \| 93 \| 26 \| 129 \| Uncharacterized protein \| Uncharacterized protein; afdb-uniprot50 \| AF-A0A4Q3U4W6-F1-MODEL\_V4 \| 1.0 \| 0.003294 \| 117 \| 0.186 \| 102 \| 69 \| 7 \| 2 \| 94 \| 25 \| 121 \| Phage tail assembly protein \| Phage tail assembly protein; afdb-uniprot50 \| AF-A0A2W5H3D6-F1-MODEL\_V4 \| 1.0 \| 0.009882 \| 117 \| 0.159 \| 94 \| 71 \| 3 \| 3 \| 92 \| 65 \| 154 \| Uncharacterized protein \| Uncharacterized protein; afdb-uniprot50 \| AF-A0A1M7ZLP0-F1-MODEL\_V4 \| 1.0 \| 0.005524 \| 116 \| 0.241 \| 91 \| 56 \| 4 \| 16 \| 93 \| 4 \| 94 \| Uncharacterized protein \| Uncharacterized protein; afdb-uniprot50 \| AF-A0A4U2PVC1-F1-MODEL\_V4 \| 1.0 \| 0.002895 \| 116 \| 0.155 \| 103 \| 74 \| 5 \| 2 \| 92 \| 8 \| 109 \| Phage tail assembly protein \| Phage tail assembly protein; afdb-uniprot50 \| AF-A0A0P7I4K8-F1-MODEL\_V4 \| 1.0 \| 0.007154 \| 115 \| 0.206 \| 97 \| 66 \| 5 \| 2 \| 92 \| 5 \| 96 \| Uncharacterized protein \| Uncharacterized protein; afdb-uniprot50 \| AF-E2CI76-F1-MODEL\_V4 \| 1.0 \| 0.005524 \| 115 \| 0.183 \| 98 \| 67 \| 5 \| 2 \| 92 \| 42 \| 133 \| Uncharacterized protein \| Uncharacterized protein; afdb-uniprot50 \| AF-A0A7S8IJ10-F1-MODEL\_V4 \| 1.0 \| 0.007631 \| 113 \| 0.18 \| 100 \| 70 \| 3 \| 2 \| 92 \| 14 \| 110 \| Uncharacterized protein \| Uncharacterized protein; afdb-uniprot50 \| AF-A0A7Z0LWV3-F1-MODEL\_V4 \| 1.0 \| 0.009264 \| 112 \| 0.148 \| 101 \| 70 \| 4 \| 1 \| 94 \| 16 \| 107 \| Phage tail assembly protein \| Phage tail assembly protein; afdb-uniprot50 \| AF-A0A1L5P7E5-F1-MODEL\_V4 \| 1.0 \| 0.009264 \| 111 \| 0.189 \| 95 \| 68 \| 4 \| 2 \| 92 \| 60 \| 149 \| Mu-like prophage FluMu protein \| Mu-like prophage FluMu protein; afdb-uniprot50 \| AF-A0A2A2IKT4-F1-MODEL\_V4 \| 1.0 \| 0.009882 \| 110 \| 0.229 \| 87 \| 55 \| 3 \| 18 \| 92 \| 2 \| 88 \| Uncharacterized protein \| Uncharacterized protein; afdb-uniprot50 \| AF-A0A2T6LZL0-F1-MODEL\_V4 \| 1.0 \| 0.008141 \| 110 \| 0.171 \| 99 \| 72 \| 3 \| 1 \| 92 \| 1 \| 96 \| Tail assembly chaperone E/41/14-like protein \| Tail assembly chaperone E/41/14-like protein; afdb-uniprot50 \| AF-A0A2V2FWG8-F1-MODEL\_V4 \| 1.0 \| 0.008141 \| 108 \| 0.185 \| 113 \| 72 \| 6 \| 1 \| 93 \| 1 \| 113 \| Uncharacterized protein \| Uncharacterized protein; afdb-uniprot50 \| AF-A0A2D9ER84-F1-MODEL\_V4 \| 1.0 \| 0.008684 \| 106 \| 0.113 \| 106 \| 79 \| 4 \| 1 \| 94 \| 8 \| 110 \| Phage tail protein \| Phage tail protein; afdb-uniprot50 \| AF-A0A137SPX7-F1-MODEL\_V4 \| 1.0 \| 0.009264 \| 106 \| 0.145 \| 110 \| 77 \| 4 \| 1 \| 93 \| 6 \| 115 \| Uncharacterized protein \| Uncharacterized protein |
| Top keywords  (threshold 1.00e-02 (evalue)) | **tail, assembly, Phage, or, E, chaperone, 14\_like, gp41, Putative, prophage** |
| Output files | ../../similar\_structures/20\_FANPEZAQ\_CDS\_0020\_afdb-proteome\_foldseek.tsv ../../similar\_structures/20\_FANPEZAQ\_CDS\_0020\_afdb-uniprot50\_foldseek.tsv ../../similar\_structures/20\_FANPEZAQ\_CDS\_0020\_merged.svg ../../similar\_structures/20\_FANPEZAQ\_CDS\_0020\_pdb\_foldseek.tsv |

  
  
  

Return to summary | Go to previous | Go to next

  


---

**Sequence/structure alignments coloring**  
Each object in the alignment figures is colored according to its E-value following this color coding:

1e-100
10
