## Supplementary material for "Completing the BASEL phage collection to unlock hidden diversity for systematic exploration of phage-host interactions": Data S2: 21.html

FANPEZAQ\_CDS\_0021


Return to summary | Go to previous | Go to next

|  |  |
| --- | --- |
| FANPEZAQ\_CDS\_0021  Page creation date: 02 Sep 2024, 12:00  Project folder: n/a  Input sequences file: Escherichia\_virus\_HeidiAbel.gb | phage gpe tail p2 tape measure chaperone hypothetical |

### Sequence information

|  |  |
| --- | --- |
| Name | FANPEZAQ\_CDS\_0021  21\_FANPEZAQ\_CDS\_0021 (pipeline id) |
| Imported annotations | Escherichia\_virus\_HeidiAbel Bas97 |
| Protein sequence | MALASHTGWQLSEIQRLRVSRFVWWLDGLPKKQN |
| Number of residues | 34 |
| Molecular weight (Da) | 4052.66 |
| Output files | ../../query\_sequences/21\_FANPEZAQ\_CDS\_0021.fasta |

### Putative domain architecture and protein family

#### Search results (HHblits)^1^

|  |  |
| --- | --- |
| Domain family databases searched | Pfam, Ncbi-cd, Cath, Phrogs |
| Results, scheme(s)  (Top layers only; threshold 1.00e-03 (evalue)) | xml version="1.0" encoding="utf-8" standalone="no"?       2024-09-02T21:08:16.703320 image/svg+xml   Matplotlib v3.7.2, https://matplotlib.org/ |
| Results, table  (E-value ≤ 1.00e-03 (evalue)) | -- |
| Top keywords  (threshold 1.00e-03 (evalue)) | -- |
| Output files | ../../domain\_architecture/21\_FANPEZAQ\_CDS\_0021\_cath.hhr ../../domain\_architecture/21\_FANPEZAQ\_CDS\_0021\_merged.svg ../../domain\_architecture/21\_FANPEZAQ\_CDS\_0021\_ncbi-cd.hhr ../../domain\_architecture/21\_FANPEZAQ\_CDS\_0021\_pfam.hhr ../../domain\_architecture/21\_FANPEZAQ\_CDS\_0021\_phrogs.hhr |

### Identical protein sequences/structures

#### Search results

|  |  |
| --- | --- |
| Protein sequence databases searched | Pdb, Swissprot, Refseq |
| Identical proteins found | -- |
| Top keywords | -- |
| Output files | -- |

### Similar protein sequences/structures

#### Sequence similarity search results (HHblits)^1^

|  |  |
| --- | --- |
| Sequence databases searched | Uniclust, Pdb70 |
| Results, scheme(s)  (Top layers only, threshold 1.00e-03 (evalue)) | xml version="1.0" encoding="utf-8" standalone="no"?       2024-09-02T21:08:38.105559 image/svg+xml   Matplotlib v3.7.2, https://matplotlib.org/ |
| Results, table(s)  (threshold 1.00e-03 (evalue)) | db \| id \| prob \| evalue \| pvalue \| score \| cols \| query \| query\_len \| template \| template\_len \| name \| description; uniclust \| UniRef100\_A0A5C2H343 \| 99.4 \| 4.6e-16 \| 9.2e-22 \| 91.7 \| 33 \| (1, 33) \| 34 \| (39, 71) \| 72 \| Uncharacterized protein \| Uncharacterized protein; uniclust \| UniRef100\_A0A060H7C1 \| 99.4 \| 2.8e-15 \| 5.7e-21 \| 82.7 \| 33 \| (1, 33) \| 34 \| (13, 45) \| 46 \| Uncharacterized protein \| Uncharacterized protein; uniclust \| UniRef100\_A0A858NPH9 \| 99.3 \| 3.6e-15 \| 7e-21 \| 78.7 \| 33 \| (1, 33) \| 34 \| (2, 34) \| 35 \| Uncharacterized protein \| Uncharacterized protein; uniclust \| UniRef100\_A0A7L8G6A2 \| 99.2 \| 1.3e-13 \| 2.5e-19 \| 89.7 \| 33 \| (1, 33) \| 34 \| (108, 140) \| 145 \| Tape measure chaperone \| Tape measure chaperone; uniclust \| UniRef100\_A0A975AA02 \| 99.0 \| 3e-12 \| 5.4e-18 \| 71.0 \| 34 \| (1, 34) \| 34 \| (12, 45) \| 45 \| GpE family phage tail protein \| GpE family phage tail protein; uniclust \| UniRef100\_A0A0P9QIX4 \| 98.8 \| 2.1e-11 \| 4.4e-17 \| 67.0 \| 26 \| (2, 27) \| 34 \| (16, 41) \| 42 \| Uncharacterized protein \| Uncharacterized protein; uniclust \| UniRef100\_A0A8S5RUP2 \| 98.5 \| 1.9e-09 \| 3.5e-15 \| 58.5 \| 31 \| (1, 31) \| 34 \| (2, 32) \| 37 \| Phage protein \| Phage protein; uniclust \| UniRef100\_UPI001F35B4CE \| 98.4 \| 2.6e-09 \| 4.8e-15 \| 61.5 \| 33 \| (1, 33) \| 34 \| (19, 51) \| 53 \| hypothetical protein \| hypothetical protein; uniclust \| UniRef100\_R9TPW5 \| 98.4 \| 2.7e-09 \| 4.9e-15 \| 62.9 \| 33 \| (2, 34) \| 34 \| (29, 61) \| 62 \| Uncharacterized protein \| Uncharacterized protein; uniclust \| UniRef100\_A0A1H4SBZ9 \| 98.1 \| 6.1e-08 \| 1.2e-13 \| 58.3 \| 26 \| (2, 27) \| 34 \| (42, 67) \| 68 \| Uncharacterized protein \| Uncharacterized protein; uniclust \| UniRef100\_A0A0C5VIR9 \| 97.8 \| 5.3e-07 \| 1e-12 \| 51.0 \| 31 \| (1, 31) \| 34 \| (6, 36) \| 44 \| GpE family phage tail protein \| GpE family phage tail protein; uniclust \| UniRef100\_A0A6I5RSZ0 \| 97.8 \| 5.8e-07 \| 1.1e-12 \| 51.0 \| 26 \| (2, 27) \| 34 \| (19, 44) \| 45 \| Uncharacterized protein \| Uncharacterized protein; uniclust \| UniRef100\_A0A1T4WVU1 \| 97.7 \| 8.6e-07 \| 1.6e-12 \| 51.6 \| 27 \| (2, 28) \| 34 \| (24, 50) \| 52 \| Phage P2 GpE \| Phage P2 GpE; uniclust \| UniRef100\_A0A023Q0E1 \| 97.7 \| 9.6e-07 \| 1.8e-12 \| 49.5 \| 29 \| (1, 29) \| 34 \| (3, 31) \| 41 \| GpE family phage tail protein \| GpE family phage tail protein; uniclust \| UniRef100\_A0A1I7J3I1 \| 97.2 \| 1.6e-05 \| 3.1e-11 \| 46.1 \| 30 \| (2, 31) \| 34 \| (15, 44) \| 48 \| Uncharacterized protein \| Uncharacterized protein; uniclust \| UniRef100\_E6VU91 \| 97.1 \| 3.8e-05 \| 6.9e-11 \| 43.5 \| 29 \| (1, 29) \| 34 \| (12, 40) \| 41 \| Uncharacterized protein \| Uncharacterized protein; uniclust \| UniRef100\_A0A024HBK0 \| 96.7 \| 0.00016 \| 3.2e-10 \| 41.2 \| 26 \| (2, 27) \| 34 \| (12, 37) \| 41 \| Uncharacterized protein \| Uncharacterized protein; uniclust \| UniRef100\_A0A450ZF00 \| 96.4 \| 0.00056 \| 1e-09 \| 38.9 \| 28 \| (3, 30) \| 34 \| (5, 32) \| 39 \| Phage P2 GpE \| Phage P2 GpE |
| Top keywords  (threshold 1.00e-03 (evalue)) | **phage, GpE, tail, P2, Tape, measure, chaperone, hypothetical** |
| Output files | ../../similar\_sequences/21\_FANPEZAQ\_CDS\_0021\_merged.svg ../../similar\_sequences/21\_FANPEZAQ\_CDS\_0021\_pdb70.a3m ../../similar\_sequences/21\_FANPEZAQ\_CDS\_0021\_pdb70.hhr ../../similar\_sequences/21\_FANPEZAQ\_CDS\_0021\_uniclust.a3m ../../similar\_sequences/21\_FANPEZAQ\_CDS\_0021\_uniclust.hhr |

#### Structure prediction (AlphaFold)^2^

|  |  |
| --- | --- |
| Stats | xml version="1.0" encoding="utf-8" standalone="no"?       2024-09-02T21:09:20.281047 image/svg+xml   Matplotlib v3.7.2, https://matplotlib.org/ |
| Predicted structure | **NGL Viewer Controls:**  - Center: *Left-Click* - Rotate: *Left-Click + Drag* - Translate: *Right-Click + Drag* - Zoom: *Shift + Left-Click + Drag* |
| Output files | ../../predicted\_structures/21\_FANPEZAQ\_CDS\_0021/features.pkl ../../predicted\_structures/21\_FANPEZAQ\_CDS\_0021/ranked\_0.pdb ../../predicted\_structures/21\_FANPEZAQ\_CDS\_0021/ranked\_0\_plots.svg ../../predicted\_structures/21\_FANPEZAQ\_CDS\_0021/result\_model\_1\_ptm\_pred\_0.pkl |

#### Structure similarity search results (Foldseek)^3^

|  |  |
| --- | --- |
| Structure databases searched | Pdb, Afdb-proteome, Afdb-uniprot50 |
| Results, scheme(s)  (Top layers only, threshold 1.00e-02 (evalue)) | xml version="1.0" encoding="utf-8" standalone="no"?       2024-09-02T21:10:44.906509 image/svg+xml   Matplotlib v3.7.2, https://matplotlib.org/ |
| Results, table  (threshold 1.00e-02 (evalue)) | -- |
| Top keywords  (threshold 1.00e-02 (evalue)) | -- |
| Output files | ../../similar\_structures/21\_FANPEZAQ\_CDS\_0021\_afdb-proteome\_foldseek.tsv ../../similar\_structures/21\_FANPEZAQ\_CDS\_0021\_afdb-uniprot50\_foldseek.tsv ../../similar\_structures/21\_FANPEZAQ\_CDS\_0021\_merged.svg ../../similar\_structures/21\_FANPEZAQ\_CDS\_0021\_pdb\_foldseek.tsv |

  
  
  

Return to summary | Go to previous | Go to next

  


---

**Sequence/structure alignments coloring**  
Each object in the alignment figures is colored according to its E-value following this color coding:

1e-100
10
