## Supplementary material for "Completing the BASEL phage collection to unlock hidden diversity for systematic exploration of phage-host interactions": Data S2: 22.html

FANPEZAQ\_CDS\_0022


Return to summary | Go to previous | Go to next

|  |  |
| --- | --- |
| FANPEZAQ\_CDS\_0022  Page creation date: 02 Sep 2024, 12:00  Project folder: n/a  Input sequences file: Escherichia\_virus\_HeidiAbel.gb | tail tape measure phage tp901 domain\_containing core region phagemin\_tail length phage\_related minor putative fragment tape\_measure |

### Sequence information

|  |  |
| --- | --- |
| Name | FANPEZAQ\_CDS\_0022  22\_FANPEZAQ\_CDS\_0022 (pipeline id) |
| Imported annotations | Escherichia\_virus\_HeidiAbel Bas97 |
| Protein sequence | MANKKLNATITIGGAVSSSLKGAFGTVKSSVEQVGAAMAKLEREQRTLTNAIQTFGRQGK NVDSLRAKYAANVAAVDKLRTATERLKRVEDARERNLAKREQYKDGIMGTIALGATIAAP IKAAMDFESTVADIKKVVDAPKGADEDTFFKAISKQVLDMSKTLPMTANEIGRIFALGGQ SGIATEDLGKFTESAVKMGVAFDVSAEQAGQSMAELRSAFGMTQDQVNTLADQMNYLDNN GSAAAKDIMEIVQRIGPLAKVAGVSAGQVAALGSTLRGMGVQNEIAATGIKNLFLTLASG DAATKKQKEVFKQLGYTSKEVSKSMQLDAEKTMTVILKQISKLPKHAQAAALTELFGKEV VGSIAPLMTRLGELQKNFDAVADRTKYAGSMQKEFEARAATTANQLILFRNQITGLGVTI GSVLLPAVNSVLQTVGPWIGKITELADAHPVVTKAIVATAGALITMRIATFAAGFAFTYL RGGALRVVGALAGARAQMALTAVSARAVGAAATVANGGLVGMASRGIPAVVTGVRAIGAA FISTGIGALITGLALGGLWIYRNWAGVKAFMTGTLQGIQQGLQPVTEKFQALWDRLGPVK TAFQWVADAAGKAWDWFTKLAEPVSYSGEELKKAGDAGQQFGKALAAGIDYVLGPIQFLI DKIIWVSDNIGALTQKAVEFKNSVSDMAGGAWQKTKDFFSNPFGSDDVPEGNTLATPLTP NNALPAPALANRGGSTYTDSSTTTIQVTQRPGENNADLAKRIADEQERRRQVRQRSMMND GVTAP |
| Number of residues | 785 |
| Molecular weight (Da) | 82722.95 |
| Output files | ../../query\_sequences/22\_FANPEZAQ\_CDS\_0022.fasta |

### Putative domain architecture and protein family

#### Search results (HHblits)^1^

|  |  |
| --- | --- |
| Domain family databases searched | Pfam, Ncbi-cd, Cath, Phrogs |
| Results, scheme(s)  (Top layers only; threshold 1.00e-03 (evalue)) | xml version="1.0" encoding="utf-8" standalone="no"?       2024-09-02T21:08:16.903637 image/svg+xml   Matplotlib v3.7.2, https://matplotlib.org/ |
| Results, table  (E-value ≤ 1.00e-03 (evalue)) | db \| id \| prob \| evalue \| pvalue \| score \| cols \| query \| query\_len \| template \| template\_len \| name \| description; pfam \| PF10145 \| 95.2 \| 9.7e-05 \| 2.8e-08 \| 57.2 \| 90 \| (206, 297) \| 785 \| (55, 146) \| 200 \| PhageMin\_Tail \| Phage-related minor tail protein; phrogs \| 28 \| 100.0 \| 4.4e-44 \| 5.6e-48 \| 424.9 \| 628 \| (1, 665) \| 785 \| (1, 648) \| 820 \| tail length tape measure protein \| tail length tape measure protein; Category: tail; p348226 VI\_02582; phrogs \| 991 \| 100.0 \| 2e-42 \| 2.5e-46 \| 400.6 \| 467 \| (107, 706) \| 785 \| (155, 628) \| 767 \| tail length tape measure protein \| tail length tape measure protein; Category: tail; p290120 VI\_00584; phrogs \| 4704 \| 100.0 \| 2.5e-39 \| 2.9e-43 \| 342.8 \| 420 \| (15, 455) \| 785 \| (31, 496) \| 576 \| tail length tape measure protein \| tail length tape measure protein; Category: tail; p73956 VI\_01292; phrogs \| 1293 \| 100.0 \| 2.6e-37 \| 3.1e-41 \| 363.6 \| 430 \| (15, 464) \| 785 \| (30, 504) \| 1391 \| tail length tape measure protein \| tail length tape measure protein; Category: tail; p160953 VI\_04898; phrogs \| 5218 \| 100.0 \| 2.1e-34 \| 2.5e-38 \| 330.9 \| 432 \| (2, 455) \| 785 \| (12, 495) \| 1296 \| NA \| NA; Category: unknown function; p362597 VI\_05060; phrogs \| 1272 \| 99.9 \| 6.9e-31 \| 8.3e-35 \| 298.3 \| 436 \| (112, 596) \| 785 \| (45, 513) \| 878 \| tail length tape measure protein \| tail length tape measure protein; Category: tail; p177841 VI\_05919; phrogs \| 5493 \| 99.9 \| 8.1e-31 \| 9.4e-35 \| 290.2 \| 324 \| (105, 444) \| 785 \| (46, 388) \| 1111 \| tail length tape measure protein \| tail length tape measure protein; Category: tail; p353434 VI\_05194; phrogs \| 1533 \| 99.9 \| 1.1e-29 \| 1.3e-33 \| 289.9 \| 267 \| (181, 461) \| 785 \| (2, 291) \| 954 \| tail length tape measure protein \| tail length tape measure protein; Category: tail; p53969 VI\_05779; phrogs \| 19074 \| 99.9 \| 2e-28 \| 2.3e-32 \| 263.2 \| 309 \| (112, 439) \| 785 \| (273, 589) \| 1750 \| NA \| NA; Category: unknown function; p33998 VI\_01128; phrogs \| 16692 \| 99.8 \| 2.3e-26 \| 2.6e-30 \| 244.2 \| 323 \| (107, 448) \| 785 \| (123, 454) \| 947 \| NA \| NA; Category: unknown function; p85627 VI\_04017; phrogs \| 794 \| 99.8 \| 3.3e-24 \| 4e-28 \| 240.7 \| 340 \| (112, 471) \| 785 \| (137, 535) \| 698 \| tail length tape measure protein \| tail length tape measure protein; Category: tail; p192014 VI\_06934; phrogs \| 5690 \| 99.7 \| 1.4e-22 \| 1.7e-26 \| 199.6 \| 138 \| (537, 681) \| 785 \| (40, 177) \| 347 \| NA \| NA; Category: unknown function; p270016 VI\_07747; phrogs \| 3654 \| 99.7 \| 1.7e-22 \| 1.9e-26 \| 210.3 \| 95 \| (2, 96) \| 785 \| (3, 97) \| 612 \| NA \| NA; Category: unknown function; p148691 VI\_02621; phrogs \| 9228 \| 99.7 \| 2e-22 \| 2.3e-26 \| 224.5 \| 435 \| (110, 590) \| 785 \| (208, 659) \| 1359 \| tail length tape measure protein \| tail length tape measure protein; Category: tail; NC\_009993\_p20; phrogs \| 4211 \| 99.7 \| 1.3e-21 \| 1.6e-25 \| 230.1 \| 318 \| (109, 447) \| 785 \| (346, 673) \| 2292 \| tail length tape measure protein \| tail length tape measure protein; Category: tail; p43110 VI\_00473; phrogs \| 5448 \| 99.6 \| 9.5e-21 \| 1.1e-24 \| 216.7 \| 347 \| (106, 460) \| 785 \| (448, 829) \| 3327 \| tail length tape measure protein \| tail length tape measure protein; Category: tail; NC\_030924\_p25; phrogs \| 19282 \| 99.6 \| 6.7e-20 \| 7.6e-24 \| 178.8 \| 310 \| (6, 324) \| 785 \| (1, 336) \| 434 \| NA \| NA; Category: unknown function; p400074 VI\_09057; phrogs \| 28420 \| 99.6 \| 1.6e-19 \| 1.8e-23 \| 183.8 \| 303 \| (113, 443) \| 785 \| (47, 350) \| 873 \| tail length tape measure protein \| tail length tape measure protein; Category: tail; p93440 VI\_05600; phrogs \| 36688 \| 99.5 \| 1.9e-19 \| 2.1e-23 \| 188.5 \| 325 \| (112, 453) \| 785 \| (62, 388) \| 1303 \| tail length tape measure protein \| tail length tape measure protein; Category: tail; KP836355\_p32; phrogs \| 5755 \| 99.5 \| 3.5e-19 \| 4e-23 \| 175.7 \| 382 \| (1, 423) \| 785 \| (1, 436) \| 441 \| tail length tape measure protein \| tail length tape measure protein; Category: tail; p325579 VI\_04535; phrogs \| 34270 \| 99.5 \| 4.1e-19 \| 4.6e-23 \| 175.3 \| 318 \| (108, 440) \| 785 \| (61, 386) \| 606 \| tail length tape measure protein \| tail length tape measure protein; Category: tail; NC\_025427\_p16; phrogs \| 14917 \| 99.5 \| 7.7e-19 \| 8.7e-23 \| 190.8 \| 306 \| (101, 424) \| 785 \| (314, 624) \| 2249 \| tail length tape measure protein \| tail length tape measure protein; Category: tail; NC\_022774\_p53; phrogs \| 37649 \| 99.4 \| 4.5e-18 \| 5.1e-22 \| 182.1 \| 317 \| (108, 444) \| 785 \| (332, 650) \| 1827 \| tail length tape measure protein \| tail length tape measure protein; Category: tail; NC\_029075\_p20; phrogs \| 36752 \| 99.4 \| 4.6e-18 \| 5.2e-22 \| 174.1 \| 326 \| (119, 471) \| 785 \| (71, 406) \| 1076 \| NA \| NA; Category: unknown function; p212730 VI\_04312; phrogs \| 10030 \| 99.3 \| 5.6e-17 \| 6.4e-21 \| 178.5 \| 352 \| (107, 465) \| 785 \| (171, 536) \| 2102 \| tail length tape measure protein \| tail length tape measure protein; Category: tail; MG757153\_p31; phrogs \| 35707 \| 99.3 \| 7.1e-17 \| 8e-21 \| 161.5 \| 324 \| (114, 443) \| 785 \| (72, 398) \| 715 \| tail length tape measure protein \| tail length tape measure protein; Category: tail; p193300 VI\_04290; phrogs \| 17727 \| 99.3 \| 7.6e-17 \| 8.6e-21 \| 155.9 \| 283 \| (277, 627) \| 785 \| (53, 341) \| 448 \| tail length tape measure protein \| tail length tape measure protein; Category: tail; p417437 VI\_07567; phrogs \| 8462 \| 99.3 \| 9.4e-17 \| 1.1e-20 \| 168.7 \| 235 \| (347, 595) \| 785 \| (171, 418) \| 671 \| tail length tape measure protein \| tail length tape measure protein; Category: tail; p424879 VI\_01041; phrogs \| 10780 \| 99.3 \| 1.1e-16 \| 1.3e-20 \| 179.7 \| 311 \| (108, 439) \| 785 \| (63, 385) \| 1884 \| tail length tape measure protein \| tail length tape measure protein; Category: tail; NC\_030942\_p18; phrogs \| 2723 \| 99.3 \| 1.7e-16 \| 2e-20 \| 168.5 \| 346 \| (111, 469) \| 785 \| (162, 608) \| 1577 \| tail length tape measure protein \| tail length tape measure protein; Category: tail; JN006062\_p23; phrogs \| 15478 \| 99.3 \| 1.8e-16 \| 2e-20 \| 154.1 \| 325 \| (286, 706) \| 785 \| (3, 341) \| 473 \| tail length tape measure protein \| tail length tape measure protein; Category: tail; NC\_019506\_p24; phrogs \| 973 \| 99.3 \| 2.2e-16 \| 2.7e-20 \| 171.3 \| 297 \| (119, 440) \| 785 \| (71, 375) \| 583 \| tail length tape measure protein \| tail length tape measure protein; Category: tail; p232472 VI\_03267; phrogs \| 38390 \| 99.2 \| 9.7e-16 \| 1.1e-19 \| 154.8 \| 294 \| (112, 423) \| 785 \| (48, 341) \| 1029 \| tail length tape measure protein \| tail length tape measure protein; Category: tail; NC\_015466\_p80; phrogs \| 30543 \| 99.2 \| 1.5e-15 \| 1.7e-19 \| 154.6 \| 325 \| (108, 432) \| 785 \| (52, 381) \| 1167 \| tail length tape measure protein \| tail length tape measure protein; Category: tail; MG711466\_p67; phrogs \| 8125 \| 99.2 \| 3.1e-15 \| 3.5e-19 \| 157.3 \| 373 \| (251, 701) \| 785 \| (210, 597) \| 662 \| tail length tape measure protein \| tail length tape measure protein; Category: tail; p114197 VI\_11397; phrogs \| 23013 \| 99.1 \| 1.1e-14 \| 1.3e-18 \| 151.2 \| 333 \| (121, 471) \| 785 \| (132, 467) \| 1379 \| tail length tape measure protein \| tail length tape measure protein; Category: tail; JF713456\_p61; phrogs \| 4200 \| 99.0 \| 2.6e-14 \| 3e-18 \| 153.0 \| 107 \| (422, 584) \| 785 \| (141, 247) \| 810 \| tail length tape measure protein \| tail length tape measure protein; Category: tail; p335142 VI\_05069; phrogs \| 339 \| 99.0 \| 3.9e-14 \| 4.7e-18 \| 163.4 \| 307 \| (108, 447) \| 785 \| (76, 387) \| 1128 \| tail length tape measure protein \| tail length tape measure protein; Category: tail; p256393 VI\_10266; phrogs \| 1068 \| 98.9 \| 1.1e-13 \| 1.3e-17 \| 152.2 \| 191 \| (17, 217) \| 785 \| (11, 208) \| 788 \| tail length tape measure protein \| tail length tape measure protein; Category: tail; p214035 VI\_06128; phrogs \| 17594 \| 98.9 \| 2.8e-13 \| 3.1e-17 \| 144.4 \| 339 \| (112, 456) \| 785 \| (166, 513) \| 1580 \| tail length tape measure protein \| tail length tape measure protein; Category: tail; MF185731\_p36; phrogs \| 3336 \| 98.8 \| 4.7e-13 \| 5.6e-17 \| 149.9 \| 313 \| (108, 595) \| 785 \| (185, 508) \| 1052 \| tail length tape measure protein \| tail length tape measure protein; Category: tail; NC\_029097\_p21; phrogs \| 26705 \| 98.8 \| 7.4e-13 \| 8.3e-17 \| 136.9 \| 298 \| (119, 443) \| 785 \| (301, 599) \| 1303 \| NA \| NA; Category: unknown function; p49245 VI\_05491; phrogs \| 58 \| 98.5 \| 1.6e-11 \| 2e-15 \| 145.7 \| 58 \| (538, 595) \| 785 \| (663, 723) \| 1263 \| tail length tape measure protein \| tail length tape measure protein; Category: tail; p145088 VI\_05474; phrogs \| 9696 \| 98.5 \| 2.2e-11 \| 2.6e-15 \| 108.6 \| 112 \| (107, 223) \| 785 \| (64, 175) \| 175 \| tail length tape measure protein \| tail length tape measure protein; Category: tail; p98305 VI\_00740; phrogs \| 3824 \| 98.5 \| 2.8e-11 \| 3.2e-15 \| 126.5 \| 192 \| (247, 458) \| 785 \| (71, 272) \| 682 \| NA \| NA; Category: unknown function; NC\_031117\_p21; phrogs \| 17321 \| 98.4 \| 4.4e-11 \| 4.9e-15 \| 123.2 \| 194 \| (108, 301) \| 785 \| (74, 269) \| 1484 \| NA \| NA; Category: unknown function; p152511 VI\_00610; phrogs \| 17114 \| 98.3 \| 1.1e-10 \| 1.3e-14 \| 106.6 \| 147 \| (538, 691) \| 785 \| (24, 170) \| 369 \| NA \| NA; Category: unknown function; p385881 VI\_10695; phrogs \| 23108 \| 98.2 \| 4.2e-10 \| 4.8e-14 \| 120.8 \| 324 \| (109, 443) \| 785 \| (307, 643) \| 1401 \| tail length tape measure protein \| tail length tape measure protein; Category: tail; NC\_025463\_p37; phrogs \| 13339 \| 98.1 \| 6.8e-10 \| 7.7e-14 \| 91.2 \| 130 \| (161, 290) \| 785 \| (2, 131) \| 133 \| tail length tape measure protein \| tail length tape measure protein; Category: tail; p142514 VI\_00726; phrogs \| 4200 \| 98.0 \| 2.2e-09 \| 2.5e-13 \| 114.5 \| 145 \| (554, 707) \| 785 \| (459, 616) \| 810 \| tail length tape measure protein \| tail length tape measure protein; Category: tail; p335142 VI\_05069; phrogs \| 3981 \| 97.9 \| 2.9e-09 \| 3.4e-13 \| 112.3 \| 48 \| (396, 444) \| 785 \| (257, 304) \| 730 \| tail length tape measure protein \| tail length tape measure protein; Category: tail; p408809 VI\_04997; phrogs \| 14441 \| 97.9 \| 3e-09 \| 3.5e-13 \| 92.0 \| 108 \| (101, 211) \| 785 \| (56, 163) \| 164 \| tail length tape measure protein \| tail length tape measure protein; Category: tail; p275139 VI\_05876; phrogs \| 4200 \| 97.9 \| 3.5e-09 \| 4e-13 \| 112.8 \| 137 \| (555, 710) \| 785 \| (394, 531) \| 810 \| tail length tape measure protein \| tail length tape measure protein; Category: tail; p335142 VI\_05069; phrogs \| 4197 \| 97.8 \| 5.6e-09 \| 6.5e-13 \| 118.6 \| 59 \| (535, 593) \| 785 \| (950, 1011) \| 1729 \| tail length tape measure protein \| tail length tape measure protein; Category: tail; p189664 VI\_00038; phrogs \| 34405 \| 97.8 \| 6.7e-09 \| 7.4e-13 \| 109.9 \| 295 \| (116, 422) \| 785 \| (162, 475) \| 2675 \| tail length tape measure protein \| tail length tape measure protein; Category: tail; MF001361\_p11; phrogs \| 4372 \| 97.6 \| 3.1e-08 \| 3.6e-12 \| 112.1 \| 104 \| (116, 219) \| 785 \| (187, 291) \| 1826 \| tail length tape measure protein \| tail length tape measure protein; Category: tail; MF467948\_p23; phrogs \| 6090 \| 97.5 \| 4.5e-08 \| 5.3e-12 \| 110.7 \| 52 \| (393, 444) \| 785 \| (640, 691) \| 1808 \| tail length tape measure protein \| tail length tape measure protein; Category: tail; MF919510\_p42; phrogs \| 32669 \| 97.5 \| 5.7e-08 \| 6.3e-12 \| 97.3 \| 141 \| (536, 678) \| 785 \| (693, 873) \| 1036 \| NA \| NA; Category: unknown function; p276907 VI\_03881; phrogs \| 27190 \| 97.5 \| 7.4e-08 \| 8.3e-12 \| 97.3 \| 128 \| (83, 217) \| 785 \| (75, 206) \| 1014 \| tail length tape measure protein \| tail length tape measure protein; Category: tail; p213007 VI\_01051; phrogs \| 1686 \| 97.3 \| 1.8e-07 \| 2.1e-11 \| 103.2 \| 57 \| (539, 595) \| 785 \| (681, 741) \| 1788 \| tail length tape measure protein \| tail length tape measure protein; Category: tail; NC\_024384\_p40; phrogs \| 34082 \| 97.2 \| 2.6e-07 \| 3e-11 \| 88.4 \| 131 \| (537, 706) \| 785 \| (290, 421) \| 529 \| tail length tape measure protein \| tail length tape measure protein; Category: tail; MH019216\_p23; phrogs \| 21157 \| 97.2 \| 2.8e-07 \| 3.2e-11 \| 93.4 \| 141 \| (419, 581) \| 785 \| (575, 719) \| 768 \| minor tail protein \| minor tail protein; Category: tail; NC\_002668\_p50; phrogs \| 1204 \| 97.2 \| 3.1e-07 \| 3.7e-11 \| 105.3 \| 126 \| (122, 271) \| 785 \| (79, 205) \| 1355 \| tail protein \| tail protein; Category: tail; p387522 VI\_00450; phrogs \| 8927 \| 97.1 \| 6.2e-07 \| 7.2e-11 \| 100.4 \| 146 \| (148, 303) \| 785 \| (237, 395) \| 1569 \| tail length tape measure protein \| tail length tape measure protein; Category: tail; KY203335\_p44; phrogs \| 16297 \| 97.0 \| 7.6e-07 \| 8.6e-11 \| 89.1 \| 108 \| (108, 221) \| 785 \| (77, 188) \| 776 \| tail length tape measure protein \| tail length tape measure protein; Category: tail; p238715 VI\_06415; phrogs \| 13137 \| 97.0 \| 7.9e-07 \| 8.8e-11 \| 86.0 \| 161 \| (112, 273) \| 785 \| (345, 526) \| 541 \| tail length tape measure protein \| tail length tape measure protein; Category: tail; p31102 VI\_05860; phrogs \| 971 \| 96.9 \| 1.1e-06 \| 1.3e-10 \| 98.5 \| 43 \| (530, 572) \| 785 \| (416, 461) \| 1164 \| endolysin \| endolysin; Category: lysis; NC\_031076\_p13; phrogs \| 29312 \| 96.9 \| 1.5e-06 \| 1.6e-10 \| 82.8 \| 242 \| (22, 278) \| 785 \| (305, 547) \| 558 \| tail length tape measure protein \| tail length tape measure protein; Category: tail; NC\_007581\_p131; phrogs \| 2335 \| 96.8 \| 1.8e-06 \| 2.1e-10 \| 97.7 \| 143 \| (61, 213) \| 785 \| (55, 201) \| 1433 \| tail length tape measure protein \| tail length tape measure protein; Category: tail; p247749 VI\_04302; phrogs \| 3519 \| 96.8 \| 1.8e-06 \| 2.2e-10 \| 88.1 \| 104 \| (111, 220) \| 785 \| (75, 182) \| 448 \| tail length tape measure protein \| tail length tape measure protein; Category: tail; p424447 VI\_08065; phrogs \| 10613 \| 96.7 \| 3.6e-06 \| 4.2e-10 \| 71.5 \| 84 \| (3, 89) \| 785 \| (2, 97) \| 138 \| tail length tape measure protein \| tail length tape measure protein; Category: tail; p365128 VI\_01223; phrogs \| 8927 \| 96.6 \| 5.4e-06 \| 6.2e-10 \| 92.8 \| 39 \| (211, 249) \| 785 \| (223, 261) \| 1569 \| tail length tape measure protein \| tail length tape measure protein; Category: tail; KY203335\_p44; phrogs \| 1535 \| 96.5 \| 6.7e-06 \| 7.9e-10 \| 92.6 \| 58 \| (533, 590) \| 785 \| (845, 905) \| 1217 \| tail length tape measure protein \| tail length tape measure protein; Category: tail; p395424 VI\_04399; phrogs \| 9388 \| 96.4 \| 1.1e-05 \| 1.3e-09 \| 81.6 \| 44 \| (535, 578) \| 785 \| (399, 442) \| 629 \| NA \| NA; Category: unknown function; p379982 VI\_11797; phrogs \| 38386 \| 96.4 \| 1.2e-05 \| 1.3e-09 \| 82.8 \| 134 \| (536, 677) \| 785 \| (771, 941) \| 1703 \| NA \| NA; Category: unknown function; p413626 VI\_12283; phrogs \| 26805 \| 96.4 \| 1.2e-05 \| 1.3e-09 \| 70.5 \| 56 \| (537, 592) \| 785 \| (167, 222) \| 253 \| NA \| NA; Category: unknown function; p228647 VI\_05149; phrogs \| 12798 \| 96.3 \| 1.4e-05 \| 1.6e-09 \| 86.0 \| 112 \| (110, 227) \| 785 \| (53, 178) \| 1052 \| tail length tape measure protein \| tail length tape measure protein; Category: tail; p167255 VI\_04008; phrogs \| 8010 \| 96.1 \| 2.9e-05 \| 3.4e-09 \| 74.1 \| 107 \| (108, 220) \| 785 \| (72, 182) \| 325 \| tail length tape measure protein \| tail length tape measure protein; Category: tail; p301250 VI\_08079; phrogs \| 38258 \| 96.0 \| 3.8e-05 \| 4.2e-09 \| 70.3 \| 63 \| (533, 595) \| 785 \| (165, 237) \| 427 \| NA \| NA; Category: unknown function; p292485 VI\_04908; phrogs \| 26804 \| 95.9 \| 4.7e-05 \| 5.2e-09 \| 76.3 \| 73 \| (398, 470) \| 785 \| (291, 363) \| 1065 \| NA \| NA; Category: unknown function; p121571 VI\_12455; phrogs \| 34923 \| 95.2 \| 0.00022 \| 2.4e-08 \| 67.4 \| 179 \| (260, 455) \| 785 \| (145, 330) \| 537 \| tail length tape measure protein \| tail length tape measure protein; Category: tail; MG675557\_p40; phrogs \| 19166 \| 95.2 \| 0.00024 \| 2.8e-08 \| 61.2 \| 111 \| (111, 227) \| 785 \| (47, 159) \| 186 \| NA \| NA; Category: unknown function; p255465 VI\_06352; phrogs \| 305 \| 95.1 \| 0.00027 \| 3.3e-08 \| 79.2 \| 69 \| (644, 712) \| 785 \| (797, 867) \| 1015 \| tail length tape measure protein \| tail length tape measure protein; Category: tail; MF975638\_p8; phrogs \| 17020 \| 95.0 \| 0.00033 \| 3.8e-08 \| 62.2 \| 25 \| (651, 675) \| 785 \| (80, 104) \| 238 \| NA \| NA; Category: unknown function; p388498 VI\_05144; phrogs \| 15002 \| 94.8 \| 0.00044 \| 5e-08 \| 56.7 \| 88 \| (1, 95) \| 785 \| (1, 119) \| 144 \| NA \| NA; Category: unknown function; p434568 VI\_05139; phrogs \| 9959 \| 94.8 \| 0.00044 \| 5.1e-08 \| 76.5 \| 93 \| (108, 206) \| 785 \| (59, 151) \| 1333 \| tail length tape measure protein \| tail length tape measure protein; Category: tail; p39219 VI\_01014; phrogs \| 7740 \| 94.5 \| 0.00064 \| 7.5e-08 \| 63.4 \| 144 \| (108, 272) \| 785 \| (72, 220) \| 255 \| tail length tape measure protein \| tail length tape measure protein; Category: tail; p336818 VI\_11619; phrogs \| 33100 \| 94.5 \| 0.00068 \| 7.5e-08 \| 50.8 \| 75 \| (148, 222) \| 785 \| (13, 87) \| 109 \| NA \| NA; Category: unknown function; p346208 VI\_06081; phrogs \| 25337 \| 94.2 \| 0.001 \| 1.2e-07 \| 68.4 \| 174 \| (119, 299) \| 785 \| (190, 372) \| 1419 \| tail length tape measure protein \| tail length tape measure protein; Category: tail; NC\_031926\_p4 |
| Top keywords  (threshold 1.00e-03 (evalue)) | **tail, length, tape, measure, p335142, VI\_05069, minor, KY203335\_p44, Phage\_related, p348226** |
| Output files | ../../domain\_architecture/22\_FANPEZAQ\_CDS\_0022\_cath.hhr ../../domain\_architecture/22\_FANPEZAQ\_CDS\_0022\_merged.svg ../../domain\_architecture/22\_FANPEZAQ\_CDS\_0022\_ncbi-cd.hhr ../../domain\_architecture/22\_FANPEZAQ\_CDS\_0022\_pfam.hhr ../../domain\_architecture/22\_FANPEZAQ\_CDS\_0022\_phrogs.hhr |

### Identical protein sequences/structures

#### Search results

|  |  |
| --- | --- |
| Protein sequence databases searched | Pdb, Swissprot, Refseq |
| Identical proteins found | -- |
| Top keywords | -- |
| Output files | -- |

### Similar protein sequences/structures

#### Sequence similarity search results (HHblits)^1^

|  |  |
| --- | --- |
| Sequence databases searched | Uniclust, Pdb70 |
| Results, scheme(s)  (Top layers only, threshold 1.00e-03 (evalue)) | xml version="1.0" encoding="utf-8" standalone="no"?       2024-09-02T21:08:38.711127 image/svg+xml   Matplotlib v3.7.2, https://matplotlib.org/ |
| Results, table(s)  (threshold 1.00e-03 (evalue)) | db \| id \| prob \| evalue \| pvalue \| score \| cols \| query \| query\_len \| template \| template\_len \| name \| description; uniclust \| UniRef100\_A0A088FV68 \| 100.0 \| 9.9e-84 \| 1.8e-89 \| 618.7 \| 770 \| (1, 785) \| 785 \| (6, 777) \| 777 \| Tail protein \| Tail protein; uniclust \| UniRef100\_A0A060GZT2 \| 100.0 \| 4.9e-79 \| 8.6e-85 \| 629.3 \| 762 \| (1, 785) \| 785 \| (66, 836) \| 925 \| Phage tail protein \| Phage tail protein; uniclust \| UniRef100\_A0A011NAN7 \| 100.0 \| 1.5e-71 \| 2.7e-77 \| 556.5 \| 756 \| (4, 782) \| 785 \| (90, 925) \| 925 \| Tail protein \| Tail protein; uniclust \| UniRef100\_A0A061JKM9 \| 100.0 \| 1.9e-70 \| 3.4e-76 \| 574.7 \| 735 \| (4, 784) \| 785 \| (79, 856) \| 1009 \| Phage tail tape measure protein domain-containing protein \| Phage tail tape measure protein domain-containing protein; uniclust \| UniRef100\_A0A126R353 \| 100.0 \| 2.4e-70 \| 4.2e-76 \| 557.8 \| 731 \| (5, 783) \| 785 \| (80, 845) \| 851 \| Putative phage-related membrane protein \| Putative phage-related membrane protein; uniclust \| UniRef100\_A0A031GY25 \| 100.0 \| 9.5e-70 \| 1.7e-75 \| 572.0 \| 743 \| (4, 783) \| 785 \| (116, 902) \| 907 \| Bacteriophage-related tail transmembrane protein \| Bacteriophage-related tail transmembrane protein; uniclust \| UniRef100\_A0A0C3DQL6 \| 100.0 \| 3.4e-68 \| 6e-74 \| 554.7 \| 562 \| (4, 589) \| 785 \| (78, 658) \| 946 \| Phage tail tape measure protein domain-containing protein \| Phage tail tape measure protein domain-containing protein; uniclust \| UniRef100\_A0A081MYF0 \| 100.0 \| 9.2e-66 \| 1.6e-71 \| 538.1 \| 741 \| (4, 784) \| 785 \| (78, 852) \| 862 \| Phage tail tape measure protein domain-containing protein \| Phage tail tape measure protein domain-containing protein; uniclust \| UniRef100\_A0A0M1HSJ1 \| 100.0 \| 1.9e-65 \| 3.3e-71 \| 536.8 \| 565 \| (1, 586) \| 785 \| (79, 654) \| 877 \| Membrane protein \| Membrane protein; uniclust \| UniRef100\_A0A0A6TUZ1 \| 100.0 \| 1e-63 \| 1.8e-69 \| 487.5 \| 611 \| (4, 633) \| 785 \| (79, 699) \| 966 \| Phage-related tail protein \| Phage-related tail protein; uniclust \| UniRef100\_A0A062I8W3 \| 100.0 \| 1.8e-62 \| 3.1e-68 \| 493.3 \| 732 \| (17, 783) \| 785 \| (89, 849) \| 851 \| Phage tail tape measure protein, TP901 family, core region (Fragment) \| Phage tail tape measure protein, TP901 family, core region (Fragment); uniclust \| UniRef100\_A0A077L0E5 \| 100.0 \| 3.5e-61 \| 6.2e-67 \| 487.1 \| 634 \| (4, 646) \| 785 \| (61, 729) \| 1047 \| Phage tail tape measure protein domain-containing protein \| Phage tail tape measure protein domain-containing protein; uniclust \| UniRef100\_A0A077NKH9 \| 100.0 \| 1.1e-60 \| 2e-66 \| 490.4 \| 650 \| (5, 674) \| 785 \| (86, 747) \| 984 \| Putative bacteriophage tail fiber protein T \| Putative bacteriophage tail fiber protein T; uniclust \| UniRef100\_A0A1G6JE81 \| 100.0 \| 5.9e-60 \| 1e-65 \| 468.8 \| 714 \| (1, 785) \| 785 \| (1, 724) \| 725 \| Phage tail tape measure protein, TP901 family, core region \| Phage tail tape measure protein, TP901 family, core region; uniclust \| UniRef100\_A0A081N7U2 \| 100.0 \| 9.9e-60 \| 1.8e-65 \| 467.2 \| 569 \| (4, 579) \| 785 \| (74, 662) \| 806 \| Phage tail tape measure protein domain-containing protein \| Phage tail tape measure protein domain-containing protein; uniclust \| UniRef100\_A0A0A1HVB3 \| 100.0 \| 1e-59 \| 1.9e-65 \| 481.5 \| 736 \| (4, 783) \| 785 \| (134, 941) \| 944 \| Phage tail tape measure protein, TP901 family \| Phage tail tape measure protein, TP901 family; uniclust \| UniRef100\_A0A1E3G7C9 \| 100.0 \| 3e-59 \| 5.4e-65 \| 461.1 \| 558 \| (4, 586) \| 785 \| (79, 644) \| 783 \| Phage tail tape measure protein \| Phage tail tape measure protein; uniclust \| UniRef100\_A0A0A2XLQ8 \| 100.0 \| 3.9e-59 \| 7e-65 \| 470.3 \| 579 \| (106, 703) \| 785 \| (60, 678) \| 811 \| Phage tail protein \| Phage tail protein; uniclust \| UniRef100\_A0A0D7V1W3 \| 100.0 \| 2.3e-58 \| 4.1e-64 \| 464.5 \| 732 \| (4, 782) \| 785 \| (78, 887) \| 887 \| Phage tail tape measure protein domain-containing protein \| Phage tail tape measure protein domain-containing protein; uniclust \| UniRef100\_A0A2P5N6P9 \| 100.0 \| 4.1e-58 \| 7.3e-64 \| 445.5 \| 531 \| (1, 588) \| 785 \| (23, 562) \| 676 \| Phage tail tape measure protein \| Phage tail tape measure protein; uniclust \| UniRef100\_A0A022PGH9 \| 100.0 \| 6.5e-58 \| 1.2e-63 \| 453.7 \| 629 \| (118, 783) \| 785 \| (208, 887) \| 889 \| Phage tail tape measure protein, TP901 family \| Phage tail tape measure protein, TP901 family; uniclust \| UniRef100\_A0A2P8QYQ4 \| 100.0 \| 3.1e-57 \| 5.5e-63 \| 461.0 \| 600 \| (63, 672) \| 785 \| (95, 715) \| 833 \| Phage tail tape measure protein \| Phage tail tape measure protein; uniclust \| UniRef100\_A0A080IDD0 \| 100.0 \| 4.4e-55 \| 7.9e-61 \| 433.4 \| 621 \| (5, 675) \| 785 \| (78, 720) \| 944 \| Phage tail tape measure protein, TP901 family, core region \| Phage tail tape measure protein, TP901 family, core region; uniclust \| UniRef100\_A0A009QNM6 \| 100.0 \| 5.2e-54 \| 9.2e-60 \| 442.3 \| 724 \| (4, 784) \| 785 \| (80, 893) \| 893 \| Putative bacteriophage tail protein gpT \| Putative bacteriophage tail protein gpT; uniclust \| UniRef100\_A0A014N9D9 \| 100.0 \| 6.3e-54 \| 1.1e-59 \| 451.1 \| 623 \| (4, 668) \| 785 \| (80, 729) \| 921 \| Tail protein (Fragment) \| Tail protein (Fragment); uniclust \| UniRef100\_A0A017HCD3 \| 100.0 \| 1.2e-53 \| 2.2e-59 \| 449.9 \| 482 \| (111, 598) \| 785 \| (175, 678) \| 941 \| Phage tail length tape-measure protein \| Phage tail length tape-measure protein; uniclust \| UniRef100\_A0A080ITW0 \| 100.0 \| 1.6e-53 \| 2.8e-59 \| 424.5 \| 655 \| (5, 674) \| 785 \| (95, 779) \| 953 \| Phage tail tape measure protein, TP901 family, core region \| Phage tail tape measure protein, TP901 family, core region; uniclust \| UniRef100\_A0A0G9K5Q8 \| 100.0 \| 3e-53 \| 5.3e-59 \| 420.1 \| 552 \| (4, 585) \| 785 \| (73, 625) \| 782 \| Tail tape measure protein \| Tail tape measure protein; uniclust \| UniRef100\_A0A0D0HTB3 \| 100.0 \| 3.8e-53 \| 6.8e-59 \| 422.4 \| 529 \| (17, 594) \| 785 \| (169, 731) \| 1047 \| Phage-related minor tail protein \| Phage-related minor tail protein; uniclust \| UniRef100\_A0A0E2H9C3 \| 100.0 \| 4e-53 \| 7.1e-59 \| 455.9 \| 471 \| (110, 626) \| 785 \| (392, 872) \| 1237 \| Phage tail tape measure protein, TP901 family, core region \| Phage tail tape measure protein, TP901 family, core region; uniclust \| UniRef100\_A0A0A0GNB0 \| 100.0 \| 7.3e-53 \| 1.3e-58 \| 438.6 \| 643 \| (20, 674) \| 785 \| (108, 773) \| 998 \| Phage tail tape measure protein domain-containing protein \| Phage tail tape measure protein domain-containing protein; uniclust \| UniRef100\_A0A167GZN6 \| 100.0 \| 8.7e-53 \| 1.6e-58 \| 429.0 \| 591 \| (61, 660) \| 785 \| (210, 821) \| 1235 \| Transglycosylase \| Transglycosylase; uniclust \| UniRef100\_A0A059V4B6 \| 100.0 \| 1.1e-52 \| 2e-58 \| 434.9 \| 670 \| (1, 702) \| 785 \| (134, 854) \| 972 \| Bacteriophage tail protein \| Bacteriophage tail protein; uniclust \| UniRef100\_A0A0Q4IFW9 \| 100.0 \| 2.6e-52 \| 4.7e-58 \| 411.3 \| 707 \| (5, 783) \| 785 \| (80, 840) \| 845 \| Phage tail tape measure protein \| Phage tail tape measure protein; uniclust \| UniRef100\_A0A0A3A3N3 \| 100.0 \| 3.8e-52 \| 6.9e-58 \| 409.4 \| 562 \| (4, 589) \| 785 \| (80, 663) \| 778 \| Tail protein (Fragment) \| Tail protein (Fragment); uniclust \| UniRef100\_A0A1M6SBV5 \| 100.0 \| 1.4e-51 \| 2.4e-57 \| 406.4 \| 460 \| (1, 465) \| 785 \| (75, 540) \| 851 \| Phage tail tape measure protein, TP901 family, core region (Fragment) \| Phage tail tape measure protein, TP901 family, core region (Fragment); uniclust \| UniRef100\_A0A075KAY1 \| 100.0 \| 1.5e-51 \| 2.6e-57 \| 420.2 \| 561 \| (1, 585) \| 785 \| (2, 574) \| 760 \| Phage tail tape measure protein, TP901 family \| Phage tail tape measure protein, TP901 family; uniclust \| UniRef100\_A0A1V3IZV8 \| 100.0 \| 3.1e-51 \| 5.5e-57 \| 403.4 \| 570 \| (4, 583) \| 785 \| (78, 688) \| 717 \| Phage tail tape measure protein \| Phage tail tape measure protein; uniclust \| UniRef100\_A0A0L1KF90 \| 100.0 \| 3.2e-51 \| 5.6e-57 \| 410.6 \| 709 \| (1, 784) \| 785 \| (1, 743) \| 746 \| Tp901 family phage tail tape measure protein \| Tp901 family phage tail tape measure protein; uniclust \| UniRef100\_A0A496N6E3 \| 100.0 \| 2.3e-50 \| 4.2e-56 \| 388.0 \| 568 \| (26, 596) \| 785 \| (26, 644) \| 844 \| Phage tail tape measure protein (Fragment) \| Phage tail tape measure protein (Fragment); uniclust \| UniRef100\_A0A0K6GTP1 \| 100.0 \| 3.5e-50 \| 6.3e-56 \| 390.0 \| 724 \| (4, 782) \| 785 \| (79, 883) \| 883 \| Phage tail tape measure protein, TP901 family, core region \| Phage tail tape measure protein, TP901 family, core region; uniclust \| UniRef100\_A0A062IV08 \| 100.0 \| 1.9e-49 \| 3.4e-55 \| 382.9 \| 554 \| (4, 584) \| 785 \| (79, 656) \| 687 \| Phage tail tape measure protein, TP901 family, core region (Fragment) \| Phage tail tape measure protein, TP901 family, core region (Fragment); uniclust \| UniRef100\_A0A059UU34 \| 100.0 \| 3.2e-49 \| 5.7e-55 \| 395.5 \| 414 \| (110, 584) \| 785 \| (122, 535) \| 729 \| Phage tail protein \| Phage tail protein; uniclust \| UniRef100\_A0A099I5C4 \| 100.0 \| 4.3e-49 \| 7.6e-55 \| 410.4 \| 551 \| (1, 594) \| 785 \| (26, 628) \| 1011 \| Phage tail length tape measure protein \| Phage tail length tape measure protein; uniclust \| UniRef100\_A0A2W6QW57 \| 100.0 \| 9.4e-49 \| 1.7e-54 \| 379.1 \| 523 \| (119, 675) \| 785 \| (196, 720) \| 936 \| Phage tail tape measure protein (Fragment) \| Phage tail tape measure protein (Fragment); uniclust \| UniRef100\_A0A095YJN6 \| 100.0 \| 1.3e-48 \| 2.4e-54 \| 389.2 \| 375 \| (109, 491) \| 785 \| (193, 567) \| 1004 \| Phage tail tape measure protein, TP901 family, core region \| Phage tail tape measure protein, TP901 family, core region; uniclust \| UniRef100\_A0A6L5WL94 \| 100.0 \| 2.8e-48 \| 5.1e-54 \| 382.3 \| 476 \| (115, 596) \| 785 \| (106, 614) \| 807 \| Phage tail tape measure protein \| Phage tail tape measure protein; uniclust \| UniRef100\_A0A484GDB7 \| 100.0 \| 6.8e-48 \| 1.2e-53 \| 370.5 \| 648 \| (4, 673) \| 785 \| (9, 728) \| 902 \| Phage tail tape measure protein \| Phage tail tape measure protein; uniclust \| UniRef100\_A0A554WXF0 \| 100.0 \| 1.8e-47 \| 3.3e-53 \| 358.2 \| 561 \| (4, 594) \| 785 \| (72, 634) \| 749 \| Phage tail tape measure protein \| Phage tail tape measure protein; uniclust \| UniRef100\_A0A5S9Q3L2 \| 100.0 \| 2.6e-47 \| 4.8e-53 \| 361.8 \| 611 \| (112, 782) \| 785 \| (173, 795) \| 795 \| Phage tail tape measure protein domain-containing protein \| Phage tail tape measure protein domain-containing protein; uniclust \| UniRef100\_A0A022PG58 \| 100.0 \| 5.5e-47 \| 9.7e-53 \| 400.4 \| 655 \| (4, 674) \| 785 \| (163, 923) \| 1097 \| Phage tail tape measure protein, TP901 family \| Phage tail tape measure protein, TP901 family; uniclust \| UniRef100\_A0A0H3ZXY1 \| 100.0 \| 5.4e-47 \| 9.7e-53 \| 382.2 \| 733 \| (4, 783) \| 785 \| (86, 881) \| 939 \| Phage tail length tape-measure protein \| Phage tail length tape-measure protein; uniclust \| UniRef100\_A0A133QGT8 \| 100.0 \| 6.3e-47 \| 1.1e-52 \| 378.4 \| 541 \| (20, 593) \| 785 \| (47, 623) \| 1124 \| Phage tail tape measure protein, TP901 family \| Phage tail tape measure protein, TP901 family; uniclust \| UniRef100\_A0A0B7D869 \| 100.0 \| 6.4e-47 \| 1.2e-52 \| 366.0 \| 524 \| (111, 695) \| 785 \| (110, 650) \| 786 \| Phage-related minor tail protein \| Phage-related minor tail protein; uniclust \| UniRef100\_A0A2J4VH89 \| 100.0 \| 2.5e-46 \| 4.5e-52 \| 356.5 \| 616 \| (5, 670) \| 785 \| (78, 715) \| 815 \| Phage tail tape measure protein (Fragment) \| Phage tail tape measure protein (Fragment); uniclust \| UniRef100\_A0A0B5A7A7 \| 100.0 \| 2.8e-46 \| 5.1e-52 \| 380.4 \| 420 \| (111, 590) \| 785 \| (158, 580) \| 994 \| Membrane protein P6 \| Membrane protein P6; uniclust \| UniRef100\_A0A1B9KNC1 \| 100.0 \| 3.5e-46 \| 6.5e-52 \| 348.5 \| 496 \| (111, 625) \| 785 \| (51, 547) \| 725 \| Phage tail tape measure protein \| Phage tail tape measure protein; uniclust \| UniRef100\_A0A1H0P6W7 \| 100.0 \| 4.9e-46 \| 8.9e-52 \| 378.7 \| 464 \| (1, 471) \| 785 \| (29, 496) \| 1223 \| Phage tail tape measure protein, TP901 family, core region \| Phage tail tape measure protein, TP901 family, core region; uniclust \| UniRef100\_A0A0M4SUP3 \| 100.0 \| 2.1e-45 \| 3.7e-51 \| 357.8 \| 478 \| (113, 596) \| 785 \| (77, 588) \| 749 \| Phage tail tape measure protein, TP901 family \| Phage tail tape measure protein, TP901 family; uniclust \| UniRef100\_A0A1H2M3U9 \| 100.0 \| 3.5e-45 \| 6.5e-51 \| 353.3 \| 486 \| (117, 617) \| 785 \| (332, 828) \| 1105 \| Phage tail tape measure protein, TP901 family, core region \| Phage tail tape measure protein, TP901 family, core region; uniclust \| UniRef100\_A0A137SPW2 \| 100.0 \| 6.8e-45 \| 1.2e-50 \| 376.3 \| 466 \| (4, 488) \| 785 \| (81, 552) \| 1180 \| Phage tail tape measure protein, TP901 family \| Phage tail tape measure protein, TP901 family; uniclust \| UniRef100\_A0A0R1KHJ8 \| 100.0 \| 1e-44 \| 1.8e-50 \| 377.2 \| 457 \| (110, 589) \| 785 \| (274, 737) \| 1166 \| Phage tail tape measure protein domain-containing protein \| Phage tail tape measure protein domain-containing protein; uniclust \| UniRef100\_A0A0N8W5Z8 \| 100.0 \| 1.5e-44 \| 2.8e-50 \| 345.1 \| 478 \| (108, 622) \| 785 \| (57, 539) \| 708 \| Phage tail tape measure protein domain-containing protein \| Phage tail tape measure protein domain-containing protein; uniclust \| UniRef100\_A0A844HR58 \| 100.0 \| 1.5e-44 \| 2.8e-50 \| 337.8 \| 522 \| (113, 667) \| 785 \| (11, 545) \| 719 \| Phage tail tape measure protein \| Phage tail tape measure protein; uniclust \| UniRef100\_A0A0B5ATH8 \| 100.0 \| 1.7e-44 \| 2.9e-50 \| 374.6 \| 543 \| (4, 593) \| 785 \| (16, 597) \| 973 \| Phage tail tape measure protein domain-containing protein \| Phage tail tape measure protein domain-containing protein; uniclust \| UniRef100\_A0A3A6WDZ8 \| 100.0 \| 2.1e-44 \| 3.8e-50 \| 355.7 \| 561 \| (1, 583) \| 785 \| (74, 661) \| 853 \| Phage tail tape measure protein \| Phage tail tape measure protein; uniclust \| UniRef100\_A0A212KXI4 \| 100.0 \| 2.5e-44 \| 4.4e-50 \| 353.0 \| 568 \| (1, 576) \| 785 \| (1, 617) \| 766 \| Phage tail tape measure protein domain-containing protein \| Phage tail tape measure protein domain-containing protein; uniclust \| UniRef100\_A0A143WZ13 \| 100.0 \| 2.9e-44 \| 5.1e-50 \| 374.1 \| 423 \| (112, 591) \| 785 \| (153, 581) \| 890 \| Phage-related minor tail protein \| Phage-related minor tail protein; uniclust \| UniRef100\_A0A268TAT5 \| 100.0 \| 3.8e-44 \| 6.7e-50 \| 362.4 \| 535 \| (115, 664) \| 785 \| (119, 703) \| 790 \| Phage tail tape measure protein \| Phage tail tape measure protein; uniclust \| UniRef100\_A0A375H2Y0 \| 100.0 \| 4.2e-44 \| 7.6e-50 \| 331.1 \| 430 \| (156, 594) \| 785 \| (4, 443) \| 585 \| Phage tail tape measure protein domain-containing protein \| Phage tail tape measure protein domain-containing protein; uniclust \| UniRef100\_A0A376SBX2 \| 100.0 \| 4.8e-44 \| 8.6e-50 \| 338.9 \| 508 \| (117, 674) \| 785 \| (19, 533) \| 630 \| Tail length tape measure protein \| Tail length tape measure protein; uniclust \| UniRef100\_A0A2X0VQT4 \| 100.0 \| 5.8e-44 \| 1.1e-49 \| 346.3 \| 572 \| (4, 624) \| 785 \| (73, 654) \| 817 \| Phage-related minor tail protein \| Phage-related minor tail protein; uniclust \| UniRef100\_A0A068ZAA7 \| 100.0 \| 7.6e-44 \| 1.4e-49 \| 356.5 \| 637 \| (5, 675) \| 785 \| (79, 753) \| 911 \| Uncharacterized protein \| Uncharacterized protein; uniclust \| UniRef100\_A0A0D0GQG2 \| 100.0 \| 8.7e-44 \| 1.6e-49 \| 366.3 \| 746 \| (4, 784) \| 785 \| (225, 1106) \| 1154 \| Uncharacterized protein \| Uncharacterized protein; uniclust \| UniRef100\_A0A061P2R0 \| 100.0 \| 2e-43 \| 3.6e-49 \| 357.6 \| 441 \| (111, 593) \| 785 \| (162, 604) \| 834 \| Phage tail length tape-measure protein \| Phage tail length tape-measure protein; uniclust \| UniRef100\_A0A069F3H5 \| 100.0 \| 2.2e-43 \| 3.9e-49 \| 363.5 \| 431 \| (111, 593) \| 785 \| (157, 590) \| 899 \| Phage tail tape measure protein domain-containing protein \| Phage tail tape measure protein domain-containing protein; uniclust \| UniRef100\_A0A0R6PDG4 \| 100.0 \| 2.5e-43 \| 4.4e-49 \| 367.7 \| 434 \| (112, 592) \| 785 \| (158, 594) \| 1290 \| Tail protein \| Tail protein; uniclust \| UniRef100\_A0A3B9Q8T3 \| 100.0 \| 3.7e-43 \| 6.6e-49 \| 341.7 \| 572 \| (4, 587) \| 785 \| (73, 656) \| 788 \| Phage tail tape measure protein \| Phage tail tape measure protein; uniclust \| UniRef100\_A0A1I7FWK0 \| 100.0 \| 3.9e-43 \| 7e-49 \| 366.6 \| 335 \| (112, 462) \| 785 \| (202, 537) \| 1046 \| Phage tail tape measure protein, TP901 family, core region \| Phage tail tape measure protein, TP901 family, core region; uniclust \| UniRef100\_A0A081RQL7 \| 100.0 \| 4.2e-43 \| 7.6e-49 \| 336.7 \| 544 \| (17, 587) \| 785 \| (88, 641) \| 647 \| Phage-related minor tail protein \| Phage-related minor tail protein; uniclust \| UniRef100\_A0A1S6TMZ4 \| 100.0 \| 6.7e-43 \| 1.2e-48 \| 331.3 \| 475 \| (109, 595) \| 785 \| (74, 576) \| 765 \| Phage tail tape measure protein, TP901 family \| Phage tail tape measure protein, TP901 family; uniclust \| UniRef100\_A0A0A8H9G3 \| 100.0 \| 7.3e-43 \| 1.3e-48 \| 331.4 \| 476 \| (115, 595) \| 785 \| (87, 575) \| 740 \| Phage tail tape measure protein, TP901 family \| Phage tail tape measure protein, TP901 family; uniclust \| UniRef100\_A0A379VM56 \| 100.0 \| 1.1e-42 \| 2e-48 \| 336.5 \| 659 \| (6, 674) \| 785 \| (80, 848) \| 1068 \| Phage tail tape measure protein \| Phage tail tape measure protein; uniclust \| UniRef100\_A0A2N1WF95 \| 100.0 \| 1.6e-42 \| 2.8e-48 \| 313.6 \| 502 \| (258, 784) \| 785 \| (4, 506) \| 507 \| Phage tail tape measure protein (Fragment) \| Phage tail tape measure protein (Fragment); uniclust \| UniRef100\_A0A317PGW3 \| 100.0 \| 2.2e-42 \| 4e-48 \| 322.8 \| 413 \| (114, 586) \| 785 \| (50, 462) \| 551 \| TP901 family phage tail tape measure protein (Fragment) \| TP901 family phage tail tape measure protein (Fragment); uniclust \| UniRef100\_A0A450W6Y7 \| 100.0 \| 3.9e-42 \| 7.1e-48 \| 327.0 \| 652 \| (12, 673) \| 785 \| (84, 737) \| 1065 \| Phage tail tape measure protein, TP901 family, core region \| Phage tail tape measure protein, TP901 family, core region; uniclust \| UniRef100\_A0A109WD97 \| 100.0 \| 4.5e-42 \| 8.1e-48 \| 351.6 \| 458 \| (3, 471) \| 785 \| (105, 626) \| 1175 \| Phage tail tape measure protein domain-containing protein \| Phage tail tape measure protein domain-containing protein; uniclust \| UniRef100\_A0A2C6C9V6 \| 100.0 \| 4.6e-42 \| 8.4e-48 \| 333.7 \| 499 \| (109, 619) \| 785 \| (174, 695) \| 855 \| Phage tail tape measure protein \| Phage tail tape measure protein; uniclust \| UniRef100\_A0A135IJM6 \| 100.0 \| 5e-42 \| 8.9e-48 \| 343.8 \| 488 \| (106, 626) \| 785 \| (64, 562) \| 933 \| Phage tail tape measure protein domain-containing protein \| Phage tail tape measure protein domain-containing protein; uniclust \| UniRef100\_A0A077K8S2 \| 100.0 \| 5.4e-42 \| 9.5e-48 \| 355.7 \| 729 \| (1, 784) \| 785 \| (82, 896) \| 899 \| Uncharacterized protein \| Uncharacterized protein; uniclust \| UniRef100\_A0A132P719 \| 100.0 \| 5.9e-42 \| 1e-47 \| 370.8 \| 351 \| (110, 472) \| 785 \| (155, 506) \| 1062 \| Phage tail protein \| Phage tail protein; uniclust \| UniRef100\_A0A358PSL6 \| 100.0 \| 6.2e-42 \| 1.1e-47 \| 337.6 \| 441 \| (111, 594) \| 785 \| (149, 595) \| 971 \| Phage tail tape measure protein \| Phage tail tape measure protein; uniclust \| UniRef100\_A0A1Q1PVX7 \| 100.0 \| 8e-42 \| 1.4e-47 \| 363.1 \| 441 \| (23, 474) \| 785 \| (385, 871) \| 2029 \| Tail tape measure protein \| Tail tape measure protein; uniclust \| UniRef100\_A0A0R2DDY7 \| 100.0 \| 8e-42 \| 1.5e-47 \| 339.5 \| 518 \| (111, 705) \| 785 \| (280, 807) \| 1044 \| Phage tail tape measure protein domain-containing protein \| Phage tail tape measure protein domain-containing protein; uniclust \| UniRef100\_A0A2S0M4J9 \| 100.0 \| 8.3e-42 \| 1.5e-47 \| 340.0 \| 469 \| (112, 595) \| 785 \| (73, 567) \| 814 \| Phage tail tape measure protein \| Phage tail tape measure protein; uniclust \| UniRef100\_A0A323UJ65 \| 100.0 \| 1.3e-41 \| 2.4e-47 \| 341.6 \| 507 \| (109, 621) \| 785 \| (54, 595) \| 891 \| Phage tail tape measure protein \| Phage tail tape measure protein; uniclust \| UniRef100\_A0A1C4CAH5 \| 100.0 \| 1.3e-41 \| 2.4e-47 \| 343.4 \| 422 \| (112, 584) \| 785 \| (225, 648) \| 1275 \| Phage tail tape measure protein \| Phage tail tape measure protein; uniclust \| UniRef100\_A0A1A9RNP4 \| 100.0 \| 1.4e-41 \| 2.5e-47 \| 344.9 \| 615 \| (2, 624) \| 785 \| (1, 660) \| 899 \| Phage tail tape measure protein \| Phage tail tape measure protein; uniclust \| UniRef100\_A0A064AK64 \| 100.0 \| 2.2e-41 \| 4e-47 \| 340.7 \| 460 \| (114, 586) \| 785 \| (263, 730) \| 951 \| Phage tail tape measure protein domain-containing protein \| Phage tail tape measure protein domain-containing protein; uniclust \| UniRef100\_A0A021X9J0 \| 100.0 \| 3e-41 \| 5.2e-47 \| 346.8 \| 499 \| (109, 621) \| 785 \| (88, 626) \| 838 \| TP901 family phage tail tape measure protein \| TP901 family phage tail tape measure protein; uniclust \| UniRef100\_A0A022PJ34 \| 100.0 \| 2.9e-41 \| 5.3e-47 \| 327.4 \| 667 \| (4, 702) \| 785 \| (22, 735) \| 840 \| Phage tail tape measure protein, TP901 family \| Phage tail tape measure protein, TP901 family; uniclust \| UniRef100\_UPI00137359FD \| 100.0 \| 3.2e-41 \| 5.8e-47 \| 327.6 \| 583 \| (2, 595) \| 785 \| (1, 632) \| 661 \| phage tail tape measure protein \| phage tail tape measure protein; uniclust \| UniRef100\_A0A136C2Y5 \| 100.0 \| 3.9e-41 \| 7e-47 \| 352.3 \| 418 \| (54, 478) \| 785 \| (211, 651) \| 1262 \| Phage tail tape measure protein \| Phage tail tape measure protein; uniclust \| UniRef100\_A0A1G8XEH5 \| 100.0 \| 4.5e-41 \| 8.1e-47 \| 337.6 \| 339 \| (257, 612) \| 785 \| (261, 623) \| 860 \| Phage-related minor tail protein \| Phage-related minor tail protein; uniclust \| UniRef100\_A0A5P0YP70 \| 100.0 \| 5.5e-41 \| 9.7e-47 \| 338.9 \| 460 \| (111, 624) \| 785 \| (75, 544) \| 762 \| Phage tail tape measure protein \| Phage tail tape measure protein; uniclust \| UniRef100\_A0A0A7RU22 \| 100.0 \| 6.1e-41 \| 1.1e-46 \| 334.7 \| 331 \| (111, 453) \| 785 \| (64, 395) \| 699 \| Tail tape measure protein \| Tail tape measure protein; uniclust \| UniRef100\_A0A060LZ15 \| 100.0 \| 2.1e-40 \| 3.8e-46 \| 357.9 \| 370 \| (110, 492) \| 785 \| (173, 547) \| 1524 \| Metalloprotease \| Metalloprotease; uniclust \| UniRef100\_A0A1T5K1D4 \| 100.0 \| 2.9e-40 \| 5.2e-46 \| 344.0 \| 427 \| (110, 575) \| 785 \| (159, 589) \| 1280 \| Phage tail tape measure protein, TP901 family, core region \| Phage tail tape measure protein, TP901 family, core region; uniclust \| UniRef100\_A0A286BYN5 \| 100.0 \| 2.9e-40 \| 5.2e-46 \| 315.2 \| 510 \| (118, 674) \| 785 \| (265, 778) \| 879 \| Phage tail tape measure protein, TP901 family, core region \| Phage tail tape measure protein, TP901 family, core region; uniclust \| UniRef100\_A0A2X4YRE2 \| 100.0 \| 3.1e-40 \| 5.6e-46 \| 313.2 \| 497 \| (112, 625) \| 785 \| (35, 594) \| 676 \| Phage protein \| Phage protein; uniclust \| UniRef100\_A0A4V2PT27 \| 100.0 \| 3.1e-40 \| 5.7e-46 \| 318.8 \| 497 \| (120, 623) \| 785 \| (71, 574) \| 1021 \| TP901 family phage tail tape measure protein \| TP901 family phage tail tape measure protein; uniclust \| UniRef100\_A0A1M5Z9G5 \| 100.0 \| 3.8e-40 \| 6.9e-46 \| 323.3 \| 617 \| (7, 657) \| 785 \| (137, 774) \| 1021 \| Phage-related minor tail protein \| Phage-related minor tail protein; uniclust \| UniRef100\_A0A0F6XYW3 \| 100.0 \| 5.2e-40 \| 9.4e-46 \| 326.6 \| 536 \| (113, 705) \| 785 \| (240, 818) \| 1005 \| Phage tail tape measure protein domain-containing protein \| Phage tail tape measure protein domain-containing protein; uniclust \| UniRef100\_A0A0D1X6H1 \| 100.0 \| 6.9e-40 \| 1.2e-45 \| 344.7 \| 429 \| (111, 591) \| 785 \| (161, 657) \| 1211 \| Phage tail tape measure protein, TP901 family, core region \| Phage tail tape measure protein, TP901 family, core region; uniclust \| UniRef100\_A0A8X6G804 \| 100.0 \| 7.9e-40 \| 1.4e-45 \| 310.8 \| 580 \| (5, 587) \| 785 \| (70, 656) \| 850 \| Phage tail tape measure protein \| Phage tail tape measure protein; uniclust \| UniRef100\_A0A1V9DAU4 \| 100.0 \| 9.2e-40 \| 1.7e-45 \| 314.8 \| 710 \| (6, 782) \| 785 \| (102, 840) \| 842 \| Phage tail tape measure protein \| Phage tail tape measure protein; uniclust \| UniRef100\_A0A0F4VRI4 \| 100.0 \| 9.6e-40 \| 1.7e-45 \| 342.3 \| 448 \| (118, 594) \| 785 \| (174, 632) \| 998 \| Phage tail length tape-measure protein \| Phage tail length tape-measure protein; uniclust \| UniRef100\_A0A1F0GHX6 \| 100.0 \| 9.6e-40 \| 1.7e-45 \| 330.8 \| 465 \| (111, 620) \| 785 \| (90, 560) \| 880 \| Phage tail tape measure protein domain-containing protein \| Phage tail tape measure protein domain-containing protein; uniclust \| UniRef100\_H9A124 \| 100.0 \| 1.3e-39 \| 2.4e-45 \| 313.6 \| 471 \| (110, 592) \| 785 \| (168, 671) \| 848 \| Tail tape measure protein \| Tail tape measure protein; uniclust \| UniRef100\_A0A6M0FWE7 \| 100.0 \| 1.6e-39 \| 2.9e-45 \| 324.6 \| 346 \| (110, 460) \| 785 \| (76, 422) \| 751 \| Phage tail tape measure protein \| Phage tail tape measure protein; uniclust \| UniRef100\_A0A174B5Q2 \| 100.0 \| 2.2e-39 \| 3.9e-45 \| 342.4 \| 443 \| (111, 595) \| 785 \| (106, 553) \| 1168 \| Phage-related minor tail protein \| Phage-related minor tail protein; uniclust \| UniRef100\_A0A1G3M6W5 \| 100.0 \| 2.2e-39 \| 4e-45 \| 334.0 \| 494 \| (110, 628) \| 785 \| (59, 569) \| 767 \| Phage tail tape measure protein \| Phage tail tape measure protein; uniclust \| UniRef100\_A0A0U4IIN6 \| 100.0 \| 2.2e-39 \| 4e-45 \| 335.9 \| 439 \| (111, 596) \| 785 \| (243, 685) \| 905 \| Putative tail tape measure protein \| Putative tail tape measure protein; uniclust \| UniRef100\_UPI0002AF4875 \| 100.0 \| 2.5e-39 \| 4.5e-45 \| 298.8 \| 563 \| (179, 782) \| 785 \| (15, 619) \| 619 \| phage tail tape measure protein \| phage tail tape measure protein; uniclust \| UniRef100\_A0A3L2QDC9 \| 100.0 \| 3.4e-39 \| 6.1e-45 \| 329.3 \| 535 \| (116, 670) \| 785 \| (112, 722) \| 978 \| Phage tail tape measure protein (Fragment) \| Phage tail tape measure protein (Fragment); uniclust \| UniRef100\_A0A0E3K1L2 \| 100.0 \| 3.5e-39 \| 6.4e-45 \| 308.6 \| 533 \| (111, 703) \| 785 \| (72, 609) \| 723 \| Phage protein \| Phage protein; uniclust \| UniRef100\_A0A411WHS8 \| 100.0 \| 5.2e-39 \| 9.4e-45 \| 316.6 \| 680 \| (2, 697) \| 785 \| (1, 717) \| 937 \| Phage tail tape measure protein \| Phage tail tape measure protein; uniclust \| UniRef100\_A0A1A9RDG3 \| 100.0 \| 5.7e-39 \| 1e-44 \| 308.4 \| 589 \| (6, 623) \| 785 \| (79, 692) \| 900 \| Phage tail tape measure protein \| Phage tail tape measure protein; uniclust \| UniRef100\_A0A1T1J9S7 \| 100.0 \| 7.2e-39 \| 1.3e-44 \| 317.6 \| 426 \| (98, 589) \| 785 \| (163, 596) \| 798 \| Phage tail tape measure protein \| Phage tail tape measure protein; uniclust \| UniRef100\_A0A1X3LEP7 \| 100.0 \| 9.9e-39 \| 1.8e-44 \| 321.2 \| 438 \| (121, 619) \| 785 \| (253, 703) \| 891 \| Phage tail tape measure protein, TP901 family, core region \| Phage tail tape measure protein, TP901 family, core region; uniclust \| UniRef100\_A0A1I4GFH8 \| 100.0 \| 1e-38 \| 1.8e-44 \| 295.4 \| 453 \| (113, 618) \| 785 \| (61, 518) \| 598 \| Phage tail tape measure protein, TP901 family, core region \| Phage tail tape measure protein, TP901 family, core region; uniclust \| UniRef100\_A0A0S4TWE8 \| 100.0 \| 2.3e-38 \| 4.1e-44 \| 295.8 \| 446 \| (279, 782) \| 785 \| (5, 487) \| 488 \| Phage-related tail transmembrane protein \| Phage-related tail transmembrane protein; uniclust \| UniRef100\_A0A173R5S4 \| 100.0 \| 2.7e-38 \| 4.8e-44 \| 327.2 \| 311 \| (115, 437) \| 785 \| (232, 543) \| 1265 \| Phage-related minor tail protein \| Phage-related minor tail protein; uniclust \| UniRef100\_A0A1C5YW05 \| 100.0 \| 2.9e-38 \| 5.2e-44 \| 310.0 \| 426 \| (111, 596) \| 785 \| (61, 490) \| 868 \| Phage-related minor tail protein \| Phage-related minor tail protein; uniclust \| UniRef100\_A0A072YR64 \| 100.0 \| 3.3e-38 \| 6.1e-44 \| 307.7 \| 443 \| (114, 584) \| 785 \| (61, 521) \| 940 \| Tail tape measure protein \| Tail tape measure protein; uniclust \| UniRef100\_UPI00068C4C6A \| 100.0 \| 3.6e-38 \| 6.7e-44 \| 297.1 \| 479 \| (110, 593) \| 785 \| (90, 588) \| 933 \| phage tail tape measure protein \| phage tail tape measure protein; uniclust \| UniRef100\_A0A2G6CRZ3 \| 100.0 \| 5.6e-38 \| 1e-43 \| 299.4 \| 556 \| (1, 567) \| 785 \| (1, 574) \| 707 \| Phage tail tape measure protein \| Phage tail tape measure protein; uniclust \| UniRef100\_UPI001EF7523E \| 100.0 \| 5.5e-38 \| 1e-43 \| 291.4 \| 528 \| (117, 647) \| 785 \| (5, 649) \| 795 \| phage tail tape measure protein \| phage tail tape measure protein; uniclust \| UniRef100\_A0A069RFS6 \| 100.0 \| 6.9e-38 \| 1.2e-43 \| 316.1 \| 439 \| (111, 593) \| 785 \| (165, 610) \| 1085 \| TP901 family phage tail tape measure protein \| TP901 family phage tail tape measure protein; uniclust \| UniRef100\_A0A064AM58 \| 100.0 \| 8.2e-38 \| 1.5e-43 \| 318.1 \| 296 \| (118, 424) \| 785 \| (194, 497) \| 925 \| Phage tail tape measure protein domain-containing protein \| Phage tail tape measure protein domain-containing protein; uniclust \| UniRef100\_A0A3S0BBD4 \| 100.0 \| 8.7e-38 \| 1.6e-43 \| 302.6 \| 461 \| (112, 597) \| 785 \| (83, 546) \| 668 \| Phage tail tape measure protein \| Phage tail tape measure protein; uniclust \| UniRef100\_A0A0H3J8I0 \| 100.0 \| 1e-37 \| 1.8e-43 \| 332.0 \| 306 \| (125, 449) \| 785 \| (252, 565) \| 1065 \| Minor tail protein \| Minor tail protein; uniclust \| UniRef100\_A0A0U2B0N6 \| 100.0 \| 1.1e-37 \| 2e-43 \| 309.2 \| 600 \| (4, 623) \| 785 \| (110, 759) \| 1243 \| Phage tail tape measure protein domain-containing protein \| Phage tail tape measure protein domain-containing protein; uniclust \| UniRef100\_A0A1H2FMR0 \| 100.0 \| 1.8e-37 \| 3.2e-43 \| 315.2 \| 478 \| (106, 594) \| 785 \| (72, 587) \| 825 \| Phage tail tape measure protein, TP901 family, core region \| Phage tail tape measure protein, TP901 family, core region; uniclust \| UniRef100\_A0A1Y4I868 \| 100.0 \| 2e-37 \| 3.6e-43 \| 320.3 \| 526 \| (6, 594) \| 785 \| (9, 594) \| 825 \| Phage tail tape measure protein (Fragment) \| Phage tail tape measure protein (Fragment); uniclust \| UniRef100\_A0A5A9EN03 \| 100.0 \| 2e-37 \| 3.7e-43 \| 288.3 \| 536 \| (77, 631) \| 785 \| (142, 684) \| 808 \| Phage tail tape measure protein (Fragment) \| Phage tail tape measure protein (Fragment); uniclust \| UniRef100\_A0A3Q8WGQ2 \| 100.0 \| 2.4e-37 \| 4.4e-43 \| 310.4 \| 544 \| (30, 601) \| 785 \| (159, 723) \| 1541 \| Phage tail tape measure protein \| Phage tail tape measure protein; uniclust \| UniRef100\_A0A0F7PRC0 \| 100.0 \| 3.3e-37 \| 5.9e-43 \| 321.3 \| 435 \| (115, 592) \| 785 \| (275, 724) \| 1436 \| Phage tail tape mesure protein \| Phage tail tape mesure protein; uniclust \| UniRef100\_A0A252F2N6 \| 100.0 \| 3.8e-37 \| 6.8e-43 \| 304.6 \| 528 \| (15, 596) \| 785 \| (117, 680) \| 922 \| Phage tail tape measure protein domain-containing protein \| Phage tail tape measure protein domain-containing protein; uniclust \| UniRef100\_A0A077RGZ1 \| 100.0 \| 4.3e-37 \| 7.6e-43 \| 339.4 \| 466 \| (110, 593) \| 785 \| (435, 918) \| 2100 \| lysostaphin \| lysostaphin; uniclust \| UniRef100\_A0A7Z0HAF9 \| 100.0 \| 4.7e-37 \| 8.5e-43 \| 310.1 \| 432 \| (111, 587) \| 785 \| (83, 521) \| 1141 \| Phage tail tape measure protein \| Phage tail tape measure protein; uniclust \| UniRef100\_A0A3A6P993 \| 100.0 \| 5.7e-37 \| 1e-42 \| 295.3 \| 572 \| (11, 622) \| 785 \| (27, 637) \| 754 \| Phage tail tape measure protein \| Phage tail tape measure protein; uniclust \| UniRef100\_A0A0Q2XJT6 \| 100.0 \| 7.4e-37 \| 1.4e-42 \| 290.1 \| 492 \| (111, 610) \| 785 \| (69, 579) \| 764 \| Phage tail tape measure protein domain-containing protein \| Phage tail tape measure protein domain-containing protein; uniclust \| UniRef100\_A0A349GR42 \| 100.0 \| 1e-36 \| 1.9e-42 \| 296.3 \| 437 \| (112, 572) \| 785 \| (188, 625) \| 826 \| Phage tail tape measure protein \| Phage tail tape measure protein; uniclust \| UniRef100\_A0A0R1H356 \| 100.0 \| 1.2e-36 \| 2e-42 \| 322.2 \| 410 \| (32, 463) \| 785 \| (186, 625) \| 1227 \| Tape measure protein \| Tape measure protein; uniclust \| UniRef100\_A0A371IQX0 \| 100.0 \| 1.2e-36 \| 2.1e-42 \| 306.6 \| 555 \| (3, 594) \| 785 \| (65, 642) \| 1121 \| Phage tail tape measure protein \| Phage tail tape measure protein; uniclust \| UniRef100\_A0A081NWQ5 \| 100.0 \| 1.3e-36 \| 2.2e-42 \| 318.0 \| 316 \| (113, 445) \| 785 \| (133, 456) \| 854 \| Phage tail tape measure protein domain-containing protein \| Phage tail tape measure protein domain-containing protein; uniclust \| UniRef100\_A0A0D1IZE2 \| 100.0 \| 1.4e-36 \| 2.6e-42 \| 327.1 \| 432 \| (111, 594) \| 785 \| (130, 570) \| 1126 \| Phage tail tape measure protein domain-containing protein \| Phage tail tape measure protein domain-containing protein; uniclust \| UniRef100\_A0A855AEA3 \| 100.0 \| 1.8e-36 \| 3.3e-42 \| 307.1 \| 456 \| (111, 623) \| 785 \| (61, 530) \| 1054 \| Phage tail tape measure protein domain-containing protein \| Phage tail tape measure protein domain-containing protein; uniclust \| UniRef100\_E1QHA5 \| 100.0 \| 2.3e-36 \| 4.1e-42 \| 293.7 \| 523 \| (62, 589) \| 785 \| (97, 633) \| 993 \| Phage tail tape measure protein, TP901 family \| Phage tail tape measure protein, TP901 family; uniclust \| UniRef100\_A0A176EZ66 \| 100.0 \| 2.5e-36 \| 4.4e-42 \| 311.2 \| 362 \| (108, 474) \| 785 \| (70, 431) \| 1055 \| Phage tail tape measure protein domain-containing protein \| Phage tail tape measure protein domain-containing protein; uniclust \| UniRef100\_A0A381EBY2 \| 100.0 \| 2.9e-36 \| 5.2e-42 \| 294.3 \| 539 \| (17, 582) \| 785 \| (93, 667) \| 874 \| Uncharacterized conserved protein \| Uncharacterized conserved protein; uniclust \| UniRef100\_A0A2X1LDV0 \| 100.0 \| 3e-36 \| 5.4e-42 \| 287.1 \| 417 \| (147, 587) \| 785 \| (60, 478) \| 492 \| Tail protein (Modular protein) \| Tail protein (Modular protein); uniclust \| UniRef100\_A0A7V7G2N6 \| 100.0 \| 3.3e-36 \| 5.9e-42 \| 287.0 \| 651 \| (1, 783) \| 785 \| (2, 729) \| 730 \| Phage tail tape measure protein \| Phage tail tape measure protein; uniclust \| UniRef100\_A0A0A2XR61 \| 100.0 \| 3.7e-36 \| 6.7e-42 \| 298.6 \| 608 \| (1, 619) \| 785 \| (110, 769) \| 1042 \| Phage tail tape measure protein domain-containing protein \| Phage tail tape measure protein domain-containing protein; uniclust \| UniRef100\_A0A2K1FR02 \| 100.0 \| 4.7e-36 \| 8.6e-42 \| 277.1 \| 452 \| (118, 589) \| 785 \| (51, 508) \| 657 \| Phage tail tape measure protein \| Phage tail tape measure protein; uniclust \| UniRef100\_A0A0M0KJB6 \| 100.0 \| 5.1e-36 \| 9e-42 \| 318.0 \| 440 \| (6, 460) \| 785 \| (35, 499) \| 1120 \| Phage tail tape measure protein domain-containing protein \| Phage tail tape measure protein domain-containing protein; uniclust \| UniRef100\_A0A510Y3F9 \| 100.0 \| 6.4e-36 \| 1.2e-41 \| 309.2 \| 335 \| (111, 457) \| 785 \| (73, 408) \| 1202 \| Phage tail tape measure protein domain-containing protein \| Phage tail tape measure protein domain-containing protein; uniclust \| UniRef100\_A0A4D6ALN7 \| 100.0 \| 7.6e-36 \| 1.4e-41 \| 286.6 \| 457 \| (111, 620) \| 785 \| (75, 547) \| 887 \| Tail length tape-measure protein \| Tail length tape-measure protein; uniclust \| UniRef100\_UPI00036738C7 \| 100.0 \| 8.2e-36 \| 1.5e-41 \| 277.6 \| 540 \| (100, 653) \| 785 \| (121, 686) \| 801 \| phage tail tape measure protein \| phage tail tape measure protein; uniclust \| UniRef100\_A0A0E0TET7 \| 100.0 \| 1e-35 \| 1.8e-41 \| 331.7 \| 492 \| (56, 592) \| 785 \| (212, 731) \| 1632 \| Tail tape measure protein TP901 core region protein \| Tail tape measure protein TP901 core region protein; uniclust \| UniRef100\_A0A1G5JIN5 \| 100.0 \| 1.2e-35 \| 2.1e-41 \| 300.9 \| 338 \| (115, 466) \| 785 \| (61, 399) \| 812 \| Phage tail tape measure protein, TP901 family, core region \| Phage tail tape measure protein, TP901 family, core region; uniclust \| UniRef100\_A0A3T0L2L7 \| 100.0 \| 1.3e-35 \| 2.4e-41 \| 277.4 \| 505 \| (70, 594) \| 785 \| (126, 634) \| 836 \| Phage tail tape measure protein \| Phage tail tape measure protein; uniclust \| UniRef100\_A0A812RFT8 \| 100.0 \| 1.4e-35 \| 2.6e-41 \| 295.3 \| 470 \| (103, 592) \| 785 \| (168, 637) \| 1893 \| site-specific DNA-methyltransferase (adenine-specific) \| site-specific DNA-methyltransferase (adenine-specific); uniclust \| UniRef100\_A0A061NKC8 \| 100.0 \| 1.6e-35 \| 2.9e-41 \| 311.9 \| 340 \| (112, 463) \| 785 \| (61, 401) \| 968 \| Phage tail length tape-measure protein \| Phage tail length tape-measure protein; uniclust \| UniRef100\_A0A345CNQ0 \| 100.0 \| 1.8e-35 \| 3.3e-41 \| 282.4 \| 587 \| (3, 600) \| 785 \| (2, 639) \| 716 \| Phage tail tape measure protein \| Phage tail tape measure protein; uniclust \| UniRef100\_A0A099WJF5 \| 100.0 \| 3e-35 \| 5.3e-41 \| 304.8 \| 326 \| (118, 470) \| 785 \| (258, 590) \| 952 \| Phage tail protein \| Phage tail protein; uniclust \| UniRef100\_A0A2R3MZN4 \| 100.0 \| 3.3e-35 \| 6e-41 \| 287.2 \| 458 \| (112, 596) \| 785 \| (202, 665) \| 911 \| Phage tail tape measure protein \| Phage tail tape measure protein; uniclust \| UniRef100\_A0A0V9UNT3 \| 100.0 \| 3.5e-35 \| 6.1e-41 \| 300.4 \| 316 \| (114, 441) \| 785 \| (70, 388) \| 809 \| Phage tail tape measure protein domain-containing protein \| Phage tail tape measure protein domain-containing protein; uniclust \| UniRef100\_A0A059N3D7 \| 100.0 \| 3.6e-35 \| 6.3e-41 \| 312.5 \| 478 \| (110, 671) \| 785 \| (242, 744) \| 920 \| Phage tail tape measure protein, TP901 family \| Phage tail tape measure protein, TP901 family; uniclust \| UniRef100\_A0A157BCK0 \| 100.0 \| 4.1e-35 \| 7.5e-41 \| 271.9 \| 450 \| (178, 674) \| 785 \| (16, 472) \| 568 \| Phage Tail Tape Measure protein \| Phage Tail Tape Measure protein; uniclust \| UniRef100\_UPI000935C447 \| 100.0 \| 4.4e-35 \| 8.1e-41 \| 272.5 \| 580 \| (4, 601) \| 785 \| (77, 677) \| 792 \| phage tail tape measure protein \| phage tail tape measure protein; uniclust \| UniRef100\_A0A085AFM7 \| 100.0 \| 5.2e-35 \| 9.3e-41 \| 300.9 \| 450 \| (21, 474) \| 785 \| (169, 624) \| 945 \| Phage tail tape measure protein domain-containing protein \| Phage tail tape measure protein domain-containing protein; uniclust \| UniRef100\_E6TVF9 \| 100.0 \| 5.7e-35 \| 1e-40 \| 292.1 \| 424 \| (111, 591) \| 785 \| (163, 595) \| 950 \| Phage tail tape measure protein, TP901 family \| Phage tail tape measure protein, TP901 family; uniclust \| UniRef100\_A0A133ZYY1 \| 100.0 \| 5.9e-35 \| 1.1e-40 \| 302.5 \| 319 \| (111, 439) \| 785 \| (60, 383) \| 1184 \| Phage tail tape measure protein, TP901 family \| Phage tail tape measure protein, TP901 family; uniclust \| UniRef100\_A0A0A0IDQ6 \| 100.0 \| 6.2e-35 \| 1.1e-40 \| 303.8 \| 423 \| (111, 595) \| 785 \| (165, 599) \| 876 \| TP901 family phage tail tape measure protein \| TP901 family phage tail tape measure protein; uniclust \| UniRef100\_A0A0F0CCS2 \| 100.0 \| 6.3e-35 \| 1.1e-40 \| 307.9 \| 351 \| (111, 473) \| 785 \| (229, 671) \| 1807 \| Phage-related minor tail protein \| Phage-related minor tail protein; uniclust \| UniRef100\_A0A024QH21 \| 100.0 \| 7.1e-35 \| 1.3e-40 \| 293.7 \| 398 \| (111, 589) \| 785 \| (50, 457) \| 763 \| Phage tail tape measure protein, TP901 family, core region \| Phage tail tape measure protein, TP901 family, core region; uniclust \| UniRef100\_A0A1K1MLR7 \| 100.0 \| 7.2e-35 \| 1.3e-40 \| 312.4 \| 322 \| (112, 445) \| 785 \| (164, 487) \| 1544 \| Phage tail tape measure protein, TP901 family, core region \| Phage tail tape measure protein, TP901 family, core region; uniclust \| UniRef100\_A0A5T0QQZ5 \| 100.0 \| 7.5e-35 \| 1.4e-40 \| 267.5 \| 455 \| (115, 574) \| 785 \| (85, 552) \| 554 \| Phage tail tape measure protein (Fragment) \| Phage tail tape measure protein (Fragment); uniclust \| UniRef100\_A0A6N7JJS5 \| 100.0 \| 8e-35 \| 1.5e-40 \| 285.6 \| 428 \| (23, 456) \| 785 \| (357, 788) \| 1494 \| Phage tail tape measure protein \| Phage tail tape measure protein; uniclust \| UniRef100\_A0A160M9X1 \| 100.0 \| 8.8e-35 \| 1.6e-40 \| 306.8 \| 442 \| (23, 475) \| 785 \| (176, 646) \| 1265 \| Phage tail tape measure protein domain-containing protein \| Phage tail tape measure protein domain-containing protein; uniclust \| UniRef100\_UPI001FF55D27 \| 100.0 \| 9.6e-35 \| 1.8e-40 \| 273.7 \| 482 \| (106, 593) \| 785 \| (221, 719) \| 902 \| phage tail tape measure protein \| phage tail tape measure protein; uniclust \| UniRef100\_A0A090IMA5 \| 100.0 \| 1.1e-34 \| 1.9e-40 \| 302.6 \| 461 \| (5, 470) \| 785 \| (136, 603) \| 920 \| Phage-related tail protein \| Phage-related tail protein; uniclust \| UniRef100\_A0A0Q7K0A6 \| 100.0 \| 1.2e-34 \| 2.1e-40 \| 302.6 \| 526 \| (118, 701) \| 785 \| (84, 621) \| 889 \| Phage tail tape measure protein domain-containing protein \| Phage tail tape measure protein domain-containing protein; uniclust \| UniRef100\_A0A0B3BNP9 \| 100.0 \| 1.3e-34 \| 2.3e-40 \| 286.3 \| 538 \| (63, 621) \| 785 \| (109, 660) \| 800 \| Phage tail tape measure protein, TP901 family \| Phage tail tape measure protein, TP901 family; uniclust \| UniRef100\_A0A0T1XR25 \| 100.0 \| 1.3e-34 \| 2.4e-40 \| 299.6 \| 331 \| (110, 461) \| 785 \| (58, 390) \| 826 \| Phage tail tape measure protein domain-containing protein \| Phage tail tape measure protein domain-containing protein; uniclust \| UniRef100\_A0A080KEC1 \| 100.0 \| 1.5e-34 \| 2.8e-40 \| 273.1 \| 481 \| (115, 647) \| 785 \| (74, 567) \| 777 \| Phage-related minor tail protein \| Phage-related minor tail protein; uniclust \| UniRef100\_A0A075X0G2 \| 100.0 \| 1.6e-34 \| 2.9e-40 \| 286.5 \| 473 \| (114, 587) \| 785 \| (74, 604) \| 778 \| Phage tail tape measure protein domain-containing protein \| Phage tail tape measure protein domain-containing protein; uniclust \| UniRef100\_A0A060PHZ7 \| 100.0 \| 2.6e-34 \| 4.7e-40 \| 308.3 \| 436 \| (23, 470) \| 785 \| (51, 514) \| 1297 \| Phage tail tape measure protein domain-containing protein \| Phage tail tape measure protein domain-containing protein; uniclust \| UniRef100\_A0A8S5PUM7 \| 100.0 \| 3.7e-34 \| 6.7e-40 \| 280.6 \| 576 \| (9, 595) \| 785 \| (79, 697) \| 1042 \| Minor tail protein (Fragment) \| Minor tail protein (Fragment); uniclust \| UniRef100\_A0A143DED3 \| 100.0 \| 3.9e-34 \| 6.9e-40 \| 277.2 \| 486 \| (1, 621) \| 785 \| (2, 496) \| 606 \| Phage tail tape measure protein \| Phage tail tape measure protein; uniclust \| UniRef100\_UPI0005E6ED2E \| 100.0 \| 5.3e-34 \| 9.6e-40 \| 270.2 \| 474 \| (4, 494) \| 785 \| (166, 659) \| 732 \| phage tail tape measure protein \| phage tail tape measure protein; uniclust \| UniRef100\_UPI0022E9545A \| 100.0 \| 6.1e-34 \| 1.1e-39 \| 268.5 \| 478 \| (103, 592) \| 785 \| (308, 786) \| 902 \| phage tail tape measure protein \| phage tail tape measure protein; uniclust \| UniRef100\_A0A367EFZ0 \| 100.0 \| 7e-34 \| 1.3e-39 \| 281.8 \| 414 \| (112, 586) \| 785 \| (70, 488) \| 857 \| Phage tail tape measure protein \| Phage tail tape measure protein; uniclust \| UniRef100\_A0A269PKX6 \| 100.0 \| 7.5e-34 \| 1.4e-39 \| 283.2 \| 454 \| (110, 590) \| 785 \| (52, 528) \| 1007 \| Phage tail tape measure protein \| Phage tail tape measure protein; uniclust \| UniRef100\_A0A073KF69 \| 100.0 \| 7.8e-34 \| 1.4e-39 \| 301.0 \| 332 \| (109, 458) \| 785 \| (71, 406) \| 1013 \| Phage tail tape measure protein domain-containing protein \| Phage tail tape measure protein domain-containing protein; uniclust \| UniRef100\_D5EFA3 \| 100.0 \| 8.1e-34 \| 1.5e-39 \| 264.9 \| 557 \| (5, 591) \| 785 \| (73, 636) \| 810 \| Phage tail tape measure protein, TP901 family \| Phage tail tape measure protein, TP901 family; uniclust \| UniRef100\_A0A076LFK6 \| 100.0 \| 9e-34 \| 1.6e-39 \| 259.3 \| 354 \| (111, 468) \| 785 \| (81, 434) \| 466 \| Phage tail length tape-measure protein \| Phage tail length tape-measure protein; uniclust \| UniRef100\_A0A0D0IGY2 \| 100.0 \| 9.3e-34 \| 1.7e-39 \| 281.4 \| 336 \| (433, 782) \| 785 \| (494, 892) \| 892 \| Phage-related minor tail protein \| Phage-related minor tail protein; uniclust \| UniRef100\_A0A0Q0F7A5 \| 100.0 \| 1.1e-33 \| 2e-39 \| 269.8 \| 513 \| (113, 691) \| 785 \| (52, 581) \| 720 \| Prophage PSSSM-04, tail length tape-measure protein \| Prophage PSSSM-04, tail length tape-measure protein; uniclust \| UniRef100\_A0A0U4NFW6 \| 100.0 \| 1.1e-33 \| 2.1e-39 \| 286.9 \| 418 \| (111, 583) \| 785 \| (213, 634) \| 1751 \| Chromosome partition protein Smc \| Chromosome partition protein Smc; uniclust \| UniRef100\_A0A090IER3 \| 100.0 \| 1.2e-33 \| 2.1e-39 \| 290.5 \| 454 \| (102, 578) \| 785 \| (44, 512) \| 747 \| Phage tail tape measure protein, TP901 family, core region,Phage-related minor tail protein \| Phage tail tape measure protein, TP901 family, core region,Phage-related minor tail protein; uniclust \| UniRef100\_A0A5B0BRS7 \| 100.0 \| 1.3e-33 \| 2.3e-39 \| 293.4 \| 421 \| (112, 581) \| 785 \| (170, 597) \| 1481 \| Phage tail tape measure protein \| Phage tail tape measure protein; uniclust \| UniRef100\_A0A1C2K2M3 \| 100.0 \| 1.3e-33 \| 2.4e-39 \| 280.2 \| 470 \| (4, 473) \| 785 \| (127, 609) \| 884 \| Phage tail tape measure protein (Fragment) \| Phage tail tape measure protein (Fragment); uniclust \| UniRef100\_A0A2T3FUX6 \| 100.0 \| 1.4e-33 \| 2.5e-39 \| 287.3 \| 448 \| (1, 464) \| 785 \| (1, 495) \| 1969 \| Phage tail tape measure protein \| Phage tail tape measure protein; uniclust \| UniRef100\_UPI00069E9770 \| 100.0 \| 1.5e-33 \| 2.7e-39 \| 260.4 \| 545 \| (117, 674) \| 785 \| (31, 653) \| 730 \| phage tail tape measure protein \| phage tail tape measure protein; uniclust \| UniRef100\_A0A1X3JZY9 \| 100.0 \| 1.6e-33 \| 2.9e-39 \| 276.2 \| 488 \| (184, 702) \| 785 \| (81, 613) \| 727 \| Putative phage tail protein \| Putative phage tail protein; uniclust \| UniRef100\_A0A1L7N5W8 \| 100.0 \| 1.9e-33 \| 3.5e-39 \| 283.5 \| 579 \| (5, 610) \| 785 \| (79, 692) \| 1199 \| Phage tail tape measure protein domain-containing protein \| Phage tail tape measure protein domain-containing protein; uniclust \| UniRef100\_A0A4Y5WA60 \| 100.0 \| 2.2e-33 \| 3.9e-39 \| 279.3 \| 526 \| (239, 782) \| 785 \| (200, 807) \| 807 \| Phage tail tape measure protein \| Phage tail tape measure protein; uniclust \| UniRef100\_A0A1V0P119 \| 100.0 \| 2.7e-33 \| 4.9e-39 \| 288.1 \| 440 \| (6, 453) \| 785 \| (58, 522) \| 1114 \| Phage tail component \| Phage tail component; uniclust \| UniRef100\_D3FXW3 \| 100.0 \| 3e-33 \| 5.4e-39 \| 269.0 \| 416 \| (111, 581) \| 785 \| (56, 475) \| 718 \| TP901 family phage tail tape measure protein \| TP901 family phage tail tape measure protein; uniclust \| UniRef100\_A0A072YD48 \| 100.0 \| 3.7e-33 \| 6.6e-39 \| 300.1 \| 341 \| (110, 465) \| 785 \| (283, 624) \| 1179 \| Phage tail tape measure protein domain-containing protein \| Phage tail tape measure protein domain-containing protein; uniclust \| UniRef100\_A0A142XX90 \| 100.0 \| 3.7e-33 \| 6.6e-39 \| 286.5 \| 349 \| (111, 474) \| 785 \| (60, 408) \| 1044 \| Phage-related minor tail protein \| Phage-related minor tail protein; uniclust \| UniRef100\_A0A369RL83 \| 100.0 \| 4e-33 \| 7.3e-39 \| 259.4 \| 459 \| (13, 477) \| 785 \| (78, 536) \| 574 \| Phage tail tape measure protein, TP901 family, core region \| Phage tail tape measure protein, TP901 family, core region; uniclust \| UniRef100\_A0A073JQH9 \| 100.0 \| 4.5e-33 \| 8.1e-39 \| 282.2 \| 488 \| (111, 627) \| 785 \| (68, 571) \| 980 \| Phage tail length tape measure protein \| Phage tail length tape measure protein; uniclust \| UniRef100\_A0A2T0BLK1 \| 100.0 \| 5e-33 \| 9e-39 \| 281.2 \| 328 \| (116, 455) \| 785 \| (128, 456) \| 1128 \| Phage-related minor tail protein \| Phage-related minor tail protein; uniclust \| UniRef100\_A0A1I4SVB4 \| 100.0 \| 5.3e-33 \| 9.5e-39 \| 287.8 \| 359 \| (98, 461) \| 785 \| (132, 490) \| 1153 \| Phage tail tape measure protein, TP901 family, core region \| Phage tail tape measure protein, TP901 family, core region; uniclust \| UniRef100\_A0A3B8TYN9 \| 100.0 \| 6e-33 \| 1.1e-38 \| 275.2 \| 452 \| (57, 596) \| 785 \| (103, 573) \| 737 \| Phage tail tape measure protein domain-containing protein \| Phage tail tape measure protein domain-containing protein; uniclust \| UniRef100\_A0A6N9Q7V1 \| 100.0 \| 6.7e-33 \| 1.2e-38 \| 274.2 \| 427 \| (112, 591) \| 785 \| (158, 596) \| 874 \| Phage tail tape measure protein \| Phage tail tape measure protein; uniclust \| UniRef100\_UPI00117E3743 \| 100.0 \| 9.3e-33 \| 1.7e-38 \| 252.0 \| 422 \| (59, 490) \| 785 \| (33, 467) \| 486 \| phage tail tape measure protein \| phage tail tape measure protein; uniclust \| UniRef100\_A0A072Y8W2 \| 100.0 \| 1.2e-32 \| 2.1e-38 \| 280.0 \| 310 \| (116, 437) \| 785 \| (96, 406) \| 1003 \| Phage tail tape measure protein \| Phage tail tape measure protein; uniclust \| UniRef100\_A0A4P9VH79 \| 100.0 \| 1.2e-32 \| 2.3e-38 \| 263.9 \| 352 \| (254, 624) \| 785 \| (159, 514) \| 665 \| Phage tail tape measure protein \| Phage tail tape measure protein; uniclust \| UniRef100\_A0A358QWZ2 \| 100.0 \| 1.3e-32 \| 2.3e-38 \| 281.2 \| 324 \| (116, 451) \| 785 \| (182, 506) \| 1247 \| Phage tail tape measure protein \| Phage tail tape measure protein; uniclust \| UniRef100\_A0A2T4MNE9 \| 100.0 \| 1.3e-32 \| 2.3e-38 \| 285.7 \| 331 \| (111, 457) \| 785 \| (137, 468) \| 1547 \| Probable transglycosylase IsaA \| Probable transglycosylase IsaA; uniclust \| UniRef100\_UPI000A7274A8 \| 100.0 \| 1.3e-32 \| 2.4e-38 \| 258.4 \| 480 \| (275, 782) \| 785 \| (7, 561) \| 561 \| phage tail tape measure protein \| phage tail tape measure protein; uniclust \| UniRef100\_A0A2D5RV15 \| 100.0 \| 1.4e-32 \| 2.5e-38 \| 267.0 \| 525 \| (109, 662) \| 785 \| (51, 590) \| 804 \| Phage tail tape measure protein \| Phage tail tape measure protein; uniclust \| UniRef100\_A0A1F9II93 \| 100.0 \| 2.2e-32 \| 4e-38 \| 263.2 \| 422 \| (337, 782) \| 785 \| (77, 567) \| 567 \| Phage tail tape measure protein (Fragment) \| Phage tail tape measure protein (Fragment); uniclust \| UniRef100\_A0A218KMB6 \| 100.0 \| 2.6e-32 \| 4.7e-38 \| 254.5 \| 377 \| (191, 587) \| 785 \| (3, 379) \| 520 \| Phage tail tape measure protein \| Phage tail tape measure protein; uniclust \| UniRef100\_UPI0009B38736 \| 100.0 \| 2.6e-32 \| 4.7e-38 \| 259.6 \| 439 \| (113, 596) \| 785 \| (96, 541) \| 612 \| phage tail tape measure protein \| phage tail tape measure protein; uniclust \| UniRef100\_A0A1S2U586 \| 99.9 \| 2.7e-32 \| 4.9e-38 \| 254.8 \| 482 \| (69, 593) \| 785 \| (34, 516) \| 654 \| Phage tail tape measure protein \| Phage tail tape measure protein; uniclust \| UniRef100\_A0A1M7N201 \| 99.9 \| 2.9e-32 \| 5.1e-38 \| 277.4 \| 414 \| (112, 585) \| 785 \| (52, 478) \| 870 \| Phage tail tape measure protein, TP901 family, core region \| Phage tail tape measure protein, TP901 family, core region; uniclust \| UniRef100\_A0A084ABK3 \| 99.9 \| 2.9e-32 \| 5.2e-38 \| 297.8 \| 341 \| (110, 460) \| 785 \| (224, 574) \| 1524 \| Phage tail length tape-measure protein \| Phage tail length tape-measure protein; uniclust \| UniRef100\_A0A0H3NQN0 \| 99.9 \| 3.2e-32 \| 5.9e-38 \| 247.4 \| 418 \| (213, 674) \| 785 \| (3, 424) \| 515 \| Phage protein \| Phage protein; uniclust \| UniRef100\_A0A7C4AJ87 \| 99.9 \| 4.2e-32 \| 7.7e-38 \| 258.0 \| 537 \| (23, 594) \| 785 \| (20, 564) \| 747 \| Phage tail tape measure protein \| Phage tail tape measure protein; uniclust \| UniRef100\_A0A031IV85 \| 99.9 \| 6.6e-32 \| 1.2e-37 \| 283.8 \| 432 \| (99, 573) \| 785 \| (48, 492) \| 856 \| Prophage PSPPH06, TP901 family tail tape measure protein \| Prophage PSPPH06, TP901 family tail tape measure protein; uniclust \| UniRef100\_A0A090YKY6 \| 99.9 \| 7.1e-32 \| 1.3e-37 \| 273.9 \| 437 \| (112, 596) \| 785 \| (103, 549) \| 1340 \| Phage tail tape measure protein, TP901 family, core region \| Phage tail tape measure protein, TP901 family, core region; uniclust \| UniRef100\_A0A098AYW7 \| 99.9 \| 9.7e-32 \| 1.7e-37 \| 272.0 \| 426 \| (1, 441) \| 785 \| (2, 430) \| 944 \| Phage-related minor tail protein \| Phage-related minor tail protein; uniclust \| UniRef100\_A0A1C2DED9 \| 99.9 \| 1.2e-31 \| 2.1e-37 \| 279.9 \| 452 \| (114, 621) \| 785 \| (52, 544) \| 778 \| Phage tail tape measure protein domain-containing protein \| Phage tail tape measure protein domain-containing protein; uniclust \| UniRef100\_A0A0H2YTM6 \| 99.9 \| 1.2e-31 \| 2.1e-37 \| 280.5 \| 456 \| (112, 582) \| 785 \| (208, 677) \| 1086 \| Phage tail tape measure protein, TP901 family \| Phage tail tape measure protein, TP901 family; uniclust \| UniRef100\_A0A0E2H1X6 \| 99.9 \| 1.4e-31 \| 2.5e-37 \| 272.4 \| 464 \| (113, 596) \| 785 \| (65, 570) \| 815 \| Phage tail tape measure protein, TP901 family, core region (Fragment) \| Phage tail tape measure protein, TP901 family, core region (Fragment); uniclust \| UniRef100\_UPI000EFC6317 \| 99.9 \| 1.4e-31 \| 2.5e-37 \| 251.9 \| 466 \| (119, 631) \| 785 \| (24, 493) \| 747 \| phage tail tape measure protein \| phage tail tape measure protein; uniclust \| UniRef100\_A0A1H3I7B8 \| 99.9 \| 1.5e-31 \| 2.6e-37 \| 268.2 \| 463 \| (111, 593) \| 785 \| (52, 555) \| 823 \| Phage-related minor tail protein \| Phage-related minor tail protein; uniclust \| UniRef100\_A0A7U5BUK8 \| 99.9 \| 1.5e-31 \| 2.6e-37 \| 268.7 \| 434 \| (124, 593) \| 785 \| (72, 512) \| 937 \| Phage tail tape measure protein domain-containing protein \| Phage tail tape measure protein domain-containing protein; uniclust \| UniRef100\_A0A0R2JK06 \| 99.9 \| 1.7e-31 \| 3.1e-37 \| 283.5 \| 341 \| (111, 465) \| 785 \| (221, 561) \| 1705 \| TP901 family phage tail tape measure protein \| TP901 family phage tail tape measure protein; uniclust \| UniRef100\_A0A497UTH2 \| 99.9 \| 1.8e-31 \| 3.2e-37 \| 253.6 \| 490 \| (109, 624) \| 785 \| (62, 574) \| 727 \| TP901 family phage tail tape measure protein \| TP901 family phage tail tape measure protein; uniclust \| UniRef100\_A0A1Q4L0K8 \| 99.9 \| 1.8e-31 \| 3.2e-37 \| 261.5 \| 327 \| (111, 449) \| 785 \| (68, 395) \| 847 \| Phage tail tape measure protein \| Phage tail tape measure protein; uniclust \| UniRef100\_A0A2E7EBE4 \| 99.9 \| 1.9e-31 \| 3.4e-37 \| 278.3 \| 353 \| (100, 457) \| 785 \| (259, 611) \| 1382 \| Phage tail tape measure protein \| Phage tail tape measure protein; uniclust \| UniRef100\_A0A150K6J4 \| 99.9 \| 1.9e-31 \| 3.4e-37 \| 273.2 \| 422 \| (117, 593) \| 785 \| (175, 605) \| 1428 \| Phage tail tape measure protein domain-containing protein \| Phage tail tape measure protein domain-containing protein; uniclust \| UniRef100\_A0A0A1GX62 \| 99.9 \| 2e-31 \| 3.5e-37 \| 293.5 \| 376 \| (110, 492) \| 785 \| (162, 547) \| 1536 \| Phage tail tape measure protein \| Phage tail tape measure protein; uniclust \| UniRef100\_A0A0U3NKE8 \| 99.9 \| 2.5e-31 \| 4.4e-37 \| 288.9 \| 337 \| (111, 461) \| 785 \| (233, 569) \| 1515 \| Phage tail length tape-measure protein \| Phage tail length tape-measure protein; uniclust \| UniRef100\_A0A1K1LB63 \| 99.9 \| 2.5e-31 \| 4.6e-37 \| 265.8 \| 327 \| (116, 446) \| 785 \| (61, 387) \| 1011 \| Phage tape measure \| Phage tape measure; uniclust \| UniRef100\_A0A256CBP9 \| 99.9 \| 2.9e-31 \| 5.2e-37 \| 248.2 \| 504 \| (22, 592) \| 785 \| (19, 524) \| 726 \| Phage tail tape measure protein \| Phage tail tape measure protein; uniclust \| UniRef100\_A0A0J1IJR7 \| 99.9 \| 3.5e-31 \| 6.1e-37 \| 285.9 \| 401 \| (53, 473) \| 785 \| (245, 663) \| 1200 \| Phage tail tape measure protein domain-containing protein \| Phage tail tape measure protein domain-containing protein; uniclust \| UniRef100\_UPI00210E7B92 \| 99.9 \| 3.3e-31 \| 6.1e-37 \| 245.7 \| 426 \| (107, 593) \| 785 \| (139, 565) \| 740 \| phage tail tape measure protein \| phage tail tape measure protein; uniclust \| UniRef100\_A0A0R3MWR6 \| 99.9 \| 4.6e-31 \| 8.3e-37 \| 265.3 \| 601 \| (2, 664) \| 785 \| (25, 733) \| 876 \| Phage tail protein \| Phage tail protein; uniclust \| UniRef100\_A0A8H9QJE6 \| 99.9 \| 4.7e-31 \| 8.3e-37 \| 277.4 \| 377 \| (85, 473) \| 785 \| (199, 578) \| 1231 \| Phage tail tape measure protein \| Phage tail tape measure protein; uniclust \| UniRef100\_A0A0F9I3U5 \| 99.9 \| 4.8e-31 \| 8.7e-37 \| 259.5 \| 472 \| (112, 593) \| 785 \| (67, 567) \| 655 \| Phage tail tape measure protein domain-containing protein \| Phage tail tape measure protein domain-containing protein; uniclust \| UniRef100\_A0A1G8PK41 \| 99.9 \| 5.4e-31 \| 9.5e-37 \| 272.4 \| 421 \| (117, 580) \| 785 \| (110, 543) \| 727 \| Phage-related minor tail protein \| Phage-related minor tail protein; uniclust \| UniRef100\_A0A0E2E9Q1 \| 99.9 \| 5.4e-31 \| 9.7e-37 \| 257.5 \| 451 \| (112, 594) \| 785 \| (67, 537) \| 653 \| Phage tail tape measure protein, TP901 family, core region \| Phage tail tape measure protein, TP901 family, core region; uniclust \| UniRef100\_A0A1W1VPP1 \| 99.9 \| 5.5e-31 \| 9.9e-37 \| 257.9 \| 447 \| (111, 596) \| 785 \| (63, 524) \| 946 \| Phage tail tape measure protein, TP901 family, core region \| Phage tail tape measure protein, TP901 family, core region; uniclust \| UniRef100\_A0A0C7QMD9 \| 99.9 \| 5.6e-31 \| 1e-36 \| 272.6 \| 349 \| (111, 474) \| 785 \| (234, 583) \| 1211 \| Phage tail tape measure protein \| Phage tail tape measure protein; uniclust \| UniRef100\_A0A078MEL8 \| 99.9 \| 7.1e-31 \| 1.3e-36 \| 269.8 \| 364 \| (63, 448) \| 785 \| (24, 391) \| 1010 \| Phage-related minor tail protein \| Phage-related minor tail protein; uniclust \| UniRef100\_A0A917G2Q0 \| 99.9 \| 6.9e-31 \| 1.3e-36 \| 251.0 \| 490 \| (1, 494) \| 785 \| (1, 494) \| 1013 \| Phage tail tape measure protein domain-containing protein \| Phage tail tape measure protein domain-containing protein; uniclust \| UniRef100\_A0A239C875 \| 99.9 \| 7.8e-31 \| 1.4e-36 \| 251.1 \| 468 \| (101, 582) \| 785 \| (137, 619) \| 800 \| Phage tail tape measure protein, TP901 family, core region \| Phage tail tape measure protein, TP901 family, core region; uniclust \| UniRef100\_A0A3C0QQX1 \| 99.9 \| 8.1e-31 \| 1.5e-36 \| 262.6 \| 452 \| (114, 595) \| 785 \| (64, 525) \| 866 \| Phage tail tape measure protein (Fragment) \| Phage tail tape measure protein (Fragment); uniclust \| UniRef100\_A0A1B9LEZ4 \| 99.9 \| 8.6e-31 \| 1.6e-36 \| 249.1 \| 509 \| (114, 632) \| 785 \| (203, 731) \| 955 \| Phage tail tape measure protein \| Phage tail tape measure protein; uniclust \| UniRef100\_A0A0S4UKB0 \| 99.9 \| 9.2e-31 \| 1.7e-36 \| 255.2 \| 729 \| (4, 784) \| 785 \| (79, 889) \| 901 \| Phage-related tail transmembrane protein \| Phage-related tail transmembrane protein; uniclust \| UniRef100\_A0A7J6YK69 \| 99.9 \| 9.4e-31 \| 1.7e-36 \| 266.0 \| 440 \| (110, 580) \| 785 \| (1938, 2378) \| 2523 \| Uncharacterized protein \| Uncharacterized protein; uniclust \| UniRef100\_A0A1Q6K3L0 \| 99.9 \| 9.7e-31 \| 1.8e-36 \| 256.5 \| 308 \| (111, 433) \| 785 \| (159, 470) \| 1048 \| Phage tail tape measure protein \| Phage tail tape measure protein; uniclust \| UniRef100\_A0A099WNM6 \| 99.9 \| 1.1e-30 \| 1.9e-36 \| 272.8 \| 337 \| (111, 460) \| 785 \| (72, 413) \| 1584 \| Phage tail tape measure protein domain-containing protein \| Phage tail tape measure protein domain-containing protein; uniclust \| UniRef100\_A0A4S3L0X3 \| 99.9 \| 1.1e-30 \| 2e-36 \| 246.1 \| 469 \| (1, 473) \| 785 \| (1, 471) \| 778 \| Phage tail tape measure protein \| Phage tail tape measure protein; uniclust \| UniRef100\_A0A0C2R458 \| 99.9 \| 1.2e-30 \| 2.1e-36 \| 296.2 \| 324 \| (111, 451) \| 785 \| (196, 523) \| 1391 \| Peptidase M23 domain-containing protein \| Peptidase M23 domain-containing protein; uniclust \| UniRef100\_A0A074LW09 \| 99.9 \| 1.4e-30 \| 2.5e-36 \| 273.3 \| 315 \| (117, 448) \| 785 \| (241, 563) \| 962 \| Phage tail tape measure protein domain-containing protein \| Phage tail tape measure protein domain-containing protein; uniclust \| UniRef100\_A0A1B8Q889 \| 99.9 \| 1.4e-30 \| 2.5e-36 \| 280.5 \| 348 \| (107, 459) \| 785 \| (245, 592) \| 1395 \| Phage tail tape measure protein \| Phage tail tape measure protein; uniclust \| UniRef100\_A0A133YZY4 \| 99.9 \| 1.5e-30 \| 2.6e-36 \| 287.0 \| 359 \| (111, 477) \| 785 \| (357, 723) \| 1470 \| Phage tail tape measure protein domain-containing protein \| Phage tail tape measure protein domain-containing protein; uniclust \| UniRef100\_A0A0G3WGN7 \| 99.9 \| 1.7e-30 \| 2.9e-36 \| 285.2 \| 321 \| (113, 444) \| 785 \| (133, 453) \| 1035 \| Tape measure protein \| Tape measure protein; uniclust \| UniRef100\_A0A090FCA3 \| 99.9 \| 1.7e-30 \| 3e-36 \| 270.4 \| 399 \| (110, 580) \| 785 \| (80, 480) \| 887 \| Phage tail tape measure protein domain-containing protein \| Phage tail tape measure protein domain-containing protein; uniclust \| UniRef100\_A0A919ZAN7 \| 99.9 \| 1.8e-30 \| 3.2e-36 \| 258.6 \| 349 \| (111, 465) \| 785 \| (151, 558) \| 1016 \| Phage tail tape measure protein domain-containing protein \| Phage tail tape measure protein domain-containing protein; uniclust \| UniRef100\_A0A0F5ARQ0 \| 99.9 \| 1.9e-30 \| 3.4e-36 \| 260.7 \| 430 \| (104, 575) \| 785 \| (50, 492) \| 806 \| Phage tail tape measure protein domain-containing protein \| Phage tail tape measure protein domain-containing protein; uniclust \| UniRef100\_A0A1H1G2L2 \| 99.9 \| 1.9e-30 \| 3.5e-36 \| 255.2 \| 458 \| (5, 472) \| 785 \| (94, 579) \| 991 \| Phage tail tape measure protein, TP901 family, core region \| Phage tail tape measure protein, TP901 family, core region; uniclust \| UniRef100\_UPI001C007827 \| 99.9 \| 2e-30 \| 3.7e-36 \| 244.5 \| 448 \| (114, 594) \| 785 \| (232, 683) \| 870 \| phage tail tape measure protein \| phage tail tape measure protein; uniclust \| UniRef100\_A0A1C6E246 \| 99.9 \| 2.2e-30 \| 4e-36 \| 250.3 \| 331 \| (111, 454) \| 785 \| (153, 483) \| 566 \| Phage-related minor tail protein \| Phage-related minor tail protein; uniclust \| UniRef100\_A0A1H0GBX4 \| 99.9 \| 2.3e-30 \| 4.2e-36 \| 265.2 \| 461 \| (5, 469) \| 785 \| (67, 566) \| 1109 \| Phage tail tape measure protein, TP901 family, core region \| Phage tail tape measure protein, TP901 family, core region; uniclust \| UniRef100\_UPI001F56DA87 \| 99.9 \| 2.5e-30 \| 4.5e-36 \| 250.4 \| 395 \| (60, 464) \| 785 \| (76, 476) \| 902 \| phage tail tape measure protein \| phage tail tape measure protein; uniclust \| UniRef100\_A0A1I5RNE0 \| 99.9 \| 3e-30 \| 5.5e-36 \| 254.9 \| 427 \| (109, 584) \| 785 \| (203, 656) \| 792 \| Phage tail tape measure protein, TP901 family, core region (Fragment) \| Phage tail tape measure protein, TP901 family, core region (Fragment); uniclust \| UniRef100\_A0A430RXV9 \| 99.9 \| 3.2e-30 \| 5.9e-36 \| 234.4 \| 354 \| (111, 469) \| 785 \| (56, 409) \| 486 \| Phage tail tape measure protein (Fragment) \| Phage tail tape measure protein (Fragment); uniclust \| UniRef100\_UPI0010102470 \| 99.9 \| 3.3e-30 \| 6e-36 \| 231.1 \| 485 \| (234, 781) \| 785 \| (4, 496) \| 497 \| phage tail tape measure protein \| phage tail tape measure protein; uniclust \| UniRef100\_A0A098F348 \| 99.9 \| 4.5e-30 \| 8e-36 \| 275.5 \| 411 \| (28, 443) \| 785 \| (57, 496) \| 1182 \| Phage tail tape measure protein, family, core region \| Phage tail tape measure protein, family, core region; uniclust \| UniRef100\_A0A2X1J0M0 \| 99.9 \| 4.5e-30 \| 8.2e-36 \| 240.7 \| 559 \| (6, 574) \| 785 \| (80, 669) \| 684 \| TP901 family phage tail tape measure protein \| TP901 family phage tail tape measure protein; uniclust \| UniRef100\_A0A1V4SUX2 \| 99.9 \| 4.8e-30 \| 8.6e-36 \| 270.8 \| 345 \| (110, 466) \| 785 \| (344, 699) \| 1239 \| Chromosome partition protein Smc \| Chromosome partition protein Smc; uniclust \| UniRef100\_A0A1C5VU61 \| 99.9 \| 4.9e-30 \| 8.8e-36 \| 258.1 \| 388 \| (51, 448) \| 785 \| (79, 486) \| 883 \| Phage-related protein \| Phage-related protein; uniclust \| UniRef100\_A0A0C3I468 \| 99.9 \| 5.5e-30 \| 1e-35 \| 271.7 \| 343 \| (111, 468) \| 785 \| (235, 578) \| 1721 \| Tail tape measure protein \| Tail tape measure protein; uniclust \| UniRef100\_UPI0021053B5F \| 99.9 \| 6.2e-30 \| 1.1e-35 \| 243.7 \| 331 \| (113, 443) \| 785 \| (31, 363) \| 967 \| phage tail tape measure protein \| phage tail tape measure protein; uniclust \| UniRef100\_A0A174TRV8 \| 99.9 \| 6.8e-30 \| 1.2e-35 \| 269.1 \| 349 \| (117, 470) \| 785 \| (202, 555) \| 1242 \| Phage-related minor tail protein \| Phage-related minor tail protein; uniclust \| UniRef100\_A0A068QV54 \| 99.9 \| 7.6e-30 \| 1.3e-35 \| 263.0 \| 555 \| (103, 702) \| 785 \| (41, 612) \| 715 \| Putative phage protein \| Putative phage protein; uniclust \| UniRef100\_A0A7W3T073 \| 99.9 \| 7.5e-30 \| 1.3e-35 \| 261.8 \| 338 \| (111, 469) \| 785 \| (68, 407) \| 1083 \| Phage tail tape measure protein \| Phage tail tape measure protein; uniclust \| UniRef100\_A0A0R2FUF3 \| 99.9 \| 7.7e-30 \| 1.4e-35 \| 270.2 \| 333 \| (111, 450) \| 785 \| (172, 514) \| 1325 \| TP901 family phage tail tape measure protein \| TP901 family phage tail tape measure protein; uniclust \| UniRef100\_A0A1I6URJ1 \| 99.9 \| 8.1e-30 \| 1.5e-35 \| 270.7 \| 333 \| (110, 456) \| 785 \| (325, 665) \| 1585 \| Phage tail tape measure protein, TP901 family, core region \| Phage tail tape measure protein, TP901 family, core region; uniclust \| UniRef100\_A0A1A7PS85 \| 99.9 \| 8.1e-30 \| 1.5e-35 \| 244.8 \| 585 \| (3, 595) \| 785 \| (2, 621) \| 696 \| Phage tail tape measure protein domain-containing protein (Fragment) \| Phage tail tape measure protein domain-containing protein (Fragment); uniclust \| UniRef100\_A0A1B9AU19 \| 99.9 \| 9.1e-30 \| 1.7e-35 \| 246.6 \| 423 \| (119, 581) \| 785 \| (222, 672) \| 899 \| Phage tail tape measure protein \| Phage tail tape measure protein; uniclust \| UniRef100\_A0A081R6K7 \| 99.9 \| 1.1e-29 \| 1.9e-35 \| 282.1 \| 355 \| (111, 469) \| 785 \| (201, 564) \| 1424 \| Phage tail tape measure protein, TP901 family, core region \| Phage tail tape measure protein, TP901 family, core region; uniclust \| UniRef100\_UPI001F397BD3 \| 99.9 \| 1.2e-29 \| 2.1e-35 \| 250.4 \| 318 \| (116, 445) \| 785 \| (163, 494) \| 914 \| phage tail tape measure protein \| phage tail tape measure protein; uniclust \| UniRef100\_A0A090I2H9 \| 99.9 \| 1.5e-29 \| 2.6e-35 \| 248.8 \| 417 \| (111, 587) \| 785 \| (84, 503) \| 665 \| Phage tail tape measure protein, TP901 family \| Phage tail tape measure protein, TP901 family; uniclust \| UniRef100\_A0A011T795 \| 99.9 \| 1.5e-29 \| 2.6e-35 \| 257.7 \| 330 \| (112, 447) \| 785 \| (81, 410) \| 753 \| Tail protein \| Tail protein; uniclust \| UniRef100\_A0A0B4BHD1 \| 99.9 \| 1.8e-29 \| 3.2e-35 \| 250.9 \| 469 \| (1, 594) \| 785 \| (3, 480) \| 851 \| Phage tail length tape measure protein \| Phage tail length tape measure protein; uniclust \| UniRef100\_A0A0E3GSM8 \| 99.9 \| 1.9e-29 \| 3.4e-35 \| 259.9 \| 321 \| (243, 609) \| 785 \| (213, 549) \| 994 \| Phage-like protein \| Phage-like protein; uniclust \| UniRef100\_A0A412CHP1 \| 99.9 \| 1.9e-29 \| 3.5e-35 \| 242.6 \| 460 \| (118, 594) \| 785 \| (67, 551) \| 835 \| Phage tail tape measure protein \| Phage tail tape measure protein; uniclust \| UniRef100\_UPI000933C153 \| 99.9 \| 2.2e-29 \| 4.1e-35 \| 244.5 \| 509 \| (103, 628) \| 785 \| (168, 710) \| 1200 \| phage tail tape measure protein \| phage tail tape measure protein; uniclust \| UniRef100\_A0A060PPH4 \| 99.9 \| 2.5e-29 \| 4.4e-35 \| 255.9 \| 333 \| (110, 457) \| 785 \| (57, 390) \| 937 \| Phage tail tape measure protein domain-containing protein \| Phage tail tape measure protein domain-containing protein; uniclust \| UniRef100\_A0A023CIR9 \| 99.9 \| 2.6e-29 \| 4.6e-35 \| 265.8 \| 423 \| (14, 451) \| 785 \| (154, 607) \| 1850 \| Bacteriophage-related protein \| Bacteriophage-related protein; uniclust \| UniRef100\_A0A2Z6DXQ1 \| 99.9 \| 2.8e-29 \| 5.1e-35 \| 249.9 \| 577 \| (6, 594) \| 785 \| (5, 652) \| 1035 \| Phage tail tape measure protein \| Phage tail tape measure protein; uniclust \| UniRef100\_A0A178HLZ5 \| 99.9 \| 2.9e-29 \| 5.3e-35 \| 239.5 \| 466 \| (115, 595) \| 785 \| (64, 555) \| 977 \| Phage tail tape measure protein domain-containing protein \| Phage tail tape measure protein domain-containing protein; uniclust \| UniRef100\_A0A143XDR8 \| 99.9 \| 3e-29 \| 5.4e-35 \| 260.9 \| 321 \| (241, 626) \| 785 \| (350, 720) \| 1063 \| Phage-related minor tail protein \| Phage-related minor tail protein; uniclust \| UniRef100\_A0A7D4BEW5 \| 99.9 \| 3.6e-29 \| 6.5e-35 \| 252.8 \| 399 \| (115, 584) \| 785 \| (54, 453) \| 805 \| Phage tail tape measure protein \| Phage tail tape measure protein; uniclust \| UniRef100\_A0A0B0SDM8 \| 99.9 \| 4.3e-29 \| 7.8e-35 \| 245.0 \| 468 \| (111, 585) \| 785 \| (58, 540) \| 966 \| Phage tail tape measure protein domain-containing protein \| Phage tail tape measure protein domain-containing protein; uniclust \| UniRef100\_A0A973FFH6 \| 99.9 \| 4.3e-29 \| 7.9e-35 \| 229.8 \| 342 \| (118, 460) \| 785 \| (78, 421) \| 672 \| Phage tail tape measure protein \| Phage tail tape measure protein; uniclust \| UniRef100\_A0A1R1MK84 \| 99.9 \| 4.7e-29 \| 8.6e-35 \| 232.7 \| 469 \| (114, 595) \| 785 \| (61, 553) \| 767 \| Phage tail tape measure protein \| Phage tail tape measure protein; uniclust \| UniRef100\_A0A1I1GTR9 \| 99.9 \| 4.9e-29 \| 8.7e-35 \| 262.6 \| 329 \| (111, 450) \| 785 \| (192, 523) \| 1274 \| Phage tail tape measure protein, TP901 family, core region \| Phage tail tape measure protein, TP901 family, core region; uniclust \| UniRef100\_A0A024Q9Y0 \| 99.9 \| 5.4e-29 \| 9.6e-35 \| 266.7 \| 325 \| (113, 447) \| 785 \| (67, 401) \| 1226 \| Phage tail tape measure protein, TP901 family, core region \| Phage tail tape measure protein, TP901 family, core region; uniclust \| UniRef100\_A0A4R3J9G1 \| 99.9 \| 5.5e-29 \| 9.9e-35 \| 236.8 \| 410 \| (111, 574) \| 785 \| (74, 493) \| 600 \| TP901 family phage tail tape measure protein \| TP901 family phage tail tape measure protein; uniclust \| UniRef100\_A0A135YQ01 \| 99.9 \| 6.4e-29 \| 1.2e-34 \| 240.4 \| 335 \| (111, 460) \| 785 \| (66, 401) \| 777 \| Phage tail tape measure protein, TP901 family \| Phage tail tape measure protein, TP901 family; uniclust \| UniRef100\_A0A072NTF3 \| 99.9 \| 6.8e-29 \| 1.2e-34 \| 272.0 \| 627 \| (4, 705) \| 785 \| (212, 894) \| 1237 \| Phage-related minor tail protein \| Phage-related minor tail protein; uniclust \| UniRef100\_UPI001CFF0720 \| 99.9 \| 7e-29 \| 1.3e-34 \| 243.6 \| 492 \| (68, 592) \| 785 \| (13, 509) \| 1008 \| phage tail tape measure protein \| phage tail tape measure protein; uniclust \| UniRef100\_A0A1E7Q036 \| 99.9 \| 7.2e-29 \| 1.3e-34 \| 241.5 \| 312 \| (255, 575) \| 785 \| (258, 578) \| 749 \| Phage tail tape measure protein \| Phage tail tape measure protein; uniclust \| UniRef100\_A0A916IVX2 \| 99.9 \| 9.2e-29 \| 1.7e-34 \| 236.4 \| 553 \| (3, 580) \| 785 \| (48, 653) \| 763 \| Phage tail tape measure protein domain-containing protein \| Phage tail tape measure protein domain-containing protein; uniclust \| UniRef100\_A0A0L0W671 \| 99.9 \| 1e-28 \| 1.8e-34 \| 269.3 \| 324 \| (111, 451) \| 785 \| (220, 546) \| 1170 \| Phage-like minor tail protein (Fragment) \| Phage-like minor tail protein (Fragment); uniclust \| UniRef100\_A0A3P6KZI8 \| 99.9 \| 1.1e-28 \| 1.9e-34 \| 228.8 \| 459 \| (5, 476) \| 785 \| (78, 558) \| 568 \| Tail fiber protein T (Tape measure) (GpT) \| Tail fiber protein T (Tape measure) (GpT); uniclust \| UniRef100\_UPI000CDD6178 \| 99.9 \| 1.3e-28 \| 2.4e-34 \| 223.3 \| 415 \| (118, 572) \| 785 \| (89, 507) \| 508 \| phage tail tape measure protein \| phage tail tape measure protein; uniclust \| UniRef100\_A0A0F9FT22 \| 99.9 \| 1.4e-28 \| 2.5e-34 \| 233.1 \| 397 \| (117, 585) \| 785 \| (56, 457) \| 578 \| Phage tail tape measure protein domain-containing protein (Fragment) \| Phage tail tape measure protein domain-containing protein (Fragment); uniclust \| UniRef100\_A0A495V9N5 \| 99.9 \| 1.4e-28 \| 2.6e-34 \| 237.2 \| 450 \| (118, 590) \| 785 \| (62, 512) \| 1090 \| TP901 family phage tail tape measure protein \| TP901 family phage tail tape measure protein; uniclust \| UniRef100\_UPI0008A95C93 \| 99.9 \| 1.4e-28 \| 2.6e-34 \| 247.6 \| 327 \| (111, 445) \| 785 \| (510, 844) \| 1366 \| phage tail tape measure protein \| phage tail tape measure protein; uniclust \| UniRef100\_A0A8T8DYB4 \| 99.9 \| 1.5e-28 \| 2.7e-34 \| 240.9 \| 442 \| (111, 577) \| 785 \| (59, 503) \| 877 \| Phage tail tape measure protein \| Phage tail tape measure protein; uniclust \| UniRef100\_A0A080IQ97 \| 99.9 \| 1.6e-28 \| 2.9e-34 \| 239.3 \| 440 \| (111, 621) \| 785 \| (68, 518) \| 633 \| Phage tail tape measure protein, TP901 family, core region \| Phage tail tape measure protein, TP901 family, core region; uniclust \| UniRef100\_A0A1S1V6R2 \| 99.9 \| 1.6e-28 \| 2.9e-34 \| 248.2 \| 420 \| (119, 592) \| 785 \| (111, 544) \| 833 \| Phage-related minor tail protein \| Phage-related minor tail protein; uniclust \| UniRef100\_A0A378E4L1 \| 99.9 \| 1.7e-28 \| 3.1e-34 \| 238.9 \| 538 \| (4, 571) \| 785 \| (78, 644) \| 690 \| Phage tail tape measure protein, family \| Phage tail tape measure protein, family; uniclust \| UniRef100\_A0A1Y5SUX3 \| 99.9 \| 1.9e-28 \| 3.6e-34 \| 239.1 \| 469 \| (115, 594) \| 785 \| (64, 554) \| 1263 \| Phage-related minor tail protein \| Phage-related minor tail protein; uniclust \| UniRef100\_A0A1G3UAI7 \| 99.9 \| 2e-28 \| 3.6e-34 \| 238.6 \| 592 \| (1, 625) \| 785 \| (1, 617) \| 850 \| Phage tail tape measure protein \| Phage tail tape measure protein; uniclust \| UniRef100\_A0A0D0FE37 \| 99.9 \| 2.1e-28 \| 3.6e-34 \| 262.3 \| 320 \| (116, 449) \| 785 \| (64, 383) \| 1025 \| Tape measure protein N-terminal domain-containing protein \| Tape measure protein N-terminal domain-containing protein; uniclust \| UniRef100\_A0A2N9Y986 \| 99.9 \| 2.2e-28 \| 3.9e-34 \| 226.1 \| 350 \| (105, 464) \| 785 \| (67, 416) \| 444 \| Phage tail tape measure protein (Fragment) \| Phage tail tape measure protein (Fragment); uniclust \| UniRef100\_A0A0U5MPA6 \| 99.9 \| 2.2e-28 \| 4e-34 \| 232.4 \| 409 \| (118, 596) \| 785 \| (60, 475) \| 598 \| Phage tail tape measure protein domain-containing protein \| Phage tail tape measure protein domain-containing protein; uniclust \| UniRef100\_A0A1H8A1R0 \| 99.9 \| 2.5e-28 \| 4.6e-34 \| 241.6 \| 463 \| (4, 469) \| 785 \| (75, 544) \| 1108 \| Phage tail tape measure protein, TP901 family, core region \| Phage tail tape measure protein, TP901 family, core region; uniclust \| UniRef100\_A0A158S093 \| 99.9 \| 3.1e-28 \| 5.5e-34 \| 238.2 \| 292 \| (398, 783) \| 785 \| (302, 605) \| 606 \| Phage tail tape measure protein \| Phage tail tape measure protein; uniclust \| UniRef100\_A0A024QBD0 \| 99.9 \| 3.1e-28 \| 5.6e-34 \| 260.5 \| 332 \| (111, 460) \| 785 \| (172, 508) \| 1287 \| Chromosome segregation protein SMC \| Chromosome segregation protein SMC; uniclust \| UniRef100\_UPI00191A810C \| 99.9 \| 3.1e-28 \| 5.6e-34 \| 209.9 \| 378 \| (2, 383) \| 785 \| (3, 380) \| 388 \| phage tail tape measure protein \| phage tail tape measure protein; uniclust \| UniRef100\_A0A1L8I8U0 \| 99.9 \| 3.3e-28 \| 5.9e-34 \| 242.0 \| 345 \| (111, 460) \| 785 \| (176, 525) \| 653 \| Phage tail tape measure protein (Fragment) \| Phage tail tape measure protein (Fragment); uniclust \| UniRef100\_UPI000AEE5402 \| 99.9 \| 3.3e-28 \| 6e-34 \| 241.7 \| 472 \| (109, 589) \| 785 \| (356, 1001) \| 1155 \| phage tail tape measure protein \| phage tail tape measure protein; uniclust \| UniRef100\_A0A4R2N8L9 \| 99.9 \| 3.3e-28 \| 6.1e-34 \| 220.7 \| 440 \| (23, 466) \| 785 \| (16, 461) \| 585 \| TP901 family phage tail tape measure protein \| TP901 family phage tail tape measure protein; uniclust \| UniRef100\_A0A069D3M4 \| 99.9 \| 3.7e-28 \| 6.6e-34 \| 263.9 \| 300 \| (111, 419) \| 785 \| (128, 439) \| 1113 \| Minor tail protein GP26-like protein \| Minor tail protein GP26-like protein; uniclust \| UniRef100\_A0A133MP93 \| 99.9 \| 3.9e-28 \| 7e-34 \| 246.1 \| 328 \| (111, 456) \| 785 \| (209, 538) \| 992 \| Phage tail tape measure protein, TP901 family \| Phage tail tape measure protein, TP901 family; uniclust \| UniRef100\_A0A0H4TEZ5 \| 99.9 \| 4.4e-28 \| 8e-34 \| 240.1 \| 327 \| (115, 446) \| 785 \| (159, 490) \| 975 \| Tail tape measure protein \| Tail tape measure protein; uniclust \| UniRef100\_A0A5H2QW59 \| 99.9 \| 4.4e-28 \| 8.1e-34 \| 234.0 \| 435 \| (117, 595) \| 785 \| (54, 496) \| 727 \| Tail length tape-measure protein \| Tail length tape-measure protein; uniclust \| UniRef100\_A0A0K1LJH9 \| 99.9 \| 4.6e-28 \| 8.2e-34 \| 247.9 \| 448 \| (1, 594) \| 785 \| (61, 521) \| 657 \| Putative tail protein \| Putative tail protein; uniclust \| UniRef100\_UPI00097D468B \| 99.9 \| 5.8e-28 \| 1.1e-33 \| 238.7 \| 461 \| (114, 583) \| 785 \| (484, 948) \| 1083 \| phage tail tape measure protein \| phage tail tape measure protein; uniclust \| UniRef100\_A0A1W7LHP8 \| 99.9 \| 5.9e-28 \| 1.1e-33 \| 259.2 \| 329 \| (111, 450) \| 785 \| (203, 531) \| 1810 \| Phage tail tape measure protein domain-containing protein \| Phage tail tape measure protein domain-containing protein; uniclust \| UniRef100\_A0A024YKU6 \| 99.9 \| 6.6e-28 \| 1.2e-33 \| 250.3 \| 414 \| (111, 593) \| 785 \| (79, 507) \| 782 \| Phage tail tape measure protein domain-containing protein (Fragment) \| Phage tail tape measure protein domain-containing protein (Fragment); uniclust \| UniRef100\_A0A386PK18 \| 99.9 \| 7.4e-28 \| 1.3e-33 \| 243.0 \| 323 \| (112, 446) \| 785 \| (48, 371) \| 871 \| Phage tail tape measure protein \| Phage tail tape measure protein; uniclust \| UniRef100\_UPI00042A37CB \| 99.9 \| 7.9e-28 \| 1.4e-33 \| 238.9 \| 568 \| (4, 595) \| 785 \| (44, 668) \| 994 \| phage tail tape measure protein \| phage tail tape measure protein; uniclust \| UniRef100\_UPI001FCE6FAE \| 99.9 \| 8.2e-28 \| 1.5e-33 \| 207.3 \| 352 \| (407, 783) \| 785 \| (17, 369) \| 371 \| hypothetical protein \| hypothetical protein; uniclust \| UniRef100\_A0A380FL26 \| 99.9 \| 1e-27 \| 1.9e-33 \| 246.6 \| 442 \| (113, 587) \| 785 \| (172, 620) \| 1292 \| Phage tail tape measure protein, TP901 family, core region \| Phage tail tape measure protein, TP901 family, core region; uniclust \| UniRef100\_A0A377U3B0 \| 99.9 \| 1.1e-27 \| 1.9e-33 \| 225.7 \| 290 \| (276, 591) \| 785 \| (43, 338) \| 555 \| Phage tail tape measure protein, family \| Phage tail tape measure protein, family; uniclust \| UniRef100\_A0A379TRM7 \| 99.9 \| 1.2e-27 \| 2.2e-33 \| 231.0 \| 672 \| (4, 703) \| 785 \| (3, 766) \| 874 \| Putative bacteriophage protein \| Putative bacteriophage protein; uniclust \| UniRef100\_A0A0D0Q2R1 \| 99.9 \| 1.2e-27 \| 2.2e-33 \| 251.0 \| 360 \| (111, 483) \| 785 \| (266, 690) \| 1291 \| Tail tape measure protein \| Tail tape measure protein; uniclust \| UniRef100\_A0A285NFD1 \| 99.9 \| 1.2e-27 \| 2.2e-33 \| 226.2 \| 349 \| (114, 471) \| 785 \| (63, 412) \| 806 \| Phage tail tape measure protein, TP901 family, core region \| Phage tail tape measure protein, TP901 family, core region; uniclust \| UniRef100\_UPI001EFBD742 \| 99.9 \| 1.3e-27 \| 2.4e-33 \| 222.3 \| 548 \| (116, 703) \| 785 \| (53, 617) \| 694 \| phage tail tape measure protein \| phage tail tape measure protein; uniclust \| UniRef100\_UPI001918F45D \| 99.9 \| 1.4e-27 \| 2.5e-33 \| 209.9 \| 239 \| (352, 594) \| 785 \| (4, 243) \| 375 \| phage tail tape measure protein \| phage tail tape measure protein; uniclust \| UniRef100\_A0A0J9EDB0 \| 99.9 \| 1.4e-27 \| 2.5e-33 \| 236.7 \| 413 \| (188, 619) \| 785 \| (130, 564) \| 1019 \| Phage tail length tape-measure protein \| Phage tail length tape-measure protein; uniclust \| UniRef100\_A0A0D0P0W2 \| 99.9 \| 1.6e-27 \| 2.9e-33 \| 233.3 \| 427 \| (102, 573) \| 785 \| (42, 480) \| 633 \| Contig\_52, whole genome shotgun sequence \| Contig\_52, whole genome shotgun sequence; uniclust \| UniRef100\_A0A2X1MWP2 \| 99.9 \| 1.6e-27 \| 2.9e-33 \| 216.1 \| 448 \| (119, 616) \| 785 \| (39, 493) \| 493 \| Phage related tail protein \| Phage related tail protein; uniclust \| UniRef100\_A0A136H2X7 \| 99.9 \| 1.8e-27 \| 3.1e-33 \| 241.1 \| 203 \| (4, 214) \| 785 \| (77, 292) \| 717 \| Tape measure protein N-terminal domain-containing protein \| Tape measure protein N-terminal domain-containing protein; uniclust \| UniRef100\_A0A0A6VGD4 \| 99.9 \| 1.8e-27 \| 3.2e-33 \| 239.5 \| 393 \| (115, 589) \| 785 \| (261, 664) \| 842 \| Phage tail tape measure protein domain-containing protein \| Phage tail tape measure protein domain-containing protein; uniclust \| UniRef100\_UPI0021BF577E \| 99.9 \| 2.1e-27 \| 3.8e-33 \| 230.2 \| 425 \| (112, 595) \| 785 \| (236, 674) \| 999 \| phage tail tape measure protein \| phage tail tape measure protein; uniclust \| UniRef100\_A0A401UU86 \| 99.9 \| 2.2e-27 \| 4.1e-33 \| 244.6 \| 428 \| (111, 585) \| 785 \| (155, 592) \| 1714 \| Phage tail tape measure protein domain-containing protein \| Phage tail tape measure protein domain-containing protein; uniclust \| UniRef100\_A0A379S656 \| 99.9 \| 2.2e-27 \| 4.1e-33 \| 219.6 \| 331 \| (196, 583) \| 785 \| (4, 334) \| 517 \| Transglycosylase \| Transglycosylase; uniclust \| UniRef100\_A0A1Q6JQ74 \| 99.9 \| 2.5e-27 \| 4.6e-33 \| 217.5 \| 453 \| (5, 472) \| 785 \| (75, 533) \| 643 \| Phage tail tape measure protein \| Phage tail tape measure protein; uniclust \| UniRef100\_A0A0R8VA05 \| 99.9 \| 3e-27 \| 5.4e-33 \| 247.3 \| 405 \| (118, 591) \| 785 \| (220, 629) \| 1657 \| Tail tape measure protein \| Tail tape measure protein; uniclust \| UniRef100\_A0A084E6E2 \| 99.9 \| 3.1e-27 \| 5.6e-33 \| 251.3 \| 322 \| (111, 444) \| 785 \| (78, 401) \| 1183 \| Phage tail tape measure protein, TP901 family, core region \| Phage tail tape measure protein, TP901 family, core region; uniclust \| UniRef100\_A0A0J0VAR5 \| 99.9 \| 3.2e-27 \| 5.7e-33 \| 241.2 \| 319 \| (205, 592) \| 785 \| (331, 658) \| 941 \| Replication protein \| Replication protein; uniclust \| UniRef100\_UPI0015D2B999 \| 99.9 \| 3.1e-27 \| 5.8e-33 \| 203.6 \| 378 \| (303, 702) \| 785 \| (7, 384) \| 385 \| phage tail tape measure protein \| phage tail tape measure protein; uniclust \| UniRef100\_A0A1B2DVD5 \| 99.9 \| 3.3e-27 \| 5.9e-33 \| 252.1 \| 330 \| (113, 462) \| 785 \| (170, 561) \| 1489 \| Phage tail tape measure protein domain-containing protein \| Phage tail tape measure protein domain-containing protein; uniclust \| UniRef100\_I0GWR9 \| 99.9 \| 3.4e-27 \| 6.2e-33 \| 225.3 \| 464 \| (120, 618) \| 785 \| (68, 546) \| 951 \| Putative tail tape measure protein \| Putative tail tape measure protein; uniclust \| UniRef100\_A0A484YXF6 \| 99.9 \| 3.6e-27 \| 6.6e-33 \| 208.3 \| 313 \| (157, 469) \| 785 \| (5, 317) \| 362 \| Phage tail tape measure protein \| Phage tail tape measure protein; uniclust \| UniRef100\_A0A235CJC1 \| 99.9 \| 3.6e-27 \| 6.6e-33 \| 232.3 \| 486 \| (147, 674) \| 785 \| (81, 584) \| 952 \| Phage tail protein \| Phage tail protein; uniclust \| UniRef100\_A0A072YJ62 \| 99.9 \| 4e-27 \| 7.2e-33 \| 237.0 \| 456 \| (111, 594) \| 785 \| (67, 643) \| 859 \| Phage tail tape measure protein \| Phage tail tape measure protein; uniclust \| UniRef100\_A0A2J0QV17 \| 99.9 \| 4.7e-27 \| 8.5e-33 \| 227.3 \| 472 \| (163, 674) \| 785 \| (12, 501) \| 602 \| Phage tail tape measure protein (Fragment) \| Phage tail tape measure protein (Fragment); uniclust \| UniRef100\_A0A1I6REX1 \| 99.9 \| 4.8e-27 \| 8.6e-33 \| 233.4 \| 328 \| (111, 456) \| 785 \| (56, 385) \| 761 \| Phage tail tape measure protein, TP901 family, core region \| Phage tail tape measure protein, TP901 family, core region; uniclust \| UniRef100\_A0A1U7M6P2 \| 99.9 \| 5e-27 \| 9e-33 \| 250.1 \| 361 \| (111, 477) \| 785 \| (194, 609) \| 1210 \| Chromosome partition protein Smc \| Chromosome partition protein Smc; uniclust \| UniRef100\_A0A5C4YDP2 \| 99.9 \| 5e-27 \| 9.2e-33 \| 215.8 \| 475 \| (116, 595) \| 785 \| (86, 575) \| 602 \| Phage tail tape measure protein (Fragment) \| Phage tail tape measure protein (Fragment); uniclust \| UniRef100\_UPI0018EADAE6 \| 99.9 \| 5.3e-27 \| 9.7e-33 \| 223.4 \| 480 \| (109, 622) \| 785 \| (50, 545) \| 705 \| phage tail tape measure protein \| phage tail tape measure protein; uniclust \| UniRef100\_A0A1Z4PN55 \| 99.9 \| 6e-27 \| 1.1e-32 \| 225.5 \| 444 \| (110, 596) \| 785 \| (53, 505) \| 775 \| Putative bacteriophage protein \| Putative bacteriophage protein; uniclust \| UniRef100\_A0A1C3H4A0 \| 99.9 \| 6.9e-27 \| 1.3e-32 \| 240.5 \| 349 \| (108, 461) \| 785 \| (342, 690) \| 1661 \| Phage tail length tape-measure protein \| Phage tail length tape-measure protein; uniclust \| UniRef100\_UPI0005878020 \| 99.9 \| 7.2e-27 \| 1.3e-32 \| 218.0 \| 529 \| (30, 600) \| 785 \| (159, 736) \| 745 \| phage tail tape measure protein \| phage tail tape measure protein; uniclust \| UniRef100\_A0A0Q1AXG2 \| 99.9 \| 7.8e-27 \| 1.4e-32 \| 248.8 \| 330 \| (112, 455) \| 785 \| (171, 509) \| 1635 \| Putative membrane protein \| Putative membrane protein; uniclust \| UniRef100\_A0A089HYG9 \| 99.9 \| 8.1e-27 \| 1.4e-32 \| 244.1 \| 308 \| (116, 447) \| 785 \| (128, 441) \| 838 \| Phage tail tape measure protein domain-containing protein \| Phage tail tape measure protein domain-containing protein; uniclust \| UniRef100\_C9MV98 \| 99.9 \| 1e-26 \| 1.9e-32 \| 212.0 \| 372 \| (110, 485) \| 785 \| (194, 565) \| 600 \| Phage tail tape measure protein, TP901 family \| Phage tail tape measure protein, TP901 family; uniclust \| UniRef100\_H0U9Y6 \| 99.9 \| 1e-26 \| 1.9e-32 \| 219.1 \| 413 \| (118, 591) \| 785 \| (82, 503) \| 654 \| Phage tail tape measure protein, TP901 family, core region \| Phage tail tape measure protein, TP901 family, core region; uniclust \| UniRef100\_A0A1M6KNC3 \| 99.9 \| 1.1e-26 \| 1.9e-32 \| 235.6 \| 367 \| (111, 484) \| 785 \| (76, 492) \| 865 \| Phage tail tape measure protein, TP901 family, core region (Fragment) \| Phage tail tape measure protein, TP901 family, core region (Fragment); uniclust \| UniRef100\_A0A0J1FKM1 \| 99.9 \| 1.2e-26 \| 2.1e-32 \| 235.5 \| 336 \| (111, 460) \| 785 \| (63, 409) \| 839 \| Phage-related minor tail protein \| Phage-related minor tail protein; uniclust \| UniRef100\_UPI001495E0A7 \| 99.9 \| 1.2e-26 \| 2.1e-32 \| 220.6 \| 325 \| (113, 449) \| 785 \| (53, 378) \| 738 \| phage tail tape measure protein \| phage tail tape measure protein; uniclust \| UniRef100\_A0A1X3ITD6 \| 99.9 \| 1.3e-26 \| 2.4e-32 \| 207.2 \| 340 \| (118, 467) \| 785 \| (43, 384) \| 403 \| Phage tail tape measure protein, TP901 family, core region (Fragment) \| Phage tail tape measure protein, TP901 family, core region (Fragment); uniclust \| UniRef100\_A0A2M7BJQ9 \| 99.9 \| 1.4e-26 \| 2.5e-32 \| 215.9 \| 352 \| (100, 456) \| 785 \| (231, 582) \| 615 \| Phage tail tape measure protein (Fragment) \| Phage tail tape measure protein (Fragment); uniclust \| UniRef100\_UPI001EF72727 \| 99.9 \| 1.5e-26 \| 2.6e-32 \| 234.4 \| 406 \| (146, 587) \| 785 \| (11, 420) \| 975 \| phage tail tape measure protein \| phage tail tape measure protein; uniclust \| UniRef100\_A0A380DUS3 \| 99.9 \| 1.4e-26 \| 2.6e-32 \| 235.8 \| 326 \| (111, 455) \| 785 \| (326, 655) \| 1604 \| TP901 family prophage L54a \| TP901 family prophage L54a; uniclust \| UniRef100\_UPI0018A2AB22 \| 99.9 \| 1.5e-26 \| 2.7e-32 \| 233.2 \| 537 \| (20, 587) \| 785 \| (241, 823) \| 1216 \| hypothetical protein \| hypothetical protein; uniclust \| UniRef100\_A0A0N7I3V5 \| 99.9 \| 1.9e-26 \| 3.3e-32 \| 236.2 \| 379 \| (121, 593) \| 785 \| (73, 459) \| 762 \| Phage tail tape measure protein domain-containing protein \| Phage tail tape measure protein domain-containing protein; uniclust \| UniRef100\_A0A1C6H3B5 \| 99.9 \| 1.9e-26 \| 3.4e-32 \| 243.1 \| 328 \| (112, 451) \| 785 \| (195, 523) \| 1278 \| Phage-related minor tail protein \| Phage-related minor tail protein; uniclust \| UniRef100\_A0A343TII8 \| 99.9 \| 1.9e-26 \| 3.5e-32 \| 233.8 \| 363 \| (114, 489) \| 785 \| (52, 419) \| 943 \| Phage tail tape measure protein \| Phage tail tape measure protein; uniclust \| UniRef100\_UPI001EFFEF77 \| 99.9 \| 1.9e-26 \| 3.5e-32 \| 234.4 \| 344 \| (111, 466) \| 785 \| (151, 495) \| 1485 \| phage tail tape measure protein \| phage tail tape measure protein; uniclust \| UniRef100\_A0A174N846 \| 99.9 \| 2e-26 \| 3.6e-32 \| 238.1 \| 425 \| (116, 596) \| 785 \| (61, 499) \| 1019 \| Phage-related minor tail protein \| Phage-related minor tail protein; uniclust \| UniRef100\_A0A377ALG4 \| 99.9 \| 2e-26 \| 3.7e-32 \| 223.6 \| 478 \| (186, 673) \| 785 \| (59, 618) \| 646 \| TP901 family phage tail tape measure protein \| TP901 family phage tail tape measure protein; uniclust \| UniRef100\_A0A0A7RUN0 \| 99.9 \| 2.1e-26 \| 3.8e-32 \| 232.6 \| 329 \| (110, 460) \| 785 \| (66, 398) \| 757 \| Tail-tape measure protein \| Tail-tape measure protein; uniclust \| UniRef100\_UPI00065279AA \| 99.9 \| 2.2e-26 \| 4e-32 \| 224.8 \| 296 \| (115, 422) \| 785 \| (94, 390) \| 779 \| phage tail tape measure protein \| phage tail tape measure protein; uniclust \| UniRef100\_A0A1H6HI37 \| 99.9 \| 2.5e-26 \| 4.6e-32 \| 223.4 \| 471 \| (5, 488) \| 785 \| (87, 590) \| 838 \| Phage tail tape measure protein, TP901 family, core region \| Phage tail tape measure protein, TP901 family, core region; uniclust \| UniRef100\_UPI00079563B0 \| 99.9 \| 2.6e-26 \| 4.7e-32 \| 221.4 \| 442 \| (111, 581) \| 785 \| (47, 505) \| 668 \| phage tail tape measure protein \| phage tail tape measure protein; uniclust \| UniRef100\_A0A1Q9TM56 \| 99.9 \| 2.6e-26 \| 4.7e-32 \| 218.3 \| 358 \| (5, 369) \| 785 \| (78, 450) \| 505 \| Phage tail tape measure protein \| Phage tail tape measure protein; uniclust \| UniRef100\_A0A3R9NYT5 \| 99.9 \| 2.6e-26 \| 4.8e-32 \| 218.4 \| 305 \| (119, 441) \| 785 \| (62, 367) \| 714 \| Phage tail tape measure protein \| Phage tail tape measure protein; uniclust \| UniRef100\_A0A848AWP4 \| 99.9 \| 2.7e-26 \| 4.8e-32 \| 223.2 \| 532 \| (112, 671) \| 785 \| (53, 656) \| 774 \| Phage tail tape measure protein domain-containing protein \| Phage tail tape measure protein domain-containing protein; uniclust \| UniRef100\_A0A0J6WUL5 \| 99.9 \| 3.1e-26 \| 5.5e-32 \| 242.0 \| 323 \| (109, 447) \| 785 \| (148, 480) \| 1003 \| Phage tail tape measure protein domain-containing protein \| Phage tail tape measure protein domain-containing protein; uniclust \| UniRef100\_UPI0003B73CF5 \| 99.9 \| 3.2e-26 \| 5.9e-32 \| 218.0 \| 542 \| (37, 586) \| 785 \| (103, 675) \| 913 \| phage tail tape measure protein \| phage tail tape measure protein; uniclust \| UniRef100\_A0A1C9C1A8 \| 99.9 \| 3.4e-26 \| 6e-32 \| 245.2 \| 291 \| (247, 592) \| 785 \| (486, 788) \| 1575 \| Phage protein \| Phage protein; uniclust \| UniRef100\_A0A4Y8RH88 \| 99.9 \| 3.3e-26 \| 6.1e-32 \| 226.2 \| 441 \| (114, 574) \| 785 \| (59, 500) \| 1437 \| Phage tail tape measure protein \| Phage tail tape measure protein; uniclust \| UniRef100\_A0A1T4WD18 \| 99.9 \| 3.7e-26 \| 6.7e-32 \| 221.4 \| 424 \| (113, 592) \| 785 \| (63, 490) \| 712 \| Phage tail tape measure protein, TP901 family, core region \| Phage tail tape measure protein, TP901 family, core region; uniclust \| UniRef100\_A0A0J1IMP2 \| 99.9 \| 4.1e-26 \| 7.3e-32 \| 246.6 \| 329 \| (111, 461) \| 785 \| (299, 641) \| 1304 \| Phage tail tape measure protein domain-containing protein \| Phage tail tape measure protein domain-containing protein; uniclust \| UniRef100\_A0A0C3M8Z2 \| 99.9 \| 4.2e-26 \| 7.5e-32 \| 246.1 \| 340 \| (110, 464) \| 785 \| (322, 715) \| 1915 \| Tail protein \| Tail protein; uniclust \| UniRef100\_E2CFK1 \| 99.9 \| 4.2e-26 \| 7.6e-32 \| 221.7 \| 462 \| (103, 593) \| 785 \| (73, 545) \| 796 \| Putative phage tail tape measure protein, core region \| Putative phage tail tape measure protein, core region; uniclust \| UniRef100\_A0A259MGX1 \| 99.9 \| 4.2e-26 \| 7.8e-32 \| 214.9 \| 454 \| (113, 585) \| 785 \| (70, 523) \| 813 \| Phage tail tape measure protein \| Phage tail tape measure protein; uniclust \| UniRef100\_UPI00039627E7 \| 99.9 \| 4.7e-26 \| 8.6e-32 \| 213.3 \| 541 \| (110, 702) \| 785 \| (77, 634) \| 764 \| phage tail tape measure protein \| phage tail tape measure protein; uniclust \| UniRef100\_C9RFS8 \| 99.9 \| 4.8e-26 \| 8.8e-32 \| 216.2 \| 289 \| (113, 421) \| 785 \| (62, 351) \| 730 \| Phage tail tape measure protein, TP901 family \| Phage tail tape measure protein, TP901 family; uniclust \| UniRef100\_A0A086F263 \| 99.9 \| 4.9e-26 \| 8.9e-32 \| 220.0 \| 563 \| (17, 591) \| 785 \| (11, 620) \| 839 \| Tail protein \| Tail protein; uniclust \| UniRef100\_UPI000949D917 \| 99.9 \| 5.3e-26 \| 9.8e-32 \| 198.5 \| 341 \| (118, 462) \| 785 \| (73, 413) \| 419 \| phage tail tape measure protein \| phage tail tape measure protein; uniclust \| UniRef100\_A0A2Z4PV63 \| 99.9 \| 5.6e-26 \| 1e-31 \| 224.0 \| 389 \| (80, 472) \| 785 \| (8, 406) \| 729 \| Phage tail tape measure protein \| Phage tail tape measure protein; uniclust \| UniRef100\_A0A0F7L6E4 \| 99.9 \| 5.9e-26 \| 1e-31 \| 236.7 \| 395 \| (111, 580) \| 785 \| (50, 448) \| 846 \| TP901 family phage tail tape measure protein \| TP901 family phage tail tape measure protein; uniclust \| UniRef100\_A0A1X0A497 \| 99.9 \| 6.2e-26 \| 1.1e-31 \| 238.9 \| 319 \| (118, 448) \| 785 \| (330, 651) \| 1468 \| Phage tail tape measure protein \| Phage tail tape measure protein; uniclust \| UniRef100\_A0A660N6E2 \| 99.9 \| 6.1e-26 \| 1.1e-31 \| 202.2 \| 347 \| (118, 468) \| 785 \| (40, 391) \| 491 \| Phage tail tape measure protein (Fragment) \| Phage tail tape measure protein (Fragment); uniclust \| UniRef100\_A0A1C6EPE4 \| 99.9 \| 6.7e-26 \| 1.2e-31 \| 243.0 \| 398 \| (4, 443) \| 785 \| (19, 416) \| 1134 \| Phage-related protein \| Phage-related protein; uniclust \| UniRef100\_A0A2S9YZW5 \| 99.9 \| 7.8e-26 \| 1.4e-31 \| 244.2 \| 334 \| (115, 464) \| 785 \| (248, 595) \| 2271 \| Phage tail tape measure protein \| Phage tail tape measure protein; uniclust \| UniRef100\_A0A7Z7J1X9 \| 99.9 \| 8.7e-26 \| 1.6e-31 \| 205.2 \| 310 \| (304, 628) \| 785 \| (4, 315) \| 577 \| Phage-related tail protein \| Phage-related tail protein; uniclust \| UniRef100\_A0A1C6F7V7 \| 99.9 \| 8.9e-26 \| 1.6e-31 \| 239.6 \| 421 \| (26, 463) \| 785 \| (123, 552) \| 1280 \| Chromosome segregation protein \| Chromosome segregation protein; uniclust \| UniRef100\_A0A071MDL4 \| 99.9 \| 9.4e-26 \| 1.7e-31 \| 223.7 \| 478 \| (240, 780) \| 785 \| (328, 850) \| 850 \| Membrane protein \| Membrane protein; uniclust \| UniRef100\_A0A0J6WRI9 \| 99.9 \| 1.1e-25 \| 2e-31 \| 219.3 \| 430 \| (115, 584) \| 785 \| (84, 518) \| 737 \| Phage tail tape measure protein domain-containing protein \| Phage tail tape measure protein domain-containing protein; uniclust \| UniRef100\_A0A3D1NR59 \| 99.9 \| 1.1e-25 \| 2e-31 \| 215.1 \| 406 \| (260, 702) \| 785 \| (86, 502) \| 686 \| Phage tail tape measure protein \| Phage tail tape measure protein; uniclust \| UniRef100\_A0A829MBP5 \| 99.9 \| 1.2e-25 \| 2.1e-31 \| 224.9 \| 454 \| (117, 591) \| 785 \| (215, 675) \| 1046 \| Phage tail tape measure protein, TP901 family, core region \| Phage tail tape measure protein, TP901 family, core region; uniclust \| UniRef100\_A0A100JLS2 \| 99.9 \| 1.2e-25 \| 2.2e-31 \| 235.2 \| 415 \| (110, 620) \| 785 \| (85, 517) \| 702 \| Phage-related minor tail protein \| Phage-related minor tail protein; uniclust \| UniRef100\_A0A2I0DVT3 \| 99.9 \| 1.2e-25 \| 2.2e-31 \| 221.9 \| 426 \| (103, 573) \| 785 \| (42, 479) \| 735 \| Phage tail tape measure protein \| Phage tail tape measure protein; uniclust \| UniRef100\_A0A0F9MG89 \| 99.9 \| 1.3e-25 \| 2.3e-31 \| 210.1 \| 327 \| (119, 457) \| 785 \| (104, 431) \| 489 \| Phage tail tape measure protein domain-containing protein (Fragment) \| Phage tail tape measure protein domain-containing protein (Fragment); uniclust \| UniRef100\_UPI001878DDDC \| 99.9 \| 1.3e-25 \| 2.3e-31 \| 206.9 \| 349 \| (115, 467) \| 785 \| (53, 401) \| 619 \| phage tail tape measure protein \| phage tail tape measure protein; uniclust \| UniRef100\_A0A2J4RFS2 \| 99.9 \| 1.3e-25 \| 2.3e-31 \| 212.4 \| 347 \| (4, 357) \| 785 \| (78, 432) \| 434 \| Phage tail tape measure protein (Fragment) \| Phage tail tape measure protein (Fragment); uniclust \| UniRef100\_A0A9D5UUJ2 \| 99.9 \| 1.3e-25 \| 2.4e-31 \| 226.7 \| 451 \| (5, 459) \| 785 \| (80, 581) \| 1509 \| Phage tail tape measure protein \| Phage tail tape measure protein; uniclust \| UniRef100\_A0A1C0TVM0 \| 99.9 \| 1.4e-25 \| 2.4e-31 \| 216.7 \| 344 \| (101, 467) \| 785 \| (43, 395) \| 486 \| Phage tail tape measure protein \| Phage tail tape measure protein; uniclust \| UniRef100\_A0A355D6E1 \| 99.9 \| 1.5e-25 \| 2.6e-31 \| 226.9 \| 343 \| (111, 464) \| 785 \| (64, 406) \| 1089 \| Phage tail tape measure protein \| Phage tail tape measure protein; uniclust \| UniRef100\_A0A081L6U1 \| 99.9 \| 1.5e-25 \| 2.7e-31 \| 248.8 \| 317 \| (117, 450) \| 785 \| (304, 628) \| 1423 \| Transglycosylase SLT domain-containing protein \| Transglycosylase SLT domain-containing protein; uniclust \| UniRef100\_A0A1V3J0L3 \| 99.9 \| 1.6e-25 \| 3e-31 \| 201.2 \| 330 \| (430, 782) \| 785 \| (3, 404) \| 404 \| Phage tail tape measure protein \| Phage tail tape measure protein; uniclust \| UniRef100\_A0A1C6E118 \| 99.9 \| 1.7e-25 \| 3.1e-31 \| 232.9 \| 327 \| (114, 440) \| 785 \| (58, 399) \| 1176 \| Phage-related protein \| Phage-related protein; uniclust \| UniRef100\_UPI002227BD23 \| 99.9 \| 1.7e-25 \| 3.1e-31 \| 214.8 \| 413 \| (117, 586) \| 785 \| (96, 517) \| 984 \| phage tail tape measure protein \| phage tail tape measure protein; uniclust \| UniRef100\_A0A4Q4AZ49 \| 99.9 \| 1.7e-25 \| 3.1e-31 \| 202.4 \| 451 \| (282, 782) \| 785 \| (7, 464) \| 464 \| Phage tail tape measure protein \| Phage tail tape measure protein; uniclust \| UniRef100\_A0A143Z8T0 \| 99.9 \| 1.9e-25 \| 3.5e-31 \| 220.3 \| 427 \| (111, 597) \| 785 \| (153, 588) \| 858 \| Phage tail tape measure protein \| Phage tail tape measure protein; uniclust \| UniRef100\_A0A1X7MGH5 \| 99.9 \| 2.1e-25 \| 3.9e-31 \| 199.4 \| 314 \| (265, 594) \| 785 \| (2, 321) \| 448 \| Phage tail tape measure protein, family, core region \| Phage tail tape measure protein, family, core region; uniclust \| UniRef100\_A0A143HC28 \| 99.9 \| 2.2e-25 \| 3.9e-31 \| 238.9 \| 373 \| (4, 444) \| 785 \| (5, 385) \| 1162 \| Phage tail tape measure protein domain-containing protein \| Phage tail tape measure protein domain-containing protein; uniclust \| UniRef100\_A0A1V0UQY8 \| 99.9 \| 2.5e-25 \| 4.6e-31 \| 222.5 \| 324 \| (110, 445) \| 785 \| (61, 385) \| 954 \| Phage tail tape measure protein \| Phage tail tape measure protein; uniclust \| UniRef100\_A0A0H3U2U8 \| 99.9 \| 2.6e-25 \| 4.8e-31 \| 216.8 \| 417 \| (118, 594) \| 785 \| (180, 605) \| 951 \| Tail length tape measure \| Tail length tape measure; uniclust \| UniRef100\_A0A1I5H9Z3 \| 99.9 \| 2.7e-25 \| 5e-31 \| 213.3 \| 471 \| (5, 485) \| 785 \| (79, 564) \| 874 \| Phage tail tape measure protein, TP901 family, core region \| Phage tail tape measure protein, TP901 family, core region; uniclust \| UniRef100\_A0A7W9YIK7 \| 99.9 \| 2.8e-25 \| 5e-31 \| 228.3 \| 451 \| (118, 589) \| 785 \| (230, 715) \| 1645 \| TP901 family phage tail tape measure protein \| TP901 family phage tail tape measure protein; uniclust \| UniRef100\_A0A0U4II47 \| 99.9 \| 3e-25 \| 5.3e-31 \| 228.2 \| 319 \| (115, 449) \| 785 \| (146, 467) \| 753 \| Tail tape measure protein \| Tail tape measure protein; uniclust \| UniRef100\_A0A099S5E7 \| 99.9 \| 3.1e-25 \| 5.4e-31 \| 234.5 \| 273 \| (393, 701) \| 785 \| (328, 638) \| 875 \| Phage-like protein \| Phage-like protein; uniclust \| UniRef100\_A0A2P5GLT3 \| 99.9 \| 3.2e-25 \| 5.8e-31 \| 212.5 \| 517 \| (124, 670) \| 785 \| (90, 646) \| 761 \| Phage tail tape measure protein \| Phage tail tape measure protein; uniclust \| UniRef100\_A0A1S1YB82 \| 99.9 \| 3.4e-25 \| 6.2e-31 \| 219.9 \| 421 \| (111, 593) \| 785 \| (50, 486) \| 784 \| Phage tail tape measure protein domain-containing protein \| Phage tail tape measure protein domain-containing protein; uniclust \| UniRef100\_A0A1E3GZT2 \| 99.9 \| 3.7e-25 \| 6.6e-31 \| 219.7 \| 332 \| (112, 462) \| 785 \| (62, 403) \| 766 \| Phage-related minor tail protein \| Phage-related minor tail protein; uniclust \| UniRef100\_A0A2T4MMQ9 \| 99.9 \| 3.8e-25 \| 7e-31 \| 208.6 \| 309 \| (112, 432) \| 785 \| (170, 479) \| 541 \| Phage tail tape measure protein (Fragment) \| Phage tail tape measure protein (Fragment); uniclust \| UniRef100\_A0A0R1GKD1 \| 99.9 \| 3.9e-25 \| 7e-31 \| 242.2 \| 347 \| (110, 466) \| 785 \| (303, 659) \| 2004 \| NlpC/P60 domain-containing protein \| NlpC/P60 domain-containing protein; uniclust \| UniRef100\_A0A1A7KG91 \| 99.9 \| 4e-25 \| 7.1e-31 \| 222.7 \| 468 \| (118, 622) \| 785 \| (67, 554) \| 793 \| Phage-related minor tail protein \| Phage-related minor tail protein; uniclust \| UniRef100\_A0A173RDD3 \| 99.9 \| 4.2e-25 \| 7.4e-31 \| 242.2 \| 323 \| (111, 444) \| 785 \| (82, 404) \| 1392 \| Phage-related protein \| Phage-related protein; uniclust \| UniRef100\_A0A4T2BU47 \| 99.9 \| 4.2e-25 \| 7.5e-31 \| 234.5 \| 327 \| (119, 457) \| 785 \| (252, 667) \| 1102 \| Phage tail tape measure protein \| Phage tail tape measure protein; uniclust \| UniRef100\_A0A4Z0GIX0 \| 99.9 \| 4.2e-25 \| 7.6e-31 \| 226.2 \| 326 \| (110, 445) \| 785 \| (272, 598) \| 1299 \| Phage tail tape measure protein (Fragment) \| Phage tail tape measure protein (Fragment); uniclust \| UniRef100\_A0A5C5TFK9 \| 99.9 \| 4.2e-25 \| 7.6e-31 \| 207.0 \| 423 \| (19, 448) \| 785 \| (47, 494) \| 535 \| Phage tail tape measure protein \| Phage tail tape measure protein; uniclust \| UniRef100\_A0A143ZV16 \| 99.9 \| 4.8e-25 \| 8.4e-31 \| 238.9 \| 321 \| (111, 443) \| 785 \| (46, 367) \| 1213 \| Phage-related minor tail protein \| Phage-related minor tail protein; uniclust \| UniRef100\_A0A239EK42 \| 99.9 \| 4.6e-25 \| 8.5e-31 \| 215.5 \| 445 \| (107, 582) \| 785 \| (63, 512) \| 1202 \| Phage tail tape measure protein, TP901 family, core region \| Phage tail tape measure protein, TP901 family, core region; uniclust \| UniRef100\_A0A291LZ62 \| 99.9 \| 4.7e-25 \| 8.6e-31 \| 206.3 \| 423 \| (4, 439) \| 785 \| (5, 437) \| 746 \| Phage tail tape measure protein \| Phage tail tape measure protein; uniclust \| UniRef100\_UPI001118EFF6 \| 99.9 \| 5.2e-25 \| 9.5e-31 \| 193.5 \| 316 \| (208, 580) \| 785 \| (3, 318) \| 377 \| phage tail tape measure protein \| phage tail tape measure protein; uniclust \| UniRef100\_A0A432QRK4 \| 99.9 \| 5.2e-25 \| 9.5e-31 \| 210.0 \| 348 \| (116, 472) \| 785 \| (9, 359) \| 908 \| Phage tail tape measure protein (Fragment) \| Phage tail tape measure protein (Fragment); uniclust \| UniRef100\_A0A7C4LLA7 \| 99.9 \| 5.3e-25 \| 9.7e-31 \| 215.0 \| 347 \| (115, 473) \| 785 \| (57, 404) \| 926 \| Phage tail tape measure protein \| Phage tail tape measure protein; uniclust \| UniRef100\_A0A3R6NIQ7 \| 99.9 \| 5.4e-25 \| 9.9e-31 \| 220.3 \| 320 \| (111, 448) \| 785 \| (111, 432) \| 1172 \| Phage tail tape measure protein \| Phage tail tape measure protein; uniclust \| UniRef100\_A0A0K1Y580 \| 99.8 \| 5.8e-25 \| 1e-30 \| 237.1 \| 433 \| (114, 594) \| 785 \| (69, 531) \| 1587 \| Phage tail tape measure protein domain-containing protein \| Phage tail tape measure protein domain-containing protein; uniclust \| UniRef100\_A0A806D8P0 \| 99.8 \| 5.8e-25 \| 1.1e-30 \| 209.0 \| 468 \| (120, 594) \| 785 \| (204, 704) \| 876 \| Phage tail tape measure protein, TP901 family \| Phage tail tape measure protein, TP901 family; uniclust \| UniRef100\_A0A7X8V2E7 \| 99.8 \| 6.9e-25 \| 1.3e-30 \| 194.7 \| 288 \| (278, 588) \| 785 \| (9, 296) \| 471 \| Phage tail tape measure protein \| Phage tail tape measure protein; uniclust \| UniRef100\_A0A2W4KBS4 \| 99.8 \| 7.1e-25 \| 1.3e-30 \| 212.3 \| 346 \| (112, 463) \| 785 \| (84, 439) \| 738 \| Phage tail tape measure protein (Fragment) \| Phage tail tape measure protein (Fragment); uniclust \| UniRef100\_A0A3N4UR92 \| 99.8 \| 7.3e-25 \| 1.3e-30 \| 217.4 \| 448 \| (121, 592) \| 785 \| (124, 616) \| 812 \| TP901 family phage tail tape measure protein \| TP901 family phage tail tape measure protein; uniclust \| UniRef100\_A0A0A0YSZ0 \| 99.8 \| 7.7e-25 \| 1.4e-30 \| 216.1 \| 409 \| (109, 572) \| 785 \| (60, 481) \| 637 \| Phage tail tape measure protein domain-containing protein \| Phage tail tape measure protein domain-containing protein; uniclust \| UniRef100\_A0A5C7JCR9 \| 99.8 \| 7.6e-25 \| 1.4e-30 \| 199.4 \| 344 \| (115, 464) \| 785 \| (85, 429) \| 580 \| Phage tail tape measure protein (Fragment) \| Phage tail tape measure protein (Fragment); uniclust \| UniRef100\_A0A8T4G949 \| 99.8 \| 8.2e-25 \| 1.5e-30 \| 208.2 \| 306 \| (117, 439) \| 785 \| (47, 353) \| 713 \| Phage tail tape measure protein \| Phage tail tape measure protein; uniclust \| UniRef100\_A0A8I1W8W3 \| 99.8 \| 8.2e-25 \| 1.5e-30 \| 210.5 \| 438 \| (24, 466) \| 785 \| (174, 617) \| 802 \| Phage tail tape measure protein \| Phage tail tape measure protein; pdb70 \| 6V8I\_CF \| 99.2 \| 9.4e-16 \| 1.3e-19 \| 163.0 \| 95 \| (155, 250) \| 785 \| (175, 269) \| 1154 \| Distal Tail Protein, gp58, Tail-Associated \| 6V8I\_CF Distal Tail Protein, gp58, Tail-Associated phage tail, tail tip, tape |
| Top keywords  (threshold 1.00e-03 (evalue)) | **tail, Phage, tape, measure, TP901, domain\_containing, core, region, Phage\_related, Fragment** |
| Output files | ../../similar\_sequences/22\_FANPEZAQ\_CDS\_0022\_merged.svg ../../similar\_sequences/22\_FANPEZAQ\_CDS\_0022\_pdb70.a3m ../../similar\_sequences/22\_FANPEZAQ\_CDS\_0022\_pdb70.hhr ../../similar\_sequences/22\_FANPEZAQ\_CDS\_0022\_uniclust.a3m ../../similar\_sequences/22\_FANPEZAQ\_CDS\_0022\_uniclust.hhr |

#### Structure prediction (AlphaFold)^2^

|  |  |
| --- | --- |
| Stats | xml version="1.0" encoding="utf-8" standalone="no"?       2024-09-02T21:09:22.305241 image/svg+xml   Matplotlib v3.7.2, https://matplotlib.org/ |
| Predicted structure | **NGL Viewer Controls:**  - Center: *Left-Click* - Rotate: *Left-Click + Drag* - Translate: *Right-Click + Drag* - Zoom: *Shift + Left-Click + Drag* |
| Output files | ../../predicted\_structures/22\_FANPEZAQ\_CDS\_0022/features.pkl ../../predicted\_structures/22\_FANPEZAQ\_CDS\_0022/ranked\_0.pdb ../../predicted\_structures/22\_FANPEZAQ\_CDS\_0022/ranked\_0\_plots.svg ../../predicted\_structures/22\_FANPEZAQ\_CDS\_0022/result\_model\_1\_ptm\_pred\_0.pkl |

#### Structure similarity search results (Foldseek)^3^

|  |  |
| --- | --- |
| Structure databases searched | Pdb, Afdb-proteome, Afdb-uniprot50 |
| Results, scheme(s)  (Top layers only, threshold 1.00e-02 (evalue)) | xml version="1.0" encoding="utf-8" standalone="no"?       2024-09-02T21:10:49.053219 image/svg+xml   Matplotlib v3.7.2, https://matplotlib.org/ |
| Results, table  (threshold 1.00e-02 (evalue)) | db \| id \| prob \| evalue \| bits \| fident \| alnlen \| mismatch \| gapopen \| qstart \| qend \| tstart \| tend \| name \| description; afdb-proteome \| AF-Q8ZMV3-F1-MODEL\_V4 \| 1.0 \| 1.278e-23 \| 829 \| 0.198 \| 920 \| 552 \| 21 \| 19 \| 785 \| 48 \| 934 \| Fels-2 prophage protein \| Fels-2 prophage protein; afdb-proteome \| AF-A0A0H3H1R5-F1-MODEL\_V4 \| 1.0 \| 3.576e-18 \| 539 \| 0.202 \| 846 \| 510 \| 25 \| 19 \| 781 \| 50 \| 813 \| Putative prophage tail length determinator \| Putative prophage tail length determinator; afdb-proteome \| AF-A0A0H3GQS8-F1-MODEL\_V4 \| 1.0 \| 3.039e-14 \| 434 \| 0.163 \| 883 \| 530 \| 30 \| 15 \| 785 \| 11 \| 796 \| Phage tail tape measure protein, family \| Phage tail tape measure protein, family; afdb-proteome \| AF-A0A5P3FW64-F1-MODEL\_V4 \| 1.0 \| 3.051e-11 \| 405 \| 0.176 \| 589 \| 352 \| 14 \| 1 \| 459 \| 1 \| 586 \| Phage tail tape measure protein \| Phage tail tape measure protein; afdb-proteome \| AF-Q2FWU3-F1-MODEL\_V4 \| 1.0 \| 1.937e-12 \| 377 \| 0.142 \| 921 \| 545 \| 39 \| 1 \| 702 \| 1 \| 895 \| Lysostaphin \| Lysostaphin; afdb-proteome \| AF-Q8ZKJ4-F1-MODEL\_V4 \| 1.0 \| 4.604e-13 \| 352 \| 0.174 \| 785 \| 510 \| 32 \| 35 \| 754 \| 3 \| 714 \| Putative phage tail protein \| Putative phage tail protein; afdb-proteome \| AF-Q2FYC8-F1-MODEL\_V4 \| 1.0 \| 4.623e-10 \| 349 \| 0.142 \| 631 \| 368 \| 19 \| 19 \| 519 \| 127 \| 714 \| Lysostaphin \| Lysostaphin; afdb-proteome \| AF-G3XD83-F1-MODEL\_V4 \| 1.0 \| 1.953e-06 \| 208 \| 0.157 \| 513 \| 341 \| 19 \| 2 \| 469 \| 1 \| 467 \| Uncharacterized protein \| Uncharacterized protein; afdb-proteome \| AF-A0A5K4F6A0-F1-MODEL\_V4 \| 0.382 \| 0.007632 \| 47 \| 0.088 \| 782 \| 393 \| 30 \| 103 \| 785 \| 14 \| 574 \| G\_PROTEIN\_RECEP\_F1\_2 domain-containing protein \| G\_PROTEIN\_RECEP\_F1\_2 domain-containing protein; afdb-uniprot50 \| AF-A0A5U1APE0-F1-MODEL\_V4 \| 1.0 \| 4.447e-25 \| 1458 \| 0.456 \| 353 \| 189 \| 1 \| 117 \| 469 \| 12 \| 361 \| Phage tail tape measure protein \| Phage tail tape measure protein; afdb-uniprot50 \| AF-A0A5U1APS5-F1-MODEL\_V4 \| 1.0 \| 1.639e-26 \| 1366 \| 0.423 \| 418 \| 216 \| 5 \| 61 \| 469 \| 1 \| 402 \| Phage tail tape measure protein \| Phage tail tape measure protein; afdb-uniprot50 \| AF-A0A761LW72-F1-MODEL\_V4 \| 1.0 \| 9.78e-26 \| 1344 \| 0.432 \| 414 \| 210 \| 5 \| 61 \| 465 \| 1 \| 398 \| Phage tail tape measure protein \| Phage tail tape measure protein; afdb-uniprot50 \| AF-A0A5Y1GPG4-F1-MODEL\_V4 \| 1.0 \| 6.507e-21 \| 1221 \| 0.496 \| 298 \| 147 \| 1 \| 133 \| 430 \| 2 \| 296 \| Phage tail tape measure protein \| Phage tail tape measure protein; afdb-uniprot50 \| AF-A0A7T7R8S8-F1-MODEL\_V4 \| 1.0 \| 2.979e-34 \| 1216 \| 0.314 \| 796 \| 504 \| 9 \| 1 \| 785 \| 1 \| 765 \| Phage tail tape measure protein \| Phage tail tape measure protein; afdb-uniprot50 \| AF-A0A450W6Y7-F1-MODEL\_V4 \| 1.0 \| 1.115e-33 \| 1146 \| 0.316 \| 791 \| 442 \| 10 \| 21 \| 755 \| 50 \| 797 \| Phage tail tape measure protein, TP901 family, core region \| Phage tail tape measure protein, TP901 family, core region; afdb-uniprot50 \| AF-A0A5T2W9F8-F1-MODEL\_V4 \| 1.0 \| 1.945e-24 \| 1138 \| 0.334 \| 577 \| 278 \| 10 \| 118 \| 687 \| 2 \| 479 \| Phage tail tape measure protein \| Phage tail tape measure protein; afdb-uniprot50 \| AF-A0A630L2F6-F1-MODEL\_V4 \| 1.0 \| 1.745e-18 \| 1128 \| 0.496 \| 266 \| 131 \| 1 \| 134 \| 399 \| 2 \| 264 \| Phage tail tape measure protein \| Phage tail tape measure protein; afdb-uniprot50 \| AF-A0A5Z1EPF8-F1-MODEL\_V4 \| 1.0 \| 9.86e-20 \| 1127 \| 0.344 \| 348 \| 219 \| 3 \| 117 \| 464 \| 12 \| 350 \| Phage tail tape measure protein \| Phage tail tape measure protein; afdb-uniprot50 \| AF-A0A4P6BLS6-F1-MODEL\_V4 \| 1.0 \| 1.278e-28 \| 1087 \| 0.309 \| 795 \| 390 \| 16 \| 18 \| 777 \| 17 \| 687 \| PhageMin\_Tail domain-containing protein \| PhageMin\_Tail domain-containing protein; afdb-uniprot50 \| AF-A0A2J7U4C6-F1-MODEL\_V4 \| 1.0 \| 1.249e-26 \| 1075 \| 0.347 \| 656 \| 337 \| 17 \| 61 \| 692 \| 1 \| 589 \| Phage tail tape measure protein \| Phage tail tape measure protein; afdb-uniprot50 \| AF-A0A5Z1ENM2-F1-MODEL\_V4 \| 1.0 \| 1.23e-18 \| 1070 \| 0.331 \| 347 \| 224 \| 4 \| 117 \| 461 \| 12 \| 352 \| Phage tail tape measure protein \| Phage tail tape measure protein; afdb-uniprot50 \| AF-A0A601P240-F1-MODEL\_V4 \| 1.0 \| 3.662e-25 \| 1069 \| 0.323 \| 631 \| 320 \| 13 \| 117 \| 729 \| 12 \| 553 \| Phage tail tape measure protein \| Phage tail tape measure protein; afdb-uniprot50 \| AF-A0A754TYG6-F1-MODEL\_V4 \| 1.0 \| 1.002e-17 \| 1045 \| 0.513 \| 251 \| 122 \| 0 \| 180 \| 430 \| 1 \| 251 \| Phage tail tape measure protein \| Phage tail tape measure protein; afdb-uniprot50 \| AF-A0A5T1M5R4-F1-MODEL\_V4 \| 1.0 \| 2.144e-19 \| 1034 \| 0.332 \| 388 \| 248 \| 4 \| 79 \| 464 \| 21 \| 399 \| Phage tail tape measure protein \| Phage tail tape measure protein; afdb-uniprot50 \| AF-F7U3Q6-F1-MODEL\_V4 \| 1.0 \| 7.336e-28 \| 1033 \| 0.269 \| 789 \| 432 \| 18 \| 63 \| 782 \| 2 \| 714 \| Family phage tail tape measure protein \| Family phage tail tape measure protein; afdb-uniprot50 \| AF-A0A560D0Q3-F1-MODEL\_V4 \| 1.0 \| 5.173e-28 \| 1015 \| 0.33 \| 727 \| 362 \| 19 \| 70 \| 752 \| 7 \| 652 \| TP901 family phage tail tape measure protein \| TP901 family phage tail tape measure protein; afdb-uniprot50 \| AF-A0A7U3GJM6-F1-MODEL\_V4 \| 1.0 \| 3.593e-20 \| 1009 \| 0.343 \| 405 \| 231 \| 7 \| 39 \| 433 \| 8 \| 387 \| Phage tail tape measure protein \| Phage tail tape measure protein; afdb-uniprot50 \| AF-A0A4R2N8L9-F1-MODEL\_V4 \| 1.0 \| 3.677e-22 \| 1007 \| 0.34 \| 487 \| 305 \| 6 \| 6 \| 484 \| 1 \| 479 \| TP901 family phage tail tape measure protein \| TP901 family phage tail tape measure protein; afdb-uniprot50 \| AF-A0A7X8PBI2-F1-MODEL\_V4 \| 1.0 \| 4.48e-29 \| 999 \| 0.284 \| 826 \| 471 \| 20 \| 1 \| 782 \| 1 \| 750 \| Phage tail tape measure protein \| Phage tail tape measure protein; afdb-uniprot50 \| AF-A0A430S197-F1-MODEL\_V4 \| 1.0 \| 5.571e-21 \| 993 \| 0.329 \| 465 \| 282 \| 8 \| 62 \| 508 \| 4 \| 456 \| Phage tail tape measure protein \| Phage tail tape measure protein; afdb-uniprot50 \| AF-A0A4P7EI80-F1-MODEL\_V4 \| 1.0 \| 3.677e-22 \| 982 \| 0.302 \| 505 \| 290 \| 7 \| 19 \| 466 \| 51 \| 550 \| Phage tail tape measure protein \| Phage tail tape measure protein; afdb-uniprot50 \| AF-A0A1M6LSY3-F1-MODEL\_V4 \| 1.0 \| 3.689e-29 \| 981 \| 0.256 \| 829 \| 467 \| 17 \| 19 \| 702 \| 45 \| 868 \| Phage tail tape measure protein, TP901 family, core region \| Phage tail tape measure protein, TP901 family, core region; afdb-uniprot50 \| AF-F5SA52-F1-MODEL\_V4 \| 1.0 \| 8.378e-16 \| 980 \| 0.554 \| 222 \| 99 \| 0 \| 155 \| 376 \| 3 \| 224 \| Tail tape measure protein \| Tail tape measure protein; afdb-uniprot50 \| AF-A0A2A2IKN4-F1-MODEL\_V4 \| 1.0 \| 2.856e-27 \| 977 \| 0.289 \| 797 \| 404 \| 20 \| 19 \| 785 \| 20 \| 683 \| Phage tail tape measure protein \| Phage tail tape measure protein; afdb-uniprot50 \| AF-A0A823TKD2-F1-MODEL\_V4 \| 1.0 \| 9.337e-22 \| 973 \| 0.267 \| 527 \| 330 \| 8 \| 29 \| 552 \| 2 \| 475 \| Phage tail tape measure protein \| Phage tail tape measure protein; afdb-uniprot50 \| AF-A0A315BPW6-F1-MODEL\_V4 \| 1.0 \| 1.283e-25 \| 963 \| 0.271 \| 789 \| 424 \| 11 \| 1 \| 785 \| 8 \| 649 \| Phage tail tape measure protein \| Phage tail tape measure protein; afdb-uniprot50 \| AF-A0A745AEG2-F1-MODEL\_V4 \| 1.0 \| 2.07e-16 \| 961 \| 0.501 \| 251 \| 122 \| 1 \| 118 \| 368 \| 33 \| 280 \| Phage tail tape measure protein \| Phage tail tape measure protein; afdb-uniprot50 \| AF-A0A4Q0GN92-F1-MODEL\_V4 \| 1.0 \| 1.298e-26 \| 961 \| 0.268 \| 823 \| 412 \| 18 \| 22 \| 782 \| 50 \| 744 \| Phage tail tape measure protein \| Phage tail tape measure protein; afdb-uniprot50 \| AF-A0A484YXF6-F1-MODEL\_V4 \| 1.0 \| 1.25e-16 \| 958 \| 0.522 \| 241 \| 115 \| 0 \| 190 \| 430 \| 2 \| 242 \| Phage tail tape measure protein \| Phage tail tape measure protein; afdb-uniprot50 \| AF-Q4EB59-F1-MODEL\_V4 \| 1.0 \| 1.799e-24 \| 957 \| 0.279 \| 636 \| 378 \| 8 \| 24 \| 619 \| 4 \| 598 \| Phage tail tape meausure protein, TP901 family, putative \| Phage tail tape meausure protein, TP901 family, putative; afdb-uniprot50 \| AF-A0A751FK49-F1-MODEL\_V4 \| 1.0 \| 5.594e-18 \| 955 \| 0.467 \| 276 \| 147 \| 0 \| 190 \| 465 \| 3 \| 278 \| Phage tail tape measure protein \| Phage tail tape measure protein; afdb-uniprot50 \| AF-G3IRG2-F1-MODEL\_V4 \| 1.0 \| 2.571e-28 \| 952 \| 0.315 \| 842 \| 423 \| 22 \| 1 \| 780 \| 1 \| 750 \| Phage tail tape measure protein, TP901 family \| Phage tail tape measure protein, TP901 family; afdb-uniprot50 \| AF-A0A8A6KE23-F1-MODEL\_V4 \| 1.0 \| 1.499e-25 \| 943 \| 0.266 \| 789 \| 481 \| 18 \| 1 \| 784 \| 1 \| 696 \| Phage tail tape measure protein \| Phage tail tape measure protein; afdb-uniprot50 \| AF-B6WUA5-F1-MODEL\_V4 \| 1.0 \| 3.311e-23 \| 940 \| 0.274 \| 677 \| 350 \| 16 \| 1 \| 672 \| 1 \| 540 \| Phage tail tape measure protein, TP901 family \| Phage tail tape measure protein, TP901 family; afdb-uniprot50 \| AF-A0A430B9P2-F1-MODEL\_V4 \| 1.0 \| 2.592e-22 \| 935 \| 0.311 \| 575 \| 322 \| 14 \| 70 \| 620 \| 2 \| 526 \| Phage tail tape measure protein \| Phage tail tape measure protein; afdb-uniprot50 \| AF-A0A2M7BJQ9-F1-MODEL\_V4 \| 1.0 \| 4.196e-20 \| 929 \| 0.272 \| 480 \| 313 \| 6 \| 19 \| 480 \| 61 \| 522 \| Phage tail tape measure protein \| Phage tail tape measure protein; afdb-uniprot50 \| AF-A0A8B0U511-F1-MODEL\_V4 \| 1.0 \| 1.35e-26 \| 927 \| 0.267 \| 821 \| 483 \| 23 \| 1 \| 785 \| 1 \| 738 \| Phage tail tape measure protein \| Phage tail tape measure protein; afdb-uniprot50 \| AF-A0A259MGX1-F1-MODEL\_V4 \| 1.0 \| 5.59e-28 \| 922 \| 0.29 \| 799 \| 443 \| 25 \| 13 \| 782 \| 1 \| 704 \| Phage tail tape measure protein \| Phage tail tape measure protein; afdb-uniprot50 \| AF-A0A846BM80-F1-MODEL\_V4 \| 1.0 \| 6.358e-19 \| 921 \| 0.325 \| 446 \| 255 \| 7 \| 117 \| 562 \| 38 \| 437 \| Phage tail tape measure protein \| Phage tail tape measure protein; afdb-uniprot50 \| AF-A0A350LX86-F1-MODEL\_V4 \| 1.0 \| 4.362e-20 \| 918 \| 0.294 \| 455 \| 296 \| 7 \| 66 \| 499 \| 15 \| 465 \| Phage tail tape measure protein \| Phage tail tape measure protein; afdb-uniprot50 \| AF-A0A844HR58-F1-MODEL\_V4 \| 1.0 \| 3.958e-25 \| 915 \| 0.271 \| 767 \| 395 \| 18 \| 116 \| 783 \| 14 \| 715 \| Phage tail tape measure protein \| Phage tail tape measure protein; afdb-uniprot50 \| AF-A0A660N6E2-F1-MODEL\_V4 \| 1.0 \| 5.359e-21 \| 913 \| 0.303 \| 530 \| 286 \| 11 \| 46 \| 539 \| 5 \| 487 \| Phage tail tape measure protein \| Phage tail tape measure protein; afdb-uniprot50 \| AF-A0A348AJ09-F1-MODEL\_V4 \| 1.0 \| 7.539e-27 \| 911 \| 0.262 \| 843 \| 486 \| 16 \| 19 \| 785 \| 11 \| 793 \| Phage-related minor tail protein \| Phage-related minor tail protein; afdb-uniprot50 \| AF-A0A2W5VBB3-F1-MODEL\_V4 \| 1.0 \| 5.972e-27 \| 911 \| 0.286 \| 839 \| 461 \| 22 \| 19 \| 783 \| 52 \| 826 \| Phage tail tape measure protein \| Phage tail tape measure protein; afdb-uniprot50 \| AF-A0A7W5RGC2-F1-MODEL\_V4 \| 1.0 \| 6.454e-27 \| 907 \| 0.284 \| 847 \| 478 \| 19 \| 1 \| 784 \| 2 \| 783 \| TP901 family phage tail tape measure protein \| TP901 family phage tail tape measure protein; afdb-uniprot50 \| AF-A0A1S9ZKE4-F1-MODEL\_V4 \| 1.0 \| 3.259e-25 \| 906 \| 0.287 \| 695 \| 432 \| 14 \| 36 \| 702 \| 2 \| 660 \| Phage tail tape measure protein \| Phage tail tape measure protein; afdb-uniprot50 \| AF-G2HXA7-F1-MODEL\_V4 \| 1.0 \| 7.168e-26 \| 903 \| 0.272 \| 799 \| 458 \| 21 \| 3 \| 785 \| 1 \| 691 \| Phage tail tape measure protein \| Phage tail tape measure protein; afdb-uniprot50 \| AF-G4CJF3-F1-MODEL\_V4 \| 1.0 \| 1.029e-26 \| 903 \| 0.255 \| 758 \| 471 \| 19 \| 19 \| 710 \| 21 \| 750 \| Phage tail tape measure protein \| Phage tail tape measure protein; afdb-uniprot50 \| AF-A0A1M3AG88-F1-MODEL\_V4 \| 1.0 \| 1.156e-26 \| 901 \| 0.26 \| 920 \| 459 \| 23 \| 13 \| 785 \| 1 \| 845 \| Phage tail tape measure protein \| Phage tail tape measure protein; afdb-uniprot50 \| AF-A0A359KC48-F1-MODEL\_V4 \| 1.0 \| 8.776e-20 \| 894 \| 0.346 \| 513 \| 266 \| 12 \| 22 \| 475 \| 53 \| 554 \| Phage tail tape measure protein \| Phage tail tape measure protein; afdb-uniprot50 \| AF-A0A5M7PDK9-F1-MODEL\_V4 \| 1.0 \| 3.272e-22 \| 891 \| 0.279 \| 644 \| 362 \| 13 \| 70 \| 706 \| 7 \| 555 \| Phage tail tape measure protein \| Phage tail tape measure protein; afdb-uniprot50 \| AF-A0A1I3JGJ4-F1-MODEL\_V4 \| 1.0 \| 1.771e-26 \| 890 \| 0.319 \| 793 \| 435 \| 21 \| 62 \| 784 \| 14 \| 771 \| Phage tail tape measure protein, TP901 family, core region \| Phage tail tape measure protein, TP901 family, core region; afdb-uniprot50 \| AF-A0A5C7R6S2-F1-MODEL\_V4 \| 1.0 \| 1.108e-19 \| 888 \| 0.313 \| 492 \| 317 \| 6 \| 1 \| 480 \| 1 \| 483 \| Phage tail tape measure protein \| Phage tail tape measure protein; afdb-uniprot50 \| AF-A0A2G0WA81-F1-MODEL\_V4 \| 1.0 \| 1.289e-22 \| 888 \| 0.272 \| 729 \| 371 \| 16 \| 71 \| 750 \| 2 \| 619 \| PhageMin\_Tail domain-containing protein \| PhageMin\_Tail domain-containing protein; afdb-uniprot50 \| AF-A0A378UD30-F1-MODEL\_V4 \| 1.0 \| 6.71e-27 \| 887 \| 0.257 \| 793 \| 466 \| 17 \| 18 \| 710 \| 20 \| 789 \| Putative phage-related tail protein \| Putative phage-related tail protein; afdb-uniprot50 \| AF-W4M094-F1-MODEL\_V4 \| 1.0 \| 5.529e-17 \| 886 \| 0.341 \| 340 \| 219 \| 3 \| 143 \| 480 \| 6 \| 342 \| PhageMin\_Tail domain-containing protein \| PhageMin\_Tail domain-containing protein; afdb-uniprot50 \| AF-A0A5A9EN03-F1-MODEL\_V4 \| 1.0 \| 4.327e-26 \| 886 \| 0.269 \| 762 \| 457 \| 14 \| 19 \| 700 \| 32 \| 773 \| Phage tail tape measure protein \| Phage tail tape measure protein; afdb-uniprot50 \| AF-K8AC29-F1-MODEL\_V4 \| 1.0 \| 5.702e-23 \| 882 \| 0.254 \| 711 \| 402 \| 19 \| 71 \| 754 \| 2 \| 611 \| Phage tail length tape-measure protein \| Phage tail length tape-measure protein; afdb-uniprot50 \| AF-A0A1V0BIB8-F1-MODEL\_V4 \| 1.0 \| 4.327e-26 \| 881 \| 0.227 \| 985 \| 507 \| 19 \| 19 \| 782 \| 28 \| 979 \| Phage tail tape measure protein \| Phage tail tape measure protein; afdb-uniprot50 \| AF-A0A2X4DBT0-F1-MODEL\_V4 \| 1.0 \| 2.684e-25 \| 880 \| 0.234 \| 891 \| 510 \| 22 \| 19 \| 780 \| 64 \| 910 \| Phage-like tail protein \| Phage-like tail protein; afdb-uniprot50 \| AF-A0A606E632-F1-MODEL\_V4 \| 1.0 \| 4.449e-15 \| 879 \| 0.493 \| 243 \| 120 \| 1 \| 115 \| 357 \| 1 \| 240 \| Phage tail tape measure protein \| Phage tail tape measure protein; afdb-uniprot50 \| AF-A0A5M7L7B0-F1-MODEL\_V4 \| 1.0 \| 7.779e-23 \| 878 \| 0.259 \| 689 \| 388 \| 17 \| 87 \| 755 \| 3 \| 588 \| Phage tail tape measure protein \| Phage tail tape measure protein; afdb-uniprot50 \| AF-A0A2U0TFG7-F1-MODEL\_V4 \| 1.0 \| 8.054e-26 \| 874 \| 0.249 \| 763 \| 449 \| 16 \| 7 \| 754 \| 1 \| 654 \| TP901 family phage tail tape measure protein \| TP901 family phage tail tape measure protein; afdb-uniprot50 \| AF-A0A423XU08-F1-MODEL\_V4 \| 1.0 \| 2.16e-23 \| 873 \| 0.257 \| 733 \| 398 \| 20 \| 74 \| 754 \| 2 \| 639 \| Phage tail tape measure protein \| Phage tail tape measure protein; afdb-uniprot50 \| AF-A0A6I2KDC7-F1-MODEL\_V4 \| 1.0 \| 1.23e-18 \| 869 \| 0.301 \| 478 \| 292 \| 10 \| 120 \| 572 \| 4 \| 464 \| Phage tail tape measure protein \| Phage tail tape measure protein; afdb-uniprot50 \| AF-A0A5C4PN41-F1-MODEL\_V4 \| 1.0 \| 1.967e-25 \| 866 \| 0.313 \| 720 \| 416 \| 14 \| 14 \| 702 \| 12 \| 683 \| Phage tail tape measure protein \| Phage tail tape measure protein; afdb-uniprot50 \| AF-A0A1M5CNY5-F1-MODEL\_V4 \| 1.0 \| 2.633e-20 \| 864 \| 0.268 \| 536 \| 345 \| 9 \| 5 \| 508 \| 2 \| 522 \| Phage tail tape measure protein, TP901 family, core region \| Phage tail tape measure protein, TP901 family, core region; afdb-uniprot50 \| AF-A0A0F9TCL8-F1-MODEL\_V4 \| 1.0 \| 8.981e-22 \| 863 \| 0.286 \| 608 \| 342 \| 11 \| 19 \| 558 \| 55 \| 638 \| PhageMin\_Tail domain-containing protein \| PhageMin\_Tail domain-containing protein; afdb-uniprot50 \| AF-A0A0X1U7S0-F1-MODEL\_V4 \| 1.0 \| 1.583e-23 \| 862 \| 0.264 \| 832 \| 393 \| 24 \| 14 \| 776 \| 1 \| 682 \| Phage-related minor tail protein \| Phage-related minor tail protein; afdb-uniprot50 \| AF-A0A259CEH9-F1-MODEL\_V4 \| 1.0 \| 6.38e-26 \| 862 \| 0.271 \| 814 \| 458 \| 15 \| 46 \| 780 \| 3 \| 760 \| Phage tail tape measure protein \| Phage tail tape measure protein; afdb-uniprot50 \| AF-A0A2N9ASI3-F1-MODEL\_V4 \| 1.0 \| 2.686e-15 \| 861 \| 0.229 \| 340 \| 254 \| 3 \| 117 \| 455 \| 138 \| 470 \| PhageMin\_Tail domain-containing protein \| PhageMin\_Tail domain-containing protein; afdb-uniprot50 \| AF-A0A1T4V251-F1-MODEL\_V4 \| 1.0 \| 4.1e-18 \| 859 \| 0.301 \| 451 \| 272 \| 10 \| 91 \| 519 \| 9 \| 438 \| Phage-related minor tail protein \| Phage-related minor tail protein; afdb-uniprot50 \| AF-A0A519ZJF7-F1-MODEL\_V4 \| 1.0 \| 5.215e-22 \| 858 \| 0.226 \| 690 \| 426 \| 16 \| 119 \| 782 \| 38 \| 645 \| Phage tail tape measure protein \| Phage tail tape measure protein; afdb-uniprot50 \| AF-A0A6N8NKN5-F1-MODEL\_V4 \| 1.0 \| 3.272e-22 \| 857 \| 0.311 \| 642 \| 299 \| 13 \| 156 \| 749 \| 2 \| 548 \| Phage tail tape measure protein \| Phage tail tape measure protein; afdb-uniprot50 \| AF-A0A0Q6ZZL4-F1-MODEL\_V4 \| 1.0 \| 7.658e-25 \| 857 \| 0.246 \| 820 \| 489 \| 20 \| 1 \| 780 \| 1 \| 731 \| PhageMin\_Tail domain-containing protein \| PhageMin\_Tail domain-containing protein; afdb-uniprot50 \| AF-A0A254UT32-F1-MODEL\_V4 \| 1.0 \| 6.066e-25 \| 856 \| 0.249 \| 857 \| 452 \| 22 \| 27 \| 785 \| 9 \| 771 \| Phage tail tape measure protein \| Phage tail tape measure protein; afdb-uniprot50 \| AF-V5SJ01-F1-MODEL\_V4 \| 1.0 \| 3.297e-26 \| 854 \| 0.258 \| 836 \| 486 \| 23 \| 13 \| 785 \| 1 \| 765 \| PhageMin\_Tail domain-containing protein \| PhageMin\_Tail domain-containing protein; afdb-uniprot50 \| AF-A0A5F1B8J0-F1-MODEL\_V4 \| 1.0 \| 4.067e-24 \| 852 \| 0.244 \| 844 \| 449 \| 15 \| 6 \| 782 \| 2 \| 723 \| Phage tail tape measure protein \| Phage tail tape measure protein; afdb-uniprot50 \| AF-A0A3P6KAY6-F1-MODEL\_V4 \| 1.0 \| 2.389e-25 \| 852 \| 0.249 \| 818 \| 463 \| 20 \| 66 \| 776 \| 5 \| 778 \| Phage-like protein \| Phage-like protein; afdb-uniprot50 \| AF-S5XQ18-F1-MODEL\_V4 \| 1.0 \| 1.283e-25 \| 852 \| 0.258 \| 877 \| 499 \| 24 \| 1 \| 784 \| 1 \| 818 \| Phage-related tail protein \| Phage-related tail protein; afdb-uniprot50 \| AF-A0A2W5DL54-F1-MODEL\_V4 \| 1.0 \| 7.452e-26 \| 852 \| 0.223 \| 941 \| 515 \| 20 \| 22 \| 776 \| 54 \| 964 \| Phage tail tape measure protein \| Phage tail tape measure protein; afdb-uniprot50 \| AF-A0A378QQC7-F1-MODEL\_V4 \| 1.0 \| 1.16e-23 \| 852 \| 0.301 \| 684 \| 388 \| 16 \| 39 \| 702 \| 2 \| 615 \| Phage-related minor tail protein \| Phage-related minor tail protein; afdb-uniprot50 \| AF-A0A521D146-F1-MODEL\_V4 \| 1.0 \| 5.835e-25 \| 850 \| 0.288 \| 790 \| 454 \| 20 \| 26 \| 759 \| 1 \| 738 \| Phage tail tape measure protein, TP901 family, core region \| Phage tail tape measure protein, TP901 family, core region; afdb-uniprot50 \| AF-A0A1Y3P9U4-F1-MODEL\_V4 \| 1.0 \| 7.281e-24 \| 845 \| 0.263 \| 759 \| 445 \| 20 \| 64 \| 755 \| 2 \| 713 \| PhageMin\_Tail domain-containing protein \| PhageMin\_Tail domain-containing protein; afdb-uniprot50 \| AF-A0A7W6A8D0-F1-MODEL\_V4 \| 1.0 \| 8.604e-25 \| 844 \| 0.241 \| 845 \| 477 \| 24 \| 22 \| 785 \| 53 \| 814 \| TP901 family phage tail tape measure protein \| TP901 family phage tail tape measure protein; afdb-uniprot50 \| AF-A0A3N1J100-F1-MODEL\_V4 \| 1.0 \| 9.446e-23 \| 840 \| 0.252 \| 757 \| 390 \| 22 \| 32 \| 754 \| 5 \| 619 \| TP901 family phage tail tape measure protein \| TP901 family phage tail tape measure protein; afdb-uniprot50 \| AF-A0A1I4C772-F1-MODEL\_V4 \| 1.0 \| 3.016e-25 \| 840 \| 0.253 \| 895 \| 476 \| 22 \| 1 \| 785 \| 1 \| 812 \| Phage tail tape measure protein, TP901 family, core region \| Phage tail tape measure protein, TP901 family, core region; afdb-uniprot50 \| AF-A0A4P8FYQ0-F1-MODEL\_V4 \| 1.0 \| 3.299e-16 \| 838 \| 0.363 \| 333 \| 205 \| 1 \| 155 \| 480 \| 3 \| 335 \| Phage tail tape measure protein \| Phage tail tape measure protein; afdb-uniprot50 \| AF-A0A2K2G0G8-F1-MODEL\_V4 \| 1.0 \| 1.684e-25 \| 833 \| 0.274 \| 849 \| 444 \| 27 \| 19 \| 782 \| 58 \| 819 \| Phage tail tape measure protein \| Phage tail tape measure protein; afdb-uniprot50 \| AF-A0A5C4LLJ5-F1-MODEL\_V4 \| 1.0 \| 1.894e-15 \| 831 \| 0.231 \| 371 \| 259 \| 5 \| 109 \| 461 \| 132 \| 494 \| Phage tail tape measure protein \| Phage tail tape measure protein; afdb-uniprot50 \| AF-A0A0Q2UDJ2-F1-MODEL\_V4 \| 1.0 \| 9.086e-23 \| 831 \| 0.231 \| 854 \| 488 \| 20 \| 37 \| 773 \| 2 \| 803 \| PhageMin\_Tail domain-containing protein \| PhageMin\_Tail domain-containing protein; afdb-uniprot50 \| AF-A0A1V3YRW6-F1-MODEL\_V4 \| 1.0 \| 1.535e-17 \| 826 \| 0.245 \| 432 \| 276 \| 8 \| 61 \| 457 \| 3 \| 419 \| Phage tail tape measure protein \| Phage tail tape measure protein; afdb-uniprot50 \| AF-A0A212KXI4-F1-MODEL\_V4 \| 1.0 \| 2.758e-24 \| 824 \| 0.258 \| 865 \| 432 \| 24 \| 1 \| 775 \| 1 \| 745 \| PhageMin\_Tail domain-containing protein \| PhageMin\_Tail domain-containing protein; afdb-uniprot50 \| AF-A0A6F9MD22-F1-MODEL\_V4 \| 1.0 \| 1.799e-24 \| 823 \| 0.244 \| 776 \| 491 \| 18 \| 55 \| 784 \| 13 \| 738 \| Phage tail tape measure protein \| Phage tail tape measure protein; afdb-uniprot50 \| AF-B9D1S6-F1-MODEL\_V4 \| 1.0 \| 5.613e-25 \| 820 \| 0.253 \| 881 \| 484 \| 25 \| 1 \| 764 \| 1 \| 824 \| Phage tail tape measure protein, TP901 family \| Phage tail tape measure protein, TP901 family; afdb-uniprot50 \| AF-A0A6P0VE51-F1-MODEL\_V4 \| 1.0 \| 1.061e-22 \| 819 \| 0.235 \| 753 \| 467 \| 16 \| 71 \| 782 \| 2 \| 686 \| Phage tail tape measure protein \| Phage tail tape measure protein; afdb-uniprot50 \| AF-A0A1W1Z477-F1-MODEL\_V4 \| 1.0 \| 5.613e-25 \| 819 \| 0.246 \| 901 \| 463 \| 26 \| 36 \| 783 \| 3 \| 840 \| Phage tail tape measure protein, TP901 family, core region \| Phage tail tape measure protein, TP901 family, core region; afdb-uniprot50 \| AF-A0A285V0T5-F1-MODEL\_V4 \| 1.0 \| 1.22e-24 \| 819 \| 0.243 \| 787 \| 460 \| 20 \| 1 \| 713 \| 1 \| 725 \| TP901 family phage tail tape measure protein \| TP901 family phage tail tape measure protein; afdb-uniprot50 \| AF-B3R3J6-F1-MODEL\_V4 \| 1.0 \| 2.271e-24 \| 815 \| 0.203 \| 926 \| 557 \| 26 \| 19 \| 782 \| 59 \| 965 \| Bacteriophage P2 tail protein GPT putative tail length determinator \| Bacteriophage P2 tail protein GPT putative tail length determinator; afdb-uniprot50 \| AF-B6IMG4-F1-MODEL\_V4 \| 1.0 \| 5.613e-25 \| 811 \| 0.235 \| 914 \| 486 \| 26 \| 63 \| 782 \| 2 \| 896 \| Phage tail tape measure protein, TP901 family, core region \| Phage tail tape measure protein, TP901 family, core region; afdb-uniprot50 \| AF-A0A017HCD3-F1-MODEL\_V4 \| 1.0 \| 2.867e-24 \| 808 \| 0.22 \| 900 \| 502 \| 23 \| 1 \| 782 \| 1 \| 818 \| Phage tail length tape-measure protein \| Phage tail length tape-measure protein; afdb-uniprot50 \| AF-A0A285P0E0-F1-MODEL\_V4 \| 1.0 \| 2.769e-21 \| 807 \| 0.268 \| 718 \| 381 \| 19 \| 71 \| 750 \| 2 \| 612 \| Phage tail tape measure protein, TP901 family, core region \| Phage tail tape measure protein, TP901 family, core region; afdb-uniprot50 \| AF-A0A836WW58-F1-MODEL\_V4 \| 1.0 \| 1.108e-19 \| 805 \| 0.294 \| 605 \| 339 \| 16 \| 36 \| 613 \| 4 \| 547 \| Phage tail tape measure protein \| Phage tail tape measure protein; afdb-uniprot50 \| AF-A0A1E3VZJ0-F1-MODEL\_V4 \| 1.0 \| 3.899e-17 \| 802 \| 0.243 \| 469 \| 326 \| 10 \| 63 \| 519 \| 2 \| 453 \| PhageMin\_Tail domain-containing protein \| PhageMin\_Tail domain-containing protein; afdb-uniprot50 \| AF-A0A378QE64-F1-MODEL\_V4 \| 1.0 \| 7.57e-24 \| 802 \| 0.262 \| 808 \| 427 \| 19 \| 39 \| 779 \| 2 \| 707 \| Phage-related minor tail protein \| Phage-related minor tail protein; afdb-uniprot50 \| AF-A0A1G8HWZ8-F1-MODEL\_V4 \| 1.0 \| 2.177e-17 \| 801 \| 0.331 \| 456 \| 274 \| 7 \| 42 \| 480 \| 168 \| 609 \| Phage tail tape measure protein, TP901 family, core region \| Phage tail tape measure protein, TP901 family, core region; afdb-uniprot50 \| AF-A0A7C3GEH9-F1-MODEL\_V4 \| 1.0 \| 7.311e-21 \| 800 \| 0.246 \| 642 \| 377 \| 18 \| 29 \| 655 \| 2 \| 551 \| Phage tail tape measure protein \| Phage tail tape measure protein; afdb-uniprot50 \| AF-A0A4R2D627-F1-MODEL\_V4 \| 1.0 \| 3.763e-24 \| 799 \| 0.251 \| 792 \| 432 \| 21 \| 3 \| 726 \| 1 \| 699 \| TP901 family phage tail tape measure protein \| TP901 family phage tail tape measure protein; afdb-uniprot50 \| AF-A0A8B2QSJ7-F1-MODEL\_V4 \| 1.0 \| 3.867e-23 \| 795 \| 0.228 \| 792 \| 484 \| 21 \| 19 \| 776 \| 12 \| 710 \| Phage tail tape measure protein \| Phage tail tape measure protein; afdb-uniprot50 \| AF-A0A386UJJ2-F1-MODEL\_V4 \| 1.0 \| 4.067e-24 \| 788 \| 0.234 \| 914 \| 483 \| 27 \| 1 \| 782 \| 1 \| 829 \| Phage tail tape measure protein \| Phage tail tape measure protein; afdb-uniprot50 \| AF-A0A6G5QFI2-F1-MODEL\_V4 \| 1.0 \| 2.427e-23 \| 784 \| 0.246 \| 817 \| 503 \| 21 \| 1 \| 764 \| 1 \| 757 \| Phage tail tape measure protein, TP901 family \| Phage tail tape measure protein, TP901 family; afdb-uniprot50 \| AF-A0A074U4R1-F1-MODEL\_V4 \| 1.0 \| 1.665e-24 \| 783 \| 0.238 \| 992 \| 495 \| 29 \| 19 \| 780 \| 51 \| 1011 \| Tail protein \| Tail protein; afdb-uniprot50 \| AF-A0A753E0J5-F1-MODEL\_V4 \| 1.0 \| 7.119e-12 \| 782 \| 0.505 \| 180 \| 89 \| 0 \| 220 \| 399 \| 1 \| 180 \| Phage tail tape measure protein \| Phage tail tape measure protein; afdb-uniprot50 \| AF-A0A135IJM6-F1-MODEL\_V4 \| 1.0 \| 2.16e-23 \| 781 \| 0.212 \| 965 \| 506 \| 19 \| 46 \| 784 \| 9 \| 945 \| PhageMin\_Tail domain-containing protein \| PhageMin\_Tail domain-containing protein; afdb-uniprot50 \| AF-A0A1H8A1R0-F1-MODEL\_V4 \| 1.0 \| 4.228e-24 \| 781 \| 0.211 \| 1063 \| 535 \| 26 \| 19 \| 779 \| 38 \| 1098 \| Phage tail tape measure protein, TP901 family, core region \| Phage tail tape measure protein, TP901 family, core region; afdb-uniprot50 \| AF-A0A2P5N6P9-F1-MODEL\_V4 \| 1.0 \| 2.738e-20 \| 780 \| 0.218 \| 772 \| 429 \| 19 \| 25 \| 780 \| 2 \| 614 \| Phage tail tape measure protein \| Phage tail tape measure protein; afdb-uniprot50 \| AF-A0A1E3G7C9-F1-MODEL\_V4 \| 1.0 \| 1.922e-23 \| 780 \| 0.24 \| 820 \| 439 \| 17 \| 22 \| 785 \| 53 \| 744 \| Phage tail tape measure protein \| Phage tail tape measure protein; afdb-uniprot50 \| AF-A0A5C7E451-F1-MODEL\_V4 \| 1.0 \| 1.021e-22 \| 779 \| 0.212 \| 775 \| 511 \| 19 \| 39 \| 779 \| 21 \| 729 \| Phage tail tape measure protein \| Phage tail tape measure protein; afdb-uniprot50 \| AF-A0A268TUE9-F1-MODEL\_V4 \| 1.0 \| 1.254e-23 \| 779 \| 0.243 \| 818 \| 498 \| 22 \| 6 \| 764 \| 1 \| 756 \| Phage tail tape measure protein \| Phage tail tape measure protein; afdb-uniprot50 \| AF-A0A239C875-F1-MODEL\_V4 \| 1.0 \| 8.407e-23 \| 778 \| 0.267 \| 804 \| 464 \| 19 \| 27 \| 779 \| 4 \| 733 \| Phage tail tape measure protein, TP901 family, core region \| Phage tail tape measure protein, TP901 family, core region; afdb-uniprot50 \| AF-A0A1M4WE74-F1-MODEL\_V4 \| 1.0 \| 5.702e-23 \| 777 \| 0.245 \| 876 \| 468 \| 21 \| 13 \| 782 \| 1 \| 789 \| Phage tail tape measure protein, TP901 family, core region \| Phage tail tape measure protein, TP901 family, core region; afdb-uniprot50 \| AF-A0A2P8QYQ4-F1-MODEL\_V4 \| 1.0 \| 1.061e-22 \| 776 \| 0.228 \| 822 \| 485 \| 20 \| 16 \| 780 \| 1 \| 729 \| Phage tail tape measure protein \| Phage tail tape measure protein; afdb-uniprot50 \| AF-A0A845ZTZ3-F1-MODEL\_V4 \| 1.0 \| 9.086e-23 \| 773 \| 0.243 \| 794 \| 478 \| 27 \| 63 \| 781 \| 2 \| 747 \| Phage tail tape measure protein \| Phage tail tape measure protein; afdb-uniprot50 \| AF-A0A349GR42-F1-MODEL\_V4 \| 1.0 \| 1.183e-18 \| 772 \| 0.274 \| 532 \| 307 \| 9 \| 5 \| 461 \| 2 \| 529 \| Phage tail tape measure protein \| Phage tail tape measure protein; afdb-uniprot50 \| AF-A0A0U1DA91-F1-MODEL\_V4 \| 1.0 \| 1.454e-19 \| 771 \| 0.251 \| 695 \| 354 \| 12 \| 132 \| 785 \| 2 \| 570 \| Phage tail tape measure protein \| Phage tail tape measure protein; afdb-uniprot50 \| AF-A0A3P4AP62-F1-MODEL\_V4 \| 1.0 \| 2.623e-23 \| 771 \| 0.242 \| 828 \| 460 \| 18 \| 81 \| 762 \| 25 \| 830 \| PhageMin\_Tail domain-containing protein \| PhageMin\_Tail domain-containing protein; afdb-uniprot50 \| AF-W4SK08-F1-MODEL\_V4 \| 1.0 \| 4.789e-18 \| 769 \| 0.359 \| 414 \| 200 \| 7 \| 19 \| 375 \| 49 \| 454 \| Phage-related tail protein \| Phage-related tail protein; afdb-uniprot50 \| AF-A0A3B0IZ82-F1-MODEL\_V4 \| 1.0 \| 4.734e-17 \| 766 \| 0.289 \| 439 \| 258 \| 7 \| 19 \| 411 \| 39 \| 469 \| PhageMin\_Tail domain-containing protein \| PhageMin\_Tail domain-containing protein; afdb-uniprot50 \| AF-G1UYT9-F1-MODEL\_V4 \| 1.0 \| 7.341e-18 \| 766 \| 0.317 \| 539 \| 311 \| 12 \| 18 \| 516 \| 184 \| 705 \| PhageMin\_Tail domain-containing protein \| PhageMin\_Tail domain-containing protein; afdb-uniprot50 \| AF-A0A4S3L0X3-F1-MODEL\_V4 \| 1.0 \| 1.608e-21 \| 766 \| 0.217 \| 763 \| 484 \| 15 \| 18 \| 708 \| 18 \| 739 \| Phage tail tape measure protein \| Phage tail tape measure protein; afdb-uniprot50 \| AF-A0A7C5VJK5-F1-MODEL\_V4 \| 1.0 \| 9.707e-22 \| 765 \| 0.258 \| 703 \| 425 \| 14 \| 62 \| 690 \| 4 \| 683 \| Phage tail tape measure protein \| Phage tail tape measure protein; afdb-uniprot50 \| AF-N6VPY0-F1-MODEL\_V4 \| 1.0 \| 1.773e-16 \| 764 \| 0.28 \| 453 \| 305 \| 7 \| 39 \| 487 \| 4 \| 439 \| Phage tail protein \| Phage tail protein; afdb-uniprot50 \| AF-A0A327Q7T9-F1-MODEL\_V4 \| 1.0 \| 3.928e-21 \| 762 \| 0.208 \| 768 \| 471 \| 22 \| 47 \| 780 \| 4 \| 668 \| TP901 family phage tail tape measure protein \| TP901 family phage tail tape measure protein; afdb-uniprot50 \| AF-A0A2Z4PV63-F1-MODEL\_V4 \| 1.0 \| 1.087e-14 \| 761 \| 0.328 \| 317 \| 209 \| 1 \| 160 \| 472 \| 5 \| 321 \| Phage tail tape measure protein \| Phage tail tape measure protein; afdb-uniprot50 \| AF-A0A161V1N0-F1-MODEL\_V4 \| 1.0 \| 5.859e-22 \| 758 \| 0.224 \| 875 \| 470 \| 22 \| 1 \| 782 \| 1 \| 759 \| Phage-related minor tail protein \| Phage-related minor tail protein; afdb-uniprot50 \| AF-A0A1B9MVQ4-F1-MODEL\_V4 \| 1.0 \| 9.446e-23 \| 757 \| 0.282 \| 764 \| 421 \| 22 \| 79 \| 756 \| 6 \| 727 \| Phage tail tape measure protein \| Phage tail tape measure protein; afdb-uniprot50 \| AF-A0A807AA81-F1-MODEL\_V4 \| 1.0 \| 7.869e-24 \| 757 \| 0.207 \| 905 \| 544 \| 23 \| 19 \| 781 \| 50 \| 922 \| Phage tail tape measure protein \| Phage tail tape measure protein; afdb-uniprot50 \| AF-A0A751FIF2-F1-MODEL\_V4 \| 1.0 \| 2.381e-18 \| 756 \| 0.322 \| 483 \| 242 \| 8 \| 193 \| 675 \| 2 \| 399 \| Phage tail tape measure protein \| Phage tail tape measure protein; afdb-uniprot50 \| AF-A0A6L7FWV9-F1-MODEL\_V4 \| 1.0 \| 1.711e-23 \| 756 \| 0.249 \| 858 \| 457 \| 27 \| 20 \| 785 \| 46 \| 808 \| Phage tail tape measure protein \| Phage tail tape measure protein; afdb-uniprot50 \| AF-A0A0K1NEY0-F1-MODEL\_V4 \| 1.0 \| 5.421e-22 \| 754 \| 0.232 \| 847 \| 532 \| 22 \| 22 \| 781 \| 53 \| 868 \| PhageMin\_Tail domain-containing protein \| PhageMin\_Tail domain-containing protein; afdb-uniprot50 \| AF-A0A7T5EN76-F1-MODEL\_V4 \| 1.0 \| 1.953e-21 \| 753 \| 0.238 \| 797 \| 470 \| 22 \| 18 \| 776 \| 18 \| 715 \| Phage tail tape measure protein \| Phage tail tape measure protein; afdb-uniprot50 \| AF-A0A2N3KSG5-F1-MODEL\_V4 \| 1.0 \| 2.494e-22 \| 752 \| 0.221 \| 809 \| 471 \| 14 \| 19 \| 772 \| 51 \| 755 \| Phage tail tape measure protein \| Phage tail tape measure protein; afdb-uniprot50 \| AF-A0A7M1L7G1-F1-MODEL\_V4 \| 1.0 \| 9.086e-23 \| 752 \| 0.22 \| 917 \| 523 \| 28 \| 19 \| 782 \| 35 \| 912 \| Phage tail tape measure protein \| Phage tail tape measure protein; afdb-uniprot50 \| AF-A0A2N9Y986-F1-MODEL\_V4 \| 1.0 \| 3.137e-15 \| 748 \| 0.254 \| 432 \| 275 \| 7 \| 16 \| 440 \| 1 \| 392 \| Phage tail tape measure protein \| Phage tail tape measure protein; afdb-uniprot50 \| AF-A0A1J5HRD0-F1-MODEL\_V4 \| 1.0 \| 4.295e-22 \| 748 \| 0.219 \| 913 \| 480 \| 25 \| 2 \| 783 \| 1 \| 811 \| Phage tail tape measure protein \| Phage tail tape measure protein; afdb-uniprot50 \| AF-A0A1T0CPH3-F1-MODEL\_V4 \| 1.0 \| 1.393e-22 \| 746 \| 0.24 \| 879 \| 485 \| 18 \| 36 \| 777 \| 2 \| 834 \| Phage tail tape measure protein \| Phage tail tape measure protein; afdb-uniprot50 \| AF-A0A6Y5D9Q6-F1-MODEL\_V4 \| 1.0 \| 7.999e-12 \| 745 \| 0.51 \| 182 \| 89 \| 0 \| 187 \| 368 \| 1 \| 182 \| Phage tail tape measure protein \| Phage tail tape measure protein; afdb-uniprot50 \| AF-A0A7Y8PBA4-F1-MODEL\_V4 \| 1.0 \| 2.334e-23 \| 745 \| 0.242 \| 808 \| 530 \| 21 \| 1 \| 780 \| 1 \| 754 \| Phage tail tape measure protein \| Phage tail tape measure protein; afdb-uniprot50 \| AF-G1USF9-F1-MODEL\_V4 \| 1.0 \| 1.289e-22 \| 744 \| 0.216 \| 863 \| 487 \| 21 \| 2 \| 777 \| 1 \| 760 \| PhageMin\_Tail domain-containing protein \| PhageMin\_Tail domain-containing protein; afdb-uniprot50 \| AF-A0A2E6Y2Q7-F1-MODEL\_V4 \| 1.0 \| 5.075e-23 \| 744 \| 0.228 \| 872 \| 526 \| 25 \| 13 \| 784 \| 2 \| 826 \| Phage tail tape measure protein \| Phage tail tape measure protein; afdb-uniprot50 \| AF-A0A076LFK6-F1-MODEL\_V4 \| 1.0 \| 2.047e-15 \| 743 \| 0.437 \| 318 \| 159 \| 6 \| 76 \| 383 \| 6 \| 313 \| Phage tail length tape-measure protein \| Phage tail length tape-measure protein; afdb-uniprot50 \| AF-A0A7W8HL33-F1-MODEL\_V4 \| 1.0 \| 5.155e-21 \| 741 \| 0.228 \| 789 \| 447 \| 24 \| 64 \| 783 \| 6 \| 701 \| TP901 family phage tail tape measure protein \| TP901 family phage tail tape measure protein; afdb-uniprot50 \| AF-L0R7N5-F1-MODEL\_V4 \| 1.0 \| 1.324e-21 \| 736 \| 0.241 \| 849 \| 453 \| 23 \| 15 \| 782 \| 14 \| 752 \| PhageMin\_Tail domain-containing protein \| PhageMin\_Tail domain-containing protein; afdb-uniprot50 \| AF-A0A6M3KKZ6-F1-MODEL\_V4 \| 1.0 \| 2.563e-21 \| 736 \| 0.197 \| 881 \| 533 \| 22 \| 19 \| 782 \| 34 \| 857 \| Putative tail protein \| Putative tail protein; afdb-uniprot50 \| AF-A0A6L5WL94-F1-MODEL\_V4 \| 1.0 \| 1.23e-18 \| 735 \| 0.251 \| 613 \| 413 \| 12 \| 25 \| 628 \| 4 \| 579 \| Phage tail tape measure protein \| Phage tail tape measure protein; afdb-uniprot50 \| AF-A0A0C1HF59-F1-MODEL\_V4 \| 1.0 \| 2.993e-21 \| 734 \| 0.248 \| 769 \| 444 \| 23 \| 80 \| 780 \| 11 \| 713 \| Phage tail tape measure protein, TP901 family \| Phage tail tape measure protein, TP901 family; afdb-uniprot50 \| AF-A0A840X1L3-F1-MODEL\_V4 \| 1.0 \| 5.636e-22 \| 734 \| 0.264 \| 799 \| 392 \| 24 \| 15 \| 712 \| 27 \| 730 \| TP901 family phage tail tape measure protein \| TP901 family phage tail tape measure protein; afdb-uniprot50 \| AF-A0A753E616-F1-MODEL\_V4 \| 1.0 \| 6.485e-14 \| 732 \| 0.43 \| 260 \| 140 \| 1 \| 220 \| 471 \| 1 \| 260 \| Phage tail tape measure protein \| Phage tail tape measure protein; afdb-uniprot50 \| AF-A0A7G5EHY0-F1-MODEL\_V4 \| 1.0 \| 5.016e-22 \| 732 \| 0.213 \| 874 \| 499 \| 23 \| 19 \| 782 \| 47 \| 841 \| Phage tail tape measure protein \| Phage tail tape measure protein; afdb-uniprot50 \| AF-A0A2T3NIG3-F1-MODEL\_V4 \| 1.0 \| 1.202e-16 \| 732 \| 0.298 \| 510 \| 305 \| 12 \| 42 \| 519 \| 167 \| 655 \| Phage tail tape measure protein \| Phage tail tape measure protein; afdb-uniprot50 \| AF-A0A660MJK7-F1-MODEL\_V4 \| 1.0 \| 2.238e-16 \| 730 \| 0.254 \| 503 \| 289 \| 7 \| 19 \| 440 \| 69 \| 566 \| Phage tail tape measure protein \| Phage tail tape measure protein; afdb-uniprot50 \| AF-A7MW86-F1-MODEL\_V4 \| 1.0 \| 7.544e-17 \| 728 \| 0.326 \| 499 \| 306 \| 7 \| 42 \| 519 \| 168 \| 657 \| PhageMin\_Tail domain-containing protein \| PhageMin\_Tail domain-containing protein; afdb-uniprot50 \| AF-A0A4P5QGR5-F1-MODEL\_V4 \| 1.0 \| 2.264e-17 \| 728 \| 0.24 \| 598 \| 346 \| 9 \| 19 \| 513 \| 60 \| 652 \| Phage tail tape measure protein \| Phage tail tape measure protein; afdb-uniprot50 \| AF-A0A0U2B0N6-F1-MODEL\_V4 \| 1.0 \| 5.215e-22 \| 728 \| 0.235 \| 837 \| 465 \| 22 \| 1 \| 712 \| 1 \| 787 \| PhageMin\_Tail domain-containing protein \| PhageMin\_Tail domain-containing protein; afdb-uniprot50 \| AF-A0A1E3XJQ2-F1-MODEL\_V4 \| 1.0 \| 9.123e-20 \| 727 \| 0.252 \| 718 \| 397 \| 20 \| 16 \| 678 \| 1 \| 633 \| Phage tail tape measure protein \| Phage tail tape measure protein; afdb-uniprot50 \| AF-A0A1I5ZGS7-F1-MODEL\_V4 \| 1.0 \| 2.03e-21 \| 725 \| 0.255 \| 762 \| 421 \| 20 \| 36 \| 733 \| 5 \| 683 \| Phage tail tape measure protein, TP901 family, core region \| Phage tail tape measure protein, TP901 family, core region; afdb-uniprot50 \| AF-A0A369YAP6-F1-MODEL\_V4 \| 1.0 \| 6.41e-13 \| 723 \| 0.329 \| 282 \| 166 \| 6 \| 122 \| 392 \| 73 \| 342 \| Phage tail tape measure protein \| Phage tail tape measure protein; afdb-uniprot50 \| AF-A0A0N0ZNR6-F1-MODEL\_V4 \| 1.0 \| 5.859e-22 \| 723 \| 0.216 \| 1000 \| 498 \| 25 \| 36 \| 783 \| 2 \| 967 \| PhageMin\_Tail domain-containing protein \| PhageMin\_Tail domain-containing protein; afdb-uniprot50 \| AF-E5Y5W4-F1-MODEL\_V4 \| 1.0 \| 5.215e-22 \| 722 \| 0.259 \| 827 \| 452 \| 23 \| 19 \| 743 \| 36 \| 803 \| Phage tail tape measure protein, TP901 family, core region \| Phage tail tape measure protein, TP901 family, core region; afdb-uniprot50 \| AF-A0A0F8XZV2-F1-MODEL\_V4 \| 1.0 \| 1.946e-14 \| 719 \| 0.236 \| 422 \| 285 \| 7 \| 46 \| 464 \| 2 \| 389 \| PhageMin\_Tail domain-containing protein \| PhageMin\_Tail domain-containing protein; afdb-uniprot50 \| AF-A0A2W5GET3-F1-MODEL\_V4 \| 1.0 \| 4.535e-20 \| 718 \| 0.198 \| 730 \| 369 \| 17 \| 110 \| 784 \| 35 \| 603 \| Phage tail tape measure protein \| Phage tail tape measure protein; afdb-uniprot50 \| AF-A0A504UBL4-F1-MODEL\_V4 \| 1.0 \| 2.281e-21 \| 718 \| 0.196 \| 860 \| 499 \| 26 \| 18 \| 785 \| 4 \| 763 \| Phage tail tape measure protein \| Phage tail tape measure protein; afdb-uniprot50 \| AF-A0A5S9Q3L2-F1-MODEL\_V4 \| 1.0 \| 6.844e-22 \| 718 \| 0.216 \| 827 \| 538 \| 17 \| 19 \| 782 \| 14 \| 793 \| PhageMin\_Tail domain-containing protein \| PhageMin\_Tail domain-containing protein; afdb-uniprot50 \| AF-A0A2C6C9V6-F1-MODEL\_V4 \| 1.0 \| 2.494e-22 \| 718 \| 0.198 \| 966 \| 567 \| 26 \| 1 \| 778 \| 5 \| 950 \| Phage tail tape measure protein \| Phage tail tape measure protein; afdb-uniprot50 \| AF-A0A0D0J8M0-F1-MODEL\_V4 \| 1.0 \| 2.169e-20 \| 714 \| 0.237 \| 788 \| 445 \| 19 \| 62 \| 780 \| 13 \| 713 \| Phage tail tape measure protein, family \| Phage tail tape measure protein, family; afdb-uniprot50 \| AF-A0A4Q8ME49-F1-MODEL\_V4 \| 1.0 \| 1.529e-20 \| 712 \| 0.237 \| 807 \| 446 \| 26 \| 47 \| 780 \| 9 \| 718 \| Phage tail tape measure protein \| Phage tail tape measure protein; afdb-uniprot50 \| AF-A0A7X6JUT3-F1-MODEL\_V4 \| 1.0 \| 1.738e-21 \| 711 \| 0.205 \| 1002 \| 504 \| 27 \| 19 \| 780 \| 27 \| 975 \| Phage tail tape measure protein \| Phage tail tape measure protein; afdb-uniprot50 \| AF-A0A5C8WLB6-F1-MODEL\_V4 \| 1.0 \| 1.188e-15 \| 709 \| 0.217 \| 474 \| 295 \| 9 \| 27 \| 444 \| 3 \| 456 \| Phage tail tape measure protein \| Phage tail tape measure protein; afdb-uniprot50 \| AF-B7AAN0-F1-MODEL\_V4 \| 1.0 \| 2.664e-21 \| 705 \| 0.224 \| 932 \| 480 \| 18 \| 39 \| 754 \| 4 \| 908 \| Phage tail tape measure protein, TP901 family \| Phage tail tape measure protein, TP901 family; afdb-uniprot50 \| AF-A0A7X6NRZ0-F1-MODEL\_V4 \| 1.0 \| 3.444e-13 \| 704 \| 0.228 \| 342 \| 241 \| 6 \| 117 \| 443 \| 6 \| 339 \| Phage tail tape measure protein \| Phage tail tape measure protein; afdb-uniprot50 \| AF-A0A5P8MUD0-F1-MODEL\_V4 \| 1.0 \| 3.99e-19 \| 704 \| 0.21 \| 808 \| 455 \| 21 \| 31 \| 785 \| 13 \| 690 \| Phage tail tape measure protein \| Phage tail tape measure protein; afdb-uniprot50 \| AF-B9NP85-F1-MODEL\_V4 \| 1.0 \| 3.928e-21 \| 704 \| 0.209 \| 917 \| 470 \| 27 \| 19 \| 785 \| 51 \| 862 \| Tail tape measure protein \| Tail tape measure protein; afdb-uniprot50 \| AF-A0A316GV47-F1-MODEL\_V4 \| 1.0 \| 3.076e-20 \| 703 \| 0.219 \| 841 \| 470 \| 23 \| 1 \| 785 \| 1 \| 710 \| TP901 family phage tail tape measure protein \| TP901 family phage tail tape measure protein; afdb-uniprot50 \| AF-A0A806J4A9-F1-MODEL\_V4 \| 1.0 \| 4.958e-21 \| 703 \| 0.193 \| 862 \| 548 \| 25 \| 28 \| 781 \| 5 \| 826 \| Bacteriophage P2 tail protein GPT putative tail length determinator \| Bacteriophage P2 tail protein GPT putative tail length determinator; afdb-uniprot50 \| AF-A0A3D3B5Z8-F1-MODEL\_V4 \| 1.0 \| 3.112e-21 \| 702 \| 0.22 \| 757 \| 480 \| 17 \| 16 \| 714 \| 1 \| 705 \| Phage tail tape measure protein \| Phage tail tape measure protein; afdb-uniprot50 \| AF-A0A840BWZ4-F1-MODEL\_V4 \| 1.0 \| 9.055e-16 \| 697 \| 0.23 \| 499 \| 317 \| 10 \| 27 \| 467 \| 6 \| 495 \| TP901 family phage tail tape measure protein \| TP901 family phage tail tape measure protein; afdb-uniprot50 \| AF-E3H9B1-F1-MODEL\_V4 \| 1.0 \| 1.511e-19 \| 697 \| 0.235 \| 714 \| 407 \| 17 \| 1 \| 633 \| 1 \| 656 \| Phage tail tape measure protein, TP901 family \| Phage tail tape measure protein, TP901 family; afdb-uniprot50 \| AF-A0A0H3ZYV9-F1-MODEL\_V4 \| 1.0 \| 2.655e-14 \| 696 \| 0.226 \| 445 \| 308 \| 10 \| 98 \| 519 \| 5 \| 436 \| Phage tail length tape-measure protein \| Phage tail length tape-measure protein; afdb-uniprot50 \| AF-A0A1H9HD03-F1-MODEL\_V4 \| 1.0 \| 1.121e-20 \| 692 \| 0.221 \| 823 \| 503 \| 23 \| 34 \| 784 \| 6 \| 762 \| Phage tail tape measure protein, TP901 family, core region \| Phage tail tape measure protein, TP901 family, core region; afdb-uniprot50 \| AF-A0A855G9Q1-F1-MODEL\_V4 \| 1.0 \| 7.033e-21 \| 692 \| 0.216 \| 837 \| 420 \| 20 \| 1 \| 785 \| 1 \| 653 \| Phage tail tape measure protein \| Phage tail tape measure protein; afdb-uniprot50 \| AF-A0A660MG03-F1-MODEL\_V4 \| 1.0 \| 1.765e-19 \| 692 \| 0.237 \| 694 \| 431 \| 13 \| 40 \| 702 \| 60 \| 685 \| Phage tail tape measure protein \| Phage tail tape measure protein; afdb-uniprot50 \| AF-A0A6B8RHZ8-F1-MODEL\_V4 \| 1.0 \| 3.456e-20 \| 688 \| 0.206 \| 769 \| 470 \| 24 \| 19 \| 713 \| 15 \| 716 \| Phage tail tape measure protein \| Phage tail tape measure protein; afdb-uniprot50 \| AF-A0A4R2MR69-F1-MODEL\_V4 \| 1.0 \| 2.633e-20 \| 688 \| 0.226 \| 844 \| 506 \| 21 \| 19 \| 784 \| 28 \| 802 \| TP901 family phage tail tape measure protein \| TP901 family phage tail tape measure protein; afdb-uniprot50 \| AF-A0A315BEH9-F1-MODEL\_V4 \| 1.0 \| 8.878e-21 \| 686 \| 0.192 \| 836 \| 526 \| 20 \| 19 \| 755 \| 45 \| 830 \| Phage tail tape measure protein \| Phage tail tape measure protein; afdb-uniprot50 \| AF-A0A1C3E538-F1-MODEL\_V4 \| 1.0 \| 6.384e-16 \| 685 \| 0.304 \| 542 \| 289 \| 12 \| 19 \| 477 \| 76 \| 612 \| Phage tail tape measure protein \| Phage tail tape measure protein; afdb-uniprot50 \| AF-A0A369Y1L1-F1-MODEL\_V4 \| 1.0 \| 2.846e-20 \| 685 \| 0.209 \| 853 \| 464 \| 25 \| 19 \| 784 \| 77 \| 805 \| Phage tail tape measure protein \| Phage tail tape measure protein; afdb-uniprot50 \| AF-A0A2G6CD52-F1-MODEL\_V4 \| 1.0 \| 1.454e-19 \| 683 \| 0.229 \| 786 \| 418 \| 23 \| 47 \| 785 \| 10 \| 654 \| Phage tail tape measure protein \| Phage tail tape measure protein; afdb-uniprot50 \| AF-A0A0F8XNY0-F1-MODEL\_V4 \| 1.0 \| 4.753e-14 \| 682 \| 0.22 \| 404 \| 281 \| 10 \| 126 \| 519 \| 2 \| 381 \| PhageMin\_Tail domain-containing protein \| PhageMin\_Tail domain-containing protein; afdb-uniprot50 \| AF-A0A1Q6PXA2-F1-MODEL\_V4 \| 1.0 \| 1.108e-19 \| 677 \| 0.223 \| 702 \| 422 \| 18 \| 79 \| 708 \| 21 \| 671 \| Phage tail tape measure protein \| Phage tail tape measure protein; afdb-uniprot50 \| AF-A0A379CMS6-F1-MODEL\_V4 \| 1.0 \| 3.607e-17 \| 674 \| 0.265 \| 598 \| 327 \| 12 \| 27 \| 538 \| 113 \| 683 \| Phage tail tape measure protein, TP901 family, core region \| Phage tail tape measure protein, TP901 family, core region; afdb-uniprot50 \| AF-A0A635X8N3-F1-MODEL\_V4 \| 1.0 \| 1.477e-17 \| 673 \| 0.276 \| 567 \| 297 \| 11 \| 214 \| 755 \| 2 \| 479 \| Phage tail tape measure protein \| Phage tail tape measure protein; afdb-uniprot50 \| AF-A0A2G6FTD7-F1-MODEL\_V4 \| 1.0 \| 9.86e-20 \| 672 \| 0.202 \| 787 \| 511 \| 15 \| 26 \| 785 \| 1 \| 697 \| Phage tail tape measure protein \| Phage tail tape measure protein; afdb-uniprot50 \| AF-A0A354YUM2-F1-MODEL\_V4 \| 1.0 \| 2.055e-12 \| 671 \| 0.228 \| 315 \| 229 \| 6 \| 109 \| 416 \| 31 \| 338 \| Phage tail tape measure protein \| Phage tail tape measure protein; afdb-uniprot50 \| AF-A0A4R5HEK1-F1-MODEL\_V4 \| 1.0 \| 3.299e-16 \| 671 \| 0.252 \| 595 \| 347 \| 9 \| 19 \| 538 \| 77 \| 648 \| Phage tail tape measure protein \| Phage tail tape measure protein; afdb-uniprot50 \| AF-A0A2J9QJ08-F1-MODEL\_V4 \| 1.0 \| 4.196e-20 \| 670 \| 0.223 \| 852 \| 479 \| 24 \| 16 \| 777 \| 1 \| 759 \| Phage tail tape measure protein \| Phage tail tape measure protein; afdb-uniprot50 \| AF-A0A1C2K2M3-F1-MODEL\_V4 \| 1.0 \| 2.418e-16 \| 668 \| 0.255 \| 587 \| 327 \| 11 \| 19 \| 514 \| 77 \| 644 \| Phage tail tape measure protein \| Phage tail tape measure protein; afdb-uniprot50 \| AF-A0A248UM23-F1-MODEL\_V4 \| 1.0 \| 8.214e-21 \| 668 \| 0.189 \| 886 \| 510 \| 23 \| 1 \| 783 \| 1 \| 780 \| Phage tail tape measure protein, TP901 family, core region \| Phage tail tape measure protein, TP901 family, core region; afdb-uniprot50 \| AF-A0A1H0GBX4-F1-MODEL\_V4 \| 1.0 \| 4.083e-21 \| 668 \| 0.192 \| 997 \| 544 \| 27 \| 3 \| 785 \| 1 \| 949 \| Phage tail tape measure protein, TP901 family, core region \| Phage tail tape measure protein, TP901 family, core region; afdb-uniprot50 \| AF-D3UG43-F1-MODEL\_V4 \| 1.0 \| 1.59e-20 \| 667 \| 0.207 \| 832 \| 509 \| 24 \| 6 \| 785 \| 1 \| 733 \| Putative phage tape-measure protein \| Putative phage tape-measure protein; afdb-uniprot50 \| AF-A0A4V2PT27-F1-MODEL\_V4 \| 1.0 \| 5.359e-21 \| 666 \| 0.224 \| 852 \| 513 \| 25 \| 16 \| 785 \| 1 \| 786 \| TP901 family phage tail tape measure protein \| TP901 family phage tail tape measure protein; afdb-uniprot50 \| AF-A0A2X3K7Y1-F1-MODEL\_V4 \| 1.0 \| 3.365e-11 \| 665 \| 0.554 \| 184 \| 82 \| 0 \| 177 \| 360 \| 4 \| 187 \| Putative phage tail tape measure protein \| Putative phage tail tape measure protein; afdb-uniprot50 \| AF-B6WRQ7-F1-MODEL\_V4 \| 1.0 \| 4.362e-20 \| 664 \| 0.218 \| 850 \| 505 \| 18 \| 19 \| 784 \| 3 \| 776 \| Phage tail tape measure protein, TP901 family \| Phage tail tape measure protein, TP901 family; afdb-uniprot50 \| AF-A0A0F2N598-F1-MODEL\_V4 \| 1.0 \| 5.236e-19 \| 664 \| 0.219 \| 737 \| 430 \| 18 \| 39 \| 750 \| 209 \| 824 \| PhageMin\_Tail domain-containing protein \| PhageMin\_Tail domain-containing protein; afdb-uniprot50 \| AF-A0A6N2Z5X0-F1-MODEL\_V4 \| 1.0 \| 3.593e-20 \| 663 \| 0.231 \| 817 \| 475 \| 26 \| 27 \| 776 \| 124 \| 854 \| Chromosome partition protein Smc \| Chromosome partition protein Smc; afdb-uniprot50 \| AF-A0A4Y9VRB4-F1-MODEL\_V4 \| 1.0 \| 7.144e-19 \| 661 \| 0.191 \| 831 \| 472 \| 21 \| 22 \| 782 \| 53 \| 753 \| Phage tail tape measure protein \| Phage tail tape measure protein; afdb-uniprot50 \| AF-A0A0N8W5Z8-F1-MODEL\_V4 \| 1.0 \| 1.415e-20 \| 660 \| 0.194 \| 835 \| 492 \| 23 \| 63 \| 779 \| 2 \| 773 \| PhageMin\_Tail domain-containing protein \| PhageMin\_Tail domain-containing protein; afdb-uniprot50 \| AF-A0A1C6E118-F1-MODEL\_V4 \| 1.0 \| 8.12e-20 \| 660 \| 0.248 \| 749 \| 436 \| 22 \| 53 \| 734 \| 9 \| 697 \| Phage-related protein \| Phage-related protein; afdb-uniprot50 \| AF-A0A1I4SVB4-F1-MODEL\_V4 \| 1.0 \| 6.432e-20 \| 660 \| 0.211 \| 857 \| 491 \| 22 \| 1 \| 716 \| 1 \| 813 \| Phage tail tape measure protein, TP901 family, core region \| Phage tail tape measure protein, TP901 family, core region; afdb-uniprot50 \| AF-A0A378QS12-F1-MODEL\_V4 \| 1.0 \| 6.687e-20 \| 659 \| 0.2 \| 798 \| 476 \| 22 \| 27 \| 701 \| 6 \| 764 \| Phage-related minor tail protein \| Phage-related minor tail protein; afdb-uniprot50 \| AF-A0A3G8M4E6-F1-MODEL\_V4 \| 1.0 \| 1.006e-14 \| 656 \| 0.232 \| 478 \| 321 \| 13 \| 18 \| 469 \| 15 \| 472 \| Phage tail tape measure protein \| Phage tail tape measure protein; afdb-uniprot50 \| AF-A0A7X3TZT8-F1-MODEL\_V4 \| 1.0 \| 3.838e-19 \| 654 \| 0.22 \| 872 \| 521 \| 25 \| 19 \| 785 \| 34 \| 851 \| Phage tail tape measure protein \| Phage tail tape measure protein; afdb-uniprot50 \| AF-A0A323TXZ9-F1-MODEL\_V4 \| 1.0 \| 1.165e-20 \| 654 \| 0.214 \| 928 \| 520 \| 24 \| 3 \| 785 \| 1 \| 864 \| Phage tail tape measure protein \| Phage tail tape measure protein; afdb-uniprot50 \| AF-A0A1I3T075-F1-MODEL\_V4 \| 1.0 \| 5.725e-20 \| 653 \| 0.209 \| 810 \| 462 \| 23 \| 1 \| 785 \| 1 \| 656 \| Phage tail tape measure protein, TP901 family, core region \| Phage tail tape measure protein, TP901 family, core region; afdb-uniprot50 \| AF-A0A1T4QE70-F1-MODEL\_V4 \| 1.0 \| 6.952e-20 \| 651 \| 0.184 \| 949 \| 514 \| 28 \| 3 \| 782 \| 1 \| 858 \| Phage tail tape measure protein, TP901 family, core region \| Phage tail tape measure protein, TP901 family, core region; afdb-uniprot50 \| AF-A0A5C7P5Y8-F1-MODEL\_V4 \| 1.0 \| 1.398e-19 \| 649 \| 0.201 \| 848 \| 412 \| 21 \| 19 \| 735 \| 28 \| 740 \| Phage tail tape measure protein \| Phage tail tape measure protein; afdb-uniprot50 \| AF-A0A0A3A3N3-F1-MODEL\_V4 \| 1.0 \| 9.9e-17 \| 648 \| 0.207 \| 627 \| 386 \| 16 \| 19 \| 570 \| 57 \| 647 \| Tail protein \| Tail protein; afdb-uniprot50 \| AF-A0A373FPP0-F1-MODEL\_V4 \| 1.0 \| 3.076e-20 \| 648 \| 0.196 \| 1035 \| 538 \| 30 \| 19 \| 782 \| 41 \| 1052 \| Phage tail tape measure protein \| Phage tail tape measure protein; afdb-uniprot50 \| AF-A0A448V531-F1-MODEL\_V4 \| 1.0 \| 6.187e-20 \| 647 \| 0.215 \| 830 \| 515 \| 21 \| 16 \| 776 \| 1 \| 762 \| Phage-related minor tail protein \| Phage-related minor tail protein; afdb-uniprot50 \| AF-A0A5N7YE88-F1-MODEL\_V4 \| 1.0 \| 5.976e-17 \| 645 \| 0.328 \| 493 \| 251 \| 10 \| 193 \| 685 \| 3 \| 415 \| Phage tail tape measure protein \| Phage tail tape measure protein; afdb-uniprot50 \| AF-A0A840DVF7-F1-MODEL\_V4 \| 1.0 \| 3.51e-18 \| 642 \| 0.232 \| 732 \| 423 \| 21 \| 16 \| 661 \| 1 \| 679 \| TP901 family phage tail tape measure protein \| TP901 family phage tail tape measure protein; afdb-uniprot50 \| AF-A0A376ZDS5-F1-MODEL\_V4 \| 1.0 \| 5.815e-18 \| 641 \| 0.189 \| 690 \| 466 \| 15 \| 118 \| 785 \| 4 \| 621 \| Tail protein (Modular protein) \| Tail protein (Modular protein); afdb-uniprot50 \| AF-A0A366CXV4-F1-MODEL\_V4 \| 1.0 \| 2.903e-15 \| 641 \| 0.251 \| 588 \| 312 \| 9 \| 20 \| 483 \| 124 \| 707 \| TP901 family phage tail tape measure protein \| TP901 family phage tail tape measure protein; afdb-uniprot50 \| AF-A0A6B8M5R6-F1-MODEL\_V4 \| 1.0 \| 2.76e-14 \| 640 \| 0.244 \| 462 \| 315 \| 10 \| 4 \| 450 \| 10 \| 452 \| Phage tail tape measure protein \| Phage tail tape measure protein; afdb-uniprot50 \| AF-G1UXP3-F1-MODEL\_V4 \| 1.0 \| 5.725e-20 \| 638 \| 0.253 \| 813 \| 472 \| 26 \| 18 \| 754 \| 28 \| 781 \| PhageMin\_Tail domain-containing protein \| PhageMin\_Tail domain-containing protein; afdb-uniprot50 \| AF-A0A2S9IP78-F1-MODEL\_V4 \| 1.0 \| 1.506e-12 \| 635 \| 0.365 \| 309 \| 185 \| 3 \| 51 \| 358 \| 2 \| 300 \| Phage tail tape measure protein \| Phage tail tape measure protein; afdb-uniprot50 \| AF-E1QHA5-F1-MODEL\_V4 \| 1.0 \| 4.362e-20 \| 633 \| 0.197 \| 1022 \| 525 \| 26 \| 19 \| 782 \| 10 \| 993 \| Phage tail tape measure protein, TP901 family \| Phage tail tape measure protein, TP901 family; afdb-uniprot50 \| AF-A0A3P3EPW8-F1-MODEL\_V4 \| 1.0 \| 1.156e-16 \| 630 \| 0.165 \| 745 \| 403 \| 20 \| 116 \| 785 \| 1 \| 601 \| Phage tail tape measure protein \| Phage tail tape measure protein; afdb-uniprot50 \| AF-A0A1E2RY73-F1-MODEL\_V4 \| 1.0 \| 1.041e-17 \| 627 \| 0.204 \| 762 \| 440 \| 28 \| 49 \| 785 \| 6 \| 625 \| Phage-related minor tail protein \| Phage-related minor tail protein; afdb-uniprot50 \| AF-A0A286BYN5-F1-MODEL\_V4 \| 1.0 \| 5.176e-18 \| 627 \| 0.196 \| 708 \| 451 \| 16 \| 109 \| 782 \| 372 \| 995 \| Phage tail tape measure protein, TP901 family, core region \| Phage tail tape measure protein, TP901 family, core region; afdb-uniprot50 \| AF-A0A2M8RTD1-F1-MODEL\_V4 \| 1.0 \| 2.228e-19 \| 625 \| 0.216 \| 785 \| 507 \| 22 \| 19 \| 734 \| 16 \| 761 \| Phage tail tape measure protein \| Phage tail tape measure protein; afdb-uniprot50 \| AF-A0A7C7D7E0-F1-MODEL\_V4 \| 1.0 \| 1.732e-14 \| 624 \| 0.215 \| 479 \| 331 \| 13 \| 21 \| 465 \| 16 \| 483 \| Phage tail tape measure protein \| Phage tail tape measure protein; afdb-uniprot50 \| AF-A0A2W4IRV0-F1-MODEL\_V4 \| 1.0 \| 8.186e-14 \| 623 \| 0.202 \| 519 \| 345 \| 13 \| 6 \| 516 \| 1 \| 458 \| Phage tail tape measure protein \| Phage tail tape measure protein; afdb-uniprot50 \| AF-A0A510KEQ7-F1-MODEL\_V4 \| 1.0 \| 6.358e-19 \| 623 \| 0.196 \| 834 \| 517 \| 22 \| 19 \| 715 \| 48 \| 864 \| Phage tail tape measure protein, TP901 family \| Phage tail tape measure protein, TP901 family; afdb-uniprot50 \| AF-A0A1L9CWC2-F1-MODEL\_V4 \| 1.0 \| 8.987e-12 \| 621 \| 0.227 \| 339 \| 245 \| 7 \| 91 \| 422 \| 3 \| 331 \| Phage tail tape measure protein \| Phage tail tape measure protein; afdb-uniprot50 \| AF-A0A2T3R1S0-F1-MODEL\_V4 \| 1.0 \| 3.854e-16 \| 619 \| 0.265 \| 594 \| 337 \| 13 \| 19 \| 519 \| 69 \| 655 \| Phage tail tape measure protein \| Phage tail tape measure protein; afdb-uniprot50 \| AF-A0A1B8Q889-F1-MODEL\_V4 \| 1.0 \| 3.99e-19 \| 617 \| 0.207 \| 795 \| 464 \| 19 \| 20 \| 704 \| 14 \| 752 \| Phage tail tape measure protein \| Phage tail tape measure protein; afdb-uniprot50 \| AF-A0A1Q8Q232-F1-MODEL\_V4 \| 1.0 \| 5.236e-19 \| 616 \| 0.177 \| 1005 \| 553 \| 26 \| 19 \| 781 \| 19 \| 991 \| Phage tail tape measure protein \| Phage tail tape measure protein; afdb-uniprot50 \| AF-A0A7W2GS65-F1-MODEL\_V4 \| 1.0 \| 2.675e-18 \| 612 \| 0.212 \| 770 \| 461 \| 19 \| 14 \| 757 \| 1 \| 650 \| Phage tail tape measure protein \| Phage tail tape measure protein; afdb-uniprot50 \| AF-A0A238HFL8-F1-MODEL\_V4 \| 1.0 \| 1.494e-18 \| 612 \| 0.175 \| 874 \| 545 \| 27 \| 24 \| 785 \| 55 \| 864 \| Phage-related minor tail protein \| Phage-related minor tail protein; afdb-uniprot50 \| AF-T4VE82-F1-MODEL\_V4 \| 1.0 \| 2.177e-17 \| 611 \| 0.192 \| 750 \| 466 \| 22 \| 64 \| 782 \| 4 \| 644 \| Phage tail tape measure protein, TP901 family, core region \| Phage tail tape measure protein, TP901 family, core region; afdb-uniprot50 \| AF-A0A2G2Q4G5-F1-MODEL\_V4 \| 1.0 \| 1.013e-18 \| 611 \| 0.21 \| 789 \| 485 \| 20 \| 2 \| 705 \| 1 \| 736 \| Phage tail tape measure protein \| Phage tail tape measure protein; afdb-uniprot50 \| AF-A0A327JMP0-F1-MODEL\_V4 \| 1.0 \| 1.17e-17 \| 605 \| 0.177 \| 872 \| 548 \| 25 \| 8 \| 783 \| 1 \| 798 \| Phage tail tape measure protein \| Phage tail tape measure protein; afdb-uniprot50 \| AF-J0Q8Z0-F1-MODEL\_V4 \| 1.0 \| 2.079e-13 \| 604 \| 0.273 \| 355 \| 242 \| 6 \| 155 \| 508 \| 3 \| 342 \| Phage tail tape measure protein, TP901 family, core region \| Phage tail tape measure protein, TP901 family, core region; afdb-uniprot50 \| AF-A0A852QWW4-F1-MODEL\_V4 \| 1.0 \| 1.006e-14 \| 604 \| 0.214 \| 518 \| 311 \| 15 \| 18 \| 526 \| 16 \| 446 \| TP901 family phage tail tape measure protein \| TP901 family phage tail tape measure protein; afdb-uniprot50 \| AF-R5BHC1-F1-MODEL\_V4 \| 1.0 \| 2.514e-16 \| 604 \| 0.227 \| 694 \| 416 \| 22 \| 132 \| 781 \| 2 \| 619 \| Tail tape measure protein TIGR01760 family \| Tail tape measure protein TIGR01760 family; afdb-uniprot50 \| AF-A0A844UUJ6-F1-MODEL\_V4 \| 1.0 \| 1.314e-17 \| 604 \| 0.187 \| 780 \| 461 \| 21 \| 129 \| 785 \| 2 \| 731 \| Phage tail tape measure protein \| Phage tail tape measure protein; afdb-uniprot50 \| AF-A0A2D5PEJ4-F1-MODEL\_V4 \| 1.0 \| 1.082e-17 \| 604 \| 0.209 \| 820 \| 391 \| 18 \| 19 \| 755 \| 74 \| 718 \| Phage tail tape measure protein \| Phage tail tape measure protein; afdb-uniprot50 \| AF-A0A485DIL8-F1-MODEL\_V4 \| 1.0 \| 9.267e-18 \| 603 \| 0.213 \| 796 \| 506 \| 21 \| 21 \| 780 \| 2 \| 713 \| Phage protein \| Phage protein; afdb-uniprot50 \| AF-A0A8B2NXI7-F1-MODEL\_V4 \| 1.0 \| 5.659e-19 \| 603 \| 0.218 \| 781 \| 439 \| 29 \| 98 \| 785 \| 27 \| 728 \| Phage tail tape measure protein \| Phage tail tape measure protein; afdb-uniprot50 \| AF-U6Q3L1-F1-MODEL\_V4 \| 1.0 \| 2.015e-17 \| 602 \| 0.214 \| 778 \| 499 \| 23 \| 1 \| 702 \| 1 \| 741 \| Phage tail tape measure protein, TP901 family, core region \| Phage tail tape measure protein, TP901 family, core region; afdb-uniprot50 \| AF-A0A1F1HUA7-F1-MODEL\_V4 \| 1.0 \| 5.116e-17 \| 601 \| 0.197 \| 679 \| 428 \| 20 \| 21 \| 628 \| 48 \| 680 \| PhageMin\_Tail domain-containing protein \| PhageMin\_Tail domain-containing protein; afdb-uniprot50 \| AF-A0A4Y6KLY6-F1-MODEL\_V4 \| 1.0 \| 2.573e-18 \| 600 \| 0.165 \| 989 \| 560 \| 24 \| 40 \| 785 \| 94 \| 1059 \| Phage tail tape measure protein \| Phage tail tape measure protein; afdb-uniprot50 \| AF-A0A2G1QM28-F1-MODEL\_V4 \| 1.0 \| 3.96e-15 \| 599 \| 0.185 \| 701 \| 405 \| 14 \| 40 \| 719 \| 20 \| 575 \| Phage tail tape measure protein \| Phage tail tape measure protein; afdb-uniprot50 \| AF-A0A2J9QPE7-F1-MODEL\_V4 \| 1.0 \| 1.398e-19 \| 599 \| 0.188 \| 906 \| 538 \| 23 \| 19 \| 782 \| 44 \| 893 \| Phage tail tape measure protein \| Phage tail tape measure protein; afdb-uniprot50 \| AF-A0A0M9GN37-F1-MODEL\_V4 \| 1.0 \| 9.267e-18 \| 597 \| 0.179 \| 779 \| 442 \| 17 \| 39 \| 785 \| 3 \| 615 \| PhageMin\_Tail domain-containing protein \| PhageMin\_Tail domain-containing protein; afdb-uniprot50 \| AF-A0A2X7FXT8-F1-MODEL\_V4 \| 1.0 \| 9.375e-19 \| 597 \| 0.187 \| 940 \| 531 \| 24 \| 23 \| 785 \| 50 \| 933 \| Tail protein (Modular protein) \| Tail protein (Modular protein); afdb-uniprot50 \| AF-A0A2P1S7S0-F1-MODEL\_V4 \| 1.0 \| 1.961e-18 \| 597 \| 0.21 \| 939 \| 521 \| 20 \| 18 \| 776 \| 108 \| 1005 \| Phage tail tape measure protein \| Phage tail tape measure protein; afdb-uniprot50 \| AF-A0A7X5J7E5-F1-MODEL\_V4 \| 1.0 \| 3.444e-13 \| 596 \| 0.2 \| 445 \| 326 \| 9 \| 13 \| 432 \| 47 \| 486 \| Phage tail tape measure protein \| Phage tail tape measure protein; afdb-uniprot50 \| AF-A0A846BMG1-F1-MODEL\_V4 \| 1.0 \| 1.432e-11 \| 595 \| 0.309 \| 317 \| 195 \| 6 \| 64 \| 371 \| 4 \| 305 \| Phage tail tape measure protein \| Phage tail tape measure protein; afdb-uniprot50 \| AF-A0A496JTX2-F1-MODEL\_V4 \| 1.0 \| 2.119e-18 \| 595 \| 0.218 \| 842 \| 448 \| 25 \| 19 \| 710 \| 34 \| 815 \| Phage tail tape measure protein \| Phage tail tape measure protein; afdb-uniprot50 \| AF-V7ZKH4-F1-MODEL\_V4 \| 1.0 \| 6.587e-12 \| 594 \| 0.197 \| 385 \| 242 \| 8 \| 117 \| 436 \| 20 \| 402 \| PhageMin\_Tail domain-containing protein \| PhageMin\_Tail domain-containing protein; afdb-uniprot50 \| AF-A0A486PRA3-F1-MODEL\_V4 \| 1.0 \| 6.046e-18 \| 594 \| 0.18 \| 925 \| 555 \| 29 \| 19 \| 785 \| 32 \| 911 \| Putative prophage tail length determinator \| Putative prophage tail length determinator; afdb-uniprot50 \| AF-A0A414PRA3-F1-MODEL\_V4 \| 1.0 \| 1.615e-18 \| 594 \| 0.209 \| 917 \| 507 \| 28 \| 27 \| 782 \| 88 \| 947 \| Phage tail tape measure protein \| Phage tail tape measure protein; afdb-uniprot50 \| AF-A0A1Q8Z201-F1-MODEL\_V4 \| 1.0 \| 2.706e-19 \| 594 \| 0.167 \| 980 \| 565 \| 27 \| 21 \| 785 \| 48 \| 991 \| Phage tail tape measure protein \| Phage tail tape measure protein; afdb-uniprot50 \| AF-A0A0R2D4C9-F1-MODEL\_V4 \| 1.0 \| 7.341e-18 \| 590 \| 0.199 \| 717 \| 486 \| 21 \| 12 \| 702 \| 141 \| 795 \| PhageMin\_Tail domain-containing protein \| PhageMin\_Tail domain-containing protein; afdb-uniprot50 \| AF-A0A286IFD9-F1-MODEL\_V4 \| 1.0 \| 2.675e-18 \| 588 \| 0.207 \| 816 \| 504 \| 27 \| 3 \| 785 \| 1 \| 706 \| TP901 family phage tail tape measure protein \| TP901 family phage tail tape measure protein; afdb-uniprot50 \| AF-A0A2V4L3Y4-F1-MODEL\_V4 \| 1.0 \| 5.815e-18 \| 588 \| 0.211 \| 902 \| 446 \| 25 \| 19 \| 783 \| 28 \| 800 \| Phage tail tape measure protein \| Phage tail tape measure protein; afdb-uniprot50 \| AF-A0A376TE73-F1-MODEL\_V4 \| 1.0 \| 3.124e-18 \| 587 \| 0.188 \| 833 \| 526 \| 25 \| 1 \| 781 \| 15 \| 749 \| Phage tail tape measure protein \| Phage tail tape measure protein; afdb-uniprot50 \| AF-A0A843BA66-F1-MODEL\_V4 \| 1.0 \| 8.575e-18 \| 585 \| 0.225 \| 695 \| 415 \| 19 \| 165 \| 780 \| 2 \| 651 \| Phage tail tape measure protein \| Phage tail tape measure protein; afdb-uniprot50 \| AF-A0A744CZM2-F1-MODEL\_V4 \| 1.0 \| 5.529e-17 \| 583 \| 0.202 \| 731 \| 447 \| 19 \| 120 \| 774 \| 26 \| 696 \| Phage tail tape measure protein \| Phage tail tape measure protein; afdb-uniprot50 \| AF-A0A414Q2W1-F1-MODEL\_V4 \| 1.0 \| 1.814e-18 \| 582 \| 0.212 \| 942 \| 504 \| 29 \| 19 \| 776 \| 81 \| 968 \| Phage tail tape measure protein \| Phage tail tape measure protein; afdb-uniprot50 \| AF-U6Q3L9-F1-MODEL\_V4 \| 1.0 \| 6.459e-17 \| 579 \| 0.22 \| 767 \| 505 \| 19 \| 1 \| 701 \| 1 \| 740 \| Phage tail tape measure protein, TP901 family, core region \| Phage tail tape measure protein, TP901 family, core region; afdb-uniprot50 \| AF-A0A826S7R5-F1-MODEL\_V4 \| 1.0 \| 8.575e-18 \| 578 \| 0.175 \| 887 \| 545 \| 20 \| 17 \| 785 \| 1 \| 818 \| Phage tail tape measure protein \| Phage tail tape measure protein; afdb-uniprot50 \| AF-A0A3L9GV37-F1-MODEL\_V4 \| 1.0 \| 2.038e-18 \| 578 \| 0.195 \| 863 \| 506 \| 27 \| 21 \| 735 \| 48 \| 869 \| Phage tail tape measure protein \| Phage tail tape measure protein; afdb-uniprot50 \| AF-A0A168UYB6-F1-MODEL\_V4 \| 1.0 \| 1.006e-14 \| 577 \| 0.202 \| 672 \| 374 \| 17 \| 117 \| 715 \| 14 \| 596 \| PhageMin\_Tail domain-containing protein \| PhageMin\_Tail domain-containing protein; afdb-uniprot50 \| AF-A0A7W6N9T0-F1-MODEL\_V4 \| 1.0 \| 4.165e-16 \| 577 \| 0.174 \| 783 \| 484 \| 18 \| 46 \| 785 \| 2 \| 664 \| TP901 family phage tail tape measure protein \| TP901 family phage tail tape measure protein; afdb-uniprot50 \| AF-A0A846VL41-F1-MODEL\_V4 \| 1.0 \| 2.936e-16 \| 577 \| 0.19 \| 820 \| 486 \| 24 \| 13 \| 784 \| 1 \| 690 \| TP901 family phage tail tape measure protein \| TP901 family phage tail tape measure protein; afdb-uniprot50 \| AF-A0A504KM96-F1-MODEL\_V4 \| 1.0 \| 1.299e-16 \| 577 \| 0.18 \| 802 \| 407 \| 23 \| 104 \| 782 \| 56 \| 729 \| Phage tail tape measure protein \| Phage tail tape measure protein; afdb-uniprot50 \| AF-A0A679GE32-F1-MODEL\_V4 \| 1.0 \| 1.421e-17 \| 576 \| 0.209 \| 773 \| 503 \| 16 \| 19 \| 702 \| 45 \| 798 \| PhageMin\_Tail domain-containing protein \| PhageMin\_Tail domain-containing protein; afdb-uniprot50 \| AF-A0A731W7G9-F1-MODEL\_V4 \| 1.0 \| 4.999e-15 \| 575 \| 0.186 \| 639 \| 427 \| 14 \| 105 \| 722 \| 7 \| 573 \| Phage tail tape measure protein \| Phage tail tape measure protein; afdb-uniprot50 \| AF-V4N3Q7-F1-MODEL\_V4 \| 1.0 \| 1.518e-16 \| 574 \| 0.199 \| 731 \| 451 \| 20 \| 89 \| 749 \| 4 \| 669 \| Tail protein \| Tail protein; afdb-uniprot50 \| AF-A0A411WMH1-F1-MODEL\_V4 \| 1.0 \| 1.314e-17 \| 574 \| 0.163 \| 930 \| 574 \| 25 \| 19 \| 781 \| 35 \| 927 \| Phage tail tape measure protein \| Phage tail tape measure protein; afdb-uniprot50 \| AF-A0A1W2EJS2-F1-MODEL\_V4 \| 1.0 \| 7.457e-16 \| 572 \| 0.155 \| 859 \| 480 \| 25 \| 1 \| 785 \| 1 \| 687 \| Phage tail tape measure protein, TP901 family, core region \| Phage tail tape measure protein, TP901 family, core region; afdb-uniprot50 \| AF-A0A2S9GTE6-F1-MODEL\_V4 \| 1.0 \| 1.17e-17 \| 572 \| 0.17 \| 919 \| 549 \| 29 \| 19 \| 780 \| 56 \| 917 \| Phage tail tape measure protein, TP901 family, core region \| Phage tail tape measure protein, TP901 family, core region; afdb-uniprot50 \| AF-A0A0T9KM18-F1-MODEL\_V4 \| 1.0 \| 3.794e-18 \| 570 \| 0.171 \| 950 \| 569 \| 26 \| 22 \| 781 \| 54 \| 974 \| Tail fiber component T of bacteriophage P2 \| Tail fiber component T of bacteriophage P2; afdb-uniprot50 \| AF-A0A432WB56-F1-MODEL\_V4 \| 1.0 \| 3.51e-18 \| 570 \| 0.182 \| 778 \| 484 \| 20 \| 19 \| 698 \| 37 \| 760 \| Phage tail tape measure protein \| Phage tail tape measure protein; afdb-uniprot50 \| AF-U7T719-F1-MODEL\_V4 \| 1.0 \| 7.061e-18 \| 568 \| 0.196 \| 913 \| 524 \| 29 \| 1 \| 778 \| 2 \| 839 \| Phage tail tape measure protein, TP901 family, core region \| Phage tail tape measure protein, TP901 family, core region; afdb-uniprot50 \| AF-A0A1N6MRW7-F1-MODEL\_V4 \| 1.0 \| 5.319e-17 \| 566 \| 0.177 \| 829 \| 538 \| 22 \| 19 \| 754 \| 45 \| 822 \| PhageMin\_Tail domain-containing protein \| PhageMin\_Tail domain-containing protein; afdb-uniprot50 \| AF-A0A375H2Y0-F1-MODEL\_V4 \| 1.0 \| 4.165e-16 \| 565 \| 0.264 \| 579 \| 316 \| 9 \| 206 \| 784 \| 7 \| 475 \| PhageMin\_Tail domain-containing protein \| PhageMin\_Tail domain-containing protein; afdb-uniprot50 \| AF-A0A1F1DK89-F1-MODEL\_V4 \| 1.0 \| 1.341e-12 \| 565 \| 0.234 \| 474 \| 308 \| 13 \| 1 \| 466 \| 1 \| 427 \| PhageMin\_Tail domain-containing protein \| PhageMin\_Tail domain-containing protein; afdb-uniprot50 \| AF-A0A7X7BBI3-F1-MODEL\_V4 \| 1.0 \| 6.166e-13 \| 564 \| 0.18 \| 516 \| 347 \| 9 \| 19 \| 463 \| 15 \| 525 \| Phage tail tape measure protein \| Phage tail tape measure protein; afdb-uniprot50 \| AF-A0A3B8S463-F1-MODEL\_V4 \| 1.0 \| 3.39e-15 \| 563 \| 0.215 \| 593 \| 359 \| 24 \| 50 \| 564 \| 4 \| 567 \| Phage tail tape measure protein \| Phage tail tape measure protein; afdb-uniprot50 \| AF-A0A7C7DFP0-F1-MODEL\_V4 \| 1.0 \| 8.153e-17 \| 563 \| 0.188 \| 802 \| 477 \| 31 \| 1 \| 750 \| 1 \| 680 \| Phage tail tape measure protein \| Phage tail tape measure protein; afdb-uniprot50 \| AF-A0A826YZU6-F1-MODEL\_V4 \| 1.0 \| 5.137e-14 \| 560 \| 0.195 \| 589 \| 374 \| 15 \| 88 \| 644 \| 5 \| 525 \| Phage tail tape measure protein \| Phage tail tape measure protein; afdb-uniprot50 \| AF-A0A3L2QDC9-F1-MODEL\_V4 \| 1.0 \| 1.299e-16 \| 560 \| 0.206 \| 737 \| 453 \| 19 \| 132 \| 785 \| 2 \| 689 \| Phage tail tape measure protein \| Phage tail tape measure protein; afdb-uniprot50 \| AF-A0A547PW98-F1-MODEL\_V4 \| 1.0 \| 7.632e-18 \| 560 \| 0.197 \| 890 \| 498 \| 25 \| 49 \| 784 \| 6 \| 832 \| Phage tail tape measure protein \| Phage tail tape measure protein; afdb-uniprot50 \| AF-A0A2E1XVX6-F1-MODEL\_V4 \| 1.0 \| 2.186e-14 \| 557 \| 0.22 \| 650 \| 386 \| 14 \| 1 \| 628 \| 1 \| 551 \| Phage tail tape measure protein \| Phage tail tape measure protein; afdb-uniprot50 \| AF-A0A0N1N416-F1-MODEL\_V4 \| 1.0 \| 5.403e-15 \| 557 \| 0.174 \| 733 \| 465 \| 17 \| 33 \| 721 \| 2 \| 638 \| PhageMin\_Tail domain-containing protein \| PhageMin\_Tail domain-containing protein; afdb-uniprot50 \| AF-A0A174F216-F1-MODEL\_V4 \| 1.0 \| 6.848e-12 \| 557 \| 0.21 \| 433 \| 321 \| 8 \| 42 \| 460 \| 199 \| 624 \| Phage tail tape measure protein, family \| Phage tail tape measure protein, family; afdb-uniprot50 \| AF-A0A6P1K7K2-F1-MODEL\_V4 \| 1.0 \| 2.177e-17 \| 553 \| 0.179 \| 945 \| 535 \| 26 \| 23 \| 785 \| 50 \| 935 \| Phage tail tape measure protein \| Phage tail tape measure protein; afdb-uniprot50 \| AF-A0A7T4T096-F1-MODEL\_V4 \| 1.0 \| 5.529e-17 \| 552 \| 0.181 \| 745 \| 535 \| 20 \| 12 \| 702 \| 169 \| 892 \| Phage tail tape measure protein \| Phage tail tape measure protein; afdb-uniprot50 \| AF-A0A0F3RNQ7-F1-MODEL\_V4 \| 1.0 \| 3.553e-09 \| 551 \| 0.317 \| 227 \| 151 \| 2 \| 132 \| 358 \| 2 \| 224 \| Phage tail tape measure protein, TP901 family, core region \| Phage tail tape measure protein, TP901 family, core region; afdb-uniprot50 \| AF-A0A2W6QW57-F1-MODEL\_V4 \| 1.0 \| 7.875e-14 \| 549 \| 0.22 \| 550 \| 379 \| 15 \| 132 \| 672 \| 2 \| 510 \| Phage tail tape measure protein \| Phage tail tape measure protein; afdb-uniprot50 \| AF-A0A6P1Q224-F1-MODEL\_V4 \| 1.0 \| 3.899e-17 \| 549 \| 0.183 \| 813 \| 513 \| 23 \| 19 \| 782 \| 13 \| 723 \| PhageMin\_Tail domain-containing protein \| PhageMin\_Tail domain-containing protein; afdb-uniprot50 \| AF-Q5D5E9-F1-MODEL\_V4 \| 1.0 \| 2.815e-09 \| 547 \| 0.345 \| 223 \| 144 \| 1 \| 116 \| 338 \| 8 \| 228 \| Prophage P2W3, tail tape measure protein \| Prophage P2W3, tail tape measure protein; afdb-uniprot50 \| AF-A0A7V5PAS2-F1-MODEL\_V4 \| 1.0 \| 5.575e-11 \| 547 \| 0.175 \| 411 \| 314 \| 8 \| 70 \| 463 \| 4 \| 406 \| Phage tail tape measure protein \| Phage tail tape measure protein; afdb-uniprot50 \| AF-A0A827RXC1-F1-MODEL\_V4 \| 1.0 \| 1.156e-16 \| 547 \| 0.193 \| 782 \| 472 \| 21 \| 120 \| 785 \| 36 \| 774 \| Phage tail tape measure protein \| Phage tail tape measure protein; afdb-uniprot50 \| AF-A8I7U7-F1-MODEL\_V4 \| 1.0 \| 4.679e-16 \| 546 \| 0.186 \| 844 \| 469 \| 25 \| 14 \| 784 \| 38 \| 736 \| Putative phage-related tail protein \| Putative phage-related tail protein; afdb-uniprot50 \| AF-A0A2D9FXU5-F1-MODEL\_V4 \| 1.0 \| 1.705e-16 \| 545 \| 0.187 \| 730 \| 454 \| 25 \| 76 \| 780 \| 18 \| 633 \| Phage tail tape measure protein \| Phage tail tape measure protein; afdb-uniprot50 \| AF-A0A3A6S8Q3-F1-MODEL\_V4 \| 1.0 \| 2.971e-17 \| 544 \| 0.198 \| 784 \| 480 \| 22 \| 21 \| 717 \| 48 \| 769 \| Phage tail tape measure protein \| Phage tail tape measure protein; afdb-uniprot50 \| AF-A0A2U1TTC3-F1-MODEL\_V4 \| 1.0 \| 1.535e-17 \| 544 \| 0.167 \| 955 \| 559 \| 24 \| 19 \| 785 \| 47 \| 952 \| Phage tail tape measure protein \| Phage tail tape measure protein; afdb-uniprot50 \| AF-A0A2T4M4G0-F1-MODEL\_V4 \| 1.0 \| 3.275e-12 \| 543 \| 0.198 \| 513 \| 348 \| 15 \| 19 \| 480 \| 324 \| 824 \| Phage tail tape measure protein \| Phage tail tape measure protein; afdb-uniprot50 \| AF-A0A1B9AKU2-F1-MODEL\_V4 \| 1.0 \| 3.338e-17 \| 543 \| 0.207 \| 760 \| 478 \| 22 \| 27 \| 702 \| 60 \| 778 \| Phage tail tape measure protein \| Phage tail tape measure protein; afdb-uniprot50 \| AF-A0A1X3ITD6-F1-MODEL\_V4 \| 1.0 \| 1.591e-10 \| 542 \| 0.174 \| 412 \| 277 \| 10 \| 36 \| 442 \| 2 \| 355 \| Phage tail tape measure protein, TP901 family, core region \| Phage tail tape measure protein, TP901 family, core region; afdb-uniprot50 \| AF-S4FQD2-F1-MODEL\_V4 \| 1.0 \| 1.193e-12 \| 542 \| 0.205 \| 443 \| 269 \| 9 \| 118 \| 479 \| 2 \| 442 \| Phage tail tape measure protein, TP901 family \| Phage tail tape measure protein, TP901 family; afdb-uniprot50 \| AF-A0A484GDB7-F1-MODEL\_V4 \| 1.0 \| 2.103e-14 \| 542 \| 0.17 \| 579 \| 436 \| 8 \| 125 \| 690 \| 2 \| 548 \| Phage tail tape measure protein \| Phage tail tape measure protein; afdb-uniprot50 \| AF-A0A4U3CH96-F1-MODEL\_V4 \| 1.0 \| 1.773e-16 \| 542 \| 0.2 \| 759 \| 464 \| 26 \| 1 \| 702 \| 1 \| 673 \| PhageMin\_Tail domain-containing protein \| PhageMin\_Tail domain-containing protein; afdb-uniprot50 \| AF-A0A2T5UQZ4-F1-MODEL\_V4 \| 1.0 \| 1.725e-17 \| 541 \| 0.199 \| 879 \| 459 \| 32 \| 1 \| 785 \| 1 \| 728 \| TP901 family phage tail tape measure protein \| TP901 family phage tail tape measure protein; afdb-uniprot50 \| AF-A0A348FYG3-F1-MODEL\_V4 \| 1.0 \| 2.152e-16 \| 539 \| 0.189 \| 761 \| 483 \| 22 \| 13 \| 750 \| 1 \| 650 \| PhageMin\_Tail domain-containing protein \| PhageMin\_Tail domain-containing protein; afdb-uniprot50 \| AF-A0A0H3H1R5-F1-MODEL\_V4 \| 1.0 \| 1.314e-17 \| 539 \| 0.202 \| 846 \| 510 \| 25 \| 19 \| 781 \| 50 \| 813 \| Putative prophage tail length determinator \| Putative prophage tail length determinator; afdb-uniprot50 \| AF-A0A078LGR8-F1-MODEL\_V4 \| 1.0 \| 5.976e-17 \| 538 \| 0.192 \| 841 \| 506 \| 28 \| 19 \| 755 \| 43 \| 813 \| Phage tail tape measure protein \| Phage tail tape measure protein; afdb-uniprot50 \| AF-A0A8B5P8D7-F1-MODEL\_V4 \| 1.0 \| 8.282e-15 \| 537 \| 0.199 \| 658 \| 422 \| 18 \| 120 \| 754 \| 12 \| 587 \| Phage tail tape measure protein \| Phage tail tape measure protein; afdb-uniprot50 \| AF-A0A126NVC8-F1-MODEL\_V4 \| 1.0 \| 7.371e-15 \| 535 \| 0.163 \| 778 \| 453 \| 21 \| 36 \| 785 \| 5 \| 612 \| PhageMin\_Tail domain-containing protein \| PhageMin\_Tail domain-containing protein; afdb-uniprot50 \| AF-A0A1C4WZH7-F1-MODEL\_V4 \| 1.0 \| 2.238e-16 \| 534 \| 0.187 \| 741 \| 466 \| 28 \| 2 \| 700 \| 1 \| 647 \| Phage tail tape measure protein, TP901 family, core region \| Phage tail tape measure protein, TP901 family, core region; afdb-uniprot50 \| AF-A0A653HUE7-F1-MODEL\_V4 \| 1.0 \| 1.299e-16 \| 534 \| 0.179 \| 915 \| 563 \| 22 \| 19 \| 785 \| 29 \| 903 \| PhageMin\_Tail domain-containing protein \| PhageMin\_Tail domain-containing protein; afdb-uniprot50 \| AF-A0A1F1C065-F1-MODEL\_V4 \| 1.0 \| 6.98e-17 \| 532 \| 0.182 \| 878 \| 494 \| 26 \| 21 \| 781 \| 48 \| 818 \| PhageMin\_Tail domain-containing protein \| PhageMin\_Tail domain-containing protein; afdb-uniprot50 \| AF-A0A3D4QR57-F1-MODEL\_V4 \| 1.0 \| 3.275e-12 \| 531 \| 0.176 \| 471 \| 312 \| 10 \| 19 \| 422 \| 34 \| 495 \| Phage tail tape measure protein \| Phage tail tape measure protein; afdb-uniprot50 \| AF-A0A6B4AIV4-F1-MODEL\_V4 \| 1.0 \| 3.679e-12 \| 530 \| 0.213 \| 482 \| 301 \| 13 \| 49 \| 464 \| 203 \| 672 \| Phage tail tape measure protein \| Phage tail tape measure protein; afdb-uniprot50 \| AF-A0A0B0SDM8-F1-MODEL\_V4 \| 1.0 \| 2.015e-17 \| 530 \| 0.224 \| 837 \| 421 \| 19 \| 152 \| 762 \| 2 \| 835 \| PhageMin\_Tail domain-containing protein \| PhageMin\_Tail domain-containing protein; afdb-uniprot50 \| AF-F9PY49-F1-MODEL\_V4 \| 1.0 \| 1.5e-15 \| 528 \| 0.198 \| 724 \| 463 \| 24 \| 19 \| 668 \| 3 \| 682 \| Phage tail tape measure protein, TP901 family \| Phage tail tape measure protein, TP901 family; afdb-uniprot50 \| AF-A0A6I6AWA7-F1-MODEL\_V4 \| 1.0 \| 4.38e-17 \| 528 \| 0.194 \| 838 \| 496 \| 25 \| 19 \| 758 \| 37 \| 793 \| Phage tail tape measure protein \| Phage tail tape measure protein; afdb-uniprot50 \| AF-A0A2D1KVM8-F1-MODEL\_V4 \| 1.0 \| 2.614e-16 \| 528 \| 0.183 \| 778 \| 524 \| 16 \| 18 \| 702 \| 122 \| 880 \| Phage tail tape measure protein \| Phage tail tape measure protein; afdb-uniprot50 \| AF-A0A268IIP2-F1-MODEL\_V4 \| 1.0 \| 5.466e-16 \| 527 \| 0.212 \| 705 \| 465 \| 19 \| 19 \| 669 \| 52 \| 719 \| Phage tail tape measure protein \| Phage tail tape measure protein; afdb-uniprot50 \| AF-A0A1Q6U7T4-F1-MODEL\_V4 \| 1.0 \| 1.864e-17 \| 526 \| 0.179 \| 858 \| 507 \| 23 \| 19 \| 781 \| 40 \| 795 \| Phage tail tape measure protein \| Phage tail tape measure protein; afdb-uniprot50 \| AF-A0A1B9NQD5-F1-MODEL\_V4 \| 1.0 \| 1.916e-16 \| 526 \| 0.16 \| 978 \| 561 \| 25 \| 19 \| 782 \| 43 \| 973 \| Phage tail tape measure protein \| Phage tail tape measure protein; afdb-uniprot50 \| AF-A0A1E4UTG7-F1-MODEL\_V4 \| 1.0 \| 3.053e-16 \| 525 \| 0.215 \| 936 \| 475 \| 26 \| 18 \| 783 \| 76 \| 921 \| Phage tail tape measure protein \| Phage tail tape measure protein; afdb-uniprot50 \| AF-A0A1W2EA98-F1-MODEL\_V4 \| 1.0 \| 3.664e-15 \| 523 \| 0.186 \| 729 \| 461 \| 20 \| 39 \| 754 \| 2 \| 611 \| Phage tail tape measure protein, TP901 family, core region \| Phage tail tape measure protein, TP901 family, core region; afdb-uniprot50 \| AF-A0A845QX85-F1-MODEL\_V4 \| 1.0 \| 2.804e-12 \| 522 \| 0.2 \| 515 \| 321 \| 13 \| 20 \| 460 \| 68 \| 565 \| Phage tail tape measure protein \| Phage tail tape measure protein; afdb-uniprot50 \| AF-A0A8B5S3L7-F1-MODEL\_V4 \| 1.0 \| 1.916e-16 \| 521 \| 0.154 \| 933 \| 591 \| 23 \| 19 \| 782 \| 22 \| 925 \| Phage tail tape measure protein \| Phage tail tape measure protein; afdb-uniprot50 \| AF-A0A2D5B2W5-F1-MODEL\_V4 \| 1.0 \| 1.752e-15 \| 520 \| 0.185 \| 729 \| 486 \| 26 \| 35 \| 700 \| 13 \| 696 \| Phage tail tape measure protein \| Phage tail tape measure protein; afdb-uniprot50 \| AF-A0A7V9PI93-F1-MODEL\_V4 \| 1.0 \| 2.583e-15 \| 520 \| 0.19 \| 721 \| 456 \| 19 \| 67 \| 702 \| 2 \| 679 \| Phage tail tape measure protein \| Phage tail tape measure protein; afdb-uniprot50 \| AF-A0A5J6LCN0-F1-MODEL\_V4 \| 1.0 \| 1.559e-15 \| 519 \| 0.168 \| 869 \| 581 \| 22 \| 19 \| 782 \| 58 \| 889 \| Phage tail tape measure protein \| Phage tail tape measure protein; afdb-uniprot50 \| AF-A0A0F9RHG1-F1-MODEL\_V4 \| 1.0 \| 1.087e-14 \| 519 \| 0.183 \| 699 \| 461 \| 27 \| 70 \| 700 \| 23 \| 679 \| PhageMin\_Tail domain-containing protein \| PhageMin\_Tail domain-containing protein; afdb-uniprot50 \| AF-A0A1H8B8F5-F1-MODEL\_V4 \| 1.0 \| 7.606e-11 \| 518 \| 0.175 \| 444 \| 290 \| 9 \| 43 \| 466 \| 17 \| 404 \| Phage tail tape measure protein, TP901 family, core region \| Phage tail tape measure protein, TP901 family, core region; afdb-uniprot50 \| AF-A0A3T0L2L7-F1-MODEL\_V4 \| 1.0 \| 3.299e-16 \| 517 \| 0.188 \| 759 \| 494 \| 26 \| 4 \| 710 \| 42 \| 730 \| Phage tail tape measure protein \| Phage tail tape measure protein; afdb-uniprot50 \| AF-A0A841GB24-F1-MODEL\_V4 \| 1.0 \| 1.801e-14 \| 516 \| 0.161 \| 773 \| 444 \| 23 \| 62 \| 781 \| 9 \| 629 \| TP901 family phage tail tape measure protein \| TP901 family phage tail tape measure protein; afdb-uniprot50 \| AF-A0A2J9H2B8-F1-MODEL\_V4 \| 1.0 \| 4.006e-16 \| 515 \| 0.178 \| 859 \| 514 \| 25 \| 22 \| 778 \| 50 \| 818 \| Phage tail tape measure protein \| Phage tail tape measure protein; afdb-uniprot50 \| AF-A0A2N6UUM1-F1-MODEL\_V4 \| 1.0 \| 2.238e-16 \| 515 \| 0.161 \| 976 \| 549 \| 33 \| 63 \| 785 \| 3 \| 961 \| Phage tail tape measure protein \| Phage tail tape measure protein; afdb-uniprot50 \| AF-A0A1M3MNY4-F1-MODEL\_V4 \| 1.0 \| 5.682e-16 \| 514 \| 0.202 \| 819 \| 511 \| 24 \| 51 \| 780 \| 2 \| 766 \| Phage tail tape measure protein \| Phage tail tape measure protein; afdb-uniprot50 \| AF-A0A1V5V4L3-F1-MODEL\_V4 \| 1.0 \| 8.059e-16 \| 514 \| 0.207 \| 742 \| 441 \| 23 \| 65 \| 716 \| 10 \| 694 \| Phage-related minor tail protein \| Phage-related minor tail protein; afdb-uniprot50 \| AF-G8PUN4-F1-MODEL\_V4 \| 1.0 \| 5.403e-15 \| 514 \| 0.182 \| 1019 \| 516 \| 31 \| 20 \| 782 \| 93 \| 1049 \| Phage-related tail protein \| Phage-related tail protein; afdb-uniprot50 \| AF-A0A0F9CXC0-F1-MODEL\_V4 \| 1.0 \| 3.484e-14 \| 513 \| 0.202 \| 607 \| 433 \| 18 \| 122 \| 702 \| 5 \| 586 \| PhageMin\_Tail domain-containing protein \| PhageMin\_Tail domain-containing protein; afdb-uniprot50 \| AF-A0A2S5N2H2-F1-MODEL\_V4 \| 1.0 \| 7.966e-15 \| 513 \| 0.163 \| 831 \| 475 \| 25 \| 4 \| 781 \| 65 \| 727 \| Phage tail tape measure protein \| Phage tail tape measure protein; afdb-uniprot50 \| AF-A0A5E9SAB7-F1-MODEL\_V4 \| 1.0 \| 9.198e-14 \| 512 \| 0.206 \| 629 \| 408 \| 20 \| 32 \| 623 \| 2 \| 576 \| Phage tail tape measure protein \| Phage tail tape measure protein; afdb-uniprot50 \| AF-A0A0M7AQ48-F1-MODEL\_V4 \| 1.0 \| 1.752e-15 \| 512 \| 0.182 \| 794 \| 458 \| 25 \| 1 \| 766 \| 1 \| 631 \| Phage tail tape measure protein, TP901 family, core region \| Phage tail tape measure protein, TP901 family, core region; afdb-uniprot50 \| AF-A0A1F4X418-F1-MODEL\_V4 \| 1.0 \| 3.722e-13 \| 511 \| 0.147 \| 727 \| 447 \| 21 \| 35 \| 750 \| 7 \| 571 \| Phage tail tape measure protein \| Phage tail tape measure protein; afdb-uniprot50 \| AF-A0A1G9I6R0-F1-MODEL\_V4 \| 1.0 \| 4.753e-14 \| 511 \| 0.192 \| 581 \| 396 \| 16 \| 27 \| 566 \| 62 \| 610 \| Phage tail tape measure protein, TP901 family, core region \| Phage tail tape measure protein, TP901 family, core region; afdb-uniprot50 \| AF-A0A6L8LYC8-F1-MODEL\_V4 \| 1.0 \| 4.626e-15 \| 511 \| 0.161 \| 871 \| 532 \| 26 \| 6 \| 785 \| 1 \| 763 \| Phage tail tape measure protein \| Phage tail tape measure protein; afdb-uniprot50 \| AF-A0A0N1LUL1-F1-MODEL\_V4 \| 1.0 \| 1.5e-15 \| 510 \| 0.172 \| 864 \| 451 \| 27 \| 19 \| 785 \| 66 \| 762 \| PhageMin\_Tail domain-containing protein \| PhageMin\_Tail domain-containing protein; afdb-uniprot50 \| AF-A0A0U5L5G9-F1-MODEL\_V4 \| 1.0 \| 4.33e-16 \| 510 \| 0.184 \| 845 \| 526 \| 22 \| 19 \| 781 \| 50 \| 812 \| Tail protein \| Tail protein; afdb-uniprot50 \| AF-A0A0F9RC03-F1-MODEL\_V4 \| 1.0 \| 4.039e-10 \| 509 \| 0.179 \| 424 \| 273 \| 10 \| 27 \| 433 \| 5 \| 370 \| PhageMin\_Tail domain-containing protein \| PhageMin\_Tail domain-containing protein; afdb-uniprot50 \| AF-A0A268U7H8-F1-MODEL\_V4 \| 1.0 \| 1.082e-17 \| 508 \| 0.197 \| 826 \| 510 \| 24 \| 2 \| 720 \| 1 \| 780 \| Phage tail tape measure protein \| Phage tail tape measure protein; afdb-uniprot50 \| AF-A0A417HJE2-F1-MODEL\_V4 \| 1.0 \| 1.916e-16 \| 508 \| 0.181 \| 844 \| 497 \| 25 \| 19 \| 702 \| 10 \| 819 \| Phage tail tape measure protein \| Phage tail tape measure protein; afdb-uniprot50 \| AF-A0A517IVI2-F1-MODEL\_V4 \| 1.0 \| 6.769e-11 \| 506 \| 0.154 \| 427 \| 317 \| 10 \| 28 \| 443 \| 10 \| 403 \| Phage tail tape measure protein \| Phage tail tape measure protein; afdb-uniprot50 \| AF-A0A1X0SVA8-F1-MODEL\_V4 \| 1.0 \| 3.351e-14 \| 506 \| 0.16 \| 761 \| 435 \| 18 \| 70 \| 785 \| 2 \| 603 \| Phage tail tape measure protein \| Phage tail tape measure protein; afdb-uniprot50 \| AF-A0A559QNE5-F1-MODEL\_V4 \| 1.0 \| 3.018e-15 \| 506 \| 0.184 \| 833 \| 551 \| 26 \| 31 \| 780 \| 164 \| 950 \| TP901 family phage tail tape measure protein \| TP901 family phage tail tape measure protein; afdb-uniprot50 \| AF-K0ZBN7-F1-MODEL\_V4 \| 1.0 \| 5.403e-15 \| 505 \| 0.183 \| 745 \| 448 \| 28 \| 1 \| 655 \| 1 \| 674 \| Phage tail tape measure protein, TP901 family, core region \| Phage tail tape measure protein, TP901 family, core region; afdb-uniprot50 \| AF-A0A2T6LZL7-F1-MODEL\_V4 \| 1.0 \| 6.384e-16 \| 505 \| 0.18 \| 809 \| 560 \| 24 \| 1 \| 785 \| 1 \| 730 \| TP901 family phage tail tape measure protein \| TP901 family phage tail tape measure protein; afdb-uniprot50 \| AF-A0A0F9T8W6-F1-MODEL\_V4 \| 1.0 \| 6.436e-10 \| 504 \| 0.17 \| 392 \| 293 \| 10 \| 86 \| 465 \| 18 \| 389 \| PhageMin\_Tail domain-containing protein \| PhageMin\_Tail domain-containing protein; afdb-uniprot50 \| AF-A0A175R8T9-F1-MODEL\_V4 \| 1.0 \| 5.617e-15 \| 504 \| 0.18 \| 782 \| 480 \| 19 \| 2 \| 702 \| 1 \| 702 \| PhageMin\_Tail domain-containing protein \| PhageMin\_Tail domain-containing protein; afdb-uniprot50 \| AF-W7CZ00-F1-MODEL\_V4 \| 1.0 \| 7.457e-16 \| 504 \| 0.2 \| 687 \| 449 \| 25 \| 20 \| 662 \| 55 \| 684 \| Tail tape measure protein \| Tail tape measure protein; afdb-uniprot50 \| AF-A0A4R8HHY1-F1-MODEL\_V4 \| 1.0 \| 3.809e-15 \| 504 \| 0.177 \| 884 \| 506 \| 26 \| 27 \| 780 \| 20 \| 811 \| TP901 family phage tail tape measure protein \| TP901 family phage tail tape measure protein; afdb-uniprot50 \| AF-A0A1R1MK84-F1-MODEL\_V4 \| 1.0 \| 7.457e-16 \| 503 \| 0.197 \| 809 \| 516 \| 27 \| 55 \| 773 \| 2 \| 766 \| Phage tail tape measure protein \| Phage tail tape measure protein; afdb-uniprot50 \| AF-A0A4R2GR49-F1-MODEL\_V4 \| 1.0 \| 5.682e-16 \| 501 \| 0.186 \| 847 \| 503 \| 26 \| 21 \| 764 \| 24 \| 787 \| TP901 family phage tail tape measure protein \| TP901 family phage tail tape measure protein; afdb-uniprot50 \| AF-A0A231UT38-F1-MODEL\_V4 \| 1.0 \| 5.617e-15 \| 501 \| 0.172 \| 765 \| 449 \| 24 \| 49 \| 785 \| 76 \| 684 \| Phage tail tape measure protein \| Phage tail tape measure protein; afdb-uniprot50 \| AF-A0A268TWS7-F1-MODEL\_V4 \| 1.0 \| 1.202e-16 \| 500 \| 0.186 \| 813 \| 527 \| 24 \| 27 \| 785 \| 16 \| 747 \| Phage tail tape measure protein \| Phage tail tape measure protein; afdb-uniprot50 \| AF-A0A1W2BBP6-F1-MODEL\_V4 \| 1.0 \| 2.903e-15 \| 500 \| 0.161 \| 799 \| 487 \| 22 \| 26 \| 785 \| 1 \| 655 \| Phage tail tape measure protein, TP901 family, core region \| Phage tail tape measure protein, TP901 family, core region; afdb-uniprot50 \| AF-A0A349HHP7-F1-MODEL\_V4 \| 1.0 \| 8.645e-12 \| 499 \| 0.175 \| 523 \| 360 \| 8 \| 12 \| 475 \| 40 \| 550 \| Phage tail tape measure protein \| Phage tail tape measure protein; afdb-uniprot50 \| AF-D5EFA3-F1-MODEL\_V4 \| 1.0 \| 1.143e-15 \| 499 \| 0.163 \| 846 \| 549 \| 25 \| 19 \| 782 \| 42 \| 810 \| Phage tail tape measure protein, TP901 family \| Phage tail tape measure protein, TP901 family; afdb-uniprot50 \| AF-A0A077L0E5-F1-MODEL\_V4 \| 1.0 \| 9.413e-16 \| 499 \| 0.177 \| 841 \| 589 \| 24 \| 19 \| 783 \| 22 \| 835 \| PhageMin\_Tail domain-containing protein \| PhageMin\_Tail domain-containing protein; afdb-uniprot50 \| AF-A0A0B1Q8A4-F1-MODEL\_V4 \| 1.0 \| 1.006e-14 \| 498 \| 0.171 \| 799 \| 501 \| 24 \| 11 \| 754 \| 1 \| 693 \| PhageMin\_Tail domain-containing protein \| PhageMin\_Tail domain-containing protein; afdb-uniprot50 \| AF-A0A4Z0DLG0-F1-MODEL\_V4 \| 1.0 \| 3.211e-17 \| 497 \| 0.201 \| 887 \| 533 \| 28 \| 1 \| 782 \| 1 \| 816 \| Phage tail tape measure protein \| Phage tail tape measure protein; afdb-uniprot50 \| AF-K8EVH7-F1-MODEL\_V4 \| 1.0 \| 1.29e-12 \| 495 \| 0.176 \| 539 \| 343 \| 15 \| 12 \| 461 \| 45 \| 571 \| Phage tail tape measure protein, TP901 family,core region \| Phage tail tape measure protein, TP901 family,core region; afdb-uniprot50 \| AF-A0A660ZYG0-F1-MODEL\_V4 \| 1.0 \| 3.261e-15 \| 495 \| 0.175 \| 736 \| 479 \| 25 \| 16 \| 689 \| 1 \| 670 \| Phage tail tape measure protein \| Phage tail tape measure protein; afdb-uniprot50 \| AF-A0A3B0B7K3-F1-MODEL\_V4 \| 1.0 \| 2.212e-15 \| 495 \| 0.151 \| 997 \| 549 \| 28 \| 1 \| 785 \| 1 \| 912 \| Phage tail tape measure protein \| Phage tail tape measure protein; afdb-uniprot50 \| AF-A0A7X3HJK4-F1-MODEL\_V4 \| 1.0 \| 4.117e-15 \| 494 \| 0.176 \| 875 \| 550 \| 29 \| 19 \| 782 \| 42 \| 856 \| Phage tail tape measure protein \| Phage tail tape measure protein; afdb-uniprot50 \| AF-A0A1Q3NZU9-F1-MODEL\_V4 \| 1.0 \| 4.773e-11 \| 493 \| 0.192 \| 453 \| 323 \| 10 \| 35 \| 474 \| 7 \| 429 \| Phage tail tape measure protein \| Phage tail tape measure protein; afdb-uniprot50 \| AF-A0A7W6BZB0-F1-MODEL\_V4 \| 1.0 \| 5.137e-14 \| 493 \| 0.177 \| 798 \| 463 \| 26 \| 1 \| 785 \| 1 \| 617 \| TP901 family phage tail tape measure protein \| TP901 family phage tail tape measure protein; afdb-uniprot50 \| AF-A0A839Y122-F1-MODEL\_V4 \| 1.0 \| 1.969e-15 \| 493 \| 0.17 \| 867 \| 491 \| 27 \| 50 \| 750 \| 2 \| 805 \| TP901 family phage tail tape measure protein \| TP901 family phage tail tape measure protein; afdb-uniprot50 \| AF-A0A642C2T3-F1-MODEL\_V4 \| 1.0 \| 7.726e-09 \| 492 \| 0.137 \| 327 \| 256 \| 10 \| 117 \| 433 \| 4 \| 314 \| Phage tail tape measure protein \| Phage tail tape measure protein; afdb-uniprot50 \| AF-A0A7W5WZ30-F1-MODEL\_V4 \| 1.0 \| 6e-14 \| 492 \| 0.191 \| 794 \| 475 \| 23 \| 1 \| 728 \| 1 \| 693 \| TP901 family phage tail tape measure protein \| TP901 family phage tail tape measure protein; afdb-uniprot50 \| AF-A0A1W1ZU85-F1-MODEL\_V4 \| 1.0 \| 4.28e-15 \| 492 \| 0.2 \| 808 \| 464 \| 25 \| 31 \| 785 \| 13 \| 691 \| Phage tail tape measure protein, TP901 family, core region \| Phage tail tape measure protein, TP901 family, core region; afdb-uniprot50 \| AF-S3XXC4-F1-MODEL\_V4 \| 1.0 \| 1.443e-15 \| 492 \| 0.182 \| 941 \| 530 \| 37 \| 19 \| 785 \| 10 \| 884 \| Phage tail tape measure protein, TP901 family, core region \| Phage tail tape measure protein, TP901 family, core region; afdb-uniprot50 \| AF-A0A1B8P3Y8-F1-MODEL\_V4 \| 1.0 \| 1.346e-09 \| 491 \| 0.182 \| 356 \| 260 \| 9 \| 77 \| 426 \| 18 \| 348 \| Phage-related minor tail protein \| Phage-related minor tail protein; afdb-uniprot50 \| AF-A0A829CHN2-F1-MODEL\_V4 \| 1.0 \| 1.489e-11 \| 491 \| 0.193 \| 516 \| 342 \| 12 \| 23 \| 480 \| 50 \| 549 \| Phage tail tape measure protein, TP901 family, core region \| Phage tail tape measure protein, TP901 family, core region; afdb-uniprot50 \| AF-A0A0Q4C350-F1-MODEL\_V4 \| 1.0 \| 6.384e-16 \| 491 \| 0.196 \| 825 \| 537 \| 29 \| 2 \| 781 \| 1 \| 744 \| PhageMin\_Tail domain-containing protein \| PhageMin\_Tail domain-containing protein; afdb-uniprot50 \| AF-A0A1S1V6R2-F1-MODEL\_V4 \| 1.0 \| 8.61e-15 \| 491 \| 0.194 \| 795 \| 487 \| 29 \| 62 \| 785 \| 8 \| 719 \| Phage-related minor tail protein \| Phage-related minor tail protein; afdb-uniprot50 \| AF-A0A1Q6JQ74-F1-MODEL\_V4 \| 1.0 \| 1.226e-11 \| 489 \| 0.205 \| 506 \| 322 \| 12 \| 19 \| 466 \| 44 \| 527 \| Phage tail tape measure protein \| Phage tail tape measure protein; afdb-uniprot50 \| AF-A0A2M8WDP6-F1-MODEL\_V4 \| 1.0 \| 1.894e-15 \| 489 \| 0.173 \| 838 \| 561 \| 29 \| 1 \| 781 \| 1 \| 763 \| Phage tail tape measure protein \| Phage tail tape measure protein; afdb-uniprot50 \| AF-A0A2W6YTE4-F1-MODEL\_V4 \| 1.0 \| 9.305e-15 \| 489 \| 0.176 \| 854 \| 504 \| 30 \| 15 \| 784 \| 13 \| 750 \| Phage tail tape measure protein \| Phage tail tape measure protein; afdb-uniprot50 \| AF-A0A1L3SQ06-F1-MODEL\_V4 \| 1.0 \| 2.023e-14 \| 488 \| 0.176 \| 767 \| 463 \| 23 \| 26 \| 723 \| 1 \| 667 \| Phage tail tape measure protein \| Phage tail tape measure protein; afdb-uniprot50 \| AF-H6U8K4-F1-MODEL\_V4 \| 1.0 \| 2.825e-16 \| 488 \| 0.221 \| 766 \| 404 \| 24 \| 166 \| 776 \| 1 \| 728 \| Phage tail tape measure protein TP901 family core region \| Phage tail tape measure protein TP901 family core region; afdb-uniprot50 \| AF-A0A291LZ62-F1-MODEL\_V4 \| 1.0 \| 1.372e-14 \| 488 \| 0.178 \| 855 \| 519 \| 24 \| 1 \| 785 \| 2 \| 742 \| Phage tail tape measure protein \| Phage tail tape measure protein; afdb-uniprot50 \| AF-A0A380JYP8-F1-MODEL\_V4 \| 1.0 \| 6e-14 \| 488 \| 0.19 \| 678 \| 443 \| 19 \| 67 \| 702 \| 15 \| 628 \| Phage protein \| Phage protein; afdb-uniprot50 \| AF-A0A4R3NZQ3-F1-MODEL\_V4 \| 1.0 \| 1.752e-15 \| 487 \| 0.205 \| 795 \| 497 \| 29 \| 39 \| 776 \| 4 \| 720 \| TP901 family phage tail tape measure protein \| TP901 family phage tail tape measure protein; afdb-uniprot50 \| AF-A0A379YC51-F1-MODEL\_V4 \| 1.0 \| 1.335e-15 \| 487 \| 0.183 \| 800 \| 499 \| 25 \| 19 \| 690 \| 48 \| 820 \| Phage tail tape measure protein, TP901 family, core region \| Phage tail tape measure protein, TP901 family, core region; afdb-uniprot50 \| AF-A0A101IPF4-F1-MODEL\_V4 \| 1.0 \| 8.447e-10 \| 486 \| 0.175 \| 358 \| 265 \| 11 \| 70 \| 422 \| 10 \| 342 \| Phage tail tape measure protein, TP901 family, core region \| Phage tail tape measure protein, TP901 family, core region; afdb-uniprot50 \| AF-A0A5R8LBZ4-F1-MODEL\_V4 \| 1.0 \| 1.32e-14 \| 486 \| 0.186 \| 741 \| 457 \| 22 \| 35 \| 711 \| 14 \| 672 \| Phage tail tape measure protein \| Phage tail tape measure protein; afdb-uniprot50 \| AF-A0A2X2BQ74-F1-MODEL\_V4 \| 1.0 \| 4.117e-15 \| 485 \| 0.168 \| 823 \| 516 \| 28 \| 81 \| 784 \| 3 \| 775 \| Phage tail fiber protein \| Phage tail fiber protein; afdb-uniprot50 \| AF-W7BV28-F1-MODEL\_V4 \| 1.0 \| 1.909e-09 \| 484 \| 0.189 \| 328 \| 206 \| 8 \| 137 \| 408 \| 4 \| 327 \| TP901 family phage tail tape measure protein \| TP901 family phage tail tape measure protein; afdb-uniprot50 \| AF-A0A6M3L5E0-F1-MODEL\_V4 \| 1.0 \| 1.512e-09 \| 484 \| 0.135 \| 392 \| 306 \| 11 \| 88 \| 469 \| 6 \| 374 \| Putative tail protein \| Putative tail protein; afdb-uniprot50 \| AF-A0A2D6X7M0-F1-MODEL\_V4 \| 1.0 \| 1.732e-14 \| 484 \| 0.188 \| 784 \| 494 \| 27 \| 66 \| 785 \| 3 \| 707 \| Phage tail tape measure protein \| Phage tail tape measure protein; afdb-uniprot50 \| AF-A0A4R7DE84-F1-MODEL\_V4 \| 1.0 \| 8.951e-15 \| 484 \| 0.179 \| 813 \| 546 \| 28 \| 1 \| 781 \| 1 \| 724 \| TP901 family phage tail tape measure protein \| TP901 family phage tail tape measure protein; afdb-uniprot50 \| AF-R2T824-F1-MODEL\_V4 \| 1.0 \| 7.457e-16 \| 483 \| 0.192 \| 775 \| 513 \| 23 \| 19 \| 702 \| 123 \| 875 \| Phage tail tape measure protein, TP901 family, core region \| Phage tail tape measure protein, TP901 family, core region; afdb-uniprot50 \| AF-A0A021VU74-F1-MODEL\_V4 \| 1.0 \| 4.449e-15 \| 482 \| 0.161 \| 977 \| 554 \| 30 \| 14 \| 785 \| 1 \| 916 \| PhageMin\_Tail domain-containing protein \| PhageMin\_Tail domain-containing protein; afdb-uniprot50 \| AF-A0A0C5XG03-F1-MODEL\_V4 \| 1.0 \| 2.212e-15 \| 482 \| 0.174 \| 913 \| 527 \| 32 \| 61 \| 785 \| 7 \| 880 \| Phage tail length tape-measure protein \| Phage tail length tape-measure protein; afdb-uniprot50 \| AF-A0A3R8SD78-F1-MODEL\_V4 \| 1.0 \| 1.969e-15 \| 482 \| 0.163 \| 915 \| 547 \| 30 \| 39 \| 765 \| 2 \| 885 \| Phage tail tape measure protein \| Phage tail tape measure protein; afdb-uniprot50 \| AF-A0A7C5G4Y5-F1-MODEL\_V4 \| 1.0 \| 4.039e-10 \| 481 \| 0.179 \| 424 \| 313 \| 14 \| 62 \| 473 \| 8 \| 408 \| Phage tail tape measure protein \| Phage tail tape measure protein; afdb-uniprot50 \| AF-A0A0Q5FLM7-F1-MODEL\_V4 \| 1.0 \| 3.622e-14 \| 481 \| 0.157 \| 923 \| 528 \| 26 \| 76 \| 785 \| 14 \| 899 \| PhageMin\_Tail domain-containing protein \| PhageMin\_Tail domain-containing protein; afdb-uniprot50 \| AF-A0A1I5W4T8-F1-MODEL\_V4 \| 1.0 \| 3.224e-14 \| 481 \| 0.222 \| 637 \| 406 \| 20 \| 109 \| 710 \| 50 \| 631 \| Phage tail tape measure protein, TP901 family, core region \| Phage tail tape measure protein, TP901 family, core region; afdb-uniprot50 \| AF-A0A6B4JNP3-F1-MODEL\_V4 \| 1.0 \| 1.673e-11 \| 481 \| 0.178 \| 488 \| 336 \| 12 \| 1 \| 459 \| 1 \| 452 \| Phage tail tape measure protein \| Phage tail tape measure protein; afdb-uniprot50 \| AF-A0A0Q5GQH4-F1-MODEL\_V4 \| 1.0 \| 7.457e-16 \| 480 \| 0.18 \| 896 \| 548 \| 24 \| 1 \| 780 \| 1 \| 825 \| PhageMin\_Tail domain-containing protein \| PhageMin\_Tail domain-containing protein; afdb-uniprot50 \| AF-A0A6G3L4W1-F1-MODEL\_V4 \| 1.0 \| 1.666e-14 \| 480 \| 0.186 \| 921 \| 515 \| 25 \| 42 \| 785 \| 167 \| 1029 \| Phage tail tape measure protein \| Phage tail tape measure protein; afdb-uniprot50 \| AF-X0RKC4-F1-MODEL\_V4 \| 1.0 \| 2.686e-15 \| 480 \| 0.183 \| 772 \| 496 \| 27 \| 27 \| 706 \| 53 \| 781 \| PhageMin\_Tail domain-containing protein \| PhageMin\_Tail domain-containing protein; afdb-uniprot50 \| AF-A0A376KHT3-F1-MODEL\_V4 \| 1.0 \| 2.554e-14 \| 479 \| 0.193 \| 697 \| 424 \| 19 \| 21 \| 639 \| 48 \| 684 \| Tail protein (Modular protein) \| Tail protein (Modular protein); afdb-uniprot50 \| AF-A0A0Q0XGG6-F1-MODEL\_V4 \| 1.0 \| 1.946e-14 \| 479 \| 0.17 \| 778 \| 508 \| 27 \| 51 \| 780 \| 9 \| 696 \| PhageMin\_Tail domain-containing protein \| PhageMin\_Tail domain-containing protein; afdb-uniprot50 \| AF-E2S7P3-F1-MODEL\_V4 \| 1.0 \| 4.626e-15 \| 478 \| 0.175 \| 808 \| 497 \| 27 \| 61 \| 750 \| 11 \| 766 \| Phage tail tape measure protein, TP901 family \| Phage tail tape measure protein, TP901 family; afdb-uniprot50 \| AF-A0A3A9ICH5-F1-MODEL\_V4 \| 1.0 \| 4.572e-14 \| 478 \| 0.176 \| 917 \| 585 \| 26 \| 3 \| 784 \| 1 \| 881 \| Phage tail tape measure protein \| Phage tail tape measure protein; afdb-uniprot50 \| AF-A0A2E3N1K5-F1-MODEL\_V4 \| 1.0 \| 1.372e-14 \| 477 \| 0.163 \| 868 \| 510 \| 30 \| 1 \| 780 \| 1 \| 739 \| Phage tail tape measure protein \| Phage tail tape measure protein; afdb-uniprot50 \| AF-A0A0X8GNA4-F1-MODEL\_V4 \| 1.0 \| 2.363e-14 \| 477 \| 0.188 \| 869 \| 546 \| 29 \| 19 \| 782 \| 49 \| 862 \| PhageMin\_Tail domain-containing protein \| PhageMin\_Tail domain-containing protein; afdb-uniprot50 \| AF-J3DZE3-F1-MODEL\_V4 \| 1.0 \| 2.869e-14 \| 476 \| 0.192 \| 862 \| 456 \| 25 \| 1 \| 783 \| 26 \| 726 \| Phage tail tape measure protein, TP901 family \| Phage tail tape measure protein, TP901 family; afdb-uniprot50 \| AF-A0A836WW60-F1-MODEL\_V4 \| 1.0 \| 7.173e-16 \| 476 \| 0.152 \| 906 \| 584 \| 31 \| 28 \| 780 \| 4 \| 878 \| Phage tail tape measure protein \| Phage tail tape measure protein; afdb-uniprot50 \| AF-A0A7X9B3Q6-F1-MODEL\_V4 \| 1.0 \| 2.848e-10 \| 475 \| 0.159 \| 421 \| 310 \| 11 \| 77 \| 489 \| 20 \| 404 \| Phage tail tape measure protein \| Phage tail tape measure protein; afdb-uniprot50 \| AF-A0A5C7JCR9-F1-MODEL\_V4 \| 1.0 \| 2.055e-12 \| 475 \| 0.174 \| 619 \| 429 \| 19 \| 47 \| 625 \| 2 \| 578 \| Phage tail tape measure protein \| Phage tail tape measure protein; afdb-uniprot50 \| AF-A0A7C4AJ87-F1-MODEL\_V4 \| 1.0 \| 2.363e-14 \| 475 \| 0.174 \| 795 \| 502 \| 25 \| 30 \| 785 \| 6 \| 684 \| Phage tail tape measure protein \| Phage tail tape measure protein; afdb-uniprot50 \| AF-R9J8Y0-F1-MODEL\_V4 \| 1.0 \| 2.023e-14 \| 475 \| 0.161 \| 683 \| 481 \| 21 \| 39 \| 702 \| 3 \| 612 \| Phage tail tape measure protein, TP901 family, core region \| Phage tail tape measure protein, TP901 family, core region; afdb-uniprot50 \| AF-K6DYV4-F1-MODEL\_V4 \| 1.0 \| 1.732e-14 \| 474 \| 0.193 \| 744 \| 452 \| 27 \| 1 \| 701 \| 1 \| 639 \| PhageMin\_Tail domain-containing protein \| PhageMin\_Tail domain-containing protein; afdb-uniprot50 \| AF-A0A377ALG4-F1-MODEL\_V4 \| 1.0 \| 3.237e-11 \| 473 \| 0.231 \| 384 \| 254 \| 7 \| 182 \| 556 \| 11 \| 362 \| TP901 family phage tail tape measure protein \| TP901 family phage tail tape measure protein; afdb-uniprot50 \| AF-A0A2N3J4A2-F1-MODEL\_V4 \| 1.0 \| 3.261e-15 \| 473 \| 0.165 \| 859 \| 530 \| 30 \| 19 \| 782 \| 46 \| 812 \| Phage tail protein \| Phage tail protein; afdb-uniprot50 \| AF-A0A6G2L5R9-F1-MODEL\_V4 \| 1.0 \| 3.664e-15 \| 473 \| 0.171 \| 852 \| 524 \| 20 \| 1 \| 679 \| 1 \| 843 \| Phage tail tape measure protein \| Phage tail tape measure protein; afdb-uniprot50 \| AF-A0A061P2R0-F1-MODEL\_V4 \| 1.0 \| 1.692e-12 \| 472 \| 0.171 \| 607 \| 393 \| 20 \| 5 \| 541 \| 2 \| 568 \| Phage tail length tape-measure protein \| Phage tail length tape-measure protein; afdb-uniprot50 \| AF-F2J629-F1-MODEL\_V4 \| 1.0 \| 9.413e-16 \| 472 \| 0.188 \| 835 \| 516 \| 32 \| 13 \| 760 \| 1 \| 760 \| Phage tail tape measure protein, TP901 family, core region \| Phage tail tape measure protein, TP901 family, core region; afdb-uniprot50 \| AF-A0A653LRG4-F1-MODEL\_V4 \| 1.0 \| 5.839e-15 \| 472 \| 0.184 \| 785 \| 504 \| 26 \| 19 \| 708 \| 41 \| 783 \| PhageMin\_Tail domain-containing protein \| PhageMin\_Tail domain-containing protein; afdb-uniprot50 \| AF-A0A6V6Y4C0-F1-MODEL\_V4 \| 1.0 \| 1.045e-14 \| 471 \| 0.176 \| 835 \| 534 \| 22 \| 49 \| 782 \| 3 \| 784 \| Phage tail tape measure protein, TP901 family \| Phage tail tape measure protein, TP901 family; afdb-uniprot50 \| AF-I3DCP6-F1-MODEL\_V4 \| 1.0 \| 4.449e-15 \| 471 \| 0.16 \| 876 \| 548 \| 26 \| 19 \| 782 \| 16 \| 815 \| Phage tail tape measure protein, TP901 family \| Phage tail tape measure protein, TP901 family; afdb-uniprot50 \| AF-A0A1B3E8J2-F1-MODEL\_V4 \| 1.0 \| 3.664e-15 \| 471 \| 0.169 \| 873 \| 554 \| 26 \| 20 \| 782 \| 36 \| 847 \| Tail tape measure protein \| Tail tape measure protein; afdb-uniprot50 \| AF-A0A852Q908-F1-MODEL\_V4 \| 1.0 \| 4.999e-15 \| 471 \| 0.184 \| 927 \| 514 \| 30 \| 21 \| 780 \| 39 \| 890 \| Phage tail tape measure protein \| Phage tail tape measure protein; afdb-uniprot50 \| AF-A0A839FTM8-F1-MODEL\_V4 \| 1.0 \| 2.363e-14 \| 471 \| 0.177 \| 753 \| 465 \| 25 \| 7 \| 708 \| 1 \| 649 \| TP901 family phage tail tape measure protein \| TP901 family phage tail tape measure protein; afdb-uniprot50 \| AF-A0A446I8U4-F1-MODEL\_V4 \| 1.0 \| 2.023e-14 \| 471 \| 0.161 \| 835 \| 521 \| 26 \| 19 \| 702 \| 74 \| 879 \| Phage tail tape measure protein \| Phage tail tape measure protein; afdb-uniprot50 \| AF-A0A7X7XZN6-F1-MODEL\_V4 \| 1.0 \| 8.61e-15 \| 470 \| 0.183 \| 756 \| 506 \| 20 \| 27 \| 708 \| 53 \| 770 \| Phage tail tape measure protein \| Phage tail tape measure protein; afdb-uniprot50 \| AF-A0A2E5BTY7-F1-MODEL\_V4 \| 1.0 \| 6.56e-15 \| 470 \| 0.177 \| 864 \| 539 \| 27 \| 18 \| 780 \| 49 \| 841 \| Phage tail tape measure protein \| Phage tail tape measure protein; afdb-uniprot50 \| AF-A0A6N6NKD3-F1-MODEL\_V4 \| 1.0 \| 6.485e-14 \| 470 \| 0.182 \| 749 \| 486 \| 23 \| 1 \| 654 \| 1 \| 717 \| Phage tail tape measure protein \| Phage tail tape measure protein; afdb-uniprot50 \| AF-A0A1E4AIA1-F1-MODEL\_V4 \| 1.0 \| 1.541e-14 \| 469 \| 0.182 \| 806 \| 484 \| 26 \| 35 \| 776 \| 8 \| 702 \| Phage tail tape measure protein \| Phage tail tape measure protein; afdb-uniprot50 \| AF-F1Z9C8-F1-MODEL\_V4 \| 1.0 \| 8.282e-15 \| 469 \| 0.173 \| 828 \| 525 \| 33 \| 1 \| 776 \| 1 \| 720 \| TP901 family phage tail tape measure protein \| TP901 family phage tail tape measure protein; afdb-uniprot50 \| AF-A0A075JTQ7-F1-MODEL\_V4 \| 1.0 \| 3.622e-14 \| 469 \| 0.151 \| 811 \| 532 \| 26 \| 69 \| 785 \| 18 \| 765 \| PhageMin\_Tail domain-containing protein \| PhageMin\_Tail domain-containing protein; afdb-uniprot50 \| AF-A0A556S942-F1-MODEL\_V4 \| 1.0 \| 8.61e-15 \| 469 \| 0.16 \| 851 \| 533 \| 17 \| 19 \| 702 \| 36 \| 871 \| Phage tail tape measure protein \| Phage tail tape measure protein; afdb-uniprot50 \| AF-A0A258KVP7-F1-MODEL\_V4 \| 1.0 \| 3.622e-14 \| 468 \| 0.18 \| 769 \| 497 \| 26 \| 96 \| 781 \| 8 \| 725 \| Phage tail tape measure protein \| Phage tail tape measure protein; afdb-uniprot50 \| AF-A0A6G6ZVQ1-F1-MODEL\_V4 \| 1.0 \| 1.483e-14 \| 468 \| 0.176 \| 906 \| 475 \| 28 \| 1 \| 783 \| 22 \| 778 \| Phage tail tape measure protein \| Phage tail tape measure protein; afdb-uniprot50 \| AF-A0A6N2ZLI6-F1-MODEL\_V4 \| 1.0 \| 1.006e-14 \| 468 \| 0.151 \| 837 \| 565 \| 22 \| 19 \| 782 \| 19 \| 782 \| Phage-related minor tail protein \| Phage-related minor tail protein; afdb-uniprot50 \| AF-S6GUR2-F1-MODEL\_V4 \| 1.0 \| 1.045e-14 \| 468 \| 0.176 \| 883 \| 546 \| 26 \| 19 \| 782 \| 57 \| 876 \| PhageMin\_Tail domain-containing protein \| PhageMin\_Tail domain-containing protein; afdb-uniprot50 \| AF-A0A1D9FNG0-F1-MODEL\_V4 \| 1.0 \| 1.85e-13 \| 467 \| 0.15 \| 773 \| 507 \| 28 \| 60 \| 783 \| 9 \| 680 \| Phage tail tape measure protein \| Phage tail tape measure protein; afdb-uniprot50 \| AF-A0A157SJY1-F1-MODEL\_V4 \| 1.0 \| 7.966e-15 \| 467 \| 0.169 \| 957 \| 572 \| 25 \| 22 \| 785 \| 53 \| 979 \| Bacteriophage protein \| Bacteriophage protein; afdb-uniprot50 \| AF-A0A072Y8W2-F1-MODEL\_V4 \| 1.0 \| 2.008e-10 \| 467 \| 0.165 \| 471 \| 304 \| 12 \| 1 \| 465 \| 102 \| 489 \| Phage tail tape measure protein \| Phage tail tape measure protein; afdb-uniprot50 \| AF-A0A2W4M4D8-F1-MODEL\_V4 \| 1.0 \| 4.591e-11 \| 466 \| 0.161 \| 507 \| 356 \| 12 \| 16 \| 510 \| 1 \| 450 \| Phage tail tape measure protein \| Phage tail tape measure protein; afdb-uniprot50 \| AF-A0A7V8U794-F1-MODEL\_V4 \| 1.0 \| 3.809e-15 \| 466 \| 0.178 \| 814 \| 550 \| 26 \| 1 \| 781 \| 1 \| 728 \| Phage tail tape measure protein \| Phage tail tape measure protein; afdb-uniprot50 \| AF-A0A7I0K7H0-F1-MODEL\_V4 \| 1.0 \| 4.626e-15 \| 466 \| 0.188 \| 813 \| 534 \| 26 \| 47 \| 782 \| 3 \| 766 \| Phage tail tape measure protein \| Phage tail tape measure protein; afdb-uniprot50 \| AF-A0A0F7LMA5-F1-MODEL\_V4 \| 1.0 \| 2.103e-14 \| 466 \| 0.153 \| 878 \| 577 \| 23 \| 19 \| 785 \| 62 \| 883 \| Tail protein \| Tail protein; afdb-uniprot50 \| AF-A0A223S641-F1-MODEL\_V4 \| 1.0 \| 1.087e-14 \| 466 \| 0.162 \| 912 \| 568 \| 32 \| 35 \| 785 \| 5 \| 881 \| Phage tail tape measure protein \| Phage tail tape measure protein; afdb-uniprot50 \| AF-A0A1V3VRH9-F1-MODEL\_V4 \| 1.0 \| 8.781e-10 \| 465 \| 0.17 \| 429 \| 314 \| 10 \| 32 \| 439 \| 4 \| 411 \| Phage tail tape measure protein \| Phage tail tape measure protein; afdb-uniprot50 \| AF-A0A0F9TFS1-F1-MODEL\_V4 \| 1.0 \| 1.879e-11 \| 464 \| 0.177 \| 541 \| 354 \| 20 \| 42 \| 572 \| 5 \| 464 \| PhageMin\_Tail domain-containing protein \| PhageMin\_Tail domain-containing protein; afdb-uniprot50 \| AF-A0A632WVI2-F1-MODEL\_V4 \| 1.0 \| 2.4e-12 \| 464 \| 0.162 \| 700 \| 455 \| 21 \| 118 \| 781 \| 31 \| 634 \| Phage tail tape measure protein \| Phage tail tape measure protein; afdb-uniprot50 \| AF-Q30VW6-F1-MODEL\_V4 \| 1.0 \| 4.753e-14 \| 464 \| 0.155 \| 859 \| 519 \| 25 \| 1 \| 785 \| 1 \| 726 \| Phage tail tape measure protein, TP901 family \| Phage tail tape measure protein, TP901 family; afdb-uniprot50 \| AF-A0A1W9UFW0-F1-MODEL\_V4 \| 1.0 \| 1.872e-14 \| 464 \| 0.183 \| 821 \| 534 \| 22 \| 61 \| 785 \| 3 \| 782 \| Phage tail tape measure protein \| Phage tail tape measure protein; afdb-uniprot50 \| AF-A0A316QC74-F1-MODEL\_V4 \| 1.0 \| 2e-13 \| 463 \| 0.154 \| 720 \| 518 \| 20 \| 19 \| 702 \| 19 \| 683 \| Phage tail tape measure protein \| Phage tail tape measure protein; afdb-uniprot50 \| AF-A0A497UTH2-F1-MODEL\_V4 \| 1.0 \| 8.847e-14 \| 461 \| 0.175 \| 771 \| 511 \| 23 \| 64 \| 776 \| 11 \| 714 \| TP901 family phage tail tape measure protein \| TP901 family phage tail tape measure protein; afdb-uniprot50 \| AF-W6N4J1-F1-MODEL\_V4 \| 1.0 \| 7.286e-14 \| 461 \| 0.16 \| 771 \| 518 \| 23 \| 75 \| 780 \| 19 \| 724 \| Phage tail length tape-measure protein \| Phage tail length tape-measure protein; afdb-uniprot50 \| AF-A0A1I4ZMH4-F1-MODEL\_V4 \| 1.0 \| 4.999e-15 \| 461 \| 0.16 \| 998 \| 585 \| 31 \| 12 \| 785 \| 175 \| 1143 \| Phage tail tape measure protein, TP901 family, core region \| Phage tail tape measure protein, TP901 family, core region; afdb-uniprot50 \| AF-A0A2U1BMI5-F1-MODEL\_V4 \| 1.0 \| 4.231e-14 \| 460 \| 0.162 \| 851 \| 563 \| 25 \| 22 \| 780 \| 53 \| 845 \| TP901 family phage tail tape measure protein \| TP901 family phage tail tape measure protein; afdb-uniprot50 \| AF-B6XAD2-F1-MODEL\_V4 \| 1.0 \| 1.074e-13 \| 460 \| 0.149 \| 898 \| 548 \| 22 \| 19 \| 784 \| 57 \| 870 \| Phage tail tape measure protein, TP901 family \| Phage tail tape measure protein, TP901 family; afdb-uniprot50 \| AF-A0A6L5FX98-F1-MODEL\_V4 \| 1.0 \| 2.161e-13 \| 459 \| 0.173 \| 776 \| 493 \| 30 \| 63 \| 781 \| 16 \| 699 \| Phage tail tape measure protein \| Phage tail tape measure protein; afdb-uniprot50 \| AF-A0A7Y4MDX7-F1-MODEL\_V4 \| 1.0 \| 5.137e-14 \| 459 \| 0.186 \| 747 \| 452 \| 27 \| 47 \| 711 \| 2 \| 674 \| Phage tail tape measure protein \| Phage tail tape measure protein; afdb-uniprot50 \| AF-A0A4U7BTH2-F1-MODEL\_V4 \| 1.0 \| 1.726e-07 \| 458 \| 0.202 \| 252 \| 195 \| 3 \| 126 \| 376 \| 5 \| 251 \| Phage tail tape measure protein \| Phage tail tape measure protein; afdb-uniprot50 \| AF-A0A081RQL7-F1-MODEL\_V4 \| 1.0 \| 5.575e-11 \| 458 \| 0.135 \| 540 \| 382 \| 12 \| 19 \| 498 \| 48 \| 562 \| Phage-related minor tail protein \| Phage-related minor tail protein; afdb-uniprot50 \| AF-A0A3S8ZPZ1-F1-MODEL\_V4 \| 1.0 \| 6.238e-14 \| 458 \| 0.158 \| 971 \| 580 \| 30 \| 19 \| 782 \| 46 \| 985 \| Phage tail tape measure protein \| Phage tail tape measure protein; afdb-uniprot50 \| AF-A0A662KJ26-F1-MODEL\_V4 \| 1.0 \| 4.848e-09 \| 456 \| 0.175 \| 398 \| 288 \| 12 \| 61 \| 441 \| 1 \| 375 \| Phage tail tape measure protein \| Phage tail tape measure protein; afdb-uniprot50 \| AF-A0A833NTL2-F1-MODEL\_V4 \| 1.0 \| 1.977e-12 \| 456 \| 0.19 \| 620 \| 387 \| 17 \| 21 \| 570 \| 48 \| 622 \| Phage tail tape measure protein \| Phage tail tape measure protein; afdb-uniprot50 \| AF-A0A256CBP9-F1-MODEL\_V4 \| 1.0 \| 1.466e-13 \| 456 \| 0.14 \| 816 \| 514 \| 24 \| 19 \| 785 \| 9 \| 685 \| Phage tail tape measure protein \| Phage tail tape measure protein; afdb-uniprot50 \| AF-A0A1I7FWK0-F1-MODEL\_V4 \| 1.0 \| 2.554e-14 \| 456 \| 0.144 \| 899 \| 585 \| 26 \| 17 \| 781 \| 3 \| 850 \| Phage tail tape measure protein, TP901 family, core region \| Phage tail tape measure protein, TP901 family, core region; afdb-uniprot50 \| AF-A0A658JFX6-F1-MODEL\_V4 \| 1.0 \| 2.063e-09 \| 455 \| 0.157 \| 419 \| 315 \| 12 \| 35 \| 441 \| 2 \| 394 \| Phage tail tape measure protein \| Phage tail tape measure protein; afdb-uniprot50 \| AF-A0A081MYF0-F1-MODEL\_V4 \| 1.0 \| 2.728e-13 \| 455 \| 0.173 \| 812 \| 471 \| 21 \| 27 \| 776 \| 60 \| 732 \| PhageMin\_Tail domain-containing protein \| PhageMin\_Tail domain-containing protein; afdb-uniprot50 \| AF-X1H190-F1-MODEL\_V4 \| 1.0 \| 1.026e-09 \| 454 \| 0.162 \| 443 \| 288 \| 13 \| 19 \| 447 \| 7 \| 380 \| PhageMin\_Tail domain-containing protein \| PhageMin\_Tail domain-containing protein; afdb-uniprot50 \| AF-X1FNH0-F1-MODEL\_V4 \| 1.0 \| 5.24e-09 \| 453 \| 0.162 \| 407 \| 303 \| 10 \| 119 \| 508 \| 2 \| 387 \| PhageMin\_Tail domain-containing protein \| PhageMin\_Tail domain-containing protein; afdb-uniprot50 \| AF-A0A1H1RSV7-F1-MODEL\_V4 \| 1.0 \| 2.554e-14 \| 453 \| 0.174 \| 888 \| 472 \| 24 \| 1 \| 783 \| 22 \| 753 \| Phage tail tape measure protein, TP901 family, core region \| Phage tail tape measure protein, TP901 family, core region; afdb-uniprot50 \| AF-A0A416DY15-F1-MODEL\_V4 \| 1.0 \| 2.554e-14 \| 453 \| 0.169 \| 909 \| 564 \| 36 \| 1 \| 785 \| 1 \| 842 \| Phage tail tape measure protein \| Phage tail tape measure protein; afdb-uniprot50 \| AF-A0A0F9L912-F1-MODEL\_V4 \| 1.0 \| 1.014e-08 \| 452 \| 0.192 \| 390 \| 276 \| 12 \| 70 \| 443 \| 4 \| 370 \| PhageMin\_Tail domain-containing protein \| PhageMin\_Tail domain-containing protein; afdb-uniprot50 \| AF-A0A1E4MVY1-F1-MODEL\_V4 \| 1.0 \| 4.572e-14 \| 452 \| 0.166 \| 812 \| 496 \| 27 \| 1 \| 784 \| 1 \| 659 \| Phage tail tape measure protein \| Phage tail tape measure protein; afdb-uniprot50 \| AF-A0A239A9Y9-F1-MODEL\_V4 \| 1.0 \| 1.117e-13 \| 452 \| 0.171 \| 907 \| 541 \| 29 \| 19 \| 785 \| 33 \| 868 \| Phage tail tape measure protein, TP901 family, core region \| Phage tail tape measure protein, TP901 family, core region; afdb-uniprot50 \| AF-A0A268TN51-F1-MODEL\_V4 \| 1.0 \| 2.728e-13 \| 451 \| 0.172 \| 804 \| 501 \| 22 \| 6 \| 764 \| 1 \| 684 \| Phage tail tape measure protein \| Phage tail tape measure protein; afdb-uniprot50 \| AF-A0A5S5CR07-F1-MODEL\_V4 \| 1.0 \| 5.403e-15 \| 451 \| 0.162 \| 1014 \| 554 \| 36 \| 1 \| 784 \| 1 \| 948 \| TP901 family phage tail tape measure protein \| TP901 family phage tail tape measure protein; afdb-uniprot50 \| AF-A0A6M3XFH2-F1-MODEL\_V4 \| 1.0 \| 2.635e-10 \| 450 \| 0.203 \| 535 \| 320 \| 20 \| 46 \| 570 \| 3 \| 441 \| Putative tail protein \| Putative tail protein; afdb-uniprot50 \| AF-A0A235B980-F1-MODEL\_V4 \| 1.0 \| 2.079e-13 \| 450 \| 0.16 \| 705 \| 485 \| 23 \| 50 \| 702 \| 2 \| 651 \| Phage tail tape measure protein \| Phage tail tape measure protein; afdb-uniprot50 \| AF-A0A7V2AMB8-F1-MODEL\_V4 \| 1.0 \| 7.009e-14 \| 450 \| 0.177 \| 739 \| 490 \| 23 \| 1 \| 676 \| 1 \| 684 \| Phage tail tape measure protein \| Phage tail tape measure protein; afdb-uniprot50 \| AF-A0A371PCG1-F1-MODEL\_V4 \| 1.0 \| 2.655e-14 \| 450 \| 0.16 \| 784 \| 506 \| 21 \| 1 \| 701 \| 27 \| 740 \| Phage tail tape measure protein \| Phage tail tape measure protein; afdb-uniprot50 \| AF-A0A840DK27-F1-MODEL\_V4 \| 1.0 \| 1.524e-13 \| 449 \| 0.17 \| 756 \| 479 \| 27 \| 1 \| 701 \| 1 \| 663 \| TP901 family phage tail tape measure protein \| TP901 family phage tail tape measure protein; afdb-uniprot50 \| AF-A0A3D3CGB4-F1-MODEL\_V4 \| 1.0 \| 1.13e-14 \| 448 \| 0.168 \| 892 \| 525 \| 35 \| 52 \| 782 \| 13 \| 848 \| Phage tail tape measure protein \| Phage tail tape measure protein; afdb-uniprot50 \| AF-A0A166IL48-F1-MODEL\_V4 \| 1.0 \| 2.728e-13 \| 448 \| 0.156 \| 839 \| 422 \| 21 \| 19 \| 755 \| 56 \| 710 \| PhageMin\_Tail domain-containing protein \| PhageMin\_Tail domain-containing protein; afdb-uniprot50 \| AF-A0A7U9XL26-F1-MODEL\_V4 \| 1.0 \| 3.224e-14 \| 448 \| 0.17 \| 761 \| 493 \| 25 \| 19 \| 702 \| 79 \| 777 \| Chromosome partition protein Smc \| Chromosome partition protein Smc; afdb-uniprot50 \| AF-A0A0F9GIW1-F1-MODEL\_V4 \| 1.0 \| 7.316e-11 \| 447 \| 0.225 \| 444 \| 287 \| 15 \| 166 \| 604 \| 2 \| 393 \| PhageMin\_Tail domain-containing protein \| PhageMin\_Tail domain-containing protein; afdb-uniprot50 \| AF-A0A1I7G2Q8-F1-MODEL\_V4 \| 1.0 \| 3.885e-10 \| 447 \| 0.156 \| 461 \| 317 \| 14 \| 27 \| 439 \| 72 \| 508 \| Phage tail tape measure protein, TP901 family, core region \| Phage tail tape measure protein, TP901 family, core region; afdb-uniprot50 \| AF-A0A6M3J046-F1-MODEL\_V4 \| 1.0 \| 9.601e-11 \| 447 \| 0.176 \| 709 \| 366 \| 26 \| 91 \| 782 \| 24 \| 531 \| Putative tail protein \| Putative tail protein; afdb-uniprot50 \| AF-A0A7Y6Z4I9-F1-MODEL\_V4 \| 1.0 \| 2.055e-12 \| 447 \| 0.173 \| 768 \| 448 \| 26 \| 17 \| 765 \| 1 \| 600 \| Phage tail tape measure protein \| Phage tail tape measure protein; afdb-uniprot50 \| AF-A0A1X3I4F2-F1-MODEL\_V4 \| 1.0 \| 5.795e-11 \| 447 \| 0.161 \| 527 \| 359 \| 14 \| 19 \| 480 \| 47 \| 555 \| Putative tail protein \| Putative tail protein; afdb-uniprot50 \| AF-A0A4S3LKS8-F1-MODEL\_V4 \| 1.0 \| 4.023e-13 \| 446 \| 0.158 \| 758 \| 503 \| 20 \| 20 \| 715 \| 41 \| 725 \| Phage tail tape measure protein \| Phage tail tape measure protein; afdb-uniprot50 \| AF-H4FBM6-F1-MODEL\_V4 \| 1.0 \| 3.484e-14 \| 446 \| 0.165 \| 847 \| 541 \| 22 \| 19 \| 785 \| 3 \| 763 \| Phage tail tape measure protein, TP901 family \| Phage tail tape measure protein, TP901 family; afdb-uniprot50 \| AF-A0A5Q2N0Q6-F1-MODEL\_V4 \| 1.0 \| 5.552e-14 \| 446 \| 0.177 \| 886 \| 579 \| 35 \| 2 \| 783 \| 1 \| 840 \| Phage tail tape measure protein, TP901 family protein \| Phage tail tape measure protein, TP901 family protein; afdb-uniprot50 \| AF-A0A410QG87-F1-MODEL\_V4 \| 1.0 \| 8.186e-14 \| 445 \| 0.149 \| 910 \| 576 \| 34 \| 19 \| 783 \| 49 \| 905 \| Phage tail tape measure protein \| Phage tail tape measure protein; afdb-uniprot50 \| AF-A0A7H0U435-F1-MODEL\_V4 \| 1.0 \| 4.069e-14 \| 445 \| 0.158 \| 917 \| 575 \| 27 \| 19 \| 783 \| 39 \| 910 \| Phage tail tape measure protein \| Phage tail tape measure protein; afdb-uniprot50 \| AF-A0A1M6EWV2-F1-MODEL\_V4 \| 1.0 \| 1.245e-09 \| 445 \| 0.191 \| 439 \| 300 \| 15 \| 52 \| 472 \| 18 \| 419 \| Phage tail tape measure protein, TP901 family, core region \| Phage tail tape measure protein, TP901 family, core region; afdb-uniprot50 \| AF-A0A024Q9Y0-F1-MODEL\_V4 \| 1.0 \| 1.117e-13 \| 445 \| 0.187 \| 698 \| 446 \| 25 \| 69 \| 702 \| 16 \| 656 \| Phage tail tape measure protein, TP901 family, core region \| Phage tail tape measure protein, TP901 family, core region; afdb-uniprot50 \| AF-A0A5M6DGJ8-F1-MODEL\_V4 \| 1.0 \| 2e-13 \| 445 \| 0.164 \| 819 \| 531 \| 26 \| 63 \| 785 \| 2 \| 762 \| Phage tail tape measure protein \| Phage tail tape measure protein; afdb-uniprot50 \| AF-A0A3D4MZY3-F1-MODEL\_V4 \| 1.0 \| 1.584e-13 \| 445 \| 0.166 \| 847 \| 506 \| 22 \| 22 \| 702 \| 50 \| 862 \| Phage tail tape measure protein \| Phage tail tape measure protein; afdb-uniprot50 \| AF-A0A0D7V1W3-F1-MODEL\_V4 \| 1.0 \| 2.336e-13 \| 444 \| 0.175 \| 861 \| 552 \| 26 \| 22 \| 780 \| 54 \| 858 \| PhageMin\_Tail domain-containing protein \| PhageMin\_Tail domain-containing protein; afdb-uniprot50 \| AF-A0A5Y1L0N9-F1-MODEL\_V4 \| 1.0 \| 6.166e-13 \| 443 \| 0.202 \| 692 \| 460 \| 23 \| 34 \| 701 \| 10 \| 633 \| Phage tail tape measure protein \| Phage tail tape measure protein; afdb-uniprot50 \| AF-A0A374E9U4-F1-MODEL\_V4 \| 1.0 \| 6.956e-10 \| 441 \| 0.197 \| 486 \| 342 \| 15 \| 13 \| 466 \| 47 \| 516 \| Phage tail tape measure protein \| Phage tail tape measure protein; afdb-uniprot50 \| AF-A0A5P9CRD9-F1-MODEL\_V4 \| 1.0 \| 1.924e-13 \| 441 \| 0.187 \| 801 \| 490 \| 23 \| 19 \| 700 \| 56 \| 814 \| Chromosome segregation protein \| Chromosome segregation protein; afdb-uniprot50 \| AF-A0A2I1IQK4-F1-MODEL\_V4 \| 1.0 \| 9.562e-14 \| 441 \| 0.171 \| 793 \| 498 \| 29 \| 6 \| 702 \| 2 \| 731 \| Phage tail tape measure protein \| Phage tail tape measure protein; afdb-uniprot50 \| AF-A0A3L7JE15-F1-MODEL\_V4 \| 1.0 \| 1.85e-13 \| 441 \| 0.156 \| 833 \| 474 \| 23 \| 19 \| 753 \| 85 \| 786 \| Phage tail tape measure protein \| Phage tail tape measure protein; afdb-uniprot50 \| AF-A0A0B1QAA3-F1-MODEL\_V4 \| 1.0 \| 1.584e-13 \| 440 \| 0.192 \| 804 \| 504 \| 26 \| 2 \| 734 \| 1 \| 729 \| PhageMin\_Tail domain-containing protein \| PhageMin\_Tail domain-containing protein; afdb-uniprot50 \| AF-A0A3S2TUB2-F1-MODEL\_V4 \| 1.0 \| 6.742e-14 \| 440 \| 0.171 \| 753 \| 502 \| 23 \| 77 \| 746 \| 19 \| 732 \| Phage tail tape measure protein \| Phage tail tape measure protein; afdb-uniprot50 \| AF-A0A425XBK8-F1-MODEL\_V4 \| 1.0 \| 9.198e-14 \| 440 \| 0.163 \| 797 \| 536 \| 28 \| 1 \| 713 \| 1 \| 750 \| Phage tail tape measure protein \| Phage tail tape measure protein; afdb-uniprot50 \| AF-A0A5F1PHV9-F1-MODEL\_V4 \| 1.0 \| 5.863e-12 \| 439 \| 0.183 \| 644 \| 399 \| 20 \| 23 \| 590 \| 50 \| 642 \| Phage tail tape measure protein \| Phage tail tape measure protein; afdb-uniprot50 \| AF-A0A841TAB5-F1-MODEL\_V4 \| 1.0 \| 4.885e-13 \| 439 \| 0.169 \| 763 \| 462 \| 23 \| 49 \| 755 \| 3 \| 649 \| Phage tail tape measure protein \| Phage tail tape measure protein; afdb-uniprot50 \| AF-A0A6L6J9S0-F1-MODEL\_V4 \| 1.0 \| 9.94e-14 \| 439 \| 0.188 \| 917 \| 515 \| 31 \| 1 \| 779 \| 2 \| 826 \| Phage tail tape measure protein \| Phage tail tape measure protein; afdb-uniprot50 \| AF-A0A0F9VTR3-F1-MODEL\_V4 \| 1.0 \| 1.584e-13 \| 439 \| 0.185 \| 783 \| 452 \| 27 \| 27 \| 775 \| 8 \| 638 \| PhageMin\_Tail domain-containing protein \| PhageMin\_Tail domain-containing protein; afdb-uniprot50 \| AF-C7MM53-F1-MODEL\_V4 \| 1.0 \| 7.875e-14 \| 439 \| 0.174 \| 740 \| 478 \| 22 \| 27 \| 701 \| 11 \| 682 \| Phage tail tape measure protein, TP901 family \| Phage tail tape measure protein, TP901 family; afdb-uniprot50 \| AF-A0A2G6CBN5-F1-MODEL\_V4 \| 1.0 \| 1.699e-09 \| 438 \| 0.175 \| 398 \| 284 \| 10 \| 19 \| 392 \| 11 \| 388 \| Phage tail tape measure protein \| Phage tail tape measure protein; afdb-uniprot50 \| AF-A0A0F9ELH6-F1-MODEL\_V4 \| 1.0 \| 1.198e-09 \| 438 \| 0.148 \| 432 \| 306 \| 15 \| 61 \| 467 \| 4 \| 398 \| PhageMin\_Tail domain-containing protein \| PhageMin\_Tail domain-containing protein; afdb-uniprot50 \| AF-T2RFP8-F1-MODEL\_V4 \| 1.0 \| 4.265e-08 \| 437 \| 0.123 \| 349 \| 266 \| 10 \| 125 \| 456 \| 2 \| 327 \| Phage tail tape measure protein, TP901 family, core region \| Phage tail tape measure protein, TP901 family, core region; afdb-uniprot50 \| AF-A0A2X3H9A1-F1-MODEL\_V4 \| 1.0 \| 1.01e-11 \| 437 \| 0.178 \| 621 \| 404 \| 17 \| 19 \| 570 \| 35 \| 618 \| Putative prophage tail length determinator \| Putative prophage tail length determinator; afdb-uniprot50 \| AF-A0A258E9Z4-F1-MODEL\_V4 \| 1.0 \| 7.575e-14 \| 437 \| 0.176 \| 759 \| 529 \| 20 \| 67 \| 781 \| 2 \| 708 \| Phage tail tape measure protein \| Phage tail tape measure protein; afdb-uniprot50 \| AF-A0A7T9D1E2-F1-MODEL\_V4 \| 1.0 \| 1.712e-13 \| 437 \| 0.159 \| 765 \| 485 \| 26 \| 118 \| 768 \| 2 \| 722 \| Phage tail tape measure protein \| Phage tail tape measure protein; afdb-uniprot50 \| AF-D8HAR8-F1-MODEL\_V4 \| 1.0 \| 5.552e-14 \| 437 \| 0.166 \| 770 \| 545 \| 29 \| 51 \| 785 \| 3 \| 710 \| Putative phage tail tape measure protein \| Putative phage tail tape measure protein; afdb-uniprot50 \| AF-A0A1C6J674-F1-MODEL\_V4 \| 1.0 \| 8.51e-14 \| 437 \| 0.16 \| 833 \| 555 \| 34 \| 49 \| 785 \| 2 \| 785 \| Phage-related minor tail protein \| Phage-related minor tail protein; afdb-uniprot50 \| AF-A0A1F8UPJ4-F1-MODEL\_V4 \| 1.0 \| 4.753e-14 \| 437 \| 0.156 \| 809 \| 520 \| 29 \| 4 \| 702 \| 6 \| 761 \| Phage tail tape measure protein \| Phage tail tape measure protein; afdb-uniprot50 \| AF-A0A7C4LLA7-F1-MODEL\_V4 \| 1.0 \| 4.753e-14 \| 437 \| 0.164 \| 768 \| 505 \| 26 \| 73 \| 755 \| 9 \| 724 \| Phage tail tape measure protein \| Phage tail tape measure protein; afdb-uniprot50 \| AF-A0A367YQV0-F1-MODEL\_V4 \| 1.0 \| 2.247e-13 \| 437 \| 0.176 \| 797 \| 464 \| 24 \| 1 \| 701 \| 38 \| 737 \| Phage tail tape measure protein \| Phage tail tape measure protein; afdb-uniprot50 \| AF-F4A0G3-F1-MODEL\_V4 \| 1.0 \| 8.51e-14 \| 436 \| 0.183 \| 830 \| 490 \| 31 \| 32 \| 782 \| 2 \| 722 \| Phage tail tape measure protein, TP901 family \| Phage tail tape measure protein, TP901 family; afdb-uniprot50 \| AF-A0A7V5P9D7-F1-MODEL\_V4 \| 1.0 \| 1.022e-12 \| 436 \| 0.163 \| 649 \| 429 \| 24 \| 77 \| 702 \| 18 \| 575 \| Phage tail tape measure protein \| Phage tail tape measure protein; afdb-uniprot50 \| AF-A0A7X1ZGF5-F1-MODEL\_V4 \| 1.0 \| 1.924e-13 \| 436 \| 0.175 \| 904 \| 507 \| 33 \| 19 \| 777 \| 56 \| 865 \| Phage tail tape measure protein \| Phage tail tape measure protein; afdb-uniprot50 \| AF-A0A1Q8Q338-F1-MODEL\_V4 \| 1.0 \| 2.336e-13 \| 436 \| 0.17 \| 743 \| 483 \| 23 \| 13 \| 702 \| 84 \| 745 \| Phage tail tape measure protein \| Phage tail tape measure protein; afdb-uniprot50 \| AF-A0A845R2K9-F1-MODEL\_V4 \| 1.0 \| 1.074e-13 \| 436 \| 0.141 \| 806 \| 540 \| 21 \| 19 \| 702 \| 42 \| 817 \| Phage tail tape measure protein \| Phage tail tape measure protein; afdb-uniprot50 \| AF-A0A2E4CSW6-F1-MODEL\_V4 \| 1.0 \| 2.103e-14 \| 435 \| 0.152 \| 844 \| 568 \| 27 \| 1 \| 781 \| 1 \| 759 \| Phage tail tape measure protein \| Phage tail tape measure protein; afdb-uniprot50 \| AF-A0A1C3K3I9-F1-MODEL\_V4 \| 1.0 \| 2.186e-14 \| 435 \| 0.17 \| 914 \| 555 \| 32 \| 7 \| 782 \| 1 \| 848 \| Phage tail length tape-measure protein \| Phage tail length tape-measure protein; afdb-uniprot50 \| AF-I0GWR9-F1-MODEL\_V4 \| 1.0 \| 2.804e-12 \| 435 \| 0.193 \| 681 \| 463 \| 18 \| 62 \| 702 \| 5 \| 638 \| Putative tail tape measure protein \| Putative tail tape measure protein; afdb-uniprot50 \| AF-A0A4Y8K9M9-F1-MODEL\_V4 \| 1.0 \| 5.552e-14 \| 435 \| 0.152 \| 997 \| 556 \| 30 \| 46 \| 784 \| 3 \| 968 \| Phage tail tape measure protein \| Phage tail tape measure protein; afdb-uniprot50 \| AF-A0A1Y5SUX3-F1-MODEL\_V4 \| 1.0 \| 8.987e-12 \| 435 \| 0.201 \| 609 \| 404 \| 19 \| 63 \| 629 \| 2 \| 569 \| Phage-related minor tail protein \| Phage-related minor tail protein; afdb-uniprot50 \| AF-A0A7B5ZAU9-F1-MODEL\_V4 \| 1.0 \| 8.781e-10 \| 434 \| 0.178 \| 476 \| 308 \| 11 \| 26 \| 434 \| 1 \| 460 \| Phage tail tape measure protein \| Phage tail tape measure protein; afdb-uniprot50 \| AF-A0A1M3PG12-F1-MODEL\_V4 \| 1.0 \| 1.666e-14 \| 433 \| 0.18 \| 844 \| 528 \| 29 \| 1 \| 782 \| 1 \| 742 \| Phage tail tape measure protein \| Phage tail tape measure protein; afdb-uniprot50 \| AF-A0A2D8YMB0-F1-MODEL\_V4 \| 1.0 \| 3.15e-12 \| 433 \| 0.187 \| 662 \| 483 \| 19 \| 62 \| 702 \| 7 \| 634 \| Phage tail tape measure protein \| Phage tail tape measure protein; afdb-uniprot50 \| AF-A0A069F3H5-F1-MODEL\_V4 \| 1.0 \| 1.712e-13 \| 433 \| 0.151 \| 812 \| 525 \| 29 \| 77 \| 778 \| 17 \| 774 \| PhageMin\_Tail domain-containing protein \| PhageMin\_Tail domain-containing protein; afdb-uniprot50 \| AF-A0A2D5HGI6-F1-MODEL\_V4 \| 1.0 \| 3.15e-12 \| 432 \| 0.187 \| 652 \| 442 \| 23 \| 1 \| 620 \| 1 \| 596 \| Phage tail tape measure protein \| Phage tail tape measure protein; afdb-uniprot50 \| AF-A0A847KSJ3-F1-MODEL\_V4 \| 1.0 \| 1.924e-13 \| 432 \| 0.173 \| 832 \| 513 \| 31 \| 1 \| 785 \| 14 \| 717 \| Phage tail tape measure protein \| Phage tail tape measure protein; afdb-uniprot50 \| AF-A0A3P1S6F0-F1-MODEL\_V4 \| 1.0 \| 1.074e-13 \| 432 \| 0.145 \| 996 \| 561 \| 30 \| 9 \| 776 \| 1 \| 933 \| Phage tail tape measure protein \| Phage tail tape measure protein; afdb-uniprot50 \| AF-A0A352TM73-F1-MODEL\_V4 \| 1.0 \| 8.186e-14 \| 432 \| 0.174 \| 791 \| 515 \| 26 \| 1 \| 702 \| 1 \| 742 \| Phage tail tape measure protein \| Phage tail tape measure protein; afdb-uniprot50 \| AF-A0A1R4FZY8-F1-MODEL\_V4 \| 1.0 \| 2.161e-13 \| 432 \| 0.181 \| 776 \| 482 \| 26 \| 1 \| 701 \| 1 \| 697 \| Phage tail length tape-measure protein \| Phage tail length tape-measure protein; afdb-uniprot50 \| AF-A0A315X605-F1-MODEL\_V4 \| 1.0 \| 8.092e-13 \| 432 \| 0.161 \| 731 \| 447 \| 22 \| 43 \| 712 \| 21 \| 646 \| Phage tail tape measure protein \| Phage tail tape measure protein; afdb-uniprot50 \| AF-A0A417T5R9-F1-MODEL\_V4 \| 1.0 \| 3.722e-13 \| 431 \| 0.167 \| 760 \| 524 \| 25 \| 19 \| 702 \| 22 \| 748 \| Phage tail tape measure protein \| Phage tail tape measure protein; afdb-uniprot50 \| AF-A0A0V0PX43-F1-MODEL\_V4 \| 1.0 \| 5.137e-14 \| 431 \| 0.177 \| 846 \| 558 \| 29 \| 36 \| 776 \| 7 \| 819 \| PhageMin\_Tail domain-containing protein \| PhageMin\_Tail domain-containing protein; afdb-uniprot50 \| AF-A0A4Q7RAQ5-F1-MODEL\_V4 \| 1.0 \| 8.412e-13 \| 431 \| 0.189 \| 744 \| 476 \| 23 \| 3 \| 719 \| 203 \| 846 \| Lambda family phage tail tape measure protein/TP901 family phage tail tape measure protein \| Lambda family phage tail tape measure protein/TP901 family phage tail tape measure protein; afdb-uniprot50 \| AF-A0A373ZD47-F1-MODEL\_V4 \| 1.0 \| 4.52e-13 \| 430 \| 0.152 \| 881 \| 571 \| 33 \| 2 \| 784 \| 1 \| 803 \| Phage tail tape measure protein \| Phage tail tape measure protein; afdb-uniprot50 \| AF-A0A558GXF9-F1-MODEL\_V4 \| 1.0 \| 6.664e-13 \| 430 \| 0.179 \| 960 \| 533 \| 32 \| 19 \| 755 \| 9 \| 936 \| Phage tail tape measure protein \| Phage tail tape measure protein; afdb-uniprot50 \| AF-A0A5J6SLX3-F1-MODEL\_V4 \| 1.0 \| 7.009e-14 \| 430 \| 0.18 \| 831 \| 465 \| 26 \| 1 \| 667 \| 1 \| 779 \| Phage tail tape measure protein \| Phage tail tape measure protein; afdb-uniprot50 \| AF-A0A5U3G3I6-F1-MODEL\_V4 \| 1.0 \| 7.518e-10 \| 429 \| 0.164 \| 524 \| 343 \| 13 \| 118 \| 626 \| 31 \| 474 \| Phage tail tape measure protein \| Phage tail tape measure protein; afdb-uniprot50 \| AF-A0A6G3HIH4-F1-MODEL\_V4 \| 1.0 \| 1.284e-15 \| 429 \| 0.215 \| 725 \| 406 \| 16 \| 211 \| 785 \| 2 \| 713 \| Phage tail tape measure protein \| Phage tail tape measure protein; afdb-uniprot50 \| AF-A0A1I2YB35-F1-MODEL\_V4 \| 1.0 \| 5.552e-14 \| 429 \| 0.162 \| 862 \| 530 \| 29 \| 62 \| 785 \| 4 \| 811 \| Phage tail tape measure protein, TP901 family, core region \| Phage tail tape measure protein, TP901 family, core region; afdb-uniprot50 \| AF-A0A7S8E854-F1-MODEL\_V4 \| 1.0 \| 2.836e-13 \| 428 \| 0.174 \| 850 \| 523 \| 32 \| 61 \| 776 \| 1 \| 805 \| Phage tail tape measure protein \| Phage tail tape measure protein; afdb-uniprot50 \| AF-A0A1C5PPL3-F1-MODEL\_V4 \| 1.0 \| 1.628e-12 \| 428 \| 0.148 \| 701 \| 510 \| 23 \| 53 \| 702 \| 2 \| 666 \| Phage-related minor tail protein \| Phage-related minor tail protein; afdb-uniprot50 \| AF-A0A5M7L9A2-F1-MODEL\_V4 \| 1.0 \| 4.182e-13 \| 428 \| 0.151 \| 816 \| 509 \| 26 \| 19 \| 755 \| 34 \| 744 \| Phage tail tape measure protein \| Phage tail tape measure protein; afdb-uniprot50 \| AF-A0A6G8J7Q0-F1-MODEL\_V4 \| 1.0 \| 4.699e-13 \| 428 \| 0.153 \| 880 \| 569 \| 26 \| 19 \| 780 \| 54 \| 875 \| Phage tail tape measure protein \| Phage tail tape measure protein; afdb-uniprot50 \| AF-A0A7W4EBY4-F1-MODEL\_V4 \| 1.0 \| 6.485e-14 \| 428 \| 0.16 \| 842 \| 525 \| 25 \| 35 \| 781 \| 2 \| 756 \| Phage tail tape measure protein \| Phage tail tape measure protein; afdb-uniprot50 \| AF-E6MGX4-F1-MODEL\_V4 \| 1.0 \| 1.41e-13 \| 427 \| 0.141 \| 907 \| 585 \| 31 \| 1 \| 785 \| 1 \| 835 \| Phage tail tape measure protein, TP901 family \| Phage tail tape measure protein, TP901 family; afdb-uniprot50 \| AF-A0A2N2CA29-F1-MODEL\_V4 \| 1.0 \| 2.428e-13 \| 427 \| 0.18 \| 761 \| 503 \| 29 \| 1 \| 702 \| 1 \| 699 \| Phage tail tape measure protein \| Phage tail tape measure protein; afdb-uniprot50 \| AF-B8FNZ4-F1-MODEL\_V4 \| 1.0 \| 9.94e-14 \| 427 \| 0.183 \| 790 \| 439 \| 25 \| 21 \| 662 \| 77 \| 808 \| Phage tail tape measure protein, TP901 family \| Phage tail tape measure protein, TP901 family; afdb-uniprot50 \| AF-A0A2G0E7P5-F1-MODEL\_V4 \| 1.0 \| 5.931e-13 \| 426 \| 0.174 \| 687 \| 398 \| 19 \| 1 \| 532 \| 1 \| 673 \| Phage tail tape measure protein \| Phage tail tape measure protein; afdb-uniprot50 \| AF-A0A6B4HS58-F1-MODEL\_V4 \| 1.0 \| 6.41e-13 \| 426 \| 0.15 \| 729 \| 501 \| 26 \| 27 \| 702 \| 61 \| 723 \| Phage tail tape measure protein \| Phage tail tape measure protein; afdb-uniprot50 \| AF-A0A239GVA7-F1-MODEL\_V4 \| 1.0 \| 3.042e-09 \| 425 \| 0.332 \| 304 \| 171 \| 5 \| 26 \| 321 \| 1 \| 280 \| Phage tail tape measure protein, TP901 family, core region \| Phage tail tape measure protein, TP901 family, core region; afdb-uniprot50 \| AF-A0A7V6Y6C8-F1-MODEL\_V4 \| 1.0 \| 2.524e-13 \| 425 \| 0.159 \| 859 \| 536 \| 32 \| 19 \| 781 \| 25 \| 793 \| Phage tail tape measure protein \| Phage tail tape measure protein; afdb-uniprot50 \| AF-A0A5E4XFP1-F1-MODEL\_V4 \| 1.0 \| 2.961e-10 \| 425 \| 0.153 \| 553 \| 380 \| 11 \| 19 \| 519 \| 46 \| 562 \| Phage tail tape measure protein \| Phage tail tape measure protein; afdb-uniprot50 \| AF-A0A838LDI5-F1-MODEL\_V4 \| 1.0 \| 1.356e-13 \| 425 \| 0.152 \| 995 \| 515 \| 29 \| 1 \| 784 \| 1 \| 877 \| Phage tail tape measure protein \| Phage tail tape measure protein; afdb-uniprot50 \| AF-A0A552V6Z6-F1-MODEL\_V4 \| 1.0 \| 1.712e-13 \| 424 \| 0.155 \| 867 \| 585 \| 30 \| 9 \| 767 \| 1 \| 827 \| Phage tail tape measure protein \| Phage tail tape measure protein; afdb-uniprot50 \| AF-A0A484ZHI6-F1-MODEL\_V4 \| 1.0 \| 1.466e-13 \| 424 \| 0.161 \| 860 \| 565 \| 26 \| 19 \| 782 \| 58 \| 856 \| Phage-related minor tail protein \| Phage-related minor tail protein; afdb-uniprot50 \| AF-A0A350TW28-F1-MODEL\_V4 \| 1.0 \| 3.58e-13 \| 424 \| 0.164 \| 789 \| 506 \| 30 \| 1 \| 709 \| 1 \| 715 \| Phage tail tape measure protein \| Phage tail tape measure protein; afdb-uniprot50 \| AF-A0A3N5VV08-F1-MODEL\_V4 \| 1.0 \| 5.863e-12 \| 423 \| 0.145 \| 640 \| 444 \| 17 \| 117 \| 702 \| 24 \| 614 \| Phage tail tape measure protein \| Phage tail tape measure protein; afdb-uniprot50 \| AF-A0A7Z9YAJ2-F1-MODEL\_V4 \| 1.0 \| 1.432e-11 \| 423 \| 0.173 \| 653 \| 445 \| 19 \| 76 \| 702 \| 5 \| 588 \| Phage tail tape measure protein \| Phage tail tape measure protein; afdb-uniprot50 \| AF-A0A4R3JH78-F1-MODEL\_V4 \| 1.0 \| 1.179e-11 \| 423 \| 0.153 \| 647 \| 469 \| 17 \| 87 \| 701 \| 23 \| 622 \| TP901 family phage tail tape measure protein \| TP901 family phage tail tape measure protein; afdb-uniprot50 \| AF-A0A4Y3W6Q9-F1-MODEL\_V4 \| 1.0 \| 3.065e-13 \| 423 \| 0.148 \| 803 \| 533 \| 23 \| 1 \| 748 \| 2 \| 708 \| PhageMin\_Tail domain-containing protein \| PhageMin\_Tail domain-containing protein; afdb-uniprot50 \| AF-A0A3A9AGY9-F1-MODEL\_V4 \| 1.0 \| 5.137e-14 \| 422 \| 0.167 \| 793 \| 513 \| 28 \| 19 \| 702 \| 24 \| 777 \| Phage tail tape measure protein \| Phage tail tape measure protein; afdb-uniprot50 \| AF-A0A1D3N8J3-F1-MODEL\_V4 \| 1.0 \| 4.52e-13 \| 422 \| 0.175 \| 759 \| 517 \| 21 \| 19 \| 702 \| 85 \| 809 \| Putative tail tape measure protein \| Putative tail tape measure protein; afdb-uniprot50 \| AF-R7FHV4-F1-MODEL\_V4 \| 1.0 \| 8.847e-14 \| 422 \| 0.162 \| 915 \| 518 \| 32 \| 4 \| 702 \| 6 \| 887 \| Phage tail tape measure protein family \| Phage tail tape measure protein family; afdb-uniprot50 \| AF-A0A2T2UAU0-F1-MODEL\_V4 \| 1.0 \| 1.954e-11 \| 421 \| 0.151 \| 725 \| 432 \| 31 \| 1 \| 714 \| 21 \| 572 \| Phage tail tape measure protein \| Phage tail tape measure protein; afdb-uniprot50 \| AF-A6TR98-F1-MODEL\_V4 \| 1.0 \| 2.161e-13 \| 421 \| 0.16 \| 873 \| 556 \| 31 \| 1 \| 779 \| 4 \| 793 \| Phage tail tape measure protein, TP901 family \| Phage tail tape measure protein, TP901 family; afdb-uniprot50 \| AF-A0A374VKX6-F1-MODEL\_V4 \| 1.0 \| 1.712e-13 \| 421 \| 0.176 \| 767 \| 503 \| 25 \| 19 \| 702 \| 22 \| 742 \| Phage tail tape measure protein \| Phage tail tape measure protein; afdb-uniprot50 \| AF-D4M9M3-F1-MODEL\_V4 \| 1.0 \| 1.524e-13 \| 421 \| 0.171 \| 917 \| 531 \| 30 \| 3 \| 780 \| 1 \| 827 \| Phage tail tape measure protein, TP901 family, core region \| Phage tail tape measure protein, TP901 family, core region; afdb-uniprot50 \| AF-A0A178HLZ5-F1-MODEL\_V4 \| 1.0 \| 1.062e-12 \| 421 \| 0.173 \| 766 \| 468 \| 20 \| 50 \| 706 \| 2 \| 711 \| PhageMin\_Tail domain-containing protein \| PhageMin\_Tail domain-containing protein; afdb-uniprot50 \| AF-A0A7C1VR42-F1-MODEL\_V4 \| 1.0 \| 6.41e-13 \| 421 \| 0.134 \| 922 \| 581 \| 27 \| 2 \| 776 \| 1 \| 852 \| Phage tail tape measure protein \| Phage tail tape measure protein; afdb-uniprot50 \| AF-A0A4E0QXR2-F1-MODEL\_V4 \| 1.0 \| 4.134e-12 \| 420 \| 0.174 \| 663 \| 425 \| 27 \| 81 \| 701 \| 25 \| 606 \| Phage tail tape measure protein \| Phage tail tape measure protein; afdb-uniprot50 \| AF-A0A7W2CT18-F1-MODEL\_V4 \| 1.0 \| 1.117e-13 \| 420 \| 0.162 \| 888 \| 543 \| 28 \| 1 \| 779 \| 1 \| 796 \| Phage tail tape measure protein \| Phage tail tape measure protein; afdb-uniprot50 \| AF-A0A5K1ITD8-F1-MODEL\_V4 \| 1.0 \| 1.628e-12 \| 420 \| 0.171 \| 730 \| 434 \| 24 \| 116 \| 785 \| 83 \| 701 \| Phage-related minor tail protein \| Phage-related minor tail protein; afdb-uniprot50 \| AF-A0A0F9IXZ9-F1-MODEL\_V4 \| 1.0 \| 1.466e-13 \| 419 \| 0.172 \| 755 \| 454 \| 31 \| 64 \| 700 \| 11 \| 712 \| PhageMin\_Tail domain-containing protein \| PhageMin\_Tail domain-containing protein; afdb-uniprot50 \| AF-A0A3M2BDP3-F1-MODEL\_V4 \| 1.0 \| 3.15e-12 \| 419 \| 0.161 \| 670 \| 449 \| 24 \| 53 \| 702 \| 33 \| 609 \| Phage tail tape measure protein \| Phage tail tape measure protein; afdb-uniprot50 \| AF-A0A542YF06-F1-MODEL\_V4 \| 1.0 \| 3.065e-13 \| 419 \| 0.174 \| 787 \| 507 \| 22 \| 10 \| 704 \| 1 \| 736 \| TP901 family phage tail tape measure protein \| TP901 family phage tail tape measure protein; afdb-uniprot50 \| AF-A0A1F4QW09-F1-MODEL\_V4 \| 1.0 \| 2.477e-08 \| 418 \| 0.17 \| 435 \| 274 \| 15 \| 14 \| 433 \| 1 \| 363 \| Phage tail tape measure protein \| Phage tail tape measure protein; afdb-uniprot50 \| AF-Q7NAC3-F1-MODEL\_V4 \| 1.0 \| 5.279e-13 \| 418 \| 0.141 \| 875 \| 576 \| 23 \| 18 \| 784 \| 6 \| 812 \| PhageMin\_Tail domain-containing protein \| PhageMin\_Tail domain-containing protein; afdb-uniprot50 \| AF-R6LX22-F1-MODEL\_V4 \| 1.0 \| 3.187e-13 \| 418 \| 0.162 \| 825 \| 500 \| 26 \| 32 \| 712 \| 8 \| 785 \| Phage tail tape measure protein TP901 family \| Phage tail tape measure protein TP901 family; afdb-uniprot50 \| AF-A0A376SKG7-F1-MODEL\_V4 \| 1.0 \| 1.245e-09 \| 417 \| 0.177 \| 456 \| 294 \| 15 \| 23 \| 417 \| 50 \| 485 \| Tail fiber component of prophage CP-933T \| Tail fiber component of prophage CP-933T; afdb-uniprot50 \| AF-A0A661DYN7-F1-MODEL\_V4 \| 1.0 \| 3.595e-10 \| 417 \| 0.146 \| 663 \| 422 \| 21 \| 46 \| 687 \| 2 \| 541 \| Phage tail tape measure protein \| Phage tail tape measure protein; afdb-uniprot50 \| AF-A0A6N9QF02-F1-MODEL\_V4 \| 1.0 \| 4.182e-13 \| 417 \| 0.168 \| 741 \| 480 \| 25 \| 1 \| 710 \| 1 \| 635 \| Phage tail tape measure protein \| Phage tail tape measure protein; afdb-uniprot50 \| AF-A0A0B3BNP9-F1-MODEL\_V4 \| 1.0 \| 5.705e-13 \| 417 \| 0.167 \| 872 \| 506 \| 31 \| 1 \| 782 \| 1 \| 742 \| Phage tail tape measure protein, TP901 family \| Phage tail tape measure protein, TP901 family; afdb-uniprot50 \| AF-A0A1M5S5P3-F1-MODEL\_V4 \| 1.0 \| 2.728e-13 \| 417 \| 0.17 \| 810 \| 483 \| 30 \| 25 \| 783 \| 16 \| 687 \| Phage tail tape measure protein, TP901 family, core region \| Phage tail tape measure protein, TP901 family, core region; afdb-uniprot50 \| AF-A0A2A2P5S7-F1-MODEL\_V4 \| 1.0 \| 1.548e-11 \| 417 \| 0.157 \| 629 \| 453 \| 19 \| 108 \| 702 \| 63 \| 648 \| Phage tail tape measure protein \| Phage tail tape measure protein; afdb-uniprot50 \| AF-W0FHT5-F1-MODEL\_V4 \| 1.0 \| 9.94e-14 \| 417 \| 0.153 \| 911 \| 529 \| 31 \| 1 \| 701 \| 1 \| 878 \| Phage tail tape measure protein, family, core region \| Phage tail tape measure protein, family, core region; afdb-uniprot50 \| AF-W7ZHM9-F1-MODEL\_V4 \| 1.0 \| 4.086e-11 \| 416 \| 0.159 \| 591 \| 401 \| 23 \| 75 \| 637 \| 7 \| 529 \| Phage tail length tape-measure protein \| Phage tail length tape-measure protein; afdb-uniprot50 \| AF-A0A2J6HSG0-F1-MODEL\_V4 \| 1.0 \| 5.019e-12 \| 416 \| 0.162 \| 701 \| 485 \| 22 \| 39 \| 696 \| 3 \| 644 \| Phage tail tape measure protein \| Phage tail tape measure protein; afdb-uniprot50 \| AF-Q896A9-F1-MODEL\_V4 \| 1.0 \| 1.062e-12 \| 415 \| 0.137 \| 801 \| 533 \| 28 \| 8 \| 779 \| 1 \| 672 \| Phage protein \| Phage protein; afdb-uniprot50 \| AF-C3X1X5-F1-MODEL\_V4 \| 1.0 \| 5.078e-13 \| 415 \| 0.159 \| 855 \| 544 \| 22 \| 19 \| 781 \| 34 \| 805 \| Phage tail tape measure protein, TP901 family, core region \| Phage tail tape measure protein, TP901 family, core region; afdb-uniprot50 \| AF-A0A1Q3R2P7-F1-MODEL\_V4 \| 1.0 \| 4.347e-13 \| 414 \| 0.2 \| 698 \| 440 \| 24 \| 148 \| 782 \| 12 \| 654 \| Phage tail tape measure protein \| Phage tail tape measure protein; afdb-uniprot50 \| AF-A0A7W9VFJ5-F1-MODEL\_V4 \| 1.0 \| 4.231e-14 \| 414 \| 0.178 \| 831 \| 536 \| 31 \| 1 \| 750 \| 2 \| 766 \| TP901 family phage tail tape measure protein \| TP901 family phage tail tape measure protein; afdb-uniprot50 \| AF-A0A644VK59-F1-MODEL\_V4 \| 1.0 \| 6.41e-13 \| 414 \| 0.143 \| 752 \| 524 \| 28 \| 27 \| 702 \| 61 \| 768 \| PhageMin\_Tail domain-containing protein \| PhageMin\_Tail domain-containing protein; afdb-uniprot50 \| AF-A0A0F9M5T9-F1-MODEL\_V4 \| 1.0 \| 2.256e-10 \| 413 \| 0.149 \| 741 \| 419 \| 22 \| 50 \| 785 \| 6 \| 539 \| PhageMin\_Tail domain-containing protein \| PhageMin\_Tail domain-containing protein; afdb-uniprot50 \| AF-A0A6I6WW05-F1-MODEL\_V4 \| 1.0 \| 7.784e-13 \| 413 \| 0.159 \| 878 \| 521 \| 30 \| 1 \| 769 \| 1 \| 770 \| Phage tail tape measure protein \| Phage tail tape measure protein; afdb-uniprot50 \| AF-A0A0H3F7C7-F1-MODEL\_V4 \| 1.0 \| 5.931e-13 \| 413 \| 0.164 \| 905 \| 525 \| 26 \| 19 \| 781 \| 49 \| 863 \| Phage tail tape measure protein, TP901 family \| Phage tail tape measure protein, TP901 family; afdb-uniprot50 \| AF-H1D5F2-F1-MODEL\_V4 \| 1.0 \| 2.524e-13 \| 412 \| 0.229 \| 627 \| 358 \| 18 \| 207 \| 785 \| 4 \| 553 \| Phage tail tape measure protein, TP901 family, core region \| Phage tail tape measure protein, TP901 family, core region; afdb-uniprot50 \| AF-A0A3R9NYT5-F1-MODEL\_V4 \| 1.0 \| 3.539e-12 \| 412 \| 0.146 \| 804 \| 499 \| 25 \| 65 \| 777 \| 5 \| 711 \| Phage tail tape measure protein \| Phage tail tape measure protein; afdb-uniprot50 \| AF-A0A124GKD2-F1-MODEL\_V4 \| 1.0 \| 3.58e-13 \| 412 \| 0.145 \| 751 \| 512 \| 23 \| 117 \| 779 \| 37 \| 745 \| PhageMin\_Tail domain-containing protein \| PhageMin\_Tail domain-containing protein; afdb-uniprot50 \| AF-A0A3D5P8N1-F1-MODEL\_V4 \| 1.0 \| 5.078e-13 \| 412 \| 0.139 \| 972 \| 595 \| 31 \| 1 \| 749 \| 1 \| 953 \| Phage tail tape measure protein \| Phage tail tape measure protein; afdb-uniprot50 \| AF-A0A7Y5FSF4-F1-MODEL\_V4 \| 1.0 \| 5.341e-14 \| 412 \| 0.179 \| 803 \| 517 \| 30 \| 19 \| 765 \| 11 \| 727 \| Phage tail tape measure protein \| Phage tail tape measure protein; afdb-uniprot50 \| AF-A0A1M5PWE6-F1-MODEL\_V4 \| 1.0 \| 8.092e-13 \| 412 \| 0.16 \| 899 \| 573 \| 29 \| 19 \| 783 \| 146 \| 996 \| Phage tail tape measure protein, TP901 family, core region \| Phage tail tape measure protein, TP901 family, core region; afdb-uniprot50 \| AF-A0A2G2JMR9-F1-MODEL\_V4 \| 1.0 \| 3.825e-12 \| 411 \| 0.155 \| 686 \| 465 \| 28 \| 63 \| 702 \| 10 \| 626 \| Phage tail tape measure protein \| Phage tail tape measure protein; afdb-uniprot50 \| AF-A0A7V6YSF2-F1-MODEL\_V4 \| 1.0 \| 6.166e-13 \| 411 \| 0.166 \| 783 \| 490 \| 26 \| 63 \| 747 \| 5 \| 722 \| Phage tail tape measure protein \| Phage tail tape measure protein; afdb-uniprot50 \| AF-A0A2W4KBI5-F1-MODEL\_V4 \| 1.0 \| 4.182e-13 \| 411 \| 0.158 \| 894 \| 551 \| 26 \| 35 \| 776 \| 2 \| 845 \| Phage tail tape measure protein \| Phage tail tape measure protein; afdb-uniprot50 \| AF-F4HB27-F1-MODEL\_V4 \| 1.0 \| 8.746e-13 \| 411 \| 0.176 \| 940 \| 529 \| 28 \| 19 \| 781 \| 77 \| 947 \| Phage-related minor tail protein \| Phage-related minor tail protein; afdb-uniprot50 \| AF-A0A1P8WB67-F1-MODEL\_V4 \| 1.0 \| 1.654e-10 \| 410 \| 0.174 \| 557 \| 373 \| 17 \| 32 \| 556 \| 4 \| 505 \| Phage tail tape measure protein, TP901 family, core region \| Phage tail tape measure protein, TP901 family, core region; afdb-uniprot50 \| AF-A0A7Y2L7P9-F1-MODEL\_V4 \| 1.0 \| 7.606e-11 \| 410 \| 0.183 \| 633 \| 368 \| 19 \| 2 \| 620 \| 1 \| 498 \| Phage tail tape measure protein \| Phage tail tape measure protein; afdb-uniprot50 \| AF-A0A1I3C2N2-F1-MODEL\_V4 \| 1.0 \| 6.095e-12 \| 410 \| 0.197 \| 593 \| 384 \| 16 \| 160 \| 702 \| 1 \| 551 \| Phage tail tape measure protein, TP901 family, core region \| Phage tail tape measure protein, TP901 family, core region; afdb-uniprot50 \| AF-A0A2A4V258-F1-MODEL\_V4 \| 1.0 \| 1.449e-12 \| 410 \| 0.164 \| 722 \| 492 \| 23 \| 46 \| 732 \| 18 \| 662 \| Phage tail tape measure protein \| Phage tail tape measure protein; afdb-uniprot50 \| AF-A0A1Q6K3L0-F1-MODEL\_V4 \| 1.0 \| 1.24e-12 \| 410 \| 0.154 \| 895 \| 562 \| 32 \| 19 \| 785 \| 81 \| 908 \| Phage tail tape measure protein \| Phage tail tape measure protein; afdb-uniprot50 \| AF-A0A1D9FP83-F1-MODEL\_V4 \| 1.0 \| 9.826e-13 \| 410 \| 0.147 \| 740 \| 486 \| 26 \| 1 \| 702 \| 1 \| 633 \| Phage tail tape measure protein \| Phage tail tape measure protein; afdb-uniprot50 \| AF-A0A4V6M1R7-F1-MODEL\_V4 \| 1.0 \| 7.487e-13 \| 409 \| 0.182 \| 784 \| 502 \| 29 \| 88 \| 784 \| 35 \| 766 \| PhageMin\_Tail domain-containing protein \| PhageMin\_Tail domain-containing protein; afdb-uniprot50 \| AF-A0A6S4UXC8-F1-MODEL\_V4 \| 1.0 \| 1.022e-12 \| 409 \| 0.158 \| 977 \| 523 \| 31 \| 19 \| 782 \| 56 \| 945 \| PhageMin\_Tail domain-containing protein \| PhageMin\_Tail domain-containing protein; afdb-uniprot50 \| AF-A0A6D2F7P3-F1-MODEL\_V4 \| 1.0 \| 1.198e-09 \| 408 \| 0.165 \| 509 \| 332 \| 13 \| 19 \| 452 \| 44 \| 534 \| Phage tail tape measure protein \| Phage tail tape measure protein; afdb-uniprot50 \| AF-A0A285M600-F1-MODEL\_V4 \| 1.0 \| 2.221e-12 \| 408 \| 0.153 \| 842 \| 522 \| 23 \| 19 \| 780 \| 71 \| 801 \| Phage tail tape measure protein, TP901 family, core region \| Phage tail tape measure protein, TP901 family, core region; afdb-uniprot50 \| AF-A0A7U6KQY0-F1-MODEL\_V4 \| 1.0 \| 3.065e-13 \| 408 \| 0.162 \| 868 \| 553 \| 23 \| 19 \| 782 \| 49 \| 846 \| PhageMin\_Tail domain-containing protein \| PhageMin\_Tail domain-containing protein; afdb-uniprot50 \| AF-A0A2U3F164-F1-MODEL\_V4 \| 1.0 \| 4.699e-13 \| 408 \| 0.153 \| 918 \| 543 \| 28 \| 19 \| 781 \| 47 \| 884 \| Phage tail tape measure protein \| Phage tail tape measure protein; afdb-uniprot50 \| AF-A0A402A530-F1-MODEL\_V4 \| 1.0 \| 5.575e-11 \| 407 \| 0.174 \| 649 \| 444 \| 21 \| 112 \| 702 \| 53 \| 667 \| PhageMin\_Tail domain-containing protein \| PhageMin\_Tail domain-containing protein; afdb-uniprot50 \| AF-J3UKH6-F1-MODEL\_V4 \| 1.0 \| 4.885e-13 \| 407 \| 0.133 \| 879 \| 586 \| 27 \| 20 \| 785 \| 2 \| 817 \| TP901 family phage tail tape measure protein \| TP901 family phage tail tape measure protein; afdb-uniprot50 \| AF-A0A2C9CPT9-F1-MODEL\_V4 \| 1.0 \| 4.828e-12 \| 407 \| 0.152 \| 741 \| 493 \| 26 \| 83 \| 780 \| 2 \| 650 \| Phage tail tape measure protein, lambda family/phage tail tape measure protein, TP901 family, core region \| Phage tail tape measure protein, lambda family/phage tail tape measure protein, TP901 family, core region; afdb-uniprot50 \| AF-A0A433KZN9-F1-MODEL\_V4 \| 1.0 \| 3.418e-09 \| 407 \| 0.181 \| 496 \| 331 \| 16 \| 19 \| 458 \| 78 \| 554 \| Phage tail tape measure protein \| Phage tail tape measure protein; afdb-uniprot50 \| AF-A0A1H0P6W7-F1-MODEL\_V4 \| 1.0 \| 5.019e-12 \| 407 \| 0.176 \| 730 \| 498 \| 18 \| 19 \| 702 \| 19 \| 690 \| Phage tail tape measure protein, TP901 family, core region \| Phage tail tape measure protein, TP901 family, core region; afdb-uniprot50 \| AF-A0A765X8R7-F1-MODEL\_V4 \| 1.0 \| 1.179e-11 \| 406 \| 0.155 \| 783 \| 506 \| 25 \| 80 \| 781 \| 3 \| 710 \| Phage tail tape measure protein \| Phage tail tape measure protein; afdb-uniprot50 \| AF-A0A150ISK2-F1-MODEL\_V4 \| 1.0 \| 3.93e-11 \| 406 \| 0.171 \| 641 \| 400 \| 24 \| 115 \| 702 \| 47 \| 609 \| Phage-related minor tail protein \| Phage-related minor tail protein; afdb-uniprot50 \| AF-A0A1Y6FRA0-F1-MODEL\_V4 \| 1.0 \| 2.221e-12 \| 406 \| 0.16 \| 814 \| 519 \| 31 \| 1 \| 702 \| 1 \| 761 \| Phage tail tape measure protein, TP901 family, core region \| Phage tail tape measure protein, TP901 family, core region; afdb-uniprot50 \| AF-Q4E9A1-F1-MODEL\_V4 \| 1.0 \| 2.42e-06 \| 405 \| 0.317 \| 195 \| 132 \| 1 \| 144 \| 338 \| 2 \| 195 \| PhageMin\_Tail domain-containing protein \| PhageMin\_Tail domain-containing protein; afdb-uniprot50 \| AF-A0A497GWY3-F1-MODEL\_V4 \| 1.0 \| 7.316e-11 \| 405 \| 0.148 \| 547 \| 384 \| 18 \| 2 \| 509 \| 1 \| 504 \| Phage tail tape measure protein \| Phage tail tape measure protein; afdb-uniprot50 \| AF-W7PY06-F1-MODEL\_V4 \| 1.0 \| 4.416e-11 \| 405 \| 0.15 \| 730 \| 492 \| 23 \| 39 \| 741 \| 2 \| 630 \| PhageMin\_Tail domain-containing protein \| PhageMin\_Tail domain-containing protein; afdb-uniprot50 \| AF-A0A2S8FUK5-F1-MODEL\_V4 \| 1.0 \| 4.086e-11 \| 405 \| 0.16 \| 711 \| 467 \| 22 \| 83 \| 774 \| 18 \| 617 \| Phage tail tape measure protein \| Phage tail tape measure protein; afdb-uniprot50 \| AF-A0A1F3BA55-F1-MODEL\_V4 \| 1.0 \| 2.195e-11 \| 405 \| 0.161 \| 695 \| 468 \| 23 \| 48 \| 703 \| 3 \| 621 \| Phage tail tape measure protein \| Phage tail tape measure protein; afdb-uniprot50 \| AF-A0A5P3FW64-F1-MODEL\_V4 \| 1.0 \| 1.121e-10 \| 405 \| 0.176 \| 589 \| 352 \| 14 \| 1 \| 459 \| 1 \| 586 \| Phage tail tape measure protein \| Phage tail tape measure protein; afdb-uniprot50 \| AF-A0A7V2ILC6-F1-MODEL\_V4 \| 1.0 \| 2.292e-08 \| 404 \| 0.146 \| 437 \| 327 \| 12 \| 39 \| 472 \| 11 \| 404 \| Phage tail tape measure protein \| Phage tail tape measure protein; afdb-uniprot50 \| AF-A0A2M8E3Z2-F1-MODEL\_V4 \| 1.0 \| 9.866e-10 \| 404 \| 0.171 \| 573 \| 378 \| 23 \| 97 \| 651 \| 9 \| 502 \| Phage tail tape measure protein \| Phage tail tape measure protein; afdb-uniprot50 \| AF-A0A523V8K2-F1-MODEL\_V4 \| 1.0 \| 3.404e-12 \| 403 \| 0.165 \| 712 \| 483 \| 21 \| 35 \| 702 \| 17 \| 660 \| Phage tail tape measure protein \| Phage tail tape measure protein; afdb-uniprot50 \| AF-A0A7W8U3E0-F1-MODEL\_V4 \| 1.0 \| 2.221e-12 \| 403 \| 0.177 \| 806 \| 545 \| 24 \| 35 \| 782 \| 231 \| 976 \| TP901 family phage tail tape measure protein \| TP901 family phage tail tape measure protein; afdb-uniprot50 \| AF-A0A3S0BBD4-F1-MODEL\_V4 \| 1.0 \| 1.212e-10 \| 402 \| 0.181 \| 720 \| 434 \| 20 \| 35 \| 716 \| 5 \| 606 \| Phage tail tape measure protein \| Phage tail tape measure protein; afdb-uniprot50 \| AF-A0A5P0YP70-F1-MODEL\_V4 \| 1.0 \| 2.495e-12 \| 402 \| 0.162 \| 796 \| 484 \| 32 \| 41 \| 767 \| 13 \| 694 \| Phage tail tape measure protein \| Phage tail tape measure protein; afdb-uniprot50 \| AF-M0EPI4-F1-MODEL\_V4 \| 1.0 \| 2.221e-12 \| 402 \| 0.192 \| 750 \| 452 \| 35 \| 63 \| 783 \| 5 \| 629 \| Phage tail tape measure protein, TP901 family \| Phage tail tape measure protein, TP901 family; afdb-uniprot50 \| AF-A0A7Z7QQC7-F1-MODEL\_V4 \| 1.0 \| 2.594e-12 \| 402 \| 0.171 \| 768 \| 508 \| 27 \| 1 \| 701 \| 1 \| 706 \| Phage tail tape measure protein, TP901 family, core region \| Phage tail tape measure protein, TP901 family, core region; afdb-uniprot50 \| AF-A0A1R0WG41-F1-MODEL\_V4 \| 1.0 \| 4.182e-13 \| 402 \| 0.156 \| 1005 \| 571 \| 31 \| 1 \| 784 \| 3 \| 951 \| Phage tail tape measure protein \| Phage tail tape measure protein; afdb-uniprot50 \| AF-A0A135YZ87-F1-MODEL\_V4 \| 1.0 \| 2.948e-13 \| 402 \| 0.183 \| 833 \| 510 \| 28 \| 19 \| 709 \| 164 \| 967 \| Phage tail tape measure protein, TP901 family \| Phage tail tape measure protein, TP901 family; afdb-uniprot50 \| AF-A0A0F9TKT3-F1-MODEL\_V4 \| 1.0 \| 3.365e-11 \| 401 \| 0.159 \| 690 \| 464 \| 25 \| 61 \| 704 \| 3 \| 622 \| PhageMin\_Tail domain-containing protein \| PhageMin\_Tail domain-containing protein; afdb-uniprot50 \| AF-K4Z9C6-F1-MODEL\_V4 \| 1.0 \| 4.885e-13 \| 401 \| 0.152 \| 806 \| 547 \| 28 \| 46 \| 776 \| 3 \| 747 \| Phage tail tape measure protein, TP901 family \| Phage tail tape measure protein, TP901 family; afdb-uniprot50 \| AF-G5S7F4-F1-MODEL\_V4 \| 1.0 \| 1.449e-12 \| 401 \| 0.16 \| 873 \| 479 \| 26 \| 1 \| 755 \| 11 \| 747 \| Phage tail tape measure protein, TP901 family \| Phage tail tape measure protein, TP901 family; afdb-uniprot50 \| AF-A0A371IQX0-F1-MODEL\_V4 \| 1.0 \| 9.826e-13 \| 401 \| 0.167 \| 762 \| 471 \| 27 \| 6 \| 701 \| 3 \| 666 \| Phage tail tape measure protein \| Phage tail tape measure protein; afdb-uniprot50 \| AF-A0A5C5XSP4-F1-MODEL\_V4 \| 1.0 \| 2.915e-12 \| 400 \| 0.16 \| 849 \| 514 \| 31 \| 6 \| 785 \| 2 \| 720 \| Phage-related minor tail protein \| Phage-related minor tail protein; afdb-uniprot50 \| AF-C8WM92-F1-MODEL\_V4 \| 1.0 \| 1.01e-11 \| 400 \| 0.199 \| 676 \| 453 \| 26 \| 61 \| 702 \| 13 \| 634 \| Phage tail tape measure protein, TP901 family \| Phage tail tape measure protein, TP901 family; afdb-uniprot50 \| AF-F3ZVD4-F1-MODEL\_V4 \| 1.0 \| 8.316e-12 \| 400 \| 0.173 \| 703 \| 474 \| 24 \| 46 \| 702 \| 2 \| 643 \| Phage tail tape measure protein, TP901 family \| Phage tail tape measure protein, TP901 family; afdb-uniprot50 \| AF-A0A1H1G2L2-F1-MODEL\_V4 \| 1.0 \| 2.495e-12 \| 400 \| 0.169 \| 908 \| 565 \| 32 \| 19 \| 785 \| 65 \| 923 \| Phage tail tape measure protein, TP901 family, core region \| Phage tail tape measure protein, TP901 family, core region; afdb-uniprot50 \| AF-A0A378AFF2-F1-MODEL\_V4 \| 1.0 \| 4.663e-09 \| 399 \| 0.172 \| 487 \| 321 \| 12 \| 20 \| 438 \| 45 \| 517 \| Putative prophage tail length determinator \| Putative prophage tail length determinator; afdb-uniprot50 \| AF-A0A5M6CYT6-F1-MODEL\_V4 \| 1.0 \| 2.466e-11 \| 399 \| 0.14 \| 776 \| 463 \| 22 \| 26 \| 777 \| 1 \| 596 \| Phage tail tape measure protein \| Phage tail tape measure protein; afdb-uniprot50 \| AF-A0A4R3J9G1-F1-MODEL\_V4 \| 1.0 \| 9.235e-11 \| 399 \| 0.154 \| 732 \| 446 \| 24 \| 36 \| 735 \| 5 \| 595 \| TP901 family phage tail tape measure protein \| TP901 family phage tail tape measure protein; afdb-uniprot50 \| AF-A0A379FY57-F1-MODEL\_V4 \| 1.0 \| 1.079e-10 \| 399 \| 0.153 \| 663 \| 430 \| 15 \| 21 \| 604 \| 39 \| 648 \| Chromosome segregation protein \| Chromosome segregation protein; afdb-uniprot50 \| AF-A0A537MFF8-F1-MODEL\_V4 \| 1.0 \| 3.065e-13 \| 399 \| 0.168 \| 891 \| 526 \| 32 \| 22 \| 781 \| 53 \| 859 \| Phage tail tape measure protein \| Phage tail tape measure protein; afdb-uniprot50 \| AF-A0A506TXK5-F1-MODEL\_V4 \| 1.0 \| 1.901e-12 \| 399 \| 0.159 \| 801 \| 506 \| 24 \| 6 \| 774 \| 1 \| 665 \| Phage tail tape measure protein \| Phage tail tape measure protein; afdb-uniprot50 \| AF-A0A354PGC8-F1-MODEL\_V4 \| 1.0 \| 8.987e-12 \| 399 \| 0.165 \| 724 \| 465 \| 23 \| 83 \| 702 \| 18 \| 705 \| Phage tail tape measure protein \| Phage tail tape measure protein; afdb-uniprot50 \| AF-A0A7J2N4A4-F1-MODEL\_V4 \| 1.0 \| 1.148e-12 \| 399 \| 0.168 \| 790 \| 477 \| 23 \| 19 \| 716 \| 207 \| 908 \| Phage tail tape measure protein \| Phage tail tape measure protein; afdb-uniprot50 \| AF-A0A268TN88-F1-MODEL\_V4 \| 1.0 \| 3.379e-08 \| 398 \| 0.164 \| 371 \| 239 \| 4 \| 27 \| 362 \| 9 \| 343 \| Phage tail tape measure protein \| Phage tail tape measure protein; afdb-uniprot50 \| AF-A0A2Z4WE31-F1-MODEL\_V4 \| 1.0 \| 1.28e-08 \| 398 \| 0.175 \| 417 \| 265 \| 13 \| 80 \| 444 \| 2 \| 391 \| Phage tail tape measure protein \| Phage tail tape measure protein; afdb-uniprot50 \| AF-A0A346QYG9-F1-MODEL\_V4 \| 1.0 \| 1.022e-12 \| 398 \| 0.199 \| 667 \| 387 \| 23 \| 166 \| 754 \| 8 \| 605 \| Phage tail tape measure protein \| Phage tail tape measure protein; afdb-uniprot50 \| AF-A0A6B5AE33-F1-MODEL\_V4 \| 1.0 \| 4.885e-13 \| 398 \| 0.151 \| 910 \| 575 \| 40 \| 1 \| 783 \| 1 \| 839 \| Phage tail tape measure protein \| Phage tail tape measure protein; afdb-uniprot50 \| AF-A0A1Y3CT02-F1-MODEL\_V4 \| 1.0 \| 6.485e-14 \| 398 \| 0.164 \| 862 \| 565 \| 28 \| 19 \| 785 \| 56 \| 856 \| Phage tail tape measure protein \| Phage tail tape measure protein; afdb-uniprot50 \| AF-A0A1I5H9Z3-F1-MODEL\_V4 \| 1.0 \| 7.487e-13 \| 398 \| 0.173 \| 861 \| 551 \| 28 \| 19 \| 785 \| 51 \| 844 \| Phage tail tape measure protein, TP901 family, core region \| Phage tail tape measure protein, TP901 family, core region; afdb-uniprot50 \| AF-A0A0P0EGD7-F1-MODEL\_V4 \| 1.0 \| 1.78e-13 \| 398 \| 0.156 \| 1005 \| 555 \| 39 \| 39 \| 785 \| 43 \| 1012 \| PhageMin\_Tail domain-containing protein \| PhageMin\_Tail domain-containing protein; afdb-uniprot50 \| AF-A0A380DV55-F1-MODEL\_V4 \| 1.0 \| 1.566e-12 \| 398 \| 0.158 \| 790 \| 534 \| 31 \| 1 \| 702 \| 1 \| 747 \| Phage tail tape measure protein, TP901 family, core region \| Phage tail tape measure protein, TP901 family, core region; afdb-uniprot50 \| AF-A0A523CMP8-F1-MODEL\_V4 \| 1.0 \| 2.995e-11 \| 397 \| 0.126 \| 829 \| 488 \| 21 \| 6 \| 785 \| 3 \| 643 \| Phage tail tape measure protein \| Phage tail tape measure protein; afdb-uniprot50 \| AF-A0A3E0KFX5-F1-MODEL\_V4 \| 1.0 \| 9.826e-13 \| 397 \| 0.192 \| 741 \| 478 \| 25 \| 35 \| 708 \| 23 \| 709 \| Phage tail tape measure protein \| Phage tail tape measure protein; afdb-uniprot50 \| AF-A0A0M6WZJ0-F1-MODEL\_V4 \| 1.0 \| 6.336e-12 \| 397 \| 0.157 \| 728 \| 524 \| 21 \| 27 \| 701 \| 62 \| 752 \| PhageMin\_Tail domain-containing protein \| PhageMin\_Tail domain-containing protein; afdb-uniprot50 \| AF-A0A0D0HTB3-F1-MODEL\_V4 \| 1.0 \| 1.977e-12 \| 397 \| 0.166 \| 900 \| 590 \| 27 \| 8 \| 776 \| 157 \| 1026 \| Phage-related minor tail protein \| Phage-related minor tail protein; afdb-uniprot50 \| AF-A0A3A9F184-F1-MODEL\_V4 \| 1.0 \| 1.739e-11 \| 397 \| 0.178 \| 667 \| 463 \| 23 \| 77 \| 701 \| 3 \| 626 \| Phage tail tape measure protein \| Phage tail tape measure protein; afdb-uniprot50 \| AF-A0A523V4W2-F1-MODEL\_V4 \| 1.0 \| 5.729e-10 \| 396 \| 0.149 \| 576 \| 385 \| 19 \| 61 \| 627 \| 1 \| 480 \| Phage tail tape measure protein \| Phage tail tape measure protein; afdb-uniprot50 \| AF-A0A0A8FI92-F1-MODEL\_V4 \| 1.0 \| 5.384e-08 \| 396 \| 0.157 \| 369 \| 280 \| 13 \| 89 \| 445 \| 97 \| 446 \| PhageMin\_Tail domain-containing protein \| PhageMin\_Tail domain-containing protein; afdb-uniprot50 \| AF-A0A6M0SSJ1-F1-MODEL\_V4 \| 1.0 \| 2.309e-12 \| 396 \| 0.149 \| 869 \| 527 \| 30 \| 1 \| 775 \| 1 \| 750 \| Phage tail tape measure protein \| Phage tail tape measure protein; afdb-uniprot50 \| AF-A0A3D4VTZ8-F1-MODEL\_V4 \| 1.0 \| 8.092e-13 \| 396 \| 0.165 \| 892 \| 552 \| 35 \| 8 \| 782 \| 8 \| 823 \| Phage tail tape measure protein \| Phage tail tape measure protein; afdb-uniprot50 \| AF-A0A7C6ZVA5-F1-MODEL\_V4 \| 1.0 \| 2.564e-11 \| 396 \| 0.164 \| 676 \| 450 \| 20 \| 46 \| 700 \| 2 \| 583 \| Phage tail tape measure protein \| Phage tail tape measure protein; afdb-uniprot50 \| AF-A0A6N9P328-F1-MODEL\_V4 \| 1.0 \| 3.976e-12 \| 396 \| 0.215 \| 705 \| 454 \| 24 \| 53 \| 704 \| 3 \| 661 \| Phage tail tape measure protein \| Phage tail tape measure protein; afdb-uniprot50 \| AF-R9MQH3-F1-MODEL\_V4 \| 1.0 \| 1.449e-12 \| 396 \| 0.155 \| 823 \| 519 \| 28 \| 1 \| 702 \| 22 \| 789 \| Phage tail tape measure protein, TP901 family, core region \| Phage tail tape measure protein, TP901 family, core region; afdb-uniprot50 \| AF-A0A354XNE6-F1-MODEL\_V4 \| 1.0 \| 1.109e-09 \| 395 \| 0.164 \| 596 \| 398 \| 16 \| 22 \| 552 \| 54 \| 614 \| Phage tail tape measure protein \| Phage tail tape measure protein; afdb-uniprot50 \| AF-A0A7X2D754-F1-MODEL\_V4 \| 1.0 \| 1.038e-10 \| 395 \| 0.15 \| 690 \| 480 \| 23 \| 42 \| 684 \| 2 \| 632 \| Phage tail tape measure protein \| Phage tail tape measure protein; afdb-uniprot50 \| AF-A0A1S7PL82-F1-MODEL\_V4 \| 1.0 \| 2.771e-11 \| 395 \| 0.173 \| 776 \| 448 \| 26 \| 2 \| 723 \| 1 \| 636 \| Putative Phage tail tape measure protein, family, core region \| Putative Phage tail tape measure protein, family, core region; afdb-uniprot50 \| AF-A0A0F9IVD7-F1-MODEL\_V4 \| 1.0 \| 1.325e-11 \| 395 \| 0.206 \| 620 \| 426 \| 26 \| 96 \| 702 \| 5 \| 571 \| PhageMin\_Tail domain-containing protein \| PhageMin\_Tail domain-containing protein; afdb-uniprot50 \| AF-C9RFS8-F1-MODEL\_V4 \| 1.0 \| 7.784e-13 \| 395 \| 0.154 \| 788 \| 514 \| 31 \| 63 \| 772 \| 17 \| 729 \| Phage tail tape measure protein, TP901 family \| Phage tail tape measure protein, TP901 family; afdb-uniprot50 \| AF-A0A839XX57-F1-MODEL\_V4 \| 1.0 \| 8.746e-13 \| 395 \| 0.156 \| 928 \| 563 \| 32 \| 2 \| 785 \| 8 \| 859 \| TP901 family phage tail tape measure protein \| TP901 family phage tail tape measure protein; afdb-uniprot50 \| AF-A0A2E0VFX7-F1-MODEL\_V4 \| 1.0 \| 6.336e-12 \| 395 \| 0.137 \| 790 \| 484 \| 27 \| 49 \| 785 \| 5 \| 649 \| Phage tail tape measure protein \| Phage tail tape measure protein; afdb-uniprot50 \| AF-A0A6N7XH90-F1-MODEL\_V4 \| 1.0 \| 7.119e-12 \| 394 \| 0.161 \| 747 \| 493 \| 29 \| 109 \| 785 \| 52 \| 734 \| Phage tail tape measure protein \| Phage tail tape measure protein; afdb-uniprot50 \| AF-A0A5S4Z6P0-F1-MODEL\_V4 \| 1.0 \| 2.697e-12 \| 394 \| 0.157 \| 787 \| 494 \| 33 \| 70 \| 784 \| 6 \| 695 \| Phage tail tape measure protein \| Phage tail tape measure protein; afdb-uniprot50 \| AF-E5Y2G1-F1-MODEL\_V4 \| 1.0 \| 1.506e-12 \| 394 \| 0.163 \| 917 \| 522 \| 30 \| 1 \| 785 \| 1 \| 803 \| Phage tail tape measure protein, TP901 family, core region \| Phage tail tape measure protein, TP901 family, core region; afdb-uniprot50 \| AF-A0A7Y0EFA1-F1-MODEL\_V4 \| 1.0 \| 1.879e-11 \| 394 \| 0.137 \| 712 \| 488 \| 18 \| 77 \| 702 \| 21 \| 692 \| Phage tail tape measure protein \| Phage tail tape measure protein; afdb-uniprot50 \| AF-A0A5K7WX33-F1-MODEL\_V4 \| 1.0 \| 8.412e-13 \| 394 \| 0.16 \| 936 \| 560 \| 27 \| 27 \| 784 \| 62 \| 949 \| PhageMin\_Tail domain-containing protein \| PhageMin\_Tail domain-containing protein; afdb-uniprot50 \| AF-A0A380FL26-F1-MODEL\_V4 \| 1.0 \| 5.863e-12 \| 394 \| 0.153 \| 731 \| 506 \| 27 \| 26 \| 702 \| 51 \| 722 \| Phage tail tape measure protein, TP901 family, core region \| Phage tail tape measure protein, TP901 family, core region; afdb-uniprot50 \| AF-A0A7X7IZG9-F1-MODEL\_V4 \| 1.0 \| 1.829e-12 \| 393 \| 0.163 \| 803 \| 507 \| 29 \| 79 \| 774 \| 22 \| 766 \| Phage tail tape measure protein \| Phage tail tape measure protein; afdb-uniprot50 \| AF-A0A2S0JRG7-F1-MODEL\_V4 \| 1.0 \| 1.148e-12 \| 393 \| 0.155 \| 798 \| 492 \| 28 \| 19 \| 778 \| 23 \| 676 \| Phage tail tape measure protein \| Phage tail tape measure protein; afdb-uniprot50 \| AF-A0A5D4RWQ3-F1-MODEL\_V4 \| 1.0 \| 8.645e-12 \| 393 \| 0.16 \| 699 \| 467 \| 22 \| 70 \| 701 \| 4 \| 649 \| Phage tail tape measure protein \| Phage tail tape measure protein; afdb-uniprot50 \| AF-A0A4Q2R8E7-F1-MODEL\_V4 \| 1.0 \| 3.114e-11 \| 393 \| 0.151 \| 839 \| 452 \| 21 \| 32 \| 755 \| 192 \| 885 \| Phage tail tape measure protein \| Phage tail tape measure protein; afdb-uniprot50 \| AF-A0A5Y5IG90-F1-MODEL\_V4 \| 1.0 \| 2.635e-10 \| 392 \| 0.284 \| 436 \| 217 \| 9 \| 250 \| 682 \| 3 \| 346 \| Phage tail tape measure protein \| Phage tail tape measure protein; afdb-uniprot50 \| AF-A0A3R6KUE5-F1-MODEL\_V4 \| 1.0 \| 2.112e-11 \| 392 \| 0.16 \| 716 \| 451 \| 23 \| 19 \| 675 \| 45 \| 669 \| Phage tail tape measure protein \| Phage tail tape measure protein; afdb-uniprot50 \| AF-A0A4R1Z735-F1-MODEL\_V4 \| 1.0 \| 3.404e-12 \| 392 \| 0.161 \| 857 \| 525 \| 24 \| 32 \| 781 \| 3 \| 772 \| TP901 family phage tail tape measure protein \| TP901 family phage tail tape measure protein; afdb-uniprot50 \| AF-A0A1Y3TQP8-F1-MODEL\_V4 \| 1.0 \| 7.037e-11 \| 392 \| 0.16 \| 637 \| 437 \| 22 \| 77 \| 702 \| 22 \| 571 \| Phage tail tape measure protein \| Phage tail tape measure protein; afdb-uniprot50 \| AF-A0A3Q9B6I7-F1-MODEL\_V4 \| 1.0 \| 9.601e-11 \| 391 \| 0.153 \| 721 \| 465 \| 29 \| 90 \| 785 \| 17 \| 616 \| Phage tail tape measure protein \| Phage tail tape measure protein; afdb-uniprot50 \| AF-S9TGE4-F1-MODEL\_V4 \| 1.0 \| 2.031e-11 \| 391 \| 0.162 \| 775 \| 459 \| 21 \| 5 \| 750 \| 2 \| 615 \| TP901 family phage tail tape measure protein \| TP901 family phage tail tape measure protein; afdb-uniprot50 \| AF-A0A846I152-F1-MODEL\_V4 \| 1.0 \| 3.404e-12 \| 391 \| 0.169 \| 736 \| 492 \| 27 \| 1 \| 701 \| 1 \| 651 \| Phage tail tape measure protein \| Phage tail tape measure protein; afdb-uniprot50 \| AF-A0A371IJP1-F1-MODEL\_V4 \| 1.0 \| 1.901e-12 \| 391 \| 0.152 \| 916 \| 529 \| 30 \| 27 \| 785 \| 130 \| 954 \| Phage tail tape measure protein \| Phage tail tape measure protein; afdb-uniprot50 \| AF-A0A0D1X6H1-F1-MODEL\_V4 \| 1.0 \| 1.566e-12 \| 391 \| 0.137 \| 970 \| 586 \| 30 \| 19 \| 784 \| 40 \| 962 \| Phage tail tape measure protein, TP901 family, core region \| Phage tail tape measure protein, TP901 family, core region; afdb-uniprot50 \| AF-V4PS02-F1-MODEL\_V4 \| 1.0 \| 1.506e-12 \| 390 \| 0.171 \| 849 \| 573 \| 27 \| 1 \| 782 \| 1 \| 785 \| PhageMin\_Tail domain-containing protein \| PhageMin\_Tail domain-containing protein; afdb-uniprot50 \| AF-A0A1C3EE79-F1-MODEL\_V4 \| 1.0 \| 1.759e-12 \| 390 \| 0.179 \| 880 \| 536 \| 24 \| 19 \| 782 \| 57 \| 866 \| Phage tail tape measure protein \| Phage tail tape measure protein; afdb-uniprot50 \| AF-A0A2A4H5A1-F1-MODEL\_V4 \| 1.0 \| 4.644e-12 \| 390 \| 0.151 \| 790 \| 515 \| 25 \| 1 \| 702 \| 1 \| 722 \| Phage tail tape measure protein \| Phage tail tape measure protein; afdb-uniprot50 \| AF-A0A1W9SV37-F1-MODEL\_V4 \| 1.0 \| 4.297e-12 \| 389 \| 0.17 \| 751 \| 502 \| 24 \| 46 \| 722 \| 5 \| 708 \| Phage tail tape measure protein \| Phage tail tape measure protein; afdb-uniprot50 \| AF-A0A1X9MF98-F1-MODEL\_V4 \| 1.0 \| 5.64e-12 \| 389 \| 0.185 \| 819 \| 504 \| 31 \| 1 \| 702 \| 1 \| 773 \| Phage-related minor tail protein \| Phage-related minor tail protein; afdb-uniprot50 \| AF-A0A6M0STF1-F1-MODEL\_V4 \| 1.0 \| 2.848e-10 \| 388 \| 0.148 \| 794 \| 445 \| 28 \| 1 \| 785 \| 1 \| 572 \| Phage tail tape measure protein \| Phage tail tape measure protein; afdb-uniprot50 \| AF-A0A4D7BBQ9-F1-MODEL\_V4 \| 1.0 \| 2.466e-11 \| 388 \| 0.155 \| 664 \| 425 \| 20 \| 116 \| 755 \| 29 \| 580 \| Phage tail tape measure protein \| Phage tail tape measure protein; afdb-uniprot50 \| AF-A0A6P0MHK5-F1-MODEL\_V4 \| 1.0 \| 4.828e-12 \| 388 \| 0.159 \| 805 \| 527 \| 29 \| 70 \| 784 \| 4 \| 748 \| Phage tail tape measure protein \| Phage tail tape measure protein; afdb-uniprot50 \| AF-A0A661DRW5-F1-MODEL\_V4 \| 1.0 \| 1.179e-11 \| 388 \| 0.151 \| 772 \| 461 \| 25 \| 1 \| 702 \| 6 \| 653 \| Phage tail tape measure protein \| Phage tail tape measure protein; afdb-uniprot50 \| AF-A0A5C6C1J2-F1-MODEL\_V4 \| 1.0 \| 1.325e-11 \| 388 \| 0.177 \| 777 \| 467 \| 25 \| 1 \| 705 \| 1 \| 677 \| Phage-related minor tail protein \| Phage-related minor tail protein; afdb-uniprot50 \| AF-G4Q995-F1-MODEL\_V4 \| 1.0 \| 5.51e-10 \| 388 \| 0.163 \| 537 \| 335 \| 17 \| 1 \| 463 \| 1 \| 497 \| Phage tail tape measure protein, TP901 family, core region domain protein \| Phage tail tape measure protein, TP901 family, core region domain protein; afdb-uniprot50 \| AF-A0A0P6WTM8-F1-MODEL\_V4 \| 1.0 \| 6.664e-13 \| 388 \| 0.168 \| 730 \| 460 \| 26 \| 62 \| 705 \| 5 \| 673 \| PhageMin\_Tail domain-containing protein \| PhageMin\_Tail domain-containing protein; afdb-uniprot50 \| AF-A0A502CNZ0-F1-MODEL\_V4 \| 1.0 \| 1.506e-12 \| 388 \| 0.163 \| 1003 \| 572 \| 34 \| 19 \| 785 \| 39 \| 1009 \| Phage tail tape measure protein \| Phage tail tape measure protein; afdb-uniprot50 \| AF-A0A417X761-F1-MODEL\_V4 \| 1.0 \| 1.455e-09 \| 387 \| 0.145 \| 618 \| 389 \| 20 \| 81 \| 662 \| 5 \| 519 \| Phage tail tape measure protein \| Phage tail tape measure protein; afdb-uniprot50 \| AF-A0A4Q0YND4-F1-MODEL\_V4 \| 1.0 \| 1.985e-09 \| 387 \| 0.179 \| 564 \| 369 \| 16 \| 19 \| 508 \| 57 \| 600 \| Phage tail tape measure protein \| Phage tail tape measure protein; afdb-uniprot50 \| AF-A0A841GQ15-F1-MODEL\_V4 \| 1.0 \| 3.825e-12 \| 387 \| 0.148 \| 739 \| 493 \| 18 \| 1 \| 701 \| 2 \| 641 \| TP901 family phage tail tape measure protein \| TP901 family phage tail tape measure protein; afdb-uniprot50 \| AF-A0A5T7NPI6-F1-MODEL\_V4 \| 1.0 \| 4.086e-11 \| 386 \| 0.17 \| 700 \| 411 \| 19 \| 19 \| 623 \| 44 \| 668 \| Phage tail tape measure protein \| Phage tail tape measure protein; afdb-uniprot50 \| AF-A0A0F9R9A9-F1-MODEL\_V4 \| 1.0 \| 5.64e-12 \| 386 \| 0.183 \| 794 \| 500 \| 29 \| 63 \| 764 \| 5 \| 741 \| PhageMin\_Tail domain-containing protein \| PhageMin\_Tail domain-containing protein; afdb-uniprot50 \| AF-A0A6L9H1L1-F1-MODEL\_V4 \| 1.0 \| 8.545e-11 \| 386 \| 0.172 \| 609 \| 415 \| 20 \| 113 \| 702 \| 1 \| 539 \| Phage tail tape measure protein \| Phage tail tape measure protein; afdb-uniprot50 \| AF-A0A4S2HCP5-F1-MODEL\_V4 \| 1.0 \| 4.644e-12 \| 386 \| 0.146 \| 825 \| 524 \| 20 \| 22 \| 702 \| 50 \| 837 \| Phage tail tape measure protein \| Phage tail tape measure protein; afdb-uniprot50 \| AF-A0A417YR53-F1-MODEL\_V4 \| 1.0 \| 5.279e-13 \| 386 \| 0.158 \| 1074 \| 560 \| 37 \| 19 \| 785 \| 131 \| 1167 \| Phage tail tape measure protein \| Phage tail tape measure protein; afdb-uniprot50 \| AF-A0A5C7LFW0-F1-MODEL\_V4 \| 1.0 \| 1.416e-10 \| 385 \| 0.131 \| 645 \| 438 \| 17 \| 70 \| 700 \| 21 \| 556 \| Phage tail tape measure protein \| Phage tail tape measure protein; afdb-uniprot50 \| AF-A0A2E8I968-F1-MODEL\_V4 \| 1.0 \| 2.666e-11 \| 385 \| 0.159 \| 791 \| 446 \| 30 \| 1 \| 750 \| 1 \| 613 \| Phage tail tape measure protein \| Phage tail tape measure protein; afdb-uniprot50 \| AF-A0A3C0IJN3-F1-MODEL\_V4 \| 1.0 \| 1.109e-09 \| 385 \| 0.178 \| 573 \| 361 \| 19 \| 19 \| 519 \| 91 \| 625 \| Phage tail tape measure protein \| Phage tail tape measure protein; afdb-uniprot50 \| AF-A0A0F9KJV0-F1-MODEL\_V4 \| 1.0 \| 6.025e-11 \| 385 \| 0.152 \| 707 \| 485 \| 24 \| 13 \| 677 \| 2 \| 635 \| PhageMin\_Tail domain-containing protein \| PhageMin\_Tail domain-containing protein; afdb-uniprot50 \| AF-A0A2N5EEH0-F1-MODEL\_V4 \| 1.0 \| 2.804e-12 \| 385 \| 0.178 \| 783 \| 459 \| 24 \| 42 \| 776 \| 9 \| 654 \| Phage tail tape measure protein \| Phage tail tape measure protein; afdb-uniprot50 \| AF-A0A7C6HAK4-F1-MODEL\_V4 \| 1.0 \| 9.235e-11 \| 385 \| 0.145 \| 661 \| 386 \| 20 \| 108 \| 754 \| 111 \| 606 \| Phage tail tape measure protein \| Phage tail tape measure protein; afdb-uniprot50 \| AF-A0A1F4MH22-F1-MODEL\_V4 \| 1.0 \| 7.606e-11 \| 385 \| 0.164 \| 640 \| 443 \| 25 \| 76 \| 702 \| 6 \| 566 \| Phage tail tape measure protein \| Phage tail tape measure protein; afdb-uniprot50 \| AF-A0A2I6QM47-F1-MODEL\_V4 \| 1.0 \| 4.134e-12 \| 385 \| 0.153 \| 919 \| 582 \| 28 \| 6 \| 785 \| 1 \| 862 \| Phage tail length tape-measure protein \| Phage tail length tape-measure protein; afdb-uniprot50 \| AF-A0A7X5L8G2-F1-MODEL\_V4 \| 1.0 \| 3.679e-12 \| 385 \| 0.158 \| 777 \| 494 \| 32 \| 2 \| 702 \| 3 \| 695 \| Phage tail tape measure protein \| Phage tail tape measure protein; afdb-uniprot50 \| AF-A0A660IYM3-F1-MODEL\_V4 \| 1.0 \| 1.466e-13 \| 385 \| 0.155 \| 876 \| 490 \| 29 \| 9 \| 650 \| 1 \| 860 \| Phage tail tape measure protein \| Phage tail tape measure protein; afdb-uniprot50 \| AF-A0A7G2KFG2-F1-MODEL\_V4 \| 1.0 \| 2.594e-12 \| 384 \| 0.233 \| 549 \| 329 \| 12 \| 212 \| 678 \| 2 \| 540 \| Phage tail tape measure protein, TP901 family, core region \| Phage tail tape measure protein, TP901 family, core region; afdb-uniprot50 \| AF-F4N710-F1-MODEL\_V4 \| 1.0 \| 4.591e-11 \| 384 \| 0.143 \| 731 \| 489 \| 23 \| 9 \| 687 \| 1 \| 646 \| PhageMin\_Tail domain-containing protein \| PhageMin\_Tail domain-containing protein; afdb-uniprot50 \| AF-A0A2G3DYB1-F1-MODEL\_V4 \| 1.0 \| 1.05e-11 \| 384 \| 0.14 \| 860 \| 541 \| 32 \| 1 \| 782 \| 1 \| 739 \| Phage tail tape measure protein \| Phage tail tape measure protein; afdb-uniprot50 \| AF-E6U1K4-F1-MODEL\_V4 \| 1.0 \| 4.248e-11 \| 384 \| 0.16 \| 680 \| 446 \| 21 \| 38 \| 702 \| 11 \| 580 \| Phage tail tape measure protein, TP901 family \| Phage tail tape measure protein, TP901 family; afdb-uniprot50 \| AF-A0A2E0SQ31-F1-MODEL\_V4 \| 1.0 \| 3.03e-12 \| 384 \| 0.151 \| 925 \| 558 \| 31 \| 19 \| 785 \| 26 \| 881 \| Phage tail tape measure protein \| Phage tail tape measure protein; afdb-uniprot50 \| AF-A0A845QYL5-F1-MODEL\_V4 \| 1.0 \| 1.362e-10 \| 384 \| 0.165 \| 669 \| 435 \| 24 \| 118 \| 708 \| 52 \| 674 \| Phage tail tape measure protein \| Phage tail tape measure protein; afdb-uniprot50 \| AF-B9CLJ5-F1-MODEL\_V4 \| 1.0 \| 1.759e-12 \| 384 \| 0.161 \| 844 \| 541 \| 24 \| 1 \| 702 \| 1 \| 819 \| Phage tail tape measure protein, TP901 family \| Phage tail tape measure protein, TP901 family; afdb-uniprot50 \| AF-A0A830K262-F1-MODEL\_V4 \| 1.0 \| 4.365e-10 \| 383 \| 0.239 \| 422 \| 283 \| 8 \| 222 \| 617 \| 1 \| 410 \| Phage tail tape measure protein \| Phage tail tape measure protein; afdb-uniprot50 \| AF-A0A1V3IZV8-F1-MODEL\_V4 \| 1.0 \| 1.399e-09 \| 383 \| 0.153 \| 614 \| 421 \| 16 \| 19 \| 555 \| 49 \| 640 \| Phage tail tape measure protein \| Phage tail tape measure protein; afdb-uniprot50 \| AF-A0A7Y9Z9G2-F1-MODEL\_V4 \| 1.0 \| 1.166e-10 \| 383 \| 0.162 \| 783 \| 484 \| 27 \| 37 \| 784 \| 2 \| 647 \| TP901 family phage tail tape measure protein \| TP901 family phage tail tape measure protein; afdb-uniprot50 \| AF-A0A0F9T832-F1-MODEL\_V4 \| 1.0 \| 9.981e-11 \| 383 \| 0.162 \| 659 \| 465 \| 19 \| 64 \| 702 \| 2 \| 593 \| PhageMin\_Tail domain-containing protein \| PhageMin\_Tail domain-containing protein; afdb-uniprot50 \| AF-R7IG02-F1-MODEL\_V4 \| 1.0 \| 2.136e-12 \| 383 \| 0.158 \| 845 \| 563 \| 34 \| 1 \| 767 \| 1 \| 775 \| Phage tail tape measure protein TP901 family core region \| Phage tail tape measure protein TP901 family core region; afdb-uniprot50 \| AF-A0A0R2KRB1-F1-MODEL\_V4 \| 1.0 \| 3.275e-12 \| 383 \| 0.159 \| 805 \| 537 \| 26 \| 19 \| 716 \| 22 \| 793 \| PhageMin\_Tail domain-containing protein \| PhageMin\_Tail domain-containing protein; afdb-uniprot50 \| AF-A0A3A6JVW7-F1-MODEL\_V4 \| 1.0 \| 2.961e-10 \| 382 \| 0.177 \| 635 \| 468 \| 21 \| 94 \| 702 \| 19 \| 624 \| Phage tail tape measure protein \| Phage tail tape measure protein; afdb-uniprot50 \| AF-A0A5C5X117-F1-MODEL\_V4 \| 1.0 \| 6.095e-12 \| 382 \| 0.164 \| 789 \| 520 \| 24 \| 63 \| 776 \| 4 \| 727 \| Phage-related minor tail protein \| Phage-related minor tail protein; afdb-uniprot50 \| AF-A0A1T4MQV6-F1-MODEL\_V4 \| 1.0 \| 8.645e-12 \| 382 \| 0.143 \| 872 \| 576 \| 29 \| 27 \| 785 \| 59 \| 872 \| Phage tail tape measure protein, TP901 family, core region \| Phage tail tape measure protein, TP901 family, core region; afdb-uniprot50 \| AF-A0A6N7TNS1-F1-MODEL\_V4 \| 1.0 \| 7.907e-11 \| 382 \| 0.176 \| 623 \| 440 \| 19 \| 117 \| 708 \| 48 \| 628 \| Phage tail tape measure protein \| Phage tail tape measure protein; afdb-uniprot50 \| AF-A0A0F9HCP7-F1-MODEL\_V4 \| 1.0 \| 9.981e-11 \| 381 \| 0.151 \| 744 \| 467 \| 31 \| 39 \| 748 \| 2 \| 614 \| PhageMin\_Tail domain-containing protein \| PhageMin\_Tail domain-containing protein; afdb-uniprot50 \| AF-A0A0A0CMA7-F1-MODEL\_V4 \| 1.0 \| 1.179e-11 \| 381 \| 0.131 \| 963 \| 588 \| 30 \| 19 \| 784 \| 42 \| 952 \| PhageMin\_Tail domain-containing protein \| PhageMin\_Tail domain-containing protein; afdb-uniprot50 \| AF-A0A235F9B5-F1-MODEL\_V4 \| 1.0 \| 1.134e-11 \| 381 \| 0.151 \| 764 \| 487 \| 30 \| 19 \| 689 \| 40 \| 734 \| Phage tail tape measure protein \| Phage tail tape measure protein; afdb-uniprot50 \| AF-A0A1E8RGF4-F1-MODEL\_V4 \| 1.0 \| 9.452e-13 \| 381 \| 0.142 \| 952 \| 520 \| 32 \| 27 \| 702 \| 172 \| 1102 \| PhageMin\_Tail domain-containing protein \| PhageMin\_Tail domain-containing protein; afdb-uniprot50 \| AF-A0A1U7PT90-F1-MODEL\_V4 \| 1.0 \| 4.468e-12 \| 380 \| 0.145 \| 795 \| 502 \| 26 \| 5 \| 702 \| 2 \| 715 \| Phage tail tape measure protein, TP901 family, core region \| Phage tail tape measure protein, TP901 family, core region; afdb-uniprot50 \| AF-E7GJW0-F1-MODEL\_V4 \| 1.0 \| 3.825e-12 \| 379 \| 0.183 \| 772 \| 492 \| 30 \| 10 \| 702 \| 10 \| 721 \| TMP repeat-containing protein \| TMP repeat-containing protein; afdb-uniprot50 \| AF-A0A1M4VAW5-F1-MODEL\_V4 \| 1.0 \| 3.976e-12 \| 378 \| 0.152 \| 976 \| 576 \| 30 \| 16 \| 776 \| 1 \| 939 \| Phage tail tape measure protein, TP901 family, core region \| Phage tail tape measure protein, TP901 family, core region; afdb-uniprot50 \| AF-A1SE57-F1-MODEL\_V4 \| 1.0 \| 6.025e-11 \| 378 \| 0.168 \| 651 \| 443 \| 24 \| 118 \| 724 \| 41 \| 636 \| Phage tail tape measure protein, TP901 family \| Phage tail tape measure protein, TP901 family; afdb-uniprot50 \| AF-A0A7W1XA84-F1-MODEL\_V4 \| 1.0 \| 2.666e-11 \| 378 \| 0.146 \| 738 \| 496 \| 20 \| 37 \| 702 \| 2 \| 677 \| Phage tail tape measure protein \| Phage tail tape measure protein; afdb-uniprot50 \| AF-A0A1H0MFG2-F1-MODEL\_V4 \| 1.0 \| 1.325e-11 \| 377 \| 0.155 \| 734 \| 452 \| 27 \| 1 \| 641 \| 1 \| 659 \| Phage tail tape measure protein, TP901 family, core region \| Phage tail tape measure protein, TP901 family, core region; afdb-uniprot50 \| AF-A5ZRU7-F1-MODEL\_V4 \| 1.0 \| 3.93e-11 \| 377 \| 0.16 \| 754 \| 510 \| 29 \| 27 \| 702 \| 78 \| 786 \| Phage tail tape measure protein, TP901 family \| Phage tail tape measure protein, TP901 family; afdb-uniprot50 \| AF-Q2FWU3-F1-MODEL\_V4 \| 1.0 \| 7.119e-12 \| 377 \| 0.142 \| 921 \| 545 \| 39 \| 1 \| 702 \| 1 \| 895 \| Lysostaphin \| Lysostaphin; afdb-uniprot50 \| AF-A0A7Y8S2G1-F1-MODEL\_V4 \| 1.0 \| 2.256e-10 \| 376 \| 0.155 \| 636 \| 421 \| 20 \| 1 \| 551 \| 1 \| 605 \| Phage tail tape measure protein \| Phage tail tape measure protein; afdb-uniprot50 \| AF-A1VPH8-F1-MODEL\_V4 \| 1.0 \| 1.166e-10 \| 376 \| 0.155 \| 655 \| 442 \| 24 \| 76 \| 702 \| 20 \| 591 \| Phage tail tape measure protein, TP901 family \| Phage tail tape measure protein, TP901 family; afdb-uniprot50 \| AF-A0A6I2HFM4-F1-MODEL\_V4 \| 1.0 \| 2.995e-11 \| 376 \| 0.16 \| 735 \| 488 \| 22 \| 50 \| 702 \| 16 \| 703 \| Phage tail tape measure protein \| Phage tail tape measure protein; afdb-uniprot50 \| AF-A0A6N8HZ99-F1-MODEL\_V4 \| 1.0 \| 2.771e-11 \| 376 \| 0.152 \| 754 \| 527 \| 22 \| 13 \| 702 \| 42 \| 747 \| Phage-related minor tail protein \| Phage-related minor tail protein; afdb-uniprot50 \| AF-A0A1Q3LGX9-F1-MODEL\_V4 \| 1.0 \| 4.644e-12 \| 376 \| 0.179 \| 741 \| 466 \| 29 \| 19 \| 707 \| 144 \| 794 \| Phage tail tape measure protein \| Phage tail tape measure protein; afdb-uniprot50 \| AF-A0A0F9FLG5-F1-MODEL\_V4 \| 1.0 \| 6.049e-08 \| 375 \| 0.167 \| 425 \| 291 \| 15 \| 16 \| 420 \| 1 \| 382 \| PhageMin\_Tail domain-containing protein \| PhageMin\_Tail domain-containing protein; afdb-uniprot50 \| AF-A0A087M271-F1-MODEL\_V4 \| 1.0 \| 6.025e-11 \| 375 \| 0.159 \| 829 \| 485 \| 30 \| 1 \| 783 \| 1 \| 663 \| PhageMin\_Tail domain-containing protein \| PhageMin\_Tail domain-containing protein; afdb-uniprot50 \| AF-K5D6H4-F1-MODEL\_V4 \| 1.0 \| 6.848e-12 \| 375 \| 0.15 \| 842 \| 529 \| 31 \| 47 \| 779 \| 4 \| 767 \| Family phage tail tape measure protein \| Family phage tail tape measure protein; afdb-uniprot50 \| AF-A0A806NJH9-F1-MODEL\_V4 \| 1.0 \| 1.325e-11 \| 375 \| 0.158 \| 752 \| 482 \| 24 \| 1 \| 702 \| 1 \| 651 \| Tail protein \| Tail protein; afdb-uniprot50 \| AF-A0A354SEK7-F1-MODEL\_V4 \| 1.0 \| 6.587e-12 \| 375 \| 0.157 \| 920 \| 515 \| 36 \| 13 \| 779 \| 2 \| 813 \| Phage tail tape measure protein \| Phage tail tape measure protein; afdb-uniprot50 \| AF-A0A7U5GYW1-F1-MODEL\_V4 \| 1.0 \| 8.645e-12 \| 375 \| 0.15 \| 808 \| 504 \| 22 \| 18 \| 785 \| 18 \| 682 \| Phage tail tape measure protein \| Phage tail tape measure protein; afdb-uniprot50 \| AF-A0A5E8P1F1-F1-MODEL\_V4 \| 1.0 \| 1.931e-10 \| 374 \| 0.173 \| 670 \| 399 \| 20 \| 19 \| 599 \| 35 \| 638 \| Phage tail tape measure protein \| Phage tail tape measure protein; afdb-uniprot50 \| AF-A0A6B4V7Z7-F1-MODEL\_V4 \| 1.0 \| 2.031e-11 \| 374 \| 0.141 \| 856 \| 539 \| 32 \| 1 \| 774 \| 1 \| 742 \| Phage tail tape measure protein \| Phage tail tape measure protein; afdb-uniprot50 \| AF-A0A2I1RIZ6-F1-MODEL\_V4 \| 1.0 \| 5.279e-13 \| 374 \| 0.179 \| 880 \| 563 \| 32 \| 19 \| 782 \| 56 \| 892 \| Phage tail tape measure protein \| Phage tail tape measure protein; afdb-uniprot50 \| AF-A0A3A9ETB4-F1-MODEL\_V4 \| 1.0 \| 5.795e-11 \| 374 \| 0.163 \| 668 \| 455 \| 19 \| 70 \| 702 \| 6 \| 604 \| Phage tail tape measure protein \| Phage tail tape measure protein; afdb-uniprot50 \| AF-A0A4U9RAU9-F1-MODEL\_V4 \| 1.0 \| 3.694e-09 \| 374 \| 0.154 \| 504 \| 332 \| 15 \| 19 \| 441 \| 70 \| 560 \| Phage tail tape measure protein, family, core region \| Phage tail tape measure protein, family, core region; afdb-uniprot50 \| AF-A0A6N9B9M4-F1-MODEL\_V4 \| 1.0 \| 6.511e-11 \| 373 \| 0.156 \| 641 \| 410 \| 22 \| 81 \| 696 \| 4 \| 538 \| Phage tail tape measure protein \| Phage tail tape measure protein; afdb-uniprot50 \| AF-A0A5M5BW70-F1-MODEL\_V4 \| 1.0 \| 1.152e-09 \| 373 \| 0.112 \| 659 \| 405 \| 16 \| 110 \| 750 \| 211 \| 707 \| Phage tail tape measure protein \| Phage tail tape measure protein; afdb-uniprot50 \| AF-A0A7J5WEE1-F1-MODEL\_V4 \| 1.0 \| 3.237e-11 \| 373 \| 0.158 \| 870 \| 512 \| 27 \| 1 \| 777 \| 1 \| 742 \| Family phage tail tape measure \| Family phage tail tape measure; afdb-uniprot50 \| AF-A0A496UIZ6-F1-MODEL\_V4 \| 1.0 \| 3.781e-11 \| 373 \| 0.142 \| 716 \| 479 \| 27 \| 28 \| 702 \| 5 \| 626 \| Phage tail tape measure protein \| Phage tail tape measure protein; afdb-uniprot50 \| AF-A0A2E2FJ28-F1-MODEL\_V4 \| 1.0 \| 5.64e-12 \| 373 \| 0.168 \| 771 \| 484 \| 30 \| 6 \| 702 \| 2 \| 689 \| Phage tail tape measure protein \| Phage tail tape measure protein; afdb-uniprot50 \| AF-A0A5B9MJG5-F1-MODEL\_V4 \| 1.0 \| 8.987e-12 \| 373 \| 0.154 \| 868 \| 524 \| 29 \| 1 \| 785 \| 1 \| 741 \| Phage-related minor tail protein \| Phage-related minor tail protein; afdb-uniprot50 \| AF-A0A535HRI6-F1-MODEL\_V4 \| 1.0 \| 6.691e-10 \| 373 \| 0.179 \| 631 \| 442 \| 20 \| 117 \| 702 \| 16 \| 615 \| Phage tail tape measure protein \| Phage tail tape measure protein; afdb-uniprot50 \| AF-A0A848EQ20-F1-MODEL\_V4 \| 1.0 \| 6.511e-11 \| 372 \| 0.154 \| 833 \| 501 \| 37 \| 19 \| 784 \| 11 \| 706 \| Phage tail tape measure protein \| Phage tail tape measure protein; afdb-uniprot50 \| AF-A0A661KKY6-F1-MODEL\_V4 \| 1.0 \| 1.226e-11 \| 372 \| 0.137 \| 821 \| 550 \| 29 \| 64 \| 785 \| 4 \| 764 \| Phage tail tape measure protein \| Phage tail tape measure protein; afdb-uniprot50 \| AF-A0A7M2B111-F1-MODEL\_V4 \| 1.0 \| 1.275e-11 \| 372 \| 0.14 \| 800 \| 535 \| 22 \| 21 \| 701 \| 83 \| 848 \| Phage tail tape measure protein \| Phage tail tape measure protein; afdb-uniprot50 \| AF-A0A0F9SUJ2-F1-MODEL\_V4 \| 1.0 \| 4.644e-12 \| 372 \| 0.167 \| 778 \| 486 \| 25 \| 32 \| 702 \| 15 \| 737 \| Uncharacterized protein \| Uncharacterized protein; afdb-uniprot50 \| AF-A0A101W526-F1-MODEL\_V4 \| 1.0 \| 7.316e-11 \| 371 \| 0.147 \| 732 \| 500 \| 24 \| 80 \| 772 \| 18 \| 664 \| PhageMin\_Tail domain-containing protein \| PhageMin\_Tail domain-containing protein; afdb-uniprot50 \| AF-A0A2R4MEE8-F1-MODEL\_V4 \| 1.0 \| 6.848e-12 \| 371 \| 0.152 \| 750 \| 484 \| 30 \| 49 \| 783 \| 20 \| 632 \| Tape measure protein \| Tape measure protein; afdb-uniprot50 \| AF-A0A1N7MQX6-F1-MODEL\_V4 \| 1.0 \| 7.037e-11 \| 371 \| 0.146 \| 717 \| 466 \| 31 \| 60 \| 702 \| 2 \| 646 \| Phage tail tape measure protein, TP901 family, core region \| Phage tail tape measure protein, TP901 family, core region; afdb-uniprot50 \| AF-A0A3N0IV62-F1-MODEL\_V4 \| 1.0 \| 4.134e-12 \| 371 \| 0.168 \| 850 \| 499 \| 32 \| 1 \| 701 \| 1 \| 791 \| Phage tail tape measure protein \| Phage tail tape measure protein; afdb-uniprot50 \| AF-W9V4X2-F1-MODEL\_V4 \| 1.0 \| 1.449e-12 \| 371 \| 0.169 \| 827 \| 476 \| 30 \| 63 \| 705 \| 6 \| 805 \| PhageMin\_Tail domain-containing protein \| PhageMin\_Tail domain-containing protein; afdb-uniprot50 \| AF-A0A3C1WR03-F1-MODEL\_V4 \| 1.0 \| 3.93e-11 \| 371 \| 0.157 \| 700 \| 482 \| 22 \| 50 \| 700 \| 7 \| 647 \| Phage tail tape measure protein \| Phage tail tape measure protein; afdb-uniprot50 \| AF-A0A7G9W8A8-F1-MODEL\_V4 \| 1.0 \| 6.095e-12 \| 371 \| 0.15 \| 757 \| 494 \| 30 \| 1 \| 667 \| 3 \| 700 \| Phage tail tape measure protein \| Phage tail tape measure protein; afdb-uniprot50 \| AF-K9EC41-F1-MODEL\_V4 \| 1.0 \| 1.038e-10 \| 371 \| 0.16 \| 661 \| 449 \| 24 \| 64 \| 702 \| 19 \| 595 \| Phage tail tape measure protein, TP901 family, core region \| Phage tail tape measure protein, TP901 family, core region; afdb-uniprot50 \| AF-A0A553KRV1-F1-MODEL\_V4 \| 1.0 \| 1.179e-11 \| 371 \| 0.176 \| 778 \| 503 \| 26 \| 18 \| 706 \| 42 \| 770 \| Phage tail tape measure protein \| Phage tail tape measure protein; afdb-uniprot50 \| AF-A0A078MEL8-F1-MODEL\_V4 \| 1.0 \| 1.166e-10 \| 371 \| 0.159 \| 669 \| 450 \| 24 \| 49 \| 702 \| 3 \| 573 \| Phage-related minor tail protein \| Phage-related minor tail protein; afdb-uniprot50 \| AF-A0A1T4QWG9-F1-MODEL\_V4 \| 1.0 \| 8.545e-11 \| 370 \| 0.136 \| 791 \| 466 \| 27 \| 32 \| 781 \| 49 \| 663 \| Phage tail tape measure protein, TP901 family, core region \| Phage tail tape measure protein, TP901 family, core region; afdb-uniprot50 \| AF-A0A7Y5QHS0-F1-MODEL\_V4 \| 1.0 \| 4.248e-11 \| 370 \| 0.171 \| 763 \| 522 \| 29 \| 74 \| 773 \| 14 \| 728 \| Phage tail tape measure protein \| Phage tail tape measure protein; afdb-uniprot50 \| AF-A0A662HX82-F1-MODEL\_V4 \| 1.0 \| 2.995e-11 \| 370 \| 0.148 \| 809 \| 527 \| 28 \| 46 \| 772 \| 8 \| 736 \| Phage tail tape measure protein \| Phage tail tape measure protein; afdb-uniprot50 \| AF-A0A350XS20-F1-MODEL\_V4 \| 1.0 \| 5.575e-11 \| 370 \| 0.148 \| 786 \| 504 \| 22 \| 1 \| 711 \| 1 \| 695 \| Phage tail tape measure protein \| Phage tail tape measure protein; afdb-uniprot50 \| AF-A0A416ZSH5-F1-MODEL\_V4 \| 1.0 \| 4.962e-11 \| 370 \| 0.182 \| 723 \| 466 \| 25 \| 17 \| 712 \| 34 \| 658 \| Phage tail tape measure protein \| Phage tail tape measure protein; afdb-uniprot50 \| AF-A0A2A6Z7M7-F1-MODEL\_V4 \| 1.0 \| 7.606e-11 \| 370 \| 0.151 \| 694 \| 409 \| 26 \| 97 \| 755 \| 90 \| 638 \| Phage tail tape measure protein \| Phage tail tape measure protein; afdb-uniprot50 \| AF-A0A846T1D6-F1-MODEL\_V4 \| 1.0 \| 3.781e-11 \| 369 \| 0.204 \| 582 \| 324 \| 19 \| 214 \| 743 \| 2 \| 496 \| Phage tail tape measure protein \| Phage tail tape measure protein; afdb-uniprot50 \| AF-H3SBJ7-F1-MODEL\_V4 \| 1.0 \| 1.378e-11 \| 369 \| 0.15 \| 955 \| 584 \| 32 \| 19 \| 785 \| 88 \| 1002 \| Tail tape measure protein TP901 core region protein \| Tail tape measure protein TP901 core region protein; afdb-uniprot50 \| AF-A0A2V3W552-F1-MODEL\_V4 \| 1.0 \| 1.808e-11 \| 369 \| 0.147 \| 781 \| 501 \| 25 \| 14 \| 685 \| 1 \| 725 \| TP901 family phage tail tape measure protein \| TP901 family phage tail tape measure protein; afdb-uniprot50 \| AF-R9LVJ9-F1-MODEL\_V4 \| 1.0 \| 3.637e-11 \| 369 \| 0.172 \| 706 \| 480 \| 22 \| 63 \| 702 \| 5 \| 671 \| Phage tail tape measure protein, TP901 family, core region \| Phage tail tape measure protein, TP901 family, core region; afdb-uniprot50 \| AF-A0A4Q2B6C1-F1-MODEL\_V4 \| 1.0 \| 2.466e-11 \| 369 \| 0.158 \| 755 \| 505 \| 27 \| 36 \| 702 \| 10 \| 721 \| Phage tail tape measure protein \| Phage tail tape measure protein; afdb-uniprot50 \| AF-A0A497MDG9-F1-MODEL\_V4 \| 1.0 \| 1.226e-11 \| 368 \| 0.152 \| 831 \| 511 \| 29 \| 28 \| 774 \| 12 \| 732 \| Phage tail tape measure protein \| Phage tail tape measure protein; afdb-uniprot50 \| AF-A0A243J227-F1-MODEL\_V4 \| 1.0 \| 9.343e-12 \| 368 \| 0.122 \| 971 \| 573 \| 34 \| 1 \| 765 \| 1 \| 897 \| Phage tail tape measure protein \| Phage tail tape measure protein; afdb-uniprot50 \| AF-A0A431JPH2-F1-MODEL\_V4 \| 1.0 \| 5.019e-12 \| 368 \| 0.198 \| 895 \| 521 \| 33 \| 9 \| 780 \| 1 \| 821 \| Phage tail tape measure protein \| Phage tail tape measure protein; afdb-uniprot50 \| AF-A0A120A0Q4-F1-MODEL\_V4 \| 1.0 \| 1.787e-10 \| 368 \| 0.106 \| 694 \| 421 \| 21 \| 109 \| 785 \| 216 \| 727 \| Phage-related tail protein \| Phage-related tail protein; afdb-uniprot50 \| AF-A0A7Z7J4C2-F1-MODEL\_V4 \| 1.0 \| 3.288e-09 \| 367 \| 0.324 \| 348 \| 166 \| 6 \| 19 \| 302 \| 39 \| 381 \| Putative enzyme \| Putative enzyme; afdb-uniprot50 \| AF-A0A1H9SQJ9-F1-MODEL\_V4 \| 1.0 \| 1.325e-11 \| 367 \| 0.15 \| 815 \| 501 \| 28 \| 104 \| 778 \| 15 \| 777 \| Phage tail tape measure protein, TP901 family, core region \| Phage tail tape measure protein, TP901 family, core region; afdb-uniprot50 \| AF-A0A2E2B3T2-F1-MODEL\_V4 \| 1.0 \| 2.031e-11 \| 367 \| 0.131 \| 834 \| 524 \| 29 \| 76 \| 785 \| 28 \| 784 \| Phage tail tape measure protein \| Phage tail tape measure protein; afdb-uniprot50 \| AF-A0A2T4Z920-F1-MODEL\_V4 \| 1.0 \| 6.025e-11 \| 367 \| 0.152 \| 729 \| 518 \| 21 \| 44 \| 702 \| 8 \| 706 \| TP901 family phage tail tape measure protein \| TP901 family phage tail tape measure protein; afdb-uniprot50 \| AF-A0A328KRU5-F1-MODEL\_V4 \| 1.0 \| 2.087e-10 \| 367 \| 0.167 \| 667 \| 461 \| 22 \| 61 \| 702 \| 13 \| 609 \| Phage tail tape measure protein \| Phage tail tape measure protein; afdb-uniprot50 \| AF-E6LKB1-F1-MODEL\_V4 \| 1.0 \| 2.666e-11 \| 367 \| 0.137 \| 692 \| 472 \| 23 \| 90 \| 748 \| 29 \| 628 \| Phage tail tape measure protein, TP901 family \| Phage tail tape measure protein, TP901 family; afdb-uniprot50 \| AF-A0A243K0L1-F1-MODEL\_V4 \| 1.0 \| 1.325e-11 \| 367 \| 0.108 \| 1037 \| 567 \| 28 \| 1 \| 785 \| 1 \| 931 \| Phage tail tape measure protein \| Phage tail tape measure protein; afdb-uniprot50 \| AF-A0A3A9I4F3-F1-MODEL\_V4 \| 1.0 \| 2.594e-12 \| 367 \| 0.165 \| 829 \| 533 \| 29 \| 18 \| 750 \| 224 \| 989 \| Phage tail tape measure protein \| Phage tail tape measure protein; afdb-uniprot50 \| AF-A0A2G8BY92-F1-MODEL\_V4 \| 1.0 \| 4.248e-11 \| 366 \| 0.153 \| 900 \| 539 \| 26 \| 22 \| 785 \| 54 \| 866 \| Phage tail tape measure protein \| Phage tail tape measure protein; afdb-uniprot50 \| AF-A0A0F9QC99-F1-MODEL\_V4 \| 1.0 \| 2.466e-11 \| 366 \| 0.143 \| 780 \| 445 \| 19 \| 50 \| 783 \| 3 \| 605 \| PhageMin\_Tail domain-containing protein \| PhageMin\_Tail domain-containing protein; afdb-uniprot50 \| AF-A0A1T1AVS0-F1-MODEL\_V4 \| 1.0 \| 4.086e-11 \| 366 \| 0.177 \| 745 \| 433 \| 30 \| 70 \| 772 \| 20 \| 626 \| Phage tail tape measure protein \| Phage tail tape measure protein; afdb-uniprot50 \| AF-A0A1I2DTC2-F1-MODEL\_V4 \| 1.0 \| 1.566e-12 \| 366 \| 0.141 \| 905 \| 562 \| 35 \| 19 \| 785 \| 25 \| 852 \| Phage tail tape measure protein, TP901 family, core region \| Phage tail tape measure protein, TP901 family, core region; afdb-uniprot50 \| AF-A0A4U1KS68-F1-MODEL\_V4 \| 1.0 \| 3.15e-12 \| 366 \| 0.15 \| 951 \| 557 \| 32 \| 1 \| 785 \| 1 \| 866 \| Phage tail tape measure protein \| Phage tail tape measure protein; afdb-uniprot50 \| AF-A0A510Y3F9-F1-MODEL\_V4 \| 1.0 \| 8.884e-11 \| 366 \| 0.154 \| 705 \| 478 \| 18 \| 59 \| 699 \| 5 \| 655 \| PhageMin\_Tail domain-containing protein \| PhageMin\_Tail domain-containing protein; afdb-uniprot50 \| AF-V1GPQ7-F1-MODEL\_V4 \| 1.0 \| 7.999e-12 \| 365 \| 0.153 \| 873 \| 521 \| 29 \| 19 \| 781 \| 54 \| 818 \| TP901 family phage tail tape measure protein \| TP901 family phage tail tape measure protein; afdb-uniprot50 \| AF-A0A0F9UQC3-F1-MODEL\_V4 \| 1.0 \| 3.637e-11 \| 365 \| 0.136 \| 799 \| 529 \| 26 \| 46 \| 780 \| 4 \| 705 \| PhageMin\_Tail domain-containing protein \| PhageMin\_Tail domain-containing protein; afdb-uniprot50 \| AF-A0A3D5P8F7-F1-MODEL\_V4 \| 1.0 \| 8.316e-12 \| 365 \| 0.15 \| 976 \| 580 \| 33 \| 1 \| 776 \| 1 \| 926 \| Phage tail tape measure protein \| Phage tail tape measure protein; afdb-uniprot50 \| AF-A0A829ZJ99-F1-MODEL\_V4 \| 1.0 \| 2.031e-11 \| 365 \| 0.144 \| 952 \| 530 \| 29 \| 19 \| 755 \| 62 \| 943 \| Chromosome partition protein Smc \| Chromosome partition protein Smc; afdb-uniprot50 \| AF-A0A0B2A639-F1-MODEL\_V4 \| 1.0 \| 6.336e-12 \| 365 \| 0.163 \| 800 \| 493 \| 28 \| 18 \| 705 \| 24 \| 758 \| PhageMin\_Tail domain-containing protein \| PhageMin\_Tail domain-containing protein; afdb-uniprot50 \| AF-A0A380ED76-F1-MODEL\_V4 \| 1.0 \| 3.993e-09 \| 365 \| 0.15 \| 591 \| 350 \| 19 \| 19 \| 492 \| 127 \| 682 \| Phage protein \| Phage protein; afdb-uniprot50 \| AF-A0A3A8WM39-F1-MODEL\_V4 \| 1.0 \| 5.425e-12 \| 364 \| 0.182 \| 716 \| 483 \| 20 \| 19 \| 703 \| 18 \| 661 \| Phage tail tape measure protein \| Phage tail tape measure protein; afdb-uniprot50 \| AF-A0A2N6UJH0-F1-MODEL\_V4 \| 1.0 \| 1.325e-11 \| 364 \| 0.16 \| 773 \| 480 \| 26 \| 10 \| 701 \| 8 \| 692 \| Phage tail tape measure protein \| Phage tail tape measure protein; afdb-uniprot50 \| AF-A0A1I4HAG3-F1-MODEL\_V4 \| 1.0 \| 2.466e-11 \| 364 \| 0.144 \| 849 \| 513 \| 32 \| 7 \| 701 \| 1 \| 789 \| Phage tail tape measure protein, TP901 family, core region \| Phage tail tape measure protein, TP901 family, core region; afdb-uniprot50 \| AF-A0A6I1NYL3-F1-MODEL\_V4 \| 1.0 \| 1.134e-11 \| 364 \| 0.173 \| 794 \| 510 \| 27 \| 19 \| 701 \| 35 \| 792 \| Phage tail tape measure protein \| Phage tail tape measure protein; afdb-uniprot50 \| AF-A0A0F9JQN3-F1-MODEL\_V4 \| 1.0 \| 4.538e-10 \| 363 \| 0.155 \| 706 \| 427 \| 22 \| 40 \| 702 \| 2 \| 580 \| PhageMin\_Tail domain-containing protein \| PhageMin\_Tail domain-containing protein; afdb-uniprot50 \| AF-A0A2W6TNQ0-F1-MODEL\_V4 \| 1.0 \| 2.635e-10 \| 363 \| 0.143 \| 803 \| 500 \| 24 \| 14 \| 785 \| 1 \| 646 \| PhageMin\_Tail domain-containing protein \| PhageMin\_Tail domain-containing protein; afdb-uniprot50 \| AF-A0A1C5VU61-F1-MODEL\_V4 \| 1.0 \| 1.858e-10 \| 363 \| 0.13 \| 772 \| 496 \| 27 \| 117 \| 784 \| 19 \| 718 \| Phage-related protein \| Phage-related protein; afdb-uniprot50 \| AF-A0A859DR59-F1-MODEL\_V4 \| 1.0 \| 2.282e-11 \| 363 \| 0.149 \| 878 \| 516 \| 28 \| 19 \| 702 \| 81 \| 921 \| Phage tail tape measure protein \| Phage tail tape measure protein; afdb-uniprot50 \| AF-A0A2B0C957-F1-MODEL\_V4 \| 1.0 \| 1.01e-11 \| 363 \| 0.172 \| 796 \| 473 \| 29 \| 6 \| 702 \| 15 \| 723 \| Phage tail tape measure protein \| Phage tail tape measure protein; afdb-uniprot50 \| AF-A0A1I0H884-F1-MODEL\_V4 \| 1.0 \| 2.282e-11 \| 363 \| 0.148 \| 857 \| 486 \| 31 \| 13 \| 702 \| 2 \| 781 \| Phage tail tape measure protein, TP901 family, core region \| Phage tail tape measure protein, TP901 family, core region; afdb-uniprot50 \| AF-A0A822P8U8-F1-MODEL\_V4 \| 1.0 \| 2.564e-11 \| 363 \| 0.142 \| 1057 \| 577 \| 41 \| 19 \| 785 \| 74 \| 1090 \| Phage tail tape measure protein \| Phage tail tape measure protein; afdb-uniprot50 \| AF-B1V2P5-F1-MODEL\_V4 \| 1.0 \| 1.548e-11 \| 363 \| 0.134 \| 1001 \| 602 \| 39 \| 21 \| 785 \| 216 \| 1187 \| Putative phage tail tape measure protein, family, core region domain protein \| Putative phage tail tape measure protein, family, core region domain protein; afdb-uniprot50 \| AF-A0A543NFS2-F1-MODEL\_V4 \| 1.0 \| 3.03e-12 \| 363 \| 0.146 \| 948 \| 533 \| 33 \| 50 \| 785 \| 2 \| 884 \| TP901 family phage tail tape measure protein \| TP901 family phage tail tape measure protein; afdb-uniprot50 \| AF-A0A7Z1FTZ1-F1-MODEL\_V4 \| 1.0 \| 1.739e-11 \| 362 \| 0.118 \| 989 \| 582 \| 30 \| 61 \| 783 \| 8 \| 972 \| Phage tail tape measure protein \| Phage tail tape measure protein; afdb-uniprot50 \| AF-A0A6N9P0N2-F1-MODEL\_V4 \| 1.0 \| 1.954e-11 \| 362 \| 0.158 \| 774 \| 464 \| 31 \| 19 \| 700 \| 44 \| 721 \| Phage tail tape measure protein \| Phage tail tape measure protein; afdb-uniprot50 \| AF-D8F206-F1-MODEL\_V4 \| 1.0 \| 6.263e-11 \| 361 \| 0.157 \| 744 \| 457 \| 29 \| 35 \| 718 \| 19 \| 652 \| Phage tail tape measure protein, TP901 family \| Phage tail tape measure protein, TP901 family; afdb-uniprot50 \| AF-A0A1Y0EH74-F1-MODEL\_V4 \| 1.0 \| 1.01e-11 \| 361 \| 0.133 \| 810 \| 520 \| 29 \| 6 \| 735 \| 1 \| 708 \| Phage-related minor tail protein \| Phage-related minor tail protein; afdb-uniprot50 \| AF-A0A7V1ZUT4-F1-MODEL\_V4 \| 1.0 \| 3.781e-11 \| 360 \| 0.152 \| 694 \| 486 \| 20 \| 61 \| 702 \| 4 \| 646 \| Phage tail tape measure protein \| Phage tail tape measure protein; afdb-uniprot50 \| AF-A0A7V6XHU3-F1-MODEL\_V4 \| 1.0 \| 1.489e-11 \| 360 \| 0.157 \| 843 \| 510 \| 37 \| 1 \| 702 \| 1 \| 783 \| Phage tail tape measure protein \| Phage tail tape measure protein; afdb-uniprot50 \| AF-A0A0R2DDY7-F1-MODEL\_V4 \| 1.0 \| 8.22e-11 \| 360 \| 0.14 \| 902 \| 530 \| 26 \| 1 \| 702 \| 1 \| 856 \| PhageMin\_Tail domain-containing protein \| PhageMin\_Tail domain-containing protein; afdb-uniprot50 \| AF-F6AX89-F1-MODEL\_V4 \| 1.0 \| 1.609e-11 \| 359 \| 0.141 \| 790 \| 500 \| 28 \| 19 \| 702 \| 33 \| 749 \| Phage tail tape measure protein, TP901 family \| Phage tail tape measure protein, TP901 family; afdb-uniprot50 \| AF-A0A3D1WP41-F1-MODEL\_V4 \| 1.0 \| 4.086e-11 \| 359 \| 0.142 \| 767 \| 515 \| 28 \| 19 \| 702 \| 91 \| 797 \| Phage tail tape measure protein \| Phage tail tape measure protein; afdb-uniprot50 \| AF-A0A497QGF2-F1-MODEL\_V4 \| 1.0 \| 8.884e-11 \| 358 \| 0.151 \| 728 \| 497 \| 22 \| 19 \| 702 \| 15 \| 665 \| Phage tail tape measure protein \| Phage tail tape measure protein; afdb-uniprot50 \| AF-A0A496UZA7-F1-MODEL\_V4 \| 1.0 \| 9.981e-11 \| 358 \| 0.155 \| 669 \| 480 \| 19 \| 81 \| 708 \| 17 \| 641 \| Phage tail tape measure protein \| Phage tail tape measure protein; afdb-uniprot50 \| AF-A0A354UB38-F1-MODEL\_V4 \| 1.0 \| 1.295e-09 \| 358 \| 0.145 \| 619 \| 435 \| 14 \| 115 \| 701 \| 109 \| 665 \| Phage tail tape measure protein \| Phage tail tape measure protein; afdb-uniprot50 \| AF-A0A1G6LVA0-F1-MODEL\_V4 \| 1.0 \| 1.609e-11 \| 358 \| 0.17 \| 810 \| 546 \| 31 \| 49 \| 785 \| 6 \| 762 \| Phage tail tape measure protein, TP901 family, core region \| Phage tail tape measure protein, TP901 family, core region; afdb-uniprot50 \| AF-A0A151BBD5-F1-MODEL\_V4 \| 1.0 \| 3.2e-10 \| 357 \| 0.157 \| 747 \| 451 \| 29 \| 19 \| 718 \| 10 \| 624 \| Phage-like protein \| Phage-like protein; afdb-uniprot50 \| AF-A0A1I6REX1-F1-MODEL\_V4 \| 1.0 \| 2.535e-10 \| 357 \| 0.143 \| 803 \| 441 \| 27 \| 6 \| 783 \| 3 \| 583 \| Phage tail tape measure protein, TP901 family, core region \| Phage tail tape measure protein, TP901 family, core region; afdb-uniprot50 \| AF-A0A6G7JLG1-F1-MODEL\_V4 \| 1.0 \| 4.086e-11 \| 357 \| 0.135 \| 859 \| 550 \| 27 \| 1 \| 783 \| 1 \| 742 \| Phage tail tape measure protein \| Phage tail tape measure protein; afdb-uniprot50 \| AF-A0A1H5Z5X2-F1-MODEL\_V4 \| 1.0 \| 5.955e-10 \| 357 \| 0.145 \| 666 \| 446 \| 24 \| 49 \| 702 \| 7 \| 561 \| Phage tail length tape-measure protein \| Phage tail length tape-measure protein; afdb-uniprot50 \| AF-A0A0F9TFD4-F1-MODEL\_V4 \| 1.0 \| 2.564e-11 \| 356 \| 0.14 \| 781 \| 501 \| 24 \| 1 \| 702 \| 1 \| 689 \| PhageMin\_Tail domain-containing protein \| PhageMin\_Tail domain-containing protein; afdb-uniprot50 \| AF-A0A098AYW7-F1-MODEL\_V4 \| 1.0 \| 4.773e-11 \| 355 \| 0.138 \| 785 \| 520 \| 24 \| 18 \| 785 \| 18 \| 662 \| Phage-related minor tail protein \| Phage-related minor tail protein; afdb-uniprot50 \| AF-A0A489F6U0-F1-MODEL\_V4 \| 1.0 \| 1.654e-10 \| 355 \| 0.157 \| 703 \| 444 \| 23 \| 67 \| 662 \| 3 \| 663 \| Phage protein, tail length tape measure protein \| Phage protein, tail length tape measure protein; afdb-uniprot50 \| AF-A0A7U6KQJ0-F1-MODEL\_V4 \| 1.0 \| 2.575e-08 \| 354 \| 0.289 \| 321 \| 207 \| 5 \| 1 \| 311 \| 1 \| 310 \| PhageMin\_Tail domain-containing protein \| PhageMin\_Tail domain-containing protein; afdb-uniprot50 \| AF-A0A354KNI6-F1-MODEL\_V4 \| 1.0 \| 1.673e-11 \| 354 \| 0.154 \| 746 \| 469 \| 26 \| 35 \| 701 \| 6 \| 668 \| Phage tail tape measure protein \| Phage tail tape measure protein; afdb-uniprot50 \| AF-A0A847STA2-F1-MODEL\_V4 \| 1.0 \| 2.771e-11 \| 354 \| 0.146 \| 791 \| 494 \| 28 \| 1 \| 703 \| 1 \| 697 \| Phage tail tape measure protein \| Phage tail tape measure protein; afdb-uniprot50 \| AF-A0A7T0D4M4-F1-MODEL\_V4 \| 1.0 \| 1.739e-11 \| 354 \| 0.161 \| 797 \| 534 \| 23 \| 19 \| 701 \| 76 \| 851 \| Phage tail tape measure protein \| Phage tail tape measure protein; afdb-uniprot50 \| AF-E0MRI5-F1-MODEL\_V4 \| 1.0 \| 7.037e-11 \| 353 \| 0.161 \| 807 \| 438 \| 29 \| 36 \| 785 \| 11 \| 635 \| Phage tail tape measure protein, family, core region \| Phage tail tape measure protein, family, core region; afdb-uniprot50 \| AF-A0A1X7HG53-F1-MODEL\_V4 \| 1.0 \| 2.17e-10 \| 353 \| 0.154 \| 731 \| 444 \| 29 \| 70 \| 702 \| 12 \| 666 \| Phage tail tape measure protein, TP901 family, core region \| Phage tail tape measure protein, TP901 family, core region; afdb-uniprot50 \| AF-A0A6M3L4K5-F1-MODEL\_V4 \| 1.0 \| 9.981e-11 \| 353 \| 0.163 \| 790 \| 443 \| 29 \| 61 \| 784 \| 1 \| 638 \| Putative tail protein \| Putative tail protein; afdb-uniprot50 \| AF-H0U9Y6-F1-MODEL\_V4 \| 1.0 \| 1.109e-09 \| 352 \| 0.144 \| 814 \| 477 \| 26 \| 16 \| 772 \| 1 \| 651 \| Phage tail tape measure protein, TP901 family, core region \| Phage tail tape measure protein, TP901 family, core region; afdb-uniprot50 \| AF-A0A1M6L0R6-F1-MODEL\_V4 \| 1.0 \| 4.468e-12 \| 352 \| 0.162 \| 737 \| 505 \| 26 \| 80 \| 772 \| 25 \| 692 \| Phage tail tape measure protein, TP901 family, core region \| Phage tail tape measure protein, TP901 family, core region; afdb-uniprot50 \| AF-A0A660UYP3-F1-MODEL\_V4 \| 1.0 \| 7.037e-11 \| 352 \| 0.139 \| 755 \| 470 \| 28 \| 109 \| 716 \| 63 \| 784 \| Phage tail tape measure protein \| Phage tail tape measure protein; afdb-uniprot50 \| AF-R7JDX5-F1-MODEL\_V4 \| 1.0 \| 5.64e-12 \| 352 \| 0.155 \| 851 \| 545 \| 27 \| 29 \| 776 \| 3 \| 782 \| Phage tail tape measure protein TP901 family \| Phage tail tape measure protein TP901 family; afdb-uniprot50 \| AF-A0A518GEF8-F1-MODEL\_V4 \| 1.0 \| 9.235e-11 \| 352 \| 0.155 \| 879 \| 530 \| 29 \| 70 \| 785 \| 8 \| 836 \| Phage-related minor tail protein \| Phage-related minor tail protein; afdb-uniprot50 \| AF-A0A2I0FHY8-F1-MODEL\_V4 \| 1.0 \| 4.248e-11 \| 352 \| 0.151 \| 919 \| 545 \| 36 \| 61 \| 785 \| 4 \| 881 \| Phage tail tape measure protein \| Phage tail tape measure protein; afdb-uniprot50 \| AF-U6SUL4-F1-MODEL\_V4 \| 1.0 \| 3.637e-11 \| 352 \| 0.138 \| 836 \| 526 \| 25 \| 3 \| 702 \| 1 \| 777 \| PhageMin\_Tail domain-containing protein \| PhageMin\_Tail domain-containing protein; afdb-uniprot50 \| AF-A0A349YIH1-F1-MODEL\_V4 \| 1.0 \| 1.416e-10 \| 352 \| 0.122 \| 1103 \| 538 \| 31 \| 19 \| 785 \| 79 \| 1087 \| Phage tail tape measure protein \| Phage tail tape measure protein; afdb-uniprot50 \| AF-A0A2E7ZRD4-F1-MODEL\_V4 \| 1.0 \| 1.152e-09 \| 351 \| 0.14 \| 706 \| 469 \| 26 \| 80 \| 713 \| 16 \| 655 \| Phage tail tape measure protein \| Phage tail tape measure protein; afdb-uniprot50 \| AF-D0GNX1-F1-MODEL\_V4 \| 1.0 \| 4.416e-11 \| 351 \| 0.137 \| 885 \| 546 \| 34 \| 2 \| 785 \| 1 \| 768 \| Phage tail tape measure protein, TP901 family \| Phage tail tape measure protein, TP901 family; afdb-uniprot50 \| AF-A0A150KS49-F1-MODEL\_V4 \| 1.0 \| 7.316e-11 \| 351 \| 0.151 \| 767 \| 501 \| 23 \| 18 \| 759 \| 18 \| 659 \| PhageMin\_Tail domain-containing protein \| PhageMin\_Tail domain-containing protein; afdb-uniprot50 \| AF-A0A3T1DR99-F1-MODEL\_V4 \| 1.0 \| 9.024e-09 \| 349 \| 0.161 \| 673 \| 423 \| 27 \| 39 \| 690 \| 2 \| 553 \| PhageMin\_Tail domain-containing protein \| PhageMin\_Tail domain-containing protein; afdb-uniprot50 \| AF-A0A512CS43-F1-MODEL\_V4 \| 1.0 \| 1.858e-10 \| 349 \| 0.157 \| 750 \| 499 \| 24 \| 1 \| 708 \| 1 \| 659 \| PhageMin\_Tail domain-containing protein \| PhageMin\_Tail domain-containing protein; afdb-uniprot50 \| AF-F2JRA3-F1-MODEL\_V4 \| 1.0 \| 1.673e-11 \| 349 \| 0.145 \| 781 \| 545 \| 27 \| 1 \| 719 \| 1 \| 720 \| Phage tail tape measure protein, TP901 family \| Phage tail tape measure protein, TP901 family; afdb-uniprot50 \| AF-A0A1B7LM07-F1-MODEL\_V4 \| 1.0 \| 1.166e-10 \| 349 \| 0.131 \| 910 \| 499 \| 34 \| 19 \| 783 \| 74 \| 836 \| PhageMin\_Tail domain-containing protein \| PhageMin\_Tail domain-containing protein; afdb-uniprot50 \| AF-A0A2E1AEH8-F1-MODEL\_V4 \| 1.0 \| 3.781e-11 \| 349 \| 0.133 \| 896 \| 544 \| 34 \| 1 \| 748 \| 9 \| 819 \| Phage tail tape measure protein \| Phage tail tape measure protein; afdb-uniprot50 \| AF-A0A847H9X0-F1-MODEL\_V4 \| 1.0 \| 3.237e-11 \| 349 \| 0.137 \| 893 \| 551 \| 37 \| 48 \| 785 \| 4 \| 831 \| Phage tail tape measure protein \| Phage tail tape measure protein; afdb-uniprot50 \| AF-G9RPT7-F1-MODEL\_V4 \| 1.0 \| 5.158e-11 \| 349 \| 0.155 \| 886 \| 525 \| 35 \| 4 \| 702 \| 6 \| 854 \| Phage tail tape measure protein, TP901 family, core region \| Phage tail tape measure protein, TP901 family, core region; afdb-uniprot50 \| AF-Q2FYC8-F1-MODEL\_V4 \| 1.0 \| 1.699e-09 \| 349 \| 0.142 \| 631 \| 368 \| 19 \| 19 \| 519 \| 127 \| 714 \| Lysostaphin \| Lysostaphin; afdb-uniprot50 \| AF-A0A7H0RK43-F1-MODEL\_V4 \| 1.0 \| 1.472e-10 \| 348 \| 0.17 \| 686 \| 423 \| 22 \| 139 \| 706 \| 36 \| 693 \| Phage tail tape measure protein \| Phage tail tape measure protein; afdb-uniprot50 \| AF-A0A2W4GQJ4-F1-MODEL\_V4 \| 1.0 \| 1.954e-11 \| 348 \| 0.157 \| 823 \| 533 \| 26 \| 19 \| 785 \| 34 \| 751 \| Phage tail tape measure protein \| Phage tail tape measure protein; afdb-uniprot50 \| AF-A0A841RZ32-F1-MODEL\_V4 \| 1.0 \| 2.708e-09 \| 348 \| 0.161 \| 594 \| 385 \| 27 \| 117 \| 702 \| 43 \| 531 \| TP901 family phage tail tape measure protein \| TP901 family phage tail tape measure protein; afdb-uniprot50 \| AF-A0A2W4GR67-F1-MODEL\_V4 \| 1.0 \| 1.739e-11 \| 348 \| 0.14 \| 942 \| 550 \| 24 \| 50 \| 785 \| 17 \| 904 \| Phage tail tape measure protein \| Phage tail tape measure protein; afdb-uniprot50 \| AF-A0A0A8B9B9-F1-MODEL\_V4 \| 1.0 \| 4.416e-11 \| 348 \| 0.135 \| 1033 \| 582 \| 33 \| 14 \| 785 \| 13 \| 994 \| PhageMin\_Tail domain-containing protein \| PhageMin\_Tail domain-containing protein; afdb-uniprot50 \| AF-A0A0J6WRI9-F1-MODEL\_V4 \| 1.0 \| 1.53e-10 \| 347 \| 0.133 \| 841 \| 535 \| 34 \| 2 \| 782 \| 1 \| 707 \| PhageMin\_Tail domain-containing protein \| PhageMin\_Tail domain-containing protein; afdb-uniprot50 \| AF-E6SKI6-F1-MODEL\_V4 \| 1.0 \| 2.848e-10 \| 347 \| 0.176 \| 718 \| 473 \| 23 \| 36 \| 708 \| 2 \| 645 \| Phage tail tape measure protein, TP901 family \| Phage tail tape measure protein, TP901 family; afdb-uniprot50 \| AF-A0A843HVN5-F1-MODEL\_V4 \| 1.0 \| 1.038e-10 \| 347 \| 0.164 \| 841 \| 492 \| 27 \| 19 \| 702 \| 119 \| 905 \| Phage tail tape measure protein \| Phage tail tape measure protein; afdb-uniprot50 \| AF-A0A3A9G8N3-F1-MODEL\_V4 \| 1.0 \| 3.163e-09 \| 346 \| 0.142 \| 686 \| 475 \| 19 \| 28 \| 685 \| 121 \| 720 \| Phage tail tape measure protein \| Phage tail tape measure protein; afdb-uniprot50 \| AF-A0A2M9KTE8-F1-MODEL\_V4 \| 1.0 \| 1.489e-11 \| 346 \| 0.163 \| 950 \| 543 \| 36 \| 1 \| 785 \| 1 \| 863 \| Phage tail tape measure protein \| Phage tail tape measure protein; afdb-uniprot50 \| AF-A0A2E1Y1Q6-F1-MODEL\_V4 \| 1.0 \| 7.606e-11 \| 346 \| 0.159 \| 783 \| 498 \| 26 \| 35 \| 706 \| 5 \| 737 \| Phage tail tape measure protein \| Phage tail tape measure protein; afdb-uniprot50 \| AF-A0A3R6GFF2-F1-MODEL\_V4 \| 1.0 \| 2.112e-11 \| 345 \| 0.158 \| 871 \| 519 \| 34 \| 1 \| 785 \| 1 \| 743 \| Phage tail tape measure protein \| Phage tail tape measure protein; afdb-uniprot50 \| AF-A0A2I5TL78-F1-MODEL\_V4 \| 1.0 \| 1.31e-10 \| 345 \| 0.154 \| 805 \| 451 \| 28 \| 1 \| 779 \| 167 \| 767 \| Phage tail tape measure protein \| Phage tail tape measure protein; afdb-uniprot50 \| AF-A0A069AZP6-F1-MODEL\_V4 \| 1.0 \| 5.099e-10 \| 345 \| 0.15 \| 719 \| 483 \| 26 \| 18 \| 668 \| 213 \| 871 \| Putative phage tail tape measure protein, family, core region domain protein \| Putative phage tail tape measure protein, family, core region domain protein; afdb-uniprot50 \| AF-A0A840PRY4-F1-MODEL\_V4 \| 1.0 \| 7.316e-11 \| 345 \| 0.14 \| 806 \| 471 \| 26 \| 18 \| 759 \| 13 \| 660 \| TP901 family phage tail tape measure protein \| TP901 family phage tail tape measure protein; afdb-uniprot50 \| AF-A0A848AWW3-F1-MODEL\_V4 \| 1.0 \| 1.416e-10 \| 345 \| 0.168 \| 734 \| 446 \| 24 \| 83 \| 681 \| 24 \| 727 \| Phage tail tape measure protein \| Phage tail tape measure protein; afdb-uniprot50 \| AF-A0A4Y3TD25-F1-MODEL\_V4 \| 1.0 \| 2.535e-10 \| 345 \| 0.136 \| 761 \| 511 \| 26 \| 117 \| 785 \| 65 \| 770 \| PhageMin\_Tail domain-containing protein \| PhageMin\_Tail domain-containing protein; afdb-uniprot50 \| AF-A0A419HC04-F1-MODEL\_V4 \| 1.0 \| 1.091e-11 \| 345 \| 0.151 \| 966 \| 532 \| 32 \| 36 \| 785 \| 10 \| 903 \| Phage tail tape measure protein \| Phage tail tape measure protein; afdb-uniprot50 \| AF-A0A6N6XWZ4-F1-MODEL\_V4 \| 1.0 \| 6.769e-11 \| 345 \| 0.162 \| 961 \| 533 \| 37 \| 19 \| 781 \| 44 \| 930 \| Phage tail tape measure protein \| Phage tail tape measure protein; afdb-uniprot50 \| AF-A0A368U1R4-F1-MODEL\_V4 \| 1.0 \| 2.848e-10 \| 345 \| 0.158 \| 718 \| 504 \| 22 \| 39 \| 705 \| 2 \| 669 \| Phage tail tape measure protein \| Phage tail tape measure protein; afdb-uniprot50 \| AF-A0A3M2MF09-F1-MODEL\_V4 \| 1.0 \| 5.795e-11 \| 345 \| 0.154 \| 887 \| 554 \| 33 \| 15 \| 785 \| 28 \| 834 \| Phage tail tape measure protein \| Phage tail tape measure protein; afdb-uniprot50 \| AF-A0A1W7LS78-F1-MODEL\_V4 \| 1.0 \| 7.316e-11 \| 345 \| 0.141 \| 892 \| 522 \| 28 \| 1 \| 710 \| 4 \| 833 \| PhageMin\_Tail domain-containing protein \| PhageMin\_Tail domain-containing protein; afdb-uniprot50 \| AF-A0A7D4GDK8-F1-MODEL\_V4 \| 1.0 \| 1.31e-10 \| 344 \| 0.161 \| 781 \| 508 \| 28 \| 1 \| 702 \| 1 \| 713 \| Phage tail tape measure protein \| Phage tail tape measure protein; afdb-uniprot50 \| AF-A0A496XFC7-F1-MODEL\_V4 \| 1.0 \| 3.595e-10 \| 343 \| 0.183 \| 724 \| 482 \| 27 \| 17 \| 686 \| 1 \| 668 \| Phage tail tape measure protein \| Phage tail tape measure protein; afdb-uniprot50 \| AF-A0A1G0RDW7-F1-MODEL\_V4 \| 1.0 \| 4.591e-11 \| 343 \| 0.143 \| 834 \| 533 \| 30 \| 61 \| 785 \| 11 \| 771 \| Phage tail tape measure protein \| Phage tail tape measure protein; afdb-uniprot50 \| AF-A0A7Y0ENB1-F1-MODEL\_V4 \| 1.0 \| 1.166e-10 \| 343 \| 0.151 \| 781 \| 462 \| 23 \| 19 \| 662 \| 79 \| 795 \| Phage tail tape measure protein \| Phage tail tape measure protein; afdb-uniprot50 \| AF-A0A143ZV16-F1-MODEL\_V4 \| 1.0 \| 7.431e-09 \| 343 \| 0.17 \| 667 \| 453 \| 19 \| 97 \| 701 \| 29 \| 656 \| Phage-related minor tail protein \| Phage-related minor tail protein; afdb-uniprot50 \| AF-D8J9U6-F1-MODEL\_V4 \| 1.0 \| 9.713e-12 \| 343 \| 0.155 \| 861 \| 476 \| 28 \| 1 \| 708 \| 8 \| 769 \| Prophage pi3 protein 14 \| Prophage pi3 protein 14; afdb-uniprot50 \| AF-A0A174ERH0-F1-MODEL\_V4 \| 1.0 \| 3.237e-11 \| 343 \| 0.157 \| 864 \| 520 \| 30 \| 21 \| 701 \| 64 \| 902 \| Phage tail tape measure protein, TP901 family \| Phage tail tape measure protein, TP901 family; afdb-uniprot50 \| AF-A0A369RL83-F1-MODEL\_V4 \| 1.0 \| 7.726e-09 \| 342 \| 0.251 \| 394 \| 225 \| 10 \| 6 \| 332 \| 1 \| 391 \| Phage tail tape measure protein, TP901 family, core region \| Phage tail tape measure protein, TP901 family, core region; afdb-uniprot50 \| AF-A0A352HSY4-F1-MODEL\_V4 \| 1.0 \| 2.345e-10 \| 342 \| 0.162 \| 714 \| 448 \| 29 \| 61 \| 727 \| 25 \| 635 \| Phage tail tape measure protein \| Phage tail tape measure protein; afdb-uniprot50 \| AF-A0A1G5EF29-F1-MODEL\_V4 \| 1.0 \| 6.691e-10 \| 342 \| 0.157 \| 724 \| 485 \| 22 \| 62 \| 702 \| 4 \| 685 \| Phage tail tape measure protein, TP901 family, core region \| Phage tail tape measure protein, TP901 family, core region; afdb-uniprot50 \| AF-A0A2K3J8S3-F1-MODEL\_V4 \| 1.0 \| 1.879e-11 \| 342 \| 0.151 \| 902 \| 559 \| 32 \| 11 \| 785 \| 10 \| 831 \| Phage tail tape measure protein \| Phage tail tape measure protein; afdb-uniprot50 \| AF-A0A285Q2V0-F1-MODEL\_V4 \| 1.0 \| 5.362e-11 \| 342 \| 0.159 \| 815 \| 466 \| 27 \| 13 \| 702 \| 1 \| 721 \| Phage tail tape measure protein, TP901 family, core region \| Phage tail tape measure protein, TP901 family, core region; afdb-uniprot50 \| AF-J4WBN7-F1-MODEL\_V4 \| 1.0 \| 7.907e-11 \| 342 \| 0.159 \| 869 \| 559 \| 24 \| 19 \| 782 \| 125 \| 926 \| Tail tape measure protein, TIGR01760 family \| Tail tape measure protein, TIGR01760 family; afdb-uniprot50 \| AF-A0A859FEZ2-F1-MODEL\_V4 \| 1.0 \| 9.49e-10 \| 342 \| 0.168 \| 688 \| 466 \| 24 \| 80 \| 702 \| 19 \| 665 \| Phage tail tape measure protein \| Phage tail tape measure protein; afdb-uniprot50 \| AF-A0A1E5JY92-F1-MODEL\_V4 \| 1.0 \| 4.718e-10 \| 342 \| 0.151 \| 800 \| 520 \| 29 \| 1 \| 700 \| 1 \| 741 \| Phage tail tape measure protein \| Phage tail tape measure protein; afdb-uniprot50 \| AF-A0A174GWB8-F1-MODEL\_V4 \| 1.0 \| 1.362e-10 \| 342 \| 0.118 \| 803 \| 486 \| 28 \| 27 \| 649 \| 72 \| 832 \| TP901 family phage tail tape measure protein \| TP901 family phage tail tape measure protein; afdb-uniprot50 \| AF-A0A1G2Z547-F1-MODEL\_V4 \| 1.0 \| 1.591e-10 \| 341 \| 0.153 \| 802 \| 521 \| 33 \| 91 \| 785 \| 38 \| 788 \| Phage tail tape measure protein \| Phage tail tape measure protein; afdb-uniprot50 \| AF-A0A524AE58-F1-MODEL\_V4 \| 1.0 \| 6.691e-10 \| 341 \| 0.142 \| 721 \| 497 \| 27 \| 60 \| 702 \| 15 \| 691 \| Phage tail tape measure protein \| Phage tail tape measure protein; afdb-uniprot50 \| AF-A0A194AHA9-F1-MODEL\_V4 \| 1.0 \| 1.591e-10 \| 341 \| 0.142 \| 742 \| 494 \| 26 \| 1 \| 702 \| 1 \| 639 \| PhageMin\_Tail domain-containing protein \| PhageMin\_Tail domain-containing protein; afdb-uniprot50 \| AF-A0A6M3IS27-F1-MODEL\_V4 \| 1.0 \| 7.518e-10 \| 339 \| 0.169 \| 761 \| 462 \| 24 \| 61 \| 722 \| 1 \| 690 \| Putative tail protein \| Putative tail protein; afdb-uniprot50 \| AF-A0A1V6CLC6-F1-MODEL\_V4 \| 1.0 \| 2.087e-10 \| 339 \| 0.135 \| 892 \| 550 \| 34 \| 1 \| 785 \| 1 \| 777 \| Phage-related minor tail protein \| Phage-related minor tail protein; afdb-uniprot50 \| AF-D7K0E2-F1-MODEL\_V4 \| 1.0 \| 5.24e-09 \| 339 \| 0.102 \| 694 \| 422 \| 20 \| 117 \| 776 \| 4 \| 530 \| Putative bacteriophage tail fiber protein T (Tape measure) \| Putative bacteriophage tail fiber protein T (Tape measure); afdb-uniprot50 \| AF-A0A6N9NRD9-F1-MODEL\_V4 \| 1.0 \| 2.063e-09 \| 339 \| 0.154 \| 694 \| 447 \| 19 \| 120 \| 779 \| 4 \| 591 \| Phage tail tape measure protein \| Phage tail tape measure protein; afdb-uniprot50 \| AF-A0A4Q5NKK3-F1-MODEL\_V4 \| 1.0 \| 1.315e-07 \| 338 \| 0.208 \| 365 \| 262 \| 9 \| 9 \| 355 \| 69 \| 424 \| Phage tail tape measure protein \| Phage tail tape measure protein; afdb-uniprot50 \| AF-A0A517QH69-F1-MODEL\_V4 \| 1.0 \| 3.458e-10 \| 338 \| 0.141 \| 807 \| 499 \| 23 \| 1 \| 782 \| 3 \| 640 \| Phage-related minor tail protein \| Phage-related minor tail protein; afdb-uniprot50 \| AF-A0A285PTB9-F1-MODEL\_V4 \| 1.0 \| 1.079e-10 \| 338 \| 0.149 \| 785 \| 496 \| 26 \| 105 \| 772 \| 41 \| 770 \| Phage tail tape measure protein \| Phage tail tape measure protein; afdb-uniprot50 \| AF-G5FA09-F1-MODEL\_V4 \| 1.0 \| 4.773e-11 \| 338 \| 0.143 \| 1021 \| 593 \| 37 \| 1 \| 784 \| 1 \| 976 \| PhageMin\_Tail domain-containing protein \| PhageMin\_Tail domain-containing protein; afdb-uniprot50 \| AF-A0A2C3G875-F1-MODEL\_V4 \| 1.0 \| 3.737e-10 \| 338 \| 0.138 \| 860 \| 526 \| 24 \| 1 \| 702 \| 1 \| 803 \| Phage tail tape measure protein \| Phage tail tape measure protein; afdb-uniprot50 \| AF-A0A0Q6RQW9-F1-MODEL\_V4 \| 1.0 \| 2.318e-09 \| 337 \| 0.14 \| 830 \| 476 \| 26 \| 32 \| 785 \| 14 \| 681 \| PhageMin\_Tail domain-containing protein \| PhageMin\_Tail domain-containing protein; afdb-uniprot50 \| AF-A0A7W3T073-F1-MODEL\_V4 \| 1.0 \| 7.518e-10 \| 337 \| 0.184 \| 725 \| 466 \| 31 \| 53 \| 702 \| 2 \| 675 \| Phage tail tape measure protein \| Phage tail tape measure protein; afdb-uniprot50 \| AF-A0A2E7QIC9-F1-MODEL\_V4 \| 1.0 \| 7.606e-11 \| 337 \| 0.161 \| 767 \| 493 \| 27 \| 28 \| 702 \| 4 \| 711 \| Phage tail tape measure protein \| Phage tail tape measure protein; afdb-uniprot50 \| AF-A0A735D9H4-F1-MODEL\_V4 \| 1.0 \| 1.489e-11 \| 336 \| 0.262 \| 525 \| 285 \| 17 \| 250 \| 755 \| 3 \| 444 \| Phage tail tape measure protein \| Phage tail tape measure protein; afdb-uniprot50 \| AF-A0A1G4U6I3-F1-MODEL\_V4 \| 1.0 \| 2.926e-09 \| 336 \| 0.166 \| 793 \| 455 \| 34 \| 5 \| 746 \| 2 \| 639 \| Phage tail tape measure protein, TP901 family, core region \| Phage tail tape measure protein, TP901 family, core region; afdb-uniprot50 \| AF-A0A060PHZ7-F1-MODEL\_V4 \| 1.0 \| 1.654e-10 \| 336 \| 0.151 \| 791 \| 476 \| 27 \| 19 \| 700 \| 21 \| 724 \| PhageMin\_Tail domain-containing protein \| PhageMin\_Tail domain-containing protein; afdb-uniprot50 \| AF-A0A847KXS4-F1-MODEL\_V4 \| 1.0 \| 9.49e-10 \| 335 \| 0.136 \| 820 \| 508 \| 26 \| 3 \| 750 \| 1 \| 692 \| Phage tail tape measure protein \| Phage tail tape measure protein; afdb-uniprot50 \| AF-A0A521F9Y7-F1-MODEL\_V4 \| 1.0 \| 1.931e-10 \| 335 \| 0.134 \| 926 \| 520 \| 33 \| 77 \| 784 \| 20 \| 881 \| Phage tail tape measure protein, TP901 family, core region \| Phage tail tape measure protein, TP901 family, core region; afdb-uniprot50 \| AF-A0A7X5KST3-F1-MODEL\_V4 \| 1.0 \| 5.469e-06 \| 335 \| 0.144 \| 375 \| 276 \| 9 \| 87 \| 437 \| 21 \| 374 \| Phage tail tape measure protein \| Phage tail tape measure protein; afdb-uniprot50 \| AF-A0A2E2MD95-F1-MODEL\_V4 \| 1.0 \| 1.265e-07 \| 334 \| 0.178 \| 455 \| 288 \| 17 \| 1 \| 439 \| 1 \| 385 \| Phage tail tape measure protein \| Phage tail tape measure protein; afdb-uniprot50 \| AF-A0A6B4SLJ1-F1-MODEL\_V4 \| 1.0 \| 6.191e-10 \| 334 \| 0.157 \| 735 \| 415 \| 31 \| 19 \| 620 \| 22 \| 684 \| Phage tail tape measure protein \| Phage tail tape measure protein; afdb-uniprot50 \| AF-A0A859FFP0-F1-MODEL\_V4 \| 1.0 \| 1.038e-10 \| 334 \| 0.178 \| 837 \| 510 \| 34 \| 62 \| 776 \| 8 \| 788 \| Phage tail tape measure protein \| Phage tail tape measure protein; afdb-uniprot50 \| AF-A0A354P661-F1-MODEL\_V4 \| 1.0 \| 1.26e-10 \| 334 \| 0.172 \| 834 \| 459 \| 29 \| 1 \| 702 \| 1 \| 734 \| Phage tail tape measure protein \| Phage tail tape measure protein; afdb-uniprot50 \| AF-A0A653R895-F1-MODEL\_V4 \| 1.0 \| 3.326e-10 \| 334 \| 0.154 \| 766 \| 462 \| 24 \| 5 \| 702 \| 2 \| 649 \| Putative Phage tail tape measure protein \| Putative Phage tail tape measure protein; afdb-uniprot50 \| AF-A0A1C3H6T6-F1-MODEL\_V4 \| 1.0 \| 5.955e-10 \| 334 \| 0.148 \| 708 \| 496 \| 26 \| 27 \| 702 \| 13 \| 645 \| Phage tail length tape-measure protein \| Phage tail length tape-measure protein; afdb-uniprot50 \| AF-R6CVJ2-F1-MODEL\_V4 \| 1.0 \| 2.635e-10 \| 334 \| 0.132 \| 847 \| 544 \| 27 \| 21 \| 709 \| 83 \| 896 \| Phage tail tape measure protein TP901 family core region \| Phage tail tape measure protein TP901 family core region; afdb-uniprot50 \| AF-A0A662GNB5-F1-MODEL\_V4 \| 1.0 \| 1.472e-10 \| 333 \| 0.168 \| 725 \| 461 \| 24 \| 117 \| 773 \| 59 \| 709 \| Phage tail tape measure protein \| Phage tail tape measure protein; afdb-uniprot50 \| AF-A0A3L8PL57-F1-MODEL\_V4 \| 1.0 \| 8.22e-11 \| 333 \| 0.155 \| 783 \| 512 \| 27 \| 91 \| 760 \| 20 \| 765 \| Phage tail tape measure protein \| Phage tail tape measure protein; afdb-uniprot50 \| AF-A0A0F9GIC6-F1-MODEL\_V4 \| 1.0 \| 4.538e-10 \| 333 \| 0.164 \| 758 \| 461 \| 23 \| 26 \| 702 \| 8 \| 673 \| PhageMin\_Tail domain-containing protein \| PhageMin\_Tail domain-containing protein; afdb-uniprot50 \| AF-R6MP60-F1-MODEL\_V4 \| 1.0 \| 1.121e-10 \| 333 \| 0.157 \| 819 \| 502 \| 28 \| 1 \| 708 \| 1 \| 742 \| Phage tail tape measure protein TP901 family \| Phage tail tape measure protein TP901 family; afdb-uniprot50 \| AF-A0A7X9BRP2-F1-MODEL\_V4 \| 1.0 \| 1.121e-10 \| 333 \| 0.139 \| 844 \| 511 \| 34 \| 1 \| 689 \| 1 \| 783 \| Phage tail tape measure protein \| Phage tail tape measure protein; afdb-uniprot50 \| AF-A0A173WSY1-F1-MODEL\_V4 \| 1.0 \| 9.601e-11 \| 332 \| 0.132 \| 799 \| 483 \| 31 \| 39 \| 784 \| 5 \| 645 \| Phage-related minor tail protein \| Phage-related minor tail protein; afdb-uniprot50 \| AF-D5RM85-F1-MODEL\_V4 \| 1.0 \| 4.718e-10 \| 331 \| 0.174 \| 717 \| 474 \| 31 \| 61 \| 702 \| 3 \| 676 \| Phage tail tape measure protein, TP901 family \| Phage tail tape measure protein, TP901 family; afdb-uniprot50 \| AF-A0A2I6S817-F1-MODEL\_V4 \| 1.0 \| 1.245e-09 \| 330 \| 0.147 \| 645 \| 453 \| 31 \| 75 \| 677 \| 6 \| 595 \| Phage tail tape measure protein \| Phage tail tape measure protein; afdb-uniprot50 \| AF-A0A5J6LAV7-F1-MODEL\_V4 \| 1.0 \| 1.591e-10 \| 330 \| 0.138 \| 797 \| 487 \| 28 \| 117 \| 784 \| 45 \| 770 \| Phage tail tape measure protein \| Phage tail tape measure protein; afdb-uniprot50 \| AF-A0A4R8I8J4-F1-MODEL\_V4 \| 1.0 \| 6.691e-10 \| 329 \| 0.136 \| 825 \| 518 \| 29 \| 70 \| 779 \| 3 \| 747 \| TP901 family phage tail tape measure protein \| TP901 family phage tail tape measure protein; afdb-uniprot50 \| AF-A0A073KF69-F1-MODEL\_V4 \| 1.0 \| 1.152e-09 \| 329 \| 0.164 \| 772 \| 459 \| 27 \| 1 \| 702 \| 1 \| 656 \| PhageMin\_Tail domain-containing protein \| PhageMin\_Tail domain-containing protein; afdb-uniprot50 \| AF-A0A661VDL1-F1-MODEL\_V4 \| 1.0 \| 2.41e-09 \| 328 \| 0.155 \| 806 \| 482 \| 30 \| 19 \| 785 \| 2 \| 647 \| Phage tail tape measure protein \| Phage tail tape measure protein; afdb-uniprot50 \| AF-A0A3E3EFA4-F1-MODEL\_V4 \| 1.0 \| 8.22e-11 \| 328 \| 0.157 \| 799 \| 510 \| 33 \| 94 \| 771 \| 18 \| 773 \| Phage tail tape measure protein \| Phage tail tape measure protein; afdb-uniprot50 \| AF-A0A3R6RKG2-F1-MODEL\_V4 \| 1.0 \| 4.962e-11 \| 328 \| 0.136 \| 955 \| 555 \| 33 \| 1 \| 767 \| 1 \| 873 \| Phage tail tape measure protein \| Phage tail tape measure protein; afdb-uniprot50 \| AF-A0A497QP70-F1-MODEL\_V4 \| 1.0 \| 3.326e-10 \| 328 \| 0.154 \| 840 \| 536 \| 34 \| 13 \| 779 \| 1 \| 738 \| Phage tail tape measure protein \| Phage tail tape measure protein; afdb-uniprot50 \| AF-A0A6B2YV45-F1-MODEL\_V4 \| 1.0 \| 3.93e-11 \| 328 \| 0.16 \| 995 \| 560 \| 30 \| 14 \| 785 \| 1 \| 942 \| Phage tail tape measure protein \| Phage tail tape measure protein; afdb-uniprot50 \| AF-U2Q4W9-F1-MODEL\_V4 \| 1.0 \| 1.985e-09 \| 328 \| 0.151 \| 674 \| 472 \| 21 \| 80 \| 702 \| 19 \| 643 \| Phage tail tape measure protein, TP901 family \| Phage tail tape measure protein, TP901 family; afdb-uniprot50 \| AF-A0A2S5M680-F1-MODEL\_V4 \| 1.0 \| 4.538e-10 \| 327 \| 0.131 \| 850 \| 468 \| 32 \| 19 \| 785 \| 25 \| 686 \| Phage tail tape measure protein \| Phage tail tape measure protein; afdb-uniprot50 \| AF-A0A6N8H0F9-F1-MODEL\_V4 \| 1.0 \| 2.848e-10 \| 327 \| 0.147 \| 730 \| 478 \| 19 \| 117 \| 770 \| 65 \| 725 \| Phage tail tape measure protein \| Phage tail tape measure protein; afdb-uniprot50 \| AF-A0A7C4CJ38-F1-MODEL\_V4 \| 1.0 \| 8.158e-07 \| 326 \| 0.137 \| 401 \| 281 \| 15 \| 64 \| 437 \| 10 \| 372 \| Phage tail tape measure protein \| Phage tail tape measure protein; afdb-uniprot50 \| AF-M0BS65-F1-MODEL\_V4 \| 1.0 \| 7.518e-10 \| 326 \| 0.177 \| 760 \| 439 \| 30 \| 6 \| 701 \| 3 \| 640 \| Phage tail tape measure protein, TP901 family \| Phage tail tape measure protein, TP901 family; afdb-uniprot50 \| AF-A0A2R3MZN4-F1-MODEL\_V4 \| 1.0 \| 3.458e-10 \| 326 \| 0.155 \| 861 \| 542 \| 34 \| 19 \| 785 \| 106 \| 874 \| Phage tail tape measure protein \| Phage tail tape measure protein; afdb-uniprot50 \| AF-A0A495R7S4-F1-MODEL\_V4 \| 1.0 \| 1.26e-10 \| 326 \| 0.141 \| 933 \| 511 \| 31 \| 81 \| 768 \| 22 \| 909 \| TP901 family phage tail tape measure protein \| TP901 family phage tail tape measure protein; afdb-uniprot50 \| AF-A0A0F3FS32-F1-MODEL\_V4 \| 1.0 \| 3.595e-10 \| 326 \| 0.162 \| 743 \| 504 \| 25 \| 19 \| 701 \| 77 \| 760 \| PhageMin\_Tail domain-containing protein \| PhageMin\_Tail domain-containing protein; afdb-uniprot50 \| AF-A0A7W3T5L2-F1-MODEL\_V4 \| 1.0 \| 8.884e-11 \| 326 \| 0.183 \| 734 \| 460 \| 31 \| 4 \| 701 \| 66 \| 695 \| Phage tail tape measure protein \| Phage tail tape measure protein; afdb-uniprot50 \| AF-A0A1I5WTC2-F1-MODEL\_V4 \| 1.0 \| 1.673e-11 \| 325 \| 0.143 \| 835 \| 515 \| 34 \| 77 \| 785 \| 18 \| 777 \| Phage tail tape measure protein, TP901 family, core region \| Phage tail tape measure protein, TP901 family, core region; afdb-uniprot50 \| AF-A0A1C6GH83-F1-MODEL\_V4 \| 1.0 \| 2.438e-10 \| 325 \| 0.118 \| 811 \| 504 \| 30 \| 1 \| 706 \| 1 \| 705 \| Phage-related minor tail protein \| Phage-related minor tail protein; afdb-uniprot50 \| AF-D3I9W8-F1-MODEL\_V4 \| 1.0 \| 2.438e-10 \| 324 \| 0.182 \| 712 \| 460 \| 32 \| 70 \| 706 \| 9 \| 673 \| Phage tail tape measure protein, TP901 family, core region \| Phage tail tape measure protein, TP901 family, core region; afdb-uniprot50 \| AF-A0A221KDC2-F1-MODEL\_V4 \| 1.0 \| 5.51e-10 \| 323 \| 0.148 \| 767 \| 446 \| 24 \| 118 \| 713 \| 46 \| 776 \| Phage tail tape measure protein \| Phage tail tape measure protein; afdb-uniprot50 \| AF-A0A402CRD8-F1-MODEL\_V4 \| 1.0 \| 2.961e-10 \| 323 \| 0.136 \| 806 \| 514 \| 31 \| 65 \| 785 \| 7 \| 715 \| PhageMin\_Tail domain-containing protein \| PhageMin\_Tail domain-containing protein; afdb-uniprot50 \| AF-A0A6L8JRL2-F1-MODEL\_V4 \| 1.0 \| 1.121e-10 \| 323 \| 0.156 \| 772 \| 500 \| 28 \| 34 \| 702 \| 4 \| 726 \| Phage tail tape measure protein \| Phage tail tape measure protein; afdb-uniprot50 \| AF-A0A1Q7RJA2-F1-MODEL\_V4 \| 1.0 \| 2.256e-10 \| 322 \| 0.155 \| 861 \| 499 \| 34 \| 97 \| 781 \| 16 \| 823 \| Phage tail tape measure protein \| Phage tail tape measure protein; afdb-uniprot50 \| AF-A0A3N2KV37-F1-MODEL\_V4 \| 1.0 \| 6.511e-11 \| 321 \| 0.139 \| 938 \| 549 \| 34 \| 1 \| 785 \| 1 \| 832 \| Phage tail tape measure protein \| Phage tail tape measure protein; afdb-uniprot50 \| AF-A0A5M8P9Z2-F1-MODEL\_V4 \| 1.0 \| 2.23e-09 \| 320 \| 0.148 \| 653 \| 438 \| 25 \| 114 \| 708 \| 42 \| 634 \| Phage tail tape measure protein \| Phage tail tape measure protein; afdb-uniprot50 \| AF-U5BPA9-F1-MODEL\_V4 \| 1.0 \| 3.84e-09 \| 320 \| 0.15 \| 705 \| 458 \| 26 \| 118 \| 781 \| 56 \| 660 \| Uncharacterized protein \| Uncharacterized protein; afdb-uniprot50 \| AF-A0A7U3G9U5-F1-MODEL\_V4 \| 1.0 \| 1.699e-09 \| 320 \| 0.138 \| 709 \| 421 \| 29 \| 118 \| 699 \| 44 \| 689 \| Phage tail tape measure protein \| Phage tail tape measure protein; afdb-uniprot50 \| AF-A0A6H1ZVM3-F1-MODEL\_V4 \| 1.0 \| 1.931e-10 \| 320 \| 0.141 \| 926 \| 532 \| 33 \| 62 \| 785 \| 5 \| 869 \| Putative tail protein \| Putative tail protein; afdb-uniprot50 \| AF-D8JAW3-F1-MODEL\_V4 \| 1.0 \| 5.51e-10 \| 319 \| 0.167 \| 662 \| 427 \| 24 \| 132 \| 702 \| 2 \| 630 \| Phage tail tape measure protein, TP901 family \| Phage tail tape measure protein, TP901 family; afdb-uniprot50 \| AF-A0A6B1XSV2-F1-MODEL\_V4 \| 1.0 \| 9.866e-10 \| 318 \| 0.133 \| 841 \| 539 \| 32 \| 12 \| 772 \| 32 \| 762 \| Phage tail tape measure protein \| Phage tail tape measure protein; afdb-uniprot50 \| AF-V7HXZ1-F1-MODEL\_V4 \| 1.0 \| 1.512e-09 \| 318 \| 0.148 \| 672 \| 462 \| 20 \| 81 \| 702 \| 3 \| 613 \| PhageMin\_Tail domain-containing protein \| PhageMin\_Tail domain-containing protein; afdb-uniprot50 \| AF-A0A7C6P3T7-F1-MODEL\_V4 \| 1.0 \| 5.729e-10 \| 317 \| 0.148 \| 803 \| 499 \| 34 \| 110 \| 779 \| 50 \| 800 \| Phage tail tape measure protein \| Phage tail tape measure protein; afdb-uniprot50 \| AF-K9DZ49-F1-MODEL\_V4 \| 1.0 \| 3.418e-09 \| 317 \| 0.142 \| 675 \| 431 \| 27 \| 90 \| 706 \| 30 \| 614 \| Phage tail tape measure protein, TP901 family, core region \| Phage tail tape measure protein, TP901 family, core region; afdb-uniprot50 \| AF-A0A2P9EK54-F1-MODEL\_V4 \| 1.0 \| 5.179e-08 \| 316 \| 0.158 \| 619 \| 404 \| 17 \| 19 \| 613 \| 142 \| 667 \| Putative phage minor tail protein \| Putative phage minor tail protein; afdb-uniprot50 \| AF-A0A7C2SUJ3-F1-MODEL\_V4 \| 1.0 \| 2.505e-09 \| 316 \| 0.137 \| 828 \| 503 \| 24 \| 53 \| 785 \| 6 \| 716 \| Phage tail tape measure protein \| Phage tail tape measure protein; afdb-uniprot50 \| AF-A0A497ITP3-F1-MODEL\_V4 \| 1.0 \| 3.885e-10 \| 316 \| 0.138 \| 854 \| 468 \| 29 \| 4 \| 677 \| 171 \| 936 \| Phage tail tape measure protein \| Phage tail tape measure protein; afdb-uniprot50 \| AF-A0A1Q6KUJ6-F1-MODEL\_V4 \| 1.0 \| 1.198e-09 \| 316 \| 0.158 \| 727 \| 447 \| 26 \| 86 \| 719 \| 25 \| 679 \| Phage tail tape measure protein \| Phage tail tape measure protein; afdb-uniprot50 \| AF-A0A5R1ZFV2-F1-MODEL\_V4 \| 1.0 \| 4.848e-09 \| 315 \| 0.145 \| 717 \| 437 \| 22 \| 18 \| 677 \| 141 \| 738 \| Phage tail tape measure protein \| Phage tail tape measure protein; afdb-uniprot50 \| AF-A0A351TP38-F1-MODEL\_V4 \| 1.0 \| 5.51e-10 \| 315 \| 0.153 \| 774 \| 512 \| 29 \| 19 \| 701 \| 92 \| 812 \| Phage tail tape measure protein \| Phage tail tape measure protein; afdb-uniprot50 \| AF-A0A3D9L500-F1-MODEL\_V4 \| 1.0 \| 1.121e-10 \| 315 \| 0.141 \| 942 \| 566 \| 33 \| 1 \| 759 \| 1 \| 882 \| TP901 family phage tail tape measure protein \| TP901 family phage tail tape measure protein; afdb-uniprot50 \| AF-A0A0F9U446-F1-MODEL\_V4 \| 1.0 \| 9.129e-10 \| 313 \| 0.163 \| 782 \| 448 \| 33 \| 6 \| 709 \| 1 \| 653 \| PhageMin\_Tail domain-containing protein \| PhageMin\_Tail domain-containing protein; afdb-uniprot50 \| AF-A0A6A7URX7-F1-MODEL\_V4 \| 1.0 \| 4.905e-10 \| 313 \| 0.131 \| 889 \| 532 \| 34 \| 43 \| 783 \| 27 \| 822 \| Phage tail tape measure protein \| Phage tail tape measure protein; afdb-uniprot50 \| AF-R6FSK8-F1-MODEL\_V4 \| 1.0 \| 7.816e-10 \| 313 \| 0.133 \| 905 \| 502 \| 32 \| 21 \| 701 \| 67 \| 912 \| Phage tail tape measure protein TP901 family core region \| Phage tail tape measure protein TP901 family core region; afdb-uniprot50 \| AF-A0A2G6ZIE4-F1-MODEL\_V4 \| 1.0 \| 4.365e-10 \| 312 \| 0.175 \| 780 \| 442 \| 31 \| 70 \| 702 \| 4 \| 729 \| TP901 family phage tail tape measure protein \| TP901 family phage tail tape measure protein; afdb-uniprot50 \| AF-A0A255T722-F1-MODEL\_V4 \| 1.0 \| 7.816e-10 \| 312 \| 0.154 \| 697 \| 456 \| 29 \| 46 \| 707 \| 27 \| 624 \| Phage tail tape measure protein \| Phage tail tape measure protein; afdb-uniprot50 \| AF-A0A0Q8PQD2-F1-MODEL\_V4 \| 1.0 \| 1.512e-09 \| 311 \| 0.148 \| 807 \| 486 \| 35 \| 80 \| 712 \| 14 \| 792 \| PhageMin\_Tail domain-containing protein \| PhageMin\_Tail domain-containing protein; afdb-uniprot50 \| AF-A0A6I4YMZ5-F1-MODEL\_V4 \| 1.0 \| 2.708e-09 \| 310 \| 0.151 \| 821 \| 506 \| 26 \| 14 \| 752 \| 81 \| 792 \| Phage tail tape measure protein \| Phage tail tape measure protein; afdb-uniprot50 \| AF-J8W777-F1-MODEL\_V4 \| 1.0 \| 4.905e-10 \| 310 \| 0.136 \| 914 \| 564 \| 32 \| 16 \| 778 \| 20 \| 858 \| Tail tape measure protein, TIGR01760 family \| Tail tape measure protein, TIGR01760 family; afdb-uniprot50 \| AF-A0A1W2GQA3-F1-MODEL\_V4 \| 1.0 \| 1.699e-09 \| 309 \| 0.151 \| 752 \| 495 \| 26 \| 45 \| 702 \| 18 \| 720 \| Phage tail tape measure protein, TP901 family, core region \| Phage tail tape measure protein, TP901 family, core region; afdb-uniprot50 \| AF-A0A1I2G2M1-F1-MODEL\_V4 \| 1.0 \| 7.148e-09 \| 309 \| 0.168 \| 694 \| 437 \| 20 \| 87 \| 702 \| 19 \| 650 \| Phage tail tape measure protein, TP901 family, core region \| Phage tail tape measure protein, TP901 family, core region; afdb-uniprot50 \| AF-A0A3C0QQX1-F1-MODEL\_V4 \| 1.0 \| 1.152e-09 \| 308 \| 0.164 \| 742 \| 471 \| 30 \| 80 \| 702 \| 15 \| 726 \| Phage tail tape measure protein \| Phage tail tape measure protein; afdb-uniprot50 \| AF-A0A6B4MZE8-F1-MODEL\_V4 \| 1.0 \| 4.039e-10 \| 308 \| 0.156 \| 822 \| 513 \| 30 \| 21 \| 702 \| 73 \| 853 \| Phage tail tape measure protein \| Phage tail tape measure protein; afdb-uniprot50 \| AF-A0A1S6GAN8-F1-MODEL\_V4 \| 1.0 \| 2.41e-09 \| 308 \| 0.148 \| 793 \| 480 \| 31 \| 51 \| 702 \| 2 \| 739 \| Phage tail tape measure protein \| Phage tail tape measure protein; afdb-uniprot50 \| AF-A0A1I4PAD3-F1-MODEL\_V4 \| 1.0 \| 1.512e-09 \| 308 \| 0.136 \| 745 \| 509 \| 30 \| 19 \| 728 \| 67 \| 711 \| Phage tail tape measure protein, TP901 family, core region \| Phage tail tape measure protein, TP901 family, core region; afdb-uniprot50 \| AF-A0A1D7XZW9-F1-MODEL\_V4 \| 1.0 \| 7.148e-09 \| 308 \| 0.169 \| 666 \| 430 \| 24 \| 117 \| 714 \| 60 \| 670 \| Phage tail length tape-measure protein \| Phage tail length tape-measure protein; afdb-uniprot50 \| AF-A0A410MCA2-F1-MODEL\_V4 \| 1.0 \| 5.955e-10 \| 307 \| 0.138 \| 801 \| 481 \| 32 \| 2 \| 748 \| 161 \| 806 \| Phage tail tape measure protein \| Phage tail tape measure protein; afdb-uniprot50 \| AF-A0A774SDM2-F1-MODEL\_V4 \| 1.0 \| 2.328e-06 \| 306 \| 0.154 \| 448 \| 340 \| 12 \| 19 \| 443 \| 142 \| 573 \| Phage tail tape measure protein \| Phage tail tape measure protein; afdb-uniprot50 \| AF-A0A4U9R711-F1-MODEL\_V4 \| 1.0 \| 2.345e-10 \| 306 \| 0.126 \| 886 \| 509 \| 31 \| 81 \| 782 \| 16 \| 820 \| Phage protein \| Phage protein; afdb-uniprot50 \| AF-F7SRZ5-F1-MODEL\_V4 \| 1.0 \| 8.125e-10 \| 306 \| 0.16 \| 850 \| 481 \| 30 \| 46 \| 702 \| 6 \| 815 \| Phage tail tape measure protein, TP901 family, core region \| Phage tail tape measure protein, TP901 family, core region; afdb-uniprot50 \| AF-B2TQU2-F1-MODEL\_V4 \| 1.0 \| 1.673e-11 \| 306 \| 0.152 \| 850 \| 509 \| 35 \| 9 \| 701 \| 7 \| 801 \| Tail tape measure protein \| Tail tape measure protein; afdb-uniprot50 \| AF-F9MRG1-F1-MODEL\_V4 \| 1.0 \| 1.054e-08 \| 305 \| 0.162 \| 790 \| 461 \| 33 \| 32 \| 782 \| 27 \| 654 \| Phage tail tape measure protein, TP901 family \| Phage tail tape measure protein, TP901 family; afdb-uniprot50 \| AF-A0A1N7LR98-F1-MODEL\_V4 \| 1.0 \| 3.885e-10 \| 305 \| 0.201 \| 799 \| 483 \| 32 \| 46 \| 785 \| 196 \| 898 \| Phage tail tape measure protein, TP901 family, core region \| Phage tail tape measure protein, TP901 family, core region; afdb-uniprot50 \| AF-A0A7J2RGM5-F1-MODEL\_V4 \| 1.0 \| 1.331e-08 \| 305 \| 0.138 \| 709 \| 479 \| 22 \| 14 \| 706 \| 1 \| 593 \| Phage tail tape measure protein \| Phage tail tape measure protein; afdb-uniprot50 \| AF-A0A414RNY6-F1-MODEL\_V4 \| 1.0 \| 5.729e-10 \| 305 \| 0.143 \| 768 \| 480 \| 30 \| 63 \| 785 \| 8 \| 642 \| Phage tail tape measure protein \| Phage tail tape measure protein; afdb-uniprot50 \| AF-A0A6M1QSR6-F1-MODEL\_V4 \| 1.0 \| 2.23e-09 \| 304 \| 0.156 \| 967 \| 542 \| 38 \| 52 \| 785 \| 9 \| 934 \| Phage tail tape measure protein \| Phage tail tape measure protein; afdb-uniprot50 \| AF-B9DJ04-F1-MODEL\_V4 \| 1.0 \| 1.66e-07 \| 302 \| 0.151 \| 587 \| 356 \| 15 \| 19 \| 483 \| 127 \| 693 \| Truncated phiSLT orf2067-like protein (Fragment 1) \| Truncated phiSLT orf2067-like protein (Fragment 1); afdb-uniprot50 \| AF-A0A6M3KH64-F1-MODEL\_V4 \| 1.0 \| 1.985e-09 \| 302 \| 0.139 \| 730 \| 512 \| 28 \| 14 \| 702 \| 14 \| 667 \| Putative tail protein \| Putative tail protein; afdb-uniprot50 \| AF-A0A1W6WWI4-F1-MODEL\_V4 \| 1.0 \| 1.367e-07 \| 302 \| 0.18 \| 621 \| 424 \| 18 \| 118 \| 702 \| 17 \| 588 \| Phage tail tape measure protein \| Phage tail tape measure protein; afdb-uniprot50 \| AF-A0A2H5YGJ4-F1-MODEL\_V4 \| 1.0 \| 2.41e-09 \| 301 \| 0.153 \| 880 \| 500 \| 38 \| 36 \| 785 \| 2 \| 766 \| PhageMin\_Tail domain-containing protein \| PhageMin\_Tail domain-containing protein; afdb-uniprot50 \| AF-A0A6M3M5N5-F1-MODEL\_V4 \| 1.0 \| 1.634e-09 \| 301 \| 0.145 \| 741 \| 491 \| 26 \| 87 \| 755 \| 27 \| 696 \| Putative tail protein \| Putative tail protein; afdb-uniprot50 \| AF-U5RUF6-F1-MODEL\_V4 \| 1.0 \| 1.572e-09 \| 301 \| 0.143 \| 869 \| 517 \| 27 \| 13 \| 702 \| 62 \| 881 \| Phage tail tape measure protein, TP901 family \| Phage tail tape measure protein, TP901 family; afdb-uniprot50 \| AF-A0A2D6FNM6-F1-MODEL\_V4 \| 1.0 \| 9.752e-09 \| 300 \| 0.146 \| 723 \| 476 \| 22 \| 34 \| 702 \| 10 \| 645 \| Phage tail tape measure protein \| Phage tail tape measure protein; afdb-uniprot50 \| AF-A0A4R4D151-F1-MODEL\_V4 \| 1.0 \| 3.042e-09 \| 299 \| 0.148 \| 776 \| 471 \| 30 \| 98 \| 785 \| 31 \| 704 \| Phage tail tape measure protein \| Phage tail tape measure protein; afdb-uniprot50 \| AF-A0A6M3J350-F1-MODEL\_V4 \| 1.0 \| 5.955e-10 \| 299 \| 0.158 \| 771 \| 503 \| 30 \| 79 \| 767 \| 22 \| 728 \| Putative tail protein \| Putative tail protein; afdb-uniprot50 \| AF-A0A518I8T5-F1-MODEL\_V4 \| 1.0 \| 9.381e-09 \| 299 \| 0.168 \| 716 \| 490 \| 25 \| 62 \| 707 \| 5 \| 684 \| Phage-related minor tail protein \| Phage-related minor tail protein; afdb-uniprot50 \| AF-A0A497SD96-F1-MODEL\_V4 \| 1.0 \| 1.699e-09 \| 298 \| 0.144 \| 876 \| 533 \| 33 \| 1 \| 772 \| 4 \| 766 \| Phage tail tape measure protein \| Phage tail tape measure protein; afdb-uniprot50 \| AF-A0A7U9SJK2-F1-MODEL\_V4 \| 1.0 \| 1.909e-09 \| 298 \| 0.16 \| 790 \| 489 \| 30 \| 117 \| 785 \| 40 \| 775 \| PhageMin\_Tail domain-containing protein \| PhageMin\_Tail domain-containing protein; afdb-uniprot50 \| AF-J9GWP7-F1-MODEL\_V4 \| 1.0 \| 3.126e-08 \| 297 \| 0.166 \| 572 \| 372 \| 19 \| 151 \| 677 \| 3 \| 514 \| Protein containing Phage tail tape measure protein TP901, core region domain protein \| Protein containing Phage tail tape measure protein TP901, core region domain protein; afdb-uniprot50 \| AF-A0A822VLY9-F1-MODEL\_V4 \| 1.0 \| 2.17e-10 \| 297 \| 0.139 \| 875 \| 547 \| 32 \| 21 \| 785 \| 69 \| 847 \| Prophage LambdaSa04, TP901 family tail tape measure protein \| Prophage LambdaSa04, TP901 family tail tape measure protein; afdb-uniprot50 \| AF-A0A1H3EM43-F1-MODEL\_V4 \| 1.0 \| 1.495e-08 \| 296 \| 0.152 \| 749 \| 497 \| 32 \| 35 \| 744 \| 11 \| 660 \| Phage tail tape measure protein, TP901 family, core region \| Phage tail tape measure protein, TP901 family, core region; afdb-uniprot50 \| AF-A0A3C1GIL4-F1-MODEL\_V4 \| 1.0 \| 7.148e-09 \| 296 \| 0.165 \| 805 \| 483 \| 32 \| 13 \| 785 \| 1 \| 648 \| Phage tail tape measure protein \| Phage tail tape measure protein; afdb-uniprot50 \| AF-A0A658JZU8-F1-MODEL\_V4 \| 1.0 \| 2.23e-09 \| 295 \| 0.154 \| 855 \| 521 \| 31 \| 19 \| 712 \| 49 \| 862 \| Phage tail tape measure protein \| Phage tail tape measure protein; afdb-uniprot50 \| AF-A0A3E5B8P9-F1-MODEL\_V4 \| 1.0 \| 2.204e-08 \| 294 \| 0.126 \| 729 \| 460 \| 25 \| 32 \| 719 \| 27 \| 619 \| Phage tail tape measure protein \| Phage tail tape measure protein; afdb-uniprot50 \| AF-A0A098YPR7-F1-MODEL\_V4 \| 1.0 \| 6.614e-09 \| 292 \| 0.164 \| 693 \| 408 \| 28 \| 41 \| 713 \| 29 \| 570 \| PhageMin\_Tail domain-containing protein \| PhageMin\_Tail domain-containing protein; afdb-uniprot50 \| AF-A0A0F9U574-F1-MODEL\_V4 \| 1.0 \| 2.145e-09 \| 291 \| 0.15 \| 753 \| 496 \| 31 \| 53 \| 706 \| 7 \| 714 \| PhageMin\_Tail domain-containing protein \| PhageMin\_Tail domain-containing protein; afdb-uniprot50 \| AF-M1ZYX2-F1-MODEL\_V4 \| 1.0 \| 3.694e-09 \| 291 \| 0.13 \| 791 \| 480 \| 23 \| 19 \| 749 \| 127 \| 769 \| Phage protein \| Phage protein; afdb-uniprot50 \| AF-A0A094IDZ6-F1-MODEL\_V4 \| 1.0 \| 1.026e-09 \| 291 \| 0.129 \| 779 \| 527 \| 30 \| 19 \| 702 \| 5 \| 726 \| Phage-related minor tail protein \| Phage-related minor tail protein; afdb-uniprot50 \| AF-A0A3D5S7X8-F1-MODEL\_V4 \| 1.0 \| 1.198e-09 \| 291 \| 0.15 \| 817 \| 545 \| 31 \| 14 \| 716 \| 38 \| 819 \| Phage tail tape measure protein \| Phage tail tape measure protein; afdb-uniprot50 \| AF-N0BGY2-F1-MODEL\_V4 \| 1.0 \| 1.766e-09 \| 291 \| 0.131 \| 951 \| 559 \| 29 \| 74 \| 785 \| 16 \| 938 \| Phage tail tape measure protein, TP901 family, core region \| Phage tail tape measure protein, TP901 family, core region; afdb-uniprot50 \| AF-G4WKD4-F1-MODEL\_V4 \| 1.0 \| 5.04e-09 \| 291 \| 0.14 \| 792 \| 504 \| 31 \| 49 \| 706 \| 7 \| 755 \| Putative tail tape measure protein \| Putative tail tape measure protein; afdb-uniprot50 \| AF-A0A1N6R4M9-F1-MODEL\_V4 \| 1.0 \| 6.614e-09 \| 290 \| 0.155 \| 747 \| 498 \| 23 \| 64 \| 747 \| 2 \| 678 \| Phage tail tape measure protein, TP901 family, core region \| Phage tail tape measure protein, TP901 family, core region; afdb-uniprot50 \| AF-A0A3N5RQ82-F1-MODEL\_V4 \| 1.0 \| 4.103e-08 \| 290 \| 0.147 \| 714 \| 480 \| 24 \| 16 \| 701 \| 1 \| 613 \| Phage tail tape measure protein \| Phage tail tape measure protein; afdb-uniprot50 \| AF-A0A6M3K6S3-F1-MODEL\_V4 \| 1.0 \| 7.431e-09 \| 290 \| 0.13 \| 868 \| 567 \| 24 \| 46 \| 784 \| 140 \| 948 \| Putative tail protein \| Putative tail protein; afdb-uniprot50 \| AF-A0A1C5SWG1-F1-MODEL\_V4 \| 1.0 \| 1.438e-08 \| 290 \| 0.154 \| 685 \| 445 \| 29 \| 81 \| 702 \| 6 \| 618 \| Phage tail tape measure protein, TP901 family, core region \| Phage tail tape measure protein, TP901 family, core region; afdb-uniprot50 \| AF-A0A7C5LDN6-F1-MODEL\_V4 \| 1.0 \| 3.163e-09 \| 289 \| 0.14 \| 900 \| 510 \| 33 \| 1 \| 772 \| 1 \| 764 \| Phage tail tape measure protein \| Phage tail tape measure protein; afdb-uniprot50 \| AF-A0A221KA30-F1-MODEL\_V4 \| 1.0 \| 1.634e-09 \| 289 \| 0.139 \| 887 \| 504 \| 28 \| 97 \| 755 \| 8 \| 862 \| Phage tail tape measure protein \| Phage tail tape measure protein; afdb-uniprot50 \| AF-A0A2E9DMI6-F1-MODEL\_V4 \| 1.0 \| 1.383e-08 \| 289 \| 0.145 \| 731 \| 435 \| 23 \| 79 \| 780 \| 22 \| 591 \| Phage tail tape measure protein \| Phage tail tape measure protein; afdb-uniprot50 \| AF-A0A3A9H1N1-F1-MODEL\_V4 \| 1.0 \| 8.481e-07 \| 288 \| 0.139 \| 486 \| 326 \| 15 \| 19 \| 442 \| 192 \| 647 \| Phage tail tape measure protein \| Phage tail tape measure protein; afdb-uniprot50 \| AF-A0A0F9RZT3-F1-MODEL\_V4 \| 1.0 \| 1.455e-09 \| 288 \| 0.166 \| 746 \| 509 \| 21 \| 19 \| 702 \| 23 \| 717 \| PhageMin\_Tail domain-containing protein \| PhageMin\_Tail domain-containing protein; afdb-uniprot50 \| AF-A0A5B0D944-F1-MODEL\_V4 \| 1.0 \| 5.795e-11 \| 286 \| 0.293 \| 538 \| 281 \| 14 \| 264 \| 753 \| 4 \| 490 \| Phage tail tape measure protein \| Phage tail tape measure protein; afdb-uniprot50 \| AF-K1ZEP0-F1-MODEL\_V4 \| 1.0 \| 1.352e-06 \| 286 \| 0.147 \| 469 \| 314 \| 13 \| 1 \| 455 \| 1 \| 397 \| PhageMin\_Tail domain-containing protein \| PhageMin\_Tail domain-containing protein; afdb-uniprot50 \| AF-A0A7K0ATB2-F1-MODEL\_V4 \| 1.0 \| 4.663e-09 \| 286 \| 0.149 \| 803 \| 497 \| 32 \| 60 \| 780 \| 24 \| 722 \| Phage tail tape measure protein \| Phage tail tape measure protein; afdb-uniprot50 \| AF-A0A6I4Z5B8-F1-MODEL\_V4 \| 1.0 \| 5.597e-08 \| 285 \| 0.128 \| 804 \| 484 \| 22 \| 14 \| 785 \| 208 \| 826 \| Phage tail tape measure protein \| Phage tail tape measure protein; afdb-uniprot50 \| AF-A0A0N9MLM3-F1-MODEL\_V4 \| 1.0 \| 7.907e-11 \| 285 \| 0.15 \| 1043 \| 538 \| 38 \| 61 \| 781 \| 7 \| 1022 \| Phage tail tape measure protein, TP901 family \| Phage tail tape measure protein, TP901 family; afdb-uniprot50 \| AF-A0A2P8EI71-F1-MODEL\_V4 \| 1.0 \| 1.096e-08 \| 285 \| 0.108 \| 883 \| 562 \| 25 \| 18 \| 709 \| 43 \| 890 \| TP901 family phage tail tape measure protein \| TP901 family phage tail tape measure protein; afdb-uniprot50 \| AF-R5IIS8-F1-MODEL\_V4 \| 1.0 \| 9.381e-09 \| 283 \| 0.131 \| 827 \| 523 \| 30 \| 19 \| 714 \| 87 \| 848 \| Phage tail tape measure protein TP901 family core region \| Phage tail tape measure protein TP901 family core region; afdb-uniprot50 \| AF-A0A085EXN0-F1-MODEL\_V4 \| 1.0 \| 1.217e-07 \| 282 \| 0.154 \| 732 \| 473 \| 28 \| 46 \| 723 \| 2 \| 641 \| Putative bacteriophage protein \| Putative bacteriophage protein; afdb-uniprot50 \| AF-A0A6M0RDA6-F1-MODEL\_V4 \| 1.0 \| 2.677e-08 \| 282 \| 0.128 \| 909 \| 527 \| 30 \| 3 \| 785 \| 7 \| 775 \| Phage tail tape measure protein \| Phage tail tape measure protein; afdb-uniprot50 \| AF-A0A7X7EMK5-F1-MODEL\_V4 \| 1.0 \| 4.486e-09 \| 282 \| 0.115 \| 939 \| 552 \| 33 \| 42 \| 770 \| 18 \| 887 \| Phage tail tape measure protein \| Phage tail tape measure protein; afdb-uniprot50 \| AF-A0A2G6ESM5-F1-MODEL\_V4 \| 1.0 \| 1.331e-08 \| 282 \| 0.129 \| 763 \| 532 \| 27 \| 19 \| 707 \| 11 \| 714 \| Phage tail tape measure protein \| Phage tail tape measure protein; afdb-uniprot50 \| AF-A0A416BKF0-F1-MODEL\_V4 \| 1.0 \| 7.346e-08 \| 280 \| 0.152 \| 682 \| 432 \| 27 \| 111 \| 686 \| 17 \| 658 \| Phage tail tape measure protein \| Phage tail tape measure protein; afdb-uniprot50 \| AF-A0A0F9MJ74-F1-MODEL\_V4 \| 1.0 \| 4.663e-09 \| 280 \| 0.145 \| 793 \| 547 \| 25 \| 1 \| 701 \| 2 \| 755 \| PhageMin\_Tail domain-containing protein \| PhageMin\_Tail domain-containing protein; afdb-uniprot50 \| AF-A0A853EHV0-F1-MODEL\_V4 \| 1.0 \| 1.31e-10 \| 279 \| 0.146 \| 893 \| 563 \| 37 \| 1 \| 746 \| 1 \| 840 \| Phage tail tape measure protein \| Phage tail tape measure protein; afdb-uniprot50 \| AF-A0A180FCQ4-F1-MODEL\_V4 \| 1.0 \| 6.049e-08 \| 279 \| 0.136 \| 654 \| 437 \| 19 \| 77 \| 719 \| 15 \| 551 \| Phage tail tape measure protein, TP901 family, core region \| Phage tail tape measure protein, TP901 family, core region; afdb-uniprot50 \| AF-A0A3D0Y8L6-F1-MODEL\_V4 \| 1.0 \| 5.04e-09 \| 279 \| 0.133 \| 891 \| 560 \| 36 \| 1 \| 784 \| 2 \| 787 \| Phage tail tape measure protein \| Phage tail tape measure protein; afdb-uniprot50 \| AF-A0A1Q7RI84-F1-MODEL\_V4 \| 1.0 \| 6.614e-09 \| 279 \| 0.16 \| 858 \| 526 \| 31 \| 21 \| 702 \| 91 \| 929 \| Phage tail tape measure protein \| Phage tail tape measure protein; afdb-uniprot50 \| AF-A0A1S7MQ78-F1-MODEL\_V4 \| 1.0 \| 9.273e-08 \| 277 \| 0.147 \| 850 \| 461 \| 28 \| 24 \| 785 \| 90 \| 763 \| PhageMin\_Tail domain-containing protein \| PhageMin\_Tail domain-containing protein; afdb-uniprot50 \| AF-A0A2U2DG04-F1-MODEL\_V4 \| 1.0 \| 1.139e-08 \| 277 \| 0.138 \| 908 \| 475 \| 33 \| 15 \| 783 \| 39 \| 777 \| PhageMin\_Tail domain-containing protein \| PhageMin\_Tail domain-containing protein; afdb-uniprot50 \| AF-A0A2K8U7C3-F1-MODEL\_V4 \| 1.0 \| 2.646e-07 \| 276 \| 0.155 \| 668 \| 454 \| 26 \| 64 \| 702 \| 7 \| 593 \| Phage tail tape measure protein \| Phage tail tape measure protein; afdb-uniprot50 \| AF-A0A661TQT9-F1-MODEL\_V4 \| 1.0 \| 8.68e-09 \| 276 \| 0.162 \| 738 \| 469 \| 31 \| 46 \| 700 \| 2 \| 673 \| Phage tail tape measure protein \| Phage tail tape measure protein; afdb-uniprot50 \| AF-A0A3A6RBH3-F1-MODEL\_V4 \| 1.0 \| 5.04e-09 \| 276 \| 0.138 \| 996 \| 576 \| 38 \| 14 \| 785 \| 141 \| 1077 \| Phage tail tape measure protein \| Phage tail tape measure protein; afdb-uniprot50 \| AF-A0A2D7ZNU8-F1-MODEL\_V4 \| 1.0 \| 3.84e-09 \| 274 \| 0.143 \| 774 \| 470 \| 26 \| 118 \| 750 \| 87 \| 808 \| Phage tail tape measure protein \| Phage tail tape measure protein; afdb-uniprot50 \| AF-A0A8B5R2V2-F1-MODEL\_V4 \| 1.0 \| 5.099e-10 \| 274 \| 0.171 \| 705 \| 450 \| 30 \| 66 \| 701 \| 3 \| 642 \| Phage tail tape measure protein \| Phage tail tape measure protein; afdb-uniprot50 \| AF-A0A2G2J0Y5-F1-MODEL\_V4 \| 1.0 \| 4.103e-08 \| 274 \| 0.147 \| 718 \| 487 \| 27 \| 39 \| 712 \| 14 \| 650 \| Phage tail tape measure protein \| Phage tail tape measure protein; afdb-uniprot50 \| AF-A0A6M3KIG7-F1-MODEL\_V4 \| 1.0 \| 4.792e-08 \| 273 \| 0.132 \| 746 \| 515 \| 28 \| 1 \| 702 \| 1 \| 657 \| Putative tail protein \| Putative tail protein; afdb-uniprot50 \| AF-E2CI78-F1-MODEL\_V4 \| 1.0 \| 3.652e-08 \| 271 \| 0.153 \| 875 \| 458 \| 31 \| 14 \| 784 \| 38 \| 733 \| Putative phage tail tape measure protein, core region \| Putative phage tail tape measure protein, core region; afdb-uniprot50 \| AF-A0A7Z8JJT8-F1-MODEL\_V4 \| 1.0 \| 1.495e-08 \| 271 \| 0.147 \| 740 \| 481 \| 28 \| 1 \| 702 \| 1 \| 628 \| Phage tail tape measure protein \| Phage tail tape measure protein; afdb-uniprot50 \| AF-A0A1W9TY32-F1-MODEL\_V4 \| 1.0 \| 1.331e-08 \| 271 \| 0.137 \| 732 \| 473 \| 28 \| 1 \| 702 \| 1 \| 603 \| Phage tail tape measure protein \| Phage tail tape measure protein; afdb-uniprot50 \| AF-A0A7J3WET8-F1-MODEL\_V4 \| 1.0 \| 1.383e-08 \| 269 \| 0.142 \| 889 \| 532 \| 35 \| 1 \| 777 \| 3 \| 772 \| Phage tail tape measure protein \| Phage tail tape measure protein; afdb-uniprot50 \| AF-D1Y710-F1-MODEL\_V4 \| 1.0 \| 5.819e-08 \| 268 \| 0.153 \| 683 \| 449 \| 21 \| 20 \| 625 \| 41 \| 670 \| Phage tail tape measure protein, TP901 family \| Phage tail tape measure protein, TP901 family; afdb-uniprot50 \| AF-A0A662BU24-F1-MODEL\_V4 \| 1.0 \| 9.381e-09 \| 268 \| 0.126 \| 792 \| 519 \| 26 \| 19 \| 704 \| 12 \| 736 \| PhageMin\_Tail domain-containing protein \| PhageMin\_Tail domain-containing protein; afdb-uniprot50 \| AF-A0A1Q6LLB6-F1-MODEL\_V4 \| 1.0 \| 3.752e-07 \| 268 \| 0.139 \| 695 \| 446 \| 24 \| 100 \| 699 \| 40 \| 677 \| Phage tail tape measure protein \| Phage tail tape measure protein; afdb-uniprot50 \| AF-A0A2E0R799-F1-MODEL\_V4 \| 1.0 \| 1.265e-07 \| 267 \| 0.228 \| 503 \| 263 \| 16 \| 13 \| 465 \| 17 \| 444 \| Phage tail tape measure protein \| Phage tail tape measure protein; afdb-uniprot50 \| AF-A0A4Y9G537-F1-MODEL\_V4 \| 1.0 \| 1.641e-06 \| 267 \| 0.205 \| 429 \| 230 \| 9 \| 18 \| 345 \| 54 \| 472 \| Phage tail tape measure protein \| Phage tail tape measure protein; afdb-uniprot50 \| AF-A0A2T7H7Y6-F1-MODEL\_V4 \| 1.0 \| 1.939e-07 \| 267 \| 0.136 \| 798 \| 468 \| 24 \| 39 \| 782 \| 75 \| 705 \| PhageMin\_Tail domain-containing protein \| PhageMin\_Tail domain-containing protein; afdb-uniprot50 \| AF-A0A4Q6Y918-F1-MODEL\_V4 \| 1.0 \| 2.12e-08 \| 266 \| 0.168 \| 794 \| 515 \| 32 \| 18 \| 772 \| 6 \| 692 \| Phage tail tape measure protein \| Phage tail tape measure protein; afdb-uniprot50 \| AF-A0A367HZ43-F1-MODEL\_V4 \| 1.0 \| 2.04e-08 \| 265 \| 0.138 \| 1094 \| 550 \| 35 \| 32 \| 785 \| 2 \| 1042 \| Phage tail tape measure protein \| Phage tail tape measure protein; afdb-uniprot50 \| AF-A0A3D2IQU7-F1-MODEL\_V4 \| 1.0 \| 3.947e-08 \| 264 \| 0.137 \| 763 \| 487 \| 26 \| 42 \| 704 \| 4 \| 695 \| Phage tail tape measure protein \| Phage tail tape measure protein; afdb-uniprot50 \| AF-A0A6I4XSS1-F1-MODEL\_V4 \| 1.0 \| 4.737e-07 \| 263 \| 0.181 \| 447 \| 253 \| 16 \| 1 \| 423 \| 2 \| 359 \| Phage tail tape measure protein \| Phage tail tape measure protein; afdb-uniprot50 \| AF-A0A412EM13-F1-MODEL\_V4 \| 1.0 \| 3.694e-09 \| 263 \| 0.142 \| 808 \| 537 \| 25 \| 1 \| 713 \| 1 \| 747 \| Phage tail tape measure protein \| Phage tail tape measure protein; afdb-uniprot50 \| AF-A0A2G2DHY8-F1-MODEL\_V4 \| 1.0 \| 3.752e-07 \| 262 \| 0.133 \| 809 \| 494 \| 26 \| 32 \| 784 \| 13 \| 670 \| Phage tail tape measure protein \| Phage tail tape measure protein; afdb-uniprot50 \| AF-A0A2P8NKI0-F1-MODEL\_V4 \| 1.0 \| 5.98e-07 \| 261 \| 0.146 \| 722 \| 457 \| 27 \| 49 \| 731 \| 6 \| 606 \| Phage tail tape measure protein \| Phage tail tape measure protein; afdb-uniprot50 \| AF-A0A2M8NE79-F1-MODEL\_V4 \| 1.0 \| 4.103e-08 \| 260 \| 0.125 \| 837 \| 526 \| 32 \| 19 \| 704 \| 28 \| 809 \| Phage tail tape measure protein \| Phage tail tape measure protein; afdb-uniprot50 \| AF-A0A6N8TKC3-F1-MODEL\_V4 \| 1.0 \| 2.096e-07 \| 259 \| 0.131 \| 838 \| 455 \| 27 \| 19 \| 783 \| 52 \| 689 \| Phage tail tape measure protein \| Phage tail tape measure protein; afdb-uniprot50 \| AF-A0A7J2HKH6-F1-MODEL\_V4 \| 1.0 \| 2.893e-08 \| 259 \| 0.129 \| 862 \| 527 \| 28 \| 109 \| 785 \| 23 \| 845 \| Phage tail tape measure protein \| Phage tail tape measure protein; afdb-uniprot50 \| AF-A0A6M3KID6-F1-MODEL\_V4 \| 1.0 \| 3.513e-08 \| 254 \| 0.122 \| 757 \| 470 \| 27 \| 31 \| 702 \| 83 \| 729 \| Putative tail protein \| Putative tail protein; afdb-uniprot50 \| AF-A0A349NKI0-F1-MODEL\_V4 \| 1.0 \| 2.265e-07 \| 252 \| 0.131 \| 593 \| 392 \| 17 \| 155 \| 702 \| 2 \| 516 \| Phage tail tape measure protein \| Phage tail tape measure protein; afdb-uniprot50 \| AF-A0A4Z0V6T5-F1-MODEL\_V4 \| 1.0 \| 4.925e-07 \| 252 \| 0.168 \| 664 \| 433 \| 25 \| 70 \| 690 \| 30 \| 617 \| Phage tail tape measure protein \| Phage tail tape measure protein; afdb-uniprot50 \| AF-A0A3E4VC51-F1-MODEL\_V4 \| 1.0 \| 1.794e-07 \| 251 \| 0.142 \| 807 \| 462 \| 34 \| 49 \| 684 \| 5 \| 752 \| Phage tail tape measure protein \| Phage tail tape measure protein; afdb-uniprot50 \| AF-A4W8K8-F1-MODEL\_V4 \| 1.0 \| 1.28e-08 \| 248 \| 0.2 \| 614 \| 354 \| 15 \| 243 \| 785 \| 2 \| 549 \| Phage tail-like protein \| Phage tail-like protein; afdb-uniprot50 \| AF-A0A662F5X9-F1-MODEL\_V4 \| 1.0 \| 5.322e-07 \| 248 \| 0.123 \| 735 \| 470 \| 25 \| 10 \| 680 \| 1 \| 624 \| Phage tail tape measure protein \| Phage tail tape measure protein; afdb-uniprot50 \| AF-A0A2W5EKK6-F1-MODEL\_V4 \| 1.0 \| 5.533e-07 \| 247 \| 0.137 \| 691 \| 389 \| 24 \| 109 \| 785 \| 200 \| 697 \| PhageMin\_Tail domain-containing protein \| PhageMin\_Tail domain-containing protein; afdb-uniprot50 \| AF-A0A1H6JMQ9-F1-MODEL\_V4 \| 1.0 \| 3.007e-08 \| 247 \| 0.14 \| 740 \| 445 \| 28 \| 50 \| 674 \| 10 \| 673 \| Phage tail tape measure protein, TP901 family, core region \| Phage tail tape measure protein, TP901 family, core region; afdb-uniprot50 \| AF-A0A662PR09-F1-MODEL\_V4 \| 1.0 \| 4.737e-07 \| 246 \| 0.148 \| 672 \| 437 \| 21 \| 66 \| 701 \| 2 \| 573 \| Phage tail tape measure protein \| Phage tail tape measure protein; afdb-uniprot50 \| AF-H1PPE3-F1-MODEL\_V4 \| 1.0 \| 1.519e-06 \| 244 \| 0.111 \| 664 \| 486 \| 22 \| 109 \| 702 \| 54 \| 683 \| Phage tail tape measure protein, TP901 family, core region \| Phage tail tape measure protein, TP901 family, core region; afdb-uniprot50 \| AF-A0A376FD60-F1-MODEL\_V4 \| 1.0 \| 3.568e-06 \| 243 \| 0.146 \| 513 \| 300 \| 14 \| 19 \| 458 \| 34 \| 481 \| Phage Tail Tape Measure protein \| Phage Tail Tape Measure protein; afdb-uniprot50 \| AF-A0A1C5ZH20-F1-MODEL\_V4 \| 1.0 \| 4.737e-07 \| 243 \| 0.156 \| 775 \| 447 \| 32 \| 103 \| 702 \| 37 \| 779 \| Phage-related protein \| Phage-related protein; afdb-uniprot50 \| AF-A0A662Q4G6-F1-MODEL\_V4 \| 1.0 \| 8.716e-06 \| 242 \| 0.138 \| 506 \| 367 \| 13 \| 10 \| 485 \| 313 \| 779 \| Phage tail tape measure protein \| Phage tail tape measure protein; afdb-uniprot50 \| AF-A0A2N6Q6V0-F1-MODEL\_V4 \| 1.0 \| 4.982e-08 \| 242 \| 0.125 \| 836 \| 503 \| 33 \| 44 \| 780 \| 29 \| 735 \| Phage tail tape measure protein \| Phage tail tape measure protein; afdb-uniprot50 \| AF-A0A212L7A0-F1-MODEL\_V4 \| 1.0 \| 5.752e-07 \| 237 \| 0.12 \| 797 \| 460 \| 31 \| 1 \| 776 \| 1 \| 577 \| PhageMin\_Tail domain-containing protein \| PhageMin\_Tail domain-containing protein; afdb-uniprot50 \| AF-A0A1Y0C490-F1-MODEL\_V4 \| 1.0 \| 5.819e-08 \| 203 \| 0.155 \| 836 \| 516 \| 32 \| 4 \| 755 \| 39 \| 768 \| Phage tail tape measure protein \| Phage tail tape measure protein; afdb-uniprot50 \| AF-A0A7X7BVE1-F1-MODEL\_V4 \| 1.0 \| 9.68e-05 \| 170 \| 0.223 \| 466 \| 289 \| 9 \| 239 \| 702 \| 3 \| 397 \| Phage tail tape measure protein \| Phage tail tape measure protein |
| Top keywords  (threshold 1.00e-02 (evalue)) | **tail, Phage, tape, measure, TP901, domain\_containing, PhageMin\_Tail, core, region, Phage\_related** |
| Output files | ../../similar\_structures/22\_FANPEZAQ\_CDS\_0022\_afdb-proteome\_foldseek.tsv ../../similar\_structures/22\_FANPEZAQ\_CDS\_0022\_afdb-uniprot50\_foldseek.tsv ../../similar\_structures/22\_FANPEZAQ\_CDS\_0022\_merged.svg ../../similar\_structures/22\_FANPEZAQ\_CDS\_0022\_pdb\_foldseek.tsv |

  
  
  

Return to summary | Go to previous | Go to next

  


---

**Sequence/structure alignments coloring**  
Each object in the alignment figures is colored according to its E-value following this color coding:

1e-100
10
