## Supplementary material for "Completing the BASEL phage collection to unlock hidden diversity for systematic exploration of phage-host interactions": Data S2: 23.html

FANPEZAQ\_CDS\_0023


Return to summary | Go to previous | Go to next

|  |  |
| --- | --- |
| FANPEZAQ\_CDS\_0023  Page creation date: 02 Sep 2024, 12:00  Project folder: n/a  Input sequences file: Escherichia\_virus\_HeidiAbel.gb | phage tail domain\_containing gpu lysm p2 u tube contractile system injection n\_terminal oxidoreductase peptidoglycan\_binding putative fragment bacteriophage hypothetical phage\_related dna fiber prophage |

### Sequence information

|  |  |
| --- | --- |
| Name | FANPEZAQ\_CDS\_0023  23\_FANPEZAQ\_CDS\_0023 (pipeline id) |
| Imported annotations | Escherichia\_virus\_HeidiAbel Bas97 |
| Protein sequence | MTSFTILTGQTNIMMMLGGFQFCILTAAYQELNRHSEYKWPSQHRFGQRPSSQFVGIGEE TITLAGIIYPEYRGGFQQVEQMRSMAGRGQPLLLVDGLGKLWGRWVIEVIDEKQSLFAAF GAPRKQEFNLQLRRFN |
| Number of residues | 136 |
| Molecular weight (Da) | 15595.79 |
| Output files | ../../query\_sequences/23\_FANPEZAQ\_CDS\_0023.fasta |

### Putative domain architecture and protein family

#### Search results (HHblits)^1^

|  |  |
| --- | --- |
| Domain family databases searched | Pfam, Ncbi-cd, Cath, Phrogs |
| Results, scheme(s)  (Top layers only; threshold 1.00e-03 (evalue)) | xml version="1.0" encoding="utf-8" standalone="no"?       2024-09-02T21:08:17.150768 image/svg+xml   Matplotlib v3.7.2, https://matplotlib.org/ |
| Results, table  (E-value ≤ 1.00e-03 (evalue)) | db \| id \| prob \| evalue \| pvalue \| score \| cols \| query \| query\_len \| template \| template\_len \| name \| description; pfam \| PF06995 \| 99.7 \| 1e-22 \| 1.8e-26 \| 126.0 \| 118 \| (18, 135) \| 136 \| (1, 118) \| 120 \| Phage\_P2\_GpU \| Phage P2 GpU; pfam \| PF19266 \| 99.5 \| 5.8e-19 \| 1.1e-22 \| 114.2 \| 112 \| (21, 135) \| 136 \| (22, 151) \| 159 \| CIS\_tube \| Contractile injection system tube protein; pfam \| PF19512 \| 99.4 \| 2.7e-17 \| 4.8e-21 \| 103.9 \| 104 \| (28, 135) \| 136 \| (4, 125) \| 137 \| DUF6046 \| Domain of unknown function (DUF6046); pfam \| PF07157 \| 96.9 \| 8.2e-07 \| 1.5e-10 \| 49.2 \| 75 \| (16, 97) \| 136 \| (5, 80) \| 86 \| DNA\_circ\_N \| DNA circularisation protein N-terminus; phrogs \| 76 \| 100.0 \| 9.5e-35 \| 1.2e-38 \| 205.3 \| 124 \| (13, 136) \| 136 \| (1, 124) \| 162 \| tail protein \| tail protein; Category: tail; p258311 VI\_01887; phrogs \| 788 \| 99.9 \| 3.8e-29 \| 4.5e-33 \| 189.2 \| 122 \| (13, 135) \| 136 \| (1, 122) \| 298 \| tail protein \| tail protein; Category: tail; p182850 VI\_05566; phrogs \| 4094 \| 99.7 \| 2.3e-23 \| 2.7e-27 \| 151.9 \| 121 \| (16, 136) \| 136 \| (167, 287) \| 293 \| NA \| NA; Category: unknown function; p422642 VI\_05571; phrogs \| 380 \| 99.6 \| 2.2e-20 \| 2.8e-24 \| 137.2 \| 119 \| (13, 136) \| 136 \| (5, 132) \| 230 \| endolysin \| endolysin; Category: lysis; NC\_005294\_p57; phrogs \| 5184 \| 99.2 \| 1.1e-15 \| 1.3e-19 \| 103.5 \| 109 \| (19, 136) \| 136 \| (16, 136) \| 145 \| NA \| NA; Category: unknown function; p243320 VI\_01361; phrogs \| 32879 \| 99.1 \| 6.3e-15 \| 7e-19 \| 103.7 \| 124 \| (11, 135) \| 136 \| (164, 291) \| 292 \| NA \| NA; Category: unknown function; p423527 VI\_06236; phrogs \| 29415 \| 98.7 \| 4.2e-12 \| 4.7e-16 \| 70.2 \| 58 \| (13, 70) \| 136 \| (1, 58) \| 59 \| NA \| NA; Category: unknown function; p408258 VI\_11010; phrogs \| 21362 \| 98.2 \| 2.9e-10 \| 3.4e-14 \| 82.8 \| 111 \| (3, 118) \| 136 \| (9, 119) \| 329 \| NA \| NA; Category: unknown function; p152991 VI\_08595; phrogs \| 23201 \| 96.2 \| 1.9e-05 \| 2.1e-09 \| 53.6 \| 105 \| (26, 136) \| 136 \| (17, 129) \| 222 \| NA \| NA; Category: unknown function; p369082 VI\_04056; phrogs \| 27169 \| 96.2 \| 2.5e-05 \| 2.8e-09 \| 43.5 \| 35 \| (101, 135) \| 136 \| (2, 36) \| 72 \| NA \| NA; Category: unknown function; p366908 VI\_01591; phrogs \| 1517 \| 96.1 \| 2.8e-05 \| 3.5e-09 \| 53.6 \| 99 \| (31, 134) \| 136 \| (22, 133) \| 194 \| endolysin \| endolysin; Category: lysis; NC\_020843\_p101; phrogs \| 817 \| 95.3 \| 0.00018 \| 2.2e-08 \| 48.1 \| 91 \| (40, 135) \| 136 \| (4, 110) \| 165 \| tail fiber protein \| tail fiber protein; Category: tail; p327551 VI\_10369 |
| Top keywords  (threshold 1.00e-03 (evalue)) | **tail, endolysin, lysis, Phage, P2, GpU, Contractile, injection, system, tube** |
| Output files | ../../domain\_architecture/23\_FANPEZAQ\_CDS\_0023\_cath.hhr ../../domain\_architecture/23\_FANPEZAQ\_CDS\_0023\_merged.svg ../../domain\_architecture/23\_FANPEZAQ\_CDS\_0023\_ncbi-cd.hhr ../../domain\_architecture/23\_FANPEZAQ\_CDS\_0023\_pfam.hhr ../../domain\_architecture/23\_FANPEZAQ\_CDS\_0023\_phrogs.hhr |

### Identical protein sequences/structures

#### Search results

|  |  |
| --- | --- |
| Protein sequence databases searched | Pdb, Swissprot, Refseq |
| Identical proteins found | -- |
| Top keywords | -- |
| Output files | -- |

### Similar protein sequences/structures

#### Sequence similarity search results (HHblits)^1^

|  |  |
| --- | --- |
| Sequence databases searched | Uniclust, Pdb70 |
| Results, scheme(s)  (Top layers only, threshold 1.00e-03 (evalue)) | xml version="1.0" encoding="utf-8" standalone="no"?       2024-09-02T21:08:39.322659 image/svg+xml   Matplotlib v3.7.2, https://matplotlib.org/ |
| Results, table(s)  (threshold 1.00e-03 (evalue)) | db \| id \| prob \| evalue \| pvalue \| score \| cols \| query \| query\_len \| template \| template\_len \| name \| description; uniclust \| UniRef100\_A0A009PIW6 \| 100.0 \| 7.9e-42 \| 1.7e-47 \| 246.7 \| 136 \| (1, 136) \| 136 \| (1, 136) \| 202 \| Phage P2 GpU family protein \| Phage P2 GpU family protein; uniclust \| UniRef100\_A0A061YJX4 \| 100.0 \| 1.9e-40 \| 4.3e-46 \| 238.6 \| 134 \| (3, 136) \| 136 \| (29, 162) \| 187 \| Phage tail protein \| Phage tail protein; uniclust \| UniRef100\_A0A011NB97 \| 100.0 \| 1.4e-38 \| 2.9e-44 \| 233.6 \| 131 \| (6, 136) \| 136 \| (24, 154) \| 235 \| Oxidoreductase \| Oxidoreductase; uniclust \| UniRef100\_A0A158E8Q2 \| 100.0 \| 3.3e-38 \| 6.7e-44 \| 238.3 \| 133 \| (4, 136) \| 136 \| (2, 134) \| 311 \| Bacteriophage tail-related protein \| Bacteriophage tail-related protein; uniclust \| UniRef100\_A0A060H3V2 \| 100.0 \| 7.7e-37 \| 1.6e-42 \| 216.2 \| 130 \| (7, 136) \| 136 \| (29, 158) \| 175 \| Tail protein \| Tail protein; uniclust \| UniRef100\_A0A022PJH1 \| 100.0 \| 9.8e-37 \| 2e-42 \| 218.3 \| 128 \| (9, 136) \| 136 \| (6, 133) \| 211 \| Phage protein U \| Phage protein U; uniclust \| UniRef100\_A0A074LRJ1 \| 100.0 \| 2.1e-35 \| 4.5e-41 \| 211.6 \| 131 \| (5, 136) \| 136 \| (21, 157) \| 188 \| Phage tail protein \| Phage tail protein; uniclust \| UniRef100\_A0A059WRP1 \| 100.0 \| 2.8e-35 \| 5.9e-41 \| 207.2 \| 125 \| (11, 136) \| 136 \| (1, 129) \| 165 \| Tail protein \| Tail protein; uniclust \| UniRef100\_A0A098B0B0 \| 100.0 \| 3.4e-34 \| 7.5e-40 \| 197.6 \| 125 \| (11, 136) \| 136 \| (1, 129) \| 138 \| Phage P2 GpU \| Phage P2 GpU; uniclust \| UniRef100\_A0A066RZT5 \| 100.0 \| 2.8e-33 \| 5.6e-39 \| 194.1 \| 132 \| (5, 136) \| 136 \| (24, 156) \| 162 \| p2 GpU family protein \| p2 GpU family protein; uniclust \| UniRef100\_A0A022PDG1 \| 100.0 \| 1.1e-32 \| 2.1e-38 \| 194.0 \| 125 \| (11, 135) \| 136 \| (6, 130) \| 190 \| Phage protein U \| Phage protein U; uniclust \| UniRef100\_A0A024L4N8 \| 100.0 \| 1.3e-32 \| 2.5e-38 \| 193.4 \| 128 \| (9, 136) \| 136 \| (38, 165) \| 206 \| Bacteriophage tail protein GpU \| Bacteriophage tail protein GpU; uniclust \| UniRef100\_A0A017HDE4 \| 99.9 \| 2.3e-32 \| 4.9e-38 \| 197.0 \| 131 \| (6, 136) \| 136 \| (27, 157) \| 194 \| Phage P2 GpU \| Phage P2 GpU; uniclust \| UniRef100\_A0A064AE81 \| 99.9 \| 3e-32 \| 6.2e-38 \| 190.2 \| 127 \| (9, 136) \| 136 \| (22, 154) \| 160 \| Phage tail protein \| Phage tail protein; uniclust \| UniRef100\_A0A4R3LHW8 \| 99.9 \| 7.4e-32 \| 1.4e-37 \| 199.4 \| 128 \| (9, 136) \| 136 \| (1, 128) \| 288 \| Phage protein U \| Phage protein U; uniclust \| UniRef100\_A0A137SQ39 \| 99.9 \| 2.2e-31 \| 4.4e-37 \| 186.3 \| 126 \| (10, 136) \| 136 \| (19, 150) \| 165 \| Phage P2 GpU \| Phage P2 GpU; uniclust \| UniRef100\_A0A134C9G6 \| 99.9 \| 2.4e-31 \| 4.9e-37 \| 185.2 \| 127 \| (9, 136) \| 136 \| (1, 133) \| 165 \| Phage tail protein (Fragment) \| Phage tail protein (Fragment); uniclust \| UniRef100\_A0A011VGA6 \| 99.9 \| 3.5e-31 \| 7.5e-37 \| 189.5 \| 131 \| (6, 136) \| 136 \| (17, 147) \| 183 \| Tail protein \| Tail protein; uniclust \| UniRef100\_A0A120KRM4 \| 99.9 \| 4.8e-31 \| 1e-36 \| 189.5 \| 125 \| (10, 135) \| 136 \| (46, 176) \| 197 \| Phage tail protein \| Phage tail protein; uniclust \| UniRef100\_A0A0G0CA59 \| 99.9 \| 1.1e-30 \| 2.3e-36 \| 185.2 \| 125 \| (11, 136) \| 136 \| (1, 125) \| 176 \| Phage protein U \| Phage protein U; uniclust \| UniRef100\_A0A348FYG2 \| 99.9 \| 3.9e-30 \| 7.7e-36 \| 179.7 \| 130 \| (7, 136) \| 136 \| (21, 150) \| 176 \| Oxidoreductase \| Oxidoreductase; uniclust \| UniRef100\_A0A022PKF1 \| 99.9 \| 5.6e-30 \| 1.2e-35 \| 197.9 \| 127 \| (9, 136) \| 136 \| (19, 145) \| 322 \| Phage protein U \| Phage protein U; uniclust \| UniRef100\_A0A098F6K3 \| 99.9 \| 7.3e-30 \| 1.5e-35 \| 186.9 \| 129 \| (7, 136) \| 136 \| (3, 137) \| 239 \| SH3 domain-containing protein \| SH3 domain-containing protein; uniclust \| UniRef100\_A0A0H3HPD4 \| 99.9 \| 1.5e-29 \| 2.9e-35 \| 176.7 \| 126 \| (11, 136) \| 136 \| (20, 145) \| 189 \| Phage-related tail protein \| Phage-related tail protein; uniclust \| UniRef100\_A0A1C0A2B2 \| 99.9 \| 1.8e-29 \| 3.6e-35 \| 186.3 \| 125 \| (11, 136) \| 136 \| (26, 150) \| 247 \| Phage tail protein \| Phage tail protein; uniclust \| UniRef100\_A0A1X3HX14 \| 99.9 \| 3.1e-29 \| 6.1e-35 \| 167.8 \| 124 \| (12, 135) \| 136 \| (5, 128) \| 131 \| Putative bacteriophage tail fiber protein U \| Putative bacteriophage tail fiber protein U; uniclust \| UniRef100\_A0A1T4WWY7 \| 99.9 \| 2.1e-28 \| 4.1e-34 \| 169.6 \| 128 \| (9, 136) \| 136 \| (26, 156) \| 169 \| Phage P2 GpU \| Phage P2 GpU; uniclust \| UniRef100\_A0A8J7IC90 \| 99.9 \| 2.4e-28 \| 4.7e-34 \| 175.1 \| 129 \| (7, 135) \| 136 \| (33, 161) \| 222 \| Phage tail protein \| Phage tail protein; uniclust \| UniRef100\_A0A4P5VNR5 \| 99.9 \| 9.2e-28 \| 1.8e-33 \| 178.9 \| 127 \| (10, 136) \| 136 \| (1, 127) \| 308 \| Phage tail protein \| Phage tail protein; uniclust \| UniRef100\_UPI0013D8B061 \| 99.9 \| 1.1e-27 \| 2e-33 \| 158.3 \| 117 \| (20, 136) \| 136 \| (17, 133) \| 139 \| phage tail protein \| phage tail protein; uniclust \| UniRef100\_A0A0E3BS28 \| 99.9 \| 1.4e-27 \| 2.9e-33 \| 182.9 \| 127 \| (9, 136) \| 136 \| (19, 145) \| 327 \| Phage P2 GpU family protein \| Phage P2 GpU family protein; uniclust \| UniRef100\_A0A3A9CQH5 \| 99.9 \| 2e-27 \| 3.9e-33 \| 165.8 \| 126 \| (10, 136) \| 136 \| (41, 172) \| 172 \| Phage tail protein \| Phage tail protein; uniclust \| UniRef100\_A0A104N430 \| 99.9 \| 4.1e-27 \| 7.9e-33 \| 169.4 \| 123 \| (13, 135) \| 136 \| (1, 124) \| 236 \| Phage tail protein \| Phage tail protein; uniclust \| UniRef100\_UPI0018EE9540 \| 99.9 \| 4.7e-27 \| 8.6e-33 \| 174.3 \| 123 \| (14, 136) \| 136 \| (1, 123) \| 363 \| phage tail protein \| phage tail protein; uniclust \| UniRef100\_A0A0D2JH39 \| 99.9 \| 7e-27 \| 1.5e-32 \| 160.7 \| 127 \| (8, 136) \| 136 \| (1, 127) \| 142 \| Uncharacterized protein \| Uncharacterized protein; uniclust \| UniRef100\_A0A0A8H9Q9 \| 99.9 \| 1e-26 \| 2.1e-32 \| 157.7 \| 122 \| (12, 135) \| 136 \| (1, 122) \| 134 \| Phage P2 GpU family protein \| Phage P2 GpU family protein; uniclust \| UniRef100\_A0A0A3TKD9 \| 99.9 \| 1.1e-26 \| 2.2e-32 \| 159.5 \| 126 \| (10, 136) \| 136 \| (1, 130) \| 157 \| Phage protein U \| Phage protein U; uniclust \| UniRef100\_A0A072R0Y2 \| 99.9 \| 3e-26 \| 6.1e-32 \| 160.3 \| 126 \| (10, 135) \| 136 \| (1, 132) \| 168 \| Uncharacterized protein \| Uncharacterized protein; uniclust \| UniRef100\_A0A143HCV2 \| 99.9 \| 4.4e-26 \| 8.7e-32 \| 167.9 \| 127 \| (10, 136) \| 136 \| (12, 143) \| 260 \| SH3b domain-containing protein \| SH3b domain-containing protein; uniclust \| UniRef100\_UPI002227D0FF \| 99.9 \| 9.2e-26 \| 1.7e-31 \| 156.0 \| 125 \| (11, 135) \| 136 \| (2, 126) \| 199 \| phage tail protein \| phage tail protein; uniclust \| UniRef100\_B6IMG5 \| 99.9 \| 9.8e-26 \| 1.9e-31 \| 155.5 \| 123 \| (13, 136) \| 136 \| (1, 123) \| 162 \| Phage P2 GpU \| Phage P2 GpU; uniclust \| UniRef100\_A0A2E1R8X5 \| 99.9 \| 1.9e-25 \| 3.4e-31 \| 151.8 \| 132 \| (5, 136) \| 136 \| (2, 133) \| 174 \| Phage tail protein \| Phage tail protein; uniclust \| UniRef100\_A0A024E920 \| 99.9 \| 3.9e-25 \| 8.1e-31 \| 171.3 \| 118 \| (19, 136) \| 136 \| (214, 331) \| 339 \| Phage tail protein \| Phage tail protein; uniclust \| UniRef100\_N9WEM7 \| 99.8 \| 5.6e-25 \| 1.1e-30 \| 152.4 \| 123 \| (13, 135) \| 136 \| (62, 184) \| 186 \| Phage P2 GpU protein \| Phage P2 GpU protein; uniclust \| UniRef100\_A0A0M1I1F2 \| 99.8 \| 1.1e-24 \| 2.1e-30 \| 146.1 \| 128 \| (9, 136) \| 136 \| (7, 134) \| 153 \| Phage tail protein \| Phage tail protein; uniclust \| UniRef100\_UPI001F442EBF \| 99.8 \| 1.9e-24 \| 3.4e-30 \| 153.9 \| 126 \| (11, 136) \| 136 \| (118, 243) \| 246 \| phage tail protein \| phage tail protein; uniclust \| UniRef100\_A0A1M7R7F9 \| 99.8 \| 5.3e-24 \| 1.1e-29 \| 161.0 \| 124 \| (12, 136) \| 136 \| (1, 124) \| 284 \| Phage protein U \| Phage protein U; uniclust \| UniRef100\_I7LIF3 \| 99.8 \| 6.5e-24 \| 1.2e-29 \| 145.5 \| 122 \| (13, 135) \| 136 \| (1, 126) \| 183 \| Phage protein U \| Phage protein U; uniclust \| UniRef100\_A0A1H5DBV8 \| 99.8 \| 1.1e-23 \| 2.2e-29 \| 151.2 \| 122 \| (13, 135) \| 136 \| (1, 122) \| 213 \| Phage protein U \| Phage protein U; uniclust \| UniRef100\_A0A239C7Y7 \| 99.8 \| 1.2e-23 \| 2.4e-29 \| 160.0 \| 125 \| (11, 136) \| 136 \| (172, 300) \| 300 \| Phage P2 GpU \| Phage P2 GpU; uniclust \| UniRef100\_A0A7Z0SJA2 \| 99.8 \| 2e-23 \| 3.8e-29 \| 153.4 \| 127 \| (10, 136) \| 136 \| (109, 235) \| 273 \| Phage tail protein \| Phage tail protein; uniclust \| UniRef100\_UPI001D0D63E4 \| 99.8 \| 2.5e-23 \| 4.6e-29 \| 141.9 \| 116 \| (20, 135) \| 136 \| (2, 117) \| 176 \| phage tail protein \| phage tail protein; uniclust \| UniRef100\_A0A4P9VIK5 \| 99.8 \| 2.6e-23 \| 4.7e-29 \| 142.0 \| 119 \| (18, 136) \| 136 \| (23, 142) \| 172 \| Phage tail protein \| Phage tail protein; uniclust \| UniRef100\_A0A1H1G2U6 \| 99.8 \| 3e-23 \| 5.9e-29 \| 139.7 \| 126 \| (10, 135) \| 136 \| (1, 127) \| 136 \| Phage tail protein \| Phage tail protein; uniclust \| UniRef100\_A0A0S2F794 \| 99.8 \| 3.2e-23 \| 6.3e-29 \| 137.2 \| 92 \| (44, 135) \| 136 \| (2, 93) \| 120 \| Phage P2 GpU family protein \| Phage P2 GpU family protein; uniclust \| UniRef100\_A0A0T7DUZ2 \| 99.8 \| 3.1e-23 \| 6.4e-29 \| 144.8 \| 132 \| (2, 135) \| 136 \| (17, 148) \| 156 \| Phage tail protein \| Phage tail protein; uniclust \| UniRef100\_A0A2I0GV19 \| 99.8 \| 3.5e-23 \| 6.8e-29 \| 134.5 \| 89 \| (48, 136) \| 136 \| (1, 89) \| 105 \| Phage tail protein \| Phage tail protein; uniclust \| UniRef100\_A0A522WZG7 \| 99.8 \| 3.7e-23 \| 7.2e-29 \| 133.0 \| 100 \| (9, 108) \| 136 \| (2, 102) \| 104 \| Phage tail protein (Fragment) \| Phage tail protein (Fragment); uniclust \| UniRef100\_A0A0C1G7H4 \| 99.8 \| 6e-23 \| 1.2e-28 \| 145.1 \| 126 \| (11, 136) \| 136 \| (1, 128) \| 164 \| Tail protein \| Tail protein; uniclust \| UniRef100\_A0A069PK85 \| 99.8 \| 7.2e-23 \| 1.4e-28 \| 153.1 \| 117 \| (20, 136) \| 136 \| (170, 286) \| 292 \| Oxidoreductase \| Oxidoreductase; uniclust \| UniRef100\_A0A1J5BJG5 \| 99.8 \| 7.4e-23 \| 1.4e-28 \| 147.9 \| 122 \| (14, 136) \| 136 \| (45, 166) \| 218 \| Phage tail protein \| Phage tail protein; uniclust \| UniRef100\_A0A1U7MFM3 \| 99.8 \| 1.1e-22 \| 2.1e-28 \| 150.0 \| 123 \| (12, 135) \| 136 \| (1, 127) \| 286 \| Phage P2 GpU \| Phage P2 GpU; uniclust \| UniRef100\_A0A345DE50 \| 99.8 \| 1.2e-22 \| 2.2e-28 \| 139.4 \| 126 \| (11, 136) \| 136 \| (36, 161) \| 182 \| Phage tail protein \| Phage tail protein; uniclust \| UniRef100\_A0A369RKP4 \| 99.8 \| 1.7e-22 \| 3.2e-28 \| 134.1 \| 114 \| (14, 136) \| 136 \| (1, 114) \| 138 \| Phage tail protein GpU \| Phage tail protein GpU; uniclust \| UniRef100\_UPI00227ED58D \| 99.8 \| 1.8e-22 \| 3.4e-28 \| 144.6 \| 126 \| (10, 135) \| 136 \| (2, 127) \| 254 \| phage tail protein \| phage tail protein; uniclust \| UniRef100\_A0A9D5ZW77 \| 99.8 \| 2.6e-22 \| 4.9e-28 \| 152.2 \| 122 \| (13, 135) \| 136 \| (1, 122) \| 361 \| Phage tail protein \| Phage tail protein; uniclust \| UniRef100\_A0A9D8DA11 \| 99.8 \| 2.9e-22 \| 5.5e-28 \| 124.4 \| 83 \| (9, 91) \| 136 \| (1, 83) \| 83 \| Phage tail protein (Fragment) \| Phage tail protein (Fragment); uniclust \| UniRef100\_A0A4Q4GV95 \| 99.8 \| 3e-22 \| 5.5e-28 \| 135.0 \| 122 \| (13, 134) \| 136 \| (1, 123) \| 149 \| Oxidoreductase \| Oxidoreductase; uniclust \| UniRef100\_UPI00027C90E7 \| 99.8 \| 4.2e-22 \| 7.9e-28 \| 157.6 \| 123 \| (13, 135) \| 136 \| (1, 123) \| 563 \| phage tail protein \| phage tail protein; uniclust \| UniRef100\_A0A0N4UTZ6 \| 99.8 \| 9.4e-22 \| 1.7e-27 \| 145.3 \| 123 \| (13, 135) \| 136 \| (1, 123) \| 326 \| Phage tail protein \| Phage tail protein; uniclust \| UniRef100\_A0A7X3TZ69 \| 99.8 \| 9.8e-22 \| 1.9e-27 \| 147.5 \| 127 \| (10, 136) \| 136 \| (31, 158) \| 327 \| Phage tail protein \| Phage tail protein; uniclust \| UniRef100\_A0A379YFH8 \| 99.8 \| 1.1e-21 \| 2.1e-27 \| 140.8 \| 125 \| (12, 136) \| 136 \| (25, 166) \| 207 \| Phage protein U \| Phage protein U; uniclust \| UniRef100\_A0A2D8Q6P1 \| 99.8 \| 1.3e-21 \| 2.4e-27 \| 148.4 \| 126 \| (11, 136) \| 136 \| (282, 407) \| 416 \| Phage tail protein \| Phage tail protein; uniclust \| UniRef100\_A0A7K0GNT9 \| 99.8 \| 1.3e-21 \| 2.5e-27 \| 133.5 \| 127 \| (10, 136) \| 136 \| (37, 163) \| 173 \| Phage tail protein \| Phage tail protein; uniclust \| UniRef100\_A0A430HF79 \| 99.7 \| 1.6e-21 \| 3e-27 \| 142.9 \| 125 \| (11, 135) \| 136 \| (2, 126) \| 302 \| Phage tail protein \| Phage tail protein; uniclust \| UniRef100\_A0A8D4LNE0 \| 99.7 \| 2.1e-21 \| 3.9e-27 \| 135.9 \| 124 \| (12, 135) \| 136 \| (5, 128) \| 208 \| Phage tail protein \| Phage tail protein; uniclust \| UniRef100\_A0A0H3GS01 \| 99.7 \| 2.2e-21 \| 4.3e-27 \| 130.8 \| 84 \| (53, 136) \| 136 \| (1, 84) \| 134 \| Putative prophage tail protein \| Putative prophage tail protein; uniclust \| UniRef100\_A0A2M7G5P3 \| 99.7 \| 2.7e-21 \| 5.5e-27 \| 147.8 \| 125 \| (11, 136) \| 136 \| (1, 125) \| 306 \| Phage tail protein \| Phage tail protein; uniclust \| UniRef100\_A0A495YAP1 \| 99.7 \| 4e-21 \| 7.8e-27 \| 134.2 \| 118 \| (19, 136) \| 136 \| (45, 162) \| 170 \| Oxidoreductase \| Oxidoreductase; uniclust \| UniRef100\_A0A1V3RQN1 \| 99.7 \| 5.3e-21 \| 1e-26 \| 139.0 \| 126 \| (9, 136) \| 136 \| (54, 179) \| 233 \| Phage tail protein \| Phage tail protein; uniclust \| UniRef100\_A0A1Q6KUC1 \| 99.7 \| 5.8e-21 \| 1.1e-26 \| 138.8 \| 127 \| (9, 136) \| 136 \| (14, 150) \| 228 \| LysM domain-containing protein \| LysM domain-containing protein; uniclust \| UniRef100\_UPI0011303F02 \| 99.7 \| 7.3e-21 \| 1.4e-26 \| 125.1 \| 118 \| (13, 131) \| 136 \| (1, 118) \| 120 \| phage tail protein \| phage tail protein; uniclust \| UniRef100\_A0A075KG62 \| 99.7 \| 7.3e-21 \| 1.4e-26 \| 134.4 \| 109 \| (28, 136) \| 136 \| (70, 182) \| 189 \| p2 GpU family protein \| p2 GpU family protein; uniclust \| UniRef100\_A0A4Q6D3J7 \| 99.7 \| 8.4e-21 \| 1.6e-26 \| 121.4 \| 85 \| (52, 136) \| 136 \| (2, 86) \| 99 \| Phage tail protein (Fragment) \| Phage tail protein (Fragment); uniclust \| UniRef100\_A0A0Q8AZW1 \| 99.7 \| 9e-21 \| 1.7e-26 \| 127.6 \| 124 \| (12, 135) \| 136 \| (1, 126) \| 141 \| Phage tail protein \| Phage tail protein; uniclust \| UniRef100\_A0A0F4NJK3 \| 99.7 \| 1.4e-20 \| 2.9e-26 \| 131.4 \| 122 \| (11, 136) \| 136 \| (10, 131) \| 150 \| Tail protein \| Tail protein; uniclust \| UniRef100\_A0A0E4G101 \| 99.7 \| 2.5e-20 \| 4.8e-26 \| 130.0 \| 118 \| (19, 136) \| 136 \| (19, 150) \| 171 \| Phage tail protein \| Phage tail protein; uniclust \| UniRef100\_A0A853I3C0 \| 99.7 \| 2.7e-20 \| 5e-26 \| 132.0 \| 117 \| (20, 136) \| 136 \| (25, 142) \| 227 \| Phage tail protein \| Phage tail protein; uniclust \| UniRef100\_A0A011NCS8 \| 99.7 \| 2.8e-20 \| 5.8e-26 \| 143.1 \| 124 \| (11, 135) \| 136 \| (1, 124) \| 313 \| Phage tail protein \| Phage tail protein; uniclust \| UniRef100\_A0A256CB29 \| 99.7 \| 3.1e-20 \| 5.9e-26 \| 125.1 \| 124 \| (12, 135) \| 136 \| (2, 125) \| 144 \| Oxidoreductase \| Oxidoreductase; uniclust \| UniRef100\_UPI0013EEEE0C \| 99.7 \| 3.4e-20 \| 6.2e-26 \| 138.3 \| 124 \| (13, 136) \| 136 \| (1, 124) \| 347 \| phage tail protein \| phage tail protein; uniclust \| UniRef100\_A0A940VSA1 \| 99.7 \| 4.2e-20 \| 7.7e-26 \| 126.4 \| 124 \| (12, 136) \| 136 \| (1, 130) \| 173 \| Uncharacterized protein \| Uncharacterized protein; uniclust \| UniRef100\_A0A0J8DFL5 \| 99.7 \| 6.1e-20 \| 1.3e-25 \| 135.3 \| 128 \| (8, 136) \| 136 \| (11, 144) \| 212 \| Uncharacterized protein \| Uncharacterized protein; uniclust \| UniRef100\_A0A4P9VGW4 \| 99.7 \| 8.3e-20 \| 1.5e-25 \| 135.1 \| 126 \| (11, 136) \| 136 \| (184, 309) \| 320 \| Phage tail protein \| Phage tail protein; uniclust \| UniRef100\_A0A1Q6LL69 \| 99.7 \| 8.4e-20 \| 1.6e-25 \| 133.2 \| 124 \| (11, 135) \| 136 \| (1, 134) \| 232 \| LysM domain-containing protein \| LysM domain-containing protein; uniclust \| UniRef100\_UPI00135C3713 \| 99.7 \| 1.3e-19 \| 2.5e-25 \| 129.6 \| 128 \| (9, 136) \| 136 \| (22, 166) \| 205 \| phage tail protein \| phage tail protein; uniclust \| UniRef100\_A0A2H4J1T3 \| 99.7 \| 1.5e-19 \| 3e-25 \| 119.5 \| 103 \| (34, 136) \| 136 \| (3, 105) \| 116 \| Tail protein \| Tail protein; uniclust \| UniRef100\_A0A8S0FTG4 \| 99.7 \| 2e-19 \| 3.9e-25 \| 117.6 \| 101 \| (36, 136) \| 136 \| (2, 104) \| 110 \| Phage tail protein \| Phage tail protein; uniclust \| UniRef100\_A0A2W5VBP0 \| 99.7 \| 2.4e-19 \| 4.3e-25 \| 122.4 \| 123 \| (13, 135) \| 136 \| (3, 125) \| 168 \| Oxidoreductase \| Oxidoreductase; uniclust \| UniRef100\_A0A8S5U2I9 \| 99.6 \| 2.4e-19 \| 4.5e-25 \| 126.9 \| 124 \| (12, 136) \| 136 \| (19, 149) \| 221 \| Uncharacterized protein \| Uncharacterized protein; uniclust \| UniRef100\_A0A0H5AI46 \| 99.6 \| 2.6e-19 \| 4.9e-25 \| 132.0 \| 118 \| (19, 136) \| 136 \| (163, 280) \| 286 \| Phage tail protein \| Phage tail protein; uniclust \| UniRef100\_A0A8J3E9E2 \| 99.6 \| 3e-19 \| 5.7e-25 \| 121.2 \| 121 \| (10, 136) \| 136 \| (1, 122) \| 140 \| Phage tail protein \| Phage tail protein; uniclust \| UniRef100\_A0A064AGR8 \| 99.6 \| 3.6e-19 \| 7.3e-25 \| 131.1 \| 123 \| (12, 135) \| 136 \| (30, 157) \| 219 \| Phage tail protein \| Phage tail protein; uniclust \| UniRef100\_A0A926ZYZ2 \| 99.6 \| 3.9e-19 \| 7.4e-25 \| 122.1 \| 124 \| (11, 135) \| 136 \| (16, 139) \| 159 \| Phage tail protein \| Phage tail protein; uniclust \| UniRef100\_A0A4Y6UBW6 \| 99.6 \| 4.5e-19 \| 8.2e-25 \| 121.9 \| 126 \| (10, 136) \| 136 \| (22, 147) \| 176 \| Phage tail protein \| Phage tail protein; uniclust \| UniRef100\_UPI00217550BA \| 99.6 \| 4.8e-19 \| 8.8e-25 \| 114.1 \| 88 \| (13, 100) \| 136 \| (1, 88) \| 114 \| phage tail protein \| phage tail protein; uniclust \| UniRef100\_A0A4P7KXL2 \| 99.6 \| 6.4e-19 \| 1.2e-24 \| 119.7 \| 122 \| (13, 135) \| 136 \| (1, 122) \| 162 \| Phage P2 GpU \| Phage P2 GpU; uniclust \| UniRef100\_A0A5T7Y1R6 \| 99.6 \| 9.7e-19 \| 1.8e-24 \| 121.9 \| 129 \| (8, 136) \| 136 \| (58, 188) \| 194 \| Phage tail protein \| Phage tail protein; uniclust \| UniRef100\_A0A0S3TZR2 \| 99.6 \| 9.3e-19 \| 1.9e-24 \| 129.7 \| 108 \| (28, 136) \| 136 \| (66, 190) \| 220 \| Contractile injection system tube protein N-terminal domain-containing protein \| Contractile injection system tube protein N-terminal domain-containing protein; uniclust \| UniRef100\_A0A0X8GMF9 \| 99.6 \| 9.9e-19 \| 1.9e-24 \| 132.4 \| 116 \| (20, 136) \| 136 \| (183, 298) \| 304 \| Phage tail protein \| Phage tail protein; uniclust \| UniRef100\_A0A6L9ABV2 \| 99.6 \| 1.3e-18 \| 2.4e-24 \| 110.5 \| 95 \| (13, 108) \| 136 \| (1, 95) \| 95 \| Phage tail protein (Fragment) \| Phage tail protein (Fragment); uniclust \| UniRef100\_A0A0E1UDG7 \| 99.6 \| 1.6e-18 \| 2.9e-24 \| 129.5 \| 118 \| (19, 136) \| 136 \| (173, 290) \| 296 \| Phage P2 GpU family protein \| Phage P2 GpU family protein; uniclust \| UniRef100\_A0A2X0VB68 \| 99.6 \| 1.6e-18 \| 3e-24 \| 118.5 \| 127 \| (9, 136) \| 136 \| (1, 135) \| 152 \| Phage protein U \| Phage protein U; uniclust \| UniRef100\_A0A095WG46 \| 99.6 \| 1.5e-18 \| 3.1e-24 \| 123.8 \| 124 \| (12, 136) \| 136 \| (1, 132) \| 174 \| Uncharacterized protein \| Uncharacterized protein; uniclust \| UniRef100\_A0A2D5PEI7 \| 99.6 \| 1.9e-18 \| 3.7e-24 \| 114.7 \| 123 \| (11, 136) \| 136 \| (1, 124) \| 125 \| Phage tail protein \| Phage tail protein; uniclust \| UniRef100\_R5BE61 \| 99.6 \| 2.7e-18 \| 4.9e-24 \| 119.5 \| 123 \| (12, 135) \| 136 \| (36, 164) \| 191 \| p2 GpU family protein \| p2 GpU family protein; uniclust \| UniRef100\_A0A377DJJ0 \| 99.6 \| 2.5e-18 \| 4.9e-24 \| 117.6 \| 90 \| (47, 136) \| 136 \| (15, 104) \| 142 \| Tail fiber protein \| Tail fiber protein; uniclust \| UniRef100\_A0A0S4RVH7 \| 99.6 \| 4.1e-18 \| 8e-24 \| 116.4 \| 123 \| (11, 135) \| 136 \| (12, 134) \| 138 \| Phage tail protein \| Phage tail protein; uniclust \| UniRef100\_A0A014P2G8 \| 99.6 \| 5.1e-18 \| 1.1e-23 \| 130.0 \| 107 \| (28, 135) \| 136 \| (63, 183) \| 266 \| Peptidoglycan-binding protein \| Peptidoglycan-binding protein; uniclust \| UniRef100\_A0A524RVY3 \| 99.6 \| 5.9e-18 \| 1.1e-23 \| 122.5 \| 125 \| (12, 136) \| 136 \| (3, 127) \| 257 \| Phage tail protein \| Phage tail protein; uniclust \| UniRef100\_A0A7V8U724 \| 99.6 \| 6.1e-18 \| 1.1e-23 \| 116.8 \| 124 \| (12, 135) \| 136 \| (55, 178) \| 179 \| Phage tail protein \| Phage tail protein; uniclust \| UniRef100\_A0A075WU95 \| 99.6 \| 9.4e-18 \| 1.8e-23 \| 118.1 \| 121 \| (12, 135) \| 136 \| (3, 124) \| 175 \| Phage tail protein \| Phage tail protein; uniclust \| UniRef100\_A0A162QMP6 \| 99.6 \| 1e-17 \| 1.9e-23 \| 121.1 \| 122 \| (13, 135) \| 136 \| (1, 128) \| 254 \| Putative peptidoglycan binding domain protein \| Putative peptidoglycan binding domain protein; uniclust \| UniRef100\_UPI001F3F7F85 \| 99.5 \| 1.3e-17 \| 2.4e-23 \| 126.4 \| 122 \| (13, 135) \| 136 \| (1, 122) \| 383 \| phage tail protein \| phage tail protein; uniclust \| UniRef100\_A0A022MNT4 \| 99.5 \| 1.1e-17 \| 2.5e-23 \| 131.4 \| 108 \| (27, 135) \| 136 \| (86, 202) \| 302 \| Peptidase M23B \| Peptidase M23B; uniclust \| UniRef100\_A0A6J4N4J5 \| 99.5 \| 1.6e-17 \| 2.9e-23 \| 114.6 \| 124 \| (11, 135) \| 136 \| (1, 124) \| 176 \| Phage protein U \| Phage protein U; uniclust \| UniRef100\_A0A0N1FE70 \| 99.5 \| 1.7e-17 \| 3.3e-23 \| 113.0 \| 127 \| (9, 135) \| 136 \| (1, 132) \| 142 \| Phage tail protein \| Phage tail protein; uniclust \| UniRef100\_UPI0000380ED2 \| 99.5 \| 1.8e-17 \| 3.4e-23 \| 108.7 \| 105 \| (3, 107) \| 136 \| (18, 122) \| 126 \| phage tail protein \| phage tail protein; uniclust \| UniRef100\_UPI00197DB023 \| 99.5 \| 2.3e-17 \| 4.2e-23 \| 112.8 \| 124 \| (11, 136) \| 136 \| (36, 165) \| 165 \| phage tail protein \| phage tail protein; uniclust \| UniRef100\_A0A0Q0VG01 \| 99.5 \| 2.2e-17 \| 4.3e-23 \| 124.6 \| 123 \| (13, 136) \| 136 \| (1, 124) \| 284 \| Phage tail protein \| Phage tail protein; uniclust \| UniRef100\_A0A085AFM6 \| 99.5 \| 2.3e-17 \| 4.5e-23 \| 117.4 \| 123 \| (12, 136) \| 136 \| (24, 147) \| 184 \| Uncharacterized protein \| Uncharacterized protein; uniclust \| UniRef100\_A0A6L9HIG7 \| 99.5 \| 2.7e-17 \| 4.9e-23 \| 113.5 \| 123 \| (14, 136) \| 136 \| (48, 176) \| 176 \| Phage tail protein \| Phage tail protein; uniclust \| UniRef100\_A0A068T9F9 \| 99.5 \| 3.1e-17 \| 6.3e-23 \| 117.3 \| 131 \| (3, 134) \| 136 \| (9, 144) \| 168 \| Phage tail protein \| Phage tail protein; uniclust \| UniRef100\_A0A285NP30 \| 99.5 \| 3.3e-17 \| 6.4e-23 \| 111.1 \| 124 \| (10, 135) \| 136 \| (3, 127) \| 134 \| Phage protein U \| Phage protein U; uniclust \| UniRef100\_A0A4S2HDA3 \| 99.5 \| 3.7e-17 \| 6.7e-23 \| 112.3 \| 124 \| (12, 136) \| 136 \| (40, 168) \| 170 \| Phage tail protein \| Phage tail protein; uniclust \| UniRef100\_A0A1R1MK76 \| 99.5 \| 3.8e-17 \| 7.1e-23 \| 107.0 \| 123 \| (13, 136) \| 136 \| (1, 124) \| 124 \| Phage tail protein \| Phage tail protein; uniclust \| UniRef100\_A0A179CYQ5 \| 99.5 \| 3.8e-17 \| 7.6e-23 \| 125.4 \| 122 \| (13, 136) \| 136 \| (1, 122) \| 316 \| Tail protein \| Tail protein; uniclust \| UniRef100\_UPI000427E38B \| 99.5 \| 4.5e-17 \| 8.5e-23 \| 111.2 \| 123 \| (12, 136) \| 136 \| (2, 124) \| 148 \| phage tail protein \| phage tail protein; uniclust \| UniRef100\_UPI000764B768 \| 99.5 \| 6e-17 \| 1.1e-22 \| 106.6 \| 123 \| (12, 135) \| 136 \| (3, 125) \| 127 \| phage tail protein \| phage tail protein; uniclust \| UniRef100\_A0A951W1P6 \| 99.5 \| 9.2e-17 \| 1.7e-22 \| 109.1 \| 123 \| (13, 136) \| 136 \| (2, 125) \| 157 \| Phage tail protein \| Phage tail protein; uniclust \| UniRef100\_UPI001ABB63DB \| 99.5 \| 1.1e-16 \| 2e-22 \| 108.5 \| 124 \| (11, 134) \| 136 \| (2, 131) \| 148 \| phage tail protein \| phage tail protein; uniclust \| UniRef100\_UPI00223F0CB2 \| 99.5 \| 1.6e-16 \| 3e-22 \| 113.7 \| 118 \| (19, 136) \| 136 \| (105, 222) \| 228 \| phage tail protein \| phage tail protein; uniclust \| UniRef100\_A0A812RFT8 \| 99.5 \| 1.8e-16 \| 3.4e-22 \| 135.6 \| 108 \| (29, 136) \| 136 \| (746, 853) \| 1893 \| site-specific DNA-methyltransferase (adenine-specific) \| site-specific DNA-methyltransferase (adenine-specific); uniclust \| UniRef100\_A0A7X7NI24 \| 99.4 \| 2.8e-16 \| 5.1e-22 \| 108.7 \| 121 \| (13, 135) \| 136 \| (1, 121) \| 176 \| Phage tail protein \| Phage tail protein; uniclust \| UniRef100\_A0A376J404 \| 99.4 \| 2.6e-16 \| 5.2e-22 \| 100.6 \| 80 \| (12, 91) \| 136 \| (2, 81) \| 85 \| GpU phage protein \| GpU phage protein; uniclust \| UniRef100\_UPI001F36ED2C \| 99.4 \| 3.1e-16 \| 5.7e-22 \| 119.0 \| 124 \| (12, 135) \| 136 \| (8, 143) \| 375 \| phage tail protein \| phage tail protein; uniclust \| UniRef100\_A0A620N5X6 \| 99.4 \| 4e-16 \| 7.3e-22 \| 121.6 \| 122 \| (13, 135) \| 136 \| (1, 122) \| 488 \| Phage protein D \| Phage protein D; uniclust \| UniRef100\_A0A1C5WMN6 \| 99.4 \| 4.1e-16 \| 7.8e-22 \| 114.3 \| 124 \| (12, 136) \| 136 \| (114, 244) \| 244 \| Bacterial SH3 domain \| Bacterial SH3 domain; uniclust \| UniRef100\_A0A3P6JH52 \| 99.4 \| 4.5e-16 \| 8.9e-22 \| 108.7 \| 124 \| (12, 136) \| 136 \| (1, 130) \| 147 \| Phage tail protein \| Phage tail protein; uniclust \| UniRef100\_UPI001A8E6AE5 \| 99.4 \| 4.9e-16 \| 9e-22 \| 118.2 \| 110 \| (27, 136) \| 136 \| (235, 344) \| 383 \| N-acetylmuramidase domain-containing protein \| N-acetylmuramidase domain-containing protein; uniclust \| UniRef100\_A0A1G3UAP7 \| 99.4 \| 8.6e-16 \| 1.7e-21 \| 103.9 \| 120 \| (13, 135) \| 136 \| (2, 121) \| 124 \| Phage tail protein \| Phage tail protein; uniclust \| UniRef100\_A0A318KUR9 \| 99.4 \| 9.9e-16 \| 1.8e-21 \| 113.7 \| 122 \| (13, 135) \| 136 \| (1, 122) \| 303 \| Phage protein U \| Phage protein U; uniclust \| UniRef100\_A0A2W5H3R5 \| 99.4 \| 1.2e-15 \| 2.2e-21 \| 104.5 \| 95 \| (16, 110) \| 136 \| (1, 95) \| 163 \| Phage tail protein \| Phage tail protein; uniclust \| UniRef100\_A0A482IL72 \| 99.4 \| 1.6e-15 \| 2.9e-21 \| 113.0 \| 123 \| (12, 136) \| 136 \| (1, 123) \| 311 \| Phage tail protein \| Phage tail protein; uniclust \| UniRef100\_A0A024YYK6 \| 99.4 \| 1.6e-15 \| 3.6e-21 \| 119.1 \| 109 \| (27, 136) \| 136 \| (58, 184) \| 297 \| Peptidase M23B \| Peptidase M23B; uniclust \| UniRef100\_A0A2W5C6U1 \| 99.4 \| 2.2e-15 \| 4.3e-21 \| 102.3 \| 120 \| (11, 135) \| 136 \| (1, 124) \| 128 \| Phage tail protein \| Phage tail protein; uniclust \| UniRef100\_UPI0005B3BCC5 \| 99.4 \| 2.4e-15 \| 4.4e-21 \| 93.0 \| 82 \| (13, 94) \| 136 \| (1, 82) \| 85 \| phage tail protein \| phage tail protein; uniclust \| UniRef100\_A0A9D1QXB7 \| 99.4 \| 2.4e-15 \| 4.5e-21 \| 112.3 \| 124 \| (11, 135) \| 136 \| (2, 125) \| 317 \| Phage tail protein \| Phage tail protein; uniclust \| UniRef100\_A0A084T293 \| 99.4 \| 2.2e-15 \| 4.5e-21 \| 117.0 \| 109 \| (27, 136) \| 136 \| (50, 193) \| 283 \| LysM domain-containing protein \| LysM domain-containing protein; uniclust \| UniRef100\_A0A9E8EL20 \| 99.4 \| 2.8e-15 \| 5.1e-21 \| 103.3 \| 122 \| (13, 135) \| 136 \| (1, 122) \| 168 \| Phage tail protein \| Phage tail protein; uniclust \| UniRef100\_UPI0005609077 \| 99.4 \| 2.8e-15 \| 5.2e-21 \| 94.8 \| 93 \| (43, 136) \| 136 \| (2, 94) \| 97 \| phage tail protein \| phage tail protein; uniclust \| UniRef100\_E2CFJ9 \| 99.4 \| 3.1e-15 \| 5.9e-21 \| 103.9 \| 123 \| (11, 135) \| 136 \| (1, 123) \| 159 \| Putative phage tail protein \| Putative phage tail protein; uniclust \| UniRef100\_A0A8S5NYM1 \| 99.3 \| 3.5e-15 \| 6.5e-21 \| 103.9 \| 124 \| (11, 135) \| 136 \| (1, 130) \| 182 \| Uncharacterized protein \| Uncharacterized protein; uniclust \| UniRef100\_A0A1I5H9R0 \| 99.3 \| 3.6e-15 \| 6.9e-21 \| 103.6 \| 124 \| (11, 134) \| 136 \| (1, 124) \| 149 \| Phage protein U \| Phage protein U; uniclust \| UniRef100\_A0A2P1VUY8 \| 99.3 \| 4.4e-15 \| 8.3e-21 \| 101.1 \| 122 \| (13, 136) \| 136 \| (2, 124) \| 143 \| Phage protein U \| Phage protein U; uniclust \| UniRef100\_A0A970G5W5 \| 99.3 \| 4.6e-15 \| 8.5e-21 \| 108.8 \| 125 \| (12, 136) \| 136 \| (1, 129) \| 270 \| Phage tail protein \| Phage tail protein; uniclust \| UniRef100\_A0A3D1NSN0 \| 99.3 \| 5.6e-15 \| 1.1e-20 \| 110.6 \| 122 \| (13, 135) \| 136 \| (1, 122) \| 280 \| Phage tail protein \| Phage tail protein; uniclust \| UniRef100\_A0A165W2G2 \| 99.3 \| 7.4e-15 \| 1.5e-20 \| 102.0 \| 124 \| (9, 135) \| 136 \| (4, 128) \| 142 \| Phage tail protein \| Phage tail protein; uniclust \| UniRef100\_UPI002094A02D \| 99.3 \| 8.5e-15 \| 1.6e-20 \| 98.4 \| 90 \| (46, 135) \| 136 \| (1, 90) \| 141 \| phage tail protein \| phage tail protein; uniclust \| UniRef100\_A0A150S1J3 \| 99.3 \| 7.9e-15 \| 1.6e-20 \| 110.0 \| 115 \| (21, 136) \| 136 \| (27, 155) \| 231 \| LysM domain-containing protein \| LysM domain-containing protein; uniclust \| UniRef100\_A0A090ILA7 \| 99.3 \| 8.8e-15 \| 1.7e-20 \| 99.3 \| 120 \| (11, 135) \| 136 \| (1, 121) \| 127 \| Phage U-like protein \| Phage U-like protein; uniclust \| UniRef100\_UPI000CF8F1C8 \| 99.3 \| 1.2e-14 \| 2.2e-20 \| 102.1 \| 123 \| (12, 135) \| 136 \| (59, 190) \| 191 \| phage tail protein \| phage tail protein; uniclust \| UniRef100\_A0A5S4YPE8 \| 99.3 \| 1.3e-14 \| 2.4e-20 \| 107.0 \| 118 \| (19, 136) \| 136 \| (126, 257) \| 279 \| Phage tail protein \| Phage tail protein; uniclust \| UniRef100\_UPI00215D9118 \| 99.3 \| 1.6e-14 \| 2.9e-20 \| 94.8 \| 109 \| (13, 122) \| 136 \| (1, 109) \| 120 \| phage tail protein \| phage tail protein; uniclust \| UniRef100\_A0A1S0V6E3 \| 99.3 \| 1.8e-14 \| 3.3e-20 \| 86.6 \| 56 \| (80, 135) \| 136 \| (1, 56) \| 66 \| Tail protein (Fragment) \| Tail protein (Fragment); uniclust \| UniRef100\_E5VKI8 \| 99.3 \| 2e-14 \| 3.6e-20 \| 99.1 \| 125 \| (11, 136) \| 136 \| (1, 131) \| 166 \| Phage P2 GpU \| Phage P2 GpU; uniclust \| UniRef100\_A0A1C6BMU7 \| 99.3 \| 2.3e-14 \| 4.3e-20 \| 96.5 \| 122 \| (13, 135) \| 136 \| (1, 129) \| 129 \| Phage-related protein \| Phage-related protein; uniclust \| UniRef100\_R7HX73 \| 99.3 \| 2.8e-14 \| 5.1e-20 \| 100.5 \| 116 \| (20, 136) \| 136 \| (71, 192) \| 193 \| Phage protein U \| Phage protein U; uniclust \| UniRef100\_UPI001270A914 \| 99.2 \| 2.9e-14 \| 5.5e-20 \| 82.9 \| 56 \| (13, 68) \| 136 \| (1, 56) \| 57 \| phage tail protein \| phage tail protein; uniclust \| UniRef100\_UPI00068D9337 \| 99.2 \| 4e-14 \| 7.3e-20 \| 105.3 \| 122 \| (13, 135) \| 136 \| (1, 122) \| 295 \| phage tail protein \| phage tail protein; uniclust \| UniRef100\_UPI0005EE32AE \| 99.2 \| 4.8e-14 \| 9e-20 \| 91.1 \| 79 \| (10, 88) \| 136 \| (21, 99) \| 100 \| phage tail protein \| phage tail protein; uniclust \| UniRef100\_A0A423PRP6 \| 99.2 \| 5.4e-14 \| 9.9e-20 \| 91.3 \| 79 \| (11, 89) \| 136 \| (31, 109) \| 111 \| Tail protein (Fragment) \| Tail protein (Fragment); uniclust \| UniRef100\_A0A2Z4RAD7 \| 99.2 \| 5.7e-14 \| 1.1e-19 \| 105.8 \| 122 \| (13, 135) \| 136 \| (1, 122) \| 258 \| Phage protein U \| Phage protein U; uniclust \| UniRef100\_A0A0D6KHC7 \| 99.2 \| 8.2e-14 \| 1.7e-19 \| 106.1 \| 112 \| (22, 135) \| 136 \| (28, 156) \| 238 \| LysM domain protein \| LysM domain protein; uniclust \| UniRef100\_A0A3G2V4J2 \| 99.2 \| 1e-13 \| 1.9e-19 \| 98.4 \| 127 \| (10, 136) \| 136 \| (1, 146) \| 196 \| Phage tail protein \| Phage tail protein; uniclust \| UniRef100\_A0A126S8J7 \| 99.2 \| 1.1e-13 \| 2.2e-19 \| 86.7 \| 67 \| (70, 136) \| 136 \| (3, 69) \| 76 \| Phage protein U \| Phage protein U; uniclust \| UniRef100\_A0A840MJS2 \| 99.2 \| 1.4e-13 \| 2.5e-19 \| 107.4 \| 123 \| (12, 135) \| 136 \| (4, 126) \| 451 \| Phage protein U \| Phage protein U; uniclust \| UniRef100\_A0A1H7YIQ2 \| 99.2 \| 1.5e-13 \| 2.9e-19 \| 99.0 \| 124 \| (12, 136) \| 136 \| (1, 134) \| 180 \| Phage P2 GpU \| Phage P2 GpU; uniclust \| UniRef100\_A0A6C1BSU7 \| 99.2 \| 1.6e-13 \| 3e-19 \| 95.0 \| 121 \| (13, 136) \| 136 \| (2, 122) \| 168 \| Phage tail protein \| Phage tail protein; uniclust \| UniRef100\_A0A7G7ZA20 \| 99.1 \| 2.5e-13 \| 4.6e-19 \| 88.7 \| 80 \| (56, 135) \| 136 \| (11, 90) \| 114 \| Phage tail protein \| Phage tail protein; uniclust \| UniRef100\_A0A2W4LKE1 \| 99.1 \| 2.7e-13 \| 5.2e-19 \| 96.1 \| 109 \| (27, 136) \| 136 \| (23, 142) \| 169 \| Peptidoglycan-binding protein (Fragment) \| Peptidoglycan-binding protein (Fragment); uniclust \| UniRef100\_A0A1T4W4X0 \| 99.1 \| 3e-13 \| 5.5e-19 \| 91.7 \| 126 \| (11, 136) \| 136 \| (1, 133) \| 145 \| Phage P2 GpU \| Phage P2 GpU; uniclust \| UniRef100\_A0A3C1WUQ3 \| 99.1 \| 3.2e-13 \| 5.8e-19 \| 90.8 \| 123 \| (13, 135) \| 136 \| (3, 136) \| 137 \| Phage tail protein \| Phage tail protein; uniclust \| UniRef100\_A0A836CGV5 \| 99.1 \| 3.3e-13 \| 6.1e-19 \| 113.2 \| 107 \| (30, 136) \| 136 \| (703, 809) \| 1085 \| Phage tail tape measure protein domain-containing protein \| Phage tail tape measure protein domain-containing protein; uniclust \| UniRef100\_A0A8J6Y7F4 \| 99.1 \| 3.6e-13 \| 6.5e-19 \| 94.4 \| 122 \| (13, 135) \| 136 \| (18, 161) \| 181 \| Phage tail protein \| Phage tail protein; uniclust \| UniRef100\_A0A062UU40 \| 99.1 \| 3.2e-13 \| 6.6e-19 \| 103.7 \| 103 \| (27, 135) \| 136 \| (49, 161) \| 258 \| Contractile injection system tube protein N-terminal domain-containing protein \| Contractile injection system tube protein N-terminal domain-containing protein; uniclust \| UniRef100\_UPI0021562D4A \| 99.1 \| 4.5e-13 \| 8.3e-19 \| 88.2 \| 86 \| (11, 96) \| 136 \| (27, 113) \| 119 \| phage tail protein \| phage tail protein; uniclust \| UniRef100\_A0A1B6BCU7 \| 99.1 \| 4.2e-13 \| 9e-19 \| 104.3 \| 113 \| (21, 135) \| 136 \| (45, 175) \| 265 \| Contractile injection system tube protein N-terminal domain-containing protein \| Contractile injection system tube protein N-terminal domain-containing protein; uniclust \| UniRef100\_A0A066U3S9 \| 99.1 \| 4.5e-13 \| 9.8e-19 \| 107.1 \| 113 \| (20, 136) \| 136 \| (66, 194) \| 325 \| Peptidase M23B \| Peptidase M23B; uniclust \| UniRef100\_A0A173SAB9 \| 99.1 \| 5.7e-13 \| 1.1e-18 \| 91.6 \| 123 \| (12, 135) \| 136 \| (29, 157) \| 158 \| Phage protein U \| Phage protein U; uniclust \| UniRef100\_UPI000B16A0A2 \| 99.1 \| 9.3e-13 \| 1.7e-18 \| 95.0 \| 122 \| (13, 135) \| 136 \| (2, 123) \| 223 \| phage tail protein \| phage tail protein; uniclust \| UniRef100\_Q602Z4 \| 99.1 \| 1.1e-12 \| 2.1e-18 \| 92.6 \| 122 \| (13, 135) \| 136 \| (2, 123) \| 189 \| Conserved domain protein \| Conserved domain protein; uniclust \| UniRef100\_A0A450YW55 \| 99.0 \| 1.5e-12 \| 2.7e-18 \| 98.2 \| 121 \| (13, 135) \| 136 \| (2, 122) \| 319 \| Phage P2 GpU \| Phage P2 GpU; uniclust \| UniRef100\_A0A3S4YZC4 \| 99.0 \| 1.7e-12 \| 3.1e-18 \| 85.6 \| 104 \| (11, 114) \| 136 \| (2, 111) \| 114 \| Phage protein U \| Phage protein U; uniclust \| UniRef100\_A0A2M8PTG2 \| 99.0 \| 1.6e-12 \| 3.2e-18 \| 96.1 \| 109 \| (27, 136) \| 136 \| (27, 143) \| 221 \| Peptidoglycan-binding protein \| Peptidoglycan-binding protein; uniclust \| UniRef100\_UPI00190F23EE \| 99.0 \| 1.8e-12 \| 3.4e-18 \| 79.2 \| 73 \| (33, 105) \| 136 \| (3, 75) \| 76 \| phage tail protein \| phage tail protein; uniclust \| UniRef100\_A0A7Z0UY93 \| 99.0 \| 2e-12 \| 3.7e-18 \| 86.6 \| 71 \| (65, 135) \| 136 \| (2, 72) \| 131 \| Phage tail protein \| Phage tail protein; uniclust \| UniRef100\_Q31HS8 \| 99.0 \| 2.7e-12 \| 4.9e-18 \| 87.1 \| 124 \| (12, 136) \| 136 \| (5, 139) \| 142 \| Phage P2 GpU family protein \| Phage P2 GpU family protein; uniclust \| UniRef100\_A0A8X6WSB0 \| 99.0 \| 2.8e-12 \| 5.2e-18 \| 96.5 \| 108 \| (28, 135) \| 136 \| (10, 117) \| 309 \| Phage late control gene d protein GpD \| Phage late control gene d protein GpD; uniclust \| UniRef100\_A0A317H5Z8 \| 99.0 \| 3.3e-12 \| 6e-18 \| 90.5 \| 125 \| (11, 135) \| 136 \| (47, 171) \| 192 \| Phage tail protein \| Phage tail protein; uniclust \| UniRef100\_UPI001F273361 \| 99.0 \| 3.4e-12 \| 6.4e-18 \| 86.9 \| 120 \| (14, 136) \| 136 \| (8, 129) \| 131 \| phage tail protein \| phage tail protein; uniclust \| UniRef100\_A0A0D6ARK4 \| 99.0 \| 4e-12 \| 7.3e-18 \| 86.9 \| 124 \| (11, 135) \| 136 \| (1, 127) \| 148 \| Phage protein \| Phage protein; uniclust \| UniRef100\_UPI0015C58154 \| 99.0 \| 4.1e-12 \| 7.5e-18 \| 79.8 \| 84 \| (13, 96) \| 136 \| (3, 86) \| 88 \| phage tail protein \| phage tail protein; uniclust \| UniRef100\_UPI0021794BC2 \| 99.0 \| 4.2e-12 \| 7.7e-18 \| 83.5 \| 88 \| (47, 135) \| 136 \| (2, 90) \| 116 \| phage tail protein \| phage tail protein; uniclust \| UniRef100\_A0A0B1YHP6 \| 98.9 \| 5e-12 \| 1.1e-17 \| 101.2 \| 119 \| (4, 133) \| 136 \| (11, 129) \| 335 \| DNA circulation N-terminal domain-containing protein \| DNA circulation N-terminal domain-containing protein; uniclust \| UniRef100\_UPI0020C15A30 \| 98.9 \| 5.8e-12 \| 1.1e-17 \| 82.4 \| 89 \| (47, 135) \| 136 \| (2, 90) \| 112 \| phage tail protein \| phage tail protein; uniclust \| UniRef100\_UPI00224967BF \| 98.9 \| 6.1e-12 \| 1.1e-17 \| 88.6 \| 107 \| (27, 133) \| 136 \| (54, 160) \| 183 \| phage tail protein \| phage tail protein; uniclust \| UniRef100\_UPI001F07EFB3 \| 98.9 \| 7.1e-12 \| 1.3e-17 \| 103.9 \| 88 \| (48, 135) \| 136 \| (722, 809) \| 847 \| phage tail tape measure protein \| phage tail tape measure protein; uniclust \| UniRef100\_UPI001FCD6EEF \| 98.9 \| 9.2e-12 \| 1.7e-17 \| 80.1 \| 86 \| (9, 94) \| 136 \| (7, 94) \| 99 \| phage tail protein \| phage tail protein; uniclust \| UniRef100\_A0A016QLM3 \| 98.9 \| 1e-11 \| 2e-17 \| 92.3 \| 109 \| (27, 135) \| 136 \| (34, 167) \| 213 \| Peptidoglycan-binding LysM \| Peptidoglycan-binding LysM; uniclust \| UniRef100\_A0A3A4NRP5 \| 98.9 \| 1.4e-11 \| 2.7e-17 \| 93.1 \| 107 \| (28, 135) \| 136 \| (23, 139) \| 292 \| Contractile injection system tube protein N-terminal domain-containing protein \| Contractile injection system tube protein N-terminal domain-containing protein; uniclust \| UniRef100\_A0A014NMA0 \| 98.9 \| 1.3e-11 \| 2.8e-17 \| 98.5 \| 107 \| (28, 135) \| 136 \| (78, 202) \| 294 \| Peptidoglycan-binding protein \| Peptidoglycan-binding protein; uniclust \| UniRef100\_A0A174T297 \| 98.9 \| 1.5e-11 \| 3e-17 \| 91.5 \| 125 \| (9, 136) \| 136 \| (10, 140) \| 195 \| Phage tail protein \| Phage tail protein; uniclust \| UniRef100\_A0A0P6Y0I8 \| 98.9 \| 1.5e-11 \| 3.3e-17 \| 98.8 \| 112 \| (21, 135) \| 136 \| (30, 154) \| 330 \| Contractile injection system tube protein N-terminal domain-containing protein \| Contractile injection system tube protein N-terminal domain-containing protein; uniclust \| UniRef100\_A0A0S4XMR6 \| 98.9 \| 1.8e-11 \| 3.5e-17 \| 82.3 \| 117 \| (13, 135) \| 136 \| (1, 117) \| 119 \| Phage tail protein \| Phage tail protein; uniclust \| UniRef100\_UPI0018835B78 \| 98.9 \| 2e-11 \| 3.7e-17 \| 78.1 \| 60 \| (12, 71) \| 136 \| (27, 86) \| 93 \| phage tail protein \| phage tail protein; uniclust \| UniRef100\_UPI000583C80F \| 98.9 \| 2e-11 \| 3.8e-17 \| 74.4 \| 60 \| (13, 73) \| 136 \| (1, 60) \| 66 \| phage tail protein \| phage tail protein; uniclust \| UniRef100\_A0A0L6JHX4 \| 98.8 \| 2.2e-11 \| 4.5e-17 \| 92.7 \| 110 \| (27, 136) \| 136 \| (52, 176) \| 233 \| Contractile injection system tube protein N-terminal domain-containing protein \| Contractile injection system tube protein N-terminal domain-containing protein; uniclust \| UniRef100\_A0A2N8GSE5 \| 98.8 \| 2.7e-11 \| 5e-17 \| 78.6 \| 76 \| (19, 94) \| 136 \| (22, 97) \| 100 \| Phage tail protein (Fragment) \| Phage tail protein (Fragment); uniclust \| UniRef100\_A0A1C0V8H7 \| 98.8 \| 3.8e-11 \| 7.1e-17 \| 83.5 \| 121 \| (13, 134) \| 136 \| (5, 125) \| 163 \| Phage tail protein \| Phage tail protein; uniclust \| UniRef100\_A0A945U371 \| 98.8 \| 3.8e-11 \| 7.1e-17 \| 85.2 \| 117 \| (18, 136) \| 136 \| (17, 157) \| 168 \| Peptidoglycan-binding protein (Fragment) \| Peptidoglycan-binding protein (Fragment); uniclust \| UniRef100\_A0A842IWC8 \| 98.8 \| 3.9e-11 \| 7.2e-17 \| 86.5 \| 124 \| (12, 136) \| 136 \| (31, 159) \| 200 \| Phage tail protein \| Phage tail protein; uniclust \| UniRef100\_A0A327Q841 \| 98.8 \| 5.7e-11 \| 1e-16 \| 88.8 \| 122 \| (13, 135) \| 136 \| (1, 122) \| 280 \| GpU protein \| GpU protein; uniclust \| UniRef100\_UPI000629E4F5 \| 98.8 \| 6.6e-11 \| 1.2e-16 \| 85.4 \| 63 \| (74, 136) \| 136 \| (113, 175) \| 212 \| phage regulatory CII family protein \| phage regulatory CII family protein; uniclust \| UniRef100\_A0A976C8U3 \| 98.8 \| 6.7e-11 \| 1.2e-16 \| 88.5 \| 123 \| (13, 136) \| 136 \| (1, 123) \| 280 \| Uncharacterized protein \| Uncharacterized protein; uniclust \| UniRef100\_A0A2G2BDF5 \| 98.8 \| 6.7e-11 \| 1.3e-16 \| 76.3 \| 59 \| (78, 136) \| 136 \| (11, 69) \| 92 \| Phage tail protein \| Phage tail protein; uniclust \| UniRef100\_A0A378TUW3 \| 98.8 \| 7.2e-11 \| 1.3e-16 \| 96.1 \| 121 \| (13, 134) \| 136 \| (1, 121) \| 606 \| Putative phage tail fiber protein \| Putative phage tail fiber protein; uniclust \| UniRef100\_A0A8S5UB81 \| 98.7 \| 8.9e-11 \| 1.6e-16 \| 80.4 \| 120 \| (13, 133) \| 136 \| (2, 130) \| 145 \| Uncharacterized protein \| Uncharacterized protein; uniclust \| UniRef100\_A0A0V7ZMG3 \| 98.7 \| 8e-11 \| 1.6e-16 \| 91.1 \| 107 \| (28, 134) \| 136 \| (66, 187) \| 256 \| Contractile injection system tube protein N-terminal domain-containing protein \| Contractile injection system tube protein N-terminal domain-containing protein; uniclust \| UniRef100\_A0A8E3ZHR9 \| 98.7 \| 8.8e-11 \| 1.7e-16 \| 85.1 \| 57 \| (13, 69) \| 136 \| (1, 57) \| 196 \| Late control gene D protein \| Late control gene D protein; uniclust \| UniRef100\_A0A3S4FEQ5 \| 98.7 \| 9e-11 \| 1.7e-16 \| 68.7 \| 56 \| (13, 68) \| 136 \| (1, 56) \| 57 \| Phage tail protein \| Phage tail protein; uniclust \| UniRef100\_A0A2S7JR76 \| 98.7 \| 9.1e-11 \| 1.7e-16 \| 86.6 \| 118 \| (16, 136) \| 136 \| (63, 180) \| 250 \| Phage tail protein \| Phage tail protein; uniclust \| UniRef100\_UPI002364701C \| 98.7 \| 1.1e-10 \| 2e-16 \| 83.3 \| 122 \| (13, 136) \| 136 \| (64, 186) \| 193 \| phage tail protein \| phage tail protein; uniclust \| UniRef100\_A0A011PSV8 \| 98.7 \| 9.4e-11 \| 2e-16 \| 91.2 \| 107 \| (28, 135) \| 136 \| (39, 160) \| 244 \| Contractile injection system tube protein N-terminal domain-containing protein \| Contractile injection system tube protein N-terminal domain-containing protein; uniclust \| UniRef100\_UPI000829C05D \| 98.7 \| 1.2e-10 \| 2.2e-16 \| 83.3 \| 122 \| (12, 135) \| 136 \| (2, 124) \| 196 \| phage tail protein \| phage tail protein; uniclust \| UniRef100\_A0A377K9Z3 \| 98.7 \| 1.3e-10 \| 2.4e-16 \| 88.6 \| 75 \| (45, 119) \| 136 \| (247, 321) \| 328 \| Putative phage tail tape measure protein \| Putative phage tail tape measure protein; uniclust \| UniRef100\_A0A1E2WLU3 \| 98.7 \| 1.3e-10 \| 2.5e-16 \| 88.9 \| 108 \| (28, 136) \| 136 \| (40, 161) \| 259 \| Contractile injection system tube protein N-terminal domain-containing protein \| Contractile injection system tube protein N-terminal domain-containing protein; uniclust \| UniRef100\_A0A0P9CJQ9 \| 98.7 \| 1.7e-10 \| 3.1e-16 \| 74.7 \| 86 \| (50, 135) \| 136 \| (2, 89) \| 101 \| Oxidoreductase (Fragment) \| Oxidoreductase (Fragment); uniclust \| UniRef100\_A0A377I4B1 \| 98.7 \| 2.1e-10 \| 3.9e-16 \| 73.6 \| 84 \| (13, 98) \| 136 \| (1, 84) \| 91 \| Putative bacteriophage V tail protein \| Putative bacteriophage V tail protein; uniclust \| UniRef100\_A0A177M7D1 \| 98.7 \| 1.9e-10 \| 4e-16 \| 95.1 \| 108 \| (27, 135) \| 136 \| (42, 161) \| 428 \| Contractile injection system tube protein N-terminal domain-containing protein \| Contractile injection system tube protein N-terminal domain-containing protein; uniclust \| UniRef100\_A0A0Q6MK11 \| 98.7 \| 1.9e-10 \| 4e-16 \| 90.9 \| 107 \| (28, 135) \| 136 \| (58, 178) \| 288 \| Contractile injection system tube protein N-terminal domain-containing protein \| Contractile injection system tube protein N-terminal domain-containing protein; uniclust \| UniRef100\_B9NM79 \| 98.6 \| 2.6e-10 \| 4.8e-16 \| 72.4 \| 75 \| (62, 136) \| 136 \| (2, 76) \| 90 \| Putative phage tail protein U \| Putative phage tail protein U; uniclust \| UniRef100\_A0A0B6D192 \| 98.6 \| 2.9e-10 \| 5.3e-16 \| 75.6 \| 116 \| (15, 136) \| 136 \| (2, 117) \| 118 \| Phage P2 GpU family protein \| Phage P2 GpU family protein; uniclust \| UniRef100\_A0A1Q7YAK2 \| 98.6 \| 2.6e-10 \| 5.5e-16 \| 95.3 \| 105 \| (27, 135) \| 136 \| (43, 156) \| 447 \| Contractile injection system tube protein N-terminal domain-containing protein \| Contractile injection system tube protein N-terminal domain-containing protein; uniclust \| UniRef100\_A0A7X6FT88 \| 98.6 \| 3.4e-10 \| 6.3e-16 \| 68.1 \| 61 \| (12, 72) \| 136 \| (1, 61) \| 66 \| Uncharacterized protein \| Uncharacterized protein; uniclust \| UniRef100\_UPI00192B53BF \| 98.6 \| 3.6e-10 \| 6.7e-16 \| 71.6 \| 62 \| (75, 136) \| 136 \| (9, 70) \| 88 \| phage tail protein \| phage tail protein; uniclust \| UniRef100\_UPI0021B6119D \| 98.6 \| 3.7e-10 \| 6.8e-16 \| 75.2 \| 115 \| (13, 135) \| 136 \| (2, 116) \| 119 \| phage tail protein \| phage tail protein; uniclust \| UniRef100\_A0A9E6URY0 \| 98.6 \| 3.7e-10 \| 6.8e-16 \| 89.2 \| 115 \| (20, 135) \| 136 \| (263, 377) \| 447 \| Phage tail protein \| Phage tail protein; uniclust \| UniRef100\_A0A1B9SFL0 \| 98.6 \| 3.3e-10 \| 7.2e-16 \| 90.0 \| 116 \| (7, 133) \| 136 \| (16, 131) \| 285 \| Phage tail protein \| Phage tail protein; uniclust \| UniRef100\_A0A0F0HZW4 \| 98.6 \| 3.4e-10 \| 7.3e-16 \| 94.3 \| 107 \| (27, 135) \| 136 \| (38, 157) \| 443 \| Contractile injection system tube protein N-terminal domain-containing protein \| Contractile injection system tube protein N-terminal domain-containing protein; uniclust \| UniRef100\_A0A0H3ZMX6 \| 98.6 \| 4.9e-10 \| 8.9e-16 \| 72.3 \| 72 \| (18, 89) \| 136 \| (17, 88) \| 98 \| Uncharacterized protein \| Uncharacterized protein; uniclust \| UniRef100\_UPI000E5BC930 \| 98.6 \| 5.4e-10 \| 1e-15 \| 72.8 \| 98 \| (13, 112) \| 136 \| (1, 98) \| 98 \| phage tail protein \| phage tail protein; uniclust \| UniRef100\_A0A1H8C8S6 \| 98.6 \| 6.5e-10 \| 1.3e-15 \| 86.8 \| 113 \| (21, 135) \| 136 \| (50, 181) \| 266 \| Contractile injection system tube protein N-terminal domain-containing protein \| Contractile injection system tube protein N-terminal domain-containing protein; uniclust \| UniRef100\_A0A536CDU8 \| 98.6 \| 7.7e-10 \| 1.4e-15 \| 77.1 \| 115 \| (18, 135) \| 136 \| (16, 149) \| 159 \| Contractile injection system tube protein N-terminal domain-containing protein (Fragment) \| Contractile injection system tube protein N-terminal domain-containing protein (Fragment); uniclust \| UniRef100\_A0A0D6P4T5 \| 98.5 \| 7e-10 \| 1.5e-15 \| 87.6 \| 119 \| (7, 132) \| 136 \| (18, 136) \| 269 \| Phage protein \| Phage protein; uniclust \| UniRef100\_A0A2X1L4N2 \| 98.5 \| 8.2e-10 \| 1.5e-15 \| 73.4 \| 69 \| (68, 136) \| 136 \| (1, 69) \| 108 \| Tail fiber protein \| Tail fiber protein; uniclust \| UniRef100\_UPI001F155EF8 \| 98.5 \| 8.5e-10 \| 1.6e-15 \| 77.9 \| 108 \| (20, 130) \| 136 \| (55, 162) \| 174 \| phage tail protein \| phage tail protein; uniclust \| UniRef100\_A0A545SBB2 \| 98.5 \| 8.8e-10 \| 1.6e-15 \| 65.0 \| 58 \| (11, 68) \| 136 \| (2, 59) \| 59 \| Oxidoreductase (Fragment) \| Oxidoreductase (Fragment); uniclust \| UniRef100\_A0A2A2HJB8 \| 98.5 \| 1e-09 \| 1.9e-15 \| 67.1 \| 48 \| (89, 136) \| 136 \| (1, 48) \| 66 \| Phage tail protein (Fragment) \| Phage tail protein (Fragment); uniclust \| UniRef100\_A0A080IDD0 \| 98.5 \| 9.8e-10 \| 1.9e-15 \| 96.6 \| 87 \| (50, 136) \| 136 \| (814, 900) \| 944 \| Phage tail tape measure protein, TP901 family, core region \| Phage tail tape measure protein, TP901 family, core region; uniclust \| UniRef100\_UPI001ABB4451 \| 98.5 \| 1.1e-09 \| 2e-15 \| 65.9 \| 62 \| (11, 72) \| 136 \| (2, 63) \| 65 \| phage tail protein \| phage tail protein; uniclust \| UniRef100\_A0A3E0DS09 \| 98.5 \| 1.1e-09 \| 2e-15 \| 70.9 \| 78 \| (11, 90) \| 136 \| (1, 79) \| 92 \| GpU protein \| GpU protein; uniclust \| UniRef100\_A0A074LNR3 \| 98.5 \| 1e-09 \| 2.1e-15 \| 82.5 \| 121 \| (5, 135) \| 136 \| (6, 126) \| 211 \| LysM domain-containing protein \| LysM domain-containing protein; uniclust \| UniRef100\_A0A661D5Z7 \| 98.5 \| 1.3e-09 \| 2.5e-15 \| 79.6 \| 113 \| (21, 135) \| 136 \| (28, 154) \| 170 \| Contractile injection system tube protein N-terminal domain-containing protein (Fragment) \| Contractile injection system tube protein N-terminal domain-containing protein (Fragment); uniclust \| UniRef100\_A0A022PIZ6 \| 98.5 \| 1.2e-09 \| 2.6e-15 \| 86.4 \| 107 \| (28, 134) \| 136 \| (39, 161) \| 281 \| Contractile injection system tube protein N-terminal domain-containing protein \| Contractile injection system tube protein N-terminal domain-containing protein; uniclust \| UniRef100\_A0A959IW21 \| 98.5 \| 1.5e-09 \| 2.9e-15 \| 76.5 \| 125 \| (8, 133) \| 136 \| (3, 148) \| 149 \| Contractile injection system tube protein N-terminal domain-containing protein (Fragment) \| Contractile injection system tube protein N-terminal domain-containing protein (Fragment); uniclust \| UniRef100\_A0A0F5IZK2 \| 98.5 \| 1.5e-09 \| 3e-15 \| 82.7 \| 107 \| (28, 135) \| 136 \| (36, 157) \| 230 \| Contractile injection system tube protein N-terminal domain-containing protein \| Contractile injection system tube protein N-terminal domain-containing protein; uniclust \| UniRef100\_UPI001BD310B7 \| 98.5 \| 1.9e-09 \| 3.4e-15 \| 73.9 \| 91 \| (18, 108) \| 136 \| (40, 139) \| 140 \| phage tail protein \| phage tail protein; uniclust \| UniRef100\_A0A011NTK7 \| 98.5 \| 1.7e-09 \| 3.5e-15 \| 86.0 \| 107 \| (27, 134) \| 136 \| (34, 161) \| 293 \| Uncharacterized protein \| Uncharacterized protein; uniclust \| UniRef100\_A0A0A7HAL1 \| 98.5 \| 1.8e-09 \| 3.8e-15 \| 83.6 \| 114 \| (20, 135) \| 136 \| (47, 164) \| 248 \| Peptidoglycan-binding protein LysM \| Peptidoglycan-binding protein LysM; uniclust \| UniRef100\_A0A812QV43 \| 98.4 \| 3.2e-09 \| 5.9e-15 \| 90.9 \| 92 \| (36, 127) \| 136 \| (530, 633) \| 1104 \| GpFI protein \| GpFI protein; uniclust \| UniRef100\_A0A656GLE8 \| 98.4 \| 4.1e-09 \| 7.5e-15 \| 64.7 \| 71 \| (35, 105) \| 136 \| (2, 72) \| 72 \| Uncharacterized protein (Fragment) \| Uncharacterized protein (Fragment); uniclust \| UniRef100\_A0A8T5Z531 \| 98.4 \| 3.9e-09 \| 7.5e-15 \| 66.5 \| 62 \| (75, 136) \| 136 \| (7, 68) \| 74 \| Phage tail protein \| Phage tail protein; uniclust \| UniRef100\_UPI0006D28F55 \| 98.4 \| 4.2e-09 \| 7.8e-15 \| 69.4 \| 99 \| (13, 111) \| 136 \| (2, 104) \| 108 \| phage tail protein \| phage tail protein; uniclust \| UniRef100\_A0A101JEW8 \| 98.4 \| 3.9e-09 \| 7.9e-15 \| 79.9 \| 108 \| (27, 135) \| 136 \| (30, 176) \| 203 \| Contractile injection system tube protein N-terminal domain-containing protein \| Contractile injection system tube protein N-terminal domain-containing protein; uniclust \| UniRef100\_UPI001E4C1914 \| 98.4 \| 4.6e-09 \| 8.4e-15 \| 66.3 \| 73 \| (63, 135) \| 136 \| (2, 74) \| 84 \| phage tail protein \| phage tail protein; uniclust \| UniRef100\_A0A0X8JTK8 \| 98.4 \| 5.2e-09 \| 9.7e-15 \| 78.9 \| 122 \| (13, 135) \| 136 \| (1, 132) \| 263 \| Phage tail protein \| Phage tail protein; uniclust \| UniRef100\_UPI000B1FC26E \| 98.4 \| 5.7e-09 \| 1e-14 \| 65.0 \| 57 \| (6, 62) \| 136 \| (21, 77) \| 77 \| phage tail protein \| phage tail protein; uniclust \| UniRef100\_A0A4V1M2L6 \| 98.4 \| 5.7e-09 \| 1e-14 \| 72.8 \| 73 \| (13, 86) \| 136 \| (1, 73) \| 155 \| Uncharacterized protein \| Uncharacterized protein; uniclust \| UniRef100\_UPI001483AD28 \| 98.3 \| 6.2e-09 \| 1.1e-14 \| 66.8 \| 50 \| (87, 136) \| 136 \| (2, 51) \| 88 \| phage tail protein \| phage tail protein; uniclust \| UniRef100\_A0A6P0M9Y5 \| 98.3 \| 8.7e-09 \| 1.6e-14 \| 69.6 \| 118 \| (12, 134) \| 136 \| (1, 122) \| 125 \| Phage tail protein \| Phage tail protein; uniclust \| UniRef100\_UPI000AA14590 \| 98.3 \| 9e-09 \| 1.7e-14 \| 70.5 \| 68 \| (48, 115) \| 136 \| (68, 135) \| 136 \| phage tail protein \| phage tail protein; uniclust \| UniRef100\_A0A011MMX4 \| 98.3 \| 9.6e-09 \| 2e-14 \| 83.5 \| 112 \| (19, 134) \| 136 \| (25, 145) \| 352 \| Contractile injection system tube protein N-terminal domain-containing protein \| Contractile injection system tube protein N-terminal domain-containing protein; uniclust \| UniRef100\_A0A2M8WDQ8 \| 98.3 \| 1.2e-08 \| 2.1e-14 \| 67.5 \| 79 \| (57, 135) \| 136 \| (29, 107) \| 108 \| Uncharacterized protein \| Uncharacterized protein; uniclust \| UniRef100\_A0A496NCJ1 \| 98.3 \| 1.2e-08 \| 2.2e-14 \| 65.5 \| 89 \| (48, 136) \| 136 \| (1, 91) \| 91 \| Phage tail protein (Fragment) \| Phage tail protein (Fragment); uniclust \| UniRef100\_A0A936B9Z8 \| 98.2 \| 1.4e-08 \| 2.8e-14 \| 80.0 \| 112 \| (22, 135) \| 136 \| (75, 215) \| 288 \| Contractile injection system tube protein N-terminal domain-containing protein \| Contractile injection system tube protein N-terminal domain-containing protein; uniclust \| UniRef100\_UPI00211C54AE \| 98.2 \| 1.6e-08 \| 2.9e-14 \| 69.4 \| 106 \| (25, 132) \| 136 \| (2, 107) \| 136 \| phage tail protein \| phage tail protein; uniclust \| UniRef100\_A0A4C4L837 \| 98.2 \| 1.7e-08 \| 3.2e-14 \| 65.8 \| 55 \| (10, 64) \| 136 \| (27, 81) \| 100 \| Uncharacterized protein \| Uncharacterized protein; uniclust \| UniRef100\_UPI001F061B49 \| 98.2 \| 2e-08 \| 3.7e-14 \| 68.8 \| 78 \| (57, 134) \| 136 \| (8, 85) \| 135 \| phage tail protein \| phage tail protein; uniclust \| UniRef100\_UPI00146D20ED \| 98.2 \| 2.1e-08 \| 3.8e-14 \| 60.4 \| 56 \| (81, 136) \| 136 \| (1, 56) \| 63 \| phage tail protein \| phage tail protein; uniclust \| UniRef100\_A0A1Q8A398 \| 98.2 \| 2.1e-08 \| 4e-14 \| 70.5 \| 103 \| (27, 135) \| 136 \| (30, 140) \| 148 \| Contractile injection system tube protein N-terminal domain-containing protein \| Contractile injection system tube protein N-terminal domain-containing protein; uniclust \| UniRef100\_A0A7S4ZV00 \| 98.2 \| 2.1e-08 \| 4.1e-14 \| 69.5 \| 112 \| (14, 125) \| 136 \| (6, 120) \| 131 \| Phage P2 GpU family protein \| Phage P2 GpU family protein; uniclust \| UniRef100\_UPI001374B094 \| 98.2 \| 2.2e-08 \| 4.1e-14 \| 68.4 \| 60 \| (76, 135) \| 136 \| (59, 118) \| 133 \| phage tail protein \| phage tail protein; uniclust \| UniRef100\_A0A0M0SE01 \| 98.2 \| 2e-08 \| 4.2e-14 \| 80.5 \| 116 \| (18, 136) \| 136 \| (49, 176) \| 296 \| Contractile injection system tube protein N-terminal domain-containing protein \| Contractile injection system tube protein N-terminal domain-containing protein; uniclust \| UniRef100\_A0A2I7R3K3 \| 98.2 \| 2.5e-08 \| 4.6e-14 \| 68.1 \| 121 \| (14, 135) \| 136 \| (6, 129) \| 131 \| Uncharacterized protein \| Uncharacterized protein; uniclust \| UniRef100\_A0A2G6EZG3 \| 98.2 \| 2.6e-08 \| 4.8e-14 \| 75.5 \| 123 \| (12, 135) \| 136 \| (31, 159) \| 246 \| Phage tail protein \| Phage tail protein; uniclust \| UniRef100\_A0A853IPJ9 \| 98.2 \| 2.7e-08 \| 4.9e-14 \| 62.8 \| 73 \| (64, 136) \| 136 \| (2, 77) \| 82 \| Phage tail protein \| Phage tail protein; uniclust \| UniRef100\_UPI0018F0E200 \| 98.1 \| 3.6e-08 \| 6.6e-14 \| 59.2 \| 57 \| (80, 136) \| 136 \| (3, 59) \| 62 \| phage tail protein \| phage tail protein; uniclust \| UniRef100\_A0A1Q7YH54 \| 98.1 \| 3.5e-08 \| 6.7e-14 \| 71.0 \| 114 \| (18, 135) \| 136 \| (17, 137) \| 157 \| Contractile injection system tube protein N-terminal domain-containing protein \| Contractile injection system tube protein N-terminal domain-containing protein; uniclust \| UniRef100\_A0A484X1E5 \| 98.1 \| 3.6e-08 \| 6.8e-14 \| 61.6 \| 60 \| (13, 72) \| 136 \| (1, 60) \| 74 \| GpU phage protein \| GpU phage protein; uniclust \| UniRef100\_A0A1V5LS04 \| 98.1 \| 3.9e-08 \| 7.5e-14 \| 72.1 \| 105 \| (28, 135) \| 136 \| (43, 161) \| 174 \| Contractile injection system tube protein N-terminal domain-containing protein \| Contractile injection system tube protein N-terminal domain-containing protein; uniclust \| UniRef100\_A0A7W0VBY2 \| 98.1 \| 4.5e-08 \| 8.2e-14 \| 67.4 \| 87 \| (49, 135) \| 136 \| (21, 127) \| 137 \| Contractile injection system tube protein N-terminal domain-containing protein \| Contractile injection system tube protein N-terminal domain-containing protein; uniclust \| UniRef100\_UPI001EE21EE9 \| 98.1 \| 4.5e-08 \| 8.2e-14 \| 63.6 \| 77 \| (13, 89) \| 136 \| (1, 80) \| 96 \| phage tail protein \| phage tail protein; uniclust \| UniRef100\_UPI0007EEB1E3 \| 98.1 \| 4.6e-08 \| 8.5e-14 \| 63.2 \| 83 \| (15, 102) \| 136 \| (2, 84) \| 88 \| phage tail protein \| phage tail protein; uniclust \| UniRef100\_UPI0020223C4F \| 98.1 \| 4.8e-08 \| 8.8e-14 \| 61.5 \| 68 \| (68, 136) \| 136 \| (3, 71) \| 80 \| phage tail protein \| phage tail protein; uniclust \| UniRef100\_A0A0F9K3Y5 \| 98.1 \| 4.6e-08 \| 9.2e-14 \| 76.6 \| 108 \| (28, 135) \| 136 \| (33, 151) \| 272 \| Tip attachment protein J domain-containing protein \| Tip attachment protein J domain-containing protein; uniclust \| UniRef100\_A0A366DKR3 \| 98.1 \| 5.3e-08 \| 9.8e-14 \| 62.6 \| 64 \| (13, 76) \| 136 \| (1, 64) \| 90 \| GpU protein \| GpU protein; uniclust \| UniRef100\_A0A099ICH7 \| 98.1 \| 4.9e-08 \| 1e-13 \| 74.2 \| 110 \| (20, 133) \| 136 \| (20, 149) \| 202 \| Contractile injection system tube protein N-terminal domain-containing protein \| Contractile injection system tube protein N-terminal domain-containing protein; uniclust \| UniRef100\_A0A947CVW6 \| 98.1 \| 5.5e-08 \| 1e-13 \| 70.1 \| 117 \| (18, 135) \| 136 \| (16, 142) \| 187 \| Contractile injection system tube protein N-terminal domain-containing protein \| Contractile injection system tube protein N-terminal domain-containing protein; uniclust \| UniRef100\_UPI001FC8D6E6 \| 98.1 \| 5.6e-08 \| 1e-13 \| 59.9 \| 51 \| (86, 136) \| 136 \| (2, 52) \| 71 \| phage tail protein \| phage tail protein; uniclust \| UniRef100\_A0A968ZAB9 \| 98.1 \| 5.7e-08 \| 1.1e-13 \| 73.2 \| 107 \| (28, 135) \| 136 \| (39, 166) \| 263 \| Protein kinase domain-containing protein \| Protein kinase domain-containing protein; uniclust \| UniRef100\_A0A6B2WT17 \| 98.1 \| 5.6e-08 \| 1.1e-13 \| 70.4 \| 114 \| (22, 136) \| 136 \| (39, 170) \| 173 \| Peptidase M23 (Fragment) \| Peptidase M23 (Fragment); uniclust \| UniRef100\_A0A928YH49 \| 98.1 \| 5.8e-08 \| 1.1e-13 \| 70.0 \| 121 \| (12, 135) \| 136 \| (2, 128) \| 166 \| Uncharacterized protein \| Uncharacterized protein; uniclust \| UniRef100\_A0A0Q9U2K6 \| 98.1 \| 5.6e-08 \| 1.2e-13 \| 82.5 \| 107 \| (27, 135) \| 136 \| (31, 151) \| 489 \| Contractile injection system tube protein N-terminal domain-containing protein \| Contractile injection system tube protein N-terminal domain-containing protein; uniclust \| UniRef100\_A0A7J0CYE3 \| 98.1 \| 6.5e-08 \| 1.2e-13 \| 74.6 \| 108 \| (27, 135) \| 136 \| (38, 163) \| 319 \| LysM domain-containing protein \| LysM domain-containing protein; uniclust \| UniRef100\_A0A7V9L357 \| 98.0 \| 9.6e-08 \| 1.8e-13 \| 70.2 \| 121 \| (14, 135) \| 136 \| (40, 183) \| 214 \| Contractile injection system tube protein N-terminal domain-containing protein \| Contractile injection system tube protein N-terminal domain-containing protein; uniclust \| UniRef100\_A0A9D1LT46 \| 98.0 \| 1e-07 \| 1.9e-13 \| 69.6 \| 122 \| (13, 135) \| 136 \| (2, 129) \| 202 \| Phage tail protein \| Phage tail protein; uniclust \| UniRef100\_UPI00202512B1 \| 98.0 \| 1.1e-07 \| 2e-13 \| 56.0 \| 47 \| (90, 136) \| 136 \| (3, 49) \| 55 \| phage tail protein \| phage tail protein; uniclust \| UniRef100\_A0A433VYQ6 \| 98.0 \| 1.1e-07 \| 2e-13 \| 70.6 \| 109 \| (28, 136) \| 136 \| (24, 157) \| 197 \| Contractile injection system tube protein N-terminal domain-containing protein \| Contractile injection system tube protein N-terminal domain-containing protein; uniclust \| UniRef100\_A0A0H3NMP3 \| 98.0 \| 1.3e-07 \| 2.4e-13 \| 59.8 \| 43 \| (94, 136) \| 136 \| (1, 43) \| 73 \| Phage-related tail protein \| Phage-related tail protein; uniclust \| UniRef100\_A0A1L6I298 \| 98.0 \| 1.3e-07 \| 2.6e-13 \| 69.8 \| 113 \| (9, 131) \| 136 \| (4, 116) \| 165 \| Uncharacterized protein \| Uncharacterized protein; uniclust \| UniRef100\_A0A2V2A0Z3 \| 98.0 \| 1.6e-07 \| 3e-13 \| 62.0 \| 85 \| (52, 136) \| 136 \| (16, 102) \| 104 \| GpU protein \| GpU protein; uniclust \| UniRef100\_A0A3M1ER04 \| 97.9 \| 1.8e-07 \| 3.4e-13 \| 65.8 \| 91 \| (27, 118) \| 136 \| (21, 130) \| 131 \| Peptigoglycan-binding protein LysM (Fragment) \| Peptigoglycan-binding protein LysM (Fragment); uniclust \| UniRef100\_A0A853IKN5 \| 97.9 \| 2.2e-07 \| 4.1e-13 \| 54.2 \| 43 \| (94, 136) \| 136 \| (2, 44) \| 52 \| Phage tail protein \| Phage tail protein; uniclust \| UniRef100\_UPI000BA48FEE \| 97.9 \| 2.3e-07 \| 4.2e-13 \| 73.6 \| 70 \| (19, 88) \| 136 \| (158, 227) \| 392 \| phage tail protein \| phage tail protein; uniclust \| UniRef100\_UPI001BD14020 \| 97.9 \| 2.4e-07 \| 4.3e-13 \| 62.7 \| 59 \| (77, 135) \| 136 \| (21, 79) \| 119 \| phage tail protein \| phage tail protein; uniclust \| UniRef100\_UPI001FE72D82 \| 97.9 \| 2.4e-07 \| 4.5e-13 \| 67.8 \| 109 \| (27, 136) \| 136 \| (24, 147) \| 204 \| hypothetical protein \| hypothetical protein; uniclust \| UniRef100\_UPI001F18FFEC \| 97.9 \| 2.9e-07 \| 5.4e-13 \| 64.7 \| 88 \| (42, 135) \| 136 \| (42, 129) \| 152 \| phage tail protein \| phage tail protein; uniclust \| UniRef100\_A0A3E4VCF7 \| 97.9 \| 3e-07 \| 5.8e-13 \| 56.8 \| 61 \| (75, 136) \| 136 \| (3, 65) \| 65 \| Uncharacterized protein \| Uncharacterized protein; uniclust \| UniRef100\_UPI001F4C87A2 \| 97.9 \| 3.2e-07 \| 5.9e-13 \| 53.3 \| 41 \| (96, 136) \| 136 \| (2, 42) \| 51 \| phage tail protein \| phage tail protein; uniclust \| UniRef100\_A0A1Z4PN67 \| 97.8 \| 3.7e-07 \| 6.9e-13 \| 63.6 \| 118 \| (13, 133) \| 136 \| (4, 127) \| 131 \| Uncharacterized protein \| Uncharacterized protein; uniclust \| UniRef100\_UPI0022B72820 \| 97.8 \| 4.2e-07 \| 7.7e-13 \| 61.8 \| 51 \| (86, 136) \| 136 \| (2, 52) \| 122 \| phage tail protein \| phage tail protein; uniclust \| UniRef100\_UPI001F173E80 \| 97.8 \| 4.5e-07 \| 8.4e-13 \| 58.5 \| 69 \| (14, 82) \| 136 \| (6, 74) \| 83 \| phage tail protein \| phage tail protein; uniclust \| UniRef100\_UPI0021C3E1B0 \| 97.8 \| 4.7e-07 \| 8.6e-13 \| 62.1 \| 61 \| (13, 74) \| 136 \| (1, 61) \| 129 \| phage tail protein \| phage tail protein; uniclust \| UniRef100\_UPI001C999792 \| 97.8 \| 4.8e-07 \| 8.9e-13 \| 59.9 \| 90 \| (11, 100) \| 136 \| (2, 91) \| 103 \| phage tail protein \| phage tail protein; uniclust \| UniRef100\_A0A0F9VF18 \| 97.8 \| 4.7e-07 \| 9.3e-13 \| 68.1 \| 109 \| (27, 135) \| 136 \| (32, 159) \| 187 \| Contractile injection system tube protein N-terminal domain-containing protein \| Contractile injection system tube protein N-terminal domain-containing protein; uniclust \| UniRef100\_A0A0H3ZVP6 \| 97.8 \| 5.5e-07 \| 1e-12 \| 55.6 \| 58 \| (78, 135) \| 136 \| (3, 60) \| 69 \| Phage tail protein \| Phage tail protein; uniclust \| UniRef100\_H1D5E7 \| 97.8 \| 5.6e-07 \| 1e-12 \| 56.8 \| 62 \| (12, 73) \| 136 \| (1, 65) \| 78 \| Uncharacterized protein \| Uncharacterized protein; uniclust \| UniRef100\_A0A935R531 \| 97.8 \| 5.7e-07 \| 1.1e-12 \| 65.0 \| 112 \| (20, 133) \| 136 \| (27, 148) \| 169 \| Contractile injection system tube protein N-terminal domain-containing protein \| Contractile injection system tube protein N-terminal domain-containing protein; uniclust \| UniRef100\_A0A4Q3K0V1 \| 97.8 \| 5.9e-07 \| 1.1e-12 \| 51.7 \| 48 \| (85, 132) \| 136 \| (1, 48) \| 48 \| Oxidoreductase (Fragment) \| Oxidoreductase (Fragment); uniclust \| UniRef100\_UPI001B7CF361 \| 97.8 \| 6e-07 \| 1.1e-12 \| 67.9 \| 109 \| (27, 136) \| 136 \| (79, 202) \| 254 \| hypothetical protein \| hypothetical protein; uniclust \| UniRef100\_A0A3S5K2M8 \| 97.8 \| 6e-07 \| 1.1e-12 \| 71.7 \| 109 \| (27, 136) \| 136 \| (35, 156) \| 408 \| Contractile injection system tube protein N-terminal domain-containing protein \| Contractile injection system tube protein N-terminal domain-containing protein; uniclust \| UniRef100\_A0A524GYW0 \| 97.8 \| 6e-07 \| 1.2e-12 \| 66.3 \| 58 \| (77, 135) \| 136 \| (13, 75) \| 159 \| LysM peptidoglycan-binding domain-containing protein \| LysM peptidoglycan-binding domain-containing protein; uniclust \| UniRef100\_A0A378MZA4 \| 97.7 \| 7.7e-07 \| 1.4e-12 \| 57.2 \| 54 \| (80, 133) \| 136 \| (4, 58) \| 86 \| Phage protein U \| Phage protein U; uniclust \| UniRef100\_UPI000AA7DEF5 \| 97.7 \| 7.7e-07 \| 1.4e-12 \| 53.6 \| 55 \| (81, 136) \| 136 \| (3, 59) \| 60 \| phage tail protein \| phage tail protein; uniclust \| UniRef100\_A0A447N5T3 \| 97.7 \| 7.8e-07 \| 1.4e-12 \| 50.4 \| 41 \| (13, 53) \| 136 \| (1, 41) \| 44 \| Gp25 \| Gp25; uniclust \| UniRef100\_A0A074LMC3 \| 97.7 \| 7.8e-07 \| 1.5e-12 \| 69.1 \| 113 \| (9, 133) \| 136 \| (1, 113) \| 243 \| LysM domain-containing protein \| LysM domain-containing protein; uniclust \| UniRef100\_UPI0009BF56FE \| 97.7 \| 8.6e-07 \| 1.6e-12 \| 69.2 \| 63 \| (11, 73) \| 136 \| (2, 64) \| 318 \| contractile injection system protein, VgrG/Pvc8 family \| contractile injection system protein, VgrG/Pvc8 family; uniclust \| UniRef100\_A0A1S6HMA4 \| 97.7 \| 8.5e-07 \| 1.6e-12 \| 64.1 \| 108 \| (28, 135) \| 136 \| (30, 154) \| 167 \| Contractile injection system tube protein N-terminal domain-containing protein \| Contractile injection system tube protein N-terminal domain-containing protein; uniclust \| UniRef100\_A0A2Z3EGZ9 \| 97.7 \| 1e-06 \| 1.8e-12 \| 63.1 \| 115 \| (20, 135) \| 136 \| (28, 151) \| 167 \| Contractile injection system tube protein N-terminal domain-containing protein \| Contractile injection system tube protein N-terminal domain-containing protein; uniclust \| UniRef100\_A0A060BT42 \| 97.7 \| 9.6e-07 \| 1.9e-12 \| 65.3 \| 80 \| (56, 136) \| 136 \| (6, 102) \| 177 \| CAZy families CBM50 protein (Fragment) \| CAZy families CBM50 protein (Fragment); uniclust \| UniRef100\_A0A174USZ0 \| 97.7 \| 9.1e-07 \| 1.9e-12 \| 63.6 \| 117 \| (10, 135) \| 136 \| (1, 118) \| 129 \| Minor capsid protein \| Minor capsid protein; uniclust \| UniRef100\_A0A073CFQ3 \| 97.7 \| 9.9e-07 \| 2e-12 \| 70.9 \| 109 \| (28, 136) \| 136 \| (118, 295) \| 314 \| Contractile injection system tube protein N-terminal domain-containing protein \| Contractile injection system tube protein N-terminal domain-containing protein; uniclust \| UniRef100\_UPI001659B0F1 \| 97.7 \| 1.1e-06 \| 2e-12 \| 58.1 \| 57 \| (79, 135) \| 136 \| (2, 58) \| 97 \| phage tail protein \| phage tail protein; uniclust \| UniRef100\_A0A2D7W1J1 \| 97.7 \| 1e-06 \| 2e-12 \| 74.9 \| 124 \| (10, 136) \| 136 \| (255, 408) \| 515 \| Contractile injection system tube protein N-terminal domain-containing protein \| Contractile injection system tube protein N-terminal domain-containing protein; uniclust \| UniRef100\_A0A061NGS6 \| 97.7 \| 9.6e-07 \| 2.1e-12 \| 70.8 \| 126 \| (5, 135) \| 136 \| (19, 156) \| 255 \| Phage-like element PBSX protein XkdP \| Phage-like element PBSX protein XkdP; uniclust \| UniRef100\_A0A2A7WUN3 \| 97.7 \| 1.2e-06 \| 2.3e-12 \| 67.3 \| 107 \| (28, 135) \| 136 \| (38, 147) \| 227 \| LysM domain-containing protein \| LysM domain-containing protein; uniclust \| UniRef100\_R6PMQ9 \| 97.7 \| 1.3e-06 \| 2.4e-12 \| 56.5 \| 68 \| (68, 135) \| 136 \| (3, 70) \| 88 \| Uncharacterized protein \| Uncharacterized protein; uniclust \| UniRef100\_S5MV70 \| 97.7 \| 1.3e-06 \| 2.4e-12 \| 52.6 \| 57 \| (68, 124) \| 136 \| (2, 58) \| 60 \| Phage tail protein \| Phage tail protein; uniclust \| UniRef100\_A0A105T8J5 \| 97.7 \| 1.3e-06 \| 2.4e-12 \| 64.1 \| 57 \| (19, 75) \| 136 \| (142, 198) \| 200 \| Phage protein U \| Phage protein U; uniclust \| UniRef100\_UPI00227354E9 \| 97.7 \| 1.3e-06 \| 2.4e-12 \| 59.7 \| 111 \| (13, 124) \| 136 \| (3, 116) \| 123 \| phage tail protein \| phage tail protein; uniclust \| UniRef100\_A0A1J0AH11 \| 97.7 \| 1.3e-06 \| 2.5e-12 \| 67.7 \| 116 \| (20, 136) \| 136 \| (17, 150) \| 263 \| Contractile injection system tube protein N-terminal domain-containing protein \| Contractile injection system tube protein N-terminal domain-containing protein; uniclust \| UniRef100\_UPI00157B406D \| 97.6 \| 1.4e-06 \| 2.6e-12 \| 51.6 \| 36 \| (100, 135) \| 136 \| (19, 54) \| 55 \| phage tail protein \| phage tail protein; uniclust \| UniRef100\_A0A846T4R7 \| 97.6 \| 1.6e-06 \| 2.8e-12 \| 53.3 \| 60 \| (13, 72) \| 136 \| (1, 60) \| 66 \| Uncharacterized protein \| Uncharacterized protein; uniclust \| UniRef100\_A0A318ENP1 \| 97.6 \| 1.7e-06 \| 3e-12 \| 56.3 \| 70 \| (14, 83) \| 136 \| (3, 77) \| 91 \| GpU protein \| GpU protein; uniclust \| UniRef100\_A0A1F3S067 \| 97.6 \| 1.7e-06 \| 3.4e-12 \| 67.4 \| 105 \| (28, 134) \| 136 \| (40, 162) \| 233 \| Contractile injection system tube protein N-terminal domain-containing protein (Fragment) \| Contractile injection system tube protein N-terminal domain-containing protein (Fragment); uniclust \| UniRef100\_A0A2N5DAB4 \| 97.6 \| 2.1e-06 \| 4.1e-12 \| 73.2 \| 108 \| (27, 135) \| 136 \| (37, 166) \| 496 \| Contractile injection system tube protein N-terminal domain-containing protein \| Contractile injection system tube protein N-terminal domain-containing protein; uniclust \| UniRef100\_UPI001F0A3F5E \| 97.6 \| 2.3e-06 \| 4.2e-12 \| 51.9 \| 39 \| (39, 77) \| 136 \| (4, 43) \| 61 \| phage tail protein \| phage tail protein; uniclust \| UniRef100\_UPI0022A7BF70 \| 97.5 \| 2.9e-06 \| 5.3e-12 \| 54.9 \| 54 \| (83, 136) \| 136 \| (4, 57) \| 87 \| phage tail protein \| phage tail protein; uniclust \| UniRef100\_A0A955R5H5 \| 97.5 \| 3e-06 \| 5.5e-12 \| 66.9 \| 116 \| (18, 135) \| 136 \| (16, 146) \| 350 \| LysM peptidoglycan-binding domain-containing protein \| LysM peptidoglycan-binding domain-containing protein; uniclust \| UniRef100\_A0A1C6JGS8 \| 97.5 \| 3.4e-06 \| 6.4e-12 \| 62.8 \| 121 \| (12, 136) \| 136 \| (2, 129) \| 201 \| Phage protein U \| Phage protein U; uniclust \| UniRef100\_A0A1M6M9A3 \| 97.5 \| 3.3e-06 \| 6.5e-12 \| 64.5 \| 114 \| (10, 133) \| 136 \| (1, 115) \| 199 \| LysM domain-containing protein \| LysM domain-containing protein; uniclust \| UniRef100\_U2F2B2 \| 97.5 \| 3.6e-06 \| 6.7e-12 \| 52.2 \| 61 \| (74, 134) \| 136 \| (6, 66) \| 69 \| Phage tail protein \| Phage tail protein; uniclust \| UniRef100\_UPI0009EDCD06 \| 97.5 \| 3.8e-06 \| 7.2e-12 \| 54.9 \| 78 \| (14, 99) \| 136 \| (1, 78) \| 85 \| phage tail protein \| phage tail protein; uniclust \| UniRef100\_A0A935P3X5 \| 97.5 \| 3.9e-06 \| 7.3e-12 \| 63.3 \| 106 \| (28, 135) \| 136 \| (56, 178) \| 238 \| Contractile injection system tube protein N-terminal domain-containing protein \| Contractile injection system tube protein N-terminal domain-containing protein; uniclust \| UniRef100\_A0A1Q3WFK6 \| 97.5 \| 3.6e-06 \| 7.4e-12 \| 70.9 \| 106 \| (28, 134) \| 136 \| (29, 147) \| 414 \| Contractile injection system tube protein N-terminal domain-containing protein \| Contractile injection system tube protein N-terminal domain-containing protein; uniclust \| UniRef100\_A0A084SKL0 \| 97.5 \| 3.6e-06 \| 7.6e-12 \| 68.9 \| 107 \| (28, 135) \| 136 \| (23, 153) \| 306 \| Contractile injection system tube protein N-terminal domain-containing protein \| Contractile injection system tube protein N-terminal domain-containing protein; uniclust \| UniRef100\_UPI001E5FDDE9 \| 97.5 \| 4.3e-06 \| 7.9e-12 \| 49.0 \| 40 \| (10, 49) \| 136 \| (8, 47) \| 51 \| phage tail protein \| phage tail protein; uniclust \| UniRef100\_A0A9D1ATR1 \| 97.5 \| 4.6e-06 \| 8.4e-12 \| 66.4 \| 113 \| (21, 135) \| 136 \| (42, 171) \| 372 \| LysM peptidoglycan-binding domain-containing protein (Fragment) \| LysM peptidoglycan-binding domain-containing protein (Fragment); uniclust \| UniRef100\_UPI001F06D85D \| 97.5 \| 4.6e-06 \| 8.4e-12 \| 54.2 \| 59 \| (76, 135) \| 136 \| (3, 61) \| 89 \| phage tail protein \| phage tail protein; uniclust \| UniRef100\_A0A1X7MDE4 \| 97.4 \| 5.2e-06 \| 9.5e-12 \| 45.9 \| 35 \| (12, 46) \| 136 \| (3, 37) \| 38 \| [weak similarity to] phage P2 GpU (Fragment) \| [weak similarity to] phage P2 GpU (Fragment); uniclust \| UniRef100\_A0A4P5W163 \| 97.4 \| 5.3e-06 \| 9.8e-12 \| 64.0 \| 115 \| (19, 135) \| 136 \| (20, 155) \| 283 \| Contractile injection system tube protein N-terminal domain-containing protein \| Contractile injection system tube protein N-terminal domain-containing protein; uniclust \| UniRef100\_A0A0Q4KRA9 \| 97.4 \| 4.9e-06 \| 1e-11 \| 66.4 \| 116 \| (17, 135) \| 136 \| (35, 215) \| 264 \| Contractile injection system tube protein N-terminal domain-containing protein \| Contractile injection system tube protein N-terminal domain-containing protein; uniclust \| UniRef100\_A0A0K9NAG5 \| 97.4 \| 5.1e-06 \| 1e-11 \| 62.4 \| 114 \| (19, 134) \| 136 \| (23, 143) \| 169 \| Contractile injection system tube protein N-terminal domain-containing protein \| Contractile injection system tube protein N-terminal domain-containing protein; uniclust \| UniRef100\_A0A7X6JBU4 \| 97.4 \| 6e-06 \| 1.1e-11 \| 54.8 \| 59 \| (78, 136) \| 136 \| (3, 64) \| 100 \| Phage tail protein \| Phage tail protein; uniclust \| UniRef100\_N8S9H2 \| 97.4 \| 6.3e-06 \| 1.2e-11 \| 47.0 \| 40 \| (13, 52) \| 136 \| (1, 40) \| 44 \| Uncharacterized protein \| Uncharacterized protein; uniclust \| UniRef100\_UPI0018EF83E4 \| 97.4 \| 7e-06 \| 1.3e-11 \| 70.6 \| 108 \| (28, 136) \| 136 \| (27, 147) \| 834 \| VgrG-related protein \| VgrG-related protein; uniclust \| UniRef100\_UPI001E41EFA3 \| 97.4 \| 7.2e-06 \| 1.3e-11 \| 49.2 \| 34 \| (102, 135) \| 136 \| (2, 35) \| 57 \| phage tail protein \| phage tail protein; uniclust \| UniRef100\_A0A941G883 \| 97.4 \| 7.4e-06 \| 1.4e-11 \| 56.1 \| 69 \| (68, 136) \| 136 \| (1, 69) \| 120 \| Phage tail protein \| Phage tail protein; uniclust \| UniRef100\_A0A014LIK0 \| 97.4 \| 7.2e-06 \| 1.5e-11 \| 67.8 \| 105 \| (28, 133) \| 136 \| (112, 250) \| 323 \| Signal peptide protein \| Signal peptide protein; uniclust \| UniRef100\_A0A3S0DNP6 \| 97.3 \| 8.6e-06 \| 1.7e-11 \| 62.3 \| 105 \| (27, 134) \| 136 \| (33, 146) \| 202 \| Contractile injection system tube protein N-terminal domain-containing protein (Fragment) \| Contractile injection system tube protein N-terminal domain-containing protein (Fragment); uniclust \| UniRef100\_A0A2W4LKG2 \| 97.3 \| 8.7e-06 \| 1.7e-11 \| 58.0 \| 45 \| (91, 136) \| 136 \| (2, 47) \| 124 \| Peptidoglycan-binding protein (Fragment) \| Peptidoglycan-binding protein (Fragment); uniclust \| UniRef100\_A0A2D4SL56 \| 97.3 \| 8.9e-06 \| 1.7e-11 \| 63.5 \| 106 \| (27, 133) \| 136 \| (39, 192) \| 245 \| Contractile injection system tube protein N-terminal domain-containing protein \| Contractile injection system tube protein N-terminal domain-containing protein; uniclust \| UniRef100\_A0A069A2Y0 \| 97.3 \| 8.5e-06 \| 1.8e-11 \| 64.9 \| 122 \| (8, 136) \| 136 \| (18, 149) \| 233 \| LysM domain protein \| LysM domain protein; uniclust \| UniRef100\_A0A173TJ12 \| 97.3 \| 1e-05 \| 1.9e-11 \| 60.7 \| 115 \| (17, 135) \| 136 \| (21, 150) \| 218 \| LysM domain-containing protein \| LysM domain-containing protein; uniclust \| UniRef100\_UPI00049A5F88 \| 97.3 \| 1.2e-05 \| 2.2e-11 \| 57.1 \| 122 \| (13, 135) \| 136 \| (1, 126) \| 150 \| phage tail protein \| phage tail protein; uniclust \| UniRef100\_A0A173R5A8 \| 97.3 \| 1.2e-05 \| 2.4e-11 \| 62.6 \| 123 \| (11, 136) \| 136 \| (1, 128) \| 227 \| LysM domain-containing protein \| LysM domain-containing protein; uniclust \| UniRef100\_A0A075RDD3 \| 97.3 \| 1.3e-05 \| 2.6e-11 \| 61.6 \| 110 \| (15, 135) \| 136 \| (3, 112) \| 198 \| LysM domain/BON superfamily protein \| LysM domain/BON superfamily protein; uniclust \| UniRef100\_UPI0013FA864C \| 97.2 \| 1.6e-05 \| 2.9e-11 \| 50.3 \| 69 \| (29, 99) \| 136 \| (5, 73) \| 74 \| phage tail protein \| phage tail protein; uniclust \| UniRef100\_UPI001BDA1730 \| 97.2 \| 1.7e-05 \| 3.1e-11 \| 52.2 \| 81 \| (54, 134) \| 136 \| (8, 88) \| 92 \| phage tail protein \| phage tail protein; uniclust \| UniRef100\_A0A1N7BSF8 \| 97.2 \| 1.6e-05 \| 3.1e-11 \| 63.1 \| 108 \| (27, 135) \| 136 \| (31, 169) \| 266 \| Contractile injection system tube protein N-terminal domain-containing protein \| Contractile injection system tube protein N-terminal domain-containing protein; uniclust \| UniRef100\_A0A7V8YCD3 \| 97.2 \| 1.7e-05 \| 3.2e-11 \| 62.0 \| 113 \| (20, 134) \| 136 \| (20, 149) \| 303 \| LysM peptidoglycan-binding domain-containing protein \| LysM peptidoglycan-binding domain-containing protein; uniclust \| UniRef100\_UPI001FE103EA \| 97.2 \| 1.8e-05 \| 3.2e-11 \| 51.6 \| 71 \| (20, 90) \| 136 \| (2, 72) \| 87 \| phage tail protein \| phage tail protein; uniclust \| UniRef100\_A0A1T4KBT6 \| 97.2 \| 1.8e-05 \| 3.7e-11 \| 61.0 \| 123 \| (1, 134) \| 136 \| (1, 123) \| 184 \| LysM domain-containing protein \| LysM domain-containing protein; uniclust \| UniRef100\_UPI001F05C40E \| 97.2 \| 2.1e-05 \| 3.8e-11 \| 60.6 \| 108 \| (28, 136) \| 136 \| (22, 147) \| 265 \| hypothetical protein \| hypothetical protein; uniclust \| UniRef100\_A0A1Z4HE74 \| 97.2 \| 2.1e-05 \| 4e-11 \| 58.2 \| 108 \| (28, 136) \| 136 \| (35, 161) \| 191 \| Contractile injection system tube protein N-terminal domain-containing protein \| Contractile injection system tube protein N-terminal domain-containing protein; uniclust \| UniRef100\_A0A1V2H6I8 \| 97.2 \| 2.4e-05 \| 4.5e-11 \| 62.2 \| 97 \| (12, 113) \| 136 \| (45, 141) \| 291 \| Phage tail protein \| Phage tail protein; uniclust \| UniRef100\_UPI002019748F \| 97.1 \| 2.6e-05 \| 4.7e-11 \| 50.1 \| 48 \| (48, 95) \| 136 \| (5, 52) \| 80 \| phage tail protein \| phage tail protein; uniclust \| UniRef100\_A0A376J1X2 \| 97.1 \| 2.6e-05 \| 4.8e-11 \| 45.5 \| 38 \| (81, 118) \| 136 \| (3, 40) \| 48 \| GpU phage protein \| GpU phage protein; uniclust \| UniRef100\_A0A0B0HFG9 \| 97.1 \| 2.7e-05 \| 4.9e-11 \| 47.1 \| 50 \| (86, 136) \| 136 \| (2, 53) \| 58 \| Uncharacterized protein \| Uncharacterized protein; uniclust \| UniRef100\_A0A0F3RPF7 \| 97.1 \| 2.7e-05 \| 5e-11 \| 53.9 \| 44 \| (93, 136) \| 136 \| (80, 123) \| 125 \| Phage tail tape measure protein, TP901 family, core region \| Phage tail tape measure protein, TP901 family, core region; uniclust \| UniRef100\_C8PII4 \| 97.1 \| 3.5e-05 \| 6.5e-11 \| 44.4 \| 40 \| (95, 134) \| 136 \| (2, 41) \| 45 \| Phage P2 GpU \| Phage P2 GpU; uniclust \| UniRef100\_A0A399IQQ4 \| 97.1 \| 3.5e-05 \| 6.7e-11 \| 58.3 \| 111 \| (12, 134) \| 136 \| (2, 112) \| 196 \| LysM peptidoglycan-binding domain-containing protein \| LysM peptidoglycan-binding domain-containing protein; uniclust \| UniRef100\_A0A090T0X2 \| 97.1 \| 3.7e-05 \| 6.8e-11 \| 58.6 \| 107 \| (28, 135) \| 136 \| (24, 150) \| 241 \| Contractile injection system tube protein N-terminal domain-containing protein \| Contractile injection system tube protein N-terminal domain-containing protein; uniclust \| UniRef100\_A0A497PS16 \| 97.1 \| 3.9e-05 \| 7.3e-11 \| 60.1 \| 107 \| (28, 135) \| 136 \| (59, 212) \| 258 \| Uncharacterized protein \| Uncharacterized protein; uniclust \| UniRef100\_E2CCF4 \| 97.1 \| 4.1e-05 \| 7.5e-11 \| 45.8 \| 38 \| (99, 136) \| 136 \| (4, 41) \| 54 \| Phage-related tail protein \| Phage-related tail protein; uniclust \| UniRef100\_A0A017RV10 \| 97.1 \| 3.7e-05 \| 7.6e-11 \| 62.7 \| 127 \| (3, 135) \| 136 \| (33, 171) \| 282 \| LysM domain-containing protein \| LysM domain-containing protein; uniclust \| UniRef100\_A0A952YFU8 \| 97.1 \| 4.2e-05 \| 7.7e-11 \| 47.8 \| 50 \| (87, 136) \| 136 \| (4, 53) \| 68 \| Phage tail protein \| Phage tail protein; uniclust \| UniRef100\_A0A3E4VD00 \| 97.1 \| 4.2e-05 \| 7.8e-11 \| 56.0 \| 62 \| (12, 73) \| 136 \| (109, 174) \| 179 \| SH3b domain-containing protein \| SH3b domain-containing protein; uniclust \| UniRef100\_A0A6B3MKD8 \| 97.0 \| 4e-05 \| 8.2e-11 \| 58.0 \| 115 \| (17, 135) \| 136 \| (32, 154) \| 165 \| Uncharacterized protein \| Uncharacterized protein; uniclust \| UniRef100\_A0A8S0FXT8 \| 97.0 \| 4.5e-05 \| 8.3e-11 \| 54.1 \| 48 \| (88, 135) \| 136 \| (3, 50) \| 144 \| Phage tail protein \| Phage tail protein; uniclust \| UniRef100\_A0A1G6LTE8 \| 97.0 \| 4.3e-05 \| 8.6e-11 \| 58.7 \| 113 \| (20, 136) \| 136 \| (19, 145) \| 186 \| Uncharacterized protein \| Uncharacterized protein; uniclust \| UniRef100\_UPI00207C8BD9 \| 97.0 \| 4.7e-05 \| 8.6e-11 \| 45.4 \| 30 \| (106, 135) \| 136 \| (1, 30) \| 53 \| phage tail protein \| phage tail protein; uniclust \| UniRef100\_UPI00202188DC \| 97.0 \| 4.8e-05 \| 8.8e-11 \| 49.7 \| 51 \| (27, 77) \| 136 \| (7, 57) \| 87 \| phage tail protein \| phage tail protein; uniclust \| UniRef100\_A0A6I0EJT0 \| 97.0 \| 5.3e-05 \| 9.8e-11 \| 44.8 \| 37 \| (100, 136) \| 136 \| (15, 51) \| 51 \| Uncharacterized protein \| Uncharacterized protein; uniclust \| UniRef100\_UPI002153A50E \| 97.0 \| 5.4e-05 \| 9.9e-11 \| 47.1 \| 64 \| (28, 91) \| 136 \| (2, 65) \| 66 \| phage tail protein \| phage tail protein; uniclust \| UniRef100\_UPI001D1340C6 \| 97.0 \| 5.4e-05 \| 1e-10 \| 56.3 \| 110 \| (12, 131) \| 136 \| (3, 112) \| 198 \| hypothetical protein \| hypothetical protein; uniclust \| UniRef100\_A0A8G2HV68 \| 97.0 \| 5.8e-05 \| 1.1e-10 \| 48.8 \| 44 \| (92, 136) \| 136 \| (8, 51) \| 79 \| Uncharacterized protein \| Uncharacterized protein; uniclust \| UniRef100\_A0A951MSF9 \| 97.0 \| 5.8e-05 \| 1.1e-10 \| 58.4 \| 107 \| (28, 135) \| 136 \| (55, 179) \| 254 \| Contractile injection system tube protein N-terminal domain-containing protein \| Contractile injection system tube protein N-terminal domain-containing protein; uniclust \| UniRef100\_A0A1Z8QW35 \| 96.9 \| 6.1e-05 \| 1.3e-10 \| 60.0 \| 108 \| (28, 135) \| 136 \| (32, 161) \| 230 \| Contractile injection system tube protein N-terminal domain-containing protein \| Contractile injection system tube protein N-terminal domain-containing protein; uniclust \| UniRef100\_E6X1M4 \| 96.9 \| 7.3e-05 \| 1.3e-10 \| 52.6 \| 117 \| (13, 135) \| 136 \| (1, 117) \| 135 \| Uncharacterized protein \| Uncharacterized protein; uniclust \| UniRef100\_A0A011PPH1 \| 96.9 \| 6.9e-05 \| 1.4e-10 \| 58.3 \| 101 \| (29, 136) \| 136 \| (13, 117) \| 197 \| Mu-like prophage DNA circulation protein \| Mu-like prophage DNA circulation protein; uniclust \| UniRef100\_A0A935YIZ6 \| 96.9 \| 8.4e-05 \| 1.5e-10 \| 54.5 \| 117 \| (17, 135) \| 136 \| (28, 147) \| 178 \| Phage tail protein \| Phage tail protein; uniclust \| UniRef100\_A0A0C1UAF4 \| 96.9 \| 7.1e-05 \| 1.6e-10 \| 60.9 \| 121 \| (10, 136) \| 136 \| (28, 157) \| 248 \| LysM domain protein \| LysM domain protein; uniclust \| UniRef100\_UPI001FE4EA06 \| 96.8 \| 0.00013 \| 2.4e-10 \| 49.7 \| 70 \| (66, 135) \| 136 \| (13, 85) \| 109 \| phage tail protein \| phage tail protein; uniclust \| UniRef100\_A0A069A1N0 \| 96.8 \| 0.00011 \| 2.5e-10 \| 61.1 \| 128 \| (5, 135) \| 136 \| (24, 165) \| 294 \| LysM domain protein \| LysM domain protein; uniclust \| UniRef100\_A0A081R180 \| 96.8 \| 0.00012 \| 2.6e-10 \| 59.6 \| 111 \| (20, 135) \| 136 \| (47, 168) \| 262 \| Phage protein \| Phage protein; uniclust \| UniRef100\_A6VY17 \| 96.8 \| 0.00014 \| 2.6e-10 \| 48.0 \| 79 \| (44, 124) \| 136 \| (2, 80) \| 90 \| Uncharacterized protein \| Uncharacterized protein; uniclust \| UniRef100\_F4BFP8 \| 96.8 \| 0.00015 \| 2.7e-10 \| 47.5 \| 67 \| (63, 135) \| 136 \| (2, 69) \| 86 \| Uncharacterized protein \| Uncharacterized protein; uniclust \| UniRef100\_A0A6I7NZD3 \| 96.8 \| 0.00015 \| 2.8e-10 \| 60.5 \| 104 \| (27, 134) \| 136 \| (42, 161) \| 393 \| Contractile injection system tube protein N-terminal domain-containing protein \| Contractile injection system tube protein N-terminal domain-containing protein; uniclust \| UniRef100\_A0A0D1XSK1 \| 96.8 \| 0.00014 \| 2.9e-10 \| 58.8 \| 102 \| (28, 135) \| 136 \| (44, 154) \| 248 \| Uncharacterized protein \| Uncharacterized protein; uniclust \| UniRef100\_A0A3M3B317 \| 96.7 \| 0.00017 \| 3.2e-10 \| 51.8 \| 55 \| (19, 73) \| 136 \| (71, 125) \| 142 \| Uncharacterized protein (Fragment) \| Uncharacterized protein (Fragment); uniclust \| UniRef100\_A0A3C0WX82 \| 96.7 \| 0.00018 \| 3.3e-10 \| 57.6 \| 108 \| (27, 135) \| 136 \| (62, 180) \| 340 \| Contractile injection system tube protein N-terminal domain-containing protein \| Contractile injection system tube protein N-terminal domain-containing protein; uniclust \| UniRef100\_A0A6J7DL88 \| 96.7 \| 0.0002 \| 3.6e-10 \| 57.0 \| 122 \| (12, 136) \| 136 \| (57, 215) \| 320 \| Unannotated protein \| Unannotated protein; uniclust \| UniRef100\_A0A6I5PEH0 \| 96.7 \| 0.0002 \| 3.6e-10 \| 57.4 \| 108 \| (28, 136) \| 136 \| (24, 150) \| 341 \| Beta/gamma crystallin 'Greek key' domain-containing protein \| Beta/gamma crystallin 'Greek key' domain-containing protein; uniclust \| UniRef100\_A0A1U7IQP6 \| 96.7 \| 0.0002 \| 3.8e-10 \| 58.3 \| 109 \| (28, 136) \| 136 \| (149, 273) \| 395 \| Contractile injection system tube protein N-terminal domain-containing protein \| Contractile injection system tube protein N-terminal domain-containing protein; uniclust \| UniRef100\_A0A853I0P3 \| 96.7 \| 0.00021 \| 4e-10 \| 55.5 \| 107 \| (28, 135) \| 136 \| (32, 150) \| 225 \| LysM peptidoglycan-binding domain-containing protein \| LysM peptidoglycan-binding domain-containing protein; uniclust \| UniRef100\_A0A6M0RCM0 \| 96.7 \| 0.00021 \| 4.1e-10 \| 54.8 \| 115 \| (11, 135) \| 136 \| (1, 115) \| 184 \| DNA-binding protein \| DNA-binding protein; uniclust \| UniRef100\_A0A061N9X2 \| 96.7 \| 0.00022 \| 4.6e-10 \| 55.7 \| 112 \| (10, 135) \| 136 \| (1, 112) \| 190 \| Uncharacterized protein \| Uncharacterized protein; uniclust \| UniRef100\_A0A377B712 \| 96.6 \| 0.00026 \| 4.8e-10 \| 45.5 \| 50 \| (41, 90) \| 136 \| (12, 61) \| 76 \| Putative phage P2 GpU family protein \| Putative phage P2 GpU family protein; uniclust \| UniRef100\_A0A0E2HQC6 \| 96.6 \| 0.00023 \| 4.8e-10 \| 58.6 \| 107 \| (20, 135) \| 136 \| (45, 164) \| 276 \| LysM domain-containing protein \| LysM domain-containing protein; uniclust \| UniRef100\_A0A072NRR3 \| 96.6 \| 0.00023 \| 4.8e-10 \| 58.3 \| 104 \| (28, 136) \| 136 \| (49, 165) \| 262 \| LysM domain-containing protein \| LysM domain-containing protein; uniclust \| UniRef100\_UPI00055A433D \| 96.6 \| 0.00026 \| 4.8e-10 \| 58.9 \| 107 \| (20, 134) \| 136 \| (23, 155) \| 456 \| hypothetical protein \| hypothetical protein; uniclust \| UniRef100\_A0A8S0FZB8 \| 96.6 \| 0.00028 \| 5.2e-10 \| 46.4 \| 42 \| (13, 55) \| 136 \| (1, 42) \| 86 \| Uncharacterized protein \| Uncharacterized protein; uniclust \| UniRef100\_UPI001F192CD3 \| 96.6 \| 0.00029 \| 5.2e-10 \| 53.8 \| 109 \| (28, 136) \| 136 \| (34, 209) \| 226 \| hypothetical protein \| hypothetical protein; uniclust \| UniRef100\_UPI001E4DB374 \| 96.6 \| 0.0003 \| 5.5e-10 \| 41.9 \| 40 \| (94, 133) \| 136 \| (3, 42) \| 51 \| phage tail protein \| phage tail protein; uniclust \| UniRef100\_UPI001E431A0C \| 96.6 \| 0.00031 \| 5.8e-10 \| 50.8 \| 110 \| (15, 135) \| 136 \| (3, 112) \| 155 \| hypothetical protein \| hypothetical protein; uniclust \| UniRef100\_UPI0022B5F671 \| 96.6 \| 0.00032 \| 5.8e-10 \| 43.2 \| 43 \| (27, 69) \| 136 \| (5, 47) \| 60 \| phage tail protein \| phage tail protein; uniclust \| UniRef100\_UPI001F2F5FEE \| 96.6 \| 0.00032 \| 5.9e-10 \| 47.1 \| 46 \| (20, 65) \| 136 \| (2, 47) \| 97 \| phage tail protein \| phage tail protein; uniclust \| UniRef100\_A0A1I5USI5 \| 96.6 \| 0.00032 \| 5.9e-10 \| 48.6 \| 115 \| (13, 135) \| 136 \| (2, 116) \| 118 \| Phage P2 GpU \| Phage P2 GpU; uniclust \| UniRef100\_A0A327JCS4 \| 96.6 \| 0.00033 \| 6.1e-10 \| 53.8 \| 104 \| (30, 135) \| 136 \| (11, 114) \| 237 \| CYTH domain-containing protein \| CYTH domain-containing protein; uniclust \| UniRef100\_A0A2T4UGD8 \| 96.6 \| 0.00033 \| 6.1e-10 \| 59.8 \| 116 \| (18, 135) \| 136 \| (410, 544) \| 622 \| LysM domain-containing protein \| LysM domain-containing protein; uniclust \| UniRef100\_UPI0008A65577 \| 96.6 \| 0.00038 \| 6.9e-10 \| 48.2 \| 48 \| (48, 95) \| 136 \| (5, 52) \| 116 \| phage tail protein \| phage tail protein; uniclust \| UniRef100\_UPI0018970A81 \| 96.5 \| 0.00038 \| 7e-10 \| 44.2 \| 53 \| (83, 135) \| 136 \| (3, 55) \| 71 \| phage tail protein \| phage tail protein; uniclust \| UniRef100\_UPI00126FC877 \| 96.5 \| 0.0004 \| 7.4e-10 \| 45.4 \| 60 \| (76, 135) \| 136 \| (3, 62) \| 83 \| phage tail protein \| phage tail protein; uniclust \| UniRef100\_UPI0019A7B555 \| 96.5 \| 0.00041 \| 7.5e-10 \| 43.9 \| 41 \| (11, 51) \| 136 \| (1, 42) \| 69 \| hypothetical protein \| hypothetical protein; uniclust \| UniRef100\_UPI001F2E8B4F \| 96.5 \| 0.00043 \| 7.9e-10 \| 41.2 \| 45 \| (91, 135) \| 136 \| (5, 49) \| 50 \| phage tail protein \| phage tail protein; uniclust \| UniRef100\_A0A0P7ZF50 \| 96.5 \| 0.00045 \| 8.2e-10 \| 46.2 \| 35 \| (100, 135) \| 136 \| (11, 45) \| 94 \| Contractile injection system tube protein N-terminal domain-containing protein \| Contractile injection system tube protein N-terminal domain-containing protein; uniclust \| UniRef100\_UPI0002DAFDF3 \| 96.5 \| 0.00052 \| 9.6e-10 \| 53.9 \| 119 \| (13, 135) \| 136 \| (2, 129) \| 280 \| hypothetical protein \| hypothetical protein; uniclust \| UniRef100\_A0A448QV13 \| 96.5 \| 0.00053 \| 9.7e-10 \| 45.8 \| 47 \| (89, 135) \| 136 \| (8, 54) \| 93 \| Phage P2 GpU \| Phage P2 GpU; uniclust \| UniRef100\_A0A432DSZ4 \| 96.5 \| 0.00049 \| 9.9e-10 \| 53.4 \| 82 \| (53, 135) \| 136 \| (16, 115) \| 189 \| LysM peptidoglycan-binding domain-containing protein \| LysM peptidoglycan-binding domain-containing protein; uniclust \| UniRef100\_A0A2E3R265 \| 96.5 \| 0.00054 \| 9.9e-10 \| 49.9 \| 106 \| (20, 129) \| 136 \| (14, 130) \| 160 \| Uncharacterized protein \| Uncharacterized protein; uniclust \| UniRef100\_A0A958XQD9 \| 96.5 \| 0.00055 \| 1e-09 \| 52.9 \| 107 \| (28, 134) \| 136 \| (41, 159) \| 233 \| Contractile injection system tube protein N-terminal domain-containing protein \| Contractile injection system tube protein N-terminal domain-containing protein; uniclust \| UniRef100\_A0A6P0YVG0 \| 96.4 \| 0.00055 \| 1e-09 \| 44.8 \| 46 \| (90, 135) \| 136 \| (10, 55) \| 83 \| Contractile injection system tube protein N-terminal domain-containing protein (Fragment) \| Contractile injection system tube protein N-terminal domain-containing protein (Fragment); uniclust \| UniRef100\_A0A1C4SHA2 \| 96.4 \| 0.00052 \| 1e-09 \| 52.9 \| 71 \| (64, 135) \| 136 \| (8, 94) \| 186 \| LysM domain-containing protein (Fragment) \| LysM domain-containing protein (Fragment); uniclust \| UniRef100\_A0A761H723 \| 96.4 \| 0.00058 \| 1.1e-09 \| 45.9 \| 48 \| (13, 61) \| 136 \| (1, 48) \| 96 \| Uncharacterized protein \| Uncharacterized protein; uniclust \| UniRef100\_A0A0D6PEJ5 \| 96.4 \| 0.00057 \| 1.1e-09 \| 55.2 \| 85 \| (28, 116) \| 136 \| (39, 123) \| 274 \| Phage protein \| Phage protein; uniclust \| UniRef100\_UPI0011AF9C52 \| 96.4 \| 0.00063 \| 1.1e-09 \| 50.9 \| 122 \| (11, 134) \| 136 \| (10, 139) \| 191 \| hypothetical protein \| hypothetical protein; uniclust \| UniRef100\_A0A1G9JFT4 \| 96.4 \| 0.0006 \| 1.2e-09 \| 54.2 \| 115 \| (11, 135) \| 136 \| (4, 121) \| 225 \| DNA circularisation protein N-terminus \| DNA circularisation protein N-terminus; uniclust \| UniRef100\_UPI000D8ED903 \| 96.4 \| 0.00065 \| 1.2e-09 \| 44.3 \| 52 \| (20, 71) \| 136 \| (16, 70) \| 81 \| hypothetical protein \| hypothetical protein; uniclust \| UniRef100\_A0A1G5H8K2 \| 96.4 \| 0.00061 \| 1.3e-09 \| 57.7 \| 59 \| (76, 135) \| 136 \| (216, 275) \| 329 \| Contractile injection system tube protein N-terminal domain-containing protein \| Contractile injection system tube protein N-terminal domain-containing protein; uniclust \| UniRef100\_A0A1Q7LVG0 \| 96.4 \| 0.00067 \| 1.3e-09 \| 56.0 \| 101 \| (28, 135) \| 136 \| (37, 141) \| 287 \| Carboxypeptidase regulatory-like domain-containing protein \| Carboxypeptidase regulatory-like domain-containing protein; uniclust \| UniRef100\_UPI002252A225 \| 96.4 \| 0.00075 \| 1.4e-09 \| 48.2 \| 52 \| (18, 69) \| 136 \| (81, 132) \| 139 \| phage tail protein \| phage tail protein; uniclust \| UniRef100\_A0A081C209 \| 96.4 \| 0.00066 \| 1.4e-09 \| 54.0 \| 103 \| (29, 136) \| 136 \| (15, 120) \| 204 \| Uncharacterized protein \| Uncharacterized protein; uniclust \| UniRef100\_A0A3M0WJW7 \| 96.4 \| 0.00075 \| 1.4e-09 \| 50.0 \| 107 \| (28, 135) \| 136 \| (25, 144) \| 156 \| Uncharacterized protein \| Uncharacterized protein; uniclust \| UniRef100\_A0A9D9KBG8 \| 96.4 \| 0.00076 \| 1.4e-09 \| 52.8 \| 108 \| (28, 135) \| 136 \| (23, 145) \| 238 \| Uncharacterized protein \| Uncharacterized protein; uniclust \| UniRef100\_A0A2E3A8K4 \| 96.3 \| 0.00086 \| 1.6e-09 \| 48.1 \| 99 \| (28, 132) \| 136 \| (27, 135) \| 137 \| Contractile injection system tube protein N-terminal domain-containing protein \| Contractile injection system tube protein N-terminal domain-containing protein; uniclust \| UniRef100\_A0A7C6PPU2 \| 96.3 \| 0.00087 \| 1.6e-09 \| 50.2 \| 109 \| (28, 136) \| 136 \| (42, 163) \| 191 \| Contractile injection system tube protein N-terminal domain-containing protein \| Contractile injection system tube protein N-terminal domain-containing protein; uniclust \| UniRef100\_A0A061NTU5 \| 96.3 \| 0.00089 \| 1.6e-09 \| 43.2 \| 61 \| (74, 135) \| 136 \| (5, 67) \| 75 \| Uncharacterized protein \| Uncharacterized protein; uniclust \| UniRef100\_A0A1C7GNG6 \| 96.3 \| 0.0008 \| 1.7e-09 \| 52.4 \| 103 \| (4, 112) \| 136 \| (1, 103) \| 177 \| LysM domain-containing protein \| LysM domain-containing protein; pdb70 \| 6J0N\_d \| 99.3 \| 5.1e-16 \| 4.4e-20 \| 109.7 \| 112 \| (23, 135) \| 136 \| (28, 155) \| 229 \| Pvc1, Pvc9, Pvc11, Pvc12, Pvc4 \| 6J0N\_d Pvc1, Pvc9, Pvc11, Pvc12, Pvc4 assembly, Photorhabdus asymbiotica, PVC, contractile; pdb70 \| 6RAO\_G \| 99.2 \| 1.1e-15 \| 9.2e-20 \| 107.8 \| 112 \| (23, 135) \| 136 \| (28, 155) \| 229 \| Afp1, Afp2, Afp3, Afp5, Afp9 \| 6RAO\_G Afp1, Afp2, Afp3, Afp5, Afp9 Anti-feeding prophage, secretion system, AFP |
| Top keywords  (threshold 1.00e-03 (evalue)) | **Phage, tail, domain\_containing, Contractile, system, injection, N\_terminal, tube, GpU, LysM** |
| Output files | ../../similar\_sequences/23\_FANPEZAQ\_CDS\_0023\_merged.svg ../../similar\_sequences/23\_FANPEZAQ\_CDS\_0023\_pdb70.a3m ../../similar\_sequences/23\_FANPEZAQ\_CDS\_0023\_pdb70.hhr ../../similar\_sequences/23\_FANPEZAQ\_CDS\_0023\_uniclust.a3m ../../similar\_sequences/23\_FANPEZAQ\_CDS\_0023\_uniclust.hhr |

#### Structure prediction (AlphaFold)^2^

|  |  |
| --- | --- |
| Stats | xml version="1.0" encoding="utf-8" standalone="no"?       2024-09-02T21:09:23.942820 image/svg+xml   Matplotlib v3.7.2, https://matplotlib.org/ |
| Predicted structure | **NGL Viewer Controls:**  - Center: *Left-Click* - Rotate: *Left-Click + Drag* - Translate: *Right-Click + Drag* - Zoom: *Shift + Left-Click + Drag* |
| Output files | ../../predicted\_structures/23\_FANPEZAQ\_CDS\_0023/features.pkl ../../predicted\_structures/23\_FANPEZAQ\_CDS\_0023/ranked\_0.pdb ../../predicted\_structures/23\_FANPEZAQ\_CDS\_0023/ranked\_0\_plots.svg ../../predicted\_structures/23\_FANPEZAQ\_CDS\_0023/result\_model\_1\_ptm\_pred\_0.pkl |

#### Structure similarity search results (Foldseek)^3^

|  |  |
| --- | --- |
| Structure databases searched | Pdb, Afdb-proteome, Afdb-uniprot50 |
| Results, scheme(s)  (Top layers only, threshold 1.00e-02 (evalue)) | xml version="1.0" encoding="utf-8" standalone="no"?       2024-09-02T21:10:52.344980 image/svg+xml   Matplotlib v3.7.2, https://matplotlib.org/ |
| Results, table  (threshold 1.00e-02 (evalue)) | db \| id \| prob \| evalue \| bits \| fident \| alnlen \| mismatch \| gapopen \| qstart \| qend \| tstart \| tend \| name \| description; pdb \| 6U5B\_H \| 1.0 \| 1.055e-11 \| 483 \| 0.378 \| 140 \| 71 \| 3 \| 13 \| 136 \| 144 \| 283 \| Ripcord PA0626 \| Ripcord PA0626; pdb \| 6J0N\_B \| 1.0 \| 2.363e-05 \| 199 \| 0.147 \| 122 \| 87 \| 6 \| 29 \| 134 \| 32 \| 152 \| Pvc7 \| Pvc7; pdb \| 7AEB\_M \| 1.0 \| 2.494e-05 \| 173 \| 0.114 \| 149 \| 106 \| 6 \| 13 \| 136 \| 8 \| 155 \| Phage tail protein \| Phage tail protein; pdb \| 7B5H\_AK \| 1.0 \| 2.928e-05 \| 169 \| 0.126 \| 158 \| 102 \| 9 \| 13 \| 136 \| 5 \| 160 \| All3321 protein \| All3321 protein; pdb \| 2X8K\_C \| 1.0 \| 0.0002495 \| 169 \| 0.097 \| 133 \| 99 \| 8 \| 13 \| 136 \| 3 \| 123 \| HYPOTHETICAL PROTEIN 19.1 \| HYPOTHETICAL PROTEIN 19.1; pdb \| 6RBK\_A \| 1.0 \| 0.000146 \| 160 \| 0.127 \| 118 \| 88 \| 4 \| 30 \| 132 \| 34 \| 151 \| Afp7 \| Afp7; pdb \| 8EON\_C \| 1.0 \| 0.0008551 \| 153 \| 0.096 \| 135 \| 110 \| 5 \| 4 \| 136 \| 5 \| 129 \| Baseplate component gp37 \| Baseplate component gp37; pdb \| 6TUI\_G \| 1.0 \| 0.0007282 \| 150 \| 0.125 \| 136 \| 95 \| 7 \| 12 \| 136 \| 6 \| 128 \| Tail tube protein Rcc01691 \| Tail tube protein Rcc01691; pdb \| 7YFZ\_B \| 1.0 \| 0.001244 \| 147 \| 0.112 \| 124 \| 98 \| 4 \| 16 \| 136 \| 1 \| 115 \| Pam3 tube initiator gp17 \| Pam3 tube initiator gp17; pdb \| 6V8I\_AC \| 1.0 \| 0.001004 \| 121 \| 0.081 \| 172 \| 100 \| 9 \| 13 \| 136 \| 3 \| 164 \| Distal Tail Protein, gp58 \| Distal Tail Protein, gp58; pdb \| 6TBA\_5A \| 1.0 \| 0.003263 \| 117 \| 0.127 \| 141 \| 96 \| 10 \| 6 \| 136 \| 1 \| 124 \| Phage major tail protein, TP901-1 family \| Phage major tail protein, TP901-1 family; pdb \| 8HDR\_1 \| 1.0 \| 0.004264 \| 109 \| 0.107 \| 140 \| 102 \| 8 \| 6 \| 134 \| 4 \| 131 \| pam3 tube protein \| pam3 tube protein; pdb \| 6V8I\_EA \| 1.0 \| 0.006204 \| 109 \| 0.094 \| 148 \| 102 \| 11 \| 7 \| 136 \| 1 \| 134 \| Major Tail Protein, gp53 \| Major Tail Protein, gp53; pdb \| 6RAO\_A \| 1.0 \| 0.00191 \| 104 \| 0.113 \| 150 \| 102 \| 9 \| 2 \| 136 \| 9 \| 142 \| Afp1 \| Afp1; pdb \| 5IV5\_U \| 1.0 \| 0.003831 \| 100 \| 0.11 \| 127 \| 78 \| 9 \| 37 \| 136 \| 41 \| 159 \| Baseplate tail-tube protein gp48 \| Baseplate tail-tube protein gp48; pdb \| 6J0F\_A \| 0.998 \| 0.005881 \| 95 \| 0.108 \| 147 \| 100 \| 9 \| 1 \| 131 \| 1 \| 132 \| Pvc1 \| Pvc1; pdb \| 6J0B\_A \| 0.998 \| 0.001313 \| 94 \| 0.124 \| 145 \| 104 \| 9 \| 2 \| 134 \| 9 \| 142 \| Pvc1 \| Pvc1; pdb \| 6V8I\_EB \| 0.998 \| 0.004499 \| 93 \| 0.121 \| 148 \| 93 \| 14 \| 10 \| 136 \| 3 \| 134 \| Major Tail Protein, gp53 \| Major Tail Protein, gp53; pdb \| 8BL4\_P \| 0.988 \| 0.006546 \| 82 \| 0.091 \| 131 \| 88 \| 10 \| 29 \| 136 \| 24 \| 146 \| Phage tail protein \| Phage tail protein; pdb \| 6RAO\_F \| 0.923 \| 0.005881 \| 69 \| 0.133 \| 142 \| 93 \| 10 \| 13 \| 136 \| 12 \| 141 \| Afp5 \| Afp5; afdb-proteome \| AF-Q8ZMV4-F1-MODEL\_V4 \| 1.0 \| 7.529e-13 \| 587 \| 0.346 \| 124 \| 81 \| 0 \| 13 \| 136 \| 1 \| 124 \| Fels-2 prophage protein \| Fels-2 prophage protein; afdb-proteome \| AF-A0A0H3GQH5-F1-MODEL\_V4 \| 1.0 \| 6.411e-13 \| 582 \| 0.338 \| 124 \| 82 \| 0 \| 13 \| 136 \| 1 \| 124 \| Putative bacteriophage tail protein \| Putative bacteriophage tail protein; afdb-proteome \| AF-G3XD65-F1-MODEL\_V4 \| 1.0 \| 7.943e-13 \| 536 \| 0.35 \| 154 \| 81 \| 5 \| 2 \| 136 \| 131 \| 284 \| Uncharacterized protein \| Uncharacterized protein; afdb-proteome \| AF-Q8ZKJ5-F1-MODEL\_V4 \| 1.0 \| 2.727e-09 \| 403 \| 0.227 \| 123 \| 94 \| 1 \| 13 \| 135 \| 1 \| 122 \| Putative methyl-accepting chemotaxis protein \| Putative methyl-accepting chemotaxis protein; afdb-proteome \| AF-A0A0H3GS01-F1-MODEL\_V4 \| 1.0 \| 5.783e-05 \| 222 \| 0.424 \| 73 \| 42 \| 0 \| 64 \| 136 \| 1 \| 73 \| Putative prophage tail protein \| Putative prophage tail protein; afdb-proteome \| AF-P71389-F1-MODEL\_V4 \| 1.0 \| 4.668e-05 \| 199 \| 0.099 \| 131 \| 106 \| 5 \| 7 \| 136 \| 1 \| 120 \| Mu-like prophage FluMu DNA circularization protein \| Mu-like prophage FluMu DNA circularization protein; afdb-proteome \| AF-A0A3P7DQ66-F1-MODEL\_V4 \| 1.0 \| 0.0002884 \| 154 \| 0.11 \| 163 \| 100 \| 10 \| 13 \| 136 \| 328 \| 484 \| Uncharacterized protein \| Uncharacterized protein; afdb-proteome \| AF-A0A0H3GKS4-F1-MODEL\_V4 \| 1.0 \| 0.000377 \| 145 \| 0.091 \| 153 \| 101 \| 10 \| 13 \| 136 \| 31 \| 174 \| Uncharacterized protein \| Uncharacterized protein; afdb-proteome \| AF-A0A0H3H550-F1-MODEL\_V4 \| 1.0 \| 0.006798 \| 131 \| 0.118 \| 110 \| 82 \| 5 \| 30 \| 136 \| 12 \| 109 \| Minor tail fiber protein M \| Minor tail fiber protein M; afdb-proteome \| AF-A0A133CKK6-F1-MODEL\_V4 \| 1.0 \| 0.003212 \| 121 \| 0.156 \| 128 \| 94 \| 7 \| 13 \| 133 \| 5 \| 125 \| Phage tail protein \| Phage tail protein; afdb-proteome \| AF-Q57791-F1-MODEL\_V4 \| 1.0 \| 0.002592 \| 121 \| 0.086 \| 127 \| 95 \| 7 \| 13 \| 136 \| 209 \| 317 \| Uncharacterized protein MJ0345 \| Uncharacterized protein MJ0345; afdb-proteome \| AF-Q8ZQ89-F1-MODEL\_V4 \| 1.0 \| 0.002592 \| 114 \| 0.086 \| 162 \| 105 \| 12 \| 1 \| 136 \| 6 \| 150 \| Gifsy-2 prophage probable major tail protein \| Gifsy-2 prophage probable major tail protein; afdb-proteome \| AF-Q2FX66-F1-MODEL\_V4 \| 1.0 \| 0.002592 \| 110 \| 0.108 \| 157 \| 82 \| 9 \| 27 \| 136 \| 20 \| 165 \| Conserved hypothetical phage protein \| Conserved hypothetical phage protein; afdb-proteome \| AF-Q2FYC9-F1-MODEL\_V4 \| 1.0 \| 0.003044 \| 109 \| 0.09 \| 132 \| 99 \| 9 \| 13 \| 133 \| 5 \| 126 \| Holin-like protein \| Holin-like protein; afdb-proteome \| AF-A0A077ZLB1-F1-MODEL\_V4 \| 1.0 \| 0.006798 \| 101 \| 0.118 \| 143 \| 96 \| 11 \| 19 \| 134 \| 23 \| 162 \| Uncharacterized protein \| Uncharacterized protein; afdb-proteome \| AF-Q2FX62-F1-MODEL\_V4 \| 0.998 \| 0.008886 \| 94 \| 0.106 \| 141 \| 90 \| 11 \| 13 \| 134 \| 12 \| 135 \| Phage structural protein, putative \| Phage structural protein, putative; afdb-uniprot50 \| AF-A0A1V2DRL6-F1-MODEL\_V4 \| 1.0 \| 2.002e-17 \| 820 \| 0.559 \| 127 \| 56 \| 0 \| 10 \| 136 \| 1 \| 127 \| Phage tail protein \| Phage tail protein; afdb-uniprot50 \| AF-A0A6M4Y8X7-F1-MODEL\_V4 \| 1.0 \| 1.304e-17 \| 816 \| 0.564 \| 131 \| 57 \| 0 \| 6 \| 136 \| 3 \| 133 \| Phage tail protein \| Phage tail protein; afdb-uniprot50 \| AF-A0A660NPZ3-F1-MODEL\_V4 \| 1.0 \| 1.053e-16 \| 794 \| 0.559 \| 127 \| 56 \| 0 \| 10 \| 136 \| 8 \| 134 \| Phage tail protein \| Phage tail protein; afdb-uniprot50 \| AF-A0A2W5L0T9-F1-MODEL\_V4 \| 1.0 \| 8.057e-17 \| 790 \| 0.58 \| 124 \| 52 \| 0 \| 13 \| 136 \| 10 \| 133 \| Uncharacterized protein \| Uncharacterized protein; afdb-uniprot50 \| AF-A0A1G6JEA6-F1-MODEL\_V4 \| 1.0 \| 6.503e-17 \| 788 \| 0.492 \| 130 \| 66 \| 0 \| 7 \| 136 \| 5 \| 134 \| Uncharacterized protein \| Uncharacterized protein; afdb-uniprot50 \| AF-A0A2Z3IG89-F1-MODEL\_V4 \| 1.0 \| 2.228e-17 \| 782 \| 0.544 \| 136 \| 59 \| 1 \| 1 \| 136 \| 1 \| 133 \| Phage tail protein \| Phage tail protein; afdb-uniprot50 \| AF-A0A315BRY9-F1-MODEL\_V4 \| 1.0 \| 6.164e-17 \| 768 \| 0.588 \| 136 \| 53 \| 1 \| 1 \| 136 \| 2 \| 134 \| Uncharacterized protein \| Uncharacterized protein; afdb-uniprot50 \| AF-A0A410UF80-F1-MODEL\_V4 \| 1.0 \| 2.914e-16 \| 767 \| 0.434 \| 129 \| 73 \| 0 \| 8 \| 136 \| 25 \| 153 \| Phage tail protein \| Phage tail protein; afdb-uniprot50 \| AF-A0A2Z6AYW2-F1-MODEL\_V4 \| 1.0 \| 2.762e-16 \| 766 \| 0.496 \| 127 \| 64 \| 0 \| 10 \| 136 \| 1 \| 127 \| P2 GpU family protein \| P2 GpU family protein; afdb-uniprot50 \| AF-A0A081N7U3-F1-MODEL\_V4 \| 1.0 \| 3.422e-16 \| 762 \| 0.417 \| 127 \| 74 \| 0 \| 10 \| 136 \| 1 \| 127 \| Tail protein \| Tail protein; afdb-uniprot50 \| AF-A0A1N6I0S3-F1-MODEL\_V4 \| 1.0 \| 5.846e-16 \| 758 \| 0.472 \| 127 \| 67 \| 0 \| 10 \| 136 \| 10 \| 136 \| Uncharacterized protein \| Uncharacterized protein; afdb-uniprot50 \| AF-A0A291LZ32-F1-MODEL\_V4 \| 1.0 \| 3.422e-16 \| 757 \| 0.425 \| 127 \| 73 \| 0 \| 10 \| 136 \| 1 \| 127 \| Phage tail protein \| Phage tail protein; afdb-uniprot50 \| AF-A0A212KMX5-F1-MODEL\_V4 \| 1.0 \| 7.641e-16 \| 756 \| 0.511 \| 127 \| 62 \| 0 \| 10 \| 136 \| 1 \| 127 \| Uncharacterized protein \| Uncharacterized protein; afdb-uniprot50 \| AF-A0A5A9EN75-F1-MODEL\_V4 \| 1.0 \| 8.505e-16 \| 756 \| 0.409 \| 127 \| 75 \| 0 \| 10 \| 136 \| 1 \| 127 \| Phage tail protein \| Phage tail protein; afdb-uniprot50 \| AF-A0A212KBL8-F1-MODEL\_V4 \| 1.0 \| 8.973e-16 \| 754 \| 0.488 \| 127 \| 65 \| 0 \| 10 \| 136 \| 1 \| 127 \| Uncharacterized protein \| Uncharacterized protein; afdb-uniprot50 \| AF-A0A4Z0WCE3-F1-MODEL\_V4 \| 1.0 \| 1.706e-15 \| 747 \| 0.451 \| 124 \| 68 \| 0 \| 13 \| 136 \| 31 \| 154 \| Phage tail protein \| Phage tail protein; afdb-uniprot50 \| AF-A0A430HF79-F1-MODEL\_V4 \| 1.0 \| 1.453e-15 \| 746 \| 0.456 \| 127 \| 69 \| 0 \| 10 \| 136 \| 1 \| 127 \| Phage tail protein \| Phage tail protein; afdb-uniprot50 \| AF-A0A4U8YGX8-F1-MODEL\_V4 \| 1.0 \| 1.305e-15 \| 739 \| 0.484 \| 126 \| 65 \| 0 \| 11 \| 136 \| 17 \| 142 \| Myoviridae gpu \| Myoviridae gpu; afdb-uniprot50 \| AF-A0A7C8M0S1-F1-MODEL\_V4 \| 1.0 \| 5.846e-16 \| 738 \| 0.462 \| 132 \| 71 \| 0 \| 5 \| 136 \| 20 \| 151 \| Uncharacterized protein \| Uncharacterized protein; afdb-uniprot50 \| AF-A0A291G0E7-F1-MODEL\_V4 \| 1.0 \| 1.706e-15 \| 737 \| 0.48 \| 127 \| 66 \| 0 \| 10 \| 136 \| 2 \| 128 \| Phage-related tail protein (GpU) \| Phage-related tail protein (GpU); afdb-uniprot50 \| AF-A0A345DE50-F1-MODEL\_V4 \| 1.0 \| 1.533e-15 \| 736 \| 0.382 \| 128 \| 79 \| 0 \| 9 \| 136 \| 34 \| 161 \| Uncharacterized protein \| Uncharacterized protein; afdb-uniprot50 \| AF-Q31Q86-F1-MODEL\_V4 \| 1.0 \| 2.619e-15 \| 736 \| 0.464 \| 127 \| 68 \| 0 \| 10 \| 136 \| 1 \| 127 \| Uncharacterized protein \| Uncharacterized protein; afdb-uniprot50 \| AF-B8GS04-F1-MODEL\_V4 \| 1.0 \| 2.114e-15 \| 730 \| 0.419 \| 124 \| 72 \| 0 \| 13 \| 136 \| 1 \| 124 \| Putative phage tail protein U \| Putative phage tail protein U; afdb-uniprot50 \| AF-A0A1S1TJY4-F1-MODEL\_V4 \| 1.0 \| 1.377e-15 \| 730 \| 0.472 \| 125 \| 66 \| 0 \| 12 \| 136 \| 2 \| 126 \| Uncharacterized protein \| Uncharacterized protein; afdb-uniprot50 \| AF-A0A1G7SAR7-F1-MODEL\_V4 \| 1.0 \| 1.706e-15 \| 727 \| 0.46 \| 128 \| 68 \| 1 \| 10 \| 136 \| 1 \| 128 \| Uncharacterized protein \| Uncharacterized protein; afdb-uniprot50 \| AF-A0A118DSW1-F1-MODEL\_V4 \| 1.0 \| 8.505e-16 \| 720 \| 0.448 \| 136 \| 74 \| 1 \| 1 \| 136 \| 1 \| 135 \| Uncharacterized protein \| Uncharacterized protein; afdb-uniprot50 \| AF-A0A8A6KDU1-F1-MODEL\_V4 \| 1.0 \| 4.475e-15 \| 718 \| 0.537 \| 121 \| 56 \| 0 \| 14 \| 134 \| 1 \| 121 \| Phage tail protein \| Phage tail protein; afdb-uniprot50 \| AF-A0A6S6Y3Q4-F1-MODEL\_V4 \| 1.0 \| 3.245e-15 \| 715 \| 0.433 \| 127 \| 72 \| 0 \| 10 \| 136 \| 2 \| 128 \| Phage tail protein \| Phage tail protein; afdb-uniprot50 \| AF-A0A1L2AZL7-F1-MODEL\_V4 \| 1.0 \| 1.305e-15 \| 713 \| 0.419 \| 136 \| 79 \| 0 \| 1 \| 136 \| 16 \| 151 \| Phage tail protein \| Phage tail protein; afdb-uniprot50 \| AF-C6BVX0-F1-MODEL\_V4 \| 1.0 \| 5.544e-15 \| 712 \| 0.395 \| 124 \| 75 \| 0 \| 13 \| 136 \| 2 \| 125 \| p2 GpU family protein \| p2 GpU family protein; afdb-uniprot50 \| AF-A0A7K0GNT9-F1-MODEL\_V4 \| 1.0 \| 6.167e-16 \| 707 \| 0.422 \| 135 \| 78 \| 0 \| 2 \| 136 \| 29 \| 163 \| Uncharacterized protein \| Uncharacterized protein; afdb-uniprot50 \| AF-K5Z1D9-F1-MODEL\_V4 \| 1.0 \| 1.706e-15 \| 707 \| 0.448 \| 136 \| 74 \| 1 \| 1 \| 136 \| 5 \| 139 \| Phage P2 GpU \| Phage P2 GpU; afdb-uniprot50 \| AF-A0A239EIZ1-F1-MODEL\_V4 \| 1.0 \| 2.915e-15 \| 705 \| 0.438 \| 130 \| 73 \| 0 \| 7 \| 136 \| 1 \| 130 \| Uncharacterized protein \| Uncharacterized protein; afdb-uniprot50 \| AF-A0A1C3ELB9-F1-MODEL\_V4 \| 1.0 \| 6.51e-15 \| 703 \| 0.491 \| 124 \| 63 \| 0 \| 13 \| 136 \| 1 \| 124 \| Phage tail protein \| Phage tail protein; afdb-uniprot50 \| AF-A0A1C3K3E1-F1-MODEL\_V4 \| 1.0 \| 6.868e-15 \| 703 \| 0.42 \| 126 \| 73 \| 0 \| 11 \| 136 \| 3 \| 128 \| Phage tail protein \| Phage tail protein; afdb-uniprot50 \| AF-A0A4P5VNR5-F1-MODEL\_V4 \| 1.0 \| 8.065e-15 \| 702 \| 0.416 \| 125 \| 73 \| 0 \| 12 \| 136 \| 2 \| 126 \| Uncharacterized protein \| Uncharacterized protein; afdb-uniprot50 \| AF-A0A6B3LC06-F1-MODEL\_V4 \| 1.0 \| 8.065e-15 \| 700 \| 0.343 \| 128 \| 83 \| 1 \| 10 \| 136 \| 1 \| 128 \| Phage tail protein \| Phage tail protein; afdb-uniprot50 \| AF-A0A3N2E1A2-F1-MODEL\_V4 \| 1.0 \| 3.81e-15 \| 700 \| 0.395 \| 134 \| 81 \| 0 \| 3 \| 136 \| 15 \| 148 \| Uncharacterized protein \| Uncharacterized protein; afdb-uniprot50 \| AF-A0A285D6T1-F1-MODEL\_V4 \| 1.0 \| 1.054e-14 \| 696 \| 0.373 \| 126 \| 79 \| 0 \| 10 \| 135 \| 1 \| 126 \| Uncharacterized protein \| Uncharacterized protein; afdb-uniprot50 \| AF-A0A348HI90-F1-MODEL\_V4 \| 1.0 \| 8.065e-15 \| 695 \| 0.425 \| 127 \| 73 \| 0 \| 10 \| 136 \| 2 \| 128 \| Phage protein U \| Phage protein U; afdb-uniprot50 \| AF-A0A3G6WB93-F1-MODEL\_V4 \| 1.0 \| 1.534e-14 \| 689 \| 0.428 \| 126 \| 72 \| 0 \| 10 \| 135 \| 1 \| 126 \| Phage tail protein \| Phage tail protein; afdb-uniprot50 \| AF-A0A2E3N1P0-F1-MODEL\_V4 \| 1.0 \| 5.254e-15 \| 686 \| 0.425 \| 127 \| 73 \| 0 \| 10 \| 136 \| 2 \| 128 \| Phage tail protein \| Phage tail protein; afdb-uniprot50 \| AF-A0A1Y2K0S7-F1-MODEL\_V4 \| 1.0 \| 5.544e-15 \| 685 \| 0.431 \| 132 \| 74 \| 1 \| 6 \| 136 \| 2 \| 133 \| Putative P2 GpU family protein \| Putative P2 GpU family protein; afdb-uniprot50 \| AF-A0A158E8Q2-F1-MODEL\_V4 \| 1.0 \| 8.977e-15 \| 683 \| 0.426 \| 129 \| 74 \| 0 \| 8 \| 136 \| 5 \| 133 \| Bacteriophage tail-related protein \| Bacteriophage tail-related protein; afdb-uniprot50 \| AF-A0A1E3G8A9-F1-MODEL\_V4 \| 1.0 \| 1.618e-14 \| 682 \| 0.401 \| 127 \| 76 \| 0 \| 10 \| 136 \| 2 \| 128 \| Uncharacterized protein \| Uncharacterized protein; afdb-uniprot50 \| AF-J7SX82-F1-MODEL\_V4 \| 1.0 \| 6.51e-15 \| 681 \| 0.401 \| 132 \| 79 \| 0 \| 5 \| 136 \| 19 \| 150 \| Uncharacterized protein \| Uncharacterized protein; afdb-uniprot50 \| AF-A0A6G8F2C6-F1-MODEL\_V4 \| 1.0 \| 4.02e-15 \| 681 \| 0.413 \| 133 \| 76 \| 1 \| 6 \| 136 \| 18 \| 150 \| Uncharacterized protein \| Uncharacterized protein; afdb-uniprot50 \| AF-A0A4R3LHW8-F1-MODEL\_V4 \| 1.0 \| 1.801e-14 \| 680 \| 0.42 \| 126 \| 73 \| 0 \| 11 \| 136 \| 3 \| 128 \| Uncharacterized protein \| Uncharacterized protein; afdb-uniprot50 \| AF-A0A349FC45-F1-MODEL\_V4 \| 1.0 \| 3.077e-14 \| 679 \| 0.33 \| 127 \| 85 \| 0 \| 10 \| 136 \| 1 \| 127 \| Phage tail protein \| Phage tail protein; afdb-uniprot50 \| AF-A0A7Y4DZ99-F1-MODEL\_V4 \| 1.0 \| 3.246e-14 \| 679 \| 0.328 \| 128 \| 85 \| 1 \| 10 \| 136 \| 1 \| 128 \| Phage tail protein \| Phage tail protein; afdb-uniprot50 \| AF-A0A4R3L5P7-F1-MODEL\_V4 \| 1.0 \| 4.721e-15 \| 679 \| 0.433 \| 136 \| 66 \| 2 \| 12 \| 136 \| 2 \| 137 \| Uncharacterized protein \| Uncharacterized protein; afdb-uniprot50 \| AF-A0A4P9VGW4-F1-MODEL\_V4 \| 1.0 \| 3.613e-14 \| 678 \| 0.37 \| 124 \| 78 \| 0 \| 13 \| 136 \| 186 \| 309 \| Uncharacterized protein \| Uncharacterized protein; afdb-uniprot50 \| AF-A0A2S2E5C8-F1-MODEL\_V4 \| 1.0 \| 2.917e-14 \| 676 \| 0.359 \| 128 \| 82 \| 0 \| 9 \| 136 \| 2 \| 129 \| Putative tape measure protein \| Putative tape measure protein; afdb-uniprot50 \| AF-A0A1X7Q475-F1-MODEL\_V4 \| 1.0 \| 1.707e-14 \| 673 \| 0.409 \| 127 \| 75 \| 0 \| 10 \| 136 \| 2 \| 128 \| Uncharacterized protein \| Uncharacterized protein; afdb-uniprot50 \| AF-A0A066RZT5-F1-MODEL\_V4 \| 1.0 \| 1.618e-14 \| 672 \| 0.359 \| 128 \| 81 \| 1 \| 10 \| 136 \| 1 \| 128 \| p2 GpU family protein \| p2 GpU family protein; afdb-uniprot50 \| AF-I3TTC4-F1-MODEL\_V4 \| 1.0 \| 5.257e-14 \| 672 \| 0.406 \| 123 \| 73 \| 0 \| 14 \| 136 \| 285 \| 407 \| Uncharacterized protein \| Uncharacterized protein; afdb-uniprot50 \| AF-A0A1Y1QYE0-F1-MODEL\_V4 \| 1.0 \| 2.231e-14 \| 670 \| 0.437 \| 128 \| 68 \| 2 \| 13 \| 136 \| 1 \| 128 \| Uncharacterized protein \| Uncharacterized protein; afdb-uniprot50 \| AF-A0A135IJT4-F1-MODEL\_V4 \| 1.0 \| 3.613e-14 \| 665 \| 0.44 \| 127 \| 71 \| 0 \| 10 \| 136 \| 2 \| 128 \| Phage tail protein \| Phage tail protein; afdb-uniprot50 \| AF-A0A145VQC9-F1-MODEL\_V4 \| 1.0 \| 6.868e-15 \| 665 \| 0.398 \| 148 \| 68 \| 2 \| 10 \| 136 \| 1 \| 148 \| Uncharacterized protein \| Uncharacterized protein; afdb-uniprot50 \| AF-A0A149SP38-F1-MODEL\_V4 \| 1.0 \| 3.812e-14 \| 662 \| 0.341 \| 129 \| 85 \| 0 \| 8 \| 136 \| 21 \| 149 \| Uncharacterized protein \| Uncharacterized protein; afdb-uniprot50 \| AF-A0A3R8L896-F1-MODEL\_V4 \| 1.0 \| 5.257e-14 \| 661 \| 0.39 \| 128 \| 77 \| 1 \| 10 \| 136 \| 1 \| 128 \| Phage tail protein \| Phage tail protein; afdb-uniprot50 \| AF-A0A4P7EJT5-F1-MODEL\_V4 \| 1.0 \| 2.354e-14 \| 661 \| 0.423 \| 130 \| 75 \| 0 \| 7 \| 136 \| 1 \| 130 \| Phage tail protein \| Phage tail protein; afdb-uniprot50 \| AF-A0A0F9VVN6-F1-MODEL\_V4 \| 1.0 \| 6.173e-14 \| 658 \| 0.475 \| 124 \| 65 \| 0 \| 13 \| 136 \| 1 \| 124 \| Uncharacterized protein \| Uncharacterized protein; afdb-uniprot50 \| AF-A0A376TF55-F1-MODEL\_V4 \| 1.0 \| 8.982e-14 \| 658 \| 0.443 \| 124 \| 69 \| 0 \| 13 \| 136 \| 1 \| 124 \| GpU phage protein \| GpU phage protein; afdb-uniprot50 \| AF-A0A554X131-F1-MODEL\_V4 \| 1.0 \| 4.983e-14 \| 657 \| 0.388 \| 126 \| 77 \| 0 \| 11 \| 136 \| 3 \| 128 \| Phage P2 GpU \| Phage P2 GpU; afdb-uniprot50 \| AF-A0A4S5DES3-F1-MODEL\_V4 \| 1.0 \| 4.022e-14 \| 655 \| 0.362 \| 127 \| 81 \| 0 \| 10 \| 136 \| 1 \| 127 \| Phage tail protein \| Phage tail protein; afdb-uniprot50 \| AF-A0A4R3LF92-F1-MODEL\_V4 \| 1.0 \| 5.546e-14 \| 654 \| 0.433 \| 127 \| 72 \| 0 \| 10 \| 136 \| 1 \| 127 \| Uncharacterized protein \| Uncharacterized protein; afdb-uniprot50 \| AF-A0A849B434-F1-MODEL\_V4 \| 1.0 \| 9.476e-14 \| 654 \| 0.411 \| 124 \| 73 \| 0 \| 13 \| 136 \| 1 \| 124 \| Oxidoreductase \| Oxidoreductase; afdb-uniprot50 \| AF-A0A554XB21-F1-MODEL\_V4 \| 1.0 \| 5.257e-14 \| 652 \| 0.433 \| 127 \| 72 \| 0 \| 10 \| 136 \| 1 \| 127 \| Phage P2 GpU \| Phage P2 GpU; afdb-uniprot50 \| AF-A0A7S8C741-F1-MODEL\_V4 \| 1.0 \| 5.257e-14 \| 650 \| 0.385 \| 127 \| 78 \| 0 \| 10 \| 136 \| 1 \| 127 \| Phage tail protein \| Phage tail protein; afdb-uniprot50 \| AF-V4P7Q6-F1-MODEL\_V4 \| 1.0 \| 1.055e-13 \| 649 \| 0.411 \| 124 \| 73 \| 0 \| 13 \| 136 \| 2 \| 125 \| Oxidoreductase \| Oxidoreductase; afdb-uniprot50 \| AF-A0A326L399-F1-MODEL\_V4 \| 1.0 \| 1.534e-13 \| 649 \| 0.336 \| 125 \| 83 \| 0 \| 12 \| 136 \| 2 \| 126 \| Oxidoreductase \| Oxidoreductase; afdb-uniprot50 \| AF-A0A6L2ZPM8-F1-MODEL\_V4 \| 1.0 \| 1.534e-13 \| 647 \| 0.427 \| 124 \| 71 \| 0 \| 13 \| 136 \| 1 \| 124 \| Tail protein \| Tail protein; afdb-uniprot50 \| AF-A0A4P9VFE7-F1-MODEL\_V4 \| 1.0 \| 1.708e-13 \| 647 \| 0.365 \| 126 \| 80 \| 0 \| 11 \| 136 \| 3 \| 128 \| Phage tail protein \| Phage tail protein; afdb-uniprot50 \| AF-A0A1N6Q3K9-F1-MODEL\_V4 \| 1.0 \| 9.476e-14 \| 646 \| 0.403 \| 124 \| 74 \| 0 \| 13 \| 136 \| 1 \| 124 \| Uncharacterized protein \| Uncharacterized protein; afdb-uniprot50 \| AF-A0A3M3TVG5-F1-MODEL\_V4 \| 1.0 \| 8.513e-14 \| 643 \| 0.381 \| 131 \| 74 \| 1 \| 13 \| 136 \| 23 \| 153 \| Uncharacterized protein \| Uncharacterized protein; afdb-uniprot50 \| AF-A0A7V8FKF8-F1-MODEL\_V4 \| 1.0 \| 3.812e-14 \| 642 \| 0.387 \| 129 \| 79 \| 0 \| 8 \| 136 \| 23 \| 151 \| Uncharacterized protein \| Uncharacterized protein; afdb-uniprot50 \| AF-A0A6L9FGD7-F1-MODEL\_V4 \| 1.0 \| 5.852e-14 \| 642 \| 0.375 \| 136 \| 84 \| 1 \| 2 \| 136 \| 17 \| 152 \| Uncharacterized protein \| Uncharacterized protein; afdb-uniprot50 \| AF-N9WEM7-F1-MODEL\_V4 \| 1.0 \| 1.174e-13 \| 642 \| 0.387 \| 129 \| 79 \| 0 \| 8 \| 136 \| 57 \| 185 \| Phage P2 GpU protein \| Phage P2 GpU protein; afdb-uniprot50 \| AF-A0A429LGV1-F1-MODEL\_V4 \| 1.0 \| 2.232e-13 \| 641 \| 0.443 \| 124 \| 69 \| 0 \| 13 \| 136 \| 1 \| 124 \| Phage tail protein \| Phage tail protein; afdb-uniprot50 \| AF-A0A524RVY3-F1-MODEL\_V4 \| 1.0 \| 1.802e-13 \| 641 \| 0.307 \| 127 \| 88 \| 0 \| 10 \| 136 \| 1 \| 127 \| Phage tail protein \| Phage tail protein; afdb-uniprot50 \| AF-A0A7W6RFB2-F1-MODEL\_V4 \| 1.0 \| 4.022e-14 \| 640 \| 0.4 \| 135 \| 73 \| 3 \| 10 \| 136 \| 1 \| 135 \| Uncharacterized protein \| Uncharacterized protein; afdb-uniprot50 \| AF-A0A1D2QSE6-F1-MODEL\_V4 \| 1.0 \| 5.257e-14 \| 640 \| 0.443 \| 133 \| 65 \| 1 \| 13 \| 136 \| 1 \| 133 \| Uncharacterized protein \| Uncharacterized protein; afdb-uniprot50 \| AF-A0A395R3N0-F1-MODEL\_V4 \| 1.0 \| 1.454e-13 \| 638 \| 0.443 \| 124 \| 69 \| 0 \| 13 \| 136 \| 1 \| 124 \| Oxidoreductase \| Oxidoreductase; afdb-uniprot50 \| AF-A0A1W1XJU9-F1-MODEL\_V4 \| 1.0 \| 5.546e-14 \| 636 \| 0.382 \| 136 \| 84 \| 0 \| 1 \| 136 \| 16 \| 151 \| Phage protein U \| Phage protein U; afdb-uniprot50 \| AF-A0A4Q0YJQ4-F1-MODEL\_V4 \| 1.0 \| 2.232e-13 \| 635 \| 0.475 \| 124 \| 65 \| 0 \| 13 \| 136 \| 1 \| 124 \| Oxidoreductase \| Oxidoreductase; afdb-uniprot50 \| AF-A0A1C6YVI2-F1-MODEL\_V4 \| 1.0 \| 9.476e-14 \| 635 \| 0.335 \| 128 \| 85 \| 0 \| 8 \| 135 \| 24 \| 151 \| Uncharacterized protein \| Uncharacterized protein; afdb-uniprot50 \| AF-B6I123-F1-MODEL\_V4 \| 1.0 \| 1.379e-13 \| 634 \| 0.346 \| 130 \| 83 \| 1 \| 9 \| 136 \| 9 \| 138 \| Uncharacterized protein \| Uncharacterized protein; afdb-uniprot50 \| AF-A0A6L8HM89-F1-MODEL\_V4 \| 1.0 \| 2.006e-13 \| 633 \| 0.404 \| 126 \| 75 \| 0 \| 11 \| 136 \| 3 \| 128 \| Phage tail protein \| Phage tail protein; afdb-uniprot50 \| AF-A0A2C9B8U0-F1-MODEL\_V4 \| 1.0 \| 1.802e-13 \| 633 \| 0.395 \| 124 \| 75 \| 0 \| 13 \| 136 \| 1 \| 124 \| Uncharacterized protein \| Uncharacterized protein; afdb-uniprot50 \| AF-A0A375FXQ5-F1-MODEL\_V4 \| 1.0 \| 4.245e-13 \| 633 \| 0.395 \| 124 \| 75 \| 0 \| 13 \| 136 \| 1 \| 124 \| Bacteriophage P2 tail protein GPU \| Bacteriophage P2 tail protein GPU; afdb-uniprot50 \| AF-A0A396RZM8-F1-MODEL\_V4 \| 1.0 \| 2.622e-13 \| 633 \| 0.435 \| 124 \| 70 \| 0 \| 13 \| 136 \| 1 \| 124 \| Oxidoreductase \| Oxidoreductase; afdb-uniprot50 \| AF-A0A1G8XD62-F1-MODEL\_V4 \| 1.0 \| 2.232e-13 \| 632 \| 0.411 \| 124 \| 73 \| 0 \| 13 \| 136 \| 1 \| 124 \| Uncharacterized protein \| Uncharacterized protein; afdb-uniprot50 \| AF-A0A1Y6CS08-F1-MODEL\_V4 \| 1.0 \| 1.055e-13 \| 631 \| 0.35 \| 134 \| 83 \| 2 \| 6 \| 136 \| 4 \| 136 \| Uncharacterized protein \| Uncharacterized protein; afdb-uniprot50 \| AF-Q1QXT0-F1-MODEL\_V4 \| 1.0 \| 3.079e-13 \| 630 \| 0.419 \| 124 \| 72 \| 0 \| 13 \| 136 \| 1 \| 124 \| Phage P2 GpU \| Phage P2 GpU; afdb-uniprot50 \| AF-C1D6Q1-F1-MODEL\_V4 \| 1.0 \| 2.006e-13 \| 629 \| 0.39 \| 128 \| 77 \| 1 \| 10 \| 136 \| 2 \| 129 \| Phage P2 GpU \| Phage P2 GpU; afdb-uniprot50 \| AF-A0A318KNF0-F1-MODEL\_V4 \| 1.0 \| 2.766e-13 \| 628 \| 0.376 \| 125 \| 77 \| 1 \| 13 \| 136 \| 2 \| 126 \| Uncharacterized protein \| Uncharacterized protein; afdb-uniprot50 \| AF-A0A1K0JN57-F1-MODEL\_V4 \| 1.0 \| 5.26e-13 \| 628 \| 0.395 \| 124 \| 75 \| 0 \| 13 \| 136 \| 1 \| 124 \| Bacteriophage P2 tail protein GPU \| Bacteriophage P2 tail protein GPU; afdb-uniprot50 \| AF-A0A0P6X2A9-F1-MODEL\_V4 \| 1.0 \| 3.079e-13 \| 626 \| 0.322 \| 127 \| 86 \| 0 \| 10 \| 136 \| 2 \| 128 \| Uncharacterized protein \| Uncharacterized protein; afdb-uniprot50 \| AF-A0A3N1XDI5-F1-MODEL\_V4 \| 1.0 \| 3.427e-13 \| 626 \| 0.443 \| 124 \| 69 \| 0 \| 13 \| 136 \| 1 \| 124 \| Uncharacterized protein \| Uncharacterized protein; afdb-uniprot50 \| AF-D4M9M2-F1-MODEL\_V4 \| 1.0 \| 6.875e-13 \| 624 \| 0.376 \| 125 \| 77 \| 1 \| 13 \| 136 \| 1 \| 125 \| Phage protein U \| Phage protein U; afdb-uniprot50 \| AF-A0A1E3G6A9-F1-MODEL\_V4 \| 1.0 \| 2.766e-13 \| 624 \| 0.447 \| 123 \| 66 \| 2 \| 14 \| 136 \| 1 \| 121 \| Uncharacterized protein \| Uncharacterized protein; afdb-uniprot50 \| AF-A0A4Y7X8T9-F1-MODEL\_V4 \| 1.0 \| 7.253e-13 \| 624 \| 0.403 \| 124 \| 74 \| 0 \| 13 \| 136 \| 1 \| 124 \| Phage tail protein \| Phage tail protein; afdb-uniprot50 \| AF-A0A1H1SML2-F1-MODEL\_V4 \| 1.0 \| 4.479e-13 \| 622 \| 0.435 \| 124 \| 70 \| 0 \| 13 \| 136 \| 1 \| 124 \| Uncharacterized protein \| Uncharacterized protein; afdb-uniprot50 \| AF-A0A7V8INR8-F1-MODEL\_V4 \| 1.0 \| 3.248e-13 \| 621 \| 0.387 \| 124 \| 76 \| 0 \| 13 \| 136 \| 1 \| 124 \| Tail formation protein phage P2 GpU \| Tail formation protein phage P2 GpU; afdb-uniprot50 \| AF-A0A1X3JWZ1-F1-MODEL\_V4 \| 1.0 \| 1.379e-13 \| 619 \| 0.34 \| 138 \| 87 \| 3 \| 1 \| 136 \| 1 \| 136 \| Tail formation protein \| Tail formation protein; afdb-uniprot50 \| AF-A0A7V8U724-F1-MODEL\_V4 \| 1.0 \| 1.307e-13 \| 619 \| 0.348 \| 132 \| 86 \| 0 \| 5 \| 136 \| 48 \| 179 \| Uncharacterized protein \| Uncharacterized protein; afdb-uniprot50 \| AF-A0A0N4UTZ6-F1-MODEL\_V4 \| 1.0 \| 7.253e-13 \| 617 \| 0.435 \| 124 \| 70 \| 0 \| 13 \| 136 \| 1 \| 124 \| Uncharacterized protein \| Uncharacterized protein; afdb-uniprot50 \| AF-A0A0H3NZQ0-F1-MODEL\_V4 \| 1.0 \| 6.875e-13 \| 614 \| 0.395 \| 124 \| 75 \| 0 \| 13 \| 136 \| 1 \| 124 \| Phage-related tail protein \| Phage-related tail protein; afdb-uniprot50 \| AF-A0A853I3C0-F1-MODEL\_V4 \| 1.0 \| 1.379e-13 \| 614 \| 0.381 \| 144 \| 79 \| 4 \| 1 \| 136 \| 1 \| 142 \| Phage tail protein \| Phage tail protein; afdb-uniprot50 \| AF-A0A2C6DLN0-F1-MODEL\_V4 \| 1.0 \| 4.985e-13 \| 613 \| 0.37 \| 124 \| 78 \| 0 \| 13 \| 136 \| 1 \| 124 \| Phage protein U \| Phage protein U; afdb-uniprot50 \| AF-A0A8A7WU58-F1-MODEL\_V4 \| 1.0 \| 7.652e-13 \| 612 \| 0.411 \| 124 \| 73 \| 0 \| 13 \| 136 \| 1 \| 124 \| Phage tail protein \| Phage tail protein; afdb-uniprot50 \| AF-A0A2N3KSK9-F1-MODEL\_V4 \| 1.0 \| 8.982e-14 \| 611 \| 0.353 \| 130 \| 83 \| 1 \| 7 \| 136 \| 1 \| 129 \| Phage tail protein \| Phage tail protein; afdb-uniprot50 \| AF-A0A2D0KY36-F1-MODEL\_V4 \| 1.0 \| 6.875e-13 \| 611 \| 0.403 \| 124 \| 74 \| 0 \| 13 \| 136 \| 1 \| 124 \| Tail assembly protein \| Tail assembly protein; afdb-uniprot50 \| AF-A0A2Z4UEV7-F1-MODEL\_V4 \| 1.0 \| 8.517e-13 \| 609 \| 0.395 \| 124 \| 75 \| 0 \| 13 \| 136 \| 1 \| 124 \| Phage P2 GpU \| Phage P2 GpU; afdb-uniprot50 \| AF-A0A0T9UNC2-F1-MODEL\_V4 \| 1.0 \| 6.177e-13 \| 609 \| 0.379 \| 124 \| 77 \| 0 \| 13 \| 136 \| 1 \| 124 \| Phage-related tail protein \| Phage-related tail protein; afdb-uniprot50 \| AF-A0A5P9F1S5-F1-MODEL\_V4 \| 1.0 \| 3.079e-13 \| 609 \| 0.311 \| 135 \| 93 \| 0 \| 2 \| 136 \| 16 \| 150 \| Phage P2 GpU \| Phage P2 GpU; afdb-uniprot50 \| AF-A0A5U3EXN1-F1-MODEL\_V4 \| 1.0 \| 1.055e-12 \| 609 \| 0.395 \| 124 \| 75 \| 0 \| 13 \| 136 \| 1 \| 124 \| Phage tail protein \| Phage tail protein; afdb-uniprot50 \| AF-A0A2W5DF00-F1-MODEL\_V4 \| 1.0 \| 8.073e-13 \| 607 \| 0.403 \| 124 \| 74 \| 0 \| 13 \| 136 \| 1 \| 124 \| Oxidoreductase \| Oxidoreductase; afdb-uniprot50 \| AF-A0A5E4XFM2-F1-MODEL\_V4 \| 1.0 \| 4.477e-14 \| 607 \| 0.394 \| 142 \| 80 \| 1 \| 1 \| 136 \| 1 \| 142 \| Oxidoreductase \| Oxidoreductase; afdb-uniprot50 \| AF-A0A5J2S1K9-F1-MODEL\_V4 \| 1.0 \| 6.875e-13 \| 607 \| 0.346 \| 124 \| 81 \| 0 \| 13 \| 136 \| 1 \| 124 \| Phage tail protein \| Phage tail protein; afdb-uniprot50 \| AF-A0A1I0YHT2-F1-MODEL\_V4 \| 1.0 \| 6.516e-13 \| 607 \| 0.387 \| 124 \| 76 \| 0 \| 13 \| 136 \| 1 \| 124 \| Uncharacterized protein \| Uncharacterized protein; afdb-uniprot50 \| AF-A0A7S5LS35-F1-MODEL\_V4 \| 1.0 \| 7.652e-13 \| 607 \| 0.358 \| 131 \| 77 \| 1 \| 13 \| 136 \| 48 \| 178 \| Phage tail protein \| Phage tail protein; afdb-uniprot50 \| AF-A0A2G3K2E9-F1-MODEL\_V4 \| 1.0 \| 1.175e-12 \| 606 \| 0.403 \| 124 \| 74 \| 0 \| 13 \| 136 \| 1 \| 124 \| Oxidoreductase \| Oxidoreductase; afdb-uniprot50 \| AF-A0A7C8LYL7-F1-MODEL\_V4 \| 1.0 \| 6.173e-14 \| 606 \| 0.383 \| 146 \| 75 \| 3 \| 6 \| 136 \| 1 \| 146 \| Uncharacterized protein \| Uncharacterized protein; afdb-uniprot50 \| AF-A0A157QP36-F1-MODEL\_V4 \| 1.0 \| 8.517e-13 \| 606 \| 0.379 \| 124 \| 77 \| 0 \| 13 \| 136 \| 1 \| 124 \| Phage protein U \| Phage protein U; afdb-uniprot50 \| AF-Q87ZN2-F1-MODEL\_V4 \| 1.0 \| 9.481e-13 \| 606 \| 0.335 \| 131 \| 80 \| 1 \| 13 \| 136 \| 118 \| 248 \| Uncharacterized protein \| Uncharacterized protein; afdb-uniprot50 \| AF-H8L2K9-F1-MODEL\_V4 \| 1.0 \| 8.982e-14 \| 606 \| 0.382 \| 136 \| 79 \| 1 \| 6 \| 136 \| 139 \| 274 \| Phage protein U \| Phage protein U; afdb-uniprot50 \| AF-A0A7X3TZ69-F1-MODEL\_V4 \| 1.0 \| 3.615e-13 \| 606 \| 0.346 \| 130 \| 84 \| 1 \| 6 \| 134 \| 24 \| 153 \| Uncharacterized protein \| Uncharacterized protein; afdb-uniprot50 \| AF-A0A329B3N7-F1-MODEL\_V4 \| 1.0 \| 6.516e-13 \| 605 \| 0.395 \| 124 \| 75 \| 0 \| 13 \| 136 \| 1 \| 124 \| Uncharacterized protein \| Uncharacterized protein; afdb-uniprot50 \| AF-A0A258E9H2-F1-MODEL\_V4 \| 1.0 \| 4.479e-13 \| 605 \| 0.308 \| 136 \| 94 \| 0 \| 1 \| 136 \| 19 \| 154 \| Uncharacterized protein \| Uncharacterized protein; afdb-uniprot50 \| AF-A0A564M9A9-F1-MODEL\_V4 \| 1.0 \| 7.652e-13 \| 605 \| 0.346 \| 124 \| 81 \| 0 \| 13 \| 136 \| 1 \| 124 \| Uncharacterized protein \| Uncharacterized protein; afdb-uniprot50 \| AF-A0A0M0P397-F1-MODEL\_V4 \| 1.0 \| 6.875e-13 \| 604 \| 0.36 \| 122 \| 78 \| 0 \| 13 \| 134 \| 1 \| 122 \| Oxidoreductase \| Oxidoreductase; afdb-uniprot50 \| AF-A0A271U8M1-F1-MODEL\_V4 \| 1.0 \| 1.307e-12 \| 604 \| 0.411 \| 124 \| 73 \| 0 \| 13 \| 136 \| 1 \| 124 \| Oxidoreductase \| Oxidoreductase; afdb-uniprot50 \| AF-A0A4Q3KFT0-F1-MODEL\_V4 \| 1.0 \| 1.455e-12 \| 604 \| 0.427 \| 124 \| 71 \| 0 \| 13 \| 136 \| 1 \| 124 \| Phage tail protein \| Phage tail protein; afdb-uniprot50 \| AF-A0A838C006-F1-MODEL\_V4 \| 1.0 \| 8.986e-13 \| 604 \| 0.346 \| 124 \| 81 \| 0 \| 13 \| 136 \| 1 \| 124 \| Phage tail protein \| Phage tail protein; afdb-uniprot50 \| AF-A0A4R0GC61-F1-MODEL\_V4 \| 1.0 \| 8.073e-13 \| 604 \| 0.354 \| 124 \| 80 \| 0 \| 13 \| 136 \| 1 \| 124 \| Phage tail protein \| Phage tail protein; afdb-uniprot50 \| AF-A0A1I5KZ84-F1-MODEL\_V4 \| 1.0 \| 7.253e-13 \| 604 \| 0.333 \| 135 \| 85 \| 1 \| 7 \| 136 \| 152 \| 286 \| Uncharacterized protein \| Uncharacterized protein; afdb-uniprot50 \| AF-A0A537MFJ2-F1-MODEL\_V4 \| 1.0 \| 1e-12 \| 603 \| 0.346 \| 124 \| 81 \| 0 \| 13 \| 136 \| 2 \| 125 \| Oxidoreductase \| Oxidoreductase; afdb-uniprot50 \| AF-A0A1M7YZC4-F1-MODEL\_V4 \| 1.0 \| 1.534e-13 \| 603 \| 0.363 \| 143 \| 83 \| 3 \| 1 \| 136 \| 5 \| 146 \| Phage P2 GpU \| Phage P2 GpU; afdb-uniprot50 \| AF-A0A239CL96-F1-MODEL\_V4 \| 1.0 \| 7.652e-13 \| 602 \| 0.393 \| 127 \| 77 \| 0 \| 10 \| 136 \| 1 \| 127 \| Uncharacterized protein \| Uncharacterized protein; afdb-uniprot50 \| AF-A0A1Q6U7S4-F1-MODEL\_V4 \| 1.0 \| 2.918e-13 \| 602 \| 0.43 \| 123 \| 70 \| 0 \| 14 \| 136 \| 1 \| 123 \| Uncharacterized protein \| Uncharacterized protein; afdb-uniprot50 \| AF-A0A3M5X4R0-F1-MODEL\_V4 \| 1.0 \| 1.113e-12 \| 601 \| 0.418 \| 117 \| 68 \| 0 \| 20 \| 136 \| 11 \| 127 \| Uncharacterized protein \| Uncharacterized protein; afdb-uniprot50 \| AF-A0A7X6X091-F1-MODEL\_V4 \| 1.0 \| 1.709e-12 \| 601 \| 0.419 \| 124 \| 72 \| 0 \| 13 \| 136 \| 1 \| 124 \| Phage tail protein \| Phage tail protein; afdb-uniprot50 \| AF-A0A5S3YY93-F1-MODEL\_V4 \| 1.0 \| 2.485e-13 \| 601 \| 0.38 \| 142 \| 76 \| 3 \| 7 \| 136 \| 3 \| 144 \| Uncharacterized protein \| Uncharacterized protein; afdb-uniprot50 \| AF-A0A0H3NJV7-F1-MODEL\_V4 \| 1.0 \| 9.481e-13 \| 601 \| 0.379 \| 124 \| 77 \| 0 \| 13 \| 136 \| 1 \| 124 \| Phage-related tail protein \| Phage-related tail protein; afdb-uniprot50 \| AF-F5S5D7-F1-MODEL\_V4 \| 1.0 \| 1e-12 \| 601 \| 0.37 \| 124 \| 78 \| 0 \| 13 \| 136 \| 2 \| 125 \| Phage P2 GpU \| Phage P2 GpU; afdb-uniprot50 \| AF-A0A3Q9W528-F1-MODEL\_V4 \| 1.0 \| 1.535e-12 \| 600 \| 0.379 \| 124 \| 77 \| 0 \| 13 \| 136 \| 1 \| 124 \| Oxidoreductase \| Oxidoreductase; afdb-uniprot50 \| AF-A0A7X0AIN6-F1-MODEL\_V4 \| 1.0 \| 1.902e-12 \| 600 \| 0.37 \| 124 \| 78 \| 0 \| 13 \| 136 \| 1 \| 124 \| Uncharacterized protein \| Uncharacterized protein; afdb-uniprot50 \| AF-A0A6P2P2T0-F1-MODEL\_V4 \| 1.0 \| 1.379e-12 \| 600 \| 0.338 \| 124 \| 82 \| 0 \| 13 \| 136 \| 1 \| 124 \| Oxidoreductase \| Oxidoreductase; afdb-uniprot50 \| AF-A0A090SIS3-F1-MODEL\_V4 \| 1.0 \| 8.073e-13 \| 599 \| 0.346 \| 127 \| 80 \| 1 \| 13 \| 136 \| 4 \| 130 \| Phage tail protein \| Phage tail protein; afdb-uniprot50 \| AF-A0A4Q3U4S0-F1-MODEL\_V4 \| 1.0 \| 1.379e-12 \| 599 \| 0.327 \| 122 \| 82 \| 0 \| 13 \| 134 \| 2 \| 123 \| Phage tail protein \| Phage tail protein; afdb-uniprot50 \| AF-A0A2D0ITE4-F1-MODEL\_V4 \| 1.0 \| 1.055e-12 \| 599 \| 0.403 \| 124 \| 74 \| 0 \| 13 \| 136 \| 1 \| 124 \| Tail assembly protein \| Tail assembly protein; afdb-uniprot50 \| AF-C3X1X4-F1-MODEL\_V4 \| 1.0 \| 1.175e-12 \| 599 \| 0.37 \| 124 \| 78 \| 0 \| 13 \| 136 \| 1 \| 124 \| Uncharacterized protein \| Uncharacterized protein; afdb-uniprot50 \| AF-A0A376RU30-F1-MODEL\_V4 \| 1.0 \| 1.055e-12 \| 599 \| 0.346 \| 124 \| 81 \| 0 \| 13 \| 136 \| 1 \| 124 \| Bacteriophage tail protein GpU \| Bacteriophage tail protein GpU; afdb-uniprot50 \| AF-G0AIR7-F1-MODEL\_V4 \| 1.0 \| 1.379e-12 \| 599 \| 0.379 \| 124 \| 77 \| 0 \| 13 \| 136 \| 1 \| 124 \| Phage P2 GpU \| Phage P2 GpU; afdb-uniprot50 \| AF-A0A1H8J1Q9-F1-MODEL\_V4 \| 1.0 \| 1.708e-13 \| 599 \| 0.352 \| 136 \| 83 \| 1 \| 6 \| 136 \| 140 \| 275 \| Uncharacterized protein \| Uncharacterized protein; afdb-uniprot50 \| AF-A0A7X3NEZ9-F1-MODEL\_V4 \| 1.0 \| 1.113e-12 \| 598 \| 0.338 \| 124 \| 82 \| 0 \| 13 \| 136 \| 1 \| 124 \| Phage tail protein \| Phage tail protein; afdb-uniprot50 \| AF-A0A519MN22-F1-MODEL\_V4 \| 1.0 \| 1.175e-12 \| 598 \| 0.379 \| 124 \| 77 \| 0 \| 13 \| 136 \| 1 \| 124 \| Phage tail protein \| Phage tail protein; afdb-uniprot50 \| AF-A0A376HJR0-F1-MODEL\_V4 \| 1.0 \| 1.709e-12 \| 598 \| 0.373 \| 123 \| 77 \| 0 \| 14 \| 136 \| 1 \| 123 \| Tail fiber protein \| Tail fiber protein; afdb-uniprot50 \| AF-A0A4Q9EHW1-F1-MODEL\_V4 \| 1.0 \| 1.307e-12 \| 598 \| 0.362 \| 124 \| 79 \| 0 \| 13 \| 136 \| 1 \| 124 \| Phage tail protein \| Phage tail protein; afdb-uniprot50 \| AF-A0A5T5B4P6-F1-MODEL\_V4 \| 1.0 \| 1.055e-12 \| 598 \| 0.346 \| 124 \| 81 \| 0 \| 13 \| 136 \| 1 \| 124 \| Phage tail protein \| Phage tail protein; afdb-uniprot50 \| AF-A0A5M7L4M4-F1-MODEL\_V4 \| 1.0 \| 1.902e-12 \| 597 \| 0.354 \| 124 \| 80 \| 0 \| 13 \| 136 \| 1 \| 124 \| Phage tail protein \| Phage tail protein; afdb-uniprot50 \| AF-A0A1B7IQW7-F1-MODEL\_V4 \| 1.0 \| 1.113e-12 \| 597 \| 0.403 \| 124 \| 74 \| 0 \| 13 \| 136 \| 1 \| 124 \| Putative phage tail protein \| Putative phage tail protein; afdb-uniprot50 \| AF-A0A6L3ZUK1-F1-MODEL\_V4 \| 1.0 \| 1.379e-12 \| 596 \| 0.39 \| 123 \| 75 \| 0 \| 14 \| 136 \| 1 \| 123 \| Oxidoreductase \| Oxidoreductase; afdb-uniprot50 \| AF-A0A7Y0I342-F1-MODEL\_V4 \| 1.0 \| 2.622e-13 \| 596 \| 0.355 \| 138 \| 86 \| 1 \| 2 \| 136 \| 14 \| 151 \| Uncharacterized protein \| Uncharacterized protein; afdb-uniprot50 \| AF-A0A0P7LSF8-F1-MODEL\_V4 \| 1.0 \| 1.379e-12 \| 596 \| 0.346 \| 124 \| 81 \| 0 \| 13 \| 136 \| 1 \| 124 \| Phage tail protein \| Phage tail protein; afdb-uniprot50 \| AF-A0A485Z843-F1-MODEL\_V4 \| 1.0 \| 1.307e-12 \| 596 \| 0.346 \| 124 \| 81 \| 0 \| 13 \| 136 \| 1 \| 124 \| Phage tail protein \| Phage tail protein; afdb-uniprot50 \| AF-A0A7W6WBR9-F1-MODEL\_V4 \| 1.0 \| 1.455e-12 \| 596 \| 0.368 \| 125 \| 77 \| 2 \| 13 \| 136 \| 2 \| 125 \| Uncharacterized protein \| Uncharacterized protein; afdb-uniprot50 \| AF-A0A702B4K5-F1-MODEL\_V4 \| 1.0 \| 2.007e-12 \| 596 \| 0.37 \| 124 \| 78 \| 0 \| 13 \| 136 \| 1 \| 124 \| Phage tail protein \| Phage tail protein; afdb-uniprot50 \| AF-A0A6L5PEK8-F1-MODEL\_V4 \| 1.0 \| 2.355e-13 \| 595 \| 0.33 \| 127 \| 84 \| 1 \| 10 \| 136 \| 1 \| 126 \| Phage tail protein \| Phage tail protein; afdb-uniprot50 \| AF-A0A250DSZ6-F1-MODEL\_V4 \| 1.0 \| 1.62e-12 \| 595 \| 0.382 \| 123 \| 76 \| 0 \| 14 \| 136 \| 1 \| 123 \| Oxidoreductase \| Oxidoreductase; afdb-uniprot50 \| AF-A0A024L4N8-F1-MODEL\_V4 \| 1.0 \| 1.379e-12 \| 595 \| 0.346 \| 124 \| 81 \| 0 \| 13 \| 136 \| 1 \| 124 \| Bacteriophage tail protein GpU \| Bacteriophage tail protein GpU; afdb-uniprot50 \| AF-A0A8A5ILR9-F1-MODEL\_V4 \| 1.0 \| 1.113e-12 \| 595 \| 0.346 \| 124 \| 81 \| 0 \| 13 \| 136 \| 1 \| 124 \| Phage tail protein \| Phage tail protein; afdb-uniprot50 \| AF-A0A2W5VBP0-F1-MODEL\_V4 \| 1.0 \| 1.62e-12 \| 595 \| 0.296 \| 125 \| 88 \| 0 \| 12 \| 136 \| 2 \| 126 \| Oxidoreductase \| Oxidoreductase; afdb-uniprot50 \| AF-A0A7Z0SJA2-F1-MODEL\_V4 \| 1.0 \| 7.652e-13 \| 595 \| 0.333 \| 135 \| 90 \| 0 \| 2 \| 136 \| 101 \| 235 \| Phage tail protein \| Phage tail protein; afdb-uniprot50 \| AF-A0A4E0GYK4-F1-MODEL\_V4 \| 1.0 \| 1.175e-12 \| 594 \| 0.346 \| 124 \| 81 \| 0 \| 13 \| 136 \| 1 \| 124 \| Phage tail protein \| Phage tail protein; afdb-uniprot50 \| AF-A0A246KFY5-F1-MODEL\_V4 \| 1.0 \| 2.007e-12 \| 593 \| 0.33 \| 124 \| 83 \| 0 \| 13 \| 136 \| 1 \| 124 \| Phage protein U \| Phage protein U; afdb-uniprot50 \| AF-A0A5U9GBD7-F1-MODEL\_V4 \| 1.0 \| 1.455e-12 \| 593 \| 0.346 \| 124 \| 81 \| 0 \| 13 \| 136 \| 1 \| 124 \| Phage tail protein \| Phage tail protein; afdb-uniprot50 \| AF-A0A495Y9Q8-F1-MODEL\_V4 \| 1.0 \| 1e-12 \| 593 \| 0.342 \| 140 \| 87 \| 1 \| 2 \| 136 \| 150 \| 289 \| Oxidoreductase \| Oxidoreductase; afdb-uniprot50 \| AF-M1PBX3-F1-MODEL\_V4 \| 1.0 \| 4.024e-13 \| 592 \| 0.403 \| 124 \| 73 \| 1 \| 13 \| 136 \| 1 \| 123 \| Phage tail protein U \| Phage tail protein U; afdb-uniprot50 \| AF-E2CJS7-F1-MODEL\_V4 \| 1.0 \| 5.26e-13 \| 592 \| 0.343 \| 131 \| 85 \| 1 \| 6 \| 136 \| 1 \| 130 \| p2 GpU family protein \| p2 GpU family protein; afdb-uniprot50 \| AF-A0A660MIA9-F1-MODEL\_V4 \| 1.0 \| 1.902e-12 \| 592 \| 0.384 \| 125 \| 75 \| 2 \| 13 \| 136 \| 1 \| 124 \| Phage tail protein \| Phage tail protein; afdb-uniprot50 \| AF-A0A2D5S2F6-F1-MODEL\_V4 \| 1.0 \| 3.814e-13 \| 591 \| 0.313 \| 134 \| 88 \| 3 \| 7 \| 136 \| 1 \| 134 \| Oxidoreductase \| Oxidoreductase; afdb-uniprot50 \| AF-A0A241WFD3-F1-MODEL\_V4 \| 1.0 \| 6.516e-13 \| 591 \| 0.392 \| 125 \| 75 \| 1 \| 13 \| 136 \| 1 \| 125 \| Oxidoreductase \| Oxidoreductase; afdb-uniprot50 \| AF-A0A1I4NEJ2-F1-MODEL\_V4 \| 1.0 \| 9.481e-13 \| 591 \| 0.306 \| 137 \| 90 \| 1 \| 5 \| 136 \| 151 \| 287 \| Uncharacterized protein \| Uncharacterized protein; afdb-uniprot50 \| AF-A0A1M3HI07-F1-MODEL\_V4 \| 1.0 \| 9.481e-13 \| 590 \| 0.384 \| 125 \| 76 \| 1 \| 13 \| 136 \| 1 \| 125 \| Uncharacterized protein \| Uncharacterized protein; afdb-uniprot50 \| AF-A0A5V0HLG2-F1-MODEL\_V4 \| 1.0 \| 1.902e-12 \| 590 \| 0.346 \| 124 \| 81 \| 0 \| 13 \| 136 \| 1 \| 124 \| Phage tail protein \| Phage tail protein; afdb-uniprot50 \| AF-A0A838CHN8-F1-MODEL\_V4 \| 1.0 \| 1.175e-12 \| 590 \| 0.321 \| 140 \| 90 \| 1 \| 2 \| 136 \| 92 \| 231 \| Phage tail protein \| Phage tail protein; afdb-uniprot50 \| AF-A0A2G2NAY4-F1-MODEL\_V4 \| 1.0 \| 7.253e-13 \| 589 \| 0.513 \| 111 \| 54 \| 0 \| 26 \| 136 \| 3 \| 113 \| Phage tail protein \| Phage tail protein; afdb-uniprot50 \| AF-A0A6C9KZW5-F1-MODEL\_V4 \| 1.0 \| 2.117e-12 \| 589 \| 0.338 \| 124 \| 82 \| 0 \| 13 \| 136 \| 1 \| 124 \| Phage tail protein \| Phage tail protein; afdb-uniprot50 \| AF-A0A2G0Q023-F1-MODEL\_V4 \| 1.0 \| 1.902e-12 \| 589 \| 0.395 \| 124 \| 75 \| 0 \| 13 \| 136 \| 1 \| 124 \| Tail assembly protein \| Tail assembly protein; afdb-uniprot50 \| AF-A0A1G5TM27-F1-MODEL\_V4 \| 1.0 \| 1.62e-12 \| 587 \| 0.416 \| 125 \| 71 \| 2 \| 13 \| 136 \| 1 \| 124 \| Uncharacterized protein \| Uncharacterized protein; afdb-uniprot50 \| AF-A0A659PR58-F1-MODEL\_V4 \| 1.0 \| 2.234e-12 \| 586 \| 0.33 \| 124 \| 83 \| 0 \| 13 \| 136 \| 1 \| 124 \| Phage tail protein \| Phage tail protein; afdb-uniprot50 \| AF-A0A1B9LPX2-F1-MODEL\_V4 \| 1.0 \| 2.234e-12 \| 586 \| 0.344 \| 122 \| 80 \| 0 \| 13 \| 134 \| 1 \| 122 \| Uncharacterized protein \| Uncharacterized protein; afdb-uniprot50 \| AF-A0A486S2L3-F1-MODEL\_V4 \| 1.0 \| 2.92e-12 \| 586 \| 0.346 \| 124 \| 81 \| 0 \| 13 \| 136 \| 1 \| 124 \| Putative bacteriophage tail protein \| Putative bacteriophage tail protein; afdb-uniprot50 \| AF-B7LNA5-F1-MODEL\_V4 \| 1.0 \| 6.875e-13 \| 586 \| 0.309 \| 126 \| 87 \| 0 \| 11 \| 136 \| 25 \| 150 \| Putative Phage-related tail protein \| Putative Phage-related tail protein; afdb-uniprot50 \| AF-D4GLP7-F1-MODEL\_V4 \| 1.0 \| 2.767e-12 \| 586 \| 0.379 \| 124 \| 77 \| 0 \| 13 \| 136 \| 1 \| 124 \| Uncharacterized protein \| Uncharacterized protein; afdb-uniprot50 \| AF-A0A833KUX0-F1-MODEL\_V4 \| 1.0 \| 2.117e-12 \| 586 \| 0.346 \| 124 \| 81 \| 0 \| 13 \| 136 \| 1 \| 124 \| Phage tail protein \| Phage tail protein; afdb-uniprot50 \| AF-A0A6P1Q0K5-F1-MODEL\_V4 \| 1.0 \| 3.617e-12 \| 585 \| 0.322 \| 124 \| 84 \| 0 \| 13 \| 136 \| 1 \| 124 \| Uncharacterized protein \| Uncharacterized protein; afdb-uniprot50 \| AF-A0A1E7WJB8-F1-MODEL\_V4 \| 1.0 \| 2.007e-12 \| 585 \| 0.377 \| 122 \| 76 \| 0 \| 13 \| 134 \| 1 \| 122 \| Phage P2 GpU \| Phage P2 GpU; afdb-uniprot50 \| AF-A0A4P0TK52-F1-MODEL\_V4 \| 1.0 \| 8.073e-13 \| 585 \| 0.381 \| 139 \| 71 \| 3 \| 13 \| 136 \| 114 \| 252 \| Putative tail formation protein \| Putative tail formation protein; afdb-uniprot50 \| AF-A0A4V2B685-F1-MODEL\_V4 \| 1.0 \| 6.875e-13 \| 584 \| 0.305 \| 134 \| 90 \| 1 \| 6 \| 136 \| 3 \| 136 \| Phage tail protein \| Phage tail protein; afdb-uniprot50 \| AF-A0A3B0TD64-F1-MODEL\_V4 \| 1.0 \| 1.239e-12 \| 584 \| 0.362 \| 127 \| 79 \| 1 \| 12 \| 136 \| 2 \| 128 \| Uncharacterized protein \| Uncharacterized protein; afdb-uniprot50 \| AF-A0A2W4T7M1-F1-MODEL\_V4 \| 1.0 \| 8.517e-13 \| 584 \| 0.354 \| 127 \| 81 \| 1 \| 10 \| 136 \| 2 \| 127 \| Phage tail protein \| Phage tail protein; afdb-uniprot50 \| AF-A0A515CWV7-F1-MODEL\_V4 \| 1.0 \| 1.709e-12 \| 584 \| 0.379 \| 124 \| 77 \| 0 \| 13 \| 136 \| 1 \| 124 \| Phage tail protein \| Phage tail protein; afdb-uniprot50 \| AF-F4VCL7-F1-MODEL\_V4 \| 1.0 \| 2.234e-12 \| 584 \| 0.346 \| 124 \| 81 \| 0 \| 13 \| 136 \| 1 \| 124 \| Putative bacteriophage tail protein \| Putative bacteriophage tail protein; afdb-uniprot50 \| AF-A0A248LHR8-F1-MODEL\_V4 \| 1.0 \| 3.25e-12 \| 582 \| 0.333 \| 126 \| 84 \| 0 \| 11 \| 136 \| 3 \| 128 \| Phage protein U \| Phage protein U; afdb-uniprot50 \| AF-A0A1I1Q5P9-F1-MODEL\_V4 \| 1.0 \| 6.516e-13 \| 582 \| 0.356 \| 143 \| 80 \| 3 \| 6 \| 136 \| 1 \| 143 \| Uncharacterized protein \| Uncharacterized protein; afdb-uniprot50 \| AF-A0A4R9NLD4-F1-MODEL\_V4 \| 1.0 \| 3.617e-12 \| 582 \| 0.346 \| 124 \| 81 \| 0 \| 13 \| 136 \| 1 \| 124 \| Phage tail protein \| Phage tail protein; afdb-uniprot50 \| AF-A0A0Q4MMS8-F1-MODEL\_V4 \| 1.0 \| 4.728e-12 \| 582 \| 0.362 \| 124 \| 79 \| 0 \| 13 \| 136 \| 1 \| 124 \| Phage tail protein \| Phage tail protein; afdb-uniprot50 \| AF-A0A4Y6UBW6-F1-MODEL\_V4 \| 1.0 \| 3.615e-13 \| 582 \| 0.294 \| 136 \| 95 \| 1 \| 1 \| 136 \| 13 \| 147 \| Phage tail protein \| Phage tail protein; afdb-uniprot50 \| AF-A0A1I0AWP0-F1-MODEL\_V4 \| 1.0 \| 2.117e-12 \| 582 \| 0.357 \| 126 \| 78 \| 2 \| 13 \| 136 \| 1 \| 125 \| Uncharacterized protein \| Uncharacterized protein; afdb-uniprot50 \| AF-A0A516SM14-F1-MODEL\_V4 \| 1.0 \| 1.055e-12 \| 582 \| 0.354 \| 144 \| 84 \| 2 \| 2 \| 136 \| 118 \| 261 \| Phage tail protein \| Phage tail protein; afdb-uniprot50 \| AF-A0A7X9X5I9-F1-MODEL\_V4 \| 1.0 \| 2.357e-12 \| 581 \| 0.36 \| 122 \| 78 \| 0 \| 13 \| 134 \| 2 \| 123 \| Phage tail protein \| Phage tail protein; afdb-uniprot50 \| AF-A0A2V3UC18-F1-MODEL\_V4 \| 1.0 \| 4.248e-12 \| 580 \| 0.299 \| 127 \| 89 \| 0 \| 10 \| 136 \| 1 \| 127 \| Uncharacterized protein \| Uncharacterized protein; afdb-uniprot50 \| AF-A0A853IGN0-F1-MODEL\_V4 \| 1.0 \| 4.026e-12 \| 580 \| 0.322 \| 127 \| 86 \| 0 \| 10 \| 136 \| 1 \| 127 \| Phage tail protein \| Phage tail protein; afdb-uniprot50 \| AF-A0A7T5QS73-F1-MODEL\_V4 \| 1.0 \| 4.026e-12 \| 580 \| 0.379 \| 124 \| 77 \| 0 \| 13 \| 136 \| 1 \| 124 \| Phage tail protein \| Phage tail protein; afdb-uniprot50 \| AF-A0A2A7V0X6-F1-MODEL\_V4 \| 1.0 \| 3.08e-12 \| 580 \| 0.395 \| 124 \| 75 \| 0 \| 13 \| 136 \| 1 \| 124 \| Oxidoreductase \| Oxidoreductase; afdb-uniprot50 \| AF-A0A0P7PMR1-F1-MODEL\_V4 \| 1.0 \| 4.481e-12 \| 580 \| 0.354 \| 124 \| 80 \| 0 \| 13 \| 136 \| 1 \| 124 \| Oxidoreductase \| Oxidoreductase; afdb-uniprot50 \| AF-A0A222EWH4-F1-MODEL\_V4 \| 1.0 \| 2.234e-12 \| 579 \| 0.362 \| 124 \| 79 \| 0 \| 13 \| 136 \| 1 \| 124 \| Oxidoreductase \| Oxidoreductase; afdb-uniprot50 \| AF-A0A0G3S980-F1-MODEL\_V4 \| 1.0 \| 4.026e-12 \| 579 \| 0.346 \| 124 \| 81 \| 0 \| 13 \| 136 \| 1 \| 124 \| Tail protein \| Tail protein; afdb-uniprot50 \| AF-A0A1I1UCM4-F1-MODEL\_V4 \| 1.0 \| 3.08e-12 \| 578 \| 0.317 \| 126 \| 84 \| 1 \| 13 \| 136 \| 3 \| 128 \| Uncharacterized protein \| Uncharacterized protein; afdb-uniprot50 \| AF-A0A356D2V6-F1-MODEL\_V4 \| 1.0 \| 5.857e-12 \| 577 \| 0.346 \| 124 \| 81 \| 0 \| 13 \| 136 \| 1 \| 124 \| Oxidoreductase \| Oxidoreductase; afdb-uniprot50 \| AF-A0A559QNF7-F1-MODEL\_V4 \| 1.0 \| 5.552e-12 \| 577 \| 0.379 \| 124 \| 77 \| 0 \| 13 \| 136 \| 1 \| 124 \| Uncharacterized protein \| Uncharacterized protein; afdb-uniprot50 \| AF-A0A5E5ANB0-F1-MODEL\_V4 \| 1.0 \| 4.481e-12 \| 577 \| 0.338 \| 124 \| 82 \| 0 \| 13 \| 136 \| 1 \| 124 \| Phage tail protein \| Phage tail protein; afdb-uniprot50 \| AF-T0QJT7-F1-MODEL\_V4 \| 1.0 \| 2.623e-12 \| 576 \| 0.37 \| 124 \| 78 \| 0 \| 13 \| 136 \| 1 \| 124 \| Uncharacterized protein \| Uncharacterized protein; afdb-uniprot50 \| AF-A0A1C7BUG8-F1-MODEL\_V4 \| 1.0 \| 1.379e-12 \| 576 \| 0.375 \| 141 \| 72 \| 3 \| 12 \| 136 \| 144 \| 284 \| Uncharacterized protein \| Uncharacterized protein; afdb-uniprot50 \| AF-A0A348FYG2-F1-MODEL\_V4 \| 1.0 \| 1.62e-12 \| 575 \| 0.325 \| 129 \| 81 \| 2 \| 12 \| 136 \| 2 \| 128 \| Oxidoreductase \| Oxidoreductase; afdb-uniprot50 \| AF-A0A3S4IAG6-F1-MODEL\_V4 \| 1.0 \| 6.18e-12 \| 575 \| 0.346 \| 124 \| 81 \| 0 \| 13 \| 136 \| 1 \| 124 \| Phage tail protein \| Phage tail protein; afdb-uniprot50 \| AF-A0A256CB29-F1-MODEL\_V4 \| 1.0 \| 4.988e-12 \| 574 \| 0.311 \| 122 \| 84 \| 0 \| 13 \| 134 \| 3 \| 124 \| Uncharacterized protein \| Uncharacterized protein; afdb-uniprot50 \| AF-A0A1Z9Z2S2-F1-MODEL\_V4 \| 1.0 \| 5.262e-12 \| 574 \| 0.352 \| 125 \| 80 \| 1 \| 13 \| 136 \| 1 \| 125 \| Oxidoreductase \| Oxidoreductase; afdb-uniprot50 \| AF-A0A212J3T6-F1-MODEL\_V4 \| 1.0 \| 4.728e-12 \| 573 \| 0.36 \| 125 \| 78 \| 2 \| 13 \| 136 \| 1 \| 124 \| Phage P2 GpU protein \| Phage P2 GpU protein; afdb-uniprot50 \| AF-A0A3D9UGE8-F1-MODEL\_V4 \| 1.0 \| 4.026e-12 \| 572 \| 0.37 \| 124 \| 78 \| 0 \| 13 \| 136 \| 1 \| 124 \| Uncharacterized protein \| Uncharacterized protein; afdb-uniprot50 \| AF-A0A0H3KJY5-F1-MODEL\_V4 \| 1.0 \| 6.52e-12 \| 571 \| 0.37 \| 124 \| 78 \| 0 \| 13 \| 136 \| 1 \| 124 \| Probable bacteriophage protein U \| Probable bacteriophage protein U; afdb-uniprot50 \| AF-A0A853X1L4-F1-MODEL\_V4 \| 1.0 \| 4.026e-12 \| 569 \| 0.357 \| 126 \| 78 \| 2 \| 13 \| 136 \| 1 \| 125 \| Oxidoreductase \| Oxidoreductase; afdb-uniprot50 \| AF-A0A516S9T9-F1-MODEL\_V4 \| 1.0 \| 3.08e-12 \| 569 \| 0.344 \| 145 \| 86 \| 2 \| 1 \| 136 \| 154 \| 298 \| Phage tail protein \| Phage tail protein; afdb-uniprot50 \| AF-A0A855M4M1-F1-MODEL\_V4 \| 1.0 \| 8.522e-12 \| 568 \| 0.338 \| 124 \| 82 \| 0 \| 13 \| 136 \| 1 \| 124 \| Phage tail protein \| Phage tail protein; afdb-uniprot50 \| AF-A0A228PZF6-F1-MODEL\_V4 \| 1.0 \| 4.026e-12 \| 567 \| 0.373 \| 126 \| 76 \| 2 \| 13 \| 136 \| 1 \| 125 \| Oxidoreductase \| Oxidoreductase; afdb-uniprot50 \| AF-A0A7X1ZI31-F1-MODEL\_V4 \| 1.0 \| 2.766e-13 \| 567 \| 0.383 \| 146 \| 75 \| 5 \| 1 \| 136 \| 16 \| 156 \| Phage tail protein \| Phage tail protein; afdb-uniprot50 \| AF-A0A4D7AYY7-F1-MODEL\_V4 \| 1.0 \| 4.481e-12 \| 566 \| 0.251 \| 127 \| 93 \| 2 \| 12 \| 136 \| 2 \| 128 \| Phage tail protein \| Phage tail protein; afdb-uniprot50 \| AF-A0A5E4YCW2-F1-MODEL\_V4 \| 1.0 \| 8.991e-12 \| 566 \| 0.338 \| 124 \| 82 \| 0 \| 13 \| 136 \| 1 \| 124 \| Phage tail protein \| Phage tail protein; afdb-uniprot50 \| AF-A0A7U3R1Q9-F1-MODEL\_V4 \| 1.0 \| 4.728e-12 \| 566 \| 0.29 \| 141 \| 94 \| 2 \| 1 \| 136 \| 164 \| 303 \| Uncharacterized protein \| Uncharacterized protein; afdb-uniprot50 \| AF-A0A7X4CLP0-F1-MODEL\_V4 \| 1.0 \| 4.026e-12 \| 564 \| 0.366 \| 131 \| 76 \| 2 \| 13 \| 136 \| 1 \| 131 \| Phage tail protein \| Phage tail protein; afdb-uniprot50 \| AF-A0A263HHG5-F1-MODEL\_V4 \| 1.0 \| 2.92e-12 \| 564 \| 0.335 \| 131 \| 86 \| 1 \| 7 \| 136 \| 1 \| 131 \| Oxidoreductase \| Oxidoreductase; afdb-uniprot50 \| AF-A0A740LUZ9-F1-MODEL\_V4 \| 1.0 \| 7.652e-13 \| 564 \| 0.313 \| 134 \| 90 \| 1 \| 5 \| 136 \| 17 \| 150 \| Phage tail protein \| Phage tail protein; afdb-uniprot50 \| AF-A0A376ZJI1-F1-MODEL\_V4 \| 1.0 \| 1.056e-11 \| 564 \| 0.314 \| 124 \| 85 \| 0 \| 13 \| 136 \| 1 \| 124 \| Tail fiber protein \| Tail fiber protein; afdb-uniprot50 \| AF-A0A2E3Q565-F1-MODEL\_V4 \| 1.0 \| 5.552e-12 \| 562 \| 0.318 \| 135 \| 85 \| 2 \| 8 \| 136 \| 178 \| 311 \| Uncharacterized protein \| Uncharacterized protein; afdb-uniprot50 \| AF-A0A7Z0EYT7-F1-MODEL\_V4 \| 1.0 \| 8.991e-12 \| 561 \| 0.346 \| 124 \| 81 \| 0 \| 13 \| 136 \| 1 \| 124 \| Uncharacterized protein \| Uncharacterized protein; afdb-uniprot50 \| AF-A0A5T7Y1R6-F1-MODEL\_V4 \| 1.0 \| 1e-12 \| 561 \| 0.272 \| 143 \| 97 \| 2 \| 1 \| 136 \| 43 \| 185 \| Phage tail protein \| Phage tail protein; afdb-uniprot50 \| AF-A0A4V5PSP7-F1-MODEL\_V4 \| 1.0 \| 5.552e-12 \| 560 \| 0.312 \| 144 \| 91 \| 2 \| 1 \| 136 \| 147 \| 290 \| Phage tail protein \| Phage tail protein; afdb-uniprot50 \| AF-D6I841-F1-MODEL\_V4 \| 1.0 \| 3.816e-12 \| 559 \| 0.31 \| 129 \| 86 \| 2 \| 11 \| 136 \| 24 \| 152 \| Predicted protein \| Predicted protein; afdb-uniprot50 \| AF-A0A4P7QXJ5-F1-MODEL\_V4 \| 1.0 \| 6.878e-12 \| 557 \| 0.36 \| 125 \| 78 \| 2 \| 13 \| 136 \| 1 \| 124 \| Phage tail protein \| Phage tail protein; afdb-uniprot50 \| AF-A0A4R6E0B6-F1-MODEL\_V4 \| 1.0 \| 3.615e-13 \| 557 \| 0.293 \| 150 \| 92 \| 3 \| 1 \| 136 \| 12 \| 161 \| Uncharacterized protein \| Uncharacterized protein; afdb-uniprot50 \| AF-A0A377Q627-F1-MODEL\_V4 \| 1.0 \| 1.001e-11 \| 556 \| 0.384 \| 125 \| 75 \| 2 \| 13 \| 136 \| 1 \| 124 \| Phage protein U \| Phage protein U; afdb-uniprot50 \| AF-A0A080KNK2-F1-MODEL\_V4 \| 1.0 \| 7.656e-12 \| 556 \| 0.32 \| 125 \| 83 \| 2 \| 13 \| 136 \| 1 \| 124 \| Phage protein U \| Phage protein U; afdb-uniprot50 \| AF-A0A2C5TNN0-F1-MODEL\_V4 \| 1.0 \| 8.522e-12 \| 556 \| 0.354 \| 124 \| 80 \| 0 \| 13 \| 136 \| 2 \| 125 \| Phage tail protein \| Phage tail protein; afdb-uniprot50 \| AF-A0A0X8GMF9-F1-MODEL\_V4 \| 1.0 \| 5.552e-12 \| 556 \| 0.321 \| 140 \| 87 \| 3 \| 2 \| 136 \| 134 \| 270 \| Uncharacterized protein \| Uncharacterized protein; afdb-uniprot50 \| AF-A0A4Q0GPA2-F1-MODEL\_V4 \| 1.0 \| 8.522e-12 \| 555 \| 0.306 \| 124 \| 86 \| 0 \| 13 \| 136 \| 1 \| 124 \| Phage tail protein \| Phage tail protein; afdb-uniprot50 \| AF-A0A516SHQ7-F1-MODEL\_V4 \| 1.0 \| 1.621e-11 \| 552 \| 0.37 \| 124 \| 77 \| 1 \| 14 \| 136 \| 1 \| 124 \| Phage tail protein \| Phage tail protein; afdb-uniprot50 \| AF-A0A024HAP1-F1-MODEL\_V4 \| 1.0 \| 1.803e-12 \| 552 \| 0.364 \| 140 \| 80 \| 2 \| 6 \| 136 \| 140 \| 279 \| Pyocin R2\_PP, tail formation protein \| Pyocin R2\_PP, tail formation protein; afdb-uniprot50 \| AF-A0A4P9VIK5-F1-MODEL\_V4 \| 1.0 \| 8.517e-13 \| 551 \| 0.381 \| 144 \| 79 \| 3 \| 1 \| 136 \| 1 \| 142 \| Uncharacterized protein \| Uncharacterized protein; afdb-uniprot50 \| AF-A0A381GQM7-F1-MODEL\_V4 \| 1.0 \| 1.114e-11 \| 550 \| 0.336 \| 122 \| 81 \| 0 \| 13 \| 134 \| 1 \| 122 \| Phage tail protein \| Phage tail protein; afdb-uniprot50 \| AF-A0A3N2E0P3-F1-MODEL\_V4 \| 1.0 \| 1e-12 \| 549 \| 0.367 \| 136 \| 77 \| 1 \| 10 \| 136 \| 1 \| 136 \| Uncharacterized protein \| Uncharacterized protein; afdb-uniprot50 \| AF-D4BTZ4-F1-MODEL\_V4 \| 1.0 \| 2.624e-11 \| 548 \| 0.362 \| 124 \| 79 \| 0 \| 13 \| 136 \| 1 \| 124 \| Phage P2 GpU \| Phage P2 GpU; afdb-uniprot50 \| AF-A0A4Y6W0D4-F1-MODEL\_V4 \| 1.0 \| 1.308e-11 \| 547 \| 0.355 \| 121 \| 78 \| 0 \| 16 \| 136 \| 2 \| 122 \| Oxidoreductase \| Oxidoreductase; afdb-uniprot50 \| AF-A0A2W6Y6Z1-F1-MODEL\_V4 \| 1.0 \| 3.251e-11 \| 546 \| 0.306 \| 124 \| 86 \| 0 \| 13 \| 136 \| 1 \| 124 \| Oxidoreductase \| Oxidoreductase; afdb-uniprot50 \| AF-A0A0Q8AZW1-F1-MODEL\_V4 \| 1.0 \| 1.621e-11 \| 546 \| 0.301 \| 126 \| 86 \| 2 \| 13 \| 136 \| 2 \| 127 \| Uncharacterized protein \| Uncharacterized protein; afdb-uniprot50 \| AF-A0A2Y0TQQ1-F1-MODEL\_V4 \| 1.0 \| 2.487e-11 \| 545 \| 0.306 \| 124 \| 86 \| 0 \| 13 \| 136 \| 1 \| 124 \| Phage tail protein \| Phage tail protein; afdb-uniprot50 \| AF-A0A1V9UC49-F1-MODEL\_V4 \| 1.0 \| 1.239e-12 \| 544 \| 0.297 \| 148 \| 91 \| 2 \| 2 \| 136 \| 147 \| 294 \| Uncharacterized protein \| Uncharacterized protein; afdb-uniprot50 \| AF-A0A369YGR3-F1-MODEL\_V4 \| 1.0 \| 1.903e-11 \| 542 \| 0.343 \| 128 \| 83 \| 1 \| 10 \| 136 \| 3 \| 130 \| Oxidoreductase \| Oxidoreductase; afdb-uniprot50 \| AF-A0A085ARU2-F1-MODEL\_V4 \| 1.0 \| 2.234e-12 \| 542 \| 0.287 \| 139 \| 95 \| 1 \| 2 \| 136 \| 17 \| 155 \| Phage tail protein \| Phage tail protein; afdb-uniprot50 \| AF-A0A7S6MLX5-F1-MODEL\_V4 \| 1.0 \| 2.008e-11 \| 536 \| 0.307 \| 140 \| 88 \| 3 \| 5 \| 136 \| 119 \| 257 \| Phage tail protein \| Phage tail protein; afdb-uniprot50 \| AF-A0A4R1TKW3-F1-MODEL\_V4 \| 1.0 \| 2.624e-11 \| 536 \| 0.303 \| 135 \| 87 \| 3 \| 8 \| 136 \| 182 \| 315 \| Uncharacterized protein \| Uncharacterized protein; afdb-uniprot50 \| AF-A0A7S8IIF9-F1-MODEL\_V4 \| 1.0 \| 3.251e-11 \| 533 \| 0.301 \| 126 \| 86 \| 1 \| 13 \| 136 \| 1 \| 126 \| Uncharacterized protein \| Uncharacterized protein; afdb-uniprot50 \| AF-A0A1C3EBL2-F1-MODEL\_V4 \| 1.0 \| 6.52e-12 \| 533 \| 0.251 \| 139 \| 101 \| 2 \| 1 \| 136 \| 5 \| 143 \| Uncharacterized protein \| Uncharacterized protein; afdb-uniprot50 \| AF-A0A1W0CDP0-F1-MODEL\_V4 \| 1.0 \| 1.38e-11 \| 533 \| 0.313 \| 137 \| 88 \| 3 \| 5 \| 136 \| 134 \| 269 \| Uncharacterized protein \| Uncharacterized protein; afdb-uniprot50 \| AF-A0A7Z1Y6H2-F1-MODEL\_V4 \| 1.0 \| 2.118e-11 \| 532 \| 0.308 \| 123 \| 85 \| 0 \| 14 \| 136 \| 1 \| 123 \| Phage tail protein \| Phage tail protein; afdb-uniprot50 \| AF-A0A1B6NQL9-F1-MODEL\_V4 \| 1.0 \| 4.25e-11 \| 531 \| 0.33 \| 124 \| 83 \| 0 \| 13 \| 136 \| 1 \| 124 \| Phage P2 GpU \| Phage P2 GpU; afdb-uniprot50 \| AF-A0A2A4XTZ4-F1-MODEL\_V4 \| 1.0 \| 3.619e-11 \| 529 \| 0.27 \| 122 \| 89 \| 0 \| 13 \| 134 \| 1 \| 122 \| Uncharacterized protein \| Uncharacterized protein; afdb-uniprot50 \| AF-A0A4R7RZR3-F1-MODEL\_V4 \| 1.0 \| 2.624e-11 \| 528 \| 0.309 \| 126 \| 85 \| 2 \| 10 \| 134 \| 1 \| 125 \| Uncharacterized protein \| Uncharacterized protein; afdb-uniprot50 \| AF-A0A0H5CXN7-F1-MODEL\_V4 \| 1.0 \| 3.43e-11 \| 528 \| 0.282 \| 124 \| 89 \| 0 \| 13 \| 136 \| 5 \| 128 \| Phage protein U \| Phage protein U; afdb-uniprot50 \| AF-A0A0K1K621-F1-MODEL\_V4 \| 1.0 \| 2.769e-11 \| 528 \| 0.309 \| 142 \| 91 \| 2 \| 2 \| 136 \| 107 \| 248 \| Uncharacterized protein \| Uncharacterized protein; afdb-uniprot50 \| AF-A0A1T4W4X0-F1-MODEL\_V4 \| 1.0 \| 8.991e-12 \| 525 \| 0.272 \| 132 \| 87 \| 4 \| 13 \| 136 \| 3 \| 133 \| Uncharacterized protein \| Uncharacterized protein; afdb-uniprot50 \| AF-A0A2X3DKM3-F1-MODEL\_V4 \| 1.0 \| 7.26e-11 \| 523 \| 0.298 \| 124 \| 87 \| 0 \| 13 \| 136 \| 1 \| 124 \| Putative bacteriophage tail protein \| Putative bacteriophage tail protein; afdb-uniprot50 \| AF-A0A2S6N2W6-F1-MODEL\_V4 \| 1.0 \| 3.818e-11 \| 522 \| 0.293 \| 126 \| 87 \| 2 \| 13 \| 136 \| 2 \| 127 \| Uncharacterized protein \| Uncharacterized protein; afdb-uniprot50 \| AF-A0A6L3Y3D4-F1-MODEL\_V4 \| 1.0 \| 2.769e-11 \| 522 \| 0.338 \| 124 \| 82 \| 0 \| 13 \| 136 \| 3 \| 126 \| Phage tail protein \| Phage tail protein; afdb-uniprot50 \| AF-A0A6B3IYA5-F1-MODEL\_V4 \| 1.0 \| 1.71e-11 \| 522 \| 0.248 \| 145 \| 92 \| 1 \| 9 \| 136 \| 22 \| 166 \| Uncharacterized protein \| Uncharacterized protein; afdb-uniprot50 \| AF-A0A425B1Z8-F1-MODEL\_V4 \| 1.0 \| 8.081e-11 \| 520 \| 0.282 \| 124 \| 89 \| 0 \| 13 \| 136 \| 1 \| 124 \| Uncharacterized protein \| Uncharacterized protein; afdb-uniprot50 \| AF-A0A317H5Z8-F1-MODEL\_V4 \| 1.0 \| 3.818e-11 \| 520 \| 0.219 \| 132 \| 103 \| 0 \| 5 \| 136 \| 41 \| 172 \| Phage tail protein \| Phage tail protein; afdb-uniprot50 \| AF-A0A844Z703-F1-MODEL\_V4 \| 1.0 \| 4.99e-11 \| 519 \| 0.371 \| 121 \| 76 \| 0 \| 16 \| 136 \| 2 \| 122 \| Oxidoreductase \| Oxidoreductase; afdb-uniprot50 \| AF-A0A2D2CYK8-F1-MODEL\_V4 \| 1.0 \| 6.183e-11 \| 518 \| 0.256 \| 125 \| 91 \| 2 \| 13 \| 136 \| 3 \| 126 \| Phage tail protein \| Phage tail protein; afdb-uniprot50 \| AF-M5JLR0-F1-MODEL\_V4 \| 1.0 \| 1.001e-10 \| 517 \| 0.241 \| 124 \| 94 \| 0 \| 13 \| 136 \| 1 \| 124 \| Fels-2 prophage protein \| Fels-2 prophage protein; afdb-uniprot50 \| AF-Q31HS8-F1-MODEL\_V4 \| 1.0 \| 8.991e-12 \| 517 \| 0.273 \| 139 \| 89 \| 2 \| 9 \| 136 \| 2 \| 139 \| Phage P2 GpU family protein \| Phage P2 GpU family protein; afdb-uniprot50 \| AF-A0A0P6W270-F1-MODEL\_V4 \| 1.0 \| 1.114e-10 \| 516 \| 0.233 \| 124 \| 94 \| 1 \| 13 \| 136 \| 1 \| 123 \| Uncharacterized protein \| Uncharacterized protein; afdb-uniprot50 \| AF-A0A3S0GWK0-F1-MODEL\_V4 \| 1.0 \| 1.536e-11 \| 516 \| 0.338 \| 127 \| 81 \| 1 \| 13 \| 136 \| 3 \| 129 \| Phage tail protein \| Phage tail protein; afdb-uniprot50 \| AF-A0A6B8KGJ5-F1-MODEL\_V4 \| 1.0 \| 3.619e-11 \| 516 \| 0.263 \| 129 \| 93 \| 1 \| 10 \| 136 \| 1 \| 129 \| Uncharacterized protein \| Uncharacterized protein; afdb-uniprot50 \| AF-A0A318TZH9-F1-MODEL\_V4 \| 1.0 \| 8.995e-11 \| 515 \| 0.217 \| 124 \| 97 \| 0 \| 13 \| 136 \| 2 \| 125 \| Uncharacterized protein \| Uncharacterized protein; afdb-uniprot50 \| AF-A0A0J5GKQ4-F1-MODEL\_V4 \| 1.0 \| 3.251e-11 \| 515 \| 0.297 \| 141 \| 92 \| 4 \| 1 \| 136 \| 104 \| 242 \| Uncharacterized protein \| Uncharacterized protein; afdb-uniprot50 \| AF-A0A7W2GS23-F1-MODEL\_V4 \| 1.0 \| 6.523e-11 \| 513 \| 0.328 \| 131 \| 84 \| 1 \| 10 \| 136 \| 1 \| 131 \| Phage tail protein \| Phage tail protein; afdb-uniprot50 \| AF-A0A143I3T8-F1-MODEL\_V4 \| 1.0 \| 1.381e-10 \| 513 \| 0.363 \| 121 \| 77 \| 0 \| 16 \| 136 \| 2 \| 122 \| Uncharacterized protein \| Uncharacterized protein; afdb-uniprot50 \| AF-A0A8B2NS16-F1-MODEL\_V4 \| 1.0 \| 1.114e-10 \| 507 \| 0.262 \| 122 \| 90 \| 0 \| 14 \| 135 \| 6 \| 127 \| Oxidoreductase \| Oxidoreductase; afdb-uniprot50 \| AF-A0A2T5UQY9-F1-MODEL\_V4 \| 1.0 \| 1.904e-10 \| 506 \| 0.33 \| 121 \| 81 \| 0 \| 16 \| 136 \| 2 \| 122 \| Uncharacterized protein \| Uncharacterized protein; afdb-uniprot50 \| AF-A0A2W7APL3-F1-MODEL\_V4 \| 1.0 \| 8.526e-11 \| 504 \| 0.268 \| 123 \| 88 \| 2 \| 13 \| 134 \| 1 \| 122 \| Phage tail protein \| Phage tail protein; afdb-uniprot50 \| AF-A0A037US03-F1-MODEL\_V4 \| 1.0 \| 1.537e-10 \| 504 \| 0.185 \| 124 \| 101 \| 0 \| 13 \| 136 \| 1 \| 124 \| Tail protein \| Tail protein; afdb-uniprot50 \| AF-A0A7X2IMF3-F1-MODEL\_V4 \| 1.0 \| 1.056e-10 \| 502 \| 0.297 \| 141 \| 87 \| 2 \| 8 \| 136 \| 158 \| 298 \| Uncharacterized protein \| Uncharacterized protein; afdb-uniprot50 \| AF-A0A847KTZ8-F1-MODEL\_V4 \| 1.0 \| 8.995e-11 \| 494 \| 0.292 \| 130 \| 92 \| 0 \| 7 \| 136 \| 10 \| 139 \| Phage tail protein \| Phage tail protein; afdb-uniprot50 \| AF-A0A6I6AWA7-F1-MODEL\_V4 \| 1.0 \| 1.904e-10 \| 494 \| 0.385 \| 109 \| 67 \| 0 \| 28 \| 136 \| 797 \| 905 \| Phage tail tape measure protein \| Phage tail tape measure protein; afdb-uniprot50 \| AF-A0A1Q8D7K3-F1-MODEL\_V4 \| 1.0 \| 2.489e-10 \| 491 \| 0.33 \| 121 \| 80 \| 1 \| 13 \| 133 \| 1 \| 120 \| Phage tail protein \| Phage tail protein; afdb-uniprot50 \| AF-A0A0D0K1T5-F1-MODEL\_V4 \| 1.0 \| 1.457e-10 \| 491 \| 0.289 \| 128 \| 88 \| 2 \| 10 \| 134 \| 2 \| 129 \| Tail protein \| Tail protein; afdb-uniprot50 \| AF-A0A6M3KNW0-F1-MODEL\_V4 \| 1.0 \| 2.236e-10 \| 489 \| 0.349 \| 126 \| 79 \| 2 \| 12 \| 136 \| 2 \| 125 \| Uncharacterized protein \| Uncharacterized protein; afdb-uniprot50 \| AF-A0A5S9QVS9-F1-MODEL\_V4 \| 1.0 \| 6.523e-11 \| 489 \| 0.257 \| 128 \| 91 \| 2 \| 13 \| 136 \| 3 \| 130 \| Uncharacterized protein \| Uncharacterized protein; afdb-uniprot50 \| AF-A0A833LH97-F1-MODEL\_V4 \| 1.0 \| 4.993e-10 \| 488 \| 0.233 \| 124 \| 95 \| 0 \| 13 \| 136 \| 1 \| 124 \| Phage tail protein \| Phage tail protein; afdb-uniprot50 \| AF-A0A212LR60-F1-MODEL\_V4 \| 1.0 \| 1.457e-10 \| 488 \| 0.293 \| 126 \| 84 \| 4 \| 13 \| 133 \| 1 \| 126 \| Phage protein U \| Phage protein U; afdb-uniprot50 \| AF-A0A0R3KUL0-F1-MODEL\_V4 \| 1.0 \| 5.863e-10 \| 487 \| 0.225 \| 124 \| 96 \| 0 \| 13 \| 136 \| 1 \| 124 \| Uncharacterized protein \| Uncharacterized protein; afdb-uniprot50 \| AF-A0A4Z1QXY8-F1-MODEL\_V4 \| 1.0 \| 4.252e-10 \| 482 \| 0.212 \| 127 \| 97 \| 1 \| 11 \| 134 \| 22 \| 148 \| Phage tail protein \| Phage tail protein; afdb-uniprot50 \| AF-A0A1M5AJI1-F1-MODEL\_V4 \| 1.0 \| 1.904e-10 \| 481 \| 0.33 \| 127 \| 78 \| 4 \| 13 \| 136 \| 3 \| 125 \| Uncharacterized protein \| Uncharacterized protein; afdb-uniprot50 \| AF-A0A165XHF3-F1-MODEL\_V4 \| 1.0 \| 8.081e-11 \| 481 \| 0.296 \| 128 \| 86 \| 3 \| 12 \| 136 \| 2 \| 128 \| Phage P2 GpU \| Phage P2 GpU; afdb-uniprot50 \| AF-A0A1S2U5I4-F1-MODEL\_V4 \| 1.0 \| 2.236e-10 \| 480 \| 0.264 \| 125 \| 89 \| 2 \| 13 \| 136 \| 1 \| 123 \| Phage tail protein \| Phage tail protein; afdb-uniprot50 \| AF-M5K5B9-F1-MODEL\_V4 \| 1.0 \| 8.53e-10 \| 480 \| 0.274 \| 124 \| 90 \| 0 \| 13 \| 136 \| 1 \| 124 \| Fels-2 prophage protein \| Fels-2 prophage protein; afdb-uniprot50 \| AF-A0A3L7AM95-F1-MODEL\_V4 \| 1.0 \| 3.82e-10 \| 480 \| 0.278 \| 122 \| 88 \| 0 \| 13 \| 134 \| 1 \| 122 \| Phage tail protein \| Phage tail protein; afdb-uniprot50 \| AF-A0A4R2GR57-F1-MODEL\_V4 \| 1.0 \| 3.253e-10 \| 479 \| 0.234 \| 132 \| 96 \| 1 \| 10 \| 136 \| 1 \| 132 \| Phage protein U \| Phage protein U; afdb-uniprot50 \| AF-A0A371WU12-F1-MODEL\_V4 \| 1.0 \| 9.495e-10 \| 479 \| 0.245 \| 122 \| 92 \| 0 \| 13 \| 134 \| 1 \| 122 \| Phage tail protein \| Phage tail protein; afdb-uniprot50 \| AF-A0A1T4WWY7-F1-MODEL\_V4 \| 1.0 \| 1.537e-10 \| 477 \| 0.323 \| 130 \| 82 \| 3 \| 13 \| 136 \| 2 \| 131 \| Uncharacterized protein \| Uncharacterized protein; afdb-uniprot50 \| AF-A0A2X4TYK3-F1-MODEL\_V4 \| 1.0 \| 4.252e-10 \| 473 \| 0.413 \| 104 \| 61 \| 0 \| 33 \| 136 \| 2 \| 105 \| Phage protein U \| Phage protein U; afdb-uniprot50 \| AF-A0A7W6WAV6-F1-MODEL\_V4 \| 1.0 \| 3.083e-10 \| 472 \| 0.261 \| 130 \| 88 \| 4 \| 13 \| 136 \| 3 \| 130 \| Uncharacterized protein \| Uncharacterized protein; afdb-uniprot50 \| AF-A0A5E7CNT0-F1-MODEL\_V4 \| 1.0 \| 5.863e-10 \| 471 \| 0.256 \| 125 \| 90 \| 2 \| 13 \| 136 \| 3 \| 125 \| Uncharacterized protein \| Uncharacterized protein; afdb-uniprot50 \| AF-A0A4Q7FRM2-F1-MODEL\_V4 \| 1.0 \| 4.252e-10 \| 471 \| 0.207 \| 135 \| 99 \| 1 \| 10 \| 136 \| 1 \| 135 \| Uncharacterized protein \| Uncharacterized protein; afdb-uniprot50 \| AF-A0A5Y6QJ23-F1-MODEL\_V4 \| 1.0 \| 6.186e-10 \| 470 \| 0.256 \| 125 \| 90 \| 2 \| 13 \| 136 \| 1 \| 123 \| Phage protein U \| Phage protein U; afdb-uniprot50 \| AF-A0A218KPW4-F1-MODEL\_V4 \| 1.0 \| 6.526e-10 \| 469 \| 0.333 \| 123 \| 77 \| 1 \| 14 \| 136 \| 1 \| 118 \| Phage tail protein \| Phage tail protein; afdb-uniprot50 \| AF-A0A5C8S9X4-F1-MODEL\_V4 \| 1.0 \| 2.119e-10 \| 469 \| 0.248 \| 141 \| 92 \| 3 \| 10 \| 136 \| 1 \| 141 \| Phage tail protein \| Phage tail protein; afdb-uniprot50 \| AF-A0A3G2V4J2-F1-MODEL\_V4 \| 1.0 \| 1.537e-10 \| 468 \| 0.23 \| 143 \| 91 \| 3 \| 13 \| 136 \| 4 \| 146 \| Uncharacterized protein \| Uncharacterized protein; afdb-uniprot50 \| AF-A0A6N6MG51-F1-MODEL\_V4 \| 1.0 \| 4.03e-10 \| 467 \| 0.255 \| 129 \| 91 \| 2 \| 13 \| 136 \| 1 \| 129 \| Phage tail protein \| Phage tail protein; afdb-uniprot50 \| AF-A0A285V255-F1-MODEL\_V4 \| 1.0 \| 1.309e-09 \| 465 \| 0.258 \| 124 \| 92 \| 0 \| 13 \| 136 \| 1 \| 124 \| Uncharacterized protein \| Uncharacterized protein; afdb-uniprot50 \| AF-A0A1G7PUT3-F1-MODEL\_V4 \| 1.0 \| 9e-10 \| 464 \| 0.258 \| 124 \| 90 \| 1 \| 13 \| 134 \| 1 \| 124 \| Phage P2 GpU \| Phage P2 GpU; afdb-uniprot50 \| AF-A0A377LS59-F1-MODEL\_V4 \| 1.0 \| 8.526e-11 \| 464 \| 0.278 \| 147 \| 95 \| 1 \| 1 \| 136 \| 19 \| 165 \| Phage P2 GpU family protein \| Phage P2 GpU family protein; afdb-uniprot50 \| AF-A0A522WZG7-F1-MODEL\_V4 \| 1.0 \| 4.486e-10 \| 462 \| 0.535 \| 99 \| 46 \| 0 \| 10 \| 108 \| 1 \| 99 \| Phage tail protein \| Phage tail protein; afdb-uniprot50 \| AF-A0A5N3PHE3-F1-MODEL\_V4 \| 1.0 \| 5.268e-10 \| 462 \| 0.232 \| 125 \| 93 \| 1 \| 13 \| 134 \| 1 \| 125 \| Uncharacterized protein \| Uncharacterized protein; afdb-uniprot50 \| AF-A0A1S2E7C7-F1-MODEL\_V4 \| 1.0 \| 1.622e-09 \| 461 \| 0.225 \| 124 \| 95 \| 1 \| 13 \| 136 \| 1 \| 123 \| Phage tail protein \| Phage tail protein; afdb-uniprot50 \| AF-A0A1W2EJV4-F1-MODEL\_V4 \| 1.0 \| 1.711e-09 \| 461 \| 0.196 \| 122 \| 98 \| 0 \| 13 \| 134 \| 1 \| 122 \| Uncharacterized protein \| Uncharacterized protein; afdb-uniprot50 \| AF-A0A4Y8RF33-F1-MODEL\_V4 \| 1.0 \| 1.905e-09 \| 460 \| 0.219 \| 123 \| 96 \| 0 \| 13 \| 135 \| 5 \| 127 \| Oxidoreductase \| Oxidoreductase; afdb-uniprot50 \| AF-A0A346QYH1-F1-MODEL\_V4 \| 1.0 \| 3.083e-10 \| 460 \| 0.31 \| 119 \| 82 \| 0 \| 18 \| 136 \| 20 \| 138 \| Oxidoreductase \| Oxidoreductase; afdb-uniprot50 \| AF-A0A485ALS4-F1-MODEL\_V4 \| 1.0 \| 1.309e-09 \| 460 \| 0.366 \| 109 \| 69 \| 0 \| 28 \| 136 \| 6 \| 114 \| Phage protein U \| Phage protein U; afdb-uniprot50 \| AF-A0A846VJH3-F1-MODEL\_V4 \| 1.0 \| 1.381e-09 \| 459 \| 0.187 \| 128 \| 100 \| 1 \| 13 \| 136 \| 4 \| 131 \| Phage protein U \| Phage protein U; afdb-uniprot50 \| AF-A0A7L5Y3W4-F1-MODEL\_V4 \| 1.0 \| 6.526e-10 \| 459 \| 0.195 \| 133 \| 98 \| 2 \| 13 \| 136 \| 4 \| 136 \| Phage tail protein \| Phage tail protein; afdb-uniprot50 \| AF-A0A0M7ANC8-F1-MODEL\_V4 \| 1.0 \| 1.905e-09 \| 459 \| 0.209 \| 124 \| 98 \| 0 \| 13 \| 136 \| 1 \| 124 \| Phage protein U \| Phage protein U; afdb-uniprot50 \| AF-A0A1H0KW29-F1-MODEL\_V4 \| 1.0 \| 1.309e-09 \| 458 \| 0.288 \| 125 \| 86 \| 2 \| 13 \| 136 \| 3 \| 125 \| Phage tail protein \| Phage tail protein; afdb-uniprot50 \| AF-A0A4Q6DUT2-F1-MODEL\_V4 \| 1.0 \| 6.186e-10 \| 457 \| 0.325 \| 123 \| 81 \| 2 \| 13 \| 134 \| 1 \| 122 \| Uncharacterized protein \| Uncharacterized protein; afdb-uniprot50 \| AF-A0A549T0T0-F1-MODEL\_V4 \| 1.0 \| 2.237e-09 \| 457 \| 0.217 \| 124 \| 97 \| 0 \| 13 \| 136 \| 1 \| 124 \| Phage tail protein \| Phage tail protein; afdb-uniprot50 \| AF-A0A0G0CA59-F1-MODEL\_V4 \| 1.0 \| 1.711e-09 \| 455 \| 0.161 \| 124 \| 103 \| 1 \| 13 \| 136 \| 1 \| 123 \| Uncharacterized protein \| Uncharacterized protein; afdb-uniprot50 \| AF-A0A6I1JKM9-F1-MODEL\_V4 \| 1.0 \| 1.905e-09 \| 454 \| 0.25 \| 124 \| 93 \| 0 \| 13 \| 136 \| 1 \| 124 \| Putative phage tail assembly protein \| Putative phage tail assembly protein; afdb-uniprot50 \| AF-A0A1L9QZK1-F1-MODEL\_V4 \| 1.0 \| 1.176e-09 \| 453 \| 0.286 \| 122 \| 85 \| 1 \| 13 \| 134 \| 1 \| 120 \| Uncharacterized protein \| Uncharacterized protein; afdb-uniprot50 \| AF-A0A1N7LQZ5-F1-MODEL\_V4 \| 1.0 \| 3.083e-10 \| 453 \| 0.281 \| 128 \| 90 \| 2 \| 10 \| 136 \| 2 \| 128 \| Phage protein U \| Phage protein U; afdb-uniprot50 \| AF-A0A369RKP4-F1-MODEL\_V4 \| 1.0 \| 5.555e-11 \| 453 \| 0.39 \| 123 \| 66 \| 3 \| 14 \| 136 \| 1 \| 114 \| Phage tail protein GpU \| Phage tail protein GpU; afdb-uniprot50 \| AF-A0A7C4AK56-F1-MODEL\_V4 \| 1.0 \| 8.085e-10 \| 452 \| 0.244 \| 127 \| 92 \| 3 \| 13 \| 136 \| 2 \| 127 \| Phage tail protein \| Phage tail protein; afdb-uniprot50 \| AF-A0A377NJ59-F1-MODEL\_V4 \| 1.0 \| 6.526e-10 \| 451 \| 0.319 \| 119 \| 78 \| 1 \| 21 \| 136 \| 11 \| 129 \| Phage protein U \| Phage protein U; afdb-uniprot50 \| AF-A0A7C4EKB5-F1-MODEL\_V4 \| 1.0 \| 3.085e-09 \| 451 \| 0.193 \| 124 \| 99 \| 1 \| 13 \| 136 \| 1 \| 123 \| Phage tail protein \| Phage tail protein; afdb-uniprot50 \| AF-A0A0M3AYT1-F1-MODEL\_V4 \| 1.0 \| 3.085e-09 \| 450 \| 0.185 \| 124 \| 101 \| 0 \| 13 \| 136 \| 1 \| 124 \| Tail protein \| Tail protein; afdb-uniprot50 \| AF-A0A1E4M031-F1-MODEL\_V4 \| 1.0 \| 3.085e-09 \| 449 \| 0.233 \| 124 \| 94 \| 1 \| 13 \| 136 \| 1 \| 123 \| Uncharacterized protein \| Uncharacterized protein; afdb-uniprot50 \| AF-A0A3S0YGQ2-F1-MODEL\_V4 \| 1.0 \| 2.489e-10 \| 449 \| 0.233 \| 137 \| 100 \| 3 \| 3 \| 136 \| 14 \| 148 \| Phage tail protein \| Phage tail protein; afdb-uniprot50 \| AF-A0A512JPP3-F1-MODEL\_V4 \| 1.0 \| 4.488e-09 \| 448 \| 0.177 \| 124 \| 102 \| 0 \| 13 \| 136 \| 1 \| 124 \| Uncharacterized protein \| Uncharacterized protein; afdb-uniprot50 \| AF-F8ESZ6-F1-MODEL\_V4 \| 1.0 \| 5.265e-11 \| 448 \| 0.255 \| 184 \| 88 \| 4 \| 1 \| 136 \| 2 \| 184 \| p2 GpU family protein \| p2 GpU family protein; afdb-uniprot50 \| AF-A0A7W9D3K1-F1-MODEL\_V4 \| 1.0 \| 1.711e-09 \| 447 \| 0.286 \| 122 \| 84 \| 2 \| 16 \| 134 \| 2 \| 123 \| Uncharacterized protein \| Uncharacterized protein; afdb-uniprot50 \| AF-A0A7X5J7J9-F1-MODEL\_V4 \| 1.0 \| 7.664e-10 \| 446 \| 0.211 \| 137 \| 102 \| 3 \| 5 \| 136 \| 2 \| 137 \| Uncharacterized protein \| Uncharacterized protein; afdb-uniprot50 \| AF-A0A379QTF1-F1-MODEL\_V4 \| 1.0 \| 2.49e-09 \| 445 \| 0.227 \| 123 \| 94 \| 1 \| 13 \| 135 \| 1 \| 122 \| P2 GpU \| P2 GpU; afdb-uniprot50 \| AF-A0A2L0IJT3-F1-MODEL\_V4 \| 1.0 \| 1.115e-09 \| 442 \| 0.415 \| 113 \| 66 \| 0 \| 24 \| 136 \| 26 \| 138 \| Phage tail protein \| Phage tail protein; afdb-uniprot50 \| AF-Q602Z4-F1-MODEL\_V4 \| 1.0 \| 2.36e-09 \| 442 \| 0.201 \| 124 \| 98 \| 1 \| 13 \| 136 \| 2 \| 124 \| Conserved domain protein \| Conserved domain protein; afdb-uniprot50 \| AF-A0A017HDE4-F1-MODEL\_V4 \| 1.0 \| 2.12e-09 \| 441 \| 0.325 \| 123 \| 79 \| 2 \| 17 \| 136 \| 3 \| 124 \| Phage P2 GpU \| Phage P2 GpU; afdb-uniprot50 \| AF-A0A847GAS6-F1-MODEL\_V4 \| 1.0 \| 4.254e-09 \| 441 \| 0.216 \| 125 \| 96 \| 2 \| 13 \| 136 \| 2 \| 125 \| Phage tail protein \| Phage tail protein; afdb-uniprot50 \| AF-E2CHK4-F1-MODEL\_V4 \| 1.0 \| 1.905e-09 \| 440 \| 0.275 \| 127 \| 88 \| 3 \| 13 \| 136 \| 1 \| 126 \| Putative phage tail protein U \| Putative phage tail protein U; afdb-uniprot50 \| AF-A0A3G8MA71-F1-MODEL\_V4 \| 1.0 \| 4.254e-09 \| 439 \| 0.157 \| 127 \| 104 \| 1 \| 13 \| 136 \| 5 \| 131 \| Uncharacterized protein \| Uncharacterized protein; afdb-uniprot50 \| AF-B6IMG5-F1-MODEL\_V4 \| 1.0 \| 2.36e-09 \| 439 \| 0.219 \| 123 \| 94 \| 1 \| 13 \| 135 \| 1 \| 121 \| Uncharacterized protein \| Uncharacterized protein; afdb-uniprot50 \| AF-A0A397LXE3-F1-MODEL\_V4 \| 1.0 \| 5.56e-09 \| 439 \| 0.241 \| 124 \| 94 \| 0 \| 13 \| 136 \| 1 \| 124 \| Uncharacterized protein \| Uncharacterized protein; afdb-uniprot50 \| AF-A0A651FVX8-F1-MODEL\_V4 \| 1.0 \| 2.12e-09 \| 438 \| 0.225 \| 124 \| 95 \| 1 \| 13 \| 136 \| 1 \| 123 \| Phage tail protein \| Phage tail protein; afdb-uniprot50 \| AF-R5HI01-F1-MODEL\_V4 \| 1.0 \| 2.771e-09 \| 438 \| 0.193 \| 124 \| 99 \| 1 \| 13 \| 136 \| 1 \| 123 \| Putative phage-related protein \| Putative phage-related protein; afdb-uniprot50 \| AF-E2CJH4-F1-MODEL\_V4 \| 1.0 \| 8.089e-09 \| 438 \| 0.217 \| 124 \| 97 \| 0 \| 13 \| 136 \| 1 \| 124 \| p2 GpU family protein \| p2 GpU family protein; afdb-uniprot50 \| AF-A0A258L8T5-F1-MODEL\_V4 \| 1.0 \| 1.176e-09 \| 438 \| 0.259 \| 135 \| 98 \| 1 \| 1 \| 135 \| 1 \| 133 \| Uncharacterized protein \| Uncharacterized protein; afdb-uniprot50 \| AF-R8AR70-F1-MODEL\_V4 \| 1.0 \| 3.822e-09 \| 436 \| 0.24 \| 125 \| 92 \| 2 \| 13 \| 136 \| 2 \| 124 \| GpU family protein \| GpU family protein; afdb-uniprot50 \| AF-A0NQA4-F1-MODEL\_V4 \| 1.0 \| 3.822e-09 \| 436 \| 0.204 \| 122 \| 97 \| 0 \| 13 \| 134 \| 1 \| 122 \| Putative bacteriophage tail protein \| Putative bacteriophage tail protein; afdb-uniprot50 \| AF-A0A7C7PSI2-F1-MODEL\_V4 \| 1.0 \| 2.771e-09 \| 436 \| 0.217 \| 124 \| 94 \| 2 \| 13 \| 136 \| 4 \| 124 \| Phage tail protein \| Phage tail protein; afdb-uniprot50 \| AF-A0A376MUH3-F1-MODEL\_V4 \| 1.0 \| 9e-10 \| 434 \| 0.285 \| 126 \| 86 \| 2 \| 13 \| 136 \| 1 \| 124 \| Tail fiber protein \| Tail fiber protein; afdb-uniprot50 \| AF-A0A806DA75-F1-MODEL\_V4 \| 1.0 \| 3.621e-10 \| 434 \| 0.255 \| 168 \| 80 \| 3 \| 13 \| 136 \| 1 \| 167 \| P2 GpU family protein \| P2 GpU family protein; afdb-uniprot50 \| AF-A0A2G1CVT3-F1-MODEL\_V4 \| 1.0 \| 5.27e-09 \| 432 \| 0.233 \| 124 \| 91 \| 2 \| 13 \| 136 \| 2 \| 121 \| Uncharacterized protein \| Uncharacterized protein; afdb-uniprot50 \| AF-G2HXA8-F1-MODEL\_V4 \| 1.0 \| 6.529e-09 \| 431 \| 0.209 \| 124 \| 94 \| 2 \| 13 \| 136 \| 1 \| 120 \| Hypothetical phage protein \| Hypothetical phage protein; afdb-uniprot50 \| AF-A0A7U9JER2-F1-MODEL\_V4 \| 1.0 \| 4.254e-09 \| 431 \| 0.336 \| 104 \| 69 \| 0 \| 33 \| 136 \| 2 \| 105 \| Oxidoreductase \| Oxidoreductase; afdb-uniprot50 \| AF-U1ASZ1-F1-MODEL\_V4 \| 1.0 \| 5.56e-09 \| 431 \| 0.225 \| 124 \| 95 \| 1 \| 13 \| 136 \| 1 \| 123 \| Phage tail protein \| Phage tail protein; afdb-uniprot50 \| AF-A0A7Y4XET2-F1-MODEL\_V4 \| 1.0 \| 2.12e-09 \| 431 \| 0.167 \| 131 \| 108 \| 1 \| 6 \| 136 \| 9 \| 138 \| Phage tail protein \| Phage tail protein; afdb-uniprot50 \| AF-A0A437NT45-F1-MODEL\_V4 \| 1.0 \| 1.623e-08 \| 430 \| 0.217 \| 124 \| 97 \| 0 \| 13 \| 136 \| 1 \| 124 \| Phage tail protein \| Phage tail protein; afdb-uniprot50 \| AF-A0A2A3D6X4-F1-MODEL\_V4 \| 1.0 \| 6.529e-09 \| 429 \| 0.209 \| 124 \| 97 \| 1 \| 13 \| 136 \| 1 \| 123 \| Phage tail protein \| Phage tail protein; afdb-uniprot50 \| AF-A0A411WI49-F1-MODEL\_V4 \| 1.0 \| 5.27e-09 \| 429 \| 0.224 \| 125 \| 96 \| 1 \| 12 \| 136 \| 2 \| 125 \| Phage tail protein \| Phage tail protein; afdb-uniprot50 \| AF-A0A6D0J052-F1-MODEL\_V4 \| 1.0 \| 7.668e-09 \| 428 \| 0.353 \| 116 \| 75 \| 0 \| 13 \| 128 \| 1 \| 116 \| Phage tail protein \| Phage tail protein; afdb-uniprot50 \| AF-L0NDK6-F1-MODEL\_V4 \| 1.0 \| 2.627e-09 \| 428 \| 0.246 \| 126 \| 90 \| 4 \| 16 \| 136 \| 2 \| 127 \| Putative Phage P2 GpU \| Putative Phage P2 GpU; afdb-uniprot50 \| AF-A0A327K1Q9-F1-MODEL\_V4 \| 1.0 \| 3.085e-09 \| 427 \| 0.265 \| 128 \| 85 \| 4 \| 13 \| 136 \| 3 \| 125 \| Uncharacterized protein \| Uncharacterized protein; afdb-uniprot50 \| AF-A0A2D5PEI7-F1-MODEL\_V4 \| 1.0 \| 2.771e-09 \| 426 \| 0.244 \| 127 \| 90 \| 5 \| 12 \| 136 \| 2 \| 124 \| Uncharacterized protein \| Uncharacterized protein; afdb-uniprot50 \| AF-A0A0Q6ZRY4-F1-MODEL\_V4 \| 1.0 \| 5.56e-09 \| 423 \| 0.23 \| 126 \| 94 \| 3 \| 13 \| 136 \| 1 \| 125 \| Uncharacterized protein \| Uncharacterized protein; afdb-uniprot50 \| AF-A0A5M8P9Q4-F1-MODEL\_V4 \| 1.0 \| 6.189e-09 \| 423 \| 0.246 \| 134 \| 93 \| 4 \| 10 \| 136 \| 1 \| 133 \| Phage tail protein \| Phage tail protein; afdb-uniprot50 \| AF-A0A4P7KXL2-F1-MODEL\_V4 \| 1.0 \| 2.924e-09 \| 423 \| 0.219 \| 123 \| 95 \| 1 \| 13 \| 135 \| 1 \| 122 \| Phage P2 GpU \| Phage P2 GpU; afdb-uniprot50 \| AF-A0A125QFM2-F1-MODEL\_V4 \| 1.0 \| 1.242e-08 \| 422 \| 0.201 \| 124 \| 98 \| 1 \| 13 \| 136 \| 1 \| 123 \| Phage P2 GpU \| Phage P2 GpU; afdb-uniprot50 \| AF-A0A7C3M160-F1-MODEL\_V4 \| 1.0 \| 1.309e-09 \| 421 \| 0.22 \| 136 \| 100 \| 4 \| 1 \| 136 \| 1 \| 130 \| Uncharacterized protein \| Uncharacterized protein; afdb-uniprot50 \| AF-A0A432QVN4-F1-MODEL\_V4 \| 1.0 \| 5.557e-10 \| 420 \| 0.27 \| 137 \| 87 \| 3 \| 13 \| 136 \| 1 \| 137 \| Uncharacterized protein \| Uncharacterized protein; afdb-uniprot50 \| AF-A0A4P5QXL4-F1-MODEL\_V4 \| 1.0 \| 1.176e-09 \| 420 \| 0.211 \| 137 \| 105 \| 2 \| 1 \| 136 \| 7 \| 141 \| Uncharacterized protein \| Uncharacterized protein; afdb-uniprot50 \| AF-A0A377PAA7-F1-MODEL\_V4 \| 1.0 \| 4.254e-09 \| 420 \| 0.227 \| 123 \| 94 \| 1 \| 13 \| 135 \| 1 \| 122 \| Phage protein U \| Phage protein U; afdb-uniprot50 \| AF-A0A221T8K1-F1-MODEL\_V4 \| 1.0 \| 4.735e-09 \| 419 \| 0.211 \| 123 \| 96 \| 1 \| 13 \| 135 \| 1 \| 122 \| Phage tail protein \| Phage tail protein; afdb-uniprot50 \| AF-A0A085AFM6-F1-MODEL\_V4 \| 1.0 \| 1.309e-09 \| 418 \| 0.206 \| 145 \| 104 \| 3 \| 1 \| 136 \| 1 \| 143 \| Uncharacterized protein \| Uncharacterized protein; afdb-uniprot50 \| AF-A0A4R6M7R2-F1-MODEL\_V4 \| 1.0 \| 1.177e-08 \| 417 \| 0.217 \| 124 \| 93 \| 2 \| 13 \| 136 \| 1 \| 120 \| Phage protein U \| Phage protein U; afdb-uniprot50 \| AF-A0A6N8TE18-F1-MODEL\_V4 \| 1.0 \| 4.032e-09 \| 417 \| 0.224 \| 129 \| 94 \| 3 \| 13 \| 136 \| 9 \| 136 \| Uncharacterized protein \| Uncharacterized protein; afdb-uniprot50 \| AF-A0A7A6ZBU5-F1-MODEL\_V4 \| 1.0 \| 8.535e-09 \| 416 \| 0.211 \| 123 \| 96 \| 1 \| 13 \| 135 \| 1 \| 122 \| Phage tail protein \| Phage tail protein; afdb-uniprot50 \| AF-A0A7X7NI24-F1-MODEL\_V4 \| 1.0 \| 2.121e-08 \| 415 \| 0.169 \| 124 \| 101 \| 1 \| 13 \| 136 \| 1 \| 122 \| Phage tail protein \| Phage tail protein; afdb-uniprot50 \| AF-A0A3B9Q8H5-F1-MODEL\_V4 \| 1.0 \| 4.488e-09 \| 414 \| 0.179 \| 134 \| 105 \| 4 \| 7 \| 136 \| 1 \| 133 \| Uncharacterized protein \| Uncharacterized protein; afdb-uniprot50 \| AF-A0A829E1U0-F1-MODEL\_V4 \| 1.0 \| 9.5e-09 \| 414 \| 0.203 \| 123 \| 97 \| 1 \| 13 \| 135 \| 1 \| 122 \| Phage P2 GpU family protein \| Phage P2 GpU family protein; afdb-uniprot50 \| AF-A0A1C3E6K8-F1-MODEL\_V4 \| 1.0 \| 4.735e-09 \| 413 \| 0.206 \| 126 \| 94 \| 4 \| 12 \| 136 \| 2 \| 122 \| Uncharacterized protein \| Uncharacterized protein; afdb-uniprot50 \| AF-A0A1M7R7F9-F1-MODEL\_V4 \| 1.0 \| 1.242e-08 \| 413 \| 0.233 \| 124 \| 94 \| 1 \| 13 \| 136 \| 1 \| 123 \| Phage protein U \| Phage protein U; afdb-uniprot50 \| AF-A0A2M7MVA7-F1-MODEL\_V4 \| 1.0 \| 2.238e-08 \| 413 \| 0.185 \| 124 \| 100 \| 1 \| 13 \| 136 \| 2 \| 124 \| Uncharacterized protein \| Uncharacterized protein; afdb-uniprot50 \| AF-A0A0B6D192-F1-MODEL\_V4 \| 1.0 \| 5.866e-09 \| 412 \| 0.186 \| 123 \| 94 \| 3 \| 14 \| 136 \| 1 \| 117 \| Phage P2 GpU family protein \| Phage P2 GpU family protein; afdb-uniprot50 \| AF-A0A8B2QSL7-F1-MODEL\_V4 \| 1.0 \| 9.004e-09 \| 412 \| 0.243 \| 123 \| 89 \| 2 \| 13 \| 134 \| 1 \| 120 \| Oxidoreductase \| Oxidoreductase; afdb-uniprot50 \| AF-A0A855HFC3-F1-MODEL\_V4 \| 1.0 \| 6.529e-09 \| 411 \| 0.466 \| 103 \| 54 \| 1 \| 8 \| 109 \| 5 \| 107 \| Phage tail protein \| Phage tail protein; afdb-uniprot50 \| AF-A0A1J5BJG5-F1-MODEL\_V4 \| 1.0 \| 9.5e-09 \| 410 \| 0.182 \| 126 \| 101 \| 2 \| 9 \| 134 \| 26 \| 149 \| Uncharacterized protein \| Uncharacterized protein; afdb-uniprot50 \| AF-A0A0F4NJK3-F1-MODEL\_V4 \| 1.0 \| 1.712e-08 \| 409 \| 0.204 \| 122 \| 93 \| 2 \| 13 \| 134 \| 1 \| 118 \| Uncharacterized protein \| Uncharacterized protein; afdb-uniprot50 \| AF-A0A7J4ETH5-F1-MODEL\_V4 \| 1.0 \| 1.177e-08 \| 409 \| 0.266 \| 124 \| 87 \| 3 \| 13 \| 136 \| 1 \| 120 \| Uncharacterized protein \| Uncharacterized protein; afdb-uniprot50 \| AF-A0A2M7G5P3-F1-MODEL\_V4 \| 1.0 \| 9.004e-09 \| 409 \| 0.169 \| 124 \| 102 \| 1 \| 12 \| 135 \| 2 \| 124 \| Uncharacterized protein \| Uncharacterized protein; afdb-uniprot50 \| AF-A0A6L5BH81-F1-MODEL\_V4 \| 1.0 \| 2.628e-08 \| 407 \| 0.169 \| 124 \| 102 \| 1 \| 13 \| 136 \| 1 \| 123 \| Uncharacterized protein \| Uncharacterized protein; afdb-uniprot50 \| AF-A0A2X2GG49-F1-MODEL\_V4 \| 1.0 \| 4.032e-09 \| 406 \| 0.25 \| 108 \| 81 \| 0 \| 28 \| 135 \| 9 \| 116 \| Phage protein U \| Phage protein U; afdb-uniprot50 \| AF-A0A4R5HEI0-F1-MODEL\_V4 \| 1.0 \| 4.254e-09 \| 406 \| 0.274 \| 124 \| 87 \| 2 \| 14 \| 136 \| 3 \| 124 \| Phage tail protein \| Phage tail protein; afdb-uniprot50 \| AF-A0A6F8NNB5-F1-MODEL\_V4 \| 1.0 \| 1.806e-09 \| 406 \| 0.266 \| 120 \| 79 \| 2 \| 26 \| 136 \| 37 \| 156 \| Bacteriophage P2 tail protein GPU \| Bacteriophage P2 tail protein GPU; afdb-uniprot50 \| AF-A0A1G3UAP7-F1-MODEL\_V4 \| 1.0 \| 9.5e-09 \| 405 \| 0.233 \| 124 \| 92 \| 1 \| 13 \| 136 \| 2 \| 122 \| Uncharacterized protein \| Uncharacterized protein; afdb-uniprot50 \| AF-A0A2T5NQ46-F1-MODEL\_V4 \| 1.0 \| 1.623e-08 \| 405 \| 0.233 \| 124 \| 94 \| 1 \| 13 \| 136 \| 1 \| 123 \| Uncharacterized protein \| Uncharacterized protein; afdb-uniprot50 \| AF-A0A5C7EXC1-F1-MODEL\_V4 \| 1.0 \| 1.538e-08 \| 405 \| 0.186 \| 123 \| 99 \| 1 \| 13 \| 135 \| 3 \| 124 \| Phage tail protein \| Phage tail protein; afdb-uniprot50 \| AF-A0A0M1VTG3-F1-MODEL\_V4 \| 1.0 \| 1.002e-08 \| 402 \| 0.257 \| 128 \| 90 \| 4 \| 13 \| 136 \| 2 \| 128 \| Uncharacterized protein \| Uncharacterized protein; afdb-uniprot50 \| AF-A0A7U9G627-F1-MODEL\_V4 \| 1.0 \| 2.238e-08 \| 402 \| 0.203 \| 123 \| 97 \| 1 \| 13 \| 135 \| 1 \| 122 \| Putative phage-related protein \| Putative phage-related protein; afdb-uniprot50 \| AF-A0A1X0TI83-F1-MODEL\_V4 \| 1.0 \| 3.256e-08 \| 402 \| 0.211 \| 123 \| 96 \| 1 \| 13 \| 135 \| 1 \| 122 \| Phage tail protein \| Phage tail protein; afdb-uniprot50 \| AF-A0A7Z1M3M8-F1-MODEL\_V4 \| 1.0 \| 1.806e-08 \| 402 \| 0.193 \| 124 \| 99 \| 1 \| 13 \| 136 \| 2 \| 124 \| Phage protein U \| Phage protein U; afdb-uniprot50 \| AF-A0A1R1MK76-F1-MODEL\_V4 \| 1.0 \| 9.004e-09 \| 399 \| 0.188 \| 127 \| 97 \| 3 \| 13 \| 136 \| 1 \| 124 \| Uncharacterized protein \| Uncharacterized protein; afdb-uniprot50 \| AF-E8KG72-F1-MODEL\_V4 \| 1.0 \| 3.435e-08 \| 398 \| 0.169 \| 124 \| 102 \| 1 \| 13 \| 136 \| 3 \| 125 \| Uncharacterized protein \| Uncharacterized protein; afdb-uniprot50 \| AF-A0A6J4N4J5-F1-MODEL\_V4 \| 1.0 \| 2.361e-08 \| 398 \| 0.201 \| 124 \| 98 \| 1 \| 13 \| 136 \| 3 \| 125 \| Uncharacterized protein \| Uncharacterized protein; afdb-uniprot50 \| AF-A0A1M7RIG2-F1-MODEL\_V4 \| 1.0 \| 3.086e-08 \| 398 \| 0.225 \| 124 \| 95 \| 1 \| 13 \| 136 \| 1 \| 123 \| Phage protein U \| Phage protein U; afdb-uniprot50 \| AF-A0A432PWE8-F1-MODEL\_V4 \| 1.0 \| 1.538e-08 \| 396 \| 0.288 \| 125 \| 79 \| 5 \| 13 \| 136 \| 1 \| 116 \| Uncharacterized protein \| Uncharacterized protein; afdb-uniprot50 \| AF-A0A379WIT7-F1-MODEL\_V4 \| 1.0 \| 1.242e-08 \| 396 \| 0.219 \| 123 \| 95 \| 1 \| 13 \| 135 \| 1 \| 122 \| Methyl-accepting chemotaxis protein \| Methyl-accepting chemotaxis protein; afdb-uniprot50 \| AF-A0A836CGV5-F1-MODEL\_V4 \| 1.0 \| 1.457e-09 \| 396 \| 0.274 \| 135 \| 89 \| 3 \| 8 \| 135 \| 676 \| 808 \| Uncharacterized protein \| Uncharacterized protein; afdb-uniprot50 \| AF-A0A1M5PY95-F1-MODEL\_V4 \| 1.0 \| 4.486e-10 \| 395 \| 0.237 \| 143 \| 89 \| 4 \| 8 \| 136 \| 14 \| 150 \| Uncharacterized protein \| Uncharacterized protein; afdb-uniprot50 \| AF-A0A1T4QF76-F1-MODEL\_V4 \| 1.0 \| 3.824e-08 \| 394 \| 0.14 \| 128 \| 105 \| 4 \| 13 \| 136 \| 2 \| 128 \| Uncharacterized protein \| Uncharacterized protein; afdb-uniprot50 \| AF-A0A1Y5SUW4-F1-MODEL\_V4 \| 1.0 \| 2.925e-08 \| 394 \| 0.169 \| 124 \| 102 \| 1 \| 13 \| 136 \| 1 \| 123 \| Phage P2 GpU \| Phage P2 GpU; afdb-uniprot50 \| AF-A0A659PNI1-F1-MODEL\_V4 \| 1.0 \| 2.361e-08 \| 393 \| 0.323 \| 102 \| 69 \| 0 \| 35 \| 136 \| 2 \| 103 \| Phage tail protein \| Phage tail protein; afdb-uniprot50 \| AF-A0A745FX58-F1-MODEL\_V4 \| 1.0 \| 4.256e-08 \| 393 \| 0.343 \| 102 \| 67 \| 0 \| 35 \| 136 \| 1 \| 102 \| Phage tail protein \| Phage tail protein; afdb-uniprot50 \| AF-A0A5S4YPE8-F1-MODEL\_V4 \| 1.0 \| 3.434e-09 \| 393 \| 0.275 \| 120 \| 78 \| 2 \| 26 \| 136 \| 138 \| 257 \| Phage tail protein \| Phage tail protein; afdb-uniprot50 \| AF-A0A5V6NIV2-F1-MODEL\_V4 \| 1.0 \| 1.458e-08 \| 392 \| 0.187 \| 128 \| 99 \| 4 \| 13 \| 136 \| 2 \| 128 \| Uncharacterized protein \| Uncharacterized protein; afdb-uniprot50 \| AF-A0A1C0V8H7-F1-MODEL\_V4 \| 1.0 \| 1.382e-08 \| 392 \| 0.193 \| 129 \| 100 \| 4 \| 10 \| 136 \| 2 \| 128 \| Uncharacterized protein \| Uncharacterized protein; afdb-uniprot50 \| AF-K1JX06-F1-MODEL\_V4 \| 1.0 \| 1.906e-08 \| 391 \| 0.171 \| 128 \| 101 \| 4 \| 13 \| 136 \| 7 \| 133 \| Uncharacterized protein \| Uncharacterized protein; afdb-uniprot50 \| AF-A0A2T7H7Y5-F1-MODEL\_V4 \| 1.0 \| 2.773e-08 \| 390 \| 0.226 \| 128 \| 91 \| 4 \| 13 \| 134 \| 3 \| 128 \| Uncharacterized protein \| Uncharacterized protein; afdb-uniprot50 \| AF-A0A7Z0SRH5-F1-MODEL\_V4 \| 1.0 \| 6.192e-08 \| 390 \| 0.217 \| 124 \| 96 \| 1 \| 13 \| 136 \| 1 \| 123 \| Phage tail protein \| Phage tail protein; afdb-uniprot50 \| AF-A0A4R5W1S1-F1-MODEL\_V4 \| 1.0 \| 3.256e-08 \| 390 \| 0.209 \| 124 \| 97 \| 1 \| 12 \| 135 \| 2 \| 124 \| Phage tail protein \| Phage tail protein; afdb-uniprot50 \| AF-A0A1U7MFM3-F1-MODEL\_V4 \| 1.0 \| 1.382e-08 \| 389 \| 0.2 \| 130 \| 95 \| 5 \| 13 \| 136 \| 2 \| 128 \| Phage P2 GpU \| Phage P2 GpU; afdb-uniprot50 \| AF-A0A2G3J6J3-F1-MODEL\_V4 \| 1.0 \| 4.256e-08 \| 389 \| 0.217 \| 124 \| 96 \| 1 \| 13 \| 136 \| 1 \| 123 \| Uncharacterized protein \| Uncharacterized protein; afdb-uniprot50 \| AF-E2CFJ9-F1-MODEL\_V4 \| 1.0 \| 3.086e-08 \| 387 \| 0.259 \| 127 \| 88 \| 2 \| 13 \| 136 \| 3 \| 126 \| Putative phage tail protein \| Putative phage tail protein; afdb-uniprot50 \| AF-A0A2D3VW39-F1-MODEL\_V4 \| 1.0 \| 3.435e-08 \| 386 \| 0.193 \| 124 \| 96 \| 2 \| 13 \| 136 \| 1 \| 120 \| Uncharacterized protein \| Uncharacterized protein; afdb-uniprot50 \| AF-A0A7V8L4R9-F1-MODEL\_V4 \| 1.0 \| 1.382e-08 \| 386 \| 0.17 \| 123 \| 101 \| 1 \| 13 \| 135 \| 1 \| 122 \| Phage P2 GpU family protein \| Phage P2 GpU family protein; afdb-uniprot50 \| AF-I7LIF3-F1-MODEL\_V4 \| 1.0 \| 2.121e-08 \| 385 \| 0.207 \| 130 \| 94 \| 4 \| 13 \| 136 \| 1 \| 127 \| Uncharacterized protein \| Uncharacterized protein; afdb-uniprot50 \| AF-A0A4V3D5Q8-F1-MODEL\_V4 \| 1.0 \| 2.121e-08 \| 385 \| 0.162 \| 123 \| 102 \| 1 \| 12 \| 134 \| 2 \| 123 \| Phage protein U \| Phage protein U; afdb-uniprot50 \| AF-Q3RD92-F1-MODEL\_V4 \| 1.0 \| 3.086e-08 \| 384 \| 0.424 \| 99 \| 57 \| 0 \| 38 \| 136 \| 1 \| 99 \| Phage P2 GpU \| Phage P2 GpU; afdb-uniprot50 \| AF-A0A1V3RQN1-F1-MODEL\_V4 \| 1.0 \| 1.002e-08 \| 384 \| 0.173 \| 127 \| 103 \| 2 \| 10 \| 136 \| 1 \| 125 \| Uncharacterized protein \| Uncharacterized protein; afdb-uniprot50 \| AF-D8IV23-F1-MODEL\_V4 \| 1.0 \| 1.906e-08 \| 384 \| 0.171 \| 134 \| 110 \| 1 \| 3 \| 136 \| 13 \| 145 \| Uncharacterized protein \| Uncharacterized protein; afdb-uniprot50 \| AF-A0A318KUR9-F1-MODEL\_V4 \| 1.0 \| 4.998e-08 \| 384 \| 0.245 \| 122 \| 91 \| 1 \| 13 \| 134 \| 1 \| 121 \| Phage protein U \| Phage protein U; afdb-uniprot50 \| AF-A0A0E4CZC6-F1-MODEL\_V4 \| 1.0 \| 7.268e-09 \| 382 \| 0.181 \| 149 \| 101 \| 6 \| 1 \| 136 \| 17 \| 157 \| Uncharacterized protein \| Uncharacterized protein; afdb-uniprot50 \| AF-A0A432PWN2-F1-MODEL\_V4 \| 1.0 \| 6.533e-08 \| 381 \| 0.233 \| 124 \| 90 \| 3 \| 13 \| 136 \| 1 \| 119 \| Uncharacterized protein \| Uncharacterized protein; afdb-uniprot50 \| AF-A0A3D1NSN0-F1-MODEL\_V4 \| 1.0 \| 1.058e-07 \| 381 \| 0.18 \| 122 \| 99 \| 1 \| 13 \| 134 \| 1 \| 121 \| Uncharacterized protein \| Uncharacterized protein; afdb-uniprot50 \| AF-A0A662Z774-F1-MODEL\_V4 \| 1.0 \| 2.773e-08 \| 380 \| 0.187 \| 128 \| 99 \| 4 \| 13 \| 136 \| 6 \| 132 \| Uncharacterized protein \| Uncharacterized protein; afdb-uniprot50 \| AF-A0A541BHI4-F1-MODEL\_V4 \| 1.0 \| 6.533e-08 \| 379 \| 0.193 \| 124 \| 98 \| 2 \| 13 \| 136 \| 2 \| 123 \| Uncharacterized protein \| Uncharacterized protein; afdb-uniprot50 \| AF-A0A1H5DBV8-F1-MODEL\_V4 \| 1.0 \| 1.058e-07 \| 379 \| 0.153 \| 124 \| 104 \| 1 \| 13 \| 136 \| 1 \| 123 \| Phage protein U \| Phage protein U; afdb-uniprot50 \| AF-A0A6I2ITM1-F1-MODEL\_V4 \| 1.0 \| 2.238e-08 \| 378 \| 0.4 \| 105 \| 63 \| 0 \| 2 \| 106 \| 17 \| 121 \| Phage tail protein \| Phage tail protein; afdb-uniprot50 \| AF-A0A1L9Q5U4-F1-MODEL\_V4 \| 1.0 \| 9.009e-08 \| 378 \| 0.178 \| 129 \| 100 \| 3 \| 13 \| 136 \| 3 \| 130 \| Uncharacterized protein \| Uncharacterized protein; afdb-uniprot50 \| AF-E0NXI7-F1-MODEL\_V4 \| 1.0 \| 1.242e-08 \| 378 \| 0.207 \| 135 \| 96 \| 5 \| 10 \| 136 \| 39 \| 170 \| Uncharacterized protein \| Uncharacterized protein; afdb-uniprot50 \| AF-A0A3G3HJJ9-F1-MODEL\_V4 \| 1.0 \| 4.737e-08 \| 377 \| 0.178 \| 123 \| 100 \| 1 \| 13 \| 135 \| 1 \| 122 \| Uncharacterized protein \| Uncharacterized protein; afdb-uniprot50 \| AF-A0A2N6FYN8-F1-MODEL\_V4 \| 1.0 \| 4.737e-08 \| 376 \| 0.208 \| 125 \| 93 \| 4 \| 13 \| 136 \| 1 \| 120 \| Uncharacterized protein \| Uncharacterized protein; afdb-uniprot50 \| AF-A0A559IZL2-F1-MODEL\_V4 \| 1.0 \| 1.31e-08 \| 376 \| 0.207 \| 135 \| 90 \| 6 \| 13 \| 136 \| 1 \| 129 \| Phage tail protein \| Phage tail protein; afdb-uniprot50 \| AF-A0A822TT98-F1-MODEL\_V4 \| 1.0 \| 1.458e-08 \| 376 \| 0.186 \| 123 \| 99 \| 1 \| 13 \| 135 \| 1 \| 122 \| Phage tail protein \| Phage tail protein; afdb-uniprot50 \| AF-A0A5M8P636-F1-MODEL\_V4 \| 1.0 \| 3.824e-08 \| 375 \| 0.147 \| 129 \| 105 \| 1 \| 13 \| 136 \| 2 \| 130 \| Uncharacterized protein \| Uncharacterized protein; afdb-uniprot50 \| AF-A0A561QKN3-F1-MODEL\_V4 \| 1.0 \| 1.538e-08 \| 374 \| 0.227 \| 132 \| 91 \| 6 \| 13 \| 136 \| 1 \| 129 \| Uncharacterized protein \| Uncharacterized protein; afdb-uniprot50 \| AF-U4T472-F1-MODEL\_V4 \| 1.0 \| 1.624e-07 \| 372 \| 0.177 \| 124 \| 101 \| 1 \| 13 \| 136 \| 1 \| 123 \| Putative methyl-accepting chemotaxis protein \| Putative methyl-accepting chemotaxis protein; afdb-uniprot50 \| AF-A0A622TXM7-F1-MODEL\_V4 \| 1.0 \| 1.623e-08 \| 371 \| 0.195 \| 128 \| 94 \| 5 \| 13 \| 134 \| 2 \| 126 \| Phage tail protein \| Phage tail protein; afdb-uniprot50 \| AF-A0A150KS36-F1-MODEL\_V4 \| 1.0 \| 2.361e-08 \| 371 \| 0.25 \| 132 \| 88 \| 6 \| 13 \| 136 \| 3 \| 131 \| SH3b domain-containing protein \| SH3b domain-containing protein; afdb-uniprot50 \| AF-E5VKI8-F1-MODEL\_V4 \| 1.0 \| 2.491e-08 \| 370 \| 0.159 \| 132 \| 100 \| 7 \| 13 \| 136 \| 3 \| 131 \| Uncharacterized protein \| Uncharacterized protein; afdb-uniprot50 \| AF-A0A2Z4RAD7-F1-MODEL\_V4 \| 1.0 \| 1.003e-07 \| 370 \| 0.184 \| 125 \| 99 \| 2 \| 13 \| 136 \| 1 \| 123 \| Uncharacterized protein \| Uncharacterized protein; afdb-uniprot50 \| AF-W4EGN0-F1-MODEL\_V4 \| 1.0 \| 1.906e-08 \| 369 \| 0.163 \| 147 \| 106 \| 6 \| 1 \| 136 \| 5 \| 145 \| Uncharacterized protein \| Uncharacterized protein; afdb-uniprot50 \| AF-A0A2W5JTV2-F1-MODEL\_V4 \| 1.0 \| 3.824e-08 \| 368 \| 0.209 \| 124 \| 94 \| 2 \| 13 \| 136 \| 2 \| 121 \| Uncharacterized protein \| Uncharacterized protein; afdb-uniprot50 \| AF-A0A0Q0VG01-F1-MODEL\_V4 \| 1.0 \| 1.459e-07 \| 368 \| 0.2 \| 125 \| 98 \| 2 \| 13 \| 136 \| 1 \| 124 \| Uncharacterized protein \| Uncharacterized protein; afdb-uniprot50 \| AF-A0A7J0BHU5-F1-MODEL\_V4 \| 1.0 \| 4.034e-08 \| 368 \| 0.165 \| 139 \| 111 \| 4 \| 2 \| 136 \| 151 \| 288 \| Uncharacterized protein \| Uncharacterized protein; afdb-uniprot50 \| AF-D2ZHP5-F1-MODEL\_V4 \| 1.0 \| 1.623e-08 \| 368 \| 0.195 \| 123 \| 98 \| 1 \| 13 \| 135 \| 1 \| 122 \| Phage P2 GpU \| Phage P2 GpU; afdb-uniprot50 \| AF-A0A7C3GV02-F1-MODEL\_V4 \| 1.0 \| 6.892e-08 \| 366 \| 0.207 \| 130 \| 96 \| 3 \| 12 \| 136 \| 2 \| 129 \| Phage tail protein \| Phage tail protein; afdb-uniprot50 \| AF-A0A3A9CQH5-F1-MODEL\_V4 \| 1.0 \| 1.806e-08 \| 366 \| 0.2 \| 145 \| 104 \| 7 \| 1 \| 136 \| 25 \| 166 \| Phage tail protein \| Phage tail protein; afdb-uniprot50 \| AF-A0A521DSD1-F1-MODEL\_V4 \| 1.0 \| 1.178e-07 \| 365 \| 0.237 \| 122 \| 91 \| 2 \| 13 \| 134 \| 1 \| 120 \| Phage protein U \| Phage protein U; afdb-uniprot50 \| AF-A0A859FBV8-F1-MODEL\_V4 \| 1.0 \| 2.491e-08 \| 365 \| 0.189 \| 132 \| 96 \| 6 \| 13 \| 136 \| 1 \| 129 \| Phage tail protein \| Phage tail protein; afdb-uniprot50 \| AF-A0A2W6YBV7-F1-MODEL\_V4 \| 1.0 \| 9.009e-08 \| 365 \| 0.185 \| 124 \| 97 \| 3 \| 13 \| 133 \| 1 \| 123 \| Uncharacterized protein \| Uncharacterized protein; afdb-uniprot50 \| AF-S7UNK7-F1-MODEL\_V4 \| 1.0 \| 4.737e-08 \| 365 \| 0.17 \| 123 \| 101 \| 1 \| 13 \| 135 \| 3 \| 124 \| p2 GpU family protein \| p2 GpU family protein; afdb-uniprot50 \| AF-A0A450WGZ2-F1-MODEL\_V4 \| 1.0 \| 5.563e-08 \| 365 \| 0.209 \| 124 \| 97 \| 1 \| 13 \| 136 \| 2 \| 124 \| Phage protein U \| Phage protein U; afdb-uniprot50 \| AF-A0A5B8JQ08-F1-MODEL\_V4 \| 1.0 \| 2.361e-08 \| 364 \| 0.204 \| 137 \| 100 \| 5 \| 7 \| 136 \| 1 \| 135 \| Phage tail protein \| Phage tail protein; afdb-uniprot50 \| AF-A0A0D6ARK4-F1-MODEL\_V4 \| 1.0 \| 2.925e-08 \| 364 \| 0.267 \| 127 \| 89 \| 2 \| 12 \| 135 \| 2 \| 127 \| Phage protein \| Phage protein; afdb-uniprot50 \| AF-A0A7J5WF49-F1-MODEL\_V4 \| 1.0 \| 3.824e-08 \| 364 \| 0.164 \| 134 \| 106 \| 5 \| 7 \| 136 \| 164 \| 295 \| Uncharacterized protein \| Uncharacterized protein; afdb-uniprot50 \| AF-A0A6N7B4J7-F1-MODEL\_V4 \| 1.0 \| 1.003e-07 \| 364 \| 0.163 \| 122 \| 100 \| 1 \| 13 \| 134 \| 2 \| 121 \| Uncharacterized protein \| Uncharacterized protein; afdb-uniprot50 \| AF-A0A1Y3SYD9-F1-MODEL\_V4 \| 1.0 \| 2.925e-08 \| 363 \| 0.21 \| 133 \| 93 \| 7 \| 13 \| 136 \| 2 \| 131 \| Uncharacterized protein \| Uncharacterized protein; afdb-uniprot50 \| AF-R5BE61-F1-MODEL\_V4 \| 1.0 \| 6.192e-08 \| 363 \| 0.181 \| 132 \| 97 \| 6 \| 11 \| 134 \| 35 \| 163 \| p2 GpU family protein \| p2 GpU family protein; afdb-uniprot50 \| AF-A0A316MLA5-F1-MODEL\_V4 \| 1.0 \| 4.49e-08 \| 359 \| 0.218 \| 142 \| 91 \| 7 \| 13 \| 136 \| 2 \| 141 \| Uncharacterized protein \| Uncharacterized protein; afdb-uniprot50 \| AF-A0A4U8YXW6-F1-MODEL\_V4 \| 1.0 \| 5.273e-08 \| 358 \| 0.201 \| 134 \| 88 \| 5 \| 14 \| 136 \| 3 \| 128 \| Myoviridae gpu \| Myoviridae gpu; afdb-uniprot50 \| AF-A0A511Z868-F1-MODEL\_V4 \| 1.0 \| 5.273e-08 \| 358 \| 0.174 \| 132 \| 98 \| 6 \| 13 \| 136 \| 1 \| 129 \| Uncharacterized protein \| Uncharacterized protein; afdb-uniprot50 \| AF-F3YY63-F1-MODEL\_V4 \| 1.0 \| 1.907e-07 \| 357 \| 0.14 \| 128 \| 105 \| 4 \| 13 \| 136 \| 2 \| 128 \| Uncharacterized protein \| Uncharacterized protein; afdb-uniprot50 \| AF-A0A2P8EI36-F1-MODEL\_V4 \| 1.0 \| 1.459e-07 \| 357 \| 0.147 \| 129 \| 103 \| 5 \| 13 \| 136 \| 2 \| 128 \| GpU protein \| GpU protein; afdb-uniprot50 \| AF-B3QTJ0-F1-MODEL\_V4 \| 1.0 \| 6.533e-08 \| 357 \| 0.201 \| 129 \| 91 \| 4 \| 13 \| 136 \| 1 \| 122 \| Uncharacterized protein \| Uncharacterized protein; afdb-uniprot50 \| AF-A0A855IT43-F1-MODEL\_V4 \| 1.0 \| 1.116e-07 \| 356 \| 0.148 \| 128 \| 99 \| 4 \| 13 \| 136 \| 2 \| 123 \| Uncharacterized protein \| Uncharacterized protein; afdb-uniprot50 \| AF-A0A416B7P4-F1-MODEL\_V4 \| 1.0 \| 5.273e-08 \| 356 \| 0.18 \| 133 \| 97 \| 7 \| 13 \| 136 \| 3 \| 132 \| Uncharacterized protein \| Uncharacterized protein; afdb-uniprot50 \| AF-A0A840MJS2-F1-MODEL\_V4 \| 1.0 \| 5.563e-08 \| 356 \| 0.211 \| 123 \| 96 \| 1 \| 13 \| 135 \| 5 \| 126 \| Phage protein U \| Phage protein U; afdb-uniprot50 \| AF-A0A620N5X6-F1-MODEL\_V4 \| 1.0 \| 3.256e-08 \| 356 \| 0.219 \| 123 \| 95 \| 1 \| 13 \| 135 \| 1 \| 122 \| Uncharacterized protein \| Uncharacterized protein; afdb-uniprot50 \| AF-A0A833UXN4-F1-MODEL\_V4 \| 1.0 \| 1.459e-07 \| 355 \| 0.32 \| 128 \| 82 \| 1 \| 2 \| 124 \| 16 \| 143 \| Uncharacterized protein \| Uncharacterized protein; afdb-uniprot50 \| AF-A0A3A9GXW5-F1-MODEL\_V4 \| 1.0 \| 3.435e-08 \| 354 \| 0.201 \| 134 \| 91 \| 6 \| 13 \| 136 \| 3 \| 130 \| Phage tail protein \| Phage tail protein; afdb-uniprot50 \| AF-A0A6D2GEA9-F1-MODEL\_V4 \| 1.0 \| 4.737e-08 \| 354 \| 0.203 \| 123 \| 97 \| 1 \| 13 \| 135 \| 1 \| 122 \| Methyl-accepting chemotaxis protein \| Methyl-accepting chemotaxis protein; afdb-uniprot50 \| AF-J5HM67-F1-MODEL\_V4 \| 1.0 \| 4.256e-08 \| 354 \| 0.2 \| 135 \| 92 \| 6 \| 13 \| 136 \| 20 \| 149 \| Phage P2 GpU \| Phage P2 GpU; afdb-uniprot50 \| AF-A0A4S2HDA3-F1-MODEL\_V4 \| 1.0 \| 4.256e-08 \| 353 \| 0.179 \| 134 \| 94 \| 6 \| 13 \| 136 \| 41 \| 168 \| Phage tail protein \| Phage tail protein; afdb-uniprot50 \| AF-E5Y5W8-F1-MODEL\_V4 \| 1.0 \| 9.504e-08 \| 352 \| 0.215 \| 130 \| 94 \| 4 \| 13 \| 136 \| 3 \| 130 \| Uncharacterized protein \| Uncharacterized protein; afdb-uniprot50 \| AF-A0A4U7J7L0-F1-MODEL\_V4 \| 1.0 \| 1.178e-07 \| 352 \| 0.155 \| 129 \| 103 \| 4 \| 13 \| 136 \| 2 \| 129 \| Phage tail protein \| Phage tail protein; afdb-uniprot50 \| AF-A0A450WK91-F1-MODEL\_V4 \| 1.0 \| 7.271e-08 \| 352 \| 0.177 \| 124 \| 101 \| 1 \| 13 \| 136 \| 2 \| 124 \| Phage P2 GpU \| Phage P2 GpU; afdb-uniprot50 \| AF-E1QHA1-F1-MODEL\_V4 \| 1.0 \| 1.624e-07 \| 351 \| 0.173 \| 127 \| 101 \| 3 \| 13 \| 136 \| 5 \| 130 \| Uncharacterized protein \| Uncharacterized protein; afdb-uniprot50 \| AF-A0A1M6GF68-F1-MODEL\_V4 \| 1.0 \| 2.238e-08 \| 351 \| 0.179 \| 134 \| 101 \| 5 \| 10 \| 136 \| 20 \| 151 \| Uncharacterized protein \| Uncharacterized protein; afdb-uniprot50 \| AF-A0A497UT05-F1-MODEL\_V4 \| 1.0 \| 1.907e-07 \| 351 \| 0.186 \| 123 \| 98 \| 2 \| 13 \| 134 \| 1 \| 122 \| Phage protein U \| Phage protein U; afdb-uniprot50 \| AF-A0A2J9QJ37-F1-MODEL\_V4 \| 1.0 \| 1.383e-07 \| 351 \| 0.188 \| 122 \| 98 \| 1 \| 14 \| 135 \| 3 \| 123 \| Phage tail protein \| Phage tail protein; afdb-uniprot50 \| AF-A0A327JMS0-F1-MODEL\_V4 \| 1.0 \| 1.003e-07 \| 350 \| 0.258 \| 112 \| 77 \| 1 \| 28 \| 133 \| 4 \| 115 \| Uncharacterized protein \| Uncharacterized protein; afdb-uniprot50 \| AF-E5WEF4-F1-MODEL\_V4 \| 1.0 \| 6.892e-08 \| 350 \| 0.204 \| 132 \| 94 \| 6 \| 13 \| 136 \| 2 \| 130 \| Uncharacterized protein \| Uncharacterized protein; afdb-uniprot50 \| AF-U2BMK3-F1-MODEL\_V4 \| 1.0 \| 3.435e-08 \| 348 \| 0.2 \| 135 \| 91 \| 6 \| 13 \| 136 \| 6 \| 134 \| Uncharacterized protein \| Uncharacterized protein; afdb-uniprot50 \| AF-A0A0P0A749-F1-MODEL\_V4 \| 1.0 \| 9.009e-08 \| 348 \| 0.198 \| 126 \| 90 \| 3 \| 13 \| 133 \| 1 \| 120 \| Uncharacterized protein \| Uncharacterized protein; afdb-uniprot50 \| AF-A0A376P7R0-F1-MODEL\_V4 \| 1.0 \| 6.896e-07 \| 347 \| 0.294 \| 112 \| 79 \| 0 \| 13 \| 124 \| 1 \| 112 \| Tail fiber protein \| Tail fiber protein; afdb-uniprot50 \| AF-A0A6L9HIG7-F1-MODEL\_V4 \| 1.0 \| 1.712e-08 \| 347 \| 0.187 \| 144 \| 102 \| 7 \| 6 \| 136 \| 35 \| 176 \| Phage tail protein \| Phage tail protein; afdb-uniprot50 \| AF-A0A4U8UH88-F1-MODEL\_V4 \| 1.0 \| 3.626e-07 \| 346 \| 0.209 \| 124 \| 95 \| 1 \| 13 \| 136 \| 2 \| 122 \| Uncharacterized protein \| Uncharacterized protein; afdb-uniprot50 \| AF-N2AGK0-F1-MODEL\_V4 \| 1.0 \| 1.311e-07 \| 345 \| 0.157 \| 133 \| 100 \| 7 \| 13 \| 136 \| 3 \| 132 \| Uncharacterized protein \| Uncharacterized protein; afdb-uniprot50 \| AF-A0A2D3WLM9-F1-MODEL\_V4 \| 1.0 \| 2.362e-07 \| 345 \| 0.206 \| 121 \| 92 \| 2 \| 13 \| 133 \| 1 \| 117 \| Uncharacterized protein \| Uncharacterized protein; afdb-uniprot50 \| AF-A0A212KXX9-F1-MODEL\_V4 \| 1.0 \| 1.539e-07 \| 344 \| 0.121 \| 132 \| 104 \| 6 \| 13 \| 136 \| 2 \| 129 \| Uncharacterized protein \| Uncharacterized protein; afdb-uniprot50 \| AF-A0A5P9CRA6-F1-MODEL\_V4 \| 1.0 \| 2.63e-07 \| 344 \| 0.16 \| 125 \| 101 \| 3 \| 13 \| 136 \| 1 \| 122 \| Phage P2 GpU \| Phage P2 GpU; afdb-uniprot50 \| AF-A0A1I7ND59-F1-MODEL\_V4 \| 1.0 \| 3.258e-07 \| 343 \| 0.25 \| 104 \| 78 \| 0 \| 33 \| 136 \| 2 \| 105 \| Uncharacterized protein \| Uncharacterized protein; afdb-uniprot50 \| AF-A0A8B3A6D8-F1-MODEL\_V4 \| 1.0 \| 1.003e-07 \| 343 \| 0.181 \| 132 \| 97 \| 6 \| 13 \| 136 \| 1 \| 129 \| DNA-packaging protein \| DNA-packaging protein; afdb-uniprot50 \| AF-E2CGG6-F1-MODEL\_V4 \| 1.0 \| 5.872e-07 \| 343 \| 0.133 \| 127 \| 107 \| 2 \| 13 \| 136 \| 2 \| 128 \| Gp45 \| Gp45; afdb-uniprot50 \| AF-A0A075WU95-F1-MODEL\_V4 \| 1.0 \| 1.311e-07 \| 343 \| 0.217 \| 124 \| 93 \| 3 \| 14 \| 136 \| 3 \| 123 \| Uncharacterized protein \| Uncharacterized protein; afdb-uniprot50 \| AF-A0A377ICM6-F1-MODEL\_V4 \| 1.0 \| 3.824e-08 \| 343 \| 0.185 \| 124 \| 99 \| 1 \| 13 \| 136 \| 1 \| 122 \| Putative bacteriophage V tail protein \| Putative bacteriophage V tail protein; afdb-uniprot50 \| AF-A0A3A9CSQ2-F1-MODEL\_V4 \| 1.0 \| 3.258e-07 \| 342 \| 0.186 \| 129 \| 99 \| 5 \| 13 \| 136 \| 2 \| 129 \| Phage tail protein \| Phage tail protein; afdb-uniprot50 \| AF-A0A1H1FLQ9-F1-MODEL\_V4 \| 1.0 \| 3.258e-07 \| 342 \| 0.2 \| 110 \| 86 \| 2 \| 28 \| 136 \| 7 \| 115 \| Uncharacterized protein \| Uncharacterized protein; afdb-uniprot50 \| AF-A0A7V6Q5B3-F1-MODEL\_V4 \| 1.0 \| 1.459e-07 \| 342 \| 0.181 \| 132 \| 97 \| 6 \| 13 \| 136 \| 2 \| 130 \| Uncharacterized protein \| Uncharacterized protein; afdb-uniprot50 \| AF-A0A327Q841-F1-MODEL\_V4 \| 1.0 \| 6.195e-07 \| 342 \| 0.137 \| 124 \| 106 \| 1 \| 13 \| 136 \| 1 \| 123 \| GpU protein \| GpU protein; afdb-uniprot50 \| AF-A0A165W2G2-F1-MODEL\_V4 \| 1.0 \| 1.807e-07 \| 341 \| 0.155 \| 135 \| 104 \| 4 \| 6 \| 136 \| 1 \| 129 \| Uncharacterized protein \| Uncharacterized protein; afdb-uniprot50 \| AF-A0A6C1BSU7-F1-MODEL\_V4 \| 1.0 \| 1.178e-07 \| 341 \| 0.161 \| 124 \| 101 \| 2 \| 13 \| 136 \| 2 \| 122 \| Phage tail protein \| Phage tail protein; afdb-uniprot50 \| AF-A0A450YW55-F1-MODEL\_V4 \| 1.0 \| 2.927e-07 \| 341 \| 0.169 \| 124 \| 101 \| 1 \| 13 \| 136 \| 2 \| 123 \| Phage P2 GpU \| Phage P2 GpU; afdb-uniprot50 \| AF-A0A385Q1W0-F1-MODEL\_V4 \| 1.0 \| 1.807e-07 \| 340 \| 0.195 \| 133 \| 95 \| 7 \| 13 \| 136 \| 3 \| 132 \| Phage tail protein \| Phage tail protein; afdb-uniprot50 \| AF-A0A413BFN5-F1-MODEL\_V4 \| 1.0 \| 1.459e-07 \| 340 \| 0.175 \| 137 \| 101 \| 7 \| 9 \| 136 \| 82 \| 215 \| Uncharacterized protein \| Uncharacterized protein; afdb-uniprot50 \| AF-A0A239C7Y7-F1-MODEL\_V4 \| 1.0 \| 4.998e-08 \| 340 \| 0.169 \| 142 \| 102 \| 6 \| 4 \| 136 \| 170 \| 304 \| Phage P2 GpU \| Phage P2 GpU; afdb-uniprot50 \| AF-A0A7C2IU81-F1-MODEL\_V4 \| 1.0 \| 7.675e-07 \| 339 \| 0.161 \| 130 \| 101 \| 3 \| 13 \| 136 \| 1 \| 128 \| Uncharacterized protein \| Uncharacterized protein; afdb-uniprot50 \| AF-A0A3M4ZPG5-F1-MODEL\_V4 \| 1.0 \| 7.675e-07 \| 338 \| 0.404 \| 84 \| 50 \| 0 \| 53 \| 136 \| 8 \| 91 \| Uncharacterized protein \| Uncharacterized protein; afdb-uniprot50 \| AF-A0A828ULQ4-F1-MODEL\_V4 \| 1.0 \| 1.624e-07 \| 338 \| 0.279 \| 118 \| 77 \| 2 \| 13 \| 130 \| 1 \| 110 \| Phage P2 GpU \| Phage P2 GpU; afdb-uniprot50 \| AF-A0A5A8F1S4-F1-MODEL\_V4 \| 1.0 \| 2.012e-07 \| 338 \| 0.183 \| 136 \| 87 \| 6 \| 13 \| 136 \| 2 \| 125 \| Uncharacterized protein \| Uncharacterized protein; afdb-uniprot50 \| AF-G4KPI6-F1-MODEL\_V4 \| 1.0 \| 1.242e-07 \| 338 \| 0.149 \| 134 \| 100 \| 5 \| 13 \| 136 \| 1 \| 130 \| Uncharacterized protein \| Uncharacterized protein; afdb-uniprot50 \| AF-A0A1M6SJI7-F1-MODEL\_V4 \| 1.0 \| 7.671e-08 \| 338 \| 0.222 \| 135 \| 88 \| 6 \| 13 \| 136 \| 2 \| 130 \| Uncharacterized protein \| Uncharacterized protein; afdb-uniprot50 \| AF-K4ZKW4-F1-MODEL\_V4 \| 1.0 \| 1.907e-07 \| 337 \| 0.204 \| 132 \| 94 \| 6 \| 13 \| 136 \| 1 \| 129 \| Phage P2 GpU \| Phage P2 GpU; afdb-uniprot50 \| AF-A0A1Q6PU11-F1-MODEL\_V4 \| 1.0 \| 1.459e-07 \| 337 \| 0.2 \| 135 \| 91 \| 5 \| 13 \| 136 \| 2 \| 130 \| Uncharacterized protein \| Uncharacterized protein; afdb-uniprot50 \| AF-A0A2A2ILD3-F1-MODEL\_V4 \| 1.0 \| 8.094e-08 \| 337 \| 0.223 \| 134 \| 89 \| 4 \| 13 \| 136 \| 2 \| 130 \| Uncharacterized protein \| Uncharacterized protein; afdb-uniprot50 \| AF-A0A173SAB9-F1-MODEL\_V4 \| 1.0 \| 1.242e-07 \| 337 \| 0.164 \| 146 \| 105 \| 6 \| 2 \| 136 \| 19 \| 158 \| Phage protein U \| Phage protein U; afdb-uniprot50 \| AF-W6THI8-F1-MODEL\_V4 \| 1.0 \| 4.258e-07 \| 336 \| 0.488 \| 84 \| 43 \| 0 \| 53 \| 136 \| 3 \| 86 \| Phage protein U \| Phage protein U; afdb-uniprot50 \| AF-A0A2N2EIQ3-F1-MODEL\_V4 \| 1.0 \| 8.539e-08 \| 336 \| 0.143 \| 132 \| 102 \| 6 \| 13 \| 136 \| 1 \| 129 \| Uncharacterized protein \| Uncharacterized protein; afdb-uniprot50 \| AF-A0A3P6KUM2-F1-MODEL\_V4 \| 1.0 \| 3.824e-08 \| 336 \| 0.171 \| 140 \| 105 \| 6 \| 5 \| 136 \| 31 \| 167 \| Phage protein U \| Phage protein U; afdb-uniprot50 \| AF-A0A7M3MAX1-F1-MODEL\_V4 \| 1.0 \| 1.003e-06 \| 335 \| 0.157 \| 127 \| 102 \| 3 \| 13 \| 136 \| 2 \| 126 \| Uncharacterized protein \| Uncharacterized protein; afdb-uniprot50 \| AF-A0A369Y010-F1-MODEL\_V4 \| 1.0 \| 2.63e-07 \| 335 \| 0.165 \| 133 \| 102 \| 3 \| 13 \| 136 \| 1 \| 133 \| Uncharacterized protein \| Uncharacterized protein; afdb-uniprot50 \| AF-A0A3S0JS18-F1-MODEL\_V4 \| 1.0 \| 4.256e-08 \| 334 \| 0.189 \| 132 \| 96 \| 6 \| 13 \| 136 \| 1 \| 129 \| Phage tail protein \| Phage tail protein; afdb-uniprot50 \| AF-A0A414ZQ39-F1-MODEL\_V4 \| 1.0 \| 5.273e-08 \| 334 \| 0.165 \| 145 \| 105 \| 5 \| 1 \| 136 \| 14 \| 151 \| Uncharacterized protein \| Uncharacterized protein; afdb-uniprot50 \| AF-A0A482IL72-F1-MODEL\_V4 \| 1.0 \| 8.539e-08 \| 334 \| 0.185 \| 124 \| 99 \| 1 \| 13 \| 136 \| 2 \| 123 \| Phage tail protein \| Phage tail protein; afdb-uniprot50 \| AF-A0A8B2E404-F1-MODEL\_V4 \| 1.0 \| 8.094e-08 \| 334 \| 0.196 \| 127 \| 94 \| 2 \| 13 \| 136 \| 1 \| 122 \| Phage tail protein \| Phage tail protein; afdb-uniprot50 \| AF-A0A6N9P7S3-F1-MODEL\_V4 \| 1.0 \| 4.258e-07 \| 333 \| 0.152 \| 131 \| 103 \| 6 \| 13 \| 136 \| 2 \| 131 \| Phage tail protein \| Phage tail protein; afdb-uniprot50 \| AF-A0A4P8G292-F1-MODEL\_V4 \| 1.0 \| 5e-07 \| 332 \| 0.201 \| 109 \| 83 \| 2 \| 29 \| 136 \| 27 \| 132 \| Uncharacterized protein \| Uncharacterized protein; afdb-uniprot50 \| AF-J0QKB2-F1-MODEL\_V4 \| 1.0 \| 1.539e-07 \| 332 \| 0.223 \| 112 \| 81 \| 2 \| 31 \| 136 \| 16 \| 127 \| Uncharacterized protein \| Uncharacterized protein; afdb-uniprot50 \| AF-A8SB39-F1-MODEL\_V4 \| 1.0 \| 2.492e-07 \| 331 \| 0.157 \| 133 \| 100 \| 6 \| 13 \| 136 \| 1 \| 130 \| Uncharacterized protein \| Uncharacterized protein; afdb-uniprot50 \| AF-A0A7U6KE04-F1-MODEL\_V4 \| 1.0 \| 4.737e-08 \| 331 \| 0.185 \| 135 \| 94 \| 5 \| 13 \| 136 \| 50 \| 179 \| Uncharacterized protein \| Uncharacterized protein; afdb-uniprot50 \| AF-A0A2N1QG54-F1-MODEL\_V4 \| 1.0 \| 1.539e-07 \| 329 \| 0.143 \| 132 \| 102 \| 6 \| 13 \| 136 \| 1 \| 129 \| Uncharacterized protein \| Uncharacterized protein; afdb-uniprot50 \| AF-A0A3A9EU93-F1-MODEL\_V4 \| 1.0 \| 1.311e-07 \| 329 \| 0.159 \| 132 \| 100 \| 6 \| 13 \| 136 \| 3 \| 131 \| Uncharacterized protein \| Uncharacterized protein; afdb-uniprot50 \| AF-A0A0C2YV44-F1-MODEL\_V4 \| 1.0 \| 1.907e-07 \| 329 \| 0.172 \| 133 \| 102 \| 4 \| 6 \| 136 \| 2 \| 128 \| Uncharacterized protein \| Uncharacterized protein; afdb-uniprot50 \| AF-A0A7X5C9Z1-F1-MODEL\_V4 \| 1.0 \| 5.563e-08 \| 329 \| 0.149 \| 134 \| 103 \| 6 \| 11 \| 136 \| 41 \| 171 \| Phage tail protein \| Phage tail protein; afdb-uniprot50 \| AF-A0A2E2N194-F1-MODEL\_V4 \| 1.0 \| 9.009e-08 \| 328 \| 0.212 \| 132 \| 89 \| 5 \| 13 \| 135 \| 1 \| 126 \| Uncharacterized protein \| Uncharacterized protein; afdb-uniprot50 \| AF-A0A2S7JR76-F1-MODEL\_V4 \| 1.0 \| 2.492e-07 \| 328 \| 0.151 \| 132 \| 109 \| 2 \| 5 \| 136 \| 52 \| 180 \| Uncharacterized protein \| Uncharacterized protein; afdb-uniprot50 \| AF-A0A3D4L0U5-F1-MODEL\_V4 \| 1.0 \| 6.195e-07 \| 327 \| 0.161 \| 130 \| 102 \| 5 \| 13 \| 136 \| 2 \| 130 \| Uncharacterized protein \| Uncharacterized protein; afdb-uniprot50 \| AF-A0A5R8Y635-F1-MODEL\_V4 \| 1.0 \| 2.63e-07 \| 327 \| 0.181 \| 127 \| 98 \| 3 \| 13 \| 136 \| 3 \| 126 \| Uncharacterized protein \| Uncharacterized protein; afdb-uniprot50 \| AF-A0A1A9G3Z3-F1-MODEL\_V4 \| 1.0 \| 7.275e-07 \| 327 \| 0.169 \| 124 \| 99 \| 3 \| 13 \| 133 \| 1 \| 123 \| Uncharacterized protein \| Uncharacterized protein; afdb-uniprot50 \| AF-A0A064AGR8-F1-MODEL\_V4 \| 1.0 \| 6.195e-07 \| 327 \| 0.154 \| 136 \| 107 \| 6 \| 7 \| 136 \| 22 \| 155 \| Uncharacterized protein \| Uncharacterized protein; afdb-uniprot50 \| AF-A0A1C0A2B2-F1-MODEL\_V4 \| 1.0 \| 8.094e-08 \| 326 \| 0.198 \| 136 \| 95 \| 4 \| 9 \| 136 \| 24 \| 153 \| Uncharacterized protein \| Uncharacterized protein; afdb-uniprot50 \| AF-A0A416GNR4-F1-MODEL\_V4 \| 1.0 \| 3.826e-07 \| 326 \| 0.184 \| 130 \| 94 \| 7 \| 16 \| 136 \| 114 \| 240 \| SH3b domain-containing protein \| SH3b domain-containing protein; afdb-uniprot50 \| AF-A0A6N9K7R1-F1-MODEL\_V4 \| 1.0 \| 2.362e-07 \| 325 \| 0.167 \| 137 \| 96 \| 7 \| 12 \| 136 \| 2 \| 132 \| Uncharacterized protein \| Uncharacterized protein; afdb-uniprot50 \| AF-A0A3C1WUQ3-F1-MODEL\_V4 \| 1.0 \| 3.626e-07 \| 325 \| 0.176 \| 136 \| 99 \| 6 \| 13 \| 136 \| 3 \| 137 \| Uncharacterized protein \| Uncharacterized protein; afdb-uniprot50 \| AF-A0A1V4IUT3-F1-MODEL\_V4 \| 1.0 \| 1.713e-07 \| 324 \| 0.181 \| 132 \| 97 \| 6 \| 13 \| 136 \| 1 \| 129 \| Phage P2 GpU \| Phage P2 GpU; afdb-uniprot50 \| AF-A0A162QMP6-F1-MODEL\_V4 \| 1.0 \| 1.116e-07 \| 324 \| 0.198 \| 131 \| 94 \| 6 \| 13 \| 135 \| 1 \| 128 \| Putative peptidoglycan binding domain protein \| Putative peptidoglycan binding domain protein; afdb-uniprot50 \| AF-A0A329UAJ4-F1-MODEL\_V4 \| 1.0 \| 9.509e-07 \| 323 \| 0.1 \| 130 \| 110 \| 5 \| 13 \| 136 \| 2 \| 130 \| Uncharacterized protein \| Uncharacterized protein; afdb-uniprot50 \| AF-A0A7K1PMP8-F1-MODEL\_V4 \| 1.0 \| 9.509e-07 \| 322 \| 0.193 \| 124 \| 96 \| 3 \| 13 \| 133 \| 1 \| 123 \| Uncharacterized protein \| Uncharacterized protein; afdb-uniprot50 \| AF-A0A5Y3VZG7-F1-MODEL\_V4 \| 1.0 \| 9.013e-07 \| 322 \| 0.184 \| 125 \| 96 \| 3 \| 13 \| 135 \| 1 \| 121 \| Uncharacterized protein \| Uncharacterized protein; afdb-uniprot50 \| AF-A0A0X1U7T4-F1-MODEL\_V4 \| 1.0 \| 8.543e-07 \| 321 \| 0.166 \| 132 \| 99 \| 6 \| 13 \| 136 \| 1 \| 129 \| Phage P2 GpU \| Phage P2 GpU; afdb-uniprot50 \| AF-A0A6N2ZLQ2-F1-MODEL\_V4 \| 1.0 \| 4.74e-07 \| 321 \| 0.192 \| 135 \| 98 \| 6 \| 10 \| 136 \| 56 \| 187 \| Phage P2 GpU \| Phage P2 GpU; afdb-uniprot50 \| AF-I0GRI8-F1-MODEL\_V4 \| 1.0 \| 2.774e-07 \| 320 \| 0.222 \| 144 \| 90 \| 7 \| 6 \| 136 \| 28 \| 162 \| Putative phage protein \| Putative phage protein; afdb-uniprot50 \| AF-A0A369RIA8-F1-MODEL\_V4 \| 1.0 \| 3.088e-07 \| 319 \| 0.321 \| 115 \| 73 \| 1 \| 14 \| 128 \| 1 \| 110 \| Phage protein U \| Phage protein U; afdb-uniprot50 \| AF-A0A7C8KPW6-F1-MODEL\_V4 \| 1.0 \| 4.74e-07 \| 319 \| 0.212 \| 132 \| 93 \| 6 \| 13 \| 136 \| 3 \| 131 \| Phage tail protein \| Phage tail protein; afdb-uniprot50 \| AF-A0A1B1NT78-F1-MODEL\_V4 \| 1.0 \| 3.437e-07 \| 319 \| 0.167 \| 137 \| 108 \| 4 \| 1 \| 136 \| 2 \| 133 \| Uncharacterized protein \| Uncharacterized protein; afdb-uniprot50 \| AF-A0A261E7R4-F1-MODEL\_V4 \| 1.0 \| 2.012e-07 \| 319 \| 0.159 \| 138 \| 112 \| 3 \| 1 \| 136 \| 14 \| 149 \| Uncharacterized protein \| Uncharacterized protein; afdb-uniprot50 \| AF-R7HX73-F1-MODEL\_V4 \| 1.0 \| 4.258e-07 \| 319 \| 0.143 \| 146 \| 109 \| 5 \| 2 \| 136 \| 52 \| 192 \| Uncharacterized protein \| Uncharacterized protein; afdb-uniprot50 \| AF-W7S4G8-F1-MODEL\_V4 \| 1.0 \| 4.036e-07 \| 319 \| 0.183 \| 131 \| 96 \| 6 \| 13 \| 135 \| 3 \| 130 \| SH3b domain-containing protein \| SH3b domain-containing protein; afdb-uniprot50 \| AF-A0A6L9GVJ5-F1-MODEL\_V4 \| 1.0 \| 4.492e-07 \| 318 \| 0.177 \| 135 \| 94 \| 6 \| 13 \| 136 \| 3 \| 131 \| Phage tail protein \| Phage tail protein; afdb-uniprot50 \| AF-R5GZY0-F1-MODEL\_V4 \| 1.0 \| 4.036e-07 \| 317 \| 0.171 \| 134 \| 96 \| 6 \| 13 \| 136 \| 2 \| 130 \| p2 GpU family protein \| p2 GpU family protein; afdb-uniprot50 \| AF-A0A1H7YIQ2-F1-MODEL\_V4 \| 1.0 \| 2.012e-07 \| 317 \| 0.125 \| 135 \| 101 \| 6 \| 13 \| 136 \| 1 \| 129 \| Phage P2 GpU \| Phage P2 GpU; afdb-uniprot50 \| AF-A0A1T4PTF8-F1-MODEL\_V4 \| 1.0 \| 1.54e-06 \| 317 \| 0.121 \| 132 \| 110 \| 4 \| 10 \| 136 \| 28 \| 158 \| Phage P2 GpU \| Phage P2 GpU; afdb-uniprot50 \| AF-A0A1Q6KUC1-F1-MODEL\_V4 \| 1.0 \| 1.907e-07 \| 317 \| 0.222 \| 135 \| 90 \| 6 \| 13 \| 135 \| 3 \| 134 \| LysM domain-containing protein \| LysM domain-containing protein; afdb-uniprot50 \| AF-A0A4Q2LYV9-F1-MODEL\_V4 \| 1.0 \| 2.927e-07 \| 316 \| 0.152 \| 131 \| 102 \| 5 \| 13 \| 136 \| 1 \| 129 \| Phage tail protein \| Phage tail protein; afdb-uniprot50 \| AF-A0A377A2A2-F1-MODEL\_V4 \| 1.0 \| 1.311e-06 \| 316 \| 0.322 \| 96 \| 65 \| 0 \| 41 \| 136 \| 1 \| 96 \| Tail fiber protein \| Tail fiber protein; afdb-uniprot50 \| AF-A0A765XB11-F1-MODEL\_V4 \| 1.0 \| 3.828e-06 \| 314 \| 0.293 \| 109 \| 77 \| 0 \| 13 \| 121 \| 1 \| 109 \| Phage tail protein \| Phage tail protein; afdb-uniprot50 \| AF-A0A4Q2B414-F1-MODEL\_V4 \| 1.0 \| 5.872e-07 \| 314 \| 0.155 \| 135 \| 97 \| 6 \| 13 \| 136 \| 1 \| 129 \| Phage tail protein \| Phage tail protein; afdb-uniprot50 \| AF-A0A5D8QDT5-F1-MODEL\_V4 \| 1.0 \| 6.195e-07 \| 314 \| 0.115 \| 130 \| 108 \| 5 \| 13 \| 136 \| 2 \| 130 \| Uncharacterized protein \| Uncharacterized protein; afdb-uniprot50 \| AF-A0A2V2FV54-F1-MODEL\_V4 \| 1.0 \| 1.003e-06 \| 313 \| 0.138 \| 130 \| 105 \| 4 \| 13 \| 136 \| 2 \| 130 \| Uncharacterized protein \| Uncharacterized protein; afdb-uniprot50 \| AF-A0A316QBZ5-F1-MODEL\_V4 \| 1.0 \| 3.258e-07 \| 313 \| 0.17 \| 135 \| 95 \| 6 \| 13 \| 136 \| 2 \| 130 \| Uncharacterized protein \| Uncharacterized protein; afdb-uniprot50 \| AF-A0A0P8WV76-F1-MODEL\_V4 \| 1.0 \| 4.036e-07 \| 312 \| 0.155 \| 135 \| 97 \| 6 \| 13 \| 136 \| 2 \| 130 \| Uncharacterized protein \| Uncharacterized protein; afdb-uniprot50 \| AF-A0A078MHP7-F1-MODEL\_V4 \| 1.0 \| 3.258e-07 \| 311 \| 0.124 \| 137 \| 99 \| 6 \| 13 \| 136 \| 1 \| 129 \| Phage P2 GpU \| Phage P2 GpU; afdb-uniprot50 \| AF-I9DAV5-F1-MODEL\_V4 \| 1.0 \| 4.036e-07 \| 311 \| 0.179 \| 134 \| 95 \| 4 \| 13 \| 136 \| 1 \| 129 \| p2 GpU family protein \| p2 GpU family protein; afdb-uniprot50 \| AF-A0A7I9SHL0-F1-MODEL\_V4 \| 1.0 \| 2.494e-06 \| 310 \| 0.252 \| 103 \| 75 \| 2 \| 33 \| 134 \| 1 \| 102 \| Phage tail protein \| Phage tail protein; afdb-uniprot50 \| AF-A0A4Q9TKB8-F1-MODEL\_V4 \| 1.0 \| 6.195e-07 \| 310 \| 0.143 \| 132 \| 102 \| 6 \| 13 \| 136 \| 1 \| 129 \| Phage tail protein \| Phage tail protein; afdb-uniprot50 \| AF-A0A496P7D4-F1-MODEL\_V4 \| 1.0 \| 2.927e-07 \| 310 \| 0.2 \| 135 \| 91 \| 6 \| 13 \| 136 \| 2 \| 130 \| Phage tail protein \| Phage tail protein; afdb-uniprot50 \| AF-R6LLD3-F1-MODEL\_V4 \| 1.0 \| 1.625e-06 \| 309 \| 0.148 \| 135 \| 98 \| 5 \| 13 \| 136 \| 2 \| 130 \| Uncharacterized protein \| Uncharacterized protein; afdb-uniprot50 \| AF-K9D590-F1-MODEL\_V4 \| 1.0 \| 4.036e-07 \| 309 \| 0.198 \| 131 \| 96 \| 5 \| 13 \| 136 \| 2 \| 130 \| Uncharacterized protein \| Uncharacterized protein; afdb-uniprot50 \| AF-A0A3R6V8J2-F1-MODEL\_V4 \| 1.0 \| 4.492e-07 \| 308 \| 0.151 \| 132 \| 101 \| 6 \| 13 \| 136 \| 1 \| 129 \| Uncharacterized protein \| Uncharacterized protein; afdb-uniprot50 \| AF-A0A7T1BPS3-F1-MODEL\_V4 \| 1.0 \| 8.543e-07 \| 308 \| 0.153 \| 124 \| 103 \| 2 \| 13 \| 136 \| 9 \| 130 \| Phage tail protein \| Phage tail protein; afdb-uniprot50 \| AF-E3HBK8-F1-MODEL\_V4 \| 1.0 \| 2.927e-07 \| 308 \| 0.165 \| 133 \| 96 \| 5 \| 13 \| 136 \| 1 \| 127 \| Uncharacterized protein \| Uncharacterized protein; afdb-uniprot50 \| AF-A0A553SNJ4-F1-MODEL\_V4 \| 1.0 \| 3.437e-07 \| 308 \| 0.201 \| 139 \| 98 \| 8 \| 6 \| 135 \| 7 \| 141 \| Phage tail protein \| Phage tail protein; afdb-uniprot50 \| AF-A0A6I2R3F1-F1-MODEL\_V4 \| 1.0 \| 6.896e-07 \| 307 \| 0.17 \| 135 \| 95 \| 6 \| 13 \| 136 \| 3 \| 131 \| Uncharacterized protein \| Uncharacterized protein; afdb-uniprot50 \| AF-D1AFB3-F1-MODEL\_V4 \| 1.0 \| 4.492e-07 \| 307 \| 0.159 \| 132 \| 98 \| 6 \| 13 \| 136 \| 2 \| 128 \| Uncharacterized protein \| Uncharacterized protein; afdb-uniprot50 \| AF-A0A198XEA5-F1-MODEL\_V4 \| 1.0 \| 3.437e-07 \| 307 \| 0.177 \| 124 \| 99 \| 3 \| 13 \| 135 \| 1 \| 122 \| Uncharacterized protein \| Uncharacterized protein; afdb-uniprot50 \| AF-A0A2X0YZL3-F1-MODEL\_V4 \| 1.0 \| 6.195e-07 \| 306 \| 0.176 \| 130 \| 98 \| 5 \| 13 \| 135 \| 3 \| 130 \| N-acetylmuramoyl-L-alanine amidase \| N-acetylmuramoyl-L-alanine amidase; afdb-uniprot50 \| AF-A0A1M6N828-F1-MODEL\_V4 \| 1.0 \| 1.117e-06 \| 305 \| 0.169 \| 130 \| 101 \| 5 \| 13 \| 136 \| 2 \| 130 \| Uncharacterized protein \| Uncharacterized protein; afdb-uniprot50 \| AF-G2FT53-F1-MODEL\_V4 \| 1.0 \| 5.276e-07 \| 305 \| 0.11 \| 136 \| 103 \| 6 \| 13 \| 136 \| 2 \| 131 \| Uncharacterized protein \| Uncharacterized protein; afdb-uniprot50 \| AF-A0A0J8DFL5-F1-MODEL\_V4 \| 1.0 \| 1.058e-06 \| 304 \| 0.143 \| 132 \| 102 \| 7 \| 13 \| 136 \| 2 \| 130 \| Uncharacterized protein \| Uncharacterized protein; afdb-uniprot50 \| AF-A0A150MJG2-F1-MODEL\_V4 \| 1.0 \| 6.896e-07 \| 304 \| 0.183 \| 131 \| 96 \| 6 \| 13 \| 135 \| 3 \| 130 \| SH3b domain-containing protein \| SH3b domain-containing protein; afdb-uniprot50 \| AF-B9NM79-F1-MODEL\_V4 \| 1.0 \| 2.631e-06 \| 303 \| 0.472 \| 74 \| 39 \| 0 \| 63 \| 136 \| 3 \| 76 \| Putative phage tail protein U \| Putative phage tail protein U; afdb-uniprot50 \| AF-A0A7G9GXJ0-F1-MODEL\_V4 \| 1.0 \| 4.495e-06 \| 303 \| 0.131 \| 129 \| 106 \| 3 \| 13 \| 136 \| 24 \| 151 \| Phage tail protein \| Phage tail protein; afdb-uniprot50 \| AF-E6LKB5-F1-MODEL\_V4 \| 1.0 \| 1.383e-06 \| 302 \| 0.143 \| 132 \| 102 \| 8 \| 13 \| 136 \| 3 \| 131 \| Uncharacterized protein \| Uncharacterized protein; afdb-uniprot50 \| AF-A0A399IPH0-F1-MODEL\_V4 \| 1.0 \| 3.089e-06 \| 302 \| 0.145 \| 131 \| 104 \| 5 \| 13 \| 136 \| 2 \| 131 \| Uncharacterized protein \| Uncharacterized protein; afdb-uniprot50 \| AF-A0A4U1VS70-F1-MODEL\_V4 \| 1.0 \| 1.311e-06 \| 300 \| 0.179 \| 134 \| 104 \| 4 \| 5 \| 136 \| 2 \| 131 \| Uncharacterized protein \| Uncharacterized protein; afdb-uniprot50 \| AF-A0A1C6BMU7-F1-MODEL\_V4 \| 1.0 \| 3.828e-06 \| 299 \| 0.178 \| 129 \| 98 \| 5 \| 13 \| 134 \| 1 \| 128 \| Uncharacterized protein \| Uncharacterized protein; afdb-uniprot50 \| AF-A0A6M0SNX6-F1-MODEL\_V4 \| 1.0 \| 8.543e-07 \| 299 \| 0.148 \| 135 \| 98 \| 5 \| 13 \| 136 \| 2 \| 130 \| Uncharacterized protein \| Uncharacterized protein; afdb-uniprot50 \| AF-A0A7C9LIZ2-F1-MODEL\_V4 \| 1.0 \| 1.311e-06 \| 297 \| 0.133 \| 135 \| 100 \| 6 \| 13 \| 136 \| 1 \| 129 \| Uncharacterized protein \| Uncharacterized protein; afdb-uniprot50 \| AF-A0A844Q469-F1-MODEL\_V4 \| 1.0 \| 7.275e-07 \| 296 \| 0.162 \| 129 \| 99 \| 5 \| 13 \| 134 \| 5 \| 131 \| Uncharacterized protein \| Uncharacterized protein; afdb-uniprot50 \| AF-A0A2D3P2W1-F1-MODEL\_V4 \| 1.0 \| 4.038e-06 \| 296 \| 0.13 \| 138 \| 113 \| 5 \| 4 \| 136 \| 15 \| 150 \| Uncharacterized protein \| Uncharacterized protein; afdb-uniprot50 \| AF-A0A3A6JMF6-F1-MODEL\_V4 \| 1.0 \| 1.625e-06 \| 295 \| 0.098 \| 132 \| 108 \| 6 \| 13 \| 136 \| 5 \| 133 \| Uncharacterized protein \| Uncharacterized protein; afdb-uniprot50 \| AF-A0A3C1Q6J4-F1-MODEL\_V4 \| 1.0 \| 2.927e-07 \| 295 \| 0.16 \| 143 \| 102 \| 6 \| 6 \| 136 \| 23 \| 159 \| Uncharacterized protein \| Uncharacterized protein; afdb-uniprot50 \| AF-A0A3G2Q678-F1-MODEL\_V4 \| 1.0 \| 7.275e-07 \| 294 \| 0.195 \| 133 \| 103 \| 3 \| 6 \| 136 \| 1 \| 131 \| Uncharacterized protein \| Uncharacterized protein; afdb-uniprot50 \| AF-A0A699YKY6-F1-MODEL\_V4 \| 1.0 \| 5.003e-06 \| 293 \| 0.428 \| 84 \| 48 \| 0 \| 53 \| 136 \| 5 \| 88 \| Uncharacterized protein \| Uncharacterized protein; afdb-uniprot50 \| AF-A0A0G3WAU0-F1-MODEL\_V4 \| 1.0 \| 4.74e-07 \| 291 \| 0.174 \| 132 \| 95 \| 6 \| 16 \| 136 \| 1 \| 129 \| Putative phage P2 GpU family protein \| Putative phage P2 GpU family protein; afdb-uniprot50 \| AF-G9PUJ3-F1-MODEL\_V4 \| 1.0 \| 1.058e-06 \| 290 \| 0.186 \| 118 \| 84 \| 3 \| 25 \| 136 \| 9 \| 120 \| Uncharacterized protein \| Uncharacterized protein; afdb-uniprot50 \| AF-G4KQ70-F1-MODEL\_V4 \| 1.0 \| 3.259e-06 \| 290 \| 0.113 \| 132 \| 106 \| 7 \| 13 \| 136 \| 2 \| 130 \| Uncharacterized protein \| Uncharacterized protein; afdb-uniprot50 \| AF-A0A2W5H3R5-F1-MODEL\_V4 \| 1.0 \| 1.243e-06 \| 289 \| 0.37 \| 100 \| 62 \| 1 \| 16 \| 114 \| 1 \| 100 \| Uncharacterized protein \| Uncharacterized protein; afdb-uniprot50 \| AF-U2PM82-F1-MODEL\_V4 \| 1.0 \| 4.038e-06 \| 289 \| 0.156 \| 134 \| 102 \| 7 \| 13 \| 136 \| 1 \| 133 \| Uncharacterized protein \| Uncharacterized protein; afdb-uniprot50 \| AF-A0A285M8K9-F1-MODEL\_V4 \| 1.0 \| 1.54e-06 \| 288 \| 0.15 \| 126 \| 101 \| 3 \| 14 \| 136 \| 4 \| 126 \| Uncharacterized protein \| Uncharacterized protein; afdb-uniprot50 \| AF-A0A379ZRU9-F1-MODEL\_V4 \| 1.0 \| 5.569e-06 \| 288 \| 0.342 \| 76 \| 50 \| 0 \| 61 \| 136 \| 51 \| 126 \| Phage tail protein \| Phage tail protein; afdb-uniprot50 \| AF-A0A2X1LFE2-F1-MODEL\_V4 \| 1.0 \| 1.312e-05 \| 287 \| 0.312 \| 96 \| 66 \| 0 \| 13 \| 108 \| 1 \| 96 \| Tail fiber protein \| Tail fiber protein; afdb-uniprot50 \| AF-A0A836RHU7-F1-MODEL\_V4 \| 1.0 \| 3.828e-06 \| 287 \| 0.135 \| 133 \| 100 \| 5 \| 13 \| 136 \| 3 \| 129 \| Uncharacterized protein \| Uncharacterized protein; afdb-uniprot50 \| AF-A0A1Q6LL69-F1-MODEL\_V4 \| 1.0 \| 2.123e-06 \| 287 \| 0.155 \| 122 \| 96 \| 4 \| 18 \| 136 \| 36 \| 153 \| LysM domain-containing protein \| LysM domain-containing protein; afdb-uniprot50 \| AF-A0A1Q6JPI8-F1-MODEL\_V4 \| 1.0 \| 1.178e-06 \| 285 \| 0.136 \| 132 \| 100 \| 5 \| 16 \| 136 \| 1 \| 129 \| Uncharacterized protein \| Uncharacterized protein; afdb-uniprot50 \| AF-A0A2X1KFU1-F1-MODEL\_V4 \| 1.0 \| 5.875e-06 \| 285 \| 0.452 \| 84 \| 46 \| 0 \| 53 \| 136 \| 20 \| 103 \| GpU phage protein \| GpU phage protein; afdb-uniprot50 \| AF-A0A3M3AZU3-F1-MODEL\_V4 \| 1.0 \| 1.715e-05 \| 284 \| 0.402 \| 72 \| 43 \| 0 \| 64 \| 135 \| 1 \| 72 \| Uncharacterized protein \| Uncharacterized protein; afdb-uniprot50 \| AF-A0A3P6KJ42-F1-MODEL\_V4 \| 1.0 \| 2.24e-06 \| 284 \| 0.149 \| 134 \| 97 \| 6 \| 13 \| 135 \| 2 \| 129 \| Uncharacterized protein \| Uncharacterized protein; afdb-uniprot50 \| AF-A0A0C1G7H4-F1-MODEL\_V4 \| 1.0 \| 6.899e-06 \| 284 \| 0.14 \| 107 \| 91 \| 1 \| 29 \| 134 \| 18 \| 124 \| Uncharacterized protein \| Uncharacterized protein; afdb-uniprot50 \| AF-A0A644SV89-F1-MODEL\_V4 \| 1.0 \| 2.631e-06 \| 283 \| 0.164 \| 134 \| 97 \| 5 \| 13 \| 136 \| 1 \| 129 \| Uncharacterized protein \| Uncharacterized protein; afdb-uniprot50 \| AF-A0A3D1M336-F1-MODEL\_V4 \| 1.0 \| 3.628e-06 \| 283 \| 0.185 \| 113 \| 87 \| 2 \| 29 \| 136 \| 84 \| 196 \| Uncharacterized protein \| Uncharacterized protein; afdb-uniprot50 \| AF-A0A2V2GP09-F1-MODEL\_V4 \| 1.0 \| 1.54e-06 \| 282 \| 0.103 \| 135 \| 104 \| 6 \| 13 \| 136 \| 2 \| 130 \| Uncharacterized protein \| Uncharacterized protein; afdb-uniprot50 \| AF-A0A827SX25-F1-MODEL\_V4 \| 1.0 \| 6.539e-06 \| 281 \| 0.355 \| 90 \| 58 \| 0 \| 47 \| 136 \| 1 \| 90 \| Phage tail protein \| Phage tail protein; afdb-uniprot50 \| AF-A0A4R4E4X3-F1-MODEL\_V4 \| 1.0 \| 2.24e-06 \| 281 \| 0.125 \| 135 \| 101 \| 6 \| 13 \| 136 \| 2 \| 130 \| Uncharacterized protein \| Uncharacterized protein; afdb-uniprot50 \| AF-A0A6D1AID6-F1-MODEL\_V4 \| 1.0 \| 1.179e-05 \| 280 \| 0.451 \| 93 \| 51 \| 0 \| 31 \| 123 \| 1 \| 93 \| Phage tail protein \| Phage tail protein; afdb-uniprot50 \| AF-A0A2G6EZG3-F1-MODEL\_V4 \| 1.0 \| 1.541e-05 \| 279 \| 0.115 \| 130 \| 108 \| 5 \| 13 \| 136 \| 32 \| 160 \| Uncharacterized protein \| Uncharacterized protein; afdb-uniprot50 \| AF-A0A7I8DHZ1-F1-MODEL\_V4 \| 1.0 \| 6.539e-06 \| 278 \| 0.139 \| 129 \| 103 \| 5 \| 13 \| 136 \| 2 \| 127 \| Uncharacterized protein \| Uncharacterized protein; afdb-uniprot50 \| AF-A0A5S9NAT8-F1-MODEL\_V4 \| 1.0 \| 1.244e-05 \| 277 \| 0.14 \| 128 \| 102 \| 4 \| 13 \| 133 \| 1 \| 127 \| Uncharacterized protein \| Uncharacterized protein; afdb-uniprot50 \| AF-A4P157-F1-MODEL\_V4 \| 1.0 \| 1.059e-05 \| 274 \| 0.319 \| 97 \| 65 \| 1 \| 9 \| 104 \| 2 \| 98 \| Probable phage-related tail protein \| Probable phage-related tail protein; afdb-uniprot50 \| AF-A0A510JEI6-F1-MODEL\_V4 \| 1.0 \| 8.102e-06 \| 271 \| 0.156 \| 134 \| 102 \| 7 \| 13 \| 136 \| 1 \| 133 \| Uncharacterized protein \| Uncharacterized protein; afdb-uniprot50 \| AF-A0A0S2F794-F1-MODEL\_V4 \| 1.0 \| 1.004e-05 \| 265 \| 0.373 \| 91 \| 55 \| 2 \| 47 \| 136 \| 2 \| 91 \| Phage P2 GpU family protein \| Phage P2 GpU family protein; afdb-uniprot50 \| AF-A0A7G7ZA20-F1-MODEL\_V4 \| 1.0 \| 2.93e-05 \| 264 \| 0.379 \| 79 \| 49 \| 0 \| 58 \| 136 \| 13 \| 91 \| Phage tail protein \| Phage tail protein; afdb-uniprot50 \| AF-A0A3C1LVB9-F1-MODEL\_V4 \| 1.0 \| 1.809e-05 \| 264 \| 0.125 \| 128 \| 102 \| 4 \| 10 \| 136 \| 1 \| 119 \| Uncharacterized protein \| Uncharacterized protein; afdb-uniprot50 \| AF-A0A842IWC8-F1-MODEL\_V4 \| 1.0 \| 1.004e-05 \| 264 \| 0.142 \| 133 \| 100 \| 5 \| 13 \| 136 \| 29 \| 156 \| Phage tail protein \| Phage tail protein; afdb-uniprot50 \| AF-A0A351VDG0-F1-MODEL\_V4 \| 1.0 \| 1.384e-05 \| 263 \| 0.121 \| 140 \| 100 \| 6 \| 13 \| 136 \| 3 \| 135 \| LysM domain-containing protein \| LysM domain-containing protein; afdb-uniprot50 \| AF-A0A7X3VGF6-F1-MODEL\_V4 \| 1.0 \| 2.365e-05 \| 261 \| 0.457 \| 70 \| 38 \| 0 \| 67 \| 136 \| 4 \| 73 \| Uncharacterized protein \| Uncharacterized protein; afdb-uniprot50 \| AF-A0A4D7AUF4-F1-MODEL\_V4 \| 1.0 \| 1.625e-05 \| 261 \| 0.178 \| 112 \| 78 \| 4 \| 33 \| 136 \| 2 \| 107 \| Phage tail protein \| Phage tail protein; afdb-uniprot50 \| AF-A0A4S3LUR7-F1-MODEL\_V4 \| 1.0 \| 4.262e-05 \| 259 \| 0.405 \| 79 \| 47 \| 0 \| 58 \| 136 \| 1 \| 79 \| Oxidoreductase \| Oxidoreductase; afdb-uniprot50 \| AF-A0A6N8HZ03-F1-MODEL\_V4 \| 1.0 \| 7.279e-06 \| 259 \| 0.119 \| 134 \| 106 \| 6 \| 13 \| 136 \| 2 \| 133 \| Uncharacterized protein \| Uncharacterized protein; afdb-uniprot50 \| AF-A0A846M0R6-F1-MODEL\_V4 \| 1.0 \| 4.497e-05 \| 258 \| 0.392 \| 79 \| 48 \| 0 \| 58 \| 136 \| 2 \| 80 \| Phage protein U \| Phage protein U; afdb-uniprot50 \| AF-A0A2B7YL04-F1-MODEL\_V4 \| 1.0 \| 2.632e-05 \| 258 \| 0.157 \| 133 \| 101 \| 7 \| 13 \| 136 \| 1 \| 131 \| Uncharacterized protein \| Uncharacterized protein; afdb-uniprot50 \| AF-A0A132MI06-F1-MODEL\_V4 \| 1.0 \| 2.123e-06 \| 258 \| 0.165 \| 145 \| 103 \| 8 \| 1 \| 135 \| 20 \| 156 \| LysM domain-containing protein \| LysM domain-containing protein; afdb-uniprot50 \| AF-A0A316S5B6-F1-MODEL\_V4 \| 1.0 \| 1.625e-05 \| 257 \| 0.205 \| 112 \| 75 \| 4 \| 33 \| 136 \| 2 \| 107 \| Uncharacterized protein \| Uncharacterized protein; afdb-uniprot50 \| AF-V5TVJ3-F1-MODEL\_V4 \| 1.0 \| 4.262e-05 \| 257 \| 0.379 \| 79 \| 49 \| 0 \| 58 \| 136 \| 2 \| 80 \| Uncharacterized protein \| Uncharacterized protein; afdb-uniprot50 \| AF-A0A6P0MS31-F1-MODEL\_V4 \| 1.0 \| 1.059e-05 \| 257 \| 0.134 \| 134 \| 102 \| 6 \| 13 \| 136 \| 3 \| 132 \| Uncharacterized protein \| Uncharacterized protein; afdb-uniprot50 \| AF-A0A1F0F4I9-F1-MODEL\_V4 \| 1.0 \| 4.742e-06 \| 257 \| 0.177 \| 141 \| 96 \| 6 \| 5 \| 136 \| 8 \| 137 \| Uncharacterized protein \| Uncharacterized protein; afdb-uniprot50 \| AF-A0A1I1D0Z6-F1-MODEL\_V4 \| 1.0 \| 3.439e-06 \| 256 \| 0.172 \| 145 \| 99 \| 7 \| 1 \| 136 \| 3 \| 135 \| Uncharacterized protein \| Uncharacterized protein; afdb-uniprot50 \| AF-A0A1F0F528-F1-MODEL\_V4 \| 1.0 \| 6.198e-06 \| 256 \| 0.172 \| 133 \| 92 \| 5 \| 13 \| 136 \| 14 \| 137 \| Uncharacterized protein \| Uncharacterized protein; afdb-uniprot50 \| AF-A0A6P0RQA4-F1-MODEL\_V4 \| 1.0 \| 6.539e-06 \| 250 \| 0.171 \| 128 \| 95 \| 5 \| 14 \| 136 \| 3 \| 124 \| Uncharacterized protein \| Uncharacterized protein; afdb-uniprot50 \| AF-A0A4R4CZH4-F1-MODEL\_V4 \| 1.0 \| 1.244e-05 \| 249 \| 0.151 \| 132 \| 101 \| 5 \| 10 \| 136 \| 2 \| 127 \| LysM domain-containing protein \| LysM domain-containing protein; afdb-uniprot50 \| AF-C4GFW6-F1-MODEL\_V4 \| 1.0 \| 4.038e-06 \| 249 \| 0.173 \| 144 \| 94 \| 7 \| 2 \| 136 \| 23 \| 150 \| Uncharacterized protein \| Uncharacterized protein; afdb-uniprot50 \| AF-A0A1C5WXV0-F1-MODEL\_V4 \| 1.0 \| 3.261e-05 \| 246 \| 0.169 \| 124 \| 92 \| 5 \| 13 \| 136 \| 3 \| 115 \| LysM domain-containing protein \| LysM domain-containing protein; afdb-uniprot50 \| AF-A0A011PPH1-F1-MODEL\_V4 \| 1.0 \| 1.625e-05 \| 245 \| 0.132 \| 128 \| 96 \| 6 \| 13 \| 136 \| 1 \| 117 \| Mu-like prophage DNA circulation protein \| Mu-like prophage DNA circulation protein; afdb-uniprot50 \| AF-A0A1Q6PWR2-F1-MODEL\_V4 \| 1.0 \| 6.201e-05 \| 245 \| 0.116 \| 129 \| 108 \| 4 \| 13 \| 136 \| 3 \| 130 \| Uncharacterized protein \| Uncharacterized protein; afdb-uniprot50 \| AF-A0A6L5BK85-F1-MODEL\_V4 \| 1.0 \| 5.281e-05 \| 243 \| 0.348 \| 89 \| 52 \| 2 \| 54 \| 136 \| 1 \| 89 \| Uncharacterized protein \| Uncharacterized protein; afdb-uniprot50 \| AF-R9L5R6-F1-MODEL\_V4 \| 1.0 \| 1.004e-05 \| 243 \| 0.107 \| 130 \| 107 \| 7 \| 13 \| 136 \| 3 \| 129 \| Uncharacterized protein \| Uncharacterized protein; afdb-uniprot50 \| AF-A0A3D4CGN3-F1-MODEL\_V4 \| 1.0 \| 5.571e-05 \| 242 \| 0.106 \| 122 \| 98 \| 5 \| 13 \| 134 \| 5 \| 115 \| Uncharacterized protein \| Uncharacterized protein; afdb-uniprot50 \| AF-N8SB57-F1-MODEL\_V4 \| 1.0 \| 0.0001461 \| 239 \| 0.364 \| 74 \| 47 \| 0 \| 63 \| 136 \| 3 \| 76 \| Uncharacterized protein \| Uncharacterized protein; afdb-uniprot50 \| AF-A0A6N2U2G7-F1-MODEL\_V4 \| 1.0 \| 4.04e-05 \| 239 \| 0.132 \| 121 \| 95 \| 5 \| 14 \| 133 \| 3 \| 114 \| Uncharacterized protein \| Uncharacterized protein; afdb-uniprot50 \| AF-A0A5Q0GYR9-F1-MODEL\_V4 \| 1.0 \| 4.26e-06 \| 239 \| 0.207 \| 140 \| 93 \| 5 \| 13 \| 136 \| 6 \| 143 \| Peptidoglycan-binding protein \| Peptidoglycan-binding protein; afdb-uniprot50 \| AF-R6UJW8-F1-MODEL\_V4 \| 1.0 \| 5.005e-05 \| 238 \| 0.112 \| 125 \| 100 \| 5 \| 10 \| 134 \| 1 \| 114 \| LysM domain-containing protein \| LysM domain-containing protein; afdb-uniprot50 \| AF-A0A358PQ27-F1-MODEL\_V4 \| 1.0 \| 6.899e-06 \| 238 \| 0.143 \| 146 \| 110 \| 10 \| 1 \| 136 \| 1 \| 141 \| Uncharacterized protein \| Uncharacterized protein; afdb-uniprot50 \| AF-A0A848D1N5-F1-MODEL\_V4 \| 1.0 \| 1.179e-05 \| 238 \| 0.144 \| 145 \| 101 \| 7 \| 10 \| 136 \| 1 \| 140 \| LysM peptidoglycan-binding domain-containing protein \| LysM peptidoglycan-binding domain-containing protein; afdb-uniprot50 \| AF-A0A6N2T955-F1-MODEL\_V4 \| 1.0 \| 1.244e-05 \| 238 \| 0.149 \| 134 \| 95 \| 7 \| 14 \| 136 \| 4 \| 129 \| Uncharacterized protein \| Uncharacterized protein; afdb-uniprot50 \| AF-A0A7G9WG84-F1-MODEL\_V4 \| 1.0 \| 6.201e-05 \| 237 \| 0.137 \| 131 \| 100 \| 6 \| 13 \| 135 \| 2 \| 127 \| Uncharacterized protein \| Uncharacterized protein; afdb-uniprot50 \| AF-A0A6N6VMR6-F1-MODEL\_V4 \| 1.0 \| 1.715e-05 \| 237 \| 0.112 \| 142 \| 104 \| 5 \| 8 \| 136 \| 7 \| 139 \| Uncharacterized protein \| Uncharacterized protein; afdb-uniprot50 \| AF-A0A1C6JGS8-F1-MODEL\_V4 \| 1.0 \| 3.829e-05 \| 237 \| 0.16 \| 131 \| 97 \| 7 \| 14 \| 136 \| 1 \| 126 \| Uncharacterized protein \| Uncharacterized protein; afdb-uniprot50 \| AF-A0A842J060-F1-MODEL\_V4 \| 1.0 \| 3.091e-05 \| 237 \| 0.128 \| 132 \| 103 \| 4 \| 13 \| 136 \| 29 \| 156 \| Phage tail protein \| Phage tail protein; afdb-uniprot50 \| AF-A0A7C6A8Q0-F1-MODEL\_V4 \| 1.0 \| 1.715e-05 \| 237 \| 0.184 \| 125 \| 88 \| 7 \| 13 \| 136 \| 1 \| 112 \| LysM domain-containing protein \| LysM domain-containing protein; afdb-uniprot50 \| AF-A0A2T6G9C8-F1-MODEL\_V4 \| 1.0 \| 2.241e-05 \| 235 \| 0.138 \| 137 \| 100 \| 4 \| 13 \| 136 \| 1 \| 132 \| Terminase \| Terminase; afdb-uniprot50 \| AF-A0A3S9SLX9-F1-MODEL\_V4 \| 1.0 \| 2.014e-05 \| 235 \| 0.165 \| 139 \| 96 \| 6 \| 7 \| 136 \| 20 \| 147 \| Uncharacterized protein \| Uncharacterized protein; afdb-uniprot50 \| AF-A0A1H8DLN7-F1-MODEL\_V4 \| 1.0 \| 5.878e-05 \| 234 \| 0.123 \| 121 \| 95 \| 5 \| 14 \| 134 \| 4 \| 113 \| DNA circularisation protein N-terminus \| DNA circularisation protein N-terminus; afdb-uniprot50 \| AF-A0A2X4V2I9-F1-MODEL\_V4 \| 1.0 \| 1.384e-05 \| 234 \| 0.145 \| 158 \| 103 \| 6 \| 2 \| 136 \| 6 \| 154 \| Uncharacterized protein \| Uncharacterized protein; afdb-uniprot50 \| AF-A0A7Z0UY93-F1-MODEL\_V4 \| 1.0 \| 0.0001716 \| 233 \| 0.356 \| 73 \| 47 \| 0 \| 64 \| 136 \| 1 \| 73 \| Phage tail protein \| Phage tail protein; afdb-uniprot50 \| AF-A0A7W0G3T0-F1-MODEL\_V4 \| 1.0 \| 2.241e-05 \| 233 \| 0.18 \| 144 \| 98 \| 9 \| 7 \| 136 \| 1 \| 138 \| Uncharacterized protein \| Uncharacterized protein; afdb-uniprot50 \| AF-A0A7X3MI16-F1-MODEL\_V4 \| 1.0 \| 5.878e-05 \| 232 \| 0.093 \| 128 \| 108 \| 6 \| 14 \| 136 \| 1 \| 125 \| Uncharacterized protein \| Uncharacterized protein; afdb-uniprot50 \| AF-A0A417HQS1-F1-MODEL\_V4 \| 1.0 \| 6.543e-05 \| 231 \| 0.122 \| 131 \| 104 \| 7 \| 13 \| 136 \| 3 \| 129 \| Uncharacterized protein \| Uncharacterized protein; afdb-uniprot50 \| AF-A9I936-F1-MODEL\_V4 \| 1.0 \| 2.365e-05 \| 231 \| 0.131 \| 145 \| 107 \| 7 \| 1 \| 136 \| 1 \| 135 \| Uncharacterized protein \| Uncharacterized protein; afdb-uniprot50 \| AF-A0A0P0RPN1-F1-MODEL\_V4 \| 1.0 \| 2.632e-05 \| 230 \| 0.129 \| 124 \| 93 \| 3 \| 28 \| 136 \| 28 \| 151 \| LysM domain \| LysM domain; afdb-uniprot50 \| AF-A0A0F5EPQ5-F1-MODEL\_V4 \| 1.0 \| 4.744e-05 \| 229 \| 0.128 \| 132 \| 97 \| 4 \| 14 \| 136 \| 15 \| 137 \| Uncharacterized protein \| Uncharacterized protein; afdb-uniprot50 \| AF-A0A2M8P9D4-F1-MODEL\_V4 \| 1.0 \| 2.632e-05 \| 229 \| 0.157 \| 159 \| 110 \| 6 \| 1 \| 136 \| 1 \| 158 \| Uncharacterized protein \| Uncharacterized protein; afdb-uniprot50 \| AF-A0A2E8DR59-F1-MODEL\_V4 \| 1.0 \| 0.0001244 \| 229 \| 0.131 \| 122 \| 98 \| 4 \| 20 \| 136 \| 80 \| 198 \| Uncharacterized protein \| Uncharacterized protein; afdb-uniprot50 \| AF-A0A174T7I7-F1-MODEL\_V4 \| 1.0 \| 3.261e-05 \| 229 \| 0.148 \| 128 \| 98 \| 7 \| 13 \| 135 \| 3 \| 124 \| Phage protein D \| Phage protein D; afdb-uniprot50 \| AF-A0A2I0GV19-F1-MODEL\_V4 \| 1.0 \| 0.0002015 \| 228 \| 0.309 \| 84 \| 58 \| 0 \| 53 \| 136 \| 2 \| 85 \| Uncharacterized protein \| Uncharacterized protein; afdb-uniprot50 \| AF-A0A562GHF5-F1-MODEL\_V4 \| 1.0 \| 3.261e-05 \| 228 \| 0.153 \| 137 \| 94 \| 7 \| 12 \| 136 \| 2 \| 128 \| Uncharacterized protein \| Uncharacterized protein; afdb-uniprot50 \| AF-A0A7X1U0D0-F1-MODEL\_V4 \| 1.0 \| 2.632e-05 \| 227 \| 0.175 \| 131 \| 90 \| 5 \| 21 \| 136 \| 19 \| 146 \| LysM peptidoglycan-binding domain-containing protein \| LysM peptidoglycan-binding domain-containing protein; afdb-uniprot50 \| AF-G1WGB9-F1-MODEL\_V4 \| 1.0 \| 1.625e-05 \| 226 \| 0.119 \| 134 \| 102 \| 9 \| 13 \| 136 \| 3 \| 130 \| Uncharacterized protein \| Uncharacterized protein; afdb-uniprot50 \| AF-A0A412CI02-F1-MODEL\_V4 \| 1.0 \| 6.201e-05 \| 226 \| 0.15 \| 146 \| 105 \| 7 \| 2 \| 136 \| 17 \| 154 \| Uncharacterized protein \| Uncharacterized protein; afdb-uniprot50 \| AF-A0A436RST7-F1-MODEL\_V4 \| 1.0 \| 0.0002778 \| 225 \| 0.232 \| 99 \| 75 \| 1 \| 13 \| 111 \| 1 \| 98 \| Phage tail protein \| Phage tail protein; afdb-uniprot50 \| AF-A0A285NP30-F1-MODEL\_V4 \| 1.0 \| 8.106e-05 \| 225 \| 0.13 \| 130 \| 100 \| 5 \| 13 \| 136 \| 1 \| 123 \| Phage protein U \| Phage protein U; afdb-uniprot50 \| AF-A0A2V4VHT5-F1-MODEL\_V4 \| 1.0 \| 2.125e-05 \| 225 \| 0.153 \| 150 \| 101 \| 9 \| 5 \| 136 \| 2 \| 143 \| LysM domain-containing protein \| LysM domain-containing protein; afdb-uniprot50 \| AF-A0A630BA39-F1-MODEL\_V4 \| 1.0 \| 0.000191 \| 224 \| 0.215 \| 88 \| 69 \| 0 \| 47 \| 134 \| 2 \| 89 \| Phage tail protein \| Phage tail protein; afdb-uniprot50 \| AF-A0A1H9SSE7-F1-MODEL\_V4 \| 1.0 \| 2.93e-05 \| 224 \| 0.148 \| 141 \| 102 \| 7 \| 10 \| 136 \| 1 \| 137 \| LysM domain-containing protein \| LysM domain-containing protein; afdb-uniprot50 \| AF-A0A125V4N9-F1-MODEL\_V4 \| 1.0 \| 3.091e-05 \| 223 \| 0.127 \| 133 \| 101 \| 4 \| 13 \| 136 \| 3 \| 129 \| Phage-like element pbsx protein xkdP \| Phage-like element pbsx protein xkdP; afdb-uniprot50 \| AF-A0A1W9V6L5-F1-MODEL\_V4 \| 1.0 \| 2.93e-05 \| 223 \| 0.115 \| 147 \| 104 \| 7 \| 6 \| 136 \| 1 \| 137 \| Uncharacterized protein \| Uncharacterized protein; afdb-uniprot50 \| AF-A0A1H7MLA5-F1-MODEL\_V4 \| 1.0 \| 4.497e-05 \| 223 \| 0.13 \| 123 \| 92 \| 4 \| 29 \| 136 \| 34 \| 156 \| LysM domain-containing protein \| LysM domain-containing protein; afdb-uniprot50 \| AF-A0A6L5WH99-F1-MODEL\_V4 \| 1.0 \| 0.0001461 \| 222 \| 0.239 \| 92 \| 70 \| 0 \| 43 \| 134 \| 3 \| 94 \| Oxidoreductase \| Oxidoreductase; afdb-uniprot50 \| AF-A0A0P9CJQ9-F1-MODEL\_V4 \| 1.0 \| 0.0001059 \| 222 \| 0.321 \| 87 \| 57 \| 1 \| 50 \| 134 \| 2 \| 88 \| Uncharacterized protein \| Uncharacterized protein; afdb-uniprot50 \| AF-A0A0H3GS01-F1-MODEL\_V4 \| 1.0 \| 0.0002126 \| 222 \| 0.424 \| 73 \| 42 \| 0 \| 64 \| 136 \| 1 \| 73 \| Putative prophage tail protein \| Putative prophage tail protein; afdb-uniprot50 \| AF-R6N2M2-F1-MODEL\_V4 \| 1.0 \| 0.0001004 \| 222 \| 0.12 \| 124 \| 98 \| 5 \| 13 \| 136 \| 4 \| 116 \| LysM domain protein \| LysM domain protein; afdb-uniprot50 \| AF-A0A2E8S0C4-F1-MODEL\_V4 \| 1.0 \| 4.497e-05 \| 222 \| 0.176 \| 153 \| 103 \| 13 \| 3 \| 136 \| 30 \| 178 \| LysM domain-containing protein \| LysM domain-containing protein; afdb-uniprot50 \| AF-A0A1Y4MDW2-F1-MODEL\_V4 \| 1.0 \| 0.0001716 \| 221 \| 0.147 \| 129 \| 103 \| 5 \| 13 \| 136 \| 3 \| 129 \| Uncharacterized protein \| Uncharacterized protein; afdb-uniprot50 \| AF-A0A143HGM4-F1-MODEL\_V4 \| 1.0 \| 5.878e-05 \| 221 \| 0.153 \| 130 \| 95 \| 7 \| 10 \| 136 \| 1 \| 118 \| LysM domain-containing protein \| LysM domain-containing protein; afdb-uniprot50 \| AF-A0A5T2A9N3-F1-MODEL\_V4 \| 1.0 \| 0.0006204 \| 220 \| 0.252 \| 107 \| 76 \| 3 \| 13 \| 118 \| 1 \| 104 \| Phage tail protein \| Phage tail protein; afdb-uniprot50 \| AF-A0A1W6WBE5-F1-MODEL\_V4 \| 1.0 \| 0.0001626 \| 220 \| 0.107 \| 130 \| 99 \| 5 \| 10 \| 136 \| 1 \| 116 \| LysM domain-containing protein \| LysM domain-containing protein; afdb-uniprot50 \| AF-A0A3S4Z3K3-F1-MODEL\_V4 \| 1.0 \| 0.0001461 \| 219 \| 0.252 \| 111 \| 77 \| 1 \| 13 \| 117 \| 4 \| 114 \| Phage protein U \| Phage protein U; afdb-uniprot50 \| AF-A0A075KG62-F1-MODEL\_V4 \| 1.0 \| 3.63e-05 \| 218 \| 0.133 \| 142 \| 105 \| 5 \| 1 \| 136 \| 45 \| 174 \| p2 GpU family protein \| p2 GpU family protein; afdb-uniprot50 \| AF-V2YJG8-F1-MODEL\_V4 \| 1.0 \| 7.282e-05 \| 218 \| 0.104 \| 125 \| 95 \| 5 \| 13 \| 134 \| 4 \| 114 \| LysM domain-containing protein \| LysM domain-containing protein; afdb-uniprot50 \| AF-A0A1R0FQC3-F1-MODEL\_V4 \| 1.0 \| 4.744e-05 \| 218 \| 0.081 \| 159 \| 116 \| 7 \| 1 \| 136 \| 13 \| 164 \| Uncharacterized protein \| Uncharacterized protein; afdb-uniprot50 \| AF-A0A562HK82-F1-MODEL\_V4 \| 1.0 \| 6.903e-05 \| 218 \| 0.102 \| 146 \| 114 \| 7 \| 2 \| 136 \| 12 \| 151 \| LysM domain-containing protein \| LysM domain-containing protein; afdb-uniprot50 \| AF-A0A1V5KAZ2-F1-MODEL\_V4 \| 1.0 \| 0.0001385 \| 217 \| 0.161 \| 130 \| 93 \| 5 \| 13 \| 136 \| 1 \| 120 \| Uncharacterized protein \| Uncharacterized protein; afdb-uniprot50 \| AF-A0A1H7XXV0-F1-MODEL\_V4 \| 1.0 \| 3.261e-05 \| 217 \| 0.107 \| 140 \| 104 \| 7 \| 6 \| 136 \| 7 \| 134 \| Uncharacterized protein \| Uncharacterized protein; afdb-uniprot50 \| AF-A0A4R6IUX9-F1-MODEL\_V4 \| 1.0 \| 3.829e-05 \| 217 \| 0.15 \| 159 \| 105 \| 6 \| 5 \| 134 \| 6 \| 163 \| LysM domain-containing protein \| LysM domain-containing protein; afdb-uniprot50 \| AF-A0A6N7W5B6-F1-MODEL\_V4 \| 1.0 \| 6.903e-05 \| 216 \| 0.169 \| 130 \| 92 \| 8 \| 13 \| 136 \| 3 \| 122 \| Uncharacterized protein \| Uncharacterized protein; afdb-uniprot50 \| AF-A0A416C480-F1-MODEL\_V4 \| 1.0 \| 8.106e-05 \| 216 \| 0.15 \| 120 \| 87 \| 4 \| 20 \| 136 \| 13 \| 120 \| Uncharacterized protein \| Uncharacterized protein; afdb-uniprot50 \| AF-A0A5B8K6Q3-F1-MODEL\_V4 \| 1.0 \| 4.744e-05 \| 216 \| 0.158 \| 139 \| 94 \| 7 \| 13 \| 136 \| 1 \| 131 \| LysM peptidoglycan-binding domain-containing protein \| LysM peptidoglycan-binding domain-containing protein; afdb-uniprot50 \| AF-A0A374UJF1-F1-MODEL\_V4 \| 1.0 \| 0.000181 \| 215 \| 0.133 \| 127 \| 95 \| 5 \| 10 \| 134 \| 1 \| 114 \| DNA\_circ\_N domain-containing protein \| DNA\_circ\_N domain-containing protein; afdb-uniprot50 \| AF-A0A3M1D9D3-F1-MODEL\_V4 \| 1.0 \| 3.44e-05 \| 215 \| 0.123 \| 154 \| 109 \| 5 \| 6 \| 136 \| 2 \| 152 \| LysM peptidoglycan-binding domain-containing protein \| LysM peptidoglycan-binding domain-containing protein; afdb-uniprot50 \| AF-A0A3N8BG80-F1-MODEL\_V4 \| 1.0 \| 7.683e-05 \| 215 \| 0.119 \| 159 \| 110 \| 8 \| 5 \| 136 \| 2 \| 157 \| Uncharacterized protein \| Uncharacterized protein; afdb-uniprot50 \| AF-A0A6I4VAC5-F1-MODEL\_V4 \| 1.0 \| 0.0002366 \| 215 \| 0.08 \| 125 \| 103 \| 4 \| 13 \| 136 \| 18 \| 131 \| DNA circulation protein \| DNA circulation protein; afdb-uniprot50 \| AF-A0A1V2YD69-F1-MODEL\_V4 \| 1.0 \| 5.571e-05 \| 214 \| 0.102 \| 137 \| 102 \| 7 \| 13 \| 136 \| 3 \| 131 \| Uncharacterized protein \| Uncharacterized protein; afdb-uniprot50 \| AF-A0A6N9NZL1-F1-MODEL\_V4 \| 1.0 \| 9.519e-05 \| 214 \| 0.105 \| 133 \| 100 \| 6 \| 14 \| 136 \| 6 \| 129 \| Uncharacterized protein \| Uncharacterized protein; afdb-uniprot50 \| AF-A0A3P3U9B4-F1-MODEL\_V4 \| 1.0 \| 5.878e-05 \| 214 \| 0.136 \| 139 \| 98 \| 7 \| 13 \| 136 \| 1 \| 132 \| LysM peptidoglycan-binding domain-containing protein \| LysM peptidoglycan-binding domain-containing protein; afdb-uniprot50 \| AF-A0A2M8WDQ8-F1-MODEL\_V4 \| 1.0 \| 4.744e-05 \| 213 \| 0.261 \| 107 \| 72 \| 1 \| 37 \| 136 \| 2 \| 108 \| Uncharacterized protein \| Uncharacterized protein; afdb-uniprot50 \| AF-A0A0S8GTR5-F1-MODEL\_V4 \| 1.0 \| 0.0001004 \| 213 \| 0.139 \| 129 \| 105 \| 5 \| 10 \| 136 \| 1 \| 125 \| Uncharacterized protein \| Uncharacterized protein; afdb-uniprot50 \| AF-A0A497RM57-F1-MODEL\_V4 \| 1.0 \| 4.04e-05 \| 213 \| 0.151 \| 132 \| 85 \| 6 \| 13 \| 136 \| 1 \| 113 \| Uncharacterized protein \| Uncharacterized protein; afdb-uniprot50 \| AF-A0A1Q6J5B4-F1-MODEL\_V4 \| 1.0 \| 0.0001244 \| 213 \| 0.106 \| 131 \| 98 \| 7 \| 10 \| 136 \| 1 \| 116 \| Uncharacterized protein \| Uncharacterized protein; afdb-uniprot50 \| AF-A0A8B5L9S7-F1-MODEL\_V4 \| 1.0 \| 8.552e-05 \| 213 \| 0.135 \| 133 \| 100 \| 4 \| 13 \| 136 \| 3 \| 129 \| LysM peptidoglycan-binding domain-containing protein \| LysM peptidoglycan-binding domain-containing protein; afdb-uniprot50 \| AF-A0A849CUH9-F1-MODEL\_V4 \| 1.0 \| 7.282e-05 \| 211 \| 0.116 \| 129 \| 99 \| 8 \| 13 \| 136 \| 3 \| 121 \| Uncharacterized protein \| Uncharacterized protein; afdb-uniprot50 \| AF-A0A327JCS4-F1-MODEL\_V4 \| 1.0 \| 0.0004747 \| 211 \| 0.173 \| 104 \| 84 \| 2 \| 31 \| 134 \| 12 \| 113 \| Uncharacterized protein \| Uncharacterized protein; afdb-uniprot50 \| AF-A0A1H1IJ14-F1-MODEL\_V4 \| 1.0 \| 9.519e-05 \| 211 \| 0.128 \| 132 \| 97 \| 6 \| 10 \| 136 \| 1 \| 119 \| Uncharacterized protein \| Uncharacterized protein; afdb-uniprot50 \| AF-A0A542AM64-F1-MODEL\_V4 \| 1.0 \| 0.0005283 \| 210 \| 0.072 \| 125 \| 101 \| 5 \| 14 \| 136 \| 5 \| 116 \| LysM domain-containing protein \| LysM domain-containing protein; afdb-uniprot50 \| AF-A0A352UR19-F1-MODEL\_V4 \| 1.0 \| 4.497e-05 \| 210 \| 0.133 \| 142 \| 100 \| 10 \| 13 \| 136 \| 2 \| 138 \| LysM domain-containing protein \| LysM domain-containing protein; afdb-uniprot50 \| AF-A0A061NGS6-F1-MODEL\_V4 \| 1.0 \| 1.312e-05 \| 210 \| 0.147 \| 136 \| 100 \| 6 \| 13 \| 136 \| 8 \| 139 \| Phage-like element PBSX protein XkdP \| Phage-like element PBSX protein XkdP; afdb-uniprot50 \| AF-A0A1Y4EYM2-F1-MODEL\_V4 \| 1.0 \| 0.0001059 \| 210 \| 0.124 \| 137 \| 105 \| 7 \| 2 \| 134 \| 53 \| 178 \| LysM domain-containing protein \| LysM domain-containing protein; afdb-uniprot50 \| AF-A0A479ZZ86-F1-MODEL\_V4 \| 1.0 \| 6.543e-05 \| 209 \| 0.134 \| 156 \| 100 \| 9 \| 7 \| 136 \| 1 \| 147 \| Uncharacterized protein \| Uncharacterized protein; afdb-uniprot50 \| AF-A0A8A7K8X4-F1-MODEL\_V4 \| 1.0 \| 0.0001385 \| 209 \| 0.104 \| 125 \| 100 \| 5 \| 14 \| 136 \| 3 \| 117 \| Uncharacterized protein \| Uncharacterized protein; afdb-uniprot50 \| AF-A0A5D8QJ58-F1-MODEL\_V4 \| 1.0 \| 6.903e-05 \| 209 \| 0.132 \| 136 \| 96 \| 9 \| 13 \| 136 \| 5 \| 130 \| Uncharacterized protein \| Uncharacterized protein; afdb-uniprot50 \| AF-C7RHB7-F1-MODEL\_V4 \| 1.0 \| 5.281e-05 \| 209 \| 0.128 \| 132 \| 101 \| 5 \| 6 \| 136 \| 1 \| 119 \| Uncharacterized protein \| Uncharacterized protein; afdb-uniprot50 \| AF-A0A423PRP6-F1-MODEL\_V4 \| 1.0 \| 0.0001385 \| 208 \| 0.465 \| 86 \| 46 \| 0 \| 5 \| 90 \| 25 \| 110 \| Tail protein \| Tail protein; afdb-uniprot50 \| AF-G9YSH0-F1-MODEL\_V4 \| 1.0 \| 0.0004265 \| 208 \| 0.1 \| 119 \| 96 \| 4 \| 16 \| 134 \| 2 \| 109 \| LysM domain protein \| LysM domain protein; afdb-uniprot50 \| AF-C0ED24-F1-MODEL\_V4 \| 1.0 \| 6.903e-05 \| 208 \| 0.086 \| 138 \| 106 \| 6 \| 13 \| 136 \| 1 \| 132 \| LysM domain protein \| LysM domain protein; afdb-uniprot50 \| AF-A0A5C7J2Y4-F1-MODEL\_V4 \| 1.0 \| 7.282e-05 \| 208 \| 0.095 \| 157 \| 112 \| 7 \| 1 \| 136 \| 3 \| 150 \| Uncharacterized protein \| Uncharacterized protein; afdb-uniprot50 \| AF-A0A3D1ACM7-F1-MODEL\_V4 \| 1.0 \| 3.091e-05 \| 208 \| 0.125 \| 128 \| 99 \| 4 \| 21 \| 136 \| 12 \| 138 \| Peptidoglycan-binding protein \| Peptidoglycan-binding protein; afdb-uniprot50 \| AF-A0A498CN55-F1-MODEL\_V4 \| 1.0 \| 0.0001461 \| 207 \| 0.114 \| 131 \| 102 \| 5 \| 13 \| 136 \| 6 \| 129 \| LysM peptidoglycan-binding domain-containing protein \| LysM peptidoglycan-binding domain-containing protein; afdb-uniprot50 \| AF-D3EHD2-F1-MODEL\_V4 \| 1.0 \| 0.0001244 \| 207 \| 0.122 \| 139 \| 102 \| 6 \| 13 \| 136 \| 1 \| 134 \| Peptidoglycan-binding lysin domain protein \| Peptidoglycan-binding lysin domain protein; afdb-uniprot50 \| AF-A0A497FQZ2-F1-MODEL\_V4 \| 1.0 \| 0.0001626 \| 206 \| 0.104 \| 124 \| 93 \| 5 \| 13 \| 134 \| 1 \| 108 \| Uncharacterized protein \| Uncharacterized protein; afdb-uniprot50 \| AF-A0A661D5Z7-F1-MODEL\_V4 \| 1.0 \| 5.571e-05 \| 206 \| 0.118 \| 152 \| 109 \| 7 \| 7 \| 136 \| 1 \| 149 \| Uncharacterized protein \| Uncharacterized protein; afdb-uniprot50 \| AF-A0A5P8WCD5-F1-MODEL\_V4 \| 1.0 \| 0.0001004 \| 206 \| 0.121 \| 157 \| 113 \| 10 \| 1 \| 136 \| 3 \| 155 \| Uncharacterized protein \| Uncharacterized protein; afdb-uniprot50 \| AF-A0A2L2NSD1-F1-MODEL\_V4 \| 1.0 \| 3.44e-05 \| 206 \| 0.195 \| 138 \| 76 \| 10 \| 21 \| 136 \| 22 \| 146 \| Uncharacterized protein \| Uncharacterized protein; afdb-uniprot50 \| AF-A0A6H2GWJ3-F1-MODEL\_V4 \| 1.0 \| 6.903e-05 \| 206 \| 0.136 \| 147 \| 97 \| 7 \| 10 \| 136 \| 1 \| 137 \| LysM peptidoglycan-binding domain-containing protein \| LysM peptidoglycan-binding domain-containing protein; afdb-uniprot50 \| AF-D4LIC3-F1-MODEL\_V4 \| 1.0 \| 0.0001541 \| 206 \| 0.108 \| 138 \| 100 \| 8 \| 13 \| 136 \| 3 \| 131 \| LysM domain \| LysM domain; afdb-uniprot50 \| AF-A0A2D5YZE5-F1-MODEL\_V4 \| 1.0 \| 9.022e-05 \| 205 \| 0.14 \| 128 \| 98 \| 8 \| 14 \| 136 \| 1 \| 121 \| Uncharacterized protein \| Uncharacterized protein; afdb-uniprot50 \| AF-A0A3D3FCD8-F1-MODEL\_V4 \| 1.0 \| 0.0001541 \| 205 \| 0.1 \| 130 \| 100 \| 6 \| 10 \| 136 \| 1 \| 116 \| DNA\_circ\_N domain-containing protein \| DNA\_circ\_N domain-containing protein; afdb-uniprot50 \| AF-L9WCV5-F1-MODEL\_V4 \| 1.0 \| 6.201e-05 \| 205 \| 0.115 \| 138 \| 110 \| 6 \| 1 \| 135 \| 1 \| 129 \| Uncharacterized protein \| Uncharacterized protein; afdb-uniprot50 \| AF-A0A0S2ZJ55-F1-MODEL\_V4 \| 1.0 \| 0.0001244 \| 205 \| 0.104 \| 134 \| 104 \| 6 \| 6 \| 136 \| 3 \| 123 \| Uncharacterized protein \| Uncharacterized protein; afdb-uniprot50 \| AF-A0A1X0SYI6-F1-MODEL\_V4 \| 1.0 \| 0.0004265 \| 204 \| 0.081 \| 123 \| 100 \| 5 \| 14 \| 136 \| 13 \| 122 \| DNA circulation family protein \| DNA circulation family protein; afdb-uniprot50 \| AF-A0A2V2E9H6-F1-MODEL\_V4 \| 1.0 \| 0.0002243 \| 204 \| 0.12 \| 125 \| 99 \| 4 \| 14 \| 136 \| 4 \| 119 \| Uncharacterized protein \| Uncharacterized protein; afdb-uniprot50 \| AF-A0A1C5SWE6-F1-MODEL\_V4 \| 1.0 \| 0.000191 \| 204 \| 0.088 \| 136 \| 106 \| 5 \| 14 \| 136 \| 4 \| 134 \| LysM domain/BON superfamily protein \| LysM domain/BON superfamily protein; afdb-uniprot50 \| AF-A0A838RHN1-F1-MODEL\_V4 \| 1.0 \| 7.683e-05 \| 204 \| 0.102 \| 147 \| 107 \| 8 \| 13 \| 136 \| 9 \| 153 \| LysM peptidoglycan-binding domain-containing protein \| LysM peptidoglycan-binding domain-containing protein; afdb-uniprot50 \| AF-A0A1H8U727-F1-MODEL\_V4 \| 1.0 \| 7.282e-05 \| 204 \| 0.112 \| 160 \| 108 \| 11 \| 3 \| 136 \| 40 \| 191 \| Uncharacterized protein \| Uncharacterized protein; afdb-uniprot50 \| AF-A0A5E6YD38-F1-MODEL\_V4 \| 1.0 \| 0.0008556 \| 203 \| 0.308 \| 94 \| 58 \| 1 \| 10 \| 96 \| 3 \| 96 \| Uncharacterized protein \| Uncharacterized protein; afdb-uniprot50 \| AF-A0A6M3M430-F1-MODEL\_V4 \| 1.0 \| 0.0001244 \| 203 \| 0.113 \| 159 \| 111 \| 7 \| 1 \| 136 \| 26 \| 177 \| Uncharacterized protein \| Uncharacterized protein; afdb-uniprot50 \| AF-A0A845W825-F1-MODEL\_V4 \| 1.0 \| 2.632e-05 \| 203 \| 0.108 \| 193 \| 114 \| 10 \| 1 \| 136 \| 20 \| 211 \| LysM peptidoglycan-binding domain-containing protein \| LysM peptidoglycan-binding domain-containing protein; afdb-uniprot50 \| AF-A0A349URD4-F1-MODEL\_V4 \| 1.0 \| 0.0001461 \| 202 \| 0.157 \| 127 \| 97 \| 7 \| 14 \| 136 \| 1 \| 121 \| Uncharacterized protein \| Uncharacterized protein; afdb-uniprot50 \| AF-A0A3A6HVN4-F1-MODEL\_V4 \| 1.0 \| 0.0002366 \| 202 \| 0.139 \| 129 \| 96 \| 4 \| 13 \| 136 \| 1 \| 119 \| Uncharacterized protein \| Uncharacterized protein; afdb-uniprot50 \| AF-A0A1C5ZE84-F1-MODEL\_V4 \| 1.0 \| 0.0004499 \| 202 \| 0.115 \| 113 \| 90 \| 4 \| 24 \| 136 \| 17 \| 119 \| Uncharacterized protein \| Uncharacterized protein; afdb-uniprot50 \| AF-A0A0R2DDQ3-F1-MODEL\_V4 \| 1.0 \| 0.0001716 \| 202 \| 0.147 \| 115 \| 81 \| 5 \| 27 \| 136 \| 234 \| 336 \| Uncharacterized protein \| Uncharacterized protein; afdb-uniprot50 \| AF-A0A2C8EVW5-F1-MODEL\_V4 \| 1.0 \| 0.001245 \| 201 \| 0.396 \| 63 \| 38 \| 0 \| 73 \| 135 \| 3 \| 65 \| Phage protein U \| Phage protein U; afdb-uniprot50 \| AF-A0A7X0IVG1-F1-MODEL\_V4 \| 1.0 \| 9.022e-05 \| 201 \| 0.196 \| 112 \| 85 \| 3 \| 28 \| 136 \| 4 \| 113 \| Uncharacterized protein \| Uncharacterized protein; afdb-uniprot50 \| AF-A0A1V5B941-F1-MODEL\_V4 \| 1.0 \| 3.829e-05 \| 201 \| 0.156 \| 134 \| 98 \| 8 \| 13 \| 136 \| 8 \| 136 \| LysM domain/BON superfamily protein \| LysM domain/BON superfamily protein; afdb-uniprot50 \| AF-A0A562STB2-F1-MODEL\_V4 \| 1.0 \| 8.552e-05 \| 201 \| 0.166 \| 120 \| 86 \| 4 \| 29 \| 134 \| 30 \| 149 \| LysM domain-containing protein \| LysM domain-containing protein; afdb-uniprot50 \| AF-A0A661PHT0-F1-MODEL\_V4 \| 1.0 \| 0.0002366 \| 201 \| 0.136 \| 154 \| 91 \| 7 \| 15 \| 136 \| 16 \| 159 \| Uncharacterized protein \| Uncharacterized protein; afdb-uniprot50 \| AF-A0A1Z5IQW4-F1-MODEL\_V4 \| 1.0 \| 0.0002243 \| 201 \| 0.095 \| 115 \| 97 \| 3 \| 27 \| 136 \| 140 \| 252 \| LysM domain protein \| LysM domain protein; afdb-uniprot50 \| AF-A0A0C1PPW5-F1-MODEL\_V4 \| 1.0 \| 0.0001626 \| 201 \| 0.068 \| 131 \| 109 \| 5 \| 11 \| 136 \| 219 \| 341 \| Muramidase (Flagellum-specific) \| Muramidase (Flagellum-specific); afdb-uniprot50 \| AF-A0A5C4LKH5-F1-MODEL\_V4 \| 1.0 \| 0.0005283 \| 200 \| 0.128 \| 109 \| 86 \| 3 \| 28 \| 136 \| 4 \| 103 \| Uncharacterized protein \| Uncharacterized protein; afdb-uniprot50 \| AF-A0A4R2CQ90-F1-MODEL\_V4 \| 1.0 \| 0.0002778 \| 200 \| 0.161 \| 136 \| 95 \| 7 \| 10 \| 136 \| 1 \| 126 \| Uncharacterized protein \| Uncharacterized protein; afdb-uniprot50 \| AF-A0A3B7MQB9-F1-MODEL\_V4 \| 1.0 \| 9.519e-05 \| 200 \| 0.111 \| 143 \| 99 \| 5 \| 21 \| 136 \| 32 \| 173 \| LysM peptidoglycan-binding domain-containing protein \| LysM peptidoglycan-binding domain-containing protein; afdb-uniprot50 \| AF-A0A7W1N1F9-F1-MODEL\_V4 \| 1.0 \| 8.106e-05 \| 200 \| 0.107 \| 168 \| 102 \| 9 \| 7 \| 136 \| 1 \| 158 \| Uncharacterized protein \| Uncharacterized protein; afdb-uniprot50 \| AF-A0A2E7P7M5-F1-MODEL\_V4 \| 1.0 \| 0.0001004 \| 200 \| 0.121 \| 140 \| 107 \| 7 \| 1 \| 136 \| 1 \| 128 \| DNA\_circ\_N domain-containing protein \| DNA\_circ\_N domain-containing protein; afdb-uniprot50 \| AF-B0PCG8-F1-MODEL\_V4 \| 1.0 \| 0.0004265 \| 199 \| 0.119 \| 126 \| 93 \| 6 \| 16 \| 136 \| 2 \| 114 \| Uncharacterized protein \| Uncharacterized protein; afdb-uniprot50 \| AF-A0A5C7QIA0-F1-MODEL\_V4 \| 1.0 \| 7.282e-05 \| 199 \| 0.143 \| 132 \| 94 \| 8 \| 14 \| 136 \| 1 \| 122 \| Uncharacterized protein \| Uncharacterized protein; afdb-uniprot50 \| AF-A0A133ZRU1-F1-MODEL\_V4 \| 1.0 \| 0.0002633 \| 199 \| 0.126 \| 134 \| 97 \| 7 \| 14 \| 136 \| 8 \| 132 \| Uncharacterized protein \| Uncharacterized protein; afdb-uniprot50 \| AF-A0A6M3XXM1-F1-MODEL\_V4 \| 1.0 \| 6.903e-05 \| 198 \| 0.106 \| 141 \| 108 \| 8 \| 6 \| 136 \| 3 \| 135 \| Uncharacterized protein \| Uncharacterized protein; afdb-uniprot50 \| AF-A0A6L9ZDI7-F1-MODEL\_V4 \| 1.0 \| 8.106e-05 \| 198 \| 0.15 \| 159 \| 107 \| 8 \| 1 \| 134 \| 1 \| 156 \| LysM domain-containing protein \| LysM domain-containing protein; afdb-uniprot50 \| AF-A0A416BNH2-F1-MODEL\_V4 \| 1.0 \| 0.0001385 \| 198 \| 0.092 \| 130 \| 105 \| 5 \| 14 \| 136 \| 4 \| 127 \| LysM peptidoglycan-binding domain-containing protein \| LysM peptidoglycan-binding domain-containing protein; afdb-uniprot50 \| AF-A0A072R0Y2-F1-MODEL\_V4 \| 1.0 \| 0.001717 \| 197 \| 0.162 \| 117 \| 92 \| 1 \| 10 \| 120 \| 1 \| 117 \| Uncharacterized protein \| Uncharacterized protein; afdb-uniprot50 \| AF-A0A6B2FIX0-F1-MODEL\_V4 \| 1.0 \| 0.0003831 \| 197 \| 0.07 \| 142 \| 107 \| 5 \| 11 \| 136 \| 9 \| 141 \| Uncharacterized protein \| Uncharacterized protein; afdb-uniprot50 \| AF-A0A357B502-F1-MODEL\_V4 \| 1.0 \| 0.0007286 \| 197 \| 0.084 \| 119 \| 98 \| 4 \| 16 \| 134 \| 2 \| 109 \| Peptidoglycan-binding protein LysM \| Peptidoglycan-binding protein LysM; afdb-uniprot50 \| AF-A0A7U0I7H4-F1-MODEL\_V4 \| 1.0 \| 0.0001179 \| 196 \| 0.158 \| 151 \| 94 \| 8 \| 13 \| 136 \| 4 \| 148 \| Uncharacterized protein \| Uncharacterized protein; afdb-uniprot50 \| AF-A0A7S9CMN8-F1-MODEL\_V4 \| 1.0 \| 0.0002931 \| 196 \| 0.115 \| 130 \| 104 \| 6 \| 10 \| 136 \| 37 \| 158 \| Uncharacterized protein \| Uncharacterized protein; afdb-uniprot50 \| AF-A0A2W6YVY0-F1-MODEL\_V4 \| 1.0 \| 0.0002778 \| 196 \| 0.1 \| 129 \| 99 \| 7 \| 11 \| 136 \| 3 \| 117 \| DNA\_circ\_N domain-containing protein \| DNA\_circ\_N domain-containing protein; afdb-uniprot50 \| AF-A0A174LM59-F1-MODEL\_V4 \| 1.0 \| 0.000181 \| 195 \| 0.124 \| 129 \| 96 \| 3 \| 20 \| 136 \| 13 \| 136 \| LysM domain/BON superfamily protein \| LysM domain/BON superfamily protein; afdb-uniprot50 \| AF-A0A412CG89-F1-MODEL\_V4 \| 1.0 \| 0.0004042 \| 195 \| 0.137 \| 153 \| 105 \| 7 \| 2 \| 136 \| 82 \| 225 \| Uncharacterized protein \| Uncharacterized protein; afdb-uniprot50 \| AF-A0A140PTD7-F1-MODEL\_V4 \| 1.0 \| 0.0002778 \| 194 \| 0.116 \| 129 \| 98 \| 6 \| 11 \| 136 \| 17 \| 132 \| Uncharacterized protein \| Uncharacterized protein; afdb-uniprot50 \| AF-M9LYN7-F1-MODEL\_V4 \| 1.0 \| 0.0004747 \| 194 \| 0.087 \| 137 \| 106 \| 7 \| 13 \| 136 \| 1 \| 131 \| LysM domain-containing protein \| LysM domain-containing protein; afdb-uniprot50 \| AF-A0A246IW45-F1-MODEL\_V4 \| 1.0 \| 3.63e-05 \| 194 \| 0.176 \| 147 \| 98 \| 6 \| 13 \| 136 \| 9 \| 155 \| Uncharacterized protein \| Uncharacterized protein; afdb-uniprot50 \| AF-A0A542ARR8-F1-MODEL\_V4 \| 1.0 \| 0.0002243 \| 194 \| 0.087 \| 137 \| 103 \| 8 \| 11 \| 136 \| 3 \| 128 \| Uncharacterized protein \| Uncharacterized protein; afdb-uniprot50 \| AF-A0A3S5K2M8-F1-MODEL\_V4 \| 1.0 \| 7.683e-05 \| 194 \| 0.104 \| 162 \| 110 \| 11 \| 1 \| 136 \| 4 \| 156 \| Uncharacterized protein \| Uncharacterized protein; afdb-uniprot50 \| AF-A0A7Z6QNR3-F1-MODEL\_V4 \| 1.0 \| 0.0002778 \| 194 \| 0.083 \| 131 \| 105 \| 7 \| 13 \| 136 \| 197 \| 319 \| LysM peptidoglycan-binding domain-containing protein \| LysM peptidoglycan-binding domain-containing protein; afdb-uniprot50 \| AF-A0A2P5GVD3-F1-MODEL\_V4 \| 1.0 \| 0.0004265 \| 193 \| 0.164 \| 134 \| 99 \| 7 \| 6 \| 136 \| 1 \| 124 \| Uncharacterized protein \| Uncharacterized protein; afdb-uniprot50 \| AF-A0A653QVV7-F1-MODEL\_V4 \| 1.0 \| 0.0001313 \| 193 \| 0.082 \| 134 \| 107 \| 7 \| 8 \| 136 \| 9 \| 131 \| Uncharacterized protein \| Uncharacterized protein; afdb-uniprot50 \| AF-A0A1C6CLA3-F1-MODEL\_V4 \| 1.0 \| 0.0001461 \| 193 \| 0.159 \| 138 \| 98 \| 8 \| 10 \| 136 \| 1 \| 131 \| LysM domain/BON superfamily protein \| LysM domain/BON superfamily protein; afdb-uniprot50 \| AF-A0A1B9AUD6-F1-MODEL\_V4 \| 1.0 \| 0.0001385 \| 193 \| 0.124 \| 137 \| 103 \| 6 \| 13 \| 136 \| 4 \| 136 \| LysM domain-containing protein \| LysM domain-containing protein; afdb-uniprot50 \| AF-R8W901-F1-MODEL\_V4 \| 1.0 \| 0.0001118 \| 193 \| 0.147 \| 142 \| 96 \| 7 \| 14 \| 136 \| 3 \| 138 \| LysM domain-containing protein \| LysM domain-containing protein; afdb-uniprot50 \| AF-A0A3N9F9V1-F1-MODEL\_V4 \| 1.0 \| 0.0003442 \| 193 \| 0.119 \| 184 \| 108 \| 7 \| 6 \| 136 \| 2 \| 184 \| Uncharacterized protein \| Uncharacterized protein; afdb-uniprot50 \| AF-J4UC17-F1-MODEL\_V4 \| 1.0 \| 0.0002243 \| 192 \| 0.095 \| 136 \| 102 \| 7 \| 14 \| 136 \| 3 \| 130 \| Uncharacterized protein \| Uncharacterized protein; afdb-uniprot50 \| AF-K1JXW6-F1-MODEL\_V4 \| 1.0 \| 0.0001004 \| 192 \| 0.122 \| 131 \| 99 \| 7 \| 13 \| 136 \| 3 \| 124 \| Uncharacterized protein \| Uncharacterized protein; afdb-uniprot50 \| AF-A0A2P5KA36-F1-MODEL\_V4 \| 1.0 \| 0.0003093 \| 192 \| 0.148 \| 141 \| 103 \| 8 \| 1 \| 136 \| 2 \| 130 \| Uncharacterized protein \| Uncharacterized protein; afdb-uniprot50 \| AF-A0A7X0Y1A5-F1-MODEL\_V4 \| 1.0 \| 0.0003632 \| 192 \| 0.094 \| 127 \| 96 \| 5 \| 14 \| 136 \| 1 \| 112 \| LysM domain-containing protein \| LysM domain-containing protein; afdb-uniprot50 \| AF-A0A373VS92-F1-MODEL\_V4 \| 1.0 \| 0.0006204 \| 192 \| 0.111 \| 135 \| 101 \| 7 \| 14 \| 136 \| 1 \| 128 \| LysM domain-containing protein \| LysM domain-containing protein; afdb-uniprot50 \| AF-A0A0N0E150-F1-MODEL\_V4 \| 1.0 \| 0.0005008 \| 192 \| 0.069 \| 130 \| 111 \| 5 \| 12 \| 136 \| 49 \| 173 \| Uncharacterized protein \| Uncharacterized protein; afdb-uniprot50 \| AF-A0A849CT25-F1-MODEL\_V4 \| 1.0 \| 0.0006204 \| 191 \| 0.207 \| 101 \| 74 \| 3 \| 40 \| 136 \| 11 \| 109 \| Phage tail protein \| Phage tail protein; afdb-uniprot50 \| AF-A0A413PG52-F1-MODEL\_V4 \| 1.0 \| 0.0003263 \| 191 \| 0.1 \| 129 \| 101 \| 8 \| 13 \| 136 \| 3 \| 121 \| Uncharacterized protein \| Uncharacterized protein; afdb-uniprot50 \| AF-A0A176XP81-F1-MODEL\_V4 \| 1.0 \| 0.0006204 \| 191 \| 0.077 \| 129 \| 110 \| 4 \| 9 \| 136 \| 11 \| 131 \| Uncharacterized protein \| Uncharacterized protein; afdb-uniprot50 \| AF-E5Y1X5-F1-MODEL\_V4 \| 1.0 \| 0.0004265 \| 191 \| 0.125 \| 144 \| 108 \| 7 \| 1 \| 136 \| 5 \| 138 \| Uncharacterized protein \| Uncharacterized protein; afdb-uniprot50 \| AF-A0A6M8UDV6-F1-MODEL\_V4 \| 1.0 \| 0.0002126 \| 191 \| 0.113 \| 158 \| 108 \| 10 \| 1 \| 136 \| 6 \| 153 \| Uncharacterized protein \| Uncharacterized protein; afdb-uniprot50 \| AF-C0Q8T6-F1-MODEL\_V4 \| 1.0 \| 6.543e-05 \| 191 \| 0.156 \| 128 \| 94 \| 4 \| 20 \| 136 \| 27 \| 151 \| LysM domain-containing protein \| LysM domain-containing protein; afdb-uniprot50 \| AF-A0A193QH76-F1-MODEL\_V4 \| 1.0 \| 0.0003831 \| 191 \| 0.132 \| 128 \| 91 \| 5 \| 24 \| 136 \| 49 \| 171 \| Uncharacterized protein \| Uncharacterized protein; afdb-uniprot50 \| AF-A0A0R2CKH2-F1-MODEL\_V4 \| 1.0 \| 0.0003263 \| 191 \| 0.089 \| 112 \| 93 \| 5 \| 27 \| 136 \| 143 \| 247 \| Uncharacterized protein \| Uncharacterized protein; afdb-uniprot50 \| AF-A0A7C6NZD4-F1-MODEL\_V4 \| 1.0 \| 0.0002778 \| 190 \| 0.076 \| 144 \| 107 \| 6 \| 13 \| 136 \| 6 \| 143 \| LysM peptidoglycan-binding domain-containing protein \| LysM peptidoglycan-binding domain-containing protein; afdb-uniprot50 \| AF-A0A350PIS1-F1-MODEL\_V4 \| 1.0 \| 0.0003442 \| 190 \| 0.092 \| 163 \| 113 \| 7 \| 5 \| 133 \| 17 \| 178 \| Uncharacterized protein \| Uncharacterized protein; afdb-uniprot50 \| AF-A0A348HFQ1-F1-MODEL\_V4 \| 1.0 \| 0.0003632 \| 190 \| 0.094 \| 127 \| 108 \| 5 \| 13 \| 136 \| 61 \| 183 \| Glutamate decarboxylase \| Glutamate decarboxylase; afdb-uniprot50 \| AF-Z9JLS4-F1-MODEL\_V4 \| 1.0 \| 0.0005283 \| 189 \| 0.328 \| 73 \| 49 \| 0 \| 64 \| 136 \| 1 \| 73 \| Uncharacterized protein \| Uncharacterized protein; afdb-uniprot50 \| AF-A0A0F9D999-F1-MODEL\_V4 \| 1.0 \| 0.0004747 \| 189 \| 0.141 \| 127 \| 99 \| 5 \| 13 \| 136 \| 1 \| 120 \| Uncharacterized protein \| Uncharacterized protein; afdb-uniprot50 \| AF-A0A7C7EU00-F1-MODEL\_V4 \| 1.0 \| 0.0005283 \| 189 \| 0.075 \| 132 \| 101 \| 6 \| 7 \| 134 \| 1 \| 115 \| LysM peptidoglycan-binding domain-containing protein \| LysM peptidoglycan-binding domain-containing protein; afdb-uniprot50 \| AF-A0A5Q4Z738-F1-MODEL\_V4 \| 1.0 \| 0.0007286 \| 189 \| 0.112 \| 116 \| 91 \| 4 \| 28 \| 136 \| 25 \| 135 \| Uncharacterized protein \| Uncharacterized protein; afdb-uniprot50 \| AF-A0A317H4C1-F1-MODEL\_V4 \| 1.0 \| 0.0002778 \| 189 \| 0.125 \| 144 \| 107 \| 9 \| 1 \| 136 \| 1 \| 133 \| DNA\_circ\_N domain-containing protein \| DNA\_circ\_N domain-containing protein; afdb-uniprot50 \| AF-A0A174KNK1-F1-MODEL\_V4 \| 1.0 \| 0.0003093 \| 189 \| 0.1 \| 140 \| 103 \| 8 \| 11 \| 136 \| 3 \| 133 \| Uncharacterized protein \| Uncharacterized protein; afdb-uniprot50 \| AF-A0A5M6IU86-F1-MODEL\_V4 \| 1.0 \| 0.0006906 \| 189 \| 0.135 \| 133 \| 101 \| 6 \| 4 \| 132 \| 18 \| 140 \| Uncharacterized protein \| Uncharacterized protein; afdb-uniprot50 \| AF-A0A178TQB9-F1-MODEL\_V4 \| 1.0 \| 0.0002243 \| 189 \| 0.136 \| 132 \| 104 \| 5 \| 13 \| 136 \| 148 \| 277 \| Uncharacterized protein \| Uncharacterized protein; afdb-uniprot50 \| AF-A0A2G5KZW3-F1-MODEL\_V4 \| 1.0 \| 0.0002633 \| 188 \| 0.074 \| 147 \| 107 \| 7 \| 13 \| 136 \| 2 \| 142 \| Uncharacterized protein \| Uncharacterized protein; afdb-uniprot50 \| AF-A0A1H9HQS4-F1-MODEL\_V4 \| 1.0 \| 0.0008556 \| 188 \| 0.09 \| 155 \| 108 \| 6 \| 6 \| 136 \| 10 \| 155 \| Uncharacterized protein \| Uncharacterized protein; afdb-uniprot50 \| AF-A0A410WT00-F1-MODEL\_V4 \| 1.0 \| 0.0003093 \| 188 \| 0.113 \| 132 \| 107 \| 6 \| 13 \| 136 \| 147 \| 276 \| Uncharacterized protein \| Uncharacterized protein; afdb-uniprot50 \| AF-A0A0N0UTD2-F1-MODEL\_V4 \| 1.0 \| 0.0004499 \| 188 \| 0.085 \| 128 \| 103 \| 7 \| 14 \| 136 \| 188 \| 306 \| Uncharacterized protein \| Uncharacterized protein; afdb-uniprot50 \| AF-A0A0G3CNH3-F1-MODEL\_V4 \| 1.0 \| 0.0004265 \| 187 \| 0.11 \| 127 \| 89 \| 4 \| 29 \| 136 \| 33 \| 154 \| Uncharacterized protein \| Uncharacterized protein; afdb-uniprot50 \| AF-A0A1Q3NUK7-F1-MODEL\_V4 \| 1.0 \| 0.0006204 \| 187 \| 0.097 \| 133 \| 104 \| 7 \| 6 \| 134 \| 1 \| 121 \| DNA\_circ\_N domain-containing protein \| DNA\_circ\_N domain-containing protein; afdb-uniprot50 \| AF-A0A763M6N7-F1-MODEL\_V4 \| 1.0 \| 0.001245 \| 187 \| 0.238 \| 88 \| 67 \| 0 \| 48 \| 135 \| 513 \| 600 \| Phage tail tape measure protein \| Phage tail tape measure protein; afdb-uniprot50 \| AF-A0A3C0RSF2-F1-MODEL\_V4 \| 1.0 \| 0.002016 \| 186 \| 0.148 \| 121 \| 88 \| 5 \| 13 \| 124 \| 3 \| 117 \| Uncharacterized protein \| Uncharacterized protein; afdb-uniprot50 \| AF-A0A133NAR5-F1-MODEL\_V4 \| 1.0 \| 0.0002015 \| 186 \| 0.145 \| 131 \| 92 \| 5 \| 11 \| 136 \| 21 \| 136 \| Uncharacterized protein \| Uncharacterized protein; afdb-uniprot50 \| AF-A0A6N1B959-F1-MODEL\_V4 \| 1.0 \| 0.0005008 \| 186 \| 0.072 \| 152 \| 110 \| 7 \| 7 \| 136 \| 1 \| 143 \| Uncharacterized protein \| Uncharacterized protein; afdb-uniprot50 \| AF-A0A1Z5IUN3-F1-MODEL\_V4 \| 1.0 \| 0.0005283 \| 186 \| 0.097 \| 134 \| 104 \| 6 \| 9 \| 136 \| 198 \| 320 \| LysM domain-containing protein \| LysM domain-containing protein; afdb-uniprot50 \| AF-A0A3C0YSN0-F1-MODEL\_V4 \| 1.0 \| 0.0002931 \| 185 \| 0.093 \| 128 \| 105 \| 4 \| 13 \| 136 \| 1 \| 121 \| Uncharacterized protein \| Uncharacterized protein; afdb-uniprot50 \| AF-A0A401X7L6-F1-MODEL\_V4 \| 1.0 \| 0.0004265 \| 185 \| 0.092 \| 130 \| 113 \| 4 \| 8 \| 136 \| 59 \| 184 \| Uncharacterized protein \| Uncharacterized protein; afdb-uniprot50 \| AF-A0A3R6PJA9-F1-MODEL\_V4 \| 1.0 \| 0.0004265 \| 185 \| 0.129 \| 139 \| 103 \| 7 \| 8 \| 136 \| 23 \| 153 \| LysM peptidoglycan-binding domain-containing protein \| LysM peptidoglycan-binding domain-containing protein; afdb-uniprot50 \| AF-R0EE86-F1-MODEL\_V4 \| 1.0 \| 0.0003093 \| 185 \| 0.12 \| 150 \| 104 \| 10 \| 4 \| 136 \| 28 \| 166 \| Mu-like prophage DNA circulation protein \| Mu-like prophage DNA circulation protein; afdb-uniprot50 \| AF-A0A2V2A0Z3-F1-MODEL\_V4 \| 1.0 \| 0.003833 \| 184 \| 0.25 \| 76 \| 55 \| 1 \| 61 \| 136 \| 29 \| 102 \| GpU protein \| GpU protein; afdb-uniprot50 \| AF-A0A317HDG5-F1-MODEL\_V4 \| 1.0 \| 0.0003263 \| 184 \| 0.117 \| 145 \| 108 \| 6 \| 1 \| 136 \| 16 \| 149 \| Uncharacterized protein \| Uncharacterized protein; afdb-uniprot50 \| AF-A0A352S729-F1-MODEL\_V4 \| 1.0 \| 0.0003831 \| 184 \| 0.107 \| 139 \| 99 \| 6 \| 13 \| 136 \| 39 \| 167 \| DNA circulation family protein \| DNA circulation family protein; afdb-uniprot50 \| AF-A0A5M6IYQ8-F1-MODEL\_V4 \| 1.0 \| 0.0007286 \| 183 \| 0.156 \| 134 \| 89 \| 8 \| 10 \| 136 \| 1 \| 117 \| Uncharacterized protein \| Uncharacterized protein; afdb-uniprot50 \| AF-A0A661VCX6-F1-MODEL\_V4 \| 1.0 \| 0.0005283 \| 183 \| 0.121 \| 123 \| 96 \| 6 \| 13 \| 133 \| 7 \| 119 \| DNA\_circ\_N domain-containing protein \| DNA\_circ\_N domain-containing protein; afdb-uniprot50 \| AF-A0A447N0F3-F1-MODEL\_V4 \| 1.0 \| 0.004044 \| 182 \| 0.378 \| 82 \| 51 \| 0 \| 13 \| 94 \| 1 \| 82 \| Gp25 \| Gp25; afdb-uniprot50 \| AF-A0A349YQ66-F1-MODEL\_V4 \| 1.0 \| 0.001118 \| 182 \| 0.088 \| 124 \| 100 \| 5 \| 14 \| 136 \| 1 \| 112 \| LysM domain-containing protein \| LysM domain-containing protein; afdb-uniprot50 \| AF-A0A2E5S6T5-F1-MODEL\_V4 \| 1.0 \| 0.0003831 \| 182 \| 0.126 \| 150 \| 108 \| 8 \| 1 \| 136 \| 6 \| 146 \| Uncharacterized protein \| Uncharacterized protein; afdb-uniprot50 \| AF-A0A4R7PWP3-F1-MODEL\_V4 \| 1.0 \| 0.001005 \| 182 \| 0.097 \| 144 \| 100 \| 5 \| 14 \| 136 \| 12 \| 146 \| Uncharacterized protein \| Uncharacterized protein; afdb-uniprot50 \| AF-A0A3B7DR28-F1-MODEL\_V4 \| 1.0 \| 0.0004499 \| 182 \| 0.119 \| 142 \| 110 \| 8 \| 1 \| 136 \| 35 \| 167 \| Uncharacterized protein \| Uncharacterized protein; afdb-uniprot50 \| AF-A0A2W4KCL7-F1-MODEL\_V4 \| 1.0 \| 0.001118 \| 182 \| 0.065 \| 137 \| 107 \| 6 \| 14 \| 136 \| 3 \| 132 \| Peptidoglycan-binding protein \| Peptidoglycan-binding protein; afdb-uniprot50 \| AF-A0A661FPN4-F1-MODEL\_V4 \| 1.0 \| 0.0002633 \| 182 \| 0.138 \| 130 \| 90 \| 4 \| 29 \| 136 \| 38 \| 167 \| Peptidoglycan-binding protein \| Peptidoglycan-binding protein; afdb-uniprot50 \| AF-A0A7C9KHI7-F1-MODEL\_V4 \| 1.0 \| 0.0003263 \| 181 \| 0.128 \| 132 \| 101 \| 6 \| 13 \| 136 \| 3 \| 128 \| Uncharacterized protein \| Uncharacterized protein; afdb-uniprot50 \| AF-A0A3D3SD07-F1-MODEL\_V4 \| 1.0 \| 0.0006204 \| 181 \| 0.107 \| 139 \| 113 \| 6 \| 1 \| 136 \| 1 \| 131 \| Uncharacterized protein \| Uncharacterized protein; afdb-uniprot50 \| AF-A0A2Z3I6W3-F1-MODEL\_V4 \| 1.0 \| 0.0002243 \| 181 \| 0.161 \| 136 \| 99 \| 8 \| 5 \| 136 \| 4 \| 128 \| Uncharacterized protein \| Uncharacterized protein; afdb-uniprot50 \| AF-A0A0M0X770-F1-MODEL\_V4 \| 1.0 \| 0.0008556 \| 181 \| 0.137 \| 131 \| 96 \| 5 \| 10 \| 136 \| 2 \| 119 \| LysM domain-containing protein \| LysM domain-containing protein; afdb-uniprot50 \| AF-A0A495RHL5-F1-MODEL\_V4 \| 1.0 \| 0.0007687 \| 180 \| 0.117 \| 205 \| 103 \| 5 \| 1 \| 136 \| 4 \| 199 \| Uncharacterized protein \| Uncharacterized protein; afdb-uniprot50 \| AF-A0A2T6LTV8-F1-MODEL\_V4 \| 1.0 \| 0.001717 \| 179 \| 0.096 \| 125 \| 102 \| 6 \| 13 \| 136 \| 5 \| 119 \| Uncharacterized protein \| Uncharacterized protein; afdb-uniprot50 \| AF-A0A2T0QI76-F1-MODEL\_V4 \| 1.0 \| 0.000811 \| 179 \| 0.094 \| 137 \| 101 \| 6 \| 14 \| 136 \| 27 \| 154 \| Uncharacterized protein \| Uncharacterized protein; afdb-uniprot50 \| AF-A0A2G3Q549-F1-MODEL\_V4 \| 1.0 \| 0.0006204 \| 179 \| 0.11 \| 127 \| 96 \| 5 \| 23 \| 136 \| 10 \| 132 \| Peptidoglycan-binding protein \| Peptidoglycan-binding protein; afdb-uniprot50 \| AF-A0A4P5VDU8-F1-MODEL\_V4 \| 1.0 \| 0.0001179 \| 179 \| 0.138 \| 130 \| 87 \| 7 \| 29 \| 134 \| 39 \| 167 \| Uncharacterized protein \| Uncharacterized protein; afdb-uniprot50 \| AF-A0A2D9CHW4-F1-MODEL\_V4 \| 1.0 \| 0.001811 \| 178 \| 0.089 \| 112 \| 94 \| 4 \| 28 \| 136 \| 28 \| 134 \| Uncharacterized protein \| Uncharacterized protein; afdb-uniprot50 \| AF-A0A2B7YZ51-F1-MODEL\_V4 \| 1.0 \| 0.0003263 \| 178 \| 0.121 \| 132 \| 100 \| 6 \| 8 \| 136 \| 5 \| 123 \| Uncharacterized protein \| Uncharacterized protein; afdb-uniprot50 \| AF-A0A1H8DHX8-F1-MODEL\_V4 \| 1.0 \| 0.0007286 \| 178 \| 0.109 \| 164 \| 106 \| 9 \| 2 \| 136 \| 57 \| 209 \| Uncharacterized protein \| Uncharacterized protein; afdb-uniprot50 \| AF-A0A0A0IRV2-F1-MODEL\_V4 \| 1.0 \| 0.0004265 \| 178 \| 0.141 \| 134 \| 87 \| 10 \| 12 \| 136 \| 2 \| 116 \| Uncharacterized protein \| Uncharacterized protein; afdb-uniprot50 \| AF-A0A6L9ABV2-F1-MODEL\_V4 \| 1.0 \| 0.002016 \| 177 \| 0.229 \| 96 \| 73 \| 1 \| 13 \| 108 \| 1 \| 95 \| Uncharacterized protein \| Uncharacterized protein; afdb-uniprot50 \| AF-A0A257VSV7-F1-MODEL\_V4 \| 1.0 \| 0.001386 \| 177 \| 0.094 \| 127 \| 102 \| 4 \| 14 \| 136 \| 18 \| 135 \| Uncharacterized protein \| Uncharacterized protein; afdb-uniprot50 \| AF-A0A5Q2VGY7-F1-MODEL\_V4 \| 1.0 \| 0.0008556 \| 177 \| 0.097 \| 164 \| 110 \| 8 \| 1 \| 136 \| 1 \| 154 \| Uncharacterized protein \| Uncharacterized protein; afdb-uniprot50 \| AF-A0A158D0R9-F1-MODEL\_V4 \| 1.0 \| 0.0006546 \| 177 \| 0.117 \| 136 \| 111 \| 5 \| 4 \| 136 \| 41 \| 170 \| Uncharacterized protein \| Uncharacterized protein; afdb-uniprot50 \| AF-A0A843GUK1-F1-MODEL\_V4 \| 1.0 \| 0.0007286 \| 176 \| 0.102 \| 136 \| 104 \| 6 \| 10 \| 136 \| 60 \| 186 \| Uncharacterized protein \| Uncharacterized protein; afdb-uniprot50 \| AF-A0A097R3F3-F1-MODEL\_V4 \| 1.0 \| 0.0006906 \| 176 \| 0.117 \| 153 \| 101 \| 9 \| 8 \| 136 \| 12 \| 154 \| Uncharacterized protein \| Uncharacterized protein; afdb-uniprot50 \| AF-A0A239MZ86-F1-MODEL\_V4 \| 1.0 \| 0.0009027 \| 175 \| 0.145 \| 137 \| 108 \| 5 \| 4 \| 136 \| 41 \| 172 \| Uncharacterized protein \| Uncharacterized protein; afdb-uniprot50 \| AF-A0A5M5NPH4-F1-MODEL\_V4 \| 1.0 \| 0.0001626 \| 175 \| 0.152 \| 157 \| 103 \| 7 \| 6 \| 136 \| 1 \| 153 \| LysM peptidoglycan-binding domain-containing protein \| LysM peptidoglycan-binding domain-containing protein; afdb-uniprot50 \| AF-A0A0R2FPR3-F1-MODEL\_V4 \| 1.0 \| 0.0003831 \| 175 \| 0.104 \| 143 \| 90 \| 8 \| 27 \| 136 \| 20 \| 157 \| Muramidase (Flagellum-specific) \| Muramidase (Flagellum-specific); afdb-uniprot50 \| AF-A0A0M7MQ27-F1-MODEL\_V4 \| 1.0 \| 0.002367 \| 174 \| 0.081 \| 110 \| 95 \| 4 \| 28 \| 136 \| 63 \| 167 \| Uncharacterized protein \| Uncharacterized protein; afdb-uniprot50 \| AF-A0A1E5QLG5-F1-MODEL\_V4 \| 1.0 \| 9.022e-05 \| 172 \| 0.174 \| 132 \| 89 \| 9 \| 20 \| 136 \| 15 \| 141 \| Uncharacterized protein \| Uncharacterized protein; afdb-uniprot50 \| AF-A0A1Y4GLM1-F1-MODEL\_V4 \| 1.0 \| 0.0002778 \| 172 \| 0.108 \| 138 \| 104 \| 10 \| 7 \| 136 \| 1 \| 127 \| Uncharacterized protein \| Uncharacterized protein; afdb-uniprot50 \| AF-R6NK38-F1-MODEL\_V4 \| 1.0 \| 0.0009523 \| 172 \| 0.094 \| 138 \| 100 \| 7 \| 14 \| 136 \| 7 \| 134 \| Uncharacterized protein \| Uncharacterized protein; afdb-uniprot50 \| AF-A0A2N9X683-F1-MODEL\_V4 \| 1.0 \| 0.001627 \| 172 \| 0.131 \| 129 \| 99 \| 7 \| 13 \| 136 \| 41 \| 161 \| Uncharacterized protein \| Uncharacterized protein; afdb-uniprot50 \| AF-A0A267S2K4-F1-MODEL\_V4 \| 1.0 \| 0.001811 \| 172 \| 0.129 \| 124 \| 87 \| 5 \| 28 \| 136 \| 48 \| 165 \| Uncharacterized protein \| Uncharacterized protein; afdb-uniprot50 \| AF-A0A7C8AGL9-F1-MODEL\_V4 \| 1.0 \| 0.0006546 \| 171 \| 0.139 \| 129 \| 97 \| 10 \| 13 \| 136 \| 1 \| 120 \| Uncharacterized protein \| Uncharacterized protein; afdb-uniprot50 \| AF-A0A661DK94-F1-MODEL\_V4 \| 1.0 \| 0.0006906 \| 171 \| 0.1 \| 129 \| 91 \| 3 \| 28 \| 136 \| 21 \| 144 \| Uncharacterized protein \| Uncharacterized protein; afdb-uniprot50 \| AF-A0A212KEY3-F1-MODEL\_V4 \| 1.0 \| 0.001313 \| 171 \| 0.136 \| 139 \| 100 \| 6 \| 2 \| 136 \| 3 \| 125 \| Uncharacterized protein \| Uncharacterized protein; afdb-uniprot50 \| AF-A0A371YK61-F1-MODEL\_V4 \| 1.0 \| 0.001313 \| 171 \| 0.097 \| 113 \| 89 \| 4 \| 28 \| 136 \| 38 \| 141 \| Uncharacterized protein \| Uncharacterized protein; afdb-uniprot50 \| AF-A0A378E5K6-F1-MODEL\_V4 \| 1.0 \| 0.006549 \| 170 \| 0.311 \| 77 \| 53 \| 0 \| 16 \| 92 \| 4 \| 80 \| Phage tail protein \| Phage tail protein; afdb-uniprot50 \| AF-A0A497FI72-F1-MODEL\_V4 \| 1.0 \| 0.0006204 \| 170 \| 0.109 \| 128 \| 103 \| 4 \| 13 \| 136 \| 1 \| 121 \| Uncharacterized protein \| Uncharacterized protein; afdb-uniprot50 \| AF-G1UUM6-F1-MODEL\_V4 \| 1.0 \| 0.003264 \| 170 \| 0.129 \| 139 \| 104 \| 7 \| 1 \| 136 \| 1 \| 125 \| Uncharacterized protein \| Uncharacterized protein; afdb-uniprot50 \| AF-A0A542BN30-F1-MODEL\_V4 \| 1.0 \| 0.002497 \| 170 \| 0.106 \| 132 \| 109 \| 6 \| 6 \| 136 \| 1 \| 124 \| Uncharacterized protein \| Uncharacterized protein; afdb-uniprot50 \| AF-A0A7R6YQJ1-F1-MODEL\_V4 \| 1.0 \| 0.002244 \| 170 \| 0.093 \| 129 \| 112 \| 3 \| 8 \| 136 \| 20 \| 143 \| Uncharacterized protein \| Uncharacterized protein; afdb-uniprot50 \| AF-A0A0B6S238-F1-MODEL\_V4 \| 1.0 \| 0.001386 \| 170 \| 0.089 \| 146 \| 101 \| 6 \| 14 \| 136 \| 15 \| 151 \| Uncharacterized protein \| Uncharacterized protein; afdb-uniprot50 \| AF-A0A2E7CJN2-F1-MODEL\_V4 \| 1.0 \| 0.0004499 \| 170 \| 0.125 \| 151 \| 93 \| 7 \| 15 \| 135 \| 6 \| 147 \| LysM domain-containing protein \| LysM domain-containing protein; afdb-uniprot50 \| AF-A0A7H9S354-F1-MODEL\_V4 \| 1.0 \| 0.00106 \| 170 \| 0.161 \| 124 \| 82 \| 5 \| 28 \| 136 \| 52 \| 168 \| Uncharacterized protein \| Uncharacterized protein; afdb-uniprot50 \| AF-A0A6N8MHG0-F1-MODEL\_V4 \| 1.0 \| 0.0009523 \| 170 \| 0.144 \| 125 \| 83 \| 7 \| 28 \| 136 \| 73 \| 189 \| Uncharacterized protein \| Uncharacterized protein; afdb-uniprot50 \| AF-A0A3D5P8C3-F1-MODEL\_V4 \| 1.0 \| 0.001462 \| 170 \| 0.057 \| 139 \| 108 \| 6 \| 14 \| 136 \| 3 \| 134 \| Uncharacterized protein \| Uncharacterized protein; afdb-uniprot50 \| AF-A0A0S8DK82-F1-MODEL\_V4 \| 1.0 \| 0.0001716 \| 169 \| 0.133 \| 165 \| 104 \| 10 \| 6 \| 136 \| 1 \| 160 \| Uncharacterized protein \| Uncharacterized protein; afdb-uniprot50 \| AF-A0A4D0QGZ7-F1-MODEL\_V4 \| 1.0 \| 0.0007286 \| 169 \| 0.079 \| 139 \| 103 \| 6 \| 14 \| 136 \| 12 \| 141 \| Uncharacterized protein \| Uncharacterized protein; afdb-uniprot50 \| AF-A0A1V3IL17-F1-MODEL\_V4 \| 1.0 \| 0.002127 \| 169 \| 0.123 \| 113 \| 88 \| 5 \| 28 \| 136 \| 54 \| 159 \| Uncharacterized protein \| Uncharacterized protein; afdb-uniprot50 \| AF-A0A1U9NH92-F1-MODEL\_V4 \| 1.0 \| 0.001811 \| 168 \| 0.079 \| 126 \| 106 \| 5 \| 14 \| 136 \| 3 \| 121 \| Uncharacterized protein \| Uncharacterized protein; afdb-uniprot50 \| AF-A0A7T9IJ30-F1-MODEL\_V4 \| 1.0 \| 0.0005283 \| 168 \| 0.148 \| 128 \| 99 \| 6 \| 14 \| 136 \| 1 \| 123 \| Uncharacterized protein \| Uncharacterized protein; afdb-uniprot50 \| AF-A0A5C7NN52-F1-MODEL\_V4 \| 1.0 \| 0.002367 \| 168 \| 0.083 \| 132 \| 103 \| 7 \| 13 \| 136 \| 2 \| 123 \| Uncharacterized protein \| Uncharacterized protein; afdb-uniprot50 \| AF-A0A4V5MR96-F1-MODEL\_V4 \| 1.0 \| 0.001542 \| 168 \| 0.068 \| 161 \| 114 \| 7 \| 1 \| 136 \| 3 \| 152 \| Uncharacterized protein \| Uncharacterized protein; afdb-uniprot50 \| AF-A0A3A9BCQ7-F1-MODEL\_V4 \| 1.0 \| 0.0006546 \| 168 \| 0.088 \| 135 \| 102 \| 8 \| 13 \| 136 \| 7 \| 131 \| LysM peptidoglycan-binding domain-containing protein \| LysM peptidoglycan-binding domain-containing protein; afdb-uniprot50 \| AF-A0A354M6D3-F1-MODEL\_V4 \| 1.0 \| 0.0003263 \| 168 \| 0.13 \| 161 \| 100 \| 9 \| 2 \| 136 \| 65 \| 211 \| Uncharacterized protein \| Uncharacterized protein; afdb-uniprot50 \| AF-A0A1H0K7P7-F1-MODEL\_V4 \| 1.0 \| 0.003264 \| 168 \| 0.1 \| 140 \| 112 \| 6 \| 1 \| 136 \| 3 \| 132 \| Uncharacterized protein \| Uncharacterized protein; afdb-uniprot50 \| AF-A0A354WTT6-F1-MODEL\_V4 \| 1.0 \| 0.002635 \| 167 \| 0.09 \| 121 \| 93 \| 4 \| 28 \| 136 \| 69 \| 184 \| Uncharacterized protein \| Uncharacterized protein; afdb-uniprot50 \| AF-A0A202DYA2-F1-MODEL\_V4 \| 1.0 \| 0.0002778 \| 167 \| 0.122 \| 131 \| 96 \| 5 \| 20 \| 134 \| 27 \| 154 \| LysM domain-containing protein \| LysM domain-containing protein; afdb-uniprot50 \| AF-A0A6N3BSE7-F1-MODEL\_V4 \| 1.0 \| 0.002244 \| 167 \| 0.088 \| 158 \| 99 \| 5 \| 15 \| 136 \| 64 \| 212 \| Uncharacterized protein \| Uncharacterized protein; afdb-uniprot50 \| AF-A0A0F9NHG4-F1-MODEL\_V4 \| 1.0 \| 0.001386 \| 166 \| 0.115 \| 139 \| 107 \| 8 \| 1 \| 132 \| 12 \| 141 \| Uncharacterized protein \| Uncharacterized protein; afdb-uniprot50 \| AF-A0A1C7W148-F1-MODEL\_V4 \| 1.0 \| 0.002933 \| 166 \| 0.106 \| 132 \| 109 \| 6 \| 6 \| 136 \| 1 \| 124 \| Uncharacterized protein \| Uncharacterized protein; afdb-uniprot50 \| AF-N6U770-F1-MODEL\_V4 \| 1.0 \| 0.00278 \| 166 \| 0.102 \| 136 \| 115 \| 5 \| 2 \| 136 \| 35 \| 164 \| Uncharacterized protein \| Uncharacterized protein; afdb-uniprot50 \| AF-A0A379DMP2-F1-MODEL\_V4 \| 1.0 \| 0.001313 \| 166 \| 0.084 \| 154 \| 100 \| 6 \| 2 \| 136 \| 24 \| 155 \| Uncharacterized protein \| Uncharacterized protein; afdb-uniprot50 \| AF-A0A7X8YZY7-F1-MODEL\_V4 \| 1.0 \| 0.0004499 \| 166 \| 0.112 \| 133 \| 103 \| 6 \| 11 \| 134 \| 149 \| 275 \| Uncharacterized protein \| Uncharacterized protein; afdb-uniprot50 \| AF-A0A3G2V8Z8-F1-MODEL\_V4 \| 1.0 \| 0.001627 \| 166 \| 0.086 \| 138 \| 107 \| 7 \| 7 \| 136 \| 38 \| 164 \| Uncharacterized protein \| Uncharacterized protein; afdb-uniprot50 \| AF-A0A494R194-F1-MODEL\_V4 \| 1.0 \| 0.003264 \| 165 \| 0.301 \| 73 \| 44 \| 1 \| 63 \| 135 \| 3 \| 68 \| Uncharacterized protein \| Uncharacterized protein; afdb-uniprot50 \| AF-A0A366MWP2-F1-MODEL\_V4 \| 1.0 \| 0.002244 \| 165 \| 0.083 \| 131 \| 103 \| 8 \| 13 \| 136 \| 3 \| 123 \| Uncharacterized protein \| Uncharacterized protein; afdb-uniprot50 \| AF-A0A2P1S0D5-F1-MODEL\_V4 \| 1.0 \| 0.001911 \| 165 \| 0.108 \| 129 \| 93 \| 8 \| 14 \| 136 \| 23 \| 135 \| Uncharacterized protein \| Uncharacterized protein; afdb-uniprot50 \| AF-A0A3D0E3C9-F1-MODEL\_V4 \| 1.0 \| 0.001811 \| 165 \| 0.129 \| 139 \| 97 \| 9 \| 13 \| 136 \| 1 \| 130 \| LysM peptidoglycan-binding domain-containing protein \| LysM peptidoglycan-binding domain-containing protein; afdb-uniprot50 \| AF-A0A060QFF2-F1-MODEL\_V4 \| 1.0 \| 0.004749 \| 165 \| 0.082 \| 170 \| 113 \| 6 \| 1 \| 136 \| 1 \| 161 \| Putative bacteriophage protein \| Putative bacteriophage protein; afdb-uniprot50 \| AF-U2M6B1-F1-MODEL\_V4 \| 1.0 \| 0.0005008 \| 165 \| 0.119 \| 168 \| 113 \| 9 \| 3 \| 136 \| 165 \| 331 \| Uncharacterized protein \| Uncharacterized protein; afdb-uniprot50 \| AF-A0A0F8XZK0-F1-MODEL\_V4 \| 1.0 \| 0.001245 \| 164 \| 0.105 \| 151 \| 108 \| 8 \| 5 \| 136 \| 3 \| 145 \| Uncharacterized protein \| Uncharacterized protein; afdb-uniprot50 \| AF-H6U8J6-F1-MODEL\_V4 \| 1.0 \| 0.006549 \| 164 \| 0.224 \| 89 \| 67 \| 2 \| 48 \| 135 \| 3 \| 90 \| GpU \| GpU; afdb-uniprot50 \| AF-A0A3M3XDC7-F1-MODEL\_V4 \| 1.0 \| 0.004749 \| 163 \| 0.095 \| 178 \| 106 \| 6 \| 1 \| 136 \| 1 \| 165 \| Uncharacterized protein \| Uncharacterized protein; afdb-uniprot50 \| AF-A0A496CXI1-F1-MODEL\_V4 \| 1.0 \| 0.0002496 \| 163 \| 0.12 \| 124 \| 87 \| 7 \| 29 \| 134 \| 30 \| 149 \| LysM domain-containing protein \| LysM domain-containing protein; afdb-uniprot50 \| AF-A0A7Y8YBS2-F1-MODEL\_V4 \| 1.0 \| 0.002933 \| 163 \| 0.046 \| 151 \| 120 \| 6 \| 4 \| 136 \| 20 \| 164 \| Uncharacterized protein \| Uncharacterized protein; afdb-uniprot50 \| AF-A0A5C7BSU6-F1-MODEL\_V4 \| 1.0 \| 0.002016 \| 162 \| 0.107 \| 130 \| 90 \| 5 \| 28 \| 136 \| 22 \| 146 \| Uncharacterized protein \| Uncharacterized protein; afdb-uniprot50 \| AF-A0A6B5QZN2-F1-MODEL\_V4 \| 1.0 \| 0.001386 \| 162 \| 0.082 \| 145 \| 105 \| 6 \| 8 \| 136 \| 9 \| 141 \| Uncharacterized protein \| Uncharacterized protein; afdb-uniprot50 \| AF-A0A528TS09-F1-MODEL\_V4 \| 1.0 \| 0.003833 \| 161 \| 0.101 \| 128 \| 110 \| 3 \| 9 \| 136 \| 21 \| 143 \| Uncharacterized protein \| Uncharacterized protein; afdb-uniprot50 \| AF-A0A661R841-F1-MODEL\_V4 \| 1.0 \| 0.001386 \| 161 \| 0.102 \| 146 \| 101 \| 7 \| 9 \| 136 \| 9 \| 142 \| Uncharacterized protein \| Uncharacterized protein; afdb-uniprot50 \| AF-A0A1X0TE28-F1-MODEL\_V4 \| 1.0 \| 0.002497 \| 161 \| 0.13 \| 130 \| 96 \| 7 \| 16 \| 136 \| 40 \| 161 \| Uncharacterized protein \| Uncharacterized protein; afdb-uniprot50 \| AF-H3K667-F1-MODEL\_V4 \| 1.0 \| 0.001118 \| 161 \| 0.069 \| 144 \| 115 \| 7 \| 2 \| 136 \| 18 \| 151 \| Uncharacterized protein \| Uncharacterized protein; afdb-uniprot50 \| AF-A0A497RKB8-F1-MODEL\_V4 \| 1.0 \| 0.002016 \| 160 \| 0.187 \| 133 \| 90 \| 7 \| 13 \| 136 \| 5 \| 128 \| Uncharacterized protein \| Uncharacterized protein; afdb-uniprot50 \| AF-I2IJR8-F1-MODEL\_V4 \| 1.0 \| 0.001717 \| 160 \| 0.109 \| 146 \| 114 \| 8 \| 1 \| 136 \| 33 \| 172 \| Uncharacterized protein \| Uncharacterized protein; afdb-uniprot50 \| AF-A0A497PXB8-F1-MODEL\_V4 \| 1.0 \| 0.001462 \| 160 \| 0.085 \| 117 \| 89 \| 5 \| 26 \| 132 \| 74 \| 182 \| Uncharacterized protein \| Uncharacterized protein; afdb-uniprot50 \| AF-A0A261SPV5-F1-MODEL\_V4 \| 1.0 \| 0.002635 \| 159 \| 0.087 \| 114 \| 90 \| 6 \| 28 \| 136 \| 6 \| 110 \| Uncharacterized protein \| Uncharacterized protein; afdb-uniprot50 \| AF-J6H051-F1-MODEL\_V4 \| 1.0 \| 0.0006204 \| 159 \| 0.15 \| 133 \| 85 \| 7 \| 13 \| 136 \| 23 \| 136 \| Uncharacterized protein \| Uncharacterized protein; afdb-uniprot50 \| AF-H0TLJ4-F1-MODEL\_V4 \| 1.0 \| 0.004501 \| 159 \| 0.12 \| 133 \| 110 \| 6 \| 5 \| 136 \| 30 \| 156 \| Uncharacterized protein \| Uncharacterized protein; afdb-uniprot50 \| AF-A0A5B0HDI0-F1-MODEL\_V4 \| 1.0 \| 0.005884 \| 159 \| 0.1 \| 109 \| 95 \| 1 \| 28 \| 136 \| 53 \| 158 \| Uncharacterized protein \| Uncharacterized protein; afdb-uniprot50 \| AF-A0A419QIV5-F1-MODEL\_V4 \| 1.0 \| 0.00118 \| 159 \| 0.168 \| 125 \| 87 \| 6 \| 13 \| 133 \| 4 \| 115 \| Uncharacterized protein \| Uncharacterized protein; afdb-uniprot50 \| AF-A0A7C5SLD3-F1-MODEL\_V4 \| 1.0 \| 0.002635 \| 158 \| 0.106 \| 131 \| 105 \| 7 \| 13 \| 136 \| 1 \| 126 \| Uncharacterized protein \| Uncharacterized protein; afdb-uniprot50 \| AF-A0A316J465-F1-MODEL\_V4 \| 1.0 \| 0.001313 \| 158 \| 0.116 \| 146 \| 106 \| 9 \| 2 \| 136 \| 22 \| 155 \| Uncharacterized protein \| Uncharacterized protein; afdb-uniprot50 \| AF-A0A3E2KNQ6-F1-MODEL\_V4 \| 1.0 \| 0.002244 \| 158 \| 0.097 \| 133 \| 105 \| 9 \| 8 \| 136 \| 25 \| 146 \| Uncharacterized protein \| Uncharacterized protein; afdb-uniprot50 \| AF-A0A261UEK8-F1-MODEL\_V4 \| 1.0 \| 0.001811 \| 158 \| 0.062 \| 129 \| 107 \| 6 \| 13 \| 136 \| 48 \| 167 \| Uncharacterized protein \| Uncharacterized protein; afdb-uniprot50 \| AF-A0A6S7DDP2-F1-MODEL\_V4 \| 1.0 \| 0.006208 \| 158 \| 0.084 \| 119 \| 102 \| 5 \| 19 \| 136 \| 55 \| 167 \| Uncharacterized protein \| Uncharacterized protein; afdb-uniprot50 \| AF-A0A2A7QBL7-F1-MODEL\_V4 \| 1.0 \| 0.002497 \| 158 \| 0.112 \| 142 \| 106 \| 10 \| 3 \| 134 \| 10 \| 141 \| Uncharacterized protein \| Uncharacterized protein; afdb-uniprot50 \| AF-A0A0S8ER01-F1-MODEL\_V4 \| 1.0 \| 0.002933 \| 158 \| 0.133 \| 127 \| 98 \| 6 \| 15 \| 136 \| 162 \| 281 \| Uncharacterized protein \| Uncharacterized protein; afdb-uniprot50 \| AF-A0A537KYH8-F1-MODEL\_V4 \| 1.0 \| 0.001245 \| 157 \| 0.116 \| 146 \| 106 \| 7 \| 6 \| 133 \| 15 \| 155 \| Uncharacterized protein \| Uncharacterized protein; afdb-uniprot50 \| AF-A0A4Q3P5X9-F1-MODEL\_V4 \| 1.0 \| 0.002016 \| 157 \| 0.1 \| 180 \| 96 \| 10 \| 14 \| 136 \| 34 \| 204 \| Uncharacterized protein \| Uncharacterized protein; afdb-uniprot50 \| AF-A0A2N0CX77-F1-MODEL\_V4 \| 1.0 \| 0.0004747 \| 156 \| 0.206 \| 121 \| 82 \| 4 \| 28 \| 136 \| 6 \| 124 \| Uncharacterized protein \| Uncharacterized protein; afdb-uniprot50 \| AF-A0A6I7P5X6-F1-MODEL\_V4 \| 1.0 \| 0.000811 \| 156 \| 0.125 \| 136 \| 86 \| 7 \| 29 \| 136 \| 34 \| 164 \| Peptidoglycan-binding protein \| Peptidoglycan-binding protein; afdb-uniprot50 \| AF-A0A7H1NU42-F1-MODEL\_V4 \| 1.0 \| 0.00278 \| 156 \| 0.086 \| 116 \| 92 \| 5 \| 26 \| 136 \| 83 \| 189 \| Uncharacterized protein \| Uncharacterized protein; afdb-uniprot50 \| AF-A0A5C7M9Y1-F1-MODEL\_V4 \| 1.0 \| 0.003633 \| 156 \| 0.1 \| 140 \| 107 \| 12 \| 7 \| 136 \| 1 \| 131 \| Uncharacterized protein \| Uncharacterized protein; afdb-uniprot50 \| AF-A0A3D0M341-F1-MODEL\_V4 \| 1.0 \| 0.003264 \| 155 \| 0.074 \| 135 \| 107 \| 7 \| 7 \| 134 \| 1 \| 124 \| Uncharacterized protein \| Uncharacterized protein; afdb-uniprot50 \| AF-A0A1Y4D3Z5-F1-MODEL\_V4 \| 1.0 \| 0.001627 \| 155 \| 0.137 \| 131 \| 95 \| 9 \| 13 \| 136 \| 12 \| 131 \| Uncharacterized protein \| Uncharacterized protein; afdb-uniprot50 \| AF-A0A6V8IG24-F1-MODEL\_V4 \| 1.0 \| 0.002933 \| 155 \| 0.074 \| 148 \| 108 \| 7 \| 13 \| 136 \| 56 \| 198 \| Uncharacterized protein \| Uncharacterized protein; afdb-uniprot50 \| AF-W7DNG3-F1-MODEL\_V4 \| 1.0 \| 0.005577 \| 155 \| 0.096 \| 114 \| 90 \| 4 \| 27 \| 136 \| 98 \| 202 \| Uncharacterized protein \| Uncharacterized protein; afdb-uniprot50 \| AF-A0A2T0BLH2-F1-MODEL\_V4 \| 1.0 \| 0.004044 \| 155 \| 0.08 \| 125 \| 108 \| 6 \| 14 \| 133 \| 3 \| 125 \| Phage tail protein \| Phage tail protein; afdb-uniprot50 \| AF-A0A1Y2SAM4-F1-MODEL\_V4 \| 1.0 \| 0.00501 \| 154 \| 0.138 \| 130 \| 95 \| 7 \| 16 \| 136 \| 37 \| 158 \| Uncharacterized protein \| Uncharacterized protein; afdb-uniprot50 \| AF-A0A561JSY6-F1-MODEL\_V4 \| 1.0 \| 0.001386 \| 154 \| 0.111 \| 144 \| 109 \| 8 \| 2 \| 136 \| 39 \| 172 \| Uncharacterized protein \| Uncharacterized protein; afdb-uniprot50 \| AF-A0A7I7AC95-F1-MODEL\_V4 \| 1.0 \| 0.004501 \| 154 \| 0.128 \| 125 \| 85 \| 8 \| 28 \| 136 \| 55 \| 171 \| Uncharacterized protein \| Uncharacterized protein; afdb-uniprot50 \| AF-A0A3S0C2K4-F1-MODEL\_V4 \| 1.0 \| 0.001313 \| 153 \| 0.146 \| 116 \| 83 \| 6 \| 27 \| 136 \| 42 \| 147 \| Uncharacterized protein \| Uncharacterized protein; afdb-uniprot50 \| AF-A0A166CB51-F1-MODEL\_V4 \| 1.0 \| 0.003094 \| 153 \| 0.055 \| 127 \| 106 \| 5 \| 14 \| 136 \| 4 \| 120 \| Putative peptidoglycan binding domain protein \| Putative peptidoglycan binding domain protein; afdb-uniprot50 \| AF-A0A1M7AAE4-F1-MODEL\_V4 \| 1.0 \| 0.0004747 \| 153 \| 0.157 \| 146 \| 89 \| 6 \| 20 \| 134 \| 35 \| 177 \| Uncharacterized protein \| Uncharacterized protein; afdb-uniprot50 \| AF-A0A840C384-F1-MODEL\_V4 \| 1.0 \| 0.00856 \| 152 \| 0.061 \| 130 \| 109 \| 4 \| 11 \| 136 \| 12 \| 132 \| Uncharacterized protein \| Uncharacterized protein; afdb-uniprot50 \| AF-A0A4P7WFQ6-F1-MODEL\_V4 \| 1.0 \| 0.0007286 \| 152 \| 0.097 \| 133 \| 94 \| 5 \| 25 \| 136 \| 69 \| 196 \| Uncharacterized protein \| Uncharacterized protein; afdb-uniprot50 \| AF-A0A2J9H268-F1-MODEL\_V4 \| 1.0 \| 0.002497 \| 152 \| 0.088 \| 159 \| 94 \| 6 \| 20 \| 136 \| 57 \| 206 \| Uncharacterized protein \| Uncharacterized protein; afdb-uniprot50 \| AF-A0A246Q0L6-F1-MODEL\_V4 \| 1.0 \| 0.007691 \| 151 \| 0.056 \| 125 \| 96 \| 5 \| 28 \| 136 \| 30 \| 148 \| Uncharacterized protein \| Uncharacterized protein; afdb-uniprot50 \| AF-A0A7H1NTS9-F1-MODEL\_V4 \| 1.0 \| 0.006549 \| 150 \| 0.041 \| 144 \| 97 \| 4 \| 28 \| 136 \| 26 \| 163 \| Uncharacterized protein \| Uncharacterized protein; afdb-uniprot50 \| AF-A0A0U5F1N5-F1-MODEL\_V4 \| 1.0 \| 0.0008556 \| 150 \| 0.073 \| 150 \| 108 \| 8 \| 13 \| 136 \| 55 \| 199 \| Burkholderia phage Bcep781 gp51 \| Burkholderia phage Bcep781 gp51; afdb-uniprot50 \| AF-M7CDH5-F1-MODEL\_V4 \| 1.0 \| 0.005286 \| 150 \| 0.116 \| 129 \| 99 \| 6 \| 16 \| 136 \| 86 \| 207 \| Uncharacterized protein \| Uncharacterized protein; afdb-uniprot50 \| AF-A0A7X5QF72-F1-MODEL\_V4 \| 1.0 \| 0.005577 \| 150 \| 0.062 \| 143 \| 108 \| 6 \| 13 \| 136 \| 319 \| 454 \| Uncharacterized protein \| Uncharacterized protein; afdb-uniprot50 \| AF-A0A0H3B1Y1-F1-MODEL\_V4 \| 1.0 \| 0.004749 \| 149 \| 0.081 \| 135 \| 107 \| 9 \| 7 \| 136 \| 1 \| 123 \| Uncharacterized protein \| Uncharacterized protein; afdb-uniprot50 \| AF-A0A1J5RYQ4-F1-MODEL\_V4 \| 1.0 \| 0.006549 \| 149 \| 0.069 \| 130 \| 114 \| 5 \| 8 \| 136 \| 35 \| 158 \| Uncharacterized protein \| Uncharacterized protein; afdb-uniprot50 \| AF-A0A420W5B5-F1-MODEL\_V4 \| 1.0 \| 0.003264 \| 149 \| 0.13 \| 130 \| 96 \| 8 \| 16 \| 136 \| 37 \| 158 \| Uncharacterized protein \| Uncharacterized protein; afdb-uniprot50 \| AF-A0A1Q8E695-F1-MODEL\_V4 \| 1.0 \| 0.004267 \| 149 \| 0.075 \| 132 \| 106 \| 8 \| 14 \| 136 \| 4 \| 128 \| Uncharacterized protein \| Uncharacterized protein; afdb-uniprot50 \| AF-A0A0C1PS50-F1-MODEL\_V4 \| 1.0 \| 0.002635 \| 149 \| 0.063 \| 142 \| 111 \| 9 \| 5 \| 136 \| 195 \| 324 \| N-acetylmuramoyl-L-alanine amidase, family 4 \| N-acetylmuramoyl-L-alanine amidase, family 4; afdb-uniprot50 \| AF-A0A1U7GIE9-F1-MODEL\_V4 \| 1.0 \| 0.003833 \| 148 \| 0.128 \| 132 \| 97 \| 7 \| 14 \| 136 \| 1 \| 123 \| Uncharacterized protein \| Uncharacterized protein; afdb-uniprot50 \| AF-A0A0D6N7K5-F1-MODEL\_V4 \| 1.0 \| 0.003094 \| 148 \| 0.076 \| 156 \| 114 \| 8 \| 2 \| 136 \| 29 \| 175 \| Uncharacterized protein \| Uncharacterized protein; afdb-uniprot50 \| AF-A0A3A1YWM1-F1-MODEL\_V4 \| 1.0 \| 0.00501 \| 148 \| 0.084 \| 166 \| 103 \| 7 \| 11 \| 136 \| 8 \| 164 \| Uncharacterized protein \| Uncharacterized protein; afdb-uniprot50 \| AF-A0A3A9FPZ3-F1-MODEL\_V4 \| 1.0 \| 0.002635 \| 148 \| 0.142 \| 140 \| 108 \| 8 \| 1 \| 132 \| 8 \| 143 \| Uncharacterized protein \| Uncharacterized protein; afdb-uniprot50 \| AF-A0A239HRH5-F1-MODEL\_V4 \| 1.0 \| 0.00856 \| 147 \| 0.1 \| 110 \| 87 \| 4 \| 30 \| 136 \| 12 \| 112 \| Phage-related protein \| Phage-related protein; afdb-uniprot50 \| AF-A0A7X9XB50-F1-MODEL\_V4 \| 1.0 \| 0.001386 \| 147 \| 0.067 \| 149 \| 111 \| 6 \| 13 \| 136 \| 3 \| 148 \| Uncharacterized protein \| Uncharacterized protein; afdb-uniprot50 \| AF-A0A4R1FNK5-F1-MODEL\_V4 \| 1.0 \| 0.003094 \| 147 \| 0.084 \| 130 \| 102 \| 6 \| 16 \| 136 \| 38 \| 159 \| Uncharacterized protein \| Uncharacterized protein; afdb-uniprot50 \| AF-A0A563VQY9-F1-MODEL\_V4 \| 1.0 \| 0.0002243 \| 147 \| 0.161 \| 130 \| 90 \| 6 \| 21 \| 136 \| 22 \| 146 \| Rhs element Vgr protein (Modular protein) \| Rhs element Vgr protein (Modular protein); afdb-uniprot50 \| AF-A0A6A7KA75-F1-MODEL\_V4 \| 1.0 \| 0.002367 \| 146 \| 0.121 \| 132 \| 94 \| 8 \| 13 \| 135 \| 2 \| 120 \| Uncharacterized protein \| Uncharacterized protein; afdb-uniprot50 \| AF-A0A4Z0RKI1-F1-MODEL\_V4 \| 1.0 \| 0.004044 \| 146 \| 0.103 \| 116 \| 93 \| 5 \| 25 \| 136 \| 12 \| 120 \| LysM domain-containing protein \| LysM domain-containing protein; afdb-uniprot50 \| AF-A0A2D8LFG3-F1-MODEL\_V4 \| 1.0 \| 0.002016 \| 144 \| 0.063 \| 141 \| 107 \| 9 \| 7 \| 136 \| 1 \| 127 \| Uncharacterized protein \| Uncharacterized protein; afdb-uniprot50 \| AF-A0A346PHI6-F1-MODEL\_V4 \| 1.0 \| 0.009031 \| 144 \| 0.131 \| 114 \| 86 \| 6 \| 29 \| 136 \| 14 \| 120 \| Phage associated protein \| Phage associated protein; afdb-uniprot50 \| AF-A0A379S622-F1-MODEL\_V4 \| 1.0 \| 0.004267 \| 143 \| 0.066 \| 150 \| 112 \| 11 \| 6 \| 136 \| 19 \| 159 \| Uncharacterized protein \| Uncharacterized protein; afdb-uniprot50 \| AF-A0A1B9Y3L0-F1-MODEL\_V4 \| 1.0 \| 0.0009027 \| 142 \| 0.104 \| 143 \| 97 \| 8 \| 7 \| 136 \| 5 \| 129 \| Uncharacterized protein \| Uncharacterized protein; afdb-uniprot50 \| AF-A0A3C1Y293-F1-MODEL\_V4 \| 1.0 \| 0.005884 \| 142 \| 0.145 \| 117 \| 86 \| 6 \| 25 \| 136 \| 11 \| 118 \| Uncharacterized protein \| Uncharacterized protein; afdb-uniprot50 \| AF-A0A1V6D5U1-F1-MODEL\_V4 \| 1.0 \| 0.009528 \| 141 \| 0.069 \| 130 \| 103 \| 11 \| 14 \| 136 \| 4 \| 122 \| Uncharacterized protein \| Uncharacterized protein; afdb-uniprot50 \| AF-A0A257QIG8-F1-MODEL\_V4 \| 1.0 \| 0.006909 \| 140 \| 0.105 \| 151 \| 108 \| 9 \| 2 \| 136 \| 31 \| 170 \| Uncharacterized protein \| Uncharacterized protein; afdb-uniprot50 \| AF-A0A654D8R3-F1-MODEL\_V4 \| 1.0 \| 0.002127 \| 139 \| 0.091 \| 142 \| 112 \| 10 \| 3 \| 136 \| 2 \| 134 \| Uncharacterized protein \| Uncharacterized protein; afdb-uniprot50 \| AF-A0A412UZK8-F1-MODEL\_V4 \| 1.0 \| 0.00118 \| 139 \| 0.137 \| 124 \| 87 \| 5 \| 18 \| 134 \| 18 \| 128 \| Uncharacterized protein \| Uncharacterized protein; afdb-uniprot50 \| AF-A0A6M3IHI8-F1-MODEL\_V4 \| 1.0 \| 0.003633 \| 139 \| 0.116 \| 112 \| 83 \| 5 \| 30 \| 136 \| 148 \| 248 \| Uncharacterized protein \| Uncharacterized protein; afdb-uniprot50 \| AF-A0A3A2J2V4-F1-MODEL\_V4 \| 1.0 \| 0.00856 \| 138 \| 0.094 \| 127 \| 97 \| 8 \| 20 \| 133 \| 39 \| 160 \| Uncharacterized protein \| Uncharacterized protein; afdb-uniprot50 \| AF-A0A1H0N722-F1-MODEL\_V4 \| 1.0 \| 0.005884 \| 138 \| 0.073 \| 123 \| 106 \| 5 \| 16 \| 134 \| 2 \| 120 \| Phage-related protein \| Phage-related protein; afdb-uniprot50 \| AF-A0A2L0WYZ1-F1-MODEL\_V4 \| 1.0 \| 0.006549 \| 137 \| 0.115 \| 130 \| 99 \| 8 \| 15 \| 136 \| 52 \| 173 \| Uncharacterized protein \| Uncharacterized protein; afdb-uniprot50 \| AF-A0A7Y7ISW7-F1-MODEL\_V4 \| 1.0 \| 0.007691 \| 137 \| 0.067 \| 163 \| 119 \| 10 \| 2 \| 136 \| 32 \| 189 \| Uncharacterized protein \| Uncharacterized protein; afdb-uniprot50 \| AF-A0A270BII5-F1-MODEL\_V4 \| 1.0 \| 0.003444 \| 136 \| 0.08 \| 137 \| 97 \| 5 \| 25 \| 136 \| 68 \| 200 \| Uncharacterized protein \| Uncharacterized protein; afdb-uniprot50 \| AF-A0A644T6N0-F1-MODEL\_V4 \| 1.0 \| 0.001245 \| 136 \| 0.129 \| 155 \| 95 \| 9 \| 12 \| 136 \| 8 \| 152 \| Uncharacterized protein \| Uncharacterized protein; afdb-uniprot50 \| AF-A0A497RPK8-F1-MODEL\_V4 \| 1.0 \| 0.009528 \| 136 \| 0.107 \| 112 \| 86 \| 4 \| 29 \| 132 \| 123 \| 228 \| Uncharacterized protein \| Uncharacterized protein; afdb-uniprot50 \| AF-A0A6L4BC15-F1-MODEL\_V4 \| 1.0 \| 0.009528 \| 135 \| 0.08 \| 112 \| 94 \| 4 \| 30 \| 136 \| 13 \| 120 \| Phage tail protein \| Phage tail protein; afdb-uniprot50 \| AF-A0A1M5UNR8-F1-MODEL\_V4 \| 1.0 \| 0.0007687 \| 135 \| 0.101 \| 158 \| 95 \| 8 \| 21 \| 136 \| 25 \| 177 \| LysM domain-containing protein \| LysM domain-containing protein; afdb-uniprot50 \| AF-A0A4Y3TK87-F1-MODEL\_V4 \| 1.0 \| 0.00729 \| 135 \| 0.071 \| 140 \| 100 \| 6 \| 23 \| 136 \| 65 \| 200 \| Uncharacterized protein \| Uncharacterized protein; afdb-uniprot50 \| AF-X8IK28-F1-MODEL\_V4 \| 1.0 \| 0.008114 \| 134 \| 0.129 \| 124 \| 96 \| 5 \| 11 \| 133 \| 17 \| 129 \| Uncharacterized protein \| Uncharacterized protein; afdb-uniprot50 \| AF-A0A7X7YWV6-F1-MODEL\_V4 \| 1.0 \| 0.009528 \| 134 \| 0.136 \| 125 \| 100 \| 5 \| 13 \| 134 \| 1 \| 120 \| Uncharacterized protein \| Uncharacterized protein; afdb-uniprot50 \| AF-A0A0F9ES18-F1-MODEL\_V4 \| 1.0 \| 0.00856 \| 134 \| 0.093 \| 139 \| 110 \| 9 \| 1 \| 133 \| 33 \| 161 \| Uncharacterized protein \| Uncharacterized protein; afdb-uniprot50 \| AF-A0A1J5KCD7-F1-MODEL\_V4 \| 1.0 \| 0.002497 \| 132 \| 0.095 \| 147 \| 99 \| 11 \| 6 \| 136 \| 1 \| 129 \| Uncharacterized protein \| Uncharacterized protein; afdb-uniprot50 \| AF-A0A0R1L1L1-F1-MODEL\_V4 \| 1.0 \| 0.007691 \| 132 \| 0.095 \| 136 \| 93 \| 10 \| 13 \| 136 \| 122 \| 239 \| Mn2+ Fe2+ transporter \| Mn2+ Fe2+ transporter; afdb-uniprot50 \| AF-B6WQ21-F1-MODEL\_V4 \| 1.0 \| 0.005884 \| 131 \| 0.069 \| 130 \| 96 \| 7 \| 19 \| 136 \| 82 \| 198 \| Uncharacterized protein \| Uncharacterized protein; afdb-uniprot50 \| AF-A0A2T4TTF0-F1-MODEL\_V4 \| 1.0 \| 0.003094 \| 130 \| 0.107 \| 139 \| 97 \| 8 \| 18 \| 136 \| 22 \| 153 \| Uncharacterized protein \| Uncharacterized protein; afdb-uniprot50 \| AF-A0A316LZT1-F1-MODEL\_V4 \| 1.0 \| 0.006909 \| 130 \| 0.105 \| 114 \| 94 \| 5 \| 25 \| 134 \| 11 \| 120 \| Phage tail protein \| Phage tail protein; afdb-uniprot50 \| AF-A0A2V7R7U0-F1-MODEL\_V4 \| 1.0 \| 0.001462 \| 130 \| 0.113 \| 150 \| 103 \| 11 \| 5 \| 134 \| 2 \| 141 \| Uncharacterized protein \| Uncharacterized protein; afdb-uniprot50 \| AF-A0A6I4YV07-F1-MODEL\_V4 \| 1.0 \| 0.002933 \| 129 \| 0.128 \| 140 \| 95 \| 12 \| 13 \| 136 \| 31 \| 159 \| Uncharacterized protein \| Uncharacterized protein; afdb-uniprot50 \| AF-A0A2W4ILC1-F1-MODEL\_V4 \| 1.0 \| 0.007691 \| 129 \| 0.101 \| 158 \| 102 \| 10 \| 1 \| 134 \| 5 \| 146 \| Uncharacterized protein \| Uncharacterized protein; afdb-uniprot50 \| AF-A0A352SX32-F1-MODEL\_V4 \| 1.0 \| 0.00856 \| 128 \| 0.075 \| 132 \| 98 \| 10 \| 13 \| 135 \| 2 \| 118 \| Uncharacterized protein \| Uncharacterized protein; afdb-uniprot50 \| AF-A0A2D6MH57-F1-MODEL\_V4 \| 1.0 \| 0.009031 \| 128 \| 0.091 \| 120 \| 91 \| 5 \| 23 \| 132 \| 118 \| 229 \| Uncharacterized protein \| Uncharacterized protein; afdb-uniprot50 \| AF-A0A2X3JWJ8-F1-MODEL\_V4 \| 1.0 \| 0.002016 \| 125 \| 0.083 \| 143 \| 95 \| 9 \| 5 \| 133 \| 15 \| 135 \| Putative DNA circulation protein \| Putative DNA circulation protein; afdb-uniprot50 \| AF-A0A497RR60-F1-MODEL\_V4 \| 1.0 \| 0.00856 \| 124 \| 0.128 \| 125 \| 81 \| 6 \| 23 \| 132 \| 144 \| 255 \| Uncharacterized protein \| Uncharacterized protein; afdb-uniprot50 \| AF-A0A1Z4HHM5-F1-MODEL\_V4 \| 1.0 \| 0.006208 \| 119 \| 0.11 \| 172 \| 90 \| 7 \| 13 \| 136 \| 6 \| 162 \| Peptidoglycan-binding LysM \| Peptidoglycan-binding LysM; afdb-uniprot50 \| AF-A0A0F9EZB2-F1-MODEL\_V4 \| 1.0 \| 0.00501 \| 115 \| 0.154 \| 136 \| 83 \| 12 \| 14 \| 133 \| 23 \| 142 \| Uncharacterized protein \| Uncharacterized protein; afdb-uniprot50 \| AF-A0A2T0YM39-F1-MODEL\_V4 \| 1.0 \| 0.005577 \| 114 \| 0.079 \| 151 \| 109 \| 9 \| 6 \| 136 \| 1 \| 141 \| Uncharacterized protein \| Uncharacterized protein; afdb-uniprot50 \| AF-G5H934-F1-MODEL\_V4 \| 1.0 \| 0.00729 \| 111 \| 0.156 \| 147 \| 93 \| 9 \| 1 \| 133 \| 6 \| 135 \| Uncharacterized protein \| Uncharacterized protein; afdb-uniprot50 \| AF-A0A136Q8M2-F1-MODEL\_V4 \| 1.0 \| 0.006208 \| 107 \| 0.095 \| 126 \| 95 \| 7 \| 14 \| 136 \| 20 \| 129 \| Uncharacterized protein \| Uncharacterized protein |
| Top keywords  (threshold 1.00e-02 (evalue)) | **Phage, tail, GpU, LysM, domain\_containing, P2, U, Oxidoreductase, Putative, peptidoglycan\_binding** |
| Output files | ../../similar\_structures/23\_FANPEZAQ\_CDS\_0023\_afdb-proteome\_foldseek.tsv ../../similar\_structures/23\_FANPEZAQ\_CDS\_0023\_afdb-uniprot50\_foldseek.tsv ../../similar\_structures/23\_FANPEZAQ\_CDS\_0023\_merged.svg ../../similar\_structures/23\_FANPEZAQ\_CDS\_0023\_pdb\_foldseek.tsv |

  
  
  

Return to summary | Go to previous | Go to next

  


---

**Sequence/structure alignments coloring**  
Each object in the alignment figures is colored according to its E-value following this color coding:

1e-100
10
