## Supplementary material for "Completing the BASEL phage collection to unlock hidden diversity for systematic exploration of phage-host interactions": Data S2: 24.html

FANPEZAQ\_CDS\_0024


Return to summary | Go to previous | Go to next

|  |  |
| --- | --- |
| FANPEZAQ\_CDS\_0024  Page creation date: 02 Sep 2024, 12:00  Project folder: n/a  Input sequences file: Escherichia\_virus\_HeidiAbel.gb | lysm domain\_containing tail phage x peptidoglycan\_binding prophage p2\_like fragment bon superfamily putative spore coat assembly morphogenetic safa associated with spovid cazy families cbm50 lectin membrane |

### Sequence information

|  |  |
| --- | --- |
| Name | FANPEZAQ\_CDS\_0024  24\_FANPEZAQ\_CDS\_0024 (pipeline id) |
| Imported annotations | Escherichia\_virus\_HeidiAbel Bas97 |
| Protein sequence | MQTYITKDGDTADYIAWKYYGNQDAGTVEALVDANKGLADRGPVLPAGLVINLPEITTPA TTQGVKLWD |
| Number of residues | 69 |
| Molecular weight (Da) | 7426.24 |
| Output files | ../../query\_sequences/24\_FANPEZAQ\_CDS\_0024.fasta |

### Putative domain architecture and protein family

#### Search results (HHblits)^1^

|  |  |
| --- | --- |
| Domain family databases searched | Pfam, Ncbi-cd, Cath, Phrogs |
| Results, scheme(s)  (Top layers only; threshold 1.00e-03 (evalue)) | xml version="1.0" encoding="utf-8" standalone="no"?       2024-09-02T21:08:17.350755 image/svg+xml   Matplotlib v3.7.2, https://matplotlib.org/ |
| Results, table  (E-value ≤ 1.00e-03 (evalue)) | db \| id \| prob \| evalue \| pvalue \| score \| cols \| query \| query\_len \| template \| template\_len \| name \| description; phrogs \| 46 \| 99.8 \| 1.4e-26 \| 1.9e-30 \| 132.6 \| 66 \| (1, 69) \| 69 \| (9, 76) \| 85 \| baseplate hub \| baseplate hub; Category: tail; p199511 VI\_11453; phrogs \| 7145 \| 98.7 \| 1.3e-12 \| 1.6e-16 \| 75.2 \| 55 \| (3, 59) \| 69 \| (38, 93) \| 97 \| baseplate wedge subunit \| baseplate wedge subunit; Category: tail; MF360958\_p187; phrogs \| 21966 \| 98.4 \| 5.5e-11 \| 6.3e-15 \| 69.9 \| 51 \| (2, 54) \| 69 \| (78, 128) \| 128 \| NA \| NA; Category: unknown function; AY855346\_p20; phrogs \| 7348 \| 98.2 \| 3.7e-10 \| 4.3e-14 \| 64.9 \| 53 \| (2, 58) \| 69 \| (5, 57) \| 104 \| endolysin \| endolysin; Category: lysis; MG428991\_p108; phrogs \| 8272 \| 96.9 \| 1.6e-06 \| 1.8e-10 \| 47.8 \| 53 \| (2, 56) \| 69 \| (38, 90) \| 93 \| tail protein \| tail protein; Category: tail; NC\_029013\_p171; phrogs \| 14367 \| 96.5 \| 6.8e-06 \| 8e-10 \| 45.8 \| 54 \| (3, 58) \| 69 \| (36, 91) \| 93 \| NA \| NA; Category: unknown function; p23038 VI\_12315; phrogs \| 975 \| 96.0 \| 3.9e-05 \| 4.8e-09 \| 48.8 \| 56 \| (2, 59) \| 69 \| (5, 72) \| 236 \| baseplate protein \| baseplate protein; Category: tail; NC\_020871\_p93; phrogs \| 380 \| 95.9 \| 4.5e-05 \| 5.9e-09 \| 47.7 \| 39 \| (2, 58) \| 69 \| (162, 203) \| 230 \| endolysin \| endolysin; Category: lysis; NC\_005294\_p57; phrogs \| 232 \| 95.0 \| 0.00032 \| 4.1e-08 \| 43.3 \| 35 \| (2, 38) \| 69 \| (49, 83) \| 209 \| baseplate wedge subunit \| baseplate wedge subunit; Category: tail; NC\_027132\_p2; phrogs \| 4033 \| 94.2 \| 0.001 \| 1.2e-07 \| 43.3 \| 51 \| (2, 57) \| 69 \| (133, 183) \| 320 \| NA \| NA; Category: unknown function; p55048 VI\_12418 |
| Top keywords  (threshold 1.00e-03 (evalue)) | **tail, baseplate, wedge, endolysin, lysis, hub, p199511, VI\_11453, MF360958\_p187, AY855346\_p20** |
| Output files | ../../domain\_architecture/24\_FANPEZAQ\_CDS\_0024\_cath.hhr ../../domain\_architecture/24\_FANPEZAQ\_CDS\_0024\_merged.svg ../../domain\_architecture/24\_FANPEZAQ\_CDS\_0024\_ncbi-cd.hhr ../../domain\_architecture/24\_FANPEZAQ\_CDS\_0024\_pfam.hhr ../../domain\_architecture/24\_FANPEZAQ\_CDS\_0024\_phrogs.hhr |

### Identical protein sequences/structures

#### Search results

|  |  |
| --- | --- |
| Protein sequence databases searched | Pdb, Swissprot, Refseq |
| Identical proteins found | -- |
| Top keywords | -- |
| Output files | -- |

### Similar protein sequences/structures

#### Sequence similarity search results (HHblits)^1^

|  |  |
| --- | --- |
| Sequence databases searched | Uniclust, Pdb70 |
| Results, scheme(s)  (Top layers only, threshold 1.00e-03 (evalue)) | xml version="1.0" encoding="utf-8" standalone="no"?       2024-09-02T21:08:39.917925 image/svg+xml   Matplotlib v3.7.2, https://matplotlib.org/ |
| Results, table(s)  (threshold 1.00e-03 (evalue)) | db \| id \| prob \| evalue \| pvalue \| score \| cols \| query \| query\_len \| template \| template\_len \| name \| description; uniclust \| UniRef100\_A0A0B4B3T0 \| 99.8 \| 7.2e-24 \| 1.4e-29 \| 113.1 \| 69 \| (1, 69) \| 69 \| (12, 80) \| 83 \| p2-like protein prophage tail protein X-like protein \| p2-like protein prophage tail protein X-like protein; uniclust \| UniRef100\_A0A009QNP1 \| 99.8 \| 8.2e-23 \| 1.8e-28 \| 112.9 \| 68 \| (1, 69) \| 69 \| (16, 83) \| 92 \| Phage Tail Protein X family protein \| Phage Tail Protein X family protein; uniclust \| UniRef100\_A0A074TD60 \| 99.8 \| 2.4e-22 \| 4.9e-28 \| 108.7 \| 68 \| (1, 69) \| 69 \| (3, 70) \| 86 \| Tail protein \| Tail protein; uniclust \| UniRef100\_A0A085ARU3 \| 99.8 \| 7.6e-22 \| 1.5e-27 \| 105.2 \| 69 \| (1, 69) \| 69 \| (7, 75) \| 78 \| Phage tail protein \| Phage tail protein; uniclust \| UniRef100\_A0A165XHI0 \| 99.8 \| 8e-22 \| 1.7e-27 \| 107.2 \| 68 \| (1, 69) \| 69 \| (15, 83) \| 85 \| Phage Tail Protein X \| Phage Tail Protein X; uniclust \| UniRef100\_A0A069CJM3 \| 99.8 \| 1.1e-21 \| 2.2e-27 \| 108.0 \| 68 \| (1, 69) \| 69 \| (10, 77) \| 95 \| Phage tail protein \| Phage tail protein; uniclust \| UniRef100\_A0A1Q8SPL8 \| 99.7 \| 1.5e-21 \| 3e-27 \| 101.8 \| 68 \| (1, 69) \| 69 \| (4, 72) \| 72 \| Phage tail protein \| Phage tail protein; uniclust \| UniRef100\_A0A060GYH3 \| 99.7 \| 2.9e-21 \| 5.9e-27 \| 101.5 \| 69 \| (1, 69) \| 69 \| (5, 73) \| 73 \| Tail protein \| Tail protein; uniclust \| UniRef100\_A0A0B6D337 \| 99.7 \| 5.9e-21 \| 1.2e-26 \| 103.4 \| 66 \| (2, 69) \| 69 \| (18, 84) \| 86 \| Phage Tail Protein X family protein \| Phage Tail Protein X family protein; uniclust \| UniRef100\_A0A069PL70 \| 99.7 \| 3e-20 \| 6.3e-26 \| 101.0 \| 66 \| (2, 69) \| 69 \| (18, 83) \| 84 \| Phage tail protein \| Phage tail protein; uniclust \| UniRef100\_A0A024E969 \| 99.7 \| 3.6e-20 \| 7.4e-26 \| 100.8 \| 67 \| (1, 69) \| 69 \| (17, 83) \| 86 \| Tail protein X-related protein \| Tail protein X-related protein; uniclust \| UniRef100\_Q94MD7 \| 99.7 \| 5e-20 \| 1e-25 \| 97.4 \| 68 \| (1, 69) \| 69 \| (7, 74) \| 74 \| Tail component protein (Fragment) \| Tail component protein (Fragment); uniclust \| UniRef100\_A0A077KTR2 \| 99.7 \| 9.6e-20 \| 2e-25 \| 96.3 \| 67 \| (1, 69) \| 69 \| (7, 73) \| 73 \| Phage tail protein \| Phage tail protein; uniclust \| UniRef100\_A0A0N1C803 \| 99.6 \| 3.1e-19 \| 6.5e-25 \| 96.3 \| 68 \| (2, 69) \| 69 \| (8, 75) \| 78 \| Phage tail protein \| Phage tail protein; uniclust \| UniRef100\_A0A0C5VGD3 \| 99.6 \| 5.8e-19 \| 1.2e-24 \| 99.8 \| 68 \| (2, 69) \| 69 \| (3, 106) \| 107 \| p2-like prophage tail protein X \| p2-like prophage tail protein X; uniclust \| UniRef100\_A0A0E4BWV6 \| 99.6 \| 1.3e-18 \| 2.6e-24 \| 93.7 \| 69 \| (1, 69) \| 69 \| (7, 79) \| 79 \| Phage tail protein \| Phage tail protein; uniclust \| UniRef100\_A0A0D8L3F8 \| 99.6 \| 1.2e-18 \| 2.6e-24 \| 93.7 \| 67 \| (2, 69) \| 69 \| (9, 77) \| 77 \| Phage tail protein \| Phage tail protein; uniclust \| UniRef100\_A0A0E3UN65 \| 99.6 \| 4.7e-18 \| 9.3e-24 \| 97.0 \| 56 \| (2, 59) \| 69 \| (59, 114) \| 120 \| LysM domain-containing protein \| LysM domain-containing protein; uniclust \| UniRef100\_A0A1R4JJ60 \| 99.5 \| 1.1e-17 \| 2.3e-23 \| 91.9 \| 56 \| (1, 58) \| 69 \| (32, 87) \| 90 \| LysM domain-containing protein \| LysM domain-containing protein; uniclust \| UniRef100\_A0A0F8ZZ29 \| 99.5 \| 1.3e-17 \| 2.7e-23 \| 88.2 \| 55 \| (2, 58) \| 69 \| (14, 69) \| 73 \| LysM domain-containing protein (Fragment) \| LysM domain-containing protein (Fragment); uniclust \| UniRef100\_A0A0A2WI51 \| 99.5 \| 1.4e-17 \| 2.7e-23 \| 96.5 \| 55 \| (3, 59) \| 69 \| (62, 116) \| 129 \| Peptidoglycan-binding lysin domain-containing protein \| Peptidoglycan-binding lysin domain-containing protein; uniclust \| UniRef100\_A0A1B8SM38 \| 99.5 \| 1.7e-17 \| 3.4e-23 \| 88.1 \| 54 \| (2, 57) \| 69 \| (18, 71) \| 75 \| LysM domain-containing protein \| LysM domain-containing protein; uniclust \| UniRef100\_A0A023XZ78 \| 99.5 \| 1.8e-17 \| 3.7e-23 \| 96.2 \| 56 \| (2, 59) \| 69 \| (65, 120) \| 131 \| LysM peptidoglycan-binding domain-containing protein \| LysM peptidoglycan-binding domain-containing protein; uniclust \| UniRef100\_A0A219Y8Z5 \| 99.5 \| 2.4e-17 \| 5e-23 \| 90.4 \| 67 \| (2, 69) \| 69 \| (17, 84) \| 88 \| Tail protein \| Tail protein; uniclust \| UniRef100\_A0A031FQW4 \| 99.5 \| 2.5e-17 \| 5.2e-23 \| 90.4 \| 69 \| (1, 69) \| 69 \| (18, 87) \| 87 \| Phage tail X \| Phage tail X; uniclust \| UniRef100\_A0A3B9ZHN5 \| 99.5 \| 2.9e-17 \| 6e-23 \| 94.0 \| 56 \| (2, 59) \| 69 \| (55, 110) \| 114 \| LysM domain-containing protein \| LysM domain-containing protein; uniclust \| UniRef100\_A0A535FPJ6 \| 99.5 \| 3.2e-17 \| 6.5e-23 \| 84.4 \| 53 \| (2, 58) \| 69 \| (7, 59) \| 61 \| LysM peptidoglycan-binding domain-containing protein \| LysM peptidoglycan-binding domain-containing protein; uniclust \| UniRef100\_A0A011MIB9 \| 99.5 \| 4.7e-17 \| 9.9e-23 \| 90.8 \| 64 \| (2, 68) \| 69 \| (17, 80) \| 95 \| Membrane protein \| Membrane protein; uniclust \| UniRef100\_A0A0C2Z8D7 \| 99.5 \| 6.1e-17 \| 1.2e-22 \| 85.2 \| 68 \| (1, 69) \| 69 \| (1, 68) \| 68 \| Phage tail protein \| Phage tail protein; uniclust \| UniRef100\_A0A0S4XN94 \| 99.5 \| 7e-17 \| 1.5e-22 \| 85.3 \| 66 \| (1, 69) \| 69 \| (2, 68) \| 68 \| Putative Phage Tail Protein X \| Putative Phage Tail Protein X; uniclust \| UniRef100\_A0A0D0S6Z4 \| 99.5 \| 8.5e-17 \| 1.7e-22 \| 93.4 \| 54 \| (2, 57) \| 69 \| (70, 123) \| 136 \| NODE\_43, whole genome shotgun sequence \| NODE\_43, whole genome shotgun sequence; uniclust \| UniRef100\_A0A1A9RBR4 \| 99.5 \| 1.1e-16 \| 2e-22 \| 84.7 \| 66 \| (3, 69) \| 69 \| (10, 77) \| 77 \| Phage tail protein \| Phage tail protein; uniclust \| UniRef100\_A0A3A9HNX8 \| 99.5 \| 1.2e-16 \| 2.3e-22 \| 83.6 \| 66 \| (1, 69) \| 69 \| (6, 71) \| 71 \| Phage tail protein \| Phage tail protein; uniclust \| UniRef100\_A0A0F9YGF8 \| 99.5 \| 1.6e-16 \| 3.3e-22 \| 93.7 \| 50 \| (2, 55) \| 69 \| (92, 141) \| 142 \| LysM domain-containing protein \| LysM domain-containing protein; uniclust \| UniRef100\_A0A0D3LEX1 \| 99.4 \| 2.3e-16 \| 4.6e-22 \| 88.3 \| 54 \| (2, 57) \| 69 \| (43, 96) \| 97 \| Putative phospholipid-binding protein,LysM domain-containing protein \| Putative phospholipid-binding protein,LysM domain-containing protein; uniclust \| UniRef100\_A0A0A8HA04 \| 99.4 \| 2.3e-16 \| 4.7e-22 \| 82.1 \| 64 \| (1, 69) \| 69 \| (2, 65) \| 65 \| Phage tail protein X family protein \| Phage tail protein X family protein; uniclust \| UniRef100\_A0A7C2W7N3 \| 99.4 \| 2.6e-16 \| 5.3e-22 \| 92.2 \| 53 \| (2, 56) \| 69 \| (82, 134) \| 135 \| LysM peptidoglycan-binding domain-containing protein \| LysM peptidoglycan-binding domain-containing protein; uniclust \| UniRef100\_A0A0D0RZI1 \| 99.4 \| 2.7e-16 \| 5.6e-22 \| 95.0 \| 54 \| (2, 57) \| 69 \| (112, 165) \| 167 \| LysM domain protein \| LysM domain protein; uniclust \| UniRef100\_A0A1I4XDP5 \| 99.4 \| 3.6e-16 \| 6.9e-22 \| 85.3 \| 55 \| (2, 58) \| 69 \| (35, 89) \| 92 \| LysM domain-containing protein \| LysM domain-containing protein; uniclust \| UniRef100\_A0A521DSA7 \| 99.4 \| 3.6e-16 \| 7.2e-22 \| 84.6 \| 66 \| (1, 69) \| 69 \| (12, 77) \| 80 \| Phage Tail Protein X \| Phage Tail Protein X; uniclust \| UniRef100\_A0A1F8QSY2 \| 99.4 \| 4.9e-16 \| 9.9e-22 \| 81.8 \| 56 \| (1, 56) \| 69 \| (11, 67) \| 68 \| LysM domain-containing protein \| LysM domain-containing protein; uniclust \| UniRef100\_A0A0Q8DXV3 \| 99.4 \| 5.6e-16 \| 1.1e-21 \| 86.5 \| 53 \| (2, 56) \| 69 \| (48, 100) \| 101 \| Peptidoglycan-binding protein \| Peptidoglycan-binding protein; uniclust \| UniRef100\_A0A3M1G446 \| 99.4 \| 6.9e-16 \| 1.3e-21 \| 78.1 \| 52 \| (2, 55) \| 69 \| (10, 61) \| 62 \| LysM peptidoglycan-binding domain-containing protein (Fragment) \| LysM peptidoglycan-binding domain-containing protein (Fragment); uniclust \| UniRef100\_A0A060BNX4 \| 99.4 \| 7.1e-16 \| 1.4e-21 \| 80.6 \| 55 \| (1, 57) \| 69 \| (11, 65) \| 68 \| CAZy families CBM50 protein \| CAZy families CBM50 protein; uniclust \| UniRef100\_A0A0G3EL81 \| 99.4 \| 6.7e-16 \| 1.4e-21 \| 100.6 \| 55 \| (2, 58) \| 69 \| (230, 284) \| 286 \| LysM domain/BON superfamily protein \| LysM domain/BON superfamily protein; uniclust \| UniRef100\_A0A1W1D095 \| 99.4 \| 9.1e-16 \| 1.8e-21 \| 90.8 \| 52 \| (1, 55) \| 69 \| (96, 147) \| 147 \| Ferric siderophore transport system, periplasmic binding protein TonB \| Ferric siderophore transport system, periplasmic binding protein TonB; uniclust \| UniRef100\_A0A0S8E6Q2 \| 99.4 \| 9.8e-16 \| 2.1e-21 \| 94.9 \| 54 \| (2, 58) \| 69 \| (123, 177) \| 191 \| LysM domain-containing protein \| LysM domain-containing protein; uniclust \| UniRef100\_A0A011UMN3 \| 99.4 \| 1e-15 \| 2.2e-21 \| 83.5 \| 67 \| (3, 69) \| 69 \| (10, 78) \| 80 \| Phage tail protein \| Phage tail protein; uniclust \| UniRef100\_A0A1I7HNC1 \| 99.4 \| 1.4e-15 \| 2.6e-21 \| 86.6 \| 66 \| (3, 69) \| 69 \| (73, 138) \| 138 \| P2-like prophage tail protein X \| P2-like prophage tail protein X; uniclust \| UniRef100\_A0A1G0ZIW6 \| 99.4 \| 1.2e-15 \| 2.6e-21 \| 94.8 \| 55 \| (2, 58) \| 69 \| (136, 190) \| 192 \| LysM domain-containing protein \| LysM domain-containing protein; uniclust \| UniRef100\_A0A0E3ZHJ8 \| 99.4 \| 1.3e-15 \| 2.7e-21 \| 87.4 \| 55 \| (2, 58) \| 69 \| (63, 118) \| 120 \| Peptidoglycan-binding protein \| Peptidoglycan-binding protein; uniclust \| UniRef100\_A0A1G1PBG2 \| 99.4 \| 1.5e-15 \| 3.1e-21 \| 85.1 \| 53 \| (2, 56) \| 69 \| (43, 96) \| 97 \| LysM domain-containing protein \| LysM domain-containing protein; uniclust \| UniRef100\_A0A290XBI2 \| 99.4 \| 1.7e-15 \| 3.3e-21 \| 82.1 \| 57 \| (1, 59) \| 69 \| (26, 82) \| 87 \| LysM domain-containing protein \| LysM domain-containing protein; uniclust \| UniRef100\_A0A1W1BHR3 \| 99.4 \| 2e-15 \| 4.1e-21 \| 99.5 \| 57 \| (1, 59) \| 69 \| (254, 310) \| 321 \| LysM domain-containing protein \| LysM domain-containing protein; uniclust \| UniRef100\_D2ZKL4 \| 99.4 \| 2.3e-15 \| 4.2e-21 \| 82.9 \| 64 \| (5, 69) \| 69 \| (46, 109) \| 109 \| Phage Tail Protein X \| Phage Tail Protein X; uniclust \| UniRef100\_A0A7C3XNE8 \| 99.4 \| 2.5e-15 \| 4.8e-21 \| 86.9 \| 50 \| (2, 55) \| 69 \| (81, 130) \| 130 \| LysM domain-containing protein \| LysM domain-containing protein; uniclust \| UniRef100\_A0A061NN19 \| 99.4 \| 2.4e-15 \| 5e-21 \| 86.7 \| 64 \| (1, 68) \| 69 \| (18, 81) \| 114 \| Phage tail protein \| Phage tail protein; uniclust \| UniRef100\_A0A6I0F174 \| 99.4 \| 2.6e-15 \| 5.1e-21 \| 86.6 \| 52 \| (2, 55) \| 69 \| (75, 126) \| 127 \| LysM peptidoglycan-binding domain-containing protein \| LysM peptidoglycan-binding domain-containing protein; uniclust \| UniRef100\_A0A085ZVS9 \| 99.4 \| 2.5e-15 \| 5.1e-21 \| 90.6 \| 55 \| (2, 58) \| 69 \| (103, 157) \| 161 \| Peptidoglycan-binding protein \| Peptidoglycan-binding protein; uniclust \| UniRef100\_A0A0B5FQ72 \| 99.3 \| 3.1e-15 \| 6.4e-21 \| 84.3 \| 53 \| (2, 58) \| 69 \| (39, 91) \| 99 \| LysM domain-containing protein \| LysM domain-containing protein; uniclust \| UniRef100\_A0A1G0XZP4 \| 99.3 \| 5e-15 \| 9.5e-21 \| 80.5 \| 53 \| (2, 56) \| 69 \| (35, 87) \| 88 \| LysM domain-containing protein \| LysM domain-containing protein; uniclust \| UniRef100\_A0A2A4RLX5 \| 99.3 \| 5.1e-15 \| 1.1e-20 \| 94.5 \| 55 \| (2, 58) \| 69 \| (179, 233) \| 246 \| LysM domain-containing protein \| LysM domain-containing protein; uniclust \| UniRef100\_UPI0011B4CCD0 \| 99.3 \| 5.9e-15 \| 1.1e-20 \| 73.3 \| 51 \| (1, 55) \| 69 \| (2, 52) \| 53 \| LysM domain-containing protein \| LysM domain-containing protein; uniclust \| UniRef100\_A0A0R2XHY5 \| 99.3 \| 5.7e-15 \| 1.2e-20 \| 96.9 \| 55 \| (2, 58) \| 69 \| (249, 303) \| 305 \| LysM domain-containing protein \| LysM domain-containing protein; uniclust \| UniRef100\_A0A098PW87 \| 99.3 \| 6.3e-15 \| 1.2e-20 \| 83.3 \| 53 \| (2, 56) \| 69 \| (60, 112) \| 113 \| LysM peptidoglycan-binding domain-containing protein \| LysM peptidoglycan-binding domain-containing protein; uniclust \| UniRef100\_A0A009H911 \| 99.3 \| 6.4e-15 \| 1.3e-20 \| 93.2 \| 56 \| (2, 59) \| 69 \| (150, 205) \| 227 \| LysM domain-containing protein \| LysM domain-containing protein; uniclust \| UniRef100\_A0A023BS42 \| 99.3 \| 7.6e-15 \| 1.5e-20 \| 90.1 \| 56 \| (2, 59) \| 69 \| (99, 154) \| 184 \| Peptidase M23B \| Peptidase M23B; uniclust \| UniRef100\_A0A060C391 \| 99.3 \| 7.9e-15 \| 1.6e-20 \| 87.4 \| 53 \| (2, 56) \| 69 \| (97, 149) \| 150 \| CAZy families CBM50 protein (Fragment) \| CAZy families CBM50 protein (Fragment); uniclust \| UniRef100\_A0A0A1FIF4 \| 99.3 \| 8.8e-15 \| 1.8e-20 \| 89.3 \| 53 \| (2, 56) \| 69 \| (128, 180) \| 182 \| BON domain protein \| BON domain protein; uniclust \| UniRef100\_A0A0D2JA12 \| 99.3 \| 8.6e-15 \| 1.8e-20 \| 80.1 \| 64 \| (1, 68) \| 69 \| (11, 74) \| 81 \| Tail protein \| Tail protein; uniclust \| UniRef100\_A0A1V5UVW7 \| 99.3 \| 9.1e-15 \| 1.8e-20 \| 81.6 \| 53 \| (3, 58) \| 69 \| (44, 96) \| 98 \| LysM domain-containing protein \| LysM domain-containing protein; uniclust \| UniRef100\_A0A3M1EEN8 \| 99.3 \| 9.3e-15 \| 1.8e-20 \| 85.5 \| 56 \| (2, 59) \| 69 \| (74, 129) \| 139 \| LysM peptidoglycan-binding domain-containing protein \| LysM peptidoglycan-binding domain-containing protein; uniclust \| UniRef100\_A0A060BU78 \| 99.3 \| 9.8e-15 \| 1.9e-20 \| 79.9 \| 53 \| (2, 56) \| 69 \| (35, 87) \| 89 \| CAZy families CBM50 protein (Fragment) \| CAZy families CBM50 protein (Fragment); uniclust \| UniRef100\_A0A1U9NM08 \| 99.3 \| 1.1e-14 \| 2.2e-20 \| 88.4 \| 55 \| (1, 57) \| 69 \| (117, 171) \| 172 \| LysM domain/BON superfamily protein \| LysM domain/BON superfamily protein; uniclust \| UniRef100\_A0A7X6LA34 \| 99.3 \| 1.2e-14 \| 2.3e-20 \| 76.7 \| 54 \| (2, 59) \| 69 \| (9, 62) \| 70 \| LysM peptidoglycan-binding domain-containing protein \| LysM peptidoglycan-binding domain-containing protein; uniclust \| UniRef100\_A0A6G5QG46 \| 99.3 \| 1.3e-14 \| 2.5e-20 \| 75.5 \| 67 \| (1, 69) \| 69 \| (2, 68) \| 68 \| Phage tail protein X family protein \| Phage tail protein X family protein; uniclust \| UniRef100\_A0A1R4HC34 \| 99.3 \| 1.4e-14 \| 2.8e-20 \| 93.1 \| 52 \| (2, 55) \| 69 \| (237, 288) \| 288 \| LysM domain-containing protein \| LysM domain-containing protein; uniclust \| UniRef100\_A0A0P6WT71 \| 99.3 \| 1.5e-14 \| 3.2e-20 \| 88.9 \| 58 \| (2, 59) \| 69 \| (113, 171) \| 177 \| LysM domain-containing protein \| LysM domain-containing protein; uniclust \| UniRef100\_E5AKR2 \| 99.3 \| 2e-14 \| 3.6e-20 \| 77.1 \| 67 \| (2, 69) \| 69 \| (23, 90) \| 90 \| Tail protein X \| Tail protein X; uniclust \| UniRef100\_A0A1D2UDF4 \| 99.3 \| 2e-14 \| 3.9e-20 \| 85.4 \| 51 \| (3, 57) \| 69 \| (95, 145) \| 146 \| Peptidoglycan-binding protein \| Peptidoglycan-binding protein; uniclust \| UniRef100\_A0A9D8L854 \| 99.3 \| 2.3e-14 \| 4.3e-20 \| 76.0 \| 66 \| (3, 69) \| 69 \| (12, 78) \| 78 \| Tail protein X \| Tail protein X; uniclust \| UniRef100\_A0A346Y079 \| 99.3 \| 2.2e-14 \| 4.5e-20 \| 89.2 \| 54 \| (1, 58) \| 69 \| (138, 191) \| 194 \| LysM domain-containing protein \| LysM domain-containing protein; uniclust \| UniRef100\_A0A087N8U9 \| 99.3 \| 2.3e-14 \| 4.5e-20 \| 77.3 \| 66 \| (3, 69) \| 69 \| (11, 79) \| 80 \| Phage tail protein \| Phage tail protein; uniclust \| UniRef100\_A0A0S8KMS1 \| 99.3 \| 2.3e-14 \| 4.6e-20 \| 83.8 \| 56 \| (2, 57) \| 69 \| (74, 130) \| 131 \| LysM domain-containing protein \| LysM domain-containing protein; uniclust \| UniRef100\_A0A0P7CPN1 \| 99.3 \| 2.3e-14 \| 4.7e-20 \| 85.2 \| 53 \| (2, 58) \| 69 \| (82, 134) \| 141 \| LysM domain-containing protein \| LysM domain-containing protein; uniclust \| UniRef100\_A0A023BZG8 \| 99.3 \| 2.5e-14 \| 5e-20 \| 86.2 \| 56 \| (2, 59) \| 69 \| (104, 159) \| 163 \| Peptidoglycan-binding protein \| Peptidoglycan-binding protein; uniclust \| UniRef100\_A0A011M151 \| 99.3 \| 2.6e-14 \| 5.3e-20 \| 88.3 \| 55 \| (3, 59) \| 69 \| (109, 163) \| 197 \| Peptidoglycan-binding protein LysM \| Peptidoglycan-binding protein LysM; uniclust \| UniRef100\_A0A068MT20 \| 99.2 \| 2.6e-14 \| 5.4e-20 \| 89.7 \| 54 \| (3, 58) \| 69 \| (144, 197) \| 208 \| LysM domain-containing protein \| LysM domain-containing protein; uniclust \| UniRef100\_A0A1Q7YIP8 \| 99.2 \| 3.1e-14 \| 5.9e-20 \| 74.6 \| 53 \| (2, 56) \| 69 \| (17, 69) \| 71 \| LysM domain-containing protein \| LysM domain-containing protein; uniclust \| UniRef100\_A0A4R5HGF7 \| 99.2 \| 3e-14 \| 6e-20 \| 75.3 \| 68 \| (2, 69) \| 69 \| (3, 70) \| 70 \| Phage tail protein \| Phage tail protein; uniclust \| UniRef100\_A0A0M9U3I1 \| 99.2 \| 3.2e-14 \| 6.1e-20 \| 77.9 \| 57 \| (3, 59) \| 69 \| (31, 88) \| 91 \| LysM domain-containing protein \| LysM domain-containing protein; uniclust \| UniRef100\_A0A0P8WJ67 \| 99.2 \| 3.1e-14 \| 6.2e-20 \| 78.8 \| 63 \| (3, 68) \| 69 \| (17, 79) \| 91 \| Phage tail protein X \| Phage tail protein X; uniclust \| UniRef100\_A0A0H5RIV1 \| 99.2 \| 3.4e-14 \| 6.4e-20 \| 71.9 \| 50 \| (2, 55) \| 69 \| (11, 60) \| 60 \| Mannose-binding lectin \| Mannose-binding lectin; uniclust \| UniRef100\_A0A0K0Y197 \| 99.2 \| 3.3e-14 \| 6.5e-20 \| 92.7 \| 55 \| (2, 58) \| 69 \| (247, 301) \| 303 \| LysM domain/BON superfamily protein \| LysM domain/BON superfamily protein; uniclust \| UniRef100\_A0A1V6GXV1 \| 99.2 \| 3.2e-14 \| 6.7e-20 \| 95.1 \| 53 \| (2, 56) \| 69 \| (310, 362) \| 362 \| LysM domain/BON superfamily protein \| LysM domain/BON superfamily protein; uniclust \| UniRef100\_A0A1G3Z8F3 \| 99.2 \| 3.7e-14 \| 7.5e-20 \| 90.9 \| 52 \| (2, 57) \| 69 \| (198, 249) \| 251 \| LysM domain-containing protein \| LysM domain-containing protein; uniclust \| UniRef100\_A0A2N3IK62 \| 99.2 \| 4.3e-14 \| 8.6e-20 \| 83.6 \| 53 \| (2, 56) \| 69 \| (83, 136) \| 137 \| LysM domain \| LysM domain; uniclust \| UniRef100\_A0A0B0VWC3 \| 99.2 \| 4.8e-14 \| 9e-20 \| 76.8 \| 64 \| (3, 69) \| 69 \| (7, 70) \| 95 \| Phage tail protein \| Phage tail protein; uniclust \| UniRef100\_A0A948S9G9 \| 99.2 \| 5.2e-14 \| 9.5e-20 \| 73.0 \| 51 \| (2, 54) \| 69 \| (23, 73) \| 73 \| LysM peptidoglycan-binding domain-containing protein \| LysM peptidoglycan-binding domain-containing protein; uniclust \| UniRef100\_A0A024K1G4 \| 99.2 \| 5.6e-14 \| 1.1e-19 \| 88.5 \| 53 \| (2, 58) \| 69 \| (174, 226) \| 237 \| Mannose-binding lectin \| Mannose-binding lectin; uniclust \| UniRef100\_A0A0Q4L1W6 \| 99.2 \| 5.7e-14 \| 1.2e-19 \| 80.4 \| 54 \| (2, 59) \| 69 \| (43, 96) \| 107 \| LysM domain-containing protein \| LysM domain-containing protein; uniclust \| UniRef100\_A0A1G0YUF0 \| 99.2 \| 6.1e-14 \| 1.3e-19 \| 91.7 \| 56 \| (2, 59) \| 69 \| (239, 294) \| 298 \| LysM domain-containing protein \| LysM domain-containing protein; uniclust \| UniRef100\_A0A2V8QG62 \| 99.2 \| 7.4e-14 \| 1.4e-19 \| 83.8 \| 54 \| (2, 57) \| 69 \| (108, 161) \| 163 \| Peptidoglycan-binding protein \| Peptidoglycan-binding protein; uniclust \| UniRef100\_A0A1G8UZV7 \| 99.2 \| 7.2e-14 \| 1.4e-19 \| 85.3 \| 52 \| (2, 55) \| 69 \| (124, 176) \| 176 \| LysM domain-containing protein \| LysM domain-containing protein; uniclust \| UniRef100\_A0A3M2CM87 \| 99.2 \| 7.4e-14 \| 1.5e-19 \| 88.7 \| 54 \| (2, 57) \| 69 \| (190, 243) \| 249 \| LysM domain-containing protein \| LysM domain-containing protein; uniclust \| UniRef100\_A0A1G1LW77 \| 99.2 \| 7.7e-14 \| 1.5e-19 \| 87.3 \| 56 \| (2, 59) \| 69 \| (158, 213) \| 218 \| LysM domain-containing protein \| LysM domain-containing protein; uniclust \| UniRef100\_A0A3M1PZI9 \| 99.2 \| 8.5e-14 \| 1.7e-19 \| 93.3 \| 52 \| (2, 57) \| 69 \| (224, 275) \| 405 \| LysM peptidoglycan-binding domain-containing protein \| LysM peptidoglycan-binding domain-containing protein; uniclust \| UniRef100\_A0A1I3PDS2 \| 99.2 \| 8.8e-14 \| 1.8e-19 \| 88.9 \| 53 \| (2, 56) \| 69 \| (123, 175) \| 249 \| LysM repeat-containing protein \| LysM repeat-containing protein; uniclust \| UniRef100\_A0A016QMV4 \| 99.2 \| 9e-14 \| 1.8e-19 \| 89.3 \| 53 \| (2, 58) \| 69 \| (214, 266) \| 273 \| Peptidoglycan-binding LysM \| Peptidoglycan-binding LysM; uniclust \| UniRef100\_A0A024YYK6 \| 99.2 \| 8.5e-14 \| 1.8e-19 \| 91.6 \| 54 \| (2, 59) \| 69 \| (201, 254) \| 297 \| Peptidase M23B \| Peptidase M23B; uniclust \| UniRef100\_A0A084T293 \| 99.2 \| 8.9e-14 \| 1.8e-19 \| 90.6 \| 54 \| (2, 59) \| 69 \| (210, 263) \| 283 \| LysM domain-containing protein \| LysM domain-containing protein; uniclust \| UniRef100\_A0A069A2Y0 \| 99.2 \| 9.5e-14 \| 2e-19 \| 88.7 \| 52 \| (2, 55) \| 69 \| (182, 233) \| 233 \| LysM domain protein \| LysM domain protein; uniclust \| UniRef100\_A0A962ECX3 \| 99.2 \| 1.1e-13 \| 2.1e-19 \| 81.3 \| 56 \| (2, 59) \| 69 \| (92, 147) \| 152 \| LysM peptidoglycan-binding domain-containing protein \| LysM peptidoglycan-binding domain-containing protein; uniclust \| UniRef100\_A0A0B1KPG3 \| 99.2 \| 1.2e-13 \| 2.3e-19 \| 72.1 \| 68 \| (2, 69) \| 69 \| (3, 72) \| 72 \| Phage tail protein \| Phage tail protein; uniclust \| UniRef100\_A0A1G1N886 \| 99.2 \| 1.2e-13 \| 2.4e-19 \| 86.6 \| 54 \| (2, 57) \| 69 \| (132, 185) \| 201 \| LysM domain-containing protein \| LysM domain-containing protein; uniclust \| UniRef100\_A0A086ML12 \| 99.2 \| 1.2e-13 \| 2.5e-19 \| 87.9 \| 55 \| (2, 58) \| 69 \| (165, 219) \| 230 \| Peptigoglycan-binding protein LysM \| Peptigoglycan-binding protein LysM; uniclust \| UniRef100\_A0A0N7LZZ4 \| 99.2 \| 1.3e-13 \| 2.6e-19 \| 86.1 \| 53 \| (2, 56) \| 69 \| (154, 206) \| 207 \| LysM domain/BON superfamily protein \| LysM domain/BON superfamily protein; uniclust \| UniRef100\_A0A4P5YH32 \| 99.2 \| 1.6e-13 \| 3.1e-19 \| 86.8 \| 52 \| (2, 57) \| 69 \| (178, 229) \| 231 \| LysM domain-containing protein \| LysM domain-containing protein; uniclust \| UniRef100\_A0A1Z9KPR5 \| 99.2 \| 1.5e-13 \| 3.1e-19 \| 87.4 \| 54 \| (3, 59) \| 69 \| (175, 228) \| 235 \| LysM domain-containing protein \| LysM domain-containing protein; uniclust \| UniRef100\_A0A2H0LH77 \| 99.2 \| 1.7e-13 \| 3.4e-19 \| 85.6 \| 55 \| (2, 58) \| 69 \| (132, 188) \| 204 \| LysM domain-containing protein \| LysM domain-containing protein; uniclust \| UniRef100\_A0A399ZZW7 \| 99.2 \| 1.7e-13 \| 3.4e-19 \| 93.3 \| 56 \| (2, 59) \| 69 \| (373, 428) \| 435 \| LysM domain-containing protein \| LysM domain-containing protein; uniclust \| UniRef100\_A0A0S7WJV5 \| 99.2 \| 1.7e-13 \| 3.5e-19 \| 82.7 \| 56 \| (3, 58) \| 69 \| (89, 146) \| 149 \| LysM domain-containing protein \| LysM domain-containing protein; uniclust \| UniRef100\_A0A1X7MBB4 \| 99.1 \| 2e-13 \| 4e-19 \| 78.1 \| 53 \| (2, 56) \| 69 \| (59, 111) \| 113 \| LysM domain/BON superfamily protein \| LysM domain/BON superfamily protein; uniclust \| UniRef100\_A0A2E8Y824 \| 99.1 \| 2.1e-13 \| 4e-19 \| 74.1 \| 53 \| (1, 55) \| 69 \| (31, 84) \| 84 \| LysM domain-containing protein \| LysM domain-containing protein; uniclust \| UniRef100\_A0A2A4RLX5 \| 99.1 \| 2e-13 \| 4e-19 \| 87.6 \| 53 \| (2, 58) \| 69 \| (105, 157) \| 246 \| LysM domain-containing protein \| LysM domain-containing protein; uniclust \| UniRef100\_A0A358MIT7 \| 99.1 \| 2.4e-13 \| 4.4e-19 \| 69.3 \| 51 \| (3, 55) \| 69 \| (14, 64) \| 65 \| LysM domain-containing protein \| LysM domain-containing protein; uniclust \| UniRef100\_A0A0H3L797 \| 99.1 \| 2.5e-13 \| 4.6e-19 \| 79.2 \| 64 \| (5, 69) \| 69 \| (93, 156) \| 156 \| Phage tail protein X \| Phage tail protein X; uniclust \| UniRef100\_A0A022MNT4 \| 99.1 \| 2.2e-13 \| 4.7e-19 \| 90.1 \| 53 \| (2, 58) \| 69 \| (221, 273) \| 302 \| Peptidase M23B \| Peptidase M23B; uniclust \| UniRef100\_A0A143HC07 \| 99.1 \| 2.4e-13 \| 4.8e-19 \| 77.5 \| 55 \| (1, 58) \| 69 \| (2, 56) \| 108 \| Phage tail protein \| Phage tail protein; uniclust \| UniRef100\_A0A0A0HQW3 \| 99.1 \| 2.5e-13 \| 5.1e-19 \| 87.1 \| 54 \| (2, 57) \| 69 \| (191, 244) \| 246 \| LysM domain protein \| LysM domain protein; uniclust \| UniRef100\_A0A6C0GK95 \| 99.1 \| 2.8e-13 \| 5.2e-19 \| 76.9 \| 55 \| (2, 58) \| 69 \| (53, 107) \| 119 \| LysM peptidoglycan-binding domain-containing protein \| LysM peptidoglycan-binding domain-containing protein; uniclust \| UniRef100\_A0A098B9N3 \| 99.1 \| 2.7e-13 \| 5.4e-19 \| 88.9 \| 53 \| (2, 58) \| 69 \| (193, 245) \| 309 \| Peptidoglycan-binding LysM \| Peptidoglycan-binding LysM; uniclust \| UniRef100\_A0A2G2AA53 \| 99.1 \| 2.8e-13 \| 5.5e-19 \| 82.8 \| 54 \| (2, 57) \| 69 \| (124, 177) \| 178 \| LysM domain-containing protein \| LysM domain-containing protein; uniclust \| UniRef100\_A0A0V0PS59 \| 99.1 \| 2.9e-13 \| 5.6e-19 \| 76.3 \| 50 \| (1, 57) \| 69 \| (3, 52) \| 103 \| Peptidoglycan-binding protein \| Peptidoglycan-binding protein; uniclust \| UniRef100\_A0A1M9YTZ0 \| 99.1 \| 3.2e-13 \| 6e-19 \| 79.7 \| 50 \| (2, 55) \| 69 \| (103, 152) \| 152 \| Putative mannose-specific lectin \| Putative mannose-specific lectin; uniclust \| UniRef100\_A0A1G1IT91 \| 99.1 \| 3.2e-13 \| 6.6e-19 \| 85.7 \| 55 \| (2, 58) \| 69 \| (156, 210) \| 223 \| LysM domain-containing protein \| LysM domain-containing protein; uniclust \| UniRef100\_A0A2V2D9M4 \| 99.1 \| 3.5e-13 \| 6.8e-19 \| 68.5 \| 49 \| (1, 56) \| 69 \| (7, 55) \| 57 \| LysM domain-containing protein \| LysM domain-containing protein; uniclust \| UniRef100\_A0A2D5WBI1 \| 99.1 \| 3.4e-13 \| 6.9e-19 \| 87.1 \| 52 \| (1, 58) \| 69 \| (139, 190) \| 266 \| LysM domain-containing protein \| LysM domain-containing protein; uniclust \| UniRef100\_A0A0N1F3Q5 \| 99.1 \| 3.5e-13 \| 6.9e-19 \| 73.0 \| 65 \| (5, 69) \| 69 \| (12, 79) \| 79 \| Phage tail protein \| Phage tail protein; uniclust \| UniRef100\_A0A0E3XT95 \| 99.1 \| 3.9e-13 \| 7.8e-19 \| 86.7 \| 54 \| (1, 58) \| 69 \| (38, 91) \| 278 \| Mannose-binding protein \| Mannose-binding protein; uniclust \| UniRef100\_A0A2W4LKG2 \| 99.1 \| 4.1e-13 \| 8e-19 \| 77.9 \| 53 \| (2, 58) \| 69 \| (66, 118) \| 124 \| Peptidoglycan-binding protein (Fragment) \| Peptidoglycan-binding protein (Fragment); uniclust \| UniRef100\_A0A9E4JKM9 \| 99.1 \| 4.6e-13 \| 8.4e-19 \| 80.3 \| 59 \| (1, 59) \| 69 \| (24, 83) \| 192 \| LysM peptidoglycan-binding domain-containing protein \| LysM peptidoglycan-binding domain-containing protein; uniclust \| UniRef100\_A0A1V6BES0 \| 99.1 \| 4.4e-13 \| 8.7e-19 \| 70.6 \| 48 \| (3, 57) \| 69 \| (15, 62) \| 67 \| Gamma-D-glutamyl-L-diamino acid endopeptidase 1 \| Gamma-D-glutamyl-L-diamino acid endopeptidase 1; uniclust \| UniRef100\_A0A7C1AWR5 \| 99.1 \| 4.9e-13 \| 9.4e-19 \| 76.3 \| 56 \| (2, 59) \| 69 \| (54, 109) \| 115 \| LysM peptidoglycan-binding domain-containing protein \| LysM peptidoglycan-binding domain-containing protein; uniclust \| UniRef100\_A0A959MSN4 \| 99.1 \| 5.1e-13 \| 9.4e-19 \| 68.5 \| 56 \| (2, 59) \| 69 \| (5, 60) \| 67 \| LysM peptidoglycan-binding domain-containing protein \| LysM peptidoglycan-binding domain-containing protein; uniclust \| UniRef100\_A0A7C5GIV3 \| 99.1 \| 5e-13 \| 9.5e-19 \| 80.0 \| 53 \| (1, 55) \| 69 \| (111, 163) \| 163 \| LysM peptidoglycan-binding domain-containing protein \| LysM peptidoglycan-binding domain-containing protein; uniclust \| UniRef100\_A0A1D7YSS0 \| 99.1 \| 4.7e-13 \| 9.6e-19 \| 83.7 \| 56 \| (2, 59) \| 69 \| (140, 195) \| 199 \| LysM domain-containing protein \| LysM domain-containing protein; uniclust \| UniRef100\_A0A849XXF2 \| 99.1 \| 5.4e-13 \| 9.9e-19 \| 69.7 \| 51 \| (2, 55) \| 69 \| (25, 75) \| 75 \| LysM peptidoglycan-binding domain-containing protein \| LysM peptidoglycan-binding domain-containing protein; uniclust \| UniRef100\_A0A3M1E5Y8 \| 99.1 \| 5.5e-13 \| 1e-18 \| 87.7 \| 55 \| (2, 58) \| 69 \| (323, 377) \| 378 \| LysM peptidoglycan-binding domain-containing protein \| LysM peptidoglycan-binding domain-containing protein; uniclust \| UniRef100\_A0A2E2NN19 \| 99.1 \| 5.6e-13 \| 1.1e-18 \| 87.5 \| 55 \| (2, 58) \| 69 \| (271, 325) \| 327 \| LysM domain-containing protein \| LysM domain-containing protein; uniclust \| UniRef100\_A0A0F9II26 \| 99.1 \| 5.8e-13 \| 1.1e-18 \| 71.0 \| 50 \| (2, 55) \| 69 \| (26, 76) \| 76 \| LysM domain-containing protein (Fragment) \| LysM domain-containing protein (Fragment); uniclust \| UniRef100\_A0A351VDG3 \| 99.1 \| 5.5e-13 \| 1.1e-18 \| 75.2 \| 56 \| (1, 59) \| 69 \| (3, 58) \| 99 \| Phage tail protein \| Phage tail protein; uniclust \| UniRef100\_A0A2A2TBS7 \| 99.1 \| 5.8e-13 \| 1.1e-18 \| 70.4 \| 49 \| (2, 56) \| 69 \| (5, 53) \| 73 \| LysM domain-containing protein \| LysM domain-containing protein; uniclust \| UniRef100\_A0A142X5D6 \| 99.1 \| 5.6e-13 \| 1.1e-18 \| 88.2 \| 54 \| (2, 57) \| 69 \| (183, 236) \| 334 \| LysM domain/BON superfamily protein \| LysM domain/BON superfamily protein; uniclust \| UniRef100\_A0A3A4QHX3 \| 99.1 \| 5.6e-13 \| 1.2e-18 \| 90.2 \| 58 \| (2, 59) \| 69 \| (210, 267) \| 387 \| LysM peptidoglycan-binding domain-containing protein \| LysM peptidoglycan-binding domain-containing protein; uniclust \| UniRef100\_A0A2D5B1A9 \| 99.1 \| 5.5e-13 \| 1.2e-18 \| 89.2 \| 53 \| (2, 58) \| 69 \| (181, 233) \| 333 \| LysM domain-containing protein \| LysM domain-containing protein; uniclust \| UniRef100\_A0A1C3E468 \| 99.1 \| 5.7e-13 \| 1.2e-18 \| 90.3 \| 52 \| (1, 54) \| 69 \| (210, 261) \| 392 \| LysM domain-containing protein \| LysM domain-containing protein; uniclust \| UniRef100\_A0A7Y2EMS1 \| 99.1 \| 6.5e-13 \| 1.2e-18 \| 72.6 \| 53 \| (2, 56) \| 69 \| (39, 91) \| 92 \| LysM peptidoglycan-binding domain-containing protein \| LysM peptidoglycan-binding domain-containing protein; uniclust \| UniRef100\_A0A074LNR3 \| 99.1 \| 6.6e-13 \| 1.3e-18 \| 82.9 \| 51 \| (1, 55) \| 69 \| (161, 211) \| 211 \| LysM domain-containing protein \| LysM domain-containing protein; uniclust \| UniRef100\_A0A937VFL9 \| 99.1 \| 7e-13 \| 1.3e-18 \| 76.1 \| 51 \| (2, 54) \| 69 \| (78, 128) \| 128 \| LysM peptidoglycan-binding domain-containing protein \| LysM peptidoglycan-binding domain-containing protein; uniclust \| UniRef100\_A0A177Q767 \| 99.1 \| 6.6e-13 \| 1.3e-18 \| 86.2 \| 55 \| (2, 58) \| 69 \| (225, 279) \| 282 \| LysM domain-containing protein \| LysM domain-containing protein; uniclust \| UniRef100\_A0A1G2XHC5 \| 99.1 \| 6.7e-13 \| 1.3e-18 \| 80.2 \| 55 \| (2, 58) \| 69 \| (106, 160) \| 164 \| LysM domain-containing protein \| LysM domain-containing protein; uniclust \| UniRef100\_A0A069DLW2 \| 99.1 \| 6.2e-13 \| 1.4e-18 \| 94.0 \| 51 \| (1, 57) \| 69 \| (25, 75) \| 570 \| LysM domain-containing protein \| LysM domain-containing protein; uniclust \| UniRef100\_A0A1H6F920 \| 99.1 \| 7e-13 \| 1.4e-18 \| 89.4 \| 53 \| (2, 56) \| 69 \| (334, 386) \| 386 \| LysM domain/BON superfamily protein \| LysM domain/BON superfamily protein; uniclust \| UniRef100\_A0A0F0HL11 \| 99.1 \| 7.6e-13 \| 1.5e-18 \| 74.4 \| 51 \| (3, 56) \| 69 \| (45, 95) \| 97 \| LysM domain-containing protein \| LysM domain-containing protein; uniclust \| UniRef100\_A0A098MA02 \| 99.1 \| 7.8e-13 \| 1.5e-18 \| 76.6 \| 49 \| (2, 57) \| 69 \| (8, 56) \| 120 \| LysM domain-containing protein \| LysM domain-containing protein; uniclust \| UniRef100\_A0A3C0A1K5 \| 99.1 \| 8.2e-13 \| 1.6e-18 \| 71.1 \| 53 \| (2, 56) \| 69 \| (26, 78) \| 80 \| LysM domain-containing protein (Fragment) \| LysM domain-containing protein (Fragment); uniclust \| UniRef100\_A0A7X7RKI7 \| 99.1 \| 8.2e-13 \| 1.6e-18 \| 83.0 \| 56 \| (2, 57) \| 69 \| (180, 235) \| 236 \| LysM peptidoglycan-binding domain-containing protein \| LysM peptidoglycan-binding domain-containing protein; uniclust \| UniRef100\_A0A0G0VNZ0 \| 99.1 \| 8e-13 \| 1.6e-18 \| 78.9 \| 51 \| (2, 56) \| 69 \| (88, 138) \| 143 \| LysM domain-containing protein \| LysM domain-containing protein; uniclust \| UniRef100\_A0A0P6YX09 \| 99.1 \| 9e-13 \| 1.6e-18 \| 67.5 \| 52 \| (2, 59) \| 69 \| (11, 62) \| 66 \| LysM domain-containing protein \| LysM domain-containing protein; uniclust \| UniRef100\_A0A1Q7A1S2 \| 99.1 \| 9.7e-13 \| 1.8e-18 \| 77.3 \| 55 \| (3, 59) \| 69 \| (24, 79) \| 150 \| LysM domain-containing protein \| LysM domain-containing protein; uniclust \| UniRef100\_A0A2D5Z103 \| 99.1 \| 9.7e-13 \| 1.8e-18 \| 78.6 \| 49 \| (2, 54) \| 69 \| (119, 167) \| 167 \| LysM domain-containing protein \| LysM domain-containing protein; uniclust \| UniRef100\_A0A066U3S9 \| 99.1 \| 8.7e-13 \| 1.9e-18 \| 87.9 \| 54 \| (2, 59) \| 69 \| (211, 264) \| 325 \| Peptidase M23B \| Peptidase M23B; uniclust \| UniRef100\_A0A1J5SZ61 \| 99.1 \| 9.4e-13 \| 1.9e-18 \| 87.7 \| 55 \| (2, 58) \| 69 \| (297, 351) \| 353 \| Chromosome partition protein Smc \| Chromosome partition protein Smc; uniclust \| UniRef100\_A0A7V3PXX5 \| 99.1 \| 1.1e-12 \| 2e-18 \| 66.4 \| 54 \| (3, 58) \| 69 \| (1, 54) \| 62 \| LysM peptidoglycan-binding domain-containing protein \| LysM peptidoglycan-binding domain-containing protein; uniclust \| UniRef100\_UPI0002EF452D \| 99.0 \| 1.1e-12 \| 2.1e-18 \| 70.3 \| 56 \| (2, 58) \| 69 \| (17, 74) \| 80 \| LysM peptidoglycan-binding domain-containing protein \| LysM peptidoglycan-binding domain-containing protein; uniclust \| UniRef100\_A0A1G6JKH2 \| 99.0 \| 1e-12 \| 2.1e-18 \| 86.9 \| 52 \| (1, 58) \| 69 \| (3, 54) \| 325 \| LysM domain-containing protein \| LysM domain-containing protein; uniclust \| UniRef100\_A0A136P411 \| 99.0 \| 1.1e-12 \| 2.1e-18 \| 68.7 \| 51 \| (2, 54) \| 69 \| (26, 76) \| 76 \| LysM domain/BON superfamily protein \| LysM domain/BON superfamily protein; uniclust \| UniRef100\_A0A2E0Y0Y8 \| 99.0 \| 1.1e-12 \| 2.2e-18 \| 85.0 \| 52 \| (2, 57) \| 69 \| (188, 239) \| 312 \| LysM domain-containing protein \| LysM domain-containing protein; uniclust \| UniRef100\_A0A520XTY3 \| 99.0 \| 1.1e-12 \| 2.2e-18 \| 81.7 \| 55 \| (2, 58) \| 69 \| (141, 195) \| 213 \| Peptidoglycan-binding protein LysM \| Peptidoglycan-binding protein LysM; uniclust \| UniRef100\_A0A356KNA6 \| 99.0 \| 1.2e-12 \| 2.3e-18 \| 70.9 \| 55 \| (2, 58) \| 69 \| (30, 84) \| 87 \| LysM domain-containing protein (Fragment) \| LysM domain-containing protein (Fragment); uniclust \| UniRef100\_A0A6J5KM05 \| 99.0 \| 1.1e-12 \| 2.3e-18 \| 73.6 \| 51 \| (3, 56) \| 69 \| (42, 92) \| 94 \| LysM domain \| LysM domain; uniclust \| UniRef100\_A0A7X1TZH6 \| 99.0 \| 1.3e-12 \| 2.3e-18 \| 66.0 \| 52 \| (3, 58) \| 69 \| (6, 57) \| 61 \| LysM peptidoglycan-binding domain-containing protein \| LysM peptidoglycan-binding domain-containing protein; uniclust \| UniRef100\_A0A014P2G8 \| 99.0 \| 1.1e-12 \| 2.4e-18 \| 85.3 \| 54 \| (2, 59) \| 69 \| (202, 255) \| 266 \| Peptidoglycan-binding protein \| Peptidoglycan-binding protein; uniclust \| UniRef100\_A0A2E2N8R8 \| 99.0 \| 1.1e-12 \| 2.4e-18 \| 87.9 \| 54 \| (2, 57) \| 69 \| (291, 344) \| 347 \| LysM domain-containing protein \| LysM domain-containing protein; uniclust \| UniRef100\_A0A1W9UYL2 \| 99.0 \| 1.3e-12 \| 2.4e-18 \| 83.9 \| 54 \| (2, 57) \| 69 \| (235, 288) \| 289 \| LysM domain-containing protein \| LysM domain-containing protein; uniclust \| UniRef100\_A0A1H2WQU9 \| 99.0 \| 1.2e-12 \| 2.4e-18 \| 85.0 \| 50 \| (3, 54) \| 69 \| (219, 268) \| 268 \| LysM domain-containing protein \| LysM domain-containing protein; uniclust \| UniRef100\_A0A1Z9QC33 \| 99.0 \| 1.2e-12 \| 2.5e-18 \| 86.2 \| 57 \| (1, 59) \| 69 \| (233, 289) \| 293 \| LysM domain-containing protein \| LysM domain-containing protein; uniclust \| UniRef100\_A0A1W1CAC6 \| 99.0 \| 1.3e-12 \| 2.5e-18 \| 71.4 \| 53 \| (1, 55) \| 69 \| (38, 90) \| 90 \| LysM domain-containing protein \| LysM domain-containing protein; uniclust \| UniRef100\_A0A1H7C4U6 \| 99.0 \| 1.3e-12 \| 2.5e-18 \| 70.8 \| 53 \| (2, 56) \| 69 \| (33, 85) \| 86 \| LysM domain-containing protein \| LysM domain-containing protein; uniclust \| UniRef100\_A0A0C2VQS2 \| 99.0 \| 1.2e-12 \| 2.5e-18 \| 81.3 \| 53 \| (2, 56) \| 69 \| (137, 189) \| 190 \| LysM domain-containing protein \| LysM domain-containing protein; uniclust \| UniRef100\_A0A6I7PWC1 \| 99.0 \| 1.3e-12 \| 2.7e-18 \| 86.8 \| 51 \| (2, 54) \| 69 \| (302, 352) \| 352 \| LysM peptidoglycan-binding domain-containing protein \| LysM peptidoglycan-binding domain-containing protein; uniclust \| UniRef100\_A0A1V4QZM9 \| 99.0 \| 1.3e-12 \| 2.7e-18 \| 85.4 \| 56 \| (2, 57) \| 69 \| (229, 284) \| 288 \| LysM domain-containing protein \| LysM domain-containing protein; uniclust \| UniRef100\_A0A151Z132 \| 99.0 \| 1.3e-12 \| 2.7e-18 \| 84.1 \| 51 \| (1, 57) \| 69 \| (38, 88) \| 243 \| LysM domain-containing protein \| LysM domain-containing protein; uniclust \| UniRef100\_A0A376KHT5 \| 99.0 \| 1.5e-12 \| 2.8e-18 \| 74.9 \| 63 \| (4, 69) \| 69 \| (3, 65) \| 139 \| Tail protein X (GpX) \| Tail protein X (GpX); uniclust \| UniRef100\_A0A348PCI9 \| 99.0 \| 1.5e-12 \| 2.9e-18 \| 85.5 \| 53 \| (2, 58) \| 69 \| (174, 226) \| 301 \| LysM domain-containing protein \| LysM domain-containing protein; uniclust \| UniRef100\_A0A3M1EYD0 \| 99.0 \| 1.5e-12 \| 3e-18 \| 78.9 \| 53 \| (2, 57) \| 69 \| (121, 174) \| 175 \| LysM peptidoglycan-binding domain-containing protein \| LysM peptidoglycan-binding domain-containing protein; uniclust \| UniRef100\_A0A1M6M9A3 \| 99.0 \| 1.5e-12 \| 3e-18 \| 80.8 \| 50 \| (2, 55) \| 69 \| (149, 198) \| 199 \| LysM domain-containing protein \| LysM domain-containing protein; uniclust \| UniRef100\_A0A0B3RRN2 \| 99.0 \| 1.7e-12 \| 3.4e-18 \| 85.0 \| 54 \| (3, 58) \| 69 \| (261, 314) \| 316 \| LysM domain protein \| LysM domain protein; uniclust \| UniRef100\_A0A0C2W9T9 \| 99.0 \| 1.9e-12 \| 3.8e-18 \| 82.3 \| 51 \| (1, 57) \| 69 \| (1, 51) \| 247 \| LysM domain-containing protein \| LysM domain-containing protein; uniclust \| UniRef100\_A0A0Q9LMA6 \| 99.0 \| 1.8e-12 \| 3.8e-18 \| 93.1 \| 49 \| (1, 55) \| 69 \| (22, 70) \| 708 \| LysM domain-containing protein \| LysM domain-containing protein; uniclust \| UniRef100\_A0A015KHE2 \| 99.0 \| 1.9e-12 \| 3.9e-18 \| 92.0 \| 50 \| (1, 56) \| 69 \| (93, 142) \| 640 \| LysM domain-containing protein \| LysM domain-containing protein; uniclust \| UniRef100\_A0A1G1NHF1 \| 99.0 \| 2.1e-12 \| 4e-18 \| 71.3 \| 54 \| (3, 58) \| 69 \| (22, 77) \| 95 \| LysM domain-containing protein \| LysM domain-containing protein; uniclust \| UniRef100\_A0A0A3HNG1 \| 99.0 \| 1.9e-12 \| 4.1e-18 \| 83.8 \| 50 \| (2, 56) \| 69 \| (194, 243) \| 244 \| LysM domain-containing protein \| LysM domain-containing protein; uniclust \| UniRef100\_A0A2E2B0B4 \| 99.0 \| 2e-12 \| 4.1e-18 \| 86.4 \| 56 \| (2, 59) \| 69 \| (297, 353) \| 362 \| LysM domain-containing protein \| LysM domain-containing protein; uniclust \| UniRef100\_A0A1G0XXQ9 \| 99.0 \| 2.2e-12 \| 4.4e-18 \| 84.0 \| 56 \| (2, 59) \| 69 \| (214, 269) \| 282 \| LysM domain-containing protein \| LysM domain-containing protein; uniclust \| UniRef100\_A0A3M1ZW81 \| 99.0 \| 2.3e-12 \| 4.5e-18 \| 75.6 \| 55 \| (2, 56) \| 69 \| (83, 138) \| 139 \| LysM peptidoglycan-binding domain-containing protein \| LysM peptidoglycan-binding domain-containing protein; uniclust \| UniRef100\_A0A177Q305 \| 99.0 \| 2.4e-12 \| 4.6e-18 \| 75.1 \| 56 \| (2, 59) \| 69 \| (75, 130) \| 134 \| LysM domain-containing protein \| LysM domain-containing protein; uniclust \| UniRef100\_A0A1P8WSI9 \| 99.0 \| 2.2e-12 \| 4.7e-18 \| 86.3 \| 56 \| (1, 58) \| 69 \| (207, 262) \| 337 \| LysM domain/BON superfamily protein \| LysM domain/BON superfamily protein; uniclust \| UniRef100\_A0A0G0CX04 \| 99.0 \| 2.4e-12 \| 4.8e-18 \| 82.3 \| 49 \| (3, 55) \| 69 \| (183, 231) \| 232 \| LysM domain-containing protein \| LysM domain-containing protein; uniclust \| UniRef100\_A0A7C4EC90 \| 99.0 \| 2.5e-12 \| 4.9e-18 \| 84.8 \| 53 \| (2, 58) \| 69 \| (193, 245) \| 332 \| LysM peptidoglycan-binding domain-containing protein \| LysM peptidoglycan-binding domain-containing protein; uniclust \| UniRef100\_A0A061NGS6 \| 99.0 \| 2.3e-12 \| 5e-18 \| 83.7 \| 56 \| (2, 59) \| 69 \| (186, 242) \| 255 \| Phage-like element PBSX protein XkdP \| Phage-like element PBSX protein XkdP; uniclust \| UniRef100\_A0A2D5B1A9 \| 99.0 \| 2.3e-12 \| 5e-18 \| 86.4 \| 53 \| (2, 58) \| 69 \| (258, 310) \| 333 \| LysM domain-containing protein \| LysM domain-containing protein; uniclust \| UniRef100\_A0A0E3UH91 \| 99.0 \| 2.5e-12 \| 5.1e-18 \| 84.7 \| 53 \| (1, 55) \| 69 \| (261, 313) \| 313 \| LysM domain-containing protein \| LysM domain-containing protein; uniclust \| UniRef100\_A0A7C1J8S6 \| 99.0 \| 2.8e-12 \| 5.1e-18 \| 67.3 \| 48 \| (3, 54) \| 69 \| (29, 76) \| 76 \| LysM peptidoglycan-binding domain-containing protein \| LysM peptidoglycan-binding domain-containing protein; uniclust \| UniRef100\_A0A7J9WBE9 \| 99.0 \| 2.6e-12 \| 5.1e-18 \| 78.3 \| 53 \| (2, 58) \| 69 \| (119, 171) \| 173 \| LysM peptidoglycan-binding domain-containing protein \| LysM peptidoglycan-binding domain-containing protein; uniclust \| UniRef100\_A0A0T7DV60 \| 99.0 \| 2.6e-12 \| 5.2e-18 \| 69.6 \| 68 \| (1, 68) \| 69 \| (5, 75) \| 77 \| Phage tail protein \| Phage tail protein; uniclust \| UniRef100\_A0A0G0HFV7 \| 99.0 \| 2.6e-12 \| 5.2e-18 \| 79.0 \| 50 \| (3, 56) \| 69 \| (131, 180) \| 182 \| D-mannose binding lectin \| D-mannose binding lectin; uniclust \| UniRef100\_A0A0P0FPI5 \| 99.0 \| 2.7e-12 \| 5.3e-18 \| 82.3 \| 55 \| (2, 58) \| 69 \| (200, 254) \| 256 \| LysM domain/BON superfamily protein \| LysM domain/BON superfamily protein; uniclust \| UniRef100\_A0A1M4LC88 \| 99.0 \| 2.9e-12 \| 5.3e-18 \| 83.0 \| 67 \| (2, 69) \| 69 \| (301, 368) \| 368 \| Phage tail tape measure protein domain-containing protein \| Phage tail tape measure protein domain-containing protein; uniclust \| UniRef100\_A0A1I2AZK8 \| 99.0 \| 2.6e-12 \| 5.4e-18 \| 88.3 \| 52 \| (1, 58) \| 69 \| (4, 55) \| 439 \| Morphogenetic protein associated with SpoVID (Fragment) \| Morphogenetic protein associated with SpoVID (Fragment); uniclust \| UniRef100\_UPI0003462B98 \| 99.0 \| 3e-12 \| 5.5e-18 \| 68.2 \| 58 \| (2, 59) \| 69 \| (4, 62) \| 83 \| LysM peptidoglycan-binding domain-containing protein \| LysM peptidoglycan-binding domain-containing protein; uniclust \| UniRef100\_A0A0F9YZQ3 \| 99.0 \| 2.7e-12 \| 5.6e-18 \| 81.6 \| 53 \| (2, 58) \| 69 \| (92, 144) \| 217 \| Cell-wall-binding phage-encoded protein \| Cell-wall-binding phage-encoded protein; uniclust \| UniRef100\_A0A0B1YCV7 \| 99.0 \| 2.8e-12 \| 5.6e-18 \| 85.8 \| 53 \| (1, 59) \| 69 \| (19, 71) \| 363 \| Morphogenetic protein associated with SpoVID \| Morphogenetic protein associated with SpoVID; uniclust \| UniRef100\_A0A0S8ICA2 \| 99.0 \| 2.9e-12 \| 5.6e-18 \| 72.2 \| 56 \| (2, 57) \| 69 \| (45, 101) \| 103 \| LysM domain-containing protein \| LysM domain-containing protein; uniclust \| UniRef100\_A0A0B8MZE7 \| 99.0 \| 3e-12 \| 5.7e-18 \| 79.4 \| 49 \| (2, 54) \| 69 \| (162, 210) \| 210 \| 5'-nucleotidase \| 5'-nucleotidase; uniclust \| UniRef100\_A0A080LZ55 \| 99.0 \| 2.8e-12 \| 5.8e-18 \| 73.5 \| 56 \| (2, 59) \| 69 \| (40, 99) \| 105 \| LysM domain-containing protein \| LysM domain-containing protein; uniclust \| UniRef100\_A0A517QRQ4 \| 99.0 \| 3.1e-12 \| 6e-18 \| 82.3 \| 52 \| (2, 56) \| 69 \| (215, 266) \| 267 \| LysM domain/BON superfamily protein \| LysM domain/BON superfamily protein; uniclust \| UniRef100\_A0A2D7F4L0 \| 99.0 \| 3.3e-12 \| 6.1e-18 \| 68.5 \| 51 \| (2, 54) \| 69 \| (32, 82) \| 82 \| Peptidoglycan-binding protein \| Peptidoglycan-binding protein; uniclust \| UniRef100\_A0A496RP82 \| 99.0 \| 3.2e-12 \| 6.2e-18 \| 76.7 \| 55 \| (2, 58) \| 69 \| (87, 141) \| 156 \| LysM peptidoglycan-binding domain-containing protein \| LysM peptidoglycan-binding domain-containing protein; uniclust \| UniRef100\_A0A1V5WI12 \| 99.0 \| 3.2e-12 \| 6.3e-18 \| 71.3 \| 52 \| (2, 56) \| 69 \| (43, 94) \| 97 \| LysM domain/BON superfamily protein \| LysM domain/BON superfamily protein; uniclust \| UniRef100\_A0A068TBM2 \| 99.0 \| 3.2e-12 \| 6.3e-18 \| 68.4 \| 68 \| (1, 68) \| 69 \| (1, 72) \| 73 \| Phage tail protein \| Phage tail protein; uniclust \| UniRef100\_A0A1F8UA20 \| 99.0 \| 3.1e-12 \| 6.5e-18 \| 88.2 \| 54 \| (2, 59) \| 69 \| (398, 451) \| 455 \| LysM domain-containing protein \| LysM domain-containing protein; uniclust \| UniRef100\_A0A1F4ZYX1 \| 99.0 \| 3.6e-12 \| 6.6e-18 \| 69.1 \| 51 \| (3, 55) \| 69 \| (42, 92) \| 92 \| LysM domain-containing protein \| LysM domain-containing protein; uniclust \| UniRef100\_A0A1G2ZRS0 \| 99.0 \| 3.1e-12 \| 6.6e-18 \| 81.9 \| 55 \| (2, 58) \| 69 \| (138, 192) \| 227 \| LysM domain-containing protein \| LysM domain-containing protein; uniclust \| UniRef100\_A0A1G9Q464 \| 99.0 \| 3.7e-12 \| 6.8e-18 \| 72.5 \| 55 \| (2, 58) \| 69 \| (42, 96) \| 128 \| LysM domain-containing protein \| LysM domain-containing protein; uniclust \| UniRef100\_A0A0C7NBC1 \| 99.0 \| 3.5e-12 \| 6.9e-18 \| 81.4 \| 50 \| (3, 58) \| 69 \| (90, 139) \| 255 \| Spore coat assembly protein SafA \| Spore coat assembly protein SafA; uniclust \| UniRef100\_A0A090ZT01 \| 99.0 \| 3.3e-12 \| 7e-18 \| 83.8 \| 54 \| (2, 57) \| 69 \| (212, 266) \| 278 \| LysM domain protein \| LysM domain protein; uniclust \| UniRef100\_A0A6L3F9N8 \| 99.0 \| 3.8e-12 \| 7.4e-18 \| 82.8 \| 52 \| (2, 55) \| 69 \| (260, 311) \| 312 \| LysM peptidoglycan-binding domain-containing protein \| LysM peptidoglycan-binding domain-containing protein; uniclust \| UniRef100\_A0A1A5XTK2 \| 99.0 \| 3.9e-12 \| 7.8e-18 \| 72.7 \| 51 \| (1, 58) \| 69 \| (1, 51) \| 107 \| Peptidoglycan-binding protein LysM \| Peptidoglycan-binding protein LysM; uniclust \| UniRef100\_A0A2E2N8R8 \| 99.0 \| 3.9e-12 \| 8.3e-18 \| 85.5 \| 52 \| (2, 55) \| 69 \| (207, 258) \| 347 \| LysM domain-containing protein \| LysM domain-containing protein; uniclust \| UniRef100\_X0VWZ0 \| 99.0 \| 4.6e-12 \| 8.5e-18 \| 76.2 \| 53 \| (2, 58) \| 69 \| (58, 110) \| 191 \| LysM domain-containing protein (Fragment) \| LysM domain-containing protein (Fragment); uniclust \| UniRef100\_A0A0F9QKI0 \| 99.0 \| 4.1e-12 \| 8.6e-18 \| 72.9 \| 53 \| (3, 58) \| 69 \| (37, 89) \| 103 \| LysM domain-containing protein \| LysM domain-containing protein; uniclust \| UniRef100\_A0A1F9QVD6 \| 99.0 \| 4.2e-12 \| 8.8e-18 \| 88.2 \| 57 \| (1, 59) \| 69 \| (428, 484) \| 487 \| LysM domain-containing protein \| LysM domain-containing protein; uniclust \| UniRef100\_A0A348PCI9 \| 99.0 \| 4.4e-12 \| 8.9e-18 \| 83.3 \| 52 \| (3, 56) \| 69 \| (248, 300) \| 301 \| LysM domain-containing protein \| LysM domain-containing protein; uniclust \| UniRef100\_A0A2A5DCA3 \| 99.0 \| 4.3e-12 \| 8.9e-18 \| 82.7 \| 53 \| (2, 58) \| 69 \| (135, 187) \| 264 \| LysM domain-containing protein \| LysM domain-containing protein; uniclust \| UniRef100\_A0A0A2SNG7 \| 99.0 \| 4.2e-12 \| 8.9e-18 \| 89.8 \| 51 \| (1, 57) \| 69 \| (31, 81) \| 565 \| Peptidoglycan-binding protein \| Peptidoglycan-binding protein; uniclust \| UniRef100\_A0A0P1EW10 \| 99.0 \| 4.5e-12 \| 9e-18 \| 69.4 \| 63 \| (7, 69) \| 69 \| (13, 77) \| 80 \| Phage Tail Protein X \| Phage Tail Protein X; uniclust \| UniRef100\_A0A1C3WM59 \| 99.0 \| 4.9e-12 \| 9.1e-18 \| 68.3 \| 67 \| (2, 69) \| 69 \| (23, 90) \| 90 \| p2-like prophage tail protein X \| p2-like prophage tail protein X; uniclust \| UniRef100\_UPI001373773E \| 99.0 \| 4.8e-12 \| 9.1e-18 \| 71.8 \| 54 \| (2, 57) \| 69 \| (60, 113) \| 114 \| LysM peptidoglycan-binding domain-containing protein \| LysM peptidoglycan-binding domain-containing protein; uniclust \| UniRef100\_A0A2E8CYE4 \| 99.0 \| 4.6e-12 \| 9.2e-18 \| 81.2 \| 55 \| (2, 58) \| 69 \| (158, 212) \| 243 \| LysM domain-containing protein \| LysM domain-containing protein; uniclust \| UniRef100\_A0A0S8DVM3 \| 98.9 \| 5.2e-12 \| 9.9e-18 \| 68.9 \| 54 \| (2, 57) \| 69 \| (21, 74) \| 86 \| LysM domain-containing protein \| LysM domain-containing protein; uniclust \| UniRef100\_A0A524HEJ6 \| 98.9 \| 5.4e-12 \| 9.9e-18 \| 67.0 \| 53 \| (2, 56) \| 69 \| (27, 80) \| 81 \| LysM domain-containing protein \| LysM domain-containing protein; uniclust \| UniRef100\_A0A9D7QCG6 \| 98.9 \| 5.5e-12 \| 1e-17 \| 63.6 \| 51 \| (2, 54) \| 69 \| (10, 60) \| 60 \| LysM peptidoglycan-binding domain-containing protein \| LysM peptidoglycan-binding domain-containing protein; uniclust \| UniRef100\_A0A0H5SYF8 \| 98.9 \| 4.6e-12 \| 1e-17 \| 89.0 \| 52 \| (2, 57) \| 69 \| (460, 511) \| 513 \| Putative membrane protein \| Putative membrane protein; uniclust \| UniRef100\_A0A1F2Y198 \| 98.9 \| 5.2e-12 \| 1e-17 \| 70.3 \| 51 \| (2, 56) \| 69 \| (37, 88) \| 89 \| LysM domain-containing protein \| LysM domain-containing protein; uniclust \| UniRef100\_A0A080KJZ3 \| 98.9 \| 5.5e-12 \| 1e-17 \| 65.7 \| 62 \| (7, 69) \| 69 \| (6, 67) \| 67 \| p2-like prophage tail protein X \| p2-like prophage tail protein X; uniclust \| UniRef100\_A0A1Q2SNZ6 \| 98.9 \| 5.4e-12 \| 1.1e-17 \| 82.9 \| 53 \| (2, 56) \| 69 \| (257, 309) \| 311 \| Peptidoglycan-binding LysM \| Peptidoglycan-binding LysM; uniclust \| UniRef100\_A0A293NIF4 \| 98.9 \| 5.2e-12 \| 1.1e-17 \| 83.7 \| 54 \| (1, 57) \| 69 \| (216, 270) \| 298 \| LysM domain-containing protein \| LysM domain-containing protein; uniclust \| UniRef100\_A0A089HPX8 \| 98.9 \| 5.4e-12 \| 1.1e-17 \| 89.4 \| 53 \| (1, 59) \| 69 \| (58, 110) \| 610 \| LysM domain-containing protein \| LysM domain-containing protein; uniclust \| UniRef100\_A0A5C5VW67 \| 98.9 \| 5.9e-12 \| 1.1e-17 \| 75.3 \| 51 \| (2, 54) \| 69 \| (106, 156) \| 156 \| LysM domain/BON superfamily protein \| LysM domain/BON superfamily protein; uniclust \| UniRef100\_A0A1C6BB91 \| 98.9 \| 5.6e-12 \| 1.1e-17 \| 86.9 \| 50 \| (2, 55) \| 69 \| (424, 473) \| 474 \| LysM domain/BON superfamily protein \| LysM domain/BON superfamily protein; uniclust \| UniRef100\_A0A0D6KHC7 \| 98.9 \| 5.5e-12 \| 1.1e-17 \| 81.0 \| 54 \| (3, 60) \| 69 \| (173, 226) \| 238 \| LysM domain protein \| LysM domain protein; uniclust \| UniRef100\_E2CFJ8 \| 98.9 \| 6e-12 \| 1.1e-17 \| 66.7 \| 57 \| (1, 59) \| 69 \| (5, 61) \| 73 \| Phage Tail Protein X \| Phage Tail Protein X; uniclust \| UniRef100\_A0A1W1BYN0 \| 98.9 \| 5.6e-12 \| 1.1e-17 \| 84.3 \| 57 \| (2, 60) \| 69 \| (236, 292) \| 346 \| LysM domain-containing protein \| LysM domain-containing protein; uniclust \| UniRef100\_A0A838EI61 \| 98.9 \| 6e-12 \| 1.1e-17 \| 72.2 \| 52 \| (2, 55) \| 69 \| (65, 117) \| 118 \| LysM peptidoglycan-binding domain-containing protein \| LysM peptidoglycan-binding domain-containing protein; uniclust \| UniRef100\_A0A2A4M5P7 \| 98.9 \| 5.6e-12 \| 1.2e-17 \| 83.6 \| 51 \| (2, 59) \| 69 \| (247, 297) \| 303 \| LysM domain-containing protein \| LysM domain-containing protein; uniclust \| UniRef100\_A0A1C0AD30 \| 98.9 \| 6.1e-12 \| 1.2e-17 \| 80.2 \| 47 \| (1, 54) \| 69 \| (17, 63) \| 231 \| LysM domain-containing protein \| LysM domain-containing protein; uniclust \| UniRef100\_A0A2A4RPQ5 \| 98.9 \| 5.9e-12 \| 1.3e-17 \| 84.9 \| 55 \| (1, 57) \| 69 \| (212, 266) \| 354 \| LysM domain-containing protein \| LysM domain-containing protein; uniclust \| UniRef100\_A0A2N9MSL1 \| 98.9 \| 6.6e-12 \| 1.3e-17 \| 71.7 \| 54 \| (2, 59) \| 69 \| (50, 103) \| 111 \| Peptidoglycan-binding LysM \| Peptidoglycan-binding LysM; uniclust \| UniRef100\_A0A1F2RWZ5 \| 98.9 \| 6.9e-12 \| 1.3e-17 \| 72.0 \| 54 \| (3, 59) \| 69 \| (9, 62) \| 122 \| LysM domain-containing protein \| LysM domain-containing protein; uniclust \| UniRef100\_A0A037ZGX1 \| 98.9 \| 6.2e-12 \| 1.3e-17 \| 82.5 \| 53 \| (3, 57) \| 69 \| (27, 79) \| 277 \| LysM domain-containing protein \| LysM domain-containing protein; uniclust \| UniRef100\_A0A288TXV0 \| 98.9 \| 7.2e-12 \| 1.3e-17 \| 62.3 \| 51 \| (2, 55) \| 69 \| (1, 53) \| 54 \| LysM domain-containing protein \| LysM domain-containing protein; uniclust \| UniRef100\_A0A0A8JME8 \| 98.9 \| 6.5e-12 \| 1.4e-17 \| 85.5 \| 50 \| (1, 56) \| 69 \| (26, 75) \| 392 \| LysM domain-containing protein \| LysM domain-containing protein; uniclust \| UniRef100\_A0A937KBK3 \| 98.9 \| 7.3e-12 \| 1.4e-17 \| 64.9 \| 54 \| (2, 57) \| 69 \| (13, 66) \| 67 \| LysM peptidoglycan-binding domain-containing protein \| LysM peptidoglycan-binding domain-containing protein; uniclust \| UniRef100\_A0A1Q3SS48 \| 98.9 \| 7.3e-12 \| 1.4e-17 \| 79.3 \| 52 \| (2, 55) \| 69 \| (201, 252) \| 252 \| LysM domain-containing protein \| LysM domain-containing protein; uniclust \| UniRef100\_A0A924W8C7 \| 98.9 \| 7.2e-12 \| 1.4e-17 \| 79.7 \| 55 \| (2, 58) \| 69 \| (200, 254) \| 256 \| LysM peptidoglycan-binding domain-containing protein \| LysM peptidoglycan-binding domain-containing protein; uniclust \| UniRef100\_A0A8J3HSV3 \| 98.9 \| 7.5e-12 \| 1.4e-17 \| 73.9 \| 50 \| (2, 55) \| 69 \| (100, 149) \| 149 \| LysM domain-containing protein \| LysM domain-containing protein; uniclust \| UniRef100\_A0A7Y7XD64 \| 98.9 \| 7.7e-12 \| 1.4e-17 \| 68.7 \| 52 \| (2, 55) \| 69 \| (42, 93) \| 93 \| LysM peptidoglycan-binding domain-containing protein \| LysM peptidoglycan-binding domain-containing protein; uniclust \| UniRef100\_A0A1H3K5G8 \| 98.9 \| 7.1e-12 \| 1.4e-17 \| 85.1 \| 53 \| (1, 59) \| 69 \| (49, 101) \| 400 \| Morphogenetic protein associated with SpoVID \| Morphogenetic protein associated with SpoVID; uniclust \| UniRef100\_A0A2E2B0B4 \| 98.9 \| 7.2e-12 \| 1.4e-17 \| 84.0 \| 53 \| (2, 58) \| 69 \| (229, 281) \| 362 \| LysM domain-containing protein \| LysM domain-containing protein; uniclust \| UniRef100\_A0A1Z9QC33 \| 98.9 \| 7e-12 \| 1.5e-17 \| 82.8 \| 54 \| (2, 58) \| 69 \| (157, 211) \| 293 \| LysM domain-containing protein \| LysM domain-containing protein; uniclust \| UniRef100\_A0A4P5YIC6 \| 98.9 \| 7.5e-12 \| 1.5e-17 \| 86.7 \| 52 \| (2, 57) \| 69 \| (346, 397) \| 510 \| LysM domain-containing protein \| LysM domain-containing protein; uniclust \| UniRef100\_A0A1G0Y4L6 \| 98.9 \| 7.8e-12 \| 1.5e-17 \| 80.3 \| 57 \| (1, 59) \| 69 \| (196, 252) \| 255 \| LysM domain-containing protein \| LysM domain-containing protein; uniclust \| UniRef100\_A0A0P1GCB9 \| 98.9 \| 8.2e-12 \| 1.7e-17 \| 78.5 \| 51 \| (3, 55) \| 69 \| (152, 202) \| 203 \| LysM domain/BON superfamily protein \| LysM domain/BON superfamily protein; uniclust \| UniRef100\_A0A9D0S2Q7 \| 98.9 \| 9e-12 \| 1.7e-17 \| 78.5 \| 51 \| (2, 54) \| 69 \| (204, 254) \| 254 \| LysM peptidoglycan-binding domain-containing protein \| LysM peptidoglycan-binding domain-containing protein; uniclust \| UniRef100\_A0A2K2TWU4 \| 98.9 \| 9.1e-12 \| 1.7e-17 \| 72.8 \| 51 \| (2, 54) \| 69 \| (87, 137) \| 137 \| LysM domain-containing protein \| LysM domain-containing protein; uniclust \| UniRef100\_A0A0K2SFQ4 \| 98.9 \| 8.9e-12 \| 1.8e-17 \| 77.6 \| 51 \| (2, 58) \| 69 \| (21, 71) \| 199 \| LysM domain-containing protein \| LysM domain-containing protein; uniclust \| UniRef100\_A0A953GLI6 \| 98.9 \| 9.5e-12 \| 1.8e-17 \| 76.2 \| 52 \| (2, 55) \| 69 \| (143, 194) \| 195 \| LysM peptidoglycan-binding domain-containing protein \| LysM peptidoglycan-binding domain-containing protein; uniclust \| UniRef100\_A0A3M2EYF6 \| 98.9 \| 9.9e-12 \| 1.8e-17 \| 64.3 \| 51 \| (2, 54) \| 69 \| (19, 69) \| 69 \| LysM peptidoglycan-binding domain-containing protein \| LysM peptidoglycan-binding domain-containing protein; uniclust \| UniRef100\_A0A135L2D6 \| 98.9 \| 8.8e-12 \| 1.8e-17 \| 82.9 \| 52 \| (1, 58) \| 69 \| (1, 52) \| 317 \| LysM domain-containing protein \| LysM domain-containing protein; uniclust \| UniRef100\_A0A0D6KAX3 \| 98.9 \| 9.9e-12 \| 1.9e-17 \| 66.0 \| 51 \| (2, 56) \| 69 \| (22, 72) \| 74 \| LysM domain protein \| LysM domain protein; uniclust \| UniRef100\_A0A524GYW0 \| 98.9 \| 9.4e-12 \| 1.9e-17 \| 75.6 \| 54 \| (2, 59) \| 69 \| (94, 147) \| 159 \| LysM peptidoglycan-binding domain-containing protein \| LysM peptidoglycan-binding domain-containing protein; uniclust \| UniRef100\_A0A956BD51 \| 98.9 \| 1e-11 \| 1.9e-17 \| 74.8 \| 53 \| (2, 58) \| 69 \| (107, 159) \| 176 \| LysM peptidoglycan-binding domain-containing protein \| LysM peptidoglycan-binding domain-containing protein; uniclust \| UniRef100\_A0A0S2W5J5 \| 98.9 \| 1e-11 \| 1.9e-17 \| 79.0 \| 53 \| (2, 58) \| 69 \| (160, 212) \| 278 \| LysM domain-containing protein \| LysM domain-containing protein; uniclust \| UniRef100\_A0A1G8DH49 \| 98.9 \| 9.8e-12 \| 2e-17 \| 82.7 \| 53 \| (1, 59) \| 69 \| (1, 53) \| 323 \| Morphogenetic protein associated with SpoVID \| Morphogenetic protein associated with SpoVID; uniclust \| UniRef100\_A0A086YNE6 \| 98.9 \| 9.6e-12 \| 2.1e-17 \| 88.7 \| 47 \| (2, 55) \| 69 \| (515, 561) \| 618 \| Peptidoglycan-binding protein \| Peptidoglycan-binding protein; uniclust \| UniRef100\_A0A7V3QFU2 \| 98.9 \| 1.1e-11 \| 2.1e-17 \| 62.2 \| 54 \| (2, 57) \| 69 \| (4, 57) \| 58 \| LysM domain-containing protein \| LysM domain-containing protein; uniclust \| UniRef100\_A0A533Z935 \| 98.9 \| 1.1e-11 \| 2.1e-17 \| 77.2 \| 55 \| (1, 59) \| 69 \| (4, 58) \| 237 \| LysM peptidoglycan-binding domain-containing protein \| LysM peptidoglycan-binding domain-containing protein; uniclust \| UniRef100\_A0A178IH20 \| 98.9 \| 1.1e-11 \| 2.1e-17 \| 88.9 \| 51 \| (2, 54) \| 69 \| (729, 779) \| 779 \| LysM domain-containing protein \| LysM domain-containing protein; uniclust \| UniRef100\_A0A1V2SEX1 \| 98.9 \| 1.1e-11 \| 2.2e-17 \| 75.0 \| 53 \| (2, 56) \| 69 \| (94, 154) \| 156 \| LysM domain-containing protein (Fragment) \| LysM domain-containing protein (Fragment); uniclust \| UniRef100\_A0A0B0ELG5 \| 98.9 \| 1.1e-11 \| 2.2e-17 \| 81.8 \| 54 \| (2, 57) \| 69 \| (260, 313) \| 315 \| LysM domain-containing protein \| LysM domain-containing protein; uniclust \| UniRef100\_A0A2A5FW86 \| 98.9 \| 1.2e-11 \| 2.4e-17 \| 79.4 \| 51 \| (1, 54) \| 69 \| (216, 266) \| 267 \| LysM domain-containing protein \| LysM domain-containing protein; uniclust \| UniRef100\_A0A2P5MCF5 \| 98.9 \| 1.2e-11 \| 2.4e-17 \| 82.1 \| 53 \| (3, 59) \| 69 \| (281, 333) \| 338 \| LysM domain-containing protein \| LysM domain-containing protein; uniclust \| UniRef100\_A0A0G0JZZ0 \| 98.9 \| 1.1e-11 \| 2.4e-17 \| 79.4 \| 53 \| (2, 58) \| 69 \| (90, 142) \| 227 \| LysM domain-containing protein \| LysM domain-containing protein; uniclust \| UniRef100\_A0A1N6VA67 \| 98.9 \| 1.2e-11 \| 2.4e-17 \| 82.6 \| 54 \| (2, 58) \| 69 \| (35, 88) \| 329 \| LysM domain-containing protein \| LysM domain-containing protein; uniclust \| UniRef100\_A0A0K9YT35 \| 98.9 \| 1.3e-11 \| 2.4e-17 \| 74.2 \| 51 \| (1, 58) \| 69 \| (27, 77) \| 157 \| Peptidoglycan-binding protein LysM \| Peptidoglycan-binding protein LysM; uniclust \| UniRef100\_A0A2N3DL64 \| 98.9 \| 1.3e-11 \| 2.5e-17 \| 64.5 \| 60 \| (6, 69) \| 69 \| (10, 70) \| 70 \| Phage tail protein \| Phage tail protein; uniclust \| UniRef100\_A0A4Y9QV55 \| 98.9 \| 1.4e-11 \| 2.5e-17 \| 71.6 \| 49 \| (2, 54) \| 69 \| (96, 144) \| 144 \| LysM domain-containing protein \| LysM domain-containing protein; uniclust \| UniRef100\_A0A7C5KF11 \| 98.9 \| 1.4e-11 \| 2.5e-17 \| 65.3 \| 53 \| (3, 58) \| 69 \| (18, 70) \| 79 \| LysM domain-containing protein \| LysM domain-containing protein; uniclust \| UniRef100\_A0A1W9WAD3 \| 98.9 \| 1.4e-11 \| 2.6e-17 \| 59.6 \| 46 \| (7, 54) \| 69 \| (2, 47) \| 47 \| LysM domain-containing protein \| LysM domain-containing protein; uniclust \| UniRef100\_A0A350UMH8 \| 98.9 \| 1.3e-11 \| 2.6e-17 \| 80.2 \| 54 \| (1, 58) \| 69 \| (11, 64) \| 297 \| LysM domain-containing protein \| LysM domain-containing protein; uniclust \| UniRef100\_A0A024QAB1 \| 98.9 \| 1.3e-11 \| 2.7e-17 \| 82.9 \| 53 \| (1, 59) \| 69 \| (29, 81) \| 347 \| Spore coat assembly protein ExsA \| Spore coat assembly protein ExsA; uniclust \| UniRef100\_A0A0S8C8Z0 \| 98.9 \| 1.3e-11 \| 2.7e-17 \| 84.3 \| 53 \| (2, 57) \| 69 \| (71, 123) \| 397 \| LysM domain-containing protein \| LysM domain-containing protein; uniclust \| UniRef100\_A0A2D5NJU9 \| 98.9 \| 1.3e-11 \| 2.8e-17 \| 80.1 \| 54 \| (1, 58) \| 69 \| (124, 177) \| 250 \| LysM domain-containing protein \| LysM domain-containing protein; uniclust \| UniRef100\_A0A081LB99 \| 98.9 \| 1.3e-11 \| 2.8e-17 \| 86.9 \| 53 \| (1, 59) \| 69 \| (14, 66) \| 538 \| SpoVID-associated morphogenetic protein \| SpoVID-associated morphogenetic protein; uniclust \| UniRef100\_A0A084SYV7 \| 98.9 \| 1.3e-11 \| 2.8e-17 \| 85.5 \| 51 \| (2, 55) \| 69 \| (72, 122) \| 435 \| Peptidoglycan-binding protein \| Peptidoglycan-binding protein; uniclust \| UniRef100\_A0A838HEQ8 \| 98.9 \| 1.5e-11 \| 2.8e-17 \| 61.6 \| 51 \| (2, 54) \| 69 \| (7, 57) \| 57 \| LysM peptidoglycan-binding domain-containing protein \| LysM peptidoglycan-binding domain-containing protein; uniclust \| UniRef100\_A0A078MBJ7 \| 98.9 \| 1.4e-11 \| 2.8e-17 \| 80.1 \| 51 \| (2, 55) \| 69 \| (205, 256) \| 257 \| LysM domain protein \| LysM domain protein; uniclust \| UniRef100\_A0A2T6CMD0 \| 98.9 \| 1.4e-11 \| 2.9e-17 \| 89.4 \| 51 \| (2, 54) \| 69 \| (820, 870) \| 870 \| LysM domain-containing protein \| LysM domain-containing protein; uniclust \| UniRef100\_A0A0A3I541 \| 98.9 \| 1.5e-11 \| 3e-17 \| 83.6 \| 50 \| (1, 56) \| 69 \| (3, 52) \| 415 \| Uncharacterized protein \| Uncharacterized protein; uniclust \| UniRef100\_A0A0G0CX04 \| 98.9 \| 1.4e-11 \| 3e-17 \| 78.9 \| 54 \| (2, 59) \| 69 \| (109, 162) \| 232 \| LysM domain-containing protein \| LysM domain-containing protein; uniclust \| UniRef100\_A0A0U1NKI6 \| 98.9 \| 1.4e-11 \| 3e-17 \| 83.1 \| 52 \| (2, 55) \| 69 \| (303, 354) \| 355 \| LysM domain/BON superfamily protein \| LysM domain/BON superfamily protein; uniclust \| UniRef100\_A0A2P5LPX3 \| 98.9 \| 1.7e-11 \| 3.2e-17 \| 69.6 \| 57 \| (2, 58) \| 69 \| (42, 98) \| 118 \| Peptidoglycan-binding protein \| Peptidoglycan-binding protein; uniclust \| UniRef100\_A0A074LMC3 \| 98.9 \| 1.6e-11 \| 3.2e-17 \| 78.2 \| 53 \| (2, 58) \| 69 \| (149, 201) \| 243 \| LysM domain-containing protein \| LysM domain-containing protein; uniclust \| UniRef100\_A0A1W9VZ66 \| 98.9 \| 1.6e-11 \| 3.2e-17 \| 81.0 \| 53 \| (2, 58) \| 69 \| (75, 127) \| 327 \| LysM domain-containing protein \| LysM domain-containing protein; uniclust \| UniRef100\_A0A379LXN4 \| 98.9 \| 1.7e-11 \| 3.2e-17 \| 69.5 \| 52 \| (2, 57) \| 69 \| (55, 106) \| 108 \| LysM domain/BON superfamily protein \| LysM domain/BON superfamily protein; uniclust \| UniRef100\_A0A2A4XSY9 \| 98.9 \| 1.7e-11 \| 3.3e-17 \| 65.9 \| 68 \| (2, 69) \| 69 \| (7, 77) \| 77 \| Phage tail protein \| Phage tail protein; uniclust \| UniRef100\_A0A1I7J0W0 \| 98.9 \| 1.7e-11 \| 3.3e-17 \| 82.3 \| 53 \| (1, 59) \| 69 \| (34, 86) \| 372 \| LysM repeat-containing protein \| LysM repeat-containing protein; uniclust \| UniRef100\_A0A0G0YVI2 \| 98.9 \| 1.8e-11 \| 3.3e-17 \| 66.3 \| 49 \| (3, 55) \| 69 \| (36, 84) \| 84 \| LysM domain-containing protein \| LysM domain-containing protein; uniclust \| UniRef100\_A0A1X1NF73 \| 98.9 \| 1.7e-11 \| 3.3e-17 \| 68.1 \| 52 \| (2, 56) \| 69 \| (40, 91) \| 92 \| LysM domain-containing protein \| LysM domain-containing protein; uniclust \| UniRef100\_A0A345ZPL7 \| 98.9 \| 1.8e-11 \| 3.4e-17 \| 71.2 \| 54 \| (2, 57) \| 69 \| (84, 137) \| 139 \| LysM peptidoglycan-binding domain-containing protein \| LysM peptidoglycan-binding domain-containing protein; uniclust \| UniRef100\_A0A0G1F008 \| 98.9 \| 1.8e-11 \| 3.5e-17 \| 63.0 \| 52 \| (3, 56) \| 69 \| (8, 59) \| 60 \| LysM domain-containing protein (Fragment) \| LysM domain-containing protein (Fragment); uniclust \| UniRef100\_A0A0X8HDA8 \| 98.9 \| 1.8e-11 \| 3.6e-17 \| 77.4 \| 51 \| (3, 55) \| 69 \| (174, 224) \| 224 \| LysM domain/BON superfamily protein \| LysM domain/BON superfamily protein; uniclust \| UniRef100\_UPI000665D35C \| 98.9 \| 2e-11 \| 3.6e-17 \| 63.4 \| 69 \| (1, 69) \| 69 \| (1, 70) \| 70 \| tail protein X \| tail protein X; uniclust \| UniRef100\_A0A097ARJ0 \| 98.9 \| 1.9e-11 \| 3.7e-17 \| 74.1 \| 49 \| (2, 56) \| 69 \| (33, 81) \| 166 \| Spore coat assembly protein SafA \| Spore coat assembly protein SafA; uniclust \| UniRef100\_A0A1F9QAA3 \| 98.9 \| 1.9e-11 \| 3.7e-17 \| 76.8 \| 54 \| (2, 58) \| 69 \| (152, 205) \| 219 \| LysM domain-containing protein \| LysM domain-containing protein; uniclust \| UniRef100\_A0A143XDP1 \| 98.9 \| 1.8e-11 \| 3.8e-17 \| 84.7 \| 52 \| (3, 58) \| 69 \| (397, 448) \| 451 \| LysM domain/BON superfamily protein \| LysM domain/BON superfamily protein; uniclust \| UniRef100\_A0A7V2K829 \| 98.9 \| 2.1e-11 \| 3.8e-17 \| 68.9 \| 57 \| (2, 60) \| 69 \| (54, 110) \| 119 \| LysM peptidoglycan-binding domain-containing protein \| LysM peptidoglycan-binding domain-containing protein; uniclust \| UniRef100\_A0A7C4EC90 \| 98.9 \| 2e-11 \| 3.9e-17 \| 80.8 \| 57 \| (2, 60) \| 69 \| (269, 326) \| 332 \| LysM peptidoglycan-binding domain-containing protein \| LysM peptidoglycan-binding domain-containing protein; uniclust \| UniRef100\_A0A2R7Z0I0 \| 98.9 \| 1.9e-11 \| 3.9e-17 \| 72.6 \| 47 \| (2, 55) \| 69 \| (85, 131) \| 132 \| LysM domain-containing protein \| LysM domain-containing protein; uniclust \| UniRef100\_A0A0A0HMA7 \| 98.9 \| 1.8e-11 \| 3.9e-17 \| 83.9 \| 55 \| (2, 58) \| 69 \| (71, 126) \| 411 \| Amino acid ABC transporter substrate-binding protein, PAAT family \| Amino acid ABC transporter substrate-binding protein, PAAT family; uniclust \| UniRef100\_A0A0A7HAL1 \| 98.8 \| 2e-11 \| 4e-17 \| 78.7 \| 54 \| (2, 57) \| 69 \| (192, 245) \| 248 \| Peptidoglycan-binding protein LysM \| Peptidoglycan-binding protein LysM; uniclust \| UniRef100\_A0A7W3WB71 \| 98.8 \| 2.2e-11 \| 4e-17 \| 61.6 \| 53 \| (2, 56) \| 69 \| (7, 59) \| 60 \| LysM peptidoglycan-binding domain-containing protein \| LysM peptidoglycan-binding domain-containing protein; uniclust \| UniRef100\_A0A1G0ZHH7 \| 98.8 \| 2e-11 \| 4.1e-17 \| 77.9 \| 48 \| (2, 56) \| 69 \| (173, 220) \| 220 \| LysM domain-containing protein \| LysM domain-containing protein; uniclust \| UniRef100\_A0A1Q7T6T4 \| 98.8 \| 2.1e-11 \| 4.1e-17 \| 65.3 \| 52 \| (2, 55) \| 69 \| (22, 74) \| 75 \| LysM domain-containing protein \| LysM domain-containing protein; uniclust \| UniRef100\_A0A1R4IUY4 \| 98.8 \| 2.1e-11 \| 4.2e-17 \| 76.3 \| 47 \| (2, 55) \| 69 \| (145, 191) \| 192 \| TolA protein \| TolA protein; uniclust \| UniRef100\_S3J5W6 \| 98.8 \| 2.3e-11 \| 4.2e-17 \| 68.8 \| 55 \| (1, 59) \| 69 \| (1, 55) \| 120 \| LysM domain-containing protein \| LysM domain-containing protein; uniclust \| UniRef100\_A0A1I3WQT9 \| 98.8 \| 2.1e-11 \| 4.3e-17 \| 84.9 \| 51 \| (1, 57) \| 69 \| (12, 62) \| 522 \| Spore coat assembly protein SafA \| Spore coat assembly protein SafA; uniclust \| UniRef100\_A0A6I0F283 \| 98.8 \| 2.2e-11 \| 4.3e-17 \| 64.7 \| 50 \| (3, 59) \| 69 \| (15, 64) \| 68 \| LysM peptidoglycan-binding domain-containing protein \| LysM peptidoglycan-binding domain-containing protein; uniclust \| UniRef100\_A0A925ALC7 \| 98.8 \| 2.3e-11 \| 4.4e-17 \| 70.8 \| 49 \| (2, 54) \| 69 \| (85, 133) \| 133 \| LysM peptidoglycan-binding domain-containing protein \| LysM peptidoglycan-binding domain-containing protein; uniclust \| UniRef100\_A0A3C0VA40 \| 98.8 \| 2.4e-11 \| 4.5e-17 \| 75.7 \| 53 \| (2, 56) \| 69 \| (185, 237) \| 238 \| LysM domain-containing protein \| LysM domain-containing protein; uniclust \| UniRef100\_A0A2H5Z0M6 \| 98.8 \| 2.2e-11 \| 4.5e-17 \| 72.6 \| 50 \| (2, 58) \| 69 \| (88, 137) \| 138 \| N-acetylmuramoyl-L-alanine amidase XlyA \| N-acetylmuramoyl-L-alanine amidase XlyA; uniclust \| UniRef100\_A0A0G0VNZ0 \| 98.8 \| 2.3e-11 \| 4.6e-17 \| 73.1 \| 53 \| (2, 58) \| 69 \| (18, 70) \| 143 \| LysM domain-containing protein \| LysM domain-containing protein; uniclust \| UniRef100\_A0A1R0F851 \| 98.8 \| 2.3e-11 \| 4.6e-17 \| 81.0 \| 56 \| (2, 59) \| 69 \| (265, 320) \| 335 \| Nucleoid-associated protein YgaU, contains BON and LysM domains \| Nucleoid-associated protein YgaU, contains BON and LysM domains; uniclust \| UniRef100\_A0A2W5H3R5 \| 98.8 \| 2.5e-11 \| 4.6e-17 \| 71.7 \| 68 \| (2, 69) \| 69 \| (95, 163) \| 163 \| Phage tail protein \| Phage tail protein; uniclust \| UniRef100\_A0A534MS82 \| 98.8 \| 2.5e-11 \| 4.6e-17 \| 71.2 \| 54 \| (2, 57) \| 69 \| (86, 139) \| 142 \| LysM peptidoglycan-binding domain-containing protein (Fragment) \| LysM peptidoglycan-binding domain-containing protein (Fragment); uniclust \| UniRef100\_A0A357KMI0 \| 98.8 \| 2.6e-11 \| 4.7e-17 \| 67.8 \| 49 \| (3, 54) \| 69 \| (61, 109) \| 110 \| LysM domain-containing protein \| LysM domain-containing protein; uniclust \| UniRef100\_A0A2D6N5F9 \| 98.8 \| 2.4e-11 \| 4.7e-17 \| 67.4 \| 55 \| (2, 58) \| 69 \| (18, 73) \| 90 \| LysM domain-containing protein \| LysM domain-containing protein; uniclust \| UniRef100\_A0A2D5XPP0 \| 98.8 \| 2.4e-11 \| 4.7e-17 \| 74.3 \| 56 \| (2, 59) \| 69 \| (98, 153) \| 165 \| LysM domain-containing protein \| LysM domain-containing protein; uniclust \| UniRef100\_A0A7C5HFW9 \| 98.8 \| 2.6e-11 \| 4.8e-17 \| 72.8 \| 56 \| (1, 58) \| 69 \| (122, 177) \| 184 \| LysM peptidoglycan-binding domain-containing protein \| LysM peptidoglycan-binding domain-containing protein; uniclust \| UniRef100\_A0A1F7R001 \| 98.8 \| 2.3e-11 \| 4.9e-17 \| 77.5 \| 52 \| (2, 59) \| 69 \| (52, 103) \| 217 \| LysM domain-containing protein \| LysM domain-containing protein; uniclust \| UniRef100\_A0A109MUW9 \| 98.8 \| 2.5e-11 \| 5e-17 \| 85.8 \| 53 \| (1, 59) \| 69 \| (11, 63) \| 595 \| Uncharacterized protein (Fragment) \| Uncharacterized protein (Fragment); uniclust \| UniRef100\_UPI00117B2ACC \| 98.8 \| 2.8e-11 \| 5.1e-17 \| 61.1 \| 53 \| (1, 59) \| 69 \| (1, 53) \| 59 \| SafA/ExsA family spore coat assembly protein \| SafA/ExsA family spore coat assembly protein; uniclust \| UniRef100\_A0A1Z9KPR5 \| 98.8 \| 2.5e-11 \| 5.2e-17 \| 77.9 \| 54 \| (2, 59) \| 69 \| (110, 163) \| 235 \| LysM domain-containing protein \| LysM domain-containing protein; uniclust \| UniRef100\_A0A202DNT9 \| 98.8 \| 2.7e-11 \| 5.2e-17 \| 81.5 \| 53 \| (2, 56) \| 69 \| (365, 417) \| 419 \| LysM domain-containing protein \| LysM domain-containing protein; uniclust \| UniRef100\_A0A8J3GFQ9 \| 98.8 \| 2.8e-11 \| 5.2e-17 \| 65.7 \| 51 \| (2, 55) \| 69 \| (40, 90) \| 91 \| LysM domain-containing protein \| LysM domain-containing protein; uniclust \| UniRef100\_A0A2T7TER6 \| 98.8 \| 2.7e-11 \| 5.3e-17 \| 67.2 \| 52 \| (2, 57) \| 69 \| (30, 83) \| 92 \| LysM domain-containing protein \| LysM domain-containing protein; uniclust \| UniRef100\_A0A2A5DCA3 \| 98.8 \| 2.5e-11 \| 5.3e-17 \| 79.3 \| 52 \| (2, 55) \| 69 \| (211, 263) \| 264 \| LysM domain-containing protein \| LysM domain-containing protein; uniclust \| UniRef100\_A0A0P6YH39 \| 98.8 \| 2.9e-11 \| 5.3e-17 \| 66.9 \| 52 \| (2, 55) \| 69 \| (52, 103) \| 103 \| LysM domain-containing protein \| LysM domain-containing protein; uniclust \| UniRef100\_A0A432QVU7 \| 98.8 \| 2.9e-11 \| 5.4e-17 \| 63.6 \| 69 \| (1, 69) \| 69 \| (1, 71) \| 75 \| Phage tail protein \| Phage tail protein; uniclust \| UniRef100\_A0A2V5Q5V7 \| 98.8 \| 2.7e-11 \| 5.4e-17 \| 83.1 \| 55 \| (2, 58) \| 69 \| (406, 460) \| 465 \| LysM domain-containing protein \| LysM domain-containing protein; uniclust \| UniRef100\_A0A120MKI8 \| 98.8 \| 2.7e-11 \| 5.4e-17 \| 82.2 \| 50 \| (2, 55) \| 69 \| (335, 384) \| 385 \| LysM domain/BON superfamily protein \| LysM domain/BON superfamily protein; uniclust \| UniRef100\_A0A2E5G5P4 \| 98.8 \| 3e-11 \| 5.5e-17 \| 70.4 \| 54 \| (2, 59) \| 69 \| (86, 139) \| 146 \| LysM domain-containing protein \| LysM domain-containing protein; uniclust \| UniRef100\_A0A961BZ49 \| 98.8 \| 3.1e-11 \| 5.7e-17 \| 73.2 \| 50 \| (2, 55) \| 69 \| (136, 185) \| 185 \| LysM peptidoglycan-binding domain-containing protein \| LysM peptidoglycan-binding domain-containing protein; uniclust \| UniRef100\_A0A7X4CNS4 \| 98.8 \| 3.2e-11 \| 5.8e-17 \| 62.7 \| 67 \| (3, 69) \| 69 \| (4, 70) \| 70 \| Phage tail protein \| Phage tail protein; uniclust \| UniRef100\_A0A353M381 \| 98.8 \| 2.9e-11 \| 5.8e-17 \| 80.3 \| 53 \| (2, 58) \| 69 \| (269, 321) \| 324 \| LysM peptidoglycan-binding domain-containing protein \| LysM peptidoglycan-binding domain-containing protein; uniclust \| UniRef100\_A0A099S5L2 \| 98.8 \| 3e-11 \| 6e-17 \| 74.3 \| 49 \| (1, 55) \| 69 \| (1, 49) \| 179 \| Peptidoglycan-binding protein \| Peptidoglycan-binding protein; uniclust \| UniRef100\_A0A7C1J0I8 \| 98.8 \| 3.2e-11 \| 6.1e-17 \| 71.2 \| 53 \| (3, 57) \| 69 \| (86, 139) \| 140 \| LysM peptidoglycan-binding domain-containing protein \| LysM peptidoglycan-binding domain-containing protein; uniclust \| UniRef100\_A0A084JKQ0 \| 98.8 \| 3e-11 \| 6.3e-17 \| 85.2 \| 54 \| (2, 59) \| 69 \| (486, 539) \| 549 \| Peptidoglycan-binding protein \| Peptidoglycan-binding protein; uniclust \| UniRef100\_A0A2E2MDG1 \| 98.8 \| 3.3e-11 \| 6.3e-17 \| 79.1 \| 50 \| (3, 56) \| 69 \| (274, 323) \| 324 \| LysM domain-containing protein \| LysM domain-containing protein; uniclust \| UniRef100\_A0A1F9NKC6 \| 98.8 \| 3.1e-11 \| 6.3e-17 \| 78.7 \| 61 \| (2, 68) \| 69 \| (116, 176) \| 263 \| Endolytic peptidoglycan transglycosylase RlpA \| Endolytic peptidoglycan transglycosylase RlpA; uniclust \| UniRef100\_A0A3M1Q3A8 \| 98.8 \| 3.2e-11 \| 6.5e-17 \| 81.5 \| 56 \| (2, 57) \| 69 \| (204, 259) \| 370 \| LysM domain-containing protein \| LysM domain-containing protein; uniclust \| UniRef100\_UPI001FC8EFCB \| 98.8 \| 3.6e-11 \| 6.6e-17 \| 62.5 \| 51 \| (3, 55) \| 69 \| (18, 68) \| 70 \| LysM peptidoglycan-binding domain-containing protein \| LysM peptidoglycan-binding domain-containing protein; uniclust \| UniRef100\_A0A064AM34 \| 98.8 \| 3.4e-11 \| 6.7e-17 \| 64.8 \| 56 \| (1, 59) \| 69 \| (9, 64) \| 75 \| Tail protein \| Tail protein; uniclust \| UniRef100\_A0A2D5NJ77 \| 98.8 \| 3.3e-11 \| 6.8e-17 \| 80.2 \| 55 \| (3, 59) \| 69 \| (254, 309) \| 314 \| LysM domain-containing protein \| LysM domain-containing protein; uniclust \| UniRef100\_A0A916G808 \| 98.8 \| 3.5e-11 \| 6.9e-17 \| 77.6 \| 54 \| (2, 59) \| 69 \| (99, 152) \| 258 \| LysM domain-containing protein \| LysM domain-containing protein; uniclust \| UniRef100\_A0A072P1Q2 \| 98.8 \| 3.3e-11 \| 6.9e-17 \| 84.2 \| 52 \| (1, 58) \| 69 \| (16, 67) \| 504 \| Spore coat assembly protein SafA \| Spore coat assembly protein SafA; uniclust \| UniRef100\_A0A0M0LN26 \| 98.8 \| 3.6e-11 \| 7e-17 \| 82.0 \| 52 \| (1, 58) \| 69 \| (15, 66) \| 456 \| LysM domain-containing protein \| LysM domain-containing protein; uniclust \| UniRef100\_A0A535BUW1 \| 98.8 \| 3.7e-11 \| 7.1e-17 \| 66.0 \| 52 \| (2, 55) \| 69 \| (38, 90) \| 90 \| LysM peptidoglycan-binding domain-containing protein \| LysM peptidoglycan-binding domain-containing protein; uniclust \| UniRef100\_A0A1G2Y5P4 \| 98.8 \| 3.5e-11 \| 7.3e-17 \| 80.9 \| 57 \| (2, 58) \| 69 \| (190, 249) \| 340 \| LysM domain-containing protein \| LysM domain-containing protein; uniclust \| UniRef100\_A0A2V8X4F0 \| 98.8 \| 3.9e-11 \| 7.3e-17 \| 64.3 \| 52 \| (2, 56) \| 69 \| (26, 77) \| 80 \| Peptidoglycan-binding protein LysM (Fragment) \| Peptidoglycan-binding protein LysM (Fragment); uniclust \| UniRef100\_A0A969XUJ1 \| 98.8 \| 4e-11 \| 7.4e-17 \| 69.3 \| 51 \| (3, 55) \| 69 \| (88, 138) \| 138 \| LysM peptidoglycan-binding domain-containing protein \| LysM peptidoglycan-binding domain-containing protein; uniclust \| UniRef100\_A0A4P5YIC6 \| 98.8 \| 3.7e-11 \| 7.5e-17 \| 83.5 \| 53 \| (2, 58) \| 69 \| (276, 328) \| 510 \| LysM domain-containing protein \| LysM domain-containing protein; uniclust \| UniRef100\_A0A533ZFK9 \| 98.8 \| 4.1e-11 \| 7.6e-17 \| 67.5 \| 54 \| (1, 58) \| 69 \| (1, 54) \| 116 \| LysM peptidoglycan-binding domain-containing protein \| LysM peptidoglycan-binding domain-containing protein; uniclust \| UniRef100\_A0A097ARJ0 \| 98.8 \| 4e-11 \| 7.8e-17 \| 72.8 \| 49 \| (2, 56) \| 69 \| (91, 139) \| 166 \| Spore coat assembly protein SafA \| Spore coat assembly protein SafA; uniclust \| UniRef100\_A0A2A5A5I2 \| 98.8 \| 4e-11 \| 7.8e-17 \| 77.8 \| 52 \| (2, 55) \| 69 \| (232, 283) \| 283 \| LysM domain-containing protein \| LysM domain-containing protein; uniclust \| UniRef100\_A0A1H0NML7 \| 98.8 \| 4.3e-11 \| 7.9e-17 \| 65.0 \| 54 \| (5, 59) \| 69 \| (4, 57) \| 91 \| P2-like prophage tail protein X \| P2-like prophage tail protein X; uniclust \| UniRef100\_A0A7C5FFV2 \| 98.8 \| 4.4e-11 \| 8.2e-17 \| 78.9 \| 56 \| (1, 58) \| 69 \| (310, 365) \| 372 \| LysM peptidoglycan-binding domain-containing protein \| LysM peptidoglycan-binding domain-containing protein; uniclust \| UniRef100\_A0A0F9YZQ3 \| 98.8 \| 4e-11 \| 8.2e-17 \| 76.6 \| 49 \| (3, 55) \| 69 \| (168, 216) \| 217 \| Cell-wall-binding phage-encoded protein \| Cell-wall-binding phage-encoded protein; uniclust \| UniRef100\_A0A085LA63 \| 98.8 \| 4.2e-11 \| 8.3e-17 \| 75.8 \| 50 \| (2, 58) \| 69 \| (138, 187) \| 215 \| LysM domain-containing protein \| LysM domain-containing protein; uniclust \| UniRef100\_A0A0P7KLW9 \| 98.8 \| 4.5e-11 \| 8.4e-17 \| 68.2 \| 55 \| (2, 58) \| 69 \| (62, 116) \| 118 \| LysM domain-containing protein \| LysM domain-containing protein; uniclust \| UniRef100\_A0A0S8HL66 \| 98.8 \| 4.3e-11 \| 8.4e-17 \| 78.9 \| 54 \| (2, 59) \| 69 \| (139, 193) \| 312 \| LysM domain-containing protein \| LysM domain-containing protein; uniclust \| UniRef100\_A0A8T3PST5 \| 98.8 \| 4.6e-11 \| 8.5e-17 \| 66.2 \| 52 \| (3, 58) \| 69 \| (49, 100) \| 103 \| LysM peptidoglycan-binding domain-containing protein \| LysM peptidoglycan-binding domain-containing protein; uniclust \| UniRef100\_A0A1A2S6J1 \| 98.8 \| 4.7e-11 \| 8.6e-17 \| 72.3 \| 53 \| (2, 58) \| 69 \| (3, 55) \| 195 \| LysM domain-containing protein \| LysM domain-containing protein; uniclust \| UniRef100\_A0A1B6BCU7 \| 98.8 \| 4.1e-11 \| 8.6e-17 \| 78.5 \| 54 \| (2, 59) \| 69 \| (194, 247) \| 265 \| Contractile injection system tube protein N-terminal domain-containing protein \| Contractile injection system tube protein N-terminal domain-containing protein; uniclust \| UniRef100\_A0A090IT45 \| 98.8 \| 4.2e-11 \| 8.6e-17 \| 64.4 \| 67 \| (1, 69) \| 69 \| (2, 69) \| 69 \| Putative phage tail X protein \| Putative phage tail X protein; uniclust \| UniRef100\_A0A0S8BGR5 \| 98.8 \| 4.2e-11 \| 8.7e-17 \| 75.2 \| 49 \| (2, 57) \| 69 \| (140, 188) \| 190 \| LysM domain-containing protein \| LysM domain-containing protein; uniclust \| UniRef100\_A0A7X7N646 \| 98.8 \| 4.8e-11 \| 8.8e-17 \| 63.8 \| 51 \| (2, 54) \| 69 \| (32, 82) \| 82 \| LysM peptidoglycan-binding domain-containing protein \| LysM peptidoglycan-binding domain-containing protein; uniclust \| UniRef100\_A0A966QMG2 \| 98.8 \| 4.8e-11 \| 8.9e-17 \| 69.3 \| 51 \| (2, 54) \| 69 \| (91, 142) \| 143 \| LysM peptidoglycan-binding domain-containing protein \| LysM peptidoglycan-binding domain-containing protein; uniclust \| UniRef100\_A0A9D8NX83 \| 98.8 \| 4.9e-11 \| 9e-17 \| 69.6 \| 54 \| (3, 56) \| 69 \| (93, 147) \| 148 \| LysM peptidoglycan-binding domain-containing protein \| LysM peptidoglycan-binding domain-containing protein; uniclust \| UniRef100\_A0A1Q9QNI0 \| 98.8 \| 4.7e-11 \| 9e-17 \| 68.1 \| 55 \| (2, 59) \| 69 \| (40, 109) \| 111 \| LysM domain-containing protein \| LysM domain-containing protein; uniclust \| UniRef100\_A0A1V6D8Z8 \| 98.8 \| 4.6e-11 \| 9.2e-17 \| 81.6 \| 53 \| (2, 58) \| 69 \| (321, 374) \| 417 \| LysM domain/BON superfamily protein \| LysM domain/BON superfamily protein; uniclust \| UniRef100\_A0A0A0J9G3 \| 98.8 \| 4.7e-11 \| 9.4e-17 \| 72.5 \| 47 \| (2, 55) \| 69 \| (109, 155) \| 156 \| LysM domain-containing protein \| LysM domain-containing protein; uniclust \| UniRef100\_A0A1F7FIL3 \| 98.8 \| 4.8e-11 \| 9.4e-17 \| 76.2 \| 50 \| (2, 55) \| 69 \| (186, 235) \| 236 \| LysM domain-containing protein \| LysM domain-containing protein; uniclust \| UniRef100\_A0A1F9Q953 \| 98.8 \| 4.7e-11 \| 9.5e-17 \| 83.6 \| 55 \| (2, 59) \| 69 \| (468, 522) \| 528 \| LysM domain-containing protein \| LysM domain-containing protein; uniclust \| UniRef100\_A0A292YEE7 \| 98.8 \| 4.9e-11 \| 9.5e-17 \| 78.9 \| 50 \| (1, 56) \| 69 \| (87, 136) \| 342 \| LysM domain-containing protein \| LysM domain-containing protein; uniclust \| UniRef100\_A0A1G6JHR3 \| 98.8 \| 4.9e-11 \| 9.6e-17 \| 74.0 \| 54 \| (3, 58) \| 69 \| (129, 182) \| 187 \| BON domain-containing protein \| BON domain-containing protein; uniclust \| UniRef100\_A0A2E8JYY7 \| 98.8 \| 4.8e-11 \| 9.7e-17 \| 77.7 \| 53 \| (2, 56) \| 69 \| (137, 189) \| 269 \| LysM domain-containing protein \| LysM domain-containing protein; uniclust \| UniRef100\_A0A017RV10 \| 98.8 \| 4.8e-11 \| 9.7e-17 \| 78.4 \| 52 \| (2, 56) \| 69 \| (203, 257) \| 282 \| LysM domain-containing protein \| LysM domain-containing protein; uniclust \| UniRef100\_A0A7X8VU95 \| 98.8 \| 5.4e-11 \| 9.9e-17 \| 65.2 \| 53 \| (2, 55) \| 69 \| (43, 95) \| 96 \| LysM peptidoglycan-binding domain-containing protein \| LysM peptidoglycan-binding domain-containing protein; uniclust \| UniRef100\_A0A918W949 \| 98.8 \| 5.5e-11 \| 1e-16 \| 70.5 \| 54 \| (2, 57) \| 69 \| (109, 162) \| 165 \| LysM domain-containing protein \| LysM domain-containing protein; uniclust \| UniRef100\_A0A024YSH0 \| 98.8 \| 5.3e-11 \| 1e-16 \| 73.9 \| 53 \| (2, 56) \| 69 \| (148, 201) \| 204 \| LysM domain-containing protein \| LysM domain-containing protein; uniclust \| UniRef100\_A0A1M7PRK3 \| 98.8 \| 5e-11 \| 1e-16 \| 74.4 \| 53 \| (2, 59) \| 69 \| (127, 179) \| 187 \| LysM domain-containing protein \| LysM domain-containing protein; uniclust \| UniRef100\_A0A2D5AB17 \| 98.8 \| 5.2e-11 \| 1e-16 \| 83.3 \| 51 \| (2, 54) \| 69 \| (497, 548) \| 548 \| LysM domain-containing protein \| LysM domain-containing protein; uniclust \| UniRef100\_A0A2E1VR95 \| 98.8 \| 5.4e-11 \| 1.1e-16 \| 76.6 \| 52 \| (3, 58) \| 69 \| (203, 254) \| 259 \| LysM domain-containing protein \| LysM domain-containing protein; uniclust \| UniRef100\_A0A3M0ZNV8 \| 98.8 \| 5.8e-11 \| 1.1e-16 \| 61.0 \| 49 \| (3, 58) \| 69 \| (10, 58) \| 65 \| LysM domain-containing protein (Fragment) \| LysM domain-containing protein (Fragment); uniclust \| UniRef100\_A0A0C2M3Y0 \| 98.8 \| 5.6e-11 \| 1.1e-16 \| 64.5 \| 50 \| (2, 57) \| 69 \| (11, 60) \| 82 \| LysM peptidoglycan-binding domain-containing protein \| LysM peptidoglycan-binding domain-containing protein; uniclust \| UniRef100\_A0A257ECJ0 \| 98.8 \| 5.9e-11 \| 1.1e-16 \| 75.4 \| 52 \| (2, 55) \| 69 \| (218, 269) \| 269 \| LysM domain-containing protein \| LysM domain-containing protein; uniclust \| UniRef100\_A0A1W9P516 \| 98.8 \| 5.9e-11 \| 1.1e-16 \| 66.4 \| 56 \| (2, 59) \| 69 \| (22, 79) \| 100 \| LysM domain-containing protein \| LysM domain-containing protein; uniclust \| UniRef100\_A0A1L9QA14 \| 98.8 \| 5.9e-11 \| 1.1e-16 \| 63.2 \| 69 \| (1, 69) \| 69 \| (1, 73) \| 74 \| Phage tail protein \| Phage tail protein; uniclust \| UniRef100\_A0A1Q6PX12 \| 98.8 \| 5.4e-11 \| 1.1e-16 \| 65.0 \| 55 \| (2, 59) \| 69 \| (7, 61) \| 76 \| Phage tail protein \| Phage tail protein; uniclust \| UniRef100\_A0A0S7ZZD8 \| 98.8 \| 5.2e-11 \| 1.1e-16 \| 82.7 \| 53 \| (2, 56) \| 69 \| (85, 137) \| 446 \| LysM domain-containing protein \| LysM domain-containing protein; uniclust \| UniRef100\_A0A081BQ49 \| 98.8 \| 5.8e-11 \| 1.1e-16 \| 82.2 \| 54 \| (1, 58) \| 69 \| (1, 54) \| 533 \| LysM domain-containing protein \| LysM domain-containing protein; uniclust \| UniRef100\_UPI0020B20EE2 \| 98.8 \| 6.1e-11 \| 1.1e-16 \| 60.4 \| 51 \| (2, 54) \| 69 \| (12, 62) \| 62 \| LysM peptidoglycan-binding domain-containing protein \| LysM peptidoglycan-binding domain-containing protein; uniclust \| UniRef100\_A0A0F8XQA1 \| 98.8 \| 6e-11 \| 1.1e-16 \| 73.3 \| 51 \| (2, 54) \| 69 \| (147, 198) \| 198 \| LysM domain-containing protein (Fragment) \| LysM domain-containing protein (Fragment); uniclust \| UniRef100\_A0A376EW91 \| 98.8 \| 6.1e-11 \| 1.1e-16 \| 59.9 \| 50 \| (2, 54) \| 69 \| (3, 52) \| 56 \| Inner membrane protein \| Inner membrane protein; uniclust \| UniRef100\_A0A1V4QZM9 \| 98.8 \| 5.6e-11 \| 1.1e-16 \| 78.2 \| 56 \| (2, 57) \| 69 \| (128, 183) \| 288 \| LysM domain-containing protein \| LysM domain-containing protein; uniclust \| UniRef100\_A0A351FFR9 \| 98.8 \| 6.3e-11 \| 1.2e-16 \| 66.9 \| 55 \| (2, 58) \| 69 \| (61, 115) \| 117 \| LysM domain-containing protein (Fragment) \| LysM domain-containing protein (Fragment); uniclust \| UniRef100\_A0A0F2PY82 \| 98.8 \| 5.8e-11 \| 1.2e-16 \| 75.3 \| 47 \| (2, 54) \| 69 \| (20, 66) \| 216 \| LysM domain-containing protein \| LysM domain-containing protein; uniclust \| UniRef100\_A0A356JRW9 \| 98.8 \| 6.7e-11 \| 1.2e-16 \| 66.5 \| 52 \| (2, 55) \| 69 \| (61, 112) \| 113 \| LysM peptidoglycan-binding domain-containing protein \| LysM peptidoglycan-binding domain-containing protein; uniclust \| UniRef100\_UPI00117BF9B8 \| 98.8 \| 6.7e-11 \| 1.2e-16 \| 63.7 \| 54 \| (2, 59) \| 69 \| (12, 65) \| 86 \| LysM peptidoglycan-binding domain-containing protein \| LysM peptidoglycan-binding domain-containing protein; uniclust \| UniRef100\_A0A7X7ENM6 \| 98.8 \| 6.6e-11 \| 1.2e-16 \| 76.4 \| 53 \| (2, 58) \| 69 \| (166, 218) \| 298 \| LysM peptidoglycan-binding domain-containing protein \| LysM peptidoglycan-binding domain-containing protein; uniclust \| UniRef100\_A0A136KIR5 \| 98.8 \| 6.5e-11 \| 1.2e-16 \| 74.3 \| 51 \| (2, 56) \| 69 \| (165, 215) \| 218 \| LysM domain/BON superfamily protein \| LysM domain/BON superfamily protein; uniclust \| UniRef100\_A0A2A2RQ54 \| 98.8 \| 6.5e-11 \| 1.2e-16 \| 74.8 \| 51 \| (2, 54) \| 69 \| (181, 231) \| 231 \| LysM domain-containing protein \| LysM domain-containing protein; uniclust \| UniRef100\_A0A060BXH1 \| 98.8 \| 6.4e-11 \| 1.3e-16 \| 73.4 \| 54 \| (2, 59) \| 69 \| (12, 65) \| 184 \| CAZy families CBM50\|CE1 protein (Fragment) \| CAZy families CBM50\|CE1 protein (Fragment); uniclust \| UniRef100\_A0A7X8YZD2 \| 98.8 \| 6.7e-11 \| 1.3e-16 \| 68.9 \| 50 \| (2, 55) \| 69 \| (82, 131) \| 132 \| LysM peptidoglycan-binding domain-containing protein \| LysM peptidoglycan-binding domain-containing protein; uniclust \| UniRef100\_A0A286RCX4 \| 98.8 \| 6.6e-11 \| 1.3e-16 \| 75.8 \| 55 \| (2, 58) \| 69 \| (165, 219) \| 261 \| Peptidoglycan-binding LysM \| Peptidoglycan-binding LysM; uniclust \| UniRef100\_A0A4Z0QZ87 \| 98.8 \| 6.5e-11 \| 1.3e-16 \| 75.2 \| 52 \| (2, 59) \| 69 \| (1, 52) \| 227 \| LysM peptidoglycan-binding domain-containing protein \| LysM peptidoglycan-binding domain-containing protein; uniclust \| UniRef100\_A0A096B6Z9 \| 98.8 \| 6.1e-11 \| 1.3e-16 \| 75.4 \| 48 \| (3, 56) \| 69 \| (155, 202) \| 204 \| LysM domain-containing protein \| LysM domain-containing protein; uniclust \| UniRef100\_A0A935WEF4 \| 98.8 \| 7.1e-11 \| 1.3e-16 \| 67.2 \| 53 \| (1, 55) \| 69 \| (70, 122) \| 123 \| LysM peptidoglycan-binding domain-containing protein \| LysM peptidoglycan-binding domain-containing protein; uniclust \| UniRef100\_A0A350XTG1 \| 98.8 \| 7.2e-11 \| 1.3e-16 \| 65.5 \| 52 \| (2, 55) \| 69 \| (52, 103) \| 104 \| LysM domain-containing protein \| LysM domain-containing protein; uniclust \| UniRef100\_A0A938DEZ6 \| 98.8 \| 7.3e-11 \| 1.3e-16 \| 64.7 \| 52 \| (2, 57) \| 69 \| (43, 94) \| 96 \| LysM peptidoglycan-binding domain-containing protein \| LysM peptidoglycan-binding domain-containing protein; uniclust \| UniRef100\_A0A7C5W2S4 \| 98.8 \| 7.3e-11 \| 1.3e-16 \| 68.4 \| 51 \| (1, 55) \| 69 \| (89, 139) \| 140 \| LysM peptidoglycan-binding domain-containing protein \| LysM peptidoglycan-binding domain-containing protein; uniclust \| UniRef100\_A0A078M1Q6 \| 98.8 \| 6.7e-11 \| 1.3e-16 \| 79.8 \| 51 \| (2, 55) \| 69 \| (314, 364) \| 364 \| Elastin-binding protein EbpS \| Elastin-binding protein EbpS; uniclust \| UniRef100\_A0A0E2ELW5 \| 98.8 \| 6.7e-11 \| 1.4e-16 \| 81.3 \| 53 \| (1, 58) \| 69 \| (14, 66) \| 427 \| LysM domain-containing protein \| LysM domain-containing protein; uniclust \| UniRef100\_A0A0J1DQ06 \| 98.8 \| 6.6e-11 \| 1.4e-16 \| 70.4 \| 55 \| (2, 59) \| 69 \| (60, 115) \| 128 \| LysM domain-containing protein \| LysM domain-containing protein; uniclust \| UniRef100\_A0A2D7GZ53 \| 98.8 \| 7.2e-11 \| 1.4e-16 \| 75.2 \| 51 \| (1, 55) \| 69 \| (185, 235) \| 236 \| LysM domain-containing protein \| LysM domain-containing protein; uniclust \| UniRef100\_A0A7X8UXT2 \| 98.8 \| 7.5e-11 \| 1.4e-16 \| 70.1 \| 51 \| (2, 54) \| 69 \| (103, 154) \| 154 \| LysM peptidoglycan-binding domain-containing protein \| LysM peptidoglycan-binding domain-containing protein; uniclust \| UniRef100\_A0A7K0PL28 \| 98.8 \| 7.1e-11 \| 1.4e-16 \| 68.1 \| 48 \| (2, 56) \| 69 \| (59, 106) \| 108 \| LysM peptidoglycan-binding domain-containing protein \| LysM peptidoglycan-binding domain-containing protein; uniclust \| UniRef100\_A0A1G7JKS9 \| 98.7 \| 7.3e-11 \| 1.4e-16 \| 79.3 \| 52 \| (1, 58) \| 69 \| (1, 52) \| 367 \| Morphogenetic protein associated with SpoVID (Fragment) \| Morphogenetic protein associated with SpoVID (Fragment); uniclust \| UniRef100\_A0A0A3IYD3 \| 98.7 \| 7.4e-11 \| 1.4e-16 \| 77.2 \| 50 \| (1, 56) \| 69 \| (1, 50) \| 298 \| Morphogenetic protein associated with SpoVID \| Morphogenetic protein associated with SpoVID; uniclust \| UniRef100\_A0A1V4MSH9 \| 98.7 \| 7.9e-11 \| 1.5e-16 \| 69.9 \| 57 \| (2, 58) \| 69 \| (106, 162) \| 166 \| LysM domain-containing protein \| LysM domain-containing protein; uniclust \| UniRef100\_A0A1N7P9I2 \| 98.7 \| 7.4e-11 \| 1.5e-16 \| 76.2 \| 52 \| (1, 58) \| 69 \| (1, 52) \| 262 \| LysM domain-containing protein \| LysM domain-containing protein; uniclust \| UniRef100\_A0A2W4JPI4 \| 98.7 \| 7.3e-11 \| 1.5e-16 \| 76.2 \| 56 \| (2, 59) \| 69 \| (174, 229) \| 243 \| LysM domain-containing protein \| LysM domain-containing protein; uniclust \| UniRef100\_A0A1V2MFZ6 \| 98.7 \| 7.6e-11 \| 1.5e-16 \| 69.2 \| 52 \| (2, 55) \| 69 \| (78, 129) \| 130 \| LysM domain-containing protein \| LysM domain-containing protein; uniclust \| UniRef100\_A0A202DMI1 \| 98.7 \| 7.6e-11 \| 1.5e-16 \| 74.9 \| 55 \| (2, 58) \| 69 \| (173, 227) \| 228 \| LysM domain-containing protein \| LysM domain-containing protein; uniclust \| UniRef100\_A0A9E7IT89 \| 98.7 \| 7.8e-11 \| 1.5e-16 \| 61.3 \| 52 \| (2, 56) \| 69 \| (8, 59) \| 61 \| LysM peptidoglycan-binding domain-containing protein \| LysM peptidoglycan-binding domain-containing protein; uniclust \| UniRef100\_A0A080LV77 \| 98.7 \| 7.1e-11 \| 1.5e-16 \| 68.0 \| 52 \| (3, 58) \| 69 \| (44, 95) \| 101 \| LysM domain-containing protein \| LysM domain-containing protein; uniclust \| UniRef100\_A0A060CH00 \| 98.7 \| 7.6e-11 \| 1.5e-16 \| 70.5 \| 48 \| (2, 56) \| 69 \| (85, 132) \| 136 \| CAZy families CE4\|CBM50 protein (Fragment) \| CAZy families CE4\|CBM50 protein (Fragment); uniclust \| UniRef100\_A0A0C1UAF4 \| 98.7 \| 7.1e-11 \| 1.5e-16 \| 77.1 \| 54 \| (2, 58) \| 69 \| (186, 239) \| 248 \| LysM domain protein \| LysM domain protein; uniclust \| UniRef100\_A0A069A1N0 \| 98.7 \| 7.3e-11 \| 1.6e-16 \| 78.5 \| 54 \| (2, 58) \| 69 \| (201, 255) \| 294 \| LysM domain protein \| LysM domain protein; uniclust \| UniRef100\_A0A0G0JZZ0 \| 98.7 \| 7.5e-11 \| 1.6e-16 \| 75.9 \| 53 \| (3, 59) \| 69 \| (165, 217) \| 227 \| LysM domain-containing protein \| LysM domain-containing protein; uniclust \| UniRef100\_A0A0J1FA30 \| 98.7 \| 8.1e-11 \| 1.6e-16 \| 61.0 \| 48 \| (1, 54) \| 69 \| (7, 54) \| 59 \| LysM domain protein \| LysM domain protein; uniclust \| UniRef100\_X0S693 \| 98.7 \| 8.6e-11 \| 1.6e-16 \| 69.3 \| 57 \| (3, 59) \| 69 \| (4, 60) \| 158 \| LysM domain-containing protein (Fragment) \| LysM domain-containing protein (Fragment); uniclust \| UniRef100\_A0A0B0I074 \| 98.7 \| 9e-11 \| 1.7e-16 \| 65.2 \| 53 \| (1, 59) \| 69 \| (1, 53) \| 104 \| Putative peptidoglycan endopeptidase LytE \| Putative peptidoglycan endopeptidase LytE; uniclust \| UniRef100\_A0A957XDI4 \| 98.7 \| 9.1e-11 \| 1.7e-16 \| 70.9 \| 54 \| (2, 57) \| 69 \| (135, 188) \| 189 \| LysM peptidoglycan-binding domain-containing protein \| LysM peptidoglycan-binding domain-containing protein; uniclust \| UniRef100\_A0A2V5KHA2 \| 98.7 \| 8.9e-11 \| 1.7e-16 \| 70.6 \| 52 \| (2, 55) \| 69 \| (114, 165) \| 166 \| LysM domain-containing protein (Fragment) \| LysM domain-containing protein (Fragment); uniclust \| UniRef100\_A0A1C4SHA2 \| 98.7 \| 8.8e-11 \| 1.7e-16 \| 72.7 \| 54 \| (2, 59) \| 69 \| (112, 165) \| 186 \| LysM domain-containing protein (Fragment) \| LysM domain-containing protein (Fragment); uniclust \| UniRef100\_A0A058ZQ48 \| 98.7 \| 8.4e-11 \| 1.8e-16 \| 73.1 \| 55 \| (3, 59) \| 69 \| (97, 151) \| 174 \| Uncharacterized protein \| Uncharacterized protein; uniclust \| UniRef100\_A0A0Q9NJ51 \| 98.7 \| 8.6e-11 \| 1.8e-16 \| 81.3 \| 52 \| (1, 58) \| 69 \| (21, 72) \| 464 \| LysM domain-containing protein \| LysM domain-containing protein; uniclust \| UniRef100\_A0A090AGM7 \| 98.7 \| 8.6e-11 \| 1.8e-16 \| 74.3 \| 42 \| (2, 46) \| 69 \| (42, 83) \| 203 \| Uncharacterized protein \| Uncharacterized protein; uniclust \| UniRef100\_A0A1F7G2S0 \| 98.7 \| 9.3e-11 \| 1.8e-16 \| 78.1 \| 54 \| (2, 57) \| 69 \| (311, 364) \| 368 \| LysM domain-containing protein \| LysM domain-containing protein; uniclust \| UniRef100\_A0A7V9DN14 \| 98.7 \| 8.9e-11 \| 1.8e-16 \| 69.6 \| 51 \| (2, 59) \| 69 \| (58, 108) \| 130 \| LysM peptidoglycan-binding domain-containing protein \| LysM peptidoglycan-binding domain-containing protein; uniclust \| UniRef100\_A0A0E2BBV5 \| 98.7 \| 9e-11 \| 1.8e-16 \| 70.4 \| 56 \| (1, 59) \| 69 \| (8, 63) \| 141 \| Phage tail protein X \| Phage tail protein X; uniclust \| UniRef100\_UPI001F499B57 \| 98.7 \| 1e-10 \| 1.9e-16 \| 62.8 \| 62 \| (1, 68) \| 69 \| (23, 84) \| 84 \| tail protein X \| tail protein X; uniclust \| UniRef100\_A0A2S5MYD0 \| 98.7 \| 1e-10 \| 1.9e-16 \| 68.1 \| 52 \| (2, 57) \| 69 \| (91, 142) \| 144 \| Heavy metal transporter (Fragment) \| Heavy metal transporter (Fragment); uniclust \| UniRef100\_A0A1Z8RYG9 \| 98.7 \| 9.1e-11 \| 1.9e-16 \| 67.2 \| 52 \| (2, 57) \| 69 \| (40, 91) \| 99 \| LysM domain-containing protein \| LysM domain-containing protein; uniclust \| UniRef100\_A0A1V6DV92 \| 98.7 \| 9.2e-11 \| 1.9e-16 \| 76.2 \| 55 \| (2, 58) \| 69 \| (192, 246) \| 248 \| LysM domain protein \| LysM domain protein; uniclust \| UniRef100\_A0A1W9R610 \| 98.7 \| 1e-10 \| 1.9e-16 \| 57.6 \| 49 \| (7, 57) \| 69 \| (2, 50) \| 51 \| LysM domain-containing protein \| LysM domain-containing protein; uniclust \| UniRef100\_A0A7C1HTJ5 \| 98.7 \| 1e-10 \| 1.9e-16 \| 70.3 \| 53 \| (2, 56) \| 69 \| (112, 164) \| 164 \| BON domain-containing protein (Fragment) \| BON domain-containing protein (Fragment); uniclust \| UniRef100\_A0A1Z9KNB0 \| 98.7 \| 1e-10 \| 2e-16 \| 78.6 \| 55 \| (1, 58) \| 69 \| (239, 293) \| 364 \| LysM domain-containing protein \| LysM domain-containing protein; uniclust \| UniRef100\_UPI001F1763FC \| 98.7 \| 1.1e-10 \| 2e-16 \| 62.2 \| 54 \| (1, 60) \| 69 \| (1, 54) \| 80 \| LysM domain-containing protein \| LysM domain-containing protein; uniclust \| UniRef100\_A0A972MN72 \| 98.7 \| 1.1e-10 \| 2e-16 \| 71.8 \| 52 \| (1, 54) \| 69 \| (168, 219) \| 219 \| LysM peptidoglycan-binding domain-containing protein \| LysM peptidoglycan-binding domain-containing protein; uniclust \| UniRef100\_A0A933KDG5 \| 98.7 \| 1.1e-10 \| 2e-16 \| 64.8 \| 55 \| (2, 58) \| 69 \| (43, 97) \| 104 \| LysM peptidoglycan-binding domain-containing protein (Fragment) \| LysM peptidoglycan-binding domain-containing protein (Fragment); uniclust \| UniRef100\_A0A935LSQ5 \| 98.7 \| 1.1e-10 \| 2e-16 \| 58.0 \| 49 \| (3, 55) \| 69 \| (2, 50) \| 51 \| LysM peptidoglycan-binding domain-containing protein \| LysM peptidoglycan-binding domain-containing protein; uniclust \| UniRef100\_A0A1R0XIF0 \| 98.7 \| 1.1e-10 \| 2.1e-16 \| 72.8 \| 52 \| (1, 58) \| 69 \| (1, 52) \| 217 \| LysM domain-containing protein (Fragment) \| LysM domain-containing protein (Fragment); uniclust \| UniRef100\_A0A098F1J4 \| 98.7 \| 1e-10 \| 2.1e-16 \| 84.8 \| 52 \| (1, 58) \| 69 \| (12, 63) \| 807 \| Spore coat assembly protein SafA \| Spore coat assembly protein SafA; uniclust \| UniRef100\_A0A7C6PRM4 \| 98.7 \| 1.1e-10 \| 2.1e-16 \| 57.9 \| 47 \| (2, 55) \| 69 \| (3, 49) \| 49 \| LysM peptidoglycan-binding domain-containing protein \| LysM peptidoglycan-binding domain-containing protein; uniclust \| UniRef100\_A0A0C9PWS5 \| 98.7 \| 1.1e-10 \| 2.1e-16 \| 78.2 \| 54 \| (1, 58) \| 69 \| (243, 296) \| 377 \| LysM domain-containing protein \| LysM domain-containing protein |
| Top keywords  (threshold 1.00e-03 (evalue)) | **LysM, domain\_containing, peptidoglycan\_binding, tail, Phage, Fragment, BON, X, superfamily, Spore** |
| Output files | ../../similar\_sequences/24\_FANPEZAQ\_CDS\_0024\_merged.svg ../../similar\_sequences/24\_FANPEZAQ\_CDS\_0024\_pdb70.a3m ../../similar\_sequences/24\_FANPEZAQ\_CDS\_0024\_pdb70.hhr ../../similar\_sequences/24\_FANPEZAQ\_CDS\_0024\_uniclust.a3m ../../similar\_sequences/24\_FANPEZAQ\_CDS\_0024\_uniclust.hhr |

#### Structure prediction (AlphaFold)^2^

|  |  |
| --- | --- |
| Stats | xml version="1.0" encoding="utf-8" standalone="no"?       2024-09-02T21:09:24.638686 image/svg+xml   Matplotlib v3.7.2, https://matplotlib.org/ |
| Predicted structure | **NGL Viewer Controls:**  - Center: *Left-Click* - Rotate: *Left-Click + Drag* - Translate: *Right-Click + Drag* - Zoom: *Shift + Left-Click + Drag* |
| Output files | ../../predicted\_structures/24\_FANPEZAQ\_CDS\_0024/features.pkl ../../predicted\_structures/24\_FANPEZAQ\_CDS\_0024/ranked\_0.pdb ../../predicted\_structures/24\_FANPEZAQ\_CDS\_0024/ranked\_0\_plots.svg ../../predicted\_structures/24\_FANPEZAQ\_CDS\_0024/result\_model\_1\_ptm\_pred\_0.pkl |

#### Structure similarity search results (Foldseek)^3^

|  |  |
| --- | --- |
| Structure databases searched | Pdb, Afdb-proteome, Afdb-uniprot50 |
| Results, scheme(s)  (Top layers only, threshold 1.00e-02 (evalue)) | xml version="1.0" encoding="utf-8" standalone="no"?       2024-09-02T21:10:52.721427 image/svg+xml   Matplotlib v3.7.2, https://matplotlib.org/ |
| Results, table  (threshold 1.00e-02 (evalue)) | db \| id \| prob \| evalue \| bits \| fident \| alnlen \| mismatch \| gapopen \| qstart \| qend \| tstart \| tend \| name \| description; pdb \| 6U5B\_6 \| 1.0 \| 0.0003522 \| 149 \| 0.436 \| 55 \| 30 \| 1 \| 2 \| 56 \| 1 \| 54 \| Glue PA0627 \| Glue PA0627; pdb \| 2LTF\_A \| 1.0 \| 0.0001977 \| 139 \| 0.397 \| 68 \| 40 \| 1 \| 2 \| 69 \| 1 \| 67 \| Tail protein X \| Tail protein X; pdb \| 2LTF\_A \| 1.0 \| 0.0001378 \| 135 \| 0.397 \| 68 \| 40 \| 1 \| 2 \| 69 \| 1 \| 67 \| Tail protein X \| Tail protein X; pdb \| 2LTF\_A \| 1.0 \| 0.0003785 \| 127 \| 0.397 \| 68 \| 40 \| 1 \| 2 \| 69 \| 1 \| 67 \| Tail protein X \| Tail protein X; afdb-proteome \| AF-A0A0H3GM11-F1-MODEL\_V4 \| 1.0 \| 6.871e-09 \| 283 \| 0.441 \| 68 \| 37 \| 1 \| 2 \| 69 \| 1 \| 67 \| Uncharacterized protein \| Uncharacterized protein; afdb-proteome \| AF-Q8ZMT5-F1-MODEL\_V4 \| 1.0 \| 1.53e-07 \| 244 \| 0.426 \| 68 \| 38 \| 1 \| 2 \| 69 \| 1 \| 67 \| Fels-2 prophage protein \| Fels-2 prophage protein; afdb-proteome \| AF-A0A0H3GWZ2-F1-MODEL\_V4 \| 1.0 \| 4.516e-07 \| 232 \| 0.397 \| 68 \| 40 \| 1 \| 2 \| 69 \| 1 \| 67 \| Putative prophage tail component protein \| Putative prophage tail component protein; afdb-proteome \| AF-Q8ZKJ6-F1-MODEL\_V4 \| 1.0 \| 0.0001938 \| 145 \| 0.354 \| 62 \| 37 \| 2 \| 2 \| 63 \| 4 \| 62 \| Putative inner membrane protein \| Putative inner membrane protein; afdb-proteome \| AF-G3XD62-F1-MODEL\_V4 \| 1.0 \| 0.0006607 \| 139 \| 0.411 \| 68 \| 38 \| 2 \| 2 \| 69 \| 3 \| 68 \| Uncharacterized protein \| Uncharacterized protein; afdb-uniprot50 \| AF-A0A212KMZ3-F1-MODEL\_V4 \| 1.0 \| 9.95e-08 \| 267 \| 0.5 \| 70 \| 32 \| 2 \| 2 \| 69 \| 3 \| 71 \| Tail protein X \| Tail protein X; afdb-uniprot50 \| AF-A0A3G6WBC5-F1-MODEL\_V4 \| 1.0 \| 6.004e-08 \| 264 \| 0.608 \| 69 \| 27 \| 0 \| 1 \| 69 \| 1 \| 69 \| Phage tail protein \| Phage tail protein; afdb-uniprot50 \| AF-A0A4R0EZY9-F1-MODEL\_V4 \| 1.0 \| 1.069e-07 \| 260 \| 0.414 \| 70 \| 39 \| 2 \| 1 \| 69 \| 1 \| 69 \| Phage tail protein \| Phage tail protein; afdb-uniprot50 \| AF-A0A1Y3CEI3-F1-MODEL\_V4 \| 1.0 \| 4.867e-07 \| 260 \| 0.478 \| 69 \| 36 \| 0 \| 1 \| 69 \| 1 \| 69 \| Uncharacterized protein \| Uncharacterized protein; afdb-uniprot50 \| AF-A0A450SYU4-F1-MODEL\_V4 \| 1.0 \| 1.427e-07 \| 260 \| 0.507 \| 69 \| 32 \| 2 \| 2 \| 69 \| 3 \| 70 \| P2-like prophage tail protein X \| P2-like prophage tail protein X; afdb-uniprot50 \| AF-A0A1G5ACY5-F1-MODEL\_V4 \| 1.0 \| 1.427e-07 \| 258 \| 0.521 \| 69 \| 32 \| 1 \| 1 \| 69 \| 1 \| 68 \| P2-like prophage tail protein X \| P2-like prophage tail protein X; afdb-uniprot50 \| AF-A0A136HGF2-F1-MODEL\_V4 \| 1.0 \| 1.328e-07 \| 258 \| 0.528 \| 70 \| 31 \| 2 \| 1 \| 69 \| 1 \| 69 \| Phage tail protein \| Phage tail protein; afdb-uniprot50 \| AF-A0A2W5H3R5-F1-MODEL\_V4 \| 1.0 \| 1.534e-07 \| 254 \| 0.521 \| 69 \| 32 \| 1 \| 2 \| 69 \| 95 \| 163 \| Uncharacterized protein \| Uncharacterized protein; afdb-uniprot50 \| AF-A0A2U1XYZ3-F1-MODEL\_V4 \| 1.0 \| 1.149e-07 \| 250 \| 0.536 \| 69 \| 31 \| 1 \| 1 \| 69 \| 1 \| 68 \| Phage tail protein \| Phage tail protein; afdb-uniprot50 \| AF-D2ZKL4-F1-MODEL\_V4 \| 1.0 \| 1.076e-06 \| 243 \| 0.455 \| 68 \| 36 \| 1 \| 2 \| 69 \| 43 \| 109 \| Phage Tail Protein X \| Phage Tail Protein X; afdb-uniprot50 \| AF-A0A4Q0YIT0-F1-MODEL\_V4 \| 1.0 \| 1.076e-06 \| 239 \| 0.5 \| 68 \| 32 \| 2 \| 2 \| 69 \| 1 \| 66 \| Phage tail protein \| Phage tail protein; afdb-uniprot50 \| AF-F4BFP7-F1-MODEL\_V4 \| 1.0 \| 6.044e-07 \| 237 \| 0.455 \| 68 \| 35 \| 1 \| 1 \| 68 \| 1 \| 66 \| Uncharacterized protein \| Uncharacterized protein; afdb-uniprot50 \| AF-A0A1Q5TNQ8-F1-MODEL\_V4 \| 1.0 \| 1.244e-06 \| 237 \| 0.455 \| 68 \| 36 \| 1 \| 2 \| 69 \| 1 \| 67 \| Tail protein X \| Tail protein X; afdb-uniprot50 \| AF-B8GS03-F1-MODEL\_V4 \| 1.0 \| 1.337e-06 \| 237 \| 0.426 \| 68 \| 37 \| 2 \| 1 \| 67 \| 1 \| 67 \| P2-like protein prophage tail protein X-like protein \| P2-like protein prophage tail protein X-like protein; afdb-uniprot50 \| AF-A0A846VV59-F1-MODEL\_V4 \| 1.0 \| 2.381e-06 \| 236 \| 0.323 \| 68 \| 46 \| 0 \| 2 \| 69 \| 7 \| 74 \| Phage tail protein \| Phage tail protein; afdb-uniprot50 \| AF-C0B391-F1-MODEL\_V4 \| 1.0 \| 5.231e-07 \| 234 \| 0.391 \| 69 \| 40 \| 2 \| 2 \| 69 \| 1 \| 68 \| Phage Tail Protein X \| Phage Tail Protein X; afdb-uniprot50 \| AF-A0A348FYG1-F1-MODEL\_V4 \| 1.0 \| 6.044e-07 \| 234 \| 0.528 \| 70 \| 31 \| 2 \| 1 \| 69 \| 1 \| 69 \| Tail protein X \| Tail protein X; afdb-uniprot50 \| AF-A0A2J0SUU1-F1-MODEL\_V4 \| 1.0 \| 9.318e-07 \| 234 \| 0.536 \| 69 \| 31 \| 1 \| 2 \| 69 \| 3 \| 71 \| Phage tail protein \| Phage tail protein; afdb-uniprot50 \| AF-A0A7U4KP86-F1-MODEL\_V4 \| 1.0 \| 9.318e-07 \| 233 \| 0.397 \| 68 \| 40 \| 1 \| 2 \| 69 \| 1 \| 67 \| Tail protein X \| Tail protein X; afdb-uniprot50 \| AF-R9VJF3-F1-MODEL\_V4 \| 1.0 \| 1.784e-06 \| 233 \| 0.426 \| 68 \| 38 \| 1 \| 2 \| 69 \| 1 \| 67 \| Tail protein X \| Tail protein X; afdb-uniprot50 \| AF-A0A5D4YFH7-F1-MODEL\_V4 \| 1.0 \| 2.061e-06 \| 233 \| 0.397 \| 68 \| 40 \| 1 \| 2 \| 69 \| 1 \| 67 \| Phage tail protein \| Phage tail protein; afdb-uniprot50 \| AF-A0A502FVN3-F1-MODEL\_V4 \| 1.0 \| 1.437e-06 \| 232 \| 0.367 \| 68 \| 42 \| 1 \| 2 \| 69 \| 1 \| 67 \| Phage tail protein \| Phage tail protein; afdb-uniprot50 \| AF-A0A239EHV7-F1-MODEL\_V4 \| 1.0 \| 4.867e-07 \| 230 \| 0.478 \| 69 \| 34 \| 1 \| 1 \| 69 \| 1 \| 67 \| P2-like prophage tail protein X \| P2-like prophage tail protein X; afdb-uniprot50 \| AF-A0A0G4QHL0-F1-MODEL\_V4 \| 1.0 \| 9.318e-07 \| 229 \| 0.391 \| 69 \| 40 \| 2 \| 2 \| 69 \| 1 \| 68 \| Phage tail protein X \| Phage tail protein X; afdb-uniprot50 \| AF-A0A2W4T7L0-F1-MODEL\_V4 \| 1.0 \| 4.899e-06 \| 228 \| 0.455 \| 68 \| 36 \| 1 \| 2 \| 69 \| 3 \| 69 \| Phage tail protein \| Phage tail protein; afdb-uniprot50 \| AF-A0A4Y6U9C4-F1-MODEL\_V4 \| 1.0 \| 1.076e-06 \| 228 \| 0.394 \| 71 \| 40 \| 1 \| 2 \| 69 \| 1 \| 71 \| Phage tail protein \| Phage tail protein; afdb-uniprot50 \| AF-C9XYT5-F1-MODEL\_V4 \| 1.0 \| 1.917e-06 \| 226 \| 0.405 \| 69 \| 38 \| 3 \| 2 \| 69 \| 1 \| 67 \| Tail protein X \| Tail protein X; afdb-uniprot50 \| AF-A0A1X7Q4C3-F1-MODEL\_V4 \| 1.0 \| 5.231e-07 \| 226 \| 0.565 \| 69 \| 27 \| 3 \| 2 \| 69 \| 3 \| 69 \| p2-like prophage tail protein X \| p2-like prophage tail protein X; afdb-uniprot50 \| AF-A0A1T4WWY3-F1-MODEL\_V4 \| 1.0 \| 2.381e-06 \| 224 \| 0.428 \| 70 \| 38 \| 2 \| 1 \| 69 \| 1 \| 69 \| p2-like prophage tail protein X \| p2-like prophage tail protein X; afdb-uniprot50 \| AF-A0A2D3TFL6-F1-MODEL\_V4 \| 1.0 \| 2.215e-06 \| 224 \| 0.406 \| 64 \| 37 \| 1 \| 2 \| 65 \| 1 \| 63 \| Phage tail protein \| Phage tail protein; afdb-uniprot50 \| AF-A0A6G8F2N9-F1-MODEL\_V4 \| 1.0 \| 4.241e-06 \| 223 \| 0.5 \| 68 \| 33 \| 1 \| 1 \| 68 \| 1 \| 67 \| Uncharacterized protein \| Uncharacterized protein; afdb-uniprot50 \| AF-A0A2X3CVI6-F1-MODEL\_V4 \| 1.0 \| 3.178e-06 \| 223 \| 0.382 \| 68 \| 41 \| 1 \| 2 \| 69 \| 1 \| 67 \| Tail component protein \| Tail component protein; afdb-uniprot50 \| AF-A0A496K2J9-F1-MODEL\_V4 \| 1.0 \| 2.381e-06 \| 221 \| 0.436 \| 71 \| 36 \| 2 \| 2 \| 69 \| 1 \| 70 \| Phage tail protein \| Phage tail protein; afdb-uniprot50 \| AF-A0A1C3WM59-F1-MODEL\_V4 \| 1.0 \| 2.559e-06 \| 220 \| 0.405 \| 69 \| 39 \| 2 \| 2 \| 69 \| 23 \| 90 \| p2-like prophage tail protein X \| p2-like prophage tail protein X; afdb-uniprot50 \| AF-A0A738YWL5-F1-MODEL\_V4 \| 1.0 \| 7.028e-06 \| 219 \| 0.367 \| 68 \| 42 \| 1 \| 2 \| 69 \| 1 \| 67 \| Phage tail protein \| Phage tail protein; afdb-uniprot50 \| AF-A0A8A5W2Y0-F1-MODEL\_V4 \| 1.0 \| 3.946e-06 \| 219 \| 0.441 \| 68 \| 37 \| 1 \| 2 \| 69 \| 3 \| 69 \| Tail protein X \| Tail protein X; afdb-uniprot50 \| AF-R9AS75-F1-MODEL\_V4 \| 1.0 \| 2.381e-06 \| 219 \| 0.376 \| 69 \| 41 \| 2 \| 2 \| 69 \| 3 \| 70 \| Uncharacterized protein \| Uncharacterized protein; afdb-uniprot50 \| AF-A0A426CLA1-F1-MODEL\_V4 \| 1.0 \| 5.266e-06 \| 218 \| 0.417 \| 67 \| 37 \| 2 \| 2 \| 68 \| 6 \| 70 \| Phage tail protein \| Phage tail protein; afdb-uniprot50 \| AF-A0A841ILH7-F1-MODEL\_V4 \| 1.0 \| 3.671e-06 \| 217 \| 0.397 \| 68 \| 40 \| 1 \| 2 \| 69 \| 1 \| 67 \| Phage tail protein X \| Phage tail protein X; afdb-uniprot50 \| AF-A0A1X3I4F6-F1-MODEL\_V4 \| 1.0 \| 5.266e-06 \| 217 \| 0.411 \| 68 \| 39 \| 1 \| 2 \| 69 \| 19 \| 85 \| Uncharacterized protein \| Uncharacterized protein; afdb-uniprot50 \| AF-S6GQ04-F1-MODEL\_V4 \| 1.0 \| 8.119e-06 \| 216 \| 0.347 \| 69 \| 43 \| 2 \| 2 \| 69 \| 1 \| 68 \| Gp8 \| Gp8; afdb-uniprot50 \| AF-A0A6M3KNH7-F1-MODEL\_V4 \| 1.0 \| 9.38e-06 \| 216 \| 0.304 \| 69 \| 48 \| 0 \| 1 \| 69 \| 1 \| 69 \| Putative tail protein \| Putative tail protein; afdb-uniprot50 \| AF-A0A432QVU7-F1-MODEL\_V4 \| 1.0 \| 2.061e-06 \| 216 \| 0.338 \| 71 \| 45 \| 1 \| 1 \| 69 \| 1 \| 71 \| Phage tail protein \| Phage tail protein; afdb-uniprot50 \| AF-A0A809H9W3-F1-MODEL\_V4 \| 1.0 \| 2.751e-06 \| 215 \| 0.397 \| 68 \| 40 \| 1 \| 2 \| 69 \| 1 \| 67 \| Phage tail protein \| Phage tail protein; afdb-uniprot50 \| AF-A0A0Q7XZV6-F1-MODEL\_V4 \| 1.0 \| 2.956e-06 \| 214 \| 0.441 \| 68 \| 36 \| 2 \| 2 \| 68 \| 3 \| 69 \| Phage tail protein \| Phage tail protein; afdb-uniprot50 \| AF-E2CFJ8-F1-MODEL\_V4 \| 1.0 \| 7.554e-06 \| 213 \| 0.43 \| 65 \| 35 \| 1 \| 1 \| 65 \| 1 \| 63 \| Phage Tail Protein X \| Phage Tail Protein X; afdb-uniprot50 \| AF-A0A6L2ZQ46-F1-MODEL\_V4 \| 1.0 \| 5.266e-06 \| 213 \| 0.428 \| 70 \| 39 \| 1 \| 1 \| 69 \| 1 \| 70 \| Tail protein X \| Tail protein X; afdb-uniprot50 \| AF-A0A6H1ZLH4-F1-MODEL\_V4 \| 1.0 \| 2.215e-06 \| 213 \| 0.38 \| 71 \| 41 \| 1 \| 2 \| 69 \| 1 \| 71 \| Putative tail protein \| Putative tail protein; afdb-uniprot50 \| AF-I7E1A5-F1-MODEL\_V4 \| 1.0 \| 2.061e-06 \| 212 \| 0.5 \| 68 \| 34 \| 0 \| 2 \| 69 \| 6 \| 73 \| Phage tail protein X \| Phage tail protein X; afdb-uniprot50 \| AF-A0A7L9REI3-F1-MODEL\_V4 \| 1.0 \| 5.66e-06 \| 211 \| 0.338 \| 68 \| 44 \| 1 \| 2 \| 69 \| 1 \| 67 \| Phage tail protein \| Phage tail protein; afdb-uniprot50 \| AF-A0A0H3L797-F1-MODEL\_V4 \| 1.0 \| 4.899e-06 \| 210 \| 0.397 \| 68 \| 40 \| 1 \| 2 \| 69 \| 90 \| 156 \| Phage tail protein X \| Phage tail protein X; afdb-uniprot50 \| AF-A0A6L9FJD9-F1-MODEL\_V4 \| 1.0 \| 3.946e-06 \| 209 \| 0.411 \| 68 \| 38 \| 1 \| 2 \| 69 \| 3 \| 68 \| Phage tail protein \| Phage tail protein; afdb-uniprot50 \| AF-A0A7Z7RKR3-F1-MODEL\_V4 \| 1.0 \| 9.38e-06 \| 209 \| 0.388 \| 67 \| 39 \| 2 \| 1 \| 66 \| 1 \| 66 \| Phage Tail Protein X \| Phage Tail Protein X; afdb-uniprot50 \| AF-A0A1H0NML7-F1-MODEL\_V4 \| 1.0 \| 1.252e-05 \| 209 \| 0.468 \| 64 \| 33 \| 1 \| 2 \| 65 \| 1 \| 63 \| P2-like prophage tail protein X \| P2-like prophage tail protein X; afdb-uniprot50 \| AF-A0A484S6Y7-F1-MODEL\_V4 \| 1.0 \| 3.671e-06 \| 208 \| 0.405 \| 69 \| 39 \| 2 \| 2 \| 69 \| 1 \| 68 \| Phage tail completion protein \| Phage tail completion protein; afdb-uniprot50 \| AF-A0A659L5Z7-F1-MODEL\_V4 \| 1.0 \| 1.446e-05 \| 208 \| 0.367 \| 68 \| 42 \| 1 \| 2 \| 69 \| 1 \| 67 \| Phage tail protein \| Phage tail protein; afdb-uniprot50 \| AF-A0A143DFK3-F1-MODEL\_V4 \| 1.0 \| 8.727e-06 \| 208 \| 0.514 \| 68 \| 32 \| 1 \| 2 \| 69 \| 3 \| 69 \| Uncharacterized protein \| Uncharacterized protein; afdb-uniprot50 \| AF-E5AKR2-F1-MODEL\_V4 \| 1.0 \| 5.266e-06 \| 207 \| 0.463 \| 69 \| 35 \| 2 \| 2 \| 69 \| 23 \| 90 \| Tail protein X \| Tail protein X; afdb-uniprot50 \| AF-A0A7K0GNQ8-F1-MODEL\_V4 \| 1.0 \| 3.415e-06 \| 206 \| 0.442 \| 70 \| 35 \| 2 \| 2 \| 69 \| 1 \| 68 \| Phage tail protein \| Phage tail protein; afdb-uniprot50 \| AF-Q31Q85-F1-MODEL\_V4 \| 1.0 \| 7.028e-06 \| 206 \| 0.434 \| 69 \| 38 \| 1 \| 1 \| 69 \| 1 \| 68 \| Uncharacterized protein \| Uncharacterized protein; afdb-uniprot50 \| AF-A0A1T4W4K7-F1-MODEL\_V4 \| 1.0 \| 1.084e-05 \| 206 \| 0.414 \| 70 \| 37 \| 3 \| 2 \| 69 \| 4 \| 71 \| p2-like prophage tail protein X \| p2-like prophage tail protein X; afdb-uniprot50 \| AF-A0A6I4T3J1-F1-MODEL\_V4 \| 1.0 \| 4.558e-06 \| 206 \| 0.478 \| 69 \| 34 \| 2 \| 2 \| 69 \| 4 \| 71 \| Phage tail protein \| Phage tail protein; afdb-uniprot50 \| AF-A0A248LIW4-F1-MODEL\_V4 \| 1.0 \| 7.028e-06 \| 205 \| 0.463 \| 69 \| 35 \| 2 \| 2 \| 69 \| 1 \| 68 \| Tail protein X \| Tail protein X; afdb-uniprot50 \| AF-A0A4R5HGF7-F1-MODEL\_V4 \| 1.0 \| 1.93e-05 \| 205 \| 0.405 \| 69 \| 41 \| 0 \| 1 \| 69 \| 1 \| 69 \| Phage tail protein \| Phage tail protein; afdb-uniprot50 \| AF-G9PUI9-F1-MODEL\_V4 \| 1.0 \| 2.075e-05 \| 205 \| 0.362 \| 58 \| 35 \| 2 \| 1 \| 57 \| 1 \| 57 \| Uncharacterized protein \| Uncharacterized protein; afdb-uniprot50 \| AF-A0A1S0V5S2-F1-MODEL\_V4 \| 1.0 \| 5.66e-06 \| 205 \| 0.492 \| 69 \| 34 \| 1 \| 2 \| 69 \| 4 \| 72 \| Tail protein \| Tail protein; afdb-uniprot50 \| AF-A0A2N0DX25-F1-MODEL\_V4 \| 1.0 \| 2.075e-05 \| 205 \| 0.369 \| 65 \| 39 \| 2 \| 2 \| 66 \| 3 \| 65 \| Phage tail protein X \| Phage tail protein X; afdb-uniprot50 \| AF-M4Z448-F1-MODEL\_V4 \| 1.0 \| 1.446e-05 \| 204 \| 0.397 \| 68 \| 40 \| 1 \| 2 \| 69 \| 3 \| 69 \| Putative phage tail protein \| Putative phage tail protein; afdb-uniprot50 \| AF-A0A616MEN0-F1-MODEL\_V4 \| 1.0 \| 1.671e-05 \| 204 \| 0.376 \| 69 \| 42 \| 1 \| 2 \| 69 \| 3 \| 71 \| Phage tail protein \| Phage tail protein; afdb-uniprot50 \| AF-A0A350LX84-F1-MODEL\_V4 \| 1.0 \| 4.899e-06 \| 204 \| 0.45 \| 71 \| 35 \| 2 \| 1 \| 69 \| 1 \| 69 \| Phage tail protein \| Phage tail protein; afdb-uniprot50 \| AF-A0A165XHI0-F1-MODEL\_V4 \| 1.0 \| 4.558e-06 \| 204 \| 0.417 \| 67 \| 38 \| 1 \| 3 \| 69 \| 12 \| 77 \| Phage Tail Protein X \| Phage Tail Protein X; afdb-uniprot50 \| AF-F4V8N9-F1-MODEL\_V4 \| 1.0 \| 1.671e-05 \| 203 \| 0.391 \| 69 \| 41 \| 1 \| 1 \| 69 \| 23 \| 90 \| Conserved domain protein \| Conserved domain protein; afdb-uniprot50 \| AF-A0A292AJG7-F1-MODEL\_V4 \| 1.0 \| 7.554e-06 \| 202 \| 0.391 \| 69 \| 39 \| 3 \| 2 \| 69 \| 1 \| 67 \| Phage tail protein \| Phage tail protein; afdb-uniprot50 \| AF-A0A3Q9W3P2-F1-MODEL\_V4 \| 1.0 \| 1.93e-05 \| 202 \| 0.382 \| 68 \| 41 \| 1 \| 2 \| 69 \| 5 \| 71 \| Phage tail protein \| Phage tail protein; afdb-uniprot50 \| AF-A0A2S6N2T4-F1-MODEL\_V4 \| 1.0 \| 1.93e-05 \| 202 \| 0.428 \| 70 \| 38 \| 2 \| 2 \| 69 \| 5 \| 74 \| Uncharacterized protein \| Uncharacterized protein; afdb-uniprot50 \| AF-A0A7W6WBC8-F1-MODEL\_V4 \| 1.0 \| 3.415e-06 \| 201 \| 0.457 \| 70 \| 34 \| 2 \| 3 \| 69 \| 15 \| 83 \| Phage tail protein X \| Phage tail protein X; afdb-uniprot50 \| AF-A0A1K0IP73-F1-MODEL\_V4 \| 1.0 \| 1.252e-05 \| 200 \| 0.47 \| 68 \| 35 \| 1 \| 2 \| 69 \| 1 \| 67 \| Tail protein X \| Tail protein X; afdb-uniprot50 \| AF-A0A2S9IC13-F1-MODEL\_V4 \| 1.0 \| 1.165e-05 \| 199 \| 0.426 \| 68 \| 38 \| 1 \| 2 \| 69 \| 1 \| 67 \| Phage tail protein \| Phage tail protein; afdb-uniprot50 \| AF-A0A541BL22-F1-MODEL\_V4 \| 1.0 \| 1.008e-05 \| 199 \| 0.4 \| 70 \| 38 \| 3 \| 2 \| 69 \| 1 \| 68 \| Phage tail protein \| Phage tail protein; afdb-uniprot50 \| AF-A0A1H9YCX3-F1-MODEL\_V4 \| 1.0 \| 6.539e-06 \| 199 \| 0.42 \| 69 \| 38 \| 2 \| 2 \| 69 \| 6 \| 73 \| P2-like prophage tail protein X \| P2-like prophage tail protein X; afdb-uniprot50 \| AF-A0A142BH73-F1-MODEL\_V4 \| 1.0 \| 1.671e-05 \| 198 \| 0.441 \| 68 \| 36 \| 1 \| 2 \| 69 \| 1 \| 66 \| Tail X family protein \| Tail X family protein; afdb-uniprot50 \| AF-C6DCA5-F1-MODEL\_V4 \| 1.0 \| 2.397e-05 \| 198 \| 0.382 \| 68 \| 41 \| 1 \| 2 \| 69 \| 1 \| 67 \| Tail X family protein \| Tail X family protein; afdb-uniprot50 \| AF-A0A4P7LCT2-F1-MODEL\_V4 \| 1.0 \| 1.446e-05 \| 198 \| 0.347 \| 69 \| 43 \| 2 \| 2 \| 69 \| 1 \| 68 \| Phage Tail Protein X \| Phage Tail Protein X; afdb-uniprot50 \| AF-A0A1S1U741-F1-MODEL\_V4 \| 1.0 \| 3.178e-06 \| 198 \| 0.43 \| 72 \| 36 \| 2 \| 2 \| 69 \| 1 \| 71 \| Uncharacterized protein \| Uncharacterized protein; afdb-uniprot50 \| AF-A0A1N6I0M6-F1-MODEL\_V4 \| 1.0 \| 1.446e-05 \| 197 \| 0.417 \| 67 \| 38 \| 1 \| 2 \| 68 \| 3 \| 68 \| p2-like prophage tail protein X \| p2-like prophage tail protein X; afdb-uniprot50 \| AF-A0A8B2TUJ1-F1-MODEL\_V4 \| 1.0 \| 2.576e-05 \| 196 \| 0.308 \| 68 \| 47 \| 0 \| 2 \| 69 \| 1 \| 68 \| Phage tail protein \| Phage tail protein; afdb-uniprot50 \| AF-A0A1I1F5Z9-F1-MODEL\_V4 \| 1.0 \| 1.165e-05 \| 196 \| 0.375 \| 72 \| 42 \| 2 \| 1 \| 69 \| 1 \| 72 \| P2-like prophage tail protein X \| P2-like prophage tail protein X; afdb-uniprot50 \| AF-A0A1X7MAH8-F1-MODEL\_V4 \| 1.0 \| 6.124e-05 \| 195 \| 0.384 \| 65 \| 39 \| 1 \| 1 \| 65 \| 1 \| 64 \| Phage tail protein X \| Phage tail protein X; afdb-uniprot50 \| AF-A0A0H5CZ07-F1-MODEL\_V4 \| 1.0 \| 3.695e-05 \| 194 \| 0.367 \| 68 \| 42 \| 1 \| 2 \| 69 \| 3 \| 69 \| Phage Tail Protein X \| Phage Tail Protein X; afdb-uniprot50 \| AF-N9SPH0-F1-MODEL\_V4 \| 1.0 \| 3.695e-05 \| 194 \| 0.371 \| 70 \| 42 \| 2 \| 1 \| 69 \| 1 \| 69 \| Uncharacterized protein \| Uncharacterized protein; afdb-uniprot50 \| AF-A0A496AHX0-F1-MODEL\_V4 \| 1.0 \| 1.252e-05 \| 193 \| 0.246 \| 69 \| 52 \| 0 \| 1 \| 69 \| 1 \| 69 \| Uncharacterized protein \| Uncharacterized protein; afdb-uniprot50 \| AF-A0A285VTB6-F1-MODEL\_V4 \| 1.0 \| 2.075e-05 \| 193 \| 0.449 \| 69 \| 36 \| 2 \| 2 \| 69 \| 3 \| 70 \| P2-like prophage tail protein X \| P2-like prophage tail protein X; afdb-uniprot50 \| AF-A0A2T5UR74-F1-MODEL\_V4 \| 1.0 \| 1.446e-05 \| 193 \| 0.449 \| 69 \| 36 \| 2 \| 2 \| 69 \| 3 \| 70 \| Phage tail protein X \| Phage tail protein X; afdb-uniprot50 \| AF-A0A1V0BI36-F1-MODEL\_V4 \| 1.0 \| 1.554e-05 \| 193 \| 0.426 \| 68 \| 38 \| 1 \| 2 \| 69 \| 4 \| 70 \| Phage tail protein \| Phage tail protein; afdb-uniprot50 \| AF-A0A149SNW1-F1-MODEL\_V4 \| 1.0 \| 1.671e-05 \| 193 \| 0.464 \| 71 \| 36 \| 2 \| 1 \| 69 \| 1 \| 71 \| Uncharacterized protein \| Uncharacterized protein; afdb-uniprot50 \| AF-A0A1A9VKI1-F1-MODEL\_V4 \| 1.0 \| 2.576e-05 \| 193 \| 0.447 \| 67 \| 36 \| 1 \| 3 \| 69 \| 342 \| 407 \| Uncharacterized protein \| Uncharacterized protein; afdb-uniprot50 \| AF-A0A367WTU8-F1-MODEL\_V4 \| 1.0 \| 1.93e-05 \| 192 \| 0.434 \| 69 \| 36 \| 2 \| 1 \| 69 \| 1 \| 66 \| Tail protein X \| Tail protein X; afdb-uniprot50 \| AF-A0A439F0D7-F1-MODEL\_V4 \| 1.0 \| 1.554e-05 \| 192 \| 0.441 \| 68 \| 37 \| 1 \| 2 \| 69 \| 1 \| 67 \| Phage tail protein \| Phage tail protein; afdb-uniprot50 \| AF-A0A167GZN0-F1-MODEL\_V4 \| 1.0 \| 3.972e-05 \| 192 \| 0.42 \| 69 \| 38 \| 2 \| 2 \| 69 \| 1 \| 68 \| Tail protein \| Tail protein; afdb-uniprot50 \| AF-A0A1X3DKC3-F1-MODEL\_V4 \| 1.0 \| 2.976e-05 \| 192 \| 0.362 \| 69 \| 42 \| 2 \| 2 \| 69 \| 6 \| 73 \| Phage tail protein \| Phage tail protein; afdb-uniprot50 \| AF-A0A376SBV1-F1-MODEL\_V4 \| 1.0 \| 0.0002245 \| 192 \| 0.428 \| 56 \| 31 \| 1 \| 2 \| 57 \| 1 \| 55 \| Phage tail protein X \| Phage tail protein X; afdb-uniprot50 \| AF-V0XX61-F1-MODEL\_V4 \| 1.0 \| 0.0002413 \| 190 \| 0.421 \| 57 \| 32 \| 1 \| 2 \| 58 \| 1 \| 56 \| Phage Tail Protein X \| Phage Tail Protein X; afdb-uniprot50 \| AF-A0A7X5THN7-F1-MODEL\_V4 \| 1.0 \| 3.438e-05 \| 190 \| 0.397 \| 68 \| 40 \| 1 \| 2 \| 69 \| 1 \| 67 \| Phage tail protein \| Phage tail protein; afdb-uniprot50 \| AF-A0A2T4JP28-F1-MODEL\_V4 \| 1.0 \| 2.075e-05 \| 188 \| 0.382 \| 68 \| 40 \| 1 \| 2 \| 69 \| 6 \| 71 \| Phage tail protein \| Phage tail protein; afdb-uniprot50 \| AF-A0A1I7HNC1-F1-MODEL\_V4 \| 1.0 \| 2.576e-05 \| 188 \| 0.426 \| 68 \| 38 \| 1 \| 2 \| 69 \| 72 \| 138 \| P2-like prophage tail protein X \| P2-like prophage tail protein X; afdb-uniprot50 \| AF-A0A840C414-F1-MODEL\_V4 \| 1.0 \| 2.075e-05 \| 187 \| 0.449 \| 69 \| 37 \| 1 \| 1 \| 69 \| 1 \| 68 \| Phage tail protein X \| Phage tail protein X; afdb-uniprot50 \| AF-A0A7Z0SIU9-F1-MODEL\_V4 \| 1.0 \| 3.972e-05 \| 187 \| 0.352 \| 68 \| 41 \| 2 \| 2 \| 68 \| 3 \| 68 \| Tail protein X \| Tail protein X; afdb-uniprot50 \| AF-L8MGI2-F1-MODEL\_V4 \| 1.0 \| 2.976e-05 \| 186 \| 0.376 \| 69 \| 42 \| 1 \| 1 \| 69 \| 1 \| 68 \| Phage-related tail protein \| Phage-related tail protein; afdb-uniprot50 \| AF-A0A7Y6Z4E9-F1-MODEL\_V4 \| 1.0 \| 0.000126 \| 185 \| 0.375 \| 64 \| 39 \| 1 \| 2 \| 65 \| 5 \| 67 \| Tail protein X \| Tail protein X; afdb-uniprot50 \| AF-A0A3S4CJZ7-F1-MODEL\_V4 \| 1.0 \| 2.976e-05 \| 184 \| 0.376 \| 69 \| 41 \| 1 \| 1 \| 69 \| 1 \| 67 \| Phage Tail Protein X \| Phage Tail Protein X; afdb-uniprot50 \| AF-G0AIP8-F1-MODEL\_V4 \| 1.0 \| 3.438e-05 \| 184 \| 0.4 \| 70 \| 39 \| 2 \| 2 \| 69 \| 1 \| 69 \| Putative phage tail protein \| Putative phage tail protein; afdb-uniprot50 \| AF-A0A1Q9PIY2-F1-MODEL\_V4 \| 1.0 \| 8.785e-05 \| 184 \| 0.344 \| 58 \| 35 \| 2 \| 1 \| 58 \| 1 \| 55 \| Uncharacterized protein \| Uncharacterized protein; afdb-uniprot50 \| AF-A0A5B9YAQ9-F1-MODEL\_V4 \| 1.0 \| 0.000126 \| 184 \| 0.306 \| 62 \| 40 \| 2 \| 2 \| 63 \| 4 \| 62 \| Phage tail protein \| Phage tail protein; afdb-uniprot50 \| AF-A0A1R4LK82-F1-MODEL\_V4 \| 1.0 \| 4.269e-05 \| 183 \| 0.42 \| 69 \| 40 \| 0 \| 1 \| 69 \| 1 \| 69 \| Phage Tail Protein X \| Phage Tail Protein X; afdb-uniprot50 \| AF-H6SQI2-F1-MODEL\_V4 \| 1.0 \| 3.199e-05 \| 183 \| 0.397 \| 68 \| 38 \| 2 \| 2 \| 69 \| 5 \| 69 \| Uncharacterized protein \| Uncharacterized protein; afdb-uniprot50 \| AF-A0A6P2ZWQ5-F1-MODEL\_V4 \| 1.0 \| 1.346e-05 \| 183 \| 0.352 \| 71 \| 42 \| 2 \| 2 \| 69 \| 1 \| 70 \| Tail X family protein \| Tail X family protein; afdb-uniprot50 \| AF-A0A3N2E0P9-F1-MODEL\_V4 \| 1.0 \| 1.084e-05 \| 183 \| 0.375 \| 72 \| 38 \| 2 \| 2 \| 69 \| 3 \| 71 \| Phage tail protein X \| Phage tail protein X; afdb-uniprot50 \| AF-A0A1R3W3I9-F1-MODEL\_V4 \| 1.0 \| 3.695e-05 \| 182 \| 0.42 \| 69 \| 38 \| 2 \| 2 \| 69 \| 1 \| 68 \| p2-like prophage tail protein X \| p2-like prophage tail protein X; afdb-uniprot50 \| AF-A0A2K2G664-F1-MODEL\_V4 \| 1.0 \| 2.576e-05 \| 182 \| 0.428 \| 70 \| 37 \| 2 \| 2 \| 69 \| 3 \| 71 \| Phage tail protein \| Phage tail protein; afdb-uniprot50 \| AF-A0A6G5QMP5-F1-MODEL\_V4 \| 1.0 \| 3.695e-05 \| 180 \| 0.352 \| 68 \| 43 \| 1 \| 2 \| 69 \| 1 \| 67 \| Phage tail protein X family protein \| Phage tail protein X family protein; afdb-uniprot50 \| AF-A0A5D0CMM3-F1-MODEL\_V4 \| 1.0 \| 0.0001456 \| 180 \| 0.254 \| 59 \| 41 \| 2 \| 2 \| 60 \| 1 \| 56 \| Phage tail protein \| Phage tail protein; afdb-uniprot50 \| AF-A0A7T8QHI0-F1-MODEL\_V4 \| 1.0 \| 3.695e-05 \| 180 \| 0.449 \| 69 \| 37 \| 1 \| 1 \| 69 \| 12 \| 79 \| Tail protein X \| Tail protein X; afdb-uniprot50 \| AF-A0A4Q8MDG8-F1-MODEL\_V4 \| 1.0 \| 3.199e-05 \| 180 \| 0.426 \| 68 \| 37 \| 2 \| 3 \| 69 \| 18 \| 84 \| Phage tail protein \| Phage tail protein; afdb-uniprot50 \| AF-A0A0J5WBN4-F1-MODEL\_V4 \| 1.0 \| 4.932e-05 \| 178 \| 0.405 \| 69 \| 39 \| 2 \| 2 \| 69 \| 1 \| 68 \| Tail protein \| Tail protein; afdb-uniprot50 \| AF-A0A7X4CNS4-F1-MODEL\_V4 \| 1.0 \| 0.0001943 \| 178 \| 0.338 \| 68 \| 45 \| 0 \| 2 \| 69 \| 3 \| 70 \| Phage tail protein \| Phage tail protein; afdb-uniprot50 \| AF-A0A2E2MZM5-F1-MODEL\_V4 \| 1.0 \| 8.785e-05 \| 178 \| 0.309 \| 71 \| 46 \| 2 \| 1 \| 69 \| 1 \| 70 \| Phage tail protein \| Phage tail protein; afdb-uniprot50 \| AF-A0A6C1BPB3-F1-MODEL\_V4 \| 1.0 \| 0.0001015 \| 177 \| 0.373 \| 67 \| 39 \| 2 \| 2 \| 68 \| 1 \| 64 \| Uncharacterized protein \| Uncharacterized protein; afdb-uniprot50 \| AF-A0A2N3KSN4-F1-MODEL\_V4 \| 1.0 \| 0.0001456 \| 177 \| 0.308 \| 68 \| 47 \| 0 \| 2 \| 69 \| 1 \| 68 \| Phage tail protein \| Phage tail protein; afdb-uniprot50 \| AF-A0A6L5PED9-F1-MODEL\_V4 \| 1.0 \| 0.0001354 \| 176 \| 0.424 \| 66 \| 37 \| 1 \| 4 \| 69 \| 1 \| 65 \| Phage tail protein \| Phage tail protein; afdb-uniprot50 \| AF-A0A5C7CK72-F1-MODEL\_V4 \| 1.0 \| 5.697e-05 \| 176 \| 0.391 \| 69 \| 39 \| 3 \| 2 \| 69 \| 1 \| 67 \| Phage tail protein \| Phage tail protein; afdb-uniprot50 \| AF-A0A7X6X0F0-F1-MODEL\_V4 \| 1.0 \| 2.576e-05 \| 176 \| 0.4 \| 70 \| 37 \| 3 \| 2 \| 69 \| 1 \| 67 \| Phage tail protein \| Phage tail protein; afdb-uniprot50 \| AF-A0A1E3G6A1-F1-MODEL\_V4 \| 1.0 \| 0.0001565 \| 176 \| 0.294 \| 68 \| 47 \| 1 \| 2 \| 69 \| 3 \| 69 \| Uncharacterized protein \| Uncharacterized protein; afdb-uniprot50 \| AF-A0A7W6DBB8-F1-MODEL\_V4 \| 1.0 \| 4.588e-05 \| 176 \| 0.485 \| 68 \| 33 \| 2 \| 3 \| 69 \| 17 \| 83 \| Phage tail protein X \| Phage tail protein X; afdb-uniprot50 \| AF-A0A4P9VJG3-F1-MODEL\_V4 \| 1.0 \| 0.0001354 \| 176 \| 0.382 \| 68 \| 39 \| 2 \| 2 \| 69 \| 38 \| 102 \| Uncharacterized protein \| Uncharacterized protein; afdb-uniprot50 \| AF-A0A249A0N5-F1-MODEL\_V4 \| 1.0 \| 2.23e-05 \| 175 \| 0.361 \| 72 \| 41 \| 3 \| 1 \| 69 \| 1 \| 70 \| Phage Tail Protein X \| Phage Tail Protein X; afdb-uniprot50 \| AF-A0A849VAT6-F1-MODEL\_V4 \| 1.0 \| 7.604e-05 \| 175 \| 0.323 \| 71 \| 44 \| 2 \| 1 \| 68 \| 1 \| 70 \| Uncharacterized protein \| Uncharacterized protein; afdb-uniprot50 \| AF-A0A498RQG7-F1-MODEL\_V4 \| 1.0 \| 4.932e-05 \| 174 \| 0.347 \| 69 \| 43 \| 2 \| 2 \| 69 \| 1 \| 68 \| Phage Tail Protein X \| Phage Tail Protein X; afdb-uniprot50 \| AF-A0A0F2RD92-F1-MODEL\_V4 \| 1.0 \| 2.976e-05 \| 174 \| 0.411 \| 68 \| 38 \| 2 \| 2 \| 69 \| 3 \| 68 \| Uncharacterized protein \| Uncharacterized protein; afdb-uniprot50 \| AF-A0A1M4LC88-F1-MODEL\_V4 \| 1.0 \| 8.173e-05 \| 174 \| 0.42 \| 69 \| 38 \| 2 \| 2 \| 69 \| 301 \| 368 \| PhageMin\_Tail domain-containing protein \| PhageMin\_Tail domain-containing protein; afdb-uniprot50 \| AF-A0A4U2PVD7-F1-MODEL\_V4 \| 1.0 \| 0.0002787 \| 173 \| 0.35 \| 57 \| 34 \| 2 \| 1 \| 57 \| 1 \| 54 \| Phage tail protein \| Phage tail protein; afdb-uniprot50 \| AF-A0A495YI39-F1-MODEL\_V4 \| 1.0 \| 7.604e-05 \| 173 \| 0.463 \| 69 \| 35 \| 2 \| 2 \| 69 \| 6 \| 73 \| Phage tail protein \| Phage tail protein; afdb-uniprot50 \| AF-A0A1A9RBR4-F1-MODEL\_V4 \| 1.0 \| 4.932e-05 \| 173 \| 0.385 \| 70 \| 39 \| 2 \| 3 \| 69 \| 10 \| 78 \| Uncharacterized protein \| Uncharacterized protein; afdb-uniprot50 \| AF-A0A854I2Y0-F1-MODEL\_V4 \| 1.0 \| 9.442e-05 \| 173 \| 0.296 \| 64 \| 42 \| 2 \| 2 \| 65 \| 1 \| 61 \| Uncharacterized protein \| Uncharacterized protein; afdb-uniprot50 \| AF-A0A1Y6HDB3-F1-MODEL\_V4 \| 1.0 \| 7.604e-05 \| 171 \| 0.414 \| 70 \| 38 \| 2 \| 2 \| 69 \| 1 \| 69 \| Phage-related tail protein \| Phage-related tail protein; afdb-uniprot50 \| AF-J8W766-F1-MODEL\_V4 \| 1.0 \| 0.0001172 \| 171 \| 0.372 \| 59 \| 32 \| 3 \| 1 \| 57 \| 1 \| 56 \| Phage tail protein X \| Phage tail protein X; afdb-uniprot50 \| AF-A0A261E6J5-F1-MODEL\_V4 \| 1.0 \| 0.0001172 \| 171 \| 0.348 \| 66 \| 41 \| 1 \| 2 \| 65 \| 6 \| 71 \| Uncharacterized protein \| Uncharacterized protein; afdb-uniprot50 \| AF-A0A2S4U2P3-F1-MODEL\_V4 \| 1.0 \| 8.173e-05 \| 170 \| 0.367 \| 68 \| 42 \| 1 \| 2 \| 69 \| 1 \| 67 \| Phage tail protein \| Phage tail protein; afdb-uniprot50 \| AF-A0A095GKT5-F1-MODEL\_V4 \| 1.0 \| 9.442e-05 \| 170 \| 0.318 \| 69 \| 45 \| 2 \| 2 \| 69 \| 1 \| 68 \| Phage tail protein \| Phage tail protein; afdb-uniprot50 \| AF-A0A545T5R5-F1-MODEL\_V4 \| 1.0 \| 0.000126 \| 170 \| 0.317 \| 104 \| 35 \| 2 \| 2 \| 69 \| 3 \| 106 \| Uncharacterized protein \| Uncharacterized protein; afdb-uniprot50 \| AF-A0A2G6EI15-F1-MODEL\_V4 \| 1.0 \| 6.582e-05 \| 169 \| 0.333 \| 69 \| 42 \| 3 \| 2 \| 69 \| 1 \| 66 \| Phage tail protein \| Phage tail protein; afdb-uniprot50 \| AF-A0A165NX17-F1-MODEL\_V4 \| 1.0 \| 0.0002787 \| 169 \| 0.333 \| 69 \| 42 \| 3 \| 2 \| 69 \| 3 \| 68 \| Phage Tail Protein X \| Phage Tail Protein X; afdb-uniprot50 \| AF-A0A2D3VR18-F1-MODEL\_V4 \| 1.0 \| 0.0001015 \| 169 \| 0.376 \| 69 \| 41 \| 2 \| 1 \| 68 \| 1 \| 68 \| Phage tail protein \| Phage tail protein; afdb-uniprot50 \| AF-A0A376WTB8-F1-MODEL\_V4 \| 1.0 \| 0.0002787 \| 169 \| 0.369 \| 65 \| 39 \| 2 \| 2 \| 65 \| 1 \| 64 \| Phage tail protein X \| Phage tail protein X; afdb-uniprot50 \| AF-A0A3A9IJM0-F1-MODEL\_V4 \| 1.0 \| 0.0001682 \| 169 \| 0.306 \| 62 \| 40 \| 2 \| 1 \| 61 \| 1 \| 60 \| Phage tail protein \| Phage tail protein; afdb-uniprot50 \| AF-A0A7X1YEJ4-F1-MODEL\_V4 \| 1.0 \| 0.0004297 \| 169 \| 0.428 \| 63 \| 35 \| 1 \| 2 \| 64 \| 3 \| 64 \| Phage tail protein \| Phage tail protein; afdb-uniprot50 \| AF-A0A6L9JTD6-F1-MODEL\_V4 \| 1.0 \| 0.0002245 \| 168 \| 0.308 \| 68 \| 45 \| 2 \| 2 \| 69 \| 1 \| 66 \| Phage tail protein \| Phage tail protein; afdb-uniprot50 \| AF-A0A2G6DPR2-F1-MODEL\_V4 \| 1.0 \| 4.588e-05 \| 168 \| 0.309 \| 71 \| 44 \| 4 \| 1 \| 69 \| 1 \| 68 \| Phage tail protein \| Phage tail protein; afdb-uniprot50 \| AF-A0A246KGZ2-F1-MODEL\_V4 \| 1.0 \| 6.124e-05 \| 168 \| 0.361 \| 72 \| 42 \| 3 \| 1 \| 69 \| 1 \| 71 \| Phage Tail Protein X \| Phage Tail Protein X; afdb-uniprot50 \| AF-A0A5Q3RUR6-F1-MODEL\_V4 \| 1.0 \| 0.0002413 \| 168 \| 0.357 \| 70 \| 41 \| 2 \| 2 \| 69 \| 5 \| 72 \| Phage tail protein \| Phage tail protein; afdb-uniprot50 \| AF-A0A1Y3SU18-F1-MODEL\_V4 \| 1.0 \| 0.0004619 \| 168 \| 0.41 \| 56 \| 30 \| 2 \| 1 \| 56 \| 1 \| 53 \| Phage tail protein \| Phage tail protein; afdb-uniprot50 \| AF-A0A376J471-F1-MODEL\_V4 \| 1.0 \| 0.0001943 \| 168 \| 0.367 \| 68 \| 42 \| 1 \| 2 \| 69 \| 24 \| 90 \| Tail protein X (GpX) \| Tail protein X (GpX); afdb-uniprot50 \| AF-A0A7X3TZ69-F1-MODEL\_V4 \| 1.0 \| 8.785e-05 \| 168 \| 0.397 \| 68 \| 37 \| 2 \| 4 \| 69 \| 259 \| 324 \| Uncharacterized protein \| Uncharacterized protein; afdb-uniprot50 \| AF-A0A1C3EE73-F1-MODEL\_V4 \| 1.0 \| 0.0001565 \| 167 \| 0.294 \| 68 \| 46 \| 2 \| 2 \| 69 \| 1 \| 66 \| Uncharacterized protein \| Uncharacterized protein; afdb-uniprot50 \| AF-A0A3G2IBA3-F1-MODEL\_V4 \| 1.0 \| 0.0001456 \| 167 \| 0.388 \| 67 \| 39 \| 1 \| 2 \| 68 \| 3 \| 67 \| Phage tail protein \| Phage tail protein; afdb-uniprot50 \| AF-A0A7Z1A3P8-F1-MODEL\_V4 \| 1.0 \| 0.0001456 \| 167 \| 0.289 \| 69 \| 45 \| 2 \| 2 \| 69 \| 3 \| 68 \| Uncharacterized protein \| Uncharacterized protein; afdb-uniprot50 \| AF-A0A1T4QDP8-F1-MODEL\_V4 \| 1.0 \| 0.0002593 \| 167 \| 0.298 \| 67 \| 44 \| 2 \| 2 \| 68 \| 3 \| 66 \| Phage Tail Protein X \| Phage Tail Protein X; afdb-uniprot50 \| AF-B7MRX8-F1-MODEL\_V4 \| 1.0 \| 0.0001943 \| 167 \| 0.318 \| 69 \| 43 \| 4 \| 2 \| 69 \| 24 \| 89 \| Tail protein X from prophage \| Tail protein X from prophage; afdb-uniprot50 \| AF-A0A0S4XN94-F1-MODEL\_V4 \| 1.0 \| 8.785e-05 \| 166 \| 0.357 \| 70 \| 41 \| 3 \| 1 \| 69 \| 1 \| 67 \| Putative Phage Tail Protein X \| Putative Phage Tail Protein X; afdb-uniprot50 \| AF-A0A1K2ALR6-F1-MODEL\_V4 \| 1.0 \| 0.0001943 \| 166 \| 0.411 \| 68 \| 38 \| 2 \| 2 \| 69 \| 3 \| 68 \| p2-like prophage tail protein X \| p2-like prophage tail protein X; afdb-uniprot50 \| AF-A0A7S8EHN3-F1-MODEL\_V4 \| 1.0 \| 0.0001808 \| 166 \| 0.417 \| 67 \| 38 \| 1 \| 3 \| 69 \| 17 \| 82 \| Phage tail protein \| Phage tail protein; afdb-uniprot50 \| AF-A0A1B8QCU9-F1-MODEL\_V4 \| 1.0 \| 0.0001565 \| 165 \| 0.323 \| 68 \| 44 \| 2 \| 2 \| 69 \| 5 \| 70 \| Uncharacterized protein \| Uncharacterized protein; afdb-uniprot50 \| AF-A0A2D5PEH8-F1-MODEL\_V4 \| 1.0 \| 0.0001354 \| 165 \| 0.294 \| 68 \| 48 \| 0 \| 2 \| 69 \| 4 \| 71 \| Phage tail protein \| Phage tail protein; afdb-uniprot50 \| AF-A0A317PI36-F1-MODEL\_V4 \| 1.0 \| 0.0002245 \| 165 \| 0.367 \| 68 \| 43 \| 0 \| 2 \| 69 \| 6 \| 73 \| Phage tail protein X \| Phage tail protein X; afdb-uniprot50 \| AF-A0A2Z4UEW3-F1-MODEL\_V4 \| 1.0 \| 8.173e-05 \| 165 \| 0.361 \| 72 \| 37 \| 3 \| 3 \| 69 \| 13 \| 80 \| Phage tail protein X \| Phage tail protein X; afdb-uniprot50 \| AF-A0A2N1QH04-F1-MODEL\_V4 \| 1.0 \| 0.0002413 \| 164 \| 0.292 \| 65 \| 43 \| 2 \| 1 \| 65 \| 1 \| 62 \| Phage tail protein \| Phage tail protein; afdb-uniprot50 \| AF-A0A3Q8DHW5-F1-MODEL\_V4 \| 1.0 \| 0.0001808 \| 163 \| 0.347 \| 69 \| 42 \| 2 \| 2 \| 68 \| 6 \| 73 \| Uncharacterized protein \| Uncharacterized protein; afdb-uniprot50 \| AF-A0A6L9HTM9-F1-MODEL\_V4 \| 1.0 \| 0.0001015 \| 163 \| 0.297 \| 74 \| 44 \| 3 \| 1 \| 69 \| 1 \| 71 \| Phage tail protein \| Phage tail protein; afdb-uniprot50 \| AF-A0A3A8YFV1-F1-MODEL\_V4 \| 1.0 \| 0.0001943 \| 163 \| 0.384 \| 65 \| 36 \| 3 \| 2 \| 65 \| 4 \| 65 \| Phage tail protein \| Phage tail protein; afdb-uniprot50 \| AF-A0A250DS69-F1-MODEL\_V4 \| 1.0 \| 0.0002245 \| 162 \| 0.362 \| 69 \| 42 \| 2 \| 2 \| 69 \| 3 \| 70 \| Phage tail protein \| Phage tail protein; afdb-uniprot50 \| AF-A0A1S1CG91-F1-MODEL\_V4 \| 1.0 \| 0.0002088 \| 162 \| 0.388 \| 72 \| 39 \| 3 \| 2 \| 69 \| 1 \| 71 \| LysM domain-containing protein \| LysM domain-containing protein; afdb-uniprot50 \| AF-A0A1G8NEF7-F1-MODEL\_V4 \| 1.0 \| 0.0001682 \| 161 \| 0.411 \| 68 \| 38 \| 1 \| 2 \| 69 \| 1 \| 66 \| P2-like prophage tail protein X \| P2-like prophage tail protein X; afdb-uniprot50 \| AF-A0A6L5YES5-F1-MODEL\_V4 \| 1.0 \| 0.0007121 \| 161 \| 0.359 \| 64 \| 40 \| 1 \| 2 \| 65 \| 7 \| 69 \| Phage tail protein \| Phage tail protein; afdb-uniprot50 \| AF-A0A806CIL3-F1-MODEL\_V4 \| 1.0 \| 0.000126 \| 161 \| 0.362 \| 69 \| 42 \| 2 \| 2 \| 69 \| 3 \| 70 \| Tail X family protein \| Tail X family protein; afdb-uniprot50 \| AF-A0A142YH45-F1-MODEL\_V4 \| 1.0 \| 0.0007654 \| 161 \| 0.345 \| 55 \| 34 \| 1 \| 2 \| 56 \| 221 \| 273 \| Phage Tail Protein X \| Phage Tail Protein X; afdb-uniprot50 \| AF-A0A066RM09-F1-MODEL\_V4 \| 1.0 \| 0.0001354 \| 160 \| 0.449 \| 69 \| 37 \| 1 \| 1 \| 69 \| 1 \| 68 \| Tail protein \| Tail protein; afdb-uniprot50 \| AF-A0A4R2MR54-F1-MODEL\_V4 \| 1.0 \| 0.0001943 \| 159 \| 0.371 \| 70 \| 40 \| 3 \| 2 \| 69 \| 1 \| 68 \| Phage tail protein X \| Phage tail protein X; afdb-uniprot50 \| AF-A0A4R3ABI5-F1-MODEL\_V4 \| 1.0 \| 0.000372 \| 159 \| 0.318 \| 69 \| 45 \| 2 \| 2 \| 69 \| 3 \| 70 \| Phage tail protein X \| Phage tail protein X; afdb-uniprot50 \| AF-A0A840FZY4-F1-MODEL\_V4 \| 1.0 \| 0.000322 \| 159 \| 0.42 \| 69 \| 38 \| 2 \| 2 \| 69 \| 3 \| 70 \| Phage tail protein X \| Phage tail protein X; afdb-uniprot50 \| AF-C2BG19-F1-MODEL\_V4 \| 1.0 \| 0.0002593 \| 159 \| 0.294 \| 68 \| 42 \| 3 \| 2 \| 69 \| 3 \| 64 \| Uncharacterized protein \| Uncharacterized protein; afdb-uniprot50 \| AF-A0A2S8YUV6-F1-MODEL\_V4 \| 1.0 \| 0.0002787 \| 158 \| 0.455 \| 68 \| 35 \| 2 \| 2 \| 69 \| 3 \| 68 \| Phage tail protein \| Phage tail protein; afdb-uniprot50 \| AF-A0A5P8MUY6-F1-MODEL\_V4 \| 1.0 \| 0.000322 \| 158 \| 0.4 \| 70 \| 40 \| 2 \| 2 \| 69 \| 6 \| 75 \| Phage tail protein \| Phage tail protein; afdb-uniprot50 \| AF-A0A5M6I273-F1-MODEL\_V4 \| 1.0 \| 0.0001682 \| 157 \| 0.385 \| 70 \| 39 \| 2 \| 1 \| 68 \| 1 \| 68 \| Phage tail protein \| Phage tail protein; afdb-uniprot50 \| AF-A0A6A8A117-F1-MODEL\_V4 \| 1.0 \| 0.0002593 \| 157 \| 0.323 \| 68 \| 43 \| 2 \| 2 \| 69 \| 3 \| 67 \| Phage tail protein \| Phage tail protein; afdb-uniprot50 \| AF-A0A397PIN3-F1-MODEL\_V4 \| 1.0 \| 0.0002088 \| 157 \| 0.42 \| 69 \| 38 \| 2 \| 2 \| 69 \| 3 \| 70 \| Phage tail protein X \| Phage tail protein X; afdb-uniprot50 \| AF-A0A7W9RBM3-F1-MODEL\_V4 \| 1.0 \| 0.0001354 \| 157 \| 0.385 \| 70 \| 40 \| 2 \| 2 \| 69 \| 4 \| 72 \| Phage tail protein X \| Phage tail protein X; afdb-uniprot50 \| AF-A0A554XC87-F1-MODEL\_V4 \| 1.0 \| 0.0006164 \| 156 \| 0.4 \| 65 \| 34 \| 3 \| 1 \| 60 \| 2 \| 66 \| Phage Tail Protein X \| Phage Tail Protein X; afdb-uniprot50 \| AF-A0A1G5MFC1-F1-MODEL\_V4 \| 1.0 \| 0.0002996 \| 156 \| 0.367 \| 68 \| 43 \| 0 \| 2 \| 69 \| 3 \| 70 \| P2-like prophage tail protein X \| P2-like prophage tail protein X; afdb-uniprot50 \| AF-A0A4Q3U2H3-F1-MODEL\_V4 \| 1.0 \| 0.0002413 \| 156 \| 0.318 \| 69 \| 45 \| 2 \| 2 \| 69 \| 9 \| 76 \| Phage tail protein \| Phage tail protein; afdb-uniprot50 \| AF-A0A6B4FAW5-F1-MODEL\_V4 \| 1.0 \| 0.0005735 \| 155 \| 0.275 \| 69 \| 46 \| 3 \| 1 \| 69 \| 1 \| 65 \| LysM domain-containing protein \| LysM domain-containing protein; afdb-uniprot50 \| AF-A0A7H1VPF4-F1-MODEL\_V4 \| 1.0 \| 0.001098 \| 155 \| 0.258 \| 62 \| 43 \| 2 \| 2 \| 63 \| 8 \| 66 \| Tail protein X \| Tail protein X; afdb-uniprot50 \| AF-A0A512IX42-F1-MODEL\_V4 \| 1.0 \| 0.0004297 \| 155 \| 0.363 \| 77 \| 37 \| 2 \| 2 \| 69 \| 4 \| 77 \| Uncharacterized protein \| Uncharacterized protein; afdb-uniprot50 \| AF-V4JIT5-F1-MODEL\_V4 \| 1.0 \| 0.0001456 \| 155 \| 0.333 \| 105 \| 32 \| 4 \| 2 \| 69 \| 3 \| 106 \| Tail protein X \| Tail protein X; afdb-uniprot50 \| AF-A0A315BEJ3-F1-MODEL\_V4 \| 1.0 \| 0.0002996 \| 154 \| 0.352 \| 68 \| 41 \| 3 \| 2 \| 69 \| 10 \| 74 \| Uncharacterized protein \| Uncharacterized protein; afdb-uniprot50 \| AF-A0A5E4XFK5-F1-MODEL\_V4 \| 1.0 \| 0.0006164 \| 154 \| 0.382 \| 68 \| 40 \| 2 \| 2 \| 69 \| 11 \| 76 \| Phage tail protein \| Phage tail protein; afdb-uniprot50 \| AF-A0A4Y6VUF0-F1-MODEL\_V4 \| 1.0 \| 0.0001808 \| 154 \| 0.375 \| 72 \| 40 \| 2 \| 2 \| 69 \| 22 \| 92 \| Phage tail protein \| Phage tail protein; afdb-uniprot50 \| AF-A0A7W0S7Q0-F1-MODEL\_V4 \| 1.0 \| 0.0007654 \| 154 \| 0.333 \| 54 \| 34 \| 1 \| 2 \| 55 \| 137 \| 188 \| Lectin \| Lectin; afdb-uniprot50 \| AF-A0A7Y7V0E2-F1-MODEL\_V4 \| 1.0 \| 0.0003998 \| 153 \| 0.357 \| 70 \| 44 \| 1 \| 1 \| 69 \| 1 \| 70 \| Tail protein X \| Tail protein X; afdb-uniprot50 \| AF-A0A653HUK6-F1-MODEL\_V4 \| 1.0 \| 0.0003461 \| 153 \| 0.333 \| 69 \| 44 \| 2 \| 2 \| 69 \| 5 \| 72 \| Phage Tail Protein X \| Phage Tail Protein X; afdb-uniprot50 \| AF-A0A0T7FB11-F1-MODEL\_V4 \| 1.0 \| 0.000372 \| 153 \| 0.356 \| 73 \| 43 \| 3 \| 1 \| 69 \| 1 \| 73 \| Uncharacterized protein \| Uncharacterized protein; afdb-uniprot50 \| AF-A0A1L9QA14-F1-MODEL\_V4 \| 1.0 \| 0.0004297 \| 153 \| 0.281 \| 71 \| 49 \| 1 \| 1 \| 69 \| 1 \| 71 \| Uncharacterized protein \| Uncharacterized protein; afdb-uniprot50 \| AF-A0A0E4BWV6-F1-MODEL\_V4 \| 1.0 \| 0.0005336 \| 153 \| 0.402 \| 72 \| 39 \| 1 \| 2 \| 69 \| 4 \| 75 \| Uncharacterized protein \| Uncharacterized protein; afdb-uniprot50 \| AF-L0NDQ6-F1-MODEL\_V4 \| 1.0 \| 0.002102 \| 153 \| 0.396 \| 58 \| 34 \| 1 \| 3 \| 60 \| 54 \| 110 \| Phage tail protein X (Modular protein) \| Phage tail protein X (Modular protein); afdb-uniprot50 \| AF-A0A812QV43-F1-MODEL\_V4 \| 1.0 \| 0.0003461 \| 153 \| 0.353 \| 65 \| 40 \| 1 \| 1 \| 65 \| 631 \| 693 \| GpFI protein \| GpFI protein; afdb-uniprot50 \| AF-A0A3A9ERJ5-F1-MODEL\_V4 \| 1.0 \| 0.0006164 \| 152 \| 0.301 \| 63 \| 41 \| 2 \| 1 \| 63 \| 1 \| 60 \| Phage tail protein \| Phage tail protein; afdb-uniprot50 \| AF-A0A833GDU7-F1-MODEL\_V4 \| 1.0 \| 0.0002787 \| 152 \| 0.376 \| 69 \| 41 \| 2 \| 2 \| 69 \| 3 \| 70 \| Phage tail protein \| Phage tail protein; afdb-uniprot50 \| AF-N9PQN2-F1-MODEL\_V4 \| 1.0 \| 0.0001456 \| 151 \| 0.347 \| 69 \| 43 \| 1 \| 1 \| 69 \| 1 \| 67 \| Uncharacterized protein \| Uncharacterized protein; afdb-uniprot50 \| AF-A0A842IYZ7-F1-MODEL\_V4 \| 1.0 \| 0.0007121 \| 151 \| 0.268 \| 67 \| 46 \| 2 \| 2 \| 68 \| 7 \| 70 \| Tail protein X \| Tail protein X; afdb-uniprot50 \| AF-R7HY28-F1-MODEL\_V4 \| 1.0 \| 0.001363 \| 151 \| 0.321 \| 56 \| 35 \| 2 \| 2 \| 57 \| 3 \| 55 \| Phage Tail Protein X \| Phage Tail Protein X; afdb-uniprot50 \| AF-A0A366DKA8-F1-MODEL\_V4 \| 1.0 \| 0.0004619 \| 151 \| 0.397 \| 68 \| 39 \| 2 \| 3 \| 69 \| 14 \| 80 \| Phage tail protein X \| Phage tail protein X; afdb-uniprot50 \| AF-A0A4P5SNE7-F1-MODEL\_V4 \| 1.0 \| 0.0005735 \| 151 \| 0.357 \| 56 \| 29 \| 3 \| 2 \| 55 \| 119 \| 169 \| Uncharacterized protein \| Uncharacterized protein; afdb-uniprot50 \| AF-A0A7W6RH01-F1-MODEL\_V4 \| 1.0 \| 0.0005336 \| 150 \| 0.385 \| 70 \| 39 \| 2 \| 1 \| 68 \| 10 \| 77 \| Phage tail protein X \| Phage tail protein X; afdb-uniprot50 \| AF-M5JVY0-F1-MODEL\_V4 \| 1.0 \| 0.0002413 \| 150 \| 0.347 \| 72 \| 43 \| 2 \| 2 \| 69 \| 8 \| 79 \| p2-like prophage tail protein X \| p2-like prophage tail protein X; afdb-uniprot50 \| AF-A0A0C5VGD3-F1-MODEL\_V4 \| 1.0 \| 0.0006164 \| 150 \| 0.33 \| 103 \| 34 \| 3 \| 2 \| 69 \| 3 \| 105 \| p2-like prophage tail protein X \| p2-like prophage tail protein X; afdb-uniprot50 \| AF-A0A2J9QNL2-F1-MODEL\_V4 \| 1.0 \| 0.0004964 \| 149 \| 0.289 \| 69 \| 46 \| 3 \| 2 \| 69 \| 1 \| 67 \| Phage tail protein \| Phage tail protein; afdb-uniprot50 \| AF-A0A7Y0MMX1-F1-MODEL\_V4 \| 1.0 \| 0.0003998 \| 149 \| 0.314 \| 70 \| 44 \| 3 \| 2 \| 69 \| 1 \| 68 \| Phage tail protein \| Phage tail protein; afdb-uniprot50 \| AF-A0A2N3D6G0-F1-MODEL\_V4 \| 1.0 \| 0.0004619 \| 149 \| 0.342 \| 70 \| 42 \| 3 \| 2 \| 69 \| 4 \| 71 \| Phage tail protein \| Phage tail protein; afdb-uniprot50 \| AF-A0A1V3DCJ7-F1-MODEL\_V4 \| 1.0 \| 0.0004619 \| 149 \| 0.375 \| 72 \| 40 \| 3 \| 2 \| 69 \| 1 \| 71 \| Phage tail protein \| Phage tail protein; afdb-uniprot50 \| AF-A0A806LPR1-F1-MODEL\_V4 \| 1.0 \| 0.001268 \| 149 \| 0.301 \| 63 \| 41 \| 2 \| 1 \| 63 \| 1 \| 60 \| Phage tail protein \| Phage tail protein; afdb-uniprot50 \| AF-A0A1T3BRC0-F1-MODEL\_V4 \| 1.0 \| 0.0002996 \| 148 \| 0.492 \| 67 \| 31 \| 2 \| 5 \| 69 \| 1 \| 66 \| p2-like prophage tail protein X \| p2-like prophage tail protein X; afdb-uniprot50 \| AF-A0A0B6D337-F1-MODEL\_V4 \| 1.0 \| 0.0006626 \| 148 \| 0.281 \| 71 \| 49 \| 2 \| 1 \| 69 \| 1 \| 71 \| Phage Tail Protein X family protein \| Phage Tail Protein X family protein; afdb-uniprot50 \| AF-A0A7L5ZYB4-F1-MODEL\_V4 \| 1.0 \| 0.0005735 \| 148 \| 0.276 \| 65 \| 44 \| 2 \| 1 \| 65 \| 1 \| 62 \| LysM domain-containing protein \| LysM domain-containing protein; afdb-uniprot50 \| AF-A0A537KXH0-F1-MODEL\_V4 \| 1.0 \| 0.0004619 \| 147 \| 0.323 \| 68 \| 45 \| 1 \| 2 \| 69 \| 1 \| 67 \| Phage tail protein \| Phage tail protein; afdb-uniprot50 \| AF-R9B495-F1-MODEL\_V4 \| 1.0 \| 0.0002413 \| 147 \| 0.362 \| 69 \| 42 \| 1 \| 1 \| 69 \| 1 \| 67 \| Uncharacterized protein \| Uncharacterized protein; afdb-uniprot50 \| AF-A0A379KDD4-F1-MODEL\_V4 \| 1.0 \| 0.0006626 \| 147 \| 0.303 \| 66 \| 45 \| 1 \| 2 \| 67 \| 1 \| 65 \| Phage Tail Protein X \| Phage Tail Protein X; afdb-uniprot50 \| AF-A0A1N7LQZ2-F1-MODEL\_V4 \| 1.0 \| 0.0005735 \| 146 \| 0.388 \| 72 \| 40 \| 3 \| 1 \| 69 \| 1 \| 71 \| p2-like prophage tail protein X \| p2-like prophage tail protein X; afdb-uniprot50 \| AF-A0A4R2T1X8-F1-MODEL\_V4 \| 1.0 \| 0.0003998 \| 146 \| 0.328 \| 70 \| 44 \| 2 \| 2 \| 69 \| 3 \| 71 \| Phage tail protein X \| Phage tail protein X; afdb-uniprot50 \| AF-A0A345DE49-F1-MODEL\_V4 \| 1.0 \| 0.0007654 \| 146 \| 0.279 \| 68 \| 47 \| 1 \| 2 \| 69 \| 3 \| 68 \| Uncharacterized protein \| Uncharacterized protein; afdb-uniprot50 \| AF-A0A529HSU4-F1-MODEL\_V4 \| 1.0 \| 0.000372 \| 146 \| 0.405 \| 69 \| 39 \| 2 \| 2 \| 69 \| 16 \| 83 \| Phage tail protein \| Phage tail protein; afdb-uniprot50 \| AF-A0A5U3G616-F1-MODEL\_V4 \| 1.0 \| 0.0005735 \| 145 \| 0.333 \| 66 \| 40 \| 3 \| 1 \| 65 \| 1 \| 63 \| Phage tail protein \| Phage tail protein; afdb-uniprot50 \| AF-A0A7U9H248-F1-MODEL\_V4 \| 1.0 \| 0.001268 \| 145 \| 0.258 \| 62 \| 43 \| 2 \| 2 \| 63 \| 6 \| 64 \| Uncharacterized protein \| Uncharacterized protein; afdb-uniprot50 \| AF-A0A363RF60-F1-MODEL\_V4 \| 1.0 \| 0.0008843 \| 144 \| 0.308 \| 68 \| 45 \| 2 \| 2 \| 69 \| 1 \| 66 \| Phage tail protein \| Phage tail protein; afdb-uniprot50 \| AF-A0A371J2Y1-F1-MODEL\_V4 \| 1.0 \| 0.0007121 \| 144 \| 0.323 \| 65 \| 41 \| 2 \| 1 \| 65 \| 1 \| 62 \| Phage tail protein \| Phage tail protein; afdb-uniprot50 \| AF-A0A0L0WAM0-F1-MODEL\_V4 \| 1.0 \| 0.002102 \| 144 \| 0.213 \| 61 \| 45 \| 2 \| 2 \| 62 \| 8 \| 65 \| Phage tail protein X \| Phage tail protein X; afdb-uniprot50 \| AF-A0A3G2VEY6-F1-MODEL\_V4 \| 1.0 \| 0.001465 \| 144 \| 0.338 \| 71 \| 43 \| 1 \| 3 \| 69 \| 7 \| 77 \| Phage tail protein \| Phage tail protein; afdb-uniprot50 \| AF-A0A2D3WQQ6-F1-MODEL\_V4 \| 1.0 \| 0.0004619 \| 143 \| 0.333 \| 69 \| 42 \| 3 \| 1 \| 68 \| 1 \| 66 \| Phage tail protein \| Phage tail protein; afdb-uniprot50 \| AF-A0A857E7Y2-F1-MODEL\_V4 \| 1.0 \| 0.0008843 \| 143 \| 0.328 \| 70 \| 43 \| 3 \| 2 \| 69 \| 1 \| 68 \| Phage tail protein \| Phage tail protein; afdb-uniprot50 \| AF-A0A7X2H1T7-F1-MODEL\_V4 \| 1.0 \| 0.001268 \| 143 \| 0.292 \| 65 \| 43 \| 2 \| 1 \| 65 \| 1 \| 62 \| Phage tail protein \| Phage tail protein; afdb-uniprot50 \| AF-E2CJS6-F1-MODEL\_V4 \| 1.0 \| 0.001098 \| 143 \| 0.328 \| 70 \| 46 \| 1 \| 1 \| 69 \| 1 \| 70 \| Putative tail protein X \| Putative tail protein X; afdb-uniprot50 \| AF-A0A7U9X278-F1-MODEL\_V4 \| 1.0 \| 0.000322 \| 143 \| 0.328 \| 64 \| 38 \| 3 \| 3 \| 64 \| 5 \| 65 \| Uncharacterized protein \| Uncharacterized protein; afdb-uniprot50 \| AF-E2CN21-F1-MODEL\_V4 \| 1.0 \| 0.0008227 \| 143 \| 0.378 \| 74 \| 40 \| 3 \| 2 \| 69 \| 5 \| 78 \| Phage Tail Protein X \| Phage Tail Protein X; afdb-uniprot50 \| AF-A0A2K3TWI8-F1-MODEL\_V4 \| 1.0 \| 0.0008227 \| 142 \| 0.323 \| 68 \| 45 \| 1 \| 2 \| 69 \| 1 \| 67 \| Phage tail protein \| Phage tail protein; afdb-uniprot50 \| AF-A0A2G6EZ32-F1-MODEL\_V4 \| 1.0 \| 0.0008227 \| 142 \| 0.25 \| 68 \| 46 \| 3 \| 2 \| 69 \| 6 \| 68 \| Phage tail protein \| Phage tail protein; afdb-uniprot50 \| AF-A0A4Q0GTV4-F1-MODEL\_V4 \| 1.0 \| 0.00118 \| 142 \| 0.246 \| 69 \| 50 \| 2 \| 2 \| 69 \| 1 \| 68 \| Phage tail protein \| Phage tail protein; afdb-uniprot50 \| AF-B6JEG3-F1-MODEL\_V4 \| 1.0 \| 0.0003998 \| 142 \| 0.394 \| 71 \| 39 \| 2 \| 2 \| 69 \| 4 \| 73 \| Putative phage tail protein \| Putative phage tail protein; afdb-uniprot50 \| AF-A0A2G2DHZ7-F1-MODEL\_V4 \| 1.0 \| 0.001956 \| 142 \| 0.394 \| 71 \| 41 \| 2 \| 1 \| 69 \| 6 \| 76 \| Phage tail protein \| Phage tail protein; afdb-uniprot50 \| AF-A0A377A283-F1-MODEL\_V4 \| 1.0 \| 0.001022 \| 142 \| 0.308 \| 68 \| 44 \| 3 \| 2 \| 69 \| 166 \| 230 \| Head completion/stabilization protein L \| Head completion/stabilization protein L; afdb-uniprot50 \| AF-A0A7C5AES2-F1-MODEL\_V4 \| 1.0 \| 0.004997 \| 142 \| 0.29 \| 55 \| 39 \| 0 \| 2 \| 56 \| 306 \| 360 \| LysM peptidoglycan-binding domain-containing protein \| LysM peptidoglycan-binding domain-containing protein; afdb-uniprot50 \| AF-A0A7I8DM73-F1-MODEL\_V4 \| 1.0 \| 0.001956 \| 141 \| 0.274 \| 62 \| 42 \| 2 \| 2 \| 63 \| 3 \| 61 \| Uncharacterized protein \| Uncharacterized protein; afdb-uniprot50 \| AF-A0A0P6VPS0-F1-MODEL\_V4 \| 1.0 \| 0.004649 \| 141 \| 0.298 \| 57 \| 39 \| 1 \| 1 \| 56 \| 1 \| 57 \| Uncharacterized protein \| Uncharacterized protein; afdb-uniprot50 \| AF-A0A6B8RNW6-F1-MODEL\_V4 \| 1.0 \| 0.004326 \| 140 \| 0.233 \| 60 \| 43 \| 2 \| 2 \| 61 \| 5 \| 61 \| LysM domain-containing protein \| LysM domain-containing protein; afdb-uniprot50 \| AF-A0A3L7AJJ2-F1-MODEL\_V4 \| 1.0 \| 0.0005735 \| 140 \| 0.371 \| 70 \| 42 \| 2 \| 2 \| 69 \| 4 \| 73 \| Phage tail protein \| Phage tail protein; afdb-uniprot50 \| AF-A0A3N2E0V9-F1-MODEL\_V4 \| 1.0 \| 0.0006626 \| 140 \| 0.287 \| 101 \| 39 \| 3 \| 2 \| 69 \| 3 \| 103 \| Phage tail protein X \| Phage tail protein X; afdb-uniprot50 \| AF-A0A2E4ZYN4-F1-MODEL\_V4 \| 1.0 \| 0.005371 \| 140 \| 0.258 \| 58 \| 43 \| 0 \| 2 \| 59 \| 192 \| 249 \| LysM domain-containing protein \| LysM domain-containing protein; afdb-uniprot50 \| AF-A0A537MKE3-F1-MODEL\_V4 \| 1.0 \| 0.001098 \| 139 \| 0.338 \| 68 \| 40 \| 3 \| 4 \| 69 \| 1 \| 65 \| Phage tail protein \| Phage tail protein; afdb-uniprot50 \| AF-A0A2P1VUX4-F1-MODEL\_V4 \| 1.0 \| 0.001268 \| 139 \| 0.304 \| 69 \| 48 \| 0 \| 1 \| 69 \| 1 \| 69 \| Phage Tail Protein X \| Phage Tail Protein X; afdb-uniprot50 \| AF-A0A1C0V8F9-F1-MODEL\_V4 \| 1.0 \| 0.001956 \| 139 \| 0.338 \| 65 \| 40 \| 2 \| 2 \| 65 \| 4 \| 66 \| Uncharacterized protein \| Uncharacterized protein; afdb-uniprot50 \| AF-A0A4Q5K7X7-F1-MODEL\_V4 \| 1.0 \| 0.001465 \| 139 \| 0.253 \| 67 \| 45 \| 2 \| 1 \| 65 \| 1 \| 64 \| Uncharacterized protein \| Uncharacterized protein; afdb-uniprot50 \| AF-A0A3G2Q4R3-F1-MODEL\_V4 \| 1.0 \| 0.00182 \| 139 \| 0.217 \| 69 \| 52 \| 1 \| 2 \| 68 \| 1 \| 69 \| Uncharacterized protein \| Uncharacterized protein; afdb-uniprot50 \| AF-A0A087N8U9-F1-MODEL\_V4 \| 1.0 \| 0.0004619 \| 139 \| 0.347 \| 72 \| 42 \| 3 \| 2 \| 69 \| 5 \| 75 \| Uncharacterized protein \| Uncharacterized protein; afdb-uniprot50 \| AF-A0A0M4SYX6-F1-MODEL\_V4 \| 1.0 \| 0.001022 \| 138 \| 0.279 \| 68 \| 48 \| 1 \| 2 \| 69 \| 8 \| 74 \| Uncharacterized protein \| Uncharacterized protein; afdb-uniprot50 \| AF-A0A7W7INH6-F1-MODEL\_V4 \| 1.0 \| 0.001465 \| 138 \| 0.277 \| 72 \| 46 \| 3 \| 1 \| 69 \| 7 \| 75 \| Phage tail protein X \| Phage tail protein X; afdb-uniprot50 \| AF-A0A6N2Z4M7-F1-MODEL\_V4 \| 1.0 \| 0.002102 \| 137 \| 0.281 \| 64 \| 43 \| 2 \| 2 \| 65 \| 3 \| 63 \| Uncharacterized protein \| Uncharacterized protein; afdb-uniprot50 \| AF-A0A2W6YQP5-F1-MODEL\_V4 \| 1.0 \| 0.001022 \| 137 \| 0.405 \| 74 \| 37 \| 3 \| 1 \| 69 \| 1 \| 72 \| Phage tail protein \| Phage tail protein; afdb-uniprot50 \| AF-A0A6P1LZV2-F1-MODEL\_V4 \| 1.0 \| 0.0008843 \| 136 \| 0.294 \| 68 \| 46 \| 1 \| 2 \| 69 \| 3 \| 68 \| Uncharacterized protein \| Uncharacterized protein; afdb-uniprot50 \| AF-E1W2C5-F1-MODEL\_V4 \| 1.0 \| 0.001022 \| 136 \| 0.267 \| 71 \| 47 \| 3 \| 2 \| 69 \| 1 \| 69 \| Putative tail synthesis protein \| Putative tail synthesis protein; afdb-uniprot50 \| AF-A0A7V8RB52-F1-MODEL\_V4 \| 1.0 \| 0.0008227 \| 135 \| 0.406 \| 64 \| 35 \| 2 \| 8 \| 69 \| 2 \| 64 \| Phage tail protein \| Phage tail protein; afdb-uniprot50 \| AF-A0A411WHQ6-F1-MODEL\_V4 \| 1.0 \| 0.001022 \| 135 \| 0.283 \| 67 \| 45 \| 2 \| 2 \| 68 \| 6 \| 69 \| Uncharacterized protein \| Uncharacterized protein; afdb-uniprot50 \| AF-A0A8B3RD81-F1-MODEL\_V4 \| 1.0 \| 0.00182 \| 133 \| 0.414 \| 70 \| 36 \| 4 \| 1 \| 68 \| 1 \| 67 \| Phage tail protein \| Phage tail protein; afdb-uniprot50 \| AF-A0A3G8M8D8-F1-MODEL\_V4 \| 1.0 \| 0.00182 \| 133 \| 0.308 \| 68 \| 47 \| 0 \| 2 \| 69 \| 1 \| 68 \| Phage tail protein \| Phage tail protein; afdb-uniprot50 \| AF-A0A318KRC1-F1-MODEL\_V4 \| 1.0 \| 0.00182 \| 133 \| 0.323 \| 71 \| 45 \| 3 \| 1 \| 69 \| 1 \| 70 \| Phage tail protein X \| Phage tail protein X; afdb-uniprot50 \| AF-A0A1G8HXC6-F1-MODEL\_V4 \| 1.0 \| 0.002806 \| 133 \| 0.25 \| 72 \| 51 \| 1 \| 1 \| 69 \| 1 \| 72 \| Phage Tail Protein X \| Phage Tail Protein X; afdb-uniprot50 \| AF-A0A842J0J6-F1-MODEL\_V4 \| 1.0 \| 0.003744 \| 132 \| 0.264 \| 68 \| 47 \| 2 \| 2 \| 69 \| 7 \| 71 \| Tail protein X \| Tail protein X; afdb-uniprot50 \| AF-B3QTI9-F1-MODEL\_V4 \| 1.0 \| 0.001465 \| 132 \| 0.271 \| 81 \| 44 \| 3 \| 2 \| 69 \| 5 \| 83 \| Putative inner membrane protein \| Putative inner membrane protein; afdb-uniprot50 \| AF-A0A064AM34-F1-MODEL\_V4 \| 1.0 \| 0.002259 \| 131 \| 0.23 \| 65 \| 47 \| 2 \| 2 \| 66 \| 10 \| 71 \| Tail protein \| Tail protein; afdb-uniprot50 \| AF-A0A3B0MEG3-F1-MODEL\_V4 \| 1.0 \| 0.001268 \| 131 \| 0.342 \| 73 \| 38 \| 3 \| 2 \| 69 \| 118 \| 185 \| Uncharacterized protein \| Uncharacterized protein; afdb-uniprot50 \| AF-A0A7V8A966-F1-MODEL\_V4 \| 1.0 \| 0.003241 \| 129 \| 0.242 \| 70 \| 49 \| 3 \| 2 \| 69 \| 1 \| 68 \| Uncharacterized protein \| Uncharacterized protein; afdb-uniprot50 \| AF-A0A1I1XSG4-F1-MODEL\_V4 \| 1.0 \| 0.0005336 \| 128 \| 0.394 \| 71 \| 35 \| 4 \| 2 \| 69 \| 8 \| 73 \| P2-like prophage tail protein X \| P2-like prophage tail protein X; afdb-uniprot50 \| AF-A0A845R965-F1-MODEL\_V4 \| 1.0 \| 0.002428 \| 127 \| 0.323 \| 65 \| 39 \| 3 \| 3 \| 67 \| 4 \| 63 \| Uncharacterized protein \| Uncharacterized protein; afdb-uniprot50 \| AF-A0A5M8P5L1-F1-MODEL\_V4 \| 1.0 \| 0.00261 \| 126 \| 0.323 \| 65 \| 38 \| 3 \| 2 \| 62 \| 4 \| 66 \| Uncharacterized protein \| Uncharacterized protein; afdb-uniprot50 \| AF-Q0FZ17-F1-MODEL\_V4 \| 1.0 \| 0.003015 \| 125 \| 0.391 \| 69 \| 38 \| 3 \| 4 \| 69 \| 2 \| 69 \| Uncharacterized protein \| Uncharacterized protein; afdb-uniprot50 \| AF-A0A3L7JDR3-F1-MODEL\_V4 \| 1.0 \| 0.004326 \| 124 \| 0.319 \| 72 \| 46 \| 1 \| 1 \| 69 \| 1 \| 72 \| Uncharacterized protein \| Uncharacterized protein; afdb-uniprot50 \| AF-A0A4Q3Q4D9-F1-MODEL\_V4 \| 1.0 \| 0.0005735 \| 124 \| 0.352 \| 71 \| 38 \| 3 \| 2 \| 69 \| 7 \| 72 \| Phage tail protein \| Phage tail protein; afdb-uniprot50 \| AF-A8SB42-F1-MODEL\_V4 \| 1.0 \| 0.0006626 \| 124 \| 0.376 \| 69 \| 34 \| 4 \| 2 \| 64 \| 10 \| 75 \| Phage Tail Protein X \| Phage Tail Protein X; afdb-uniprot50 \| AF-A0A1Y4BIR9-F1-MODEL\_V4 \| 1.0 \| 0.004326 \| 124 \| 0.281 \| 64 \| 43 \| 2 \| 2 \| 65 \| 11 \| 71 \| Uncharacterized protein \| Uncharacterized protein; afdb-uniprot50 \| AF-A0A7C3CH29-F1-MODEL\_V4 \| 1.0 \| 0.008281 \| 124 \| 0.315 \| 57 \| 38 \| 1 \| 1 \| 56 \| 270 \| 326 \| LysM peptidoglycan-binding domain-containing protein \| LysM peptidoglycan-binding domain-containing protein; afdb-uniprot50 \| AF-M5JRD9-F1-MODEL\_V4 \| 1.0 \| 0.004997 \| 123 \| 0.314 \| 70 \| 47 \| 1 \| 1 \| 69 \| 1 \| 70 \| Tail X family protein \| Tail X family protein; afdb-uniprot50 \| AF-A0A318EKS8-F1-MODEL\_V4 \| 1.0 \| 0.004025 \| 123 \| 0.312 \| 64 \| 41 \| 2 \| 1 \| 64 \| 18 \| 78 \| Uncharacterized protein \| Uncharacterized protein; afdb-uniprot50 \| AF-A0A643EZQ1-F1-MODEL\_V4 \| 1.0 \| 0.004997 \| 122 \| 0.3 \| 70 \| 48 \| 1 \| 1 \| 69 \| 1 \| 70 \| Phage tail protein \| Phage tail protein; afdb-uniprot50 \| AF-A0A6N9P0A8-F1-MODEL\_V4 \| 1.0 \| 0.001956 \| 121 \| 0.303 \| 66 \| 43 \| 2 \| 1 \| 65 \| 1 \| 64 \| Phage tail protein \| Phage tail protein; afdb-uniprot50 \| AF-A0A7C2KV08-F1-MODEL\_V4 \| 1.0 \| 0.008281 \| 120 \| 0.292 \| 65 \| 45 \| 1 \| 2 \| 66 \| 185 \| 248 \| LysM peptidoglycan-binding domain-containing protein \| LysM peptidoglycan-binding domain-containing protein; afdb-uniprot50 \| AF-A0A285MC51-F1-MODEL\_V4 \| 1.0 \| 0.004997 \| 119 \| 0.271 \| 70 \| 47 \| 3 \| 2 \| 69 \| 3 \| 70 \| P2-like prophage tail protein X \| P2-like prophage tail protein X; afdb-uniprot50 \| AF-A0A7Y3K1T7-F1-MODEL\_V4 \| 1.0 \| 0.007169 \| 119 \| 0.271 \| 70 \| 48 \| 2 \| 2 \| 69 \| 149 \| 217 \| LysM peptidoglycan-binding domain-containing protein \| LysM peptidoglycan-binding domain-containing protein; afdb-uniprot50 \| AF-A0A2K2G0B1-F1-MODEL\_V4 \| 1.0 \| 0.008901 \| 118 \| 0.281 \| 71 \| 47 \| 3 \| 2 \| 69 \| 1 \| 70 \| Phage tail protein \| Phage tail protein; afdb-uniprot50 \| AF-A0A258L1B2-F1-MODEL\_V4 \| 1.0 \| 0.003744 \| 116 \| 0.342 \| 70 \| 43 \| 2 \| 2 \| 68 \| 6 \| 75 \| Phage tail protein \| Phage tail protein; afdb-uniprot50 \| AF-A0A329U822-F1-MODEL\_V4 \| 1.0 \| 0.009567 \| 115 \| 0.296 \| 64 \| 40 \| 3 \| 2 \| 65 \| 3 \| 61 \| Phage tail protein \| Phage tail protein; afdb-uniprot50 \| AF-A0A4V2V312-F1-MODEL\_V4 \| 1.0 \| 0.003015 \| 115 \| 0.341 \| 79 \| 41 \| 4 \| 2 \| 69 \| 7 \| 85 \| Phage tail protein X \| Phage tail protein X; afdb-uniprot50 \| AF-A0A7C6DLC9-F1-MODEL\_V4 \| 1.0 \| 0.007169 \| 115 \| 0.234 \| 64 \| 45 \| 1 \| 2 \| 61 \| 203 \| 266 \| LysM peptidoglycan-binding domain-containing protein \| LysM peptidoglycan-binding domain-containing protein; afdb-uniprot50 \| AF-A0A7V4P1T4-F1-MODEL\_V4 \| 1.0 \| 0.007705 \| 115 \| 0.323 \| 68 \| 46 \| 0 \| 2 \| 69 \| 207 \| 274 \| LysM peptidoglycan-binding domain-containing protein \| LysM peptidoglycan-binding domain-containing protein; afdb-uniprot50 \| AF-A0A4R2GR23-F1-MODEL\_V4 \| 1.0 \| 0.005773 \| 113 \| 0.346 \| 75 \| 40 \| 5 \| 2 \| 69 \| 1 \| 73 \| Phage tail protein X \| Phage tail protein X; afdb-uniprot50 \| AF-A0A414D8J5-F1-MODEL\_V4 \| 1.0 \| 0.003241 \| 113 \| 0.257 \| 66 \| 38 \| 3 \| 2 \| 59 \| 8 \| 70 \| LysM domain-containing protein \| LysM domain-containing protein; afdb-uniprot50 \| AF-A0A1V5X535-F1-MODEL\_V4 \| 1.0 \| 0.008901 \| 113 \| 0.296 \| 64 \| 43 \| 1 \| 2 \| 65 \| 113 \| 174 \| LysM domain/BON superfamily protein \| LysM domain/BON superfamily protein; afdb-uniprot50 \| AF-A0A7W6IJI9-F1-MODEL\_V4 \| 1.0 \| 0.002259 \| 112 \| 0.356 \| 73 \| 42 \| 2 \| 1 \| 69 \| 1 \| 72 \| Phage tail protein X \| Phage tail protein X; afdb-uniprot50 \| AF-F4HB38-F1-MODEL\_V4 \| 1.0 \| 0.004025 \| 112 \| 0.271 \| 70 \| 47 \| 3 \| 2 \| 69 \| 8 \| 75 \| Phage Tail Protein X \| Phage Tail Protein X; afdb-uniprot50 \| AF-A0A2W6Z0D8-F1-MODEL\_V4 \| 1.0 \| 0.003241 \| 112 \| 0.369 \| 73 \| 41 \| 4 \| 2 \| 69 \| 4 \| 76 \| Phage tail protein \| Phage tail protein; afdb-uniprot50 \| AF-A0A0T6ZD76-F1-MODEL\_V4 \| 1.0 \| 0.005371 \| 112 \| 0.22 \| 77 \| 48 \| 3 \| 2 \| 69 \| 1 \| 74 \| Uncharacterized protein \| Uncharacterized protein; afdb-uniprot50 \| AF-A0A1C3E6J5-F1-MODEL\_V4 \| 1.0 \| 0.007705 \| 111 \| 0.217 \| 69 \| 51 \| 2 \| 2 \| 69 \| 1 \| 67 \| Uncharacterized protein \| Uncharacterized protein; afdb-uniprot50 \| AF-A0A7C3XER5-F1-MODEL\_V4 \| 1.0 \| 0.008901 \| 111 \| 0.313 \| 67 \| 41 \| 2 \| 2 \| 63 \| 183 \| 249 \| LysM peptidoglycan-binding domain-containing protein \| LysM peptidoglycan-binding domain-containing protein; afdb-uniprot50 \| AF-A0A371WU87-F1-MODEL\_V4 \| 1.0 \| 0.005371 \| 107 \| 0.281 \| 71 \| 48 \| 2 \| 2 \| 69 \| 4 \| 74 \| Phage tail protein \| Phage tail protein; afdb-uniprot50 \| AF-A0A6I1JS86-F1-MODEL\_V4 \| 1.0 \| 0.004997 \| 103 \| 0.324 \| 74 \| 45 \| 3 \| 1 \| 69 \| 1 \| 74 \| Phage tail protein \| Phage tail protein; afdb-uniprot50 \| AF-A0A174T8M1-F1-MODEL\_V4 \| 0.998 \| 0.006205 \| 94 \| 0.217 \| 69 \| 41 \| 4 \| 1 \| 59 \| 1 \| 66 \| LysM domain-containing protein \| LysM domain-containing protein; afdb-uniprot50 \| AF-A0A256CAT5-F1-MODEL\_V4 \| 0.995 \| 0.006669 \| 88 \| 0.295 \| 71 \| 45 \| 3 \| 1 \| 69 \| 1 \| 68 \| Uncharacterized protein \| Uncharacterized protein |
| Top keywords  (threshold 1.00e-02 (evalue)) | **tail, Phage, X, prophage, P2\_like, domain\_containing, LysM, Putative, peptidoglycan\_binding, component** |
| Output files | ../../similar\_structures/24\_FANPEZAQ\_CDS\_0024\_afdb-proteome\_foldseek.tsv ../../similar\_structures/24\_FANPEZAQ\_CDS\_0024\_afdb-uniprot50\_foldseek.tsv ../../similar\_structures/24\_FANPEZAQ\_CDS\_0024\_merged.svg ../../similar\_structures/24\_FANPEZAQ\_CDS\_0024\_pdb\_foldseek.tsv |

  
  
  

Return to summary | Go to previous | Go to next

  


---

**Sequence/structure alignments coloring**  
Each object in the alignment figures is colored according to its E-value following this color coding:

1e-100
10
