## Supplementary material for "Completing the BASEL phage collection to unlock hidden diversity for systematic exploration of phage-host interactions": Data S2: 25.html

FANPEZAQ\_CDS\_0025


Return to summary | Go to previous | Go to next

|  |  |
| --- | --- |
| FANPEZAQ\_CDS\_0025  Page creation date: 02 Sep 2024, 12:00  Project folder: n/a  Input sequences file: Escherichia\_virus\_HeidiAbel.gb | d phage late control type secretion vi system vgr vgrg rhs domain\_containing tip tail element fragment gene ob\_fold gp5 iv phage\_base\_v gpd baseplate putative assembly prophage vgrg\_related in kda tssi phage\_related vgrg1 bacteriophage beta impa muso2 and fold conserved lysm roll hub structural spike escherichia coli regulatory engineered yes organism\_taxid |

### Sequence information

|  |  |
| --- | --- |
| Name | FANPEZAQ\_CDS\_0025  25\_FANPEZAQ\_CDS\_0025 (pipeline id) |
| Imported annotations | Escherichia\_virus\_HeidiAbel Bas97 |
| Protein sequence | MPETGAELELFLGYDNWTQRMGLFICDEVELAGWPGTMTIRARAATYDKSKGGKTDLQTQ KSRSWTKGTKFGDMVKKIASEHGMESAVSESMAKITLPHTDQADESDLNLLLRLAKKYDG VVKPSGGKITVTKRGEGKTASGEELPTITLTAQDCTAFRLTSQKRENAGTVVAYYQMTKK AARNEVKVGEGEPVKRIGKYYPTQAMALDAAKAELAKRQRNQVTVAVTLPGDPSIAAECK LVLVGFRDGIDGEWLITRADHRLDDNGYSCDVEATKPNSEEEPKVEVIERKKPQESDPPD DDGPGHVIEEK |
| Number of residues | 311 |
| Molecular weight (Da) | 34138.16 |
| Output files | ../../query\_sequences/25\_FANPEZAQ\_CDS\_0025.fasta |

### Putative domain architecture and protein family

#### Search results (HHblits)^1^

|  |  |
| --- | --- |
| Domain family databases searched | Pfam, Ncbi-cd, Cath, Phrogs |
| Results, scheme(s)  (Top layers only; threshold 1.00e-03 (evalue)) | xml version="1.0" encoding="utf-8" standalone="no"?       2024-09-02T21:08:17.614933 image/svg+xml   Matplotlib v3.7.2, https://matplotlib.org/ |
| Results, table  (E-value ≤ 1.00e-03 (evalue)) | db \| id \| prob \| evalue \| pvalue \| score \| cols \| query \| query\_len \| template \| template\_len \| name \| description; pfam \| PF05954 \| 99.4 \| 7.3e-18 \| 1.2e-21 \| 135.2 \| 250 \| (3, 266) \| 311 \| (27, 297) \| 298 \| Phage\_GPD \| Phage tail baseplate hub (GPD); pfam \| PF14594 \| 98.9 \| 6e-14 \| 1e-17 \| 114.4 \| 244 \| (20, 275) \| 311 \| (54, 343) \| 343 \| Sipho\_Gp37 \| Siphovirus ReqiPepy6 Gp37-like protein; pfam \| PF06605 \| 98.7 \| 1.9e-12 \| 3.2e-16 \| 101.0 \| 205 \| (62, 278) \| 311 \| (12, 253) \| 263 \| Prophage\_tail \| Prophage endopeptidase tail; pfam \| PF13550 \| 97.9 \| 3.2e-09 \| 5.3e-13 \| 75.0 \| 146 \| (104, 260) \| 311 \| (3, 163) \| 164 \| Phage-tail\_3 \| Putative phage tail protein; pfam \| PF16467 \| 97.8 \| 6.9e-09 \| 1.2e-12 \| 67.6 \| 74 \| (201, 276) \| 311 \| (24, 97) \| 102 \| DUF5048 \| Domain of unknown function (DUF5048); cath \| 1wruA01 \| 98.5 \| 1.3e-11 \| 2.2e-15 \| 102.5 \| 64 \| (218, 281) \| 311 \| (107, 177) \| 177 \| 43 kda tail protein \| CATHCODE: 2.30.300.10 NAME: 43 kda tail protein. Chain: a. Synonym: gpp, gene product 44. Engineered: yes SOURCE: Enterobacteria phage mu. Organism\_taxid: 10677. Gene: 44. Expressed in: escherichia coli. Expression\_system\_taxid: 562. CLASS: Mainly Beta, ARCH: Roll, TOPOL: Phage tail proteins - horseshoe like beta roll fold , HOMOL: Baseplate protein-like domain - beta roll fold; cath \| 3cddA01 \| 98.3 \| 8.7e-11 \| 1.5e-14 \| 97.4 \| 59 \| (225, 283) \| 311 \| (121, 185) \| 187 \| Prophage muso2, 43 kda tail protein \| CATHCODE: 2.30.300.10 NAME: Prophage muso2, 43 kda tail protein. Chain: a, b, c, d, e, f. Engineered: yes SOURCE: Shewanella oneidensis. Organism\_taxid: 211586. Strain: mr-1. Gene: so\_2699. Expressed in: escherichia coli bl21(de3). Expression\_system\_taxid: 469008. CLASS: Mainly Beta, ARCH: Roll, TOPOL: Phage tail proteins - horseshoe like beta roll fold , HOMOL: Baseplate protein-like domain - beta roll fold; cath \| 3d37B01 \| 98.2 \| 2.5e-10 \| 4.3e-14 \| 93.2 \| 61 \| (221, 281) \| 311 \| (122, 188) \| 188 \| Tail protein, 43 kda \| CATHCODE: 2.30.300.10 NAME: Tail protein, 43 kda. Chain: a, b. Engineered: yes SOURCE: Neisseria meningitidis mc58. Organism\_taxid: 122586. Strain: mc58 / serogroup b. Gene: gi:7226348, nmb1110. Expressed in: escherichia coli. CLASS: Mainly Beta, ARCH: Roll, TOPOL: Phage tail proteins - horseshoe like beta roll fold , HOMOL: Baseplate protein-like domain - beta roll fold; cath \| 4uhvA02 \| 97.7 \| 1.2e-08 \| 2.3e-12 \| 69.6 \| 78 \| (58, 136) \| 311 \| (8, 90) \| 92 \| Vgrg1, valine-glycine repeat protein g1 \| CATHCODE: 3.55.50.10 NAME: Vgrg1, valine-glycine repeat protein g1. Chain: a, b. Engineered: yes SOURCE: Pseudomonas aeruginosa. Organism\_taxid: 208964. Strain: pao1. Expressed in: escherichia coli. Expression\_system\_taxid: 469008. CLASS: Alpha Beta, ARCH: 3-Layer(bab) Sandwich, TOPOL: Phage tail protein beta-alpha-beta fold, HOMOL: Baseplate protein-like domains; cath \| 1wruA02 \| 97.6 \| 1.4e-08 \| 2.6e-12 \| 70.6 \| 71 \| (61, 132) \| 311 \| (13, 87) \| 88 \| 43 kda tail protein \| CATHCODE: 3.55.50.10 NAME: 43 kda tail protein. Chain: a. Synonym: gpp, gene product 44. Engineered: yes SOURCE: Enterobacteria phage mu. Organism\_taxid: 10677. Gene: 44. Expressed in: escherichia coli. Expression\_system\_taxid: 562. CLASS: Alpha Beta, ARCH: 3-Layer(bab) Sandwich, TOPOL: Phage tail protein beta-alpha-beta fold, HOMOL: Baseplate protein-like domains; cath \| 3d37A02 \| 97.5 \| 3.5e-08 \| 6.6e-12 \| 66.8 \| 71 \| (61, 132) \| 311 \| (11, 84) \| 85 \| Tail protein, 43 kda \| CATHCODE: 3.55.50.10 NAME: Tail protein, 43 kda. Chain: a, b. Engineered: yes SOURCE: Neisseria meningitidis mc58. Organism\_taxid: 122586. Strain: mc58 / serogroup b. Gene: gi:7226348, nmb1110. Expressed in: escherichia coli. CLASS: Alpha Beta, ARCH: 3-Layer(bab) Sandwich, TOPOL: Phage tail protein beta-alpha-beta fold, HOMOL: Baseplate protein-like domains; cath \| 2p5zX02 \| 97.4 \| 8.1e-08 \| 1.6e-11 \| 65.6 \| 76 \| (59, 135) \| 311 \| (9, 94) \| 97 \| Type vi secretion system component \| CATHCODE: 3.55.50.10 NAME: Type vi secretion system component. Chain: x. Fragment: n-terminal domain. Engineered: yes SOURCE: Escherichia coli o6. Organism\_taxid: 217992. Strain: o6:h1, cft073, upec. Atcc: 700928. Gene: c3393. Expressed in: escherichia coli bl21(de3). Expression\_system\_taxid: 469008. CLASS: Alpha Beta, ARCH: 3-Layer(bab) Sandwich, TOPOL: Phage tail protein beta-alpha-beta fold, HOMOL: Baseplate protein-like domains; phrogs \| 52 \| 100.0 \| 4.8e-40 \| 5.8e-44 \| 290.6 \| 275 \| (1, 283) \| 311 \| (52, 328) \| 333 \| tail protein \| tail protein; Category: tail; KY290949\_p37; phrogs \| 177 \| 99.9 \| 1e-33 \| 1.2e-37 \| 252.2 \| 269 \| (3, 293) \| 311 \| (49, 352) \| 374 \| baseplate hub \| baseplate hub; Category: tail; p166740 VI\_07350; phrogs \| 400 \| 99.9 \| 2.6e-28 \| 3.1e-32 \| 212.7 \| 250 \| (2, 276) \| 311 \| (50, 322) \| 324 \| tail protein \| tail protein; Category: tail; p235698 VI\_00931; phrogs \| 1887 \| 99.5 \| 3.2e-19 \| 3.8e-23 \| 154.6 \| 258 \| (5, 274) \| 311 \| (23, 312) \| 316 \| tail protein \| tail protein; Category: tail; NC\_022766\_p29; phrogs \| 4930 \| 99.5 \| 5.5e-19 \| 6.3e-23 \| 146.7 \| 255 \| (3, 278) \| 311 \| (50, 337) \| 363 \| baseplate hub \| baseplate hub; Category: tail; p87959 VI\_12426; phrogs \| 11191 \| 99.3 \| 2.2e-16 \| 2.5e-20 \| 121.9 \| 159 \| (3, 182) \| 311 \| (52, 221) \| 262 \| NA \| NA; Category: unknown function; p213285 VI\_11403; phrogs \| 196 \| 99.0 \| 2.3e-14 \| 2.8e-18 \| 124.3 \| 214 \| (4, 277) \| 311 \| (102, 331) \| 333 \| baseplate hub \| baseplate hub; Category: tail; NC\_020843\_p99; phrogs \| 1390 \| 98.7 \| 2.4e-12 \| 2.8e-16 \| 106.8 \| 243 \| (4, 277) \| 311 \| (70, 336) \| 338 \| tail protein \| tail protein; Category: tail; p10114 VI\_10453; phrogs \| 4456 \| 98.6 \| 5.4e-12 \| 6e-16 \| 91.5 \| 107 \| (1, 117) \| 311 \| (57, 172) \| 173 \| NA \| NA; Category: unknown function; p151864 VI\_00910; phrogs \| 20686 \| 98.6 \| 1.1e-11 \| 1.3e-15 \| 82.1 \| 80 \| (198, 278) \| 311 \| (12, 92) \| 97 \| NA \| NA; Category: unknown function; p199509 VI\_11453; phrogs \| 12925 \| 98.3 \| 1.1e-10 \| 1.3e-14 \| 77.6 \| 87 \| (192, 278) \| 311 \| (6, 94) \| 102 \| tail protein \| tail protein; Category: tail; p409900 VI\_07763; phrogs \| 350 \| 97.1 \| 4.3e-07 \| 5.2e-11 \| 79.5 \| 80 \| (204, 283) \| 311 \| (324, 409) \| 412 \| tail protein \| tail protein; Category: tail; MF448340\_p356; phrogs \| 9932 \| 97.1 \| 5.2e-07 \| 5.8e-11 \| 63.2 \| 96 \| (191, 286) \| 311 \| (12, 120) \| 148 \| NA \| NA; Category: unknown function; p278086 VI\_09755; phrogs \| 33062 \| 96.4 \| 1.1e-05 \| 1.2e-09 \| 45.1 \| 47 \| (230, 277) \| 311 \| (2, 49) \| 54 \| NA \| NA; Category: unknown function; p376724 VI\_10043; phrogs \| 24827 \| 96.2 \| 2.1e-05 \| 2.4e-09 \| 60.3 \| 72 \| (208, 279) \| 311 \| (204, 286) \| 288 \| tail protein \| tail protein; Category: tail; NC\_019443\_p15; phrogs \| 6616 \| 95.9 \| 5e-05 \| 5.7e-09 \| 68.3 \| 222 \| (18, 265) \| 311 \| (121, 354) \| 873 \| tail protein \| tail protein; Category: tail; p153366 VI\_00729; phrogs \| 21793 \| 95.9 \| 5.3e-05 \| 6e-09 \| 61.8 \| 150 \| (103, 264) \| 311 \| (301, 465) \| 476 \| NA \| NA; Category: unknown function; NC\_021327\_p131; phrogs \| 33423 \| 94.7 \| 0.00052 \| 5.8e-08 \| 51.0 \| 100 \| (4, 117) \| 311 \| (52, 157) \| 265 \| NA \| NA; Category: unknown function; p240160 VI\_01320; phrogs \| 5313 \| 94.6 \| 0.00062 \| 7.3e-08 \| 63.0 \| 209 \| (69, 286) \| 311 \| (571, 805) \| 903 \| NA \| NA; Category: unknown function; p230345 VI\_04092; phrogs \| 13879 \| 94.3 \| 0.00092 \| 1.1e-07 \| 57.2 \| 224 \| (3, 246) \| 311 \| (252, 489) \| 519 \| NA \| NA; Category: unknown function; p308556 VI\_00914; phrogs \| 326 \| 94.2 \| 0.0009 \| 1.1e-07 \| 58.4 \| 187 \| (69, 264) \| 311 \| (107, 317) \| 486 \| tail protein with endopeptidase domain \| tail protein with endopeptidase domain; Category: other; p133814 VI\_05051 |
| Top keywords  (threshold 1.00e-03 (evalue)) | **tail, Beta, Phage, Baseplate, fold, roll, Gene, escherichia, coli, Engineered** |
| Output files | ../../domain\_architecture/25\_FANPEZAQ\_CDS\_0025\_cath.hhr ../../domain\_architecture/25\_FANPEZAQ\_CDS\_0025\_merged.svg ../../domain\_architecture/25\_FANPEZAQ\_CDS\_0025\_ncbi-cd.hhr ../../domain\_architecture/25\_FANPEZAQ\_CDS\_0025\_pfam.hhr ../../domain\_architecture/25\_FANPEZAQ\_CDS\_0025\_phrogs.hhr |

### Identical protein sequences/structures

#### Search results

|  |  |
| --- | --- |
| Protein sequence databases searched | Pdb, Swissprot, Refseq |
| Identical proteins found | -- |
| Top keywords | -- |
| Output files | -- |

### Similar protein sequences/structures

#### Sequence similarity search results (HHblits)^1^

|  |  |
| --- | --- |
| Sequence databases searched | Uniclust, Pdb70 |
| Results, scheme(s)  (Top layers only, threshold 1.00e-03 (evalue)) | xml version="1.0" encoding="utf-8" standalone="no"?       2024-09-02T21:08:40.558087 image/svg+xml   Matplotlib v3.7.2, https://matplotlib.org/ |
| Results, table(s)  (threshold 1.00e-03 (evalue)) | db \| id \| prob \| evalue \| pvalue \| score \| cols \| query \| query\_len \| template \| template\_len \| name \| description; uniclust \| UniRef100\_A0A0S8GYZ7 \| 100.0 \| 4.5e-41 \| 8.2e-47 \| 294.5 \| 290 \| (4, 307) \| 311 \| (143, 471) \| 781 \| Type VI secretion protein ImpA (Fragment) \| Type VI secretion protein ImpA (Fragment); uniclust \| UniRef100\_A0A166E304 \| 100.0 \| 9.2e-41 \| 1.7e-46 \| 278.7 \| 287 \| (4, 305) \| 311 \| (62, 380) \| 520 \| Phage-related baseplate assembly protein \| Phage-related baseplate assembly protein; uniclust \| UniRef100\_A0A062V571 \| 100.0 \| 2.5e-40 \| 4.6e-46 \| 280.6 \| 268 \| (2, 285) \| 311 \| (84, 378) \| 510 \| Phage protein D \| Phage protein D; uniclust \| UniRef100\_A0A074LMF2 \| 100.0 \| 6e-40 \| 1.1e-45 \| 277.6 \| 265 \| (3, 283) \| 311 \| (99, 389) \| 421 \| Phage late control D family protein \| Phage late control D family protein; uniclust \| UniRef100\_A0A011P577 \| 100.0 \| 8e-40 \| 1.5e-45 \| 279.3 \| 273 \| (1, 280) \| 311 \| (89, 368) \| 445 \| Tail protein \| Tail protein; uniclust \| UniRef100\_A0A017T2T3 \| 100.0 \| 1.1e-39 \| 2e-45 \| 297.7 \| 289 \| (4, 306) \| 311 \| (93, 422) \| 905 \| VgrG protein \| VgrG protein; uniclust \| UniRef100\_A0A021XHC5 \| 100.0 \| 1.5e-39 \| 2.8e-45 \| 299.4 \| 289 \| (4, 306) \| 311 \| (168, 495) \| 1005 \| Gp5/Type VI secretion system Vgr protein OB-fold domain-containing protein \| Gp5/Type VI secretion system Vgr protein OB-fold domain-containing protein; uniclust \| UniRef100\_A0A022FQY4 \| 100.0 \| 3.1e-39 \| 5.6e-45 \| 298.6 \| 290 \| (4, 306) \| 311 \| (158, 483) \| 1025 \| ImpA family type VI secretion-associated protein \| ImpA family type VI secretion-associated protein; uniclust \| UniRef100\_A0A1I4TS69 \| 100.0 \| 7.1e-39 \| 1.3e-44 \| 282.9 \| 287 \| (4, 304) \| 311 \| (70, 392) \| 713 \| Type VI secretion system secreted protein VgrG \| Type VI secretion system secreted protein VgrG; uniclust \| UniRef100\_A0A1W0CDX9 \| 100.0 \| 1.9e-38 \| 3.5e-44 \| 279.7 \| 288 \| (4, 305) \| 311 \| (65, 395) \| 944 \| Gp5/Type VI secretion system Vgr protein OB-fold domain-containing protein \| Gp5/Type VI secretion system Vgr protein OB-fold domain-containing protein; uniclust \| UniRef100\_A0A010RV36 \| 100.0 \| 3.8e-38 \| 6.9e-44 \| 286.3 \| 288 \| (4, 305) \| 311 \| (182, 508) \| 910 \| Type IV secretion protein Rhs \| Type IV secretion protein Rhs; uniclust \| UniRef100\_A0A011MPL4 \| 100.0 \| 7.8e-38 \| 1.4e-43 \| 277.8 \| 288 \| (4, 305) \| 311 \| (94, 421) \| 797 \| Type VI secretion system Vgr family protein \| Type VI secretion system Vgr family protein; uniclust \| UniRef100\_A0A071LQ37 \| 100.0 \| 7.8e-38 \| 1.4e-43 \| 285.2 \| 289 \| (3, 305) \| 311 \| (134, 464) \| 1175 \| Type IV secretion protein Rhs \| Type IV secretion protein Rhs; uniclust \| UniRef100\_A0A017HSZ5 \| 100.0 \| 1.3e-37 \| 2.5e-43 \| 280.2 \| 289 \| (4, 306) \| 311 \| (192, 520) \| 1037 \| VgrG protein \| VgrG protein; uniclust \| UniRef100\_A0A1H0PUY2 \| 100.0 \| 2.2e-37 \| 3.9e-43 \| 266.0 \| 288 \| (3, 305) \| 311 \| (59, 397) \| 619 \| Rhs element Vgr protein \| Rhs element Vgr protein; uniclust \| UniRef100\_A0A017SWS8 \| 100.0 \| 2.3e-37 \| 4.2e-43 \| 288.6 \| 289 \| (4, 307) \| 311 \| (139, 471) \| 957 \| VgrG protein \| VgrG protein; uniclust \| UniRef100\_A0A0S8A627 \| 100.0 \| 3.3e-37 \| 6.1e-43 \| 252.4 \| 291 \| (3, 307) \| 311 \| (56, 387) \| 431 \| Type VI secretion protein ImpA (Fragment) \| Type VI secretion protein ImpA (Fragment); uniclust \| UniRef100\_A0A0K1E835 \| 100.0 \| 3.4e-37 \| 6.3e-43 \| 275.4 \| 284 \| (4, 303) \| 311 \| (199, 522) \| 1030 \| Gp5/Type VI secretion system Vgr protein OB-fold domain-containing protein \| Gp5/Type VI secretion system Vgr protein OB-fold domain-containing protein; uniclust \| UniRef100\_A0A011NCB8 \| 100.0 \| 7.6e-37 \| 1.4e-42 \| 261.9 \| 289 \| (1, 304) \| 311 \| (119, 422) \| 496 \| Regulatory protein \| Regulatory protein; uniclust \| UniRef100\_A0A075K3F1 \| 100.0 \| 8.3e-37 \| 1.5e-42 \| 273.1 \| 291 \| (4, 309) \| 311 \| (183, 520) \| 918 \| Type IV secretion protein Rhs \| Type IV secretion protein Rhs; uniclust \| UniRef100\_A0A081P6T1 \| 100.0 \| 8.5e-37 \| 1.6e-42 \| 243.3 \| 260 \| (3, 278) \| 311 \| (91, 372) \| 374 \| Phage protein D \| Phage protein D; uniclust \| UniRef100\_A0A1E7J776 \| 100.0 \| 1.3e-36 \| 2.3e-42 \| 248.7 \| 268 \| (3, 282) \| 311 \| (80, 383) \| 395 \| Phage late control D family protein \| Phage late control D family protein; uniclust \| UniRef100\_A0A014PYH8 \| 100.0 \| 1.3e-36 \| 2.4e-42 \| 263.6 \| 277 \| (1, 284) \| 311 \| (169, 469) \| 561 \| Late control protein D \| Late control protein D; uniclust \| UniRef100\_A0A2E5HMJ2 \| 100.0 \| 1.5e-36 \| 2.7e-42 \| 248.8 \| 264 \| (2, 280) \| 311 \| (64, 366) \| 374 \| Phage late control D family protein \| Phage late control D family protein; uniclust \| UniRef100\_A0A023XMA1 \| 100.0 \| 1.5e-36 \| 2.8e-42 \| 271.9 \| 286 \| (3, 303) \| 311 \| (141, 470) \| 845 \| Type VI secretion protein, VrgS/VgrG family \| Type VI secretion protein, VrgS/VgrG family; uniclust \| UniRef100\_A0A2U2N2P6 \| 100.0 \| 1.7e-36 \| 3e-42 \| 265.2 \| 295 \| (4, 310) \| 311 \| (113, 446) \| 873 \| Type VI secretion system tip protein VgrG \| Type VI secretion system tip protein VgrG; uniclust \| UniRef100\_A0A0E1SGK6 \| 100.0 \| 1.8e-36 \| 3.3e-42 \| 259.8 \| 288 \| (4, 305) \| 311 \| (70, 399) \| 741 \| Rhs element Vgr protein \| Rhs element Vgr protein; uniclust \| UniRef100\_A0A0N1N1R4 \| 100.0 \| 4.6e-36 \| 8.4e-42 \| 246.0 \| 275 \| (1, 282) \| 311 \| (67, 349) \| 394 \| Late control protein \| Late control protein; uniclust \| UniRef100\_A0A017T4K1 \| 100.0 \| 4.7e-36 \| 8.5e-42 \| 274.3 \| 286 \| (4, 304) \| 311 \| (197, 518) \| 895 \| VgrG protein \| VgrG protein; uniclust \| UniRef100\_A0A016XIQ1 \| 100.0 \| 4.7e-36 \| 8.6e-42 \| 250.5 \| 273 \| (1, 280) \| 311 \| (92, 367) \| 458 \| Phage late control protein \| Phage late control protein; uniclust \| UniRef100\_A0A060H9H7 \| 100.0 \| 1e-35 \| 1.9e-41 \| 234.5 \| 267 \| (11, 284) \| 311 \| (2, 270) \| 292 \| Phage tail protein \| Phage tail protein; uniclust \| UniRef100\_A0A0F7PH30 \| 100.0 \| 1.5e-35 \| 2.8e-41 \| 242.6 \| 289 \| (4, 306) \| 311 \| (59, 389) \| 485 \| Rhs element Vgr protein \| Rhs element Vgr protein; uniclust \| UniRef100\_A0A0S8CC80 \| 100.0 \| 2.7e-35 \| 4.9e-41 \| 251.9 \| 261 \| (2, 278) \| 311 \| (192, 482) \| 728 \| Gp5/Type VI secretion system Vgr protein OB-fold domain-containing protein \| Gp5/Type VI secretion system Vgr protein OB-fold domain-containing protein; uniclust \| UniRef100\_A0A0P8A270 \| 100.0 \| 4e-35 \| 7.4e-41 \| 236.6 \| 260 \| (3, 278) \| 311 \| (65, 349) \| 448 \| Phage late controlD protein (GPD) \| Phage late controlD protein (GPD); uniclust \| UniRef100\_A0A0C5VFD9 \| 100.0 \| 5.2e-35 \| 9.6e-41 \| 232.9 \| 268 \| (1, 279) \| 311 \| (69, 342) \| 350 \| Phage protein D \| Phage protein D; uniclust \| UniRef100\_A0A016QM46 \| 100.0 \| 7.6e-35 \| 1.4e-40 \| 265.6 \| 265 \| (3, 278) \| 311 \| (240, 532) \| 899 \| Rhs element Vgr protein \| Rhs element Vgr protein; uniclust \| UniRef100\_A0A011PNN2 \| 100.0 \| 9.9e-35 \| 1.8e-40 \| 242.0 \| 263 \| (3, 281) \| 311 \| (88, 377) \| 383 \| Phage protein D \| Phage protein D; uniclust \| UniRef100\_A0A010SW75 \| 100.0 \| 1.5e-34 \| 2.7e-40 \| 263.0 \| 285 \| (4, 305) \| 311 \| (147, 477) \| 1128 \| Type VI secretion protein ImpA \| Type VI secretion protein ImpA; uniclust \| UniRef100\_A0A162LXF1 \| 100.0 \| 2.2e-34 \| 4e-40 \| 250.7 \| 287 \| (4, 305) \| 311 \| (70, 395) \| 919 \| Gp5/Type VI secretion system Vgr protein OB-fold domain-containing protein \| Gp5/Type VI secretion system Vgr protein OB-fold domain-containing protein; uniclust \| UniRef100\_A0A068QS18 \| 100.0 \| 2.7e-34 \| 5e-40 \| 252.1 \| 287 \| (4, 305) \| 311 \| (76, 405) \| 832 \| SURP motif domain-containing protein \| SURP motif domain-containing protein; uniclust \| UniRef100\_A0A2E2NRL3 \| 100.0 \| 2.8e-34 \| 5.1e-40 \| 229.9 \| 264 \| (3, 283) \| 311 \| (63, 349) \| 370 \| Phage late control D family protein \| Phage late control D family protein; uniclust \| UniRef100\_A0A0A2WGR2 \| 100.0 \| 3.4e-34 \| 6.3e-40 \| 241.3 \| 265 \| (3, 281) \| 311 \| (99, 402) \| 415 \| Phage protein D \| Phage protein D; uniclust \| UniRef100\_A0A061Q576 \| 100.0 \| 3.8e-34 \| 6.9e-40 \| 252.9 \| 291 \| (4, 308) \| 311 \| (72, 399) \| 848 \| VgrG protein \| VgrG protein; uniclust \| UniRef100\_A0A0T5NZY3 \| 100.0 \| 3.8e-34 \| 7e-40 \| 246.9 \| 289 \| (2, 308) \| 311 \| (75, 386) \| 591 \| Gp5/Type VI secretion system Vgr protein OB-fold domain-containing protein \| Gp5/Type VI secretion system Vgr protein OB-fold domain-containing protein; uniclust \| UniRef100\_A0A0C1ITH4 \| 100.0 \| 5.4e-34 \| 9.9e-40 \| 231.2 \| 263 \| (2, 279) \| 311 \| (103, 376) \| 414 \| Phage protein D \| Phage protein D; uniclust \| UniRef100\_A0A022PIB0 \| 100.0 \| 8.1e-34 \| 1.5e-39 \| 233.1 \| 275 \| (1, 282) \| 311 \| (69, 384) \| 479 \| Phage protein D \| Phage protein D; uniclust \| UniRef100\_A0A0A8H9K0 \| 100.0 \| 9.8e-34 \| 1.8e-39 \| 231.7 \| 265 \| (2, 277) \| 311 \| (107, 376) \| 381 \| Phage protein D \| Phage protein D; uniclust \| UniRef100\_A0A023PRE5 \| 100.0 \| 1.2e-33 \| 2.1e-39 \| 258.5 \| 288 \| (6, 309) \| 311 \| (104, 432) \| 1093 \| Gp5/Type VI secretion system Vgr protein OB-fold domain-containing protein \| Gp5/Type VI secretion system Vgr protein OB-fold domain-containing protein; uniclust \| UniRef100\_A0A084A131 \| 100.0 \| 1.2e-33 \| 2.1e-39 \| 244.4 \| 288 \| (4, 304) \| 311 \| (62, 390) \| 809 \| Baseplate hub subunit and tail lysozyme \| Baseplate hub subunit and tail lysozyme; uniclust \| UniRef100\_A0A080M5H4 \| 100.0 \| 1.3e-33 \| 2.4e-39 \| 217.1 \| 245 \| (20, 279) \| 311 \| (4, 272) \| 275 \| Tail protein \| Tail protein; uniclust \| UniRef100\_A0A2S5R6X2 \| 100.0 \| 1.4e-33 \| 2.5e-39 \| 216.6 \| 250 \| (22, 278) \| 311 \| (2, 251) \| 260 \| Phage late control D protein (GPD) \| Phage late control D protein (GPD); uniclust \| UniRef100\_A0A1W9R245 \| 100.0 \| 1.6e-33 \| 2.9e-39 \| 235.0 \| 286 \| (3, 306) \| 311 \| (72, 380) \| 431 \| Type IV secretion protein Rhs (Fragment) \| Type IV secretion protein Rhs (Fragment); uniclust \| UniRef100\_A0A1F5AGG1 \| 100.0 \| 1.7e-33 \| 3.2e-39 \| 255.3 \| 270 \| (6, 289) \| 311 \| (72, 381) \| 834 \| Gp5/Type VI secretion system Vgr protein OB-fold domain-containing protein \| Gp5/Type VI secretion system Vgr protein OB-fold domain-containing protein; uniclust \| UniRef100\_A0A178ILA2 \| 100.0 \| 1.8e-33 \| 3.2e-39 \| 244.5 \| 288 \| (4, 305) \| 311 \| (58, 392) \| 883 \| Gp5/Type VI secretion system Vgr protein OB-fold domain-containing protein \| Gp5/Type VI secretion system Vgr protein OB-fold domain-containing protein; uniclust \| UniRef100\_A0A063B487 \| 100.0 \| 1.9e-33 \| 3.4e-39 \| 217.7 \| 266 \| (20, 295) \| 311 \| (3, 274) \| 300 \| Phage late control D family protein \| Phage late control D family protein; uniclust \| UniRef100\_A0A0F9ANU3 \| 100.0 \| 2.2e-33 \| 4e-39 \| 241.6 \| 285 \| (5, 306) \| 311 \| (131, 456) \| 708 \| Gp5/Type VI secretion system Vgr protein OB-fold domain-containing protein (Fragment) \| Gp5/Type VI secretion system Vgr protein OB-fold domain-containing protein (Fragment); uniclust \| UniRef100\_A0A2T5PAE8 \| 100.0 \| 2.2e-33 \| 4e-39 \| 232.7 \| 287 \| (4, 304) \| 311 \| (108, 434) \| 606 \| Type VI secretion system tip protein VgrG (Fragment) \| Type VI secretion system tip protein VgrG (Fragment); uniclust \| UniRef100\_A0A0B5KGS8 \| 100.0 \| 2.6e-33 \| 4.8e-39 \| 244.0 \| 291 \| (4, 307) \| 311 \| (259, 591) \| 848 \| ImpA family type VI secretion-associated protein \| ImpA family type VI secretion-associated protein; uniclust \| UniRef100\_A0A022FWW3 \| 100.0 \| 3.7e-33 \| 6.8e-39 \| 260.4 \| 288 \| (4, 305) \| 311 \| (143, 472) \| 1100 \| Type IV secretion protein Rhs \| Type IV secretion protein Rhs; uniclust \| UniRef100\_A0A257YB17 \| 100.0 \| 4.6e-33 \| 8.4e-39 \| 226.9 \| 290 \| (4, 307) \| 311 \| (65, 392) \| 401 \| Gp5/Type VI secretion system Vgr protein OB-fold domain-containing protein (Fragment) \| Gp5/Type VI secretion system Vgr protein OB-fold domain-containing protein (Fragment); uniclust \| UniRef100\_A0A0G3Q9L4 \| 100.0 \| 5.1e-33 \| 9.4e-39 \| 243.2 \| 291 \| (4, 305) \| 311 \| (65, 395) \| 915 \| ImpA family type VI secretion-associated protein \| ImpA family type VI secretion-associated protein; uniclust \| UniRef100\_A0A014LZB8 \| 100.0 \| 5.6e-33 \| 1e-38 \| 248.0 \| 289 \| (2, 308) \| 311 \| (88, 399) \| 640 \| Type VI secretion protein VgrG \| Type VI secretion protein VgrG; uniclust \| UniRef100\_A0A2M6UWZ5 \| 100.0 \| 9.9e-33 \| 1.8e-38 \| 217.1 \| 268 \| (1, 283) \| 311 \| (56, 329) \| 373 \| Late control protein (Fragment) \| Late control protein (Fragment); uniclust \| UniRef100\_A0A1H8RSK6 \| 100.0 \| 1.4e-32 \| 2.6e-38 \| 215.2 \| 267 \| (1, 278) \| 311 \| (53, 320) \| 325 \| Phage protein D \| Phage protein D; uniclust \| UniRef100\_A0A093RA64 \| 100.0 \| 1.5e-32 \| 2.8e-38 \| 225.8 \| 285 \| (7, 308) \| 311 \| (144, 469) \| 516 \| Type IV secretion protein Rhs (Fragment) \| Type IV secretion protein Rhs (Fragment); uniclust \| UniRef100\_A0A1H2FLK2 \| 100.0 \| 1.9e-32 \| 3.5e-38 \| 214.3 \| 266 \| (1, 282) \| 311 \| (58, 328) \| 335 \| Phage protein D \| Phage protein D; uniclust \| UniRef100\_A0A0A7JME1 \| 100.0 \| 1.9e-32 \| 3.5e-38 \| 212.9 \| 270 \| (1, 280) \| 311 \| (51, 327) \| 354 \| Phage late control protein \| Phage late control protein; uniclust \| UniRef100\_A0A066RIQ8 \| 100.0 \| 2.3e-32 \| 4.3e-38 \| 237.6 \| 290 \| (3, 307) \| 311 \| (57, 380) \| 666 \| Type IV secretion protein Rhs \| Type IV secretion protein Rhs; uniclust \| UniRef100\_A0A2V0QT28 \| 100.0 \| 2.6e-32 \| 4.8e-38 \| 227.5 \| 291 \| (4, 307) \| 311 \| (83, 415) \| 528 \| Uncharacterized conserved protein \| Uncharacterized conserved protein; uniclust \| UniRef100\_A0A0C2D2R7 \| 99.9 \| 3e-32 \| 5.5e-38 \| 236.8 \| 291 \| (4, 308) \| 311 \| (53, 384) \| 900 \| Putative VGR-RELATED protein \| Putative VGR-RELATED protein; uniclust \| UniRef100\_A0A1N6KVX8 \| 99.9 \| 3.5e-32 \| 6.4e-38 \| 246.6 \| 286 \| (4, 305) \| 311 \| (69, 421) \| 1049 \| Rhs element Vgr protein \| Rhs element Vgr protein; uniclust \| UniRef100\_A0A084A0B4 \| 99.9 \| 4e-32 \| 7.4e-38 \| 235.0 \| 285 \| (3, 301) \| 311 \| (53, 380) \| 880 \| Type VI secretion system Vgr family protein \| Type VI secretion system Vgr family protein; uniclust \| UniRef100\_A0A162L1Y1 \| 99.9 \| 6e-32 \| 1.1e-37 \| 235.8 \| 288 \| (3, 306) \| 311 \| (73, 399) \| 702 \| Gp5/Type VI secretion system Vgr protein OB-fold domain-containing protein \| Gp5/Type VI secretion system Vgr protein OB-fold domain-containing protein; uniclust \| UniRef100\_A0A3B9Q8K7 \| 99.9 \| 7.8e-32 \| 1.4e-37 \| 212.8 \| 268 \| (2, 280) \| 311 \| (64, 377) \| 378 \| Late control D family protein \| Late control D family protein; uniclust \| UniRef100\_A0A023D7D4 \| 99.9 \| 8.8e-32 \| 1.6e-37 \| 233.3 \| 287 \| (4, 305) \| 311 \| (79, 402) \| 675 \| Secretion system Type VI Rhs element Vgr \| Secretion system Type VI Rhs element Vgr; uniclust \| UniRef100\_A0A017T2T0 \| 99.9 \| 1e-31 \| 1.9e-37 \| 238.7 \| 273 \| (4, 288) \| 311 \| (127, 436) \| 662 \| VgrG protein \| VgrG protein; uniclust \| UniRef100\_A0A0C1GDZ6 \| 99.9 \| 1.4e-31 \| 2.6e-37 \| 235.9 \| 288 \| (4, 304) \| 311 \| (64, 392) \| 871 \| Type IV secretion protein Rhs \| Type IV secretion protein Rhs; uniclust \| UniRef100\_A0A072N5Y6 \| 99.9 \| 1.5e-31 \| 2.7e-37 \| 233.6 \| 285 \| (4, 305) \| 311 \| (94, 422) \| 761 \| VgrG protein \| VgrG protein; uniclust \| UniRef100\_A0A0N1LTT8 \| 99.9 \| 1.5e-31 \| 2.7e-37 \| 219.2 \| 274 \| (1, 281) \| 311 \| (64, 345) \| 373 \| Late control protein \| Late control protein; uniclust \| UniRef100\_A0A074VFM2 \| 99.9 \| 1.6e-31 \| 3e-37 \| 208.0 \| 272 \| (25, 310) \| 311 \| (3, 291) \| 300 \| Phage protein D (Fragment) \| Phage protein D (Fragment); uniclust \| UniRef100\_A0A4U1GXL6 \| 99.9 \| 1.7e-31 \| 3e-37 \| 228.6 \| 286 \| (5, 305) \| 311 \| (83, 405) \| 858 \| Type VI secretion system tip protein VgrG \| Type VI secretion system tip protein VgrG; uniclust \| UniRef100\_A0A2D3T999 \| 99.9 \| 1.7e-31 \| 3.2e-37 \| 208.3 \| 254 \| (21, 281) \| 311 \| (3, 258) \| 292 \| Phage tail protein \| Phage tail protein; uniclust \| UniRef100\_A0A099EW46 \| 99.9 \| 1.8e-31 \| 3.3e-37 \| 233.0 \| 283 \| (4, 301) \| 311 \| (75, 399) \| 762 \| Type IV secretion protein Rhs \| Type IV secretion protein Rhs; uniclust \| UniRef100\_A0A1E4HSI3 \| 99.9 \| 2.2e-31 \| 3.9e-37 \| 227.3 \| 289 \| (4, 305) \| 311 \| (66, 393) \| 642 \| Gp5/Type VI secretion system Vgr protein OB-fold domain-containing protein \| Gp5/Type VI secretion system Vgr protein OB-fold domain-containing protein; uniclust \| UniRef100\_A0A175QC54 \| 99.9 \| 2.4e-31 \| 4.5e-37 \| 213.6 \| 291 \| (4, 305) \| 311 \| (88, 418) \| 470 \| Type IV secretion protein Rhs (Fragment) \| Type IV secretion protein Rhs (Fragment); uniclust \| UniRef100\_A0A0X8EV94 \| 99.9 \| 3.4e-31 \| 6.1e-37 \| 230.3 \| 292 \| (4, 309) \| 311 \| (62, 393) \| 708 \| Gp5/Type VI secretion system Vgr protein OB-fold domain-containing protein \| Gp5/Type VI secretion system Vgr protein OB-fold domain-containing protein; uniclust \| UniRef100\_A0A017SVJ8 \| 99.9 \| 3.6e-31 \| 6.6e-37 \| 240.2 \| 288 \| (4, 305) \| 311 \| (137, 478) \| 967 \| VgrG protein \| VgrG protein; uniclust \| UniRef100\_A0A087NA94 \| 99.9 \| 3.6e-31 \| 6.7e-37 \| 210.9 \| 272 \| (1, 279) \| 311 \| (81, 382) \| 398 \| Uncharacterized protein \| Uncharacterized protein; uniclust \| UniRef100\_A0A1V6ENZ6 \| 99.9 \| 4.1e-31 \| 7.5e-37 \| 215.4 \| 244 \| (4, 261) \| 311 \| (84, 353) \| 383 \| Phage-related baseplate assembly protein \| Phage-related baseplate assembly protein; uniclust \| UniRef100\_A0A6N2ELR8 \| 99.9 \| 4.8e-31 \| 8.7e-37 \| 225.4 \| 279 \| (17, 308) \| 311 \| (82, 397) \| 828 \| Type VI secretion system tip protein VgrG (Fragment) \| Type VI secretion system tip protein VgrG (Fragment); uniclust \| UniRef100\_A0A1L3I5I6 \| 99.9 \| 5.6e-31 \| 1e-36 \| 204.0 \| 271 \| (1, 279) \| 311 \| (54, 325) \| 349 \| Putative phage late control gene D protein \| Putative phage late control gene D protein; uniclust \| UniRef100\_A0A377AJP2 \| 99.9 \| 5.8e-31 \| 1.1e-36 \| 212.6 \| 245 \| (26, 283) \| 311 \| (3, 263) \| 295 \| Phage late control D family protein \| Phage late control D family protein; uniclust \| UniRef100\_A0A023PRF0 \| 99.9 \| 6.1e-31 \| 1.1e-36 \| 219.4 \| 276 \| (17, 308) \| 311 \| (86, 400) \| 562 \| VgrG protein \| VgrG protein; uniclust \| UniRef100\_A0A064AHQ8 \| 99.9 \| 6.3e-31 \| 1.2e-36 \| 213.4 \| 268 \| (1, 282) \| 311 \| (70, 352) \| 356 \| Late control protein D \| Late control protein D; uniclust \| UniRef100\_A0A432PWE3 \| 99.9 \| 6.8e-31 \| 1.3e-36 \| 204.2 \| 262 \| (1, 276) \| 311 \| (59, 326) \| 330 \| Phage late control D family protein \| Phage late control D family protein; uniclust \| UniRef100\_A0A1G4UZT6 \| 99.9 \| 7.1e-31 \| 1.3e-36 \| 214.5 \| 243 \| (57, 305) \| 311 \| (20, 299) \| 427 \| Rhs element Vgr protein (Fragment) \| Rhs element Vgr protein (Fragment); uniclust \| UniRef100\_A0A0E3VX40 \| 99.9 \| 1.5e-30 \| 2.7e-36 \| 214.2 \| 275 \| (2, 284) \| 311 \| (75, 360) \| 419 \| Uncharacterized protein \| Uncharacterized protein; uniclust \| UniRef100\_A0A072N097 \| 99.9 \| 1.5e-30 \| 2.8e-36 \| 229.1 \| 274 \| (17, 307) \| 311 \| (200, 511) \| 1019 \| Lysozyme \| Lysozyme; uniclust \| UniRef100\_A0A399WGD3 \| 99.9 \| 1.8e-30 \| 3.3e-36 \| 229.6 \| 287 \| (4, 305) \| 311 \| (62, 411) \| 1055 \| Uncharacterized protein \| Uncharacterized protein; uniclust \| UniRef100\_A0A932J5G0 \| 99.9 \| 2.2e-30 \| 4.1e-36 \| 211.1 \| 289 \| (4, 306) \| 311 \| (70, 402) \| 500 \| Type VI secretion system tip protein VgrG \| Type VI secretion system tip protein VgrG; uniclust \| UniRef100\_A0A063BIY2 \| 99.9 \| 2.4e-30 \| 4.4e-36 \| 218.9 \| 280 \| (17, 309) \| 311 \| (126, 457) \| 622 \| Type VI secretion system Vgr family protein (Fragment) \| Type VI secretion system Vgr family protein (Fragment); uniclust \| UniRef100\_A0A017STW5 \| 99.9 \| 2.8e-30 \| 5.2e-36 \| 235.4 \| 290 \| (3, 307) \| 311 \| (192, 528) \| 951 \| VgrG protein \| VgrG protein; uniclust \| UniRef100\_A0A0N0VWG2 \| 99.9 \| 3.1e-30 \| 5.6e-36 \| 220.6 \| 272 \| (17, 305) \| 311 \| (69, 383) \| 777 \| Rhs element Vgr protein \| Rhs element Vgr protein; uniclust \| UniRef100\_A0A1Q3H876 \| 99.9 \| 3.4e-30 \| 6.2e-36 \| 223.9 \| 291 \| (4, 308) \| 311 \| (61, 395) \| 1047 \| Gp5/Type VI secretion system Vgr protein OB-fold domain-containing protein \| Gp5/Type VI secretion system Vgr protein OB-fold domain-containing protein; uniclust \| UniRef100\_A0A0B1XX70 \| 99.9 \| 3.7e-30 \| 6.7e-36 \| 224.5 \| 266 \| (2, 283) \| 311 \| (68, 357) \| 589 \| Type VI secretion protein VgrG \| Type VI secretion protein VgrG; uniclust \| UniRef100\_A0A066PR67 \| 99.9 \| 4e-30 \| 7.3e-36 \| 223.7 \| 288 \| (4, 304) \| 311 \| (77, 407) \| 673 \| Putative type VI secretion system protein VgrGB \| Putative type VI secretion system protein VgrGB; uniclust \| UniRef100\_A0A085WRZ9 \| 99.9 \| 4.6e-30 \| 8.4e-36 \| 221.9 \| 271 \| (4, 289) \| 311 \| (169, 478) \| 618 \| VgrG protein \| VgrG protein; uniclust \| UniRef100\_A0A1H4BCX2 \| 99.9 \| 4.8e-30 \| 8.7e-36 \| 219.8 \| 289 \| (2, 308) \| 311 \| (70, 382) \| 583 \| Rhs element Vgr protein \| Rhs element Vgr protein; uniclust \| UniRef100\_A0A085ZBA7 \| 99.9 \| 4.9e-30 \| 8.9e-36 \| 236.0 \| 281 \| (4, 305) \| 311 \| (170, 487) \| 858 \| Rhs element Vgr protein \| Rhs element Vgr protein; uniclust \| UniRef100\_A0A0B6XFU3 \| 99.9 \| 5.3e-30 \| 9.7e-36 \| 225.6 \| 277 \| (17, 310) \| 311 \| (171, 484) \| 836 \| Gp5/Type VI secretion system Vgr protein OB-fold domain-containing protein \| Gp5/Type VI secretion system Vgr protein OB-fold domain-containing protein; uniclust \| UniRef100\_A0A0Q4EHZ4 \| 99.9 \| 5.3e-30 \| 9.8e-36 \| 217.7 \| 294 \| (1, 308) \| 311 \| (49, 380) \| 676 \| ImpA family type VI secretion-associated protein \| ImpA family type VI secretion-associated protein; uniclust \| UniRef100\_A0A3N5LHC6 \| 99.9 \| 5.5e-30 \| 1e-35 \| 210.3 \| 260 \| (3, 277) \| 311 \| (90, 378) \| 394 \| Phage late control D family protein \| Phage late control D family protein; uniclust \| UniRef100\_A0A1E7HX75 \| 99.9 \| 5.9e-30 \| 1.1e-35 \| 229.0 \| 263 \| (17, 291) \| 311 \| (69, 374) \| 831 \| Rhs element Vgr protein \| Rhs element Vgr protein; uniclust \| UniRef100\_A0A1R1JJ15 \| 99.9 \| 6.3e-30 \| 1.1e-35 \| 223.0 \| 276 \| (17, 304) \| 311 \| (121, 442) \| 903 \| Type IV secretion protein Rhs \| Type IV secretion protein Rhs; uniclust \| UniRef100\_A0A009Q2V4 \| 99.9 \| 7.5e-30 \| 1.4e-35 \| 214.6 \| 269 \| (1, 277) \| 311 \| (96, 370) \| 585 \| Putative phage late control D protein \| Putative phage late control D protein; uniclust \| UniRef100\_A0A1E4J654 \| 99.9 \| 7.6e-30 \| 1.4e-35 \| 220.0 \| 290 \| (4, 305) \| 311 \| (49, 391) \| 1057 \| Type IV secretion protein Rhs \| Type IV secretion protein Rhs; uniclust \| UniRef100\_A0A1N6X884 \| 99.9 \| 8.3e-30 \| 1.5e-35 \| 201.9 \| 262 \| (3, 280) \| 311 \| (62, 350) \| 351 \| Phage protein D \| Phage protein D; uniclust \| UniRef100\_A0A177Q3K8 \| 99.9 \| 9.3e-30 \| 1.7e-35 \| 219.7 \| 286 \| (4, 307) \| 311 \| (65, 419) \| 570 \| Gp5/Type VI secretion system Vgr protein OB-fold domain-containing protein \| Gp5/Type VI secretion system Vgr protein OB-fold domain-containing protein; uniclust \| UniRef100\_A0A090G3M4 \| 99.9 \| 9.4e-30 \| 1.7e-35 \| 213.7 \| 276 \| (1, 285) \| 311 \| (129, 417) \| 463 \| Late control D family protein \| Late control D family protein; uniclust \| UniRef100\_A0A023Q190 \| 99.9 \| 9.5e-30 \| 1.7e-35 \| 228.0 \| 267 \| (4, 286) \| 311 \| (127, 430) \| 766 \| Type IV secretion protein Rhs \| Type IV secretion protein Rhs; uniclust \| UniRef100\_A0A022PJC1 \| 99.9 \| 1.2e-29 \| 2.1e-35 \| 222.3 \| 273 \| (17, 306) \| 311 \| (210, 521) \| 860 \| Rhs element Vgr protein \| Rhs element Vgr protein; uniclust \| UniRef100\_A0A5M7LZ08 \| 99.9 \| 1.2e-29 \| 2.2e-35 \| 197.4 \| 272 \| (4, 286) \| 311 \| (54, 362) \| 374 \| Type VI secretion system tip protein VgrG (Fragment) \| Type VI secretion system tip protein VgrG (Fragment); uniclust \| UniRef100\_A0A538CK92 \| 99.9 \| 1.3e-29 \| 2.3e-35 \| 194.6 \| 229 \| (35, 279) \| 311 \| (9, 261) \| 266 \| Phage late control D family protein \| Phage late control D family protein; uniclust \| UniRef100\_A0A1C6AKC2 \| 99.9 \| 1.3e-29 \| 2.3e-35 \| 208.8 \| 257 \| (3, 277) \| 311 \| (98, 373) \| 375 \| Phage protein D \| Phage protein D; uniclust \| UniRef100\_A0A017TAU5 \| 99.9 \| 1.3e-29 \| 2.4e-35 \| 221.8 \| 290 \| (4, 304) \| 311 \| (63, 381) \| 749 \| VgrG protein \| VgrG protein; uniclust \| UniRef100\_A0A061PST6 \| 99.9 \| 1.6e-29 \| 2.9e-35 \| 222.4 \| 286 \| (3, 304) \| 311 \| (70, 393) \| 676 \| VgrG protein \| VgrG protein; uniclust \| UniRef100\_A0A088UGI5 \| 99.9 \| 1.6e-29 \| 3e-35 \| 232.7 \| 291 \| (4, 306) \| 311 \| (80, 417) \| 2022 \| Rhs element Vgr family protein \| Rhs element Vgr family protein; uniclust \| UniRef100\_A0A069BDU2 \| 99.9 \| 1.8e-29 \| 3.3e-35 \| 220.9 \| 287 \| (4, 304) \| 311 \| (59, 387) \| 874 \| Type IV secretion protein Rhs \| Type IV secretion protein Rhs; uniclust \| UniRef100\_A0A0H2LRL4 \| 99.9 \| 2.1e-29 \| 3.8e-35 \| 236.7 \| 291 \| (4, 305) \| 311 \| (72, 402) \| 2113 \| Putative deoxyribonuclease RhsC \| Putative deoxyribonuclease RhsC; uniclust \| UniRef100\_A0A1G3DZZ5 \| 99.9 \| 2.3e-29 \| 4.3e-35 \| 204.7 \| 275 \| (17, 305) \| 311 \| (68, 364) \| 517 \| Type IV secretion protein Rhs (Fragment) \| Type IV secretion protein Rhs (Fragment); uniclust \| UniRef100\_A0A2G6L1C0 \| 99.9 \| 2.5e-29 \| 4.6e-35 \| 204.6 \| 275 \| (17, 306) \| 311 \| (70, 390) \| 481 \| Type VI secretion system tip protein VgrG (Fragment) \| Type VI secretion system tip protein VgrG (Fragment); uniclust \| UniRef100\_A0A4Q6CNI6 \| 99.9 \| 2.5e-29 \| 4.6e-35 \| 195.0 \| 265 \| (2, 276) \| 311 \| (51, 315) \| 332 \| Phage late control D family protein \| Phage late control D family protein; uniclust \| UniRef100\_A0A0E3V6K5 \| 99.9 \| 2.8e-29 \| 5.1e-35 \| 221.3 \| 282 \| (4, 305) \| 311 \| (60, 381) \| 610 \| Gp5/Type VI secretion system Vgr protein OB-fold domain-containing protein \| Gp5/Type VI secretion system Vgr protein OB-fold domain-containing protein; uniclust \| UniRef100\_A0A356TQN0 \| 99.9 \| 2.8e-29 \| 5.2e-35 \| 220.4 \| 293 \| (3, 308) \| 311 \| (62, 390) \| 856 \| Gp5/Type VI secretion system Vgr protein OB-fold domain-containing protein \| Gp5/Type VI secretion system Vgr protein OB-fold domain-containing protein; uniclust \| UniRef100\_A0A0Q4RTT5 \| 99.9 \| 3.2e-29 \| 5.9e-35 \| 211.5 \| 287 \| (4, 304) \| 311 \| (49, 384) \| 848 \| Type IV secretion protein Rhs \| Type IV secretion protein Rhs; uniclust \| UniRef100\_A0A0K8MDT9 \| 99.9 \| 3.3e-29 \| 6e-35 \| 219.9 \| 287 \| (4, 304) \| 311 \| (243, 572) \| 1067 \| Phage-related baseplate assembly protein \| Phage-related baseplate assembly protein; uniclust \| UniRef100\_A0A086ELE6 \| 99.9 \| 3.4e-29 \| 6.2e-35 \| 217.6 \| 258 \| (38, 310) \| 311 \| (98, 391) \| 470 \| Type VI secretion system tip protein VgrG (Fragment) \| Type VI secretion system tip protein VgrG (Fragment); uniclust \| UniRef100\_A0A017TBW0 \| 99.9 \| 3.7e-29 \| 6.8e-35 \| 220.6 \| 257 \| (37, 304) \| 311 \| (91, 375) \| 825 \| VgrG protein \| VgrG protein; uniclust \| UniRef100\_A0A077PRC6 \| 99.9 \| 3.7e-29 \| 6.8e-35 \| 213.4 \| 274 \| (17, 307) \| 311 \| (90, 406) \| 664 \| Putative VrgG protein \| Putative VrgG protein; uniclust \| UniRef100\_A0A085GIL1 \| 99.9 \| 4.1e-29 \| 7.5e-35 \| 219.3 \| 274 \| (17, 304) \| 311 \| (101, 410) \| 1073 \| Gp5/Type VI secretion system Vgr protein OB-fold domain-containing protein \| Gp5/Type VI secretion system Vgr protein OB-fold domain-containing protein; uniclust \| UniRef100\_A0A017T830 \| 99.9 \| 7.2e-29 \| 1.3e-34 \| 194.6 \| 248 \| (53, 306) \| 311 \| (39, 329) \| 367 \| VgrG protein \| VgrG protein; uniclust \| UniRef100\_A0A0G3A0D9 \| 99.9 \| 7.7e-29 \| 1.4e-34 \| 216.5 \| 260 \| (4, 280) \| 311 \| (94, 374) \| 568 \| Phage baseplate assembly protein V \| Phage baseplate assembly protein V; uniclust \| UniRef100\_A0A356KF96 \| 99.9 \| 7.9e-29 \| 1.5e-34 \| 201.4 \| 264 \| (3, 282) \| 311 \| (68, 360) \| 454 \| Phage late control D family protein \| Phage late control D family protein; uniclust \| UniRef100\_A0A090JZ97 \| 99.9 \| 8.2e-29 \| 1.5e-34 \| 211.9 \| 274 \| (18, 307) \| 311 \| (81, 393) \| 742 \| VgrG protein \| VgrG protein; uniclust \| UniRef100\_A0A1H1G3E9 \| 99.9 \| 9.5e-29 \| 1.7e-34 \| 192.7 \| 270 \| (1, 277) \| 311 \| (57, 331) \| 333 \| Phage protein D \| Phage protein D; uniclust \| UniRef100\_A0A0A6D7Z2 \| 99.9 \| 1e-28 \| 1.9e-34 \| 203.1 \| 269 \| (20, 305) \| 311 \| (95, 409) \| 550 \| Type IV secretion protein Rhs (Fragment) \| Type IV secretion protein Rhs (Fragment); uniclust \| UniRef100\_A0A1V1VC84 \| 99.9 \| 1.1e-28 \| 2.1e-34 \| 198.0 \| 277 \| (17, 307) \| 311 \| (79, 392) \| 455 \| Type VI secretion system tip protein VgrG \| Type VI secretion system tip protein VgrG; uniclust \| UniRef100\_A0A0D0GR35 \| 99.9 \| 1.2e-28 \| 2.2e-34 \| 198.2 \| 272 \| (1, 279) \| 311 \| (120, 416) \| 428 \| Phage late control D family protein \| Phage late control D family protein; uniclust \| UniRef100\_A0A1C5CUP6 \| 99.9 \| 1.2e-28 \| 2.2e-34 \| 203.4 \| 264 \| (4, 277) \| 311 \| (62, 356) \| 521 \| Uncharacterized conserved protein, implicated in type VI secretion and phage assembly (Fragment) \| Uncharacterized conserved protein, implicated in type VI secretion and phage assembly (Fragment); uniclust \| UniRef100\_A0A0S9QJG9 \| 99.9 \| 1.7e-28 \| 3.1e-34 \| 214.8 \| 240 \| (58, 305) \| 311 \| (2, 277) \| 679 \| Gp5/Type VI secretion system Vgr protein OB-fold domain-containing protein (Fragment) \| Gp5/Type VI secretion system Vgr protein OB-fold domain-containing protein (Fragment); uniclust \| UniRef100\_A0A0B5KGT4 \| 99.9 \| 2e-28 \| 3.7e-34 \| 204.2 \| 288 \| (4, 305) \| 311 \| (48, 376) \| 644 \| Rhs element Vgr protein \| Rhs element Vgr protein; uniclust \| UniRef100\_A0A2K9EFG9 \| 99.9 \| 2.1e-28 \| 3.9e-34 \| 212.0 \| 283 \| (4, 301) \| 311 \| (64, 388) \| 726 \| Type VI secretion system tip protein VgrG \| Type VI secretion system tip protein VgrG; uniclust \| UniRef100\_A0A0F0H101 \| 99.9 \| 2.2e-28 \| 3.9e-34 \| 215.2 \| 263 \| (3, 279) \| 311 \| (85, 403) \| 653 \| Gp5/Type VI secretion system Vgr protein OB-fold domain-containing protein \| Gp5/Type VI secretion system Vgr protein OB-fold domain-containing protein; uniclust \| UniRef100\_A0A0F2S4F5 \| 99.9 \| 2.4e-28 \| 4.3e-34 \| 209.2 \| 287 \| (4, 304) \| 311 \| (59, 384) \| 742 \| Gp5/Type VI secretion system Vgr protein OB-fold domain-containing protein \| Gp5/Type VI secretion system Vgr protein OB-fold domain-containing protein; uniclust \| UniRef100\_A0A174TRQ2 \| 99.9 \| 2.4e-28 \| 4.4e-34 \| 205.8 \| 256 \| (17, 285) \| 311 \| (191, 457) \| 530 \| Phage protein D \| Phage protein D; uniclust \| UniRef100\_A0A1I3SYP4 \| 99.9 \| 2.5e-28 \| 4.6e-34 \| 191.2 \| 270 \| (1, 277) \| 311 \| (52, 328) \| 381 \| Phage protein D \| Phage protein D; uniclust \| UniRef100\_A0A2J7TZ74 \| 99.9 \| 2.5e-28 \| 4.7e-34 \| 212.9 \| 272 \| (18, 307) \| 311 \| (148, 471) \| 978 \| Gp5/Type VI secretion system Vgr protein OB-fold domain-containing protein \| Gp5/Type VI secretion system Vgr protein OB-fold domain-containing protein; uniclust \| UniRef100\_A0A0C1KLU7 \| 99.9 \| 2.7e-28 \| 5e-34 \| 211.3 \| 288 \| (4, 304) \| 311 \| (88, 417) \| 969 \| Type IV secretion protein Rhs \| Type IV secretion protein Rhs; uniclust \| UniRef100\_A0A1F9MHR6 \| 99.9 \| 2.7e-28 \| 5e-34 \| 201.4 \| 232 \| (36, 283) \| 311 \| (97, 364) \| 446 \| Late control protein D \| Late control protein D; uniclust \| UniRef100\_A0A060I7P9 \| 99.9 \| 2.8e-28 \| 5.1e-34 \| 216.6 \| 287 \| (5, 304) \| 311 \| (83, 414) \| 930 \| Type VI secretion system Rhs element Vgr family protein \| Type VI secretion system Rhs element Vgr family protein; uniclust \| UniRef100\_A0A143XPS7 \| 99.9 \| 3.5e-28 \| 6.4e-34 \| 206.5 \| 268 \| (1, 282) \| 311 \| (98, 373) \| 483 \| Phage late control gene D protein (GPD) \| Phage late control gene D protein (GPD); uniclust \| UniRef100\_A0A2A2HZ20 \| 99.9 \| 3.6e-28 \| 6.6e-34 \| 201.5 \| 285 \| (6, 306) \| 311 \| (42, 367) \| 607 \| Type IV secretion protein Rhs (Fragment) \| Type IV secretion protein Rhs (Fragment); uniclust \| UniRef100\_A0A090SM11 \| 99.9 \| 3.9e-28 \| 7.2e-34 \| 208.3 \| 242 \| (56, 305) \| 311 \| (9, 293) \| 644 \| Probable vgr related protein \| Probable vgr related protein; uniclust \| UniRef100\_A0A0M0SQ28 \| 99.9 \| 4e-28 \| 7.3e-34 \| 203.5 \| 260 \| (2, 278) \| 311 \| (64, 349) \| 572 \| Gp5/Type VI secretion system Vgr protein OB-fold domain-containing protein \| Gp5/Type VI secretion system Vgr protein OB-fold domain-containing protein; uniclust \| UniRef100\_A0A1S8DCS2 \| 99.9 \| 4.7e-28 \| 8.6e-34 \| 191.2 \| 253 \| (37, 304) \| 311 \| (14, 309) \| 414 \| Type VI secretion system tip protein VgrG (Fragment) \| Type VI secretion system tip protein VgrG (Fragment); uniclust \| UniRef100\_A0A031FRL6 \| 99.9 \| 5.4e-28 \| 9.9e-34 \| 194.6 \| 272 \| (1, 280) \| 311 \| (59, 345) \| 355 \| Phage protein D-like protein \| Phage protein D-like protein; uniclust \| UniRef100\_A0A947C8Q0 \| 99.9 \| 5.8e-28 \| 1.1e-33 \| 194.0 \| 291 \| (4, 307) \| 311 \| (57, 397) \| 449 \| Type VI secretion system tip protein VgrG (Fragment) \| Type VI secretion system tip protein VgrG (Fragment); uniclust \| UniRef100\_A0A133ZYU8 \| 99.9 \| 7.6e-28 \| 1.4e-33 \| 201.2 \| 267 \| (1, 281) \| 311 \| (83, 360) \| 445 \| Phage late control D protein \| Phage late control D protein; uniclust \| UniRef100\_A0A0C1ZDR4 \| 99.9 \| 7.8e-28 \| 1.4e-33 \| 220.0 \| 290 \| (4, 305) \| 311 \| (278, 613) \| 980 \| VgrG protein \| VgrG protein; uniclust \| UniRef100\_A0A2A5C5S4 \| 99.9 \| 8e-28 \| 1.5e-33 \| 209.8 \| 288 \| (5, 305) \| 311 \| (58, 382) \| 816 \| Gp5/Type VI secretion system Vgr protein OB-fold domain-containing protein \| Gp5/Type VI secretion system Vgr protein OB-fold domain-containing protein; uniclust \| UniRef100\_A0A349VLS7 \| 99.9 \| 8.1e-28 \| 1.5e-33 \| 189.7 \| 211 \| (4, 227) \| 311 \| (57, 294) \| 300 \| Type VI secretion system tip protein VgrG (Fragment) \| Type VI secretion system tip protein VgrG (Fragment); uniclust \| UniRef100\_A0A212RQ57 \| 99.9 \| 8.8e-28 \| 1.6e-33 \| 206.3 \| 294 \| (2, 305) \| 311 \| (76, 445) \| 695 \| Type VI secretion system secreted protein VgrG \| Type VI secretion system secreted protein VgrG; uniclust \| UniRef100\_A0A016A3P0 \| 99.9 \| 1e-27 \| 1.9e-33 \| 216.3 \| 282 \| (4, 305) \| 311 \| (100, 418) \| 721 \| Phage late control D family protein \| Phage late control D family protein; uniclust \| UniRef100\_A0A193QJR7 \| 99.9 \| 1.1e-27 \| 1.9e-33 \| 188.1 \| 271 \| (1, 278) \| 311 \| (63, 360) \| 374 \| Phage late control gene D protein (GPD) \| Phage late control gene D protein (GPD); uniclust \| UniRef100\_A0A4U1HBY7 \| 99.9 \| 1.3e-27 \| 2.3e-33 \| 192.2 \| 289 \| (4, 305) \| 311 \| (52, 377) \| 501 \| Type VI secretion system tip protein VgrG \| Type VI secretion system tip protein VgrG; uniclust \| UniRef100\_A0A0Q8PZU1 \| 99.9 \| 1.4e-27 \| 2.5e-33 \| 207.8 \| 290 \| (4, 305) \| 311 \| (294, 629) \| 937 \| Gp5/Type VI secretion system Vgr protein OB-fold domain-containing protein \| Gp5/Type VI secretion system Vgr protein OB-fold domain-containing protein; uniclust \| UniRef100\_A0A2N1PDD4 \| 99.9 \| 1.5e-27 \| 2.7e-33 \| 210.4 \| 281 \| (3, 304) \| 311 \| (108, 425) \| 648 \| Gp5/Type VI secretion system Vgr protein OB-fold domain-containing protein \| Gp5/Type VI secretion system Vgr protein OB-fold domain-containing protein; uniclust \| UniRef100\_A0A0P9T8U5 \| 99.9 \| 1.7e-27 \| 3.2e-33 \| 203.3 \| 289 \| (4, 305) \| 311 \| (71, 401) \| 729 \| Rhs element Vgr protein \| Rhs element Vgr protein; uniclust \| UniRef100\_A0A081J2H0 \| 99.9 \| 2e-27 \| 3.7e-33 \| 184.7 \| 246 \| (4, 263) \| 311 \| (49, 322) \| 331 \| Type VI secretion system tip protein VgrG (Fragment) \| Type VI secretion system tip protein VgrG (Fragment); uniclust \| UniRef100\_A0A0E3KUE2 \| 99.9 \| 2.1e-27 \| 3.9e-33 \| 196.6 \| 269 \| (3, 285) \| 311 \| (76, 365) \| 574 \| VgrG protein \| VgrG protein; uniclust \| UniRef100\_UPI000573CFEF \| 99.9 \| 2.2e-27 \| 4.1e-33 \| 191.4 \| 289 \| (4, 305) \| 311 \| (48, 377) \| 477 \| type VI secretion system tip protein TssI/VgrG \| type VI secretion system tip protein TssI/VgrG; uniclust \| UniRef100\_A0A0K9N9X5 \| 99.9 \| 2.7e-27 \| 4.9e-33 \| 197.6 \| 264 \| (3, 280) \| 311 \| (112, 411) \| 430 \| Phage late control gene D protein (GPD) \| Phage late control gene D protein (GPD); uniclust \| UniRef100\_A0A524PWE2 \| 99.9 \| 2.7e-27 \| 5e-33 \| 177.2 \| 244 \| (17, 278) \| 311 \| (8, 274) \| 277 \| Phage late control D family protein (Fragment) \| Phage late control D family protein (Fragment); uniclust \| UniRef100\_A0A246IXC8 \| 99.9 \| 2.7e-27 \| 5e-33 \| 202.2 \| 289 \| (4, 305) \| 311 \| (72, 398) \| 942 \| Gp5/Type VI secretion system Vgr protein OB-fold domain-containing protein \| Gp5/Type VI secretion system Vgr protein OB-fold domain-containing protein; uniclust \| UniRef100\_A0A085G1R1 \| 99.9 \| 2.8e-27 \| 5.1e-33 \| 209.3 \| 270 \| (17, 305) \| 311 \| (116, 427) \| 925 \| VgrG family protein \| VgrG family protein; uniclust \| UniRef100\_A0A255HM11 \| 99.9 \| 2.9e-27 \| 5.3e-33 \| 216.6 \| 267 \| (7, 288) \| 311 \| (95, 407) \| 1043 \| Gp5/Type VI secretion system Vgr protein OB-fold domain-containing protein \| Gp5/Type VI secretion system Vgr protein OB-fold domain-containing protein; uniclust \| UniRef100\_A0A0B3BQR0 \| 99.9 \| 3e-27 \| 5.5e-33 \| 212.4 \| 283 \| (6, 307) \| 311 \| (107, 433) \| 1205 \| Type VI secretion system secreted protein VgrG \| Type VI secretion system secreted protein VgrG; uniclust \| UniRef100\_A0A0H3K4B3 \| 99.9 \| 3.1e-27 \| 5.7e-33 \| 188.5 \| 271 \| (1, 278) \| 311 \| (52, 329) \| 480 \| Phage late control gene D protein GPD \| Phage late control gene D protein GPD; uniclust \| UniRef100\_A0A1W6LMB9 \| 99.9 \| 3.5e-27 \| 6.5e-33 \| 206.3 \| 289 \| (4, 305) \| 311 \| (55, 380) \| 1193 \| Type VI secretion system Vgr family protein \| Type VI secretion system Vgr family protein; uniclust \| UniRef100\_A0A0C2EHH0 \| 99.9 \| 4e-27 \| 7.4e-33 \| 199.0 \| 271 \| (18, 305) \| 311 \| (101, 415) \| 608 \| VgrG protein \| VgrG protein; uniclust \| UniRef100\_A0A0D8FQF9 \| 99.9 \| 5.2e-27 \| 9.4e-33 \| 206.6 \| 259 \| (3, 280) \| 311 \| (122, 401) \| 645 \| Phage-like baseplate assembly protein \| Phage-like baseplate assembly protein; uniclust \| UniRef100\_A0A0Q8LLQ7 \| 99.9 \| 6.2e-27 \| 1.1e-32 \| 198.8 \| 263 \| (3, 283) \| 311 \| (75, 364) \| 601 \| Gp5/Type VI secretion system Vgr protein OB-fold domain-containing protein \| Gp5/Type VI secretion system Vgr protein OB-fold domain-containing protein; uniclust \| UniRef100\_A0A0Q4S1L3 \| 99.9 \| 6.7e-27 \| 1.2e-32 \| 200.1 \| 287 \| (5, 305) \| 311 \| (50, 388) \| 770 \| ImpA family type VI secretion-associated protein (Fragment) \| ImpA family type VI secretion-associated protein (Fragment); uniclust \| UniRef100\_A0A031FNF9 \| 99.9 \| 8.6e-27 \| 1.6e-32 \| 211.3 \| 279 \| (17, 309) \| 311 \| (106, 424) \| 1055 \| Rhs element Vgr protein \| Rhs element Vgr protein; uniclust \| UniRef100\_A0A017T1P1 \| 99.9 \| 9e-27 \| 1.6e-32 \| 213.9 \| 290 \| (5, 306) \| 311 \| (66, 387) \| 1090 \| VgrG protein \| VgrG protein; uniclust \| UniRef100\_A0A268TJ33 \| 99.9 \| 9.6e-27 \| 1.8e-32 \| 185.4 \| 256 \| (1, 277) \| 311 \| (53, 308) \| 314 \| Phage tail protein \| Phage tail protein; uniclust \| UniRef100\_A0A017TAS0 \| 99.9 \| 1e-26 \| 1.9e-32 \| 207.1 \| 289 \| (4, 303) \| 311 \| (89, 410) \| 890 \| VgrG protein \| VgrG protein; uniclust \| UniRef100\_A0A518AUG2 \| 99.9 \| 1.1e-26 \| 1.9e-32 \| 209.9 \| 288 \| (4, 305) \| 311 \| (90, 451) \| 1273 \| Phage-related baseplate assembly protein \| Phage-related baseplate assembly protein; uniclust \| UniRef100\_A0A0J6KFU2 \| 99.9 \| 1.2e-26 \| 2.1e-32 \| 211.9 \| 286 \| (4, 304) \| 311 \| (111, 470) \| 1143 \| Type VI secretion system protein \| Type VI secretion system protein; uniclust \| UniRef100\_A0A0D0K4C1 \| 99.9 \| 1.4e-26 \| 2.6e-32 \| 198.7 \| 290 \| (4, 305) \| 311 \| (133, 468) \| 722 \| Type VI secretion system tip protein VgrG (Fragment) \| Type VI secretion system tip protein VgrG (Fragment); uniclust \| UniRef100\_A0A3D9MY61 \| 99.9 \| 1.7e-26 \| 3e-32 \| 213.2 \| 278 \| (17, 308) \| 311 \| (121, 452) \| 1218 \| Rhs element Vgr protein \| Rhs element Vgr protein; uniclust \| UniRef100\_A0A374MIX7 \| 99.9 \| 1.8e-26 \| 3.4e-32 \| 195.5 \| 280 \| (3, 304) \| 311 \| (62, 373) \| 476 \| Type VI secretion system tip protein VgrG (Fragment) \| Type VI secretion system tip protein VgrG (Fragment); uniclust \| UniRef100\_A0A2K1QDW4 \| 99.9 \| 1.9e-26 \| 3.5e-32 \| 193.9 \| 291 \| (3, 305) \| 311 \| (160, 485) \| 752 \| Type VI secretion system tip protein VgrG (Fragment) \| Type VI secretion system tip protein VgrG (Fragment); uniclust \| UniRef100\_A0A150Q779 \| 99.9 \| 2.4e-26 \| 4.4e-32 \| 204.9 \| 269 \| (5, 287) \| 311 \| (75, 361) \| 1028 \| Gp5/Type VI secretion system Vgr protein OB-fold domain-containing protein \| Gp5/Type VI secretion system Vgr protein OB-fold domain-containing protein; uniclust \| UniRef100\_A0A023Y2K3 \| 99.9 \| 2.4e-26 \| 4.4e-32 \| 212.7 \| 288 \| (3, 304) \| 311 \| (143, 522) \| 1156 \| Type IV secretion protein Rhs \| Type IV secretion protein Rhs; uniclust \| UniRef100\_A0A3S1JAZ2 \| 99.9 \| 2.5e-26 \| 4.5e-32 \| 184.9 \| 288 \| (4, 305) \| 311 \| (52, 378) \| 502 \| Type VI secretion system tip protein VgrG (Fragment) \| Type VI secretion system tip protein VgrG (Fragment); uniclust \| UniRef100\_A0A085EST2 \| 99.9 \| 2.8e-26 \| 5.2e-32 \| 211.8 \| 284 \| (4, 303) \| 311 \| (107, 449) \| 1280 \| Type VI secretion system Vgr family protein \| Type VI secretion system Vgr family protein; uniclust \| UniRef100\_A0A963HZS7 \| 99.9 \| 2.8e-26 \| 5.2e-32 \| 185.5 \| 268 \| (4, 285) \| 311 \| (51, 355) \| 524 \| Type VI secretion system tip protein VgrG \| Type VI secretion system tip protein VgrG; uniclust \| UniRef100\_A0A0B4BFG3 \| 99.9 \| 2.9e-26 \| 5.3e-32 \| 200.8 \| 287 \| (4, 305) \| 311 \| (77, 402) \| 769 \| Gp5/Type VI secretion system Vgr protein OB-fold domain-containing protein \| Gp5/Type VI secretion system Vgr protein OB-fold domain-containing protein; uniclust \| UniRef100\_A0A094QB96 \| 99.9 \| 3e-26 \| 5.5e-32 \| 189.8 \| 259 \| (2, 279) \| 311 \| (59, 348) \| 596 \| Gp5/Type VI secretion system Vgr protein OB-fold domain-containing protein \| Gp5/Type VI secretion system Vgr protein OB-fold domain-containing protein; uniclust \| UniRef100\_A0A077KP81 \| 99.9 \| 3.1e-26 \| 5.7e-32 \| 213.8 \| 281 \| (4, 304) \| 311 \| (132, 450) \| 1149 \| Gp5/Type VI secretion system Vgr protein OB-fold domain-containing protein \| Gp5/Type VI secretion system Vgr protein OB-fold domain-containing protein; uniclust \| UniRef100\_A0A327JI00 \| 99.9 \| 3.5e-26 \| 6.4e-32 \| 175.6 \| 261 \| (2, 276) \| 311 \| (59, 325) \| 328 \| Phage late control D family protein \| Phage late control D family protein; uniclust \| UniRef100\_A0A022FT39 \| 99.9 \| 3.5e-26 \| 6.4e-32 \| 205.8 \| 277 \| (17, 305) \| 311 \| (135, 469) \| 1045 \| Type IV secretion protein Rhs \| Type IV secretion protein Rhs; uniclust \| UniRef100\_A0A228HLH2 \| 99.9 \| 4e-26 \| 7.2e-32 \| 210.4 \| 285 \| (3, 305) \| 311 \| (69, 398) \| 1515 \| Gp5/Type VI secretion system Vgr protein OB-fold domain-containing protein \| Gp5/Type VI secretion system Vgr protein OB-fold domain-containing protein; uniclust \| UniRef100\_A0A0E1SC86 \| 99.9 \| 4.3e-26 \| 7.9e-32 \| 207.0 \| 258 \| (18, 288) \| 311 \| (96, 398) \| 1057 \| Rhs element Vgr protein \| Rhs element Vgr protein; uniclust \| UniRef100\_A0A0A1FJE6 \| 99.9 \| 4.7e-26 \| 8.7e-32 \| 198.9 \| 275 \| (17, 305) \| 311 \| (237, 549) \| 1075 \| VgrG protein \| VgrG protein; uniclust \| UniRef100\_A0A0C2HQP1 \| 99.9 \| 5.3e-26 \| 9.8e-32 \| 186.8 \| 258 \| (2, 281) \| 311 \| (67, 347) \| 361 \| Phage protein D \| Phage protein D; uniclust \| UniRef100\_A0A1I4YEC9 \| 99.9 \| 5.8e-26 \| 1.1e-31 \| 211.0 \| 284 \| (5, 303) \| 311 \| (67, 440) \| 873 \| Rhs element Vgr protein (Fragment) \| Rhs element Vgr protein (Fragment); uniclust \| UniRef100\_A0A6L4A988 \| 99.9 \| 5.9e-26 \| 1.1e-31 \| 184.7 \| 209 \| (61, 278) \| 311 \| (3, 237) \| 487 \| VgrG-related protein (Fragment) \| VgrG-related protein (Fragment); uniclust \| UniRef100\_A0A016CPP4 \| 99.9 \| 6.6e-26 \| 1.2e-31 \| 200.0 \| 280 \| (4, 304) \| 311 \| (91, 401) \| 745 \| Phage-related baseplate assembly family protein \| Phage-related baseplate assembly family protein; uniclust \| UniRef100\_A0A084DLK1 \| 99.9 \| 7e-26 \| 1.3e-31 \| 210.8 \| 279 \| (17, 307) \| 311 \| (127, 458) \| 1317 \| Type IV secretion protein Rhs \| Type IV secretion protein Rhs; uniclust \| UniRef100\_A0A3M1LWC1 \| 99.9 \| 9.3e-26 \| 1.7e-31 \| 189.8 \| 292 \| (5, 309) \| 311 \| (59, 387) \| 681 \| Type VI secretion system tip protein VgrG (Fragment) \| Type VI secretion system tip protein VgrG (Fragment); uniclust \| UniRef100\_A0A0P9ZJ72 \| 99.9 \| 9.9e-26 \| 1.8e-31 \| 185.0 \| 271 \| (18, 305) \| 311 \| (83, 399) \| 602 \| Rhs element Vgr protein (Fragment) \| Rhs element Vgr protein (Fragment); uniclust \| UniRef100\_A0A6A7XZM2 \| 99.9 \| 1.2e-25 \| 2.1e-31 \| 202.9 \| 275 \| (4, 289) \| 311 \| (144, 467) \| 942 \| Type VI secretion system tip protein VgrG \| Type VI secretion system tip protein VgrG; uniclust \| UniRef100\_A0A198GE76 \| 99.9 \| 1.2e-25 \| 2.2e-31 \| 192.4 \| 284 \| (4, 305) \| 311 \| (67, 387) \| 619 \| VgrG family protein \| VgrG family protein; uniclust \| UniRef100\_UPI000AA40F38 \| 99.9 \| 1.4e-25 \| 2.5e-31 \| 176.6 \| 276 \| (17, 304) \| 311 \| (16, 326) \| 382 \| type VI secretion system tip protein TssI/VgrG \| type VI secretion system tip protein TssI/VgrG; uniclust \| UniRef100\_A0A1I7HLI2 \| 99.9 \| 1.5e-25 \| 2.8e-31 \| 202.5 \| 267 \| (4, 287) \| 311 \| (68, 386) \| 863 \| Type VI secretion system secreted protein VgrG \| Type VI secretion system secreted protein VgrG; uniclust \| UniRef100\_A0A0M2S027 \| 99.9 \| 1.5e-25 \| 2.8e-31 \| 186.2 \| 262 \| (3, 276) \| 311 \| (61, 380) \| 380 \| Phage protein D \| Phage protein D; uniclust \| UniRef100\_A0A015YA37 \| 99.9 \| 1.6e-25 \| 2.9e-31 \| 199.4 \| 279 \| (4, 304) \| 311 \| (86, 403) \| 651 \| Phage late control D family protein \| Phage late control D family protein; uniclust \| UniRef100\_A0A7V1ZWM7 \| 99.9 \| 1.6e-25 \| 3e-31 \| 180.3 \| 267 \| (3, 285) \| 311 \| (71, 364) \| 596 \| Phage late control D family protein \| Phage late control D family protein; uniclust \| UniRef100\_A0A2M8S304 \| 99.9 \| 1.8e-25 \| 3.3e-31 \| 196.3 \| 241 \| (58, 306) \| 311 \| (86, 362) \| 579 \| Type VI secretion system tip protein VgrG (Fragment) \| Type VI secretion system tip protein VgrG (Fragment); uniclust \| UniRef100\_A0A2A4Y9Q6 \| 99.9 \| 1.9e-25 \| 3.4e-31 \| 203.2 \| 260 \| (17, 289) \| 311 \| (73, 377) \| 1037 \| Gp5/Type VI secretion system Vgr protein OB-fold domain-containing protein \| Gp5/Type VI secretion system Vgr protein OB-fold domain-containing protein; uniclust \| UniRef100\_A0A014MDT4 \| 99.9 \| 2.2e-25 \| 4e-31 \| 176.6 \| 263 \| (3, 277) \| 311 \| (79, 394) \| 401 \| Type IV secretion protein Rhs \| Type IV secretion protein Rhs; uniclust \| UniRef100\_A0A2N8DM59 \| 99.9 \| 2.2e-25 \| 4.1e-31 \| 185.9 \| 254 \| (37, 305) \| 311 \| (88, 384) \| 680 \| Type VI secretion system tip protein VgrG (Fragment) \| Type VI secretion system tip protein VgrG (Fragment); uniclust \| UniRef100\_A0A7G8BMN3 \| 99.9 \| 2.5e-25 \| 4.6e-31 \| 191.1 \| 293 \| (4, 308) \| 311 \| (56, 402) \| 986 \| Type VI secretion system tip protein VgrG \| Type VI secretion system tip protein VgrG; uniclust \| UniRef100\_A0A0A2ECB3 \| 99.9 \| 2.6e-25 \| 4.8e-31 \| 199.7 \| 282 \| (3, 304) \| 311 \| (163, 476) \| 733 \| Gp5/Type VI secretion system Vgr protein OB-fold domain-containing protein \| Gp5/Type VI secretion system Vgr protein OB-fold domain-containing protein; uniclust \| UniRef100\_A0A0D0R540 \| 99.9 \| 2.7e-25 \| 5e-31 \| 196.9 \| 287 \| (4, 304) \| 311 \| (144, 515) \| 939 \| VgrG protein \| VgrG protein; uniclust \| UniRef100\_A0A0M6XR59 \| 99.9 \| 2.7e-25 \| 5e-31 \| 181.3 \| 284 \| (4, 301) \| 311 \| (57, 384) \| 468 \| Type VI secretion system Vgr family protein \| Type VI secretion system Vgr family protein; uniclust \| UniRef100\_A0A380U1Y3 \| 99.9 \| 2.8e-25 \| 5.1e-31 \| 189.5 \| 274 \| (17, 305) \| 311 \| (146, 456) \| 778 \| Uncharacterized protein conserved in bacteria \| Uncharacterized protein conserved in bacteria; uniclust \| UniRef100\_A0A0T9NGE2 \| 99.9 \| 4e-25 \| 7.4e-31 \| 184.9 \| 287 \| (4, 304) \| 311 \| (56, 383) \| 690 \| Rhs element Vgr protein \| Rhs element Vgr protein; uniclust \| UniRef100\_A0A933X3D3 \| 99.9 \| 4.1e-25 \| 7.6e-31 \| 189.4 \| 278 \| (16, 307) \| 311 \| (69, 385) \| 1211 \| Type VI secretion system tip protein VgrG \| Type VI secretion system tip protein VgrG; uniclust \| UniRef100\_A0A085WSA7 \| 99.9 \| 4.5e-25 \| 8.2e-31 \| 188.4 \| 273 \| (17, 305) \| 311 \| (75, 373) \| 1140 \| VgrG protein \| VgrG protein; uniclust \| UniRef100\_A0A077MYI5 \| 99.9 \| 4.7e-25 \| 8.6e-31 \| 196.6 \| 290 \| (4, 310) \| 311 \| (153, 475) \| 942 \| Gp5/Type VI secretion system Vgr protein OB-fold domain-containing protein \| Gp5/Type VI secretion system Vgr protein OB-fold domain-containing protein; uniclust \| UniRef100\_A0A0N0UY75 \| 99.9 \| 5e-25 \| 9.2e-31 \| 186.0 \| 264 \| (3, 283) \| 311 \| (71, 358) \| 595 \| Gp5/Type VI secretion system Vgr protein OB-fold domain-containing protein \| Gp5/Type VI secretion system Vgr protein OB-fold domain-containing protein; uniclust \| UniRef100\_A0A2E8S0G6 \| 99.8 \| 5.7e-25 \| 1.1e-30 \| 181.7 \| 268 \| (3, 283) \| 311 \| (75, 393) \| 407 \| Phage late control D family protein \| Phage late control D family protein; uniclust \| UniRef100\_A0A068QXL9 \| 99.8 \| 5.9e-25 \| 1.1e-30 \| 179.8 \| 273 \| (1, 281) \| 311 \| (75, 480) \| 498 \| Putative phage protein (D protein) (Modular protein) \| Putative phage protein (D protein) (Modular protein); uniclust \| UniRef100\_A0A0X8HC04 \| 99.8 \| 6.2e-25 \| 1.1e-30 \| 192.5 \| 274 \| (17, 306) \| 311 \| (72, 400) \| 1032 \| Electron transfer flavoprotein subunit alpha \| Electron transfer flavoprotein subunit alpha; uniclust \| UniRef100\_A0A090AED5 \| 99.8 \| 6.5e-25 \| 1.2e-30 \| 187.1 \| 262 \| (3, 280) \| 311 \| (69, 371) \| 418 \| Phage protein D \| Phage protein D; uniclust \| UniRef100\_A0A1G0FFT8 \| 99.8 \| 8.3e-25 \| 1.5e-30 \| 180.9 \| 281 \| (4, 310) \| 311 \| (57, 361) \| 542 \| Gp5/Type VI secretion system Vgr protein OB-fold domain-containing protein \| Gp5/Type VI secretion system Vgr protein OB-fold domain-containing protein; uniclust \| UniRef100\_A0A1B9SFQ1 \| 99.8 \| 1.1e-24 \| 2.1e-30 \| 178.9 \| 268 \| (17, 300) \| 311 \| (82, 377) \| 394 \| Phage tail protein \| Phage tail protein; uniclust \| UniRef100\_A0A0B5FHG4 \| 99.8 \| 1.2e-24 \| 2.2e-30 \| 174.7 \| 270 \| (2, 288) \| 311 \| (61, 390) \| 432 \| Phage protein D-like protein \| Phage protein D-like protein; uniclust \| UniRef100\_A0A178J7K0 \| 99.8 \| 1.3e-24 \| 2.3e-30 \| 194.3 \| 285 \| (7, 307) \| 311 \| (65, 390) \| 1022 \| Type IV secretion protein Rhs \| Type IV secretion protein Rhs; uniclust \| UniRef100\_A0A257MWB5 \| 99.8 \| 1.3e-24 \| 2.3e-30 \| 174.8 \| 289 \| (4, 304) \| 311 \| (55, 403) \| 462 \| Rhs-family protein \| Rhs-family protein; uniclust \| UniRef100\_A0A1X0TCA5 \| 99.8 \| 1.3e-24 \| 2.3e-30 \| 182.3 \| 291 \| (2, 305) \| 311 \| (222, 548) \| 823 \| Type VI secretion protein Vgr \| Type VI secretion protein Vgr; uniclust \| UniRef100\_A0A099PCF2 \| 99.8 \| 1.3e-24 \| 2.4e-30 \| 177.7 \| 260 \| (3, 277) \| 311 \| (89, 387) \| 391 \| Tip attachment protein J domain-containing protein \| Tip attachment protein J domain-containing protein; uniclust \| UniRef100\_A0A1I1H522 \| 99.8 \| 1.5e-24 \| 2.7e-30 \| 199.4 \| 268 \| (5, 286) \| 311 \| (62, 369) \| 950 \| Rhs element Vgr protein \| Rhs element Vgr protein; uniclust \| UniRef100\_A0A0K9L316 \| 99.8 \| 1.5e-24 \| 2.7e-30 \| 170.0 \| 262 \| (1, 276) \| 311 \| (59, 335) \| 393 \| Phage late control gene D protein \| Phage late control gene D protein; uniclust \| UniRef100\_A0A063BL00 \| 99.8 \| 1.8e-24 \| 3.2e-30 \| 194.3 \| 270 \| (17, 302) \| 311 \| (136, 437) \| 985 \| Rhs element Vgr protein \| Rhs element Vgr protein; uniclust \| UniRef100\_A0A1B8H0M3 \| 99.8 \| 2.1e-24 \| 3.8e-30 \| 188.7 \| 289 \| (5, 310) \| 311 \| (107, 426) \| 644 \| Type VI secretion protein (Fragment) \| Type VI secretion protein (Fragment); uniclust \| UniRef100\_A0A0W8JA15 \| 99.8 \| 2.2e-24 \| 4e-30 \| 190.5 \| 276 \| (18, 307) \| 311 \| (90, 402) \| 1057 \| VgrG protein \| VgrG protein; uniclust \| UniRef100\_A0A0X3APK5 \| 99.8 \| 2.3e-24 \| 4.1e-30 \| 193.3 \| 279 \| (4, 304) \| 311 \| (103, 423) \| 813 \| Uncharacterized conserved protein, implicated in type VI secretion and phage assembly \| Uncharacterized conserved protein, implicated in type VI secretion and phage assembly; uniclust \| UniRef100\_A0A3M1QV85 \| 99.8 \| 2.7e-24 \| 4.9e-30 \| 168.1 \| 275 \| (17, 307) \| 311 \| (117, 442) \| 446 \| Type VI secretion system tip protein VgrG (Fragment) \| Type VI secretion system tip protein VgrG (Fragment); uniclust \| UniRef100\_A0A1Q7UPE7 \| 99.8 \| 2.8e-24 \| 5.1e-30 \| 171.7 \| 263 \| (3, 280) \| 311 \| (61, 349) \| 437 \| Phage late control D family protein (Fragment) \| Phage late control D family protein (Fragment); uniclust \| UniRef100\_A0A097QZD7 \| 99.8 \| 3e-24 \| 5.5e-30 \| 183.5 \| 269 \| (17, 304) \| 311 \| (111, 421) \| 749 \| Type IV secretion protein Rhs \| Type IV secretion protein Rhs; uniclust \| UniRef100\_A0A2A6PZI3 \| 99.8 \| 3.2e-24 \| 5.9e-30 \| 179.8 \| 281 \| (6, 305) \| 311 \| (113, 435) \| 637 \| Type VI secretion system tip protein VgrG (Fragment) \| Type VI secretion system tip protein VgrG (Fragment); uniclust \| UniRef100\_A0A022PDU6 \| 99.8 \| 3.4e-24 \| 6.1e-30 \| 188.4 \| 258 \| (3, 275) \| 311 \| (73, 356) \| 575 \| Gp5/Type VI secretion system Vgr protein OB-fold domain-containing protein \| Gp5/Type VI secretion system Vgr protein OB-fold domain-containing protein; uniclust \| UniRef100\_A0A0M2WHW5 \| 99.8 \| 4.2e-24 \| 7.6e-30 \| 194.8 \| 287 \| (4, 305) \| 311 \| (77, 457) \| 1005 \| Phage-like baseplate assembly protein \| Phage-like baseplate assembly protein; uniclust \| UniRef100\_A0A518AYV3 \| 99.8 \| 4.9e-24 \| 9e-30 \| 193.7 \| 284 \| (4, 303) \| 311 \| (106, 451) \| 787 \| Phage-related baseplate assembly protein \| Phage-related baseplate assembly protein; uniclust \| UniRef100\_A0A074M4V0 \| 99.8 \| 5.3e-24 \| 9.7e-30 \| 173.3 \| 255 \| (3, 277) \| 311 \| (53, 334) \| 366 \| Uncharacterized protein \| Uncharacterized protein; uniclust \| UniRef100\_A0A0S4X111 \| 99.8 \| 5.8e-24 \| 1.1e-29 \| 186.6 \| 289 \| (4, 304) \| 311 \| (127, 510) \| 899 \| Vgr-related protein (Fragment) \| Vgr-related protein (Fragment); uniclust \| UniRef100\_A0A940QC34 \| 99.8 \| 6.6e-24 \| 1.2e-29 \| 163.6 \| 260 \| (3, 277) \| 311 \| (68, 350) \| 352 \| Uncharacterized protein \| Uncharacterized protein; uniclust \| UniRef100\_A0A085U800 \| 99.8 \| 6.8e-24 \| 1.3e-29 \| 163.5 \| 240 \| (36, 282) \| 311 \| (11, 274) \| 287 \| Gene D protein \| Gene D protein; uniclust \| UniRef100\_A0A3M1TEK0 \| 99.8 \| 7.5e-24 \| 1.4e-29 \| 168.4 \| 266 \| (2, 283) \| 311 \| (104, 395) \| 516 \| Phage late control D family protein \| Phage late control D family protein; uniclust \| UniRef100\_A0A4U1HBW3 \| 99.8 \| 7.7e-24 \| 1.4e-29 \| 179.1 \| 287 \| (4, 305) \| 311 \| (68, 407) \| 545 \| Type VI secretion system tip protein VgrG \| Type VI secretion system tip protein VgrG; uniclust \| UniRef100\_A0A522CBE1 \| 99.8 \| 8.6e-24 \| 1.6e-29 \| 164.0 \| 245 \| (5, 266) \| 311 \| (55, 339) \| 356 \| Type VI secretion system tip protein VgrG (Fragment) \| Type VI secretion system tip protein VgrG (Fragment); uniclust \| UniRef100\_A0A0Q6V477 \| 99.8 \| 8.7e-24 \| 1.6e-29 \| 175.2 \| 262 \| (3, 277) \| 311 \| (67, 388) \| 417 \| Phage tail protein \| Phage tail protein; uniclust \| UniRef100\_A0A376EWE4 \| 99.8 \| 1e-23 \| 1.8e-29 \| 168.9 \| 290 \| (1, 304) \| 311 \| (74, 380) \| 550 \| Gene D protein \| Gene D protein; uniclust \| UniRef100\_A0A090SLR1 \| 99.8 \| 1e-23 \| 1.9e-29 \| 165.7 \| 128 \| (3, 139) \| 311 \| (62, 198) \| 266 \| VgrG protein \| VgrG protein; uniclust \| UniRef100\_I7IJM6 \| 99.8 \| 1.1e-23 \| 2e-29 \| 190.1 \| 289 \| (4, 305) \| 311 \| (72, 402) \| 1638 \| RHs family protein (Fragment) \| RHs family protein (Fragment); uniclust \| UniRef100\_A0A066RRK7 \| 99.8 \| 1.2e-23 \| 2.1e-29 \| 190.7 \| 276 \| (17, 305) \| 311 \| (68, 388) \| 1616 \| Gp5/Type VI secretion system Vgr protein OB-fold domain-containing protein \| Gp5/Type VI secretion system Vgr protein OB-fold domain-containing protein; uniclust \| UniRef100\_A0A3N4GAW7 \| 99.8 \| 1.2e-23 \| 2.1e-29 \| 177.4 \| 287 \| (4, 304) \| 311 \| (63, 392) \| 763 \| Type VI secretion system tip protein VgrG \| Type VI secretion system tip protein VgrG; uniclust \| UniRef100\_A0A356T4I1 \| 99.8 \| 1.2e-23 \| 2.1e-29 \| 172.4 \| 257 \| (17, 286) \| 311 \| (23, 316) \| 500 \| Type VI secretion system tip protein VgrG (Fragment) \| Type VI secretion system tip protein VgrG (Fragment); uniclust \| UniRef100\_A0A126ZFA3 \| 99.8 \| 1.2e-23 \| 2.2e-29 \| 172.9 \| 269 \| (3, 285) \| 311 \| (79, 422) \| 466 \| Type VI secretion protein \| Type VI secretion protein; uniclust \| UniRef100\_A0A653V686 \| 99.8 \| 1.2e-23 \| 2.2e-29 \| 175.9 \| 278 \| (4, 302) \| 311 \| (82, 408) \| 636 \| Gp5/Type VI secretion system Vgr protein OB-fold domain-containing protein \| Gp5/Type VI secretion system Vgr protein OB-fold domain-containing protein; uniclust \| UniRef100\_A0A1I3FF82 \| 99.8 \| 1.3e-23 \| 2.3e-29 \| 161.0 \| 252 \| (37, 301) \| 311 \| (21, 317) \| 318 \| Type VI secretion system secreted protein VgrG \| Type VI secretion system secreted protein VgrG; uniclust \| UniRef100\_UPI000A45C3D4 \| 99.8 \| 1.3e-23 \| 2.4e-29 \| 152.5 \| 223 \| (58, 285) \| 311 \| (4, 244) \| 251 \| type VI secretion system tip protein TssI/VgrG \| type VI secretion system tip protein TssI/VgrG; uniclust \| UniRef100\_A0A369UJ95 \| 99.8 \| 1.4e-23 \| 2.6e-29 \| 184.3 \| 287 \| (4, 303) \| 311 \| (62, 393) \| 1177 \| Type VI secretion system tip protein VgrG \| Type VI secretion system tip protein VgrG; uniclust \| UniRef100\_A0A519EAL2 \| 99.8 \| 1.6e-23 \| 3e-29 \| 173.8 \| 282 \| (4, 302) \| 311 \| (57, 376) \| 568 \| Type VI secretion system tip protein VgrG (Fragment) \| Type VI secretion system tip protein VgrG (Fragment); uniclust \| UniRef100\_A0A084ZAN1 \| 99.8 \| 1.7e-23 \| 3.2e-29 \| 172.9 \| 269 \| (17, 305) \| 311 \| (70, 380) \| 583 \| Rhs element Vgr family protein (Fragment) \| Rhs element Vgr family protein (Fragment); uniclust \| UniRef100\_A0A177QF44 \| 99.8 \| 1.8e-23 \| 3.3e-29 \| 178.4 \| 285 \| (4, 304) \| 311 \| (56, 390) \| 872 \| Gp5/Type VI secretion system Vgr protein OB-fold domain-containing protein \| Gp5/Type VI secretion system Vgr protein OB-fold domain-containing protein; uniclust \| UniRef100\_B8KYT8 \| 99.8 \| 1.9e-23 \| 3.6e-29 \| 166.7 \| 248 \| (4, 265) \| 311 \| (67, 348) \| 387 \| Type VI secretion system Vgr family protein \| Type VI secretion system Vgr family protein; uniclust \| UniRef100\_A0A1H3JED8 \| 99.8 \| 2e-23 \| 3.7e-29 \| 179.8 \| 260 \| (3, 278) \| 311 \| (74, 363) \| 567 \| Rhs element Vgr protein \| Rhs element Vgr protein; uniclust \| UniRef100\_A0A2D9TST2 \| 99.8 \| 2e-23 \| 3.7e-29 \| 183.4 \| 275 \| (17, 306) \| 311 \| (69, 382) \| 1081 \| Gp5/Type VI secretion system Vgr protein OB-fold domain-containing protein \| Gp5/Type VI secretion system Vgr protein OB-fold domain-containing protein; uniclust \| UniRef100\_A0A077FEP7 \| 99.8 \| 2.2e-23 \| 4.1e-29 \| 170.5 \| 272 \| (19, 306) \| 311 \| (69, 382) \| 531 \| Rhs element Vgr protein \| Rhs element Vgr protein; uniclust \| UniRef100\_A0A380U204 \| 99.8 \| 2.2e-23 \| 4.1e-29 \| 182.2 \| 274 \| (17, 305) \| 311 \| (215, 525) \| 1051 \| Extracellular phospholipase A1 \| Extracellular phospholipase A1; uniclust \| UniRef100\_A0A4Y9T3M5 \| 99.8 \| 2.5e-23 \| 4.6e-29 \| 174.7 \| 279 \| (17, 310) \| 311 \| (66, 390) \| 898 \| Type VI secretion system tip protein VgrG \| Type VI secretion system tip protein VgrG; uniclust \| UniRef100\_A0A0F9V267 \| 99.8 \| 2.6e-23 \| 4.9e-29 \| 169.5 \| 274 \| (17, 307) \| 311 \| (73, 395) \| 511 \| Gp5/Type VI secretion system Vgr protein OB-fold domain-containing protein \| Gp5/Type VI secretion system Vgr protein OB-fold domain-containing protein; uniclust \| UniRef100\_A0A0U2P820 \| 99.8 \| 2.9e-23 \| 5.2e-29 \| 179.5 \| 254 \| (17, 285) \| 311 \| (85, 366) \| 692 \| Gp5/Type VI secretion system Vgr protein OB-fold domain-containing protein \| Gp5/Type VI secretion system Vgr protein OB-fold domain-containing protein; uniclust \| UniRef100\_A0A953GUI7 \| 99.8 \| 2.9e-23 \| 5.4e-29 \| 175.1 \| 286 \| (4, 301) \| 311 \| (50, 375) \| 942 \| Type VI secretion system tip protein VgrG \| Type VI secretion system tip protein VgrG; uniclust \| UniRef100\_A0A2L1UY19 \| 99.8 \| 3e-23 \| 5.6e-29 \| 166.3 \| 230 \| (17, 264) \| 311 \| (98, 355) \| 390 \| Type IV secretion protein Rhs \| Type IV secretion protein Rhs; uniclust \| UniRef100\_A0A078M1E5 \| 99.8 \| 3.6e-23 \| 6.6e-29 \| 163.9 \| 214 \| (19, 248) \| 311 \| (99, 336) \| 341 \| Rhs element Vgr protein \| Rhs element Vgr protein; uniclust \| UniRef100\_A0A071M6W2 \| 99.8 \| 4e-23 \| 7.4e-29 \| 191.8 \| 288 \| (4, 306) \| 311 \| (156, 536) \| 1273 \| Type IV secretion protein Rhs \| Type IV secretion protein Rhs; uniclust \| UniRef100\_A0A0D0GEX9 \| 99.8 \| 4.1e-23 \| 7.5e-29 \| 185.5 \| 254 \| (17, 285) \| 311 \| (172, 456) \| 1050 \| Rhs element Vgr family protein \| Rhs element Vgr family protein; uniclust \| UniRef100\_A0A3B8ISA8 \| 99.8 \| 4.3e-23 \| 7.8e-29 \| 167.0 \| 213 \| (2, 230) \| 311 \| (73, 308) \| 311 \| Rhs element Vgr protein (Fragment) \| Rhs element Vgr protein (Fragment); uniclust \| UniRef100\_A0A1V5B929 \| 99.8 \| 4.3e-23 \| 7.9e-29 \| 160.6 \| 259 \| (2, 277) \| 311 \| (61, 350) \| 355 \| Phage late control gene D protein (GPD) \| Phage late control gene D protein (GPD); uniclust \| UniRef100\_A0A024E5Z2 \| 99.8 \| 4.4e-23 \| 8.1e-29 \| 178.4 \| 279 \| (6, 303) \| 311 \| (73, 383) \| 626 \| Putative Rhs element Vgr protein \| Putative Rhs element Vgr protein; uniclust \| UniRef100\_A0A519M875 \| 99.8 \| 4.7e-23 \| 8.6e-29 \| 166.4 \| 259 \| (4, 283) \| 311 \| (64, 353) \| 407 \| Type VI secretion system tip protein VgrG (Fragment) \| Type VI secretion system tip protein VgrG (Fragment); uniclust \| UniRef100\_UPI00027C90E7 \| 99.8 \| 4.7e-23 \| 8.6e-29 \| 168.9 \| 270 \| (1, 277) \| 311 \| (243, 544) \| 563 \| phage tail protein \| phage tail protein; uniclust \| UniRef100\_A0A1Z8A3M6 \| 99.8 \| 4.8e-23 \| 8.8e-29 \| 177.2 \| 259 \| (37, 308) \| 311 \| (90, 387) \| 1207 \| Gp5/Type VI secretion system Vgr protein OB-fold domain-containing protein \| Gp5/Type VI secretion system Vgr protein OB-fold domain-containing protein; uniclust \| UniRef100\_A0A828PCY0 \| 99.8 \| 5.4e-23 \| 9.9e-29 \| 175.2 \| 292 \| (2, 305) \| 311 \| (159, 485) \| 945 \| Type VI secretion system tip protein VgrG \| Type VI secretion system tip protein VgrG; uniclust \| UniRef100\_A0A2G0VNE3 \| 99.8 \| 5.5e-23 \| 1e-28 \| 156.3 \| 124 \| (3, 137) \| 311 \| (72, 206) \| 213 \| Type VI secretion system tip protein VgrG \| Type VI secretion system tip protein VgrG; uniclust \| UniRef100\_UPI0020A7DE59 \| 99.8 \| 6e-23 \| 1.1e-28 \| 165.5 \| 252 \| (17, 281) \| 311 \| (71, 355) \| 579 \| type VI secretion system tip protein TssI/VgrG \| type VI secretion system tip protein TssI/VgrG; uniclust \| UniRef100\_A0A066PIW0 \| 99.8 \| 6e-23 \| 1.1e-28 \| 173.4 \| 247 \| (17, 280) \| 311 \| (111, 380) \| 397 \| Phage late control protein D \| Phage late control protein D; uniclust \| UniRef100\_A0A165W2E7 \| 99.8 \| 6.4e-23 \| 1.2e-28 \| 163.6 \| 266 \| (1, 278) \| 311 \| (50, 322) \| 324 \| Late control protein D \| Late control protein D; uniclust \| UniRef100\_A0A0J6NA85 \| 99.8 \| 6.6e-23 \| 1.2e-28 \| 175.2 \| 285 \| (4, 304) \| 311 \| (69, 424) \| 752 \| Type VI secretion system protein (Fragment) \| Type VI secretion system protein (Fragment); uniclust \| UniRef100\_A0A2N2R031 \| 99.8 \| 6.6e-23 \| 1.2e-28 \| 170.2 \| 265 \| (3, 283) \| 311 \| (86, 407) \| 454 \| Type VI secretion system tip protein VgrG (Fragment) \| Type VI secretion system tip protein VgrG (Fragment); uniclust \| UniRef100\_A0A417HIZ6 \| 99.8 \| 6.7e-23 \| 1.2e-28 \| 162.1 \| 259 \| (3, 276) \| 311 \| (77, 364) \| 367 \| Phage late control D family protein \| Phage late control D family protein; uniclust \| UniRef100\_A0A5N4FUG3 \| 99.8 \| 7.4e-23 \| 1.4e-28 \| 166.5 \| 258 \| (38, 307) \| 311 \| (98, 389) \| 514 \| Type VI secretion system tip protein VgrG (Fragment) \| Type VI secretion system tip protein VgrG (Fragment); uniclust \| UniRef100\_A0A849ZEB7 \| 99.8 \| 7.4e-23 \| 1.4e-28 \| 169.4 \| 290 \| (4, 308) \| 311 \| (84, 503) \| 625 \| Type VI secretion system tip protein VgrG (Fragment) \| Type VI secretion system tip protein VgrG (Fragment); uniclust \| UniRef100\_A0A071LW85 \| 99.8 \| 8e-23 \| 1.5e-28 \| 186.0 \| 250 \| (17, 279) \| 311 \| (142, 438) \| 982 \| Type IV secretion protein Rhs \| Type IV secretion protein Rhs; uniclust \| UniRef100\_A0A158K0S5 \| 99.8 \| 8e-23 \| 1.5e-28 \| 181.6 \| 253 \| (17, 284) \| 311 \| (96, 379) \| 863 \| Rhs element Vgr protein \| Rhs element Vgr protein; uniclust \| UniRef100\_A0A0E3ZX09 \| 99.8 \| 8.2e-23 \| 1.5e-28 \| 172.8 \| 264 \| (4, 285) \| 311 \| (61, 342) \| 576 \| Gp5/Type VI secretion system Vgr protein OB-fold domain-containing protein \| Gp5/Type VI secretion system Vgr protein OB-fold domain-containing protein; uniclust \| UniRef100\_A0A147GLC0 \| 99.8 \| 8.3e-23 \| 1.5e-28 \| 165.9 \| 239 \| (17, 267) \| 311 \| (118, 389) \| 398 \| Type VI secretion protein ImpA (Fragment) \| Type VI secretion protein ImpA (Fragment); uniclust \| UniRef100\_A0A010SYM9 \| 99.8 \| 8.6e-23 \| 1.6e-28 \| 172.5 \| 258 \| (7, 279) \| 311 \| (126, 419) \| 431 \| Type IV secretion protein Rhs \| Type IV secretion protein Rhs; uniclust \| UniRef100\_A0A069PJ07 \| 99.8 \| 8.9e-23 \| 1.6e-28 \| 187.6 \| 253 \| (18, 285) \| 311 \| (145, 424) \| 988 \| Type IV secretion protein Rhs \| Type IV secretion protein Rhs; uniclust \| UniRef100\_A0A0G3BGP6 \| 99.8 \| 8.9e-23 \| 1.6e-28 \| 179.4 \| 290 \| (5, 306) \| 311 \| (89, 420) \| 1041 \| Type VI secretion system tip protein VgrG \| Type VI secretion system tip protein VgrG; uniclust \| UniRef100\_A0A7W0Y6A5 \| 99.8 \| 9.5e-23 \| 1.7e-28 \| 158.4 \| 262 \| (3, 277) \| 311 \| (54, 351) \| 356 \| Phage late control D family protein \| Phage late control D family protein; uniclust \| UniRef100\_A0A080M199 \| 99.8 \| 1e-22 \| 1.8e-28 \| 169.3 \| 262 \| (6, 282) \| 311 \| (79, 370) \| 398 \| Phage protein D \| Phage protein D; uniclust \| UniRef100\_A0A1U7N292 \| 99.8 \| 1e-22 \| 1.9e-28 \| 165.1 \| 250 \| (3, 277) \| 311 \| (66, 337) \| 347 \| Phage protein D \| Phage protein D; uniclust \| UniRef100\_UPI00056E02FB \| 99.8 \| 1.1e-22 \| 2e-28 \| 157.4 \| 246 \| (4, 262) \| 311 \| (50, 329) \| 340 \| type VI secretion system tip protein TssI/VgrG \| type VI secretion system tip protein TssI/VgrG; uniclust \| UniRef100\_A0A954DSN9 \| 99.8 \| 1.1e-22 \| 2.1e-28 \| 171.4 \| 285 \| (4, 305) \| 311 \| (276, 607) \| 923 \| Type VI secretion system tip protein VgrG \| Type VI secretion system tip protein VgrG; uniclust \| UniRef100\_A0A1H4H7U1 \| 99.8 \| 1.4e-22 \| 2.6e-28 \| 177.7 \| 277 \| (4, 300) \| 311 \| (94, 406) \| 643 \| Uncharacterized conserved protein, implicated in type VI secretion and phage assembly \| Uncharacterized conserved protein, implicated in type VI secretion and phage assembly; uniclust \| UniRef100\_A0A0J6RCQ0 \| 99.8 \| 1.4e-22 \| 2.6e-28 \| 173.8 \| 275 \| (17, 304) \| 311 \| (85, 399) \| 647 \| Gp5/Type VI secretion system Vgr protein OB-fold domain-containing protein (Fragment) \| Gp5/Type VI secretion system Vgr protein OB-fold domain-containing protein (Fragment); uniclust \| UniRef100\_A0A9D5K2R1 \| 99.8 \| 1.5e-22 \| 2.8e-28 \| 149.0 \| 225 \| (58, 286) \| 311 \| (3, 249) \| 275 \| Type VI secretion system tip protein VgrG \| Type VI secretion system tip protein VgrG; uniclust \| UniRef100\_A0A0A0EXH8 \| 99.8 \| 1.6e-22 \| 2.9e-28 \| 174.0 \| 284 \| (2, 304) \| 311 \| (67, 404) \| 668 \| Gp5/Type VI secretion system Vgr protein OB-fold domain-containing protein (Fragment) \| Gp5/Type VI secretion system Vgr protein OB-fold domain-containing protein (Fragment); uniclust \| UniRef100\_A0A2X1MAT3 \| 99.8 \| 1.6e-22 \| 2.9e-28 \| 163.4 \| 174 \| (1, 181) \| 311 \| (63, 237) \| 280 \| Phage protein D \| Phage protein D; uniclust \| UniRef100\_UPI0003D2A615 \| 99.8 \| 1.7e-22 \| 3.2e-28 \| 156.5 \| 255 \| (1, 262) \| 311 \| (12, 309) \| 310 \| contractile injection system protein, VgrG/Pvc8 family \| contractile injection system protein, VgrG/Pvc8 family; uniclust \| UniRef100\_A0A1A9VKH9 \| 99.8 \| 1.8e-22 \| 3.3e-28 \| 158.9 \| 268 \| (7, 281) \| 311 \| (49, 317) \| 462 \| Uncharacterized protein \| Uncharacterized protein; uniclust \| UniRef100\_A0A0B1XTW2 \| 99.8 \| 1.8e-22 \| 3.3e-28 \| 177.6 \| 287 \| (3, 303) \| 311 \| (98, 442) \| 1005 \| Type VI secretion protein (Fragment) \| Type VI secretion protein (Fragment); uniclust \| UniRef100\_A0A1X3D1R8 \| 99.8 \| 1.9e-22 \| 3.4e-28 \| 169.3 \| 274 \| (17, 303) \| 311 \| (94, 399) \| 534 \| Gp5/Type VI secretion system Vgr protein OB-fold domain-containing protein (Fragment) \| Gp5/Type VI secretion system Vgr protein OB-fold domain-containing protein (Fragment); uniclust \| UniRef100\_UPI0003807C1F \| 99.8 \| 2e-22 \| 3.7e-28 \| 179.5 \| 280 \| (17, 307) \| 311 \| (68, 386) \| 1399 \| type VI secretion system tip protein TssI/VgrG \| type VI secretion system tip protein TssI/VgrG; uniclust \| UniRef100\_A0A1H4SWS9 \| 99.8 \| 2.1e-22 \| 3.9e-28 \| 174.9 \| 254 \| (38, 304) \| 311 \| (89, 380) \| 1019 \| Rhs element Vgr protein \| Rhs element Vgr protein; uniclust \| UniRef100\_A0A061NEY3 \| 99.8 \| 2.3e-22 \| 4.1e-28 \| 172.2 \| 252 \| (3, 278) \| 311 \| (79, 346) \| 432 \| Phage-like element PBSX protein XkdQ \| Phage-like element PBSX protein XkdQ; uniclust \| UniRef100\_A0A534PM04 \| 99.8 \| 2.3e-22 \| 4.3e-28 \| 163.6 \| 258 \| (6, 278) \| 311 \| (53, 336) \| 565 \| Phage late control D family protein \| Phage late control D family protein; uniclust \| UniRef100\_A0A8X6GUF6 \| 99.8 \| 2.4e-22 \| 4.3e-28 \| 160.7 \| 264 \| (8, 278) \| 311 \| (73, 337) \| 529 \| Phage tail protein \| Phage tail protein; uniclust \| UniRef100\_A0A017T573 \| 99.8 \| 2.6e-22 \| 4.8e-28 \| 182.4 \| 273 \| (17, 304) \| 311 \| (95, 421) \| 956 \| VgrG protein \| VgrG protein; uniclust \| UniRef100\_UPI000AD481B1 \| 99.8 \| 2.8e-22 \| 5.1e-28 \| 149.0 \| 207 \| (58, 268) \| 311 \| (8, 235) \| 251 \| type VI secretion system tip protein VgrG \| type VI secretion system tip protein VgrG; uniclust \| UniRef100\_A0A085ULB1 \| 99.8 \| 2.8e-22 \| 5.1e-28 \| 152.2 \| 201 \| (56, 265) \| 311 \| (27, 250) \| 263 \| Type IV secretion protein Rhs (Fragment) \| Type IV secretion protein Rhs (Fragment); uniclust \| UniRef100\_A0A250KKR6 \| 99.8 \| 3e-22 \| 5.6e-28 \| 168.5 \| 279 \| (4, 304) \| 311 \| (59, 369) \| 593 \| Type IV secretion protein Rhs \| Type IV secretion protein Rhs; uniclust \| UniRef100\_A0A0N0D4A4 \| 99.8 \| 3.1e-22 \| 5.7e-28 \| 156.3 \| 167 \| (2, 183) \| 311 \| (67, 250) \| 301 \| Rhs element Vgr protein, subgroup (Fragment) \| Rhs element Vgr protein, subgroup (Fragment); uniclust \| UniRef100\_A0A071M8E6 \| 99.8 \| 3.2e-22 \| 5.9e-28 \| 183.1 \| 285 \| (4, 306) \| 311 \| (122, 491) \| 1175 \| Rhs element Vgr protein \| Rhs element Vgr protein; uniclust \| UniRef100\_A0A136Q991 \| 99.8 \| 3.2e-22 \| 5.9e-28 \| 156.3 \| 211 \| (1, 221) \| 311 \| (55, 272) \| 292 \| Phage late control protein (Fragment) \| Phage late control protein (Fragment); uniclust \| UniRef100\_A0A380U1Z0 \| 99.8 \| 3.6e-22 \| 6.7e-28 \| 181.9 \| 273 \| (17, 305) \| 311 \| (199, 507) \| 2004 \| Cell wall-associated polypeptide CWBP200 \| Cell wall-associated polypeptide CWBP200; uniclust \| UniRef100\_A0A105DDV7 \| 99.8 \| 4.6e-22 \| 8.5e-28 \| 184.0 \| 272 \| (17, 303) \| 311 \| (130, 434) \| 1296 \| Type IV secretion protein Rhs \| Type IV secretion protein Rhs; uniclust \| UniRef100\_A0A069NIW4 \| 99.8 \| 5.5e-22 \| 1e-27 \| 172.3 \| 276 \| (17, 304) \| 311 \| (103, 430) \| 789 \| Type VI secretion protein ImpA (Fragment) \| Type VI secretion protein ImpA (Fragment); uniclust \| UniRef100\_A0A0T7DUV3 \| 99.8 \| 5.6e-22 \| 1e-27 \| 161.6 \| 258 \| (1, 277) \| 311 \| (69, 342) \| 344 \| Putative bacteriophage regulatory protein \| Putative bacteriophage regulatory protein; uniclust \| UniRef100\_A0A0D0MWE3 \| 99.8 \| 5.6e-22 \| 1e-27 \| 168.6 \| 285 \| (4, 303) \| 311 \| (96, 442) \| 636 \| Gp5/Type VI secretion system Vgr protein OB-fold domain-containing protein (Fragment) \| Gp5/Type VI secretion system Vgr protein OB-fold domain-containing protein (Fragment); uniclust \| UniRef100\_A0A0H2LUU0 \| 99.8 \| 5.7e-22 \| 1e-27 \| 178.7 \| 236 \| (17, 265) \| 311 \| (72, 336) \| 1189 \| Phage-related baseplate assembly protein \| Phage-related baseplate assembly protein; uniclust \| UniRef100\_A0A1Q7M720 \| 99.8 \| 6e-22 \| 1.1e-27 \| 159.6 \| 259 \| (3, 281) \| 311 \| (88, 365) \| 448 \| Phage late control D family protein \| Phage late control D family protein; uniclust \| UniRef100\_A0A354P3G0 \| 99.8 \| 6.4e-22 \| 1.2e-27 \| 175.1 \| 270 \| (17, 301) \| 311 \| (101, 446) \| 944 \| Gp5/Type VI secretion system Vgr protein OB-fold domain-containing protein \| Gp5/Type VI secretion system Vgr protein OB-fold domain-containing protein; uniclust \| UniRef100\_A0A2D4SL60 \| 99.8 \| 6.6e-22 \| 1.2e-27 \| 158.0 \| 263 \| (3, 282) \| 311 \| (61, 347) \| 355 \| Phage late control D family protein \| Phage late control D family protein; uniclust \| UniRef100\_A0A011TTI3 \| 99.8 \| 7.1e-22 \| 1.3e-27 \| 161.3 \| 274 \| (1, 287) \| 311 \| (68, 353) \| 397 \| Late control D family protein \| Late control D family protein; uniclust \| UniRef100\_UPI00129B7A46 \| 99.8 \| 7.4e-22 \| 1.4e-27 \| 151.9 \| 232 \| (17, 265) \| 311 \| (10, 271) \| 286 \| type VI secretion system tip protein TssI/VgrG \| type VI secretion system tip protein TssI/VgrG; uniclust \| UniRef100\_A0A023Y2E8 \| 99.8 \| 7.8e-22 \| 1.4e-27 \| 184.5 \| 288 \| (4, 303) \| 311 \| (140, 520) \| 1157 \| Type IV secretion protein Rhs \| Type IV secretion protein Rhs; uniclust \| UniRef100\_A0A0S3TZS7 \| 99.8 \| 8e-22 \| 1.5e-27 \| 166.9 \| 244 \| (18, 278) \| 311 \| (117, 394) \| 810 \| Rhs element Vgr protein \| Rhs element Vgr protein; uniclust \| UniRef100\_A0A4U1HWQ4 \| 99.8 \| 9e-22 \| 1.6e-27 \| 165.4 \| 256 \| (37, 307) \| 311 \| (92, 376) \| 643 \| Type VI secretion system tip protein VgrG \| Type VI secretion system tip protein VgrG; uniclust \| UniRef100\_A0A0Q8WLP0 \| 99.8 \| 9.3e-22 \| 1.7e-27 \| 165.8 \| 276 \| (17, 307) \| 311 \| (76, 392) \| 558 \| Type VI secretion protein VgrG \| Type VI secretion protein VgrG; uniclust \| UniRef100\_A0A1Y5GX49 \| 99.8 \| 9.5e-22 \| 1.7e-27 \| 177.3 \| 276 \| (17, 307) \| 311 \| (73, 385) \| 1309 \| Type VI secretion system tip protein VgrG \| Type VI secretion system tip protein VgrG; uniclust \| UniRef100\_A0A518AZS0 \| 99.8 \| 9.8e-22 \| 1.8e-27 \| 174.6 \| 283 \| (4, 302) \| 311 \| (59, 408) \| 1054 \| Phage-related baseplate assembly protein \| Phage-related baseplate assembly protein; uniclust \| UniRef100\_A0A081C212 \| 99.8 \| 1.1e-21 \| 2e-27 \| 161.9 \| 255 \| (3, 279) \| 311 \| (84, 361) \| 371 \| Uncharacterized protein \| Uncharacterized protein; uniclust \| UniRef100\_F2NUD2 \| 99.8 \| 1.1e-21 \| 2e-27 \| 165.0 \| 259 \| (3, 277) \| 311 \| (63, 348) \| 619 \| Uncharacterized protein \| Uncharacterized protein; uniclust \| UniRef100\_A0A0X8C270 \| 99.8 \| 1.1e-21 \| 2e-27 \| 182.0 \| 277 \| (4, 302) \| 311 \| (101, 412) \| 1104 \| Gp5/Type VI secretion system Vgr protein OB-fold domain-containing protein \| Gp5/Type VI secretion system Vgr protein OB-fold domain-containing protein; uniclust \| UniRef100\_A0A080IYI6 \| 99.8 \| 1.1e-21 \| 2.1e-27 \| 162.1 \| 291 \| (2, 304) \| 311 \| (154, 479) \| 646 \| Rhs element Vgr family protein \| Rhs element Vgr family protein; uniclust \| UniRef100\_A0A4R4ILG7 \| 99.8 \| 1.2e-21 \| 2.3e-27 \| 158.0 \| 256 \| (37, 305) \| 311 \| (112, 401) \| 496 \| Type VI secretion system tip protein VgrG (Fragment) \| Type VI secretion system tip protein VgrG (Fragment); uniclust \| UniRef100\_A0A849Z2N6 \| 99.7 \| 1.4e-21 \| 2.6e-27 \| 158.1 \| 257 \| (36, 304) \| 311 \| (92, 375) \| 492 \| Type VI secretion system tip protein VgrG (Fragment) \| Type VI secretion system tip protein VgrG (Fragment); uniclust \| UniRef100\_A0A0J8DB12 \| 99.7 \| 1.4e-21 \| 2.6e-27 \| 161.8 \| 253 \| (2, 281) \| 311 \| (71, 327) \| 330 \| Uncharacterized protein \| Uncharacterized protein; uniclust \| UniRef100\_A0A2V8RQE1 \| 99.7 \| 1.5e-21 \| 2.7e-27 \| 151.8 \| 247 \| (4, 265) \| 311 \| (62, 354) \| 360 \| Type VI secretion system tip protein VgrG (Fragment) \| Type VI secretion system tip protein VgrG (Fragment); uniclust \| UniRef100\_UPI001ED99487 \| 99.7 \| 1.5e-21 \| 2.7e-27 \| 170.6 \| 287 \| (4, 305) \| 311 \| (759, 1086) \| 1451 \| type VI secretion system tip protein TssI/VgrG \| type VI secretion system tip protein TssI/VgrG; uniclust \| UniRef100\_UPI001ED99487 \| 99.7 \| 1.5e-21 \| 2.7e-27 \| 170.6 \| 287 \| (4, 305) \| 311 \| (197, 524) \| 1451 \| type VI secretion system tip protein TssI/VgrG \| type VI secretion system tip protein TssI/VgrG; uniclust \| UniRef100\_A0A023Y2A2 \| 99.7 \| 1.5e-21 \| 2.7e-27 \| 177.8 \| 284 \| (2, 303) \| 311 \| (279, 612) \| 1207 \| Type IV secretion protein Rhs \| Type IV secretion protein Rhs; uniclust \| UniRef100\_A0A2E7ADA3 \| 99.7 \| 1.5e-21 \| 2.8e-27 \| 168.6 \| 275 \| (16, 303) \| 311 \| (67, 374) \| 975 \| Gp5/Type VI secretion system Vgr protein OB-fold domain-containing protein \| Gp5/Type VI secretion system Vgr protein OB-fold domain-containing protein; uniclust \| UniRef100\_A0A2D6B585 \| 99.7 \| 1.6e-21 \| 3e-27 \| 164.6 \| 287 \| (5, 303) \| 311 \| (266, 607) \| 917 \| Type VI secretion system tip protein VgrG \| Type VI secretion system tip protein VgrG; uniclust \| UniRef100\_A0A150PRS2 \| 99.7 \| 1.6e-21 \| 3e-27 \| 169.3 \| 288 \| (4, 305) \| 311 \| (58, 382) \| 732 \| Gp5/Type VI secretion system Vgr protein OB-fold domain-containing protein (Fragment) \| Gp5/Type VI secretion system Vgr protein OB-fold domain-containing protein (Fragment); uniclust \| UniRef100\_A0A1H0U3P0 \| 99.7 \| 1.6e-21 \| 3e-27 \| 143.2 \| 123 \| (4, 135) \| 311 \| (50, 180) \| 204 \| Rhs element Vgr protein (Fragment) \| Rhs element Vgr protein (Fragment); uniclust \| UniRef100\_A0A2X3JV88 \| 99.7 \| 1.6e-21 \| 3e-27 \| 150.5 \| 252 \| (37, 305) \| 311 \| (31, 321) \| 337 \| Rhs element Vgr protein \| Rhs element Vgr protein; uniclust \| UniRef100\_A0A068T9R7 \| 99.7 \| 1.7e-21 \| 3.1e-27 \| 154.8 \| 272 \| (2, 281) \| 311 \| (58, 339) \| 355 \| Phage late control D family protein \| Phage late control D family protein; uniclust \| UniRef100\_A0A016XHX9 \| 99.7 \| 1.7e-21 \| 3.2e-27 \| 169.5 \| 264 \| (3, 282) \| 311 \| (89, 375) \| 432 \| Baseplate protein \| Baseplate protein; uniclust \| UniRef100\_A0A2A4HGW6 \| 99.7 \| 1.8e-21 \| 3.4e-27 \| 153.1 \| 250 \| (17, 283) \| 311 \| (68, 357) \| 370 \| Type VI secretion system tip protein VgrG (Fragment) \| Type VI secretion system tip protein VgrG (Fragment); uniclust \| UniRef100\_UPI0018EF83E4 \| 99.7 \| 2e-21 \| 3.7e-27 \| 162.7 \| 257 \| (3, 277) \| 311 \| (262, 556) \| 834 \| VgrG-related protein \| VgrG-related protein; uniclust \| UniRef100\_A0A010RVC8 \| 99.7 \| 2.1e-21 \| 3.9e-27 \| 175.4 \| 274 \| (18, 303) \| 311 \| (83, 417) \| 709 \| Type IV secretion protein Rhs \| Type IV secretion protein Rhs; uniclust \| UniRef100\_A0A7J6YLS0 \| 99.7 \| 2.2e-21 \| 4e-27 \| 162.6 \| 270 \| (7, 283) \| 311 \| (73, 343) \| 756 \| phospholipase A2 \| phospholipase A2; uniclust \| UniRef100\_A0A351G4S7 \| 99.7 \| 2.3e-21 \| 4.3e-27 \| 149.0 \| 247 \| (17, 278) \| 311 \| (85, 362) \| 374 \| Phage late control D family protein \| Phage late control D family protein; uniclust \| UniRef100\_A0A024L6E0 \| 99.7 \| 2.7e-21 \| 4.9e-27 \| 164.1 \| 260 \| (17, 308) \| 311 \| (81, 383) \| 723 \| Rhs element Vgr protein \| Rhs element Vgr protein; uniclust \| UniRef100\_A0A3B0ZLP7 \| 99.7 \| 3e-21 \| 5.5e-27 \| 170.5 \| 274 \| (16, 303) \| 311 \| (81, 392) \| 1041 \| Gp5/Type VI secretion system Vgr protein OB-fold domain-containing protein \| Gp5/Type VI secretion system Vgr protein OB-fold domain-containing protein; uniclust \| UniRef100\_A0A0F2QXK9 \| 99.7 \| 3.1e-21 \| 5.7e-27 \| 152.0 \| 213 \| (19, 248) \| 311 \| (85, 321) \| 347 \| Type IV secretion protein Rhs (Fragment) \| Type IV secretion protein Rhs (Fragment); uniclust \| UniRef100\_A0A1Q8BNB2 \| 99.7 \| 3.1e-21 \| 5.8e-27 \| 160.0 \| 258 \| (6, 278) \| 311 \| (64, 347) \| 627 \| Gp5/Type VI secretion system Vgr protein OB-fold domain-containing protein \| Gp5/Type VI secretion system Vgr protein OB-fold domain-containing protein; uniclust \| UniRef100\_A0A074V5I6 \| 99.7 \| 3.9e-21 \| 7.1e-27 \| 167.4 \| 273 \| (19, 307) \| 311 \| (72, 420) \| 847 \| Gp5/Type VI secretion system Vgr protein OB-fold domain-containing protein \| Gp5/Type VI secretion system Vgr protein OB-fold domain-containing protein; uniclust \| UniRef100\_A0A7V8K0W3 \| 99.7 \| 4e-21 \| 7.3e-27 \| 167.1 \| 277 \| (17, 305) \| 311 \| (91, 401) \| 1332 \| Actin cross-linking toxin VgrG1 \| Actin cross-linking toxin VgrG1; uniclust \| UniRef100\_A0A349NY43 \| 99.7 \| 4e-21 \| 7.3e-27 \| 147.8 \| 240 \| (2, 257) \| 311 \| (60, 326) \| 328 \| Rhs element Vgr protein (Fragment) \| Rhs element Vgr protein (Fragment); uniclust \| UniRef100\_UPI001FCF2336 \| 99.7 \| 4.2e-21 \| 7.6e-27 \| 165.4 \| 292 \| (2, 305) \| 311 \| (591, 917) \| 1169 \| type VI secretion system membrane subunit TssM \| type VI secretion system membrane subunit TssM; uniclust \| UniRef100\_A0A661NEM7 \| 99.7 \| 4.3e-21 \| 7.9e-27 \| 157.4 \| 278 \| (17, 308) \| 311 \| (69, 376) \| 561 \| Type VI secretion system tip protein VgrG (Fragment) \| Type VI secretion system tip protein VgrG (Fragment); uniclust \| UniRef100\_A0A0Q7MJZ9 \| 99.7 \| 5e-21 \| 9.2e-27 \| 159.6 \| 266 \| (4, 283) \| 311 \| (60, 360) \| 372 \| Phage tail protein \| Phage tail protein; uniclust \| UniRef100\_A0A126UXB1 \| 99.7 \| 5.3e-21 \| 9.7e-27 \| 161.4 \| 208 \| (94, 305) \| 311 \| (12, 253) \| 540 \| Gp5/Type VI secretion system Vgr protein OB-fold domain-containing protein \| Gp5/Type VI secretion system Vgr protein OB-fold domain-containing protein; uniclust \| UniRef100\_A0A318UXY0 \| 99.7 \| 5.4e-21 \| 9.9e-27 \| 167.5 \| 274 \| (19, 306) \| 311 \| (79, 400) \| 1054 \| Rhs element Vgr protein (Fragment) \| Rhs element Vgr protein (Fragment); uniclust \| UniRef100\_A0A1E4W345 \| 99.7 \| 5.7e-21 \| 1e-26 \| 162.5 \| 208 \| (96, 306) \| 311 \| (2, 243) \| 580 \| Type VI secretion protein ImpA (Fragment) \| Type VI secretion protein ImpA (Fragment); uniclust \| UniRef100\_A0A059USW8 \| 99.7 \| 6.5e-21 \| 1.2e-26 \| 168.0 \| 279 \| (17, 307) \| 311 \| (92, 426) \| 855 \| Rhs element Vgr protein \| Rhs element Vgr protein; uniclust \| UniRef100\_A0A7Y4J6W0 \| 99.7 \| 6.8e-21 \| 1.3e-26 \| 137.1 \| 109 \| (17, 134) \| 311 \| (12, 126) \| 185 \| Type VI secretion system tip protein VgrG (Fragment) \| Type VI secretion system tip protein VgrG (Fragment); uniclust \| UniRef100\_A0A1H2PKZ2 \| 99.7 \| 7e-21 \| 1.3e-26 \| 160.3 \| 258 \| (17, 288) \| 311 \| (77, 378) \| 609 \| Type VI secretion system secreted protein VgrG \| Type VI secretion system secreted protein VgrG; uniclust \| UniRef100\_A0A017TA00 \| 99.7 \| 7.4e-21 \| 1.4e-26 \| 168.0 \| 276 \| (17, 309) \| 311 \| (68, 391) \| 945 \| VgrG protein \| VgrG protein; uniclust \| UniRef100\_A0A1Y2SPT5 \| 99.7 \| 7.7e-21 \| 1.4e-26 \| 155.8 \| 277 \| (17, 310) \| 311 \| (74, 384) \| 520 \| Rhs element Vgr family protein \| Rhs element Vgr family protein; uniclust \| UniRef100\_A0A356TFU1 \| 99.7 \| 8.2e-21 \| 1.5e-26 \| 164.5 \| 291 \| (5, 305) \| 311 \| (250, 581) \| 934 \| Gp5/Type VI secretion system Vgr protein OB-fold domain-containing protein \| Gp5/Type VI secretion system Vgr protein OB-fold domain-containing protein; uniclust \| UniRef100\_A0A1E4LS02 \| 99.7 \| 9.4e-21 \| 1.7e-26 \| 149.1 \| 260 \| (18, 284) \| 311 \| (2, 269) \| 317 \| Late control protein (Fragment) \| Late control protein (Fragment); uniclust \| UniRef100\_A0A4U7JJ27 \| 99.7 \| 9.8e-21 \| 1.8e-26 \| 154.8 \| 262 \| (3, 279) \| 311 \| (91, 377) \| 639 \| Gp5/Type VI secretion system Vgr protein OB-fold domain-containing protein \| Gp5/Type VI secretion system Vgr protein OB-fold domain-containing protein; uniclust \| UniRef100\_A0A086CZD6 \| 99.7 \| 1.1e-20 \| 2e-26 \| 166.3 \| 286 \| (5, 306) \| 311 \| (82, 450) \| 859 \| Type IV secretion protein Rhs (Fragment) \| Type IV secretion protein Rhs (Fragment); uniclust \| UniRef100\_A0A158G169 \| 99.7 \| 1.1e-20 \| 2e-26 \| 156.1 \| 268 \| (19, 302) \| 311 \| (101, 400) \| 486 \| Rhs element Vgr protein \| Rhs element Vgr protein; uniclust \| UniRef100\_A0A563DMM3 \| 99.7 \| 1.1e-20 \| 2.1e-26 \| 153.5 \| 257 \| (4, 284) \| 311 \| (98, 398) \| 453 \| Uncharacterized protein (Fragment) \| Uncharacterized protein (Fragment); uniclust \| UniRef100\_A0A7S7SKC8 \| 99.7 \| 1.1e-20 \| 2.1e-26 \| 160.0 \| 274 \| (17, 302) \| 311 \| (72, 384) \| 937 \| Type VI secretion system tip protein VgrG \| Type VI secretion system tip protein VgrG; uniclust \| UniRef100\_A0A376JF19 \| 99.7 \| 1.2e-20 \| 2.2e-26 \| 149.1 \| 172 \| (1, 185) \| 311 \| (74, 246) \| 295 \| Phage late control D family protein \| Phage late control D family protein; uniclust \| UniRef100\_A0A812QV43 \| 99.7 \| 1.2e-20 \| 2.2e-26 \| 162.0 \| 266 \| (1, 273) \| 311 \| (734, 1005) \| 1104 \| GpFI protein \| GpFI protein; uniclust \| UniRef100\_UPI00142415E8 \| 99.7 \| 1.3e-20 \| 2.3e-26 \| 159.2 \| 290 \| (5, 307) \| 311 \| (59, 399) \| 901 \| type VI secretion system tip protein TssI/VgrG \| type VI secretion system tip protein TssI/VgrG; uniclust \| UniRef100\_UPI00079AD8BF \| 99.7 \| 1.3e-20 \| 2.5e-26 \| 154.7 \| 270 \| (1, 277) \| 311 \| (362, 658) \| 665 \| phage major capsid protein, P2 family \| phage major capsid protein, P2 family; uniclust \| UniRef100\_A0A0N8P2U1 \| 99.7 \| 1.4e-20 \| 2.5e-26 \| 147.5 \| 179 \| (1, 186) \| 311 \| (59, 238) \| 267 \| Uncharacterized protein (Fragment) \| Uncharacterized protein (Fragment); uniclust \| UniRef100\_A0A3M1GYX0 \| 99.7 \| 1.6e-20 \| 2.9e-26 \| 145.4 \| 265 \| (3, 282) \| 311 \| (60, 356) \| 389 \| Phage late control D family protein (Fragment) \| Phage late control D family protein (Fragment); uniclust \| UniRef100\_UPI000B4AB36B \| 99.7 \| 1.7e-20 \| 3.1e-26 \| 153.2 \| 286 \| (4, 305) \| 311 \| (125, 489) \| 625 \| type VI secretion system tip protein TssI/VgrG \| type VI secretion system tip protein TssI/VgrG; uniclust \| UniRef100\_UPI0005AAC8F8 \| 99.7 \| 1.8e-20 \| 3.3e-26 \| 150.1 \| 256 \| (18, 287) \| 311 \| (52, 334) \| 432 \| type VI secretion system tip protein VgrG \| type VI secretion system tip protein VgrG; uniclust \| UniRef100\_A0A109PY16 \| 99.7 \| 1.9e-20 \| 3.4e-26 \| 155.4 \| 254 \| (4, 277) \| 311 \| (94, 379) \| 493 \| Gp5/Type VI secretion system Vgr protein OB-fold domain-containing protein \| Gp5/Type VI secretion system Vgr protein OB-fold domain-containing protein; uniclust \| UniRef100\_UPI000AEA10C1 \| 99.7 \| 2e-20 \| 3.6e-26 \| 137.5 \| 209 \| (53, 268) \| 311 \| (6, 240) \| 255 \| type VI secretion system tip protein TssI/VgrG \| type VI secretion system tip protein TssI/VgrG; uniclust \| UniRef100\_UPI0009F9D285 \| 99.7 \| 2e-20 \| 3.7e-26 \| 163.8 \| 276 \| (17, 305) \| 311 \| (67, 377) \| 1451 \| type VI secretion system tip protein TssI/VgrG \| type VI secretion system tip protein TssI/VgrG; uniclust \| UniRef100\_A0A379AXR7 \| 99.7 \| 2.1e-20 \| 3.8e-26 \| 158.7 \| 272 \| (17, 305) \| 311 \| (80, 397) \| 723 \| Uncharacterized protein conserved in bacteria \| Uncharacterized protein conserved in bacteria; uniclust \| UniRef100\_A0A840P1G0 \| 99.7 \| 2.1e-20 \| 3.8e-26 \| 155.9 \| 265 \| (1, 280) \| 311 \| (499, 769) \| 775 \| Phage protein D \| Phage protein D; uniclust \| UniRef100\_A0A451AF14 \| 99.7 \| 2.1e-20 \| 3.8e-26 \| 137.1 \| 202 \| (22, 230) \| 311 \| (2, 208) \| 211 \| Phage late control gene D protein (GPD) \| Phage late control gene D protein (GPD); uniclust \| UniRef100\_A0A930IWN4 \| 99.7 \| 2.1e-20 \| 3.9e-26 \| 148.4 \| 277 \| (5, 302) \| 311 \| (60, 369) \| 398 \| Uncharacterized protein \| Uncharacterized protein; uniclust \| UniRef100\_A0A4U9VH05 \| 99.7 \| 2.2e-20 \| 4e-26 \| 153.1 \| 277 \| (17, 310) \| 311 \| (69, 378) \| 473 \| Uncharacterized protein conserved in bacteria \| Uncharacterized protein conserved in bacteria; uniclust \| UniRef100\_A0A0J5KJT2 \| 99.7 \| 2.3e-20 \| 4.3e-26 \| 159.9 \| 235 \| (65, 305) \| 311 \| (1, 269) \| 787 \| Type IV secretion protein Rhs (Fragment) \| Type IV secretion protein Rhs (Fragment); uniclust \| UniRef100\_UPI0003754FB8 \| 99.7 \| 2.4e-20 \| 4.4e-26 \| 147.5 \| 239 \| (5, 265) \| 311 \| (62, 330) \| 335 \| contractile injection system protein, VgrG/Pvc8 family \| contractile injection system protein, VgrG/Pvc8 family; uniclust \| UniRef100\_A0A0U1PQF4 \| 99.7 \| 2.5e-20 \| 4.5e-26 \| 166.7 \| 249 \| (18, 281) \| 311 \| (93, 379) \| 930 \| Type IV secretion protein Rhs (Fragment) \| Type IV secretion protein Rhs (Fragment); uniclust \| UniRef100\_A0A4Q1SII8 \| 99.7 \| 2.5e-20 \| 4.6e-26 \| 160.0 \| 261 \| (4, 285) \| 311 \| (101, 390) \| 596 \| Type VI secretion system tip protein VgrG \| Type VI secretion system tip protein VgrG; uniclust \| UniRef100\_A0A7K0IVA3 \| 99.7 \| 2.5e-20 \| 4.7e-26 \| 155.9 \| 282 \| (7, 305) \| 311 \| (80, 400) \| 804 \| Type VI secretion system tip protein VgrG \| Type VI secretion system tip protein VgrG; uniclust \| UniRef100\_A0A0Q9QUG3 \| 99.7 \| 2.6e-20 \| 4.7e-26 \| 152.1 \| 286 \| (2, 304) \| 311 \| (86, 428) \| 474 \| Uncharacterized protein \| Uncharacterized protein; uniclust \| UniRef100\_A0A916CQA2 \| 99.7 \| 2.6e-20 \| 4.8e-26 \| 159.5 \| 273 \| (17, 304) \| 311 \| (81, 393) \| 949 \| Actin cross-linking toxin VgrG1 \| Actin cross-linking toxin VgrG1; uniclust \| UniRef100\_A0A3B0Z0G3 \| 99.7 \| 2.7e-20 \| 4.9e-26 \| 166.6 \| 266 \| (4, 284) \| 311 \| (64, 376) \| 913 \| VgrG protein \| VgrG protein; uniclust \| UniRef100\_A0A1Q7URD9 \| 99.7 \| 2.8e-20 \| 5.1e-26 \| 142.1 \| 257 \| (6, 277) \| 311 \| (58, 340) \| 350 \| Phage late control D family protein (Fragment) \| Phage late control D family protein (Fragment); uniclust \| UniRef100\_A0A956DL55 \| 99.7 \| 3.2e-20 \| 5.8e-26 \| 143.1 \| 245 \| (57, 305) \| 311 \| (104, 370) \| 374 \| Type VI secretion system tip protein VgrG (Fragment) \| Type VI secretion system tip protein VgrG (Fragment); uniclust \| UniRef100\_A0A376W2Q8 \| 99.7 \| 3.2e-20 \| 5.9e-26 \| 141.2 \| 228 \| (1, 235) \| 311 \| (13, 271) \| 273 \| Late control gene D protein from prophage \| Late control gene D protein from prophage; uniclust \| UniRef100\_UPI00124E8932 \| 99.7 \| 3.3e-20 \| 6.1e-26 \| 161.6 \| 288 \| (5, 304) \| 311 \| (209, 542) \| 999 \| type VI secretion system tip protein TssI/VgrG \| type VI secretion system tip protein TssI/VgrG; uniclust \| UniRef100\_UPI0006494E4F \| 99.7 \| 3.3e-20 \| 6.1e-26 \| 137.1 \| 229 \| (4, 243) \| 311 \| (3, 258) \| 259 \| type VI secretion system tip protein VgrG \| type VI secretion system tip protein VgrG; uniclust \| UniRef100\_A0A1E5AAS2 \| 99.7 \| 3.5e-20 \| 6.5e-26 \| 152.9 \| 161 \| (4, 184) \| 311 \| (95, 263) \| 332 \| Uncharacterized protein \| Uncharacterized protein; uniclust \| UniRef100\_A0A285CU56 \| 99.7 \| 3.7e-20 \| 6.7e-26 \| 155.9 \| 219 \| (80, 308) \| 311 \| (2, 240) \| 456 \| Gp5/Type VI secretion system Vgr protein OB-fold domain-containing protein \| Gp5/Type VI secretion system Vgr protein OB-fold domain-containing protein; uniclust \| UniRef100\_E2CFJ7 \| 99.7 \| 4.2e-20 \| 7.8e-26 \| 143.6 \| 272 \| (2, 281) \| 311 \| (51, 331) \| 335 \| Phage late control D protein (GPD) \| Phage late control D protein (GPD); uniclust \| UniRef100\_A0A8S5UBA3 \| 99.7 \| 4.4e-20 \| 8.1e-26 \| 151.9 \| 262 \| (2, 277) \| 311 \| (61, 344) \| 668 \| Tail protein \| Tail protein; uniclust \| UniRef100\_A0A9D8CM58 \| 99.7 \| 4.4e-20 \| 8.1e-26 \| 160.7 \| 289 \| (5, 304) \| 311 \| (704, 1015) \| 1314 \| Type VI secretion system tip protein VgrG \| Type VI secretion system tip protein VgrG; uniclust \| UniRef100\_A0A257DQE2 \| 99.7 \| 4.4e-20 \| 8.1e-26 \| 157.8 \| 278 \| (17, 308) \| 311 \| (78, 395) \| 710 \| Type VI secretion protein VgrG (Fragment) \| Type VI secretion protein VgrG (Fragment); uniclust \| UniRef100\_A0A2S6N506 \| 99.7 \| 4.5e-20 \| 8.2e-26 \| 142.6 \| 243 \| (6, 265) \| 311 \| (58, 329) \| 331 \| Gp5/Type VI secretion system Vgr protein OB-fold domain-containing protein \| Gp5/Type VI secretion system Vgr protein OB-fold domain-containing protein; uniclust \| UniRef100\_UPI00207C1DAF \| 99.7 \| 4.6e-20 \| 8.5e-26 \| 159.5 \| 285 \| (7, 308) \| 311 \| (58, 388) \| 1198 \| type VI secretion system tip protein TssI/VgrG \| type VI secretion system tip protein TssI/VgrG; uniclust \| UniRef100\_A0A1I2UCB8 \| 99.7 \| 4.7e-20 \| 8.6e-26 \| 162.9 \| 291 \| (4, 306) \| 311 \| (70, 407) \| 1076 \| Rhs element Vgr protein \| Rhs element Vgr protein; uniclust \| UniRef100\_A0A010SJZ7 \| 99.7 \| 4.9e-20 \| 8.9e-26 \| 160.0 \| 220 \| (18, 303) \| 311 \| (107, 346) \| 548 \| Type VI secretion protein \| Type VI secretion protein; uniclust \| UniRef100\_A0A1V4Z8I8 \| 99.7 \| 4.9e-20 \| 9e-26 \| 158.7 \| 251 \| (3, 277) \| 311 \| (54, 323) \| 707 \| Chromosome segregation protein \| Chromosome segregation protein; uniclust \| UniRef100\_A0A1G6Z7R3 \| 99.7 \| 4.9e-20 \| 9.1e-26 \| 162.1 \| 274 \| (3, 301) \| 311 \| (61, 361) \| 627 \| Uncharacterized conserved protein, implicated in type VI secretion and phage assembly \| Uncharacterized conserved protein, implicated in type VI secretion and phage assembly; uniclust \| UniRef100\_A0A0K2RMD0 \| 99.7 \| 5.4e-20 \| 9.8e-26 \| 142.0 \| 247 \| (17, 280) \| 311 \| (8, 274) \| 318 \| Rhs element Vgr protein \| Rhs element Vgr protein; uniclust \| UniRef100\_A0A850B9U5 \| 99.7 \| 5.7e-20 \| 1.1e-25 \| 161.0 \| 284 \| (5, 303) \| 311 \| (70, 408) \| 1439 \| Type VI secretion system tip protein VgrG \| Type VI secretion system tip protein VgrG; uniclust \| UniRef100\_A0A0B4BF20 \| 99.7 \| 5.9e-20 \| 1.1e-25 \| 149.1 \| 275 \| (17, 304) \| 311 \| (83, 397) \| 484 \| Gp5/Type VI secretion system Vgr protein OB-fold domain-containing protein (Fragment) \| Gp5/Type VI secretion system Vgr protein OB-fold domain-containing protein (Fragment); uniclust \| UniRef100\_A0A3Q9JH54 \| 99.7 \| 6.1e-20 \| 1.1e-25 \| 157.3 \| 245 \| (56, 307) \| 311 \| (102, 410) \| 912 \| Type VI secretion system tip protein VgrG \| Type VI secretion system tip protein VgrG; uniclust \| UniRef100\_A0A1H4XF09 \| 99.7 \| 6.3e-20 \| 1.2e-25 \| 155.2 \| 252 \| (37, 304) \| 311 \| (293, 588) \| 900 \| Type VI secretion system secreted protein VgrG \| Type VI secretion system secreted protein VgrG; uniclust \| UniRef100\_A0A542MBF0 \| 99.7 \| 6.3e-20 \| 1.2e-25 \| 160.9 \| 290 \| (3, 304) \| 311 \| (106, 486) \| 908 \| Rhs element Vgr protein \| Rhs element Vgr protein; uniclust \| UniRef100\_A0A518B2T4 \| 99.7 \| 6.3e-20 \| 1.2e-25 \| 153.8 \| 272 \| (17, 305) \| 311 \| (72, 409) \| 812 \| Phage-related baseplate assembly protein \| Phage-related baseplate assembly protein; uniclust \| UniRef100\_A0A074LLZ7 \| 99.7 \| 6.4e-20 \| 1.2e-25 \| 149.0 \| 263 \| (7, 279) \| 311 \| (80, 375) \| 385 \| Uncharacterized protein \| Uncharacterized protein; uniclust \| UniRef100\_A0A1W2F9M8 \| 99.7 \| 6.5e-20 \| 1.2e-25 \| 147.1 \| 250 \| (17, 281) \| 311 \| (90, 382) \| 389 \| Rhs element Vgr protein (Fragment) \| Rhs element Vgr protein (Fragment); uniclust \| UniRef100\_A0A023WVR6 \| 99.7 \| 6.5e-20 \| 1.2e-25 \| 156.0 \| 265 \| (3, 282) \| 311 \| (78, 363) \| 435 \| Baseplate protein \| Baseplate protein; uniclust \| UniRef100\_A0A518B4Q1 \| 99.7 \| 6.5e-20 \| 1.2e-25 \| 156.4 \| 282 \| (6, 304) \| 311 \| (59, 413) \| 995 \| Phage-related baseplate assembly protein \| Phage-related baseplate assembly protein; uniclust \| UniRef100\_A0A4P9VTF7 \| 99.7 \| 7e-20 \| 1.3e-25 \| 162.6 \| 276 \| (17, 306) \| 311 \| (69, 386) \| 1114 \| Type VI secretion system tip protein VgrG \| Type VI secretion system tip protein VgrG; uniclust \| UniRef100\_A0A1Q9WIR8 \| 99.7 \| 7.1e-20 \| 1.3e-25 \| 140.8 \| 248 \| (53, 305) \| 311 \| (12, 298) \| 365 \| Type IV secretion protein Rhs (Fragment) \| Type IV secretion protein Rhs (Fragment); uniclust \| UniRef100\_A0A2J4Y4X1 \| 99.7 \| 7.3e-20 \| 1.3e-25 \| 153.6 \| 248 \| (17, 279) \| 311 \| (91, 375) \| 515 \| Type VI secretion system tip protein VgrG (Fragment) \| Type VI secretion system tip protein VgrG (Fragment); uniclust \| UniRef100\_A0A084ZJJ8 \| 99.7 \| 7.7e-20 \| 1.4e-25 \| 147.1 \| 274 \| (8, 310) \| 311 \| (115, 429) \| 430 \| VgrG family protein \| VgrG family protein; uniclust \| UniRef100\_A0A150Q280 \| 99.7 \| 7.9e-20 \| 1.5e-25 \| 149.9 \| 259 \| (2, 282) \| 311 \| (53, 321) \| 329 \| Uncharacterized protein \| Uncharacterized protein; uniclust \| UniRef100\_A0A1V3NAA1 \| 99.7 \| 8.3e-20 \| 1.5e-25 \| 154.3 \| 255 \| (17, 286) \| 311 \| (67, 358) \| 641 \| Gp5/Type VI secretion system Vgr protein OB-fold domain-containing protein (Fragment) \| Gp5/Type VI secretion system Vgr protein OB-fold domain-containing protein (Fragment); uniclust \| UniRef100\_A0A4P9VIU8 \| 99.7 \| 8.4e-20 \| 1.5e-25 \| 161.4 \| 285 \| (7, 307) \| 311 \| (58, 383) \| 1432 \| Type VI secretion system tip protein VgrG \| Type VI secretion system tip protein VgrG; uniclust \| UniRef100\_A0A8J4ZPK6 \| 99.7 \| 8.9e-20 \| 1.6e-25 \| 164.3 \| 259 \| (38, 308) \| 311 \| (1038, 1334) \| 2288 \| Protein ClpV1 \| Protein ClpV1; uniclust \| UniRef100\_A0A084DHQ4 \| 99.7 \| 9.3e-20 \| 1.7e-25 \| 163.5 \| 287 \| (4, 305) \| 311 \| (116, 500) \| 978 \| Type IV secretion protein Rhs \| Type IV secretion protein Rhs; uniclust \| UniRef100\_A0A3M2UZC3 \| 99.7 \| 9.5e-20 \| 1.7e-25 \| 144.4 \| 226 \| (3, 241) \| 311 \| (87, 340) \| 341 \| Type VI secretion system Vgr protein \| Type VI secretion system Vgr protein; uniclust \| UniRef100\_A0A2T3WF36 \| 99.7 \| 1e-19 \| 1.8e-25 \| 152.1 \| 246 \| (3, 264) \| 311 \| (73, 358) \| 466 \| Type VI secretion system tip protein VgrG (Fragment) \| Type VI secretion system tip protein VgrG (Fragment); uniclust \| UniRef100\_A0A1T2CG57 \| 99.7 \| 1e-19 \| 1.9e-25 \| 132.8 \| 165 \| (109, 279) \| 311 \| (2, 166) \| 173 \| Phage late control D family protein (Fragment) \| Phage late control D family protein (Fragment); uniclust \| UniRef100\_UPI001FE95B91 \| 99.7 \| 1e-19 \| 1.9e-25 \| 153.5 \| 289 \| (4, 304) \| 311 \| (54, 382) \| 675 \| type VI secretion system tip protein TssI/VgrG \| type VI secretion system tip protein TssI/VgrG; uniclust \| UniRef100\_A0A518AY46 \| 99.7 \| 1.1e-19 \| 1.9e-25 \| 157.9 \| 286 \| (4, 305) \| 311 \| (57, 413) \| 1249 \| Filamentous hemagglutinin \| Filamentous hemagglutinin; uniclust \| UniRef100\_A0A1V2GC00 \| 99.7 \| 1.2e-19 \| 2.1e-25 \| 143.7 \| 271 \| (1, 278) \| 311 \| (62, 350) \| 367 \| Late control protein \| Late control protein; uniclust \| UniRef100\_A0A0S4UVJ0 \| 99.7 \| 1.2e-19 \| 2.2e-25 \| 158.4 \| 285 \| (4, 306) \| 311 \| (107, 476) \| 822 \| Gp5/Type VI secretion system Vgr protein OB-fold domain-containing protein (Fragment) \| Gp5/Type VI secretion system Vgr protein OB-fold domain-containing protein (Fragment); uniclust \| UniRef100\_A0A2M8B170 \| 99.7 \| 1.3e-19 \| 2.3e-25 \| 134.0 \| 126 \| (4, 138) \| 311 \| (60, 196) \| 202 \| Type VI secretion system tip protein VgrG (Fragment) \| Type VI secretion system tip protein VgrG (Fragment); uniclust \| UniRef100\_A0A812RFT8 \| 99.7 \| 1.3e-19 \| 2.3e-25 \| 161.7 \| 268 \| (4, 278) \| 311 \| (1045, 1314) \| 1893 \| site-specific DNA-methyltransferase (adenine-specific) \| site-specific DNA-methyltransferase (adenine-specific); uniclust \| UniRef100\_A0A1G9SJN9 \| 99.7 \| 1.3e-19 \| 2.4e-25 \| 167.0 \| 275 \| (17, 308) \| 311 \| (68, 399) \| 1956 \| Rhs element Vgr protein \| Rhs element Vgr protein; uniclust \| UniRef100\_A0A1B3JDP2 \| 99.7 \| 1.3e-19 \| 2.5e-25 \| 166.8 \| 274 \| (17, 307) \| 311 \| (199, 515) \| 2127 \| Uncharacterized protein \| Uncharacterized protein; uniclust \| UniRef100\_A0A1M5NRK7 \| 99.7 \| 1.4e-19 \| 2.5e-25 \| 147.0 \| 257 \| (16, 290) \| 311 \| (73, 348) \| 365 \| Phage protein D \| Phage protein D; uniclust \| UniRef100\_UPI0013A681D6 \| 99.7 \| 1.4e-19 \| 2.5e-25 \| 150.9 \| 273 \| (17, 307) \| 311 \| (198, 509) \| 650 \| type VI secretion system tip protein TssI/VgrG \| type VI secretion system tip protein TssI/VgrG; uniclust \| UniRef100\_A0A1Q7JQQ3 \| 99.7 \| 1.4e-19 \| 2.5e-25 \| 163.8 \| 264 \| (2, 281) \| 311 \| (61, 350) \| 1718 \| Methyl-accepting transducer domain-containing protein \| Methyl-accepting transducer domain-containing protein; uniclust \| UniRef100\_A0A0A1YX04 \| 99.7 \| 1.4e-19 \| 2.6e-25 \| 154.3 \| 247 \| (17, 278) \| 311 \| (105, 393) \| 579 \| Type IV secretion protein Rhs (Fragment) \| Type IV secretion protein Rhs (Fragment); uniclust \| UniRef100\_A0A8J4ZXS2 \| 99.7 \| 1.5e-19 \| 2.7e-25 \| 165.0 \| 259 \| (38, 308) \| 311 \| (1011, 1307) \| 3044 \| Protein ClpV1 \| Protein ClpV1; uniclust \| UniRef100\_A0A3N7E1V0 \| 99.7 \| 1.6e-19 \| 2.9e-25 \| 145.4 \| 235 \| (17, 263) \| 311 \| (101, 383) \| 415 \| Type VI secretion system tip protein VgrG (Fragment) \| Type VI secretion system tip protein VgrG (Fragment); uniclust \| UniRef100\_A0A853I7P2 \| 99.7 \| 1.7e-19 \| 3e-25 \| 137.5 \| 187 \| (1, 196) \| 311 \| (55, 241) \| 248 \| Uncharacterized protein (Fragment) \| Uncharacterized protein (Fragment); uniclust \| UniRef100\_A0A017RU14 \| 99.7 \| 1.7e-19 \| 3e-25 \| 154.0 \| 252 \| (3, 277) \| 311 \| (85, 354) \| 396 \| Uncharacterized protein \| Uncharacterized protein; uniclust \| UniRef100\_A0A0D6BIP0 \| 99.7 \| 1.8e-19 \| 3.2e-25 \| 150.4 \| 219 \| (55, 304) \| 311 \| (114, 369) \| 539 \| ImpA family type VI secretion-associated protein \| ImpA family type VI secretion-associated protein; uniclust \| UniRef100\_A0A285JXF0 \| 99.7 \| 1.8e-19 \| 3.3e-25 \| 146.3 \| 262 \| (9, 277) \| 311 \| (176, 481) \| 494 \| Gp5/Type VI secretion system Vgr protein OB-fold domain-containing protein \| Gp5/Type VI secretion system Vgr protein OB-fold domain-containing protein; uniclust \| UniRef100\_UPI000A3B321B \| 99.7 \| 1.8e-19 \| 3.4e-25 \| 150.1 \| 233 \| (68, 305) \| 311 \| (1, 267) \| 751 \| type VI secretion system tip protein TssI/VgrG \| type VI secretion system tip protein TssI/VgrG; pdb70 \| 4MTK\_A \| 99.9 \| 8.3e-29 \| 6.2e-33 \| 229.3 \| 263 \| (3, 278) \| 311 \| (51, 350) \| 643 \| VgrG1 \| 4MTK\_A VgrG1 Beta-barrel, OB-fold, Beta-helix, Type VI HET: SO4, TAM; pdb70 \| 4MTK\_B \| 99.9 \| 8.3e-29 \| 6.2e-33 \| 229.3 \| 263 \| (3, 278) \| 311 \| (51, 350) \| 643 \| VgrG1 \| 4MTK\_B VgrG1 Beta-barrel, OB-fold, Beta-helix, Type VI HET: TAM, SO4; pdb70 \| 4MTK\_C \| 99.9 \| 8.3e-29 \| 6.2e-33 \| 229.3 \| 263 \| (3, 278) \| 311 \| (51, 350) \| 643 \| VgrG1 \| 4MTK\_C VgrG1 Beta-barrel, OB-fold, Beta-helix, Type VI HET: TAM, SO4; pdb70 \| 4MTK\_D \| 99.9 \| 8.3e-29 \| 6.2e-33 \| 229.3 \| 263 \| (3, 278) \| 311 \| (51, 350) \| 643 \| VgrG1 \| 4MTK\_D VgrG1 Beta-barrel, OB-fold, Beta-helix, Type VI HET: SO4, TAM; pdb70 \| 4MTK\_E \| 99.9 \| 8.3e-29 \| 6.2e-33 \| 229.3 \| 263 \| (3, 278) \| 311 \| (51, 350) \| 643 \| VgrG1 \| 4MTK\_E VgrG1 Beta-barrel, OB-fold, Beta-helix, Type VI HET: SO4, TAM; pdb70 \| 4MTK\_F \| 99.9 \| 8.3e-29 \| 6.2e-33 \| 229.3 \| 263 \| (3, 278) \| 311 \| (51, 350) \| 643 \| VgrG1 \| 4MTK\_F VgrG1 Beta-barrel, OB-fold, Beta-helix, Type VI HET: SO4, TAM; pdb70 \| 4UHV\_A \| 99.9 \| 1e-28 \| 7.8e-33 \| 229.0 \| 263 \| (3, 278) \| 311 \| (51, 350) \| 651 \| VGRG1, VALINE-GLYCINE REPEAT PROTEIN G1 \| 4UHV\_A VGRG1, VALINE-GLYCINE REPEAT PROTEIN G1 STRUCTURAL PROTEIN, VGRG1, VIRULENCE, TOXIN HET: NA, MSE, CL; pdb70 \| 4UHV\_B \| 99.9 \| 1e-28 \| 7.8e-33 \| 229.0 \| 263 \| (3, 278) \| 311 \| (51, 350) \| 651 \| VGRG1, VALINE-GLYCINE REPEAT PROTEIN G1 \| 4UHV\_B VGRG1, VALINE-GLYCINE REPEAT PROTEIN G1 STRUCTURAL PROTEIN, VGRG1, VIRULENCE, TOXIN HET: CL, NA, MSE; pdb70 \| 2P5Z\_X \| 99.8 \| 1.3e-25 \| 9.6e-30 \| 201.6 \| 261 \| (3, 278) \| 311 \| (61, 368) \| 491 \| Type VI secretion system component \| 2P5Z\_X Type VI secretion system component structural genomics, UNKNOWN FUNCTION, PSI-2; pdb70 \| 6RBK\_C \| 99.8 \| 1.9e-25 \| 1.4e-29 \| 202.2 \| 256 \| (4, 277) \| 311 \| (59, 343) \| 529 \| Afp1, Afp2, Afp3, Afp16 \| 6RBK\_C Afp1, Afp2, Afp3, Afp16 Anti-feeding prophage, secretion system, AFP; pdb70 \| 6J0M\_C \| 99.8 \| 2.1e-25 \| 1.6e-29 \| 202.3 \| 259 \| (3, 274) \| 311 \| (61, 348) \| 538 \| Pvc8 \| 6J0M\_C Pvc8 assembly, Photorhabdus asymbiotica, PVC, contractile; pdb70 \| 1WRU\_A \| 99.7 \| 4.4e-23 \| 3.3e-27 \| 178.9 \| 257 \| (2, 279) \| 311 \| (48, 338) \| 379 \| 43 kDa tail protein \| 1WRU\_A 43 kDa tail protein BACTERIOPHAGE MU, BASEPLATE, GENE PRODUCT; pdb70 \| 3D37\_A \| 99.7 \| 2.3e-22 \| 1.7e-26 \| 174.4 \| 259 \| (2, 279) \| 311 \| (53, 340) \| 381 \| Tail protein, 43 kDa \| 3D37\_A Tail protein, 43 kDa tail protein, structural genomics, PSI; pdb70 \| 3D37\_B \| 99.7 \| 2.3e-22 \| 1.7e-26 \| 174.4 \| 259 \| (2, 279) \| 311 \| (53, 340) \| 381 \| Tail protein, 43 kDa \| 3D37\_B Tail protein, 43 kDa tail protein, structural genomics, PSI; pdb70 \| 3CDD\_F \| 99.7 \| 1.4e-21 \| 1e-25 \| 168.0 \| 259 \| (3, 279) \| 311 \| (58, 343) \| 361 \| Prophage MuSo2, 43 kDa tail \| 3CDD\_F Prophage MuSo2, 43 kDa tail MuSo2, Shewanella oneidensis MR-1, Structural HET: MSE; pdb70 \| 3CDD\_D \| 99.7 \| 1.4e-21 \| 1e-25 \| 168.0 \| 259 \| (3, 279) \| 311 \| (58, 343) \| 361 \| Prophage MuSo2, 43 kDa tail \| 3CDD\_D Prophage MuSo2, 43 kDa tail MuSo2, Shewanella oneidensis MR-1, Structural |
| Top keywords  (threshold 1.00e-03 (evalue)) | **Type, secretion, VI, system, VgrG, Vgr, Fragment, Rhs, Phage, tip** |
| Output files | ../../similar\_sequences/25\_FANPEZAQ\_CDS\_0025\_merged.svg ../../similar\_sequences/25\_FANPEZAQ\_CDS\_0025\_pdb70.a3m ../../similar\_sequences/25\_FANPEZAQ\_CDS\_0025\_pdb70.hhr ../../similar\_sequences/25\_FANPEZAQ\_CDS\_0025\_uniclust.a3m ../../similar\_sequences/25\_FANPEZAQ\_CDS\_0025\_uniclust.hhr |

#### Structure prediction (AlphaFold)^2^

|  |  |
| --- | --- |
| Stats | xml version="1.0" encoding="utf-8" standalone="no"?       2024-09-02T21:09:25.673745 image/svg+xml   Matplotlib v3.7.2, https://matplotlib.org/ |
| Predicted structure | **NGL Viewer Controls:**  - Center: *Left-Click* - Rotate: *Left-Click + Drag* - Translate: *Right-Click + Drag* - Zoom: *Shift + Left-Click + Drag* |
| Output files | ../../predicted\_structures/25\_FANPEZAQ\_CDS\_0025/features.pkl ../../predicted\_structures/25\_FANPEZAQ\_CDS\_0025/ranked\_0.pdb ../../predicted\_structures/25\_FANPEZAQ\_CDS\_0025/ranked\_0\_plots.svg ../../predicted\_structures/25\_FANPEZAQ\_CDS\_0025/result\_model\_1\_ptm\_pred\_0.pkl |

#### Structure similarity search results (Foldseek)^3^

|  |  |
| --- | --- |
| Structure databases searched | Pdb, Afdb-proteome, Afdb-uniprot50 |
| Results, scheme(s)  (Top layers only, threshold 1.00e-02 (evalue)) | xml version="1.0" encoding="utf-8" standalone="no"?       2024-09-02T21:10:56.958834 image/svg+xml   Matplotlib v3.7.2, https://matplotlib.org/ |
| Results, table  (threshold 1.00e-02 (evalue)) | db \| id \| prob \| evalue \| bits \| fident \| alnlen \| mismatch \| gapopen \| qstart \| qend \| tstart \| tend \| name \| description; pdb \| 7B5H\_AJ \| 1.0 \| 2.032e-14 \| 435 \| 0.169 \| 307 \| 208 \| 12 \| 7 \| 284 \| 70 \| 358 \| All3320 protein \| All3320 protein; pdb \| 7AEF\_Q \| 1.0 \| 2.392e-13 \| 408 \| 0.164 \| 297 \| 198 \| 12 \| 7 \| 275 \| 74 \| 348 \| Phosphoserine phosphatase SerB \| Phosphoserine phosphatase SerB; pdb \| 6U5H\_C \| 1.0 \| 2.866e-14 \| 395 \| 0.261 \| 298 \| 164 \| 17 \| 1 \| 275 \| 51 \| 315 \| Probable bacteriophage protein Pyocin R2 \| Probable bacteriophage protein Pyocin R2; pdb \| 6J0M\_A \| 1.0 \| 1.239e-10 \| 298 \| 0.129 \| 308 \| 214 \| 17 \| 7 \| 279 \| 64 \| 352 \| Pvc8 \| Pvc8; pdb \| 3CDD\_B \| 1.0 \| 3.129e-11 \| 287 \| 0.146 \| 280 \| 193 \| 17 \| 19 \| 277 \| 66 \| 320 \| Prophage MuSo2, 43 kDa tail protein \| Prophage MuSo2, 43 kDa tail protein; pdb \| 3CDD\_C \| 1.0 \| 2.79e-11 \| 283 \| 0.134 \| 313 \| 207 \| 17 \| 7 \| 277 \| 34 \| 324 \| Prophage MuSo2, 43 kDa tail protein \| Prophage MuSo2, 43 kDa tail protein; pdb \| 3CDD\_D \| 1.0 \| 4.949e-11 \| 269 \| 0.141 \| 317 \| 204 \| 20 \| 7 \| 277 \| 33 \| 327 \| Prophage MuSo2, 43 kDa tail protein \| Prophage MuSo2, 43 kDa tail protein; pdb \| 3D37\_A \| 1.0 \| 4.631e-10 \| 267 \| 0.134 \| 283 \| 200 \| 17 \| 19 \| 276 \| 54 \| 316 \| Tail protein, 43 kDa \| Tail protein, 43 kDa; pdb \| 3CDD\_A \| 1.0 \| 1.747e-10 \| 266 \| 0.167 \| 281 \| 184 \| 17 \| 19 \| 277 \| 67 \| 319 \| Prophage MuSo2, 43 kDa tail protein \| Prophage MuSo2, 43 kDa tail protein; pdb \| 3CDD\_F \| 1.0 \| 6.593e-11 \| 258 \| 0.146 \| 313 \| 200 \| 18 \| 7 \| 277 \| 32 \| 319 \| Prophage MuSo2, 43 kDa tail protein \| Prophage MuSo2, 43 kDa tail protein; pdb \| 3CDD\_E \| 1.0 \| 6.169e-10 \| 256 \| 0.148 \| 283 \| 184 \| 18 \| 19 \| 277 \| 69 \| 318 \| Prophage MuSo2, 43 kDa tail protein \| Prophage MuSo2, 43 kDa tail protein; pdb \| 3D37\_B \| 1.0 \| 3.864e-09 \| 245 \| 0.125 \| 287 \| 190 \| 17 \| 19 \| 276 \| 52 \| 306 \| Tail protein, 43 kDa \| Tail protein, 43 kDa; pdb \| 8GRA\_G \| 1.0 \| 6.919e-10 \| 242 \| 0.116 \| 317 \| 187 \| 24 \| 17 \| 279 \| 75 \| 352 \| Type VI secretion system spike protein VgrG \| Type VI secretion system spike protein VgrG; pdb \| 7Q5P\_B \| 1.0 \| 4.092e-09 \| 239 \| 0.1 \| 320 \| 216 \| 24 \| 5 \| 278 \| 49 \| 342 \| Type VI secretion protein VgrG \| Type VI secretion protein VgrG; pdb \| 6RBK\_C \| 1.0 \| 3.415e-08 \| 223 \| 0.125 \| 302 \| 203 \| 17 \| 7 \| 282 \| 58 \| 324 \| Afp8 \| Afp8; pdb \| 6H3N\_B \| 1.0 \| 9.05e-08 \| 218 \| 0.117 \| 314 \| 210 \| 24 \| 5 \| 277 \| 49 \| 336 \| VgrG1 \| VgrG1; pdb \| 7YFZ\_V \| 1.0 \| 1.503e-06 \| 185 \| 0.093 \| 289 \| 167 \| 17 \| 8 \| 276 \| 2 \| 215 \| Pam3 hub gp19 \| Pam3 hub gp19; pdb \| 5IV5\_YD \| 1.0 \| 6.733e-07 \| 177 \| 0.096 \| 322 \| 224 \| 20 \| 7 \| 289 \| 68 \| 361 \| Baseplate hub protein gp27 \| Baseplate hub protein gp27; pdb \| 2Z6B\_D \| 1.0 \| 1.195e-06 \| 168 \| 0.09 \| 309 \| 206 \| 19 \| 16 \| 281 \| 78 \| 354 \| Baseplate structural protein Gp27 \| Baseplate structural protein Gp27; pdb \| 3GS9\_A \| 1.0 \| 1.065e-06 \| 165 \| 0.095 \| 304 \| 207 \| 19 \| 7 \| 275 \| 37 \| 307 \| Protein gp18 \| Protein gp18; pdb \| 1WTH\_D \| 1.0 \| 5.618e-06 \| 157 \| 0.089 \| 313 \| 215 \| 17 \| 16 \| 291 \| 78 \| 357 \| Baseplate structural protein Gp27 \| Baseplate structural protein Gp27; pdb \| 4UHV\_A \| 1.0 \| 1.118e-05 \| 149 \| 0.103 \| 328 \| 216 \| 24 \| 4 \| 281 \| 52 \| 351 \| VGRG1, VALINE-GLYCINE REPEAT PROTEIN G1 \| VGRG1, VALINE-GLYCINE REPEAT PROTEIN G1; pdb \| 7ZN2\_C \| 1.0 \| 7.483e-06 \| 144 \| 0.126 \| 229 \| 138 \| 18 \| 89 \| 281 \| 519 \| 721 \| Tail tube protein \| Tail tube protein; pdb \| 8EON\_L \| 1.0 \| 7.853e-05 \| 140 \| 0.124 \| 290 \| 157 \| 17 \| 7 \| 279 \| 78 \| 287 \| Baseplate hub gp41 \| Baseplate hub gp41; pdb \| 6V8I\_BE \| 1.0 \| 0.000521 \| 109 \| 0.095 \| 304 \| 198 \| 21 \| 22 \| 280 \| 63 \| 334 \| Tail-Associated Lysin, gp59 \| Tail-Associated Lysin, gp59; pdb \| 2P5Z\_X \| 1.0 \| 0.0002334 \| 105 \| 0.112 \| 302 \| 168 \| 24 \| 9 \| 280 \| 49 \| 280 \| Type VI secretion system component \| Type VI secretion system component; pdb \| 4V5I\_A0 \| 0.999 \| 1.769e-05 \| 99 \| 0.063 \| 332 \| 219 \| 26 \| 8 \| 281 \| 75 \| 372 \| ORF16 \| ORF16; afdb-proteome \| AF-G3XCU8-F1-MODEL\_V4 \| 1.0 \| 9.522e-25 \| 896 \| 0.29 \| 282 \| 183 \| 5 \| 1 \| 275 \| 51 \| 322 \| Uncharacterized protein \| Uncharacterized protein; afdb-proteome \| AF-Q8ZMV5-F1-MODEL\_V4 \| 1.0 \| 1.912e-25 \| 875 \| 0.291 \| 309 \| 184 \| 8 \| 1 \| 281 \| 63 \| 364 \| Fels-2 prophage protein \| Fels-2 prophage protein; afdb-proteome \| AF-A0A0H3GX06-F1-MODEL\_V4 \| 1.0 \| 8.017e-25 \| 869 \| 0.301 \| 332 \| 180 \| 8 \| 1 \| 288 \| 63 \| 386 \| Putative prophage tail protein \| Putative prophage tail protein; afdb-proteome \| AF-A0A0H3GM37-F1-MODEL\_V4 \| 1.0 \| 2.125e-24 \| 868 \| 0.289 \| 307 \| 182 \| 9 \| 1 \| 279 \| 63 \| 361 \| Late control gene D protein (GpD) \| Late control gene D protein (GpD); afdb-proteome \| AF-Q8ZKJ7-F1-MODEL\_V4 \| 1.0 \| 3.213e-19 \| 652 \| 0.214 \| 289 \| 200 \| 9 \| 2 \| 277 \| 68 \| 342 \| Putative cytoplasmic protein \| Putative cytoplasmic protein; afdb-proteome \| AF-Q32HG2-F1-MODEL\_V4 \| 1.0 \| 1.923e-13 \| 430 \| 0.301 \| 196 \| 126 \| 4 \| 1 \| 187 \| 30 \| 223 \| Uncharacterized protein \| Uncharacterized protein; afdb-proteome \| AF-Q9I1A6-F1-MODEL\_V4 \| 1.0 \| 3.517e-10 \| 280 \| 0.129 \| 316 \| 211 \| 19 \| 4 \| 283 \| 54 \| 341 \| Phage\_base\_V domain-containing protein \| Phage\_base\_V domain-containing protein; afdb-proteome \| AF-Q8ZRL1-F1-MODEL\_V4 \| 1.0 \| 2.471e-09 \| 256 \| 0.124 \| 313 \| 211 \| 24 \| 7 \| 277 \| 159 \| 450 \| Putative cytoplasmic protein \| Putative cytoplasmic protein; afdb-proteome \| AF-Q9I741-F1-MODEL\_V4 \| 1.0 \| 2.448e-08 \| 240 \| 0.112 \| 321 \| 213 \| 21 \| 5 \| 277 \| 53 \| 349 \| Type VI secretion system spike protein VgrG1a \| Type VI secretion system spike protein VgrG1a; afdb-proteome \| AF-Q9I6M7-F1-MODEL\_V4 \| 1.0 \| 7.778e-09 \| 240 \| 0.135 \| 325 \| 196 \| 22 \| 17 \| 283 \| 62 \| 359 \| Type VI secretion system spike protein VgrG2b \| Type VI secretion system spike protein VgrG2b; afdb-proteome \| AF-Q9I737-F1-MODEL\_V4 \| 1.0 \| 5.159e-08 \| 229 \| 0.118 \| 321 \| 211 \| 21 \| 5 \| 277 \| 56 \| 352 \| Type VI secretion system spike protein VgrG1b \| Type VI secretion system spike protein VgrG1b; afdb-proteome \| AF-Q9I3K1-F1-MODEL\_V4 \| 1.0 \| 1.461e-08 \| 222 \| 0.129 \| 316 \| 192 \| 21 \| 23 \| 283 \| 72 \| 359 \| Type VI secretion system spike protein VgrG2a \| Type VI secretion system spike protein VgrG2a; afdb-proteome \| AF-Q9HYC3-F1-MODEL\_V4 \| 1.0 \| 4.384e-09 \| 210 \| 0.152 \| 342 \| 196 \| 24 \| 7 \| 276 \| 39 \| 358 \| Type VI secretion system spike protein VgrG4b \| Type VI secretion system spike protein VgrG4b; afdb-proteome \| AF-Q9I0F3-F1-MODEL\_V4 \| 1.0 \| 1.72e-07 \| 208 \| 0.131 \| 319 \| 209 \| 25 \| 5 \| 277 \| 93 \| 389 \| Type VI secretion system spike protein VgrG1c \| Type VI secretion system spike protein VgrG1c; afdb-proteome \| AF-Q9HYU9-F1-MODEL\_V4 \| 1.0 \| 1.23e-08 \| 202 \| 0.159 \| 338 \| 198 \| 24 \| 7 \| 276 \| 39 \| 358 \| Phage\_base\_V domain-containing protein \| Phage\_base\_V domain-containing protein; afdb-proteome \| AF-Q5F989-F1-MODEL\_V4 \| 1.0 \| 1.689e-05 \| 161 \| 0.131 \| 213 \| 145 \| 16 \| 95 \| 276 \| 215 \| 418 \| Uncharacterized protein \| Uncharacterized protein; afdb-proteome \| AF-Q2FX67-F1-MODEL\_V4 \| 1.0 \| 5.682e-06 \| 151 \| 0.12 \| 298 \| 201 \| 19 \| 20 \| 280 \| 62 \| 335 \| Phi ETA orf 55-like protein \| Phi ETA orf 55-like protein; afdb-proteome \| AF-Q2FYD0-F1-MODEL\_V4 \| 1.0 \| 0.0001121 \| 121 \| 0.076 \| 434 \| 205 \| 24 \| 22 \| 280 \| 68 \| 480 \| SLT orf 527-like protein \| SLT orf 527-like protein; afdb-uniprot50 \| AF-Q3RCU8-F1-MODEL\_V4 \| 1.0 \| 3.926e-43 \| 1669 \| 0.438 \| 303 \| 163 \| 2 \| 1 \| 297 \| 1 \| 302 \| Phage late control D \| Phage late control D; afdb-uniprot50 \| AF-Q87CI2-F1-MODEL\_V4 \| 1.0 \| 6.459e-41 \| 1576 \| 0.442 \| 303 \| 162 \| 2 \| 1 \| 297 \| 55 \| 356 \| Phage-related tail protein \| Phage-related tail protein; afdb-uniprot50 \| AF-A0A2J0SR82-F1-MODEL\_V4 \| 1.0 \| 4.806e-40 \| 1538 \| 0.486 \| 296 \| 146 \| 1 \| 1 \| 290 \| 57 \| 352 \| Phage tail protein \| Phage tail protein; afdb-uniprot50 \| AF-K5ZNX9-F1-MODEL\_V4 \| 1.0 \| 2.956e-37 \| 1478 \| 0.491 \| 281 \| 142 \| 1 \| 1 \| 280 \| 57 \| 337 \| Uncharacterized protein \| Uncharacterized protein; afdb-uniprot50 \| AF-A0A158E854-F1-MODEL\_V4 \| 1.0 \| 5.882e-37 \| 1378 \| 0.513 \| 292 \| 136 \| 1 \| 2 \| 287 \| 58 \| 349 \| Late control D family protein \| Late control D family protein; afdb-uniprot50 \| AF-A0A1H2ZGX1-F1-MODEL\_V4 \| 1.0 \| 5.724e-34 \| 1337 \| 0.431 \| 276 \| 157 \| 0 \| 1 \| 276 \| 58 \| 333 \| Uncharacterized protein \| Uncharacterized protein; afdb-uniprot50 \| AF-A0A1N6R0G0-F1-MODEL\_V4 \| 1.0 \| 1.206e-33 \| 1312 \| 0.41 \| 280 \| 157 \| 3 \| 1 \| 279 \| 1 \| 273 \| Uncharacterized protein \| Uncharacterized protein; afdb-uniprot50 \| AF-A0A2G2N728-F1-MODEL\_V4 \| 1.0 \| 5.673e-33 \| 1297 \| 0.417 \| 278 \| 162 \| 0 \| 1 \| 278 \| 55 \| 332 \| Phage tail protein \| Phage tail protein; afdb-uniprot50 \| AF-A0A5J5GRJ3-F1-MODEL\_V4 \| 1.0 \| 5.357e-33 \| 1293 \| 0.415 \| 255 \| 149 \| 0 \| 22 \| 276 \| 2 \| 256 \| Late control protein D \| Late control protein D; afdb-uniprot50 \| AF-A0A4Q7BES0-F1-MODEL\_V4 \| 1.0 \| 5.673e-33 \| 1275 \| 0.457 \| 280 \| 145 \| 2 \| 1 \| 279 \| 54 \| 327 \| Phage tail protein \| Phage tail protein; afdb-uniprot50 \| AF-A0A2N1ANJ8-F1-MODEL\_V4 \| 1.0 \| 6.062e-34 \| 1271 \| 0.445 \| 285 \| 152 \| 2 \| 1 \| 279 \| 55 \| 339 \| Phage tail protein \| Phage tail protein; afdb-uniprot50 \| AF-A0A0P8YPJ6-F1-MODEL\_V4 \| 1.0 \| 1.702e-33 \| 1266 \| 0.361 \| 277 \| 177 \| 0 \| 2 \| 278 \| 8 \| 284 \| Uncharacterized protein \| Uncharacterized protein; afdb-uniprot50 \| AF-A0A7Z1WDF0-F1-MODEL\_V4 \| 1.0 \| 2.668e-32 \| 1209 \| 0.459 \| 272 \| 140 \| 2 \| 12 \| 282 \| 1 \| 266 \| Phage tail protein \| Phage tail protein; afdb-uniprot50 \| AF-A0A149SNT1-F1-MODEL\_V4 \| 1.0 \| 1.129e-32 \| 1203 \| 0.366 \| 322 \| 189 \| 5 \| 1 \| 311 \| 55 \| 372 \| Uncharacterized protein \| Uncharacterized protein; afdb-uniprot50 \| AF-A0A2Z6AZ21-F1-MODEL\_V4 \| 1.0 \| 2.644e-31 \| 1200 \| 0.436 \| 277 \| 153 \| 3 \| 1 \| 276 \| 56 \| 330 \| Late control D family protein \| Late control D family protein; afdb-uniprot50 \| AF-A0A145VT91-F1-MODEL\_V4 \| 1.0 \| 9.974e-32 \| 1187 \| 0.43 \| 279 \| 159 \| 0 \| 1 \| 279 \| 55 \| 333 \| Uncharacterized protein \| Uncharacterized protein; afdb-uniprot50 \| AF-A0A143DDT4-F1-MODEL\_V4 \| 1.0 \| 5.26e-31 \| 1179 \| 0.414 \| 285 \| 159 \| 3 \| 1 \| 279 \| 55 \| 337 \| Uncharacterized protein \| Uncharacterized protein; afdb-uniprot50 \| AF-A0A844HSX7-F1-MODEL\_V4 \| 1.0 \| 4.469e-32 \| 1172 \| 0.388 \| 286 \| 168 \| 1 \| 1 \| 279 \| 58 \| 343 \| Late control protein D \| Late control protein D; afdb-uniprot50 \| AF-A0A7S8C754-F1-MODEL\_V4 \| 1.0 \| 1.463e-29 \| 1106 \| 0.389 \| 280 \| 163 \| 3 \| 1 \| 279 \| 51 \| 323 \| Phage late control D family protein \| Phage late control D family protein; afdb-uniprot50 \| AF-A0A1H9YCN2-F1-MODEL\_V4 \| 1.0 \| 1.305e-29 \| 1091 \| 0.377 \| 278 \| 172 \| 1 \| 1 \| 278 \| 55 \| 331 \| Uncharacterized protein \| Uncharacterized protein; afdb-uniprot50 \| AF-A0A1Z5H6V1-F1-MODEL\_V4 \| 1.0 \| 6.881e-29 \| 1088 \| 0.397 \| 279 \| 160 \| 3 \| 1 \| 278 \| 51 \| 322 \| Uncharacterized protein \| Uncharacterized protein; afdb-uniprot50 \| AF-A0A2G6CS06-F1-MODEL\_V4 \| 1.0 \| 7.717e-29 \| 1082 \| 0.376 \| 279 \| 165 \| 5 \| 1 \| 277 \| 52 \| 323 \| Uncharacterized protein \| Uncharacterized protein; afdb-uniprot50 \| AF-A0A2W4TBH4-F1-MODEL\_V4 \| 1.0 \| 1.369e-28 \| 1080 \| 0.358 \| 282 \| 171 \| 4 \| 1 \| 279 \| 55 \| 329 \| Uncharacterized protein \| Uncharacterized protein; afdb-uniprot50 \| AF-B7RNP1-F1-MODEL\_V4 \| 1.0 \| 1.738e-29 \| 1080 \| 0.377 \| 278 \| 173 \| 0 \| 1 \| 278 \| 58 \| 335 \| Phage late control D \| Phage late control D; afdb-uniprot50 \| AF-A0A4R3LEZ2-F1-MODEL\_V4 \| 1.0 \| 1.949e-29 \| 1065 \| 0.361 \| 296 \| 165 \| 6 \| 1 \| 281 \| 55 \| 341 \| Uncharacterized protein \| Uncharacterized protein; afdb-uniprot50 \| AF-A0A0J1K2H0-F1-MODEL\_V4 \| 1.0 \| 3.056e-28 \| 1064 \| 0.371 \| 283 \| 171 \| 4 \| 1 \| 279 \| 54 \| 333 \| Tail protein \| Tail protein; afdb-uniprot50 \| AF-A8U2S0-F1-MODEL\_V4 \| 1.0 \| 1.382e-29 \| 1063 \| 0.323 \| 284 \| 176 \| 5 \| 1 \| 276 \| 85 \| 360 \| Late control gene D protein \| Late control gene D protein; afdb-uniprot50 \| AF-A0A4V1IMX3-F1-MODEL\_V4 \| 1.0 \| 5.421e-28 \| 1051 \| 0.35 \| 277 \| 172 \| 3 \| 1 \| 276 \| 50 \| 319 \| Phage late control D family protein \| Phage late control D family protein; afdb-uniprot50 \| AF-A0A7C8M556-F1-MODEL\_V4 \| 1.0 \| 4.878e-29 \| 1044 \| 0.409 \| 281 \| 161 \| 3 \| 1 \| 279 \| 53 \| 330 \| Uncharacterized protein \| Uncharacterized protein; afdb-uniprot50 \| AF-A0A1X7JLX1-F1-MODEL\_V4 \| 1.0 \| 2.315e-29 \| 1037 \| 0.375 \| 280 \| 161 \| 5 \| 1 \| 275 \| 56 \| 326 \| Uncharacterized protein \| Uncharacterized protein; afdb-uniprot50 \| AF-A0A167H3I8-F1-MODEL\_V4 \| 1.0 \| 6.819e-28 \| 1032 \| 0.361 \| 282 \| 170 \| 3 \| 1 \| 279 \| 51 \| 325 \| Uncharacterized protein \| Uncharacterized protein; afdb-uniprot50 \| AF-B2I805-F1-MODEL\_V4 \| 1.0 \| 3.843e-28 \| 1030 \| 0.4 \| 225 \| 129 \| 1 \| 79 \| 297 \| 2 \| 226 \| Late control D family protein \| Late control D family protein; afdb-uniprot50 \| AF-A0A2U1XZ03-F1-MODEL\_V4 \| 1.0 \| 1.437e-27 \| 1023 \| 0.373 \| 284 \| 168 \| 5 \| 1 \| 281 \| 55 \| 331 \| Uncharacterized protein \| Uncharacterized protein; afdb-uniprot50 \| AF-A0A3M6B5Q4-F1-MODEL\_V4 \| 1.0 \| 1.707e-27 \| 1019 \| 0.31 \| 283 \| 176 \| 6 \| 1 \| 275 \| 19 \| 290 \| Prophage PSPPH02, late control protein D protein \| Prophage PSPPH02, late control protein D protein; afdb-uniprot50 \| AF-A0A0P9S9Z1-F1-MODEL\_V4 \| 1.0 \| 2.147e-27 \| 1016 \| 0.307 \| 283 \| 177 \| 6 \| 1 \| 275 \| 19 \| 290 \| Prophage PSPPH02, late control protein D protein \| Prophage PSPPH02, late control protein D protein; afdb-uniprot50 \| AF-A0A329B6J9-F1-MODEL\_V4 \| 1.0 \| 1.437e-27 \| 1015 \| 0.315 \| 285 \| 181 \| 7 \| 2 \| 280 \| 16 \| 292 \| Uncharacterized protein \| Uncharacterized protein; afdb-uniprot50 \| AF-A0A3N2E0R5-F1-MODEL\_V4 \| 1.0 \| 2.549e-27 \| 1015 \| 0.341 \| 278 \| 178 \| 3 \| 1 \| 276 \| 53 \| 327 \| Uncharacterized protein \| Uncharacterized protein; afdb-uniprot50 \| AF-A0A5E4XF48-F1-MODEL\_V4 \| 1.0 \| 1.612e-27 \| 1015 \| 0.348 \| 281 \| 177 \| 2 \| 1 \| 280 \| 51 \| 326 \| Late control protein \| Late control protein; afdb-uniprot50 \| AF-A0A210BC43-F1-MODEL\_V4 \| 1.0 \| 7.648e-28 \| 1014 \| 0.245 \| 310 \| 200 \| 8 \| 1 \| 284 \| 8 \| 309 \| Late control protein \| Late control protein; afdb-uniprot50 \| AF-A0A4D9YTW0-F1-MODEL\_V4 \| 1.0 \| 1.221e-28 \| 1014 \| 0.286 \| 332 \| 181 \| 12 \| 2 \| 288 \| 14 \| 334 \| Phage late control D family protein \| Phage late control D family protein; afdb-uniprot50 \| AF-A0A5A9EMZ8-F1-MODEL\_V4 \| 1.0 \| 2.912e-29 \| 1013 \| 0.349 \| 312 \| 178 \| 7 \| 1 \| 304 \| 97 \| 391 \| Uncharacterized protein \| Uncharacterized protein; afdb-uniprot50 \| AF-A0A3S0U918-F1-MODEL\_V4 \| 1.0 \| 1.807e-27 \| 1009 \| 0.357 \| 277 \| 170 \| 3 \| 1 \| 276 \| 51 \| 320 \| Phage tail protein \| Phage tail protein; afdb-uniprot50 \| AF-A0A4R1PBW7-F1-MODEL\_V4 \| 1.0 \| 4.79e-27 \| 1004 \| 0.353 \| 286 \| 171 \| 7 \| 1 \| 280 \| 41 \| 318 \| Uncharacterized protein \| Uncharacterized protein; afdb-uniprot50 \| AF-A0A6G8F2E2-F1-MODEL\_V4 \| 1.0 \| 7.579e-27 \| 999 \| 0.347 \| 282 \| 177 \| 5 \| 1 \| 278 \| 51 \| 329 \| Phage late control protein \| Phage late control protein; afdb-uniprot50 \| AF-A0A2T5J1F0-F1-MODEL\_V4 \| 1.0 \| 1.009e-26 \| 999 \| 0.364 \| 280 \| 167 \| 6 \| 2 \| 277 \| 56 \| 328 \| Uncharacterized protein \| Uncharacterized protein; afdb-uniprot50 \| AF-A0A5S9Q3J3-F1-MODEL\_V4 \| 1.0 \| 9.532e-27 \| 997 \| 0.368 \| 285 \| 167 \| 5 \| 1 \| 279 \| 51 \| 328 \| Uncharacterized protein \| Uncharacterized protein; afdb-uniprot50 \| AF-A0A5J6LDM7-F1-MODEL\_V4 \| 1.0 \| 3.997e-26 \| 994 \| 0.307 \| 276 \| 177 \| 7 \| 8 \| 278 \| 1 \| 267 \| Phage late control D family protein \| Phage late control D family protein; afdb-uniprot50 \| AF-A0A496N5Y9-F1-MODEL\_V4 \| 1.0 \| 3.236e-28 \| 994 \| 0.28 \| 332 \| 181 \| 10 \| 1 \| 282 \| 15 \| 338 \| Phage late control D family protein \| Phage late control D family protein; afdb-uniprot50 \| AF-A0A7Z0N080-F1-MODEL\_V4 \| 1.0 \| 1.522e-27 \| 990 \| 0.363 \| 278 \| 167 \| 6 \| 1 \| 276 \| 51 \| 320 \| Uncharacterized protein \| Uncharacterized protein; afdb-uniprot50 \| AF-A0A7K0GN29-F1-MODEL\_V4 \| 1.0 \| 3.178e-26 \| 989 \| 0.403 \| 238 \| 133 \| 4 \| 39 \| 274 \| 2 \| 232 \| Phage late control D family protein \| Phage late control D family protein; afdb-uniprot50 \| AF-A0A1H7EQQ5-F1-MODEL\_V4 \| 1.0 \| 1.597e-26 \| 989 \| 0.348 \| 287 \| 175 \| 7 \| 1 \| 282 \| 54 \| 333 \| Uncharacterized protein \| Uncharacterized protein; afdb-uniprot50 \| AF-A0A809Z5I0-F1-MODEL\_V4 \| 1.0 \| 1.258e-25 \| 988 \| 0.306 \| 281 \| 187 \| 3 \| 1 \| 280 \| 72 \| 345 \| Regulator of late gene expression \| Regulator of late gene expression; afdb-uniprot50 \| AF-A0A1Y6CY31-F1-MODEL\_V4 \| 1.0 \| 1.691e-26 \| 985 \| 0.329 \| 279 \| 177 \| 5 \| 1 \| 279 \| 51 \| 319 \| Uncharacterized protein \| Uncharacterized protein; afdb-uniprot50 \| AF-A0A362XU25-F1-MODEL\_V4 \| 1.0 \| 2.526e-26 \| 985 \| 0.341 \| 281 \| 175 \| 5 \| 2 \| 279 \| 55 \| 328 \| Uncharacterized protein \| Uncharacterized protein; afdb-uniprot50 \| AF-A0A7W6CNX6-F1-MODEL\_V4 \| 1.0 \| 1.508e-26 \| 977 \| 0.293 \| 286 \| 183 \| 8 \| 1 \| 276 \| 7 \| 283 \| Uncharacterized protein \| Uncharacterized protein; afdb-uniprot50 \| AF-E5AKN5-F1-MODEL\_V4 \| 1.0 \| 1.188e-25 \| 976 \| 0.322 \| 285 \| 181 \| 6 \| 1 \| 280 \| 61 \| 338 \| Gene D protein \| Gene D protein; afdb-uniprot50 \| AF-A0A2E3N1J0-F1-MODEL\_V4 \| 1.0 \| 5.324e-26 \| 975 \| 0.31 \| 287 \| 183 \| 5 \| 1 \| 279 \| 51 \| 330 \| Uncharacterized protein \| Uncharacterized protein; afdb-uniprot50 \| AF-A0A4R2PFS0-F1-MODEL\_V4 \| 1.0 \| 1.791e-26 \| 975 \| 0.314 \| 286 \| 181 \| 5 \| 1 \| 279 \| 55 \| 332 \| Uncharacterized protein \| Uncharacterized protein; afdb-uniprot50 \| AF-A0A4V3GTL6-F1-MODEL\_V4 \| 1.0 \| 2.407e-27 \| 974 \| 0.332 \| 283 \| 175 \| 7 \| 10 \| 288 \| 1 \| 273 \| Uncharacterized protein \| Uncharacterized protein; afdb-uniprot50 \| AF-A0A8A7WWT4-F1-MODEL\_V4 \| 1.0 \| 3.001e-26 \| 973 \| 0.347 \| 282 \| 172 \| 8 \| 2 \| 279 \| 55 \| 328 \| Phage late control D family protein \| Phage late control D family protein; afdb-uniprot50 \| AF-A0A4Q3Z0D3-F1-MODEL\_V4 \| 1.0 \| 2.676e-26 \| 971 \| 0.35 \| 280 \| 181 \| 1 \| 2 \| 281 \| 59 \| 337 \| Late control protein D \| Late control protein D; afdb-uniprot50 \| AF-A0A318KN30-F1-MODEL\_V4 \| 1.0 \| 7.156e-27 \| 968 \| 0.333 \| 300 \| 173 \| 8 \| 1 \| 280 \| 64 \| 356 \| Uncharacterized protein \| Uncharacterized protein; afdb-uniprot50 \| AF-A0A6L2ZP70-F1-MODEL\_V4 \| 1.0 \| 1.597e-26 \| 966 \| 0.346 \| 283 \| 168 \| 7 \| 1 \| 276 \| 52 \| 324 \| Phage late control gene D protein \| Phage late control gene D protein; afdb-uniprot50 \| AF-A0A2V3Y0H4-F1-MODEL\_V4 \| 1.0 \| 2.676e-26 \| 966 \| 0.296 \| 283 \| 183 \| 7 \| 1 \| 275 \| 60 \| 334 \| Uncharacterized protein \| Uncharacterized protein; afdb-uniprot50 \| AF-A0A368VC74-F1-MODEL\_V4 \| 1.0 \| 7.953e-26 \| 965 \| 0.334 \| 281 \| 177 \| 5 \| 2 \| 279 \| 55 \| 328 \| Uncharacterized protein \| Uncharacterized protein; afdb-uniprot50 \| AF-A0A4U8YMH7-F1-MODEL\_V4 \| 1.0 \| 7.953e-26 \| 964 \| 0.306 \| 284 \| 186 \| 6 \| 1 \| 281 \| 51 \| 326 \| Phage late control gene d protein (Gpd) \| Phage late control gene d protein (Gpd); afdb-uniprot50 \| AF-L8MFF9-F1-MODEL\_V4 \| 1.0 \| 8.92e-26 \| 964 \| 0.352 \| 284 \| 172 \| 6 \| 2 \| 281 \| 57 \| 332 \| Phage late control D \| Phage late control D; afdb-uniprot50 \| AF-A0A3N1QKI9-F1-MODEL\_V4 \| 1.0 \| 1.122e-25 \| 963 \| 0.345 \| 281 \| 174 \| 6 \| 2 \| 279 \| 55 \| 328 \| Uncharacterized protein \| Uncharacterized protein; afdb-uniprot50 \| AF-A0A519ZAP9-F1-MODEL\_V4 \| 1.0 \| 2.009e-26 \| 961 \| 0.325 \| 286 \| 167 \| 8 \| 7 \| 275 \| 2 \| 278 \| Uncharacterized protein \| Uncharacterized protein; afdb-uniprot50 \| AF-A0A2D8Q6U7-F1-MODEL\_V4 \| 1.0 \| 2.526e-26 \| 961 \| 0.339 \| 286 \| 174 \| 6 \| 1 \| 280 \| 51 \| 327 \| Phage tail protein \| Phage tail protein; afdb-uniprot50 \| AF-A0A4Y6UAT9-F1-MODEL\_V4 \| 1.0 \| 2.253e-26 \| 961 \| 0.301 \| 312 \| 184 \| 6 \| 1 \| 278 \| 81 \| 392 \| Uncharacterized protein \| Uncharacterized protein; afdb-uniprot50 \| AF-A0A256CB22-F1-MODEL\_V4 \| 1.0 \| 6.323e-26 \| 957 \| 0.342 \| 283 \| 176 \| 5 \| 1 \| 281 \| 56 \| 330 \| Uncharacterized protein \| Uncharacterized protein; afdb-uniprot50 \| AF-A0A166Z710-F1-MODEL\_V4 \| 1.0 \| 4.483e-26 \| 956 \| 0.267 \| 307 \| 198 \| 8 \| 1 \| 290 \| 49 \| 345 \| Late control protein D \| Late control protein D; afdb-uniprot50 \| AF-A0A1A9VKH9-F1-MODEL\_V4 \| 1.0 \| 1.508e-26 \| 953 \| 0.359 \| 270 \| 165 \| 3 \| 7 \| 275 \| 49 \| 311 \| Ankyrin\_rpt-contain\_dom domain-containing protein \| Ankyrin\_rpt-contain\_dom domain-containing protein; afdb-uniprot50 \| AF-A0A7W4WU07-F1-MODEL\_V4 \| 1.0 \| 5.324e-26 \| 952 \| 0.309 \| 284 \| 182 \| 10 \| 2 \| 279 \| 22 \| 297 \| Phage protein D \| Phage protein D; afdb-uniprot50 \| AF-A0A6L7X9K2-F1-MODEL\_V4 \| 1.0 \| 5.276e-25 \| 951 \| 0.377 \| 225 \| 137 \| 3 \| 57 \| 279 \| 5 \| 228 \| Uncharacterized protein \| Uncharacterized protein; afdb-uniprot50 \| AF-A0A5K6V8S0-F1-MODEL\_V4 \| 1.0 \| 9.532e-27 \| 950 \| 0.353 \| 283 \| 170 \| 4 \| 1 \| 276 \| 53 \| 329 \| Late control protein \| Late control protein; afdb-uniprot50 \| AF-A0A7W6E035-F1-MODEL\_V4 \| 1.0 \| 1.991e-25 \| 949 \| 0.322 \| 276 \| 180 \| 2 \| 1 \| 276 \| 56 \| 324 \| Uncharacterized protein \| Uncharacterized protein; afdb-uniprot50 \| AF-A0A713UWD1-F1-MODEL\_V4 \| 1.0 \| 3.365e-26 \| 948 \| 0.3 \| 313 \| 179 \| 11 \| 1 \| 282 \| 18 \| 321 \| Phage late control D family protein \| Phage late control D family protein; afdb-uniprot50 \| AF-A0A1I1UD34-F1-MODEL\_V4 \| 1.0 \| 5.324e-26 \| 948 \| 0.292 \| 287 \| 190 \| 9 \| 1 \| 279 \| 50 \| 331 \| Uncharacterized protein \| Uncharacterized protein; afdb-uniprot50 \| AF-A0A5M6IGE8-F1-MODEL\_V4 \| 1.0 \| 4.483e-26 \| 946 \| 0.317 \| 287 \| 178 \| 6 \| 1 \| 277 \| 55 \| 333 \| Late control protein D \| Late control protein D; afdb-uniprot50 \| AF-A0A840UI14-F1-MODEL\_V4 \| 1.0 \| 2.364e-25 \| 945 \| 0.327 \| 281 \| 179 \| 5 \| 2 \| 279 \| 55 \| 328 \| Uncharacterized protein \| Uncharacterized protein; afdb-uniprot50 \| AF-A0A3G2VC57-F1-MODEL\_V4 \| 1.0 \| 3.178e-26 \| 942 \| 0.237 \| 299 \| 204 \| 8 \| 1 \| 282 \| 10 \| 301 \| Uncharacterized protein \| Uncharacterized protein; afdb-uniprot50 \| AF-A0A3T0N1H3-F1-MODEL\_V4 \| 1.0 \| 3.997e-26 \| 941 \| 0.325 \| 280 \| 179 \| 6 \| 1 \| 278 \| 51 \| 322 \| Phage tail protein \| Phage tail protein; afdb-uniprot50 \| AF-A0A1H8RSK6-F1-MODEL\_V4 \| 1.0 \| 6.323e-26 \| 941 \| 0.313 \| 287 \| 180 \| 6 \| 1 \| 279 \| 51 \| 328 \| Uncharacterized protein \| Uncharacterized protein; afdb-uniprot50 \| AF-A0A519MN03-F1-MODEL\_V4 \| 1.0 \| 1e-25 \| 940 \| 0.371 \| 285 \| 167 \| 7 \| 2 \| 282 \| 65 \| 341 \| Phage late control D family protein \| Phage late control D family protein; afdb-uniprot50 \| AF-A0A2E3M0S7-F1-MODEL\_V4 \| 1.0 \| 1.009e-26 \| 939 \| 0.329 \| 282 \| 179 \| 5 \| 1 \| 280 \| 63 \| 336 \| Phage tail protein \| Phage tail protein; afdb-uniprot50 \| AF-A0A3G2FX49-F1-MODEL\_V4 \| 1.0 \| 8.423e-26 \| 937 \| 0.311 \| 260 \| 170 \| 4 \| 22 \| 279 \| 5 \| 257 \| Phage late control D family protein \| Phage late control D family protein; afdb-uniprot50 \| AF-A0A381EBY3-F1-MODEL\_V4 \| 1.0 \| 2.232e-25 \| 937 \| 0.312 \| 282 \| 184 \| 5 \| 1 \| 279 \| 49 \| 323 \| Phage protein D \| Phage protein D; afdb-uniprot50 \| AF-A0A847KNG7-F1-MODEL\_V4 \| 1.0 \| 1.88e-25 \| 937 \| 0.263 \| 288 \| 196 \| 7 \| 1 \| 280 \| 51 \| 330 \| Phage late control D family protein \| Phage late control D family protein; afdb-uniprot50 \| AF-A0A1X7NEM5-F1-MODEL\_V4 \| 1.0 \| 9.446e-26 \| 936 \| 0.332 \| 280 \| 176 \| 6 \| 1 \| 278 \| 49 \| 319 \| Uncharacterized protein \| Uncharacterized protein; afdb-uniprot50 \| AF-W6TE35-F1-MODEL\_V4 \| 1.0 \| 3.809e-27 \| 936 \| 0.33 \| 312 \| 191 \| 4 \| 1 \| 311 \| 51 \| 345 \| Gene D protein \| Gene D protein; afdb-uniprot50 \| AF-A0A3E0X405-F1-MODEL\_V4 \| 1.0 \| 3.335e-25 \| 932 \| 0.323 \| 281 \| 180 \| 6 \| 1 \| 278 \| 59 \| 332 \| Late control protein \| Late control protein; afdb-uniprot50 \| AF-G3IRG5-F1-MODEL\_V4 \| 1.0 \| 2.504e-25 \| 931 \| 0.313 \| 281 \| 178 \| 9 \| 1 \| 275 \| 51 \| 322 \| Late control D family protein \| Late control D family protein; afdb-uniprot50 \| AF-A0A1G7PWP3-F1-MODEL\_V4 \| 1.0 \| 5.027e-26 \| 931 \| 0.259 \| 293 \| 197 \| 8 \| 1 \| 282 \| 50 \| 333 \| Uncharacterized protein \| Uncharacterized protein; afdb-uniprot50 \| AF-A0A2E2MXP4-F1-MODEL\_V4 \| 1.0 \| 3.365e-26 \| 930 \| 0.305 \| 314 \| 199 \| 8 \| 1 \| 311 \| 67 \| 364 \| Late control protein \| Late control protein; afdb-uniprot50 \| AF-A0A315BKV5-F1-MODEL\_V4 \| 1.0 \| 3.001e-26 \| 928 \| 0.302 \| 324 \| 198 \| 9 \| 2 \| 310 \| 61 \| 371 \| Uncharacterized protein \| Uncharacterized protein; afdb-uniprot50 \| AF-A0A5P9F186-F1-MODEL\_V4 \| 1.0 \| 7.028e-25 \| 927 \| 0.316 \| 281 \| 181 \| 5 \| 1 \| 278 \| 50 \| 322 \| Phage late control gene D protein (GPD) \| Phage late control gene D protein (GPD); afdb-uniprot50 \| AF-A0A235EW76-F1-MODEL\_V4 \| 1.0 \| 3.149e-25 \| 927 \| 0.316 \| 297 \| 178 \| 6 \| 1 \| 279 \| 54 \| 343 \| Late control protein \| Late control protein; afdb-uniprot50 \| AF-A0A3S4CKJ0-F1-MODEL\_V4 \| 1.0 \| 3.149e-25 \| 925 \| 0.307 \| 280 \| 184 \| 5 \| 2 \| 279 \| 52 \| 323 \| Phage late control gene D protein (GPD) \| Phage late control gene D protein (GPD); afdb-uniprot50 \| AF-A0A2T6F1B4-F1-MODEL\_V4 \| 1.0 \| 2.651e-25 \| 925 \| 0.266 \| 304 \| 205 \| 7 \| 1 \| 296 \| 54 \| 347 \| Late control protein \| Late control protein; afdb-uniprot50 \| AF-A0A0Q3BFB6-F1-MODEL\_V4 \| 1.0 \| 7.028e-25 \| 923 \| 0.311 \| 279 \| 182 \| 5 \| 1 \| 277 \| 51 \| 321 \| Uncharacterized protein \| Uncharacterized protein; afdb-uniprot50 \| AF-A0A7W6RE50-F1-MODEL\_V4 \| 1.0 \| 3.149e-25 \| 922 \| 0.278 \| 294 \| 197 \| 10 \| 1 \| 288 \| 61 \| 345 \| Uncharacterized protein \| Uncharacterized protein; afdb-uniprot50 \| AF-A0A2N1WF73-F1-MODEL\_V4 \| 1.0 \| 5.917e-25 \| 921 \| 0.332 \| 283 \| 179 \| 6 \| 1 \| 280 \| 61 \| 336 \| Late control protein \| Late control protein; afdb-uniprot50 \| AF-C3X1X3-F1-MODEL\_V4 \| 1.0 \| 1.568e-24 \| 921 \| 0.262 \| 290 \| 201 \| 6 \| 1 \| 284 \| 55 \| 337 \| Uncharacterized protein \| Uncharacterized protein; afdb-uniprot50 \| AF-A0A1X7Q5G6-F1-MODEL\_V4 \| 1.0 \| 6.696e-26 \| 921 \| 0.296 \| 293 \| 183 \| 6 \| 1 \| 278 \| 56 \| 340 \| Uncharacterized protein \| Uncharacterized protein; afdb-uniprot50 \| AF-A0A7Z0UXS5-F1-MODEL\_V4 \| 1.0 \| 1.676e-25 \| 921 \| 0.295 \| 318 \| 208 \| 9 \| 1 \| 311 \| 98 \| 406 \| Uncharacterized protein \| Uncharacterized protein; afdb-uniprot50 \| AF-A0A0Q2Y5J5-F1-MODEL\_V4 \| 1.0 \| 4.442e-25 \| 920 \| 0.255 \| 297 \| 204 \| 7 \| 2 \| 290 \| 51 \| 338 \| Uncharacterized protein \| Uncharacterized protein; afdb-uniprot50 \| AF-A0A096FN45-F1-MODEL\_V4 \| 1.0 \| 4.442e-25 \| 920 \| 0.267 \| 310 \| 195 \| 7 \| 2 \| 287 \| 49 \| 350 \| Tail protein \| Tail protein; afdb-uniprot50 \| AF-A0A2E5BTI9-F1-MODEL\_V4 \| 1.0 \| 3.961e-25 \| 918 \| 0.29 \| 286 \| 189 \| 7 \| 1 \| 279 \| 51 \| 329 \| Late control protein \| Late control protein; afdb-uniprot50 \| AF-A0A5B0VXU9-F1-MODEL\_V4 \| 1.0 \| 3.961e-25 \| 918 \| 0.273 \| 296 \| 191 \| 7 \| 1 \| 279 \| 63 \| 351 \| Phage late control D family protein \| Phage late control D family protein; afdb-uniprot50 \| AF-A0A089PYY8-F1-MODEL\_V4 \| 1.0 \| 1.059e-25 \| 918 \| 0.304 \| 305 \| 198 \| 6 \| 2 \| 295 \| 54 \| 355 \| Uncharacterized protein \| Uncharacterized protein; afdb-uniprot50 \| AF-A0A2N7P6U8-F1-MODEL\_V4 \| 1.0 \| 1.112e-24 \| 917 \| 0.307 \| 280 \| 184 \| 5 \| 1 \| 278 \| 52 \| 323 \| Uncharacterized protein \| Uncharacterized protein; afdb-uniprot50 \| AF-A0A1G7D4E3-F1-MODEL\_V4 \| 1.0 \| 6.636e-25 \| 917 \| 0.287 \| 289 \| 194 \| 7 \| 1 \| 284 \| 51 \| 332 \| Uncharacterized protein \| Uncharacterized protein; afdb-uniprot50 \| AF-A0A8B2NZ52-F1-MODEL\_V4 \| 1.0 \| 6.266e-25 \| 916 \| 0.292 \| 284 \| 184 \| 8 \| 2 \| 277 \| 54 \| 328 \| Late control protein D \| Late control protein D; afdb-uniprot50 \| AF-A0A6L9FMV3-F1-MODEL\_V4 \| 1.0 \| 4.937e-24 \| 916 \| 0.291 \| 285 \| 194 \| 6 \| 1 \| 279 \| 54 \| 336 \| Late control protein D \| Late control protein D; afdb-uniprot50 \| AF-A0A2S8R677-F1-MODEL\_V4 \| 1.0 \| 2.974e-25 \| 916 \| 0.283 \| 307 \| 184 \| 10 \| 1 \| 280 \| 62 \| 359 \| Late control protein D \| Late control protein D; afdb-uniprot50 \| AF-A0A1D2QS63-F1-MODEL\_V4 \| 1.0 \| 7.092e-26 \| 914 \| 0.304 \| 292 \| 178 \| 8 \| 1 \| 280 \| 49 \| 327 \| Uncharacterized protein \| Uncharacterized protein; afdb-uniprot50 \| AF-A0A345DE48-F1-MODEL\_V4 \| 1.0 \| 5.971e-26 \| 914 \| 0.353 \| 283 \| 169 \| 8 \| 1 \| 277 \| 49 \| 323 \| Uncharacterized protein \| Uncharacterized protein; afdb-uniprot50 \| AF-A0A328X5L9-F1-MODEL\_V4 \| 1.0 \| 7.028e-25 \| 911 \| 0.324 \| 287 \| 177 \| 8 \| 2 \| 280 \| 64 \| 341 \| Uncharacterized protein \| Uncharacterized protein; afdb-uniprot50 \| AF-A0A545T5X0-F1-MODEL\_V4 \| 1.0 \| 2.089e-24 \| 908 \| 0.279 \| 283 \| 189 \| 5 \| 1 \| 279 \| 74 \| 345 \| Uncharacterized protein \| Uncharacterized protein; afdb-uniprot50 \| AF-A0A1W1Z5V8-F1-MODEL\_V4 \| 1.0 \| 8.839e-25 \| 906 \| 0.267 \| 284 \| 194 \| 8 \| 2 \| 278 \| 51 \| 327 \| Uncharacterized protein \| Uncharacterized protein; afdb-uniprot50 \| AF-A0A1N6Q3I2-F1-MODEL\_V4 \| 1.0 \| 5.537e-24 \| 905 \| 0.301 \| 282 \| 185 \| 6 \| 1 \| 278 \| 55 \| 328 \| Uncharacterized protein \| Uncharacterized protein; afdb-uniprot50 \| AF-A0A251ZIV1-F1-MODEL\_V4 \| 1.0 \| 8.423e-26 \| 904 \| 0.259 \| 308 \| 201 \| 8 \| 1 \| 288 \| 71 \| 371 \| Uncharacterized protein \| Uncharacterized protein; afdb-uniprot50 \| AF-A0A143DG31-F1-MODEL\_V4 \| 1.0 \| 4.662e-24 \| 903 \| 0.29 \| 282 \| 187 \| 8 \| 1 \| 276 \| 50 \| 324 \| Uncharacterized protein \| Uncharacterized protein; afdb-uniprot50 \| AF-A0A1E7Q062-F1-MODEL\_V4 \| 1.0 \| 2.628e-24 \| 902 \| 0.362 \| 281 \| 169 \| 5 \| 2 \| 279 \| 55 \| 328 \| Uncharacterized protein \| Uncharacterized protein; afdb-uniprot50 \| AF-A0A7Z1UDY9-F1-MODEL\_V4 \| 1.0 \| 2.783e-24 \| 902 \| 0.223 \| 326 \| 208 \| 8 \| 1 \| 289 \| 67 \| 384 \| Uncharacterized protein \| Uncharacterized protein; afdb-uniprot50 \| AF-A0A6H1S4A5-F1-MODEL\_V4 \| 1.0 \| 2.947e-24 \| 902 \| 0.225 \| 328 \| 207 \| 8 \| 1 \| 289 \| 69 \| 388 \| Late control D family protein \| Late control D family protein; afdb-uniprot50 \| AF-A0A348HI91-F1-MODEL\_V4 \| 1.0 \| 5.587e-25 \| 901 \| 0.31 \| 283 \| 183 \| 8 \| 1 \| 279 \| 55 \| 329 \| Phage protein D \| Phage protein D; afdb-uniprot50 \| AF-A0A350LX83-F1-MODEL\_V4 \| 1.0 \| 2.108e-25 \| 900 \| 0.314 \| 283 \| 184 \| 6 \| 1 \| 281 \| 51 \| 325 \| Phage tail protein \| Phage tail protein; afdb-uniprot50 \| AF-A0A1E3G722-F1-MODEL\_V4 \| 1.0 \| 6.696e-26 \| 899 \| 0.345 \| 278 \| 175 \| 2 \| 2 \| 279 \| 54 \| 324 \| Uncharacterized protein \| Uncharacterized protein; afdb-uniprot50 \| AF-A0A1M4WEC4-F1-MODEL\_V4 \| 1.0 \| 1.398e-24 \| 897 \| 0.269 \| 286 \| 194 \| 9 \| 1 \| 278 \| 51 \| 329 \| Uncharacterized protein \| Uncharacterized protein; afdb-uniprot50 \| AF-G3XCU8-F1-MODEL\_V4 \| 1.0 \| 3.5e-24 \| 896 \| 0.29 \| 282 \| 183 \| 5 \| 1 \| 275 \| 51 \| 322 \| Uncharacterized protein \| Uncharacterized protein; afdb-uniprot50 \| AF-A0A1Y2K0X3-F1-MODEL\_V4 \| 1.0 \| 1.424e-26 \| 896 \| 0.342 \| 283 \| 176 \| 6 \| 1 \| 280 \| 51 \| 326 \| Uncharacterized protein \| Uncharacterized protein; afdb-uniprot50 \| AF-W0HLP1-F1-MODEL\_V4 \| 1.0 \| 4.704e-25 \| 896 \| 0.293 \| 300 \| 182 \| 7 \| 1 \| 277 \| 64 \| 356 \| Late control gene D protein \| Late control gene D protein; afdb-uniprot50 \| AF-A0A1B4NNN3-F1-MODEL\_V4 \| 1.0 \| 7.51e-26 \| 896 \| 0.311 \| 318 \| 183 \| 9 \| 2 \| 291 \| 74 \| 383 \| Late control protein D \| Late control protein D; afdb-uniprot50 \| AF-A0A840P1G0-F1-MODEL\_V4 \| 1.0 \| 2.232e-25 \| 896 \| 0.269 \| 289 \| 188 \| 10 \| 1 \| 281 \| 499 \| 772 \| Phage protein D \| Phage protein D; afdb-uniprot50 \| AF-C5BEQ4-F1-MODEL\_V4 \| 1.0 \| 9.913e-25 \| 895 \| 0.311 \| 299 \| 179 \| 8 \| 1 \| 280 \| 59 \| 349 \| Phage late control gene D protein (GPD) \| Phage late control gene D protein (GPD); afdb-uniprot50 \| AF-A0A6D0JLG2-F1-MODEL\_V4 \| 1.0 \| 5.537e-24 \| 895 \| 0.284 \| 288 \| 192 \| 7 \| 1 \| 277 \| 56 \| 340 \| Late control protein \| Late control protein; afdb-uniprot50 \| AF-C6C6Z8-F1-MODEL\_V4 \| 1.0 \| 1.247e-24 \| 895 \| 0.22 \| 326 \| 208 \| 7 \| 2 \| 289 \| 77 \| 394 \| Late control D family protein \| Late control D family protein; afdb-uniprot50 \| AF-A0A166F5B7-F1-MODEL\_V4 \| 1.0 \| 2.089e-24 \| 894 \| 0.269 \| 289 \| 196 \| 8 \| 1 \| 281 \| 51 \| 332 \| Phage late control gene D protein (GPD) \| Phage late control gene D protein (GPD); afdb-uniprot50 \| AF-A0A212J3N4-F1-MODEL\_V4 \| 1.0 \| 8.839e-25 \| 893 \| 0.295 \| 284 \| 188 \| 7 \| 2 \| 280 \| 52 \| 328 \| Gene D protein \| Gene D protein; afdb-uniprot50 \| AF-A0A7Z1Y516-F1-MODEL\_V4 \| 1.0 \| 2.783e-24 \| 892 \| 0.272 \| 286 \| 192 \| 7 \| 2 \| 275 \| 61 \| 342 \| Late control protein D \| Late control protein D; afdb-uniprot50 \| AF-N6VTB1-F1-MODEL\_V4 \| 1.0 \| 3.564e-26 \| 892 \| 0.282 \| 340 \| 169 \| 7 \| 10 \| 282 \| 1 \| 332 \| Phage late control protein D \| Phage late control protein D; afdb-uniprot50 \| AF-A0A4U3FAV9-F1-MODEL\_V4 \| 1.0 \| 2.089e-24 \| 892 \| 0.263 \| 323 \| 205 \| 8 \| 1 \| 298 \| 63 \| 377 \| Uncharacterized protein \| Uncharacterized protein; afdb-uniprot50 \| AF-A0A844YYZ9-F1-MODEL\_V4 \| 1.0 \| 4.402e-24 \| 889 \| 0.277 \| 288 \| 192 \| 7 \| 1 \| 279 \| 11 \| 291 \| Phage late control D family protein \| Phage late control D family protein; afdb-uniprot50 \| AF-A0A1N6M6E7-F1-MODEL\_V4 \| 1.0 \| 4.195e-25 \| 889 \| 0.272 \| 294 \| 195 \| 9 \| 1 \| 281 \| 33 \| 320 \| Phage late control gene D protein (GPD) \| Phage late control gene D protein (GPD); afdb-uniprot50 \| AF-A0A2D2C160-F1-MODEL\_V4 \| 1.0 \| 1.398e-24 \| 889 \| 0.267 \| 284 \| 194 \| 7 \| 2 \| 278 \| 51 \| 327 \| Late control protein D \| Late control protein D; afdb-uniprot50 \| AF-A0A7Y6Z4E7-F1-MODEL\_V4 \| 1.0 \| 2.089e-24 \| 889 \| 0.273 \| 285 \| 193 \| 7 \| 1 \| 279 \| 51 \| 327 \| Uncharacterized protein \| Uncharacterized protein; afdb-uniprot50 \| AF-S6I325-F1-MODEL\_V4 \| 1.0 \| 3.706e-24 \| 889 \| 0.283 \| 293 \| 192 \| 7 \| 1 \| 282 \| 54 \| 339 \| Uncharacterized protein \| Uncharacterized protein; afdb-uniprot50 \| AF-A0A0A3ZBQ9-F1-MODEL\_V4 \| 1.0 \| 1.481e-24 \| 889 \| 0.264 \| 306 \| 192 \| 6 \| 1 \| 280 \| 58 \| 356 \| Late control protein D \| Late control protein D; afdb-uniprot50 \| AF-A0A0S2F7D3-F1-MODEL\_V4 \| 1.0 \| 5.228e-24 \| 888 \| 0.333 \| 282 \| 176 \| 8 \| 2 \| 279 \| 31 \| 304 \| Phage late control gene D family protein \| Phage late control gene D family protein; afdb-uniprot50 \| AF-A0A2G6RW27-F1-MODEL\_V4 \| 1.0 \| 6.21e-24 \| 888 \| 0.285 \| 242 \| 168 \| 4 \| 57 \| 296 \| 10 \| 248 \| Uncharacterized protein \| Uncharacterized protein; afdb-uniprot50 \| AF-A0A2S5TZV2-F1-MODEL\_V4 \| 1.0 \| 1.568e-24 \| 887 \| 0.265 \| 282 \| 194 \| 7 \| 2 \| 277 \| 52 \| 326 \| Uncharacterized protein \| Uncharacterized protein; afdb-uniprot50 \| AF-A0A3M2YE85-F1-MODEL\_V4 \| 1.0 \| 1.661e-24 \| 887 \| 0.28 \| 282 \| 186 \| 6 \| 1 \| 275 \| 85 \| 356 \| Tail protein D \| Tail protein D; afdb-uniprot50 \| AF-A0A6G8AI05-F1-MODEL\_V4 \| 1.0 \| 4.442e-25 \| 886 \| 0.284 \| 313 \| 206 \| 6 \| 2 \| 311 \| 65 \| 362 \| Phage late control D family protein \| Phage late control D family protein; afdb-uniprot50 \| AF-A0A378YM62-F1-MODEL\_V4 \| 1.0 \| 2.364e-25 \| 886 \| 0.311 \| 315 \| 171 \| 10 \| 1 \| 279 \| 73 \| 377 \| Phage protein D \| Phage protein D; afdb-uniprot50 \| AF-D6JBF4-F1-MODEL\_V4 \| 1.0 \| 4.937e-24 \| 886 \| 0.244 \| 311 \| 203 \| 7 \| 1 \| 286 \| 110 \| 413 \| Uncharacterized protein \| Uncharacterized protein; afdb-uniprot50 \| AF-A0A199YKM6-F1-MODEL\_V4 \| 1.0 \| 8.839e-25 \| 884 \| 0.308 \| 279 \| 183 \| 5 \| 1 \| 276 \| 51 \| 322 \| Uncharacterized protein \| Uncharacterized protein; afdb-uniprot50 \| AF-A0A212KMW2-F1-MODEL\_V4 \| 1.0 \| 4.402e-24 \| 884 \| 0.341 \| 287 \| 173 \| 9 \| 1 \| 278 \| 51 \| 330 \| Late control D family protein \| Late control D family protein; afdb-uniprot50 \| AF-A0A7W7KUK6-F1-MODEL\_V4 \| 1.0 \| 3.121e-24 \| 883 \| 0.347 \| 279 \| 171 \| 7 \| 2 \| 276 \| 55 \| 326 \| Uncharacterized protein \| Uncharacterized protein; afdb-uniprot50 \| AF-A0A2G6EX41-F1-MODEL\_V4 \| 1.0 \| 1.167e-23 \| 882 \| 0.315 \| 282 \| 183 \| 5 \| 2 \| 280 \| 65 \| 339 \| Late control protein D \| Late control protein D; afdb-uniprot50 \| AF-A0A4R3V673-F1-MODEL\_V4 \| 1.0 \| 2.089e-24 \| 882 \| 0.27 \| 299 \| 197 \| 10 \| 2 \| 291 \| 60 \| 346 \| Uncharacterized protein \| Uncharacterized protein; afdb-uniprot50 \| AF-A0A7C8G8Q1-F1-MODEL\_V4 \| 1.0 \| 3.5e-24 \| 882 \| 0.261 \| 310 \| 199 \| 8 \| 1 \| 288 \| 63 \| 364 \| Phage late control D family protein \| Phage late control D family protein; afdb-uniprot50 \| AF-A0A6N3HNE7-F1-MODEL\_V4 \| 1.0 \| 5.228e-24 \| 882 \| 0.25 \| 311 \| 201 \| 6 \| 1 \| 286 \| 109 \| 412 \| Phage late control gene D protein (GPD) \| Phage late control gene D protein (GPD); afdb-uniprot50 \| AF-A0A423PQL7-F1-MODEL\_V4 \| 1.0 \| 1e-25 \| 881 \| 0.299 \| 284 \| 182 \| 6 \| 1 \| 281 \| 29 \| 298 \| Uncharacterized protein \| Uncharacterized protein; afdb-uniprot50 \| AF-A0A3R9DQ60-F1-MODEL\_V4 \| 1.0 \| 5.864e-24 \| 881 \| 0.309 \| 284 \| 183 \| 7 \| 1 \| 277 \| 52 \| 329 \| Phage late control D family protein \| Phage late control D family protein; afdb-uniprot50 \| AF-A0A0T9KDK3-F1-MODEL\_V4 \| 1.0 \| 3.961e-25 \| 880 \| 0.254 \| 310 \| 195 \| 8 \| 1 \| 282 \| 130 \| 431 \| Late control gene D protein \| Late control gene D protein; afdb-uniprot50 \| AF-A0A559QNE2-F1-MODEL\_V4 \| 1.0 \| 1.386e-23 \| 878 \| 0.303 \| 280 \| 185 \| 6 \| 1 \| 277 \| 56 \| 328 \| Uncharacterized protein \| Uncharacterized protein; afdb-uniprot50 \| AF-A0A455W3W8-F1-MODEL\_V4 \| 1.0 \| 2.481e-24 \| 876 \| 0.255 \| 321 \| 189 \| 9 \| 1 \| 279 \| 63 \| 375 \| Regulator of late gene expression \| Regulator of late gene expression; afdb-uniprot50 \| AF-Q8ZMV5-F1-MODEL\_V4 \| 1.0 \| 7.028e-25 \| 875 \| 0.291 \| 309 \| 184 \| 8 \| 1 \| 281 \| 63 \| 364 \| Fels-2 prophage protein \| Fels-2 prophage protein; afdb-uniprot50 \| AF-A0A3A8E5R3-F1-MODEL\_V4 \| 1.0 \| 1.167e-23 \| 874 \| 0.318 \| 283 \| 181 \| 7 \| 2 \| 279 \| 54 \| 329 \| Uncharacterized protein \| Uncharacterized protein; afdb-uniprot50 \| AF-A0A1S1NEL7-F1-MODEL\_V4 \| 1.0 \| 4.157e-24 \| 874 \| 0.267 \| 288 \| 191 \| 6 \| 1 \| 276 \| 54 \| 333 \| Phage tail protein \| Phage tail protein; afdb-uniprot50 \| AF-A0A5V6NKW8-F1-MODEL\_V4 \| 1.0 \| 4.662e-24 \| 873 \| 0.278 \| 284 \| 195 \| 3 \| 1 \| 275 \| 61 \| 343 \| Late control protein D \| Late control protein D; afdb-uniprot50 \| AF-A0A6L3Z4U3-F1-MODEL\_V4 \| 1.0 \| 9.361e-25 \| 872 \| 0.269 \| 319 \| 204 \| 9 \| 1 \| 311 \| 52 \| 349 \| Late control protein D \| Late control protein D; afdb-uniprot50 \| AF-A0A198GQJ0-F1-MODEL\_V4 \| 1.0 \| 2.628e-24 \| 871 \| 0.264 \| 306 \| 192 \| 6 \| 1 \| 280 \| 61 \| 359 \| Phage late control gene D protein \| Phage late control gene D protein; afdb-uniprot50 \| AF-A0A5U9S5Z9-F1-MODEL\_V4 \| 1.0 \| 4.402e-24 \| 871 \| 0.278 \| 327 \| 183 \| 8 \| 1 \| 282 \| 163 \| 481 \| Phage late control D family protein \| Phage late control D family protein; afdb-uniprot50 \| AF-A0A221DKQ4-F1-MODEL\_V4 \| 1.0 \| 5.864e-24 \| 868 \| 0.266 \| 330 \| 193 \| 10 \| 1 \| 290 \| 63 \| 383 \| Uncharacterized protein \| Uncharacterized protein; afdb-uniprot50 \| AF-A0A516W9U2-F1-MODEL\_V4 \| 1.0 \| 1.646e-23 \| 867 \| 0.306 \| 284 \| 186 \| 6 \| 1 \| 280 \| 59 \| 335 \| Phage late control D family protein \| Phage late control D family protein; afdb-uniprot50 \| AF-A0A1C3EBM6-F1-MODEL\_V4 \| 1.0 \| 1.236e-23 \| 866 \| 0.301 \| 279 \| 187 \| 3 \| 1 \| 278 \| 52 \| 323 \| Uncharacterized protein \| Uncharacterized protein; afdb-uniprot50 \| AF-B5JYF8-F1-MODEL\_V4 \| 1.0 \| 6.21e-24 \| 866 \| 0.297 \| 279 \| 186 \| 6 \| 2 \| 277 \| 57 \| 328 \| Phage late control D \| Phage late control D; afdb-uniprot50 \| AF-A0A269PKZ6-F1-MODEL\_V4 \| 1.0 \| 5.864e-24 \| 866 \| 0.281 \| 291 \| 190 \| 6 \| 1 \| 279 \| 53 \| 336 \| Uncharacterized protein \| Uncharacterized protein; afdb-uniprot50 \| AF-A0A7S6S5Y4-F1-MODEL\_V4 \| 1.0 \| 5.228e-24 \| 866 \| 0.271 \| 287 \| 190 \| 7 \| 1 \| 278 \| 51 \| 327 \| Late control protein \| Late control protein; afdb-uniprot50 \| AF-A0A4Q6CNI6-F1-MODEL\_V4 \| 1.0 \| 1.646e-23 \| 865 \| 0.249 \| 277 \| 198 \| 5 \| 1 \| 277 \| 13 \| 279 \| Uncharacterized protein \| Uncharacterized protein; afdb-uniprot50 \| AF-A0A381AML1-F1-MODEL\_V4 \| 1.0 \| 2.192e-23 \| 865 \| 0.275 \| 287 \| 191 \| 7 \| 2 \| 279 \| 35 \| 313 \| Phage protein D \| Phage protein D; afdb-uniprot50 \| AF-A0A1C7L3E6-F1-MODEL\_V4 \| 1.0 \| 1.554e-23 \| 865 \| 0.283 \| 282 \| 192 \| 5 \| 1 \| 279 \| 59 \| 333 \| Phage late control protein \| Phage late control protein; afdb-uniprot50 \| AF-A0A4S1DI58-F1-MODEL\_V4 \| 1.0 \| 1.167e-23 \| 865 \| 0.326 \| 282 \| 175 \| 9 \| 1 \| 275 \| 69 \| 342 \| DNA primase \| DNA primase; afdb-uniprot50 \| AF-A0A0Q5LSP6-F1-MODEL\_V4 \| 1.0 \| 3.925e-24 \| 864 \| 0.264 \| 314 \| 208 \| 6 \| 1 \| 311 \| 61 \| 354 \| Late control protein \| Late control protein; afdb-uniprot50 \| AF-A0A329B2U7-F1-MODEL\_V4 \| 1.0 \| 3.706e-24 \| 863 \| 0.303 \| 264 \| 171 \| 6 \| 22 \| 280 \| 4 \| 259 \| Uncharacterized protein \| Uncharacterized protein; afdb-uniprot50 \| AF-A0A1M7YYR7-F1-MODEL\_V4 \| 1.0 \| 2.628e-24 \| 863 \| 0.278 \| 294 \| 185 \| 6 \| 1 \| 274 \| 51 \| 337 \| Phage late control gene D protein (GPD) \| Phage late control gene D protein (GPD); afdb-uniprot50 \| AF-Q1QKT4-F1-MODEL\_V4 \| 1.0 \| 5.864e-24 \| 862 \| 0.27 \| 288 \| 194 \| 8 \| 2 \| 281 \| 53 \| 332 \| Phage late control D \| Phage late control D; afdb-uniprot50 \| AF-A0A1E3G781-F1-MODEL\_V4 \| 1.0 \| 1.167e-23 \| 861 \| 0.255 \| 290 \| 198 \| 8 \| 1 \| 279 \| 56 \| 338 \| Uncharacterized protein \| Uncharacterized protein; afdb-uniprot50 \| AF-A0A5S9P2X8-F1-MODEL\_V4 \| 1.0 \| 3.121e-24 \| 860 \| 0.25 \| 287 \| 200 \| 7 \| 1 \| 281 \| 52 \| 329 \| Uncharacterized protein \| Uncharacterized protein; afdb-uniprot50 \| AF-A0A292AJG1-F1-MODEL\_V4 \| 1.0 \| 9.361e-25 \| 860 \| 0.263 \| 322 \| 189 \| 11 \| 1 \| 288 \| 70 \| 377 \| Late control protein D \| Late control protein D; afdb-uniprot50 \| AF-A0A395R2S1-F1-MODEL\_V4 \| 1.0 \| 1.177e-24 \| 860 \| 0.28 \| 317 \| 197 \| 6 \| 2 \| 294 \| 59 \| 368 \| Late control protein \| Late control protein; afdb-uniprot50 \| AF-A0A2A2B6A0-F1-MODEL\_V4 \| 1.0 \| 4.402e-24 \| 859 \| 0.255 \| 294 \| 196 \| 9 \| 1 \| 278 \| 33 \| 319 \| Uncharacterized protein \| Uncharacterized protein; afdb-uniprot50 \| AF-E2CJS5-F1-MODEL\_V4 \| 1.0 \| 7.442e-25 \| 859 \| 0.271 \| 335 \| 201 \| 10 \| 1 \| 311 \| 54 \| 369 \| Late control D protein \| Late control D protein; afdb-uniprot50 \| AF-F4BFP6-F1-MODEL\_V4 \| 1.0 \| 8.839e-25 \| 859 \| 0.249 \| 317 \| 220 \| 6 \| 2 \| 311 \| 53 \| 358 \| Phage-related tail protein \| Phage-related tail protein; afdb-uniprot50 \| AF-A0A1H0NNX3-F1-MODEL\_V4 \| 1.0 \| 6.636e-25 \| 859 \| 0.292 \| 339 \| 195 \| 8 \| 1 \| 310 \| 69 \| 391 \| Uncharacterized protein \| Uncharacterized protein; afdb-uniprot50 \| AF-E2CN20-F1-MODEL\_V4 \| 1.0 \| 3.5e-24 \| 858 \| 0.327 \| 281 \| 177 \| 7 \| 2 \| 279 \| 56 \| 327 \| Putative bacteriophage late control D protein \| Putative bacteriophage late control D protein; afdb-uniprot50 \| AF-A0A812QV43-F1-MODEL\_V4 \| 1.0 \| 1.398e-24 \| 858 \| 0.313 \| 287 \| 182 \| 7 \| 1 \| 280 \| 734 \| 1012 \| GpFI protein \| GpFI protein; afdb-uniprot50 \| AF-A0A829E6A7-F1-MODEL\_V4 \| 1.0 \| 2.192e-23 \| 857 \| 0.252 \| 297 \| 208 \| 9 \| 2 \| 292 \| 60 \| 348 \| Uncharacterized protein \| Uncharacterized protein; afdb-uniprot50 \| AF-A0A1T4W4M5-F1-MODEL\_V4 \| 1.0 \| 2.628e-24 \| 856 \| 0.3 \| 290 \| 182 \| 9 \| 1 \| 278 \| 51 \| 331 \| Uncharacterized protein \| Uncharacterized protein; afdb-uniprot50 \| AF-A0A1S1TJX6-F1-MODEL\_V4 \| 1.0 \| 4.704e-25 \| 856 \| 0.261 \| 333 \| 206 \| 9 \| 1 \| 309 \| 62 \| 378 \| Uncharacterized protein \| Uncharacterized protein; afdb-uniprot50 \| AF-A0A2W5XH22-F1-MODEL\_V4 \| 1.0 \| 3.093e-23 \| 853 \| 0.295 \| 281 \| 185 \| 7 \| 2 \| 279 \| 55 \| 325 \| Late control protein \| Late control protein; afdb-uniprot50 \| AF-E3G761-F1-MODEL\_V4 \| 1.0 \| 2.459e-23 \| 850 \| 0.241 \| 306 \| 206 \| 10 \| 1 \| 290 \| 63 \| 358 \| Putative prophage tail protein \| Putative prophage tail protein; afdb-uniprot50 \| AF-A0A4Q0GTT7-F1-MODEL\_V4 \| 1.0 \| 3.468e-23 \| 850 \| 0.27 \| 288 \| 193 \| 8 \| 2 \| 279 \| 78 \| 358 \| Phage late control D family protein \| Phage late control D family protein; afdb-uniprot50 \| AF-A0A515D4Y2-F1-MODEL\_V4 \| 1.0 \| 6.964e-24 \| 850 \| 0.285 \| 301 \| 182 \| 7 \| 1 \| 275 \| 154 \| 447 \| Phage late control D family protein \| Phage late control D family protein; afdb-uniprot50 \| AF-A0A2U9U220-F1-MODEL\_V4 \| 1.0 \| 3.468e-23 \| 849 \| 0.235 \| 293 \| 202 \| 8 \| 1 \| 279 \| 56 \| 340 \| Uncharacterized protein \| Uncharacterized protein; afdb-uniprot50 \| AF-A0A841IGU3-F1-MODEL\_V4 \| 1.0 \| 1.743e-23 \| 848 \| 0.237 \| 333 \| 202 \| 11 \| 1 \| 290 \| 30 \| 353 \| Uncharacterized protein \| Uncharacterized protein; afdb-uniprot50 \| AF-A0A377KBH9-F1-MODEL\_V4 \| 1.0 \| 2.604e-23 \| 848 \| 0.29 \| 310 \| 184 \| 8 \| 1 \| 282 \| 117 \| 418 \| Late control protein D protein \| Late control protein D protein; afdb-uniprot50 \| AF-A0A853HRL3-F1-MODEL\_V4 \| 1.0 \| 6.266e-25 \| 845 \| 0.286 \| 297 \| 177 \| 6 \| 1 \| 278 \| 53 \| 333 \| Late control protein \| Late control protein; afdb-uniprot50 \| AF-A0A5S3YZL2-F1-MODEL\_V4 \| 1.0 \| 4.892e-23 \| 845 \| 0.251 \| 290 \| 199 \| 5 \| 1 \| 279 \| 55 \| 337 \| Phage tail protein \| Phage tail protein; afdb-uniprot50 \| AF-A0A4D7AWL8-F1-MODEL\_V4 \| 1.0 \| 1.454e-22 \| 845 \| 0.244 \| 286 \| 202 \| 7 \| 1 \| 279 \| 67 \| 345 \| Uncharacterized protein \| Uncharacterized protein; afdb-uniprot50 \| AF-A0A1B6NQN7-F1-MODEL\_V4 \| 1.0 \| 2.192e-23 \| 844 \| 0.265 \| 290 \| 191 \| 8 \| 2 \| 276 \| 52 \| 334 \| Phage late control D family protein \| Phage late control D family protein; afdb-uniprot50 \| AF-N9R5A0-F1-MODEL\_V4 \| 1.0 \| 1.236e-23 \| 844 \| 0.254 \| 307 \| 209 \| 12 \| 1 \| 299 \| 73 \| 367 \| Uncharacterized protein \| Uncharacterized protein; afdb-uniprot50 \| AF-A0A7X2ILW0-F1-MODEL\_V4 \| 1.0 \| 4.362e-23 \| 843 \| 0.281 \| 281 \| 186 \| 6 \| 1 \| 275 \| 51 \| 321 \| Late control protein \| Late control protein; afdb-uniprot50 \| AF-A0A827Q2Z4-F1-MODEL\_V4 \| 1.0 \| 9.276e-24 \| 842 \| 0.292 \| 267 \| 168 \| 7 \| 24 \| 274 \| 2 \| 263 \| Late control protein \| Late control protein; afdb-uniprot50 \| AF-A0A379CYH0-F1-MODEL\_V4 \| 1.0 \| 1.468e-23 \| 842 \| 0.245 \| 318 \| 199 \| 11 \| 1 \| 288 \| 81 \| 387 \| Phage protein D \| Phage protein D; afdb-uniprot50 \| AF-A0A2V3Q6R4-F1-MODEL\_V4 \| 1.0 \| 1.468e-23 \| 841 \| 0.253 \| 328 \| 199 \| 8 \| 1 \| 291 \| 65 \| 383 \| Uncharacterized protein \| Uncharacterized protein; afdb-uniprot50 \| AF-A0A4Z0WCL9-F1-MODEL\_V4 \| 1.0 \| 1.04e-23 \| 840 \| 0.28 \| 278 \| 182 \| 8 \| 1 \| 275 \| 53 \| 315 \| Uncharacterized protein \| Uncharacterized protein; afdb-uniprot50 \| AF-A0A4Q3Z459-F1-MODEL\_V4 \| 1.0 \| 3.89e-23 \| 840 \| 0.273 \| 278 \| 191 \| 7 \| 1 \| 275 \| 51 \| 320 \| Phage tail protein \| Phage tail protein; afdb-uniprot50 \| AF-A0A348FYG0-F1-MODEL\_V4 \| 1.0 \| 2.628e-24 \| 839 \| 0.286 \| 311 \| 199 \| 4 \| 2 \| 311 \| 52 \| 340 \| Phage late control protein \| Phage late control protein; afdb-uniprot50 \| AF-A0A1S1WTC3-F1-MODEL\_V4 \| 1.0 \| 3.5e-24 \| 839 \| 0.29 \| 317 \| 180 \| 11 \| 1 \| 280 \| 67 \| 375 \| Phage tail protein \| Phage tail protein; afdb-uniprot50 \| AF-A0A6C8XZE7-F1-MODEL\_V4 \| 1.0 \| 3.093e-23 \| 839 \| 0.247 \| 311 \| 202 \| 6 \| 1 \| 286 \| 77 \| 380 \| Phage late control D family protein \| Phage late control D family protein; afdb-uniprot50 \| AF-A0A7Z1M7H1-F1-MODEL\_V4 \| 1.0 \| 2.481e-24 \| 838 \| 0.249 \| 305 \| 204 \| 8 \| 2 \| 291 \| 52 \| 346 \| Uncharacterized protein \| Uncharacterized protein; afdb-uniprot50 \| AF-A0A524RW11-F1-MODEL\_V4 \| 1.0 \| 1.468e-23 \| 838 \| 0.281 \| 291 \| 196 \| 7 \| 7 \| 290 \| 68 \| 352 \| Uncharacterized protein \| Uncharacterized protein; afdb-uniprot50 \| AF-A0A3R8K6F9-F1-MODEL\_V4 \| 1.0 \| 3.5e-24 \| 838 \| 0.318 \| 298 \| 179 \| 7 \| 1 \| 278 \| 61 \| 354 \| Uncharacterized protein \| Uncharacterized protein; afdb-uniprot50 \| AF-A0A6L3Y0N7-F1-MODEL\_V4 \| 1.0 \| 1.568e-24 \| 837 \| 0.265 \| 305 \| 195 \| 9 \| 21 \| 308 \| 3 \| 295 \| Late control protein D \| Late control protein D; afdb-uniprot50 \| AF-A0A2A4XT50-F1-MODEL\_V4 \| 1.0 \| 3.275e-23 \| 837 \| 0.281 \| 274 \| 182 \| 6 \| 9 \| 277 \| 55 \| 318 \| Uncharacterized protein \| Uncharacterized protein; afdb-uniprot50 \| AF-A0A7S8IIV4-F1-MODEL\_V4 \| 1.0 \| 9.735e-23 \| 837 \| 0.26 \| 284 \| 198 \| 6 \| 1 \| 278 \| 52 \| 329 \| Late control protein D \| Late control protein D; afdb-uniprot50 \| AF-A0A2S8JPK2-F1-MODEL\_V4 \| 1.0 \| 2.604e-23 \| 837 \| 0.295 \| 288 \| 186 \| 8 \| 1 \| 274 \| 62 \| 346 \| Late control protein \| Late control protein; afdb-uniprot50 \| AF-N9UTV2-F1-MODEL\_V4 \| 1.0 \| 5.811e-23 \| 836 \| 0.243 \| 288 \| 202 \| 6 \| 1 \| 279 \| 72 \| 352 \| Phage-related tail protein \| Phage-related tail protein; afdb-uniprot50 \| AF-A0A836MBB7-F1-MODEL\_V4 \| 1.0 \| 1.398e-24 \| 836 \| 0.292 \| 314 \| 178 \| 9 \| 1 \| 277 \| 72 \| 378 \| Tail protein \| Tail protein; afdb-uniprot50 \| AF-C1DBI8-F1-MODEL\_V4 \| 1.0 \| 5.537e-24 \| 836 \| 0.287 \| 316 \| 182 \| 9 \| 2 \| 281 \| 89 \| 397 \| Tail protein D \| Tail protein D; afdb-uniprot50 \| AF-A0A1W9H416-F1-MODEL\_V4 \| 1.0 \| 4.892e-23 \| 834 \| 0.284 \| 288 \| 183 \| 10 \| 1 \| 275 \| 57 \| 334 \| Uncharacterized protein \| Uncharacterized protein; afdb-uniprot50 \| AF-A0A2T5TZJ6-F1-MODEL\_V4 \| 1.0 \| 1.743e-23 \| 834 \| 0.299 \| 284 \| 184 \| 6 \| 1 \| 276 \| 66 \| 342 \| Uncharacterized protein \| Uncharacterized protein; afdb-uniprot50 \| AF-A0A432WBJ7-F1-MODEL\_V4 \| 1.0 \| 1.846e-23 \| 834 \| 0.287 \| 292 \| 187 \| 8 \| 1 \| 279 \| 59 \| 342 \| Late control protein D \| Late control protein D; afdb-uniprot50 \| AF-A0A516SAU9-F1-MODEL\_V4 \| 1.0 \| 7.309e-23 \| 833 \| 0.255 \| 286 \| 195 \| 6 \| 1 \| 278 \| 51 \| 326 \| Late control protein \| Late control protein; afdb-uniprot50 \| AF-A0A2S6N2T0-F1-MODEL\_V4 \| 1.0 \| 2.192e-23 \| 832 \| 0.261 \| 294 \| 200 \| 8 \| 1 \| 286 \| 51 \| 335 \| Uncharacterized protein \| Uncharacterized protein; afdb-uniprot50 \| AF-A0A326L1B4-F1-MODEL\_V4 \| 1.0 \| 7.309e-23 \| 832 \| 0.277 \| 299 \| 191 \| 6 \| 2 \| 281 \| 56 \| 348 \| Late control protein \| Late control protein; afdb-uniprot50 \| AF-A0A846VID6-F1-MODEL\_V4 \| 1.0 \| 3.89e-23 \| 831 \| 0.25 \| 287 \| 200 \| 6 \| 1 \| 279 \| 56 \| 335 \| Uncharacterized protein \| Uncharacterized protein; afdb-uniprot50 \| AF-A0A380ALZ0-F1-MODEL\_V4 \| 1.0 \| 8.271e-24 \| 830 \| 0.279 \| 286 \| 191 \| 7 \| 2 \| 274 \| 62 \| 345 \| Phage protein D \| Phage protein D; afdb-uniprot50 \| AF-A0A484GCS1-F1-MODEL\_V4 \| 1.0 \| 4.62e-23 \| 829 \| 0.265 \| 298 \| 189 \| 8 \| 2 \| 279 \| 53 \| 340 \| Phage late control D family protein \| Phage late control D family protein; afdb-uniprot50 \| AF-A0A1Z9Z2K2-F1-MODEL\_V4 \| 1.0 \| 1.373e-22 \| 829 \| 0.286 \| 283 \| 187 \| 10 \| 2 \| 277 \| 71 \| 345 \| Uncharacterized protein \| Uncharacterized protein; afdb-uniprot50 \| AF-A0A2G6IRT2-F1-MODEL\_V4 \| 1.0 \| 8.839e-25 \| 827 \| 0.288 \| 326 \| 205 \| 9 \| 1 \| 309 \| 54 \| 369 \| Uncharacterized protein \| Uncharacterized protein; afdb-uniprot50 \| AF-A0A4R5EG26-F1-MODEL\_V4 \| 1.0 \| 3.673e-23 \| 826 \| 0.234 \| 286 \| 204 \| 8 \| 1 \| 279 \| 52 \| 329 \| Late control protein D \| Late control protein D; afdb-uniprot50 \| AF-A0A7X4GD43-F1-MODEL\_V4 \| 1.0 \| 5.811e-23 \| 826 \| 0.271 \| 284 \| 190 \| 7 \| 1 \| 275 \| 59 \| 334 \| Phage late control D family protein \| Phage late control D family protein; afdb-uniprot50 \| AF-A0A222EYQ7-F1-MODEL\_V4 \| 1.0 \| 6.901e-23 \| 825 \| 0.291 \| 285 \| 184 \| 7 \| 2 \| 275 \| 38 \| 315 \| Late control protein \| Late control protein; afdb-uniprot50 \| AF-A0A3G4VA69-F1-MODEL\_V4 \| 1.0 \| 3.065e-22 \| 825 \| 0.27 \| 281 \| 194 \| 6 \| 1 \| 277 \| 51 \| 324 \| Uncharacterized protein \| Uncharacterized protein; afdb-uniprot50 \| AF-A0A7T9VKR8-F1-MODEL\_V4 \| 1.0 \| 1.04e-23 \| 825 \| 0.293 \| 303 \| 194 \| 9 \| 1 \| 297 \| 53 \| 341 \| Uncharacterized protein \| Uncharacterized protein; afdb-uniprot50 \| AF-A0A506V2R9-F1-MODEL\_V4 \| 1.0 \| 1.955e-23 \| 825 \| 0.248 \| 326 \| 197 \| 12 \| 2 \| 290 \| 66 \| 380 \| Phage late control D family protein \| Phage late control D family protein; afdb-uniprot50 \| AF-A0A840E0Y3-F1-MODEL\_V4 \| 1.0 \| 5.181e-23 \| 824 \| 0.271 \| 273 \| 178 \| 9 \| 12 \| 278 \| 1 \| 258 \| Uncharacterized protein \| Uncharacterized protein; afdb-uniprot50 \| AF-A0A1G5AD05-F1-MODEL\_V4 \| 1.0 \| 6.964e-24 \| 824 \| 0.343 \| 282 \| 172 \| 7 \| 1 \| 279 \| 51 \| 322 \| Uncharacterized protein \| Uncharacterized protein; afdb-uniprot50 \| AF-A0A2V3UNG3-F1-MODEL\_V4 \| 1.0 \| 1.454e-22 \| 823 \| 0.254 \| 287 \| 200 \| 6 \| 1 \| 281 \| 51 \| 329 \| Uncharacterized protein \| Uncharacterized protein; afdb-uniprot50 \| AF-V5ZC41-F1-MODEL\_V4 \| 1.0 \| 5.181e-23 \| 823 \| 0.262 \| 308 \| 190 \| 9 \| 1 \| 279 \| 63 \| 362 \| Late control gene D protein GpD \| Late control gene D protein GpD; afdb-uniprot50 \| AF-A0A839IXE6-F1-MODEL\_V4 \| 1.0 \| 6.21e-24 \| 822 \| 0.291 \| 288 \| 187 \| 9 \| 1 \| 277 \| 53 \| 334 \| Phage late control D family protein \| Phage late control D family protein; afdb-uniprot50 \| AF-A0A833GCM2-F1-MODEL\_V4 \| 1.0 \| 3.275e-23 \| 822 \| 0.293 \| 303 \| 184 \| 7 \| 1 \| 281 \| 51 \| 345 \| Uncharacterized protein \| Uncharacterized protein; afdb-uniprot50 \| AF-A0A379CBI1-F1-MODEL\_V4 \| 1.0 \| 4.119e-23 \| 820 \| 0.288 \| 305 \| 184 \| 8 \| 1 \| 279 \| 68 \| 365 \| Phage protein D \| Phage protein D; afdb-uniprot50 \| AF-A0A1Y2S8A5-F1-MODEL\_V4 \| 1.0 \| 2.07e-23 \| 819 \| 0.258 \| 333 \| 188 \| 8 \| 1 \| 282 \| 75 \| 399 \| Phage protein \| Phage protein; afdb-uniprot50 \| AF-A0A085A808-F1-MODEL\_V4 \| 1.0 \| 2.459e-23 \| 818 \| 0.253 \| 272 \| 187 \| 8 \| 22 \| 282 \| 5 \| 271 \| Phage late control gene D protein \| Phage late control gene D protein; afdb-uniprot50 \| AF-A0A1Y1QXI4-F1-MODEL\_V4 \| 1.0 \| 6.154e-23 \| 814 \| 0.299 \| 284 \| 185 \| 7 \| 2 \| 279 \| 52 \| 327 \| Uncharacterized protein \| Uncharacterized protein; afdb-uniprot50 \| AF-A0A3B0MDS7-F1-MODEL\_V4 \| 1.0 \| 3.437e-22 \| 813 \| 0.262 \| 282 \| 196 \| 6 \| 1 \| 279 \| 51 \| 323 \| Uncharacterized protein \| Uncharacterized protein; afdb-uniprot50 \| AF-A0A2K3TWD5-F1-MODEL\_V4 \| 1.0 \| 4.119e-23 \| 813 \| 0.291 \| 312 \| 178 \| 8 \| 1 \| 277 \| 62 \| 365 \| Late control protein D \| Late control protein D; afdb-uniprot50 \| AF-A0A0C5VFD9-F1-MODEL\_V4 \| 1.0 \| 1.031e-22 \| 811 \| 0.252 \| 285 \| 193 \| 5 \| 1 \| 276 \| 37 \| 310 \| Phage protein D \| Phage protein D; afdb-uniprot50 \| AF-A0A2M8RTD2-F1-MODEL\_V4 \| 1.0 \| 1.236e-23 \| 811 \| 0.26 \| 323 \| 186 \| 9 \| 1 \| 277 \| 66 \| 381 \| Phage tail protein \| Phage tail protein; afdb-uniprot50 \| AF-A0A7X6FQ70-F1-MODEL\_V4 \| 1.0 \| 4.119e-23 \| 810 \| 0.239 \| 317 \| 217 \| 7 \| 1 \| 311 \| 27 \| 325 \| Late control protein D \| Late control protein D; afdb-uniprot50 \| AF-A0A2P1VV04-F1-MODEL\_V4 \| 1.0 \| 3.275e-23 \| 809 \| 0.298 \| 288 \| 184 \| 10 \| 1 \| 276 \| 53 \| 334 \| Phage tail protein \| Phage tail protein; afdb-uniprot50 \| AF-J0ZBK7-F1-MODEL\_V4 \| 1.0 \| 9.824e-24 \| 808 \| 0.274 \| 342 \| 173 \| 7 \| 1 \| 275 \| 56 \| 389 \| Uncharacterized protein \| Uncharacterized protein; afdb-uniprot50 \| AF-A0A2N3KSJ2-F1-MODEL\_V4 \| 1.0 \| 4.119e-23 \| 807 \| 0.318 \| 289 \| 179 \| 10 \| 2 \| 281 \| 54 \| 333 \| Uncharacterized protein \| Uncharacterized protein; afdb-uniprot50 \| AF-A0A522JN25-F1-MODEL\_V4 \| 1.0 \| 3.065e-22 \| 807 \| 0.311 \| 295 \| 181 \| 9 \| 1 \| 280 \| 69 \| 356 \| Phage late control D family protein \| Phage late control D family protein; afdb-uniprot50 \| AF-A0A1N6MRZ2-F1-MODEL\_V4 \| 1.0 \| 5.487e-23 \| 807 \| 0.237 \| 312 \| 205 \| 7 \| 1 \| 286 \| 86 \| 390 \| Late control gene D protein \| Late control gene D protein; afdb-uniprot50 \| AF-A0A4Q0YKX8-F1-MODEL\_V4 \| 1.0 \| 9.735e-23 \| 802 \| 0.297 \| 279 \| 185 \| 7 \| 1 \| 275 \| 53 \| 324 \| Late control protein \| Late control protein; afdb-uniprot50 \| AF-A0A6G0ZW07-F1-MODEL\_V4 \| 1.0 \| 4.805e-21 \| 801 \| 0.322 \| 220 \| 145 \| 4 \| 60 \| 277 \| 2 \| 219 \| Uncharacterized protein \| Uncharacterized protein; afdb-uniprot50 \| AF-A0A2W6YRE0-F1-MODEL\_V4 \| 1.0 \| 7.74e-23 \| 801 \| 0.277 \| 295 \| 193 \| 9 \| 1 \| 282 \| 61 \| 348 \| Late control protein \| Late control protein; afdb-uniprot50 \| AF-A0A8A6BLQ2-F1-MODEL\_V4 \| 1.0 \| 2.301e-22 \| 800 \| 0.271 \| 287 \| 191 \| 5 \| 1 \| 276 \| 54 \| 333 \| Phage tail protein \| Phage tail protein; afdb-uniprot50 \| AF-A0A5S9R5J4-F1-MODEL\_V4 \| 1.0 \| 2.415e-21 \| 799 \| 0.245 \| 224 \| 165 \| 3 \| 57 \| 277 \| 1 \| 223 \| Uncharacterized protein \| Uncharacterized protein; afdb-uniprot50 \| AF-A0A5N8AC16-F1-MODEL\_V4 \| 1.0 \| 4.848e-22 \| 799 \| 0.281 \| 288 \| 189 \| 9 \| 2 \| 278 \| 53 \| 333 \| Phage late control D family protein \| Phage late control D family protein; afdb-uniprot50 \| AF-A0A3B0T6D5-F1-MODEL\_V4 \| 1.0 \| 3.673e-23 \| 798 \| 0.336 \| 285 \| 157 \| 8 \| 1 \| 279 \| 51 \| 309 \| Uncharacterized protein \| Uncharacterized protein; afdb-uniprot50 \| AF-A0A2S2E755-F1-MODEL\_V4 \| 1.0 \| 8.602e-22 \| 796 \| 0.243 \| 283 \| 200 \| 8 \| 1 \| 277 \| 50 \| 324 \| Putative baseplate hub protein \| Putative baseplate hub protein; afdb-uniprot50 \| AF-A0A258L5C4-F1-MODEL\_V4 \| 1.0 \| 2.58e-22 \| 795 \| 0.241 \| 286 \| 201 \| 7 \| 1 \| 277 \| 47 \| 325 \| Uncharacterized protein \| Uncharacterized protein; afdb-uniprot50 \| AF-A0A4Q6D1D6-F1-MODEL\_V4 \| 1.0 \| 1.361e-21 \| 792 \| 0.246 \| 280 \| 199 \| 5 \| 1 \| 279 \| 50 \| 318 \| Uncharacterized protein \| Uncharacterized protein; afdb-uniprot50 \| AF-B8GS02-F1-MODEL\_V4 \| 1.0 \| 1.224e-22 \| 792 \| 0.292 \| 287 \| 179 \| 8 \| 1 \| 276 \| 51 \| 324 \| Phage-related tail protein \| Phage-related tail protein; afdb-uniprot50 \| AF-A0A6B8KJK1-F1-MODEL\_V4 \| 1.0 \| 6.839e-22 \| 792 \| 0.261 \| 287 \| 196 \| 8 \| 2 \| 279 \| 51 \| 330 \| Uncharacterized protein \| Uncharacterized protein; afdb-uniprot50 \| AF-F7VG01-F1-MODEL\_V4 \| 1.0 \| 1.373e-22 \| 791 \| 0.266 \| 270 \| 173 \| 6 \| 41 \| 292 \| 2 \| 264 \| Phage late control D family protein \| Phage late control D family protein; afdb-uniprot50 \| AF-A0A432QTK6-F1-MODEL\_V4 \| 1.0 \| 7.67e-22 \| 791 \| 0.286 \| 279 \| 183 \| 6 \| 1 \| 276 \| 51 \| 316 \| Uncharacterized protein \| Uncharacterized protein; afdb-uniprot50 \| AF-A0A537MFG1-F1-MODEL\_V4 \| 1.0 \| 2.733e-22 \| 789 \| 0.269 \| 304 \| 186 \| 9 \| 2 \| 276 \| 57 \| 353 \| Phage late control D family protein \| Phage late control D family protein; afdb-uniprot50 \| AF-A0A1I3SYP4-F1-MODEL\_V4 \| 1.0 \| 6.517e-23 \| 788 \| 0.24 \| 320 \| 220 \| 10 \| 1 \| 311 \| 52 \| 357 \| Uncharacterized protein \| Uncharacterized protein; afdb-uniprot50 \| AF-A0A1M5PXU2-F1-MODEL\_V4 \| 1.0 \| 1.082e-21 \| 787 \| 0.237 \| 286 \| 200 \| 9 \| 1 \| 277 \| 56 \| 332 \| Phage protein D \| Phage protein D; afdb-uniprot50 \| AF-A0A7W6WMD7-F1-MODEL\_V4 \| 1.0 \| 5.437e-22 \| 786 \| 0.256 \| 292 \| 195 \| 7 \| 1 \| 281 \| 52 \| 332 \| Uncharacterized protein \| Uncharacterized protein; afdb-uniprot50 \| AF-A0A258E9L1-F1-MODEL\_V4 \| 1.0 \| 1.022e-21 \| 784 \| 0.278 \| 284 \| 189 \| 6 \| 1 \| 275 \| 54 \| 330 \| Uncharacterized protein \| Uncharacterized protein; afdb-uniprot50 \| AF-A0A7X3TYP5-F1-MODEL\_V4 \| 1.0 \| 1.727e-22 \| 783 \| 0.278 \| 291 \| 189 \| 9 \| 2 \| 279 \| 53 \| 335 \| Uncharacterized protein \| Uncharacterized protein; afdb-uniprot50 \| AF-J1JNX9-F1-MODEL\_V4 \| 1.0 \| 1.454e-22 \| 782 \| 0.272 \| 283 \| 185 \| 10 \| 1 \| 277 \| 43 \| 310 \| Uncharacterized protein \| Uncharacterized protein; afdb-uniprot50 \| AF-A0A4Y7X954-F1-MODEL\_V4 \| 1.0 \| 9.193e-23 \| 782 \| 0.251 \| 326 \| 203 \| 12 \| 1 \| 294 \| 73 \| 389 \| DNA primase \| DNA primase; afdb-uniprot50 \| AF-A0A5C8S7D7-F1-MODEL\_V4 \| 1.0 \| 1.082e-21 \| 780 \| 0.221 \| 293 \| 201 \| 9 \| 1 \| 277 \| 56 \| 337 \| Uncharacterized protein \| Uncharacterized protein; afdb-uniprot50 \| AF-A0A6L9HG12-F1-MODEL\_V4 \| 1.0 \| 3.245e-22 \| 775 \| 0.181 \| 319 \| 238 \| 10 \| 2 \| 311 \| 1 \| 305 \| Uncharacterized protein \| Uncharacterized protein; afdb-uniprot50 \| AF-A0A0Q8AN74-F1-MODEL\_V4 \| 1.0 \| 5.811e-23 \| 775 \| 0.242 \| 297 \| 203 \| 6 \| 18 \| 311 \| 84 \| 361 \| Uncharacterized protein \| Uncharacterized protein; afdb-uniprot50 \| AF-A0A1C3HKM4-F1-MODEL\_V4 \| 1.0 \| 2.051e-22 \| 773 \| 0.229 \| 335 \| 207 \| 9 \| 2 \| 310 \| 64 \| 373 \| Phage late control gene D protein (GPD) \| Phage late control gene D protein (GPD); afdb-uniprot50 \| AF-A0A7G5CTF9-F1-MODEL\_V4 \| 1.0 \| 6.098e-22 \| 765 \| 0.22 \| 331 \| 207 \| 9 \| 1 \| 288 \| 62 \| 384 \| Phage late control D family protein \| Phage late control D family protein; afdb-uniprot50 \| AF-A0A2N3DW94-F1-MODEL\_V4 \| 1.0 \| 5.042e-20 \| 764 \| 0.251 \| 223 \| 163 \| 4 \| 60 \| 279 \| 2 \| 223 \| Late control protein \| Late control protein; afdb-uniprot50 \| AF-A0A852Y1I1-F1-MODEL\_V4 \| 1.0 \| 2.134e-20 \| 764 \| 0.257 \| 284 \| 195 \| 7 \| 1 \| 276 \| 55 \| 330 \| Uncharacterized protein \| Uncharacterized protein; afdb-uniprot50 \| AF-A0A554XC54-F1-MODEL\_V4 \| 1.0 \| 5.758e-22 \| 764 \| 0.286 \| 290 \| 190 \| 7 \| 1 \| 280 \| 52 \| 334 \| Phage late control D protein (GPD) \| Phage late control D protein (GPD); afdb-uniprot50 \| AF-A0A7X3ZHS0-F1-MODEL\_V4 \| 1.0 \| 8.123e-22 \| 762 \| 0.26 \| 280 \| 195 \| 6 \| 1 \| 276 \| 51 \| 322 \| Uncharacterized protein \| Uncharacterized protein; afdb-uniprot50 \| AF-A0A806CG79-F1-MODEL\_V4 \| 1.0 \| 3.82e-21 \| 762 \| 0.218 \| 288 \| 208 \| 9 \| 1 \| 282 \| 54 \| 330 \| Late control D family protein \| Late control D family protein; afdb-uniprot50 \| AF-A0A317PFJ8-F1-MODEL\_V4 \| 1.0 \| 4.805e-21 \| 762 \| 0.212 \| 292 \| 209 \| 9 \| 1 \| 279 \| 55 \| 338 \| Uncharacterized protein \| Uncharacterized protein; afdb-uniprot50 \| AF-A0A512IX34-F1-MODEL\_V4 \| 1.0 \| 2.733e-22 \| 759 \| 0.225 \| 337 \| 225 \| 11 \| 1 \| 311 \| 56 \| 382 \| Bacteriophage late control gene D protein \| Bacteriophage late control gene D protein; afdb-uniprot50 \| AF-A0A1D2QMU3-F1-MODEL\_V4 \| 1.0 \| 2.393e-20 \| 758 \| 0.242 \| 231 \| 166 \| 4 \| 57 \| 281 \| 5 \| 232 \| Uncharacterized protein \| Uncharacterized protein; afdb-uniprot50 \| AF-A0A1Y1R1L0-F1-MODEL\_V4 \| 1.0 \| 4.848e-22 \| 758 \| 0.261 \| 271 \| 184 \| 6 \| 13 \| 278 \| 3 \| 262 \| Uncharacterized protein \| Uncharacterized protein; afdb-uniprot50 \| AF-A0A5V4G7N6-F1-MODEL\_V4 \| 1.0 \| 1.454e-22 \| 758 \| 0.285 \| 298 \| 162 \| 7 \| 32 \| 286 \| 1 \| 290 \| Phage late control D family protein \| Phage late control D family protein; afdb-uniprot50 \| AF-A0A1N7LR18-F1-MODEL\_V4 \| 1.0 \| 6.266e-25 \| 757 \| 0.306 \| 284 \| 184 \| 5 \| 1 \| 279 \| 52 \| 327 \| Uncharacterized protein \| Uncharacterized protein; afdb-uniprot50 \| AF-A0A4Q8MFD5-F1-MODEL\_V4 \| 1.0 \| 4.537e-21 \| 757 \| 0.256 \| 285 \| 196 \| 6 \| 1 \| 277 \| 52 \| 328 \| Uncharacterized protein \| Uncharacterized protein; afdb-uniprot50 \| AF-J8V8B9-F1-MODEL\_V4 \| 1.0 \| 9.11e-22 \| 757 \| 0.28 \| 285 \| 178 \| 10 \| 1 \| 276 \| 51 \| 317 \| Tail protein D \| Tail protein D; afdb-uniprot50 \| AF-V4JM50-F1-MODEL\_V4 \| 1.0 \| 3.89e-23 \| 755 \| 0.292 \| 284 \| 183 \| 7 \| 1 \| 278 \| 63 \| 334 \| Uncharacterized protein \| Uncharacterized protein; afdb-uniprot50 \| AF-A0A1Y6HD93-F1-MODEL\_V4 \| 1.0 \| 3.855e-22 \| 754 \| 0.319 \| 285 \| 178 \| 8 \| 30 \| 311 \| 2 \| 273 \| Bacteriophage P2 gpD protein \| Bacteriophage P2 gpD protein; afdb-uniprot50 \| AF-A0A2Z6GCV6-F1-MODEL\_V4 \| 1.0 \| 1.92e-21 \| 752 \| 0.221 \| 398 \| 185 \| 8 \| 1 \| 280 \| 54 \| 444 \| Late control D family protein \| Late control D family protein; afdb-uniprot50 \| AF-F8KPK8-F1-MODEL\_V4 \| 1.0 \| 9.11e-22 \| 751 \| 0.222 \| 283 \| 200 \| 12 \| 1 \| 281 \| 45 \| 309 \| Phage late control D family protein \| Phage late control D family protein; afdb-uniprot50 \| AF-A0A285M2K7-F1-MODEL\_V4 \| 1.0 \| 1.512e-20 \| 751 \| 0.239 \| 284 \| 201 \| 8 \| 2 \| 278 \| 54 \| 329 \| Phage late control gene D protein (GPD) \| Phage late control gene D protein (GPD); afdb-uniprot50 \| AF-A0A496KC99-F1-MODEL\_V4 \| 1.0 \| 3.855e-22 \| 750 \| 0.237 \| 358 \| 204 \| 9 \| 1 \| 308 \| 85 \| 423 \| Late control protein \| Late control protein; afdb-uniprot50 \| AF-J0Q1E2-F1-MODEL\_V4 \| 1.0 \| 1.349e-20 \| 749 \| 0.281 \| 224 \| 149 \| 5 \| 57 \| 277 \| 70 \| 284 \| Uncharacterized protein \| Uncharacterized protein; afdb-uniprot50 \| AF-A0A6N8TBC0-F1-MODEL\_V4 \| 1.0 \| 3.82e-21 \| 749 \| 0.236 \| 288 \| 202 \| 11 \| 2 \| 279 \| 54 \| 333 \| Uncharacterized protein \| Uncharacterized protein; afdb-uniprot50 \| AF-A0A2G9WV33-F1-MODEL\_V4 \| 1.0 \| 2.033e-21 \| 749 \| 0.244 \| 307 \| 211 \| 9 \| 1 \| 296 \| 55 \| 351 \| Uncharacterized protein \| Uncharacterized protein; afdb-uniprot50 \| AF-W1J5W8-F1-MODEL\_V4 \| 1.0 \| 6.458e-22 \| 749 \| 0.256 \| 312 \| 187 \| 10 \| 1 \| 280 \| 106 \| 404 \| Late control gene D protein \| Late control gene D protein; afdb-uniprot50 \| AF-A0A4Y5WA89-F1-MODEL\_V4 \| 1.0 \| 2.173e-22 \| 749 \| 0.265 \| 324 \| 194 \| 10 \| 1 \| 292 \| 77 \| 388 \| Late control protein \| Late control protein; afdb-uniprot50 \| AF-A0A2N2S1D8-F1-MODEL\_V4 \| 1.0 \| 1.441e-21 \| 745 \| 0.245 \| 391 \| 177 \| 10 \| 2 \| 281 \| 187 \| 570 \| Uncharacterized protein \| Uncharacterized protein; afdb-uniprot50 \| AF-A0A2V4DTS8-F1-MODEL\_V4 \| 1.0 \| 4.537e-21 \| 744 \| 0.209 \| 329 \| 207 \| 15 \| 1 \| 288 \| 56 \| 372 \| Uncharacterized protein \| Uncharacterized protein; afdb-uniprot50 \| AF-J0Q3C9-F1-MODEL\_V4 \| 1.0 \| 1.631e-22 \| 744 \| 0.266 \| 338 \| 174 \| 7 \| 1 \| 272 \| 56 \| 385 \| Uncharacterized protein \| Uncharacterized protein; afdb-uniprot50 \| AF-A0A7I0BEW8-F1-MODEL\_V4 \| 1.0 \| 2.015e-20 \| 740 \| 0.26 \| 230 \| 156 \| 6 \| 57 \| 278 \| 15 \| 238 \| Late control protein \| Late control protein; afdb-uniprot50 \| AF-E2CCF5-F1-MODEL\_V4 \| 1.0 \| 2.033e-21 \| 740 \| 0.203 \| 330 \| 222 \| 10 \| 1 \| 311 \| 51 \| 358 \| Phage late control D protein (GPD) \| Phage late control D protein (GPD); afdb-uniprot50 \| AF-A0A1G6QR90-F1-MODEL\_V4 \| 1.0 \| 8.602e-22 \| 739 \| 0.272 \| 264 \| 164 \| 5 \| 57 \| 294 \| 5 \| 266 \| Uncharacterized protein \| Uncharacterized protein; afdb-uniprot50 \| AF-A0A2N5EEH1-F1-MODEL\_V4 \| 1.0 \| 3.406e-21 \| 738 \| 0.231 \| 302 \| 200 \| 8 \| 1 \| 277 \| 62 \| 356 \| Late control protein D \| Late control protein D; afdb-uniprot50 \| AF-A0A327JNY9-F1-MODEL\_V4 \| 1.0 \| 3.574e-20 \| 737 \| 0.241 \| 290 \| 200 \| 11 \| 2 \| 281 \| 60 \| 339 \| Uncharacterized protein \| Uncharacterized protein; afdb-uniprot50 \| AF-A0A3G2IAI4-F1-MODEL\_V4 \| 1.0 \| 5.864e-24 \| 737 \| 0.273 \| 303 \| 190 \| 8 \| 1 \| 276 \| 59 \| 358 \| Uncharacterized protein \| Uncharacterized protein; afdb-uniprot50 \| AF-A0A1X3DKM2-F1-MODEL\_V4 \| 1.0 \| 4.848e-22 \| 736 \| 0.244 \| 331 \| 191 \| 8 \| 1 \| 280 \| 74 \| 396 \| Late control protein \| Late control protein; afdb-uniprot50 \| AF-E3PAT4-F1-MODEL\_V4 \| 1.0 \| 3.187e-20 \| 733 \| 0.241 \| 248 \| 163 \| 4 \| 63 \| 286 \| 1 \| 247 \| Putative phage late gene regulator \| Putative phage late gene regulator; afdb-uniprot50 \| AF-A0A4R3NVK0-F1-MODEL\_V4 \| 1.0 \| 1.349e-20 \| 731 \| 0.229 \| 283 \| 195 \| 9 \| 1 \| 277 \| 52 \| 317 \| Uncharacterized protein \| Uncharacterized protein; afdb-uniprot50 \| AF-A0A5N3PH75-F1-MODEL\_V4 \| 1.0 \| 1.349e-20 \| 731 \| 0.243 \| 292 \| 199 \| 11 \| 1 \| 279 \| 54 \| 336 \| Late control protein \| Late control protein; afdb-uniprot50 \| AF-A0A844H206-F1-MODEL\_V4 \| 1.0 \| 3.64e-22 \| 729 \| 0.267 \| 318 \| 208 \| 7 \| 1 \| 310 \| 51 \| 351 \| Phage tail protein \| Phage tail protein; afdb-uniprot50 \| AF-A0A366DKG6-F1-MODEL\_V4 \| 1.0 \| 7.177e-21 \| 726 \| 0.238 \| 319 \| 223 \| 6 \| 1 \| 309 \| 52 \| 360 \| Phage protein D \| Phage protein D; afdb-uniprot50 \| AF-E2CFJ7-F1-MODEL\_V4 \| 1.0 \| 1.003e-19 \| 725 \| 0.243 \| 287 \| 200 \| 10 \| 1 \| 278 \| 50 \| 328 \| Phage late control D protein (GPD) \| Phage late control D protein (GPD); afdb-uniprot50 \| AF-Q31Q84-F1-MODEL\_V4 \| 1.0 \| 2.415e-21 \| 725 \| 0.27 \| 285 \| 188 \| 8 \| 1 \| 275 \| 52 \| 326 \| Phage late control gene D protein GPD \| Phage late control gene D protein GPD; afdb-uniprot50 \| AF-D4M9M1-F1-MODEL\_V4 \| 1.0 \| 6.777e-21 \| 724 \| 0.277 \| 285 \| 183 \| 10 \| 1 \| 279 \| 54 \| 321 \| Phage protein D \| Phage protein D; afdb-uniprot50 \| AF-A0A175R4Y2-F1-MODEL\_V4 \| 1.0 \| 1.92e-21 \| 724 \| 0.25 \| 324 \| 222 \| 12 \| 1 \| 311 \| 51 \| 366 \| Uncharacterized protein \| Uncharacterized protein; afdb-uniprot50 \| AF-A0A6L8M6A2-F1-MODEL\_V4 \| 1.0 \| 3.375e-20 \| 722 \| 0.244 \| 278 \| 201 \| 6 \| 1 \| 277 \| 47 \| 316 \| Uncharacterized protein \| Uncharacterized protein; afdb-uniprot50 \| AF-A0A2C9D686-F1-MODEL\_V4 \| 1.0 \| 5.655e-20 \| 721 \| 0.228 \| 289 \| 205 \| 10 \| 1 \| 279 \| 55 \| 335 \| Tail protein \| Tail protein; afdb-uniprot50 \| AF-A0A2D0L424-F1-MODEL\_V4 \| 1.0 \| 3.01e-20 \| 720 \| 0.188 \| 414 \| 193 \| 8 \| 1 \| 279 \| 75 \| 480 \| Phage protein \| Phage protein; afdb-uniprot50 \| AF-U3BZ25-F1-MODEL\_V4 \| 1.0 \| 2.015e-20 \| 719 \| 0.25 \| 283 \| 199 \| 8 \| 1 \| 279 \| 47 \| 320 \| Uncharacterized protein \| Uncharacterized protein; afdb-uniprot50 \| AF-A0A3N2E0P8-F1-MODEL\_V4 \| 1.0 \| 2.259e-20 \| 716 \| 0.236 \| 288 \| 194 \| 10 \| 1 \| 277 \| 75 \| 347 \| Phage protein D \| Phage protein D; afdb-uniprot50 \| AF-A0A451AIR0-F1-MODEL\_V4 \| 1.0 \| 4.245e-20 \| 715 \| 0.377 \| 244 \| 143 \| 4 \| 1 \| 242 \| 51 \| 287 \| Uncharacterized protein \| Uncharacterized protein; afdb-uniprot50 \| AF-A0A068T9R7-F1-MODEL\_V4 \| 1.0 \| 8.602e-22 \| 715 \| 0.236 \| 296 \| 201 \| 14 \| 1 \| 281 \| 53 \| 338 \| Phage late control D family protein \| Phage late control D family protein; afdb-uniprot50 \| AF-A0A3M4MW84-F1-MODEL\_V4 \| 1.0 \| 1.885e-19 \| 714 \| 0.295 \| 210 \| 139 \| 2 \| 72 \| 275 \| 2 \| 208 \| Uncharacterized protein \| Uncharacterized protein; afdb-uniprot50 \| AF-A0A2W6ZLW1-F1-MODEL\_V4 \| 1.0 \| 9.56e-21 \| 714 \| 0.236 \| 326 \| 223 \| 12 \| 1 \| 311 \| 54 \| 368 \| Late control protein \| Late control protein; afdb-uniprot50 \| AF-A0A7W2GRY8-F1-MODEL\_V4 \| 1.0 \| 1.146e-21 \| 713 \| 0.275 \| 283 \| 191 \| 8 \| 1 \| 277 \| 51 \| 325 \| Late control protein \| Late control protein; afdb-uniprot50 \| AF-A0A7L5XWR6-F1-MODEL\_V4 \| 1.0 \| 1.135e-20 \| 712 \| 0.323 \| 229 \| 148 \| 2 \| 84 \| 311 \| 1 \| 223 \| Uncharacterized protein \| Uncharacterized protein; afdb-uniprot50 \| AF-A0A679JNK0-F1-MODEL\_V4 \| 1.0 \| 3.01e-20 \| 712 \| 0.216 \| 296 \| 200 \| 12 \| 1 \| 279 \| 57 \| 337 \| Uncharacterized protein \| Uncharacterized protein; afdb-uniprot50 \| AF-A0A4Y9VRL2-F1-MODEL\_V4 \| 1.0 \| 4.045e-21 \| 712 \| 0.238 \| 385 \| 180 \| 13 \| 2 \| 281 \| 58 \| 434 \| Uncharacterized protein \| Uncharacterized protein; afdb-uniprot50 \| AF-A0A379TSJ9-F1-MODEL\_V4 \| 1.0 \| 1.146e-21 \| 708 \| 0.223 \| 331 \| 199 \| 13 \| 5 \| 292 \| 2 \| 317 \| Putative bacteriophage regulatory protein \| Putative bacteriophage regulatory protein; afdb-uniprot50 \| AF-Q0FZ18-F1-MODEL\_V4 \| 1.0 \| 1.512e-20 \| 708 \| 0.189 \| 291 \| 211 \| 13 \| 1 \| 275 \| 55 \| 336 \| Phage-related tail protein \| Phage-related tail protein; afdb-uniprot50 \| AF-A0A0S4XLQ7-F1-MODEL\_V4 \| 1.0 \| 2.816e-19 \| 703 \| 0.266 \| 278 \| 186 \| 9 \| 1 \| 275 \| 50 \| 312 \| Putative phage tail protein D \| Putative phage tail protein D; afdb-uniprot50 \| AF-U2FA85-F1-MODEL\_V4 \| 1.0 \| 4.761e-20 \| 702 \| 0.245 \| 285 \| 193 \| 11 \| 1 \| 275 \| 50 \| 322 \| Tail protein D, putative \| Tail protein D, putative; afdb-uniprot50 \| AF-A0A6I1K313-F1-MODEL\_V4 \| 1.0 \| 6.716e-20 \| 700 \| 0.232 \| 292 \| 203 \| 13 \| 1 \| 280 \| 54 \| 336 \| Phage late control D family protein \| Phage late control D family protein; afdb-uniprot50 \| AF-A0A3T0L2M0-F1-MODEL\_V4 \| 1.0 \| 7.532e-20 \| 698 \| 0.242 \| 276 \| 190 \| 9 \| 1 \| 275 \| 52 \| 309 \| Phage tail protein \| Phage tail protein; afdb-uniprot50 \| AF-E6X1M2-F1-MODEL\_V4 \| 1.0 \| 7.532e-20 \| 697 \| 0.246 \| 272 \| 183 \| 9 \| 7 \| 277 \| 50 \| 300 \| Phage tail protein D \| Phage tail protein D; afdb-uniprot50 \| AF-A0A165XHG8-F1-MODEL\_V4 \| 1.0 \| 5.34e-20 \| 697 \| 0.231 \| 294 \| 201 \| 9 \| 1 \| 277 \| 51 \| 336 \| Phage late control gene D protein (GPD) \| Phage late control gene D protein (GPD); afdb-uniprot50 \| AF-A0A1T4WXQ6-F1-MODEL\_V4 \| 1.0 \| 4.009e-20 \| 695 \| 0.241 \| 277 \| 191 \| 8 \| 7 \| 275 \| 57 \| 322 \| Uncharacterized protein \| Uncharacterized protein; afdb-uniprot50 \| AF-A0A7Y3Y9A7-F1-MODEL\_V4 \| 1.0 \| 6.342e-20 \| 691 \| 0.23 \| 282 \| 203 \| 8 \| 2 \| 279 \| 46 \| 317 \| Uncharacterized protein \| Uncharacterized protein; afdb-uniprot50 \| AF-A0A0B6CUQ7-F1-MODEL\_V4 \| 1.0 \| 7.601e-21 \| 689 \| 0.25 \| 284 \| 185 \| 8 \| 1 \| 276 \| 52 \| 315 \| Phage late control D family protein \| Phage late control D family protein; afdb-uniprot50 \| AF-A0A651FN49-F1-MODEL\_V4 \| 1.0 \| 2.816e-19 \| 687 \| 0.212 \| 292 \| 210 \| 9 \| 1 \| 280 \| 56 \| 339 \| Late control D family protein \| Late control D family protein; afdb-uniprot50 \| AF-A0A6L5YCW9-F1-MODEL\_V4 \| 1.0 \| 5.292e-19 \| 685 \| 0.235 \| 289 \| 199 \| 10 \| 1 \| 281 \| 54 \| 328 \| Uncharacterized protein \| Uncharacterized protein; afdb-uniprot50 \| AF-D2U2W5-F1-MODEL\_V4 \| 1.0 \| 2.534e-20 \| 685 \| 0.217 \| 299 \| 198 \| 10 \| 2 \| 278 \| 38 \| 322 \| Phage transcriptional regulator \| Phage transcriptional regulator; afdb-uniprot50 \| AF-A0A179SHY2-F1-MODEL\_V4 \| 1.0 \| 3.345e-19 \| 684 \| 0.221 \| 289 \| 204 \| 10 \| 1 \| 275 \| 55 \| 336 \| Uncharacterized protein \| Uncharacterized protein; afdb-uniprot50 \| AF-A0A063B487-F1-MODEL\_V4 \| 1.0 \| 9.389e-19 \| 681 \| 0.268 \| 212 \| 142 \| 4 \| 72 \| 275 \| 2 \| 208 \| Uncharacterized protein \| Uncharacterized protein; afdb-uniprot50 \| AF-A0A1I7E360-F1-MODEL\_V4 \| 1.0 \| 5.34e-20 \| 681 \| 0.273 \| 256 \| 146 \| 4 \| 72 \| 289 \| 265 \| 518 \| Phage tail tape measure protein, TP901 family, core region \| Phage tail tape measure protein, TP901 family, core region; afdb-uniprot50 \| AF-A0A7X6WN95-F1-MODEL\_V4 \| 1.0 \| 1.403e-18 \| 680 \| 0.293 \| 252 \| 169 \| 5 \| 2 \| 251 \| 56 \| 300 \| Phage late control D family protein \| Phage late control D family protein; afdb-uniprot50 \| AF-A0A8B4S4Z3-F1-MODEL\_V4 \| 1.0 \| 3.542e-19 \| 678 \| 0.266 \| 255 \| 155 \| 8 \| 26 \| 277 \| 2 \| 227 \| Phage protein D \| Phage protein D; afdb-uniprot50 \| AF-A0A1I5USE1-F1-MODEL\_V4 \| 1.0 \| 1.273e-20 \| 678 \| 0.305 \| 278 \| 167 \| 15 \| 1 \| 276 \| 43 \| 296 \| Uncharacterized protein \| Uncharacterized protein; afdb-uniprot50 \| AF-A0A7X5L5E7-F1-MODEL\_V4 \| 1.0 \| 1.499e-19 \| 675 \| 0.179 \| 324 \| 235 \| 14 \| 1 \| 311 \| 68 \| 373 \| Uncharacterized protein \| Uncharacterized protein; afdb-uniprot50 \| AF-A0A7X3T0M9-F1-MODEL\_V4 \| 1.0 \| 3.973e-19 \| 674 \| 0.275 \| 269 \| 161 \| 6 \| 1 \| 243 \| 63 \| 323 \| Phage late control D family protein \| Phage late control D family protein; afdb-uniprot50 \| AF-B6JEG4-F1-MODEL\_V4 \| 1.0 \| 4.997e-19 \| 674 \| 0.215 \| 292 \| 209 \| 11 \| 1 \| 279 \| 52 \| 336 \| Putative phage late control protein D \| Putative phage late control protein D; afdb-uniprot50 \| AF-A0A109LFY2-F1-MODEL\_V4 \| 1.0 \| 3.345e-19 \| 672 \| 0.247 \| 251 \| 172 \| 7 \| 39 \| 279 \| 1 \| 244 \| Phage late control protein D protein (GPD) \| Phage late control protein D protein (GPD); afdb-uniprot50 \| AF-A0A7Y4XEN4-F1-MODEL\_V4 \| 1.0 \| 4.997e-19 \| 672 \| 0.208 \| 288 \| 204 \| 10 \| 2 \| 279 \| 59 \| 332 \| Cro/Cl family transcriptional regulator \| Cro/Cl family transcriptional regulator; afdb-uniprot50 \| AF-A0A4Y8P2F6-F1-MODEL\_V4 \| 1.0 \| 2.766e-17 \| 670 \| 0.432 \| 164 \| 93 \| 0 \| 2 \| 165 \| 60 \| 223 \| Phage late control D family protein \| Phage late control D family protein; afdb-uniprot50 \| AF-A0A4P8G0Y2-F1-MODEL\_V4 \| 1.0 \| 1.192e-19 \| 669 \| 0.234 \| 294 \| 203 \| 10 \| 1 \| 279 \| 51 \| 337 \| Phage late control D family protein \| Phage late control D family protein; afdb-uniprot50 \| AF-A0A435G0Q0-F1-MODEL\_V4 \| 1.0 \| 1.403e-18 \| 667 \| 0.24 \| 245 \| 172 \| 7 \| 41 \| 279 \| 3 \| 239 \| Uncharacterized protein \| Uncharacterized protein; afdb-uniprot50 \| AF-A0A4R7WMC5-F1-MODEL\_V4 \| 1.0 \| 1.587e-19 \| 667 \| 0.183 \| 322 \| 239 \| 10 \| 1 \| 311 \| 56 \| 364 \| Uncharacterized protein \| Uncharacterized protein; afdb-uniprot50 \| AF-E5VKJ0-F1-MODEL\_V4 \| 1.0 \| 2.982e-19 \| 664 \| 0.212 \| 292 \| 207 \| 10 \| 1 \| 284 \| 66 \| 342 \| Uncharacterized protein \| Uncharacterized protein; afdb-uniprot50 \| AF-A0A285V0M3-F1-MODEL\_V4 \| 1.0 \| 6.285e-19 \| 662 \| 0.215 \| 288 \| 212 \| 8 \| 1 \| 279 \| 53 \| 335 \| Uncharacterized protein \| Uncharacterized protein; afdb-uniprot50 \| AF-C6LFY9-F1-MODEL\_V4 \| 1.0 \| 1.181e-18 \| 657 \| 0.188 \| 265 \| 197 \| 9 \| 18 \| 279 \| 89 \| 338 \| Uncharacterized protein \| Uncharacterized protein; afdb-uniprot50 \| AF-A0A4R2GH06-F1-MODEL\_V4 \| 1.0 \| 1.053e-18 \| 657 \| 0.222 \| 296 \| 198 \| 11 \| 1 \| 280 \| 53 \| 332 \| Uncharacterized protein \| Uncharacterized protein; afdb-uniprot50 \| AF-A0A1E4LS02-F1-MODEL\_V4 \| 1.0 \| 1.499e-19 \| 656 \| 0.209 \| 291 \| 196 \| 12 \| 26 \| 299 \| 5 \| 278 \| Uncharacterized protein \| Uncharacterized protein; afdb-uniprot50 \| AF-A0A1A9WZ35-F1-MODEL\_V4 \| 1.0 \| 2.982e-19 \| 656 \| 0.201 \| 352 \| 229 \| 11 \| 2 \| 311 \| 278 \| 619 \| Uncharacterized protein \| Uncharacterized protein; afdb-uniprot50 \| AF-A0A0J6NA89-F1-MODEL\_V4 \| 1.0 \| 1.115e-18 \| 655 \| 0.243 \| 292 \| 193 \| 10 \| 2 \| 279 \| 64 \| 341 \| Bacteriophage regulatory protein \| Bacteriophage regulatory protein; afdb-uniprot50 \| AF-A0A2A2HM81-F1-MODEL\_V4 \| 1.0 \| 2.982e-19 \| 654 \| 0.222 \| 301 \| 194 \| 10 \| 2 \| 278 \| 67 \| 351 \| Phage protein D \| Phage protein D; afdb-uniprot50 \| AF-A0A2G1CU53-F1-MODEL\_V4 \| 1.0 \| 1.063e-19 \| 652 \| 0.248 \| 278 \| 193 \| 9 \| 1 \| 275 \| 48 \| 312 \| Phage tail protein \| Phage tail protein; afdb-uniprot50 \| AF-A0A3L7ALJ4-F1-MODEL\_V4 \| 1.0 \| 4.415e-18 \| 652 \| 0.201 \| 288 \| 211 \| 9 \| 1 \| 277 \| 51 \| 330 \| Uncharacterized protein \| Uncharacterized protein; afdb-uniprot50 \| AF-Q8ZKJ7-F1-MODEL\_V4 \| 1.0 \| 1.181e-18 \| 652 \| 0.214 \| 289 \| 200 \| 9 \| 2 \| 277 \| 68 \| 342 \| Putative cytoplasmic protein \| Putative cytoplasmic protein; afdb-uniprot50 \| AF-A0A022PIB0-F1-MODEL\_V4 \| 1.0 \| 1.764e-18 \| 651 \| 0.285 \| 263 \| 155 \| 8 \| 1 \| 238 \| 14 \| 268 \| Phage protein D \| Phage protein D; afdb-uniprot50 \| AF-A0A1G3UAZ7-F1-MODEL\_V4 \| 1.0 \| 7.464e-19 \| 651 \| 0.228 \| 280 \| 196 \| 10 \| 1 \| 275 \| 50 \| 314 \| Uncharacterized protein \| Uncharacterized protein; afdb-uniprot50 \| AF-A0A5R8Y6A7-F1-MODEL\_V4 \| 1.0 \| 1.262e-19 \| 651 \| 0.215 \| 316 \| 223 \| 15 \| 9 \| 309 \| 61 \| 366 \| Uncharacterized protein \| Uncharacterized protein; afdb-uniprot50 \| AF-A0A6C8XQL5-F1-MODEL\_V4 \| 1.0 \| 6.716e-20 \| 651 \| 0.201 \| 328 \| 211 \| 15 \| 2 \| 292 \| 74 \| 387 \| Phage protein D \| Phage protein D; afdb-uniprot50 \| AF-A0A7W4RH00-F1-MODEL\_V4 \| 1.0 \| 9.304e-18 \| 649 \| 0.311 \| 202 \| 133 \| 6 \| 84 \| 280 \| 1 \| 201 \| Phage protein D \| Phage protein D; afdb-uniprot50 \| AF-A0A3R8K9T6-F1-MODEL\_V4 \| 1.0 \| 2.982e-19 \| 649 \| 0.227 \| 308 \| 204 \| 11 \| 2 \| 292 \| 75 \| 365 \| Phage late control D family protein \| Phage late control D family protein; afdb-uniprot50 \| AF-A0A6C1BNV6-F1-MODEL\_V4 \| 1.0 \| 1.485e-18 \| 647 \| 0.196 \| 285 \| 205 \| 12 \| 2 \| 280 \| 64 \| 330 \| Uncharacterized protein \| Uncharacterized protein; afdb-uniprot50 \| AF-A0A090G3M4-F1-MODEL\_V4 \| 1.0 \| 3.51e-18 \| 647 \| 0.221 \| 294 \| 210 \| 10 \| 1 \| 284 \| 52 \| 336 \| Uncharacterized protein \| Uncharacterized protein; afdb-uniprot50 \| AF-A0A5W6TUF8-F1-MODEL\_V4 \| 1.0 \| 4.952e-18 \| 646 \| 0.245 \| 281 \| 180 \| 7 \| 1 \| 256 \| 66 \| 339 \| Phage late control D family protein \| Phage late control D family protein; afdb-uniprot50 \| AF-A0A087KKM1-F1-MODEL\_V4 \| 1.0 \| 7.532e-20 \| 646 \| 0.238 \| 327 \| 198 \| 13 \| 2 \| 292 \| 75 \| 386 \| Bacteriophage regulatory protein \| Bacteriophage regulatory protein; afdb-uniprot50 \| AF-A0A7X5I0X9-F1-MODEL\_V4 \| 1.0 \| 8.865e-19 \| 645 \| 0.192 \| 301 \| 213 \| 11 \| 15 \| 310 \| 78 \| 353 \| LysM peptidoglycan-binding domain-containing protein \| LysM peptidoglycan-binding domain-containing protein; afdb-uniprot50 \| AF-A0A1B9JGB6-F1-MODEL\_V4 \| 1.0 \| 1.053e-18 \| 644 \| 0.175 \| 400 \| 198 \| 9 \| 2 \| 277 \| 57 \| 448 \| Uncharacterized protein \| Uncharacterized protein; afdb-uniprot50 \| AF-A0A1H0W5E2-F1-MODEL\_V4 \| 1.0 \| 7.834e-18 \| 641 \| 0.267 \| 239 \| 155 \| 5 \| 72 \| 308 \| 2 \| 222 \| Phage late control gene D protein (GPD) \| Phage late control gene D protein (GPD); afdb-uniprot50 \| AF-A0A6P0VER2-F1-MODEL\_V4 \| 1.0 \| 1.666e-18 \| 641 \| 0.21 \| 299 \| 199 \| 14 \| 1 \| 284 \| 9 \| 285 \| Uncharacterized protein \| Uncharacterized protein; afdb-uniprot50 \| AF-A0A743PFW0-F1-MODEL\_V4 \| 1.0 \| 8.946e-20 \| 641 \| 0.223 \| 327 \| 203 \| 12 \| 2 \| 292 \| 64 \| 375 \| Phage protein D \| Phage protein D; afdb-uniprot50 \| AF-A0A7W3CBM1-F1-MODEL\_V4 \| 1.0 \| 1.78e-19 \| 641 \| 0.212 \| 329 \| 206 \| 14 \| 2 \| 292 \| 75 \| 388 \| Phage late control D family protein \| Phage late control D family protein; afdb-uniprot50 \| AF-A0A7U6KDT6-F1-MODEL\_V4 \| 1.0 \| 2.791e-18 \| 639 \| 0.193 \| 264 \| 200 \| 9 \| 17 \| 279 \| 83 \| 334 \| Putative bacteriophage regulatory protein \| Putative bacteriophage regulatory protein; afdb-uniprot50 \| AF-A0A1M3AKH3-F1-MODEL\_V4 \| 1.0 \| 7.464e-19 \| 639 \| 0.213 \| 323 \| 219 \| 13 \| 1 \| 311 \| 53 \| 352 \| Late control D family protein \| Late control D family protein; afdb-uniprot50 \| AF-A0A1C6ENZ2-F1-MODEL\_V4 \| 1.0 \| 5.881e-18 \| 639 \| 0.223 \| 264 \| 191 \| 7 \| 17 \| 279 \| 83 \| 333 \| LysM domain/BON superfamily protein \| LysM domain/BON superfamily protein; afdb-uniprot50 \| AF-A0A2A3D6W1-F1-MODEL\_V4 \| 1.0 \| 5.553e-18 \| 638 \| 0.203 \| 295 \| 214 \| 11 \| 1 \| 284 \| 52 \| 336 \| Uncharacterized protein \| Uncharacterized protein; afdb-uniprot50 \| AF-A0A7X5CGI8-F1-MODEL\_V4 \| 1.0 \| 6.228e-18 \| 638 \| 0.16 \| 299 \| 226 \| 11 \| 1 \| 289 \| 58 \| 341 \| LysM peptidoglycan-binding domain-containing protein \| LysM peptidoglycan-binding domain-containing protein; afdb-uniprot50 \| AF-A0A0R3KWX4-F1-MODEL\_V4 \| 1.0 \| 8.371e-19 \| 637 \| 0.222 \| 328 \| 220 \| 14 \| 1 \| 311 \| 55 \| 364 \| Uncharacterized protein \| Uncharacterized protein; afdb-uniprot50 \| AF-A0A1J5I2G1-F1-MODEL\_V4 \| 1.0 \| 1.666e-18 \| 636 \| 0.18 \| 294 \| 209 \| 13 \| 1 \| 277 \| 54 \| 332 \| Uncharacterized protein \| Uncharacterized protein; afdb-uniprot50 \| AF-A0A2D3T999-F1-MODEL\_V4 \| 1.0 \| 3.285e-17 \| 635 \| 0.339 \| 165 \| 109 \| 0 \| 113 \| 277 \| 3 \| 167 \| Uncharacterized protein \| Uncharacterized protein; afdb-uniprot50 \| AF-A0A6L5TD89-F1-MODEL\_V4 \| 1.0 \| 5.604e-19 \| 634 \| 0.216 \| 328 \| 217 \| 14 \| 1 \| 311 \| 57 \| 361 \| Uncharacterized protein \| Uncharacterized protein; afdb-uniprot50 \| AF-A0A1B4SFB1-F1-MODEL\_V4 \| 1.0 \| 1.324e-18 \| 632 \| 0.19 \| 341 \| 215 \| 11 \| 2 \| 311 \| 73 \| 383 \| Cro/Cl family transcriptional regulator \| Cro/Cl family transcriptional regulator; afdb-uniprot50 \| AF-A0A1M3BPS0-F1-MODEL\_V4 \| 1.0 \| 6.596e-18 \| 631 \| 0.223 \| 282 \| 195 \| 10 \| 7 \| 272 \| 51 \| 324 \| Uncharacterized protein \| Uncharacterized protein; afdb-uniprot50 \| AF-A8SB41-F1-MODEL\_V4 \| 1.0 \| 7.834e-18 \| 631 \| 0.188 \| 260 \| 199 \| 6 \| 18 \| 276 \| 82 \| 330 \| Uncharacterized protein \| Uncharacterized protein; afdb-uniprot50 \| AF-R7JNW8-F1-MODEL\_V4 \| 1.0 \| 1.181e-18 \| 630 \| 0.216 \| 291 \| 202 \| 10 \| 2 \| 279 \| 59 \| 336 \| Phage late control gene D protein (GPD) putative \| Phage late control gene D protein (GPD) putative; afdb-uniprot50 \| AF-A0A0P6VKL9-F1-MODEL\_V4 \| 1.0 \| 1.39e-17 \| 629 \| 0.205 \| 282 \| 206 \| 8 \| 9 \| 279 \| 72 \| 346 \| Uncharacterized protein \| Uncharacterized protein; afdb-uniprot50 \| AF-A0A370X1T7-F1-MODEL\_V4 \| 1.0 \| 8.296e-18 \| 628 \| 0.304 \| 240 \| 149 \| 5 \| 74 \| 310 \| 38 \| 262 \| Uncharacterized protein \| Uncharacterized protein; afdb-uniprot50 \| AF-A0A371J384-F1-MODEL\_V4 \| 1.0 \| 4.718e-19 \| 628 \| 0.184 \| 314 \| 213 \| 15 \| 2 \| 296 \| 61 \| 350 \| LysM peptidoglycan-binding domain-containing protein \| LysM peptidoglycan-binding domain-containing protein; afdb-uniprot50 \| AF-A0A161VRP1-F1-MODEL\_V4 \| 1.0 \| 5.935e-19 \| 627 \| 0.185 \| 297 \| 204 \| 13 \| 2 \| 279 \| 19 \| 296 \| Late control D family protein \| Late control D family protein; afdb-uniprot50 \| AF-A0A1M7LUC3-F1-MODEL\_V4 \| 1.0 \| 8.296e-18 \| 627 \| 0.197 \| 289 \| 207 \| 12 \| 2 \| 283 \| 58 \| 328 \| LysM domain-containing protein \| LysM domain-containing protein; afdb-uniprot50 \| AF-A0A2V2GIE6-F1-MODEL\_V4 \| 1.0 \| 1.105e-17 \| 627 \| 0.208 \| 264 \| 195 \| 7 \| 18 \| 278 \| 79 \| 331 \| Uncharacterized protein \| Uncharacterized protein; afdb-uniprot50 \| AF-A0A2U1CFJ9-F1-MODEL\_V4 \| 1.0 \| 1.666e-18 \| 625 \| 0.175 \| 308 \| 226 \| 13 \| 16 \| 311 \| 185 \| 476 \| Uncharacterized protein \| Uncharacterized protein; afdb-uniprot50 \| AF-A0A4U8UEB9-F1-MODEL\_V4 \| 1.0 \| 1.181e-18 \| 624 \| 0.265 \| 279 \| 182 \| 10 \| 1 \| 276 \| 50 \| 308 \| Phage tail protein \| Phage tail protein; afdb-uniprot50 \| AF-A0A0Q7A3Q8-F1-MODEL\_V4 \| 1.0 \| 8.785e-18 \| 624 \| 0.202 \| 271 \| 197 \| 9 \| 20 \| 279 \| 69 \| 331 \| Uncharacterized protein \| Uncharacterized protein; afdb-uniprot50 \| AF-C1D832-F1-MODEL\_V4 \| 1.0 \| 1.764e-18 \| 624 \| 0.23 \| 299 \| 194 \| 10 \| 2 \| 278 \| 64 \| 348 \| Probable bacteriophage regulatory protein \| Probable bacteriophage regulatory protein; afdb-uniprot50 \| AF-A0A3E2T6D6-F1-MODEL\_V4 \| 1.0 \| 1.17e-17 \| 623 \| 0.209 \| 287 \| 202 \| 11 \| 2 \| 278 \| 58 \| 329 \| Uncharacterized protein \| Uncharacterized protein; afdb-uniprot50 \| AF-A0A4D7AV46-F1-MODEL\_V4 \| 1.0 \| 2.466e-17 \| 622 \| 0.211 \| 284 \| 202 \| 10 \| 2 \| 276 \| 62 \| 332 \| Late control protein D \| Late control protein D; afdb-uniprot50 \| AF-A0A2U8VQW8-F1-MODEL\_V4 \| 1.0 \| 5.553e-18 \| 622 \| 0.217 \| 303 \| 205 \| 10 \| 1 \| 280 \| 52 \| 345 \| Uncharacterized protein \| Uncharacterized protein; afdb-uniprot50 \| AF-A0A2U0TFH0-F1-MODEL\_V4 \| 1.0 \| 6.656e-19 \| 620 \| 0.188 \| 302 \| 220 \| 10 \| 20 \| 311 \| 72 \| 358 \| Uncharacterized protein \| Uncharacterized protein; afdb-uniprot50 \| AF-A0A1R1MKJ1-F1-MODEL\_V4 \| 1.0 \| 9.943e-19 \| 619 \| 0.246 \| 288 \| 188 \| 14 \| 2 \| 276 \| 59 \| 330 \| Uncharacterized protein \| Uncharacterized protein; afdb-uniprot50 \| AF-A0A385Q3K1-F1-MODEL\_V4 \| 1.0 \| 6.656e-19 \| 619 \| 0.181 \| 297 \| 216 \| 8 \| 16 \| 310 \| 72 \| 343 \| LysM peptidoglycan-binding domain-containing protein \| LysM peptidoglycan-binding domain-containing protein; afdb-uniprot50 \| AF-A0A1S7Q5S5-F1-MODEL\_V4 \| 1.0 \| 1.651e-17 \| 617 \| 0.208 \| 283 \| 205 \| 10 \| 9 \| 280 \| 117 \| 391 \| Phage late control D family protein \| Phage late control D family protein; afdb-uniprot50 \| AF-A0A4R6MB46-F1-MODEL\_V4 \| 1.0 \| 6.985e-18 \| 616 \| 0.2 \| 279 \| 198 \| 8 \| 1 \| 279 \| 50 \| 303 \| Uncharacterized protein \| Uncharacterized protein; afdb-uniprot50 \| AF-A0A1X3HWQ9-F1-MODEL\_V4 \| 1.0 \| 3.479e-17 \| 616 \| 0.25 \| 264 \| 173 \| 6 \| 1 \| 246 \| 68 \| 324 \| Bacteriophage late protein control protein D \| Bacteriophage late protein control protein D; afdb-uniprot50 \| AF-A0A285M8H1-F1-MODEL\_V4 \| 1.0 \| 1.764e-18 \| 616 \| 0.194 \| 304 \| 218 \| 11 \| 18 \| 311 \| 71 \| 357 \| Uncharacterized protein \| Uncharacterized protein; afdb-uniprot50 \| AF-A0A451AF14-F1-MODEL\_V4 \| 1.0 \| 2.199e-17 \| 615 \| 0.416 \| 197 \| 108 \| 2 \| 22 \| 218 \| 2 \| 191 \| Phage late control gene D protein (GPD) \| Phage late control gene D protein (GPD); afdb-uniprot50 \| AF-A0A163TCJ4-F1-MODEL\_V4 \| 1.0 \| 1.403e-18 \| 614 \| 0.209 \| 272 \| 185 \| 11 \| 25 \| 279 \| 1 \| 259 \| Late control D family protein \| Late control D family protein; afdb-uniprot50 \| AF-A0A268U3E7-F1-MODEL\_V4 \| 1.0 \| 3.51e-18 \| 614 \| 0.2 \| 274 \| 198 \| 9 \| 1 \| 274 \| 48 \| 300 \| Uncharacterized protein \| Uncharacterized protein; afdb-uniprot50 \| AF-A0A2S9IK07-F1-MODEL\_V4 \| 1.0 \| 1.472e-17 \| 614 \| 0.21 \| 295 \| 203 \| 11 \| 1 \| 279 \| 53 \| 333 \| Late control D family protein \| Late control D family protein; afdb-uniprot50 \| AF-A0A0H3ZXP3-F1-MODEL\_V4 \| 1.0 \| 3.102e-17 \| 612 \| 0.245 \| 200 \| 147 \| 3 \| 82 \| 278 \| 4 \| 202 \| Phage protein D \| Phage protein D; afdb-uniprot50 \| AF-A0A037UNC0-F1-MODEL\_V4 \| 1.0 \| 5.553e-18 \| 612 \| 0.245 \| 273 \| 186 \| 10 \| 20 \| 279 \| 68 \| 333 \| Late control protein D \| Late control protein D; afdb-uniprot50 \| AF-A0A2T7H7Y0-F1-MODEL\_V4 \| 1.0 \| 1.485e-18 \| 611 \| 0.217 \| 294 \| 206 \| 9 \| 1 \| 279 \| 54 \| 338 \| Uncharacterized protein \| Uncharacterized protein; afdb-uniprot50 \| AF-A0A3P6KIP4-F1-MODEL\_V4 \| 1.0 \| 8.296e-18 \| 611 \| 0.184 \| 293 \| 210 \| 10 \| 2 \| 281 \| 86 \| 362 \| Phage protein D \| Phage protein D; afdb-uniprot50 \| AF-A0A1B9LF16-F1-MODEL\_V4 \| 1.0 \| 2.114e-19 \| 608 \| 0.253 \| 308 \| 193 \| 10 \| 1 \| 281 \| 56 \| 353 \| Uncharacterized protein \| Uncharacterized protein; afdb-uniprot50 \| AF-A0A6P0J3I4-F1-MODEL\_V4 \| 1.0 \| 2.488e-18 \| 607 \| 0.167 \| 304 \| 210 \| 12 \| 3 \| 279 \| 20 \| 307 \| Phage late control D family protein \| Phage late control D family protein; afdb-uniprot50 \| AF-A0A3D0U3C5-F1-MODEL\_V4 \| 1.0 \| 9.943e-19 \| 607 \| 0.206 \| 305 \| 190 \| 14 \| 2 \| 278 \| 26 \| 306 \| Uncharacterized protein \| Uncharacterized protein; afdb-uniprot50 \| AF-A0A212S2I3-F1-MODEL\_V4 \| 1.0 \| 9.853e-18 \| 606 \| 0.243 \| 267 \| 186 \| 8 \| 22 \| 279 \| 77 \| 336 \| Uncharacterized protein \| Uncharacterized protein; afdb-uniprot50 \| AF-W4SDY7-F1-MODEL\_V4 \| 1.0 \| 2.588e-16 \| 604 \| 0.331 \| 172 \| 112 \| 3 \| 109 \| 277 \| 2 \| 173 \| Uncharacterized protein \| Uncharacterized protein; afdb-uniprot50 \| AF-A0A6I0CE52-F1-MODEL\_V4 \| 1.0 \| 5.828e-17 \| 603 \| 0.206 \| 295 \| 202 \| 11 \| 2 \| 279 \| 64 \| 343 \| Late control protein D \| Late control protein D; afdb-uniprot50 \| AF-A0A6L2ZNF4-F1-MODEL\_V4 \| 1.0 \| 3.479e-17 \| 602 \| 0.28 \| 203 \| 128 \| 3 \| 95 \| 279 \| 4 \| 206 \| Late control gene D protein \| Late control gene D protein; afdb-uniprot50 \| AF-A0A7C2ARE1-F1-MODEL\_V4 \| 1.0 \| 1.239e-17 \| 602 \| 0.222 \| 283 \| 194 \| 13 \| 3 \| 276 \| 65 \| 330 \| Uncharacterized protein \| Uncharacterized protein; afdb-uniprot50 \| AF-A0A376EWE4-F1-MODEL\_V4 \| 1.0 \| 2.488e-18 \| 601 \| 0.233 \| 330 \| 215 \| 13 \| 2 \| 311 \| 75 \| 386 \| Gene D protein \| Gene D protein; afdb-uniprot50 \| AF-A0A3A6K9Q8-F1-MODEL\_V4 \| 1.0 \| 1.472e-17 \| 600 \| 0.179 \| 268 \| 197 \| 11 \| 15 \| 276 \| 81 \| 331 \| Preprotein translocase subunit TatB \| Preprotein translocase subunit TatB; afdb-uniprot50 \| AF-E2CJH8-F1-MODEL\_V4 \| 1.0 \| 1.651e-17 \| 600 \| 0.211 \| 274 \| 194 \| 12 \| 19 \| 279 \| 70 \| 334 \| Late control D family protein \| Late control D family protein; afdb-uniprot50 \| AF-A0NQA6-F1-MODEL\_V4 \| 1.0 \| 6.985e-18 \| 595 \| 0.214 \| 280 \| 186 \| 12 \| 19 \| 279 \| 67 \| 331 \| Uncharacterized protein \| Uncharacterized protein; afdb-uniprot50 \| AF-M5K2K4-F1-MODEL\_V4 \| 1.0 \| 1.651e-17 \| 595 \| 0.22 \| 327 \| 224 \| 15 \| 1 \| 311 \| 53 \| 364 \| Phage late control D family protein \| Phage late control D family protein; afdb-uniprot50 \| AF-A0A3A9CQQ7-F1-MODEL\_V4 \| 1.0 \| 1.651e-17 \| 595 \| 0.182 \| 307 \| 217 \| 14 \| 1 \| 297 \| 175 \| 457 \| LysM peptidoglycan-binding domain-containing protein \| LysM peptidoglycan-binding domain-containing protein; afdb-uniprot50 \| AF-A0A3D2IA26-F1-MODEL\_V4 \| 1.0 \| 4.634e-17 \| 592 \| 0.195 \| 332 \| 224 \| 19 \| 1 \| 311 \| 58 \| 367 \| Uncharacterized protein \| Uncharacterized protein; afdb-uniprot50 \| AF-A0A7X2NKR7-F1-MODEL\_V4 \| 1.0 \| 1.545e-16 \| 592 \| 0.194 \| 293 \| 204 \| 10 \| 1 \| 280 \| 57 \| 330 \| LysM peptidoglycan-binding domain-containing protein \| LysM peptidoglycan-binding domain-containing protein; afdb-uniprot50 \| AF-A0A1G6EJ79-F1-MODEL\_V4 \| 1.0 \| 2.929e-17 \| 591 \| 0.219 \| 292 \| 200 \| 13 \| 1 \| 279 \| 53 \| 329 \| Uncharacterized protein \| Uncharacterized protein; afdb-uniprot50 \| AF-A0A0X1U7U0-F1-MODEL\_V4 \| 1.0 \| 1.16e-16 \| 590 \| 0.178 \| 269 \| 199 \| 9 \| 15 \| 276 \| 74 \| 327 \| Phage late control protein D \| Phage late control protein D; afdb-uniprot50 \| AF-A0A5R9GK92-F1-MODEL\_V4 \| 1.0 \| 2.35e-18 \| 588 \| 0.222 \| 306 \| 186 \| 10 \| 4 \| 279 \| 83 \| 366 \| Phage late control D family protein \| Phage late control D family protein; afdb-uniprot50 \| AF-A0A318LI70-F1-MODEL\_V4 \| 1.0 \| 6.228e-18 \| 587 \| 0.233 \| 334 \| 215 \| 14 \| 2 \| 311 \| 62 \| 378 \| Uncharacterized protein \| Uncharacterized protein; afdb-uniprot50 \| AF-A0A2D2CYN5-F1-MODEL\_V4 \| 1.0 \| 4.131e-17 \| 585 \| 0.193 \| 300 \| 212 \| 12 \| 2 \| 279 \| 54 \| 345 \| Uncharacterized protein \| Uncharacterized protein; afdb-uniprot50 \| AF-A0A317H3Z9-F1-MODEL\_V4 \| 1.0 \| 1.39e-17 \| 581 \| 0.167 \| 347 \| 236 \| 15 \| 1 \| 311 \| 84 \| 413 \| Uncharacterized protein \| Uncharacterized protein; afdb-uniprot50 \| AF-A0A434TY84-F1-MODEL\_V4 \| 1.0 \| 5.244e-18 \| 580 \| 0.213 \| 295 \| 201 \| 12 \| 26 \| 310 \| 8 \| 281 \| Uncharacterized protein \| Uncharacterized protein; afdb-uniprot50 \| AF-A0A327KRF2-F1-MODEL\_V4 \| 1.0 \| 9.676e-16 \| 574 \| 0.301 \| 189 \| 128 \| 3 \| 93 \| 277 \| 1 \| 189 \| Uncharacterized protein \| Uncharacterized protein; afdb-uniprot50 \| AF-A0A7W6PUG3-F1-MODEL\_V4 \| 1.0 \| 9.22e-17 \| 574 \| 0.203 \| 275 \| 188 \| 12 \| 19 \| 277 \| 82 \| 341 \| Uncharacterized protein \| Uncharacterized protein; afdb-uniprot50 \| AF-A0A5E8H666-F1-MODEL\_V4 \| 1.0 \| 7.397e-18 \| 572 \| 0.21 \| 309 \| 207 \| 12 \| 20 \| 311 \| 69 \| 357 \| Phage protein D \| Phage protein D; afdb-uniprot50 \| AF-A0A0D6KHQ0-F1-MODEL\_V4 \| 1.0 \| 4.907e-17 \| 571 \| 0.153 \| 299 \| 211 \| 11 \| 7 \| 279 \| 73 \| 355 \| Bacteriophage protein D \| Bacteriophage protein D; afdb-uniprot50 \| AF-A0A2I8Q7T7-F1-MODEL\_V4 \| 1.0 \| 1.16e-16 \| 571 \| 0.231 \| 272 \| 178 \| 7 \| 2 \| 249 \| 81 \| 345 \| Uncharacterized protein \| Uncharacterized protein; afdb-uniprot50 \| AF-A0A7B9FRA4-F1-MODEL\_V4 \| 1.0 \| 1.472e-17 \| 570 \| 0.268 \| 246 \| 134 \| 7 \| 89 \| 291 \| 1 \| 243 \| Phage late control D family protein \| Phage late control D family protein; afdb-uniprot50 \| AF-A0A2C9D5M1-F1-MODEL\_V4 \| 1.0 \| 5.197e-17 \| 569 \| 0.2 \| 279 \| 192 \| 12 \| 20 \| 279 \| 68 \| 334 \| Tail protein \| Tail protein; afdb-uniprot50 \| AF-A0A445ZUU9-F1-MODEL\_V4 \| 1.0 \| 1.312e-17 \| 569 \| 0.256 \| 277 \| 172 \| 11 \| 1 \| 261 \| 51 \| 309 \| Uncharacterized protein \| Uncharacterized protein; afdb-uniprot50 \| AF-A0A2X2S5N7-F1-MODEL\_V4 \| 1.0 \| 6.536e-17 \| 568 \| 0.236 \| 250 \| 158 \| 6 \| 79 \| 308 \| 2 \| 238 \| Phage tail protein \| Phage tail protein; afdb-uniprot50 \| AF-A0A833AM78-F1-MODEL\_V4 \| 1.0 \| 1.835e-16 \| 568 \| 0.198 \| 247 \| 164 \| 8 \| 57 \| 278 \| 9 \| 246 \| Uncharacterized protein \| Uncharacterized protein; afdb-uniprot50 \| AF-A0A3B8Q684-F1-MODEL\_V4 \| 1.0 \| 5.503e-17 \| 568 \| 0.185 \| 297 \| 201 \| 11 \| 7 \| 278 \| 77 \| 357 \| Uncharacterized protein \| Uncharacterized protein; afdb-uniprot50 \| AF-A0A0C1ITH4-F1-MODEL\_V4 \| 1.0 \| 1.545e-16 \| 565 \| 0.225 \| 288 \| 198 \| 10 \| 3 \| 278 \| 60 \| 334 \| Uncharacterized protein \| Uncharacterized protein; afdb-uniprot50 \| AF-Q602Z6-F1-MODEL\_V4 \| 1.0 \| 4.863e-16 \| 565 \| 0.219 \| 287 \| 198 \| 11 \| 2 \| 279 \| 71 \| 340 \| Putative prophage MuMc02, late control gene D protein \| Putative prophage MuMc02, late control gene D protein; afdb-uniprot50 \| AF-A0A1I3I3R8-F1-MODEL\_V4 \| 1.0 \| 2.588e-16 \| 565 \| 0.199 \| 301 \| 201 \| 10 \| 3 \| 278 \| 70 \| 355 \| Phage protein D \| Phage protein D; afdb-uniprot50 \| AF-A0A2S5R6X2-F1-MODEL\_V4 \| 1.0 \| 3.046e-15 \| 562 \| 0.353 \| 178 \| 115 \| 0 \| 75 \| 252 \| 2 \| 179 \| Phage late control D protein (GPD) \| Phage late control D protein (GPD); afdb-uniprot50 \| AF-F3YY66-F1-MODEL\_V4 \| 1.0 \| 7.763e-17 \| 561 \| 0.237 \| 278 \| 183 \| 11 \| 15 \| 279 \| 76 \| 337 \| Late control D family protein \| Late control D family protein; afdb-uniprot50 \| AF-A0A377YUK4-F1-MODEL\_V4 \| 1.0 \| 5.15e-16 \| 558 \| 0.261 \| 210 \| 130 \| 4 \| 95 \| 279 \| 1 \| 210 \| Late control gene D protein \| Late control gene D protein; afdb-uniprot50 \| AF-A0A832EBM3-F1-MODEL\_V4 \| 1.0 \| 2.741e-16 \| 558 \| 0.212 \| 296 \| 191 \| 11 \| 9 \| 278 \| 82 \| 361 \| Uncharacterized protein \| Uncharacterized protein; afdb-uniprot50 \| AF-A0A6G5QFX8-F1-MODEL\_V4 \| 1.0 \| 3.074e-16 \| 557 \| 0.172 \| 255 \| 195 \| 7 \| 20 \| 274 \| 67 \| 305 \| Phage protein D \| Phage protein D; afdb-uniprot50 \| AF-A0A0S9PYR5-F1-MODEL\_V4 \| 1.0 \| 3.447e-16 \| 557 \| 0.169 \| 295 \| 215 \| 12 \| 1 \| 278 \| 52 \| 333 \| Uncharacterized protein \| Uncharacterized protein; afdb-uniprot50 \| AF-A0A411WI36-F1-MODEL\_V4 \| 1.0 \| 1.651e-17 \| 557 \| 0.201 \| 333 \| 213 \| 15 \| 2 \| 311 \| 63 \| 365 \| Late control protein D \| Late control protein D; afdb-uniprot50 \| AF-A0A0C2UPU9-F1-MODEL\_V4 \| 1.0 \| 1.251e-18 \| 556 \| 0.252 \| 277 \| 179 \| 10 \| 7 \| 279 \| 49 \| 301 \| Uncharacterized protein \| Uncharacterized protein; afdb-uniprot50 \| AF-H1CIQ5-F1-MODEL\_V4 \| 1.0 \| 3.074e-16 \| 556 \| 0.182 \| 324 \| 225 \| 17 \| 1 \| 308 \| 58 \| 357 \| Uncharacterized protein \| Uncharacterized protein; afdb-uniprot50 \| AF-A0A7Y5KGH0-F1-MODEL\_V4 \| 1.0 \| 1.651e-17 \| 555 \| 0.175 \| 313 \| 204 \| 12 \| 3 \| 282 \| 5 \| 296 \| Phage late control D family protein \| Phage late control D family protein; afdb-uniprot50 \| AF-U3TRV7-F1-MODEL\_V4 \| 1.0 \| 4.131e-17 \| 554 \| 0.23 \| 260 \| 163 \| 10 \| 59 \| 294 \| 3 \| 249 \| Uncharacterized protein \| Uncharacterized protein; afdb-uniprot50 \| AF-A0A1W2EJU3-F1-MODEL\_V4 \| 1.0 \| 1.377e-16 \| 554 \| 0.165 \| 278 \| 206 \| 12 \| 20 \| 278 \| 67 \| 337 \| Uncharacterized protein \| Uncharacterized protein; afdb-uniprot50 \| AF-A0A2N9MSM3-F1-MODEL\_V4 \| 1.0 \| 6.536e-17 \| 554 \| 0.174 \| 304 \| 196 \| 12 \| 9 \| 279 \| 75 \| 356 \| Uncharacterized protein \| Uncharacterized protein; afdb-uniprot50 \| AF-A0A6I3SN83-F1-MODEL\_V4 \| 1.0 \| 1.545e-16 \| 552 \| 0.161 \| 297 \| 202 \| 9 \| 9 \| 279 \| 92 \| 367 \| Uncharacterized protein \| Uncharacterized protein; afdb-uniprot50 \| AF-A0A285MDK0-F1-MODEL\_V4 \| 1.0 \| 2.308e-16 \| 551 \| 0.196 \| 275 \| 191 \| 10 \| 20 \| 279 \| 68 \| 327 \| Uncharacterized protein \| Uncharacterized protein; afdb-uniprot50 \| AF-A0A3M2ZMQ4-F1-MODEL\_V4 \| 1.0 \| 8.628e-16 \| 550 \| 0.297 \| 232 \| 146 \| 5 \| 1 \| 225 \| 53 \| 274 \| Tail protein D \| Tail protein D; afdb-uniprot50 \| AF-A0A2V4UUX4-F1-MODEL\_V4 \| 1.0 \| 2.219e-18 \| 550 \| 0.272 \| 279 \| 165 \| 8 \| 12 \| 280 \| 68 \| 318 \| Phage baseplate assembly protein V \| Phage baseplate assembly protein V; afdb-uniprot50 \| AF-A0A6H2NTK7-F1-MODEL\_V4 \| 1.0 \| 7.693e-16 \| 549 \| 0.3 \| 213 \| 141 \| 4 \| 7 \| 218 \| 49 \| 254 \| Phage tail protein \| Phage tail protein; afdb-uniprot50 \| AF-A0A6P0MTJ9-F1-MODEL\_V4 \| 1.0 \| 1.301e-16 \| 549 \| 0.213 \| 295 \| 192 \| 14 \| 4 \| 281 \| 62 \| 333 \| Uncharacterized protein \| Uncharacterized protein; afdb-uniprot50 \| AF-A0A1C6BMG6-F1-MODEL\_V4 \| 1.0 \| 1.095e-16 \| 549 \| 0.19 \| 283 \| 195 \| 13 \| 7 \| 278 \| 73 \| 332 \| Phage protein D \| Phage protein D; afdb-uniprot50 \| AF-A0A6L5BH82-F1-MODEL\_V4 \| 1.0 \| 1.545e-16 \| 547 \| 0.191 \| 323 \| 223 \| 11 \| 2 \| 308 \| 68 \| 368 \| Uncharacterized protein \| Uncharacterized protein; afdb-uniprot50 \| AF-A0A6P0L7N5-F1-MODEL\_V4 \| 1.0 \| 3.074e-16 \| 546 \| 0.166 \| 301 \| 206 \| 13 \| 7 \| 279 \| 73 \| 356 \| Phage late control D family protein \| Phage late control D family protein; afdb-uniprot50 \| AF-A0A3M1TEK0-F1-MODEL\_V4 \| 1.0 \| 1.459e-16 \| 546 \| 0.184 \| 331 \| 219 \| 15 \| 7 \| 308 \| 109 \| 417 \| Phage late control D family protein \| Phage late control D family protein; afdb-uniprot50 \| AF-A0A7X6ACD1-F1-MODEL\_V4 \| 1.0 \| 1.377e-16 \| 544 \| 0.166 \| 306 \| 210 \| 14 \| 3 \| 279 \| 44 \| 333 \| Uncharacterized protein \| Uncharacterized protein; afdb-uniprot50 \| AF-A0A5B9EBV9-F1-MODEL\_V4 \| 1.0 \| 6.477e-16 \| 544 \| 0.187 \| 299 \| 201 \| 10 \| 7 \| 279 \| 74 \| 356 \| Phage late control D family protein \| Phage late control D family protein; afdb-uniprot50 \| AF-A0A329TWB6-F1-MODEL\_V4 \| 1.0 \| 1.034e-16 \| 544 \| 0.19 \| 309 \| 204 \| 13 \| 2 \| 282 \| 65 \| 355 \| Preprotein translocase subunit TatB \| Preprotein translocase subunit TatB; afdb-uniprot50 \| AF-A0A838CFT9-F1-MODEL\_V4 \| 1.0 \| 4.592e-16 \| 543 \| 0.235 \| 208 \| 134 \| 3 \| 113 \| 307 \| 2 \| 197 \| Late control protein \| Late control protein; afdb-uniprot50 \| AF-A0A534NNF3-F1-MODEL\_V4 \| 1.0 \| 7.763e-17 \| 543 \| 0.196 \| 320 \| 201 \| 13 \| 7 \| 304 \| 80 \| 365 \| Phage late control D family protein \| Phage late control D family protein; afdb-uniprot50 \| AF-A0A534TDB4-F1-MODEL\_V4 \| 1.0 \| 5.828e-17 \| 543 \| 0.181 \| 331 \| 217 \| 15 \| 7 \| 311 \| 51 \| 353 \| Uncharacterized protein \| Uncharacterized protein; afdb-uniprot50 \| AF-A0A2J4V7J1-F1-MODEL\_V4 \| 1.0 \| 7.693e-16 \| 541 \| 0.266 \| 229 \| 126 \| 3 \| 95 \| 281 \| 2 \| 230 \| Uncharacterized protein \| Uncharacterized protein; afdb-uniprot50 \| AF-A0A4R1J9U4-F1-MODEL\_V4 \| 1.0 \| 4.592e-16 \| 541 \| 0.144 \| 298 \| 209 \| 11 \| 7 \| 279 \| 69 \| 345 \| Phage protein D \| Phage protein D; afdb-uniprot50 \| AF-A0A2T6G5A6-F1-MODEL\_V4 \| 1.0 \| 4.907e-17 \| 540 \| 0.185 \| 296 \| 199 \| 9 \| 7 \| 278 \| 84 \| 361 \| Uncharacterized protein \| Uncharacterized protein; afdb-uniprot50 \| AF-A0A0P9CMZ8-F1-MODEL\_V4 \| 1.0 \| 2.444e-16 \| 539 \| 0.247 \| 218 \| 141 \| 5 \| 102 \| 311 \| 2 \| 204 \| Uncharacterized protein \| Uncharacterized protein; afdb-uniprot50 \| AF-A0A0Q2UDB3-F1-MODEL\_V4 \| 1.0 \| 5.775e-16 \| 539 \| 0.211 \| 289 \| 197 \| 13 \| 2 \| 276 \| 54 \| 325 \| Uncharacterized protein \| Uncharacterized protein; afdb-uniprot50 \| AF-A0A5M3Y7N1-F1-MODEL\_V4 \| 1.0 \| 6.477e-16 \| 539 \| 0.166 \| 300 \| 205 \| 11 \| 7 \| 278 \| 58 \| 340 \| Mlr6558 protein \| Mlr6558 protein; afdb-uniprot50 \| AF-A0A7V1ZWM7-F1-MODEL\_V4 \| 1.0 \| 1.377e-16 \| 539 \| 0.218 \| 325 \| 199 \| 15 \| 9 \| 305 \| 77 \| 374 \| Phage late control D family protein \| Phage late control D family protein; afdb-uniprot50 \| AF-A0A6M0FC52-F1-MODEL\_V4 \| 1.0 \| 5.454e-16 \| 538 \| 0.167 \| 299 \| 204 \| 14 \| 9 \| 279 \| 76 \| 357 \| Phage late control D family protein \| Phage late control D family protein; afdb-uniprot50 \| AF-A0A6I7P4G6-F1-MODEL\_V4 \| 1.0 \| 2.058e-16 \| 538 \| 0.194 \| 308 \| 193 \| 13 \| 3 \| 278 \| 73 \| 357 \| Phage late control D family protein \| Phage late control D family protein; afdb-uniprot50 \| AF-A0A850BD04-F1-MODEL\_V4 \| 1.0 \| 2.741e-16 \| 537 \| 0.188 \| 302 \| 196 \| 13 \| 7 \| 279 \| 68 \| 349 \| Phage late control D family protein \| Phage late control D family protein; afdb-uniprot50 \| AF-A0A7W6JTC3-F1-MODEL\_V4 \| 1.0 \| 3.255e-16 \| 537 \| 0.196 \| 300 \| 191 \| 12 \| 9 \| 278 \| 77 \| 356 \| Phage protein D \| Phage protein D; afdb-uniprot50 \| AF-A0A1V4SNX6-F1-MODEL\_V4 \| 1.0 \| 4.863e-16 \| 535 \| 0.18 \| 304 \| 195 \| 13 \| 7 \| 279 \| 66 \| 346 \| Phage late control protein Gpd \| Phage late control protein Gpd; afdb-uniprot50 \| AF-A0A534WF19-F1-MODEL\_V4 \| 1.0 \| 3.447e-16 \| 535 \| 0.161 \| 298 \| 204 \| 14 \| 9 \| 278 \| 67 \| 346 \| Phage late control D family protein \| Phage late control D family protein; afdb-uniprot50 \| AF-L9Z2P0-F1-MODEL\_V4 \| 1.0 \| 2.287e-15 \| 534 \| 0.164 \| 285 \| 218 \| 9 \| 3 \| 279 \| 37 \| 309 \| Phage protein D \| Phage protein D; afdb-uniprot50 \| AF-A0A7Y3X8C2-F1-MODEL\_V4 \| 1.0 \| 5.15e-16 \| 534 \| 0.173 \| 300 \| 203 \| 14 \| 7 \| 279 \| 79 \| 360 \| Phage late control D family protein \| Phage late control D family protein; afdb-uniprot50 \| AF-A0A7X5DYJ1-F1-MODEL\_V4 \| 1.0 \| 1.621e-15 \| 533 \| 0.164 \| 274 \| 212 \| 9 \| 17 \| 287 \| 80 \| 339 \| Uncharacterized protein \| Uncharacterized protein; afdb-uniprot50 \| AF-A0A2E2G3Z9-F1-MODEL\_V4 \| 1.0 \| 2.308e-16 \| 532 \| 0.178 \| 303 \| 194 \| 16 \| 9 \| 281 \| 73 \| 350 \| Uncharacterized protein \| Uncharacterized protein; afdb-uniprot50 \| AF-A0A2V8IMH5-F1-MODEL\_V4 \| 1.0 \| 7.693e-16 \| 532 \| 0.154 \| 304 \| 202 \| 12 \| 9 \| 279 \| 72 \| 353 \| Uncharacterized protein \| Uncharacterized protein; afdb-uniprot50 \| AF-A0A6M0AY93-F1-MODEL\_V4 \| 1.0 \| 8.628e-16 \| 531 \| 0.169 \| 283 \| 191 \| 12 \| 24 \| 279 \| 6 \| 271 \| Phage late control D family protein \| Phage late control D family protein; afdb-uniprot50 \| AF-A0A524PWE2-F1-MODEL\_V4 \| 1.0 \| 1.217e-15 \| 531 \| 0.163 \| 281 \| 193 \| 9 \| 24 \| 279 \| 11 \| 274 \| Phage late control D family protein \| Phage late control D family protein; afdb-uniprot50 \| AF-A0A1V4VPA0-F1-MODEL\_V4 \| 1.0 \| 3.447e-16 \| 531 \| 0.161 \| 303 \| 201 \| 10 \| 9 \| 279 \| 78 \| 359 \| Phage late control gene D protein (GPD) \| Phage late control gene D protein (GPD); afdb-uniprot50 \| AF-A0A7X3GEI3-F1-MODEL\_V4 \| 1.0 \| 7.264e-16 \| 531 \| 0.185 \| 302 \| 201 \| 11 \| 7 \| 279 \| 76 \| 361 \| Uncharacterized protein \| Uncharacterized protein; afdb-uniprot50 \| AF-A0A0Q6TBZ7-F1-MODEL\_V4 \| 1.0 \| 1.025e-15 \| 531 \| 0.186 \| 305 \| 199 \| 12 \| 3 \| 278 \| 76 \| 360 \| Uncharacterized protein \| Uncharacterized protein; afdb-uniprot50 \| AF-A0A1R1C2A7-F1-MODEL\_V4 \| 1.0 \| 2.179e-16 \| 531 \| 0.177 \| 299 \| 196 \| 11 \| 9 \| 279 \| 88 \| 364 \| Uncharacterized protein \| Uncharacterized protein; afdb-uniprot50 \| AF-A0A6L5TFX2-F1-MODEL\_V4 \| 1.0 \| 1.717e-15 \| 529 \| 0.191 \| 293 \| 209 \| 13 \| 1 \| 281 \| 57 \| 333 \| Preprotein translocase subunit TatB \| Preprotein translocase subunit TatB; afdb-uniprot50 \| AF-A0A1L6M343-F1-MODEL\_V4 \| 1.0 \| 1.085e-15 \| 528 \| 0.167 \| 299 \| 204 \| 12 \| 7 \| 279 \| 76 \| 355 \| VgrG protein \| VgrG protein; afdb-uniprot50 \| AF-A0A4R1J4F9-F1-MODEL\_V4 \| 1.0 \| 6.859e-16 \| 528 \| 0.165 \| 303 \| 204 \| 14 \| 7 \| 279 \| 73 \| 356 \| Phage protein D \| Phage protein D; afdb-uniprot50 \| AF-A0A1E7J776-F1-MODEL\_V4 \| 1.0 \| 3.447e-16 \| 528 \| 0.16 \| 305 \| 198 \| 13 \| 9 \| 279 \| 80 \| 360 \| Uncharacterized protein \| Uncharacterized protein; afdb-uniprot50 \| AF-A0A534XWT7-F1-MODEL\_V4 \| 1.0 \| 7.693e-16 \| 528 \| 0.161 \| 297 \| 205 \| 13 \| 9 \| 278 \| 67 \| 346 \| Uncharacterized protein \| Uncharacterized protein; afdb-uniprot50 \| AF-A0A1C0TMK1-F1-MODEL\_V4 \| 1.0 \| 1.785e-13 \| 527 \| 0.239 \| 146 \| 111 \| 0 \| 134 \| 279 \| 13 \| 158 \| Phage tail protein \| Phage tail protein; afdb-uniprot50 \| AF-A0A1I5AWE8-F1-MODEL\_V4 \| 1.0 \| 1.445e-15 \| 524 \| 0.243 \| 234 \| 127 \| 3 \| 99 \| 282 \| 2 \| 235 \| Uncharacterized protein \| Uncharacterized protein; afdb-uniprot50 \| AF-A0A7V6P996-F1-MODEL\_V4 \| 1.0 \| 2.444e-16 \| 524 \| 0.18 \| 294 \| 201 \| 12 \| 1 \| 279 \| 55 \| 323 \| Late control D family protein \| Late control D family protein; afdb-uniprot50 \| AF-A0A4Q7FRN2-F1-MODEL\_V4 \| 1.0 \| 1.217e-15 \| 524 \| 0.163 \| 288 \| 208 \| 11 \| 18 \| 281 \| 118 \| 396 \| Uncharacterized protein \| Uncharacterized protein; afdb-uniprot50 \| AF-A0A6L6VHW7-F1-MODEL\_V4 \| 1.0 \| 5.723e-15 \| 521 \| 0.212 \| 207 \| 155 \| 5 \| 79 \| 279 \| 9 \| 213 \| Uncharacterized protein \| Uncharacterized protein; afdb-uniprot50 \| AF-A0A0L6JHZ3-F1-MODEL\_V4 \| 1.0 \| 2.308e-16 \| 519 \| 0.189 \| 301 \| 191 \| 11 \| 9 \| 279 \| 87 \| 364 \| Uncharacterized protein \| Uncharacterized protein; afdb-uniprot50 \| AF-A0A3A4NZJ3-F1-MODEL\_V4 \| 1.0 \| 1.925e-15 \| 519 \| 0.191 \| 303 \| 197 \| 13 \| 7 \| 277 \| 63 \| 349 \| Phage late control D family protein \| Phage late control D family protein; afdb-uniprot50 \| AF-A0A3N5R5L3-F1-MODEL\_V4 \| 1.0 \| 2.588e-16 \| 519 \| 0.174 \| 310 \| 207 \| 14 \| 3 \| 279 \| 6 \| 299 \| Type IV secretion protein Rhs \| Type IV secretion protein Rhs; afdb-uniprot50 \| AF-A0A840BJW7-F1-MODEL\_V4 \| 1.0 \| 6.859e-16 \| 518 \| 0.168 \| 297 \| 201 \| 13 \| 9 \| 278 \| 72 \| 349 \| Phage protein D \| Phage protein D; afdb-uniprot50 \| AF-A0A535MWT5-F1-MODEL\_V4 \| 1.0 \| 1.025e-15 \| 518 \| 0.177 \| 327 \| 212 \| 13 \| 9 \| 311 \| 60 \| 353 \| Uncharacterized protein \| Uncharacterized protein; afdb-uniprot50 \| AF-A0A1L6M143-F1-MODEL\_V4 \| 1.0 \| 1.445e-15 \| 517 \| 0.165 \| 303 \| 202 \| 12 \| 7 \| 279 \| 82 \| 363 \| Phage protein D \| Phage protein D; afdb-uniprot50 \| AF-A0A328FDU1-F1-MODEL\_V4 \| 1.0 \| 1.818e-15 \| 517 \| 0.163 \| 305 \| 199 \| 12 \| 7 \| 278 \| 80 \| 361 \| Uncharacterized protein \| Uncharacterized protein; afdb-uniprot50 \| AF-A0A7W6N992-F1-MODEL\_V4 \| 1.0 \| 3.255e-16 \| 516 \| 0.205 \| 326 \| 224 \| 16 \| 2 \| 311 \| 52 \| 358 \| Uncharacterized protein \| Uncharacterized protein; afdb-uniprot50 \| AF-A0A7X5KSP1-F1-MODEL\_V4 \| 1.0 \| 1.085e-15 \| 516 \| 0.212 \| 273 \| 184 \| 12 \| 17 \| 280 \| 83 \| 333 \| Uncharacterized protein \| Uncharacterized protein; afdb-uniprot50 \| AF-A0A1U7IXV3-F1-MODEL\_V4 \| 1.0 \| 2.039e-15 \| 515 \| 0.151 \| 303 \| 206 \| 11 \| 7 \| 279 \| 73 \| 354 \| Uncharacterized protein \| Uncharacterized protein; afdb-uniprot50 \| AF-S4XY39-F1-MODEL\_V4 \| 1.0 \| 2.422e-15 \| 515 \| 0.177 \| 305 \| 199 \| 11 \| 9 \| 277 \| 69 \| 357 \| Uncharacterized protein \| Uncharacterized protein; afdb-uniprot50 \| AF-A0A2R5EJX7-F1-MODEL\_V4 \| 1.0 \| 8.147e-16 \| 515 \| 0.194 \| 298 \| 193 \| 12 \| 7 \| 277 \| 80 \| 357 \| Uncharacterized protein \| Uncharacterized protein; afdb-uniprot50 \| AF-A0A6J4XBG3-F1-MODEL\_V4 \| 1.0 \| 7.264e-16 \| 515 \| 0.197 \| 309 \| 184 \| 13 \| 7 \| 277 \| 76 \| 358 \| Uncharacterized protein \| Uncharacterized protein; afdb-uniprot50 \| AF-A0A0Q9LKQ7-F1-MODEL\_V4 \| 1.0 \| 7.693e-16 \| 515 \| 0.182 \| 296 \| 198 \| 11 \| 9 \| 279 \| 86 \| 362 \| Uncharacterized protein \| Uncharacterized protein; afdb-uniprot50 \| AF-A0A1H9D136-F1-MODEL\_V4 \| 1.0 \| 2.876e-15 \| 515 \| 0.191 \| 303 \| 190 \| 12 \| 9 \| 278 \| 78 \| 358 \| Phage late control gene D protein (GPD) \| Phage late control gene D protein (GPD); afdb-uniprot50 \| AF-A0A6L6WH01-F1-MODEL\_V4 \| 1.0 \| 6.477e-16 \| 514 \| 0.444 \| 144 \| 80 \| 0 \| 133 \| 276 \| 3 \| 146 \| Uncharacterized protein \| Uncharacterized protein; afdb-uniprot50 \| AF-F0B7N5-F1-MODEL\_V4 \| 1.0 \| 4.509e-14 \| 513 \| 0.343 \| 192 \| 124 \| 2 \| 75 \| 264 \| 2 \| 193 \| Phage protein D \| Phage protein D; afdb-uniprot50 \| AF-A0A848YEL6-F1-MODEL\_V4 \| 1.0 \| 1.717e-15 \| 513 \| 0.192 \| 302 \| 194 \| 12 \| 7 \| 278 \| 82 \| 363 \| Phage late control D family protein \| Phage late control D family protein; afdb-uniprot50 \| AF-A0A534UFN3-F1-MODEL\_V4 \| 1.0 \| 1.289e-15 \| 513 \| 0.189 \| 322 \| 213 \| 11 \| 9 \| 307 \| 67 \| 363 \| Uncharacterized protein \| Uncharacterized protein; afdb-uniprot50 \| AF-A0A534VJ89-F1-MODEL\_V4 \| 1.0 \| 5.454e-16 \| 513 \| 0.166 \| 313 \| 207 \| 15 \| 9 \| 293 \| 60 \| 346 \| Phage late control D family protein \| Phage late control D family protein; afdb-uniprot50 \| AF-A0A833GXP7-F1-MODEL\_V4 \| 1.0 \| 8.073e-15 \| 512 \| 0.153 \| 300 \| 211 \| 10 \| 3 \| 277 \| 76 \| 357 \| Phage late control D family protein \| Phage late control D family protein; afdb-uniprot50 \| AF-A0A520K035-F1-MODEL\_V4 \| 1.0 \| 4.297e-15 \| 511 \| 0.162 \| 308 \| 208 \| 13 \| 3 \| 279 \| 71 \| 359 \| Phage late control D family protein \| Phage late control D family protein; afdb-uniprot50 \| AF-A0A1Q8A378-F1-MODEL\_V4 \| 1.0 \| 6.477e-16 \| 510 \| 0.185 \| 307 \| 199 \| 15 \| 3 \| 279 \| 69 \| 354 \| Uncharacterized protein \| Uncharacterized protein; afdb-uniprot50 \| AF-A0A5S9ITQ6-F1-MODEL\_V4 \| 1.0 \| 3.046e-15 \| 509 \| 0.165 \| 303 \| 200 \| 11 \| 7 \| 279 \| 65 \| 344 \| Uncharacterized protein \| Uncharacterized protein; afdb-uniprot50 \| AF-A0A2E0L150-F1-MODEL\_V4 \| 1.0 \| 1.531e-15 \| 509 \| 0.162 \| 307 \| 201 \| 13 \| 7 \| 279 \| 66 \| 350 \| Phage\_base\_V domain-containing protein \| Phage\_base\_V domain-containing protein; afdb-uniprot50 \| AF-A0A165RMG4-F1-MODEL\_V4 \| 1.0 \| 4.297e-15 \| 508 \| 0.173 \| 277 \| 200 \| 11 \| 20 \| 279 \| 69 \| 333 \| Phage late control gene D protein (GPD) \| Phage late control gene D protein (GPD); afdb-uniprot50 \| AF-A0A1V4W1U0-F1-MODEL\_V4 \| 1.0 \| 4.336e-16 \| 508 \| 0.193 \| 299 \| 192 \| 11 \| 9 \| 279 \| 88 \| 365 \| Phage late control gene D protein (GPD) \| Phage late control gene D protein (GPD); afdb-uniprot50 \| AF-A0A401HTD7-F1-MODEL\_V4 \| 1.0 \| 3.618e-15 \| 508 \| 0.164 \| 298 \| 206 \| 11 \| 7 \| 279 \| 87 \| 366 \| Uncharacterized protein \| Uncharacterized protein; afdb-uniprot50 \| AF-A0A0L8VEK1-F1-MODEL\_V4 \| 1.0 \| 1.531e-15 \| 508 \| 0.163 \| 300 \| 205 \| 12 \| 3 \| 275 \| 38 \| 318 \| Phage\_base\_V domain-containing protein \| Phage\_base\_V domain-containing protein; afdb-uniprot50 \| AF-A0A285M2Q7-F1-MODEL\_V4 \| 1.0 \| 2.716e-15 \| 507 \| 0.213 \| 286 \| 200 \| 13 \| 2 \| 276 \| 59 \| 330 \| Uncharacterized protein \| Uncharacterized protein; afdb-uniprot50 \| AF-A0A2N2D1G3-F1-MODEL\_V4 \| 1.0 \| 1.621e-15 \| 507 \| 0.187 \| 298 \| 198 \| 12 \| 7 \| 279 \| 88 \| 366 \| Uncharacterized protein \| Uncharacterized protein; afdb-uniprot50 \| AF-A0A661RNA4-F1-MODEL\_V4 \| 1.0 \| 2.422e-15 \| 505 \| 0.195 \| 302 \| 198 \| 14 \| 7 \| 279 \| 73 \| 358 \| Uncharacterized protein \| Uncharacterized protein; afdb-uniprot50 \| AF-A0A1Y3RUL2-F1-MODEL\_V4 \| 1.0 \| 6.797e-15 \| 505 \| 0.144 \| 269 \| 206 \| 12 \| 57 \| 311 \| 15 \| 273 \| Uncharacterized protein \| Uncharacterized protein; afdb-uniprot50 \| AF-A0A7V6UDV8-F1-MODEL\_V4 \| 1.0 \| 5.404e-15 \| 503 \| 0.189 \| 280 \| 198 \| 15 \| 2 \| 276 \| 48 \| 303 \| Uncharacterized protein \| Uncharacterized protein; afdb-uniprot50 \| AF-A0A6P1IRG3-F1-MODEL\_V4 \| 1.0 \| 8.628e-16 \| 503 \| 0.168 \| 302 \| 197 \| 14 \| 9 \| 278 \| 79 \| 358 \| Phage late control D family protein \| Phage late control D family protein; afdb-uniprot50 \| AF-G8PLB4-F1-MODEL\_V4 \| 1.0 \| 1.925e-15 \| 503 \| 0.213 \| 323 \| 226 \| 12 \| 2 \| 308 \| 58 \| 368 \| Bacteriophage regulatory protein \| Bacteriophage regulatory protein; afdb-uniprot50 \| AF-A0A376W2Q8-F1-MODEL\_V4 \| 1.0 \| 8.55e-15 \| 502 \| 0.228 \| 223 \| 145 \| 5 \| 1 \| 203 \| 11 \| 226 \| Late control gene D protein from prophage \| Late control gene D protein from prophage; afdb-uniprot50 \| AF-A0A3N5XNR8-F1-MODEL\_V4 \| 1.0 \| 2.716e-15 \| 502 \| 0.136 \| 307 \| 207 \| 11 \| 7 \| 279 \| 68 \| 350 \| Phage late control D family protein \| Phage late control D family protein; afdb-uniprot50 \| AF-A0A7W7MBI9-F1-MODEL\_V4 \| 1.0 \| 3.046e-15 \| 500 \| 0.17 \| 300 \| 200 \| 10 \| 9 \| 278 \| 69 \| 349 \| Phage protein D \| Phage protein D; afdb-uniprot50 \| AF-U2LWR3-F1-MODEL\_V4 \| 1.0 \| 1.149e-15 \| 498 \| 0.12 \| 315 \| 241 \| 11 \| 7 \| 311 \| 64 \| 352 \| Uncharacterized protein \| Uncharacterized protein; afdb-uniprot50 \| AF-A0A0E3KUE2-F1-MODEL\_V4 \| 1.0 \| 6.061e-15 \| 497 \| 0.179 \| 295 \| 204 \| 12 \| 7 \| 279 \| 78 \| 356 \| VgrG protein \| VgrG protein; afdb-uniprot50 \| AF-A0A4R4G0N1-F1-MODEL\_V4 \| 1.0 \| 8.973e-14 \| 494 \| 0.273 \| 219 \| 140 \| 6 \| 10 \| 220 \| 2 \| 209 \| Late control protein \| Late control protein; afdb-uniprot50 \| AF-A0A3M9Z207-F1-MODEL\_V4 \| 1.0 \| 1.217e-15 \| 494 \| 0.155 \| 309 \| 199 \| 16 \| 7 \| 279 \| 82 \| 364 \| Phage late control D family protein \| Phage late control D family protein; afdb-uniprot50 \| AF-A0A3M1T127-F1-MODEL\_V4 \| 1.0 \| 4.55e-15 \| 493 \| 0.17 \| 334 \| 221 \| 14 \| 7 \| 309 \| 72 \| 380 \| Phage late control D family protein \| Phage late control D family protein; afdb-uniprot50 \| AF-A0A4C2EPK7-F1-MODEL\_V4 \| 1.0 \| 1.075e-14 \| 492 \| 0.171 \| 285 \| 204 \| 12 \| 7 \| 277 \| 77 \| 343 \| Uncharacterized protein \| Uncharacterized protein; afdb-uniprot50 \| AF-A0A7Y5L630-F1-MODEL\_V4 \| 1.0 \| 4.297e-15 \| 492 \| 0.173 \| 305 \| 194 \| 12 \| 9 \| 279 \| 73 \| 353 \| Phage late control D family protein \| Phage late control D family protein; afdb-uniprot50 \| AF-F2NYF5-F1-MODEL\_V4 \| 1.0 \| 2.876e-15 \| 492 \| 0.148 \| 309 \| 208 \| 16 \| 7 \| 282 \| 66 \| 352 \| Uncharacterized protein \| Uncharacterized protein; afdb-uniprot50 \| AF-G2G245-F1-MODEL\_V4 \| 1.0 \| 4.258e-14 \| 491 \| 0.199 \| 246 \| 166 \| 6 \| 57 \| 279 \| 1 \| 238 \| Uncharacterized protein \| Uncharacterized protein; afdb-uniprot50 \| AF-A0A5U2MRG6-F1-MODEL\_V4 \| 1.0 \| 8.706e-17 \| 491 \| 0.168 \| 302 \| 197 \| 12 \| 7 \| 277 \| 2 \| 280 \| Phage late control D family protein \| Phage late control D family protein; afdb-uniprot50 \| AF-A0A4R2R9S9-F1-MODEL\_V4 \| 1.0 \| 2.039e-15 \| 490 \| 0.157 \| 299 \| 208 \| 15 \| 9 \| 279 \| 88 \| 370 \| Uncharacterized protein \| Uncharacterized protein; afdb-uniprot50 \| AF-A0A081N302-F1-MODEL\_V4 \| 1.0 \| 2.039e-15 \| 487 \| 0.378 \| 169 \| 103 \| 1 \| 113 \| 279 \| 1 \| 169 \| Uncharacterized protein \| Uncharacterized protein; afdb-uniprot50 \| AF-A0A0F4NJB2-F1-MODEL\_V4 \| 1.0 \| 3.651e-16 \| 487 \| 0.207 \| 284 \| 191 \| 11 \| 1 \| 279 \| 53 \| 307 \| Uncharacterized protein \| Uncharacterized protein; afdb-uniprot50 \| AF-A0A080M5H4-F1-MODEL\_V4 \| 1.0 \| 2.691e-14 \| 486 \| 0.181 \| 248 \| 169 \| 10 \| 57 \| 279 \| 10 \| 248 \| Tail protein \| Tail protein; afdb-uniprot50 \| AF-A0A0V7ZK90-F1-MODEL\_V4 \| 1.0 \| 5.103e-15 \| 486 \| 0.156 \| 325 \| 219 \| 12 \| 7 \| 290 \| 77 \| 387 \| Uncharacterized protein \| Uncharacterized protein; afdb-uniprot50 \| AF-A0A5B8ST50-F1-MODEL\_V4 \| 1.0 \| 1.769e-12 \| 485 \| 0.309 \| 139 \| 94 \| 2 \| 145 \| 281 \| 1 \| 139 \| Uncharacterized protein \| Uncharacterized protein; afdb-uniprot50 \| AF-A0A521P0G8-F1-MODEL\_V4 \| 1.0 \| 3.585e-14 \| 485 \| 0.265 \| 181 \| 128 \| 3 \| 112 \| 290 \| 2 \| 179 \| Uncharacterized protein \| Uncharacterized protein; afdb-uniprot50 \| AF-A0A545T2R6-F1-MODEL\_V4 \| 1.0 \| 6.419e-15 \| 485 \| 0.162 \| 296 \| 200 \| 15 \| 9 \| 278 \| 72 \| 345 \| Uncharacterized protein \| Uncharacterized protein; afdb-uniprot50 \| AF-A0A410ULA3-F1-MODEL\_V4 \| 1.0 \| 3.797e-14 \| 484 \| 0.149 \| 302 \| 215 \| 14 \| 3 \| 279 \| 65 \| 349 \| Phage late control D family protein \| Phage late control D family protein; afdb-uniprot50 \| AF-A0A1F8N4S5-F1-MODEL\_V4 \| 1.0 \| 1.149e-15 \| 484 \| 0.211 \| 307 \| 187 \| 12 \| 7 \| 281 \| 66 \| 349 \| Phage\_base\_V domain-containing protein \| Phage\_base\_V domain-containing protein; afdb-uniprot50 \| AF-F2NYH1-F1-MODEL\_V4 \| 1.0 \| 3.416e-15 \| 484 \| 0.142 \| 301 \| 209 \| 14 \| 7 \| 277 \| 66 \| 347 \| Uncharacterized protein \| Uncharacterized protein; afdb-uniprot50 \| AF-G4KQ66-F1-MODEL\_V4 \| 1.0 \| 1.353e-14 \| 483 \| 0.149 \| 274 \| 203 \| 11 \| 9 \| 277 \| 53 \| 301 \| Uncharacterized protein \| Uncharacterized protein; afdb-uniprot50 \| AF-A0A1I6IQH4-F1-MODEL\_V4 \| 1.0 \| 5.103e-15 \| 483 \| 0.157 \| 292 \| 208 \| 14 \| 7 \| 279 \| 77 \| 349 \| Uncharacterized protein \| Uncharacterized protein; afdb-uniprot50 \| AF-A0A534PEG0-F1-MODEL\_V4 \| 1.0 \| 7.199e-15 \| 483 \| 0.175 \| 324 \| 208 \| 13 \| 9 \| 310 \| 69 \| 355 \| Phage late control D family protein \| Phage late control D family protein; afdb-uniprot50 \| AF-A0A3B9YXC3-F1-MODEL\_V4 \| 1.0 \| 7.623e-15 \| 481 \| 0.175 \| 308 \| 197 \| 15 \| 7 \| 279 \| 83 \| 368 \| Uncharacterized protein \| Uncharacterized protein; afdb-uniprot50 \| AF-A0A7C6PS95-F1-MODEL\_V4 \| 1.0 \| 9.589e-15 \| 481 \| 0.183 \| 310 \| 197 \| 14 \| 3 \| 278 \| 62 \| 349 \| Phage late control D family protein \| Phage late control D family protein; afdb-uniprot50 \| AF-A0A660RRK5-F1-MODEL\_V4 \| 1.0 \| 1.277e-14 \| 481 \| 0.164 \| 304 \| 207 \| 14 \| 7 \| 281 \| 65 \| 350 \| Type IV secretion protein Rhs \| Type IV secretion protein Rhs; afdb-uniprot50 \| AF-A0A376YJ37-F1-MODEL\_V4 \| 1.0 \| 4.021e-14 \| 480 \| 0.235 \| 242 \| 153 \| 6 \| 1 \| 218 \| 29 \| 262 \| Gene late control D protein \| Gene late control D protein; afdb-uniprot50 \| AF-A0A2N6CT05-F1-MODEL\_V4 \| 1.0 \| 1.908e-14 \| 480 \| 0.153 \| 319 \| 209 \| 12 \| 7 \| 284 \| 82 \| 380 \| Type IV secretion protein Rhs \| Type IV secretion protein Rhs; afdb-uniprot50 \| AF-A0A6G3WNF9-F1-MODEL\_V4 \| 1.0 \| 7.623e-15 \| 480 \| 0.197 \| 319 \| 199 \| 17 \| 3 \| 281 \| 3 \| 304 \| VgrG-related protein \| VgrG-related protein; afdb-uniprot50 \| AF-A0A0Q4KJV3-F1-MODEL\_V4 \| 1.0 \| 2.021e-14 \| 479 \| 0.188 \| 297 \| 201 \| 12 \| 7 \| 279 \| 63 \| 343 \| Uncharacterized protein \| Uncharacterized protein; afdb-uniprot50 \| AF-B9NUV5-F1-MODEL\_V4 \| 1.0 \| 2.14e-14 \| 479 \| 0.176 \| 301 \| 204 \| 11 \| 7 \| 279 \| 85 \| 369 \| Uncharacterized protein \| Uncharacterized protein; afdb-uniprot50 \| AF-A0A7V3SCS7-F1-MODEL\_V4 \| 1.0 \| 3.416e-15 \| 479 \| 0.162 \| 308 \| 197 \| 12 \| 9 \| 281 \| 67 \| 348 \| Type IV secretion protein Rhs \| Type IV secretion protein Rhs; afdb-uniprot50 \| AF-A0A661I686-F1-MODEL\_V4 \| 1.0 \| 3.018e-14 \| 478 \| 0.172 \| 301 \| 202 \| 12 \| 7 \| 278 \| 60 \| 342 \| Uncharacterized protein \| Uncharacterized protein; afdb-uniprot50 \| AF-A0A853R2R6-F1-MODEL\_V4 \| 1.0 \| 4.469e-13 \| 477 \| 0.333 \| 198 \| 129 \| 3 \| 1 \| 197 \| 55 \| 250 \| Late control protein D \| Late control protein D; afdb-uniprot50 \| AF-A0A2R6EXY7-F1-MODEL\_V4 \| 1.0 \| 9.054e-15 \| 477 \| 0.174 \| 287 \| 202 \| 13 \| 7 \| 279 \| 72 \| 337 \| Uncharacterized protein \| Uncharacterized protein; afdb-uniprot50 \| AF-A0A6N8BHD5-F1-MODEL\_V4 \| 1.0 \| 2.287e-15 \| 476 \| 0.164 \| 304 \| 205 \| 9 \| 7 \| 279 \| 79 \| 364 \| Phage late control D family protein \| Phage late control D family protein; afdb-uniprot50 \| AF-F2NUD6-F1-MODEL\_V4 \| 1.0 \| 5.103e-15 \| 476 \| 0.152 \| 309 \| 207 \| 15 \| 7 \| 282 \| 66 \| 352 \| Uncharacterized protein \| Uncharacterized protein; afdb-uniprot50 \| AF-A0A4R4G4D4-F1-MODEL\_V4 \| 1.0 \| 5.672e-14 \| 475 \| 0.28 \| 228 \| 145 \| 8 \| 1 \| 220 \| 51 \| 267 \| Late control protein \| Late control protein; afdb-uniprot50 \| AF-A0A4P7HPH6-F1-MODEL\_V4 \| 1.0 \| 6.061e-15 \| 474 \| 0.156 \| 314 \| 204 \| 11 \| 9 \| 280 \| 82 \| 376 \| Phage late control D family protein \| Phage late control D family protein; afdb-uniprot50 \| AF-X5J8E7-F1-MODEL\_V4 \| 1.0 \| 1.517e-14 \| 474 \| 0.145 \| 296 \| 205 \| 12 \| 9 \| 276 \| 70 \| 345 \| Rhs element Vgr protein \| Rhs element Vgr protein; afdb-uniprot50 \| AF-A0A6S7B3U2-F1-MODEL\_V4 \| 1.0 \| 5.404e-15 \| 473 \| 0.149 \| 308 \| 213 \| 15 \| 7 \| 278 \| 81 \| 375 \| Uncharacterized protein \| Uncharacterized protein; afdb-uniprot50 \| AF-A0A356KG57-F1-MODEL\_V4 \| 1.0 \| 1.015e-14 \| 473 \| 0.165 \| 344 \| 223 \| 19 \| 3 \| 311 \| 61 \| 375 \| Uncharacterized protein \| Uncharacterized protein; afdb-uniprot50 \| AF-A0A418VPR5-F1-MODEL\_V4 \| 1.0 \| 1.075e-14 \| 472 \| 0.154 \| 291 \| 201 \| 10 \| 16 \| 279 \| 77 \| 349 \| Phage late control D family protein \| Phage late control D family protein; afdb-uniprot50 \| AF-A0A3A0AIX8-F1-MODEL\_V4 \| 1.0 \| 2.266e-14 \| 472 \| 0.163 \| 306 \| 201 \| 13 \| 7 \| 279 \| 65 \| 348 \| Type IV secretion protein Rhs \| Type IV secretion protein Rhs; afdb-uniprot50 \| AF-A0A5N8XB31-F1-MODEL\_V4 \| 1.0 \| 8.073e-15 \| 472 \| 0.146 \| 307 \| 209 \| 12 \| 9 \| 284 \| 72 \| 356 \| Phage\_base\_V domain-containing protein \| Phage\_base\_V domain-containing protein; afdb-uniprot50 \| AF-A0A3A8NI17-F1-MODEL\_V4 \| 1.0 \| 9.589e-15 \| 471 \| 0.176 \| 301 \| 194 \| 14 \| 9 \| 279 \| 63 \| 339 \| Phage late control D family protein \| Phage late control D family protein; afdb-uniprot50 \| AF-A0A7C7R7X2-F1-MODEL\_V4 \| 1.0 \| 4.775e-14 \| 471 \| 0.173 \| 305 \| 199 \| 12 \| 3 \| 275 \| 59 \| 342 \| Phage late control D family protein \| Phage late control D family protein; afdb-uniprot50 \| AF-A0A3M1T848-F1-MODEL\_V4 \| 1.0 \| 2.021e-14 \| 471 \| 0.172 \| 325 \| 206 \| 15 \| 7 \| 297 \| 70 \| 365 \| Phage late control D family protein \| Phage late control D family protein; afdb-uniprot50 \| AF-A0A7K0GTP6-F1-MODEL\_V4 \| 1.0 \| 2.4e-14 \| 471 \| 0.17 \| 293 \| 203 \| 12 \| 7 \| 276 \| 64 \| 339 \| Type VI secretion system tip protein VgrG \| Type VI secretion system tip protein VgrG; afdb-uniprot50 \| AF-A0A255Z5N1-F1-MODEL\_V4 \| 1.0 \| 2.14e-14 \| 471 \| 0.182 \| 301 \| 196 \| 11 \| 7 \| 279 \| 69 \| 347 \| Type IV secretion protein Rhs \| Type IV secretion protein Rhs; afdb-uniprot50 \| AF-A0A1H4FYY6-F1-MODEL\_V4 \| 1.0 \| 2.159e-15 \| 470 \| 0.167 \| 322 \| 195 \| 15 \| 7 \| 277 \| 93 \| 392 \| Phage protein D \| Phage protein D; afdb-uniprot50 \| AF-A0A7Y2SMN3-F1-MODEL\_V4 \| 1.0 \| 3.797e-14 \| 470 \| 0.177 \| 298 \| 192 \| 10 \| 7 \| 273 \| 64 \| 339 \| VgrG-related protein \| VgrG-related protein; afdb-uniprot50 \| AF-A0A442UMS5-F1-MODEL\_V4 \| 1.0 \| 1.606e-14 \| 469 \| 0.166 \| 306 \| 210 \| 12 \| 3 \| 279 \| 78 \| 367 \| Phage late control D family protein \| Phage late control D family protein; afdb-uniprot50 \| AF-A0A3G6TJU6-F1-MODEL\_V4 \| 1.0 \| 8.55e-15 \| 469 \| 0.124 \| 305 \| 214 \| 12 \| 7 \| 281 \| 72 \| 353 \| Type VI secretion system tip protein VgrG \| Type VI secretion system tip protein VgrG; afdb-uniprot50 \| AF-L8JQG5-F1-MODEL\_V4 \| 1.0 \| 1.206e-14 \| 468 \| 0.179 \| 295 \| 200 \| 11 \| 7 \| 275 \| 72 \| 350 \| VgrG protein \| VgrG protein; afdb-uniprot50 \| AF-A0A497BPI8-F1-MODEL\_V4 \| 1.0 \| 1.353e-14 \| 468 \| 0.194 \| 303 \| 188 \| 15 \| 9 \| 278 \| 66 \| 345 \| Phage\_base\_V domain-containing protein \| Phage\_base\_V domain-containing protein; afdb-uniprot50 \| AF-A8U2N1-F1-MODEL\_V4 \| 1.0 \| 2.85e-14 \| 467 \| 0.303 \| 227 \| 140 \| 7 \| 1 \| 218 \| 57 \| 274 \| Late control gene D protein \| Late control gene D protein; afdb-uniprot50 \| AF-A0A7W7V134-F1-MODEL\_V4 \| 1.0 \| 8e-14 \| 467 \| 0.17 \| 305 \| 208 \| 14 \| 3 \| 279 \| 65 \| 352 \| Phage protein D \| Phage protein D; afdb-uniprot50 \| AF-A0A1L6M2U5-F1-MODEL\_V4 \| 1.0 \| 2.4e-14 \| 467 \| 0.155 \| 309 \| 201 \| 14 \| 3 \| 278 \| 109 \| 390 \| Phage protein D \| Phage protein D; afdb-uniprot50 \| AF-A0A0S2FQJ5-F1-MODEL\_V4 \| 1.0 \| 1.802e-14 \| 466 \| 0.147 \| 292 \| 194 \| 15 \| 20 \| 278 \| 82 \| 351 \| Phage late control gene D family protein \| Phage late control gene D family protein; afdb-uniprot50 \| AF-A0A7J0CY33-F1-MODEL\_V4 \| 1.0 \| 3.385e-14 \| 466 \| 0.208 \| 312 \| 194 \| 17 \| 3 \| 276 \| 18 \| 314 \| Uncharacterized protein \| Uncharacterized protein; afdb-uniprot50 \| AF-A0A353VJI0-F1-MODEL\_V4 \| 1.0 \| 1.802e-14 \| 466 \| 0.174 \| 293 \| 203 \| 13 \| 7 \| 277 \| 72 \| 347 \| Rhs element Vgr protein \| Rhs element Vgr protein; afdb-uniprot50 \| AF-A0A7Y4QYQ7-F1-MODEL\_V4 \| 1.0 \| 2.266e-14 \| 466 \| 0.156 \| 301 \| 198 \| 13 \| 7 \| 276 \| 72 \| 347 \| Type VI secretion system tip protein VgrG \| Type VI secretion system tip protein VgrG; afdb-uniprot50 \| AF-A0A6I5QAM8-F1-MODEL\_V4 \| 1.0 \| 7.555e-14 \| 465 \| 0.165 \| 296 \| 204 \| 12 \| 7 \| 277 \| 74 \| 351 \| Type VI secretion system tip protein VgrG \| Type VI secretion system tip protein VgrG; afdb-uniprot50 \| AF-Q08VZ1-F1-MODEL\_V4 \| 1.0 \| 1.925e-15 \| 463 \| 0.152 \| 375 \| 222 \| 15 \| 9 \| 307 \| 198 \| 552 \| Uncharacterized protein \| Uncharacterized protein; afdb-uniprot50 \| AF-A0A7X6GUB8-F1-MODEL\_V4 \| 1.0 \| 3.618e-15 \| 462 \| 0.145 \| 282 \| 220 \| 9 \| 39 \| 311 \| 1 \| 270 \| Uncharacterized protein \| Uncharacterized protein; afdb-uniprot50 \| AF-A0A3A0A9Z2-F1-MODEL\_V4 \| 1.0 \| 8e-14 \| 462 \| 0.161 \| 298 \| 208 \| 10 \| 7 \| 279 \| 76 \| 356 \| Rhs element Vgr protein \| Rhs element Vgr protein; afdb-uniprot50 \| AF-A0A5B8ATZ1-F1-MODEL\_V4 \| 1.0 \| 1.353e-14 \| 460 \| 0.155 \| 348 \| 198 \| 16 \| 7 \| 277 \| 63 \| 391 \| Uncharacterized protein \| Uncharacterized protein; afdb-uniprot50 \| AF-A0A2S6NI59-F1-MODEL\_V4 \| 1.0 \| 3.618e-15 \| 459 \| 0.137 \| 305 \| 205 \| 13 \| 7 \| 276 \| 19 \| 300 \| Uncharacterized protein \| Uncharacterized protein; afdb-uniprot50 \| AF-A0A1N6X884-F1-MODEL\_V4 \| 1.0 \| 4.258e-14 \| 459 \| 0.136 \| 307 \| 207 \| 14 \| 7 \| 279 \| 66 \| 348 \| Phage protein D \| Phage protein D; afdb-uniprot50 \| AF-A0A6N7YBH3-F1-MODEL\_V4 \| 1.0 \| 3.018e-14 \| 459 \| 0.196 \| 301 \| 194 \| 14 \| 7 \| 279 \| 67 \| 347 \| VgrG-related protein \| VgrG-related protein; afdb-uniprot50 \| AF-A0A4U7JJ27-F1-MODEL\_V4 \| 1.0 \| 1.701e-14 \| 459 \| 0.149 \| 301 \| 204 \| 16 \| 7 \| 276 \| 95 \| 374 \| Uncharacterized protein \| Uncharacterized protein; afdb-uniprot50 \| AF-A0A7T8BBA2-F1-MODEL\_V4 \| 1.0 \| 1.006e-13 \| 458 \| 0.11 \| 299 \| 221 \| 12 \| 7 \| 277 \| 65 \| 346 \| Phage late control D family protein \| Phage late control D family protein; afdb-uniprot50 \| AF-A0A3M1GYX0-F1-MODEL\_V4 \| 1.0 \| 1.802e-14 \| 458 \| 0.146 \| 335 \| 232 \| 16 \| 9 \| 311 \| 66 \| 378 \| Phage late control D family protein \| Phage late control D family protein; afdb-uniprot50 \| AF-A0A4Z0M9I4-F1-MODEL\_V4 \| 1.0 \| 1.701e-14 \| 458 \| 0.149 \| 321 \| 210 \| 12 \| 7 \| 280 \| 112 \| 416 \| Phage late control D family protein \| Phage late control D family protein; afdb-uniprot50 \| AF-A0A497AT53-F1-MODEL\_V4 \| 1.0 \| 4.509e-14 \| 458 \| 0.171 \| 291 \| 199 \| 11 \| 17 \| 281 \| 78 \| 352 \| Type IV secretion protein Rhs \| Type IV secretion protein Rhs; afdb-uniprot50 \| AF-A0A7W1S8H0-F1-MODEL\_V4 \| 1.0 \| 1.908e-14 \| 458 \| 0.156 \| 306 \| 208 \| 15 \| 7 \| 281 \| 67 \| 353 \| VgrG-related protein \| VgrG-related protein; afdb-uniprot50 \| AF-A0A318JR32-F1-MODEL\_V4 \| 1.0 \| 9.331e-12 \| 457 \| 0.293 \| 150 \| 103 \| 3 \| 135 \| 281 \| 4 \| 153 \| Late control gene D protein (GPD) \| Late control gene D protein (GPD); afdb-uniprot50 \| AF-J0ZTM2-F1-MODEL\_V4 \| 1.0 \| 2.496e-12 \| 455 \| 0.31 \| 145 \| 92 \| 3 \| 145 \| 281 \| 12 \| 156 \| Uncharacterized protein \| Uncharacterized protein; afdb-uniprot50 \| AF-A0A7Y2WZB6-F1-MODEL\_V4 \| 1.0 \| 1.139e-14 \| 455 \| 0.175 \| 290 \| 185 \| 13 \| 20 \| 281 \| 72 \| 335 \| VgrG-related protein \| VgrG-related protein; afdb-uniprot50 \| AF-A0A418QD23-F1-MODEL\_V4 \| 1.0 \| 1.266e-13 \| 453 \| 0.166 \| 313 \| 202 \| 13 \| 3 \| 281 \| 42 \| 329 \| Phage\_base\_V domain-containing protein \| Phage\_base\_V domain-containing protein; afdb-uniprot50 \| AF-A0A2S6I9B9-F1-MODEL\_V4 \| 1.0 \| 1.006e-13 \| 453 \| 0.195 \| 292 \| 196 \| 11 \| 7 \| 275 \| 76 \| 351 \| Rhs element Vgr protein \| Rhs element Vgr protein; afdb-uniprot50 \| AF-A0A1T2CG57-F1-MODEL\_V4 \| 1.0 \| 5.57e-12 \| 452 \| 0.362 \| 127 \| 81 \| 0 \| 153 \| 279 \| 3 \| 129 \| Uncharacterized protein \| Uncharacterized protein; afdb-uniprot50 \| AF-A0A317N0I9-F1-MODEL\_V4 \| 1.0 \| 1.503e-13 \| 452 \| 0.18 \| 294 \| 191 \| 12 \| 9 \| 278 \| 70 \| 337 \| Phage protein D \| Phage protein D; afdb-uniprot50 \| AF-A0A511MZG9-F1-MODEL\_V4 \| 1.0 \| 5.672e-14 \| 452 \| 0.191 \| 293 \| 181 \| 12 \| 19 \| 278 \| 75 \| 344 \| Uncharacterized protein \| Uncharacterized protein; afdb-uniprot50 \| AF-A0A098LIX2-F1-MODEL\_V4 \| 1.0 \| 2.667e-13 \| 452 \| 0.133 \| 293 \| 213 \| 9 \| 7 \| 276 \| 74 \| 348 \| Rhs element Vgr protein \| Rhs element Vgr protein; afdb-uniprot50 \| AF-A0A377A6L9-F1-MODEL\_V4 \| 1.0 \| 9.972e-13 \| 451 \| 0.232 \| 219 \| 144 \| 3 \| 68 \| 262 \| 2 \| 220 \| Putative phage late gene regulator \| Putative phage late gene regulator; afdb-uniprot50 \| AF-A0A7V1N7G8-F1-MODEL\_V4 \| 1.0 \| 8.55e-15 \| 451 \| 0.157 \| 324 \| 196 \| 14 \| 24 \| 285 \| 12 \| 320 \| Phage late control D family protein \| Phage late control D family protein; afdb-uniprot50 \| AF-A0A1M7TZB9-F1-MODEL\_V4 \| 1.0 \| 6.361e-14 \| 451 \| 0.186 \| 295 \| 197 \| 16 \| 9 \| 277 \| 82 \| 359 \| Phage protein D \| Phage protein D; afdb-uniprot50 \| AF-A0A1G9EX68-F1-MODEL\_V4 \| 1.0 \| 3.018e-14 \| 451 \| 0.163 \| 331 \| 196 \| 15 \| 7 \| 276 \| 62 \| 372 \| Phage protein D \| Phage protein D; afdb-uniprot50 \| AF-Q3A0I0-F1-MODEL\_V4 \| 1.0 \| 3.197e-14 \| 451 \| 0.18 \| 293 \| 196 \| 16 \| 9 \| 281 \| 62 \| 330 \| Phage protein D and tail spike protein, putative \| Phage protein D and tail spike protein, putative; afdb-uniprot50 \| AF-R5H1V4-F1-MODEL\_V4 \| 1.0 \| 2.691e-14 \| 450 \| 0.145 \| 302 \| 208 \| 14 \| 7 \| 279 \| 72 \| 352 \| Uncharacterized protein \| Uncharacterized protein; afdb-uniprot50 \| AF-A0A7C3CDK3-F1-MODEL\_V4 \| 1.0 \| 1.34e-13 \| 450 \| 0.174 \| 293 \| 196 \| 12 \| 17 \| 281 \| 76 \| 350 \| Phage late control D family protein \| Phage late control D family protein; afdb-uniprot50 \| AF-A0A158HQY1-F1-MODEL\_V4 \| 1.0 \| 8e-14 \| 449 \| 0.148 \| 331 \| 206 \| 14 \| 7 \| 277 \| 64 \| 378 \| Phage late control gene D protein (GPD) \| Phage late control gene D protein (GPD); afdb-uniprot50 \| AF-N1V1U4-F1-MODEL\_V4 \| 1.0 \| 9.589e-15 \| 449 \| 0.179 \| 296 \| 202 \| 14 \| 1 \| 273 \| 1 \| 278 \| Phage\_base\_V domain-containing protein \| Phage\_base\_V domain-containing protein; afdb-uniprot50 \| AF-A0A2G5L080-F1-MODEL\_V4 \| 1.0 \| 1.066e-13 \| 449 \| 0.134 \| 305 \| 208 \| 11 \| 3 \| 276 \| 68 \| 347 \| Phage\_base\_V domain-containing protein \| Phage\_base\_V domain-containing protein; afdb-uniprot50 \| AF-A0A7X4YHH4-F1-MODEL\_V4 \| 1.0 \| 2.991e-13 \| 448 \| 0.123 \| 291 \| 202 \| 12 \| 20 \| 278 \| 84 \| 353 \| Uncharacterized protein \| Uncharacterized protein; afdb-uniprot50 \| AF-A0A7J4PP46-F1-MODEL\_V4 \| 1.0 \| 8.473e-14 \| 448 \| 0.134 \| 304 \| 210 \| 14 \| 7 \| 277 \| 73 \| 356 \| Phage late control D family protein \| Phage late control D family protein; afdb-uniprot50 \| AF-A0A1W9R245-F1-MODEL\_V4 \| 1.0 \| 1.686e-13 \| 448 \| 0.139 \| 293 \| 211 \| 11 \| 7 \| 275 \| 75 \| 350 \| Type IV secretion protein Rhs \| Type IV secretion protein Rhs; afdb-uniprot50 \| AF-A0A6G6GPZ0-F1-MODEL\_V4 \| 1.0 \| 2.002e-13 \| 448 \| 0.155 \| 302 \| 212 \| 12 \| 3 \| 279 \| 68 \| 351 \| Type VI secretion system tip protein VgrG \| Type VI secretion system tip protein VgrG; afdb-uniprot50 \| AF-A0A850P3R3-F1-MODEL\_V4 \| 1.0 \| 9.589e-15 \| 447 \| 0.135 \| 303 \| 221 \| 14 \| 7 \| 277 \| 27 \| 320 \| Uncharacterized protein \| Uncharacterized protein; afdb-uniprot50 \| AF-A0A7U7FD77-F1-MODEL\_V4 \| 1.0 \| 5.197e-17 \| 447 \| 0.185 \| 312 \| 161 \| 10 \| 26 \| 288 \| 155 \| 422 \| Putative phage-related protein \| Putative phage-related protein; afdb-uniprot50 \| AF-A0A0F9U2B2-F1-MODEL\_V4 \| 1.0 \| 4.509e-14 \| 447 \| 0.138 \| 389 \| 232 \| 16 \| 7 \| 311 \| 90 \| 459 \| Uncharacterized protein \| Uncharacterized protein; afdb-uniprot50 \| AF-A0A620N5X6-F1-MODEL\_V4 \| 1.0 \| 3.762e-13 \| 447 \| 0.196 \| 219 \| 153 \| 7 \| 74 \| 278 \| 275 \| 484 \| Uncharacterized protein \| Uncharacterized protein; afdb-uniprot50 \| AF-A0A2A3CLV9-F1-MODEL\_V4 \| 1.0 \| 3.553e-13 \| 446 \| 0.216 \| 250 \| 136 \| 9 \| 37 \| 279 \| 1 \| 197 \| Uncharacterized protein \| Uncharacterized protein; afdb-uniprot50 \| AF-U1KQI4-F1-MODEL\_V4 \| 1.0 \| 2.002e-13 \| 446 \| 0.177 \| 299 \| 202 \| 15 \| 3 \| 276 \| 59 \| 338 \| Rhs element Vgr protein \| Rhs element Vgr protein; afdb-uniprot50 \| AF-W7VHJ1-F1-MODEL\_V4 \| 1.0 \| 3.197e-14 \| 445 \| 0.178 \| 269 \| 184 \| 12 \| 26 \| 273 \| 6 \| 258 \| Phage protein \| Phage protein; afdb-uniprot50 \| AF-A0A1M5CLP3-F1-MODEL\_V4 \| 1.0 \| 1.206e-14 \| 445 \| 0.151 \| 304 \| 210 \| 12 \| 3 \| 276 \| 74 \| 359 \| Phage protein D \| Phage protein D; afdb-uniprot50 \| AF-A0A0F2NZ57-F1-MODEL\_V4 \| 1.0 \| 5.356e-14 \| 445 \| 0.173 \| 306 \| 199 \| 13 \| 7 \| 282 \| 72 \| 353 \| Phage\_base\_V domain-containing protein \| Phage\_base\_V domain-containing protein; afdb-uniprot50 \| AF-A0A1Q3SKN4-F1-MODEL\_V4 \| 1.0 \| 7.134e-14 \| 445 \| 0.173 \| 305 \| 199 \| 12 \| 9 \| 281 \| 71 \| 354 \| Phage\_base\_V domain-containing protein \| Phage\_base\_V domain-containing protein; afdb-uniprot50 \| AF-A0A7Y5SFG2-F1-MODEL\_V4 \| 1.0 \| 1.066e-13 \| 444 \| 0.156 \| 313 \| 207 \| 15 \| 7 \| 281 \| 65 \| 358 \| VgrG-related protein \| VgrG-related protein; afdb-uniprot50 \| AF-A0A202DY36-F1-MODEL\_V4 \| 1.0 \| 1.891e-13 \| 444 \| 0.163 \| 299 \| 206 \| 14 \| 7 \| 280 \| 68 \| 347 \| Phage\_base\_V domain-containing protein \| Phage\_base\_V domain-containing protein; afdb-uniprot50 \| AF-A0A826JQU7-F1-MODEL\_V4 \| 1.0 \| 3.553e-13 \| 443 \| 0.218 \| 211 \| 142 \| 6 \| 84 \| 278 \| 2 \| 205 \| Phage late control D family protein \| Phage late control D family protein; afdb-uniprot50 \| AF-X5R3B7-F1-MODEL\_V4 \| 1.0 \| 6.006e-14 \| 443 \| 0.157 \| 305 \| 201 \| 16 \| 14 \| 279 \| 87 \| 374 \| Uncharacterized protein \| Uncharacterized protein; afdb-uniprot50 \| AF-A0A7V5YZF1-F1-MODEL\_V4 \| 1.0 \| 7.555e-14 \| 443 \| 0.155 \| 308 \| 206 \| 16 \| 7 \| 281 \| 65 \| 351 \| Type IV secretion protein Rhs \| Type IV secretion protein Rhs; afdb-uniprot50 \| AF-A0A4R1AAX1-F1-MODEL\_V4 \| 1.0 \| 4.775e-14 \| 443 \| 0.14 \| 292 \| 208 \| 15 \| 9 \| 276 \| 74 \| 346 \| Type VI secretion system tip protein VgrG \| Type VI secretion system tip protein VgrG; afdb-uniprot50 \| AF-A0A3C2AA05-F1-MODEL\_V4 \| 1.0 \| 8.473e-14 \| 443 \| 0.158 \| 296 \| 204 \| 12 \| 7 \| 276 \| 71 \| 347 \| Phage\_base\_V domain-containing protein \| Phage\_base\_V domain-containing protein; afdb-uniprot50 \| AF-A0A3M1GZV7-F1-MODEL\_V4 \| 1.0 \| 3.385e-14 \| 442 \| 0.169 \| 295 \| 197 \| 14 \| 15 \| 280 \| 73 \| 348 \| Phage late control D family protein \| Phage late control D family protein; afdb-uniprot50 \| AF-A0A2P5MZM2-F1-MODEL\_V4 \| 1.0 \| 2.667e-13 \| 441 \| 0.156 \| 288 \| 200 \| 14 \| 15 \| 278 \| 76 \| 344 \| Uncharacterized protein \| Uncharacterized protein; afdb-uniprot50 \| AF-A0A850ABE1-F1-MODEL\_V4 \| 1.0 \| 3.168e-13 \| 441 \| 0.159 \| 307 \| 207 \| 15 \| 7 \| 279 \| 78 \| 367 \| Phage late control D family protein \| Phage late control D family protein; afdb-uniprot50 \| AF-A0A7C3CCR0-F1-MODEL\_V4 \| 1.0 \| 1.34e-13 \| 441 \| 0.133 \| 299 \| 208 \| 11 \| 7 \| 276 \| 74 \| 350 \| Type VI secretion system tip protein VgrG \| Type VI secretion system tip protein VgrG; afdb-uniprot50 \| AF-A0A7V8CYH2-F1-MODEL\_V4 \| 1.0 \| 1.34e-13 \| 441 \| 0.185 \| 285 \| 187 \| 14 \| 20 \| 279 \| 74 \| 338 \| VgrG-related protein \| VgrG-related protein; afdb-uniprot50 \| AF-A0A497TMB9-F1-MODEL\_V4 \| 1.0 \| 5.011e-13 \| 441 \| 0.17 \| 294 \| 202 \| 12 \| 7 \| 276 \| 75 \| 350 \| Rhs element Vgr protein \| Rhs element Vgr protein; afdb-uniprot50 \| AF-A0A767SPK6-F1-MODEL\_V4 \| 1.0 \| 3.948e-12 \| 440 \| 0.342 \| 181 \| 109 \| 4 \| 1 \| 179 \| 24 \| 196 \| Phage late control D family protein \| Phage late control D family protein; afdb-uniprot50 \| AF-J2D8Z2-F1-MODEL\_V4 \| 1.0 \| 1.503e-13 \| 440 \| 0.148 \| 310 \| 197 \| 14 \| 9 \| 278 \| 76 \| 358 \| Phage protein D \| Phage protein D; afdb-uniprot50 \| AF-A0A2V8S145-F1-MODEL\_V4 \| 1.0 \| 9.054e-15 \| 440 \| 0.156 \| 346 \| 203 \| 14 \| 3 \| 281 \| 38 \| 361 \| Uncharacterized protein \| Uncharacterized protein; afdb-uniprot50 \| AF-A0A447N5J0-F1-MODEL\_V4 \| 1.0 \| 1.108e-11 \| 439 \| 0.347 \| 161 \| 95 \| 4 \| 8 \| 166 \| 2 \| 154 \| Phage late control gene D \| Phage late control gene D; afdb-uniprot50 \| AF-A0A5C1IFS9-F1-MODEL\_V4 \| 1.0 \| 4.22e-13 \| 439 \| 0.123 \| 292 \| 216 \| 11 \| 9 \| 276 \| 77 \| 352 \| Type VI secretion system tip protein VgrG \| Type VI secretion system tip protein VgrG; afdb-uniprot50 \| AF-A0A2A5AR63-F1-MODEL\_V4 \| 1.0 \| 1.891e-13 \| 439 \| 0.172 \| 301 \| 192 \| 14 \| 7 \| 275 \| 72 \| 347 \| Type VI secretion protein VgrG \| Type VI secretion protein VgrG; afdb-uniprot50 \| AF-Q09DF8-F1-MODEL\_V4 \| 1.0 \| 1.503e-13 \| 439 \| 0.175 \| 302 \| 202 \| 14 \| 7 \| 282 \| 73 \| 353 \| Rhs element Vgr protein \| Rhs element Vgr protein; afdb-uniprot50 \| AF-A0A853I7P2-F1-MODEL\_V4 \| 1.0 \| 2.121e-13 \| 438 \| 0.341 \| 205 \| 118 \| 5 \| 1 \| 203 \| 49 \| 238 \| Uncharacterized protein \| Uncharacterized protein; afdb-uniprot50 \| AF-A0A1M4WE38-F1-MODEL\_V4 \| 1.0 \| 1.066e-13 \| 438 \| 0.15 \| 299 \| 202 \| 13 \| 7 \| 276 \| 75 \| 350 \| Rhs element Vgr protein \| Rhs element Vgr protein; afdb-uniprot50 \| AF-A0A315XAN7-F1-MODEL\_V4 \| 1.0 \| 2.991e-13 \| 438 \| 0.155 \| 303 \| 210 \| 12 \| 3 \| 276 \| 67 \| 352 \| Phage\_base\_V domain-containing protein \| Phage\_base\_V domain-containing protein; afdb-uniprot50 \| AF-A0A2M9BS42-F1-MODEL\_V4 \| 1.0 \| 6.361e-14 \| 438 \| 0.18 \| 300 \| 193 \| 14 \| 7 \| 276 \| 68 \| 344 \| Rhs element Vgr protein \| Rhs element Vgr protein; afdb-uniprot50 \| AF-A0A8B5FWM5-F1-MODEL\_V4 \| 1.0 \| 4.181e-12 \| 437 \| 0.338 \| 195 \| 119 \| 4 \| 1 \| 193 \| 63 \| 249 \| Uncharacterized protein \| Uncharacterized protein; afdb-uniprot50 \| AF-A0A7V8YCD2-F1-MODEL\_V4 \| 1.0 \| 2.85e-14 \| 437 \| 0.133 \| 353 \| 226 \| 13 \| 7 \| 311 \| 47 \| 367 \| Phage late control D family protein \| Phage late control D family protein; afdb-uniprot50 \| AF-A0A0F9SYY8-F1-MODEL\_V4 \| 1.0 \| 5.057e-14 \| 437 \| 0.192 \| 306 \| 188 \| 16 \| 9 \| 278 \| 67 \| 349 \| Uncharacterized protein \| Uncharacterized protein; afdb-uniprot50 \| AF-A0A5K7S6F5-F1-MODEL\_V4 \| 1.0 \| 9.502e-14 \| 437 \| 0.163 \| 300 \| 202 \| 14 \| 7 \| 278 \| 74 \| 352 \| VgrG protein \| VgrG protein; afdb-uniprot50 \| AF-A0A244EQM1-F1-MODEL\_V4 \| 1.0 \| 2.667e-13 \| 437 \| 0.177 \| 293 \| 201 \| 13 \| 7 \| 276 \| 69 \| 344 \| Type IV secretion protein Rhs \| Type IV secretion protein Rhs; afdb-uniprot50 \| AF-B9M7I2-F1-MODEL\_V4 \| 1.0 \| 1.184e-12 \| 436 \| 0.148 \| 296 \| 211 \| 10 \| 3 \| 274 \| 45 \| 323 \| Phage protein D and tail spike protein, putative \| Phage protein D and tail spike protein, putative; afdb-uniprot50 \| AF-A0A2E7CJ66-F1-MODEL\_V4 \| 1.0 \| 1.419e-13 \| 435 \| 0.164 \| 291 \| 189 \| 14 \| 22 \| 278 \| 72 \| 342 \| Uncharacterized protein \| Uncharacterized protein; afdb-uniprot50 \| AF-A0A852ZLG6-F1-MODEL\_V4 \| 1.0 \| 3.168e-13 \| 435 \| 0.193 \| 300 \| 201 \| 10 \| 7 \| 279 \| 85 \| 370 \| Uncharacterized protein \| Uncharacterized protein; afdb-uniprot50 \| AF-A0A0K2RMD0-F1-MODEL\_V4 \| 1.0 \| 2.021e-14 \| 434 \| 0.175 \| 296 \| 193 \| 15 \| 12 \| 280 \| 1 \| 272 \| Rhs element Vgr protein \| Rhs element Vgr protein; afdb-uniprot50 \| AF-A0A535WQL9-F1-MODEL\_V4 \| 1.0 \| 2.246e-13 \| 434 \| 0.18 \| 311 \| 193 \| 12 \| 7 \| 275 \| 66 \| 356 \| Phage late control D family protein \| Phage late control D family protein; afdb-uniprot50 \| AF-A0A5N7Z8R8-F1-MODEL\_V4 \| 1.0 \| 1.592e-13 \| 434 \| 0.162 \| 296 \| 202 \| 16 \| 7 \| 276 \| 75 \| 350 \| Type VI secretion system tip protein VgrG \| Type VI secretion system tip protein VgrG; afdb-uniprot50 \| AF-A0A7C3NZA9-F1-MODEL\_V4 \| 1.0 \| 8.473e-14 \| 434 \| 0.173 \| 311 \| 196 \| 12 \| 7 \| 281 \| 67 \| 352 \| Phage late control D family protein \| Phage late control D family protein; afdb-uniprot50 \| AF-A0A3D2C425-F1-MODEL\_V4 \| 1.0 \| 1.195e-13 \| 433 \| 0.175 \| 290 \| 187 \| 13 \| 19 \| 278 \| 74 \| 341 \| Uncharacterized protein \| Uncharacterized protein; afdb-uniprot50 \| AF-A0A1M6GFP2-F1-MODEL\_V4 \| 1.0 \| 2.002e-13 \| 433 \| 0.201 \| 283 \| 191 \| 15 \| 1 \| 273 \| 52 \| 309 \| Uncharacterized conserved protein, implicated in type VI secretion and phage assembly \| Uncharacterized conserved protein, implicated in type VI secretion and phage assembly; afdb-uniprot50 \| AF-A0A2E6F191-F1-MODEL\_V4 \| 1.0 \| 4.732e-13 \| 433 \| 0.136 \| 293 \| 213 \| 10 \| 7 \| 276 \| 64 \| 339 \| Phage\_base\_V domain-containing protein \| Phage\_base\_V domain-containing protein; afdb-uniprot50 \| AF-A0A3M1XG71-F1-MODEL\_V4 \| 1.0 \| 6.361e-14 \| 433 \| 0.182 \| 302 \| 196 \| 15 \| 12 \| 281 \| 79 \| 361 \| Phage\_base\_V domain-containing protein \| Phage\_base\_V domain-containing protein; afdb-uniprot50 \| AF-A0A420WUX9-F1-MODEL\_V4 \| 1.0 \| 5.15e-16 \| 432 \| 0.331 \| 178 \| 102 \| 1 \| 118 \| 278 \| 2 \| 179 \| Late control gene D protein (GPD) \| Late control gene D protein (GPD); afdb-uniprot50 \| AF-A0A538CK92-F1-MODEL\_V4 \| 1.0 \| 2.519e-13 \| 432 \| 0.176 \| 261 \| 168 \| 9 \| 62 \| 278 \| 16 \| 273 \| Phage late control D family protein \| Phage late control D family protein; afdb-uniprot50 \| AF-A0A5R9JEF4-F1-MODEL\_V4 \| 1.0 \| 4.509e-14 \| 432 \| 0.149 \| 315 \| 205 \| 13 \| 7 \| 279 \| 38 \| 331 \| Uncharacterized protein \| Uncharacterized protein; afdb-uniprot50 \| AF-A0A0C1QXT6-F1-MODEL\_V4 \| 1.0 \| 2.002e-13 \| 432 \| 0.187 \| 288 \| 189 \| 14 \| 19 \| 278 \| 88 \| 358 \| Uncharacterized protein \| Uncharacterized protein; afdb-uniprot50 \| AF-A0A849TPI3-F1-MODEL\_V4 \| 1.0 \| 1.701e-14 \| 432 \| 0.162 \| 332 \| 217 \| 14 \| 9 \| 311 \| 62 \| 361 \| Uncharacterized protein \| Uncharacterized protein; afdb-uniprot50 \| AF-A0A1I0S6T2-F1-MODEL\_V4 \| 1.0 \| 4.22e-13 \| 432 \| 0.136 \| 300 \| 214 \| 15 \| 7 \| 279 \| 75 \| 356 \| Rhs element Vgr protein \| Rhs element Vgr protein; afdb-uniprot50 \| AF-A0A2K8QSP8-F1-MODEL\_V4 \| 1.0 \| 2.246e-13 \| 432 \| 0.153 \| 299 \| 209 \| 14 \| 7 \| 279 \| 72 \| 352 \| Rhs element Vgr protein \| Rhs element Vgr protein; afdb-uniprot50 \| AF-A0A1Q3QSK3-F1-MODEL\_V4 \| 1.0 \| 5.356e-14 \| 431 \| 0.169 \| 330 \| 198 \| 16 \| 7 \| 281 \| 98 \| 406 \| Uncharacterized protein \| Uncharacterized protein; afdb-uniprot50 \| AF-A0A3C2A1R6-F1-MODEL\_V4 \| 1.0 \| 8e-14 \| 431 \| 0.144 \| 305 \| 202 \| 14 \| 7 \| 278 \| 71 \| 349 \| Rhs element Vgr protein \| Rhs element Vgr protein; afdb-uniprot50 \| AF-A0A399D1A6-F1-MODEL\_V4 \| 1.0 \| 2.121e-13 \| 431 \| 0.159 \| 294 \| 202 \| 14 \| 7 \| 275 \| 75 \| 348 \| Type VI secretion system tip protein VgrG \| Type VI secretion system tip protein VgrG; afdb-uniprot50 \| AF-A0A1H8J908-F1-MODEL\_V4 \| 1.0 \| 7.069e-13 \| 431 \| 0.152 \| 295 \| 207 \| 12 \| 7 \| 276 \| 73 \| 349 \| Rhs element Vgr protein \| Rhs element Vgr protein; afdb-uniprot50 \| AF-Q32HG2-F1-MODEL\_V4 \| 1.0 \| 7.069e-13 \| 430 \| 0.301 \| 196 \| 126 \| 4 \| 1 \| 187 \| 30 \| 223 \| Uncharacterized protein \| Uncharacterized protein; afdb-uniprot50 \| AF-K5YW23-F1-MODEL\_V4 \| 1.0 \| 1.503e-13 \| 430 \| 0.169 \| 289 \| 196 \| 11 \| 7 \| 276 \| 59 \| 322 \| Uncharacterized protein \| Uncharacterized protein; afdb-uniprot50 \| AF-A0A7V2RMU7-F1-MODEL\_V4 \| 1.0 \| 3.355e-13 \| 430 \| 0.142 \| 301 \| 204 \| 12 \| 7 \| 276 \| 76 \| 353 \| Phage late control D family protein \| Phage late control D family protein; afdb-uniprot50 \| AF-A0A7S9UGH2-F1-MODEL\_V4 \| 1.0 \| 7.134e-14 \| 430 \| 0.17 \| 328 \| 201 \| 15 \| 7 \| 281 \| 97 \| 406 \| Phage late control D family protein \| Phage late control D family protein; afdb-uniprot50 \| AF-A0A4R2IXT5-F1-MODEL\_V4 \| 1.0 \| 8.473e-14 \| 430 \| 0.188 \| 308 \| 202 \| 14 \| 7 \| 281 \| 64 \| 356 \| Phage protein D \| Phage protein D; afdb-uniprot50 \| AF-A0A6C0GDX1-F1-MODEL\_V4 \| 1.0 \| 3.984e-13 \| 430 \| 0.157 \| 298 \| 203 \| 12 \| 7 \| 276 \| 72 \| 349 \| Type VI secretion system tip protein VgrG \| Type VI secretion system tip protein VgrG; afdb-uniprot50 \| AF-A0A0Q0VUT6-F1-MODEL\_V4 \| 1.0 \| 1.686e-13 \| 429 \| 0.174 \| 298 \| 195 \| 12 \| 11 \| 277 \| 77 \| 354 \| Uncharacterized protein \| Uncharacterized protein; afdb-uniprot50 \| AF-A0A3N5Q711-F1-MODEL\_V4 \| 1.0 \| 6.361e-14 \| 429 \| 0.195 \| 292 \| 183 \| 16 \| 20 \| 279 \| 67 \| 338 \| Phage late control D family protein \| Phage late control D family protein; afdb-uniprot50 \| AF-A0A2N2JMI8-F1-MODEL\_V4 \| 1.0 \| 3.762e-13 \| 429 \| 0.142 \| 330 \| 212 \| 15 \| 3 \| 279 \| 64 \| 375 \| Uncharacterized protein \| Uncharacterized protein; afdb-uniprot50 \| AF-D0LLS6-F1-MODEL\_V4 \| 1.0 \| 3.797e-14 \| 429 \| 0.173 \| 300 \| 198 \| 11 \| 22 \| 287 \| 85 \| 368 \| Putative phage late control gene D protein \| Putative phage late control gene D protein; afdb-uniprot50 \| AF-H8GWZ7-F1-MODEL\_V4 \| 1.0 \| 7.928e-13 \| 429 \| 0.14 \| 300 \| 211 \| 15 \| 7 \| 281 \| 67 \| 344 \| Rhs element Vgr protein \| Rhs element Vgr protein; afdb-uniprot50 \| AF-A0A136KQ05-F1-MODEL\_V4 \| 1.0 \| 8.473e-14 \| 429 \| 0.162 \| 296 \| 194 \| 16 \| 16 \| 279 \| 76 \| 349 \| Phage late controlD protein (GPD) \| Phage late controlD protein (GPD); afdb-uniprot50 \| AF-A0A5N5ZAG1-F1-MODEL\_V4 \| 1.0 \| 5.952e-13 \| 429 \| 0.136 \| 294 \| 211 \| 12 \| 7 \| 276 \| 76 \| 350 \| Type VI secretion system tip protein VgrG \| Type VI secretion system tip protein VgrG; afdb-uniprot50 \| AF-A0A1Q3MS08-F1-MODEL\_V4 \| 1.0 \| 5.62e-13 \| 429 \| 0.15 \| 305 \| 212 \| 14 \| 7 \| 281 \| 64 \| 351 \| Phage\_base\_V domain-containing protein \| Phage\_base\_V domain-containing protein; afdb-uniprot50 \| AF-A0A4Y9QX69-F1-MODEL\_V4 \| 1.0 \| 6.303e-13 \| 428 \| 0.105 \| 302 \| 223 \| 13 \| 7 \| 281 \| 70 \| 351 \| Type VI secretion system tip protein VgrG \| Type VI secretion system tip protein VgrG; afdb-uniprot50 \| AF-U2J4H7-F1-MODEL\_V4 \| 1.0 \| 4.469e-13 \| 428 \| 0.153 \| 307 \| 210 \| 12 \| 7 \| 282 \| 61 \| 348 \| Phage\_base\_V domain-containing protein \| Phage\_base\_V domain-containing protein; afdb-uniprot50 \| AF-A0A1F8LMF4-F1-MODEL\_V4 \| 1.0 \| 8.973e-14 \| 428 \| 0.198 \| 313 \| 194 \| 14 \| 7 \| 281 \| 65 \| 358 \| Phage\_base\_V domain-containing protein \| Phage\_base\_V domain-containing protein; afdb-uniprot50 \| AF-A0A4R5H677-F1-MODEL\_V4 \| 1.0 \| 2.14e-14 \| 427 \| 0.213 \| 314 \| 195 \| 14 \| 3 \| 278 \| 86 \| 385 \| Phage late control D family protein \| Phage late control D family protein; afdb-uniprot50 \| AF-A0A7I0NT04-F1-MODEL\_V4 \| 1.0 \| 1.34e-13 \| 427 \| 0.174 \| 332 \| 197 \| 16 \| 7 \| 278 \| 87 \| 401 \| Phage late control D family protein \| Phage late control D family protein; afdb-uniprot50 \| AF-A0A1P8MVG2-F1-MODEL\_V4 \| 1.0 \| 9.972e-13 \| 427 \| 0.152 \| 322 \| 217 \| 15 \| 7 \| 280 \| 115 \| 428 \| Uncharacterized protein \| Uncharacterized protein; afdb-uniprot50 \| AF-A0A3D0UKR5-F1-MODEL\_V4 \| 1.0 \| 3.762e-13 \| 427 \| 0.178 \| 292 \| 194 \| 12 \| 17 \| 281 \| 76 \| 348 \| Phage\_base\_V domain-containing protein \| Phage\_base\_V domain-containing protein; afdb-uniprot50 \| AF-A0A2N6N316-F1-MODEL\_V4 \| 1.0 \| 3.168e-13 \| 427 \| 0.168 \| 297 \| 205 \| 12 \| 7 \| 278 \| 75 \| 354 \| Type IV secretion protein Rhs \| Type IV secretion protein Rhs; afdb-uniprot50 \| AF-A0A4S3LP71-F1-MODEL\_V4 \| 1.0 \| 7.069e-13 \| 426 \| 0.268 \| 246 \| 133 \| 6 \| 1 \| 207 \| 63 \| 300 \| Phage late control D family protein \| Phage late control D family protein; afdb-uniprot50 \| AF-B0P8P7-F1-MODEL\_V4 \| 1.0 \| 2.121e-13 \| 426 \| 0.151 \| 298 \| 210 \| 14 \| 7 \| 278 \| 96 \| 376 \| Uncharacterized protein \| Uncharacterized protein; afdb-uniprot50 \| AF-A0A2T4X8Z9-F1-MODEL\_V4 \| 1.0 \| 8.892e-13 \| 426 \| 0.145 \| 295 \| 210 \| 11 \| 9 \| 279 \| 76 \| 352 \| Rhs element Vgr protein \| Rhs element Vgr protein; afdb-uniprot50 \| AF-A0A1G9ZZY8-F1-MODEL\_V4 \| 1.0 \| 3.762e-13 \| 426 \| 0.145 \| 296 \| 208 \| 16 \| 7 \| 276 \| 74 \| 350 \| Rhs element Vgr protein \| Rhs element Vgr protein; afdb-uniprot50 \| AF-A0A402CWS6-F1-MODEL\_V4 \| 1.0 \| 6.675e-13 \| 426 \| 0.131 \| 304 \| 216 \| 14 \| 9 \| 281 \| 67 \| 353 \| Phage\_base\_V domain-containing protein \| Phage\_base\_V domain-containing protein; afdb-uniprot50 \| AF-A0A6I3ME80-F1-MODEL\_V4 \| 1.0 \| 7.134e-14 \| 425 \| 0.159 \| 332 \| 196 \| 15 \| 9 \| 276 \| 67 \| 379 \| Uncharacterized protein \| Uncharacterized protein; afdb-uniprot50 \| AF-A0A553KYW3-F1-MODEL\_V4 \| 1.0 \| 7.928e-13 \| 425 \| 0.132 \| 301 \| 206 \| 14 \| 7 \| 276 \| 75 \| 351 \| Type VI secretion system tip protein VgrG \| Type VI secretion system tip protein VgrG; afdb-uniprot50 \| AF-A0A1I0RS25-F1-MODEL\_V4 \| 1.0 \| 3.355e-13 \| 425 \| 0.175 \| 297 \| 204 \| 12 \| 7 \| 279 \| 72 \| 351 \| Rhs element Vgr protein \| Rhs element Vgr protein; afdb-uniprot50 \| AF-A0A1V5B929-F1-MODEL\_V4 \| 1.0 \| 5.011e-13 \| 424 \| 0.187 \| 304 \| 195 \| 13 \| 7 \| 277 \| 66 \| 350 \| Phage late control gene D protein (GPD) \| Phage late control gene D protein (GPD); afdb-uniprot50 \| AF-A0A0A8WVU1-F1-MODEL\_V4 \| 1.0 \| 2.991e-13 \| 424 \| 0.151 \| 311 \| 202 \| 15 \| 7 \| 279 \| 76 \| 362 \| Phage late control gene D protein \| Phage late control gene D protein; afdb-uniprot50 \| AF-A0A7Z7BGZ7-F1-MODEL\_V4 \| 1.0 \| 9.502e-14 \| 424 \| 0.176 \| 317 \| 199 \| 16 \| 7 \| 279 \| 82 \| 380 \| Uncharacterized protein \| Uncharacterized protein; afdb-uniprot50 \| AF-A0A7J5ERL7-F1-MODEL\_V4 \| 1.0 \| 4.021e-14 \| 424 \| 0.157 \| 331 \| 207 \| 14 \| 22 \| 311 \| 81 \| 380 \| Phage late control D family protein \| Phage late control D family protein; afdb-uniprot50 \| AF-A0A1M5XUS7-F1-MODEL\_V4 \| 1.0 \| 3.168e-13 \| 424 \| 0.165 \| 303 \| 206 \| 15 \| 7 \| 282 \| 71 \| 353 \| Rhs element Vgr protein \| Rhs element Vgr protein; afdb-uniprot50 \| AF-A0A3N4MGZ2-F1-MODEL\_V4 \| 1.0 \| 2.825e-13 \| 424 \| 0.155 \| 295 \| 199 \| 15 \| 7 \| 275 \| 75 \| 345 \| Type VI secretion system tip protein VgrG \| Type VI secretion system tip protein VgrG; afdb-uniprot50 \| AF-A0A497IHJ3-F1-MODEL\_V4 \| 1.0 \| 1.891e-13 \| 423 \| 0.173 \| 306 \| 198 \| 15 \| 9 \| 279 \| 75 \| 360 \| Uncharacterized protein \| Uncharacterized protein; afdb-uniprot50 \| AF-A0A1B6BCT6-F1-MODEL\_V4 \| 1.0 \| 1.254e-12 \| 423 \| 0.186 \| 301 \| 199 \| 16 \| 7 \| 279 \| 82 \| 364 \| Phage protein D \| Phage protein D; afdb-uniprot50 \| AF-A0A1I4DGB1-F1-MODEL\_V4 \| 1.0 \| 7.555e-14 \| 423 \| 0.169 \| 318 \| 203 \| 17 \| 7 \| 280 \| 89 \| 389 \| Phage protein D \| Phage protein D; afdb-uniprot50 \| AF-A0A2E8S0G6-F1-MODEL\_V4 \| 1.0 \| 1.328e-12 \| 423 \| 0.118 \| 362 \| 236 \| 19 \| 3 \| 309 \| 74 \| 407 \| Uncharacterized protein \| Uncharacterized protein; afdb-uniprot50 \| AF-A0A6L9YV50-F1-MODEL\_V4 \| 1.0 \| 1.686e-13 \| 423 \| 0.153 \| 300 \| 210 \| 12 \| 8 \| 279 \| 1 \| 284 \| Type VI secretion system tip protein VgrG \| Type VI secretion system tip protein VgrG; afdb-uniprot50 \| AF-A0A5S3QJL7-F1-MODEL\_V4 \| 1.0 \| 5.011e-13 \| 423 \| 0.124 \| 297 \| 210 \| 11 \| 9 \| 277 \| 77 \| 351 \| Type VI secretion system tip protein VgrG \| Type VI secretion system tip protein VgrG; afdb-uniprot50 \| AF-A0A5J6P870-F1-MODEL\_V4 \| 1.0 \| 2.002e-13 \| 422 \| 0.157 \| 299 \| 198 \| 13 \| 9 \| 278 \| 73 \| 346 \| Uncharacterized protein \| Uncharacterized protein; afdb-uniprot50 \| AF-A0A3B8IAA2-F1-MODEL\_V4 \| 1.0 \| 8.473e-14 \| 422 \| 0.144 \| 305 \| 202 \| 15 \| 7 \| 278 \| 71 \| 349 \| Rhs element Vgr protein \| Rhs element Vgr protein; afdb-uniprot50 \| AF-A0A368NIY7-F1-MODEL\_V4 \| 1.0 \| 1.407e-12 \| 422 \| 0.146 \| 300 \| 211 \| 12 \| 3 \| 276 \| 67 \| 347 \| Type VI secretion system tip protein VgrG \| Type VI secretion system tip protein VgrG; afdb-uniprot50 \| AF-A0A317I459-F1-MODEL\_V4 \| 1.0 \| 2.4e-14 \| 421 \| 0.138 \| 333 \| 204 \| 17 \| 7 \| 279 \| 86 \| 395 \| Uncharacterized protein \| Uncharacterized protein; afdb-uniprot50 \| AF-A0A1A1YQV5-F1-MODEL\_V4 \| 1.0 \| 1.066e-13 \| 421 \| 0.148 \| 317 \| 208 \| 15 \| 7 \| 278 \| 97 \| 396 \| Uncharacterized protein \| Uncharacterized protein; afdb-uniprot50 \| AF-A0A1M7AA40-F1-MODEL\_V4 \| 1.0 \| 1.328e-12 \| 421 \| 0.132 \| 294 \| 214 \| 12 \| 7 \| 276 \| 81 \| 357 \| Rhs element Vgr protein \| Rhs element Vgr protein; afdb-uniprot50 \| AF-A0A1H9K9T5-F1-MODEL\_V4 \| 1.0 \| 1.891e-13 \| 421 \| 0.157 \| 304 \| 202 \| 17 \| 9 \| 282 \| 74 \| 353 \| Rhs element Vgr protein \| Rhs element Vgr protein; afdb-uniprot50 \| AF-A0A4R5VFK6-F1-MODEL\_V4 \| 1.0 \| 9.416e-13 \| 421 \| 0.152 \| 296 \| 206 \| 14 \| 9 \| 278 \| 76 \| 352 \| Type VI secretion system tip protein VgrG \| Type VI secretion system tip protein VgrG; afdb-uniprot50 \| AF-A0A1J5Q762-F1-MODEL\_V4 \| 1.0 \| 1.266e-13 \| 420 \| 0.132 \| 295 \| 206 \| 12 \| 9 \| 276 \| 65 \| 336 \| Phage late control protein Gpd \| Phage late control protein Gpd; afdb-uniprot50 \| AF-A0A846AR29-F1-MODEL\_V4 \| 1.0 \| 8.396e-13 \| 420 \| 0.155 \| 303 \| 206 \| 15 \| 3 \| 275 \| 38 \| 320 \| Type VI secretion system tip protein VgrG \| Type VI secretion system tip protein VgrG; afdb-uniprot50 \| AF-A0A5M8QXA8-F1-MODEL\_V4 \| 1.0 \| 9.972e-13 \| 420 \| 0.156 \| 306 \| 208 \| 11 \| 3 \| 279 \| 62 \| 346 \| Type VI secretion system tip protein VgrG \| Type VI secretion system tip protein VgrG; afdb-uniprot50 \| AF-A0A844MNW0-F1-MODEL\_V4 \| 1.0 \| 1.254e-12 \| 420 \| 0.145 \| 295 \| 209 \| 12 \| 7 \| 276 \| 74 \| 350 \| Type VI secretion system tip protein VgrG \| Type VI secretion system tip protein VgrG; afdb-uniprot50 \| AF-A0A660LA17-F1-MODEL\_V4 \| 1.0 \| 2.121e-13 \| 419 \| 0.198 \| 303 \| 205 \| 13 \| 3 \| 281 \| 66 \| 354 \| Uncharacterized protein involved in type VI secretion and phage assembly \| Uncharacterized protein involved in type VI secretion and phage assembly; afdb-uniprot50 \| AF-A0A7C6ZWP8-F1-MODEL\_V4 \| 1.0 \| 1.328e-12 \| 419 \| 0.168 \| 297 \| 194 \| 12 \| 16 \| 281 \| 75 \| 349 \| VgrG-related protein \| VgrG-related protein; afdb-uniprot50 \| AF-A0A645AVJ7-F1-MODEL\_V4 \| 1.0 \| 6.736e-14 \| 418 \| 0.174 \| 316 \| 197 \| 19 \| 3 \| 280 \| 36 \| 325 \| Uncharacterized protein \| Uncharacterized protein; afdb-uniprot50 \| AF-A0A2W2B7E0-F1-MODEL\_V4 \| 1.0 \| 2.121e-13 \| 418 \| 0.177 \| 293 \| 198 \| 12 \| 9 \| 277 \| 61 \| 334 \| Type IV secretion protein Rhs \| Type IV secretion protein Rhs; afdb-uniprot50 \| AF-A0A7U8YTK7-F1-MODEL\_V4 \| 1.0 \| 2.991e-13 \| 418 \| 0.141 \| 311 \| 205 \| 16 \| 8 \| 276 \| 1 \| 291 \| Rhs element Vgr protein \| Rhs element Vgr protein; afdb-uniprot50 \| AF-A0A369QEE2-F1-MODEL\_V4 \| 1.0 \| 1.328e-12 \| 418 \| 0.172 \| 301 \| 203 \| 16 \| 7 \| 279 \| 78 \| 360 \| Phage\_base\_V domain-containing protein \| Phage\_base\_V domain-containing protein; afdb-uniprot50 \| AF-A0A4U9HGJ7-F1-MODEL\_V4 \| 1.0 \| 4.732e-13 \| 416 \| 0.271 \| 217 \| 141 \| 4 \| 1 \| 204 \| 59 \| 271 \| Phage protein D \| Phage protein D; afdb-uniprot50 \| AF-A0A1I1ZGB1-F1-MODEL\_V4 \| 1.0 \| 2.496e-12 \| 416 \| 0.146 \| 293 \| 206 \| 12 \| 9 \| 276 \| 77 \| 350 \| Rhs element Vgr protein \| Rhs element Vgr protein; afdb-uniprot50 \| AF-A0A6A8Q9P6-F1-MODEL\_V4 \| 1.0 \| 1.769e-12 \| 416 \| 0.148 \| 296 \| 204 \| 11 \| 7 \| 275 \| 73 \| 347 \| Type VI secretion system tip protein VgrG \| Type VI secretion system tip protein VgrG; afdb-uniprot50 \| AF-A0A6P1ATK7-F1-MODEL\_V4 \| 1.0 \| 8.732e-11 \| 415 \| 0.515 \| 126 \| 61 \| 0 \| 1 \| 126 \| 45 \| 170 \| Phage tail protein \| Phage tail protein; afdb-uniprot50 \| AF-A0A378MZ86-F1-MODEL\_V4 \| 1.0 \| 1.656e-11 \| 415 \| 0.331 \| 172 \| 102 \| 5 \| 1 \| 166 \| 14 \| 178 \| Phage protein D \| Phage protein D; afdb-uniprot50 \| AF-A0A2X1MAT3-F1-MODEL\_V4 \| 1.0 \| 1.108e-11 \| 415 \| 0.343 \| 172 \| 103 \| 4 \| 10 \| 179 \| 1 \| 164 \| Phage protein D \| Phage protein D; afdb-uniprot50 \| AF-A0A4V1UKC6-F1-MODEL\_V4 \| 1.0 \| 1.769e-12 \| 415 \| 0.148 \| 243 \| 171 \| 12 \| 57 \| 275 \| 7 \| 237 \| Type VI secretion system tip protein VgrG \| Type VI secretion system tip protein VgrG; afdb-uniprot50 \| AF-A0A2P7THA8-F1-MODEL\_V4 \| 1.0 \| 4.469e-13 \| 415 \| 0.152 \| 296 \| 204 \| 11 \| 9 \| 277 \| 68 \| 343 \| Type VI secretion system tip protein VgrG \| Type VI secretion system tip protein VgrG; afdb-uniprot50 \| AF-A0A375AES2-F1-MODEL\_V4 \| 1.0 \| 7.069e-13 \| 415 \| 0.148 \| 296 \| 210 \| 14 \| 7 \| 277 \| 72 \| 350 \| VgrG protein \| VgrG protein; afdb-uniprot50 \| AF-N1ZL07-F1-MODEL\_V4 \| 1.0 \| 1.066e-13 \| 414 \| 0.181 \| 309 \| 196 \| 13 \| 9 \| 279 \| 84 \| 373 \| Uncharacterized protein \| Uncharacterized protein; afdb-uniprot50 \| AF-G5IB76-F1-MODEL\_V4 \| 1.0 \| 3.553e-13 \| 414 \| 0.143 \| 313 \| 218 \| 12 \| 2 \| 281 \| 80 \| 375 \| Uncharacterized protein \| Uncharacterized protein; afdb-uniprot50 \| AF-A0A7Y6QLZ6-F1-MODEL\_V4 \| 1.0 \| 3.018e-14 \| 414 \| 0.144 \| 312 \| 213 \| 14 \| 7 \| 278 \| 99 \| 396 \| Phage late control D family protein \| Phage late control D family protein; afdb-uniprot50 \| AF-A0A2N5Z8J6-F1-MODEL\_V4 \| 1.0 \| 4.469e-13 \| 414 \| 0.162 \| 295 \| 203 \| 14 \| 7 \| 277 \| 70 \| 344 \| Rhs element Vgr protein \| Rhs element Vgr protein; afdb-uniprot50 \| AF-A8LSA3-F1-MODEL\_V4 \| 1.0 \| 9.416e-13 \| 414 \| 0.181 \| 297 \| 193 \| 15 \| 7 \| 275 \| 72 \| 346 \| Phage\_base\_V domain-containing protein \| Phage\_base\_V domain-containing protein; afdb-uniprot50 \| AF-A0A2N6A013-F1-MODEL\_V4 \| 1.0 \| 1.407e-12 \| 414 \| 0.141 \| 297 \| 212 \| 12 \| 7 \| 278 \| 73 \| 351 \| Rhs element Vgr protein \| Rhs element Vgr protein; afdb-uniprot50 \| AF-A0A349D7K0-F1-MODEL\_V4 \| 1.0 \| 2.378e-13 \| 414 \| 0.157 \| 298 \| 203 \| 14 \| 7 \| 275 \| 72 \| 350 \| Rhs element Vgr protein \| Rhs element Vgr protein; afdb-uniprot50 \| AF-A0A0N0UY75-F1-MODEL\_V4 \| 1.0 \| 1.769e-12 \| 414 \| 0.132 \| 294 \| 213 \| 11 \| 7 \| 275 \| 75 \| 351 \| Phage\_base\_V domain-containing protein \| Phage\_base\_V domain-containing protein; afdb-uniprot50 \| AF-A0A351UYP7-F1-MODEL\_V4 \| 1.0 \| 7.928e-13 \| 413 \| 0.145 \| 302 \| 207 \| 13 \| 7 \| 279 \| 75 \| 354 \| Rhs element Vgr protein \| Rhs element Vgr protein; afdb-uniprot50 \| AF-A0A651HNM5-F1-MODEL\_V4 \| 1.0 \| 1.517e-14 \| 413 \| 0.156 \| 313 \| 213 \| 15 \| 3 \| 279 \| 88 \| 385 \| Phage late control D family protein \| Phage late control D family protein; afdb-uniprot50 \| AF-A0A538EV62-F1-MODEL\_V4 \| 1.0 \| 3.355e-13 \| 413 \| 0.16 \| 275 \| 179 \| 11 \| 32 \| 275 \| 58 \| 311 \| Phage\_base\_V domain-containing protein \| Phage\_base\_V domain-containing protein; afdb-uniprot50 \| AF-A0A810LDR6-F1-MODEL\_V4 \| 1.0 \| 1.006e-13 \| 412 \| 0.18 \| 326 \| 192 \| 17 \| 3 \| 279 \| 64 \| 363 \| Uncharacterized protein \| Uncharacterized protein; afdb-uniprot50 \| AF-A0A0M6ZMU0-F1-MODEL\_V4 \| 1.0 \| 7.486e-13 \| 412 \| 0.169 \| 295 \| 200 \| 12 \| 7 \| 276 \| 71 \| 345 \| Rhs element Vgr protein \| Rhs element Vgr protein; afdb-uniprot50 \| AF-A0A369WSX9-F1-MODEL\_V4 \| 1.0 \| 3.948e-12 \| 412 \| 0.15 \| 300 \| 210 \| 11 \| 3 \| 274 \| 79 \| 361 \| Type VI secretion system tip protein VgrG \| Type VI secretion system tip protein VgrG; afdb-uniprot50 \| AF-A0A7A6RA84-F1-MODEL\_V4 \| 1.0 \| 1.232e-10 \| 411 \| 0.365 \| 167 \| 96 \| 4 \| 1 \| 165 \| 63 \| 221 \| Phage late control D family protein \| Phage late control D family protein; afdb-uniprot50 \| AF-A0A417HIZ6-F1-MODEL\_V4 \| 1.0 \| 1.592e-13 \| 411 \| 0.186 \| 306 \| 197 \| 14 \| 7 \| 279 \| 80 \| 366 \| Uncharacterized protein \| Uncharacterized protein; afdb-uniprot50 \| AF-A0A8A6KDY8-F1-MODEL\_V4 \| 1.0 \| 2.85e-14 \| 411 \| 0.166 \| 324 \| 200 \| 16 \| 7 \| 277 \| 89 \| 395 \| Phage late control D family protein \| Phage late control D family protein; afdb-uniprot50 \| AF-D1JAM5-F1-MODEL\_V4 \| 1.0 \| 4.732e-13 \| 411 \| 0.177 \| 326 \| 223 \| 14 \| 7 \| 302 \| 74 \| 384 \| Phage\_base\_V domain-containing protein \| Phage\_base\_V domain-containing protein; afdb-uniprot50 \| AF-A0A2L2NSD3-F1-MODEL\_V4 \| 1.0 \| 9.972e-13 \| 411 \| 0.138 \| 311 \| 214 \| 13 \| 7 \| 281 \| 98 \| 390 \| Rhs element Vgr protein \| Rhs element Vgr protein; afdb-uniprot50 \| AF-A0A381E6W6-F1-MODEL\_V4 \| 1.0 \| 8.32e-12 \| 410 \| 0.213 \| 183 \| 134 \| 5 \| 92 \| 274 \| 1 \| 173 \| Phage protein D \| Phage protein D; afdb-uniprot50 \| AF-A0A512ISE5-F1-MODEL\_V4 \| 1.0 \| 1.066e-13 \| 410 \| 0.189 \| 311 \| 203 \| 15 \| 7 \| 286 \| 60 \| 352 \| Uncharacterized protein \| Uncharacterized protein; afdb-uniprot50 \| AF-A0A7X5W735-F1-MODEL\_V4 \| 1.0 \| 6.736e-14 \| 410 \| 0.179 \| 289 \| 186 \| 15 \| 23 \| 279 \| 88 \| 357 \| Uncharacterized protein \| Uncharacterized protein; afdb-uniprot50 \| AF-A0A0A2EIY0-F1-MODEL\_V4 \| 1.0 \| 6.675e-13 \| 410 \| 0.114 \| 287 \| 209 \| 11 \| 19 \| 279 \| 84 \| 351 \| Phage\_base\_V domain-containing protein \| Phage\_base\_V domain-containing protein; afdb-uniprot50 \| AF-A0A4U1I5U2-F1-MODEL\_V4 \| 1.0 \| 8.892e-13 \| 410 \| 0.136 \| 301 \| 205 \| 14 \| 7 \| 276 \| 72 \| 348 \| Type VI secretion system tip protein VgrG \| Type VI secretion system tip protein VgrG; afdb-uniprot50 \| AF-A0A2D0N1B1-F1-MODEL\_V4 \| 1.0 \| 5.011e-13 \| 410 \| 0.157 \| 304 \| 195 \| 16 \| 7 \| 276 \| 69 \| 345 \| Phage\_base\_V domain-containing protein \| Phage\_base\_V domain-containing protein; afdb-uniprot50 \| AF-A0A4R7HI80-F1-MODEL\_V4 \| 1.0 \| 4.22e-13 \| 410 \| 0.188 \| 307 \| 206 \| 15 \| 7 \| 281 \| 64 \| 359 \| Uncharacterized protein involved in type VI secretion and phage assembly \| Uncharacterized protein involved in type VI secretion and phage assembly; afdb-uniprot50 \| AF-A0A6G9ZEV2-F1-MODEL\_V4 \| 1.0 \| 5.952e-13 \| 410 \| 0.15 \| 300 \| 200 \| 13 \| 15 \| 277 \| 190 \| 471 \| VgrG-related protein \| VgrG-related protein; afdb-uniprot50 \| AF-A0A2A3WJY1-F1-MODEL\_V4 \| 1.0 \| 9.331e-12 \| 409 \| 0.304 \| 194 \| 127 \| 3 \| 1 \| 188 \| 56 \| 247 \| Late control protein \| Late control protein; afdb-uniprot50 \| AF-A0A7L4Q7F5-F1-MODEL\_V4 \| 1.0 \| 1.49e-12 \| 409 \| 0.174 \| 310 \| 201 \| 14 \| 7 \| 278 \| 73 \| 365 \| Phage late control D family protein \| Phage late control D family protein; afdb-uniprot50 \| AF-C7XD81-F1-MODEL\_V4 \| 1.0 \| 4.69e-12 \| 409 \| 0.175 \| 239 \| 169 \| 7 \| 57 \| 276 \| 7 \| 236 \| Rhs element Vgr protein \| Rhs element Vgr protein; afdb-uniprot50 \| AF-A0A1H3K869-F1-MODEL\_V4 \| 1.0 \| 5.57e-12 \| 409 \| 0.166 \| 301 \| 203 \| 12 \| 9 \| 281 \| 66 \| 346 \| Phage protein D \| Phage protein D; afdb-uniprot50 \| AF-A0A563VQY9-F1-MODEL\_V4 \| 1.0 \| 2.964e-12 \| 408 \| 0.12 \| 299 \| 209 \| 10 \| 15 \| 281 \| 270 \| 546 \| Rhs element Vgr protein (Modular protein) \| Rhs element Vgr protein (Modular protein); afdb-uniprot50 \| AF-A0A7X8ICU0-F1-MODEL\_V4 \| 1.0 \| 6.361e-14 \| 407 \| 0.171 \| 314 \| 196 \| 18 \| 7 \| 279 \| 83 \| 373 \| Phage late control D family protein \| Phage late control D family protein; afdb-uniprot50 \| AF-A0A7V2INA6-F1-MODEL\_V4 \| 1.0 \| 1.984e-12 \| 407 \| 0.156 \| 344 \| 209 \| 18 \| 7 \| 287 \| 64 \| 389 \| Phage late control D family protein \| Phage late control D family protein; afdb-uniprot50 \| AF-A0A4Y8ABY2-F1-MODEL\_V4 \| 1.0 \| 6.675e-13 \| 407 \| 0.134 \| 289 \| 206 \| 12 \| 13 \| 275 \| 79 \| 349 \| Rhs element Vgr protein \| Rhs element Vgr protein; afdb-uniprot50 \| AF-A0A495EIK4-F1-MODEL\_V4 \| 1.0 \| 8.396e-13 \| 407 \| 0.147 \| 299 \| 203 \| 14 \| 7 \| 276 \| 75 \| 350 \| Rhs element Vgr protein \| Rhs element Vgr protein; afdb-uniprot50 \| AF-A0A7W9EDQ9-F1-MODEL\_V4 \| 1.0 \| 8.811e-12 \| 406 \| 0.38 \| 189 \| 109 \| 3 \| 2 \| 189 \| 55 \| 236 \| Uncharacterized protein \| Uncharacterized protein; afdb-uniprot50 \| AF-A0A3P6KJ21-F1-MODEL\_V4 \| 1.0 \| 5.259e-12 \| 406 \| 0.162 \| 289 \| 206 \| 12 \| 7 \| 278 \| 103 \| 372 \| Phage protein D \| Phage protein D; afdb-uniprot50 \| AF-A0A402CKJ8-F1-MODEL\_V4 \| 1.0 \| 7.555e-14 \| 406 \| 0.161 \| 310 \| 208 \| 12 \| 7 \| 278 \| 78 \| 373 \| Phage protein D \| Phage protein D; afdb-uniprot50 \| AF-A0A5M6DLW7-F1-MODEL\_V4 \| 1.0 \| 1.874e-12 \| 406 \| 0.149 \| 295 \| 205 \| 15 \| 7 \| 275 \| 75 \| 349 \| Type VI secretion system tip protein VgrG \| Type VI secretion system tip protein VgrG; afdb-uniprot50 \| AF-A0A412YNE6-F1-MODEL\_V4 \| 1.0 \| 1.118e-12 \| 406 \| 0.143 \| 300 \| 204 \| 11 \| 7 \| 275 \| 73 \| 350 \| Type VI secretion system tip protein VgrG \| Type VI secretion system tip protein VgrG; afdb-uniprot50 \| AF-A0A258B3I5-F1-MODEL\_V4 \| 1.0 \| 1.407e-12 \| 406 \| 0.16 \| 299 \| 204 \| 12 \| 9 \| 279 \| 75 \| 354 \| Phage\_base\_V domain-containing protein \| Phage\_base\_V domain-containing protein; afdb-uniprot50 \| AF-A0A316QJQ2-F1-MODEL\_V4 \| 1.0 \| 3.728e-12 \| 405 \| 0.176 \| 300 \| 203 \| 14 \| 7 \| 279 \| 83 \| 365 \| Uncharacterized protein \| Uncharacterized protein; afdb-uniprot50 \| AF-A0A3A0B1P0-F1-MODEL\_V4 \| 1.0 \| 2.4e-14 \| 405 \| 0.18 \| 316 \| 201 \| 12 \| 9 \| 280 \| 83 \| 384 \| Uncharacterized protein \| Uncharacterized protein; afdb-uniprot50 \| AF-A0A7Y2DHA3-F1-MODEL\_V4 \| 1.0 \| 9.416e-13 \| 405 \| 0.132 \| 303 \| 210 \| 11 \| 7 \| 279 \| 75 \| 354 \| Type VI secretion system tip protein VgrG \| Type VI secretion system tip protein VgrG; afdb-uniprot50 \| AF-A0A838RL76-F1-MODEL\_V4 \| 1.0 \| 1.056e-12 \| 405 \| 0.168 \| 297 \| 193 \| 15 \| 19 \| 281 \| 77 \| 353 \| VgrG-related protein \| VgrG-related protein; afdb-uniprot50 \| AF-A0A538PYD2-F1-MODEL\_V4 \| 1.0 \| 1.195e-13 \| 404 \| 0.173 \| 328 \| 210 \| 13 \| 22 \| 309 \| 83 \| 389 \| Phage late control D family protein \| Phage late control D family protein; afdb-uniprot50 \| AF-A0A364Y3U2-F1-MODEL\_V4 \| 1.0 \| 1.874e-12 \| 404 \| 0.137 \| 305 \| 206 \| 13 \| 7 \| 279 \| 69 \| 348 \| Rhs element Vgr protein \| Rhs element Vgr protein; afdb-uniprot50 \| AF-A0A447R078-F1-MODEL\_V4 \| 1.0 \| 8.732e-11 \| 403 \| 0.333 \| 168 \| 102 \| 4 \| 1 \| 166 \| 25 \| 184 \| Phage late control gene D \| Phage late control gene D; afdb-uniprot50 \| AF-A0A3R6R790-F1-MODEL\_V4 \| 1.0 \| 1.056e-12 \| 403 \| 0.151 \| 303 \| 197 \| 16 \| 7 \| 275 \| 85 \| 361 \| Uncharacterized protein \| Uncharacterized protein; afdb-uniprot50 \| AF-A0A2E2FL30-F1-MODEL\_V4 \| 1.0 \| 1.254e-12 \| 403 \| 0.158 \| 315 \| 201 \| 16 \| 7 \| 278 \| 57 \| 350 \| Uncharacterized protein \| Uncharacterized protein; afdb-uniprot50 \| AF-A0A2K8QX00-F1-MODEL\_V4 \| 1.0 \| 5.952e-13 \| 403 \| 0.13 \| 329 \| 213 \| 18 \| 7 \| 278 \| 77 \| 389 \| Phage control protein (GPD) \| Phage control protein (GPD); afdb-uniprot50 \| AF-A0A1G3LKX5-F1-MODEL\_V4 \| 1.0 \| 8.892e-13 \| 403 \| 0.177 \| 355 \| 210 \| 21 \| 7 \| 311 \| 67 \| 389 \| Uncharacterized protein \| Uncharacterized protein; afdb-uniprot50 \| AF-A0A841BZE2-F1-MODEL\_V4 \| 1.0 \| 1.34e-13 \| 403 \| 0.165 \| 308 \| 198 \| 16 \| 7 \| 279 \| 33 \| 316 \| Phage protein D/phage baseplate assembly protein gpV \| Phage protein D/phage baseplate assembly protein gpV; afdb-uniprot50 \| AF-A0A2T0WPJ0-F1-MODEL\_V4 \| 1.0 \| 7.069e-13 \| 402 \| 0.14 \| 307 \| 209 \| 13 \| 7 \| 279 \| 72 \| 357 \| Phage protein D \| Phage protein D; afdb-uniprot50 \| AF-A0A1I5CAJ2-F1-MODEL\_V4 \| 1.0 \| 4.732e-13 \| 400 \| 0.153 \| 312 \| 206 \| 16 \| 7 \| 278 \| 82 \| 375 \| Uncharacterized protein \| Uncharacterized protein; afdb-uniprot50 \| AF-A0A4Q6ANH2-F1-MODEL\_V4 \| 1.0 \| 5.57e-12 \| 400 \| 0.148 \| 296 \| 208 \| 14 \| 7 \| 276 \| 75 \| 352 \| Type VI secretion system tip protein VgrG \| Type VI secretion system tip protein VgrG; afdb-uniprot50 \| AF-A0A6N3K7N7-F1-MODEL\_V4 \| 1.0 \| 1.407e-12 \| 400 \| 0.159 \| 301 \| 202 \| 14 \| 9 \| 281 \| 57 \| 334 \| Type IV secretion protein Rhs \| Type IV secretion protein Rhs; afdb-uniprot50 \| AF-A0A1Q7S8W7-F1-MODEL\_V4 \| 1.0 \| 1.328e-12 \| 400 \| 0.146 \| 315 \| 219 \| 15 \| 2 \| 281 \| 37 \| 336 \| Phage\_base\_V domain-containing protein \| Phage\_base\_V domain-containing protein; afdb-uniprot50 \| AF-A0A6I7PB73-F1-MODEL\_V4 \| 1.0 \| 3.521e-12 \| 400 \| 0.131 \| 296 \| 214 \| 13 \| 7 \| 276 \| 71 \| 349 \| Type VI secretion system tip protein VgrG \| Type VI secretion system tip protein VgrG; afdb-uniprot50 \| AF-A0A7H5F1M2-F1-MODEL\_V4 \| 1.0 \| 2.225e-12 \| 400 \| 0.163 \| 293 \| 204 \| 12 \| 7 \| 275 \| 74 \| 349 \| Type VI secretion system tip protein VgrG \| Type VI secretion system tip protein VgrG; afdb-uniprot50 \| AF-A0A484HHC7-F1-MODEL\_V4 \| 1.0 \| 9.972e-13 \| 400 \| 0.157 \| 298 \| 211 \| 14 \| 7 \| 276 \| 66 \| 351 \| Phage\_base\_V domain-containing protein \| Phage\_base\_V domain-containing protein; afdb-uniprot50 \| AF-A0A5P9HZY4-F1-MODEL\_V4 \| 1.0 \| 4.181e-12 \| 399 \| 0.151 \| 298 \| 204 \| 12 \| 7 \| 278 \| 72 \| 346 \| Phage late control gene D protein (GPD) \| Phage late control gene D protein (GPD); afdb-uniprot50 \| AF-A0A1L3I302-F1-MODEL\_V4 \| 1.0 \| 8.396e-13 \| 398 \| 0.261 \| 176 \| 124 \| 3 \| 137 \| 310 \| 45 \| 216 \| Phage late control gene D protein (GPD) \| Phage late control gene D protein (GPD); afdb-uniprot50 \| AF-A0A7C1JGV2-F1-MODEL\_V4 \| 1.0 \| 7.069e-13 \| 398 \| 0.171 \| 297 \| 193 \| 15 \| 16 \| 281 \| 78 \| 352 \| Phage\_base\_V domain-containing protein \| Phage\_base\_V domain-containing protein; afdb-uniprot50 \| AF-A0A1E5G2C5-F1-MODEL\_V4 \| 1.0 \| 3.553e-13 \| 397 \| 0.174 \| 310 \| 191 \| 17 \| 9 \| 279 \| 88 \| 371 \| Uncharacterized protein \| Uncharacterized protein; afdb-uniprot50 \| AF-A0A480AGP3-F1-MODEL\_V4 \| 1.0 \| 8.892e-13 \| 397 \| 0.158 \| 303 \| 195 \| 17 \| 7 \| 275 \| 63 \| 339 \| Phage\_base\_V domain-containing protein \| Phage\_base\_V domain-containing protein; afdb-uniprot50 \| AF-A0A2M8Q5J0-F1-MODEL\_V4 \| 1.0 \| 8.396e-13 \| 397 \| 0.19 \| 309 \| 194 \| 18 \| 7 \| 281 \| 67 \| 353 \| Type IV secretion protein Rhs \| Type IV secretion protein Rhs; afdb-uniprot50 \| AF-A0A2V2LGK4-F1-MODEL\_V4 \| 1.0 \| 6.615e-12 \| 396 \| 0.147 \| 291 \| 211 \| 13 \| 7 \| 279 \| 71 \| 342 \| Phage\_base\_V domain-containing protein \| Phage\_base\_V domain-containing protein; afdb-uniprot50 \| AF-A0A1N6HBQ1-F1-MODEL\_V4 \| 1.0 \| 3.948e-12 \| 396 \| 0.132 \| 295 \| 213 \| 14 \| 7 \| 276 \| 74 \| 350 \| Rhs element Vgr protein \| Rhs element Vgr protein; afdb-uniprot50 \| AF-A0A512BIY2-F1-MODEL\_V4 \| 1.0 \| 1.476e-11 \| 395 \| 0.16 \| 292 \| 205 \| 10 \| 9 \| 276 \| 76 \| 351 \| Type IV secretion protein Rhs \| Type IV secretion protein Rhs; afdb-uniprot50 \| AF-A0A4R1YY35-F1-MODEL\_V4 \| 1.0 \| 6.675e-13 \| 395 \| 0.162 \| 301 \| 200 \| 19 \| 7 \| 277 \| 72 \| 350 \| Phage\_base\_V domain-containing protein \| Phage\_base\_V domain-containing protein; afdb-uniprot50 \| AF-A0A3D9UDY1-F1-MODEL\_V4 \| 1.0 \| 5.519e-11 \| 394 \| 0.276 \| 195 \| 130 \| 5 \| 1 \| 193 \| 75 \| 260 \| Late control gene D protein (GPD) \| Late control gene D protein (GPD); afdb-uniprot50 \| AF-A0A7X9I2D4-F1-MODEL\_V4 \| 1.0 \| 1.753e-11 \| 394 \| 0.237 \| 257 \| 148 \| 6 \| 1 \| 217 \| 63 \| 311 \| Phage tail protein \| Phage tail protein; afdb-uniprot50 \| AF-A0A424H668-F1-MODEL\_V4 \| 1.0 \| 5.898e-12 \| 394 \| 0.126 \| 301 \| 213 \| 13 \| 7 \| 275 \| 75 \| 357 \| Phage\_base\_V domain-containing protein \| Phage\_base\_V domain-containing protein; afdb-uniprot50 \| AF-A0A1I3Q7R5-F1-MODEL\_V4 \| 1.0 \| 2.357e-12 \| 393 \| 0.138 \| 297 \| 208 \| 18 \| 7 \| 276 \| 77 \| 352 \| Rhs element Vgr protein \| Rhs element Vgr protein; afdb-uniprot50 \| AF-A0A368JR84-F1-MODEL\_V4 \| 1.0 \| 3.324e-12 \| 393 \| 0.163 \| 299 \| 199 \| 13 \| 7 \| 276 \| 76 \| 352 \| Type VI secretion system tip protein VgrG \| Type VI secretion system tip protein VgrG; afdb-uniprot50 \| AF-A0A2A2EVG0-F1-MODEL\_V4 \| 1.0 \| 7.419e-12 \| 392 \| 0.424 \| 125 \| 72 \| 0 \| 155 \| 279 \| 2 \| 126 \| Uncharacterized protein \| Uncharacterized protein; afdb-uniprot50 \| AF-A0A257Q6I6-F1-MODEL\_V4 \| 1.0 \| 1.769e-12 \| 392 \| 0.134 \| 290 \| 192 \| 11 \| 14 \| 276 \| 68 \| 325 \| Uncharacterized protein \| Uncharacterized protein; afdb-uniprot50 \| AF-A0A369ASP5-F1-MODEL\_V4 \| 1.0 \| 3.521e-12 \| 392 \| 0.154 \| 297 \| 201 \| 14 \| 7 \| 275 \| 91 \| 365 \| Phage protein D \| Phage protein D; afdb-uniprot50 \| AF-A0A644Z3M6-F1-MODEL\_V4 \| 1.0 \| 1.184e-12 \| 392 \| 0.138 \| 310 \| 213 \| 13 \| 9 \| 281 \| 83 \| 375 \| Uncharacterized protein \| Uncharacterized protein; afdb-uniprot50 \| AF-A0A1V0Q0E9-F1-MODEL\_V4 \| 1.0 \| 1.129e-13 \| 392 \| 0.156 \| 320 \| 205 \| 14 \| 9 \| 279 \| 94 \| 397 \| Phage protein D \| Phage protein D; afdb-uniprot50 \| AF-A0A2S6I8V1-F1-MODEL\_V4 \| 1.0 \| 5.259e-12 \| 392 \| 0.14 \| 298 \| 205 \| 13 \| 7 \| 276 \| 76 \| 350 \| Rhs element Vgr protein \| Rhs element Vgr protein; afdb-uniprot50 \| AF-A0A3M1YH05-F1-MODEL\_V4 \| 1.0 \| 5.307e-13 \| 392 \| 0.151 \| 303 \| 204 \| 14 \| 7 \| 279 \| 72 \| 351 \| Type VI secretion system tip protein VgrG \| Type VI secretion system tip protein VgrG; afdb-uniprot50 \| AF-A0A1T5N4P7-F1-MODEL\_V4 \| 1.0 \| 4.428e-12 \| 392 \| 0.116 \| 293 \| 220 \| 11 \| 7 \| 276 \| 79 \| 355 \| Rhs element Vgr protein \| Rhs element Vgr protein; afdb-uniprot50 \| AF-A0A165X1V4-F1-MODEL\_V4 \| 1.0 \| 2.225e-12 \| 391 \| 0.191 \| 219 \| 145 \| 7 \| 112 \| 311 \| 4 \| 209 \| Phage late control gene D protein (GPD) \| Phage late control gene D protein (GPD); afdb-uniprot50 \| AF-A0A5S9QSE6-F1-MODEL\_V4 \| 1.0 \| 7.419e-12 \| 391 \| 0.133 \| 292 \| 211 \| 10 \| 7 \| 275 \| 72 \| 344 \| Phage\_base\_V domain-containing protein \| Phage\_base\_V domain-containing protein; afdb-uniprot50 \| AF-A0A2D0MYA0-F1-MODEL\_V4 \| 1.0 \| 2.101e-12 \| 391 \| 0.165 \| 302 \| 199 \| 17 \| 7 \| 278 \| 76 \| 354 \| Type IV secretion protein Rhs \| Type IV secretion protein Rhs; afdb-uniprot50 \| AF-A0A559QPW3-F1-MODEL\_V4 \| 1.0 \| 1.328e-12 \| 391 \| 0.155 \| 302 \| 203 \| 13 \| 7 \| 275 \| 72 \| 354 \| Rhs element Vgr protein \| Rhs element Vgr protein; afdb-uniprot50 \| AF-A0A850NG49-F1-MODEL\_V4 \| 1.0 \| 7.486e-13 \| 390 \| 0.138 \| 297 \| 211 \| 13 \| 7 \| 276 \| 61 \| 339 \| Uncharacterized protein \| Uncharacterized protein; afdb-uniprot50 \| AF-R5JY02-F1-MODEL\_V4 \| 1.0 \| 6.247e-12 \| 390 \| 0.167 \| 311 \| 203 \| 13 \| 7 \| 278 \| 84 \| 377 \| Uncharacterized protein \| Uncharacterized protein; afdb-uniprot50 \| AF-A0A3P1CVH5-F1-MODEL\_V4 \| 1.0 \| 3.728e-12 \| 390 \| 0.132 \| 303 \| 209 \| 12 \| 9 \| 278 \| 74 \| 355 \| Type VI secretion system tip protein VgrG \| Type VI secretion system tip protein VgrG; afdb-uniprot50 \| AF-F2NUE0-F1-MODEL\_V4 \| 1.0 \| 3.521e-12 \| 390 \| 0.155 \| 257 \| 174 \| 12 \| 61 \| 290 \| 149 \| 389 \| Uncharacterized protein \| Uncharacterized protein; afdb-uniprot50 \| AF-A0A2E7URL9-F1-MODEL\_V4 \| 1.0 \| 1.056e-12 \| 389 \| 0.132 \| 303 \| 204 \| 15 \| 9 \| 278 \| 36 \| 312 \| Uncharacterized protein \| Uncharacterized protein; afdb-uniprot50 \| AF-A0A7U9SDK9-F1-MODEL\_V4 \| 1.0 \| 7.928e-13 \| 389 \| 0.13 \| 315 \| 212 \| 16 \| 9 \| 281 \| 83 \| 377 \| Uncharacterized protein \| Uncharacterized protein; afdb-uniprot50 \| AF-A0A257KZG6-F1-MODEL\_V4 \| 1.0 \| 6.247e-12 \| 389 \| 0.136 \| 294 \| 204 \| 14 \| 9 \| 274 \| 75 \| 346 \| Type IV secretion protein Rhs \| Type IV secretion protein Rhs; afdb-uniprot50 \| AF-A0A2D8HSQ5-F1-MODEL\_V4 \| 1.0 \| 9.331e-12 \| 389 \| 0.134 \| 297 \| 217 \| 16 \| 1 \| 281 \| 53 \| 325 \| Phage\_base\_V domain-containing protein \| Phage\_base\_V domain-containing protein; afdb-uniprot50 \| AF-A0A136LBS7-F1-MODEL\_V4 \| 1.0 \| 4.428e-12 \| 389 \| 0.119 \| 302 \| 212 \| 13 \| 7 \| 276 \| 78 \| 357 \| Rhs element Vgr protein \| Rhs element Vgr protein; afdb-uniprot50 \| AF-A0A853IAA6-F1-MODEL\_V4 \| 1.0 \| 1.769e-12 \| 388 \| 0.148 \| 316 \| 212 \| 15 \| 3 \| 278 \| 61 \| 359 \| Uncharacterized protein \| Uncharacterized protein; afdb-uniprot50 \| AF-A0A6C2CEP6-F1-MODEL\_V4 \| 1.0 \| 1.174e-11 \| 388 \| 0.153 \| 299 \| 210 \| 13 \| 7 \| 281 \| 73 \| 352 \| Type VI secretion system tip protein VgrG \| Type VI secretion system tip protein VgrG; afdb-uniprot50 \| AF-A0A518BPM5-F1-MODEL\_V4 \| 1.0 \| 1.874e-12 \| 388 \| 0.165 \| 315 \| 189 \| 14 \| 9 \| 281 \| 73 \| 355 \| Phage late control gene D protein (GPD) \| Phage late control gene D protein (GPD); afdb-uniprot50 \| AF-J0P5E3-F1-MODEL\_V4 \| 1.0 \| 2.799e-12 \| 388 \| 0.137 \| 299 \| 207 \| 17 \| 7 \| 276 \| 73 \| 349 \| Rhs element Vgr protein \| Rhs element Vgr protein; afdb-uniprot50 \| AF-A0A2N0KBG2-F1-MODEL\_V4 \| 1.0 \| 2.496e-12 \| 387 \| 0.143 \| 285 \| 211 \| 11 \| 7 \| 275 \| 88 \| 355 \| Uncharacterized protein \| Uncharacterized protein; afdb-uniprot50 \| AF-A0A1Y3U421-F1-MODEL\_V4 \| 1.0 \| 1.49e-12 \| 387 \| 0.166 \| 295 \| 197 \| 16 \| 7 \| 275 \| 85 \| 356 \| Uncharacterized protein \| Uncharacterized protein; afdb-uniprot50 \| AF-A0A1Y3SD88-F1-MODEL\_V4 \| 1.0 \| 3.521e-12 \| 387 \| 0.168 \| 309 \| 202 \| 16 \| 3 \| 279 \| 79 \| 364 \| Uncharacterized protein \| Uncharacterized protein; afdb-uniprot50 \| AF-A0A7Y9PD51-F1-MODEL\_V4 \| 1.0 \| 2.378e-13 \| 387 \| 0.181 \| 342 \| 209 \| 20 \| 7 \| 311 \| 51 \| 358 \| Phage protein D \| Phage protein D; afdb-uniprot50 \| AF-A0A3D1AC35-F1-MODEL\_V4 \| 1.0 \| 4.966e-12 \| 387 \| 0.138 \| 295 \| 218 \| 12 \| 7 \| 279 \| 73 \| 353 \| Rhs element Vgr protein \| Rhs element Vgr protein; afdb-uniprot50 \| AF-A0A379YH32-F1-MODEL\_V4 \| 1.0 \| 2.335e-11 \| 386 \| 0.309 \| 194 \| 125 \| 4 \| 1 \| 193 \| 63 \| 248 \| Phage protein D \| Phage protein D; afdb-uniprot50 \| AF-A0A7G9WG86-F1-MODEL\_V4 \| 1.0 \| 2.496e-12 \| 386 \| 0.137 \| 269 \| 191 \| 16 \| 20 \| 277 \| 64 \| 302 \| Phage late control D family protein \| Phage late control D family protein; afdb-uniprot50 \| AF-A0A3S0U1E7-F1-MODEL\_V4 \| 1.0 \| 3.728e-12 \| 386 \| 0.149 \| 335 \| 212 \| 16 \| 7 \| 291 \| 59 \| 370 \| Uncharacterized protein \| Uncharacterized protein; afdb-uniprot50 \| AF-A0A2N5YLC6-F1-MODEL\_V4 \| 1.0 \| 2.082e-11 \| 386 \| 0.129 \| 293 \| 213 \| 12 \| 7 \| 275 \| 69 \| 343 \| Rhs element Vgr protein \| Rhs element Vgr protein; afdb-uniprot50 \| AF-A0A4P5VE00-F1-MODEL\_V4 \| 1.0 \| 2.643e-12 \| 386 \| 0.126 \| 293 \| 205 \| 14 \| 13 \| 276 \| 72 \| 342 \| Type IV secretion protein Rhs \| Type IV secretion protein Rhs; afdb-uniprot50 \| AF-A0A257QUN5-F1-MODEL\_V4 \| 1.0 \| 1.184e-12 \| 385 \| 0.148 \| 282 \| 187 \| 12 \| 19 \| 275 \| 82 \| 335 \| Uncharacterized protein \| Uncharacterized protein; afdb-uniprot50 \| AF-A0A3D4KWM2-F1-MODEL\_V4 \| 1.0 \| 3.139e-12 \| 385 \| 0.126 \| 325 \| 209 \| 14 \| 7 \| 275 \| 83 \| 388 \| Uncharacterized protein \| Uncharacterized protein; afdb-uniprot50 \| AF-A0A291INC8-F1-MODEL\_V4 \| 1.0 \| 5.259e-12 \| 385 \| 0.167 \| 287 \| 196 \| 14 \| 18 \| 281 \| 72 \| 338 \| Baseplate assembly protein V \| Baseplate assembly protein V; afdb-uniprot50 \| AF-C7QK16-F1-MODEL\_V4 \| 1.0 \| 4.69e-12 \| 385 \| 0.163 \| 311 \| 207 \| 14 \| 3 \| 276 \| 60 \| 354 \| Rhs element Vgr protein \| Rhs element Vgr protein; afdb-uniprot50 \| AF-A0A371WSZ2-F1-MODEL\_V4 \| 1.0 \| 8e-14 \| 384 \| 0.153 \| 346 \| 215 \| 18 \| 7 \| 310 \| 61 \| 370 \| Uncharacterized protein \| Uncharacterized protein; afdb-uniprot50 \| AF-A0A661IT63-F1-MODEL\_V4 \| 1.0 \| 1.407e-12 \| 384 \| 0.133 \| 314 \| 207 \| 16 \| 7 \| 277 \| 94 \| 385 \| Uncharacterized protein \| Uncharacterized protein; afdb-uniprot50 \| AF-A0A4Y6PVJ4-F1-MODEL\_V4 \| 1.0 \| 7.069e-13 \| 384 \| 0.141 \| 339 \| 201 \| 17 \| 9 \| 278 \| 79 \| 396 \| Phage late control D family protein \| Phage late control D family protein; afdb-uniprot50 \| AF-A0A1M4ZE13-F1-MODEL\_V4 \| 1.0 \| 1.108e-11 \| 384 \| 0.126 \| 293 \| 218 \| 14 \| 7 \| 279 \| 76 \| 350 \| Uncharacterized conserved protein, implicated in type VI secretion and phage assembly \| Uncharacterized conserved protein, implicated in type VI secretion and phage assembly; afdb-uniprot50 \| AF-A0A1G7A3Q8-F1-MODEL\_V4 \| 1.0 \| 6.247e-12 \| 384 \| 0.138 \| 304 \| 199 \| 18 \| 7 \| 275 \| 76 \| 351 \| Rhs element Vgr protein \| Rhs element Vgr protein; afdb-uniprot50 \| AF-A0A6D1ICX5-F1-MODEL\_V4 \| 1.0 \| 3.324e-12 \| 384 \| 0.179 \| 301 \| 211 \| 12 \| 9 \| 281 \| 66 \| 358 \| Type IV secretion protein Rhs \| Type IV secretion protein Rhs; afdb-uniprot50 \| AF-A0A7C6P4H4-F1-MODEL\_V4 \| 1.0 \| 2.799e-12 \| 383 \| 0.134 \| 298 \| 205 \| 16 \| 3 \| 274 \| 82 \| 352 \| Phage late control D family protein \| Phage late control D family protein; afdb-uniprot50 \| AF-A0A1Q3UZH9-F1-MODEL\_V4 \| 1.0 \| 3.762e-13 \| 383 \| 0.138 \| 332 \| 202 \| 16 \| 7 \| 290 \| 61 \| 356 \| Uncharacterized protein \| Uncharacterized protein; afdb-uniprot50 \| AF-A0A7Z9KYN7-F1-MODEL\_V4 \| 1.0 \| 1.476e-11 \| 382 \| 0.136 \| 330 \| 216 \| 15 \| 7 \| 279 \| 77 \| 394 \| Uncharacterized protein \| Uncharacterized protein; afdb-uniprot50 \| AF-A0A4Q3UH64-F1-MODEL\_V4 \| 1.0 \| 2.378e-13 \| 382 \| 0.163 \| 311 \| 210 \| 12 \| 7 \| 277 \| 116 \| 416 \| Phage late control D family protein \| Phage late control D family protein; afdb-uniprot50 \| AF-A0A4Y8NQP5-F1-MODEL\_V4 \| 1.0 \| 9.416e-13 \| 382 \| 0.142 \| 322 \| 199 \| 13 \| 18 \| 278 \| 106 \| 411 \| Uncharacterized protein \| Uncharacterized protein; afdb-uniprot50 \| AF-A0A536PPK3-F1-MODEL\_V4 \| 1.0 \| 2.643e-12 \| 382 \| 0.136 \| 308 \| 216 \| 16 \| 8 \| 281 \| 1 \| 292 \| VgrG-related protein \| VgrG-related protein; afdb-uniprot50 \| AF-A0A553FM29-F1-MODEL\_V4 \| 1.0 \| 7.928e-13 \| 382 \| 0.131 \| 311 \| 205 \| 14 \| 7 \| 282 \| 72 \| 352 \| Type VI secretion system tip protein VgrG \| Type VI secretion system tip protein VgrG; afdb-uniprot50 \| AF-A0A1B9SFQ1-F1-MODEL\_V4 \| 1.0 \| 2.667e-13 \| 381 \| 0.17 \| 311 \| 192 \| 16 \| 9 \| 282 \| 62 \| 343 \| Uncharacterized protein \| Uncharacterized protein; afdb-uniprot50 \| AF-A0A3D2CCS9-F1-MODEL\_V4 \| 1.0 \| 1.984e-12 \| 381 \| 0.139 \| 301 \| 204 \| 15 \| 9 \| 278 \| 55 \| 331 \| Uncharacterized protein \| Uncharacterized protein; afdb-uniprot50 \| AF-A0A136LD44-F1-MODEL\_V4 \| 1.0 \| 6.247e-12 \| 381 \| 0.146 \| 313 \| 208 \| 16 \| 7 \| 281 \| 70 \| 361 \| Rhs element Vgr protein \| Rhs element Vgr protein; afdb-uniprot50 \| AF-A0A0E3SC16-F1-MODEL\_V4 \| 1.0 \| 8.32e-12 \| 380 \| 0.132 \| 286 \| 201 \| 11 \| 22 \| 277 \| 90 \| 358 \| Phage protein D \| Phage protein D; afdb-uniprot50 \| AF-A0A4R5WB95-F1-MODEL\_V4 \| 1.0 \| 2.225e-12 \| 380 \| 0.175 \| 291 \| 193 \| 14 \| 9 \| 273 \| 63 \| 332 \| Type IV secretion protein Rhs \| Type IV secretion protein Rhs; afdb-uniprot50 \| AF-A0A2S6GL78-F1-MODEL\_V4 \| 1.0 \| 7.006e-12 \| 380 \| 0.147 \| 305 \| 207 \| 13 \| 7 \| 279 \| 63 \| 346 \| Rhs element Vgr protein \| Rhs element Vgr protein; afdb-uniprot50 \| AF-A0A1W2GZA8-F1-MODEL\_V4 \| 1.0 \| 6.247e-12 \| 380 \| 0.156 \| 300 \| 198 \| 15 \| 7 \| 276 \| 76 \| 350 \| Rhs element Vgr protein \| Rhs element Vgr protein; afdb-uniprot50 \| AF-A0A7G8V716-F1-MODEL\_V4 \| 1.0 \| 8.32e-12 \| 380 \| 0.135 \| 296 \| 211 \| 15 \| 7 \| 276 \| 74 \| 350 \| Type VI secretion system tip protein VgrG \| Type VI secretion system tip protein VgrG; afdb-uniprot50 \| AF-A0A1T5DNZ3-F1-MODEL\_V4 \| 1.0 \| 3.139e-12 \| 379 \| 0.144 \| 311 \| 212 \| 18 \| 3 \| 279 \| 83 \| 373 \| Phage protein D \| Phage protein D; afdb-uniprot50 \| AF-A0A7Z7N0R5-F1-MODEL\_V4 \| 1.0 \| 2.002e-13 \| 379 \| 0.157 \| 331 \| 202 \| 14 \| 9 \| 281 \| 102 \| 413 \| Phage protein D \| Phage protein D; afdb-uniprot50 \| AF-A0A150S1I8-F1-MODEL\_V4 \| 1.0 \| 3.948e-12 \| 379 \| 0.134 \| 380 \| 207 \| 15 \| 4 \| 278 \| 88 \| 450 \| Uncharacterized protein \| Uncharacterized protein; afdb-uniprot50 \| AF-A0A7Y0MDW7-F1-MODEL\_V4 \| 1.0 \| 1.984e-12 \| 379 \| 0.17 \| 310 \| 204 \| 15 \| 1 \| 282 \| 59 \| 343 \| Type IV secretion protein Rhs \| Type IV secretion protein Rhs; afdb-uniprot50 \| AF-A0A1Z9NC79-F1-MODEL\_V4 \| 1.0 \| 4.69e-12 \| 379 \| 0.141 \| 310 \| 206 \| 14 \| 7 \| 280 \| 73 \| 358 \| Phage\_base\_V domain-containing protein \| Phage\_base\_V domain-containing protein; afdb-uniprot50 \| AF-A0A377AID4-F1-MODEL\_V4 \| 1.0 \| 8.732e-11 \| 378 \| 0.263 \| 171 \| 101 \| 3 \| 113 \| 258 \| 2 \| 172 \| Late control protein D protein \| Late control protein D protein; afdb-uniprot50 \| AF-A0A377AGI3-F1-MODEL\_V4 \| 1.0 \| 2.082e-11 \| 378 \| 0.247 \| 230 \| 158 \| 4 \| 1 \| 222 \| 49 \| 271 \| Gene late control D protein \| Gene late control D protein; afdb-uniprot50 \| AF-A0A2T7QF39-F1-MODEL\_V4 \| 1.0 \| 6.247e-12 \| 378 \| 0.137 \| 321 \| 207 \| 14 \| 7 \| 276 \| 78 \| 379 \| Phage\_base\_V domain-containing protein \| Phage\_base\_V domain-containing protein; afdb-uniprot50 \| AF-A0A856MT28-F1-MODEL\_V4 \| 1.0 \| 2.938e-11 \| 378 \| 0.122 \| 295 \| 213 \| 12 \| 15 \| 281 \| 98 \| 374 \| Type IV secretion protein Rhs \| Type IV secretion protein Rhs; afdb-uniprot50 \| AF-A0A5C9AP84-F1-MODEL\_V4 \| 1.0 \| 5.165e-10 \| 377 \| 0.35 \| 151 \| 90 \| 3 \| 1 \| 150 \| 63 \| 206 \| Phage late control D family protein \| Phage late control D family protein; afdb-uniprot50 \| AF-A0A0P8A270-F1-MODEL\_V4 \| 1.0 \| 7.006e-12 \| 377 \| 0.166 \| 307 \| 212 \| 14 \| 3 \| 281 \| 69 \| 359 \| Phage late controlD protein (GPD) \| Phage late controlD protein (GPD); afdb-uniprot50 \| AF-A0A8A3HJ98-F1-MODEL\_V4 \| 1.0 \| 2.082e-11 \| 377 \| 0.191 \| 245 \| 165 \| 12 \| 63 \| 278 \| 2 \| 242 \| VgrG-related protein \| VgrG-related protein; afdb-uniprot50 \| AF-A0A2A2HU16-F1-MODEL\_V4 \| 1.0 \| 7.419e-12 \| 377 \| 0.163 \| 312 \| 199 \| 11 \| 7 \| 281 \| 63 \| 349 \| Phage\_base\_V domain-containing protein \| Phage\_base\_V domain-containing protein; afdb-uniprot50 \| AF-A0A6P0W330-F1-MODEL\_V4 \| 1.0 \| 1.243e-11 \| 377 \| 0.138 \| 267 \| 200 \| 13 \| 20 \| 275 \| 93 \| 340 \| Phage\_base\_V domain-containing protein \| Phage\_base\_V domain-containing protein; afdb-uniprot50 \| AF-B6XAD0-F1-MODEL\_V4 \| 1.0 \| 7.286e-10 \| 376 \| 0.299 \| 167 \| 107 \| 4 \| 1 \| 165 \| 59 \| 217 \| Phage late control gene D protein (GPD) \| Phage late control gene D protein (GPD); afdb-uniprot50 \| AF-W8ZT94-F1-MODEL\_V4 \| 1.0 \| 2.774e-11 \| 376 \| 0.295 \| 196 \| 127 \| 3 \| 1 \| 187 \| 62 \| 255 \| Uncharacterized protein \| Uncharacterized protein; afdb-uniprot50 \| AF-A0A7Y5NVM8-F1-MODEL\_V4 \| 1.0 \| 3.728e-12 \| 376 \| 0.177 \| 304 \| 181 \| 15 \| 23 \| 278 \| 79 \| 361 \| Uncharacterized protein \| Uncharacterized protein; afdb-uniprot50 \| AF-A0A6L6J115-F1-MODEL\_V4 \| 1.0 \| 3.457e-10 \| 375 \| 0.329 \| 167 \| 104 \| 3 \| 1 \| 166 \| 51 \| 210 \| Phage tail protein \| Phage tail protein; afdb-uniprot50 \| AF-A0A840VPW3-F1-MODEL\_V4 \| 1.0 \| 9.416e-13 \| 375 \| 0.138 \| 289 \| 190 \| 10 \| 16 \| 278 \| 66 \| 321 \| Phage protein D \| Phage protein D; afdb-uniprot50 \| AF-A0A1S6FJC9-F1-MODEL\_V4 \| 1.0 \| 2.357e-12 \| 375 \| 0.136 \| 307 \| 204 \| 14 \| 7 \| 277 \| 72 \| 353 \| Phage\_base\_V domain-containing protein \| Phage\_base\_V domain-containing protein; afdb-uniprot50 \| AF-A0A530QSH5-F1-MODEL\_V4 \| 1.0 \| 3.762e-13 \| 374 \| 0.246 \| 239 \| 116 \| 7 \| 133 \| 311 \| 3 \| 237 \| Uncharacterized protein \| Uncharacterized protein; afdb-uniprot50 \| AF-A0A349E0J1-F1-MODEL\_V4 \| 1.0 \| 1.243e-11 \| 374 \| 0.166 \| 307 \| 204 \| 15 \| 7 \| 280 \| 61 \| 348 \| Phage\_base\_V domain-containing protein \| Phage\_base\_V domain-containing protein; afdb-uniprot50 \| AF-M4Z9U0-F1-MODEL\_V4 \| 1.0 \| 3.948e-12 \| 374 \| 0.185 \| 308 \| 202 \| 17 \| 7 \| 284 \| 65 \| 353 \| Phage\_base\_V domain-containing protein \| Phage\_base\_V domain-containing protein; afdb-uniprot50 \| AF-A0A135IJV0-F1-MODEL\_V4 \| 1.0 \| 3.878e-10 \| 373 \| 0.292 \| 123 \| 85 \| 2 \| 157 \| 278 \| 3 \| 124 \| Phage tail protein \| Phage tail protein; afdb-uniprot50 \| AF-A0A377Z1K9-F1-MODEL\_V4 \| 1.0 \| 5.165e-10 \| 373 \| 0.359 \| 167 \| 97 \| 4 \| 1 \| 165 \| 63 \| 221 \| Putative prophage tail protein \| Putative prophage tail protein; afdb-uniprot50 \| AF-A0A1I0CAS2-F1-MODEL\_V4 \| 1.0 \| 1.769e-12 \| 373 \| 0.123 \| 292 \| 217 \| 13 \| 7 \| 279 \| 86 \| 357 \| Phage protein D \| Phage protein D; afdb-uniprot50 \| AF-A0A101WDJ3-F1-MODEL\_V4 \| 1.0 \| 1.563e-11 \| 373 \| 0.164 \| 298 \| 205 \| 14 \| 7 \| 278 \| 84 \| 363 \| Uncharacterized protein \| Uncharacterized protein; afdb-uniprot50 \| AF-A0A536GUG3-F1-MODEL\_V4 \| 1.0 \| 8.811e-12 \| 373 \| 0.146 \| 294 \| 202 \| 15 \| 17 \| 276 \| 74 \| 352 \| VgrG-related protein \| VgrG-related protein; afdb-uniprot50 \| AF-A0A349BRR9-F1-MODEL\_V4 \| 1.0 \| 5.259e-12 \| 372 \| 0.122 \| 302 \| 214 \| 16 \| 7 \| 275 \| 76 \| 359 \| Uncharacterized protein \| Uncharacterized protein; afdb-uniprot50 \| AF-J3UA61-F1-MODEL\_V4 \| 1.0 \| 1.769e-12 \| 372 \| 0.12 \| 332 \| 247 \| 18 \| 7 \| 311 \| 103 \| 416 \| Uncharacterized protein \| Uncharacterized protein; afdb-uniprot50 \| AF-A0A7V7PRG0-F1-MODEL\_V4 \| 1.0 \| 1.394e-11 \| 372 \| 0.166 \| 282 \| 184 \| 15 \| 20 \| 273 \| 69 \| 327 \| Phage\_base\_V domain-containing protein \| Phage\_base\_V domain-containing protein; afdb-uniprot50 \| AF-A0A6G3WKK2-F1-MODEL\_V4 \| 1.0 \| 1.316e-11 \| 372 \| 0.173 \| 253 \| 175 \| 11 \| 57 \| 281 \| 7 \| 253 \| VgrG-related protein \| VgrG-related protein; afdb-uniprot50 \| AF-A0A1X4H071-F1-MODEL\_V4 \| 1.0 \| 2.643e-12 \| 372 \| 0.15 \| 306 \| 205 \| 18 \| 14 \| 280 \| 73 \| 362 \| Phage\_base\_V domain-containing protein \| Phage\_base\_V domain-containing protein; afdb-uniprot50 \| AF-A0A257NMK5-F1-MODEL\_V4 \| 1.0 \| 1.874e-12 \| 369 \| 0.132 \| 302 \| 200 \| 13 \| 7 \| 279 \| 60 \| 328 \| Uncharacterized protein \| Uncharacterized protein; afdb-uniprot50 \| AF-A0A1G5HB84-F1-MODEL\_V4 \| 1.0 \| 3.139e-12 \| 369 \| 0.137 \| 341 \| 209 \| 16 \| 7 \| 281 \| 85 \| 406 \| Uncharacterized protein \| Uncharacterized protein; afdb-uniprot50 \| AF-A0A2E1UZT3-F1-MODEL\_V4 \| 1.0 \| 6.247e-12 \| 368 \| 0.147 \| 311 \| 202 \| 16 \| 7 \| 279 \| 75 \| 360 \| Phage\_base\_V domain-containing protein \| Phage\_base\_V domain-containing protein; afdb-uniprot50 \| AF-A0A3D2XI45-F1-MODEL\_V4 \| 1.0 \| 3.111e-11 \| 368 \| 0.15 \| 305 \| 208 \| 15 \| 7 \| 279 \| 73 \| 358 \| Rhs element Vgr protein \| Rhs element Vgr protein; afdb-uniprot50 \| AF-A0A1I3BRU1-F1-MODEL\_V4 \| 1.0 \| 3.553e-13 \| 367 \| 0.165 \| 326 \| 197 \| 16 \| 7 \| 278 \| 101 \| 405 \| Phage protein D \| Phage protein D; afdb-uniprot50 \| AF-A0A1Q7VC12-F1-MODEL\_V4 \| 1.0 \| 1.118e-12 \| 366 \| 0.166 \| 319 \| 205 \| 13 \| 7 \| 281 \| 87 \| 388 \| Uncharacterized protein \| Uncharacterized protein; afdb-uniprot50 \| AF-A0A850I9N9-F1-MODEL\_V4 \| 1.0 \| 4.428e-12 \| 365 \| 0.154 \| 318 \| 202 \| 17 \| 7 \| 279 \| 84 \| 379 \| Phage late control D family protein \| Phage late control D family protein; afdb-uniprot50 \| AF-A0A7X1E8L6-F1-MODEL\_V4 \| 1.0 \| 3.913e-11 \| 365 \| 0.129 \| 332 \| 220 \| 14 \| 20 \| 311 \| 75 \| 377 \| Uncharacterized protein \| Uncharacterized protein; afdb-uniprot50 \| AF-A0A150KDD5-F1-MODEL\_V4 \| 1.0 \| 1.266e-13 \| 364 \| 0.182 \| 313 \| 209 \| 16 \| 1 \| 296 \| 13 \| 295 \| Uncharacterized protein \| Uncharacterized protein; afdb-uniprot50 \| AF-A0A6N2W373-F1-MODEL\_V4 \| 1.0 \| 4.428e-12 \| 363 \| 0.141 \| 311 \| 203 \| 18 \| 2 \| 275 \| 67 \| 350 \| Uncharacterized protein \| Uncharacterized protein; afdb-uniprot50 \| AF-A0A7Y5K2K5-F1-MODEL\_V4 \| 1.0 \| 3.324e-12 \| 363 \| 0.146 \| 328 \| 207 \| 19 \| 18 \| 284 \| 147 \| 462 \| Uncharacterized protein \| Uncharacterized protein; afdb-uniprot50 \| AF-A0A7X0CIA8-F1-MODEL\_V4 \| 1.0 \| 2.774e-11 \| 363 \| 0.131 \| 304 \| 215 \| 16 \| 7 \| 282 \| 63 \| 345 \| Rhs element Vgr protein \| Rhs element Vgr protein; afdb-uniprot50 \| AF-G2ZT36-F1-MODEL\_V4 \| 1.0 \| 3.521e-12 \| 362 \| 0.157 \| 317 \| 205 \| 14 \| 7 \| 280 \| 71 \| 368 \| Uncharacterized protein \| Uncharacterized protein; afdb-uniprot50 \| AF-A0A7Y5LNK5-F1-MODEL\_V4 \| 1.0 \| 2.774e-11 \| 362 \| 0.133 \| 329 \| 210 \| 15 \| 7 \| 278 \| 76 \| 386 \| Uncharacterized protein \| Uncharacterized protein; afdb-uniprot50 \| AF-A0A6P1MD19-F1-MODEL\_V4 \| 1.0 \| 1.316e-11 \| 361 \| 0.145 \| 295 \| 208 \| 12 \| 3 \| 274 \| 83 \| 356 \| Uncharacterized protein \| Uncharacterized protein; afdb-uniprot50 \| AF-A0A4R1HQU8-F1-MODEL\_V4 \| 1.0 \| 1.857e-11 \| 361 \| 0.18 \| 304 \| 201 \| 10 \| 7 \| 281 \| 63 \| 347 \| Phage protein D \| Phage protein D; afdb-uniprot50 \| AF-A0A3A9G426-F1-MODEL\_V4 \| 1.0 \| 5.212e-11 \| 360 \| 0.145 \| 309 \| 206 \| 18 \| 2 \| 276 \| 74 \| 358 \| Phage late control D family protein \| Phage late control D family protein; afdb-uniprot50 \| AF-A0A760MUK4-F1-MODEL\_V4 \| 1.0 \| 3.489e-11 \| 357 \| 0.24 \| 229 \| 142 \| 6 \| 1 \| 205 \| 72 \| 292 \| Phage late control D family protein \| Phage late control D family protein; afdb-uniprot50 \| AF-A0A2C8EXW9-F1-MODEL\_V4 \| 1.0 \| 1.088e-09 \| 356 \| 0.353 \| 167 \| 100 \| 3 \| 1 \| 166 \| 77 \| 236 \| Phage protein D \| Phage protein D; afdb-uniprot50 \| AF-A0A149USP9-F1-MODEL\_V4 \| 1.0 \| 1.966e-11 \| 356 \| 0.112 \| 303 \| 213 \| 11 \| 17 \| 281 \| 106 \| 390 \| Uncharacterized protein \| Uncharacterized protein; afdb-uniprot50 \| AF-A0A1S9CQS2-F1-MODEL\_V4 \| 1.0 \| 3.728e-12 \| 356 \| 0.096 \| 354 \| 244 \| 18 \| 7 \| 308 \| 105 \| 434 \| NlpC/P60 domain-containing protein \| NlpC/P60 domain-containing protein; afdb-uniprot50 \| AF-A0A6C8RYY1-F1-MODEL\_V4 \| 1.0 \| 1.221e-09 \| 354 \| 0.378 \| 140 \| 78 \| 3 \| 27 \| 165 \| 1 \| 132 \| Phage late control D family protein \| Phage late control D family protein; afdb-uniprot50 \| AF-A0A6M8HQ44-F1-MODEL\_V4 \| 1.0 \| 4.966e-12 \| 354 \| 0.129 \| 308 \| 213 \| 15 \| 7 \| 277 \| 55 \| 344 \| Uncharacterized protein \| Uncharacterized protein; afdb-uniprot50 \| AF-A0A3S0CSF5-F1-MODEL\_V4 \| 1.0 \| 3.913e-11 \| 354 \| 0.152 \| 328 \| 216 \| 17 \| 3 \| 285 \| 63 \| 373 \| Uncharacterized protein \| Uncharacterized protein; afdb-uniprot50 \| AF-A0A2E7UTL5-F1-MODEL\_V4 \| 1.0 \| 8.32e-12 \| 353 \| 0.131 \| 304 \| 203 \| 14 \| 9 \| 278 \| 55 \| 331 \| Uncharacterized protein \| Uncharacterized protein; afdb-uniprot50 \| AF-A0A6S5WYG3-F1-MODEL\_V4 \| 1.0 \| 3.913e-11 \| 352 \| 0.312 \| 205 \| 127 \| 5 \| 1 \| 204 \| 75 \| 266 \| Uncharacterized protein \| Uncharacterized protein; afdb-uniprot50 \| AF-A0A1Y4S3J0-F1-MODEL\_V4 \| 1.0 \| 2.335e-11 \| 352 \| 0.147 \| 291 \| 212 \| 13 \| 3 \| 275 \| 79 \| 351 \| Uncharacterized protein \| Uncharacterized protein; afdb-uniprot50 \| AF-A0A357YYD2-F1-MODEL\_V4 \| 1.0 \| 2.619e-11 \| 352 \| 0.149 \| 294 \| 195 \| 16 \| 9 \| 273 \| 64 \| 331 \| Phage\_base\_V domain-containing protein \| Phage\_base\_V domain-containing protein; afdb-uniprot50 \| AF-A0A836CGV5-F1-MODEL\_V4 \| 1.0 \| 2.596e-10 \| 352 \| 0.269 \| 189 \| 129 \| 4 \| 1 \| 187 \| 895 \| 1076 \| Uncharacterized protein \| Uncharacterized protein; afdb-uniprot50 \| AF-A0A257SB31-F1-MODEL\_V4 \| 1.0 \| 2.473e-11 \| 351 \| 0.154 \| 285 \| 198 \| 10 \| 12 \| 276 \| 71 \| 332 \| Uncharacterized protein \| Uncharacterized protein; afdb-uniprot50 \| AF-A0A7X5INZ1-F1-MODEL\_V4 \| 1.0 \| 1.476e-11 \| 350 \| 0.131 \| 304 \| 201 \| 13 \| 9 \| 276 \| 78 \| 354 \| Uncharacterized protein \| Uncharacterized protein; afdb-uniprot50 \| AF-A0A3N8BFJ3-F1-MODEL\_V4 \| 1.0 \| 3.695e-11 \| 350 \| 0.162 \| 308 \| 207 \| 13 \| 3 \| 279 \| 74 \| 361 \| Phage\_base\_V domain-containing protein \| Phage\_base\_V domain-containing protein; afdb-uniprot50 \| AF-F3IS59-F1-MODEL\_V4 \| 1.0 \| 3.235e-09 \| 349 \| 0.246 \| 126 \| 89 \| 1 \| 156 \| 275 \| 1 \| 126 \| Tail protein D \| Tail protein D; afdb-uniprot50 \| AF-A0A1I0IWR4-F1-MODEL\_V4 \| 1.0 \| 4.144e-11 \| 348 \| 0.105 \| 293 \| 224 \| 16 \| 3 \| 275 \| 78 \| 352 \| Phage protein D \| Phage protein D; afdb-uniprot50 \| AF-A0A1Y4T7B9-F1-MODEL\_V4 \| 1.0 \| 2.335e-11 \| 348 \| 0.161 \| 315 \| 197 \| 17 \| 3 \| 279 \| 75 \| 360 \| Uncharacterized protein \| Uncharacterized protein; afdb-uniprot50 \| AF-A0A192A527-F1-MODEL\_V4 \| 1.0 \| 8.32e-12 \| 348 \| 0.165 \| 314 \| 197 \| 14 \| 7 \| 279 \| 70 \| 359 \| Uncharacterized protein \| Uncharacterized protein; afdb-uniprot50 \| AF-A0A0F9MJ86-F1-MODEL\_V4 \| 1.0 \| 1.966e-11 \| 347 \| 0.152 \| 335 \| 206 \| 17 \| 7 \| 281 \| 78 \| 394 \| Uncharacterized protein \| Uncharacterized protein; afdb-uniprot50 \| AF-A0A2X4X699-F1-MODEL\_V4 \| 1.0 \| 3.426e-09 \| 346 \| 0.321 \| 168 \| 105 \| 4 \| 1 \| 166 \| 63 \| 223 \| Phage protein D \| Phage protein D; afdb-uniprot50 \| AF-A0A090SGD1-F1-MODEL\_V4 \| 1.0 \| 1.037e-10 \| 345 \| 0.271 \| 140 \| 99 \| 2 \| 145 \| 281 \| 15 \| 154 \| Uncharacterized protein \| Uncharacterized protein; afdb-uniprot50 \| AF-A0A1C6AKC2-F1-MODEL\_V4 \| 1.0 \| 9.331e-12 \| 345 \| 0.149 \| 295 \| 199 \| 17 \| 9 \| 275 \| 91 \| 361 \| Phage protein D \| Phage protein D; afdb-uniprot50 \| AF-A0A7W4CKY1-F1-MODEL\_V4 \| 1.0 \| 7.857e-12 \| 345 \| 0.174 \| 316 \| 193 \| 14 \| 7 \| 278 \| 42 \| 333 \| Phage\_base\_V domain-containing protein \| Phage\_base\_V domain-containing protein; afdb-uniprot50 \| AF-A0A4Q9KNI5-F1-MODEL\_V4 \| 1.0 \| 9.416e-13 \| 344 \| 0.158 \| 329 \| 194 \| 21 \| 9 \| 281 \| 87 \| 388 \| Phage late control D family protein \| Phage late control D family protein; afdb-uniprot50 \| AF-A0A1R3VFD6-F1-MODEL\_V4 \| 1.0 \| 6.247e-12 \| 344 \| 0.137 \| 319 \| 215 \| 14 \| 7 \| 278 \| 82 \| 387 \| Phage protein D-like protein \| Phage protein D-like protein; afdb-uniprot50 \| AF-A0A6G2IEN4-F1-MODEL\_V4 \| 1.0 \| 2.082e-11 \| 344 \| 0.145 \| 310 \| 201 \| 15 \| 9 \| 281 \| 79 \| 361 \| Phage\_base\_V domain-containing protein \| Phage\_base\_V domain-containing protein; afdb-uniprot50 \| AF-A0A4D7ATE7-F1-MODEL\_V4 \| 1.0 \| 6.438e-09 \| 343 \| 0.169 \| 136 \| 110 \| 3 \| 143 \| 278 \| 11 \| 143 \| Uncharacterized protein \| Uncharacterized protein; afdb-uniprot50 \| AF-A4X0M8-F1-MODEL\_V4 \| 1.0 \| 1.476e-11 \| 343 \| 0.168 \| 320 \| 200 \| 18 \| 3 \| 276 \| 89 \| 388 \| Uncharacterized protein \| Uncharacterized protein; afdb-uniprot50 \| AF-Q479Z3-F1-MODEL\_V4 \| 1.0 \| 1.641e-10 \| 343 \| 0.142 \| 294 \| 202 \| 14 \| 9 \| 274 \| 75 \| 346 \| Rhs element Vgr protein \| Rhs element Vgr protein; afdb-uniprot50 \| AF-A0A7G2TQH6-F1-MODEL\_V4 \| 1.0 \| 5.519e-11 \| 342 \| 0.15 \| 305 \| 192 \| 15 \| 7 \| 275 \| 72 \| 345 \| Type IV secretion protein Rhs \| Type IV secretion protein Rhs; afdb-uniprot50 \| AF-A0A849VCJ4-F1-MODEL\_V4 \| 1.0 \| 3.294e-11 \| 342 \| 0.151 \| 310 \| 201 \| 15 \| 7 \| 279 \| 63 \| 347 \| Uncharacterized protein \| Uncharacterized protein; afdb-uniprot50 \| AF-R7UKH8-F1-MODEL\_V4 \| 1.0 \| 4.069e-09 \| 339 \| 0.283 \| 166 \| 107 \| 6 \| 1 \| 164 \| 48 \| 203 \| Uncharacterized protein \| Uncharacterized protein; afdb-uniprot50 \| AF-A0A0D6MQM0-F1-MODEL\_V4 \| 1.0 \| 1.163e-10 \| 338 \| 0.114 \| 296 \| 210 \| 12 \| 13 \| 277 \| 76 \| 350 \| Uncharacterized protein \| Uncharacterized protein; afdb-uniprot50 \| AF-A0A6N8HUA9-F1-MODEL\_V4 \| 1.0 \| 9.247e-11 \| 337 \| 0.14 \| 305 \| 204 \| 16 \| 9 \| 276 \| 81 \| 364 \| Uncharacterized protein \| Uncharacterized protein; afdb-uniprot50 \| AF-A0A1D9H202-F1-MODEL\_V4 \| 1.0 \| 9.331e-12 \| 337 \| 0.164 \| 323 \| 194 \| 14 \| 7 \| 278 \| 69 \| 366 \| Uncharacterized protein \| Uncharacterized protein; afdb-uniprot50 \| AF-A0A0F9NFA5-F1-MODEL\_V4 \| 1.0 \| 1.563e-11 \| 336 \| 0.137 \| 392 \| 209 \| 17 \| 7 \| 279 \| 70 \| 451 \| Uncharacterized protein \| Uncharacterized protein; afdb-uniprot50 \| AF-A0A401Z4L8-F1-MODEL\_V4 \| 1.0 \| 4.921e-11 \| 336 \| 0.128 \| 319 \| 209 \| 16 \| 13 \| 280 \| 82 \| 382 \| Type IV secretion protein Rhs \| Type IV secretion protein Rhs; afdb-uniprot50 \| AF-A8T9K3-F1-MODEL\_V4 \| 1.0 \| 2.314e-10 \| 335 \| 0.116 \| 301 \| 215 \| 16 \| 7 \| 275 \| 233 \| 514 \| Phage\_base\_V domain-containing protein \| Phage\_base\_V domain-containing protein; afdb-uniprot50 \| AF-A0A5U6FP48-F1-MODEL\_V4 \| 1.0 \| 3.235e-09 \| 333 \| 0.267 \| 168 \| 113 \| 4 \| 1 \| 165 \| 66 \| 226 \| Late control protein \| Late control protein; afdb-uniprot50 \| AF-A0A074LLZ7-F1-MODEL\_V4 \| 1.0 \| 7.069e-13 \| 333 \| 0.145 \| 317 \| 200 \| 12 \| 7 \| 279 \| 79 \| 368 \| Uncharacterized protein \| Uncharacterized protein; afdb-uniprot50 \| AF-A1ZC10-F1-MODEL\_V4 \| 1.0 \| 2.335e-11 \| 332 \| 0.165 \| 321 \| 203 \| 19 \| 7 \| 280 \| 62 \| 364 \| Rhs element Vgr protein \| Rhs element Vgr protein; afdb-uniprot50 \| AF-A0A2Z5ZLK9-F1-MODEL\_V4 \| 1.0 \| 8.245e-11 \| 329 \| 0.138 \| 317 \| 206 \| 15 \| 7 \| 279 \| 69 \| 362 \| Phage late control protein D \| Phage late control protein D; afdb-uniprot50 \| AF-C0BXW3-F1-MODEL\_V4 \| 1.0 \| 1.232e-10 \| 329 \| 0.137 \| 312 \| 212 \| 13 \| 9 \| 281 \| 84 \| 377 \| Uncharacterized protein \| Uncharacterized protein; afdb-uniprot50 \| AF-A0A5Q4DFT5-F1-MODEL\_V4 \| 1.0 \| 1.463e-10 \| 329 \| 0.179 \| 335 \| 196 \| 16 \| 7 \| 276 \| 61 \| 381 \| Phage late control D family protein \| Phage late control D family protein; afdb-uniprot50 \| AF-A0A2N0F3R2-F1-MODEL\_V4 \| 1.0 \| 9.882e-12 \| 329 \| 0.142 \| 336 \| 221 \| 18 \| 7 \| 297 \| 111 \| 424 \| Uncharacterized protein \| Uncharacterized protein; afdb-uniprot50 \| AF-A0A430AW93-F1-MODEL\_V4 \| 1.0 \| 8.732e-11 \| 329 \| 0.137 \| 312 \| 215 \| 16 \| 7 \| 277 \| 107 \| 405 \| Flagellin \| Flagellin; afdb-uniprot50 \| AF-A0A329MVT4-F1-MODEL\_V4 \| 1.0 \| 5.212e-11 \| 329 \| 0.122 \| 310 \| 211 \| 20 \| 7 \| 279 \| 466 \| 751 \| Uncharacterized protein \| Uncharacterized protein; afdb-uniprot50 \| AF-A0A2R7SMS6-F1-MODEL\_V4 \| 1.0 \| 5.165e-10 \| 328 \| 0.204 \| 181 \| 118 \| 4 \| 148 \| 311 \| 23 \| 194 \| Late control protein D \| Late control protein D; afdb-uniprot50 \| AF-A0A0F9U2F1-F1-MODEL\_V4 \| 1.0 \| 6.942e-11 \| 328 \| 0.165 \| 302 \| 187 \| 17 \| 22 \| 281 \| 105 \| 383 \| Uncharacterized protein \| Uncharacterized protein; afdb-uniprot50 \| AF-A0A7Y5J5A2-F1-MODEL\_V4 \| 1.0 \| 3.083e-10 \| 328 \| 0.116 \| 325 \| 212 \| 14 \| 7 \| 279 \| 79 \| 380 \| Type VI secretion system tip protein VgrG \| Type VI secretion system tip protein VgrG; afdb-uniprot50 \| AF-A0A1H5RLG9-F1-MODEL\_V4 \| 1.0 \| 2.473e-11 \| 325 \| 0.131 \| 251 \| 175 \| 10 \| 83 \| 311 \| 295 \| 524 \| Uncharacterized protein \| Uncharacterized protein; afdb-uniprot50 \| AF-G2JAY7-F1-MODEL\_V4 \| 1.0 \| 3.265e-10 \| 324 \| 0.14 \| 321 \| 203 \| 13 \| 7 \| 280 \| 74 \| 368 \| Uncharacterized protein \| Uncharacterized protein; afdb-uniprot50 \| AF-A0A3A5IJU8-F1-MODEL\_V4 \| 1.0 \| 3.695e-11 \| 324 \| 0.128 \| 334 \| 218 \| 17 \| 7 \| 304 \| 115 \| 411 \| Uncharacterized protein \| Uncharacterized protein; afdb-uniprot50 \| AF-A0A827NCX0-F1-MODEL\_V4 \| 1.0 \| 2.724e-09 \| 323 \| 0.25 \| 168 \| 117 \| 3 \| 27 \| 193 \| 2 \| 161 \| Phage late control D family protein \| Phage late control D family protein; afdb-uniprot50 \| AF-A0A7C3BT38-F1-MODEL\_V4 \| 1.0 \| 1.028e-09 \| 322 \| 0.16 \| 225 \| 154 \| 7 \| 83 \| 281 \| 4 \| 219 \| VgrG-related protein \| VgrG-related protein; afdb-uniprot50 \| AF-A0A352UL14-F1-MODEL\_V4 \| 1.0 \| 6.555e-11 \| 321 \| 0.125 \| 286 \| 198 \| 17 \| 9 \| 274 \| 10 \| 263 \| Uncharacterized protein \| Uncharacterized protein; afdb-uniprot50 \| AF-A0A497Y796-F1-MODEL\_V4 \| 1.0 \| 6.496e-10 \| 320 \| 0.11 \| 290 \| 212 \| 16 \| 13 \| 275 \| 72 \| 342 \| Phage\_base\_V domain-containing protein \| Phage\_base\_V domain-containing protein; afdb-uniprot50 \| AF-A0A7C5MPP8-F1-MODEL\_V4 \| 1.0 \| 4.921e-11 \| 318 \| 0.133 \| 321 \| 215 \| 14 \| 7 \| 279 \| 68 \| 373 \| Phage\_base\_V domain-containing protein \| Phage\_base\_V domain-containing protein; afdb-uniprot50 \| AF-A0A4Y7RGG6-F1-MODEL\_V4 \| 1.0 \| 5.519e-11 \| 317 \| 0.15 \| 339 \| 197 \| 20 \| 7 \| 274 \| 59 \| 377 \| Phage late control gene D protein (GPD) \| Phage late control gene D protein (GPD); afdb-uniprot50 \| AF-A0A7Y0GUL4-F1-MODEL\_V4 \| 1.0 \| 2.619e-11 \| 315 \| 0.158 \| 322 \| 200 \| 16 \| 7 \| 280 \| 69 \| 367 \| Uncharacterized protein \| Uncharacterized protein; afdb-uniprot50 \| AF-A0A2N2FIC8-F1-MODEL\_V4 \| 1.0 \| 4.388e-11 \| 315 \| 0.14 \| 334 \| 208 \| 18 \| 7 \| 288 \| 69 \| 375 \| Uncharacterized protein \| Uncharacterized protein; afdb-uniprot50 \| AF-A0A371J250-F1-MODEL\_V4 \| 1.0 \| 2.273e-08 \| 314 \| 0.153 \| 124 \| 103 \| 2 \| 156 \| 279 \| 2 \| 123 \| LysM peptidoglycan-binding domain-containing protein \| LysM peptidoglycan-binding domain-containing protein; afdb-uniprot50 \| AF-G4Q993-F1-MODEL\_V4 \| 1.0 \| 6.19e-11 \| 313 \| 0.126 \| 315 \| 203 \| 16 \| 1 \| 282 \| 1 \| 276 \| Uncharacterized protein \| Uncharacterized protein; afdb-uniprot50 \| AF-A0A7Z1V5Y3-F1-MODEL\_V4 \| 1.0 \| 2.619e-11 \| 313 \| 0.134 \| 402 \| 213 \| 16 \| 13 \| 311 \| 86 \| 455 \| Uncharacterized protein \| Uncharacterized protein; afdb-uniprot50 \| AF-U2X5L0-F1-MODEL\_V4 \| 1.0 \| 9.793e-11 \| 312 \| 0.128 \| 287 \| 207 \| 16 \| 10 \| 275 \| 2 \| 266 \| Phage protein \| Phage protein; afdb-uniprot50 \| AF-A0A654C107-F1-MODEL\_V4 \| 1.0 \| 3.913e-11 \| 312 \| 0.106 \| 319 \| 216 \| 18 \| 7 \| 286 \| 103 \| 391 \| Uncharacterized protein \| Uncharacterized protein; afdb-uniprot50 \| AF-A0A2V2RGV6-F1-MODEL\_V4 \| 1.0 \| 3.055e-09 \| 311 \| 0.181 \| 160 \| 107 \| 7 \| 144 \| 281 \| 21 \| 178 \| Uncharacterized protein \| Uncharacterized protein; afdb-uniprot50 \| AF-A0A060QI63-F1-MODEL\_V4 \| 1.0 \| 1.098e-10 \| 310 \| 0.138 \| 304 \| 197 \| 19 \| 13 \| 279 \| 10 \| 285 \| Uncharacterized protein \| Uncharacterized protein; afdb-uniprot50 \| AF-A0A4D7QFP1-F1-MODEL\_V4 \| 1.0 \| 1.381e-10 \| 310 \| 0.108 \| 305 \| 225 \| 17 \| 7 \| 279 \| 21 \| 310 \| Uncharacterized protein \| Uncharacterized protein; afdb-uniprot50 \| AF-A0A433X014-F1-MODEL\_V4 \| 1.0 \| 1.549e-10 \| 308 \| 0.136 \| 300 \| 196 \| 15 \| 18 \| 279 \| 81 \| 355 \| Uncharacterized protein \| Uncharacterized protein; afdb-uniprot50 \| AF-A0A6P0SVX6-F1-MODEL\_V4 \| 1.0 \| 2.407e-08 \| 305 \| 0.161 \| 155 \| 107 \| 4 \| 148 \| 279 \| 14 \| 168 \| Uncharacterized protein \| Uncharacterized protein; afdb-uniprot50 \| AF-A4N701-F1-MODEL\_V4 \| 1.0 \| 1.174e-11 \| 305 \| 0.251 \| 282 \| 156 \| 9 \| 2 \| 281 \| 67 \| 295 \| Bacteriophage P2 gpD protein \| Bacteriophage P2 gpD protein; afdb-uniprot50 \| AF-A0A3D5LUE2-F1-MODEL\_V4 \| 1.0 \| 1.293e-09 \| 304 \| 0.202 \| 168 \| 116 \| 5 \| 148 \| 309 \| 9 \| 164 \| LysM domain-containing protein \| LysM domain-containing protein; afdb-uniprot50 \| AF-A0A5E4KK76-F1-MODEL\_V4 \| 1.0 \| 2.146e-08 \| 302 \| 0.17 \| 164 \| 108 \| 5 \| 141 \| 278 \| 13 \| 174 \| Uncharacterized protein \| Uncharacterized protein; afdb-uniprot50 \| AF-A0A011PC04-F1-MODEL\_V4 \| 1.0 \| 2.859e-08 \| 300 \| 0.135 \| 155 \| 111 \| 5 \| 146 \| 277 \| 10 \| 164 \| Phage protein D \| Phage protein D; afdb-uniprot50 \| AF-A0A511BLX4-F1-MODEL\_V4 \| 1.0 \| 1.641e-10 \| 298 \| 0.114 \| 305 \| 200 \| 13 \| 17 \| 279 \| 80 \| 356 \| Uncharacterized protein \| Uncharacterized protein; afdb-uniprot50 \| AF-A0A843SHV9-F1-MODEL\_V4 \| 1.0 \| 2.451e-10 \| 298 \| 0.171 \| 320 \| 196 \| 16 \| 7 \| 279 \| 69 \| 366 \| Uncharacterized protein \| Uncharacterized protein; afdb-uniprot50 \| AF-A0A0C2UIQ9-F1-MODEL\_V4 \| 1.0 \| 4.106e-10 \| 298 \| 0.107 \| 298 \| 221 \| 13 \| 7 \| 281 \| 74 \| 349 \| Phage-like element PBSX protein xkdQ \| Phage-like element PBSX protein xkdQ; afdb-uniprot50 \| AF-A0A089QIV9-F1-MODEL\_V4 \| 1.0 \| 2.911e-10 \| 298 \| 0.131 \| 297 \| 207 \| 19 \| 18 \| 278 \| 123 \| 404 \| Putative flagellin-like protein \| Putative flagellin-like protein; afdb-uniprot50 \| AF-A0A0V0PZL6-F1-MODEL\_V4 \| 1.0 \| 3.596e-08 \| 296 \| 0.28 \| 157 \| 106 \| 2 \| 37 \| 193 \| 3 \| 152 \| Uncharacterized protein \| Uncharacterized protein; afdb-uniprot50 \| AF-A0A1H9SQV1-F1-MODEL\_V4 \| 1.0 \| 1.037e-10 \| 296 \| 0.137 \| 335 \| 237 \| 16 \| 1 \| 311 \| 48 \| 354 \| Uncharacterized protein \| Uncharacterized protein; afdb-uniprot50 \| AF-A0A257QRW0-F1-MODEL\_V4 \| 1.0 \| 4.349e-10 \| 296 \| 0.145 \| 317 \| 198 \| 14 \| 7 \| 273 \| 71 \| 364 \| Uncharacterized protein \| Uncharacterized protein; afdb-uniprot50 \| AF-A0A376FE78-F1-MODEL\_V4 \| 1.0 \| 2.724e-09 \| 295 \| 0.195 \| 179 \| 107 \| 5 \| 150 \| 292 \| 1 \| 178 \| Gene D protein \| Gene D protein; afdb-uniprot50 \| AF-A0A3G2ECV7-F1-MODEL\_V4 \| 1.0 \| 1.641e-10 \| 294 \| 0.157 \| 323 \| 197 \| 16 \| 7 \| 280 \| 69 \| 365 \| Uncharacterized protein \| Uncharacterized protein; afdb-uniprot50 \| AF-R5RWZ2-F1-MODEL\_V4 \| 1.0 \| 7.286e-10 \| 291 \| 0.136 \| 285 \| 197 \| 17 \| 11 \| 276 \| 55 \| 309 \| Uncharacterized protein \| Uncharacterized protein; afdb-uniprot50 \| AF-A0A510KG70-F1-MODEL\_V4 \| 1.0 \| 8.732e-11 \| 289 \| 0.155 \| 327 \| 212 \| 14 \| 9 \| 311 \| 47 \| 333 \| Uncharacterized protein \| Uncharacterized protein; afdb-uniprot50 \| AF-A0A5R8Q954-F1-MODEL\_V4 \| 1.0 \| 4.106e-10 \| 287 \| 0.141 \| 303 \| 203 \| 16 \| 7 \| 275 \| 35 \| 314 \| Uncharacterized protein \| Uncharacterized protein; afdb-uniprot50 \| AF-A0A4P9VGZ3-F1-MODEL\_V4 \| 1.0 \| 1.21e-08 \| 286 \| 0.361 \| 141 \| 83 \| 3 \| 26 \| 166 \| 6 \| 139 \| Uncharacterized protein \| Uncharacterized protein; afdb-uniprot50 \| AF-A0A1M4LET7-F1-MODEL\_V4 \| 1.0 \| 3.055e-09 \| 285 \| 0.212 \| 174 \| 111 \| 5 \| 160 \| 311 \| 50 \| 219 \| Uncharacterized protein \| Uncharacterized protein; afdb-uniprot50 \| AF-A0A536FPD7-F1-MODEL\_V4 \| 1.0 \| 3.628e-09 \| 284 \| 0.154 \| 233 \| 165 \| 8 \| 76 \| 281 \| 2 \| 229 \| VgrG-related protein \| VgrG-related protein; afdb-uniprot50 \| AF-R0KRW8-F1-MODEL\_V4 \| 1.0 \| 1.132e-07 \| 283 \| 0.311 \| 138 \| 86 \| 3 \| 37 \| 173 \| 4 \| 133 \| Gene D protein \| Gene D protein; afdb-uniprot50 \| AF-A0A556RSP6-F1-MODEL\_V4 \| 1.0 \| 5.42e-09 \| 282 \| 0.239 \| 163 \| 99 \| 6 \| 1 \| 146 \| 14 \| 168 \| Uncharacterized protein \| Uncharacterized protein; afdb-uniprot50 \| AF-A0A7C5ITK4-F1-MODEL\_V4 \| 1.0 \| 4.523e-08 \| 281 \| 0.161 \| 192 \| 131 \| 6 \| 106 \| 275 \| 4 \| 187 \| Type VI secretion system tip protein VgrG \| Type VI secretion system tip protein VgrG; afdb-uniprot50 \| AF-A0A522CNZ2-F1-MODEL\_V4 \| 1.0 \| 1.626e-09 \| 279 \| 0.143 \| 279 \| 204 \| 17 \| 9 \| 279 \| 52 \| 303 \| Uncharacterized protein \| Uncharacterized protein; afdb-uniprot50 \| AF-A0A2C7A780-F1-MODEL\_V4 \| 1.0 \| 2.749e-10 \| 278 \| 0.134 \| 431 \| 222 \| 14 \| 7 \| 311 \| 69 \| 474 \| Uncharacterized protein \| Uncharacterized protein; afdb-uniprot50 \| AF-A0A372IVE0-F1-MODEL\_V4 \| 1.0 \| 1.706e-08 \| 277 \| 0.198 \| 207 \| 138 \| 7 \| 102 \| 281 \| 10 \| 215 \| Type IV secretion protein Rhs \| Type IV secretion protein Rhs; afdb-uniprot50 \| AF-A0A2S0JG92-F1-MODEL\_V4 \| 1.0 \| 4.605e-10 \| 274 \| 0.135 \| 273 \| 193 \| 12 \| 20 \| 275 \| 64 \| 310 \| Uncharacterized protein \| Uncharacterized protein; afdb-uniprot50 \| AF-A0A7Z0PR13-F1-MODEL\_V4 \| 1.0 \| 1.641e-10 \| 274 \| 0.111 \| 313 \| 228 \| 18 \| 9 \| 300 \| 60 \| 343 \| Uncharacterized protein \| Uncharacterized protein; afdb-uniprot50 \| AF-R5KQT4-F1-MODEL\_V4 \| 1.0 \| 2.596e-10 \| 273 \| 0.128 \| 311 \| 204 \| 19 \| 9 \| 293 \| 49 \| 318 \| Uncharacterized protein \| Uncharacterized protein; afdb-uniprot50 \| AF-A0A520HRV1-F1-MODEL\_V4 \| 1.0 \| 3.027e-08 \| 273 \| 0.135 \| 244 \| 173 \| 12 \| 65 \| 282 \| 2 \| 233 \| Phage\_base\_V domain-containing protein \| Phage\_base\_V domain-containing protein; afdb-uniprot50 \| AF-B9NXS0-F1-MODEL\_V4 \| 1.0 \| 1.931e-09 \| 272 \| 0.309 \| 126 \| 83 \| 3 \| 157 \| 281 \| 5 \| 127 \| Phage tail protein D \| Phage tail protein D; afdb-uniprot50 \| AF-A0A7U6BTU5-F1-MODEL\_V4 \| 1.0 \| 5.47e-10 \| 272 \| 0.137 \| 328 \| 228 \| 16 \| 1 \| 311 \| 53 \| 342 \| NlpC/P60 domain-containing protein \| NlpC/P60 domain-containing protein; afdb-uniprot50 \| AF-A0A6L4A988-F1-MODEL\_V4 \| 1.0 \| 3.206e-08 \| 271 \| 0.16 \| 218 \| 148 \| 7 \| 84 \| 278 \| 2 \| 207 \| VgrG-related protein \| VgrG-related protein; afdb-uniprot50 \| AF-A0A3E2B4M2-F1-MODEL\_V4 \| 1.0 \| 2.294e-09 \| 270 \| 0.125 \| 296 \| 196 \| 17 \| 9 \| 277 \| 53 \| 312 \| Uncharacterized protein \| Uncharacterized protein; afdb-uniprot50 \| AF-A0A535AFH3-F1-MODEL\_V4 \| 1.0 \| 7.22e-09 \| 270 \| 0.145 \| 227 \| 155 \| 9 \| 85 \| 281 \| 2 \| 219 \| VgrG-related protein \| VgrG-related protein; afdb-uniprot50 \| AF-A0A0P9G0N8-F1-MODEL\_V4 \| 1.0 \| 3.177e-07 \| 269 \| 0.187 \| 144 \| 99 \| 2 \| 153 \| 278 \| 1 \| 144 \| Uncharacterized protein \| Uncharacterized protein; afdb-uniprot50 \| AF-A0A1C2Y2A6-F1-MODEL\_V4 \| 1.0 \| 1.722e-09 \| 269 \| 0.142 \| 288 \| 191 \| 16 \| 20 \| 276 \| 66 \| 328 \| Uncharacterized protein \| Uncharacterized protein; afdb-uniprot50 \| AF-U2C6T1-F1-MODEL\_V4 \| 1.0 \| 1.437e-08 \| 266 \| 0.148 \| 235 \| 161 \| 13 \| 61 \| 277 \| 18 \| 231 \| Uncharacterized protein \| Uncharacterized protein; afdb-uniprot50 \| AF-A0A1C5QMY9-F1-MODEL\_V4 \| 1.0 \| 1.079e-08 \| 265 \| 0.113 \| 290 \| 202 \| 19 \| 11 \| 277 \| 55 \| 312 \| Mu-like prophage tail protein gpP \| Mu-like prophage tail protein gpP; afdb-uniprot50 \| AF-A0A6B3GF16-F1-MODEL\_V4 \| 1.0 \| 6.38e-08 \| 263 \| 0.176 \| 210 \| 130 \| 10 \| 106 \| 280 \| 7 \| 208 \| VgrG-related protein \| VgrG-related protein; afdb-uniprot50 \| AF-A0A3B8Q5W1-F1-MODEL\_V4 \| 1.0 \| 5.688e-08 \| 263 \| 0.101 \| 177 \| 138 \| 5 \| 106 \| 275 \| 1 \| 163 \| Rhs element Vgr protein \| Rhs element Vgr protein; afdb-uniprot50 \| AF-Q67SN3-F1-MODEL\_V4 \| 1.0 \| 3.206e-08 \| 262 \| 0.092 \| 388 \| 221 \| 22 \| 7 \| 279 \| 309 \| 680 \| Uncharacterized protein \| Uncharacterized protein; afdb-uniprot50 \| AF-A0A3P1WM06-F1-MODEL\_V4 \| 1.0 \| 8.838e-06 \| 261 \| 0.277 \| 90 \| 64 \| 1 \| 57 \| 146 \| 12 \| 100 \| Phage late control D family protein \| Phage late control D family protein; afdb-uniprot50 \| AF-A0A1W9WK55-F1-MODEL\_V4 \| 1.0 \| 9.081e-09 \| 261 \| 0.147 \| 305 \| 174 \| 17 \| 9 \| 281 \| 259 \| 509 \| Phage\_base\_V domain-containing protein \| Phage\_base\_V domain-containing protein; afdb-uniprot50 \| AF-A0A2G6AV19-F1-MODEL\_V4 \| 1.0 \| 4.605e-10 \| 259 \| 0.131 \| 305 \| 209 \| 19 \| 19 \| 300 \| 75 \| 346 \| NlpC/P60 domain-containing protein \| NlpC/P60 domain-containing protein; afdb-uniprot50 \| AF-A0A1F7WZV6-F1-MODEL\_V4 \| 1.0 \| 1.21e-08 \| 259 \| 0.117 \| 212 \| 151 \| 9 \| 105 \| 288 \| 862 \| 1065 \| Uncharacterized protein \| Uncharacterized protein; afdb-uniprot50 \| AF-A0A523ZZ81-F1-MODEL\_V4 \| 1.0 \| 6.695e-07 \| 253 \| 0.13 \| 161 \| 116 \| 5 \| 145 \| 281 \| 16 \| 176 \| Phage\_base\_V domain-containing protein \| Phage\_base\_V domain-containing protein; afdb-uniprot50 \| AF-W4M5L5-F1-MODEL\_V4 \| 1.0 \| 2.232e-06 \| 252 \| 0.154 \| 149 \| 103 \| 2 \| 154 \| 279 \| 4 \| 152 \| Uncharacterized protein \| Uncharacterized protein; afdb-uniprot50 \| AF-A0A2G2BDD8-F1-MODEL\_V4 \| 1.0 \| 2.294e-09 \| 250 \| 0.352 \| 119 \| 77 \| 0 \| 161 \| 279 \| 62 \| 180 \| Uncharacterized protein \| Uncharacterized protein; afdb-uniprot50 \| AF-A0A539E6J3-F1-MODEL\_V4 \| 1.0 \| 1.344e-07 \| 250 \| 0.131 \| 183 \| 131 \| 9 \| 114 \| 275 \| 4 \| 179 \| Rhs element Vgr protein \| Rhs element Vgr protein; afdb-uniprot50 \| AF-A0A3C1JKA1-F1-MODEL\_V4 \| 1.0 \| 9.53e-08 \| 250 \| 0.143 \| 195 \| 128 \| 9 \| 114 \| 279 \| 2 \| 186 \| Phage\_base\_V domain-containing protein \| Phage\_base\_V domain-containing protein; afdb-uniprot50 \| AF-A0A0P0E7V7-F1-MODEL\_V4 \| 1.0 \| 2.859e-08 \| 249 \| 0.103 \| 289 \| 213 \| 16 \| 18 \| 279 \| 345 \| 614 \| Uncharacterized protein \| Uncharacterized protein; afdb-uniprot50 \| AF-A0A7G9B2C8-F1-MODEL\_V4 \| 1.0 \| 1.088e-09 \| 248 \| 0.155 \| 290 \| 186 \| 18 \| 11 \| 276 \| 55 \| 309 \| Uncharacterized protein \| Uncharacterized protein; afdb-uniprot50 \| AF-A0A859FGK9-F1-MODEL\_V4 \| 1.0 \| 3.055e-09 \| 247 \| 0.145 \| 316 \| 216 \| 18 \| 17 \| 311 \| 72 \| 354 \| Uncharacterized protein \| Uncharacterized protein; afdb-uniprot50 \| AF-A0A220MH41-F1-MODEL\_V4 \| 1.0 \| 5.072e-08 \| 246 \| 0.108 \| 285 \| 202 \| 16 \| 19 \| 276 \| 65 \| 324 \| Uncharacterized protein \| Uncharacterized protein; afdb-uniprot50 \| AF-D4CX49-F1-MODEL\_V4 \| 1.0 \| 1.931e-09 \| 246 \| 0.155 \| 334 \| 217 \| 18 \| 1 \| 310 \| 42 \| 334 \| Uncharacterized protein \| Uncharacterized protein; afdb-uniprot50 \| AF-A0A318XLJ8-F1-MODEL\_V4 \| 1.0 \| 5.072e-08 \| 245 \| 0.091 \| 296 \| 213 \| 19 \| 18 \| 279 \| 99 \| 372 \| Uncharacterized protein \| Uncharacterized protein; afdb-uniprot50 \| AF-A0A6I1I493-F1-MODEL\_V4 \| 1.0 \| 1.199e-07 \| 244 \| 0.328 \| 146 \| 89 \| 3 \| 22 \| 166 \| 2 \| 139 \| Uncharacterized protein \| Uncharacterized protein; afdb-uniprot50 \| AF-A0A352X1L2-F1-MODEL\_V4 \| 1.0 \| 4.271e-08 \| 244 \| 0.129 \| 293 \| 192 \| 25 \| 14 \| 279 \| 53 \| 309 \| Uncharacterized protein \| Uncharacterized protein; afdb-uniprot50 \| AF-J1H9W8-F1-MODEL\_V4 \| 1.0 \| 8.498e-08 \| 242 \| 0.14 \| 220 \| 162 \| 9 \| 73 \| 275 \| 2 \| 211 \| Uncharacterized protein \| Uncharacterized protein; afdb-uniprot50 \| AF-A0A519GJE0-F1-MODEL\_V4 \| 1.0 \| 2.833e-07 \| 241 \| 0.171 \| 198 \| 133 \| 9 \| 106 \| 280 \| 4 \| 193 \| Type VI secretion system tip protein VgrG \| Type VI secretion system tip protein VgrG; afdb-uniprot50 \| AF-A0A2E4F7X2-F1-MODEL\_V4 \| 1.0 \| 5.026e-07 \| 239 \| 0.143 \| 174 \| 116 \| 5 \| 141 \| 281 \| 3 \| 176 \| Uncharacterized protein \| Uncharacterized protein; afdb-uniprot50 \| AF-A0A3A4U4Y7-F1-MODEL\_V4 \| 1.0 \| 3.808e-08 \| 238 \| 0.16 \| 268 \| 165 \| 12 \| 37 \| 281 \| 446 \| 676 \| Uncharacterized protein \| Uncharacterized protein; afdb-uniprot50 \| AF-A0A7Z0MMM6-F1-MODEL\_V4 \| 1.0 \| 2.127e-07 \| 235 \| 0.117 \| 179 \| 134 \| 7 \| 110 \| 276 \| 26 \| 192 \| Uncharacterized protein \| Uncharacterized protein; afdb-uniprot50 \| AF-A0A837LG98-F1-MODEL\_V4 \| 1.0 \| 6.438e-09 \| 233 \| 0.134 \| 312 \| 193 \| 20 \| 23 \| 278 \| 74 \| 364 \| Type IV secretion protein Rhs \| Type IV secretion protein Rhs; afdb-uniprot50 \| AF-A0A0L6JT74-F1-MODEL\_V4 \| 1.0 \| 3.206e-08 \| 233 \| 0.102 \| 381 \| 213 \| 18 \| 11 \| 279 \| 451 \| 814 \| Uncharacterized protein \| Uncharacterized protein; afdb-uniprot50 \| AF-A0A2E4HHK0-F1-MODEL\_V4 \| 1.0 \| 1.775e-06 \| 232 \| 0.168 \| 154 \| 104 \| 6 \| 144 \| 275 \| 7 \| 158 \| Phage\_base\_V domain-containing protein \| Phage\_base\_V domain-containing protein; afdb-uniprot50 \| AF-A0A0P9YAT2-F1-MODEL\_V4 \| 1.0 \| 8.838e-06 \| 231 \| 0.263 \| 110 \| 75 \| 2 \| 172 \| 275 \| 2 \| 111 \| Tail protein D \| Tail protein D; afdb-uniprot50 \| AF-A0A845L7G9-F1-MODEL\_V4 \| 1.0 \| 6.818e-09 \| 229 \| 0.118 \| 305 \| 205 \| 17 \| 7 \| 274 \| 44 \| 321 \| Uncharacterized protein \| Uncharacterized protein; afdb-uniprot50 \| AF-A0A1H9HAT7-F1-MODEL\_V4 \| 1.0 \| 2.027e-08 \| 228 \| 0.111 \| 313 \| 207 \| 18 \| 3 \| 276 \| 26 \| 306 \| Uncharacterized protein \| Uncharacterized protein; afdb-uniprot50 \| AF-A0A061P2C1-F1-MODEL\_V4 \| 1.0 \| 3.365e-07 \| 222 \| 0.101 \| 305 \| 215 \| 21 \| 8 \| 275 \| 28 \| 310 \| Uncharacterized protein \| Uncharacterized protein; afdb-uniprot50 \| AF-A0A829CIG2-F1-MODEL\_V4 \| 1.0 \| 8.345e-06 \| 221 \| 0.272 \| 125 \| 86 \| 4 \| 63 \| 185 \| 1 \| 122 \| Phage late control D family protein \| Phage late control D family protein; afdb-uniprot50 \| AF-A0A2V7V433-F1-MODEL\_V4 \| 1.0 \| 4.271e-08 \| 214 \| 0.18 \| 205 \| 124 \| 8 \| 105 \| 277 \| 1 \| 193 \| Phage\_base\_V domain-containing protein \| Phage\_base\_V domain-containing protein; afdb-uniprot50 \| AF-A0A833G3K2-F1-MODEL\_V4 \| 1.0 \| 2.089e-05 \| 209 \| 0.248 \| 137 \| 92 \| 6 \| 20 \| 152 \| 66 \| 195 \| Late control D family protein \| Late control D family protein; afdb-uniprot50 \| AF-A0A265Q3H3-F1-MODEL\_V4 \| 1.0 \| 2.675e-07 \| 201 \| 0.14 \| 242 \| 151 \| 13 \| 69 \| 275 \| 1 \| 220 \| Uncharacterized protein \| Uncharacterized protein; afdb-uniprot50 \| AF-A0A4U9KC87-F1-MODEL\_V4 \| 1.0 \| 1.879e-06 \| 201 \| 0.139 \| 165 \| 108 \| 12 \| 144 \| 282 \| 17 \| 173 \| Phage protein D \| Phage protein D; afdb-uniprot50 \| AF-A0A8B4Q938-F1-MODEL\_V4 \| 1.0 \| 1.508e-07 \| 200 \| 0.109 \| 319 \| 201 \| 17 \| 7 \| 276 \| 35 \| 319 \| Uncharacterized protein \| Uncharacterized protein; afdb-uniprot50 \| AF-A0A2S5HQZ7-F1-MODEL\_V4 \| 1.0 \| 9.444e-07 \| 199 \| 0.118 \| 363 \| 205 \| 23 \| 9 \| 278 \| 808 \| 1148 \| Peptidase M23 \| Peptidase M23; afdb-uniprot50 \| AF-A0A2L2X7P9-F1-MODEL\_V4 \| 1.0 \| 1.188e-06 \| 198 \| 0.123 \| 218 \| 157 \| 12 \| 88 \| 278 \| 821 \| 1031 \| Peptidase, M23/M37 family \| Peptidase, M23/M37 family; afdb-uniprot50 \| AF-A0A2G2BI84-F1-MODEL\_V4 \| 1.0 \| 6.265e-06 \| 195 \| 0.38 \| 92 \| 54 \| 1 \| 191 \| 279 \| 25 \| 116 \| Uncharacterized protein \| Uncharacterized protein; afdb-uniprot50 \| AF-A0A3D0XCR8-F1-MODEL\_V4 \| 1.0 \| 3.773e-07 \| 193 \| 0.114 \| 324 \| 191 \| 26 \| 7 \| 279 \| 34 \| 312 \| Uncharacterized protein \| Uncharacterized protein; afdb-uniprot50 \| AF-A0A7H0YH63-F1-MODEL\_V4 \| 1.0 \| 1.582e-06 \| 192 \| 0.098 \| 355 \| 213 \| 19 \| 9 \| 279 \| 687 \| 1018 \| M23 family metallopeptidase \| M23 family metallopeptidase; afdb-uniprot50 \| AF-A0A3C1CUM3-F1-MODEL\_V4 \| 1.0 \| 3.334e-06 \| 189 \| 0.114 \| 289 \| 204 \| 24 \| 9 \| 276 \| 49 \| 306 \| Uncharacterized protein \| Uncharacterized protein; afdb-uniprot50 \| AF-A0A7M2GS28-F1-MODEL\_V4 \| 1.0 \| 1e-06 \| 187 \| 0.169 \| 165 \| 108 \| 8 \| 147 \| 284 \| 157 \| 319 \| Uncharacterized protein \| Uncharacterized protein; afdb-uniprot50 \| AF-A0A2R8BS21-F1-MODEL\_V4 \| 1.0 \| 0.0002603 \| 177 \| 0.268 \| 93 \| 68 \| 0 \| 189 \| 281 \| 11 \| 103 \| Uncharacterized protein \| Uncharacterized protein; afdb-uniprot50 \| AF-A0A662L4J5-F1-MODEL\_V4 \| 1.0 \| 0.0001167 \| 176 \| 0.131 \| 137 \| 93 \| 4 \| 166 \| 278 \| 1 \| 135 \| Type IV secretion protein Rhs \| Type IV secretion protein Rhs; afdb-uniprot50 \| AF-A0A0F9MNU2-F1-MODEL\_V4 \| 1.0 \| 5.863e-05 \| 172 \| 0.126 \| 182 \| 136 \| 7 \| 108 \| 275 \| 610 \| 782 \| Uncharacterized protein \| Uncharacterized protein; afdb-uniprot50 \| AF-A0A1I3WTN1-F1-MODEL\_V4 \| 1.0 \| 0.0002458 \| 164 \| 0.149 \| 154 \| 108 \| 3 \| 146 \| 277 \| 83 \| 235 \| Phage\_base\_V domain-containing protein \| Phage\_base\_V domain-containing protein; afdb-uniprot50 \| AF-A0A431HHS2-F1-MODEL\_V4 \| 1.0 \| 1.759e-05 \| 161 \| 0.095 \| 325 \| 204 \| 21 \| 22 \| 275 \| 728 \| 1033 \| Uncharacterized protein \| Uncharacterized protein; afdb-uniprot50 \| AF-A0A6P0J1U6-F1-MODEL\_V4 \| 1.0 \| 0.0009733 \| 145 \| 0.141 \| 134 \| 87 \| 3 \| 171 \| 279 \| 2 \| 132 \| Uncharacterized protein \| Uncharacterized protein; afdb-uniprot50 \| AF-A0A316PEN3-F1-MODEL\_V4 \| 1.0 \| 6.575e-05 \| 128 \| 0.126 \| 142 \| 106 \| 5 \| 153 \| 291 \| 340 \| 466 \| Uncharacterized protein \| Uncharacterized protein; afdb-uniprot50 \| AF-A0A7C1Y5F9-F1-MODEL\_V4 \| 1.0 \| 0.0002757 \| 125 \| 0.152 \| 197 \| 117 \| 6 \| 157 \| 311 \| 2 \| 190 \| Uncharacterized protein \| Uncharacterized protein; afdb-uniprot50 \| AF-A0A0F9L0Y6-F1-MODEL\_V4 \| 1.0 \| 0.001937 \| 118 \| 0.117 \| 170 \| 123 \| 7 \| 147 \| 302 \| 12 \| 168 \| Uncharacterized protein \| Uncharacterized protein; afdb-uniprot50 \| AF-A0A1V5V4Y2-F1-MODEL\_V4 \| 1.0 \| 0.006838 \| 115 \| 0.215 \| 93 \| 65 \| 4 \| 224 \| 311 \| 74 \| 163 \| Uncharacterized protein \| Uncharacterized protein; afdb-uniprot50 \| AF-A0A0F9QPT5-F1-MODEL\_V4 \| 1.0 \| 0.001373 \| 110 \| 0.078 \| 395 \| 169 \| 16 \| 59 \| 275 \| 13 \| 390 \| Uncharacterized protein \| Uncharacterized protein; afdb-uniprot50 \| AF-A0A1V5LRQ3-F1-MODEL\_V4 \| 1.0 \| 0.005133 \| 101 \| 0.185 \| 124 \| 94 \| 5 \| 191 \| 311 \| 6 \| 125 \| Uncharacterized protein \| Uncharacterized protein |
| Top keywords  (threshold 1.00e-02 (evalue)) | **D, Phage, late, control, Type, secretion, Rhs, domain\_containing, tail, gene** |
| Output files | ../../similar\_structures/25\_FANPEZAQ\_CDS\_0025\_afdb-proteome\_foldseek.tsv ../../similar\_structures/25\_FANPEZAQ\_CDS\_0025\_afdb-uniprot50\_foldseek.tsv ../../similar\_structures/25\_FANPEZAQ\_CDS\_0025\_merged.svg ../../similar\_structures/25\_FANPEZAQ\_CDS\_0025\_pdb\_foldseek.tsv |

  
  
  

Return to summary | Go to previous | Go to next

  


---

**Sequence/structure alignments coloring**  
Each object in the alignment figures is colored according to its E-value following this color coding:

1e-100
10
