## Supplementary material for "Completing the BASEL phage collection to unlock hidden diversity for systematic exploration of phage-host interactions": Data S2: 27.html

FANPEZAQ\_CDS\_0027


Return to summary | Go to previous | Go to next

|  |  |
| --- | --- |
| FANPEZAQ\_CDS\_0027  Page creation date: 02 Sep 2024, 12:00  Project folder: n/a  Input sequences file: Escherichia\_virus\_HeidiAbel.gb | hypothetical domain\_containing duf551 duf6378 morphogenetic fragment restriction alleviation lar ky705409\_p32 nc\_031918\_p6 mg676466\_p62 nc\_004333\_p63 mf153391\_p29 he956707\_p19 sav\_like kow hnh endonuclease duf3310 cupin duf3085 datp dgtp diphosphohydrolase n\_terminal duf4326 jacalin\_type lectin 3'\_phosphatase 5'\_polynucleotide kinase phage duf2203 |

### Sequence information

|  |  |
| --- | --- |
| Name | FANPEZAQ\_CDS\_0027  27\_FANPEZAQ\_CDS\_0027 (pipeline id) |
| Imported annotations | Escherichia\_virus\_HeidiAbel Bas97 |
| Protein sequence | MNTSNKTLLQILVEELPKRGGWPKGAVKIHQDHDGETWAWLNHDGTERLFKLDQVAKNAR QFAKDETEENMVTRDQYESALAEKEHPLPWDEKNTDADGWISWPGGECPVDGDAIVEVKY RKPNPYQFNNDRAGDFTWSHDGFDGDIIAYRLPHPHEVKTAGIDVQRDAVKSNDAQLEAD LNDCIGQPQSNDYARDEILYELANFAATTKTSLDIGVATEIAIWLSGKGYRKQ |
| Number of residues | 233 |
| Molecular weight (Da) | 26353.72 |
| Output files | ../../query\_sequences/27\_FANPEZAQ\_CDS\_0027.fasta |

### Putative domain architecture and protein family

#### Search results (HHblits)^1^

|  |  |
| --- | --- |
| Domain family databases searched | Pfam, Ncbi-cd, Cath, Phrogs |
| Results, scheme(s)  (Top layers only; threshold 1.00e-03 (evalue)) | xml version="1.0" encoding="utf-8" standalone="no"?       2024-09-02T21:08:18.031237 image/svg+xml   Matplotlib v3.7.2, https://matplotlib.org/ |
| Results, table  (E-value ≤ 1.00e-03 (evalue)) | db \| id \| prob \| evalue \| pvalue \| score \| cols \| query \| query\_len \| template \| template\_len \| name \| description; phrogs \| 26870 \| 100.0 \| 2.9e-80 \| 3.4e-84 \| 531.8 \| 218 \| (1, 233) \| 233 \| (1, 233) \| 233 \| NA \| NA; Category: unknown function; KY705409\_p32; phrogs \| 7798 \| 99.9 \| 4.6e-34 \| 5.7e-38 \| 244.2 \| 107 \| (5, 126) \| 233 \| (10, 125) \| 211 \| NA \| NA; Category: unknown function; NC\_031918\_p6; phrogs \| 9174 \| 99.9 \| 7.4e-34 \| 9.2e-38 \| 226.9 \| 104 \| (92, 217) \| 233 \| (5, 115) \| 124 \| NA \| NA; Category: unknown function; MG676466\_p62; phrogs \| 16943 \| 99.5 \| 2.4e-18 \| 2.7e-22 \| 139.8 \| 74 \| (80, 156) \| 233 \| (92, 165) \| 166 \| NA \| NA; Category: unknown function; NC\_004333\_p63; phrogs \| 34383 \| 96.6 \| 4.9e-06 \| 5.5e-10 \| 60.3 \| 34 \| (6, 39) \| 233 \| (1, 34) \| 85 \| NA \| NA; Category: unknown function; MF153391\_p29; phrogs \| 13874 \| 95.5 \| 0.00013 \| 1.6e-08 \| 52.0 \| 57 \| (100, 157) \| 233 \| (4, 69) \| 75 \| NA \| NA; Category: unknown function; HE956707\_p19 |
| Top keywords  (threshold 1.00e-03 (evalue)) | **KY705409\_p32, NC\_031918\_p6, MG676466\_p62, NC\_004333\_p63, MF153391\_p29, HE956707\_p19** |
| Output files | ../../domain\_architecture/27\_FANPEZAQ\_CDS\_0027\_cath.hhr ../../domain\_architecture/27\_FANPEZAQ\_CDS\_0027\_merged.svg ../../domain\_architecture/27\_FANPEZAQ\_CDS\_0027\_ncbi-cd.hhr ../../domain\_architecture/27\_FANPEZAQ\_CDS\_0027\_pfam.hhr ../../domain\_architecture/27\_FANPEZAQ\_CDS\_0027\_phrogs.hhr |

### Identical protein sequences/structures

#### Search results

|  |  |
| --- | --- |
| Protein sequence databases searched | Pdb, Swissprot, Refseq |
| Identical proteins found | -- |
| Top keywords | -- |
| Output files | -- |

### Similar protein sequences/structures

#### Sequence similarity search results (HHblits)^1^

|  |  |
| --- | --- |
| Sequence databases searched | Uniclust, Pdb70 |
| Results, scheme(s)  (Top layers only, threshold 1.00e-03 (evalue)) | xml version="1.0" encoding="utf-8" standalone="no"?       2024-09-02T21:08:40.951344 image/svg+xml   Matplotlib v3.7.2, https://matplotlib.org/ |
| Results, table(s)  (threshold 1.00e-03 (evalue)) | db \| id \| prob \| evalue \| pvalue \| score \| cols \| query \| query\_len \| template \| template\_len \| name \| description; uniclust \| UniRef100\_A0A193GZ03 \| 100.0 \| 1.6e-69 \| 3.3e-75 \| 467.6 \| 217 \| (1, 233) \| 233 \| (1, 230) \| 230 \| Uncharacterized protein \| Uncharacterized protein; uniclust \| UniRef100\_A0A088FV72 \| 100.0 \| 1.4e-53 \| 2.6e-59 \| 347.9 \| 141 \| (93, 233) \| 233 \| (19, 159) \| 159 \| Uncharacterized protein \| Uncharacterized protein; uniclust \| UniRef100\_A0A291AXI6 \| 100.0 \| 1.8e-50 \| 4.8e-56 \| 368.0 \| 159 \| (5, 191) \| 233 \| (4, 166) \| 274 \| Uncharacterized protein \| Uncharacterized protein; uniclust \| UniRef100\_A0A173GC52 \| 99.9 \| 4.3e-31 \| 1.3e-36 \| 234.7 \| 101 \| (1, 123) \| 233 \| (7, 108) \| 201 \| Uncharacterized protein \| Uncharacterized protein; uniclust \| UniRef100\_A0A7D7FL43 \| 99.9 \| 1.3e-30 \| 2.4e-36 \| 209.9 \| 122 \| (6, 153) \| 233 \| (1, 131) \| 131 \| Uncharacterized protein \| Uncharacterized protein; uniclust \| UniRef100\_A0A1X1C2R2 \| 99.8 \| 4.6e-25 \| 1.1e-30 \| 184.9 \| 88 \| (69, 162) \| 233 \| (24, 112) \| 132 \| Uncharacterized protein \| Uncharacterized protein; uniclust \| UniRef100\_A0A1I9SES2 \| 99.8 \| 6.6e-23 \| 1.6e-28 \| 180.1 \| 95 \| (6, 124) \| 233 \| (1, 99) \| 185 \| Uncharacterized protein \| Uncharacterized protein; uniclust \| UniRef100\_UPI000A1DC278 \| 99.7 \| 5.3e-21 \| 9.6e-27 \| 156.3 \| 101 \| (6, 124) \| 233 \| (1, 106) \| 133 \| hypothetical protein \| hypothetical protein; uniclust \| UniRef100\_UPI000F510781 \| 99.6 \| 1.3e-18 \| 2.5e-24 \| 135.8 \| 86 \| (6, 119) \| 233 \| (2, 87) \| 95 \| hypothetical protein \| hypothetical protein; uniclust \| UniRef100\_A0A6G5Y9E3 \| 99.5 \| 2.2e-17 \| 4.1e-23 \| 141.5 \| 89 \| (75, 190) \| 233 \| (3, 91) \| 181 \| Uncharacterized protein \| Uncharacterized protein; uniclust \| UniRef100\_A0A513ZYB6 \| 99.5 \| 2.6e-17 \| 4.7e-23 \| 136.9 \| 85 \| (76, 162) \| 233 \| (54, 141) \| 143 \| Uncharacterized protein \| Uncharacterized protein; uniclust \| UniRef100\_A0A653TFN3 \| 99.4 \| 3.7e-16 \| 7.7e-22 \| 138.6 \| 85 \| (76, 162) \| 233 \| (113, 198) \| 200 \| Uncharacterized protein \| Uncharacterized protein; uniclust \| UniRef100\_UPI0005C660F9 \| 99.4 \| 4.2e-16 \| 8.5e-22 \| 123.1 \| 82 \| (3, 87) \| 233 \| (1, 86) \| 91 \| hypothetical protein \| hypothetical protein; uniclust \| UniRef100\_J3DEC6 \| 99.4 \| 4.3e-16 \| 8.7e-22 \| 133.0 \| 86 \| (71, 162) \| 233 \| (27, 149) \| 151 \| DUF551 domain-containing protein \| DUF551 domain-containing protein; uniclust \| UniRef100\_A0A2E7P804 \| 99.4 \| 6.7e-16 \| 1.5e-21 \| 122.7 \| 65 \| (96, 162) \| 233 \| (12, 82) \| 87 \| Uncharacterized protein \| Uncharacterized protein; uniclust \| UniRef100\_A0A3B7DLK8 \| 99.4 \| 6.9e-16 \| 1.8e-21 \| 119.9 \| 62 \| (93, 156) \| 233 \| (4, 71) \| 74 \| Uncharacterized protein \| Uncharacterized protein; uniclust \| UniRef100\_UPI000D50E511 \| 99.4 \| 2.8e-15 \| 5.1e-21 \| 139.1 \| 116 \| (95, 216) \| 233 \| (120, 249) \| 348 \| hypothetical protein \| hypothetical protein; uniclust \| UniRef100\_A0A5J6TWV0 \| 99.3 \| 9.6e-15 \| 1.8e-20 \| 129.9 \| 190 \| (11, 221) \| 233 \| (9, 211) \| 231 \| Uncharacterized protein \| Uncharacterized protein; uniclust \| UniRef100\_A0A411H6I8 \| 99.3 \| 2.1e-14 \| 4.2e-20 \| 133.1 \| 97 \| (94, 216) \| 233 \| (63, 176) \| 280 \| DUF551 domain-containing protein \| DUF551 domain-containing protein; uniclust \| UniRef100\_A0A6J5LK44 \| 99.3 \| 2.4e-14 \| 4.6e-20 \| 122.3 \| 94 \| (97, 205) \| 233 \| (5, 102) \| 155 \| SaV-like \| SaV-like; uniclust \| UniRef100\_A0A7Y7SSS1 \| 99.2 \| 4.9e-14 \| 9.5e-20 \| 123.4 \| 140 \| (5, 158) \| 233 \| (32, 181) \| 183 \| Uncharacterized protein \| Uncharacterized protein; uniclust \| UniRef100\_A0A6G5YAH6 \| 99.2 \| 5.3e-14 \| 9.8e-20 \| 116.6 \| 62 \| (92, 155) \| 233 \| (70, 131) \| 131 \| Uncharacterized protein \| Uncharacterized protein; uniclust \| UniRef100\_A0A2S1PDQ1 \| 99.2 \| 5.5e-14 \| 1e-19 \| 132.1 \| 150 \| (11, 172) \| 233 \| (44, 215) \| 347 \| KOW domain-containing protein \| KOW domain-containing protein; uniclust \| UniRef100\_UPI0018DF4812 \| 99.2 \| 6.3e-14 \| 1.2e-19 \| 118.6 \| 97 \| (98, 206) \| 233 \| (2, 100) \| 144 \| hypothetical protein \| hypothetical protein; uniclust \| UniRef100\_A0A6A6K281 \| 99.2 \| 7.7e-14 \| 1.4e-19 \| 125.2 \| 93 \| (5, 123) \| 233 \| (65, 158) \| 244 \| Uncharacterized protein \| Uncharacterized protein; uniclust \| UniRef100\_A0A2E7P1H3 \| 99.2 \| 8.7e-14 \| 1.6e-19 \| 109.2 \| 62 \| (95, 157) \| 233 \| (29, 90) \| 91 \| Uncharacterized protein \| Uncharacterized protein; uniclust \| UniRef100\_UPI000681C15B \| 99.2 \| 1e-13 \| 2.3e-19 \| 124.3 \| 92 \| (6, 124) \| 233 \| (1, 92) \| 190 \| hypothetical protein \| hypothetical protein; uniclust \| UniRef100\_UPI0023312D73 \| 99.2 \| 1.8e-13 \| 3.3e-19 \| 110.8 \| 75 \| (6, 81) \| 233 \| (1, 83) \| 110 \| hypothetical protein \| hypothetical protein; uniclust \| UniRef100\_A0A6G5Y6M2 \| 99.1 \| 2.3e-13 \| 4.2e-19 \| 116.6 \| 62 \| (92, 155) \| 233 \| (104, 165) \| 165 \| Uncharacterized protein \| Uncharacterized protein; uniclust \| UniRef100\_A0A6G5Y600 \| 99.1 \| 3.6e-13 \| 6.6e-19 \| 125.9 \| 140 \| (6, 159) \| 233 \| (1, 162) \| 351 \| Uncharacterized protein \| Uncharacterized protein; uniclust \| UniRef100\_UPI00065FE3AD \| 99.1 \| 4.4e-13 \| 9e-19 \| 118.1 \| 71 \| (85, 158) \| 233 \| (107, 177) \| 179 \| hypothetical protein \| hypothetical protein; uniclust \| UniRef100\_A0A0E2NX12 \| 99.1 \| 4.9e-13 \| 1.1e-18 \| 101.5 \| 58 \| (96, 156) \| 233 \| (6, 63) \| 65 \| Uncharacterized protein \| Uncharacterized protein; uniclust \| UniRef100\_A0A1X3M1B6 \| 99.1 \| 1e-12 \| 2e-18 \| 122.2 \| 105 \| (5, 113) \| 233 \| (55, 171) \| 287 \| Morphogenetic protein \| Morphogenetic protein; uniclust \| UniRef100\_A0A5B9N4V6 \| 99.0 \| 1.8e-12 \| 3.4e-18 \| 119.8 \| 134 \| (8, 158) \| 233 \| (22, 160) \| 307 \| Uncharacterized protein \| Uncharacterized protein; uniclust \| UniRef100\_A0A127KN71 \| 99.0 \| 2.4e-12 \| 5.3e-18 \| 110.0 \| 61 \| (100, 162) \| 233 \| (69, 131) \| 133 \| Uncharacterized protein \| Uncharacterized protein; uniclust \| UniRef100\_A0A950H461 \| 99.0 \| 3e-12 \| 5.5e-18 \| 110.0 \| 60 \| (95, 156) \| 233 \| (98, 157) \| 163 \| Uncharacterized protein \| Uncharacterized protein; uniclust \| UniRef100\_A0A1X1D6V7 \| 99.0 \| 3.9e-12 \| 7.2e-18 \| 100.6 \| 76 \| (6, 81) \| 233 \| (1, 89) \| 92 \| DUF551 domain-containing protein \| DUF551 domain-containing protein; uniclust \| UniRef100\_UPI001D1510C6 \| 99.0 \| 4e-12 \| 7.4e-18 \| 98.5 \| 71 \| (6, 76) \| 233 \| (1, 81) \| 81 \| hypothetical protein \| hypothetical protein; uniclust \| UniRef100\_A0A2R3UAP6 \| 98.9 \| 5.1e-12 \| 1.1e-17 \| 118.5 \| 73 \| (6, 81) \| 233 \| (1, 82) \| 269 \| Uncharacterized protein \| Uncharacterized protein; uniclust \| UniRef100\_UPI001CCC15E2 \| 98.9 \| 7.8e-12 \| 1.4e-17 \| 94.0 \| 56 \| (95, 154) \| 233 \| (12, 67) \| 67 \| hypothetical protein \| hypothetical protein; uniclust \| UniRef100\_A0A127KN71 \| 98.9 \| 6.8e-12 \| 1.5e-17 \| 107.4 \| 57 \| (94, 153) \| 233 \| (7, 68) \| 133 \| Uncharacterized protein \| Uncharacterized protein; uniclust \| UniRef100\_UPI0003EB40DC \| 98.9 \| 8.5e-12 \| 1.6e-17 \| 95.6 \| 73 \| (6, 85) \| 233 \| (1, 73) \| 75 \| hypothetical protein \| hypothetical protein; uniclust \| UniRef100\_A0A1U9HYR5 \| 98.9 \| 1.3e-11 \| 2.4e-17 \| 97.0 \| 77 \| (6, 84) \| 233 \| (1, 82) \| 85 \| Uncharacterized protein \| Uncharacterized protein; uniclust \| UniRef100\_UPI000FE13C13 \| 98.8 \| 2.9e-11 \| 5.7e-17 \| 96.0 \| 61 \| (96, 158) \| 233 \| (23, 83) \| 85 \| hypothetical protein \| hypothetical protein; uniclust \| UniRef100\_A0A5C8J0V6 \| 98.8 \| 3.6e-11 \| 6.6e-17 \| 103.4 \| 63 \| (98, 162) \| 233 \| (8, 70) \| 159 \| Uncharacterized protein \| Uncharacterized protein; uniclust \| UniRef100\_A0A6G5Y1Q1 \| 98.8 \| 4.9e-11 \| 9.2e-17 \| 92.3 \| 36 \| (5, 40) \| 233 \| (2, 37) \| 76 \| Uncharacterized protein \| Uncharacterized protein; uniclust \| UniRef100\_UPI00226A3416 \| 98.8 \| 6.3e-11 \| 1.2e-16 \| 99.7 \| 60 \| (95, 156) \| 233 \| (73, 132) \| 135 \| hypothetical protein \| hypothetical protein; uniclust \| UniRef100\_UPI00102A3018 \| 98.8 \| 6.6e-11 \| 1.2e-16 \| 87.8 \| 34 \| (6, 39) \| 233 \| (1, 34) \| 61 \| hypothetical protein \| hypothetical protein; uniclust \| UniRef100\_A0A949XH11 \| 98.7 \| 8e-11 \| 1.5e-16 \| 97.4 \| 66 \| (91, 157) \| 233 \| (47, 114) \| 119 \| Uncharacterized protein \| Uncharacterized protein; uniclust \| UniRef100\_UPI001E79528C \| 98.7 \| 8.2e-11 \| 1.5e-16 \| 108.9 \| 68 \| (6, 80) \| 233 \| (1, 72) \| 291 \| HNH endonuclease \| HNH endonuclease; uniclust \| UniRef100\_A0A6G5Y609 \| 98.7 \| 9.6e-11 \| 1.8e-16 \| 96.0 \| 66 \| (88, 154) \| 233 \| (41, 111) \| 112 \| Uncharacterized protein \| Uncharacterized protein; uniclust \| UniRef100\_A0A2X5UEE0 \| 98.7 \| 9.6e-11 \| 1.8e-16 \| 107.4 \| 81 \| (5, 87) \| 233 \| (51, 140) \| 259 \| Uncharacterized protein \| Uncharacterized protein; uniclust \| UniRef100\_UPI001B343D12 \| 98.7 \| 9.8e-11 \| 1.8e-16 \| 102.6 \| 64 \| (96, 162) \| 233 \| (73, 136) \| 182 \| hypothetical protein \| hypothetical protein; uniclust \| UniRef100\_A0A345MLB9 \| 98.7 \| 1e-10 \| 1.9e-16 \| 100.8 \| 62 \| (6, 85) \| 233 \| (1, 63) \| 160 \| DUF6378 domain-containing protein \| DUF6378 domain-containing protein; uniclust \| UniRef100\_A0A7X2IEN7 \| 98.7 \| 2e-10 \| 3.7e-16 \| 97.0 \| 63 \| (95, 157) \| 233 \| (72, 135) \| 136 \| DUF3310 domain-containing protein \| DUF3310 domain-containing protein; uniclust \| UniRef100\_UPI001E45F460 \| 98.6 \| 2.7e-10 \| 5e-16 \| 105.9 \| 86 \| (6, 121) \| 233 \| (1, 86) \| 296 \| hypothetical protein \| hypothetical protein; uniclust \| UniRef100\_A0A2C9DUX5 \| 98.6 \| 3.7e-10 \| 7.9e-16 \| 92.8 \| 64 \| (95, 159) \| 233 \| (21, 90) \| 99 \| Uncharacterized protein \| Uncharacterized protein; uniclust \| UniRef100\_UPI0013C3000B \| 98.6 \| 4.8e-10 \| 9.4e-16 \| 86.6 \| 60 \| (96, 157) \| 233 \| (4, 67) \| 71 \| hypothetical protein \| hypothetical protein; uniclust \| UniRef100\_A0A246JRV2 \| 98.6 \| 6.1e-10 \| 1.2e-15 \| 86.5 \| 62 \| (97, 158) \| 233 \| (4, 71) \| 73 \| Uncharacterized protein \| Uncharacterized protein; uniclust \| UniRef100\_A0A2E5T5S2 \| 98.5 \| 9.7e-10 \| 1.9e-15 \| 92.7 \| 61 \| (96, 157) \| 233 \| (52, 112) \| 125 \| Uncharacterized protein \| Uncharacterized protein; uniclust \| UniRef100\_A0A1H8H2L7 \| 98.5 \| 8e-10 \| 1.9e-15 \| 93.8 \| 66 \| (97, 162) \| 233 \| (8, 73) \| 114 \| Uncharacterized protein \| Uncharacterized protein; uniclust \| UniRef100\_A0A2S1GM15 \| 98.5 \| 1.1e-09 \| 2e-15 \| 91.4 \| 64 \| (91, 156) \| 233 \| (40, 103) \| 122 \| Uncharacterized protein \| Uncharacterized protein; uniclust \| UniRef100\_UPI0019821726 \| 98.5 \| 1.3e-09 \| 2.3e-15 \| 96.5 \| 58 \| (95, 155) \| 233 \| (129, 187) \| 187 \| hypothetical protein \| hypothetical protein; uniclust \| UniRef100\_UPI000DE54B3E \| 98.5 \| 1.4e-09 \| 2.5e-15 \| 99.7 \| 61 \| (96, 158) \| 233 \| (185, 247) \| 249 \| hypothetical protein \| hypothetical protein; uniclust \| UniRef100\_A0A1E4GP42 \| 98.5 \| 1.4e-09 \| 2.6e-15 \| 92.3 \| 57 \| (96, 154) \| 233 \| (82, 138) \| 138 \| Uncharacterized protein \| Uncharacterized protein; uniclust \| UniRef100\_A0A1B0VDS6 \| 98.5 \| 1.5e-09 \| 2.8e-15 \| 103.9 \| 81 \| (5, 87) \| 233 \| (51, 140) \| 381 \| Morphogenetic protein \| Morphogenetic protein; uniclust \| UniRef100\_A0A386KMJ4 \| 98.5 \| 1.6e-09 \| 3.1e-15 \| 95.3 \| 59 \| (95, 155) \| 233 \| (109, 167) \| 168 \| Uncharacterized protein \| Uncharacterized protein; uniclust \| UniRef100\_A0A8E7FPP9 \| 98.5 \| 1.7e-09 \| 3.2e-15 \| 97.7 \| 96 \| (6, 119) \| 233 \| (1, 111) \| 201 \| Uncharacterized protein \| Uncharacterized protein; uniclust \| UniRef100\_A0A074JFI6 \| 98.4 \| 1.9e-09 \| 4.3e-15 \| 91.3 \| 66 \| (96, 162) \| 233 \| (12, 84) \| 114 \| Uncharacterized protein \| Uncharacterized protein; uniclust \| UniRef100\_UPI00106EAB0B \| 98.4 \| 2.2e-09 \| 4.4e-15 \| 85.6 \| 60 \| (96, 157) \| 233 \| (17, 83) \| 84 \| hypothetical protein \| hypothetical protein; uniclust \| UniRef100\_UPI001C915A73 \| 98.4 \| 2.7e-09 \| 5e-15 \| 85.9 \| 78 \| (69, 154) \| 233 \| (8, 94) \| 95 \| hypothetical protein \| hypothetical protein; uniclust \| UniRef100\_UPI00096A5A9B \| 98.4 \| 2.7e-09 \| 5e-15 \| 95.5 \| 57 \| (95, 154) \| 233 \| (34, 93) \| 204 \| hypothetical protein \| hypothetical protein; uniclust \| UniRef100\_UPI00143CFDF2 \| 98.4 \| 2.7e-09 \| 5e-15 \| 89.6 \| 57 \| (96, 154) \| 233 \| (65, 123) \| 126 \| hypothetical protein \| hypothetical protein; uniclust \| UniRef100\_A0A0Q8RJ07 \| 98.4 \| 2.7e-09 \| 5.2e-15 \| 87.5 \| 64 \| (96, 160) \| 233 \| (9, 79) \| 99 \| Uncharacterized protein \| Uncharacterized protein; uniclust \| UniRef100\_A0A386KLU1 \| 98.4 \| 2.8e-09 \| 5.7e-15 \| 92.7 \| 58 \| (96, 155) \| 233 \| (81, 140) \| 141 \| Uncharacterized protein \| Uncharacterized protein; uniclust \| UniRef100\_A0A962YGS3 \| 98.4 \| 3.9e-09 \| 7.2e-15 \| 94.7 \| 61 \| (96, 158) \| 233 \| (109, 169) \| 206 \| Uncharacterized protein (Fragment) \| Uncharacterized protein (Fragment); uniclust \| UniRef100\_A0A941I2F7 \| 98.4 \| 4e-09 \| 7.3e-15 \| 89.9 \| 70 \| (84, 154) \| 233 \| (64, 139) \| 139 \| Uncharacterized protein \| Uncharacterized protein; uniclust \| UniRef100\_A0A6G5YBR4 \| 98.4 \| 4.1e-09 \| 7.5e-15 \| 86.0 \| 74 \| (4, 84) \| 233 \| (23, 97) \| 103 \| Uncharacterized protein \| Uncharacterized protein; uniclust \| UniRef100\_A0A1I3FVE1 \| 98.4 \| 5.3e-09 \| 9.7e-15 \| 89.0 \| 126 \| (7, 156) \| 233 \| (5, 136) \| 137 \| Uncharacterized protein \| Uncharacterized protein; uniclust \| UniRef100\_A0A482MK44 \| 98.4 \| 5.2e-09 \| 1e-14 \| 93.3 \| 60 \| (92, 154) \| 233 \| (114, 174) \| 174 \| Uncharacterized protein \| Uncharacterized protein; uniclust \| UniRef100\_A0A0C3RLK6 \| 98.3 \| 5.1e-09 \| 1.1e-14 \| 82.3 \| 59 \| (97, 157) \| 233 \| (4, 70) \| 73 \| Uncharacterized protein \| Uncharacterized protein; uniclust \| UniRef100\_A0A4R5PK78 \| 98.3 \| 5.6e-09 \| 1.1e-14 \| 80.3 \| 56 \| (97, 154) \| 233 \| (6, 65) \| 67 \| Cupin \| Cupin; uniclust \| UniRef100\_A0A6G5YPA9 \| 98.3 \| 7.7e-09 \| 1.4e-14 \| 96.0 \| 137 \| (6, 157) \| 233 \| (1, 155) \| 271 \| Uncharacterized protein \| Uncharacterized protein; uniclust \| UniRef100\_A0A931HEC0 \| 98.3 \| 7.5e-09 \| 1.5e-14 \| 81.8 \| 60 \| (95, 157) \| 233 \| (8, 71) \| 79 \| Uncharacterized protein \| Uncharacterized protein; uniclust \| UniRef100\_UPI000969E4E3 \| 98.3 \| 1.1e-08 \| 2.1e-14 \| 92.9 \| 74 \| (68, 155) \| 233 \| (129, 203) \| 205 \| hypothetical protein \| hypothetical protein; uniclust \| UniRef100\_A0A385EED3 \| 98.3 \| 1.3e-08 \| 2.5e-14 \| 87.2 \| 58 \| (97, 156) \| 233 \| (69, 137) \| 138 \| Uncharacterized protein \| Uncharacterized protein; uniclust \| UniRef100\_A0A6H1ZBT8 \| 98.2 \| 1.6e-08 \| 2.9e-14 \| 87.8 \| 68 \| (75, 155) \| 233 \| (82, 154) \| 154 \| Uncharacterized protein \| Uncharacterized protein; uniclust \| UniRef100\_UPI00221E5426 \| 98.2 \| 1.7e-08 \| 3.2e-14 \| 93.7 \| 62 \| (96, 159) \| 233 \| (82, 143) \| 267 \| hypothetical protein \| hypothetical protein; uniclust \| UniRef100\_A0A2D7XBX7 \| 98.1 \| 4.6e-08 \| 8.5e-14 \| 82.9 \| 59 \| (95, 154) \| 233 \| (70, 128) \| 128 \| DUF3085 domain-containing protein \| DUF3085 domain-containing protein; uniclust \| UniRef100\_A0A6G5YJZ6 \| 98.1 \| 5e-08 \| 9.2e-14 \| 86.8 \| 72 \| (97, 172) \| 233 \| (18, 101) \| 183 \| dATP/dGTP diphosphohydrolase N-terminal domain-containing protein \| dATP/dGTP diphosphohydrolase N-terminal domain-containing protein; uniclust \| UniRef100\_A0A0B3RIM0 \| 98.1 \| 4.1e-08 \| 9.2e-14 \| 81.7 \| 64 \| (96, 160) \| 233 \| (10, 82) \| 98 \| Uncharacterized protein \| Uncharacterized protein; uniclust \| UniRef100\_A0A368TTC8 \| 98.1 \| 5.8e-08 \| 1.1e-13 \| 78.2 \| 53 \| (98, 153) \| 233 \| (38, 90) \| 91 \| Uncharacterized protein \| Uncharacterized protein; uniclust \| UniRef100\_A0A0R3Q2Y3 \| 98.1 \| 6.3e-08 \| 1.2e-13 \| 79.9 \| 53 \| (96, 153) \| 233 \| (53, 105) \| 106 \| Uncharacterized protein \| Uncharacterized protein; uniclust \| UniRef100\_A0A516MMQ1 \| 98.1 \| 7.6e-08 \| 1.4e-13 \| 88.6 \| 62 \| (94, 155) \| 233 \| (171, 233) \| 239 \| Uncharacterized protein \| Uncharacterized protein; uniclust \| UniRef100\_A0A345BNB0 \| 98.0 \| 1.2e-07 \| 2.2e-13 \| 82.4 \| 81 \| (6, 90) \| 233 \| (1, 82) \| 151 \| Uncharacterized protein \| Uncharacterized protein; uniclust \| UniRef100\_A0A481W720 \| 98.0 \| 1.3e-07 \| 2.4e-13 \| 86.7 \| 63 \| (96, 160) \| 233 \| (100, 163) \| 229 \| Uncharacterized protein \| Uncharacterized protein; uniclust \| UniRef100\_A0A1H7CNT9 \| 98.0 \| 9.6e-08 \| 2.5e-13 \| 81.7 \| 65 \| (96, 160) \| 233 \| (11, 87) \| 111 \| Uncharacterized protein \| Uncharacterized protein; uniclust \| UniRef100\_A0A4Q2SVH6 \| 98.0 \| 1.4e-07 \| 2.5e-13 \| 71.3 \| 55 \| (99, 156) \| 233 \| (2, 56) \| 61 \| Uncharacterized protein \| Uncharacterized protein; uniclust \| UniRef100\_A0A0F5PG93 \| 98.0 \| 1.5e-07 \| 3.1e-13 \| 74.9 \| 58 \| (96, 155) \| 233 \| (11, 73) \| 76 \| Uncharacterized protein \| Uncharacterized protein; uniclust \| UniRef100\_A0A7S5R7Q8 \| 97.9 \| 2.1e-07 \| 3.8e-13 \| 71.0 \| 56 \| (98, 155) \| 233 \| (2, 61) \| 64 \| Uncharacterized protein \| Uncharacterized protein; uniclust \| UniRef100\_A0A246JRV1 \| 97.9 \| 2.2e-07 \| 4.2e-13 \| 75.7 \| 61 \| (96, 156) \| 233 \| (16, 87) \| 90 \| Uncharacterized protein \| Uncharacterized protein; uniclust \| UniRef100\_UPI00192B0E5B \| 97.9 \| 2.3e-07 \| 4.3e-13 \| 75.9 \| 44 \| (96, 141) \| 233 \| (20, 65) \| 98 \| hypothetical protein \| hypothetical protein; uniclust \| UniRef100\_N0DPA9 \| 97.9 \| 3.3e-07 \| 6.1e-13 \| 81.6 \| 60 \| (96, 156) \| 233 \| (65, 124) \| 176 \| Uncharacterized protein \| Uncharacterized protein; uniclust \| UniRef100\_A0A516MMP7 \| 97.9 \| 3.5e-07 \| 6.4e-13 \| 82.3 \| 89 \| (6, 124) \| 233 \| (2, 93) \| 191 \| Uncharacterized protein \| Uncharacterized protein; uniclust \| UniRef100\_A0A0F9BXC9 \| 97.8 \| 3.7e-07 \| 6.8e-13 \| 81.5 \| 65 \| (96, 162) \| 233 \| (49, 115) \| 179 \| Uncharacterized protein (Fragment) \| Uncharacterized protein (Fragment); uniclust \| UniRef100\_UPI0013B66D81 \| 97.8 \| 4.2e-07 \| 7.7e-13 \| 67.4 \| 32 \| (6, 37) \| 233 \| (1, 32) \| 54 \| hypothetical protein \| hypothetical protein; uniclust \| UniRef100\_A0A2D2W544 \| 97.8 \| 4e-07 \| 7.8e-13 \| 81.0 \| 59 \| (97, 156) \| 233 \| (78, 136) \| 160 \| Uncharacterized protein \| Uncharacterized protein; uniclust \| UniRef100\_A0A7W5S4A8 \| 97.8 \| 5e-07 \| 9.2e-13 \| 73.7 \| 61 \| (97, 160) \| 233 \| (9, 73) \| 95 \| Uncharacterized protein \| Uncharacterized protein; uniclust \| UniRef100\_A0A1T4SRW2 \| 97.8 \| 6.3e-07 \| 1.2e-12 \| 67.0 \| 54 \| (98, 154) \| 233 \| (2, 55) \| 56 \| Uncharacterized protein \| Uncharacterized protein; uniclust \| UniRef100\_A0A2W5IUS1 \| 97.7 \| 7.8e-07 \| 1.4e-12 \| 68.5 \| 57 \| (97, 155) \| 233 \| (2, 63) \| 66 \| Uncharacterized protein \| Uncharacterized protein; uniclust \| UniRef100\_A0A431IDW0 \| 97.7 \| 8.5e-07 \| 1.6e-12 \| 83.2 \| 61 \| (96, 156) \| 233 \| (96, 170) \| 264 \| Uncharacterized protein \| Uncharacterized protein; uniclust \| UniRef100\_A0A2E6JZ26 \| 97.7 \| 9.2e-07 \| 1.7e-12 \| 79.4 \| 58 \| (96, 155) \| 233 \| (119, 179) \| 183 \| Uncharacterized protein \| Uncharacterized protein; uniclust \| UniRef100\_A0A976S1A0 \| 97.7 \| 9.8e-07 \| 1.8e-12 \| 76.1 \| 80 \| (69, 157) \| 233 \| (44, 134) \| 136 \| Uncharacterized protein \| Uncharacterized protein; uniclust \| UniRef100\_UPI001F11253C \| 97.7 \| 1.2e-06 \| 2.2e-12 \| 73.3 \| 28 \| (5, 32) \| 233 \| (46, 73) \| 108 \| hypothetical protein \| hypothetical protein; uniclust \| UniRef100\_A0A431RAX2 \| 97.7 \| 1.2e-06 \| 2.3e-12 \| 67.8 \| 58 \| (98, 155) \| 233 \| (5, 68) \| 68 \| Uncharacterized protein \| Uncharacterized protein; uniclust \| UniRef100\_UPI002283A885 \| 97.7 \| 1.3e-06 \| 2.4e-12 \| 73.9 \| 68 \| (6, 83) \| 233 \| (5, 73) \| 117 \| hypothetical protein \| hypothetical protein; uniclust \| UniRef100\_W0ADT7 \| 97.6 \| 1.6e-06 \| 2.9e-12 \| 74.9 \| 71 \| (73, 154) \| 233 \| (57, 133) \| 135 \| Uncharacterized protein \| Uncharacterized protein; uniclust \| UniRef100\_UPI0008AB9E75 \| 97.6 \| 1.6e-06 \| 2.9e-12 \| 75.2 \| 54 \| (96, 152) \| 233 \| (82, 138) \| 139 \| DUF4326 domain-containing protein \| DUF4326 domain-containing protein; uniclust \| UniRef100\_A0A4S0XR95 \| 97.6 \| 1.9e-06 \| 3.6e-12 \| 68.2 \| 63 \| (95, 157) \| 233 \| (4, 70) \| 77 \| Uncharacterized protein \| Uncharacterized protein; uniclust \| UniRef100\_UPI002091CB86 \| 97.6 \| 2.2e-06 \| 4.1e-12 \| 77.0 \| 68 \| (87, 157) \| 233 \| (91, 166) \| 168 \| hypothetical protein \| hypothetical protein; uniclust \| UniRef100\_UPI00111C8B24 \| 97.6 \| 2.3e-06 \| 4.2e-12 \| 71.1 \| 72 \| (6, 80) \| 233 \| (1, 73) \| 103 \| hypothetical protein \| hypothetical protein; uniclust \| UniRef100\_A0A0A6CX03 \| 97.5 \| 2.5e-06 \| 4.9e-12 \| 69.1 \| 58 \| (96, 153) \| 233 \| (15, 81) \| 82 \| Uncharacterized protein \| Uncharacterized protein; uniclust \| UniRef100\_A0A7S5SKF1 \| 97.5 \| 3e-06 \| 5.5e-12 \| 82.2 \| 88 \| (19, 124) \| 233 \| (12, 110) \| 347 \| Uncharacterized protein \| Uncharacterized protein; uniclust \| UniRef100\_A0A6S7EY07 \| 97.5 \| 3.1e-06 \| 5.7e-12 \| 65.5 \| 51 \| (102, 156) \| 233 \| (11, 61) \| 66 \| Uncharacterized protein \| Uncharacterized protein; uniclust \| UniRef100\_A0A3S9U7R6 \| 97.5 \| 3.2e-06 \| 5.9e-12 \| 79.4 \| 65 \| (96, 162) \| 233 \| (128, 192) \| 258 \| DUF6378 domain-containing protein \| DUF6378 domain-containing protein; uniclust \| UniRef100\_A0A179BTQ8 \| 97.5 \| 3.2e-06 \| 6.4e-12 \| 67.9 \| 57 \| (97, 155) \| 233 \| (5, 75) \| 77 \| Uncharacterized protein \| Uncharacterized protein; uniclust \| UniRef100\_A0A291GC54 \| 97.5 \| 2.7e-06 \| 7e-12 \| 72.3 \| 66 \| (96, 162) \| 233 \| (9, 84) \| 101 \| Uncharacterized protein \| Uncharacterized protein; uniclust \| UniRef100\_UPI001294A8BB \| 97.4 \| 5.2e-06 \| 9.6e-12 \| 66.0 \| 56 \| (98, 155) \| 233 \| (12, 67) \| 77 \| hypothetical protein \| hypothetical protein; uniclust \| UniRef100\_UPI000A11260D \| 97.4 \| 5.6e-06 \| 1e-11 \| 80.2 \| 76 \| (7, 85) \| 233 \| (136, 223) \| 333 \| hypothetical protein \| hypothetical protein; uniclust \| UniRef100\_A0A839YDB8 \| 97.4 \| 6.2e-06 \| 1.2e-11 \| 65.1 \| 58 \| (97, 156) \| 233 \| (5, 65) \| 70 \| Uncharacterized protein \| Uncharacterized protein; uniclust \| UniRef100\_A0A5C0ZYP1 \| 97.4 \| 7.8e-06 \| 1.4e-11 \| 84.0 \| 69 \| (96, 166) \| 233 \| (421, 502) \| 630 \| Uncharacterized protein \| Uncharacterized protein; uniclust \| UniRef100\_A0A1P8DJC3 \| 97.3 \| 9.4e-06 \| 1.7e-11 \| 75.7 \| 55 \| (97, 154) \| 233 \| (170, 227) \| 232 \| Uncharacterized protein \| Uncharacterized protein; uniclust \| UniRef100\_A0A6I1NN18 \| 97.3 \| 1.1e-05 \| 2e-11 \| 73.0 \| 63 \| (96, 160) \| 233 \| (112, 175) \| 181 \| Uncharacterized protein \| Uncharacterized protein; uniclust \| UniRef100\_A0A1Z3U571 \| 97.3 \| 1.2e-05 \| 2.2e-11 \| 77.0 \| 64 \| (96, 162) \| 233 \| (71, 138) \| 292 \| Uncharacterized protein \| Uncharacterized protein; uniclust \| UniRef100\_UPI0011D19E68 \| 97.3 \| 1.1e-05 \| 2.3e-11 \| 74.1 \| 40 \| (68, 123) \| 233 \| (47, 86) \| 178 \| hypothetical protein \| hypothetical protein; uniclust \| UniRef100\_U5N140 \| 97.3 \| 1.4e-05 \| 2.5e-11 \| 59.2 \| 42 \| (110, 154) \| 233 \| (9, 50) \| 50 \| Uncharacterized protein \| Uncharacterized protein; uniclust \| UniRef100\_UPI0008D812AA \| 97.3 \| 1.4e-05 \| 2.5e-11 \| 68.5 \| 63 \| (97, 159) \| 233 \| (12, 80) \| 121 \| hypothetical protein \| hypothetical protein; uniclust \| UniRef100\_A0A2E2D777 \| 97.2 \| 1.3e-05 \| 2.8e-11 \| 75.0 \| 57 \| (96, 157) \| 233 \| (38, 95) \| 199 \| Uncharacterized protein \| Uncharacterized protein; uniclust \| UniRef100\_A0A6J5RMM4 \| 97.2 \| 1.6e-05 \| 2.9e-11 \| 73.6 \| 56 \| (96, 154) \| 233 \| (160, 215) \| 215 \| DUF6378 domain-containing protein \| DUF6378 domain-containing protein; uniclust \| UniRef100\_UPI0011408B34 \| 97.2 \| 1.9e-05 \| 3.4e-11 \| 64.2 \| 55 \| (103, 161) \| 233 \| (9, 63) \| 85 \| hypothetical protein \| hypothetical protein; uniclust \| UniRef100\_A0A5D9C913 \| 97.2 \| 2e-05 \| 3.9e-11 \| 62.2 \| 56 \| (96, 154) \| 233 \| (4, 65) \| 67 \| Uncharacterized protein \| Uncharacterized protein; uniclust \| UniRef100\_A0A7S5R948 \| 97.2 \| 2.1e-05 \| 4.1e-11 \| 69.5 \| 55 \| (96, 153) \| 233 \| (83, 142) \| 142 \| Uncharacterized protein \| Uncharacterized protein; uniclust \| UniRef100\_UPI001CFF8498 \| 97.1 \| 3.1e-05 \| 5.7e-11 \| 69.2 \| 60 \| (98, 157) \| 233 \| (11, 73) \| 161 \| hypothetical protein \| hypothetical protein; uniclust \| UniRef100\_A0A5C7PPU1 \| 97.1 \| 3.1e-05 \| 5.8e-11 \| 77.5 \| 95 \| (86, 184) \| 233 \| (245, 340) \| 438 \| Uncharacterized protein \| Uncharacterized protein; uniclust \| UniRef100\_A0A2E0UI72 \| 97.1 \| 3.6e-05 \| 6.5e-11 \| 70.9 \| 98 \| (96, 200) \| 233 \| (62, 166) \| 201 \| Jacalin-type lectin domain-containing protein \| Jacalin-type lectin domain-containing protein; uniclust \| UniRef100\_A0A0C3NUH1 \| 97.1 \| 3.6e-05 \| 6.6e-11 \| 64.9 \| 55 \| (98, 154) \| 233 \| (42, 102) \| 106 \| Uncharacterized protein \| Uncharacterized protein; uniclust \| UniRef100\_A0A7L5BCL0 \| 97.1 \| 4.2e-05 \| 7.6e-11 \| 65.8 \| 58 \| (98, 157) \| 233 \| (61, 121) \| 121 \| Restriction alleviation protein, Lar family \| Restriction alleviation protein, Lar family; uniclust \| UniRef100\_A0A3P3ZQD0 \| 97.0 \| 5.6e-05 \| 1e-10 \| 68.9 \| 53 \| (96, 154) \| 233 \| (123, 175) \| 183 \| Uncharacterized protein \| Uncharacterized protein; uniclust \| UniRef100\_A0A385IKG2 \| 97.0 \| 5.8e-05 \| 1.1e-10 \| 73.1 \| 65 \| (92, 158) \| 233 \| (63, 129) \| 277 \| 3'-phosphatase, 5'-polynucleotide kinase \| 3'-phosphatase, 5'-polynucleotide kinase; uniclust \| UniRef100\_A0A379CP65 \| 96.9 \| 7.7e-05 \| 1.5e-10 \| 69.6 \| 149 \| (15, 215) \| 233 \| (17, 172) \| 199 \| Uncharacterized protein \| Uncharacterized protein; uniclust \| UniRef100\_UPI00070EF9DC \| 96.9 \| 8.6e-05 \| 1.6e-10 \| 66.9 \| 65 \| (98, 162) \| 233 \| (3, 68) \| 165 \| hypothetical protein \| hypothetical protein; uniclust \| UniRef100\_A0A3B7DND1 \| 96.9 \| 8.6e-05 \| 1.6e-10 \| 69.5 \| 59 \| (93, 153) \| 233 \| (4, 64) \| 222 \| Uncharacterized protein \| Uncharacterized protein; uniclust \| UniRef100\_UPI002022F4A1 \| 96.8 \| 0.00013 \| 2.3e-10 \| 74.4 \| 60 \| (96, 156) \| 233 \| (267, 329) \| 487 \| hypothetical protein \| hypothetical protein; uniclust \| UniRef100\_B5AX45 \| 96.8 \| 0.00013 \| 2.4e-10 \| 72.1 \| 61 \| (96, 160) \| 233 \| (140, 200) \| 353 \| Uncharacterized protein \| Uncharacterized protein; uniclust \| UniRef100\_UPI001C8B0C5D \| 96.8 \| 0.00014 \| 2.5e-10 \| 66.3 \| 68 \| (6, 86) \| 233 \| (1, 68) \| 176 \| hypothetical protein \| hypothetical protein; uniclust \| UniRef100\_A0A0F6TI71 \| 96.8 \| 0.00015 \| 2.7e-10 \| 63.8 \| 52 \| (96, 151) \| 233 \| (74, 130) \| 132 \| Phage protein \| Phage protein; uniclust \| UniRef100\_UPI000DF92DE2 \| 96.8 \| 0.00015 \| 2.7e-10 \| 56.7 \| 52 \| (101, 154) \| 233 \| (3, 61) \| 64 \| hypothetical protein \| hypothetical protein; uniclust \| UniRef100\_A0A350UCK3 \| 96.8 \| 0.00016 \| 3e-10 \| 64.0 \| 61 \| (98, 161) \| 233 \| (2, 63) \| 142 \| Uncharacterized protein \| Uncharacterized protein; uniclust \| UniRef100\_A0A516MIH6 \| 96.7 \| 0.00019 \| 3.5e-10 \| 53.7 \| 48 \| (102, 152) \| 233 \| (2, 49) \| 50 \| Uncharacterized protein \| Uncharacterized protein; uniclust \| UniRef100\_A0A259NIN2 \| 96.7 \| 0.0002 \| 3.7e-10 \| 68.3 \| 93 \| (62, 162) \| 233 \| (39, 131) \| 252 \| Uncharacterized protein \| Uncharacterized protein; uniclust \| UniRef100\_A0A2D9H0Z1 \| 96.7 \| 0.00023 \| 4.3e-10 \| 62.5 \| 56 \| (98, 156) \| 233 \| (2, 59) \| 133 \| Uncharacterized protein \| Uncharacterized protein; uniclust \| UniRef100\_A0A2L1KEG8 \| 96.6 \| 0.00027 \| 5e-10 \| 61.7 \| 56 \| (98, 156) \| 233 \| (55, 119) \| 126 \| Uncharacterized protein \| Uncharacterized protein; uniclust \| UniRef100\_A0A931MMN7 \| 96.6 \| 0.00029 \| 5.3e-10 \| 58.6 \| 54 \| (96, 152) \| 233 \| (33, 90) \| 91 \| Uncharacterized protein \| Uncharacterized protein; uniclust \| UniRef100\_UPI002302FDAA \| 96.6 \| 0.00029 \| 5.3e-10 \| 66.6 \| 63 \| (95, 160) \| 233 \| (36, 99) \| 230 \| hypothetical protein \| hypothetical protein; uniclust \| UniRef100\_UPI001124F1A2 \| 96.6 \| 0.00033 \| 6.1e-10 \| 61.4 \| 67 \| (87, 153) \| 233 \| (59, 129) \| 129 \| hypothetical protein \| hypothetical protein; uniclust \| UniRef100\_A0A5C7LQ10 \| 96.6 \| 0.00034 \| 6.2e-10 \| 64.1 \| 61 \| (96, 156) \| 233 \| (108, 171) \| 177 \| DUF2203 family protein \| DUF2203 family protein; uniclust \| UniRef100\_A0A6A6K0Z2 \| 96.5 \| 0.00044 \| 8e-10 \| 62.4 \| 38 \| (66, 118) \| 233 \| (29, 66) \| 156 \| Uncharacterized protein \| Uncharacterized protein; uniclust \| UniRef100\_A0A7S5QYL1 \| 96.5 \| 0.00051 \| 9.4e-10 \| 55.6 \| 59 \| (98, 158) \| 233 \| (2, 69) \| 76 \| Uncharacterized protein \| Uncharacterized protein; uniclust \| UniRef100\_A0A2D6BGZ8 \| 96.4 \| 0.00059 \| 1.1e-09 \| 57.5 \| 55 \| (98, 154) \| 233 \| (16, 82) \| 94 \| Uncharacterized protein \| Uncharacterized protein; uniclust \| UniRef100\_A0A021XBI4 \| 96.4 \| 0.00065 \| 1.2e-09 \| 52.7 \| 52 \| (99, 155) \| 233 \| (2, 57) \| 59 \| Uncharacterized protein \| Uncharacterized protein; uniclust \| UniRef100\_A0A1V0M647 \| 96.3 \| 0.00085 \| 1.6e-09 \| 61.3 \| 56 \| (98, 156) \| 233 \| (98, 162) \| 169 \| Uncharacterized protein \| Uncharacterized protein; uniclust \| UniRef100\_W0A8K8 \| 96.3 \| 0.00092 \| 1.7e-09 \| 64.9 \| 57 \| (96, 157) \| 233 \| (82, 139) \| 274 \| Restriction alleviation protein Lar \| Restriction alleviation protein Lar |
| Top keywords  (threshold 1.00e-03 (evalue)) | **hypothetical, domain\_containing, DUF551, DUF6378, Morphogenetic, Fragment, Restriction, alleviation, Lar, SaV\_like** |
| Output files | ../../similar\_sequences/27\_FANPEZAQ\_CDS\_0027\_merged.svg ../../similar\_sequences/27\_FANPEZAQ\_CDS\_0027\_pdb70.a3m ../../similar\_sequences/27\_FANPEZAQ\_CDS\_0027\_pdb70.hhr ../../similar\_sequences/27\_FANPEZAQ\_CDS\_0027\_uniclust.a3m ../../similar\_sequences/27\_FANPEZAQ\_CDS\_0027\_uniclust.hhr |

#### Structure prediction (AlphaFold)^2^

|  |  |
| --- | --- |
| Stats | xml version="1.0" encoding="utf-8" standalone="no"?       2024-09-02T21:09:27.189806 image/svg+xml   Matplotlib v3.7.2, https://matplotlib.org/ |
| Predicted structure | **NGL Viewer Controls:**  - Center: *Left-Click* - Rotate: *Left-Click + Drag* - Translate: *Right-Click + Drag* - Zoom: *Shift + Left-Click + Drag* |
| Output files | ../../predicted\_structures/27\_FANPEZAQ\_CDS\_0027/features.pkl ../../predicted\_structures/27\_FANPEZAQ\_CDS\_0027/ranked\_0.pdb ../../predicted\_structures/27\_FANPEZAQ\_CDS\_0027/ranked\_0\_plots.svg ../../predicted\_structures/27\_FANPEZAQ\_CDS\_0027/result\_model\_1\_ptm\_pred\_0.pkl |

#### Structure similarity search results (Foldseek)^3^

|  |  |
| --- | --- |
| Structure databases searched | Pdb, Afdb-proteome, Afdb-uniprot50 |
| Results, scheme(s)  (Top layers only, threshold 1.00e-02 (evalue)) | xml version="1.0" encoding="utf-8" standalone="no"?       2024-09-02T21:10:57.290511 image/svg+xml   Matplotlib v3.7.2, https://matplotlib.org/ |
| Results, table  (threshold 1.00e-02 (evalue)) | db \| id \| prob \| evalue \| bits \| fident \| alnlen \| mismatch \| gapopen \| qstart \| qend \| tstart \| tend \| name \| description; afdb-uniprot50 \| AF-A0A8B2FBR4-F1-MODEL\_V4 \| 1.0 \| 7.831e-07 \| 146 \| 0.197 \| 203 \| 111 \| 11 \| 6 \| 186 \| 2 \| 174 \| Uncharacterized protein \| Uncharacterized protein; afdb-uniprot50 \| AF-A0A0F9BXC9-F1-MODEL\_V4 \| 1.0 \| 7.324e-05 \| 120 \| 0.174 \| 166 \| 111 \| 6 \| 78 \| 230 \| 27 \| 179 \| Uncharacterized protein \| Uncharacterized protein; afdb-uniprot50 \| AF-A0A259NIN2-F1-MODEL\_V4 \| 1.0 \| 0.0004499 \| 112 \| 0.179 \| 162 \| 98 \| 6 \| 72 \| 233 \| 49 \| 175 \| Uncharacterized protein \| Uncharacterized protein; afdb-uniprot50 \| AF-A0A7X5MK68-F1-MODEL\_V4 \| 1.0 \| 4.086e-05 \| 104 \| 0.229 \| 170 \| 78 \| 10 \| 9 \| 158 \| 57 \| 193 \| Uncharacterized protein \| Uncharacterized protein |
| Top keywords  (threshold 1.00e-02 (evalue)) | -- |
| Output files | ../../similar\_structures/27\_FANPEZAQ\_CDS\_0027\_afdb-proteome\_foldseek.tsv ../../similar\_structures/27\_FANPEZAQ\_CDS\_0027\_afdb-uniprot50\_foldseek.tsv ../../similar\_structures/27\_FANPEZAQ\_CDS\_0027\_merged.svg ../../similar\_structures/27\_FANPEZAQ\_CDS\_0027\_pdb\_foldseek.tsv |

  
  
  

Return to summary | Go to previous | Go to next

  


---

**Sequence/structure alignments coloring**  
Each object in the alignment figures is colored according to its E-value following this color coding:

1e-100
10
