## Supplementary material for "Completing the BASEL phage collection to unlock hidden diversity for systematic exploration of phage-host interactions": Data S2: 28.html

FANPEZAQ\_CDS\_0028


Return to summary | Go to previous | Go to next

|  |  |
| --- | --- |
| FANPEZAQ\_CDS\_0028  Page creation date: 02 Sep 2024, 12:00  Project folder: n/a  Input sequences file: Escherichia\_virus\_HeidiAbel.gb | parb partition spo0j domain\_containing repb partitioning chromosome sulfiredoxin chromosome\_partitioning parb\_like plasmid probable fragment putative and dna nuclease dna\_binding stage sporulation j segregation transcriptional regulator hth nucleoid occlusion rna nucleotide metabolism n\_terminal cro c1\_type |

### Sequence information

|  |  |
| --- | --- |
| Name | FANPEZAQ\_CDS\_0028  28\_FANPEZAQ\_CDS\_0028 (pipeline id) |
| Imported annotations | Escherichia\_virus\_HeidiAbel Bas97 |
| Protein sequence | MATIGQLYKAGATEAKPRKVFFVPLAELYVEDGYNIRDVDQAHVEEFRDAYLAGEYVPPL TVEVTQQGIKVIDGHHRFAGAKMAAETGLELRLEVNDITGTSEADKIALMVTSSQGKPLE PLERAKAYARLKAQGWTNDEIARKVKRSPSDVANMLALAECPEPIKDMVRGGQMSYVTAT ELTRQHGTEAEAVAAQALEEAKAAGKDKVTKKFTEPKAPKAPKAKPEAIQPPFGKDKLTR LADLAVQIELVHSDDRLEDAEPDESMTLRAPAGVVREFMALVADYVGGVPE |
| Number of residues | 291 |
| Molecular weight (Da) | 31696.74 |
| Output files | ../../query\_sequences/28\_FANPEZAQ\_CDS\_0028.fasta |

### Putative domain architecture and protein family

#### Search results (HHblits)^1^

|  |  |
| --- | --- |
| Domain family databases searched | Pfam, Ncbi-cd, Cath, Phrogs |
| Results, scheme(s)  (Top layers only; threshold 1.00e-03 (evalue)) | xml version="1.0" encoding="utf-8" standalone="no"?       2024-09-02T21:08:18.303612 image/svg+xml   Matplotlib v3.7.2, https://matplotlib.org/ |
| Results, table  (E-value ≤ 1.00e-03 (evalue)) | db \| id \| prob \| evalue \| pvalue \| score \| cols \| query \| query\_len \| template \| template\_len \| name \| description; pfam \| PF20188 \| 98.6 \| 3.8e-12 \| 7.9e-16 \| 107.2 \| 113 \| (31, 147) \| 291 \| (3, 118) \| 242 \| DUF6551 \| Family of unknown function (DUF6551); pfam \| PF08857 \| 98.4 \| 4.1e-11 \| 8.1e-15 \| 95.6 \| 90 \| (57, 151) \| 291 \| (38, 138) \| 153 \| ParBc\_2 \| Putative ParB-like nuclease; pfam \| PF14072 \| 97.9 \| 2.2e-09 \| 4.6e-13 \| 93.6 \| 94 \| (28, 124) \| 291 \| (34, 159) \| 337 \| DndB \| DNA-sulfur modification-associated; pfam \| PF07506 \| 96.9 \| 6.9e-07 \| 1.4e-10 \| 69.5 \| 73 \| (116, 188) \| 291 \| (2, 76) \| 171 \| RepB \| RepB plasmid partitioning protein; pfam \| PF02195 \| 96.8 \| 1e-06 \| 2.3e-10 \| 57.8 \| 70 \| (23, 95) \| 291 \| (3, 74) \| 91 \| ParBc \| ParB/Sulfiredoxin domain; pfam \| PF18090 \| 95.8 \| 3.5e-05 \| 7.8e-09 \| 47.8 \| 69 \| (119, 187) \| 291 \| (1, 72) \| 75 \| SoPB\_HTH \| Centromere-binding protein HTH domain; pfam \| PF08535 \| 95.3 \| 0.0001 \| 2.2e-08 \| 48.9 \| 56 \| (134, 189) \| 291 \| (2, 58) \| 92 \| KorB \| KorB domain; ncbi-cd \| cd16396 \| 97.9 \| 1.7e-09 \| 4.6e-13 \| 70.8 \| 78 \| (19, 96) \| 291 \| (2, 80) \| 95 \| Noc\_N \| cd16396 Noc\_N; nucleoid occlusion protein, N-terminal domain, and related domains of the ParB partitioning protein family.; ncbi-cd \| cd16393 \| 97.9 \| 2.3e-09 \| 6.3e-13 \| 70.1 \| 76 \| (21, 96) \| 291 \| (2, 78) \| 97 \| SPO0J\_N \| cd16393 SPO0J\_N; Thermus thermophilus stage 0 sporulation protein J-like N-terminal domain, ParB family member.; ncbi-cd \| cd16390 \| 97.8 \| 2.6e-09 \| 6.5e-13 \| 82.1 \| 71 \| (21, 96) \| 291 \| (2, 84) \| 162 \| ParB\_N\_Srx\_like \| cd16390 ParB\_N\_Srx\_like; uncharacterized family distantly related to the N-terminal domain of the ParB/Srx superfamily.; ncbi-cd \| cd16402 \| 97.5 \| 2e-08 \| 5.3e-12 \| 64.9 \| 67 \| (25, 96) \| 291 \| (2, 68) \| 87 \| ParB\_N\_like\_MT \| cd16402 ParB\_N\_like\_MT; ParB N-terminal-like domain, some attached to C-terminal S-adenosylmethionine-dependent methyltransferase domain.; ncbi-cd \| cd16397 \| 97.3 \| 6e-08 \| 1.6e-11 \| 64.5 \| 77 \| (19, 96) \| 291 \| (4, 87) \| 100 \| IbrB\_like \| cd16397 IbrB\_like; immunoglobulin-binding regulator IbrB activates eib genes. IbrB (along with IbrA) activates immunoglobulin-binding eib genes in Escherichia coli.; ncbi-cd \| cd16403 \| 97.3 \| 6.1e-08 \| 1.6e-11 \| 62.8 \| 68 \| (25, 96) \| 291 \| (2, 69) \| 88 \| ParB\_N\_like\_MT \| cd16403 ParB\_N\_like\_MT; ParB N-terminal-like domain, some attached to C-terminal S-adenosylmethionine-dependent methyltransferase.; ncbi-cd \| cd16408 \| 97.3 \| 6.1e-08 \| 1.7e-11 \| 61.6 \| 58 \| (39, 96) \| 291 \| (12, 70) \| 84 \| ParB\_N\_like \| cd16408 ParB\_N\_like; ParB N-terminal, parA -binding, -like domain of bacterial and plasmid parABS partitioning systems.; ncbi-cd \| cd16405 \| 97.3 \| 6.8e-08 \| 1.8e-11 \| 62.9 \| 72 \| (24, 96) \| 291 \| (2, 78) \| 91 \| RepB\_like\_N \| cd16405 RepB\_like\_N; plasmid segregation replication protein B like protein, N-terminal domain. RepB, found on plasmids and secondary chromosomes, works along with repA in directing plasmid segregation, and has been shown in Rhizobium etli to require the parS centromere-like sequence for full transcriptional repression of the repABC operon, inducing plasmid incompatibility.; ncbi-cd \| cd16410 \| 97.3 \| 7.5e-08 \| 2e-11 \| 62.1 \| 65 \| (40, 111) \| 291 \| (13, 77) \| 80 \| ParB\_N\_like \| cd16410 ParB\_N\_like; ParB N-terminal, parA-binding, -like domain of bacterial and plasmid parABS partitioning systems.; ncbi-cd \| cd16407 \| 97.2 \| 1.1e-07 \| 2.9e-11 \| 60.6 \| 70 \| (27, 96) \| 291 \| (3, 73) \| 86 \| ParB\_N\_like \| cd16407 ParB\_N\_like; ParB N-terminal, parA-binding, -like domain of bacterial and plasmid parABS partitioning systems.; ncbi-cd \| cd16398 \| 97.1 \| 2.4e-07 \| 6.5e-11 \| 60.3 \| 70 \| (27, 96) \| 291 \| (3, 75) \| 91 \| KorB\_N\_like \| cd16398 KorB\_N\_like; ParB-like partition protein of low copy number plasmid RK2, N-terminal domain and related domains.; ncbi-cd \| cd16388 \| 97.0 \| 3.3e-07 \| 9.2e-11 \| 56.2 \| 71 \| (21, 96) \| 291 \| (2, 74) \| 77 \| SbnI\_like\_N \| cd16388 SbnI\_like\_N; N-terminal domain of transcriptional regulators similar to SbnI. Siderophore staphylobactin biosynthesis protein SbnI of Staphylococcus aureus is a ParB/Spo0J like protein required for the expression of genes in the sbn operon, which is responsible for staphyloferrin B (SB) biosynthesis.; ncbi-cd \| cd16411 \| 97.0 \| 4.4e-07 \| 1.2e-10 \| 58.2 \| 70 \| (24, 96) \| 291 \| (2, 76) \| 90 \| ParB\_N\_like \| cd16411 ParB\_N\_like; ParB N-terminal, parA -binding, domain of bacterial and plasmid parABS partitioning systems.; ncbi-cd \| cd16395 \| 96.9 \| 4.9e-07 \| 1.3e-10 \| 58.8 \| 73 \| (21, 96) \| 291 \| (2, 80) \| 90 \| Srx \| cd16395 Srx; Sulfiredoxin reactivates peroxiredoxins after oxidative inactivation. Sulfiredoxin reduces and thereby re-activates 2-cys peroxiredoxins.; ncbi-cd \| cd16409 \| 96.9 \| 4.9e-07 \| 1.3e-10 \| 55.5 \| 56 \| (41, 96) \| 291 \| (3, 59) \| 74 \| ParB\_N\_like \| cd16409 ParB\_N\_like; ParB N-terminal-like domain of bacterial and plasmid parABS partitioning systems.; ncbi-cd \| cd16404 \| 96.9 \| 5.4e-07 \| 1.4e-10 \| 55.7 \| 55 \| (39, 96) \| 291 \| (12, 66) \| 69 \| pNOB8\_ParB\_N\_like \| cd16404 pNOB8\_ParB\_N\_like; pNOB8 ParB-like N-terminal domain, plasmid partitioning system protein domain. archaeal pNOB8 ParB acts in a plasmid partitioning system made up of 3 parts: AspA, ParA motor protein, and ParB, which links ParA to the protein-DNA superhelix.; ncbi-cd \| cd16394 \| 96.9 \| 7e-07 \| 1.9e-10 \| 54.3 \| 55 \| (39, 94) \| 291 \| (12, 66) \| 67 \| sopB\_N \| cd16394 sopB\_N; N-terminal domain of sopB protein, which promotes proper partitioning of F1 plasmid.; ncbi-cd \| cd16400 \| 96.7 \| 1.6e-06 \| 4.4e-10 \| 53.2 \| 67 \| (23, 96) \| 291 \| (2, 69) \| 72 \| ParB\_Srx\_like\_nuclease \| cd16400 ParB\_Srx\_like\_nuclease; ParB/Srx\_like nuclease and putative transcriptional regulators related to SbnI. This family contains a Pyrococcus Furiosus enzyme reported to have DNA nuclease activity and resembles the N-terminal domain of ParB proteins of the parABS bacterial chromosome partitioning system.; ncbi-cd \| cd16401 \| 96.6 \| 2e-06 \| 5.4e-10 \| 54.4 \| 56 \| (39, 96) \| 291 \| (12, 67) \| 85 \| ParB\_N\_like\_MT \| cd16401 ParB\_N\_like\_MT; ParB N-terminal-like domain, some attached to C-terminal S-adenosylmethionine-dependent methyltransferase domain.; ncbi-cd \| cd16406 \| 96.5 \| 3e-06 \| 8e-10 \| 53.7 \| 55 \| (41, 95) \| 291 \| (3, 66) \| 82 \| ParB\_N\_like \| cd16406 ParB\_N\_like; ParB N-terminal, parA-binding, -like domain of bacterial and plasmid parABS partitioning systems.; ncbi-cd \| cd16387 \| 96.3 \| 5.5e-06 \| 1.5e-09 \| 46.5 \| 50 \| (42, 91) \| 291 \| (3, 53) \| 54 \| ParB\_N\_Srx \| cd16387 ParB\_N\_Srx; ParB N-terminal domain and sulfiredoxin protein-related families. The ParB N-terminal domain/Sulfiredoxin (Srx) superfamily contains proteins with diverse activities.; ncbi-cd \| cd16844 \| 96.2 \| 1e-05 \| 2.7e-09 \| 46.5 \| 51 \| (41, 94) \| 291 \| (3, 53) \| 54 \| ParB\_N\_like\_MT \| cd16844 ParB\_N\_like\_MT; ParB N-terminal-like domain, some attached to C-terminal S-adenosylmethionine-dependent methyltransferase domain.; ncbi-cd \| cd16392 \| 96.1 \| 1.1e-05 \| 3.1e-09 \| 48.8 \| 56 \| (40, 95) \| 291 \| (10, 69) \| 72 \| toxin-ParB \| cd16392 toxin-ParB; toxin domain of the ParB/Srx superfamily. toxin domain with similarity to the N-terminal domain of ParB, a DNA-binding component of the prokaryotic parABS partitioning system and related proteins.; cath \| 2hwjA01 \| 98.3 \| 8.8e-11 \| 1.4e-14 \| 92.2 \| 92 \| (16, 112) \| 291 \| (5, 105) \| 134 \| Hypothetical protein atu1540 \| CATHCODE: 3.90.1530.10 NAME: Hypothetical protein atu1540. Chain: a, b, c, d, e, f. Synonym: agr\_c\_2837p. Engineered: yes SOURCE: Agrobacterium tumefaciens str. C58. Organism\_taxid: 176299. Strain: c58 / atcc 33970. Expressed in: escherichia coli. Expression\_system\_taxid: 562. CLASS: Alpha Beta, ARCH: Alpha-Beta Complex, TOPOL: Conserved hypothetical protein from pyrococcus furiosus pfu- 392566-001, ParB domain, HOMOL: Conserved hypothetical protein from pyrococcus furiosus pfu- 392566-001, ParB domain; cath \| 1vz0C02 \| 97.8 \| 5.4e-09 \| 9.5e-13 \| 73.9 \| 84 \| (106, 189) \| 291 \| (6, 89) \| 113 \| Chromosome-partitioning protein spo0j \| CATHCODE: 1.10.10.2830 NAME: Chromosome-partitioning protein spo0j. Chain: a, b, c, d, e, f, g, h. Fragment: DNA-binding domain, residues 1-222. Engineered: yes SOURCE: Thermus thermophilus hb27. Organism\_taxid: 262724. Atcc: baa-163d. Gene: spo0c, parb, tt\_c1604. Expressed in: escherichia coli bl21(de3). Expression\_system\_taxid: 469008. Expression\_system\_variant: c41 CLASS: Mainly Alpha, ARCH: Orthogonal Bundle, TOPOL: Arc Repressor Mutant, subunit A, HOMOL: Arc Repressor Mutant, subunit A; cath \| 3mkyP00 \| 97.7 \| 9.5e-09 \| 1.6e-12 \| 75.8 \| 73 \| (118, 190) \| 291 \| (1, 76) \| 114 \| Dna (5'- d(\*cp\*tp\*gp\*gp\*gp\*ap\*cp\*cp\*ap\*tp\*gp\*gp\*tp\*cp\*cp\*cp\*ap\*g)-3') \| CATHCODE: 1.10.10.2830 NAME: Dna (5'- d(\*cp\*tp\*gp\*gp\*gp\*ap\*cp\*cp\*ap\*tp\*gp\*gp\*tp\*cp\*cp\*cp\*ap\*g)-3').Chain: u, t. Engineered: yes. Protein sopb. Chain: b, p. Fragment: unp residues 155 to 323. Synonym: plasmid partition protein b. Engineered: yes SOURCE: Yes. Other\_details: the DNA was chemically synthesized.. CLASS: Mainly Alpha, ARCH: Orthogonal Bundle, TOPOL: Arc Repressor Mutant, subunit A, HOMOL: Arc Repressor Mutant, subunit A; cath \| 1xw3A01 \| 97.5 \| 5.2e-08 \| 8.8e-12 \| 67.8 \| 74 \| (19, 95) \| 291 \| (3, 82) \| 96 \| Sulfiredoxin \| CATHCODE: 3.90.1530.10 NAME: Sulfiredoxin. Chain: a. Fragment: residues 32-137. Engineered: yes SOURCE: Homo sapiens. Human. Organism\_taxid: 9606. Expressed in: escherichia coli. Expression\_system\_taxid: 562. CLASS: Alpha Beta, ARCH: Alpha-Beta Complex, TOPOL: Conserved hypothetical protein from pyrococcus furiosus pfu- 392566-001, ParB domain, HOMOL: Conserved hypothetical protein from pyrococcus furiosus pfu- 392566-001, ParB domain; cath \| 1vk1A01 \| 97.2 \| 2.2e-07 \| 3.7e-11 \| 67.0 \| 74 \| (17, 95) \| 291 \| (15, 91) \| 112 \| Conserved hypothetical protein \| CATHCODE: 3.90.1530.10 NAME: Conserved hypothetical protein. Chain: a. Engineered: yes SOURCE: Pyrococcus furiosus. Organism\_taxid: 2261. Expressed in: escherichia coli. Expression\_system\_taxid: 562 CLASS: Alpha Beta, ARCH: Alpha-Beta Complex, TOPOL: Conserved hypothetical protein from pyrococcus furiosus pfu- 392566-001, ParB domain, HOMOL: Conserved hypothetical protein from pyrococcus furiosus pfu- 392566-001, ParB domain; cath \| 1vz0A01 \| 96.8 \| 1.5e-06 \| 2.6e-10 \| 54.1 \| 56 \| (38, 93) \| 291 \| (5, 60) \| 63 \| Chromosome-partitioning protein spo0j \| CATHCODE: 3.90.1530.30 NAME: Chromosome-partitioning protein spo0j. Chain: a, b, c, d, e, f, g, h. Fragment: DNA-binding domain, residues 1-222. Engineered: yes SOURCE: Thermus thermophilus hb27. Organism\_taxid: 262724. Atcc: baa-163d. Gene: spo0c, parb, tt\_c1604. Expressed in: escherichia coli bl21(de3). Expression\_system\_taxid: 469008. Expression\_system\_variant: c41 CLASS: Alpha Beta, ARCH: Alpha-Beta Complex, TOPOL: Conserved hypothetical protein from pyrococcus furiosus pfu- 392566-001, ParB domain, HOMOL: Conserved hypothetical protein from pyrococcus furiosus pfu- 392566-001, ParB domain; phrogs \| 1033 \| 100.0 \| 8.5e-40 \| 1.1e-43 \| 293.7 \| 270 \| (1, 286) \| 291 \| (3, 283) \| 288 \| ParB-like partition protein \| ParB-like partition protein; Category: DNA, RNA and nucleotide metabolism; NC\_031091\_p46; phrogs \| 4715 \| 99.8 \| 1.5e-25 \| 1.8e-29 \| 194.6 \| 176 \| (16, 196) \| 291 \| (3, 215) \| 257 \| ParB-like partition protein \| ParB-like partition protein; Category: DNA, RNA and nucleotide metabolism; p121521 VI\_12455; phrogs \| 620 \| 99.8 \| 7.8e-24 \| 9.7e-28 \| 177.3 \| 134 \| (13, 161) \| 291 \| (19, 152) \| 191 \| ParB-like partition protein \| ParB-like partition protein; Category: DNA, RNA and nucleotide metabolism; MF189178\_p69; phrogs \| 6218 \| 99.7 \| 4.8e-23 \| 5.7e-27 \| 189.8 \| 164 \| (19, 189) \| 291 \| (2, 173) \| 502 \| ParB-like partition protein \| ParB-like partition protein; Category: DNA, RNA and nucleotide metabolism; KJ829260\_p91; phrogs \| 2159 \| 99.7 \| 1.9e-22 \| 2.3e-26 \| 167.2 \| 147 \| (14, 161) \| 291 \| (48, 198) \| 214 \| ParB-like partition protein \| ParB-like partition protein; Category: DNA, RNA and nucleotide metabolism; p9193 VI\_02947; phrogs \| 498 \| 99.7 \| 4.5e-22 \| 5.5e-26 \| 183.2 \| 130 \| (15, 150) \| 291 \| (7, 145) \| 419 \| DNA methyltransferase \| DNA methyltransferase; Category: other; p48104 VI\_12205; phrogs \| 3951 \| 99.6 \| 1.5e-20 \| 1.7e-24 \| 161.8 \| 175 \| (16, 198) \| 291 \| (6, 185) \| 300 \| NA \| NA; Category: unknown function; p296951 VI\_02154; phrogs \| 3264 \| 99.6 \| 2.2e-20 \| 2.8e-24 \| 162.8 \| 132 \| (15, 154) \| 291 \| (7, 141) \| 272 \| ParB-like partition protein \| ParB-like partition protein; Category: DNA, RNA and nucleotide metabolism; p313310 VI\_04191; phrogs \| 15396 \| 99.6 \| 4.9e-20 \| 5.5e-24 \| 145.5 \| 132 \| (15, 158) \| 291 \| (1, 135) \| 201 \| ParB-like partition nuclease \| ParB-like partition nuclease; Category: DNA, RNA and nucleotide metabolism; NC\_024369\_p1; phrogs \| 18726 \| 99.6 \| 6.9e-20 \| 7.7e-24 \| 162.8 \| 181 \| (18, 200) \| 291 \| (1, 215) \| 572 \| NA \| NA; Category: unknown function; p234010 VI\_05492; phrogs \| 8071 \| 99.6 \| 7.4e-20 \| 8.4e-24 \| 159.7 \| 129 \| (15, 159) \| 291 \| (5, 133) \| 394 \| ParB-like partition protein \| ParB-like partition protein; Category: DNA, RNA and nucleotide metabolism; KU886268\_p74; phrogs \| 4272 \| 99.6 \| 1.2e-19 \| 1.4e-23 \| 166.0 \| 173 \| (17, 189) \| 291 \| (294, 474) \| 658 \| NA \| NA; Category: unknown function; p55719 VI\_12435; phrogs \| 723 \| 99.5 \| 1.5e-18 \| 1.8e-22 \| 145.9 \| 133 \| (15, 161) \| 291 \| (14, 149) \| 219 \| ParB-like partition protein \| ParB-like partition protein; Category: DNA, RNA and nucleotide metabolism; NC\_031253\_p1; phrogs \| 18056 \| 99.4 \| 4.4e-18 \| 5e-22 \| 121.1 \| 77 \| (18, 103) \| 291 \| (1, 77) \| 94 \| ParB-like partition nuclease \| ParB-like partition nuclease; Category: DNA, RNA and nucleotide metabolism; NC\_007924\_p29; phrogs \| 1146 \| 99.3 \| 2.3e-16 \| 2.8e-20 \| 138.7 \| 167 \| (15, 188) \| 291 \| (42, 219) \| 333 \| ParB-like partition protein \| ParB-like partition protein; Category: DNA, RNA and nucleotide metabolism; p43547 VI\_12111; phrogs \| 26387 \| 99.3 \| 3e-16 \| 3.4e-20 \| 127.6 \| 148 \| (16, 170) \| 291 \| (6, 162) \| 252 \| ParB-like partition nuclease \| ParB-like partition nuclease; Category: DNA, RNA and nucleotide metabolism; NC\_031231\_p92; phrogs \| 15982 \| 99.3 \| 3.3e-16 \| 3.8e-20 \| 137.6 \| 178 \| (16, 196) \| 291 \| (25, 226) \| 452 \| NA \| NA; Category: unknown function; p176212 VI\_01150; phrogs \| 9353 \| 99.2 \| 7.5e-16 \| 8.8e-20 \| 129.5 \| 123 \| (17, 142) \| 291 \| (8, 136) \| 246 \| NA \| NA; Category: unknown function; p407464 VI\_01025; phrogs \| 12047 \| 99.2 \| 1.1e-15 \| 1.3e-19 \| 131.9 \| 157 \| (22, 188) \| 291 \| (2, 188) \| 359 \| ParB-like partition nuclease \| ParB-like partition nuclease; Category: DNA, RNA and nucleotide metabolism; NC\_007581\_p185; phrogs \| 7783 \| 99.1 \| 1.5e-14 \| 1.7e-18 \| 131.1 \| 186 \| (17, 208) \| 291 \| (33, 224) \| 669 \| NA \| NA; Category: unknown function; p235076 VI\_06411; phrogs \| 3694 \| 99.0 \| 1.9e-14 \| 2.2e-18 \| 126.7 \| 147 \| (15, 163) \| 291 \| (10, 201) \| 375 \| NA \| NA; Category: unknown function; p237513 VI\_09544; phrogs \| 27290 \| 99.0 \| 7.5e-14 \| 8.4e-18 \| 121.9 \| 172 \| (15, 189) \| 291 \| (25, 214) \| 517 \| NA \| NA; Category: unknown function; p222055 VI\_01195; phrogs \| 7405 \| 98.7 \| 3.6e-12 \| 4.2e-16 \| 105.9 \| 111 \| (16, 132) \| 291 \| (21, 134) \| 249 \| NA \| NA; Category: unknown function; p151533 VI\_12464; phrogs \| 8071 \| 98.5 \| 2.4e-11 \| 2.7e-15 \| 105.3 \| 122 \| (15, 147) \| 291 \| (210, 336) \| 394 \| ParB-like partition protein \| ParB-like partition protein; Category: DNA, RNA and nucleotide metabolism; KU886268\_p74; phrogs \| 5475 \| 98.4 \| 4.4e-11 \| 5e-15 \| 106.3 \| 98 \| (20, 125) \| 291 \| (2, 100) \| 519 \| NA \| NA; Category: unknown function; p62317 VI\_01138; phrogs \| 32069 \| 98.3 \| 1.6e-10 \| 1.8e-14 \| 95.4 \| 171 \| (17, 188) \| 291 \| (39, 221) \| 348 \| ParB-like partition protein \| ParB-like partition protein; Category: DNA, RNA and nucleotide metabolism; NC\_019713\_p89; phrogs \| 18017 \| 98.1 \| 9.6e-10 \| 1.1e-13 \| 87.7 \| 164 \| (16, 189) \| 291 \| (3, 211) \| 228 \| NA \| NA; Category: unknown function; p240016 VI\_08189; phrogs \| 8947 \| 98.1 \| 1.2e-09 \| 1.3e-13 \| 86.5 \| 130 \| (18, 156) \| 291 \| (10, 146) \| 216 \| NA \| NA; Category: unknown function; p43781 VI\_12381; phrogs \| 6392 \| 97.8 \| 1.1e-08 \| 1.3e-12 \| 82.7 \| 134 \| (16, 158) \| 291 \| (6, 146) \| 265 \| ParB-like partition protein \| ParB-like partition protein; Category: DNA, RNA and nucleotide metabolism; KX589269\_p1; phrogs \| 26377 \| 97.7 \| 1.7e-08 \| 1.9e-12 \| 82.0 \| 99 \| (43, 148) \| 291 \| (29, 128) \| 326 \| ParB-like partition protein \| ParB-like partition protein; Category: DNA, RNA and nucleotide metabolism; NC\_007409\_p70; phrogs \| 17439 \| 97.6 \| 3.6e-08 \| 4.1e-12 \| 78.4 \| 135 \| (26, 167) \| 291 \| (22, 160) \| 226 \| NA \| NA; Category: unknown function; p97272 VI\_12440; phrogs \| 21315 \| 97.4 \| 8.7e-08 \| 9.8e-12 \| 82.3 \| 88 \| (39, 131) \| 291 \| (27, 117) \| 450 \| NA \| NA; Category: unknown function; p108440 VI\_04298; phrogs \| 5931 \| 96.9 \| 1.2e-06 \| 1.4e-10 \| 75.6 \| 75 \| (34, 117) \| 291 \| (46, 123) \| 303 \| NA \| NA; Category: unknown function; NC\_023604\_p41; phrogs \| 16545 \| 96.9 \| 1.6e-06 \| 1.8e-10 \| 63.4 \| 134 \| (21, 160) \| 291 \| (2, 136) \| 154 \| ParB-like partition protein \| ParB-like partition protein; Category: DNA, RNA and nucleotide metabolism; KC821630\_p51; phrogs \| 31451 \| 96.8 \| 1.9e-06 \| 2.1e-10 \| 68.8 \| 139 \| (42, 193) \| 291 \| (32, 175) \| 298 \| NA \| NA; Category: unknown function; p274462 VI\_03342; phrogs \| 2859 \| 96.8 \| 1.8e-06 \| 2.2e-10 \| 66.7 \| 71 \| (19, 100) \| 291 \| (53, 124) \| 149 \| NA \| NA; Category: unknown function; KM209228\_p98; phrogs \| 12305 \| 96.7 \| 2.7e-06 \| 3.1e-10 \| 69.7 \| 62 \| (54, 127) \| 291 \| (40, 102) \| 265 \| ParB-like partition protein \| ParB-like partition protein; Category: DNA, RNA and nucleotide metabolism; KY065505\_p11; phrogs \| 17622 \| 96.6 \| 5.7e-06 \| 6.4e-10 \| 63.6 \| 85 \| (26, 115) \| 291 \| (15, 104) \| 218 \| NA \| NA; Category: unknown function; p234013 VI\_05492; phrogs \| 21912 \| 96.5 \| 6.4e-06 \| 7.2e-10 \| 61.3 \| 56 \| (42, 101) \| 291 \| (52, 119) \| 158 \| DNA binding protein \| DNA binding protein; Category: DNA, RNA and nucleotide metabolism; MH045566\_p52; phrogs \| 26754 \| 96.5 \| 6.7e-06 \| 7.5e-10 \| 62.6 \| 77 \| (17, 102) \| 291 \| (67, 144) \| 207 \| ParB-like partition nuclease \| ParB-like partition nuclease; Category: DNA, RNA and nucleotide metabolism; NC\_011421\_p55; phrogs \| 26392 \| 96.4 \| 1.3e-05 \| 1.4e-09 \| 61.8 \| 81 \| (40, 131) \| 291 \| (4, 85) \| 231 \| NA \| NA; Category: unknown function; KF005319\_p27; phrogs \| 27360 \| 96.3 \| 1.5e-05 \| 1.7e-09 \| 64.4 \| 83 \| (47, 131) \| 291 \| (1, 89) \| 280 \| NA \| NA; Category: unknown function; NC\_006268\_p50; phrogs \| 26063 \| 96.0 \| 3.5e-05 \| 4e-09 \| 62.3 \| 136 \| (21, 160) \| 291 \| (6, 161) \| 334 \| NA \| NA; Category: unknown function; p11818 VI\_06587; phrogs \| 26588 \| 95.9 \| 5e-05 \| 5.6e-09 \| 67.4 \| 76 \| (17, 101) \| 291 \| (97, 173) \| 932 \| ParB-like partition nuclease \| ParB-like partition nuclease; Category: DNA, RNA and nucleotide metabolism; NC\_025423\_p216; phrogs \| 35656 \| 95.5 \| 0.00014 \| 1.6e-08 \| 61.1 \| 140 \| (50, 189) \| 291 \| (69, 238) \| 432 \| NA \| NA; Category: unknown function; KU234533\_p86; phrogs \| 30771 \| 94.9 \| 0.00038 \| 4.3e-08 \| 60.3 \| 107 \| (19, 126) \| 291 \| (40, 161) \| 549 \| NA \| NA; Category: unknown function; p308958 VI\_06640; phrogs \| 17442 \| 94.7 \| 0.00051 \| 5.7e-08 \| 50.9 \| 97 \| (13, 116) \| 291 \| (69, 165) \| 185 \| NA \| NA; Category: unknown function; p407489 VI\_05339 |
| Top keywords  (threshold 1.00e-03 (evalue)) | **and, ParB, DNA, ParB\_like, partition, RNA, nucleotide, metabolism, N\_terminal, plasmid** |
| Output files | ../../domain\_architecture/28\_FANPEZAQ\_CDS\_0028\_cath.hhr ../../domain\_architecture/28\_FANPEZAQ\_CDS\_0028\_merged.svg ../../domain\_architecture/28\_FANPEZAQ\_CDS\_0028\_ncbi-cd.hhr ../../domain\_architecture/28\_FANPEZAQ\_CDS\_0028\_pfam.hhr ../../domain\_architecture/28\_FANPEZAQ\_CDS\_0028\_phrogs.hhr |

### Identical protein sequences/structures

#### Search results

|  |  |
| --- | --- |
| Protein sequence databases searched | Pdb, Swissprot, Refseq |
| Identical proteins found | -- |
| Top keywords | -- |
| Output files | -- |

### Similar protein sequences/structures

#### Sequence similarity search results (HHblits)^1^

|  |  |
| --- | --- |
| Sequence databases searched | Uniclust, Pdb70 |
| Results, scheme(s)  (Top layers only, threshold 1.00e-03 (evalue)) | xml version="1.0" encoding="utf-8" standalone="no"?       2024-09-02T21:08:41.732191 image/svg+xml   Matplotlib v3.7.2, https://matplotlib.org/ |
| Results, table(s)  (threshold 1.00e-03 (evalue)) | db \| id \| prob \| evalue \| pvalue \| score \| cols \| query \| query\_len \| template \| template\_len \| name \| description; uniclust \| UniRef100\_A0A071M4B4 \| 100.0 \| 4.4e-41 \| 8.9e-47 \| 295.9 \| 270 \| (1, 288) \| 291 \| (40, 315) \| 388 \| Chromosome partitioning protein ParB \| Chromosome partitioning protein ParB; uniclust \| UniRef100\_A0A076G7J3 \| 100.0 \| 7.3e-40 \| 1.4e-45 \| 282.0 \| 260 \| (1, 290) \| 291 \| (19, 282) \| 452 \| ParB/Sulfiredoxin domain-containing protein \| ParB/Sulfiredoxin domain-containing protein; uniclust \| UniRef100\_A0A0F3IEX4 \| 100.0 \| 8.8e-38 \| 1.7e-43 \| 249.2 \| 170 \| (17, 192) \| 291 \| (33, 205) \| 231 \| Probable chromosome-partitioning protein ParB (Fragment) \| Probable chromosome-partitioning protein ParB (Fragment); uniclust \| UniRef100\_A0A0J1DM72 \| 100.0 \| 4.8e-37 \| 9.2e-43 \| 261.2 \| 179 \| (16, 200) \| 291 \| (86, 268) \| 389 \| HTH cro/C1-type domain-containing protein \| HTH cro/C1-type domain-containing protein; uniclust \| UniRef100\_A0A0Q8XC42 \| 100.0 \| 5.2e-37 \| 9.8e-43 \| 244.9 \| 266 \| (1, 285) \| 291 \| (13, 281) \| 282 \| ParB/Sulfiredoxin domain-containing protein \| ParB/Sulfiredoxin domain-containing protein; uniclust \| UniRef100\_A0A073JT65 \| 100.0 \| 1e-36 \| 2.1e-42 \| 264.7 \| 177 \| (19, 201) \| 291 \| (65, 245) \| 352 \| Nucleoid occlusion protein \| Nucleoid occlusion protein; uniclust \| UniRef100\_A0A011VWW9 \| 100.0 \| 1.3e-36 \| 2.7e-42 \| 263.8 \| 177 \| (18, 200) \| 291 \| (77, 258) \| 349 \| Stage 0 sporulation protein J \| Stage 0 sporulation protein J; uniclust \| UniRef100\_A0A0Q4N015 \| 100.0 \| 1.4e-36 \| 2.8e-42 \| 264.4 \| 266 \| (1, 290) \| 291 \| (19, 287) \| 512 \| Chromosome partitioning protein ParB \| Chromosome partitioning protein ParB; uniclust \| UniRef100\_A0A0G1XWI0 \| 100.0 \| 1.6e-36 \| 3.2e-42 \| 256.5 \| 178 \| (18, 201) \| 291 \| (43, 224) \| 311 \| ParB-like protein partition protein \| ParB-like protein partition protein; uniclust \| UniRef100\_A0A0G3EIV5 \| 100.0 \| 1.6e-36 \| 3.3e-42 \| 262.9 \| 176 \| (18, 199) \| 291 \| (75, 254) \| 355 \| Putative chromosome-partitioning protein ParB \| Putative chromosome-partitioning protein ParB; uniclust \| UniRef100\_A0A0G1TMM8 \| 100.0 \| 1.9e-36 \| 3.7e-42 \| 245.8 \| 170 \| (15, 190) \| 291 \| (29, 201) \| 248 \| ParB-like protein partition protein \| ParB-like protein partition protein; uniclust \| UniRef100\_A0A097AUS5 \| 100.0 \| 2.7e-36 \| 5.3e-42 \| 251.7 \| 178 \| (17, 200) \| 291 \| (51, 233) \| 313 \| Nucleoid occlusion protein Noc \| Nucleoid occlusion protein Noc; uniclust \| UniRef100\_A0A088SZT7 \| 100.0 \| 3.6e-36 \| 7.3e-42 \| 256.6 \| 178 \| (18, 201) \| 291 \| (42, 223) \| 313 \| Chromosome-partitioning protein Spo0J \| Chromosome-partitioning protein Spo0J; uniclust \| UniRef100\_A0A0C2D3E5 \| 100.0 \| 3.9e-36 \| 7.8e-42 \| 265.3 \| 176 \| (18, 199) \| 291 \| (127, 306) \| 405 \| Chromosome (Plasmid) partitioning protein ParB \| Chromosome (Plasmid) partitioning protein ParB; uniclust \| UniRef100\_A0A010PSR0 \| 100.0 \| 4.2e-36 \| 8.7e-42 \| 268.1 \| 177 \| (18, 200) \| 291 \| (96, 276) \| 408 \| Chromosome partitioning protein ParB \| Chromosome partitioning protein ParB; uniclust \| UniRef100\_A0A257WQE5 \| 100.0 \| 4.8e-36 \| 9.4e-42 \| 239.7 \| 169 \| (17, 191) \| 291 \| (45, 217) \| 229 \| Chromosome partitioning protein ParB (Fragment) \| Chromosome partitioning protein ParB (Fragment); uniclust \| UniRef100\_A0A0D0QNA4 \| 100.0 \| 7.3e-36 \| 1.4e-41 \| 252.3 \| 178 \| (18, 201) \| 291 \| (73, 254) \| 339 \| Chromosome (Plasmid) partitioning protein ParB / Stage 0 sporulation protein J \| Chromosome (Plasmid) partitioning protein ParB / Stage 0 sporulation protein J; uniclust \| UniRef100\_A0A0G0M240 \| 100.0 \| 8.9e-36 \| 1.8e-41 \| 252.3 \| 178 \| (18, 201) \| 291 \| (51, 233) \| 318 \| Chromosome segregation DNA-binding protein \| Chromosome segregation DNA-binding protein; uniclust \| UniRef100\_A0A094S1U1 \| 100.0 \| 1.4e-35 \| 2.7e-41 \| 238.1 \| 262 \| (1, 288) \| 291 \| (20, 282) \| 306 \| DNA-binding protein \| DNA-binding protein; uniclust \| UniRef100\_A0A011VV16 \| 100.0 \| 2e-35 \| 4.1e-41 \| 260.0 \| 178 \| (18, 201) \| 291 \| (92, 274) \| 401 \| Chromosome partitioning protein ParB \| Chromosome partitioning protein ParB; uniclust \| UniRef100\_A0A010RRU4 \| 100.0 \| 2.3e-35 \| 4.7e-41 \| 267.7 \| 177 \| (18, 200) \| 291 \| (186, 366) \| 515 \| Probable chromosome-partitioning protein ParB \| Probable chromosome-partitioning protein ParB; uniclust \| UniRef100\_A0A0D5LM02 \| 100.0 \| 2.6e-35 \| 4.9e-41 \| 245.5 \| 174 \| (19, 200) \| 291 \| (60, 239) \| 325 \| Chromosome partitioning protein ParB \| Chromosome partitioning protein ParB; uniclust \| UniRef100\_A0A0A8B5V6 \| 100.0 \| 2.9e-35 \| 5.7e-41 \| 259.2 \| 166 \| (19, 190) \| 291 \| (181, 350) \| 463 \| Chromosome partitioning protein ParB \| Chromosome partitioning protein ParB; uniclust \| UniRef100\_A0A009GHB1 \| 100.0 \| 3.3e-35 \| 6.5e-41 \| 249.2 \| 179 \| (18, 202) \| 291 \| (67, 249) \| 342 \| Probable chromosome-partitioning protein ParB \| Probable chromosome-partitioning protein ParB; uniclust \| UniRef100\_A0A1I1P9F0 \| 100.0 \| 3.5e-35 \| 7e-41 \| 246.6 \| 215 \| (1, 216) \| 291 \| (18, 233) \| 299 \| ParB/RepB/Spo0J family partition protein \| ParB/RepB/Spo0J family partition protein; uniclust \| UniRef100\_A0A1G9Q6I4 \| 100.0 \| 3.7e-35 \| 7.2e-41 \| 244.2 \| 181 \| (14, 199) \| 291 \| (24, 209) \| 316 \| Chromosome segregation DNA-binding protein \| Chromosome segregation DNA-binding protein; uniclust \| UniRef100\_A0A014P5P6 \| 100.0 \| 3.7e-35 \| 7.6e-41 \| 265.9 \| 176 \| (19, 200) \| 291 \| (153, 333) \| 490 \| Chromosome partitioning protein ParB \| Chromosome partitioning protein ParB; uniclust \| UniRef100\_A0A1Q6R1A4 \| 100.0 \| 4.3e-35 \| 8.3e-41 \| 241.5 \| 178 \| (18, 201) \| 291 \| (34, 215) \| 316 \| HTH cro/C1-type domain-containing protein \| HTH cro/C1-type domain-containing protein; uniclust \| UniRef100\_A0A1G3VWE4 \| 100.0 \| 4.4e-35 \| 8.4e-41 \| 241.0 \| 172 \| (16, 193) \| 291 \| (45, 219) \| 306 \| ParB/Sulfiredoxin domain-containing protein \| ParB/Sulfiredoxin domain-containing protein; uniclust \| UniRef100\_A0A011SW19 \| 100.0 \| 5.6e-35 \| 1.1e-40 \| 257.5 \| 173 \| (19, 199) \| 291 \| (84, 262) \| 416 \| Chromosome partitioning protein ParB \| Chromosome partitioning protein ParB; uniclust \| UniRef100\_A0A1B1AIB8 \| 100.0 \| 1.2e-34 \| 2.2e-40 \| 249.5 \| 174 \| (19, 200) \| 291 \| (92, 271) \| 432 \| ParB/Sulfiredoxin domain-containing protein \| ParB/Sulfiredoxin domain-containing protein; uniclust \| UniRef100\_A0A081DF57 \| 100.0 \| 1.9e-34 \| 3.7e-40 \| 239.8 \| 176 \| (19, 200) \| 291 \| (57, 237) \| 313 \| Chromosome (Plasmid) partitioning protein ParB \| Chromosome (Plasmid) partitioning protein ParB; uniclust \| UniRef100\_A0A1F6CPK3 \| 100.0 \| 3.4e-34 \| 6.7e-40 \| 243.5 \| 181 \| (13, 199) \| 291 \| (16, 203) \| 323 \| ParB/Sulfiredoxin domain-containing protein \| ParB/Sulfiredoxin domain-containing protein; uniclust \| UniRef100\_A0A094QAV1 \| 100.0 \| 4e-34 \| 7.9e-40 \| 248.1 \| 177 \| (19, 200) \| 291 \| (61, 241) \| 402 \| Chromosome partitioning protein ParB \| Chromosome partitioning protein ParB; uniclust \| UniRef100\_A0A0G0YQH3 \| 100.0 \| 4.1e-34 \| 8.3e-40 \| 250.7 \| 180 \| (16, 200) \| 291 \| (122, 306) \| 390 \| ParB-like protein partition protein \| ParB-like protein partition protein; uniclust \| UniRef100\_A0A3B8LAV2 \| 100.0 \| 7.7e-34 \| 1.5e-39 \| 241.6 \| 176 \| (18, 199) \| 291 \| (55, 234) \| 332 \| Chromosome partitioning protein ParB \| Chromosome partitioning protein ParB; uniclust \| UniRef100\_A0A2D9PNL7 \| 100.0 \| 8.3e-34 \| 1.6e-39 \| 236.7 \| 181 \| (15, 201) \| 291 \| (44, 227) \| 326 \| Chromosome partitioning protein ParB \| Chromosome partitioning protein ParB; uniclust \| UniRef100\_A0A117LBP9 \| 100.0 \| 1.2e-33 \| 2.3e-39 \| 231.6 \| 178 \| (18, 201) \| 291 \| (35, 216) \| 299 \| Chromosome-partitioning protein ParB \| Chromosome-partitioning protein ParB; uniclust \| UniRef100\_A0A081B6M9 \| 100.0 \| 1.3e-33 \| 2.5e-39 \| 238.0 \| 166 \| (17, 188) \| 291 \| (65, 236) \| 346 \| ParB-like partition protein \| ParB-like partition protein; uniclust \| UniRef100\_A0A142X869 \| 100.0 \| 1.3e-33 \| 2.6e-39 \| 243.6 \| 169 \| (18, 191) \| 291 \| (50, 224) \| 331 \| Chromosome-partitioning protein Spo0J \| Chromosome-partitioning protein Spo0J; uniclust \| UniRef100\_A0A1C3K881 \| 100.0 \| 1.4e-33 \| 2.7e-39 \| 234.3 \| 212 \| (1, 215) \| 291 \| (7, 223) \| 316 \| Predicted transcriptional regulators \| Predicted transcriptional regulators; uniclust \| UniRef100\_A0A1Q6QWS0 \| 100.0 \| 1.5e-33 \| 2.9e-39 \| 248.5 \| 178 \| (18, 201) \| 291 \| (20, 201) \| 434 \| HTH cro/C1-type domain-containing protein \| HTH cro/C1-type domain-containing protein; uniclust \| UniRef100\_A0A1W9QJJ8 \| 100.0 \| 1.5e-33 \| 3e-39 \| 234.3 \| 170 \| (16, 191) \| 291 \| (56, 229) \| 311 \| ParB/Sulfiredoxin domain-containing protein (Fragment) \| ParB/Sulfiredoxin domain-containing protein (Fragment); uniclust \| UniRef100\_A0A1F5N9U7 \| 100.0 \| 1.6e-33 \| 3.2e-39 \| 238.8 \| 181 \| (15, 201) \| 291 \| (61, 246) \| 333 \| ParB/Sulfiredoxin domain-containing protein \| ParB/Sulfiredoxin domain-containing protein; uniclust \| UniRef100\_A0A0F2IXB0 \| 100.0 \| 1.9e-33 \| 3.7e-39 \| 229.3 \| 180 \| (16, 201) \| 291 \| (19, 203) \| 283 \| Stage 0 sporulation protein J \| Stage 0 sporulation protein J; uniclust \| UniRef100\_A0A101FLX1 \| 100.0 \| 1.9e-33 \| 3.8e-39 \| 235.7 \| 183 \| (13, 201) \| 291 \| (31, 216) \| 319 \| ParB-like partition protein \| ParB-like partition protein; uniclust \| UniRef100\_A0A0F9YZP5 \| 100.0 \| 2.1e-33 \| 4e-39 \| 238.9 \| 166 \| (18, 189) \| 291 \| (33, 201) \| 352 \| ParB-like protein partition protein \| ParB-like protein partition protein; uniclust \| UniRef100\_A0A074JJ97 \| 100.0 \| 2.7e-33 \| 5.2e-39 \| 242.8 \| 164 \| (18, 187) \| 291 \| (43, 212) \| 480 \| DNA-binding protein \| DNA-binding protein; uniclust \| UniRef100\_A0A1F2X3R4 \| 100.0 \| 2.9e-33 \| 5.7e-39 \| 237.7 \| 180 \| (14, 199) \| 291 \| (29, 211) \| 319 \| ParB/Sulfiredoxin domain-containing protein \| ParB/Sulfiredoxin domain-containing protein; uniclust \| UniRef100\_A0A024EM96 \| 100.0 \| 3.1e-33 \| 5.9e-39 \| 240.3 \| 256 \| (14, 287) \| 291 \| (78, 357) \| 397 \| ParB/Sulfiredoxin domain-containing protein \| ParB/Sulfiredoxin domain-containing protein; uniclust \| UniRef100\_A0A0G0GAF4 \| 100.0 \| 7.5e-33 \| 1.5e-38 \| 238.3 \| 178 \| (18, 200) \| 291 \| (103, 292) \| 387 \| ParB-like protein partition protein \| ParB-like protein partition protein; uniclust \| UniRef100\_A0A077ZKX5 \| 100.0 \| 9.6e-33 \| 1.9e-38 \| 247.8 \| 179 \| (16, 200) \| 291 \| (279, 463) \| 547 \| CbiA and ParBc and KorB domain containing protein \| CbiA and ParBc and KorB domain containing protein; uniclust \| UniRef100\_A0A062WZP7 \| 100.0 \| 1.4e-32 \| 2.6e-38 \| 240.0 \| 178 \| (18, 200) \| 291 \| (136, 317) \| 472 \| Chromosome segregation DNA-binding protein \| Chromosome segregation DNA-binding protein; uniclust \| UniRef100\_A0A0X7A5Y3 \| 100.0 \| 2.2e-32 \| 4.3e-38 \| 227.7 \| 249 \| (14, 283) \| 291 \| (30, 286) \| 305 \| ParB/Sulfiredoxin domain-containing protein \| ParB/Sulfiredoxin domain-containing protein; uniclust \| UniRef100\_A0A0F8XGC3 \| 99.9 \| 2.5e-32 \| 4.9e-38 \| 238.1 \| 176 \| (13, 193) \| 291 \| (65, 242) \| 379 \| ParB/Sulfiredoxin domain-containing protein \| ParB/Sulfiredoxin domain-containing protein; uniclust \| UniRef100\_A0A212AQY6 \| 99.9 \| 2.7e-32 \| 5.2e-38 \| 215.6 \| 171 \| (16, 192) \| 291 \| (46, 221) \| 231 \| ParB/Sulfiredoxin domain-containing protein \| ParB/Sulfiredoxin domain-containing protein; uniclust \| UniRef100\_A0A0F9ICC8 \| 99.9 \| 2.7e-32 \| 5.5e-38 \| 232.8 \| 179 \| (16, 200) \| 291 \| (24, 209) \| 306 \| ParB/Sulfiredoxin domain-containing protein (Fragment) \| ParB/Sulfiredoxin domain-containing protein (Fragment); uniclust \| UniRef100\_A0A024BX61 \| 99.9 \| 2.9e-32 \| 5.6e-38 \| 242.7 \| 178 \| (18, 200) \| 291 \| (77, 267) \| 498 \| Chromosome partitioning protein ParB \| Chromosome partitioning protein ParB; uniclust \| UniRef100\_A0A1V4VEC1 \| 99.9 \| 3.6e-32 \| 7e-38 \| 229.3 \| 180 \| (16, 200) \| 291 \| (65, 251) \| 328 \| Chromosome-partitioning protein Spo0J \| Chromosome-partitioning protein Spo0J; uniclust \| UniRef100\_A0A0M7DV75 \| 99.9 \| 4.7e-32 \| 8.9e-38 \| 228.5 \| 269 \| (1, 286) \| 291 \| (8, 300) \| 396 \| ParB/RepB/Spo0J family partition protein \| ParB/RepB/Spo0J family partition protein; uniclust \| UniRef100\_A0A257V0R5 \| 99.9 \| 7.2e-32 \| 1.4e-37 \| 213.8 \| 175 \| (16, 195) \| 291 \| (36, 214) \| 237 \| ParB/Sulfiredoxin domain-containing protein \| ParB/Sulfiredoxin domain-containing protein; uniclust \| UniRef100\_A0A066UG41 \| 99.9 \| 9.6e-32 \| 1.9e-37 \| 232.3 \| 194 \| (17, 216) \| 291 \| (123, 329) \| 403 \| Probable chromosome-partitioning protein ParB \| Probable chromosome-partitioning protein ParB; uniclust \| UniRef100\_A0A1C5QNA2 \| 99.9 \| 1.2e-31 \| 2.4e-37 \| 222.9 \| 178 \| (17, 201) \| 291 \| (6, 188) \| 302 \| Nucleoid occlusion protein \| Nucleoid occlusion protein; uniclust \| UniRef100\_A0A011TZQ8 \| 99.9 \| 1.8e-31 \| 3.5e-37 \| 239.8 \| 173 \| (17, 194) \| 291 \| (212, 388) \| 538 \| Chromosome partitioning protein ParB \| Chromosome partitioning protein ParB; uniclust \| UniRef100\_A0A929SNF7 \| 99.9 \| 1.8e-31 \| 3.6e-37 \| 243.5 \| 178 \| (18, 201) \| 291 \| (5, 186) \| 584 \| ParB/RepB/Spo0J family partition protein \| ParB/RepB/Spo0J family partition protein; uniclust \| UniRef100\_A0A1A9KGS4 \| 99.9 \| 2e-31 \| 3.7e-37 \| 217.7 \| 273 \| (2, 290) \| 291 \| (71, 354) \| 357 \| ParB/Sulfiredoxin domain-containing protein \| ParB/Sulfiredoxin domain-containing protein; uniclust \| UniRef100\_A0A2H0T7Y9 \| 99.9 \| 2.3e-31 \| 4.4e-37 \| 206.5 \| 160 \| (5, 170) \| 291 \| (15, 177) \| 178 \| Chromosome partitioning protein ParB (Fragment) \| Chromosome partitioning protein ParB (Fragment); uniclust \| UniRef100\_A0A068VSG1 \| 99.9 \| 2.6e-31 \| 4.9e-37 \| 234.7 \| 179 \| (17, 200) \| 291 \| (243, 425) \| 533 \| Chromosome partitioning protein \| Chromosome partitioning protein; uniclust \| UniRef100\_A0A3A0CR42 \| 99.9 \| 2.5e-31 \| 5e-37 \| 225.4 \| 170 \| (15, 190) \| 291 \| (47, 220) \| 333 \| HTH cro/C1-type domain-containing protein \| HTH cro/C1-type domain-containing protein; uniclust \| UniRef100\_A0A060DLG4 \| 99.9 \| 2.7e-31 \| 5.5e-37 \| 233.8 \| 175 \| (16, 198) \| 291 \| (76, 258) \| 389 \| ParB/Sulfiredoxin domain-containing protein \| ParB/Sulfiredoxin domain-containing protein; uniclust \| UniRef100\_A0A022KT50 \| 99.9 \| 3.1e-31 \| 6.2e-37 \| 242.2 \| 176 \| (15, 196) \| 291 \| (302, 484) \| 593 \| Chromosome partitioning protein ParB \| Chromosome partitioning protein ParB; uniclust \| UniRef100\_A0A2J0LBM3 \| 99.9 \| 9e-31 \| 1.7e-36 \| 212.2 \| 166 \| (17, 188) \| 291 \| (57, 225) \| 269 \| Chromosome partitioning protein ParB (Fragment) \| Chromosome partitioning protein ParB (Fragment); uniclust \| UniRef100\_A0A096HPB0 \| 99.9 \| 1.3e-30 \| 2.6e-36 \| 222.2 \| 272 \| (3, 286) \| 291 \| (17, 321) \| 353 \| Chromosome partitioning protein ParB \| Chromosome partitioning protein ParB; uniclust \| UniRef100\_A0A022PNW0 \| 99.9 \| 1.6e-30 \| 3e-36 \| 227.1 \| 269 \| (13, 288) \| 291 \| (90, 371) \| 486 \| ParB/Sulfiredoxin domain-containing protein \| ParB/Sulfiredoxin domain-containing protein; uniclust \| UniRef100\_A0A075GB43 \| 99.9 \| 2e-30 \| 3.9e-36 \| 222.7 \| 178 \| (17, 200) \| 291 \| (48, 229) \| 359 \| Transcriptional regulator (ParB, spo0J) \| Transcriptional regulator (ParB, spo0J); uniclust \| UniRef100\_A0A0F9U0H1 \| 99.9 \| 2.3e-30 \| 4.7e-36 \| 227.9 \| 203 \| (14, 216) \| 291 \| (72, 285) \| 392 \| ParB/Sulfiredoxin domain-containing protein \| ParB/Sulfiredoxin domain-containing protein; uniclust \| UniRef100\_A0A930FAM5 \| 99.9 \| 2.8e-30 \| 5.4e-36 \| 194.7 \| 160 \| (18, 183) \| 291 \| (4, 166) \| 167 \| ParB/RepB/Spo0J family partition protein (Fragment) \| ParB/RepB/Spo0J family partition protein (Fragment); uniclust \| UniRef100\_A0A233HUD9 \| 99.9 \| 3.8e-30 \| 7.4e-36 \| 219.6 \| 264 \| (13, 287) \| 291 \| (50, 321) \| 373 \| Repressor KorB domain-containing protein \| Repressor KorB domain-containing protein; uniclust \| UniRef100\_A0A382LW29 \| 99.9 \| 4e-30 \| 7.5e-36 \| 194.4 \| 152 \| (15, 172) \| 291 \| (19, 174) \| 175 \| ParB/Sulfiredoxin domain-containing protein (Fragment) \| ParB/Sulfiredoxin domain-containing protein (Fragment); uniclust \| UniRef100\_A0A7X9GCQ7 \| 99.9 \| 5.2e-30 \| 9.9e-36 \| 213.2 \| 178 \| (18, 201) \| 291 \| (8, 189) \| 343 \| ParB/RepB/Spo0J family partition protein \| ParB/RepB/Spo0J family partition protein; uniclust \| UniRef100\_A0A4V1URX4 \| 99.9 \| 6.3e-30 \| 1.2e-35 \| 214.1 \| 173 \| (20, 197) \| 291 \| (49, 227) \| 318 \| ParB/RepB/Spo0J family partition protein \| ParB/RepB/Spo0J family partition protein; uniclust \| UniRef100\_A0A021VUN1 \| 99.9 \| 7e-30 \| 1.4e-35 \| 213.3 \| 167 \| (15, 187) \| 291 \| (25, 192) \| 297 \| ParB/Sulfiredoxin domain-containing protein \| ParB/Sulfiredoxin domain-containing protein; uniclust \| UniRef100\_A0A060DS69 \| 99.9 \| 7.4e-30 \| 1.5e-35 \| 222.6 \| 173 \| (18, 197) \| 291 \| (89, 267) \| 361 \| Chromosome partitioning protein \| Chromosome partitioning protein; uniclust \| UniRef100\_A0A3L7VX77 \| 99.9 \| 7.8e-30 \| 1.5e-35 \| 200.8 \| 165 \| (21, 191) \| 291 \| (44, 213) \| 216 \| ParB/RepB/Spo0J family partition protein (Fragment) \| ParB/RepB/Spo0J family partition protein (Fragment); uniclust \| UniRef100\_A0A0L6JYD4 \| 99.9 \| 9.1e-30 \| 1.8e-35 \| 211.5 \| 193 \| (14, 214) \| 291 \| (31, 229) \| 302 \| ParB/Sulfiredoxin domain-containing protein \| ParB/Sulfiredoxin domain-containing protein; uniclust \| UniRef100\_A0A0T5ZL74 \| 99.9 \| 9.4e-30 \| 1.8e-35 \| 215.1 \| 173 \| (16, 194) \| 291 \| (67, 243) \| 379 \| Chromosome partitioning protein ParB, chromosome partitioning protein, ParB family \| Chromosome partitioning protein ParB, chromosome partitioning protein, ParB family; uniclust \| UniRef100\_A0A1F4I2D7 \| 99.9 \| 9.9e-30 \| 1.9e-35 \| 216.0 \| 165 \| (18, 187) \| 291 \| (48, 223) \| 344 \| ParB/Sulfiredoxin domain-containing protein \| ParB/Sulfiredoxin domain-containing protein; uniclust \| UniRef100\_A0A2G2FUH5 \| 99.9 \| 1.2e-29 \| 2.3e-35 \| 214.2 \| 171 \| (16, 192) \| 291 \| (36, 211) \| 342 \| Chromosome partitioning protein ParB \| Chromosome partitioning protein ParB; uniclust \| UniRef100\_A0A0F8XDG1 \| 99.9 \| 1.5e-29 \| 2.9e-35 \| 213.4 \| 173 \| (15, 193) \| 291 \| (10, 185) \| 287 \| ParB/Spo0J HTH domain-containing protein (Fragment) \| ParB/Spo0J HTH domain-containing protein (Fragment); uniclust \| UniRef100\_A0A139NDU2 \| 99.9 \| 1.6e-29 \| 3e-35 \| 194.0 \| 178 \| (18, 201) \| 291 \| (9, 189) \| 209 \| Chromosome (Plasmid) partitioning protein ParB / Stage 0 sporulation protein J \| Chromosome (Plasmid) partitioning protein ParB / Stage 0 sporulation protein J; uniclust \| UniRef100\_A0A432HCS4 \| 99.9 \| 1.8e-29 \| 3.3e-35 \| 206.8 \| 168 \| (16, 188) \| 291 \| (59, 229) \| 325 \| Chromosome partitioning protein ParB \| Chromosome partitioning protein ParB; uniclust \| UniRef100\_A0A090I4P8 \| 99.9 \| 1.7e-29 \| 3.4e-35 \| 219.0 \| 180 \| (17, 202) \| 291 \| (30, 214) \| 392 \| ParB/Sulfiredoxin domain-containing protein \| ParB/Sulfiredoxin domain-containing protein; uniclust \| UniRef100\_A0A088C411 \| 99.9 \| 1.9e-29 \| 3.6e-35 \| 201.1 \| 196 \| (13, 214) \| 291 \| (20, 216) \| 262 \| ParB-like protein \| ParB-like protein; uniclust \| UniRef100\_A0A0C1V2T0 \| 99.9 \| 2e-29 \| 3.8e-35 \| 221.3 \| 180 \| (1, 189) \| 291 \| (2, 184) \| 540 \| ParB/RepB/Spo0J family partition protein \| ParB/RepB/Spo0J family partition protein; uniclust \| UniRef100\_A0A086DUB0 \| 99.9 \| 2.1e-29 \| 4.1e-35 \| 208.4 \| 172 \| (16, 192) \| 291 \| (111, 288) \| 295 \| Chromosome partitioning protein ParB (Fragment) \| Chromosome partitioning protein ParB (Fragment); uniclust \| UniRef100\_A0A3G8F6G3 \| 99.9 \| 2.1e-29 \| 4.2e-35 \| 219.0 \| 173 \| (17, 195) \| 291 \| (34, 210) \| 361 \| Chromosome (Plasmid) partitioning protein \| Chromosome (Plasmid) partitioning protein; uniclust \| UniRef100\_A0A086D7K6 \| 99.9 \| 2.6e-29 \| 5.1e-35 \| 211.1 \| 180 \| (17, 202) \| 291 \| (49, 232) \| 332 \| Probable chromosome-partitioning protein ParB (Fragment) \| Probable chromosome-partitioning protein ParB (Fragment); uniclust \| UniRef100\_D1B9D9 \| 99.9 \| 3e-29 \| 5.6e-35 \| 209.0 \| 178 \| (17, 200) \| 291 \| (96, 276) \| 368 \| ParB-like partition protein \| ParB-like partition protein; uniclust \| UniRef100\_A0A062Y0Q3 \| 99.9 \| 2.8e-29 \| 5.6e-35 \| 216.8 \| 167 \| (17, 191) \| 291 \| (65, 236) \| 354 \| ParB/Sulfiredoxin domain-containing protein \| ParB/Sulfiredoxin domain-containing protein; uniclust \| UniRef100\_A0A1F9PB14 \| 99.9 \| 3.2e-29 \| 6.1e-35 \| 197.7 \| 170 \| (14, 188) \| 291 \| (53, 225) \| 258 \| Chromosome partitioning protein ParB (Fragment) \| Chromosome partitioning protein ParB (Fragment); uniclust \| UniRef100\_A0A2G4ISH6 \| 99.9 \| 3.6e-29 \| 6.9e-35 \| 213.9 \| 165 \| (18, 188) \| 291 \| (40, 210) \| 432 \| Chromosome partitioning protein ParB \| Chromosome partitioning protein ParB; uniclust \| UniRef100\_A0A1P8WKH2 \| 99.9 \| 3.7e-29 \| 7.2e-35 \| 209.3 \| 171 \| (17, 191) \| 291 \| (35, 208) \| 322 \| Putative chromosome-partitioning protein ParB \| Putative chromosome-partitioning protein ParB; uniclust \| UniRef100\_A0A0F9CE03 \| 99.9 \| 4e-29 \| 7.9e-35 \| 208.7 \| 173 \| (15, 193) \| 291 \| (11, 186) \| 275 \| ParB/Sulfiredoxin domain-containing protein \| ParB/Sulfiredoxin domain-containing protein; uniclust \| UniRef100\_A0A0F9EZV6 \| 99.9 \| 4.6e-29 \| 9e-35 \| 223.9 \| 177 \| (15, 197) \| 291 \| (10, 192) \| 536 \| ParB/Sulfiredoxin domain-containing protein \| ParB/Sulfiredoxin domain-containing protein; uniclust \| UniRef100\_A0A095WUL4 \| 99.9 \| 5.5e-29 \| 1.1e-34 \| 222.1 \| 175 \| (16, 195) \| 291 \| (194, 387) \| 560 \| Chromosome partitioning protein ParB \| Chromosome partitioning protein ParB; uniclust \| UniRef100\_A0A0F2RAT7 \| 99.9 \| 5.5e-29 \| 1.1e-34 \| 212.0 \| 174 \| (18, 197) \| 291 \| (64, 241) \| 332 \| ParB/Sulfiredoxin domain-containing protein \| ParB/Sulfiredoxin domain-containing protein; uniclust \| UniRef100\_A0A0G1ET61 \| 99.9 \| 6.4e-29 \| 1.3e-34 \| 219.6 \| 180 \| (15, 199) \| 291 \| (38, 232) \| 479 \| ParB-like protein partition protein \| ParB-like protein partition protein; uniclust \| UniRef100\_A0A2A5EJ42 \| 99.9 \| 7.7e-29 \| 1.5e-34 \| 191.7 \| 132 \| (19, 156) \| 291 \| (38, 175) \| 175 \| Chromosome partitioning protein ParB (Fragment) \| Chromosome partitioning protein ParB (Fragment); uniclust \| UniRef100\_A0A7R8WWZ7 \| 99.9 \| 8.6e-29 \| 1.6e-34 \| 191.0 \| 166 \| (23, 194) \| 291 \| (6, 175) \| 209 \| ParB/Sulfiredoxin domain-containing protein (Fragment) \| ParB/Sulfiredoxin domain-containing protein (Fragment); uniclust \| UniRef100\_A0A2U3KH09 \| 99.9 \| 1e-28 \| 1.9e-34 \| 211.5 \| 169 \| (15, 189) \| 291 \| (147, 320) \| 421 \| Chromosome segregation DNA-binding protein \| Chromosome segregation DNA-binding protein; uniclust \| UniRef100\_UPI0004B65802 \| 99.9 \| 1.2e-28 \| 2.3e-34 \| 197.3 \| 269 \| (2, 289) \| 291 \| (4, 286) \| 289 \| ParB N-terminal domain-containing protein \| ParB N-terminal domain-containing protein; uniclust \| UniRef100\_A0A078MLS8 \| 99.9 \| 1.3e-28 \| 2.4e-34 \| 209.0 \| 176 \| (17, 198) \| 291 \| (123, 305) \| 405 \| Putative chromosome-partitioning protein ParB \| Putative chromosome-partitioning protein ParB; uniclust \| UniRef100\_A0A085FMH6 \| 99.9 \| 1.3e-28 \| 2.6e-34 \| 211.3 \| 203 \| (14, 216) \| 291 \| (66, 280) \| 357 \| Uncharacterized protein \| Uncharacterized protein; uniclust \| UniRef100\_A0A352SVJ6 \| 99.9 \| 1.7e-28 \| 3.1e-34 \| 185.7 \| 162 \| (13, 180) \| 291 \| (18, 183) \| 183 \| Nucleoid occlusion protein (Fragment) \| Nucleoid occlusion protein (Fragment); uniclust \| UniRef100\_A0A3M0WWG3 \| 99.9 \| 1.9e-28 \| 3.6e-34 \| 188.3 \| 133 \| (16, 154) \| 291 \| (28, 164) \| 165 \| ParB/RepB/Spo0J family partition protein (Fragment) \| ParB/RepB/Spo0J family partition protein (Fragment); uniclust \| UniRef100\_A0A5C6FSQ3 \| 99.9 \| 1.9e-28 \| 3.7e-34 \| 200.2 \| 180 \| (16, 201) \| 291 \| (45, 231) \| 308 \| Nucleoid occlusion protein \| Nucleoid occlusion protein; uniclust \| UniRef100\_A0A517R8W8 \| 99.9 \| 2.1e-28 \| 4.1e-34 \| 202.9 \| 178 \| (18, 199) \| 291 \| (48, 231) \| 308 \| Putative chromosome-partitioning protein ParB \| Putative chromosome-partitioning protein ParB; uniclust \| UniRef100\_A0A1V4R7Y3 \| 99.9 \| 2.4e-28 \| 4.6e-34 \| 204.8 \| 164 \| (22, 191) \| 291 \| (41, 210) \| 298 \| ParB/Sulfiredoxin domain-containing protein \| ParB/Sulfiredoxin domain-containing protein; uniclust \| UniRef100\_A0A524ABT4 \| 99.9 \| 2.4e-28 \| 4.6e-34 \| 201.3 \| 167 \| (18, 190) \| 291 \| (2, 171) \| 271 \| ParB/RepB/Spo0J family partition protein \| ParB/RepB/Spo0J family partition protein; uniclust \| UniRef100\_A0A2N8GU19 \| 99.9 \| 3.5e-28 \| 6.5e-34 \| 187.1 \| 177 \| (16, 198) \| 291 \| (39, 220) \| 223 \| Nucleoid occlusion protein (Fragment) \| Nucleoid occlusion protein (Fragment); uniclust \| UniRef100\_A0A4P7A9C7 \| 99.9 \| 4.2e-28 \| 7.9e-34 \| 200.2 \| 212 \| (3, 217) \| 291 \| (6, 225) \| 319 \| ParB family chromosome partitioning protein \| ParB family chromosome partitioning protein; uniclust \| UniRef100\_A0A059W6T3 \| 99.9 \| 4.2e-28 \| 8e-34 \| 207.6 \| 175 \| (19, 198) \| 291 \| (33, 212) \| 415 \| Chromosome partitioning protein ParB \| Chromosome partitioning protein ParB; uniclust \| UniRef100\_A0A0R2P7U3 \| 99.9 \| 4.6e-28 \| 8.7e-34 \| 198.2 \| 179 \| (16, 199) \| 291 \| (52, 233) \| 313 \| Probable chromosome-partitioning protein ParB \| Probable chromosome-partitioning protein ParB; uniclust \| UniRef100\_A0A2D4RST1 \| 99.9 \| 5e-28 \| 9.4e-34 \| 194.5 \| 172 \| (16, 193) \| 291 \| (11, 189) \| 291 \| Chromosome partitioning protein ParB \| Chromosome partitioning protein ParB; uniclust \| UniRef100\_A0A0F9CXE9 \| 99.9 \| 4.7e-28 \| 9.4e-34 \| 207.9 \| 170 \| (17, 192) \| 291 \| (29, 201) \| 317 \| ParB/Sulfiredoxin domain-containing protein \| ParB/Sulfiredoxin domain-containing protein; uniclust \| UniRef100\_A0A0M9UL29 \| 99.9 \| 5.7e-28 \| 1.1e-33 \| 218.2 \| 170 \| (15, 190) \| 291 \| (401, 578) \| 679 \| Chromosome (Plasmid) partitioning protein ParB \| Chromosome (Plasmid) partitioning protein ParB; uniclust \| UniRef100\_UPI0004B4361E \| 99.9 \| 5.9e-28 \| 1.1e-33 \| 200.5 \| 185 \| (18, 211) \| 291 \| (84, 274) \| 338 \| ParB/RepB/Spo0J family partition protein \| ParB/RepB/Spo0J family partition protein; uniclust \| UniRef100\_A0A402D6S3 \| 99.9 \| 6e-28 \| 1.1e-33 \| 208.1 \| 169 \| (15, 189) \| 291 \| (72, 248) \| 474 \| Chromosome partitioning protein ParB \| Chromosome partitioning protein ParB; uniclust \| UniRef100\_A0A497B792 \| 99.9 \| 7.9e-28 \| 1.5e-33 \| 199.3 \| 170 \| (16, 190) \| 291 \| (32, 203) \| 314 \| ParB/Sulfiredoxin domain-containing protein \| ParB/Sulfiredoxin domain-containing protein; uniclust \| UniRef100\_A0A1F8TYH4 \| 99.9 \| 1e-27 \| 2e-33 \| 198.8 \| 189 \| (3, 199) \| 291 \| (8, 204) \| 279 \| Chromosome partitioning protein ParB \| Chromosome partitioning protein ParB; uniclust \| UniRef100\_A0A0J5KM24 \| 99.9 \| 1.2e-27 \| 2.2e-33 \| 196.3 \| 256 \| (2, 286) \| 291 \| (4, 265) \| 367 \| Chromosome partitioning protein ParB \| Chromosome partitioning protein ParB; uniclust \| UniRef100\_A0A2E6VPY2 \| 99.9 \| 1.8e-27 \| 3.3e-33 \| 195.9 \| 168 \| (17, 189) \| 291 \| (39, 211) \| 370 \| Chromosome partitioning protein ParB \| Chromosome partitioning protein ParB; uniclust \| UniRef100\_A0A009RW17 \| 99.9 \| 1.6e-27 \| 3.3e-33 \| 209.9 \| 161 \| (21, 187) \| 291 \| (67, 231) \| 362 \| ParB/RepB/Spo0J family partition domain protein \| ParB/RepB/Spo0J family partition domain protein; uniclust \| UniRef100\_A0A1E3ZL90 \| 99.9 \| 1.9e-27 \| 3.7e-33 \| 220.6 \| 180 \| (15, 200) \| 291 \| (82, 268) \| 726 \| ParB/Sulfiredoxin domain-containing protein \| ParB/Sulfiredoxin domain-containing protein; uniclust \| UniRef100\_A0A060D5B2 \| 99.9 \| 2.1e-27 \| 4.1e-33 \| 199.5 \| 182 \| (2, 184) \| 291 \| (12, 208) \| 296 \| Uncharacterized protein \| Uncharacterized protein; uniclust \| UniRef100\_A0A2E3TXI9 \| 99.9 \| 2.3e-27 \| 4.3e-33 \| 192.2 \| 164 \| (18, 187) \| 291 \| (35, 201) \| 295 \| Chromosome partitioning protein ParB \| Chromosome partitioning protein ParB; uniclust \| UniRef100\_A0A171DQU0 \| 99.9 \| 2.6e-27 \| 5e-33 \| 198.9 \| 178 \| (17, 199) \| 291 \| (41, 234) \| 313 \| Chromosome partitioning protein ParB \| Chromosome partitioning protein ParB; uniclust \| UniRef100\_A0A015USM3 \| 99.9 \| 2.5e-27 \| 5e-33 \| 221.2 \| 178 \| (17, 200) \| 291 \| (103, 284) \| 724 \| ParB/RepB/Spo0J family partition domain protein \| ParB/RepB/Spo0J family partition domain protein; uniclust \| UniRef100\_A0A2N7QGC9 \| 99.9 \| 2.6e-27 \| 5.1e-33 \| 186.2 \| 134 \| (18, 157) \| 291 \| (54, 190) \| 190 \| ParB/Sulfiredoxin domain-containing protein \| ParB/Sulfiredoxin domain-containing protein; uniclust \| UniRef100\_A0A023WZP1 \| 99.9 \| 2.6e-27 \| 5.2e-33 \| 205.4 \| 166 \| (18, 189) \| 291 \| (77, 249) \| 349 \| Partitioning protein \| Partitioning protein; uniclust \| UniRef100\_A0A938R653 \| 99.9 \| 3.3e-27 \| 6.2e-33 \| 197.2 \| 176 \| (17, 198) \| 291 \| (60, 238) \| 361 \| ParB/RepB/Spo0J family partition protein \| ParB/RepB/Spo0J family partition protein; uniclust \| UniRef100\_A0A0F5MQM1 \| 99.9 \| 3.8e-27 \| 7.6e-33 \| 198.2 \| 173 \| (17, 195) \| 291 \| (40, 219) \| 284 \| Probable chromosome-partitioning protein ParB \| Probable chromosome-partitioning protein ParB; uniclust \| UniRef100\_A0A1F2QAI7 \| 99.9 \| 4.2e-27 \| 8.1e-33 \| 185.6 \| 164 \| (20, 190) \| 291 \| (32, 199) \| 221 \| ParB/Sulfiredoxin domain-containing protein (Fragment) \| ParB/Sulfiredoxin domain-containing protein (Fragment); uniclust \| UniRef100\_A0A1G2ESN9 \| 99.9 \| 4.2e-27 \| 8.2e-33 \| 199.8 \| 182 \| (13, 199) \| 291 \| (36, 228) \| 339 \| ParB/Sulfiredoxin domain-containing protein \| ParB/Sulfiredoxin domain-containing protein; uniclust \| UniRef100\_A0A083UPG1 \| 99.9 \| 5.2e-27 \| 1e-32 \| 203.7 \| 178 \| (17, 200) \| 291 \| (91, 275) \| 361 \| Partitioning protein \| Partitioning protein; uniclust \| UniRef100\_A0A0F9HNR6 \| 99.9 \| 5.6e-27 \| 1.1e-32 \| 183.0 \| 128 \| (17, 150) \| 291 \| (56, 187) \| 187 \| ParB/Sulfiredoxin domain-containing protein (Fragment) \| ParB/Sulfiredoxin domain-containing protein (Fragment); uniclust \| UniRef100\_A0A0F9MK30 \| 99.9 \| 5.8e-27 \| 1.1e-32 \| 193.8 \| 171 \| (14, 190) \| 291 \| (4, 177) \| 276 \| ParB/Sulfiredoxin domain-containing protein \| ParB/Sulfiredoxin domain-containing protein; uniclust \| UniRef100\_A0A0A8WV70 \| 99.9 \| 5.8e-27 \| 1.2e-32 \| 201.9 \| 168 \| (13, 188) \| 291 \| (55, 226) \| 325 \| Putative chromosome-partitioning protein ParB \| Putative chromosome-partitioning protein ParB; uniclust \| UniRef100\_A0A1F8L508 \| 99.9 \| 7.9e-27 \| 1.5e-32 \| 187.4 \| 179 \| (17, 201) \| 291 \| (28, 208) \| 292 \| ParB/Sulfiredoxin domain-containing protein \| ParB/Sulfiredoxin domain-containing protein; uniclust \| UniRef100\_A0A0G1NIH2 \| 99.9 \| 8.2e-27 \| 1.5e-32 \| 200.1 \| 182 \| (17, 202) \| 291 \| (178, 363) \| 463 \| ParB-like protein partition protein \| ParB-like protein partition protein; uniclust \| UniRef100\_A0A0U5F961 \| 99.9 \| 9.5e-27 \| 1.8e-32 \| 196.6 \| 212 \| (3, 216) \| 291 \| (19, 236) \| 395 \| FtsK gamma domain-containing protein \| FtsK gamma domain-containing protein; uniclust \| UniRef100\_A0A142XAM1 \| 99.9 \| 1e-26 \| 2e-32 \| 194.5 \| 173 \| (16, 198) \| 291 \| (15, 192) \| 283 \| Putative chromosome-partitioning protein ParB \| Putative chromosome-partitioning protein ParB; uniclust \| UniRef100\_A0A072MWG2 \| 99.9 \| 1.1e-26 \| 2.1e-32 \| 208.4 \| 175 \| (15, 195) \| 291 \| (232, 434) \| 546 \| Chromosome partitioning protein ParB \| Chromosome partitioning protein ParB; uniclust \| UniRef100\_A0A2H5YTQ1 \| 99.9 \| 1.1e-26 \| 2.2e-32 \| 198.9 \| 177 \| (19, 200) \| 291 \| (53, 234) \| 370 \| Putative chromosome-partitioning protein ParB \| Putative chromosome-partitioning protein ParB; uniclust \| UniRef100\_A0A3D3EDT8 \| 99.9 \| 1.1e-26 \| 2.2e-32 \| 182.2 \| 144 \| (18, 167) \| 291 \| (52, 201) \| 206 \| Chromosome partitioning protein ParB (Fragment) \| Chromosome partitioning protein ParB (Fragment); uniclust \| UniRef100\_A0A1G0G5V5 \| 99.9 \| 1.2e-26 \| 2.3e-32 \| 192.7 \| 176 \| (17, 199) \| 291 \| (22, 204) \| 325 \| Probable chromosome-partitioning protein ParB \| Probable chromosome-partitioning protein ParB; uniclust \| UniRef100\_A0A081XJM3 \| 99.9 \| 1.2e-26 \| 2.4e-32 \| 203.7 \| 177 \| (20, 201) \| 291 \| (58, 252) \| 408 \| Plasmid partitioning protein \| Plasmid partitioning protein; uniclust \| UniRef100\_A0A0Q6YVV5 \| 99.9 \| 1.6e-26 \| 3e-32 \| 198.2 \| 180 \| (17, 200) \| 291 \| (59, 243) \| 414 \| ParB/Sulfiredoxin domain-containing protein \| ParB/Sulfiredoxin domain-containing protein; uniclust \| UniRef100\_A0A3A0DHT8 \| 99.9 \| 1.8e-26 \| 3.3e-32 \| 192.4 \| 176 \| (17, 198) \| 291 \| (61, 241) \| 400 \| Chromosome partitioning protein ParB \| Chromosome partitioning protein ParB; uniclust \| UniRef100\_A0A1Y4U457 \| 99.9 \| 1.8e-26 \| 3.5e-32 \| 185.9 \| 177 \| (17, 199) \| 291 \| (72, 251) \| 275 \| Chromosome partitioning protein ParB (Fragment) \| Chromosome partitioning protein ParB (Fragment); uniclust \| UniRef100\_A0A0P6YU31 \| 99.9 \| 2e-26 \| 3.8e-32 \| 199.1 \| 179 \| (16, 200) \| 291 \| (46, 236) \| 405 \| Chromosome partitioning protein ParB \| Chromosome partitioning protein ParB; uniclust \| UniRef100\_A0A1A5PA14 \| 99.9 \| 3.6e-26 \| 6.9e-32 \| 197.1 \| 176 \| (22, 202) \| 291 \| (58, 250) \| 390 \| ParB/Sulfiredoxin domain-containing protein \| ParB/Sulfiredoxin domain-containing protein; uniclust \| UniRef100\_A0A517P8X0 \| 99.9 \| 3.8e-26 \| 7.1e-32 \| 191.6 \| 177 \| (19, 200) \| 291 \| (56, 234) \| 407 \| Chromosome-partitioning protein Spo0J \| Chromosome-partitioning protein Spo0J; uniclust \| UniRef100\_A0A1F4ZSL5 \| 99.9 \| 3.7e-26 \| 7.3e-32 \| 211.1 \| 174 \| (14, 190) \| 291 \| (32, 214) \| 610 \| ParB/Sulfiredoxin domain-containing protein \| ParB/Sulfiredoxin domain-containing protein; uniclust \| UniRef100\_A0A2S1XUK5 \| 99.9 \| 3.8e-26 \| 7.5e-32 \| 195.0 \| 173 \| (17, 195) \| 291 \| (52, 229) \| 333 \| Chromosome partitioning protein \| Chromosome partitioning protein; uniclust \| UniRef100\_A0A356EUL3 \| 99.9 \| 4.7e-26 \| 8.7e-32 \| 191.9 \| 170 \| (16, 191) \| 291 \| (159, 331) \| 442 \| ParB/Sulfiredoxin domain-containing protein \| ParB/Sulfiredoxin domain-containing protein; uniclust \| UniRef100\_A0A928HHV2 \| 99.9 \| 4.7e-26 \| 8.8e-32 \| 192.5 \| 166 \| (17, 188) \| 291 \| (172, 340) \| 445 \| ParB/RepB/Spo0J family partition protein \| ParB/RepB/Spo0J family partition protein; uniclust \| UniRef100\_A0A0F5ILC4 \| 99.9 \| 4.7e-26 \| 8.9e-32 \| 184.5 \| 178 \| (17, 200) \| 291 \| (11, 194) \| 268 \| ParB-like partition protein \| ParB-like partition protein; uniclust \| UniRef100\_A0A0Q5I357 \| 99.9 \| 4.7e-26 \| 9.2e-32 \| 195.1 \| 175 \| (20, 200) \| 291 \| (36, 241) \| 337 \| ParB/Sulfiredoxin domain-containing protein \| ParB/Sulfiredoxin domain-containing protein; uniclust \| UniRef100\_A0A432MEU0 \| 99.9 \| 4.9e-26 \| 9.3e-32 \| 190.3 \| 169 \| (19, 191) \| 291 \| (58, 231) \| 365 \| ParB/RepB/Spo0J family partition protein \| ParB/RepB/Spo0J family partition protein; uniclust \| UniRef100\_A0A015M8C0 \| 99.9 \| 5.3e-26 \| 1.1e-31 \| 203.5 \| 176 \| (17, 197) \| 291 \| (79, 274) \| 424 \| Plasmid stablization protein ParB \| Plasmid stablization protein ParB; uniclust \| UniRef100\_A0A087ABQ3 \| 99.9 \| 5.3e-26 \| 1.1e-31 \| 207.8 \| 167 \| (16, 188) \| 291 \| (26, 195) \| 529 \| Chromosome partitioning protein parB \| Chromosome partitioning protein parB; uniclust \| UniRef100\_A0A095TLQ3 \| 99.9 \| 5.5e-26 \| 1.1e-31 \| 201.7 \| 174 \| (17, 198) \| 291 \| (40, 222) \| 415 \| ParB-like partitioning protein \| ParB-like partitioning protein; uniclust \| UniRef100\_UPI0004152AE1 \| 99.9 \| 5.8e-26 \| 1.1e-31 \| 198.7 \| 178 \| (19, 202) \| 291 \| (290, 471) \| 573 \| ParB/RepB/Spo0J family partition protein \| ParB/RepB/Spo0J family partition protein; uniclust \| UniRef100\_A0A077XU49 \| 99.9 \| 6.1e-26 \| 1.2e-31 \| 210.4 \| 181 \| (17, 203) \| 291 \| (60, 243) \| 659 \| Putative ParB-like partition protein \| Putative ParB-like partition protein; uniclust \| UniRef100\_A0A1G1ZRB4 \| 99.9 \| 6.7e-26 \| 1.3e-31 \| 189.1 \| 174 \| (15, 193) \| 291 \| (13, 197) \| 366 \| ParB/Sulfiredoxin domain-containing protein \| ParB/Sulfiredoxin domain-containing protein; uniclust \| UniRef100\_A0A1C6R5S8 \| 99.9 \| 7.4e-26 \| 1.4e-31 \| 202.9 \| 172 \| (15, 193) \| 291 \| (103, 292) \| 538 \| ParB/RepB/Spo0J family partition protein \| ParB/RepB/Spo0J family partition protein; uniclust \| UniRef100\_A0A024YNU6 \| 99.9 \| 7.6e-26 \| 1.4e-31 \| 193.0 \| 180 \| (16, 200) \| 291 \| (19, 204) \| 402 \| Chromosome partitioning protein \| Chromosome partitioning protein; uniclust \| UniRef100\_A0A963MDU3 \| 99.9 \| 8.9e-26 \| 1.6e-31 \| 164.8 \| 150 \| (32, 186) \| 291 \| (3, 155) \| 170 \| ParB/RepB/Spo0J family partition protein (Fragment) \| ParB/RepB/Spo0J family partition protein (Fragment); uniclust \| UniRef100\_A0A1H6C3X3 \| 99.9 \| 8.5e-26 \| 1.7e-31 \| 202.7 \| 186 \| (14, 206) \| 291 \| (26, 218) \| 555 \| Chromosome partitioning protein, ParB family \| Chromosome partitioning protein, ParB family; uniclust \| UniRef100\_A0A1F5DHD6 \| 99.9 \| 8.6e-26 \| 1.7e-31 \| 193.4 \| 180 \| (15, 200) \| 291 \| (15, 212) \| 368 \| ParB/Sulfiredoxin domain-containing protein \| ParB/Sulfiredoxin domain-containing protein; uniclust \| UniRef100\_A0A1H6A853 \| 99.9 \| 9.7e-26 \| 1.8e-31 \| 182.0 \| 165 \| (15, 185) \| 291 \| (14, 182) \| 273 \| Chromosome partitioning protein, ParB family \| Chromosome partitioning protein, ParB family; uniclust \| UniRef100\_A0A0U3N394 \| 99.9 \| 1.1e-25 \| 2.1e-31 \| 208.4 \| 182 \| (17, 200) \| 291 \| (81, 269) \| 675 \| ParB/Sulfiredoxin domain-containing protein \| ParB/Sulfiredoxin domain-containing protein; uniclust \| UniRef100\_A0A1E7NI96 \| 99.9 \| 1.2e-25 \| 2.2e-31 \| 179.8 \| 180 \| (13, 198) \| 291 \| (24, 206) \| 279 \| Chromosome partitioning protein ParB \| Chromosome partitioning protein ParB; uniclust \| UniRef100\_A0A073CBP1 \| 99.9 \| 1.3e-25 \| 2.5e-31 \| 201.1 \| 173 \| (21, 199) \| 291 \| (95, 306) \| 444 \| ParB \| ParB; uniclust \| UniRef100\_A0A1N7ST09 \| 99.9 \| 1.5e-25 \| 2.9e-31 \| 186.8 \| 202 \| (16, 217) \| 291 \| (41, 251) \| 347 \| ParB-like partition protein \| ParB-like partition protein; uniclust \| UniRef100\_A0A085FM87 \| 99.9 \| 1.5e-25 \| 3e-31 \| 195.6 \| 165 \| (15, 185) \| 291 \| (72, 243) \| 354 \| ParB-like partition protein \| ParB-like partition protein; uniclust \| UniRef100\_A0A170PJZ6 \| 99.9 \| 1.6e-25 \| 3e-31 \| 193.1 \| 180 \| (15, 198) \| 291 \| (63, 266) \| 410 \| Putative ParB-like partition protein \| Putative ParB-like partition protein; uniclust \| UniRef100\_A0A2E5HJM8 \| 99.9 \| 1.7e-25 \| 3.1e-31 \| 185.0 \| 179 \| (16, 199) \| 291 \| (52, 233) \| 382 \| Chromosome partitioning protein ParB \| Chromosome partitioning protein ParB; uniclust \| UniRef100\_A0A014MAM2 \| 99.9 \| 1.6e-25 \| 3.2e-31 \| 216.0 \| 177 \| (16, 198) \| 291 \| (118, 314) \| 921 \| ParB/Sulfiredoxin domain-containing protein \| ParB/Sulfiredoxin domain-containing protein; uniclust \| UniRef100\_A0A0G1YN89 \| 99.9 \| 1.9e-25 \| 3.6e-31 \| 183.7 \| 177 \| (18, 199) \| 291 \| (2, 191) \| 315 \| ParB-like protein partition protein \| ParB-like protein partition protein; uniclust \| UniRef100\_A0A0D5A6I8 \| 99.9 \| 1.8e-25 \| 3.6e-31 \| 205.7 \| 166 \| (18, 189) \| 291 \| (109, 279) \| 583 \| Chromosome partitioning protein ParB \| Chromosome partitioning protein ParB; uniclust \| UniRef100\_A0A0C1R7E8 \| 99.9 \| 1.8e-25 \| 3.6e-31 \| 192.7 \| 177 \| (19, 201) \| 291 \| (45, 260) \| 342 \| Plasmid partitioning protein ParB \| Plasmid partitioning protein ParB; uniclust \| UniRef100\_A0A086PDT6 \| 99.9 \| 1.9e-25 \| 3.7e-31 \| 187.3 \| 163 \| (15, 185) \| 291 \| (23, 190) \| 326 \| ParB-like protein \| ParB-like protein; uniclust \| UniRef100\_A0A022MM69 \| 99.9 \| 1.9e-25 \| 3.7e-31 \| 196.7 \| 174 \| (19, 198) \| 291 \| (54, 246) \| 404 \| ParB/Sulfiredoxin domain-containing protein \| ParB/Sulfiredoxin domain-containing protein; uniclust \| UniRef100\_A0A015YUD5 \| 99.9 \| 2e-25 \| 3.8e-31 \| 179.5 \| 198 \| (15, 215) \| 291 \| (6, 214) \| 273 \| ParB/RepB/Spo0J family partition domain protein \| ParB/RepB/Spo0J family partition domain protein; uniclust \| UniRef100\_A0A0D5YU03 \| 99.9 \| 2.2e-25 \| 4.2e-31 \| 199.2 \| 176 \| (16, 198) \| 291 \| (31, 208) \| 600 \| Putative partitioning protein \| Putative partitioning protein; uniclust \| UniRef100\_A0A071MC08 \| 99.9 \| 2.3e-25 \| 4.4e-31 \| 198.6 \| 234 \| (13, 247) \| 291 \| (77, 327) \| 558 \| Pyridoxal phosphate biosynthetic protein PdxJ \| Pyridoxal phosphate biosynthetic protein PdxJ; uniclust \| UniRef100\_A0A2D2LWZ4 \| 99.9 \| 2.7e-25 \| 5.1e-31 \| 183.7 \| 166 \| (18, 191) \| 291 \| (26, 196) \| 321 \| ParB/Sulfiredoxin domain-containing protein \| ParB/Sulfiredoxin domain-containing protein; uniclust \| UniRef100\_A0A1B8RYV7 \| 99.9 \| 2.7e-25 \| 5.1e-31 \| 187.9 \| 152 \| (16, 172) \| 291 \| (34, 187) \| 347 \| ParB/Sulfiredoxin domain-containing protein \| ParB/Sulfiredoxin domain-containing protein; uniclust \| UniRef100\_A0A0D6KEL4 \| 99.9 \| 2.6e-25 \| 5.2e-31 \| 191.5 \| 176 \| (20, 200) \| 291 \| (74, 287) \| 362 \| Partitioning protein ParB \| Partitioning protein ParB; uniclust \| UniRef100\_E0NJQ3 \| 99.9 \| 3.2e-25 \| 6e-31 \| 181.5 \| 168 \| (16, 189) \| 291 \| (91, 261) \| 346 \| ParB-like protein \| ParB-like protein; uniclust \| UniRef100\_A0A086P5Z5 \| 99.9 \| 3.1e-25 \| 6e-31 \| 192.4 \| 165 \| (17, 188) \| 291 \| (48, 215) \| 368 \| Chromosome partitioning protein ParB \| Chromosome partitioning protein ParB; uniclust \| UniRef100\_A0A4Y8ZAY6 \| 99.9 \| 3.3e-25 \| 6.1e-31 \| 160.7 \| 149 \| (23, 176) \| 291 \| (1, 153) \| 155 \| ParB/RepB/Spo0J family partition protein (Fragment) \| ParB/RepB/Spo0J family partition protein (Fragment); uniclust \| UniRef100\_A0A1G3URH6 \| 99.9 \| 3.3e-25 \| 6.4e-31 \| 187.0 \| 166 \| (15, 188) \| 291 \| (31, 201) \| 311 \| ParB/Sulfiredoxin domain-containing protein \| ParB/Sulfiredoxin domain-containing protein; uniclust \| UniRef100\_A0A1C7HFJ6 \| 99.9 \| 3.3e-25 \| 6.6e-31 \| 192.9 \| 168 \| (17, 189) \| 291 \| (25, 196) \| 388 \| ParB/Sulfiredoxin domain-containing protein \| ParB/Sulfiredoxin domain-containing protein; uniclust \| UniRef100\_A0A069RGP5 \| 99.9 \| 3.5e-25 \| 6.9e-31 \| 187.0 \| 238 \| (18, 289) \| 291 \| (42, 298) \| 304 \| ParB-like protein \| ParB-like protein; uniclust \| UniRef100\_A0A096XNS1 \| 99.9 \| 3.7e-25 \| 7.5e-31 \| 198.5 \| 166 \| (17, 190) \| 291 \| (35, 205) \| 436 \| ParB-like partition protein \| ParB-like partition protein; uniclust \| UniRef100\_A0A0K2SP45 \| 99.9 \| 4e-25 \| 7.9e-31 \| 190.1 \| 153 \| (14, 172) \| 291 \| (11, 165) \| 341 \| Chromosome partitioning protein ParB \| Chromosome partitioning protein ParB; uniclust \| UniRef100\_A0A140L4M6 \| 99.9 \| 4.2e-25 \| 7.9e-31 \| 186.4 \| 174 \| (18, 197) \| 291 \| (8, 197) \| 358 \| Chromosome-partitioning protein Spo0J \| Chromosome-partitioning protein Spo0J; uniclust \| UniRef100\_A0A139KWY8 \| 99.9 \| 4.2e-25 \| 8e-31 \| 200.9 \| 179 \| (18, 199) \| 291 \| (2, 215) \| 698 \| Chromosome partitioning protein ParB \| Chromosome partitioning protein ParB; uniclust \| UniRef100\_A0A8S9ARD5 \| 99.9 \| 5.2e-25 \| 9.9e-31 \| 184.3 \| 175 \| (16, 195) \| 291 \| (50, 236) \| 373 \| ParB/RepB/Spo0J family partition protein \| ParB/RepB/Spo0J family partition protein; uniclust \| UniRef100\_A0A011N723 \| 99.8 \| 5.3e-25 \| 1.1e-30 \| 196.5 \| 164 \| (18, 189) \| 291 \| (78, 246) \| 410 \| Transcriptional repressor protein KorB \| Transcriptional repressor protein KorB; uniclust \| UniRef100\_A0A2K0XC05 \| 99.8 \| 5.7e-25 \| 1.1e-30 \| 198.0 \| 177 \| (14, 196) \| 291 \| (8, 191) \| 623 \| ParB/Sulfiredoxin domain-containing protein \| ParB/Sulfiredoxin domain-containing protein; uniclust \| UniRef100\_A0A011W170 \| 99.8 \| 5.6e-25 \| 1.1e-30 \| 205.1 \| 168 \| (15, 189) \| 291 \| (117, 291) \| 608 \| Chromosome partitioning protein ParB \| Chromosome partitioning protein ParB; uniclust \| UniRef100\_A0A1D2R6B9 \| 99.8 \| 6e-25 \| 1.2e-30 \| 182.6 \| 176 \| (17, 197) \| 291 \| (9, 194) \| 281 \| ParB/Sulfiredoxin domain-containing protein \| ParB/Sulfiredoxin domain-containing protein; uniclust \| UniRef100\_A0A3S0DWZ9 \| 99.8 \| 7e-25 \| 1.3e-30 \| 160.8 \| 126 \| (15, 146) \| 291 \| (17, 145) \| 145 \| ParB/RepB/Spo0J family partition protein (Fragment) \| ParB/RepB/Spo0J family partition protein (Fragment); uniclust \| UniRef100\_A0A1G6LR52 \| 99.8 \| 6.7e-25 \| 1.3e-30 \| 200.7 \| 166 \| (16, 187) \| 291 \| (12, 181) \| 540 \| ParB/RepB/Spo0J family partition protein \| ParB/RepB/Spo0J family partition protein; uniclust \| UniRef100\_A0A0C5BR18 \| 99.8 \| 6.8e-25 \| 1.4e-30 \| 184.2 \| 162 \| (21, 187) \| 291 \| (15, 178) \| 281 \| ParB/Sulfiredoxin domain-containing protein \| ParB/Sulfiredoxin domain-containing protein; uniclust \| UniRef100\_A0A0F9W5T8 \| 99.8 \| 7.3e-25 \| 1.4e-30 \| 177.0 \| 165 \| (21, 189) \| 291 \| (2, 171) \| 296 \| ParB/Sulfiredoxin domain-containing protein \| ParB/Sulfiredoxin domain-containing protein; uniclust \| UniRef100\_A0A076ZED2 \| 99.8 \| 7.7e-25 \| 1.5e-30 \| 187.6 \| 191 \| (2, 197) \| 291 \| (42, 255) \| 345 \| ParB-like protein \| ParB-like protein; uniclust \| UniRef100\_A0A1W9HD66 \| 99.8 \| 8.5e-25 \| 1.6e-30 \| 182.8 \| 165 \| (18, 188) \| 291 \| (102, 272) \| 365 \| ParB/Sulfiredoxin domain-containing protein \| ParB/Sulfiredoxin domain-containing protein; uniclust \| UniRef100\_A0A1K1LLY6 \| 99.8 \| 9.6e-25 \| 1.9e-30 \| 183.9 \| 226 \| (19, 250) \| 291 \| (45, 291) \| 323 \| Chromosome partitioning protein, ParB family \| Chromosome partitioning protein, ParB family; uniclust \| UniRef100\_A0A069IHU9 \| 99.8 \| 1.1e-24 \| 2.2e-30 \| 173.0 \| 179 \| (17, 202) \| 291 \| (8, 192) \| 218 \| ParB/Sulfiredoxin domain-containing protein (Fragment) \| ParB/Sulfiredoxin domain-containing protein (Fragment); uniclust \| UniRef100\_A0A061M5E0 \| 99.8 \| 1.2e-24 \| 2.3e-30 \| 188.0 \| 163 \| (17, 186) \| 291 \| (35, 199) \| 349 \| Transcriptional repressor protein korB \| Transcriptional repressor protein korB; uniclust \| UniRef100\_A0A0C2LAE5 \| 99.8 \| 1.2e-24 \| 2.3e-30 \| 186.9 \| 179 \| (16, 200) \| 291 \| (39, 257) \| 338 \| ParB/RepB/Spo0J family partition protein \| ParB/RepB/Spo0J family partition protein; uniclust \| UniRef100\_A0A075JJA6 \| 99.8 \| 1.2e-24 \| 2.4e-30 \| 201.0 \| 166 \| (18, 189) \| 291 \| (84, 267) \| 552 \| HTH cro/C1-type domain-containing protein \| HTH cro/C1-type domain-containing protein; uniclust \| UniRef100\_A0A0F8X7Y3 \| 99.8 \| 1.3e-24 \| 2.5e-30 \| 169.8 \| 144 \| (18, 166) \| 291 \| (54, 204) \| 206 \| ParB/Sulfiredoxin domain-containing protein (Fragment) \| ParB/Sulfiredoxin domain-containing protein (Fragment); uniclust \| UniRef100\_A0A069P132 \| 99.8 \| 1.3e-24 \| 2.7e-30 \| 208.3 \| 181 \| (14, 201) \| 291 \| (111, 298) \| 811 \| Chromosome partitioning protein ParB \| Chromosome partitioning protein ParB; uniclust \| UniRef100\_A0A0C5K7X7 \| 99.8 \| 1.4e-24 \| 2.8e-30 \| 191.4 \| 219 \| (17, 244) \| 291 \| (104, 333) \| 386 \| ParB/Sulfiredoxin domain-containing protein \| ParB/Sulfiredoxin domain-containing protein; uniclust \| UniRef100\_A0A523DAK4 \| 99.8 \| 1.8e-24 \| 3.4e-30 \| 190.3 \| 179 \| (15, 199) \| 291 \| (14, 203) \| 553 \| ParB/RepB/Spo0J family partition protein \| ParB/RepB/Spo0J family partition protein; uniclust \| UniRef100\_A0A4V2UP75 \| 99.8 \| 1.8e-24 \| 3.5e-30 \| 190.9 \| 167 \| (17, 189) \| 291 \| (81, 253) \| 606 \| ParB/RepB/Spo0J family partition protein \| ParB/RepB/Spo0J family partition protein; uniclust \| UniRef100\_A0A1T1B7P7 \| 99.8 \| 1.8e-24 \| 3.5e-30 \| 184.3 \| 178 \| (17, 199) \| 291 \| (47, 256) \| 352 \| ParB/Sulfiredoxin domain-containing protein \| ParB/Sulfiredoxin domain-containing protein; uniclust \| UniRef100\_A0A016XIJ2 \| 99.8 \| 1.8e-24 \| 3.5e-30 \| 195.8 \| 169 \| (16, 190) \| 291 \| (2, 185) \| 612 \| Chromosome partitioning protein ParB \| Chromosome partitioning protein ParB; uniclust \| UniRef100\_A0A0F9LDT1 \| 99.8 \| 2e-24 \| 3.8e-30 \| 181.5 \| 176 \| (18, 197) \| 291 \| (71, 256) \| 326 \| ParB/Sulfiredoxin domain-containing protein \| ParB/Sulfiredoxin domain-containing protein; uniclust \| UniRef100\_A0A015W8Q1 \| 99.8 \| 2.1e-24 \| 3.9e-30 \| 194.8 \| 177 \| (17, 200) \| 291 \| (67, 249) \| 691 \| ParB/RepB/Spo0J family partition domain protein \| ParB/RepB/Spo0J family partition domain protein; uniclust \| UniRef100\_A0A1H5Z312 \| 99.8 \| 2.2e-24 \| 4.1e-30 \| 173.7 \| 174 \| (19, 198) \| 291 \| (9, 188) \| 264 \| ParB-like nuclease \| ParB-like nuclease; uniclust \| UniRef100\_A0A1I6NRN6 \| 99.8 \| 2.2e-24 \| 4.4e-30 \| 201.4 \| 176 \| (18, 199) \| 291 \| (107, 285) \| 681 \| ParB/RepB/Spo0J family partition protein \| ParB/RepB/Spo0J family partition protein; uniclust \| UniRef100\_A0A1V4W4D9 \| 99.8 \| 2.4e-24 \| 4.5e-30 \| 196.2 \| 171 \| (24, 201) \| 291 \| (30, 207) \| 789 \| Chromosome-partitioning protein Spo0J \| Chromosome-partitioning protein Spo0J; uniclust \| UniRef100\_A0A0F8W8Q8 \| 99.8 \| 2.3e-24 \| 4.5e-30 \| 180.2 \| 169 \| (16, 188) \| 291 \| (29, 208) \| 313 \| ParB/Sulfiredoxin domain-containing protein (Fragment) \| ParB/Sulfiredoxin domain-containing protein (Fragment); uniclust \| UniRef100\_A0A0A8F238 \| 99.8 \| 2.3e-24 \| 4.5e-30 \| 183.4 \| 174 \| (18, 197) \| 291 \| (42, 234) \| 349 \| Putative plasmid partitioning protein, parb2 \| Putative plasmid partitioning protein, parb2; uniclust \| UniRef100\_A0A174EVK5 \| 99.8 \| 2.3e-24 \| 4.6e-30 \| 186.7 \| 168 \| (15, 189) \| 291 \| (3, 182) \| 342 \| Chromosome-partitioning protein parB \| Chromosome-partitioning protein parB; uniclust \| UniRef100\_A0A254TB70 \| 99.8 \| 2.5e-24 \| 4.9e-30 \| 202.3 \| 173 \| (20, 198) \| 291 \| (56, 238) \| 748 \| ParB/Sulfiredoxin domain-containing protein \| ParB/Sulfiredoxin domain-containing protein; uniclust \| UniRef100\_A0A143XPK3 \| 99.8 \| 2.6e-24 \| 5e-30 \| 180.6 \| 179 \| (15, 198) \| 291 \| (38, 238) \| 325 \| Chromosome-partitioning protein Spo0J \| Chromosome-partitioning protein Spo0J; uniclust \| UniRef100\_A0A0G1AJS6 \| 99.8 \| 2.6e-24 \| 5e-30 \| 184.1 \| 186 \| (17, 207) \| 291 \| (31, 222) \| 336 \| Chromosome segregation DNA-binding protein \| Chromosome segregation DNA-binding protein; uniclust \| UniRef100\_A0A0D6QE72 \| 99.8 \| 3e-24 \| 5.8e-30 \| 196.1 \| 179 \| (14, 198) \| 291 \| (40, 221) \| 596 \| Putative chromosome-partitioning protein ParB \| Putative chromosome-partitioning protein ParB; uniclust \| UniRef100\_A0A1M5YKM5 \| 99.8 \| 4e-24 \| 7.6e-30 \| 180.1 \| 267 \| (5, 285) \| 291 \| (55, 336) \| 348 \| Chromosome partitioning protein, ParB family \| Chromosome partitioning protein, ParB family; uniclust \| UniRef100\_A0A1Q3SMN2 \| 99.8 \| 4.1e-24 \| 8e-30 \| 185.3 \| 167 \| (18, 192) \| 291 \| (56, 228) \| 375 \| ParB/Sulfiredoxin domain-containing protein \| ParB/Sulfiredoxin domain-containing protein; uniclust \| UniRef100\_X1CJ30 \| 99.8 \| 4.8e-24 \| 8.8e-30 \| 164.9 \| 170 \| (14, 189) \| 291 \| (41, 213) \| 248 \| ParB/Sulfiredoxin domain-containing protein (Fragment) \| ParB/Sulfiredoxin domain-containing protein (Fragment); uniclust \| UniRef100\_A0A0G1WPA1 \| 99.8 \| 5e-24 \| 9.8e-30 \| 180.6 \| 184 \| (13, 200) \| 291 \| (28, 220) \| 324 \| ParB/Sulfiredoxin domain-containing protein \| ParB/Sulfiredoxin domain-containing protein; uniclust \| UniRef100\_A0A212L1T9 \| 99.8 \| 5.3e-24 \| 1e-29 \| 177.2 \| 183 \| (16, 201) \| 291 \| (34, 229) \| 322 \| Putative Nucleoid occlusion protein \| Putative Nucleoid occlusion protein; uniclust \| UniRef100\_A0A9E2BEM2 \| 99.8 \| 5.5e-24 \| 1e-29 \| 172.1 \| 166 \| (17, 188) \| 291 \| (41, 209) \| 302 \| Chromosome-partitioning protein ParB \| Chromosome-partitioning protein ParB; uniclust \| UniRef100\_A0A0F8ZIF1 \| 99.8 \| 5.7e-24 \| 1.1e-29 \| 176.2 \| 189 \| (15, 207) \| 291 \| (18, 216) \| 298 \| ParB/Sulfiredoxin domain-containing protein \| ParB/Sulfiredoxin domain-containing protein; uniclust \| UniRef100\_A0A5N5XVG2 \| 99.8 \| 6.6e-24 \| 1.3e-29 \| 174.6 \| 171 \| (18, 193) \| 291 \| (6, 194) \| 296 \| ParB/RepB/Spo0J family partition protein \| ParB/RepB/Spo0J family partition protein; uniclust \| UniRef100\_A0A069PKE4 \| 99.8 \| 7.2e-24 \| 1.4e-29 \| 181.9 \| 204 \| (13, 216) \| 291 \| (65, 278) \| 404 \| Transcriptional regulator \| Transcriptional regulator; uniclust \| UniRef100\_A0A967V135 \| 99.8 \| 7.9e-24 \| 1.5e-29 \| 174.7 \| 164 \| (19, 188) \| 291 \| (25, 192) \| 416 \| ParB/RepB/Spo0J family partition protein \| ParB/RepB/Spo0J family partition protein; uniclust \| UniRef100\_A0A1I4DE87 \| 99.8 \| 7.7e-24 \| 1.5e-29 \| 180.7 \| 166 \| (18, 189) \| 291 \| (143, 312) \| 399 \| ParB/RepB/Spo0J family partition protein \| ParB/RepB/Spo0J family partition protein; uniclust \| UniRef100\_A0A3D1LXR1 \| 99.8 \| 8e-24 \| 1.5e-29 \| 178.2 \| 175 \| (17, 195) \| 291 \| (39, 224) \| 313 \| Chromosome partitioning protein ParB \| Chromosome partitioning protein ParB; uniclust \| UniRef100\_A0A3A5APM2 \| 99.8 \| 8.4e-24 \| 1.6e-29 \| 175.1 \| 171 \| (15, 192) \| 291 \| (77, 251) \| 381 \| ParB/RepB/Spo0J family partition protein \| ParB/RepB/Spo0J family partition protein; uniclust \| UniRef100\_A0A142KA42 \| 99.8 \| 9.1e-24 \| 1.8e-29 \| 190.6 \| 170 \| (13, 188) \| 291 \| (17, 190) \| 527 \| ParB-like nuclease domain protein \| ParB-like nuclease domain protein; uniclust \| UniRef100\_A0A2N2QZ62 \| 99.8 \| 9.4e-24 \| 1.8e-29 \| 171.3 \| 177 \| (6, 189) \| 291 \| (30, 209) \| 271 \| Chromosome partitioning protein ParB (Fragment) \| Chromosome partitioning protein ParB (Fragment); uniclust \| UniRef100\_A0A318SDS1 \| 99.8 \| 1e-23 \| 2e-29 \| 177.8 \| 175 \| (18, 197) \| 291 \| (29, 232) \| 316 \| ParB family protein \| ParB family protein; uniclust \| UniRef100\_A0A1F3KBJ9 \| 99.8 \| 1e-23 \| 2e-29 \| 190.3 \| 177 \| (17, 199) \| 291 \| (5, 189) \| 583 \| ParB/Sulfiredoxin domain-containing protein \| ParB/Sulfiredoxin domain-containing protein; uniclust \| UniRef100\_A0A1E3S303 \| 99.8 \| 1e-23 \| 2e-29 \| 185.8 \| 169 \| (14, 188) \| 291 \| (20, 193) \| 487 \| ParB/Sulfiredoxin domain-containing protein \| ParB/Sulfiredoxin domain-containing protein; uniclust \| UniRef100\_A0A0F7VPB0 \| 99.8 \| 1e-23 \| 2e-29 \| 187.8 \| 181 \| (13, 199) \| 291 \| (40, 240) \| 409 \| ParB \| ParB; uniclust \| UniRef100\_A0A351H5H0 \| 99.8 \| 1.1e-23 \| 2.1e-29 \| 173.2 \| 182 \| (16, 202) \| 291 \| (34, 218) \| 311 \| Nucleoid occlusion protein \| Nucleoid occlusion protein; uniclust \| UniRef100\_A0A959ZJQ8 \| 99.8 \| 1.1e-23 \| 2.1e-29 \| 185.4 \| 159 \| (17, 179) \| 291 \| (11, 172) \| 544 \| ParB/RepB/Spo0J family partition protein \| ParB/RepB/Spo0J family partition protein; uniclust \| UniRef100\_UPI002072E589 \| 99.8 \| 1.2e-23 \| 2.3e-29 \| 178.4 \| 162 \| (21, 188) \| 291 \| (3, 176) \| 428 \| ParB/RepB/Spo0J family partition protein \| ParB/RepB/Spo0J family partition protein; uniclust \| UniRef100\_A0A222WSL8 \| 99.8 \| 1.3e-23 \| 2.4e-29 \| 176.0 \| 160 \| (22, 187) \| 291 \| (4, 165) \| 380 \| ParB/Sulfiredoxin domain-containing protein \| ParB/Sulfiredoxin domain-containing protein; uniclust \| UniRef100\_A0A4Y6UB58 \| 99.8 \| 1.3e-23 \| 2.4e-29 \| 171.3 \| 163 \| (20, 188) \| 291 \| (17, 204) \| 323 \| ParB/RepB/Spo0J family partition protein \| ParB/RepB/Spo0J family partition protein; uniclust \| UniRef100\_A0A1C6HC15 \| 99.8 \| 1.6e-23 \| 2.9e-29 \| 179.7 \| 173 \| (21, 199) \| 291 \| (290, 466) \| 549 \| Probable chromosome-partitioning protein parB \| Probable chromosome-partitioning protein parB; uniclust \| UniRef100\_A0A0C1MVS1 \| 99.8 \| 1.5e-23 \| 3e-29 \| 187.7 \| 204 \| (17, 246) \| 291 \| (79, 338) \| 413 \| Plasmid partitioning protein ParB \| Plasmid partitioning protein ParB; uniclust \| UniRef100\_A0A3A4U2T7 \| 99.8 \| 1.8e-23 \| 3.4e-29 \| 174.8 \| 172 \| (13, 190) \| 291 \| (74, 248) \| 407 \| ParB/RepB/Spo0J family partition protein \| ParB/RepB/Spo0J family partition protein; uniclust \| UniRef100\_A0A064CDN1 \| 99.8 \| 1.8e-23 \| 3.5e-29 \| 177.5 \| 220 \| (18, 248) \| 291 \| (54, 308) \| 326 \| Peptide transporter \| Peptide transporter; uniclust \| UniRef100\_A0A086ZNU6 \| 99.8 \| 1.9e-23 \| 3.7e-29 \| 185.5 \| 168 \| (15, 188) \| 291 \| (80, 248) \| 548 \| ParB-like nuclease domain \| ParB-like nuclease domain; uniclust \| UniRef100\_A0A352BEU1 \| 99.8 \| 2.2e-23 \| 4.1e-29 \| 154.9 \| 147 \| (17, 168) \| 291 \| (11, 174) \| 175 \| ParB/Sulfiredoxin domain-containing protein \| ParB/Sulfiredoxin domain-containing protein; uniclust \| UniRef100\_A0A1U7J7S0 \| 99.8 \| 2.2e-23 \| 4.5e-29 \| 180.5 \| 176 \| (17, 198) \| 291 \| (44, 256) \| 339 \| Chromosome partitioning protein ParB \| Chromosome partitioning protein ParB; uniclust \| UniRef100\_A0A016QJQ7 \| 99.8 \| 2.3e-23 \| 4.5e-29 \| 179.9 \| 177 \| (19, 201) \| 291 \| (44, 253) \| 361 \| ParB/Sulfiredoxin domain-containing protein \| ParB/Sulfiredoxin domain-containing protein; uniclust \| UniRef100\_A0A2X1TFX5 \| 99.8 \| 2.4e-23 \| 4.5e-29 \| 160.8 \| 162 \| (54, 216) \| 291 \| (3, 166) \| 211 \| ParB/RepB/Spo0J family partition protein \| ParB/RepB/Spo0J family partition protein; uniclust \| UniRef100\_A0A317EEA9 \| 99.8 \| 2.4e-23 \| 4.6e-29 \| 186.4 \| 182 \| (15, 198) \| 291 \| (115, 303) \| 601 \| ParB/Sulfiredoxin domain-containing protein \| ParB/Sulfiredoxin domain-containing protein; uniclust \| UniRef100\_A0A016QT58 \| 99.8 \| 2.4e-23 \| 4.6e-29 \| 178.5 \| 172 \| (21, 199) \| 291 \| (59, 259) \| 346 \| ParB-like partition protein \| ParB-like partition protein; uniclust \| UniRef100\_A0A5M8SQ23 \| 99.8 \| 2.6e-23 \| 4.8e-29 \| 167.8 \| 185 \| (15, 205) \| 291 \| (6, 196) \| 286 \| ParB/RepB/Spo0J family partition protein \| ParB/RepB/Spo0J family partition protein; uniclust \| UniRef100\_A0A173SU81 \| 99.8 \| 2.5e-23 \| 4.9e-29 \| 177.5 \| 179 \| (15, 198) \| 291 \| (43, 242) \| 351 \| Probable chromosome-partitioning protein parB \| Probable chromosome-partitioning protein parB; uniclust \| UniRef100\_A0A060ULH7 \| 99.8 \| 2.5e-23 \| 4.9e-29 \| 189.1 \| 175 \| (18, 200) \| 291 \| (30, 208) \| 583 \| ParB/Sulfiredoxin domain-containing protein \| ParB/Sulfiredoxin domain-containing protein; uniclust \| UniRef100\_A0A0M0BQB3 \| 99.8 \| 2.7e-23 \| 5.1e-29 \| 165.4 \| 178 \| (20, 201) \| 291 \| (4, 184) \| 249 \| ParB/Sulfiredoxin domain-containing protein \| ParB/Sulfiredoxin domain-containing protein; uniclust \| UniRef100\_A0A1H4K415 \| 99.8 \| 2.8e-23 \| 5.3e-29 \| 185.1 \| 180 \| (16, 201) \| 291 \| (19, 201) \| 578 \| Chromosome partitioning protein, ParB family \| Chromosome partitioning protein, ParB family; uniclust \| UniRef100\_A0A2L2WSX3 \| 99.8 \| 3.2e-23 \| 6.1e-29 \| 180.8 \| 179 \| (15, 199) \| 291 \| (10, 193) \| 515 \| Chromosome partitioning protein, ParB family \| Chromosome partitioning protein, ParB family; uniclust \| UniRef100\_A0A922VHK8 \| 99.8 \| 3.4e-23 \| 6.3e-29 \| 165.4 \| 175 \| (18, 197) \| 291 \| (27, 203) \| 313 \| Chromosome partitioning protein ParB \| Chromosome partitioning protein ParB; uniclust \| UniRef100\_A0A1H8AXE2 \| 99.8 \| 3.2e-23 \| 6.5e-29 \| 178.8 \| 176 \| (17, 197) \| 291 \| (42, 227) \| 326 \| Chromosome partitioning protein, ParB family \| Chromosome partitioning protein, ParB family; uniclust \| UniRef100\_A0A2E5S2I9 \| 99.8 \| 3.4e-23 \| 6.5e-29 \| 170.7 \| 168 \| (15, 188) \| 291 \| (19, 189) \| 284 \| Chromosome partitioning protein ParB \| Chromosome partitioning protein ParB; uniclust \| UniRef100\_A0A660VCE9 \| 99.8 \| 3.6e-23 \| 6.6e-29 \| 167.2 \| 162 \| (14, 181) \| 291 \| (112, 275) \| 344 \| HTH cro/C1-type domain-containing protein \| HTH cro/C1-type domain-containing protein; uniclust \| UniRef100\_A0A2D9HGI4 \| 99.8 \| 3.6e-23 \| 6.6e-29 \| 167.1 \| 179 \| (15, 198) \| 291 \| (51, 233) \| 318 \| ParB/Sulfiredoxin domain-containing protein \| ParB/Sulfiredoxin domain-containing protein; uniclust \| UniRef100\_A0A3D3LYG7 \| 99.8 \| 3.6e-23 \| 7e-29 \| 182.2 \| 169 \| (14, 188) \| 291 \| (8, 182) \| 487 \| ParB/Sulfiredoxin domain-containing protein \| ParB/Sulfiredoxin domain-containing protein; uniclust \| UniRef100\_A0A011NAJ4 \| 99.8 \| 4.4e-23 \| 8.9e-29 \| 182.5 \| 165 \| (18, 187) \| 291 \| (82, 255) \| 373 \| Chromosome-partitioning protein ParB \| Chromosome-partitioning protein ParB; uniclust \| UniRef100\_A0A1G8EV61 \| 99.8 \| 4.8e-23 \| 9.3e-29 \| 177.0 \| 156 \| (21, 182) \| 291 \| (88, 250) \| 352 \| Chromosome partitioning protein, ParB family \| Chromosome partitioning protein, ParB family; uniclust \| UniRef100\_C3NJE7 \| 99.8 \| 5.2e-23 \| 9.6e-29 \| 152.5 \| 160 \| (22, 187) \| 291 \| (2, 167) \| 183 \| ParB-like partition protein \| ParB-like partition protein; uniclust \| UniRef100\_A0A2H9LMS7 \| 99.8 \| 5e-23 \| 9.7e-29 \| 169.0 \| 164 \| (19, 187) \| 291 \| (11, 179) \| 265 \| ParB/Sulfiredoxin domain-containing protein \| ParB/Sulfiredoxin domain-containing protein; uniclust \| UniRef100\_A0A1Q3SGT3 \| 99.8 \| 5e-23 \| 9.8e-29 \| 174.0 \| 137 \| (18, 160) \| 291 \| (55, 197) \| 296 \| ParB/Sulfiredoxin domain-containing protein \| ParB/Sulfiredoxin domain-containing protein; uniclust \| UniRef100\_A0A517T448 \| 99.8 \| 5.6e-23 \| 1e-28 \| 170.1 \| 175 \| (19, 199) \| 291 \| (48, 225) \| 421 \| Putative chromosome-partitioning protein ParB \| Putative chromosome-partitioning protein ParB; uniclust \| UniRef100\_UPI000698FBAD \| 99.8 \| 5.5e-23 \| 1e-28 \| 177.9 \| 162 \| (20, 187) \| 291 \| (3, 166) \| 497 \| ParB/RepB/Spo0J family partition protein \| ParB/RepB/Spo0J family partition protein; uniclust \| UniRef100\_A0A938L9U1 \| 99.8 \| 5.9e-23 \| 1.1e-28 \| 171.8 \| 164 \| (19, 187) \| 291 \| (197, 363) \| 466 \| ParB/RepB/Spo0J family partition protein \| ParB/RepB/Spo0J family partition protein; uniclust \| UniRef100\_A0A3M1FTJ6 \| 99.8 \| 6.1e-23 \| 1.1e-28 \| 172.6 \| 163 \| (20, 188) \| 291 \| (227, 392) \| 487 \| ParB/RepB/Spo0J family partition protein \| ParB/RepB/Spo0J family partition protein; uniclust \| UniRef100\_A0A1Q3T2X0 \| 99.8 \| 5.9e-23 \| 1.1e-28 \| 177.3 \| 151 \| (13, 168) \| 291 \| (31, 186) \| 380 \| ParB/Sulfiredoxin domain-containing protein \| ParB/Sulfiredoxin domain-containing protein; uniclust \| UniRef100\_A0A1T4V321 \| 99.8 \| 6.3e-23 \| 1.2e-28 \| 176.7 \| 168 \| (15, 187) \| 291 \| (44, 213) \| 561 \| Probable chromosome-partitioning protein ParB \| Probable chromosome-partitioning protein ParB; uniclust \| UniRef100\_A0A2N5KH80 \| 99.8 \| 6.3e-23 \| 1.2e-28 \| 175.0 \| 168 \| (17, 191) \| 291 \| (34, 206) \| 325 \| ParB/Sulfiredoxin domain-containing protein \| ParB/Sulfiredoxin domain-containing protein; uniclust \| UniRef100\_A0A354BGG0 \| 99.8 \| 6.6e-23 \| 1.3e-28 \| 172.7 \| 166 \| (16, 188) \| 291 \| (87, 257) \| 367 \| ParB/Sulfiredoxin domain-containing protein \| ParB/Sulfiredoxin domain-containing protein; uniclust \| UniRef100\_A0A1V4MDQ5 \| 99.8 \| 6.4e-23 \| 1.3e-28 \| 188.3 \| 174 \| (18, 199) \| 291 \| (64, 243) \| 593 \| site-specific DNA-methyltransferase (cytosine-N(4)-specific) \| site-specific DNA-methyltransferase (cytosine-N(4)-specific); uniclust \| UniRef100\_A0A2V2GRE7 \| 99.8 \| 6.6e-23 \| 1.3e-28 \| 172.6 \| 182 \| (15, 201) \| 291 \| (19, 207) \| 327 \| ParB/Sulfiredoxin domain-containing protein \| ParB/Sulfiredoxin domain-containing protein; uniclust \| UniRef100\_A0A069P7S0 \| 99.8 \| 6.6e-23 \| 1.3e-28 \| 175.2 \| 217 \| (16, 242) \| 291 \| (64, 289) \| 335 \| Chromosome partitioning protein ParB \| Chromosome partitioning protein ParB; uniclust \| UniRef100\_A0A166QSF2 \| 99.8 \| 8.5e-23 \| 1.6e-28 \| 165.7 \| 166 \| (18, 189) \| 291 \| (28, 197) \| 290 \| Probable chromosome-partitioning protein ParB \| Probable chromosome-partitioning protein ParB; uniclust \| UniRef100\_UPI00069DE912 \| 99.8 \| 8.7e-23 \| 1.6e-28 \| 171.7 \| 168 \| (16, 187) \| 291 \| (42, 214) \| 372 \| ParB/RepB/Spo0J family partition protein \| ParB/RepB/Spo0J family partition protein; uniclust \| UniRef100\_A0A060A0B8 \| 99.8 \| 8.4e-23 \| 1.7e-28 \| 183.1 \| 169 \| (17, 188) \| 291 \| (61, 248) \| 483 \| Chromosome (Plasmid) partitioning protein ParB / Stage 0 sporulation protein J \| Chromosome (Plasmid) partitioning protein ParB / Stage 0 sporulation protein J; uniclust \| UniRef100\_A0A8J6TJ42 \| 99.8 \| 9.3e-23 \| 1.8e-28 \| 183.7 \| 179 \| (15, 202) \| 291 \| (168, 352) \| 692 \| ParB/RepB/Spo0J family partition protein \| ParB/RepB/Spo0J family partition protein; uniclust \| UniRef100\_A0A535C431 \| 99.8 \| 9.8e-23 \| 1.8e-28 \| 168.0 \| 167 \| (17, 189) \| 291 \| (134, 303) \| 406 \| ParB/RepB/Spo0J family partition protein \| ParB/RepB/Spo0J family partition protein; uniclust \| UniRef100\_A0A1Q9PA47 \| 99.8 \| 1e-22 \| 1.9e-28 \| 172.6 \| 143 \| (18, 166) \| 291 \| (3, 153) \| 338 \| Putative chromosome-partitioning protein ParB \| Putative chromosome-partitioning protein ParB; uniclust \| UniRef100\_A0A0F9TED6 \| 99.8 \| 1.1e-22 \| 2e-28 \| 174.8 \| 167 \| (18, 189) \| 291 \| (31, 202) \| 430 \| ParB/Sulfiredoxin domain-containing protein \| ParB/Sulfiredoxin domain-containing protein; uniclust \| UniRef100\_A0A6M5YN73 \| 99.8 \| 1.1e-22 \| 2.1e-28 \| 166.3 \| 165 \| (20, 189) \| 291 \| (51, 221) \| 378 \| Chromosome (Plasmid) partitioning protein ParB \| Chromosome (Plasmid) partitioning protein ParB; uniclust \| UniRef100\_A0A0Q6VIP4 \| 99.8 \| 1.1e-22 \| 2.1e-28 \| 188.1 \| 172 \| (20, 197) \| 291 \| (40, 220) \| 649 \| ParB/Sulfiredoxin domain-containing protein \| ParB/Sulfiredoxin domain-containing protein; uniclust \| UniRef100\_A0A084CND8 \| 99.8 \| 1.1e-22 \| 2.1e-28 \| 175.4 \| 168 \| (17, 187) \| 291 \| (53, 229) \| 358 \| Nucleoid occlusion protein \| Nucleoid occlusion protein; uniclust \| UniRef100\_A0A0N1JNZ4 \| 99.8 \| 1.1e-22 \| 2.2e-28 \| 170.2 \| 164 \| (18, 188) \| 291 \| (71, 241) \| 337 \| Partitioning protein \| Partitioning protein; uniclust \| UniRef100\_A0A4Q3NWG3 \| 99.8 \| 1.2e-22 \| 2.2e-28 \| 170.6 \| 167 \| (17, 188) \| 291 \| (36, 204) \| 480 \| Aldehyde dehydrogenase family protein \| Aldehyde dehydrogenase family protein; uniclust \| UniRef100\_A0A1F4NGN6 \| 99.8 \| 1.1e-22 \| 2.2e-28 \| 176.3 \| 168 \| (17, 190) \| 291 \| (63, 235) \| 329 \| HTH cro/C1-type domain-containing protein \| HTH cro/C1-type domain-containing protein; uniclust \| UniRef100\_A0A1K2ICX5 \| 99.8 \| 1.2e-22 \| 2.2e-28 \| 169.1 \| 175 \| (16, 197) \| 291 \| (30, 206) \| 369 \| Chromosome partitioning protein, ParB family \| Chromosome partitioning protein, ParB family; uniclust \| UniRef100\_A0A0Q7DD19 \| 99.8 \| 1.1e-22 \| 2.2e-28 \| 174.8 \| 176 \| (15, 199) \| 291 \| (30, 210) \| 354 \| ParB/Sulfiredoxin domain-containing protein \| ParB/Sulfiredoxin domain-containing protein; uniclust \| UniRef100\_A0A373ZZS3 \| 99.8 \| 1.2e-22 \| 2.2e-28 \| 186.3 \| 173 \| (20, 198) \| 291 \| (60, 236) \| 738 \| ParB/RepB/Spo0J family partition protein \| ParB/RepB/Spo0J family partition protein; uniclust \| UniRef100\_A0A0A8WL90 \| 99.8 \| 1.2e-22 \| 2.3e-28 \| 163.8 \| 170 \| (16, 188) \| 291 \| (9, 186) \| 229 \| ParB/Sulfiredoxin domain-containing protein \| ParB/Sulfiredoxin domain-containing protein; uniclust \| UniRef100\_A0A1F8SA75 \| 99.8 \| 1.4e-22 \| 2.5e-28 \| 159.7 \| 168 \| (16, 189) \| 291 \| (35, 206) \| 245 \| ParB/Sulfiredoxin domain-containing protein (Fragment) \| ParB/Sulfiredoxin domain-containing protein (Fragment); uniclust \| UniRef100\_A0A0E2B209 \| 99.8 \| 1.4e-22 \| 2.7e-28 \| 160.9 \| 160 \| (18, 183) \| 291 \| (21, 184) \| 236 \| ParB-like protein \| ParB-like protein; uniclust \| UniRef100\_A0A7C6QZE1 \| 99.8 \| 1.6e-22 \| 2.9e-28 \| 170.8 \| 164 \| (17, 186) \| 291 \| (33, 198) \| 502 \| ParB/RepB/Spo0J family partition protein \| ParB/RepB/Spo0J family partition protein; uniclust \| UniRef100\_A0A662TCT7 \| 99.8 \| 1.6e-22 \| 2.9e-28 \| 163.6 \| 171 \| (17, 193) \| 291 \| (5, 178) \| 293 \| ParB/Sulfiredoxin domain-containing protein \| ParB/Sulfiredoxin domain-containing protein; uniclust \| UniRef100\_A0A1G2SN81 \| 99.8 \| 1.7e-22 \| 3.1e-28 \| 168.6 \| 175 \| (18, 197) \| 291 \| (11, 190) \| 346 \| ParB/Sulfiredoxin domain-containing protein \| ParB/Sulfiredoxin domain-containing protein; uniclust \| UniRef100\_A0A2T5IQH8 \| 99.8 \| 1.7e-22 \| 3.2e-28 \| 164.7 \| 180 \| (17, 200) \| 291 \| (45, 231) \| 313 \| ParB/RepB/Spo0J family partition protein \| ParB/RepB/Spo0J family partition protein; uniclust \| UniRef100\_A0A084SGC0 \| 99.8 \| 1.7e-22 \| 3.2e-28 \| 148.8 \| 117 \| (17, 139) \| 291 \| (11, 131) \| 134 \| ParB/Sulfiredoxin domain-containing protein (Fragment) \| ParB/Sulfiredoxin domain-containing protein (Fragment); uniclust \| UniRef100\_A0A1N0EX50 \| 99.8 \| 1.7e-22 \| 3.3e-28 \| 155.7 \| 145 \| (39, 188) \| 291 \| (2, 149) \| 184 \| Chromosome partitioning protein ParB \| Chromosome partitioning protein ParB; uniclust \| UniRef100\_A0A069I5N8 \| 99.8 \| 1.7e-22 \| 3.4e-28 \| 197.0 \| 178 \| (17, 197) \| 291 \| (231, 414) \| 944 \| ParB/Sulfiredoxin domain-containing protein \| ParB/Sulfiredoxin domain-containing protein; uniclust \| UniRef100\_A0A395X326 \| 99.8 \| 1.9e-22 \| 3.6e-28 \| 163.2 \| 179 \| (14, 197) \| 291 \| (27, 225) \| 272 \| ParB/RepB/Spo0J family partition protein (Fragment) \| ParB/RepB/Spo0J family partition protein (Fragment); uniclust \| UniRef100\_A0A0F6L5S3 \| 99.8 \| 1.9e-22 \| 3.7e-28 \| 180.2 \| 147 \| (19, 172) \| 291 \| (75, 227) \| 448 \| ParB/RepB/Spo0J family partition domain protein \| ParB/RepB/Spo0J family partition domain protein; uniclust \| UniRef100\_A0A349FWC2 \| 99.8 \| 2e-22 \| 3.8e-28 \| 151.5 \| 118 \| (14, 137) \| 291 \| (32, 152) \| 154 \| ParB/Sulfiredoxin domain-containing protein \| ParB/Sulfiredoxin domain-containing protein; uniclust \| UniRef100\_A0A1Q3SIH1 \| 99.8 \| 2e-22 \| 3.8e-28 \| 165.4 \| 165 \| (18, 189) \| 291 \| (38, 206) \| 299 \| ParB/Sulfiredoxin domain-containing protein \| ParB/Sulfiredoxin domain-containing protein; uniclust \| UniRef100\_A0A0Q8BHN4 \| 99.8 \| 2e-22 \| 3.8e-28 \| 174.6 \| 161 \| (22, 190) \| 291 \| (36, 211) \| 419 \| ParB/Sulfiredoxin domain-containing protein \| ParB/Sulfiredoxin domain-containing protein; uniclust \| UniRef100\_A6W8V9 \| 99.8 \| 2e-22 \| 3.8e-28 \| 169.8 \| 164 \| (16, 185) \| 291 \| (80, 245) \| 398 \| ParB-like partition protein \| ParB-like partition protein; uniclust \| UniRef100\_A0A3D2QDE7 \| 99.8 \| 2e-22 \| 3.8e-28 \| 149.6 \| 140 \| (54, 198) \| 291 \| (2, 145) \| 153 \| Nucleoid occlusion protein (Fragment) \| Nucleoid occlusion protein (Fragment); uniclust \| UniRef100\_A0A258BLJ1 \| 99.8 \| 2e-22 \| 3.9e-28 \| 168.7 \| 177 \| (18, 196) \| 291 \| (18, 202) \| 330 \| Chromosome partitioning protein ParB (Fragment) \| Chromosome partitioning protein ParB (Fragment); uniclust \| UniRef100\_A0A7C6XDD3 \| 99.8 \| 2.1e-22 \| 4e-28 \| 166.1 \| 175 \| (20, 200) \| 291 \| (41, 219) \| 344 \| ParB/RepB/Spo0J family partition protein \| ParB/RepB/Spo0J family partition protein; uniclust \| UniRef100\_A0A1C0B7H8 \| 99.8 \| 2.2e-22 \| 4.2e-28 \| 170.3 \| 177 \| (14, 198) \| 291 \| (26, 209) \| 336 \| Chromosome-partitioning protein Spo0J \| Chromosome-partitioning protein Spo0J; uniclust \| UniRef100\_A0A0D6KAG1 \| 99.8 \| 2.2e-22 \| 4.2e-28 \| 171.7 \| 174 \| (21, 199) \| 291 \| (117, 328) \| 402 \| Partitioning protein ParB \| Partitioning protein ParB; uniclust \| UniRef100\_UPI001BCFECD9 \| 99.8 \| 2.5e-22 \| 4.5e-28 \| 162.8 \| 268 \| (14, 288) \| 291 \| (42, 315) \| 348 \| hypothetical protein \| hypothetical protein; uniclust \| UniRef100\_A0A1V5NPZ0 \| 99.8 \| 2.5e-22 \| 4.6e-28 \| 158.4 \| 171 \| (21, 198) \| 291 \| (3, 177) \| 248 \| Chromosome-partitioning protein Spo0J \| Chromosome-partitioning protein Spo0J; uniclust \| UniRef100\_A0A064A7Z1 \| 99.8 \| 2.3e-22 \| 4.7e-28 \| 180.6 \| 173 \| (17, 193) \| 291 \| (66, 278) \| 412 \| Chromosome partitioning protein ParB \| Chromosome partitioning protein ParB; uniclust \| UniRef100\_A0A2A6U4Q8 \| 99.8 \| 2.8e-22 \| 5.1e-28 \| 153.6 \| 165 \| (29, 199) \| 291 \| (2, 169) \| 227 \| Chromosome partitioning protein ParB (Fragment) \| Chromosome partitioning protein ParB (Fragment); uniclust \| UniRef100\_A0A1I9YVG4 \| 99.8 \| 2.6e-22 \| 5.1e-28 \| 173.5 \| 192 \| (15, 214) \| 291 \| (23, 232) \| 385 \| ParB/RepB/Spo0J family partition protein \| ParB/RepB/Spo0J family partition protein; uniclust \| UniRef100\_A0A059G147 \| 99.8 \| 2.7e-22 \| 5.1e-28 \| 179.3 \| 182 \| (13, 195) \| 291 \| (82, 268) \| 555 \| Partitioning protein-like protein (Fragment) \| Partitioning protein-like protein (Fragment); uniclust \| UniRef100\_A0A0L0M783 \| 99.8 \| 2.8e-22 \| 5.6e-28 \| 174.2 \| 177 \| (20, 201) \| 291 \| (22, 203) \| 361 \| Chromosome (Plasmid) partitioning protein ParB \| Chromosome (Plasmid) partitioning protein ParB; uniclust \| UniRef100\_A0A165R355 \| 99.8 \| 3.1e-22 \| 5.7e-28 \| 178.7 \| 170 \| (17, 192) \| 291 \| (126, 298) \| 734 \| ParB/Sulfiredoxin domain-containing protein (Fragment) \| ParB/Sulfiredoxin domain-containing protein (Fragment); uniclust \| UniRef100\_A0A097ENY6 \| 99.8 \| 3.1e-22 \| 5.8e-28 \| 164.2 \| 169 \| (14, 188) \| 291 \| (40, 213) \| 305 \| Chromosome partitioning protein ParB \| Chromosome partitioning protein ParB; uniclust \| UniRef100\_A0A0F5JTH4 \| 99.8 \| 3e-22 \| 5.9e-28 \| 173.7 \| 256 \| (19, 289) \| 291 \| (82, 352) \| 377 \| ParB/Sulfiredoxin domain-containing protein \| ParB/Sulfiredoxin domain-containing protein; uniclust \| UniRef100\_A0A0F9LWS5 \| 99.8 \| 3.4e-22 \| 6.3e-28 \| 154.7 \| 166 \| (18, 189) \| 291 \| (7, 175) \| 202 \| ParB/Sulfiredoxin domain-containing protein (Fragment) \| ParB/Sulfiredoxin domain-containing protein (Fragment); uniclust \| UniRef100\_A0A1F9UGF1 \| 99.8 \| 3.5e-22 \| 6.4e-28 \| 161.0 \| 167 \| (17, 189) \| 291 \| (8, 183) \| 332 \| ParB/Sulfiredoxin domain-containing protein \| ParB/Sulfiredoxin domain-containing protein; uniclust \| UniRef100\_A0A4V2P9K0 \| 99.8 \| 3.5e-22 \| 6.6e-28 \| 178.5 \| 179 \| (18, 201) \| 291 \| (70, 269) \| 577 \| ParB/RepB/Spo0J family partition protein \| ParB/RepB/Spo0J family partition protein; uniclust \| UniRef100\_A0A1N6KBF8 \| 99.8 \| 3.5e-22 \| 6.8e-28 \| 180.4 \| 179 \| (18, 201) \| 291 \| (26, 206) \| 630 \| ParB/RepB/Spo0J family partition protein \| ParB/RepB/Spo0J family partition protein; uniclust \| UniRef100\_UPI0002BEB314 \| 99.8 \| 3.6e-22 \| 6.8e-28 \| 169.2 \| 173 \| (15, 193) \| 291 \| (47, 226) \| 387 \| ParB/RepB/Spo0J family partition protein \| ParB/RepB/Spo0J family partition protein; uniclust \| UniRef100\_A0A1P8WH68 \| 99.8 \| 3.5e-22 \| 6.9e-28 \| 168.0 \| 178 \| (15, 198) \| 291 \| (18, 201) \| 291 \| Putative chromosome-partitioning protein ParB \| Putative chromosome-partitioning protein ParB; uniclust \| UniRef100\_A0A3S3RBF0 \| 99.8 \| 3.9e-22 \| 7.4e-28 \| 148.1 \| 123 \| (15, 142) \| 291 \| (29, 153) \| 155 \| ParB/RepB/Spo0J family partition protein \| ParB/RepB/Spo0J family partition protein; uniclust \| UniRef100\_A0A7X7KEX9 \| 99.8 \| 4.2e-22 \| 7.7e-28 \| 159.0 \| 162 \| (17, 184) \| 291 \| (142, 306) \| 308 \| ParB/RepB/Spo0J family partition protein (Fragment) \| ParB/RepB/Spo0J family partition protein (Fragment); uniclust \| UniRef100\_A0A066YG78 \| 99.8 \| 4.1e-22 \| 7.9e-28 \| 173.4 \| 174 \| (20, 202) \| 291 \| (58, 251) \| 376 \| ParB/Sulfiredoxin domain-containing protein \| ParB/Sulfiredoxin domain-containing protein; uniclust \| UniRef100\_A0A3G9IH11 \| 99.8 \| 4.2e-22 \| 8e-28 \| 167.0 \| 166 \| (18, 188) \| 291 \| (106, 273) \| 359 \| ParB/Sulfiredoxin domain-containing protein \| ParB/Sulfiredoxin domain-containing protein; uniclust \| UniRef100\_A0A7W1GI18 \| 99.8 \| 4.3e-22 \| 8e-28 \| 156.1 \| 172 \| (14, 190) \| 291 \| (11, 190) \| 231 \| ParB/RepB/Spo0J family partition protein \| ParB/RepB/Spo0J family partition protein; uniclust \| UniRef100\_B6AP60 \| 99.8 \| 4.5e-22 \| 8.4e-28 \| 162.4 \| 177 \| (17, 199) \| 291 \| (29, 209) \| 319 \| ParB-like partition protein \| ParB-like partition protein; uniclust \| UniRef100\_A0A0G0QGC9 \| 99.8 \| 4.6e-22 \| 8.4e-28 \| 162.0 \| 182 \| (13, 199) \| 291 \| (94, 278) \| 361 \| ParB-like protein partition protein \| ParB-like protein partition protein; uniclust \| UniRef100\_A0A225D4E0 \| 99.8 \| 4.9e-22 \| 9.4e-28 \| 162.6 \| 183 \| (1, 188) \| 291 \| (1, 187) \| 266 \| ParB/Sulfiredoxin domain-containing protein \| ParB/Sulfiredoxin domain-containing protein; uniclust \| UniRef100\_A0A2D9Y5I6 \| 99.8 \| 5.6e-22 \| 1e-27 \| 172.9 \| 177 \| (17, 198) \| 291 \| (6, 197) \| 579 \| ParB/Sulfiredoxin domain-containing protein \| ParB/Sulfiredoxin domain-containing protein; uniclust \| UniRef100\_A0A0W7YQU1 \| 99.8 \| 5.6e-22 \| 1.1e-27 \| 174.9 \| 180 \| (17, 199) \| 291 \| (38, 234) \| 446 \| ParB/Sulfiredoxin domain-containing protein \| ParB/Sulfiredoxin domain-containing protein; uniclust \| UniRef100\_A0A0J0YDN7 \| 99.8 \| 5.7e-22 \| 1.1e-27 \| 173.1 \| 245 \| (17, 290) \| 291 \| (62, 321) \| 351 \| ParB/Sulfiredoxin domain-containing protein \| ParB/Sulfiredoxin domain-containing protein; uniclust \| UniRef100\_A0A3B9QB56 \| 99.8 \| 6e-22 \| 1.1e-27 \| 164.4 \| 176 \| (17, 199) \| 291 \| (28, 207) \| 310 \| ParB/Spo0J HTH domain-containing protein \| ParB/Spo0J HTH domain-containing protein; uniclust \| UniRef100\_A0A2D5W999 \| 99.8 \| 6.4e-22 \| 1.2e-27 \| 165.8 \| 168 \| (18, 190) \| 291 \| (150, 323) \| 419 \| Chromosome partitioning protein ParB \| Chromosome partitioning protein ParB; uniclust \| UniRef100\_A0A2R3UAM1 \| 99.8 \| 6.5e-22 \| 1.2e-27 \| 161.5 \| 213 \| (3, 216) \| 291 \| (4, 225) \| 305 \| Endonuclease \| Endonuclease; uniclust \| UniRef100\_A0A0G1Q7M5 \| 99.8 \| 6.8e-22 \| 1.2e-27 \| 150.7 \| 169 \| (15, 188) \| 291 \| (31, 203) \| 218 \| Stage 0 sporulation protein J \| Stage 0 sporulation protein J; uniclust \| UniRef100\_A0A1T1HAZ1 \| 99.8 \| 6.7e-22 \| 1.3e-27 \| 164.3 \| 168 \| (15, 188) \| 291 \| (32, 204) \| 344 \| ParB/Sulfiredoxin domain-containing protein \| ParB/Sulfiredoxin domain-containing protein; uniclust \| UniRef100\_A0A259CJI7 \| 99.8 \| 6.8e-22 \| 1.3e-27 \| 162.2 \| 157 \| (13, 175) \| 291 \| (47, 209) \| 299 \| ParB/Sulfiredoxin domain-containing protein \| ParB/Sulfiredoxin domain-containing protein; uniclust \| UniRef100\_UPI000B35E396 \| 99.8 \| 7.6e-22 \| 1.4e-27 \| 151.0 \| 147 \| (17, 168) \| 291 \| (31, 180) \| 224 \| ParB/RepB/Spo0J family partition protein \| ParB/RepB/Spo0J family partition protein; uniclust \| UniRef100\_A0A1C6F112 \| 99.8 \| 7.3e-22 \| 1.4e-27 \| 165.8 \| 168 \| (21, 193) \| 291 \| (33, 211) \| 321 \| Nucleoid occlusion protein \| Nucleoid occlusion protein; uniclust \| UniRef100\_A0A7Y9T2W1 \| 99.8 \| 7.6e-22 \| 1.4e-27 \| 161.3 \| 178 \| (17, 200) \| 291 \| (4, 184) \| 326 \| ParB/RepB/Spo0J family partition protein \| ParB/RepB/Spo0J family partition protein; uniclust \| UniRef100\_A0A2I7QW51 \| 99.8 \| 7.7e-22 \| 1.4e-27 \| 159.0 \| 271 \| (2, 288) \| 291 \| (6, 282) \| 286 \| Uncharacterized protein \| Uncharacterized protein; uniclust \| UniRef100\_A0A1G7A3Z4 \| 99.8 \| 7.7e-22 \| 1.5e-27 \| 169.2 \| 170 \| (16, 190) \| 291 \| (51, 223) \| 388 \| Chromosome partitioning protein, ParB family \| Chromosome partitioning protein, ParB family; uniclust \| UniRef100\_A0A0P9EKV8 \| 99.8 \| 7.9e-22 \| 1.5e-27 \| 166.9 \| 158 \| (21, 187) \| 291 \| (3, 162) \| 389 \| Nucleoid occlusion protein \| Nucleoid occlusion protein; uniclust \| UniRef100\_A0A1M6KVA2 \| 99.8 \| 8.8e-22 \| 1.7e-27 \| 161.8 \| 162 \| (20, 189) \| 291 \| (34, 200) \| 304 \| ParB/RepB/Spo0J family partition protein \| ParB/RepB/Spo0J family partition protein; uniclust \| UniRef100\_F3AXU5 \| 99.8 \| 8.9e-22 \| 1.7e-27 \| 152.4 \| 145 \| (17, 169) \| 291 \| (3, 154) \| 199 \| ParB/Sulfiredoxin domain-containing protein \| ParB/Sulfiredoxin domain-containing protein; uniclust \| UniRef100\_A0A1Q3SL05 \| 99.8 \| 8.4e-22 \| 1.7e-27 \| 173.1 \| 178 \| (16, 199) \| 291 \| (45, 231) \| 373 \| ParB/Sulfiredoxin domain-containing protein \| ParB/Sulfiredoxin domain-containing protein; uniclust \| UniRef100\_A0A2E2NA70 \| 99.8 \| 9.1e-22 \| 1.7e-27 \| 165.0 \| 167 \| (16, 188) \| 291 \| (198, 368) \| 466 \| HTH cro/C1-type domain-containing protein \| HTH cro/C1-type domain-containing protein; uniclust \| UniRef100\_A0A1V2SIP1 \| 99.8 \| 9.2e-22 \| 1.7e-27 \| 162.6 \| 161 \| (20, 186) \| 291 \| (3, 165) \| 409 \| ParB/Sulfiredoxin domain-containing protein \| ParB/Sulfiredoxin domain-containing protein; uniclust \| UniRef100\_A0A087C1L2 \| 99.8 \| 9.4e-22 \| 1.8e-27 \| 177.3 \| 172 \| (17, 194) \| 291 \| (386, 604) \| 710 \| Chromosome partitioning protein ParB \| Chromosome partitioning protein ParB; uniclust \| UniRef100\_A0A1V6MIC0 \| 99.8 \| 9.4e-22 \| 1.8e-27 \| 159.3 \| 175 \| (18, 199) \| 291 \| (39, 231) \| 263 \| Plasmid partitioning protein (Fragment) \| Plasmid partitioning protein (Fragment); uniclust \| UniRef100\_A0A1C5G7E9 \| 99.8 \| 9.4e-22 \| 1.8e-27 \| 180.5 \| 163 \| (20, 187) \| 291 \| (58, 223) \| 580 \| Chromosome partitioning protein, ParB family \| Chromosome partitioning protein, ParB family; uniclust \| UniRef100\_A0A031HLV3 \| 99.8 \| 9.7e-22 \| 1.9e-27 \| 184.6 \| 171 \| (22, 197) \| 291 \| (115, 302) \| 792 \| ParB-like partition protein \| ParB-like partition protein; uniclust \| UniRef100\_A0A1G5CHR0 \| 99.8 \| 1.2e-21 \| 2.2e-27 \| 162.4 \| 165 \| (17, 189) \| 291 \| (31, 197) \| 354 \| Chromosome partitioning protein, ParB family \| Chromosome partitioning protein, ParB family; uniclust \| UniRef100\_UPI001607B1A3 \| 99.8 \| 1.2e-21 \| 2.2e-27 \| 152.4 \| 195 \| (69, 288) \| 291 \| (2, 197) \| 209 \| chromosome partitioning protein ParB \| chromosome partitioning protein ParB; uniclust \| UniRef100\_A0A2L2NV26 \| 99.8 \| 1.1e-21 \| 2.2e-27 \| 167.3 \| 164 \| (16, 184) \| 291 \| (16, 184) \| 291 \| Chromosome/plasmid partitioning protein ParB \| Chromosome/plasmid partitioning protein ParB; uniclust \| UniRef100\_A0A800ETG4 \| 99.8 \| 1.2e-21 \| 2.3e-27 \| 154.0 \| 152 \| (14, 171) \| 291 \| (31, 185) \| 273 \| ParB/RepB/Spo0J family partition protein (Fragment) \| ParB/RepB/Spo0J family partition protein (Fragment); uniclust \| UniRef100\_A0A7V9IA21 \| 99.8 \| 1.2e-21 \| 2.3e-27 \| 164.4 \| 182 \| (13, 200) \| 291 \| (32, 220) \| 411 \| ParB/RepB/Spo0J family partition protein \| ParB/RepB/Spo0J family partition protein; uniclust \| UniRef100\_A0A7X1ZX84 \| 99.8 \| 1.3e-21 \| 2.3e-27 \| 158.8 \| 169 \| (15, 189) \| 291 \| (31, 202) \| 305 \| ParB/RepB/Spo0J family partition protein \| ParB/RepB/Spo0J family partition protein; uniclust \| UniRef100\_A0A098MHM3 \| 99.8 \| 1.3e-21 \| 2.4e-27 \| 160.2 \| 154 \| (15, 174) \| 291 \| (19, 176) \| 285 \| Plasmid partitioning protein ParB \| Plasmid partitioning protein ParB; uniclust \| UniRef100\_A0A1C5TTA9 \| 99.8 \| 1.3e-21 \| 2.4e-27 \| 161.1 \| 178 \| (15, 197) \| 291 \| (43, 240) \| 289 \| Chromosome-partitioning protein parB \| Chromosome-partitioning protein parB; uniclust \| UniRef100\_A0A077FQM9 \| 99.7 \| 1.3e-21 \| 2.5e-27 \| 171.9 \| 166 \| (17, 188) \| 291 \| (71, 244) \| 360 \| Probable chromosome-partitioning protein ParB \| Probable chromosome-partitioning protein ParB; uniclust \| UniRef100\_A0A173WSZ4 \| 99.7 \| 1.3e-21 \| 2.6e-27 \| 156.8 \| 179 \| (13, 196) \| 291 \| (27, 226) \| 231 \| Nucleoid occlusion protein \| Nucleoid occlusion protein; uniclust \| UniRef100\_A0A0M4DV65 \| 99.7 \| 1.4e-21 \| 2.6e-27 \| 165.7 \| 176 \| (20, 199) \| 291 \| (36, 235) \| 376 \| ParB/Sulfiredoxin domain-containing protein \| ParB/Sulfiredoxin domain-containing protein; uniclust \| UniRef100\_A0A0B3BFF3 \| 99.7 \| 1.3e-21 \| 2.7e-27 \| 178.8 \| 171 \| (20, 197) \| 291 \| (35, 235) \| 489 \| Putative transcriptional regulator \| Putative transcriptional regulator; uniclust \| UniRef100\_A0A2M7TEW5 \| 99.7 \| 1.5e-21 \| 2.8e-27 \| 149.1 \| 165 \| (18, 188) \| 291 \| (4, 171) \| 182 \| ParB/Sulfiredoxin domain-containing protein \| ParB/Sulfiredoxin domain-containing protein; uniclust \| UniRef100\_A0A0P9H9C4 \| 99.7 \| 1.5e-21 \| 2.8e-27 \| 147.9 \| 164 \| (19, 188) \| 291 \| (2, 176) \| 188 \| Chromosome partitioning protein ParB (Fragment) \| Chromosome partitioning protein ParB (Fragment); uniclust \| UniRef100\_A0A133XIV0 \| 99.7 \| 1.5e-21 \| 2.8e-27 \| 164.9 \| 166 \| (17, 187) \| 291 \| (43, 215) \| 341 \| Chromosome partitioning protein ParB \| Chromosome partitioning protein ParB; uniclust \| UniRef100\_A0A519FZE1 \| 99.7 \| 1.5e-21 \| 2.9e-27 \| 176.2 \| 171 \| (17, 193) \| 291 \| (7, 185) \| 686 \| ParB/RepB/Spo0J family partition protein \| ParB/RepB/Spo0J family partition protein; uniclust \| UniRef100\_A0A953F209 \| 99.7 \| 1.6e-21 \| 3e-27 \| 149.0 \| 163 \| (17, 185) \| 291 \| (32, 204) \| 221 \| ParB/RepB/Spo0J family partition protein (Fragment) \| ParB/RepB/Spo0J family partition protein (Fragment); uniclust \| UniRef100\_A0A0T7G344 \| 99.7 \| 1.6e-21 \| 3e-27 \| 180.1 \| 181 \| (15, 197) \| 291 \| (33, 220) \| 667 \| ParB domain protein nuclease (Fragment) \| ParB domain protein nuclease (Fragment); uniclust \| UniRef100\_A0A7C5GQE9 \| 99.7 \| 1.6e-21 \| 3e-27 \| 160.0 \| 169 \| (15, 189) \| 291 \| (33, 209) \| 291 \| ParB/RepB/Spo0J family partition protein \| ParB/RepB/Spo0J family partition protein; uniclust \| UniRef100\_A0A2M7E1N5 \| 99.7 \| 1.6e-21 \| 3e-27 \| 161.1 \| 165 \| (15, 185) \| 291 \| (8, 186) \| 278 \| ParB/Sulfiredoxin domain-containing protein \| ParB/Sulfiredoxin domain-containing protein; uniclust \| UniRef100\_A0A512MCK4 \| 99.7 \| 1.6e-21 \| 3.1e-27 \| 158.5 \| 177 \| (14, 196) \| 291 \| (26, 207) \| 288 \| ParB/Sulfiredoxin domain-containing protein \| ParB/Sulfiredoxin domain-containing protein; uniclust \| UniRef100\_A0A5S4ZSV0 \| 99.7 \| 1.7e-21 \| 3.2e-27 \| 170.0 \| 164 \| (18, 187) \| 291 \| (2, 172) \| 638 \| ParB/RepB/Spo0J family partition protein \| ParB/RepB/Spo0J family partition protein; uniclust \| UniRef100\_A0A0H3ZQX6 \| 99.7 \| 1.7e-21 \| 3.2e-27 \| 165.1 \| 152 \| (17, 177) \| 291 \| (31, 186) \| 319 \| Chromosome (Plasmid) partitioning protein ParB \| Chromosome (Plasmid) partitioning protein ParB; uniclust \| UniRef100\_A0A1X7DWC0 \| 99.7 \| 1.7e-21 \| 3.4e-27 \| 167.0 \| 179 \| (16, 197) \| 291 \| (6, 193) \| 370 \| Chromosome partitioning protein, ParB family (Fragment) \| Chromosome partitioning protein, ParB family (Fragment); uniclust \| UniRef100\_A0A0N1NWJ0 \| 99.7 \| 1.8e-21 \| 3.4e-27 \| 183.2 \| 174 \| (20, 198) \| 291 \| (675, 854) \| 1019 \| ParB-like partition protein \| ParB-like partition protein; uniclust \| UniRef100\_A0A0C2AKN6 \| 99.7 \| 1.8e-21 \| 3.5e-27 \| 172.9 \| 168 \| (15, 188) \| 291 \| (32, 221) \| 476 \| ParB/Sulfiredoxin domain-containing protein \| ParB/Sulfiredoxin domain-containing protein; uniclust \| UniRef100\_A0A177NYB8 \| 99.7 \| 1.9e-21 \| 3.7e-27 \| 183.8 \| 177 \| (18, 196) \| 291 \| (7, 190) \| 726 \| ParB/Sulfiredoxin domain-containing protein \| ParB/Sulfiredoxin domain-containing protein; uniclust \| UniRef100\_A0A1C4K463 \| 99.7 \| 1.9e-21 \| 3.7e-27 \| 170.0 \| 174 \| (21, 199) \| 291 \| (76, 272) \| 431 \| ParB/RepB/Spo0J family partition protein \| ParB/RepB/Spo0J family partition protein; uniclust \| UniRef100\_A0A1F9UE43 \| 99.7 \| 2.1e-21 \| 4e-27 \| 161.9 \| 167 \| (18, 189) \| 291 \| (13, 183) \| 369 \| ParB/Sulfiredoxin domain-containing protein \| ParB/Sulfiredoxin domain-containing protein; uniclust \| UniRef100\_A0A537LQR2 \| 99.7 \| 2.2e-21 \| 4e-27 \| 149.5 \| 145 \| (19, 169) \| 291 \| (85, 232) \| 234 \| ParB/RepB/Spo0J family partition protein (Fragment) \| ParB/RepB/Spo0J family partition protein (Fragment); uniclust \| UniRef100\_A0A352Z5M4 \| 99.7 \| 2.1e-21 \| 4.1e-27 \| 160.9 \| 181 \| (14, 199) \| 291 \| (17, 200) \| 309 \| ParB/Sulfiredoxin domain-containing protein \| ParB/Sulfiredoxin domain-containing protein; uniclust \| UniRef100\_A0A497FIQ1 \| 99.7 \| 2.2e-21 \| 4.2e-27 \| 165.4 \| 165 \| (18, 188) \| 291 \| (3, 170) \| 348 \| ParB/Sulfiredoxin domain-containing protein \| ParB/Sulfiredoxin domain-containing protein; uniclust \| UniRef100\_UPI00041C85F7 \| 99.7 \| 2.4e-21 \| 4.4e-27 \| 159.9 \| 169 \| (16, 189) \| 291 \| (24, 198) \| 400 \| ParB/RepB/Spo0J family partition protein \| ParB/RepB/Spo0J family partition protein; uniclust \| UniRef100\_A0A257NIY2 \| 99.7 \| 2.4e-21 \| 4.5e-27 \| 145.5 \| 152 \| (43, 198) \| 291 \| (2, 156) \| 180 \| ParB/Sulfiredoxin domain-containing protein (Fragment) \| ParB/Sulfiredoxin domain-containing protein (Fragment); uniclust \| UniRef100\_A0A225DDB5 \| 99.7 \| 2.4e-21 \| 4.5e-27 \| 161.8 \| 179 \| (17, 201) \| 291 \| (44, 233) \| 312 \| Chromosome (Plasmid) partitioning protein ParB / Stage 0 sporulation protein J \| Chromosome (Plasmid) partitioning protein ParB / Stage 0 sporulation protein J; uniclust \| UniRef100\_A0A075MDE3 \| 99.7 \| 2.4e-21 \| 4.5e-27 \| 149.0 \| 145 \| (20, 170) \| 291 \| (36, 183) \| 188 \| Chromosome (Plasmid) partitioning protein ParB/ Stage 0 sporulation protein J \| Chromosome (Plasmid) partitioning protein ParB/ Stage 0 sporulation protein J; uniclust \| UniRef100\_A0A6N6T5G8 \| 99.7 \| 2.4e-21 \| 4.5e-27 \| 156.7 \| 176 \| (18, 194) \| 291 \| (5, 186) \| 270 \| ParB/RepB/Spo0J family partition protein \| ParB/RepB/Spo0J family partition protein; uniclust \| UniRef100\_A0A3A4Z2I3 \| 99.7 \| 2.5e-21 \| 4.7e-27 \| 158.3 \| 164 \| (19, 187) \| 291 \| (38, 206) \| 313 \| ParB/RepB/Spo0J family partition protein \| ParB/RepB/Spo0J family partition protein; uniclust \| UniRef100\_A0A950B1H5 \| 99.7 \| 2.6e-21 \| 4.8e-27 \| 156.9 \| 165 \| (20, 189) \| 291 \| (50, 218) \| 345 \| ParB/RepB/Spo0J family partition protein \| ParB/RepB/Spo0J family partition protein; uniclust \| UniRef100\_A0A176EXX9 \| 99.7 \| 2.4e-21 \| 4.8e-27 \| 170.2 \| 223 \| (13, 247) \| 291 \| (29, 266) \| 330 \| ParB/Sulfiredoxin domain-containing protein \| ParB/Sulfiredoxin domain-containing protein; uniclust \| UniRef100\_A0A0S8I9V8 \| 99.7 \| 2.6e-21 \| 4.9e-27 \| 160.2 \| 168 \| (18, 186) \| 291 \| (5, 178) \| 291 \| Chromosome partitioning protein ParB \| Chromosome partitioning protein ParB; uniclust \| UniRef100\_A0A952V660 \| 99.7 \| 2.7e-21 \| 5.1e-27 \| 172.3 \| 180 \| (14, 199) \| 291 \| (25, 221) \| 581 \| ParB/RepB/Spo0J family partition protein \| ParB/RepB/Spo0J family partition protein; uniclust \| UniRef100\_A0A1F8MEB9 \| 99.7 \| 2.8e-21 \| 5.3e-27 \| 156.1 \| 176 \| (21, 201) \| 291 \| (3, 183) \| 259 \| ParB/Sulfiredoxin domain-containing protein \| ParB/Sulfiredoxin domain-containing protein; uniclust \| UniRef100\_A0A7X6ND40 \| 99.7 \| 2.9e-21 \| 5.3e-27 \| 168.9 \| 179 \| (16, 199) \| 291 \| (430, 620) \| 706 \| ParB/RepB/Spo0J family partition protein \| ParB/RepB/Spo0J family partition protein; uniclust \| UniRef100\_A0A977UBZ6 \| 99.7 \| 3e-21 \| 5.4e-27 \| 145.0 \| 162 \| (23, 189) \| 291 \| (5, 171) \| 196 \| ParB/RepB/Spo0J family partition protein \| ParB/RepB/Spo0J family partition protein; uniclust \| UniRef100\_A0A537PWK3 \| 99.7 \| 3e-21 \| 5.5e-27 \| 145.1 \| 144 \| (39, 187) \| 291 \| (4, 152) \| 197 \| ParB/RepB/Spo0J family partition protein (Fragment) \| ParB/RepB/Spo0J family partition protein (Fragment); uniclust \| UniRef100\_A0A0F9L3S6 \| 99.7 \| 3e-21 \| 5.7e-27 \| 167.9 \| 180 \| (14, 198) \| 291 \| (49, 237) \| 403 \| ParB/Sulfiredoxin domain-containing protein \| ParB/Sulfiredoxin domain-containing protein; uniclust \| UniRef100\_A0A7V9GCS1 \| 99.7 \| 3.1e-21 \| 5.8e-27 \| 149.5 \| 167 \| (16, 189) \| 291 \| (27, 197) \| 230 \| ParB/RepB/Spo0J family partition protein (Fragment) \| ParB/RepB/Spo0J family partition protein (Fragment); uniclust \| UniRef100\_A0A0N8Q4D0 \| 99.7 \| 3e-21 \| 5.9e-27 \| 180.4 \| 176 \| (21, 201) \| 291 \| (63, 243) \| 631 \| Probable chromosome-partitioning protein ParB \| Probable chromosome-partitioning protein ParB; uniclust \| UniRef100\_A0A0F4I4G6 \| 99.7 \| 3.2e-21 \| 5.9e-27 \| 150.2 \| 164 \| (30, 198) \| 291 \| (4, 184) \| 221 \| ParB/Sulfiredoxin domain-containing protein (Fragment) \| ParB/Sulfiredoxin domain-containing protein (Fragment); uniclust \| UniRef100\_A0A5Q4ZYV4 \| 99.7 \| 3.1e-21 \| 6e-27 \| 165.3 \| 172 \| (16, 188) \| 291 \| (32, 208) \| 340 \| ParB/Sulfiredoxin domain-containing protein \| ParB/Sulfiredoxin domain-containing protein; uniclust \| UniRef100\_R7M4B8 \| 99.7 \| 3.3e-21 \| 6.1e-27 \| 160.9 \| 174 \| (20, 199) \| 291 \| (7, 184) \| 444 \| ParB-like partition protein \| ParB-like partition protein; uniclust \| UniRef100\_A0A6L7GCN8 \| 99.7 \| 3.2e-21 \| 6.1e-27 \| 161.7 \| 181 \| (1, 187) \| 291 \| (1, 184) \| 304 \| ParB/RepB/Spo0J family partition protein \| ParB/RepB/Spo0J family partition protein; uniclust \| UniRef100\_A0A1V2H5Y6 \| 99.7 \| 3.1e-21 \| 6.1e-27 \| 171.4 \| 165 \| (17, 187) \| 291 \| (132, 304) \| 419 \| Chromosome partitioning protein ParB \| Chromosome partitioning protein ParB; uniclust \| UniRef100\_A0A0Q4MYY7 \| 99.7 \| 3.6e-21 \| 7e-27 \| 162.3 \| 168 \| (15, 188) \| 291 \| (35, 205) \| 324 \| ParB/Sulfiredoxin domain-containing protein \| ParB/Sulfiredoxin domain-containing protein; uniclust \| UniRef100\_UPI00216A9ACB \| 99.7 \| 3.8e-21 \| 7e-27 \| 139.6 \| 139 \| (21, 165) \| 291 \| (3, 153) \| 157 \| ParB/RepB/Spo0J family partition protein \| ParB/RepB/Spo0J family partition protein; uniclust \| UniRef100\_E7H2H0 \| 99.7 \| 3.9e-21 \| 7.3e-27 \| 163.3 \| 167 \| (16, 190) \| 291 \| (26, 200) \| 388 \| ParB/Sulfiredoxin domain-containing protein \| ParB/Sulfiredoxin domain-containing protein; uniclust \| UniRef100\_A0A936A9A9 \| 99.7 \| 4e-21 \| 7.4e-27 \| 161.3 \| 170 \| (17, 191) \| 291 \| (33, 205) \| 467 \| Probable chromosome-partitioning protein ParB \| Probable chromosome-partitioning protein ParB; uniclust \| UniRef100\_A0A060HQA9 \| 99.7 \| 3.9e-21 \| 7.4e-27 \| 156.0 \| 172 \| (18, 195) \| 291 \| (3, 182) \| 257 \| ParB/Sulfiredoxin domain-containing protein \| ParB/Sulfiredoxin domain-containing protein; uniclust \| UniRef100\_A0A073IWS9 \| 99.7 \| 4e-21 \| 8e-27 \| 182.2 \| 178 \| (19, 198) \| 291 \| (62, 247) \| 704 \| ParB/Sulfiredoxin domain-containing protein \| ParB/Sulfiredoxin domain-containing protein; uniclust \| UniRef100\_UPI00126055D0 \| 99.7 \| 4.4e-21 \| 8.1e-27 \| 149.1 \| 166 \| (18, 189) \| 291 \| (49, 220) \| 248 \| ParB/RepB/Spo0J family partition protein \| ParB/RepB/Spo0J family partition protein; uniclust \| UniRef100\_A0A5F0LUW4 \| 99.7 \| 4.2e-21 \| 8.1e-27 \| 176.4 \| 166 \| (18, 189) \| 291 \| (37, 212) \| 703 \| ParB/RepB/Spo0J family partition protein \| ParB/RepB/Spo0J family partition protein; uniclust \| UniRef100\_A0A1C5P2R5 \| 99.7 \| 4.4e-21 \| 8.3e-27 \| 159.0 \| 176 \| (14, 194) \| 291 \| (36, 220) \| 309 \| Probable chromosome-partitioning protein parB \| Probable chromosome-partitioning protein parB; uniclust \| UniRef100\_UPI001F0133BB \| 99.7 \| 4.6e-21 \| 8.5e-27 \| 154.7 \| 167 \| (15, 187) \| 291 \| (29, 198) \| 332 \| ParB/RepB/Spo0J family partition protein \| ParB/RepB/Spo0J family partition protein; uniclust \| UniRef100\_A0A962QQD6 \| 99.7 \| 4.5e-21 \| 8.6e-27 \| 161.1 \| 149 \| (19, 173) \| 291 \| (23, 176) \| 342 \| ParB/RepB/Spo0J family partition protein \| ParB/RepB/Spo0J family partition protein; uniclust \| UniRef100\_A0A920G1W5 \| 99.7 \| 5.1e-21 \| 9.4e-27 \| 158.9 \| 175 \| (20, 199) \| 291 \| (171, 348) \| 422 \| Probable chromosome-partitioning protein ParB \| Probable chromosome-partitioning protein ParB; uniclust \| UniRef100\_A0A840CS83 \| 99.7 \| 5.2e-21 \| 1e-26 \| 176.9 \| 177 \| (14, 196) \| 291 \| (294, 473) \| 832 \| ParB/RepB/Spo0J family partition protein \| ParB/RepB/Spo0J family partition protein; uniclust \| UniRef100\_A0A021VU08 \| 99.7 \| 5.3e-21 \| 1e-26 \| 170.6 \| 176 \| (15, 196) \| 291 \| (40, 219) \| 533 \| Chromosomal partitioning protein ParB \| Chromosomal partitioning protein ParB; uniclust \| UniRef100\_A0A011TXK3 \| 99.7 \| 5.1e-21 \| 1e-26 \| 167.8 \| 186 \| (14, 206) \| 291 \| (20, 210) \| 362 \| Plasmid stablization protein ParB \| Plasmid stablization protein ParB; uniclust \| UniRef100\_A0A2T2VQX6 \| 99.7 \| 5.2e-21 \| 1e-26 \| 156.9 \| 131 \| (66, 201) \| 291 \| (6, 139) \| 228 \| ParB/Spo0J HTH domain-containing protein (Fragment) \| ParB/Spo0J HTH domain-containing protein (Fragment); uniclust \| UniRef100\_A0A1J9PXM3 \| 99.7 \| 5.5e-21 \| 1e-26 \| 163.3 \| 235 \| (13, 247) \| 291 \| (27, 290) \| 453 \| Pyridoxal phosphate biosynthetic protein PdxJ \| Pyridoxal phosphate biosynthetic protein PdxJ; uniclust \| UniRef100\_A0A3M2CVE6 \| 99.7 \| 5.6e-21 \| 1e-26 \| 156.5 \| 163 \| (21, 190) \| 291 \| (31, 197) \| 327 \| ParB/RepB/Spo0J family partition protein \| ParB/RepB/Spo0J family partition protein; uniclust \| UniRef100\_A0A173ZK65 \| 99.7 \| 5.5e-21 \| 1e-26 \| 172.6 \| 176 \| (17, 197) \| 291 \| (331, 531) \| 660 \| Probable chromosome-partitioning protein parB \| Probable chromosome-partitioning protein parB; uniclust \| UniRef100\_A0A246BRZ4 \| 99.7 \| 5.5e-21 \| 1.1e-26 \| 162.3 \| 175 \| (16, 198) \| 291 \| (32, 233) \| 315 \| ParB/Sulfiredoxin domain-containing protein \| ParB/Sulfiredoxin domain-containing protein; uniclust \| UniRef100\_A0A0F9U9W1 \| 99.7 \| 5.7e-21 \| 1.1e-26 \| 163.3 \| 155 \| (15, 176) \| 291 \| (35, 193) \| 383 \| ParB/Sulfiredoxin domain-containing protein \| ParB/Sulfiredoxin domain-containing protein; uniclust \| UniRef100\_A0A011Q0A6 \| 99.7 \| 5.5e-21 \| 1.1e-26 \| 188.3 \| 179 \| (16, 197) \| 291 \| (179, 363) \| 924 \| Plasmid partitioning protein \| Plasmid partitioning protein; uniclust \| UniRef100\_A0A352YU83 \| 99.7 \| 6e-21 \| 1.1e-26 \| 159.1 \| 166 \| (16, 187) \| 291 \| (33, 210) \| 321 \| ParB/Sulfiredoxin domain-containing protein \| ParB/Sulfiredoxin domain-containing protein; uniclust \| UniRef100\_A0A7C3VM87 \| 99.7 \| 6.3e-21 \| 1.2e-26 \| 153.5 \| 162 \| (18, 184) \| 291 \| (5, 176) \| 323 \| ParB/RepB/Spo0J family partition protein \| ParB/RepB/Spo0J family partition protein; uniclust \| UniRef100\_A0A1G0T9I7 \| 99.7 \| 6.3e-21 \| 1.2e-26 \| 160.2 \| 176 \| (19, 199) \| 291 \| (44, 224) \| 466 \| ParB/Sulfiredoxin domain-containing protein \| ParB/Sulfiredoxin domain-containing protein; uniclust \| UniRef100\_A0A916DA64 \| 99.7 \| 6.1e-21 \| 1.2e-26 \| 159.4 \| 171 \| (19, 199) \| 291 \| (42, 213) \| 333 \| Chromosome-partitioning protein Spo0J \| Chromosome-partitioning protein Spo0J; uniclust \| UniRef100\_A0A0A7PKV5 \| 99.7 \| 6e-21 \| 1.2e-26 \| 169.5 \| 166 \| (13, 183) \| 291 \| (45, 220) \| 372 \| ParB-like partition protein \| ParB-like partition protein; uniclust \| UniRef100\_A0A357EIG5 \| 99.7 \| 6.6e-21 \| 1.2e-26 \| 152.6 \| 165 \| (18, 188) \| 291 \| (20, 190) \| 281 \| ParB/Sulfiredoxin domain-containing protein (Fragment) \| ParB/Sulfiredoxin domain-containing protein (Fragment); uniclust \| UniRef100\_A0A920UHB7 \| 99.7 \| 6.8e-21 \| 1.3e-26 \| 144.4 \| 152 \| (15, 173) \| 291 \| (10, 168) \| 208 \| ParB/Sulfiredoxin domain-containing protein \| ParB/Sulfiredoxin domain-containing protein; uniclust \| UniRef100\_A0A1F3T0T4 \| 99.7 \| 7.1e-21 \| 1.3e-26 \| 157.5 \| 173 \| (18, 196) \| 291 \| (49, 226) \| 308 \| ParB/Sulfiredoxin domain-containing protein \| ParB/Sulfiredoxin domain-containing protein; uniclust \| UniRef100\_A0A2E9QWR4 \| 99.7 \| 6.9e-21 \| 1.3e-26 \| 163.4 \| 181 \| (14, 200) \| 291 \| (73, 261) \| 349 \| ParB/Sulfiredoxin domain-containing protein \| ParB/Sulfiredoxin domain-containing protein; uniclust \| UniRef100\_A0A1I7FCB2 \| 99.7 \| 7e-21 \| 1.4e-26 \| 165.6 \| 170 \| (18, 194) \| 291 \| (42, 221) \| 384 \| ParB/RepB/Spo0J family partition protein \| ParB/RepB/Spo0J family partition protein; uniclust \| UniRef100\_A0A523SL81 \| 99.7 \| 7.5e-21 \| 1.4e-26 \| 169.9 \| 175 \| (17, 198) \| 291 \| (9, 186) \| 623 \| ParB/RepB/Spo0J family partition protein \| ParB/RepB/Spo0J family partition protein; uniclust \| UniRef100\_A0A934DLA7 \| 99.7 \| 7.7e-21 \| 1.4e-26 \| 156.3 \| 175 \| (15, 196) \| 291 \| (29, 206) \| 386 \| ParB/RepB/Spo0J family partition protein \| ParB/RepB/Spo0J family partition protein; uniclust \| UniRef100\_A0A1F6FNT4 \| 99.7 \| 7.6e-21 \| 1.4e-26 \| 168.6 \| 178 \| (16, 199) \| 291 \| (8, 194) \| 627 \| ParB/Sulfiredoxin domain-containing protein \| ParB/Sulfiredoxin domain-containing protein; uniclust \| UniRef100\_A0A0P8BD48 \| 99.7 \| 7.3e-21 \| 1.4e-26 \| 164.7 \| 180 \| (16, 200) \| 291 \| (45, 266) \| 349 \| Chromosome partitioning protein, ParB family \| Chromosome partitioning protein, ParB family; uniclust \| UniRef100\_X1ICS2 \| 99.7 \| 7.7e-21 \| 1.5e-26 \| 147.0 \| 174 \| (22, 199) \| 291 \| (8, 185) \| 202 \| ParB/Sulfiredoxin domain-containing protein (Fragment) \| ParB/Sulfiredoxin domain-containing protein (Fragment); uniclust \| UniRef100\_A0A7Y4RZN6 \| 99.7 \| 8e-21 \| 1.5e-26 \| 159.6 \| 167 \| (14, 188) \| 291 \| (31, 202) \| 332 \| ParB/RepB/Spo0J family partition protein \| ParB/RepB/Spo0J family partition protein; uniclust \| UniRef100\_A0A068YQ66 \| 99.7 \| 8.1e-21 \| 1.6e-26 \| 164.0 \| 169 \| (14, 189) \| 291 \| (57, 226) \| 381 \| Chromosome (Plasmid) partitioning protein ParB / Stage 0 sporulation protein J \| Chromosome (Plasmid) partitioning protein ParB / Stage 0 sporulation protein J; uniclust \| UniRef100\_A0A0Q5HQC1 \| 99.7 \| 8.3e-21 \| 1.6e-26 \| 165.0 \| 166 \| (17, 190) \| 291 \| (29, 205) \| 372 \| ParB/Sulfiredoxin domain-containing protein \| ParB/Sulfiredoxin domain-containing protein; uniclust \| UniRef100\_A0A662E073 \| 99.7 \| 8.9e-21 \| 1.6e-26 \| 150.6 \| 173 \| (19, 197) \| 291 \| (2, 177) \| 290 \| ParB/Sulfiredoxin domain-containing protein \| ParB/Sulfiredoxin domain-containing protein; uniclust \| UniRef100\_A0A940CRG9 \| 99.7 \| 9.7e-21 \| 1.8e-26 \| 162.0 \| 162 \| (19, 185) \| 291 \| (3, 172) \| 555 \| ParB/RepB/Spo0J family partition protein \| ParB/RepB/Spo0J family partition protein; uniclust \| UniRef100\_A0A0L6J3Z5 \| 99.7 \| 9.1e-21 \| 1.8e-26 \| 178.3 \| 178 \| (19, 198) \| 291 \| (114, 304) \| 786 \| ParB/Sulfiredoxin domain-containing protein \| ParB/Sulfiredoxin domain-containing protein; uniclust \| UniRef100\_A0A011U8K2 \| 99.7 \| 8.9e-21 \| 1.8e-26 \| 185.1 \| 180 \| (15, 197) \| 291 \| (144, 336) \| 961 \| DNA-binding protein \| DNA-binding protein; uniclust \| UniRef100\_A0A5R8QHF2 \| 99.7 \| 1.1e-20 \| 1.9e-26 \| 164.8 \| 168 \| (19, 192) \| 291 \| (12, 183) \| 675 \| ParB/RepB/Spo0J family partition protein \| ParB/RepB/Spo0J family partition protein; uniclust \| UniRef100\_UPI001A972BC8 \| 99.7 \| 1e-20 \| 1.9e-26 \| 156.8 \| 164 \| (19, 188) \| 291 \| (52, 226) \| 367 \| ParB/RepB/Spo0J family partition protein \| ParB/RepB/Spo0J family partition protein; uniclust \| UniRef100\_A0A0C2R4G1 \| 99.7 \| 1e-20 \| 2e-26 \| 163.8 \| 152 \| (16, 172) \| 291 \| (4, 161) \| 406 \| ParB/Sulfiredoxin domain-containing protein \| ParB/Sulfiredoxin domain-containing protein; uniclust \| UniRef100\_A0A1V5UI04 \| 99.7 \| 1.1e-20 \| 2.1e-26 \| 157.8 \| 181 \| (15, 200) \| 291 \| (35, 217) \| 283 \| Putative chromosome-partitioning protein ParB \| Putative chromosome-partitioning protein ParB; uniclust \| UniRef100\_A0A067YD33 \| 99.7 \| 1.1e-20 \| 2.1e-26 \| 162.3 \| 143 \| (17, 165) \| 291 \| (45, 192) \| 326 \| Chromosome-partitioning protein ParB \| Chromosome-partitioning protein ParB; uniclust \| UniRef100\_A0A6G8FUF2 \| 99.7 \| 1.2e-20 \| 2.2e-26 \| 152.9 \| 166 \| (14, 185) \| 291 \| (73, 240) \| 330 \| ParB/RepB/Spo0J family partition protein \| ParB/RepB/Spo0J family partition protein; uniclust \| UniRef100\_A0A016QQ86 \| 99.7 \| 1.2e-20 \| 2.3e-26 \| 160.9 \| 180 \| (17, 201) \| 291 \| (29, 238) \| 313 \| ParB-like partition protein \| ParB-like partition protein; uniclust \| UniRef100\_A0A6N7Z139 \| 99.7 \| 1.2e-20 \| 2.3e-26 \| 150.7 \| 167 \| (17, 188) \| 291 \| (32, 205) \| 305 \| ParB/RepB/Spo0J family partition protein \| ParB/RepB/Spo0J family partition protein; pdb70 \| 6SDK\_B \| 99.8 \| 1.3e-25 \| 1.2e-29 \| 190.6 \| 176 \| (18, 199) \| 291 \| (2, 181) \| 199 \| Stage 0 sporulation protein J \| 6SDK\_B Stage 0 sporulation protein J ParB, Chromosome segregation, Chromosome organization HET: MSE, CDP; pdb70 \| 6SDK\_C \| 99.8 \| 1.3e-25 \| 1.2e-29 \| 190.6 \| 176 \| (18, 199) \| 291 \| (2, 181) \| 199 \| Stage 0 sporulation protein J \| 6SDK\_C Stage 0 sporulation protein J ParB, Chromosome segregation, Chromosome organization HET: MSE, CDP; pdb70 \| 1VZ0\_D \| 99.8 \| 2.7e-25 \| 2.5e-29 \| 193.9 \| 178 \| (16, 199) \| 291 \| (19, 198) \| 230 \| CHROMOSOME PARTITIONING PROTEIN PARB \| 1VZ0\_D CHROMOSOME PARTITIONING PROTEIN PARB NUCLEAR PROTEIN, CHROMOSOME SEGREGATION, DNA-BINDING HET: CO; pdb70 \| 4UMK\_A \| 99.8 \| 2e-24 \| 1.8e-28 \| 189.5 \| 179 \| (14, 198) \| 291 \| (27, 208) \| 240 \| PROBABLE CHROMOSOME-PARTITIONING PROTEIN PARB \| 4UMK\_A PROBABLE CHROMOSOME-PARTITIONING PROTEIN PARB DNA BINDING PROTEIN-DNA COMPLEX HET: SO4; pdb70 \| 4UMK\_C \| 99.8 \| 2e-24 \| 1.8e-28 \| 189.5 \| 179 \| (14, 198) \| 291 \| (27, 208) \| 240 \| PROBABLE CHROMOSOME-PARTITIONING PROTEIN PARB \| 4UMK\_C PROBABLE CHROMOSOME-PARTITIONING PROTEIN PARB DNA BINDING PROTEIN-DNA COMPLEX HET: SO4; pdb70 \| 5K5A\_B \| 99.7 \| 3.1e-22 \| 2.8e-26 \| 181.6 \| 166 \| (14, 188) \| 291 \| (5, 230) \| 300 \| ParB domain protein nuclease \| 5K5A\_B ParB domain protein nuclease ParB, pNOB8, partition, DNA segregation HET: MSE; pdb70 \| 5K5D\_C \| 99.7 \| 6.7e-22 \| 6e-26 \| 179.4 \| 166 \| (14, 188) \| 291 \| (5, 230) \| 300 \| ParB domain protein nuclease \| 5K5D\_C ParB domain protein nuclease ParB-N, pnob8, partition, HYDROLASE HET: CIT, MSE; pdb70 \| 6RYK\_A \| 99.6 \| 7.8e-20 \| 7.1e-24 \| 159.2 \| 159 \| (14, 184) \| 291 \| (28, 189) \| 229 \| ParB-like nuclease domain protein \| 6RYK\_A ParB-like nuclease domain protein ParABS, cytoskeleton, bactofilin, CTP, CELL HET: CTP; pdb70 \| 6KY4\_A \| 99.0 \| 6.9e-14 \| 6.5e-18 \| 106.9 \| 87 \| (17, 109) \| 291 \| (16, 102) \| 112 \| Sulfiredoxin, chloroplastic/mitochondrial (E.C.1.8.98.2) \| 6KY4\_A Sulfiredoxin, chloroplastic/mitochondrial (E.C.1.8.98.2) sulfiredoxin, cysteine, sulfinic acid, peroxiredoxin HET: PO4, ADP; pdb70 \| 1R71\_B \| 99.0 \| 1e-13 \| 9.5e-18 \| 115.1 \| 103 \| (77, 187) \| 291 \| (1, 105) \| 178 \| Transcriptional repressor protein korB/DNA Complex \| 1R71\_B Transcriptional repressor protein korB/DNA Complex IncP, plasmid partitioning, protein-DNA complex; pdb70 \| 1XW3\_A \| 98.8 \| 5.7e-13 \| 5.4e-17 \| 101.2 \| 91 \| (8, 106) \| 291 \| (2, 98) \| 110 \| SULFIREDOXIN \| 1XW3\_A SULFIREDOXIN retroreduction, sulfinic acid, peroxiredoxin, OXIDOREDUCTASE; pdb70 \| 3CYI\_A \| 98.8 \| 1.2e-12 \| 1.1e-16 \| 99.2 \| 91 \| (11, 109) \| 291 \| (5, 101) \| 110 \| Sulfiredoxin-1 (E.C.1.8.98.2) \| 3CYI\_A Sulfiredoxin-1 (E.C.1.8.98.2) Cys sulfinic acid reduction, Antioxidant HET: ATP; pdb70 \| 2B6F\_A \| 98.8 \| 1.4e-12 \| 1.3e-16 \| 100.9 \| 81 \| (12, 97) \| 291 \| (17, 103) \| 121 \| Sulfiredoxin (E.C.1.18.-.-) \| 2B6F\_A Sulfiredoxin (E.C.1.18.-.-) PARB DOMAIN FOLD, OXIDOREDUCTASE HET: ATP, MG; pdb70 \| 1VK1\_A \| 98.4 \| 9e-11 \| 8.3e-15 \| 101.2 \| 91 \| (17, 117) \| 291 \| (15, 109) \| 242 \| PROTEIN \| 1VK1\_A PROTEIN reductive methylation, dimethyl lysine, Pyrococcus; pdb70 \| 5X0J\_A \| 98.1 \| 7e-10 \| 6.5e-14 \| 95.4 \| 91 \| (17, 117) \| 291 \| (15, 109) \| 242 \| Free serine kinase \| 5X0J\_A Free serine kinase Thermococcus kodakarensis, cysteine biosynthesis, TRANSFERASE HET: SEP, AMP; pdb70 \| 6NR6\_B \| 98.1 \| 8.8e-10 \| 8.1e-14 \| 95.8 \| 90 \| (17, 118) \| 291 \| (5, 97) \| 254 \| Bifunctional transcriptional regulator/O-phospho-L-serine synthase SbnI \| 6NR6\_B Bifunctional transcriptional regulator/O-phospho-L-serine synthase SbnI free serine kinase, siderophore, ParB/Srx HET: ADP; pdb70 \| 3MKW\_B \| 98.0 \| 2.2e-09 \| 2e-13 \| 84.9 \| 81 \| (110, 190) \| 291 \| (15, 98) \| 138 \| PROTEIN/DNA Complex \| 3MKW\_B PROTEIN/DNA Complex segregation, sopB, F plasmid, centromere HET: SO4; pdb70 \| 5UJE\_A \| 98.0 \| 2.2e-09 \| 2.1e-13 \| 92.3 \| 75 \| (16, 97) \| 291 \| (6, 82) \| 242 \| SbnI protein \| 5UJE\_A SbnI protein staphyloferrin B, heme, regulator, siderophore HET: GOL, MSE; pdb70 \| 3MKY\_B \| 98.0 \| 3.3e-09 \| 3e-13 \| 87.7 \| 83 \| (108, 190) \| 291 \| (12, 98) \| 189 \| PROTEIN/DNA Complex \| 3MKY\_B PROTEIN/DNA Complex partition, F plasmid, sopB, centromere HET: SO4; pdb70 \| 3MKY\_P \| 97.9 \| 4.9e-09 \| 4.6e-13 \| 86.6 \| 81 \| (110, 190) \| 291 \| (15, 98) \| 189 \| PROTEIN/DNA Complex \| 3MKY\_P PROTEIN/DNA Complex partition, F plasmid, sopB, centromere HET: SO4; pdb70 \| 3MKZ\_A \| 97.7 \| 2e-08 \| 1.9e-12 \| 76.7 \| 80 \| (115, 194) \| 291 \| (3, 85) \| 121 \| Protein sopB \| 3MKZ\_A Protein sopB partition, SopB, F plasmid, centromere; pdb70 \| 3MKZ\_N \| 97.7 \| 2e-08 \| 1.9e-12 \| 76.7 \| 80 \| (115, 194) \| 291 \| (3, 85) \| 121 \| Protein sopB \| 3MKZ\_N Protein sopB partition, SopB, F plasmid, centromere; pdb70 \| 2HWJ\_A \| 97.6 \| 5.1e-08 \| 4.4e-12 \| 82.2 \| 76 \| (15, 97) \| 291 \| (4, 89) \| 205 \| Hypothetical protein Atu1540 \| 2HWJ\_A Hypothetical protein Atu1540 Agrobacterium tumefaciens, Hypothetical protein, Structural; pdb70 \| 2HWJ\_C \| 97.6 \| 5.1e-08 \| 4.4e-12 \| 82.2 \| 76 \| (15, 97) \| 291 \| (4, 89) \| 205 \| Hypothetical protein Atu1540 \| 2HWJ\_C Hypothetical protein Atu1540 Agrobacterium tumefaciens, Hypothetical protein, Structural HET: MSE; pdb70 \| 1ZX4\_B \| 97.0 \| 1.2e-06 \| 1.2e-10 \| 71.3 \| 74 \| (115, 188) \| 291 \| (4, 78) \| 192 \| Plasmid Partition par B protein/DNA \| 1ZX4\_B Plasmid Partition par B protein/DNA partition P1 plasmid HET: CIT, MSE; pdb70 \| 2NTZ\_A \| 96.6 \| 6.1e-06 \| 5.8e-10 \| 66.8 \| 73 \| (115, 187) \| 291 \| (4, 77) \| 192 \| ParB/DNA Complex \| 2NTZ\_A ParB/DNA Complex partition, segregation, parb, para, CELL |
| Top keywords  (threshold 1.00e-03 (evalue)) | **ParB, domain\_containing, Sulfiredoxin, partition, partitioning, Chromosome, Spo0J, RepB, Fragment, chromosome\_partitioning** |
| Output files | ../../similar\_sequences/28\_FANPEZAQ\_CDS\_0028\_merged.svg ../../similar\_sequences/28\_FANPEZAQ\_CDS\_0028\_pdb70.a3m ../../similar\_sequences/28\_FANPEZAQ\_CDS\_0028\_pdb70.hhr ../../similar\_sequences/28\_FANPEZAQ\_CDS\_0028\_uniclust.a3m ../../similar\_sequences/28\_FANPEZAQ\_CDS\_0028\_uniclust.hhr |

#### Structure prediction (AlphaFold)^2^

|  |  |
| --- | --- |
| Stats | xml version="1.0" encoding="utf-8" standalone="no"?       2024-09-02T21:09:28.200035 image/svg+xml   Matplotlib v3.7.2, https://matplotlib.org/ |
| Predicted structure | **NGL Viewer Controls:**  - Center: *Left-Click* - Rotate: *Left-Click + Drag* - Translate: *Right-Click + Drag* - Zoom: *Shift + Left-Click + Drag* |
| Output files | ../../predicted\_structures/28\_FANPEZAQ\_CDS\_0028/features.pkl ../../predicted\_structures/28\_FANPEZAQ\_CDS\_0028/ranked\_0.pdb ../../predicted\_structures/28\_FANPEZAQ\_CDS\_0028/ranked\_0\_plots.svg ../../predicted\_structures/28\_FANPEZAQ\_CDS\_0028/result\_model\_1\_ptm\_pred\_0.pkl |

#### Structure similarity search results (Foldseek)^3^

|  |  |
| --- | --- |
| Structure databases searched | Pdb, Afdb-proteome, Afdb-uniprot50 |
| Results, scheme(s)  (Top layers only, threshold 1.00e-02 (evalue)) | xml version="1.0" encoding="utf-8" standalone="no"?       2024-09-02T21:11:01.550994 image/svg+xml   Matplotlib v3.7.2, https://matplotlib.org/ |
| Results, table  (threshold 1.00e-02 (evalue)) | db \| id \| prob \| evalue \| bits \| fident \| alnlen \| mismatch \| gapopen \| qstart \| qend \| tstart \| tend \| name \| description; pdb \| 6T1F\_D \| 1.0 \| 1.275e-08 \| 286 \| 0.182 \| 186 \| 134 \| 9 \| 21 \| 201 \| 2 \| 174 \| Chromosome-partitioning protein ParB \| Chromosome-partitioning protein ParB; pdb \| 7NFU\_A \| 1.0 \| 2.384e-09 \| 279 \| 0.159 \| 219 \| 161 \| 13 \| 4 \| 215 \| 9 \| 211 \| Nucleoid occlusion protein \| Nucleoid occlusion protein; pdb \| 7NFU\_B \| 1.0 \| 6.663e-09 \| 276 \| 0.178 \| 202 \| 148 \| 12 \| 19 \| 215 \| 15 \| 203 \| Nucleoid occlusion protein \| Nucleoid occlusion protein; pdb \| 7BNK\_B \| 1.0 \| 9.502e-10 \| 262 \| 0.236 \| 186 \| 132 \| 8 \| 19 \| 200 \| 1 \| 180 \| ParB family protein \| ParB family protein; pdb \| 7BNR\_A \| 1.0 \| 3.675e-09 \| 249 \| 0.227 \| 185 \| 135 \| 6 \| 19 \| 200 \| 1 \| 180 \| ParB family protein \| ParB family protein; pdb \| 1VZ0\_C \| 1.0 \| 8.943e-08 \| 243 \| 0.257 \| 171 \| 113 \| 5 \| 41 \| 203 \| 22 \| 186 \| Chromosome-partitioning protein Spo0J \| Chromosome-partitioning protein Spo0J; pdb \| 6SDK\_B \| 1.0 \| 3.298e-09 \| 232 \| 0.262 \| 187 \| 126 \| 8 \| 19 \| 200 \| 3 \| 182 \| Stage 0 sporulation protein J \| Stage 0 sporulation protein J; pdb \| 7BM8\_B \| 1.0 \| 1.084e-08 \| 231 \| 0.202 \| 183 \| 125 \| 8 \| 18 \| 195 \| 1 \| 167 \| Chromosome-partitioning protein ParB \| Chromosome-partitioning protein ParB; pdb \| 7O0N\_B \| 1.0 \| 1.027e-08 \| 224 \| 0.232 \| 185 \| 130 \| 7 \| 21 \| 200 \| 2 \| 179 \| ParB family protein \| ParB family protein; pdb \| 7NG0\_A \| 1.0 \| 9.73e-09 \| 224 \| 0.167 \| 185 \| 144 \| 8 \| 18 \| 198 \| 1 \| 179 \| Nucleoid occlusion protein \| Nucleoid occlusion protein; pdb \| 6T1F\_C \| 1.0 \| 5.941e-07 \| 197 \| 0.192 \| 187 \| 137 \| 8 \| 21 \| 201 \| 2 \| 180 \| Chromosome-partitioning protein ParB \| Chromosome-partitioning protein ParB; pdb \| 7OL9\_A \| 1.0 \| 1.11e-07 \| 194 \| 0.166 \| 186 \| 145 \| 7 \| 21 \| 201 \| 2 \| 182 \| Nucleoid occlusion protein \| Nucleoid occlusion protein; pdb \| 6T1F\_A \| 1.0 \| 1.137e-06 \| 192 \| 0.187 \| 187 \| 134 \| 8 \| 21 \| 201 \| 2 \| 176 \| Chromosome-partitioning protein ParB \| Chromosome-partitioning protein ParB; pdb \| 6KY4\_A \| 1.0 \| 0.0002409 \| 176 \| 0.178 \| 84 \| 64 \| 2 \| 18 \| 101 \| 1 \| 79 \| Sulfiredoxin, chloroplastic/mitochondrial \| Sulfiredoxin, chloroplastic/mitochondrial; pdb \| 6RYK\_B \| 1.0 \| 0.0002992 \| 136 \| 0.142 \| 133 \| 102 \| 6 \| 19 \| 145 \| 5 \| 131 \| ParB-like nuclease domain protein \| ParB-like nuclease domain protein; pdb \| 5K5D\_B \| 1.0 \| 0.0001941 \| 125 \| 0.131 \| 244 \| 155 \| 16 \| 17 \| 214 \| 1 \| 233 \| ParB domain protein nuclease \| ParB domain protein nuclease; pdb \| 6S6H\_A \| 1.0 \| 0.008564 \| 123 \| 0.258 \| 120 \| 79 \| 3 \| 107 \| 225 \| 6 \| 116 \| Chromosome-partitioning protein ParB \| Chromosome-partitioning protein ParB; pdb \| 5X0G\_A \| 1.0 \| 0.006897 \| 118 \| 0.206 \| 92 \| 61 \| 6 \| 16 \| 104 \| 12 \| 94 \| Free serine kinase \| Free serine kinase; pdb \| 6S6H\_B \| 1.0 \| 0.00619 \| 117 \| 0.248 \| 141 \| 93 \| 4 \| 107 \| 243 \| 6 \| 137 \| Chromosome-partitioning protein ParB \| Chromosome-partitioning protein ParB; pdb \| 5X0F\_A \| 1.0 \| 0.008564 \| 116 \| 0.206 \| 92 \| 61 \| 6 \| 16 \| 104 \| 13 \| 95 \| Free serine kinase \| Free serine kinase; pdb \| 4UMK\_B \| 1.0 \| 0.000638 \| 115 \| 0.204 \| 166 \| 123 \| 6 \| 31 \| 194 \| 2 \| 160 \| PROBABLE CHROMOSOME-PARTITIONING PROTEIN PARB \| PROBABLE CHROMOSOME-PARTITIONING PROTEIN PARB; pdb \| 5K5A\_B \| 1.0 \| 0.0002992 \| 113 \| 0.135 \| 236 \| 141 \| 17 \| 15 \| 206 \| 2 \| 218 \| ParB domain protein nuclease \| ParB domain protein nuclease; pdb \| 3HY2\_Y \| 1.0 \| 0.002749 \| 110 \| 0.2 \| 110 \| 73 \| 6 \| 16 \| 119 \| 1 \| 101 \| Sulfiredoxin-1 \| Sulfiredoxin-1; pdb \| 5K5D\_A \| 1.0 \| 0.0009318 \| 106 \| 0.128 \| 233 \| 138 \| 17 \| 17 \| 205 \| 1 \| 212 \| ParB domain protein nuclease \| ParB domain protein nuclease; pdb \| 4UMK\_A \| 1.0 \| 0.004015 \| 104 \| 0.189 \| 164 \| 122 \| 4 \| 40 \| 200 \| 1 \| 156 \| PROBABLE CHROMOSOME-PARTITIONING PROTEIN PARB \| PROBABLE CHROMOSOME-PARTITIONING PROTEIN PARB; pdb \| 2B6F\_A \| 0.999 \| 0.008564 \| 96 \| 0.216 \| 97 \| 61 \| 7 \| 11 \| 98 \| 14 \| 104 \| Sulfiredoxin \| Sulfiredoxin; pdb \| 1VK1\_A \| 0.995 \| 0.0003714 \| 88 \| 0.15 \| 212 \| 102 \| 15 \| 16 \| 160 \| 4 \| 204 \| Conserved hypothetical protein \| Conserved hypothetical protein; pdb \| 5X0K\_A \| 0.99 \| 0.006534 \| 83 \| 0.136 \| 168 \| 88 \| 10 \| 16 \| 135 \| 12 \| 170 \| Free serine kinase \| Free serine kinase; pdb \| 5X0B\_A \| 0.961 \| 0.003064 \| 74 \| 0.133 \| 224 \| 105 \| 14 \| 16 \| 172 \| 13 \| 214 \| Free serine kinase \| Free serine kinase; afdb-proteome \| AF-A0A077ZKX5-F1-MODEL\_V4 \| 1.0 \| 4.292e-13 \| 349 \| 0.201 \| 307 \| 199 \| 15 \| 4 \| 291 \| 250 \| 529 \| CbiA and ParBc and KorB domain containing protein \| CbiA and ParBc and KorB domain containing protein; afdb-proteome \| AF-Q9HT12-F1-MODEL\_V4 \| 1.0 \| 1.2e-12 \| 343 \| 0.205 \| 297 \| 196 \| 15 \| 1 \| 282 \| 18 \| 289 \| Probable chromosome-partitioning protein ParB \| Probable chromosome-partitioning protein ParB; afdb-proteome \| AF-A0A132Z8J9-F1-MODEL\_V4 \| 1.0 \| 8.215e-13 \| 334 \| 0.201 \| 302 \| 189 \| 15 \| 5 \| 282 \| 16 \| 289 \| Chromosome partitioning protein ParB \| Chromosome partitioning protein ParB; afdb-proteome \| AF-P77174-F1-MODEL\_V4 \| 1.0 \| 2.393e-08 \| 316 \| 0.185 \| 140 \| 108 \| 5 \| 20 \| 155 \| 50 \| 187 \| Uncharacterized protein YbdM \| Uncharacterized protein YbdM; afdb-proteome \| AF-Q8DMW1-F1-MODEL\_V4 \| 1.0 \| 2.92e-11 \| 308 \| 0.174 \| 270 \| 196 \| 10 \| 19 \| 282 \| 3 \| 251 \| Chromosome segregation protein \| Chromosome segregation protein; afdb-proteome \| AF-Q5F4Z9-F1-MODEL\_V4 \| 1.0 \| 1.61e-11 \| 304 \| 0.194 \| 303 \| 194 \| 14 \| 7 \| 291 \| 15 \| 285 \| Chromosome partitioning protein ParB \| Chromosome partitioning protein ParB; afdb-proteome \| AF-A0A0H3GZZ3-F1-MODEL\_V4 \| 1.0 \| 6.227e-11 \| 294 \| 0.199 \| 286 \| 178 \| 14 \| 1 \| 272 \| 6 \| 254 \| ParB family protein \| ParB family protein; afdb-proteome \| AF-Q2G118-F1-MODEL\_V4 \| 1.0 \| 2.482e-11 \| 290 \| 0.187 \| 310 \| 199 \| 14 \| 1 \| 287 \| 1 \| 280 \| Chromosome partioning protein, ParB family, putative \| Chromosome partioning protein, ParB family, putative; afdb-proteome \| AF-Q8ZR25-F1-MODEL\_V4 \| 1.0 \| 1.351e-07 \| 287 \| 0.157 \| 140 \| 112 \| 5 \| 20 \| 155 \| 46 \| 183 \| Putative transcriptional regulator \| Putative transcriptional regulator; afdb-proteome \| AF-Q9I1Y9-F1-MODEL\_V4 \| 1.0 \| 2.321e-07 \| 267 \| 0.2 \| 150 \| 113 \| 5 \| 20 \| 163 \| 51 \| 199 \| ParB domain-containing protein \| ParB domain-containing protein; afdb-proteome \| AF-K0EKP4-F1-MODEL\_V4 \| 1.0 \| 3.987e-07 \| 266 \| 0.202 \| 148 \| 104 \| 6 \| 17 \| 160 \| 18 \| 155 \| StrR-like regulatory protein \| StrR-like regulatory protein; afdb-proteome \| AF-A0A0H3GPU5-F1-MODEL\_V4 \| 1.0 \| 6.147e-07 \| 264 \| 0.201 \| 149 \| 108 \| 7 \| 20 \| 160 \| 46 \| 191 \| ParB domain-containing protein \| ParB domain-containing protein; afdb-proteome \| AF-Q2FUQ5-F1-MODEL\_V4 \| 1.0 \| 3.332e-10 \| 260 \| 0.184 \| 287 \| 188 \| 17 \| 4 \| 271 \| 18 \| 277 \| ParB domain-containing protein \| ParB domain-containing protein; afdb-proteome \| AF-O25758-F1-MODEL\_V4 \| 1.0 \| 1.94e-10 \| 260 \| 0.189 \| 296 \| 204 \| 13 \| 1 \| 282 \| 15 \| 288 \| Probable chromosome-partitioning protein ParB \| Probable chromosome-partitioning protein ParB; afdb-proteome \| AF-Q93GP5-F1-MODEL\_V4 \| 1.0 \| 1.47e-08 \| 258 \| 0.18 \| 277 \| 192 \| 13 \| 19 \| 283 \| 37 \| 290 \| Putative ParB-like nuclease domain \| Putative ParB-like nuclease domain; afdb-proteome \| AF-Q9PJ25-F1-MODEL\_V4 \| 1.0 \| 1.36e-09 \| 254 \| 0.136 \| 272 \| 201 \| 13 \| 19 \| 282 \| 30 \| 275 \| Probable chromosome-partitioning protein ParB \| Probable chromosome-partitioning protein ParB; afdb-proteome \| AF-K0F0D7-F1-MODEL\_V4 \| 1.0 \| 1.461e-06 \| 243 \| 0.135 \| 192 \| 154 \| 7 \| 14 \| 198 \| 42 \| 228 \| Putative Chromosome partitioning protein \| Putative Chromosome partitioning protein; afdb-proteome \| AF-Q50201-F1-MODEL\_V4 \| 1.0 \| 3.712e-10 \| 231 \| 0.159 \| 314 \| 216 \| 14 \| 1 \| 285 \| 37 \| 331 \| Probable chromosome-partitioning protein ParB \| Probable chromosome-partitioning protein ParB; afdb-proteome \| AF-P9WIJ9-F1-MODEL\_V4 \| 1.0 \| 8.358e-10 \| 220 \| 0.182 \| 302 \| 194 \| 13 \| 19 \| 291 \| 59 \| 336 \| Probable chromosome-partitioning protein ParB \| Probable chromosome-partitioning protein ParB; afdb-proteome \| AF-K0F2G5-F1-MODEL\_V4 \| 1.0 \| 8.823e-10 \| 215 \| 0.184 \| 309 \| 195 \| 17 \| 10 \| 282 \| 30 \| 317 \| Stage 0 sporulation protein J, parB family protein \| Stage 0 sporulation protein J, parB family protein; afdb-proteome \| AF-A0A0H3GXN9-F1-MODEL\_V4 \| 1.0 \| 6.488e-07 \| 201 \| 0.133 \| 270 \| 171 \| 14 \| 3 \| 214 \| 1 \| 265 \| Uncharacterized protein \| Uncharacterized protein; afdb-proteome \| AF-P76068-F1-MODEL\_V4 \| 1.0 \| 0.0001375 \| 178 \| 0.228 \| 92 \| 63 \| 4 \| 15 \| 106 \| 2 \| 85 \| Uncharacterized protein YnaK \| Uncharacterized protein YnaK; afdb-proteome \| AF-A0A0H3GWE9-F1-MODEL\_V4 \| 1.0 \| 6.147e-07 \| 175 \| 0.18 \| 222 \| 165 \| 10 \| 2 \| 214 \| 18 \| 231 \| DNA-binding protein \| DNA-binding protein; afdb-proteome \| AF-Q45GC6-F1-MODEL\_V4 \| 1.0 \| 0.0002778 \| 156 \| 0.229 \| 87 \| 62 \| 3 \| 13 \| 98 \| 62 \| 144 \| Sulfiredoxin \| Sulfiredoxin; afdb-proteome \| AF-Q8GY89-F1-MODEL\_V4 \| 1.0 \| 0.0003449 \| 152 \| 0.191 \| 89 \| 67 \| 3 \| 11 \| 98 \| 31 \| 115 \| Sulfiredoxin, chloroplastic/mitochondrial \| Sulfiredoxin, chloroplastic/mitochondrial; afdb-proteome \| AF-I1NDB8-F1-MODEL\_V4 \| 1.0 \| 0.0009641 \| 148 \| 0.17 \| 88 \| 66 \| 4 \| 13 \| 98 \| 34 \| 116 \| Sulfiredoxin \| Sulfiredoxin; afdb-proteome \| AF-Q32IW8-F1-MODEL\_V4 \| 1.0 \| 0.0001169 \| 148 \| 0.183 \| 136 \| 89 \| 5 \| 20 \| 152 \| 50 \| 166 \| ParB domain-containing protein \| ParB domain-containing protein; afdb-proteome \| AF-Q93GQ7-F1-MODEL\_V4 \| 1.0 \| 2.586e-07 \| 147 \| 0.143 \| 313 \| 184 \| 16 \| 20 \| 286 \| 44 \| 318 \| Plasmid partition protein B \| Plasmid partition protein B; afdb-proteome \| AF-Q327G3-F1-MODEL\_V4 \| 1.0 \| 7.632e-07 \| 146 \| 0.147 \| 298 \| 195 \| 16 \| 20 \| 283 \| 44 \| 316 \| Plasmid segregation protein \| Plasmid segregation protein; afdb-proteome \| AF-A0A0H3H577-F1-MODEL\_V4 \| 1.0 \| 2.07e-05 \| 139 \| 0.121 \| 272 \| 183 \| 14 \| 19 \| 287 \| 14 \| 232 \| ParB domain-containing protein \| ParB domain-containing protein; afdb-proteome \| AF-A0A0H3GZ74-F1-MODEL\_V4 \| 1.0 \| 5.823e-07 \| 139 \| 0.153 \| 306 \| 189 \| 15 \| 20 \| 286 \| 43 \| 317 \| Plasmid partition protein B \| Plasmid partition protein B; afdb-proteome \| AF-Q326T4-F1-MODEL\_V4 \| 1.0 \| 2.252e-06 \| 127 \| 0.141 \| 296 \| 187 \| 16 \| 26 \| 287 \| 41 \| 303 \| VirB \| VirB; afdb-proteome \| AF-A0A0H3GY80-F1-MODEL\_V4 \| 1.0 \| 3.368e-05 \| 123 \| 0.156 \| 287 \| 166 \| 15 \| 15 \| 291 \| 2 \| 222 \| ParB domain-containing protein \| ParB domain-containing protein; afdb-proteome \| AF-X8FLN4-F1-MODEL\_V4 \| 1.0 \| 0.001334 \| 122 \| 0.262 \| 122 \| 76 \| 7 \| 63 \| 176 \| 2 \| 117 \| ParB/RepB/Spo0J family partition domain protein \| ParB/RepB/Spo0J family partition domain protein; afdb-proteome \| AF-Q9D975-F1-MODEL\_V4 \| 1.0 \| 0.007533 \| 110 \| 0.214 \| 107 \| 69 \| 6 \| 19 \| 119 \| 39 \| 136 \| Sulfiredoxin-1 \| Sulfiredoxin-1; afdb-proteome \| AF-Q9BYN0-F1-MODEL\_V4 \| 1.0 \| 0.007136 \| 104 \| 0.198 \| 116 \| 74 \| 7 \| 10 \| 119 \| 35 \| 137 \| Sulfiredoxin-1 \| Sulfiredoxin-1; afdb-proteome \| AF-Q9VX10-F1-MODEL\_V4 \| 0.999 \| 0.008861 \| 98 \| 0.149 \| 127 \| 84 \| 7 \| 8 \| 119 \| 44 \| 161 \| Putative sulfiredoxin \| Putative sulfiredoxin; afdb-proteome \| AF-K0F507-F1-MODEL\_V4 \| 0.997 \| 0.0004282 \| 91 \| 0.128 \| 367 \| 205 \| 17 \| 4 \| 285 \| 18 \| 354 \| Uncharacterized protein \| Uncharacterized protein; afdb-proteome \| AF-Q54RQ8-F1-MODEL\_V4 \| 0.993 \| 0.004887 \| 86 \| 0.122 \| 179 \| 115 \| 9 \| 19 \| 169 \| 9 \| 173 \| BEACH domain-containing protein lvsE \| BEACH domain-containing protein lvsE; afdb-proteome \| AF-Q7TP44-F1-MODEL\_V4 \| 0.967 \| 0.004154 \| 75 \| 0.167 \| 215 \| 120 \| 9 \| 19 \| 195 \| 39 \| 232 \| Sulfiredoxin \| Sulfiredoxin; afdb-uniprot50 \| AF-A0A3S4JMI3-F1-MODEL\_V4 \| 1.0 \| 2.427e-28 \| 1052 \| 0.453 \| 291 \| 130 \| 5 \| 1 \| 287 \| 1 \| 266 \| ParB/RepB/Spo0J family partition protein \| ParB/RepB/Spo0J family partition protein; afdb-uniprot50 \| AF-A0A0P0QGS8-F1-MODEL\_V4 \| 1.0 \| 7.16e-28 \| 1028 \| 0.46 \| 291 \| 128 \| 5 \| 1 \| 287 \| 1 \| 266 \| DNA-binding protein \| DNA-binding protein; afdb-uniprot50 \| AF-A0A198GH96-F1-MODEL\_V4 \| 1.0 \| 8.171e-27 \| 980 \| 0.435 \| 289 \| 134 \| 5 \| 1 \| 287 \| 1 \| 262 \| Uncharacterized protein \| Uncharacterized protein; afdb-uniprot50 \| AF-A0A318NRJ4-F1-MODEL\_V4 \| 1.0 \| 1.612e-27 \| 979 \| 0.443 \| 291 \| 133 \| 5 \| 1 \| 287 \| 1 \| 266 \| DNA-binding protein \| DNA-binding protein; afdb-uniprot50 \| AF-A0A2S9I472-F1-MODEL\_V4 \| 1.0 \| 2.545e-26 \| 946 \| 0.461 \| 297 \| 125 \| 7 \| 1 \| 291 \| 1 \| 268 \| Chromosome partitioning protein ParB \| Chromosome partitioning protein ParB; afdb-uniprot50 \| AF-A0A437T2I0-F1-MODEL\_V4 \| 1.0 \| 1.691e-25 \| 898 \| 0.433 \| 307 \| 134 \| 8 \| 1 \| 289 \| 1 \| 285 \| DUF550 domain-containing protein \| DUF550 domain-containing protein; afdb-uniprot50 \| AF-A0A209B1U4-F1-MODEL\_V4 \| 1.0 \| 4.989e-25 \| 897 \| 0.419 \| 293 \| 139 \| 7 \| 1 \| 287 \| 1 \| 268 \| ParB domain-containing protein \| ParB domain-containing protein; afdb-uniprot50 \| AF-A0A377XPM9-F1-MODEL\_V4 \| 1.0 \| 1.63e-22 \| 892 \| 0.477 \| 249 \| 110 \| 4 \| 1 \| 245 \| 1 \| 233 \| ParB/RepB/Spo0J family partition protein \| ParB/RepB/Spo0J family partition protein; afdb-uniprot50 \| AF-A0A484A9Q6-F1-MODEL\_V4 \| 1.0 \| 1.773e-23 \| 872 \| 0.433 \| 284 \| 124 \| 7 \| 1 \| 280 \| 1 \| 251 \| DNA-binding plasmid partitioning protein \| DNA-binding plasmid partitioning protein; afdb-uniprot50 \| AF-K8C8K5-F1-MODEL\_V4 \| 1.0 \| 4.695e-23 \| 870 \| 0.459 \| 259 \| 107 \| 4 \| 23 \| 280 \| 1 \| 227 \| Predicted transcriptional regulators \| Predicted transcriptional regulators; afdb-uniprot50 \| AF-V5U266-F1-MODEL\_V4 \| 1.0 \| 8.065e-23 \| 859 \| 0.463 \| 259 \| 106 \| 5 \| 23 \| 280 \| 1 \| 227 \| Uncharacterized protein \| Uncharacterized protein; afdb-uniprot50 \| AF-A0A561G4M7-F1-MODEL\_V4 \| 1.0 \| 8.458e-21 \| 750 \| 0.335 \| 274 \| 170 \| 6 \| 1 \| 271 \| 8 \| 272 \| ParB/RepB/Spo0J family partition protein \| ParB/RepB/Spo0J family partition protein; afdb-uniprot50 \| AF-A0A378DHA4-F1-MODEL\_V4 \| 1.0 \| 1.128e-17 \| 731 \| 0.561 \| 171 \| 70 \| 3 \| 1 \| 167 \| 1 \| 170 \| ParB/RepB/Spo0J family partition protein \| ParB/RepB/Spo0J family partition protein; afdb-uniprot50 \| AF-A0A5E7QKL3-F1-MODEL\_V4 \| 1.0 \| 6.609e-20 \| 700 \| 0.254 \| 302 \| 187 \| 8 \| 1 \| 285 \| 2 \| 282 \| ParB domain-containing protein \| ParB domain-containing protein; afdb-uniprot50 \| AF-A0A6N8BW42-F1-MODEL\_V4 \| 1.0 \| 4.746e-18 \| 691 \| 0.329 \| 237 \| 143 \| 4 \| 14 \| 246 \| 9 \| 233 \| Uncharacterized protein \| Uncharacterized protein; afdb-uniprot50 \| AF-A0A2A2GPN7-F1-MODEL\_V4 \| 1.0 \| 2.01e-20 \| 686 \| 0.274 \| 302 \| 187 \| 12 \| 1 \| 287 \| 38 \| 322 \| Uncharacterized protein \| Uncharacterized protein; afdb-uniprot50 \| AF-A0A7W9JWW8-F1-MODEL\_V4 \| 1.0 \| 5.323e-20 \| 679 \| 0.278 \| 305 \| 171 \| 9 \| 1 \| 282 \| 3 \| 281 \| ParB-like chromosome segregation protein Spo0J \| ParB-like chromosome segregation protein Spo0J; afdb-uniprot50 \| AF-A0A2L1KUX8-F1-MODEL\_V4 \| 1.0 \| 3.271e-20 \| 678 \| 0.285 \| 301 \| 181 \| 11 \| 4 \| 287 \| 40 \| 323 \| Uncharacterized protein \| Uncharacterized protein; afdb-uniprot50 \| AF-A0A1I1VMS0-F1-MODEL\_V4 \| 1.0 \| 4.061e-20 \| 674 \| 0.298 \| 291 \| 184 \| 8 \| 1 \| 287 \| 7 \| 281 \| Chromosome partitioning protein, ParB family \| Chromosome partitioning protein, ParB family; afdb-uniprot50 \| AF-A0A847BHV1-F1-MODEL\_V4 \| 1.0 \| 2.698e-19 \| 673 \| 0.27 \| 307 \| 185 \| 11 \| 1 \| 291 \| 5 \| 288 \| Uncharacterized protein \| Uncharacterized protein; afdb-uniprot50 \| AF-A0A411X2Y8-F1-MODEL\_V4 \| 1.0 \| 3.453e-20 \| 667 \| 0.303 \| 297 \| 175 \| 8 \| 8 \| 291 \| 31 \| 308 \| Uncharacterized protein \| Uncharacterized protein; afdb-uniprot50 \| AF-A9BYB8-F1-MODEL\_V4 \| 1.0 \| 3.536e-19 \| 661 \| 0.321 \| 271 \| 167 \| 6 \| 1 \| 259 \| 38 \| 303 \| Uncharacterized protein \| Uncharacterized protein; afdb-uniprot50 \| AF-A0A1B8TI45-F1-MODEL\_V4 \| 1.0 \| 7.493e-17 \| 660 \| 0.324 \| 228 \| 135 \| 7 \| 1 \| 213 \| 24 \| 247 \| Uncharacterized protein \| Uncharacterized protein; afdb-uniprot50 \| AF-A0A2W5FW74-F1-MODEL\_V4 \| 1.0 \| 4.635e-19 \| 657 \| 0.286 \| 286 \| 176 \| 9 \| 3 \| 273 \| 4 \| 276 \| ParB domain-containing protein \| ParB domain-containing protein; afdb-uniprot50 \| AF-A0A0E3BYE5-F1-MODEL\_V4 \| 1.0 \| 7.542e-19 \| 652 \| 0.286 \| 304 \| 181 \| 11 \| 1 \| 287 \| 9 \| 293 \| Uncharacterized protein \| Uncharacterized protein; afdb-uniprot50 \| AF-A0A2Z3I5V1-F1-MODEL\_V4 \| 1.0 \| 3.35e-19 \| 649 \| 0.289 \| 311 \| 184 \| 11 \| 1 \| 287 \| 1 \| 298 \| Uncharacterized protein \| Uncharacterized protein; afdb-uniprot50 \| AF-A0A7W7JCN3-F1-MODEL\_V4 \| 1.0 \| 6.412e-19 \| 646 \| 0.263 \| 304 \| 189 \| 9 \| 1 \| 289 \| 3 \| 286 \| ParB-like chromosome segregation protein Spo0J \| ParB-like chromosome segregation protein Spo0J; afdb-uniprot50 \| AF-A0A193GNC7-F1-MODEL\_V4 \| 1.0 \| 2.294e-19 \| 642 \| 0.289 \| 287 \| 175 \| 10 \| 16 \| 288 \| 25 \| 296 \| Uncharacterized protein \| Uncharacterized protein; afdb-uniprot50 \| AF-A0A5P8XR09-F1-MODEL\_V4 \| 1.0 \| 3.94e-19 \| 641 \| 0.305 \| 298 \| 176 \| 10 \| 3 \| 282 \| 4 \| 288 \| ParB domain-containing protein \| ParB domain-containing protein; afdb-uniprot50 \| AF-A0A0S4PBI2-F1-MODEL\_V4 \| 1.0 \| 2.987e-17 \| 635 \| 0.285 \| 252 \| 157 \| 5 \| 1 \| 246 \| 18 \| 252 \| Phage protein \| Phage protein; afdb-uniprot50 \| AF-A0A5K7YEB8-F1-MODEL\_V4 \| 1.0 \| 7.542e-19 \| 635 \| 0.313 \| 281 \| 174 \| 10 \| 16 \| 287 \| 61 \| 331 \| Uncharacterized protein \| Uncharacterized protein; afdb-uniprot50 \| AF-A0A7X0PCX7-F1-MODEL\_V4 \| 1.0 \| 2.917e-18 \| 630 \| 0.271 \| 280 \| 179 \| 8 \| 1 \| 268 \| 1 \| 267 \| ParB-like chromosome segregation protein Spo0J \| ParB-like chromosome segregation protein Spo0J; afdb-uniprot50 \| AF-A0A484VAU1-F1-MODEL\_V4 \| 1.0 \| 2.618e-18 \| 621 \| 0.282 \| 294 \| 186 \| 8 \| 3 \| 287 \| 9 \| 286 \| Predicted transcriptional regulators \| Predicted transcriptional regulators; afdb-uniprot50 \| AF-A0A556AID6-F1-MODEL\_V4 \| 1.0 \| 3.513e-17 \| 618 \| 0.281 \| 259 \| 165 \| 9 \| 1 \| 246 \| 16 \| 266 \| ParB/RepB/Spo0J family partition protein \| ParB/RepB/Spo0J family partition protein; afdb-uniprot50 \| AF-A0A7U3GA52-F1-MODEL\_V4 \| 1.0 \| 1.938e-17 \| 617 \| 0.258 \| 298 \| 172 \| 9 \| 1 \| 289 \| 11 \| 268 \| ParB/RepB/Spo0J family partition protein \| ParB/RepB/Spo0J family partition protein; afdb-uniprot50 \| AF-A0A6N4J675-F1-MODEL\_V4 \| 1.0 \| 3.079e-18 \| 617 \| 0.276 \| 300 \| 188 \| 10 \| 1 \| 288 \| 2 \| 284 \| ParB domain-containing protein \| ParB domain-containing protein; afdb-uniprot50 \| AF-A0A4P7L9Q5-F1-MODEL\_V4 \| 1.0 \| 3.733e-19 \| 617 \| 0.281 \| 295 \| 181 \| 11 \| 7 \| 287 \| 45 \| 322 \| Uncharacterized protein \| Uncharacterized protein; afdb-uniprot50 \| AF-A0A6M1CKP2-F1-MODEL\_V4 \| 1.0 \| 1.35e-14 \| 608 \| 0.57 \| 149 \| 59 \| 3 \| 1 \| 145 \| 1 \| 148 \| DNA-binding protein \| DNA-binding protein; afdb-uniprot50 \| AF-A0A0P9Z862-F1-MODEL\_V4 \| 1.0 \| 1.012e-17 \| 607 \| 0.273 \| 300 \| 190 \| 12 \| 2 \| 287 \| 3 \| 288 \| Carbon storage regulator CsrA \| Carbon storage regulator CsrA; afdb-uniprot50 \| AF-A0A1H2PPM0-F1-MODEL\_V4 \| 1.0 \| 4.232e-16 \| 605 \| 0.266 \| 248 \| 167 \| 7 \| 7 \| 244 \| 25 \| 267 \| Chromosome segregation protein Spo0J, contains ParB-like nuclease domain \| Chromosome segregation protein Spo0J, contains ParB-like nuclease domain; afdb-uniprot50 \| AF-A0A5N7WBC4-F1-MODEL\_V4 \| 1.0 \| 4.035e-18 \| 604 \| 0.289 \| 294 \| 174 \| 10 \| 11 \| 287 \| 14 \| 289 \| Ftsk\_gamma domain-containing protein \| Ftsk\_gamma domain-containing protein; afdb-uniprot50 \| AF-A0A4Q2V971-F1-MODEL\_V4 \| 1.0 \| 1.128e-17 \| 602 \| 0.262 \| 293 \| 187 \| 9 \| 6 \| 287 \| 16 \| 290 \| Uncharacterized protein \| Uncharacterized protein; afdb-uniprot50 \| AF-A0A1Q9BSC2-F1-MODEL\_V4 \| 1.0 \| 7.961e-19 \| 598 \| 0.302 \| 298 \| 189 \| 11 \| 4 \| 287 \| 32 \| 324 \| Uncharacterized protein \| Uncharacterized protein; afdb-uniprot50 \| AF-N2IPQ2-F1-MODEL\_V4 \| 1.0 \| 1.647e-17 \| 596 \| 0.291 \| 278 \| 167 \| 8 \| 1 \| 265 \| 1 \| 261 \| Uncharacterized protein \| Uncharacterized protein; afdb-uniprot50 \| AF-A0A261UUL3-F1-MODEL\_V4 \| 1.0 \| 1.012e-17 \| 594 \| 0.238 \| 306 \| 189 \| 12 \| 2 \| 288 \| 9 \| 289 \| Uncharacterized protein \| Uncharacterized protein; afdb-uniprot50 \| AF-A0A2R7NKF8-F1-MODEL\_V4 \| 1.0 \| 1.739e-17 \| 592 \| 0.263 \| 300 \| 177 \| 10 \| 8 \| 287 \| 15 \| 290 \| KorB domain-containing protein \| KorB domain-containing protein; afdb-uniprot50 \| AF-A0A1C7LIL5-F1-MODEL\_V4 \| 1.0 \| 1.588e-14 \| 591 \| 0.317 \| 205 \| 129 \| 5 \| 5 \| 202 \| 22 \| 222 \| Uncharacterized protein \| Uncharacterized protein; afdb-uniprot50 \| AF-A1VVC1-F1-MODEL\_V4 \| 1.0 \| 5.416e-17 \| 591 \| 0.268 \| 264 \| 164 \| 10 \| 1 \| 246 \| 33 \| 285 \| Uncharacterized protein \| Uncharacterized protein; afdb-uniprot50 \| AF-A0A2C9D2B9-F1-MODEL\_V4 \| 1.0 \| 2.681e-17 \| 590 \| 0.27 \| 300 \| 186 \| 10 \| 1 \| 291 \| 1 \| 276 \| Plasmid partitioning protein \| Plasmid partitioning protein; afdb-uniprot50 \| AF-A0A2S9P286-F1-MODEL\_V4 \| 1.0 \| 1.835e-17 \| 588 \| 0.256 \| 300 \| 193 \| 12 \| 1 \| 284 \| 9 \| 294 \| Chromosome partitioning protein ParB \| Chromosome partitioning protein ParB; afdb-uniprot50 \| AF-B7XGN8-F1-MODEL\_V4 \| 1.0 \| 4.746e-18 \| 588 \| 0.249 \| 309 \| 188 \| 10 \| 1 \| 286 \| 7 \| 294 \| KorB domain-containing protein \| KorB domain-containing protein; afdb-uniprot50 \| AF-A0A846VCZ1-F1-MODEL\_V4 \| 1.0 \| 5.717e-17 \| 585 \| 0.238 \| 289 \| 195 \| 8 \| 1 \| 282 \| 2 \| 272 \| ParB/RepB/Spo0J family partition protein \| ParB/RepB/Spo0J family partition protein; afdb-uniprot50 \| AF-A0A426FNN8-F1-MODEL\_V4 \| 1.0 \| 2.83e-17 \| 582 \| 0.22 \| 313 \| 196 \| 11 \| 3 \| 287 \| 34 \| 326 \| ParB domain-containing protein \| ParB domain-containing protein; afdb-uniprot50 \| AF-A0A6P1IN57-F1-MODEL\_V4 \| 1.0 \| 5.131e-17 \| 582 \| 0.257 \| 299 \| 182 \| 10 \| 1 \| 287 \| 54 \| 324 \| Uncharacterized protein \| Uncharacterized protein; afdb-uniprot50 \| AF-A0A7Y8KX42-F1-MODEL\_V4 \| 1.0 \| 4.232e-16 \| 581 \| 0.278 \| 273 \| 169 \| 8 \| 1 \| 247 \| 22 \| 292 \| Uncharacterized protein \| Uncharacterized protein; afdb-uniprot50 \| AF-A0A2D2DTI9-F1-MODEL\_V4 \| 1.0 \| 4.861e-17 \| 580 \| 0.268 \| 287 \| 168 \| 9 \| 11 \| 291 \| 5 \| 255 \| Uncharacterized protein \| Uncharacterized protein; afdb-uniprot50 \| AF-A0A158EA19-F1-MODEL\_V4 \| 1.0 \| 1.835e-17 \| 578 \| 0.266 \| 311 \| 184 \| 12 \| 1 \| 287 \| 5 \| 295 \| Nucleoid occlusion protein \| Nucleoid occlusion protein; afdb-uniprot50 \| AF-A0A4V1RYX4-F1-MODEL\_V4 \| 1.0 \| 3.513e-17 \| 576 \| 0.266 \| 300 \| 183 \| 10 \| 5 \| 287 \| 18 \| 297 \| Uncharacterized protein \| Uncharacterized protein; afdb-uniprot50 \| AF-A0A6J5FJG2-F1-MODEL\_V4 \| 1.0 \| 8.813e-17 \| 575 \| 0.254 \| 311 \| 187 \| 11 \| 1 \| 282 \| 9 \| 303 \| ParB domain-containing protein \| ParB domain-containing protein; afdb-uniprot50 \| AF-W7XAG2-F1-MODEL\_V4 \| 1.0 \| 2.539e-17 \| 574 \| 0.268 \| 313 \| 188 \| 14 \| 1 \| 291 \| 10 \| 303 \| Uncharacterized protein \| Uncharacterized protein; afdb-uniprot50 \| AF-A0A3A6QPU2-F1-MODEL\_V4 \| 1.0 \| 1.287e-16 \| 572 \| 0.179 \| 351 \| 201 \| 11 \| 1 \| 287 \| 20 \| 347 \| Uncharacterized protein \| Uncharacterized protein; afdb-uniprot50 \| AF-A0A1H9NHG0-F1-MODEL\_V4 \| 1.0 \| 3.709e-17 \| 571 \| 0.26 \| 292 \| 188 \| 10 \| 9 \| 289 \| 20 \| 294 \| ParB/RepB/Spo0J family partition protein \| ParB/RepB/Spo0J family partition protein; afdb-uniprot50 \| AF-A0A1E5X7A6-F1-MODEL\_V4 \| 1.0 \| 5.416e-17 \| 571 \| 0.268 \| 324 \| 196 \| 11 \| 1 \| 291 \| 5 \| 320 \| ParB domain-containing protein \| ParB domain-containing protein; afdb-uniprot50 \| AF-Q5NWX7-F1-MODEL\_V4 \| 1.0 \| 3.798e-16 \| 570 \| 0.305 \| 259 \| 156 \| 9 \| 1 \| 246 \| 15 \| 262 \| Uncharacterized protein \| Uncharacterized protein; afdb-uniprot50 \| AF-A0A5E4SIJ2-F1-MODEL\_V4 \| 1.0 \| 6.724e-17 \| 569 \| 0.236 \| 317 \| 193 \| 11 \| 1 \| 291 \| 4 \| 297 \| Uncharacterized protein \| Uncharacterized protein; afdb-uniprot50 \| AF-A0A5C7NQX4-F1-MODEL\_V4 \| 1.0 \| 1.55e-15 \| 567 \| 0.237 \| 257 \| 173 \| 9 \| 1 \| 246 \| 20 \| 264 \| Uncharacterized protein \| Uncharacterized protein; afdb-uniprot50 \| AF-A0A427FDH5-F1-MODEL\_V4 \| 1.0 \| 3.915e-17 \| 566 \| 0.261 \| 298 \| 185 \| 10 \| 8 \| 282 \| 18 \| 303 \| Uncharacterized protein \| Uncharacterized protein; afdb-uniprot50 \| AF-A0A3S7HQT3-F1-MODEL\_V4 \| 1.0 \| 4.132e-17 \| 566 \| 0.27 \| 310 \| 187 \| 11 \| 5 \| 287 \| 43 \| 340 \| Uncharacterized protein \| Uncharacterized protein; afdb-uniprot50 \| AF-A0A6G8CZU1-F1-MODEL\_V4 \| 1.0 \| 2.539e-17 \| 565 \| 0.274 \| 288 \| 180 \| 11 \| 14 \| 287 \| 16 \| 288 \| Uncharacterized protein \| Uncharacterized protein; afdb-uniprot50 \| AF-A0A6N8DQB4-F1-MODEL\_V4 \| 1.0 \| 6.329e-15 \| 563 \| 0.236 \| 254 \| 166 \| 7 \| 1 \| 246 \| 1 \| 234 \| Uncharacterized protein \| Uncharacterized protein; afdb-uniprot50 \| AF-A0A443KCC4-F1-MODEL\_V4 \| 1.0 \| 1.88e-16 \| 557 \| 0.273 \| 304 \| 177 \| 12 \| 1 \| 291 \| 1 \| 273 \| ParB/RepB/Spo0J family partition protein \| ParB/RepB/Spo0J family partition protein; afdb-uniprot50 \| AF-A0A0F2QVW0-F1-MODEL\_V4 \| 1.0 \| 1.824e-15 \| 557 \| 0.251 \| 258 \| 169 \| 9 \| 1 \| 246 \| 9 \| 254 \| Uncharacterized protein \| Uncharacterized protein; afdb-uniprot50 \| AF-A0A1B8TLL9-F1-MODEL\_V4 \| 1.0 \| 1.514e-16 \| 557 \| 0.212 \| 324 \| 199 \| 11 \| 1 \| 290 \| 21 \| 322 \| ParB domain-containing protein \| ParB domain-containing protein; afdb-uniprot50 \| AF-A0A2W4VYK9-F1-MODEL\_V4 \| 1.0 \| 9.82e-17 \| 555 \| 0.264 \| 318 \| 185 \| 11 \| 1 \| 272 \| 29 \| 343 \| Uncharacterized protein \| Uncharacterized protein; afdb-uniprot50 \| AF-A0A1B8HMK7-F1-MODEL\_V4 \| 1.0 \| 1.327e-17 \| 555 \| 0.299 \| 304 \| 183 \| 14 \| 8 \| 287 \| 44 \| 341 \| Uncharacterized protein \| Uncharacterized protein; afdb-uniprot50 \| AF-A0A7V8FVR6-F1-MODEL\_V4 \| 1.0 \| 5.131e-17 \| 553 \| 0.279 \| 304 \| 178 \| 12 \| 3 \| 287 \| 6 \| 287 \| Uncharacterized protein \| Uncharacterized protein; afdb-uniprot50 \| AF-A0A502TIG2-F1-MODEL\_V4 \| 1.0 \| 6.524e-16 \| 552 \| 0.251 \| 282 \| 179 \| 9 \| 11 \| 282 \| 10 \| 269 \| Uncharacterized protein \| Uncharacterized protein; afdb-uniprot50 \| AF-A0A022PNW0-F1-MODEL\_V4 \| 1.0 \| 7.909e-17 \| 552 \| 0.29 \| 286 \| 184 \| 11 \| 16 \| 287 \| 60 \| 340 \| Uncharacterized protein \| Uncharacterized protein; afdb-uniprot50 \| AF-A0A7Z1BBA0-F1-MODEL\_V4 \| 1.0 \| 1.219e-16 \| 550 \| 0.264 \| 302 \| 185 \| 11 \| 1 \| 291 \| 39 \| 314 \| Uncharacterized protein \| Uncharacterized protein; afdb-uniprot50 \| AF-A0A1N7P1X5-F1-MODEL\_V4 \| 1.0 \| 4.978e-16 \| 548 \| 0.225 \| 297 \| 204 \| 11 \| 1 \| 282 \| 21 \| 306 \| Uncharacterized protein \| Uncharacterized protein; afdb-uniprot50 \| AF-A0A659AP85-F1-MODEL\_V4 \| 1.0 \| 1.868e-14 \| 545 \| 0.311 \| 228 \| 143 \| 8 \| 2 \| 217 \| 11 \| 236 \| Uncharacterized protein \| Uncharacterized protein; afdb-uniprot50 \| AF-A0A1V0M6I1-F1-MODEL\_V4 \| 1.0 \| 2.898e-16 \| 545 \| 0.277 \| 285 \| 176 \| 10 \| 16 \| 287 \| 86 \| 353 \| Uncharacterized protein \| Uncharacterized protein; afdb-uniprot50 \| AF-A0A4R3V4Q9-F1-MODEL\_V4 \| 1.0 \| 7.269e-16 \| 545 \| 0.275 \| 279 \| 167 \| 10 \| 1 \| 247 \| 14 \| 289 \| ParB-like chromosome segregation protein Spo0J \| ParB-like chromosome segregation protein Spo0J; afdb-uniprot50 \| AF-A0A1H6NM10-F1-MODEL\_V4 \| 1.0 \| 1.094e-16 \| 544 \| 0.231 \| 315 \| 194 \| 12 \| 1 \| 288 \| 1 \| 294 \| Uncharacterized protein \| Uncharacterized protein; afdb-uniprot50 \| AF-A0A2X1TFX5-F1-MODEL\_V4 \| 1.0 \| 7.673e-16 \| 543 \| 0.425 \| 242 \| 103 \| 7 \| 53 \| 291 \| 2 \| 210 \| ParB/RepB/Spo0J family partition protein \| ParB/RepB/Spo0J family partition protein; afdb-uniprot50 \| AF-A0A5Q6PET4-F1-MODEL\_V4 \| 1.0 \| 7.493e-17 \| 543 \| 0.258 \| 302 \| 189 \| 10 \| 2 \| 287 \| 11 \| 293 \| Uncharacterized protein \| Uncharacterized protein; afdb-uniprot50 \| AF-A0A069PKE4-F1-MODEL\_V4 \| 1.0 \| 4.978e-16 \| 543 \| 0.259 \| 308 \| 186 \| 12 \| 1 \| 291 \| 15 \| 297 \| Transcriptional regulator \| Transcriptional regulator; afdb-uniprot50 \| AF-A0A2N5BQD3-F1-MODEL\_V4 \| 1.0 \| 8.349e-17 \| 541 \| 0.27 \| 299 \| 184 \| 14 \| 7 \| 287 \| 35 \| 317 \| Uncharacterized protein \| Uncharacterized protein; afdb-uniprot50 \| AF-A0A7Y1B9U3-F1-MODEL\_V4 \| 1.0 \| 4.132e-17 \| 541 \| 0.242 \| 363 \| 180 \| 13 \| 3 \| 291 \| 15 \| 356 \| Chromosome partitioning protein ParB \| Chromosome partitioning protein ParB; afdb-uniprot50 \| AF-A0A4Q2L955-F1-MODEL\_V4 \| 1.0 \| 2.464e-16 \| 540 \| 0.217 \| 317 \| 200 \| 12 \| 1 \| 291 \| 1 \| 295 \| Uncharacterized protein \| Uncharacterized protein; afdb-uniprot50 \| AF-A0A7Y3JE79-F1-MODEL\_V4 \| 1.0 \| 4.575e-15 \| 540 \| 0.26 \| 257 \| 169 \| 7 \| 1 \| 246 \| 18 \| 264 \| ParB N-terminal domain-containing protein \| ParB N-terminal domain-containing protein; afdb-uniprot50 \| AF-A0A840VDW9-F1-MODEL\_V4 \| 1.0 \| 5.546e-16 \| 540 \| 0.245 \| 309 \| 193 \| 9 \| 1 \| 287 \| 1 \| 291 \| Uncharacterized protein \| Uncharacterized protein; afdb-uniprot50 \| AF-A0A2K4L1R5-F1-MODEL\_V4 \| 1.0 \| 6.524e-16 \| 535 \| 0.192 \| 332 \| 201 \| 10 \| 7 \| 291 \| 32 \| 343 \| Uncharacterized protein \| Uncharacterized protein; afdb-uniprot50 \| AF-A0A2N8QQS3-F1-MODEL\_V4 \| 1.0 \| 5.546e-16 \| 534 \| 0.243 \| 304 \| 192 \| 13 \| 1 \| 291 \| 28 \| 306 \| Uncharacterized protein \| Uncharacterized protein; afdb-uniprot50 \| AF-A0A157RM33-F1-MODEL\_V4 \| 1.0 \| 8.55e-16 \| 534 \| 0.267 \| 277 \| 170 \| 10 \| 1 \| 247 \| 13 \| 286 \| Protein of uncharacterized function (DUF550) \| Protein of uncharacterized function (DUF550); afdb-uniprot50 \| AF-A0A522ZXA9-F1-MODEL\_V4 \| 1.0 \| 6.886e-16 \| 531 \| 0.24 \| 304 \| 186 \| 11 \| 1 \| 291 \| 18 \| 289 \| Uncharacterized protein \| Uncharacterized protein; afdb-uniprot50 \| AF-A0A1H2PMU7-F1-MODEL\_V4 \| 1.0 \| 9.026e-16 \| 530 \| 0.243 \| 287 \| 176 \| 11 \| 1 \| 271 \| 18 \| 279 \| ParB-like nuclease domain-containing protein \| ParB-like nuclease domain-containing protein; afdb-uniprot50 \| AF-A0A178LKV7-F1-MODEL\_V4 \| 1.0 \| 1.006e-15 \| 530 \| 0.211 \| 327 \| 198 \| 11 \| 5 \| 291 \| 23 \| 329 \| Uncharacterized protein \| Uncharacterized protein; afdb-uniprot50 \| AF-A0A6M5HTZ0-F1-MODEL\_V4 \| 1.0 \| 8.1e-16 \| 529 \| 0.21 \| 295 \| 198 \| 10 \| 1 \| 287 \| 1 \| 268 \| Chromosome partitioning protein ParB \| Chromosome partitioning protein ParB; afdb-uniprot50 \| AF-A0A3D9P4K5-F1-MODEL\_V4 \| 1.0 \| 8.349e-17 \| 529 \| 0.247 \| 311 \| 199 \| 12 \| 2 \| 287 \| 6 \| 306 \| Uncharacterized protein \| Uncharacterized protein; afdb-uniprot50 \| AF-A0A6L3Y803-F1-MODEL\_V4 \| 1.0 \| 7.222e-14 \| 526 \| 0.244 \| 249 \| 165 \| 7 \| 2 \| 244 \| 4 \| 235 \| Chromosome partitioning protein ParB \| Chromosome partitioning protein ParB; afdb-uniprot50 \| AF-A0A502HT17-F1-MODEL\_V4 \| 1.0 \| 4.978e-16 \| 526 \| 0.221 \| 303 \| 198 \| 10 \| 2 \| 288 \| 5 \| 285 \| ParB domain-containing protein \| ParB domain-containing protein; afdb-uniprot50 \| AF-A0A3N6N757-F1-MODEL\_V4 \| 1.0 \| 7.269e-16 \| 525 \| 0.28 \| 324 \| 179 \| 11 \| 1 \| 283 \| 15 \| 325 \| Uncharacterized protein \| Uncharacterized protein; afdb-uniprot50 \| AF-A0A857JBI8-F1-MODEL\_V4 \| 1.0 \| 1.469e-15 \| 524 \| 0.231 \| 303 \| 187 \| 10 \| 1 \| 289 \| 1 \| 271 \| Uncharacterized protein \| Uncharacterized protein; afdb-uniprot50 \| AF-A0A0F5LX83-F1-MODEL\_V4 \| 1.0 \| 2.523e-15 \| 524 \| 0.226 \| 291 \| 190 \| 9 \| 2 \| 286 \| 4 \| 265 \| Chromosome partitioning protein ParB \| Chromosome partitioning protein ParB; afdb-uniprot50 \| AF-A0A0Q8XC42-F1-MODEL\_V4 \| 1.0 \| 5.346e-13 \| 523 \| 0.31 \| 177 \| 113 \| 5 \| 21 \| 190 \| 1 \| 175 \| Uncharacterized protein \| Uncharacterized protein; afdb-uniprot50 \| AF-A0A601DCY2-F1-MODEL\_V4 \| 1.0 \| 3.889e-15 \| 523 \| 0.232 \| 293 \| 188 \| 10 \| 2 \| 287 \| 4 \| 266 \| Chromosome partitioning protein ParB \| Chromosome partitioning protein ParB; afdb-uniprot50 \| AF-A0A4V3UTC2-F1-MODEL\_V4 \| 1.0 \| 4.106e-15 \| 520 \| 0.204 \| 298 \| 201 \| 9 \| 1 \| 287 \| 15 \| 287 \| ParB domain-containing protein \| ParB domain-containing protein; afdb-uniprot50 \| AF-A0A6I4KMK4-F1-MODEL\_V4 \| 1.0 \| 1.287e-16 \| 519 \| 0.307 \| 299 \| 158 \| 9 \| 15 \| 286 \| 1 \| 277 \| ParB/RepB/Spo0J family partition protein \| ParB/RepB/Spo0J family partition protein; afdb-uniprot50 \| AF-A0A1J5QET2-F1-MODEL\_V4 \| 1.0 \| 1.824e-15 \| 517 \| 0.25 \| 304 \| 190 \| 12 \| 1 \| 291 \| 36 \| 314 \| Uncharacterized protein \| Uncharacterized protein; afdb-uniprot50 \| AF-A0A072TPP5-F1-MODEL\_V4 \| 1.0 \| 5.546e-16 \| 517 \| 0.271 \| 280 \| 170 \| 10 \| 23 \| 291 \| 1 \| 257 \| Curli production assembly/transport component CsgG \| Curli production assembly/transport component CsgG; afdb-uniprot50 \| AF-A4JUB9-F1-MODEL\_V4 \| 1.0 \| 2.811e-15 \| 516 \| 0.223 \| 295 \| 195 \| 9 \| 1 \| 283 \| 23 \| 295 \| Uncharacterized protein \| Uncharacterized protein; afdb-uniprot50 \| AF-U7GET7-F1-MODEL\_V4 \| 1.0 \| 1.469e-15 \| 514 \| 0.244 \| 307 \| 194 \| 12 \| 1 \| 282 \| 1 \| 294 \| Uncharacterized protein \| Uncharacterized protein; afdb-uniprot50 \| AF-A0A5C7KRJ6-F1-MODEL\_V4 \| 1.0 \| 2.25e-13 \| 514 \| 0.288 \| 218 \| 142 \| 8 \| 2 \| 207 \| 22 \| 238 \| Uncharacterized protein \| Uncharacterized protein; afdb-uniprot50 \| AF-A0A4R3Z3N6-F1-MODEL\_V4 \| 1.0 \| 2.663e-15 \| 513 \| 0.246 \| 292 \| 181 \| 9 \| 2 \| 287 \| 32 \| 290 \| Uncharacterized protein \| Uncharacterized protein; afdb-uniprot50 \| AF-A0A837FD39-F1-MODEL\_V4 \| 1.0 \| 4.467e-16 \| 509 \| 0.314 \| 312 \| 176 \| 10 \| 1 \| 291 \| 1 \| 295 \| Chromosome partitioning protein ParB \| Chromosome partitioning protein ParB; afdb-uniprot50 \| AF-A0A850QVK8-F1-MODEL\_V4 \| 1.0 \| 9.303e-17 \| 509 \| 0.259 \| 335 \| 185 \| 13 \| 2 \| 287 \| 23 \| 343 \| Uncharacterized protein \| Uncharacterized protein; afdb-uniprot50 \| AF-A0A0P0RBG0-F1-MODEL\_V4 \| 1.0 \| 5.51e-14 \| 509 \| 0.252 \| 257 \| 175 \| 8 \| 1 \| 247 \| 4 \| 253 \| Transcriptional regulator \| Transcriptional regulator; afdb-uniprot50 \| AF-A0A373FB27-F1-MODEL\_V4 \| 1.0 \| 3.598e-16 \| 508 \| 0.272 \| 294 \| 167 \| 9 \| 28 \| 291 \| 2 \| 278 \| Chromosome partitioning protein ParB \| Chromosome partitioning protein ParB; afdb-uniprot50 \| AF-A0A0E3BGD4-F1-MODEL\_V4 \| 1.0 \| 4.467e-16 \| 508 \| 0.267 \| 329 \| 184 \| 11 \| 2 \| 291 \| 4 \| 314 \| Uncharacterized protein \| Uncharacterized protein; afdb-uniprot50 \| AF-A0A7Z7BHH9-F1-MODEL\_V4 \| 1.0 \| 1.183e-15 \| 507 \| 0.2 \| 315 \| 204 \| 12 \| 1 \| 287 \| 2 \| 296 \| ParB-like nuclease domain-containing protein \| ParB-like nuclease domain-containing protein; afdb-uniprot50 \| AF-A0A509JFZ1-F1-MODEL\_V4 \| 1.0 \| 1.425e-14 \| 506 \| 0.269 \| 267 \| 164 \| 12 \| 1 \| 245 \| 32 \| 289 \| Uncharacterized protein \| Uncharacterized protein; afdb-uniprot50 \| AF-A0A1G0NK38-F1-MODEL\_V4 \| 1.0 \| 3.773e-14 \| 505 \| 0.238 \| 260 \| 174 \| 10 \| 1 \| 246 \| 15 \| 264 \| Uncharacterized protein \| Uncharacterized protein; afdb-uniprot50 \| AF-A0A096HPB0-F1-MODEL\_V4 \| 1.0 \| 3.798e-16 \| 499 \| 0.25 \| 312 \| 175 \| 11 \| 20 \| 291 \| 6 \| 298 \| Uncharacterized protein \| Uncharacterized protein; afdb-uniprot50 \| AF-A0A4R1MTX3-F1-MODEL\_V4 \| 1.0 \| 1.087e-14 \| 496 \| 0.238 \| 294 \| 185 \| 11 \| 1 \| 282 \| 15 \| 281 \| Uncharacterized protein \| Uncharacterized protein; afdb-uniprot50 \| AF-A0A494XEZ0-F1-MODEL\_V4 \| 1.0 \| 8.295e-15 \| 495 \| 0.233 \| 291 \| 188 \| 10 \| 1 \| 281 \| 17 \| 282 \| Transcriptional regulator \| Transcriptional regulator; afdb-uniprot50 \| AF-A0A7W9ADE5-F1-MODEL\_V4 \| 1.0 \| 4.945e-14 \| 493 \| 0.258 \| 263 \| 167 \| 11 \| 1 \| 246 \| 26 \| 277 \| ParB-like chromosome segregation protein Spo0J \| ParB-like chromosome segregation protein Spo0J; afdb-uniprot50 \| AF-A0A4V3Z014-F1-MODEL\_V4 \| 1.0 \| 4.334e-15 \| 492 \| 0.217 \| 358 \| 190 \| 14 \| 1 \| 291 \| 20 \| 354 \| Uncharacterized protein \| Uncharacterized protein; afdb-uniprot50 \| AF-A0A4Q3X4M1-F1-MODEL\_V4 \| 1.0 \| 1.959e-12 \| 491 \| 0.252 \| 198 \| 135 \| 6 \| 1 \| 187 \| 24 \| 219 \| Uncharacterized protein \| Uncharacterized protein; afdb-uniprot50 \| AF-A0A4R5I757-F1-MODEL\_V4 \| 1.0 \| 6.482e-14 \| 491 \| 0.229 \| 253 \| 169 \| 9 \| 3 \| 246 \| 10 \| 245 \| Uncharacterized protein \| Uncharacterized protein; afdb-uniprot50 \| AF-A0A240UI40-F1-MODEL\_V4 \| 1.0 \| 1.425e-14 \| 489 \| 0.258 \| 298 \| 191 \| 10 \| 2 \| 272 \| 30 \| 324 \| Uncharacterized protein \| Uncharacterized protein; afdb-uniprot50 \| AF-A0A4V5T687-F1-MODEL\_V4 \| 1.0 \| 1.279e-14 \| 487 \| 0.209 \| 296 \| 196 \| 10 \| 2 \| 287 \| 3 \| 270 \| Chromosome partitioning protein ParB \| Chromosome partitioning protein ParB; afdb-uniprot50 \| AF-A0A7U2LLI1-F1-MODEL\_V4 \| 1.0 \| 1.971e-14 \| 487 \| 0.225 \| 302 \| 195 \| 12 \| 1 \| 290 \| 55 \| 329 \| Uncharacterized protein \| Uncharacterized protein; afdb-uniprot50 \| AF-A0A2T4MZN0-F1-MODEL\_V4 \| 1.0 \| 2.081e-14 \| 486 \| 0.194 \| 339 \| 200 \| 11 \| 1 \| 287 \| 19 \| 336 \| Uncharacterized protein \| Uncharacterized protein; afdb-uniprot50 \| AF-A0A1E3FN09-F1-MODEL\_V4 \| 1.0 \| 4.106e-15 \| 485 \| 0.249 \| 321 \| 192 \| 12 \| 1 \| 291 \| 1 \| 302 \| Uncharacterized protein \| Uncharacterized protein; afdb-uniprot50 \| AF-A0A1H9NKE0-F1-MODEL\_V4 \| 1.0 \| 1.769e-14 \| 483 \| 0.233 \| 300 \| 195 \| 12 \| 2 \| 282 \| 4 \| 287 \| Uncharacterized protein \| Uncharacterized protein; afdb-uniprot50 \| AF-A0A6G9QRE5-F1-MODEL\_V4 \| 1.0 \| 2.464e-16 \| 483 \| 0.246 \| 317 \| 189 \| 12 \| 3 \| 291 \| 4 \| 298 \| Uncharacterized protein \| Uncharacterized protein; afdb-uniprot50 \| AF-A0A2E3EM85-F1-MODEL\_V4 \| 1.0 \| 8.295e-15 \| 480 \| 0.228 \| 298 \| 187 \| 11 \| 10 \| 287 \| 4 \| 278 \| ParB domain-containing protein \| ParB domain-containing protein; afdb-uniprot50 \| AF-A0A370N771-F1-MODEL\_V4 \| 1.0 \| 2.319e-14 \| 480 \| 0.215 \| 306 \| 197 \| 12 \| 1 \| 289 \| 15 \| 294 \| Transcriptional regulator \| Transcriptional regulator; afdb-uniprot50 \| AF-A0A7X6KYA8-F1-MODEL\_V4 \| 1.0 \| 9.404e-12 \| 479 \| 0.302 \| 172 \| 113 \| 5 \| 18 \| 188 \| 1 \| 166 \| ParB/RepB/Spo0J family partition protein \| ParB/RepB/Spo0J family partition protein; afdb-uniprot50 \| AF-A0A3D9EDD3-F1-MODEL\_V4 \| 1.0 \| 2.081e-14 \| 479 \| 0.21 \| 290 \| 184 \| 8 \| 5 \| 289 \| 23 \| 272 \| ParB/RepB/Spo0J family partition protein \| ParB/RepB/Spo0J family partition protein; afdb-uniprot50 \| AF-A0A5E4SS88-F1-MODEL\_V4 \| 1.0 \| 1.113e-13 \| 479 \| 0.222 \| 310 \| 197 \| 10 \| 1 \| 287 \| 16 \| 304 \| Chromosome-partitioning protein Spo0J \| Chromosome-partitioning protein Spo0J; afdb-uniprot50 \| AF-A0A106DRD1-F1-MODEL\_V4 \| 1.0 \| 9.243e-15 \| 479 \| 0.242 \| 293 \| 189 \| 12 \| 13 \| 287 \| 10 \| 287 \| ParB domain-containing protein \| ParB domain-containing protein; afdb-uniprot50 \| AF-A0A0F9U0H1-F1-MODEL\_V4 \| 1.0 \| 3.208e-14 \| 478 \| 0.238 \| 302 \| 190 \| 13 \| 2 \| 283 \| 20 \| 301 \| Uncharacterized protein \| Uncharacterized protein; afdb-uniprot50 \| AF-A0A4Q2ZCP1-F1-MODEL\_V4 \| 1.0 \| 8.909e-12 \| 475 \| 0.245 \| 208 \| 144 \| 8 \| 9 \| 205 \| 31 \| 236 \| ParB domain-containing protein \| ParB domain-containing protein; afdb-uniprot50 \| AF-A0A7Z0F9Z9-F1-MODEL\_V4 \| 1.0 \| 3.386e-14 \| 475 \| 0.225 \| 311 \| 194 \| 14 \| 1 \| 291 \| 15 \| 298 \| ParB domain-containing protein \| ParB domain-containing protein; afdb-uniprot50 \| AF-A0A522FGG9-F1-MODEL\_V4 \| 1.0 \| 3.983e-14 \| 471 \| 0.259 \| 301 \| 187 \| 11 \| 2 \| 291 \| 28 \| 303 \| ParB/RepB/Spo0J family partition protein \| ParB/RepB/Spo0J family partition protein; afdb-uniprot50 \| AF-A0A7Z7NRH8-F1-MODEL\_V4 \| 1.0 \| 1.868e-14 \| 471 \| 0.212 \| 292 \| 204 \| 10 \| 3 \| 282 \| 10 \| 287 \| Uncharacterized protein \| Uncharacterized protein; afdb-uniprot50 \| AF-A0A7U6D4Y3-F1-MODEL\_V4 \| 1.0 \| 4.334e-15 \| 469 \| 0.234 \| 316 \| 182 \| 12 \| 1 \| 282 \| 1 \| 290 \| Putative transcriptional regulator \| Putative transcriptional regulator; afdb-uniprot50 \| AF-A0A7U9K830-F1-MODEL\_V4 \| 1.0 \| 1.676e-14 \| 468 \| 0.237 \| 295 \| 156 \| 11 \| 1 \| 279 \| 9 \| 250 \| Uncharacterized protein \| Uncharacterized protein; afdb-uniprot50 \| AF-A0A4Z0IUQ2-F1-MODEL\_V4 \| 1.0 \| 1.148e-14 \| 468 \| 0.208 \| 321 \| 193 \| 13 \| 2 \| 282 \| 5 \| 304 \| KorB domain-containing protein \| KorB domain-containing protein; afdb-uniprot50 \| AF-H0BRY7-F1-MODEL\_V4 \| 1.0 \| 1.31e-13 \| 465 \| 0.25 \| 291 \| 174 \| 13 \| 15 \| 291 \| 1 \| 261 \| Uncharacterized protein \| Uncharacterized protein; afdb-uniprot50 \| AF-A0A7S6Z1Q0-F1-MODEL\_V4 \| 1.0 \| 5.064e-13 \| 464 \| 0.239 \| 259 \| 170 \| 12 \| 1 \| 246 \| 16 \| 260 \| ParB N-terminal domain-containing protein \| ParB N-terminal domain-containing protein; afdb-uniprot50 \| AF-A0A085FMH6-F1-MODEL\_V4 \| 1.0 \| 4.685e-14 \| 464 \| 0.214 \| 321 \| 198 \| 13 \| 1 \| 291 \| 17 \| 313 \| Uncharacterized protein \| Uncharacterized protein; afdb-uniprot50 \| AF-A0A377Q615-F1-MODEL\_V4 \| 1.0 \| 1.912e-13 \| 461 \| 0.218 \| 321 \| 190 \| 15 \| 1 \| 291 \| 13 \| 302 \| ParB/RepB/Spo0J family partition protein \| ParB/RepB/Spo0J family partition protein; afdb-uniprot50 \| AF-A4JWD7-F1-MODEL\_V4 \| 1.0 \| 3.039e-14 \| 458 \| 0.203 \| 305 \| 201 \| 9 \| 7 \| 291 \| 11 \| 293 \| Uncharacterized protein \| Uncharacterized protein; afdb-uniprot50 \| AF-A0A7X3HRP0-F1-MODEL\_V4 \| 1.0 \| 6.482e-14 \| 456 \| 0.207 \| 313 \| 206 \| 11 \| 1 \| 291 \| 19 \| 311 \| Uncharacterized protein \| Uncharacterized protein; afdb-uniprot50 \| AF-A0A5K1NGU7-F1-MODEL\_V4 \| 1.0 \| 1.912e-13 \| 455 \| 0.192 \| 302 \| 207 \| 10 \| 3 \| 291 \| 32 \| 309 \| Uncharacterized protein \| Uncharacterized protein; afdb-uniprot50 \| AF-A0A3D9VR34-F1-MODEL\_V4 \| 1.0 \| 2.507e-13 \| 452 \| 0.181 \| 298 \| 201 \| 11 \| 1 \| 282 \| 47 \| 317 \| Uncharacterized protein \| Uncharacterized protein; afdb-uniprot50 \| AF-D3VM21-F1-MODEL\_V4 \| 1.0 \| 1.31e-13 \| 449 \| 0.221 \| 294 \| 158 \| 10 \| 1 \| 278 \| 7 \| 245 \| Uncharacterized protein \| Uncharacterized protein; afdb-uniprot50 \| AF-A0A2E8AAK1-F1-MODEL\_V4 \| 1.0 \| 7.222e-14 \| 448 \| 0.267 \| 292 \| 191 \| 10 \| 13 \| 287 \| 21 \| 306 \| Uncharacterized protein \| Uncharacterized protein; afdb-uniprot50 \| AF-A0A2D7YR92-F1-MODEL\_V4 \| 1.0 \| 1.382e-13 \| 445 \| 0.214 \| 317 \| 188 \| 13 \| 1 \| 272 \| 19 \| 319 \| Uncharacterized protein \| Uncharacterized protein; afdb-uniprot50 \| AF-A0A1D3KAB3-F1-MODEL\_V4 \| 1.0 \| 1.626e-13 \| 444 \| 0.216 \| 282 \| 181 \| 11 \| 16 \| 282 \| 51 \| 307 \| Uncharacterized protein \| Uncharacterized protein; afdb-uniprot50 \| AF-A0A645D0H4-F1-MODEL\_V4 \| 1.0 \| 1.382e-13 \| 440 \| 0.24 \| 275 \| 140 \| 11 \| 21 \| 279 \| 1 \| 222 \| Uncharacterized protein \| Uncharacterized protein; afdb-uniprot50 \| AF-A0A6A6K0T1-F1-MODEL\_V4 \| 1.0 \| 1.062e-15 \| 439 \| 0.277 \| 285 \| 130 \| 7 \| 1 \| 282 \| 1 \| 212 \| Uncharacterized protein \| Uncharacterized protein; afdb-uniprot50 \| AF-A0A077M210-F1-MODEL\_V4 \| 1.0 \| 2.359e-11 \| 438 \| 0.241 \| 186 \| 126 \| 5 \| 1 \| 185 \| 1 \| 172 \| ParB domain-containing protein \| ParB domain-containing protein; afdb-uniprot50 \| AF-A0A2D0K822-F1-MODEL\_V4 \| 1.0 \| 2.375e-13 \| 438 \| 0.206 \| 300 \| 161 \| 12 \| 1 \| 280 \| 6 \| 248 \| Chromosome partitioning protein ParB \| Chromosome partitioning protein ParB; afdb-uniprot50 \| AF-A0A2S4KFT6-F1-MODEL\_V4 \| 1.0 \| 1.055e-13 \| 438 \| 0.233 \| 304 \| 195 \| 13 \| 1 \| 291 \| 40 \| 318 \| Uncharacterized protein \| Uncharacterized protein; afdb-uniprot50 \| AF-A0A021VUN1-F1-MODEL\_V4 \| 1.0 \| 9.124e-11 \| 433 \| 0.289 \| 169 \| 113 \| 5 \| 18 \| 185 \| 1 \| 163 \| ParB domain-containing protein \| ParB domain-containing protein; afdb-uniprot50 \| AF-A0A1E3LIY1-F1-MODEL\_V4 \| 1.0 \| 7.396e-13 \| 433 \| 0.211 \| 308 \| 198 \| 13 \| 1 \| 282 \| 16 \| 304 \| Uncharacterized protein \| Uncharacterized protein; afdb-uniprot50 \| AF-A0A220WPE6-F1-MODEL\_V4 \| 1.0 \| 7.007e-13 \| 432 \| 0.24 \| 295 \| 181 \| 14 \| 9 \| 282 \| 250 \| 522 \| Uncharacterized protein \| Uncharacterized protein; afdb-uniprot50 \| AF-A0A6H1ZXB1-F1-MODEL\_V4 \| 1.0 \| 1.665e-12 \| 429 \| 0.167 \| 280 \| 179 \| 8 \| 15 \| 285 \| 9 \| 243 \| ParB domain-containing protein \| ParB domain-containing protein; afdb-uniprot50 \| AF-A0A6N6T5G8-F1-MODEL\_V4 \| 1.0 \| 3.932e-10 \| 428 \| 0.224 \| 183 \| 137 \| 4 \| 18 \| 196 \| 1 \| 182 \| ParB/RepB/Spo0J family partition protein \| ParB/RepB/Spo0J family partition protein; afdb-uniprot50 \| AF-A0A2A3UEJ5-F1-MODEL\_V4 \| 1.0 \| 8.241e-13 \| 421 \| 0.188 \| 286 \| 194 \| 12 \| 13 \| 288 \| 11 \| 268 \| ParB domain-containing protein \| ParB domain-containing protein; afdb-uniprot50 \| AF-A0A1G3YGW4-F1-MODEL\_V4 \| 1.0 \| 2.929e-11 \| 418 \| 0.221 \| 190 \| 137 \| 6 \| 14 \| 199 \| 6 \| 188 \| ParB domain-containing protein \| ParB domain-containing protein; afdb-uniprot50 \| AF-A0A1G5PGC5-F1-MODEL\_V4 \| 1.0 \| 1.812e-13 \| 417 \| 0.211 \| 303 \| 179 \| 9 \| 22 \| 287 \| 40 \| 319 \| Chromosome partitioning protein, ParB family \| Chromosome partitioning protein, ParB family; afdb-uniprot50 \| AF-A0A542ZE30-F1-MODEL\_V4 \| 1.0 \| 1.946e-10 \| 416 \| 0.291 \| 168 \| 112 \| 4 \| 20 \| 185 \| 3 \| 165 \| ParB/RepB/Spo0J family partition protein \| ParB/RepB/Spo0J family partition protein; afdb-uniprot50 \| AF-A0A373FCW8-F1-MODEL\_V4 \| 1.0 \| 1.204e-12 \| 415 \| 0.225 \| 293 \| 187 \| 12 \| 13 \| 290 \| 69 \| 336 \| Uncharacterized protein \| Uncharacterized protein; afdb-uniprot50 \| AF-A0A420RV90-F1-MODEL\_V4 \| 1.0 \| 2.567e-12 \| 413 \| 0.189 \| 290 \| 197 \| 10 \| 9 \| 283 \| 36 \| 302 \| PAPS\_reduct domain-containing protein \| PAPS\_reduct domain-containing protein; afdb-uniprot50 \| AF-A0A729D027-F1-MODEL\_V4 \| 1.0 \| 3.724e-10 \| 412 \| 0.205 \| 190 \| 139 \| 2 \| 19 \| 196 \| 7 \| 196 \| Uncharacterized protein \| Uncharacterized protein; afdb-uniprot50 \| AF-A0A5R2ATN3-F1-MODEL\_V4 \| 1.0 \| 2.842e-10 \| 407 \| 0.216 \| 171 \| 123 \| 5 \| 13 \| 172 \| 241 \| 411 \| Chromosome partitioning protein ParB \| Chromosome partitioning protein ParB; afdb-uniprot50 \| AF-A0A6N7YJC7-F1-MODEL\_V4 \| 1.0 \| 4.881e-10 \| 406 \| 0.237 \| 194 \| 136 \| 8 \| 19 \| 204 \| 68 \| 257 \| ParB/RepB/Spo0J family partition protein \| ParB/RepB/Spo0J family partition protein; afdb-uniprot50 \| AF-A0A6L5X784-F1-MODEL\_V4 \| 1.0 \| 1.654e-10 \| 404 \| 0.188 \| 207 \| 148 \| 8 \| 18 \| 214 \| 1 \| 197 \| ParB/RepB/Spo0J family partition protein \| ParB/RepB/Spo0J family partition protein; afdb-uniprot50 \| AF-A0A644X8U2-F1-MODEL\_V4 \| 1.0 \| 1.406e-10 \| 404 \| 0.202 \| 188 \| 140 \| 6 \| 13 \| 197 \| 1 \| 181 \| Nucleoid occlusion protein \| Nucleoid occlusion protein; afdb-uniprot50 \| AF-A0A7X2BNE3-F1-MODEL\_V4 \| 1.0 \| 2.182e-12 \| 395 \| 0.182 \| 345 \| 204 \| 17 \| 2 \| 291 \| 22 \| 343 \| Uncharacterized protein \| Uncharacterized protein; afdb-uniprot50 \| AF-A0A4R8EP77-F1-MODEL\_V4 \| 1.0 \| 7.396e-13 \| 395 \| 0.18 \| 315 \| 210 \| 14 \| 1 \| 282 \| 26 \| 325 \| Uncharacterized protein \| Uncharacterized protein; afdb-uniprot50 \| AF-Q9LCY0-F1-MODEL\_V4 \| 1.0 \| 2.235e-11 \| 389 \| 0.231 \| 203 \| 147 \| 5 \| 1 \| 201 \| 5 \| 200 \| Putative partitioning protein \| Putative partitioning protein; afdb-uniprot50 \| AF-X1RK19-F1-MODEL\_V4 \| 1.0 \| 1.041e-09 \| 388 \| 0.2 \| 190 \| 141 \| 7 \| 18 \| 200 \| 1 \| 186 \| ParB domain-containing protein \| ParB domain-containing protein; afdb-uniprot50 \| AF-V0UPH2-F1-MODEL\_V4 \| 1.0 \| 1.341e-12 \| 388 \| 0.226 \| 296 \| 193 \| 13 \| 19 \| 287 \| 19 \| 305 \| Uncharacterized protein \| Uncharacterized protein; afdb-uniprot50 \| AF-A0A1F7U682-F1-MODEL\_V4 \| 1.0 \| 2.117e-11 \| 387 \| 0.209 \| 243 \| 172 \| 12 \| 14 \| 246 \| 15 \| 247 \| ParB domain-containing protein \| ParB domain-containing protein; afdb-uniprot50 \| AF-A0A2M9YIY9-F1-MODEL\_V4 \| 1.0 \| 9.863e-10 \| 387 \| 0.21 \| 185 \| 131 \| 6 \| 13 \| 182 \| 527 \| 711 \| Chromosome partitioning protein ParB \| Chromosome partitioning protein ParB; afdb-uniprot50 \| AF-A0A1E3LMZ5-F1-MODEL\_V4 \| 1.0 \| 9.183e-13 \| 386 \| 0.211 \| 335 \| 192 \| 13 \| 4 \| 288 \| 1 \| 313 \| Uncharacterized protein \| Uncharacterized protein; afdb-uniprot50 \| AF-A0A5I0PMP9-F1-MODEL\_V4 \| 1.0 \| 1.855e-12 \| 386 \| 0.233 \| 295 \| 191 \| 10 \| 19 \| 287 \| 55 \| 340 \| DNA-binding protein \| DNA-binding protein; afdb-uniprot50 \| AF-A0A7Z0PSX1-F1-MODEL\_V4 \| 1.0 \| 1.16e-09 \| 385 \| 0.218 \| 169 \| 124 \| 6 \| 18 \| 184 \| 1 \| 163 \| ParB/RepB/Spo0J family partition protein \| ParB/RepB/Spo0J family partition protein; afdb-uniprot50 \| AF-A0A1E3Y3Z3-F1-MODEL\_V4 \| 1.0 \| 9.343e-10 \| 384 \| 0.164 \| 182 \| 146 \| 3 \| 17 \| 197 \| 1 \| 177 \| ParB domain-containing protein \| ParB domain-containing protein; afdb-uniprot50 \| AF-A0A6M3IM54-F1-MODEL\_V4 \| 1.0 \| 4.25e-09 \| 383 \| 0.155 \| 186 \| 149 \| 4 \| 18 \| 199 \| 1 \| 182 \| ParB domain-containing protein \| ParB domain-containing protein; afdb-uniprot50 \| AF-A0A062XQP4-F1-MODEL\_V4 \| 1.0 \| 5.064e-13 \| 382 \| 0.212 \| 297 \| 199 \| 11 \| 1 \| 287 \| 3 \| 274 \| HTH cro/C1-type domain-containing protein \| HTH cro/C1-type domain-containing protein; afdb-uniprot50 \| AF-A0A522PSB4-F1-MODEL\_V4 \| 1.0 \| 6.061e-10 \| 382 \| 0.207 \| 198 \| 140 \| 8 \| 13 \| 199 \| 6 \| 197 \| ParB/RepB/Spo0J family partition protein \| ParB/RepB/Spo0J family partition protein; afdb-uniprot50 \| AF-A0A811H1P2-F1-MODEL\_V4 \| 1.0 \| 8.44e-12 \| 381 \| 0.246 \| 256 \| 149 \| 6 \| 53 \| 287 \| 2 \| 234 \| Uncharacterized protein \| Uncharacterized protein; afdb-uniprot50 \| AF-A0A522RG87-F1-MODEL\_V4 \| 1.0 \| 3.092e-11 \| 381 \| 0.225 \| 200 \| 140 \| 9 \| 5 \| 200 \| 22 \| 210 \| ParB/RepB/Spo0J family partition protein \| ParB/RepB/Spo0J family partition protein; afdb-uniprot50 \| AF-A0A6L6XV94-F1-MODEL\_V4 \| 1.0 \| 5.311e-11 \| 380 \| 0.205 \| 200 \| 132 \| 8 \| 19 \| 210 \| 63 \| 243 \| ParB/RepB/Spo0J family partition protein \| ParB/RepB/Spo0J family partition protein; afdb-uniprot50 \| AF-A0A2A2M1Z6-F1-MODEL\_V4 \| 1.0 \| 9.863e-10 \| 379 \| 0.205 \| 185 \| 137 \| 6 \| 18 \| 194 \| 1 \| 183 \| ParB domain-containing protein \| ParB domain-containing protein; afdb-uniprot50 \| AF-A0A098EFW5-F1-MODEL\_V4 \| 1.0 \| 4.306e-13 \| 379 \| 0.175 \| 297 \| 209 \| 14 \| 1 \| 282 \| 1 \| 276 \| Chromosome-partitioning protein Spo0J \| Chromosome-partitioning protein Spo0J; afdb-uniprot50 \| AF-A0A1G2AS51-F1-MODEL\_V4 \| 1.0 \| 1.099e-09 \| 379 \| 0.216 \| 189 \| 134 \| 8 \| 19 \| 200 \| 20 \| 201 \| ParB domain-containing protein \| ParB domain-containing protein; afdb-uniprot50 \| AF-A0A4R8JJU0-F1-MODEL\_V4 \| 1.0 \| 2.054e-10 \| 379 \| 0.177 \| 203 \| 152 \| 8 \| 19 \| 214 \| 8 \| 202 \| ParB/RepB/Spo0J family partition protein \| ParB/RepB/Spo0J family partition protein; afdb-uniprot50 \| AF-A0A512RS84-F1-MODEL\_V4 \| 1.0 \| 1.52e-09 \| 379 \| 0.16 \| 181 \| 145 \| 5 \| 13 \| 192 \| 28 \| 202 \| ParB domain-containing protein \| ParB domain-containing protein; afdb-uniprot50 \| AF-E2S7S5-F1-MODEL\_V4 \| 1.0 \| 2.842e-10 \| 377 \| 0.265 \| 196 \| 124 \| 8 \| 13 \| 202 \| 1 \| 182 \| ParB-like protein \| ParB-like protein; afdb-uniprot50 \| AF-A0A6H1Z9T6-F1-MODEL\_V4 \| 1.0 \| 1.16e-09 \| 377 \| 0.163 \| 202 \| 150 \| 8 \| 19 \| 214 \| 6 \| 194 \| ParB domain-containing protein \| ParB domain-containing protein; afdb-uniprot50 \| AF-A0A1M3EWF3-F1-MODEL\_V4 \| 1.0 \| 2.006e-11 \| 377 \| 0.226 \| 238 \| 142 \| 13 \| 19 \| 246 \| 33 \| 238 \| ParB domain-containing protein \| ParB domain-containing protein; afdb-uniprot50 \| AF-Q141V7-F1-MODEL\_V4 \| 1.0 \| 9.863e-10 \| 377 \| 0.211 \| 184 \| 134 \| 6 \| 19 \| 197 \| 81 \| 258 \| ParB family protein \| ParB family protein; afdb-uniprot50 \| AF-A0A2V9TQV9-F1-MODEL\_V4 \| 1.0 \| 1.758e-12 \| 376 \| 0.192 \| 296 \| 211 \| 14 \| 2 \| 282 \| 3 \| 285 \| Chromosome partitioning protein ParB \| Chromosome partitioning protein ParB; afdb-uniprot50 \| AF-A0A6G9HIL3-F1-MODEL\_V4 \| 1.0 \| 8.909e-12 \| 376 \| 0.243 \| 275 \| 177 \| 9 \| 19 \| 267 \| 55 \| 324 \| Putative transcriptional regulator \| Putative transcriptional regulator; afdb-uniprot50 \| AF-A0A1Q3THK1-F1-MODEL\_V4 \| 1.0 \| 3.343e-10 \| 375 \| 0.172 \| 185 \| 133 \| 8 \| 19 \| 189 \| 35 \| 213 \| ParB domain-containing protein \| ParB domain-containing protein; afdb-uniprot50 \| AF-A0A1H2IE39-F1-MODEL\_V4 \| 1.0 \| 2.289e-10 \| 374 \| 0.237 \| 236 \| 138 \| 10 \| 20 \| 246 \| 26 \| 228 \| ParB-like nuclease domain-containing protein \| ParB-like nuclease domain-containing protein; afdb-uniprot50 \| AF-Q72H91-F1-MODEL\_V4 \| 1.0 \| 2.71e-12 \| 372 \| 0.182 \| 290 \| 201 \| 10 \| 1 \| 282 \| 4 \| 265 \| Chromosome-partitioning protein Spo0J \| Chromosome-partitioning protein Spo0J; afdb-uniprot50 \| AF-A0A6P1V5I1-F1-MODEL\_V4 \| 1.0 \| 7.175e-12 \| 372 \| 0.186 \| 333 \| 210 \| 14 \| 1 \| 287 \| 10 \| 327 \| DNA-binding protein \| DNA-binding protein; afdb-uniprot50 \| AF-A0A1F4FSB8-F1-MODEL\_V4 \| 1.0 \| 2.344e-09 \| 371 \| 0.232 \| 185 \| 127 \| 9 \| 19 \| 198 \| 1 \| 175 \| BZIP domain-containing protein \| BZIP domain-containing protein; afdb-uniprot50 \| AF-A0A239A1N1-F1-MODEL\_V4 \| 1.0 \| 3.724e-10 \| 371 \| 0.213 \| 183 \| 130 \| 6 \| 19 \| 194 \| 17 \| 192 \| ParB/RepB/Spo0J family partition protein \| ParB/RepB/Spo0J family partition protein; afdb-uniprot50 \| AF-A0A5N9VV60-F1-MODEL\_V4 \| 1.0 \| 5.88e-09 \| 370 \| 0.202 \| 178 \| 130 \| 2 \| 26 \| 191 \| 2 \| 179 \| Uncharacterized protein \| Uncharacterized protein; afdb-uniprot50 \| AF-A0A2A7RHV9-F1-MODEL\_V4 \| 1.0 \| 4.306e-13 \| 370 \| 0.193 \| 310 \| 203 \| 16 \| 1 \| 291 \| 13 \| 294 \| Chromosome partitioning protein ParB \| Chromosome partitioning protein ParB; afdb-uniprot50 \| AF-A0A1F2QTS0-F1-MODEL\_V4 \| 1.0 \| 8.241e-13 \| 369 \| 0.171 \| 286 \| 203 \| 11 \| 13 \| 282 \| 38 \| 305 \| ParB domain-containing protein \| ParB domain-containing protein; afdb-uniprot50 \| AF-A0A5Q4ZYV4-F1-MODEL\_V4 \| 1.0 \| 3.637e-11 \| 369 \| 0.191 \| 297 \| 185 \| 12 \| 20 \| 282 \| 32 \| 307 \| Uncharacterized protein \| Uncharacterized protein; afdb-uniprot50 \| AF-A0A2P9H4Z5-F1-MODEL\_V4 \| 1.0 \| 3.423e-09 \| 369 \| 0.243 \| 185 \| 128 \| 8 \| 19 \| 197 \| 10 \| 188 \| Putative partition protein \| Putative partition protein; afdb-uniprot50 \| AF-A0A3D2NSA5-F1-MODEL\_V4 \| 1.0 \| 5.153e-10 \| 368 \| 0.316 \| 183 \| 117 \| 4 \| 71 \| 246 \| 8 \| 189 \| Uncharacterized protein \| Uncharacterized protein; afdb-uniprot50 \| AF-A0A2E4W5V6-F1-MODEL\_V4 \| 1.0 \| 1.946e-10 \| 368 \| 0.176 \| 244 \| 177 \| 11 \| 13 \| 247 \| 2 \| 230 \| ParB domain-containing protein \| ParB domain-containing protein; afdb-uniprot50 \| AF-A0A378MEE2-F1-MODEL\_V4 \| 1.0 \| 8.188e-11 \| 367 \| 0.227 \| 211 \| 146 \| 10 \| 1 \| 204 \| 1 \| 201 \| Probable chromosome-partitioning protein parB \| Probable chromosome-partitioning protein parB; afdb-uniprot50 \| AF-A0A6G7Z243-F1-MODEL\_V4 \| 1.0 \| 7.129e-10 \| 367 \| 0.267 \| 191 \| 125 \| 7 \| 17 \| 198 \| 1 \| 185 \| ParB/RepB/Spo0J family partition protein \| ParB/RepB/Spo0J family partition protein; afdb-uniprot50 \| AF-A0A4S2FYS1-F1-MODEL\_V4 \| 1.0 \| 4.486e-09 \| 367 \| 0.167 \| 185 \| 144 \| 6 \| 19 \| 198 \| 14 \| 193 \| ParB/RepB/Spo0J family partition protein \| ParB/RepB/Spo0J family partition protein; afdb-uniprot50 \| AF-A0A1J5D517-F1-MODEL\_V4 \| 1.0 \| 1.416e-12 \| 366 \| 0.204 \| 299 \| 196 \| 12 \| 1 \| 282 \| 7 \| 280 \| ParB domain-containing protein \| ParB domain-containing protein; afdb-uniprot50 \| AF-A0A2V3UB25-F1-MODEL\_V4 \| 1.0 \| 7.525e-10 \| 366 \| 0.175 \| 188 \| 149 \| 5 \| 19 \| 202 \| 1 \| 186 \| ParB family chromosome partitioning protein \| ParB family chromosome partitioning protein; afdb-uniprot50 \| AF-E8V178-F1-MODEL\_V4 \| 1.0 \| 1.993e-09 \| 366 \| 0.221 \| 185 \| 132 \| 7 \| 19 \| 197 \| 10 \| 188 \| ParB-like partition protein \| ParB-like partition protein; afdb-uniprot50 \| AF-W7DNX5-F1-MODEL\_V4 \| 1.0 \| 4.624e-10 \| 363 \| 0.204 \| 205 \| 153 \| 8 \| 1 \| 201 \| 4 \| 202 \| ParB family chromosome partition protein \| ParB family chromosome partition protein; afdb-uniprot50 \| AF-A0A1F4T8B3-F1-MODEL\_V4 \| 1.0 \| 1.577e-12 \| 363 \| 0.169 \| 289 \| 210 \| 10 \| 4 \| 282 \| 16 \| 284 \| ParB domain-containing protein \| ParB domain-containing protein; afdb-uniprot50 \| AF-A0A7W2GL23-F1-MODEL\_V4 \| 1.0 \| 2.948e-13 \| 363 \| 0.193 \| 310 \| 200 \| 15 \| 1 \| 287 \| 5 \| 287 \| ParB/RepB/Spo0J family partition protein \| ParB/RepB/Spo0J family partition protein; afdb-uniprot50 \| AF-A0A3B8Q0U2-F1-MODEL\_V4 \| 1.0 \| 2.304e-12 \| 363 \| 0.205 \| 292 \| 195 \| 14 \| 12 \| 291 \| 28 \| 294 \| Chromosome partitioning protein ParB \| Chromosome partitioning protein ParB; afdb-uniprot50 \| AF-A0A1G2EPD7-F1-MODEL\_V4 \| 1.0 \| 1.788e-09 \| 362 \| 0.169 \| 213 \| 158 \| 9 \| 13 \| 214 \| 11 \| 215 \| ParB domain-containing protein \| ParB domain-containing protein; afdb-uniprot50 \| AF-A0A1M3PL20-F1-MODEL\_V4 \| 1.0 \| 3.243e-09 \| 362 \| 0.201 \| 194 \| 146 \| 8 \| 19 \| 205 \| 1 \| 192 \| ParB domain-containing protein \| ParB domain-containing protein; afdb-uniprot50 \| AF-M6UQM9-F1-MODEL\_V4 \| 1.0 \| 3.724e-10 \| 362 \| 0.196 \| 219 \| 138 \| 8 \| 13 \| 193 \| 352 \| 570 \| ParB-like protein \| ParB-like protein; afdb-uniprot50 \| AF-A0A7X6YYW6-F1-MODEL\_V4 \| 1.0 \| 2.692e-10 \| 361 \| 0.22 \| 177 \| 129 \| 6 \| 19 \| 192 \| 35 \| 205 \| ParB/RepB/Spo0J family partition protein \| ParB/RepB/Spo0J family partition protein; afdb-uniprot50 \| AF-A0A2H0SUV7-F1-MODEL\_V4 \| 1.0 \| 1.946e-10 \| 361 \| 0.225 \| 191 \| 138 \| 8 \| 15 \| 201 \| 2 \| 186 \| ParB domain-containing protein \| ParB domain-containing protein; afdb-uniprot50 \| AF-A0A327KRU4-F1-MODEL\_V4 \| 1.0 \| 4.409e-12 \| 361 \| 0.22 \| 281 \| 175 \| 13 \| 21 \| 282 \| 5 \| 260 \| Chromosome partitioning protein ParB \| Chromosome partitioning protein ParB; afdb-uniprot50 \| AF-A0A1I4DE87-F1-MODEL\_V4 \| 1.0 \| 9.631e-11 \| 361 \| 0.272 \| 202 \| 123 \| 7 \| 11 \| 205 \| 124 \| 308 \| ParB/RepB/Spo0J family partition protein \| ParB/RepB/Spo0J family partition protein; afdb-uniprot50 \| AF-A0A357IVD1-F1-MODEL\_V4 \| 1.0 \| 1.16e-09 \| 361 \| 0.207 \| 212 \| 146 \| 9 \| 13 \| 203 \| 155 \| 365 \| Site-specific DNA-methyltransferase (adenine-specific) \| Site-specific DNA-methyltransferase (adenine-specific); afdb-uniprot50 \| AF-E9S8W9-F1-MODEL\_V4 \| 1.0 \| 6.753e-10 \| 360 \| 0.187 \| 203 \| 146 \| 7 \| 18 \| 212 \| 31 \| 222 \| ParB-like protein \| ParB-like protein; afdb-uniprot50 \| AF-A0A7U6FS42-F1-MODEL\_V4 \| 1.0 \| 2.117e-11 \| 360 \| 0.201 \| 318 \| 203 \| 11 \| 5 \| 291 \| 20 \| 317 \| Uncharacterized protein \| Uncharacterized protein; afdb-uniprot50 \| AF-A0A7Y9UCD1-F1-MODEL\_V4 \| 1.0 \| 1.705e-11 \| 360 \| 0.226 \| 278 \| 169 \| 15 \| 1 \| 263 \| 38 \| 284 \| ParB/RepB/Spo0J family partition protein \| ParB/RepB/Spo0J family partition protein; afdb-uniprot50 \| AF-A0A3M1TME7-F1-MODEL\_V4 \| 1.0 \| 1.023e-12 \| 359 \| 0.213 \| 295 \| 186 \| 14 \| 14 \| 291 \| 31 \| 296 \| ParB/RepB/Spo0J family partition protein \| ParB/RepB/Spo0J family partition protein; afdb-uniprot50 \| AF-A0A849PIM8-F1-MODEL\_V4 \| 1.0 \| 2.55e-10 \| 359 \| 0.189 \| 216 \| 143 \| 10 \| 13 \| 205 \| 1 \| 207 \| ParB/RepB/Spo0J family partition protein \| ParB/RepB/Spo0J family partition protein; afdb-uniprot50 \| AF-A0A7C6QC06-F1-MODEL\_V4 \| 1.0 \| 2.757e-09 \| 359 \| 0.182 \| 192 \| 145 \| 7 \| 13 \| 198 \| 29 \| 214 \| ParB/RepB/Spo0J family partition protein \| ParB/RepB/Spo0J family partition protein; afdb-uniprot50 \| AF-A0A839IZR4-F1-MODEL\_V4 \| 1.0 \| 7.083e-08 \| 358 \| 0.38 \| 100 \| 59 \| 2 \| 1 \| 97 \| 1 \| 100 \| Uncharacterized protein \| Uncharacterized protein; afdb-uniprot50 \| AF-A0A380DKU1-F1-MODEL\_V4 \| 1.0 \| 1.843e-10 \| 358 \| 0.234 \| 205 \| 145 \| 8 \| 1 \| 200 \| 1 \| 198 \| ParB family partitioning protein \| ParB family partitioning protein; afdb-uniprot50 \| AF-A0A1H6A853-F1-MODEL\_V4 \| 1.0 \| 7.129e-10 \| 358 \| 0.221 \| 190 \| 136 \| 7 \| 14 \| 198 \| 13 \| 195 \| Chromosome partitioning protein, ParB family \| Chromosome partitioning protein, ParB family; afdb-uniprot50 \| AF-A0A1G5IQC4-F1-MODEL\_V4 \| 1.0 \| 1.16e-09 \| 358 \| 0.158 \| 208 \| 162 \| 6 \| 1 \| 202 \| 11 \| 211 \| ParB/RepB/Spo0J family partition protein \| ParB/RepB/Spo0J family partition protein; afdb-uniprot50 \| AF-A0A7X6P274-F1-MODEL\_V4 \| 1.0 \| 3.724e-10 \| 357 \| 0.213 \| 248 \| 156 \| 11 \| 19 \| 265 \| 1 \| 210 \| ParB/RepB/Spo0J family partition protein \| ParB/RepB/Spo0J family partition protein; afdb-uniprot50 \| AF-A0A511N9H7-F1-MODEL\_V4 \| 1.0 \| 1.615e-11 \| 357 \| 0.186 \| 274 \| 184 \| 11 \| 21 \| 282 \| 2 \| 248 \| Putative chromosome 1-partitioning protein ParB \| Putative chromosome 1-partitioning protein ParB; afdb-uniprot50 \| AF-A0A381KQP5-F1-MODEL\_V4 \| 1.0 \| 2.929e-11 \| 357 \| 0.16 \| 355 \| 205 \| 11 \| 19 \| 291 \| 8 \| 351 \| ParB/RepB/Spo0J family partition protein \| ParB/RepB/Spo0J family partition protein; afdb-uniprot50 \| AF-A0A1Q3PK79-F1-MODEL\_V4 \| 1.0 \| 5.311e-11 \| 357 \| 0.232 \| 232 \| 150 \| 10 \| 1 \| 218 \| 1 \| 218 \| ParB domain-containing protein \| ParB domain-containing protein; afdb-uniprot50 \| AF-A0A521U4H3-F1-MODEL\_V4 \| 1.0 \| 9.064e-09 \| 357 \| 0.185 \| 183 \| 138 \| 8 \| 19 \| 197 \| 13 \| 188 \| ParB/RepB/Spo0J family partition protein \| ParB/RepB/Spo0J family partition protein; afdb-uniprot50 \| AF-A0A3M1FTJ6-F1-MODEL\_V4 \| 1.0 \| 1.14e-12 \| 356 \| 0.205 \| 282 \| 197 \| 11 \| 14 \| 282 \| 218 \| 485 \| ParB/RepB/Spo0J family partition protein \| ParB/RepB/Spo0J family partition protein; afdb-uniprot50 \| AF-R5J9W3-F1-MODEL\_V4 \| 1.0 \| 3.724e-10 \| 354 \| 0.183 \| 191 \| 148 \| 6 \| 13 \| 200 \| 2 \| 187 \| Nucleoid occlusion protein \| Nucleoid occlusion protein; afdb-uniprot50 \| AF-A0A1G3IP66-F1-MODEL\_V4 \| 1.0 \| 4.079e-13 \| 352 \| 0.201 \| 318 \| 195 \| 16 \| 1 \| 291 \| 6 \| 291 \| Chromosome partitioning protein ParB \| Chromosome partitioning protein ParB; afdb-uniprot50 \| AF-A0A1M6XZ26-F1-MODEL\_V4 \| 1.0 \| 1.694e-09 \| 352 \| 0.227 \| 189 \| 129 \| 11 \| 19 \| 198 \| 12 \| 192 \| ParB/RepB/Spo0J family partition protein \| ParB/RepB/Spo0J family partition protein; afdb-uniprot50 \| AF-A0A7C3C974-F1-MODEL\_V4 \| 1.0 \| 9.631e-11 \| 351 \| 0.234 \| 235 \| 169 \| 6 \| 1 \| 230 \| 5 \| 233 \| ParB/RepB/Spo0J family partition protein \| ParB/RepB/Spo0J family partition protein; afdb-uniprot50 \| AF-A0A2E9KZY7-F1-MODEL\_V4 \| 1.0 \| 2.359e-11 \| 351 \| 0.171 \| 285 \| 204 \| 10 \| 15 \| 282 \| 2 \| 271 \| Chromosome partitioning protein ParB \| Chromosome partitioning protein ParB; afdb-uniprot50 \| AF-A0A516P4G1-F1-MODEL\_V4 \| 1.0 \| 1.099e-09 \| 351 \| 0.172 \| 191 \| 140 \| 10 \| 19 \| 202 \| 3 \| 182 \| ParB/RepB/Spo0J family partition protein \| ParB/RepB/Spo0J family partition protein; afdb-uniprot50 \| AF-A0A536RK66-F1-MODEL\_V4 \| 1.0 \| 2.104e-09 \| 350 \| 0.201 \| 174 \| 130 \| 6 \| 1 \| 172 \| 24 \| 190 \| ParB/RepB/Spo0J family partition protein \| ParB/RepB/Spo0J family partition protein; afdb-uniprot50 \| AF-A0A1G0HZB1-F1-MODEL\_V4 \| 1.0 \| 7.175e-12 \| 350 \| 0.195 \| 296 \| 207 \| 13 \| 5 \| 291 \| 19 \| 292 \| Probable chromosome-partitioning protein ParB \| Probable chromosome-partitioning protein ParB; afdb-uniprot50 \| AF-A0A3D5A0J3-F1-MODEL\_V4 \| 1.0 \| 9.183e-13 \| 350 \| 0.191 \| 298 \| 212 \| 12 \| 1 \| 282 \| 23 \| 307 \| Stage 0 sporulation protein J \| Stage 0 sporulation protein J; afdb-uniprot50 \| AF-A0A497JKX0-F1-MODEL\_V4 \| 1.0 \| 1.16e-09 \| 350 \| 0.205 \| 200 \| 147 \| 6 \| 4 \| 200 \| 59 \| 249 \| HTH cro/C1-type domain-containing protein \| HTH cro/C1-type domain-containing protein; afdb-uniprot50 \| AF-A0A7C5RVI1-F1-MODEL\_V4 \| 1.0 \| 1.373e-11 \| 349 \| 0.195 \| 261 \| 184 \| 9 \| 15 \| 271 \| 17 \| 255 \| ParB/RepB/Spo0J family partition protein \| ParB/RepB/Spo0J family partition protein; afdb-uniprot50 \| AF-A0A7V2YAS1-F1-MODEL\_V4 \| 1.0 \| 6.44e-12 \| 349 \| 0.187 \| 293 \| 209 \| 12 \| 7 \| 291 \| 11 \| 282 \| ParB/RepB/Spo0J family partition protein \| ParB/RepB/Spo0J family partition protein; afdb-uniprot50 \| AF-A0A524MV92-F1-MODEL\_V4 \| 1.0 \| 2.22e-09 \| 349 \| 0.217 \| 193 \| 135 \| 8 \| 1 \| 184 \| 39 \| 224 \| ParB/RepB/Spo0J family partition protein \| ParB/RepB/Spo0J family partition protein; afdb-uniprot50 \| AF-A0A495IX89-F1-MODEL\_V4 \| 1.0 \| 4.381e-10 \| 349 \| 0.192 \| 203 \| 142 \| 8 \| 3 \| 189 \| 23 \| 219 \| ParB/RepB/Spo0J family partition protein \| ParB/RepB/Spo0J family partition protein; afdb-uniprot50 \| AF-A0A847JDU6-F1-MODEL\_V4 \| 1.0 \| 6.44e-12 \| 348 \| 0.218 \| 293 \| 185 \| 14 \| 13 \| 288 \| 8 \| 273 \| ParB/RepB/Spo0J family partition protein \| ParB/RepB/Spo0J family partition protein; afdb-uniprot50 \| AF-E5C648-F1-MODEL\_V4 \| 1.0 \| 1.993e-09 \| 348 \| 0.192 \| 192 \| 139 \| 9 \| 14 \| 199 \| 30 \| 211 \| ParB-like partition protein \| ParB-like partition protein; afdb-uniprot50 \| AF-A0A356HZ39-F1-MODEL\_V4 \| 1.0 \| 5.475e-12 \| 347 \| 0.15 \| 293 \| 216 \| 12 \| 2 \| 282 \| 8 \| 279 \| ParB domain-containing protein \| ParB domain-containing protein; afdb-uniprot50 \| AF-A0A846QGQ9-F1-MODEL\_V4 \| 1.0 \| 1.27e-12 \| 347 \| 0.188 \| 307 \| 212 \| 11 \| 4 \| 291 \| 12 \| 300 \| ParB family chromosome partitioning protein \| ParB family chromosome partitioning protein; afdb-uniprot50 \| AF-A0A519M6E1-F1-MODEL\_V4 \| 1.0 \| 6.753e-10 \| 346 \| 0.212 \| 212 \| 151 \| 7 \| 10 \| 214 \| 14 \| 216 \| ParB/RepB/Spo0J family partition protein \| ParB/RepB/Spo0J family partition protein; afdb-uniprot50 \| AF-A0A1E4LQW4-F1-MODEL\_V4 \| 1.0 \| 1.301e-11 \| 346 \| 0.221 \| 289 \| 200 \| 12 \| 2 \| 282 \| 1 \| 272 \| ParB domain-containing protein \| ParB domain-containing protein; afdb-uniprot50 \| AF-A0A430RCV0-F1-MODEL\_V4 \| 1.0 \| 1.406e-10 \| 346 \| 0.266 \| 218 \| 125 \| 9 \| 19 \| 205 \| 8 \| 221 \| ParB domain-containing protein \| ParB domain-containing protein; afdb-uniprot50 \| AF-A0A562SHY2-F1-MODEL\_V4 \| 1.0 \| 9.863e-10 \| 346 \| 0.202 \| 198 \| 142 \| 9 \| 19 \| 210 \| 1 \| 188 \| ParB/RepB/Spo0J family partition protein \| ParB/RepB/Spo0J family partition protein; afdb-uniprot50 \| AF-A0A4Y9IUR5-F1-MODEL\_V4 \| 1.0 \| 9.343e-10 \| 346 \| 0.189 \| 206 \| 152 \| 8 \| 18 \| 214 \| 1 \| 200 \| ParB/RepB/Spo0J family partition protein \| ParB/RepB/Spo0J family partition protein; afdb-uniprot50 \| AF-A0A661BEU3-F1-MODEL\_V4 \| 1.0 \| 5.742e-10 \| 345 \| 0.212 \| 174 \| 131 \| 4 \| 13 \| 184 \| 15 \| 184 \| ParB domain-containing protein \| ParB domain-containing protein; afdb-uniprot50 \| AF-A0A511FBI8-F1-MODEL\_V4 \| 1.0 \| 5.311e-11 \| 345 \| 0.225 \| 280 \| 171 \| 16 \| 18 \| 287 \| 1 \| 244 \| ParB family chromosome partitioning protein \| ParB family chromosome partitioning protein; afdb-uniprot50 \| AF-A0A0U1QTP7-F1-MODEL\_V4 \| 1.0 \| 2.117e-11 \| 345 \| 0.193 \| 330 \| 218 \| 12 \| 5 \| 287 \| 17 \| 345 \| DNA-binding protein homolog \| DNA-binding protein homolog; afdb-uniprot50 \| AF-A0A2E3VV26-F1-MODEL\_V4 \| 1.0 \| 8.851e-10 \| 345 \| 0.175 \| 216 \| 161 \| 9 \| 1 \| 214 \| 17 \| 217 \| Partitioning protein \| Partitioning protein; afdb-uniprot50 \| AF-A0A512ICY0-F1-MODEL\_V4 \| 1.0 \| 2.359e-11 \| 345 \| 0.176 \| 317 \| 207 \| 18 \| 1 \| 291 \| 320 \| 608 \| Chromosome partitioning protein ParB \| Chromosome partitioning protein ParB; afdb-uniprot50 \| AF-A0A520FL95-F1-MODEL\_V4 \| 1.0 \| 7.525e-10 \| 344 \| 0.217 \| 170 \| 125 \| 5 \| 18 \| 183 \| 1 \| 166 \| Chromosome partitioning protein ParB \| Chromosome partitioning protein ParB; afdb-uniprot50 \| AF-A0A637WJU4-F1-MODEL\_V4 \| 1.0 \| 1.125e-08 \| 344 \| 0.229 \| 187 \| 128 \| 4 \| 1 \| 172 \| 34 \| 219 \| DNA-binding protein \| DNA-binding protein; afdb-uniprot50 \| AF-A0A1M3DP83-F1-MODEL\_V4 \| 1.0 \| 4.25e-09 \| 344 \| 0.189 \| 179 \| 124 \| 7 \| 19 \| 182 \| 3 \| 175 \| ParB domain-containing protein \| ParB domain-containing protein; afdb-uniprot50 \| AF-A0A3D5I6M5-F1-MODEL\_V4 \| 1.0 \| 5.88e-09 \| 343 \| 0.216 \| 189 \| 130 \| 10 \| 23 \| 205 \| 1 \| 177 \| Chromosome partitioning protein ParB \| Chromosome partitioning protein ParB; afdb-uniprot50 \| AF-A0A726LUF6-F1-MODEL\_V4 \| 1.0 \| 1.262e-10 \| 343 \| 0.172 \| 273 \| 185 \| 6 \| 19 \| 253 \| 7 \| 276 \| Uncharacterized protein \| Uncharacterized protein; afdb-uniprot50 \| AF-A0A2V7X017-F1-MODEL\_V4 \| 1.0 \| 2.49e-11 \| 343 \| 0.18 \| 311 \| 205 \| 16 \| 1 \| 282 \| 120 \| 409 \| ParB domain-containing protein \| ParB domain-containing protein; afdb-uniprot50 \| AF-W4RPT4-F1-MODEL\_V4 \| 1.0 \| 1.406e-10 \| 342 \| 0.217 \| 253 \| 179 \| 12 \| 1 \| 246 \| 1 \| 241 \| Chromosome (Plasmid) partitioning protein ParB \| Chromosome (Plasmid) partitioning protein ParB; afdb-uniprot50 \| AF-A0A4V2B1M2-F1-MODEL\_V4 \| 1.0 \| 1.225e-09 \| 342 \| 0.197 \| 197 \| 149 \| 6 \| 10 \| 202 \| 34 \| 225 \| ParB/RepB/Spo0J family partition protein \| ParB/RepB/Spo0J family partition protein; afdb-uniprot50 \| AF-A0A5B8Z9Y2-F1-MODEL\_V4 \| 1.0 \| 2.629e-11 \| 342 \| 0.201 \| 298 \| 200 \| 16 \| 1 \| 282 \| 1 \| 276 \| ParB/RepB/Spo0J family partition protein \| ParB/RepB/Spo0J family partition protein; afdb-uniprot50 \| AF-A0A829R838-F1-MODEL\_V4 \| 1.0 \| 7.574e-12 \| 342 \| 0.187 \| 298 \| 204 \| 15 \| 1 \| 282 \| 1 \| 276 \| ParB family chromosome partioning protein \| ParB family chromosome partioning protein; afdb-uniprot50 \| AF-A0A4Z1A271-F1-MODEL\_V4 \| 1.0 \| 3e-10 \| 342 \| 0.18 \| 233 \| 155 \| 7 \| 5 \| 203 \| 350 \| 580 \| Chromosome partitioning protein ParB \| Chromosome partitioning protein ParB; afdb-uniprot50 \| AF-A0A3M1MVL3-F1-MODEL\_V4 \| 1.0 \| 3.072e-09 \| 341 \| 0.2 \| 195 \| 143 \| 8 \| 13 \| 201 \| 30 \| 217 \| ParB/RepB/Spo0J family partition protein \| ParB/RepB/Spo0J family partition protein; afdb-uniprot50 \| AF-A0A7C4YD08-F1-MODEL\_V4 \| 1.0 \| 3.957e-12 \| 341 \| 0.18 \| 288 \| 198 \| 14 \| 1 \| 282 \| 3 \| 258 \| ParB/RepB/Spo0J family partition protein \| ParB/RepB/Spo0J family partition protein; afdb-uniprot50 \| AF-A0A4R9H704-F1-MODEL\_V4 \| 1.0 \| 3.724e-10 \| 341 \| 0.178 \| 224 \| 146 \| 8 \| 13 \| 200 \| 404 \| 625 \| Chromosome partitioning protein ParB \| Chromosome partitioning protein ParB; afdb-uniprot50 \| AF-K1AA06-F1-MODEL\_V4 \| 1.0 \| 7.944e-10 \| 340 \| 0.228 \| 166 \| 121 \| 5 \| 19 \| 182 \| 3 \| 163 \| Chromosome partitioning protein \| Chromosome partitioning protein; afdb-uniprot50 \| AF-A0A7C6LJS3-F1-MODEL\_V4 \| 1.0 \| 5.186e-12 \| 340 \| 0.2 \| 310 \| 206 \| 14 \| 1 \| 291 \| 4 \| 290 \| ParB/RepB/Spo0J family partition protein \| ParB/RepB/Spo0J family partition protein; afdb-uniprot50 \| AF-A0A7W9ZEF2-F1-MODEL\_V4 \| 1.0 \| 9.927e-12 \| 340 \| 0.212 \| 287 \| 190 \| 14 \| 13 \| 282 \| 32 \| 299 \| ParB family chromosome partitioning protein \| ParB family chromosome partitioning protein; afdb-uniprot50 \| AF-A0A069JDM1-F1-MODEL\_V4 \| 1.0 \| 5.607e-11 \| 340 \| 0.225 \| 275 \| 171 \| 14 \| 13 \| 263 \| 8 \| 264 \| ParB domain-containing protein \| ParB domain-containing protein; afdb-uniprot50 \| AF-R5VX17-F1-MODEL\_V4 \| 1.0 \| 4.736e-09 \| 340 \| 0.155 \| 212 \| 151 \| 8 \| 19 \| 210 \| 3 \| 206 \| ParB-like partition protein \| ParB-like partition protein; afdb-uniprot50 \| AF-A0A7V6Y4G7-F1-MODEL\_V4 \| 1.0 \| 6.44e-12 \| 339 \| 0.183 \| 295 \| 194 \| 13 \| 9 \| 282 \| 10 \| 278 \| ParB/RepB/Spo0J family partition protein \| ParB/RepB/Spo0J family partition protein; afdb-uniprot50 \| AF-A0A3A9SIV7-F1-MODEL\_V4 \| 1.0 \| 6.797e-12 \| 339 \| 0.166 \| 283 \| 208 \| 11 \| 12 \| 282 \| 34 \| 300 \| ParB/RepB/Spo0J family partition protein \| ParB/RepB/Spo0J family partition protein; afdb-uniprot50 \| AF-A0A2V9YQC4-F1-MODEL\_V4 \| 1.0 \| 1.106e-11 \| 339 \| 0.163 \| 294 \| 207 \| 14 \| 13 \| 291 \| 49 \| 318 \| Chromosome partitioning protein ParB \| Chromosome partitioning protein ParB; afdb-uniprot50 \| AF-A0A363TPC3-F1-MODEL\_V4 \| 1.0 \| 4.913e-12 \| 338 \| 0.18 \| 299 \| 212 \| 10 \| 1 \| 289 \| 10 \| 285 \| ParB domain-containing protein \| ParB domain-containing protein; afdb-uniprot50 \| AF-A0A7X8E5Z8-F1-MODEL\_V4 \| 1.0 \| 2.117e-11 \| 338 \| 0.164 \| 286 \| 206 \| 11 \| 9 \| 282 \| 21 \| 285 \| ParB/RepB/Spo0J family partition protein \| ParB/RepB/Spo0J family partition protein; afdb-uniprot50 \| AF-A0A5J6WBF0-F1-MODEL\_V4 \| 1.0 \| 1.654e-10 \| 337 \| 0.155 \| 283 \| 203 \| 11 \| 19 \| 291 \| 4 \| 260 \| ParB/RepB/Spo0J family partition protein \| ParB/RepB/Spo0J family partition protein; afdb-uniprot50 \| AF-A0A318EAH7-F1-MODEL\_V4 \| 1.0 \| 4.767e-11 \| 337 \| 0.171 \| 298 \| 194 \| 11 \| 13 \| 291 \| 3 \| 266 \| Probable chromosome-partitioning protein ParB \| Probable chromosome-partitioning protein ParB; afdb-uniprot50 \| AF-A0A509J6R8-F1-MODEL\_V4 \| 1.0 \| 1.45e-11 \| 337 \| 0.179 \| 296 \| 192 \| 19 \| 13 \| 282 \| 6 \| 276 \| Probable chromosome-partitioning protein ParB \| Probable chromosome-partitioning protein ParB; afdb-uniprot50 \| AF-H5SEN9-F1-MODEL\_V4 \| 1.0 \| 1.233e-11 \| 337 \| 0.192 \| 302 \| 212 \| 10 \| 1 \| 290 \| 5 \| 286 \| Chromosome partitioning protein, ParB family \| Chromosome partitioning protein, ParB family; afdb-uniprot50 \| AF-A0A2E4E0Y8-F1-MODEL\_V4 \| 1.0 \| 1.048e-11 \| 337 \| 0.175 \| 314 \| 213 \| 17 \| 1 \| 291 \| 37 \| 327 \| Probable chromosome-partitioning protein ParB \| Probable chromosome-partitioning protein ParB; afdb-uniprot50 \| AF-A0A7K2V6D7-F1-MODEL\_V4 \| 1.0 \| 2.359e-11 \| 337 \| 0.204 \| 294 \| 183 \| 15 \| 13 \| 268 \| 7 \| 287 \| ParB/RepB/Spo0J family partition protein \| ParB/RepB/Spo0J family partition protein; afdb-uniprot50 \| AF-A0A7D7UY71-F1-MODEL\_V4 \| 1.0 \| 4.881e-10 \| 337 \| 0.238 \| 235 \| 145 \| 13 \| 19 \| 247 \| 8 \| 214 \| ParB/RepB/Spo0J family partition protein \| ParB/RepB/Spo0J family partition protein; afdb-uniprot50 \| AF-A0A016BNU9-F1-MODEL\_V4 \| 1.0 \| 7.301e-09 \| 337 \| 0.198 \| 191 \| 140 \| 9 \| 13 \| 196 \| 7 \| 191 \| ParB/RepB/Spo0J family partition domain protein \| ParB/RepB/Spo0J family partition domain protein; afdb-uniprot50 \| AF-A0A2D8R2Z7-F1-MODEL\_V4 \| 1.0 \| 7.757e-11 \| 337 \| 0.19 \| 304 \| 195 \| 15 \| 1 \| 287 \| 1 \| 270 \| ParB domain-containing protein \| ParB domain-containing protein; afdb-uniprot50 \| AF-A0A0Q6Z6C7-F1-MODEL\_V4 \| 1.0 \| 1.9e-11 \| 336 \| 0.209 \| 287 \| 189 \| 15 \| 19 \| 291 \| 15 \| 277 \| Probable chromosome-partitioning protein ParB \| Probable chromosome-partitioning protein ParB; afdb-uniprot50 \| AF-A0A3M1JT40-F1-MODEL\_V4 \| 1.0 \| 6.797e-12 \| 336 \| 0.187 \| 282 \| 196 \| 10 \| 14 \| 282 \| 22 \| 283 \| ParB/RepB/Spo0J family partition protein \| ParB/RepB/Spo0J family partition protein; afdb-uniprot50 \| AF-A0A3M7LGN8-F1-MODEL\_V4 \| 1.0 \| 2.104e-09 \| 336 \| 0.167 \| 203 \| 145 \| 9 \| 19 \| 204 \| 13 \| 208 \| ParB/RepB/Spo0J family partition protein \| ParB/RepB/Spo0J family partition protein; afdb-uniprot50 \| AF-A0A5V0QUK1-F1-MODEL\_V4 \| 1.0 \| 1.365e-09 \| 335 \| 0.221 \| 208 \| 141 \| 11 \| 1 \| 195 \| 1 \| 200 \| ParB/RepB/Spo0J family partition protein \| ParB/RepB/Spo0J family partition protein; afdb-uniprot50 \| AF-A0A4P0XZV3-F1-MODEL\_V4 \| 1.0 \| 1.605e-09 \| 335 \| 0.182 \| 219 \| 156 \| 10 \| 3 \| 205 \| 1 \| 212 \| PRTRC system ParB family protein \| PRTRC system ParB family protein; afdb-uniprot50 \| AF-A0A3N1JZ34-F1-MODEL\_V4 \| 1.0 \| 3.072e-09 \| 335 \| 0.182 \| 192 \| 144 \| 7 \| 13 \| 197 \| 3 \| 188 \| ParB/RepB/Spo0J family partition protein \| ParB/RepB/Spo0J family partition protein; afdb-uniprot50 \| AF-A0A2K0XC05-F1-MODEL\_V4 \| 1.0 \| 1.735e-08 \| 335 \| 0.161 \| 204 \| 157 \| 9 \| 19 \| 214 \| 15 \| 212 \| ParB domain-containing protein \| ParB domain-containing protein; afdb-uniprot50 \| AF-A0A0M0BQB3-F1-MODEL\_V4 \| 1.0 \| 3e-10 \| 334 \| 0.207 \| 284 \| 171 \| 12 \| 19 \| 291 \| 3 \| 243 \| ParB domain-containing protein \| ParB domain-containing protein; afdb-uniprot50 \| AF-A0A1F2VSB4-F1-MODEL\_V4 \| 1.0 \| 3.264e-11 \| 334 \| 0.179 \| 295 \| 204 \| 14 \| 8 \| 291 \| 9 \| 276 \| Probable chromosome-partitioning protein ParB \| Probable chromosome-partitioning protein ParB; afdb-uniprot50 \| AF-A0A6L6YJM1-F1-MODEL\_V4 \| 1.0 \| 1.332e-10 \| 334 \| 0.149 \| 275 \| 186 \| 15 \| 1 \| 253 \| 19 \| 267 \| ParB/RepB/Spo0J family partition protein \| ParB/RepB/Spo0J family partition protein; afdb-uniprot50 \| AF-A0A7K3KR56-F1-MODEL\_V4 \| 1.0 \| 3.613e-09 \| 334 \| 0.165 \| 206 \| 157 \| 8 \| 18 \| 214 \| 1 \| 200 \| ParB/RepB/Spo0J family partition protein \| ParB/RepB/Spo0J family partition protein; afdb-uniprot50 \| AF-A0A7W1RZL1-F1-MODEL\_V4 \| 1.0 \| 5.742e-10 \| 333 \| 0.25 \| 172 \| 123 \| 4 \| 13 \| 182 \| 22 \| 189 \| ParB/RepB/Spo0J family partition protein \| ParB/RepB/Spo0J family partition protein; afdb-uniprot50 \| AF-C8WZ31-F1-MODEL\_V4 \| 1.0 \| 3.551e-12 \| 333 \| 0.172 \| 325 \| 209 \| 15 \| 1 \| 291 \| 7 \| 305 \| ParB-like partition protein \| ParB-like partition protein; afdb-uniprot50 \| AF-A0A2L2XH21-F1-MODEL\_V4 \| 1.0 \| 1.53e-11 \| 333 \| 0.216 \| 291 \| 192 \| 16 \| 16 \| 287 \| 1 \| 274 \| Chromosome partitioning protein ParB \| Chromosome partitioning protein ParB; afdb-uniprot50 \| AF-A0A3E0KLR5-F1-MODEL\_V4 \| 1.0 \| 7.757e-11 \| 332 \| 0.184 \| 288 \| 193 \| 11 \| 20 \| 291 \| 26 \| 287 \| Chromosome partitioning protein ParB \| Chromosome partitioning protein ParB; afdb-uniprot50 \| AF-A0A358G3B0-F1-MODEL\_V4 \| 1.0 \| 9.343e-10 \| 332 \| 0.185 \| 221 \| 158 \| 10 \| 1 \| 214 \| 12 \| 217 \| Chromosome partitioning protein ParB \| Chromosome partitioning protein ParB; afdb-uniprot50 \| AF-A0A7C7K384-F1-MODEL\_V4 \| 1.0 \| 4.026e-09 \| 332 \| 0.155 \| 206 \| 162 \| 7 \| 1 \| 200 \| 11 \| 210 \| PRTRC system ParB family protein \| PRTRC system ParB family protein; afdb-uniprot50 \| AF-A0A382GYA7-F1-MODEL\_V4 \| 1.0 \| 1.066e-08 \| 331 \| 0.202 \| 168 \| 124 \| 6 \| 20 \| 182 \| 14 \| 176 \| ParB domain-containing protein \| ParB domain-containing protein; afdb-uniprot50 \| AF-A0A4D7QF63-F1-MODEL\_V4 \| 1.0 \| 1.373e-11 \| 331 \| 0.191 \| 308 \| 192 \| 14 \| 4 \| 288 \| 18 \| 291 \| ParB/RepB/Spo0J family partition protein \| ParB/RepB/Spo0J family partition protein; afdb-uniprot50 \| AF-A0A521WSH2-F1-MODEL\_V4 \| 1.0 \| 7.348e-11 \| 331 \| 0.15 \| 300 \| 206 \| 15 \| 13 \| 291 \| 26 \| 297 \| ParB/RepB/Spo0J family partition protein \| ParB/RepB/Spo0J family partition protein; afdb-uniprot50 \| AF-A0A1G3HSG6-F1-MODEL\_V4 \| 1.0 \| 4.177e-12 \| 331 \| 0.199 \| 301 \| 201 \| 12 \| 1 \| 282 \| 6 \| 285 \| Chromosome partitioning protein ParB \| Chromosome partitioning protein ParB; afdb-uniprot50 \| AF-A0A6D1VXK9-F1-MODEL\_V4 \| 1.0 \| 5.88e-09 \| 331 \| 0.18 \| 194 \| 147 \| 9 \| 13 \| 201 \| 5 \| 191 \| Chromosome partitioning protein ParB \| Chromosome partitioning protein ParB; afdb-uniprot50 \| AF-A0A1N7S8P1-F1-MODEL\_V4 \| 1.0 \| 1.066e-08 \| 331 \| 0.207 \| 202 \| 138 \| 7 \| 1 \| 197 \| 1 \| 185 \| ParB family protein \| ParB family protein; afdb-uniprot50 \| AF-A0A6L2LGI1-F1-MODEL\_V4 \| 1.0 \| 2.55e-10 \| 331 \| 0.192 \| 250 \| 157 \| 9 \| 12 \| 251 \| 312 \| 526 \| ParB domain-containing protein \| ParB domain-containing protein; afdb-uniprot50 \| AF-A0A382I217-F1-MODEL\_V4 \| 1.0 \| 1.45e-11 \| 330 \| 0.194 \| 278 \| 185 \| 12 \| 4 \| 269 \| 16 \| 266 \| ParB domain-containing protein \| ParB domain-containing protein; afdb-uniprot50 \| AF-A0A497SN10-F1-MODEL\_V4 \| 1.0 \| 6.71e-08 \| 330 \| 0.21 \| 171 \| 122 \| 8 \| 21 \| 184 \| 1 \| 165 \| ParB domain-containing protein \| ParB domain-containing protein; afdb-uniprot50 \| AF-A0A4Y3KAD6-F1-MODEL\_V4 \| 1.0 \| 9.124e-11 \| 330 \| 0.169 \| 313 \| 195 \| 11 \| 1 \| 290 \| 13 \| 283 \| Chromosome partitioning protein ParB \| Chromosome partitioning protein ParB; afdb-uniprot50 \| AF-K9H3Q4-F1-MODEL\_V4 \| 1.0 \| 9.404e-12 \| 330 \| 0.192 \| 307 \| 203 \| 13 \| 2 \| 291 \| 38 \| 316 \| Chromosome (Plasmid) partitioning protein ParB \| Chromosome (Plasmid) partitioning protein ParB; afdb-uniprot50 \| AF-A0A2H1J509-F1-MODEL\_V4 \| 1.0 \| 3.839e-11 \| 330 \| 0.17 \| 300 \| 189 \| 16 \| 12 \| 282 \| 250 \| 518 \| Chromosome partitioning protein, ParB family \| Chromosome partitioning protein, ParB family; afdb-uniprot50 \| AF-A0A6F9ZNA4-F1-MODEL\_V4 \| 1.0 \| 6.207e-09 \| 329 \| 0.162 \| 197 \| 146 \| 8 \| 1 \| 189 \| 2 \| 187 \| Chromosome-partitioning protein Spo0J \| Chromosome-partitioning protein Spo0J; afdb-uniprot50 \| AF-K5DPP1-F1-MODEL\_V4 \| 1.0 \| 1.324e-08 \| 328 \| 0.197 \| 187 \| 141 \| 7 \| 13 \| 192 \| 1 \| 185 \| Transcriptional regulator \| Transcriptional regulator; afdb-uniprot50 \| AF-A0A7C6WVD0-F1-MODEL\_V4 \| 1.0 \| 2.006e-11 \| 328 \| 0.181 \| 292 \| 187 \| 12 \| 13 \| 282 \| 2 \| 263 \| ParB/RepB/Spo0J family partition protein \| ParB/RepB/Spo0J family partition protein; afdb-uniprot50 \| AF-A0A7R8B2R3-F1-MODEL\_V4 \| 1.0 \| 6.797e-12 \| 328 \| 0.224 \| 289 \| 190 \| 12 \| 12 \| 287 \| 42 \| 309 \| Chromosome partitioning protein ParB \| Chromosome partitioning protein ParB; afdb-uniprot50 \| AF-A0A2V9VKL2-F1-MODEL\_V4 \| 1.0 \| 2.006e-11 \| 328 \| 0.2 \| 300 \| 202 \| 14 \| 3 \| 282 \| 116 \| 397 \| Chromosome partitioning protein ParB \| Chromosome partitioning protein ParB; afdb-uniprot50 \| AF-A0A0F9I9R4-F1-MODEL\_V4 \| 1.0 \| 5.032e-11 \| 328 \| 0.188 \| 286 \| 153 \| 12 \| 14 \| 283 \| 9 \| 231 \| ParB domain-containing protein \| ParB domain-containing protein; afdb-uniprot50 \| AF-A0A7V7CWH9-F1-MODEL\_V4 \| 1.0 \| 6.398e-10 \| 327 \| 0.257 \| 198 \| 129 \| 8 \| 13 \| 205 \| 38 \| 222 \| ParB/RepB/Spo0J family partition protein \| ParB/RepB/Spo0J family partition protein; afdb-uniprot50 \| AF-A0A2W4SXL5-F1-MODEL\_V4 \| 1.0 \| 9.927e-12 \| 327 \| 0.208 \| 298 \| 178 \| 13 \| 13 \| 282 \| 6 \| 273 \| Chromosome partitioning protein ParB \| Chromosome partitioning protein ParB; afdb-uniprot50 \| AF-A0A2V6T438-F1-MODEL\_V4 \| 1.0 \| 7.574e-12 \| 327 \| 0.207 \| 289 \| 190 \| 13 \| 15 \| 291 \| 19 \| 280 \| ParB domain-containing protein \| ParB domain-containing protein; afdb-uniprot50 \| AF-A0A7D7ZNA8-F1-MODEL\_V4 \| 1.0 \| 2.182e-12 \| 327 \| 0.211 \| 298 \| 184 \| 13 \| 13 \| 290 \| 33 \| 299 \| Probable chromosome-partitioning protein ParB \| Probable chromosome-partitioning protein ParB; afdb-uniprot50 \| AF-A0A2J0MU83-F1-MODEL\_V4 \| 1.0 \| 5.779e-12 \| 327 \| 0.178 \| 292 \| 207 \| 12 \| 1 \| 282 \| 33 \| 301 \| Chromosome partitioning protein ParB \| Chromosome partitioning protein ParB; afdb-uniprot50 \| AF-A0A2E8PUX0-F1-MODEL\_V4 \| 1.0 \| 7.129e-10 \| 327 \| 0.167 \| 275 \| 182 \| 13 \| 13 \| 246 \| 155 \| 423 \| ParB domain-containing protein \| ParB domain-containing protein; afdb-uniprot50 \| AF-A0A7U4TW42-F1-MODEL\_V4 \| 1.0 \| 6.398e-10 \| 326 \| 0.282 \| 198 \| 120 \| 4 \| 96 \| 282 \| 5 \| 191 \| Uncharacterized protein \| Uncharacterized protein; afdb-uniprot50 \| AF-A0A4P1KK94-F1-MODEL\_V4 \| 1.0 \| 8.851e-10 \| 326 \| 0.253 \| 189 \| 129 \| 6 \| 12 \| 195 \| 28 \| 209 \| Chromosome-partitioning protein parB \| Chromosome-partitioning protein parB; afdb-uniprot50 \| AF-A0A2D5ZN56-F1-MODEL\_V4 \| 1.0 \| 7.995e-12 \| 326 \| 0.182 \| 291 \| 207 \| 11 \| 1 \| 282 \| 8 \| 276 \| ParB domain-containing protein \| ParB domain-containing protein; afdb-uniprot50 \| AF-A0A7G9A8K7-F1-MODEL\_V4 \| 1.0 \| 9.124e-11 \| 326 \| 0.23 \| 265 \| 174 \| 10 \| 41 \| 287 \| 14 \| 266 \| Putative transcriptional regulators \| Putative transcriptional regulators; afdb-uniprot50 \| AF-A0A537KSY3-F1-MODEL\_V4 \| 1.0 \| 4.053e-11 \| 326 \| 0.202 \| 306 \| 204 \| 15 \| 4 \| 291 \| 15 \| 298 \| ParB/RepB/Spo0J family partition protein \| ParB/RepB/Spo0J family partition protein; afdb-uniprot50 \| AF-A0A137SW11-F1-MODEL\_V4 \| 1.0 \| 2.117e-11 \| 325 \| 0.196 \| 300 \| 206 \| 14 \| 4 \| 291 \| 19 \| 295 \| Putative stage 0 sporulation protein J \| Putative stage 0 sporulation protein J; afdb-uniprot50 \| AF-A0A4Q7IXJ8-F1-MODEL\_V4 \| 1.0 \| 1.168e-11 \| 325 \| 0.146 \| 300 \| 218 \| 12 \| 7 \| 291 \| 23 \| 299 \| ParB/RepB/Spo0J family partition protein \| ParB/RepB/Spo0J family partition protein; afdb-uniprot50 \| AF-A0A3M2BDT7-F1-MODEL\_V4 \| 1.0 \| 2.929e-11 \| 325 \| 0.18 \| 294 \| 203 \| 16 \| 13 \| 291 \| 47 \| 317 \| ParB/RepB/Spo0J family partition protein \| ParB/RepB/Spo0J family partition protein; afdb-uniprot50 \| AF-A0A7K0PKF7-F1-MODEL\_V4 \| 1.0 \| 1.406e-10 \| 325 \| 0.183 \| 343 \| 206 \| 17 \| 1 \| 287 \| 1 \| 325 \| ParB/RepB/Spo0J family partition protein \| ParB/RepB/Spo0J family partition protein; afdb-uniprot50 \| AF-A0A2V8F9Y2-F1-MODEL\_V4 \| 1.0 \| 6.552e-09 \| 324 \| 0.256 \| 164 \| 113 \| 6 \| 14 \| 172 \| 18 \| 177 \| Stage 0 sporulation protein J \| Stage 0 sporulation protein J; afdb-uniprot50 \| AF-A0A2S5UTJ6-F1-MODEL\_V4 \| 1.0 \| 9.343e-10 \| 324 \| 0.195 \| 205 \| 131 \| 10 \| 13 \| 193 \| 15 \| 209 \| ParB domain-containing protein \| ParB domain-containing protein; afdb-uniprot50 \| AF-A0A839QSA6-F1-MODEL\_V4 \| 1.0 \| 3.092e-11 \| 324 \| 0.182 \| 285 \| 191 \| 16 \| 19 \| 283 \| 78 \| 340 \| ParB family chromosome partitioning protein \| ParB family chromosome partitioning protein; afdb-uniprot50 \| AF-A0A4R6WAD4-F1-MODEL\_V4 \| 1.0 \| 2.612e-09 \| 324 \| 0.155 \| 187 \| 138 \| 6 \| 19 \| 191 \| 11 \| 191 \| ParB/RepB/Spo0J family partition protein \| ParB/RepB/Spo0J family partition protein; afdb-uniprot50 \| AF-A0A2G8CD03-F1-MODEL\_V4 \| 1.0 \| 4.352e-08 \| 323 \| 0.149 \| 194 \| 151 \| 7 \| 19 \| 205 \| 6 \| 192 \| Chromosome partitioning protein ParB \| Chromosome partitioning protein ParB; afdb-uniprot50 \| AF-A0A1T4NQJ2-F1-MODEL\_V4 \| 1.0 \| 2.775e-11 \| 323 \| 0.186 \| 274 \| 200 \| 11 \| 19 \| 282 \| 37 \| 297 \| Chromosome partitioning protein, ParB family \| Chromosome partitioning protein, ParB family; afdb-uniprot50 \| AF-A0A518BT11-F1-MODEL\_V4 \| 1.0 \| 5.918e-11 \| 323 \| 0.192 \| 306 \| 198 \| 14 \| 10 \| 283 \| 10 \| 298 \| Putative chromosome-partitioning protein ParB \| Putative chromosome-partitioning protein ParB; afdb-uniprot50 \| AF-A0A1N7JF37-F1-MODEL\_V4 \| 1.0 \| 5.475e-12 \| 323 \| 0.189 \| 332 \| 199 \| 19 \| 1 \| 291 \| 6 \| 308 \| ParB/RepB/Spo0J family partition protein \| ParB/RepB/Spo0J family partition protein; afdb-uniprot50 \| AF-A0A496S0Y3-F1-MODEL\_V4 \| 1.0 \| 6.247e-11 \| 322 \| 0.17 \| 299 \| 203 \| 11 \| 1 \| 282 \| 5 \| 275 \| ParB domain-containing protein \| ParB domain-containing protein; afdb-uniprot50 \| AF-A0A7X8DA56-F1-MODEL\_V4 \| 1.0 \| 7.574e-12 \| 322 \| 0.182 \| 301 \| 207 \| 15 \| 1 \| 288 \| 5 \| 279 \| ParB/RepB/Spo0J family partition protein \| ParB/RepB/Spo0J family partition protein; afdb-uniprot50 \| AF-A0A3R7RVA8-F1-MODEL\_V4 \| 1.0 \| 8.643e-11 \| 322 \| 0.162 \| 289 \| 209 \| 12 \| 5 \| 282 \| 14 \| 280 \| ParB/RepB/Spo0J family partition protein \| ParB/RepB/Spo0J family partition protein; afdb-uniprot50 \| AF-A0A522RQE6-F1-MODEL\_V4 \| 1.0 \| 2.775e-11 \| 322 \| 0.176 \| 306 \| 202 \| 17 \| 1 \| 282 \| 6 \| 285 \| ParB/RepB/Spo0J family partition protein \| ParB/RepB/Spo0J family partition protein; afdb-uniprot50 \| AF-A0A3B9TXJ6-F1-MODEL\_V4 \| 1.0 \| 3.613e-09 \| 321 \| 0.217 \| 175 \| 125 \| 6 \| 14 \| 182 \| 75 \| 243 \| ParB domain-containing protein \| ParB domain-containing protein; afdb-uniprot50 \| AF-A0A1F5PE80-F1-MODEL\_V4 \| 1.0 \| 1.373e-11 \| 321 \| 0.202 \| 287 \| 194 \| 14 \| 14 \| 283 \| 7 \| 275 \| ParB domain-containing protein \| ParB domain-containing protein; afdb-uniprot50 \| AF-A0A2G9ZDA8-F1-MODEL\_V4 \| 1.0 \| 3.445e-11 \| 321 \| 0.202 \| 287 \| 200 \| 11 \| 3 \| 282 \| 13 \| 277 \| Chromosome partitioning protein ParB \| Chromosome partitioning protein ParB; afdb-uniprot50 \| AF-A0A0F9XAK5-F1-MODEL\_V4 \| 1.0 \| 2.168e-10 \| 321 \| 0.186 \| 301 \| 202 \| 15 \| 5 \| 291 \| 11 \| 282 \| Uncharacterized protein \| Uncharacterized protein; afdb-uniprot50 \| AF-A0A2A4LKK3-F1-MODEL\_V4 \| 1.0 \| 1.615e-11 \| 321 \| 0.19 \| 310 \| 197 \| 16 \| 4 \| 288 \| 18 \| 298 \| Chromosome partitioning protein ParB \| Chromosome partitioning protein ParB; afdb-uniprot50 \| AF-A0A7H8KV52-F1-MODEL\_V4 \| 1.0 \| 2.629e-11 \| 321 \| 0.208 \| 307 \| 212 \| 17 \| 1 \| 291 \| 60 \| 351 \| ParB/RepB/Spo0J family partition protein \| ParB/RepB/Spo0J family partition protein; afdb-uniprot50 \| AF-A0A4R7C1F4-F1-MODEL\_V4 \| 1.0 \| 1.946e-10 \| 321 \| 0.183 \| 289 \| 191 \| 11 \| 13 \| 287 \| 80 \| 337 \| ParB/RepB/Spo0J family partition protein \| ParB/RepB/Spo0J family partition protein; afdb-uniprot50 \| AF-A0A7W8YCJ4-F1-MODEL\_V4 \| 1.0 \| 3.839e-11 \| 321 \| 0.185 \| 286 \| 189 \| 15 \| 19 \| 283 \| 359 \| 621 \| ParB/RepB/Spo0J family partition protein \| ParB/RepB/Spo0J family partition protein; afdb-uniprot50 \| AF-A0A1V3YZ69-F1-MODEL\_V4 \| 1.0 \| 4.736e-09 \| 320 \| 0.205 \| 219 \| 149 \| 11 \| 1 \| 203 \| 1 \| 210 \| Uncharacterized protein \| Uncharacterized protein; afdb-uniprot50 \| AF-A0A1G6F7D8-F1-MODEL\_V4 \| 1.0 \| 2.235e-11 \| 320 \| 0.184 \| 309 \| 204 \| 11 \| 1 \| 282 \| 7 \| 294 \| Chromosome partitioning protein, ParB family \| Chromosome partitioning protein, ParB family; afdb-uniprot50 \| AF-U7UNL4-F1-MODEL\_V4 \| 1.0 \| 2.117e-11 \| 320 \| 0.156 \| 287 \| 197 \| 12 \| 19 \| 291 \| 60 \| 315 \| ParB-like protein \| ParB-like protein; afdb-uniprot50 \| AF-A0A7X6XF92-F1-MODEL\_V4 \| 1.0 \| 1.406e-10 \| 320 \| 0.185 \| 275 \| 186 \| 10 \| 13 \| 271 \| 1 \| 253 \| ParB/RepB/Spo0J family partition protein \| ParB/RepB/Spo0J family partition protein; afdb-uniprot50 \| AF-A0A0F9TED6-F1-MODEL\_V4 \| 1.0 \| 4.999e-09 \| 320 \| 0.208 \| 211 \| 145 \| 10 \| 2 \| 200 \| 10 \| 210 \| ParB domain-containing protein \| ParB domain-containing protein; afdb-uniprot50 \| AF-A0A383DD87-F1-MODEL\_V4 \| 1.0 \| 1.788e-09 \| 319 \| 0.228 \| 184 \| 123 \| 8 \| 1 \| 176 \| 1 \| 173 \| ParB domain-containing protein \| ParB domain-containing protein; afdb-uniprot50 \| AF-A0A527D7P3-F1-MODEL\_V4 \| 1.0 \| 4.352e-08 \| 319 \| 0.22 \| 186 \| 133 \| 7 \| 21 \| 201 \| 36 \| 214 \| ParB/RepB/Spo0J family partition protein \| ParB/RepB/Spo0J family partition protein; afdb-uniprot50 \| AF-A0A3A4PA16-F1-MODEL\_V4 \| 1.0 \| 4.767e-11 \| 319 \| 0.146 \| 307 \| 224 \| 12 \| 1 \| 290 \| 5 \| 290 \| ParB/RepB/Spo0J family partition protein \| ParB/RepB/Spo0J family partition protein; afdb-uniprot50 \| AF-A0A1F2SQJ3-F1-MODEL\_V4 \| 1.0 \| 2.117e-11 \| 319 \| 0.201 \| 302 \| 206 \| 15 \| 1 \| 287 \| 17 \| 298 \| ParB domain-containing protein \| ParB domain-containing protein; afdb-uniprot50 \| AF-A0A7C3CPY4-F1-MODEL\_V4 \| 1.0 \| 6.398e-10 \| 319 \| 0.216 \| 217 \| 144 \| 10 \| 8 \| 205 \| 45 \| 254 \| ParB/RepB/Spo0J family partition protein \| ParB/RepB/Spo0J family partition protein; afdb-uniprot50 \| AF-A0A4R2PPN9-F1-MODEL\_V4 \| 1.0 \| 6.595e-11 \| 319 \| 0.219 \| 315 \| 189 \| 15 \| 9 \| 290 \| 29 \| 319 \| ParB family chromosome partitioning protein \| ParB family chromosome partitioning protein; afdb-uniprot50 \| AF-E5Y7U8-F1-MODEL\_V4 \| 1.0 \| 5.779e-12 \| 319 \| 0.19 \| 310 \| 191 \| 16 \| 13 \| 291 \| 37 \| 317 \| ParB-like partition protein \| ParB-like partition protein; afdb-uniprot50 \| AF-A0A542JTD4-F1-MODEL\_V4 \| 1.0 \| 1.831e-08 \| 319 \| 0.215 \| 204 \| 140 \| 11 \| 19 \| 215 \| 150 \| 340 \| ParB family chromosome partitioning protein \| ParB family chromosome partitioning protein; afdb-uniprot50 \| AF-A0A7W7FP95-F1-MODEL\_V4 \| 1.0 \| 4.624e-10 \| 319 \| 0.192 \| 285 \| 198 \| 15 \| 4 \| 280 \| 51 \| 311 \| ParB/RepB/Spo0J family partition protein \| ParB/RepB/Spo0J family partition protein; afdb-uniprot50 \| AF-A0A7Y7ILR4-F1-MODEL\_V4 \| 1.0 \| 2.168e-10 \| 319 \| 0.187 \| 283 \| 179 \| 13 \| 19 \| 284 \| 3 \| 251 \| ParB/RepB/Spo0J family partition protein \| ParB/RepB/Spo0J family partition protein; afdb-uniprot50 \| AF-A0A5D0XVA2-F1-MODEL\_V4 \| 1.0 \| 6.961e-11 \| 319 \| 0.17 \| 287 \| 190 \| 16 \| 19 \| 282 \| 221 \| 482 \| ParB/RepB/Spo0J family partition protein \| ParB/RepB/Spo0J family partition protein; afdb-uniprot50 \| AF-A0A3N9WYB0-F1-MODEL\_V4 \| 1.0 \| 1.831e-08 \| 319 \| 0.224 \| 196 \| 137 \| 9 \| 10 \| 199 \| 21 \| 207 \| ParB domain-containing protein \| ParB domain-containing protein; afdb-uniprot50 \| AF-A0A238XIP6-F1-MODEL\_V4 \| 1.0 \| 4.736e-09 \| 319 \| 0.153 \| 247 \| 184 \| 10 \| 5 \| 247 \| 72 \| 297 \| Chromosome partitioning protein, ParB family \| Chromosome partitioning protein, ParB family; afdb-uniprot50 \| AF-A0A0F9T9X7-F1-MODEL\_V4 \| 1.0 \| 1.788e-09 \| 318 \| 0.188 \| 270 \| 171 \| 12 \| 22 \| 282 \| 16 \| 246 \| ParB domain-containing protein \| ParB domain-containing protein; afdb-uniprot50 \| AF-R6QZ87-F1-MODEL\_V4 \| 1.0 \| 1.373e-11 \| 318 \| 0.167 \| 298 \| 208 \| 11 \| 1 \| 282 \| 21 \| 294 \| ParB partition protein \| ParB partition protein; afdb-uniprot50 \| AF-A0A2N5IVP5-F1-MODEL\_V4 \| 1.0 \| 8.587e-09 \| 318 \| 0.24 \| 204 \| 128 \| 12 \| 19 \| 205 \| 5 \| 198 \| ParB-like protein \| ParB-like protein; afdb-uniprot50 \| AF-A0A7R9L9W8-F1-MODEL\_V4 \| 1.0 \| 2.416e-10 \| 318 \| 0.153 \| 286 \| 202 \| 11 \| 19 \| 291 \| 282 \| 540 \| Hypothetical protein \| Hypothetical protein; afdb-uniprot50 \| AF-A0A7C5XEJ0-F1-MODEL\_V4 \| 1.0 \| 3.166e-10 \| 317 \| 0.186 \| 284 \| 198 \| 11 \| 1 \| 271 \| 2 \| 265 \| ParB/RepB/Spo0J family partition protein \| ParB/RepB/Spo0J family partition protein; afdb-uniprot50 \| AF-A0A7G9GAZ6-F1-MODEL\_V4 \| 1.0 \| 2.359e-11 \| 317 \| 0.172 \| 290 \| 211 \| 10 \| 4 \| 282 \| 38 \| 309 \| ParB/RepB/Spo0J family partition protein \| ParB/RepB/Spo0J family partition protein; afdb-uniprot50 \| AF-A0A2D8HHY6-F1-MODEL\_V4 \| 1.0 \| 2.117e-11 \| 317 \| 0.198 \| 302 \| 209 \| 10 \| 7 \| 291 \| 22 \| 307 \| Chromosome partitioning protein ParB \| Chromosome partitioning protein ParB; afdb-uniprot50 \| AF-A0A2E2FIX1-F1-MODEL\_V4 \| 1.0 \| 5.404e-08 \| 317 \| 0.147 \| 183 \| 146 \| 6 \| 19 \| 198 \| 3 \| 178 \| ParB domain-containing protein \| ParB domain-containing protein; afdb-uniprot50 \| AF-A0A7K1TL10-F1-MODEL\_V4 \| 1.0 \| 9.568e-09 \| 317 \| 0.175 \| 188 \| 139 \| 7 \| 19 \| 200 \| 23 \| 200 \| ParB/RepB/Spo0J family partition protein \| ParB/RepB/Spo0J family partition protein; afdb-uniprot50 \| AF-Q11ZU5-F1-MODEL\_V4 \| 1.0 \| 1.644e-08 \| 317 \| 0.2 \| 165 \| 123 \| 5 \| 13 \| 169 \| 4 \| 167 \| ParB family protein \| ParB family protein; afdb-uniprot50 \| AF-A0A7T3CHN4-F1-MODEL\_V4 \| 1.0 \| 3.839e-11 \| 317 \| 0.173 \| 289 \| 197 \| 13 \| 19 \| 287 \| 307 \| 573 \| ParB/RepB/Spo0J family partition protein \| ParB/RepB/Spo0J family partition protein; afdb-uniprot50 \| AF-A0A368UB33-F1-MODEL\_V4 \| 1.0 \| 2.823e-08 \| 317 \| 0.169 \| 189 \| 144 \| 7 \| 19 \| 200 \| 10 \| 192 \| PRTRC system ParB family protein \| PRTRC system ParB family protein; afdb-uniprot50 \| AF-A0A434ELE8-F1-MODEL\_V4 \| 1.0 \| 6.552e-09 \| 316 \| 0.219 \| 191 \| 132 \| 8 \| 2 \| 180 \| 10 \| 195 \| ParB/RepB/Spo0J family partition protein \| ParB/RepB/Spo0J family partition protein; afdb-uniprot50 \| AF-A0A2V8CRA1-F1-MODEL\_V4 \| 1.0 \| 1.041e-09 \| 316 \| 0.225 \| 217 \| 152 \| 6 \| 1 \| 204 \| 4 \| 217 \| Chromosome partitioning protein ParB \| Chromosome partitioning protein ParB; afdb-uniprot50 \| AF-A0A552UA07-F1-MODEL\_V4 \| 1.0 \| 5.311e-11 \| 316 \| 0.232 \| 301 \| 187 \| 16 \| 4 \| 291 \| 9 \| 278 \| ParB/RepB/Spo0J family partition protein \| ParB/RepB/Spo0J family partition protein; afdb-uniprot50 \| AF-A0A494TNC1-F1-MODEL\_V4 \| 1.0 \| 3.839e-11 \| 316 \| 0.159 \| 308 \| 216 \| 13 \| 1 \| 291 \| 12 \| 293 \| ParB/RepB/Spo0J family partition protein \| ParB/RepB/Spo0J family partition protein; afdb-uniprot50 \| AF-A0A497RGL2-F1-MODEL\_V4 \| 1.0 \| 1.475e-08 \| 316 \| 0.194 \| 185 \| 143 \| 4 \| 19 \| 201 \| 6 \| 186 \| ParB domain-containing protein \| ParB domain-containing protein; afdb-uniprot50 \| AF-A0A7K3LT60-F1-MODEL\_V4 \| 1.0 \| 1.293e-09 \| 316 \| 0.213 \| 244 \| 153 \| 15 \| 19 \| 251 \| 5 \| 220 \| ParB/RepB/Spo0J family partition protein \| ParB/RepB/Spo0J family partition protein; afdb-uniprot50 \| AF-A0A4Z0V345-F1-MODEL\_V4 \| 1.0 \| 1.254e-08 \| 316 \| 0.186 \| 209 \| 137 \| 9 \| 18 \| 195 \| 1 \| 207 \| ParB domain-containing protein \| ParB domain-containing protein; afdb-uniprot50 \| AF-A0A0R2F0I0-F1-MODEL\_V4 \| 1.0 \| 6.961e-11 \| 315 \| 0.198 \| 292 \| 185 \| 16 \| 19 \| 291 \| 1 \| 262 \| Chromosome partitioning protein ParB \| Chromosome partitioning protein ParB; afdb-uniprot50 \| AF-A0A7V2T8H7-F1-MODEL\_V4 \| 1.0 \| 3.637e-11 \| 315 \| 0.215 \| 283 \| 191 \| 11 \| 11 \| 283 \| 24 \| 285 \| ParB/RepB/Spo0J family partition protein \| ParB/RepB/Spo0J family partition protein; afdb-uniprot50 \| AF-A0A7V6W467-F1-MODEL\_V4 \| 1.0 \| 2.49e-11 \| 315 \| 0.161 \| 304 \| 209 \| 14 \| 1 \| 282 \| 9 \| 288 \| ParB/RepB/Spo0J family partition protein \| ParB/RepB/Spo0J family partition protein; afdb-uniprot50 \| AF-A0A3M0ZST4-F1-MODEL\_V4 \| 1.0 \| 1.196e-10 \| 315 \| 0.177 \| 288 \| 188 \| 13 \| 20 \| 282 \| 31 \| 294 \| ParB/RepB/Spo0J family partition protein \| ParB/RepB/Spo0J family partition protein; afdb-uniprot50 \| AF-A0A355T8A7-F1-MODEL\_V4 \| 1.0 \| 1.048e-11 \| 315 \| 0.177 \| 309 \| 206 \| 14 \| 1 \| 282 \| 10 \| 297 \| Chromosome partitioning protein ParB \| Chromosome partitioning protein ParB; afdb-uniprot50 \| AF-A0A518C0J3-F1-MODEL\_V4 \| 1.0 \| 2.629e-11 \| 315 \| 0.209 \| 296 \| 195 \| 13 \| 12 \| 291 \| 36 \| 308 \| Putative chromosome-partitioning protein ParB \| Putative chromosome-partitioning protein ParB; afdb-uniprot50 \| AF-A0A0S8F0I7-F1-MODEL\_V4 \| 1.0 \| 3.906e-08 \| 315 \| 0.18 \| 183 \| 136 \| 8 \| 20 \| 195 \| 64 \| 239 \| ParB domain-containing protein \| ParB domain-containing protein; afdb-uniprot50 \| AF-A0A5F2BGZ2-F1-MODEL\_V4 \| 1.0 \| 1.073e-10 \| 315 \| 0.161 \| 321 \| 193 \| 14 \| 13 \| 283 \| 409 \| 703 \| Chromosome partitioning protein ParB \| Chromosome partitioning protein ParB; afdb-uniprot50 \| AF-A0A7C1AMD4-F1-MODEL\_V4 \| 1.0 \| 1.788e-09 \| 314 \| 0.226 \| 190 \| 127 \| 7 \| 19 \| 206 \| 3 \| 174 \| ParB domain-containing protein \| ParB domain-containing protein; afdb-uniprot50 \| AF-A0A7X7X790-F1-MODEL\_V4 \| 1.0 \| 7.757e-11 \| 314 \| 0.185 \| 285 \| 205 \| 10 \| 7 \| 283 \| 11 \| 276 \| ParB/RepB/Spo0J family partition protein \| ParB/RepB/Spo0J family partition protein; afdb-uniprot50 \| AF-A0A2R4MAE9-F1-MODEL\_V4 \| 1.0 \| 3.092e-11 \| 314 \| 0.18 \| 304 \| 209 \| 12 \| 1 \| 283 \| 7 \| 291 \| Chromosome-partitioning protein ParB \| Chromosome-partitioning protein ParB; afdb-uniprot50 \| AF-A0A0Q7JFS1-F1-MODEL\_V4 \| 1.0 \| 3.264e-11 \| 314 \| 0.212 \| 311 \| 192 \| 19 \| 9 \| 291 \| 84 \| 369 \| Chromosome partitioning protein ParB \| Chromosome partitioning protein ParB; afdb-uniprot50 \| AF-A0A542ZDY3-F1-MODEL\_V4 \| 1.0 \| 1.073e-10 \| 314 \| 0.204 \| 294 \| 180 \| 14 \| 17 \| 290 \| 53 \| 312 \| ParB family chromosome partitioning protein \| ParB family chromosome partitioning protein; afdb-uniprot50 \| AF-A0A165DRH5-F1-MODEL\_V4 \| 1.0 \| 1.406e-10 \| 314 \| 0.183 \| 295 \| 189 \| 17 \| 19 \| 291 \| 297 \| 561 \| Chromosome partitioning protein ParB \| Chromosome partitioning protein ParB; afdb-uniprot50 \| AF-A0A1C2AW71-F1-MODEL\_V4 \| 1.0 \| 9.064e-09 \| 313 \| 0.19 \| 200 \| 148 \| 7 \| 7 \| 200 \| 15 \| 206 \| Plasmid partitioning protein ParB \| Plasmid partitioning protein ParB; afdb-uniprot50 \| AF-A0A1F4YHQ1-F1-MODEL\_V4 \| 1.0 \| 2.86e-12 \| 313 \| 0.17 \| 294 \| 211 \| 10 \| 4 \| 285 \| 9 \| 281 \| ParB domain-containing protein \| ParB domain-containing protein; afdb-uniprot50 \| AF-A0A2A4X9S6-F1-MODEL\_V4 \| 1.0 \| 4.881e-10 \| 313 \| 0.153 \| 306 \| 215 \| 15 \| 1 \| 282 \| 7 \| 292 \| Probable chromosome-partitioning protein ParB \| Probable chromosome-partitioning protein ParB; afdb-uniprot50 \| AF-F4GVT0-F1-MODEL\_V4 \| 1.0 \| 2.49e-11 \| 313 \| 0.196 \| 300 \| 202 \| 15 \| 4 \| 282 \| 19 \| 300 \| Chromosome partitioning protein \| Chromosome partitioning protein; afdb-uniprot50 \| AF-A0A543INT7-F1-MODEL\_V4 \| 1.0 \| 6.961e-11 \| 313 \| 0.192 \| 311 \| 201 \| 16 \| 1 \| 291 \| 41 \| 321 \| Chromosome segregation DNA-binding protein \| Chromosome segregation DNA-binding protein; afdb-uniprot50 \| AF-A0A2V8TIC9-F1-MODEL\_V4 \| 1.0 \| 1.705e-11 \| 313 \| 0.183 \| 295 \| 204 \| 13 \| 1 \| 283 \| 57 \| 326 \| Chromosome partitioning protein ParB \| Chromosome partitioning protein ParB; afdb-uniprot50 \| AF-A0A4Q3BDE3-F1-MODEL\_V4 \| 1.0 \| 5.404e-08 \| 313 \| 0.134 \| 201 \| 162 \| 7 \| 19 \| 214 \| 12 \| 205 \| ParB/RepB/Spo0J family partition protein \| ParB/RepB/Spo0J family partition protein; afdb-uniprot50 \| AF-A0A1C9T8A9-F1-MODEL\_V4 \| 1.0 \| 4.381e-10 \| 313 \| 0.15 \| 339 \| 210 \| 11 \| 19 \| 291 \| 7 \| 333 \| DNA-binding plasmid partitioning protein \| DNA-binding plasmid partitioning protein; afdb-uniprot50 \| AF-A0A519Z998-F1-MODEL\_V4 \| 1.0 \| 3.814e-09 \| 313 \| 0.204 \| 205 \| 141 \| 8 \| 19 \| 214 \| 61 \| 252 \| ParB/RepB/Spo0J family partition protein \| ParB/RepB/Spo0J family partition protein; afdb-uniprot50 \| AF-A0A1Y0I2T9-F1-MODEL\_V4 \| 1.0 \| 3.637e-11 \| 313 \| 0.208 \| 312 \| 205 \| 15 \| 1 \| 291 \| 74 \| 364 \| Chromosome partitioning protein ParB \| Chromosome partitioning protein ParB; afdb-uniprot50 \| AF-A0A7H2BDL5-F1-MODEL\_V4 \| 1.0 \| 2.054e-10 \| 313 \| 0.166 \| 312 \| 217 \| 17 \| 2 \| 291 \| 144 \| 434 \| ParB/RepB/Spo0J family partition protein \| ParB/RepB/Spo0J family partition protein; afdb-uniprot50 \| AF-A0A6C0GBQ2-F1-MODEL\_V4 \| 1.0 \| 4.736e-09 \| 313 \| 0.197 \| 182 \| 133 \| 7 \| 16 \| 192 \| 1 \| 174 \| ParB/RepB/Spo0J family partition protein \| ParB/RepB/Spo0J family partition protein; afdb-uniprot50 \| AF-A0A1U7GS74-F1-MODEL\_V4 \| 1.0 \| 1.262e-10 \| 312 \| 0.171 \| 285 \| 189 \| 14 \| 10 \| 282 \| 38 \| 287 \| ParB domain-containing protein \| ParB domain-containing protein; afdb-uniprot50 \| AF-A0A194AGK5-F1-MODEL\_V4 \| 1.0 \| 1.9e-11 \| 312 \| 0.178 \| 314 \| 221 \| 13 \| 1 \| 291 \| 5 \| 304 \| Chromosome partitioning protein ParB \| Chromosome partitioning protein ParB; afdb-uniprot50 \| AF-A0A5C7TZB4-F1-MODEL\_V4 \| 1.0 \| 1.843e-10 \| 312 \| 0.151 \| 290 \| 208 \| 13 \| 19 \| 282 \| 55 \| 332 \| ParB/RepB/Spo0J family partition protein \| ParB/RepB/Spo0J family partition protein; afdb-uniprot50 \| AF-A0A2N0J7B4-F1-MODEL\_V4 \| 1.0 \| 7.476e-08 \| 312 \| 0.165 \| 224 \| 149 \| 7 \| 13 \| 205 \| 28 \| 244 \| ParB/RepB/Spo0J family partition protein \| ParB/RepB/Spo0J family partition protein; afdb-uniprot50 \| AF-A0A416VSQ0-F1-MODEL\_V4 \| 1.0 \| 1.735e-08 \| 312 \| 0.202 \| 183 \| 135 \| 6 \| 19 \| 196 \| 4 \| 180 \| ParB/RepB/Spo0J family partition protein \| ParB/RepB/Spo0J family partition protein; afdb-uniprot50 \| AF-A0A661EP20-F1-MODEL\_V4 \| 1.0 \| 2.359e-11 \| 311 \| 0.173 \| 276 \| 197 \| 10 \| 19 \| 282 \| 8 \| 264 \| Probable chromosome-partitioning protein ParB \| Probable chromosome-partitioning protein ParB; afdb-uniprot50 \| AF-A0A1H0EQE3-F1-MODEL\_V4 \| 1.0 \| 6.595e-11 \| 311 \| 0.166 \| 283 \| 210 \| 10 \| 7 \| 284 \| 14 \| 275 \| Chromosome partitioning protein, ParB family \| Chromosome partitioning protein, ParB family; afdb-uniprot50 \| AF-A0A2E9PTD0-F1-MODEL\_V4 \| 1.0 \| 3.445e-11 \| 311 \| 0.202 \| 306 \| 197 \| 14 \| 1 \| 286 \| 4 \| 282 \| Stage 0 sporulation protein J \| Stage 0 sporulation protein J; afdb-uniprot50 \| AF-A0A0G2BIR2-F1-MODEL\_V4 \| 1.0 \| 5.607e-11 \| 311 \| 0.202 \| 296 \| 174 \| 12 \| 12 \| 281 \| 5 \| 264 \| Chromosome partitioning protein gp35 \| Chromosome partitioning protein gp35; afdb-uniprot50 \| AF-A0A1G1YI34-F1-MODEL\_V4 \| 1.0 \| 8.188e-11 \| 311 \| 0.176 \| 289 \| 205 \| 11 \| 4 \| 282 \| 25 \| 290 \| ParB domain-containing protein \| ParB domain-containing protein; afdb-uniprot50 \| AF-A0A5Q4DNZ7-F1-MODEL\_V4 \| 1.0 \| 3.445e-11 \| 311 \| 0.182 \| 296 \| 207 \| 14 \| 5 \| 290 \| 19 \| 289 \| ParB/RepB/Spo0J family partition protein \| ParB/RepB/Spo0J family partition protein; afdb-uniprot50 \| AF-A0A2D7MGH1-F1-MODEL\_V4 \| 1.0 \| 2.359e-11 \| 311 \| 0.169 \| 307 \| 216 \| 13 \| 1 \| 282 \| 1 \| 293 \| DNA-binding protein \| DNA-binding protein; afdb-uniprot50 \| AF-A0A841QGG9-F1-MODEL\_V4 \| 1.0 \| 2.117e-11 \| 311 \| 0.201 \| 298 \| 197 \| 12 \| 7 \| 283 \| 22 \| 299 \| ParB family chromosome partitioning protein \| ParB family chromosome partitioning protein; afdb-uniprot50 \| AF-A0A1F5DSG9-F1-MODEL\_V4 \| 1.0 \| 6.552e-09 \| 311 \| 0.182 \| 208 \| 145 \| 9 \| 18 \| 206 \| 1 \| 202 \| ParB domain-containing protein \| ParB domain-containing protein; afdb-uniprot50 \| AF-A0A1G7MC72-F1-MODEL\_V4 \| 1.0 \| 7.348e-11 \| 311 \| 0.142 \| 364 \| 196 \| 14 \| 19 \| 287 \| 28 \| 370 \| ParB/RepB/Spo0J family partition protein \| ParB/RepB/Spo0J family partition protein; afdb-uniprot50 \| AF-A0A261F4A2-F1-MODEL\_V4 \| 1.0 \| 4.881e-10 \| 311 \| 0.216 \| 277 \| 185 \| 14 \| 19 \| 282 \| 238 \| 495 \| ParB-like partition protein \| ParB-like partition protein; afdb-uniprot50 \| AF-A0A556CDC8-F1-MODEL\_V4 \| 1.0 \| 3.724e-10 \| 311 \| 0.162 \| 301 \| 201 \| 17 \| 13 \| 291 \| 319 \| 590 \| ParB/RepB/Spo0J family partition protein \| ParB/RepB/Spo0J family partition protein; afdb-uniprot50 \| AF-A0A3D9DRE9-F1-MODEL\_V4 \| 1.0 \| 4.486e-09 \| 311 \| 0.183 \| 196 \| 136 \| 8 \| 18 \| 193 \| 1 \| 192 \| ParB family chromosome partitioning protein \| ParB family chromosome partitioning protein; afdb-uniprot50 \| AF-A0A5A9X8I3-F1-MODEL\_V4 \| 1.0 \| 1.605e-09 \| 310 \| 0.185 \| 297 \| 179 \| 14 \| 12 \| 290 \| 6 \| 257 \| Uncharacterized protein \| Uncharacterized protein; afdb-uniprot50 \| AF-A0A7X9L0Y1-F1-MODEL\_V4 \| 1.0 \| 6.247e-11 \| 310 \| 0.164 \| 280 \| 190 \| 14 \| 20 \| 282 \| 25 \| 277 \| ParB/RepB/Spo0J family partition protein \| ParB/RepB/Spo0J family partition protein; afdb-uniprot50 \| AF-A0A4Y6PLY5-F1-MODEL\_V4 \| 1.0 \| 1.023e-12 \| 310 \| 0.216 \| 309 \| 184 \| 14 \| 2 \| 282 \| 15 \| 293 \| ParB/RepB/Spo0J family partition protein \| ParB/RepB/Spo0J family partition protein; afdb-uniprot50 \| AF-A0A7C7RWA4-F1-MODEL\_V4 \| 1.0 \| 4.624e-10 \| 310 \| 0.201 \| 253 \| 163 \| 14 \| 19 \| 244 \| 23 \| 263 \| ParB/RepB/Spo0J family partition protein \| ParB/RepB/Spo0J family partition protein; afdb-uniprot50 \| AF-A0A2D6G3C5-F1-MODEL\_V4 \| 1.0 \| 1.45e-11 \| 310 \| 0.189 \| 312 \| 210 \| 14 \| 1 \| 291 \| 7 \| 296 \| Chromosome partitioning protein ParB \| Chromosome partitioning protein ParB; afdb-uniprot50 \| AF-A0A1V4WRS0-F1-MODEL\_V4 \| 1.0 \| 2.823e-08 \| 310 \| 0.176 \| 170 \| 128 \| 5 \| 14 \| 176 \| 32 \| 196 \| Chromosome-partitioning protein Spo0J \| Chromosome-partitioning protein Spo0J; afdb-uniprot50 \| AF-A0A2N1Q4Q7-F1-MODEL\_V4 \| 1.0 \| 6.207e-09 \| 309 \| 0.212 \| 174 \| 127 \| 7 \| 3 \| 172 \| 10 \| 177 \| ParB domain-containing protein \| ParB domain-containing protein; afdb-uniprot50 \| AF-A0A1F4X477-F1-MODEL\_V4 \| 1.0 \| 1.373e-11 \| 309 \| 0.169 \| 284 \| 197 \| 12 \| 14 \| 282 \| 1 \| 260 \| Chromosome partitioning protein ParB \| Chromosome partitioning protein ParB; afdb-uniprot50 \| AF-A0A2H6HTM0-F1-MODEL\_V4 \| 1.0 \| 5.918e-11 \| 309 \| 0.175 \| 302 \| 207 \| 11 \| 1 \| 282 \| 6 \| 285 \| Chromosome-partitioning protein Spo0J \| Chromosome-partitioning protein Spo0J; afdb-uniprot50 \| AF-A0A227JND4-F1-MODEL\_V4 \| 1.0 \| 1.373e-11 \| 309 \| 0.182 \| 312 \| 199 \| 14 \| 1 \| 282 \| 1 \| 286 \| Chromosome partitioning protein ParB \| Chromosome partitioning protein ParB; afdb-uniprot50 \| AF-A0A2N1V271-F1-MODEL\_V4 \| 1.0 \| 7.348e-11 \| 309 \| 0.149 \| 288 \| 206 \| 11 \| 13 \| 282 \| 28 \| 294 \| ParB domain-containing protein \| ParB domain-containing protein; afdb-uniprot50 \| AF-U7D5A5-F1-MODEL\_V4 \| 1.0 \| 6.961e-11 \| 309 \| 0.158 \| 302 \| 220 \| 10 \| 1 \| 287 \| 10 \| 292 \| ParB-like partition protein \| ParB-like partition protein; afdb-uniprot50 \| AF-A0A1E4BNQ1-F1-MODEL\_V4 \| 1.0 \| 2.692e-10 \| 309 \| 0.201 \| 263 \| 170 \| 15 \| 13 \| 265 \| 3 \| 235 \| ParB domain-containing protein \| ParB domain-containing protein; afdb-uniprot50 \| AF-A0A4R3KGR8-F1-MODEL\_V4 \| 1.0 \| 1.373e-11 \| 309 \| 0.174 \| 309 \| 211 \| 15 \| 4 \| 291 \| 13 \| 298 \| ParB family chromosome partitioning protein \| ParB family chromosome partitioning protein; afdb-uniprot50 \| AF-B8GW30-F1-MODEL\_V4 \| 1.0 \| 8.909e-12 \| 309 \| 0.174 \| 310 \| 207 \| 13 \| 4 \| 289 \| 20 \| 304 \| Chromosome-partitioning protein ParB \| Chromosome-partitioning protein ParB; afdb-uniprot50 \| AF-A0A3C1SR32-F1-MODEL\_V4 \| 1.0 \| 1.017e-10 \| 309 \| 0.164 \| 298 \| 214 \| 13 \| 7 \| 291 \| 32 \| 307 \| Probable chromosome-partitioning protein ParB \| Probable chromosome-partitioning protein ParB; afdb-uniprot50 \| AF-A0A7C1ED27-F1-MODEL\_V4 \| 1.0 \| 2.49e-11 \| 309 \| 0.161 \| 310 \| 208 \| 12 \| 6 \| 282 \| 22 \| 312 \| ParB/RepB/Spo0J family partition protein \| ParB/RepB/Spo0J family partition protein; afdb-uniprot50 \| AF-A0A3D5NI57-F1-MODEL\_V4 \| 1.0 \| 4.486e-09 \| 309 \| 0.158 \| 259 \| 201 \| 9 \| 13 \| 265 \| 23 \| 270 \| ParB domain-containing protein \| ParB domain-containing protein; afdb-uniprot50 \| AF-A0A4V2UP75-F1-MODEL\_V4 \| 1.0 \| 6.961e-11 \| 309 \| 0.188 \| 318 \| 209 \| 15 \| 1 \| 291 \| 7 \| 302 \| ParB/RepB/Spo0J family partition protein \| ParB/RepB/Spo0J family partition protein; afdb-uniprot50 \| AF-A0A5C4X7V6-F1-MODEL\_V4 \| 1.0 \| 1.746e-10 \| 309 \| 0.166 \| 312 \| 210 \| 17 \| 1 \| 291 \| 323 \| 605 \| ParB/RepB/Spo0J family partition protein \| ParB/RepB/Spo0J family partition protein; afdb-uniprot50 \| AF-A0A4Y4D3V1-F1-MODEL\_V4 \| 1.0 \| 3.528e-10 \| 309 \| 0.173 \| 306 \| 196 \| 17 \| 2 \| 282 \| 360 \| 633 \| Chromosome partitioning protein ParB \| Chromosome partitioning protein ParB; afdb-uniprot50 \| AF-A0A352D877-F1-MODEL\_V4 \| 1.0 \| 9.568e-09 \| 308 \| 0.222 \| 180 \| 132 \| 5 \| 7 \| 182 \| 13 \| 188 \| ParB domain-containing protein \| ParB domain-containing protein; afdb-uniprot50 \| AF-Q8DMW1-F1-MODEL\_V4 \| 1.0 \| 1.073e-10 \| 308 \| 0.174 \| 270 \| 196 \| 10 \| 19 \| 282 \| 3 \| 251 \| Chromosome segregation protein \| Chromosome segregation protein; afdb-uniprot50 \| AF-A0A1F5JUW6-F1-MODEL\_V4 \| 1.0 \| 1.017e-10 \| 308 \| 0.202 \| 292 \| 191 \| 12 \| 19 \| 287 \| 4 \| 276 \| ParB domain-containing protein \| ParB domain-containing protein; afdb-uniprot50 \| AF-A0A1F2XIK5-F1-MODEL\_V4 \| 1.0 \| 2.49e-11 \| 308 \| 0.187 \| 304 \| 197 \| 17 \| 1 \| 282 \| 5 \| 280 \| ParB domain-containing protein \| ParB domain-containing protein; afdb-uniprot50 \| AF-A0A2D2D129-F1-MODEL\_V4 \| 1.0 \| 8.188e-11 \| 308 \| 0.191 \| 319 \| 197 \| 17 \| 1 \| 288 \| 1 \| 289 \| Chromosome partitioning protein ParB \| Chromosome partitioning protein ParB; afdb-uniprot50 \| AF-F7SA52-F1-MODEL\_V4 \| 1.0 \| 5.607e-11 \| 308 \| 0.23 \| 299 \| 191 \| 14 \| 4 \| 289 \| 20 \| 292 \| ParB-like partition protein \| ParB-like partition protein; afdb-uniprot50 \| AF-A0A7W0M8C8-F1-MODEL\_V4 \| 1.0 \| 3.166e-10 \| 308 \| 0.158 \| 297 \| 205 \| 13 \| 1 \| 287 \| 30 \| 291 \| ParB/RepB/Spo0J family partition protein \| ParB/RepB/Spo0J family partition protein; afdb-uniprot50 \| AF-A0A3C0V9C7-F1-MODEL\_V4 \| 1.0 \| 1.332e-10 \| 308 \| 0.191 \| 297 \| 203 \| 14 \| 9 \| 291 \| 52 \| 325 \| HTH cro/C1-type domain-containing protein \| HTH cro/C1-type domain-containing protein; afdb-uniprot50 \| AF-A0A7X7P058-F1-MODEL\_V4 \| 1.0 \| 2.49e-11 \| 308 \| 0.21 \| 294 \| 192 \| 15 \| 13 \| 282 \| 47 \| 324 \| ParB/RepB/Spo0J family partition protein \| ParB/RepB/Spo0J family partition protein; afdb-uniprot50 \| AF-A0A512D7U3-F1-MODEL\_V4 \| 1.0 \| 5.918e-11 \| 308 \| 0.211 \| 298 \| 196 \| 14 \| 14 \| 291 \| 56 \| 334 \| Chromosome partitioning protein ParB \| Chromosome partitioning protein ParB; afdb-uniprot50 \| AF-A0A7X5KQ06-F1-MODEL\_V4 \| 1.0 \| 3.146e-08 \| 308 \| 0.191 \| 198 \| 146 \| 9 \| 5 \| 196 \| 19 \| 208 \| ParB/RepB/Spo0J family partition protein \| ParB/RepB/Spo0J family partition protein; afdb-uniprot50 \| AF-A0A286GTP8-F1-MODEL\_V4 \| 1.0 \| 6.961e-11 \| 308 \| 0.18 \| 310 \| 208 \| 15 \| 2 \| 291 \| 34 \| 317 \| Chromosome partitioning protein, ParB family \| Chromosome partitioning protein, ParB family; afdb-uniprot50 \| AF-A0A540WI34-F1-MODEL\_V4 \| 1.0 \| 1.644e-08 \| 308 \| 0.182 \| 197 \| 138 \| 8 \| 13 \| 193 \| 5 \| 194 \| ParB/RepB/Spo0J family partition protein \| ParB/RepB/Spo0J family partition protein; afdb-uniprot50 \| AF-A0A7V3ZSZ4-F1-MODEL\_V4 \| 1.0 \| 2.168e-10 \| 307 \| 0.177 \| 299 \| 202 \| 13 \| 1 \| 291 \| 2 \| 264 \| ParB/RepB/Spo0J family partition protein \| ParB/RepB/Spo0J family partition protein; afdb-uniprot50 \| AF-O74066-F1-MODEL\_V4 \| 1.0 \| 1.788e-09 \| 307 \| 0.21 \| 290 \| 176 \| 15 \| 13 \| 289 \| 3 \| 252 \| ParB domain-containing protein \| ParB domain-containing protein; afdb-uniprot50 \| AF-A0A7U4ZBY4-F1-MODEL\_V4 \| 1.0 \| 1.654e-10 \| 307 \| 0.202 \| 282 \| 191 \| 14 \| 15 \| 283 \| 16 \| 276 \| Chromosome-partitioning protein ParB \| Chromosome-partitioning protein ParB; afdb-uniprot50 \| AF-S4NKJ4-F1-MODEL\_V4 \| 1.0 \| 5.439e-10 \| 307 \| 0.187 \| 283 \| 179 \| 13 \| 1 \| 246 \| 1 \| 269 \| ParB-like partition protein \| ParB-like partition protein; afdb-uniprot50 \| AF-A0A4Y8I9C7-F1-MODEL\_V4 \| 1.0 \| 1.017e-10 \| 307 \| 0.186 \| 295 \| 209 \| 10 \| 1 \| 282 \| 7 \| 283 \| ParB/RepB/Spo0J family partition protein \| ParB/RepB/Spo0J family partition protein; afdb-uniprot50 \| AF-A0A7X9AHL4-F1-MODEL\_V4 \| 1.0 \| 2.49e-11 \| 307 \| 0.18 \| 305 \| 201 \| 14 \| 3 \| 282 \| 9 \| 289 \| ParB/RepB/Spo0J family partition protein \| ParB/RepB/Spo0J family partition protein; afdb-uniprot50 \| AF-A0A258L6L1-F1-MODEL\_V4 \| 1.0 \| 2.117e-11 \| 307 \| 0.189 \| 296 \| 198 \| 15 \| 13 \| 291 \| 26 \| 296 \| Chromosome partitioning protein ParB \| Chromosome partitioning protein ParB; afdb-uniprot50 \| AF-A0A6M8HVA6-F1-MODEL\_V4 \| 1.0 \| 8.188e-11 \| 307 \| 0.182 \| 307 \| 210 \| 13 \| 1 \| 282 \| 6 \| 296 \| ParB/RepB/Spo0J family partition protein \| ParB/RepB/Spo0J family partition protein; afdb-uniprot50 \| AF-H6SS19-F1-MODEL\_V4 \| 1.0 \| 8.188e-11 \| 307 \| 0.193 \| 300 \| 199 \| 16 \| 7 \| 291 \| 73 \| 344 \| Chromosome segregation DNA-binding protein \| Chromosome segregation DNA-binding protein; afdb-uniprot50 \| AF-A0A7D7PZW5-F1-MODEL\_V4 \| 1.0 \| 5.742e-10 \| 307 \| 0.176 \| 295 \| 190 \| 17 \| 14 \| 282 \| 92 \| 359 \| Putative chromosome-partitioning protein ParB \| Putative chromosome-partitioning protein ParB; afdb-uniprot50 \| AF-A0A2D5W999-F1-MODEL\_V4 \| 1.0 \| 7.757e-11 \| 307 \| 0.184 \| 298 \| 211 \| 13 \| 1 \| 283 \| 134 \| 414 \| Chromosome partitioning protein ParB \| Chromosome partitioning protein ParB; afdb-uniprot50 \| AF-A0A239V883-F1-MODEL\_V4 \| 1.0 \| 2.842e-10 \| 307 \| 0.176 \| 289 \| 197 \| 14 \| 14 \| 282 \| 151 \| 418 \| Probable chromosome-partitioning protein parB \| Probable chromosome-partitioning protein parB; afdb-uniprot50 \| AF-Q4JSC7-F1-MODEL\_V4 \| 1.0 \| 7.129e-10 \| 307 \| 0.163 \| 287 \| 195 \| 13 \| 19 \| 282 \| 239 \| 503 \| Chromosome partitioning protein ParB \| Chromosome partitioning protein ParB; afdb-uniprot50 \| AF-A0A517SER9-F1-MODEL\_V4 \| 1.0 \| 4.999e-09 \| 307 \| 0.183 \| 218 \| 160 \| 9 \| 5 \| 218 \| 121 \| 324 \| Chromosome-partitioning protein Spo0J \| Chromosome-partitioning protein Spo0J; afdb-uniprot50 \| AF-A0A382NUU8-F1-MODEL\_V4 \| 1.0 \| 1.933e-08 \| 306 \| 0.206 \| 199 \| 144 \| 8 \| 7 \| 199 \| 13 \| 203 \| ParB domain-containing protein \| ParB domain-containing protein; afdb-uniprot50 \| AF-A0A1F7BL00-F1-MODEL\_V4 \| 1.0 \| 2.823e-08 \| 306 \| 0.179 \| 184 \| 129 \| 9 \| 23 \| 205 \| 2 \| 164 \| ParB domain-containing protein \| ParB domain-containing protein; afdb-uniprot50 \| AF-A0A838NRV8-F1-MODEL\_V4 \| 1.0 \| 1.53e-11 \| 306 \| 0.17 \| 288 \| 206 \| 14 \| 13 \| 282 \| 25 \| 297 \| ParB/RepB/Spo0J family partition protein \| ParB/RepB/Spo0J family partition protein; afdb-uniprot50 \| AF-A0A497Y0S2-F1-MODEL\_V4 \| 1.0 \| 3.724e-10 \| 306 \| 0.172 \| 301 \| 211 \| 14 \| 1 \| 282 \| 39 \| 320 \| ParB family chromosome partitioning protein \| ParB family chromosome partitioning protein; afdb-uniprot50 \| AF-A0A2E2LB55-F1-MODEL\_V4 \| 1.0 \| 1.735e-08 \| 306 \| 0.201 \| 203 \| 143 \| 11 \| 5 \| 201 \| 70 \| 259 \| Chromosome partitioning protein \| Chromosome partitioning protein; afdb-uniprot50 \| AF-A1VV97-F1-MODEL\_V4 \| 1.0 \| 4.594e-08 \| 306 \| 0.158 \| 164 \| 131 \| 6 \| 13 \| 172 \| 5 \| 165 \| ParB family protein \| ParB family protein; afdb-uniprot50 \| AF-A0A2E0DRV3-F1-MODEL\_V4 \| 1.0 \| 1.01e-08 \| 305 \| 0.206 \| 174 \| 128 \| 5 \| 14 \| 182 \| 27 \| 195 \| Chromosome partitioning protein ParB \| Chromosome partitioning protein ParB; afdb-uniprot50 \| AF-A0A3A4PV03-F1-MODEL\_V4 \| 1.0 \| 1.843e-10 \| 305 \| 0.173 \| 294 \| 217 \| 11 \| 1 \| 289 \| 7 \| 279 \| ParB/RepB/Spo0J family partition protein \| ParB/RepB/Spo0J family partition protein; afdb-uniprot50 \| AF-A0A7V6VL57-F1-MODEL\_V4 \| 1.0 \| 9.568e-09 \| 305 \| 0.235 \| 195 \| 132 \| 10 \| 12 \| 195 \| 94 \| 282 \| ParB/RepB/Spo0J family partition protein \| ParB/RepB/Spo0J family partition protein; afdb-uniprot50 \| AF-A0A7W1G0B4-F1-MODEL\_V4 \| 1.0 \| 9.124e-11 \| 305 \| 0.185 \| 291 \| 194 \| 13 \| 13 \| 282 \| 17 \| 285 \| ParB/RepB/Spo0J family partition protein \| ParB/RepB/Spo0J family partition protein; afdb-uniprot50 \| AF-A0A1V5DMX0-F1-MODEL\_V4 \| 1.0 \| 3.613e-09 \| 305 \| 0.187 \| 203 \| 141 \| 8 \| 13 \| 206 \| 2 \| 189 \| Chromosome-partitioning protein Spo0J \| Chromosome-partitioning protein Spo0J; afdb-uniprot50 \| AF-A0A1F6CPK3-F1-MODEL\_V4 \| 1.0 \| 2.168e-10 \| 305 \| 0.176 \| 283 \| 196 \| 14 \| 19 \| 286 \| 9 \| 269 \| ParB domain-containing protein \| ParB domain-containing protein; afdb-uniprot50 \| AF-A0A7U7GA97-F1-MODEL\_V4 \| 1.0 \| 7.348e-11 \| 305 \| 0.163 \| 311 \| 210 \| 15 \| 2 \| 287 \| 21 \| 306 \| Probable chromosome-partitioning protein ParB \| Probable chromosome-partitioning protein ParB; afdb-uniprot50 \| AF-A0A261QDL4-F1-MODEL\_V4 \| 1.0 \| 3.321e-08 \| 305 \| 0.164 \| 188 \| 145 \| 8 \| 19 \| 200 \| 5 \| 186 \| ParB domain-containing protein \| ParB domain-containing protein; afdb-uniprot50 \| AF-A0A2D6BI86-F1-MODEL\_V4 \| 1.0 \| 3.839e-11 \| 305 \| 0.14 \| 298 \| 212 \| 12 \| 13 \| 291 \| 53 \| 325 \| Chromosome partitioning protein ParB \| Chromosome partitioning protein ParB; afdb-uniprot50 \| AF-A0A495QZL2-F1-MODEL\_V4 \| 1.0 \| 1.406e-10 \| 305 \| 0.179 \| 296 \| 194 \| 14 \| 19 \| 291 \| 57 \| 326 \| ParB family chromosome partitioning protein \| ParB family chromosome partitioning protein; afdb-uniprot50 \| AF-A0A7X3NCE7-F1-MODEL\_V4 \| 1.0 \| 3.724e-10 \| 305 \| 0.179 \| 351 \| 204 \| 13 \| 22 \| 291 \| 38 \| 385 \| Uncharacterized protein \| Uncharacterized protein; afdb-uniprot50 \| AF-A0A448FNI8-F1-MODEL\_V4 \| 1.0 \| 2.98e-08 \| 304 \| 0.187 \| 155 \| 117 \| 7 \| 5 \| 156 \| 11 \| 159 \| ParB-like partition protein \| ParB-like partition protein; afdb-uniprot50 \| AF-A0A2Z3HZR4-F1-MODEL\_V4 \| 1.0 \| 1.233e-11 \| 304 \| 0.192 \| 312 \| 191 \| 14 \| 1 \| 282 \| 4 \| 284 \| Chromosome partitioning protein ParB \| Chromosome partitioning protein ParB; afdb-uniprot50 \| AF-A0A496NGH7-F1-MODEL\_V4 \| 1.0 \| 1.746e-10 \| 304 \| 0.158 \| 291 \| 206 \| 13 \| 14 \| 282 \| 48 \| 321 \| Probable chromosome-partitioning protein ParB \| Probable chromosome-partitioning protein ParB; afdb-uniprot50 \| AF-A0A4P6F2X0-F1-MODEL\_V4 \| 1.0 \| 3.445e-11 \| 304 \| 0.19 \| 320 \| 199 \| 17 \| 1 \| 291 \| 60 \| 348 \| ParB/RepB/Spo0J family partition protein \| ParB/RepB/Spo0J family partition protein; afdb-uniprot50 \| AF-A0A6N9H888-F1-MODEL\_V4 \| 1.0 \| 5.439e-10 \| 304 \| 0.167 \| 299 \| 201 \| 15 \| 14 \| 291 \| 172 \| 443 \| ParB/RepB/Spo0J family partition protein \| ParB/RepB/Spo0J family partition protein; afdb-uniprot50 \| AF-E6JYG5-F1-MODEL\_V4 \| 1.0 \| 5.439e-10 \| 304 \| 0.189 \| 291 \| 195 \| 14 \| 14 \| 285 \| 248 \| 516 \| ParB-like protein \| ParB-like protein; afdb-uniprot50 \| AF-A0A1I3C721-F1-MODEL\_V4 \| 1.0 \| 7.757e-11 \| 303 \| 0.172 \| 278 \| 207 \| 11 \| 13 \| 282 \| 29 \| 291 \| Chromosome partitioning protein, ParB family \| Chromosome partitioning protein, ParB family; afdb-uniprot50 \| AF-A0A1G2HXV3-F1-MODEL\_V4 \| 1.0 \| 3.505e-08 \| 303 \| 0.204 \| 161 \| 116 \| 6 \| 14 \| 169 \| 19 \| 172 \| ParB domain-containing protein \| ParB domain-containing protein; afdb-uniprot50 \| AF-A0A7X6H7L1-F1-MODEL\_V4 \| 1.0 \| 6.398e-10 \| 303 \| 0.177 \| 288 \| 187 \| 17 \| 19 \| 282 \| 87 \| 348 \| ParB/RepB/Spo0J family partition protein \| ParB/RepB/Spo0J family partition protein; afdb-uniprot50 \| AF-A0A2J9SJJ5-F1-MODEL\_V4 \| 1.0 \| 2.612e-09 \| 303 \| 0.162 \| 271 \| 182 \| 11 \| 11 \| 272 \| 23 \| 257 \| ParB domain-containing protein \| ParB domain-containing protein; afdb-uniprot50 \| AF-A0A4P6EPV7-F1-MODEL\_V4 \| 1.0 \| 2.054e-10 \| 303 \| 0.194 \| 313 \| 197 \| 15 \| 1 \| 282 \| 3 \| 291 \| ParB/RepB/Spo0J family partition protein \| ParB/RepB/Spo0J family partition protein; afdb-uniprot50 \| AF-A0A139DP90-F1-MODEL\_V4 \| 1.0 \| 1.831e-08 \| 303 \| 0.168 \| 196 \| 153 \| 6 \| 10 \| 200 \| 6 \| 196 \| Probable chromosome-partitioning protein ParB \| Probable chromosome-partitioning protein ParB; afdb-uniprot50 \| AF-A0A1F8UGV0-F1-MODEL\_V4 \| 1.0 \| 4.278e-11 \| 302 \| 0.211 \| 303 \| 198 \| 13 \| 2 \| 288 \| 4 \| 281 \| Chromosome partitioning protein ParB \| Chromosome partitioning protein ParB; afdb-uniprot50 \| AF-A0A1C5TMF3-F1-MODEL\_V4 \| 1.0 \| 4.516e-11 \| 302 \| 0.186 \| 284 \| 205 \| 12 \| 7 \| 282 \| 32 \| 297 \| Probable chromosome-partitioning protein parB \| Probable chromosome-partitioning protein parB; afdb-uniprot50 \| AF-A0A0S2KEG7-F1-MODEL\_V4 \| 1.0 \| 1.293e-09 \| 302 \| 0.169 \| 266 \| 182 \| 15 \| 12 \| 263 \| 9 \| 249 \| ParB-like partitioning protein \| ParB-like partitioning protein; afdb-uniprot50 \| AF-A0A7L9BNL5-F1-MODEL\_V4 \| 1.0 \| 3.343e-10 \| 302 \| 0.165 \| 296 \| 206 \| 12 \| 11 \| 282 \| 35 \| 313 \| ParB/RepB/Spo0J family partition protein \| ParB/RepB/Spo0J family partition protein; afdb-uniprot50 \| AF-A0A520Q7U8-F1-MODEL\_V4 \| 1.0 \| 2.006e-11 \| 302 \| 0.163 \| 300 \| 197 \| 13 \| 13 \| 282 \| 63 \| 338 \| ParB/RepB/Spo0J family partition protein \| ParB/RepB/Spo0J family partition protein; afdb-uniprot50 \| AF-A0A0K2RK52-F1-MODEL\_V4 \| 1.0 \| 3.343e-10 \| 302 \| 0.169 \| 313 \| 196 \| 20 \| 10 \| 291 \| 66 \| 345 \| Probable chromosome-partitioning protein ParB \| Probable chromosome-partitioning protein ParB; afdb-uniprot50 \| AF-A0A523V2R2-F1-MODEL\_V4 \| 1.0 \| 9.343e-10 \| 302 \| 0.149 \| 321 \| 205 \| 17 \| 13 \| 291 \| 4 \| 298 \| ParB/RepB/Spo0J family partition protein \| ParB/RepB/Spo0J family partition protein; afdb-uniprot50 \| AF-I8ARF6-F1-MODEL\_V4 \| 1.0 \| 1.843e-10 \| 302 \| 0.178 \| 292 \| 196 \| 15 \| 19 \| 291 \| 96 \| 362 \| ParB-like protein \| ParB-like protein; afdb-uniprot50 \| AF-A0A2G4ISH6-F1-MODEL\_V4 \| 1.0 \| 2.359e-11 \| 302 \| 0.19 \| 310 \| 207 \| 12 \| 1 \| 290 \| 8 \| 293 \| Chromosome partitioning protein ParB \| Chromosome partitioning protein ParB; afdb-uniprot50 \| AF-A0A4Y1XBM5-F1-MODEL\_V4 \| 1.0 \| 5.404e-08 \| 302 \| 0.169 \| 189 \| 147 \| 7 \| 1 \| 184 \| 6 \| 189 \| Chromosome partitioning protein ParB \| Chromosome partitioning protein ParB; afdb-uniprot50 \| AF-M6VZ53-F1-MODEL\_V4 \| 1.0 \| 1.843e-10 \| 302 \| 0.158 \| 334 \| 185 \| 16 \| 13 \| 283 \| 351 \| 651 \| ParB-like protein \| ParB-like protein; afdb-uniprot50 \| AF-A0A150PXB6-F1-MODEL\_V4 \| 1.0 \| 6.398e-10 \| 301 \| 0.243 \| 242 \| 152 \| 10 \| 13 \| 230 \| 29 \| 263 \| Chromosome partitioning protein ParB \| Chromosome partitioning protein ParB; afdb-uniprot50 \| AF-A0A7C7VJH1-F1-MODEL\_V4 \| 1.0 \| 8.643e-11 \| 301 \| 0.183 \| 294 \| 208 \| 13 \| 1 \| 282 \| 8 \| 281 \| ParB/RepB/Spo0J family partition protein \| ParB/RepB/Spo0J family partition protein; afdb-uniprot50 \| AF-A0A2V8FIU9-F1-MODEL\_V4 \| 1.0 \| 2.235e-11 \| 301 \| 0.176 \| 311 \| 197 \| 13 \| 1 \| 282 \| 4 \| 284 \| Chromosome partitioning protein ParB \| Chromosome partitioning protein ParB; afdb-uniprot50 \| AF-A0A7C2JKK4-F1-MODEL\_V4 \| 1.0 \| 5.311e-11 \| 301 \| 0.197 \| 289 \| 183 \| 15 \| 15 \| 282 \| 28 \| 288 \| ParB/RepB/Spo0J family partition protein \| ParB/RepB/Spo0J family partition protein; afdb-uniprot50 \| AF-A0A7T9F7V5-F1-MODEL\_V4 \| 1.0 \| 2.55e-10 \| 301 \| 0.149 \| 295 \| 216 \| 11 \| 13 \| 291 \| 104 \| 379 \| ParB/RepB/Spo0J family partition protein \| ParB/RepB/Spo0J family partition protein; afdb-uniprot50 \| AF-A0A350AES6-F1-MODEL\_V4 \| 1.0 \| 3.505e-08 \| 300 \| 0.185 \| 156 \| 117 \| 5 \| 13 \| 163 \| 11 \| 161 \| Chromosome partitioning protein ParB \| Chromosome partitioning protein ParB; afdb-uniprot50 \| AF-A0A523CVJ9-F1-MODEL\_V4 \| 1.0 \| 1.654e-10 \| 300 \| 0.163 \| 306 \| 210 \| 13 \| 1 \| 291 \| 5 \| 279 \| ParB/RepB/Spo0J family partition protein \| ParB/RepB/Spo0J family partition protein; afdb-uniprot50 \| AF-A0A7R9RA11-F1-MODEL\_V4 \| 1.0 \| 1.017e-10 \| 300 \| 0.187 \| 294 \| 204 \| 12 \| 1 \| 282 \| 3 \| 273 \| Stage 0 sporulation protein J \| Stage 0 sporulation protein J; afdb-uniprot50 \| AF-A0A2H1J8P3-F1-MODEL\_V4 \| 1.0 \| 3.166e-10 \| 300 \| 0.174 \| 315 \| 202 \| 18 \| 2 \| 291 \| 38 \| 319 \| Chromosome partitioning protein, ParB family \| Chromosome partitioning protein, ParB family; afdb-uniprot50 \| AF-A0A6C1SLD8-F1-MODEL\_V4 \| 1.0 \| 7.348e-11 \| 300 \| 0.182 \| 291 \| 186 \| 12 \| 20 \| 291 \| 35 \| 292 \| ParB/RepB/Spo0J family partition protein \| ParB/RepB/Spo0J family partition protein; afdb-uniprot50 \| AF-A0A7X6L5B1-F1-MODEL\_V4 \| 1.0 \| 5.153e-10 \| 300 \| 0.17 \| 316 \| 209 \| 15 \| 1 \| 291 \| 49 \| 336 \| ParB/RepB/Spo0J family partition protein \| ParB/RepB/Spo0J family partition protein; afdb-uniprot50 \| AF-A0A845MLR6-F1-MODEL\_V4 \| 1.0 \| 2.929e-11 \| 300 \| 0.203 \| 300 \| 194 \| 17 \| 7 \| 282 \| 22 \| 300 \| ParB/RepB/Spo0J family partition protein \| ParB/RepB/Spo0J family partition protein; afdb-uniprot50 \| AF-A0A838LF52-F1-MODEL\_V4 \| 1.0 \| 1.567e-10 \| 299 \| 0.19 \| 288 \| 180 \| 16 \| 13 \| 291 \| 1 \| 244 \| ParB/RepB/Spo0J family partition protein \| ParB/RepB/Spo0J family partition protein; afdb-uniprot50 \| AF-A0A520JRG8-F1-MODEL\_V4 \| 1.0 \| 1.644e-08 \| 299 \| 0.212 \| 188 \| 137 \| 6 \| 18 \| 201 \| 1 \| 181 \| Chromosome partitioning protein ParB \| Chromosome partitioning protein ParB; afdb-uniprot50 \| AF-A0A4R8DHE2-F1-MODEL\_V4 \| 1.0 \| 6.753e-10 \| 299 \| 0.146 \| 308 \| 194 \| 17 \| 13 \| 287 \| 28 \| 299 \| ParB/RepB/Spo0J family partition protein \| ParB/RepB/Spo0J family partition protein; afdb-uniprot50 \| AF-A0A382X411-F1-MODEL\_V4 \| 1.0 \| 3.613e-09 \| 298 \| 0.19 \| 199 \| 144 \| 8 \| 15 \| 205 \| 2 \| 191 \| HTH cro/C1-type domain-containing protein \| HTH cro/C1-type domain-containing protein; afdb-uniprot50 \| AF-A0A357ZK67-F1-MODEL\_V4 \| 1.0 \| 2.474e-09 \| 298 \| 0.227 \| 211 \| 141 \| 8 \| 7 \| 200 \| 33 \| 238 \| ParB domain-containing protein \| ParB domain-containing protein; afdb-uniprot50 \| AF-A0A1Y1RFK6-F1-MODEL\_V4 \| 1.0 \| 1.406e-10 \| 298 \| 0.172 \| 301 \| 217 \| 11 \| 1 \| 290 \| 13 \| 292 \| ParB domain-containing protein \| ParB domain-containing protein; afdb-uniprot50 \| AF-A0A1M3IXE2-F1-MODEL\_V4 \| 1.0 \| 1.843e-10 \| 298 \| 0.196 \| 295 \| 195 \| 15 \| 14 \| 290 \| 27 \| 297 \| ParB domain-containing protein \| ParB domain-containing protein; afdb-uniprot50 \| AF-A0A1I0F2H7-F1-MODEL\_V4 \| 1.0 \| 4.053e-11 \| 298 \| 0.182 \| 301 \| 213 \| 11 \| 5 \| 290 \| 22 \| 304 \| Chromosome partitioning protein, ParB family \| Chromosome partitioning protein, ParB family; afdb-uniprot50 \| AF-A0A1F5AZ83-F1-MODEL\_V4 \| 1.0 \| 1.153e-07 \| 298 \| 0.157 \| 197 \| 151 \| 10 \| 18 \| 206 \| 24 \| 213 \| ParB domain-containing protein \| ParB domain-containing protein; afdb-uniprot50 \| AF-C6HTU9-F1-MODEL\_V4 \| 1.0 \| 5.439e-10 \| 298 \| 0.194 \| 262 \| 175 \| 12 \| 11 \| 265 \| 21 \| 253 \| ParB-like partition protein \| ParB-like partition protein; afdb-uniprot50 \| AF-A0A512H3L9-F1-MODEL\_V4 \| 1.0 \| 1.946e-10 \| 298 \| 0.193 \| 294 \| 207 \| 13 \| 7 \| 291 \| 49 \| 321 \| Chromosome partitioning protein ParB \| Chromosome partitioning protein ParB; afdb-uniprot50 \| AF-A0A7K1A1N3-F1-MODEL\_V4 \| 1.0 \| 2.55e-10 \| 298 \| 0.201 \| 248 \| 143 \| 12 \| 13 \| 247 \| 1 \| 206 \| ParB/RepB/Spo0J family partition protein \| ParB/RepB/Spo0J family partition protein; afdb-uniprot50 \| AF-A0A1G2VKP5-F1-MODEL\_V4 \| 1.0 \| 6.753e-10 \| 297 \| 0.209 \| 196 \| 139 \| 7 \| 14 \| 200 \| 5 \| 193 \| ParB domain-containing protein \| ParB domain-containing protein; afdb-uniprot50 \| AF-A0A1X7KAE4-F1-MODEL\_V4 \| 1.0 \| 1.843e-10 \| 297 \| 0.189 \| 306 \| 201 \| 13 \| 1 \| 290 \| 4 \| 278 \| Chromosome partitioning protein, ParB family \| Chromosome partitioning protein, ParB family; afdb-uniprot50 \| AF-A0A3D1U3G5-F1-MODEL\_V4 \| 1.0 \| 4.15e-10 \| 297 \| 0.198 \| 287 \| 199 \| 12 \| 1 \| 277 \| 12 \| 277 \| Stage 0 sporulation protein J \| Stage 0 sporulation protein J; afdb-uniprot50 \| AF-E4TJE6-F1-MODEL\_V4 \| 1.0 \| 1.196e-10 \| 297 \| 0.183 \| 294 \| 193 \| 13 \| 7 \| 282 \| 11 \| 275 \| ParB-like partition protein \| ParB-like partition protein; afdb-uniprot50 \| AF-A0A2G6LB17-F1-MODEL\_V4 \| 1.0 \| 6.247e-11 \| 297 \| 0.181 \| 297 \| 197 \| 15 \| 7 \| 282 \| 10 \| 281 \| Probable chromosome-partitioning protein ParB \| Probable chromosome-partitioning protein ParB; afdb-uniprot50 \| AF-A0A0L0M783-F1-MODEL\_V4 \| 1.0 \| 9.283e-08 \| 297 \| 0.218 \| 183 \| 136 \| 5 \| 22 \| 201 \| 6 \| 184 \| Chromosome (Plasmid) partitioning protein ParB \| Chromosome (Plasmid) partitioning protein ParB; afdb-uniprot50 \| AF-A0A2S6TU51-F1-MODEL\_V4 \| 1.0 \| 3.343e-10 \| 297 \| 0.152 \| 315 \| 215 \| 15 \| 1 \| 291 \| 9 \| 295 \| Chromosome-partitioning protein ParB \| Chromosome-partitioning protein ParB; afdb-uniprot50 \| AF-A0A2W5A523-F1-MODEL\_V4 \| 1.0 \| 1.53e-11 \| 297 \| 0.178 \| 330 \| 203 \| 17 \| 1 \| 291 \| 4 \| 304 \| Chromosome partitioning protein ParB \| Chromosome partitioning protein ParB; afdb-uniprot50 \| AF-A0A318B644-F1-MODEL\_V4 \| 1.0 \| 1.993e-09 \| 296 \| 0.2 \| 215 \| 150 \| 8 \| 1 \| 201 \| 7 \| 213 \| Chromosome partitioning protein ParB \| Chromosome partitioning protein ParB; afdb-uniprot50 \| AF-A0A2N1PW21-F1-MODEL\_V4 \| 1.0 \| 3.423e-09 \| 296 \| 0.21 \| 204 \| 149 \| 7 \| 3 \| 200 \| 9 \| 206 \| Chromosome partitioning protein ParB \| Chromosome partitioning protein ParB; afdb-uniprot50 \| AF-A0A1N6LGW4-F1-MODEL\_V4 \| 1.0 \| 3.166e-10 \| 296 \| 0.162 \| 302 \| 190 \| 12 \| 3 \| 282 \| 29 \| 289 \| Chromosome partitioning protein, ParB family \| Chromosome partitioning protein, ParB family; afdb-uniprot50 \| AF-A0A2S6S2D6-F1-MODEL\_V4 \| 1.0 \| 2.692e-10 \| 296 \| 0.18 \| 315 \| 208 \| 15 \| 2 \| 290 \| 8 \| 298 \| Chromosome-partitioning protein ParB \| Chromosome-partitioning protein ParB; afdb-uniprot50 \| AF-A0A1Z9K2S5-F1-MODEL\_V4 \| 1.0 \| 1.196e-10 \| 296 \| 0.177 \| 293 \| 190 \| 14 \| 13 \| 282 \| 36 \| 300 \| Probable chromosome-partitioning protein ParB \| Probable chromosome-partitioning protein ParB; afdb-uniprot50 \| AF-A0A3M0Y0V7-F1-MODEL\_V4 \| 1.0 \| 1.017e-10 \| 296 \| 0.192 \| 301 \| 199 \| 13 \| 13 \| 291 \| 34 \| 312 \| ParB/RepB/Spo0J family partition protein \| ParB/RepB/Spo0J family partition protein; afdb-uniprot50 \| AF-A0A6N7YJS6-F1-MODEL\_V4 \| 1.0 \| 9.863e-10 \| 296 \| 0.195 \| 323 \| 192 \| 18 \| 13 \| 287 \| 7 \| 309 \| ParB/RepB/Spo0J family partition protein \| ParB/RepB/Spo0J family partition protein; afdb-uniprot50 \| AF-A0A095YDD6-F1-MODEL\_V4 \| 1.0 \| 5.742e-10 \| 296 \| 0.162 \| 314 \| 218 \| 17 \| 1 \| 291 \| 223 \| 514 \| ParB domain-containing protein \| ParB domain-containing protein; afdb-uniprot50 \| AF-A0A1V9G4E3-F1-MODEL\_V4 \| 1.0 \| 5.404e-08 \| 296 \| 0.141 \| 184 \| 147 \| 6 \| 5 \| 182 \| 9 \| 187 \| ParB domain-containing protein \| ParB domain-containing protein; afdb-uniprot50 \| AF-A0A2D8FML5-F1-MODEL\_V4 \| 1.0 \| 1.073e-10 \| 295 \| 0.225 \| 279 \| 172 \| 16 \| 23 \| 282 \| 1 \| 254 \| Chromosome partitioning protein ParB \| Chromosome partitioning protein ParB; afdb-uniprot50 \| AF-A0A6P4HEC8-F1-MODEL\_V4 \| 1.0 \| 1.993e-09 \| 295 \| 0.218 \| 252 \| 161 \| 12 \| 13 \| 246 \| 33 \| 266 \| chromosome-partitioning protein ParB-like \| chromosome-partitioning protein ParB-like; afdb-uniprot50 \| AF-A0A660TB83-F1-MODEL\_V4 \| 1.0 \| 7.995e-12 \| 295 \| 0.164 \| 291 \| 214 \| 11 \| 4 \| 282 \| 13 \| 286 \| Chromosome partitioning protein ParB \| Chromosome partitioning protein ParB; afdb-uniprot50 \| AF-A0A2E6RNQ1-F1-MODEL\_V4 \| 1.0 \| 4.767e-11 \| 295 \| 0.154 \| 298 \| 210 \| 14 \| 4 \| 291 \| 26 \| 291 \| Chromosome partitioning protein ParB \| Chromosome partitioning protein ParB; afdb-uniprot50 \| AF-A0A3D0RBB3-F1-MODEL\_V4 \| 1.0 \| 4.881e-10 \| 295 \| 0.172 \| 279 \| 195 \| 14 \| 20 \| 282 \| 38 \| 296 \| Chromosome partitioning protein ParB \| Chromosome partitioning protein ParB; afdb-uniprot50 \| AF-A0A7T8SHN7-F1-MODEL\_V4 \| 1.0 \| 2.416e-10 \| 295 \| 0.209 \| 311 \| 187 \| 14 \| 4 \| 282 \| 54 \| 337 \| ParB/RepB/Spo0J family partition protein \| ParB/RepB/Spo0J family partition protein; afdb-uniprot50 \| AF-A0A7C2KI98-F1-MODEL\_V4 \| 1.0 \| 1.511e-07 \| 295 \| 0.172 \| 174 \| 136 \| 6 \| 4 \| 172 \| 24 \| 194 \| ParB/RepB/Spo0J family partition protein \| ParB/RepB/Spo0J family partition protein; afdb-uniprot50 \| AF-A0A2E5HJM8-F1-MODEL\_V4 \| 1.0 \| 2.117e-11 \| 295 \| 0.195 \| 373 \| 182 \| 17 \| 13 \| 288 \| 19 \| 370 \| Chromosome partitioning protein ParB \| Chromosome partitioning protein ParB; afdb-uniprot50 \| AF-A0A199YNH2-F1-MODEL\_V4 \| 1.0 \| 1.262e-10 \| 295 \| 0.161 \| 315 \| 212 \| 15 \| 1 \| 282 \| 6 \| 301 \| Chromosome partitioning protein ParB \| Chromosome partitioning protein ParB; afdb-uniprot50 \| AF-A0A2V2E4M2-F1-MODEL\_V4 \| 1.0 \| 1.293e-09 \| 294 \| 0.17 \| 276 \| 186 \| 14 \| 13 \| 269 \| 2 \| 253 \| ParB domain-containing protein \| ParB domain-containing protein; afdb-uniprot50 \| AF-A0A800F760-F1-MODEL\_V4 \| 1.0 \| 1.196e-10 \| 294 \| 0.18 \| 271 \| 193 \| 11 \| 23 \| 282 \| 2 \| 254 \| ParB/RepB/Spo0J family partition protein \| ParB/RepB/Spo0J family partition protein; afdb-uniprot50 \| AF-A0A4R8MPW6-F1-MODEL\_V4 \| 1.0 \| 8.851e-10 \| 294 \| 0.18 \| 261 \| 174 \| 11 \| 4 \| 246 \| 11 \| 249 \| ParB/RepB/Spo0J family partition protein \| ParB/RepB/Spo0J family partition protein; afdb-uniprot50 \| AF-A0A1V2YCR6-F1-MODEL\_V4 \| 1.0 \| 2.168e-10 \| 294 \| 0.138 \| 297 \| 212 \| 13 \| 4 \| 282 \| 10 \| 280 \| ParB domain-containing protein \| ParB domain-containing protein; afdb-uniprot50 \| AF-A0A1V5J2I1-F1-MODEL\_V4 \| 1.0 \| 3.839e-11 \| 294 \| 0.182 \| 302 \| 207 \| 15 \| 2 \| 289 \| 14 \| 289 \| Putative chromosome-partitioning protein ParB \| Putative chromosome-partitioning protein ParB; afdb-uniprot50 \| AF-A0A7G8T7A8-F1-MODEL\_V4 \| 1.0 \| 6.061e-10 \| 294 \| 0.198 \| 303 \| 207 \| 14 \| 2 \| 291 \| 12 \| 291 \| ParB/RepB/Spo0J family partition protein \| ParB/RepB/Spo0J family partition protein; afdb-uniprot50 \| AF-A0A7C8LUY7-F1-MODEL\_V4 \| 1.0 \| 7.574e-12 \| 294 \| 0.199 \| 311 \| 194 \| 17 \| 7 \| 291 \| 15 \| 296 \| Stage 0 sporulation protein J \| Stage 0 sporulation protein J; afdb-uniprot50 \| AF-A0A2E3KDB6-F1-MODEL\_V4 \| 1.0 \| 9.124e-11 \| 294 \| 0.178 \| 302 \| 202 \| 15 \| 1 \| 282 \| 20 \| 295 \| Chromosome partitioning protein ParB \| Chromosome partitioning protein ParB; afdb-uniprot50 \| AF-A0A1U7GAU7-F1-MODEL\_V4 \| 1.0 \| 1.485e-10 \| 294 \| 0.175 \| 297 \| 189 \| 15 \| 9 \| 287 \| 43 \| 301 \| HTH cro/C1-type domain-containing protein \| HTH cro/C1-type domain-containing protein; afdb-uniprot50 \| AF-A0A2M7B686-F1-MODEL\_V4 \| 1.0 \| 2.55e-10 \| 294 \| 0.172 \| 284 \| 201 \| 11 \| 19 \| 282 \| 49 \| 318 \| ParB domain-containing protein \| ParB domain-containing protein; afdb-uniprot50 \| AF-A0A1Z8NWP4-F1-MODEL\_V4 \| 1.0 \| 5.311e-11 \| 294 \| 0.179 \| 301 \| 211 \| 13 \| 7 \| 291 \| 28 \| 308 \| ParB domain-containing protein \| ParB domain-containing protein; afdb-uniprot50 \| AF-A0A2E0P9T0-F1-MODEL\_V4 \| 1.0 \| 9.343e-10 \| 294 \| 0.179 \| 279 \| 197 \| 17 \| 20 \| 284 \| 81 \| 341 \| Chromosome partitioning protein ParB \| Chromosome partitioning protein ParB; afdb-uniprot50 \| AF-A0A2I1YC93-F1-MODEL\_V4 \| 1.0 \| 2.55e-10 \| 294 \| 0.163 \| 293 \| 202 \| 15 \| 13 \| 282 \| 105 \| 377 \| Chromosome partitioning protein ParB \| Chromosome partitioning protein ParB; afdb-uniprot50 \| AF-A0A6L9G7V0-F1-MODEL\_V4 \| 1.0 \| 1.694e-09 \| 294 \| 0.158 \| 302 \| 211 \| 14 \| 1 \| 282 \| 98 \| 376 \| ParB/RepB/Spo0J family partition protein \| ParB/RepB/Spo0J family partition protein; afdb-uniprot50 \| AF-A0A7W8TTU9-F1-MODEL\_V4 \| 1.0 \| 2.55e-10 \| 294 \| 0.184 \| 293 \| 194 \| 16 \| 13 \| 283 \| 325 \| 594 \| ParB family chromosome partitioning protein \| ParB family chromosome partitioning protein; afdb-uniprot50 \| AF-S5LZF8-F1-MODEL\_V4 \| 1.0 \| 8.643e-11 \| 293 \| 0.146 \| 273 \| 196 \| 10 \| 19 \| 282 \| 3 \| 247 \| Chromosome partitioning protein, DNA-binding protein \| Chromosome partitioning protein, DNA-binding protein; afdb-uniprot50 \| AF-A0A512BQ88-F1-MODEL\_V4 \| 1.0 \| 3.932e-10 \| 293 \| 0.201 \| 283 \| 182 \| 10 \| 18 \| 282 \| 19 \| 275 \| Chromosome partitioning protein ParB \| Chromosome partitioning protein ParB; afdb-uniprot50 \| AF-A0A7C7X6R7-F1-MODEL\_V4 \| 1.0 \| 6.398e-10 \| 293 \| 0.151 \| 290 \| 207 \| 12 \| 7 \| 282 \| 14 \| 278 \| ParB/RepB/Spo0J family partition protein \| ParB/RepB/Spo0J family partition protein; afdb-uniprot50 \| AF-A0A1Q6RN12-F1-MODEL\_V4 \| 1.0 \| 2.168e-10 \| 293 \| 0.177 \| 288 \| 201 \| 14 \| 7 \| 282 \| 15 \| 278 \| HTH cro/C1-type domain-containing protein \| HTH cro/C1-type domain-containing protein; afdb-uniprot50 \| AF-A0A2E0EKM2-F1-MODEL\_V4 \| 1.0 \| 2.842e-10 \| 293 \| 0.182 \| 318 \| 204 \| 14 \| 4 \| 287 \| 18 \| 313 \| Probable chromosome-partitioning protein ParB \| Probable chromosome-partitioning protein ParB; afdb-uniprot50 \| AF-A0A3D5E876-F1-MODEL\_V4 \| 1.0 \| 6.595e-11 \| 293 \| 0.164 \| 279 \| 195 \| 13 \| 19 \| 282 \| 59 \| 314 \| Chromosome partitioning protein ParB \| Chromosome partitioning protein ParB; afdb-uniprot50 \| AF-A0A7V2JJV2-F1-MODEL\_V4 \| 1.0 \| 2.054e-10 \| 293 \| 0.167 \| 304 \| 207 \| 11 \| 1 \| 282 \| 42 \| 321 \| ParB/RepB/Spo0J family partition protein \| ParB/RepB/Spo0J family partition protein; afdb-uniprot50 \| AF-A0A1G6LR52-F1-MODEL\_V4 \| 1.0 \| 9.064e-09 \| 293 \| 0.228 \| 197 \| 130 \| 9 \| 13 \| 195 \| 3 \| 191 \| ParB/RepB/Spo0J family partition protein \| ParB/RepB/Spo0J family partition protein; afdb-uniprot50 \| AF-A0A7X7TY23-F1-MODEL\_V4 \| 1.0 \| 7.892e-08 \| 293 \| 0.207 \| 154 \| 114 \| 4 \| 13 \| 162 \| 1 \| 150 \| ParB N-terminal domain-containing protein \| ParB N-terminal domain-containing protein; afdb-uniprot50 \| AF-A0A1F7M1P9-F1-MODEL\_V4 \| 1.0 \| 3.092e-11 \| 292 \| 0.177 \| 309 \| 197 \| 13 \| 1 \| 282 \| 3 \| 281 \| HTH cro/C1-type domain-containing protein \| HTH cro/C1-type domain-containing protein; afdb-uniprot50 \| AF-A0A651ECM7-F1-MODEL\_V4 \| 1.0 \| 2.289e-10 \| 292 \| 0.21 \| 304 \| 201 \| 12 \| 1 \| 291 \| 11 \| 288 \| ParB/RepB/Spo0J family partition protein \| ParB/RepB/Spo0J family partition protein; afdb-uniprot50 \| AF-A0A3D2CD18-F1-MODEL\_V4 \| 1.0 \| 1.406e-10 \| 292 \| 0.19 \| 283 \| 194 \| 14 \| 14 \| 282 \| 29 \| 290 \| ParB domain-containing protein \| ParB domain-containing protein; afdb-uniprot50 \| AF-A0A847AE66-F1-MODEL\_V4 \| 1.0 \| 1.301e-11 \| 292 \| 0.193 \| 295 \| 204 \| 14 \| 4 \| 282 \| 20 \| 296 \| ParB/RepB/Spo0J family partition protein \| ParB/RepB/Spo0J family partition protein; afdb-uniprot50 \| AF-A0A1C5SVQ1-F1-MODEL\_V4 \| 1.0 \| 6.247e-11 \| 292 \| 0.181 \| 292 \| 213 \| 12 \| 2 \| 282 \| 22 \| 298 \| Probable chromosome-partitioning protein parB \| Probable chromosome-partitioning protein parB; afdb-uniprot50 \| AF-A0A2W5PW74-F1-MODEL\_V4 \| 1.0 \| 3.839e-11 \| 292 \| 0.165 \| 308 \| 215 \| 12 \| 1 \| 291 \| 19 \| 301 \| Chromosome partitioning protein ParB \| Chromosome partitioning protein ParB; afdb-uniprot50 \| AF-A0A024YNU6-F1-MODEL\_V4 \| 1.0 \| 4.486e-09 \| 292 \| 0.161 \| 254 \| 172 \| 11 \| 19 \| 246 \| 3 \| 241 \| Chromosome partitioning protein \| Chromosome partitioning protein; afdb-uniprot50 \| AF-A0A7C3M1B3-F1-MODEL\_V4 \| 1.0 \| 1.225e-09 \| 292 \| 0.183 \| 283 \| 197 \| 13 \| 19 \| 282 \| 10 \| 277 \| ParB/RepB/Spo0J family partition protein \| ParB/RepB/Spo0J family partition protein; afdb-uniprot50 \| AF-A0A133UNQ5-F1-MODEL\_V4 \| 1.0 \| 2.612e-09 \| 292 \| 0.161 \| 304 \| 204 \| 14 \| 14 \| 282 \| 4 \| 291 \| ParB domain-containing protein \| ParB domain-containing protein; afdb-uniprot50 \| AF-A0A7X2FTF1-F1-MODEL\_V4 \| 1.0 \| 8.385e-10 \| 292 \| 0.174 \| 310 \| 198 \| 16 \| 2 \| 280 \| 14 \| 296 \| ParB/RepB/Spo0J family partition protein \| ParB/RepB/Spo0J family partition protein; afdb-uniprot50 \| AF-A0A1V5LF66-F1-MODEL\_V4 \| 1.0 \| 1.332e-10 \| 292 \| 0.183 \| 305 \| 205 \| 15 \| 1 \| 282 \| 7 \| 290 \| Chromosome-partitioning protein ParB \| Chromosome-partitioning protein ParB; afdb-uniprot50 \| AF-A0A2D6EYC8-F1-MODEL\_V4 \| 1.0 \| 3.724e-10 \| 292 \| 0.177 \| 305 \| 204 \| 14 \| 13 \| 291 \| 240 \| 523 \| ParB domain-containing protein \| ParB domain-containing protein; afdb-uniprot50 \| AF-A0A0F9MIX9-F1-MODEL\_V4 \| 1.0 \| 3.321e-08 \| 292 \| 0.253 \| 158 \| 102 \| 8 \| 19 \| 168 \| 5 \| 154 \| ParB domain-containing protein \| ParB domain-containing protein; afdb-uniprot50 \| AF-A0A3C1FMD8-F1-MODEL\_V4 \| 1.0 \| 3.072e-09 \| 291 \| 0.219 \| 187 \| 125 \| 7 \| 1 \| 172 \| 7 \| 187 \| Chromosome partitioning protein ParB \| Chromosome partitioning protein ParB; afdb-uniprot50 \| AF-A0A5C7QJK5-F1-MODEL\_V4 \| 1.0 \| 4.85e-08 \| 291 \| 0.15 \| 279 \| 189 \| 13 \| 19 \| 291 \| 1 \| 237 \| ParB/RepB/Spo0J family partition protein \| ParB/RepB/Spo0J family partition protein; afdb-uniprot50 \| AF-A0A1G0IPH5-F1-MODEL\_V4 \| 1.0 \| 7.944e-10 \| 291 \| 0.158 \| 296 \| 218 \| 12 \| 3 \| 291 \| 6 \| 277 \| Probable chromosome-partitioning protein ParB \| Probable chromosome-partitioning protein ParB; afdb-uniprot50 \| AF-A0A485M023-F1-MODEL\_V4 \| 1.0 \| 2.235e-11 \| 291 \| 0.186 \| 306 \| 192 \| 13 \| 8 \| 291 \| 14 \| 284 \| Chromosome-partitioning protein Spo0J \| Chromosome-partitioning protein Spo0J; afdb-uniprot50 \| AF-A0A2Z5R1P5-F1-MODEL\_V4 \| 1.0 \| 2.692e-10 \| 291 \| 0.17 \| 287 \| 190 \| 15 \| 19 \| 282 \| 39 \| 300 \| Chromosome (Plasmid) partitioning protein ParB \| Chromosome (Plasmid) partitioning protein ParB; afdb-uniprot50 \| AF-A0A2H5V7K5-F1-MODEL\_V4 \| 1.0 \| 6.398e-10 \| 291 \| 0.147 \| 291 \| 195 \| 12 \| 19 \| 287 \| 37 \| 296 \| Putative chromosome-partitioning protein ParB \| Putative chromosome-partitioning protein ParB; afdb-uniprot50 \| AF-A0A4P7WB27-F1-MODEL\_V4 \| 1.0 \| 9.863e-10 \| 291 \| 0.171 \| 304 \| 208 \| 14 \| 3 \| 282 \| 28 \| 311 \| ParB/RepB/Spo0J family partition protein \| ParB/RepB/Spo0J family partition protein; afdb-uniprot50 \| AF-A0A535XXB1-F1-MODEL\_V4 \| 1.0 \| 8.851e-10 \| 291 \| 0.203 \| 305 \| 190 \| 17 \| 5 \| 287 \| 28 \| 301 \| ParB/RepB/Spo0J family partition protein \| ParB/RepB/Spo0J family partition protein; afdb-uniprot50 \| AF-A0A142YPG3-F1-MODEL\_V4 \| 1.0 \| 3.528e-10 \| 291 \| 0.181 \| 309 \| 196 \| 16 \| 2 \| 291 \| 39 \| 309 \| Chromosome-partitioning protein Spo0J \| Chromosome-partitioning protein Spo0J; afdb-uniprot50 \| AF-A0A366IM11-F1-MODEL\_V4 \| 1.0 \| 7.944e-10 \| 291 \| 0.173 \| 294 \| 194 \| 17 \| 11 \| 282 \| 58 \| 324 \| ParB family chromosome partitioning protein \| ParB family chromosome partitioning protein; afdb-uniprot50 \| AF-A0A7W7T048-F1-MODEL\_V4 \| 1.0 \| 2.739e-07 \| 291 \| 0.195 \| 189 \| 141 \| 6 \| 19 \| 202 \| 8 \| 190 \| ParB family chromosome partitioning protein \| ParB family chromosome partitioning protein; afdb-uniprot50 \| AF-A0A4R5AUM8-F1-MODEL\_V4 \| 1.0 \| 8.385e-10 \| 290 \| 0.189 \| 274 \| 176 \| 12 \| 19 \| 287 \| 3 \| 235 \| ParB/RepB/Spo0J family partition protein \| ParB/RepB/Spo0J family partition protein; afdb-uniprot50 \| AF-A0A358UY81-F1-MODEL\_V4 \| 1.0 \| 7.129e-10 \| 290 \| 0.172 \| 290 \| 203 \| 11 \| 13 \| 287 \| 32 \| 299 \| Chromosome partitioning protein ParB \| Chromosome partitioning protein ParB; afdb-uniprot50 \| AF-R5E743-F1-MODEL\_V4 \| 1.0 \| 6.247e-11 \| 290 \| 0.156 \| 300 \| 212 \| 13 \| 1 \| 282 \| 29 \| 305 \| ParB-like protein \| ParB-like protein; afdb-uniprot50 \| AF-R6FU05-F1-MODEL\_V4 \| 1.0 \| 4.767e-11 \| 290 \| 0.161 \| 291 \| 218 \| 11 \| 4 \| 284 \| 36 \| 310 \| ParB-like protein \| ParB-like protein; afdb-uniprot50 \| AF-A0A521ZYP4-F1-MODEL\_V4 \| 1.0 \| 6.961e-11 \| 290 \| 0.177 \| 293 \| 206 \| 15 \| 10 \| 291 \| 31 \| 299 \| ParB/RepB/Spo0J family partition protein \| ParB/RepB/Spo0J family partition protein; afdb-uniprot50 \| AF-A0A662SF16-F1-MODEL\_V4 \| 1.0 \| 1.225e-09 \| 290 \| 0.169 \| 289 \| 194 \| 13 \| 18 \| 287 \| 1 \| 262 \| ParB domain-containing protein \| ParB domain-containing protein; afdb-uniprot50 \| AF-A0A1M3HAQ9-F1-MODEL\_V4 \| 1.0 \| 1.557e-08 \| 290 \| 0.158 \| 195 \| 144 \| 8 \| 8 \| 189 \| 28 \| 215 \| ParB domain-containing protein \| ParB domain-containing protein; afdb-uniprot50 \| AF-A0A355SEW4-F1-MODEL\_V4 \| 1.0 \| 5.918e-11 \| 289 \| 0.168 \| 303 \| 206 \| 12 \| 1 \| 282 \| 5 \| 282 \| Chromosome partitioning protein ParB \| Chromosome partitioning protein ParB; afdb-uniprot50 \| AF-A0A0S9NZS1-F1-MODEL\_V4 \| 1.0 \| 1.225e-09 \| 289 \| 0.182 \| 313 \| 211 \| 14 \| 1 \| 291 \| 9 \| 298 \| Chromosome partitioning protein ParB \| Chromosome partitioning protein ParB; afdb-uniprot50 \| AF-A0A3R6TEA5-F1-MODEL\_V4 \| 1.0 \| 3.445e-11 \| 289 \| 0.165 \| 290 \| 221 \| 10 \| 1 \| 282 \| 30 \| 306 \| ParB/RepB/Spo0J family partition protein \| ParB/RepB/Spo0J family partition protein; afdb-uniprot50 \| AF-A0A1T4T4U9-F1-MODEL\_V4 \| 1.0 \| 2.55e-10 \| 289 \| 0.211 \| 308 \| 196 \| 13 \| 10 \| 287 \| 30 \| 320 \| Chromosome segregation DNA-binding protein \| Chromosome segregation DNA-binding protein; afdb-uniprot50 \| AF-A0A3N1GT42-F1-MODEL\_V4 \| 1.0 \| 2.757e-09 \| 289 \| 0.168 \| 249 \| 173 \| 9 \| 19 \| 246 \| 6 \| 241 \| ParB/RepB/Spo0J family partition protein \| ParB/RepB/Spo0J family partition protein; afdb-uniprot50 \| AF-A0A4P8SIY6-F1-MODEL\_V4 \| 1.0 \| 9.863e-10 \| 289 \| 0.167 \| 298 \| 196 \| 14 \| 1 \| 272 \| 37 \| 308 \| Chromosome partitioning protein ParB \| Chromosome partitioning protein ParB; afdb-uniprot50 \| AF-A0A7U6Y6C4-F1-MODEL\_V4 \| 1.0 \| 6.398e-10 \| 289 \| 0.18 \| 327 \| 199 \| 17 \| 1 \| 291 \| 10 \| 303 \| Chromosome-partitioning protein ParB \| Chromosome-partitioning protein ParB; afdb-uniprot50 \| AF-A0A0U3PNC0-F1-MODEL\_V4 \| 1.0 \| 1.188e-08 \| 289 \| 0.176 \| 261 \| 178 \| 11 \| 19 \| 261 \| 4 \| 245 \| ParB domain-containing protein \| ParB domain-containing protein; afdb-uniprot50 \| AF-A0A021VXB0-F1-MODEL\_V4 \| 1.0 \| 1.397e-08 \| 288 \| 0.208 \| 269 \| 158 \| 10 \| 19 \| 282 \| 3 \| 221 \| ParB domain-containing protein \| ParB domain-containing protein; afdb-uniprot50 \| AF-A0A3M1V3E9-F1-MODEL\_V4 \| 1.0 \| 2.289e-10 \| 288 \| 0.204 \| 293 \| 190 \| 15 \| 14 \| 291 \| 4 \| 268 \| Probable chromosome-partitioning protein ParB \| Probable chromosome-partitioning protein ParB; afdb-uniprot50 \| AF-A0A2E8QXZ0-F1-MODEL\_V4 \| 1.0 \| 5.032e-11 \| 288 \| 0.221 \| 307 \| 186 \| 15 \| 1 \| 282 \| 15 \| 293 \| Chromosome partitioning protein ParB \| Chromosome partitioning protein ParB; afdb-uniprot50 \| AF-A0A1E3G6T5-F1-MODEL\_V4 \| 1.0 \| 2.289e-10 \| 288 \| 0.163 \| 288 \| 182 \| 15 \| 1 \| 272 \| 4 \| 248 \| ParB domain-containing protein \| ParB domain-containing protein; afdb-uniprot50 \| AF-A0A2E2QKV5-F1-MODEL\_V4 \| 1.0 \| 1.406e-10 \| 288 \| 0.209 \| 286 \| 177 \| 15 \| 20 \| 282 \| 35 \| 294 \| Chromosome partitioning protein ParB \| Chromosome partitioning protein ParB; afdb-uniprot50 \| AF-R5LI34-F1-MODEL\_V4 \| 1.0 \| 1.52e-09 \| 288 \| 0.161 \| 272 \| 180 \| 14 \| 13 \| 271 \| 2 \| 238 \| Nucleoid occlusion protein \| Nucleoid occlusion protein; afdb-uniprot50 \| AF-A0A7Y7A845-F1-MODEL\_V4 \| 1.0 \| 1.933e-08 \| 287 \| 0.222 \| 166 \| 117 \| 6 \| 13 \| 172 \| 32 \| 191 \| ParB/RepB/Spo0J family partition protein \| ParB/RepB/Spo0J family partition protein; afdb-uniprot50 \| AF-A0A2H0CD87-F1-MODEL\_V4 \| 1.0 \| 2.49e-11 \| 287 \| 0.148 \| 309 \| 196 \| 14 \| 1 \| 282 \| 19 \| 287 \| ParB domain-containing protein \| ParB domain-containing protein; afdb-uniprot50 \| AF-A0A521H3H6-F1-MODEL\_V4 \| 1.0 \| 1.567e-10 \| 287 \| 0.179 \| 317 \| 186 \| 14 \| 5 \| 282 \| 28 \| 309 \| ParB/RepB/Spo0J family partition protein \| ParB/RepB/Spo0J family partition protein; afdb-uniprot50 \| AF-A0A2Z3J328-F1-MODEL\_V4 \| 1.0 \| 1.485e-10 \| 287 \| 0.208 \| 293 \| 182 \| 12 \| 8 \| 282 \| 62 \| 322 \| Probable chromosome-partitioning protein ParB \| Probable chromosome-partitioning protein ParB; afdb-uniprot50 \| AF-A0A2D5RV49-F1-MODEL\_V4 \| 1.0 \| 8.851e-10 \| 287 \| 0.178 \| 297 \| 184 \| 14 \| 1 \| 272 \| 7 \| 268 \| Chromosome partitioning protein ParB \| Chromosome partitioning protein ParB; afdb-uniprot50 \| AF-A0A259MJE9-F1-MODEL\_V4 \| 1.0 \| 2.534e-08 \| 287 \| 0.183 \| 185 \| 132 \| 6 \| 13 \| 182 \| 74 \| 254 \| HTH cro/C1-type domain-containing protein \| HTH cro/C1-type domain-containing protein; afdb-uniprot50 \| AF-A0A0F9GST2-F1-MODEL\_V4 \| 1.0 \| 3.243e-09 \| 287 \| 0.149 \| 315 \| 207 \| 18 \| 4 \| 287 \| 1 \| 285 \| ParB domain-containing protein \| ParB domain-containing protein; afdb-uniprot50 \| AF-A0A2E9CXQ8-F1-MODEL\_V4 \| 1.0 \| 1.188e-08 \| 286 \| 0.18 \| 211 \| 157 \| 7 \| 1 \| 205 \| 57 \| 257 \| Probable chromosome-partitioning protein ParB \| Probable chromosome-partitioning protein ParB; afdb-uniprot50 \| AF-A0A2S8IL38-F1-MODEL\_V4 \| 1.0 \| 1.605e-09 \| 286 \| 0.179 \| 301 \| 187 \| 16 \| 1 \| 271 \| 1 \| 271 \| Uncharacterized protein \| Uncharacterized protein; afdb-uniprot50 \| AF-A0A2E6W466-F1-MODEL\_V4 \| 1.0 \| 3.528e-10 \| 286 \| 0.171 \| 298 \| 209 \| 14 \| 2 \| 282 \| 5 \| 281 \| Chromosome partitioning protein ParB \| Chromosome partitioning protein ParB; afdb-uniprot50 \| AF-Q1NN18-F1-MODEL\_V4 \| 1.0 \| 3.839e-11 \| 286 \| 0.187 \| 309 \| 197 \| 11 \| 1 \| 282 \| 6 \| 287 \| ParB-like partition protein \| ParB-like partition protein; afdb-uniprot50 \| AF-A0A7D7ZD79-F1-MODEL\_V4 \| 1.0 \| 2.842e-10 \| 286 \| 0.15 \| 306 \| 224 \| 12 \| 2 \| 290 \| 6 \| 292 \| ParB/RepB/Spo0J family partition protein \| ParB/RepB/Spo0J family partition protein; afdb-uniprot50 \| AF-A0A1Y5RLZ8-F1-MODEL\_V4 \| 1.0 \| 1.406e-10 \| 286 \| 0.189 \| 290 \| 190 \| 14 \| 21 \| 291 \| 36 \| 299 \| Chromosome-partitioning protein ParB \| Chromosome-partitioning protein ParB; afdb-uniprot50 \| AF-A0A1F9UGF1-F1-MODEL\_V4 \| 1.0 \| 9.343e-10 \| 286 \| 0.209 \| 306 \| 191 \| 15 \| 13 \| 291 \| 4 \| 285 \| ParB domain-containing protein \| ParB domain-containing protein; afdb-uniprot50 \| AF-A0A3A1YS35-F1-MODEL\_V4 \| 1.0 \| 5.153e-10 \| 286 \| 0.138 \| 311 \| 220 \| 17 \| 1 \| 291 \| 58 \| 340 \| ParB domain-containing protein \| ParB domain-containing protein; afdb-uniprot50 \| AF-A0A7W9SVT8-F1-MODEL\_V4 \| 1.0 \| 5.439e-10 \| 286 \| 0.197 \| 274 \| 183 \| 11 \| 18 \| 265 \| 1 \| 263 \| ParB/RepB/Spo0J family partition protein \| ParB/RepB/Spo0J family partition protein; afdb-uniprot50 \| AF-A0A0A8ESX7-F1-MODEL\_V4 \| 1.0 \| 6.552e-09 \| 286 \| 0.137 \| 277 \| 183 \| 11 \| 19 \| 287 \| 1 \| 229 \| Chromosome segregation DNA-binding protein \| Chromosome segregation DNA-binding protein; afdb-uniprot50 \| AF-A0A3E0W7R3-F1-MODEL\_V4 \| 1.0 \| 2.823e-08 \| 286 \| 0.187 \| 213 \| 147 \| 11 \| 1 \| 204 \| 21 \| 216 \| ParB domain-containing protein \| ParB domain-containing protein; afdb-uniprot50 \| AF-A0A2D6XAJ0-F1-MODEL\_V4 \| 1.0 \| 2.344e-09 \| 285 \| 0.176 \| 306 \| 180 \| 14 \| 16 \| 291 \| 1 \| 264 \| ParB domain-containing protein \| ParB domain-containing protein; afdb-uniprot50 \| AF-A0A6N7AYY8-F1-MODEL\_V4 \| 1.0 \| 3.343e-10 \| 285 \| 0.155 \| 295 \| 208 \| 13 \| 6 \| 283 \| 11 \| 281 \| Chromosome partitioning protein ParB family \| Chromosome partitioning protein ParB family; afdb-uniprot50 \| AF-A0A2D7IRV4-F1-MODEL\_V4 \| 1.0 \| 7.944e-10 \| 285 \| 0.139 \| 294 \| 229 \| 12 \| 1 \| 282 \| 7 \| 288 \| DNA-binding protein \| DNA-binding protein; afdb-uniprot50 \| AF-A0A358ICQ6-F1-MODEL\_V4 \| 1.0 \| 3e-10 \| 285 \| 0.168 \| 297 \| 209 \| 12 \| 5 \| 287 \| 18 \| 290 \| ParB domain-containing protein \| ParB domain-containing protein; afdb-uniprot50 \| AF-A0A6B1A4N4-F1-MODEL\_V4 \| 1.0 \| 4.053e-11 \| 285 \| 0.212 \| 296 \| 206 \| 11 \| 9 \| 291 \| 21 \| 302 \| ParB/RepB/Spo0J family partition protein \| ParB/RepB/Spo0J family partition protein; afdb-uniprot50 \| AF-A0A5C7UBH5-F1-MODEL\_V4 \| 1.0 \| 2.534e-08 \| 285 \| 0.229 \| 179 \| 124 \| 8 \| 13 \| 183 \| 2 \| 174 \| PRTRC system ParB family protein \| PRTRC system ParB family protein; afdb-uniprot50 \| AF-A0A1B9RA54-F1-MODEL\_V4 \| 1.0 \| 5.439e-10 \| 285 \| 0.161 \| 321 \| 188 \| 16 \| 13 \| 272 \| 1 \| 301 \| ParB domain-containing protein \| ParB domain-containing protein; afdb-uniprot50 \| AF-D7ATR0-F1-MODEL\_V4 \| 1.0 \| 6.961e-11 \| 284 \| 0.164 \| 310 \| 203 \| 16 \| 1 \| 282 \| 1 \| 282 \| ParB-like partition protein \| ParB-like partition protein; afdb-uniprot50 \| AF-A0A2K5ARA4-F1-MODEL\_V4 \| 1.0 \| 4.381e-10 \| 284 \| 0.152 \| 308 \| 202 \| 15 \| 3 \| 287 \| 1 \| 272 \| Chromosome-partitioning protein Spo0J \| Chromosome-partitioning protein Spo0J; afdb-uniprot50 \| AF-A0A2E9E3N7-F1-MODEL\_V4 \| 1.0 \| 5.607e-11 \| 284 \| 0.165 \| 314 \| 196 \| 16 \| 1 \| 282 \| 6 \| 285 \| Chromosome partitioning protein ParB \| Chromosome partitioning protein ParB; afdb-uniprot50 \| AF-A0A0D0QNA4-F1-MODEL\_V4 \| 1.0 \| 4.278e-11 \| 284 \| 0.145 \| 296 \| 197 \| 11 \| 13 \| 282 \| 28 \| 293 \| TWA4\_scaffold00001, whole genome shotgun sequence \| TWA4\_scaffold00001, whole genome shotgun sequence; afdb-uniprot50 \| AF-L0DRM7-F1-MODEL\_V4 \| 1.0 \| 2.55e-10 \| 284 \| 0.182 \| 296 \| 197 \| 14 \| 1 \| 287 \| 37 \| 296 \| ParB-like partition protein \| ParB-like partition protein; afdb-uniprot50 \| AF-R5JVN8-F1-MODEL\_V4 \| 1.0 \| 1.485e-10 \| 284 \| 0.174 \| 281 \| 203 \| 11 \| 14 \| 282 \| 52 \| 315 \| Stage 0 sporulation protein J \| Stage 0 sporulation protein J; afdb-uniprot50 \| AF-L7LEH1-F1-MODEL\_V4 \| 1.0 \| 3.613e-09 \| 284 \| 0.171 \| 304 \| 176 \| 14 \| 20 \| 272 \| 4 \| 282 \| ParB domain-containing protein \| ParB domain-containing protein; afdb-uniprot50 \| AF-A0A3M2CZK8-F1-MODEL\_V4 \| 1.0 \| 7.574e-12 \| 283 \| 0.161 \| 309 \| 198 \| 12 \| 1 \| 282 \| 4 \| 278 \| ParB/RepB/Spo0J family partition protein \| ParB/RepB/Spo0J family partition protein; afdb-uniprot50 \| AF-A0A1Q2HRV3-F1-MODEL\_V4 \| 1.0 \| 7.348e-11 \| 283 \| 0.164 \| 310 \| 213 \| 9 \| 2 \| 291 \| 1 \| 284 \| Putative chromosome-partitioning protein ParB \| Putative chromosome-partitioning protein ParB; afdb-uniprot50 \| AF-A0A373LA92-F1-MODEL\_V4 \| 1.0 \| 9.343e-10 \| 283 \| 0.178 \| 285 \| 190 \| 13 \| 12 \| 272 \| 19 \| 283 \| ParB/RepB/Spo0J family partition protein \| ParB/RepB/Spo0J family partition protein; afdb-uniprot50 \| AF-A0A3A6JNC2-F1-MODEL\_V4 \| 1.0 \| 3.814e-09 \| 283 \| 0.18 \| 277 \| 188 \| 15 \| 3 \| 246 \| 22 \| 292 \| ParB/RepB/Spo0J family partition protein \| ParB/RepB/Spo0J family partition protein; afdb-uniprot50 \| AF-A0A1G0G5V5-F1-MODEL\_V4 \| 1.0 \| 2.344e-09 \| 283 \| 0.192 \| 260 \| 171 \| 13 \| 19 \| 271 \| 15 \| 242 \| Probable chromosome-partitioning protein ParB \| Probable chromosome-partitioning protein ParB; afdb-uniprot50 \| AF-A0A1I2HRP5-F1-MODEL\_V4 \| 1.0 \| 1.041e-09 \| 283 \| 0.207 \| 313 \| 200 \| 19 \| 1 \| 291 \| 73 \| 359 \| Chromosome segregation DNA-binding protein \| Chromosome segregation DNA-binding protein; afdb-uniprot50 \| AF-A0A518GYN7-F1-MODEL\_V4 \| 1.0 \| 3.814e-09 \| 283 \| 0.171 \| 286 \| 199 \| 8 \| 11 \| 287 \| 74 \| 330 \| Putative chromosome-partitioning protein ParB \| Putative chromosome-partitioning protein ParB; afdb-uniprot50 \| AF-A0A1V9VAD5-F1-MODEL\_V4 \| 1.0 \| 2.823e-08 \| 282 \| 0.169 \| 177 \| 135 \| 6 \| 1 \| 169 \| 8 \| 180 \| ParB domain-containing protein \| ParB domain-containing protein; afdb-uniprot50 \| AF-A0A7X9CR71-F1-MODEL\_V4 \| 1.0 \| 6.595e-11 \| 282 \| 0.179 \| 290 \| 194 \| 13 \| 13 \| 282 \| 16 \| 281 \| ParB/RepB/Spo0J family partition protein \| ParB/RepB/Spo0J family partition protein; afdb-uniprot50 \| AF-A0A847NG60-F1-MODEL\_V4 \| 1.0 \| 2.49e-11 \| 282 \| 0.185 \| 281 \| 192 \| 12 \| 14 \| 282 \| 25 \| 280 \| ParB/RepB/Spo0J family partition protein \| ParB/RepB/Spo0J family partition protein; afdb-uniprot50 \| AF-A0A2N6AKC7-F1-MODEL\_V4 \| 1.0 \| 7.944e-10 \| 282 \| 0.155 \| 289 \| 206 \| 12 \| 8 \| 282 \| 24 \| 288 \| Chromosome partitioning protein ParB \| Chromosome partitioning protein ParB; afdb-uniprot50 \| AF-A0A0Q7MU60-F1-MODEL\_V4 \| 1.0 \| 7.129e-10 \| 282 \| 0.189 \| 311 \| 193 \| 16 \| 13 \| 291 \| 59 \| 342 \| ParB domain-containing protein \| ParB domain-containing protein; afdb-uniprot50 \| AF-A0A517VMS6-F1-MODEL\_V4 \| 1.0 \| 2.757e-09 \| 282 \| 0.167 \| 293 \| 193 \| 17 \| 12 \| 281 \| 3 \| 267 \| Nucleoid occlusion protein \| Nucleoid occlusion protein; afdb-uniprot50 \| AF-A0A3B9JIH2-F1-MODEL\_V4 \| 1.0 \| 1.017e-10 \| 281 \| 0.18 \| 300 \| 205 \| 13 \| 1 \| 282 \| 4 \| 280 \| Chromosome partitioning protein ParB \| Chromosome partitioning protein ParB; afdb-uniprot50 \| AF-A0A059FMC4-F1-MODEL\_V4 \| 1.0 \| 5.607e-11 \| 281 \| 0.199 \| 296 \| 192 \| 11 \| 14 \| 291 \| 17 \| 285 \| Chromosome partitioning protein ParB \| Chromosome partitioning protein ParB; afdb-uniprot50 \| AF-A0A6N8V9B5-F1-MODEL\_V4 \| 1.0 \| 3e-10 \| 281 \| 0.165 \| 314 \| 201 \| 15 \| 1 \| 282 \| 11 \| 295 \| ParB/RepB/Spo0J family partition protein \| ParB/RepB/Spo0J family partition protein; afdb-uniprot50 \| AF-A0A4Z0KDG3-F1-MODEL\_V4 \| 1.0 \| 2.474e-09 \| 281 \| 0.168 \| 302 \| 207 \| 18 \| 1 \| 282 \| 26 \| 303 \| ParB/RepB/Spo0J family partition protein \| ParB/RepB/Spo0J family partition protein; afdb-uniprot50 \| AF-A0A5U1PXS1-F1-MODEL\_V4 \| 1.0 \| 4.381e-10 \| 281 \| 0.203 \| 285 \| 178 \| 14 \| 1 \| 272 \| 6 \| 254 \| ParB/RepB/Spo0J family partition protein \| ParB/RepB/Spo0J family partition protein; afdb-uniprot50 \| AF-A0A411H8J1-F1-MODEL\_V4 \| 1.0 \| 3.072e-09 \| 281 \| 0.17 \| 310 \| 190 \| 17 \| 3 \| 290 \| 8 \| 272 \| ParB/RepB/Spo0J family partition protein \| ParB/RepB/Spo0J family partition protein; afdb-uniprot50 \| AF-A0A7H2BGE9-F1-MODEL\_V4 \| 1.0 \| 1.788e-09 \| 281 \| 0.186 \| 268 \| 171 \| 14 \| 19 \| 265 \| 9 \| 250 \| ParB/RepB/Spo0J family partition protein \| ParB/RepB/Spo0J family partition protein; afdb-uniprot50 \| AF-M4NG45-F1-MODEL\_V4 \| 1.0 \| 1.217e-07 \| 281 \| 0.175 \| 188 \| 144 \| 6 \| 19 \| 200 \| 14 \| 196 \| Probable chromosome-partitioning protein ParB \| Probable chromosome-partitioning protein ParB; afdb-uniprot50 \| AF-A0A4P7D5W3-F1-MODEL\_V4 \| 1.0 \| 2.274e-08 \| 280 \| 0.181 \| 204 \| 152 \| 8 \| 13 \| 205 \| 6 \| 205 \| ParB/RepB/Spo0J family partition protein \| ParB/RepB/Spo0J family partition protein; afdb-uniprot50 \| AF-A0A2S6QJ87-F1-MODEL\_V4 \| 1.0 \| 1.133e-10 \| 280 \| 0.184 \| 315 \| 201 \| 14 \| 1 \| 290 \| 6 \| 289 \| Chromosome-partitioning protein ParB \| Chromosome-partitioning protein ParB; afdb-uniprot50 \| AF-A0A2H9T523-F1-MODEL\_V4 \| 1.0 \| 7.525e-10 \| 280 \| 0.179 \| 301 \| 197 \| 17 \| 5 \| 273 \| 8 \| 290 \| Putative chromosome-partitioning protein ParB \| Putative chromosome-partitioning protein ParB; afdb-uniprot50 \| AF-A0A1F5FFJ9-F1-MODEL\_V4 \| 1.0 \| 8.385e-10 \| 280 \| 0.189 \| 311 \| 204 \| 13 \| 7 \| 290 \| 18 \| 307 \| HTH cro/C1-type domain-containing protein \| HTH cro/C1-type domain-containing protein; afdb-uniprot50 \| AF-A0A2A3XKZ8-F1-MODEL\_V4 \| 1.0 \| 1.099e-09 \| 280 \| 0.173 \| 288 \| 190 \| 17 \| 19 \| 283 \| 184 \| 446 \| ParB domain-containing protein \| ParB domain-containing protein; afdb-uniprot50 \| AF-A0A4P9VE58-F1-MODEL\_V4 \| 1.0 \| 4.486e-09 \| 279 \| 0.16 \| 249 \| 176 \| 10 \| 19 \| 246 \| 6 \| 242 \| ParB/RepB/Spo0J family partition protein \| ParB/RepB/Spo0J family partition protein; afdb-uniprot50 \| AF-A0A1F2Z098-F1-MODEL\_V4 \| 1.0 \| 5.439e-10 \| 279 \| 0.155 \| 303 \| 212 \| 13 \| 13 \| 291 \| 46 \| 328 \| ParB domain-containing protein \| ParB domain-containing protein; afdb-uniprot50 \| AF-A0A559UHF1-F1-MODEL\_V4 \| 1.0 \| 5.277e-09 \| 279 \| 0.17 \| 294 \| 204 \| 15 \| 8 \| 287 \| 44 \| 311 \| ParB family chromosome partitioning protein \| ParB family chromosome partitioning protein; afdb-uniprot50 \| AF-A0A5F0M8I8-F1-MODEL\_V4 \| 1.0 \| 4.026e-09 \| 279 \| 0.165 \| 290 \| 207 \| 14 \| 11 \| 282 \| 211 \| 483 \| ParB/RepB/Spo0J family partition protein \| ParB/RepB/Spo0J family partition protein; afdb-uniprot50 \| AF-A0A7C7PFL2-F1-MODEL\_V4 \| 1.0 \| 4.999e-09 \| 278 \| 0.223 \| 215 \| 148 \| 9 \| 1 \| 201 \| 1 \| 210 \| ParB/RepB/Spo0J family partition protein \| ParB/RepB/Spo0J family partition protein; afdb-uniprot50 \| AF-A0A3N5NMG6-F1-MODEL\_V4 \| 1.0 \| 4.881e-10 \| 278 \| 0.2 \| 279 \| 170 \| 11 \| 3 \| 256 \| 10 \| 260 \| ParB/RepB/Spo0J family partition protein \| ParB/RepB/Spo0J family partition protein; afdb-uniprot50 \| AF-A0A7X3YB26-F1-MODEL\_V4 \| 1.0 \| 1.225e-09 \| 278 \| 0.185 \| 286 \| 186 \| 14 \| 13 \| 282 \| 24 \| 278 \| ParB/RepB/Spo0J family partition protein \| ParB/RepB/Spo0J family partition protein; afdb-uniprot50 \| AF-A0A7C7A9V0-F1-MODEL\_V4 \| 1.0 \| 5.032e-11 \| 278 \| 0.179 \| 317 \| 197 \| 16 \| 1 \| 288 \| 4 \| 286 \| ParB/RepB/Spo0J family partition protein \| ParB/RepB/Spo0J family partition protein; afdb-uniprot50 \| AF-A0A7X5HWV5-F1-MODEL\_V4 \| 1.0 \| 4.767e-11 \| 278 \| 0.185 \| 308 \| 195 \| 13 \| 9 \| 291 \| 21 \| 297 \| ParB/RepB/Spo0J family partition protein \| ParB/RepB/Spo0J family partition protein; afdb-uniprot50 \| AF-A0A539DA23-F1-MODEL\_V4 \| 1.0 \| 2.006e-11 \| 278 \| 0.211 \| 298 \| 181 \| 12 \| 14 \| 288 \| 51 \| 317 \| Chromosome partitioning protein ParB \| Chromosome partitioning protein ParB; afdb-uniprot50 \| AF-A0A7K1LKD1-F1-MODEL\_V4 \| 1.0 \| 6.753e-10 \| 278 \| 0.183 \| 299 \| 201 \| 16 \| 13 \| 289 \| 105 \| 382 \| ParB/RepB/Spo0J family partition protein \| ParB/RepB/Spo0J family partition protein; afdb-uniprot50 \| AF-A0A827LWJ5-F1-MODEL\_V4 \| 1.0 \| 6.916e-09 \| 278 \| 0.188 \| 302 \| 201 \| 15 \| 1 \| 286 \| 22 \| 295 \| ParB/RepB/Spo0J family partition protein \| ParB/RepB/Spo0J family partition protein; afdb-uniprot50 \| AF-A0A366KCA2-F1-MODEL\_V4 \| 1.0 \| 2.22e-09 \| 278 \| 0.214 \| 247 \| 149 \| 14 \| 12 \| 246 \| 6 \| 219 \| ParB domain-containing protein \| ParB domain-containing protein; afdb-uniprot50 \| AF-A0A538S316-F1-MODEL\_V4 \| 1.0 \| 4.85e-08 \| 278 \| 0.216 \| 217 \| 136 \| 10 \| 19 \| 214 \| 184 \| 387 \| ParB domain-containing protein \| ParB domain-containing protein; afdb-uniprot50 \| AF-A0A7C6QZE1-F1-MODEL\_V4 \| 1.0 \| 4.624e-10 \| 278 \| 0.159 \| 301 \| 187 \| 15 \| 13 \| 281 \| 23 \| 289 \| ParB/RepB/Spo0J family partition protein \| ParB/RepB/Spo0J family partition protein; afdb-uniprot50 \| AF-A0A3S0HPY2-F1-MODEL\_V4 \| 1.0 \| 8.331e-08 \| 277 \| 0.184 \| 206 \| 148 \| 8 \| 18 \| 214 \| 1 \| 195 \| ParB/RepB/Spo0J family partition protein \| ParB/RepB/Spo0J family partition protein; afdb-uniprot50 \| AF-A0A2E6Z228-F1-MODEL\_V4 \| 1.0 \| 2.458e-07 \| 277 \| 0.134 \| 178 \| 146 \| 5 \| 19 \| 192 \| 6 \| 179 \| ParB domain-containing protein \| ParB domain-containing protein; afdb-uniprot50 \| AF-A0A7C6P963-F1-MODEL\_V4 \| 1.0 \| 7.476e-08 \| 277 \| 0.216 \| 157 \| 112 \| 5 \| 18 \| 170 \| 1 \| 150 \| ParB/RepB/Spo0J family partition protein \| ParB/RepB/Spo0J family partition protein; afdb-uniprot50 \| AF-A0A3D5L091-F1-MODEL\_V4 \| 1.0 \| 6.753e-10 \| 277 \| 0.151 \| 290 \| 188 \| 13 \| 20 \| 282 \| 55 \| 313 \| Chromosome partitioning protein ParB \| Chromosome partitioning protein ParB; afdb-uniprot50 \| AF-A0A2G6DSE2-F1-MODEL\_V4 \| 1.0 \| 1.406e-10 \| 277 \| 0.164 \| 280 \| 193 \| 13 \| 19 \| 282 \| 71 \| 325 \| Chromosome partitioning protein ParB \| Chromosome partitioning protein ParB; afdb-uniprot50 \| AF-A0A375JD00-F1-MODEL\_V4 \| 1.0 \| 3.613e-09 \| 277 \| 0.18 \| 305 \| 184 \| 16 \| 9 \| 272 \| 6 \| 285 \| ParB domain-containing protein \| ParB domain-containing protein; afdb-uniprot50 \| AF-A0A5C5TNX6-F1-MODEL\_V4 \| 1.0 \| 2.117e-11 \| 277 \| 0.221 \| 316 \| 186 \| 19 \| 13 \| 282 \| 36 \| 337 \| ParB/RepB/Spo0J family partition protein \| ParB/RepB/Spo0J family partition protein; afdb-uniprot50 \| AF-A0A6D1V7U6-F1-MODEL\_V4 \| 1.0 \| 7.476e-08 \| 277 \| 0.174 \| 223 \| 143 \| 9 \| 16 \| 205 \| 3 \| 217 \| ParB domain-containing protein \| ParB domain-containing protein; afdb-uniprot50 \| AF-A0A7W6KP23-F1-MODEL\_V4 \| 1.0 \| 2.739e-07 \| 276 \| 0.173 \| 190 \| 144 \| 8 \| 19 \| 199 \| 5 \| 190 \| ParB family chromosome partitioning protein \| ParB family chromosome partitioning protein; afdb-uniprot50 \| AF-A0A7C2K3S6-F1-MODEL\_V4 \| 1.0 \| 5.57e-09 \| 276 \| 0.156 \| 300 \| 201 \| 13 \| 18 \| 291 \| 4 \| 277 \| Probable chromosome-partitioning protein ParB \| Probable chromosome-partitioning protein ParB; afdb-uniprot50 \| AF-A0A3D3IUN0-F1-MODEL\_V4 \| 1.0 \| 1.654e-10 \| 276 \| 0.165 \| 315 \| 207 \| 15 \| 4 \| 288 \| 37 \| 325 \| Chromosome partitioning protein ParB \| Chromosome partitioning protein ParB; afdb-uniprot50 \| AF-D9Y7S8-F1-MODEL\_V4 \| 1.0 \| 1.788e-09 \| 276 \| 0.147 \| 285 \| 197 \| 11 \| 2 \| 272 \| 5 \| 257 \| ParB-like protein \| ParB-like protein; afdb-uniprot50 \| AF-R6M2U8-F1-MODEL\_V4 \| 1.0 \| 9.799e-08 \| 276 \| 0.203 \| 201 \| 139 \| 10 \| 4 \| 193 \| 12 \| 202 \| ParB-like partition protein \| ParB-like partition protein; afdb-uniprot50 \| AF-A0A3M1I3Z7-F1-MODEL\_V4 \| 1.0 \| 4.15e-10 \| 275 \| 0.22 \| 299 \| 188 \| 15 \| 1 \| 282 \| 4 \| 274 \| ParB/RepB/Spo0J family partition protein \| ParB/RepB/Spo0J family partition protein; afdb-uniprot50 \| AF-A0A2A4QUW5-F1-MODEL\_V4 \| 1.0 \| 4.381e-10 \| 275 \| 0.171 \| 320 \| 208 \| 15 \| 1 \| 291 \| 22 \| 313 \| Chromosome partitioning protein ParB \| Chromosome partitioning protein ParB; afdb-uniprot50 \| AF-A0A5C8PGP3-F1-MODEL\_V4 \| 1.0 \| 2.629e-11 \| 275 \| 0.227 \| 317 \| 184 \| 18 \| 13 \| 282 \| 42 \| 344 \| ParB/RepB/Spo0J family partition protein \| ParB/RepB/Spo0J family partition protein; afdb-uniprot50 \| AF-A0A221Q2J2-F1-MODEL\_V4 \| 1.0 \| 2.91e-09 \| 275 \| 0.151 \| 316 \| 195 \| 13 \| 19 \| 282 \| 4 \| 298 \| ParB domain-containing protein \| ParB domain-containing protein; afdb-uniprot50 \| AF-A0A2G9Y3U1-F1-MODEL\_V4 \| 1.0 \| 2.55e-10 \| 274 \| 0.148 \| 303 \| 207 \| 14 \| 1 \| 282 \| 2 \| 274 \| Chromosome partitioning protein ParB \| Chromosome partitioning protein ParB; afdb-uniprot50 \| AF-A0A2A4U4T4-F1-MODEL\_V4 \| 1.0 \| 2.842e-10 \| 274 \| 0.195 \| 302 \| 198 \| 14 \| 4 \| 282 \| 8 \| 287 \| Chromosome partitioning protein ParB \| Chromosome partitioning protein ParB; afdb-uniprot50 \| AF-A0A845Q5L7-F1-MODEL\_V4 \| 1.0 \| 6.398e-10 \| 274 \| 0.14 \| 314 \| 207 \| 14 \| 1 \| 282 \| 8 \| 290 \| ParB/RepB/Spo0J family partition protein \| ParB/RepB/Spo0J family partition protein; afdb-uniprot50 \| AF-A0A4Q5VAH7-F1-MODEL\_V4 \| 1.0 \| 4.736e-09 \| 274 \| 0.152 \| 288 \| 213 \| 15 \| 19 \| 291 \| 4 \| 275 \| Uncharacterized protein \| Uncharacterized protein; afdb-uniprot50 \| AF-A0A5C7QL10-F1-MODEL\_V4 \| 1.0 \| 3.814e-09 \| 274 \| 0.156 \| 256 \| 169 \| 12 \| 19 \| 247 \| 1 \| 236 \| ParB/RepB/Spo0J family partition protein \| ParB/RepB/Spo0J family partition protein; afdb-uniprot50 \| AF-A0A075KIZ2-F1-MODEL\_V4 \| 1.0 \| 1.746e-10 \| 274 \| 0.159 \| 294 \| 219 \| 12 \| 1 \| 282 \| 8 \| 285 \| ParB-like partition protein \| ParB-like partition protein; afdb-uniprot50 \| AF-G2DG78-F1-MODEL\_V4 \| 1.0 \| 4.624e-10 \| 274 \| 0.172 \| 290 \| 198 \| 14 \| 19 \| 291 \| 40 \| 304 \| Probable chromosome-partitioning protein ParB \| Probable chromosome-partitioning protein ParB; afdb-uniprot50 \| AF-A0A561EN29-F1-MODEL\_V4 \| 1.0 \| 2.041e-08 \| 274 \| 0.192 \| 250 \| 169 \| 11 \| 19 \| 246 \| 1 \| 239 \| ParB family chromosome partitioning protein \| ParB family chromosome partitioning protein; afdb-uniprot50 \| AF-A0A2X3KGI8-F1-MODEL\_V4 \| 1.0 \| 1.254e-08 \| 273 \| 0.18 \| 238 \| 151 \| 10 \| 19 \| 216 \| 35 \| 268 \| Probable chromosome-partitioning protein parB \| Probable chromosome-partitioning protein parB; afdb-uniprot50 \| AF-A0A0G0LW51-F1-MODEL\_V4 \| 1.0 \| 1.946e-10 \| 273 \| 0.198 \| 287 \| 201 \| 11 \| 3 \| 282 \| 13 \| 277 \| Chromosome (Plasmid) partitioning protein ParB / Stage 0 sporulation protein J \| Chromosome (Plasmid) partitioning protein ParB / Stage 0 sporulation protein J; afdb-uniprot50 \| AF-I8R461-F1-MODEL\_V4 \| 1.0 \| 6.595e-11 \| 273 \| 0.186 \| 317 \| 195 \| 16 \| 1 \| 288 \| 4 \| 286 \| ParB-like partition protein \| ParB-like partition protein; afdb-uniprot50 \| AF-A0A2M8QGU9-F1-MODEL\_V4 \| 1.0 \| 1.332e-10 \| 273 \| 0.203 \| 314 \| 201 \| 18 \| 1 \| 290 \| 4 \| 292 \| Stage 0 sporulation protein J \| Stage 0 sporulation protein J; afdb-uniprot50 \| AF-A0A3B9WD89-F1-MODEL\_V4 \| 1.0 \| 5.88e-09 \| 273 \| 0.167 \| 299 \| 194 \| 16 \| 12 \| 282 \| 54 \| 325 \| Chromosome partitioning protein ParB \| Chromosome partitioning protein ParB; afdb-uniprot50 \| AF-A0A1H3WS16-F1-MODEL\_V4 \| 1.0 \| 3.932e-10 \| 273 \| 0.175 \| 291 \| 194 \| 15 \| 13 \| 282 \| 79 \| 344 \| Chromosome partitioning protein, ParB family \| Chromosome partitioning protein, ParB family; afdb-uniprot50 \| AF-A0A1C3E499-F1-MODEL\_V4 \| 1.0 \| 1.475e-08 \| 273 \| 0.245 \| 220 \| 125 \| 12 \| 1 \| 195 \| 1 \| 204 \| ParB domain-containing protein \| ParB domain-containing protein; afdb-uniprot50 \| AF-A0A1T4M6I0-F1-MODEL\_V4 \| 1.0 \| 1.735e-08 \| 272 \| 0.134 \| 275 \| 196 \| 13 \| 19 \| 272 \| 3 \| 256 \| Chromosome partitioning protein, ParB family \| Chromosome partitioning protein, ParB family; afdb-uniprot50 \| AF-A0A2M7AKN5-F1-MODEL\_V4 \| 1.0 \| 6.44e-12 \| 272 \| 0.206 \| 296 \| 200 \| 11 \| 1 \| 282 \| 3 \| 277 \| ParB domain-containing protein \| ParB domain-containing protein; afdb-uniprot50 \| AF-A0A3N5J5N6-F1-MODEL\_V4 \| 1.0 \| 7.757e-11 \| 272 \| 0.198 \| 313 \| 190 \| 16 \| 1 \| 282 \| 4 \| 286 \| ParB/RepB/Spo0J family partition protein \| ParB/RepB/Spo0J family partition protein; afdb-uniprot50 \| AF-A0A7V8DCV6-F1-MODEL\_V4 \| 1.0 \| 6.398e-10 \| 272 \| 0.141 \| 310 \| 215 \| 14 \| 1 \| 282 \| 2 \| 288 \| Chromosome partitioning protein ParB family \| Chromosome partitioning protein ParB family; afdb-uniprot50 \| AF-A0A1M6HWY9-F1-MODEL\_V4 \| 1.0 \| 4.85e-08 \| 272 \| 0.192 \| 213 \| 139 \| 11 \| 13 \| 214 \| 73 \| 263 \| ParB/RepB/Spo0J family partition protein \| ParB/RepB/Spo0J family partition protein; afdb-uniprot50 \| AF-X0TEE3-F1-MODEL\_V4 \| 1.0 \| 6.509e-07 \| 271 \| 0.189 \| 158 \| 112 \| 9 \| 18 \| 168 \| 1 \| 149 \| ParB domain-containing protein \| ParB domain-containing protein; afdb-uniprot50 \| AF-T0YE96-F1-MODEL\_V4 \| 1.0 \| 4.428e-05 \| 271 \| 0.373 \| 91 \| 57 \| 0 \| 123 \| 213 \| 2 \| 92 \| Uncharacterized protein \| Uncharacterized protein; afdb-uniprot50 \| AF-A0A1F8H6F6-F1-MODEL\_V4 \| 1.0 \| 1.397e-08 \| 271 \| 0.154 \| 291 \| 198 \| 14 \| 13 \| 282 \| 8 \| 271 \| ParB domain-containing protein \| ParB domain-containing protein; afdb-uniprot50 \| AF-A0A4R6UCT0-F1-MODEL\_V4 \| 1.0 \| 6.753e-10 \| 271 \| 0.207 \| 308 \| 186 \| 15 \| 22 \| 289 \| 36 \| 325 \| ParB family chromosome partitioning protein \| ParB family chromosome partitioning protein; afdb-uniprot50 \| AF-A0A8B0SSF4-F1-MODEL\_V4 \| 1.0 \| 3.814e-09 \| 271 \| 0.186 \| 279 \| 171 \| 14 \| 5 \| 267 \| 16 \| 254 \| ParB/RepB/Spo0J family partition protein \| ParB/RepB/Spo0J family partition protein; afdb-uniprot50 \| AF-A0A0D6P2K6-F1-MODEL\_V4 \| 1.0 \| 2.757e-09 \| 270 \| 0.2 \| 264 \| 180 \| 10 \| 26 \| 282 \| 2 \| 241 \| Chromosome partitioning nuclease protein ParB \| Chromosome partitioning nuclease protein ParB; afdb-uniprot50 \| AF-A0A1Z9RIE5-F1-MODEL\_V4 \| 1.0 \| 8.643e-11 \| 270 \| 0.168 \| 285 \| 199 \| 13 \| 13 \| 282 \| 23 \| 284 \| Probable chromosome-partitioning protein ParB \| Probable chromosome-partitioning protein ParB; afdb-uniprot50 \| AF-A0A3M1XFH8-F1-MODEL\_V4 \| 1.0 \| 2.55e-10 \| 270 \| 0.202 \| 296 \| 196 \| 15 \| 12 \| 290 \| 20 \| 292 \| ParB/RepB/Spo0J family partition protein \| ParB/RepB/Spo0J family partition protein; afdb-uniprot50 \| AF-A0A1Y4DG68-F1-MODEL\_V4 \| 1.0 \| 3.423e-09 \| 270 \| 0.152 \| 296 \| 210 \| 13 \| 8 \| 287 \| 26 \| 296 \| ParB domain-containing protein \| ParB domain-containing protein; afdb-uniprot50 \| AF-A0A7X1ZX84-F1-MODEL\_V4 \| 1.0 \| 9.343e-10 \| 270 \| 0.174 \| 310 \| 200 \| 14 \| 1 \| 291 \| 5 \| 277 \| ParB/RepB/Spo0J family partition protein \| ParB/RepB/Spo0J family partition protein; afdb-uniprot50 \| AF-A0A2M6X6D0-F1-MODEL\_V4 \| 1.0 \| 1.946e-10 \| 270 \| 0.161 \| 278 \| 212 \| 10 \| 14 \| 282 \| 59 \| 324 \| Chromosome partitioning protein \| Chromosome partitioning protein; afdb-uniprot50 \| AF-A0A370GYF5-F1-MODEL\_V4 \| 1.0 \| 1.993e-09 \| 270 \| 0.176 \| 300 \| 191 \| 16 \| 13 \| 282 \| 55 \| 328 \| Chromosome segregation DNA-binding protein \| Chromosome segregation DNA-binding protein; afdb-uniprot50 \| AF-A0A1Q8LIG9-F1-MODEL\_V4 \| 1.0 \| 3.401e-07 \| 270 \| 0.194 \| 216 \| 146 \| 10 \| 2 \| 196 \| 16 \| 224 \| Chromosome (Plasmid) partitioning protein ParB / Stage 0 sporulation protein J \| Chromosome (Plasmid) partitioning protein ParB / Stage 0 sporulation protein J; afdb-uniprot50 \| AF-A0A1Q8LHK6-F1-MODEL\_V4 \| 1.0 \| 1.475e-08 \| 270 \| 0.235 \| 242 \| 134 \| 10 \| 10 \| 205 \| 50 \| 286 \| Chromosome (Plasmid) partitioning protein ParB / Stage 0 sporulation protein J \| Chromosome (Plasmid) partitioning protein ParB / Stage 0 sporulation protein J; afdb-uniprot50 \| AF-A0A7V4UP31-F1-MODEL\_V4 \| 1.0 \| 9.799e-08 \| 269 \| 0.218 \| 165 \| 120 \| 5 \| 2 \| 163 \| 5 \| 163 \| ParB/RepB/Spo0J family partition protein \| ParB/RepB/Spo0J family partition protein; afdb-uniprot50 \| AF-A0A6L7X7V6-F1-MODEL\_V4 \| 1.0 \| 8.188e-11 \| 269 \| 0.209 \| 282 \| 195 \| 9 \| 21 \| 291 \| 2 \| 266 \| ParB/RepB/Spo0J family partition protein \| ParB/RepB/Spo0J family partition protein; afdb-uniprot50 \| AF-A0A357IP86-F1-MODEL\_V4 \| 1.0 \| 7.301e-09 \| 269 \| 0.161 \| 298 \| 201 \| 12 \| 1 \| 282 \| 11 \| 275 \| ParB domain-containing protein \| ParB domain-containing protein; afdb-uniprot50 \| AF-A0A2A1ZTU4-F1-MODEL\_V4 \| 1.0 \| 2.474e-09 \| 269 \| 0.155 \| 295 \| 211 \| 14 \| 1 \| 283 \| 6 \| 274 \| ParB domain-containing protein \| ParB domain-containing protein; afdb-uniprot50 \| AF-R7KQ80-F1-MODEL\_V4 \| 1.0 \| 1.099e-09 \| 269 \| 0.143 \| 306 \| 232 \| 15 \| 1 \| 291 \| 1 \| 291 \| ParB domain-containing protein \| ParB domain-containing protein; afdb-uniprot50 \| AF-A0A1G2U0D6-F1-MODEL\_V4 \| 1.0 \| 3.072e-09 \| 269 \| 0.147 \| 291 \| 196 \| 12 \| 19 \| 282 \| 34 \| 299 \| ParB domain-containing protein \| ParB domain-containing protein; afdb-uniprot50 \| AF-A0A1Y6CZ00-F1-MODEL\_V4 \| 1.0 \| 1.475e-08 \| 269 \| 0.196 \| 264 \| 180 \| 14 \| 6 \| 247 \| 45 \| 298 \| ParB-like nuclease domain-containing protein \| ParB-like nuclease domain-containing protein; afdb-uniprot50 \| AF-A0A1Y3M0Q7-F1-MODEL\_V4 \| 1.0 \| 1.788e-09 \| 269 \| 0.178 \| 285 \| 190 \| 14 \| 19 \| 282 \| 96 \| 357 \| Chromosome partitioning protein ParB \| Chromosome partitioning protein ParB; afdb-uniprot50 \| AF-A0A7Z7VNP5-F1-MODEL\_V4 \| 1.0 \| 2.757e-09 \| 269 \| 0.15 \| 280 \| 192 \| 13 \| 1 \| 272 \| 33 \| 274 \| ParB/RepB/Spo0J family partition protein \| ParB/RepB/Spo0J family partition protein; afdb-uniprot50 \| AF-A0A087D6M2-F1-MODEL\_V4 \| 1.0 \| 8.135e-09 \| 269 \| 0.181 \| 297 \| 190 \| 20 \| 13 \| 290 \| 3 \| 265 \| ParB domain-containing protein \| ParB domain-containing protein; afdb-uniprot50 \| AF-A0A239PGG5-F1-MODEL\_V4 \| 1.0 \| 1.188e-08 \| 269 \| 0.171 \| 285 \| 171 \| 12 \| 1 \| 246 \| 9 \| 267 \| Chromosome partitioning protein, ParB family \| Chromosome partitioning protein, ParB family; afdb-uniprot50 \| AF-A0A257WQE5-F1-MODEL\_V4 \| 1.0 \| 2.154e-08 \| 268 \| 0.213 \| 173 \| 119 \| 7 \| 1 \| 162 \| 1 \| 167 \| Chromosome partitioning protein ParB \| Chromosome partitioning protein ParB; afdb-uniprot50 \| AF-A0A3A0A5C3-F1-MODEL\_V4 \| 1.0 \| 1.356e-07 \| 268 \| 0.2 \| 170 \| 126 \| 6 \| 5 \| 169 \| 27 \| 191 \| Stage 0 sporulation protein J \| Stage 0 sporulation protein J; afdb-uniprot50 \| AF-A0A4Y6UB58-F1-MODEL\_V4 \| 1.0 \| 6.061e-10 \| 268 \| 0.194 \| 308 \| 190 \| 16 \| 14 \| 291 \| 10 \| 289 \| ParB/RepB/Spo0J family partition protein \| ParB/RepB/Spo0J family partition protein; afdb-uniprot50 \| AF-A0A1W6N531-F1-MODEL\_V4 \| 1.0 \| 1.788e-09 \| 268 \| 0.186 \| 305 \| 215 \| 16 \| 1 \| 291 \| 7 \| 292 \| ParB domain-containing protein \| ParB domain-containing protein; afdb-uniprot50 \| AF-A0A7Y3NTN7-F1-MODEL\_V4 \| 1.0 \| 4.516e-11 \| 268 \| 0.146 \| 294 \| 196 \| 11 \| 16 \| 282 \| 50 \| 315 \| ParB/RepB/Spo0J family partition protein \| ParB/RepB/Spo0J family partition protein; afdb-uniprot50 \| AF-A0A1S9CBB3-F1-MODEL\_V4 \| 1.0 \| 1.567e-10 \| 267 \| 0.15 \| 293 \| 191 \| 10 \| 19 \| 281 \| 28 \| 292 \| ParB domain-containing protein \| ParB domain-containing protein; afdb-uniprot50 \| AF-A0A7V8DGJ8-F1-MODEL\_V4 \| 1.0 \| 5.153e-10 \| 267 \| 0.186 \| 300 \| 199 \| 14 \| 15 \| 291 \| 22 \| 299 \| Chromosome partitioning protein ParB family \| Chromosome partitioning protein ParB family; afdb-uniprot50 \| AF-A0A2D7FLY9-F1-MODEL\_V4 \| 1.0 \| 6.247e-11 \| 267 \| 0.2 \| 304 \| 191 \| 13 \| 13 \| 291 \| 70 \| 346 \| Chromosome partitioning protein ParB \| Chromosome partitioning protein ParB; afdb-uniprot50 \| AF-A0A1V5R4B4-F1-MODEL\_V4 \| 1.0 \| 1.654e-10 \| 266 \| 0.204 \| 313 \| 192 \| 14 \| 1 \| 291 \| 5 \| 282 \| Putative chromosome-partitioning protein ParB \| Putative chromosome-partitioning protein ParB; afdb-uniprot50 \| AF-A0A7C2F5M6-F1-MODEL\_V4 \| 1.0 \| 4.278e-11 \| 266 \| 0.164 \| 310 \| 200 \| 13 \| 1 \| 282 \| 5 \| 283 \| ParB/RepB/Spo0J family partition protein \| ParB/RepB/Spo0J family partition protein; afdb-uniprot50 \| AF-E6NHU5-F1-MODEL\_V4 \| 1.0 \| 8.851e-10 \| 266 \| 0.191 \| 297 \| 202 \| 15 \| 1 \| 282 \| 15 \| 288 \| Plasmid replication-partition related protein \| Plasmid replication-partition related protein; afdb-uniprot50 \| AF-A0A1H8CWV8-F1-MODEL\_V4 \| 1.0 \| 3.814e-09 \| 266 \| 0.139 \| 329 \| 186 \| 16 \| 13 \| 287 \| 22 \| 307 \| Chromosome partitioning protein, ParB family \| Chromosome partitioning protein, ParB family; afdb-uniprot50 \| AF-A0A7Y8ME42-F1-MODEL\_V4 \| 1.0 \| 2.289e-10 \| 266 \| 0.182 \| 307 \| 195 \| 13 \| 13 \| 290 \| 60 \| 339 \| ParB/RepB/Spo0J family partition protein \| ParB/RepB/Spo0J family partition protein; afdb-uniprot50 \| AF-A0A3A6KCT7-F1-MODEL\_V4 \| 1.0 \| 1.017e-10 \| 266 \| 0.168 \| 291 \| 217 \| 11 \| 1 \| 282 \| 118 \| 392 \| ParB/RepB/Spo0J family partition protein \| ParB/RepB/Spo0J family partition protein; afdb-uniprot50 \| AF-A0A7C6TI73-F1-MODEL\_V4 \| 1.0 \| 2.206e-07 \| 266 \| 0.2 \| 170 \| 123 \| 7 \| 14 \| 176 \| 383 \| 546 \| ParB/RepB/Spo0J family partition protein \| ParB/RepB/Spo0J family partition protein; afdb-uniprot50 \| AF-B4RER5-F1-MODEL\_V4 \| 1.0 \| 1.397e-08 \| 266 \| 0.189 \| 296 \| 201 \| 14 \| 1 \| 282 \| 52 \| 322 \| Predicted transcriptional regulator \| Predicted transcriptional regulator; afdb-uniprot50 \| AF-M4S4L0-F1-MODEL\_V4 \| 1.0 \| 1.324e-08 \| 265 \| 0.191 \| 261 \| 171 \| 12 \| 13 \| 267 \| 2 \| 228 \| ParB family protein \| ParB family protein; afdb-uniprot50 \| AF-A0A1M6YGD4-F1-MODEL\_V4 \| 1.0 \| 7.301e-09 \| 265 \| 0.187 \| 278 \| 175 \| 15 \| 13 \| 282 \| 2 \| 236 \| Chromosome partitioning protein, ParB family \| Chromosome partitioning protein, ParB family; afdb-uniprot50 \| AF-A0A1V4AMK9-F1-MODEL\_V4 \| 1.0 \| 1.099e-09 \| 265 \| 0.18 \| 305 \| 221 \| 13 \| 1 \| 291 \| 5 \| 294 \| HTH cro/C1-type domain-containing protein \| HTH cro/C1-type domain-containing protein; afdb-uniprot50 \| AF-A0A7C1FDR3-F1-MODEL\_V4 \| 1.0 \| 1.133e-10 \| 265 \| 0.201 \| 302 \| 193 \| 14 \| 12 \| 291 \| 31 \| 306 \| ParB/RepB/Spo0J family partition protein \| ParB/RepB/Spo0J family partition protein; afdb-uniprot50 \| AF-A0A235F0V7-F1-MODEL\_V4 \| 1.0 \| 3.423e-09 \| 265 \| 0.166 \| 312 \| 211 \| 17 \| 2 \| 291 \| 39 \| 323 \| ParB domain-containing protein \| ParB domain-containing protein; afdb-uniprot50 \| AF-A0A2W4UM90-F1-MODEL\_V4 \| 1.0 \| 1.694e-09 \| 265 \| 0.167 \| 311 \| 191 \| 20 \| 14 \| 291 \| 67 \| 342 \| Chromosome partitioning protein ParB \| Chromosome partitioning protein ParB; afdb-uniprot50 \| AF-A0A0N8Q4D0-F1-MODEL\_V4 \| 1.0 \| 2.206e-07 \| 265 \| 0.135 \| 177 \| 141 \| 7 \| 13 \| 183 \| 3 \| 173 \| Probable chromosome-partitioning protein ParB \| Probable chromosome-partitioning protein ParB; afdb-uniprot50 \| AF-C1AAS4-F1-MODEL\_V4 \| 1.0 \| 3.613e-09 \| 265 \| 0.164 \| 291 \| 187 \| 15 \| 14 \| 284 \| 27 \| 281 \| ParB-like partition protein family protein \| ParB-like partition protein family protein; afdb-uniprot50 \| AF-A0A355U2X2-F1-MODEL\_V4 \| 1.0 \| 1.066e-08 \| 264 \| 0.167 \| 281 \| 197 \| 13 \| 4 \| 280 \| 23 \| 270 \| Chromosome partitioning protein ParB \| Chromosome partitioning protein ParB; afdb-uniprot50 \| AF-I9WSL1-F1-MODEL\_V4 \| 1.0 \| 3.932e-10 \| 264 \| 0.166 \| 306 \| 199 \| 14 \| 1 \| 282 \| 15 \| 288 \| Transcriptional regulator involved in chromosome partitioning ParB \| Transcriptional regulator involved in chromosome partitioning ParB; afdb-uniprot50 \| AF-E2SKW7-F1-MODEL\_V4 \| 1.0 \| 4.457e-07 \| 264 \| 0.13 \| 199 \| 150 \| 6 \| 20 \| 200 \| 25 \| 218 \| ParB-like protein \| ParB-like protein; afdb-uniprot50 \| AF-A0A1Z9AP12-F1-MODEL\_V4 \| 1.0 \| 2.612e-09 \| 264 \| 0.184 \| 288 \| 194 \| 16 \| 3 \| 272 \| 8 \| 272 \| HTH cro/C1-type domain-containing protein \| HTH cro/C1-type domain-containing protein; afdb-uniprot50 \| AF-A0A364VD40-F1-MODEL\_V4 \| 1.0 \| 3.072e-09 \| 264 \| 0.186 \| 295 \| 193 \| 15 \| 13 \| 282 \| 165 \| 437 \| Chromosome partitioning protein ParB \| Chromosome partitioning protein ParB; afdb-uniprot50 \| AF-A0A2E0EQ09-F1-MODEL\_V4 \| 1.0 \| 4.736e-09 \| 263 \| 0.162 \| 302 \| 198 \| 14 \| 4 \| 282 \| 12 \| 281 \| Probable chromosome-partitioning protein ParB \| Probable chromosome-partitioning protein ParB; afdb-uniprot50 \| AF-A0A0Q6PTA8-F1-MODEL\_V4 \| 1.0 \| 9.343e-10 \| 263 \| 0.182 \| 329 \| 202 \| 17 \| 1 \| 291 \| 9 \| 308 \| Chromosome partitioning protein ParB \| Chromosome partitioning protein ParB; afdb-uniprot50 \| AF-A0A554R345-F1-MODEL\_V4 \| 1.0 \| 2.344e-09 \| 263 \| 0.188 \| 323 \| 183 \| 18 \| 13 \| 291 \| 2 \| 289 \| ParB/RepB/Spo0J family partition protein \| ParB/RepB/Spo0J family partition protein; afdb-uniprot50 \| AF-A0A0D5A6I8-F1-MODEL\_V4 \| 1.0 \| 1.092e-07 \| 263 \| 0.176 \| 227 \| 146 \| 11 \| 13 \| 205 \| 9 \| 228 \| Chromosome partitioning protein ParB \| Chromosome partitioning protein ParB; afdb-uniprot50 \| AF-A0A3D3EDT8-F1-MODEL\_V4 \| 1.0 \| 2.891e-07 \| 262 \| 0.198 \| 161 \| 111 \| 7 \| 11 \| 158 \| 20 \| 175 \| Chromosome partitioning protein ParB \| Chromosome partitioning protein ParB; afdb-uniprot50 \| AF-A0A512HN72-F1-MODEL\_V4 \| 1.0 \| 1.511e-07 \| 262 \| 0.16 \| 199 \| 145 \| 9 \| 18 \| 205 \| 1 \| 188 \| ParB domain-containing protein \| ParB domain-containing protein; afdb-uniprot50 \| AF-A0A496G6Q8-F1-MODEL\_V4 \| 1.0 \| 2.612e-09 \| 262 \| 0.172 \| 296 \| 203 \| 15 \| 4 \| 282 \| 18 \| 288 \| Chromosome partitioning protein ParB \| Chromosome partitioning protein ParB; afdb-uniprot50 \| AF-B8D0M7-F1-MODEL\_V4 \| 1.0 \| 3.423e-09 \| 262 \| 0.115 \| 268 \| 186 \| 11 \| 18 \| 282 \| 1 \| 220 \| ParB-like partition protein \| ParB-like partition protein; afdb-uniprot50 \| AF-Q120X6-F1-MODEL\_V4 \| 1.0 \| 2.474e-09 \| 262 \| 0.16 \| 312 \| 193 \| 17 \| 14 \| 291 \| 56 \| 332 \| ParB family protein \| ParB family protein; afdb-uniprot50 \| AF-A0A2E2Q773-F1-MODEL\_V4 \| 1.0 \| 4.736e-09 \| 262 \| 0.137 \| 327 \| 188 \| 20 \| 12 \| 271 \| 16 \| 315 \| Nuclease \| Nuclease; afdb-uniprot50 \| AF-A0A538CC36-F1-MODEL\_V4 \| 1.0 \| 2.22e-09 \| 261 \| 0.198 \| 212 \| 156 \| 6 \| 1 \| 205 \| 3 \| 207 \| ParB/RepB/Spo0J family partition protein \| ParB/RepB/Spo0J family partition protein; afdb-uniprot50 \| AF-A0A7W4KCA4-F1-MODEL\_V4 \| 1.0 \| 4.624e-10 \| 261 \| 0.162 \| 302 \| 192 \| 14 \| 12 \| 288 \| 32 \| 297 \| ParB/RepB/Spo0J family partition protein \| ParB/RepB/Spo0J family partition protein; afdb-uniprot50 \| AF-A0A3E0VBU9-F1-MODEL\_V4 \| 1.0 \| 1.557e-08 \| 261 \| 0.154 \| 266 \| 167 \| 15 \| 19 \| 257 \| 30 \| 264 \| ParB domain-containing protein \| ParB domain-containing protein; afdb-uniprot50 \| AF-A0A3L7PQ84-F1-MODEL\_V4 \| 1.0 \| 7.525e-10 \| 261 \| 0.152 \| 302 \| 213 \| 15 \| 13 \| 290 \| 108 \| 390 \| ParB/RepB/Spo0J family partition protein \| ParB/RepB/Spo0J family partition protein; afdb-uniprot50 \| AF-A0A382M7S9-F1-MODEL\_V4 \| 1.0 \| 5.57e-09 \| 260 \| 0.213 \| 187 \| 137 \| 6 \| 19 \| 201 \| 33 \| 213 \| ParB domain-containing protein \| ParB domain-containing protein; afdb-uniprot50 \| AF-A0A836SUD4-F1-MODEL\_V4 \| 1.0 \| 3.814e-09 \| 260 \| 0.157 \| 267 \| 184 \| 13 \| 20 \| 269 \| 30 \| 272 \| ParB/RepB/Spo0J family partition protein \| ParB/RepB/Spo0J family partition protein; afdb-uniprot50 \| AF-H5Y0W1-F1-MODEL\_V4 \| 1.0 \| 3.637e-11 \| 260 \| 0.14 \| 306 \| 221 \| 13 \| 1 \| 291 \| 3 \| 281 \| ParB-like partition protein \| ParB-like partition protein; afdb-uniprot50 \| AF-A0A0F6VZF3-F1-MODEL\_V4 \| 1.0 \| 1.254e-08 \| 260 \| 0.181 \| 292 \| 178 \| 13 \| 19 \| 291 \| 3 \| 252 \| Partition protein, phage-associated \| Partition protein, phage-associated; afdb-uniprot50 \| AF-A0A4Q1SR21-F1-MODEL\_V4 \| 1.0 \| 1.18e-06 \| 260 \| 0.2 \| 155 \| 108 \| 7 \| 19 \| 166 \| 3 \| 148 \| ParB domain-containing protein \| ParB domain-containing protein; afdb-uniprot50 \| AF-A0A662F1F4-F1-MODEL\_V4 \| 1.0 \| 5.404e-08 \| 260 \| 0.154 \| 278 \| 192 \| 12 \| 1 \| 246 \| 377 \| 643 \| Uncharacterized protein \| Uncharacterized protein; afdb-uniprot50 \| AF-A0A497AWF1-F1-MODEL\_V4 \| 1.0 \| 4.123e-08 \| 259 \| 0.163 \| 269 \| 187 \| 13 \| 1 \| 263 \| 7 \| 243 \| ParB domain-containing protein \| ParB domain-containing protein; afdb-uniprot50 \| AF-A0A1F6GQ66-F1-MODEL\_V4 \| 1.0 \| 2.344e-09 \| 259 \| 0.187 \| 278 \| 191 \| 15 \| 20 \| 282 \| 30 \| 287 \| HTH cro/C1-type domain-containing protein \| HTH cro/C1-type domain-containing protein; afdb-uniprot50 \| AF-A0A2A6SDX0-F1-MODEL\_V4 \| 1.0 \| 1.293e-09 \| 259 \| 0.172 \| 296 \| 209 \| 12 \| 1 \| 282 \| 15 \| 288 \| Chromosome partitioning protein ParB \| Chromosome partitioning protein ParB; afdb-uniprot50 \| AF-A0A293UGH0-F1-MODEL\_V4 \| 1.0 \| 7.525e-10 \| 259 \| 0.169 \| 295 \| 211 \| 12 \| 1 \| 282 \| 15 \| 288 \| Chromosome partitioning protein ParB \| Chromosome partitioning protein ParB; afdb-uniprot50 \| AF-A0A174SUC7-F1-MODEL\_V4 \| 1.0 \| 6.061e-10 \| 259 \| 0.141 \| 303 \| 196 \| 16 \| 8 \| 282 \| 22 \| 288 \| Chromosome-partitioning protein parB \| Chromosome-partitioning protein parB; afdb-uniprot50 \| AF-A0A3S0DKL5-F1-MODEL\_V4 \| 1.0 \| 6.207e-09 \| 259 \| 0.173 \| 311 \| 189 \| 17 \| 13 \| 287 \| 190 \| 468 \| ParB/RepB/Spo0J family partition protein \| ParB/RepB/Spo0J family partition protein; afdb-uniprot50 \| AF-A0A4Q6E257-F1-MODEL\_V4 \| 1.0 \| 3.166e-10 \| 258 \| 0.18 \| 315 \| 192 \| 14 \| 1 \| 282 \| 9 \| 290 \| ParB/RepB/Spo0J family partition protein \| ParB/RepB/Spo0J family partition protein; afdb-uniprot50 \| AF-A0A1F6RNA7-F1-MODEL\_V4 \| 1.0 \| 3.423e-09 \| 258 \| 0.162 \| 264 \| 181 \| 11 \| 7 \| 246 \| 23 \| 270 \| ParB domain-containing protein \| ParB domain-containing protein; afdb-uniprot50 \| AF-A0A1D8AU14-F1-MODEL\_V4 \| 1.0 \| 7.656e-07 \| 258 \| 0.18 \| 150 \| 105 \| 4 \| 20 \| 156 \| 81 \| 225 \| ParB-like nuclease domain protein \| ParB-like nuclease domain protein; afdb-uniprot50 \| AF-A0A353CWW4-F1-MODEL\_V4 \| 1.0 \| 7.129e-10 \| 258 \| 0.184 \| 304 \| 205 \| 16 \| 13 \| 291 \| 167 \| 452 \| ParB domain-containing protein \| ParB domain-containing protein; afdb-uniprot50 \| AF-A0A5C7LNT5-F1-MODEL\_V4 \| 1.0 \| 1.921e-06 \| 257 \| 0.152 \| 144 \| 115 \| 5 \| 20 \| 160 \| 27 \| 166 \| Chromosome partitioning protein ParB \| Chromosome partitioning protein ParB; afdb-uniprot50 \| AF-A0A2M7AKL3-F1-MODEL\_V4 \| 1.0 \| 4.278e-11 \| 257 \| 0.208 \| 307 \| 192 \| 17 \| 7 \| 291 \| 6 \| 283 \| ParB domain-containing protein \| ParB domain-containing protein; afdb-uniprot50 \| AF-A0A133UR66-F1-MODEL\_V4 \| 1.0 \| 6.022e-08 \| 257 \| 0.142 \| 295 \| 215 \| 12 \| 4 \| 289 \| 20 \| 285 \| ParB domain-containing protein \| ParB domain-containing protein; afdb-uniprot50 \| AF-A0A2T6WAW2-F1-MODEL\_V4 \| 1.0 \| 6.398e-10 \| 257 \| 0.189 \| 301 \| 198 \| 16 \| 1 \| 282 \| 15 \| 288 \| Chromosome partitioning protein ParB \| Chromosome partitioning protein ParB; afdb-uniprot50 \| AF-A0A7X7RYK0-F1-MODEL\_V4 \| 1.0 \| 1.694e-09 \| 257 \| 0.132 \| 303 \| 212 \| 13 \| 19 \| 291 \| 44 \| 325 \| ParB/RepB/Spo0J family partition protein \| ParB/RepB/Spo0J family partition protein; afdb-uniprot50 \| AF-A0A1F0CX37-F1-MODEL\_V4 \| 1.0 \| 7.944e-10 \| 257 \| 0.183 \| 311 \| 200 \| 13 \| 13 \| 290 \| 67 \| 356 \| ParB domain-containing protein \| ParB domain-containing protein; afdb-uniprot50 \| AF-A0A2N2T789-F1-MODEL\_V4 \| 1.0 \| 1.888e-09 \| 256 \| 0.205 \| 273 \| 171 \| 13 \| 39 \| 287 \| 1 \| 251 \| Chromosome partitioning protein ParB \| Chromosome partitioning protein ParB; afdb-uniprot50 \| AF-A0A7C1ZJF9-F1-MODEL\_V4 \| 1.0 \| 7.944e-10 \| 256 \| 0.199 \| 281 \| 178 \| 12 \| 21 \| 287 \| 2 \| 249 \| ParB/RepB/Spo0J family partition protein \| ParB/RepB/Spo0J family partition protein; afdb-uniprot50 \| AF-A0A0R1QC79-F1-MODEL\_V4 \| 1.0 \| 5.88e-09 \| 256 \| 0.181 \| 287 \| 190 \| 15 \| 13 \| 271 \| 12 \| 281 \| Chromosome partitioning protein, DNA-binding protein \| Chromosome partitioning protein, DNA-binding protein; afdb-uniprot50 \| AF-A0A2E8TD00-F1-MODEL\_V4 \| 1.0 \| 9.863e-10 \| 256 \| 0.154 \| 305 \| 213 \| 12 \| 4 \| 289 \| 11 \| 289 \| ParB domain-containing protein \| ParB domain-containing protein; afdb-uniprot50 \| AF-A0A2D7NEY1-F1-MODEL\_V4 \| 1.0 \| 8.135e-09 \| 256 \| 0.16 \| 280 \| 202 \| 13 \| 20 \| 282 \| 23 \| 286 \| Chromosome partitioning protein ParB \| Chromosome partitioning protein ParB; afdb-uniprot50 \| AF-A0A4Q5N154-F1-MODEL\_V4 \| 1.0 \| 7.301e-09 \| 256 \| 0.147 \| 319 \| 193 \| 15 \| 12 \| 271 \| 7 \| 305 \| Nuclease \| Nuclease; afdb-uniprot50 \| AF-A0A3D4V4B5-F1-MODEL\_V4 \| 1.0 \| 7.706e-09 \| 256 \| 0.164 \| 292 \| 190 \| 15 \| 12 \| 284 \| 143 \| 399 \| ParB/RepB/Spo0J family partition protein \| ParB/RepB/Spo0J family partition protein; afdb-uniprot50 \| AF-A0A5E7N365-F1-MODEL\_V4 \| 1.0 \| 2.739e-07 \| 255 \| 0.261 \| 191 \| 116 \| 5 \| 98 \| 287 \| 7 \| 173 \| Uncharacterized protein \| Uncharacterized protein; afdb-uniprot50 \| AF-A0A496RX12-F1-MODEL\_V4 \| 1.0 \| 1.125e-08 \| 255 \| 0.123 \| 275 \| 214 \| 12 \| 19 \| 282 \| 12 \| 270 \| Uncharacterized protein \| Uncharacterized protein; afdb-uniprot50 \| AF-A0A4R1KCC4-F1-MODEL\_V4 \| 1.0 \| 1.233e-11 \| 255 \| 0.197 \| 298 \| 198 \| 13 \| 1 \| 282 \| 3 \| 275 \| ParB family chromosome partitioning protein \| ParB family chromosome partitioning protein; afdb-uniprot50 \| AF-A0A134BLW4-F1-MODEL\_V4 \| 1.0 \| 6.398e-10 \| 255 \| 0.16 \| 287 \| 210 \| 10 \| 13 \| 282 \| 23 \| 295 \| Putative stage 0 sporulation protein J \| Putative stage 0 sporulation protein J; afdb-uniprot50 \| AF-A0A7W1TPW5-F1-MODEL\_V4 \| 1.0 \| 3.423e-09 \| 255 \| 0.173 \| 289 \| 194 \| 13 \| 7 \| 286 \| 28 \| 280 \| ParB/RepB/Spo0J family partition protein \| ParB/RepB/Spo0J family partition protein; afdb-uniprot50 \| AF-A0A3M2F058-F1-MODEL\_V4 \| 1.0 \| 5.153e-10 \| 255 \| 0.178 \| 303 \| 183 \| 16 \| 14 \| 282 \| 57 \| 327 \| ParB/RepB/Spo0J family partition protein \| ParB/RepB/Spo0J family partition protein; afdb-uniprot50 \| AF-A0A2M7JUR1-F1-MODEL\_V4 \| 1.0 \| 2.041e-08 \| 255 \| 0.189 \| 258 \| 158 \| 12 \| 19 \| 246 \| 1 \| 237 \| ParB domain-containing protein \| ParB domain-containing protein; afdb-uniprot50 \| AF-A0A850QHF8-F1-MODEL\_V4 \| 1.0 \| 2.274e-08 \| 255 \| 0.119 \| 317 \| 209 \| 16 \| 4 \| 287 \| 59 \| 338 \| ParB/RepB/Spo0J family partition protein \| ParB/RepB/Spo0J family partition protein; afdb-uniprot50 \| AF-A0A1C3XLZ9-F1-MODEL\_V4 \| 1.0 \| 1.475e-08 \| 255 \| 0.153 \| 287 \| 201 \| 13 \| 18 \| 291 \| 1 \| 258 \| ParB/RepB/Spo0J family partition protein \| ParB/RepB/Spo0J family partition protein; afdb-uniprot50 \| AF-A0A1F1V577-F1-MODEL\_V4 \| 1.0 \| 1.066e-08 \| 255 \| 0.175 \| 296 \| 181 \| 15 \| 19 \| 282 \| 156 \| 420 \| ParB domain-containing protein \| ParB domain-containing protein; afdb-uniprot50 \| AF-A0A1G3M1J8-F1-MODEL\_V4 \| 1.0 \| 6.509e-07 \| 254 \| 0.161 \| 161 \| 123 \| 7 \| 13 \| 168 \| 17 \| 170 \| ParB domain-containing protein \| ParB domain-containing protein; afdb-uniprot50 \| AF-A0A349XYG3-F1-MODEL\_V4 \| 1.0 \| 2.154e-08 \| 254 \| 0.166 \| 289 \| 194 \| 15 \| 18 \| 291 \| 1 \| 257 \| Plasmid partitioning protein \| Plasmid partitioning protein; afdb-uniprot50 \| AF-A0A0F9RH44-F1-MODEL\_V4 \| 1.0 \| 1.315e-06 \| 254 \| 0.171 \| 163 \| 124 \| 5 \| 18 \| 173 \| 1 \| 159 \| Uncharacterized protein \| Uncharacterized protein; afdb-uniprot50 \| AF-A0A1G5EUA4-F1-MODEL\_V4 \| 1.0 \| 9.568e-09 \| 254 \| 0.175 \| 285 \| 186 \| 15 \| 1 \| 265 \| 9 \| 264 \| Chromosome partitioning protein, ParB family \| Chromosome partitioning protein, ParB family; afdb-uniprot50 \| AF-A0A853ERN6-F1-MODEL\_V4 \| 1.0 \| 1.993e-09 \| 254 \| 0.179 \| 346 \| 168 \| 19 \| 19 \| 291 \| 3 \| 305 \| ParB/RepB/Spo0J family partition protein \| ParB/RepB/Spo0J family partition protein; afdb-uniprot50 \| AF-A0A5S4SWL2-F1-MODEL\_V4 \| 1.0 \| 1.993e-09 \| 254 \| 0.177 \| 282 \| 193 \| 13 \| 1 \| 272 \| 4 \| 256 \| ParB/RepB/Spo0J family partition protein \| ParB/RepB/Spo0J family partition protein; afdb-uniprot50 \| AF-A0A536HIP7-F1-MODEL\_V4 \| 1.0 \| 2.274e-08 \| 253 \| 0.188 \| 186 \| 143 \| 4 \| 21 \| 202 \| 1 \| 182 \| ParB/RepB/Spo0J family partition protein \| ParB/RepB/Spo0J family partition protein; afdb-uniprot50 \| AF-A0A846M3I8-F1-MODEL\_V4 \| 1.0 \| 1.694e-09 \| 253 \| 0.176 \| 266 \| 199 \| 10 \| 23 \| 282 \| 2 \| 253 \| ParB family chromosome partitioning protein \| ParB family chromosome partitioning protein; afdb-uniprot50 \| AF-A0A5C0UI65-F1-MODEL\_V4 \| 1.0 \| 1.254e-08 \| 253 \| 0.126 \| 292 \| 188 \| 12 \| 13 \| 287 \| 31 \| 272 \| Probable chromosome-partitioning protein ParB \| Probable chromosome-partitioning protein ParB; afdb-uniprot50 \| AF-A0A134AFD9-F1-MODEL\_V4 \| 1.0 \| 2.675e-08 \| 253 \| 0.124 \| 273 \| 202 \| 11 \| 1 \| 246 \| 4 \| 266 \| ParB-like protein \| ParB-like protein; afdb-uniprot50 \| AF-A0A139JRJ0-F1-MODEL\_V4 \| 1.0 \| 3.528e-10 \| 253 \| 0.182 \| 279 \| 193 \| 10 \| 1 \| 271 \| 6 \| 257 \| ParB-like protein \| ParB-like protein; afdb-uniprot50 \| AF-A0A3N4RJM6-F1-MODEL\_V4 \| 1.0 \| 1.485e-10 \| 253 \| 0.195 \| 307 \| 209 \| 17 \| 5 \| 291 \| 61 \| 349 \| ParB family chromosome partitioning protein \| ParB family chromosome partitioning protein; afdb-uniprot50 \| AF-A0A1H0AXS4-F1-MODEL\_V4 \| 1.0 \| 8.794e-08 \| 252 \| 0.166 \| 276 \| 188 \| 13 \| 18 \| 281 \| 1 \| 246 \| ParB/RepB/Spo0J family partition protein \| ParB/RepB/Spo0J family partition protein; afdb-uniprot50 \| AF-A0A2V7WSH1-F1-MODEL\_V4 \| 1.0 \| 1.946e-10 \| 251 \| 0.236 \| 254 \| 172 \| 11 \| 13 \| 256 \| 18 \| 259 \| Chromosome partitioning protein ParB \| Chromosome partitioning protein ParB; afdb-uniprot50 \| AF-A0A1W9SAC9-F1-MODEL\_V4 \| 1.0 \| 1.993e-09 \| 251 \| 0.161 \| 298 \| 204 \| 15 \| 9 \| 280 \| 20 \| 297 \| Probable chromosome-partitioning protein ParB \| Probable chromosome-partitioning protein ParB; afdb-uniprot50 \| AF-A0A7C6PVT3-F1-MODEL\_V4 \| 1.0 \| 7.944e-10 \| 251 \| 0.18 \| 311 \| 194 \| 14 \| 2 \| 275 \| 1 \| 287 \| ParB/RepB/Spo0J family partition protein \| ParB/RepB/Spo0J family partition protein; afdb-uniprot50 \| AF-A0A2N2BDG8-F1-MODEL\_V4 \| 1.0 \| 3.906e-08 \| 251 \| 0.146 \| 226 \| 150 \| 4 \| 13 \| 199 \| 19 \| 240 \| Chromosome partitioning protein ParB \| Chromosome partitioning protein ParB; afdb-uniprot50 \| AF-A0A1Q3TKN1-F1-MODEL\_V4 \| 1.0 \| 1.735e-08 \| 251 \| 0.177 \| 281 \| 180 \| 15 \| 19 \| 283 \| 44 \| 289 \| ParB domain-containing protein \| ParB domain-containing protein; afdb-uniprot50 \| AF-A0A1R1LSW7-F1-MODEL\_V4 \| 1.0 \| 5.57e-09 \| 251 \| 0.189 \| 327 \| 164 \| 14 \| 1 \| 273 \| 1 \| 280 \| ParB domain-containing protein \| ParB domain-containing protein; afdb-uniprot50 \| AF-A0A7X7NNE2-F1-MODEL\_V4 \| 1.0 \| 1.324e-08 \| 251 \| 0.18 \| 255 \| 160 \| 10 \| 13 \| 247 \| 50 \| 275 \| ParB/RepB/Spo0J family partition protein \| ParB/RepB/Spo0J family partition protein; afdb-uniprot50 \| AF-A0A0F9PBG4-F1-MODEL\_V4 \| 1.0 \| 1.284e-07 \| 250 \| 0.133 \| 233 \| 178 \| 8 \| 18 \| 245 \| 1 \| 214 \| ParB domain-containing protein \| ParB domain-containing protein; afdb-uniprot50 \| AF-A0A800BNS5-F1-MODEL\_V4 \| 1.0 \| 5.842e-07 \| 249 \| 0.225 \| 155 \| 107 \| 8 \| 1 \| 148 \| 4 \| 152 \| ParB/RepB/Spo0J family partition protein \| ParB/RepB/Spo0J family partition protein; afdb-uniprot50 \| AF-A0A6M0L040-F1-MODEL\_V4 \| 1.0 \| 6.356e-08 \| 249 \| 0.175 \| 205 \| 156 \| 9 \| 4 \| 200 \| 14 \| 213 \| ParB/RepB/Spo0J family partition protein \| ParB/RepB/Spo0J family partition protein; afdb-uniprot50 \| AF-A0A4Y4W985-F1-MODEL\_V4 \| 1.0 \| 8.385e-10 \| 249 \| 0.157 \| 304 \| 204 \| 12 \| 1 \| 282 \| 15 \| 288 \| Chromosome partitioning protein ParB \| Chromosome partitioning protein ParB; afdb-uniprot50 \| AF-A0A3D8IE22-F1-MODEL\_V4 \| 1.0 \| 1.188e-08 \| 249 \| 0.142 \| 308 \| 210 \| 15 \| 4 \| 282 \| 18 \| 300 \| Chromosome partitioning protein ParB \| Chromosome partitioning protein ParB; afdb-uniprot50 \| AF-A0A2H1L4V2-F1-MODEL\_V4 \| 1.0 \| 2.344e-09 \| 249 \| 0.189 \| 316 \| 203 \| 17 \| 1 \| 291 \| 27 \| 314 \| Chromosome partitioning protein, ParB family \| Chromosome partitioning protein, ParB family; afdb-uniprot50 \| AF-A0A430DXP8-F1-MODEL\_V4 \| 1.0 \| 2.534e-08 \| 249 \| 0.142 \| 295 \| 204 \| 14 \| 19 \| 290 \| 51 \| 319 \| ParB domain-containing protein \| ParB domain-containing protein; afdb-uniprot50 \| AF-A0A2E4QU58-F1-MODEL\_V4 \| 1.0 \| 2.842e-10 \| 249 \| 0.16 \| 317 \| 206 \| 15 \| 2 \| 291 \| 57 \| 340 \| Chromosome partitioning protein ParB \| Chromosome partitioning protein ParB; afdb-uniprot50 \| AF-A0A7C3VNC3-F1-MODEL\_V4 \| 1.0 \| 1.724e-06 \| 249 \| 0.182 \| 159 \| 119 \| 6 \| 14 \| 167 \| 39 \| 191 \| ParB domain-containing protein \| ParB domain-containing protein; afdb-uniprot50 \| AF-A0A7W6T8F3-F1-MODEL\_V4 \| 1.0 \| 7.706e-09 \| 249 \| 0.146 \| 301 \| 187 \| 17 \| 18 \| 291 \| 1 \| 258 \| ParB family chromosome partitioning protein \| ParB family chromosome partitioning protein; afdb-uniprot50 \| AF-A0A7W4IY87-F1-MODEL\_V4 \| 1.0 \| 9.343e-10 \| 248 \| 0.178 \| 292 \| 186 \| 13 \| 14 \| 283 \| 34 \| 293 \| ParB/RepB/Spo0J family partition protein \| ParB/RepB/Spo0J family partition protein; afdb-uniprot50 \| AF-A0A318SDS1-F1-MODEL\_V4 \| 1.0 \| 8.135e-09 \| 248 \| 0.2 \| 274 \| 158 \| 15 \| 13 \| 245 \| 24 \| 277 \| ParB family protein \| ParB family protein; afdb-uniprot50 \| AF-A0A3C1FVK0-F1-MODEL\_V4 \| 1.0 \| 7.706e-09 \| 248 \| 0.144 \| 304 \| 203 \| 16 \| 6 \| 287 \| 30 \| 298 \| ParB domain-containing protein \| ParB domain-containing protein; afdb-uniprot50 \| AF-A0A7K1PX19-F1-MODEL\_V4 \| 1.0 \| 1.831e-08 \| 248 \| 0.168 \| 290 \| 193 \| 14 \| 18 \| 291 \| 1 \| 258 \| ParB/RepB/Spo0J family partition protein \| ParB/RepB/Spo0J family partition protein; afdb-uniprot50 \| AF-K9D005-F1-MODEL\_V4 \| 1.0 \| 9.799e-08 \| 248 \| 0.157 \| 279 \| 186 \| 14 \| 18 \| 284 \| 1 \| 242 \| ParB-like partition protein \| ParB-like partition protein; afdb-uniprot50 \| AF-A0A318KWI1-F1-MODEL\_V4 \| 1.0 \| 7.476e-08 \| 247 \| 0.261 \| 199 \| 118 \| 9 \| 104 \| 291 \| 10 \| 190 \| KorB domain-containing protein \| KorB domain-containing protein; afdb-uniprot50 \| AF-A0A646LUJ1-F1-MODEL\_V4 \| 1.0 \| 2.4e-08 \| 247 \| 0.127 \| 275 \| 199 \| 12 \| 19 \| 282 \| 7 \| 251 \| ParB/RepB/Spo0J family partition protein \| ParB/RepB/Spo0J family partition protein; afdb-uniprot50 \| AF-K9F001-F1-MODEL\_V4 \| 1.0 \| 9.568e-09 \| 247 \| 0.143 \| 328 \| 210 \| 15 \| 4 \| 291 \| 17 \| 313 \| ParB-like partition protein \| ParB-like partition protein; afdb-uniprot50 \| AF-A0A1Q3T1A0-F1-MODEL\_V4 \| 1.0 \| 1.788e-09 \| 247 \| 0.177 \| 316 \| 204 \| 20 \| 5 \| 289 \| 32 \| 322 \| ParB domain-containing protein \| ParB domain-containing protein; afdb-uniprot50 \| AF-A0A3E0V9I7-F1-MODEL\_V4 \| 1.0 \| 4.352e-08 \| 247 \| 0.149 \| 281 \| 176 \| 15 \| 4 \| 257 \| 60 \| 304 \| ParB domain-containing protein \| ParB domain-containing protein; afdb-uniprot50 \| AF-A0A2H9LML8-F1-MODEL\_V4 \| 1.0 \| 1.98e-07 \| 246 \| 0.13 \| 300 \| 168 \| 18 \| 16 \| 291 \| 1 \| 231 \| ParB domain-containing protein \| ParB domain-containing protein; afdb-uniprot50 \| AF-A0A2T2QZM2-F1-MODEL\_V4 \| 1.0 \| 4.881e-10 \| 246 \| 0.196 \| 290 \| 190 \| 13 \| 13 \| 284 \| 34 \| 298 \| ParB domain-containing protein \| ParB domain-containing protein; afdb-uniprot50 \| AF-A0A135IPZ6-F1-MODEL\_V4 \| 1.0 \| 3.146e-08 \| 246 \| 0.193 \| 186 \| 136 \| 7 \| 18 \| 196 \| 1 \| 179 \| ParB domain-containing protein \| ParB domain-containing protein; afdb-uniprot50 \| AF-A0A3A6MUC0-F1-MODEL\_V4 \| 1.0 \| 2.385e-06 \| 246 \| 0.151 \| 185 \| 143 \| 9 \| 19 \| 193 \| 49 \| 229 \| ParB/RepB/Spo0J family partition protein \| ParB/RepB/Spo0J family partition protein; afdb-uniprot50 \| AF-A0A521TB26-F1-MODEL\_V4 \| 1.0 \| 1.777e-07 \| 245 \| 0.129 \| 271 \| 172 \| 12 \| 19 \| 282 \| 7 \| 220 \| ParB/RepB/Spo0J family partition protein \| ParB/RepB/Spo0J family partition protein; afdb-uniprot50 \| AF-A0A077FQM9-F1-MODEL\_V4 \| 1.0 \| 1.188e-08 \| 245 \| 0.155 \| 308 \| 180 \| 17 \| 5 \| 278 \| 62 \| 323 \| Probable chromosome-partitioning protein ParB \| Probable chromosome-partitioning protein ParB; afdb-uniprot50 \| AF-A0A432MEU0-F1-MODEL\_V4 \| 1.0 \| 2.416e-10 \| 245 \| 0.167 \| 329 \| 210 \| 13 \| 1 \| 291 \| 37 \| 339 \| ParB/RepB/Spo0J family partition protein \| ParB/RepB/Spo0J family partition protein; afdb-uniprot50 \| AF-A0A3D3Q403-F1-MODEL\_V4 \| 1.0 \| 1.066e-08 \| 244 \| 0.175 \| 194 \| 150 \| 6 \| 14 \| 205 \| 17 \| 202 \| Chromosome partitioning protein ParB \| Chromosome partitioning protein ParB; afdb-uniprot50 \| AF-A0A2E8Y707-F1-MODEL\_V4 \| 1.0 \| 4.881e-10 \| 244 \| 0.198 \| 277 \| 192 \| 11 \| 19 \| 284 \| 21 \| 278 \| ParB domain-containing protein \| ParB domain-containing protein; afdb-uniprot50 \| AF-A0A316RSR3-F1-MODEL\_V4 \| 1.0 \| 5.607e-11 \| 244 \| 0.193 \| 316 \| 195 \| 15 \| 1 \| 284 \| 5 \| 292 \| Stage 0 sporulation protein J \| Stage 0 sporulation protein J; afdb-uniprot50 \| AF-A0A2M9YJ10-F1-MODEL\_V4 \| 1.0 \| 1.254e-08 \| 244 \| 0.171 \| 268 \| 185 \| 16 \| 1 \| 246 \| 32 \| 284 \| Chromophore lyase \| Chromophore lyase; afdb-uniprot50 \| AF-A0A1I1EQZ5-F1-MODEL\_V4 \| 1.0 \| 1.324e-08 \| 244 \| 0.197 \| 192 \| 139 \| 8 \| 13 \| 195 \| 19 \| 204 \| ParB/RepB/Spo0J family partition protein \| ParB/RepB/Spo0J family partition protein; afdb-uniprot50 \| AF-A0A0D5MBW0-F1-MODEL\_V4 \| 1.0 \| 6.552e-09 \| 244 \| 0.167 \| 298 \| 195 \| 16 \| 4 \| 271 \| 15 \| 289 \| ParB-like partition protein \| ParB-like partition protein; afdb-uniprot50 \| AF-A0A7X6YTY0-F1-MODEL\_V4 \| 1.0 \| 3.637e-11 \| 243 \| 0.176 \| 301 \| 208 \| 18 \| 3 \| 282 \| 27 \| 308 \| ParB/RepB/Spo0J family partition protein \| ParB/RepB/Spo0J family partition protein; afdb-uniprot50 \| AF-F7UI03-F1-MODEL\_V4 \| 1.0 \| 3.7e-08 \| 243 \| 0.15 \| 305 \| 211 \| 16 \| 1 \| 291 \| 75 \| 345 \| Chromosome partitioning protein \| Chromosome partitioning protein; afdb-uniprot50 \| AF-A0A0C1N062-F1-MODEL\_V4 \| 1.0 \| 2.274e-08 \| 242 \| 0.13 \| 292 \| 185 \| 16 \| 13 \| 287 \| 26 \| 265 \| Probable chromosome-partitioning protein ParB \| Probable chromosome-partitioning protein ParB; afdb-uniprot50 \| AF-A0A7C6ZXT2-F1-MODEL\_V4 \| 1.0 \| 1.644e-08 \| 242 \| 0.183 \| 273 \| 185 \| 10 \| 13 \| 271 \| 6 \| 254 \| ParB/RepB/Spo0J family partition protein \| ParB/RepB/Spo0J family partition protein; afdb-uniprot50 \| AF-A0A2D7KGV3-F1-MODEL\_V4 \| 1.0 \| 6.753e-10 \| 242 \| 0.169 \| 307 \| 201 \| 13 \| 3 \| 282 \| 11 \| 290 \| Chromosome partitioning protein ParB \| Chromosome partitioning protein ParB; afdb-uniprot50 \| AF-A0A066YW29-F1-MODEL\_V4 \| 1.0 \| 1.406e-10 \| 242 \| 0.211 \| 312 \| 202 \| 17 \| 1 \| 290 \| 21 \| 310 \| Plasmid partitioning protein ParB \| Plasmid partitioning protein ParB; afdb-uniprot50 \| AF-A0A1Y3TUR7-F1-MODEL\_V4 \| 1.0 \| 1.431e-07 \| 242 \| 0.107 \| 260 \| 191 \| 9 \| 20 \| 243 \| 24 \| 278 \| ParB domain-containing protein \| ParB domain-containing protein; afdb-uniprot50 \| AF-A0A7V4MZE4-F1-MODEL\_V4 \| 1.0 \| 8.851e-10 \| 242 \| 0.149 \| 314 \| 203 \| 14 \| 19 \| 291 \| 27 \| 317 \| ParB/RepB/Spo0J family partition protein \| ParB/RepB/Spo0J family partition protein; afdb-uniprot50 \| AF-A0A5K1NR86-F1-MODEL\_V4 \| 1.0 \| 1.324e-08 \| 242 \| 0.156 \| 307 \| 198 \| 13 \| 1 \| 273 \| 6 \| 285 \| ParB domain-containing protein \| ParB domain-containing protein; afdb-uniprot50 \| AF-A0A3E2TLN0-F1-MODEL\_V4 \| 1.0 \| 1.225e-09 \| 241 \| 0.161 \| 298 \| 207 \| 12 \| 4 \| 283 \| 38 \| 310 \| ParB/RepB/Spo0J family partition protein \| ParB/RepB/Spo0J family partition protein; afdb-uniprot50 \| AF-A0A0K2SP45-F1-MODEL\_V4 \| 1.0 \| 4.486e-09 \| 241 \| 0.148 \| 282 \| 180 \| 12 \| 19 \| 291 \| 3 \| 233 \| Chromosome partitioning protein ParB \| Chromosome partitioning protein ParB; afdb-uniprot50 \| AF-A0A3G9J1Q0-F1-MODEL\_V4 \| 1.0 \| 1.017e-10 \| 241 \| 0.204 \| 289 \| 192 \| 13 \| 19 \| 291 \| 75 \| 341 \| Chromosome partitioning protein ParB \| Chromosome partitioning protein ParB; afdb-uniprot50 \| AF-A0A517ZB25-F1-MODEL\_V4 \| 1.0 \| 1.98e-07 \| 241 \| 0.209 \| 196 \| 123 \| 12 \| 19 \| 206 \| 4 \| 175 \| DNA methylase \| DNA methylase; afdb-uniprot50 \| AF-A0A348VT13-F1-MODEL\_V4 \| 1.0 \| 4.25e-09 \| 240 \| 0.159 \| 182 \| 136 \| 5 \| 18 \| 195 \| 1 \| 169 \| Chromosome partitioning protein ParB \| Chromosome partitioning protein ParB; afdb-uniprot50 \| AF-A0A6L7GCN8-F1-MODEL\_V4 \| 1.0 \| 6.552e-09 \| 240 \| 0.229 \| 222 \| 157 \| 11 \| 19 \| 234 \| 17 \| 230 \| ParB/RepB/Spo0J family partition protein \| ParB/RepB/Spo0J family partition protein; afdb-uniprot50 \| AF-A0A0W0ZRH2-F1-MODEL\_V4 \| 1.0 \| 7.129e-10 \| 240 \| 0.182 \| 285 \| 198 \| 12 \| 13 \| 282 \| 13 \| 277 \| Probable chromosome-partitioning protein ParB \| Probable chromosome-partitioning protein ParB; afdb-uniprot50 \| AF-A0A2M7R6K7-F1-MODEL\_V4 \| 1.0 \| 1.293e-09 \| 240 \| 0.197 \| 284 \| 197 \| 13 \| 13 \| 284 \| 24 \| 288 \| ParB domain-containing protein \| ParB domain-containing protein; afdb-uniprot50 \| AF-A0A850MS07-F1-MODEL\_V4 \| 1.0 \| 1.683e-07 \| 240 \| 0.214 \| 233 \| 153 \| 11 \| 19 \| 246 \| 3 \| 210 \| ParB/RepB/Spo0J family partition protein \| ParB/RepB/Spo0J family partition protein; afdb-uniprot50 \| AF-A0A1G9MR58-F1-MODEL\_V4 \| 1.0 \| 7.656e-07 \| 240 \| 0.115 \| 217 \| 165 \| 11 \| 2 \| 204 \| 26 \| 229 \| Chromosome partitioning protein, ParB family \| Chromosome partitioning protein, ParB family; afdb-uniprot50 \| AF-A0A7R8P8J7-F1-MODEL\_V4 \| 1.0 \| 3.222e-07 \| 240 \| 0.175 \| 228 \| 140 \| 13 \| 13 \| 205 \| 3 \| 217 \| ParB family chromosome partitioning protein \| ParB family chromosome partitioning protein; afdb-uniprot50 \| AF-A0A729SM52-F1-MODEL\_V4 \| 1.0 \| 2.757e-09 \| 239 \| 0.2 \| 225 \| 151 \| 11 \| 1 \| 214 \| 6 \| 212 \| ParB/RepB/Spo0J family partition protein \| ParB/RepB/Spo0J family partition protein; afdb-uniprot50 \| AF-A0A536BIE7-F1-MODEL\_V4 \| 1.0 \| 1.254e-08 \| 239 \| 0.2 \| 210 \| 154 \| 7 \| 3 \| 205 \| 15 \| 217 \| ParB/RepB/Spo0J family partition protein \| ParB/RepB/Spo0J family partition protein; afdb-uniprot50 \| AF-A0A377TSX5-F1-MODEL\_V4 \| 1.0 \| 1.284e-07 \| 239 \| 0.172 \| 266 \| 162 \| 13 \| 37 \| 290 \| 36 \| 255 \| PRTRC system ParB family protein \| PRTRC system ParB family protein; afdb-uniprot50 \| AF-A0A0F5MQM1-F1-MODEL\_V4 \| 1.0 \| 7.892e-08 \| 239 \| 0.148 \| 310 \| 191 \| 13 \| 1 \| 289 \| 21 \| 278 \| Probable chromosome-partitioning protein ParB \| Probable chromosome-partitioning protein ParB; afdb-uniprot50 \| AF-A0A6H3NI83-F1-MODEL\_V4 \| 1.0 \| 1.735e-08 \| 239 \| 0.164 \| 267 \| 188 \| 12 \| 7 \| 246 \| 13 \| 271 \| ParB/RepB/Spo0J family partition protein \| ParB/RepB/Spo0J family partition protein; afdb-uniprot50 \| AF-A0A3A8HK35-F1-MODEL\_V4 \| 1.0 \| 4.352e-08 \| 239 \| 0.153 \| 312 \| 183 \| 15 \| 1 \| 286 \| 6 \| 262 \| ParB/RepB/Spo0J family partition protein \| ParB/RepB/Spo0J family partition protein; afdb-uniprot50 \| AF-A0A1V6A6D7-F1-MODEL\_V4 \| 1.0 \| 2.235e-11 \| 239 \| 0.166 \| 295 \| 208 \| 12 \| 5 \| 282 \| 18 \| 291 \| Putative chromosome-partitioning protein ParB \| Putative chromosome-partitioning protein ParB; afdb-uniprot50 \| AF-A0A2S9K0I3-F1-MODEL\_V4 \| 1.0 \| 1.217e-07 \| 239 \| 0.169 \| 306 \| 190 \| 18 \| 13 \| 291 \| 34 \| 302 \| Chromosome partitioning protein ParB \| Chromosome partitioning protein ParB; afdb-uniprot50 \| AF-C2GFK6-F1-MODEL\_V4 \| 1.0 \| 4.624e-10 \| 239 \| 0.164 \| 285 \| 190 \| 13 \| 20 \| 282 \| 76 \| 334 \| ParB-like protein \| ParB-like protein; afdb-uniprot50 \| AF-A0A2H9SV72-F1-MODEL\_V4 \| 1.0 \| 1.746e-10 \| 238 \| 0.188 \| 302 \| 197 \| 15 \| 1 \| 282 \| 1 \| 274 \| Chromosome partitioning protein \| Chromosome partitioning protein; afdb-uniprot50 \| AF-A0A538PBW6-F1-MODEL\_V4 \| 1.0 \| 1.118e-06 \| 238 \| 0.171 \| 221 \| 144 \| 10 \| 12 \| 201 \| 7 \| 219 \| ParB/RepB/Spo0J family partition protein \| ParB/RepB/Spo0J family partition protein; afdb-uniprot50 \| AF-A0A6L7U0D4-F1-MODEL\_V4 \| 1.0 \| 7.944e-10 \| 238 \| 0.183 \| 321 \| 206 \| 15 \| 1 \| 289 \| 11 \| 307 \| ParB/RepB/Spo0J family partition protein \| ParB/RepB/Spo0J family partition protein; afdb-uniprot50 \| AF-A0A0R2KRI4-F1-MODEL\_V4 \| 1.0 \| 3.814e-09 \| 238 \| 0.2 \| 290 \| 177 \| 16 \| 13 \| 271 \| 15 \| 280 \| Chromosome partitioning protein, DNA-binding protein \| Chromosome partitioning protein, DNA-binding protein; afdb-uniprot50 \| AF-A0A3D4JLM1-F1-MODEL\_V4 \| 1.0 \| 6.207e-09 \| 238 \| 0.151 \| 311 \| 194 \| 11 \| 1 \| 246 \| 1 \| 306 \| Chromosome partitioning protein ParB \| Chromosome partitioning protein ParB; afdb-uniprot50 \| AF-A0A345J9F5-F1-MODEL\_V4 \| 1.0 \| 4.352e-08 \| 238 \| 0.15 \| 280 \| 188 \| 15 \| 7 \| 273 \| 65 \| 307 \| Partitioning protein, ParB family \| Partitioning protein, ParB family; afdb-uniprot50 \| AF-A0A2N3GCV6-F1-MODEL\_V4 \| 1.0 \| 2.98e-08 \| 238 \| 0.186 \| 193 \| 145 \| 8 \| 14 \| 199 \| 190 \| 377 \| ParB domain-containing protein \| ParB domain-containing protein; afdb-uniprot50 \| AF-A0A4Q3IEN1-F1-MODEL\_V4 \| 1.0 \| 2.534e-08 \| 237 \| 0.177 \| 186 \| 142 \| 6 \| 4 \| 182 \| 9 \| 190 \| ParB/RepB/Spo0J family partition protein \| ParB/RepB/Spo0J family partition protein; afdb-uniprot50 \| AF-A0A1U7GP86-F1-MODEL\_V4 \| 1.0 \| 5.842e-07 \| 237 \| 0.159 \| 257 \| 162 \| 13 \| 23 \| 246 \| 2 \| 237 \| ParB domain-containing protein \| ParB domain-containing protein; afdb-uniprot50 \| AF-A0A2H0NAZ6-F1-MODEL\_V4 \| 1.0 \| 4.85e-08 \| 236 \| 0.139 \| 294 \| 198 \| 14 \| 13 \| 287 \| 1 \| 258 \| ParB domain-containing protein \| ParB domain-containing protein; afdb-uniprot50 \| AF-A0A535EWL3-F1-MODEL\_V4 \| 1.0 \| 1.356e-07 \| 236 \| 0.142 \| 287 \| 188 \| 12 \| 7 \| 271 \| 55 \| 305 \| ParB/RepB/Spo0J family partition protein \| ParB/RepB/Spo0J family partition protein; afdb-uniprot50 \| AF-A0A523SL81-F1-MODEL\_V4 \| 1.0 \| 2.385e-06 \| 236 \| 0.167 \| 191 \| 143 \| 10 \| 19 \| 202 \| 4 \| 185 \| ParB/RepB/Spo0J family partition protein \| ParB/RepB/Spo0J family partition protein; afdb-uniprot50 \| AF-A0A1W2CT08-F1-MODEL\_V4 \| 1.0 \| 3.423e-09 \| 236 \| 0.16 \| 343 \| 198 \| 14 \| 18 \| 287 \| 1 \| 326 \| Chromosome partitioning protein, ParB family \| Chromosome partitioning protein, ParB family; afdb-uniprot50 \| AF-A0A7C7L2N7-F1-MODEL\_V4 \| 1.0 \| 1.921e-06 \| 235 \| 0.217 \| 138 \| 99 \| 6 \| 9 \| 142 \| 20 \| 152 \| ParB/RepB/Spo0J family partition protein \| ParB/RepB/Spo0J family partition protein; afdb-uniprot50 \| AF-A0A3D1XDE0-F1-MODEL\_V4 \| 1.0 \| 9.124e-11 \| 235 \| 0.184 \| 298 \| 214 \| 11 \| 3 \| 287 \| 10 \| 291 \| ParB domain-containing protein \| ParB domain-containing protein; afdb-uniprot50 \| AF-A0A6M3M0X8-F1-MODEL\_V4 \| 1.0 \| 3.505e-08 \| 235 \| 0.159 \| 308 \| 207 \| 16 \| 13 \| 291 \| 2 \| 286 \| KorB domain-containing protein \| KorB domain-containing protein; afdb-uniprot50 \| AF-D7BJI1-F1-MODEL\_V4 \| 1.0 \| 9.006e-07 \| 235 \| 0.185 \| 216 \| 142 \| 12 \| 19 \| 205 \| 1 \| 211 \| ParB-like partition protein \| ParB-like partition protein; afdb-uniprot50 \| AF-A0A3D4YD69-F1-MODEL\_V4 \| 1.0 \| 2.154e-08 \| 234 \| 0.125 \| 294 \| 187 \| 17 \| 13 \| 287 \| 30 \| 272 \| ParB domain-containing protein \| ParB domain-containing protein; afdb-uniprot50 \| AF-A0A3C0QI21-F1-MODEL\_V4 \| 1.0 \| 3.166e-10 \| 234 \| 0.129 \| 301 \| 217 \| 16 \| 1 \| 282 \| 14 \| 288 \| Chromosome partitioning protein ParB \| Chromosome partitioning protein ParB; afdb-uniprot50 \| AF-A0A2W1B217-F1-MODEL\_V4 \| 1.0 \| 3.505e-08 \| 233 \| 0.132 \| 265 \| 179 \| 9 \| 13 \| 273 \| 6 \| 223 \| Chromosome segregation protein Spo0J, contains ParB-like nuclease domain \| Chromosome segregation protein Spo0J, contains ParB-like nuclease domain; afdb-uniprot50 \| AF-A0A848FHG5-F1-MODEL\_V4 \| 1.0 \| 3.906e-08 \| 233 \| 0.144 \| 311 \| 208 \| 20 \| 1 \| 288 \| 64 \| 339 \| ParB/RepB/Spo0J family partition protein \| ParB/RepB/Spo0J family partition protein; afdb-uniprot50 \| AF-A0A7X6ZRM7-F1-MODEL\_V4 \| 1.0 \| 4.624e-10 \| 233 \| 0.182 \| 290 \| 197 \| 15 \| 13 \| 282 \| 82 \| 351 \| ParB/RepB/Spo0J family partition protein \| ParB/RepB/Spo0J family partition protein; afdb-uniprot50 \| AF-A0A3D1WJI5-F1-MODEL\_V4 \| 1.0 \| 6.356e-08 \| 232 \| 0.178 \| 269 \| 171 \| 16 \| 19 \| 270 \| 3 \| 238 \| Chromosome partitioning protein ParB \| Chromosome partitioning protein ParB; afdb-uniprot50 \| AF-A0A1C6C1D8-F1-MODEL\_V4 \| 1.0 \| 2.675e-08 \| 232 \| 0.134 \| 335 \| 218 \| 15 \| 1 \| 291 \| 1 \| 307 \| Probable chromosome-partitioning protein parB \| Probable chromosome-partitioning protein parB; afdb-uniprot50 \| AF-A0A0E2D6A4-F1-MODEL\_V4 \| 1.0 \| 5.119e-08 \| 232 \| 0.147 \| 271 \| 200 \| 11 \| 1 \| 247 \| 29 \| 292 \| ParB-like protein \| ParB-like protein; afdb-uniprot50 \| AF-B6WUQ8-F1-MODEL\_V4 \| 1.0 \| 9.343e-10 \| 232 \| 0.177 \| 310 \| 198 \| 14 \| 14 \| 291 \| 34 \| 318 \| ParB-like protein \| ParB-like protein; afdb-uniprot50 \| AF-A0A523SL66-F1-MODEL\_V4 \| 1.0 \| 9.283e-08 \| 232 \| 0.172 \| 267 \| 172 \| 10 \| 21 \| 262 \| 1 \| 243 \| ParB/RepB/Spo0J family partition protein \| ParB/RepB/Spo0J family partition protein; afdb-uniprot50 \| AF-A0A0C2AKN6-F1-MODEL\_V4 \| 1.0 \| 1.153e-07 \| 232 \| 0.165 \| 308 \| 196 \| 13 \| 1 \| 272 \| 4 \| 286 \| ParB domain-containing protein \| ParB domain-containing protein; afdb-uniprot50 \| AF-A0A3D4YDN4-F1-MODEL\_V4 \| 1.0 \| 5.704e-08 \| 231 \| 0.133 \| 306 \| 171 \| 14 \| 1 \| 275 \| 13 \| 255 \| ParB domain-containing protein \| ParB domain-containing protein; afdb-uniprot50 \| AF-A0A3C0WTF4-F1-MODEL\_V4 \| 1.0 \| 8.794e-08 \| 231 \| 0.159 \| 251 \| 177 \| 12 \| 21 \| 246 \| 1 \| 242 \| ParB domain-containing protein \| ParB domain-containing protein; afdb-uniprot50 \| AF-A0A7C2RDB0-F1-MODEL\_V4 \| 1.0 \| 1.397e-08 \| 231 \| 0.151 \| 284 \| 195 \| 11 \| 15 \| 287 \| 40 \| 288 \| ParB/RepB/Spo0J family partition protein \| ParB/RepB/Spo0J family partition protein; afdb-uniprot50 \| AF-A0A2K8WTV7-F1-MODEL\_V4 \| 1.0 \| 1.595e-07 \| 231 \| 0.157 \| 229 \| 149 \| 10 \| 10 \| 200 \| 16 \| 238 \| ParB-like partitioning protein \| ParB-like partitioning protein; afdb-uniprot50 \| AF-A0A0F6IBY4-F1-MODEL\_V4 \| 1.0 \| 1.644e-08 \| 231 \| 0.128 \| 295 \| 204 \| 15 \| 7 \| 287 \| 38 \| 293 \| ParB-like protein \| ParB-like protein; afdb-uniprot50 \| AF-A0A3P3DPP2-F1-MODEL\_V4 \| 1.0 \| 2.98e-08 \| 231 \| 0.148 \| 297 \| 196 \| 14 \| 19 \| 287 \| 52 \| 319 \| Plasmid partitioning protein RepB \| Plasmid partitioning protein RepB; afdb-uniprot50 \| AF-A0A6P2FYS8-F1-MODEL\_V4 \| 1.0 \| 3.789e-07 \| 231 \| 0.137 \| 283 \| 200 \| 16 \| 20 \| 282 \| 66 \| 324 \| ParB/RepB/Spo0J family partition protein \| ParB/RepB/Spo0J family partition protein; afdb-uniprot50 \| AF-A0A523VK37-F1-MODEL\_V4 \| 1.0 \| 1.465e-06 \| 231 \| 0.148 \| 283 \| 149 \| 8 \| 8 \| 200 \| 250 \| 530 \| MPN domain-containing protein \| MPN domain-containing protein; afdb-uniprot50 \| AF-A0A7U9T1Z3-F1-MODEL\_V4 \| 1.0 \| 1.431e-07 \| 230 \| 0.17 \| 240 \| 152 \| 9 \| 13 \| 214 \| 23 \| 253 \| Chromosome-partitioning protein Spo0J \| Chromosome-partitioning protein Spo0J; afdb-uniprot50 \| AF-A0A6H1Z970-F1-MODEL\_V4 \| 1.0 \| 3.401e-07 \| 230 \| 0.198 \| 196 \| 125 \| 11 \| 19 \| 206 \| 4 \| 175 \| Putative methyltransferase \| Putative methyltransferase; afdb-uniprot50 \| AF-A0A8B2NJQ1-F1-MODEL\_V4 \| 1.0 \| 1.876e-07 \| 230 \| 0.193 \| 181 \| 142 \| 4 \| 18 \| 196 \| 1 \| 179 \| Chromosome partitioning protein ParB \| Chromosome partitioning protein ParB; afdb-uniprot50 \| AF-X1DCG4-F1-MODEL\_V4 \| 1.0 \| 4.594e-08 \| 229 \| 0.149 \| 174 \| 140 \| 5 \| 13 \| 182 \| 21 \| 190 \| ParB domain-containing protein \| ParB domain-containing protein; afdb-uniprot50 \| AF-A0A7X7UCM7-F1-MODEL\_V4 \| 1.0 \| 4.486e-09 \| 229 \| 0.18 \| 321 \| 191 \| 16 \| 1 \| 282 \| 5 \| 292 \| ParB/RepB/Spo0J family partition protein \| ParB/RepB/Spo0J family partition protein; afdb-uniprot50 \| AF-A0A7I7Y4I0-F1-MODEL\_V4 \| 1.0 \| 6.753e-10 \| 229 \| 0.203 \| 304 \| 192 \| 15 \| 13 \| 290 \| 17 \| 296 \| Chromosome partitioning protein ParB \| Chromosome partitioning protein ParB; afdb-uniprot50 \| AF-A0A4U9YID6-F1-MODEL\_V4 \| 1.0 \| 3.789e-07 \| 229 \| 0.123 \| 234 \| 165 \| 7 \| 4 \| 202 \| 13 \| 241 \| ParB-like nuclease protein \| ParB-like nuclease protein; afdb-uniprot50 \| AF-A0A7W9PDS6-F1-MODEL\_V4 \| 1.0 \| 1.041e-09 \| 229 \| 0.183 \| 299 \| 188 \| 16 \| 14 \| 282 \| 57 \| 329 \| ParB family chromosome partitioning protein \| ParB family chromosome partitioning protein; afdb-uniprot50 \| AF-A0A6I7P420-F1-MODEL\_V4 \| 1.0 \| 2.329e-07 \| 229 \| 0.164 \| 323 \| 202 \| 16 \| 1 \| 286 \| 2 \| 293 \| ParB/RepB/Spo0J family partition protein \| ParB/RepB/Spo0J family partition protein; afdb-uniprot50 \| AF-A0A496Q7M5-F1-MODEL\_V4 \| 1.0 \| 2.028e-06 \| 228 \| 0.169 \| 230 \| 163 \| 11 \| 27 \| 247 \| 7 \| 217 \| ParB domain-containing protein \| ParB domain-containing protein; afdb-uniprot50 \| AF-A0A4Q3UFB4-F1-MODEL\_V4 \| 1.0 \| 4.123e-08 \| 228 \| 0.157 \| 305 \| 199 \| 18 \| 13 \| 290 \| 18 \| 291 \| ParB/RepB/Spo0J family partition protein \| ParB/RepB/Spo0J family partition protein; afdb-uniprot50 \| AF-A0A2N2TMN6-F1-MODEL\_V4 \| 1.0 \| 1.595e-07 \| 228 \| 0.155 \| 309 \| 199 \| 18 \| 13 \| 291 \| 20 \| 296 \| Chromosome partitioning protein ParB \| Chromosome partitioning protein ParB; afdb-uniprot50 \| AF-A0A1Y4UN77-F1-MODEL\_V4 \| 1.0 \| 3.445e-11 \| 227 \| 0.169 \| 306 \| 195 \| 14 \| 1 \| 282 \| 1 \| 271 \| Chromosome partitioning protein ParB \| Chromosome partitioning protein ParB; afdb-uniprot50 \| AF-A0A523VV34-F1-MODEL\_V4 \| 1.0 \| 3.321e-08 \| 227 \| 0.17 \| 281 \| 170 \| 11 \| 18 \| 258 \| 1 \| 258 \| ParB/RepB/Spo0J family partition protein \| ParB/RepB/Spo0J family partition protein; afdb-uniprot50 \| AF-A0A6N7YJT5-F1-MODEL\_V4 \| 1.0 \| 3.166e-10 \| 227 \| 0.174 \| 316 \| 213 \| 15 \| 1 \| 291 \| 80 \| 372 \| ParB/RepB/Spo0J family partition protein \| ParB/RepB/Spo0J family partition protein; afdb-uniprot50 \| AF-A0A326UQP0-F1-MODEL\_V4 \| 1.0 \| 6.356e-08 \| 227 \| 0.194 \| 304 \| 210 \| 10 \| 5 \| 287 \| 35 \| 324 \| ParB/RepB/Spo0J family partition protein \| ParB/RepB/Spo0J family partition protein; afdb-uniprot50 \| AF-A0A3D8P5H7-F1-MODEL\_V4 \| 1.0 \| 2.154e-08 \| 227 \| 0.185 \| 308 \| 171 \| 16 \| 13 \| 287 \| 6 \| 266 \| ParB domain-containing protein \| ParB domain-containing protein; afdb-uniprot50 \| AF-A0A1G1X644-F1-MODEL\_V4 \| 1.0 \| 2.344e-09 \| 226 \| 0.162 \| 308 \| 201 \| 16 \| 11 \| 291 \| 21 \| 298 \| ParB domain-containing protein \| ParB domain-containing protein; afdb-uniprot50 \| AF-B4RIU1-F1-MODEL\_V4 \| 1.0 \| 4.123e-08 \| 226 \| 0.233 \| 201 \| 137 \| 10 \| 3 \| 195 \| 9 \| 200 \| ParB-like partition protein \| ParB-like partition protein; afdb-uniprot50 \| AF-A0A1F5NRR0-F1-MODEL\_V4 \| 1.0 \| 1.993e-09 \| 226 \| 0.207 \| 280 \| 183 \| 12 \| 20 \| 282 \| 59 \| 316 \| ParB domain-containing protein \| ParB domain-containing protein; afdb-uniprot50 \| AF-K6TXH8-F1-MODEL\_V4 \| 1.0 \| 4.123e-08 \| 226 \| 0.157 \| 317 \| 189 \| 15 \| 1 \| 246 \| 6 \| 315 \| Putative transcriptional regulator \| Putative transcriptional regulator; afdb-uniprot50 \| AF-A0A5B8VFP4-F1-MODEL\_V4 \| 1.0 \| 1.324e-08 \| 226 \| 0.101 \| 385 \| 208 \| 17 \| 13 \| 287 \| 23 \| 379 \| ParB/RepB/Spo0J family partition protein \| ParB/RepB/Spo0J family partition protein; afdb-uniprot50 \| AF-A0A1R3U4M9-F1-MODEL\_V4 \| 1.0 \| 4.705e-07 \| 226 \| 0.151 \| 283 \| 191 \| 14 \| 18 \| 284 \| 1 \| 250 \| Putative chromosome-partitioning protein ParB \| Putative chromosome-partitioning protein ParB; afdb-uniprot50 \| AF-A0A1Q6TFJ3-F1-MODEL\_V4 \| 1.0 \| 3.146e-08 \| 225 \| 0.131 \| 320 \| 199 \| 14 \| 1 \| 291 \| 1 \| 270 \| ParB domain-containing protein \| ParB domain-containing protein; afdb-uniprot50 \| AF-A0A1T1DHW5-F1-MODEL\_V4 \| 1.0 \| 2.961e-06 \| 225 \| 0.128 \| 226 \| 174 \| 10 \| 1 \| 214 \| 22 \| 236 \| Chromosome partitioning protein ParB \| Chromosome partitioning protein ParB; afdb-uniprot50 \| AF-A0A1C6C4L8-F1-MODEL\_V4 \| 1.0 \| 3.401e-07 \| 225 \| 0.163 \| 232 \| 154 \| 9 \| 4 \| 202 \| 196 \| 420 \| Probable chromosome-partitioning protein parB \| Probable chromosome-partitioning protein parB; afdb-uniprot50 \| AF-A0A212AB45-F1-MODEL\_V4 \| 1.0 \| 3.146e-08 \| 224 \| 0.217 \| 216 \| 144 \| 8 \| 13 \| 224 \| 7 \| 201 \| ParB domain-containing protein \| ParB domain-containing protein; afdb-uniprot50 \| AF-A0A7Y7A5E4-F1-MODEL\_V4 \| 1.0 \| 5.704e-08 \| 224 \| 0.213 \| 253 \| 161 \| 14 \| 47 \| 282 \| 2 \| 233 \| ParB/RepB/Spo0J family partition protein \| ParB/RepB/Spo0J family partition protein; afdb-uniprot50 \| AF-R5KEP9-F1-MODEL\_V4 \| 1.0 \| 2.154e-08 \| 224 \| 0.129 \| 331 \| 198 \| 13 \| 1 \| 246 \| 1 \| 326 \| ParB domain protein nuclease \| ParB domain protein nuclease; afdb-uniprot50 \| AF-A0A1Q6QJG1-F1-MODEL\_V4 \| 1.0 \| 3.906e-08 \| 223 \| 0.158 \| 322 \| 195 \| 14 \| 13 \| 287 \| 26 \| 318 \| ParB domain-containing protein \| ParB domain-containing protein; afdb-uniprot50 \| AF-A0A4V5NLE5-F1-MODEL\_V4 \| 1.0 \| 1.98e-07 \| 223 \| 0.131 \| 312 \| 224 \| 15 \| 1 \| 286 \| 32 \| 322 \| Plasmid partitioning protein RepB \| Plasmid partitioning protein RepB; afdb-uniprot50 \| AF-A0A853IEB4-F1-MODEL\_V4 \| 1.0 \| 9.568e-09 \| 223 \| 0.155 \| 327 \| 209 \| 19 \| 1 \| 291 \| 1 \| 296 \| ParB/RepB/Spo0J family partition protein \| ParB/RepB/Spo0J family partition protein; afdb-uniprot50 \| AF-A0A1Q3TEL5-F1-MODEL\_V4 \| 1.0 \| 1.475e-08 \| 222 \| 0.192 \| 317 \| 177 \| 17 \| 1 \| 272 \| 1 \| 283 \| ParB domain-containing protein \| ParB domain-containing protein; afdb-uniprot50 \| AF-A0A1F5DHD6-F1-MODEL\_V4 \| 1.0 \| 1.092e-07 \| 222 \| 0.143 \| 278 \| 184 \| 15 \| 19 \| 261 \| 19 \| 277 \| ParB domain-containing protein \| ParB domain-containing protein; afdb-uniprot50 \| AF-A0A0H5Q929-F1-MODEL\_V4 \| 1.0 \| 5.153e-10 \| 221 \| 0.166 \| 301 \| 211 \| 14 \| 1 \| 282 \| 6 \| 285 \| ParB domain-containing protein \| ParB domain-containing protein; afdb-uniprot50 \| AF-A0A1F9VR61-F1-MODEL\_V4 \| 1.0 \| 1.225e-09 \| 221 \| 0.206 \| 296 \| 199 \| 14 \| 13 \| 291 \| 22 \| 298 \| ParB domain-containing protein \| ParB domain-containing protein; afdb-uniprot50 \| AF-A0A127CRA7-F1-MODEL\_V4 \| 1.0 \| 8.794e-08 \| 221 \| 0.151 \| 291 \| 197 \| 15 \| 15 \| 287 \| 1 \| 259 \| ParB domain-containing protein \| ParB domain-containing protein; afdb-uniprot50 \| AF-A0A535EHP5-F1-MODEL\_V4 \| 1.0 \| 2.274e-08 \| 220 \| 0.172 \| 308 \| 205 \| 13 \| 1 \| 282 \| 1 \| 284 \| ParB/RepB/Spo0J family partition protein \| ParB/RepB/Spo0J family partition protein; afdb-uniprot50 \| AF-A0A0D0PKE9-F1-MODEL\_V4 \| 1.0 \| 3.423e-09 \| 220 \| 0.169 \| 301 \| 219 \| 11 \| 7 \| 291 \| 64 \| 349 \| Plasmid partitioning protein ParB \| Plasmid partitioning protein ParB; afdb-uniprot50 \| AF-Q5U9L6-F1-MODEL\_V4 \| 1.0 \| 2.578e-05 \| 220 \| 0.168 \| 148 \| 110 \| 8 \| 56 \| 196 \| 1 \| 142 \| Putative ParB partitioning protein \| Putative ParB partitioning protein; afdb-uniprot50 \| AF-A0A800CKI2-F1-MODEL\_V4 \| 1.0 \| 3.126e-06 \| 220 \| 0.145 \| 233 \| 145 \| 9 \| 19 \| 199 \| 1 \| 231 \| Uncharacterized protein \| Uncharacterized protein; afdb-uniprot50 \| AF-A0A3L6ZQY3-F1-MODEL\_V4 \| 1.0 \| 2.041e-08 \| 219 \| 0.19 \| 231 \| 145 \| 11 \| 1 \| 196 \| 26 \| 249 \| ParB/RepB/Spo0J family partition protein \| ParB/RepB/Spo0J family partition protein; afdb-uniprot50 \| AF-A0A259Q3N7-F1-MODEL\_V4 \| 1.0 \| 3.789e-07 \| 219 \| 0.148 \| 277 \| 198 \| 12 \| 1 \| 265 \| 15 \| 265 \| ParB domain-containing protein \| ParB domain-containing protein; afdb-uniprot50 \| AF-E1M761-F1-MODEL\_V4 \| 1.0 \| 2.595e-07 \| 218 \| 0.19 \| 231 \| 164 \| 10 \| 56 \| 282 \| 1 \| 212 \| Stage 0 sporulation protein J \| Stage 0 sporulation protein J; afdb-uniprot50 \| AF-A0A2R4G3E4-F1-MODEL\_V4 \| 1.0 \| 6.356e-08 \| 218 \| 0.142 \| 316 \| 184 \| 16 \| 13 \| 286 \| 15 \| 285 \| ParB domain-containing protein \| ParB domain-containing protein; afdb-uniprot50 \| AF-A0A2N4X5C0-F1-MODEL\_V4 \| 1.0 \| 6.167e-07 \| 218 \| 0.188 \| 244 \| 162 \| 12 \| 23 \| 260 \| 2 \| 215 \| Chromosome partitioning protein ParB \| Chromosome partitioning protein ParB; afdb-uniprot50 \| AF-A0A109E1J8-F1-MODEL\_V4 \| 1.0 \| 6.167e-07 \| 218 \| 0.145 \| 302 \| 209 \| 17 \| 13 \| 291 \| 65 \| 340 \| ParB domain-containing protein \| ParB domain-containing protein; afdb-uniprot50 \| AF-A0A1G3BTE6-F1-MODEL\_V4 \| 1.0 \| 1.465e-06 \| 217 \| 0.111 \| 288 \| 205 \| 12 \| 15 \| 290 \| 1 \| 249 \| ParB domain-containing protein \| ParB domain-containing protein; afdb-uniprot50 \| AF-A0A3A8ZTF9-F1-MODEL\_V4 \| 1.0 \| 3.052e-07 \| 216 \| 0.158 \| 239 \| 146 \| 11 \| 13 \| 210 \| 2 \| 226 \| Chromosome partitioning protein ParB \| Chromosome partitioning protein ParB; afdb-uniprot50 \| AF-F0RQA6-F1-MODEL\_V4 \| 1.0 \| 7.253e-07 \| 216 \| 0.158 \| 233 \| 153 \| 7 \| 4 \| 199 \| 10 \| 236 \| ParB-like partition protein \| ParB-like partition protein; afdb-uniprot50 \| AF-A0A125DMD0-F1-MODEL\_V4 \| 1.0 \| 3.321e-08 \| 216 \| 0.15 \| 318 \| 190 \| 17 \| 5 \| 282 \| 57 \| 334 \| Chromosome partitioning protein ParB \| Chromosome partitioning protein ParB; afdb-uniprot50 \| AF-A0A7W9RWB0-F1-MODEL\_V4 \| 1.0 \| 4.736e-09 \| 216 \| 0.17 \| 281 \| 191 \| 16 \| 1 \| 272 \| 1 \| 248 \| ParB family chromosome partitioning protein \| ParB family chromosome partitioning protein; afdb-uniprot50 \| AF-A0A1V6GT79-F1-MODEL\_V4 \| 1.0 \| 8.794e-08 \| 216 \| 0.168 \| 296 \| 170 \| 16 \| 19 \| 282 \| 3 \| 254 \| Methyltransferase \| Methyltransferase; afdb-uniprot50 \| AF-A0A6C2U9R7-F1-MODEL\_V4 \| 1.0 \| 1.595e-07 \| 215 \| 0.141 \| 276 \| 188 \| 10 \| 18 \| 287 \| 12 \| 244 \| Chromosome-partitioning protein Spo0J \| Chromosome-partitioning protein Spo0J; afdb-uniprot50 \| AF-A0A0R2QCK2-F1-MODEL\_V4 \| 1.0 \| 4.624e-10 \| 215 \| 0.165 \| 314 \| 190 \| 14 \| 13 \| 284 \| 34 \| 317 \| ParB domain-containing protein \| ParB domain-containing protein; afdb-uniprot50 \| AF-W7X354-F1-MODEL\_V4 \| 1.0 \| 1.431e-07 \| 215 \| 0.149 \| 314 \| 194 \| 20 \| 13 \| 290 \| 57 \| 333 \| ParB/RepB/Spo0J family partition protein \| ParB/RepB/Spo0J family partition protein; afdb-uniprot50 \| AF-A0A355VXV9-F1-MODEL\_V4 \| 1.0 \| 1.511e-07 \| 214 \| 0.126 \| 300 \| 190 \| 15 \| 1 \| 287 \| 24 \| 264 \| ParB domain-containing protein \| ParB domain-containing protein; afdb-uniprot50 \| AF-A0A836VUS9-F1-MODEL\_V4 \| 1.0 \| 3.676e-06 \| 213 \| 0.182 \| 170 \| 126 \| 5 \| 2 \| 169 \| 14 \| 172 \| ParB/RepB/Spo0J family partition protein \| ParB/RepB/Spo0J family partition protein; afdb-uniprot50 \| AF-A0A0F9FPY9-F1-MODEL\_V4 \| 1.0 \| 3.401e-07 \| 213 \| 0.157 \| 280 \| 178 \| 13 \| 16 \| 287 \| 10 \| 239 \| ParB domain-containing protein \| ParB domain-containing protein; afdb-uniprot50 \| AF-A0A7X6X5M1-F1-MODEL\_V4 \| 1.0 \| 1.066e-08 \| 213 \| 0.194 \| 262 \| 182 \| 10 \| 3 \| 246 \| 23 \| 273 \| ParB/RepB/Spo0J family partition protein \| ParB/RepB/Spo0J family partition protein; afdb-uniprot50 \| AF-A0A4R5TV04-F1-MODEL\_V4 \| 1.0 \| 2.91e-09 \| 213 \| 0.189 \| 291 \| 194 \| 13 \| 13 \| 283 \| 49 \| 317 \| ParB/RepB/Spo0J family partition protein \| ParB/RepB/Spo0J family partition protein; afdb-uniprot50 \| AF-A0A2S0L073-F1-MODEL\_V4 \| 1.0 \| 3.881e-06 \| 213 \| 0.175 \| 188 \| 134 \| 7 \| 21 \| 203 \| 2 \| 173 \| ParB domain-containing protein \| ParB domain-containing protein; afdb-uniprot50 \| AF-A0A660M752-F1-MODEL\_V4 \| 1.0 \| 8.331e-08 \| 212 \| 0.222 \| 171 \| 123 \| 6 \| 3 \| 169 \| 20 \| 184 \| ParB/RepB/Spo0J family partition protein \| ParB/RepB/Spo0J family partition protein; afdb-uniprot50 \| AF-A0A378AKT6-F1-MODEL\_V4 \| 1.0 \| 5.086e-06 \| 211 \| 0.173 \| 260 \| 159 \| 12 \| 44 \| 290 \| 9 \| 225 \| PRTRC system ParB family protein \| PRTRC system ParB family protein; afdb-uniprot50 \| AF-A0A2R7NXB0-F1-MODEL\_V4 \| 1.0 \| 2.739e-07 \| 211 \| 0.19 \| 284 \| 182 \| 17 \| 1 \| 258 \| 62 \| 323 \| Chromosome partitioning protein ParB \| Chromosome partitioning protein ParB; afdb-uniprot50 \| AF-A0A246BKX2-F1-MODEL\_V4 \| 1.0 \| 1.034e-07 \| 210 \| 0.159 \| 320 \| 183 \| 11 \| 7 \| 282 \| 15 \| 292 \| ParB domain-containing protein \| ParB domain-containing protein; afdb-uniprot50 \| AF-D6TUM3-F1-MODEL\_V4 \| 1.0 \| 2.274e-08 \| 210 \| 0.187 \| 336 \| 186 \| 17 \| 14 \| 287 \| 42 \| 352 \| ParB-like partition protein \| ParB-like partition protein; afdb-uniprot50 \| AF-A0A160UJD6-F1-MODEL\_V4 \| 1.0 \| 6.509e-07 \| 209 \| 0.179 \| 178 \| 136 \| 5 \| 1 \| 172 \| 21 \| 194 \| Chromosome partitioning protein CDS \| Chromosome partitioning protein CDS; afdb-uniprot50 \| AF-A0A5E4I9I8-F1-MODEL\_V4 \| 1.0 \| 6.509e-07 \| 209 \| 0.14 \| 271 \| 178 \| 16 \| 21 \| 255 \| 2 \| 253 \| Nucleoid occlusion protein \| Nucleoid occlusion protein; afdb-uniprot50 \| AF-A0A257C4G3-F1-MODEL\_V4 \| 1.0 \| 4.352e-08 \| 209 \| 0.17 \| 276 \| 185 \| 11 \| 11 \| 273 \| 22 \| 266 \| ParB domain-containing protein \| ParB domain-containing protein; afdb-uniprot50 \| AF-A0A0D6T7F5-F1-MODEL\_V4 \| 1.0 \| 8.082e-07 \| 209 \| 0.205 \| 185 \| 138 \| 7 \| 19 \| 200 \| 4 \| 182 \| ParB domain-containing protein \| ParB domain-containing protein; afdb-uniprot50 \| AF-A0A0G1ITF5-F1-MODEL\_V4 \| 1.0 \| 4.705e-07 \| 208 \| 0.172 \| 220 \| 142 \| 10 \| 13 \| 211 \| 1 \| 201 \| ParB domain-containing protein \| ParB domain-containing protein; afdb-uniprot50 \| AF-A0A2E3SMI7-F1-MODEL\_V4 \| 1.0 \| 3.126e-06 \| 208 \| 0.147 \| 257 \| 184 \| 13 \| 13 \| 246 \| 1 \| 245 \| ParB domain-containing protein \| ParB domain-containing protein; afdb-uniprot50 \| AF-A0A845DAT5-F1-MODEL\_V4 \| 1.0 \| 2.757e-09 \| 208 \| 0.135 \| 317 \| 209 \| 14 \| 13 \| 288 \| 17 \| 309 \| ParB/RepB/Spo0J family partition protein \| ParB/RepB/Spo0J family partition protein; afdb-uniprot50 \| AF-X1SKH0-F1-MODEL\_V4 \| 1.0 \| 9.283e-08 \| 208 \| 0.18 \| 266 \| 170 \| 13 \| 22 \| 255 \| 3 \| 252 \| Uncharacterized protein \| Uncharacterized protein; afdb-uniprot50 \| AF-A0A1F5P5H6-F1-MODEL\_V4 \| 1.0 \| 1.324e-08 \| 208 \| 0.175 \| 302 \| 191 \| 16 \| 14 \| 287 \| 64 \| 335 \| ParB domain-containing protein \| ParB domain-containing protein; afdb-uniprot50 \| AF-A0A2H0QRG0-F1-MODEL\_V4 \| 1.0 \| 5.404e-08 \| 207 \| 0.106 \| 311 \| 203 \| 18 \| 10 \| 287 \| 24 \| 292 \| ParB domain-containing protein \| ParB domain-containing protein; afdb-uniprot50 \| AF-A0A443KCM7-F1-MODEL\_V4 \| 1.0 \| 3.59e-07 \| 205 \| 0.127 \| 290 \| 203 \| 14 \| 20 \| 287 \| 42 \| 303 \| Plasmid partitioning protein RepB \| Plasmid partitioning protein RepB; afdb-uniprot50 \| AF-A0A2A2GEW1-F1-MODEL\_V4 \| 1.0 \| 2.458e-07 \| 205 \| 0.147 \| 299 \| 196 \| 18 \| 19 \| 287 \| 60 \| 329 \| Plasmid partitioning protein RepB \| Plasmid partitioning protein RepB; afdb-uniprot50 \| AF-Q21QH4-F1-MODEL\_V4 \| 1.0 \| 2.329e-07 \| 205 \| 0.161 \| 279 \| 170 \| 14 \| 13 \| 284 \| 48 \| 269 \| ParB-like partition proteins \| ParB-like partition proteins; afdb-uniprot50 \| AF-A0A6N8VJ22-F1-MODEL\_V4 \| 1.0 \| 2.739e-07 \| 204 \| 0.185 \| 178 \| 130 \| 5 \| 1 \| 167 \| 15 \| 188 \| ParB/RepB/Spo0J family partition protein \| ParB/RepB/Spo0J family partition protein; afdb-uniprot50 \| AF-A0A2E7CGY7-F1-MODEL\_V4 \| 1.0 \| 1.98e-07 \| 204 \| 0.131 \| 296 \| 217 \| 11 \| 1 \| 291 \| 1 \| 261 \| HTH cro/C1-type domain-containing protein \| HTH cro/C1-type domain-containing protein; afdb-uniprot50 \| AF-A0A258WWT9-F1-MODEL\_V4 \| 1.0 \| 7.892e-08 \| 204 \| 0.132 \| 271 \| 200 \| 11 \| 19 \| 282 \| 37 \| 279 \| ParB domain-containing protein \| ParB domain-containing protein; afdb-uniprot50 \| AF-A0A539CYY2-F1-MODEL\_V4 \| 1.0 \| 6.167e-07 \| 203 \| 0.175 \| 291 \| 189 \| 11 \| 1 \| 272 \| 4 \| 262 \| ParB family protein \| ParB family protein; afdb-uniprot50 \| AF-A0A2H0TB13-F1-MODEL\_V4 \| 1.0 \| 8.385e-10 \| 203 \| 0.166 \| 289 \| 203 \| 11 \| 9 \| 282 \| 17 \| 282 \| Chromosome partitioning protein ParB \| Chromosome partitioning protein ParB; afdb-uniprot50 \| AF-A0A2D8EI10-F1-MODEL\_V4 \| 1.0 \| 9.006e-07 \| 203 \| 0.189 \| 290 \| 185 \| 15 \| 13 \| 287 \| 6 \| 260 \| ParB domain-containing protein \| ParB domain-containing protein; afdb-uniprot50 \| AF-K1MKW0-F1-MODEL\_V4 \| 1.0 \| 3.032e-05 \| 201 \| 0.122 \| 171 \| 132 \| 8 \| 3 \| 160 \| 10 \| 175 \| ParB domain-containing protein \| ParB domain-containing protein; afdb-uniprot50 \| AF-A0A2V9LAX5-F1-MODEL\_V4 \| 1.0 \| 0.0001052 \| 200 \| 0.192 \| 114 \| 87 \| 3 \| 1 \| 114 \| 5 \| 113 \| ParB domain-containing protein \| ParB domain-containing protein; afdb-uniprot50 \| AF-A0A1B2YUD7-F1-MODEL\_V4 \| 1.0 \| 0.0001908 \| 200 \| 0.137 \| 116 \| 92 \| 5 \| 14 \| 127 \| 26 \| 135 \| Chromosome (Plasmid) partitioning protein ParB \| Chromosome (Plasmid) partitioning protein ParB; afdb-uniprot50 \| AF-G5JF03-F1-MODEL\_V4 \| 1.0 \| 2.329e-07 \| 200 \| 0.146 \| 287 \| 170 \| 15 \| 12 \| 243 \| 36 \| 302 \| ParB-like protein-partition protein \| ParB-like protein-partition protein; afdb-uniprot50 \| AF-C5B6N2-F1-MODEL\_V4 \| 1.0 \| 1.465e-06 \| 200 \| 0.219 \| 187 \| 135 \| 8 \| 13 \| 193 \| 24 \| 205 \| Putative plasmid stabilization protein \| Putative plasmid stabilization protein; afdb-uniprot50 \| AF-A0A3R6DKU2-F1-MODEL\_V4 \| 1.0 \| 4.457e-07 \| 199 \| 0.113 \| 317 \| 200 \| 14 \| 12 \| 278 \| 18 \| 303 \| Chromosome partitioning protein ParB \| Chromosome partitioning protein ParB; afdb-uniprot50 \| AF-A0A239PFQ7-F1-MODEL\_V4 \| 1.0 \| 1.059e-06 \| 199 \| 0.153 \| 287 \| 173 \| 15 \| 14 \| 267 \| 16 \| 265 \| Chromosome partitioning protein, ParB family \| Chromosome partitioning protein, ParB family; afdb-uniprot50 \| AF-H4F7S0-F1-MODEL\_V4 \| 1.0 \| 3.401e-07 \| 198 \| 0.163 \| 305 \| 179 \| 18 \| 16 \| 287 \| 1 \| 262 \| Plasmid partitioning protein RepB \| Plasmid partitioning protein RepB; afdb-uniprot50 \| AF-A0A0W7WJQ7-F1-MODEL\_V4 \| 1.0 \| 5.243e-07 \| 196 \| 0.174 \| 310 \| 199 \| 18 \| 7 \| 291 \| 44 \| 321 \| ParB domain-containing protein \| ParB domain-containing protein; afdb-uniprot50 \| AF-A0A2E9QLE2-F1-MODEL\_V4 \| 1.0 \| 1.82e-06 \| 195 \| 0.108 \| 304 \| 209 \| 14 \| 1 \| 282 \| 26 \| 289 \| ParB domain-containing protein \| ParB domain-containing protein; afdb-uniprot50 \| AF-A0A379PPU0-F1-MODEL\_V4 \| 1.0 \| 1.315e-06 \| 195 \| 0.126 \| 300 \| 204 \| 18 \| 13 \| 287 \| 61 \| 327 \| Chromosome partitioning protein ParB \| Chromosome partitioning protein ParB; afdb-uniprot50 \| AF-A0A0N9UTQ1-F1-MODEL\_V4 \| 1.0 \| 1.388e-06 \| 195 \| 0.176 \| 187 \| 143 \| 6 \| 18 \| 200 \| 1 \| 180 \| ParB domain-containing protein \| ParB domain-containing protein; afdb-uniprot50 \| AF-A0A2N2LVB4-F1-MODEL\_V4 \| 1.0 \| 1.595e-07 \| 193 \| 0.164 \| 256 \| 187 \| 11 \| 21 \| 262 \| 1 \| 243 \| ParB domain-containing protein \| ParB domain-containing protein; afdb-uniprot50 \| AF-A0A842ZP40-F1-MODEL\_V4 \| 1.0 \| 4e-07 \| 193 \| 0.163 \| 306 \| 188 \| 17 \| 5 \| 290 \| 25 \| 282 \| Uncharacterized protein \| Uncharacterized protein; afdb-uniprot50 \| AF-A0A1B6BH88-F1-MODEL\_V4 \| 1.0 \| 9.445e-05 \| 192 \| 0.202 \| 143 \| 99 \| 6 \| 13 \| 144 \| 24 \| 162 \| Plasmid partitioning protein ParB \| Plasmid partitioning protein ParB; afdb-uniprot50 \| AF-A0A4R8HXN8-F1-MODEL\_V4 \| 1.0 \| 6.315e-06 \| 192 \| 0.218 \| 160 \| 119 \| 3 \| 19 \| 175 \| 5 \| 161 \| ParB family chromosome partitioning protein \| ParB family chromosome partitioning protein; afdb-uniprot50 \| AF-A0A523SL68-F1-MODEL\_V4 \| 1.0 \| 1.724e-06 \| 191 \| 0.134 \| 289 \| 178 \| 14 \| 13 \| 291 \| 4 \| 230 \| ParB/RepB/Spo0J family partition protein \| ParB/RepB/Spo0J family partition protein; afdb-uniprot50 \| AF-A0A519DTN3-F1-MODEL\_V4 \| 1.0 \| 9.006e-07 \| 191 \| 0.171 \| 280 \| 188 \| 12 \| 8 \| 271 \| 59 \| 310 \| ParB/RepB/Spo0J family partition protein \| ParB/RepB/Spo0J family partition protein; afdb-uniprot50 \| AF-F7JID4-F1-MODEL\_V4 \| 1.0 \| 3.201e-05 \| 189 \| 0.219 \| 155 \| 104 \| 8 \| 7 \| 148 \| 15 \| 165 \| ParB domain-containing protein \| ParB domain-containing protein; afdb-uniprot50 \| AF-L0A5U5-F1-MODEL\_V4 \| 1.0 \| 9.006e-07 \| 189 \| 0.163 \| 306 \| 193 \| 16 \| 14 \| 287 \| 52 \| 326 \| ParB-like partition protein \| ParB-like partition protein; afdb-uniprot50 \| AF-B9L4Q1-F1-MODEL\_V4 \| 1.0 \| 1.993e-09 \| 189 \| 0.172 \| 337 \| 207 \| 16 \| 2 \| 291 \| 15 \| 326 \| Stage 0 sporulation protein J \| Stage 0 sporulation protein J; afdb-uniprot50 \| AF-A0A0A3XRA4-F1-MODEL\_V4 \| 1.0 \| 2.274e-08 \| 189 \| 0.169 \| 277 \| 192 \| 10 \| 19 \| 281 \| 1 \| 253 \| Transcriptional regulator \| Transcriptional regulator; afdb-uniprot50 \| AF-X1JNN1-F1-MODEL\_V4 \| 1.0 \| 0.000407 \| 188 \| 0.111 \| 134 \| 111 \| 6 \| 19 \| 146 \| 11 \| 142 \| ParB domain-containing protein \| ParB domain-containing protein; afdb-uniprot50 \| AF-A0A350T1Y3-F1-MODEL\_V4 \| 1.0 \| 1.831e-08 \| 188 \| 0.154 \| 297 \| 214 \| 13 \| 1 \| 287 \| 10 \| 279 \| Stage 0 sporulation protein J \| Stage 0 sporulation protein J; afdb-uniprot50 \| AF-A0A3E4UB54-F1-MODEL\_V4 \| 1.0 \| 1.118e-06 \| 188 \| 0.136 \| 345 \| 192 \| 13 \| 14 \| 282 \| 50 \| 364 \| Chromosome partitioning protein ParB \| Chromosome partitioning protein ParB; afdb-uniprot50 \| AF-Q892Z6-F1-MODEL\_V4 \| 1.0 \| 1.276e-05 \| 187 \| 0.135 \| 199 \| 122 \| 9 \| 1 \| 156 \| 6 \| 197 \| Chromosome partitioning parB family protein \| Chromosome partitioning parB family protein; afdb-uniprot50 \| AF-A0A1G0JK78-F1-MODEL\_V4 \| 1.0 \| 1.547e-06 \| 186 \| 0.137 \| 204 \| 156 \| 9 \| 19 \| 214 \| 1 \| 192 \| ParB domain-containing protein \| ParB domain-containing protein; afdb-uniprot50 \| AF-A0A845UHA6-F1-MODEL\_V4 \| 1.0 \| 1.431e-07 \| 185 \| 0.126 \| 365 \| 216 \| 17 \| 1 \| 287 \| 38 \| 377 \| ParB/RepB/Spo0J family partition protein \| ParB/RepB/Spo0J family partition protein; afdb-uniprot50 \| AF-A0A7V8X4E6-F1-MODEL\_V4 \| 1.0 \| 3.299e-06 \| 184 \| 0.16 \| 168 \| 123 \| 6 \| 9 \| 163 \| 50 \| 212 \| ParB/RepB/Spo0J family partition protein \| ParB/RepB/Spo0J family partition protein; afdb-uniprot50 \| AF-A0A3A0AS42-F1-MODEL\_V4 \| 1.0 \| 1.066e-08 \| 184 \| 0.148 \| 329 \| 197 \| 15 \| 1 \| 287 \| 12 \| 299 \| Stage 0 sporulation protein J \| Stage 0 sporulation protein J; afdb-uniprot50 \| AF-A0A2N0BFB6-F1-MODEL\_V4 \| 1.0 \| 0.0002787 \| 183 \| 0.15 \| 113 \| 86 \| 3 \| 13 \| 119 \| 20 \| 128 \| Chromosome partitioning protein ParB \| Chromosome partitioning protein ParB; afdb-uniprot50 \| AF-A0A0F9K647-F1-MODEL\_V4 \| 1.0 \| 1.028e-05 \| 183 \| 0.114 \| 315 \| 210 \| 13 \| 27 \| 291 \| 26 \| 321 \| Uncharacterized protein \| Uncharacterized protein; afdb-uniprot50 \| AF-A0A0B6RLI6-F1-MODEL\_V4 \| 1.0 \| 6.71e-08 \| 182 \| 0.177 \| 310 \| 162 \| 14 \| 3 \| 291 \| 1 \| 238 \| ParB-like nuclease \| ParB-like nuclease; afdb-uniprot50 \| AF-A0A352R4S9-F1-MODEL\_V4 \| 1.0 \| 3.222e-07 \| 182 \| 0.161 \| 285 \| 185 \| 11 \| 14 \| 288 \| 60 \| 300 \| ParB domain-containing protein \| ParB domain-containing protein; afdb-uniprot50 \| AF-W7B412-F1-MODEL\_V4 \| 1.0 \| 2.192e-05 \| 181 \| 0.172 \| 162 \| 111 \| 9 \| 1 \| 156 \| 1 \| 145 \| Putative Partition protein ParB-like protein \| Putative Partition protein ParB-like protein; afdb-uniprot50 \| AF-A0A535YXC4-F1-MODEL\_V4 \| 1.0 \| 9.064e-09 \| 181 \| 0.177 \| 298 \| 186 \| 13 \| 13 \| 280 \| 23 \| 291 \| ParB/RepB/Spo0J family partition protein \| ParB/RepB/Spo0J family partition protein; afdb-uniprot50 \| AF-A0A2D9IDC7-F1-MODEL\_V4 \| 1.0 \| 4.85e-08 \| 179 \| 0.166 \| 294 \| 191 \| 17 \| 18 \| 291 \| 1 \| 260 \| ParB domain-containing protein \| ParB domain-containing protein; afdb-uniprot50 \| AF-A0A840SV32-F1-MODEL\_V4 \| 1.0 \| 7.083e-08 \| 178 \| 0.16 \| 361 \| 184 \| 15 \| 19 \| 290 \| 3 \| 333 \| ParB family chromosome partitioning protein \| ParB family chromosome partitioning protein; afdb-uniprot50 \| AF-A0A373Q515-F1-MODEL\_V4 \| 1.0 \| 0.0001537 \| 177 \| 0.173 \| 144 \| 111 \| 4 \| 1 \| 140 \| 1 \| 140 \| ParB/RepB/Spo0J family partition protein \| ParB/RepB/Spo0J family partition protein; afdb-uniprot50 \| AF-A0A3D9TAG4-F1-MODEL\_V4 \| 1.0 \| 8.082e-07 \| 176 \| 0.187 \| 203 \| 143 \| 11 \| 13 \| 200 \| 81 \| 276 \| ParB/RepB/Spo0J family partition protein \| ParB/RepB/Spo0J family partition protein; afdb-uniprot50 \| AF-A0A1I3QL22-F1-MODEL\_V4 \| 1.0 \| 2.206e-07 \| 176 \| 0.198 \| 272 \| 181 \| 11 \| 19 \| 280 \| 4 \| 248 \| ParB family protein \| ParB family protein; afdb-uniprot50 \| AF-A0A355VWD7-F1-MODEL\_V4 \| 1.0 \| 1.059e-06 \| 175 \| 0.123 \| 283 \| 178 \| 14 \| 19 \| 287 \| 34 \| 260 \| ParB domain-containing protein \| ParB domain-containing protein; afdb-uniprot50 \| AF-A0A1U9XHD1-F1-MODEL\_V4 \| 1.0 \| 4.565e-06 \| 175 \| 0.156 \| 281 \| 182 \| 11 \| 60 \| 291 \| 2 \| 276 \| Uncharacterized protein \| Uncharacterized protein; afdb-uniprot50 \| AF-A0A0N1HEN7-F1-MODEL\_V4 \| 1.0 \| 1.633e-06 \| 173 \| 0.169 \| 213 \| 144 \| 10 \| 19 \| 206 \| 5 \| 209 \| Uncharacterized protein \| Uncharacterized protein; afdb-uniprot50 \| AF-Q1ZJV4-F1-MODEL\_V4 \| 1.0 \| 9.506e-07 \| 173 \| 0.168 \| 303 \| 188 \| 18 \| 20 \| 287 \| 76 \| 349 \| Partitioning protein B \| Partitioning protein B; afdb-uniprot50 \| AF-A0A7V9N3J7-F1-MODEL\_V4 \| 1.0 \| 1.284e-07 \| 173 \| 0.173 \| 328 \| 200 \| 14 \| 9 \| 290 \| 24 \| 326 \| ParB/RepB/Spo0J family partition protein \| ParB/RepB/Spo0J family partition protein; afdb-uniprot50 \| AF-A0A3B8MU94-F1-MODEL\_V4 \| 1.0 \| 0.0004296 \| 170 \| 0.194 \| 113 \| 78 \| 7 \| 1 \| 106 \| 1 \| 107 \| Chromosome partitioning protein ParB \| Chromosome partitioning protein ParB; afdb-uniprot50 \| AF-A0A0Q8LGQ4-F1-MODEL\_V4 \| 1.0 \| 9.506e-07 \| 169 \| 0.176 \| 215 \| 154 \| 8 \| 4 \| 199 \| 8 \| 218 \| ParB domain-containing protein \| ParB domain-containing protein; afdb-uniprot50 \| AF-A0A564Q3X4-F1-MODEL\_V4 \| 1.0 \| 1.863e-05 \| 167 \| 0.095 \| 272 \| 166 \| 11 \| 18 \| 247 \| 1 \| 234 \| Nucleoid occlusion protein \| Nucleoid occlusion protein; afdb-uniprot50 \| AF-A0A527TY97-F1-MODEL\_V4 \| 1.0 \| 3.401e-07 \| 167 \| 0.172 \| 284 \| 192 \| 13 \| 13 \| 282 \| 1 \| 255 \| ParB/RepB/Spo0J family partition protein \| ParB/RepB/Spo0J family partition protein; afdb-uniprot50 \| AF-A0A2A4MGC7-F1-MODEL\_V4 \| 1.0 \| 1.092e-07 \| 166 \| 0.132 \| 354 \| 207 \| 15 \| 13 \| 287 \| 3 \| 335 \| ParB domain-containing protein \| ParB domain-containing protein; afdb-uniprot50 \| AF-E9SG08-F1-MODEL\_V4 \| 1.0 \| 5.209e-05 \| 165 \| 0.165 \| 169 \| 114 \| 6 \| 1 \| 160 \| 1 \| 151 \| ParB-like protein \| ParB-like protein; afdb-uniprot50 \| AF-A0A5C7ERN6-F1-MODEL\_V4 \| 1.0 \| 2.873e-05 \| 165 \| 0.169 \| 213 \| 147 \| 9 \| 18 \| 205 \| 1 \| 208 \| ParB domain-containing protein \| ParB domain-containing protein; afdb-uniprot50 \| AF-A0A7V9VJT7-F1-MODEL\_V4 \| 1.0 \| 1.876e-07 \| 163 \| 0.137 \| 320 \| 204 \| 14 \| 8 \| 291 \| 49 \| 332 \| ParB/RepB/Spo0J family partition protein \| ParB/RepB/Spo0J family partition protein; afdb-uniprot50 \| AF-A0A2N1TR59-F1-MODEL\_V4 \| 1.0 \| 6.666e-06 \| 163 \| 0.164 \| 291 \| 173 \| 12 \| 12 \| 271 \| 74 \| 325 \| Uncharacterized protein \| Uncharacterized protein; afdb-uniprot50 \| AF-A0A351AWR7-F1-MODEL\_V4 \| 1.0 \| 0.0005631 \| 162 \| 0.174 \| 109 \| 82 \| 4 \| 1 \| 107 \| 3 \| 105 \| Chromosome partitioning protein ParB \| Chromosome partitioning protein ParB; afdb-uniprot50 \| AF-A0A1W9WV54-F1-MODEL\_V4 \| 1.0 \| 3.881e-06 \| 160 \| 0.141 \| 283 \| 163 \| 17 \| 8 \| 282 \| 15 \| 225 \| ParB domain-containing protein \| ParB domain-containing protein; afdb-uniprot50 \| AF-F5YI81-F1-MODEL\_V4 \| 1.0 \| 6.71e-08 \| 160 \| 0.154 \| 317 \| 199 \| 15 \| 1 \| 287 \| 1 \| 278 \| ParB domain-containing protein \| ParB domain-containing protein; afdb-uniprot50 \| AF-A0A2W5PVF2-F1-MODEL\_V4 \| 1.0 \| 8.082e-07 \| 159 \| 0.159 \| 358 \| 187 \| 17 \| 20 \| 291 \| 14 \| 343 \| ParB domain-containing protein \| ParB domain-containing protein; afdb-uniprot50 \| AF-A0A2W6A9B4-F1-MODEL\_V4 \| 1.0 \| 1.82e-06 \| 158 \| 0.106 \| 310 \| 219 \| 13 \| 2 \| 275 \| 1 \| 288 \| ParB domain-containing protein \| ParB domain-containing protein; afdb-uniprot50 \| AF-A0A7V8WTB0-F1-MODEL\_V4 \| 1.0 \| 2.329e-07 \| 157 \| 0.16 \| 323 \| 196 \| 13 \| 7 \| 291 \| 42 \| 327 \| ParB/RepB/Spo0J family partition protein \| ParB/RepB/Spo0J family partition protein; afdb-uniprot50 \| AF-A0A8B5XDA0-F1-MODEL\_V4 \| 1.0 \| 3.401e-07 \| 156 \| 0.124 \| 321 \| 218 \| 18 \| 8 \| 282 \| 14 \| 317 \| ParB/RepB/Spo0J family partition protein \| ParB/RepB/Spo0J family partition protein; afdb-uniprot50 \| AF-A0A1S8D0Y6-F1-MODEL\_V4 \| 1.0 \| 4.705e-07 \| 156 \| 0.14 \| 291 \| 200 \| 17 \| 18 \| 291 \| 1 \| 258 \| ParB domain-containing protein \| ParB domain-containing protein; afdb-uniprot50 \| AF-A0A7Z1UWI7-F1-MODEL\_V4 \| 1.0 \| 3.222e-07 \| 156 \| 0.148 \| 289 \| 198 \| 14 \| 18 \| 290 \| 1 \| 257 \| ParB domain-containing protein \| ParB domain-containing protein; afdb-uniprot50 \| AF-A0A1M5FTR8-F1-MODEL\_V4 \| 1.0 \| 3.881e-06 \| 155 \| 0.158 \| 265 \| 188 \| 13 \| 29 \| 281 \| 1 \| 242 \| ParB/RepB/Spo0J family partition protein \| ParB/RepB/Spo0J family partition protein; afdb-uniprot50 \| AF-A0A5C7YCL1-F1-MODEL\_V4 \| 1.0 \| 2.314e-05 \| 154 \| 0.209 \| 229 \| 147 \| 10 \| 71 \| 282 \| 12 \| 223 \| ParB/RepB/Spo0J family partition protein \| ParB/RepB/Spo0J family partition protein; afdb-uniprot50 \| AF-J7QGM7-F1-MODEL\_V4 \| 1.0 \| 1.388e-06 \| 154 \| 0.174 \| 293 \| 191 \| 13 \| 13 \| 283 \| 1 \| 264 \| ParB domain protein nuclease \| ParB domain protein nuclease; afdb-uniprot50 \| AF-A0A370KZ30-F1-MODEL\_V4 \| 1.0 \| 4.705e-07 \| 154 \| 0.174 \| 356 \| 191 \| 17 \| 13 \| 287 \| 57 \| 390 \| ParB/RepB/Spo0J family partition protein \| ParB/RepB/Spo0J family partition protein; afdb-uniprot50 \| AF-A0A1M4X5Z2-F1-MODEL\_V4 \| 1.0 \| 1.145e-05 \| 152 \| 0.147 \| 278 \| 179 \| 10 \| 19 \| 273 \| 20 \| 262 \| ParB-like nuclease domain-containing protein \| ParB-like nuclease domain-containing protein; afdb-uniprot50 \| AF-A0A534ZDR6-F1-MODEL\_V4 \| 1.0 \| 0.001338 \| 150 \| 0.201 \| 114 \| 86 \| 3 \| 1 \| 114 \| 9 \| 117 \| ParB/RepB/Spo0J family partition protein \| ParB/RepB/Spo0J family partition protein; afdb-uniprot50 \| AF-A0A7Y4NL96-F1-MODEL\_V4 \| 1.0 \| 0.002561 \| 145 \| 0.261 \| 130 \| 74 \| 2 \| 136 \| 265 \| 1 \| 108 \| Uncharacterized protein \| Uncharacterized protein; afdb-uniprot50 \| AF-A0A6P1J9Y2-F1-MODEL\_V4 \| 1.0 \| 0.0002369 \| 145 \| 0.196 \| 204 \| 134 \| 5 \| 98 \| 272 \| 1 \| 203 \| Uncharacterized protein \| Uncharacterized protein; afdb-uniprot50 \| AF-A0A246JU63-F1-MODEL\_V4 \| 1.0 \| 4.324e-06 \| 145 \| 0.172 \| 279 \| 182 \| 15 \| 17 \| 280 \| 76 \| 320 \| ParB domain-containing protein \| ParB domain-containing protein; afdb-uniprot50 \| AF-A0A4Q2LMW1-F1-MODEL\_V4 \| 1.0 \| 9.445e-05 \| 144 \| 0.222 \| 207 \| 131 \| 9 \| 98 \| 288 \| 7 \| 199 \| ParB/RepB/Spo0J family partition protein \| ParB/RepB/Spo0J family partition protein; afdb-uniprot50 \| AF-A0A2H9SQB1-F1-MODEL\_V4 \| 1.0 \| 2.259e-06 \| 138 \| 0.169 \| 289 \| 181 \| 15 \| 1 \| 265 \| 3 \| 256 \| Chromosome partitioning protein ParB \| Chromosome partitioning protein ParB; afdb-uniprot50 \| AF-A0A5C8ILI9-F1-MODEL\_V4 \| 1.0 \| 7.841e-06 \| 134 \| 0.185 \| 296 \| 176 \| 18 \| 7 \| 272 \| 19 \| 279 \| ParB/RepB/Spo0J family partition protein \| ParB/RepB/Spo0J family partition protein; afdb-uniprot50 \| AF-A0A2J0L660-F1-MODEL\_V4 \| 1.0 \| 0.0004535 \| 131 \| 0.175 \| 233 \| 170 \| 10 \| 60 \| 282 \| 1 \| 221 \| Uncharacterized protein \| Uncharacterized protein; afdb-uniprot50 \| AF-A0A3B8V5W7-F1-MODEL\_V4 \| 1.0 \| 0.002854 \| 126 \| 0.179 \| 184 \| 132 \| 5 \| 107 \| 282 \| 18 \| 190 \| HTH\_ParB domain-containing protein \| HTH\_ParB domain-containing protein; afdb-uniprot50 \| AF-A0A2T5GRZ6-F1-MODEL\_V4 \| 1.0 \| 0.0003278 \| 119 \| 0.132 \| 257 \| 181 \| 11 \| 37 \| 287 \| 464 \| 684 \| ParB-like nuclease family protein \| ParB-like nuclease family protein |
| Top keywords  (threshold 1.00e-02 (evalue)) | **ParB, partition, Spo0J, RepB, domain\_containing, partitioning, Chromosome, chromosome\_partitioning, ParB\_like, Probable** |
| Output files | ../../similar\_structures/28\_FANPEZAQ\_CDS\_0028\_afdb-proteome\_foldseek.tsv ../../similar\_structures/28\_FANPEZAQ\_CDS\_0028\_afdb-uniprot50\_foldseek.tsv ../../similar\_structures/28\_FANPEZAQ\_CDS\_0028\_merged.svg ../../similar\_structures/28\_FANPEZAQ\_CDS\_0028\_pdb\_foldseek.tsv |

  
  
  

Return to summary | Go to previous | Go to next

  


---

**Sequence/structure alignments coloring**  
Each object in the alignment figures is colored according to its E-value following this color coding:

1e-100
10
