## Supplementary material for "Completing the BASEL phage collection to unlock hidden diversity for systematic exploration of phage-host interactions": Data S2: 29.html

FANPEZAQ\_CDS\_0029


Return to summary | Go to previous | Go to next

|  |  |
| --- | --- |
| FANPEZAQ\_CDS\_0029  Page creation date: 02 Sep 2024, 12:00  Project folder: n/a  Input sequences file: Escherichia\_virus\_HeidiAbel.gb | hypothetical domain\_containing phage periplasmic kow eef2k atpase duf2158 fragment p143733 vi\_10070 nc\_018850\_p11 mh020244\_p60 carboxypeptidase regulatory\_like nitrogen fixation nifz duf2171 snoal\_like prpf4b peptidylprolyl isomerase duf5641 stress\_induced\_phosphoprotein j |

### Sequence information

|  |  |
| --- | --- |
| Name | FANPEZAQ\_CDS\_0029  29\_FANPEZAQ\_CDS\_0029 (pipeline id) |
| Imported annotations | Escherichia\_virus\_HeidiAbel Bas97 |
| Protein sequence | MKGELKAGGLALVYGLRVRPEGNGKCVRLIELASYRQSELLAMALGEPVERWWRVYGLLP GNEVAQVRAQNLLPIDGDDQFLDSATGEGLNRLAEQFGFKPAELTA |
| Number of residues | 106 |
| Molecular weight (Da) | 11614.18 |
| Output files | ../../query\_sequences/29\_FANPEZAQ\_CDS\_0029.fasta |

### Putative domain architecture and protein family

#### Search results (HHblits)^1^

|  |  |
| --- | --- |
| Domain family databases searched | Pfam, Ncbi-cd, Cath, Phrogs |
| Results, scheme(s)  (Top layers only; threshold 1.00e-03 (evalue)) | xml version="1.0" encoding="utf-8" standalone="no"?       2024-09-02T21:08:18.544717 image/svg+xml   Matplotlib v3.7.2, https://matplotlib.org/ |
| Results, table  (E-value ≤ 1.00e-03 (evalue)) | db \| id \| prob \| evalue \| pvalue \| score \| cols \| query \| query\_len \| template \| template\_len \| name \| description; phrogs \| 2529 \| 99.9 \| 4.2e-29 \| 5.7e-33 \| 170.3 \| 82 \| (1, 84) \| 106 \| (1, 93) \| 101 \| NA \| NA; Category: unknown function; p143733 VI\_10070; phrogs \| 28276 \| 99.6 \| 7.2e-20 \| 8.2e-24 \| 123.4 \| 85 \| (2, 91) \| 106 \| (4, 102) \| 104 \| NA \| NA; Category: unknown function; NC\_018850\_p11; phrogs \| 6513 \| 94.3 \| 0.00087 \| 1e-07 \| 40.3 \| 52 \| (1, 57) \| 106 \| (1, 53) \| 63 \| NA \| NA; Category: unknown function; MH020244\_p60 |
| Top keywords  (threshold 1.00e-03 (evalue)) | **p143733, VI\_10070, NC\_018850\_p11, MH020244\_p60** |
| Output files | ../../domain\_architecture/29\_FANPEZAQ\_CDS\_0029\_cath.hhr ../../domain\_architecture/29\_FANPEZAQ\_CDS\_0029\_merged.svg ../../domain\_architecture/29\_FANPEZAQ\_CDS\_0029\_ncbi-cd.hhr ../../domain\_architecture/29\_FANPEZAQ\_CDS\_0029\_pfam.hhr ../../domain\_architecture/29\_FANPEZAQ\_CDS\_0029\_phrogs.hhr |

### Identical protein sequences/structures

#### Search results

|  |  |
| --- | --- |
| Protein sequence databases searched | Pdb, Swissprot, Refseq |
| Identical proteins found | -- |
| Top keywords | -- |
| Output files | -- |

### Similar protein sequences/structures

#### Sequence similarity search results (HHblits)^1^

|  |  |
| --- | --- |
| Sequence databases searched | Uniclust, Pdb70 |
| Results, scheme(s)  (Top layers only, threshold 1.00e-03 (evalue)) | xml version="1.0" encoding="utf-8" standalone="no"?       2024-09-02T21:08:41.986776 image/svg+xml   Matplotlib v3.7.2, https://matplotlib.org/ |
| Results, table(s)  (threshold 1.00e-03 (evalue)) | db \| id \| prob \| evalue \| pvalue \| score \| cols \| query \| query\_len \| template \| template\_len \| name \| description; uniclust \| UniRef100\_A0A0D9A4N1 \| 100.0 \| 1.1e-35 \| 2.8e-41 \| 206.2 \| 94 \| (1, 105) \| 106 \| (7, 112) \| 117 \| Uncharacterized protein \| Uncharacterized protein; uniclust \| UniRef100\_A0A010SYJ8 \| 100.0 \| 5.5e-34 \| 1.4e-39 \| 197.5 \| 94 \| (1, 105) \| 106 \| (15, 114) \| 114 \| Uncharacterized protein \| Uncharacterized protein; uniclust \| UniRef100\_A0A0Q8YUL3 \| 99.9 \| 3.6e-32 \| 8.2e-38 \| 184.2 \| 94 \| (1, 105) \| 106 \| (1, 101) \| 101 \| KOW domain-containing protein \| KOW domain-containing protein; uniclust \| UniRef100\_A0A1C7BVV0 \| 99.9 \| 6e-31 \| 1.3e-36 \| 177.3 \| 90 \| (1, 103) \| 106 \| (3, 98) \| 98 \| Phage protein \| Phage protein; uniclust \| UniRef100\_A0A1X7M6R0 \| 99.9 \| 4.5e-27 \| 1.1e-32 \| 161.4 \| 91 \| (1, 102) \| 106 \| (2, 104) \| 104 \| Carboxypeptidase regulatory-like domain-containing protein \| Carboxypeptidase regulatory-like domain-containing protein; uniclust \| UniRef100\_A0A088FQM9 \| 99.8 \| 6.2e-23 \| 1.4e-28 \| 140.8 \| 96 \| (1, 106) \| 106 \| (1, 105) \| 105 \| ATPase \| ATPase; uniclust \| UniRef100\_A0A3G7U449 \| 99.8 \| 3.7e-22 \| 7.6e-28 \| 136.4 \| 90 \| (1, 100) \| 106 \| (1, 102) \| 107 \| Phage protein \| Phage protein; uniclust \| UniRef100\_A0A5E7ADD6 \| 99.8 \| 4e-22 \| 8.1e-28 \| 130.7 \| 83 \| (1, 103) \| 106 \| (1, 84) \| 84 \| Uncharacterized protein \| Uncharacterized protein; uniclust \| UniRef100\_A0A1G8V272 \| 99.8 \| 8.4e-22 \| 1.7e-27 \| 131.4 \| 89 \| (1, 102) \| 106 \| (1, 91) \| 91 \| Uncharacterized protein \| Uncharacterized protein; uniclust \| UniRef100\_A0A225TZ33 \| 99.7 \| 2.3e-21 \| 4.8e-27 \| 132.3 \| 92 \| (1, 102) \| 106 \| (1, 102) \| 102 \| Uncharacterized protein \| Uncharacterized protein; uniclust \| UniRef100\_A0A022G1Y5 \| 99.7 \| 3.9e-21 \| 9.9e-27 \| 130.9 \| 70 \| (1, 81) \| 106 \| (1, 83) \| 94 \| Uncharacterized protein \| Uncharacterized protein; uniclust \| UniRef100\_A0A2X3KFA9 \| 99.7 \| 1.9e-20 \| 3.6e-26 \| 133.3 \| 91 \| (2, 103) \| 106 \| (1, 106) \| 149 \| Uncharacterized protein \| Uncharacterized protein; uniclust \| UniRef100\_A0A144RG41 \| 99.7 \| 2.5e-20 \| 5.1e-26 \| 126.6 \| 90 \| (2, 102) \| 106 \| (1, 102) \| 102 \| Uncharacterized protein \| Uncharacterized protein; uniclust \| UniRef100\_A0A2Z5A9T9 \| 99.7 \| 8.9e-20 \| 1.9e-25 \| 125.7 \| 91 \| (1, 102) \| 106 \| (1, 93) \| 108 \| Uncharacterized protein \| Uncharacterized protein; uniclust \| UniRef100\_A0A132DRQ2 \| 99.6 \| 2.8e-19 \| 6.7e-25 \| 124.4 \| 82 \| (1, 102) \| 106 \| (1, 95) \| 108 \| Uncharacterized protein \| Uncharacterized protein; uniclust \| UniRef100\_A0A104MV53 \| 99.6 \| 1.7e-18 \| 4.5e-24 \| 126.5 \| 75 \| (1, 81) \| 106 \| (10, 97) \| 145 \| Uncharacterized protein \| Uncharacterized protein; uniclust \| UniRef100\_A0A0A8RLE3 \| 99.6 \| 3e-18 \| 5.6e-24 \| 113.0 \| 89 \| (1, 103) \| 106 \| (1, 90) \| 90 \| Uncharacterized protein \| Uncharacterized protein; uniclust \| UniRef100\_A0A071LWX4 \| 99.6 \| 5.5e-18 \| 1.1e-23 \| 121.6 \| 87 \| (2, 99) \| 106 \| (26, 125) \| 146 \| Periplasmic protein \| Periplasmic protein; uniclust \| UniRef100\_A0A0N0JSM0 \| 99.4 \| 1.2e-15 \| 2.9e-21 \| 108.8 \| 96 \| (2, 104) \| 106 \| (9, 113) \| 120 \| Uncharacterized protein \| Uncharacterized protein; uniclust \| UniRef100\_A0A483MTB8 \| 99.4 \| 2.9e-15 \| 5.6e-21 \| 102.1 \| 90 \| (2, 102) \| 106 \| (1, 104) \| 104 \| Phage protein \| Phage protein; uniclust \| UniRef100\_A0A0J6LF42 \| 99.3 \| 9.5e-15 \| 1.9e-20 \| 101.5 \| 81 \| (1, 81) \| 106 \| (1, 91) \| 114 \| Uncharacterized protein \| Uncharacterized protein; uniclust \| UniRef100\_UPI001C60F1CF \| 99.3 \| 1.6e-14 \| 2.9e-20 \| 97.7 \| 86 \| (2, 98) \| 106 \| (1, 97) \| 102 \| hypothetical protein \| hypothetical protein; uniclust \| UniRef100\_A0A062ISY1 \| 99.3 \| 1.9e-14 \| 4.5e-20 \| 100.2 \| 75 \| (1, 81) \| 106 \| (1, 82) \| 102 \| Phage protein \| Phage protein; uniclust \| UniRef100\_UPI0008FB2880 \| 99.2 \| 5.4e-14 \| 1e-19 \| 97.5 \| 88 \| (2, 104) \| 106 \| (1, 89) \| 120 \| hypothetical protein \| hypothetical protein; uniclust \| UniRef100\_UPI001FC9648A \| 99.2 \| 8.3e-14 \| 1.5e-19 \| 88.3 \| 66 \| (23, 101) \| 106 \| (3, 68) \| 68 \| hypothetical protein \| hypothetical protein; uniclust \| UniRef100\_UPI00200A17EF \| 99.2 \| 8.4e-14 \| 1.5e-19 \| 87.6 \| 65 \| (1, 103) \| 106 \| (1, 65) \| 65 \| hypothetical protein \| hypothetical protein; uniclust \| UniRef100\_A0A0U3TCB6 \| 99.2 \| 1.1e-13 \| 2e-19 \| 95.3 \| 89 \| (1, 100) \| 106 \| (1, 106) \| 108 \| Uncharacterized protein \| Uncharacterized protein; uniclust \| UniRef100\_A0A1C4DD36 \| 99.2 \| 1.7e-13 \| 3.3e-19 \| 99.6 \| 79 \| (2, 81) \| 106 \| (52, 143) \| 152 \| Periplasmic protein \| Periplasmic protein; uniclust \| UniRef100\_A0A103BH94 \| 99.2 \| 1.5e-13 \| 3.5e-19 \| 99.1 \| 64 \| (3, 81) \| 106 \| (18, 94) \| 126 \| Nitrogen fixation protein NifZ \| Nitrogen fixation protein NifZ; uniclust \| UniRef100\_A0A109EH43 \| 99.1 \| 4.8e-13 \| 1.1e-18 \| 94.8 \| 64 \| (4, 81) \| 106 \| (11, 87) \| 114 \| Uncharacterized protein \| Uncharacterized protein; uniclust \| UniRef100\_A0A7Y9IXU9 \| 99.0 \| 2.3e-12 \| 4.5e-18 \| 89.3 \| 63 \| (3, 78) \| 106 \| (1, 76) \| 106 \| Uncharacterized protein \| Uncharacterized protein; uniclust \| UniRef100\_UPI00209847FF \| 99.0 \| 2.6e-12 \| 4.7e-18 \| 85.3 \| 77 \| (2, 81) \| 106 \| (1, 82) \| 87 \| hypothetical protein \| hypothetical protein; uniclust \| UniRef100\_A0A1B4LB38 \| 98.8 \| 3.7e-11 \| 8.3e-17 \| 84.6 \| 84 \| (3, 104) \| 106 \| (1, 94) \| 104 \| DUF2158 domain-containing protein \| DUF2158 domain-containing protein; uniclust \| UniRef100\_A0A1E4FXU0 \| 98.7 \| 1.3e-10 \| 2.9e-16 \| 81.5 \| 69 \| (1, 78) \| 106 \| (14, 83) \| 101 \| Uncharacterized protein \| Uncharacterized protein; uniclust \| UniRef100\_UPI0009AC47EE \| 98.7 \| 2e-10 \| 3.8e-16 \| 74.9 \| 58 \| (2, 60) \| 106 \| (1, 67) \| 74 \| hypothetical protein \| hypothetical protein; uniclust \| UniRef100\_UPI0011B2021C \| 98.7 \| 2.3e-10 \| 4.3e-16 \| 80.4 \| 81 \| (1, 81) \| 106 \| (1, 96) \| 121 \| hypothetical protein \| hypothetical protein; uniclust \| UniRef100\_UPI00059FEC15 \| 98.6 \| 3e-10 \| 5.4e-16 \| 76.8 \| 71 \| (2, 78) \| 106 \| (1, 78) \| 93 \| hypothetical protein \| hypothetical protein; uniclust \| UniRef100\_UPI00203A5C56 \| 98.6 \| 4.7e-10 \| 8.6e-16 \| 73.9 \| 45 \| (49, 103) \| 106 \| (21, 79) \| 79 \| hypothetical protein \| hypothetical protein; uniclust \| UniRef100\_A0A639VA07 \| 98.5 \| 8.1e-10 \| 1.5e-15 \| 69.7 \| 55 \| (1, 56) \| 106 \| (1, 58) \| 61 \| Uncharacterized protein \| Uncharacterized protein; uniclust \| UniRef100\_A0A3G3H4G5 \| 98.5 \| 8.6e-10 \| 1.7e-15 \| 78.1 \| 62 \| (3, 78) \| 106 \| (5, 79) \| 113 \| Uncharacterized protein \| Uncharacterized protein; uniclust \| UniRef100\_UPI0013D61B36 \| 98.5 \| 9.3e-10 \| 1.7e-15 \| 73.9 \| 66 \| (1, 81) \| 106 \| (1, 78) \| 88 \| hypothetical protein \| hypothetical protein; uniclust \| UniRef100\_A0A839T1F5 \| 98.5 \| 9.4e-10 \| 1.7e-15 \| 74.7 \| 75 \| (1, 78) \| 106 \| (1, 78) \| 94 \| Uncharacterized protein \| Uncharacterized protein; uniclust \| UniRef100\_A0A2W7MJX6 \| 98.5 \| 1.4e-09 \| 2.6e-15 \| 74.9 \| 75 \| (1, 78) \| 106 \| (1, 76) \| 96 \| Uncharacterized protein \| Uncharacterized protein; uniclust \| UniRef100\_UPI0004890263 \| 98.4 \| 2.1e-09 \| 4.2e-15 \| 75.1 \| 76 \| (3, 78) \| 106 \| (1, 91) \| 103 \| hypothetical protein \| hypothetical protein; uniclust \| UniRef100\_A0A068YI59 \| 98.4 \| 4.3e-09 \| 8.7e-15 \| 73.1 \| 71 \| (3, 81) \| 106 \| (1, 78) \| 95 \| Uncharacterized protein \| Uncharacterized protein; uniclust \| UniRef100\_A0A859WEJ4 \| 98.3 \| 8.6e-09 \| 1.6e-14 \| 71.4 \| 83 \| (2, 101) \| 106 \| (1, 96) \| 97 \| Uncharacterized protein \| Uncharacterized protein; uniclust \| UniRef100\_A0A7R8MJT1 \| 98.3 \| 1.3e-08 \| 2.4e-14 \| 71.3 \| 77 \| (2, 78) \| 106 \| (1, 88) \| 111 \| ATPase \| ATPase; uniclust \| UniRef100\_UPI001556132D \| 98.3 \| 1.4e-08 \| 2.6e-14 \| 69.7 \| 75 \| (1, 81) \| 106 \| (1, 85) \| 98 \| hypothetical protein \| hypothetical protein; uniclust \| UniRef100\_A0A1B4C0H0 \| 98.2 \| 1.4e-08 \| 3.1e-14 \| 73.0 \| 62 \| (4, 81) \| 106 \| (12, 87) \| 110 \| Uncharacterized protein \| Uncharacterized protein; uniclust \| UniRef100\_A0A6D2G8R2 \| 98.2 \| 1.9e-08 \| 3.5e-14 \| 73.9 \| 60 \| (2, 62) \| 106 \| (1, 69) \| 154 \| Periplasmic protein \| Periplasmic protein; uniclust \| UniRef100\_UPI00052A5558 \| 98.2 \| 2.4e-08 \| 4.5e-14 \| 69.7 \| 65 \| (3, 79) \| 106 \| (4, 83) \| 103 \| hypothetical protein \| hypothetical protein; uniclust \| UniRef100\_A0A096FBG0 \| 98.2 \| 3.1e-08 \| 5.8e-14 \| 69.3 \| 75 \| (1, 78) \| 106 \| (1, 85) \| 103 \| DUF2171 domain-containing protein \| DUF2171 domain-containing protein; uniclust \| UniRef100\_A0A2X3IPC5 \| 98.1 \| 3.6e-08 \| 6.6e-14 \| 65.7 \| 60 \| (22, 81) \| 106 \| (2, 71) \| 80 \| Periplasmic protein \| Periplasmic protein; uniclust \| UniRef100\_A0A1Y0EPI5 \| 98.1 \| 4e-08 \| 7.4e-14 \| 71.7 \| 77 \| (1, 78) \| 106 \| (34, 120) \| 144 \| Uncharacterized protein \| Uncharacterized protein; uniclust \| UniRef100\_UPI001C0BF18E \| 98.1 \| 4.3e-08 \| 7.9e-14 \| 68.1 \| 72 \| (3, 78) \| 106 \| (1, 85) \| 103 \| hypothetical protein \| hypothetical protein; uniclust \| UniRef100\_A0A3Q9D7Q0 \| 98.1 \| 4.5e-08 \| 8.3e-14 \| 63.8 \| 53 \| (1, 57) \| 106 \| (1, 59) \| 70 \| Uncharacterized protein \| Uncharacterized protein; uniclust \| UniRef100\_A0A7U0GBP7 \| 98.1 \| 4.9e-08 \| 9.8e-14 \| 72.7 \| 79 \| (2, 81) \| 106 \| (1, 115) \| 146 \| Uncharacterized protein \| Uncharacterized protein; uniclust \| UniRef100\_A0A1H3ELA5 \| 98.1 \| 5.8e-08 \| 1.1e-13 \| 60.1 \| 25 \| (2, 27) \| 106 \| (4, 28) \| 52 \| Uncharacterized protein \| Uncharacterized protein; uniclust \| UniRef100\_A0A7H4NPC2 \| 98.0 \| 1.1e-07 \| 2e-13 \| 61.7 \| 47 \| (46, 102) \| 106 \| (11, 67) \| 67 \| Uncharacterized protein \| Uncharacterized protein; uniclust \| UniRef100\_A0A4U9HTB8 \| 98.0 \| 1.3e-07 \| 2.4e-13 \| 55.8 \| 36 \| (2, 38) \| 106 \| (1, 39) \| 40 \| Uncharacterized protein \| Uncharacterized protein; uniclust \| UniRef100\_A0A0U4IIC9 \| 98.0 \| 1.3e-07 \| 2.4e-13 \| 47.6 \| 18 \| (73, 100) \| 106 \| (1, 18) \| 19 \| Uncharacterized protein \| Uncharacterized protein; uniclust \| UniRef100\_A0A5P8D787 \| 98.0 \| 1.5e-07 \| 2.9e-13 \| 66.1 \| 77 \| (1, 78) \| 106 \| (1, 86) \| 103 \| Uncharacterized protein \| Uncharacterized protein; uniclust \| UniRef100\_A0A2G9C6E2 \| 97.9 \| 3.4e-07 \| 6.3e-13 \| 63.5 \| 77 \| (3, 81) \| 106 \| (1, 83) \| 99 \| Uncharacterized protein \| Uncharacterized protein; uniclust \| UniRef100\_A0A965ESD1 \| 97.8 \| 3.5e-07 \| 6.7e-13 \| 65.8 \| 66 \| (1, 78) \| 106 \| (13, 98) \| 117 \| Uncharacterized protein (Fragment) \| Uncharacterized protein (Fragment); uniclust \| UniRef100\_A0A142IE47 \| 97.8 \| 4.7e-07 \| 9.2e-13 \| 66.2 \| 89 \| (4, 103) \| 106 \| (15, 125) \| 125 \| Uncharacterized protein \| Uncharacterized protein; uniclust \| UniRef100\_X1N3C4 \| 97.8 \| 5.5e-07 \| 1e-12 \| 55.2 \| 26 \| (67, 102) \| 106 \| (22, 47) \| 48 \| Uncharacterized protein (Fragment) \| Uncharacterized protein (Fragment); uniclust \| UniRef100\_UPI00115D64C9 \| 97.8 \| 6e-07 \| 1.1e-12 \| 59.5 \| 33 \| (2, 35) \| 106 \| (1, 33) \| 72 \| hypothetical protein \| hypothetical protein; uniclust \| UniRef100\_A0A0K8P477 \| 97.7 \| 7.5e-07 \| 1.5e-12 \| 64.7 \| 75 \| (3, 81) \| 106 \| (10, 97) \| 117 \| Uncharacterized protein \| Uncharacterized protein; uniclust \| UniRef100\_A0A519G502 \| 97.7 \| 8.1e-07 \| 1.6e-12 \| 62.7 \| 70 \| (1, 81) \| 106 \| (1, 82) \| 97 \| KOW domain-containing protein \| KOW domain-containing protein; uniclust \| UniRef100\_A0A6J5LA11 \| 97.7 \| 8.9e-07 \| 1.6e-12 \| 60.1 \| 74 \| (5, 99) \| 106 \| (2, 83) \| 85 \| Uncharacterized protein \| Uncharacterized protein; uniclust \| UniRef100\_A0A239N1G3 \| 97.7 \| 1.3e-06 \| 2.4e-12 \| 51.9 \| 25 \| (67, 101) \| 106 \| (15, 39) \| 39 \| Uncharacterized protein \| Uncharacterized protein; uniclust \| UniRef100\_UPI0011BDCDA8 \| 97.7 \| 1.3e-06 \| 2.5e-12 \| 61.2 \| 75 \| (2, 77) \| 106 \| (1, 87) \| 103 \| hypothetical protein \| hypothetical protein; uniclust \| UniRef100\_A0A516KWJ6 \| 97.6 \| 1.3e-06 \| 2.8e-12 \| 61.9 \| 85 \| (1, 102) \| 106 \| (2, 92) \| 97 \| Uncharacterized protein \| Uncharacterized protein; uniclust \| UniRef100\_A0A6I6SLJ2 \| 97.6 \| 1.7e-06 \| 3.3e-12 \| 61.4 \| 76 \| (1, 81) \| 106 \| (1, 86) \| 100 \| Uncharacterized protein \| Uncharacterized protein; uniclust \| UniRef100\_A0A244E847 \| 97.6 \| 1.7e-06 \| 3.5e-12 \| 62.2 \| 65 \| (1, 78) \| 106 \| (1, 72) \| 108 \| Uncharacterized protein \| Uncharacterized protein; uniclust \| UniRef100\_UPI0009E75DDD \| 97.6 \| 2.3e-06 \| 4.4e-12 \| 63.8 \| 60 \| (3, 78) \| 106 \| (26, 98) \| 141 \| hypothetical protein \| hypothetical protein; uniclust \| UniRef100\_A0A0K6IDV6 \| 97.5 \| 2.6e-06 \| 5.1e-12 \| 66.2 \| 67 \| (3, 78) \| 106 \| (1, 78) \| 185 \| Uncharacterized protein \| Uncharacterized protein; uniclust \| UniRef100\_UPI0015E516B4 \| 97.5 \| 3e-06 \| 5.6e-12 \| 53.0 \| 42 \| (2, 44) \| 106 \| (1, 42) \| 52 \| hypothetical protein \| hypothetical protein; uniclust \| UniRef100\_UPI0019579779 \| 97.5 \| 3.4e-06 \| 6.3e-12 \| 56.2 \| 32 \| (2, 34) \| 106 \| (1, 32) \| 74 \| hypothetical protein \| hypothetical protein; uniclust \| UniRef100\_A0A158D246 \| 97.5 \| 3.7e-06 \| 7.1e-12 \| 58.3 \| 64 \| (3, 78) \| 106 \| (1, 79) \| 88 \| Uncharacterized protein \| Uncharacterized protein; uniclust \| UniRef100\_A0A7D5XW54 \| 97.5 \| 3.8e-06 \| 7.2e-12 \| 62.8 \| 79 \| (1, 81) \| 106 \| (1, 91) \| 149 \| Uncharacterized protein \| Uncharacterized protein; uniclust \| UniRef100\_UPI000E2FD569 \| 97.5 \| 4e-06 \| 7.3e-12 \| 55.2 \| 48 \| (3, 60) \| 106 \| (1, 48) \| 69 \| hypothetical protein \| hypothetical protein; uniclust \| UniRef100\_A0A2M7ZW18 \| 97.5 \| 4.1e-06 \| 7.6e-12 \| 59.7 \| 77 \| (1, 78) \| 106 \| (1, 88) \| 112 \| Uncharacterized protein \| Uncharacterized protein; uniclust \| UniRef100\_A0A0G3BHA1 \| 97.5 \| 4.2e-06 \| 7.8e-12 \| 60.5 \| 69 \| (3, 78) \| 106 \| (1, 81) \| 123 \| Uncharacterized protein \| Uncharacterized protein; uniclust \| UniRef100\_A0A1W2AP14 \| 97.5 \| 4.3e-06 \| 8.1e-12 \| 63.3 \| 97 \| (3, 105) \| 106 \| (4, 128) \| 163 \| Uncharacterized protein \| Uncharacterized protein; uniclust \| UniRef100\_UPI001C462278 \| 97.5 \| 4.5e-06 \| 8.3e-12 \| 61.2 \| 67 \| (1, 78) \| 106 \| (1, 86) \| 136 \| hypothetical protein \| hypothetical protein; uniclust \| UniRef100\_UPI00187D4FC2 \| 97.4 \| 5.6e-06 \| 1e-11 \| 64.5 \| 60 \| (3, 78) \| 106 \| (4, 75) \| 212 \| hypothetical protein \| hypothetical protein; uniclust \| UniRef100\_A0A1S5R1F1 \| 97.4 \| 6.8e-06 \| 1.2e-11 \| 58.8 \| 86 \| (3, 100) \| 106 \| (6, 105) \| 113 \| Uncharacterized protein \| Uncharacterized protein; uniclust \| UniRef100\_UPI001F19C64C \| 97.4 \| 7.4e-06 \| 1.4e-11 \| 52.6 \| 44 \| (3, 60) \| 106 \| (9, 52) \| 59 \| hypothetical protein \| hypothetical protein; uniclust \| UniRef100\_A0A1H5SET1 \| 97.4 \| 7.9e-06 \| 1.4e-11 \| 59.1 \| 81 \| (1, 81) \| 106 \| (1, 90) \| 122 \| Uncharacterized protein \| Uncharacterized protein; uniclust \| UniRef100\_A0A2T7SZR4 \| 97.3 \| 8.9e-06 \| 1.6e-11 \| 59.0 \| 70 \| (3, 78) \| 106 \| (1, 80) \| 123 \| Uncharacterized protein \| Uncharacterized protein; uniclust \| UniRef100\_UPI0016401755 \| 97.3 \| 9.9e-06 \| 1.8e-11 \| 56.7 \| 65 \| (3, 81) \| 106 \| (5, 83) \| 96 \| hypothetical protein \| hypothetical protein; uniclust \| UniRef100\_UPI0006AC61BC \| 97.3 \| 1e-05 \| 2e-11 \| 56.4 \| 65 \| (3, 78) \| 106 \| (1, 80) \| 92 \| hypothetical protein \| hypothetical protein; uniclust \| UniRef100\_A0A1T1ANT1 \| 97.3 \| 1.2e-05 \| 2.2e-11 \| 56.5 \| 72 \| (3, 81) \| 106 \| (1, 87) \| 100 \| Uncharacterized protein \| Uncharacterized protein; uniclust \| UniRef100\_A0A873WNJ2 \| 97.3 \| 1.3e-05 \| 2.4e-11 \| 60.3 \| 80 \| (2, 81) \| 106 \| (1, 109) \| 146 \| Uncharacterized protein \| Uncharacterized protein; uniclust \| UniRef100\_A0A257FV18 \| 97.3 \| 1.3e-05 \| 2.5e-11 \| 58.0 \| 69 \| (1, 78) \| 106 \| (12, 100) \| 122 \| KOW domain-containing protein \| KOW domain-containing protein; uniclust \| UniRef100\_UPI0018CA9FA7 \| 97.2 \| 1.7e-05 \| 3.1e-11 \| 54.8 \| 61 \| (2, 78) \| 106 \| (1, 72) \| 89 \| hypothetical protein \| hypothetical protein; uniclust \| UniRef100\_A0A965KUK1 \| 97.2 \| 1.9e-05 \| 3.8e-11 \| 57.9 \| 69 \| (3, 77) \| 106 \| (8, 91) \| 115 \| Uncharacterized protein \| Uncharacterized protein; uniclust \| UniRef100\_UPI000A5B9EFB \| 97.2 \| 2.3e-05 \| 4.2e-11 \| 58.7 \| 45 \| (3, 59) \| 106 \| (1, 45) \| 151 \| hypothetical protein \| hypothetical protein; uniclust \| UniRef100\_A0A0Q0XEJ4 \| 97.2 \| 2.2e-05 \| 4.6e-11 \| 57.4 \| 70 \| (1, 81) \| 106 \| (18, 96) \| 109 \| Uncharacterized protein \| Uncharacterized protein; uniclust \| UniRef100\_UPI002150BB13 \| 97.1 \| 2.6e-05 \| 4.9e-11 \| 59.9 \| 79 \| (2, 104) \| 106 \| (3, 94) \| 182 \| hypothetical protein \| hypothetical protein; uniclust \| UniRef100\_A0A060BAN8 \| 97.1 \| 2.5e-05 \| 4.9e-11 \| 55.2 \| 70 \| (1, 78) \| 106 \| (1, 71) \| 91 \| DUF2158 domain-containing protein \| DUF2158 domain-containing protein; uniclust \| UniRef100\_A0A6G5Y4H1 \| 97.1 \| 3.9e-05 \| 7.2e-11 \| 52.2 \| 66 \| (1, 81) \| 106 \| (1, 69) \| 80 \| Uncharacterized protein \| Uncharacterized protein; uniclust \| UniRef100\_A0A315DX93 \| 97.0 \| 5.8e-05 \| 1.1e-10 \| 53.0 \| 78 \| (3, 81) \| 106 \| (1, 87) \| 95 \| Uncharacterized protein \| Uncharacterized protein; uniclust \| UniRef100\_A0A1W2BP56 \| 97.0 \| 5.5e-05 \| 1.1e-10 \| 56.4 \| 88 \| (2, 90) \| 106 \| (1, 101) \| 127 \| Uncharacterized protein \| Uncharacterized protein; uniclust \| UniRef100\_UPI001E552E10 \| 97.0 \| 6.1e-05 \| 1.1e-10 \| 59.1 \| 33 \| (3, 36) \| 106 \| (121, 153) \| 207 \| hypothetical protein \| hypothetical protein; uniclust \| UniRef100\_A0A924KGG3 \| 97.0 \| 6.2e-05 \| 1.1e-10 \| 53.7 \| 80 \| (1, 81) \| 106 \| (1, 88) \| 105 \| Uncharacterized protein \| Uncharacterized protein; uniclust \| UniRef100\_A0A4R8FRD1 \| 97.0 \| 6.4e-05 \| 1.2e-10 \| 53.0 \| 72 \| (2, 78) \| 106 \| (3, 76) \| 92 \| Uncharacterized protein \| Uncharacterized protein; uniclust \| UniRef100\_A0A653R688 \| 96.9 \| 9.4e-05 \| 1.8e-10 \| 53.4 \| 71 \| (2, 78) \| 106 \| (1, 88) \| 106 \| Uncharacterized protein \| Uncharacterized protein; uniclust \| UniRef100\_A0A1M7R589 \| 96.8 \| 0.00011 \| 2.1e-10 \| 53.2 \| 70 \| (3, 78) \| 106 \| (1, 87) \| 115 \| Uncharacterized protein \| Uncharacterized protein; uniclust \| UniRef100\_A0A7U4PBC5 \| 96.8 \| 0.00011 \| 2.1e-10 \| 53.3 \| 33 \| (3, 36) \| 106 \| (9, 41) \| 107 \| Uncharacterized protein \| Uncharacterized protein; uniclust \| UniRef100\_A0A071MBJ7 \| 96.8 \| 0.00014 \| 2.5e-10 \| 51.6 \| 65 \| (3, 81) \| 106 \| (6, 83) \| 99 \| Uncharacterized protein \| Uncharacterized protein; uniclust \| UniRef100\_UPI0011AE5093 \| 96.8 \| 0.00014 \| 2.6e-10 \| 52.7 \| 31 \| (1, 32) \| 106 \| (1, 33) \| 115 \| hypothetical protein \| hypothetical protein; uniclust \| UniRef100\_A0A3D8K3M3 \| 96.8 \| 0.00014 \| 2.9e-10 \| 53.9 \| 66 \| (3, 78) \| 106 \| (1, 78) \| 114 \| Uncharacterized protein \| Uncharacterized protein; uniclust \| UniRef100\_A0A828SWE6 \| 96.8 \| 0.00016 \| 3e-10 \| 50.0 \| 71 \| (2, 78) \| 106 \| (1, 79) \| 83 \| Uncharacterized protein \| Uncharacterized protein; uniclust \| UniRef100\_UPI0012F7E7BF \| 96.7 \| 0.00021 \| 3.9e-10 \| 52.2 \| 74 \| (1, 77) \| 106 \| (1, 80) \| 119 \| hypothetical protein \| hypothetical protein; uniclust \| UniRef100\_UPI0012AFD91F \| 96.7 \| 0.00023 \| 4.2e-10 \| 46.9 \| 33 \| (2, 35) \| 106 \| (1, 36) \| 63 \| hypothetical protein \| hypothetical protein; uniclust \| UniRef100\_UPI002150BB13 \| 96.7 \| 0.00024 \| 4.3e-10 \| 55.2 \| 65 \| (3, 81) \| 106 \| (94, 171) \| 182 \| hypothetical protein \| hypothetical protein; uniclust \| UniRef100\_UPI000667E3A6 \| 96.6 \| 0.00032 \| 5.9e-10 \| 47.5 \| 44 \| (3, 60) \| 106 \| (4, 47) \| 73 \| hypothetical protein \| hypothetical protein; uniclust \| UniRef100\_A0A2T7SZN9 \| 96.6 \| 0.00033 \| 6e-10 \| 51.1 \| 30 \| (3, 33) \| 106 \| (21, 52) \| 116 \| Uncharacterized protein \| Uncharacterized protein; uniclust \| UniRef100\_A0A1G3GYN4 \| 96.5 \| 0.00036 \| 7.4e-10 \| 50.8 \| 33 \| (2, 36) \| 106 \| (1, 33) \| 100 \| Uncharacterized protein \| Uncharacterized protein; uniclust \| UniRef100\_A0A5P8D3D0 \| 96.5 \| 0.0004 \| 7.4e-10 \| 49.2 \| 68 \| (2, 78) \| 106 \| (1, 80) \| 96 \| Uncharacterized protein \| Uncharacterized protein; uniclust \| UniRef100\_C5A7S6 \| 96.5 \| 0.00039 \| 7.5e-10 \| 52.0 \| 70 \| (3, 78) \| 106 \| (11, 83) \| 127 \| Uncharacterized protein \| Uncharacterized protein; uniclust \| UniRef100\_UPI000539086C \| 96.5 \| 0.00042 \| 8.2e-10 \| 50.8 \| 76 \| (3, 79) \| 106 \| (8, 96) \| 111 \| hypothetical protein \| hypothetical protein; uniclust \| UniRef100\_UPI0021F1A385 \| 96.5 \| 0.00048 \| 8.8e-10 \| 45.6 \| 35 \| (1, 37) \| 106 \| (1, 35) \| 64 \| hypothetical protein \| hypothetical protein; uniclust \| UniRef100\_UPI0011CD8128 \| 96.5 \| 0.00048 \| 8.9e-10 \| 51.9 \| 32 \| (3, 35) \| 106 \| (52, 83) \| 144 \| hypothetical protein \| hypothetical protein; uniclust \| UniRef100\_UPI001D17D500 \| 96.5 \| 0.0005 \| 9.5e-10 \| 48.0 \| 83 \| (2, 102) \| 106 \| (1, 85) \| 85 \| hypothetical protein \| hypothetical protein; uniclust \| UniRef100\_A0A431L7L8 \| 96.5 \| 0.00054 \| 1e-09 \| 50.0 \| 74 \| (1, 78) \| 106 \| (1, 88) \| 115 \| Uncharacterized protein \| Uncharacterized protein; uniclust \| UniRef100\_A0A4Y5TPK8 \| 96.4 \| 0.00059 \| 1.1e-09 \| 50.9 \| 77 \| (2, 81) \| 106 \| (1, 116) \| 129 \| Uncharacterized protein \| Uncharacterized protein; uniclust \| UniRef100\_A0A315E1P7 \| 96.4 \| 0.00063 \| 1.2e-09 \| 55.7 \| 31 \| (3, 34) \| 106 \| (4, 34) \| 272 \| Uncharacterized protein \| Uncharacterized protein; uniclust \| UniRef100\_A0A0F9SEI4 \| 96.4 \| 0.00065 \| 1.2e-09 \| 48.1 \| 64 \| (3, 79) \| 106 \| (1, 68) \| 94 \| SnoaL-like domain-containing protein \| SnoaL-like domain-containing protein; uniclust \| UniRef100\_A0A6S7CP95 \| 96.4 \| 0.00067 \| 1.2e-09 \| 48.9 \| 63 \| (3, 81) \| 106 \| (1, 73) \| 105 \| Uncharacterized protein \| Uncharacterized protein; uniclust \| UniRef100\_UPI0005CA7972 \| 96.4 \| 0.0007 \| 1.3e-09 \| 47.5 \| 67 \| (2, 78) \| 106 \| (1, 68) \| 89 \| hypothetical protein \| hypothetical protein; uniclust \| UniRef100\_A0A1D2SK25 \| 96.3 \| 0.0008 \| 1.5e-09 \| 48.1 \| 71 \| (3, 77) \| 106 \| (1, 71) \| 100 \| Uncharacterized protein \| Uncharacterized protein; uniclust \| UniRef100\_UPI0013FDE139 \| 96.3 \| 0.00082 \| 1.5e-09 \| 43.0 \| 26 \| (67, 102) \| 106 \| (27, 52) \| 52 \| hypothetical protein \| hypothetical protein |
| Top keywords  (threshold 1.00e-03 (evalue)) | **hypothetical, domain\_containing, Phage, Periplasmic, KOW, ATPase, DUF2158, Fragment, Carboxypeptidase, regulatory\_like** |
| Output files | ../../similar\_sequences/29\_FANPEZAQ\_CDS\_0029\_merged.svg ../../similar\_sequences/29\_FANPEZAQ\_CDS\_0029\_pdb70.a3m ../../similar\_sequences/29\_FANPEZAQ\_CDS\_0029\_pdb70.hhr ../../similar\_sequences/29\_FANPEZAQ\_CDS\_0029\_uniclust.a3m ../../similar\_sequences/29\_FANPEZAQ\_CDS\_0029\_uniclust.hhr |

#### Structure prediction (AlphaFold)^2^

|  |  |
| --- | --- |
| Stats | xml version="1.0" encoding="utf-8" standalone="no"?       2024-09-02T21:09:29.303587 image/svg+xml   Matplotlib v3.7.2, https://matplotlib.org/ |
| Predicted structure | **NGL Viewer Controls:**  - Center: *Left-Click* - Rotate: *Left-Click + Drag* - Translate: *Right-Click + Drag* - Zoom: *Shift + Left-Click + Drag* |
| Output files | ../../predicted\_structures/29\_FANPEZAQ\_CDS\_0029/features.pkl ../../predicted\_structures/29\_FANPEZAQ\_CDS\_0029/ranked\_0.pdb ../../predicted\_structures/29\_FANPEZAQ\_CDS\_0029/ranked\_0\_plots.svg ../../predicted\_structures/29\_FANPEZAQ\_CDS\_0029/result\_model\_1\_ptm\_pred\_0.pkl |

#### Structure similarity search results (Foldseek)^3^

|  |  |
| --- | --- |
| Structure databases searched | Pdb, Afdb-proteome, Afdb-uniprot50 |
| Results, scheme(s)  (Top layers only, threshold 1.00e-02 (evalue)) | xml version="1.0" encoding="utf-8" standalone="no"?       2024-09-02T21:11:01.797539 image/svg+xml   Matplotlib v3.7.2, https://matplotlib.org/ |
| Results, table  (threshold 1.00e-02 (evalue)) | db \| id \| prob \| evalue \| bits \| fident \| alnlen \| mismatch \| gapopen \| qstart \| qend \| tstart \| tend \| name \| description; afdb-uniprot50 \| AF-A0A7S1FJM8-F1-MODEL\_V4 \| 1.0 \| 0.002565 \| 147 \| 0.269 \| 78 \| 41 \| 3 \| 1 \| 78 \| 35 \| 96 \| Hypothetical protein \| Hypothetical protein; afdb-uniprot50 \| AF-A0A3M8RD56-F1-MODEL\_V4 \| 1.0 \| 0.0004608 \| 144 \| 0.235 \| 89 \| 53 \| 3 \| 1 \| 86 \| 1 \| 77 \| Uncharacterized protein \| Uncharacterized protein; afdb-uniprot50 \| AF-A0A7S2FYS6-F1-MODEL\_V4 \| 1.0 \| 0.005353 \| 139 \| 0.256 \| 78 \| 43 \| 3 \| 3 \| 80 \| 8 \| 70 \| Hypothetical protein \| Hypothetical protein; afdb-uniprot50 \| AF-A0A7S2C143-F1-MODEL\_V4 \| 1.0 \| 0.005691 \| 139 \| 0.194 \| 77 \| 47 \| 2 \| 1 \| 77 \| 42 \| 103 \| Hypothetical protein \| Hypothetical protein; afdb-uniprot50 \| AF-A0A7S1FKI3-F1-MODEL\_V4 \| 1.0 \| 0.007273 \| 139 \| 0.233 \| 77 \| 47 \| 2 \| 1 \| 77 \| 89 \| 153 \| Hypothetical protein \| Hypothetical protein; afdb-uniprot50 \| AF-A0A812UX06-F1-MODEL\_V4 \| 1.0 \| 0.009882 \| 138 \| 0.216 \| 83 \| 48 \| 4 \| 1 \| 82 \| 1 \| 67 \| Hypothetical protein \| Hypothetical protein; afdb-uniprot50 \| AF-A0A812LB95-F1-MODEL\_V4 \| 1.0 \| 0.001087 \| 138 \| 0.257 \| 97 \| 55 \| 5 \| 2 \| 96 \| 196 \| 277 \| Hypothetical protein \| Hypothetical protein; afdb-uniprot50 \| AF-A0A0G4HT18-F1-MODEL\_V4 \| 1.0 \| 0.003278 \| 138 \| 0.246 \| 77 \| 43 \| 2 \| 1 \| 77 \| 1 \| 62 \| Uncharacterized protein \| Uncharacterized protein; afdb-uniprot50 \| AF-A0A7S2QCT9-F1-MODEL\_V4 \| 1.0 \| 0.001229 \| 137 \| 0.206 \| 97 \| 59 \| 3 \| 4 \| 98 \| 6 \| 86 \| Hypothetical protein \| Hypothetical protein; afdb-uniprot50 \| AF-A0A7S1WPA2-F1-MODEL\_V4 \| 1.0 \| 0.008742 \| 136 \| 0.21 \| 76 \| 45 \| 3 \| 3 \| 77 \| 2 \| 63 \| Hypothetical protein \| Hypothetical protein; afdb-uniprot50 \| AF-A0A7S1LFS6-F1-MODEL\_V4 \| 1.0 \| 0.001477 \| 132 \| 0.269 \| 89 \| 47 \| 4 \| 1 \| 87 \| 103 \| 175 \| Hypothetical protein \| Hypothetical protein; afdb-uniprot50 \| AF-A0A0G4G8E2-F1-MODEL\_V4 \| 1.0 \| 0.006051 \| 132 \| 0.22 \| 86 \| 50 \| 3 \| 1 \| 85 \| 1 \| 70 \| Uncharacterized protein \| Uncharacterized protein; afdb-uniprot50 \| AF-A0A813A0V9-F1-MODEL\_V4 \| 1.0 \| 0.0008 \| 129 \| 0.218 \| 96 \| 59 \| 2 \| 5 \| 98 \| 94 \| 175 \| Eef2k protein \| Eef2k protein; afdb-uniprot50 \| AF-A0A7S4BX12-F1-MODEL\_V4 \| 1.0 \| 0.002727 \| 129 \| 0.238 \| 88 \| 49 \| 4 \| 1 \| 85 \| 214 \| 286 \| Hypothetical protein \| Hypothetical protein; afdb-uniprot50 \| AF-A0A7S1B199-F1-MODEL\_V4 \| 1.0 \| 0.006051 \| 128 \| 0.244 \| 86 \| 50 \| 3 \| 2 \| 87 \| 130 \| 200 \| Hypothetical protein \| Hypothetical protein; afdb-uniprot50 \| AF-A0A812ZDA0-F1-MODEL\_V4 \| 1.0 \| 0.001156 \| 127 \| 0.22 \| 100 \| 62 \| 3 \| 1 \| 98 \| 97 \| 182 \| Eef2k protein \| Eef2k protein; afdb-uniprot50 \| AF-A0A3D9EXJ8-F1-MODEL\_V4 \| 1.0 \| 0.005353 \| 126 \| 0.247 \| 89 \| 43 \| 3 \| 3 \| 78 \| 2 \| 79 \| Uncharacterized protein \| Uncharacterized protein; afdb-uniprot50 \| AF-A0A813JW72-F1-MODEL\_V4 \| 1.0 \| 0.002565 \| 125 \| 0.241 \| 112 \| 62 \| 5 \| 1 \| 101 \| 76 \| 175 \| Hypothetical protein \| Hypothetical protein; afdb-uniprot50 \| AF-A0A813I3Y3-F1-MODEL\_V4 \| 1.0 \| 0.006051 \| 125 \| 0.302 \| 86 \| 41 \| 4 \| 2 \| 87 \| 62 \| 128 \| Hypothetical protein \| Hypothetical protein; afdb-uniprot50 \| AF-A0A7S0BCM2-F1-MODEL\_V4 \| 1.0 \| 0.008742 \| 125 \| 0.255 \| 86 \| 49 \| 2 \| 2 \| 87 \| 17 \| 87 \| Hypothetical protein \| Hypothetical protein; afdb-uniprot50 \| AF-A0A813CZE1-F1-MODEL\_V4 \| 1.0 \| 0.004735 \| 124 \| 0.257 \| 97 \| 55 \| 5 \| 2 \| 96 \| 269 \| 350 \| Hypothetical protein \| Hypothetical protein; afdb-uniprot50 \| AF-A0A812TPH0-F1-MODEL\_V4 \| 1.0 \| 0.001775 \| 123 \| 0.226 \| 97 \| 59 \| 2 \| 4 \| 98 \| 64 \| 146 \| Eef2k protein \| Eef2k protein; afdb-uniprot50 \| AF-A0A0M0JX70-F1-MODEL\_V4 \| 1.0 \| 0.001888 \| 122 \| 0.166 \| 102 \| 70 \| 3 \| 5 \| 106 \| 21 \| 107 \| Uncharacterized protein \| Uncharacterized protein; afdb-uniprot50 \| AF-A0A0D9A4N1-F1-MODEL\_V4 \| 1.0 \| 0.005034 \| 120 \| 0.268 \| 108 \| 49 \| 4 \| 1 \| 88 \| 1 \| 98 \| Uncharacterized protein \| Uncharacterized protein; afdb-uniprot50 \| AF-A0A7S3X5C1-F1-MODEL\_V4 \| 1.0 \| 0.0009044 \| 120 \| 0.225 \| 111 \| 57 \| 5 \| 3 \| 98 \| 6 \| 102 \| Hypothetical protein \| Hypothetical protein; afdb-uniprot50 \| AF-A0A7S1J6S3-F1-MODEL\_V4 \| 1.0 \| 0.006434 \| 120 \| 0.188 \| 85 \| 53 \| 3 \| 3 \| 87 \| 10 \| 78 \| Hypothetical protein \| Hypothetical protein; afdb-uniprot50 \| AF-A0A813JBM0-F1-MODEL\_V4 \| 1.0 \| 0.008222 \| 120 \| 0.235 \| 85 \| 48 \| 4 \| 3 \| 86 \| 284 \| 352 \| Hypothetical protein \| Hypothetical protein; afdb-uniprot50 \| AF-A0A812RL46-F1-MODEL\_V4 \| 1.0 \| 0.007273 \| 120 \| 0.257 \| 97 \| 55 \| 5 \| 2 \| 96 \| 481 \| 562 \| Hypothetical protein \| Hypothetical protein; afdb-uniprot50 \| AF-A0A7S0HQB5-F1-MODEL\_V4 \| 1.0 \| 0.006434 \| 119 \| 0.233 \| 90 \| 49 \| 4 \| 2 \| 87 \| 100 \| 173 \| Hypothetical protein \| Hypothetical protein; afdb-uniprot50 \| AF-A0A812MRQ4-F1-MODEL\_V4 \| 1.0 \| 0.007733 \| 119 \| 0.22 \| 86 \| 53 \| 2 \| 1 \| 86 \| 186 \| 257 \| Hypothetical protein \| Hypothetical protein; afdb-uniprot50 \| AF-A0A7S0FC98-F1-MODEL\_V4 \| 1.0 \| 0.009294 \| 117 \| 0.255 \| 86 \| 46 \| 3 \| 5 \| 88 \| 120 \| 189 \| Hypothetical protein \| Hypothetical protein; afdb-uniprot50 \| AF-A0A0F9XCY2-F1-MODEL\_V4 \| 1.0 \| 0.003485 \| 116 \| 0.153 \| 104 \| 71 \| 3 \| 3 \| 106 \| 2 \| 88 \| Uncharacterized protein \| Uncharacterized protein; afdb-uniprot50 \| AF-A0A7S2C3M0-F1-MODEL\_V4 \| 1.0 \| 0.003083 \| 116 \| 0.223 \| 103 \| 58 \| 4 \| 4 \| 106 \| 51 \| 131 \| Hypothetical protein \| Hypothetical protein; afdb-uniprot50 \| AF-A0A7S3WU97-F1-MODEL\_V4 \| 1.0 \| 0.008742 \| 116 \| 0.206 \| 87 \| 53 \| 3 \| 1 \| 87 \| 1 \| 71 \| Hypothetical protein \| Hypothetical protein; afdb-uniprot50 \| AF-A0A813GKT4-F1-MODEL\_V4 \| 1.0 \| 0.001229 \| 114 \| 0.225 \| 102 \| 55 \| 4 \| 2 \| 103 \| 74 \| 151 \| Hypothetical protein \| Hypothetical protein; afdb-uniprot50 \| AF-A0A7S1WSB6-F1-MODEL\_V4 \| 1.0 \| 0.00684 \| 111 \| 0.209 \| 105 \| 63 \| 5 \| 4 \| 106 \| 1 \| 87 \| Hypothetical protein \| Hypothetical protein; afdb-uniprot50 \| AF-A0A813E516-F1-MODEL\_V4 \| 1.0 \| 0.006051 \| 111 \| 0.295 \| 105 \| 52 \| 5 \| 2 \| 106 \| 59 \| 141 \| Hypothetical protein \| Hypothetical protein; afdb-uniprot50 \| AF-R1DKV3-F1-MODEL\_V4 \| 1.0 \| 0.003705 \| 111 \| 0.217 \| 101 \| 57 \| 4 \| 5 \| 98 \| 66 \| 151 \| Uncharacterized protein \| Uncharacterized protein; afdb-uniprot50 \| AF-A0A7S4HC21-F1-MODEL\_V4 \| 1.0 \| 0.006434 \| 111 \| 0.196 \| 102 \| 66 \| 4 \| 5 \| 105 \| 13 \| 99 \| Hypothetical protein \| Hypothetical protein; afdb-uniprot50 \| AF-A0A7S2GV41-F1-MODEL\_V4 \| 1.0 \| 0.009882 \| 110 \| 0.145 \| 103 \| 59 \| 3 \| 4 \| 106 \| 3 \| 76 \| Hypothetical protein \| Hypothetical protein; afdb-uniprot50 \| AF-A0A812IG67-F1-MODEL\_V4 \| 1.0 \| 0.009294 \| 110 \| 0.203 \| 103 \| 56 \| 6 \| 2 \| 103 \| 100 \| 177 \| Hypothetical protein \| Hypothetical protein; afdb-uniprot50 \| AF-A0A7S0KF53-F1-MODEL\_V4 \| 1.0 \| 0.006051 \| 110 \| 0.15 \| 106 \| 68 \| 5 \| 1 \| 106 \| 10 \| 93 \| Hypothetical protein \| Hypothetical protein; afdb-uniprot50 \| AF-A0A812NLF2-F1-MODEL\_V4 \| 1.0 \| 0.007733 \| 110 \| 0.242 \| 103 \| 59 \| 4 \| 1 \| 102 \| 260 \| 344 \| PRPF4B protein \| PRPF4B protein; afdb-uniprot50 \| AF-A0A1Q9ET22-F1-MODEL\_V4 \| 1.0 \| 0.007733 \| 110 \| 0.215 \| 102 \| 64 \| 2 \| 1 \| 102 \| 560 \| 645 \| Peptidylprolyl isomerase \| Peptidylprolyl isomerase; afdb-uniprot50 \| AF-A0A7S1EZX2-F1-MODEL\_V4 \| 1.0 \| 0.007273 \| 109 \| 0.247 \| 113 \| 52 \| 5 \| 1 \| 98 \| 43 \| 137 \| Hypothetical protein \| Hypothetical protein; afdb-uniprot50 \| AF-A0A7S3BEY9-F1-MODEL\_V4 \| 1.0 \| 0.004735 \| 108 \| 0.266 \| 105 \| 59 \| 4 \| 2 \| 106 \| 9 \| 95 \| Hypothetical protein \| Hypothetical protein; afdb-uniprot50 \| AF-A0A7S3TII7-F1-MODEL\_V4 \| 1.0 \| 0.008222 \| 107 \| 0.207 \| 111 \| 66 \| 6 \| 3 \| 106 \| 2 \| 97 \| Hypothetical protein \| Hypothetical protein; afdb-uniprot50 \| AF-E9GT83-F1-MODEL\_V4 \| 1.0 \| 0.004453 \| 106 \| 0.2 \| 90 \| 59 \| 4 \| 1 \| 88 \| 339 \| 417 \| DUF5641 domain-containing protein \| DUF5641 domain-containing protein; afdb-uniprot50 \| AF-A0A7S4VFS2-F1-MODEL\_V4 \| 1.0 \| 0.009294 \| 104 \| 0.231 \| 95 \| 59 \| 4 \| 4 \| 98 \| 361 \| 441 \| Hypothetical protein \| Hypothetical protein; afdb-uniprot50 \| AF-A0A7S0FA90-F1-MODEL\_V4 \| 1.0 \| 0.007273 \| 104 \| 0.191 \| 89 \| 55 \| 3 \| 1 \| 87 \| 173 \| 246 \| Hypothetical protein \| Hypothetical protein; afdb-uniprot50 \| AF-R1B7D7-F1-MODEL\_V4 \| 1.0 \| 0.005691 \| 103 \| 0.185 \| 124 \| 61 \| 3 \| 3 \| 104 \| 170 \| 275 \| Uncharacterized protein \| Uncharacterized protein; afdb-uniprot50 \| AF-A0A2V4FDJ0-F1-MODEL\_V4 \| 1.0 \| 0.003278 \| 102 \| 0.247 \| 121 \| 57 \| 9 \| 1 \| 102 \| 1 \| 106 \| Uncharacterized protein \| Uncharacterized protein; afdb-uniprot50 \| AF-A0A7S0F8T3-F1-MODEL\_V4 \| 1.0 \| 0.008742 \| 100 \| 0.166 \| 114 \| 63 \| 6 \| 4 \| 101 \| 16 \| 113 \| Hypothetical protein \| Hypothetical protein; afdb-uniprot50 \| AF-A0A7S1TQQ4-F1-MODEL\_V4 \| 0.999 \| 0.008742 \| 99 \| 0.175 \| 108 \| 68 \| 4 \| 2 \| 103 \| 21 \| 113 \| Hypothetical protein \| Hypothetical protein; afdb-uniprot50 \| AF-A0A1Q9EN27-F1-MODEL\_V4 \| 0.999 \| 0.009294 \| 99 \| 0.202 \| 99 \| 63 \| 5 \| 1 \| 99 \| 603 \| 685 \| Stress-induced-phosphoprotein 1 \| Stress-induced-phosphoprotein 1; afdb-uniprot50 \| AF-A0A7S4RQG6-F1-MODEL\_V4 \| 0.999 \| 0.00684 \| 97 \| 0.22 \| 109 \| 63 \| 4 \| 1 \| 106 \| 70 \| 159 \| Hypothetical protein \| Hypothetical protein; afdb-uniprot50 \| AF-K8F462-F1-MODEL\_V4 \| 0.998 \| 0.00684 \| 95 \| 0.186 \| 107 \| 69 \| 5 \| 1 \| 106 \| 13 \| 102 \| Uncharacterized protein \| Uncharacterized protein; afdb-uniprot50 \| AF-A0A812KFV9-F1-MODEL\_V4 \| 0.998 \| 0.009294 \| 94 \| 0.191 \| 120 \| 61 \| 4 \| 5 \| 106 \| 307 \| 408 \| Hypothetical protein \| Hypothetical protein; afdb-uniprot50 \| AF-A0A6V7Y6U1-F1-MODEL\_V4 \| 0.997 \| 0.007733 \| 91 \| 0.166 \| 108 \| 75 \| 5 \| 2 \| 106 \| 49 \| 144 \| Hypothetical protein \| Hypothetical protein; afdb-uniprot50 \| AF-K8EAH1-F1-MODEL\_V4 \| 0.991 \| 0.009294 \| 84 \| 0.168 \| 113 \| 73 \| 3 \| 3 \| 106 \| 2 \| 102 \| J domain-containing protein \| J domain-containing protein |
| Top keywords  (threshold 1.00e-02 (evalue)) | **Hypothetical, Eef2k, domain\_containing, PRPF4B, Peptidylprolyl, isomerase, DUF5641, Stress\_induced\_phosphoprotein, J** |
| Output files | ../../similar\_structures/29\_FANPEZAQ\_CDS\_0029\_afdb-proteome\_foldseek.tsv ../../similar\_structures/29\_FANPEZAQ\_CDS\_0029\_afdb-uniprot50\_foldseek.tsv ../../similar\_structures/29\_FANPEZAQ\_CDS\_0029\_merged.svg ../../similar\_structures/29\_FANPEZAQ\_CDS\_0029\_pdb\_foldseek.tsv |

  
  
  

Return to summary | Go to previous | Go to next

  


---

**Sequence/structure alignments coloring**  
Each object in the alignment figures is colored according to its E-value following this color coding:

1e-100
10
