## Supplementary material for "Completing the BASEL phage collection to unlock hidden diversity for systematic exploration of phage-host interactions": Data S2: 32.html

FANPEZAQ\_CDS\_0032


Return to summary | Go to previous | Go to next

|  |  |
| --- | --- |
| FANPEZAQ\_CDS\_0032  Page creation date: 02 Sep 2024, 12:00  Project folder: n/a  Input sequences file: Escherichia\_virus\_HeidiAbel.gb | integrase recombinase domain\_containing tyr site\_specific phage duf4102 prophage tyrosine\_type tyrosine xerd putative dna cp4\_57 xerc catalytic dna\_binding complex a tp ap and fragment yes integration recombination c\_terminal int cps\_53 arm\_type engineered core\_binding cb inta site protein\_dna binding structural in excision alpha bundle bacteriophage transposase mainly arch orthogonal hpi genomics gp |

### Sequence information

|  |  |
| --- | --- |
| Name | FANPEZAQ\_CDS\_0032  32\_FANPEZAQ\_CDS\_0032 (pipeline id) |
| Imported annotations | Escherichia\_virus\_HeidiAbel Bas97 |
| Protein sequence | MLTERQIQAAMRAVTSETVLNDGAAGRGSGSLRLRIRAGAKGPNATWQAVWWNAGKQTSK ALGRYPDLSLADARRKYETEVRDVLTAGRDPNAVVAMAESPTIEKLFQAYIQYLRDKETA TTHTSEHLLLTGKYNAADLLGRNTIASTIEPADIANVLAKGVKRGARRTTDMQRTAMMAA FNWAMKSTHDYTQENRMDWGLKYNPVAAVPRDQAANRTLDRNLSAEEMKHVWDNAPEQSG DVMRLVMCTGQRVIEVMRIAGADVDLKARLWTMPARKTKGRKHTHMVPLTTQAVDILTTL IDLYGDGYLFPARAGAKGDIIGIPSVSRGASRMAGIKPFTPRDLRRTWKSRAGDAGIDRF TRDLIQQHAQTDTGSKHYDRFDYLPQMREAMNKWEIWLRNAIA |
| Number of residues | 403 |
| Molecular weight (Da) | 45042.87 |
| Output files | ../../query\_sequences/32\_FANPEZAQ\_CDS\_0032.fasta |

### Putative domain architecture and protein family

#### Search results (HHblits)^1^

|  |  |
| --- | --- |
| Domain family databases searched | Pfam, Ncbi-cd, Cath, Phrogs |
| Results, scheme(s)  (Top layers only; threshold 1.00e-03 (evalue)) | xml version="1.0" encoding="utf-8" standalone="no"?       2024-09-02T21:08:19.283941 image/svg+xml   Matplotlib v3.7.2, https://matplotlib.org/ |
| Results, table  (E-value ≤ 1.00e-03 (evalue)) | db \| id \| prob \| evalue \| pvalue \| score \| cols \| query \| query\_len \| template \| template\_len \| name \| description; pfam \| PF00589 \| 99.1 \| 8.5e-15 \| 2e-18 \| 116.3 \| 159 \| (221, 383) \| 403 \| (1, 171) \| 173 \| Phage\_integrase \| Phage integrase family; pfam \| PF16795 \| 98.7 \| 1.5e-12 \| 3.5e-16 \| 104.9 \| 153 \| (221, 396) \| 403 \| (2, 170) \| 171 \| Phage\_integr\_3 \| Archaeal phage integrase; pfam \| PF12835 \| 98.1 \| 4.3e-10 \| 1e-13 \| 85.4 \| 126 \| (217, 350) \| 403 \| (8, 148) \| 149 \| Integrase\_1 \| Integrase; pfam \| PF11917 \| 97.5 \| 2.9e-08 \| 6.4e-12 \| 92.1 \| 68 \| (315, 382) \| 403 \| (174, 243) \| 418 \| DUF3435 \| Protein of unknown function (DUF3435); pfam \| PF16787 \| 96.8 \| 9.7e-07 \| 2.1e-10 \| 78.3 \| 142 \| (238, 383) \| 403 \| (29, 222) \| 313 \| NDC10\_II \| Centromere DNA-binding protein complex CBF3 subunit, domain 2; ncbi-cd \| cd01193 \| 99.1 \| 2e-15 \| 5e-19 \| 120.1 \| 158 \| (219, 380) \| 403 \| (2, 175) \| 176 \| INT\_IntI\_C \| cd01193 INT\_IntI\_C; Integron integrase and similar protiens, C-terminal catalytic domain. Integron integrases mediate site-specific DNA recombination between a proximal primary site (attI) and a secondary target site (attC) found within mobile gene cassettes encoding resistance or virulence factors.; ncbi-cd \| cd01197 \| 99.1 \| 2.5e-15 \| 6e-19 \| 121.1 \| 163 \| (220, 386) \| 403 \| (4, 178) \| 181 \| INT\_FimBE\_like \| cd01197 INT\_FimBE\_like; FimB and FimE and related proteins, integrase/recombinases. This CD includes proteins similar to E.; ncbi-cd \| cd00797 \| 99.0 \| 9.1e-15 \| 2.2e-18 \| 120.0 \| 163 \| (223, 389) \| 403 \| (1, 194) \| 198 \| INT\_RitB\_C\_like \| cd00797 INT\_RitB\_C\_like; C-terminal catalytic domain of recombinase RitB, a component of the recombinase trio.; ncbi-cd \| cd01191 \| 99.0 \| 9.8e-15 \| 2.3e-18 \| 119.1 \| 164 \| (222, 392) \| 403 \| (1, 172) \| 176 \| INT\_C\_like\_2 \| cd01191 INT\_C\_like\_2; Uncharacterized site-specific tyrosine recombinase, C-terminal catalytic domain.; ncbi-cd \| cd01188 \| 99.0 \| 1.3e-14 \| 3.2e-18 \| 116.7 \| 161 \| (224, 388) \| 403 \| (1, 176) \| 179 \| INT\_RitA\_C\_like \| cd01188 INT\_RitA\_C\_like; C-terminal catalytic domain of recombinase RitA, a component of the recombinase trio.; ncbi-cd \| cd00801 \| 99.0 \| 1.7e-14 \| 4.1e-18 \| 115.9 \| 171 \| (224, 397) \| 403 \| (1, 179) \| 180 \| INT\_P4\_C \| cd00801 INT\_P4\_C; Bacteriophage P4 integrase, C-terminal catalytic domain. P4-like integrases are found in temperate bacteriophages, integrative plasmids, pathogenicity and symbiosis islands, and other mobile genetic elements.; ncbi-cd \| cd01192 \| 99.0 \| 1.7e-14 \| 4.2e-18 \| 115.9 \| 163 \| (220, 387) \| 403 \| (4, 176) \| 178 \| INT\_C\_like\_3 \| cd01192 INT\_C\_like\_3; Uncharacterized site-specific tyrosine recombinase, C-terminal catalytic domain.; ncbi-cd \| cd00796 \| 99.0 \| 2.8e-14 \| 6.7e-18 \| 111.9 \| 155 \| (221, 380) \| 403 \| (3, 162) \| 162 \| INT\_Rci\_Hp1\_C \| cd00796 INT\_Rci\_Hp1\_C; Shufflon-specific DNA recombinase Rci and Bacteriophage Hp1\_like integrase, C-terminal catalytic domain.; ncbi-cd \| cd01196 \| 99.0 \| 3.2e-14 \| 7.7e-18 \| 115.1 \| 161 \| (224, 387) \| 403 \| (1, 180) \| 183 \| INT\_C\_like\_6 \| cd01196 INT\_C\_like\_6; Uncharacterized site-specific tyrosine recombinase, C-terminal catalytic domain.; ncbi-cd \| cd01182 \| 99.0 \| 3.4e-14 \| 8.2e-18 \| 113.7 \| 162 \| (223, 388) \| 403 \| (1, 181) \| 186 \| INT\_RitC\_C\_like \| cd01182 INT\_RitC\_C\_like; C-terminal catalytic domain of recombinase RitC, a component of the recombinase trio.; ncbi-cd \| cd01195 \| 99.0 \| 4e-14 \| 9.5e-18 \| 113.2 \| 154 \| (224, 381) \| 403 \| (1, 168) \| 170 \| INT\_C\_like\_5 \| cd01195 INT\_C\_like\_5; Uncharacterized site-specific tyrosine recombinase, C-terminal catalytic domain.; ncbi-cd \| cd01186 \| 98.9 \| 4.5e-14 \| 1.1e-17 \| 112.4 \| 163 \| (222, 387) \| 403 \| (1, 182) \| 184 \| INT\_tnpA\_C\_Tn554 \| cd01186 INT\_tnpA\_C\_Tn554; Putative Transposase A from transposon Tn554, C-terminal catalytic domain. This family includes putative Transposase A from transposon Tn554.; ncbi-cd \| cd00798 \| 98.8 \| 2e-13 \| 4.8e-17 \| 107.2 \| 155 \| (225, 384) \| 403 \| (1, 170) \| 172 \| INT\_XerDC\_C \| cd00798 INT\_XerDC\_C; XerD and XerC integrases, C-terminal catalytic domains. XerDC-like integrases are involved in the site-specific integration and excision of lysogenic bacteriophage genomes, transposition of conjugative transposons, termination of chromosomal replication, and stable plasmid inheritance.; ncbi-cd \| cd01194 \| 98.8 \| 2e-13 \| 4.8e-17 \| 108.1 \| 155 \| (223, 379) \| 403 \| (1, 173) \| 174 \| INT\_C\_like\_4 \| cd01194 INT\_C\_like\_4; Uncharacterized site-specific tyrosine recombinase, C-terminal catalytic domain.; ncbi-cd \| cd00800 \| 98.8 \| 2.7e-13 \| 6.5e-17 \| 107.6 \| 152 \| (226, 384) \| 403 \| (1, 158) \| 161 \| INT\_Lambda\_C \| cd00800 INT\_Lambda\_C; C-terminal catalytic domain of Lambda integrase, a tyrosine-based site-specific recombinase.; ncbi-cd \| cd00799 \| 98.8 \| 4e-13 \| 9.7e-17 \| 108.7 \| 150 \| (238, 389) \| 403 \| (17, 184) \| 188 \| INT\_Cre\_C \| cd00799 INT\_Cre\_C; C-terminal catalytic domain of Cre recombinase (also called integrase). Cre-like recombinases are tyrosine based site specific recombinases.; ncbi-cd \| cd01185 \| 98.7 \| 7e-13 \| 1.7e-16 \| 104.8 \| 152 \| (226, 384) \| 403 \| (1, 159) \| 161 \| INTN1\_C\_like \| cd01185 INTN1\_C\_like; Integrase IntN1 of Bacteroides mobilizable transposon NBU1 and similar proteins, C-terminal catalytic domain.; ncbi-cd \| cd01184 \| 98.7 \| 8.6e-13 \| 2.1e-16 \| 104.1 \| 156 \| (223, 379) \| 403 \| (1, 179) \| 180 \| INT\_C\_like\_1 \| cd01184 INT\_C\_like\_1; Uncharacterized site-specific tyrosine recombinase, C-terminal catalytic domain.; ncbi-cd \| cd01190 \| 98.7 \| 1.1e-12 \| 2.6e-16 \| 101.9 \| 146 \| (224, 378) \| 403 \| (1, 149) \| 150 \| INT\_StrepXerD\_C\_like \| cd01190 INT\_StrepXerD\_C\_like; Putative XerD in Streptococcus pneumonia and similar proteins, C-terminal catalytic domain.; ncbi-cd \| cd00397 \| 98.6 \| 5.1e-12 \| 1.2e-15 \| 98.2 \| 139 \| (238, 379) \| 403 \| (18, 167) \| 167 \| DNA\_BRE\_C \| cd00397 DNA\_BRE\_C; DNA breaking-rejoining enzymes, C-terminal catalytic domain. The DNA breaking-rejoining enzyme superfamily includes type IB topoisomerases and tyrosine based site-specific recombinases (integrases) that share the same fold in their catalytic domain containing conserved active site residues.; ncbi-cd \| cd01189 \| 97.9 \| 1.6e-09 \| 3.8e-13 \| 82.7 \| 131 \| (225, 379) \| 403 \| (1, 146) \| 147 \| INT\_ICEBs1\_C\_like \| cd01189 INT\_ICEBs1\_C\_like; C-terminal catalytic domain of integrases from bacterial phages and conjugate transposons.; ncbi-cd \| cd11602 \| 97.8 \| 3.3e-09 \| 7.8e-13 \| 96.9 \| 163 \| (220, 387) \| 403 \| (127, 338) \| 413 \| Ndc10 \| cd11602 Ndc10; Ndc10 component of the yeast centromere-binding factor 3. Ndc10 is a multidomain protein conserved in Saccharomycotina that interacts with kinetochore components.; ncbi-cd \| cd01187 \| 97.7 \| 8.8e-09 \| 2.1e-12 \| 78.7 \| 116 \| (239, 379) \| 403 \| (15, 141) \| 142 \| INT\_tnpB\_C\_Tn554 \| cd01187 INT\_tnpB\_C\_Tn554; Putative Transposase B from transposon Tn554, C-terminal catalytic domain. This family includes putative Transposase B from transposon Tn554.; cath \| 4a8eA02 \| 99.2 \| 5.9e-16 \| 1.1e-19 \| 128.4 \| 179 \| (216, 400) \| 403 \| (3, 187) \| 197 \| Probable tyrosine recombinase xerc-like \| CATHCODE: 1.10.443.10 NAME: Probable tyrosine recombinase xerc-like. Chain: a. Synonym: xer a. Engineered: yes SOURCE: Pyrococcus abyssi. Organism\_taxid: 29292. Strain: ge5 / orsay. Expressed in: escherichia coli. Expression\_system\_taxid: 469008. CLASS: Mainly Alpha, ARCH: Orthogonal Bundle, TOPOL: hpI Integrase; Chain A, HOMOL: Intergrase catalytic core; cath \| 2a3vA02 \| 99.2 \| 8.2e-16 \| 1.5e-19 \| 127.4 \| 166 \| (215, 384) \| 403 \| (4, 214) \| 217 \| Dna (31-mer) \| CATHCODE: 1.10.443.10 NAME: Dna (31-mer). Chain: e, g. Engineered: yes. Dna (34-mer). Chain: f, h.Engineered: yes. Site-specific recombinase inti4. Chain: a, b, c, d. Engineered: yes SOURCE: Yes. Other\_details: sythetic construct. Yes. Other\_details: sythetic construct. CLASS: Mainly Alpha, ARCH: Orthogonal Bundle, TOPOL: hpI Integrase; Chain A, HOMOL: Intergrase catalytic core; cath \| 3nkhA00 \| 99.2 \| 1.2e-15 \| 2.1e-19 \| 127.8 \| 179 \| (219, 398) \| 403 \| (2, 222) \| 224 \| Integrase \| CATHCODE: 1.10.443.10 NAME: Integrase. Chain: a, b. Fragment: residues 187-406. Engineered: yes SOURCE: Staphylococcus aureus subsp. Aureus usa300\_tch1516. Organism\_taxid: 451516. Strain: strain usa300 / tch1516. Gene: usa300hou\_0850. Expressed in: escherichia coli. Expression\_system\_taxid: 511693. CLASS: Mainly Alpha, ARCH: Orthogonal Bundle, TOPOL: hpI Integrase; Chain A, HOMOL: Intergrase catalytic core; cath \| 1f44A01 \| 99.1 \| 9.3e-15 \| 1.7e-18 \| 119.7 \| 164 \| (219, 384) \| 403 \| (2, 199) \| 202 \| Dna (5'- d(\*tp\*ap\*tp\*ap\*ap\*cp\*tp\*tp\*cp\*gp\*tp\*ap\*tp\*ap\*gp\*c)-3') \| CATHCODE: 1.10.443.10 NAME: Dna (5'- d(\*tp\*ap\*tp\*ap\*ap\*cp\*tp\*tp\*cp\*gp\*tp\*ap\*tp\*ap\*gp\*c)-3'). Chain: m. Engineered: yes. Dna (5'- d(\*ap\*tp\*ap\*tp\*gp\*cp\*tp\*ap\*tp\*ap\*cp\*gp\*ap\*ap\*gp\*tp\*tp\*ap\*t)-3'). Chain: n. Engineered: yes. Cre recombinase. Chain: a. Synonym: recombinase - phage p1. Engineered: yes. Mutation: yes SOURCE: Yes. Yes. CLASS: Mainly Alpha, ARCH: Orthogonal Bundle, TOPOL: hpI Integrase; Chain A, HOMOL: Intergrase catalytic core; cath \| 1ae9A00 \| 99.0 \| 1.4e-14 \| 2.5e-18 \| 118.8 \| 158 \| (220, 384) \| 403 \| (1, 172) \| 179 \| Lambda integrase \| CATHCODE: 1.10.443.10 NAME: Lambda integrase. Chain: a, b. Fragment: catalytic domain. Engineered:yes. Mutation: yes SOURCE: Enterobacteria phage lambda. Organism\_taxid: 10710. Expressed in: escherichia coli. Expression\_system\_taxid: 562 CLASS: Mainly Alpha, ARCH: Orthogonal Bundle, TOPOL: hpI Integrase; Chain A, HOMOL: Intergrase catalytic core; cath \| 1aihA00 \| 98.8 \| 4.6e-13 \| 8.2e-17 \| 108.7 \| 154 \| (218, 393) \| 403 \| (2, 162) \| 170 \| Hp1 integrase \| CATHCODE: 1.10.443.10 NAME: Hp1 integrase. Chain: a, b, c, d. Fragment: catalytic domain, residues168 - 337. Engineered: yes SOURCE: Haemophilus phage hp1. Organism\_taxid: 10690. Strain: hp1c1. Cell\_line: haemophilus influenzae l10. Gene: genbank u24159. Expressed in: escherichia coli. Expression\_system\_taxid: 562. CLASS: Mainly Alpha, ARCH: Orthogonal Bundle, TOPOL: hpI Integrase; Chain A, HOMOL: Intergrase catalytic core; cath \| 3uxuA00 \| 98.3 \| 1.1e-10 \| 2e-14 \| 93.7 \| 140 \| (219, 384) \| 403 \| (3, 148) \| 169 \| Probable integrase \| CATHCODE: 1.10.443.10 NAME: Probable integrase. Chain: a. Fragment: catalytic domain (176-334). Engineered: yes. Mutation: yes SOURCE: Sulfolobus virus 1. Organism\_taxid: 244589. Strain: sulfolobus solfataricus. Gene: d335. Expressed in: escherichia coli. Expression\_system\_taxid: 469008. CLASS: Mainly Alpha, ARCH: Orthogonal Bundle, TOPOL: hpI Integrase; Chain A, HOMOL: Intergrase catalytic core; cath \| 4dwpA02 \| 97.9 \| 3.8e-09 \| 6.1e-13 \| 93.9 \| 163 \| (220, 384) \| 403 \| (25, 217) \| 227 \| Protelomerase \| CATHCODE: 1.10.443.30 NAME: Protelomerase. Chain: a. Engineered: yes. Dna (5'-d(\*tp\*tp\*ap\*cp\*ap\*ap\*tp\*ap\*ap\*cp\*ap\*ap\*tp\*ap\*t)- 3'). Chain: c. Engineered: yes. Dna (5'- d(\*cp\*ap\*tp\*gp\*ap\*tp\*ap\*tp\*tp\*gp\*tp\*tp\*ap\*tp\*tp\*gp\*tp\*ap\*a)-3'). Chain: d. Engineered: yes SOURCE: Agrobacterium tumefaciens. Organism\_taxid: 176299. Strain: c58 / atcc 33970. Gene: tela, atu2523. Expressed in: escherichia coli. Expression\_system\_taxid: 562. Yes. CLASS: Mainly Alpha, ARCH: Orthogonal Bundle, TOPOL: hpI Integrase; Chain A, HOMOL: Telomere resolvase; cath \| 2v6eA03 \| 97.8 \| 5.2e-09 \| 8.6e-13 \| 92.1 \| 152 \| (225, 382) \| 403 \| (32, 222) \| 225 \| Protelemorase \| CATHCODE: 1.10.443.30 NAME: Protelemorase. Chain: a, b. Fragment: c-terminally truncated active resolvase, residues 1-538. Synonym: protelomerase telk538. Engineered: yes. Telrl. Chain: c, e. Other\_details: target site for telk. Telrl. Chain: d, f. Other\_details: target site for telk SOURCE: Klebsiella phage phiko2. Organism\_taxid: 255431. Expressed in: escherichia coli. Expression\_system\_taxid: 511693. CLASS: Mainly Alpha, ARCH: Orthogonal Bundle, TOPOL: hpI Integrase; Chain A, HOMOL: Telomere resolvase; phrogs \| 7244 \| 100.0 \| 2.4e-35 \| 2.8e-39 \| 264.4 \| 222 \| (41, 295) \| 403 \| (17, 250) \| 269 \| integrase \| integrase; Category: integration and excision; p76383 VI\_09148; phrogs \| 3423 \| 99.9 \| 1.5e-29 \| 1.7e-33 \| 221.8 \| 102 \| (1, 113) \| 403 \| (1, 109) \| 217 \| NA \| NA; Category: unknown function; p166161 VI\_03799; phrogs \| 216 \| 99.9 \| 2.9e-28 \| 3.9e-32 \| 228.8 \| 211 \| (168, 395) \| 403 \| (2, 250) \| 254 \| integrase \| integrase; Category: integration and excision; p381219 VI\_06799; phrogs \| 1 \| 99.8 \| 1.7e-23 \| 2.4e-27 \| 206.0 \| 265 \| (99, 392) \| 403 \| (61, 373) \| 379 \| integrase \| integrase; Category: integration and excision; p428256 VI\_04636; phrogs \| 5127 \| 99.4 \| 1.6e-17 \| 1.9e-21 \| 156.4 \| 207 \| (147, 401) \| 403 \| (127, 339) \| 340 \| integrase \| integrase; Category: integration and excision; NC\_005361\_p7; phrogs \| 15424 \| 99.4 \| 1.8e-17 \| 2.1e-21 \| 136.5 \| 131 \| (42, 187) \| 403 \| (23, 169) \| 179 \| NA \| NA; Category: unknown function; p337988 VI\_03719; phrogs \| 7685 \| 98.5 \| 1.3e-11 \| 1.6e-15 \| 94.8 \| 85 \| (305, 395) \| 403 \| (5, 95) \| 99 \| integrase \| integrase; Category: integration and excision; p430610 VI\_04687; phrogs \| 7269 \| 98.0 \| 1.7e-09 \| 1.9e-13 \| 82.5 \| 67 \| (324, 393) \| 403 \| (17, 86) \| 105 \| integrase \| integrase; Category: integration and excision; p209079 VI\_03929; phrogs \| 17497 \| 97.9 \| 5.4e-09 \| 6.2e-13 \| 83.7 \| 97 \| (303, 402) \| 403 \| (12, 117) \| 145 \| NA \| NA; Category: unknown function; p9119 VI\_09989; phrogs \| 22184 \| 97.6 \| 4.2e-08 \| 4.8e-12 \| 83.9 \| 161 \| (97, 273) \| 403 \| (65, 243) \| 256 \| NA \| NA; Category: unknown function; p65886 VI\_01893; phrogs \| 23271 \| 97.2 \| 2.9e-07 \| 3.3e-11 \| 64.0 \| 44 \| (338, 382) \| 403 \| (8, 51) \| 67 \| NA \| NA; Category: unknown function; p201349 VI\_04413; phrogs \| 24775 \| 97.0 \| 8.4e-07 \| 9.5e-11 \| 65.7 \| 65 \| (337, 402) \| 403 \| (26, 90) \| 101 \| NA \| NA; Category: unknown function; p271771 VI\_09564; phrogs \| 35951 \| 96.5 \| 7.5e-06 \| 8.3e-10 \| 59.3 \| 56 \| (337, 394) \| 403 \| (19, 74) \| 97 \| NA \| NA; Category: unknown function; p215618 VI\_07463; phrogs \| 34557 \| 96.5 \| 7.7e-06 \| 8.5e-10 \| 63.9 \| 72 \| (306, 382) \| 403 \| (68, 142) \| 158 \| integrase \| integrase; Category: integration and excision; p391360 VI\_04907; phrogs \| 24633 \| 96.0 \| 3.6e-05 \| 4e-09 \| 66.8 \| 140 \| (237, 380) \| 403 \| (92, 258) \| 367 \| NA \| NA; Category: unknown function; p434924 VI\_10927; phrogs \| 37709 \| 94.9 \| 0.00039 \| 4.3e-08 \| 51.4 \| 83 \| (169, 265) \| 403 \| (20, 103) \| 116 \| NA \| NA; Category: unknown function; p376656 VI\_08697; phrogs \| 25190 \| 94.7 \| 0.00055 \| 6.1e-08 \| 50.0 \| 50 \| (337, 387) \| 403 \| (45, 94) \| 109 \| integrase \| integrase; Category: integration and excision; p65883 VI\_01893; phrogs \| 9208 \| 94.4 \| 0.00083 \| 9.4e-08 \| 56.2 \| 124 \| (238, 382) \| 403 \| (105, 230) \| 241 \| integrase \| integrase; Category: integration and excision; KX607102\_p66 |
| Top keywords  (threshold 1.00e-03 (evalue)) | **integrase, catalytic, a, tp, ap, and, recombinase, yes, C\_terminal, Engineered** |
| Output files | ../../domain\_architecture/32\_FANPEZAQ\_CDS\_0032\_cath.hhr ../../domain\_architecture/32\_FANPEZAQ\_CDS\_0032\_merged.svg ../../domain\_architecture/32\_FANPEZAQ\_CDS\_0032\_ncbi-cd.hhr ../../domain\_architecture/32\_FANPEZAQ\_CDS\_0032\_pfam.hhr ../../domain\_architecture/32\_FANPEZAQ\_CDS\_0032\_phrogs.hhr |

### Identical protein sequences/structures

#### Search results

|  |  |
| --- | --- |
| Protein sequence databases searched | Pdb, Swissprot, Refseq |
| Identical proteins found | -- |
| Top keywords | -- |
| Output files | -- |

### Similar protein sequences/structures

#### Sequence similarity search results (HHblits)^1^

|  |  |
| --- | --- |
| Sequence databases searched | Uniclust, Pdb70 |
| Results, scheme(s)  (Top layers only, threshold 1.00e-03 (evalue)) | xml version="1.0" encoding="utf-8" standalone="no"?       2024-09-02T21:08:42.912040 image/svg+xml   Matplotlib v3.7.2, https://matplotlib.org/ |
| Results, table(s)  (threshold 1.00e-03 (evalue)) | db \| id \| prob \| evalue \| pvalue \| score \| cols \| query \| query\_len \| template \| template\_len \| name \| description; uniclust \| UniRef100\_A0A023WX11 \| 100.0 \| 2.1e-70 \| 3.7e-76 \| 482.4 \| 373 \| (1, 402) \| 403 \| (94, 487) \| 563 \| Integrase \| Integrase; uniclust \| UniRef100\_A0A078KVJ2 \| 100.0 \| 6.9e-69 \| 1.2e-74 \| 450.5 \| 372 \| (1, 402) \| 403 \| (35, 422) \| 486 \| Prophage CP4-57 integrase \| Prophage CP4-57 integrase; uniclust \| UniRef100\_A0A009HH03 \| 100.0 \| 1.9e-66 \| 3.3e-72 \| 450.0 \| 374 \| (1, 402) \| 403 \| (70, 458) \| 525 \| Phage integrase family protein \| Phage integrase family protein; uniclust \| UniRef100\_A0A010RTI3 \| 100.0 \| 3e-66 \| 5.2e-72 \| 464.3 \| 374 \| (1, 402) \| 403 \| (92, 480) \| 605 \| Integrase \| Integrase; uniclust \| UniRef100\_A0A095BCC0 \| 100.0 \| 1.1e-65 \| 2e-71 \| 429.1 \| 373 \| (1, 402) \| 403 \| (5, 401) \| 420 \| Tyr recombinase domain-containing protein \| Tyr recombinase domain-containing protein; uniclust \| UniRef100\_A0A062VJ35 \| 100.0 \| 1.9e-65 \| 3.3e-71 \| 447.7 \| 376 \| (1, 402) \| 403 \| (72, 483) \| 578 \| Integrase family protein \| Integrase family protein; uniclust \| UniRef100\_A0A023WM80 \| 100.0 \| 3.6e-65 \| 6.3e-71 \| 430.8 \| 374 \| (1, 402) \| 403 \| (25, 413) \| 477 \| Integrase \| Integrase; uniclust \| UniRef100\_A0A009ITW1 \| 100.0 \| 8.3e-65 \| 1.5e-70 \| 447.3 \| 374 \| (1, 402) \| 403 \| (98, 489) \| 595 \| Phage integrase family protein \| Phage integrase family protein; uniclust \| UniRef100\_A0A031JRT9 \| 100.0 \| 1e-64 \| 1.7e-70 \| 451.6 \| 373 \| (1, 402) \| 403 \| (69, 462) \| 531 \| Integrase family protein \| Integrase family protein; uniclust \| UniRef100\_A0A0F7PI78 \| 100.0 \| 1.5e-64 \| 2.6e-70 \| 432.9 \| 367 \| (1, 402) \| 403 \| (64, 439) \| 472 \| Site-specific recombinase XerD \| Site-specific recombinase XerD; uniclust \| UniRef100\_A0A009GCG0 \| 100.0 \| 2.1e-64 \| 3.8e-70 \| 449.4 \| 372 \| (1, 401) \| 403 \| (80, 466) \| 655 \| Phage integrase family protein \| Phage integrase family protein; uniclust \| UniRef100\_A0A3N5UFD5 \| 100.0 \| 2.3e-64 \| 4e-70 \| 415.6 \| 372 \| (1, 402) \| 403 \| (31, 416) \| 439 \| DUF4102 domain-containing protein (Fragment) \| DUF4102 domain-containing protein (Fragment); uniclust \| UniRef100\_A0A060HVV6 \| 100.0 \| 3.9e-64 \| 6.8e-70 \| 429.9 \| 372 \| (1, 401) \| 403 \| (32, 439) \| 507 \| Phage integrase/recombinase family protein \| Phage integrase/recombinase family protein; uniclust \| UniRef100\_A0A009Q4Z9 \| 100.0 \| 4.3e-64 \| 7.5e-70 \| 445.6 \| 374 \| (1, 402) \| 403 \| (102, 493) \| 604 \| Phage integrase family protein \| Phage integrase family protein; uniclust \| UniRef100\_A0A0J1CXV2 \| 100.0 \| 8.6e-64 \| 1.5e-69 \| 412.4 \| 373 \| (1, 401) \| 403 \| (20, 409) \| 435 \| Integrase \| Integrase; uniclust \| UniRef100\_A0A010T4C0 \| 100.0 \| 1.8e-63 \| 3.1e-69 \| 441.0 \| 374 \| (1, 402) \| 403 \| (80, 470) \| 595 \| Integrase \| Integrase; uniclust \| UniRef100\_A0A1I4E0U7 \| 100.0 \| 2.5e-63 \| 4.4e-69 \| 425.7 \| 377 \| (1, 403) \| 403 \| (67, 460) \| 493 \| Phage integrase family protein \| Phage integrase family protein; uniclust \| UniRef100\_A0A069F866 \| 100.0 \| 5.1e-63 \| 9.1e-69 \| 427.2 \| 391 \| (1, 402) \| 403 \| (68, 483) \| 551 \| DUF4102 domain-containing protein \| DUF4102 domain-containing protein; uniclust \| UniRef100\_A0A011NNI0 \| 100.0 \| 6.1e-63 \| 1.1e-68 \| 426.9 \| 374 \| (1, 402) \| 403 \| (54, 448) \| 530 \| Prophage CP4-57 integrase \| Prophage CP4-57 integrase; uniclust \| UniRef100\_A0A095S8W4 \| 100.0 \| 1.5e-62 \| 2.7e-68 \| 420.2 \| 373 \| (1, 401) \| 403 \| (42, 434) \| 517 \| DNA integration/recombination/inversion protein \| DNA integration/recombination/inversion protein; uniclust \| UniRef100\_A0A011P849 \| 100.0 \| 2.2e-62 \| 3.9e-68 \| 427.6 \| 366 \| (1, 400) \| 403 \| (58, 437) \| 542 \| Tyrosine recombinase XerC \| Tyrosine recombinase XerC; uniclust \| UniRef100\_A0A1Y6D1R1 \| 100.0 \| 8.4e-62 \| 1.5e-67 \| 396.6 \| 375 \| (1, 402) \| 403 \| (6, 404) \| 422 \| Integrase \| Integrase; uniclust \| UniRef100\_A0A0A8WLC3 \| 100.0 \| 1.4e-61 \| 2.5e-67 \| 411.3 \| 364 \| (1, 402) \| 403 \| (53, 440) \| 506 \| Integrase \| Integrase; uniclust \| UniRef100\_A0A095AQU7 \| 100.0 \| 1.5e-61 \| 2.6e-67 \| 427.9 \| 377 \| (1, 403) \| 403 \| (96, 497) \| 559 \| Phage integrase \| Phage integrase; uniclust \| UniRef100\_A0A086ME39 \| 100.0 \| 1.7e-61 \| 3e-67 \| 424.3 \| 371 \| (1, 402) \| 403 \| (59, 443) \| 524 \| Tyr recombinase domain-containing protein \| Tyr recombinase domain-containing protein; uniclust \| UniRef100\_A0A0F3QL50 \| 100.0 \| 1.8e-61 \| 3.3e-67 \| 405.8 \| 363 \| (1, 398) \| 403 \| (24, 396) \| 469 \| Phage integrase family protein \| Phage integrase family protein; uniclust \| UniRef100\_A0A084TAT8 \| 100.0 \| 2.2e-61 \| 4e-67 \| 406.5 \| 375 \| (1, 401) \| 403 \| (34, 429) \| 526 \| Integrase \| Integrase; uniclust \| UniRef100\_A0A0B0Q1I0 \| 100.0 \| 2.3e-61 \| 4e-67 \| 409.6 \| 371 \| (1, 402) \| 403 \| (22, 409) \| 437 \| Integrase \| Integrase; uniclust \| UniRef100\_A0A011PLM2 \| 100.0 \| 5.6e-61 \| 9.8e-67 \| 413.0 \| 365 \| (1, 398) \| 403 \| (41, 415) \| 471 \| Putative prophage CPS-53 integrase \| Putative prophage CPS-53 integrase; uniclust \| UniRef100\_A0A1E5SVX7 \| 100.0 \| 1.1e-60 \| 1.9e-66 \| 406.5 \| 376 \| (1, 402) \| 403 \| (42, 440) \| 502 \| Tyr recombinase domain-containing protein \| Tyr recombinase domain-containing protein; uniclust \| UniRef100\_A0A0C3I7B0 \| 100.0 \| 1.8e-60 \| 3.3e-66 \| 399.4 \| 374 \| (1, 401) \| 403 \| (59, 459) \| 532 \| Tyr recombinase domain-containing protein \| Tyr recombinase domain-containing protein; uniclust \| UniRef100\_A0A023WV33 \| 100.0 \| 1.9e-60 \| 3.4e-66 \| 408.8 \| 388 \| (1, 402) \| 403 \| (41, 441) \| 489 \| Integrase \| Integrase; uniclust \| UniRef100\_A0A0D7NHD3 \| 100.0 \| 2.5e-60 \| 4.5e-66 \| 408.9 \| 372 \| (1, 402) \| 403 \| (72, 479) \| 529 \| Tyr recombinase domain-containing protein \| Tyr recombinase domain-containing protein; uniclust \| UniRef100\_A0A077N6L9 \| 100.0 \| 4.2e-60 \| 7.5e-66 \| 403.6 \| 372 \| (1, 402) \| 403 \| (69, 455) \| 487 \| Phage integrase family protein \| Phage integrase family protein; uniclust \| UniRef100\_A0A0A8TPH3 \| 100.0 \| 4.3e-60 \| 7.5e-66 \| 399.4 \| 356 \| (6, 399) \| 403 \| (26, 393) \| 439 \| Phage integrase family protein \| Phage integrase family protein; uniclust \| UniRef100\_A0A060NVI1 \| 100.0 \| 4.7e-60 \| 8.4e-66 \| 407.2 \| 374 \| (1, 402) \| 403 \| (62, 453) \| 528 \| Integrase \| Integrase; uniclust \| UniRef100\_A0A084EX67 \| 100.0 \| 5.2e-60 \| 9.3e-66 \| 399.5 \| 374 \| (1, 401) \| 403 \| (28, 420) \| 482 \| Prophage CP4-57 integrase \| Prophage CP4-57 integrase; uniclust \| UniRef100\_A0A013SVQ0 \| 100.0 \| 6.7e-60 \| 1.2e-65 \| 411.2 \| 374 \| (1, 401) \| 403 \| (55, 458) \| 529 \| Phage integrase family protein \| Phage integrase family protein; uniclust \| UniRef100\_A0A059DPB2 \| 100.0 \| 6.9e-60 \| 1.2e-65 \| 392.8 \| 366 \| (1, 398) \| 403 \| (11, 415) \| 454 \| Tyr recombinase domain-containing protein \| Tyr recombinase domain-containing protein; uniclust \| UniRef100\_A0A011PKW5 \| 100.0 \| 7.2e-60 \| 1.3e-65 \| 424.5 \| 364 \| (1, 402) \| 403 \| (156, 536) \| 651 \| Tyrosine recombinase XerD \| Tyrosine recombinase XerD; uniclust \| UniRef100\_A0A1H3N2F0 \| 100.0 \| 9.1e-60 \| 1.6e-65 \| 395.4 \| 364 \| (2, 398) \| 403 \| (6, 381) \| 421 \| Site-specific recombinase XerD \| Site-specific recombinase XerD; uniclust \| UniRef100\_A0A0B1JQG0 \| 100.0 \| 1.1e-59 \| 1.9e-65 \| 386.0 \| 366 \| (1, 400) \| 403 \| (20, 405) \| 421 \| DUF4102 domain-containing protein \| DUF4102 domain-containing protein; uniclust \| UniRef100\_A0A066PM44 \| 100.0 \| 1.5e-59 \| 2.6e-65 \| 398.4 \| 374 \| (1, 402) \| 403 \| (45, 433) \| 450 \| Integrase Int \| Integrase Int; uniclust \| UniRef100\_A0A0N8TYL7 \| 100.0 \| 4e-59 \| 7.2e-65 \| 391.8 \| 373 \| (1, 399) \| 403 \| (11, 413) \| 458 \| Tyrosine recombinase XerD \| Tyrosine recombinase XerD; uniclust \| UniRef100\_A0A076LR81 \| 100.0 \| 4.2e-59 \| 7.4e-65 \| 402.1 \| 375 \| (2, 402) \| 403 \| (90, 497) \| 525 \| Prophage integrase \| Prophage integrase; uniclust \| UniRef100\_A0A009Z8S4 \| 100.0 \| 5e-59 \| 8.9e-65 \| 388.2 \| 373 \| (1, 401) \| 403 \| (6, 406) \| 468 \| Phage integrase family protein \| Phage integrase family protein; uniclust \| UniRef100\_A0A0C1EFS8 \| 100.0 \| 6.2e-59 \| 1.1e-64 \| 393.8 \| 360 \| (1, 402) \| 403 \| (23, 389) \| 460 \| Integrase \| Integrase; uniclust \| UniRef100\_A0A010YUZ4 \| 100.0 \| 6.4e-59 \| 1.1e-64 \| 412.0 \| 364 \| (1, 398) \| 403 \| (65, 469) \| 525 \| Site-specific recombinase XerD \| Site-specific recombinase XerD; uniclust \| UniRef100\_A0A011U6N9 \| 100.0 \| 9.9e-59 \| 1.7e-64 \| 409.9 \| 365 \| (1, 401) \| 403 \| (116, 512) \| 613 \| Integrase \| Integrase; uniclust \| UniRef100\_A0A0A2YVI4 \| 100.0 \| 1.8e-58 \| 3.2e-64 \| 377.5 \| 373 \| (1, 401) \| 403 \| (7, 396) \| 436 \| Preprotein translocase \| Preprotein translocase; uniclust \| UniRef100\_A0A1T4VSE3 \| 100.0 \| 1.8e-58 \| 3.2e-64 \| 390.3 \| 401 \| (1, 402) \| 403 \| (35, 447) \| 473 \| Phage integrase family protein \| Phage integrase family protein; uniclust \| UniRef100\_A0A084EN63 \| 100.0 \| 1.9e-58 \| 3.3e-64 \| 397.8 \| 373 \| (1, 402) \| 403 \| (65, 468) \| 532 \| Phage integrase \| Phage integrase; uniclust \| UniRef100\_A0A090T816 \| 100.0 \| 2e-58 \| 3.5e-64 \| 385.7 \| 373 \| (1, 401) \| 403 \| (16, 405) \| 431 \| Phage integrase \| Phage integrase; uniclust \| UniRef100\_A0A1Q8YBY4 \| 100.0 \| 2.4e-58 \| 4.2e-64 \| 387.3 \| 372 \| (1, 402) \| 403 \| (7, 420) \| 468 \| Phage integrase domain protein \| Phage integrase domain protein; uniclust \| UniRef100\_A0A011PJQ8 \| 100.0 \| 3e-58 \| 5.2e-64 \| 407.5 \| 373 \| (1, 402) \| 403 \| (104, 525) \| 621 \| Prophage CP4-57 integrase \| Prophage CP4-57 integrase; uniclust \| UniRef100\_A0A0C1H4X8 \| 100.0 \| 7.1e-58 \| 1.3e-63 \| 377.3 \| 371 \| (1, 402) \| 403 \| (2, 390) \| 420 \| Integrase \| Integrase; uniclust \| UniRef100\_A0A0A2I197 \| 100.0 \| 8.8e-58 \| 1.6e-63 \| 370.8 \| 373 \| (1, 401) \| 403 \| (7, 395) \| 421 \| Integrase \| Integrase; uniclust \| UniRef100\_A0A009HMB5 \| 100.0 \| 9.8e-58 \| 1.7e-63 \| 395.9 \| 373 \| (1, 401) \| 403 \| (99, 502) \| 587 \| Integrase \| Integrase; uniclust \| UniRef100\_A0A094ZP55 \| 100.0 \| 1e-57 \| 1.8e-63 \| 383.9 \| 370 \| (1, 402) \| 403 \| (29, 414) \| 444 \| Phage integrase family protein \| Phage integrase family protein; uniclust \| UniRef100\_A0A011Q0B5 \| 100.0 \| 1e-57 \| 1.8e-63 \| 385.6 \| 371 \| (1, 401) \| 403 \| (30, 425) \| 479 \| Prophage CP4-57 integrase \| Prophage CP4-57 integrase; uniclust \| UniRef100\_A0A2E3D5G4 \| 100.0 \| 1.2e-57 \| 2.1e-63 \| 385.2 \| 371 \| (2, 402) \| 403 \| (52, 446) \| 463 \| Tyr recombinase domain-containing protein \| Tyr recombinase domain-containing protein; uniclust \| UniRef100\_A0A062GL24 \| 100.0 \| 1.7e-57 \| 3e-63 \| 364.2 \| 372 \| (3, 402) \| 403 \| (50, 437) \| 457 \| Phage integrase family protein \| Phage integrase family protein; uniclust \| UniRef100\_A0A075MIC5 \| 100.0 \| 1.9e-57 \| 3.4e-63 \| 386.4 \| 362 \| (1, 401) \| 403 \| (24, 408) \| 451 \| Integrase \| Integrase; uniclust \| UniRef100\_A0A077NDE6 \| 100.0 \| 6e-57 \| 1.1e-62 \| 378.0 \| 385 \| (1, 402) \| 403 \| (29, 430) \| 468 \| Tyr recombinase domain-containing protein \| Tyr recombinase domain-containing protein; uniclust \| UniRef100\_A0A0F9AQZ4 \| 100.0 \| 6.7e-57 \| 1.2e-62 \| 371.0 \| 374 \| (1, 402) \| 403 \| (40, 443) \| 451 \| Core-binding (CB) domain-containing protein (Fragment) \| Core-binding (CB) domain-containing protein (Fragment); uniclust \| UniRef100\_A0A081RJF2 \| 100.0 \| 8.8e-57 \| 1.6e-62 \| 384.3 \| 361 \| (1, 399) \| 403 \| (68, 438) \| 515 \| Phage integrase \| Phage integrase; uniclust \| UniRef100\_A0A083USI6 \| 100.0 \| 1e-56 \| 1.8e-62 \| 375.7 \| 374 \| (1, 401) \| 403 \| (38, 440) \| 567 \| Integrase \| Integrase; uniclust \| UniRef100\_A0A011URX8 \| 100.0 \| 1.5e-56 \| 2.7e-62 \| 388.3 \| 367 \| (1, 402) \| 403 \| (67, 473) \| 582 \| Integrase \| Integrase; uniclust \| UniRef100\_A0A009GC36 \| 100.0 \| 1.9e-56 \| 3.3e-62 \| 386.4 \| 372 \| (1, 402) \| 403 \| (54, 451) \| 508 \| Phage integrase family protein \| Phage integrase family protein; uniclust \| UniRef100\_A0A013XU66 \| 100.0 \| 2e-56 \| 3.6e-62 \| 381.5 \| 365 \| (1, 402) \| 403 \| (37, 414) \| 468 \| Tyr recombinase domain-containing protein \| Tyr recombinase domain-containing protein; uniclust \| UniRef100\_A0A3B9NVS3 \| 100.0 \| 2.6e-56 \| 4.5e-62 \| 371.3 \| 340 \| (1, 402) \| 403 \| (21, 366) \| 386 \| Tyr recombinase domain-containing protein \| Tyr recombinase domain-containing protein; uniclust \| UniRef100\_A0A1G4SA28 \| 100.0 \| 5.6e-56 \| 1e-61 \| 364.6 \| 373 \| (1, 401) \| 403 \| (7, 397) \| 441 \| Integrase \| Integrase; uniclust \| UniRef100\_A0A085JZ21 \| 100.0 \| 5.7e-56 \| 1e-61 \| 372.4 \| 372 \| (1, 401) \| 403 \| (55, 443) \| 470 \| Integrase (Fragment) \| Integrase (Fragment); uniclust \| UniRef100\_A0A087M879 \| 100.0 \| 6.6e-56 \| 1.2e-61 \| 376.8 \| 372 \| (1, 402) \| 403 \| (53, 454) \| 471 \| Tyr recombinase domain-containing protein \| Tyr recombinase domain-containing protein; uniclust \| UniRef100\_A0A017HRW7 \| 100.0 \| 1.3e-55 \| 2.4e-61 \| 381.8 \| 365 \| (1, 400) \| 403 \| (83, 461) \| 525 \| Site-specific recombinase, phage integrase family \| Site-specific recombinase, phage integrase family; uniclust \| UniRef100\_A0A2H5VUU6 \| 100.0 \| 1.7e-55 \| 3.1e-61 \| 352.2 \| 361 \| (1, 396) \| 403 \| (13, 383) \| 396 \| Prophage integrase IntA \| Prophage integrase IntA; uniclust \| UniRef100\_A0A162TJ54 \| 100.0 \| 2.4e-55 \| 4.4e-61 \| 357.2 \| 366 \| (1, 401) \| 403 \| (25, 401) \| 424 \| Integrase \| Integrase; uniclust \| UniRef100\_A0A059UX83 \| 100.0 \| 4.8e-55 \| 8.7e-61 \| 364.6 \| 370 \| (1, 401) \| 403 \| (35, 427) \| 476 \| Integrase \| Integrase; uniclust \| UniRef100\_A0A0B0EHE1 \| 100.0 \| 6e-55 \| 1e-60 \| 379.7 \| 328 \| (33, 398) \| 403 \| (60, 391) \| 471 \| Site-specific tyrosine recombinase \| Site-specific tyrosine recombinase; uniclust \| UniRef100\_A0A010JLF0 \| 100.0 \| 6.2e-55 \| 1.1e-60 \| 385.4 \| 368 \| (1, 400) \| 403 \| (81, 470) \| 577 \| Phage integrase \| Phage integrase; uniclust \| UniRef100\_A0A009QLU8 \| 100.0 \| 6.5e-55 \| 1.1e-60 \| 382.3 \| 374 \| (1, 402) \| 403 \| (93, 481) \| 539 \| Phage integrase family protein \| Phage integrase family protein; uniclust \| UniRef100\_A0A034T1P2 \| 100.0 \| 1.5e-54 \| 2.8e-60 \| 347.1 \| 373 \| (1, 401) \| 403 \| (2, 389) \| 401 \| Integrase family protein \| Integrase family protein; uniclust \| UniRef100\_A0A2N1AM94 \| 100.0 \| 1.6e-54 \| 2.9e-60 \| 347.6 \| 371 \| (1, 401) \| 403 \| (7, 395) \| 420 \| Integrase \| Integrase; uniclust \| UniRef100\_A0A132DDL7 \| 100.0 \| 1.9e-54 \| 3.4e-60 \| 366.0 \| 322 \| (44, 396) \| 403 \| (48, 376) \| 425 \| Site-specific recombinase XerD \| Site-specific recombinase XerD; uniclust \| UniRef100\_A0A011P551 \| 100.0 \| 3.1e-54 \| 5.5e-60 \| 381.8 \| 367 \| (1, 400) \| 403 \| (110, 499) \| 587 \| Prophage CP4-57 integrase \| Prophage CP4-57 integrase; uniclust \| UniRef100\_A0A0J7XSS8 \| 100.0 \| 3.5e-54 \| 6.2e-60 \| 369.3 \| 365 \| (1, 400) \| 403 \| (55, 432) \| 523 \| Integrase \| Integrase; uniclust \| UniRef100\_A0A1E5Q3Z9 \| 100.0 \| 4e-54 \| 7e-60 \| 363.7 \| 357 \| (1, 399) \| 403 \| (13, 380) \| 413 \| Integrase \| Integrase; uniclust \| UniRef100\_A0A068R0Z1 \| 100.0 \| 4.4e-54 \| 7.8e-60 \| 361.8 \| 369 \| (1, 402) \| 403 \| (50, 440) \| 475 \| Putative prophage terminase, large subunit \| Putative prophage terminase, large subunit; uniclust \| UniRef100\_A0A0L1KDE5 \| 100.0 \| 5.2e-54 \| 9.3e-60 \| 363.0 \| 372 \| (1, 402) \| 403 \| (35, 437) \| 474 \| Integrase family protein \| Integrase family protein; uniclust \| UniRef100\_A0A0G3XIB2 \| 100.0 \| 6.9e-54 \| 1.3e-59 \| 342.1 \| 373 \| (1, 402) \| 403 \| (1, 395) \| 428 \| Integrase \| Integrase; uniclust \| UniRef100\_A0A059E4G9 \| 100.0 \| 1.4e-53 \| 2.4e-59 \| 369.2 \| 367 \| (1, 399) \| 403 \| (74, 463) \| 511 \| Tyr recombinase domain-containing protein \| Tyr recombinase domain-containing protein; uniclust \| UniRef100\_A0A0B4WVT1 \| 100.0 \| 1.8e-53 \| 3.3e-59 \| 348.4 \| 372 \| (1, 402) \| 403 \| (35, 428) \| 433 \| Integrase/recombinase domain-containing protein \| Integrase/recombinase domain-containing protein; uniclust \| UniRef100\_A0A094YPM0 \| 100.0 \| 1.9e-53 \| 3.4e-59 \| 365.7 \| 319 \| (45, 398) \| 403 \| (94, 418) \| 472 \| Site-specific recombinase, phage integrase family \| Site-specific recombinase, phage integrase family; uniclust \| UniRef100\_A0A0C1ZFP6 \| 100.0 \| 2e-53 \| 3.5e-59 \| 353.9 \| 362 \| (1, 400) \| 403 \| (3, 379) \| 414 \| Integrase \| Integrase; uniclust \| UniRef100\_A0A6M0JVR3 \| 100.0 \| 2.4e-53 \| 4.3e-59 \| 346.3 \| 382 \| (1, 403) \| 403 \| (6, 427) \| 462 \| Integrase arm-type DNA-binding domain-containing protein \| Integrase arm-type DNA-binding domain-containing protein; uniclust \| UniRef100\_A0A0F3GSB0 \| 100.0 \| 2.9e-53 \| 5.2e-59 \| 360.6 \| 326 \| (33, 395) \| 403 \| (47, 378) \| 410 \| Phage integrase family protein \| Phage integrase family protein; uniclust \| UniRef100\_A0A010RDT3 \| 100.0 \| 5.9e-53 \| 1.1e-58 \| 358.6 \| 368 \| (1, 400) \| 403 \| (79, 474) \| 516 \| Integrase \| Integrase; uniclust \| UniRef100\_A0A0T5ZW92 \| 100.0 \| 6.1e-53 \| 1.1e-58 \| 337.7 \| 328 \| (44, 398) \| 403 \| (16, 349) \| 367 \| Putative phage integrase \| Putative phage integrase; uniclust \| UniRef100\_A0A060VRP1 \| 100.0 \| 6.7e-53 \| 1.2e-58 \| 351.8 \| 373 \| (1, 401) \| 403 \| (33, 436) \| 473 \| Phage integrase family site-specific recombinase \| Phage integrase family site-specific recombinase; uniclust \| UniRef100\_A0A0M9TXC0 \| 100.0 \| 9e-53 \| 1.6e-58 \| 343.1 \| 367 \| (1, 402) \| 403 \| (5, 398) \| 422 \| Site-specific recombinase, phage integrase family \| Site-specific recombinase, phage integrase family; uniclust \| UniRef100\_A0A037ZN86 \| 100.0 \| 9.6e-53 \| 1.7e-58 \| 354.4 \| 371 \| (1, 402) \| 403 \| (23, 404) \| 462 \| Integrase \| Integrase; uniclust \| UniRef100\_A0A068QUH7 \| 100.0 \| 1.1e-52 \| 2e-58 \| 356.3 \| 370 \| (1, 402) \| 403 \| (78, 465) \| 544 \| Integrase family protein \| Integrase family protein; uniclust \| UniRef100\_A0A0B5KFG7 \| 100.0 \| 1.6e-52 \| 2.9e-58 \| 342.9 \| 369 \| (1, 397) \| 403 \| (31, 416) \| 433 \| Integrase \| Integrase; uniclust \| UniRef100\_A0A1G7GLR9 \| 100.0 \| 1.7e-52 \| 3e-58 \| 348.1 \| 377 \| (2, 402) \| 403 \| (16, 449) \| 466 \| Site-specific recombinase XerD \| Site-specific recombinase XerD; uniclust \| UniRef100\_A0A078KST6 \| 100.0 \| 1.9e-52 \| 3.3e-58 \| 342.8 \| 376 \| (1, 402) \| 403 \| (6, 394) \| 426 \| Integrase \| Integrase; uniclust \| UniRef100\_A0A0E4BJQ1 \| 100.0 \| 1.9e-52 \| 3.4e-58 \| 341.0 \| 361 \| (1, 402) \| 403 \| (18, 393) \| 415 \| Site-specific integrase/recombinase \| Site-specific integrase/recombinase; uniclust \| UniRef100\_A0A017HLQ6 \| 100.0 \| 2.1e-52 \| 3.7e-58 \| 344.1 \| 362 \| (1, 402) \| 403 \| (39, 408) \| 427 \| Putative phage-like integrase \| Putative phage-like integrase; uniclust \| UniRef100\_A0A0M4T696 \| 100.0 \| 2.6e-52 \| 4.7e-58 \| 356.1 \| 376 \| (2, 402) \| 403 \| (65, 465) \| 510 \| Integrase \| Integrase; uniclust \| UniRef100\_A0A1V0GPP3 \| 100.0 \| 7.9e-52 \| 1.4e-57 \| 349.7 \| 374 \| (1, 401) \| 403 \| (7, 394) \| 445 \| DUF4102 domain-containing protein \| DUF4102 domain-containing protein; uniclust \| UniRef100\_A0A061JXB1 \| 100.0 \| 8e-52 \| 1.4e-57 \| 349.6 \| 371 \| (1, 401) \| 403 \| (77, 474) \| 511 \| DUF4102 domain-containing protein \| DUF4102 domain-containing protein; uniclust \| UniRef100\_A0A1H1GCP0 \| 100.0 \| 9.1e-52 \| 1.6e-57 \| 339.1 \| 376 \| (1, 401) \| 403 \| (38, 451) \| 462 \| Integrase \| Integrase; uniclust \| UniRef100\_A0A010INU5 \| 100.0 \| 9.9e-52 \| 1.8e-57 \| 345.4 \| 373 \| (1, 402) \| 403 \| (37, 454) \| 479 \| Phage integrase family protein \| Phage integrase family protein; uniclust \| UniRef100\_A0A010RYA1 \| 100.0 \| 1.3e-51 \| 2.3e-57 \| 355.7 \| 375 \| (1, 401) \| 403 \| (85, 492) \| 744 \| Integrase \| Integrase; uniclust \| UniRef100\_A0A0Q3PQJ9 \| 100.0 \| 1.5e-51 \| 2.7e-57 \| 351.1 \| 371 \| (1, 402) \| 403 \| (57, 442) \| 469 \| Tyr recombinase domain-containing protein \| Tyr recombinase domain-containing protein; uniclust \| UniRef100\_A0A1V4AHU6 \| 100.0 \| 1.7e-51 \| 3e-57 \| 343.4 \| 371 \| (1, 400) \| 403 \| (11, 414) \| 433 \| Integrase \| Integrase; uniclust \| UniRef100\_A0A0F9J930 \| 100.0 \| 2e-51 \| 3.5e-57 \| 342.5 \| 325 \| (43, 398) \| 403 \| (38, 367) \| 419 \| Tyr recombinase domain-containing protein \| Tyr recombinase domain-containing protein; uniclust \| UniRef100\_A0A0F9K6W3 \| 100.0 \| 2e-51 \| 3.6e-57 \| 345.3 \| 322 \| (46, 397) \| 403 \| (42, 382) \| 415 \| Tyr recombinase domain-containing protein \| Tyr recombinase domain-containing protein; uniclust \| UniRef100\_A0A058ZJX6 \| 100.0 \| 2.3e-51 \| 4.1e-57 \| 337.5 \| 352 \| (18, 400) \| 403 \| (46, 420) \| 456 \| Integrase family protein \| Integrase family protein; uniclust \| UniRef100\_A0A1L8CP99 \| 100.0 \| 2.4e-51 \| 4.3e-57 \| 351.0 \| 365 \| (1, 401) \| 403 \| (20, 398) \| 490 \| Uncharacterized protein \| Uncharacterized protein; uniclust \| UniRef100\_A0A097EN29 \| 100.0 \| 2.8e-51 \| 4.9e-57 \| 341.0 \| 369 \| (1, 402) \| 403 \| (7, 403) \| 459 \| Tyr recombinase domain-containing protein \| Tyr recombinase domain-containing protein; uniclust \| UniRef100\_A0A077NQI5 \| 100.0 \| 3.6e-51 \| 6.5e-57 \| 332.5 \| 373 \| (1, 402) \| 403 \| (2, 391) \| 407 \| Integrase family protein \| Integrase family protein; uniclust \| UniRef100\_A0A0D0LPS2 \| 100.0 \| 4.4e-51 \| 7.9e-57 \| 336.9 \| 360 \| (2, 395) \| 403 \| (40, 406) \| 451 \| Tyr recombinase domain-containing protein \| Tyr recombinase domain-containing protein; uniclust \| UniRef100\_A0A013XV28 \| 100.0 \| 4.8e-51 \| 8.5e-57 \| 344.5 \| 379 \| (1, 401) \| 403 \| (43, 442) \| 473 \| Integrase \| Integrase; uniclust \| UniRef100\_A0A1I9YDF8 \| 100.0 \| 5.2e-51 \| 9.2e-57 \| 339.1 \| 362 \| (3, 395) \| 403 \| (9, 418) \| 434 \| Site-specific integrase \| Site-specific integrase; uniclust \| UniRef100\_A0A101VET0 \| 100.0 \| 7.7e-51 \| 1.4e-56 \| 338.8 \| 366 \| (1, 402) \| 403 \| (12, 418) \| 484 \| Tyr recombinase domain-containing protein \| Tyr recombinase domain-containing protein; uniclust \| UniRef100\_A0A084TFE9 \| 100.0 \| 7.9e-51 \| 1.4e-56 \| 333.3 \| 367 \| (1, 401) \| 403 \| (31, 411) \| 425 \| Tyr recombinase domain-containing protein \| Tyr recombinase domain-containing protein; uniclust \| UniRef100\_A0A1Y5D4R3 \| 100.0 \| 8.4e-51 \| 1.5e-56 \| 338.6 \| 369 \| (1, 402) \| 403 \| (23, 410) \| 470 \| Tyr recombinase domain-containing protein \| Tyr recombinase domain-containing protein; uniclust \| UniRef100\_A0A0F9DQS8 \| 100.0 \| 9.4e-51 \| 1.6e-56 \| 355.4 \| 330 \| (43, 398) \| 403 \| (40, 399) \| 495 \| Tyr recombinase domain-containing protein \| Tyr recombinase domain-containing protein; uniclust \| UniRef100\_A0A090RAS3 \| 100.0 \| 9.6e-51 \| 1.7e-56 \| 329.9 \| 371 \| (2, 401) \| 403 \| (10, 409) \| 454 \| Tyr recombinase domain-containing protein \| Tyr recombinase domain-containing protein; uniclust \| UniRef100\_A0A069PA44 \| 100.0 \| 1.1e-50 \| 2e-56 \| 349.1 \| 378 \| (1, 402) \| 403 \| (60, 516) \| 572 \| Integrase \| Integrase; uniclust \| UniRef100\_A0A0R2SAA9 \| 100.0 \| 1.2e-50 \| 2.2e-56 \| 326.8 \| 362 \| (1, 394) \| 403 \| (5, 377) \| 385 \| Tyr recombinase domain-containing protein \| Tyr recombinase domain-containing protein; uniclust \| UniRef100\_A0A1H8KG10 \| 100.0 \| 1.3e-50 \| 2.4e-56 \| 329.0 \| 374 \| (1, 401) \| 403 \| (4, 395) \| 453 \| Integrase \| Integrase; uniclust \| UniRef100\_A0A1M5SXE7 \| 100.0 \| 1.5e-50 \| 2.7e-56 \| 318.8 \| 376 \| (2, 402) \| 403 \| (7, 406) \| 422 \| Site-specific recombinase XerD \| Site-specific recombinase XerD; uniclust \| UniRef100\_A0A0H2V028 \| 100.0 \| 1.7e-50 \| 3e-56 \| 327.0 \| 370 \| (1, 402) \| 403 \| (7, 391) \| 409 \| Putative integrase of prophage CP-933C \| Putative integrase of prophage CP-933C; uniclust \| UniRef100\_A0A011SXD2 \| 100.0 \| 2.5e-50 \| 4.4e-56 \| 348.2 \| 363 \| (1, 400) \| 403 \| (61, 440) \| 517 \| DUF4102 domain-containing protein \| DUF4102 domain-containing protein; uniclust \| UniRef100\_A0A1R3WVC8 \| 100.0 \| 2.8e-50 \| 5e-56 \| 328.9 \| 361 \| (1, 398) \| 403 \| (17, 393) \| 408 \| Integrase \| Integrase; uniclust \| UniRef100\_A0A0A2WI49 \| 100.0 \| 3.6e-50 \| 6.3e-56 \| 349.5 \| 321 \| (43, 395) \| 403 \| (38, 368) \| 432 \| Integrase \| Integrase; uniclust \| UniRef100\_A0A0S7XRQ7 \| 100.0 \| 4e-50 \| 7.2e-56 \| 334.4 \| 334 \| (30, 396) \| 403 \| (100, 439) \| 475 \| Tyr recombinase domain-containing protein (Fragment) \| Tyr recombinase domain-containing protein (Fragment); uniclust \| UniRef100\_A0A068Z6C2 \| 100.0 \| 4.5e-50 \| 8.1e-56 \| 324.5 \| 370 \| (2, 402) \| 403 \| (3, 386) \| 399 \| Site-specific integrase \| Site-specific integrase; uniclust \| UniRef100\_A0A060UZS8 \| 100.0 \| 4.8e-50 \| 8.5e-56 \| 347.0 \| 322 \| (43, 398) \| 403 \| (67, 406) \| 493 \| Integrase \| Integrase; uniclust \| UniRef100\_A0A076HE32 \| 100.0 \| 5e-50 \| 9e-56 \| 329.7 \| 378 \| (1, 400) \| 403 \| (22, 437) \| 445 \| Tyr recombinase domain-containing protein \| Tyr recombinase domain-containing protein; uniclust \| UniRef100\_A0A061P7F4 \| 100.0 \| 5.7e-50 \| 9.9e-56 \| 352.0 \| 326 \| (46, 399) \| 403 \| (51, 409) \| 494 \| Integrase \| Integrase; uniclust \| UniRef100\_A0A0D7E4E3 \| 100.0 \| 6.4e-50 \| 1.1e-55 \| 339.2 \| 367 \| (1, 400) \| 403 \| (46, 435) \| 451 \| Integrase DNA-binding domain-containing protein (Fragment) \| Integrase DNA-binding domain-containing protein (Fragment); uniclust \| UniRef100\_A0A059G1M6 \| 100.0 \| 6.7e-50 \| 1.2e-55 \| 318.9 \| 374 \| (1, 402) \| 403 \| (21, 416) \| 436 \| Integrase family protein (Fragment) \| Integrase family protein (Fragment); uniclust \| UniRef100\_A0A2K4WBA5 \| 100.0 \| 7.4e-50 \| 1.3e-55 \| 316.7 \| 367 \| (1, 400) \| 403 \| (2, 398) \| 401 \| Integrase \| Integrase; uniclust \| UniRef100\_A0A1E8FE80 \| 100.0 \| 8.8e-50 \| 1.6e-55 \| 329.7 \| 364 \| (1, 394) \| 403 \| (59, 441) \| 483 \| Integrase \| Integrase; uniclust \| UniRef100\_A0A0H3LRV4 \| 100.0 \| 9.7e-50 \| 1.7e-55 \| 333.4 \| 377 \| (1, 401) \| 403 \| (27, 430) \| 455 \| Phage-related integrase \| Phage-related integrase; uniclust \| UniRef100\_A0A0D1EAI8 \| 100.0 \| 1.4e-49 \| 2.4e-55 \| 334.0 \| 352 \| (19, 400) \| 403 \| (30, 403) \| 434 \| IntA protein \| IntA protein; uniclust \| UniRef100\_A0A0Q5CV90 \| 100.0 \| 2e-49 \| 3.6e-55 \| 332.6 \| 378 \| (1, 402) \| 403 \| (30, 488) \| 522 \| Integrase \| Integrase; uniclust \| UniRef100\_A0A2R3F0U9 \| 100.0 \| 2.3e-49 \| 4.1e-55 \| 320.7 \| 390 \| (1, 401) \| 403 \| (47, 465) \| 480 \| Integrase \| Integrase; uniclust \| UniRef100\_A0A0B0EHF7 \| 100.0 \| 2.4e-49 \| 4.2e-55 \| 329.5 \| 319 \| (45, 395) \| 403 \| (11, 333) \| 389 \| Phage integrase \| Phage integrase; uniclust \| UniRef100\_A0A009HKV8 \| 100.0 \| 2.5e-49 \| 4.4e-55 \| 323.9 \| 370 \| (1, 401) \| 403 \| (2, 398) \| 417 \| Phage integrase family protein \| Phage integrase family protein; uniclust \| UniRef100\_A0A0P9N4F7 \| 100.0 \| 5.2e-49 \| 9.5e-55 \| 311.5 \| 372 \| (1, 401) \| 403 \| (21, 406) \| 415 \| Integrase (Fragment) \| Integrase (Fragment); uniclust \| UniRef100\_A0A0G0N6A5 \| 100.0 \| 5.5e-49 \| 9.9e-55 \| 309.9 \| 322 \| (45, 398) \| 403 \| (8, 338) \| 351 \| Site-specific tyrosine recombinase \| Site-specific tyrosine recombinase; uniclust \| UniRef100\_A0A191UGE2 \| 100.0 \| 5.8e-49 \| 1.1e-54 \| 320.3 \| 373 \| (1, 400) \| 403 \| (36, 430) \| 450 \| Integrase \| Integrase; uniclust \| UniRef100\_A0A0D6IMZ0 \| 100.0 \| 6.2e-49 \| 1.1e-54 \| 309.6 \| 370 \| (2, 401) \| 403 \| (10, 401) \| 420 \| Prophage CP4-57 integrase \| Prophage CP4-57 integrase; uniclust \| UniRef100\_A0A0P1E854 \| 100.0 \| 7.1e-49 \| 1.3e-54 \| 332.6 \| 379 \| (2, 401) \| 403 \| (86, 477) \| 495 \| Prophage CP4-57 integrase \| Prophage CP4-57 integrase; uniclust \| UniRef100\_A0A345RBQ0 \| 100.0 \| 7.2e-49 \| 1.3e-54 \| 326.2 \| 367 \| (2, 402) \| 403 \| (37, 425) \| 441 \| Tyr recombinase domain-containing protein \| Tyr recombinase domain-containing protein; uniclust \| UniRef100\_A0A1M3BHB6 \| 100.0 \| 7.7e-49 \| 1.4e-54 \| 331.8 \| 326 \| (28, 382) \| 403 \| (66, 421) \| 477 \| Site-specific integrase \| Site-specific integrase; uniclust \| UniRef100\_A0A0T6BPG3 \| 100.0 \| 8.8e-49 \| 1.5e-54 \| 335.5 \| 329 \| (44, 400) \| 403 \| (34, 402) \| 434 \| Integrase \| Integrase; uniclust \| UniRef100\_A0A0B3S3P6 \| 100.0 \| 9.4e-49 \| 1.7e-54 \| 337.2 \| 366 \| (1, 400) \| 403 \| (57, 441) \| 479 \| Putative integrase \| Putative integrase; uniclust \| UniRef100\_A0A016QR91 \| 100.0 \| 1.1e-48 \| 1.9e-54 \| 355.0 \| 330 \| (43, 400) \| 403 \| (111, 474) \| 581 \| Site-specific integrase \| Site-specific integrase; uniclust \| UniRef100\_A0A3M1JS39 \| 100.0 \| 1.2e-48 \| 2.1e-54 \| 304.6 \| 367 \| (1, 402) \| 403 \| (11, 389) \| 391 \| Site-specific integrase \| Site-specific integrase; uniclust \| UniRef100\_A0A0M2U8J5 \| 100.0 \| 1.4e-48 \| 2.5e-54 \| 330.8 \| 328 \| (43, 399) \| 403 \| (62, 424) \| 465 \| Integrase \| Integrase; uniclust \| UniRef100\_A0A068YXB4 \| 100.0 \| 1.5e-48 \| 2.7e-54 \| 338.8 \| 373 \| (1, 401) \| 403 \| (76, 493) \| 585 \| Site-specific integrase \| Site-specific integrase; uniclust \| UniRef100\_A0A0B5FFN2 \| 100.0 \| 1.6e-48 \| 2.8e-54 \| 340.1 \| 344 \| (30, 398) \| 403 \| (32, 411) \| 452 \| Tyr recombinase domain-containing protein \| Tyr recombinase domain-containing protein; uniclust \| UniRef100\_A0A0K1UCX2 \| 100.0 \| 1.6e-48 \| 2.8e-54 \| 324.3 \| 376 \| (1, 402) \| 403 \| (8, 443) \| 460 \| Integrase \| Integrase; uniclust \| UniRef100\_A0A0A1MDV4 \| 100.0 \| 1.8e-48 \| 3.1e-54 \| 328.9 \| 327 \| (44, 399) \| 403 \| (25, 388) \| 416 \| Transposase from transposon Tn916 \| Transposase from transposon Tn916; uniclust \| UniRef100\_A0A0N1C1K5 \| 100.0 \| 1.7e-48 \| 3.1e-54 \| 317.5 \| 372 \| (1, 402) \| 403 \| (19, 421) \| 428 \| Tyr recombinase domain-containing protein \| Tyr recombinase domain-containing protein; uniclust \| UniRef100\_A0A0S8KDU2 \| 100.0 \| 2.1e-48 \| 3.8e-54 \| 326.8 \| 319 \| (46, 398) \| 403 \| (50, 383) \| 410 \| Integrase \| Integrase; uniclust \| UniRef100\_A0A136KDK1 \| 100.0 \| 2.5e-48 \| 4.5e-54 \| 326.4 \| 324 \| (46, 398) \| 403 \| (47, 393) \| 424 \| Putative integrase \| Putative integrase; uniclust \| UniRef100\_A0A0J6ZLF5 \| 100.0 \| 2.6e-48 \| 4.5e-54 \| 340.9 \| 328 \| (43, 400) \| 403 \| (57, 416) \| 475 \| Integrase \| Integrase; uniclust \| UniRef100\_A0A0E9MN93 \| 100.0 \| 2.7e-48 \| 4.8e-54 \| 317.4 \| 368 \| (1, 402) \| 403 \| (3, 383) \| 395 \| Putative recombinase \| Putative recombinase; uniclust \| UniRef100\_A0A179D3R0 \| 100.0 \| 3.3e-48 \| 5.8e-54 \| 317.9 \| 307 \| (43, 381) \| 403 \| (32, 348) \| 385 \| Phage integrase \| Phage integrase; uniclust \| UniRef100\_A0A096BD50 \| 100.0 \| 4.1e-48 \| 7.2e-54 \| 338.7 \| 326 \| (44, 399) \| 403 \| (86, 453) \| 498 \| Tyr recombinase domain-containing protein \| Tyr recombinase domain-containing protein; uniclust \| UniRef100\_A0A0F2QSG4 \| 100.0 \| 4.5e-48 \| 7.9e-54 \| 324.0 \| 339 \| (31, 397) \| 403 \| (24, 391) \| 435 \| Integrase \| Integrase; uniclust \| UniRef100\_A0A0S4KXJ1 \| 100.0 \| 5.1e-48 \| 9e-54 \| 318.8 \| 320 \| (44, 400) \| 403 \| (31, 360) \| 399 \| Putative Integrase family protein \| Putative Integrase family protein; uniclust \| UniRef100\_A0A0S2JJG5 \| 100.0 \| 5.5e-48 \| 9.6e-54 \| 324.1 \| 355 \| (2, 395) \| 403 \| (11, 379) \| 398 \| Integrase \| Integrase; uniclust \| UniRef100\_A0A0F8XP09 \| 100.0 \| 6e-48 \| 1.1e-53 \| 333.6 \| 331 \| (43, 398) \| 403 \| (123, 480) \| 515 \| Tyr recombinase domain-containing protein (Fragment) \| Tyr recombinase domain-containing protein (Fragment); uniclust \| UniRef100\_A0A061NSW1 \| 100.0 \| 6.3e-48 \| 1.1e-53 \| 340.8 \| 328 \| (46, 401) \| 403 \| (65, 427) \| 478 \| Integrase \| Integrase; uniclust \| UniRef100\_A0A0C2YL66 \| 100.0 \| 6.7e-48 \| 1.2e-53 \| 321.4 \| 367 \| (1, 401) \| 403 \| (5, 409) \| 440 \| Integrase \| Integrase; uniclust \| UniRef100\_A0A2M6ZPB3 \| 100.0 \| 7.1e-48 \| 1.3e-53 \| 320.4 \| 324 \| (44, 399) \| 403 \| (60, 396) \| 414 \| Site-specific integrase \| Site-specific integrase; uniclust \| UniRef100\_A0A285ZA31 \| 100.0 \| 7.6e-48 \| 1.3e-53 \| 313.4 \| 323 \| (61, 402) \| 403 \| (3, 341) \| 359 \| Integrase \| Integrase; uniclust \| UniRef100\_A0A083UQ90 \| 100.0 \| 7.7e-48 \| 1.4e-53 \| 315.4 \| 336 \| (15, 393) \| 403 \| (24, 372) \| 388 \| Site-specific integrase \| Site-specific integrase; uniclust \| UniRef100\_A0A0A1YGP1 \| 100.0 \| 8.1e-48 \| 1.4e-53 \| 323.4 \| 362 \| (1, 402) \| 403 \| (20, 397) \| 434 \| Integrase \| Integrase; uniclust \| UniRef100\_A0A7X4ILM6 \| 100.0 \| 1e-47 \| 1.8e-53 \| 317.9 \| 350 \| (16, 401) \| 403 \| (59, 416) \| 433 \| Tyrosine-type recombinase/integrase (Fragment) \| Tyrosine-type recombinase/integrase (Fragment); uniclust \| UniRef100\_A0A1S1V944 \| 100.0 \| 1.8e-47 \| 3.2e-53 \| 310.8 \| 328 \| (44, 399) \| 403 \| (19, 374) \| 377 \| Transposase \| Transposase; uniclust \| UniRef100\_A0A3C1G4V3 \| 100.0 \| 1.9e-47 \| 3.4e-53 \| 308.1 \| 325 \| (45, 399) \| 403 \| (15, 360) \| 385 \| Site-specific integrase \| Site-specific integrase; uniclust \| UniRef100\_A0A0N0U8F3 \| 100.0 \| 2.5e-47 \| 4.3e-53 \| 326.9 \| 327 \| (43, 398) \| 403 \| (36, 394) \| 420 \| Putative prophage phiRv2 integrase \| Putative prophage phiRv2 integrase; uniclust \| UniRef100\_A0A0Q4X9F3 \| 100.0 \| 2.5e-47 \| 4.5e-53 \| 308.1 \| 362 \| (1, 398) \| 403 \| (10, 410) \| 428 \| Integrase \| Integrase; uniclust \| UniRef100\_A0A1C7HF54 \| 100.0 \| 3.5e-47 \| 6.2e-53 \| 327.5 \| 372 \| (2, 401) \| 403 \| (35, 466) \| 486 \| Core-binding (CB) domain-containing protein \| Core-binding (CB) domain-containing protein; uniclust \| UniRef100\_A0A017TBT8 \| 100.0 \| 3.9e-47 \| 6.8e-53 \| 328.9 \| 337 \| (32, 395) \| 403 \| (41, 393) \| 461 \| Integrase \| Integrase; uniclust \| UniRef100\_A0A098RMQ0 \| 100.0 \| 5e-47 \| 8.8e-53 \| 331.1 \| 330 \| (43, 400) \| 403 \| (91, 461) \| 505 \| Integrase \| Integrase; uniclust \| UniRef100\_A0A066RIY1 \| 100.0 \| 5.2e-47 \| 9.3e-53 \| 311.4 \| 368 \| (1, 400) \| 403 \| (19, 412) \| 435 \| Tyr recombinase domain-containing protein \| Tyr recombinase domain-containing protein; uniclust \| UniRef100\_A0A089WLN7 \| 100.0 \| 6.3e-47 \| 1.1e-52 \| 314.0 \| 316 \| (43, 397) \| 403 \| (19, 353) \| 406 \| Recombinase XerD \| Recombinase XerD; uniclust \| UniRef100\_A0A1G7YW28 \| 100.0 \| 7.5e-47 \| 1.3e-52 \| 314.7 \| 326 \| (44, 399) \| 403 \| (19, 377) \| 426 \| Phage integrase family protein \| Phage integrase family protein; uniclust \| UniRef100\_A0A0F5RAY4 \| 100.0 \| 8.2e-47 \| 1.4e-52 \| 320.1 \| 323 \| (45, 397) \| 403 \| (31, 386) \| 418 \| Integrase \| Integrase; uniclust \| UniRef100\_A0A023X4M9 \| 100.0 \| 8.3e-47 \| 1.4e-52 \| 341.8 \| 326 \| (45, 399) \| 403 \| (90, 451) \| 615 \| Phage integrase family \| Phage integrase family; uniclust \| UniRef100\_A0A095TK40 \| 100.0 \| 8.6e-47 \| 1.5e-52 \| 325.6 \| 368 \| (1, 397) \| 403 \| (24, 452) \| 482 \| Phage integrase \| Phage integrase; uniclust \| UniRef100\_A0A072N9D5 \| 100.0 \| 8.6e-47 \| 1.5e-52 \| 321.0 \| 333 \| (35, 398) \| 403 \| (20, 382) \| 430 \| Integrase \| Integrase; uniclust \| UniRef100\_A0A1F5UR10 \| 100.0 \| 9.7e-47 \| 1.7e-52 \| 309.9 \| 329 \| (45, 399) \| 403 \| (17, 361) \| 367 \| Tyr recombinase domain-containing protein \| Tyr recombinase domain-containing protein; uniclust \| UniRef100\_A0A0S2W7S6 \| 100.0 \| 9.8e-47 \| 1.7e-52 \| 321.1 \| 326 \| (45, 399) \| 403 \| (76, 436) \| 523 \| Prophage LambdaBa04, site-specific recombinase, phage integrase family \| Prophage LambdaBa04, site-specific recombinase, phage integrase family; uniclust \| UniRef100\_A0A382NU96 \| 100.0 \| 9.8e-47 \| 1.8e-52 \| 289.8 \| 271 \| (1, 297) \| 403 \| (5, 287) \| 288 \| Core-binding (CB) domain-containing protein (Fragment) \| Core-binding (CB) domain-containing protein (Fragment); uniclust \| UniRef100\_A0A2D7A771 \| 100.0 \| 1.1e-46 \| 1.9e-52 \| 306.7 \| 288 \| (1, 313) \| 403 \| (17, 311) \| 332 \| Integrase \| Integrase; uniclust \| UniRef100\_A0A1C3E882 \| 100.0 \| 1.2e-46 \| 2.1e-52 \| 305.1 \| 371 \| (2, 401) \| 403 \| (9, 405) \| 417 \| Tyr recombinase domain-containing protein \| Tyr recombinase domain-containing protein; uniclust \| UniRef100\_A0A6L4B4K9 \| 100.0 \| 1.2e-46 \| 2.3e-52 \| 306.6 \| 335 \| (31, 396) \| 403 \| (104, 450) \| 469 \| Tyrosine-type recombinase/integrase \| Tyrosine-type recombinase/integrase; uniclust \| UniRef100\_A0A0A0FAJ7 \| 100.0 \| 1.4e-46 \| 2.5e-52 \| 304.4 \| 349 \| (2, 382) \| 403 \| (95, 458) \| 479 \| Uncharacterized protein \| Uncharacterized protein; uniclust \| UniRef100\_A0A3S3C496 \| 100.0 \| 1.4e-46 \| 2.5e-52 \| 307.4 \| 369 \| (2, 402) \| 403 \| (35, 416) \| 444 \| Site-specific integrase \| Site-specific integrase; uniclust \| UniRef100\_A0A024PA33 \| 100.0 \| 1.6e-46 \| 2.8e-52 \| 330.1 \| 338 \| (33, 400) \| 403 \| (52, 441) \| 494 \| Tyrosine recombinase XerC \| Tyrosine recombinase XerC; uniclust \| UniRef100\_A0A064AD33 \| 100.0 \| 1.7e-46 \| 2.9e-52 \| 331.1 \| 329 \| (43, 400) \| 403 \| (78, 452) \| 528 \| Integrase \| Integrase; uniclust \| UniRef100\_A0A096AZ84 \| 100.0 \| 1.7e-46 \| 3e-52 \| 313.0 \| 324 \| (44, 400) \| 403 \| (32, 382) \| 414 \| Tyr recombinase domain-containing protein \| Tyr recombinase domain-containing protein; uniclust \| UniRef100\_A0A0J9BJZ1 \| 100.0 \| 2e-46 \| 3.5e-52 \| 327.0 \| 327 \| (44, 400) \| 403 \| (60, 429) \| 465 \| Tyr recombinase domain-containing protein \| Tyr recombinase domain-containing protein; uniclust \| UniRef100\_A0A072NST7 \| 100.0 \| 2e-46 \| 3.6e-52 \| 322.6 \| 343 \| (26, 400) \| 403 \| (64, 458) \| 482 \| Site-specific recombinase XerD \| Site-specific recombinase XerD; uniclust \| UniRef100\_A0A0J9D0K7 \| 100.0 \| 2.3e-46 \| 4.1e-52 \| 309.2 \| 351 \| (30, 403) \| 403 \| (38, 431) \| 444 \| Tyrosine-type recombinase/integrase \| Tyrosine-type recombinase/integrase; uniclust \| UniRef100\_A0A1Q7BDQ6 \| 100.0 \| 2.3e-46 \| 4.2e-52 \| 296.9 \| 282 \| (100, 402) \| 403 \| (25, 313) \| 332 \| Tyr recombinase domain-containing protein \| Tyr recombinase domain-containing protein; uniclust \| UniRef100\_A0A0D7QFG1 \| 100.0 \| 2.5e-46 \| 4.4e-52 \| 304.5 \| 370 \| (1, 402) \| 403 \| (2, 424) \| 437 \| Tyr recombinase domain-containing protein \| Tyr recombinase domain-containing protein; uniclust \| UniRef100\_A0A0A0FSB0 \| 100.0 \| 2.6e-46 \| 4.6e-52 \| 304.3 \| 342 \| (2, 382) \| 403 \| (49, 398) \| 440 \| Uncharacterized protein \| Uncharacterized protein; uniclust \| UniRef100\_A0A0F3M6B8 \| 100.0 \| 2.7e-46 \| 4.9e-52 \| 296.7 \| 364 \| (1, 398) \| 403 \| (9, 395) \| 408 \| Integrase DNA-binding domain-containing protein \| Integrase DNA-binding domain-containing protein; uniclust \| UniRef100\_A0A0B5NC73 \| 100.0 \| 2.9e-46 \| 5e-52 \| 327.1 \| 325 \| (46, 399) \| 403 \| (78, 435) \| 478 \| Site-specific integrase \| Site-specific integrase; uniclust \| UniRef100\_A0A098BVB0 \| 100.0 \| 2.9e-46 \| 5.1e-52 \| 322.9 \| 326 \| (43, 398) \| 403 \| (55, 402) \| 470 \| Putative Site-specific recombinase XerD \| Putative Site-specific recombinase XerD; uniclust \| UniRef100\_A0A1F3SLV7 \| 100.0 \| 3e-46 \| 5.3e-52 \| 315.8 \| 305 \| (45, 382) \| 403 \| (49, 373) \| 434 \| Tyr recombinase domain-containing protein \| Tyr recombinase domain-containing protein; uniclust \| UniRef100\_A0A0H4IW46 \| 100.0 \| 3.1e-46 \| 5.5e-52 \| 302.4 \| 376 \| (1, 400) \| 403 \| (6, 410) \| 444 \| Integrase \| Integrase; uniclust \| UniRef100\_A0A087QF00 \| 100.0 \| 4.3e-46 \| 7.5e-52 \| 318.6 \| 331 \| (43, 400) \| 403 \| (55, 426) \| 467 \| Toxin-antitoxin system, toxin component, PIN family \| Toxin-antitoxin system, toxin component, PIN family; uniclust \| UniRef100\_A0A0A2WPT7 \| 100.0 \| 5.1e-46 \| 9e-52 \| 310.5 \| 313 \| (56, 396) \| 403 \| (14, 354) \| 408 \| Integrase \| Integrase; uniclust \| UniRef100\_A0A0B7IVU8 \| 100.0 \| 5.8e-46 \| 1e-51 \| 306.5 \| 368 \| (2, 402) \| 403 \| (21, 401) \| 418 \| Tyr recombinase domain-containing protein \| Tyr recombinase domain-containing protein; uniclust \| UniRef100\_A0A142WZZ4 \| 100.0 \| 6.3e-46 \| 1.1e-51 \| 316.4 \| 323 \| (44, 398) \| 403 \| (44, 391) \| 436 \| Putative prophage phiRv2 integrase \| Putative prophage phiRv2 integrase; uniclust \| UniRef100\_A0A0D8IVH4 \| 100.0 \| 6.6e-46 \| 1.2e-51 \| 313.0 \| 325 \| (44, 398) \| 403 \| (32, 392) \| 442 \| Integrase \| Integrase; uniclust \| UniRef100\_A0A1V0DCV2 \| 100.0 \| 9.5e-46 \| 1.7e-51 \| 313.3 \| 320 \| (44, 391) \| 403 \| (31, 388) \| 407 \| Tyr recombinase domain-containing protein \| Tyr recombinase domain-containing protein; uniclust \| UniRef100\_A0A1X7BYL9 \| 100.0 \| 1.1e-45 \| 2e-51 \| 295.1 \| 316 \| (58, 401) \| 403 \| (10, 340) \| 364 \| Tyrosine recombinase XerD \| Tyrosine recombinase XerD; uniclust \| UniRef100\_A0A024EK23 \| 100.0 \| 1.6e-45 \| 2.8e-51 \| 294.3 \| 373 \| (1, 401) \| 403 \| (8, 410) \| 439 \| Tyr recombinase domain-containing protein \| Tyr recombinase domain-containing protein; uniclust \| UniRef100\_A0A059NZ30 \| 100.0 \| 1.6e-45 \| 2.8e-51 \| 322.2 \| 333 \| (43, 402) \| 403 \| (81, 461) \| 483 \| Integrase \| Integrase; uniclust \| UniRef100\_A0A049DW49 \| 100.0 \| 1.6e-45 \| 2.9e-51 \| 314.4 \| 322 \| (43, 394) \| 403 \| (43, 385) \| 433 \| Tyr recombinase domain-containing protein \| Tyr recombinase domain-containing protein; uniclust \| UniRef100\_A0A084TFD8 \| 100.0 \| 1.7e-45 \| 3.1e-51 \| 307.0 \| 357 \| (1, 402) \| 403 \| (22, 394) \| 525 \| Site-specific recombinase \| Site-specific recombinase; uniclust \| UniRef100\_A0A522WA23 \| 100.0 \| 2.1e-45 \| 3.7e-51 \| 291.7 \| 281 \| (100, 402) \| 403 \| (14, 313) \| 328 \| Tyrosine-type recombinase/integrase \| Tyrosine-type recombinase/integrase; uniclust \| UniRef100\_A0A085U0R6 \| 100.0 \| 2.2e-45 \| 3.8e-51 \| 313.5 \| 382 \| (1, 401) \| 403 \| (26, 463) \| 495 \| Phage integrase \| Phage integrase; uniclust \| UniRef100\_A0A3D1TX51 \| 100.0 \| 2.2e-45 \| 3.9e-51 \| 302.7 \| 318 \| (31, 381) \| 403 \| (19, 357) \| 402 \| Site-specific integrase \| Site-specific integrase; uniclust \| UniRef100\_A0A096FMR1 \| 100.0 \| 3.2e-45 \| 5.9e-51 \| 291.8 \| 375 \| (1, 402) \| 403 \| (1, 419) \| 443 \| Integrase \| Integrase; uniclust \| UniRef100\_A0A099T6K6 \| 100.0 \| 3.6e-45 \| 6.4e-51 \| 299.3 \| 320 \| (54, 398) \| 403 \| (28, 364) \| 384 \| Integrase \| Integrase; uniclust \| UniRef100\_A0A0F5ERZ1 \| 100.0 \| 3.9e-45 \| 7.1e-51 \| 287.0 \| 372 \| (1, 402) \| 403 \| (6, 399) \| 425 \| Phage integrase family site-specific recombinase \| Phage integrase family site-specific recombinase; uniclust \| UniRef100\_A0A1U9UUM4 \| 100.0 \| 4.1e-45 \| 7.3e-51 \| 294.3 \| 352 \| (2, 382) \| 403 \| (6, 381) \| 401 \| Site-specific integrase \| Site-specific integrase; uniclust \| UniRef100\_A0A011PK30 \| 100.0 \| 5e-45 \| 8.8e-51 \| 323.0 \| 387 \| (1, 402) \| 403 \| (48, 478) \| 514 \| Phage integrase family protein \| Phage integrase family protein; uniclust \| UniRef100\_A0A086P927 \| 100.0 \| 4.9e-45 \| 8.9e-51 \| 290.5 \| 373 \| (1, 402) \| 403 \| (1, 392) \| 476 \| Integrase \| Integrase; uniclust \| UniRef100\_A0A0D8IYN9 \| 100.0 \| 5.2e-45 \| 9.2e-51 \| 310.7 \| 317 \| (45, 392) \| 403 \| (84, 431) \| 468 \| Tyr recombinase domain-containing protein \| Tyr recombinase domain-containing protein; uniclust \| UniRef100\_A0A0P1EBU4 \| 100.0 \| 5.6e-45 \| 1e-50 \| 309.3 \| 371 \| (2, 402) \| 403 \| (31, 432) \| 464 \| Site-specific recombinase XerD \| Site-specific recombinase XerD; uniclust \| UniRef100\_A0A059DPD4 \| 100.0 \| 6.3e-45 \| 1.1e-50 \| 300.5 \| 379 \| (2, 402) \| 403 \| (14, 413) \| 433 \| Tyr recombinase domain-containing protein \| Tyr recombinase domain-containing protein; uniclust \| UniRef100\_A0A1G6SUD2 \| 100.0 \| 6.5e-45 \| 1.1e-50 \| 316.0 \| 326 \| (45, 399) \| 403 \| (47, 410) \| 478 \| Integrase \| Integrase; uniclust \| UniRef100\_A0A3D6EJP8 \| 100.0 \| 6.4e-45 \| 1.2e-50 \| 293.6 \| 372 \| (2, 402) \| 403 \| (19, 430) \| 455 \| Tyr recombinase domain-containing protein \| Tyr recombinase domain-containing protein; uniclust \| UniRef100\_A0A011QXA4 \| 100.0 \| 7.5e-45 \| 1.3e-50 \| 320.2 \| 330 \| (43, 401) \| 403 \| (60, 430) \| 500 \| Integrase \| Integrase; uniclust \| UniRef100\_A0A0F8YR70 \| 100.0 \| 7.6e-45 \| 1.3e-50 \| 309.9 \| 323 \| (43, 398) \| 403 \| (28, 366) \| 462 \| Tyr recombinase domain-containing protein (Fragment) \| Tyr recombinase domain-containing protein (Fragment); uniclust \| UniRef100\_A0A061NW99 \| 100.0 \| 8.5e-45 \| 1.5e-50 \| 304.9 \| 331 \| (43, 402) \| 403 \| (21, 391) \| 417 \| Phage integrase \| Phage integrase; uniclust \| UniRef100\_A0A399RGX3 \| 100.0 \| 8.6e-45 \| 1.5e-50 \| 293.0 \| 333 \| (31, 400) \| 403 \| (3, 346) \| 386 \| Site-specific integrase \| Site-specific integrase; uniclust \| UniRef100\_A0A060DST5 \| 100.0 \| 1.1e-44 \| 2e-50 \| 304.6 \| 314 \| (45, 392) \| 403 \| (38, 357) \| 398 \| Site-specific integrase \| Site-specific integrase; uniclust \| UniRef100\_A0A177PVT5 \| 100.0 \| 1.2e-44 \| 2.1e-50 \| 311.7 \| 327 \| (31, 383) \| 403 \| (53, 437) \| 488 \| Integrase \| Integrase; uniclust \| UniRef100\_A0A2E8GAL2 \| 100.0 \| 1.3e-44 \| 2.2e-50 \| 297.2 \| 335 \| (31, 397) \| 403 \| (32, 371) \| 422 \| Site-specific integrase \| Site-specific integrase; uniclust \| UniRef100\_A0A421K355 \| 100.0 \| 1.6e-44 \| 2.9e-50 \| 285.2 \| 373 \| (1, 402) \| 403 \| (8, 388) \| 401 \| Tyr recombinase domain-containing protein \| Tyr recombinase domain-containing protein; uniclust \| UniRef100\_A0A1L7AEE4 \| 100.0 \| 1.6e-44 \| 2.9e-50 \| 303.6 \| 376 \| (2, 403) \| 403 \| (22, 437) \| 480 \| Tyr recombinase domain-containing protein \| Tyr recombinase domain-containing protein; uniclust \| UniRef100\_A0A081K581 \| 100.0 \| 1.9e-44 \| 3.5e-50 \| 298.5 \| 372 \| (1, 401) \| 403 \| (10, 426) \| 477 \| Tyr recombinase domain-containing protein \| Tyr recombinase domain-containing protein; uniclust \| UniRef100\_A0A1F6PNS7 \| 100.0 \| 2.1e-44 \| 3.7e-50 \| 297.8 \| 332 \| (32, 394) \| 403 \| (34, 387) \| 407 \| Tyr recombinase domain-containing protein \| Tyr recombinase domain-containing protein; uniclust \| UniRef100\_A0A0C9PF50 \| 100.0 \| 2.7e-44 \| 4.9e-50 \| 281.6 \| 243 \| (136, 400) \| 403 \| (2, 247) \| 286 \| Site-specific recombinase \| Site-specific recombinase; uniclust \| UniRef100\_A0A0K2JBH0 \| 100.0 \| 3e-44 \| 5.2e-50 \| 313.7 \| 322 \| (44, 398) \| 403 \| (31, 386) \| 429 \| Integrase \| Integrase; uniclust \| UniRef100\_A0A062WVW9 \| 100.0 \| 3.2e-44 \| 5.5e-50 \| 318.4 \| 321 \| (44, 393) \| 403 \| (77, 419) \| 493 \| Site-specific recombinase XerD \| Site-specific recombinase XerD; uniclust \| UniRef100\_A0A090IT75 \| 100.0 \| 4e-44 \| 7.2e-50 \| 297.4 \| 373 \| (1, 401) \| 403 \| (75, 471) \| 502 \| Putative phage integrase \| Putative phage integrase; uniclust \| UniRef100\_A0A1M3P5Q7 \| 100.0 \| 4.1e-44 \| 7.2e-50 \| 292.7 \| 324 \| (1, 354) \| 403 \| (5, 346) \| 367 \| Tyr recombinase domain-containing protein \| Tyr recombinase domain-containing protein; uniclust \| UniRef100\_A0A011V5Q7 \| 100.0 \| 4.2e-44 \| 7.5e-50 \| 297.2 \| 330 \| (45, 399) \| 403 \| (17, 393) \| 436 \| Recombinase \| Recombinase; uniclust \| UniRef100\_A0A031LMT3 \| 100.0 \| 4.3e-44 \| 7.7e-50 \| 302.3 \| 374 \| (1, 400) \| 403 \| (20, 428) \| 460 \| Integrase \| Integrase; uniclust \| UniRef100\_A0A101VH43 \| 100.0 \| 5e-44 \| 8.8e-50 \| 283.9 \| 245 \| (139, 402) \| 403 \| (19, 272) \| 294 \| Tyr recombinase domain-containing protein \| Tyr recombinase domain-containing protein; uniclust \| UniRef100\_A0A024L7S6 \| 100.0 \| 5e-44 \| 9e-50 \| 285.6 \| 373 \| (1, 401) \| 403 \| (13, 413) \| 447 \| DUF4102 domain-containing protein \| DUF4102 domain-containing protein; uniclust \| UniRef100\_A0A090P338 \| 100.0 \| 5.2e-44 \| 9.1e-50 \| 298.5 \| 311 \| (45, 394) \| 403 \| (16, 351) \| 384 \| Integrase \| Integrase; uniclust \| UniRef100\_A0A0W1DLE8 \| 100.0 \| 5.3e-44 \| 9.6e-50 \| 292.7 \| 371 \| (2, 401) \| 403 \| (12, 429) \| 459 \| Integrase \| Integrase; uniclust \| UniRef100\_A0A166RGF6 \| 100.0 \| 5.7e-44 \| 1e-49 \| 301.6 \| 329 \| (45, 399) \| 403 \| (51, 411) \| 442 \| Site-specific recombinase XerD \| Site-specific recombinase XerD; uniclust \| UniRef100\_A0A0M2NN08 \| 100.0 \| 7e-44 \| 1.2e-49 \| 303.1 \| 311 \| (45, 382) \| 403 \| (32, 383) \| 432 \| Prophage LambdaBa04, site-specific recombinase, phage integrase family \| Prophage LambdaBa04, site-specific recombinase, phage integrase family; uniclust \| UniRef100\_A0A015UQW6 \| 100.0 \| 7.9e-44 \| 1.4e-49 \| 314.6 \| 324 \| (44, 398) \| 403 \| (76, 430) \| 492 \| Phage integrase family protein \| Phage integrase family protein; uniclust \| UniRef100\_A0A086PDB9 \| 100.0 \| 7.8e-44 \| 1.4e-49 \| 298.0 \| 356 \| (1, 399) \| 403 \| (26, 394) \| 428 \| Integrase family protein \| Integrase family protein; uniclust \| UniRef100\_A0A086ZVN3 \| 100.0 \| 8.4e-44 \| 1.5e-49 \| 304.0 \| 323 \| (44, 396) \| 403 \| (89, 439) \| 495 \| Integrase/recombinase \| Integrase/recombinase; uniclust \| UniRef100\_A0A1V5MLF3 \| 100.0 \| 9e-44 \| 1.6e-49 \| 288.9 \| 319 \| (44, 400) \| 403 \| (14, 341) \| 345 \| Tyrosine recombinase XerC \| Tyrosine recombinase XerC; uniclust \| UniRef100\_A0A2V8JLK3 \| 100.0 \| 1.1e-43 \| 1.9e-49 \| 296.1 \| 332 \| (33, 398) \| 403 \| (36, 389) \| 438 \| Tyr recombinase domain-containing protein \| Tyr recombinase domain-containing protein; uniclust \| UniRef100\_A0A1M3IUQ0 \| 100.0 \| 1.1e-43 \| 2e-49 \| 281.0 \| 323 \| (43, 399) \| 403 \| (12, 350) \| 360 \| Tyr recombinase domain-containing protein \| Tyr recombinase domain-containing protein; uniclust \| UniRef100\_A0A0J8SS38 \| 100.0 \| 1.2e-43 \| 2.1e-49 \| 284.4 \| 343 \| (1, 381) \| 403 \| (28, 388) \| 414 \| Site-specific recombinase \| Site-specific recombinase; uniclust \| UniRef100\_A0A024P888 \| 100.0 \| 1.2e-43 \| 2.1e-49 \| 310.4 \| 329 \| (43, 398) \| 403 \| (36, 421) \| 487 \| Integrase \| Integrase; uniclust \| UniRef100\_A0A024JYX4 \| 100.0 \| 1.3e-43 \| 2.3e-49 \| 302.8 \| 329 \| (43, 400) \| 403 \| (28, 392) \| 431 \| Phage integrase family protein \| Phage integrase family protein; uniclust \| UniRef100\_A0A061M0A6 \| 100.0 \| 1.4e-43 \| 2.5e-49 \| 302.7 \| 307 \| (44, 382) \| 403 \| (29, 368) \| 458 \| Integrase \| Integrase; uniclust \| UniRef100\_A0A143X2R8 \| 100.0 \| 1.5e-43 \| 2.7e-49 \| 303.3 \| 327 \| (45, 400) \| 403 \| (44, 412) \| 486 \| Site-specific tyrosine recombinase XerC \| Site-specific tyrosine recombinase XerC; uniclust \| UniRef100\_A0A069AE41 \| 100.0 \| 1.5e-43 \| 2.7e-49 \| 311.7 \| 329 \| (44, 399) \| 403 \| (78, 457) \| 542 \| Prophage lambdaba04, site-specific recombinase, phage integrase family \| Prophage lambdaba04, site-specific recombinase, phage integrase family; uniclust \| UniRef100\_A0A147J3P5 \| 100.0 \| 1.7e-43 \| 3e-49 \| 294.5 \| 346 \| (31, 400) \| 403 \| (51, 409) \| 436 \| Integrase (Fragment) \| Integrase (Fragment); uniclust \| UniRef100\_A0A075KBY8 \| 100.0 \| 1.7e-43 \| 3e-49 \| 300.5 \| 339 \| (32, 400) \| 403 \| (8, 391) \| 405 \| Integrase family protein \| Integrase family protein; uniclust \| UniRef100\_A0A096CRB0 \| 100.0 \| 1.9e-43 \| 3.4e-49 \| 291.8 \| 287 \| (98, 402) \| 403 \| (94, 411) \| 490 \| Tyr recombinase domain-containing protein \| Tyr recombinase domain-containing protein; uniclust \| UniRef100\_A0A009S4D1 \| 100.0 \| 2.1e-43 \| 3.8e-49 \| 291.2 \| 372 \| (1, 400) \| 403 \| (5, 402) \| 442 \| Phage integrase family protein \| Phage integrase family protein; uniclust \| UniRef100\_A0A1H8A220 \| 100.0 \| 2.2e-43 \| 3.9e-49 \| 300.3 \| 315 \| (45, 399) \| 403 \| (67, 403) \| 426 \| Site-specific recombinase XerD \| Site-specific recombinase XerD; uniclust \| UniRef100\_A0A014L1K2 \| 100.0 \| 2.3e-43 \| 4.2e-49 \| 298.1 \| 326 \| (44, 399) \| 403 \| (80, 437) \| 472 \| Integrase \| Integrase; uniclust \| UniRef100\_A0A928E6Y9 \| 100.0 \| 2.6e-43 \| 4.7e-49 \| 283.1 \| 366 \| (1, 398) \| 403 \| (11, 403) \| 409 \| Site-specific integrase \| Site-specific integrase; uniclust \| UniRef100\_A0A1J0KSW7 \| 100.0 \| 2.8e-43 \| 5e-49 \| 285.9 \| 286 \| (99, 401) \| 403 \| (9, 308) \| 321 \| Phage integrase family protein \| Phage integrase family protein; uniclust \| UniRef100\_A0A2D9L8W3 \| 100.0 \| 2.9e-43 \| 5.2e-49 \| 281.6 \| 342 \| (2, 382) \| 403 \| (12, 376) \| 390 \| Integrase \| Integrase; uniclust \| UniRef100\_A0A0K2H1N5 \| 100.0 \| 3.7e-43 \| 6.5e-49 \| 290.3 \| 323 \| (43, 395) \| 403 \| (24, 368) \| 403 \| Integrase \| Integrase; uniclust \| UniRef100\_A0A0X8VBK9 \| 100.0 \| 3.7e-43 \| 6.6e-49 \| 290.1 \| 315 \| (44, 395) \| 403 \| (7, 346) \| 371 \| Tyrosine recombinase XerD \| Tyrosine recombinase XerD; uniclust \| UniRef100\_A0A098AZG8 \| 100.0 \| 4e-43 \| 7.1e-49 \| 295.4 \| 319 \| (54, 401) \| 403 \| (30, 380) \| 418 \| Integrase protein \| Integrase protein; uniclust \| UniRef100\_A0A011U459 \| 100.0 \| 4.5e-43 \| 7.9e-49 \| 309.4 \| 321 \| (45, 395) \| 403 \| (102, 442) \| 562 \| Integrase \| Integrase; uniclust \| UniRef100\_A0A059N5D1 \| 100.0 \| 4.5e-43 \| 8e-49 \| 300.6 \| 332 \| (43, 401) \| 403 \| (33, 403) \| 443 \| Site-specific recombinase, phage integrase family \| Site-specific recombinase, phage integrase family; uniclust \| UniRef100\_A0A0M7K4N7 \| 100.0 \| 4.6e-43 \| 8.2e-49 \| 287.6 \| 286 \| (99, 402) \| 403 \| (49, 361) \| 404 \| Prophage CP4-57 integrase \| Prophage CP4-57 integrase; uniclust \| UniRef100\_A0A0F3MLM1 \| 100.0 \| 4.8e-43 \| 8.6e-49 \| 279.9 \| 286 \| (84, 400) \| 403 \| (7, 299) \| 325 \| Phage integrase family protein \| Phage integrase family protein; uniclust \| UniRef100\_A0A059KUY7 \| 100.0 \| 4.9e-43 \| 8.6e-49 \| 306.8 \| 368 \| (1, 400) \| 403 \| (43, 425) \| 507 \| Integrase \| Integrase; uniclust \| UniRef100\_A0A0A8JYV4 \| 100.0 \| 4.9e-43 \| 8.7e-49 \| 299.7 \| 325 \| (43, 398) \| 403 \| (53, 421) \| 460 \| Integrase \| Integrase; uniclust \| UniRef100\_A0A4D7DB28 \| 100.0 \| 5e-43 \| 9.1e-49 \| 280.8 \| 372 \| (2, 402) \| 403 \| (10, 396) \| 415 \| Tyr recombinase domain-containing protein \| Tyr recombinase domain-containing protein; uniclust \| UniRef100\_A0A1D7YT86 \| 100.0 \| 5.2e-43 \| 9.2e-49 \| 279.9 \| 247 \| (125, 398) \| 403 \| (10, 264) \| 289 \| Tyr recombinase domain-containing protein \| Tyr recombinase domain-containing protein; uniclust \| UniRef100\_A0A073IUB0 \| 100.0 \| 5.2e-43 \| 9.3e-49 \| 292.7 \| 314 \| (35, 382) \| 403 \| (16, 387) \| 422 \| Integrase \| Integrase; uniclust \| UniRef100\_A0A0B1ZRK3 \| 100.0 \| 5.3e-43 \| 9.5e-49 \| 295.0 \| 363 \| (2, 401) \| 403 \| (60, 448) \| 539 \| Tyr recombinase domain-containing protein \| Tyr recombinase domain-containing protein; uniclust \| UniRef100\_A0A095YNC6 \| 100.0 \| 5.8e-43 \| 1e-48 \| 298.1 \| 326 \| (31, 396) \| 403 \| (67, 422) \| 454 \| Tyr recombinase domain-containing protein \| Tyr recombinase domain-containing protein; uniclust \| UniRef100\_A0A0F6AIS4 \| 100.0 \| 5.9e-43 \| 1e-48 \| 289.4 \| 264 \| (122, 402) \| 403 \| (33, 311) \| 358 \| Tyr recombinase domain-containing protein \| Tyr recombinase domain-containing protein; uniclust \| UniRef100\_A0A1G7JFK2 \| 100.0 \| 6.3e-43 \| 1.1e-48 \| 292.2 \| 322 \| (31, 396) \| 403 \| (57, 422) \| 440 \| Site-specific recombinase XerD \| Site-specific recombinase XerD; uniclust \| UniRef100\_A0A1V6AYS9 \| 100.0 \| 6.7e-43 \| 1.2e-48 \| 287.0 \| 320 \| (43, 398) \| 403 \| (52, 384) \| 410 \| Prophage phiRv2 integrase \| Prophage phiRv2 integrase; uniclust \| UniRef100\_A0A096A971 \| 100.0 \| 7.5e-43 \| 1.3e-48 \| 296.1 \| 317 \| (32, 381) \| 403 \| (33, 398) \| 438 \| Tyr recombinase domain-containing protein \| Tyr recombinase domain-containing protein; uniclust \| UniRef100\_A0A136KB30 \| 100.0 \| 7.6e-43 \| 1.3e-48 \| 294.8 \| 304 \| (44, 382) \| 403 \| (35, 369) \| 416 \| Phage integrase \| Phage integrase; uniclust \| UniRef100\_A0A0M9ZP03 \| 100.0 \| 8.1e-43 \| 1.4e-48 \| 290.5 \| 343 \| (32, 399) \| 403 \| (18, 393) \| 419 \| Tyr recombinase domain-containing protein \| Tyr recombinase domain-containing protein; uniclust \| UniRef100\_A0A062WVF1 \| 100.0 \| 8.6e-43 \| 1.5e-48 \| 297.9 \| 326 \| (44, 399) \| 403 \| (40, 425) \| 528 \| Site-specific recombinase XerD \| Site-specific recombinase XerD; uniclust \| UniRef100\_A0A069CWW0 \| 100.0 \| 9.6e-43 \| 1.7e-48 \| 298.0 \| 329 \| (43, 399) \| 403 \| (30, 397) \| 420 \| Phage integrase family protein (Fragment) \| Phage integrase family protein (Fragment); uniclust \| UniRef100\_A0A089ICH8 \| 100.0 \| 9.5e-43 \| 1.7e-48 \| 292.1 \| 336 \| (35, 398) \| 403 \| (50, 438) \| 461 \| Integrase \| Integrase; uniclust \| UniRef100\_A0A068F1P9 \| 100.0 \| 9.6e-43 \| 1.7e-48 \| 299.5 \| 308 \| (44, 381) \| 403 \| (130, 463) \| 544 \| Integrase \| Integrase; uniclust \| UniRef100\_A0A126QRX5 \| 100.0 \| 1.1e-42 \| 1.9e-48 \| 288.9 \| 328 \| (43, 399) \| 403 \| (76, 416) \| 475 \| Tyr recombinase domain-containing protein \| Tyr recombinase domain-containing protein; uniclust \| UniRef100\_A0A099RM23 \| 100.0 \| 1.2e-42 \| 2.1e-48 \| 286.4 \| 317 \| (34, 382) \| 403 \| (17, 359) \| 385 \| Tyr recombinase domain-containing protein \| Tyr recombinase domain-containing protein; uniclust \| UniRef100\_A0A1J5I4Z9 \| 100.0 \| 1.2e-42 \| 2.2e-48 \| 288.9 \| 324 \| (46, 401) \| 403 \| (37, 381) \| 408 \| Tyr recombinase domain-containing protein \| Tyr recombinase domain-containing protein; uniclust \| UniRef100\_A0A0L6J7P5 \| 100.0 \| 1.2e-42 \| 2.2e-48 \| 299.8 \| 341 \| (32, 400) \| 403 \| (40, 426) \| 493 \| Integrase \| Integrase; uniclust \| UniRef100\_A0A095ZM98 \| 100.0 \| 1.2e-42 \| 2.2e-48 \| 296.1 \| 316 \| (47, 395) \| 403 \| (57, 403) \| 445 \| Tyr recombinase domain-containing protein \| Tyr recombinase domain-containing protein; uniclust \| UniRef100\_A0A0M2S0L9 \| 100.0 \| 1.3e-42 \| 2.2e-48 \| 294.6 \| 324 \| (26, 382) \| 403 \| (18, 364) \| 418 \| Tyr recombinase domain-containing protein \| Tyr recombinase domain-containing protein; uniclust \| UniRef100\_UPI001488B3D3 \| 100.0 \| 1.2e-42 \| 2.2e-48 \| 271.8 \| 360 \| (1, 401) \| 403 \| (1, 370) \| 379 \| tyrosine-type recombinase/integrase \| tyrosine-type recombinase/integrase; uniclust \| UniRef100\_A0A2T5MEE6 \| 100.0 \| 1.3e-42 \| 2.3e-48 \| 283.4 \| 369 \| (6, 399) \| 403 \| (10, 396) \| 437 \| Tyr recombinase domain-containing protein \| Tyr recombinase domain-containing protein; uniclust \| UniRef100\_A0A096B432 \| 100.0 \| 1.3e-42 \| 2.3e-48 \| 292.4 \| 306 \| (69, 397) \| 403 \| (61, 395) \| 562 \| Tyr recombinase domain-containing protein \| Tyr recombinase domain-containing protein; uniclust \| UniRef100\_A0A0Q5I7C0 \| 100.0 \| 1.6e-42 \| 2.8e-48 \| 287.8 \| 333 \| (34, 397) \| 403 \| (19, 387) \| 433 \| Integrase \| Integrase; uniclust \| UniRef100\_A0A059T666 \| 100.0 \| 1.6e-42 \| 2.9e-48 \| 298.3 \| 328 \| (43, 398) \| 403 \| (60, 431) \| 464 \| Integrase \| Integrase; uniclust \| UniRef100\_A0A024H3U4 \| 100.0 \| 1.7e-42 \| 2.9e-48 \| 295.6 \| 327 \| (43, 397) \| 403 \| (37, 395) \| 432 \| Phage integrase family protein \| Phage integrase family protein; uniclust \| UniRef100\_A0A0X8JIH3 \| 100.0 \| 1.8e-42 \| 3.2e-48 \| 283.5 \| 278 \| (100, 399) \| 403 \| (35, 318) \| 386 \| Integrase \| Integrase; uniclust \| UniRef100\_A0A081RCZ7 \| 100.0 \| 2e-42 \| 3.5e-48 \| 282.5 \| 281 \| (100, 402) \| 403 \| (4, 300) \| 328 \| Phage integrase \| Phage integrase; uniclust \| UniRef100\_A0A1E7ILK3 \| 100.0 \| 2e-42 \| 3.6e-48 \| 279.1 \| 346 \| (31, 402) \| 403 \| (35, 412) \| 434 \| Tyr recombinase domain-containing protein \| Tyr recombinase domain-containing protein; uniclust \| UniRef100\_A0A0H5D4Z8 \| 100.0 \| 2.1e-42 \| 3.8e-48 \| 290.6 \| 383 \| (1, 402) \| 403 \| (12, 451) \| 486 \| Prophage CP4-57 integrase \| Prophage CP4-57 integrase; uniclust \| UniRef100\_A0A0F2NFY3 \| 100.0 \| 2.2e-42 \| 3.9e-48 \| 283.5 \| 324 \| (44, 396) \| 403 \| (74, 410) \| 459 \| Integrase \| Integrase; uniclust \| UniRef100\_A0A3M5BZX2 \| 100.0 \| 2.9e-42 \| 5.2e-48 \| 272.9 \| 277 \| (99, 395) \| 403 \| (20, 309) \| 347 \| Integrase protein \| Integrase protein; uniclust \| UniRef100\_A0A4P5VS75 \| 100.0 \| 3.4e-42 \| 6.1e-48 \| 286.3 \| 310 \| (44, 382) \| 403 \| (24, 349) \| 415 \| Integrase \| Integrase; uniclust \| UniRef100\_A0A0H2XPX9 \| 100.0 \| 3.3e-42 \| 6.1e-48 \| 271.4 \| 372 \| (1, 401) \| 403 \| (26, 415) \| 431 \| Phage integrase \| Phage integrase; uniclust \| UniRef100\_A0A0D0JR95 \| 100.0 \| 3.5e-42 \| 6.2e-48 \| 282.0 \| 286 \| (100, 402) \| 403 \| (51, 347) \| 368 \| Integrase \| Integrase; uniclust \| UniRef100\_A0A081RFF4 \| 100.0 \| 3.7e-42 \| 6.7e-48 \| 278.5 \| 286 \| (99, 401) \| 403 \| (18, 314) \| 371 \| Integrase \| Integrase; uniclust \| UniRef100\_A0A7T4N282 \| 100.0 \| 3.8e-42 \| 6.9e-48 \| 279.7 \| 374 \| (1, 401) \| 403 \| (2, 399) \| 584 \| Integrase arm-type DNA-binding domain-containing protein \| Integrase arm-type DNA-binding domain-containing protein; uniclust \| UniRef100\_A0A1V5M8V7 \| 100.0 \| 4.2e-42 \| 7.5e-48 \| 280.6 \| 311 \| (45, 381) \| 403 \| (35, 361) \| 414 \| Putative prophage phiRv2 integrase \| Putative prophage phiRv2 integrase; uniclust \| UniRef100\_A0A174EDH2 \| 100.0 \| 4.4e-42 \| 7.8e-48 \| 284.9 \| 326 \| (43, 398) \| 403 \| (49, 409) \| 420 \| Integrase \| Integrase; uniclust \| UniRef100\_A0A1C3XG07 \| 100.0 \| 4.3e-42 \| 7.8e-48 \| 281.4 \| 373 \| (2, 402) \| 403 \| (42, 438) \| 454 \| Integrase \| Integrase; uniclust \| UniRef100\_A0A1H6ZZ71 \| 100.0 \| 4.3e-42 \| 7.9e-48 \| 279.5 \| 372 \| (2, 400) \| 403 \| (67, 466) \| 506 \| Integrase \| Integrase; uniclust \| UniRef100\_A0A089HSZ3 \| 100.0 \| 4.5e-42 \| 8e-48 \| 294.8 \| 319 \| (45, 384) \| 403 \| (60, 422) \| 490 \| Integrase \| Integrase; uniclust \| UniRef100\_A0A059N3T8 \| 100.0 \| 4.7e-42 \| 8.2e-48 \| 301.0 \| 325 \| (44, 397) \| 403 \| (93, 474) \| 519 \| Site-specific recombinase, phage integrase family \| Site-specific recombinase, phage integrase family; uniclust \| UniRef100\_A0A537CBF8 \| 100.0 \| 4.8e-42 \| 8.7e-48 \| 283.4 \| 389 \| (1, 401) \| 403 \| (8, 437) \| 443 \| DUF4102 domain-containing protein (Fragment) \| DUF4102 domain-containing protein (Fragment); uniclust \| UniRef100\_A0A550JG93 \| 100.0 \| 4.9e-42 \| 8.9e-48 \| 275.3 \| 334 \| (44, 402) \| 403 \| (30, 419) \| 434 \| Site-specific integrase \| Site-specific integrase; uniclust \| UniRef100\_UPI00187E1C08 \| 100.0 \| 5e-42 \| 9.2e-48 \| 275.4 \| 366 \| (1, 397) \| 403 \| (4, 381) \| 571 \| site-specific integrase \| site-specific integrase; uniclust \| UniRef100\_A0A3N5J444 \| 100.0 \| 6e-42 \| 1.1e-47 \| 270.5 \| 333 \| (31, 394) \| 403 \| (3, 356) \| 366 \| Site-specific integrase \| Site-specific integrase; uniclust \| UniRef100\_A0A1G6JWN1 \| 100.0 \| 6.5e-42 \| 1.2e-47 \| 281.8 \| 369 \| (1, 400) \| 403 \| (75, 468) \| 473 \| Integrase DNA-binding domain-containing protein \| Integrase DNA-binding domain-containing protein; uniclust \| UniRef100\_A0A011NT00 \| 100.0 \| 7.4e-42 \| 1.3e-47 \| 295.5 \| 373 \| (1, 400) \| 403 \| (130, 537) \| 594 \| Site-specific tyrosine recombinase XerC \| Site-specific tyrosine recombinase XerC; uniclust \| UniRef100\_A0A0Q5R873 \| 100.0 \| 7.8e-42 \| 1.4e-47 \| 279.6 \| 366 \| (3, 402) \| 403 \| (15, 418) \| 435 \| Core-binding (CB) domain-containing protein \| Core-binding (CB) domain-containing protein; uniclust \| UniRef100\_A0A173G9H3 \| 100.0 \| 8.5e-42 \| 1.5e-47 \| 283.6 \| 338 \| (27, 400) \| 403 \| (15, 375) \| 393 \| Uncharacterized protein \| Uncharacterized protein; uniclust \| UniRef100\_A0A1F2SPC9 \| 100.0 \| 9e-42 \| 1.6e-47 \| 285.6 \| 323 \| (45, 400) \| 403 \| (41, 372) \| 424 \| Tyr recombinase domain-containing protein \| Tyr recombinase domain-containing protein; uniclust \| UniRef100\_A0A061NKM5 \| 100.0 \| 1.1e-41 \| 1.9e-47 \| 283.0 \| 276 \| (99, 395) \| 403 \| (50, 339) \| 353 \| Integrase \| Integrase; uniclust \| UniRef100\_A0A3B7DI48 \| 100.0 \| 1.1e-41 \| 1.9e-47 \| 275.9 \| 251 \| (134, 402) \| 403 \| (9, 277) \| 307 \| Tyr recombinase domain-containing protein \| Tyr recombinase domain-containing protein; uniclust \| UniRef100\_A0A096B2D4 \| 100.0 \| 1.1e-41 \| 2e-47 \| 301.3 \| 333 \| (45, 398) \| 403 \| (91, 471) \| 570 \| Tyr recombinase domain-containing protein \| Tyr recombinase domain-containing protein; uniclust \| UniRef100\_A0A3D1VDB8 \| 100.0 \| 1.1e-41 \| 2e-47 \| 280.0 \| 327 \| (54, 399) \| 403 \| (30, 404) \| 407 \| Tyr recombinase domain-containing protein \| Tyr recombinase domain-containing protein; uniclust \| UniRef100\_A0A1L9BA63 \| 100.0 \| 1.3e-41 \| 2.3e-47 \| 290.7 \| 330 \| (34, 395) \| 403 \| (41, 422) \| 502 \| Tyr recombinase domain-containing protein \| Tyr recombinase domain-containing protein; uniclust \| UniRef100\_A0A024YX43 \| 100.0 \| 1.3e-41 \| 2.3e-47 \| 295.0 \| 327 \| (44, 399) \| 403 \| (58, 415) \| 584 \| Integrase \| Integrase; uniclust \| UniRef100\_A0A0F9J7V5 \| 100.0 \| 1.3e-41 \| 2.4e-47 \| 287.3 \| 332 \| (33, 394) \| 403 \| (20, 372) \| 410 \| Tyr recombinase domain-containing protein \| Tyr recombinase domain-containing protein; uniclust \| UniRef100\_A0A1D8AFA3 \| 100.0 \| 1.3e-41 \| 2.4e-47 \| 277.0 \| 353 \| (1, 398) \| 403 \| (44, 411) \| 436 \| Integrase \| Integrase; uniclust \| UniRef100\_A0A1I4ZYE7 \| 100.0 \| 1.3e-41 \| 2.4e-47 \| 274.8 \| 286 \| (1, 313) \| 403 \| (30, 330) \| 339 \| Integrase DNA-binding domain-containing protein \| Integrase DNA-binding domain-containing protein; uniclust \| UniRef100\_A0A059N0G3 \| 100.0 \| 1.4e-41 \| 2.5e-47 \| 286.9 \| 336 \| (43, 401) \| 403 \| (30, 421) \| 454 \| Site-specific recombinase, phage integrase family \| Site-specific recombinase, phage integrase family; uniclust \| UniRef100\_A0A0C4WKA9 \| 100.0 \| 1.4e-41 \| 2.6e-47 \| 294.4 \| 386 \| (1, 402) \| 403 \| (66, 504) \| 539 \| Phage integrase family protein \| Phage integrase family protein; uniclust \| UniRef100\_A0A081NWM4 \| 100.0 \| 1.5e-41 \| 2.7e-47 \| 288.2 \| 308 \| (45, 382) \| 403 \| (83, 421) \| 466 \| Integrase \| Integrase; uniclust \| UniRef100\_A0A0N0K2E9 \| 100.0 \| 1.5e-41 \| 2.8e-47 \| 276.3 \| 371 \| (1, 400) \| 403 \| (2, 394) \| 435 \| Tyr recombinase domain-containing protein (Fragment) \| Tyr recombinase domain-containing protein (Fragment); uniclust \| UniRef100\_A0A023CQB7 \| 100.0 \| 1.7e-41 \| 2.9e-47 \| 294.5 \| 325 \| (45, 399) \| 403 \| (75, 439) \| 467 \| Prophage LambdaBa04, site-specific recombinase, phage integrase family protein \| Prophage LambdaBa04, site-specific recombinase, phage integrase family protein; uniclust \| UniRef100\_A0A2V2FPB6 \| 100.0 \| 1.7e-41 \| 3e-47 \| 280.1 \| 314 \| (54, 398) \| 403 \| (23, 372) \| 404 \| Site-specific integrase \| Site-specific integrase; uniclust \| UniRef100\_A0A059G3I2 \| 100.0 \| 1.9e-41 \| 3.3e-47 \| 283.9 \| 376 \| (1, 402) \| 403 \| (25, 441) \| 460 \| Phage integrase \| Phage integrase; uniclust \| UniRef100\_A0A0U4EAX9 \| 100.0 \| 2e-41 \| 3.4e-47 \| 296.9 \| 322 \| (44, 396) \| 403 \| (42, 393) \| 441 \| Integrase \| Integrase; uniclust \| UniRef100\_Q1K3L0 \| 100.0 \| 1.9e-41 \| 3.4e-47 \| 267.9 \| 327 \| (30, 382) \| 403 \| (39, 382) \| 417 \| Phage integrase \| Phage integrase; uniclust \| UniRef100\_A0A0E8XLM2 \| 100.0 \| 1.9e-41 \| 3.4e-47 \| 267.4 \| 374 \| (1, 401) \| 403 \| (23, 424) \| 445 \| Phage integrase family site specific recombinase \| Phage integrase family site specific recombinase; uniclust \| UniRef100\_A0A090GDZ2 \| 100.0 \| 1.9e-41 \| 3.4e-47 \| 276.4 \| 347 \| (2, 381) \| 403 \| (27, 388) \| 428 \| Putative Phage integrase \| Putative Phage integrase; uniclust \| UniRef100\_A0A1M6M0L3 \| 100.0 \| 2e-41 \| 3.5e-47 \| 273.7 \| 282 \| (1, 312) \| 403 \| (16, 308) \| 358 \| Phage integrase family protein \| Phage integrase family protein; uniclust \| UniRef100\_A0A069AU07 \| 100.0 \| 2.3e-41 \| 4e-47 \| 286.7 \| 325 \| (44, 398) \| 403 \| (42, 406) \| 440 \| Integrase \| Integrase; uniclust \| UniRef100\_A0A0M2UWX7 \| 100.0 \| 2.6e-41 \| 4.6e-47 \| 271.6 \| 243 \| (136, 397) \| 403 \| (10, 257) \| 313 \| Site-specific tyrosine recombinase (Fragment) \| Site-specific tyrosine recombinase (Fragment); uniclust \| UniRef100\_A0A6L4AW27 \| 100.0 \| 2.8e-41 \| 5.2e-47 \| 274.5 \| 340 \| (26, 399) \| 403 \| (70, 421) \| 470 \| Site-specific integrase \| Site-specific integrase; uniclust \| UniRef100\_A0A1Y3R4K1 \| 100.0 \| 2.9e-41 \| 5.2e-47 \| 276.7 \| 281 \| (100, 399) \| 403 \| (93, 402) \| 411 \| Tyr recombinase domain-containing protein \| Tyr recombinase domain-containing protein; uniclust \| UniRef100\_A0A2W5XPE9 \| 100.0 \| 3.4e-41 \| 6e-47 \| 281.5 \| 316 \| (55, 398) \| 403 \| (23, 364) \| 392 \| Site-specific integrase \| Site-specific integrase; uniclust \| UniRef100\_A0A5C8B1S2 \| 100.0 \| 3.3e-41 \| 6e-47 \| 271.6 \| 364 \| (2, 402) \| 403 \| (9, 389) \| 408 \| DUF4102 domain-containing protein \| DUF4102 domain-containing protein; uniclust \| UniRef100\_A0A1H5C2S2 \| 100.0 \| 3.4e-41 \| 6.2e-47 \| 271.3 \| 371 \| (2, 402) \| 403 \| (30, 412) \| 421 \| Tyr recombinase domain-containing protein \| Tyr recombinase domain-containing protein; uniclust \| UniRef100\_A0A0F9D313 \| 100.0 \| 3.6e-41 \| 6.3e-47 \| 288.1 \| 313 \| (55, 398) \| 403 \| (69, 407) \| 454 \| Tyr recombinase domain-containing protein (Fragment) \| Tyr recombinase domain-containing protein (Fragment); uniclust \| UniRef100\_A0A1C9V0P7 \| 100.0 \| 3.8e-41 \| 6.7e-47 \| 294.0 \| 323 \| (43, 400) \| 403 \| (118, 451) \| 539 \| Integrase family protein \| Integrase family protein; uniclust \| UniRef100\_A0A093UD42 \| 100.0 \| 3.9e-41 \| 7.1e-47 \| 270.6 \| 373 \| (1, 401) \| 403 \| (30, 430) \| 521 \| Integrase \| Integrase; uniclust \| UniRef100\_A0A1M6EU03 \| 100.0 \| 3.9e-41 \| 7.1e-47 \| 268.7 \| 309 \| (43, 394) \| 403 \| (14, 333) \| 347 \| Site-specific recombinase XerD \| Site-specific recombinase XerD; uniclust \| UniRef100\_A0A0H3ZTY6 \| 100.0 \| 4.2e-41 \| 7.6e-47 \| 269.5 \| 361 \| (4, 401) \| 403 \| (26, 407) \| 410 \| Phage integrase \| Phage integrase; uniclust \| UniRef100\_A0A080N279 \| 100.0 \| 4.3e-41 \| 7.7e-47 \| 290.8 \| 323 \| (44, 395) \| 403 \| (96, 459) \| 540 \| Phage integrase family protein \| Phage integrase family protein; uniclust \| UniRef100\_A0A0J1B6R9 \| 100.0 \| 4.4e-41 \| 7.8e-47 \| 288.7 \| 312 \| (43, 393) \| 403 \| (60, 379) \| 436 \| Site-specific recombinase XerD \| Site-specific recombinase XerD; uniclust \| UniRef100\_A0A1V6JU51 \| 100.0 \| 4.5e-41 \| 8e-47 \| 287.6 \| 275 \| (100, 395) \| 403 \| (113, 413) \| 455 \| Tyrosine recombinase XerC \| Tyrosine recombinase XerC; uniclust \| UniRef100\_A0A1C7HEN5 \| 100.0 \| 4.5e-41 \| 8.1e-47 \| 280.8 \| 369 \| (2, 402) \| 403 \| (8, 441) \| 453 \| Site-specific recombinase \| Site-specific recombinase; uniclust \| UniRef100\_A0A127Q2R3 \| 100.0 \| 4.5e-41 \| 8.1e-47 \| 263.6 \| 365 \| (1, 401) \| 403 \| (2, 377) \| 388 \| Phage integrase family protein \| Phage integrase family protein; uniclust \| UniRef100\_A0A2V3TQD5 \| 100.0 \| 4.7e-41 \| 8.4e-47 \| 281.0 \| 325 \| (31, 382) \| 403 \| (33, 397) \| 438 \| Site-specific recombinase XerD \| Site-specific recombinase XerD; uniclust \| UniRef100\_A0A1Q8AQL2 \| 100.0 \| 4.8e-41 \| 8.6e-47 \| 266.2 \| 245 \| (144, 402) \| 403 \| (31, 289) \| 344 \| Tyr recombinase domain-containing protein \| Tyr recombinase domain-containing protein; uniclust \| UniRef100\_A0A2E0W7N1 \| 100.0 \| 5.6e-41 \| 9.9e-47 \| 279.7 \| 319 \| (44, 395) \| 403 \| (27, 364) \| 379 \| Tyr recombinase domain-containing protein \| Tyr recombinase domain-containing protein; uniclust \| UniRef100\_A0A017H9M4 \| 100.0 \| 5.8e-41 \| 1e-46 \| 290.8 \| 323 \| (32, 382) \| 403 \| (52, 417) \| 470 \| Phage integrase \| Phage integrase; uniclust \| UniRef100\_A0A0B0EHS4 \| 100.0 \| 5.7e-41 \| 1e-46 \| 267.4 \| 322 \| (44, 395) \| 403 \| (7, 339) \| 364 \| Putative phage integrase \| Putative phage integrase; uniclust \| UniRef100\_A0A011Q867 \| 100.0 \| 6.2e-41 \| 1.1e-46 \| 286.1 \| 350 \| (29, 402) \| 403 \| (48, 445) \| 505 \| Phage integrase family protein \| Phage integrase family protein; uniclust \| UniRef100\_A0A011UHL6 \| 100.0 \| 6.3e-41 \| 1.1e-46 \| 287.1 \| 327 \| (46, 399) \| 403 \| (82, 526) \| 558 \| Integrase \| Integrase; uniclust \| UniRef100\_A0A679JWT2 \| 100.0 \| 6.2e-41 \| 1.1e-46 \| 271.2 \| 355 \| (28, 402) \| 403 \| (26, 414) \| 429 \| Prophage integrase IntA \| Prophage integrase IntA; uniclust \| UniRef100\_A0A2N5X4M8 \| 100.0 \| 6.2e-41 \| 1.1e-46 \| 266.8 \| 380 \| (2, 400) \| 403 \| (10, 424) \| 426 \| Tyr recombinase domain-containing protein \| Tyr recombinase domain-containing protein; uniclust \| UniRef100\_A0A1F2TFY9 \| 100.0 \| 6.4e-41 \| 1.1e-46 \| 273.9 \| 286 \| (99, 401) \| 403 \| (27, 324) \| 400 \| Tyr recombinase domain-containing protein \| Tyr recombinase domain-containing protein; uniclust \| UniRef100\_A0A1A0XD97 \| 100.0 \| 6.7e-41 \| 1.2e-46 \| 278.6 \| 327 \| (45, 399) \| 403 \| (18, 380) \| 436 \| Tyr recombinase domain-containing protein \| Tyr recombinase domain-containing protein; uniclust \| UniRef100\_A0A0F3M5Y8 \| 100.0 \| 6.6e-41 \| 1.2e-46 \| 265.4 \| 352 \| (1, 382) \| 403 \| (10, 396) \| 447 \| Phage integrase family protein \| Phage integrase family protein; uniclust \| UniRef100\_A0A084JMC9 \| 100.0 \| 6.9e-41 \| 1.2e-46 \| 284.0 \| 319 \| (45, 394) \| 403 \| (53, 417) \| 441 \| Tyr recombinase domain-containing protein \| Tyr recombinase domain-containing protein; uniclust \| UniRef100\_A0A1H3NLR1 \| 100.0 \| 8e-41 \| 1.4e-46 \| 283.9 \| 339 \| (1, 382) \| 403 \| (29, 389) \| 476 \| Site-specific recombinase XerD \| Site-specific recombinase XerD; uniclust \| UniRef100\_A0A098Y3T2 \| 100.0 \| 8.3e-41 \| 1.5e-46 \| 281.3 \| 313 \| (43, 382) \| 403 \| (48, 394) \| 440 \| Integrase \| Integrase; uniclust \| UniRef100\_A0A0G3V0I4 \| 100.0 \| 8.3e-41 \| 1.5e-46 \| 276.6 \| 311 \| (45, 382) \| 403 \| (21, 376) \| 411 \| Site-specific recombinase XerD \| Site-specific recombinase XerD; uniclust \| UniRef100\_A0A0Q6W4T8 \| 100.0 \| 8.4e-41 \| 1.5e-46 \| 269.3 \| 285 \| (1, 313) \| 403 \| (15, 315) \| 336 \| Tyr recombinase domain-containing protein \| Tyr recombinase domain-containing protein; uniclust \| UniRef100\_A0A060DJC5 \| 100.0 \| 9.3e-41 \| 1.6e-46 \| 296.3 \| 351 \| (6, 394) \| 403 \| (77, 468) \| 520 \| Tyr recombinase domain-containing protein \| Tyr recombinase domain-containing protein; uniclust \| UniRef100\_A0A4P7CHC9 \| 100.0 \| 9e-41 \| 1.6e-46 \| 266.7 \| 373 \| (1, 401) \| 403 \| (8, 404) \| 407 \| DUF4102 domain-containing protein \| DUF4102 domain-containing protein; uniclust \| UniRef100\_A0A6J4TB91 \| 100.0 \| 9.2e-41 \| 1.7e-46 \| 273.3 \| 321 \| (32, 382) \| 403 \| (9, 358) \| 407 \| Integrase \| Integrase; uniclust \| UniRef100\_A0A133KEU4 \| 100.0 \| 9.6e-41 \| 1.7e-46 \| 276.4 \| 314 \| (44, 396) \| 403 \| (9, 348) \| 367 \| Site-specific recombinase, phage integrase family \| Site-specific recombinase, phage integrase family; uniclust \| UniRef100\_A0A2E5C3J8 \| 100.0 \| 9.4e-41 \| 1.7e-46 \| 262.9 \| 371 \| (3, 402) \| 403 \| (6, 404) \| 418 \| Integrase \| Integrase; uniclust \| UniRef100\_A0A0A8HUI6 \| 100.0 \| 1e-40 \| 1.8e-46 \| 279.0 \| 370 \| (1, 402) \| 403 \| (36, 442) \| 454 \| Site-specific recombinase, phage integrase family (DUF4102 domain) \| Site-specific recombinase, phage integrase family (DUF4102 domain); uniclust \| UniRef100\_A0A068F5B1 \| 100.0 \| 1e-40 \| 1.8e-46 \| 285.0 \| 320 \| (43, 398) \| 403 \| (124, 467) \| 556 \| Tyrosine integrase \| Tyrosine integrase; uniclust \| UniRef100\_A0A1Y3UD86 \| 100.0 \| 1e-40 \| 1.9e-46 \| 281.8 \| 332 \| (45, 398) \| 403 \| (51, 439) \| 506 \| Site-specific integrase \| Site-specific integrase; uniclust \| UniRef100\_A0A0B5BC66 \| 100.0 \| 1.1e-40 \| 1.9e-46 \| 280.4 \| 322 \| (32, 394) \| 403 \| (10, 342) \| 381 \| Integrase \| Integrase; uniclust \| UniRef100\_A0A1Q6YTJ6 \| 100.0 \| 1.1e-40 \| 2e-46 \| 277.9 \| 331 \| (44, 399) \| 403 \| (84, 442) \| 463 \| Tyr recombinase domain-containing protein \| Tyr recombinase domain-containing protein; uniclust \| UniRef100\_A0A017HIM5 \| 100.0 \| 1.2e-40 \| 2.2e-46 \| 271.7 \| 349 \| (30, 400) \| 403 \| (8, 387) \| 416 \| Putative P4-family integrase \| Putative P4-family integrase; uniclust \| UniRef100\_A0A174NPS2 \| 100.0 \| 1.3e-40 \| 2.2e-46 \| 284.7 \| 340 \| (32, 399) \| 403 \| (34, 467) \| 480 \| Site-specific recombinase XerD \| Site-specific recombinase XerD; uniclust \| UniRef100\_A0A011UCH0 \| 100.0 \| 1.3e-40 \| 2.3e-46 \| 287.6 \| 365 \| (1, 401) \| 403 \| (40, 428) \| 534 \| Integrase \| Integrase; uniclust \| UniRef100\_A0A2D5PMG0 \| 100.0 \| 1.3e-40 \| 2.4e-46 \| 275.5 \| 307 \| (67, 402) \| 403 \| (41, 356) \| 376 \| Tyr recombinase domain-containing protein \| Tyr recombinase domain-containing protein; uniclust \| UniRef100\_A0A0F9PLM6 \| 100.0 \| 1.4e-40 \| 2.4e-46 \| 269.1 \| 311 \| (44, 382) \| 403 \| (18, 340) \| 355 \| Tyr recombinase domain-containing protein \| Tyr recombinase domain-containing protein; uniclust \| UniRef100\_A0A0C1GP24 \| 100.0 \| 1.4e-40 \| 2.4e-46 \| 271.5 \| 318 \| (31, 369) \| 403 \| (8, 342) \| 360 \| Integrase \| Integrase; uniclust \| UniRef100\_A0A0P1HC62 \| 100.0 \| 1.4e-40 \| 2.4e-46 \| 266.1 \| 365 \| (1, 401) \| 403 \| (10, 390) \| 400 \| Putative prophage CPS-53 integrase \| Putative prophage CPS-53 integrase; uniclust \| UniRef100\_A0A1M3BIR1 \| 100.0 \| 1.5e-40 \| 2.6e-46 \| 286.9 \| 317 \| (43, 381) \| 403 \| (30, 409) \| 540 \| Site-specific integrase \| Site-specific integrase; uniclust \| UniRef100\_A0A3R8YPI9 \| 100.0 \| 1.5e-40 \| 2.8e-46 \| 265.1 \| 373 \| (2, 401) \| 403 \| (15, 423) \| 443 \| Site-specific integrase \| Site-specific integrase; uniclust \| UniRef100\_A0A076MHG8 \| 100.0 \| 1.7e-40 \| 3e-46 \| 284.2 \| 333 \| (35, 399) \| 403 \| (63, 488) \| 539 \| Integrase family protein \| Integrase family protein; uniclust \| UniRef100\_A0A0U5I7C2 \| 100.0 \| 1.7e-40 \| 3e-46 \| 262.9 \| 369 \| (1, 399) \| 403 \| (5, 395) \| 407 \| Putative phage integrase family protein \| Putative phage integrase family protein; uniclust \| UniRef100\_A0A829Y7U8 \| 100.0 \| 1.7e-40 \| 3.1e-46 \| 271.3 \| 349 \| (17, 397) \| 403 \| (24, 380) \| 549 \| Tyr recombinase domain-containing protein \| Tyr recombinase domain-containing protein; uniclust \| UniRef100\_A0A651GHW1 \| 100.0 \| 1.8e-40 \| 3.2e-46 \| 275.8 \| 338 \| (28, 397) \| 403 \| (32, 418) \| 449 \| Site-specific integrase \| Site-specific integrase; uniclust \| UniRef100\_A0A1H7D2D6 \| 100.0 \| 1.8e-40 \| 3.3e-46 \| 274.2 \| 336 \| (43, 402) \| 403 \| (19, 407) \| 424 \| Site-specific recombinase XerD \| Site-specific recombinase XerD; uniclust \| UniRef100\_A0A090YSV9 \| 100.0 \| 1.9e-40 \| 3.3e-46 \| 272.2 \| 281 \| (101, 400) \| 403 \| (1, 317) \| 331 \| Phage integrase family protein \| Phage integrase family protein; uniclust \| UniRef100\_A0A1F9TZH4 \| 100.0 \| 1.9e-40 \| 3.5e-46 \| 261.1 \| 315 \| (44, 398) \| 403 \| (8, 328) \| 365 \| Tyr recombinase domain-containing protein \| Tyr recombinase domain-containing protein; uniclust \| UniRef100\_A0A2V9I892 \| 100.0 \| 2e-40 \| 3.5e-46 \| 264.6 \| 267 \| (99, 383) \| 403 \| (16, 306) \| 346 \| Tyr recombinase domain-containing protein (Fragment) \| Tyr recombinase domain-containing protein (Fragment); uniclust \| UniRef100\_A0A016QKK8 \| 100.0 \| 2e-40 \| 3.5e-46 \| 279.7 \| 326 \| (46, 400) \| 403 \| (105, 491) \| 523 \| Phage integrase \| Phage integrase; uniclust \| UniRef100\_A0A011P559 \| 100.0 \| 2.1e-40 \| 3.6e-46 \| 286.3 \| 298 \| (43, 382) \| 403 \| (81, 395) \| 440 \| Integrase \| Integrase; uniclust \| UniRef100\_A0A0D6EVV0 \| 100.0 \| 2.1e-40 \| 3.8e-46 \| 274.2 \| 346 \| (31, 399) \| 403 \| (12, 394) \| 417 \| Putative Phage related integrase \| Putative Phage related integrase; uniclust \| UniRef100\_A0A090KHJ9 \| 100.0 \| 2.1e-40 \| 3.8e-46 \| 281.4 \| 332 \| (43, 399) \| 403 \| (33, 407) \| 428 \| Uncultured bacterium genome assembly Metasoil\_fosmids\_resub \| Uncultured bacterium genome assembly Metasoil\_fosmids\_resub; uniclust \| UniRef100\_A0A024QGB3 \| 100.0 \| 2.2e-40 \| 3.8e-46 \| 284.7 \| 322 \| (54, 401) \| 403 \| (51, 412) \| 445 \| Tyrosine recombinase XerC \| Tyrosine recombinase XerC; uniclust \| UniRef100\_A0A0C1U0L8 \| 100.0 \| 2.2e-40 \| 3.9e-46 \| 273.5 \| 369 \| (1, 402) \| 403 \| (9, 411) \| 430 \| Tyr recombinase domain-containing protein \| Tyr recombinase domain-containing protein; uniclust \| UniRef100\_A0A7H1RNH4 \| 100.0 \| 2.4e-40 \| 4.4e-46 \| 270.6 \| 356 \| (2, 382) \| 403 \| (33, 471) \| 500 \| Site-specific integrase \| Site-specific integrase; uniclust \| UniRef100\_A0A069CWN5 \| 100.0 \| 2.7e-40 \| 4.7e-46 \| 282.3 \| 320 \| (32, 382) \| 403 \| (25, 377) \| 438 \| Phage integrase \| Phage integrase; uniclust \| UniRef100\_A0A0N0UZK9 \| 100.0 \| 2.8e-40 \| 5e-46 \| 265.6 \| 307 \| (69, 401) \| 403 \| (43, 376) \| 389 \| Phage integrase \| Phage integrase; uniclust \| UniRef100\_A0A061NY33 \| 100.0 \| 2.9e-40 \| 5.1e-46 \| 279.1 \| 317 \| (44, 396) \| 403 \| (51, 377) \| 397 \| DNA integration/recombination/invertion protein \| DNA integration/recombination/invertion protein; uniclust \| UniRef100\_A0A238H9E1 \| 100.0 \| 3e-40 \| 5.3e-46 \| 265.5 \| 357 \| (19, 401) \| 403 \| (20, 391) \| 413 \| Phage-related integrase \| Phage-related integrase; uniclust \| UniRef100\_A0A011V4P1 \| 100.0 \| 3e-40 \| 5.4e-46 \| 277.6 \| 327 \| (44, 400) \| 403 \| (41, 426) \| 546 \| Integrase \| Integrase; uniclust \| UniRef100\_A0A023XKK8 \| 100.0 \| 3.1e-40 \| 5.5e-46 \| 276.5 \| 373 \| (2, 403) \| 403 \| (67, 499) \| 542 \| Site-specific integrase/recombinase \| Site-specific integrase/recombinase; uniclust \| UniRef100\_A0A1I3LKL9 \| 100.0 \| 3.4e-40 \| 6.1e-46 \| 271.7 \| 307 \| (44, 382) \| 403 \| (77, 419) \| 459 \| Site-specific recombinase XerD \| Site-specific recombinase XerD; uniclust \| UniRef100\_A0A0T6W442 \| 100.0 \| 3.6e-40 \| 6.4e-46 \| 282.8 \| 379 \| (2, 400) \| 403 \| (32, 452) \| 492 \| Integrase \| Integrase; uniclust \| UniRef100\_A0A1I6MIL3 \| 100.0 \| 3.6e-40 \| 6.5e-46 \| 268.9 \| 341 \| (32, 398) \| 403 \| (18, 379) \| 397 \| Site-specific recombinase XerD \| Site-specific recombinase XerD; uniclust \| UniRef100\_A0A1H6QXH7 \| 100.0 \| 3.9e-40 \| 7e-46 \| 268.1 \| 299 \| (1, 332) \| 403 \| (33, 343) \| 353 \| Integrase DNA-binding domain-containing protein \| Integrase DNA-binding domain-containing protein; uniclust \| UniRef100\_A0A0B2JJ45 \| 100.0 \| 4e-40 \| 7.1e-46 \| 274.7 \| 319 \| (44, 394) \| 403 \| (39, 406) \| 444 \| Integrase \| Integrase; uniclust \| UniRef100\_A0A073J3R0 \| 100.0 \| 4.1e-40 \| 7.5e-46 \| 259.0 \| 357 \| (1, 400) \| 403 \| (7, 384) \| 400 \| Tyr recombinase domain-containing protein \| Tyr recombinase domain-containing protein; uniclust \| UniRef100\_A0A022PDC8 \| 100.0 \| 4.3e-40 \| 7.6e-46 \| 267.1 \| 286 \| (1, 313) \| 403 \| (28, 325) \| 339 \| Integrase DNA-binding domain-containing protein \| Integrase DNA-binding domain-containing protein; uniclust \| UniRef100\_A0A024Q9W8 \| 100.0 \| 4.5e-40 \| 7.8e-46 \| 283.4 \| 272 \| (99, 393) \| 403 \| (65, 354) \| 390 \| Tyrosine recombinase XerD \| Tyrosine recombinase XerD; uniclust \| UniRef100\_A0A1J5D474 \| 100.0 \| 4.6e-40 \| 8.1e-46 \| 284.0 \| 327 \| (43, 398) \| 403 \| (53, 414) \| 449 \| Tyr recombinase domain-containing protein \| Tyr recombinase domain-containing protein; uniclust \| UniRef100\_A0A9D1WD27 \| 100.0 \| 4.5e-40 \| 8.2e-46 \| 262.1 \| 364 \| (1, 401) \| 403 \| (8, 382) \| 392 \| Tyrosine-type recombinase/integrase \| Tyrosine-type recombinase/integrase; uniclust \| UniRef100\_A0A355B445 \| 100.0 \| 4.7e-40 \| 8.3e-46 \| 264.2 \| 292 \| (1, 313) \| 403 \| (9, 308) \| 337 \| Tyr recombinase domain-containing protein (Fragment) \| Tyr recombinase domain-containing protein (Fragment); uniclust \| UniRef100\_A0A0A0DDR9 \| 100.0 \| 4.7e-40 \| 8.4e-46 \| 267.1 \| 314 \| (45, 396) \| 403 \| (9, 331) \| 368 \| Tyr recombinase domain-containing protein \| Tyr recombinase domain-containing protein; uniclust \| UniRef100\_A0A090KIS3 \| 100.0 \| 4.8e-40 \| 8.5e-46 \| 266.4 \| 277 \| (101, 401) \| 403 \| (1, 307) \| 328 \| Uncultured bacterium genome assembly Metasoil\_fosmids\_resub \| Uncultured bacterium genome assembly Metasoil\_fosmids\_resub; uniclust \| UniRef100\_A0A022N8Z9 \| 100.0 \| 4.9e-40 \| 8.6e-46 \| 288.4 \| 317 \| (34, 381) \| 403 \| (80, 432) \| 480 \| Integrase \| Integrase; uniclust \| UniRef100\_A0A0B8N5H0 \| 100.0 \| 4.8e-40 \| 8.6e-46 \| 267.2 \| 310 \| (43, 381) \| 403 \| (11, 331) \| 386 \| Integrase \| Integrase; uniclust \| UniRef100\_A0A1C6BGX5 \| 100.0 \| 5e-40 \| 9e-46 \| 271.7 \| 330 \| (31, 395) \| 403 \| (22, 389) \| 414 \| Integrase \| Integrase; uniclust \| UniRef100\_A0A021VV25 \| 100.0 \| 5.2e-40 \| 9.3e-46 \| 270.0 \| 338 \| (27, 396) \| 403 \| (15, 375) \| 473 \| Integrase \| Integrase; uniclust \| UniRef100\_A0A286GPU9 \| 100.0 \| 5.9e-40 \| 1.1e-45 \| 270.4 \| 373 \| (1, 401) \| 403 \| (8, 411) \| 441 \| Tyr recombinase domain-containing protein \| Tyr recombinase domain-containing protein; uniclust \| UniRef100\_A0A1E3WRL1 \| 100.0 \| 6e-40 \| 1.1e-45 \| 261.5 \| 360 \| (2, 396) \| 403 \| (9, 381) \| 385 \| Tyrosine recombinase XerC \| Tyrosine recombinase XerC; uniclust \| UniRef100\_A0A021VW42 \| 100.0 \| 6.1e-40 \| 1.1e-45 \| 274.2 \| 323 \| (43, 393) \| 403 \| (55, 409) \| 437 \| Integrase \| Integrase; uniclust \| UniRef100\_A0A075M047 \| 100.0 \| 6.4e-40 \| 1.1e-45 \| 274.5 \| 304 \| (44, 381) \| 403 \| (20, 336) \| 397 \| Integrase \| Integrase; uniclust \| UniRef100\_D5V8K7 \| 100.0 \| 6.6e-40 \| 1.2e-45 \| 262.3 \| 277 \| (108, 402) \| 403 \| (3, 287) \| 326 \| Phage integrase \| Phage integrase; uniclust \| UniRef100\_A0A087MIL0 \| 100.0 \| 6.7e-40 \| 1.2e-45 \| 280.1 \| 382 \| (1, 400) \| 403 \| (29, 448) \| 507 \| Tyr recombinase domain-containing protein \| Tyr recombinase domain-containing protein; uniclust \| UniRef100\_A0A1W2A448 \| 100.0 \| 7e-40 \| 1.3e-45 \| 267.6 \| 303 \| (45, 382) \| 403 \| (33, 351) \| 419 \| Site-specific recombinase XerD \| Site-specific recombinase XerD; uniclust \| UniRef100\_A0A1G3JV15 \| 100.0 \| 7.1e-40 \| 1.3e-45 \| 266.0 \| 274 \| (100, 401) \| 403 \| (5, 288) \| 317 \| Tyr recombinase domain-containing protein \| Tyr recombinase domain-containing protein; uniclust \| UniRef100\_A0A023D6X8 \| 100.0 \| 7.3e-40 \| 1.3e-45 \| 283.9 \| 310 \| (45, 394) \| 403 \| (81, 398) \| 459 \| Prophage integrase \| Prophage integrase; uniclust \| UniRef100\_A0A096D865 \| 100.0 \| 7.3e-40 \| 1.3e-45 \| 274.4 \| 280 \| (102, 400) \| 403 \| (2, 313) \| 343 \| Tyr recombinase domain-containing protein \| Tyr recombinase domain-containing protein; uniclust \| UniRef100\_A0A141RCN1 \| 100.0 \| 7.2e-40 \| 1.3e-45 \| 259.8 \| 285 \| (100, 401) \| 403 \| (24, 318) \| 335 \| Phage integrase family (Fragment) \| Phage integrase family (Fragment); uniclust \| UniRef100\_A0A1Q6RVP7 \| 100.0 \| 7.8e-40 \| 1.4e-45 \| 270.5 \| 278 \| (99, 395) \| 403 \| (24, 307) \| 340 \| Tyr recombinase domain-containing protein \| Tyr recombinase domain-containing protein; uniclust \| UniRef100\_A0A1Q3QU81 \| 100.0 \| 8.4e-40 \| 1.5e-45 \| 266.5 \| 319 \| (56, 399) \| 403 \| (39, 374) \| 415 \| Tyr recombinase domain-containing protein \| Tyr recombinase domain-containing protein; uniclust \| UniRef100\_A0A073KMD6 \| 100.0 \| 8.9e-40 \| 1.6e-45 \| 276.0 \| 280 \| (101, 400) \| 403 \| (65, 372) \| 394 \| Integrase \| Integrase; uniclust \| UniRef100\_A0A1F3ST71 \| 100.0 \| 9.1e-40 \| 1.6e-45 \| 272.8 \| 330 \| (31, 394) \| 403 \| (28, 377) \| 403 \| Tyr recombinase domain-containing protein \| Tyr recombinase domain-containing protein; uniclust \| UniRef100\_A0A239MDX8 \| 100.0 \| 9.3e-40 \| 1.6e-45 \| 263.9 \| 273 \| (1, 300) \| 403 \| (12, 296) \| 307 \| Phage integrase family protein \| Phage integrase family protein; uniclust \| UniRef100\_A0A1C6BYJ1 \| 100.0 \| 9.6e-40 \| 1.7e-45 \| 270.6 \| 325 \| (44, 398) \| 403 \| (60, 430) \| 519 \| Integrase \| Integrase; uniclust \| UniRef100\_A0A095YZI0 \| 100.0 \| 1e-39 \| 1.8e-45 \| 278.7 \| 330 \| (44, 396) \| 403 \| (48, 422) \| 454 \| Tyr recombinase domain-containing protein \| Tyr recombinase domain-containing protein; uniclust \| UniRef100\_A0A059WLJ1 \| 100.0 \| 1.1e-39 \| 2e-45 \| 282.0 \| 333 \| (31, 394) \| 403 \| (68, 421) \| 449 \| Integrase family site specific recombinase \| Integrase family site specific recombinase; uniclust \| UniRef100\_A0A0A0HGU1 \| 100.0 \| 1.1e-39 \| 2e-45 \| 278.9 \| 307 \| (43, 383) \| 403 \| (55, 389) \| 468 \| Site-specific recombinase XerD \| Site-specific recombinase XerD; uniclust \| UniRef100\_A0A0Q4J458 \| 100.0 \| 1.2e-39 \| 2.1e-45 \| 265.3 \| 310 \| (44, 394) \| 403 \| (43, 357) \| 374 \| Tyr recombinase domain-containing protein \| Tyr recombinase domain-containing protein; uniclust \| UniRef100\_A0A023LBH6 \| 100.0 \| 1.2e-39 \| 2.2e-45 \| 280.2 \| 313 \| (44, 394) \| 403 \| (79, 403) \| 480 \| Integrase \| Integrase; uniclust \| UniRef100\_A0A177RBV1 \| 100.0 \| 1.3e-39 \| 2.3e-45 \| 270.1 \| 329 \| (45, 395) \| 403 \| (72, 417) \| 441 \| Tyr recombinase domain-containing protein \| Tyr recombinase domain-containing protein; uniclust \| UniRef100\_A0A021X9S7 \| 100.0 \| 1.4e-39 \| 2.4e-45 \| 283.0 \| 302 \| (45, 382) \| 403 \| (96, 407) \| 515 \| Site-specific recombinase XerD \| Site-specific recombinase XerD; uniclust \| UniRef100\_A0A060QBM0 \| 100.0 \| 1.4e-39 \| 2.5e-45 \| 268.3 \| 311 \| (45, 395) \| 403 \| (18, 340) \| 374 \| Integrase \| Integrase; uniclust \| UniRef100\_A0A059UQM9 \| 100.0 \| 1.4e-39 \| 2.6e-45 \| 275.1 \| 366 \| (1, 401) \| 403 \| (25, 408) \| 473 \| Phage integrase \| Phage integrase; uniclust \| UniRef100\_A0A009G861 \| 100.0 \| 1.5e-39 \| 2.6e-45 \| 285.2 \| 309 \| (32, 382) \| 403 \| (76, 394) \| 477 \| Phage integrase family protein \| Phage integrase family protein; uniclust \| UniRef100\_A0A1M5XD08 \| 100.0 \| 1.6e-39 \| 2.9e-45 \| 265.1 \| 330 \| (44, 399) \| 403 \| (8, 360) \| 462 \| Site-specific recombinase XerD \| Site-specific recombinase XerD; uniclust \| UniRef100\_A0A2E8DB97 \| 100.0 \| 1.6e-39 \| 2.9e-45 \| 269.3 \| 331 \| (32, 397) \| 403 \| (59, 405) \| 438 \| Tyr recombinase domain-containing protein \| Tyr recombinase domain-containing protein; uniclust \| UniRef100\_A0A068SP06 \| 100.0 \| 1.7e-39 \| 2.9e-45 \| 287.0 \| 322 \| (32, 398) \| 403 \| (25, 364) \| 507 \| Integrative genetic element Ppu40, integrase \| Integrative genetic element Ppu40, integrase; uniclust \| UniRef100\_A0A0T9M937 \| 100.0 \| 1.6e-39 \| 2.9e-45 \| 254.2 \| 311 \| (54, 393) \| 403 \| (2, 328) \| 345 \| Prophage CP4-57 integrase \| Prophage CP4-57 integrase; uniclust \| UniRef100\_A0A096CRU8 \| 100.0 \| 1.7e-39 \| 3e-45 \| 265.0 \| 311 \| (44, 394) \| 403 \| (32, 350) \| 351 \| Tyr recombinase domain-containing protein \| Tyr recombinase domain-containing protein; uniclust \| UniRef100\_A0A085K0B1 \| 100.0 \| 1.7e-39 \| 3.1e-45 \| 260.8 \| 285 \| (99, 403) \| 403 \| (20, 317) \| 342 \| Tyr recombinase domain-containing protein \| Tyr recombinase domain-containing protein; uniclust \| UniRef100\_A0A0D4CL96 \| 100.0 \| 1.7e-39 \| 3.1e-45 \| 274.0 \| 337 \| (32, 400) \| 403 \| (29, 391) \| 413 \| Integrase \| Integrase; pdb70 \| 1Z1B\_A \| 100.0 \| 3.4e-34 \| 2.4e-38 \| 255.7 \| 325 \| (27, 388) \| 403 \| (11, 351) \| 356 \| PROTEIN/DNA Complex \| 1Z1B\_A PROTEIN/DNA Complex PROTEIN-DNA COMPLEX, DNA BINDING PROTEIN-DNA HET: PTR; pdb70 \| 5J0N\_H \| 99.9 \| 1.1e-33 \| 7.9e-38 \| 252.4 \| 308 \| (49, 383) \| 403 \| (24, 347) \| 356 \| Integrase, Integration host factor subunit \| 5J0N\_H Integrase, Integration host factor subunit bacteriophage lambda, excision, site-specific recombination; pdb70 \| 1Z1G\_C \| 99.9 \| 1.5e-33 \| 1.1e-37 \| 251.4 \| 320 \| (27, 382) \| 403 \| (11, 346) \| 356 \| PROTEIN/DNA Complex \| 1Z1G\_C PROTEIN/DNA Complex Protein-DNA complex, DNA BINDING PROTEIN-DNA HET: MSE; pdb70 \| 1Z19\_B \| 99.9 \| 6.4e-29 \| 4.5e-33 \| 213.5 \| 263 \| (101, 388) \| 403 \| (1, 278) \| 283 \| Integrase/DNA Complex \| 1Z19\_B Integrase/DNA Complex PROTEIN-DNA COMPLEX, DNA BINDING PROTEIN-DNA; pdb70 \| 1P7D\_A \| 99.9 \| 6.4e-29 \| 4.5e-33 \| 213.5 \| 265 \| (101, 390) \| 403 \| (1, 280) \| 283 \| Lambda Integrase/DNA Complex \| 1P7D\_A Lambda Integrase/DNA Complex PROTEIN-DNA COMPLEX, DNA Binding Protein-DNA HET: PTR; pdb70 \| 5VFZ\_A \| 99.9 \| 3.3e-28 \| 2.4e-32 \| 213.1 \| 273 \| (98, 400) \| 403 \| (14, 290) \| 318 \| Gp33 \| 5VFZ\_A Gp33 Bacteriophage, Brujita, DNA-binding, Integrase, DNA HET: ACT, GOL; pdb70 \| 1A0P\_A \| 99.9 \| 1.3e-27 \| 9.2e-32 \| 205.8 \| 264 \| (106, 390) \| 403 \| (5, 288) \| 290 \| SITE-SPECIFIC RECOMBINASE XERD \| 1A0P\_A SITE-SPECIFIC RECOMBINASE XERD XERD, RECOMBINASE, DNA BINDING, DNA; pdb70 \| 6EN1\_B \| 99.9 \| 1.4e-27 \| 1e-31 \| 208.3 \| 271 \| (101, 394) \| 403 \| (2, 315) \| 317 \| Int protein/DNA Complex \| 6EN1\_B Int protein/DNA Complex transposase protein - DNA complex HET: PEG; pdb70 \| 5HXY\_A \| 99.9 \| 1.6e-27 \| 1.1e-31 \| 208.7 \| 269 \| (98, 394) \| 403 \| (34, 313) \| 317 \| Site-specific integrase/recombinase XerD related protein \| 5HXY\_A Site-specific integrase/recombinase XerD related protein recombinase, XerA, RECOMBINATION HET: PO4, MSE; pdb70 \| 6EMY\_A \| 99.9 \| 4.8e-27 \| 3.4e-31 \| 204.8 \| 272 \| (101, 394) \| 403 \| (2, 315) \| 317 \| Int protein/DNA Complex \| 6EMY\_A Int protein/DNA Complex transposase protein-DNA complex, tyrosine recombinase; pdb70 \| 2A3V\_D \| 99.9 \| 5.2e-27 \| 3.6e-31 \| 205.1 \| 267 \| (102, 395) \| 403 \| (4, 318) \| 320 \| site-specific recombinase IntI4/DNA Complex \| 2A3V\_D site-specific recombinase IntI4/DNA Complex Protein-DNA complex, RECOMBINATION; pdb70 \| 5C6K\_B \| 99.9 \| 1e-26 \| 7.2e-31 \| 200.9 \| 262 \| (98, 394) \| 403 \| (11, 274) \| 292 \| Integrase \| 5C6K\_B Integrase Integrase, tyrosine recombinase, integration, site-specific; pdb70 \| 3MGV\_D \| 99.9 \| 1.1e-26 \| 7.6e-31 \| 204.9 \| 282 \| (98, 395) \| 403 \| (15, 340) \| 343 \| Recombinase cre \| 3MGV\_D Recombinase cre CRE-LOXP, TRANSITION STATE, ISOMERASE-DNA complex; pdb70 \| 4A8E\_A \| 99.8 \| 2.6e-26 \| 1.9e-30 \| 198.3 \| 263 \| (100, 394) \| 403 \| (6, 276) \| 292 \| PROBABLE TYROSINE RECOMBINASE XERC-LIKE \| 4A8E\_A PROBABLE TYROSINE RECOMBINASE XERC-LIKE CELL CYCLE, CHROMOSOME DIMER RESOLUTION HET: SO4; pdb70 \| 1XO0\_A \| 99.8 \| 4.1e-26 \| 2.9e-30 \| 199.4 \| 274 \| (106, 395) \| 403 \| (4, 321) \| 324 \| RECOMBINASE CRE/DNA Complex \| 1XO0\_A RECOMBINASE CRE/DNA Complex CRE RECOMBINASE, HOLLIDAY JUNCTION, RECOMBINATION; pdb70 \| 4CRX\_B \| 99.8 \| 5.6e-26 \| 3.9e-30 \| 198.3 \| 273 \| (106, 394) \| 403 \| (4, 320) \| 322 \| CRE RECOMBINASE/DNA COMPLEX \| 4CRX\_B CRE RECOMBINASE/DNA COMPLEX CRE RECOMBINASE, DNA BENDING, RECOMBINATION; pdb70 \| 5JJV\_B \| 99.8 \| 1.3e-25 \| 8.9e-30 \| 200.0 \| 271 \| (105, 395) \| 403 \| (56, 361) \| 363 \| Tyrosine recombinase XerH/DNA Complex \| 5JJV\_B Tyrosine recombinase XerH/DNA Complex Xer, tyrosine recombinase, site-specific recombinase HET: GOL; pdb70 \| 5JK0\_C \| 99.8 \| 1.3e-25 \| 8.9e-30 \| 200.0 \| 271 \| (105, 395) \| 403 \| (56, 361) \| 363 \| Tyrosine recombinase XerH/DNA Complex \| 5JK0\_C Tyrosine recombinase XerH/DNA Complex Xer, tyrosine recombinase, site-specific recombinase HET: GOL, EDO; pdb70 \| 4E0P\_A \| 99.7 \| 4.3e-21 \| 3e-25 \| 175.7 \| 223 \| (150, 383) \| 403 \| (166, 430) \| 462 \| Protelomerase \| 4E0P\_A Protelomerase PROTELEMORASE, DNA BINDING PROTEIN-DNA complex HET: DT, EDO, PTR; pdb70 \| 4E0Z\_A \| 99.7 \| 4.4e-21 \| 3.1e-25 \| 175.8 \| 220 \| (150, 383) \| 403 \| (166, 430) \| 462 \| Protelomerase \| 4E0Z\_A Protelomerase PROTELEMORASE, DNA BINDING PROTEIN-DNA complex HET: GOL, PTR, DT; pdb70 \| 4DWP\_A \| 99.7 \| 5e-21 \| 3.5e-25 \| 175.3 \| 223 \| (150, 383) \| 403 \| (166, 430) \| 462 \| Protelomerase, DNA \| 4DWP\_A Protelomerase, DNA PROTELEMORASE, DNA BINDING PROTEIN-DNA complex HET: TMP; pdb70 \| 4F41\_A \| 99.6 \| 5.4e-20 \| 3.8e-24 \| 160.5 \| 244 \| (121, 383) \| 403 \| (24, 309) \| 320 \| Protelomerase/DNA complex \| 4F41\_A Protelomerase/DNA complex RECOMBINATION-DNA complex; pdb70 \| 4F43\_A \| 99.6 \| 5.4e-20 \| 3.8e-24 \| 160.5 \| 244 \| (121, 383) \| 403 \| (24, 309) \| 320 \| Protelomerase/DNA complex \| 4F43\_A Protelomerase/DNA complex RECOMBINATION-DNA complex; pdb70 \| 5DCF\_A \| 99.5 \| 1.6e-18 \| 1.1e-22 \| 147.5 \| 172 \| (221, 398) \| 403 \| (2, 189) \| 275 \| Tyrosine recombinase XerD,DNA translocase FtsK \| 5DCF\_A Tyrosine recombinase XerD,DNA translocase FtsK Recombination; pdb70 \| 1AE9\_A \| 99.5 \| 2.2e-18 \| 1.6e-22 \| 136.3 \| 156 \| (221, 383) \| 403 \| (2, 171) \| 179 \| LAMBDA INTEGRASE \| 1AE9\_A LAMBDA INTEGRASE DNA RECOMBINATION, INTEGRASE, SITE-SPECIFIC RECOMBINATION; pdb70 \| 3NKH\_A \| 99.5 \| 2.4e-18 \| 1.7e-22 \| 143.6 \| 177 \| (218, 398) \| 403 \| (21, 242) \| 244 \| Integrase \| 3NKH\_A Integrase alpha-fold, MRSA protein, Structural Genomics HET: MRD, SO4, MPD, MSE; pdb70 \| 3NKH\_B \| 99.5 \| 2.4e-18 \| 1.7e-22 \| 143.6 \| 177 \| (218, 398) \| 403 \| (21, 242) \| 244 \| Integrase \| 3NKH\_B Integrase alpha-fold, MRSA protein, Structural Genomics HET: SO4, MRD, MPD; pdb70 \| 1AE9\_B \| 99.5 \| 2.5e-18 \| 1.8e-22 \| 135.9 \| 156 \| (221, 383) \| 403 \| (2, 171) \| 179 \| LAMBDA INTEGRASE \| 1AE9\_B LAMBDA INTEGRASE DNA RECOMBINATION, INTEGRASE, SITE-SPECIFIC RECOMBINATION; pdb70 \| 5DOR\_A \| 99.4 \| 1.3e-17 \| 9.2e-22 \| 131.8 \| 156 \| (221, 393) \| 403 \| (2, 157) \| 176 \| Integrase \| 5DOR\_A Integrase tyrosine recombinase, integrase, hydrolase; pdb70 \| 5DOR\_D \| 99.4 \| 1.3e-17 \| 9.2e-22 \| 131.8 \| 156 \| (221, 393) \| 403 \| (2, 157) \| 176 \| Integrase \| 5DOR\_D Integrase tyrosine recombinase, integrase, hydrolase; pdb70 \| 1AIH\_A \| 99.3 \| 1.1e-16 \| 7.7e-21 \| 125.4 \| 150 \| (220, 391) \| 403 \| (4, 160) \| 170 \| HP1 INTEGRASE \| 1AIH\_A HP1 INTEGRASE DNA INTEGRATION, RECOMBINATION HET: SO4; pdb70 \| 1AIH\_C \| 99.3 \| 1.1e-16 \| 7.7e-21 \| 125.4 \| 150 \| (220, 391) \| 403 \| (4, 160) \| 170 \| HP1 INTEGRASE \| 1AIH\_C HP1 INTEGRASE DNA INTEGRATION, RECOMBINATION HET: SO4; pdb70 \| 3VCF\_A \| 99.0 \| 1.4e-13 \| 9.6e-18 \| 106.6 \| 152 \| (220, 398) \| 403 \| (4, 160) \| 163 \| Probable integrase \| 3VCF\_A Probable integrase catalyzes site-specific integration, RECOMBINATION; pdb70 \| 6GSA\_E \| 98.9 \| 2.5e-13 \| 1.8e-17 \| 126.5 \| 257 \| (119, 392) \| 403 \| (39, 409) \| 562 \| Centromere DNA-binding protein complex CBF3 \| 6GSA\_E Centromere DNA-binding protein complex CBF3 Centromere, CDEIII-binding, LRR domain, DNA; pdb70 \| 3UXU\_A \| 98.9 \| 4.6e-13 \| 3.2e-17 \| 104.3 \| 131 \| (220, 382) \| 403 \| (4, 146) \| 169 \| Probable integrase \| 3UXU\_A Probable integrase SSV1, Archaea, Archaeal virus, hyperthermophilic; pdb70 \| 4ACO\_A \| 98.8 \| 8.3e-13 \| 5.9e-17 \| 128.0 \| 261 \| (119, 396) \| 403 \| (38, 412) \| 956 \| CENTROMERE DNA-BINDING PROTEIN COMPLEX CBF3 \| 4ACO\_A CENTROMERE DNA-BINDING PROTEIN COMPLEX CBF3 DNA BINDING PROTEIN; pdb70 \| 6GYP\_E \| 98.8 \| 8.3e-13 \| 5.9e-17 \| 128.0 \| 261 \| (119, 396) \| 403 \| (38, 412) \| 956 \| Centromere DNA-binding protein complex CBF3 \| 6GYP\_E Centromere DNA-binding protein complex CBF3 Complex, CELL CYCLE HET: ARG, PHE, THR; pdb70 \| 3JTZ\_A \| 98.0 \| 3.6e-09 \| 2.6e-13 \| 71.5 \| 59 \| (30, 92) \| 403 \| (24, 82) \| 88 \| Integrase \| 3JTZ\_A Integrase four stranded beta-sheet, DNA BINDING; pdb70 \| 3JU0\_A \| 97.9 \| 4.8e-09 \| 3.4e-13 \| 74.0 \| 81 \| (29, 114) \| 403 \| (23, 103) \| 108 \| Phage integrase \| 3JU0\_A Phage integrase four stranded beta-sheet, DNA BINDING; pdb70 \| 3T79\_A \| 97.8 \| 1.2e-08 \| 8.7e-13 \| 90.0 \| 238 \| (119, 383) \| 403 \| (12, 303) \| 402 \| PROTEIN/DNA Complex \| 3T79\_A PROTEIN/DNA Complex DNA recombinase, DNA binding, DNA; pdb70 \| 3SQI\_A \| 97.5 \| 7.1e-08 \| 5e-12 \| 87.5 \| 238 \| (119, 383) \| 403 \| (12, 303) \| 534 \| KLLA0E03807p \| 3SQI\_A KLLA0E03807p DNA recombinase, DNA binding, DNA; pdb70 \| 2OXO\_A \| 97.3 \| 2.3e-07 \| 1.6e-11 \| 63.7 \| 85 \| (102, 192) \| 403 \| (2, 87) \| 103 \| Integrase \| 2OXO\_A Integrase DNA-binding protein, four-helix bundle, DNA; pdb70 \| 3LYS\_A \| 97.3 \| 2.5e-07 \| 1.8e-11 \| 65.1 \| 88 \| (99, 192) \| 403 \| (4, 92) \| 112 \| Prophage pi2 protein 01, integrase \| 3LYS\_A Prophage pi2 protein 01, integrase helical N-terminal domain, Structural Genomics HET: MSE; pdb70 \| 3LYS\_D \| 97.3 \| 2.5e-07 \| 1.8e-11 \| 65.1 \| 88 \| (99, 192) \| 403 \| (4, 92) \| 112 \| Prophage pi2 protein 01, integrase \| 3LYS\_D Prophage pi2 protein 01, integrase helical N-terminal domain, Structural Genomics; pdb70 \| 2KKV\_A \| 97.3 \| 2.9e-07 \| 2.1e-11 \| 65.7 \| 94 \| (99, 209) \| 403 \| (5, 100) \| 121 \| Integrase \| 2KKV\_A Integrase Protein structure, PSI, NESGC, Structural; pdb70 \| 2KHQ\_A \| 97.3 \| 3.8e-07 \| 2.7e-11 \| 63.8 \| 94 \| (101, 209) \| 403 \| (2, 96) \| 110 \| Integrase \| 2KHQ\_A Integrase all-alpha, Structural Genomics, PSI-2, Protein; pdb70 \| 2KEY\_A \| 97.2 \| 6.6e-07 \| 4.6e-11 \| 62.8 \| 88 \| (98, 192) \| 403 \| (5, 95) \| 112 \| Putative phage integrase \| 2KEY\_A Putative phage integrase Protein structure, PSI, NESG, STRUCTURAL; pdb70 \| 2KJ8\_A \| 97.1 \| 7.1e-07 \| 5e-11 \| 63.5 \| 95 \| (99, 209) \| 403 \| (3, 99) \| 118 \| Putative prophage CPS-53 integrase \| 2KJ8\_A Putative prophage CPS-53 integrase integrase, intS, intC, yfdB, CPS-53; pdb70 \| 2KIW\_A \| 97.1 \| 8.8e-07 \| 6.2e-11 \| 61.9 \| 92 \| (102, 209) \| 403 \| (1, 93) \| 111 \| Int protein \| 2KIW\_A Int protein alpha, Structural Genomics, PSI-2, Protein; pdb70 \| 2KOB\_A \| 96.9 \| 2e-06 \| 1.4e-10 \| 59.8 \| 92 \| (101, 209) \| 403 \| (2, 94) \| 108 \| Uncharacterized protein \| 2KOB\_A Uncharacterized protein alpha beta, Structural Genomics, PSI-2; pdb70 \| 2KKP\_A \| 96.9 \| 2.1e-06 \| 1.5e-10 \| 60.6 \| 96 \| (99, 209) \| 403 \| (5, 103) \| 117 \| Phage integrase \| 2KKP\_A Phage integrase SAM-like domain, alpha-helical bundle, Structural; pdb70 \| 2KD1\_A \| 96.9 \| 2.4e-06 \| 1.7e-10 \| 60.3 \| 97 \| (99, 210) \| 403 \| (4, 102) \| 118 \| DNA integration/recombination/invertion protein \| 2KD1\_A DNA integration/recombination/invertion protein PROTEIN STRUCTURE INITIATIVE, STRUCTURAL GENOMICS; pdb70 \| 3NRW\_A \| 96.8 \| 3.1e-06 \| 2.2e-10 \| 59.5 \| 96 \| (99, 209) \| 403 \| (5, 104) \| 117 \| Phage integrase/site-specific recombinase \| 3NRW\_A Phage integrase/site-specific recombinase alpha-helical domain, Structural Genomics, PSI-2; pdb70 \| 2KJ9\_A \| 96.8 \| 3.7e-06 \| 2.6e-10 \| 59.5 \| 95 \| (98, 209) \| 403 \| (8, 104) \| 118 \| Integrase \| 2KJ9\_A Integrase integrase, DNA\_BRE\_C superfamily, bacteriophage P4; pdb70 \| 2KJ5\_A \| 96.7 \| 4.9e-06 \| 3.5e-10 \| 58.5 \| 97 \| (99, 210) \| 403 \| (5, 101) \| 116 \| Phage integrase \| 2KJ5\_A Phage integrase phage integrase, GFT NMR, PSI-2; pdb70 \| 2KHV\_A \| 96.6 \| 9.7e-06 \| 6.9e-10 \| 55.8 \| 84 \| (102, 192) \| 403 \| (2, 87) \| 106 \| Phage integrase \| 2KHV\_A Phage integrase solution structure, GFT, NESG, STRUCTURAL |
| Top keywords  (threshold 1.00e-03 (evalue)) | **Integrase, recombinase, domain\_containing, Tyr, Site\_specific, Phage, DNA, XerD, complex, Prophage** |
| Output files | ../../similar\_sequences/32\_FANPEZAQ\_CDS\_0032\_merged.svg ../../similar\_sequences/32\_FANPEZAQ\_CDS\_0032\_pdb70.a3m ../../similar\_sequences/32\_FANPEZAQ\_CDS\_0032\_pdb70.hhr ../../similar\_sequences/32\_FANPEZAQ\_CDS\_0032\_uniclust.a3m ../../similar\_sequences/32\_FANPEZAQ\_CDS\_0032\_uniclust.hhr |

#### Structure prediction (AlphaFold)^2^

|  |  |
| --- | --- |
| Stats | xml version="1.0" encoding="utf-8" standalone="no"?       2024-09-02T21:09:31.882498 image/svg+xml   Matplotlib v3.7.2, https://matplotlib.org/ |
| Predicted structure | **NGL Viewer Controls:**  - Center: *Left-Click* - Rotate: *Left-Click + Drag* - Translate: *Right-Click + Drag* - Zoom: *Shift + Left-Click + Drag* |
| Output files | ../../predicted\_structures/32\_FANPEZAQ\_CDS\_0032/features.pkl ../../predicted\_structures/32\_FANPEZAQ\_CDS\_0032/ranked\_0.pdb ../../predicted\_structures/32\_FANPEZAQ\_CDS\_0032/ranked\_0\_plots.svg ../../predicted\_structures/32\_FANPEZAQ\_CDS\_0032/result\_model\_1\_ptm\_pred\_0.pkl |

#### Structure similarity search results (Foldseek)^3^

|  |  |
| --- | --- |
| Structure databases searched | Pdb, Afdb-proteome, Afdb-uniprot50 |
| Results, scheme(s)  (Top layers only, threshold 1.00e-02 (evalue)) | xml version="1.0" encoding="utf-8" standalone="no"?       2024-09-02T21:11:06.448704 image/svg+xml   Matplotlib v3.7.2, https://matplotlib.org/ |
| Results, table  (threshold 1.00e-02 (evalue)) | db \| id \| prob \| evalue \| bits \| fident \| alnlen \| mismatch \| gapopen \| qstart \| qend \| tstart \| tend \| name \| description; pdb \| 1Z19\_A \| 1.0 \| 8.811e-08 \| 288 \| 0.16 \| 305 \| 199 \| 17 \| 101 \| 382 \| 1 \| 271 \| Integrase \| Integrase; pdb \| 5VFZ\_A \| 1.0 \| 8.943e-09 \| 277 \| 0.189 \| 316 \| 199 \| 17 \| 107 \| 403 \| 9 \| 286 \| Gp33 \| Gp33; pdb \| 1Z1G\_A \| 1.0 \| 6.942e-08 \| 256 \| 0.165 \| 362 \| 240 \| 19 \| 49 \| 382 \| 17 \| 344 \| Integrase \| Integrase; pdb \| 5J0N\_E \| 1.0 \| 1.538e-06 \| 249 \| 0.166 \| 306 \| 199 \| 16 \| 99 \| 382 \| 1 \| 272 \| Integrase \| Integrase; pdb \| 6EMY\_A \| 1.0 \| 2.637e-07 \| 248 \| 0.14 \| 341 \| 215 \| 18 \| 101 \| 392 \| 1 \| 312 \| Int protein \| Int protein; pdb \| 4A8E\_A \| 1.0 \| 2.765e-07 \| 246 \| 0.167 \| 311 \| 202 \| 15 \| 104 \| 403 \| 2 \| 266 \| PROBABLE TYROSINE RECOMBINASE XERC-LIKE \| PROBABLE TYROSINE RECOMBINASE XERC-LIKE; pdb \| 1Z1B\_A \| 1.0 \| 7.281e-08 \| 244 \| 0.157 \| 380 \| 243 \| 21 \| 36 \| 382 \| 9 \| 344 \| Integrase \| Integrase; pdb \| 6EN0\_A \| 1.0 \| 4.247e-07 \| 238 \| 0.154 \| 331 \| 212 \| 17 \| 100 \| 389 \| 1 \| 304 \| Int protein \| Int protein; pdb \| 1Z19\_B \| 1.0 \| 1.952e-06 \| 237 \| 0.163 \| 311 \| 198 \| 18 \| 101 \| 382 \| 1 \| 278 \| Integrase \| Integrase; pdb \| 2A3V\_A \| 1.0 \| 1.398e-06 \| 234 \| 0.15 \| 333 \| 209 \| 15 \| 105 \| 386 \| 3 \| 312 \| site-specific recombinase IntI4 \| site-specific recombinase IntI4; pdb \| 6EN2\_B \| 1.0 \| 5.928e-07 \| 234 \| 0.132 \| 339 \| 222 \| 17 \| 100 \| 392 \| 1 \| 313 \| Int protein \| Int protein; pdb \| 5J0N\_F \| 1.0 \| 1.981e-07 \| 232 \| 0.156 \| 390 \| 236 \| 23 \| 36 \| 391 \| 16 \| 346 \| Integrase \| Integrase; pdb \| 1Z1G\_B \| 1.0 \| 2.765e-07 \| 231 \| 0.151 \| 363 \| 236 \| 16 \| 49 \| 382 \| 15 \| 334 \| Integrase \| Integrase; pdb \| 2A3V\_C \| 1.0 \| 2.857e-06 \| 225 \| 0.15 \| 325 \| 204 \| 14 \| 107 \| 382 \| 6 \| 307 \| site-specific recombinase IntI4 \| site-specific recombinase IntI4; pdb \| 6EN1\_B \| 1.0 \| 1.398e-06 \| 223 \| 0.149 \| 341 \| 215 \| 18 \| 100 \| 392 \| 1 \| 314 \| Int protein \| Int protein; pdb \| 6EMY\_B \| 1.0 \| 8.276e-07 \| 219 \| 0.132 \| 340 \| 220 \| 20 \| 101 \| 392 \| 1 \| 313 \| Int protein \| Int protein; pdb \| 1P7D\_B \| 1.0 \| 6.126e-06 \| 218 \| 0.164 \| 310 \| 199 \| 16 \| 101 \| 382 \| 1 \| 278 \| Integrase \| Integrase; pdb \| 6EN0\_B \| 1.0 \| 1.05e-06 \| 218 \| 0.144 \| 338 \| 213 \| 15 \| 101 \| 392 \| 1 \| 308 \| Int protein \| Int protein; pdb \| 5HXY\_E \| 1.0 \| 4.454e-07 \| 209 \| 0.154 \| 318 \| 201 \| 16 \| 103 \| 400 \| 2 \| 271 \| Tyrosine recombinase XerA \| Tyrosine recombinase XerA; pdb \| 5HXY\_B \| 1.0 \| 5.389e-07 \| 205 \| 0.154 \| 318 \| 196 \| 17 \| 103 \| 400 \| 2 \| 266 \| Tyrosine recombinase XerA \| Tyrosine recombinase XerA; pdb \| 5HXY\_D \| 1.0 \| 1.398e-06 \| 203 \| 0.182 \| 301 \| 196 \| 16 \| 105 \| 389 \| 2 \| 268 \| Tyrosine recombinase XerA \| Tyrosine recombinase XerA; pdb \| 2A3V\_B \| 1.0 \| 8.969e-06 \| 201 \| 0.157 \| 318 \| 196 \| 14 \| 107 \| 375 \| 6 \| 300 \| site-specific recombinase IntI4 \| site-specific recombinase IntI4; pdb \| 5HXY\_C \| 1.0 \| 7.524e-07 \| 195 \| 0.176 \| 317 \| 188 \| 18 \| 105 \| 400 \| 4 \| 268 \| Tyrosine recombinase XerA \| Tyrosine recombinase XerA; pdb \| 5HXY\_F \| 1.0 \| 3.297e-06 \| 185 \| 0.184 \| 319 \| 181 \| 23 \| 105 \| 400 \| 2 \| 264 \| Tyrosine recombinase XerA \| Tyrosine recombinase XerA; pdb \| 5JJV\_A \| 1.0 \| 4.323e-05 \| 171 \| 0.175 \| 279 \| 187 \| 17 \| 143 \| 395 \| 91 \| 352 \| Tyrosine recombinase XerH \| Tyrosine recombinase XerH; pdb \| 3RMP\_C \| 1.0 \| 0.005077 \| 163 \| 0.284 \| 88 \| 48 \| 7 \| 1 \| 87 \| 1 \| 74 \| CP4-like integrase \| CP4-like integrase; pdb \| 5JK0\_D \| 1.0 \| 1.138e-05 \| 163 \| 0.158 \| 322 \| 203 \| 20 \| 89 \| 370 \| 36 \| 329 \| Tyrosine recombinase XerH \| Tyrosine recombinase XerH; pdb \| 5JK0\_A \| 1.0 \| 4.755e-05 \| 156 \| 0.172 \| 336 \| 221 \| 20 \| 90 \| 395 \| 37 \| 345 \| Tyrosine recombinase XerH \| Tyrosine recombinase XerH; pdb \| 5C6K\_A \| 1.0 \| 0.003637 \| 154 \| 0.169 \| 153 \| 108 \| 6 \| 220 \| 370 \| 1 \| 136 \| Integrase \| Integrase; pdb \| 3RMP\_A \| 1.0 \| 0.005857 \| 153 \| 0.28 \| 89 \| 47 \| 7 \| 1 \| 86 \| 1 \| 75 \| CP4-like integrase \| CP4-like integrase; pdb \| 3JU0\_A \| 1.0 \| 0.003306 \| 153 \| 0.322 \| 93 \| 52 \| 5 \| 1 \| 93 \| 1 \| 82 \| Phage integrase \| Phage integrase; pdb \| 5C6K\_B \| 1.0 \| 0.006143 \| 138 \| 0.16 \| 175 \| 114 \| 7 \| 221 \| 393 \| 1 \| 144 \| Integrase \| Integrase; pdb \| 5JJV\_B \| 1.0 \| 0.0001121 \| 136 \| 0.165 \| 350 \| 216 \| 19 \| 70 \| 395 \| 58 \| 355 \| Tyrosine recombinase XerH \| Tyrosine recombinase XerH; pdb \| 3NKH\_A \| 1.0 \| 0.006143 \| 105 \| 0.122 \| 220 \| 136 \| 13 \| 222 \| 400 \| 5 \| 208 \| Integrase \| Integrase; pdb \| 3NKH\_B \| 1.0 \| 0.008177 \| 105 \| 0.14 \| 221 \| 139 \| 10 \| 222 \| 400 \| 6 \| 217 \| Integrase \| Integrase; afdb-proteome \| AF-A0A0H3GN77-F1-MODEL\_V4 \| 1.0 \| 3.998e-20 \| 714 \| 0.231 \| 419 \| 273 \| 17 \| 2 \| 402 \| 3 \| 390 \| Putative site specific integrase \| Putative site specific integrase; afdb-proteome \| AF-Q32GN0-F1-MODEL\_V4 \| 1.0 \| 1.645e-18 \| 657 \| 0.225 \| 421 \| 276 \| 18 \| 1 \| 402 \| 15 \| 404 \| Integrase \| Integrase; afdb-proteome \| AF-P37326-F1-MODEL\_V4 \| 1.0 \| 2.527e-18 \| 643 \| 0.218 \| 417 \| 280 \| 18 \| 1 \| 403 \| 1 \| 385 \| Prophage integrase IntS \| Prophage integrase IntS; afdb-proteome \| AF-P32053-F1-MODEL\_V4 \| 1.0 \| 3.012e-17 \| 610 \| 0.195 \| 420 \| 289 \| 17 \| 2 \| 402 \| 8 \| 397 \| Prophage integrase IntA \| Prophage integrase IntA; afdb-proteome \| AF-Q4AAY5-F1-MODEL\_V4 \| 1.0 \| 5.417e-18 \| 603 \| 0.187 \| 432 \| 284 \| 22 \| 2 \| 403 \| 3 \| 397 \| Gifsy-1 prophage protein \| Gifsy-1 prophage protein; afdb-proteome \| AF-A0A0H3GY91-F1-MODEL\_V4 \| 1.0 \| 1.384e-16 \| 568 \| 0.191 \| 424 \| 282 \| 21 \| 2 \| 402 \| 8 \| 393 \| Integrase family protein \| Integrase family protein; afdb-proteome \| AF-A0A0H3GWB4-F1-MODEL\_V4 \| 1.0 \| 3.591e-16 \| 560 \| 0.183 \| 420 \| 290 \| 19 \| 2 \| 402 \| 3 \| 388 \| Tyr recombinase domain-containing protein \| Tyr recombinase domain-containing protein; afdb-proteome \| AF-P76542-F1-MODEL\_V4 \| 1.0 \| 1.384e-16 \| 555 \| 0.203 \| 422 \| 280 \| 23 \| 2 \| 402 \| 6 \| 392 \| Prophage integrase IntZ \| Prophage integrase IntZ; afdb-proteome \| AF-A0A0H3GIF3-F1-MODEL\_V4 \| 1.0 \| 1.127e-15 \| 546 \| 0.183 \| 437 \| 289 \| 21 \| 2 \| 403 \| 3 \| 406 \| Putative P4-type integrase \| Putative P4-type integrase; afdb-proteome \| AF-Q5F9D7-F1-MODEL\_V4 \| 1.0 \| 1.2e-16 \| 540 \| 0.177 \| 429 \| 283 \| 22 \| 2 \| 402 \| 8 \| 394 \| Integrase \| Integrase; afdb-proteome \| AF-A0A0H3GS99-F1-MODEL\_V4 \| 1.0 \| 2.967e-16 \| 526 \| 0.181 \| 436 \| 288 \| 25 \| 2 \| 402 \| 3 \| 404 \| Integrase \| Integrase; afdb-proteome \| AF-P39347-F1-MODEL\_V4 \| 1.0 \| 4.938e-15 \| 496 \| 0.192 \| 400 \| 266 \| 18 \| 36 \| 402 \| 3 \| 378 \| Putative protein IntB \| Putative protein IntB; afdb-proteome \| AF-P71298-F1-MODEL\_V4 \| 1.0 \| 5.179e-15 \| 490 \| 0.192 \| 451 \| 286 \| 24 \| 2 \| 402 \| 33 \| 455 \| Prophage integrase IntF \| Prophage integrase IntF; afdb-proteome \| AF-Q8ZMR4-F1-MODEL\_V4 \| 1.0 \| 6.792e-14 \| 460 \| 0.193 \| 377 \| 247 \| 20 \| 2 \| 354 \| 8 \| 351 \| Fels-2 prophage protein \| Fels-2 prophage protein; afdb-proteome \| AF-A0A0H3H506-F1-MODEL\_V4 \| 1.0 \| 3.495e-11 \| 460 \| 0.184 \| 320 \| 226 \| 12 \| 100 \| 402 \| 39 \| 340 \| Integrase \| Integrase; afdb-proteome \| AF-A0A0H4IW46-F1-MODEL\_V4 \| 1.0 \| 4.639e-14 \| 420 \| 0.224 \| 366 \| 227 \| 19 \| 2 \| 341 \| 7 \| 341 \| Integrase \| Integrase; afdb-proteome \| AF-Q9HXQ6-F1-MODEL\_V4 \| 1.0 \| 7.382e-10 \| 372 \| 0.181 \| 319 \| 208 \| 18 \| 101 \| 393 \| 2 \| 293 \| Tyrosine recombinase XerD \| Tyrosine recombinase XerD; afdb-proteome \| AF-Q32BV7-F1-MODEL\_V4 \| 1.0 \| 1.915e-09 \| 369 \| 0.159 \| 319 \| 211 \| 20 \| 103 \| 393 \| 4 \| 293 \| Tyrosine recombinase XerD \| Tyrosine recombinase XerD; afdb-proteome \| AF-P0A8P8-F1-MODEL\_V4 \| 1.0 \| 2.549e-09 \| 368 \| 0.166 \| 319 \| 209 \| 19 \| 103 \| 393 \| 4 \| 293 \| Tyrosine recombinase XerD \| Tyrosine recombinase XerD; afdb-proteome \| AF-Q9CBU0-F1-MODEL\_V4 \| 1.0 \| 9.368e-10 \| 346 \| 0.18 \| 327 \| 198 \| 15 \| 106 \| 402 \| 3 \| 289 \| Tyrosine recombinase XerC \| Tyrosine recombinase XerC; afdb-proteome \| AF-Q2FWP2-F1-MODEL\_V4 \| 1.0 \| 2.985e-10 \| 345 \| 0.171 \| 374 \| 248 \| 16 \| 46 \| 400 \| 10 \| 340 \| Integrase \| Integrase; afdb-proteome \| AF-P9WF35-F1-MODEL\_V4 \| 1.0 \| 1.509e-09 \| 336 \| 0.182 \| 328 \| 196 \| 16 \| 106 \| 402 \| 4 \| 290 \| Tyrosine recombinase XerC \| Tyrosine recombinase XerC; afdb-proteome \| AF-P0A8P6-F1-MODEL\_V4 \| 1.0 \| 8e-09 \| 335 \| 0.18 \| 310 \| 212 \| 15 \| 101 \| 390 \| 2 \| 289 \| Tyrosine recombinase XerC \| Tyrosine recombinase XerC; afdb-proteome \| AF-Q8ZQI9-F1-MODEL\_V4 \| 1.0 \| 1.266e-10 \| 334 \| 0.154 \| 428 \| 257 \| 21 \| 36 \| 395 \| 19 \| 409 \| Putative Fels-1 prophage integrase \| Putative Fels-1 prophage integrase; afdb-proteome \| AF-Q329Y7-F1-MODEL\_V4 \| 1.0 \| 1.015e-08 \| 331 \| 0.183 \| 310 \| 211 \| 15 \| 101 \| 390 \| 2 \| 289 \| Tyrosine recombinase XerC \| Tyrosine recombinase XerC; afdb-proteome \| AF-K0ETD7-F1-MODEL\_V4 \| 1.0 \| 1.288e-08 \| 328 \| 0.175 \| 337 \| 205 \| 20 \| 103 \| 393 \| 2 \| 311 \| Tyrosine recombinase XerD \| Tyrosine recombinase XerD; afdb-proteome \| AF-P76056-F1-MODEL\_V4 \| 1.0 \| 2.587e-10 \| 327 \| 0.151 \| 442 \| 275 \| 21 \| 25 \| 401 \| 2 \| 408 \| Prophage integrase IntR \| Prophage integrase IntR; afdb-proteome \| AF-A0A1Y3JR42-F1-MODEL\_V4 \| 1.0 \| 6.711e-10 \| 320 \| 0.133 \| 412 \| 267 \| 20 \| 34 \| 402 \| 3 \| 367 \| Site-specific integrase \| Site-specific integrase; afdb-proteome \| AF-Q5F975-F1-MODEL\_V4 \| 1.0 \| 2.549e-09 \| 316 \| 0.155 \| 387 \| 231 \| 21 \| 34 \| 395 \| 2 \| 317 \| Integrase \| Integrase; afdb-proteome \| AF-A0A132P7M2-F1-MODEL\_V4 \| 1.0 \| 1.247e-09 \| 312 \| 0.144 \| 409 \| 263 \| 22 \| 32 \| 400 \| 2 \| 363 \| Integrase \| Integrase; afdb-proteome \| AF-P24218-F1-MODEL\_V4 \| 1.0 \| 1.826e-09 \| 310 \| 0.166 \| 396 \| 244 \| 17 \| 46 \| 402 \| 9 \| 357 \| Prophage integrase IntD \| Prophage integrase IntD; afdb-proteome \| AF-K0F374-F1-MODEL\_V4 \| 1.0 \| 3.038e-08 \| 309 \| 0.165 \| 332 \| 205 \| 16 \| 103 \| 402 \| 8 \| 299 \| Tyrosine recombinase XerC \| Tyrosine recombinase XerC; afdb-proteome \| AF-Q8ZQB0-F1-MODEL\_V4 \| 1.0 \| 8.932e-10 \| 309 \| 0.168 \| 428 \| 253 \| 20 \| 36 \| 396 \| 10 \| 401 \| Gifsy-2 prophage integrase \| Gifsy-2 prophage integrase; afdb-proteome \| AF-P55888-F1-MODEL\_V4 \| 1.0 \| 3.505e-08 \| 308 \| 0.172 \| 308 \| 213 \| 17 \| 103 \| 390 \| 6 \| 291 \| Tyrosine recombinase XerC \| Tyrosine recombinase XerC; afdb-proteome \| AF-A0A0H3GH57-F1-MODEL\_V4 \| 1.0 \| 4.893e-08 \| 306 \| 0.171 \| 314 \| 213 \| 17 \| 99 \| 390 \| 3 \| 291 \| Tyrosine recombinase XerC \| Tyrosine recombinase XerC; afdb-proteome \| AF-Q2FZ30-F1-MODEL\_V4 \| 1.0 \| 3.342e-08 \| 303 \| 0.185 \| 313 \| 203 \| 13 \| 106 \| 390 \| 4 \| 292 \| Tyrosine recombinase XerC \| Tyrosine recombinase XerC; afdb-proteome \| AF-P76168-F1-MODEL\_V4 \| 1.0 \| 5.73e-09 \| 301 \| 0.163 \| 386 \| 220 \| 17 \| 75 \| 401 \| 33 \| 374 \| Putative defective protein IntQ \| Putative defective protein IntQ; afdb-proteome \| AF-P46495-F1-MODEL\_V4 \| 1.0 \| 2.43e-09 \| 300 \| 0.159 \| 364 \| 224 \| 20 \| 34 \| 370 \| 2 \| 310 \| Putative integrase/recombinase HI\_1572 \| Putative integrase/recombinase HI\_1572; afdb-proteome \| AF-A0A4U3SYR4-F1-MODEL\_V4 \| 1.0 \| 8.8e-09 \| 290 \| 0.155 \| 405 \| 261 \| 29 \| 36 \| 393 \| 7 \| 377 \| Site-specific integrase \| Site-specific integrase; afdb-proteome \| AF-A0A0H3GP78-F1-MODEL\_V4 \| 1.0 \| 3.084e-09 \| 288 \| 0.156 \| 410 \| 248 \| 18 \| 36 \| 402 \| 3 \| 357 \| Site-specific recombinase, phage integrase family \| Site-specific recombinase, phage integrase family; afdb-proteome \| AF-Q32DW2-F1-MODEL\_V4 \| 1.0 \| 8.8e-09 \| 287 \| 0.163 \| 368 \| 239 \| 21 \| 48 \| 382 \| 17 \| 348 \| Integrase \| Integrase; afdb-proteome \| AF-X8FIQ7-F1-MODEL\_V4 \| 1.0 \| 2.804e-09 \| 284 \| 0.188 \| 419 \| 253 \| 28 \| 22 \| 402 \| 2 \| 371 \| Phage integrase family protein \| Phage integrase family protein; afdb-proteome \| AF-Q8ZMR5-F1-MODEL\_V4 \| 1.0 \| 1.351e-08 \| 283 \| 0.167 \| 389 \| 226 \| 23 \| 36 \| 397 \| 3 \| 320 \| Fels-2 prophage protein \| Fels-2 prophage protein; afdb-proteome \| AF-Q5FAI3-F1-MODEL\_V4 \| 1.0 \| 1.269e-07 \| 277 \| 0.174 \| 321 \| 203 \| 16 \| 103 \| 389 \| 2 \| 294 \| Tyrosine recombinase XerC \| Tyrosine recombinase XerC; afdb-proteome \| AF-Q8ZNX5-F1-MODEL\_V4 \| 1.0 \| 2.854e-07 \| 269 \| 0.17 \| 317 \| 210 \| 21 \| 93 \| 383 \| 6 \| 295 \| Putative phage integrase protein \| Putative phage integrase protein; afdb-proteome \| AF-A0A133CSM1-F1-MODEL\_V4 \| 1.0 \| 2.075e-08 \| 264 \| 0.15 \| 411 \| 261 \| 22 \| 36 \| 401 \| 7 \| 374 \| Site-specific integrase \| Site-specific integrase; afdb-proteome \| AF-P75969-F1-MODEL\_V4 \| 1.0 \| 2.511e-08 \| 262 \| 0.179 \| 384 \| 243 \| 22 \| 34 \| 379 \| 15 \| 364 \| Prophage integrase IntE \| Prophage integrase IntE; afdb-proteome \| AF-A0A132Z757-F1-MODEL\_V4 \| 1.0 \| 1.117e-08 \| 262 \| 0.159 \| 415 \| 242 \| 22 \| 36 \| 394 \| 17 \| 380 \| Tyrosine-type recombinase/integrase \| Tyrosine-type recombinase/integrase; afdb-proteome \| AF-A0A132ZA79-F1-MODEL\_V4 \| 1.0 \| 5.645e-08 \| 261 \| 0.137 \| 416 \| 277 \| 21 \| 34 \| 400 \| 3 \| 385 \| Recombinase \| Recombinase; afdb-proteome \| AF-Q2FY83-F1-MODEL\_V4 \| 1.0 \| 2.762e-08 \| 257 \| 0.107 \| 426 \| 271 \| 21 \| 34 \| 396 \| 3 \| 382 \| Bacteriophage integrase \| Bacteriophage integrase; afdb-proteome \| AF-A0A133CJD6-F1-MODEL\_V4 \| 1.0 \| 2.897e-08 \| 254 \| 0.127 \| 409 \| 263 \| 20 \| 34 \| 400 \| 2 \| 358 \| Site-specific integrase \| Site-specific integrase; afdb-proteome \| AF-A0A0H3GTY0-F1-MODEL\_V4 \| 1.0 \| 3.139e-07 \| 249 \| 0.143 \| 390 \| 236 \| 25 \| 36 \| 397 \| 3 \| 322 \| Gp27 \| Gp27; afdb-proteome \| AF-Q2FX17-F1-MODEL\_V4 \| 1.0 \| 1e-07 \| 248 \| 0.145 \| 384 \| 242 \| 27 \| 46 \| 392 \| 11 \| 345 \| Integrase, phage family, putative \| Integrase, phage family, putative; afdb-proteome \| AF-P9WMB3-F1-MODEL\_V4 \| 1.0 \| 1.269e-07 \| 248 \| 0.142 \| 385 \| 261 \| 17 \| 34 \| 391 \| 13 \| 355 \| Putative prophage phiRv2 integrase \| Putative prophage phiRv2 integrase; afdb-proteome \| AF-A0A5P3G0X5-F1-MODEL\_V4 \| 1.0 \| 1.049e-07 \| 230 \| 0.128 \| 445 \| 270 \| 29 \| 14 \| 400 \| 1 \| 385 \| Site-specific integrase \| Site-specific integrase; afdb-proteome \| AF-A0A132PA96-F1-MODEL\_V4 \| 1.0 \| 4.383e-07 \| 226 \| 0.126 \| 364 \| 214 \| 20 \| 97 \| 401 \| 17 \| 335 \| Integrase \| Integrase; afdb-proteome \| AF-A0A133CEM3-F1-MODEL\_V4 \| 1.0 \| 2.359e-07 \| 225 \| 0.148 \| 391 \| 247 \| 22 \| 56 \| 400 \| 23 \| 373 \| Site-specific integrase \| Site-specific integrase; afdb-proteome \| AF-C0H5A8-F1-MODEL\_V4 \| 1.0 \| 2.812e-06 \| 223 \| 0.146 \| 328 \| 222 \| 14 \| 99 \| 402 \| 188 \| 481 \| Tyrosine recombinase \| Tyrosine recombinase; afdb-proteome \| AF-A0A0H3GQP3-F1-MODEL\_V4 \| 1.0 \| 2.949e-06 \| 222 \| 0.155 \| 340 \| 223 \| 21 \| 66 \| 386 \| 8 \| 302 \| Integrase family protein \| Integrase family protein; afdb-proteome \| AF-A0A077ZHS4-F1-MODEL\_V4 \| 1.0 \| 6.027e-06 \| 216 \| 0.18 \| 266 \| 150 \| 12 \| 175 \| 401 \| 2 \| 238 \| MAPEG and Phage integrase and DUF72 domain contai ning protein \| MAPEG and Phage integrase and DUF72 domain contai ning protein; afdb-proteome \| AF-Q8ZMP1-F1-MODEL\_V4 \| 1.0 \| 9.708e-06 \| 205 \| 0.23 \| 200 \| 124 \| 10 \| 2 \| 186 \| 8 \| 192 \| Putative integrase \| Putative integrase; afdb-proteome \| AF-Q57813-F1-MODEL\_V4 \| 1.0 \| 1.421e-05 \| 195 \| 0.113 \| 334 \| 212 \| 14 \| 103 \| 400 \| 28 \| 313 \| Probable integrase/recombinase protein MJ0367 \| Probable integrase/recombinase protein MJ0367; afdb-proteome \| AF-A0A0H3GZJ0-F1-MODEL\_V4 \| 1.0 \| 2.641e-05 \| 185 \| 0.131 \| 349 \| 212 \| 13 \| 99 \| 399 \| 12 \| 317 \| Class I integron integrase \| Class I integron integrase; afdb-proteome \| AF-Q327R4-F1-MODEL\_V4 \| 1.0 \| 0.0003002 \| 159 \| 0.276 \| 123 \| 71 \| 6 \| 2 \| 117 \| 3 \| 114 \| Arm-DNA-bind\_3 domain-containing protein \| Arm-DNA-bind\_3 domain-containing protein; afdb-proteome \| AF-A0A132Z1U1-F1-MODEL\_V4 \| 1.0 \| 0.001196 \| 159 \| 0.145 \| 172 \| 120 \| 9 \| 227 \| 392 \| 16 \| 166 \| Integrase \| Integrase; afdb-proteome \| AF-O25386-F1-MODEL\_V4 \| 1.0 \| 2.641e-05 \| 158 \| 0.146 \| 355 \| 224 \| 19 \| 70 \| 401 \| 57 \| 355 \| Tyrosine recombinase XerH \| Tyrosine recombinase XerH; afdb-proteome \| AF-A0A0H3GVE8-F1-MODEL\_V4 \| 1.0 \| 0.0002481 \| 156 \| 0.155 \| 303 \| 170 \| 16 \| 137 \| 400 \| 7 \| 262 \| Phage integrase \| Phage integrase; afdb-proteome \| AF-P0ADH5-F1-MODEL\_V4 \| 1.0 \| 0.002118 \| 147 \| 0.185 \| 199 \| 127 \| 12 \| 210 \| 381 \| 1 \| 191 \| Type 1 fimbriae regulatory protein FimB \| Type 1 fimbriae regulatory protein FimB; afdb-proteome \| AF-Q8ZK19-F1-MODEL\_V4 \| 1.0 \| 0.004541 \| 140 \| 0.245 \| 106 \| 60 \| 7 \| 2 \| 101 \| 3 \| 94 \| Putative integrase \| Putative integrase; afdb-proteome \| AF-Q7CQX8-F1-MODEL\_V4 \| 1.0 \| 0.006974 \| 132 \| 0.162 \| 191 \| 126 \| 10 \| 219 \| 381 \| 1 \| 185 \| Putative phage integrase \| Putative phage integrase; afdb-uniprot50 \| AF-A0A2Z3I9J6-F1-MODEL\_V4 \| 1.0 \| 5.152e-36 \| 1388 \| 0.495 \| 414 \| 183 \| 10 \| 1 \| 402 \| 1 \| 400 \| Uncharacterized protein \| Uncharacterized protein; afdb-uniprot50 \| AF-A0A7W4YVC1-F1-MODEL\_V4 \| 1.0 \| 2.454e-31 \| 1266 \| 0.39 \| 422 \| 222 \| 13 \| 8 \| 402 \| 6 \| 419 \| Integrase \| Integrase; afdb-uniprot50 \| AF-V4NZZ0-F1-MODEL\_V4 \| 1.0 \| 1.656e-27 \| 1102 \| 0.298 \| 415 \| 263 \| 14 \| 2 \| 402 \| 14 \| 414 \| Integrase \| Integrase; afdb-uniprot50 \| AF-B0SV31-F1-MODEL\_V4 \| 1.0 \| 4.796e-28 \| 1092 \| 0.308 \| 415 \| 259 \| 13 \| 2 \| 402 \| 4 \| 404 \| Integrase family protein \| Integrase family protein; afdb-uniprot50 \| AF-A0A1I3YI60-F1-MODEL\_V4 \| 1.0 \| 1.078e-27 \| 1090 \| 0.306 \| 430 \| 258 \| 14 \| 1 \| 402 \| 1 \| 418 \| Tyr recombinase domain-containing protein \| Tyr recombinase domain-containing protein; afdb-uniprot50 \| AF-A0A4T2A6U0-F1-MODEL\_V4 \| 1.0 \| 7.255e-27 \| 1066 \| 0.289 \| 428 \| 268 \| 11 \| 1 \| 402 \| 2 \| 419 \| Site-specific integrase \| Site-specific integrase; afdb-uniprot50 \| AF-A0A1Q8T9Y1-F1-MODEL\_V4 \| 1.0 \| 4.295e-27 \| 1029 \| 0.301 \| 425 \| 259 \| 16 \| 1 \| 402 \| 1 \| 410 \| Tyr recombinase domain-containing protein \| Tyr recombinase domain-containing protein; afdb-uniprot50 \| AF-A0A4R8F7U9-F1-MODEL\_V4 \| 1.0 \| 2.101e-27 \| 1023 \| 0.296 \| 425 \| 261 \| 15 \| 1 \| 403 \| 1 \| 409 \| Phage integrase family protein \| Phage integrase family protein; afdb-uniprot50 \| AF-A0A1Y6D6E5-F1-MODEL\_V4 \| 1.0 \| 2.277e-26 \| 1017 \| 0.301 \| 418 \| 259 \| 17 \| 2 \| 402 \| 4 \| 405 \| Integrase \| Integrase; afdb-uniprot50 \| AF-A0A1T4VSE3-F1-MODEL\_V4 \| 1.0 \| 9.514e-26 \| 996 \| 0.275 \| 414 \| 264 \| 9 \| 15 \| 402 \| 1 \| 404 \| Phage integrase family protein \| Phage integrase family protein; afdb-uniprot50 \| AF-Q6LTT6-F1-MODEL\_V4 \| 1.0 \| 9.514e-26 \| 977 \| 0.259 \| 443 \| 273 \| 16 \| 1 \| 402 \| 1 \| 429 \| Hypothetical integrase \| Hypothetical integrase; afdb-uniprot50 \| AF-A0A4Q5KHX9-F1-MODEL\_V4 \| 1.0 \| 1.685e-25 \| 966 \| 0.283 \| 437 \| 264 \| 17 \| 1 \| 402 \| 1 \| 423 \| DUF4102 domain-containing protein \| DUF4102 domain-containing protein; afdb-uniprot50 \| AF-C6BWF0-F1-MODEL\_V4 \| 1.0 \| 1.916e-24 \| 952 \| 0.278 \| 410 \| 261 \| 14 \| 17 \| 403 \| 2 \| 399 \| Integrase family protein \| Integrase family protein; afdb-uniprot50 \| AF-A0A379FVR2-F1-MODEL\_V4 \| 1.0 \| 1.945e-25 \| 949 \| 0.264 \| 434 \| 272 \| 16 \| 2 \| 402 \| 3 \| 422 \| Prophage CP4-57 integrase \| Prophage CP4-57 integrase; afdb-uniprot50 \| AF-A0A6J6YBA8-F1-MODEL\_V4 \| 1.0 \| 5.29e-25 \| 948 \| 0.289 \| 407 \| 264 \| 13 \| 2 \| 401 \| 3 \| 391 \| Unannotated protein \| Unannotated protein; afdb-uniprot50 \| AF-A0A1T4XEC3-F1-MODEL\_V4 \| 1.0 \| 6.714e-25 \| 947 \| 0.272 \| 433 \| 266 \| 16 \| 2 \| 402 \| 4 \| 419 \| Phage integrase family protein \| Phage integrase family protein; afdb-uniprot50 \| AF-W3VAI2-F1-MODEL\_V4 \| 1.0 \| 6.714e-25 \| 922 \| 0.255 \| 435 \| 271 \| 17 \| 2 \| 400 \| 5 \| 422 \| Integrase \| Integrase; afdb-uniprot50 \| AF-E6VX36-F1-MODEL\_V4 \| 1.0 \| 4.969e-24 \| 920 \| 0.274 \| 411 \| 263 \| 15 \| 18 \| 402 \| 1 \| 402 \| Integrase family protein \| Integrase family protein; afdb-uniprot50 \| AF-A0A7Y4C939-F1-MODEL\_V4 \| 1.0 \| 2.942e-24 \| 915 \| 0.287 \| 432 \| 262 \| 15 \| 1 \| 400 \| 1 \| 418 \| Tyrosine-type recombinase/integrase \| Tyrosine-type recombinase/integrase; afdb-uniprot50 \| AF-A0A7U5HNH5-F1-MODEL\_V4 \| 1.0 \| 4.518e-24 \| 905 \| 0.286 \| 423 \| 269 \| 16 \| 1 \| 403 \| 1 \| 410 \| Putative phage integrase \| Putative phage integrase; afdb-uniprot50 \| AF-I3TMM1-F1-MODEL\_V4 \| 1.0 \| 1.397e-22 \| 895 \| 0.31 \| 370 \| 221 \| 14 \| 49 \| 402 \| 2 \| 353 \| Integrase family protein \| Integrase family protein; afdb-uniprot50 \| AF-A0A1I4Q7X5-F1-MODEL\_V4 \| 1.0 \| 1.001e-22 \| 871 \| 0.228 \| 437 \| 290 \| 15 \| 1 \| 402 \| 14 \| 438 \| Uncharacterized protein \| Uncharacterized protein; afdb-uniprot50 \| AF-A0A369C3G4-F1-MODEL\_V4 \| 1.0 \| 4.244e-23 \| 853 \| 0.265 \| 425 \| 256 \| 17 \| 1 \| 402 \| 7 \| 398 \| Site-specific recombinase XerD \| Site-specific recombinase XerD; afdb-uniprot50 \| AF-A0A2P6JM49-F1-MODEL\_V4 \| 1.0 \| 3.188e-23 \| 853 \| 0.268 \| 444 \| 269 \| 15 \| 1 \| 402 \| 3 \| 432 \| Integrase \| Integrase; afdb-uniprot50 \| AF-A0A837NCG4-F1-MODEL\_V4 \| 1.0 \| 2.855e-22 \| 846 \| 0.231 \| 428 \| 288 \| 17 \| 1 \| 401 \| 5 \| 418 \| Uncharacterized protein \| Uncharacterized protein; afdb-uniprot50 \| AF-A0A5C8S084-F1-MODEL\_V4 \| 1.0 \| 4.385e-22 \| 843 \| 0.238 \| 415 \| 272 \| 16 \| 1 \| 402 \| 5 \| 388 \| Tyrosine-type recombinase/integrase \| Tyrosine-type recombinase/integrase; afdb-uniprot50 \| AF-Q15Y37-F1-MODEL\_V4 \| 1.0 \| 1.084e-21 \| 828 \| 0.232 \| 421 \| 294 \| 17 \| 1 \| 403 \| 1 \| 410 \| Phage integrase \| Phage integrase; afdb-uniprot50 \| AF-Q0A9S4-F1-MODEL\_V4 \| 1.0 \| 7.407e-22 \| 824 \| 0.257 \| 443 \| 275 \| 18 \| 2 \| 402 \| 6 \| 436 \| Phage integrase family protein \| Phage integrase family protein; afdb-uniprot50 \| AF-A0A3D1IYE7-F1-MODEL\_V4 \| 1.0 \| 8.57e-19 \| 809 \| 0.326 \| 315 \| 193 \| 7 \| 100 \| 402 \| 11 \| 318 \| Tyr recombinase domain-containing protein \| Tyr recombinase domain-containing protein; afdb-uniprot50 \| AF-A0A142BFK5-F1-MODEL\_V4 \| 1.0 \| 3.245e-21 \| 805 \| 0.23 \| 452 \| 285 \| 15 \| 1 \| 402 \| 3 \| 441 \| Uncharacterized protein \| Uncharacterized protein; afdb-uniprot50 \| AF-A0A848W2S4-F1-MODEL\_V4 \| 1.0 \| 6.42e-22 \| 803 \| 0.264 \| 416 \| 256 \| 20 \| 3 \| 403 \| 10 \| 390 \| Tyrosine-type recombinase/integrase \| Tyrosine-type recombinase/integrase; afdb-uniprot50 \| AF-A0A7T4WGI3-F1-MODEL\_V4 \| 1.0 \| 1.121e-20 \| 792 \| 0.237 \| 413 \| 267 \| 19 \| 9 \| 403 \| 6 \| 388 \| Tyrosine-type recombinase/integrase \| Tyrosine-type recombinase/integrase; afdb-uniprot50 \| AF-A0A7G3ZYV6-F1-MODEL\_V4 \| 1.0 \| 2.438e-21 \| 788 \| 0.248 \| 426 \| 268 \| 19 \| 1 \| 402 \| 2 \| 399 \| Site-specific integrase \| Site-specific integrase; afdb-uniprot50 \| AF-A0A423PYC9-F1-MODEL\_V4 \| 1.0 \| 4.531e-21 \| 783 \| 0.229 \| 423 \| 271 \| 16 \| 2 \| 402 \| 6 \| 395 \| Uncharacterized protein \| Uncharacterized protein; afdb-uniprot50 \| AF-A0A850S105-F1-MODEL\_V4 \| 1.0 \| 6.325e-21 \| 782 \| 0.233 \| 433 \| 267 \| 21 \| 1 \| 403 \| 7 \| 404 \| Tyrosine-type recombinase/integrase \| Tyrosine-type recombinase/integrase; afdb-uniprot50 \| AF-A0A1M6HN29-F1-MODEL\_V4 \| 1.0 \| 3.744e-21 \| 781 \| 0.234 \| 426 \| 265 \| 20 \| 2 \| 403 \| 3 \| 391 \| Site-specific recombinase XerD \| Site-specific recombinase XerD; afdb-uniprot50 \| AF-A0A537RYP4-F1-MODEL\_V4 \| 1.0 \| 1.514e-21 \| 780 \| 0.254 \| 437 \| 257 \| 20 \| 2 \| 402 \| 4 \| 407 \| DUF4102 domain-containing protein \| DUF4102 domain-containing protein; afdb-uniprot50 \| AF-A0A059G9S3-F1-MODEL\_V4 \| 1.0 \| 3.049e-20 \| 775 \| 0.251 \| 417 \| 275 \| 18 \| 2 \| 402 \| 6 \| 401 \| Integrase family protein \| Integrase family protein; afdb-uniprot50 \| AF-A0A4T0UP22-F1-MODEL\_V4 \| 1.0 \| 3.689e-20 \| 774 \| 0.221 \| 415 \| 280 \| 19 \| 2 \| 402 \| 8 \| 393 \| DUF4102 domain-containing protein \| DUF4102 domain-containing protein; afdb-uniprot50 \| AF-A0A2D6KBJ9-F1-MODEL\_V4 \| 1.0 \| 2.642e-20 \| 773 \| 0.265 \| 388 \| 247 \| 16 \| 32 \| 403 \| 12 \| 377 \| Uncharacterized protein \| Uncharacterized protein; afdb-uniprot50 \| AF-A0A4R2L208-F1-MODEL\_V4 \| 1.0 \| 2.438e-21 \| 765 \| 0.238 \| 428 \| 265 \| 20 \| 2 \| 402 \| 7 \| 400 \| Integrase \| Integrase; afdb-uniprot50 \| AF-A0A2P9HI64-F1-MODEL\_V4 \| 1.0 \| 6.03e-21 \| 762 \| 0.243 \| 419 \| 268 \| 20 \| 2 \| 403 \| 36 \| 422 \| Site-specific recombinase, phage integrase family \| Site-specific recombinase, phage integrase family; afdb-uniprot50 \| AF-A0A504KCG3-F1-MODEL\_V4 \| 1.0 \| 1.019e-20 \| 761 \| 0.234 \| 426 \| 274 \| 19 \| 2 \| 402 \| 4 \| 402 \| DUF4102 domain-containing protein \| DUF4102 domain-containing protein; afdb-uniprot50 \| AF-Q0EWR9-F1-MODEL\_V4 \| 1.0 \| 1.491e-20 \| 758 \| 0.233 \| 419 \| 274 \| 18 \| 2 \| 403 \| 3 \| 391 \| Integrative genetic element Gsu32, integrase \| Integrative genetic element Gsu32, integrase; afdb-uniprot50 \| AF-A0A495GPM7-F1-MODEL\_V4 \| 1.0 \| 9.26e-21 \| 755 \| 0.247 \| 424 \| 266 \| 19 \| 2 \| 403 \| 4 \| 396 \| Integrase \| Integrase; afdb-uniprot50 \| AF-A0A5Q0M768-F1-MODEL\_V4 \| 1.0 \| 2.813e-21 \| 753 \| 0.243 \| 436 \| 263 \| 21 \| 1 \| 403 \| 3 \| 404 \| Tyrosine-type recombinase/integrase \| Tyrosine-type recombinase/integrase; afdb-uniprot50 \| AF-A0A2E1XW16-F1-MODEL\_V4 \| 1.0 \| 2.184e-20 \| 751 \| 0.243 \| 415 \| 270 \| 18 \| 2 \| 402 \| 6 \| 390 \| Tyr recombinase domain-containing protein \| Tyr recombinase domain-containing protein; afdb-uniprot50 \| AF-A0A0A1F5C1-F1-MODEL\_V4 \| 1.0 \| 2.519e-20 \| 749 \| 0.238 \| 415 \| 271 \| 17 \| 3 \| 402 \| 5 \| 389 \| Integrase \| Integrase; afdb-uniprot50 \| AF-Q2SP99-F1-MODEL\_V4 \| 1.0 \| 1.158e-19 \| 741 \| 0.223 \| 426 \| 280 \| 19 \| 2 \| 403 \| 4 \| 402 \| Integrase \| Integrase; afdb-uniprot50 \| AF-A0A0S8AFG9-F1-MODEL\_V4 \| 1.0 \| 3.689e-20 \| 740 \| 0.243 \| 419 \| 264 \| 19 \| 1 \| 402 \| 2 \| 384 \| Integrase \| Integrase; afdb-uniprot50 \| AF-A0A7C3WGF3-F1-MODEL\_V4 \| 1.0 \| 2.519e-20 \| 740 \| 0.237 \| 433 \| 266 \| 22 \| 2 \| 402 \| 6 \| 406 \| Site-specific integrase \| Site-specific integrase; afdb-uniprot50 \| AF-A0A7V2T5X5-F1-MODEL\_V4 \| 1.0 \| 1.865e-19 \| 732 \| 0.251 \| 418 \| 269 \| 16 \| 2 \| 402 \| 14 \| 404 \| Site-specific integrase \| Site-specific integrase; afdb-uniprot50 \| AF-A0A2N7M5E9-F1-MODEL\_V4 \| 1.0 \| 8.699e-20 \| 731 \| 0.229 \| 431 \| 273 \| 22 \| 2 \| 402 \| 4 \| 405 \| Integrase \| Integrase; afdb-uniprot50 \| AF-A7K117-F1-MODEL\_V4 \| 1.0 \| 3.15e-19 \| 730 \| 0.211 \| 431 \| 281 \| 17 \| 1 \| 401 \| 3 \| 404 \| Site-specific recombinase, phage integrase family protein \| Site-specific recombinase, phage integrase family protein; afdb-uniprot50 \| AF-U2H964-F1-MODEL\_V4 \| 1.0 \| 4.463e-20 \| 730 \| 0.232 \| 421 \| 273 \| 18 \| 2 \| 402 \| 66 \| 456 \| Integrase \| Integrase; afdb-uniprot50 \| AF-M9RL77-F1-MODEL\_V4 \| 1.0 \| 1.004e-19 \| 729 \| 0.226 \| 410 \| 272 \| 18 \| 2 \| 402 \| 9 \| 382 \| Putative phage integrase \| Putative phage integrase; afdb-uniprot50 \| AF-A0A849TNM8-F1-MODEL\_V4 \| 1.0 \| 1.053e-19 \| 729 \| 0.22 \| 408 \| 276 \| 18 \| 7 \| 402 \| 12 \| 389 \| Tyrosine-type recombinase/integrase \| Tyrosine-type recombinase/integrase; afdb-uniprot50 \| AF-A0A1I3R028-F1-MODEL\_V4 \| 1.0 \| 7.189e-20 \| 728 \| 0.245 \| 423 \| 262 \| 23 \| 1 \| 403 \| 1 \| 386 \| Uncharacterized protein \| Uncharacterized protein; afdb-uniprot50 \| AF-A0A0W1A9G8-F1-MODEL\_V4 \| 1.0 \| 1.695e-19 \| 727 \| 0.199 \| 417 \| 291 \| 14 \| 1 \| 402 \| 1 \| 389 \| Integrase \| Integrase; afdb-uniprot50 \| AF-A0A454JJL7-F1-MODEL\_V4 \| 1.0 \| 1.274e-19 \| 727 \| 0.241 \| 419 \| 269 \| 17 \| 1 \| 402 \| 1 \| 387 \| DUF4102 domain-containing protein \| DUF4102 domain-containing protein; afdb-uniprot50 \| AF-A0A7W2BWR4-F1-MODEL\_V4 \| 1.0 \| 1.616e-19 \| 725 \| 0.233 \| 424 \| 269 \| 22 \| 2 \| 402 \| 3 \| 393 \| Tyrosine-type recombinase/integrase \| Tyrosine-type recombinase/integrase; afdb-uniprot50 \| AF-A0A1G5PTY7-F1-MODEL\_V4 \| 1.0 \| 2.082e-20 \| 725 \| 0.22 \| 454 \| 282 \| 19 \| 1 \| 398 \| 3 \| 440 \| Phage integrase family protein \| Phage integrase family protein; afdb-uniprot50 \| AF-A0A6I2IUJ7-F1-MODEL\_V4 \| 1.0 \| 1.336e-19 \| 722 \| 0.208 \| 423 \| 283 \| 18 \| 1 \| 402 \| 1 \| 392 \| Tyrosine-type recombinase/integrase \| Tyrosine-type recombinase/integrase; afdb-uniprot50 \| AF-A0A523JSY9-F1-MODEL\_V4 \| 1.0 \| 1.469e-19 \| 722 \| 0.233 \| 416 \| 267 \| 21 \| 2 \| 401 \| 5 \| 384 \| Site-specific integrase \| Site-specific integrase; afdb-uniprot50 \| AF-A0A7Y7NB12-F1-MODEL\_V4 \| 1.0 \| 2.151e-19 \| 722 \| 0.237 \| 437 \| 263 \| 19 \| 1 \| 402 \| 7 \| 408 \| Integrase arm-type DNA-binding domain-containing protein \| Integrase arm-type DNA-binding domain-containing protein; afdb-uniprot50 \| AF-A0A2T5NNU0-F1-MODEL\_V4 \| 1.0 \| 8.699e-20 \| 720 \| 0.201 \| 417 \| 288 \| 21 \| 1 \| 402 \| 7 \| 393 \| Integrase \| Integrase; afdb-uniprot50 \| AF-A0A2H4W5G4-F1-MODEL\_V4 \| 1.0 \| 5.941e-20 \| 718 \| 0.227 \| 453 \| 285 \| 19 \| 1 \| 402 \| 10 \| 448 \| Tyr recombinase domain-containing protein \| Tyr recombinase domain-containing protein; afdb-uniprot50 \| AF-A0A5C7LRP0-F1-MODEL\_V4 \| 1.0 \| 2.864e-19 \| 715 \| 0.228 \| 428 \| 270 \| 21 \| 2 \| 402 \| 27 \| 421 \| DUF4102 domain-containing protein \| DUF4102 domain-containing protein; afdb-uniprot50 \| AF-A0A1X7AJC6-F1-MODEL\_V4 \| 1.0 \| 2.256e-19 \| 713 \| 0.248 \| 450 \| 276 \| 15 \| 1 \| 402 \| 3 \| 438 \| Phage integrase family protein \| Phage integrase family protein; afdb-uniprot50 \| AF-A0A0W0U9B0-F1-MODEL\_V4 \| 1.0 \| 2.367e-19 \| 712 \| 0.215 \| 418 \| 282 \| 17 \| 1 \| 402 \| 4 \| 391 \| Phage-related integrase \| Phage-related integrase; afdb-uniprot50 \| AF-A0A7C3KWL2-F1-MODEL\_V4 \| 1.0 \| 7.082e-19 \| 712 \| 0.228 \| 433 \| 264 \| 22 \| 2 \| 402 \| 6 \| 400 \| DUF4102 domain-containing protein \| DUF4102 domain-containing protein; afdb-uniprot50 \| AF-A0A7V2S709-F1-MODEL\_V4 \| 1.0 \| 4.397e-19 \| 710 \| 0.198 \| 413 \| 281 \| 20 \| 2 \| 402 \| 3 \| 377 \| DUF4102 domain-containing protein \| DUF4102 domain-containing protein; afdb-uniprot50 \| AF-A0A850T1C9-F1-MODEL\_V4 \| 1.0 \| 7.791e-19 \| 710 \| 0.236 \| 427 \| 272 \| 23 \| 2 \| 401 \| 6 \| 405 \| Integrase family protein \| Integrase family protein; afdb-uniprot50 \| AF-A0A2Z6G8D0-F1-MODEL\_V4 \| 1.0 \| 2.256e-19 \| 710 \| 0.215 \| 418 \| 279 \| 18 \| 2 \| 402 \| 33 \| 418 \| Prophage integrase IntA \| Prophage integrase IntA; afdb-uniprot50 \| AF-A0A2S7FLE3-F1-MODEL\_V4 \| 1.0 \| 4.256e-20 \| 708 \| 0.199 \| 421 \| 281 \| 20 \| 2 \| 401 \| 3 \| 388 \| DUF4102 domain-containing protein \| DUF4102 domain-containing protein; afdb-uniprot50 \| AF-A0A1Q3UZH6-F1-MODEL\_V4 \| 1.0 \| 3.634e-19 \| 708 \| 0.252 \| 419 \| 270 \| 18 \| 2 \| 403 \| 6 \| 398 \| Uncharacterized protein \| Uncharacterized protein; afdb-uniprot50 \| AF-Q5P090-F1-MODEL\_V4 \| 1.0 \| 6.854e-20 \| 708 \| 0.235 \| 421 \| 271 \| 18 \| 1 \| 402 \| 2 \| 390 \| Integrase \| Integrase; afdb-uniprot50 \| AF-A0A7C1UI09-F1-MODEL\_V4 \| 1.0 \| 2.256e-19 \| 708 \| 0.239 \| 413 \| 262 \| 19 \| 9 \| 402 \| 18 \| 397 \| Site-specific integrase \| Site-specific integrase; afdb-uniprot50 \| AF-V6F355-F1-MODEL\_V4 \| 1.0 \| 2.151e-19 \| 707 \| 0.222 \| 422 \| 278 \| 19 \| 2 \| 402 \| 8 \| 400 \| Putative prophage CPZ-55 integrase \| Putative prophage CPZ-55 integrase; afdb-uniprot50 \| AF-A0A2N6AWZ1-F1-MODEL\_V4 \| 1.0 \| 3.465e-19 \| 706 \| 0.243 \| 410 \| 269 \| 18 \| 2 \| 400 \| 6 \| 385 \| Integrase \| Integrase; afdb-uniprot50 \| AF-A0A2S5M9V6-F1-MODEL\_V4 \| 1.0 \| 2.184e-20 \| 706 \| 0.255 \| 411 \| 263 \| 15 \| 8 \| 402 \| 8 \| 391 \| Integrase \| Integrase; afdb-uniprot50 \| AF-A0A7C1NNI4-F1-MODEL\_V4 \| 1.0 \| 4.91e-20 \| 706 \| 0.226 \| 460 \| 280 \| 19 \| 1 \| 402 \| 13 \| 454 \| Integrase \| Integrase; afdb-uniprot50 \| AF-A0A7X9IKM6-F1-MODEL\_V4 \| 1.0 \| 2.864e-19 \| 702 \| 0.22 \| 409 \| 276 \| 20 \| 8 \| 402 \| 8 \| 387 \| Tyrosine-type recombinase/integrase \| Tyrosine-type recombinase/integrase; afdb-uniprot50 \| AF-A0A1T4XZ14-F1-MODEL\_V4 \| 1.0 \| 3.304e-19 \| 701 \| 0.234 \| 413 \| 271 \| 16 \| 2 \| 402 \| 3 \| 382 \| Integrase \| Integrase; afdb-uniprot50 \| AF-A0A1E7D9T2-F1-MODEL\_V4 \| 1.0 \| 2.445e-18 \| 701 \| 0.202 \| 430 \| 288 \| 14 \| 1 \| 403 \| 3 \| 404 \| Phage integrase \| Phage integrase; afdb-uniprot50 \| AF-A0A1Q3IXG3-F1-MODEL\_V4 \| 1.0 \| 3.465e-19 \| 701 \| 0.227 \| 426 \| 270 \| 23 \| 2 \| 403 \| 3 \| 393 \| Integrase \| Integrase; afdb-uniprot50 \| AF-A0A4R1J419-F1-MODEL\_V4 \| 1.0 \| 1.469e-19 \| 701 \| 0.213 \| 422 \| 280 \| 20 \| 2 \| 402 \| 3 \| 393 \| Site-specific recombinase XerD \| Site-specific recombinase XerD; afdb-uniprot50 \| AF-A0A4T2A4G2-F1-MODEL\_V4 \| 1.0 \| 2.331e-18 \| 701 \| 0.213 \| 422 \| 280 \| 21 \| 2 \| 402 \| 3 \| 393 \| DUF4102 domain-containing protein \| DUF4102 domain-containing protein; afdb-uniprot50 \| AF-A0A1Q3NZN8-F1-MODEL\_V4 \| 1.0 \| 1.158e-19 \| 701 \| 0.262 \| 426 \| 257 \| 20 \| 2 \| 403 \| 6 \| 398 \| Tyr recombinase domain-containing protein \| Tyr recombinase domain-containing protein; afdb-uniprot50 \| AF-A0A522J360-F1-MODEL\_V4 \| 1.0 \| 7.428e-19 \| 700 \| 0.228 \| 433 \| 272 \| 22 \| 3 \| 402 \| 19 \| 422 \| Site-specific integrase \| Site-specific integrase; afdb-uniprot50 \| AF-A0A5M8FH85-F1-MODEL\_V4 \| 1.0 \| 4.91e-20 \| 700 \| 0.239 \| 464 \| 271 \| 22 \| 1 \| 403 \| 1 \| 443 \| Integrase family protein \| Integrase family protein; afdb-uniprot50 \| AF-A0A5Y1WL89-F1-MODEL\_V4 \| 1.0 \| 8.171e-19 \| 699 \| 0.2 \| 420 \| 282 \| 21 \| 2 \| 402 \| 3 \| 387 \| Tyrosine-type recombinase/integrase \| Tyrosine-type recombinase/integrase; afdb-uniprot50 \| AF-H8YW01-F1-MODEL\_V4 \| 1.0 \| 1.274e-19 \| 699 \| 0.222 \| 462 \| 277 \| 21 \| 1 \| 401 \| 1 \| 441 \| Phage integrase family protein \| Phage integrase family protein; afdb-uniprot50 \| AF-A0A127F2R1-F1-MODEL\_V4 \| 1.0 \| 2.864e-19 \| 698 \| 0.222 \| 423 \| 276 \| 19 \| 1 \| 402 \| 1 \| 391 \| Symbiosis island integrase \| Symbiosis island integrase; afdb-uniprot50 \| AF-A0A837V9C3-F1-MODEL\_V4 \| 1.0 \| 3.634e-19 \| 697 \| 0.219 \| 419 \| 280 \| 17 \| 2 \| 402 \| 3 \| 392 \| Integrase \| Integrase; afdb-uniprot50 \| AF-E6W6Y4-F1-MODEL\_V4 \| 1.0 \| 1.38e-18 \| 695 \| 0.224 \| 423 \| 272 \| 20 \| 2 \| 402 \| 8 \| 396 \| Integrase family protein \| Integrase family protein; afdb-uniprot50 \| AF-A0A7X0ALK7-F1-MODEL\_V4 \| 1.0 \| 5.401e-20 \| 695 \| 0.215 \| 423 \| 278 \| 18 \| 1 \| 402 \| 1 \| 390 \| Integrase \| Integrase; afdb-uniprot50 \| AF-A0A3D2H5V9-F1-MODEL\_V4 \| 1.0 \| 1.865e-19 \| 695 \| 0.226 \| 428 \| 274 \| 17 \| 2 \| 402 \| 4 \| 401 \| Integrase \| Integrase; afdb-uniprot50 \| AF-A0A378IX19-F1-MODEL\_V4 \| 1.0 \| 1.316e-18 \| 695 \| 0.198 \| 418 \| 289 \| 17 \| 1 \| 402 \| 4 \| 391 \| Phage-related integrase \| Phage-related integrase; afdb-uniprot50 \| AF-A0A1I5DK39-F1-MODEL\_V4 \| 1.0 \| 1.274e-19 \| 695 \| 0.216 \| 424 \| 275 \| 18 \| 2 \| 402 \| 98 \| 487 \| Integrase \| Integrase; afdb-uniprot50 \| AF-A0A529Y462-F1-MODEL\_V4 \| 1.0 \| 6.438e-19 \| 694 \| 0.294 \| 302 \| 192 \| 9 \| 2 \| 293 \| 4 \| 294 \| Site-specific integrase \| Site-specific integrase; afdb-uniprot50 \| AF-A0A2S9KTH3-F1-MODEL\_V4 \| 1.0 \| 4.193e-19 \| 693 \| 0.216 \| 420 \| 280 \| 19 \| 2 \| 402 \| 3 \| 392 \| Integrase \| Integrase; afdb-uniprot50 \| AF-A0A4V3WAS3-F1-MODEL\_V4 \| 1.0 \| 3.998e-19 \| 693 \| 0.212 \| 432 \| 281 \| 20 \| 1 \| 402 \| 6 \| 408 \| DUF4102 domain-containing protein \| DUF4102 domain-containing protein; afdb-uniprot50 \| AF-A0A2N2FX08-F1-MODEL\_V4 \| 1.0 \| 8.988e-19 \| 692 \| 0.223 \| 443 \| 265 \| 23 \| 1 \| 402 \| 2 \| 406 \| Integrase \| Integrase; afdb-uniprot50 \| AF-A0A0P7Z5I7-F1-MODEL\_V4 \| 1.0 \| 4.612e-19 \| 690 \| 0.232 \| 425 \| 274 \| 20 \| 2 \| 402 \| 6 \| 402 \| Integrase \| Integrase; afdb-uniprot50 \| AF-A0A7C3PY40-F1-MODEL\_V4 \| 1.0 \| 2.331e-18 \| 689 \| 0.214 \| 447 \| 267 \| 28 \| 2 \| 402 \| 6 \| 414 \| DUF4102 domain-containing protein \| DUF4102 domain-containing protein; afdb-uniprot50 \| AF-A0A395JJS3-F1-MODEL\_V4 \| 1.0 \| 1.837e-18 \| 688 \| 0.207 \| 414 \| 280 \| 16 \| 1 \| 398 \| 1 \| 382 \| Integrase \| Integrase; afdb-uniprot50 \| AF-A0A1Z8Q0M2-F1-MODEL\_V4 \| 1.0 \| 1.448e-18 \| 686 \| 0.186 \| 419 \| 294 \| 18 \| 2 \| 402 \| 3 \| 392 \| Tyr recombinase domain-containing protein \| Tyr recombinase domain-containing protein; afdb-uniprot50 \| AF-A0A377P7I5-F1-MODEL\_V4 \| 1.0 \| 3.465e-19 \| 685 \| 0.219 \| 419 \| 278 \| 18 \| 2 \| 402 \| 3 \| 390 \| Prophage CPS-53 integrase \| Prophage CPS-53 integrase; afdb-uniprot50 \| AF-A0A2N4XL17-F1-MODEL\_V4 \| 1.0 \| 7.428e-19 \| 685 \| 0.214 \| 420 \| 280 \| 17 \| 2 \| 402 \| 3 \| 391 \| Integrase \| Integrase; afdb-uniprot50 \| AF-A0A239PYP0-F1-MODEL\_V4 \| 1.0 \| 3.634e-19 \| 685 \| 0.258 \| 425 \| 265 \| 20 \| 1 \| 402 \| 1 \| 398 \| Integrase \| Integrase; afdb-uniprot50 \| AF-A0A840V7L5-F1-MODEL\_V4 \| 1.0 \| 3.998e-19 \| 685 \| 0.213 \| 427 \| 280 \| 18 \| 1 \| 402 \| 31 \| 426 \| Integrase \| Integrase; afdb-uniprot50 \| AF-A0A2V7TR44-F1-MODEL\_V4 \| 1.0 \| 4.193e-19 \| 684 \| 0.23 \| 417 \| 273 \| 21 \| 2 \| 402 \| 6 \| 390 \| Uncharacterized protein \| Uncharacterized protein; afdb-uniprot50 \| AF-Q5P707-F1-MODEL\_V4 \| 1.0 \| 3.812e-19 \| 684 \| 0.235 \| 420 \| 271 \| 21 \| 1 \| 402 \| 1 \| 388 \| Phage-related integrase \| Phage-related integrase; afdb-uniprot50 \| AF-A0A845M1N2-F1-MODEL\_V4 \| 1.0 \| 1.67e-18 \| 683 \| 0.212 \| 418 \| 280 \| 19 \| 2 \| 402 \| 3 \| 388 \| Tyrosine-type recombinase/integrase \| Tyrosine-type recombinase/integrase; afdb-uniprot50 \| AF-A0A143PQ67-F1-MODEL\_V4 \| 1.0 \| 1.088e-18 \| 681 \| 0.258 \| 441 \| 249 \| 23 \| 2 \| 402 \| 5 \| 407 \| Prophage CP4-57 integrase \| Prophage CP4-57 integrase; afdb-uniprot50 \| AF-S9TLS3-F1-MODEL\_V4 \| 1.0 \| 7.791e-19 \| 680 \| 0.261 \| 421 \| 261 \| 19 \| 2 \| 403 \| 8 \| 397 \| Integrase \| Integrase; afdb-uniprot50 \| AF-A0A1Y2J7Q5-F1-MODEL\_V4 \| 1.0 \| 2.297e-17 \| 680 \| 0.191 \| 423 \| 287 \| 21 \| 2 \| 402 \| 9 \| 398 \| Tyr recombinase domain-containing protein \| Tyr recombinase domain-containing protein; afdb-uniprot50 \| AF-A0A317EG89-F1-MODEL\_V4 \| 1.0 \| 4.397e-19 \| 679 \| 0.222 \| 423 \| 274 \| 20 \| 2 \| 402 \| 9 \| 398 \| Integrase \| Integrase; afdb-uniprot50 \| AF-A0A1H5IA46-F1-MODEL\_V4 \| 1.0 \| 4.193e-19 \| 679 \| 0.217 \| 428 \| 279 \| 20 \| 2 \| 402 \| 5 \| 403 \| Site-specific recombinase XerD \| Site-specific recombinase XerD; afdb-uniprot50 \| AF-A0A290T5I9-F1-MODEL\_V4 \| 1.0 \| 1.255e-18 \| 678 \| 0.211 \| 420 \| 284 \| 16 \| 1 \| 401 \| 1 \| 392 \| Tyr recombinase domain-containing protein \| Tyr recombinase domain-containing protein; afdb-uniprot50 \| AF-A0A7V7N0U3-F1-MODEL\_V4 \| 1.0 \| 1.518e-18 \| 678 \| 0.196 \| 412 \| 285 \| 16 \| 9 \| 402 \| 6 \| 389 \| DUF4102 domain-containing protein \| DUF4102 domain-containing protein; afdb-uniprot50 \| AF-A0A286IFF7-F1-MODEL\_V4 \| 1.0 \| 2.367e-19 \| 678 \| 0.227 \| 436 \| 268 \| 19 \| 2 \| 403 \| 6 \| 406 \| Site-specific recombinase XerD \| Site-specific recombinase XerD; afdb-uniprot50 \| AF-A0A2E0QXG3-F1-MODEL\_V4 \| 1.0 \| 8.171e-19 \| 678 \| 0.22 \| 418 \| 278 \| 19 \| 2 \| 401 \| 23 \| 410 \| Integrase \| Integrase; afdb-uniprot50 \| AF-A0A2D8HMA2-F1-MODEL\_V4 \| 1.0 \| 8.57e-19 \| 676 \| 0.232 \| 421 \| 279 \| 17 \| 1 \| 402 \| 9 \| 404 \| Integrase \| Integrase; afdb-uniprot50 \| AF-A0A522KQT0-F1-MODEL\_V4 \| 1.0 \| 3.255e-18 \| 676 \| 0.216 \| 435 \| 278 \| 19 \| 1 \| 402 \| 13 \| 417 \| DUF4102 domain-containing protein \| DUF4102 domain-containing protein; afdb-uniprot50 \| AF-A0A7C8HSY9-F1-MODEL\_V4 \| 1.0 \| 4.837e-19 \| 675 \| 0.212 \| 419 \| 279 \| 19 \| 2 \| 403 \| 3 \| 387 \| Prophage integrase IntA \| Prophage integrase IntA; afdb-uniprot50 \| AF-A0A4P0Z7G3-F1-MODEL\_V4 \| 1.0 \| 6.653e-18 \| 675 \| 0.209 \| 429 \| 278 \| 18 \| 2 \| 402 \| 8 \| 403 \| Phage integrase \| Phage integrase; afdb-uniprot50 \| AF-A0A365QGV4-F1-MODEL\_V4 \| 1.0 \| 3.304e-19 \| 675 \| 0.219 \| 441 \| 272 \| 23 \| 2 \| 403 \| 7 \| 414 \| Integrase \| Integrase; afdb-uniprot50 \| AF-A0A3S0XF73-F1-MODEL\_V4 \| 1.0 \| 8.594e-16 \| 674 \| 0.276 \| 329 \| 200 \| 12 \| 92 \| 402 \| 6 \| 314 \| Site-specific integrase \| Site-specific integrase; afdb-uniprot50 \| AF-A0A1G0ESB5-F1-MODEL\_V4 \| 1.0 \| 6.139e-19 \| 674 \| 0.24 \| 428 \| 264 \| 20 \| 2 \| 402 \| 3 \| 396 \| Integrase \| Integrase; afdb-uniprot50 \| AF-A0A1M7YL42-F1-MODEL\_V4 \| 1.0 \| 1.196e-18 \| 673 \| 0.207 \| 425 \| 283 \| 17 \| 2 \| 402 \| 4 \| 398 \| Integrase \| Integrase; afdb-uniprot50 \| AF-A0A235EH93-F1-MODEL\_V4 \| 1.0 \| 6.753e-19 \| 672 \| 0.216 \| 421 \| 278 \| 19 \| 2 \| 403 \| 3 \| 390 \| Integrase \| Integrase; afdb-uniprot50 \| AF-A0A847R4Z8-F1-MODEL\_V4 \| 1.0 \| 1.837e-18 \| 671 \| 0.178 \| 419 \| 297 \| 17 \| 1 \| 403 \| 1 \| 388 \| Integrase arm-type DNA-binding domain-containing protein \| Integrase arm-type DNA-binding domain-containing protein; afdb-uniprot50 \| AF-A0A1S8CFL3-F1-MODEL\_V4 \| 1.0 \| 1.518e-18 \| 671 \| 0.214 \| 419 \| 280 \| 17 \| 2 \| 402 \| 3 \| 390 \| Integrase \| Integrase; afdb-uniprot50 \| AF-A0A2V8IYL9-F1-MODEL\_V4 \| 1.0 \| 6.343e-18 \| 671 \| 0.214 \| 429 \| 280 \| 21 \| 2 \| 402 \| 6 \| 405 \| Uncharacterized protein \| Uncharacterized protein; afdb-uniprot50 \| AF-A0A4P7R7H7-F1-MODEL\_V4 \| 1.0 \| 1.752e-18 \| 671 \| 0.222 \| 418 \| 280 \| 19 \| 2 \| 402 \| 3 \| 392 \| DUF4102 domain-containing protein \| DUF4102 domain-containing protein; afdb-uniprot50 \| AF-A0A1P8UJ48-F1-MODEL\_V4 \| 1.0 \| 3.103e-18 \| 670 \| 0.213 \| 435 \| 279 \| 18 \| 2 \| 402 \| 7 \| 412 \| Uncharacterized protein \| Uncharacterized protein; afdb-uniprot50 \| AF-A0A3B0YJF4-F1-MODEL\_V4 \| 1.0 \| 1.67e-18 \| 669 \| 0.205 \| 419 \| 282 \| 19 \| 2 \| 403 \| 10 \| 394 \| Integrase \| Integrase; afdb-uniprot50 \| AF-Q31UY1-F1-MODEL\_V4 \| 1.0 \| 3.255e-18 \| 669 \| 0.205 \| 428 \| 287 \| 18 \| 2 \| 402 \| 80 \| 481 \| Putative integrase \| Putative integrase; afdb-uniprot50 \| AF-A0A329B445-F1-MODEL\_V4 \| 1.0 \| 1.448e-18 \| 668 \| 0.223 \| 420 \| 276 \| 18 \| 1 \| 402 \| 1 \| 388 \| Integrase \| Integrase; afdb-uniprot50 \| AF-A0A2N6DE38-F1-MODEL\_V4 \| 1.0 \| 1.592e-18 \| 668 \| 0.201 \| 422 \| 284 \| 20 \| 2 \| 402 \| 3 \| 392 \| Integrase \| Integrase; afdb-uniprot50 \| AF-A0A7Y6D5N6-F1-MODEL\_V4 \| 1.0 \| 9.887e-19 \| 668 \| 0.217 \| 418 \| 280 \| 17 \| 2 \| 402 \| 3 \| 390 \| Integrase arm-type DNA-binding domain-containing protein \| Integrase arm-type DNA-binding domain-containing protein; afdb-uniprot50 \| AF-A0A1H4U7X0-F1-MODEL\_V4 \| 1.0 \| 6.753e-19 \| 668 \| 0.212 \| 465 \| 293 \| 19 \| 1 \| 402 \| 2 \| 456 \| Phage integrase family protein \| Phage integrase family protein; afdb-uniprot50 \| AF-A0A1C3H4D2-F1-MODEL\_V4 \| 1.0 \| 1.67e-18 \| 667 \| 0.208 \| 423 \| 278 \| 20 \| 1 \| 402 \| 1 \| 387 \| Integrase \| Integrase; afdb-uniprot50 \| AF-A0A2S5NSX3-F1-MODEL\_V4 \| 1.0 \| 2.821e-18 \| 667 \| 0.223 \| 420 \| 280 \| 20 \| 2 \| 402 \| 3 \| 395 \| Integrase \| Integrase; afdb-uniprot50 \| AF-A6QCY9-F1-MODEL\_V4 \| 1.0 \| 1.124e-17 \| 667 \| 0.219 \| 424 \| 279 \| 19 \| 2 \| 402 \| 8 \| 402 \| Site-specific recombinase, phage integrase family \| Site-specific recombinase, phage integrase family; afdb-uniprot50 \| AF-A0A1Q3RET7-F1-MODEL\_V4 \| 1.0 \| 3.103e-18 \| 666 \| 0.221 \| 424 \| 275 \| 20 \| 2 \| 402 \| 3 \| 394 \| Integrase \| Integrase; afdb-uniprot50 \| AF-A0A2P5IL45-F1-MODEL\_V4 \| 1.0 \| 1.67e-18 \| 666 \| 0.218 \| 425 \| 275 \| 20 \| 2 \| 403 \| 3 \| 393 \| Integrase \| Integrase; afdb-uniprot50 \| AF-W7WZL9-F1-MODEL\_V4 \| 1.0 \| 8.171e-19 \| 666 \| 0.227 \| 413 \| 264 \| 18 \| 20 \| 402 \| 50 \| 437 \| Prophage CP4-57 integrase \| Prophage CP4-57 integrase; afdb-uniprot50 \| AF-A0A6N7GTQ0-F1-MODEL\_V4 \| 1.0 \| 5.766e-18 \| 665 \| 0.244 \| 417 \| 269 \| 17 \| 2 \| 402 \| 30 \| 416 \| Tyrosine-type recombinase/integrase \| Tyrosine-type recombinase/integrase; afdb-uniprot50 \| AF-A0A443UGA5-F1-MODEL\_V4 \| 1.0 \| 7.082e-19 \| 664 \| 0.201 \| 432 \| 282 \| 15 \| 2 \| 402 \| 4 \| 403 \| Site-specific integrase \| Site-specific integrase; afdb-uniprot50 \| AF-A0A0J1CJU1-F1-MODEL\_V4 \| 1.0 \| 2.959e-18 \| 664 \| 0.222 \| 423 \| 275 \| 19 \| 1 \| 402 \| 2 \| 391 \| Integrase \| Integrase; afdb-uniprot50 \| AF-C0N6F7-F1-MODEL\_V4 \| 1.0 \| 2.565e-18 \| 663 \| 0.189 \| 427 \| 289 \| 20 \| 2 \| 402 \| 3 \| 398 \| Site-specific recombinase, phage integrase family protein \| Site-specific recombinase, phage integrase family protein; afdb-uniprot50 \| AF-A0A2N9L337-F1-MODEL\_V4 \| 1.0 \| 9.887e-19 \| 663 \| 0.189 \| 422 \| 290 \| 19 \| 2 \| 402 \| 3 \| 393 \| Phage integrase \| Phage integrase; afdb-uniprot50 \| AF-W7WT06-F1-MODEL\_V4 \| 1.0 \| 6.653e-18 \| 662 \| 0.207 \| 380 \| 257 \| 16 \| 44 \| 403 \| 6 \| 361 \| Prophage CP4-57 integrase \| Prophage CP4-57 integrase; afdb-uniprot50 \| AF-A0A509IV90-F1-MODEL\_V4 \| 1.0 \| 2.297e-17 \| 662 \| 0.211 \| 383 \| 263 \| 17 \| 36 \| 402 \| 2 \| 361 \| Integrase \| Integrase; afdb-uniprot50 \| AF-A0A550FGH2-F1-MODEL\_V4 \| 1.0 \| 2.223e-18 \| 662 \| 0.209 \| 420 \| 286 \| 14 \| 2 \| 402 \| 8 \| 400 \| DUF4102 domain-containing protein \| DUF4102 domain-containing protein; afdb-uniprot50 \| AF-A0A5W0NCW6-F1-MODEL\_V4 \| 1.0 \| 9.74e-18 \| 661 \| 0.235 \| 417 \| 274 \| 17 \| 2 \| 402 \| 3 \| 390 \| Tyrosine-type recombinase/integrase \| Tyrosine-type recombinase/integrase; afdb-uniprot50 \| AF-A0A1G3KD23-F1-MODEL\_V4 \| 1.0 \| 1.255e-18 \| 661 \| 0.217 \| 427 \| 275 \| 23 \| 2 \| 402 \| 3 \| 396 \| Integrase \| Integrase; afdb-uniprot50 \| AF-A0A6C1P3F8-F1-MODEL\_V4 \| 1.0 \| 3.103e-18 \| 661 \| 0.227 \| 409 \| 277 \| 16 \| 9 \| 403 \| 9 \| 392 \| Site-specific integrase \| Site-specific integrase; afdb-uniprot50 \| AF-A0A2E0A4M5-F1-MODEL\_V4 \| 1.0 \| 3.414e-18 \| 660 \| 0.232 \| 418 \| 269 \| 18 \| 2 \| 402 \| 3 \| 385 \| Integrase \| Integrase; afdb-uniprot50 \| AF-A0A2N8S9G8-F1-MODEL\_V4 \| 1.0 \| 1.518e-18 \| 660 \| 0.23 \| 451 \| 272 \| 21 \| 11 \| 402 \| 2 \| 436 \| Integrase \| Integrase; afdb-uniprot50 \| AF-A0A7W4PUN0-F1-MODEL\_V4 \| 1.0 \| 2.12e-18 \| 659 \| 0.22 \| 421 \| 275 \| 18 \| 2 \| 402 \| 3 \| 390 \| Tyrosine-type recombinase/integrase \| Tyrosine-type recombinase/integrase; afdb-uniprot50 \| AF-A0A2D6TQU3-F1-MODEL\_V4 \| 1.0 \| 1.927e-18 \| 659 \| 0.207 \| 425 \| 281 \| 19 \| 2 \| 403 \| 3 \| 394 \| Integrase \| Integrase; afdb-uniprot50 \| AF-A0A1T4YJI4-F1-MODEL\_V4 \| 1.0 \| 6.048e-18 \| 659 \| 0.222 \| 423 \| 277 \| 19 \| 2 \| 402 \| 3 \| 395 \| Integrase \| Integrase; afdb-uniprot50 \| AF-A0A3G8GZZ2-F1-MODEL\_V4 \| 1.0 \| 3.938e-18 \| 659 \| 0.207 \| 419 \| 286 \| 16 \| 2 \| 402 \| 8 \| 398 \| DUF4102 domain-containing protein \| DUF4102 domain-containing protein; afdb-uniprot50 \| AF-A0A256GC42-F1-MODEL\_V4 \| 1.0 \| 2.738e-16 \| 658 \| 0.243 \| 349 \| 228 \| 15 \| 69 \| 403 \| 2 \| 328 \| Phage integrase family protein \| Phage integrase family protein; afdb-uniprot50 \| AF-A0A7C7W234-F1-MODEL\_V4 \| 1.0 \| 1.141e-18 \| 658 \| 0.239 \| 410 \| 250 \| 19 \| 1 \| 379 \| 2 \| 380 \| Site-specific integrase \| Site-specific integrase; afdb-uniprot50 \| AF-A0A6M1TNW1-F1-MODEL\_V4 \| 1.0 \| 1.022e-17 \| 657 \| 0.224 \| 436 \| 276 \| 23 \| 2 \| 402 \| 6 \| 414 \| Tyrosine-type recombinase/integrase \| Tyrosine-type recombinase/integrase; afdb-uniprot50 \| AF-A0A1B4YTI6-F1-MODEL\_V4 \| 1.0 \| 6.653e-18 \| 657 \| 0.21 \| 423 \| 276 \| 20 \| 2 \| 402 \| 8 \| 394 \| Symbiosis island integrase \| Symbiosis island integrase; afdb-uniprot50 \| AF-A0A4Q6XBY6-F1-MODEL\_V4 \| 1.0 \| 9.887e-19 \| 656 \| 0.225 \| 430 \| 273 \| 23 \| 1 \| 403 \| 7 \| 403 \| DUF4102 domain-containing protein \| DUF4102 domain-containing protein; afdb-uniprot50 \| AF-A0A396S6Q7-F1-MODEL\_V4 \| 1.0 \| 4.332e-18 \| 656 \| 0.209 \| 454 \| 291 \| 18 \| 1 \| 402 \| 3 \| 440 \| Integrase \| Integrase; afdb-uniprot50 \| AF-A0A376BJV7-F1-MODEL\_V4 \| 1.0 \| 2.021e-18 \| 655 \| 0.222 \| 422 \| 275 \| 17 \| 1 \| 402 \| 1 \| 389 \| Prophage CPS-53 integrase \| Prophage CPS-53 integrase; afdb-uniprot50 \| AF-A0A850H6I7-F1-MODEL\_V4 \| 1.0 \| 8.05e-18 \| 654 \| 0.207 \| 415 \| 274 \| 19 \| 1 \| 402 \| 1 \| 373 \| Tyrosine-type recombinase/integrase \| Tyrosine-type recombinase/integrase; afdb-uniprot50 \| AF-A0A2G8BY28-F1-MODEL\_V4 \| 1.0 \| 4.332e-18 \| 654 \| 0.201 \| 417 \| 287 \| 18 \| 1 \| 401 \| 1 \| 387 \| Integrase \| Integrase; afdb-uniprot50 \| AF-A0A3S4N3C8-F1-MODEL\_V4 \| 1.0 \| 8.05e-18 \| 654 \| 0.213 \| 416 \| 286 \| 17 \| 2 \| 403 \| 3 \| 391 \| Prophage CPS-53 integrase \| Prophage CPS-53 integrase; afdb-uniprot50 \| AF-A0A3A6TZY1-F1-MODEL\_V4 \| 1.0 \| 3.255e-18 \| 653 \| 0.214 \| 414 \| 277 \| 18 \| 1 \| 399 \| 1 \| 381 \| DUF4102 domain-containing protein \| DUF4102 domain-containing protein; afdb-uniprot50 \| AF-A0A827TY01-F1-MODEL\_V4 \| 1.0 \| 1.296e-17 \| 653 \| 0.212 \| 418 \| 281 \| 17 \| 1 \| 403 \| 1 \| 385 \| Prophage integrase IntS \| Prophage integrase IntS; afdb-uniprot50 \| AF-A0A5U9KLB9-F1-MODEL\_V4 \| 1.0 \| 8.57e-19 \| 653 \| 0.208 \| 423 \| 281 \| 20 \| 1 \| 403 \| 1 \| 389 \| DUF4102 domain-containing protein \| DUF4102 domain-containing protein; afdb-uniprot50 \| AF-A0A317T8S8-F1-MODEL\_V4 \| 1.0 \| 4.765e-18 \| 653 \| 0.211 \| 420 \| 284 \| 20 \| 2 \| 403 \| 8 \| 398 \| Integrase \| Integrase; afdb-uniprot50 \| AF-A0A1V2BER2-F1-MODEL\_V4 \| 1.0 \| 4.544e-18 \| 653 \| 0.21 \| 423 \| 280 \| 20 \| 2 \| 402 \| 8 \| 398 \| Tyr recombinase domain-containing protein \| Tyr recombinase domain-containing protein; afdb-uniprot50 \| AF-A0A345DDB6-F1-MODEL\_V4 \| 1.0 \| 8.855e-18 \| 653 \| 0.2 \| 424 \| 285 \| 19 \| 1 \| 402 \| 6 \| 397 \| Prophage integrase IntA \| Prophage integrase IntA; afdb-uniprot50 \| AF-A0A3V2YQD8-F1-MODEL\_V4 \| 1.0 \| 2.565e-18 \| 652 \| 0.204 \| 421 \| 285 \| 19 \| 1 \| 403 \| 1 \| 389 \| DUF4102 domain-containing protein \| DUF4102 domain-containing protein; afdb-uniprot50 \| AF-A0A6N0IKJ5-F1-MODEL\_V4 \| 1.0 \| 4.998e-18 \| 652 \| 0.207 \| 420 \| 282 \| 17 \| 2 \| 402 \| 3 \| 390 \| Tyrosine-type recombinase/integrase \| Tyrosine-type recombinase/integrase; afdb-uniprot50 \| AF-A0A5U6SL20-F1-MODEL\_V4 \| 1.0 \| 1.088e-18 \| 652 \| 0.213 \| 431 \| 270 \| 20 \| 2 \| 403 \| 3 \| 393 \| DUF4102 domain-containing protein \| DUF4102 domain-containing protein; afdb-uniprot50 \| AF-A0A1J5TKJ8-F1-MODEL\_V4 \| 1.0 \| 3.414e-18 \| 652 \| 0.21 \| 428 \| 279 \| 19 \| 1 \| 402 \| 8 \| 402 \| Prophage CP4-57 integrase \| Prophage CP4-57 integrase; afdb-uniprot50 \| AF-A0A1M7DHR6-F1-MODEL\_V4 \| 1.0 \| 1.196e-18 \| 652 \| 0.205 \| 423 \| 283 \| 18 \| 2 \| 402 \| 28 \| 419 \| Integrase \| Integrase; afdb-uniprot50 \| AF-A0A841JT67-F1-MODEL\_V4 \| 1.0 \| 1.022e-17 \| 651 \| 0.191 \| 422 \| 290 \| 16 \| 1 \| 402 \| 2 \| 392 \| Integrase \| Integrase; afdb-uniprot50 \| AF-A0A839LID3-F1-MODEL\_V4 \| 1.0 \| 6.977e-18 \| 650 \| 0.221 \| 424 \| 275 \| 19 \| 1 \| 402 \| 7 \| 397 \| Tyrosine-type recombinase/integrase \| Tyrosine-type recombinase/integrase; afdb-uniprot50 \| AF-A0A7S7UC04-F1-MODEL\_V4 \| 1.0 \| 9.74e-18 \| 650 \| 0.189 \| 427 \| 288 \| 19 \| 2 \| 402 \| 3 \| 397 \| Uncharacterized protein \| Uncharacterized protein; afdb-uniprot50 \| AF-A0A851GY06-F1-MODEL\_V4 \| 1.0 \| 3.938e-18 \| 650 \| 0.224 \| 427 \| 280 \| 17 \| 2 \| 402 \| 50 \| 451 \| Tyrosine-type recombinase/integrase \| Tyrosine-type recombinase/integrase; afdb-uniprot50 \| AF-A0A257G8F0-F1-MODEL\_V4 \| 1.0 \| 9.74e-18 \| 649 \| 0.208 \| 426 \| 277 \| 21 \| 2 \| 402 \| 3 \| 393 \| Integrase \| Integrase; afdb-uniprot50 \| AF-A0A7C1MG73-F1-MODEL\_V4 \| 1.0 \| 2.223e-18 \| 649 \| 0.196 \| 422 \| 283 \| 19 \| 1 \| 402 \| 3 \| 388 \| Site-specific integrase \| Site-specific integrase; afdb-uniprot50 \| AF-A0A1I2A527-F1-MODEL\_V4 \| 1.0 \| 2.821e-18 \| 649 \| 0.21 \| 422 \| 280 \| 21 \| 1 \| 402 \| 1 \| 389 \| Integrase \| Integrase; afdb-uniprot50 \| AF-A0A158DHE5-F1-MODEL\_V4 \| 1.0 \| 4.998e-18 \| 649 \| 0.22 \| 421 \| 274 \| 21 \| 2 \| 402 \| 8 \| 394 \| Integrase \| Integrase; afdb-uniprot50 \| AF-A0A059FPK9-F1-MODEL\_V4 \| 1.0 \| 1.898e-17 \| 648 \| 0.227 \| 426 \| 269 \| 24 \| 2 \| 402 \| 3 \| 393 \| Phage integrase family site specific recombinase \| Phage integrase family site specific recombinase; afdb-uniprot50 \| AF-E5Y508-F1-MODEL\_V4 \| 1.0 \| 4.998e-18 \| 648 \| 0.199 \| 422 \| 280 \| 19 \| 9 \| 403 \| 6 \| 396 \| Uncharacterized protein \| Uncharacterized protein; afdb-uniprot50 \| AF-A0A381EDH6-F1-MODEL\_V4 \| 1.0 \| 2.223e-18 \| 648 \| 0.224 \| 419 \| 275 \| 15 \| 1 \| 402 \| 1 \| 386 \| Prophage CPS-53 integrase \| Prophage CPS-53 integrase; afdb-uniprot50 \| AF-A0A437QDT5-F1-MODEL\_V4 \| 1.0 \| 4.131e-18 \| 648 \| 0.224 \| 419 \| 276 \| 19 \| 2 \| 402 \| 3 \| 390 \| DUF4102 domain-containing protein \| DUF4102 domain-containing protein; afdb-uniprot50 \| AF-A0A1I7I1V6-F1-MODEL\_V4 \| 1.0 \| 7.675e-18 \| 648 \| 0.215 \| 441 \| 279 \| 21 \| 15 \| 402 \| 1 \| 427 \| Core-binding (CB) domain-containing protein \| Core-binding (CB) domain-containing protein; afdb-uniprot50 \| AF-A0A1L3ZVP2-F1-MODEL\_V4 \| 1.0 \| 4.544e-18 \| 647 \| 0.221 \| 419 \| 262 \| 21 \| 2 \| 403 \| 4 \| 375 \| Uncharacterized protein \| Uncharacterized protein; afdb-uniprot50 \| AF-A0A0F2QKR9-F1-MODEL\_V4 \| 1.0 \| 4.998e-18 \| 647 \| 0.192 \| 415 \| 285 \| 17 \| 13 \| 402 \| 5 \| 394 \| Tyr recombinase domain-containing protein \| Tyr recombinase domain-containing protein; afdb-uniprot50 \| AF-A0A1W1BZB0-F1-MODEL\_V4 \| 1.0 \| 3.58e-18 \| 647 \| 0.183 \| 426 \| 293 \| 18 \| 2 \| 402 \| 8 \| 403 \| Integrase \| Integrase; afdb-uniprot50 \| AF-A0A212JFC8-F1-MODEL\_V4 \| 1.0 \| 7.675e-18 \| 647 \| 0.213 \| 421 \| 288 \| 19 \| 1 \| 402 \| 5 \| 401 \| Site-specific recombinase, phage integrase family \| Site-specific recombinase, phage integrase family; afdb-uniprot50 \| AF-H1Y3L2-F1-MODEL\_V4 \| 1.0 \| 1.726e-17 \| 647 \| 0.209 \| 415 \| 286 \| 17 \| 2 \| 402 \| 3 \| 389 \| Integrase family protein \| Integrase family protein; afdb-uniprot50 \| AF-A0A1F0DZT6-F1-MODEL\_V4 \| 1.0 \| 7.318e-18 \| 646 \| 0.219 \| 419 \| 272 \| 20 \| 1 \| 400 \| 1 \| 383 \| Tyr recombinase domain-containing protein \| Tyr recombinase domain-containing protein; afdb-uniprot50 \| AF-A0A2D6EM10-F1-MODEL\_V4 \| 1.0 \| 1.36e-17 \| 646 \| 0.206 \| 426 \| 283 \| 21 \| 2 \| 403 \| 3 \| 397 \| Integrase \| Integrase; afdb-uniprot50 \| AF-A0A2X1WMX6-F1-MODEL\_V4 \| 1.0 \| 1.81e-17 \| 646 \| 0.217 \| 423 \| 268 \| 19 \| 1 \| 402 \| 5 \| 385 \| Prophage CP4-57 integrase \| Prophage CP4-57 integrase; afdb-uniprot50 \| AF-A0A1G0XRY8-F1-MODEL\_V4 \| 1.0 \| 1.7e-16 \| 645 \| 0.215 \| 358 \| 247 \| 13 \| 60 \| 402 \| 1 \| 339 \| Integrase \| Integrase; afdb-uniprot50 \| AF-A0A846UIU9-F1-MODEL\_V4 \| 1.0 \| 1.36e-17 \| 645 \| 0.214 \| 397 \| 266 \| 18 \| 25 \| 402 \| 4 \| 373 \| Integrase \| Integrase; afdb-uniprot50 \| AF-A0A353D6A0-F1-MODEL\_V4 \| 1.0 \| 5.242e-18 \| 645 \| 0.211 \| 425 \| 279 \| 21 \| 1 \| 403 \| 11 \| 401 \| Tyr recombinase domain-containing protein \| Tyr recombinase domain-containing protein; afdb-uniprot50 \| AF-I2DN67-F1-MODEL\_V4 \| 1.0 \| 1.898e-17 \| 644 \| 0.213 \| 421 \| 279 \| 21 \| 2 \| 403 \| 3 \| 390 \| Integrase \| Integrase; afdb-uniprot50 \| AF-A0A486XD46-F1-MODEL\_V4 \| 1.0 \| 1.296e-17 \| 644 \| 0.191 \| 423 \| 286 \| 19 \| 2 \| 402 \| 8 \| 396 \| Prophage CP4-57 integrase \| Prophage CP4-57 integrase; afdb-uniprot50 \| AF-A0A507WKG0-F1-MODEL\_V4 \| 1.0 \| 4.765e-18 \| 644 \| 0.208 \| 423 \| 284 \| 18 \| 2 \| 402 \| 3 \| 396 \| DUF4102 domain-containing protein \| DUF4102 domain-containing protein; afdb-uniprot50 \| AF-A0A845SLH3-F1-MODEL\_V4 \| 1.0 \| 8.855e-18 \| 644 \| 0.215 \| 422 \| 279 \| 20 \| 2 \| 402 \| 8 \| 398 \| Tyrosine-type recombinase/integrase \| Tyrosine-type recombinase/integrase; afdb-uniprot50 \| AF-P37326-F1-MODEL\_V4 \| 1.0 \| 9.287e-18 \| 643 \| 0.218 \| 417 \| 280 \| 18 \| 1 \| 403 \| 1 \| 385 \| Prophage integrase IntS \| Prophage integrase IntS; afdb-uniprot50 \| AF-A0A0K8NTH3-F1-MODEL\_V4 \| 1.0 \| 4.998e-18 \| 643 \| 0.208 \| 393 \| 263 \| 17 \| 36 \| 403 \| 3 \| 372 \| Phage integrase \| Phage integrase; afdb-uniprot50 \| AF-A0A2D0JZY9-F1-MODEL\_V4 \| 1.0 \| 8.05e-18 \| 643 \| 0.207 \| 429 \| 273 \| 23 \| 2 \| 402 \| 3 \| 392 \| Integrase \| Integrase; afdb-uniprot50 \| AF-A0A290XAM6-F1-MODEL\_V4 \| 1.0 \| 7.791e-19 \| 643 \| 0.226 \| 429 \| 269 \| 20 \| 1 \| 402 \| 1 \| 393 \| Integrase \| Integrase; afdb-uniprot50 \| AF-A0A523L477-F1-MODEL\_V4 \| 1.0 \| 1.569e-17 \| 643 \| 0.174 \| 419 \| 293 \| 21 \| 2 \| 402 \| 6 \| 389 \| DUF4102 domain-containing protein \| DUF4102 domain-containing protein; afdb-uniprot50 \| AF-A0A0J6KK27-F1-MODEL\_V4 \| 1.0 \| 2.021e-18 \| 643 \| 0.202 \| 430 \| 283 \| 18 \| 2 \| 403 \| 3 \| 400 \| Tyr recombinase domain-containing protein \| Tyr recombinase domain-containing protein; afdb-uniprot50 \| AF-A0A3M9XVK5-F1-MODEL\_V4 \| 1.0 \| 3.812e-19 \| 643 \| 0.24 \| 415 \| 266 \| 18 \| 1 \| 398 \| 17 \| 399 \| Site-specific integrase \| Site-specific integrase; afdb-uniprot50 \| AF-A0A5V2R899-F1-MODEL\_V4 \| 1.0 \| 1.236e-17 \| 643 \| 0.196 \| 417 \| 284 \| 16 \| 9 \| 402 \| 11 \| 399 \| Tyrosine-type recombinase/integrase \| Tyrosine-type recombinase/integrase; afdb-uniprot50 \| AF-A0A2C8ZKJ8-F1-MODEL\_V4 \| 1.0 \| 1.022e-17 \| 643 \| 0.233 \| 415 \| 270 \| 23 \| 9 \| 402 \| 75 \| 462 \| Integrase \| Integrase; afdb-uniprot50 \| AF-A0A389ML60-F1-MODEL\_V4 \| 1.0 \| 9.74e-18 \| 642 \| 0.221 \| 415 \| 270 \| 22 \| 2 \| 401 \| 13 \| 389 \| Integrase \| Integrase; afdb-uniprot50 \| AF-B8J055-F1-MODEL\_V4 \| 1.0 \| 2.69e-18 \| 641 \| 0.204 \| 421 \| 281 \| 20 \| 2 \| 402 \| 7 \| 393 \| Integrase family protein \| Integrase family protein; afdb-uniprot50 \| AF-A0A1F0DV58-F1-MODEL\_V4 \| 1.0 \| 3.255e-18 \| 641 \| 0.214 \| 425 \| 277 \| 19 \| 2 \| 402 \| 3 \| 394 \| Integrase \| Integrase; afdb-uniprot50 \| AF-A0A1Q8SPE1-F1-MODEL\_V4 \| 1.0 \| 4.131e-18 \| 640 \| 0.212 \| 418 \| 281 \| 18 \| 2 \| 401 \| 3 \| 390 \| Tyr recombinase domain-containing protein \| Tyr recombinase domain-containing protein; afdb-uniprot50 \| AF-A0A7R6PGN4-F1-MODEL\_V4 \| 1.0 \| 4.998e-18 \| 640 \| 0.191 \| 423 \| 290 \| 20 \| 1 \| 402 \| 1 \| 392 \| Phage integrase family protein \| Phage integrase family protein; afdb-uniprot50 \| AF-A0A7W0GKX6-F1-MODEL\_V4 \| 1.0 \| 7.675e-18 \| 640 \| 0.208 \| 422 \| 283 \| 18 \| 2 \| 402 \| 6 \| 397 \| Tyrosine-type recombinase/integrase \| Tyrosine-type recombinase/integrase; afdb-uniprot50 \| AF-A0A5P3MQF5-F1-MODEL\_V4 \| 1.0 \| 7.318e-18 \| 640 \| 0.232 \| 421 \| 271 \| 19 \| 2 \| 402 \| 3 \| 391 \| DUF4102 domain-containing protein \| DUF4102 domain-containing protein; afdb-uniprot50 \| AF-A0A3N1JQV1-F1-MODEL\_V4 \| 1.0 \| 8.855e-18 \| 640 \| 0.185 \| 420 \| 295 \| 18 \| 1 \| 402 \| 50 \| 440 \| Integrase \| Integrase; afdb-uniprot50 \| AF-A0A6N7EZ98-F1-MODEL\_V4 \| 1.0 \| 2.19e-17 \| 639 \| 0.185 \| 415 \| 292 \| 19 \| 2 \| 402 \| 3 \| 385 \| Tyrosine-type recombinase/integrase \| Tyrosine-type recombinase/integrase; afdb-uniprot50 \| AF-A0A2E1TM79-F1-MODEL\_V4 \| 1.0 \| 1.898e-17 \| 639 \| 0.2 \| 420 \| 285 \| 20 \| 2 \| 403 \| 3 \| 389 \| Integrase \| Integrase; afdb-uniprot50 \| AF-W2UIP2-F1-MODEL\_V4 \| 1.0 \| 4.131e-18 \| 638 \| 0.189 \| 427 \| 287 \| 21 \| 2 \| 402 \| 3 \| 396 \| Prophage CP4-57 integrase \| Prophage CP4-57 integrase; afdb-uniprot50 \| AF-A0A5F1HJD6-F1-MODEL\_V4 \| 1.0 \| 3.88e-17 \| 638 \| 0.215 \| 422 \| 279 \| 20 \| 2 \| 402 \| 8 \| 398 \| Integrase \| Integrase; afdb-uniprot50 \| AF-A0A1C3E882-F1-MODEL\_V4 \| 1.0 \| 1.726e-17 \| 637 \| 0.194 \| 421 \| 290 \| 16 \| 2 \| 403 \| 3 \| 393 \| Tyr recombinase domain-containing protein \| Tyr recombinase domain-containing protein; afdb-uniprot50 \| AF-A0A6L3SXM2-F1-MODEL\_V4 \| 1.0 \| 1.569e-17 \| 637 \| 0.215 \| 418 \| 275 \| 18 \| 2 \| 402 \| 13 \| 394 \| Tyrosine-type recombinase/integrase \| Tyrosine-type recombinase/integrase; afdb-uniprot50 \| AF-A0A5C7R8L9-F1-MODEL\_V4 \| 1.0 \| 1.426e-17 \| 637 \| 0.193 \| 429 \| 284 \| 20 \| 1 \| 402 \| 4 \| 397 \| DUF4102 domain-containing protein \| DUF4102 domain-containing protein; afdb-uniprot50 \| AF-A0A4Q6X2P3-F1-MODEL\_V4 \| 1.0 \| 6.653e-18 \| 637 \| 0.215 \| 432 \| 278 \| 20 \| 2 \| 403 \| 12 \| 412 \| DUF4102 domain-containing protein \| DUF4102 domain-containing protein; afdb-uniprot50 \| AF-A0A2T7U8C2-F1-MODEL\_V4 \| 1.0 \| 6.977e-18 \| 636 \| 0.213 \| 421 \| 278 \| 20 \| 2 \| 402 \| 3 \| 390 \| Integrase \| Integrase; afdb-uniprot50 \| AF-A0A4C3NQ15-F1-MODEL\_V4 \| 1.0 \| 2.915e-17 \| 636 \| 0.213 \| 422 \| 280 \| 19 \| 2 \| 402 \| 8 \| 398 \| DUF4102 domain-containing protein \| DUF4102 domain-containing protein; afdb-uniprot50 \| AF-A0A7Y0GZP7-F1-MODEL\_V4 \| 1.0 \| 1.071e-17 \| 636 \| 0.215 \| 426 \| 279 \| 18 \| 2 \| 402 \| 9 \| 404 \| Integrase arm-type DNA-binding domain-containing protein \| Integrase arm-type DNA-binding domain-containing protein; afdb-uniprot50 \| AF-A0A5F0WL98-F1-MODEL\_V4 \| 1.0 \| 4.332e-18 \| 635 \| 0.216 \| 420 \| 276 \| 18 \| 2 \| 402 \| 3 \| 388 \| DUF4102 domain-containing protein \| DUF4102 domain-containing protein; afdb-uniprot50 \| AF-A0A848HR69-F1-MODEL\_V4 \| 1.0 \| 2.223e-18 \| 635 \| 0.227 \| 422 \| 273 \| 17 \| 2 \| 402 \| 3 \| 392 \| Tyrosine-type recombinase/integrase \| Tyrosine-type recombinase/integrase; afdb-uniprot50 \| AF-A0A5H7WS47-F1-MODEL\_V4 \| 1.0 \| 1.179e-17 \| 635 \| 0.207 \| 419 \| 283 \| 17 \| 2 \| 402 \| 3 \| 390 \| DUF4102 domain-containing protein \| DUF4102 domain-containing protein; afdb-uniprot50 \| AF-A0A1I7NE20-F1-MODEL\_V4 \| 1.0 \| 1.496e-17 \| 635 \| 0.226 \| 419 \| 273 \| 19 \| 2 \| 402 \| 5 \| 390 \| Site-specific recombinase XerD \| Site-specific recombinase XerD; afdb-uniprot50 \| AF-A0A3M5ZHM3-F1-MODEL\_V4 \| 1.0 \| 6.874e-17 \| 635 \| 0.21 \| 422 \| 282 \| 19 \| 2 \| 403 \| 81 \| 471 \| Integrase \| Integrase; afdb-uniprot50 \| AF-A0A376D7E0-F1-MODEL\_V4 \| 1.0 \| 2.779e-17 \| 635 \| 0.206 \| 422 \| 283 \| 18 \| 2 \| 402 \| 8 \| 398 \| Integrase from prophage \| Integrase from prophage; afdb-uniprot50 \| AF-A0A6H2DRV2-F1-MODEL\_V4 \| 1.0 \| 1.179e-17 \| 634 \| 0.238 \| 411 \| 266 \| 17 \| 2 \| 403 \| 6 \| 378 \| Tyrosine-type recombinase/integrase \| Tyrosine-type recombinase/integrase; afdb-uniprot50 \| AF-A0A0M0T6P3-F1-MODEL\_V4 \| 1.0 \| 3.527e-17 \| 634 \| 0.183 \| 425 \| 288 \| 21 \| 2 \| 402 \| 3 \| 392 \| Uncharacterized protein \| Uncharacterized protein; afdb-uniprot50 \| AF-A0A1B3M9C7-F1-MODEL\_V4 \| 1.0 \| 2.297e-17 \| 634 \| 0.209 \| 424 \| 279 \| 18 \| 2 \| 402 \| 3 \| 393 \| Phage integrase family protein \| Phage integrase family protein; afdb-uniprot50 \| AF-A0A1N6DLZ0-F1-MODEL\_V4 \| 1.0 \| 2.565e-18 \| 634 \| 0.18 \| 422 \| 291 \| 17 \| 2 \| 402 \| 3 \| 390 \| Integrase \| Integrase; afdb-uniprot50 \| AF-A0A2N2UYZ7-F1-MODEL\_V4 \| 1.0 \| 1.179e-17 \| 634 \| 0.216 \| 425 \| 275 \| 21 \| 2 \| 402 \| 8 \| 398 \| Integrase \| Integrase; afdb-uniprot50 \| AF-A0A7D6SXA8-F1-MODEL\_V4 \| 1.0 \| 1.726e-17 \| 634 \| 0.192 \| 420 \| 290 \| 18 \| 2 \| 402 \| 8 \| 397 \| Tyrosine-type recombinase/integrase \| Tyrosine-type recombinase/integrase; afdb-uniprot50 \| AF-A0A2R5EJF6-F1-MODEL\_V4 \| 1.0 \| 2.565e-18 \| 634 \| 0.224 \| 446 \| 263 \| 23 \| 1 \| 402 \| 1 \| 407 \| Integrase \| Integrase; afdb-uniprot50 \| AF-A0A4Q6X281-F1-MODEL\_V4 \| 1.0 \| 1.569e-17 \| 633 \| 0.225 \| 416 \| 272 \| 23 \| 9 \| 402 \| 11 \| 398 \| DUF4102 domain-containing protein \| DUF4102 domain-containing protein; afdb-uniprot50 \| AF-A0A822P193-F1-MODEL\_V4 \| 1.0 \| 5.242e-18 \| 632 \| 0.22 \| 418 \| 272 \| 20 \| 1 \| 398 \| 1 \| 384 \| Putative prophage CPS-53 integrase CPS-53 (KpLE1) prophage \| Putative prophage CPS-53 integrase CPS-53 (KpLE1) prophage; afdb-uniprot50 \| AF-A0A4V4RAT5-F1-MODEL\_V4 \| 1.0 \| 2.489e-16 \| 632 \| 0.19 \| 421 \| 294 \| 20 \| 2 \| 403 \| 3 \| 395 \| DUF4102 domain-containing protein \| DUF4102 domain-containing protein; afdb-uniprot50 \| AF-A0A3A9I0N3-F1-MODEL\_V4 \| 1.0 \| 6.977e-18 \| 632 \| 0.214 \| 425 \| 275 \| 21 \| 2 \| 402 \| 16 \| 405 \| DUF4102 domain-containing protein \| DUF4102 domain-containing protein; afdb-uniprot50 \| AF-A0A6G6WPD0-F1-MODEL\_V4 \| 1.0 \| 6.977e-18 \| 632 \| 0.2 \| 430 \| 284 \| 22 \| 1 \| 402 \| 5 \| 402 \| Integrase arm-type DNA-binding domain-containing protein \| Integrase arm-type DNA-binding domain-containing protein; afdb-uniprot50 \| AF-A0A5P2H4F2-F1-MODEL\_V4 \| 1.0 \| 5.498e-18 \| 632 \| 0.197 \| 425 \| 273 \| 17 \| 1 \| 398 \| 24 \| 407 \| Tyrosine-type recombinase/integrase \| Tyrosine-type recombinase/integrase; afdb-uniprot50 \| AF-A0A2W0G8P2-F1-MODEL\_V4 \| 1.0 \| 3.699e-17 \| 631 \| 0.196 \| 417 \| 289 \| 18 \| 1 \| 403 \| 1 \| 385 \| DUF4102 domain-containing protein \| DUF4102 domain-containing protein; afdb-uniprot50 \| AF-A0A2D8CKN1-F1-MODEL\_V4 \| 1.0 \| 3.414e-18 \| 631 \| 0.211 \| 421 \| 276 \| 20 \| 2 \| 402 \| 5 \| 389 \| Uncharacterized protein \| Uncharacterized protein; afdb-uniprot50 \| AF-A0A534HSB6-F1-MODEL\_V4 \| 1.0 \| 3.938e-18 \| 631 \| 0.212 \| 418 \| 287 \| 18 \| 1 \| 402 \| 2 \| 393 \| DUF4102 domain-containing protein \| DUF4102 domain-containing protein; afdb-uniprot50 \| AF-A0A5C7LA00-F1-MODEL\_V4 \| 1.0 \| 3.755e-18 \| 631 \| 0.206 \| 432 \| 278 \| 18 \| 1 \| 402 \| 1 \| 397 \| DUF4102 domain-containing protein \| DUF4102 domain-containing protein; afdb-uniprot50 \| AF-A0A522LW18-F1-MODEL\_V4 \| 1.0 \| 1.022e-17 \| 631 \| 0.216 \| 402 \| 270 \| 19 \| 14 \| 402 \| 26 \| 395 \| Site-specific integrase \| Site-specific integrase; afdb-uniprot50 \| AF-A0A4Q7G062-F1-MODEL\_V4 \| 1.0 \| 1.296e-17 \| 631 \| 0.236 \| 431 \| 264 \| 21 \| 2 \| 403 \| 5 \| 399 \| Tyr recombinase domain-containing protein \| Tyr recombinase domain-containing protein; afdb-uniprot50 \| AF-A0A7I7D2M7-F1-MODEL\_V4 \| 1.0 \| 2.19e-17 \| 630 \| 0.203 \| 417 \| 286 \| 18 \| 1 \| 403 \| 1 \| 385 \| Prophage integrase IntS \| Prophage integrase IntS; afdb-uniprot50 \| AF-A0A3R0Q2U4-F1-MODEL\_V4 \| 1.0 \| 4.131e-18 \| 630 \| 0.212 \| 423 \| 279 \| 20 \| 1 \| 403 \| 1 \| 389 \| DUF4102 domain-containing protein \| DUF4102 domain-containing protein; afdb-uniprot50 \| AF-A0A0F2R7F3-F1-MODEL\_V4 \| 1.0 \| 6.653e-18 \| 630 \| 0.192 \| 425 \| 285 \| 21 \| 2 \| 402 \| 3 \| 393 \| Integrase \| Integrase; afdb-uniprot50 \| AF-A0A2N1ZXX0-F1-MODEL\_V4 \| 1.0 \| 1.645e-17 \| 630 \| 0.194 \| 417 \| 287 \| 18 \| 2 \| 402 \| 3 \| 386 \| Integrase \| Integrase; afdb-uniprot50 \| AF-A0A3G2N436-F1-MODEL\_V4 \| 1.0 \| 1.022e-17 \| 630 \| 0.238 \| 411 \| 257 \| 20 \| 14 \| 402 \| 81 \| 457 \| DUF4102 domain-containing protein \| DUF4102 domain-containing protein; afdb-uniprot50 \| AF-A0A4R1IAG8-F1-MODEL\_V4 \| 1.0 \| 1.426e-17 \| 629 \| 0.209 \| 425 \| 278 \| 19 \| 2 \| 402 \| 3 \| 393 \| Integrase \| Integrase; afdb-uniprot50 \| AF-A0A7D6P8E2-F1-MODEL\_V4 \| 1.0 \| 2.779e-17 \| 629 \| 0.183 \| 420 \| 294 \| 18 \| 2 \| 402 \| 8 \| 397 \| Integrase \| Integrase; afdb-uniprot50 \| AF-U7GEQ5-F1-MODEL\_V4 \| 1.0 \| 7.675e-18 \| 629 \| 0.2 \| 430 \| 276 \| 20 \| 2 \| 403 \| 3 \| 392 \| Uncharacterized protein \| Uncharacterized protein; afdb-uniprot50 \| AF-A0A7H1M906-F1-MODEL\_V4 \| 1.0 \| 2.19e-17 \| 629 \| 0.214 \| 419 \| 278 \| 20 \| 2 \| 401 \| 3 \| 389 \| Phage integrase family protein \| Phage integrase family protein; afdb-uniprot50 \| AF-A0A2D4TBE4-F1-MODEL\_V4 \| 1.0 \| 2.69e-18 \| 629 \| 0.201 \| 437 \| 289 \| 20 \| 2 \| 402 \| 10 \| 422 \| Uncharacterized protein \| Uncharacterized protein; afdb-uniprot50 \| AF-A0A6I2IYN0-F1-MODEL\_V4 \| 1.0 \| 9.15e-17 \| 628 \| 0.205 \| 413 \| 290 \| 18 \| 1 \| 403 \| 1 \| 385 \| Tyrosine-type recombinase/integrase \| Tyrosine-type recombinase/integrase; afdb-uniprot50 \| AF-D3PB12-F1-MODEL\_V4 \| 1.0 \| 1.071e-17 \| 628 \| 0.208 \| 418 \| 275 \| 22 \| 2 \| 402 \| 4 \| 382 \| Phage integrase \| Phage integrase; afdb-uniprot50 \| AF-A0A376M5B4-F1-MODEL\_V4 \| 1.0 \| 2.409e-17 \| 628 \| 0.208 \| 422 \| 282 \| 18 \| 2 \| 402 \| 8 \| 398 \| Integrase from prophage \| Integrase from prophage; afdb-uniprot50 \| AF-A0A1H8LKS2-F1-MODEL\_V4 \| 1.0 \| 1.496e-17 \| 627 \| 0.194 \| 427 \| 282 \| 21 \| 2 \| 402 \| 3 \| 393 \| Integrase \| Integrase; afdb-uniprot50 \| AF-A0A1A7PDT9-F1-MODEL\_V4 \| 1.0 \| 1.056e-16 \| 627 \| 0.19 \| 420 \| 291 \| 17 \| 1 \| 401 \| 7 \| 396 \| Tyr recombinase domain-containing protein \| Tyr recombinase domain-containing protein; afdb-uniprot50 \| AF-A0A560K1J7-F1-MODEL\_V4 \| 1.0 \| 1.898e-17 \| 627 \| 0.209 \| 420 \| 281 \| 20 \| 2 \| 402 \| 3 \| 390 \| Integrase \| Integrase; afdb-uniprot50 \| AF-A0A7U1D0Y1-F1-MODEL\_V4 \| 1.0 \| 1.296e-17 \| 627 \| 0.197 \| 441 \| 290 \| 20 \| 1 \| 402 \| 3 \| 418 \| Tyrosine-type recombinase/integrase \| Tyrosine-type recombinase/integrase; afdb-uniprot50 \| AF-A0A830ZYW7-F1-MODEL\_V4 \| 1.0 \| 3.645e-16 \| 626 \| 0.223 \| 381 \| 249 \| 15 \| 50 \| 402 \| 3 \| 364 \| Integrase \| Integrase; afdb-uniprot50 \| AF-A0A806JFH6-F1-MODEL\_V4 \| 1.0 \| 3.363e-17 \| 626 \| 0.206 \| 427 \| 282 \| 20 \| 2 \| 402 \| 8 \| 403 \| Integrase \| Integrase; afdb-uniprot50 \| AF-A0A1E4KI51-F1-MODEL\_V4 \| 1.0 \| 2.915e-17 \| 625 \| 0.195 \| 420 \| 291 \| 19 \| 1 \| 402 \| 3 \| 393 \| Tyr recombinase domain-containing protein \| Tyr recombinase domain-containing protein; afdb-uniprot50 \| AF-A0A811GGD4-F1-MODEL\_V4 \| 1.0 \| 2.19e-17 \| 625 \| 0.193 \| 423 \| 293 \| 18 \| 1 \| 403 \| 9 \| 403 \| Prophage CPS-53 integrase \| Prophage CPS-53 integrase; afdb-uniprot50 \| AF-A0A0H4P8E5-F1-MODEL\_V4 \| 1.0 \| 4.625e-16 \| 624 \| 0.244 \| 360 \| 233 \| 14 \| 67 \| 402 \| 30 \| 374 \| Integrase family protein \| Integrase family protein; afdb-uniprot50 \| AF-A0A6L5XMX1-F1-MODEL\_V4 \| 1.0 \| 2.527e-17 \| 624 \| 0.194 \| 422 \| 289 \| 20 \| 1 \| 402 \| 1 \| 391 \| DUF4102 domain-containing protein \| DUF4102 domain-containing protein; afdb-uniprot50 \| AF-A0A257HDF3-F1-MODEL\_V4 \| 1.0 \| 2.19e-17 \| 623 \| 0.215 \| 426 \| 275 \| 24 \| 2 \| 402 \| 3 \| 394 \| Integrase \| Integrase; afdb-uniprot50 \| AF-A0A5B0T1R7-F1-MODEL\_V4 \| 1.0 \| 5.416e-17 \| 623 \| 0.208 \| 423 \| 283 \| 19 \| 2 \| 403 \| 10 \| 401 \| Tyrosine-type recombinase/integrase \| Tyrosine-type recombinase/integrase; afdb-uniprot50 \| AF-A0A7M1AWK8-F1-MODEL\_V4 \| 1.0 \| 1.898e-17 \| 623 \| 0.183 \| 431 \| 295 \| 21 \| 2 \| 402 \| 8 \| 411 \| DUF4102 domain-containing protein \| DUF4102 domain-containing protein; afdb-uniprot50 \| AF-A0A657B792-F1-MODEL\_V4 \| 1.0 \| 8.724e-17 \| 623 \| 0.21 \| 422 \| 281 \| 20 \| 2 \| 402 \| 8 \| 398 \| Tyr recombinase domain-containing protein \| Tyr recombinase domain-containing protein; afdb-uniprot50 \| AF-A0A356X5P7-F1-MODEL\_V4 \| 1.0 \| 1.36e-17 \| 623 \| 0.21 \| 428 \| 287 \| 17 \| 2 \| 403 \| 7 \| 409 \| Uncharacterized protein \| Uncharacterized protein; afdb-uniprot50 \| AF-A0A7X4GIX6-F1-MODEL\_V4 \| 1.0 \| 1.569e-17 \| 623 \| 0.186 \| 450 \| 289 \| 20 \| 1 \| 403 \| 7 \| 426 \| Tyrosine-type recombinase/integrase \| Tyrosine-type recombinase/integrase; afdb-uniprot50 \| AF-A0A0A2WLL9-F1-MODEL\_V4 \| 1.0 \| 5.164e-17 \| 622 \| 0.208 \| 417 \| 284 \| 19 \| 2 \| 403 \| 8 \| 393 \| Phage integrase \| Phage integrase; afdb-uniprot50 \| AF-A0A2G2DS92-F1-MODEL\_V4 \| 1.0 \| 3.88e-17 \| 622 \| 0.218 \| 434 \| 277 \| 23 \| 2 \| 398 \| 8 \| 416 \| Integrase \| Integrase; afdb-uniprot50 \| AF-A0A2V8UYS4-F1-MODEL\_V4 \| 1.0 \| 2.229e-15 \| 621 \| 0.244 \| 348 \| 215 \| 16 \| 73 \| 402 \| 2 \| 319 \| Integrase \| Integrase; afdb-uniprot50 \| AF-A0A0M3TU49-F1-MODEL\_V4 \| 1.0 \| 6.554e-17 \| 621 \| 0.183 \| 415 \| 296 \| 18 \| 2 \| 403 \| 6 \| 390 \| Tyr recombinase domain-containing protein \| Tyr recombinase domain-containing protein; afdb-uniprot50 \| AF-A0A2L0VMP1-F1-MODEL\_V4 \| 1.0 \| 2.779e-17 \| 621 \| 0.213 \| 427 \| 273 \| 23 \| 2 \| 402 \| 3 \| 392 \| Integrase \| Integrase; afdb-uniprot50 \| AF-A0A175VIU2-F1-MODEL\_V4 \| 1.0 \| 4.924e-17 \| 621 \| 0.207 \| 424 \| 282 \| 21 \| 2 \| 402 \| 6 \| 398 \| Integrase \| Integrase; afdb-uniprot50 \| AF-X5NZA3-F1-MODEL\_V4 \| 1.0 \| 1.296e-17 \| 621 \| 0.211 \| 406 \| 268 \| 19 \| 18 \| 403 \| 1 \| 374 \| Integrase \| Integrase; afdb-uniprot50 \| AF-A0A1N6J5W7-F1-MODEL\_V4 \| 1.0 \| 1.277e-16 \| 620 \| 0.19 \| 421 \| 292 \| 19 \| 2 \| 402 \| 3 \| 394 \| Integrase \| Integrase; afdb-uniprot50 \| AF-Q5E028-F1-MODEL\_V4 \| 1.0 \| 2.915e-17 \| 620 \| 0.24 \| 424 \| 265 \| 23 \| 2 \| 401 \| 8 \| 398 \| Phage family integrase \| Phage family integrase; afdb-uniprot50 \| AF-A0A380QSD2-F1-MODEL\_V4 \| 1.0 \| 1.34e-16 \| 620 \| 0.196 \| 423 \| 283 \| 18 \| 2 \| 398 \| 8 \| 399 \| Putative prophage integrase \| Putative prophage integrase; afdb-uniprot50 \| AF-A0A198XHQ6-F1-MODEL\_V4 \| 1.0 \| 4.924e-17 \| 619 \| 0.197 \| 421 \| 271 \| 20 \| 1 \| 403 \| 1 \| 372 \| Prophage CP4-like integrase \| Prophage CP4-like integrase; afdb-uniprot50 \| AF-A3UDD1-F1-MODEL\_V4 \| 1.0 \| 1.81e-17 \| 619 \| 0.169 \| 426 \| 293 \| 18 \| 2 \| 402 \| 3 \| 392 \| Symbiosis island integrase \| Symbiosis island integrase; afdb-uniprot50 \| AF-A0A2G1AS07-F1-MODEL\_V4 \| 1.0 \| 8.318e-17 \| 619 \| 0.226 \| 420 \| 276 \| 19 \| 2 \| 402 \| 8 \| 397 \| Integrase \| Integrase; afdb-uniprot50 \| AF-A0A0D0PRG3-F1-MODEL\_V4 \| 1.0 \| 5.164e-17 \| 619 \| 0.182 \| 427 \| 297 \| 15 \| 4 \| 402 \| 8 \| 410 \| Contig\_52, whole genome shotgun sequence \| Contig\_52, whole genome shotgun sequence; afdb-uniprot50 \| AF-A0A382GF79-F1-MODEL\_V4 \| 1.0 \| 3.057e-17 \| 618 \| 0.199 \| 427 \| 284 \| 25 \| 1 \| 402 \| 1 \| 394 \| Tyr recombinase domain-containing protein \| Tyr recombinase domain-containing protein; afdb-uniprot50 \| AF-A0A328TT78-F1-MODEL\_V4 \| 1.0 \| 1.991e-17 \| 617 \| 0.212 \| 419 \| 280 \| 18 \| 1 \| 403 \| 1 \| 385 \| Putative prophage CPS-53 integrase \| Putative prophage CPS-53 integrase; afdb-uniprot50 \| AF-A0A3D1LLU3-F1-MODEL\_V4 \| 1.0 \| 1.898e-17 \| 617 \| 0.195 \| 424 \| 287 \| 19 \| 2 \| 402 \| 8 \| 400 \| Integrase \| Integrase; afdb-uniprot50 \| AF-A0A0Q0CJV1-F1-MODEL\_V4 \| 1.0 \| 1.474e-16 \| 617 \| 0.21 \| 422 \| 279 \| 22 \| 2 \| 402 \| 39 \| 427 \| Integrase \| Integrase; afdb-uniprot50 \| AF-A0A2D8ER74-F1-MODEL\_V4 \| 1.0 \| 8.318e-17 \| 616 \| 0.204 \| 426 \| 280 \| 21 \| 2 \| 402 \| 3 \| 394 \| Integrase \| Integrase; afdb-uniprot50 \| AF-A0A858BC33-F1-MODEL\_V4 \| 1.0 \| 1.161e-16 \| 616 \| 0.216 \| 421 \| 279 \| 21 \| 2 \| 402 \| 8 \| 397 \| DUF4102 domain-containing protein \| DUF4102 domain-containing protein; afdb-uniprot50 \| AF-A0A246JIN4-F1-MODEL\_V4 \| 1.0 \| 8.443e-18 \| 616 \| 0.221 \| 425 \| 272 \| 20 \| 2 \| 402 \| 3 \| 392 \| Integrase \| Integrase; afdb-uniprot50 \| AF-A0A7D7VIT3-F1-MODEL\_V4 \| 1.0 \| 6.249e-17 \| 616 \| 0.204 \| 421 \| 286 \| 18 \| 1 \| 402 \| 2 \| 392 \| Tyrosine-type recombinase/integrase \| Tyrosine-type recombinase/integrase; afdb-uniprot50 \| AF-A0A3P1XZM4-F1-MODEL\_V4 \| 1.0 \| 3.699e-17 \| 616 \| 0.208 \| 427 \| 279 \| 22 \| 2 \| 402 \| 14 \| 407 \| DUF4102 domain-containing protein \| DUF4102 domain-containing protein; afdb-uniprot50 \| AF-A0A520PEP9-F1-MODEL\_V4 \| 1.0 \| 1.296e-17 \| 616 \| 0.214 \| 448 \| 275 \| 19 \| 1 \| 402 \| 11 \| 427 \| DUF4102 domain-containing protein \| DUF4102 domain-containing protein; afdb-uniprot50 \| AF-C9M6R5-F1-MODEL\_V4 \| 1.0 \| 1.991e-17 \| 615 \| 0.199 \| 421 \| 279 \| 18 \| 1 \| 402 \| 1 \| 382 \| Site-specific recombinase, phage integrase family \| Site-specific recombinase, phage integrase family; afdb-uniprot50 \| AF-A0A345H8S3-F1-MODEL\_V4 \| 1.0 \| 1.961e-16 \| 615 \| 0.179 \| 423 \| 294 \| 21 \| 2 \| 400 \| 3 \| 396 \| DUF4102 domain-containing protein \| DUF4102 domain-containing protein; afdb-uniprot50 \| AF-A0A6G4WGV4-F1-MODEL\_V4 \| 1.0 \| 2.65e-17 \| 615 \| 0.235 \| 434 \| 268 \| 23 \| 1 \| 402 \| 4 \| 405 \| Integrase family protein \| Integrase family protein; afdb-uniprot50 \| AF-A0A426QL33-F1-MODEL\_V4 \| 1.0 \| 3.88e-17 \| 614 \| 0.185 \| 421 \| 295 \| 16 \| 2 \| 402 \| 14 \| 406 \| DUF4102 domain-containing protein \| DUF4102 domain-containing protein; afdb-uniprot50 \| AF-A0A0X8JKZ6-F1-MODEL\_V4 \| 1.0 \| 1.726e-17 \| 614 \| 0.186 \| 428 \| 282 \| 19 \| 2 \| 402 \| 3 \| 391 \| Integrase \| Integrase; afdb-uniprot50 \| AF-A0A8B4S2H0-F1-MODEL\_V4 \| 1.0 \| 3.057e-17 \| 614 \| 0.204 \| 446 \| 284 \| 22 \| 1 \| 402 \| 1 \| 419 \| Prophage CPS-53 integrase \| Prophage CPS-53 integrase; afdb-uniprot50 \| AF-A0A1I7M1P0-F1-MODEL\_V4 \| 1.0 \| 9.287e-18 \| 614 \| 0.217 \| 432 \| 272 \| 20 \| 2 \| 402 \| 6 \| 402 \| Site-specific recombinase XerD \| Site-specific recombinase XerD; afdb-uniprot50 \| AF-A0A3D3CKM6-F1-MODEL\_V4 \| 1.0 \| 9.15e-17 \| 613 \| 0.242 \| 408 \| 260 \| 19 \| 1 \| 398 \| 1 \| 369 \| Integrase \| Integrase; afdb-uniprot50 \| AF-A0A4U2UK76-F1-MODEL\_V4 \| 1.0 \| 8.318e-17 \| 613 \| 0.202 \| 420 \| 286 \| 18 \| 2 \| 402 \| 8 \| 397 \| DUF4102 domain-containing protein \| DUF4102 domain-containing protein; afdb-uniprot50 \| AF-A0A4R1VLQ5-F1-MODEL\_V4 \| 1.0 \| 9.287e-18 \| 613 \| 0.204 \| 426 \| 284 \| 20 \| 2 \| 402 \| 5 \| 400 \| Integrase \| Integrase; afdb-uniprot50 \| AF-A0A2U2CQD6-F1-MODEL\_V4 \| 1.0 \| 1.36e-17 \| 613 \| 0.185 \| 431 \| 282 \| 19 \| 2 \| 401 \| 33 \| 425 \| Integrase \| Integrase; afdb-uniprot50 \| AF-A0A254N809-F1-MODEL\_V4 \| 1.0 \| 2.331e-18 \| 613 \| 0.191 \| 465 \| 278 \| 22 \| 2 \| 402 \| 3 \| 433 \| Integrase \| Integrase; afdb-uniprot50 \| AF-A0A6L5XJW2-F1-MODEL\_V4 \| 1.0 \| 3.88e-17 \| 613 \| 0.198 \| 424 \| 284 \| 21 \| 2 \| 402 \| 3 \| 393 \| DUF4102 domain-containing protein \| DUF4102 domain-containing protein; afdb-uniprot50 \| AF-A0A3T7XM81-F1-MODEL\_V4 \| 1.0 \| 9.15e-17 \| 612 \| 0.197 \| 420 \| 286 \| 18 \| 2 \| 401 \| 8 \| 396 \| DUF4102 domain-containing protein \| DUF4102 domain-containing protein; afdb-uniprot50 \| AF-A0A7C8AXU5-F1-MODEL\_V4 \| 1.0 \| 5.416e-17 \| 612 \| 0.227 \| 427 \| 270 \| 20 \| 2 \| 402 \| 8 \| 400 \| Integrase arm-type DNA-binding domain-containing protein \| Integrase arm-type DNA-binding domain-containing protein; afdb-uniprot50 \| AF-A0A0X3TJA3-F1-MODEL\_V4 \| 1.0 \| 3.527e-17 \| 612 \| 0.184 \| 444 \| 290 \| 17 \| 2 \| 402 \| 3 \| 417 \| Uncharacterized protein \| Uncharacterized protein; afdb-uniprot50 \| AF-A0A317CJS7-F1-MODEL\_V4 \| 1.0 \| 3.057e-17 \| 611 \| 0.193 \| 424 \| 286 \| 19 \| 2 \| 402 \| 5 \| 395 \| Integrase \| Integrase; afdb-uniprot50 \| AF-A0A1B1NW11-F1-MODEL\_V4 \| 1.0 \| 1.87e-16 \| 611 \| 0.222 \| 427 \| 272 \| 20 \| 1 \| 402 \| 7 \| 398 \| Prophage CP4-57 integrase \| Prophage CP4-57 integrase; afdb-uniprot50 \| AF-A0A2S5QMZ1-F1-MODEL\_V4 \| 1.0 \| 6.874e-17 \| 610 \| 0.199 \| 416 \| 280 \| 21 \| 1 \| 403 \| 1 \| 376 \| Integrase \| Integrase; afdb-uniprot50 \| AF-A0A560D0T2-F1-MODEL\_V4 \| 1.0 \| 1.296e-17 \| 610 \| 0.22 \| 439 \| 265 \| 23 \| 1 \| 401 \| 5 \| 404 \| Site-specific recombinase XerD \| Site-specific recombinase XerD; afdb-uniprot50 \| AF-A0A2U8P8P1-F1-MODEL\_V4 \| 1.0 \| 1.991e-17 \| 610 \| 0.211 \| 434 \| 278 \| 24 \| 2 \| 403 \| 5 \| 406 \| DUF4102 domain-containing protein \| DUF4102 domain-containing protein; afdb-uniprot50 \| AF-A0A661G1N2-F1-MODEL\_V4 \| 1.0 \| 2.65e-17 \| 610 \| 0.188 \| 451 \| 284 \| 25 \| 2 \| 402 \| 3 \| 421 \| Integrase \| Integrase; afdb-uniprot50 \| AF-A0A095S8W4-F1-MODEL\_V4 \| 1.0 \| 3.207e-17 \| 609 \| 0.228 \| 421 \| 274 \| 20 \| 2 \| 402 \| 6 \| 395 \| DNA integration/recombination/inversion protein \| DNA integration/recombination/inversion protein; afdb-uniprot50 \| AF-A0A2U2CNI2-F1-MODEL\_V4 \| 1.0 \| 5.242e-18 \| 609 \| 0.237 \| 426 \| 267 \| 23 \| 2 \| 403 \| 11 \| 402 \| Uncharacterized protein \| Uncharacterized protein; afdb-uniprot50 \| AF-A0A379HBU4-F1-MODEL\_V4 \| 1.0 \| 4.268e-17 \| 609 \| 0.205 \| 423 \| 287 \| 17 \| 2 \| 402 \| 3 \| 398 \| Prophage CP4-57 integrase \| Prophage CP4-57 integrase; afdb-uniprot50 \| AF-A0A7X8YJ22-F1-MODEL\_V4 \| 1.0 \| 1.056e-16 \| 609 \| 0.216 \| 425 \| 274 \| 21 \| 2 \| 402 \| 8 \| 397 \| Tyrosine-type recombinase/integrase \| Tyrosine-type recombinase/integrase; afdb-uniprot50 \| AF-A0A066UH65-F1-MODEL\_V4 \| 1.0 \| 5.958e-17 \| 608 \| 0.181 \| 418 \| 300 \| 17 \| 2 \| 402 \| 3 \| 395 \| Phage integrase \| Phage integrase; afdb-uniprot50 \| AF-A0A7W9B908-F1-MODEL\_V4 \| 1.0 \| 3.363e-17 \| 608 \| 0.208 \| 427 \| 278 \| 22 \| 1 \| 402 \| 1 \| 392 \| Integrase \| Integrase; afdb-uniprot50 \| AF-A0A2G0Q224-F1-MODEL\_V4 \| 1.0 \| 1.783e-16 \| 607 \| 0.178 \| 426 \| 289 \| 22 \| 2 \| 402 \| 3 \| 392 \| Integrase \| Integrase; afdb-uniprot50 \| AF-A0A380TVV2-F1-MODEL\_V4 \| 1.0 \| 7.561e-17 \| 607 \| 0.203 \| 427 \| 282 \| 22 \| 2 \| 401 \| 8 \| 403 \| Prophage CP4-57 integrase \| Prophage CP4-57 integrase; afdb-uniprot50 \| AF-W2UZJ4-F1-MODEL\_V4 \| 1.0 \| 2.779e-17 \| 607 \| 0.193 \| 419 \| 288 \| 15 \| 2 \| 402 \| 14 \| 400 \| p4-like integrase \| p4-like integrase; afdb-uniprot50 \| AF-A0A1G9U9P8-F1-MODEL\_V4 \| 1.0 \| 1.991e-17 \| 607 \| 0.203 \| 422 \| 283 \| 17 \| 2 \| 402 \| 3 \| 392 \| Integrase \| Integrase; afdb-uniprot50 \| AF-A0A1M7T7F7-F1-MODEL\_V4 \| 1.0 \| 6.554e-17 \| 606 \| 0.175 \| 417 \| 294 \| 19 \| 2 \| 402 \| 3 \| 385 \| Integrase \| Integrase; afdb-uniprot50 \| AF-A0A3Q9JNA5-F1-MODEL\_V4 \| 1.0 \| 1.056e-16 \| 606 \| 0.211 \| 407 \| 277 \| 19 \| 2 \| 394 \| 8 \| 384 \| DUF4102 domain-containing protein \| DUF4102 domain-containing protein; afdb-uniprot50 \| AF-A0A7Y8SU33-F1-MODEL\_V4 \| 1.0 \| 2.409e-17 \| 606 \| 0.218 \| 426 \| 276 \| 19 \| 2 \| 402 \| 1 \| 394 \| Tyrosine-type recombinase/integrase \| Tyrosine-type recombinase/integrase; afdb-uniprot50 \| AF-A0A259IRK1-F1-MODEL\_V4 \| 1.0 \| 9.596e-17 \| 606 \| 0.185 \| 431 \| 291 \| 21 \| 2 \| 402 \| 3 \| 403 \| Tyr recombinase domain-containing protein \| Tyr recombinase domain-containing protein; afdb-uniprot50 \| AF-A0A7H9BH93-F1-MODEL\_V4 \| 1.0 \| 4.069e-17 \| 605 \| 0.202 \| 429 \| 281 \| 20 \| 1 \| 402 \| 15 \| 409 \| Tyrosine-type recombinase/integrase \| Tyrosine-type recombinase/integrase; afdb-uniprot50 \| AF-A0A2Z2HFA7-F1-MODEL\_V4 \| 1.0 \| 2.158e-16 \| 605 \| 0.186 \| 423 \| 288 \| 21 \| 2 \| 401 \| 43 \| 432 \| Tyr recombinase domain-containing protein \| Tyr recombinase domain-containing protein; afdb-uniprot50 \| AF-A0A6G5R766-F1-MODEL\_V4 \| 1.0 \| 2.527e-17 \| 604 \| 0.178 \| 436 \| 293 \| 18 \| 1 \| 403 \| 4 \| 407 \| Site-specific recombinase, phage integrase family (DUF4102 domain) \| Site-specific recombinase, phage integrase family (DUF4102 domain); afdb-uniprot50 \| AF-A0A509YI24-F1-MODEL\_V4 \| 1.0 \| 1.006e-16 \| 604 \| 0.185 \| 416 \| 292 \| 18 \| 2 \| 400 \| 23 \| 408 \| Integrase \| Integrase; afdb-uniprot50 \| AF-A0A4P7REU5-F1-MODEL\_V4 \| 1.0 \| 1.81e-17 \| 604 \| 0.221 \| 429 \| 271 \| 19 \| 1 \| 402 \| 7 \| 399 \| DUF4102 domain-containing protein \| DUF4102 domain-containing protein; afdb-uniprot50 \| AF-A0A6M8UC91-F1-MODEL\_V4 \| 1.0 \| 5.681e-17 \| 603 \| 0.225 \| 417 \| 264 \| 20 \| 2 \| 391 \| 8 \| 392 \| Integrase \| Integrase; afdb-uniprot50 \| AF-A0A363CZK6-F1-MODEL\_V4 \| 1.0 \| 2.872e-16 \| 603 \| 0.165 \| 428 \| 302 \| 18 \| 2 \| 403 \| 8 \| 406 \| Uncharacterized protein \| Uncharacterized protein; afdb-uniprot50 \| AF-A0A0F2P2B9-F1-MODEL\_V4 \| 1.0 \| 4.924e-17 \| 602 \| 0.189 \| 428 \| 291 \| 18 \| 2 \| 402 \| 3 \| 401 \| Integrase \| Integrase; afdb-uniprot50 \| AF-A0A1G3TPX5-F1-MODEL\_V4 \| 1.0 \| 8.724e-17 \| 602 \| 0.204 \| 420 \| 289 \| 19 \| 2 \| 402 \| 8 \| 401 \| Integrase \| Integrase; afdb-uniprot50 \| AF-A0A1I3RK68-F1-MODEL\_V4 \| 1.0 \| 8.318e-17 \| 602 \| 0.196 \| 427 \| 281 \| 21 \| 2 \| 402 \| 3 \| 393 \| Integrase \| Integrase; afdb-uniprot50 \| AF-A0A1X7JHB3-F1-MODEL\_V4 \| 1.0 \| 1.474e-16 \| 601 \| 0.186 \| 419 \| 281 \| 24 \| 2 \| 402 \| 1 \| 377 \| Integrase \| Integrase; afdb-uniprot50 \| AF-A0A8B0UBD6-F1-MODEL\_V4 \| 1.0 \| 4.009e-16 \| 601 \| 0.198 \| 414 \| 283 \| 19 \| 2 \| 402 \| 3 \| 380 \| Tyrosine-type recombinase/integrase \| Tyrosine-type recombinase/integrase; afdb-uniprot50 \| AF-A0A239LDZ8-F1-MODEL\_V4 \| 1.0 \| 1.36e-17 \| 601 \| 0.205 \| 424 \| 281 \| 19 \| 2 \| 402 \| 3 \| 393 \| Integrase \| Integrase; afdb-uniprot50 \| AF-A0A7W4WUX0-F1-MODEL\_V4 \| 1.0 \| 4.476e-17 \| 601 \| 0.189 \| 449 \| 287 \| 22 \| 1 \| 402 \| 1 \| 419 \| Integrase \| Integrase; afdb-uniprot50 \| AF-A0A2G4ASH1-F1-MODEL\_V4 \| 1.0 \| 1.161e-16 \| 600 \| 0.202 \| 415 \| 286 \| 17 \| 2 \| 402 \| 3 \| 386 \| Putative prophage CPS-53 integrase \| Putative prophage CPS-53 integrase; afdb-uniprot50 \| AF-A0A5J5FS73-F1-MODEL\_V4 \| 1.0 \| 5.416e-17 \| 600 \| 0.195 \| 425 \| 286 \| 18 \| 1 \| 402 \| 7 \| 398 \| Tyrosine-type recombinase/integrase \| Tyrosine-type recombinase/integrase; afdb-uniprot50 \| AF-A0A1S8D336-F1-MODEL\_V4 \| 1.0 \| 1.898e-17 \| 600 \| 0.209 \| 434 \| 273 \| 22 \| 2 \| 403 \| 1 \| 396 \| Uncharacterized protein \| Uncharacterized protein; afdb-uniprot50 \| AF-A0A1E7WYI8-F1-MODEL\_V4 \| 1.0 \| 4.069e-17 \| 600 \| 0.203 \| 456 \| 280 \| 21 \| 2 \| 403 \| 4 \| 430 \| Prophage CP4-57 integrase \| Prophage CP4-57 integrase; afdb-uniprot50 \| AF-A0A212LBW6-F1-MODEL\_V4 \| 1.0 \| 1.991e-17 \| 600 \| 0.192 \| 442 \| 290 \| 17 \| 2 \| 402 \| 4 \| 419 \| Putative Integrase family protein \| Putative Integrase family protein; afdb-uniprot50 \| AF-A0A5C8TMJ0-F1-MODEL\_V4 \| 1.0 \| 5.681e-17 \| 599 \| 0.22 \| 421 \| 268 \| 22 \| 2 \| 401 \| 18 \| 399 \| Tyrosine-type recombinase/integrase \| Tyrosine-type recombinase/integrase; afdb-uniprot50 \| AF-D0X1Z0-F1-MODEL\_V4 \| 1.0 \| 1.783e-16 \| 599 \| 0.213 \| 422 \| 279 \| 21 \| 2 \| 402 \| 4 \| 393 \| Tyr recombinase domain-containing protein \| Tyr recombinase domain-containing protein; afdb-uniprot50 \| AF-A0A5B0GL81-F1-MODEL\_V4 \| 1.0 \| 2.65e-17 \| 599 \| 0.214 \| 428 \| 273 \| 22 \| 2 \| 402 \| 8 \| 399 \| Tyrosine-type recombinase/integrase \| Tyrosine-type recombinase/integrase; afdb-uniprot50 \| AF-A0A7C1QT18-F1-MODEL\_V4 \| 1.0 \| 1.7e-16 \| 599 \| 0.195 \| 461 \| 288 \| 22 \| 1 \| 402 \| 1 \| 437 \| Tyr recombinase domain-containing protein \| Tyr recombinase domain-containing protein; afdb-uniprot50 \| AF-A0A6L9MH98-F1-MODEL\_V4 \| 1.0 \| 6.874e-17 \| 598 \| 0.198 \| 424 \| 284 \| 18 \| 1 \| 402 \| 3 \| 392 \| Tyrosine-type recombinase/integrase \| Tyrosine-type recombinase/integrase; afdb-uniprot50 \| AF-A0A2G6HRL6-F1-MODEL\_V4 \| 1.0 \| 1.107e-16 \| 598 \| 0.204 \| 435 \| 276 \| 23 \| 2 \| 402 \| 5 \| 403 \| Integrase \| Integrase; afdb-uniprot50 \| AF-H3RBK3-F1-MODEL\_V4 \| 1.0 \| 1.34e-16 \| 597 \| 0.201 \| 426 \| 279 \| 19 \| 2 \| 402 \| 8 \| 397 \| CP4-57 family phage integrase \| CP4-57 family phage integrase; afdb-uniprot50 \| AF-A0A3S0V2Q9-F1-MODEL\_V4 \| 1.0 \| 1.898e-17 \| 597 \| 0.209 \| 439 \| 276 \| 19 \| 2 \| 402 \| 6 \| 411 \| DUF4102 domain-containing protein \| DUF4102 domain-containing protein; afdb-uniprot50 \| AF-A0A1Y0HJN4-F1-MODEL\_V4 \| 1.0 \| 2.738e-16 \| 596 \| 0.169 \| 425 \| 300 \| 22 \| 1 \| 402 \| 5 \| 399 \| Prophage integrase IntA \| Prophage integrase IntA; afdb-uniprot50 \| AF-A0A0N0IHF8-F1-MODEL\_V4 \| 1.0 \| 1.071e-17 \| 596 \| 0.173 \| 433 \| 291 \| 19 \| 2 \| 402 \| 9 \| 406 \| Tyr recombinase domain-containing protein \| Tyr recombinase domain-containing protein; afdb-uniprot50 \| AF-A0A432ZBV6-F1-MODEL\_V4 \| 1.0 \| 3.159e-16 \| 596 \| 0.208 \| 422 \| 281 \| 19 \| 2 \| 402 \| 16 \| 405 \| Integrase \| Integrase; afdb-uniprot50 \| AF-A0A2T0XJR5-F1-MODEL\_V4 \| 1.0 \| 8.443e-18 \| 596 \| 0.201 \| 431 \| 277 \| 21 \| 1 \| 402 \| 1 \| 393 \| Integrase \| Integrase; afdb-uniprot50 \| AF-H0Q156-F1-MODEL\_V4 \| 1.0 \| 4.924e-17 \| 596 \| 0.213 \| 427 \| 279 \| 21 \| 2 \| 402 \| 28 \| 423 \| Uncharacterized protein \| Uncharacterized protein; afdb-uniprot50 \| AF-A0A2E6A368-F1-MODEL\_V4 \| 1.0 \| 2.611e-16 \| 595 \| 0.21 \| 422 \| 279 \| 20 \| 1 \| 402 \| 6 \| 393 \| Integrase \| Integrase; afdb-uniprot50 \| AF-A0A7M3B6Q3-F1-MODEL\_V4 \| 1.0 \| 7.21e-17 \| 595 \| 0.208 \| 426 \| 275 \| 21 \| 2 \| 402 \| 11 \| 399 \| Integrase \| Integrase; afdb-uniprot50 \| AF-A0A845UVE8-F1-MODEL\_V4 \| 1.0 \| 1.496e-17 \| 595 \| 0.222 \| 450 \| 280 \| 21 \| 1 \| 403 \| 10 \| 436 \| Uncharacterized protein \| Uncharacterized protein; afdb-uniprot50 \| AF-A0A482ITC2-F1-MODEL\_V4 \| 1.0 \| 3.363e-17 \| 593 \| 0.217 \| 469 \| 263 \| 25 \| 2 \| 402 \| 4 \| 436 \| DUF4102 domain-containing protein \| DUF4102 domain-containing protein; afdb-uniprot50 \| AF-A0A5S4SE16-F1-MODEL\_V4 \| 1.0 \| 7.931e-17 \| 593 \| 0.204 \| 446 \| 284 \| 21 \| 1 \| 402 \| 1 \| 419 \| Tyrosine-type recombinase/integrase \| Tyrosine-type recombinase/integrase; afdb-uniprot50 \| AF-D6VB17-F1-MODEL\_V4 \| 1.0 \| 2.297e-17 \| 593 \| 0.211 \| 421 \| 271 \| 17 \| 2 \| 401 \| 64 \| 444 \| Integrase family protein \| Integrase family protein; afdb-uniprot50 \| AF-A0A0K1QM45-F1-MODEL\_V4 \| 1.0 \| 5.088e-16 \| 592 \| 0.218 \| 411 \| 267 \| 22 \| 2 \| 391 \| 3 \| 380 \| Integrase \| Integrase; afdb-uniprot50 \| AF-A0A560WXP0-F1-MODEL\_V4 \| 1.0 \| 1.056e-16 \| 592 \| 0.207 \| 429 \| 283 \| 23 \| 2 \| 402 \| 11 \| 410 \| Integrase \| Integrase; afdb-uniprot50 \| AF-A0A1H6GE73-F1-MODEL\_V4 \| 1.0 \| 1.006e-16 \| 592 \| 0.224 \| 414 \| 262 \| 20 \| 2 \| 391 \| 3 \| 381 \| Integrase \| Integrase; afdb-uniprot50 \| AF-A0A2U0T8C2-F1-MODEL\_V4 \| 1.0 \| 4.41e-16 \| 591 \| 0.198 \| 424 \| 288 \| 20 \| 2 \| 402 \| 8 \| 402 \| Integrase \| Integrase; afdb-uniprot50 \| AF-A0A6S6TC66-F1-MODEL\_V4 \| 1.0 \| 1.546e-16 \| 591 \| 0.188 \| 429 \| 290 \| 20 \| 2 \| 402 \| 9 \| 407 \| Integrase family protein \| Integrase family protein; afdb-uniprot50 \| AF-A0A4R6Y5X0-F1-MODEL\_V4 \| 1.0 \| 2.057e-16 \| 591 \| 0.179 \| 429 \| 289 \| 20 \| 1 \| 402 \| 1 \| 393 \| Integrase \| Integrase; afdb-uniprot50 \| AF-A0A4R1HYV8-F1-MODEL\_V4 \| 1.0 \| 7.21e-17 \| 590 \| 0.219 \| 433 \| 273 \| 20 \| 2 \| 403 \| 3 \| 401 \| Site-specific recombinase XerD \| Site-specific recombinase XerD; afdb-uniprot50 \| AF-A0A5U3D0P1-F1-MODEL\_V4 \| 1.0 \| 7.561e-17 \| 590 \| 0.213 \| 449 \| 275 \| 25 \| 2 \| 403 \| 10 \| 427 \| Tyrosine-type recombinase/integrase \| Tyrosine-type recombinase/integrase; afdb-uniprot50 \| AF-A0A2U0ZIK5-F1-MODEL\_V4 \| 1.0 \| 1.546e-16 \| 589 \| 0.211 \| 431 \| 274 \| 25 \| 2 \| 403 \| 3 \| 396 \| Integrase \| Integrase; afdb-uniprot50 \| AF-A0A836Q5F6-F1-MODEL\_V4 \| 1.0 \| 4.41e-16 \| 589 \| 0.178 \| 421 \| 297 \| 21 \| 2 \| 402 \| 4 \| 395 \| DUF4102 domain-containing protein \| DUF4102 domain-containing protein; afdb-uniprot50 \| AF-A0A5E9PHW5-F1-MODEL\_V4 \| 1.0 \| 2.057e-16 \| 589 \| 0.179 \| 423 \| 300 \| 18 \| 1 \| 403 \| 1 \| 396 \| Site-specific integrase \| Site-specific integrase; afdb-uniprot50 \| AF-A0A4R6FNZ7-F1-MODEL\_V4 \| 1.0 \| 1.34e-16 \| 589 \| 0.192 \| 421 \| 280 \| 21 \| 9 \| 402 \| 7 \| 394 \| Integrase \| Integrase; afdb-uniprot50 \| AF-A0A5N0TDP5-F1-MODEL\_V4 \| 1.0 \| 1.161e-16 \| 589 \| 0.201 \| 421 \| 285 \| 18 \| 2 \| 402 \| 8 \| 397 \| Tyrosine-type recombinase/integrase \| Tyrosine-type recombinase/integrase; afdb-uniprot50 \| AF-A0A7Z9HGD8-F1-MODEL\_V4 \| 1.0 \| 7.561e-17 \| 589 \| 0.2 \| 430 \| 284 \| 20 \| 2 \| 402 \| 9 \| 407 \| DUF4102 domain-containing protein \| DUF4102 domain-containing protein; afdb-uniprot50 \| AF-A0A1G4SA28-F1-MODEL\_V4 \| 1.0 \| 4.851e-16 \| 589 \| 0.169 \| 426 \| 293 \| 20 \| 2 \| 402 \| 3 \| 392 \| Integrase \| Integrase; afdb-uniprot50 \| AF-A0A839GCA6-F1-MODEL\_V4 \| 1.0 \| 8.318e-17 \| 589 \| 0.197 \| 426 \| 283 \| 21 \| 2 \| 403 \| 3 \| 393 \| Integrase \| Integrase; afdb-uniprot50 \| AF-A0A2N2SIH5-F1-MODEL\_V4 \| 1.0 \| 1.165e-13 \| 588 \| 0.237 \| 311 \| 206 \| 11 \| 102 \| 402 \| 10 \| 299 \| Integrase \| Integrase; afdb-uniprot50 \| AF-A0A562IZK1-F1-MODEL\_V4 \| 1.0 \| 4.779e-15 \| 588 \| 0.233 \| 355 \| 232 \| 17 \| 78 \| 402 \| 19 \| 363 \| Phage integrase family protein \| Phage integrase family protein; afdb-uniprot50 \| AF-A0A4R5TYE0-F1-MODEL\_V4 \| 1.0 \| 8.318e-17 \| 588 \| 0.196 \| 418 \| 292 \| 20 \| 1 \| 402 \| 1 \| 390 \| DUF4102 domain-containing protein \| DUF4102 domain-containing protein; afdb-uniprot50 \| AF-A0A4D7WTQ9-F1-MODEL\_V4 \| 1.0 \| 8.724e-17 \| 588 \| 0.207 \| 428 \| 277 \| 19 \| 2 \| 403 \| 6 \| 397 \| DUF4102 domain-containing protein \| DUF4102 domain-containing protein; afdb-uniprot50 \| AF-A0A811B9I7-F1-MODEL\_V4 \| 1.0 \| 7.931e-17 \| 588 \| 0.177 \| 423 \| 294 \| 18 \| 2 \| 402 \| 5 \| 395 \| Integrase \| Integrase; afdb-uniprot50 \| AF-A0A2S5TDL1-F1-MODEL\_V4 \| 1.0 \| 3.475e-16 \| 588 \| 0.21 \| 423 \| 281 \| 19 \| 2 \| 402 \| 7 \| 398 \| Integrase \| Integrase; afdb-uniprot50 \| AF-A0A5S9PCI4-F1-MODEL\_V4 \| 1.0 \| 1.783e-16 \| 587 \| 0.172 \| 423 \| 297 \| 18 \| 1 \| 402 \| 1 \| 391 \| Prophage integrase IntA \| Prophage integrase IntA; afdb-uniprot50 \| AF-A0A096FMR1-F1-MODEL\_V4 \| 1.0 \| 1.81e-17 \| 586 \| 0.189 \| 454 \| 281 \| 21 \| 1 \| 402 \| 1 \| 419 \| Integrase \| Integrase; afdb-uniprot50 \| AF-A0A069E4V0-F1-MODEL\_V4 \| 1.0 \| 1.107e-16 \| 585 \| 0.216 \| 412 \| 271 \| 17 \| 9 \| 400 \| 16 \| 395 \| Phage integrase \| Phage integrase; afdb-uniprot50 \| AF-A0A0M0FY12-F1-MODEL\_V4 \| 1.0 \| 2.452e-15 \| 585 \| 0.223 \| 362 \| 234 \| 13 \| 71 \| 402 \| 47 \| 391 \| Integrase \| Integrase; afdb-uniprot50 \| AF-A0A1W6MW71-F1-MODEL\_V4 \| 1.0 \| 1.961e-16 \| 585 \| 0.197 \| 436 \| 282 \| 21 \| 2 \| 402 \| 10 \| 412 \| Uncharacterized protein \| Uncharacterized protein; afdb-uniprot50 \| AF-A0A368V6W8-F1-MODEL\_V4 \| 1.0 \| 4.205e-16 \| 585 \| 0.193 \| 423 \| 287 \| 22 \| 2 \| 402 \| 11 \| 401 \| Integrase \| Integrase; afdb-uniprot50 \| AF-A0A844WU91-F1-MODEL\_V4 \| 1.0 \| 7.813e-16 \| 584 \| 0.173 \| 421 \| 299 \| 17 \| 2 \| 402 \| 8 \| 399 \| Tyrosine-type recombinase/integrase \| Tyrosine-type recombinase/integrase; afdb-uniprot50 \| AF-A0A441UN45-F1-MODEL\_V4 \| 1.0 \| 1.124e-17 \| 584 \| 0.206 \| 432 \| 284 \| 17 \| 2 \| 402 \| 6 \| 409 \| DUF4102 domain-containing protein \| DUF4102 domain-containing protein; afdb-uniprot50 \| AF-A0A5E7JF77-F1-MODEL\_V4 \| 1.0 \| 9.15e-17 \| 584 \| 0.204 \| 450 \| 292 \| 19 \| 1 \| 400 \| 3 \| 436 \| Core-binding (CB) domain-containing protein \| Core-binding (CB) domain-containing protein; afdb-uniprot50 \| AF-A0A0P1ELQ8-F1-MODEL\_V4 \| 1.0 \| 6.249e-17 \| 584 \| 0.215 \| 436 \| 279 \| 23 \| 2 \| 401 \| 52 \| 460 \| Prophage CP4-57 integrase \| Prophage CP4-57 integrase; afdb-uniprot50 \| AF-A0A292AK98-F1-MODEL\_V4 \| 1.0 \| 5.681e-17 \| 583 \| 0.217 \| 436 \| 260 \| 22 \| 18 \| 403 \| 6 \| 410 \| Integrase \| Integrase; afdb-uniprot50 \| AF-A0A5P9HCV6-F1-MODEL\_V4 \| 1.0 \| 2.738e-16 \| 583 \| 0.19 \| 430 \| 281 \| 24 \| 2 \| 402 \| 11 \| 402 \| Prophage CPS-53 integrase \| Prophage CPS-53 integrase; afdb-uniprot50 \| AF-A0A1Q3K5N4-F1-MODEL\_V4 \| 1.0 \| 2.779e-17 \| 582 \| 0.23 \| 407 \| 265 \| 17 \| 2 \| 402 \| 3 \| 367 \| Tyr recombinase domain-containing protein \| Tyr recombinase domain-containing protein; afdb-uniprot50 \| AF-K8PFU5-F1-MODEL\_V4 \| 1.0 \| 5.416e-17 \| 582 \| 0.218 \| 425 \| 280 \| 19 \| 2 \| 403 \| 14 \| 409 \| Tyr recombinase domain-containing protein \| Tyr recombinase domain-containing protein; afdb-uniprot50 \| AF-A0A3B0W1B8-F1-MODEL\_V4 \| 1.0 \| 1.006e-16 \| 582 \| 0.186 \| 440 \| 288 \| 23 \| 1 \| 401 \| 3 \| 411 \| Phage integrase \| Phage integrase; afdb-uniprot50 \| AF-A0A0P7AIM3-F1-MODEL\_V4 \| 1.0 \| 1.961e-16 \| 581 \| 0.185 \| 415 \| 289 \| 20 \| 2 \| 403 \| 9 \| 387 \| Tyr recombinase domain-containing protein \| Tyr recombinase domain-containing protein; afdb-uniprot50 \| AF-S6CXQ8-F1-MODEL\_V4 \| 1.0 \| 1.056e-16 \| 581 \| 0.213 \| 435 \| 272 \| 19 \| 2 \| 402 \| 6 \| 404 \| Phage integrase family protein \| Phage integrase family protein; afdb-uniprot50 \| AF-A0A4Q4B2S9-F1-MODEL\_V4 \| 1.0 \| 5.597e-16 \| 581 \| 0.224 \| 428 \| 271 \| 22 \| 2 \| 402 \| 3 \| 396 \| DUF4102 domain-containing protein \| DUF4102 domain-containing protein; afdb-uniprot50 \| AF-A0A348UTP6-F1-MODEL\_V4 \| 1.0 \| 1.815e-14 \| 580 \| 0.236 \| 326 \| 213 \| 13 \| 106 \| 402 \| 3 \| 321 \| Integrase \| Integrase; afdb-uniprot50 \| AF-A0A7H4NYI2-F1-MODEL\_V4 \| 1.0 \| 1.32e-15 \| 580 \| 0.195 \| 388 \| 270 \| 15 \| 36 \| 403 \| 39 \| 404 \| Phage integrase \| Phage integrase; afdb-uniprot50 \| AF-A0A7W6S310-F1-MODEL\_V4 \| 1.0 \| 4.625e-16 \| 578 \| 0.222 \| 432 \| 270 \| 20 \| 2 \| 401 \| 7 \| 404 \| Integrase \| Integrase; afdb-uniprot50 \| AF-A0A6S5KU75-F1-MODEL\_V4 \| 1.0 \| 7.449e-16 \| 577 \| 0.218 \| 417 \| 264 \| 20 \| 1 \| 394 \| 1 \| 378 \| Integrase \| Integrase; afdb-uniprot50 \| AF-A0A345DAL3-F1-MODEL\_V4 \| 1.0 \| 6.457e-16 \| 577 \| 0.19 \| 440 \| 292 \| 22 \| 2 \| 402 \| 6 \| 420 \| Prophage integrase IntA \| Prophage integrase IntA; afdb-uniprot50 \| AF-A0A1M6UHH4-F1-MODEL\_V4 \| 1.0 \| 1.961e-16 \| 577 \| 0.191 \| 423 \| 292 \| 20 \| 8 \| 400 \| 15 \| 417 \| Arm-DNA-bind\_3 domain-containing protein \| Arm-DNA-bind\_3 domain-containing protein; afdb-uniprot50 \| AF-N9NA50-F1-MODEL\_V4 \| 1.0 \| 1.546e-16 \| 576 \| 0.194 \| 432 \| 285 \| 23 \| 2 \| 403 \| 6 \| 404 \| Tyr recombinase domain-containing protein \| Tyr recombinase domain-containing protein; afdb-uniprot50 \| AF-C3X4R8-F1-MODEL\_V4 \| 1.0 \| 9.15e-17 \| 576 \| 0.175 \| 432 \| 293 \| 20 \| 2 \| 401 \| 8 \| 408 \| Tyr recombinase domain-containing protein \| Tyr recombinase domain-containing protein; afdb-uniprot50 \| AF-E0M0I1-F1-MODEL\_V4 \| 1.0 \| 1.056e-16 \| 575 \| 0.194 \| 416 \| 289 \| 20 \| 2 \| 402 \| 3 \| 387 \| Integrase family protein \| Integrase family protein; afdb-uniprot50 \| AF-A0A7U2PLX0-F1-MODEL\_V4 \| 1.0 \| 8.443e-18 \| 575 \| 0.193 \| 435 \| 273 \| 20 \| 2 \| 403 \| 3 \| 392 \| Tyrosine-type recombinase/integrase \| Tyrosine-type recombinase/integrase; afdb-uniprot50 \| AF-A0A840MZH1-F1-MODEL\_V4 \| 1.0 \| 5.87e-16 \| 575 \| 0.183 \| 424 \| 286 \| 18 \| 2 \| 398 \| 10 \| 400 \| Integrase \| Integrase; afdb-uniprot50 \| AF-A0A6G8CS84-F1-MODEL\_V4 \| 1.0 \| 4.625e-16 \| 575 \| 0.202 \| 425 \| 283 \| 21 \| 2 \| 403 \| 11 \| 402 \| Tyrosine-type recombinase/integrase \| Tyrosine-type recombinase/integrase; afdb-uniprot50 \| AF-A0A5E7WTN4-F1-MODEL\_V4 \| 1.0 \| 5.179e-14 \| 574 \| 0.216 \| 360 \| 245 \| 15 \| 60 \| 402 \| 2 \| 341 \| Prophage integrase IntS \| Prophage integrase IntS; afdb-uniprot50 \| AF-H3RIA1-F1-MODEL\_V4 \| 1.0 \| 2.967e-15 \| 574 \| 0.208 \| 378 \| 252 \| 15 \| 53 \| 402 \| 1 \| 359 \| Site-specific recombinase, phage integrase family \| Site-specific recombinase, phage integrase family; afdb-uniprot50 \| AF-A0A2K8UAW1-F1-MODEL\_V4 \| 1.0 \| 9.014e-16 \| 574 \| 0.207 \| 404 \| 267 \| 17 \| 25 \| 400 \| 4 \| 382 \| Uncharacterized protein \| Uncharacterized protein; afdb-uniprot50 \| AF-A0A2K8L2Y2-F1-MODEL\_V4 \| 1.0 \| 2.572e-15 \| 574 \| 0.167 \| 424 \| 295 \| 22 \| 2 \| 402 \| 3 \| 391 \| Integrase \| Integrase; afdb-uniprot50 \| AF-A0A398BNA6-F1-MODEL\_V4 \| 1.0 \| 1.474e-16 \| 574 \| 0.232 \| 425 \| 261 \| 23 \| 1 \| 398 \| 1 \| 387 \| DUF4102 domain-containing protein \| DUF4102 domain-containing protein; afdb-uniprot50 \| AF-A0A3G2HYM0-F1-MODEL\_V4 \| 1.0 \| 1.218e-16 \| 574 \| 0.187 \| 443 \| 290 \| 23 \| 2 \| 402 \| 7 \| 421 \| DUF4102 domain-containing protein \| DUF4102 domain-containing protein; afdb-uniprot50 \| AF-A0A2E6KN51-F1-MODEL\_V4 \| 1.0 \| 4.851e-16 \| 573 \| 0.179 \| 423 \| 285 \| 17 \| 9 \| 402 \| 6 \| 395 \| Integrase \| Integrase; afdb-uniprot50 \| AF-A0A1L3EZE4-F1-MODEL\_V4 \| 1.0 \| 1.161e-16 \| 573 \| 0.2 \| 435 \| 282 \| 18 \| 2 \| 400 \| 7 \| 411 \| Uncharacterized protein \| Uncharacterized protein; afdb-uniprot50 \| AF-A0A7W5ZZ75-F1-MODEL\_V4 \| 1.0 \| 4.069e-17 \| 573 \| 0.18 \| 432 \| 283 \| 21 \| 2 \| 402 \| 58 \| 449 \| Integrase \| Integrase; afdb-uniprot50 \| AF-A0A6N8EHY5-F1-MODEL\_V4 \| 1.0 \| 3.057e-17 \| 572 \| 0.229 \| 406 \| 243 \| 21 \| 2 \| 367 \| 6 \| 381 \| DUF4102 domain-containing protein \| DUF4102 domain-containing protein; afdb-uniprot50 \| AF-A0A559NFV8-F1-MODEL\_V4 \| 1.0 \| 7.449e-16 \| 571 \| 0.189 \| 422 \| 278 \| 20 \| 1 \| 400 \| 2 \| 381 \| Site-specific recombinase XerD \| Site-specific recombinase XerD; afdb-uniprot50 \| AF-A0A7V2M8D0-F1-MODEL\_V4 \| 1.0 \| 2.158e-16 \| 571 \| 0.216 \| 439 \| 273 \| 22 \| 1 \| 400 \| 5 \| 411 \| DUF4102 domain-containing protein \| DUF4102 domain-containing protein; afdb-uniprot50 \| AF-A0A6F8T1B2-F1-MODEL\_V4 \| 1.0 \| 3.159e-16 \| 570 \| 0.19 \| 414 \| 296 \| 18 \| 1 \| 403 \| 25 \| 410 \| Integrase \| Integrase; afdb-uniprot50 \| AF-A0A158IH21-F1-MODEL\_V4 \| 1.0 \| 3.313e-16 \| 570 \| 0.193 \| 455 \| 282 \| 23 \| 1 \| 403 \| 6 \| 427 \| Phage integrase family protein \| Phage integrase family protein; afdb-uniprot50 \| AF-A0A257V1X1-F1-MODEL\_V4 \| 1.0 \| 7.103e-16 \| 569 \| 0.21 \| 423 \| 271 \| 17 \| 2 \| 398 \| 9 \| 394 \| Uncharacterized protein \| Uncharacterized protein; afdb-uniprot50 \| AF-A0A4Y3HQF9-F1-MODEL\_V4 \| 1.0 \| 7.813e-16 \| 568 \| 0.215 \| 417 \| 279 \| 19 \| 2 \| 403 \| 8 \| 391 \| Integrase \| Integrase; afdb-uniprot50 \| AF-A0A4R0NXE3-F1-MODEL\_V4 \| 1.0 \| 1.161e-16 \| 568 \| 0.219 \| 491 \| 264 \| 25 \| 2 \| 402 \| 6 \| 467 \| DUF4102 domain-containing protein \| DUF4102 domain-containing protein; afdb-uniprot50 \| AF-A0A1Q7BDQ6-F1-MODEL\_V4 \| 1.0 \| 6.267e-14 \| 567 \| 0.227 \| 352 \| 225 \| 11 \| 58 \| 402 \| 2 \| 313 \| Tyr recombinase domain-containing protein \| Tyr recombinase domain-containing protein; afdb-uniprot50 \| AF-D6ZZC9-F1-MODEL\_V4 \| 1.0 \| 2.158e-16 \| 567 \| 0.201 \| 422 \| 281 \| 22 \| 1 \| 401 \| 5 \| 391 \| Integrase family protein \| Integrase family protein; afdb-uniprot50 \| AF-A0A440JSK6-F1-MODEL\_V4 \| 1.0 \| 1.81e-17 \| 567 \| 0.226 \| 432 \| 264 \| 24 \| 2 \| 403 \| 5 \| 396 \| DUF4102 domain-containing protein \| DUF4102 domain-containing protein; afdb-uniprot50 \| AF-A0A3G8M944-F1-MODEL\_V4 \| 1.0 \| 4.851e-16 \| 567 \| 0.204 \| 431 \| 279 \| 22 \| 2 \| 403 \| 4 \| 399 \| Site-specific integrase \| Site-specific integrase; afdb-uniprot50 \| AF-A0A269PHV3-F1-MODEL\_V4 \| 1.0 \| 5.336e-16 \| 567 \| 0.204 \| 425 \| 280 \| 20 \| 2 \| 402 \| 8 \| 398 \| Integrase \| Integrase; afdb-uniprot50 \| AF-A0A7V2T2N7-F1-MODEL\_V4 \| 1.0 \| 1.523e-15 \| 567 \| 0.193 \| 429 \| 285 \| 20 \| 2 \| 402 \| 3 \| 398 \| DUF4102 domain-containing protein \| DUF4102 domain-containing protein; afdb-uniprot50 \| AF-A0A3S9XF91-F1-MODEL\_V4 \| 1.0 \| 9.915e-16 \| 567 \| 0.172 \| 422 \| 294 \| 20 \| 2 \| 402 \| 11 \| 398 \| DUF4102 domain-containing protein \| DUF4102 domain-containing protein; afdb-uniprot50 \| AF-A0A2E2ZXV7-F1-MODEL\_V4 \| 1.0 \| 4.779e-15 \| 566 \| 0.193 \| 382 \| 257 \| 15 \| 47 \| 402 \| 1 \| 357 \| Integrase \| Integrase; afdb-uniprot50 \| AF-A0A5R1LZU8-F1-MODEL\_V4 \| 1.0 \| 1.87e-16 \| 566 \| 0.198 \| 423 \| 277 \| 20 \| 2 \| 403 \| 4 \| 385 \| Uncharacterized protein \| Uncharacterized protein; afdb-uniprot50 \| AF-A0A1V0KP21-F1-MODEL\_V4 \| 1.0 \| 5.336e-16 \| 566 \| 0.192 \| 427 \| 283 \| 21 \| 2 \| 402 \| 8 \| 398 \| Integrase \| Integrase; afdb-uniprot50 \| AF-G4QGW8-F1-MODEL\_V4 \| 1.0 \| 2.738e-16 \| 566 \| 0.172 \| 424 \| 301 \| 19 \| 2 \| 402 \| 3 \| 399 \| DNA integration/recombination/inversion protein \| DNA integration/recombination/inversion protein; afdb-uniprot50 \| AF-A0A0Q6E4W7-F1-MODEL\_V4 \| 1.0 \| 3.475e-16 \| 566 \| 0.205 \| 443 \| 289 \| 21 \| 2 \| 402 \| 13 \| 434 \| Core-binding (CB) domain-containing protein \| Core-binding (CB) domain-containing protein; afdb-uniprot50 \| AF-A0A2E0QAK3-F1-MODEL\_V4 \| 1.0 \| 7.103e-16 \| 565 \| 0.194 \| 426 \| 283 \| 20 \| 2 \| 402 \| 8 \| 398 \| Integrase \| Integrase; afdb-uniprot50 \| AF-A0A1F9LW06-F1-MODEL\_V4 \| 1.0 \| 1.474e-16 \| 565 \| 0.204 \| 425 \| 277 \| 21 \| 2 \| 402 \| 20 \| 407 \| Uncharacterized protein \| Uncharacterized protein; afdb-uniprot50 \| AF-A0A7C7GTQ2-F1-MODEL\_V4 \| 1.0 \| 5.87e-16 \| 565 \| 0.193 \| 424 \| 288 \| 20 \| 2 \| 402 \| 21 \| 413 \| DUF4102 domain-containing protein \| DUF4102 domain-containing protein; afdb-uniprot50 \| AF-A0A0F9YIN6-F1-MODEL\_V4 \| 1.0 \| 1.546e-16 \| 564 \| 0.219 \| 424 \| 276 \| 20 \| 1 \| 402 \| 6 \| 396 \| Uncharacterized protein \| Uncharacterized protein; afdb-uniprot50 \| AF-A0A2D5IDG5-F1-MODEL\_V4 \| 1.0 \| 1.384e-15 \| 564 \| 0.168 \| 439 \| 296 \| 17 \| 2 \| 401 \| 1 \| 409 \| Integrase \| Integrase; afdb-uniprot50 \| AF-A0A6L7JZA5-F1-MODEL\_V4 \| 1.0 \| 5.597e-16 \| 563 \| 0.214 \| 419 \| 271 \| 21 \| 1 \| 400 \| 3 \| 382 \| DUF4102 domain-containing protein \| DUF4102 domain-containing protein; afdb-uniprot50 \| AF-A0A657B0N1-F1-MODEL\_V4 \| 1.0 \| 1.258e-15 \| 563 \| 0.195 \| 425 \| 287 \| 21 \| 2 \| 402 \| 7 \| 400 \| Uncharacterized protein \| Uncharacterized protein; afdb-uniprot50 \| AF-A0A5Y6ETH5-F1-MODEL\_V4 \| 1.0 \| 2.158e-16 \| 563 \| 0.217 \| 450 \| 268 \| 26 \| 2 \| 401 \| 10 \| 425 \| Tyrosine-type recombinase/integrase \| Tyrosine-type recombinase/integrase; afdb-uniprot50 \| AF-A0A534HXP5-F1-MODEL\_V4 \| 1.0 \| 8.194e-16 \| 563 \| 0.195 \| 410 \| 275 \| 19 \| 1 \| 391 \| 10 \| 383 \| DUF4102 domain-containing protein \| DUF4102 domain-containing protein; afdb-uniprot50 \| AF-A0A7X6JDQ3-F1-MODEL\_V4 \| 1.0 \| 3.012e-16 \| 562 \| 0.229 \| 418 \| 272 \| 20 \| 10 \| 403 \| 2 \| 393 \| Tyrosine-type recombinase/integrase \| Tyrosine-type recombinase/integrase; afdb-uniprot50 \| AF-A0A550KFW2-F1-MODEL\_V4 \| 1.0 \| 7.449e-16 \| 562 \| 0.205 \| 428 \| 278 \| 24 \| 2 \| 403 \| 8 \| 399 \| DUF4102 domain-containing protein \| DUF4102 domain-containing protein; afdb-uniprot50 \| AF-A0A850S1N2-F1-MODEL\_V4 \| 1.0 \| 9.454e-16 \| 562 \| 0.183 \| 447 \| 291 \| 18 \| 1 \| 402 \| 2 \| 419 \| Tyrosine-type recombinase/integrase \| Tyrosine-type recombinase/integrase; afdb-uniprot50 \| AF-A0A242NUI6-F1-MODEL\_V4 \| 1.0 \| 3.012e-16 \| 562 \| 0.193 \| 445 \| 288 \| 18 \| 1 \| 402 \| 1 \| 417 \| Tyr recombinase domain-containing protein \| Tyr recombinase domain-containing protein; afdb-uniprot50 \| AF-A0A523H6V4-F1-MODEL\_V4 \| 1.0 \| 1.405e-16 \| 561 \| 0.199 \| 416 \| 274 \| 18 \| 2 \| 394 \| 15 \| 394 \| DUF4102 domain-containing protein \| DUF4102 domain-containing protein; afdb-uniprot50 \| AF-A0A3E0DQX6-F1-MODEL\_V4 \| 1.0 \| 8.594e-16 \| 561 \| 0.194 \| 426 \| 283 \| 20 \| 2 \| 402 \| 7 \| 397 \| Integrase \| Integrase; afdb-uniprot50 \| AF-A0A2V3US64-F1-MODEL\_V4 \| 1.0 \| 7.561e-17 \| 561 \| 0.202 \| 445 \| 277 \| 24 \| 1 \| 403 \| 1 \| 409 \| Site-specific recombinase XerD \| Site-specific recombinase XerD; afdb-uniprot50 \| AF-D2TBY5-F1-MODEL\_V4 \| 1.0 \| 3.834e-13 \| 560 \| 0.185 \| 323 \| 229 \| 14 \| 96 \| 402 \| 3 \| 307 \| Int protein \| Int protein; afdb-uniprot50 \| AF-A0A842J5Y3-F1-MODEL\_V4 \| 1.0 \| 1.597e-15 \| 560 \| 0.183 \| 420 \| 289 \| 23 \| 2 \| 400 \| 8 \| 394 \| Tyrosine-type recombinase/integrase \| Tyrosine-type recombinase/integrase; afdb-uniprot50 \| AF-Q083S5-F1-MODEL\_V4 \| 1.0 \| 3.012e-16 \| 560 \| 0.195 \| 419 \| 288 \| 18 \| 2 \| 402 \| 3 \| 390 \| Phage integrase family protein \| Phage integrase family protein; afdb-uniprot50 \| AF-A0A7W5GR49-F1-MODEL\_V4 \| 1.0 \| 4.205e-16 \| 560 \| 0.203 \| 422 \| 281 \| 20 \| 14 \| 398 \| 1 \| 404 \| Integrase \| Integrase; afdb-uniprot50 \| AF-A0A5P9HXM8-F1-MODEL\_V4 \| 1.0 \| 2.263e-16 \| 560 \| 0.216 \| 411 \| 282 \| 19 \| 2 \| 400 \| 6 \| 388 \| Prophage CP4-57 integrase \| Prophage CP4-57 integrase; afdb-uniprot50 \| AF-A0A5S3UGE8-F1-MODEL\_V4 \| 1.0 \| 2.738e-16 \| 559 \| 0.183 \| 419 \| 295 \| 19 \| 2 \| 398 \| 3 \| 396 \| Tyr recombinase domain-containing protein \| Tyr recombinase domain-containing protein; afdb-uniprot50 \| AF-A0A4R6Y425-F1-MODEL\_V4 \| 1.0 \| 3.313e-16 \| 559 \| 0.173 \| 455 \| 290 \| 19 \| 1 \| 402 \| 4 \| 425 \| Integrase \| Integrase; afdb-uniprot50 \| AF-A4F5J8-F1-MODEL\_V4 \| 1.0 \| 2.263e-16 \| 558 \| 0.191 \| 417 \| 275 \| 22 \| 2 \| 389 \| 3 \| 386 \| Uncharacterized protein \| Uncharacterized protein; afdb-uniprot50 \| AF-A0A4Y6I9E4-F1-MODEL\_V4 \| 1.0 \| 2.158e-16 \| 558 \| 0.17 \| 427 \| 298 \| 20 \| 2 \| 403 \| 3 \| 398 \| DUF4102 domain-containing protein \| DUF4102 domain-containing protein; afdb-uniprot50 \| AF-A0A3S0MUP2-F1-MODEL\_V4 \| 1.0 \| 2.373e-16 \| 558 \| 0.192 \| 411 \| 291 \| 18 \| 2 \| 402 \| 7 \| 386 \| Site-specific integrase \| Site-specific integrase; afdb-uniprot50 \| AF-A0A6C9E150-F1-MODEL\_V4 \| 1.0 \| 1.621e-16 \| 558 \| 0.193 \| 434 \| 284 \| 21 \| 2 \| 402 \| 3 \| 403 \| Tyrosine-type recombinase/integrase \| Tyrosine-type recombinase/integrase; afdb-uniprot50 \| AF-A0A369TCW8-F1-MODEL\_V4 \| 1.0 \| 8.594e-16 \| 558 \| 0.23 \| 425 \| 259 \| 18 \| 13 \| 402 \| 24 \| 415 \| DUF4102 domain-containing protein \| DUF4102 domain-containing protein; afdb-uniprot50 \| AF-A0A1S9PNP5-F1-MODEL\_V4 \| 1.0 \| 1.32e-15 \| 558 \| 0.19 \| 425 \| 285 \| 20 \| 2 \| 402 \| 3 \| 392 \| Tyr recombinase domain-containing protein \| Tyr recombinase domain-containing protein; afdb-uniprot50 \| AF-A0A2W7KZ90-F1-MODEL\_V4 \| 1.0 \| 2.618e-13 \| 557 \| 0.2 \| 315 \| 223 \| 12 \| 99 \| 402 \| 12 \| 308 \| Integrase \| Integrase; afdb-uniprot50 \| AF-A0A2D7FTU9-F1-MODEL\_V4 \| 1.0 \| 2.872e-16 \| 557 \| 0.176 \| 425 \| 290 \| 18 \| 1 \| 401 \| 5 \| 393 \| Integrase \| Integrase; afdb-uniprot50 \| AF-A0A8A8H8X0-F1-MODEL\_V4 \| 1.0 \| 9.014e-16 \| 557 \| 0.189 \| 422 \| 285 \| 21 \| 1 \| 402 \| 1 \| 385 \| Tyrosine-type recombinase/integrase \| Tyrosine-type recombinase/integrase; afdb-uniprot50 \| AF-A0A7H8PYI9-F1-MODEL\_V4 \| 1.0 \| 1.384e-15 \| 557 \| 0.198 \| 433 \| 279 \| 22 \| 2 \| 402 \| 8 \| 404 \| Integrase arm-type DNA-binding domain-containing protein \| Integrase arm-type DNA-binding domain-containing protein; afdb-uniprot50 \| AF-A0A562ZNZ3-F1-MODEL\_V4 \| 1.0 \| 5.336e-16 \| 557 \| 0.241 \| 434 \| 275 \| 22 \| 2 \| 403 \| 17 \| 428 \| Integrase family protein \| Integrase family protein; afdb-uniprot50 \| AF-A0A3R0ZT27-F1-MODEL\_V4 \| 1.0 \| 2.229e-15 \| 556 \| 0.188 \| 408 \| 268 \| 19 \| 2 \| 380 \| 26 \| 399 \| DUF4102 domain-containing protein \| DUF4102 domain-containing protein; afdb-uniprot50 \| AF-A0A1M7UU15-F1-MODEL\_V4 \| 1.0 \| 1.523e-15 \| 556 \| 0.167 \| 431 \| 301 \| 23 \| 2 \| 402 \| 3 \| 405 \| Arm-DNA-bind\_3 domain-containing protein \| Arm-DNA-bind\_3 domain-containing protein; afdb-uniprot50 \| AF-A0A2K4MK02-F1-MODEL\_V4 \| 1.0 \| 3.485e-13 \| 555 \| 0.192 \| 328 \| 227 \| 13 \| 93 \| 403 \| 3 \| 309 \| Integrase \| Integrase; afdb-uniprot50 \| AF-A0A1J5IS85-F1-MODEL\_V4 \| 1.0 \| 2.452e-15 \| 555 \| 0.199 \| 416 \| 272 \| 20 \| 2 \| 391 \| 3 \| 383 \| Integrase \| Integrase; afdb-uniprot50 \| AF-A0A4Q5VI67-F1-MODEL\_V4 \| 1.0 \| 1.091e-15 \| 555 \| 0.189 \| 439 \| 287 \| 27 \| 2 \| 402 \| 8 \| 415 \| DUF4102 domain-containing protein \| DUF4102 domain-containing protein; afdb-uniprot50 \| AF-A0A0K0WKY4-F1-MODEL\_V4 \| 1.0 \| 3.645e-16 \| 554 \| 0.19 \| 405 \| 270 \| 18 \| 8 \| 402 \| 9 \| 365 \| Tyr recombinase domain-containing protein \| Tyr recombinase domain-containing protein; afdb-uniprot50 \| AF-A0A6S6UED4-F1-MODEL\_V4 \| 1.0 \| 1.243e-11 \| 552 \| 0.334 \| 203 \| 121 \| 5 \| 209 \| 403 \| 2 \| 198 \| Phage integrase family protein \| Phage integrase family protein; afdb-uniprot50 \| AF-A0A3B7L947-F1-MODEL\_V4 \| 1.0 \| 1.597e-15 \| 552 \| 0.199 \| 426 \| 285 \| 18 \| 1 \| 402 \| 3 \| 396 \| DUF4102 domain-containing protein \| DUF4102 domain-containing protein; afdb-uniprot50 \| AF-A0A8B6QQ55-F1-MODEL\_V4 \| 1.0 \| 2.304e-14 \| 551 \| 0.21 \| 361 \| 244 \| 15 \| 63 \| 402 \| 2 \| 342 \| Tyrosine-type recombinase/integrase \| Tyrosine-type recombinase/integrase; afdb-uniprot50 \| AF-A0A1Z8RU83-F1-MODEL\_V4 \| 1.0 \| 1.32e-15 \| 551 \| 0.194 \| 427 \| 274 \| 19 \| 2 \| 398 \| 4 \| 390 \| Uncharacterized protein \| Uncharacterized protein; afdb-uniprot50 \| AF-A0A2W4RA98-F1-MODEL\_V4 \| 1.0 \| 2.452e-15 \| 551 \| 0.183 \| 419 \| 283 \| 22 \| 3 \| 403 \| 6 \| 383 \| Integrase \| Integrase; afdb-uniprot50 \| AF-A0A2N4WW21-F1-MODEL\_V4 \| 1.0 \| 1.621e-16 \| 551 \| 0.22 \| 450 \| 271 \| 24 \| 1 \| 403 \| 1 \| 417 \| Integrase \| Integrase; afdb-uniprot50 \| AF-A0A560K4G8-F1-MODEL\_V4 \| 1.0 \| 2.489e-16 \| 550 \| 0.215 \| 417 \| 267 \| 20 \| 2 \| 401 \| 6 \| 379 \| Site-specific recombinase XerD \| Site-specific recombinase XerD; afdb-uniprot50 \| AF-A0A2A3LHG3-F1-MODEL\_V4 \| 1.0 \| 5.088e-16 \| 550 \| 0.201 \| 426 \| 278 \| 21 \| 2 \| 392 \| 6 \| 404 \| Uncharacterized protein \| Uncharacterized protein; afdb-uniprot50 \| AF-A0A370K4W5-F1-MODEL\_V4 \| 1.0 \| 2.611e-16 \| 550 \| 0.224 \| 445 \| 279 \| 23 \| 2 \| 398 \| 3 \| 429 \| Uncharacterized protein \| Uncharacterized protein; afdb-uniprot50 \| AF-A0A562R5J7-F1-MODEL\_V4 \| 1.0 \| 8.467e-15 \| 549 \| 0.22 \| 376 \| 250 \| 17 \| 48 \| 402 \| 2 \| 355 \| Integrase \| Integrase; afdb-uniprot50 \| AF-A0A369IHL0-F1-MODEL\_V4 \| 1.0 \| 5.87e-16 \| 549 \| 0.178 \| 426 \| 287 \| 22 \| 2 \| 401 \| 3 \| 391 \| DUF4102 domain-containing protein \| DUF4102 domain-containing protein; afdb-uniprot50 \| AF-A0A7D4SJ89-F1-MODEL\_V4 \| 1.0 \| 1.384e-15 \| 549 \| 0.189 \| 422 \| 290 \| 21 \| 1 \| 402 \| 1 \| 390 \| Tyrosine-type recombinase/integrase \| Tyrosine-type recombinase/integrase; afdb-uniprot50 \| AF-A0A1T0CMB0-F1-MODEL\_V4 \| 1.0 \| 1.757e-15 \| 549 \| 0.166 \| 426 \| 302 \| 19 \| 1 \| 402 \| 1 \| 397 \| Tyr recombinase domain-containing protein \| Tyr recombinase domain-containing protein; afdb-uniprot50 \| AF-A0A4P7BRM8-F1-MODEL\_V4 \| 1.0 \| 9.454e-16 \| 548 \| 0.179 \| 445 \| 294 \| 22 \| 2 \| 403 \| 14 \| 430 \| DUF4102 domain-containing protein \| DUF4102 domain-containing protein; afdb-uniprot50 \| AF-A0A258KZS7-F1-MODEL\_V4 \| 1.0 \| 6.156e-16 \| 547 \| 0.214 \| 411 \| 252 \| 15 \| 46 \| 402 \| 2 \| 395 \| Uncharacterized protein \| Uncharacterized protein; afdb-uniprot50 \| AF-A0A7M1LFV5-F1-MODEL\_V4 \| 1.0 \| 2.452e-15 \| 546 \| 0.185 \| 426 \| 284 \| 21 \| 2 \| 402 \| 3 \| 390 \| Tyrosine-type recombinase/integrase \| Tyrosine-type recombinase/integrase; afdb-uniprot50 \| AF-A0A5C4PWQ3-F1-MODEL\_V4 \| 1.0 \| 1.757e-15 \| 546 \| 0.201 \| 421 \| 278 \| 20 \| 9 \| 401 \| 11 \| 401 \| DUF4102 domain-containing protein \| DUF4102 domain-containing protein; afdb-uniprot50 \| AF-A0A840JLT4-F1-MODEL\_V4 \| 1.0 \| 9.454e-16 \| 546 \| 0.194 \| 427 \| 284 \| 21 \| 2 \| 402 \| 8 \| 400 \| Integrase \| Integrase; afdb-uniprot50 \| AF-A0A2D9M000-F1-MODEL\_V4 \| 1.0 \| 3.012e-16 \| 546 \| 0.175 \| 439 \| 282 \| 21 \| 2 \| 402 \| 9 \| 405 \| Integrase \| Integrase; afdb-uniprot50 \| AF-A0A430DFY4-F1-MODEL\_V4 \| 1.0 \| 4.205e-16 \| 546 \| 0.203 \| 447 \| 275 \| 24 \| 1 \| 400 \| 1 \| 413 \| DUF4102 domain-containing protein \| DUF4102 domain-containing protein; afdb-uniprot50 \| AF-A0A2A2EZC5-F1-MODEL\_V4 \| 1.0 \| 3.423e-15 \| 545 \| 0.185 \| 420 \| 282 \| 18 \| 9 \| 402 \| 11 \| 396 \| Integrase \| Integrase; afdb-uniprot50 \| AF-A0A0M4L5J0-F1-MODEL\_V4 \| 1.0 \| 5.012e-15 \| 544 \| 0.171 \| 437 \| 285 \| 23 \| 1 \| 402 \| 2 \| 396 \| Tyr recombinase domain-containing protein \| Tyr recombinase domain-containing protein; afdb-uniprot50 \| AF-A0A1H8G0Y4-F1-MODEL\_V4 \| 1.0 \| 1.384e-15 \| 544 \| 0.196 \| 418 \| 280 \| 22 \| 2 \| 403 \| 8 \| 385 \| Integrase \| Integrase; afdb-uniprot50 \| AF-Q31FR8-F1-MODEL\_V4 \| 1.0 \| 2.196e-14 \| 543 \| 0.211 \| 425 \| 275 \| 21 \| 2 \| 402 \| 5 \| 393 \| Tyrosine recombinase \| Tyrosine recombinase; afdb-uniprot50 \| AF-A0A369ZF09-F1-MODEL\_V4 \| 1.0 \| 1.842e-15 \| 543 \| 0.174 \| 424 \| 290 \| 16 \| 2 \| 402 \| 8 \| 394 \| DUF4102 domain-containing protein \| DUF4102 domain-containing protein; afdb-uniprot50 \| AF-A0A4P7RZE8-F1-MODEL\_V4 \| 1.0 \| 7.103e-16 \| 543 \| 0.179 \| 428 \| 283 \| 21 \| 2 \| 401 \| 7 \| 394 \| DUF4102 domain-containing protein \| DUF4102 domain-containing protein; afdb-uniprot50 \| AF-A0A0C5WK64-F1-MODEL\_V4 \| 1.0 \| 9.915e-16 \| 543 \| 0.184 \| 450 \| 283 \| 25 \| 2 \| 402 \| 10 \| 424 \| Putative prophage integrase \| Putative prophage integrase; afdb-uniprot50 \| AF-A0A4Z0BRC7-F1-MODEL\_V4 \| 1.0 \| 3.645e-16 \| 543 \| 0.198 \| 439 \| 289 \| 21 \| 1 \| 398 \| 2 \| 418 \| Integrase \| Integrase; afdb-uniprot50 \| AF-R9AZ04-F1-MODEL\_V4 \| 1.0 \| 1.364e-14 \| 542 \| 0.201 \| 372 \| 244 \| 18 \| 50 \| 402 \| 3 \| 340 \| Uncharacterized protein \| Uncharacterized protein; afdb-uniprot50 \| AF-F3WK33-F1-MODEL\_V4 \| 1.0 \| 1.452e-15 \| 542 \| 0.211 \| 383 \| 258 \| 17 \| 1 \| 370 \| 1 \| 352 \| Putative prophage CPS-53 integrase \| Putative prophage CPS-53 integrase; afdb-uniprot50 \| AF-Q1N825-F1-MODEL\_V4 \| 1.0 \| 1.24e-14 \| 542 \| 0.217 \| 382 \| 244 \| 17 \| 50 \| 402 \| 2 \| 357 \| Symbiosis island integrase \| Symbiosis island integrase; afdb-uniprot50 \| AF-G9EN62-F1-MODEL\_V4 \| 1.0 \| 3.591e-15 \| 542 \| 0.211 \| 411 \| 288 \| 17 \| 1 \| 400 \| 1 \| 386 \| Uncharacterized protein \| Uncharacterized protein; afdb-uniprot50 \| AF-A0A1A5PTB8-F1-MODEL\_V4 \| 1.0 \| 1.7e-16 \| 542 \| 0.18 \| 415 \| 283 \| 21 \| 9 \| 402 \| 9 \| 387 \| Tyr recombinase domain-containing protein \| Tyr recombinase domain-containing protein; afdb-uniprot50 \| AF-A0A5C5GCA0-F1-MODEL\_V4 \| 1.0 \| 2.338e-15 \| 542 \| 0.202 \| 404 \| 278 \| 17 \| 9 \| 398 \| 44 \| 417 \| DUF4102 domain-containing protein \| DUF4102 domain-containing protein; afdb-uniprot50 \| AF-D5RQ66-F1-MODEL\_V4 \| 1.0 \| 4.009e-16 \| 542 \| 0.202 \| 444 \| 284 \| 22 \| 9 \| 403 \| 7 \| 429 \| Site-specific recombinase, phage integrase family \| Site-specific recombinase, phage integrase family; afdb-uniprot50 \| AF-A0A845U6K0-F1-MODEL\_V4 \| 1.0 \| 3.159e-16 \| 541 \| 0.192 \| 416 \| 290 \| 20 \| 1 \| 403 \| 6 \| 388 \| Tyrosine-type recombinase/integrase \| Tyrosine-type recombinase/integrase; afdb-uniprot50 \| AF-A0A503K063-F1-MODEL\_V4 \| 1.0 \| 1.32e-15 \| 541 \| 0.158 \| 429 \| 292 \| 22 \| 2 \| 402 \| 6 \| 393 \| DUF4102 domain-containing protein \| DUF4102 domain-containing protein; afdb-uniprot50 \| AF-A0A7H5AGS0-F1-MODEL\_V4 \| 1.0 \| 2.38e-13 \| 540 \| 0.193 \| 336 \| 236 \| 16 \| 82 \| 402 \| 7 \| 322 \| Phage integrase \| Phage integrase; afdb-uniprot50 \| AF-A0A381F1W7-F1-MODEL\_V4 \| 1.0 \| 3.475e-16 \| 540 \| 0.193 \| 398 \| 274 \| 22 \| 1 \| 382 \| 7 \| 373 \| Prophage CP4-57 integrase \| Prophage CP4-57 integrase; afdb-uniprot50 \| AF-A0A2N2RRY9-F1-MODEL\_V4 \| 1.0 \| 1.32e-15 \| 540 \| 0.198 \| 403 \| 268 \| 20 \| 34 \| 403 \| 25 \| 405 \| Integrase \| Integrase; afdb-uniprot50 \| AF-A0A7G5KXN3-F1-MODEL\_V4 \| 1.0 \| 3.766e-15 \| 540 \| 0.167 \| 431 \| 295 \| 19 \| 2 \| 402 \| 3 \| 399 \| Tyrosine-type recombinase/integrase \| Tyrosine-type recombinase/integrase; afdb-uniprot50 \| AF-Q8EJS4-F1-MODEL\_V4 \| 1.0 \| 1.144e-15 \| 540 \| 0.18 \| 437 \| 290 \| 22 \| 2 \| 402 \| 7 \| 411 \| Integrase bacteriophage P4 family \| Integrase bacteriophage P4 family; afdb-uniprot50 \| AF-A0A239M372-F1-MODEL\_V4 \| 1.0 \| 2.229e-15 \| 540 \| 0.2 \| 429 \| 284 \| 23 \| 2 \| 402 \| 49 \| 446 \| Integrase \| Integrase; afdb-uniprot50 \| AF-A0A1E2UTE1-F1-MODEL\_V4 \| 1.0 \| 3.159e-16 \| 539 \| 0.201 \| 421 \| 286 \| 19 \| 1 \| 400 \| 1 \| 392 \| Integrase \| Integrase; afdb-uniprot50 \| AF-A0A1I5H7Y3-F1-MODEL\_V4 \| 1.0 \| 1.04e-15 \| 538 \| 0.219 \| 423 \| 269 \| 21 \| 2 \| 403 \| 4 \| 386 \| Integrase \| Integrase; afdb-uniprot50 \| AF-A0A7S9REC9-F1-MODEL\_V4 \| 1.0 \| 3.423e-15 \| 538 \| 0.194 \| 421 \| 283 \| 21 \| 2 \| 400 \| 8 \| 394 \| Tyrosine-type recombinase/integrase \| Tyrosine-type recombinase/integrase; afdb-uniprot50 \| AF-A0A0D6MPY2-F1-MODEL\_V4 \| 1.0 \| 2.698e-15 \| 538 \| 0.224 \| 428 \| 264 \| 21 \| 2 \| 399 \| 9 \| 398 \| Phage integrase \| Phage integrase; afdb-uniprot50 \| AF-A0A6I4Z264-F1-MODEL\_V4 \| 1.0 \| 2.829e-15 \| 538 \| 0.169 \| 430 \| 293 \| 18 \| 2 \| 401 \| 18 \| 413 \| Tyrosine-type recombinase/integrase \| Tyrosine-type recombinase/integrase; afdb-uniprot50 \| AF-A0A3A1PIJ8-F1-MODEL\_V4 \| 1.0 \| 8.88e-15 \| 538 \| 0.171 \| 415 \| 293 \| 22 \| 9 \| 401 \| 23 \| 408 \| DUF4102 domain-containing protein \| DUF4102 domain-containing protein; afdb-uniprot50 \| AF-A0A2W4X528-F1-MODEL\_V4 \| 1.0 \| 1.144e-15 \| 537 \| 0.205 \| 400 \| 252 \| 20 \| 2 \| 373 \| 3 \| 364 \| Integrase \| Integrase; afdb-uniprot50 \| AF-A0A378NE62-F1-MODEL\_V4 \| 1.0 \| 4.779e-15 \| 537 \| 0.205 \| 419 \| 275 \| 18 \| 2 \| 397 \| 8 \| 391 \| Prophage CP4-57 integrase \| Prophage CP4-57 integrase; afdb-uniprot50 \| AF-A0A4P5WYK2-F1-MODEL\_V4 \| 1.0 \| 3.645e-16 \| 537 \| 0.231 \| 414 \| 273 \| 20 \| 2 \| 402 \| 18 \| 399 \| Integrase \| Integrase; afdb-uniprot50 \| AF-A0A2S5PFB0-F1-MODEL\_V4 \| 1.0 \| 5.87e-16 \| 536 \| 0.199 \| 417 \| 275 \| 17 \| 9 \| 400 \| 16 \| 398 \| Integrase \| Integrase; afdb-uniprot50 \| AF-A0A1T0A3C6-F1-MODEL\_V4 \| 1.0 \| 1.24e-14 \| 536 \| 0.186 \| 423 \| 292 \| 17 \| 2 \| 403 \| 7 \| 398 \| Prophage CP4-57 integrase \| Prophage CP4-57 integrase; afdb-uniprot50 \| AF-A0A1G8PZJ9-F1-MODEL\_V4 \| 1.0 \| 2.338e-15 \| 536 \| 0.195 \| 424 \| 284 \| 17 \| 2 \| 402 \| 8 \| 397 \| Integrase \| Integrase; afdb-uniprot50 \| AF-A0A7Y0E2C5-F1-MODEL\_V4 \| 1.0 \| 5.336e-16 \| 536 \| 0.201 \| 422 \| 276 \| 21 \| 2 \| 398 \| 8 \| 393 \| Tyrosine-type recombinase/integrase \| Tyrosine-type recombinase/integrase; afdb-uniprot50 \| AF-A0A431PNN7-F1-MODEL\_V4 \| 1.0 \| 7.813e-16 \| 536 \| 0.206 \| 431 \| 273 \| 20 \| 1 \| 398 \| 8 \| 402 \| Site-specific integrase \| Site-specific integrase; afdb-uniprot50 \| AF-A0A349YDR9-F1-MODEL\_V4 \| 1.0 \| 7.339e-15 \| 536 \| 0.195 \| 441 \| 280 \| 21 \| 2 \| 398 \| 8 \| 417 \| Integrase \| Integrase; afdb-uniprot50 \| AF-A4A7S2-F1-MODEL\_V4 \| 1.0 \| 6.997e-15 \| 535 \| 0.2 \| 435 \| 282 \| 23 \| 2 \| 403 \| 8 \| 409 \| Integrase \| Integrase; afdb-uniprot50 \| AF-A0A0T6W442-F1-MODEL\_V4 \| 1.0 \| 3.766e-15 \| 535 \| 0.192 \| 394 \| 282 \| 17 \| 29 \| 400 \| 32 \| 411 \| Integrase \| Integrase; afdb-uniprot50 \| AF-A0A5C7RHY4-F1-MODEL\_V4 \| 1.0 \| 1.409e-13 \| 534 \| 0.207 \| 361 \| 254 \| 12 \| 56 \| 402 \| 11 \| 353 \| DUF4102 domain-containing protein \| DUF4102 domain-containing protein; afdb-uniprot50 \| AF-A0A2N4XR17-F1-MODEL\_V4 \| 1.0 \| 3.475e-16 \| 534 \| 0.212 \| 423 \| 270 \| 18 \| 2 \| 398 \| 7 \| 392 \| Integrase \| Integrase; afdb-uniprot50 \| AF-A0A241WFB4-F1-MODEL\_V4 \| 1.0 \| 1.025e-14 \| 534 \| 0.201 \| 422 \| 275 \| 20 \| 9 \| 402 \| 14 \| 401 \| Integrase \| Integrase; afdb-uniprot50 \| AF-G2LET3-F1-MODEL\_V4 \| 1.0 \| 4.345e-15 \| 533 \| 0.202 \| 370 \| 263 \| 15 \| 44 \| 400 \| 9 \| 359 \| Integrase \| Integrase; afdb-uniprot50 \| AF-A0A345QRY7-F1-MODEL\_V4 \| 1.0 \| 1.258e-15 \| 533 \| 0.196 \| 428 \| 279 \| 24 \| 2 \| 403 \| 9 \| 397 \| Integrase \| Integrase; afdb-uniprot50 \| AF-A0A841S252-F1-MODEL\_V4 \| 1.0 \| 8.073e-15 \| 533 \| 0.183 \| 436 \| 286 \| 23 \| 2 \| 402 \| 13 \| 413 \| Integrase \| Integrase; afdb-uniprot50 \| AF-A0A6M4GUT6-F1-MODEL\_V4 \| 1.0 \| 1.384e-15 \| 532 \| 0.181 \| 436 \| 285 \| 23 \| 2 \| 401 \| 4 \| 403 \| Prophage integrase IntA \| Prophage integrase IntA; afdb-uniprot50 \| AF-A0A2N1U9J3-F1-MODEL\_V4 \| 1.0 \| 2.027e-15 \| 532 \| 0.194 \| 422 \| 289 \| 19 \| 2 \| 402 \| 3 \| 394 \| Integrase \| Integrase; afdb-uniprot50 \| AF-A0A1E4GTV1-F1-MODEL\_V4 \| 1.0 \| 2.338e-15 \| 532 \| 0.207 \| 420 \| 276 \| 19 \| 2 \| 398 \| 9 \| 394 \| Uncharacterized protein \| Uncharacterized protein; afdb-uniprot50 \| AF-A0A430DCH7-F1-MODEL\_V4 \| 1.0 \| 1.04e-15 \| 532 \| 0.198 \| 448 \| 286 \| 23 \| 2 \| 402 \| 7 \| 428 \| Tyr recombinase domain-containing protein \| Tyr recombinase domain-containing protein; afdb-uniprot50 \| AF-A0A1B8PU39-F1-MODEL\_V4 \| 1.0 \| 4.142e-15 \| 531 \| 0.198 \| 424 \| 284 \| 22 \| 2 \| 401 \| 8 \| 399 \| Tyr recombinase domain-containing protein \| Tyr recombinase domain-containing protein; afdb-uniprot50 \| AF-A0A6I4V2B2-F1-MODEL\_V4 \| 1.0 \| 3.591e-15 \| 531 \| 0.184 \| 423 \| 285 \| 19 \| 2 \| 401 \| 8 \| 393 \| Tyrosine-type recombinase/integrase \| Tyrosine-type recombinase/integrase; afdb-uniprot50 \| AF-A0A7S7ALF1-F1-MODEL\_V4 \| 1.0 \| 8.467e-15 \| 531 \| 0.152 \| 426 \| 301 \| 20 \| 8 \| 402 \| 3 \| 399 \| Tyrosine-type recombinase/integrase \| Tyrosine-type recombinase/integrase; afdb-uniprot50 \| AF-A0A3S5DFZ5-F1-MODEL\_V4 \| 1.0 \| 1.731e-14 \| 530 \| 0.203 \| 374 \| 254 \| 15 \| 50 \| 403 \| 2 \| 351 \| Tyrosine recombinase xerD \| Tyrosine recombinase xerD; afdb-uniprot50 \| AF-A0A3B0XZC6-F1-MODEL\_V4 \| 1.0 \| 3.264e-15 \| 530 \| 0.188 \| 424 \| 282 \| 20 \| 2 \| 402 \| 4 \| 388 \| Tyr recombinase domain-containing protein \| Tyr recombinase domain-containing protein; afdb-uniprot50 \| AF-A0A7V8XVF4-F1-MODEL\_V4 \| 1.0 \| 6.691e-12 \| 529 \| 0.286 \| 251 \| 159 \| 9 \| 156 \| 402 \| 3 \| 237 \| Site-specific integrase \| Site-specific integrase; afdb-uniprot50 \| AF-A0A833P2C7-F1-MODEL\_V4 \| 1.0 \| 1.523e-15 \| 529 \| 0.196 \| 433 \| 283 \| 23 \| 1 \| 403 \| 13 \| 410 \| Integrase family \| Integrase family; afdb-uniprot50 \| AF-A0A2A6L6G2-F1-MODEL\_V4 \| 1.0 \| 1.65e-14 \| 529 \| 0.206 \| 441 \| 286 \| 23 \| 2 \| 402 \| 9 \| 425 \| Integrase \| Integrase; afdb-uniprot50 \| AF-A0A6M7UIT5-F1-MODEL\_V4 \| 1.0 \| 4.009e-16 \| 529 \| 0.185 \| 415 \| 281 \| 22 \| 9 \| 402 \| 64 \| 442 \| DUF4102 domain-containing protein \| DUF4102 domain-containing protein; afdb-uniprot50 \| AF-A0A831RFM1-F1-MODEL\_V4 \| 1.0 \| 5.257e-15 \| 528 \| 0.249 \| 401 \| 239 \| 24 \| 2 \| 367 \| 3 \| 376 \| DUF4102 domain-containing protein \| DUF4102 domain-containing protein; afdb-uniprot50 \| AF-A0A7H9F2Z4-F1-MODEL\_V4 \| 1.0 \| 5.336e-16 \| 528 \| 0.225 \| 417 \| 269 \| 19 \| 2 \| 402 \| 3 \| 381 \| Tyrosine-type recombinase/integrase \| Tyrosine-type recombinase/integrase; afdb-uniprot50 \| AF-A0A7W4FUN8-F1-MODEL\_V4 \| 1.0 \| 6.361e-15 \| 528 \| 0.169 \| 437 \| 292 \| 18 \| 2 \| 402 \| 3 \| 404 \| Site-specific integrase \| Site-specific integrase; afdb-uniprot50 \| AF-A0A2A4Y533-F1-MODEL\_V4 \| 1.0 \| 2.698e-15 \| 528 \| 0.153 \| 469 \| 298 \| 24 \| 1 \| 402 \| 10 \| 446 \| Tyr recombinase domain-containing protein \| Tyr recombinase domain-containing protein; afdb-uniprot50 \| AF-A0A528IPC3-F1-MODEL\_V4 \| 1.0 \| 3.891e-14 \| 527 \| 0.29 \| 227 \| 150 \| 6 \| 1 \| 224 \| 3 \| 221 \| Site-specific integrase \| Site-specific integrase; afdb-uniprot50 \| AF-A0A258Q6T4-F1-MODEL\_V4 \| 1.0 \| 2.829e-15 \| 527 \| 0.224 \| 375 \| 238 \| 19 \| 1 \| 354 \| 1 \| 343 \| Integrase \| Integrase; afdb-uniprot50 \| AF-A0A7S6S5Y9-F1-MODEL\_V4 \| 1.0 \| 1.075e-14 \| 527 \| 0.187 \| 415 \| 275 \| 20 \| 9 \| 400 \| 11 \| 386 \| Integrase arm-type DNA-binding domain-containing protein \| Integrase arm-type DNA-binding domain-containing protein; afdb-uniprot50 \| AF-V8FUN5-F1-MODEL\_V4 \| 1.0 \| 1.091e-15 \| 527 \| 0.191 \| 422 \| 272 \| 19 \| 2 \| 402 \| 3 \| 376 \| Tyr recombinase domain-containing protein \| Tyr recombinase domain-containing protein; afdb-uniprot50 \| AF-A0A3E0Q8B8-F1-MODEL\_V4 \| 1.0 \| 2.738e-16 \| 527 \| 0.231 \| 406 \| 265 \| 18 \| 9 \| 401 \| 8 \| 379 \| DUF4102 domain-containing protein \| DUF4102 domain-containing protein; afdb-uniprot50 \| AF-A0A661F4U9-F1-MODEL\_V4 \| 1.0 \| 2.158e-16 \| 526 \| 0.211 \| 411 \| 273 \| 17 \| 1 \| 392 \| 10 \| 388 \| Tyr recombinase domain-containing protein \| Tyr recombinase domain-containing protein; afdb-uniprot50 \| AF-A0A4P7WFD3-F1-MODEL\_V4 \| 1.0 \| 8.467e-15 \| 526 \| 0.189 \| 422 \| 296 \| 18 \| 2 \| 403 \| 9 \| 404 \| DUF4102 domain-containing protein \| DUF4102 domain-containing protein; afdb-uniprot50 \| AF-A0A7X4K6Q4-F1-MODEL\_V4 \| 1.0 \| 6.457e-16 \| 526 \| 0.224 \| 424 \| 256 \| 23 \| 13 \| 400 \| 5 \| 391 \| Tyrosine-type recombinase/integrase \| Tyrosine-type recombinase/integrase; afdb-uniprot50 \| AF-A0A800GIQ3-F1-MODEL\_V4 \| 1.0 \| 3.112e-15 \| 526 \| 0.153 \| 416 \| 308 \| 21 \| 2 \| 398 \| 1 \| 391 \| Site-specific integrase \| Site-specific integrase; afdb-uniprot50 \| AF-A0A1D8UTH6-F1-MODEL\_V4 \| 1.0 \| 4.779e-15 \| 525 \| 0.187 \| 426 \| 270 \| 20 \| 15 \| 400 \| 1 \| 390 \| Tyr recombinase domain-containing protein \| Tyr recombinase domain-containing protein; afdb-uniprot50 \| AF-A0A158AZQ9-F1-MODEL\_V4 \| 1.0 \| 2.452e-15 \| 525 \| 0.189 \| 432 \| 276 \| 23 \| 9 \| 398 \| 11 \| 410 \| Phage integrase \| Phage integrase; afdb-uniprot50 \| AF-A0A1X1PI96-F1-MODEL\_V4 \| 1.0 \| 7.813e-16 \| 525 \| 0.2 \| 443 \| 281 \| 19 \| 2 \| 398 \| 3 \| 418 \| Prophage integrase IntS \| Prophage integrase IntS; afdb-uniprot50 \| AF-A0A2W4K9L1-F1-MODEL\_V4 \| 1.0 \| 8.073e-15 \| 525 \| 0.187 \| 427 \| 293 \| 20 \| 2 \| 402 \| 14 \| 412 \| Integrase \| Integrase; afdb-uniprot50 \| AF-A0A7W7E8P4-F1-MODEL\_V4 \| 1.0 \| 7.471e-13 \| 524 \| 0.186 \| 349 \| 249 \| 15 \| 69 \| 402 \| 3 \| 331 \| Integrase \| Integrase; afdb-uniprot50 \| AF-A0A6B8KHW6-F1-MODEL\_V4 \| 1.0 \| 1.24e-14 \| 524 \| 0.204 \| 410 \| 276 \| 22 \| 2 \| 403 \| 8 \| 375 \| Tyrosine-type recombinase/integrase \| Tyrosine-type recombinase/integrase; afdb-uniprot50 \| AF-A0A241VD59-F1-MODEL\_V4 \| 1.0 \| 1.675e-15 \| 524 \| 0.167 \| 425 \| 303 \| 19 \| 1 \| 402 \| 9 \| 405 \| Tyr recombinase domain-containing protein \| Tyr recombinase domain-containing protein; afdb-uniprot50 \| AF-A0A1W9HT15-F1-MODEL\_V4 \| 1.0 \| 4.779e-15 \| 524 \| 0.164 \| 419 \| 291 \| 20 \| 2 \| 398 \| 13 \| 394 \| Core-binding (CB) domain-containing protein \| Core-binding (CB) domain-containing protein; afdb-uniprot50 \| AF-A0A7Y2KR74-F1-MODEL\_V4 \| 1.0 \| 1.091e-15 \| 524 \| 0.206 \| 427 \| 280 \| 19 \| 2 \| 398 \| 8 \| 405 \| Tyrosine-type recombinase/integrase \| Tyrosine-type recombinase/integrase; afdb-uniprot50 \| AF-A0A5S4W293-F1-MODEL\_V4 \| 1.0 \| 9.314e-15 \| 524 \| 0.198 \| 428 \| 285 \| 23 \| 2 \| 402 \| 19 \| 415 \| DUF4102 domain-containing protein \| DUF4102 domain-containing protein; afdb-uniprot50 \| AF-A0A141RCN1-F1-MODEL\_V4 \| 1.0 \| 1.324e-12 \| 523 \| 0.178 \| 313 \| 225 \| 10 \| 102 \| 402 \| 6 \| 298 \| Phage integrase family \| Phage integrase family; afdb-uniprot50 \| AF-A0A7Y9SXB0-F1-MODEL\_V4 \| 1.0 \| 9.176e-14 \| 523 \| 0.216 \| 374 \| 244 \| 16 \| 53 \| 402 \| 1 \| 349 \| Integrase \| Integrase; afdb-uniprot50 \| AF-Q3SUE5-F1-MODEL\_V4 \| 1.0 \| 1.144e-15 \| 523 \| 0.211 \| 420 \| 270 \| 23 \| 2 \| 400 \| 5 \| 384 \| Phage integrase \| Phage integrase; afdb-uniprot50 \| AF-B1YRM4-F1-MODEL\_V4 \| 1.0 \| 6.792e-13 \| 522 \| 0.237 \| 278 \| 186 \| 9 \| 136 \| 402 \| 6 \| 268 \| Integrase family protein \| Integrase family protein; afdb-uniprot50 \| AF-A0A7Z1LPX5-F1-MODEL\_V4 \| 1.0 \| 4.779e-15 \| 522 \| 0.205 \| 424 \| 273 \| 21 \| 2 \| 403 \| 8 \| 389 \| Integrase \| Integrase; afdb-uniprot50 \| AF-A0A2M8QRZ4-F1-MODEL\_V4 \| 1.0 \| 6.997e-15 \| 522 \| 0.206 \| 430 \| 275 \| 24 \| 2 \| 403 \| 4 \| 395 \| Tyr recombinase domain-containing protein \| Tyr recombinase domain-containing protein; afdb-uniprot50 \| AF-A0A196QT34-F1-MODEL\_V4 \| 1.0 \| 6.457e-16 \| 522 \| 0.191 \| 448 \| 282 \| 22 \| 2 \| 400 \| 8 \| 424 \| Tyr recombinase domain-containing protein \| Tyr recombinase domain-containing protein; afdb-uniprot50 \| AF-A0A836Q3M7-F1-MODEL\_V4 \| 1.0 \| 9.769e-15 \| 522 \| 0.182 \| 423 \| 292 \| 20 \| 2 \| 402 \| 94 \| 484 \| DUF4102 domain-containing protein \| DUF4102 domain-containing protein; afdb-uniprot50 \| AF-B1FYI2-F1-MODEL\_V4 \| 1.0 \| 6.38e-12 \| 521 \| 0.21 \| 314 \| 222 \| 15 \| 99 \| 403 \| 4 \| 300 \| Integrase family protein \| Integrase family protein; afdb-uniprot50 \| AF-A0A424MZS9-F1-MODEL\_V4 \| 1.0 \| 4.142e-15 \| 521 \| 0.186 \| 428 \| 290 \| 23 \| 1 \| 403 \| 1 \| 395 \| DUF4102 domain-containing protein \| DUF4102 domain-containing protein; afdb-uniprot50 \| AF-A0A3Z6QG17-F1-MODEL\_V4 \| 1.0 \| 3.423e-15 \| 521 \| 0.165 \| 434 \| 297 \| 23 \| 2 \| 402 \| 3 \| 404 \| DUF4102 domain-containing protein \| DUF4102 domain-containing protein; afdb-uniprot50 \| AF-A0A1G3CT50-F1-MODEL\_V4 \| 1.0 \| 2.126e-15 \| 521 \| 0.187 \| 426 \| 290 \| 18 \| 2 \| 403 \| 8 \| 401 \| Uncharacterized protein \| Uncharacterized protein; afdb-uniprot50 \| AF-A0A258D734-F1-MODEL\_V4 \| 1.0 \| 1.997e-14 \| 520 \| 0.188 \| 440 \| 279 \| 20 \| 2 \| 398 \| 8 \| 412 \| Integrase \| Integrase; afdb-uniprot50 \| AF-A0A7X5Y2R4-F1-MODEL\_V4 \| 1.0 \| 3.274e-12 \| 519 \| 0.196 \| 301 \| 198 \| 15 \| 116 \| 402 \| 6 \| 276 \| Integrase \| Integrase; afdb-uniprot50 \| AF-A0A2T3P7B2-F1-MODEL\_V4 \| 1.0 \| 1.842e-15 \| 519 \| 0.194 \| 411 \| 282 \| 18 \| 2 \| 398 \| 32 \| 407 \| Tyr recombinase domain-containing protein \| Tyr recombinase domain-containing protein; afdb-uniprot50 \| AF-A0A378IJ47-F1-MODEL\_V4 \| 1.0 \| 2.345e-12 \| 518 \| 0.223 \| 269 \| 183 \| 9 \| 146 \| 402 \| 3 \| 257 \| Integrase \| Integrase; afdb-uniprot50 \| AF-K2EIJ5-F1-MODEL\_V4 \| 1.0 \| 1.091e-15 \| 518 \| 0.197 \| 416 \| 269 \| 25 \| 2 \| 401 \| 4 \| 370 \| Tyr recombinase domain-containing protein \| Tyr recombinase domain-containing protein; afdb-uniprot50 \| AF-A0A522G4K5-F1-MODEL\_V4 \| 1.0 \| 1.573e-14 \| 518 \| 0.215 \| 418 \| 279 \| 21 \| 2 \| 403 \| 5 \| 389 \| Site-specific integrase \| Site-specific integrase; afdb-uniprot50 \| AF-A0A1N7FGQ2-F1-MODEL\_V4 \| 1.0 \| 1.258e-15 \| 518 \| 0.195 \| 419 \| 285 \| 19 \| 1 \| 402 \| 5 \| 388 \| Phage integrase family protein \| Phage integrase family protein; afdb-uniprot50 \| AF-A0A6P1RCR7-F1-MODEL\_V4 \| 1.0 \| 1.24e-14 \| 517 \| 0.192 \| 421 \| 281 \| 20 \| 9 \| 403 \| 12 \| 399 \| DUF4102 domain-containing protein \| DUF4102 domain-containing protein; afdb-uniprot50 \| AF-A0A7Z8ZRH8-F1-MODEL\_V4 \| 1.0 \| 8.194e-16 \| 517 \| 0.183 \| 435 \| 271 \| 22 \| 2 \| 402 \| 3 \| 387 \| Phage integrase family site specific recombinase \| Phage integrase family site specific recombinase; afdb-uniprot50 \| AF-A0A1Y1SIP4-F1-MODEL\_V4 \| 1.0 \| 3.264e-15 \| 516 \| 0.196 \| 428 \| 273 \| 21 \| 2 \| 398 \| 4 \| 391 \| Integrase family protein \| Integrase family protein; afdb-uniprot50 \| AF-A0A7Z0APY4-F1-MODEL\_V4 \| 1.0 \| 1.523e-15 \| 516 \| 0.194 \| 426 \| 282 \| 19 \| 9 \| 403 \| 3 \| 398 \| Integrase \| Integrase; afdb-uniprot50 \| AF-A0A6L4AZ65-F1-MODEL\_V4 \| 1.0 \| 3.264e-15 \| 515 \| 0.197 \| 405 \| 276 \| 18 \| 2 \| 392 \| 9 \| 378 \| Tyrosine-type recombinase/integrase \| Tyrosine-type recombinase/integrase; afdb-uniprot50 \| AF-A0A271K6L1-F1-MODEL\_V4 \| 1.0 \| 8.467e-15 \| 514 \| 0.204 \| 421 \| 274 \| 23 \| 2 \| 403 \| 8 \| 386 \| Uncharacterized protein \| Uncharacterized protein; afdb-uniprot50 \| AF-A0A6N7GUR2-F1-MODEL\_V4 \| 1.0 \| 2.967e-15 \| 514 \| 0.182 \| 432 \| 287 \| 18 \| 2 \| 403 \| 8 \| 403 \| DUF4102 domain-containing protein \| DUF4102 domain-containing protein; afdb-uniprot50 \| AF-A0A1G8B2A4-F1-MODEL\_V4 \| 1.0 \| 2.967e-15 \| 513 \| 0.202 \| 414 \| 267 \| 23 \| 2 \| 401 \| 4 \| 368 \| Site-specific recombinase XerD \| Site-specific recombinase XerD; afdb-uniprot50 \| AF-A0A7W9S1I0-F1-MODEL\_V4 \| 1.0 \| 2.787e-14 \| 513 \| 0.179 \| 418 \| 286 \| 19 \| 2 \| 398 \| 5 \| 386 \| Integrase \| Integrase; afdb-uniprot50 \| AF-A0A0G9KC29-F1-MODEL\_V4 \| 1.0 \| 1.009e-13 \| 513 \| 0.165 \| 436 \| 300 \| 18 \| 1 \| 403 \| 5 \| 409 \| Integrase \| Integrase; afdb-uniprot50 \| AF-A0A7L8RUW8-F1-MODEL\_V4 \| 1.0 \| 3.95e-15 \| 513 \| 0.193 \| 435 \| 286 \| 22 \| 1 \| 402 \| 5 \| 407 \| Tyrosine-type recombinase/integrase \| Tyrosine-type recombinase/integrase; afdb-uniprot50 \| AF-A0A2P8FFM7-F1-MODEL\_V4 \| 1.0 \| 1.147e-12 \| 512 \| 0.201 \| 317 \| 215 \| 13 \| 102 \| 402 \| 7 \| 301 \| Integrase \| Integrase; afdb-uniprot50 \| AF-A0A6N9BY28-F1-MODEL\_V4 \| 1.0 \| 1.815e-14 \| 512 \| 0.215 \| 390 \| 256 \| 20 \| 38 \| 403 \| 2 \| 365 \| Tyrosine-type recombinase/integrase \| Tyrosine-type recombinase/integrase; afdb-uniprot50 \| AF-A0A178L9N4-F1-MODEL\_V4 \| 1.0 \| 5.87e-16 \| 512 \| 0.204 \| 415 \| 276 \| 18 \| 8 \| 400 \| 2 \| 384 \| Uncharacterized protein \| Uncharacterized protein; afdb-uniprot50 \| AF-A0A4V1V182-F1-MODEL\_V4 \| 1.0 \| 1.573e-14 \| 512 \| 0.193 \| 423 \| 281 \| 24 \| 1 \| 401 \| 5 \| 389 \| Site-specific integrase \| Site-specific integrase; afdb-uniprot50 \| AF-A0A0Q7VZE2-F1-MODEL\_V4 \| 1.0 \| 3.834e-13 \| 511 \| 0.213 \| 333 \| 216 \| 15 \| 92 \| 402 \| 7 \| 315 \| Uncharacterized protein \| Uncharacterized protein; afdb-uniprot50 \| AF-A0A7Y9WLG8-F1-MODEL\_V4 \| 1.0 \| 1.815e-14 \| 511 \| 0.179 \| 413 \| 274 \| 23 \| 2 \| 400 \| 7 \| 368 \| Integrase \| Integrase; afdb-uniprot50 \| AF-C6XJP3-F1-MODEL\_V4 \| 1.0 \| 8.467e-15 \| 511 \| 0.179 \| 423 \| 283 \| 22 \| 2 \| 398 \| 9 \| 393 \| Integrase family protein \| Integrase family protein; afdb-uniprot50 \| AF-A0A7X8TU71-F1-MODEL\_V4 \| 1.0 \| 1.5e-14 \| 511 \| 0.174 \| 425 \| 292 \| 21 \| 2 \| 403 \| 3 \| 391 \| Site-specific integrase \| Site-specific integrase; afdb-uniprot50 \| AF-A0A6H3SY94-F1-MODEL\_V4 \| 1.0 \| 4.489e-14 \| 511 \| 0.18 \| 415 \| 281 \| 20 \| 14 \| 402 \| 22 \| 403 \| Phage integrase \| Phage integrase; afdb-uniprot50 \| AF-A0A7G8Q4D4-F1-MODEL\_V4 \| 1.0 \| 4.557e-15 \| 511 \| 0.206 \| 459 \| 260 \| 25 \| 8 \| 402 \| 8 \| 426 \| Site-specific integrase \| Site-specific integrase; afdb-uniprot50 \| AF-A0A2A4AWM1-F1-MODEL\_V4 \| 1.0 \| 6.997e-15 \| 510 \| 0.162 \| 430 \| 292 \| 20 \| 2 \| 402 \| 11 \| 401 \| Integrase \| Integrase; afdb-uniprot50 \| AF-A0A3B9UHJ3-F1-MODEL\_V4 \| 1.0 \| 3.121e-12 \| 509 \| 0.189 \| 332 \| 224 \| 14 \| 90 \| 402 \| 1 \| 306 \| Integrase \| Integrase; afdb-uniprot50 \| AF-A0A077QQF8-F1-MODEL\_V4 \| 1.0 \| 3.121e-12 \| 509 \| 0.192 \| 327 \| 235 \| 10 \| 86 \| 402 \| 9 \| 316 \| Integrase \| Integrase; afdb-uniprot50 \| AF-A0A4Q7BLR8-F1-MODEL\_V4 \| 1.0 \| 2.658e-14 \| 509 \| 0.182 \| 416 \| 289 \| 20 \| 2 \| 401 \| 12 \| 392 \| Site-specific integrase \| Site-specific integrase; afdb-uniprot50 \| AF-A0A142W0M6-F1-MODEL\_V4 \| 1.0 \| 5.783e-15 \| 509 \| 0.179 \| 435 \| 280 \| 23 \| 8 \| 400 \| 7 \| 406 \| Uncharacterized protein \| Uncharacterized protein; afdb-uniprot50 \| AF-A0A443L961-F1-MODEL\_V4 \| 1.0 \| 2.094e-14 \| 509 \| 0.188 \| 419 \| 283 \| 20 \| 1 \| 400 \| 3 \| 383 \| DUF4102 domain-containing protein \| DUF4102 domain-containing protein; afdb-uniprot50 \| AF-A0A6L6YGQ9-F1-MODEL\_V4 \| 1.0 \| 3.066e-14 \| 507 \| 0.176 \| 420 \| 287 \| 21 \| 1 \| 398 \| 1 \| 383 \| Tyrosine-type recombinase/integrase \| Tyrosine-type recombinase/integrase; afdb-uniprot50 \| AF-A0A2I8Q3E1-F1-MODEL\_V4 \| 1.0 \| 2.658e-14 \| 507 \| 0.209 \| 430 \| 281 \| 21 \| 1 \| 403 \| 3 \| 400 \| Integrase \| Integrase; afdb-uniprot50 \| AF-A0A1W9JA93-F1-MODEL\_V4 \| 1.0 \| 1.3e-14 \| 507 \| 0.181 \| 425 \| 292 \| 18 \| 16 \| 402 \| 8 \| 414 \| Tyr recombinase domain-containing protein \| Tyr recombinase domain-containing protein; afdb-uniprot50 \| AF-A0A6A4RG79-F1-MODEL\_V4 \| 1.0 \| 7.339e-15 \| 506 \| 0.198 \| 439 \| 273 \| 23 \| 2 \| 402 \| 8 \| 405 \| DUF4102 domain-containing protein \| DUF4102 domain-containing protein; afdb-uniprot50 \| AF-A0A1H0NXL1-F1-MODEL\_V4 \| 1.0 \| 1.573e-14 \| 506 \| 0.226 \| 419 \| 245 \| 18 \| 36 \| 398 \| 2 \| 397 \| Uncharacterized protein \| Uncharacterized protein; afdb-uniprot50 \| AF-A0A560FA05-F1-MODEL\_V4 \| 1.0 \| 5.783e-15 \| 506 \| 0.221 \| 393 \| 250 \| 21 \| 2 \| 371 \| 3 \| 362 \| Integrase \| Integrase; afdb-uniprot50 \| AF-A0A1A7QBR9-F1-MODEL\_V4 \| 1.0 \| 1.904e-14 \| 505 \| 0.172 \| 417 \| 294 \| 21 \| 8 \| 402 \| 6 \| 393 \| Tyr recombinase domain-containing protein \| Tyr recombinase domain-containing protein; afdb-uniprot50 \| AF-A0A4R0BFY7-F1-MODEL\_V4 \| 1.0 \| 4.938e-14 \| 505 \| 0.178 \| 437 \| 304 \| 21 \| 1 \| 402 \| 6 \| 422 \| DUF4102 domain-containing protein \| DUF4102 domain-containing protein; afdb-uniprot50 \| AF-A0A7H9JPV7-F1-MODEL\_V4 \| 1.0 \| 1.075e-14 \| 505 \| 0.179 \| 434 \| 291 \| 22 \| 2 \| 402 \| 3 \| 404 \| Integrase arm-type DNA-binding domain-containing protein \| Integrase arm-type DNA-binding domain-containing protein; afdb-uniprot50 \| AF-A0A4Q3ZIF4-F1-MODEL\_V4 \| 1.0 \| 6.38e-12 \| 504 \| 0.198 \| 312 \| 221 \| 13 \| 102 \| 402 \| 18 \| 311 \| Integrase \| Integrase; afdb-uniprot50 \| AF-A0A6N3R5Y9-F1-MODEL\_V4 \| 1.0 \| 9.481e-13 \| 504 \| 0.207 \| 361 \| 248 \| 14 \| 60 \| 402 \| 2 \| 342 \| Prophage CP4-57 integrase \| Prophage CP4-57 integrase; afdb-uniprot50 \| AF-A0A2X3G9U9-F1-MODEL\_V4 \| 1.0 \| 5.975e-14 \| 502 \| 0.198 \| 377 \| 247 \| 21 \| 2 \| 358 \| 3 \| 344 \| Putative phage integrase \| Putative phage integrase; afdb-uniprot50 \| AF-A0A3S1ZPH6-F1-MODEL\_V4 \| 1.0 \| 4.28e-14 \| 502 \| 0.195 \| 430 \| 260 \| 24 \| 2 \| 403 \| 8 \| 379 \| Site-specific integrase \| Site-specific integrase; afdb-uniprot50 \| AF-A0A3N0U659-F1-MODEL\_V4 \| 1.0 \| 9.944e-13 \| 501 \| 0.192 \| 358 \| 252 \| 12 \| 62 \| 402 \| 2 \| 339 \| DUF4102 domain-containing protein \| DUF4102 domain-containing protein; afdb-uniprot50 \| AF-M9RKD3-F1-MODEL\_V4 \| 1.0 \| 7.583e-14 \| 501 \| 0.19 \| 425 \| 284 \| 23 \| 1 \| 398 \| 4 \| 395 \| p4 family integrase \| p4 family integrase; afdb-uniprot50 \| AF-A0A0E2PF44-F1-MODEL\_V4 \| 1.0 \| 9.454e-16 \| 501 \| 0.245 \| 412 \| 249 \| 22 \| 2 \| 393 \| 22 \| 391 \| Uncharacterized protein \| Uncharacterized protein; afdb-uniprot50 \| AF-A0A2T5I0T1-F1-MODEL\_V4 \| 1.0 \| 1.573e-14 \| 501 \| 0.184 \| 422 \| 288 \| 17 \| 2 \| 402 \| 8 \| 394 \| Integrase \| Integrase; afdb-uniprot50 \| AF-A0A7W2BK59-F1-MODEL\_V4 \| 1.0 \| 1.675e-15 \| 500 \| 0.213 \| 421 \| 266 \| 23 \| 2 \| 402 \| 5 \| 380 \| Tyrosine-type recombinase/integrase \| Tyrosine-type recombinase/integrase; afdb-uniprot50 \| AF-A0A4D7QHQ2-F1-MODEL\_V4 \| 1.0 \| 1.997e-14 \| 500 \| 0.182 \| 432 \| 283 \| 21 \| 2 \| 403 \| 8 \| 399 \| DUF4102 domain-containing protein \| DUF4102 domain-containing protein; afdb-uniprot50 \| AF-A0A0Q4C992-F1-MODEL\_V4 \| 1.0 \| 1.24e-14 \| 500 \| 0.171 \| 432 \| 287 \| 23 \| 9 \| 403 \| 7 \| 404 \| Tyr recombinase domain-containing protein \| Tyr recombinase domain-containing protein; afdb-uniprot50 \| AF-A0A2A2F2K7-F1-MODEL\_V4 \| 1.0 \| 1.009e-13 \| 500 \| 0.156 \| 422 \| 303 \| 16 \| 2 \| 402 \| 8 \| 397 \| Integrase \| Integrase; afdb-uniprot50 \| AF-A0A4Q3Y2Z4-F1-MODEL\_V4 \| 1.0 \| 1.075e-14 \| 500 \| 0.193 \| 435 \| 278 \| 23 \| 2 \| 400 \| 68 \| 465 \| Site-specific integrase \| Site-specific integrase; afdb-uniprot50 \| AF-A0A5C9APT4-F1-MODEL\_V4 \| 1.0 \| 7.719e-12 \| 499 \| 0.187 \| 315 \| 222 \| 11 \| 102 \| 403 \| 5 \| 298 \| Prophage integrase IntS \| Prophage integrase IntS; afdb-uniprot50 \| AF-A0A517SMG1-F1-MODEL\_V4 \| 1.0 \| 1.452e-15 \| 499 \| 0.201 \| 411 \| 284 \| 15 \| 2 \| 401 \| 9 \| 386 \| Prophage CPS-53 integrase \| Prophage CPS-53 integrase; afdb-uniprot50 \| AF-A0A380T874-F1-MODEL\_V4 \| 1.0 \| 1.364e-14 \| 499 \| 0.177 \| 450 \| 284 \| 22 \| 2 \| 403 \| 7 \| 418 \| Integrase \| Integrase; afdb-uniprot50 \| AF-A0A0S3PXE2-F1-MODEL\_V4 \| 1.0 \| 3.216e-14 \| 498 \| 0.215 \| 380 \| 264 \| 18 \| 36 \| 403 \| 3 \| 360 \| Putative prophage CPS-53 integrase \| Putative prophage CPS-53 integrase; afdb-uniprot50 \| AF-A0A1I7D5K5-F1-MODEL\_V4 \| 1.0 \| 2.618e-13 \| 498 \| 0.201 \| 372 \| 247 \| 17 \| 53 \| 400 \| 7 \| 352 \| Site-specific recombinase XerD \| Site-specific recombinase XerD; afdb-uniprot50 \| AF-A0A2G6I7H5-F1-MODEL\_V4 \| 1.0 \| 3.264e-15 \| 498 \| 0.19 \| 430 \| 258 \| 23 \| 2 \| 389 \| 7 \| 388 \| Integrase \| Integrase; afdb-uniprot50 \| AF-A0A7C4E6T6-F1-MODEL\_V4 \| 1.0 \| 1.43e-14 \| 498 \| 0.183 \| 441 \| 285 \| 21 \| 9 \| 402 \| 14 \| 426 \| Site-specific integrase \| Site-specific integrase; afdb-uniprot50 \| AF-A0A0Q8QF83-F1-MODEL\_V4 \| 1.0 \| 7.471e-13 \| 497 \| 0.19 \| 331 \| 237 \| 14 \| 81 \| 400 \| 8 \| 318 \| Uncharacterized protein \| Uncharacterized protein; afdb-uniprot50 \| AF-A0A1X6ZV10-F1-MODEL\_V4 \| 1.0 \| 2.967e-15 \| 497 \| 0.2 \| 440 \| 271 \| 24 \| 1 \| 396 \| 29 \| 431 \| Putative prophage CPS-53 integrase \| Putative prophage CPS-53 integrase; afdb-uniprot50 \| AF-A0A826ZPT6-F1-MODEL\_V4 \| 1.0 \| 1.655e-11 \| 496 \| 0.238 \| 264 \| 177 \| 8 \| 145 \| 399 \| 5 \| 253 \| Site-specific integrase \| Site-specific integrase; afdb-uniprot50 \| AF-A0A6G7ZKC8-F1-MODEL\_V4 \| 1.0 \| 4.081e-14 \| 496 \| 0.185 \| 420 \| 291 \| 20 \| 2 \| 403 \| 5 \| 391 \| Tyrosine-type recombinase/integrase \| Tyrosine-type recombinase/integrase; afdb-uniprot50 \| AF-P39347-F1-MODEL\_V4 \| 1.0 \| 1.815e-14 \| 496 \| 0.192 \| 400 \| 266 \| 18 \| 36 \| 402 \| 3 \| 378 \| Putative protein IntB \| Putative protein IntB; afdb-uniprot50 \| AF-A0A090MGL5-F1-MODEL\_V4 \| 1.0 \| 1.182e-14 \| 496 \| 0.177 \| 440 \| 282 \| 24 \| 2 \| 402 \| 6 \| 404 \| Uncultured bacterium genome assembly Metasoil\_fosmids\_resub \| Uncultured bacterium genome assembly Metasoil\_fosmids\_resub; afdb-uniprot50 \| AF-A0A2D8M202-F1-MODEL\_V4 \| 1.0 \| 1.075e-14 \| 496 \| 0.181 \| 424 \| 286 \| 21 \| 2 \| 401 \| 7 \| 393 \| Uncharacterized protein \| Uncharacterized protein; afdb-uniprot50 \| AF-V4NAD2-F1-MODEL\_V4 \| 1.0 \| 4.217e-13 \| 495 \| 0.182 \| 367 \| 251 \| 17 \| 60 \| 402 \| 4 \| 345 \| Integrase \| Integrase; afdb-uniprot50 \| AF-A0A7H1NS61-F1-MODEL\_V4 \| 1.0 \| 3.71e-14 \| 495 \| 0.172 \| 423 \| 286 \| 24 \| 9 \| 400 \| 12 \| 401 \| Prophage integrase IntA \| Prophage integrase IntA; afdb-uniprot50 \| AF-A0A7Z2J7L8-F1-MODEL\_V4 \| 1.0 \| 1.43e-14 \| 495 \| 0.176 \| 431 \| 281 \| 20 \| 2 \| 402 \| 7 \| 393 \| Tyrosine-type recombinase/integrase \| Tyrosine-type recombinase/integrase; afdb-uniprot50 \| AF-A0A836V8A1-F1-MODEL\_V4 \| 1.0 \| 4.142e-15 \| 494 \| 0.181 \| 418 \| 275 \| 19 \| 2 \| 402 \| 7 \| 374 \| Site-specific integrase \| Site-specific integrase; afdb-uniprot50 \| AF-A0A1N7RJQ7-F1-MODEL\_V4 \| 1.0 \| 2.416e-14 \| 494 \| 0.175 \| 421 \| 292 \| 21 \| 2 \| 403 \| 45 \| 429 \| Integrase family protein \| Integrase family protein; afdb-uniprot50 \| AF-A0A354Q9H2-F1-MODEL\_V4 \| 1.0 \| 1.078e-11 \| 492 \| 0.166 \| 325 \| 237 \| 12 \| 92 \| 402 \| 3 \| 307 \| Integrase \| Integrase; afdb-uniprot50 \| AF-A0A238H9E1-F1-MODEL\_V4 \| 1.0 \| 4.57e-12 \| 492 \| 0.196 \| 330 \| 233 \| 14 \| 85 \| 402 \| 8 \| 317 \| Phage-related integrase \| Phage-related integrase; afdb-uniprot50 \| AF-A0A562Y0T8-F1-MODEL\_V4 \| 1.0 \| 1.788e-13 \| 492 \| 0.154 \| 420 \| 311 \| 20 \| 1 \| 402 \| 3 \| 396 \| Tyrosine-type recombinase/integrase \| Tyrosine-type recombinase/integrase; afdb-uniprot50 \| AF-E2CJQ7-F1-MODEL\_V4 \| 1.0 \| 6.361e-15 \| 491 \| 0.196 \| 418 \| 274 \| 26 \| 7 \| 403 \| 2 \| 378 \| Integrase family protein \| Integrase family protein; afdb-uniprot50 \| AF-R8Y101-F1-MODEL\_V4 \| 1.0 \| 1.343e-13 \| 491 \| 0.184 \| 417 \| 275 \| 22 \| 15 \| 402 \| 24 \| 404 \| Tyr recombinase domain-containing protein \| Tyr recombinase domain-containing protein; afdb-uniprot50 \| AF-A0A2W7C4Z9-F1-MODEL\_V4 \| 1.0 \| 2.829e-15 \| 491 \| 0.202 \| 440 \| 270 \| 23 \| 2 \| 398 \| 27 \| 428 \| Tyr recombinase domain-containing protein \| Tyr recombinase domain-containing protein; afdb-uniprot50 \| AF-A0A372BUY1-F1-MODEL\_V4 \| 1.0 \| 1.967e-13 \| 490 \| 0.218 \| 371 \| 242 \| 15 \| 68 \| 403 \| 9 \| 366 \| Integrase \| Integrase; afdb-uniprot50 \| AF-A0A848EHP0-F1-MODEL\_V4 \| 1.0 \| 1.3e-14 \| 489 \| 0.213 \| 426 \| 274 \| 22 \| 2 \| 399 \| 4 \| 396 \| Integrase arm-type DNA-binding domain-containing protein \| Integrase arm-type DNA-binding domain-containing protein; afdb-uniprot50 \| AF-A0A0X3TES6-F1-MODEL\_V4 \| 1.0 \| 6.267e-14 \| 489 \| 0.177 \| 416 \| 303 \| 18 \| 1 \| 402 \| 5 \| 395 \| Tyr recombinase domain-containing protein \| Tyr recombinase domain-containing protein; afdb-uniprot50 \| AF-A0A0P9UR54-F1-MODEL\_V4 \| 1.0 \| 6.361e-15 \| 488 \| 0.231 \| 401 \| 250 \| 21 \| 1 \| 377 \| 1 \| 367 \| Site-specific recombinase, phage integrase family protein \| Site-specific recombinase, phage integrase family protein; afdb-uniprot50 \| AF-A0A4D7QK21-F1-MODEL\_V4 \| 1.0 \| 6.361e-15 \| 488 \| 0.212 \| 438 \| 260 \| 26 \| 2 \| 401 \| 8 \| 398 \| DUF4102 domain-containing protein \| DUF4102 domain-containing protein; afdb-uniprot50 \| AF-A0A6I2A9Q0-F1-MODEL\_V4 \| 1.0 \| 3.121e-12 \| 487 \| 0.198 \| 322 \| 223 \| 14 \| 99 \| 403 \| 2 \| 305 \| Prophage integrase \| Prophage integrase; afdb-uniprot50 \| AF-K2JTR5-F1-MODEL\_V4 \| 1.0 \| 1.452e-15 \| 487 \| 0.229 \| 423 \| 260 \| 22 \| 2 \| 401 \| 5 \| 384 \| Integrase family protein \| Integrase family protein; afdb-uniprot50 \| AF-A0A6D0Y5S2-F1-MODEL\_V4 \| 1.0 \| 9.769e-15 \| 487 \| 0.197 \| 431 \| 279 \| 23 \| 2 \| 402 \| 8 \| 401 \| Integrase arm-type DNA-binding domain-containing protein \| Integrase arm-type DNA-binding domain-containing protein; afdb-uniprot50 \| AF-A0A1X9SW21-F1-MODEL\_V4 \| 1.0 \| 2.534e-14 \| 486 \| 0.165 \| 422 \| 285 \| 26 \| 2 \| 402 \| 6 \| 381 \| Site-specific recombinase, phage integrase family (DUF4102 domain) \| Site-specific recombinase, phage integrase family (DUF4102 domain); afdb-uniprot50 \| AF-A0A839UE18-F1-MODEL\_V4 \| 1.0 \| 2.304e-14 \| 486 \| 0.196 \| 412 \| 282 \| 15 \| 2 \| 398 \| 25 \| 402 \| Integrase \| Integrase; afdb-uniprot50 \| AF-A0A6N4CPC3-F1-MODEL\_V4 \| 1.0 \| 2.002e-11 \| 485 \| 0.195 \| 281 \| 199 \| 9 \| 137 \| 403 \| 11 \| 278 \| Uncharacterized protein \| Uncharacterized protein; afdb-uniprot50 \| AF-A0A350NYS9-F1-MODEL\_V4 \| 1.0 \| 2.203e-11 \| 485 \| 0.18 \| 310 \| 213 \| 10 \| 117 \| 401 \| 4 \| 297 \| Integrase \| Integrase; afdb-uniprot50 \| AF-N8XS28-F1-MODEL\_V4 \| 1.0 \| 7.23e-14 \| 485 \| 0.153 \| 403 \| 290 \| 22 \| 14 \| 400 \| 6 \| 373 \| Tyr recombinase domain-containing protein \| Tyr recombinase domain-containing protein; afdb-uniprot50 \| AF-A0A6I2IYY5-F1-MODEL\_V4 \| 1.0 \| 5.975e-14 \| 485 \| 0.184 \| 412 \| 271 \| 22 \| 9 \| 400 \| 12 \| 378 \| Tyrosine-type recombinase/integrase \| Tyrosine-type recombinase/integrase; afdb-uniprot50 \| AF-A0A060HK82-F1-MODEL\_V4 \| 1.0 \| 2.196e-14 \| 484 \| 0.202 \| 355 \| 241 \| 18 \| 34 \| 368 \| 7 \| 339 \| Tyr recombinase domain-containing protein \| Tyr recombinase domain-containing protein; afdb-uniprot50 \| AF-Q129B4-F1-MODEL\_V4 \| 1.0 \| 8.749e-14 \| 484 \| 0.215 \| 413 \| 281 \| 20 \| 2 \| 403 \| 7 \| 387 \| Phage integrase \| Phage integrase; afdb-uniprot50 \| AF-K2BD66-F1-MODEL\_V4 \| 1.0 \| 1.43e-14 \| 483 \| 0.197 \| 394 \| 239 \| 22 \| 2 \| 356 \| 7 \| 362 \| Tyr recombinase domain-containing protein \| Tyr recombinase domain-containing protein; afdb-uniprot50 \| AF-A0A6B8JJH7-F1-MODEL\_V4 \| 1.0 \| 2.196e-14 \| 483 \| 0.226 \| 411 \| 259 \| 20 \| 37 \| 398 \| 2 \| 402 \| Integrase \| Integrase; afdb-uniprot50 \| AF-A0A522YGD1-F1-MODEL\_V4 \| 1.0 \| 6.792e-13 \| 482 \| 0.193 \| 362 \| 242 \| 16 \| 44 \| 380 \| 5 \| 341 \| DUF4102 domain-containing protein \| DUF4102 domain-containing protein; afdb-uniprot50 \| AF-A0A560MBT4-F1-MODEL\_V4 \| 1.0 \| 2.416e-14 \| 482 \| 0.17 \| 423 \| 288 \| 21 \| 9 \| 402 \| 11 \| 399 \| Integrase \| Integrase; afdb-uniprot50 \| AF-A0A433ZTM7-F1-MODEL\_V4 \| 1.0 \| 1.343e-13 \| 481 \| 0.206 \| 353 \| 235 \| 16 \| 1 \| 335 \| 1 \| 326 \| Integrase \| Integrase; afdb-uniprot50 \| AF-A0A5C7QE83-F1-MODEL\_V4 \| 1.0 \| 1.938e-12 \| 481 \| 0.196 \| 325 \| 211 \| 14 \| 90 \| 402 \| 64 \| 350 \| Integrase \| Integrase; afdb-uniprot50 \| AF-A0A2W7FQS8-F1-MODEL\_V4 \| 1.0 \| 9.944e-13 \| 480 \| 0.227 \| 361 \| 244 \| 16 \| 60 \| 401 \| 1 \| 345 \| Integrase \| Integrase; afdb-uniprot50 \| AF-A0A127K4V2-F1-MODEL\_V4 \| 1.0 \| 1.904e-14 \| 480 \| 0.189 \| 432 \| 294 \| 23 \| 2 \| 403 \| 5 \| 410 \| Uncharacterized protein \| Uncharacterized protein; afdb-uniprot50 \| AF-A0A2W5Q3A9-F1-MODEL\_V4 \| 1.0 \| 1.815e-14 \| 480 \| 0.205 \| 453 \| 272 \| 27 \| 1 \| 402 \| 7 \| 422 \| Integrase \| Integrase; afdb-uniprot50 \| AF-A0A379NCF1-F1-MODEL\_V4 \| 1.0 \| 2.063e-13 \| 480 \| 0.2 \| 474 \| 278 \| 27 \| 2 \| 398 \| 8 \| 457 \| Prophage CP4-57 integrase \| Prophage CP4-57 integrase; afdb-uniprot50 \| AF-A0A7Y9VNF8-F1-MODEL\_V4 \| 1.0 \| 8.88e-15 \| 480 \| 0.194 \| 436 \| 277 \| 21 \| 2 \| 400 \| 8 \| 406 \| Integrase \| Integrase; afdb-uniprot50 \| AF-A0A3D1P0H0-F1-MODEL\_V4 \| 1.0 \| 5.432e-14 \| 478 \| 0.193 \| 392 \| 253 \| 20 \| 2 \| 366 \| 3 \| 358 \| Integrase \| Integrase; afdb-uniprot50 \| AF-Q6ALK6-F1-MODEL\_V4 \| 1.0 \| 3.216e-14 \| 478 \| 0.174 \| 418 \| 282 \| 19 \| 2 \| 402 \| 8 \| 379 \| Related to integrase \| Related to integrase; afdb-uniprot50 \| AF-A0A0H4IWA8-F1-MODEL\_V4 \| 1.0 \| 1.221e-13 \| 478 \| 0.161 \| 452 \| 293 \| 25 \| 2 \| 402 \| 3 \| 419 \| Core-binding (CB) domain-containing protein \| Core-binding (CB) domain-containing protein; afdb-uniprot50 \| AF-A0A176ZCC3-F1-MODEL\_V4 \| 1.0 \| 2.164e-13 \| 478 \| 0.203 \| 442 \| 247 \| 19 \| 44 \| 402 \| 6 \| 425 \| Tyr recombinase domain-containing protein \| Tyr recombinase domain-containing protein; afdb-uniprot50 \| AF-A0A6I1J6L9-F1-MODEL\_V4 \| 1.0 \| 3.548e-11 \| 477 \| 0.182 \| 317 \| 217 \| 14 \| 104 \| 402 \| 2 \| 294 \| Putative integrase \| Putative integrase; afdb-uniprot50 \| AF-A0A4U1BG56-F1-MODEL\_V4 \| 1.0 \| 1.025e-14 \| 477 \| 0.2 \| 429 \| 282 \| 21 \| 3 \| 400 \| 13 \| 411 \| DUF4102 domain-containing protein \| DUF4102 domain-containing protein; afdb-uniprot50 \| AF-A0A368KM35-F1-MODEL\_V4 \| 1.0 \| 4.639e-13 \| 476 \| 0.199 \| 361 \| 239 \| 16 \| 46 \| 401 \| 17 \| 332 \| Integrase \| Integrase; afdb-uniprot50 \| AF-G1UQY9-F1-MODEL\_V4 \| 1.0 \| 1.523e-15 \| 476 \| 0.19 \| 404 \| 257 \| 15 \| 1 \| 372 \| 1 \| 366 \| Uncharacterized protein \| Uncharacterized protein; afdb-uniprot50 \| AF-A0A5J4E109-F1-MODEL\_V4 \| 1.0 \| 3.373e-14 \| 476 \| 0.203 \| 432 \| 276 \| 23 \| 2 \| 402 \| 6 \| 400 \| Tyr recombinase domain-containing protein \| Tyr recombinase domain-containing protein; afdb-uniprot50 \| AF-X0UCK1-F1-MODEL\_V4 \| 1.0 \| 2.345e-12 \| 475 \| 0.241 \| 298 \| 192 \| 14 \| 39 \| 320 \| 2 \| 281 \| Tyr recombinase domain-containing protein \| Tyr recombinase domain-containing protein; afdb-uniprot50 \| AF-A0A3G9G2H1-F1-MODEL\_V4 \| 1.0 \| 6.38e-12 \| 475 \| 0.224 \| 325 \| 214 \| 15 \| 93 \| 403 \| 2 \| 302 \| Putative plasmid stabilization protein \| Putative plasmid stabilization protein; afdb-uniprot50 \| AF-A0A7V8FSQ5-F1-MODEL\_V4 \| 1.0 \| 9.944e-13 \| 474 \| 0.219 \| 387 \| 242 \| 19 \| 50 \| 402 \| 3 \| 363 \| Prophage integrase IntA \| Prophage integrase IntA; afdb-uniprot50 \| AF-A0A3N5Y2X2-F1-MODEL\_V4 \| 1.0 \| 3.71e-14 \| 474 \| 0.181 \| 429 \| 286 \| 22 \| 2 \| 403 \| 8 \| 398 \| DUF4102 domain-containing protein \| DUF4102 domain-containing protein; afdb-uniprot50 \| AF-A0A009KR62-F1-MODEL\_V4 \| 1.0 \| 1.55e-13 \| 474 \| 0.16 \| 411 \| 289 \| 21 \| 14 \| 402 \| 22 \| 398 \| Phage integrase family protein \| Phage integrase family protein; afdb-uniprot50 \| AF-A0A1I1WIT5-F1-MODEL\_V4 \| 1.0 \| 5.783e-15 \| 474 \| 0.199 \| 467 \| 278 \| 25 \| 2 \| 403 \| 10 \| 445 \| Integrase \| Integrase; afdb-uniprot50 \| AF-A0A7Y9NF75-F1-MODEL\_V4 \| 1.0 \| 8.492e-12 \| 473 \| 0.22 \| 327 \| 227 \| 14 \| 93 \| 398 \| 2 \| 321 \| Integrase \| Integrase; afdb-uniprot50 \| AF-A0A0F2NU29-F1-MODEL\_V4 \| 1.0 \| 7.583e-14 \| 473 \| 0.197 \| 405 \| 275 \| 19 \| 1 \| 395 \| 5 \| 369 \| Uncharacterized protein \| Uncharacterized protein; afdb-uniprot50 \| AF-A0A3M4E1T2-F1-MODEL\_V4 \| 1.0 \| 1.009e-13 \| 472 \| 0.177 \| 422 \| 286 \| 21 \| 2 \| 401 \| 8 \| 390 \| Tyr recombinase domain-containing protein \| Tyr recombinase domain-containing protein; afdb-uniprot50 \| AF-A0A1W1V7M5-F1-MODEL\_V4 \| 1.0 \| 5.194e-11 \| 471 \| 0.175 \| 320 \| 218 \| 14 \| 101 \| 402 \| 1 \| 292 \| Integrase \| Integrase; afdb-uniprot50 \| AF-A0A657BBC9-F1-MODEL\_V4 \| 1.0 \| 3.075e-11 \| 470 \| 0.182 \| 350 \| 240 \| 14 \| 65 \| 402 \| 3 \| 318 \| Tyr recombinase domain-containing protein \| Tyr recombinase domain-containing protein; afdb-uniprot50 \| AF-A0A316G5Z1-F1-MODEL\_V4 \| 1.0 \| 5.179e-14 \| 470 \| 0.182 \| 399 \| 278 \| 19 \| 20 \| 400 \| 2 \| 370 \| Integrase \| Integrase; afdb-uniprot50 \| AF-A0A5C7UIE7-F1-MODEL\_V4 \| 1.0 \| 4.938e-14 \| 469 \| 0.157 \| 431 \| 290 \| 17 \| 2 \| 398 \| 8 \| 399 \| Site-specific integrase \| Site-specific integrase; afdb-uniprot50 \| AF-A0A3E1BRB2-F1-MODEL\_V4 \| 1.0 \| 3.95e-15 \| 469 \| 0.193 \| 449 \| 271 \| 22 \| 1 \| 398 \| 7 \| 415 \| Tyr recombinase domain-containing protein \| Tyr recombinase domain-containing protein; afdb-uniprot50 \| AF-A0A2E7GWJ5-F1-MODEL\_V4 \| 1.0 \| 1.705e-13 \| 468 \| 0.165 \| 418 \| 295 \| 20 \| 2 \| 400 \| 8 \| 390 \| Uncharacterized protein \| Uncharacterized protein; afdb-uniprot50 \| AF-A0A292AEJ5-F1-MODEL\_V4 \| 1.0 \| 1.904e-14 \| 468 \| 0.193 \| 434 \| 273 \| 27 \| 1 \| 402 \| 1 \| 389 \| Tyr recombinase domain-containing protein \| Tyr recombinase domain-containing protein; afdb-uniprot50 \| AF-A0A842HU53-F1-MODEL\_V4 \| 1.0 \| 2.38e-13 \| 466 \| 0.169 \| 430 \| 292 \| 20 \| 2 \| 403 \| 4 \| 396 \| Tyrosine-type recombinase/integrase \| Tyrosine-type recombinase/integrase; afdb-uniprot50 \| AF-A0A1J5E277-F1-MODEL\_V4 \| 1.0 \| 3.891e-14 \| 465 \| 0.205 \| 409 \| 293 \| 17 \| 2 \| 401 \| 3 \| 388 \| Tyr recombinase domain-containing protein \| Tyr recombinase domain-containing protein; afdb-uniprot50 \| AF-A0A2W4L7D4-F1-MODEL\_V4 \| 1.0 \| 8.749e-14 \| 465 \| 0.201 \| 412 \| 275 \| 19 \| 1 \| 401 \| 66 \| 434 \| Integrase \| Integrase; afdb-uniprot50 \| AF-A0A1G6IJD1-F1-MODEL\_V4 \| 1.0 \| 3.323e-13 \| 464 \| 0.193 \| 387 \| 264 \| 19 \| 37 \| 402 \| 2 \| 361 \| Integrase \| Integrase; afdb-uniprot50 \| AF-A0A844XW65-F1-MODEL\_V4 \| 1.0 \| 5.975e-14 \| 464 \| 0.178 \| 432 \| 283 \| 22 \| 1 \| 402 \| 1 \| 390 \| Integrase arm-type DNA-binding domain-containing protein \| Integrase arm-type DNA-binding domain-containing protein; afdb-uniprot50 \| AF-A0A2P2EDV4-F1-MODEL\_V4 \| 1.0 \| 1.478e-13 \| 464 \| 0.171 \| 456 \| 283 \| 23 \| 2 \| 400 \| 8 \| 425 \| Tyrosine recombinase XerD \| Tyrosine recombinase XerD; afdb-uniprot50 \| AF-A0A2E7GWC7-F1-MODEL\_V4 \| 1.0 \| 1.997e-14 \| 464 \| 0.213 \| 436 \| 275 \| 26 \| 2 \| 401 \| 8 \| 411 \| Tyr recombinase domain-containing protein \| Tyr recombinase domain-containing protein; afdb-uniprot50 \| AF-H5WK80-F1-MODEL\_V4 \| 1.0 \| 3.537e-14 \| 464 \| 0.187 \| 464 \| 283 \| 23 \| 2 \| 403 \| 7 \| 438 \| Site-specific recombinase XerD \| Site-specific recombinase XerD; afdb-uniprot50 \| AF-A0A382QZ10-F1-MODEL\_V4 \| 1.0 \| 5.194e-11 \| 463 \| 0.16 \| 330 \| 242 \| 13 \| 86 \| 401 \| 9 \| 317 \| Uncharacterized protein \| Uncharacterized protein; afdb-uniprot50 \| AF-A0A7R7GHM2-F1-MODEL\_V4 \| 1.0 \| 7.492e-10 \| 461 \| 0.225 \| 200 \| 145 \| 4 \| 212 \| 403 \| 3 \| 200 \| Uncharacterized protein \| Uncharacterized protein; afdb-uniprot50 \| AF-A0A198XBH7-F1-MODEL\_V4 \| 1.0 \| 6.914e-11 \| 460 \| 0.208 \| 317 \| 215 \| 16 \| 101 \| 402 \| 2 \| 297 \| Integrase \| Integrase; afdb-uniprot50 \| AF-A0A0B8ZHW5-F1-MODEL\_V4 \| 1.0 \| 6.573e-14 \| 460 \| 0.172 \| 465 \| 275 \| 23 \| 2 \| 398 \| 9 \| 431 \| Integrase family protein \| Integrase family protein; afdb-uniprot50 \| AF-A0A6N8K839-F1-MODEL\_V4 \| 1.0 \| 7.251e-11 \| 458 \| 0.19 \| 294 \| 198 \| 11 \| 123 \| 402 \| 2 \| 269 \| Tyrosine-type recombinase/integrase \| Tyrosine-type recombinase/integrase; afdb-uniprot50 \| AF-A5UG79-F1-MODEL\_V4 \| 1.0 \| 4.793e-12 \| 458 \| 0.2 \| 365 \| 242 \| 14 \| 60 \| 401 \| 1 \| 338 \| Prophage integrase \| Prophage integrase; afdb-uniprot50 \| AF-A0A4P6L4N5-F1-MODEL\_V4 \| 1.0 \| 2.618e-13 \| 458 \| 0.193 \| 428 \| 284 \| 23 \| 2 \| 403 \| 9 \| 401 \| Site-specific integrase \| Site-specific integrase; afdb-uniprot50 \| AF-A0A165RXR2-F1-MODEL\_V4 \| 1.0 \| 7.339e-15 \| 458 \| 0.177 \| 439 \| 267 \| 23 \| 2 \| 402 \| 7 \| 389 \| Putative prophage CPS-53 integrase \| Putative prophage CPS-53 integrase; afdb-uniprot50 \| AF-A0A7C2SE25-F1-MODEL\_V4 \| 1.0 \| 5.352e-13 \| 457 \| 0.179 \| 456 \| 288 \| 23 \| 2 \| 398 \| 10 \| 438 \| DUF4102 domain-containing protein \| DUF4102 domain-containing protein; afdb-uniprot50 \| AF-A0A825BDC2-F1-MODEL\_V4 \| 1.0 \| 4.423e-13 \| 456 \| 0.179 \| 413 \| 268 \| 22 \| 7 \| 403 \| 3 \| 360 \| Uncharacterized protein \| Uncharacterized protein; afdb-uniprot50 \| AF-A0A7Y3Y9M4-F1-MODEL\_V4 \| 1.0 \| 3.021e-13 \| 456 \| 0.18 \| 422 \| 286 \| 22 \| 2 \| 403 \| 8 \| 389 \| Tyrosine-type recombinase/integrase \| Tyrosine-type recombinase/integrase; afdb-uniprot50 \| AF-A0A376VWA4-F1-MODEL\_V4 \| 1.0 \| 1.881e-10 \| 455 \| 0.203 \| 260 \| 179 \| 8 \| 158 \| 402 \| 2 \| 248 \| Integrase \| Integrase; afdb-uniprot50 \| AF-A0A0N0WLQ3-F1-MODEL\_V4 \| 1.0 \| 1.736e-11 \| 455 \| 0.21 \| 275 \| 182 \| 11 \| 141 \| 400 \| 9 \| 263 \| Site-specific recombinase \| Site-specific recombinase; afdb-uniprot50 \| AF-A0A859IFR7-F1-MODEL\_V4 \| 1.0 \| 4.217e-13 \| 455 \| 0.201 \| 382 \| 248 \| 21 \| 2 \| 357 \| 8 \| 358 \| Tyrosine-type recombinase/integrase \| Tyrosine-type recombinase/integrase; afdb-uniprot50 \| AF-A0A1G6PQW0-F1-MODEL\_V4 \| 1.0 \| 1.554e-10 \| 453 \| 0.192 \| 291 \| 204 \| 11 \| 123 \| 403 \| 5 \| 274 \| Integrase \| Integrase; afdb-uniprot50 \| AF-A0A2E7FT18-F1-MODEL\_V4 \| 1.0 \| 6.894e-14 \| 453 \| 0.189 \| 380 \| 250 \| 16 \| 2 \| 354 \| 8 \| 356 \| Integrase \| Integrase; afdb-uniprot50 \| AF-A0A286E6R0-F1-MODEL\_V4 \| 1.0 \| 2.888e-10 \| 452 \| 0.22 \| 268 \| 185 \| 8 \| 145 \| 403 \| 11 \| 263 \| Phage integrase family protein \| Phage integrase family protein; afdb-uniprot50 \| AF-A0A536TMG5-F1-MODEL\_V4 \| 1.0 \| 7.251e-11 \| 452 \| 0.207 \| 270 \| 186 \| 8 \| 145 \| 402 \| 5 \| 258 \| Site-specific integrase \| Site-specific integrase; afdb-uniprot50 \| AF-A0A3D8K2G3-F1-MODEL\_V4 \| 1.0 \| 5.993e-11 \| 452 \| 0.172 \| 290 \| 207 \| 12 \| 131 \| 403 \| 34 \| 307 \| Integrase \| Integrase; afdb-uniprot50 \| AF-A0A2R7RSS5-F1-MODEL\_V4 \| 1.0 \| 2.063e-13 \| 452 \| 0.199 \| 426 \| 271 \| 22 \| 2 \| 402 \| 8 \| 388 \| Integrase \| Integrase; afdb-uniprot50 \| AF-A0A2D5FF62-F1-MODEL\_V4 \| 1.0 \| 1.059e-13 \| 452 \| 0.159 \| 458 \| 298 \| 22 \| 1 \| 402 \| 7 \| 433 \| Integrase \| Integrase; afdb-uniprot50 \| AF-A0A1X3KZ03-F1-MODEL\_V4 \| 1.0 \| 4.093e-11 \| 451 \| 0.197 \| 293 \| 197 \| 11 \| 123 \| 402 \| 19 \| 286 \| Site-specific recombinase, phage integrase family \| Site-specific recombinase, phage integrase family; afdb-uniprot50 \| AF-A0A2N7I5S9-F1-MODEL\_V4 \| 1.0 \| 3.601e-12 \| 451 \| 0.171 \| 368 \| 257 \| 16 \| 60 \| 402 \| 1 \| 345 \| Integrase \| Integrase; afdb-uniprot50 \| AF-A0A7W8J9T4-F1-MODEL\_V4 \| 1.0 \| 1.225e-10 \| 450 \| 0.188 \| 276 \| 201 \| 7 \| 136 \| 402 \| 8 \| 269 \| Integrase \| Integrase; afdb-uniprot50 \| AF-A0A1J0S1K6-F1-MODEL\_V4 \| 1.0 \| 5.613e-13 \| 450 \| 0.166 \| 420 \| 297 \| 21 \| 2 \| 403 \| 6 \| 390 \| Tyr recombinase domain-containing protein \| Tyr recombinase domain-containing protein; afdb-uniprot50 \| AF-N8Q3U9-F1-MODEL\_V4 \| 1.0 \| 2.1e-11 \| 449 \| 0.187 \| 362 \| 254 \| 16 \| 60 \| 403 \| 1 \| 340 \| Tyr recombinase domain-containing protein \| Tyr recombinase domain-containing protein; afdb-uniprot50 \| AF-A0A1G3UJM0-F1-MODEL\_V4 \| 1.0 \| 3.168e-13 \| 448 \| 0.21 \| 346 \| 229 \| 16 \| 2 \| 330 \| 8 \| 326 \| Integrase \| Integrase; afdb-uniprot50 \| AF-A0A1D7NNW2-F1-MODEL\_V4 \| 1.0 \| 2.196e-14 \| 447 \| 0.216 \| 416 \| 239 \| 24 \| 1 \| 377 \| 1 \| 368 \| Arm-DNA-bind\_3 domain-containing protein \| Arm-DNA-bind\_3 domain-containing protein; afdb-uniprot50 \| AF-A0A3N5UFD5-F1-MODEL\_V4 \| 1.0 \| 8.342e-14 \| 446 \| 0.246 \| 357 \| 211 \| 20 \| 1 \| 340 \| 29 \| 344 \| DUF4102 domain-containing protein \| DUF4102 domain-containing protein; afdb-uniprot50 \| AF-A0A178MZZ2-F1-MODEL\_V4 \| 1.0 \| 1.876e-13 \| 446 \| 0.191 \| 417 \| 269 \| 24 \| 2 \| 401 \| 5 \| 370 \| Uncharacterized protein \| Uncharacterized protein; afdb-uniprot50 \| AF-K7A9W5-F1-MODEL\_V4 \| 1.0 \| 4.021e-13 \| 445 \| 0.182 \| 421 \| 284 \| 23 \| 1 \| 401 \| 7 \| 387 \| Tyr recombinase domain-containing protein \| Tyr recombinase domain-containing protein; afdb-uniprot50 \| AF-A0A653HY48-F1-MODEL\_V4 \| 1.0 \| 1.482e-10 \| 443 \| 0.155 \| 321 \| 227 \| 14 \| 101 \| 402 \| 1 \| 296 \| Prophage CPS-53 integrase \| Prophage CPS-53 integrase; afdb-uniprot50 \| AF-A0A5F0WJ90-F1-MODEL\_V4 \| 1.0 \| 7.836e-13 \| 442 \| 0.176 \| 424 \| 286 \| 22 \| 1 \| 401 \| 7 \| 390 \| Site-specific integrase \| Site-specific integrase; afdb-uniprot50 \| AF-A0A6J4EDZ9-F1-MODEL\_V4 \| 1.0 \| 1.262e-12 \| 442 \| 0.2 \| 425 \| 275 \| 20 \| 1 \| 401 \| 9 \| 392 \| Recombinase \| Recombinase; afdb-uniprot50 \| AF-A0A833JLG8-F1-MODEL\_V4 \| 1.0 \| 1.967e-13 \| 442 \| 0.17 \| 441 \| 278 \| 23 \| 16 \| 403 \| 34 \| 439 \| Uncharacterized protein \| Uncharacterized protein; afdb-uniprot50 \| AF-A0A534I5Z6-F1-MODEL\_V4 \| 1.0 \| 4.952e-11 \| 440 \| 0.216 \| 300 \| 198 \| 11 \| 119 \| 403 \| 12 \| 289 \| Site-specific integrase \| Site-specific integrase; afdb-uniprot50 \| AF-A0A5A7MRW9-F1-MODEL\_V4 \| 1.0 \| 2.932e-11 \| 440 \| 0.223 \| 367 \| 242 \| 17 \| 60 \| 403 \| 1 \| 347 \| Phage integrase \| Phage integrase; afdb-uniprot50 \| AF-A0A5J6R848-F1-MODEL\_V4 \| 1.0 \| 3.021e-13 \| 439 \| 0.174 \| 430 \| 279 \| 21 \| 1 \| 401 \| 7 \| 389 \| Site-specific integrase \| Site-specific integrase; afdb-uniprot50 \| AF-A0A3B9VKD5-F1-MODEL\_V4 \| 1.0 \| 4.021e-13 \| 439 \| 0.174 \| 492 \| 272 \| 27 \| 1 \| 392 \| 5 \| 462 \| Tyr recombinase domain-containing protein \| Tyr recombinase domain-containing protein; afdb-uniprot50 \| AF-A0A1L9GM55-F1-MODEL\_V4 \| 1.0 \| 5.8e-12 \| 438 \| 0.204 \| 387 \| 238 \| 18 \| 60 \| 402 \| 1 \| 361 \| Uncharacterized protein \| Uncharacterized protein; afdb-uniprot50 \| AF-A0A3M3DIU2-F1-MODEL\_V4 \| 1.0 \| 4.021e-13 \| 438 \| 0.208 \| 427 \| 264 \| 23 \| 2 \| 401 \| 7 \| 386 \| Putative Phage integrase \| Putative Phage integrase; afdb-uniprot50 \| AF-A0A2P7NYP5-F1-MODEL\_V4 \| 1.0 \| 1.243e-11 \| 437 \| 0.18 \| 361 \| 254 \| 14 \| 62 \| 402 \| 4 \| 342 \| Integrase \| Integrase; afdb-uniprot50 \| AF-A0A523H760-F1-MODEL\_V4 \| 1.0 \| 3.323e-13 \| 436 \| 0.206 \| 359 \| 229 \| 18 \| 2 \| 337 \| 9 \| 334 \| DUF4102 domain-containing protein \| DUF4102 domain-containing protein; afdb-uniprot50 \| AF-A0A2E7FR99-F1-MODEL\_V4 \| 1.0 \| 2.423e-11 \| 436 \| 0.151 \| 369 \| 247 \| 16 \| 74 \| 402 \| 14 \| 356 \| Tyr recombinase domain-containing protein \| Tyr recombinase domain-containing protein; afdb-uniprot50 \| AF-A0A5I4RF45-F1-MODEL\_V4 \| 1.0 \| 3.021e-13 \| 436 \| 0.165 \| 405 \| 266 \| 23 \| 2 \| 370 \| 3 \| 371 \| DUF4102 domain-containing protein \| DUF4102 domain-containing protein; afdb-uniprot50 \| AF-A0A5P9CRG7-F1-MODEL\_V4 \| 1.0 \| 2.236e-12 \| 436 \| 0.163 \| 416 \| 299 \| 19 \| 2 \| 401 \| 11 \| 393 \| Integrase \| Integrase; afdb-uniprot50 \| AF-I2Q1B1-F1-MODEL\_V4 \| 1.0 \| 1.55e-13 \| 436 \| 0.192 \| 411 \| 263 \| 22 \| 17 \| 391 \| 2 \| 379 \| Site-specific recombinase XerD \| Site-specific recombinase XerD; afdb-uniprot50 \| AF-A0A0A7EEY8-F1-MODEL\_V4 \| 1.0 \| 4.423e-13 \| 436 \| 0.181 \| 425 \| 284 \| 22 \| 2 \| 402 \| 7 \| 391 \| Tyr recombinase domain-containing protein \| Tyr recombinase domain-containing protein; afdb-uniprot50 \| AF-A0A5C8NMZ4-F1-MODEL\_V4 \| 1.0 \| 6.476e-13 \| 436 \| 0.18 \| 437 \| 278 \| 23 \| 9 \| 402 \| 10 \| 409 \| DUF4102 domain-containing protein \| DUF4102 domain-containing protein; afdb-uniprot50 \| AF-B8KHE9-F1-MODEL\_V4 \| 1.0 \| 4.639e-13 \| 436 \| 0.199 \| 426 \| 272 \| 22 \| 9 \| 402 \| 20 \| 408 \| Site-specific recombinase, phage integrase family \| Site-specific recombinase, phage integrase family; afdb-uniprot50 \| AF-A0A6P3BW65-F1-MODEL\_V4 \| 1.0 \| 1.602e-12 \| 433 \| 0.206 \| 344 \| 225 \| 15 \| 2 \| 326 \| 3 \| 317 \| Integrase \| Integrase; afdb-uniprot50 \| AF-A0A0W0U2X8-F1-MODEL\_V4 \| 1.0 \| 4.722e-11 \| 433 \| 0.176 \| 329 \| 242 \| 13 \| 82 \| 398 \| 4 \| 315 \| Integrase \| Integrase; afdb-uniprot50 \| AF-A0A1I2F649-F1-MODEL\_V4 \| 1.0 \| 4.865e-13 \| 433 \| 0.161 \| 470 \| 290 \| 22 \| 2 \| 402 \| 7 \| 441 \| Tyr recombinase domain-containing protein \| Tyr recombinase domain-containing protein; afdb-uniprot50 \| AF-A0A375BMV2-F1-MODEL\_V4 \| 1.0 \| 3.666e-10 \| 431 \| 0.195 \| 292 \| 204 \| 13 \| 102 \| 381 \| 16 \| 288 \| Phage integrase \| Phage integrase; afdb-uniprot50 \| AF-A0A3D0RCF0-F1-MODEL\_V4 \| 1.0 \| 3.834e-13 \| 431 \| 0.201 \| 388 \| 261 \| 22 \| 2 \| 372 \| 19 \| 374 \| Integrase \| Integrase; afdb-uniprot50 \| AF-A0A2N5Y8G1-F1-MODEL\_V4 \| 1.0 \| 2.496e-13 \| 431 \| 0.193 \| 419 \| 283 \| 25 \| 2 \| 402 \| 51 \| 432 \| Site-specific integrase \| Site-specific integrase; afdb-uniprot50 \| AF-A0A7C4EE33-F1-MODEL\_V4 \| 1.0 \| 1.262e-12 \| 431 \| 0.151 \| 450 \| 299 \| 23 \| 2 \| 402 \| 25 \| 440 \| DUF4102 domain-containing protein \| DUF4102 domain-containing protein; afdb-uniprot50 \| AF-A0A257SZ13-F1-MODEL\_V4 \| 1.0 \| 1.531e-09 \| 430 \| 0.26 \| 192 \| 133 \| 7 \| 216 \| 402 \| 6 \| 193 \| Tyr recombinase domain-containing protein \| Tyr recombinase domain-containing protein; afdb-uniprot50 \| AF-A0A8B4ZLY0-F1-MODEL\_V4 \| 1.0 \| 1.388e-12 \| 429 \| 0.179 \| 430 \| 275 \| 23 \| 2 \| 401 \| 10 \| 391 \| Site-specific integrase \| Site-specific integrase; afdb-uniprot50 \| AF-A0A1G4ZAT7-F1-MODEL\_V4 \| 1.0 \| 4.57e-12 \| 428 \| 0.167 \| 364 \| 239 \| 14 \| 52 \| 402 \| 19 \| 331 \| Integrase \| Integrase; afdb-uniprot50 \| AF-A0A1U9SCV8-F1-MODEL\_V4 \| 1.0 \| 1.046e-09 \| 427 \| 0.196 \| 259 \| 182 \| 8 \| 155 \| 403 \| 2 \| 244 \| Integrase \| Integrase; afdb-uniprot50 \| AF-A0A7J0BQK4-F1-MODEL\_V4 \| 1.0 \| 3.177e-10 \| 427 \| 0.157 \| 312 \| 225 \| 13 \| 107 \| 402 \| 2 \| 291 \| Tyr recombinase domain-containing protein \| Tyr recombinase domain-containing protein; afdb-uniprot50 \| AF-A0A3B9B7X0-F1-MODEL\_V4 \| 1.0 \| 6.792e-13 \| 427 \| 0.21 \| 342 \| 216 \| 17 \| 2 \| 321 \| 3 \| 312 \| Integrase \| Integrase; afdb-uniprot50 \| AF-A0A845MU45-F1-MODEL\_V4 \| 1.0 \| 7.449e-16 \| 427 \| 0.185 \| 420 \| 290 \| 22 \| 2 \| 402 \| 3 \| 389 \| DUF4102 domain-containing protein \| DUF4102 domain-containing protein; afdb-uniprot50 \| AF-F4MXM0-F1-MODEL\_V4 \| 1.0 \| 8.644e-10 \| 426 \| 0.236 \| 258 \| 170 \| 9 \| 153 \| 401 \| 2 \| 241 \| Integrase \| Integrase; afdb-uniprot50 \| AF-A0A6N7AFU2-F1-MODEL\_V4 \| 1.0 \| 9.04e-13 \| 426 \| 0.178 \| 415 \| 300 \| 16 \| 2 \| 403 \| 3 \| 389 \| Phage integrase family protein \| Phage integrase family protein; afdb-uniprot50 \| AF-A0A518KCN4-F1-MODEL\_V4 \| 1.0 \| 1.078e-11 \| 424 \| 0.175 \| 365 \| 241 \| 14 \| 46 \| 401 \| 14 \| 327 \| Site-specific tyrosine recombinase XerC \| Site-specific tyrosine recombinase XerC; afdb-uniprot50 \| AF-A0A5C7W994-F1-MODEL\_V4 \| 1.0 \| 3.845e-10 \| 423 \| 0.174 \| 298 \| 210 \| 9 \| 115 \| 403 \| 2 \| 272 \| Integrase \| Integrase; afdb-uniprot50 \| AF-A0A212KHW4-F1-MODEL\_V4 \| 1.0 \| 2.138e-09 \| 423 \| 0.169 \| 318 \| 230 \| 12 \| 98 \| 402 \| 2 \| 298 \| Phage integrase family protein \| Phage integrase family protein; afdb-uniprot50 \| AF-A0A090IF55-F1-MODEL\_V4 \| 1.0 \| 6.174e-13 \| 422 \| 0.182 \| 427 \| 268 \| 26 \| 2 \| 400 \| 8 \| 381 \| Putative phage integrase \| Putative phage integrase; afdb-uniprot50 \| AF-A0A512AL48-F1-MODEL\_V4 \| 1.0 \| 7.492e-10 \| 421 \| 0.203 \| 280 \| 193 \| 12 \| 136 \| 402 \| 5 \| 267 \| Tyr recombinase domain-containing protein \| Tyr recombinase domain-containing protein; afdb-uniprot50 \| AF-G4A9X4-F1-MODEL\_V4 \| 1.0 \| 9.973e-10 \| 420 \| 0.207 \| 256 \| 175 \| 10 \| 159 \| 403 \| 2 \| 240 \| Prophage CP4-57 integrase \| Prophage CP4-57 integrase; afdb-uniprot50 \| AF-A0A1X0SVT1-F1-MODEL\_V4 \| 1.0 \| 5.993e-11 \| 420 \| 0.194 \| 318 \| 219 \| 16 \| 99 \| 401 \| 9 \| 304 \| Integrase family protein \| Integrase family protein; afdb-uniprot50 \| AF-A0A0H4IW46-F1-MODEL\_V4 \| 1.0 \| 1.705e-13 \| 420 \| 0.224 \| 366 \| 227 \| 19 \| 2 \| 341 \| 7 \| 341 \| Integrase \| Integrase; afdb-uniprot50 \| AF-A0A2X4XHJ1-F1-MODEL\_V4 \| 1.0 \| 5.53e-12 \| 420 \| 0.195 \| 389 \| 243 \| 20 \| 2 \| 356 \| 3 \| 355 \| Prophage CP4-57 integrase \| Prophage CP4-57 integrase; afdb-uniprot50 \| AF-A0A1W7QVM2-F1-MODEL\_V4 \| 1.0 \| 1.043e-12 \| 420 \| 0.141 \| 444 \| 298 \| 23 \| 2 \| 402 \| 10 \| 413 \| Site-specific recombinase XerD \| Site-specific recombinase XerD; afdb-uniprot50 \| AF-A0A259Q3Y0-F1-MODEL\_V4 \| 1.0 \| 6.192e-10 \| 419 \| 0.173 \| 317 \| 223 \| 13 \| 102 \| 402 \| 1 \| 294 \| Integrase \| Integrase; afdb-uniprot50 \| AF-A0A6P4G483-F1-MODEL\_V4 \| 1.0 \| 4.435e-10 \| 418 \| 0.227 \| 299 \| 193 \| 13 \| 116 \| 398 \| 8 \| 284 \| prophage CP4-57 integrase-like \| prophage CP4-57 integrase-like; afdb-uniprot50 \| AF-A0A0F7HAR5-F1-MODEL\_V4 \| 1.0 \| 1.281e-13 \| 418 \| 0.196 \| 422 \| 270 \| 23 \| 2 \| 402 \| 3 \| 376 \| Arm-DNA-bind\_3 domain-containing protein \| Arm-DNA-bind\_3 domain-containing protein; afdb-uniprot50 \| AF-A0A5E4S4Y3-F1-MODEL\_V4 \| 1.0 \| 1.012e-10 \| 417 \| 0.199 \| 371 \| 247 \| 16 \| 60 \| 403 \| 1 \| 348 \| Integrase \| Integrase; afdb-uniprot50 \| AF-A0A2N6Q9F2-F1-MODEL\_V4 \| 1.0 \| 1.046e-09 \| 416 \| 0.176 \| 267 \| 190 \| 7 \| 149 \| 402 \| 1 \| 250 \| Tyr recombinase domain-containing protein \| Tyr recombinase domain-containing protein; afdb-uniprot50 \| AF-A0A2R7MYK0-F1-MODEL\_V4 \| 1.0 \| 2.352e-09 \| 415 \| 0.156 \| 314 \| 234 \| 11 \| 100 \| 402 \| 24 \| 317 \| Integrase \| Integrase; afdb-uniprot50 \| AF-A0A5P3AFX6-F1-MODEL\_V4 \| 1.0 \| 9.066e-10 \| 415 \| 0.175 \| 313 \| 229 \| 12 \| 99 \| 400 \| 10 \| 304 \| Prophage CP4-57 integrase \| Prophage CP4-57 integrase; afdb-uniprot50 \| AF-A0A4D9X583-F1-MODEL\_V4 \| 1.0 \| 3.323e-13 \| 415 \| 0.208 \| 378 \| 231 \| 21 \| 2 \| 346 \| 3 \| 345 \| DUF4102 domain-containing protein \| DUF4102 domain-containing protein; afdb-uniprot50 \| AF-A0A2L2XGH8-F1-MODEL\_V4 \| 1.0 \| 2.846e-09 \| 414 \| 0.176 \| 306 \| 216 \| 14 \| 101 \| 393 \| 1 \| 283 \| Tyrosine recombinase xerD \| Tyrosine recombinase xerD; afdb-uniprot50 \| AF-A0A2K8W1H0-F1-MODEL\_V4 \| 1.0 \| 2.88e-13 \| 414 \| 0.197 \| 380 \| 233 \| 20 \| 2 \| 346 \| 3 \| 345 \| Prophage P4 integrase \| Prophage P4 integrase; afdb-uniprot50 \| AF-A0A350JMT7-F1-MODEL\_V4 \| 1.0 \| 3.601e-12 \| 414 \| 0.162 \| 436 \| 296 \| 25 \| 2 \| 402 \| 8 \| 409 \| Tyr recombinase domain-containing protein \| Tyr recombinase domain-containing protein; afdb-uniprot50 \| AF-A0A4U8Z4V2-F1-MODEL\_V4 \| 1.0 \| 7.858e-10 \| 413 \| 0.222 \| 265 \| 177 \| 9 \| 149 \| 402 \| 1 \| 247 \| Tyr recombinase domain-containing protein \| Tyr recombinase domain-containing protein; afdb-uniprot50 \| AF-A0A2E4NJY1-F1-MODEL\_V4 \| 1.0 \| 7.858e-10 \| 413 \| 0.188 \| 276 \| 197 \| 9 \| 137 \| 402 \| 6 \| 264 \| Integrase \| Integrase; afdb-uniprot50 \| AF-A0A1W2A448-F1-MODEL\_V4 \| 1.0 \| 2.837e-12 \| 413 \| 0.194 \| 375 \| 236 \| 20 \| 45 \| 402 \| 33 \| 358 \| Site-specific recombinase XerD \| Site-specific recombinase XerD; afdb-uniprot50 \| AF-A0A1H4W6T1-F1-MODEL\_V4 \| 1.0 \| 6.285e-11 \| 412 \| 0.201 \| 343 \| 233 \| 16 \| 80 \| 403 \| 2 \| 322 \| Integrase \| Integrase; afdb-uniprot50 \| AF-A0A7G8BGC1-F1-MODEL\_V4 \| 1.0 \| 1.392e-09 \| 411 \| 0.188 \| 265 \| 188 \| 12 \| 101 \| 356 \| 12 \| 258 \| Tyrosine-type recombinase/integrase \| Tyrosine-type recombinase/integrase; afdb-uniprot50 \| AF-A0A399RGX3-F1-MODEL\_V4 \| 1.0 \| 2.626e-10 \| 411 \| 0.205 \| 312 \| 204 \| 15 \| 99 \| 401 \| 22 \| 298 \| Site-specific integrase \| Site-specific integrase; afdb-uniprot50 \| AF-A0A2D9RK50-F1-MODEL\_V4 \| 1.0 \| 6.592e-11 \| 411 \| 0.183 \| 366 \| 251 \| 19 \| 60 \| 402 \| 1 \| 341 \| Integrase \| Integrase; afdb-uniprot50 \| AF-A0A5P9EZI4-F1-MODEL\_V4 \| 1.0 \| 1.602e-12 \| 411 \| 0.17 \| 398 \| 260 \| 20 \| 1 \| 375 \| 2 \| 352 \| Prophage CP4-57 integrase \| Prophage CP4-57 integrase; afdb-uniprot50 \| AF-A0A523WE57-F1-MODEL\_V4 \| 1.0 \| 1.094e-12 \| 411 \| 0.181 \| 424 \| 291 \| 21 \| 2 \| 402 \| 70 \| 460 \| Uncharacterized protein \| Uncharacterized protein; afdb-uniprot50 \| AF-A0A436CCZ6-F1-MODEL\_V4 \| 1.0 \| 2.466e-09 \| 409 \| 0.184 \| 271 \| 192 \| 11 \| 146 \| 402 \| 9 \| 264 \| Integrase \| Integrase; afdb-uniprot50 \| AF-A0A8B3J0Y8-F1-MODEL\_V4 \| 1.0 \| 1.554e-10 \| 409 \| 0.198 \| 318 \| 219 \| 12 \| 99 \| 402 \| 4 \| 299 \| Integrase \| Integrase; afdb-uniprot50 \| AF-B1JDT1-F1-MODEL\_V4 \| 1.0 \| 2.236e-12 \| 409 \| 0.169 \| 396 \| 264 \| 18 \| 34 \| 401 \| 37 \| 395 \| Integrase family protein \| Integrase family protein; afdb-uniprot50 \| AF-A0A722ZTZ9-F1-MODEL\_V4 \| 1.0 \| 1.456e-12 \| 407 \| 0.18 \| 387 \| 254 \| 21 \| 2 \| 357 \| 3 \| 357 \| DUF4102 domain-containing protein \| DUF4102 domain-containing protein; afdb-uniprot50 \| AF-A0A6L2ZPK9-F1-MODEL\_V4 \| 1.0 \| 6.811e-10 \| 406 \| 0.15 \| 345 \| 249 \| 12 \| 69 \| 402 \| 6 \| 317 \| Integrase \| Integrase; afdb-uniprot50 \| AF-A0A7Y4U7C5-F1-MODEL\_V4 \| 1.0 \| 2.795e-11 \| 406 \| 0.168 \| 373 \| 255 \| 15 \| 36 \| 401 \| 3 \| 327 \| Site-specific integrase \| Site-specific integrase; afdb-uniprot50 \| AF-A0A0T5ZW92-F1-MODEL\_V4 \| 1.0 \| 4.229e-10 \| 405 \| 0.214 \| 308 \| 217 \| 13 \| 97 \| 400 \| 35 \| 321 \| Putative phage integrase \| Putative phage integrase; afdb-uniprot50 \| AF-A0A2J8HSE2-F1-MODEL\_V4 \| 1.0 \| 5.8e-12 \| 405 \| 0.161 \| 422 \| 295 \| 20 \| 2 \| 402 \| 44 \| 427 \| Tyr recombinase domain-containing protein \| Tyr recombinase domain-containing protein; afdb-uniprot50 \| AF-A0A0Q7TMR2-F1-MODEL\_V4 \| 1.0 \| 2.276e-10 \| 404 \| 0.202 \| 360 \| 233 \| 19 \| 69 \| 398 \| 2 \| 337 \| Tyr recombinase domain-containing protein \| Tyr recombinase domain-containing protein; afdb-uniprot50 \| AF-A0A7U0PNP7-F1-MODEL\_V4 \| 1.0 \| 1.078e-11 \| 404 \| 0.153 \| 423 \| 309 \| 20 \| 2 \| 402 \| 6 \| 401 \| Integrase family protein \| Integrase family protein; afdb-uniprot50 \| AF-A0A2N8M7R9-F1-MODEL\_V4 \| 1.0 \| 1.848e-12 \| 403 \| 0.169 \| 414 \| 273 \| 17 \| 9 \| 389 \| 14 \| 389 \| Integrase \| Integrase; afdb-uniprot50 \| AF-A0A090SWG2-F1-MODEL\_V4 \| 1.0 \| 1.046e-09 \| 402 \| 0.175 \| 347 \| 249 \| 15 \| 74 \| 402 \| 4 \| 331 \| Phage integrase \| Phage integrase; afdb-uniprot50 \| AF-A0A537S885-F1-MODEL\_V4 \| 1.0 \| 3.121e-12 \| 402 \| 0.17 \| 429 \| 288 \| 17 \| 2 \| 390 \| 10 \| 410 \| Site-specific integrase \| Site-specific integrase; afdb-uniprot50 \| AF-A0A225DKJ3-F1-MODEL\_V4 \| 1.0 \| 7.251e-11 \| 401 \| 0.188 \| 366 \| 239 \| 16 \| 49 \| 401 \| 22 \| 342 \| Mobile element protein \| Mobile element protein; afdb-uniprot50 \| AF-A0A257LMN8-F1-MODEL\_V4 \| 1.0 \| 2.795e-11 \| 401 \| 0.204 \| 377 \| 234 \| 20 \| 69 \| 403 \| 17 \| 369 \| Integrase \| Integrase; afdb-uniprot50 \| AF-A0A6G6J4R1-F1-MODEL\_V4 \| 1.0 \| 2.705e-12 \| 401 \| 0.168 \| 438 \| 272 \| 24 \| 1 \| 401 \| 9 \| 391 \| Site-specific integrase \| Site-specific integrase; afdb-uniprot50 \| AF-A0A0M2V4F1-F1-MODEL\_V4 \| 1.0 \| 5.53e-12 \| 400 \| 0.166 \| 414 \| 281 \| 20 \| 34 \| 402 \| 9 \| 403 \| Tyr recombinase domain-containing protein \| Tyr recombinase domain-containing protein; afdb-uniprot50 \| AF-L8MDS0-F1-MODEL\_V4 \| 1.0 \| 1.66e-08 \| 399 \| 0.278 \| 187 \| 125 \| 6 \| 223 \| 402 \| 1 \| 184 \| Prophage PSPPH01, site-specific recombinase, phage integrase family \| Prophage PSPPH01, site-specific recombinase, phage integrase family; afdb-uniprot50 \| AF-A0A5B9QPY5-F1-MODEL\_V4 \| 1.0 \| 4.722e-11 \| 399 \| 0.183 \| 366 \| 235 \| 18 \| 46 \| 401 \| 16 \| 327 \| Tyrosine recombinase XerC \| Tyrosine recombinase XerC; afdb-uniprot50 \| AF-A0A0W0U4L5-F1-MODEL\_V4 \| 1.0 \| 3.495e-10 \| 398 \| 0.206 \| 296 \| 206 \| 10 \| 112 \| 400 \| 10 \| 283 \| Integrase \| Integrase; afdb-uniprot50 \| AF-A0A510U2Z7-F1-MODEL\_V4 \| 1.0 \| 1.909e-11 \| 398 \| 0.172 \| 395 \| 266 \| 20 \| 34 \| 402 \| 35 \| 394 \| Recombinase \| Recombinase; afdb-uniprot50 \| AF-A0A656BVP0-F1-MODEL\_V4 \| 1.0 \| 3.283e-09 \| 395 \| 0.209 \| 262 \| 178 \| 9 \| 154 \| 403 \| 3 \| 247 \| Prophage Sf6-like integrase \| Prophage Sf6-like integrase; afdb-uniprot50 \| AF-A0A248XRU4-F1-MODEL\_V4 \| 1.0 \| 1.392e-09 \| 395 \| 0.182 \| 279 \| 205 \| 8 \| 130 \| 402 \| 1 \| 262 \| Integrase \| Integrase; afdb-uniprot50 \| AF-A0A853R082-F1-MODEL\_V4 \| 1.0 \| 1.909e-11 \| 395 \| 0.148 \| 405 \| 277 \| 20 \| 8 \| 389 \| 9 \| 368 \| Uncharacterized protein \| Uncharacterized protein; afdb-uniprot50 \| AF-A0A2Z4R7D1-F1-MODEL\_V4 \| 1.0 \| 2.846e-09 \| 394 \| 0.163 \| 281 \| 207 \| 11 \| 133 \| 401 \| 9 \| 273 \| Tyr recombinase domain-containing protein \| Tyr recombinase domain-containing protein; afdb-uniprot50 \| AF-A0A378MUR8-F1-MODEL\_V4 \| 1.0 \| 1.207e-09 \| 392 \| 0.197 \| 269 \| 186 \| 10 \| 146 \| 402 \| 3 \| 253 \| Prophage CP4-57 integrase \| Prophage CP4-57 integrase; afdb-uniprot50 \| AF-A0A7W1APG2-F1-MODEL\_V4 \| 1.0 \| 2.673e-08 \| 391 \| 0.216 \| 203 \| 150 \| 6 \| 207 \| 402 \| 2 \| 202 \| Site-specific integrase \| Site-specific integrase; afdb-uniprot50 \| AF-A0A192D4V7-F1-MODEL\_V4 \| 1.0 \| 1.685e-09 \| 390 \| 0.224 \| 250 \| 164 \| 12 \| 137 \| 373 \| 3 \| 235 \| Uncharacterized protein \| Uncharacterized protein; afdb-uniprot50 \| AF-A0A2A6Q2D7-F1-MODEL\_V4 \| 1.0 \| 5.041e-09 \| 389 \| 0.224 \| 236 \| 157 \| 10 \| 177 \| 402 \| 2 \| 221 \| Integrase \| Integrase; afdb-uniprot50 \| AF-A0A1H6ZMK8-F1-MODEL\_V4 \| 1.0 \| 6.38e-12 \| 389 \| 0.179 \| 430 \| 277 \| 22 \| 2 \| 401 \| 12 \| 395 \| Integrase \| Integrase; afdb-uniprot50 \| AF-A0A7W0ZY73-F1-MODEL\_V4 \| 1.0 \| 4.093e-11 \| 389 \| 0.128 \| 484 \| 312 \| 30 \| 2 \| 403 \| 5 \| 460 \| Tyr recombinase domain-containing protein \| Tyr recombinase domain-containing protein; afdb-uniprot50 \| AF-A0A7T9ZF29-F1-MODEL\_V4 \| 1.0 \| 2.985e-09 \| 388 \| 0.18 \| 310 \| 209 \| 14 \| 108 \| 403 \| 7 \| 285 \| Tyrosine-type recombinase/integrase \| Tyrosine-type recombinase/integrase; afdb-uniprot50 \| AF-A0A1G2ZM93-F1-MODEL\_V4 \| 1.0 \| 1.973e-10 \| 388 \| 0.197 \| 390 \| 247 \| 16 \| 18 \| 403 \| 4 \| 331 \| Tyr recombinase domain-containing protein \| Tyr recombinase domain-containing protein; afdb-uniprot50 \| AF-I4CE33-F1-MODEL\_V4 \| 1.0 \| 5.993e-11 \| 387 \| 0.154 \| 394 \| 276 \| 19 \| 34 \| 393 \| 2 \| 372 \| Site-specific recombinase XerD \| Site-specific recombinase XerD; afdb-uniprot50 \| AF-A0A5E5BYC9-F1-MODEL\_V4 \| 1.0 \| 2.352e-09 \| 386 \| 0.242 \| 231 \| 154 \| 8 \| 182 \| 401 \| 3 \| 223 \| Integrase \| Integrase; afdb-uniprot50 \| AF-A0A5N9VWT1-F1-MODEL\_V4 \| 1.0 \| 1.71e-10 \| 385 \| 0.176 \| 306 \| 213 \| 18 \| 2 \| 296 \| 9 \| 286 \| DUF4102 domain-containing protein \| DUF4102 domain-containing protein; afdb-uniprot50 \| AF-A0A378AUB5-F1-MODEL\_V4 \| 1.0 \| 2.352e-09 \| 384 \| 0.176 \| 295 \| 215 \| 10 \| 64 \| 347 \| 11 \| 288 \| Integrase \| Integrase; afdb-uniprot50 \| AF-A0A845AEC5-F1-MODEL\_V4 \| 1.0 \| 6.792e-13 \| 383 \| 0.187 \| 411 \| 216 \| 17 \| 2 \| 401 \| 3 \| 306 \| Integrase arm-type DNA-binding domain-containing protein \| Integrase arm-type DNA-binding domain-containing protein; afdb-uniprot50 \| AF-A0A7W5E2G6-F1-MODEL\_V4 \| 1.0 \| 3.029e-10 \| 382 \| 0.173 \| 357 \| 239 \| 16 \| 46 \| 393 \| 15 \| 324 \| Integrase \| Integrase; afdb-uniprot50 \| AF-A0A1F5UBR0-F1-MODEL\_V4 \| 1.0 \| 5.714e-11 \| 382 \| 0.158 \| 392 \| 261 \| 16 \| 18 \| 400 \| 2 \| 333 \| Uncharacterized protein \| Uncharacterized protein; afdb-uniprot50 \| AF-A0A485GR16-F1-MODEL\_V4 \| 1.0 \| 1.853e-09 \| 380 \| 0.206 \| 286 \| 184 \| 9 \| 146 \| 402 \| 11 \| 282 \| Integrase family protein \| Integrase family protein; afdb-uniprot50 \| AF-A0A1X9T2A8-F1-MODEL\_V4 \| 1.0 \| 3.283e-09 \| 379 \| 0.157 \| 324 \| 236 \| 15 \| 90 \| 398 \| 4 \| 305 \| Site-specific recombinase, phage integrase family \| Site-specific recombinase, phage integrase family; afdb-uniprot50 \| AF-A0A1I4QUM7-F1-MODEL\_V4 \| 1.0 \| 4.722e-11 \| 379 \| 0.199 \| 321 \| 211 \| 14 \| 1 \| 303 \| 6 \| 298 \| Phage integrase family protein \| Phage integrase family protein; afdb-uniprot50 \| AF-A0A564WGE9-F1-MODEL\_V4 \| 1.0 \| 2.665e-11 \| 379 \| 0.179 \| 441 \| 277 \| 20 \| 1 \| 402 \| 9 \| 403 \| Integrase family protein \| Integrase family protein; afdb-uniprot50 \| AF-A0A560UXU6-F1-MODEL\_V4 \| 1.0 \| 6.934e-08 \| 377 \| 0.252 \| 190 \| 132 \| 6 \| 220 \| 402 \| 1 \| 187 \| Phage integrase family protein \| Phage integrase family protein; afdb-uniprot50 \| AF-A0A257H209-F1-MODEL\_V4 \| 1.0 \| 6.811e-10 \| 377 \| 0.167 \| 358 \| 250 \| 15 \| 67 \| 399 \| 9 \| 343 \| Uncharacterized protein \| Uncharacterized protein; afdb-uniprot50 \| AF-A0A6A4TZN2-F1-MODEL\_V4 \| 1.0 \| 1.685e-09 \| 377 \| 0.198 \| 332 \| 210 \| 17 \| 99 \| 401 \| 29 \| 333 \| DNA integration/recombination/inversion protein \| DNA integration/recombination/inversion protein; afdb-uniprot50 \| AF-A0A655W7E3-F1-MODEL\_V4 \| 1.0 \| 8.39e-08 \| 376 \| 0.211 \| 189 \| 141 \| 4 \| 220 \| 402 \| 1 \| 187 \| Phage integrase \| Phage integrase; afdb-uniprot50 \| AF-A0A4Q3Z742-F1-MODEL\_V4 \| 1.0 \| 2.317e-08 \| 376 \| 0.248 \| 197 \| 129 \| 6 \| 217 \| 400 \| 1 \| 191 \| Tyr recombinase domain-containing protein \| Tyr recombinase domain-containing protein; afdb-uniprot50 \| AF-A0A5J6QNY1-F1-MODEL\_V4 \| 1.0 \| 3.788e-09 \| 376 \| 0.178 \| 319 \| 221 \| 14 \| 102 \| 401 \| 8 \| 304 \| Site-specific integrase \| Site-specific integrase; afdb-uniprot50 \| AF-A0A5B9QG65-F1-MODEL\_V4 \| 1.0 \| 7.492e-10 \| 376 \| 0.169 \| 348 \| 248 \| 12 \| 46 \| 389 \| 12 \| 322 \| Tyrosine recombinase XerD \| Tyrosine recombinase XerD; afdb-uniprot50 \| AF-A0A7W2L6W8-F1-MODEL\_V4 \| 1.0 \| 2.352e-09 \| 376 \| 0.19 \| 326 \| 218 \| 14 \| 99 \| 402 \| 13 \| 314 \| Site-specific integrase \| Site-specific integrase; afdb-uniprot50 \| AF-A0A257RB84-F1-MODEL\_V4 \| 1.0 \| 4.093e-11 \| 375 \| 0.167 \| 442 \| 289 \| 18 \| 2 \| 403 \| 10 \| 412 \| Tyr recombinase domain-containing protein \| Tyr recombinase domain-containing protein; afdb-uniprot50 \| AF-A0A3C1G9D8-F1-MODEL\_V4 \| 1.0 \| 3.972e-09 \| 374 \| 0.167 \| 304 \| 211 \| 16 \| 101 \| 391 \| 13 \| 287 \| Uncharacterized protein \| Uncharacterized protein; afdb-uniprot50 \| AF-A0A1E3M116-F1-MODEL\_V4 \| 1.0 \| 1.151e-09 \| 374 \| 0.186 \| 322 \| 213 \| 16 \| 101 \| 401 \| 1 \| 294 \| Uncharacterized protein \| Uncharacterized protein; afdb-uniprot50 \| AF-S5MWJ7-F1-MODEL\_V4 \| 1.0 \| 2.31e-11 \| 372 \| 0.226 \| 335 \| 215 \| 17 \| 1 \| 322 \| 7 \| 310 \| Integrase \| Integrase; afdb-uniprot50 \| AF-A0A1B6VTD4-F1-MODEL\_V4 \| 1.0 \| 6.303e-08 \| 371 \| 0.22 \| 200 \| 142 \| 5 \| 216 \| 402 \| 18 \| 216 \| Tyr recombinase domain-containing protein \| Tyr recombinase domain-containing protein; afdb-uniprot50 \| AF-A0A533QJ01-F1-MODEL\_V4 \| 1.0 \| 2.069e-10 \| 371 \| 0.19 \| 400 \| 250 \| 21 \| 34 \| 403 \| 4 \| 359 \| Putative defective protein IntQ \| Putative defective protein IntQ; afdb-uniprot50 \| AF-A0A3C7VZS0-F1-MODEL\_V4 \| 1.0 \| 1.486e-07 \| 370 \| 0.23 \| 182 \| 133 \| 5 \| 225 \| 403 \| 21 \| 198 \| Tyr recombinase domain-containing protein \| Tyr recombinase domain-containing protein; afdb-uniprot50 \| AF-A0A0S8GP47-F1-MODEL\_V4 \| 1.0 \| 6.494e-10 \| 370 \| 0.168 \| 380 \| 263 \| 18 \| 30 \| 393 \| 2 \| 344 \| Uncharacterized protein \| Uncharacterized protein; afdb-uniprot50 \| AF-A0A7Z7RKM6-F1-MODEL\_V4 \| 1.0 \| 1.439e-08 \| 369 \| 0.195 \| 245 \| 166 \| 8 \| 174 \| 402 \| 2 \| 231 \| Prophage CP4-57 integrase \| Prophage CP4-57 integrase; afdb-uniprot50 \| AF-A0A0Y0K400-F1-MODEL\_V4 \| 1.0 \| 8.218e-13 \| 369 \| 0.222 \| 382 \| 226 \| 23 \| 2 \| 354 \| 7 \| 346 \| Integrase family protein \| Integrase family protein; afdb-uniprot50 \| AF-A0A5P3VU28-F1-MODEL\_V4 \| 1.0 \| 2.31e-11 \| 369 \| 0.176 \| 413 \| 271 \| 21 \| 9 \| 396 \| 13 \| 381 \| Site-specific integrase \| Site-specific integrase; afdb-uniprot50 \| AF-A0A518HSZ1-F1-MODEL\_V4 \| 1.0 \| 7.605e-11 \| 368 \| 0.156 \| 429 \| 237 \| 20 \| 46 \| 403 \| 15 \| 389 \| Site-specific tyrosine recombinase XerC \| Site-specific tyrosine recombinase XerC; afdb-uniprot50 \| AF-Q1NEP9-F1-MODEL\_V4 \| 1.0 \| 2.038e-09 \| 367 \| 0.17 \| 310 \| 205 \| 12 \| 108 \| 402 \| 6 \| 278 \| Symbiosis island integrase \| Symbiosis island integrase; afdb-uniprot50 \| AF-A0A5E4XII2-F1-MODEL\_V4 \| 1.0 \| 5.714e-11 \| 367 \| 0.173 \| 358 \| 237 \| 18 \| 15 \| 346 \| 1 \| 325 \| Integrase \| Integrase; afdb-uniprot50 \| AF-A0A655PDZ5-F1-MODEL\_V4 \| 1.0 \| 1.417e-07 \| 366 \| 0.211 \| 189 \| 141 \| 4 \| 221 \| 403 \| 11 \| 197 \| Phage integrase \| Phage integrase; afdb-uniprot50 \| AF-A0A354XKQ8-F1-MODEL\_V4 \| 1.0 \| 8.799e-08 \| 365 \| 0.247 \| 182 \| 127 \| 5 \| 225 \| 401 \| 25 \| 201 \| Integrase \| Integrase; afdb-uniprot50 \| AF-A0A7V8QD16-F1-MODEL\_V4 \| 1.0 \| 2.236e-12 \| 365 \| 0.198 \| 358 \| 217 \| 17 \| 2 \| 321 \| 3 \| 328 \| Integrase arm-type DNA-binding domain-containing protein \| Integrase arm-type DNA-binding domain-containing protein; afdb-uniprot50 \| AF-A0A377DPL2-F1-MODEL\_V4 \| 1.0 \| 1.559e-07 \| 364 \| 0.196 \| 188 \| 142 \| 6 \| 222 \| 402 \| 3 \| 188 \| Phage integrase \| Phage integrase; afdb-uniprot50 \| AF-A0A509BEV6-F1-MODEL\_V4 \| 1.0 \| 1.486e-07 \| 364 \| 0.181 \| 193 \| 150 \| 5 \| 216 \| 402 \| 4 \| 194 \| Integrase family protein \| Integrase family protein; afdb-uniprot50 \| AF-D4C538-F1-MODEL\_V4 \| 1.0 \| 1.715e-07 \| 364 \| 0.186 \| 193 \| 149 \| 5 \| 216 \| 402 \| 26 \| 216 \| Putative prophage CP4-57 integrase \| Putative prophage CP4-57 integrase; afdb-uniprot50 \| AF-A0A0M2DWI1-F1-MODEL\_V4 \| 1.0 \| 8.516e-09 \| 364 \| 0.226 \| 234 \| 158 \| 9 \| 178 \| 402 \| 2 \| 221 \| Prophage CP4-57 integrase \| Prophage CP4-57 integrase; afdb-uniprot50 \| AF-A0A694V927-F1-MODEL\_V4 \| 1.0 \| 2.1e-11 \| 364 \| 0.169 \| 324 \| 230 \| 12 \| 1 \| 310 \| 3 \| 301 \| DUF4102 domain-containing protein \| DUF4102 domain-containing protein; afdb-uniprot50 \| AF-A0A2W5DWS9-F1-MODEL\_V4 \| 1.0 \| 1.741e-08 \| 363 \| 0.171 \| 263 \| 190 \| 10 \| 152 \| 402 \| 2 \| 248 \| Integrase \| Integrase; afdb-uniprot50 \| AF-A0A212I7A4-F1-MODEL\_V4 \| 1.0 \| 1.635e-07 \| 362 \| 0.196 \| 193 \| 147 \| 5 \| 216 \| 402 \| 26 \| 216 \| Integrase CP4-57 prophage \| Integrase CP4-57 prophage; afdb-uniprot50 \| AF-A0A3M3AIK9-F1-MODEL\_V4 \| 1.0 \| 1.247e-08 \| 362 \| 0.194 \| 262 \| 182 \| 9 \| 153 \| 402 \| 2 \| 246 \| Site-specific recombinase, phage integrase protein \| Site-specific recombinase, phage integrase protein; afdb-uniprot50 \| AF-A0A1G0VC22-F1-MODEL\_V4 \| 1.0 \| 3.666e-10 \| 362 \| 0.134 \| 380 \| 256 \| 19 \| 36 \| 400 \| 3 \| 324 \| Uncharacterized protein \| Uncharacterized protein; afdb-uniprot50 \| AF-A0A848CG60-F1-MODEL\_V4 \| 1.0 \| 1.015e-07 \| 361 \| 0.179 \| 206 \| 156 \| 8 \| 207 \| 403 \| 3 \| 204 \| Site-specific integrase \| Site-specific integrase; afdb-uniprot50 \| AF-A0A1Q3YLC3-F1-MODEL\_V4 \| 1.0 \| 6.914e-11 \| 361 \| 0.217 \| 303 \| 193 \| 16 \| 2 \| 290 \| 3 \| 275 \| Integrase \| Integrase; afdb-uniprot50 \| AF-A0A518K3T6-F1-MODEL\_V4 \| 1.0 \| 8.242e-10 \| 361 \| 0.185 \| 382 \| 260 \| 17 \| 23 \| 402 \| 2 \| 334 \| Tyrosine recombinase XerD \| Tyrosine recombinase XerD; afdb-uniprot50 \| AF-D1AHL5-F1-MODEL\_V4 \| 1.0 \| 7.144e-10 \| 361 \| 0.161 \| 389 \| 260 \| 19 \| 17 \| 393 \| 8 \| 342 \| Integrase family protein \| Integrase family protein; afdb-uniprot50 \| AF-A0A397QTS5-F1-MODEL\_V4 \| 1.0 \| 1.63e-10 \| 360 \| 0.148 \| 398 \| 272 \| 17 \| 17 \| 396 \| 11 \| 359 \| Site-specific recombinase XerD \| Site-specific recombinase XerD; afdb-uniprot50 \| AF-A0A1M6IDQ9-F1-MODEL\_V4 \| 1.0 \| 3.721e-11 \| 360 \| 0.161 \| 408 \| 279 \| 19 \| 7 \| 392 \| 8 \| 374 \| Site-specific recombinase XerD \| Site-specific recombinase XerD; afdb-uniprot50 \| AF-D8FA13-F1-MODEL\_V4 \| 1.0 \| 1.881e-10 \| 359 \| 0.172 \| 383 \| 262 \| 15 \| 34 \| 402 \| 3 \| 344 \| Site-specific recombinase, phage integrase family \| Site-specific recombinase, phage integrase family; afdb-uniprot50 \| AF-A0A432UDR0-F1-MODEL\_V4 \| 1.0 \| 2.17e-10 \| 358 \| 0.178 \| 375 \| 245 \| 16 \| 42 \| 402 \| 30 \| 355 \| Tyr recombinase domain-containing protein \| Tyr recombinase domain-containing protein; afdb-uniprot50 \| AF-G0ERT2-F1-MODEL\_V4 \| 1.0 \| 2.283e-07 \| 357 \| 0.197 \| 192 \| 143 \| 4 \| 220 \| 402 \| 20 \| 209 \| Prophage CP4-57 integrase IntA \| Prophage CP4-57 integrase IntA; afdb-uniprot50 \| AF-A0A5C9CD61-F1-MODEL\_V4 \| 1.0 \| 1.606e-09 \| 357 \| 0.178 \| 359 \| 240 \| 18 \| 46 \| 393 \| 15 \| 329 \| Uncharacterized protein \| Uncharacterized protein; afdb-uniprot50 \| AF-A0A847DD10-F1-MODEL\_V4 \| 1.0 \| 1.114e-10 \| 356 \| 0.186 \| 369 \| 267 \| 18 \| 45 \| 402 \| 36 \| 382 \| Tyrosine-type recombinase/integrase \| Tyrosine-type recombinase/integrase; afdb-uniprot50 \| AF-A0A6I6VWS2-F1-MODEL\_V4 \| 1.0 \| 6.192e-10 \| 355 \| 0.257 \| 299 \| 183 \| 14 \| 55 \| 334 \| 9 \| 287 \| Uncharacterized protein \| Uncharacterized protein; afdb-uniprot50 \| AF-A0A2V8AVC8-F1-MODEL\_V4 \| 1.0 \| 4.652e-10 \| 355 \| 0.199 \| 371 \| 246 \| 18 \| 46 \| 393 \| 19 \| 361 \| Uncharacterized protein \| Uncharacterized protein; afdb-uniprot50 \| AF-A0A3R9ZJ42-F1-MODEL\_V4 \| 1.0 \| 6.934e-08 \| 354 \| 0.169 \| 248 \| 180 \| 8 \| 164 \| 401 \| 2 \| 233 \| Integrase \| Integrase; afdb-uniprot50 \| AF-A0A6P2CVZ9-F1-MODEL\_V4 \| 1.0 \| 2.008e-08 \| 354 \| 0.205 \| 307 \| 207 \| 15 \| 101 \| 400 \| 1 \| 277 \| Uncharacterized protein \| Uncharacterized protein; afdb-uniprot50 \| AF-A0A5Y2E3L5-F1-MODEL\_V4 \| 1.0 \| 2.283e-07 \| 353 \| 0.223 \| 192 \| 133 \| 8 \| 221 \| 402 \| 1 \| 186 \| Integrase \| Integrase; afdb-uniprot50 \| AF-A0A395JEX9-F1-MODEL\_V4 \| 1.0 \| 3.234e-08 \| 353 \| 0.207 \| 241 \| 165 \| 8 \| 172 \| 402 \| 11 \| 235 \| Integrase \| Integrase; afdb-uniprot50 \| AF-A0A1Y6KVZ1-F1-MODEL\_V4 \| 1.0 \| 1.978e-07 \| 351 \| 0.244 \| 196 \| 137 \| 6 \| 216 \| 402 \| 25 \| 218 \| Prophage CP4-57 integrase \| Prophage CP4-57 integrase; afdb-uniprot50 \| AF-A0A2D4WFI9-F1-MODEL\_V4 \| 1.0 \| 1.439e-08 \| 351 \| 0.177 \| 322 \| 218 \| 14 \| 102 \| 401 \| 3 \| 299 \| Integrase \| Integrase; afdb-uniprot50 \| AF-A0A6C0TPK5-F1-MODEL\_V4 \| 1.0 \| 7.976e-11 \| 351 \| 0.211 \| 336 \| 212 \| 15 \| 1 \| 297 \| 8 \| 329 \| Integrase \| Integrase; afdb-uniprot50 \| AF-A0A376RHC7-F1-MODEL\_V4 \| 1.0 \| 1.185e-11 \| 351 \| 0.186 \| 375 \| 238 \| 22 \| 1 \| 341 \| 2 \| 343 \| Integrase \| Integrase; afdb-uniprot50 \| AF-A0A142WXX8-F1-MODEL\_V4 \| 1.0 \| 1.285e-10 \| 351 \| 0.16 \| 400 \| 281 \| 16 \| 1 \| 393 \| 17 \| 368 \| Site-specific tyrosine recombinase XerC \| Site-specific tyrosine recombinase XerC; afdb-uniprot50 \| AF-A0A1F7FFN7-F1-MODEL\_V4 \| 1.0 \| 1.881e-10 \| 350 \| 0.172 \| 383 \| 247 \| 20 \| 34 \| 395 \| 2 \| 335 \| Tyr recombinase domain-containing protein \| Tyr recombinase domain-containing protein; afdb-uniprot50 \| AF-A0A5C9C4Q8-F1-MODEL\_V4 \| 1.0 \| 5.041e-09 \| 349 \| 0.156 \| 376 \| 259 \| 14 \| 34 \| 400 \| 5 \| 331 \| Tyrosine recombinase XerC \| Tyrosine recombinase XerC; afdb-uniprot50 \| AF-A0A2N2IB33-F1-MODEL\_V4 \| 1.0 \| 9.973e-10 \| 349 \| 0.165 \| 374 \| 276 \| 15 \| 34 \| 401 \| 2 \| 345 \| Tyr recombinase domain-containing protein \| Tyr recombinase domain-containing protein; afdb-uniprot50 \| AF-A0A7U9F051-F1-MODEL\_V4 \| 1.0 \| 1.081e-08 \| 348 \| 0.204 \| 289 \| 185 \| 15 \| 137 \| 402 \| 12 \| 278 \| Tyr recombinase domain-containing protein \| Tyr recombinase domain-containing protein; afdb-uniprot50 \| AF-A0A2G6DGF4-F1-MODEL\_V4 \| 1.0 \| 1.685e-09 \| 348 \| 0.143 \| 356 \| 269 \| 16 \| 46 \| 396 \| 12 \| 336 \| Tyr recombinase domain-containing protein \| Tyr recombinase domain-containing protein; afdb-uniprot50 \| AF-A0A6C2YUQ7-F1-MODEL\_V4 \| 1.0 \| 1.531e-09 \| 348 \| 0.155 \| 361 \| 261 \| 13 \| 46 \| 393 \| 15 \| 344 \| Uncharacterized protein \| Uncharacterized protein; afdb-uniprot50 \| AF-A0A6L7K450-F1-MODEL\_V4 \| 1.0 \| 3.505e-07 \| 347 \| 0.209 \| 186 \| 138 \| 6 \| 221 \| 399 \| 9 \| 192 \| Site-specific integrase \| Site-specific integrase; afdb-uniprot50 \| AF-A0A261WKU5-F1-MODEL\_V4 \| 1.0 \| 4.044e-07 \| 347 \| 0.233 \| 184 \| 130 \| 6 \| 222 \| 398 \| 35 \| 214 \| Integrase \| Integrase; afdb-uniprot50 \| AF-A0A5S9ISB9-F1-MODEL\_V4 \| 1.0 \| 2.888e-10 \| 347 \| 0.155 \| 392 \| 243 \| 19 \| 46 \| 400 \| 8 \| 348 \| Recombinase \| Recombinase; afdb-uniprot50 \| AF-A0A7Y9UX65-F1-MODEL\_V4 \| 1.0 \| 5.904e-10 \| 347 \| 0.165 \| 406 \| 269 \| 24 \| 29 \| 401 \| 19 \| 387 \| Integrase \| Integrase; afdb-uniprot50 \| AF-A0A2T4J5A4-F1-MODEL\_V4 \| 1.0 \| 2.352e-09 \| 346 \| 0.165 \| 350 \| 235 \| 16 \| 54 \| 378 \| 3 \| 320 \| Integrase \| Integrase; afdb-uniprot50 \| AF-A0A2S8GSK0-F1-MODEL\_V4 \| 1.0 \| 7.492e-10 \| 346 \| 0.183 \| 354 \| 242 \| 16 \| 48 \| 392 \| 15 \| 330 \| Uncharacterized protein \| Uncharacterized protein; afdb-uniprot50 \| AF-A0A142XGY2-F1-MODEL\_V4 \| 1.0 \| 1.767e-09 \| 346 \| 0.172 \| 360 \| 256 \| 12 \| 46 \| 400 \| 15 \| 337 \| Site-specific tyrosine recombinase XerC \| Site-specific tyrosine recombinase XerC; afdb-uniprot50 \| AF-A0A7V9DZ64-F1-MODEL\_V4 \| 1.0 \| 1.826e-08 \| 345 \| 0.19 \| 309 \| 207 \| 12 \| 94 \| 388 \| 5 \| 284 \| Site-specific integrase \| Site-specific integrase; afdb-uniprot50 \| AF-A0A1Y3TSY9-F1-MODEL\_V4 \| 1.0 \| 3.495e-10 \| 345 \| 0.157 \| 399 \| 252 \| 24 \| 27 \| 393 \| 5 \| 351 \| Site-specific integrase \| Site-specific integrase; afdb-uniprot50 \| AF-A0A849MCQ5-F1-MODEL\_V4 \| 1.0 \| 1.606e-09 \| 344 \| 0.173 \| 374 \| 259 \| 17 \| 34 \| 400 \| 4 \| 334 \| Site-specific integrase \| Site-specific integrase; afdb-uniprot50 \| AF-A0A839ZCX8-F1-MODEL\_V4 \| 1.0 \| 3.186e-07 \| 344 \| 0.216 \| 180 \| 130 \| 5 \| 225 \| 398 \| 134 \| 308 \| Uncharacterized protein \| Uncharacterized protein; afdb-uniprot50 \| AF-A0A3E0NP80-F1-MODEL\_V4 \| 1.0 \| 5.993e-11 \| 344 \| 0.16 \| 423 \| 282 \| 23 \| 8 \| 395 \| 7 \| 391 \| Site-specific integrase \| Site-specific integrase; afdb-uniprot50 \| AF-A0A1I2MLX2-F1-MODEL\_V4 \| 1.0 \| 6.611e-08 \| 343 \| 0.186 \| 241 \| 167 \| 10 \| 175 \| 402 \| 2 \| 226 \| Phage integrase family protein \| Phage integrase family protein; afdb-uniprot50 \| AF-A0A7Z0MUD3-F1-MODEL\_V4 \| 1.0 \| 1.372e-08 \| 343 \| 0.187 \| 331 \| 227 \| 14 \| 99 \| 402 \| 14 \| 329 \| Tyrosine-type recombinase/integrase \| Tyrosine-type recombinase/integrase; afdb-uniprot50 \| AF-A0A829YLK3-F1-MODEL\_V4 \| 1.0 \| 7.164e-07 \| 342 \| 0.207 \| 193 \| 143 \| 7 \| 218 \| 402 \| 29 \| 219 \| Uncharacterized protein \| Uncharacterized protein; afdb-uniprot50 \| AF-A0A382MCM1-F1-MODEL\_V4 \| 1.0 \| 8.39e-08 \| 342 \| 0.176 \| 250 \| 181 \| 9 \| 164 \| 401 \| 2 \| 238 \| Tyr recombinase domain-containing protein \| Tyr recombinase domain-containing protein; afdb-uniprot50 \| AF-A0A1I4UQB1-F1-MODEL\_V4 \| 1.0 \| 2.673e-08 \| 342 \| 0.183 \| 261 \| 186 \| 10 \| 154 \| 402 \| 2 \| 247 \| Integrase \| Integrase; afdb-uniprot50 \| AF-A0A6L7CNV4-F1-MODEL\_V4 \| 1.0 \| 3.342e-07 \| 341 \| 0.222 \| 189 \| 138 \| 5 \| 222 \| 403 \| 8 \| 194 \| Prophage integrase IntS \| Prophage integrase IntS; afdb-uniprot50 \| AF-A0A0F2IVU2-F1-MODEL\_V4 \| 1.0 \| 1.606e-09 \| 341 \| 0.155 \| 360 \| 256 \| 16 \| 46 \| 391 \| 9 \| 334 \| Phage integrase \| Phage integrase; afdb-uniprot50 \| AF-A0A6N8JJ97-F1-MODEL\_V4 \| 1.0 \| 6.01e-08 \| 340 \| 0.206 \| 242 \| 163 \| 9 \| 174 \| 402 \| 11 \| 236 \| Tyrosine-type recombinase/integrase \| Tyrosine-type recombinase/integrase; afdb-uniprot50 \| AF-A0A3A0B0M4-F1-MODEL\_V4 \| 1.0 \| 4.105e-08 \| 340 \| 0.174 \| 315 \| 211 \| 15 \| 102 \| 401 \| 7 \| 287 \| Uncharacterized protein \| Uncharacterized protein; afdb-uniprot50 \| AF-A0A2N1RVN7-F1-MODEL\_V4 \| 1.0 \| 1.63e-10 \| 338 \| 0.157 \| 380 \| 250 \| 19 \| 44 \| 392 \| 16 \| 356 \| Tyr recombinase domain-containing protein \| Tyr recombinase domain-containing protein; afdb-uniprot50 \| AF-A0A823MIL1-F1-MODEL\_V4 \| 1.0 \| 1.799e-07 \| 337 \| 0.209 \| 205 \| 146 \| 8 \| 208 \| 401 \| 2 \| 201 \| Site-specific integrase \| Site-specific integrase; afdb-uniprot50 \| AF-A0A2U0ZBZ1-F1-MODEL\_V4 \| 1.0 \| 1.21e-06 \| 336 \| 0.235 \| 187 \| 135 \| 6 \| 222 \| 402 \| 3 \| 187 \| Phage integrase family protein \| Phage integrase family protein; afdb-uniprot50 \| AF-A0A828G9E6-F1-MODEL\_V4 \| 1.0 \| 1.531e-09 \| 336 \| 0.148 \| 404 \| 281 \| 18 \| 36 \| 393 \| 9 \| 395 \| Site-specific integrase \| Site-specific integrase; afdb-uniprot50 \| AF-A0A6F8PSE3-F1-MODEL\_V4 \| 1.0 \| 7.514e-07 \| 335 \| 0.18 \| 210 \| 161 \| 6 \| 202 \| 402 \| 5 \| 212 \| Tyr recombinase domain-containing protein \| Tyr recombinase domain-containing protein; afdb-uniprot50 \| AF-A0A2Z3H9V6-F1-MODEL\_V4 \| 1.0 \| 1.767e-09 \| 333 \| 0.172 \| 370 \| 241 \| 17 \| 46 \| 400 \| 15 \| 334 \| Uncharacterized protein \| Uncharacterized protein; afdb-uniprot50 \| AF-A0A517XN73-F1-MODEL\_V4 \| 1.0 \| 4.879e-10 \| 333 \| 0.16 \| 411 \| 252 \| 19 \| 46 \| 403 \| 15 \| 385 \| Site-specific tyrosine recombinase XerC \| Site-specific tyrosine recombinase XerC; afdb-uniprot50 \| AF-A7MTK6-F1-MODEL\_V4 \| 1.0 \| 4.967e-08 \| 332 \| 0.158 \| 302 \| 215 \| 13 \| 115 \| 402 \| 3 \| 279 \| Uncharacterized protein \| Uncharacterized protein; afdb-uniprot50 \| AF-A0A6G4KUC7-F1-MODEL\_V4 \| 1.0 \| 3.332e-10 \| 332 \| 0.205 \| 311 \| 188 \| 17 \| 44 \| 321 \| 5 \| 289 \| Integrase arm-type DNA-binding domain-containing protein \| Integrase arm-type DNA-binding domain-containing protein; afdb-uniprot50 \| AF-A0A3C0FAP9-F1-MODEL\_V4 \| 1.0 \| 5.546e-09 \| 332 \| 0.186 \| 316 \| 206 \| 14 \| 107 \| 396 \| 10 \| 300 \| Uncharacterized protein \| Uncharacterized protein; afdb-uniprot50 \| AF-A0A285R8I8-F1-MODEL\_V4 \| 1.0 \| 2.673e-08 \| 331 \| 0.161 \| 316 \| 212 \| 17 \| 101 \| 394 \| 3 \| 287 \| Integrase/recombinase XerD \| Integrase/recombinase XerD; afdb-uniprot50 \| AF-A0A2Z6ETG2-F1-MODEL\_V4 \| 1.0 \| 3.845e-10 \| 331 \| 0.191 \| 308 \| 202 \| 16 \| 2 \| 291 \| 3 \| 281 \| Integrase \| Integrase; afdb-uniprot50 \| AF-A0A139K4Z7-F1-MODEL\_V4 \| 1.0 \| 3.084e-08 \| 331 \| 0.184 \| 347 \| 242 \| 16 \| 65 \| 398 \| 3 \| 321 \| Site-specific recombinase, phage integrase family \| Site-specific recombinase, phage integrase family; afdb-uniprot50 \| AF-A0A6L8HXX8-F1-MODEL\_V4 \| 1.0 \| 1.715e-07 \| 330 \| 0.16 \| 317 \| 221 \| 17 \| 105 \| 403 \| 2 \| 291 \| Tyrosine-type recombinase/integrase \| Tyrosine-type recombinase/integrase; afdb-uniprot50 \| AF-A0A7V3D6C0-F1-MODEL\_V4 \| 1.0 \| 7.742e-09 \| 330 \| 0.157 \| 363 \| 248 \| 12 \| 46 \| 402 \| 34 \| 344 \| Tyr recombinase domain-containing protein \| Tyr recombinase domain-containing protein; afdb-uniprot50 \| AF-A0A4Q3C5E2-F1-MODEL\_V4 \| 1.0 \| 6.934e-08 \| 329 \| 0.171 \| 221 \| 171 \| 6 \| 189 \| 401 \| 4 \| 220 \| Integrase \| Integrase; afdb-uniprot50 \| AF-A0A522JXQ1-F1-MODEL\_V4 \| 1.0 \| 1.71e-10 \| 329 \| 0.186 \| 348 \| 219 \| 18 \| 1 \| 313 \| 1 \| 319 \| DUF4102 domain-containing protein \| DUF4102 domain-containing protein; afdb-uniprot50 \| AF-A0A7W2QM99-F1-MODEL\_V4 \| 1.0 \| 1.015e-07 \| 328 \| 0.21 \| 266 \| 178 \| 9 \| 152 \| 402 \| 2 \| 250 \| Site-specific integrase \| Site-specific integrase; afdb-uniprot50 \| AF-A0A737IDC5-F1-MODEL\_V4 \| 1.0 \| 1.881e-10 \| 328 \| 0.176 \| 328 \| 209 \| 17 \| 2 \| 300 \| 3 \| 298 \| DUF4102 domain-containing protein \| DUF4102 domain-containing protein; afdb-uniprot50 \| AF-A0A7W1JY51-F1-MODEL\_V4 \| 1.0 \| 8.644e-10 \| 328 \| 0.172 \| 401 \| 264 \| 18 \| 30 \| 403 \| 2 \| 361 \| Tyrosine-type recombinase/integrase \| Tyrosine-type recombinase/integrase; afdb-uniprot50 \| AF-A0A537X7F3-F1-MODEL\_V4 \| 1.0 \| 1.559e-07 \| 327 \| 0.183 \| 240 \| 170 \| 8 \| 174 \| 402 \| 7 \| 231 \| Site-specific integrase \| Site-specific integrase; afdb-uniprot50 \| AF-A0A447KKV0-F1-MODEL\_V4 \| 1.0 \| 8.799e-08 \| 327 \| 0.173 \| 265 \| 185 \| 11 \| 154 \| 402 \| 3 \| 249 \| Site-specific tyrosine recombinase XerC \| Site-specific tyrosine recombinase XerC; afdb-uniprot50 \| AF-A0A2G6C9G2-F1-MODEL\_V4 \| 1.0 \| 1.331e-06 \| 325 \| 0.222 \| 166 \| 123 \| 4 \| 238 \| 400 \| 5 \| 167 \| Integrase \| Integrase; afdb-uniprot50 \| AF-A0A7C9MK69-F1-MODEL\_V4 \| 1.0 \| 9.973e-10 \| 325 \| 0.171 \| 378 \| 258 \| 18 \| 34 \| 400 \| 2 \| 335 \| Tyrosine-type recombinase/integrase \| Tyrosine-type recombinase/integrase; afdb-uniprot50 \| AF-A0A533BAT2-F1-MODEL\_V4 \| 1.0 \| 5.816e-09 \| 325 \| 0.176 \| 386 \| 268 \| 19 \| 36 \| 401 \| 3 \| 358 \| Uncharacterized protein \| Uncharacterized protein; afdb-uniprot50 \| AF-A0A7X9AF86-F1-MODEL\_V4 \| 1.0 \| 8.39e-08 \| 324 \| 0.16 \| 318 \| 216 \| 15 \| 103 \| 393 \| 14 \| 307 \| Tyrosine recombinase XerC \| Tyrosine recombinase XerC; afdb-uniprot50 \| AF-A0A132HFJ0-F1-MODEL\_V4 \| 1.0 \| 7.251e-11 \| 324 \| 0.175 \| 411 \| 277 \| 18 \| 8 \| 392 \| 11 \| 385 \| Tyrosine recombinase XerC \| Tyrosine recombinase XerC; afdb-uniprot50 \| AF-A0A285Z377-F1-MODEL\_V4 \| 1.0 \| 8.366e-11 \| 324 \| 0.185 \| 443 \| 283 \| 25 \| 1 \| 395 \| 1 \| 413 \| Phage integrase family protein \| Phage integrase family protein; afdb-uniprot50 \| AF-A0A521UXZ2-F1-MODEL\_V4 \| 1.0 \| 1.372e-08 \| 323 \| 0.167 \| 329 \| 223 \| 17 \| 90 \| 397 \| 70 \| 368 \| Uncharacterized protein \| Uncharacterized protein; afdb-uniprot50 \| AF-A0A3M1TIK8-F1-MODEL\_V4 \| 1.0 \| 3.332e-10 \| 321 \| 0.255 \| 247 \| 146 \| 8 \| 1 \| 219 \| 9 \| 245 \| DUF4102 domain-containing protein \| DUF4102 domain-containing protein; afdb-uniprot50 \| AF-N9BVU9-F1-MODEL\_V4 \| 1.0 \| 2.283e-07 \| 321 \| 0.184 \| 309 \| 207 \| 15 \| 101 \| 402 \| 1 \| 271 \| Tyr recombinase domain-containing protein \| Tyr recombinase domain-containing protein; afdb-uniprot50 \| AF-A0A807ZPL9-F1-MODEL\_V4 \| 1.0 \| 2.17e-10 \| 320 \| 0.155 \| 353 \| 231 \| 18 \| 2 \| 319 \| 37 \| 357 \| Uncharacterized protein \| Uncharacterized protein; afdb-uniprot50 \| AF-A0A519K9Q9-F1-MODEL\_V4 \| 1.0 \| 5.117e-10 \| 320 \| 0.14 \| 405 \| 263 \| 21 \| 47 \| 400 \| 20 \| 390 \| Tyr recombinase domain-containing protein \| Tyr recombinase domain-containing protein; afdb-uniprot50 \| AF-A0A2W5FLA2-F1-MODEL\_V4 \| 1.0 \| 2.511e-07 \| 319 \| 0.201 \| 238 \| 165 \| 8 \| 177 \| 402 \| 2 \| 226 \| Integrase \| Integrase; afdb-uniprot50 \| AF-M9R7N3-F1-MODEL\_V4 \| 1.0 \| 3.234e-08 \| 318 \| 0.211 \| 289 \| 175 \| 14 \| 136 \| 396 \| 6 \| 269 \| DNA integration/recombination/inversion protein \| DNA integration/recombination/inversion protein; afdb-uniprot50 \| AF-A0A7X5VWX1-F1-MODEL\_V4 \| 1.0 \| 2.038e-09 \| 318 \| 0.157 \| 382 \| 274 \| 17 \| 34 \| 403 \| 3 \| 348 \| Tyrosine-type recombinase/integrase \| Tyrosine-type recombinase/integrase; afdb-uniprot50 \| AF-A0A3D2VB80-F1-MODEL\_V4 \| 1.0 \| 6.398e-09 \| 318 \| 0.183 \| 392 \| 248 \| 19 \| 17 \| 386 \| 10 \| 351 \| Tyr recombinase domain-containing protein \| Tyr recombinase domain-containing protein; afdb-uniprot50 \| AF-A0A382I9H7-F1-MODEL\_V4 \| 1.0 \| 1.396e-06 \| 317 \| 0.204 \| 191 \| 140 \| 8 \| 221 \| 402 \| 1 \| 188 \| Tyr recombinase domain-containing protein \| Tyr recombinase domain-containing protein; afdb-uniprot50 \| AF-A0A6L8LZX1-F1-MODEL\_V4 \| 1.0 \| 1.799e-07 \| 317 \| 0.158 \| 278 \| 202 \| 10 \| 137 \| 401 \| 3 \| 261 \| Tyrosine-type recombinase/integrase \| Tyrosine-type recombinase/integrase; afdb-uniprot50 \| AF-A0A177R241-F1-MODEL\_V4 \| 1.0 \| 1.03e-08 \| 317 \| 0.185 \| 366 \| 236 \| 15 \| 46 \| 403 \| 15 \| 326 \| Uncharacterized protein \| Uncharacterized protein; afdb-uniprot50 \| AF-U7GQ01-F1-MODEL\_V4 \| 1.0 \| 2.144e-06 \| 316 \| 0.234 \| 162 \| 120 \| 4 \| 241 \| 400 \| 3 \| 162 \| Tyr recombinase domain-containing protein \| Tyr recombinase domain-containing protein; afdb-uniprot50 \| AF-A0A3D1EUH0-F1-MODEL\_V4 \| 1.0 \| 9.368e-09 \| 316 \| 0.158 \| 384 \| 256 \| 20 \| 32 \| 401 \| 14 \| 344 \| Uncharacterized protein \| Uncharacterized protein; afdb-uniprot50 \| AF-A0A534XE27-F1-MODEL\_V4 \| 1.0 \| 6.398e-09 \| 316 \| 0.151 \| 382 \| 278 \| 16 \| 25 \| 399 \| 3 \| 345 \| Site-specific integrase \| Site-specific integrase; afdb-uniprot50 \| AF-A0A526YPQ5-F1-MODEL\_V4 \| 1.0 \| 2.812e-05 \| 315 \| 0.308 \| 94 \| 61 \| 2 \| 2 \| 92 \| 4 \| 96 \| DUF4102 domain-containing protein \| DUF4102 domain-containing protein; afdb-uniprot50 \| AF-A0A7K4GUA7-F1-MODEL\_V4 \| 1.0 \| 4.105e-08 \| 314 \| 0.147 \| 319 \| 240 \| 11 \| 99 \| 403 \| 2 \| 302 \| Tyrosine-type recombinase/integrase \| Tyrosine-type recombinase/integrase; afdb-uniprot50 \| AF-A0A146GF21-F1-MODEL\_V4 \| 1.0 \| 7.742e-09 \| 314 \| 0.131 \| 379 \| 264 \| 17 \| 15 \| 377 \| 1 \| 330 \| Site-specific recombinase XerC \| Site-specific recombinase XerC; afdb-uniprot50 \| AF-A0A0Q7Z9S4-F1-MODEL\_V4 \| 1.0 \| 8.644e-10 \| 313 \| 0.142 \| 414 \| 280 \| 18 \| 4 \| 378 \| 12 \| 389 \| Tyr recombinase domain-containing protein \| Tyr recombinase domain-containing protein; afdb-uniprot50 \| AF-G2I0W8-F1-MODEL\_V4 \| 1.0 \| 1.046e-09 \| 312 \| 0.181 \| 391 \| 233 \| 25 \| 36 \| 403 \| 7 \| 333 \| Phage DNA recombinase \| Phage DNA recombinase; afdb-uniprot50 \| AF-A0A179RXE5-F1-MODEL\_V4 \| 1.0 \| 1.826e-08 \| 312 \| 0.178 \| 381 \| 250 \| 15 \| 49 \| 390 \| 2 \| 358 \| Integrase \| Integrase; afdb-uniprot50 \| AF-A0A2V3ZGJ6-F1-MODEL\_V4 \| 1.0 \| 6.285e-11 \| 311 \| 0.156 \| 415 \| 248 \| 21 \| 9 \| 354 \| 11 \| 392 \| Uncharacterized protein \| Uncharacterized protein; afdb-uniprot50 \| AF-K0DW31-F1-MODEL\_V4 \| 1.0 \| 4.229e-10 \| 311 \| 0.167 \| 419 \| 276 \| 23 \| 1 \| 396 \| 70 \| 438 \| Phage integrase family protein \| Phage integrase family protein; afdb-uniprot50 \| AF-A0A0S8DIS8-F1-MODEL\_V4 \| 1.0 \| 5.041e-09 \| 309 \| 0.15 \| 365 \| 254 \| 16 \| 46 \| 400 \| 9 \| 327 \| Uncharacterized protein \| Uncharacterized protein; afdb-uniprot50 \| AF-A8T4I4-F1-MODEL\_V4 \| 1.0 \| 2.43e-08 \| 305 \| 0.193 \| 351 \| 186 \| 12 \| 62 \| 402 \| 1 \| 264 \| DNA integration/recombination/invertion protein \| DNA integration/recombination/invertion protein; afdb-uniprot50 \| AF-N9FYI4-F1-MODEL\_V4 \| 1.0 \| 6.831e-07 \| 304 \| 0.178 \| 235 \| 172 \| 9 \| 174 \| 401 \| 9 \| 229 \| Tyr recombinase domain-containing protein \| Tyr recombinase domain-containing protein; afdb-uniprot50 \| AF-A0A2I8S603-F1-MODEL\_V4 \| 1.0 \| 1.635e-07 \| 304 \| 0.166 \| 253 \| 168 \| 13 \| 171 \| 402 \| 10 \| 240 \| Tyr recombinase domain-containing protein \| Tyr recombinase domain-containing protein; afdb-uniprot50 \| AF-A0A433WR77-F1-MODEL\_V4 \| 1.0 \| 1.288e-07 \| 304 \| 0.179 \| 295 \| 190 \| 13 \| 119 \| 403 \| 14 \| 266 \| Uncharacterized protein \| Uncharacterized protein; afdb-uniprot50 \| AF-A0A3M2QIT5-F1-MODEL\_V4 \| 1.0 \| 1.03e-08 \| 304 \| 0.164 \| 384 \| 258 \| 18 \| 36 \| 400 \| 5 \| 344 \| Site-specific integrase \| Site-specific integrase; afdb-uniprot50 \| AF-A0A285GAL8-F1-MODEL\_V4 \| 1.0 \| 1.308e-08 \| 303 \| 0.174 \| 356 \| 229 \| 19 \| 48 \| 386 \| 67 \| 374 \| Site-specific recombinase XerD \| Site-specific recombinase XerD; afdb-uniprot50 \| AF-A0A7C6W9D4-F1-MODEL\_V4 \| 1.0 \| 2.008e-08 \| 302 \| 0.15 \| 351 \| 232 \| 16 \| 48 \| 386 \| 62 \| 358 \| Tyrosine-type recombinase/integrase \| Tyrosine-type recombinase/integrase; afdb-uniprot50 \| AF-A0A223MSG3-F1-MODEL\_V4 \| 1.0 \| 1.582e-08 \| 301 \| 0.17 \| 364 \| 251 \| 19 \| 45 \| 393 \| 21 \| 348 \| Tyr recombinase domain-containing protein \| Tyr recombinase domain-containing protein; afdb-uniprot50 \| AF-A0A1F6LJM3-F1-MODEL\_V4 \| 1.0 \| 2.713e-09 \| 301 \| 0.17 \| 387 \| 260 \| 20 \| 36 \| 401 \| 6 \| 352 \| Uncharacterized protein \| Uncharacterized protein; afdb-uniprot50 \| AF-A0A7X6WJT0-F1-MODEL\_V4 \| 1.0 \| 1.66e-08 \| 300 \| 0.13 \| 375 \| 254 \| 22 \| 45 \| 393 \| 23 \| 351 \| Tyrosine-type recombinase/integrase \| Tyrosine-type recombinase/integrase; afdb-uniprot50 \| AF-A0A382T058-F1-MODEL\_V4 \| 1.0 \| 2.176e-07 \| 297 \| 0.191 \| 324 \| 212 \| 17 \| 69 \| 371 \| 2 \| 296 \| Tyr recombinase domain-containing protein \| Tyr recombinase domain-containing protein; afdb-uniprot50 \| AF-A0A7U8TD11-F1-MODEL\_V4 \| 1.0 \| 2.713e-09 \| 296 \| 0.208 \| 316 \| 188 \| 16 \| 2 \| 288 \| 3 \| 285 \| Integrase arm-type DNA-binding domain-containing protein \| Integrase arm-type DNA-binding domain-containing protein; afdb-uniprot50 \| AF-A0A538PJK5-F1-MODEL\_V4 \| 1.0 \| 2.713e-09 \| 296 \| 0.144 \| 423 \| 259 \| 25 \| 14 \| 402 \| 1 \| 354 \| Uncharacterized protein \| Uncharacterized protein; afdb-uniprot50 \| AF-A0A5B9Q349-F1-MODEL\_V4 \| 1.0 \| 3.084e-08 \| 296 \| 0.158 \| 409 \| 234 \| 16 \| 46 \| 387 \| 15 \| 380 \| Site-specific tyrosine recombinase XerC \| Site-specific tyrosine recombinase XerC; afdb-uniprot50 \| AF-A0A518HJQ9-F1-MODEL\_V4 \| 1.0 \| 2.762e-07 \| 294 \| 0.162 \| 301 \| 214 \| 17 \| 96 \| 386 \| 2 \| 274 \| Tyrosine recombinase XerD \| Tyrosine recombinase XerD; afdb-uniprot50 \| AF-A0A521U8H4-F1-MODEL\_V4 \| 1.0 \| 1.065e-07 \| 294 \| 0.172 \| 336 \| 223 \| 20 \| 90 \| 403 \| 24 \| 326 \| Uncharacterized protein \| Uncharacterized protein; afdb-uniprot50 \| AF-A0A5Q4H1X5-F1-MODEL\_V4 \| 1.0 \| 1.065e-07 \| 294 \| 0.186 \| 290 \| 190 \| 16 \| 111 \| 382 \| 2 \| 263 \| Integrase \| Integrase; afdb-uniprot50 \| AF-A0A6M4YDB8-F1-MODEL\_V4 \| 1.0 \| 5.303e-06 \| 291 \| 0.187 \| 176 \| 131 \| 6 \| 231 \| 400 \| 2 \| 171 \| Tyr recombinase domain-containing protein \| Tyr recombinase domain-containing protein; afdb-uniprot50 \| AF-A0A2W1JGV9-F1-MODEL\_V4 \| 1.0 \| 1.21e-06 \| 291 \| 0.19 \| 205 \| 142 \| 6 \| 197 \| 395 \| 1 \| 187 \| Tyrosine recombinase XerC \| Tyrosine recombinase XerC; afdb-uniprot50 \| AF-A0A359DB79-F1-MODEL\_V4 \| 1.0 \| 9.229e-08 \| 291 \| 0.208 \| 249 \| 161 \| 10 \| 169 \| 402 \| 45 \| 272 \| Tyr recombinase domain-containing protein \| Tyr recombinase domain-containing protein; afdb-uniprot50 \| AF-K8ZZH4-F1-MODEL\_V4 \| 1.0 \| 7.403e-06 \| 290 \| 0.195 \| 184 \| 137 \| 7 \| 225 \| 403 \| 22 \| 199 \| Site-specific recombinase, phage integrase family \| Site-specific recombinase, phage integrase family; afdb-uniprot50 \| AF-A0A518G3G5-F1-MODEL\_V4 \| 1.0 \| 2.283e-07 \| 289 \| 0.175 \| 297 \| 197 \| 16 \| 105 \| 382 \| 10 \| 277 \| Tyrosine recombinase XerD \| Tyrosine recombinase XerD; afdb-uniprot50 \| AF-A0A4P7BT87-F1-MODEL\_V4 \| 1.0 \| 1.189e-08 \| 288 \| 0.18 \| 271 \| 175 \| 13 \| 2 \| 256 \| 3 \| 242 \| DUF4102 domain-containing protein \| DUF4102 domain-containing protein; afdb-uniprot50 \| AF-A0A3S4GT40-F1-MODEL\_V4 \| 1.0 \| 1.308e-08 \| 288 \| 0.199 \| 291 \| 182 \| 15 \| 2 \| 274 \| 11 \| 268 \| Integrase \| Integrase; afdb-uniprot50 \| AF-A0A3D5W661-F1-MODEL\_V4 \| 1.0 \| 9.679e-08 \| 287 \| 0.18 \| 326 \| 193 \| 12 \| 92 \| 401 \| 6 \| 273 \| Tyr recombinase domain-containing protein \| Tyr recombinase domain-containing protein; afdb-uniprot50 \| AF-A0A375A8W5-F1-MODEL\_V4 \| 1.0 \| 2.713e-09 \| 287 \| 0.186 \| 321 \| 196 \| 18 \| 2 \| 289 \| 25 \| 313 \| Mobile element protein \| Mobile element protein; afdb-uniprot50 \| AF-A0A3N5K3C0-F1-MODEL\_V4 \| 1.0 \| 1.081e-08 \| 287 \| 0.15 \| 404 \| 241 \| 22 \| 17 \| 398 \| 2 \| 325 \| Site-specific integrase \| Site-specific integrase; afdb-uniprot50 \| AF-A0A0P1I3C9-F1-MODEL\_V4 \| 1.0 \| 1.853e-09 \| 284 \| 0.151 \| 408 \| 279 \| 21 \| 1 \| 378 \| 3 \| 373 \| Uncharacterized protein \| Uncharacterized protein; afdb-uniprot50 \| AF-A0A7G7EQR9-F1-MODEL\_V4 \| 1.0 \| 3.342e-07 \| 283 \| 0.158 \| 322 \| 230 \| 14 \| 99 \| 403 \| 26 \| 323 \| Tyrosine-type recombinase/integrase \| Tyrosine-type recombinase/integrase; afdb-uniprot50 \| AF-A0A1I4UQD3-F1-MODEL\_V4 \| 1.0 \| 4.178e-06 \| 281 \| 0.177 \| 180 \| 139 \| 5 \| 228 \| 402 \| 2 \| 177 \| Phage integrase family protein \| Phage integrase family protein; afdb-uniprot50 \| AF-A0A6L9LYB1-F1-MODEL\_V4 \| 1.0 \| 1.331e-06 \| 279 \| 0.168 \| 326 \| 205 \| 15 \| 100 \| 402 \| 6 \| 288 \| Tyrosine-type recombinase/integrase \| Tyrosine-type recombinase/integrase; afdb-uniprot50 \| AF-A0A1Y5TZF1-F1-MODEL\_V4 \| 1.0 \| 1.509e-08 \| 279 \| 0.183 \| 386 \| 255 \| 20 \| 46 \| 401 \| 17 \| 372 \| Tyrosine recombinase XerC \| Tyrosine recombinase XerC; afdb-uniprot50 \| AF-A0A0F2Q7Y2-F1-MODEL\_V4 \| 1.0 \| 2.854e-06 \| 278 \| 0.165 \| 236 \| 166 \| 8 \| 169 \| 394 \| 10 \| 224 \| Tyr recombinase domain-containing protein \| Tyr recombinase domain-containing protein; afdb-uniprot50 \| AF-A0A3B0WP27-F1-MODEL\_V4 \| 1.0 \| 3.443e-09 \| 278 \| 0.135 \| 413 \| 290 \| 23 \| 3 \| 393 \| 8 \| 375 \| Tyr recombinase domain-containing protein \| Tyr recombinase domain-containing protein; afdb-uniprot50 \| AF-A0A5M6HN79-F1-MODEL\_V4 \| 1.0 \| 7.742e-09 \| 277 \| 0.147 \| 460 \| 273 \| 22 \| 36 \| 402 \| 19 \| 452 \| Tyrosine-type recombinase/integrase \| Tyrosine-type recombinase/integrase; afdb-uniprot50 \| AF-A0A7I9NUR5-F1-MODEL\_V4 \| 1.0 \| 5.303e-06 \| 274 \| 0.205 \| 190 \| 132 \| 7 \| 231 \| 403 \| 3 \| 190 \| Tyrosine-type recombinase/integrase \| Tyrosine-type recombinase/integrase; afdb-uniprot50 \| AF-A0A6G7ZA35-F1-MODEL\_V4 \| 1.0 \| 4.515e-08 \| 274 \| 0.143 \| 370 \| 252 \| 25 \| 36 \| 387 \| 3 \| 325 \| Site-specific integrase \| Site-specific integrase; afdb-uniprot50 \| AF-A0A3C2D4R1-F1-MODEL\_V4 \| 1.0 \| 4.967e-08 \| 273 \| 0.124 \| 427 \| 277 \| 20 \| 14 \| 402 \| 8 \| 375 \| Uncharacterized protein \| Uncharacterized protein; afdb-uniprot50 \| AF-A0A2J7Y8I7-F1-MODEL\_V4 \| 1.0 \| 1.587e-05 \| 272 \| 0.222 \| 166 \| 124 \| 3 \| 241 \| 402 \| 2 \| 166 \| Prophage CP4-57 integrase \| Prophage CP4-57 integrase; afdb-uniprot50 \| AF-A0A523IFE9-F1-MODEL\_V4 \| 1.0 \| 1.858e-06 \| 272 \| 0.208 \| 235 \| 166 \| 7 \| 169 \| 400 \| 4 \| 221 \| Site-specific integrase \| Site-specific integrase; afdb-uniprot50 \| AF-A0A7C4XZS9-F1-MODEL\_V4 \| 1.0 \| 2.144e-06 \| 271 \| 0.26 \| 192 \| 121 \| 10 \| 195 \| 376 \| 2 \| 182 \| Site-specific integrase \| Site-specific integrase; afdb-uniprot50 \| AF-A0A2V7QVN5-F1-MODEL\_V4 \| 1.0 \| 3.622e-06 \| 271 \| 0.15 \| 318 \| 212 \| 12 \| 100 \| 403 \| 1 \| 274 \| Tyr recombinase domain-containing protein \| Tyr recombinase domain-containing protein; afdb-uniprot50 \| AF-A0A2E9PKL6-F1-MODEL\_V4 \| 1.0 \| 2.594e-06 \| 270 \| 0.19 \| 236 \| 158 \| 10 \| 178 \| 401 \| 2 \| 216 \| Tyr recombinase domain-containing protein \| Tyr recombinase domain-containing protein; afdb-uniprot50 \| AF-A0A520LCH2-F1-MODEL\_V4 \| 1.0 \| 3.731e-08 \| 270 \| 0.154 \| 435 \| 263 \| 23 \| 13 \| 393 \| 4 \| 387 \| Core-binding (CB) domain-containing protein \| Core-binding (CB) domain-containing protein; afdb-uniprot50 \| AF-A0A0P7LNL0-F1-MODEL\_V4 \| 1.0 \| 2.897e-07 \| 269 \| 0.162 \| 259 \| 177 \| 14 \| 9 \| 252 \| 12 \| 245 \| Integrase \| Integrase; afdb-uniprot50 \| AF-A0A2M6ZPB3-F1-MODEL\_V4 \| 1.0 \| 2.713e-09 \| 269 \| 0.158 \| 397 \| 251 \| 26 \| 37 \| 402 \| 29 \| 373 \| Site-specific integrase \| Site-specific integrase; afdb-uniprot50 \| AF-A0A223MSP1-F1-MODEL\_V4 \| 1.0 \| 1.049e-06 \| 268 \| 0.161 \| 266 \| 177 \| 13 \| 146 \| 398 \| 10 \| 242 \| Tyr recombinase domain-containing protein \| Tyr recombinase domain-containing protein; afdb-uniprot50 \| AF-A0A562IZE3-F1-MODEL\_V4 \| 1.0 \| 1.417e-07 \| 261 \| 0.205 \| 268 \| 170 \| 12 \| 1 \| 237 \| 8 \| 263 \| Uncharacterized protein \| Uncharacterized protein; afdb-uniprot50 \| AF-A0A1Y4HE01-F1-MODEL\_V4 \| 1.0 \| 1.015e-07 \| 261 \| 0.153 \| 411 \| 259 \| 20 \| 1 \| 393 \| 52 \| 391 \| Tyr recombinase domain-containing protein \| Tyr recombinase domain-containing protein; afdb-uniprot50 \| AF-A0A1E4X750-F1-MODEL\_V4 \| 1.0 \| 1.509e-08 \| 261 \| 0.148 \| 430 \| 263 \| 25 \| 46 \| 403 \| 10 \| 408 \| Tyr recombinase domain-containing protein \| Tyr recombinase domain-containing protein; afdb-uniprot50 \| AF-A0A8B3HTI2-F1-MODEL\_V4 \| 1.0 \| 1.417e-07 \| 259 \| 0.221 \| 235 \| 139 \| 12 \| 2 \| 224 \| 3 \| 205 \| DUF4102 domain-containing protein \| DUF4102 domain-containing protein; afdb-uniprot50 \| AF-A0A1G7SLZ0-F1-MODEL\_V4 \| 1.0 \| 9.092e-07 \| 259 \| 0.198 \| 232 \| 144 \| 13 \| 181 \| 402 \| 2 \| 201 \| Phage integrase family protein \| Phage integrase family protein; afdb-uniprot50 \| AF-A0A259CZR1-F1-MODEL\_V4 \| 1.0 \| 5.383e-07 \| 259 \| 0.193 \| 243 \| 156 \| 13 \| 171 \| 402 \| 28 \| 241 \| Integrase \| Integrase; afdb-uniprot50 \| AF-A0A156H2Y6-F1-MODEL\_V4 \| 1.0 \| 7.627e-08 \| 258 \| 0.204 \| 274 \| 175 \| 13 \| 2 \| 261 \| 3 \| 247 \| Phage integrase family protein \| Phage integrase family protein; afdb-uniprot50 \| AF-W2TX06-F1-MODEL\_V4 \| 1.0 \| 3.139e-06 \| 258 \| 0.175 \| 239 \| 164 \| 10 \| 169 \| 396 \| 4 \| 220 \| Site-specific recombinase, phage integrase family \| Site-specific recombinase, phage integrase family; afdb-uniprot50 \| AF-A0A0H3I6J4-F1-MODEL\_V4 \| 1.0 \| 6.303e-08 \| 257 \| 0.188 \| 292 \| 189 \| 16 \| 2 \| 276 \| 3 \| 263 \| Prophage P4 integrase \| Prophage P4 integrase; afdb-uniprot50 \| AF-A0A8A3NBE6-F1-MODEL\_V4 \| 1.0 \| 2.317e-08 \| 257 \| 0.185 \| 394 \| 239 \| 17 \| 2 \| 339 \| 26 \| 393 \| Integrase family protein \| Integrase family protein; afdb-uniprot50 \| AF-A0A150J908-F1-MODEL\_V4 \| 1.0 \| 3.292e-06 \| 256 \| 0.148 \| 290 \| 200 \| 13 \| 107 \| 386 \| 2 \| 254 \| Putative tyrosine recombinase XerC-like protein \| Putative tyrosine recombinase XerC-like protein; afdb-uniprot50 \| AF-A0A836V0C8-F1-MODEL\_V4 \| 1.0 \| 5.383e-07 \| 256 \| 0.143 \| 293 \| 207 \| 15 \| 2 \| 279 \| 9 \| 272 \| DUF4102 domain-containing protein \| DUF4102 domain-containing protein; afdb-uniprot50 \| AF-A0A828R5W3-F1-MODEL\_V4 \| 1.0 \| 3.038e-07 \| 255 \| 0.217 \| 235 \| 140 \| 12 \| 2 \| 224 \| 3 \| 205 \| Integrase domain protein \| Integrase domain protein; afdb-uniprot50 \| AF-A0A5F1DSR0-F1-MODEL\_V4 \| 1.0 \| 5.73e-08 \| 254 \| 0.195 \| 292 \| 182 \| 16 \| 2 \| 274 \| 3 \| 260 \| Integrase \| Integrase; afdb-uniprot50 \| AF-A0A0C1H784-F1-MODEL\_V4 \| 1.0 \| 3.402e-05 \| 252 \| 0.159 \| 169 \| 130 \| 6 \| 221 \| 382 \| 39 \| 202 \| Virulence plasmid integrase pGP8-D \| Virulence plasmid integrase pGP8-D; afdb-uniprot50 \| AF-A0A4R7C7L2-F1-MODEL\_V4 \| 1.0 \| 7.272e-08 \| 252 \| 0.161 \| 341 \| 216 \| 19 \| 2 \| 317 \| 8 \| 303 \| Integrase \| Integrase; afdb-uniprot50 \| AF-A0A7X1NZC6-F1-MODEL\_V4 \| 1.0 \| 3.676e-07 \| 252 \| 0.148 \| 376 \| 252 \| 17 \| 49 \| 401 \| 11 \| 341 \| Tyrosine-type recombinase/integrase \| Tyrosine-type recombinase/integrase; afdb-uniprot50 \| AF-A0A2E1ZBG2-F1-MODEL\_V4 \| 1.0 \| 1.065e-07 \| 252 \| 0.145 \| 448 \| 281 \| 21 \| 18 \| 400 \| 2 \| 412 \| Tyr recombinase domain-containing protein \| Tyr recombinase domain-containing protein; afdb-uniprot50 \| AF-A0A845H3L3-F1-MODEL\_V4 \| 1.0 \| 1.171e-07 \| 251 \| 0.207 \| 227 \| 145 \| 11 \| 19 \| 235 \| 23 \| 224 \| Integrase arm-type DNA-binding domain-containing protein \| Integrase arm-type DNA-binding domain-containing protein; afdb-uniprot50 \| AF-A0A109BW84-F1-MODEL\_V4 \| 1.0 \| 5.562e-06 \| 249 \| 0.173 \| 254 \| 182 \| 10 \| 148 \| 392 \| 4 \| 238 \| Integrase \| Integrase; afdb-uniprot50 \| AF-A0A518I3J3-F1-MODEL\_V4 \| 1.0 \| 1.171e-07 \| 249 \| 0.17 \| 370 \| 235 \| 19 \| 46 \| 402 \| 15 \| 325 \| Site-specific tyrosine recombinase XerC \| Site-specific tyrosine recombinase XerC; afdb-uniprot50 \| AF-A0A2A3TB72-F1-MODEL\_V4 \| 1.0 \| 1.331e-06 \| 248 \| 0.115 \| 345 \| 244 \| 19 \| 74 \| 398 \| 4 \| 307 \| Tyr recombinase domain-containing protein \| Tyr recombinase domain-containing protein; afdb-uniprot50 \| AF-V7IKX7-F1-MODEL\_V4 \| 1.0 \| 2.511e-07 \| 244 \| 0.181 \| 281 \| 181 \| 14 \| 2 \| 265 \| 3 \| 251 \| Site-specific recombinase, phage integrase family \| Site-specific recombinase, phage integrase family; afdb-uniprot50 \| AF-A0A0F9MB95-F1-MODEL\_V4 \| 1.0 \| 5.747e-05 \| 243 \| 0.166 \| 180 \| 136 \| 5 \| 221 \| 393 \| 11 \| 183 \| Tyr recombinase domain-containing protein \| Tyr recombinase domain-containing protein; afdb-uniprot50 \| AF-A0A376MQW7-F1-MODEL\_V4 \| 1.0 \| 1.065e-07 \| 239 \| 0.221 \| 235 \| 139 \| 12 \| 2 \| 224 \| 3 \| 205 \| Phage integrase family site specific recombinase \| Phage integrase family site specific recombinase; afdb-uniprot50 \| AF-A0A327LWG9-F1-MODEL\_V4 \| 1.0 \| 2.215e-05 \| 238 \| 0.204 \| 186 \| 126 \| 7 \| 217 \| 382 \| 5 \| 188 \| Tyr recombinase domain-containing protein \| Tyr recombinase domain-containing protein; afdb-uniprot50 \| AF-A0A7U9B7Q9-F1-MODEL\_V4 \| 1.0 \| 2.762e-07 \| 237 \| 0.206 \| 233 \| 141 \| 13 \| 2 \| 224 \| 3 \| 201 \| Int \| Int; afdb-uniprot50 \| AF-A0A5C9B685-F1-MODEL\_V4 \| 1.0 \| 3.505e-07 \| 237 \| 0.174 \| 343 \| 216 \| 17 \| 46 \| 386 \| 15 \| 292 \| Core-binding (CB) domain-containing protein \| Core-binding (CB) domain-containing protein; afdb-uniprot50 \| AF-A0A4R3UKN9-F1-MODEL\_V4 \| 1.0 \| 9.092e-07 \| 236 \| 0.182 \| 236 \| 155 \| 13 \| 2 \| 227 \| 8 \| 215 \| Uncharacterized protein DUF4102 \| Uncharacterized protein DUF4102; afdb-uniprot50 \| AF-A0A0M1I116-F1-MODEL\_V4 \| 1.0 \| 1.92e-05 \| 234 \| 0.2 \| 260 \| 156 \| 12 \| 149 \| 402 \| 9 \| 222 \| Tyr recombinase domain-containing protein \| Tyr recombinase domain-containing protein; afdb-uniprot50 \| AF-A0A7U4R041-F1-MODEL\_V4 \| 1.0 \| 1.033e-05 \| 234 \| 0.168 \| 273 \| 187 \| 10 \| 144 \| 403 \| 10 \| 255 \| Tyr recombinase domain-containing protein \| Tyr recombinase domain-containing protein; afdb-uniprot50 \| AF-A0A3E0ILK8-F1-MODEL\_V4 \| 1.0 \| 1.587e-05 \| 234 \| 0.141 \| 282 \| 205 \| 10 \| 138 \| 403 \| 9 \| 269 \| Site-specific integrase \| Site-specific integrase; afdb-uniprot50 \| AF-A0A750P3X5-F1-MODEL\_V4 \| 1.0 \| 1.664e-05 \| 233 \| 0.191 \| 193 \| 130 \| 11 \| 2 \| 186 \| 3 \| 177 \| DUF4102 domain-containing protein \| DUF4102 domain-containing protein; afdb-uniprot50 \| AF-A0A5M6ILP2-F1-MODEL\_V4 \| 1.0 \| 1.831e-05 \| 232 \| 0.227 \| 180 \| 128 \| 6 \| 223 \| 397 \| 85 \| 258 \| Site-specific integrase \| Site-specific integrase; afdb-uniprot50 \| AF-A0A352UWS7-F1-MODEL\_V4 \| 1.0 \| 1.886e-07 \| 231 \| 0.217 \| 230 \| 142 \| 11 \| 2 \| 224 \| 1 \| 199 \| Core-binding (CB) domain-containing protein \| Core-binding (CB) domain-containing protein; afdb-uniprot50 \| AF-A0A7C1YJJ2-F1-MODEL\_V4 \| 1.0 \| 3.676e-07 \| 231 \| 0.199 \| 251 \| 161 \| 14 \| 2 \| 246 \| 9 \| 225 \| DUF4102 domain-containing protein \| DUF4102 domain-containing protein; afdb-uniprot50 \| AF-A0A4S5BUU1-F1-MODEL\_V4 \| 1.0 \| 2.283e-07 \| 231 \| 0.16 \| 412 \| 214 \| 21 \| 9 \| 402 \| 17 \| 314 \| Site-specific integrase \| Site-specific integrase; afdb-uniprot50 \| AF-A0A4R3L278-F1-MODEL\_V4 \| 1.0 \| 6.831e-07 \| 230 \| 0.185 \| 221 \| 145 \| 12 \| 2 \| 216 \| 3 \| 194 \| Uncharacterized protein DUF4102 \| Uncharacterized protein DUF4102; afdb-uniprot50 \| AF-A0A7G1LGL0-F1-MODEL\_V4 \| 1.0 \| 1.978e-07 \| 230 \| 0.137 \| 458 \| 272 \| 24 \| 36 \| 402 \| 4 \| 429 \| Uncharacterized protein \| Uncharacterized protein; afdb-uniprot50 \| AF-A0A7J4TFH4-F1-MODEL\_V4 \| 1.0 \| 3.292e-06 \| 229 \| 0.156 \| 332 \| 205 \| 15 \| 83 \| 400 \| 7 \| 277 \| Tyrosine-type recombinase/integrase \| Tyrosine-type recombinase/integrase; afdb-uniprot50 \| AF-A0A0X3UBE4-F1-MODEL\_V4 \| 1.0 \| 1.92e-05 \| 226 \| 0.132 \| 294 \| 184 \| 13 \| 146 \| 393 \| 11 \| 279 \| Tyr recombinase domain-containing protein \| Tyr recombinase domain-containing protein; afdb-uniprot50 \| AF-A0A3L5H774-F1-MODEL\_V4 \| 1.0 \| 7.272e-08 \| 225 \| 0.188 \| 335 \| 191 \| 21 \| 2 \| 290 \| 3 \| 302 \| ISAs1 family transposase \| ISAs1 family transposase; afdb-uniprot50 \| AF-A0A376SEM2-F1-MODEL\_V4 \| 1.0 \| 3.292e-06 \| 223 \| 0.173 \| 253 \| 177 \| 11 \| 36 \| 275 \| 3 \| 236 \| Integrase \| Integrase; afdb-uniprot50 \| AF-U3TY72-F1-MODEL\_V4 \| 1.0 \| 1.1e-06 \| 221 \| 0.179 \| 273 \| 175 \| 15 \| 2 \| 256 \| 3 \| 244 \| Integrase \| Integrase; afdb-uniprot50 \| AF-A0A450YSF1-F1-MODEL\_V4 \| 1.0 \| 5.833e-06 \| 220 \| 0.152 \| 361 \| 222 \| 18 \| 60 \| 395 \| 3 \| 304 \| Site-specific recombinase XerD \| Site-specific recombinase XerD; afdb-uniprot50 \| AF-A0A1X3L5U9-F1-MODEL\_V4 \| 1.0 \| 2.358e-06 \| 217 \| 0.18 \| 199 \| 133 \| 9 \| 34 \| 224 \| 2 \| 178 \| Integrase \| Integrase; afdb-uniprot50 \| AF-A0A7W1CIH9-F1-MODEL\_V4 \| 1.0 \| 1.137e-05 \| 215 \| 0.147 \| 339 \| 226 \| 17 \| 80 \| 382 \| 3 \| 314 \| Site-specific integrase \| Site-specific integrase; afdb-uniprot50 \| AF-A0A7X2D969-F1-MODEL\_V4 \| 1.0 \| 0.000164 \| 214 \| 0.203 \| 192 \| 144 \| 7 \| 207 \| 392 \| 10 \| 198 \| Tyrosine-type recombinase/integrase \| Tyrosine-type recombinase/integrase; afdb-uniprot50 \| AF-U9Z089-F1-MODEL\_V4 \| 1.0 \| 5.303e-06 \| 213 \| 0.198 \| 227 \| 142 \| 11 \| 36 \| 245 \| 3 \| 206 \| Core-binding (CB) domain-containing protein \| Core-binding (CB) domain-containing protein; afdb-uniprot50 \| AF-A0A267T8A9-F1-MODEL\_V4 \| 1.0 \| 0.0003196 \| 211 \| 0.19 \| 173 \| 128 \| 7 \| 207 \| 373 \| 18 \| 184 \| Recombinase XerC \| Recombinase XerC; afdb-uniprot50 \| AF-A0A7X5NGS5-F1-MODEL\_V4 \| 1.0 \| 8.022e-05 \| 210 \| 0.151 \| 244 \| 174 \| 11 \| 174 \| 403 \| 14 \| 238 \| Tyr recombinase domain-containing protein \| Tyr recombinase domain-containing protein; afdb-uniprot50 \| AF-A0A7X7MFU5-F1-MODEL\_V4 \| 1.0 \| 0.0001175 \| 209 \| 0.194 \| 190 \| 133 \| 10 \| 207 \| 386 \| 9 \| 188 \| Tyrosine-type recombinase/integrase \| Tyrosine-type recombinase/integrase; afdb-uniprot50 \| AF-Q46428-F1-MODEL\_V4 \| 1.0 \| 2.437e-05 \| 209 \| 0.15 \| 279 \| 199 \| 14 \| 99 \| 358 \| 2 \| 261 \| Tyr recombinase domain-containing protein \| Tyr recombinase domain-containing protein; afdb-uniprot50 \| AF-A0A377M8M8-F1-MODEL\_V4 \| 1.0 \| 4.382e-06 \| 208 \| 0.15 \| 232 \| 157 \| 12 \| 9 \| 224 \| 12 \| 219 \| Putative integrase \| Putative integrase; afdb-uniprot50 \| AF-A6FQ69-F1-MODEL\_V4 \| 1.0 \| 6.21e-07 \| 207 \| 0.153 \| 431 \| 250 \| 22 \| 46 \| 393 \| 11 \| 409 \| Phage integrase \| Phage integrase; afdb-uniprot50 \| AF-V0AM31-F1-MODEL\_V4 \| 1.0 \| 7.058e-06 \| 205 \| 0.168 \| 231 \| 151 \| 12 \| 34 \| 246 \| 2 \| 209 \| Core-binding (CB) domain-containing protein \| Core-binding (CB) domain-containing protein; afdb-uniprot50 \| AF-A0A4U6M1J7-F1-MODEL\_V4 \| 1.0 \| 3.622e-06 \| 203 \| 0.158 \| 234 \| 151 \| 14 \| 2 \| 224 \| 3 \| 201 \| DUF4102 domain-containing protein \| DUF4102 domain-containing protein; afdb-uniprot50 \| AF-A0A2J4P481-F1-MODEL\_V4 \| 1.0 \| 0.0001355 \| 201 \| 0.172 \| 185 \| 118 \| 8 \| 174 \| 340 \| 8 \| 175 \| Integrase \| Integrase; afdb-uniprot50 \| AF-A0A7Z7L533-F1-MODEL\_V4 \| 1.0 \| 2.474e-06 \| 201 \| 0.216 \| 194 \| 126 \| 9 \| 2 \| 187 \| 3 \| 178 \| Phage integrase family site specific recombinase \| Phage integrase family site specific recombinase; afdb-uniprot50 \| AF-A0A702LGL0-F1-MODEL\_V4 \| 1.0 \| 4.821e-06 \| 200 \| 0.182 \| 236 \| 154 \| 15 \| 2 \| 224 \| 3 \| 212 \| DUF4102 domain-containing protein \| DUF4102 domain-containing protein; afdb-uniprot50 \| AF-A0A381GQ17-F1-MODEL\_V4 \| 1.0 \| 1.858e-06 \| 198 \| 0.193 \| 238 \| 147 \| 14 \| 2 \| 224 \| 7 \| 214 \| Integrase \| Integrase; afdb-uniprot50 \| AF-A0A3R7DQE1-F1-MODEL\_V4 \| 1.0 \| 1.831e-05 \| 197 \| 0.138 \| 303 \| 195 \| 11 \| 106 \| 394 \| 34 \| 284 \| Site-specific integrase \| Site-specific integrase; afdb-uniprot50 \| AF-P0CE20-F1-MODEL\_V4 \| 1.0 \| 2.324e-05 \| 194 \| 0.141 \| 347 \| 220 \| 21 \| 77 \| 393 \| 16 \| 314 \| Virulence plasmid integrase pGP8-D \| Virulence plasmid integrase pGP8-D; afdb-uniprot50 \| AF-A0A7U0UV57-F1-MODEL\_V4 \| 1.0 \| 5.303e-06 \| 191 \| 0.119 \| 377 \| 224 \| 20 \| 48 \| 393 \| 54 \| 353 \| IS200/IS605 family transposase \| IS200/IS605 family transposase; afdb-uniprot50 \| AF-A0A7T8FLC7-F1-MODEL\_V4 \| 1.0 \| 8.541e-06 \| 189 \| 0.165 \| 230 \| 154 \| 12 \| 2 \| 224 \| 24 \| 222 \| Integrase arm-type DNA-binding domain-containing protein \| Integrase arm-type DNA-binding domain-containing protein; afdb-uniprot50 \| AF-I0QS20-F1-MODEL\_V4 \| 1.0 \| 9.536e-07 \| 188 \| 0.17 \| 335 \| 211 \| 23 \| 2 \| 322 \| 3 \| 284 \| Integrase family protein \| Integrase family protein; afdb-uniprot50 \| AF-A0A1Y4HE69-F1-MODEL\_V4 \| 1.0 \| 0.00154 \| 187 \| 0.196 \| 188 \| 131 \| 9 \| 216 \| 393 \| 3 \| 180 \| Tyr recombinase domain-containing protein \| Tyr recombinase domain-containing protein; afdb-uniprot50 \| AF-A0A1M6T5P9-F1-MODEL\_V4 \| 1.0 \| 0.001335 \| 187 \| 0.192 \| 182 \| 142 \| 4 \| 222 \| 401 \| 11 \| 189 \| Phage integrase family protein \| Phage integrase family protein; afdb-uniprot50 \| AF-A0A2D2DAJ7-F1-MODEL\_V4 \| 1.0 \| 0.0001804 \| 185 \| 0.127 \| 244 \| 174 \| 12 \| 147 \| 370 \| 7 \| 231 \| Integrase \| Integrase; afdb-uniprot50 \| AF-A0A509CWS0-F1-MODEL\_V4 \| 1.0 \| 7.293e-05 \| 184 \| 0.184 \| 201 \| 144 \| 9 \| 2 \| 197 \| 3 \| 188 \| Bacteriophage integrase \| Bacteriophage integrase; afdb-uniprot50 \| AF-A0A2V8LGT1-F1-MODEL\_V4 \| 1.0 \| 0.0005398 \| 182 \| 0.166 \| 228 \| 165 \| 8 \| 177 \| 396 \| 2 \| 212 \| Tyr recombinase domain-containing protein \| Tyr recombinase domain-containing protein; afdb-uniprot50 \| AF-A0A450Z0R5-F1-MODEL\_V4 \| 1.0 \| 0.0001804 \| 177 \| 0.139 \| 286 \| 195 \| 11 \| 125 \| 395 \| 2 \| 251 \| Site-specific recombinase XerD \| Site-specific recombinase XerD; afdb-uniprot50 \| AF-A0A3N5YUU7-F1-MODEL\_V4 \| 1.0 \| 0.0001355 \| 176 \| 0.122 \| 293 \| 193 \| 20 \| 146 \| 402 \| 12 \| 276 \| Site-specific integrase \| Site-specific integrase; afdb-uniprot50 \| AF-Q46257-F1-MODEL\_V4 \| 1.0 \| 0.0002401 \| 173 \| 0.113 \| 300 \| 209 \| 15 \| 101 \| 371 \| 14 \| 285 \| Virulence plasmid integrase pGP7-D \| Virulence plasmid integrase pGP7-D; afdb-uniprot50 \| AF-A0A2D4THJ6-F1-MODEL\_V4 \| 1.0 \| 0.003463 \| 168 \| 0.207 \| 169 \| 128 \| 5 \| 221 \| 386 \| 12 \| 177 \| Tyr recombinase domain-containing protein \| Tyr recombinase domain-containing protein; afdb-uniprot50 \| AF-A0A538PKB2-F1-MODEL\_V4 \| 1.0 \| 0.001955 \| 165 \| 0.202 \| 202 \| 137 \| 10 \| 190 \| 382 \| 3 \| 189 \| Tyr recombinase domain-containing protein \| Tyr recombinase domain-containing protein; afdb-uniprot50 \| AF-A0A1F6ZB97-F1-MODEL\_V4 \| 1.0 \| 7.293e-05 \| 164 \| 0.101 \| 363 \| 233 \| 24 \| 104 \| 400 \| 8 \| 343 \| Tyr recombinase domain-containing protein \| Tyr recombinase domain-containing protein; afdb-uniprot50 \| AF-A0A1T5DBV7-F1-MODEL\_V4 \| 1.0 \| 7.649e-05 \| 157 \| 0.151 \| 331 \| 219 \| 17 \| 74 \| 389 \| 9 \| 292 \| Integrase/recombinase XerC \| Integrase/recombinase XerC; afdb-uniprot50 \| AF-A0A0E2IVB2-F1-MODEL\_V4 \| 1.0 \| 0.003463 \| 153 \| 0.127 \| 172 \| 127 \| 6 \| 228 \| 382 \| 3 \| 168 \| Phage integrase family protein \| Phage integrase family protein; afdb-uniprot50 \| AF-A0A4Z0R2I2-F1-MODEL\_V4 \| 1.0 \| 0.00461 \| 149 \| 0.115 \| 182 \| 133 \| 4 \| 221 \| 396 \| 15 \| 174 \| Tyr recombinase domain-containing protein \| Tyr recombinase domain-containing protein; afdb-uniprot50 \| AF-A0A2S9IKS5-F1-MODEL\_V4 \| 1.0 \| 0.003809 \| 147 \| 0.185 \| 200 \| 137 \| 9 \| 212 \| 400 \| 7 \| 191 \| Integrase \| Integrase; afdb-uniprot50 \| AF-A0A0T9T9N2-F1-MODEL\_V4 \| 1.0 \| 0.004395 \| 137 \| 0.134 \| 171 \| 119 \| 7 \| 50 \| 211 \| 20 \| 170 \| Integrase \| Integrase |
| Top keywords  (threshold 1.00e-02 (evalue)) | **Integrase, recombinase, domain\_containing, Tyr, Site\_specific, DUF4102, Tyrosine\_type, Phage, Prophage, Tyrosine** |
| Output files | ../../similar\_structures/32\_FANPEZAQ\_CDS\_0032\_afdb-proteome\_foldseek.tsv ../../similar\_structures/32\_FANPEZAQ\_CDS\_0032\_afdb-uniprot50\_foldseek.tsv ../../similar\_structures/32\_FANPEZAQ\_CDS\_0032\_merged.svg ../../similar\_structures/32\_FANPEZAQ\_CDS\_0032\_pdb\_foldseek.tsv |

  
  
  

Return to summary | Go to previous | Go to next

  


---

**Sequence/structure alignments coloring**  
Each object in the alignment figures is colored according to its E-value following this color coding:

1e-100
10
