## Supplementary material for "Completing the BASEL phage collection to unlock hidden diversity for systematic exploration of phage-host interactions": Data S2: 33.html

FANPEZAQ\_CDS\_0033

Return to summary | Go to previous | Go to next

|  |
| --- |
| FANPEZAQ\_CDS\_0033  Page creation date: 02 Sep 2024, 12:00  Project folder: n/a  Input sequences file: Escherichia\_virus\_HeidiAbel.gb |

### Sequence information

|  |  |
| --- | --- |
| Name | FANPEZAQ\_CDS\_0033  33\_FANPEZAQ\_CDS\_0033 (pipeline id) |
| Imported annotations |  |
| Protein sequence | MTKAATNDQRKTPPKRGFVSRDAVMRDTDHLPYWNHLSGKFQSVRMSHCDALSRCEVTPW GLEVFAKAHHGACRLRLRRGAPPPSA |
| Number of residues | 86 |
| Molecular weight (Da) | 9712.04 |
| Output files | ../../query\_sequences/33\_FANPEZAQ\_CDS\_0033.fasta |

### Putative domain architecture and protein family

#### Search results (HHblits)^1^

|  |  |
| --- | --- |
| Domain family databases searched | Pfam, Ncbi-cd, Cath, Phrogs |
| Results, scheme(s)  (Top layers only; threshold 1.00e-03 (evalue)) | xml version="1.0" encoding="utf-8" standalone="no"?       2024-09-02T21:08:19.530960 image/svg+xml   Matplotlib v3.7.2, https://matplotlib.org/ |
| Results, table  (E-value ≤ 1.00e-03 (evalue)) | -- |
| Top keywords  (threshold 1.00e-03 (evalue)) | -- |
| Output files | ../../domain\_architecture/33\_FANPEZAQ\_CDS\_0033\_cath.hhr ../../domain\_architecture/33\_FANPEZAQ\_CDS\_0033\_merged.svg ../../domain\_architecture/33\_FANPEZAQ\_CDS\_0033\_ncbi-cd.hhr ../../domain\_architecture/33\_FANPEZAQ\_CDS\_0033\_pfam.hhr ../../domain\_architecture/33\_FANPEZAQ\_CDS\_0033\_phrogs.hhr |

### Identical protein sequences/structures

#### Search results

|  |  |
| --- | --- |
| Protein sequence databases searched | Pdb, Swissprot, Refseq |
| Identical proteins found | -- |
| Top keywords | -- |
| Output files | -- |

### Similar protein sequences/structures

#### Sequence similarity search results (HHblits)^1^

|  |  |
| --- | --- |
| Sequence databases searched | Uniclust, Pdb70 |
| Results, scheme(s)  (Top layers only, threshold 1.00e-03 (evalue)) | xml version="1.0" encoding="utf-8" standalone="no"?       2024-09-02T21:08:43.130366 image/svg+xml   Matplotlib v3.7.2, https://matplotlib.org/ |
| Results, table(s)  (threshold 1.00e-03 (evalue)) | -- |
| Top keywords  (threshold 1.00e-03 (evalue)) | -- |
| Output files | ../../similar\_sequences/33\_FANPEZAQ\_CDS\_0033\_merged.svg ../../similar\_sequences/33\_FANPEZAQ\_CDS\_0033\_pdb70.a3m ../../similar\_sequences/33\_FANPEZAQ\_CDS\_0033\_pdb70.hhr ../../similar\_sequences/33\_FANPEZAQ\_CDS\_0033\_uniclust.a3m ../../similar\_sequences/33\_FANPEZAQ\_CDS\_0033\_uniclust.hhr |

#### Structure prediction (AlphaFold)^2^

|  |  |
| --- | --- |
| Stats | xml version="1.0" encoding="utf-8" standalone="no"?       2024-09-02T21:09:32.852536 image/svg+xml   Matplotlib v3.7.2, https://matplotlib.org/ |
| Predicted structure | **NGL Viewer Controls:**  - Center: *Left-Click* - Rotate: *Left-Click + Drag* - Translate: *Right-Click + Drag* - Zoom: *Shift + Left-Click + Drag* |
| Output files | ../../predicted\_structures/33\_FANPEZAQ\_CDS\_0033/features.pkl ../../predicted\_structures/33\_FANPEZAQ\_CDS\_0033/ranked\_0.pdb ../../predicted\_structures/33\_FANPEZAQ\_CDS\_0033/ranked\_0\_plots.svg ../../predicted\_structures/33\_FANPEZAQ\_CDS\_0033/result\_model\_1\_ptm\_pred\_0.pkl |

#### Structure similarity search results (Foldseek)^3^

|  |  |
| --- | --- |
| Structure databases searched | Pdb, Afdb-proteome, Afdb-uniprot50 |
| Results, scheme(s)  (Top layers only, threshold 1.00e-02 (evalue)) | xml version="1.0" encoding="utf-8" standalone="no"?       2024-09-02T21:11:06.714357 image/svg+xml   Matplotlib v3.7.2, https://matplotlib.org/ |
| Results, table  (threshold 1.00e-02 (evalue)) | -- |
| Top keywords  (threshold 1.00e-02 (evalue)) | -- |
| Output files | ../../similar\_structures/33\_FANPEZAQ\_CDS\_0033\_afdb-proteome\_foldseek.tsv ../../similar\_structures/33\_FANPEZAQ\_CDS\_0033\_afdb-uniprot50\_foldseek.tsv ../../similar\_structures/33\_FANPEZAQ\_CDS\_0033\_merged.svg ../../similar\_structures/33\_FANPEZAQ\_CDS\_0033\_pdb\_foldseek.tsv |

  
  
  

Return to summary | Go to previous | Go to next

  

---

**Sequence/structure alignments coloring**  
Each object in the alignment figures is colored according to its E-value following this color coding:

1e-100
10
