## Supplementary material for "Completing the BASEL phage collection to unlock hidden diversity for systematic exploration of phage-host interactions": Data S2: 34.html

FANPEZAQ\_CDS\_0034


Return to summary | Go to previous | Go to next

|  |  |
| --- | --- |
| FANPEZAQ\_CDS\_0034  Page creation date: 02 Sep 2024, 12:00  Project folder: n/a  Input sequences file: Escherichia\_virus\_HeidiAbel.gb | sprt\_like domain\_containing fragment sprt acidic repeat\_containing metalloprotease transcription elongation hypothetical zinc\_dependent spartan putative protein\_like with at the n terminus zinc isoform protease hydrolase peptidase m48 metalloproteinase mpr hmg box\_containing metallopeptidase c19g7 minigluzincin capsid |

### Sequence information

|  |  |
| --- | --- |
| Name | FANPEZAQ\_CDS\_0034  34\_FANPEZAQ\_CDS\_0034 (pipeline id) |
| Imported annotations |  |
| Protein sequence | MDREGWLNALAAKFAPRFKEFGYEMPKFRVSVGFCSSGARSSTAAQCWHSKCSSDNTFEI FIMPDQVEPYLVANHLWHELTHATVGFDCGHKGAFATVCKAVGLNGPMTATTPGEKFKEY VKPFLDELGPMPHAKLTFDRGMSIKVPRLRIGLDEGGDDGEGEEIEVAPVGGASTAKPKQ TTRLKKCECQECGYTVRVTQKWLEVGPPHCPEHGAMTPEGDDESSDE |
| Number of residues | 227 |
| Molecular weight (Da) | 24902.92 |
| Output files | ../../query\_sequences/34\_FANPEZAQ\_CDS\_0034.fasta |

### Putative domain architecture and protein family

#### Search results (HHblits)^1^

|  |  |
| --- | --- |
| Domain family databases searched | Pfam, Ncbi-cd, Cath, Phrogs |
| Results, scheme(s)  (Top layers only; threshold 1.00e-03 (evalue)) | xml version="1.0" encoding="utf-8" standalone="no"?       2024-09-02T21:08:19.725489 image/svg+xml   Matplotlib v3.7.2, https://matplotlib.org/ |
| Results, table  (E-value ≤ 1.00e-03 (evalue)) | db \| id \| prob \| evalue \| pvalue \| score \| cols \| query \| query\_len \| template \| template\_len \| name \| description; phrogs \| 6361 \| 100.0 \| 2.6e-52 \| 3.4e-56 \| 356.6 \| 185 \| (1, 220) \| 227 \| (17, 204) \| 204 \| NA \| NA; Category: unknown function; MG670586\_p45; phrogs \| 1006 \| 99.8 \| 1.5e-23 \| 1.8e-27 \| 168.7 \| 75 \| (29, 112) \| 227 \| (29, 106) \| 154 \| NA \| NA; Category: unknown function; JQ512844\_p79; phrogs \| 10756 \| 99.7 \| 2.5e-21 \| 3e-25 \| 166.4 \| 109 \| (2, 111) \| 227 \| (4, 122) \| 258 \| NA \| NA; Category: unknown function; NC\_019538\_p220; phrogs \| 3768 \| 99.5 \| 3.1e-19 \| 3.8e-23 \| 148.6 \| 73 \| (29, 111) \| 227 \| (33, 108) \| 192 \| NA \| NA; Category: unknown function; NC\_029000\_p177; phrogs \| 6445 \| 99.4 \| 4.2e-18 \| 5e-22 \| 140.2 \| 82 \| (29, 116) \| 227 \| (31, 120) \| 191 \| NA \| NA; Category: unknown function; NC\_019401\_p436; phrogs \| 9897 \| 99.4 \| 5.8e-18 \| 7e-22 \| 144.8 \| 105 \| (2, 111) \| 227 \| (34, 157) \| 248 \| NA \| NA; Category: unknown function; NC\_029009\_p209; phrogs \| 27319 \| 99.1 \| 4.5e-15 \| 5e-19 \| 121.9 \| 108 \| (2, 113) \| 227 \| (44, 159) \| 220 \| NA \| NA; Category: unknown function; NC\_021327\_p65; phrogs \| 10840 \| 99.0 \| 5.5e-14 \| 6.3e-18 \| 116.8 \| 76 \| (29, 112) \| 227 \| (40, 118) \| 223 \| NA \| NA; Category: unknown function; NC\_028683\_p287; phrogs \| 24857 \| 98.3 \| 1.4e-10 \| 1.6e-14 \| 92.2 \| 72 \| (31, 111) \| 227 \| (32, 108) \| 171 \| NA \| NA; Category: unknown function; AP018399\_p45; phrogs \| 32550 \| 96.4 \| 1.2e-05 \| 1.3e-09 \| 60.1 \| 103 \| (2, 107) \| 227 \| (5, 127) \| 131 \| NA \| NA; Category: unknown function; NC\_015251\_p422; phrogs \| 1233 \| 95.0 \| 0.00029 \| 4e-08 \| 51.5 \| 41 \| (40, 87) \| 227 \| (32, 72) \| 108 \| NA \| NA; Category: unknown function; p367521 VI\_04587; phrogs \| 37767 \| 94.2 \| 0.00097 \| 1.1e-07 \| 57.9 \| 68 \| (40, 107) \| 227 \| (117, 202) \| 442 \| NA \| NA; Category: unknown function; NC\_017972\_p199 |
| Top keywords  (threshold 1.00e-03 (evalue)) | **MG670586\_p45, JQ512844\_p79, NC\_019538\_p220, NC\_029000\_p177, NC\_019401\_p436, NC\_029009\_p209, NC\_021327\_p65, NC\_028683\_p287, AP018399\_p45, NC\_015251\_p422** |
| Output files | ../../domain\_architecture/34\_FANPEZAQ\_CDS\_0034\_cath.hhr ../../domain\_architecture/34\_FANPEZAQ\_CDS\_0034\_merged.svg ../../domain\_architecture/34\_FANPEZAQ\_CDS\_0034\_ncbi-cd.hhr ../../domain\_architecture/34\_FANPEZAQ\_CDS\_0034\_pfam.hhr ../../domain\_architecture/34\_FANPEZAQ\_CDS\_0034\_phrogs.hhr |

### Identical protein sequences/structures

#### Search results

|  |  |
| --- | --- |
| Protein sequence databases searched | Pdb, Swissprot, Refseq |
| Identical proteins found | Refseq  - QIG65622.1: Hypothetical protein (Salmonella phage PT1) |
| Top keywords | **Hypothetical** |
| Output files | ../../identical\_sequences/34\_FANPEZAQ\_CDS\_0034\_refseq.fasta |

### Similar protein sequences/structures

#### Sequence similarity search results (HHblits)^1^

|  |  |
| --- | --- |
| Sequence databases searched | Uniclust, Pdb70 |
| Results, scheme(s)  (Top layers only, threshold 1.00e-03 (evalue)) | xml version="1.0" encoding="utf-8" standalone="no"?       2024-09-02T21:08:43.710066 image/svg+xml   Matplotlib v3.7.2, https://matplotlib.org/ |
| Results, table(s)  (threshold 1.00e-03 (evalue)) | db \| id \| prob \| evalue \| pvalue \| score \| cols \| query \| query\_len \| template \| template\_len \| name \| description; uniclust \| UniRef100\_A0A024GXC7 \| 100.0 \| 7.8e-50 \| 1.8e-55 \| 344.1 \| 190 \| (1, 225) \| 227 \| (40, 229) \| 259 \| Uncharacterized protein \| Uncharacterized protein; uniclust \| UniRef100\_A0A031JAB2 \| 100.0 \| 4.7e-49 \| 9.8e-55 \| 337.6 \| 193 \| (1, 225) \| 227 \| (50, 251) \| 293 \| Transcription elongation protein SprT \| Transcription elongation protein SprT; uniclust \| UniRef100\_A0A088FRV0 \| 100.0 \| 7.8e-49 \| 1.7e-54 \| 339.2 \| 193 \| (1, 226) \| 227 \| (54, 248) \| 280 \| SprT-like domain-containing protein \| SprT-like domain-containing protein; uniclust \| UniRef100\_A0A2D0W8X1 \| 100.0 \| 3.8e-39 \| 8e-45 \| 269.1 \| 189 \| (1, 227) \| 227 \| (23, 214) \| 218 \| Putative transcription elongation protein \| Putative transcription elongation protein; uniclust \| UniRef100\_UPI0011A7918A \| 100.0 \| 1.9e-37 \| 3.7e-43 \| 264.5 \| 204 \| (1, 214) \| 227 \| (93, 303) \| 309 \| SprT family zinc-dependent metalloprotease \| SprT family zinc-dependent metalloprotease; uniclust \| UniRef100\_A0A1C4U4K3 \| 100.0 \| 7.5e-34 \| 1.4e-39 \| 250.2 \| 187 \| (2, 219) \| 227 \| (256, 446) \| 448 \| SprT-like family protein \| SprT-like family protein; uniclust \| UniRef100\_A0A1C6ST94 \| 100.0 \| 6.4e-33 \| 1.2e-38 \| 241.8 \| 186 \| (1, 218) \| 227 \| (247, 436) \| 439 \| SprT-like family protein \| SprT-like family protein; uniclust \| UniRef100\_A0A090Y2V6 \| 99.9 \| 4.1e-32 \| 9.4e-38 \| 241.0 \| 189 \| (6, 226) \| 227 \| (51, 269) \| 298 \| SprT-like family protein \| SprT-like family protein; uniclust \| UniRef100\_A0A2C9WX41 \| 99.9 \| 7.9e-31 \| 1.5e-36 \| 219.1 \| 217 \| (1, 220) \| 227 \| (1, 219) \| 288 \| SprT domain-containing protein \| SprT domain-containing protein; uniclust \| UniRef100\_A0A257FUX5 \| 99.9 \| 4.9e-30 \| 9.4e-36 \| 203.6 \| 165 \| (1, 201) \| 227 \| (5, 169) \| 171 \| Uncharacterized protein \| Uncharacterized protein; uniclust \| UniRef100\_A0A498AIE9 \| 99.9 \| 9.7e-30 \| 1.8e-35 \| 216.8 \| 186 \| (1, 218) \| 227 \| (153, 340) \| 342 \| SprT-like family protein \| SprT-like family protein; uniclust \| UniRef100\_A0A143ZU24 \| 99.9 \| 2.3e-29 \| 4.9e-35 \| 216.3 \| 177 \| (7, 222) \| 227 \| (22, 223) \| 250 \| SprT-like family protein \| SprT-like family protein; uniclust \| UniRef100\_A0A158S5S8 \| 99.9 \| 9.9e-28 \| 1.9e-33 \| 192.0 \| 132 \| (1, 132) \| 227 \| (4, 141) \| 173 \| Transcription elongation protein SprT \| Transcription elongation protein SprT; uniclust \| UniRef100\_A0A014MPW4 \| 99.9 \| 1.8e-27 \| 4.1e-33 \| 214.6 \| 190 \| (9, 224) \| 227 \| (77, 287) \| 336 \| Uncharacterized protein \| Uncharacterized protein; uniclust \| UniRef100\_A0A973MLF9 \| 99.9 \| 2.4e-27 \| 4.4e-33 \| 203.5 \| 211 \| (1, 220) \| 227 \| (6, 222) \| 356 \| Uncharacterized protein \| Uncharacterized protein; uniclust \| UniRef100\_A0A0F9EIW7 \| 99.9 \| 3.4e-27 \| 7.1e-33 \| 186.6 \| 128 \| (67, 226) \| 227 \| (11, 138) \| 143 \| SprT-like domain-containing protein (Fragment) \| SprT-like domain-containing protein (Fragment); uniclust \| UniRef100\_A0A0F9B2V9 \| 99.9 \| 7.2e-27 \| 1.4e-32 \| 187.4 \| 188 \| (1, 226) \| 227 \| (1, 191) \| 192 \| SprT-like domain-containing protein (Fragment) \| SprT-like domain-containing protein (Fragment); uniclust \| UniRef100\_A0A661MH10 \| 99.9 \| 2.1e-26 \| 3.8e-32 \| 190.7 \| 188 \| (1, 218) \| 227 \| (30, 239) \| 256 \| Uncharacterized protein \| Uncharacterized protein; uniclust \| UniRef100\_A0A4Q2J0L1 \| 99.9 \| 4e-26 \| 7.3e-32 \| 192.2 \| 187 \| (1, 218) \| 227 \| (102, 292) \| 295 \| Uncharacterized protein \| Uncharacterized protein; uniclust \| UniRef100\_UPI0015E46A98 \| 99.9 \| 4.4e-26 \| 8.4e-32 \| 181.2 \| 139 \| (1, 139) \| 227 \| (12, 150) \| 169 \| hypothetical protein \| hypothetical protein; uniclust \| UniRef100\_A0A1F6D4Y3 \| 99.9 \| 9.9e-26 \| 1.9e-31 \| 184.5 \| 181 \| (1, 223) \| 227 \| (9, 198) \| 207 \| SprT-like domain-containing protein \| SprT-like domain-containing protein; uniclust \| UniRef100\_UPI001F48BFF7 \| 99.9 \| 4.6e-25 \| 8.4e-31 \| 181.7 \| 183 \| (2, 220) \| 227 \| (56, 240) \| 242 \| hypothetical protein \| hypothetical protein; uniclust \| UniRef100\_A0A0D8B6L3 \| 99.8 \| 8.5e-25 \| 1.8e-30 \| 195.1 \| 195 \| (8, 224) \| 227 \| (45, 271) \| 326 \| SprT-like family \| SprT-like family; uniclust \| UniRef100\_A0A1E7JES4 \| 99.8 \| 1.1e-24 \| 2.5e-30 \| 193.3 \| 171 \| (5, 221) \| 227 \| (47, 227) \| 288 \| SprT-like domain-containing protein \| SprT-like domain-containing protein; uniclust \| UniRef100\_A0A315DXF0 \| 99.8 \| 3.3e-24 \| 6.9e-30 \| 175.2 \| 136 \| (3, 138) \| 227 \| (21, 161) \| 178 \| Uncharacterized protein \| Uncharacterized protein; uniclust \| UniRef100\_D4Z6N5 \| 99.8 \| 1.1e-23 \| 2e-29 \| 161.8 \| 126 \| (1, 126) \| 227 \| (4, 135) \| 136 \| Transcription elongation protein SprT \| Transcription elongation protein SprT; uniclust \| UniRef100\_A0A920EZT5 \| 99.8 \| 8.9e-23 \| 1.7e-28 \| 155.3 \| 108 \| (2, 109) \| 227 \| (9, 116) \| 123 \| Uncharacterized protein \| Uncharacterized protein; uniclust \| UniRef100\_A0A010RP43 \| 99.8 \| 1.1e-22 \| 2.8e-28 \| 188.1 \| 201 \| (7, 224) \| 227 \| (77, 318) \| 356 \| Zinc metalloprotease \| Zinc metalloprotease; uniclust \| UniRef100\_A0A090SJ74 \| 99.8 \| 2.3e-22 \| 4.9e-28 \| 176.4 \| 203 \| (8, 223) \| 227 \| (16, 266) \| 291 \| Zinc metalloproteinase Mpr protein \| Zinc metalloproteinase Mpr protein; uniclust \| UniRef100\_A0A6G4A0P7 \| 99.8 \| 3.8e-22 \| 7.5e-28 \| 168.8 \| 191 \| (9, 221) \| 227 \| (5, 225) \| 233 \| SprT-like domain-containing protein \| SprT-like domain-containing protein; uniclust \| UniRef100\_A0A1C6IQ91 \| 99.8 \| 9.4e-22 \| 1.8e-27 \| 165.3 \| 178 \| (3, 222) \| 227 \| (3, 206) \| 236 \| SprT-like family \| SprT-like family; uniclust \| UniRef100\_A0A1C5HCK4 \| 99.7 \| 6.2e-21 \| 1.2e-26 \| 171.0 \| 172 \| (15, 222) \| 227 \| (39, 235) \| 382 \| SprT-like family protein \| SprT-like family protein; uniclust \| UniRef100\_A0A644ZG81 \| 99.7 \| 7.6e-21 \| 1.6e-26 \| 156.4 \| 135 \| (57, 223) \| 227 \| (11, 165) \| 171 \| Uncharacterized protein \| Uncharacterized protein; uniclust \| UniRef100\_A0A0A0HYB9 \| 99.7 \| 1.2e-20 \| 2.7e-26 \| 165.5 \| 178 \| (7, 221) \| 227 \| (12, 212) \| 241 \| SprT-like domain-containing protein \| SprT-like domain-containing protein; uniclust \| UniRef100\_A0A4R0JGF1 \| 99.7 \| 3.4e-20 \| 6.6e-26 \| 163.6 \| 168 \| (17, 222) \| 227 \| (163, 353) \| 355 \| SprT-like family protein \| SprT-like family protein; uniclust \| UniRef100\_A0A0D7CF42 \| 99.7 \| 3.2e-20 \| 7.2e-26 \| 161.6 \| 182 \| (5, 224) \| 227 \| (28, 233) \| 237 \| SprT-like domain-containing protein \| SprT-like domain-containing protein; uniclust \| UniRef100\_A0A401QRK3 \| 99.7 \| 7.5e-20 \| 1.6e-25 \| 160.8 \| 213 \| (9, 226) \| 227 \| (28, 288) \| 290 \| SprT-like domain-containing protein \| SprT-like domain-containing protein; uniclust \| UniRef100\_A0A3N5NIA4 \| 99.6 \| 7.2e-19 \| 1.3e-24 \| 136.0 \| 123 \| (2, 126) \| 227 \| (17, 140) \| 142 \| Uncharacterized protein (Fragment) \| Uncharacterized protein (Fragment); uniclust \| UniRef100\_A0A259FRY7 \| 99.6 \| 8.8e-18 \| 1.6e-23 \| 121.8 \| 86 \| (2, 87) \| 227 \| (6, 91) \| 93 \| SprT domain-containing protein (Fragment) \| SprT domain-containing protein (Fragment); uniclust \| UniRef100\_UPI002354798D \| 99.5 \| 1.3e-17 \| 2.4e-23 \| 126.7 \| 111 \| (1, 113) \| 227 \| (2, 118) \| 124 \| SprT-like domain-containing protein \| SprT-like domain-containing protein; uniclust \| UniRef100\_A0A9E2ZVF6 \| 99.5 \| 6.9e-17 \| 1.3e-22 \| 138.1 \| 170 \| (16, 220) \| 227 \| (102, 290) \| 292 \| J domain-containing protein \| J domain-containing protein; uniclust \| UniRef100\_UPI002280EFAF \| 99.5 \| 8.9e-17 \| 1.6e-22 \| 139.4 \| 187 \| (2, 219) \| 227 \| (109, 329) \| 331 \| SprT-like domain-containing protein \| SprT-like domain-containing protein; uniclust \| UniRef100\_D0ZHK0 \| 99.5 \| 1.5e-16 \| 2.8e-22 \| 138.7 \| 98 \| (12, 114) \| 227 \| (119, 233) \| 347 \| Putative zinc metalloproteinase Mpr protein \| Putative zinc metalloproteinase Mpr protein; uniclust \| UniRef100\_A0A0F9G8P3 \| 99.5 \| 1.6e-16 \| 3.3e-22 \| 119.7 \| 83 \| (108, 222) \| 227 \| (1, 84) \| 98 \| Uncharacterized protein \| Uncharacterized protein; uniclust \| UniRef100\_A0A0S3TZF6 \| 99.4 \| 3e-16 \| 6.4e-22 \| 134.7 \| 169 \| (7, 222) \| 227 \| (10, 200) \| 211 \| SprT-like domain-containing protein \| SprT-like domain-containing protein; uniclust \| UniRef100\_A0A011SPG8 \| 99.4 \| 3.1e-16 \| 6.6e-22 \| 135.6 \| 96 \| (29, 139) \| 227 \| (52, 150) \| 232 \| SprT-like domain-containing protein \| SprT-like domain-containing protein; uniclust \| UniRef100\_UPI002162A182 \| 99.4 \| 4.1e-16 \| 7.6e-22 \| 135.1 \| 168 \| (7, 212) \| 227 \| (5, 202) \| 325 \| SprT-like domain-containing protein \| SprT-like domain-containing protein; uniclust \| UniRef100\_UPI0013579F19 \| 99.4 \| 8.2e-16 \| 1.5e-21 \| 141.5 \| 187 \| (7, 223) \| 227 \| (5, 214) \| 593 \| SprT-like domain-containing protein \| SprT-like domain-containing protein; uniclust \| UniRef100\_A0A7X6P1E1 \| 99.4 \| 8.7e-16 \| 1.7e-21 \| 138.7 \| 192 \| (13, 221) \| 227 \| (25, 236) \| 420 \| SprT-like domain-containing protein \| SprT-like domain-containing protein; uniclust \| UniRef100\_UPI002108714C \| 99.4 \| 1.2e-15 \| 2.2e-21 \| 130.2 \| 169 \| (7, 220) \| 227 \| (85, 278) \| 279 \| SprT-like domain-containing protein \| SprT-like domain-containing protein; uniclust \| UniRef100\_A0A068F4D8 \| 99.4 \| 1.6e-15 \| 3.7e-21 \| 135.6 \| 73 \| (29, 110) \| 227 \| (79, 154) \| 268 \| SprT-like protease \| SprT-like protease; uniclust \| UniRef100\_A0A970D5F1 \| 99.4 \| 1.9e-15 \| 3.8e-21 \| 126.8 \| 177 \| (7, 224) \| 227 \| (14, 214) \| 215 \| SprT-like domain-containing protein \| SprT-like domain-containing protein; uniclust \| UniRef100\_A0A451CCI8 \| 99.4 \| 2.3e-15 \| 4.6e-21 \| 124.7 \| 58 \| (57, 114) \| 227 \| (18, 83) \| 183 \| SprT-like family protein \| SprT-like family protein; uniclust \| UniRef100\_A0A3D0Z8I8 \| 99.4 \| 2.8e-15 \| 5.2e-21 \| 129.9 \| 170 \| (7, 220) \| 227 \| (118, 316) \| 317 \| SprT-like domain-containing protein \| SprT-like domain-containing protein; uniclust \| UniRef100\_A0A6H9YBG4 \| 99.4 \| 3e-15 \| 5.6e-21 \| 129.4 \| 100 \| (10, 114) \| 227 \| (10, 126) \| 311 \| SprT family zinc-dependent metalloprotease \| SprT family zinc-dependent metalloprotease; uniclust \| UniRef100\_A0A1G7EMH9 \| 99.4 \| 2.8e-15 \| 5.9e-21 \| 126.1 \| 108 \| (7, 122) \| 227 \| (23, 143) \| 190 \| SprT-like family protein \| SprT-like family protein; uniclust \| UniRef100\_A0A0Q8UQ63 \| 99.3 \| 3.3e-15 \| 6.1e-21 \| 119.1 \| 132 \| (2, 136) \| 227 \| (19, 151) \| 165 \| SprT-like domain-containing protein \| SprT-like domain-containing protein; uniclust \| UniRef100\_A0A292IIQ0 \| 99.3 \| 3.4e-15 \| 6.8e-21 \| 129.4 \| 174 \| (7, 222) \| 227 \| (17, 213) \| 248 \| SprT-like domain-containing protein \| SprT-like domain-containing protein; uniclust \| UniRef100\_A0A0F0KVC3 \| 99.3 \| 4.6e-15 \| 1e-20 \| 133.2 \| 160 \| (41, 223) \| 227 \| (72, 266) \| 292 \| SprT-like family protein \| SprT-like family protein; uniclust \| UniRef100\_A0A966KII7 \| 99.3 \| 6.4e-15 \| 1.2e-20 \| 118.7 \| 114 \| (72, 221) \| 227 \| (13, 126) \| 157 \| Uncharacterized protein \| Uncharacterized protein; uniclust \| UniRef100\_A0A6P0HX66 \| 99.3 \| 6.8e-15 \| 1.3e-20 \| 122.7 \| 161 \| (1, 201) \| 227 \| (33, 209) \| 228 \| SprT family zinc-dependent metalloprotease \| SprT family zinc-dependent metalloprotease; uniclust \| UniRef100\_A0A1H9XF20 \| 99.3 \| 7.8e-15 \| 1.6e-20 \| 120.5 \| 114 \| (7, 123) \| 227 \| (7, 147) \| 169 \| SprT-like family protein (Fragment) \| SprT-like family protein (Fragment); uniclust \| UniRef100\_A0A068NQF1 \| 99.3 \| 9.1e-15 \| 2.1e-20 \| 126.1 \| 79 \| (24, 112) \| 227 \| (41, 125) \| 200 \| Acidic repeat-containing protein \| Acidic repeat-containing protein; uniclust \| UniRef100\_A0A0F9ATJ3 \| 99.3 \| 1.9e-14 \| 3.5e-20 \| 120.9 \| 199 \| (2, 220) \| 227 \| (4, 215) \| 217 \| SprT-like domain-containing protein (Fragment) \| SprT-like domain-containing protein (Fragment); uniclust \| UniRef100\_A5EQM9 \| 99.3 \| 2.2e-14 \| 3.9e-20 \| 123.7 \| 169 \| (22, 215) \| 227 \| (100, 281) \| 297 \| SprT-like domain-containing protein \| SprT-like domain-containing protein; uniclust \| UniRef100\_A0A1F3SVB2 \| 99.3 \| 1.8e-14 \| 4.3e-20 \| 124.9 \| 83 \| (14, 105) \| 227 \| (28, 120) \| 198 \| SprT-like domain-containing protein \| SprT-like domain-containing protein; uniclust \| UniRef100\_A0A095WTQ5 \| 99.3 \| 2.1e-14 \| 4.8e-20 \| 127.0 \| 101 \| (15, 134) \| 227 \| (50, 153) \| 242 \| Transcription elongation protein SprT \| Transcription elongation protein SprT; uniclust \| UniRef100\_A0A0K2SH02 \| 99.2 \| 3.4e-14 \| 7.5e-20 \| 121.0 \| 86 \| (4, 102) \| 227 \| (17, 105) \| 188 \| Metal-dependent hydrolase \| Metal-dependent hydrolase; uniclust \| UniRef100\_A0A073IWH6 \| 99.2 \| 3.8e-14 \| 7.6e-20 \| 124.2 \| 185 \| (7, 224) \| 227 \| (22, 221) \| 267 \| SprT-like domain-containing protein \| SprT-like domain-containing protein; uniclust \| UniRef100\_A0A0K2B1L4 \| 99.2 \| 4.8e-14 \| 9.7e-20 \| 117.1 \| 144 \| (42, 220) \| 227 \| (11, 178) \| 179 \| Uncharacterized protein \| Uncharacterized protein; uniclust \| UniRef100\_A0A4P7ZT04 \| 99.2 \| 5.4e-14 \| 1.1e-19 \| 127.5 \| 167 \| (5, 216) \| 227 \| (24, 212) \| 368 \| SprT-like domain-containing protein \| SprT-like domain-containing protein; uniclust \| UniRef100\_A0A146GBG5 \| 99.2 \| 5.8e-14 \| 1.3e-19 \| 127.2 \| 86 \| (16, 109) \| 227 \| (114, 207) \| 298 \| SprT-like family protein \| SprT-like family protein; uniclust \| UniRef100\_UPI0003B474FE \| 99.2 \| 9.9e-14 \| 1.8e-19 \| 114.8 \| 58 \| (57, 114) \| 227 \| (18, 84) \| 209 \| SprT-like domain-containing protein \| SprT-like domain-containing protein; uniclust \| UniRef100\_A0A5Q2W9L6 \| 99.2 \| 9.3e-14 \| 1.8e-19 \| 118.7 \| 192 \| (1, 220) \| 227 \| (10, 221) \| 224 \| SprT-like family protein \| SprT-like family protein; uniclust \| UniRef100\_A0A7Y5RTJ4 \| 99.2 \| 1e-13 \| 1.9e-19 \| 119.7 \| 167 \| (5, 219) \| 227 \| (102, 278) \| 294 \| SprT-like domain-containing protein \| SprT-like domain-containing protein; uniclust \| UniRef100\_A0A2E4W4M1 \| 99.2 \| 9.8e-14 \| 1.9e-19 \| 123.4 \| 94 \| (15, 108) \| 227 \| (59, 165) \| 326 \| SprT-like domain-containing protein \| SprT-like domain-containing protein; uniclust \| UniRef100\_A0A1V3BVK3 \| 99.2 \| 1e-13 \| 1.9e-19 \| 120.1 \| 211 \| (11, 223) \| 227 \| (28, 278) \| 294 \| SprT-like domain-containing protein \| SprT-like domain-containing protein; uniclust \| UniRef100\_A0A2D5NF12 \| 99.2 \| 8.9e-14 \| 2.2e-19 \| 123.0 \| 77 \| (25, 109) \| 227 \| (55, 139) \| 222 \| SprT-like domain-containing protein \| SprT-like domain-containing protein; uniclust \| UniRef100\_A0A0B5FXD9 \| 99.2 \| 9.8e-14 \| 2.2e-19 \| 131.7 \| 118 \| (10, 132) \| 227 \| (74, 229) \| 421 \| SprT-like domain-containing protein \| SprT-like domain-containing protein; uniclust \| UniRef100\_A0A174V594 \| 99.2 \| 1.2e-13 \| 2.8e-19 \| 120.1 \| 92 \| (5, 103) \| 227 \| (32, 126) \| 208 \| SprT domain-containing protein \| SprT domain-containing protein; uniclust \| UniRef100\_A0A7J9XW55 \| 99.2 \| 1.8e-13 \| 3.2e-19 \| 109.7 \| 122 \| (68, 223) \| 227 \| (6, 142) \| 164 \| SprT domain-containing protein \| SprT domain-containing protein; uniclust \| UniRef100\_A0A318K5J9 \| 99.1 \| 3.6e-13 \| 7.4e-19 \| 114.5 \| 120 \| (10, 129) \| 227 \| (32, 170) \| 197 \| SprT-like family protein \| SprT-like family protein; uniclust \| UniRef100\_A0A0F8YGA8 \| 99.1 \| 4e-13 \| 7.7e-19 \| 109.1 \| 130 \| (68, 216) \| 227 \| (8, 147) \| 163 \| SprT-like domain-containing protein (Fragment) \| SprT-like domain-containing protein (Fragment); uniclust \| UniRef100\_A0A358QWY1 \| 99.1 \| 4.3e-13 \| 7.9e-19 \| 110.2 \| 162 \| (45, 220) \| 227 \| (5, 195) \| 196 \| SprT-like domain-containing protein \| SprT-like domain-containing protein; uniclust \| UniRef100\_A0A354EFJ4 \| 99.1 \| 4.2e-13 \| 9.2e-19 \| 119.6 \| 98 \| (2, 112) \| 227 \| (93, 193) \| 259 \| SprT-like domain-containing protein \| SprT-like domain-containing protein; uniclust \| UniRef100\_A0A6B1ALR6 \| 99.1 \| 6.7e-13 \| 1.4e-18 \| 118.9 \| 96 \| (3, 111) \| 227 \| (98, 196) \| 298 \| SprT family zinc-dependent metalloprotease \| SprT family zinc-dependent metalloprotease; uniclust \| UniRef100\_A0A1L3GQR5 \| 99.1 \| 6.5e-13 \| 1.4e-18 \| 122.2 \| 94 \| (6, 112) \| 227 \| (170, 266) \| 324 \| SprT-like domain-containing protein \| SprT-like domain-containing protein; uniclust \| UniRef100\_A0A0A0J0D7 \| 99.1 \| 8.8e-13 \| 1.9e-18 \| 119.3 \| 88 \| (11, 111) \| 227 \| (39, 129) \| 294 \| SprT-like domain-containing protein \| SprT-like domain-containing protein; uniclust \| UniRef100\_A0A2W4I2B1 \| 99.1 \| 9.9e-13 \| 2e-18 \| 110.8 \| 78 \| (24, 112) \| 227 \| (18, 98) \| 183 \| SprT domain-containing protein \| SprT domain-containing protein; uniclust \| UniRef100\_A0A2N1P657 \| 99.0 \| 1.2e-12 \| 2.7e-18 \| 115.0 \| 88 \| (4, 107) \| 227 \| (61, 150) \| 221 \| SprT-like domain-containing protein \| SprT-like domain-containing protein; uniclust \| UniRef100\_A0A516MH30 \| 99.0 \| 1.5e-12 \| 2.8e-18 \| 108.2 \| 169 \| (19, 223) \| 227 \| (20, 208) \| 211 \| SprT-like domain-containing protein \| SprT-like domain-containing protein; uniclust \| UniRef100\_A0A8R1E689 \| 99.0 \| 2.1e-12 \| 3.8e-18 \| 116.0 \| 99 \| (11, 114) \| 227 \| (8, 123) \| 408 \| SprT-like domain-containing protein \| SprT-like domain-containing protein; uniclust \| UniRef100\_A0A2D6AHS8 \| 99.0 \| 2.3e-12 \| 4.7e-18 \| 110.9 \| 88 \| (23, 122) \| 227 \| (72, 167) \| 218 \| SprT-like domain-containing protein \| SprT-like domain-containing protein; uniclust \| UniRef100\_A0A140DWM3 \| 99.0 \| 2.2e-12 \| 5e-18 \| 111.8 \| 89 \| (7, 102) \| 227 \| (12, 105) \| 199 \| SprT-like domain-containing protein \| SprT-like domain-containing protein; uniclust \| UniRef100\_A0A2A4WS63 \| 99.0 \| 2.4e-12 \| 5.4e-18 \| 112.7 \| 94 \| (10, 112) \| 227 \| (36, 136) \| 211 \| SprT-like domain-containing protein \| SprT-like domain-containing protein; uniclust \| UniRef100\_A0A933N799 \| 99.0 \| 3e-12 \| 5.6e-18 \| 108.2 \| 182 \| (20, 218) \| 227 \| (30, 235) \| 237 \| SprT-like domain-containing protein \| SprT-like domain-containing protein; uniclust \| UniRef100\_A0A060JGM0 \| 99.0 \| 3.3e-12 \| 6.6e-18 \| 110.4 \| 96 \| (28, 139) \| 227 \| (93, 192) \| 232 \| SprT-like family \| SprT-like family; uniclust \| UniRef100\_A0A6A8MVU3 \| 99.0 \| 3.3e-12 \| 6.8e-18 \| 116.2 \| 97 \| (28, 139) \| 227 \| (203, 302) \| 337 \| SprT-like domain-containing protein \| SprT-like domain-containing protein; uniclust \| UniRef100\_A0A1U7GSA4 \| 99.0 \| 3.2e-12 \| 7e-18 \| 112.6 \| 76 \| (28, 112) \| 227 \| (43, 121) \| 236 \| SprT-like domain-containing protein \| SprT-like domain-containing protein; uniclust \| UniRef100\_A0A9D1JYK7 \| 99.0 \| 4e-12 \| 8.2e-18 \| 98.6 \| 96 \| (90, 220) \| 227 \| (5, 106) \| 107 \| Uncharacterized protein \| Uncharacterized protein; uniclust \| UniRef100\_A0A955QNM8 \| 98.9 \| 5.8e-12 \| 1.1e-17 \| 98.5 \| 107 \| (23, 129) \| 227 \| (15, 130) \| 135 \| SprT-like domain-containing protein (Fragment) \| SprT-like domain-containing protein (Fragment); uniclust \| UniRef100\_A0A076YKR6 \| 98.9 \| 6.2e-12 \| 1.3e-17 \| 108.1 \| 74 \| (29, 110) \| 227 \| (48, 124) \| 205 \| SprT-like domain-containing protein \| SprT-like domain-containing protein; uniclust \| UniRef100\_A0A0K2GJ75 \| 98.9 \| 7e-12 \| 1.5e-17 \| 113.3 \| 150 \| (6, 220) \| 227 \| (31, 199) \| 282 \| SprT-like domain-containing protein \| SprT-like domain-containing protein; uniclust \| UniRef100\_A0A1W9W5N7 \| 98.9 \| 7.4e-12 \| 1.5e-17 \| 107.6 \| 111 \| (8, 126) \| 227 \| (16, 154) \| 207 \| SprT-like domain-containing protein \| SprT-like domain-containing protein; uniclust \| UniRef100\_A0A2K9UZU6 \| 98.9 \| 1.1e-11 \| 2.2e-17 \| 102.4 \| 80 \| (23, 107) \| 227 \| (47, 139) \| 153 \| Zinc metalloproteinase Mpr protein \| Zinc metalloproteinase Mpr protein; uniclust \| UniRef100\_A0A950FD64 \| 98.9 \| 1.3e-11 \| 2.5e-17 \| 94.8 \| 98 \| (86, 220) \| 227 \| (2, 109) \| 119 \| Uncharacterized protein \| Uncharacterized protein; uniclust \| UniRef100\_A0A2U9QIZ2 \| 98.9 \| 1.6e-11 \| 2.9e-17 \| 102.0 \| 109 \| (8, 122) \| 227 \| (6, 134) \| 197 \| Zinc metalloproteinase Mpr protein \| Zinc metalloproteinase Mpr protein; uniclust \| UniRef100\_A0A8R1E603 \| 98.9 \| 1.9e-11 \| 3.6e-17 \| 117.3 \| 99 \| (11, 114) \| 227 \| (8, 123) \| 833 \| SprT-like domain-containing protein \| SprT-like domain-containing protein; uniclust \| UniRef100\_A0A0I9NBQ0 \| 98.9 \| 1.8e-11 \| 3.8e-17 \| 120.0 \| 130 \| (23, 220) \| 227 \| (371, 507) \| 590 \| Bm7449, isoform a \| Bm7449, isoform a; uniclust \| UniRef100\_A0A139TQJ4 \| 98.8 \| 2.1e-11 \| 4.3e-17 \| 107.6 \| 75 \| (28, 111) \| 227 \| (93, 172) \| 244 \| SprT-like domain-containing protein \| SprT-like domain-containing protein; uniclust \| UniRef100\_A0A1F6LM96 \| 98.8 \| 2.6e-11 \| 4.7e-17 \| 112.0 \| 195 \| (7, 219) \| 227 \| (295, 512) \| 514 \| SprT-like domain-containing protein \| SprT-like domain-containing protein; uniclust \| UniRef100\_A0A232F4E3 \| 98.8 \| 2.8e-11 \| 6.2e-17 \| 125.2 \| 151 \| (2, 219) \| 227 \| (756, 917) \| 995 \| SprT-like domain-containing protein \| SprT-like domain-containing protein; uniclust \| UniRef100\_A0A0B5A5G0 \| 98.8 \| 3.3e-11 \| 6.6e-17 \| 102.1 \| 92 \| (28, 136) \| 227 \| (66, 160) \| 190 \| SprT-like protease \| SprT-like protease; uniclust \| UniRef100\_A0A372ZJH5 \| 98.8 \| 3.9e-11 \| 7.3e-17 \| 107.4 \| 181 \| (16, 222) \| 227 \| (22, 244) \| 356 \| Uncharacterized protein \| Uncharacterized protein; uniclust \| UniRef100\_A0A6J8E1A8 \| 98.8 \| 3.9e-11 \| 8.5e-17 \| 124.0 \| 136 \| (23, 222) \| 227 \| (870, 1013) \| 1074 \| Acidic repeat-containing protein \| Acidic repeat-containing protein; uniclust \| UniRef100\_A0A653DTS8 \| 98.8 \| 4.2e-11 \| 8.9e-17 \| 121.9 \| 150 \| (2, 219) \| 227 \| (721, 882) \| 945 \| SprT-like domain-containing protein \| SprT-like domain-containing protein; uniclust \| UniRef100\_A0A2L1K6I3 \| 98.8 \| 5e-11 \| 9.3e-17 \| 101.5 \| 101 \| (7, 115) \| 227 \| (24, 138) \| 230 \| Zinc metalloproteinase Mpr protein \| Zinc metalloproteinase Mpr protein; uniclust \| UniRef100\_A0A2D5Z0A3 \| 98.8 \| 4.1e-11 \| 9.6e-17 \| 106.5 \| 93 \| (7, 108) \| 227 \| (39, 143) \| 221 \| SprT-like domain-containing protein \| SprT-like domain-containing protein; uniclust \| UniRef100\_A0A1B6C646 \| 98.8 \| 4.3e-11 \| 9.9e-17 \| 121.9 \| 133 \| (23, 220) \| 227 \| (526, 669) \| 763 \| SprT-like domain-containing protein \| SprT-like domain-containing protein; uniclust \| UniRef100\_A0A0S7ZXW9 \| 98.8 \| 4.7e-11 \| 1e-16 \| 109.4 \| 93 \| (10, 108) \| 227 \| (181, 285) \| 309 \| SprT-like domain-containing protein \| SprT-like domain-containing protein; uniclust \| UniRef100\_A0A026WDD5 \| 98.8 \| 5.1e-11 \| 1.2e-16 \| 123.7 \| 132 \| (23, 219) \| 227 \| (773, 915) \| 1001 \| Acidic repeat-containing protein \| Acidic repeat-containing protein; uniclust \| UniRef100\_UPI00111BAF9D \| 98.8 \| 6.3e-11 \| 1.2e-16 \| 89.0 \| 73 \| (57, 129) \| 227 \| (12, 93) \| 96 \| SprT-like domain-containing protein \| SprT-like domain-containing protein; uniclust \| UniRef100\_A0A2H8TPB7 \| 98.8 \| 6.1e-11 \| 1.3e-16 \| 118.9 \| 133 \| (23, 220) \| 227 \| (467, 610) \| 690 \| Acidic repeat-containing protein \| Acidic repeat-containing protein; uniclust \| UniRef100\_A0A2D7KWL3 \| 98.8 \| 6.8e-11 \| 1.4e-16 \| 100.4 \| 75 \| (29, 112) \| 227 \| (62, 139) \| 195 \| SprT domain-containing protein \| SprT domain-containing protein; uniclust \| UniRef100\_A0A016TL32 \| 98.8 \| 6.8e-11 \| 1.4e-16 \| 115.0 \| 133 \| (23, 222) \| 227 \| (363, 503) \| 591 \| SprT-like domain-containing protein \| SprT-like domain-containing protein; uniclust \| UniRef100\_A0A517XQ89 \| 98.7 \| 8.8e-11 \| 1.7e-16 \| 100.7 \| 79 \| (28, 111) \| 227 \| (89, 170) \| 225 \| SprT-like family protein \| SprT-like family protein; uniclust \| UniRef100\_A0A7M7RG74 \| 98.7 \| 8.7e-11 \| 1.8e-16 \| 118.5 \| 81 \| (23, 106) \| 227 \| (642, 729) \| 841 \| SprT-like domain-containing protein \| SprT-like domain-containing protein; uniclust \| UniRef100\_A0A143BLS1 \| 98.7 \| 9.9e-11 \| 2e-16 \| 107.2 \| 92 \| (11, 108) \| 227 \| (180, 282) \| 323 \| SprT-like domain-containing protein \| SprT-like domain-containing protein; uniclust \| UniRef100\_A0A0S9KEU5 \| 98.7 \| 1e-10 \| 2.1e-16 \| 103.9 \| 79 \| (25, 112) \| 227 \| (19, 109) \| 275 \| SprT-like domain-containing protein \| SprT-like domain-containing protein; uniclust \| UniRef100\_UPI000C6CB064 \| 98.7 \| 1.1e-10 \| 2.2e-16 \| 111.9 \| 133 \| (23, 221) \| 227 \| (294, 433) \| 494 \| acidic repeat-containing protein-like isoform X8 \| acidic repeat-containing protein-like isoform X8; uniclust \| UniRef100\_A0A0F8YXQ5 \| 98.7 \| 1.7e-10 \| 3.1e-16 \| 85.7 \| 92 \| (95, 224) \| 227 \| (1, 92) \| 94 \| Uncharacterized protein (Fragment) \| Uncharacterized protein (Fragment); uniclust \| UniRef100\_A0A023W5I8 \| 98.7 \| 1.6e-10 \| 3.1e-16 \| 97.6 \| 75 \| (29, 112) \| 227 \| (32, 109) \| 197 \| SprT-like protease \| SprT-like protease; uniclust \| UniRef100\_A0A2J7R5L2 \| 98.7 \| 1.9e-10 \| 3.8e-16 \| 110.1 \| 132 \| (23, 220) \| 227 \| (320, 461) \| 531 \| SprT-like domain-containing protein \| SprT-like domain-containing protein; uniclust \| UniRef100\_A0A7V9QDL4 \| 98.7 \| 1.8e-10 \| 3.9e-16 \| 101.2 \| 89 \| (11, 109) \| 227 \| (119, 218) \| 227 \| SprT-like domain-containing protein \| SprT-like domain-containing protein; uniclust \| UniRef100\_A0A158PCN7 \| 98.7 \| 2.2e-10 \| 4.1e-16 \| 94.5 \| 128 \| (23, 219) \| 227 \| (8, 143) \| 186 \| SprT-like domain-containing protein \| SprT-like domain-containing protein; uniclust \| UniRef100\_A0A0X3PXD0 \| 98.7 \| 2.1e-10 \| 4.2e-16 \| 112.0 \| 77 \| (23, 102) \| 227 \| (355, 445) \| 610 \| Acidic repeat-containing protein \| Acidic repeat-containing protein; uniclust \| UniRef100\_A0A315CPA0 \| 98.7 \| 2.3e-10 \| 4.3e-16 \| 102.2 \| 132 \| (3, 134) \| 227 \| (217, 352) \| 357 \| Bacteriophage T5 Orf172 DNA-binding domain-containing protein \| Bacteriophage T5 Orf172 DNA-binding domain-containing protein; uniclust \| UniRef100\_A0A1I7TER3 \| 98.7 \| 2.2e-10 \| 4.5e-16 \| 109.7 \| 92 \| (2, 102) \| 227 \| (294, 392) \| 539 \| SprT-like domain-containing protein \| SprT-like domain-containing protein; uniclust \| UniRef100\_A0A2W0DS13 \| 98.7 \| 2.2e-10 \| 4.5e-16 \| 89.0 \| 70 \| (29, 107) \| 227 \| (22, 94) \| 102 \| M48 family peptidase (Fragment) \| M48 family peptidase (Fragment); uniclust \| UniRef100\_A0A517SCM0 \| 98.7 \| 2.4e-10 \| 4.5e-16 \| 98.5 \| 180 \| (5, 224) \| 227 \| (6, 216) \| 229 \| SprT-like family protein \| SprT-like family protein; uniclust \| UniRef100\_A0A374MXY9 \| 98.7 \| 2.5e-10 \| 4.7e-16 \| 88.5 \| 95 \| (5, 103) \| 227 \| (8, 113) \| 113 \| SprT-like family protein \| SprT-like family protein; uniclust \| UniRef100\_A0A0K2UAN4 \| 98.6 \| 2.4e-10 \| 4.8e-16 \| 113.5 \| 147 \| (10, 220) \| 227 \| (571, 728) \| 791 \| Putative LOC100745142 [Bombus impatiens] \| Putative LOC100745142 [Bombus impatiens]; uniclust \| UniRef100\_A0A087Y402 \| 98.6 \| 2.2e-10 \| 4.8e-16 \| 115.8 \| 134 \| (23, 220) \| 227 \| (470, 610) \| 734 \| Acidic repeat-containing protein-like \| Acidic repeat-containing protein-like; uniclust \| UniRef100\_UPI001E52AEAC \| 98.6 \| 2.7e-10 \| 5e-16 \| 92.3 \| 132 \| (61, 219) \| 227 \| (2, 153) \| 155 \| hypothetical protein \| hypothetical protein; uniclust \| UniRef100\_UPI001A9BFEBE \| 98.6 \| 2.6e-10 \| 5e-16 \| 105.5 \| 100 \| (10, 114) \| 227 \| (142, 258) \| 398 \| SprT-like domain-containing protein \| SprT-like domain-containing protein; uniclust \| UniRef100\_A0A2A9EM23 \| 98.6 \| 2.9e-10 \| 5.7e-16 \| 105.1 \| 74 \| (29, 111) \| 227 \| (93, 169) \| 383 \| SprT-like family protein \| SprT-like family protein; uniclust \| UniRef100\_A0A1S7NI76 \| 98.6 \| 3.2e-10 \| 6.2e-16 \| 92.5 \| 125 \| (80, 224) \| 227 \| (1, 131) \| 153 \| Uncharacterized protein \| Uncharacterized protein; uniclust \| UniRef100\_A0A960IJM6 \| 98.6 \| 3.1e-10 \| 6.2e-16 \| 96.2 \| 77 \| (28, 112) \| 227 \| (50, 128) \| 191 \| SprT-like domain-containing protein \| SprT-like domain-containing protein; uniclust \| UniRef100\_A0A2E7W4W3 \| 98.6 \| 4.5e-10 \| 8.2e-16 \| 96.2 \| 139 \| (42, 217) \| 227 \| (77, 232) \| 244 \| SprT-like domain-containing protein \| SprT-like domain-containing protein; uniclust \| UniRef100\_A0A430JN74 \| 98.6 \| 4.7e-10 \| 9.3e-16 \| 90.7 \| 82 \| (14, 108) \| 227 \| (10, 94) \| 136 \| SprT domain-containing protein \| SprT domain-containing protein; uniclust \| UniRef100\_A0A0D8XAM0 \| 98.6 \| 5.5e-10 \| 1e-15 \| 101.3 \| 140 \| (11, 219) \| 227 \| (214, 363) \| 403 \| SprT-like domain-containing protein \| SprT-like domain-containing protein; uniclust \| UniRef100\_A0A076GXD3 \| 98.6 \| 4.9e-10 \| 1e-15 \| 98.1 \| 90 \| (14, 104) \| 227 \| (28, 133) \| 224 \| SprT-like domain-containing protein \| SprT-like domain-containing protein; uniclust \| UniRef100\_A0A2D3TGT7 \| 98.6 \| 5.6e-10 \| 1e-15 \| 90.7 \| 47 \| (68, 114) \| 227 \| (10, 62) \| 154 \| SprT-like domain-containing protein \| SprT-like domain-containing protein; uniclust \| UniRef100\_A0A0S7MAB5 \| 98.6 \| 4.7e-10 \| 1.1e-15 \| 107.4 \| 97 \| (2, 104) \| 227 \| (149, 254) \| 404 \| ACRC (Fragment) \| ACRC (Fragment); uniclust \| UniRef100\_A0A2E4EZM8 \| 98.6 \| 5.4e-10 \| 1.2e-15 \| 96.6 \| 85 \| (14, 111) \| 227 \| (20, 107) \| 192 \| SprT-like domain-containing protein \| SprT-like domain-containing protein; uniclust \| UniRef100\_A0A1E4ZTF4 \| 98.6 \| 5.8e-10 \| 1.2e-15 \| 100.4 \| 88 \| (17, 110) \| 227 \| (13, 109) \| 297 \| SprT-like domain-containing protein \| SprT-like domain-containing protein; uniclust \| UniRef100\_A0A3B8MXE4 \| 98.6 \| 7.3e-10 \| 1.3e-15 \| 85.0 \| 74 \| (29, 111) \| 227 \| (23, 99) \| 112 \| SprT domain-containing protein (Fragment) \| SprT domain-containing protein (Fragment); uniclust \| UniRef100\_A0A6L8C1X2 \| 98.5 \| 7.6e-10 \| 1.5e-15 \| 95.8 \| 93 \| (5, 110) \| 227 \| (135, 230) \| 235 \| SprT-like domain-containing protein \| SprT-like domain-containing protein; uniclust \| UniRef100\_UPI0013AEF1FA \| 98.5 \| 9e-10 \| 1.7e-15 \| 85.0 \| 66 \| (42, 107) \| 227 \| (11, 88) \| 116 \| SprT-like domain-containing protein \| SprT-like domain-containing protein; uniclust \| UniRef100\_K1PVW6 \| 98.5 \| 8.7e-10 \| 1.7e-15 \| 95.3 \| 135 \| (23, 221) \| 227 \| (27, 169) \| 227 \| Acidic repeat-containing protein \| Acidic repeat-containing protein; uniclust \| UniRef100\_UPI00156DE97E \| 98.5 \| 1e-09 \| 1.9e-15 \| 77.6 \| 63 \| (2, 64) \| 227 \| (5, 67) \| 70 \| hypothetical protein \| hypothetical protein; uniclust \| UniRef100\_A0A0C2QPB3 \| 98.5 \| 1.1e-09 \| 2.2e-15 \| 96.2 \| 171 \| (4, 224) \| 227 \| (30, 222) \| 229 \| SprT family zinc-dependent metalloprotease \| SprT family zinc-dependent metalloprotease; uniclust \| UniRef100\_A0A410T805 \| 98.5 \| 1.2e-09 \| 2.4e-15 \| 83.8 \| 68 \| (28, 104) \| 227 \| (29, 99) \| 103 \| SprT-like domain-containing protein \| SprT-like domain-containing protein; uniclust \| UniRef100\_A0A2E9YH35 \| 98.5 \| 1.4e-09 \| 2.6e-15 \| 84.5 \| 85 \| (17, 101) \| 227 \| (24, 121) \| 121 \| SprT-like domain-containing protein (Fragment) \| SprT-like domain-containing protein (Fragment); uniclust \| UniRef100\_A0A6I3DIT0 \| 98.5 \| 1.6e-09 \| 2.9e-15 \| 80.7 \| 70 \| (30, 108) \| 227 \| (10, 82) \| 93 \| M48 family peptidase (Fragment) \| M48 family peptidase (Fragment); uniclust \| UniRef100\_A0A0A0Q2I3 \| 98.5 \| 1.4e-09 \| 2.9e-15 \| 95.8 \| 59 \| (42, 107) \| 227 \| (58, 119) \| 227 \| SprT-like domain-containing protein \| SprT-like domain-containing protein; uniclust \| UniRef100\_A0A010SRF5 \| 98.5 \| 1.3e-09 \| 3.2e-15 \| 100.1 \| 79 \| (23, 112) \| 227 \| (86, 173) \| 266 \| Protein SprT \| Protein SprT; uniclust \| UniRef100\_UPI001F3D9D2C \| 98.5 \| 1.8e-09 \| 3.3e-15 \| 80.7 \| 45 \| (175, 219) \| 227 \| (50, 94) \| 95 \| hypothetical protein \| hypothetical protein; uniclust \| UniRef100\_UPI00201D51DF \| 98.5 \| 1.8e-09 \| 3.3e-15 \| 83.9 \| 91 \| (7, 105) \| 227 \| (17, 120) \| 120 \| SprT-like domain-containing protein \| SprT-like domain-containing protein; uniclust \| UniRef100\_A0A368G165 \| 98.5 \| 2e-09 \| 3.6e-15 \| 96.3 \| 144 \| (10, 220) \| 227 \| (115, 266) \| 342 \| SprT-like domain-containing protein \| SprT-like domain-containing protein; uniclust \| UniRef100\_A0A096MLG6 \| 98.4 \| 2.1e-09 \| 4.3e-15 \| 107.8 \| 99 \| (2, 106) \| 227 \| (580, 687) \| 810 \| Germ cell nuclear acidic peptidase \| Germ cell nuclear acidic peptidase; uniclust \| UniRef100\_A0A139AMY7 \| 98.4 \| 2.2e-09 \| 4.6e-15 \| 93.5 \| 77 \| (22, 101) \| 227 \| (28, 114) \| 200 \| SprT-like domain-containing protein (Fragment) \| SprT-like domain-containing protein (Fragment); uniclust \| UniRef100\_A0A1Q3UR33 \| 98.4 \| 2.9e-09 \| 5.3e-15 \| 91.1 \| 186 \| (4, 223) \| 227 \| (12, 211) \| 234 \| Uncharacterized protein \| Uncharacterized protein; uniclust \| UniRef100\_A0A3C0P523 \| 98.4 \| 3e-09 \| 5.6e-15 \| 78.7 \| 86 \| (101, 222) \| 227 \| (1, 86) \| 89 \| Uncharacterized protein (Fragment) \| Uncharacterized protein (Fragment); uniclust \| UniRef100\_A0A0S7Z910 \| 98.4 \| 2.7e-09 \| 5.8e-15 \| 97.8 \| 84 \| (23, 110) \| 227 \| (146, 237) \| 286 \| SprT-like domain-containing protein \| SprT-like domain-containing protein; uniclust \| UniRef100\_A7RFH1 \| 98.4 \| 3.8e-09 \| 7.3e-15 \| 89.9 \| 79 \| (24, 105) \| 227 \| (41, 123) \| 206 \| SprT-like domain-containing protein (Fragment) \| SprT-like domain-containing protein (Fragment); uniclust \| UniRef100\_A0A0K8SMS5 \| 98.4 \| 3.9e-09 \| 7.8e-15 \| 93.0 \| 132 \| (23, 220) \| 227 \| (11, 154) \| 235 \| SprT-like domain-containing protein \| SprT-like domain-containing protein; uniclust \| UniRef100\_UPI001F36BFBF \| 98.4 \| 4.6e-09 \| 8.5e-15 \| 90.1 \| 53 \| (62, 114) \| 227 \| (2, 63) \| 239 \| SprT-like domain-containing protein \| SprT-like domain-containing protein; uniclust \| UniRef100\_UPI001FE9CB55 \| 98.4 \| 4.7e-09 \| 8.6e-15 \| 87.6 \| 84 \| (29, 121) \| 227 \| (19, 105) \| 193 \| SprT-like domain-containing protein \| SprT-like domain-containing protein; uniclust \| UniRef100\_A0A962Q7G0 \| 98.4 \| 4.8e-09 \| 8.8e-15 \| 88.9 \| 131 \| (56, 219) \| 227 \| (62, 214) \| 217 \| SprT-like domain-containing protein \| SprT-like domain-containing protein; uniclust \| UniRef100\_UPI001364B730 \| 98.4 \| 5.4e-09 \| 9.8e-15 \| 86.4 \| 140 \| (43, 220) \| 227 \| (21, 177) \| 179 \| SprT family zinc-dependent metalloprotease \| SprT family zinc-dependent metalloprotease; uniclust \| UniRef100\_A0A6N9B6I9 \| 98.4 \| 5.4e-09 \| 9.9e-15 \| 76.0 \| 63 \| (42, 110) \| 227 \| (11, 76) \| 80 \| SprT domain-containing protein (Fragment) \| SprT domain-containing protein (Fragment); uniclust \| UniRef100\_A0A0F9F6D4 \| 98.4 \| 5.7e-09 \| 1e-14 \| 87.1 \| 130 \| (2, 134) \| 227 \| (24, 163) \| 192 \| SprT-like domain-containing protein (Fragment) \| SprT-like domain-containing protein (Fragment); uniclust \| UniRef100\_UPI00055C0F9E \| 98.4 \| 5.7e-09 \| 1.1e-14 \| 78.7 \| 65 \| (1, 66) \| 227 \| (17, 81) \| 99 \| hypothetical protein \| hypothetical protein; uniclust \| UniRef100\_A0A2B4RRR3 \| 98.3 \| 5.5e-09 \| 1.1e-14 \| 104.6 \| 77 \| (23, 102) \| 227 \| (691, 772) \| 883 \| HMG box-containing protein C19G7.04 \| HMG box-containing protein C19G7.04; uniclust \| UniRef100\_A0A225DJ34 \| 98.3 \| 5.8e-09 \| 1.1e-14 \| 92.9 \| 163 \| (23, 220) \| 227 \| (106, 287) \| 288 \| SprT-like domain-containing protein \| SprT-like domain-containing protein; uniclust \| UniRef100\_UPI0006B0C5A3 \| 98.3 \| 6e-09 \| 1.2e-14 \| 104.6 \| 134 \| (22, 220) \| 227 \| (764, 908) \| 972 \| mucin-4-like \| mucin-4-like; uniclust \| UniRef100\_A0A0F9MGL9 \| 98.3 \| 6.3e-09 \| 1.2e-14 \| 88.5 \| 148 \| (28, 219) \| 227 \| (50, 220) \| 222 \| SprT-like domain-containing protein \| SprT-like domain-containing protein; uniclust \| UniRef100\_A0A369RWZ3 \| 98.3 \| 6.1e-09 \| 1.2e-14 \| 102.6 \| 155 \| (2, 223) \| 227 \| (495, 655) \| 707 \| Acidic repeat-containing protein \| Acidic repeat-containing protein; uniclust \| UniRef100\_T2M7F9 \| 98.3 \| 6.2e-09 \| 1.2e-14 \| 102.3 \| 95 \| (3, 103) \| 227 \| (422, 522) \| 643 \| Acidic repeat-containing protein \| Acidic repeat-containing protein; uniclust \| UniRef100\_A0A063ZKH3 \| 98.3 \| 5.7e-09 \| 1.3e-14 \| 94.1 \| 97 \| (3, 105) \| 227 \| (60, 164) \| 238 \| SprT-like domain-containing protein \| SprT-like domain-containing protein; uniclust \| UniRef100\_A0A1H6LC21 \| 98.3 \| 6.6e-09 \| 1.3e-14 \| 89.6 \| 74 \| (28, 110) \| 227 \| (52, 134) \| 207 \| Sprt-like family \| Sprt-like family; uniclust \| UniRef100\_A0A7M2WWZ9 \| 98.3 \| 6.9e-09 \| 1.4e-14 \| 89.2 \| 75 \| (28, 112) \| 227 \| (34, 111) \| 193 \| SprT-like domain-containing protein \| SprT-like domain-containing protein; uniclust \| UniRef100\_A0A0N7ZDR3 \| 98.3 \| 6.7e-09 \| 1.4e-14 \| 109.9 \| 134 \| (23, 220) \| 227 \| (1151, 1292) \| 1372 \| SprT-like domain-containing protein \| SprT-like domain-containing protein; uniclust \| UniRef100\_A0A8F4NUB2 \| 98.3 \| 7.4e-09 \| 1.4e-14 \| 83.9 \| 56 \| (68, 123) \| 227 \| (34, 95) \| 141 \| Zinc metalloproteinase Mpr protein \| Zinc metalloproteinase Mpr protein; uniclust \| UniRef100\_A0A9D4QSU8 \| 98.3 \| 7.5e-09 \| 1.5e-14 \| 107.3 \| 134 \| (23, 220) \| 227 \| (1093, 1233) \| 1297 \| Uncharacterized protein \| Uncharacterized protein; uniclust \| UniRef100\_UPI0011194AF0 \| 98.3 \| 8.3e-09 \| 1.5e-14 \| 78.7 \| 76 \| (41, 116) \| 227 \| (5, 93) \| 103 \| SprT-like domain-containing protein \| SprT-like domain-containing protein; uniclust \| UniRef100\_A0A2E0ELX0 \| 98.3 \| 8.6e-09 \| 1.7e-14 \| 87.9 \| 73 \| (29, 110) \| 227 \| (46, 121) \| 186 \| SprT-like domain-containing protein \| SprT-like domain-containing protein; uniclust \| UniRef100\_UPI001C69A0ED \| 98.3 \| 9.9e-09 \| 1.8e-14 \| 81.1 \| 74 \| (42, 115) \| 227 \| (15, 99) \| 131 \| SprT-like domain-containing protein \| SprT-like domain-containing protein; uniclust \| UniRef100\_A0A386ZLW0 \| 98.3 \| 1e-08 \| 2e-14 \| 77.3 \| 46 \| (178, 224) \| 227 \| (38, 83) \| 90 \| Uncharacterized protein \| Uncharacterized protein; uniclust \| UniRef100\_A0A6A7FXS4 \| 98.3 \| 1.1e-08 \| 2.2e-14 \| 105.0 \| 152 \| (2, 220) \| 227 \| (1148, 1308) \| 1364 \| Acidic repeat-containing protein-like \| Acidic repeat-containing protein-like; uniclust \| UniRef100\_A0A3E4PBP9 \| 98.3 \| 1.1e-08 \| 2.2e-14 \| 88.7 \| 123 \| (55, 219) \| 227 \| (49, 187) \| 193 \| SprT-like family protein \| SprT-like family protein; uniclust \| UniRef100\_A0A444UUE3 \| 98.3 \| 1.1e-08 \| 2.2e-14 \| 90.2 \| 81 \| (22, 105) \| 227 \| (25, 113) \| 225 \| Acidic repeat-containing protein \| Acidic repeat-containing protein; uniclust \| UniRef100\_A0A942C240 \| 98.3 \| 1.3e-08 \| 2.4e-14 \| 88.2 \| 73 \| (29, 110) \| 227 \| (21, 96) \| 254 \| SprT-like domain-containing protein \| SprT-like domain-containing protein; uniclust \| UniRef100\_A0A090L5F7 \| 98.3 \| 1.3e-08 \| 2.5e-14 \| 97.9 \| 130 \| (23, 219) \| 227 \| (328, 463) \| 495 \| SprT-like domain-containing protein \| SprT-like domain-containing protein; uniclust \| UniRef100\_UPI0023581BAD \| 98.3 \| 1.4e-08 \| 2.6e-14 \| 84.4 \| 58 \| (57, 114) \| 227 \| (27, 93) \| 183 \| SprT-like domain-containing protein \| SprT-like domain-containing protein; uniclust \| UniRef100\_A0A3D4EUY1 \| 98.3 \| 1.4e-08 \| 2.6e-14 \| 79.8 \| 50 \| (59, 108) \| 227 \| (6, 63) \| 126 \| SprT-like domain-containing protein (Fragment) \| SprT-like domain-containing protein (Fragment); uniclust \| UniRef100\_UPI00190F8D28 \| 98.3 \| 1.4e-08 \| 2.6e-14 \| 81.5 \| 64 \| (57, 120) \| 227 \| (5, 77) \| 133 \| SprT-like domain-containing protein \| SprT-like domain-containing protein; uniclust \| UniRef100\_A0A0R3T453 \| 98.3 \| 1.4e-08 \| 2.6e-14 \| 102.6 \| 153 \| (1, 219) \| 227 \| (391, 557) \| 1025 \| Lipoyl synthase, mitochondrial \| Lipoyl synthase, mitochondrial; uniclust \| UniRef100\_A0A1Y6FUX7 \| 98.2 \| 1.5e-08 \| 2.7e-14 \| 72.0 \| 50 \| (1, 50) \| 227 \| (4, 54) \| 69 \| Uncharacterized protein \| Uncharacterized protein; uniclust \| UniRef100\_A0A7Y4SVY5 \| 98.2 \| 1.6e-08 \| 2.9e-14 \| 86.7 \| 168 \| (23, 222) \| 227 \| (29, 226) \| 230 \| SprT-like domain-containing protein \| SprT-like domain-containing protein; uniclust \| UniRef100\_A0A2W6D4Y0 \| 98.2 \| 1.8e-08 \| 3.2e-14 \| 84.3 \| 90 \| (18, 107) \| 227 \| (40, 147) \| 191 \| SprT-like domain-containing protein (Fragment) \| SprT-like domain-containing protein (Fragment); uniclust \| UniRef100\_A0A011NB53 \| 98.2 \| 1.5e-08 \| 3.5e-14 \| 92.7 \| 74 \| (30, 112) \| 227 \| (98, 179) \| 256 \| Protein SprT \| Protein SprT; uniclust \| UniRef100\_A0A523HVA6 \| 98.2 \| 1.9e-08 \| 3.5e-14 \| 89.6 \| 87 \| (23, 121) \| 227 \| (171, 265) \| 317 \| DUF45 domain-containing protein \| DUF45 domain-containing protein; uniclust \| UniRef100\_A0A2A5DH80 \| 98.2 \| 1.7e-08 \| 3.5e-14 \| 85.5 \| 81 \| (28, 116) \| 227 \| (44, 131) \| 160 \| SprT-like domain-containing protein \| SprT-like domain-containing protein; uniclust \| UniRef100\_A0A4Z0WHF5 \| 98.2 \| 2e-08 \| 3.7e-14 \| 84.0 \| 78 \| (24, 112) \| 227 \| (39, 119) \| 191 \| M48 family peptidase \| M48 family peptidase; uniclust \| UniRef100\_A0A4Y7LNE8 \| 98.2 \| 1.8e-08 \| 3.7e-14 \| 99.8 \| 94 \| (2, 101) \| 227 \| (366, 469) \| 593 \| EOG090X0464 \| EOG090X0464; uniclust \| UniRef100\_A0A060YC73 \| 98.2 \| 1.9e-08 \| 3.8e-14 \| 98.0 \| 80 \| (23, 105) \| 227 \| (349, 435) \| 540 \| SprT-like domain-containing protein \| SprT-like domain-containing protein; uniclust \| UniRef100\_A0A519H8W8 \| 98.2 \| 2.3e-08 \| 4.2e-14 \| 72.8 \| 65 \| (68, 138) \| 227 \| (12, 76) \| 79 \| SprT-like domain-containing protein \| SprT-like domain-containing protein; uniclust \| UniRef100\_A0A091CV90 \| 98.2 \| 2.1e-08 \| 4.3e-14 \| 99.4 \| 99 \| (2, 106) \| 227 \| (397, 504) \| 658 \| Acidic repeat-containing protein \| Acidic repeat-containing protein; uniclust \| UniRef100\_A0A0F9VUA0 \| 98.2 \| 2.3e-08 \| 4.3e-14 \| 91.3 \| 145 \| (37, 222) \| 227 \| (41, 208) \| 396 \| SprT-like domain-containing protein \| SprT-like domain-containing protein; uniclust \| UniRef100\_A0A965PII6 \| 98.2 \| 2.5e-08 \| 4.6e-14 \| 85.4 \| 97 \| (10, 108) \| 227 \| (9, 121) \| 228 \| SprT-like domain-containing protein \| SprT-like domain-containing protein; uniclust \| UniRef100\_A0A1B6FR93 \| 98.2 \| 2.4e-08 \| 4.7e-14 \| 89.8 \| 132 \| (23, 219) \| 227 \| (76, 217) \| 292 \| SprT-like domain-containing protein \| SprT-like domain-containing protein; uniclust \| UniRef100\_A0A158QQ23 \| 98.2 \| 2.8e-08 \| 5.2e-14 \| 88.5 \| 131 \| (23, 220) \| 227 \| (90, 226) \| 314 \| SprT-like domain-containing protein \| SprT-like domain-containing protein; uniclust \| UniRef100\_A0A0P7TZE5 \| 98.2 \| 2.6e-08 \| 5.6e-14 \| 100.2 \| 98 \| (2, 105) \| 227 \| (438, 543) \| 664 \| Acidic repeat containing-like (Fragment) \| Acidic repeat containing-like (Fragment); uniclust \| UniRef100\_A0A2W4S5I9 \| 98.2 \| 3.1e-08 \| 5.7e-14 \| 88.3 \| 102 \| (5, 111) \| 227 \| (120, 228) \| 314 \| SprT-like domain-containing protein \| SprT-like domain-containing protein; uniclust \| UniRef100\_A0A0C9VRQ7 \| 98.2 \| 2.9e-08 \| 5.9e-14 \| 96.9 \| 95 \| (2, 102) \| 227 \| (290, 392) \| 522 \| SprT-like domain-containing protein \| SprT-like domain-containing protein; uniclust \| UniRef100\_A0A1Y3SUR1 \| 98.1 \| 3.1e-08 \| 6.7e-14 \| 81.6 \| 94 \| (5, 102) \| 227 \| (10, 103) \| 129 \| Peptidase M48 domain-containing protein \| Peptidase M48 domain-containing protein; uniclust \| UniRef100\_A0A1W9WRE7 \| 98.1 \| 3.2e-08 \| 6.7e-14 \| 84.0 \| 94 \| (6, 110) \| 227 \| (27, 129) \| 159 \| SprT-like domain-containing protein (Fragment) \| SprT-like domain-containing protein (Fragment); uniclust \| UniRef100\_A0A0S2W4E2 \| 98.1 \| 3.7e-08 \| 6.7e-14 \| 71.5 \| 44 \| (177, 224) \| 227 \| (21, 64) \| 77 \| Uncharacterized protein \| Uncharacterized protein; uniclust \| UniRef100\_A0A8S1EWA4 \| 98.1 \| 3.9e-08 \| 7.2e-14 \| 90.3 \| 75 \| (23, 102) \| 227 \| (239, 319) \| 411 \| SprT-like domain-containing protein \| SprT-like domain-containing protein; uniclust \| UniRef100\_A0A7V9QTE1 \| 98.1 \| 4.2e-08 \| 7.7e-14 \| 82.8 \| 73 \| (29, 110) \| 227 \| (8, 83) \| 201 \| SprT-like domain-containing protein \| SprT-like domain-containing protein; uniclust \| UniRef100\_A0A815EWT2 \| 98.1 \| 4.1e-08 \| 7.9e-14 \| 91.4 \| 144 \| (10, 220) \| 227 \| (174, 326) \| 398 \| SprT-like domain-containing protein \| SprT-like domain-containing protein; uniclust \| UniRef100\_A0A967HJD9 \| 98.1 \| 4.4e-08 \| 8.1e-14 \| 83.2 \| 92 \| (13, 108) \| 227 \| (97, 200) \| 211 \| SprT family zinc-dependent metalloprotease \| SprT family zinc-dependent metalloprotease; uniclust \| UniRef100\_A0A017SUS6 \| 98.1 \| 3.5e-08 \| 8.8e-14 \| 97.8 \| 96 \| (4, 110) \| 227 \| (82, 186) \| 456 \| DUF2786 domain-containing protein \| DUF2786 domain-containing protein; uniclust \| UniRef100\_A0A6I3FXI3 \| 98.1 \| 4.9e-08 \| 9e-14 \| 85.3 \| 95 \| (29, 138) \| 227 \| (136, 233) \| 263 \| SprT-like domain-containing protein \| SprT-like domain-containing protein; uniclust \| UniRef100\_A0A4X2L320 \| 98.1 \| 4.7e-08 \| 9.1e-14 \| 91.4 \| 82 \| (22, 106) \| 227 \| (191, 281) \| 401 \| SprT-like domain-containing protein \| SprT-like domain-containing protein; uniclust \| UniRef100\_UPI001BEB5868 \| 98.1 \| 4.9e-08 \| 9.2e-14 \| 78.7 \| 79 \| (7, 88) \| 227 \| (21, 111) \| 136 \| SprT-like domain-containing protein \| SprT-like domain-containing protein; uniclust \| UniRef100\_A0A2V5M5G1 \| 98.1 \| 4.9e-08 \| 9.3e-14 \| 86.1 \| 86 \| (17, 110) \| 227 \| (85, 178) \| 256 \| SprT-like domain-containing protein \| SprT-like domain-containing protein; uniclust \| UniRef100\_A0A084VEQ8 \| 98.1 \| 4.4e-08 \| 9.7e-14 \| 100.9 \| 78 \| (24, 105) \| 227 \| (139, 225) \| 772 \| Protein with SprT-like domain at the N terminus \| Protein with SprT-like domain at the N terminus; uniclust \| UniRef100\_A0A061D4E3 \| 98.1 \| 4.8e-08 \| 9.8e-14 \| 87.6 \| 86 \| (14, 105) \| 227 \| (52, 151) \| 248 \| SPRT domain containg protein \| SPRT domain containg protein; uniclust \| UniRef100\_A0A1Z8XXQ1 \| 98.1 \| 5.3e-08 \| 1e-13 \| 85.5 \| 72 \| (29, 108) \| 227 \| (89, 168) \| 243 \| SprT-like domain-containing protein \| SprT-like domain-containing protein; uniclust \| UniRef100\_A0A813R6P1 \| 98.1 \| 5.4e-08 \| 1.1e-13 \| 99.1 \| 131 \| (23, 220) \| 227 \| (758, 895) \| 965 \| SprT-like domain-containing protein \| SprT-like domain-containing protein; uniclust \| UniRef100\_A0A060S9M2 \| 98.1 \| 5.1e-08 \| 1.1e-13 \| 99.1 \| 96 \| (4, 102) \| 227 \| (390, 497) \| 663 \| SprT-like domain-containing protein \| SprT-like domain-containing protein; uniclust \| UniRef100\_A0A0N4YL49 \| 98.1 \| 6.1e-08 \| 1.2e-13 \| 82.5 \| 89 \| (69, 220) \| 227 \| (9, 104) \| 191 \| Putative nuclear protein (inferred by orthology to a S. mansoni protein) \| Putative nuclear protein (inferred by orthology to a S. mansoni protein); uniclust \| UniRef100\_A0A932U3F1 \| 98.1 \| 6.5e-08 \| 1.2e-13 \| 80.1 \| 64 \| (42, 111) \| 227 \| (41, 110) \| 174 \| SprT-like domain-containing protein \| SprT-like domain-containing protein; uniclust \| UniRef100\_A0A2E4W6R4 \| 98.1 \| 6.3e-08 \| 1.2e-13 \| 82.8 \| 93 \| (8, 111) \| 227 \| (12, 111) \| 195 \| SprT-like domain-containing protein \| SprT-like domain-containing protein; uniclust \| UniRef100\_A0A564Y0D1 \| 98.1 \| 6.4e-08 \| 1.2e-13 \| 91.6 \| 97 \| (1, 102) \| 227 \| (240, 350) \| 481 \| SprT-like domain-containing protein \| SprT-like domain-containing protein; uniclust \| UniRef100\_A0A8J3FY24 \| 98.1 \| 6.9e-08 \| 1.3e-13 \| 82.7 \| 171 \| (15, 221) \| 227 \| (18, 210) \| 222 \| SprT-like domain-containing protein \| SprT-like domain-containing protein; uniclust \| UniRef100\_UPI0020C4C488 \| 98.1 \| 6.9e-08 \| 1.3e-13 \| 80.1 \| 135 \| (58, 223) \| 227 \| (15, 167) \| 176 \| hypothetical protein \| hypothetical protein; uniclust \| UniRef100\_A0A6A0HDZ2 \| 98.1 \| 6.7e-08 \| 1.3e-13 \| 95.7 \| 134 \| (23, 221) \| 227 \| (520, 662) \| 740 \| SprT-like domain-containing protein \| SprT-like domain-containing protein; uniclust \| UniRef100\_UPI0002659249 \| 98.1 \| 7.5e-08 \| 1.4e-13 \| 86.5 \| 71 \| (29, 101) \| 227 \| (140, 216) \| 331 \| acidic repeat-containing protein \| acidic repeat-containing protein; uniclust \| UniRef100\_A0A147E8E1 \| 98.1 \| 7.4e-08 \| 1.4e-13 \| 73.3 \| 63 \| (26, 97) \| 227 \| (29, 94) \| 95 \| SprT-like domain-containing protein (Fragment) \| SprT-like domain-containing protein (Fragment); uniclust \| UniRef100\_A0A8R1U967 \| 98.0 \| 7.8e-08 \| 1.4e-13 \| 93.7 \| 135 \| (23, 223) \| 227 \| (518, 658) \| 776 \| SprT-like domain-containing protein \| SprT-like domain-containing protein; uniclust \| UniRef100\_A0A067RTN9 \| 98.0 \| 6e-08 \| 1.4e-13 \| 100.5 \| 77 \| (23, 105) \| 227 \| (145, 230) \| 721 \| Protein with SprT-like domain at the N terminus \| Protein with SprT-like domain at the N terminus; uniclust \| UniRef100\_A0A131Z5L9 \| 98.0 \| 6.4e-08 \| 1.5e-13 \| 98.6 \| 91 \| (8, 104) \| 227 \| (113, 217) \| 646 \| Protein with SprT-like domain at the N terminus \| Protein with SprT-like domain at the N terminus; uniclust \| UniRef100\_A0A067R6Y3 \| 98.0 \| 8e-08 \| 1.5e-13 \| 95.5 \| 134 \| (23, 220) \| 227 \| (665, 808) \| 875 \| Acidic repeat-containing protein \| Acidic repeat-containing protein; uniclust \| UniRef100\_UPI001D0C490B \| 98.0 \| 8.4e-08 \| 1.5e-13 \| 75.4 \| 83 \| (42, 124) \| 227 \| (9, 105) \| 122 \| SprT-like domain-containing protein \| SprT-like domain-containing protein; uniclust \| UniRef100\_A0A3P6QM62 \| 98.0 \| 8.5e-08 \| 1.6e-13 \| 77.1 \| 89 \| (8, 102) \| 227 \| (14, 109) \| 141 \| SprT-like domain-containing protein \| SprT-like domain-containing protein; uniclust \| UniRef100\_A0A2V2LDH4 \| 98.0 \| 8.6e-08 \| 1.6e-13 \| 92.6 \| 101 \| (10, 115) \| 227 \| (527, 644) \| 698 \| SprT-like domain-containing protein \| SprT-like domain-containing protein; uniclust \| UniRef100\_A0A947HSE2 \| 98.0 \| 8.8e-08 \| 1.6e-13 \| 80.6 \| 115 \| (12, 130) \| 227 \| (14, 150) \| 194 \| Uncharacterized protein (Fragment) \| Uncharacterized protein (Fragment); uniclust \| UniRef100\_A0A660QXE7 \| 98.0 \| 8.5e-08 \| 1.6e-13 \| 73.3 \| 43 \| (69, 111) \| 227 \| (4, 46) \| 95 \| SprT domain-containing protein (Fragment) \| SprT domain-containing protein (Fragment); uniclust \| UniRef100\_A0A8B8D7V9 \| 98.0 \| 8.9e-08 \| 1.6e-13 \| 89.1 \| 135 \| (23, 221) \| 227 \| (259, 401) \| 461 \| HMG box-containing protein C19G7.04-like \| HMG box-containing protein C19G7.04-like; uniclust \| UniRef100\_UPI00186B3928 \| 98.0 \| 9.2e-08 \| 1.7e-13 \| 85.6 \| 83 \| (22, 107) \| 227 \| (129, 216) \| 318 \| acidic repeat-containing protein \| acidic repeat-containing protein; uniclust \| UniRef100\_A0A8R1DRL3 \| 98.0 \| 1e-07 \| 1.9e-13 \| 95.6 \| 92 \| (1, 101) \| 227 \| (927, 1025) \| 1163 \| Tr-type G domain-containing protein \| Tr-type G domain-containing protein; uniclust \| UniRef100\_A0A250G0W3 \| 98.0 \| 1.1e-07 \| 2e-13 \| 76.9 \| 59 \| (42, 106) \| 227 \| (62, 126) \| 132 \| SprT-like domain-containing protein \| SprT-like domain-containing protein; uniclust \| UniRef100\_A0A0N4U7B3 \| 98.0 \| 1.1e-07 \| 2.1e-13 \| 85.7 \| 127 \| (29, 221) \| 227 \| (104, 236) \| 302 \| SprT-like domain-containing protein \| SprT-like domain-containing protein; uniclust \| UniRef100\_A0A1S3GEE3 \| 98.0 \| 1.1e-07 \| 2.3e-13 \| 87.9 \| 96 \| (5, 106) \| 227 \| (94, 198) \| 311 \| Acidic repeat-containing protein \| Acidic repeat-containing protein; uniclust \| UniRef100\_A0A069QD30 \| 98.0 \| 9.6e-08 \| 2.3e-13 \| 88.9 \| 90 \| (12, 106) \| 227 \| (41, 147) \| 272 \| SprT-like domain-containing protein \| SprT-like domain-containing protein; uniclust \| UniRef100\_A0A8T0DKP9 \| 98.0 \| 1.3e-07 \| 2.3e-13 \| 85.1 \| 78 \| (22, 102) \| 227 \| (74, 165) \| 329 \| SprT-like domain-containing protein \| SprT-like domain-containing protein; uniclust \| UniRef100\_UPI0008F98D6C \| 98.0 \| 1.3e-07 \| 2.4e-13 \| 95.3 \| 76 \| (23, 101) \| 227 \| (1008, 1093) \| 1216 \| uncharacterized protein LOC109037607 \| uncharacterized protein LOC109037607; uniclust \| UniRef100\_A0A813SQV5 \| 98.0 \| 1.2e-07 \| 2.4e-13 \| 91.8 \| 87 \| (9, 100) \| 227 \| (341, 435) \| 566 \| SprT-like domain-containing protein \| SprT-like domain-containing protein; uniclust \| UniRef100\_A0A210QVD4 \| 98.0 \| 1.1e-07 \| 2.5e-13 \| 97.4 \| 86 \| (13, 104) \| 227 \| (67, 166) \| 721 \| SprT-like domain-containing protein Spartan \| SprT-like domain-containing protein Spartan; uniclust \| UniRef100\_A0A2V0P5I1 \| 98.0 \| 1.2e-07 \| 2.6e-13 \| 94.6 \| 71 \| (29, 103) \| 227 \| (118, 196) \| 569 \| Protein with SprT-like domain at the N terminus \| Protein with SprT-like domain at the N terminus; uniclust \| UniRef100\_UPI001E46D32E \| 98.0 \| 1.4e-07 \| 2.7e-13 \| 71.9 \| 73 \| (7, 86) \| 227 \| (14, 93) \| 94 \| SprT-like domain-containing protein \| SprT-like domain-containing protein; uniclust \| UniRef100\_UPI0001CBADE3 \| 98.0 \| 1.5e-07 \| 2.8e-13 \| 88.4 \| 80 \| (23, 105) \| 227 \| (299, 387) \| 499 \| acidic repeat-containing protein-like \| acidic repeat-containing protein-like; uniclust \| UniRef100\_UPI001E3CB0E9 \| 98.0 \| 1.6e-07 \| 2.9e-13 \| 72.0 \| 66 \| (42, 107) \| 227 \| (8, 89) \| 103 \| SprT-like domain-containing protein \| SprT-like domain-containing protein; uniclust \| UniRef100\_A0A6L8CSP2 \| 98.0 \| 1.6e-07 \| 3e-13 \| 85.1 \| 92 \| (4, 108) \| 227 \| (165, 259) \| 350 \| SprT-like domain-containing protein \| SprT-like domain-containing protein; uniclust \| UniRef100\_A0A2A4JTQ0 \| 97.9 \| 1.6e-07 \| 3.1e-13 \| 92.7 \| 103 \| (57, 220) \| 227 \| (427, 535) \| 608 \| SprT-like domain-containing protein \| SprT-like domain-containing protein; uniclust \| UniRef100\_A0A095C996 \| 97.9 \| 1.5e-07 \| 3.1e-13 \| 96.0 \| 97 \| (4, 102) \| 227 \| (522, 635) \| 772 \| SprT-like domain-containing protein \| SprT-like domain-containing protein; uniclust \| UniRef100\_A0A800A1Z7 \| 97.9 \| 1.8e-07 \| 3.3e-13 \| 79.3 \| 65 \| (57, 121) \| 227 \| (116, 189) \| 201 \| DUF35 domain-containing protein (Fragment) \| DUF35 domain-containing protein (Fragment); uniclust \| UniRef100\_A0A059LKE0 \| 97.9 \| 1.6e-07 \| 3.3e-13 \| 91.7 \| 76 \| (23, 102) \| 227 \| (195, 283) \| 458 \| SprT-like family protein \| SprT-like family protein; uniclust \| UniRef100\_UPI00145EA0DD \| 97.9 \| 1.9e-07 \| 3.4e-13 \| 74.4 \| 71 \| (30, 109) \| 227 \| (31, 107) \| 130 \| SprT-like domain-containing protein \| SprT-like domain-containing protein; uniclust \| UniRef100\_UPI001D095371 \| 97.9 \| 2.1e-07 \| 3.9e-13 \| 94.6 \| 134 \| (23, 220) \| 227 \| (1174, 1317) \| 1385 \| dentin sialophosphoprotein-like \| dentin sialophosphoprotein-like; uniclust \| UniRef100\_A0A2P6WC38 \| 97.9 \| 2.2e-07 \| 4.1e-13 \| 79.5 \| 81 \| (42, 134) \| 227 \| (93, 176) \| 215 \| SprT-like domain-containing protein \| SprT-like domain-containing protein; uniclust \| UniRef100\_A0A2R8FEQ8 \| 97.9 \| 2.4e-07 \| 4.5e-13 \| 80.7 \| 76 \| (25, 110) \| 227 \| (27, 105) \| 247 \| Zinc metallopeptidase SprT family \| Zinc metallopeptidase SprT family; uniclust \| UniRef100\_A0A4Q6ELT1 \| 97.9 \| 2.5e-07 \| 4.8e-13 \| 80.5 \| 91 \| (12, 106) \| 227 \| (6, 110) \| 222 \| M48 family peptidase \| M48 family peptidase; uniclust \| UniRef100\_A0A0R2KSK1 \| 97.9 \| 2.3e-07 \| 4.8e-13 \| 81.1 \| 102 \| (8, 114) \| 227 \| (25, 134) \| 190 \| SprT-like domain-containing protein \| SprT-like domain-containing protein; uniclust \| UniRef100\_A0A7Y3I8T8 \| 97.9 \| 2.6e-07 \| 4.8e-13 \| 71.9 \| 68 \| (41, 108) \| 227 \| (30, 104) \| 112 \| SprT-like domain-containing protein \| SprT-like domain-containing protein; uniclust \| UniRef100\_A0A085M4A3 \| 97.9 \| 2.5e-07 \| 4.8e-13 \| 89.3 \| 144 \| (11, 219) \| 227 \| (83, 236) \| 554 \| Alpha-carbonic anhydrase domain-containing protein \| Alpha-carbonic anhydrase domain-containing protein; uniclust \| UniRef100\_A0A068WCX1 \| 97.9 \| 2.6e-07 \| 4.9e-13 \| 90.6 \| 97 \| (1, 102) \| 227 \| (407, 517) \| 650 \| Acidic repeat-containing protein \| Acidic repeat-containing protein; uniclust \| UniRef100\_A0A2C9KYW2 \| 97.9 \| 2.7e-07 \| 4.9e-13 \| 91.3 \| 91 \| (10, 103) \| 227 \| (684, 781) \| 899 \| SprT-like domain-containing protein \| SprT-like domain-containing protein; uniclust \| UniRef100\_A0A1M5NTB3 \| 97.9 \| 2.2e-07 \| 4.9e-13 \| 77.3 \| 88 \| (6, 102) \| 227 \| (24, 120) \| 132 \| SprT-like family protein \| SprT-like family protein; uniclust \| UniRef100\_Q0A8D8 \| 97.9 \| 2.4e-07 \| 4.9e-13 \| 81.7 \| 64 \| (41, 110) \| 227 \| (83, 154) \| 216 \| SprT-like domain-containing protein \| SprT-like domain-containing protein; uniclust \| UniRef100\_A0A0C2N032 \| 97.9 \| 2.3e-07 \| 4.9e-13 \| 86.7 \| 75 \| (23, 103) \| 227 \| (87, 170) \| 312 \| SprT-like domain-containing protein Spartan \| SprT-like domain-containing protein Spartan; uniclust \| UniRef100\_A0A0A1X4E5 \| 97.9 \| 2.4e-07 \| 5.1e-13 \| 98.2 \| 89 \| (10, 103) \| 227 \| (883, 977) \| 1114 \| Acidic repeat-containing protein \| Acidic repeat-containing protein; uniclust \| UniRef100\_A0A1V9X7L4 \| 97.9 \| 2.7e-07 \| 5.2e-13 \| 90.6 \| 122 \| (30, 220) \| 227 \| (506, 637) \| 644 \| HMG box-containing protein C19G7.04-like \| HMG box-containing protein C19G7.04-like; uniclust \| UniRef100\_A0A8J1XJQ0 \| 97.9 \| 2.9e-07 \| 5.3e-13 \| 95.3 \| 134 \| (23, 220) \| 227 \| (1687, 1828) \| 1885 \| Ofus.G033133 protein \| Ofus.G033133 protein; uniclust \| UniRef100\_A0A4Y7MT59 \| 97.9 \| 2.9e-07 \| 5.4e-13 \| 93.3 \| 94 \| (2, 101) \| 227 \| (854, 957) \| 1080 \| EOG090X0464 \| EOG090X0464; uniclust \| UniRef100\_A0A6P3R813 \| 97.9 \| 2.7e-07 \| 5.8e-13 \| 94.2 \| 81 \| (23, 106) \| 227 \| (348, 436) \| 702 \| Acidic repeat-containing protein isoform X4 \| Acidic repeat-containing protein isoform X4; uniclust \| UniRef100\_A0A0D0DSM7 \| 97.9 \| 2.7e-07 \| 5.8e-13 \| 90.1 \| 95 \| (3, 103) \| 227 \| (222, 324) \| 459 \| SprT-like domain-containing protein \| SprT-like domain-containing protein; uniclust \| UniRef100\_A0A095Y7F4 \| 97.9 \| 2.7e-07 \| 6.2e-13 \| 85.3 \| 94 \| (5, 105) \| 227 \| (36, 140) \| 261 \| SprT-like domain-containing protein \| SprT-like domain-containing protein; uniclust \| UniRef100\_A0A0D2G9M8 \| 97.9 \| 3.3e-07 \| 6.2e-13 \| 75.8 \| 91 \| (10, 106) \| 227 \| (10, 108) \| 168 \| SprT-like domain-containing protein \| SprT-like domain-containing protein; uniclust \| UniRef100\_A0A2V2CUH8 \| 97.9 \| 3e-07 \| 6.2e-13 \| 79.5 \| 93 \| (12, 110) \| 227 \| (7, 108) \| 177 \| SprT-like domain-containing protein \| SprT-like domain-containing protein; uniclust \| UniRef100\_UPI001E63D35C \| 97.9 \| 3.4e-07 \| 6.3e-13 \| 71.0 \| 44 \| (176, 219) \| 227 \| (57, 100) \| 110 \| hypothetical protein \| hypothetical protein; uniclust \| UniRef100\_A0A193QM59 \| 97.8 \| 3.2e-07 \| 6.5e-13 \| 78.9 \| 64 \| (42, 111) \| 227 \| (54, 125) \| 180 \| Protein SprT \| Protein SprT; uniclust \| UniRef100\_A0A016TZM6 \| 97.8 \| 3e-07 \| 6.5e-13 \| 92.5 \| 76 \| (23, 104) \| 227 \| (91, 175) \| 566 \| SprT-like domain-containing protein \| SprT-like domain-containing protein; uniclust \| UniRef100\_A0A3L7P4X0 \| 97.8 \| 3.5e-07 \| 7e-13 \| 79.4 \| 76 \| (31, 112) \| 227 \| (48, 132) \| 192 \| SprT family protein \| SprT family protein; uniclust \| UniRef100\_A0A948QZG9 \| 97.8 \| 3.8e-07 \| 7.2e-13 \| 81.4 \| 106 \| (8, 121) \| 227 \| (119, 237) \| 266 \| SprT-like domain-containing protein \| SprT-like domain-containing protein; uniclust \| UniRef100\_A0A7J6MCN6 \| 97.8 \| 3.8e-07 \| 7.2e-13 \| 86.6 \| 72 \| (23, 101) \| 227 \| (324, 401) \| 463 \| SprT-like domain-containing protein \| SprT-like domain-containing protein; uniclust \| UniRef100\_A0A183PK73 \| 97.8 \| 4.1e-07 \| 7.4e-13 \| 79.4 \| 135 \| (22, 220) \| 227 \| (30, 178) \| 247 \| SprT-like domain-containing protein \| SprT-like domain-containing protein; uniclust \| UniRef100\_UPI001CA819BC \| 97.8 \| 3.8e-07 \| 7.6e-13 \| 81.9 \| 88 \| (8, 105) \| 227 \| (72, 168) \| 251 \| DNA-dependent metalloprotease dvc-1-like isoform X1 \| DNA-dependent metalloprotease dvc-1-like isoform X1; uniclust \| UniRef100\_A0A515MIG7 \| 97.8 \| 4.1e-07 \| 7.7e-13 \| 81.1 \| 113 \| (14, 137) \| 227 \| (127, 242) \| 271 \| SprT-like domain-containing protein \| SprT-like domain-containing protein; uniclust \| UniRef100\_A0A0C2X8A7 \| 97.8 \| 3.9e-07 \| 8e-13 \| 87.3 \| 96 \| (2, 102) \| 227 \| (163, 265) \| 421 \| SprT-like domain-containing protein \| SprT-like domain-containing protein; uniclust \| UniRef100\_UPI00138FF197 \| 97.8 \| 4.7e-07 \| 8.6e-13 \| 89.6 \| 80 \| (23, 105) \| 227 \| (294, 378) \| 877 \| uncharacterized protein LOC5522005 isoform X1 \| uncharacterized protein LOC5522005 isoform X1; uniclust \| UniRef100\_A0A1G6XTM0 \| 97.8 \| 4.4e-07 \| 8.7e-13 \| 83.9 \| 99 \| (9, 109) \| 227 \| (7, 118) \| 325 \| SprT-like family protein \| SprT-like family protein; uniclust \| UniRef100\_A0A2E7CJ87 \| 97.8 \| 4.8e-07 \| 8.8e-13 \| 75.5 \| 93 \| (14, 110) \| 227 \| (6, 107) \| 176 \| SprT-like domain-containing protein \| SprT-like domain-containing protein; uniclust \| UniRef100\_A0A0B0EPM6 \| 97.8 \| 3.9e-07 \| 9.5e-13 \| 91.3 \| 65 \| (42, 112) \| 227 \| (121, 194) \| 487 \| SprT-like family protein \| SprT-like family protein; uniclust \| UniRef100\_A0A090RVQ2 \| 97.8 \| 4.6e-07 \| 9.7e-13 \| 76.4 \| 66 \| (41, 112) \| 227 \| (6, 79) \| 146 \| Protein sprT \| Protein sprT; uniclust \| UniRef100\_A0A852Z1F4 \| 97.8 \| 5.3e-07 \| 9.7e-13 \| 71.1 \| 78 \| (121, 213) \| 227 \| (38, 115) \| 120 \| C2H2-type domain-containing protein \| C2H2-type domain-containing protein; uniclust \| UniRef100\_A0A1G8BVJ5 \| 97.8 \| 5.5e-07 \| 1e-12 \| 78.8 \| 81 \| (23, 113) \| 227 \| (20, 104) \| 252 \| SprT-like family protein \| SprT-like family protein; uniclust \| UniRef100\_A0A938L5N4 \| 97.8 \| 5.5e-07 \| 1e-12 \| 73.9 \| 76 \| (22, 109) \| 227 \| (23, 105) \| 156 \| SprT family zinc-dependent metalloprotease (Fragment) \| SprT family zinc-dependent metalloprotease (Fragment); uniclust \| UniRef100\_A0A2V7WKY5 \| 97.8 \| 5.2e-07 \| 1e-12 \| 77.3 \| 86 \| (7, 108) \| 227 \| (29, 118) \| 185 \| SprT-like domain-containing protein \| SprT-like domain-containing protein; uniclust \| UniRef100\_A0A166KJM6 \| 97.8 \| 4.9e-07 \| 1e-12 \| 88.4 \| 93 \| (3, 102) \| 227 \| (246, 347) \| 464 \| SprT-like domain-containing protein \| SprT-like domain-containing protein; uniclust \| UniRef100\_UPI000C776800 \| 97.8 \| 5.9e-07 \| 1.1e-12 \| 91.1 \| 77 \| (23, 102) \| 227 \| (1034, 1120) \| 1237 \| uncharacterized protein LOC111695131 \| uncharacterized protein LOC111695131; uniclust \| UniRef100\_A0A096NXZ5 \| 97.8 \| 5.5e-07 \| 1.1e-12 \| 84.7 \| 86 \| (14, 105) \| 227 \| (88, 187) \| 376 \| SprT-like N-terminal domain \| SprT-like N-terminal domain; uniclust \| UniRef100\_A0A163K0V2 \| 97.8 \| 5.3e-07 \| 1.1e-12 \| 87.0 \| 86 \| (13, 105) \| 227 \| (184, 283) \| 411 \| SprT-like domain-containing protein \| SprT-like domain-containing protein; uniclust \| UniRef100\_A0A183LV10 \| 97.8 \| 6.1e-07 \| 1.2e-12 \| 87.8 \| 77 \| (23, 102) \| 227 \| (410, 500) \| 627 \| SprT-like domain-containing protein \| SprT-like domain-containing protein; uniclust \| UniRef100\_A0A0S4TDI0 \| 97.8 \| 5.7e-07 \| 1.2e-12 \| 83.3 \| 76 \| (23, 105) \| 227 \| (85, 169) \| 286 \| SprT-like domain-containing protein (Fragment) \| SprT-like domain-containing protein (Fragment); uniclust \| UniRef100\_A0A384DU94 \| 97.8 \| 6.2e-07 \| 1.2e-12 \| 86.7 \| 82 \| (22, 106) \| 227 \| (298, 387) \| 497 \| Acidic repeat-containing protein \| Acidic repeat-containing protein; uniclust \| UniRef100\_A0A857ZTK7 \| 97.8 \| 6.8e-07 \| 1.2e-12 \| 71.0 \| 41 \| (68, 108) \| 227 \| (21, 64) \| 125 \| SprT-like domain-containing protein \| SprT-like domain-containing protein; uniclust \| UniRef100\_A0A4R9A316 \| 97.8 \| 7.1e-07 \| 1.3e-12 \| 71.9 \| 83 \| (25, 116) \| 227 \| (19, 104) \| 137 \| M48 family peptidase \| M48 family peptidase; uniclust \| UniRef100\_UPI001E2C7D93 \| 97.7 \| 7.2e-07 \| 1.3e-12 \| 76.9 \| 67 \| (42, 108) \| 227 \| (11, 98) \| 223 \| hypothetical protein \| hypothetical protein; uniclust \| UniRef100\_A0A1F3AAF6 \| 97.7 \| 5.6e-07 \| 1.3e-12 \| 74.3 \| 86 \| (7, 102) \| 227 \| (11, 105) \| 117 \| Uncharacterized protein \| Uncharacterized protein; uniclust \| UniRef100\_A0A927AK43 \| 97.7 \| 7.8e-07 \| 1.4e-12 \| 76.4 \| 66 \| (42, 107) \| 227 \| (41, 120) \| 215 \| Uncharacterized protein \| Uncharacterized protein; uniclust \| UniRef100\_A0A015LDH0 \| 97.7 \| 7.3e-07 \| 1.4e-12 \| 83.9 \| 86 \| (13, 105) \| 227 \| (69, 168) \| 366 \| SprT-like domain-containing protein \| SprT-like domain-containing protein; uniclust \| UniRef100\_A0A8S1E164 \| 97.7 \| 7.9e-07 \| 1.4e-12 \| 88.2 \| 73 \| (28, 102) \| 227 \| (699, 781) \| 886 \| SprT-like domain-containing protein \| SprT-like domain-containing protein; uniclust \| UniRef100\_A0A6A4WPN6 \| 97.7 \| 8.1e-07 \| 1.5e-12 \| 89.0 \| 143 \| (15, 221) \| 227 \| (784, 943) \| 1016 \| Acidic repeat-containing protein \| Acidic repeat-containing protein; uniclust \| UniRef100\_A0A1I7SSN3 \| 97.7 \| 8e-07 \| 1.5e-12 \| 90.0 \| 86 \| (12, 102) \| 227 \| (792, 884) \| 964 \| SprT-like domain-containing protein \| SprT-like domain-containing protein; uniclust \| UniRef100\_A0A023W584 \| 97.7 \| 8.3e-07 \| 1.5e-12 \| 76.5 \| 60 \| (42, 107) \| 227 \| (59, 137) \| 221 \| SprT-like domain-containing protein \| SprT-like domain-containing protein; uniclust \| UniRef100\_A0A5R8KJF2 \| 97.7 \| 8.4e-07 \| 1.5e-12 \| 79.4 \| 85 \| (15, 107) \| 227 \| (142, 233) \| 299 \| SprT family zinc-dependent metalloprotease \| SprT family zinc-dependent metalloprotease; uniclust \| UniRef100\_A0A7W1MVA6 \| 97.7 \| 9.1e-07 \| 1.7e-12 \| 77.5 \| 72 \| (29, 108) \| 227 \| (85, 164) \| 235 \| SprT-like domain-containing protein \| SprT-like domain-containing protein; uniclust \| UniRef100\_A0A060WHD9 \| 97.7 \| 7.8e-07 \| 1.7e-12 \| 91.5 \| 84 \| (14, 103) \| 227 \| (165, 262) \| 709 \| DNA-dependent metalloprotease SPRTN \| DNA-dependent metalloprotease SPRTN; uniclust \| UniRef100\_A0A1Y1LCH4 \| 97.7 \| 8e-07 \| 1.7e-12 \| 88.9 \| 87 \| (13, 105) \| 227 \| (138, 238) \| 547 \| Protein with SprT-like domain at the N terminus \| Protein with SprT-like domain at the N terminus; uniclust \| UniRef100\_A0A522YTE6 \| 97.7 \| 9.4e-07 \| 1.7e-12 \| 70.9 \| 66 \| (59, 124) \| 227 \| (60, 127) \| 133 \| Uncharacterized protein \| Uncharacterized protein; uniclust \| UniRef100\_A0A7J7JVI6 \| 97.7 \| 9.8e-07 \| 1.9e-12 \| 74.1 \| 77 \| (23, 105) \| 227 \| (36, 121) \| 165 \| SprT-like domain-containing protein \| SprT-like domain-containing protein; uniclust \| UniRef100\_A0A026WP05 \| 97.7 \| 8.4e-07 \| 1.9e-12 \| 91.1 \| 73 \| (30, 106) \| 227 \| (96, 177) \| 672 \| Protein with SprT-like domain at the N terminus (Fragment) \| Protein with SprT-like domain at the N terminus (Fragment); uniclust \| UniRef100\_A0A8T6V780 \| 97.7 \| 1e-06 \| 1.9e-12 \| 68.9 \| 80 \| (24, 114) \| 227 \| (19, 104) \| 113 \| SprT-like domain-containing protein \| SprT-like domain-containing protein; uniclust \| UniRef100\_A0A154NZ86 \| 97.7 \| 1e-06 \| 2e-12 \| 91.4 \| 134 \| (23, 220) \| 227 \| (1195, 1338) \| 1410 \| Neuropeptides capa receptor \| Neuropeptides capa receptor; uniclust \| UniRef100\_A0A075A2A6 \| 97.7 \| 1.1e-06 \| 2e-12 \| 88.5 \| 77 \| (23, 102) \| 227 \| (836, 926) \| 1080 \| SprT-like domain-containing protein \| SprT-like domain-containing protein; uniclust \| UniRef100\_A0A1X7VLW4 \| 97.7 \| 1.1e-06 \| 2.1e-12 \| 84.4 \| 75 \| (23, 100) \| 227 \| (453, 533) \| 593 \| SprT-like domain-containing protein \| SprT-like domain-containing protein; uniclust \| UniRef100\_A0A1J1I8Q7 \| 97.7 \| 1.1e-06 \| 2.1e-12 \| 92.4 \| 75 \| (29, 105) \| 227 \| (945, 1023) \| 1139 \| CLUMA\_CG009862, isoform A \| CLUMA\_CG009862, isoform A; uniclust \| UniRef100\_UPI0002CC5A87 \| 97.7 \| 1.2e-06 \| 2.2e-12 \| 72.0 \| 101 \| (9, 112) \| 227 \| (6, 125) \| 155 \| SprT-like domain-containing protein \| SprT-like domain-containing protein; uniclust \| UniRef100\_A0A024U161 \| 97.7 \| 1.1e-06 \| 2.2e-12 \| 83.2 \| 72 \| (29, 105) \| 227 \| (87, 167) \| 365 \| SprT-like domain-containing protein \| SprT-like domain-containing protein; uniclust \| UniRef100\_A0A6J1SGE2 \| 97.7 \| 1.1e-06 \| 2.2e-12 \| 89.0 \| 133 \| (23, 220) \| 227 \| (748, 890) \| 956 \| Germ cell nuclear acidic protein-like isoform X3 \| Germ cell nuclear acidic protein-like isoform X3; uniclust \| UniRef100\_A0A5A8CGL8 \| 97.7 \| 1.2e-06 \| 2.2e-12 \| 77.8 \| 89 \| (8, 106) \| 227 \| (38, 136) \| 265 \| SprT-like domain-containing protein \| SprT-like domain-containing protein; uniclust \| UniRef100\_A0A7J6QCS3 \| 97.7 \| 1.3e-06 \| 2.4e-12 \| 74.7 \| 84 \| (8, 98) \| 227 \| (49, 143) \| 205 \| SprT-like domain-containing protein (Fragment) \| SprT-like domain-containing protein (Fragment); uniclust \| UniRef100\_A0A6N9DWF4 \| 97.7 \| 1.3e-06 \| 2.4e-12 \| 75.5 \| 77 \| (22, 102) \| 227 \| (29, 122) \| 218 \| SprT family zinc-dependent metalloprotease \| SprT family zinc-dependent metalloprotease; uniclust \| UniRef100\_A0A1V5RWR0 \| 97.7 \| 1.3e-06 \| 2.4e-12 \| 73.5 \| 63 \| (41, 104) \| 227 \| (37, 106) \| 183 \| SprT-like family protein \| SprT-like family protein; uniclust \| UniRef100\_A0A6P8HBJ2 \| 97.6 \| 1.4e-06 \| 2.6e-12 \| 83.9 \| 82 \| (23, 107) \| 227 \| (292, 378) \| 495 \| Acidic repeat-containing protein-like \| Acidic repeat-containing protein-like; uniclust \| UniRef100\_A0A7J6LXX1 \| 97.6 \| 1.3e-06 \| 2.6e-12 \| 92.4 \| 72 \| (23, 101) \| 227 \| (982, 1059) \| 1207 \| JmjC domain-containing protein \| JmjC domain-containing protein; uniclust \| UniRef100\_A0A8T1MBU3 \| 97.6 \| 1.5e-06 \| 2.7e-12 \| 85.5 \| 77 \| (23, 102) \| 227 \| (131, 221) \| 770 \| lipoyl synthase (Fragment) \| lipoyl synthase (Fragment); uniclust \| UniRef100\_A0A353P8A1 \| 97.6 \| 1.5e-06 \| 2.8e-12 \| 77.2 \| 67 \| (41, 107) \| 227 \| (112, 196) \| 278 \| SprT-like domain-containing protein \| SprT-like domain-containing protein; uniclust \| UniRef100\_A0A0N5CJR3 \| 97.6 \| 1.5e-06 \| 2.8e-12 \| 75.8 \| 64 \| (33, 101) \| 227 \| (60, 129) \| 240 \| SprT-like domain-containing protein \| SprT-like domain-containing protein; uniclust \| UniRef100\_A0A965RUJ9 \| 97.6 \| 1.6e-06 \| 2.9e-12 \| 67.9 \| 72 \| (23, 105) \| 227 \| (38, 112) \| 113 \| Uncharacterized protein \| Uncharacterized protein; uniclust \| UniRef100\_A0A2W1U3I7 \| 97.6 \| 1.6e-06 \| 3.1e-12 \| 80.1 \| 77 \| (25, 111) \| 227 \| (18, 97) \| 340 \| SprT-like domain-containing protein \| SprT-like domain-containing protein; uniclust \| UniRef100\_UPI001F5BC89C \| 97.6 \| 1.8e-06 \| 3.2e-12 \| 65.3 \| 73 \| (7, 81) \| 227 \| (4, 88) \| 91 \| hypothetical protein \| hypothetical protein; uniclust \| UniRef100\_UPI000289E565 \| 97.6 \| 1.8e-06 \| 3.3e-12 \| 68.7 \| 79 \| (7, 87) \| 227 \| (4, 94) \| 125 \| SprT-like domain-containing protein \| SprT-like domain-containing protein; uniclust \| UniRef100\_A0A162V0U9 \| 97.6 \| 1.7e-06 \| 3.4e-12 \| 80.9 \| 72 \| (29, 105) \| 227 \| (115, 195) \| 325 \| SprT-like domain-containing protein \| SprT-like domain-containing protein; uniclust \| UniRef100\_A0A4Y2D086 \| 97.6 \| 1.8e-06 \| 3.5e-12 \| 78.8 \| 99 \| (3, 107) \| 227 \| (128, 234) \| 292 \| Acidic repeat-containing protein \| Acidic repeat-containing protein; uniclust \| UniRef100\_A0A925ICF3 \| 97.6 \| 2.1e-06 \| 3.9e-12 \| 74.8 \| 88 \| (29, 134) \| 227 \| (109, 199) \| 235 \| SprT-like domain-containing protein \| SprT-like domain-containing protein; uniclust \| UniRef100\_A0A1I8A746 \| 97.6 \| 2.2e-06 \| 4e-12 \| 85.4 \| 77 \| (23, 102) \| 227 \| (645, 727) \| 894 \| SprT-like domain-containing protein \| SprT-like domain-containing protein; uniclust \| UniRef100\_A0A7U9DY43 \| 97.6 \| 2.2e-06 \| 4e-12 \| 56.3 \| 35 \| (186, 221) \| 227 \| (2, 36) \| 43 \| Uncharacterized protein \| Uncharacterized protein; uniclust \| UniRef100\_UPI00201F22D7 \| 97.6 \| 2.2e-06 \| 4.1e-12 \| 88.2 \| 132 \| (23, 221) \| 227 \| (1274, 1410) \| 1470 \| dentin sialophosphoprotein-like \| dentin sialophosphoprotein-like; uniclust \| UniRef100\_UPI00193CA4CB \| 97.6 \| 2.3e-06 \| 4.2e-12 \| 88.5 \| 134 \| (23, 220) \| 227 \| (1367, 1509) \| 1604 \| uncharacterized protein LOC111060193 isoform X2 \| uncharacterized protein LOC111060193 isoform X2; uniclust \| UniRef100\_A0A7S0X6E5 \| 97.6 \| 2.1e-06 \| 4.2e-12 \| 84.4 \| 77 \| (23, 102) \| 227 \| (319, 405) \| 519 \| SprT-like domain-containing protein (Fragment) \| SprT-like domain-containing protein (Fragment); uniclust \| UniRef100\_A0A0L0DN51 \| 97.6 \| 2.4e-06 \| 4.4e-12 \| 81.8 \| 93 \| (3, 101) \| 227 \| (346, 445) \| 551 \| HMG box-containing protein \| HMG box-containing protein; uniclust \| UniRef100\_A0A2P6TE25 \| 97.6 \| 2.3e-06 \| 4.6e-12 \| 80.1 \| 77 \| (23, 105) \| 227 \| (32, 116) \| 329 \| SprT-like domain-containing Spartan \| SprT-like domain-containing Spartan; uniclust \| UniRef100\_A0A034V6Q8 \| 97.6 \| 2.4e-06 \| 4.6e-12 \| 79.5 \| 76 \| (29, 106) \| 227 \| (106, 190) \| 364 \| SprT-like domain-containing protein Spartan (Fragment) \| SprT-like domain-containing protein Spartan (Fragment); uniclust \| UniRef100\_A0A0C3RSD2 \| 97.6 \| 2.3e-06 \| 4.6e-12 \| 78.2 \| 94 \| (3, 102) \| 227 \| (45, 146) \| 263 \| SprT-like domain-containing protein (Fragment) \| SprT-like domain-containing protein (Fragment); uniclust \| UniRef100\_A0A956X7Z6 \| 97.5 \| 2.6e-06 \| 4.8e-12 \| 72.9 \| 124 \| (68, 220) \| 227 \| (52, 191) \| 205 \| Uncharacterized protein \| Uncharacterized protein; uniclust \| UniRef100\_A0A080K9R0 \| 97.5 \| 2.3e-06 \| 4.9e-12 \| 70.5 \| 70 \| (22, 102) \| 227 \| (34, 110) \| 123 \| SprT-like family (Fragment) \| SprT-like family (Fragment); uniclust \| UniRef100\_A0A6L7XKZ6 \| 97.5 \| 2.6e-06 \| 4.9e-12 \| 75.5 \| 65 \| (42, 112) \| 227 \| (98, 165) \| 254 \| SprT-like domain-containing protein \| SprT-like domain-containing protein; uniclust \| UniRef100\_A0A164WWE9 \| 97.5 \| 2.5e-06 \| 5e-12 \| 85.1 \| 94 \| (2, 101) \| 227 \| (366, 471) \| 595 \| Acidic repeat-containing-like protein \| Acidic repeat-containing-like protein; uniclust \| UniRef100\_A0A2S5R2G5 \| 97.5 \| 2.7e-06 \| 5e-12 \| 71.1 \| 65 \| (42, 112) \| 227 \| (45, 122) \| 172 \| SprT-like domain-containing protein \| SprT-like domain-containing protein; uniclust \| UniRef100\_A0A0D4DB50 \| 97.5 \| 2.7e-06 \| 5.2e-12 \| 77.1 \| 83 \| (24, 115) \| 227 \| (25, 113) \| 272 \| SprT-like protein \| SprT-like protein; uniclust \| UniRef100\_A0A822CXB0 \| 97.5 \| 2.9e-06 \| 5.3e-12 \| 68.6 \| 68 \| (31, 103) \| 227 \| (3, 79) \| 137 \| SprT-like domain-containing protein \| SprT-like domain-containing protein; uniclust \| UniRef100\_A0A433C399 \| 97.5 \| 3e-06 \| 5.6e-12 \| 67.9 \| 75 \| (25, 108) \| 227 \| (18, 102) \| 120 \| Uncharacterized protein \| Uncharacterized protein; uniclust \| UniRef100\_A0A1B6DXM7 \| 97.5 \| 3.2e-06 \| 5.9e-12 \| 80.6 \| 80 \| (23, 105) \| 227 \| (315, 404) \| 524 \| SprT-like domain-containing protein \| SprT-like domain-containing protein; uniclust \| UniRef100\_A0A2V2RIV7 \| 97.5 \| 3.3e-06 \| 6e-12 \| 72.0 \| 82 \| (13, 107) \| 227 \| (47, 135) \| 196 \| SprT-like domain-containing protein \| SprT-like domain-containing protein; uniclust \| UniRef100\_UPI001EEAE6CF \| 97.5 \| 3.3e-06 \| 6.1e-12 \| 85.7 \| 80 \| (23, 105) \| 227 \| (931, 1019) \| 1144 \| uncharacterized protein LOC124357666 \| uncharacterized protein LOC124357666; uniclust \| UniRef100\_A0A1C5Q9L4 \| 97.5 \| 3.3e-06 \| 6.4e-12 \| 75.1 \| 91 \| (5, 102) \| 227 \| (38, 131) \| 247 \| SprT-like family \| SprT-like family; uniclust \| UniRef100\_A0A4Y8K0Q1 \| 97.5 \| 3.5e-06 \| 6.5e-12 \| 72.8 \| 79 \| (27, 114) \| 227 \| (8, 89) \| 218 \| Uncharacterized protein \| Uncharacterized protein; uniclust \| UniRef100\_A0A233HV64 \| 97.5 \| 2.9e-06 \| 6.7e-12 \| 75.6 \| 50 \| (58, 107) \| 227 \| (58, 110) \| 189 \| SprT-like domain-containing protein \| SprT-like domain-containing protein; uniclust \| UniRef100\_A0A843HBC1 \| 97.5 \| 3.7e-06 \| 6.8e-12 \| 75.5 \| 48 \| (60, 107) \| 227 \| (115, 170) \| 293 \| SprT-like domain-containing protein \| SprT-like domain-containing protein; uniclust \| UniRef100\_A0A2D5VXK3 \| 97.5 \| 3.7e-06 \| 6.9e-12 \| 69.1 \| 94 \| (7, 108) \| 227 \| (30, 131) \| 152 \| SprT-like domain-containing protein \| SprT-like domain-containing protein; uniclust \| UniRef100\_A0A2S8IY47 \| 97.5 \| 3.8e-06 \| 7.1e-12 \| 62.6 \| 44 \| (176, 219) \| 227 \| (30, 73) \| 83 \| Uncharacterized protein \| Uncharacterized protein; uniclust \| UniRef100\_A0A6V7K544 \| 97.5 \| 3.8e-06 \| 7.1e-12 \| 65.2 \| 71 \| (30, 104) \| 227 \| (2, 81) \| 101 \| SprT-like domain-containing protein (Fragment) \| SprT-like domain-containing protein (Fragment); uniclust \| UniRef100\_A0A077Z859 \| 97.5 \| 3.6e-06 \| 7.4e-12 \| 81.1 \| 78 \| (23, 106) \| 227 \| (70, 156) \| 394 \| SprT-like domain containing protein \| SprT-like domain containing protein; uniclust \| UniRef100\_UPI0012E23595 \| 97.5 \| 4.2e-06 \| 7.7e-12 \| 74.6 \| 76 \| (29, 112) \| 227 \| (70, 148) \| 277 \| hypothetical protein \| hypothetical protein; uniclust \| UniRef100\_A0A4D6E533 \| 97.5 \| 4.3e-06 \| 7.9e-12 \| 73.6 \| 77 \| (25, 111) \| 227 \| (21, 100) \| 250 \| SprT-like domain-containing protein \| SprT-like domain-containing protein; uniclust \| UniRef100\_A0A131XCL1 \| 97.5 \| 4.1e-06 \| 7.9e-12 \| 81.2 \| 84 \| (14, 103) \| 227 \| (52, 149) \| 506 \| SprT-like domain-containing protein (Fragment) \| SprT-like domain-containing protein (Fragment); uniclust \| UniRef100\_F2UXT2 \| 97.5 \| 4.3e-06 \| 8e-12 \| 72.9 \| 70 \| (30, 108) \| 227 \| (69, 141) \| 231 \| SprT-like domain-containing protein \| SprT-like domain-containing protein; uniclust \| UniRef100\_A0A6J4F9I4 \| 97.5 \| 4.4e-06 \| 8e-12 \| 64.3 \| 40 \| (180, 219) \| 227 \| (8, 48) \| 99 \| Transposase \| Transposase; uniclust \| UniRef100\_A0A8S1DZ30 \| 97.5 \| 4.4e-06 \| 8.2e-12 \| 87.2 \| 73 \| (28, 102) \| 227 \| (188, 270) \| 1809 \| DH domain-containing protein \| DH domain-containing protein; uniclust \| UniRef100\_A0A2W5X165 \| 97.5 \| 4.6e-06 \| 8.5e-12 \| 76.0 \| 93 \| (4, 106) \| 227 \| (158, 255) \| 333 \| SprT-like domain-containing protein \| SprT-like domain-containing protein; uniclust \| UniRef100\_A0A0P9DLN8 \| 97.5 \| 4.3e-06 \| 8.8e-12 \| 68.3 \| 104 \| (90, 222) \| 227 \| (2, 114) \| 117 \| Uncharacterized protein (Fragment) \| Uncharacterized protein (Fragment); uniclust \| UniRef100\_A0A1S3H5D5 \| 97.4 \| 4.9e-06 \| 9.1e-12 \| 84.4 \| 80 \| (23, 105) \| 227 \| (915, 1003) \| 1114 \| Uncharacterized protein LOC106152248 \| Uncharacterized protein LOC106152248; uniclust \| UniRef100\_A0A138ZXI7 \| 97.4 \| 4.6e-06 \| 9.2e-12 \| 80.2 \| 72 \| (29, 105) \| 227 \| (139, 219) \| 425 \| Protein with SprT-like domain at the N terminus \| Protein with SprT-like domain at the N terminus; uniclust \| UniRef100\_Q8EWL3 \| 97.4 \| 5e-06 \| 9.2e-12 \| 73.5 \| 116 \| (68, 222) \| 227 \| (87, 217) \| 256 \| SprT-like domain-containing protein \| SprT-like domain-containing protein; uniclust \| UniRef100\_UPI0018851632 \| 97.4 \| 5.3e-06 \| 9.7e-12 \| 77.4 \| 135 \| (22, 221) \| 227 \| (197, 341) \| 410 \| acidic repeat-containing protein-like \| acidic repeat-containing protein-like; uniclust \| UniRef100\_A0A0C2BZ93 \| 97.4 \| 5.1e-06 \| 9.9e-12 \| 68.5 \| 75 \| (23, 103) \| 227 \| (33, 116) \| 137 \| SprT-like domain-containing protein \| SprT-like domain-containing protein; uniclust \| UniRef100\_A0A1F5UZX3 \| 97.4 \| 5.7e-06 \| 1e-11 \| 79.0 \| 86 \| (12, 105) \| 227 \| (350, 448) \| 516 \| Phospho-N-acetylmuramoyl-pentapeptide-transferase (Fragment) \| Phospho-N-acetylmuramoyl-pentapeptide-transferase (Fragment); uniclust \| UniRef100\_W6U0M4 \| 97.4 \| 5.7e-06 \| 1.1e-11 \| 82.4 \| 97 \| (1, 102) \| 227 \| (406, 516) \| 858 \| SprT-like domain-containing protein \| SprT-like domain-containing protein; uniclust \| UniRef100\_A0A3P6HE70 \| 97.4 \| 5.7e-06 \| 1.1e-11 \| 73.1 \| 97 \| (1, 103) \| 227 \| (6, 117) \| 255 \| SprT-like domain-containing protein \| SprT-like domain-containing protein; uniclust \| UniRef100\_A0A226D0T9 \| 97.4 \| 5.8e-06 \| 1.1e-11 \| 76.4 \| 71 \| (29, 103) \| 227 \| (50, 131) \| 376 \| SprT-like domain-containing protein Spartan \| SprT-like domain-containing protein Spartan; uniclust \| UniRef100\_A0A146HCT3 \| 97.4 \| 5.5e-06 \| 1.1e-11 \| 81.6 \| 82 \| (23, 106) \| 227 \| (301, 390) \| 568 \| SprT-like domain-containing protein \| SprT-like domain-containing protein; uniclust \| UniRef100\_A0A219YCL2 \| 97.4 \| 5.2e-06 \| 1.1e-11 \| 73.3 \| 71 \| (30, 110) \| 227 \| (39, 112) \| 197 \| SprT-like domain-containing protein \| SprT-like domain-containing protein; uniclust \| UniRef100\_A0A353B0T0 \| 97.4 \| 5.9e-06 \| 1.1e-11 \| 68.4 \| 69 \| (42, 116) \| 227 \| (49, 118) \| 157 \| SprT-like domain-containing protein \| SprT-like domain-containing protein; uniclust \| UniRef100\_A0A7J6ZA20 \| 97.4 \| 5.8e-06 \| 1.1e-11 \| 86.5 \| 133 \| (23, 220) \| 227 \| (1254, 1397) \| 1457 \| SprT-like domain-containing protein \| SprT-like domain-containing protein; uniclust \| UniRef100\_A0A0D6LND9 \| 97.4 \| 5.9e-06 \| 1.1e-11 \| 77.8 \| 80 \| (16, 101) \| 227 \| (264, 349) \| 447 \| SprT-like domain-containing protein \| SprT-like domain-containing protein; uniclust \| UniRef100\_A0A160TLN6 \| 97.4 \| 6e-06 \| 1.1e-11 \| 56.1 \| 29 \| (184, 212) \| 227 \| (1, 30) \| 50 \| Uncharacterized protein \| Uncharacterized protein; uniclust \| UniRef100\_A0A5S6QT35 \| 97.4 \| 6e-06 \| 1.1e-11 \| 71.8 \| 92 \| (9, 103) \| 227 \| (69, 169) \| 224 \| SprT-like domain-containing protein \| SprT-like domain-containing protein; uniclust \| UniRef100\_UPI0015AA2B96 \| 97.4 \| 6.4e-06 \| 1.2e-11 \| 68.7 \| 42 \| (73, 114) \| 227 \| (2, 49) \| 166 \| hypothetical protein \| hypothetical protein; uniclust \| UniRef100\_UPI000A922881 \| 97.4 \| 6.7e-06 \| 1.2e-11 \| 73.4 \| 56 \| (57, 112) \| 227 \| (106, 164) \| 277 \| SprT-like domain-containing protein \| SprT-like domain-containing protein; uniclust \| UniRef100\_A0A147BGG7 \| 97.4 \| 6.9e-06 \| 1.3e-11 \| 68.0 \| 66 \| (29, 99) \| 227 \| (60, 130) \| 157 \| Putative sprt-like metalloprotease (Fragment) \| Putative sprt-like metalloprotease (Fragment); uniclust \| UniRef100\_UPI000D7345FA \| 97.4 \| 6.9e-06 \| 1.3e-11 \| 74.4 \| 90 \| (9, 100) \| 227 \| (65, 160) \| 310 \| HMG box-containing protein C19G7.04-like \| HMG box-containing protein C19G7.04-like; uniclust \| UniRef100\_A0A2N1WAY1 \| 97.4 \| 7.1e-06 \| 1.3e-11 \| 67.6 \| 66 \| (42, 110) \| 227 \| (65, 137) \| 152 \| SprT-like domain-containing protein (Fragment) \| SprT-like domain-containing protein (Fragment); uniclust \| UniRef100\_A0A067TZ07 \| 97.4 \| 6.5e-06 \| 1.3e-11 \| 81.2 \| 93 \| (11, 106) \| 227 \| (254, 358) \| 477 \| SprT-like domain-containing protein \| SprT-like domain-containing protein; uniclust \| UniRef100\_A0A956U136 \| 97.4 \| 7.5e-06 \| 1.4e-11 \| 64.8 \| 65 \| (42, 112) \| 227 \| (9, 79) \| 116 \| SprT-like domain-containing protein (Fragment) \| SprT-like domain-containing protein (Fragment); uniclust \| UniRef100\_A0A7Y1ZPK4 \| 97.4 \| 7.4e-06 \| 1.4e-11 \| 72.3 \| 74 \| (29, 110) \| 227 \| (70, 151) \| 225 \| SprT-like domain-containing protein \| SprT-like domain-containing protein; uniclust \| UniRef100\_A0A915GBT6 \| 97.4 \| 7.6e-06 \| 1.4e-11 \| 71.9 \| 144 \| (16, 222) \| 227 \| (30, 182) \| 242 \| SprT-like domain-containing protein \| SprT-like domain-containing protein; uniclust \| UniRef100\_A0A947MTM0 \| 97.4 \| 8e-06 \| 1.5e-11 \| 55.3 \| 40 \| (179, 219) \| 227 \| (8, 47) \| 49 \| Uncharacterized protein \| Uncharacterized protein; uniclust \| UniRef100\_A0A7W1KMR8 \| 97.4 \| 7.8e-06 \| 1.5e-11 \| 67.2 \| 83 \| (23, 111) \| 227 \| (35, 124) \| 139 \| SprT-like domain-containing protein \| SprT-like domain-containing protein; uniclust \| UniRef100\_A0A1G8CW72 \| 97.4 \| 8.2e-06 \| 1.5e-11 \| 70.9 \| 62 \| (42, 109) \| 227 \| (68, 132) \| 222 \| SprT-like family protein \| SprT-like family protein; uniclust \| UniRef100\_A0A6L7STI3 \| 97.4 \| 8.3e-06 \| 1.5e-11 \| 55.9 \| 32 \| (68, 99) \| 227 \| (20, 51) \| 52 \| SprT family zinc-dependent metalloprotease \| SprT family zinc-dependent metalloprotease; uniclust \| UniRef100\_A0A1D2MB96 \| 97.4 \| 7.8e-06 \| 1.5e-11 \| 78.2 \| 75 \| (23, 103) \| 227 \| (78, 161) \| 400 \| SprT-like domain-containing protein Spartan (Fragment) \| SprT-like domain-containing protein Spartan (Fragment); uniclust \| UniRef100\_F0F968 \| 97.4 \| 7.7e-06 \| 1.5e-11 \| 73.3 \| 97 \| (4, 108) \| 227 \| (7, 115) \| 224 \| SprT-like domain-containing protein \| SprT-like domain-containing protein; uniclust \| UniRef100\_A0A067P3B9 \| 97.4 \| 8.2e-06 \| 1.6e-11 \| 72.7 \| 50 \| (57, 106) \| 227 \| (86, 141) \| 242 \| SprT-like domain-containing protein (Fragment) \| SprT-like domain-containing protein (Fragment); uniclust \| UniRef100\_A0A820RHD1 \| 97.4 \| 8.6e-06 \| 1.6e-11 \| 78.7 \| 68 \| (29, 100) \| 227 \| (376, 449) \| 578 \| SprT-like domain-containing protein \| SprT-like domain-containing protein; uniclust \| UniRef100\_A0A5Q3L5Z6 \| 97.3 \| 8.7e-06 \| 1.6e-11 \| 69.5 \| 87 \| (23, 118) \| 227 \| (24, 118) \| 193 \| SprT domain-containing protein \| SprT domain-containing protein; uniclust \| UniRef100\_A0A0L8I9K9 \| 97.3 \| 8.8e-06 \| 1.6e-11 \| 79.5 \| 134 \| (29, 224) \| 227 \| (457, 601) \| 658 \| SprT-like domain-containing protein \| SprT-like domain-containing protein; uniclust \| UniRef100\_A0A182IL28 \| 97.3 \| 8.7e-06 \| 1.7e-11 \| 80.2 \| 75 \| (29, 105) \| 227 \| (95, 178) \| 575 \| Protein with SprT-like domain at the N terminus \| Protein with SprT-like domain at the N terminus; uniclust \| UniRef100\_A0A484AUX1 \| 97.3 \| 8.7e-06 \| 1.7e-11 \| 77.8 \| 74 \| (28, 103) \| 227 \| (205, 283) \| 412 \| SprT-like domain-containing protein \| SprT-like domain-containing protein; uniclust \| UniRef100\_UPI001CC56835 \| 97.3 \| 9.3e-06 \| 1.7e-11 \| 71.9 \| 78 \| (22, 102) \| 227 \| (46, 132) \| 256 \| acidic repeat-containing protein-like \| acidic repeat-containing protein-like; uniclust \| UniRef100\_UPI001BB153D4 \| 97.3 \| 9.3e-06 \| 1.8e-11 \| 81.6 \| 95 \| (3, 103) \| 227 \| (513, 616) \| 730 \| acidic repeat-containing protein-like \| acidic repeat-containing protein-like; uniclust \| UniRef100\_UPI0018D5A09C \| 97.3 \| 9.8e-06 \| 1.8e-11 \| 62.4 \| 44 \| (71, 114) \| 227 \| (2, 45) \| 98 \| SprT-like domain-containing protein \| SprT-like domain-containing protein; uniclust \| UniRef100\_A0A7J8EKG4 \| 97.3 \| 9.6e-06 \| 1.8e-11 \| 75.6 \| 81 \| (23, 106) \| 227 \| (44, 132) \| 367 \| SprT-like domain-containing protein \| SprT-like domain-containing protein; uniclust \| UniRef100\_A0A2V5RCE3 \| 97.3 \| 1e-05 \| 1.8e-11 \| 67.6 \| 73 \| (28, 108) \| 227 \| (57, 136) \| 165 \| SprT-like domain-containing protein \| SprT-like domain-containing protein; uniclust \| UniRef100\_UPI000BF750D2 \| 97.3 \| 1e-05 \| 1.8e-11 \| 77.2 \| 69 \| (29, 99) \| 227 \| (358, 432) \| 497 \| acidic repeat-containing protein-like isoform X2 \| acidic repeat-containing protein-like isoform X2; uniclust \| UniRef100\_M9V2A0 \| 97.3 \| 1e-05 \| 1.9e-11 \| 60.8 \| 57 \| (29, 93) \| 227 \| (25, 84) \| 85 \| Putative SprT zinc metallopeptidase \| Putative SprT zinc metallopeptidase; uniclust \| UniRef100\_A0A0P4W780 \| 97.3 \| 9.3e-06 \| 1.9e-11 \| 80.3 \| 87 \| (14, 104) \| 227 \| (38, 139) \| 519 \| Protein with SprT-like domain at the N terminus \| Protein with SprT-like domain at the N terminus; uniclust \| UniRef100\_A0A090N044 \| 97.3 \| 9.9e-06 \| 1.9e-11 \| 77.0 \| 76 \| (23, 104) \| 227 \| (33, 117) \| 375 \| SprT-like domain-containing protein \| SprT-like domain-containing protein; uniclust \| UniRef100\_A0A7S4L4D1 \| 97.3 \| 1e-05 \| 2e-11 \| 73.0 \| 79 \| (23, 107) \| 227 \| (30, 117) \| 258 \| SprT-like domain-containing protein (Fragment) \| SprT-like domain-containing protein (Fragment); uniclust \| UniRef100\_A0A068WV36 \| 97.3 \| 9.2e-06 \| 2e-11 \| 79.4 \| 78 \| (28, 107) \| 227 \| (90, 183) \| 387 \| Protein with SprT-like domain at the N terminus \| Protein with SprT-like domain at the N terminus; uniclust \| UniRef100\_A0A497HZ04 \| 97.3 \| 9.8e-06 \| 2e-11 \| 66.1 \| 46 \| (56, 102) \| 227 \| (54, 100) \| 114 \| Uncharacterized protein \| Uncharacterized protein; uniclust \| UniRef100\_A0A6I8TF76 \| 97.3 \| 1.1e-05 \| 2.1e-11 \| 78.3 \| 71 \| (28, 100) \| 227 \| (361, 435) \| 557 \| Uncharacterized protein \| Uncharacterized protein; uniclust \| UniRef100\_A0A3M7QTT4 \| 97.3 \| 1.2e-05 \| 2.1e-11 \| 73.8 \| 78 \| (23, 105) \| 227 \| (109, 191) \| 338 \| Acidic repeat-containing (Fragment) \| Acidic repeat-containing (Fragment); uniclust \| UniRef100\_A0A803J7Z6 \| 97.3 \| 1.1e-05 \| 2.1e-11 \| 72.1 \| 78 \| (22, 105) \| 227 \| (82, 166) \| 235 \| SprT-like domain-containing protein \| SprT-like domain-containing protein; uniclust \| UniRef100\_A0A965MWP3 \| 97.3 \| 1.2e-05 \| 2.1e-11 \| 64.2 \| 53 \| (59, 112) \| 227 \| (4, 59) \| 121 \| SprT-like domain-containing protein \| SprT-like domain-containing protein; uniclust \| UniRef100\_A0A350I387 \| 97.3 \| 1.2e-05 \| 2.1e-11 \| 64.1 \| 60 \| (42, 107) \| 227 \| (47, 114) \| 120 \| SprT-like domain-containing protein \| SprT-like domain-containing protein; uniclust \| UniRef100\_A0A2H9ZV33 \| 97.3 \| 1e-05 \| 2.2e-11 \| 80.2 \| 71 \| (32, 106) \| 227 \| (87, 166) \| 459 \| SprT-like domain-containing protein \| SprT-like domain-containing protein; uniclust \| UniRef100\_A0A224XN60 \| 97.3 \| 1.1e-05 \| 2.2e-11 \| 68.8 \| 83 \| (14, 103) \| 227 \| (69, 165) \| 168 \| SprT-like domain-containing protein (Fragment) \| SprT-like domain-containing protein (Fragment); uniclust \| UniRef100\_A0A8J1IYR3 \| 97.3 \| 1e-05 \| 2.2e-11 \| 74.3 \| 89 \| (8, 104) \| 227 \| (56, 151) \| 246 \| SprT-like domain-containing protein Spartan \| SprT-like domain-containing protein Spartan; uniclust \| UniRef100\_B4CZ15 \| 97.3 \| 1.2e-05 \| 2.2e-11 \| 77.8 \| 73 \| (28, 108) \| 227 \| (437, 517) \| 582 \| Aminotransferase class V \| Aminotransferase class V; uniclust \| UniRef100\_A0A2R5LIW9 \| 97.3 \| 1.2e-05 \| 2.2e-11 \| 70.6 \| 132 \| (24, 222) \| 227 \| (98, 234) \| 238 \| Putative sprt-like metalloprotease (Fragment) \| Putative sprt-like metalloprotease (Fragment); uniclust \| UniRef100\_A0A0C3CV02 \| 97.3 \| 1.1e-05 \| 2.3e-11 \| 75.9 \| 89 \| (15, 106) \| 227 \| (114, 214) \| 332 \| SprT-like domain-containing protein \| SprT-like domain-containing protein; uniclust \| UniRef100\_A0A2E2K3M5 \| 97.3 \| 1.3e-05 \| 2.4e-11 \| 69.7 \| 63 \| (42, 110) \| 227 \| (50, 119) \| 198 \| SprT-like domain-containing protein \| SprT-like domain-containing protein; uniclust \| UniRef100\_UPI0003592C47 \| 97.3 \| 1.3e-05 \| 2.4e-11 \| 82.7 \| 133 \| (24, 220) \| 227 \| (1156, 1293) \| 1358 \| uncharacterized protein LOC101846459 \| uncharacterized protein LOC101846459; uniclust \| UniRef100\_A0A1Y4R9A6 \| 97.3 \| 1.2e-05 \| 2.4e-11 \| 70.3 \| 95 \| (4, 104) \| 227 \| (13, 111) \| 186 \| SprT-like domain-containing protein \| SprT-like domain-containing protein; uniclust \| UniRef100\_A0A183A629 \| 97.3 \| 1.3e-05 \| 2.5e-11 \| 71.1 \| 43 \| (57, 99) \| 227 \| (35, 82) \| 239 \| SprT-like domain-containing protein \| SprT-like domain-containing protein; uniclust \| UniRef100\_A0A3M7P3R5 \| 97.3 \| 1.3e-05 \| 2.5e-11 \| 74.8 \| 91 \| (9, 105) \| 227 \| (55, 159) \| 348 \| SprT-like domain-containing Spartan \| SprT-like domain-containing Spartan; uniclust \| UniRef100\_A0A1Z8WTI9 \| 97.3 \| 1.4e-05 \| 2.5e-11 \| 66.0 \| 83 \| (16, 109) \| 227 \| (41, 142) \| 151 \| SprT-like domain-containing protein \| SprT-like domain-containing protein; uniclust \| UniRef100\_R4TMW7 \| 97.3 \| 1.4e-05 \| 2.6e-11 \| 70.9 \| 44 \| (68, 111) \| 227 \| (128, 175) \| 256 \| SprT-like domain-containing protein \| SprT-like domain-containing protein; uniclust \| UniRef100\_A0A010RHC0 \| 97.3 \| 1.2e-05 \| 2.6e-11 \| 83.9 \| 73 \| (29, 103) \| 227 \| (444, 536) \| 687 \| SprT-like domain-containing protein \| SprT-like domain-containing protein; uniclust \| UniRef100\_A0A7V9Q7G0 \| 97.3 \| 1.4e-05 \| 2.6e-11 \| 68.6 \| 93 \| (7, 112) \| 227 \| (24, 123) \| 176 \| SprT-like domain-containing protein \| SprT-like domain-containing protein; uniclust \| UniRef100\_UPI001AC329C2 \| 97.3 \| 1.4e-05 \| 2.6e-11 \| 71.0 \| 39 \| (68, 106) \| 227 \| (218, 256) \| 262 \| hypothetical protein \| hypothetical protein; uniclust \| UniRef100\_A0A1E1XDU2 \| 97.3 \| 1.4e-05 \| 2.7e-11 \| 68.9 \| 69 \| (28, 101) \| 227 \| (91, 164) \| 206 \| Putative sprt-like metalloprotease (Fragment) \| Putative sprt-like metalloprotease (Fragment); uniclust \| UniRef100\_A0A7D9HBJ1 \| 97.3 \| 1.4e-05 \| 2.7e-11 \| 81.5 \| 80 \| (22, 105) \| 227 \| (708, 792) \| 909 \| SprT-like domain-containing protein \| SprT-like domain-containing protein; uniclust \| UniRef100\_A0A6P7TLT7 \| 97.3 \| 1.5e-05 \| 2.7e-11 \| 80.4 \| 135 \| (28, 224) \| 227 \| (735, 880) \| 937 \| Acidic repeat-containing protein-like \| Acidic repeat-containing protein-like; uniclust \| UniRef100\_A0A248SJR1 \| 97.3 \| 1.5e-05 \| 2.7e-11 \| 69.3 \| 74 \| (42, 121) \| 227 \| (60, 139) \| 218 \| SprT-like domain-containing protein \| SprT-like domain-containing protein; uniclust \| UniRef100\_A0A2R5L845 \| 97.3 \| 1.4e-05 \| 2.7e-11 \| 80.4 \| 94 \| (7, 106) \| 227 \| (58, 165) \| 605 \| Protein with SprT-like domain at the N terminus (Fragment) \| Protein with SprT-like domain at the N terminus (Fragment); uniclust \| UniRef100\_A0A3C1Z6Q3 \| 97.2 \| 1.6e-05 \| 2.8e-11 \| 70.9 \| 80 \| (22, 105) \| 227 \| (78, 177) \| 264 \| SprT-like domain-containing protein \| SprT-like domain-containing protein; uniclust \| UniRef100\_A0A3E0KDZ8 \| 97.2 \| 1.6e-05 \| 2.9e-11 \| 68.4 \| 67 \| (25, 102) \| 227 \| (71, 140) \| 202 \| SprT-like domain-containing protein \| SprT-like domain-containing protein; uniclust \| UniRef100\_A0A1V4KYG6 \| 97.2 \| 1.6e-05 \| 3.1e-11 \| 84.5 \| 97 \| (3, 105) \| 227 \| (1349, 1454) \| 1566 \| UDP-N-acetylglucosamine--peptide N-acetylglucosaminyltransferase 110 kDa subunit \| UDP-N-acetylglucosamine--peptide N-acetylglucosaminyltransferase 110 kDa subunit; uniclust \| UniRef100\_A0A521H7S3 \| 97.2 \| 1.7e-05 \| 3.2e-11 \| 69.1 \| 92 \| (17, 111) \| 227 \| (75, 173) \| 222 \| SprT-like domain-containing protein \| SprT-like domain-containing protein; uniclust \| UniRef100\_UPI00167373B6 \| 97.2 \| 1.8e-05 \| 3.2e-11 \| 67.6 \| 52 \| (59, 110) \| 227 \| (3, 65) \| 189 \| hypothetical protein \| hypothetical protein; uniclust \| UniRef100\_A0A7S0GYB2 \| 97.2 \| 1.7e-05 \| 3.2e-11 \| 72.0 \| 79 \| (29, 110) \| 227 \| (114, 195) \| 254 \| SprT-like domain-containing protein \| SprT-like domain-containing protein; uniclust \| UniRef100\_A0A6P1LIC7 \| 97.2 \| 1.8e-05 \| 3.3e-11 \| 71.1 \| 144 \| (42, 220) \| 227 \| (85, 251) \| 280 \| SprT-like domain-containing protein \| SprT-like domain-containing protein; uniclust \| UniRef100\_A0A0B2W3V2 \| 97.2 \| 1.8e-05 \| 3.4e-11 \| 66.1 \| 39 \| (185, 223) \| 227 \| (65, 103) \| 163 \| Acidic repeat-containing protein \| Acidic repeat-containing protein; uniclust \| UniRef100\_A0A087R6W8 \| 97.2 \| 1.8e-05 \| 3.4e-11 \| 79.5 \| 71 \| (29, 103) \| 227 \| (192, 271) \| 683 \| SprT-like domain-containing protein Spartan (Fragment) \| SprT-like domain-containing protein Spartan (Fragment); uniclust \| UniRef100\_A0A0H2STH3 \| 97.2 \| 1.9e-05 \| 3.5e-11 \| 72.6 \| 97 \| (1, 102) \| 227 \| (81, 184) \| 344 \| SprT-like domain-containing protein \| SprT-like domain-containing protein; uniclust \| UniRef100\_A0A0F9VPM0 \| 97.2 \| 1.8e-05 \| 3.7e-11 \| 68.6 \| 68 \| (29, 102) \| 227 \| (33, 108) \| 175 \| SprT-like domain-containing protein \| SprT-like domain-containing protein; uniclust \| UniRef100\_A0A6M2D328 \| 97.2 \| 1.9e-05 \| 3.7e-11 \| 70.8 \| 89 \| (2, 99) \| 227 \| (128, 222) \| 255 \| Putative sprt-like metalloprotease ovary overexpressed \| Putative sprt-like metalloprotease ovary overexpressed; uniclust \| UniRef100\_A0A1Q1N996 \| 97.2 \| 2e-05 \| 3.9e-11 \| 68.7 \| 65 \| (42, 112) \| 227 \| (44, 114) \| 196 \| SprT-like domain-containing protein \| SprT-like domain-containing protein; uniclust \| UniRef100\_A0A5A8CG24 \| 97.2 \| 2.1e-05 \| 3.9e-11 \| 72.7 \| 72 \| (29, 106) \| 227 \| (204, 285) \| 361 \| DHFR domain-containing protein \| DHFR domain-containing protein; uniclust \| UniRef100\_A0A822IAK2 \| 97.2 \| 2.1e-05 \| 3.9e-11 \| 67.2 \| 73 \| (23, 102) \| 227 \| (64, 142) \| 174 \| (mimic poison frog) hypothetical protein (Fragment) \| (mimic poison frog) hypothetical protein (Fragment); uniclust \| UniRef100\_A0A8D8EVU5 \| 97.2 \| 2.2e-05 \| 4e-11 \| 70.7 \| 73 \| (30, 104) \| 227 \| (46, 127) \| 285 \| SprT-like domain-containing protein Spartan (Fragment) \| SprT-like domain-containing protein Spartan (Fragment); uniclust \| UniRef100\_A0A5J4WFS3 \| 97.2 \| 2.1e-05 \| 4.1e-11 \| 73.5 \| 96 \| (2, 102) \| 227 \| (150, 257) \| 317 \| Putative sprt family metallopeptidase \| Putative sprt family metallopeptidase; uniclust \| UniRef100\_A0A0C2NES1 \| 97.2 \| 1.9e-05 \| 4.2e-11 \| 75.0 \| 84 \| (22, 112) \| 227 \| (31, 127) \| 311 \| SprT-like domain-containing protein \| SprT-like domain-containing protein; pdb70 \| 4QHF\_A \| 98.8 \| 6.1e-13 \| 6.4e-17 \| 98.9 \| 89 \| (1, 104) \| 227 \| (1, 93) \| 110 \| Uncharacterized protein MJ1213 \| 4QHF\_A Uncharacterized protein MJ1213 Minigluzincin, Proteolytic enzyme, HYDROLASE; pdb70 \| 4QHJ\_B \| 98.8 \| 7.5e-13 \| 7.9e-17 \| 98.3 \| 89 \| (1, 104) \| 227 \| (1, 93) \| 110 \| Uncharacterized protein MJ1213 \| 4QHJ\_B Uncharacterized protein MJ1213 Minigluzincin, Proteolytic enzyme, HYDROLASE HET: ACT; pdb70 \| 6MDX\_A \| 98.5 \| 2.5e-11 \| 2.6e-15 \| 97.7 \| 93 \| (3, 105) \| 227 \| (14, 115) \| 185 \| SprT-like domain-containing protein Spartan/DNA Complex \| 6MDX\_A SprT-like domain-containing protein Spartan/DNA Complex DPC repair, protease, DNA BINDING HET: MSE, PGE, FLC, MLZ; pdb70 \| 6MDW\_A \| 98.4 \| 7.1e-11 \| 7.3e-15 \| 96.0 \| 95 \| (3, 105) \| 227 \| (21, 124) \| 194 \| SprT-like domain-containing protein Spartan \| 6MDW\_A SprT-like domain-containing protein Spartan DPC repair protease, DNA BINDING HET: ADP, FLC, MLZ; pdb70 \| 4JIX\_A \| 98.3 \| 1.5e-10 \| 1.6e-14 \| 84.7 \| 86 \| (4, 105) \| 227 \| (9, 97) \| 112 \| Uncharacterized protein MJ0123 \| 4JIX\_A Uncharacterized protein MJ0123 Hydrolase, Metallopeptidase Zymogen, Minigluzincin HET: GOL, SO4; pdb70 \| 4JIU\_A \| 98.1 \| 1.2e-09 \| 1.3e-13 \| 79.4 \| 82 \| (7, 106) \| 227 \| (2, 87) \| 105 \| Putative uncharacterized protein \| 4JIU\_A Putative uncharacterized protein Hydrolase, Metallopeptidase Zymogen, Minigluzincin; pdb70 \| 5XBV\_A \| 97.9 \| 4.2e-09 \| 3.9e-13 \| 83.6 \| 56 \| (41, 105) \| 227 \| (77, 139) \| 149 \| Wss1p \| 5XBV\_A Wss1p protease, HYDROLASE; pdb70 \| 5XBN\_A \| 97.9 \| 4.4e-09 \| 4.1e-13 \| 83.4 \| 56 \| (41, 105) \| 227 \| (77, 139) \| 148 \| Wss1p \| 5XBN\_A Wss1p protease, HYDROLASE HET: MSE; pdb70 \| 5JIG\_A \| 97.5 \| 5.8e-08 \| 5.6e-12 \| 74.5 \| 55 \| (42, 105) \| 227 \| (61, 121) \| 127 \| Ubiquitin and WLM domain-containing metalloprotease \| 5JIG\_A Ubiquitin and WLM domain-containing metalloprotease Metalloprotease, DNA-Repair, Endoprotease, Regulation, hydrolase; pdb70 \| 3CQB\_A \| 95.8 \| 7.4e-05 \| 8.3e-09 \| 52.1 \| 68 \| (5, 85) \| 227 \| (26, 96) \| 107 \| Probable protease htpX homolog (E.C.3.4.24.-) \| 3CQB\_A Probable protease htpX homolog (E.C.3.4.24.-) Heat shock protein HtpX domain HET: EDO; pdb70 \| 6NCL\_c2 \| 95.3 \| 0.00029 \| 2.3e-08 \| 57.0 \| 46 \| (60, 105) \| 227 \| (83, 136) \| 181 \| PBCV-1 capsid \| 6NCL\_c2 PBCV-1 capsid tape-measure protein, minor capsid proteins; pdb70 \| 6NCL\_c3 \| 95.3 \| 0.00029 \| 2.3e-08 \| 57.0 \| 46 \| (60, 105) \| 227 \| (83, 136) \| 181 \| PBCV-1 capsid \| 6NCL\_c3 PBCV-1 capsid tape-measure protein, minor capsid proteins |
| Top keywords  (threshold 1.00e-03 (evalue)) | **SprT\_like, domain\_containing, Fragment, Acidic, SprT, repeat\_containing, metalloprotease, hypothetical, Spartan, Putative** |
| Output files | ../../similar\_sequences/34\_FANPEZAQ\_CDS\_0034\_merged.svg ../../similar\_sequences/34\_FANPEZAQ\_CDS\_0034\_pdb70.a3m ../../similar\_sequences/34\_FANPEZAQ\_CDS\_0034\_pdb70.hhr ../../similar\_sequences/34\_FANPEZAQ\_CDS\_0034\_uniclust.a3m ../../similar\_sequences/34\_FANPEZAQ\_CDS\_0034\_uniclust.hhr |

#### Structure prediction (AlphaFold)^2^

|  |  |
| --- | --- |
| Stats | xml version="1.0" encoding="utf-8" standalone="no"?       2024-09-02T21:09:33.674706 image/svg+xml   Matplotlib v3.7.2, https://matplotlib.org/ |
| Predicted structure | **NGL Viewer Controls:**  - Center: *Left-Click* - Rotate: *Left-Click + Drag* - Translate: *Right-Click + Drag* - Zoom: *Shift + Left-Click + Drag* |
| Output files | ../../predicted\_structures/34\_FANPEZAQ\_CDS\_0034/features.pkl ../../predicted\_structures/34\_FANPEZAQ\_CDS\_0034/ranked\_0.pdb ../../predicted\_structures/34\_FANPEZAQ\_CDS\_0034/ranked\_0\_plots.svg ../../predicted\_structures/34\_FANPEZAQ\_CDS\_0034/result\_model\_1\_ptm\_pred\_0.pkl |

#### Structure similarity search results (Foldseek)^3^

|  |  |
| --- | --- |
| Structure databases searched | Pdb, Afdb-proteome, Afdb-uniprot50 |
| Results, scheme(s)  (Top layers only, threshold 1.00e-02 (evalue)) | xml version="1.0" encoding="utf-8" standalone="no"?       2024-09-02T21:11:06.896100 image/svg+xml   Matplotlib v3.7.2, https://matplotlib.org/ |
| Results, table  (threshold 1.00e-02 (evalue)) | db \| id \| prob \| evalue \| bits \| fident \| alnlen \| mismatch \| gapopen \| qstart \| qend \| tstart \| tend \| name \| description; afdb-uniprot50 \| AF-A0A2W5L353-F1-MODEL\_V4 \| 1.0 \| 1.243e-23 \| 760 \| 0.497 \| 231 \| 101 \| 4 \| 1 \| 224 \| 1 \| 223 \| Transcription elongation protein SprT \| Transcription elongation protein SprT; afdb-uniprot50 \| AF-A0A3N8XMK1-F1-MODEL\_V4 \| 1.0 \| 2.833e-21 \| 701 \| 0.43 \| 230 \| 119 \| 5 \| 1 \| 220 \| 12 \| 239 \| Transcription elongation protein SprT \| Transcription elongation protein SprT; afdb-uniprot50 \| AF-A0A149SP00-F1-MODEL\_V4 \| 1.0 \| 3.791e-20 \| 674 \| 0.432 \| 208 \| 104 \| 4 \| 15 \| 218 \| 2 \| 199 \| Uncharacterized protein \| Uncharacterized protein; afdb-uniprot50 \| AF-S9SWB4-F1-MODEL\_V4 \| 1.0 \| 2.096e-21 \| 670 \| 0.431 \| 234 \| 105 \| 6 \| 1 \| 226 \| 36 \| 249 \| Transcription elongation protein SprT \| Transcription elongation protein SprT; afdb-uniprot50 \| AF-A0A1B0Z2A4-F1-MODEL\_V4 \| 1.0 \| 4.776e-19 \| 669 \| 0.392 \| 227 \| 94 \| 6 \| 1 \| 220 \| 1 \| 190 \| Uncharacterized protein \| Uncharacterized protein; afdb-uniprot50 \| AF-A0A1V2BSP3-F1-MODEL\_V4 \| 1.0 \| 2.46e-19 \| 648 \| 0.404 \| 225 \| 98 \| 3 \| 2 \| 221 \| 5 \| 198 \| SprT-like domain-containing protein \| SprT-like domain-containing protein; afdb-uniprot50 \| AF-A0A1L3ZVD3-F1-MODEL\_V4 \| 1.0 \| 7.738e-19 \| 641 \| 0.375 \| 224 \| 102 \| 5 \| 1 \| 218 \| 5 \| 196 \| SprT-like domain-containing protein \| SprT-like domain-containing protein; afdb-uniprot50 \| AF-A0A494WBN0-F1-MODEL\_V4 \| 1.0 \| 4.277e-20 \| 614 \| 0.42 \| 240 \| 93 \| 8 \| 2 \| 227 \| 21 \| 228 \| Transcription elongation protein SprT \| Transcription elongation protein SprT; afdb-uniprot50 \| AF-A0A349QHC6-F1-MODEL\_V4 \| 1.0 \| 1.318e-17 \| 594 \| 0.325 \| 221 \| 111 \| 5 \| 1 \| 219 \| 14 \| 198 \| Transcription elongation protein SprT \| Transcription elongation protein SprT; afdb-uniprot50 \| AF-A0A7G7MAJ5-F1-MODEL\_V4 \| 1.0 \| 2.292e-18 \| 591 \| 0.409 \| 222 \| 92 \| 6 \| 2 \| 217 \| 5 \| 193 \| SprT-like domain-containing protein \| SprT-like domain-containing protein; afdb-uniprot50 \| AF-A0A537PHM0-F1-MODEL\_V4 \| 1.0 \| 6.079e-19 \| 586 \| 0.381 \| 223 \| 99 \| 6 \| 1 \| 218 \| 54 \| 242 \| Transcription elongation protein SprT \| Transcription elongation protein SprT; afdb-uniprot50 \| AF-A0A6N6S9L2-F1-MODEL\_V4 \| 1.0 \| 2.113e-16 \| 568 \| 0.333 \| 222 \| 109 \| 4 \| 1 \| 218 \| 1 \| 187 \| Transcription elongation protein SprT \| Transcription elongation protein SprT; afdb-uniprot50 \| AF-A0A7W4YW64-F1-MODEL\_V4 \| 1.0 \| 7.285e-19 \| 549 \| 0.392 \| 237 \| 109 \| 6 \| 8 \| 227 \| 2 \| 220 \| Uncharacterized protein \| Uncharacterized protein; afdb-uniprot50 \| AF-A0A1Q8Y9H4-F1-MODEL\_V4 \| 1.0 \| 4.146e-17 \| 546 \| 0.373 \| 225 \| 122 \| 7 \| 15 \| 227 \| 2 \| 219 \| Uncharacterized protein \| Uncharacterized protein; afdb-uniprot50 \| AF-A0A257NL89-F1-MODEL\_V4 \| 1.0 \| 1.596e-18 \| 543 \| 0.371 \| 234 \| 109 \| 6 \| 1 \| 227 \| 8 \| 210 \| Uncharacterized protein \| Uncharacterized protein; afdb-uniprot50 \| AF-A0A7Y4QDV3-F1-MODEL\_V4 \| 1.0 \| 3.675e-17 \| 537 \| 0.326 \| 239 \| 113 \| 6 \| 1 \| 227 \| 1 \| 203 \| Uncharacterized protein \| Uncharacterized protein; afdb-uniprot50 \| AF-A0A562K4X6-F1-MODEL\_V4 \| 1.0 \| 7.135e-17 \| 537 \| 0.347 \| 239 \| 109 \| 9 \| 3 \| 227 \| 12 \| 217 \| Uncharacterized protein \| Uncharacterized protein; afdb-uniprot50 \| AF-A0A2M8QJE3-F1-MODEL\_V4 \| 1.0 \| 3.903e-17 \| 535 \| 0.39 \| 223 \| 95 \| 6 \| 14 \| 227 \| 6 \| 196 \| Transcription elongation protein SprT \| Transcription elongation protein SprT; afdb-uniprot50 \| AF-A0A4V0X9F2-F1-MODEL\_V4 \| 1.0 \| 9.547e-16 \| 534 \| 0.318 \| 220 \| 109 \| 6 \| 1 \| 216 \| 4 \| 186 \| Uncharacterized protein \| Uncharacterized protein; afdb-uniprot50 \| AF-A0A7X4GG86-F1-MODEL\_V4 \| 1.0 \| 2.01e-17 \| 526 \| 0.357 \| 224 \| 104 \| 5 \| 1 \| 218 \| 27 \| 216 \| Transcription elongation protein SprT \| Transcription elongation protein SprT; afdb-uniprot50 \| AF-A0A0U4H3G1-F1-MODEL\_V4 \| 1.0 \| 3.863e-16 \| 516 \| 0.36 \| 222 \| 104 \| 6 \| 1 \| 218 \| 8 \| 195 \| Transcription elongation protein SprT \| Transcription elongation protein SprT; afdb-uniprot50 \| AF-A0A419HBS0-F1-MODEL\_V4 \| 1.0 \| 3.637e-16 \| 516 \| 0.351 \| 222 \| 123 \| 7 \| 1 \| 218 \| 6 \| 210 \| Uncharacterized protein \| Uncharacterized protein; afdb-uniprot50 \| AF-A0A382AGK6-F1-MODEL\_V4 \| 1.0 \| 7.967e-16 \| 513 \| 0.297 \| 222 \| 119 \| 3 \| 1 \| 221 \| 1 \| 186 \| SprT-like domain-containing protein \| SprT-like domain-containing protein; afdb-uniprot50 \| AF-A0A158SQJ2-F1-MODEL\_V4 \| 1.0 \| 2.113e-16 \| 507 \| 0.331 \| 241 \| 114 \| 9 \| 1 \| 227 \| 28 \| 235 \| Uncharacterized protein \| Uncharacterized protein; afdb-uniprot50 \| AF-A0A498AIE9-F1-MODEL\_V4 \| 1.0 \| 2.69e-16 \| 502 \| 0.343 \| 221 \| 110 \| 5 \| 1 \| 218 \| 153 \| 341 \| SprT-like family protein \| SprT-like family protein; afdb-uniprot50 \| AF-A0A5B8DFB0-F1-MODEL\_V4 \| 1.0 \| 1.371e-15 \| 490 \| 0.324 \| 222 \| 108 \| 6 \| 1 \| 218 \| 5 \| 188 \| SprT family zinc-dependent metalloprotease \| SprT family zinc-dependent metalloprotease; afdb-uniprot50 \| AF-I6APB7-F1-MODEL\_V4 \| 1.0 \| 1.643e-15 \| 490 \| 0.308 \| 230 \| 118 \| 8 \| 2 \| 227 \| 24 \| 216 \| SprT-like family \| SprT-like family; afdb-uniprot50 \| AF-A0A7W7AN23-F1-MODEL\_V4 \| 1.0 \| 1.343e-13 \| 488 \| 0.422 \| 161 \| 83 \| 4 \| 1 \| 153 \| 4 \| 162 \| SprT-like domain-containing protein \| SprT-like domain-containing protein; afdb-uniprot50 \| AF-A0A7K3ZFI4-F1-MODEL\_V4 \| 1.0 \| 6.579e-15 \| 488 \| 0.285 \| 228 \| 120 \| 6 \| 1 \| 222 \| 10 \| 200 \| PH domain-containing protein \| PH domain-containing protein; afdb-uniprot50 \| AF-A0A7W7A6F7-F1-MODEL\_V4 \| 1.0 \| 2.245e-16 \| 474 \| 0.34 \| 241 \| 105 \| 8 \| 1 \| 226 \| 4 \| 205 \| Uncharacterized protein \| Uncharacterized protein; afdb-uniprot50 \| AF-A0A315D311-F1-MODEL\_V4 \| 1.0 \| 1.132e-14 \| 470 \| 0.29 \| 220 \| 118 \| 4 \| 1 \| 218 \| 5 \| 188 \| SprT-like domain-containing protein \| SprT-like domain-containing protein; afdb-uniprot50 \| AF-A0A3D5EPJ0-F1-MODEL\_V4 \| 1.0 \| 4.269e-14 \| 464 \| 0.301 \| 222 \| 113 \| 6 \| 2 \| 218 \| 19 \| 203 \| Transcription elongation protein SprT \| Transcription elongation protein SprT; afdb-uniprot50 \| AF-A0A4P5RR10-F1-MODEL\_V4 \| 1.0 \| 2.36e-15 \| 464 \| 0.303 \| 221 \| 113 \| 5 \| 1 \| 218 \| 22 \| 204 \| SprT-like domain-containing protein \| SprT-like domain-containing protein; afdb-uniprot50 \| AF-A0A1E1F840-F1-MODEL\_V4 \| 1.0 \| 3.223e-16 \| 461 \| 0.411 \| 214 \| 85 \| 7 \| 23 \| 227 \| 24 \| 205 \| Uncharacterized protein \| Uncharacterized protein; afdb-uniprot50 \| AF-K9CK51-F1-MODEL\_V4 \| 1.0 \| 7.423e-15 \| 453 \| 0.434 \| 205 \| 75 \| 7 \| 32 \| 227 \| 3 \| 175 \| Uncharacterized protein \| Uncharacterized protein; afdb-uniprot50 \| AF-A0A212SZK8-F1-MODEL\_V4 \| 1.0 \| 3.783e-14 \| 451 \| 0.271 \| 228 \| 129 \| 3 \| 1 \| 227 \| 10 \| 201 \| SprT-like family protein \| SprT-like family protein; afdb-uniprot50 \| AF-A0A2D6X6V1-F1-MODEL\_V4 \| 1.0 \| 2.198e-14 \| 445 \| 0.293 \| 232 \| 123 \| 5 \| 1 \| 227 \| 2 \| 197 \| Uncharacterized protein \| Uncharacterized protein; afdb-uniprot50 \| AF-A0A520VTD7-F1-MODEL\_V4 \| 1.0 \| 2.972e-14 \| 445 \| 0.275 \| 225 \| 124 \| 3 \| 1 \| 220 \| 29 \| 219 \| M48 family peptidase \| M48 family peptidase; afdb-uniprot50 \| AF-A0A1C6ST94-F1-MODEL\_V4 \| 1.0 \| 4.866e-15 \| 445 \| 0.342 \| 222 \| 107 \| 7 \| 3 \| 218 \| 249 \| 437 \| SprT-like family protein \| SprT-like family protein; afdb-uniprot50 \| AF-A0A3N7GVC1-F1-MODEL\_V4 \| 1.0 \| 6.511e-14 \| 443 \| 0.301 \| 222 \| 117 \| 8 \| 2 \| 218 \| 6 \| 194 \| Uncharacterized protein \| Uncharacterized protein; afdb-uniprot50 \| AF-A0A257FUX5-F1-MODEL\_V4 \| 1.0 \| 1.778e-11 \| 440 \| 0.351 \| 148 \| 95 \| 1 \| 1 \| 147 \| 5 \| 152 \| SprT-like domain-containing protein \| SprT-like domain-containing protein; afdb-uniprot50 \| AF-A0A5E7XVZ1-F1-MODEL\_V4 \| 1.0 \| 4.816e-14 \| 435 \| 0.355 \| 194 \| 110 \| 5 \| 2 \| 184 \| 5 \| 194 \| Transcription elongation protein SprT \| Transcription elongation protein SprT; afdb-uniprot50 \| AF-A0A5J6UEE7-F1-MODEL\_V4 \| 1.0 \| 2.048e-13 \| 432 \| 0.279 \| 236 \| 114 \| 7 \| 1 \| 219 \| 3 \| 199 \| SprT family zinc-dependent metalloprotease \| SprT family zinc-dependent metalloprotease; afdb-uniprot50 \| AF-A0A1C4U4K3-F1-MODEL\_V4 \| 1.0 \| 4.581e-15 \| 432 \| 0.36 \| 222 \| 103 \| 7 \| 3 \| 218 \| 182 \| 370 \| SprT-like family protein \| SprT-like family protein; afdb-uniprot50 \| AF-A0A0F9KWU5-F1-MODEL\_V4 \| 1.0 \| 9.829e-13 \| 431 \| 0.27 \| 222 \| 120 \| 5 \| 1 \| 218 \| 1 \| 184 \| SprT-like domain-containing protein \| SprT-like domain-containing protein; afdb-uniprot50 \| AF-D6TL66-F1-MODEL\_V4 \| 1.0 \| 1.515e-13 \| 430 \| 0.276 \| 221 \| 116 \| 6 \| 3 \| 218 \| 12 \| 193 \| Uncharacterized protein \| Uncharacterized protein; afdb-uniprot50 \| AF-A0A4Q2J0L1-F1-MODEL\_V4 \| 1.0 \| 2.335e-14 \| 430 \| 0.357 \| 221 \| 106 \| 5 \| 3 \| 218 \| 104 \| 293 \| Uncharacterized protein \| Uncharacterized protein; afdb-uniprot50 \| AF-A0A661CLR8-F1-MODEL\_V4 \| 1.0 \| 2.769e-13 \| 427 \| 0.263 \| 228 \| 120 \| 8 \| 2 \| 218 \| 6 \| 196 \| SprT-like domain-containing protein \| SprT-like domain-containing protein; afdb-uniprot50 \| AF-A0A849MYV3-F1-MODEL\_V4 \| 1.0 \| 3.353e-14 \| 427 \| 0.292 \| 239 \| 129 \| 9 \| 1 \| 226 \| 13 \| 224 \| Uncharacterized protein \| Uncharacterized protein; afdb-uniprot50 \| AF-A0A2W5Y178-F1-MODEL\_V4 \| 1.0 \| 4.816e-14 \| 421 \| 0.321 \| 233 \| 134 \| 11 \| 1 \| 227 \| 10 \| 224 \| SprT-like domain-containing protein \| SprT-like domain-containing protein; afdb-uniprot50 \| AF-A0A661MH10-F1-MODEL\_V4 \| 1.0 \| 1.343e-13 \| 417 \| 0.276 \| 257 \| 124 \| 12 \| 1 \| 227 \| 30 \| 254 \| SprT-like domain-containing protein \| SprT-like domain-containing protein; afdb-uniprot50 \| AF-A0A2W6B5M7-F1-MODEL\_V4 \| 1.0 \| 5.831e-15 \| 410 \| 0.336 \| 223 \| 113 \| 7 \| 11 \| 227 \| 2 \| 195 \| Transcription elongation protein SprT \| Transcription elongation protein SprT; afdb-uniprot50 \| AF-A0A2D6XC71-F1-MODEL\_V4 \| 1.0 \| 3.667e-11 \| 405 \| 0.329 \| 155 \| 99 \| 3 \| 2 \| 153 \| 6 \| 158 \| Transcription elongation protein SprT \| Transcription elongation protein SprT; afdb-uniprot50 \| AF-A0A3N5NIA4-F1-MODEL\_V4 \| 1.0 \| 6.634e-10 \| 403 \| 0.32 \| 128 \| 84 \| 2 \| 1 \| 127 \| 16 \| 141 \| Uncharacterized protein \| Uncharacterized protein; afdb-uniprot50 \| AF-A0A0F9B2V9-F1-MODEL\_V4 \| 1.0 \| 5.594e-11 \| 402 \| 0.327 \| 159 \| 104 \| 1 \| 1 \| 156 \| 1 \| 159 \| Uncharacterized protein \| Uncharacterized protein; afdb-uniprot50 \| AF-A0A2E4HFW3-F1-MODEL\_V4 \| 1.0 \| 2.429e-12 \| 396 \| 0.241 \| 228 \| 125 \| 7 \| 2 \| 218 \| 8 \| 198 \| Uncharacterized protein \| Uncharacterized protein; afdb-uniprot50 \| AF-A0A0Q7CQI4-F1-MODEL\_V4 \| 1.0 \| 2.997e-09 \| 353 \| 0.318 \| 138 \| 86 \| 5 \| 1 \| 135 \| 18 \| 150 \| Uncharacterized protein \| Uncharacterized protein; afdb-uniprot50 \| AF-A0A160TKS1-F1-MODEL\_V4 \| 1.0 \| 5.819e-09 \| 305 \| 0.355 \| 163 \| 76 \| 7 \| 2 \| 153 \| 5 \| 149 \| SprT-like domain-containing protein \| SprT-like domain-containing protein; afdb-uniprot50 \| AF-A0A315CPA0-F1-MODEL\_V4 \| 1.0 \| 2.45e-07 \| 296 \| 0.241 \| 141 \| 103 \| 2 \| 2 \| 138 \| 216 \| 356 \| Uncharacterized protein \| Uncharacterized protein; afdb-uniprot50 \| AF-A0A1F6D4Y3-F1-MODEL\_V4 \| 1.0 \| 4.856e-09 \| 295 \| 0.179 \| 239 \| 143 \| 9 \| 1 \| 227 \| 9 \| 206 \| Uncharacterized protein \| Uncharacterized protein; afdb-uniprot50 \| AF-A0A315EGI5-F1-MODEL\_V4 \| 1.0 \| 1.945e-08 \| 284 \| 0.289 \| 159 \| 98 \| 4 \| 2 \| 150 \| 20 \| 173 \| SprT-like domain-containing protein \| SprT-like domain-containing protein; afdb-uniprot50 \| AF-A0A144J432-F1-MODEL\_V4 \| 1.0 \| 7.95e-10 \| 283 \| 0.404 \| 163 \| 62 \| 5 \| 68 \| 227 \| 1 \| 131 \| Transcription elongation protein \| Transcription elongation protein; afdb-uniprot50 \| AF-A0A315DXF0-F1-MODEL\_V4 \| 1.0 \| 2.475e-08 \| 283 \| 0.296 \| 162 \| 102 \| 6 \| 3 \| 155 \| 22 \| 180 \| SprT-like domain-containing protein \| SprT-like domain-containing protein; afdb-uniprot50 \| AF-A0A126REC0-F1-MODEL\_V4 \| 1.0 \| 7.18e-06 \| 243 \| 0.421 \| 95 \| 49 \| 2 \| 2 \| 90 \| 5 \| 99 \| Uncharacterized protein \| Uncharacterized protein; afdb-uniprot50 \| AF-A0A259FRY7-F1-MODEL\_V4 \| 1.0 \| 3.054e-05 \| 223 \| 0.395 \| 81 \| 49 \| 0 \| 3 \| 83 \| 7 \| 87 \| SprT domain-containing protein \| SprT domain-containing protein; afdb-uniprot50 \| AF-A0A4S2U167-F1-MODEL\_V4 \| 1.0 \| 4.216e-07 \| 178 \| 0.211 \| 251 \| 151 \| 15 \| 3 \| 217 \| 32 \| 271 \| Uncharacterized protein \| Uncharacterized protein; afdb-uniprot50 \| AF-A0A1F6M955-F1-MODEL\_V4 \| 1.0 \| 6.83e-07 \| 170 \| 0.175 \| 245 \| 151 \| 17 \| 1 \| 217 \| 289 \| 510 \| SprT-like domain-containing protein \| SprT-like domain-containing protein; afdb-uniprot50 \| AF-A0A7Y6IZN4-F1-MODEL\_V4 \| 1.0 \| 1.326e-06 \| 169 \| 0.194 \| 257 \| 165 \| 13 \| 3 \| 227 \| 8 \| 254 \| Uncharacterized protein \| Uncharacterized protein; afdb-uniprot50 \| AF-A0A7D4X9Y0-F1-MODEL\_V4 \| 1.0 \| 2.424e-06 \| 163 \| 0.172 \| 266 \| 168 \| 15 \| 4 \| 227 \| 28 \| 283 \| Uncharacterized protein \| Uncharacterized protein; afdb-uniprot50 \| AF-A0A6G9F5T1-F1-MODEL\_V4 \| 1.0 \| 2.424e-06 \| 153 \| 0.175 \| 262 \| 158 \| 18 \| 1 \| 227 \| 2 \| 240 \| Uncharacterized protein \| Uncharacterized protein; afdb-uniprot50 \| AF-A0A429FBE9-F1-MODEL\_V4 \| 1.0 \| 7.626e-06 \| 149 \| 0.159 \| 269 \| 167 \| 17 \| 3 \| 227 \| 12 \| 265 \| Uncharacterized protein \| Uncharacterized protein; afdb-uniprot50 \| AF-A0A1I1XSQ6-F1-MODEL\_V4 \| 1.0 \| 3.445e-05 \| 136 \| 0.171 \| 239 \| 167 \| 13 \| 1 \| 217 \| 17 \| 246 \| Uncharacterized protein \| Uncharacterized protein; afdb-uniprot50 \| AF-A0A2S8IY47-F1-MODEL\_V4 \| 1.0 \| 0.0007934 \| 129 \| 0.411 \| 85 \| 44 \| 4 \| 146 \| 227 \| 2 \| 83 \| Uncharacterized protein \| Uncharacterized protein; afdb-uniprot50 \| AF-A0A2I1DJ04-F1-MODEL\_V4 \| 1.0 \| 0.0005868 \| 111 \| 0.126 \| 301 \| 167 \| 17 \| 4 \| 218 \| 8 \| 298 \| Uncharacterized protein \| Uncharacterized protein; afdb-uniprot50 \| AF-A0A429A0F0-F1-MODEL\_V4 \| 1.0 \| 0.000321 \| 108 \| 0.191 \| 230 \| 135 \| 13 \| 31 \| 226 \| 2 \| 214 \| Uncharacterized protein \| Uncharacterized protein; afdb-uniprot50 \| AF-A0A179BPY3-F1-MODEL\_V4 \| 1.0 \| 0.003807 \| 107 \| 0.12 \| 315 \| 173 \| 13 \| 5 \| 227 \| 26 \| 328 \| SprT-like domain-containing protein \| SprT-like domain-containing protein; afdb-uniprot50 \| AF-Q9AEZ9-F1-MODEL\_V4 \| 0.999 \| 0.0001756 \| 98 \| 0.166 \| 228 \| 145 \| 13 \| 40 \| 227 \| 15 \| 237 \| Uncharacterized protein \| Uncharacterized protein; afdb-uniprot50 \| AF-A0A2N8TS96-F1-MODEL\_V4 \| 0.992 \| 0.003807 \| 85 \| 0.184 \| 228 \| 140 \| 14 \| 34 \| 222 \| 1 \| 221 \| Uncharacterized protein \| Uncharacterized protein |
| Top keywords  (threshold 1.00e-02 (evalue)) | **SprT\_like, domain\_containing, SprT, Transcription, elongation, zinc\_dependent, metalloprotease, PH, M48, peptidase** |
| Output files | ../../similar\_structures/34\_FANPEZAQ\_CDS\_0034\_afdb-proteome\_foldseek.tsv ../../similar\_structures/34\_FANPEZAQ\_CDS\_0034\_afdb-uniprot50\_foldseek.tsv ../../similar\_structures/34\_FANPEZAQ\_CDS\_0034\_merged.svg ../../similar\_structures/34\_FANPEZAQ\_CDS\_0034\_pdb\_foldseek.tsv |

  
  
  

Return to summary | Go to previous | Go to next

  


---

**Sequence/structure alignments coloring**  
Each object in the alignment figures is colored according to its E-value following this color coding:

1e-100
10
