## Supplementary material for "Completing the BASEL phage collection to unlock hidden diversity for systematic exploration of phage-host interactions": Data S2: 35.html

FANPEZAQ\_CDS\_0035


Return to summary | Go to previous | Go to next

|  |  |
| --- | --- |
| FANPEZAQ\_CDS\_0035  Page creation date: 02 Sep 2024, 12:00  Project folder: n/a  Input sequences file: Escherichia\_virus\_HeidiAbel.gb | single\_stranded dna\_binding binding dna single\_strand replication beta a primosomal n complex mitochondrial fold yes ob fragment single engineered rna nucleic acid\_binding proteins organism\_taxid barrel coli mainly arch dihydrolipoamide acetyltransferase e2p ssb escherichia expressed in expression\_system\_taxid editing b rim1 mp18 gene c rna\_editing hypothetical strand strain and antibody putative stranded synonym |

### Sequence information

|  |  |
| --- | --- |
| Name | FANPEZAQ\_CDS\_0035  35\_FANPEZAQ\_CDS\_0035 (pipeline id) |
| Imported annotations |  |
| Protein sequence | MAGSLNKVEIIGNLGNDPEFRTLPNGNGVAQLSIATSETWKDKQTGEKKEKTEWHRVVIF GKLAEIASQYLRKGSKVFIEGALQTRKWQDQAGVDRYTTEIVVGMNGRLLMLSAANDNQA GGQRQQQQPGGGQQFSGGGQQRPPAQGAGNEPPMDFDDDIPF |
| Number of residues | 162 |
| Molecular weight (Da) | 17644.45 |
| Output files | ../../query\_sequences/35\_FANPEZAQ\_CDS\_0035.fasta |

### Putative domain architecture and protein family

#### Search results (HHblits)^1^

|  |  |
| --- | --- |
| Domain family databases searched | Pfam, Ncbi-cd, Cath, Phrogs |
| Results, scheme(s)  (Top layers only; threshold 1.00e-03 (evalue)) | xml version="1.0" encoding="utf-8" standalone="no"?       2024-09-02T21:08:19.988308 image/svg+xml   Matplotlib v3.7.2, https://matplotlib.org/ |
| Results, table  (E-value ≤ 1.00e-03 (evalue)) | db \| id \| prob \| evalue \| pvalue \| score \| cols \| query \| query\_len \| template \| template\_len \| name \| description; pfam \| PF00436 \| 99.3 \| 7.9e-17 \| 1.6e-20 \| 105.1 \| 94 \| (5, 105) \| 162 \| (1, 98) \| 102 \| SSB \| Single-strand binding protein family; pfam \| PF11506 \| 99.2 \| 2.8e-16 \| 5.7e-20 \| 101.1 \| 92 \| (4, 108) \| 162 \| (1, 92) \| 94 \| DUF3217 \| Protein of unknown function (DUF3217); ncbi-cd \| cd04496 \| 99.2 \| 4.2e-16 \| 9.4e-20 \| 100.9 \| 95 \| (8, 104) \| 162 \| (1, 95) \| 100 \| SSB\_OBF \| cd04496 SSB\_OBF; SSB\_OBF: A subfamily of OB folds similar to the OB fold of ssDNA-binding protein (SSB).; cath \| 2cwaA01 \| 99.6 \| 4.6e-20 \| 6.9e-24 \| 123.0 \| 112 \| (1, 114) \| 162 \| (1, 112) \| 126 \| Single-strand binding protein \| CATHCODE: 2.40.50.140 NAME: Single-strand binding protein. Chain: a. Synonym: ssb, helix-destabilizing protein, single-stranded DNA binding protein. Engineered: yes SOURCE: Thermus thermophilus. Organism\_taxid: 300852. Strain: hb8. Expressed in: escherichia coli. Expression\_system\_taxid: 562. CLASS: Mainly Beta, ARCH: Beta Barrel, TOPOL: OB fold (Dihydrolipoamide Acetyltransferase, E2P), HOMOL: Nucleic acid-binding proteins; cath \| 3ulpD00 \| 99.5 \| 4.5e-19 \| 6.8e-23 \| 118.0 \| 114 \| (1, 114) \| 162 \| (2, 122) \| 124 \| Single-strand binding protein \| CATHCODE: 2.40.50.140 NAME: Single-strand binding protein. Chain: a, b, c, d. Fragment: unp residues 77-200. Engineered: yes. Dna (35-mer). Chain: r, q. Engineered: yes SOURCE: Plasmodium falciparum. Organism\_taxid: 5833. Gene: pfe0435c. Expressed in: escherichia coli. Expression\_system\_taxid: 562. Expression\_system\_vector\_type: pet21a. CLASS: Mainly Beta, ARCH: Beta Barrel, TOPOL: OB fold (Dihydrolipoamide Acetyltransferase, E2P), HOMOL: Nucleic acid-binding proteins; cath \| 1se8A02 \| 99.5 \| 4.6e-19 \| 7e-23 \| 116.8 \| 108 \| (1, 110) \| 162 \| (9, 116) \| 117 \| Single-strand binding protein \| CATHCODE: 2.40.50.140 NAME: Single-strand binding protein. Chain: a. Synonym: ssb, helix-destabilizing protein. Engineered: yes SOURCE: Deinococcus radiodurans. Organism\_taxid: 1299. Gene: ssb, dr0099. Expressed in: escherichia coli. Expression\_system\_taxid: 562. CLASS: Mainly Beta, ARCH: Beta Barrel, TOPOL: OB fold (Dihydrolipoamide Acetyltransferase, E2P), HOMOL: Nucleic acid-binding proteins; cath \| 1ue6D00 \| 99.5 \| 5e-19 \| 7.5e-23 \| 116.9 \| 114 \| (2, 115) \| 162 \| (1, 115) \| 119 \| Single-strand binding protein \| CATHCODE: 2.40.50.140 NAME: Single-strand binding protein. Chain: a, b, c, d. Synonym: single-stranded DNA-binding protein. Engineered: yes SOURCE: Mycobacterium tuberculosis. Organism\_taxid: 1773. Expressed in: escherichia coli. Expression\_system\_taxid: 562. Expression\_system\_vector\_type: plasmid. CLASS: Mainly Beta, ARCH: Beta Barrel, TOPOL: OB fold (Dihydrolipoamide Acetyltransferase, E2P), HOMOL: Nucleic acid-binding proteins; cath \| 5odnC00 \| 99.5 \| 5.9e-19 \| 8.9e-23 \| 114.6 \| 108 \| (1, 109) \| 162 \| (3, 110) \| 110 \| Single-stranded DNA-binding protein \| CATHCODE: 2.40.50.140 NAME: Single-stranded DNA-binding protein. Chain: a, b, c, d, e, f, g, h. Synonym: ssb. Engineered: yes. Dna (5'-d(p\*tp\*tp\*tp\*t)-3'). Chain: i, m,j, k, n, q. Engineered: yes. Dna (5'-d(\*tp\*tp\*t)-3'). Chain: p, l. Engineered: yes SOURCE: Salinibacter ruber (strain dsm 13855 / m31). Organism\_taxid: 309807. Gene: ssb, sru\_0523. Expressed in: escherichia coli. Expression\_system\_taxid: 562. Yes. Organism\_scientific: salinibacter ruber. Organism\_taxid: 146919. CLASS: Mainly Beta, ARCH: Beta Barrel, TOPOL: OB fold (Dihydrolipoamide Acetyltransferase, E2P), HOMOL: Nucleic acid-binding proteins; cath \| 3k7uC00 \| 99.5 \| 6.3e-19 \| 9.5e-23 \| 116.5 \| 110 \| (1, 112) \| 162 \| (2, 117) \| 118 \| Antibody \| CATHCODE: 2.40.50.140 NAME: Antibody. Chain: a. Engineered: yes. Mp18 RNA editing complex protein.Chain: c. Engineered: yes SOURCE: Lama glama. Organism\_taxid: 9844. Expressed in: escherichia coli. Expression\_system\_taxid: 562. CLASS: Mainly Beta, ARCH: Beta Barrel, TOPOL: OB fold (Dihydrolipoamide Acetyltransferase, E2P), HOMOL: Nucleic acid-binding proteins; cath \| 4damC00 \| 99.5 \| 1.4e-18 \| 2e-22 \| 117.8 \| 114 \| (1, 116) \| 162 \| (11, 125) \| 128 \| Single-stranded DNA-binding protein 1 \| CATHCODE: 2.40.50.140 NAME: Single-stranded DNA-binding protein 1. Chain: a, b, c, d, e, f, g, h, i, j, k, l. Fragment: unp residues 1-116. Synonym: ssb 1, helix-destabilizing protein 1. Engineered: yes. Mutation: yes SOURCE: Streptomyces coelicolor. Organism\_taxid: 1902. Strain: m145. Gene: scc61a.04c, sco2683, ssb1. Expressed in: escherichia coli. Expression\_system\_taxid: 562. CLASS: Mainly Beta, ARCH: Beta Barrel, TOPOL: OB fold (Dihydrolipoamide Acetyltransferase, E2P), HOMOL: Nucleic acid-binding proteins; cath \| 2vw9B00 \| 99.5 \| 2.2e-18 \| 3.4e-22 \| 110.9 \| 105 \| (4, 109) \| 162 \| (1, 105) \| 106 \| Single-stranded DNA binding protein \| CATHCODE: 2.40.50.140 NAME: Single-stranded DNA binding protein. Chain: a, b. Fragment: residues 1-134. Synonym: ssb, helix-destabilizing protein. Engineered: yes. Poly-dt. Chain: c. Engineered: yes SOURCE: Helicobacter pylori. Organism\_taxid: 85962. Strain: 26695. Expressed in: escherichia coli. Expression\_system\_taxid: 562. CLASS: Mainly Beta, ARCH: Beta Barrel, TOPOL: OB fold (Dihydrolipoamide Acetyltransferase, E2P), HOMOL: Nucleic acid-binding proteins; cath \| 3kojA00 \| 99.4 \| 5.9e-18 \| 8.8e-22 \| 110.4 \| 100 \| (1, 107) \| 162 \| (7, 107) \| 108 \| Uncharacterized protein ycf41 \| CATHCODE: 2.40.50.140 NAME: Uncharacterized protein ycf41. Chain: a, b. Fragment: ssb domain. Engineered: yes. Mutation: yes SOURCE: Synechococcus elongatus pcc 6301. Anacystis nidulans. Organism\_taxid: 269084. Strain: atcc 27144/pcc 6301 /saug 1402/1. Gene: syc1425\_d, ycf41. Expressed in: escherichia coli bl21(de3). Expression\_system\_taxid: 469008. CLASS: Mainly Beta, ARCH: Beta Barrel, TOPOL: OB fold (Dihydrolipoamide Acetyltransferase, E2P), HOMOL: Nucleic acid-binding proteins; cath \| 5yyuB00 \| 99.4 \| 1.1e-17 \| 1.6e-21 \| 109.8 \| 110 \| (4, 116) \| 162 \| (1, 110) \| 112 \| Single-stranded DNA-binding protein \| CATHCODE: 2.40.50.140 NAME: Single-stranded DNA-binding protein. Chain: a, b, c, d. Synonym: ssb. Engineered: yes SOURCE: Staphylococcus aureus subsp. Aureus ed98. Organism\_taxid: 681288. Expressed in: escherichia coli bl21. Expression\_system\_taxid: 511693. CLASS: Mainly Beta, ARCH: Beta Barrel, TOPOL: OB fold (Dihydrolipoamide Acetyltransferase, E2P), HOMOL: Nucleic acid-binding proteins; cath \| 5gqoA00 \| 99.4 \| 1.4e-17 \| 2.2e-21 \| 109.6 \| 110 \| (5, 116) \| 162 \| (1, 111) \| 117 \| Single-stranded DNA-binding protein \| CATHCODE: 2.40.50.140 NAME: Single-stranded DNA-binding protein. Chain: a, b. Engineered: yes SOURCE: Mycobacterium smegmatis (strain atcc 700084 / mc(2)155). Organism\_taxid: 246196. Strain: atcc 700084 / mc(2)155. Gene: msmeg\_4701. Expressed in: escherichia coli. Expression\_system\_taxid: 511693. CLASS: Mainly Beta, ARCH: Beta Barrel, TOPOL: OB fold (Dihydrolipoamide Acetyltransferase, E2P), HOMOL: Nucleic acid-binding proteins; cath \| 1txyA00 \| 99.4 \| 2.8e-17 \| 4.2e-21 \| 105.8 \| 103 \| (4, 112) \| 162 \| (1, 103) \| 104 \| Primosomal replication protein n \| CATHCODE: 2.40.50.140 NAME: Primosomal replication protein n. Chain: a, b. Engineered: yes SOURCE: Escherichia coli. Organism\_taxid: 562. Gene: prib, b4201. Expressed in: escherichia coli. Expression\_system\_taxid: 562 CLASS: Mainly Beta, ARCH: Beta Barrel, TOPOL: OB fold (Dihydrolipoamide Acetyltransferase, E2P), HOMOL: Nucleic acid-binding proteins; cath \| 3k8aB00 \| 99.3 \| 4.9e-17 \| 7.4e-21 \| 104.4 \| 100 \| (1, 110) \| 162 \| (3, 102) \| 103 \| Putative primosomal replication protein \| CATHCODE: 2.40.50.140 NAME: Putative primosomal replication protein. Chain: a, b. Engineered: yes SOURCE: Neisseria gonorrhoeae fa 1090. Organism\_taxid: 242231. Strain: fa1090.Gene: ngo0582, prib. Expressed in: escherichia coli. Expression\_system\_taxid: 562. CLASS: Mainly Beta, ARCH: Beta Barrel, TOPOL: OB fold (Dihydrolipoamide Acetyltransferase, E2P), HOMOL: Nucleic acid-binding proteins; cath \| 3ullB00 \| 99.3 \| 5.3e-17 \| 8e-21 \| 108.9 \| 114 \| (1, 114) \| 162 \| (10, 129) \| 132 \| Dna binding protein \| CATHCODE: 2.40.50.140 NAME: Dna binding protein. Chain: a, b SOURCE: Homo sapiens. Human. Organism\_taxid: 9606. Organelle: mitochondria CLASS: Mainly Beta, ARCH: Beta Barrel, TOPOL: OB fold (Dihydrolipoamide Acetyltransferase, E2P), HOMOL: Nucleic acid-binding proteins; cath \| 4gs3A00 \| 99.3 \| 8.8e-17 \| 1.3e-20 \| 104.4 \| 99 \| (1, 110) \| 162 \| (7, 106) \| 107 \| Single-stranded DNA-binding protein \| CATHCODE: 2.40.50.140 NAME: Single-stranded DNA-binding protein. Chain: a. Fragment: unp residues 1-104. Engineered: yes SOURCE: Thermoanaerobacter tengcongensis. Organism\_taxid: 273068. Strain: dsm 15242 / jcm 11007 / nbrc 100824 / mb4. Gene: ssb, tte0829. Expressed in: escherichia coli. Expression\_system\_taxid: 469008. CLASS: Mainly Beta, ARCH: Beta Barrel, TOPOL: OB fold (Dihydrolipoamide Acetyltransferase, E2P), HOMOL: Nucleic acid-binding proteins; cath \| 3dm4A00 \| 99.2 \| 3.9e-16 \| 5.9e-20 \| 99.2 \| 97 \| (5, 110) \| 162 \| (1, 97) \| 98 \| Primosomal replication protein n \| CATHCODE: 2.40.50.140 NAME: Primosomal replication protein n. Chain: a, b. Engineered: yes SOURCE: Bordetella pertussis. Organism\_taxid: 520. Gene: prib, bp2795 CLASS: Mainly Beta, ARCH: Beta Barrel, TOPOL: OB fold (Dihydrolipoamide Acetyltransferase, E2P), HOMOL: Nucleic acid-binding proteins; cath \| 3en2A00 \| 99.2 \| 2.5e-15 \| 3.6e-19 \| 96.8 \| 86 \| (4, 92) \| 162 \| (1, 86) \| 101 \| Probable primosomal replication protein n \| CATHCODE: 2.40.50.140 NAME: Probable primosomal replication protein n. Chain: a. Engineered: yes. Mutation: yes SOURCE: Ralstonia solanacearum. Pseudomonas solanacearum. Organism\_taxid: 305.Expressed in: escherichia coli. CLASS: Mainly Beta, ARCH: Beta Barrel, TOPOL: OB fold (Dihydrolipoamide Acetyltransferase, E2P), HOMOL: Nucleic acid-binding proteins; cath \| 4dkaC00 \| 99.1 \| 3.2e-15 \| 4.7e-19 \| 97.1 \| 90 \| (3, 102) \| 162 \| (1, 92) \| 105 \| Single domain antibody vhh \| CATHCODE: 2.40.50.140 NAME: Single domain antibody vhh. Chain: a, b. Engineered: yes. Rna-editing complex protein mp81. Chain: c, d. Engineered: yes. Mutation: yes SOURCE: Lama glama. Llama. Organism\_taxid: 9844. Expressed in: escherichia coli. Expression\_system\_taxid: 469008. CLASS: Mainly Beta, ARCH: Beta Barrel, TOPOL: OB fold (Dihydrolipoamide Acetyltransferase, E2P), HOMOL: Nucleic acid-binding proteins; cath \| 2hqlA01 \| 99.1 \| 9.7e-15 \| 1.5e-18 \| 92.6 \| 93 \| (1, 106) \| 162 \| (4, 96) \| 97 \| Hypothetical protein mg376 homolog \| CATHCODE: 2.40.50.140 NAME: Hypothetical protein mg376 homolog. Chain: a, b, c, d, e, f. Synonym: g12\_orf104, single-stranded DNA-binding protein. Engineered: yes SOURCE: Mycoplasma pneumoniae. Organism\_taxid: 2104. Gene: mpn554, mp288. Expressed in: escherichia coli. Expression\_system\_taxid: 562. CLASS: Mainly Beta, ARCH: Beta Barrel, TOPOL: OB fold (Dihydrolipoamide Acetyltransferase, E2P), HOMOL: Nucleic acid-binding proteins; phrogs \| 44 \| 100.0 \| 6.1e-35 \| 8.9e-39 \| 215.9 \| 92 \| (5, 106) \| 162 \| (1, 92) \| 121 \| single strand DNA binding protein \| single strand DNA binding protein; Category: DNA, RNA and nucleotide metabolism; p256451 VI\_01358; phrogs \| 18154 \| 99.8 \| 2e-23 \| 2.4e-27 \| 151.2 \| 79 \| (73, 162) \| 162 \| (28, 113) \| 113 \| single strand DNA binding protein \| single strand DNA binding protein; Category: DNA, RNA and nucleotide metabolism; MG641885\_p29; phrogs \| 25370 \| 98.8 \| 5e-13 \| 5.7e-17 \| 81.7 \| 41 \| (4, 44) \| 162 \| (1, 41) \| 45 \| NA \| NA; Category: unknown function; p432599 VI\_04995; phrogs \| 7364 \| 98.6 \| 4.3e-12 \| 4.9e-16 \| 94.8 \| 25 \| (138, 162) \| 162 \| (134, 158) \| 158 \| single strand DNA binding protein \| single strand DNA binding protein; Category: DNA, RNA and nucleotide metabolism; NC\_005891\_p29; phrogs \| 26861 \| 98.1 \| 1.1e-09 \| 1.2e-13 \| 85.9 \| 78 \| (5, 91) \| 162 \| (134, 211) \| 242 \| NA \| NA; Category: unknown function; p359533 VI\_01260; phrogs \| 12631 \| 97.5 \| 6e-08 \| 6.7e-12 \| 63.9 \| 39 \| (2, 40) \| 162 \| (27, 65) \| 78 \| single strand DNA binding protein \| single strand DNA binding protein; Category: DNA, RNA and nucleotide metabolism; p232911 VI\_05461; phrogs \| 27667 \| 96.1 \| 2.9e-05 \| 3.2e-09 \| 48.5 \| 31 \| (77, 107) \| 162 \| (2, 32) \| 61 \| single strand DNA binding protein \| single strand DNA binding protein; Category: DNA, RNA and nucleotide metabolism; NC\_004584\_p19; phrogs \| 4477 \| 95.9 \| 4.4e-05 \| 5.2e-09 \| 63.5 \| 14 \| (149, 162) \| 162 \| (395, 408) \| 408 \| NA \| NA; Category: unknown function; NC\_020201\_p2 |
| Top keywords  (threshold 1.00e-03 (evalue)) | **Beta, yes, Engineered, OB, fold, a, Organism\_taxid, Mainly, ARCH, Barrel** |
| Output files | ../../domain\_architecture/35\_FANPEZAQ\_CDS\_0035\_cath.hhr ../../domain\_architecture/35\_FANPEZAQ\_CDS\_0035\_merged.svg ../../domain\_architecture/35\_FANPEZAQ\_CDS\_0035\_ncbi-cd.hhr ../../domain\_architecture/35\_FANPEZAQ\_CDS\_0035\_pfam.hhr ../../domain\_architecture/35\_FANPEZAQ\_CDS\_0035\_phrogs.hhr |

### Identical protein sequences/structures

#### Search results

|  |  |
| --- | --- |
| Protein sequence databases searched | Pdb, Swissprot, Refseq |
| Identical proteins found | -- |
| Top keywords | -- |
| Output files | -- |

### Similar protein sequences/structures

#### Sequence similarity search results (HHblits)^1^

|  |  |
| --- | --- |
| Sequence databases searched | Uniclust, Pdb70 |
| Results, scheme(s)  (Top layers only, threshold 1.00e-03 (evalue)) | xml version="1.0" encoding="utf-8" standalone="no"?       2024-09-02T21:08:44.373315 image/svg+xml   Matplotlib v3.7.2, https://matplotlib.org/ |
| Results, table(s)  (threshold 1.00e-03 (evalue)) | db \| id \| prob \| evalue \| pvalue \| score \| cols \| query \| query\_len \| template \| template\_len \| name \| description; uniclust \| UniRef100\_A0A098S7E1 \| 100.0 \| 5.3e-45 \| 1e-50 \| 241.9 \| 111 \| (1, 115) \| 162 \| (1, 111) \| 157 \| Single-stranded DNA-binding protein \| Single-stranded DNA-binding protein; uniclust \| UniRef100\_A0A1E7IFR0 \| 100.0 \| 1.2e-43 \| 2.2e-49 \| 234.2 \| 116 \| (2, 117) \| 162 \| (3, 118) \| 183 \| Single-stranded DNA-binding protein \| Single-stranded DNA-binding protein; uniclust \| UniRef100\_A0A0H4JBR5 \| 100.0 \| 2.4e-43 \| 4.7e-49 \| 234.4 \| 111 \| (1, 115) \| 162 \| (3, 113) \| 163 \| Single-stranded DNA-binding protein \| Single-stranded DNA-binding protein; uniclust \| UniRef100\_A0A011NXA6 \| 100.0 \| 3.4e-43 \| 6.4e-49 \| 248.0 \| 112 \| (1, 115) \| 162 \| (41, 152) \| 272 \| Single-stranded DNA-binding protein \| Single-stranded DNA-binding protein; uniclust \| UniRef100\_A0A0K1FF79 \| 100.0 \| 5.7e-43 \| 1.1e-48 \| 238.0 \| 113 \| (1, 117) \| 162 \| (13, 125) \| 180 \| Single-stranded DNA-binding protein \| Single-stranded DNA-binding protein; uniclust \| UniRef100\_A0A017T184 \| 100.0 \| 1e-42 \| 2e-48 \| 241.2 \| 111 \| (1, 115) \| 162 \| (18, 128) \| 209 \| Single-stranded DNA-binding protein \| Single-stranded DNA-binding protein; uniclust \| UniRef100\_A0A060R893 \| 100.0 \| 1.6e-42 \| 3.1e-48 \| 229.8 \| 115 \| (1, 118) \| 162 \| (1, 115) \| 171 \| Single-stranded DNA-binding protein \| Single-stranded DNA-binding protein; uniclust \| UniRef100\_A0A136KBY6 \| 100.0 \| 1.9e-42 \| 3.5e-48 \| 228.4 \| 112 \| (1, 116) \| 162 \| (24, 135) \| 174 \| Single-stranded DNA-binding protein \| Single-stranded DNA-binding protein; uniclust \| UniRef100\_A0A074LI87 \| 100.0 \| 2e-42 \| 3.9e-48 \| 231.5 \| 114 \| (1, 118) \| 162 \| (18, 131) \| 172 \| Single-stranded DNA-binding protein \| Single-stranded DNA-binding protein; uniclust \| UniRef100\_A0A0F9YE88 \| 100.0 \| 4e-42 \| 7.7e-48 \| 234.1 \| 110 \| (1, 114) \| 162 \| (30, 139) \| 198 \| Single-stranded DNA-binding protein \| Single-stranded DNA-binding protein; uniclust \| UniRef100\_A0A023WX09 \| 100.0 \| 4.7e-42 \| 9.1e-48 \| 238.5 \| 112 \| (1, 116) \| 162 \| (30, 141) \| 215 \| Single-stranded DNA-binding protein \| Single-stranded DNA-binding protein; uniclust \| UniRef100\_A0A069DGL3 \| 100.0 \| 4.9e-42 \| 9.6e-48 \| 237.2 \| 112 \| (1, 116) \| 162 \| (19, 130) \| 203 \| Single-stranded DNA-binding protein \| Single-stranded DNA-binding protein; uniclust \| UniRef100\_A0A011P4T6 \| 100.0 \| 5.3e-42 \| 1e-47 \| 238.6 \| 115 \| (1, 115) \| 162 \| (26, 140) \| 225 \| Single-stranded DNA-binding protein \| Single-stranded DNA-binding protein; uniclust \| UniRef100\_A0A0C5J9Y9 \| 100.0 \| 1.3e-41 \| 2.5e-47 \| 233.2 \| 113 \| (1, 116) \| 162 \| (40, 152) \| 238 \| Single-stranded DNA-binding protein \| Single-stranded DNA-binding protein; uniclust \| UniRef100\_A0A0G0LTD2 \| 100.0 \| 2e-41 \| 3.8e-47 \| 223.1 \| 112 \| (1, 116) \| 162 \| (5, 116) \| 159 \| Single-stranded DNA-binding protein \| Single-stranded DNA-binding protein; uniclust \| UniRef100\_A0A2E1E3C2 \| 100.0 \| 2.4e-41 \| 4.6e-47 \| 226.5 \| 113 \| (1, 118) \| 162 \| (17, 129) \| 172 \| Single-stranded DNA-binding protein \| Single-stranded DNA-binding protein; uniclust \| UniRef100\_A0A011MH48 \| 100.0 \| 3e-41 \| 5.9e-47 \| 237.6 \| 112 \| (1, 115) \| 162 \| (28, 139) \| 213 \| Single-stranded DNA-binding protein \| Single-stranded DNA-binding protein; uniclust \| UniRef100\_A0A015RX92 \| 100.0 \| 3.3e-41 \| 6.3e-47 \| 227.7 \| 111 \| (1, 115) \| 162 \| (12, 123) \| 186 \| Single-stranded DNA-binding protein \| Single-stranded DNA-binding protein; uniclust \| UniRef100\_A0A084SEV2 \| 100.0 \| 8.5e-41 \| 1.7e-46 \| 235.1 \| 112 \| (1, 116) \| 162 \| (36, 147) \| 215 \| Single-stranded DNA-binding protein \| Single-stranded DNA-binding protein; uniclust \| UniRef100\_A0A0D8J7K4 \| 100.0 \| 9e-41 \| 1.7e-46 \| 217.4 \| 117 \| (1, 121) \| 162 \| (1, 117) \| 154 \| Single-stranded DNA-binding protein \| Single-stranded DNA-binding protein; uniclust \| UniRef100\_A0A066TQG3 \| 100.0 \| 1.1e-40 \| 2.2e-46 \| 221.5 \| 111 \| (3, 117) \| 162 \| (5, 115) \| 177 \| Single-stranded DNA-binding protein \| Single-stranded DNA-binding protein; uniclust \| UniRef100\_A0A023E151 \| 100.0 \| 1.5e-40 \| 2.9e-46 \| 226.1 \| 114 \| (1, 118) \| 162 \| (4, 117) \| 195 \| Single-stranded DNA-binding protein \| Single-stranded DNA-binding protein; uniclust \| UniRef100\_A0A0W0U4V5 \| 100.0 \| 1.8e-40 \| 3.4e-46 \| 221.9 \| 111 \| (1, 115) \| 162 \| (14, 124) \| 173 \| Single-stranded DNA-binding protein \| Single-stranded DNA-binding protein; uniclust \| UniRef100\_A0A060B5W2 \| 100.0 \| 2.4e-40 \| 4.5e-46 \| 229.2 \| 116 \| (1, 116) \| 162 \| (34, 149) \| 240 \| Single-stranded DNA-binding protein \| Single-stranded DNA-binding protein; uniclust \| UniRef100\_A0A097EQ76 \| 100.0 \| 3.7e-40 \| 7e-46 \| 217.4 \| 110 \| (4, 116) \| 162 \| (5, 114) \| 172 \| Single-stranded DNA-binding protein \| Single-stranded DNA-binding protein; uniclust \| UniRef100\_A0A087M3H5 \| 100.0 \| 4e-40 \| 7.6e-46 \| 226.6 \| 113 \| (1, 117) \| 162 \| (2, 114) \| 222 \| Single-stranded DNA-binding protein \| Single-stranded DNA-binding protein; uniclust \| UniRef100\_A0A087MC81 \| 100.0 \| 6.1e-40 \| 1.2e-45 \| 229.6 \| 112 \| (1, 118) \| 162 \| (36, 147) \| 211 \| Single-stranded DNA-binding protein \| Single-stranded DNA-binding protein; uniclust \| UniRef100\_A0A0N1KPD2 \| 100.0 \| 6.6e-40 \| 1.3e-45 \| 220.6 \| 111 \| (1, 115) \| 162 \| (23, 133) \| 183 \| Single-stranded DNA-binding protein (Fragment) \| Single-stranded DNA-binding protein (Fragment); uniclust \| UniRef100\_A0A1A9RXR2 \| 100.0 \| 7.7e-40 \| 1.5e-45 \| 215.5 \| 114 \| (1, 119) \| 162 \| (2, 115) \| 156 \| Single-stranded DNA-binding protein \| Single-stranded DNA-binding protein; uniclust \| UniRef100\_A0A0G1DJJ4 \| 100.0 \| 1.2e-39 \| 2.3e-45 \| 216.4 \| 116 \| (1, 120) \| 162 \| (2, 117) \| 168 \| Single-stranded DNA-binding protein \| Single-stranded DNA-binding protein; uniclust \| UniRef100\_A0A0G1KT99 \| 100.0 \| 1.3e-39 \| 2.4e-45 \| 225.1 \| 110 \| (1, 113) \| 162 \| (23, 132) \| 207 \| Single-stranded DNA-binding protein \| Single-stranded DNA-binding protein; uniclust \| UniRef100\_A0A072Y7C9 \| 100.0 \| 1.5e-39 \| 2.9e-45 \| 224.6 \| 111 \| (1, 117) \| 162 \| (15, 125) \| 197 \| Single-stranded DNA-binding protein \| Single-stranded DNA-binding protein; uniclust \| UniRef100\_A0A143XWD3 \| 100.0 \| 1.7e-39 \| 3.3e-45 \| 222.1 \| 111 \| (1, 118) \| 162 \| (1, 113) \| 192 \| Single-stranded DNA-binding protein \| Single-stranded DNA-binding protein; uniclust \| UniRef100\_A0A023CPK7 \| 100.0 \| 1.8e-39 \| 3.5e-45 \| 221.0 \| 108 \| (2, 116) \| 162 \| (18, 125) \| 175 \| Single-stranded DNA-binding protein \| Single-stranded DNA-binding protein; uniclust \| UniRef100\_A0A014P655 \| 100.0 \| 2.9e-39 \| 5.5e-45 \| 225.1 \| 114 \| (1, 117) \| 162 \| (41, 154) \| 270 \| Single-stranded DNA-binding protein \| Single-stranded DNA-binding protein; uniclust \| UniRef100\_A0A062XTB0 \| 100.0 \| 3.1e-39 \| 5.9e-45 \| 222.6 \| 112 \| (1, 116) \| 162 \| (16, 127) \| 203 \| Single-stranded DNA-binding protein \| Single-stranded DNA-binding protein; uniclust \| UniRef100\_A0A0D2JCW1 \| 100.0 \| 3.2e-39 \| 6.1e-45 \| 220.9 \| 110 \| (1, 114) \| 162 \| (6, 115) \| 204 \| Single-stranded DNA-binding protein \| Single-stranded DNA-binding protein; uniclust \| UniRef100\_A0A1E4NJ61 \| 100.0 \| 3.7e-39 \| 6.9e-45 \| 209.7 \| 116 \| (1, 119) \| 162 \| (1, 116) \| 166 \| Single-stranded DNA-binding protein \| Single-stranded DNA-binding protein; uniclust \| UniRef100\_A0A0N8NST2 \| 100.0 \| 4.2e-39 \| 8e-45 \| 212.8 \| 108 \| (4, 117) \| 162 \| (1, 108) \| 163 \| Single-stranded DNA-binding protein \| Single-stranded DNA-binding protein; uniclust \| UniRef100\_A0A0C7NPU9 \| 100.0 \| 4.7e-39 \| 9.1e-45 \| 217.7 \| 111 \| (4, 118) \| 162 \| (40, 150) \| 187 \| Single-stranded DNA-binding protein \| Single-stranded DNA-binding protein; uniclust \| UniRef100\_A0A023WZ22 \| 100.0 \| 4.8e-39 \| 9.5e-45 \| 228.6 \| 111 \| (1, 115) \| 162 \| (27, 137) \| 224 \| Single-stranded DNA-binding protein \| Single-stranded DNA-binding protein; uniclust \| UniRef100\_A0A011TCV2 \| 100.0 \| 5.3e-39 \| 1e-44 \| 224.0 \| 115 \| (1, 115) \| 162 \| (34, 150) \| 215 \| Single-stranded DNA-binding protein \| Single-stranded DNA-binding protein; uniclust \| UniRef100\_A0A011V2J4 \| 100.0 \| 5.7e-39 \| 1.1e-44 \| 223.9 \| 110 \| (2, 117) \| 162 \| (23, 132) \| 213 \| Single-stranded DNA-binding protein \| Single-stranded DNA-binding protein; uniclust \| UniRef100\_A0A022FZQ1 \| 100.0 \| 6.2e-39 \| 1.2e-44 \| 223.6 \| 112 \| (1, 115) \| 162 \| (23, 134) \| 254 \| Single-stranded DNA-binding protein \| Single-stranded DNA-binding protein; uniclust \| UniRef100\_A0A023BY02 \| 100.0 \| 7.1e-39 \| 1.4e-44 \| 223.6 \| 114 \| (1, 117) \| 162 \| (29, 142) \| 217 \| Single-stranded DNA-binding protein \| Single-stranded DNA-binding protein; uniclust \| UniRef100\_A0A095BFI2 \| 100.0 \| 7.3e-39 \| 1.4e-44 \| 211.4 \| 114 \| (1, 117) \| 162 \| (6, 120) \| 172 \| Single-stranded DNA-binding protein \| Single-stranded DNA-binding protein; uniclust \| UniRef100\_A0A1F7RRQ4 \| 100.0 \| 7.3e-39 \| 1.4e-44 \| 216.1 \| 113 \| (2, 118) \| 162 \| (29, 141) \| 185 \| Single-stranded DNA-binding protein \| Single-stranded DNA-binding protein; uniclust \| UniRef100\_A0A0M5JTF8 \| 100.0 \| 9.9e-39 \| 1.9e-44 \| 213.8 \| 116 \| (3, 118) \| 162 \| (4, 119) \| 192 \| Single-stranded DNA-binding protein \| Single-stranded DNA-binding protein; uniclust \| UniRef100\_A0A088T2F7 \| 100.0 \| 1.2e-38 \| 2.4e-44 \| 221.7 \| 111 \| (1, 115) \| 162 \| (51, 161) \| 212 \| Single-stranded DNA-binding protein \| Single-stranded DNA-binding protein; uniclust \| UniRef100\_A0A1F3BMS9 \| 100.0 \| 1.3e-38 \| 2.5e-44 \| 208.2 \| 112 \| (1, 116) \| 162 \| (1, 112) \| 150 \| Single-stranded DNA-binding protein \| Single-stranded DNA-binding protein; uniclust \| UniRef100\_A0A0A8B1Q7 \| 100.0 \| 1.3e-38 \| 2.5e-44 \| 212.7 \| 112 \| (1, 116) \| 162 \| (7, 118) \| 175 \| Single-stranded DNA-binding protein \| Single-stranded DNA-binding protein; uniclust \| UniRef100\_A0A098R1J5 \| 100.0 \| 1.8e-38 \| 3.4e-44 \| 212.7 \| 112 \| (1, 118) \| 162 \| (2, 113) \| 160 \| Single-stranded DNA-binding protein \| Single-stranded DNA-binding protein; uniclust \| UniRef100\_A0A014M3K2 \| 100.0 \| 1.9e-38 \| 3.5e-44 \| 209.9 \| 115 \| (3, 117) \| 162 \| (4, 118) \| 193 \| Single-stranded DNA-binding protein \| Single-stranded DNA-binding protein; uniclust \| UniRef100\_A0A010YT66 \| 100.0 \| 1.8e-38 \| 3.6e-44 \| 217.6 \| 110 \| (1, 114) \| 162 \| (15, 125) \| 177 \| Single-stranded DNA-binding protein \| Single-stranded DNA-binding protein; uniclust \| UniRef100\_A0A1B8PKK7 \| 100.0 \| 2.1e-38 \| 4e-44 \| 213.1 \| 115 \| (1, 119) \| 162 \| (1, 115) \| 187 \| Single-stranded DNA-binding protein \| Single-stranded DNA-binding protein; uniclust \| UniRef100\_A0A015YND2 \| 100.0 \| 2.4e-38 \| 4.6e-44 \| 216.3 \| 115 \| (1, 118) \| 162 \| (29, 144) \| 188 \| Single-stranded DNA-binding protein \| Single-stranded DNA-binding protein; uniclust \| UniRef100\_A0A064AII8 \| 100.0 \| 2.3e-38 \| 4.6e-44 \| 223.6 \| 111 \| (1, 117) \| 162 \| (27, 137) \| 228 \| Single-stranded DNA-binding protein \| Single-stranded DNA-binding protein; uniclust \| UniRef100\_A0A0A0X2I0 \| 100.0 \| 2.8e-38 \| 5.4e-44 \| 214.4 \| 110 \| (1, 116) \| 162 \| (31, 140) \| 187 \| Single-stranded DNA-binding protein \| Single-stranded DNA-binding protein; uniclust \| UniRef100\_A0A091AL92 \| 100.0 \| 2.8e-38 \| 5.5e-44 \| 216.9 \| 113 \| (1, 118) \| 162 \| (15, 127) \| 181 \| Single-stranded DNA-binding protein \| Single-stranded DNA-binding protein; uniclust \| UniRef100\_A0A009PIV2 \| 100.0 \| 2.8e-38 \| 5.5e-44 \| 220.1 \| 112 \| (1, 115) \| 162 \| (26, 137) \| 202 \| Single-stranded DNA-binding protein \| Single-stranded DNA-binding protein; uniclust \| UniRef100\_A0A094W5Q5 \| 100.0 \| 3.4e-38 \| 6.5e-44 \| 211.2 \| 110 \| (1, 115) \| 162 \| (19, 128) \| 175 \| Single-stranded DNA-binding protein \| Single-stranded DNA-binding protein; uniclust \| UniRef100\_A0A0B0EP20 \| 100.0 \| 3.6e-38 \| 7.2e-44 \| 215.2 \| 110 \| (2, 117) \| 162 \| (22, 131) \| 170 \| Single-stranded DNA-binding protein \| Single-stranded DNA-binding protein; uniclust \| UniRef100\_A0A016XFS3 \| 100.0 \| 3.9e-38 \| 7.8e-44 \| 222.2 \| 112 \| (1, 116) \| 162 \| (36, 147) \| 217 \| Single-stranded DNA-binding protein \| Single-stranded DNA-binding protein; uniclust \| UniRef100\_A0A0A1DMG2 \| 100.0 \| 4.5e-38 \| 8.8e-44 \| 215.8 \| 112 \| (1, 116) \| 162 \| (11, 122) \| 191 \| Single-stranded DNA-binding protein \| Single-stranded DNA-binding protein; uniclust \| UniRef100\_A0A068NVF8 \| 100.0 \| 6.6e-38 \| 1.3e-43 \| 205.5 \| 112 \| (2, 119) \| 162 \| (1, 112) \| 146 \| Single-stranded DNA-binding protein \| Single-stranded DNA-binding protein; uniclust \| UniRef100\_A0A2D3R7S5 \| 100.0 \| 8.1e-38 \| 1.5e-43 \| 205.7 \| 117 \| (1, 121) \| 162 \| (1, 117) \| 170 \| Single-stranded DNA-binding protein \| Single-stranded DNA-binding protein; uniclust \| UniRef100\_A0A0F3GPA7 \| 100.0 \| 9.3e-38 \| 1.8e-43 \| 211.8 \| 109 \| (1, 113) \| 162 \| (14, 122) \| 170 \| Single-stranded DNA-binding protein \| Single-stranded DNA-binding protein; uniclust \| UniRef100\_A0A077MG65 \| 100.0 \| 1.2e-37 \| 2.3e-43 \| 208.7 \| 112 \| (2, 116) \| 162 \| (12, 123) \| 195 \| Single-stranded DNA-binding protein \| Single-stranded DNA-binding protein; uniclust \| UniRef100\_A0A096B1L2 \| 100.0 \| 1.2e-37 \| 2.3e-43 \| 205.9 \| 112 \| (2, 118) \| 162 \| (11, 122) \| 180 \| Single-stranded DNA-binding protein \| Single-stranded DNA-binding protein; uniclust \| UniRef100\_A0A072Y3A3 \| 100.0 \| 1.4e-37 \| 2.7e-43 \| 210.0 \| 111 \| (1, 117) \| 162 \| (7, 117) \| 154 \| Single-stranded DNA-binding protein \| Single-stranded DNA-binding protein; uniclust \| UniRef100\_A0A0G0BWT4 \| 100.0 \| 1.5e-37 \| 2.8e-43 \| 208.2 \| 109 \| (2, 114) \| 162 \| (11, 119) \| 177 \| Single-stranded DNA-binding protein \| Single-stranded DNA-binding protein; uniclust \| UniRef100\_A0A1B1IQN2 \| 100.0 \| 1.5e-37 \| 2.9e-43 \| 206.3 \| 111 \| (1, 116) \| 162 \| (7, 117) \| 153 \| Single-stranded DNA-binding protein \| Single-stranded DNA-binding protein; uniclust \| UniRef100\_A0A2W0AHC9 \| 100.0 \| 2.3e-37 \| 4.4e-43 \| 201.9 \| 115 \| (1, 118) \| 162 \| (1, 115) \| 162 \| Single-stranded DNA-binding protein \| Single-stranded DNA-binding protein; uniclust \| UniRef100\_A0A1Y5D6G7 \| 100.0 \| 2.8e-37 \| 5.2e-43 \| 207.3 \| 114 \| (2, 119) \| 162 \| (21, 134) \| 207 \| Single-stranded DNA-binding protein \| Single-stranded DNA-binding protein; uniclust \| UniRef100\_A0A0S8KNV4 \| 100.0 \| 3e-37 \| 5.9e-43 \| 207.3 \| 112 \| (1, 116) \| 162 \| (24, 135) \| 162 \| Single-stranded DNA-binding protein \| Single-stranded DNA-binding protein; uniclust \| UniRef100\_A0A0G2Q8L7 \| 100.0 \| 3.4e-37 \| 6.5e-43 \| 209.4 \| 117 \| (1, 117) \| 162 \| (33, 151) \| 200 \| Single-stranded DNA-binding protein \| Single-stranded DNA-binding protein; uniclust \| UniRef100\_A0A0H5QQE0 \| 100.0 \| 3.3e-37 \| 6.5e-43 \| 208.0 \| 110 \| (1, 117) \| 162 \| (5, 114) \| 170 \| Single-stranded DNA-binding protein \| Single-stranded DNA-binding protein; uniclust \| UniRef100\_A0A023BVC0 \| 100.0 \| 4.6e-37 \| 9e-43 \| 215.9 \| 112 \| (1, 116) \| 162 \| (32, 143) \| 210 \| Single-stranded DNA-binding protein \| Single-stranded DNA-binding protein; uniclust \| UniRef100\_A0A011VB38 \| 100.0 \| 5.6e-37 \| 1.1e-42 \| 217.0 \| 115 \| (1, 115) \| 162 \| (49, 165) \| 266 \| Single-stranded DNA-binding protein \| Single-stranded DNA-binding protein; uniclust \| UniRef100\_A0A0E3UIY2 \| 100.0 \| 6.3e-37 \| 1.2e-42 \| 205.1 \| 110 \| (2, 115) \| 162 \| (1, 111) \| 183 \| Single-stranded DNA-binding protein \| Single-stranded DNA-binding protein; uniclust \| UniRef100\_A0A1F7X0M2 \| 100.0 \| 7.6e-37 \| 1.4e-42 \| 202.2 \| 113 \| (1, 117) \| 162 \| (1, 113) \| 176 \| Single-stranded DNA-binding protein \| Single-stranded DNA-binding protein; uniclust \| UniRef100\_A0A084SGN0 \| 100.0 \| 1.1e-36 \| 2.1e-42 \| 193.4 \| 111 \| (1, 115) \| 162 \| (4, 114) \| 127 \| Single-stranded DNA-binding protein (Fragment) \| Single-stranded DNA-binding protein (Fragment); uniclust \| UniRef100\_A0A0A7P4Y4 \| 100.0 \| 1.4e-36 \| 2.7e-42 \| 208.3 \| 110 \| (1, 115) \| 162 \| (18, 127) \| 191 \| Single-stranded DNA-binding protein \| Single-stranded DNA-binding protein; uniclust \| UniRef100\_A0A0W0WBB9 \| 100.0 \| 1.6e-36 \| 3.1e-42 \| 202.6 \| 109 \| (1, 114) \| 162 \| (5, 113) \| 154 \| Single-stranded DNA-binding protein \| Single-stranded DNA-binding protein; uniclust \| UniRef100\_A0A084JIF5 \| 100.0 \| 1.7e-36 \| 3.4e-42 \| 208.4 \| 109 \| (1, 117) \| 162 \| (16, 127) \| 183 \| Single-stranded DNA-binding protein \| Single-stranded DNA-binding protein; uniclust \| UniRef100\_A0A0S8EKW5 \| 100.0 \| 2.9e-36 \| 5.6e-42 \| 199.6 \| 109 \| (3, 115) \| 162 \| (7, 115) \| 175 \| Single-stranded DNA-binding protein \| Single-stranded DNA-binding protein; uniclust \| UniRef100\_A0A0B6WUT2 \| 100.0 \| 3.2e-36 \| 6.2e-42 \| 201.7 \| 114 \| (2, 119) \| 162 \| (17, 130) \| 172 \| Single-stranded DNA-binding protein \| Single-stranded DNA-binding protein; uniclust \| UniRef100\_A0A0A8WRT0 \| 100.0 \| 3.9e-36 \| 7.5e-42 \| 199.5 \| 112 \| (2, 118) \| 162 \| (25, 136) \| 162 \| Single-stranded DNA-binding protein \| Single-stranded DNA-binding protein; uniclust \| UniRef100\_A0A011RQU6 \| 100.0 \| 4.9e-36 \| 9.4e-42 \| 209.2 \| 112 \| (1, 118) \| 162 \| (15, 126) \| 241 \| Single-stranded DNA-binding protein \| Single-stranded DNA-binding protein; uniclust \| UniRef100\_A0A1V5HJ78 \| 100.0 \| 8.8e-36 \| 1.7e-41 \| 195.9 \| 109 \| (1, 113) \| 162 \| (13, 121) \| 159 \| Single-stranded DNA-binding protein \| Single-stranded DNA-binding protein; uniclust \| UniRef100\_A0A0M8K5Q7 \| 100.0 \| 8.8e-36 \| 1.7e-41 \| 202.8 \| 111 \| (1, 115) \| 162 \| (29, 139) \| 201 \| Single-stranded DNA-binding protein \| Single-stranded DNA-binding protein; uniclust \| UniRef100\_A0A0G1IEC5 \| 100.0 \| 1.1e-35 \| 2.2e-41 \| 206.5 \| 111 \| (1, 116) \| 162 \| (47, 157) \| 200 \| Single-stranded DNA-binding protein \| Single-stranded DNA-binding protein; uniclust \| UniRef100\_A0A1T4V337 \| 100.0 \| 1.2e-35 \| 2.3e-41 \| 205.8 \| 112 \| (1, 115) \| 162 \| (1, 112) \| 218 \| Single-stranded DNA-binding protein \| Single-stranded DNA-binding protein; uniclust \| UniRef100\_A0A081UYA0 \| 100.0 \| 2.3e-35 \| 4.3e-41 \| 204.8 \| 114 \| (3, 116) \| 162 \| (70, 183) \| 281 \| Single-stranded DNA-binding protein \| Single-stranded DNA-binding protein; uniclust \| UniRef100\_A0A014MER4 \| 100.0 \| 2.7e-35 \| 5.3e-41 \| 194.9 \| 104 \| (3, 116) \| 162 \| (1, 104) \| 150 \| Single-stranded DNA-binding protein \| Single-stranded DNA-binding protein; uniclust \| UniRef100\_A0A0P6Y529 \| 100.0 \| 3.8e-35 \| 7.2e-41 \| 194.6 \| 112 \| (1, 116) \| 162 \| (23, 134) \| 160 \| Single-stranded DNA-binding protein \| Single-stranded DNA-binding protein; uniclust \| UniRef100\_A0A0A7LLS2 \| 100.0 \| 3.9e-35 \| 7.3e-41 \| 190.1 \| 114 \| (1, 118) \| 162 \| (6, 119) \| 148 \| Single-stranded DNA-binding protein \| Single-stranded DNA-binding protein; uniclust \| UniRef100\_A0A014LL70 \| 100.0 \| 3.9e-35 \| 7.6e-41 \| 210.0 \| 110 \| (1, 113) \| 162 \| (44, 154) \| 253 \| Single-stranded DNA-binding protein \| Single-stranded DNA-binding protein; uniclust \| UniRef100\_A0A010RG06 \| 100.0 \| 4.3e-35 \| 8e-41 \| 196.8 \| 116 \| (1, 117) \| 162 \| (7, 122) \| 218 \| Single-stranded DNA-binding protein \| Single-stranded DNA-binding protein; uniclust \| UniRef100\_A0A1V5DNT6 \| 100.0 \| 4.5e-35 \| 8.8e-41 \| 194.3 \| 113 \| (1, 118) \| 162 \| (3, 115) \| 152 \| Single-stranded DNA-binding protein \| Single-stranded DNA-binding protein; uniclust \| UniRef100\_A0A095Z4E9 \| 100.0 \| 4.8e-35 \| 9.2e-41 \| 193.0 \| 116 \| (2, 120) \| 162 \| (1, 120) \| 170 \| Single-stranded DNA-binding protein \| Single-stranded DNA-binding protein; uniclust \| UniRef100\_A0A1G0XIK8 \| 100.0 \| 5.9e-35 \| 1.1e-40 \| 198.8 \| 113 \| (1, 118) \| 162 \| (1, 114) \| 201 \| Single-stranded DNA-binding protein \| Single-stranded DNA-binding protein; uniclust \| UniRef100\_A0A101WK01 \| 100.0 \| 7e-35 \| 1.4e-40 \| 191.2 \| 112 \| (2, 119) \| 162 \| (1, 112) \| 144 \| Single-stranded DNA-binding protein \| Single-stranded DNA-binding protein; uniclust \| UniRef100\_A0A0M4CSQ1 \| 100.0 \| 7.7e-35 \| 1.5e-40 \| 195.7 \| 113 \| (1, 115) \| 162 \| (12, 124) \| 178 \| Single-stranded DNA-binding protein \| Single-stranded DNA-binding protein; uniclust \| UniRef100\_A0A0P9EXD2 \| 100.0 \| 7.9e-35 \| 1.5e-40 \| 182.0 \| 108 \| (1, 117) \| 162 \| (1, 108) \| 118 \| Single-stranded DNA-binding protein (Fragment) \| Single-stranded DNA-binding protein (Fragment); uniclust \| UniRef100\_A0A1Z9MN17 \| 100.0 \| 9.4e-35 \| 1.8e-40 \| 193.7 \| 101 \| (2, 104) \| 162 \| (1, 101) \| 157 \| Single-stranded DNA-binding protein \| Single-stranded DNA-binding protein; uniclust \| UniRef100\_A0A161QJ15 \| 100.0 \| 1.4e-34 \| 2.7e-40 \| 191.8 \| 109 \| (2, 114) \| 162 \| (6, 114) \| 171 \| Single-stranded DNA-binding protein \| Single-stranded DNA-binding protein; uniclust \| UniRef100\_A0A017RY15 \| 100.0 \| 1.5e-34 \| 3e-40 \| 197.5 \| 107 \| (1, 116) \| 162 \| (29, 135) \| 179 \| Single-stranded DNA-binding protein \| Single-stranded DNA-binding protein; uniclust \| UniRef100\_A0A136KKK6 \| 100.0 \| 1.6e-34 \| 3.1e-40 \| 188.8 \| 112 \| (1, 117) \| 162 \| (2, 113) \| 137 \| Single-stranded DNA-binding protein \| Single-stranded DNA-binding protein; uniclust \| UniRef100\_A0A0J1BEB9 \| 100.0 \| 1.9e-34 \| 3.5e-40 \| 193.4 \| 113 \| (1, 118) \| 162 \| (36, 148) \| 198 \| Single-stranded DNA-binding protein \| Single-stranded DNA-binding protein; uniclust \| UniRef100\_A0A0F9YYB5 \| 100.0 \| 3.1e-34 \| 6e-40 \| 199.4 \| 110 \| (3, 116) \| 162 \| (30, 139) \| 223 \| Single-stranded DNA-binding protein \| Single-stranded DNA-binding protein; uniclust \| UniRef100\_A0A0P0P3Y9 \| 100.0 \| 4.1e-34 \| 7.8e-40 \| 190.1 \| 115 \| (1, 115) \| 162 \| (2, 118) \| 195 \| Single-stranded DNA-binding protein \| Single-stranded DNA-binding protein; uniclust \| UniRef100\_A0A1F5RVU2 \| 100.0 \| 4.4e-34 \| 8.5e-40 \| 190.2 \| 108 \| (3, 114) \| 162 \| (27, 134) \| 169 \| Single-stranded DNA-binding protein \| Single-stranded DNA-binding protein; uniclust \| UniRef100\_A0A0A2E6F1 \| 100.0 \| 4.7e-34 \| 9e-40 \| 189.0 \| 114 \| (1, 118) \| 162 \| (16, 129) \| 171 \| Single-stranded DNA-binding protein \| Single-stranded DNA-binding protein; uniclust \| UniRef100\_A0A1B0XUD7 \| 100.0 \| 7.2e-34 \| 1.4e-39 \| 188.1 \| 113 \| (2, 118) \| 162 \| (10, 122) \| 145 \| Single-stranded DNA-binding protein \| Single-stranded DNA-binding protein; uniclust \| UniRef100\_A0A134CMY2 \| 100.0 \| 8.2e-34 \| 1.5e-39 \| 186.3 \| 103 \| (1, 104) \| 162 \| (1, 103) \| 168 \| Single-stranded DNA-binding protein \| Single-stranded DNA-binding protein; uniclust \| UniRef100\_A0A173R6J3 \| 100.0 \| 8.6e-34 \| 1.7e-39 \| 183.0 \| 106 \| (1, 115) \| 162 \| (4, 109) \| 127 \| Single-stranded DNA-binding protein \| Single-stranded DNA-binding protein; uniclust \| UniRef100\_A0A0M1LCB6 \| 100.0 \| 1.1e-33 \| 2e-39 \| 188.7 \| 106 \| (1, 115) \| 162 \| (1, 106) \| 159 \| Single-stranded DNA-binding protein \| Single-stranded DNA-binding protein; uniclust \| UniRef100\_A0A133N082 \| 100.0 \| 1.1e-33 \| 2.1e-39 \| 188.2 \| 111 \| (4, 118) \| 162 \| (1, 115) \| 172 \| Single-stranded DNA-binding protein \| Single-stranded DNA-binding protein; uniclust \| UniRef100\_A0A094P9Y5 \| 100.0 \| 1.3e-33 \| 2.6e-39 \| 192.2 \| 107 \| (1, 107) \| 162 \| (20, 126) \| 185 \| Single-stranded DNA-binding protein \| Single-stranded DNA-binding protein; uniclust \| UniRef100\_A0A031WI19 \| 100.0 \| 1.5e-33 \| 2.9e-39 \| 182.7 \| 110 \| (2, 117) \| 162 \| (7, 117) \| 151 \| Single-stranded DNA-binding protein \| Single-stranded DNA-binding protein; uniclust \| UniRef100\_A0A1Y3UI13 \| 100.0 \| 2.3e-33 \| 4.5e-39 \| 186.9 \| 113 \| (1, 118) \| 162 \| (1, 113) \| 158 \| Single-stranded DNA-binding protein \| Single-stranded DNA-binding protein; uniclust \| UniRef100\_A0A1F3DCZ5 \| 100.0 \| 2.4e-33 \| 4.5e-39 \| 187.9 \| 113 \| (3, 119) \| 162 \| (1, 113) \| 182 \| Single-stranded DNA-binding protein \| Single-stranded DNA-binding protein; uniclust \| UniRef100\_A0A059E0M9 \| 100.0 \| 2.4e-33 \| 4.5e-39 \| 184.3 \| 113 \| (1, 118) \| 162 \| (11, 123) \| 158 \| Single-stranded DNA-binding protein \| Single-stranded DNA-binding protein; uniclust \| UniRef100\_A0A068EXU3 \| 100.0 \| 3.2e-33 \| 6.1e-39 \| 188.8 \| 107 \| (2, 116) \| 162 \| (18, 124) \| 169 \| Single-stranded DNA-binding protein \| Single-stranded DNA-binding protein; uniclust \| UniRef100\_A0A0V8CZ75 \| 100.0 \| 3.4e-33 \| 6.5e-39 \| 186.5 \| 108 \| (3, 119) \| 162 \| (6, 113) \| 162 \| tRNA-binding protein \| tRNA-binding protein; uniclust \| UniRef100\_A0A0G0XSM8 \| 100.0 \| 3.9e-33 \| 7.2e-39 \| 182.6 \| 108 \| (2, 114) \| 162 \| (1, 108) \| 180 \| Single-stranded DNA-binding protein \| Single-stranded DNA-binding protein; uniclust \| UniRef100\_A0A075MAN7 \| 100.0 \| 4.1e-33 \| 7.7e-39 \| 184.6 \| 116 \| (3, 119) \| 162 \| (4, 119) \| 189 \| Single-stranded DNA-binding protein \| Single-stranded DNA-binding protein; uniclust \| UniRef100\_A0A0F0CQF3 \| 100.0 \| 4.9e-33 \| 9.3e-39 \| 189.9 \| 110 \| (1, 114) \| 162 \| (30, 139) \| 197 \| Single-stranded DNA-binding protein \| Single-stranded DNA-binding protein; uniclust \| UniRef100\_A0A077UH19 \| 100.0 \| 5e-33 \| 9.7e-39 \| 187.0 \| 112 \| (2, 119) \| 162 \| (10, 121) \| 166 \| Single-stranded DNA-binding protein \| Single-stranded DNA-binding protein; uniclust \| UniRef100\_A0A6B1E277 \| 100.0 \| 5.4e-33 \| 1e-38 \| 181.2 \| 112 \| (2, 117) \| 162 \| (1, 112) \| 164 \| Single-stranded DNA-binding protein \| Single-stranded DNA-binding protein; uniclust \| UniRef100\_A0A011RMD1 \| 100.0 \| 5.6e-33 \| 1.1e-38 \| 201.5 \| 110 \| (1, 116) \| 162 \| (46, 155) \| 262 \| Single-stranded DNA-binding protein \| Single-stranded DNA-binding protein; uniclust \| UniRef100\_A0A0M2HGC3 \| 100.0 \| 6.8e-33 \| 1.3e-38 \| 189.1 \| 111 \| (2, 112) \| 162 \| (20, 133) \| 177 \| Single-stranded DNA-binding protein \| Single-stranded DNA-binding protein; uniclust \| UniRef100\_A0A133Z2U9 \| 100.0 \| 7.9e-33 \| 1.5e-38 \| 185.0 \| 110 \| (3, 115) \| 162 \| (10, 120) \| 165 \| Single-stranded DNA-binding protein \| Single-stranded DNA-binding protein; uniclust \| UniRef100\_A0A024C9G7 \| 100.0 \| 9.8e-33 \| 1.9e-38 \| 188.4 \| 113 \| (2, 118) \| 162 \| (1, 113) \| 199 \| Single-stranded DNA-binding protein \| Single-stranded DNA-binding protein; uniclust \| UniRef100\_A0A0Q6NU73 \| 100.0 \| 1.3e-32 \| 2.4e-38 \| 179.8 \| 103 \| (1, 104) \| 162 \| (1, 104) \| 160 \| Single-stranded DNA-binding protein \| Single-stranded DNA-binding protein; uniclust \| UniRef100\_A0A0P6WWD0 \| 100.0 \| 1.5e-32 \| 3e-38 \| 183.1 \| 100 \| (2, 115) \| 162 \| (4, 103) \| 148 \| Single-stranded DNA-binding protein \| Single-stranded DNA-binding protein; uniclust \| UniRef100\_A0A013VT29 \| 100.0 \| 1.7e-32 \| 3.1e-38 \| 177.2 \| 116 \| (1, 116) \| 162 \| (3, 120) \| 159 \| Single-stranded DNA-binding protein (Fragment) \| Single-stranded DNA-binding protein (Fragment); uniclust \| UniRef100\_A0A0J8YFX1 \| 100.0 \| 1.6e-32 \| 3.1e-38 \| 183.0 \| 101 \| (2, 111) \| 162 \| (12, 112) \| 174 \| Single-stranded DNA-binding protein \| Single-stranded DNA-binding protein; uniclust \| UniRef100\_A0A1I0ACB5 \| 100.0 \| 1.9e-32 \| 3.6e-38 \| 177.3 \| 106 \| (1, 118) \| 162 \| (1, 106) \| 125 \| Single-strand DNA-binding protein \| Single-strand DNA-binding protein; uniclust \| UniRef100\_A0A2S5CMA6 \| 100.0 \| 2.1e-32 \| 3.8e-38 \| 182.7 \| 113 \| (2, 117) \| 162 \| (49, 161) \| 216 \| Single-stranded DNA-binding protein \| Single-stranded DNA-binding protein; uniclust \| UniRef100\_A0A235BXR5 \| 100.0 \| 2.4e-32 \| 4.6e-38 \| 176.9 \| 110 \| (2, 114) \| 162 \| (6, 115) \| 156 \| Single-stranded DNA-binding protein \| Single-stranded DNA-binding protein; uniclust \| UniRef100\_A0A1B9LQJ7 \| 99.9 \| 3.5e-32 \| 6.7e-38 \| 178.3 \| 146 \| (1, 162) \| 162 \| (1, 146) \| 161 \| Single-stranded DNA-binding protein \| Single-stranded DNA-binding protein; uniclust \| UniRef100\_A0A091BEV2 \| 99.9 \| 4.1e-32 \| 7.7e-38 \| 177.6 \| 111 \| (2, 115) \| 162 \| (3, 113) \| 175 \| Single-stranded DNA-binding protein (Fragment) \| Single-stranded DNA-binding protein (Fragment); uniclust \| UniRef100\_A0A2E4F7T1 \| 99.9 \| 5.2e-32 \| 9.6e-38 \| 178.3 \| 111 \| (1, 115) \| 162 \| (1, 111) \| 192 \| Single-stranded DNA-binding protein \| Single-stranded DNA-binding protein; uniclust \| UniRef100\_A0A136Q8P1 \| 99.9 \| 5.3e-32 \| 1e-37 \| 181.0 \| 106 \| (2, 114) \| 162 \| (30, 135) \| 178 \| Single-stranded DNA-binding protein \| Single-stranded DNA-binding protein; uniclust \| UniRef100\_A0A2V2DG25 \| 99.9 \| 6.2e-32 \| 1.2e-37 \| 179.7 \| 106 \| (5, 120) \| 162 \| (1, 106) \| 182 \| Single-stranded DNA-binding protein \| Single-stranded DNA-binding protein; uniclust \| UniRef100\_A0A061JJT5 \| 99.9 \| 6.1e-32 \| 1.2e-37 \| 186.7 \| 111 \| (2, 118) \| 162 \| (26, 137) \| 205 \| Single-stranded DNA-binding protein \| Single-stranded DNA-binding protein; uniclust \| UniRef100\_A0A0A0X352 \| 99.9 \| 6.7e-32 \| 1.3e-37 \| 180.6 \| 109 \| (2, 116) \| 162 \| (17, 125) \| 170 \| Single-stranded DNA-binding protein \| Single-stranded DNA-binding protein; uniclust \| UniRef100\_A0A072Z0E3 \| 99.9 \| 8.1e-32 \| 1.5e-37 \| 168.9 \| 111 \| (2, 118) \| 162 \| (1, 112) \| 120 \| Single-stranded DNA-binding protein \| Single-stranded DNA-binding protein; uniclust \| UniRef100\_A0A1F5BJ96 \| 99.9 \| 8.9e-32 \| 1.7e-37 \| 171.7 \| 110 \| (2, 117) \| 162 \| (7, 116) \| 131 \| Single-stranded DNA-binding protein \| Single-stranded DNA-binding protein; uniclust \| UniRef100\_A0A1B3AYF2 \| 99.9 \| 9e-32 \| 1.8e-37 \| 182.0 \| 108 \| (2, 112) \| 162 \| (1, 110) \| 165 \| SsDNA binding protein \| SsDNA binding protein; uniclust \| UniRef100\_A0A1F3XSW4 \| 99.9 \| 9.3e-32 \| 1.8e-37 \| 176.2 \| 113 \| (2, 118) \| 162 \| (14, 126) \| 149 \| Single-stranded DNA-binding protein \| Single-stranded DNA-binding protein; uniclust \| UniRef100\_A0A136L6G9 \| 99.9 \| 1.1e-31 \| 2e-37 \| 168.1 \| 113 \| (1, 117) \| 162 \| (1, 113) \| 123 \| Single-stranded DNA-binding protein \| Single-stranded DNA-binding protein; uniclust \| UniRef100\_A0A068MX81 \| 99.9 \| 1.4e-31 \| 2.7e-37 \| 182.4 \| 108 \| (1, 117) \| 162 \| (27, 135) \| 174 \| Single-stranded DNA-binding protein \| Single-stranded DNA-binding protein; uniclust \| UniRef100\_A0A7C3ZBM8 \| 99.9 \| 1.4e-31 \| 2.7e-37 \| 177.2 \| 111 \| (2, 116) \| 162 \| (3, 113) \| 176 \| Single-stranded DNA-binding protein \| Single-stranded DNA-binding protein; uniclust \| UniRef100\_A0A0P0DYN1 \| 99.9 \| 1.6e-31 \| 3.1e-37 \| 181.7 \| 106 \| (2, 107) \| 162 \| (24, 130) \| 187 \| Single-stranded DNA-binding protein \| Single-stranded DNA-binding protein; uniclust \| UniRef100\_A0A0E4FY13 \| 99.9 \| 1.7e-31 \| 3.2e-37 \| 180.5 \| 110 \| (1, 113) \| 162 \| (1, 111) \| 215 \| Single-stranded DNA-binding protein \| Single-stranded DNA-binding protein; uniclust \| UniRef100\_A0A0A6PYF7 \| 99.9 \| 2.2e-31 \| 4.2e-37 \| 172.9 \| 110 \| (2, 117) \| 162 \| (12, 122) \| 147 \| Single-stranded DNA-binding protein \| Single-stranded DNA-binding protein; uniclust \| UniRef100\_A0A086PBI0 \| 99.9 \| 2.2e-31 \| 4.3e-37 \| 179.0 \| 116 \| (1, 117) \| 162 \| (19, 136) \| 161 \| Single-stranded DNA-binding protein \| Single-stranded DNA-binding protein; uniclust \| UniRef100\_A0A0D5M2L7 \| 99.9 \| 2.6e-31 \| 4.9e-37 \| 160.4 \| 95 \| (1, 95) \| 162 \| (2, 96) \| 98 \| Single-stranded DNA-binding protein \| Single-stranded DNA-binding protein; uniclust \| UniRef100\_A0A0C5EQ65 \| 99.9 \| 3.1e-31 \| 5.7e-37 \| 178.8 \| 117 \| (4, 120) \| 162 \| (52, 168) \| 227 \| Single-stranded DNA-binding protein \| Single-stranded DNA-binding protein; uniclust \| UniRef100\_A0A0F9ENP5 \| 99.9 \| 3.1e-31 \| 6e-37 \| 175.2 \| 114 \| (1, 116) \| 162 \| (12, 128) \| 160 \| Single-stranded DNA-binding protein \| Single-stranded DNA-binding protein; uniclust \| UniRef100\_A0A063KS07 \| 99.9 \| 3.6e-31 \| 6.8e-37 \| 181.9 \| 112 \| (4, 116) \| 162 \| (7, 118) \| 239 \| Single-stranded DNA-binding protein \| Single-stranded DNA-binding protein; uniclust \| UniRef100\_A0A8S5LK82 \| 99.9 \| 4.6e-31 \| 8.8e-37 \| 170.9 \| 107 \| (2, 112) \| 162 \| (1, 107) \| 145 \| Single strand binding protein \| Single strand binding protein; uniclust \| UniRef100\_A0A0K2H331 \| 99.9 \| 5.1e-31 \| 1e-36 \| 176.2 \| 111 \| (2, 117) \| 162 \| (1, 111) \| 149 \| Single-stranded DNA-binding protein \| Single-stranded DNA-binding protein; uniclust \| UniRef100\_A0A0D6IC91 \| 99.9 \| 6.5e-31 \| 1.2e-36 \| 174.1 \| 114 \| (2, 118) \| 162 \| (1, 114) \| 191 \| Single-stranded DNA-binding protein \| Single-stranded DNA-binding protein; uniclust \| UniRef100\_A0A016QTA0 \| 99.9 \| 6.9e-31 \| 1.3e-36 \| 193.4 \| 111 \| (2, 117) \| 162 \| (147, 257) \| 309 \| Single-stranded DNA-binding protein \| Single-stranded DNA-binding protein; uniclust \| UniRef100\_A0A1V5FBC2 \| 99.9 \| 7.3e-31 \| 1.4e-36 \| 176.1 \| 112 \| (2, 117) \| 162 \| (5, 117) \| 191 \| Single-stranded DNA-binding protein \| Single-stranded DNA-binding protein; uniclust \| UniRef100\_A0A0K8MB88 \| 99.9 \| 8.7e-31 \| 1.7e-36 \| 175.3 \| 113 \| (1, 117) \| 162 \| (31, 146) \| 176 \| Single-stranded DNA-binding protein \| Single-stranded DNA-binding protein; uniclust \| UniRef100\_A0A0B8NGF8 \| 99.9 \| 9.5e-31 \| 1.8e-36 \| 182.3 \| 111 \| (1, 111) \| 162 \| (23, 135) \| 218 \| Single-stranded DNA-binding protein \| Single-stranded DNA-binding protein; uniclust \| UniRef100\_A0A2I7N694 \| 99.9 \| 1e-30 \| 1.9e-36 \| 169.2 \| 111 \| (1, 120) \| 162 \| (1, 111) \| 148 \| Single-stranded DNA-binding protein \| Single-stranded DNA-binding protein; uniclust \| UniRef100\_A0A022L9F6 \| 99.9 \| 1e-30 \| 2e-36 \| 187.6 \| 107 \| (1, 107) \| 162 \| (52, 159) \| 240 \| Single-stranded DNA-binding protein \| Single-stranded DNA-binding protein; uniclust \| UniRef100\_A0A2W5CU06 \| 99.9 \| 1.2e-30 \| 2.3e-36 \| 170.3 \| 107 \| (1, 107) \| 162 \| (1, 108) \| 167 \| Single-stranded DNA-binding protein \| Single-stranded DNA-binding protein; uniclust \| UniRef100\_A0A0N0JSL7 \| 99.9 \| 1.3e-30 \| 2.4e-36 \| 176.0 \| 106 \| (2, 116) \| 162 \| (27, 133) \| 164 \| Single-stranded DNA-binding protein \| Single-stranded DNA-binding protein; uniclust \| UniRef100\_A0A127SX29 \| 99.9 \| 1.4e-30 \| 2.7e-36 \| 167.5 \| 111 \| (3, 116) \| 162 \| (1, 111) \| 133 \| Single-stranded DNA-binding protein \| Single-stranded DNA-binding protein; uniclust \| UniRef100\_A0A0A7G012 \| 99.9 \| 1.5e-30 \| 3e-36 \| 169.6 \| 109 \| (1, 118) \| 162 \| (2, 110) \| 133 \| Single-stranded DNA-binding protein \| Single-stranded DNA-binding protein; uniclust \| UniRef100\_A0A133Y031 \| 99.9 \| 2.2e-30 \| 4.2e-36 \| 169.6 \| 113 \| (1, 117) \| 162 \| (20, 132) \| 140 \| Single-stranded DNA-binding protein \| Single-stranded DNA-binding protein; uniclust \| UniRef100\_A0A081PBG3 \| 99.9 \| 2.2e-30 \| 4.3e-36 \| 173.3 \| 114 \| (1, 118) \| 162 \| (11, 124) \| 155 \| Single-stranded DNA-binding protein \| Single-stranded DNA-binding protein; uniclust \| UniRef100\_A0A0F6X1X7 \| 99.9 \| 2.6e-30 \| 4.9e-36 \| 174.2 \| 107 \| (4, 118) \| 162 \| (1, 107) \| 177 \| Single-stranded DNA-binding protein \| Single-stranded DNA-binding protein; uniclust \| UniRef100\_A0A011TE40 \| 99.9 \| 2.8e-30 \| 5.3e-36 \| 171.0 \| 111 \| (3, 117) \| 162 \| (26, 143) \| 172 \| Single-stranded DNA-binding protein \| Single-stranded DNA-binding protein; uniclust \| UniRef100\_A0A0C2APC6 \| 99.9 \| 3.3e-30 \| 6.2e-36 \| 172.2 \| 105 \| (2, 107) \| 162 \| (28, 133) \| 194 \| Single-stranded DNA-binding protein \| Single-stranded DNA-binding protein; uniclust \| UniRef100\_A0A0D5YK57 \| 99.9 \| 4e-30 \| 7.6e-36 \| 153.9 \| 85 \| (1, 86) \| 162 \| (2, 86) \| 87 \| Single-strand binding protein \| Single-strand binding protein; uniclust \| UniRef100\_A0A0G4E5Q2 \| 99.9 \| 4.3e-30 \| 7.8e-36 \| 171.2 \| 118 \| (1, 119) \| 162 \| (61, 178) \| 233 \| Single-stranded DNA-binding protein \| Single-stranded DNA-binding protein; uniclust \| UniRef100\_A0A1V5ICE4 \| 99.9 \| 4.2e-30 \| 7.9e-36 \| 160.3 \| 105 \| (4, 117) \| 162 \| (1, 112) \| 129 \| Single-stranded DNA-binding protein \| Single-stranded DNA-binding protein; uniclust \| UniRef100\_A0A0F7L5K1 \| 99.9 \| 4.6e-30 \| 8.7e-36 \| 171.7 \| 107 \| (3, 119) \| 162 \| (1, 108) \| 174 \| Single-strand DNA-binding protein \| Single-strand DNA-binding protein; uniclust \| UniRef100\_A0A1G3M634 \| 99.9 \| 4.9e-30 \| 9.5e-36 \| 170.8 \| 110 \| (1, 114) \| 162 \| (1, 110) \| 158 \| Single-stranded DNA-binding protein \| Single-stranded DNA-binding protein; uniclust \| UniRef100\_A0A220SXH4 \| 99.9 \| 6.7e-30 \| 1.2e-35 \| 175.0 \| 113 \| (4, 117) \| 162 \| (18, 130) \| 261 \| Single-stranded DNA-binding protein \| Single-stranded DNA-binding protein; uniclust \| UniRef100\_A0A2W1TN85 \| 99.9 \| 6.9e-30 \| 1.3e-35 \| 174.7 \| 106 \| (1, 106) \| 162 \| (1, 107) \| 199 \| Single-stranded DNA-binding protein \| Single-stranded DNA-binding protein; uniclust \| UniRef100\_A0A2S8SU52 \| 99.9 \| 7e-30 \| 1.3e-35 \| 175.7 \| 111 \| (2, 118) \| 162 \| (4, 114) \| 220 \| Single-stranded DNA-binding protein \| Single-stranded DNA-binding protein; uniclust \| UniRef100\_A0A1G9EW49 \| 99.9 \| 7.4e-30 \| 1.4e-35 \| 170.4 \| 106 \| (2, 107) \| 162 \| (1, 111) \| 168 \| Single-stranded DNA-binding protein \| Single-stranded DNA-binding protein; uniclust \| UniRef100\_A0A078KHX1 \| 99.9 \| 8.6e-30 \| 1.6e-35 \| 165.3 \| 112 \| (1, 115) \| 162 \| (3, 114) \| 144 \| Single-stranded DNA-binding protein \| Single-stranded DNA-binding protein; uniclust \| UniRef100\_A0A1B0XTI5 \| 99.9 \| 8.6e-30 \| 1.7e-35 \| 169.9 \| 103 \| (1, 104) \| 162 \| (1, 104) \| 151 \| Primosome PriB/single-strand DNA-binding protein \| Primosome PriB/single-strand DNA-binding protein; uniclust \| UniRef100\_A0A0G0GYK1 \| 99.9 \| 9.5e-30 \| 1.8e-35 \| 169.9 \| 112 \| (3, 118) \| 162 \| (5, 116) \| 176 \| Single-stranded DNA-binding protein \| Single-stranded DNA-binding protein; uniclust \| UniRef100\_A0A2E1Q3N5 \| 99.9 \| 1.2e-29 \| 2.4e-35 \| 166.2 \| 112 \| (2, 118) \| 162 \| (1, 112) \| 145 \| Single-stranded DNA-binding protein \| Single-stranded DNA-binding protein; uniclust \| UniRef100\_A0A3D5Q9W8 \| 99.9 \| 1.3e-29 \| 2.5e-35 \| 153.7 \| 89 \| (2, 91) \| 162 \| (7, 95) \| 98 \| Single-stranded DNA-binding protein (Fragment) \| Single-stranded DNA-binding protein (Fragment); uniclust \| UniRef100\_A0A1D7QSF8 \| 99.9 \| 1.4e-29 \| 2.6e-35 \| 177.2 \| 109 \| (4, 118) \| 162 \| (1, 109) \| 227 \| Single-stranded DNA-binding protein \| Single-stranded DNA-binding protein; uniclust \| UniRef100\_A0A142XSZ2 \| 99.9 \| 1.4e-29 \| 2.7e-35 \| 171.4 \| 114 \| (2, 118) \| 162 \| (39, 158) \| 205 \| Single-stranded DNA-binding protein \| Single-stranded DNA-binding protein; uniclust \| UniRef100\_A0A199YTL7 \| 99.9 \| 1.5e-29 \| 2.8e-35 \| 153.0 \| 91 \| (3, 96) \| 162 \| (1, 91) \| 98 \| Single-stranded DNA-binding protein \| Single-stranded DNA-binding protein; uniclust \| UniRef100\_A0A5Q2N011 \| 99.9 \| 2.1e-29 \| 3.8e-35 \| 166.4 \| 110 \| (4, 117) \| 162 \| (1, 113) \| 193 \| Single-stranded DNA-binding protein \| Single-stranded DNA-binding protein; uniclust \| UniRef100\_A0A538CVG0 \| 99.9 \| 2.5e-29 \| 4.6e-35 \| 170.6 \| 113 \| (3, 118) \| 162 \| (75, 187) \| 233 \| Single-stranded DNA-binding protein \| Single-stranded DNA-binding protein; uniclust \| UniRef100\_A0A285MVQ1 \| 99.9 \| 3e-29 \| 5.7e-35 \| 166.0 \| 111 \| (1, 115) \| 162 \| (1, 111) \| 163 \| Single-stranded DNA-binding protein \| Single-stranded DNA-binding protein; uniclust \| UniRef100\_A0A1V5ZP08 \| 99.9 \| 3.1e-29 \| 5.8e-35 \| 150.5 \| 87 \| (1, 88) \| 162 \| (2, 88) \| 92 \| Single-stranded DNA-binding protein \| Single-stranded DNA-binding protein; uniclust \| UniRef100\_A0A132HW80 \| 99.9 \| 3.5e-29 \| 6.6e-35 \| 161.1 \| 111 \| (2, 117) \| 162 \| (1, 111) \| 150 \| Single-stranded DNA-binding protein \| Single-stranded DNA-binding protein; uniclust \| UniRef100\_A0A1H4XXA5 \| 99.9 \| 3.5e-29 \| 6.7e-35 \| 161.6 \| 111 \| (2, 118) \| 162 \| (1, 111) \| 133 \| Single-stranded DNA-binding protein \| Single-stranded DNA-binding protein; uniclust \| UniRef100\_A0A2D9X095 \| 99.9 \| 4.7e-29 \| 9e-35 \| 162.0 \| 109 \| (1, 118) \| 162 \| (6, 116) \| 141 \| Single-stranded DNA-binding protein \| Single-stranded DNA-binding protein; uniclust \| UniRef100\_A0A157SR46 \| 99.9 \| 6.4e-29 \| 1.2e-34 \| 162.0 \| 110 \| (1, 116) \| 162 \| (18, 127) \| 148 \| Single-stranded DNA-binding protein \| Single-stranded DNA-binding protein; uniclust \| UniRef100\_A0A0C1NEJ2 \| 99.9 \| 6.4e-29 \| 1.2e-34 \| 168.2 \| 106 \| (1, 115) \| 162 \| (27, 133) \| 190 \| Single-stranded DNA-binding protein \| Single-stranded DNA-binding protein; uniclust \| UniRef100\_A0A127EFR2 \| 99.9 \| 7.4e-29 \| 1.4e-34 \| 159.6 \| 109 \| (5, 119) \| 162 \| (1, 110) \| 133 \| Single-stranded DNA-binding protein \| Single-stranded DNA-binding protein; uniclust \| UniRef100\_A0A432EQF9 \| 99.9 \| 7.8e-29 \| 1.5e-34 \| 165.5 \| 110 \| (2, 115) \| 162 \| (7, 117) \| 186 \| Single-stranded DNA-binding protein \| Single-stranded DNA-binding protein; uniclust \| UniRef100\_A0A1G6BP95 \| 99.9 \| 8.4e-29 \| 1.6e-34 \| 159.2 \| 107 \| (4, 117) \| 162 \| (1, 107) \| 139 \| Single-stranded DNA-binding protein \| Single-stranded DNA-binding protein; uniclust \| UniRef100\_A0A061K5I9 \| 99.9 \| 8.9e-29 \| 1.7e-34 \| 171.0 \| 115 \| (4, 119) \| 162 \| (44, 158) \| 268 \| Single-stranded DNA-binding protein \| Single-stranded DNA-binding protein; uniclust \| UniRef100\_A0A0A7I9E0 \| 99.9 \| 9.5e-29 \| 1.8e-34 \| 164.2 \| 102 \| (2, 104) \| 162 \| (7, 109) \| 165 \| Single-stranded DNA-binding protein \| Single-stranded DNA-binding protein; uniclust \| UniRef100\_A0A2P4V2X1 \| 99.9 \| 1.2e-28 \| 2.2e-34 \| 147.5 \| 89 \| (1, 89) \| 162 \| (2, 90) \| 90 \| Single-stranded DNA-binding protein (Fragment) \| Single-stranded DNA-binding protein (Fragment); uniclust \| UniRef100\_A0A0B8QWW0 \| 99.9 \| 1.3e-28 \| 2.5e-34 \| 163.6 \| 106 \| (5, 117) \| 162 \| (9, 120) \| 169 \| Single-stranded DNA-binding protein \| Single-stranded DNA-binding protein; uniclust \| UniRef100\_A0A081UAD3 \| 99.9 \| 1.6e-28 \| 3e-34 \| 165.4 \| 108 \| (1, 115) \| 162 \| (33, 140) \| 174 \| Single-stranded DNA-binding protein \| Single-stranded DNA-binding protein; uniclust \| UniRef100\_A0A968VPK9 \| 99.9 \| 1.8e-28 \| 3.3e-34 \| 161.8 \| 113 \| (1, 117) \| 162 \| (1, 114) \| 214 \| Single-stranded DNA-binding protein \| Single-stranded DNA-binding protein; uniclust \| UniRef100\_A0A090CZ19 \| 99.9 \| 1.7e-28 \| 3.4e-34 \| 164.6 \| 103 \| (3, 114) \| 162 \| (12, 115) \| 168 \| Single-stranded DNA-binding protein \| Single-stranded DNA-binding protein; uniclust \| UniRef100\_A0A1G9SMN4 \| 99.9 \| 1.9e-28 \| 3.6e-34 \| 164.9 \| 108 \| (4, 117) \| 162 \| (1, 108) \| 177 \| Single-stranded DNA-binding protein \| Single-stranded DNA-binding protein; uniclust \| UniRef100\_A0A1S2MBF9 \| 99.9 \| 1.9e-28 \| 3.7e-34 \| 162.9 \| 110 \| (3, 118) \| 162 \| (20, 129) \| 153 \| Single-stranded DNA-binding protein \| Single-stranded DNA-binding protein; uniclust \| UniRef100\_A0A086Z1J5 \| 99.9 \| 1.9e-28 \| 3.8e-34 \| 162.8 \| 111 \| (3, 117) \| 162 \| (1, 112) \| 151 \| Single-stranded DNA-binding protein \| Single-stranded DNA-binding protein; uniclust \| UniRef100\_A0A0N1D9D7 \| 99.9 \| 2.1e-28 \| 4.1e-34 \| 165.3 \| 113 \| (2, 118) \| 162 \| (1, 115) \| 161 \| Single-stranded DNA-binding protein \| Single-stranded DNA-binding protein; uniclust \| UniRef100\_A0A0G1W755 \| 99.9 \| 2.2e-28 \| 4.2e-34 \| 159.2 \| 113 \| (1, 117) \| 162 \| (40, 152) \| 162 \| Single-stranded DNA-binding protein \| Single-stranded DNA-binding protein; uniclust \| UniRef100\_A0A1Q6UZU3 \| 99.9 \| 2.3e-28 \| 4.5e-34 \| 157.5 \| 109 \| (4, 118) \| 162 \| (2, 110) \| 137 \| Single-stranded DNA-binding protein \| Single-stranded DNA-binding protein; uniclust \| UniRef100\_A0A1F9CN51 \| 99.9 \| 2.6e-28 \| 4.9e-34 \| 171.6 \| 112 \| (2, 117) \| 162 \| (127, 238) \| 265 \| Single-stranded DNA-binding protein \| Single-stranded DNA-binding protein; uniclust \| UniRef100\_A0A022LTY2 \| 99.9 \| 2.6e-28 \| 5e-34 \| 172.6 \| 113 \| (1, 113) \| 162 \| (30, 144) \| 272 \| Single-stranded DNA-binding protein \| Single-stranded DNA-binding protein; uniclust \| UniRef100\_A0A5C8BC14 \| 99.9 \| 2.8e-28 \| 5.3e-34 \| 157.9 \| 110 \| (2, 118) \| 162 \| (8, 117) \| 135 \| Single-stranded DNA-binding protein \| Single-stranded DNA-binding protein; uniclust \| UniRef100\_A0A1B9F675 \| 99.9 \| 3.8e-28 \| 7.1e-34 \| 160.1 \| 111 \| (1, 116) \| 162 \| (81, 191) \| 212 \| Single-stranded DNA-binding protein \| Single-stranded DNA-binding protein; uniclust \| UniRef100\_A0A2V7U681 \| 99.9 \| 4.3e-28 \| 8e-34 \| 158.3 \| 102 \| (2, 104) \| 162 \| (1, 102) \| 166 \| Single-stranded DNA-binding protein \| Single-stranded DNA-binding protein; uniclust \| UniRef100\_A0A0N1KRF3 \| 99.9 \| 6.3e-28 \| 1.2e-33 \| 151.2 \| 111 \| (1, 114) \| 162 \| (4, 114) \| 120 \| Single-stranded DNA-binding protein (Fragment) \| Single-stranded DNA-binding protein (Fragment); uniclust \| UniRef100\_A0A2Z4R767 \| 99.9 \| 6.4e-28 \| 1.2e-33 \| 156.6 \| 113 \| (2, 118) \| 162 \| (3, 115) \| 169 \| Single-stranded DNA-binding protein \| Single-stranded DNA-binding protein; uniclust \| UniRef100\_A0A095Y5H8 \| 99.9 \| 6.5e-28 \| 1.3e-33 \| 161.5 \| 110 \| (3, 118) \| 162 \| (1, 110) \| 161 \| Single-stranded DNA-binding protein \| Single-stranded DNA-binding protein; uniclust \| UniRef100\_A0A1F2Q8N9 \| 99.9 \| 6.7e-28 \| 1.3e-33 \| 154.9 \| 113 \| (2, 117) \| 162 \| (7, 119) \| 142 \| Single-stranded DNA-binding protein \| Single-stranded DNA-binding protein; uniclust \| UniRef100\_A0A022KT57 \| 99.9 \| 6.8e-28 \| 1.3e-33 \| 167.5 \| 107 \| (1, 107) \| 162 \| (15, 122) \| 226 \| Single-stranded DNA-binding protein \| Single-stranded DNA-binding protein; uniclust \| UniRef100\_A0A0X8KMA9 \| 99.9 \| 7.3e-28 \| 1.4e-33 \| 162.7 \| 112 \| (4, 121) \| 162 \| (1, 112) \| 193 \| Single-stranded DNA-binding protein \| Single-stranded DNA-binding protein; uniclust \| UniRef100\_A0A1B3X7C3 \| 99.9 \| 8.3e-28 \| 1.6e-33 \| 157.1 \| 108 \| (5, 118) \| 162 \| (1, 108) \| 152 \| Single-stranded DNA-binding protein \| Single-stranded DNA-binding protein; uniclust \| UniRef100\_A0A1R4GZN9 \| 99.9 \| 8.7e-28 \| 1.6e-33 \| 153.4 \| 106 \| (4, 118) \| 162 \| (1, 107) \| 147 \| Single-stranded DNA-binding protein \| Single-stranded DNA-binding protein; uniclust \| UniRef100\_A0A0B5D8I7 \| 99.9 \| 8.7e-28 \| 1.7e-33 \| 168.7 \| 107 \| (1, 107) \| 162 \| (46, 153) \| 218 \| Single-stranded DNA-binding protein \| Single-stranded DNA-binding protein; uniclust \| UniRef100\_A0A1F9CN51 \| 99.9 \| 9.2e-28 \| 1.7e-33 \| 168.9 \| 109 \| (1, 113) \| 162 \| (1, 109) \| 265 \| Single-stranded DNA-binding protein \| Single-stranded DNA-binding protein; uniclust \| UniRef100\_A0A0A2VXI7 \| 99.9 \| 1e-27 \| 1.9e-33 \| 172.1 \| 159 \| (3, 161) \| 162 \| (4, 175) \| 425 \| Single-stranded DNA-binding protein \| Single-stranded DNA-binding protein; uniclust \| UniRef100\_A0A942N2T9 \| 99.9 \| 1.2e-27 \| 2.1e-33 \| 155.2 \| 112 \| (3, 118) \| 162 \| (52, 163) \| 188 \| Single-stranded DNA-binding protein \| Single-stranded DNA-binding protein; uniclust \| UniRef100\_A0A1C6R7A7 \| 99.9 \| 1.6e-27 \| 3e-33 \| 157.1 \| 105 \| (3, 107) \| 162 \| (1, 106) \| 155 \| Single-stranded DNA-binding protein \| Single-stranded DNA-binding protein; uniclust \| UniRef100\_A0A5D4T3F4 \| 99.9 \| 1.7e-27 \| 3.1e-33 \| 156.8 \| 111 \| (4, 120) \| 162 \| (1, 111) \| 184 \| Single-stranded DNA-binding protein \| Single-stranded DNA-binding protein; uniclust \| UniRef100\_A0A0A0RT45 \| 99.9 \| 2e-27 \| 3.7e-33 \| 152.7 \| 107 \| (4, 117) \| 162 \| (1, 107) \| 157 \| Single-stranded DNA-binding protein \| Single-stranded DNA-binding protein; uniclust \| UniRef100\_A0A077ELF4 \| 99.9 \| 2e-27 \| 3.7e-33 \| 151.0 \| 115 \| (1, 119) \| 162 \| (14, 128) \| 138 \| Single-stranded DNA-binding protein \| Single-stranded DNA-binding protein; uniclust \| UniRef100\_A0A945T384 \| 99.9 \| 2e-27 \| 3.8e-33 \| 150.8 \| 110 \| (4, 117) \| 162 \| (1, 110) \| 149 \| Single-stranded DNA-binding protein \| Single-stranded DNA-binding protein; uniclust \| UniRef100\_A0A1T4VV37 \| 99.9 \| 2.4e-27 \| 4.5e-33 \| 161.4 \| 103 \| (1, 104) \| 162 \| (1, 103) \| 226 \| Single-stranded DNA-binding protein \| Single-stranded DNA-binding protein; uniclust \| UniRef100\_A0A1C6LRJ7 \| 99.9 \| 2.5e-27 \| 4.7e-33 \| 159.3 \| 107 \| (5, 115) \| 162 \| (1, 107) \| 177 \| Single-stranded DNA-binding protein \| Single-stranded DNA-binding protein; uniclust \| UniRef100\_A0A8T6YXL0 \| 99.9 \| 2.6e-27 \| 4.8e-33 \| 146.7 \| 110 \| (3, 116) \| 162 \| (1, 110) \| 125 \| Single-stranded DNA-binding protein \| Single-stranded DNA-binding protein; uniclust \| UniRef100\_A0A141RPF2 \| 99.9 \| 2.7e-27 \| 5e-33 \| 152.3 \| 126 \| (37, 162) \| 162 \| (2, 130) \| 145 \| Single-stranded DNA-binding protein \| Single-stranded DNA-binding protein; uniclust \| UniRef100\_A0A1F2WYY6 \| 99.9 \| 2.7e-27 \| 5e-33 \| 151.7 \| 104 \| (3, 118) \| 162 \| (22, 125) \| 145 \| Single-stranded DNA-binding protein \| Single-stranded DNA-binding protein; uniclust \| UniRef100\_A0A078KY52 \| 99.9 \| 3.3e-27 \| 6.3e-33 \| 151.5 \| 110 \| (1, 116) \| 162 \| (1, 110) \| 141 \| Single-stranded DNA-binding protein \| Single-stranded DNA-binding protein; uniclust \| UniRef100\_A0A0K1HH46 \| 99.9 \| 4.3e-27 \| 8e-33 \| 155.1 \| 112 \| (3, 118) \| 162 \| (1, 112) \| 174 \| Single-stranded DNA-binding protein \| Single-stranded DNA-binding protein; uniclust \| UniRef100\_A0A014MGY3 \| 99.9 \| 4.5e-27 \| 8.9e-33 \| 162.2 \| 111 \| (2, 118) \| 162 \| (3, 113) \| 184 \| Single-stranded DNA-binding protein \| Single-stranded DNA-binding protein; uniclust \| UniRef100\_A0A259R8S6 \| 99.9 \| 5e-27 \| 9.5e-33 \| 152.3 \| 73 \| (42, 117) \| 162 \| (5, 77) \| 143 \| Single-stranded DNA-binding protein (Fragment) \| Single-stranded DNA-binding protein (Fragment); uniclust \| UniRef100\_A0A3D1TXE1 \| 99.9 \| 5.5e-27 \| 1e-32 \| 150.1 \| 101 \| (4, 117) \| 162 \| (28, 128) \| 138 \| Single-stranded DNA-binding protein \| Single-stranded DNA-binding protein; uniclust \| UniRef100\_A0A9D7S067 \| 99.9 \| 5.7e-27 \| 1e-32 \| 164.4 \| 112 \| (1, 115) \| 162 \| (1, 112) \| 342 \| Single-stranded DNA-binding protein \| Single-stranded DNA-binding protein; uniclust \| UniRef100\_A0A173TLD0 \| 99.9 \| 1.1e-26 \| 2.1e-32 \| 153.5 \| 105 \| (4, 117) \| 162 \| (1, 105) \| 177 \| Single-stranded DNA-binding protein \| Single-stranded DNA-binding protein; uniclust \| UniRef100\_A0A0K2JFD7 \| 99.9 \| 1.1e-26 \| 2.2e-32 \| 152.4 \| 106 \| (3, 120) \| 162 \| (1, 106) \| 164 \| Single-stranded DNA-binding protein \| Single-stranded DNA-binding protein; uniclust \| UniRef100\_A0A0C1I8N4 \| 99.9 \| 1.2e-26 \| 2.3e-32 \| 159.7 \| 109 \| (2, 116) \| 162 \| (24, 132) \| 180 \| Single-stranded DNA-binding protein \| Single-stranded DNA-binding protein; uniclust \| UniRef100\_A0A077XQ49 \| 99.9 \| 1.3e-26 \| 2.5e-32 \| 145.1 \| 111 \| (1, 115) \| 162 \| (1, 111) \| 131 \| Single-stranded DNA-binding protein \| Single-stranded DNA-binding protein; uniclust \| UniRef100\_A0A059XWD8 \| 99.9 \| 1.3e-26 \| 2.6e-32 \| 160.5 \| 109 \| (2, 118) \| 162 \| (3, 111) \| 188 \| Single-stranded DNA-binding protein \| Single-stranded DNA-binding protein; uniclust \| UniRef100\_A0A1E9Q8Q7 \| 99.9 \| 1.7e-26 \| 3.3e-32 \| 157.2 \| 106 \| (1, 107) \| 162 \| (1, 112) \| 186 \| Single-stranded DNA-binding protein \| Single-stranded DNA-binding protein; uniclust \| UniRef100\_A0A0R2CM34 \| 99.9 \| 1.8e-26 \| 3.4e-32 \| 146.3 \| 111 \| (4, 120) \| 162 \| (1, 111) \| 149 \| Single-stranded DNA-binding protein \| Single-stranded DNA-binding protein; uniclust \| UniRef100\_A0A523YRH3 \| 99.9 \| 2.1e-26 \| 3.8e-32 \| 143.0 \| 108 \| (2, 115) \| 162 \| (1, 108) \| 140 \| Single-stranded DNA-binding protein \| Single-stranded DNA-binding protein; uniclust \| UniRef100\_A0A1C7HIC5 \| 99.9 \| 2.1e-26 \| 3.9e-32 \| 157.5 \| 112 \| (2, 117) \| 162 \| (1, 112) \| 226 \| Single-stranded DNA-binding protein \| Single-stranded DNA-binding protein; uniclust \| UniRef100\_A0A160LK02 \| 99.9 \| 2.1e-26 \| 4e-32 \| 154.9 \| 101 \| (4, 119) \| 162 \| (1, 101) \| 182 \| Single-stranded DNA-binding protein \| Single-stranded DNA-binding protein; uniclust \| UniRef100\_A0A0K1F1H1 \| 99.9 \| 2.2e-26 \| 4.1e-32 \| 146.5 \| 110 \| (3, 115) \| 162 \| (1, 111) \| 159 \| Single-stranded DNA-binding protein \| Single-stranded DNA-binding protein; uniclust \| UniRef100\_A0A2D3LPR9 \| 99.9 \| 2.4e-26 \| 4.5e-32 \| 147.9 \| 107 \| (5, 115) \| 162 \| (1, 107) \| 129 \| Single-stranded DNA-binding protein \| Single-stranded DNA-binding protein; uniclust \| UniRef100\_A0A0U2Q367 \| 99.9 \| 2.5e-26 \| 4.6e-32 \| 163.3 \| 114 \| (4, 117) \| 162 \| (169, 282) \| 344 \| Single-stranded DNA-binding protein \| Single-stranded DNA-binding protein; uniclust \| UniRef100\_A0A0A8X393 \| 99.9 \| 2.7e-26 \| 4.9e-32 \| 153.4 \| 109 \| (3, 117) \| 162 \| (69, 177) \| 208 \| Single-stranded DNA-binding protein \| Single-stranded DNA-binding protein; uniclust \| UniRef100\_A0A238ZN22 \| 99.9 \| 2.9e-26 \| 5.4e-32 \| 138.9 \| 102 \| (1, 103) \| 162 \| (1, 103) \| 105 \| Single-stranded DNA-binding protein \| Single-stranded DNA-binding protein; uniclust \| UniRef100\_A0A1V6FE91 \| 99.9 \| 2.9e-26 \| 5.4e-32 \| 158.5 \| 111 \| (2, 116) \| 162 \| (1, 112) \| 249 \| Single-stranded DNA-binding protein \| Single-stranded DNA-binding protein; uniclust \| UniRef100\_A0A0S1SZJ6 \| 99.9 \| 2.9e-26 \| 5.5e-32 \| 150.4 \| 109 \| (1, 117) \| 162 \| (16, 126) \| 141 \| Single-stranded DNA-binding protein \| Single-stranded DNA-binding protein; uniclust \| UniRef100\_A0A1Y4GBS3 \| 99.9 \| 3.1e-26 \| 5.7e-32 \| 148.9 \| 113 \| (3, 120) \| 162 \| (39, 151) \| 189 \| Single-stranded DNA-binding protein \| Single-stranded DNA-binding protein; uniclust \| UniRef100\_A0A7V3PS15 \| 99.9 \| 3.2e-26 \| 6.1e-32 \| 145.2 \| 103 \| (5, 118) \| 162 \| (1, 103) \| 122 \| Single-stranded DNA-binding protein \| Single-stranded DNA-binding protein; uniclust \| UniRef100\_A0A401FYS3 \| 99.9 \| 3.6e-26 \| 6.7e-32 \| 160.8 \| 114 \| (2, 118) \| 162 \| (1, 114) \| 347 \| Single-stranded DNA-binding protein \| Single-stranded DNA-binding protein; uniclust \| UniRef100\_A0A1G6X935 \| 99.9 \| 3.6e-26 \| 6.7e-32 \| 151.7 \| 113 \| (1, 117) \| 162 \| (67, 179) \| 184 \| Single-stranded DNA-binding protein \| Single-stranded DNA-binding protein; uniclust \| UniRef100\_A0A7Z9G7J1 \| 99.9 \| 3.8e-26 \| 7.1e-32 \| 160.2 \| 114 \| (3, 120) \| 162 \| (1, 114) \| 339 \| Single-stranded DNA-binding protein (Fragment) \| Single-stranded DNA-binding protein (Fragment); uniclust \| UniRef100\_A0A0G1VV93 \| 99.9 \| 3.8e-26 \| 7.1e-32 \| 153.4 \| 111 \| (2, 116) \| 162 \| (57, 167) \| 194 \| Single-stranded DNA-binding protein \| Single-stranded DNA-binding protein; uniclust \| UniRef100\_A0A1G9Q5V4 \| 99.9 \| 4.1e-26 \| 7.7e-32 \| 146.9 \| 99 \| (3, 116) \| 162 \| (4, 102) \| 145 \| Single-stranded DNA-binding protein \| Single-stranded DNA-binding protein; uniclust \| UniRef100\_A0A0M4QA89 \| 99.9 \| 4.2e-26 \| 7.9e-32 \| 156.3 \| 104 \| (3, 106) \| 162 \| (35, 139) \| 210 \| Single-stranded DNA-binding protein \| Single-stranded DNA-binding protein; uniclust \| UniRef100\_A0A9D9DB79 \| 99.9 \| 4.3e-26 \| 8.1e-32 \| 159.8 \| 112 \| (1, 115) \| 162 \| (1, 112) \| 268 \| Single-stranded DNA-binding protein \| Single-stranded DNA-binding protein; uniclust \| UniRef100\_A0A010SZ24 \| 99.9 \| 4.8e-26 \| 9.4e-32 \| 163.3 \| 111 \| (1, 115) \| 162 \| (31, 147) \| 237 \| Single-stranded DNA-binding protein \| Single-stranded DNA-binding protein; uniclust \| UniRef100\_A0A132MZK2 \| 99.9 \| 5.6e-26 \| 1.1e-31 \| 148.4 \| 112 \| (1, 112) \| 162 \| (1, 114) \| 159 \| Single-stranded DNA-binding protein \| Single-stranded DNA-binding protein; uniclust \| UniRef100\_UPI001CCB6140 \| 99.9 \| 6.8e-26 \| 1.2e-31 \| 144.4 \| 113 \| (2, 117) \| 162 \| (1, 113) \| 165 \| single-stranded DNA-binding protein \| single-stranded DNA-binding protein; uniclust \| UniRef100\_A0A7C7W444 \| 99.9 \| 7.3e-26 \| 1.3e-31 \| 151.4 \| 114 \| (2, 118) \| 162 \| (1, 114) \| 231 \| Single-stranded DNA-binding protein \| Single-stranded DNA-binding protein; uniclust \| UniRef100\_A0A920JQB4 \| 99.9 \| 1.1e-25 \| 2.1e-31 \| 147.6 \| 114 \| (3, 119) \| 162 \| (55, 168) \| 200 \| Single-stranded DNA-binding protein \| Single-stranded DNA-binding protein; uniclust \| UniRef100\_A0A1M5RIB4 \| 99.9 \| 1.1e-25 \| 2.1e-31 \| 142.0 \| 110 \| (3, 119) \| 162 \| (1, 111) \| 135 \| Single-stranded DNA-binding protein \| Single-stranded DNA-binding protein; uniclust \| UniRef100\_A0A1C6AGN4 \| 99.9 \| 1.2e-25 \| 2.2e-31 \| 149.9 \| 106 \| (5, 117) \| 162 \| (1, 106) \| 180 \| Single-stranded DNA-binding protein \| Single-stranded DNA-binding protein; uniclust \| UniRef100\_A0A0F0LRE6 \| 99.9 \| 1.3e-25 \| 2.5e-31 \| 159.5 \| 106 \| (2, 108) \| 162 \| (35, 141) \| 234 \| Single-stranded DNA-binding protein \| Single-stranded DNA-binding protein; uniclust \| UniRef100\_A0A2D6E675 \| 99.9 \| 1.5e-25 \| 2.8e-31 \| 145.4 \| 100 \| (4, 114) \| 162 \| (1, 100) \| 140 \| Single-stranded DNA-binding protein \| Single-stranded DNA-binding protein; uniclust \| UniRef100\_A0A2N1TAT9 \| 99.9 \| 1.5e-25 \| 2.9e-31 \| 148.4 \| 112 \| (2, 117) \| 162 \| (1, 112) \| 197 \| Single-stranded DNA-binding protein \| Single-stranded DNA-binding protein; uniclust \| UniRef100\_A0A1H9QB91 \| 99.9 \| 1.5e-25 \| 2.9e-31 \| 145.3 \| 103 \| (2, 104) \| 162 \| (12, 115) \| 155 \| Single-stranded DNA-binding protein (Fragment) \| Single-stranded DNA-binding protein (Fragment); uniclust \| UniRef100\_A0A8H9CAI2 \| 99.9 \| 1.6e-25 \| 3e-31 \| 141.1 \| 117 \| (1, 118) \| 162 \| (1, 118) \| 153 \| Single-stranded DNA-binding protein \| Single-stranded DNA-binding protein; uniclust \| UniRef100\_A0A350T707 \| 99.9 \| 1.6e-25 \| 3e-31 \| 143.7 \| 112 \| (1, 116) \| 162 \| (1, 112) \| 173 \| Single-stranded DNA-binding protein (Fragment) \| Single-stranded DNA-binding protein (Fragment); uniclust \| UniRef100\_A0A077KLD4 \| 99.9 \| 1.6e-25 \| 3.1e-31 \| 152.3 \| 100 \| (9, 114) \| 162 \| (30, 129) \| 173 \| Single-stranded DNA-binding protein \| Single-stranded DNA-binding protein; uniclust \| UniRef100\_A0A1C6B8W9 \| 99.9 \| 1.7e-25 \| 3.2e-31 \| 147.2 \| 108 \| (3, 115) \| 162 \| (64, 171) \| 205 \| Single-stranded DNA-binding protein \| Single-stranded DNA-binding protein; uniclust \| UniRef100\_A0A931S483 \| 99.9 \| 1.9e-25 \| 3.5e-31 \| 141.2 \| 110 \| (2, 115) \| 162 \| (1, 110) \| 146 \| Single-stranded DNA-binding protein \| Single-stranded DNA-binding protein; uniclust \| UniRef100\_A0A0D4C1Z6 \| 99.9 \| 2e-25 \| 3.8e-31 \| 148.4 \| 100 \| (3, 104) \| 162 \| (19, 119) \| 167 \| Single-stranded DNA-binding protein \| Single-stranded DNA-binding protein; uniclust \| UniRef100\_A0A317ZLK3 \| 99.9 \| 2.1e-25 \| 4e-31 \| 144.6 \| 105 \| (2, 116) \| 162 \| (11, 116) \| 140 \| Single-stranded DNA-binding protein \| Single-stranded DNA-binding protein; uniclust \| UniRef100\_A0A087D5W7 \| 99.9 \| 2.2e-25 \| 4.2e-31 \| 160.6 \| 107 \| (1, 107) \| 162 \| (14, 121) \| 236 \| Single-stranded DNA-binding protein \| Single-stranded DNA-binding protein; uniclust \| UniRef100\_A0A5C7PTQ0 \| 99.9 \| 2.5e-25 \| 4.7e-31 \| 145.6 \| 111 \| (4, 118) \| 162 \| (1, 111) \| 165 \| Single-stranded DNA-binding protein \| Single-stranded DNA-binding protein; uniclust \| UniRef100\_A0A379VN86 \| 99.9 \| 2.6e-25 \| 4.7e-31 \| 151.4 \| 112 \| (4, 115) \| 162 \| (5, 116) \| 263 \| Single-stranded DNA-binding protein \| Single-stranded DNA-binding protein; uniclust \| UniRef100\_A0A3L7TTE1 \| 99.9 \| 3.3e-25 \| 6e-31 \| 148.0 \| 113 \| (2, 118) \| 162 \| (66, 178) \| 227 \| Single-stranded DNA-binding protein \| Single-stranded DNA-binding protein; uniclust \| UniRef100\_A0A084F1M5 \| 99.9 \| 3.2e-25 \| 6e-31 \| 149.9 \| 107 \| (4, 118) \| 162 \| (1, 108) \| 192 \| Single-stranded DNA-binding protein \| Single-stranded DNA-binding protein; uniclust \| UniRef100\_A0A2H0VXD5 \| 99.9 \| 3.7e-25 \| 7e-31 \| 147.1 \| 106 \| (1, 115) \| 162 \| (20, 126) \| 185 \| Single-stranded DNA-binding protein \| Single-stranded DNA-binding protein; uniclust \| UniRef100\_A0A9D2B152 \| 99.9 \| 3.8e-25 \| 7.1e-31 \| 157.5 \| 110 \| (1, 113) \| 162 \| (17, 126) \| 311 \| Single-stranded DNA-binding protein \| Single-stranded DNA-binding protein; uniclust \| UniRef100\_A0A1I7YM23 \| 99.9 \| 4.2e-25 \| 7.7e-31 \| 159.5 \| 119 \| (1, 120) \| 162 \| (271, 389) \| 434 \| Single-stranded DNA-binding protein \| Single-stranded DNA-binding protein; uniclust \| UniRef100\_A0A6J5R9U9 \| 99.9 \| 4.3e-25 \| 8.1e-31 \| 144.5 \| 109 \| (3, 118) \| 162 \| (22, 130) \| 153 \| Single-stranded DNA-binding protein \| Single-stranded DNA-binding protein; uniclust \| UniRef100\_A0A174F184 \| 99.9 \| 4.4e-25 \| 8.4e-31 \| 155.4 \| 109 \| (3, 118) \| 162 \| (19, 130) \| 254 \| Single-stranded DNA-binding protein \| Single-stranded DNA-binding protein; uniclust \| UniRef100\_A0A011NB88 \| 99.9 \| 4.7e-25 \| 8.8e-31 \| 139.9 \| 109 \| (5, 116) \| 162 \| (1, 109) \| 147 \| Single-stranded DNA-binding protein \| Single-stranded DNA-binding protein; uniclust \| UniRef100\_A0A0E3XA19 \| 99.9 \| 4.5e-25 \| 8.8e-31 \| 158.0 \| 106 \| (1, 107) \| 162 \| (32, 138) \| 237 \| DNA binding protein \| DNA binding protein; uniclust \| UniRef100\_A0A087D6Y5 \| 99.9 \| 4.8e-25 \| 9.2e-31 \| 146.9 \| 105 \| (2, 107) \| 162 \| (1, 109) \| 167 \| Single-stranded DNA-binding protein \| Single-stranded DNA-binding protein; uniclust \| UniRef100\_A0A0Q8L5D1 \| 99.9 \| 5.1e-25 \| 9.7e-31 \| 144.3 \| 108 \| (1, 116) \| 162 \| (2, 110) \| 143 \| Helix-destabilizing protein \| Helix-destabilizing protein; uniclust \| UniRef100\_A0A061LY12 \| 99.8 \| 5.4e-25 \| 1.1e-30 \| 160.2 \| 107 \| (1, 108) \| 162 \| (40, 147) \| 264 \| Single-stranded DNA-binding protein \| Single-stranded DNA-binding protein; uniclust \| UniRef100\_A0A140DWS2 \| 99.8 \| 5.5e-25 \| 1.1e-30 \| 144.6 \| 111 \| (1, 118) \| 162 \| (2, 112) \| 123 \| Single-stranded DNA-binding protein \| Single-stranded DNA-binding protein; uniclust \| UniRef100\_A0A0L0HLE3 \| 99.8 \| 5.8e-25 \| 1.1e-30 \| 145.4 \| 108 \| (4, 115) \| 162 \| (32, 139) \| 158 \| Single-strand binding protein \| Single-strand binding protein; uniclust \| UniRef100\_A0A1H7BM52 \| 99.8 \| 6.5e-25 \| 1.2e-30 \| 155.4 \| 112 \| (2, 113) \| 162 \| (131, 244) \| 306 \| Single-stranded DNA-binding protein \| Single-stranded DNA-binding protein; uniclust \| UniRef100\_A0A4Q0D392 \| 99.8 \| 6.6e-25 \| 1.2e-30 \| 148.2 \| 104 \| (4, 113) \| 162 \| (1, 104) \| 184 \| Single-stranded DNA-binding protein \| Single-stranded DNA-binding protein; uniclust \| UniRef100\_A0A075V2L9 \| 99.8 \| 6.5e-25 \| 1.3e-30 \| 149.2 \| 106 \| (1, 106) \| 162 \| (12, 118) \| 170 \| Single-stranded DNA-binding protein \| Single-stranded DNA-binding protein; uniclust \| UniRef100\_A0A2N2PZR6 \| 99.8 \| 7.1e-25 \| 1.3e-30 \| 138.7 \| 112 \| (2, 117) \| 162 \| (37, 148) \| 156 \| Single-stranded DNA-binding protein \| Single-stranded DNA-binding protein; uniclust \| UniRef100\_A0A0Q6VAK4 \| 99.8 \| 7.1e-25 \| 1.4e-30 \| 147.9 \| 107 \| (1, 107) \| 162 \| (1, 108) \| 163 \| Single-stranded DNA-binding protein \| Single-stranded DNA-binding protein; uniclust \| UniRef100\_V7ZAW8 \| 99.8 \| 8.2e-25 \| 1.5e-30 \| 148.9 \| 113 \| (1, 117) \| 162 \| (87, 199) \| 261 \| Single-stranded DNA-binding protein \| Single-stranded DNA-binding protein; uniclust \| UniRef100\_A0A7Z8Y919 \| 99.8 \| 8.8e-25 \| 1.7e-30 \| 147.4 \| 102 \| (3, 104) \| 162 \| (1, 104) \| 183 \| Single-stranded DNA-binding protein \| Single-stranded DNA-binding protein; uniclust \| UniRef100\_A0A1N6NNQ4 \| 99.8 \| 9.4e-25 \| 1.7e-30 \| 144.0 \| 113 \| (1, 117) \| 162 \| (45, 157) \| 207 \| Single-stranded DNA-binding protein \| Single-stranded DNA-binding protein; uniclust \| UniRef100\_A0A1V5QGS9 \| 99.8 \| 9.6e-25 \| 1.8e-30 \| 139.5 \| 106 \| (3, 117) \| 162 \| (1, 106) \| 140 \| Single-stranded DNA-binding protein \| Single-stranded DNA-binding protein; uniclust \| UniRef100\_A0A2E8QTK5 \| 99.8 \| 1e-24 \| 1.9e-30 \| 146.1 \| 110 \| (1, 115) \| 162 \| (73, 182) \| 215 \| Single-stranded DNA-binding protein \| Single-stranded DNA-binding protein; uniclust \| UniRef100\_A0A0M4MLW5 \| 99.8 \| 1e-24 \| 1.9e-30 \| 137.2 \| 103 \| (4, 107) \| 162 \| (1, 105) \| 128 \| Single-stranded DNA-binding protein \| Single-stranded DNA-binding protein; uniclust \| UniRef100\_A0A1H0IKV7 \| 99.8 \| 1e-24 \| 2e-30 \| 144.6 \| 106 \| (2, 107) \| 162 \| (1, 107) \| 154 \| Single-stranded DNA-binding protein \| Single-stranded DNA-binding protein; uniclust \| UniRef100\_A0A090VXM6 \| 99.8 \| 1.1e-24 \| 2e-30 \| 132.7 \| 85 \| (1, 86) \| 162 \| (1, 85) \| 95 \| Single-stranded DNA-binding protein \| Single-stranded DNA-binding protein; uniclust \| UniRef100\_A0A8S5LHZ5 \| 99.8 \| 1.1e-24 \| 2e-30 \| 147.4 \| 105 \| (3, 118) \| 162 \| (100, 204) \| 232 \| Single strand binding protein \| Single strand binding protein; uniclust \| UniRef100\_A0A1G7H7E7 \| 99.8 \| 1.1e-24 \| 2e-30 \| 145.3 \| 106 \| (5, 118) \| 162 \| (1, 106) \| 188 \| Single-stranded DNA-binding protein \| Single-stranded DNA-binding protein; uniclust \| UniRef100\_A0A1T4KFM7 \| 99.8 \| 1.2e-24 \| 2.3e-30 \| 152.9 \| 109 \| (3, 119) \| 162 \| (1, 110) \| 228 \| Single-stranded DNA-binding protein \| Single-stranded DNA-binding protein; uniclust \| UniRef100\_A0A086N2V0 \| 99.8 \| 1.3e-24 \| 2.5e-30 \| 154.1 \| 107 \| (1, 108) \| 162 \| (35, 142) \| 224 \| Single-stranded DNA-binding protein \| Single-stranded DNA-binding protein; uniclust \| UniRef100\_A0A0D8L8J0 \| 99.8 \| 1.6e-24 \| 3e-30 \| 130.9 \| 92 \| (2, 93) \| 162 \| (3, 94) \| 104 \| Single-stranded DNA-binding protein (Fragment) \| Single-stranded DNA-binding protein (Fragment); uniclust \| UniRef100\_A4J7S4 \| 99.8 \| 1.7e-24 \| 3.2e-30 \| 146.0 \| 106 \| (4, 119) \| 162 \| (1, 107) \| 225 \| Single-stranded DNA-binding protein \| Single-stranded DNA-binding protein; uniclust \| UniRef100\_A0A6L7UXF6 \| 99.8 \| 1.9e-24 \| 3.5e-30 \| 142.8 \| 107 \| (1, 115) \| 162 \| (59, 165) \| 191 \| Single-stranded DNA-binding protein \| Single-stranded DNA-binding protein; uniclust \| UniRef100\_A0A3M1VW93 \| 99.8 \| 2e-24 \| 3.7e-30 \| 143.9 \| 111 \| (2, 116) \| 162 \| (89, 199) \| 222 \| Single-stranded DNA-binding protein \| Single-stranded DNA-binding protein; uniclust \| UniRef100\_A0A2U8BT43 \| 99.8 \| 2.1e-24 \| 3.9e-30 \| 138.0 \| 113 \| (1, 113) \| 162 \| (1, 115) \| 137 \| Single-stranded DNA-binding protein \| Single-stranded DNA-binding protein; uniclust \| UniRef100\_A0A0A3IX47 \| 99.8 \| 2.1e-24 \| 4e-30 \| 139.4 \| 110 \| (2, 117) \| 162 \| (9, 118) \| 154 \| Single-stranded DNA-binding protein \| Single-stranded DNA-binding protein; uniclust \| UniRef100\_A0A060JLA2 \| 99.8 \| 2.3e-24 \| 4.4e-30 \| 147.3 \| 106 \| (1, 107) \| 162 \| (4, 110) \| 177 \| Single-stranded DNA-binding protein \| Single-stranded DNA-binding protein; uniclust \| UniRef100\_A0A024LSH6 \| 99.8 \| 2.4e-24 \| 4.5e-30 \| 140.6 \| 114 \| (2, 115) \| 162 \| (34, 149) \| 183 \| Single-stranded DNA-binding protein \| Single-stranded DNA-binding protein; uniclust \| UniRef100\_A0A2N5PKE3 \| 99.8 \| 2.8e-24 \| 5.2e-30 \| 135.4 \| 105 \| (5, 117) \| 162 \| (1, 106) \| 136 \| Single-stranded DNA-binding protein \| Single-stranded DNA-binding protein; uniclust \| UniRef100\_A0A2E3Q240 \| 99.8 \| 2.9e-24 \| 5.5e-30 \| 138.0 \| 89 \| (20, 118) \| 162 \| (4, 93) \| 132 \| Single-stranded DNA-binding protein (Fragment) \| Single-stranded DNA-binding protein (Fragment); uniclust \| UniRef100\_UPI00232B6A13 \| 99.8 \| 3.1e-24 \| 5.7e-30 \| 148.3 \| 111 \| (2, 117) \| 162 \| (79, 189) \| 292 \| single-stranded DNA-binding protein \| single-stranded DNA-binding protein; uniclust \| UniRef100\_A0A4Q4WRP0 \| 99.8 \| 3.1e-24 \| 5.7e-30 \| 144.6 \| 110 \| (2, 115) \| 162 \| (112, 221) \| 241 \| Single-stranded DNA-binding protein \| Single-stranded DNA-binding protein; uniclust \| UniRef100\_A0A021VMQ7 \| 99.8 \| 3e-24 \| 5.8e-30 \| 157.0 \| 107 \| (1, 107) \| 162 \| (45, 152) \| 278 \| Single-stranded DNA-binding protein \| Single-stranded DNA-binding protein; uniclust \| UniRef100\_UPI000AD85BCB \| 99.8 \| 3.3e-24 \| 6.1e-30 \| 150.8 \| 103 \| (5, 107) \| 162 \| (88, 191) \| 255 \| single-stranded DNA-binding protein \| single-stranded DNA-binding protein; uniclust \| UniRef100\_A0A066YRZ6 \| 99.8 \| 3.2e-24 \| 6.2e-30 \| 156.2 \| 106 \| (1, 107) \| 162 \| (44, 150) \| 269 \| Single-stranded DNA-binding protein \| Single-stranded DNA-binding protein; uniclust \| UniRef100\_A0A1G6H2Z0 \| 99.8 \| 3.5e-24 \| 6.7e-30 \| 150.1 \| 104 \| (2, 106) \| 162 \| (3, 108) \| 213 \| Single-stranded DNA-binding protein \| Single-stranded DNA-binding protein; uniclust \| UniRef100\_A0A1H4CFD5 \| 99.8 \| 3.8e-24 \| 7.2e-30 \| 146.7 \| 105 \| (3, 107) \| 162 \| (1, 106) \| 194 \| Single-stranded DNA-binding protein \| Single-stranded DNA-binding protein; uniclust \| UniRef100\_A0A077XMJ6 \| 99.8 \| 4.3e-24 \| 8.2e-30 \| 133.5 \| 65 \| (52, 119) \| 162 \| (2, 66) \| 105 \| Single-stranded DNA-binding protein \| Single-stranded DNA-binding protein; uniclust \| UniRef100\_A0A3L7X4Y5 \| 99.8 \| 4.5e-24 \| 8.2e-30 \| 140.7 \| 111 \| (3, 117) \| 162 \| (68, 179) \| 204 \| Single-stranded DNA-binding protein \| Single-stranded DNA-binding protein; uniclust \| UniRef100\_A0A516L280 \| 99.8 \| 4.5e-24 \| 8.4e-30 \| 135.5 \| 101 \| (2, 104) \| 162 \| (1, 101) \| 140 \| Putative single-stranded DNA-binding protein \| Putative single-stranded DNA-binding protein; uniclust \| UniRef100\_A0A011SNI5 \| 99.8 \| 4.4e-24 \| 8.7e-30 \| 156.4 \| 111 \| (1, 112) \| 162 \| (41, 152) \| 264 \| Single-stranded DNA-binding protein \| Single-stranded DNA-binding protein; uniclust \| UniRef100\_A0A1B8PQN4 \| 99.8 \| 5.2e-24 \| 9.7e-30 \| 139.9 \| 107 \| (2, 112) \| 162 \| (3, 109) \| 179 \| Single-stranded DNA-binding protein \| Single-stranded DNA-binding protein; uniclust \| UniRef100\_A0A239FC50 \| 99.8 \| 5.5e-24 \| 1e-29 \| 137.1 \| 104 \| (1, 104) \| 162 \| (1, 105) \| 138 \| Single-stranded DNA-binding protein \| Single-stranded DNA-binding protein; uniclust \| UniRef100\_A0A2W6AP70 \| 99.8 \| 5.5e-24 \| 1e-29 \| 142.8 \| 109 \| (3, 118) \| 162 \| (44, 152) \| 175 \| Single-stranded DNA-binding protein \| Single-stranded DNA-binding protein; uniclust \| UniRef100\_A0A0G1ZRH2 \| 99.8 \| 5.5e-24 \| 1.1e-29 \| 137.0 \| 80 \| (31, 114) \| 162 \| (3, 82) \| 128 \| Single-stranded DNA-binding protein \| Single-stranded DNA-binding protein; uniclust \| UniRef100\_A0A0I9UJK8 \| 99.8 \| 5.5e-24 \| 1.1e-29 \| 140.9 \| 105 \| (3, 117) \| 162 \| (12, 116) \| 150 \| Single-stranded DNA-binding protein \| Single-stranded DNA-binding protein; uniclust \| UniRef100\_A0A0D5A1F4 \| 99.8 \| 6.4e-24 \| 1.2e-29 \| 137.2 \| 117 \| (1, 117) \| 162 \| (1, 120) \| 178 \| Single-stranded DNA-binding protein \| Single-stranded DNA-binding protein; uniclust \| UniRef100\_A0A5K7ZGM1 \| 99.8 \| 7.4e-24 \| 1.4e-29 \| 147.5 \| 104 \| (4, 119) \| 162 \| (113, 216) \| 239 \| Single-stranded DNA-binding protein \| Single-stranded DNA-binding protein; uniclust \| UniRef100\_A0A1Y4CWR4 \| 99.8 \| 8e-24 \| 1.5e-29 \| 135.0 \| 111 \| (4, 118) \| 162 \| (1, 111) \| 163 \| Single-stranded DNA-binding protein \| Single-stranded DNA-binding protein; uniclust \| UniRef100\_UPI002022FF3E \| 99.8 \| 8.6e-24 \| 1.6e-29 \| 134.1 \| 100 \| (2, 104) \| 162 \| (1, 101) \| 157 \| single-stranded DNA-binding protein \| single-stranded DNA-binding protein; uniclust \| UniRef100\_A0A174BNI7 \| 99.8 \| 9.6e-24 \| 1.8e-29 \| 144.2 \| 109 \| (3, 118) \| 162 \| (5, 114) \| 222 \| Single-stranded DNA-binding protein \| Single-stranded DNA-binding protein; uniclust \| UniRef100\_A0A1R4KH11 \| 99.8 \| 9.5e-24 \| 1.8e-29 \| 147.0 \| 109 \| (3, 111) \| 162 \| (2, 113) \| 230 \| Single-stranded DNA-binding protein \| Single-stranded DNA-binding protein; uniclust \| UniRef100\_A0A432GXJ5 \| 99.8 \| 1e-23 \| 1.8e-29 \| 141.7 \| 113 \| (2, 117) \| 162 \| (81, 194) \| 234 \| Single-stranded DNA-binding protein \| Single-stranded DNA-binding protein; uniclust \| UniRef100\_A0A143ZUS2 \| 99.8 \| 9.5e-24 \| 1.8e-29 \| 142.3 \| 107 \| (3, 118) \| 162 \| (20, 127) \| 159 \| Single-stranded DNA-binding protein \| Single-stranded DNA-binding protein; uniclust \| UniRef100\_A0A1M3LAQ8 \| 99.8 \| 9.7e-24 \| 1.9e-29 \| 139.2 \| 111 \| (2, 118) \| 162 \| (7, 117) \| 141 \| Single-stranded DNA-binding protein \| Single-stranded DNA-binding protein; uniclust \| UniRef100\_A0A1E8Y6D8 \| 99.8 \| 1e-23 \| 1.9e-29 \| 138.6 \| 108 \| (5, 118) \| 162 \| (1, 108) \| 145 \| Single-stranded DNA-binding protein \| Single-stranded DNA-binding protein; uniclust \| UniRef100\_A0A516LMM7 \| 99.8 \| 1.1e-23 \| 2e-29 \| 134.8 \| 109 \| (2, 114) \| 162 \| (1, 111) \| 145 \| Single-stranded DNA-binding protein \| Single-stranded DNA-binding protein; uniclust \| UniRef100\_A0A173S9Z8 \| 99.8 \| 1.2e-23 \| 2.2e-29 \| 138.2 \| 108 \| (1, 115) \| 162 \| (27, 136) \| 183 \| Single-stranded DNA-binding protein \| Single-stranded DNA-binding protein; uniclust \| UniRef100\_A0A069A419 \| 99.8 \| 1.2e-23 \| 2.2e-29 \| 133.5 \| 106 \| (5, 116) \| 162 \| (1, 107) \| 136 \| Putative phage single-strand DNA-binding protein \| Putative phage single-strand DNA-binding protein; uniclust \| UniRef100\_A0A2N6T258 \| 99.8 \| 1.2e-23 \| 2.3e-29 \| 134.8 \| 108 \| (2, 113) \| 162 \| (1, 109) \| 141 \| Single-stranded DNA-binding protein \| Single-stranded DNA-binding protein; uniclust \| UniRef100\_A0A1W0D608 \| 99.8 \| 1.3e-23 \| 2.3e-29 \| 136.2 \| 103 \| (5, 116) \| 162 \| (1, 103) \| 170 \| Single-stranded DNA-binding protein \| Single-stranded DNA-binding protein; uniclust \| UniRef100\_A0A547EAB1 \| 99.8 \| 1.4e-23 \| 2.5e-29 \| 147.1 \| 113 \| (2, 117) \| 162 \| (1, 113) \| 324 \| Single-stranded DNA-binding protein \| Single-stranded DNA-binding protein; uniclust \| UniRef100\_A0A6L6LUW6 \| 99.8 \| 1.5e-23 \| 2.7e-29 \| 142.4 \| 108 \| (4, 115) \| 162 \| (105, 213) \| 253 \| Single-stranded DNA-binding protein \| Single-stranded DNA-binding protein; uniclust \| UniRef100\_A0A348YKY9 \| 99.8 \| 1.7e-23 \| 3.1e-29 \| 137.6 \| 110 \| (2, 115) \| 162 \| (1, 110) \| 199 \| Single-stranded DNA-binding protein \| Single-stranded DNA-binding protein; uniclust \| UniRef100\_A0A0F2KWS5 \| 99.8 \| 1.7e-23 \| 3.1e-29 \| 139.6 \| 104 \| (4, 116) \| 162 \| (1, 104) \| 180 \| Single-stranded DNA-binding protein \| Single-stranded DNA-binding protein; uniclust \| UniRef100\_A0A192A7M0 \| 99.8 \| 1.7e-23 \| 3.2e-29 \| 135.0 \| 112 \| (2, 117) \| 162 \| (1, 112) \| 157 \| Single-stranded DNA-binding protein \| Single-stranded DNA-binding protein; uniclust \| UniRef100\_A0A2P6F917 \| 99.8 \| 1.9e-23 \| 3.6e-29 \| 123.1 \| 80 \| (4, 90) \| 162 \| (1, 81) \| 83 \| Single-stranded DNA-binding protein ssb \| Single-stranded DNA-binding protein ssb; uniclust \| UniRef100\_A0A1I2DXE7 \| 99.8 \| 1.9e-23 \| 3.7e-29 \| 144.7 \| 106 \| (1, 107) \| 162 \| (1, 107) \| 186 \| Single-stranded DNA-binding protein \| Single-stranded DNA-binding protein; uniclust \| UniRef100\_A0A1D3K7U2 \| 99.8 \| 2e-23 \| 3.7e-29 \| 141.1 \| 111 \| (4, 116) \| 162 \| (22, 133) \| 204 \| Single-stranded DNA-binding protein \| Single-stranded DNA-binding protein; uniclust \| UniRef100\_UPI000CD26B27 \| 99.8 \| 2e-23 \| 3.8e-29 \| 133.5 \| 115 \| (1, 118) \| 162 \| (1, 115) \| 148 \| single-stranded DNA-binding protein \| single-stranded DNA-binding protein; uniclust \| UniRef100\_A0A2E1ZIW0 \| 99.8 \| 2.2e-23 \| 4.1e-29 \| 130.2 \| 66 \| (49, 118) \| 162 \| (12, 77) \| 107 \| Single-stranded DNA-binding protein \| Single-stranded DNA-binding protein; uniclust \| UniRef100\_A0A0F0GWP1 \| 99.8 \| 2.4e-23 \| 4.6e-29 \| 145.1 \| 106 \| (1, 107) \| 162 \| (15, 121) \| 205 \| Single-stranded DNA-binding protein \| Single-stranded DNA-binding protein; uniclust \| UniRef100\_A0A088FAZ2 \| 99.8 \| 2.6e-23 \| 4.9e-29 \| 134.9 \| 103 \| (3, 116) \| 162 \| (1, 103) \| 142 \| Single-stranded DNA binding protein \| Single-stranded DNA binding protein; uniclust \| UniRef100\_A0A2E5LYI8 \| 99.8 \| 2.7e-23 \| 4.9e-29 \| 144.9 \| 114 \| (2, 118) \| 162 \| (1, 114) \| 310 \| Multifunctional fusion protein \| Multifunctional fusion protein; uniclust \| UniRef100\_A0A7J6YTU1 \| 99.8 \| 2.8e-23 \| 5.2e-29 \| 138.4 \| 113 \| (1, 116) \| 162 \| (1, 115) \| 219 \| BON domain-containing protein \| BON domain-containing protein; uniclust \| UniRef100\_A0A968VRQ2 \| 99.8 \| 2.9e-23 \| 5.4e-29 \| 139.2 \| 114 \| (1, 118) \| 162 \| (1, 114) \| 230 \| Single-stranded DNA-binding protein (Fragment) \| Single-stranded DNA-binding protein (Fragment); uniclust \| UniRef100\_A6DQM0 \| 99.8 \| 3.2e-23 \| 5.9e-29 \| 138.3 \| 111 \| (2, 116) \| 162 \| (1, 112) \| 208 \| Single-stranded DNA-binding protein \| Single-stranded DNA-binding protein; uniclust \| UniRef100\_A0A7V1C2I3 \| 99.8 \| 3.2e-23 \| 6e-29 \| 140.1 \| 111 \| (2, 116) \| 162 \| (99, 209) \| 244 \| Single-stranded DNA-binding protein \| Single-stranded DNA-binding protein; uniclust \| UniRef100\_A0A0F4L723 \| 99.8 \| 3.2e-23 \| 6e-29 \| 130.1 \| 105 \| (2, 116) \| 162 \| (5, 110) \| 122 \| Single-stranded DNA-binding protein \| Single-stranded DNA-binding protein; uniclust \| UniRef100\_A0A7X9LYH1 \| 99.8 \| 3.5e-23 \| 6.5e-29 \| 141.7 \| 109 \| (4, 118) \| 162 \| (1, 109) \| 245 \| Single-stranded DNA-binding protein \| Single-stranded DNA-binding protein; uniclust \| UniRef100\_A0A535NDE5 \| 99.8 \| 3.6e-23 \| 6.7e-29 \| 133.3 \| 112 \| (3, 117) \| 162 \| (39, 151) \| 157 \| Single-stranded DNA-binding protein \| Single-stranded DNA-binding protein; uniclust \| UniRef100\_A0A078MU66 \| 99.8 \| 3.5e-23 \| 6.7e-29 \| 147.6 \| 111 \| (2, 113) \| 162 \| (13, 124) \| 227 \| Single-stranded DNA-binding protein \| Single-stranded DNA-binding protein; uniclust \| UniRef100\_A0A212Q4T3 \| 99.8 \| 3.7e-23 \| 6.9e-29 \| 133.4 \| 98 \| (3, 114) \| 162 \| (1, 98) \| 136 \| Single-stranded DNA-binding protein \| Single-stranded DNA-binding protein; uniclust \| UniRef100\_A0A1F3Y0G0 \| 99.8 \| 3.7e-23 \| 7.1e-29 \| 136.8 \| 112 \| (2, 116) \| 162 \| (12, 124) \| 149 \| Single-stranded DNA-binding protein \| Single-stranded DNA-binding protein; uniclust \| UniRef100\_A0A1R4G6W5 \| 99.8 \| 3.8e-23 \| 7.4e-29 \| 146.3 \| 104 \| (3, 107) \| 162 \| (21, 125) \| 234 \| Single-stranded DNA-binding protein \| Single-stranded DNA-binding protein; uniclust \| UniRef100\_A0A031H3M6 \| 99.8 \| 4e-23 \| 7.6e-29 \| 141.8 \| 103 \| (4, 107) \| 162 \| (1, 104) \| 203 \| Single-stranded DNA-binding protein \| Single-stranded DNA-binding protein; uniclust \| UniRef100\_A0A0Q2TWN2 \| 99.8 \| 4.1e-23 \| 7.7e-29 \| 129.8 \| 111 \| (3, 117) \| 162 \| (1, 112) \| 126 \| Single-stranded DNA-binding protein \| Single-stranded DNA-binding protein; uniclust \| UniRef100\_A0A075MHM8 \| 99.8 \| 4.4e-23 \| 8.1e-29 \| 142.5 \| 111 \| (4, 115) \| 162 \| (82, 192) \| 287 \| Single-stranded DNA-binding protein \| Single-stranded DNA-binding protein; uniclust \| UniRef100\_A0A241XSG0 \| 99.8 \| 4.5e-23 \| 8.3e-29 \| 133.2 \| 112 \| (1, 116) \| 162 \| (1, 112) \| 162 \| Single-stranded DNA-binding protein \| Single-stranded DNA-binding protein; uniclust \| UniRef100\_A0A023EH54 \| 99.8 \| 4.4e-23 \| 8.3e-29 \| 140.4 \| 109 \| (3, 116) \| 162 \| (67, 176) \| 194 \| Putative single-stranded dna-binding protein (Fragment) \| Putative single-stranded dna-binding protein (Fragment); uniclust \| UniRef100\_A0A955A0C7 \| 99.8 \| 4.8e-23 \| 8.9e-29 \| 120.4 \| 89 \| (2, 91) \| 162 \| (1, 89) \| 89 \| Single-stranded DNA-binding protein (Fragment) \| Single-stranded DNA-binding protein (Fragment); uniclust \| UniRef100\_A0A193GL42 \| 99.8 \| 4.9e-23 \| 9.1e-29 \| 134.0 \| 114 \| (1, 117) \| 162 \| (1, 122) \| 148 \| Single-stranded DNA-binding protein \| Single-stranded DNA-binding protein; uniclust \| UniRef100\_UPI001556A47A \| 99.8 \| 5.9e-23 \| 1.1e-28 \| 135.3 \| 110 \| (5, 118) \| 162 \| (1, 112) \| 169 \| single-stranded DNA-binding protein \| single-stranded DNA-binding protein; uniclust \| UniRef100\_A0A6J5JVU9 \| 99.8 \| 6.4e-23 \| 1.2e-28 \| 133.3 \| 113 \| (1, 117) \| 162 \| (2, 114) \| 152 \| Single-stranded DNA-binding protein \| Single-stranded DNA-binding protein; uniclust \| UniRef100\_A0A218QUP3 \| 99.8 \| 6.5e-23 \| 1.2e-28 \| 127.0 \| 106 \| (2, 117) \| 162 \| (3, 109) \| 113 \| Single-stranded DNA-binding protein \| Single-stranded DNA-binding protein; uniclust \| UniRef100\_A0A0S4SVH8 \| 99.8 \| 6.4e-23 \| 1.2e-28 \| 131.1 \| 102 \| (4, 115) \| 162 \| (1, 102) \| 126 \| Single-stranded DNA-binding protein \| Single-stranded DNA-binding protein; uniclust \| UniRef100\_A0A939ATV5 \| 99.8 \| 7.3e-23 \| 1.3e-28 \| 137.1 \| 113 \| (2, 118) \| 162 \| (107, 219) \| 226 \| Single-stranded DNA-binding protein \| Single-stranded DNA-binding protein; uniclust \| UniRef100\_A0A0Q4S0Q6 \| 99.8 \| 7.5e-23 \| 1.5e-28 \| 139.1 \| 100 \| (9, 113) \| 162 \| (36, 135) \| 165 \| Single-stranded DNA-binding protein \| Single-stranded DNA-binding protein; uniclust \| UniRef100\_A0A076NYD0 \| 99.8 \| 8.1e-23 \| 1.5e-28 \| 132.0 \| 110 \| (1, 114) \| 162 \| (52, 161) \| 162 \| Single-stranded DNA-binding protein \| Single-stranded DNA-binding protein; uniclust \| UniRef100\_A0A256VI30 \| 99.8 \| 8.1e-23 \| 1.5e-28 \| 125.8 \| 93 \| (5, 104) \| 162 \| (1, 93) \| 107 \| Single-stranded DNA-binding protein \| Single-stranded DNA-binding protein; uniclust \| UniRef100\_A0A0Q6GEB3 \| 99.8 \| 8.4e-23 \| 1.6e-28 \| 142.8 \| 112 \| (3, 114) \| 162 \| (38, 151) \| 218 \| Single-stranded DNA-binding protein \| Single-stranded DNA-binding protein; uniclust \| UniRef100\_A0A662ZJD1 \| 99.8 \| 9.9e-23 \| 1.8e-28 \| 141.3 \| 112 \| (2, 117) \| 162 \| (3, 115) \| 265 \| Single-stranded DNA-binding protein \| Single-stranded DNA-binding protein; uniclust \| UniRef100\_A0A2I6TCS3 \| 99.8 \| 9.9e-23 \| 1.8e-28 \| 126.1 \| 112 \| (2, 117) \| 162 \| (1, 112) \| 124 \| Single-stranded DNA-binding protein \| Single-stranded DNA-binding protein; uniclust \| UniRef100\_A0A024H220 \| 99.8 \| 1e-22 \| 1.9e-28 \| 141.1 \| 106 \| (2, 108) \| 162 \| (5, 111) \| 213 \| Single-stranded DNA-binding protein \| Single-stranded DNA-binding protein; uniclust \| UniRef100\_A0A6N7R1D9 \| 99.8 \| 1.1e-22 \| 2.1e-28 \| 147.5 \| 100 \| (3, 105) \| 162 \| (238, 337) \| 395 \| Single-stranded DNA-binding protein \| Single-stranded DNA-binding protein; uniclust \| UniRef100\_A0A1G2BSC3 \| 99.8 \| 1.3e-22 \| 2.3e-28 \| 131.2 \| 111 \| (2, 115) \| 162 \| (39, 149) \| 176 \| Single-stranded DNA-binding protein \| Single-stranded DNA-binding protein; uniclust \| UniRef100\_A0A1I0QCF5 \| 99.8 \| 1.2e-22 \| 2.3e-28 \| 135.1 \| 107 \| (5, 117) \| 162 \| (43, 149) \| 175 \| Single-stranded DNA-binding protein \| Single-stranded DNA-binding protein; uniclust \| UniRef100\_A0A2D7A136 \| 99.8 \| 1.3e-22 \| 2.5e-28 \| 135.7 \| 106 \| (2, 115) \| 162 \| (1, 106) \| 163 \| Single-stranded DNA-binding protein \| Single-stranded DNA-binding protein; uniclust \| UniRef100\_A0A1G6YCT7 \| 99.8 \| 1.4e-22 \| 2.6e-28 \| 141.0 \| 101 \| (1, 104) \| 162 \| (26, 128) \| 202 \| Single-strand DNA-binding protein \| Single-strand DNA-binding protein; uniclust \| UniRef100\_A0A3D0NZB0 \| 99.8 \| 1.4e-22 \| 2.7e-28 \| 127.9 \| 83 \| (28, 117) \| 162 \| (2, 84) \| 117 \| Single-stranded DNA-binding protein (Fragment) \| Single-stranded DNA-binding protein (Fragment); uniclust \| UniRef100\_A0A7V8JJS2 \| 99.8 \| 1.5e-22 \| 2.8e-28 \| 134.4 \| 112 \| (1, 115) \| 162 \| (1, 112) \| 212 \| Single-stranded DNA-binding protein \| Single-stranded DNA-binding protein; uniclust \| UniRef100\_A0A1C5RGB2 \| 99.8 \| 1.5e-22 \| 2.8e-28 \| 128.9 \| 110 \| (2, 118) \| 162 \| (7, 116) \| 123 \| Single-stranded DNA-binding protein \| Single-stranded DNA-binding protein; uniclust \| UniRef100\_A0A016QTA0 \| 99.8 \| 1.5e-22 \| 3e-28 \| 150.6 \| 109 \| (1, 113) \| 162 \| (23, 131) \| 309 \| Single-stranded DNA-binding protein \| Single-stranded DNA-binding protein; uniclust \| UniRef100\_A0A3D2SXS3 \| 99.8 \| 1.6e-22 \| 3e-28 \| 131.8 \| 111 \| (1, 115) \| 162 \| (43, 153) \| 186 \| Single-stranded DNA-binding protein \| Single-stranded DNA-binding protein; uniclust \| UniRef100\_A0A2D8B5S5 \| 99.8 \| 1.8e-22 \| 3.3e-28 \| 124.3 \| 105 \| (4, 116) \| 162 \| (1, 105) \| 118 \| Single-stranded DNA-binding protein \| Single-stranded DNA-binding protein; uniclust \| UniRef100\_L7W1E2 \| 99.8 \| 1.8e-22 \| 3.4e-28 \| 139.2 \| 113 \| (2, 118) \| 162 \| (144, 257) \| 282 \| Single-stranded DNA-binding protein \| Single-stranded DNA-binding protein; uniclust \| UniRef100\_A0A1H3N9P1 \| 99.8 \| 2e-22 \| 3.8e-28 \| 133.6 \| 112 \| (1, 115) \| 162 \| (11, 122) \| 174 \| Single-stranded DNA-binding protein \| Single-stranded DNA-binding protein; uniclust \| UniRef100\_A0A6N9JJD1 \| 99.8 \| 2.1e-22 \| 3.8e-28 \| 134.6 \| 111 \| (1, 118) \| 162 \| (1, 113) \| 180 \| Single-stranded DNA-binding protein \| Single-stranded DNA-binding protein; uniclust \| UniRef100\_A0A0F5H0G2 \| 99.8 \| 2.1e-22 \| 3.9e-28 \| 139.7 \| 108 \| (3, 118) \| 162 \| (1, 108) \| 210 \| Single-stranded DNA-binding protein \| Single-stranded DNA-binding protein; uniclust \| UniRef100\_A0A2T2WWQ9 \| 99.8 \| 2.5e-22 \| 4.8e-28 \| 124.3 \| 83 \| (30, 118) \| 162 \| (2, 84) \| 111 \| Single-stranded DNA-binding protein \| Single-stranded DNA-binding protein; uniclust \| UniRef100\_A0A143XLA3 \| 99.8 \| 2.6e-22 \| 4.9e-28 \| 132.7 \| 107 \| (3, 118) \| 162 \| (8, 117) \| 157 \| Single-stranded DNA-binding protein \| Single-stranded DNA-binding protein; uniclust \| UniRef100\_A0A6N6KQM9 \| 99.8 \| 2.8e-22 \| 5.1e-28 \| 130.8 \| 111 \| (3, 117) \| 162 \| (44, 161) \| 186 \| Single-stranded DNA-binding protein \| Single-stranded DNA-binding protein; uniclust \| UniRef100\_A0A1N6N149 \| 99.8 \| 2.7e-22 \| 5.1e-28 \| 130.2 \| 77 \| (40, 119) \| 162 \| (1, 82) \| 136 \| Single-stranded DNA-binding protein \| Single-stranded DNA-binding protein; uniclust \| UniRef100\_A0A070A9C6 \| 99.8 \| 2.8e-22 \| 5.3e-28 \| 131.5 \| 115 \| (1, 115) \| 162 \| (2, 119) \| 157 \| Single-stranded DNA-binding protein \| Single-stranded DNA-binding protein; uniclust \| UniRef100\_A0A7X2TPA8 \| 99.8 \| 2.9e-22 \| 5.5e-28 \| 132.2 \| 107 \| (2, 112) \| 162 \| (1, 109) \| 145 \| Single-stranded DNA-binding protein \| Single-stranded DNA-binding protein; uniclust \| UniRef100\_A0A2E4UVR0 \| 99.8 \| 3e-22 \| 5.7e-28 \| 129.5 \| 113 \| (2, 118) \| 162 \| (9, 121) \| 154 \| Single-stranded DNA-binding protein \| Single-stranded DNA-binding protein; uniclust \| UniRef100\_A0A0F9MDI7 \| 99.8 \| 3.2e-22 \| 5.9e-28 \| 129.2 \| 108 \| (2, 113) \| 162 \| (66, 173) \| 174 \| Single-stranded DNA-binding protein \| Single-stranded DNA-binding protein; uniclust \| UniRef100\_A0A0F9AA01 \| 99.8 \| 3.2e-22 \| 6e-28 \| 128.2 \| 99 \| (1, 110) \| 162 \| (1, 99) \| 127 \| Single-stranded DNA-binding protein \| Single-stranded DNA-binding protein; uniclust \| UniRef100\_UPI0013D7D236 \| 99.8 \| 3.3e-22 \| 6.2e-28 \| 118.1 \| 90 \| (23, 115) \| 162 \| (2, 91) \| 93 \| single-stranded DNA-binding protein \| single-stranded DNA-binding protein; uniclust \| UniRef100\_A0A966R2Y3 \| 99.8 \| 3.4e-22 \| 6.3e-28 \| 125.4 \| 110 \| (1, 115) \| 162 \| (31, 140) \| 144 \| Single-stranded DNA-binding protein \| Single-stranded DNA-binding protein; uniclust \| UniRef100\_A0A2D9CDS7 \| 99.8 \| 4.2e-22 \| 8e-28 \| 129.6 \| 106 \| (4, 118) \| 162 \| (1, 107) \| 138 \| Single-stranded DNA-binding protein \| Single-stranded DNA-binding protein; uniclust \| UniRef100\_A0A4R0XKR1 \| 99.8 \| 4.4e-22 \| 8.2e-28 \| 133.1 \| 107 \| (3, 119) \| 162 \| (15, 121) \| 183 \| Single-stranded DNA-binding protein \| Single-stranded DNA-binding protein; uniclust \| UniRef100\_A0A0B7BQE9 \| 99.8 \| 4.7e-22 \| 8.8e-28 \| 132.3 \| 110 \| (2, 116) \| 162 \| (44, 154) \| 165 \| Single-stranded DNA-binding protein \| Single-stranded DNA-binding protein; uniclust \| UniRef100\_A0A383B706 \| 99.8 \| 5.9e-22 \| 1.1e-27 \| 123.3 \| 104 \| (1, 104) \| 162 \| (29, 133) \| 136 \| Single-stranded DNA-binding protein (Fragment) \| Single-stranded DNA-binding protein (Fragment); uniclust \| UniRef100\_A0A1C9WY15 \| 99.8 \| 5.8e-22 \| 1.1e-27 \| 136.2 \| 108 \| (4, 111) \| 162 \| (6, 116) \| 181 \| Single-stranded DNA-binding protein \| Single-stranded DNA-binding protein; uniclust \| UniRef100\_A0A1D8ETT1 \| 99.8 \| 6.2e-22 \| 1.2e-27 \| 115.2 \| 84 \| (4, 88) \| 162 \| (1, 85) \| 85 \| Single-stranded DNA binding domain protein \| Single-stranded DNA binding domain protein; uniclust \| UniRef100\_A0A2E0R4C7 \| 99.8 \| 6.7e-22 \| 1.2e-27 \| 113.9 \| 79 \| (5, 89) \| 162 \| (1, 79) \| 79 \| Single-stranded DNA-binding protein (Fragment) \| Single-stranded DNA-binding protein (Fragment); uniclust \| UniRef100\_A0A7T0KD10 \| 99.8 \| 6.8e-22 \| 1.3e-27 \| 129.1 \| 112 \| (4, 118) \| 162 \| (1, 117) \| 166 \| Single-stranded DNA-binding protein \| Single-stranded DNA-binding protein; uniclust \| UniRef100\_A0A4Q3ILA5 \| 99.8 \| 7.8e-22 \| 1.5e-27 \| 117.3 \| 85 \| (1, 85) \| 162 \| (1, 85) \| 93 \| Single-stranded DNA-binding protein (Fragment) \| Single-stranded DNA-binding protein (Fragment); uniclust \| UniRef100\_A0A022LPA2 \| 99.8 \| 7.6e-22 \| 1.5e-27 \| 137.4 \| 107 \| (1, 107) \| 162 \| (1, 108) \| 204 \| Single-stranded DNA-binding protein \| Single-stranded DNA-binding protein; uniclust \| UniRef100\_A0A2S8PDV4 \| 99.8 \| 8.3e-22 \| 1.5e-27 \| 146.7 \| 110 \| (3, 118) \| 162 \| (400, 509) \| 559 \| Single-stranded DNA-binding protein \| Single-stranded DNA-binding protein; uniclust \| UniRef100\_A0A6P1ZCH8 \| 99.8 \| 8.8e-22 \| 1.6e-27 \| 131.0 \| 110 \| (2, 115) \| 162 \| (75, 185) \| 213 \| Single-stranded DNA-binding protein \| Single-stranded DNA-binding protein; uniclust \| UniRef100\_A0A3M4E9N4 \| 99.8 \| 8.8e-22 \| 1.6e-27 \| 126.1 \| 97 \| (1, 98) \| 162 \| (67, 163) \| 164 \| Single-stranded DNA-binding protein (Fragment) \| Single-stranded DNA-binding protein (Fragment); uniclust \| UniRef100\_A0A034W0Y3 \| 99.8 \| 9.5e-22 \| 1.8e-27 \| 135.7 \| 108 \| (3, 115) \| 162 \| (82, 190) \| 205 \| Single-stranded DNA-binding protein, mitochondrial \| Single-stranded DNA-binding protein, mitochondrial; uniclust \| UniRef100\_A0A1M3GUQ3 \| 99.8 \| 9.6e-22 \| 1.8e-27 \| 133.0 \| 113 \| (4, 117) \| 162 \| (1, 115) \| 186 \| Single-stranded DNA-binding protein \| Single-stranded DNA-binding protein; uniclust \| UniRef100\_A0A935RAW8 \| 99.8 \| 1e-21 \| 1.9e-27 \| 131.4 \| 109 \| (4, 115) \| 162 \| (1, 109) \| 222 \| Single-stranded DNA-binding protein \| Single-stranded DNA-binding protein; uniclust \| UniRef100\_A0A292IIP5 \| 99.8 \| 1e-21 \| 1.9e-27 \| 134.4 \| 104 \| (5, 118) \| 162 \| (1, 104) \| 187 \| Single-stranded DNA-binding protein \| Single-stranded DNA-binding protein; uniclust \| UniRef100\_A0A923AJ74 \| 99.8 \| 1.1e-21 \| 2e-27 \| 123.6 \| 83 \| (34, 120) \| 162 \| (1, 83) \| 122 \| Single-stranded DNA-binding protein \| Single-stranded DNA-binding protein; uniclust \| UniRef100\_A0A026W545 \| 99.8 \| 1.1e-21 \| 2e-27 \| 129.9 \| 108 \| (3, 115) \| 162 \| (38, 146) \| 163 \| Single-stranded DNA-binding protein \| Single-stranded DNA-binding protein; uniclust \| UniRef100\_A0A928CL58 \| 99.8 \| 1.1e-21 \| 2.1e-27 \| 126.9 \| 106 \| (2, 116) \| 162 \| (28, 133) \| 175 \| Single-stranded DNA-binding protein \| Single-stranded DNA-binding protein; uniclust \| UniRef100\_A0A068SFW7 \| 99.8 \| 1.3e-21 \| 2.4e-27 \| 126.9 \| 108 \| (3, 115) \| 162 \| (27, 134) \| 149 \| Single-stranded dna-binding protein \| Single-stranded dna-binding protein; uniclust \| UniRef100\_UPI000A9471B9 \| 99.7 \| 1.3e-21 \| 2.5e-27 \| 112.9 \| 77 \| (4, 83) \| 162 \| (1, 78) \| 78 \| single-stranded DNA-binding protein \| single-stranded DNA-binding protein; uniclust \| UniRef100\_A0A1F0Z846 \| 99.7 \| 1.4e-21 \| 2.6e-27 \| 127.7 \| 100 \| (3, 104) \| 162 \| (1, 101) \| 141 \| Single-stranded DNA-binding protein \| Single-stranded DNA-binding protein; uniclust \| UniRef100\_A0A845H8A1 \| 99.7 \| 1.4e-21 \| 2.6e-27 \| 131.9 \| 105 \| (11, 118) \| 162 \| (94, 199) \| 236 \| Single-stranded DNA-binding protein \| Single-stranded DNA-binding protein; uniclust \| UniRef100\_A0A059VWJ1 \| 99.7 \| 1.4e-21 \| 2.8e-27 \| 146.5 \| 107 \| (1, 107) \| 162 \| (50, 158) \| 336 \| Single-stranded DNA-binding protein \| Single-stranded DNA-binding protein; uniclust \| UniRef100\_A0A2H0QGF7 \| 99.7 \| 1.5e-21 \| 2.8e-27 \| 120.4 \| 94 \| (2, 104) \| 162 \| (1, 94) \| 102 \| Single-stranded DNA-binding protein \| Single-stranded DNA-binding protein; uniclust \| UniRef100\_A0A137PND5 \| 99.7 \| 1.5e-21 \| 2.8e-27 \| 130.3 \| 109 \| (4, 119) \| 162 \| (16, 124) \| 187 \| Single-stranded DNA-binding protein \| Single-stranded DNA-binding protein; uniclust \| UniRef100\_A0A8J4ZKB2 \| 99.7 \| 1.6e-21 \| 2.9e-27 \| 152.9 \| 121 \| (1, 121) \| 162 \| (1, 123) \| 1036 \| DNA topoisomerase (ATP-hydrolyzing) \| DNA topoisomerase (ATP-hydrolyzing); uniclust \| UniRef100\_A0A6J5KSU9 \| 99.7 \| 1.6e-21 \| 3e-27 \| 127.7 \| 112 \| (2, 114) \| 162 \| (1, 113) \| 145 \| Ssb Single-stranded DNA-binding protein \| Ssb Single-stranded DNA-binding protein; uniclust \| UniRef100\_UPI001B42314A \| 99.7 \| 1.7e-21 \| 3.1e-27 \| 125.5 \| 103 \| (2, 104) \| 162 \| (4, 106) \| 153 \| single-stranded DNA-binding protein \| single-stranded DNA-binding protein; uniclust \| UniRef100\_A0A7C3X101 \| 99.7 \| 1.7e-21 \| 3.2e-27 \| 125.4 \| 108 \| (4, 115) \| 162 \| (8, 116) \| 169 \| Single-stranded DNA-binding protein \| Single-stranded DNA-binding protein; uniclust \| UniRef100\_A0A2U3LGW3 \| 99.7 \| 1.8e-21 \| 3.3e-27 \| 122.5 \| 107 \| (3, 117) \| 162 \| (6, 112) \| 128 \| Single-stranded DNA-binding protein \| Single-stranded DNA-binding protein; uniclust \| UniRef100\_A0A1Q6YS66 \| 99.7 \| 1.9e-21 \| 3.5e-27 \| 129.0 \| 108 \| (4, 115) \| 162 \| (82, 189) \| 207 \| Single-stranded DNA-binding protein \| Single-stranded DNA-binding protein; uniclust \| UniRef100\_A0A952LKT7 \| 99.7 \| 2e-21 \| 3.6e-27 \| 124.6 \| 110 \| (4, 117) \| 162 \| (1, 111) \| 164 \| Single-stranded DNA-binding protein \| Single-stranded DNA-binding protein; uniclust \| UniRef100\_A0A0E3B7S8 \| 99.7 \| 2e-21 \| 3.7e-27 \| 125.2 \| 110 \| (1, 114) \| 162 \| (1, 111) \| 143 \| Single-stranded DNA-binding protein \| Single-stranded DNA-binding protein; uniclust \| UniRef100\_A0A1G8T5L8 \| 99.7 \| 2e-21 \| 3.7e-27 \| 139.1 \| 105 \| (3, 107) \| 162 \| (42, 147) \| 265 \| Single-stranded DNA-binding protein \| Single-stranded DNA-binding protein; uniclust \| UniRef100\_A0A086Z1D6 \| 99.7 \| 2.1e-21 \| 3.9e-27 \| 138.5 \| 106 \| (1, 107) \| 162 \| (39, 145) \| 271 \| Single-stranded DNA-binding protein \| Single-stranded DNA-binding protein; uniclust \| UniRef100\_A0A0F0KXD3 \| 99.7 \| 2.1e-21 \| 4e-27 \| 130.4 \| 101 \| (3, 104) \| 162 \| (7, 108) \| 166 \| Single-stranded DNA-binding protein \| Single-stranded DNA-binding protein; uniclust \| UniRef100\_A0A5J6Z6V0 \| 99.7 \| 2.2e-21 \| 4.1e-27 \| 131.1 \| 103 \| (3, 106) \| 162 \| (1, 103) \| 188 \| Single-stranded DNA-binding protein \| Single-stranded DNA-binding protein; uniclust \| UniRef100\_A0A095YG51 \| 99.7 \| 2.4e-21 \| 4.6e-27 \| 138.7 \| 106 \| (1, 106) \| 162 \| (50, 156) \| 241 \| Single-stranded DNA-binding protein \| Single-stranded DNA-binding protein; uniclust \| UniRef100\_A0A0K2GF50 \| 99.7 \| 2.5e-21 \| 4.6e-27 \| 124.9 \| 107 \| (4, 115) \| 162 \| (32, 138) \| 144 \| Single-stranded DNA-binding protein \| Single-stranded DNA-binding protein; uniclust \| UniRef100\_G1DI79 \| 99.7 \| 2.5e-21 \| 4.7e-27 \| 128.0 \| 98 \| (7, 105) \| 162 \| (63, 161) \| 203 \| Single-stranded DNA binding protein \| Single-stranded DNA binding protein; uniclust \| UniRef100\_A0A2S1PPN8 \| 99.7 \| 2.7e-21 \| 5.1e-27 \| 132.4 \| 111 \| (4, 115) \| 162 \| (5, 115) \| 229 \| Single-stranded DNA-binding protein \| Single-stranded DNA-binding protein; uniclust \| UniRef100\_A0A443SA52 \| 99.7 \| 2.8e-21 \| 5.3e-27 \| 129.2 \| 108 \| (4, 117) \| 162 \| (40, 148) \| 165 \| Single-stranded DNA-binding protein-like isoform X2 \| Single-stranded DNA-binding protein-like isoform X2; uniclust \| UniRef100\_A0A074U9W4 \| 99.7 \| 2.9e-21 \| 5.4e-27 \| 133.2 \| 116 \| (1, 116) \| 162 \| (1, 118) \| 278 \| Single-stranded DNA-binding protein \| Single-stranded DNA-binding protein; uniclust \| UniRef100\_UPI001E2AA1AE \| 99.7 \| 2.9e-21 \| 5.4e-27 \| 125.5 \| 107 \| (3, 118) \| 162 \| (41, 148) \| 179 \| single-stranded DNA-binding protein \| single-stranded DNA-binding protein; uniclust \| UniRef100\_A0A126ZWG1 \| 99.7 \| 3.1e-21 \| 5.8e-27 \| 133.8 \| 101 \| (6, 106) \| 162 \| (95, 196) \| 254 \| Single-stranded DNA-binding protein \| Single-stranded DNA-binding protein; uniclust \| UniRef100\_A0A0K2RXI4 \| 99.7 \| 3.2e-21 \| 5.9e-27 \| 130.8 \| 104 \| (4, 108) \| 162 \| (1, 105) \| 191 \| Single-stranded DNA-binding protein \| Single-stranded DNA-binding protein; uniclust \| UniRef100\_A0A0B5D4L6 \| 99.7 \| 3.2e-21 \| 6.1e-27 \| 137.1 \| 103 \| (2, 104) \| 162 \| (31, 135) \| 242 \| Single-stranded DNA-binding protein \| Single-stranded DNA-binding protein; uniclust \| UniRef100\_A0A1I4QA87 \| 99.7 \| 3.3e-21 \| 6.1e-27 \| 131.9 \| 103 \| (5, 117) \| 162 \| (1, 103) \| 222 \| Single-stranded DNA-binding protein \| Single-stranded DNA-binding protein; uniclust \| UniRef100\_A0A0L7B6A5 \| 99.7 \| 3.3e-21 \| 6.3e-27 \| 130.7 \| 103 \| (2, 104) \| 162 \| (1, 106) \| 192 \| Single-stranded DNA-binding protein \| Single-stranded DNA-binding protein; uniclust \| UniRef100\_A0A971KRS6 \| 99.7 \| 3.4e-21 \| 6.3e-27 \| 118.9 \| 111 \| (2, 116) \| 162 \| (1, 111) \| 129 \| Single-stranded DNA-binding protein \| Single-stranded DNA-binding protein; uniclust \| UniRef100\_A0A7C6LEL5 \| 99.7 \| 3.6e-21 \| 6.6e-27 \| 135.2 \| 107 \| (4, 117) \| 162 \| (1, 107) \| 321 \| Single-stranded DNA-binding protein \| Single-stranded DNA-binding protein; uniclust \| UniRef100\_A0A7Z9QKL3 \| 99.7 \| 3.6e-21 \| 6.6e-27 \| 123.0 \| 101 \| (3, 112) \| 162 \| (36, 136) \| 160 \| Single-stranded DNA-binding protein \| Single-stranded DNA-binding protein; uniclust \| UniRef100\_A0A2D9WLM2 \| 99.7 \| 3.6e-21 \| 6.6e-27 \| 138.3 \| 106 \| (3, 113) \| 162 \| (210, 315) \| 347 \| Single-stranded DNA-binding protein (Fragment) \| Single-stranded DNA-binding protein (Fragment); pdb70 \| 1EQQ\_B \| 100.0 \| 5.5e-34 \| 4.4e-38 \| 201.5 \| 160 \| (2, 162) \| 162 \| (3, 178) \| 178 \| SINGLE STRANDED DNA BINDING PROTEIN/RNA \| 1EQQ\_B SINGLE STRANDED DNA BINDING PROTEIN/RNA BETA BARREL, PROTEIN-DNA COMPLEX, REPLICATION-RNA; pdb70 \| 4MZ9\_D \| 99.9 \| 9e-34 \| 7.2e-38 \| 200.4 \| 160 \| (2, 162) \| 162 \| (3, 178) \| 178 \| Single-stranded DNA-binding protein \| 4MZ9\_D Single-stranded DNA-binding protein Single strand DNA-binding domain, SSB; pdb70 \| 5ODN\_B \| 99.9 \| 1.8e-30 \| 1.5e-34 \| 185.8 \| 117 \| (1, 121) \| 162 \| (29, 145) \| 196 \| Single-stranded DNA-binding protein/DNA Complex \| 5ODN\_B Single-stranded DNA-binding protein/DNA Complex Single-Strand Binding protein, DNA BINDING; pdb70 \| 5ODN\_F \| 99.9 \| 1.8e-30 \| 1.5e-34 \| 185.8 \| 117 \| (1, 121) \| 162 \| (29, 145) \| 196 \| Single-stranded DNA-binding protein/DNA Complex \| 5ODN\_F Single-stranded DNA-binding protein/DNA Complex Single-Strand Binding protein, DNA BINDING; pdb70 \| 5ODP\_A \| 99.9 \| 3.6e-30 \| 2.9e-34 \| 184.2 \| 118 \| (1, 122) \| 162 \| (29, 146) \| 196 \| Single-stranded DNA-binding protein/DNA Complex \| 5ODP\_A Single-stranded DNA-binding protein/DNA Complex Single-Strand Binding protein, DNA BINDING; pdb70 \| 3PGZ\_B \| 99.9 \| 6.3e-28 \| 5e-32 \| 172.1 \| 118 \| (1, 121) \| 162 \| (22, 144) \| 193 \| Single-stranded DNA-binding protein \| 3PGZ\_B Single-stranded DNA-binding protein SSGCID, BARTONELLA HENSELAE, SINGLE-STRAND BINDING; pdb70 \| 5GQO\_A \| 99.9 \| 7.3e-28 \| 5.9e-32 \| 171.4 \| 111 \| (1, 116) \| 162 \| (21, 132) \| 190 \| Single-stranded DNA-binding protein \| 5GQO\_A Single-stranded DNA-binding protein OB fold, inter-subunit clamp, Paralogous; pdb70 \| 5GQO\_B \| 99.9 \| 7.3e-28 \| 5.9e-32 \| 171.4 \| 111 \| (1, 116) \| 162 \| (21, 132) \| 190 \| Single-stranded DNA-binding protein \| 5GQO\_B Single-stranded DNA-binding protein OB fold, inter-subunit clamp, Paralogous; pdb70 \| 1UE1\_B \| 99.9 \| 2.9e-27 \| 2.3e-31 \| 164.6 \| 111 \| (2, 115) \| 162 \| (1, 112) \| 164 \| Single-strand binding protein \| 1UE1\_B Single-strand binding protein OLIGONUCLEOTIDE BINDING FOLD, DNA-BINDING PROTEIN; pdb70 \| 3TQY\_A \| 99.9 \| 7.7e-27 \| 6.1e-31 \| 161.5 \| 118 \| (1, 121) \| 162 \| (4, 121) \| 158 \| Single-stranded DNA-binding protein (E.C.2.7.7.6) \| 3TQY\_A Single-stranded DNA-binding protein (E.C.2.7.7.6) DNA replication, TRANSFERASE HET: MSE; pdb70 \| 3TQY\_C \| 99.9 \| 7.7e-27 \| 6.1e-31 \| 161.5 \| 118 \| (1, 121) \| 162 \| (4, 121) \| 158 \| Single-stranded DNA-binding protein (E.C.2.7.7.6) \| 3TQY\_C Single-stranded DNA-binding protein (E.C.2.7.7.6) DNA replication, TRANSFERASE HET: MSE; pdb70 \| 1Z9F\_A \| 99.8 \| 3.4e-26 \| 2.7e-30 \| 157.4 \| 116 \| (1, 120) \| 162 \| (12, 127) \| 153 \| Single-strand binding protein \| 1Z9F\_A Single-strand binding protein TM0604, single stranded DNA-binding protein; pdb70 \| 3AFP\_A \| 99.8 \| 5.7e-26 \| 4.6e-30 \| 158.6 \| 114 \| (2, 118) \| 162 \| (1, 115) \| 168 \| Single-stranded DNA-binding protein \| 3AFP\_A Single-stranded DNA-binding protein OB-fold, quaternary structure and stability; pdb70 \| 1QVC\_A \| 99.8 \| 6.6e-26 \| 5.3e-30 \| 154.5 \| 122 \| (1, 122) \| 162 \| (1, 122) \| 145 \| E.COLI SINGLE STRANDED DNA BINDING \| 1QVC\_A E.COLI SINGLE STRANDED DNA BINDING BETA-BARREL, SINGLE STRANDED DNA BINDING; pdb70 \| 1QVC\_B \| 99.8 \| 6.6e-26 \| 5.3e-30 \| 154.5 \| 122 \| (1, 122) \| 162 \| (1, 122) \| 145 \| E.COLI SINGLE STRANDED DNA BINDING \| 1QVC\_B E.COLI SINGLE STRANDED DNA BINDING BETA-BARREL, SINGLE STRANDED DNA BINDING; pdb70 \| 1SE8\_A \| 99.8 \| 7.3e-26 \| 5.9e-30 \| 170.8 \| 115 \| (1, 120) \| 162 \| (125, 239) \| 301 \| Single-strand binding protein \| 1SE8\_A Single-strand binding protein Single-strand binding protein, DNA BINDING; pdb70 \| 3UDG\_C \| 99.8 \| 7.3e-26 \| 5.9e-30 \| 170.8 \| 115 \| (1, 120) \| 162 \| (125, 239) \| 301 \| Single-stranded DNA-binding protein/DNA complex \| 3UDG\_C Single-stranded DNA-binding protein/DNA complex SSB, OB fold, beta-barrel, single-stranded HET: TMP; pdb70 \| 3EIV\_B \| 99.8 \| 9.3e-25 \| 7.3e-29 \| 155.8 \| 114 \| (2, 118) \| 162 \| (1, 115) \| 199 \| Crystal Structure of Single-stranded DNA-binding \| 3EIV\_B Crystal Structure of Single-stranded DNA-binding Single-stranded DNA-binding protein, Streptomyces Coelicolor; pdb70 \| 1KAW\_A \| 99.8 \| 9.4e-25 \| 7.6e-29 \| 147.0 \| 120 \| (1, 120) \| 162 \| (1, 120) \| 135 \| SINGLE-STRANDED DNA BINDING PROTEIN \| 1KAW\_A SINGLE-STRANDED DNA BINDING PROTEIN DNA-BINDING PROTEIN, SINGLE STRANDED DNA; pdb70 \| 6IRQ\_C \| 99.8 \| 2.6e-24 \| 2.1e-28 \| 142.1 \| 112 \| (1, 115) \| 162 \| (1, 112) \| 121 \| Single-stranded DNA-binding protein/DNA Complex \| 6IRQ\_C Single-stranded DNA-binding protein/DNA Complex single-strand DNA binding protein, DNA; pdb70 \| 3LGJ\_B \| 99.8 \| 4.3e-24 \| 3.4e-28 \| 149.0 \| 116 \| (2, 120) \| 162 \| (20, 140) \| 169 \| Single-stranded DNA-binding protein \| 3LGJ\_B Single-stranded DNA-binding protein NIAID, Cat Scratch Fever, Rochalimaea; pdb70 \| 2FXQ\_A \| 99.8 \| 1.1e-23 \| 9.1e-28 \| 155.8 \| 111 \| (2, 120) \| 162 \| (125, 235) \| 264 \| Single-strand binding protein \| 2FXQ\_A Single-strand binding protein 5 strand beta-sheet extended loops; pdb70 \| 1EYG\_A \| 99.8 \| 1.2e-23 \| 9.6e-28 \| 137.8 \| 113 \| (1, 113) \| 162 \| (2, 114) \| 116 \| SINGLE-STRAND DNA-BINDING PROTEIN/DNA COMPLEX \| 1EYG\_A SINGLE-STRAND DNA-BINDING PROTEIN/DNA COMPLEX PROTEIN-DNA COMPLEX OB fold; pdb70 \| 3ULP\_C \| 99.8 \| 2.6e-23 \| 2.1e-27 \| 137.7 \| 116 \| (1, 116) \| 162 \| (2, 121) \| 124 \| Single-strand binding protein/DNA complex \| 3ULP\_C Single-strand binding protein/DNA complex OB-fold, DNA binding, single-stranded DNA; pdb70 \| 3ULP\_D \| 99.8 \| 2.6e-23 \| 2.1e-27 \| 137.7 \| 116 \| (1, 116) \| 162 \| (2, 121) \| 124 \| Single-strand binding protein/DNA complex \| 3ULP\_D Single-strand binding protein/DNA complex OB-fold, DNA binding, single-stranded DNA; pdb70 \| 3A5U\_A \| 99.8 \| 2.7e-23 \| 2.2e-27 \| 138.7 \| 114 \| (2, 118) \| 162 \| (1, 115) \| 130 \| Single-stranded DNA-binding protein/DNA complex \| 3A5U\_A Single-stranded DNA-binding protein/DNA complex DNA binding protein, DNA damage; pdb70 \| 3A5U\_B \| 99.8 \| 2.7e-23 \| 2.2e-27 \| 138.7 \| 114 \| (2, 118) \| 162 \| (1, 115) \| 130 \| Single-stranded DNA-binding protein/DNA complex \| 3A5U\_B Single-stranded DNA-binding protein/DNA complex DNA binding protein, DNA damage; pdb70 \| 2VW9\_A \| 99.7 \| 4.6e-23 \| 3.7e-27 \| 138.4 \| 113 \| (4, 120) \| 162 \| (1, 113) \| 134 \| SINGLE-STRANDED DNA BINDING PROTEIN \| 2VW9\_A SINGLE-STRANDED DNA BINDING PROTEIN DNA REPLICATION, SINGLE-STRANDED DNA, SINGLE-STRANDED; pdb70 \| 2VW9\_B \| 99.7 \| 4.6e-23 \| 3.7e-27 \| 138.4 \| 113 \| (4, 120) \| 162 \| (1, 113) \| 134 \| SINGLE-STRANDED DNA BINDING PROTEIN \| 2VW9\_B SINGLE-STRANDED DNA BINDING PROTEIN DNA REPLICATION, SINGLE-STRANDED DNA, SINGLE-STRANDED; pdb70 \| 6BHW\_B \| 99.7 \| 4.6e-23 \| 3.8e-27 \| 135.6 \| 113 \| (1, 119) \| 162 \| (1, 113) \| 119 \| Single-stranded DNA-binding protein A \| 6BHW\_B Single-stranded DNA-binding protein A Single-stranded DNA binding protein, DNA HET: EDO, PEG; pdb70 \| 6BHW\_C \| 99.7 \| 4.6e-23 \| 3.8e-27 \| 135.6 \| 113 \| (1, 119) \| 162 \| (1, 113) \| 119 \| Single-stranded DNA-binding protein A \| 6BHW\_C Single-stranded DNA-binding protein A Single-stranded DNA binding protein, DNA HET: PEG, EDO; pdb70 \| 3VDY\_A \| 99.7 \| 1.1e-22 \| 8.8e-27 \| 132.9 \| 113 \| (1, 119) \| 162 \| (1, 113) \| 116 \| Single-stranded DNA-binding protein ssbB/DNA complex \| 3VDY\_A Single-stranded DNA-binding protein ssbB/DNA complex OB fold, single-strand DNA binding; pdb70 \| 4DAM\_A \| 99.7 \| 1.6e-22 \| 1.3e-26 \| 134.8 \| 111 \| (1, 116) \| 162 \| (11, 122) \| 128 \| Single-stranded DNA-binding protein 1 \| 4DAM\_A Single-stranded DNA-binding protein 1 OB-fold, DNA-binding, single-stranded DNA, DNA; pdb70 \| 4DAM\_F \| 99.7 \| 1.6e-22 \| 1.3e-26 \| 134.8 \| 111 \| (1, 116) \| 162 \| (11, 122) \| 128 \| Single-stranded DNA-binding protein 1 \| 4DAM\_F Single-stranded DNA-binding protein 1 OB-fold, DNA-binding, single-stranded DNA, DNA; pdb70 \| 6RUP\_B \| 99.7 \| 1.8e-22 \| 1.4e-26 \| 137.0 \| 116 \| (1, 119) \| 162 \| (18, 139) \| 142 \| Single-stranded DNA-binding protein, mitochondrial, SER-SER-SER-SER \| 6RUP\_B Single-stranded DNA-binding protein, mitochondrial, SER-SER-SER-SER Single-stranded DNA binding protein, mitochondria; pdb70 \| 1S3O\_A \| 99.7 \| 2.4e-22 \| 2e-26 \| 134.5 \| 115 \| (1, 118) \| 162 \| (10, 130) \| 132 \| Single-stranded DNA-binding protein, mitochondrial \| 1S3O\_A Single-stranded DNA-binding protein, mitochondrial OB fold, DNA BINDING PROTEIN; pdb70 \| 5YYU\_A \| 99.7 \| 3.8e-22 \| 3.1e-26 \| 129.9 \| 107 \| (4, 116) \| 162 \| (1, 107) \| 112 \| Single-stranded DNA-binding protein \| 5YYU\_A Single-stranded DNA-binding protein single-strand DNA binding protein, Staphylococcus; pdb70 \| 6CQO\_D \| 99.7 \| 6.5e-22 \| 5.3e-26 \| 129.8 \| 107 \| (3, 120) \| 162 \| (1, 108) \| 119 \| Single-stranded DNA-binding protein RIM1, mitochondrial \| 6CQO\_D Single-stranded DNA-binding protein RIM1, mitochondrial Mitochondrial single-stranded DNA binding proteins HET: MSE; pdb70 \| 2CCZ\_A \| 99.7 \| 7.1e-22 \| 5.8e-26 \| 130.8 \| 110 \| (1, 119) \| 162 \| (4, 113) \| 123 \| PRIMOSOMAL REPLICATION PROTEIN N \| 2CCZ\_A PRIMOSOMAL REPLICATION PROTEIN N DNA/REPLICATION, PRIMOSOME, PRIB, DNA REPLICATION; pdb70 \| 5XGT\_A \| 99.7 \| 8.5e-22 \| 6.9e-26 \| 128.0 \| 105 \| (4, 114) \| 162 \| (1, 105) \| 111 \| Single-stranded DNA-binding protein \| 5XGT\_A Single-stranded DNA-binding protein single-strand DNA binding protein, SsbA; pdb70 \| 5WQV\_B \| 99.7 \| 1.9e-21 \| 1.5e-25 \| 124.5 \| 103 \| (4, 115) \| 162 \| (1, 103) \| 104 \| Primosomal replication protein N \| 5WQV\_B Primosomal replication protein N DNA binding protein, Bacterial DNA; pdb70 \| 3KOJ\_A \| 99.7 \| 1.9e-21 \| 1.5e-25 \| 125.7 \| 100 \| (1, 110) \| 162 \| (7, 107) \| 108 \| uncharacterized protein ycf41 \| 3KOJ\_A uncharacterized protein ycf41 single-strand binding protein family, PF00436 HET: MSE; pdb70 \| 3KOJ\_B \| 99.7 \| 1.9e-21 \| 1.5e-25 \| 125.7 \| 100 \| (1, 110) \| 162 \| (7, 107) \| 108 \| uncharacterized protein ycf41 \| 3KOJ\_B uncharacterized protein ycf41 single-strand binding protein family, PF00436; pdb70 \| 3K8A\_B \| 99.7 \| 2.7e-21 \| 2.2e-25 \| 123.7 \| 96 \| (1, 101) \| 162 \| (3, 98) \| 103 \| Putative primosomal replication protein \| 3K8A\_B Putative primosomal replication protein beta-barrel, OB-fold, DNA binding protein; pdb70 \| 2IHF\_A \| 99.7 \| 2.9e-21 \| 2.3e-25 \| 140.7 \| 110 \| (1, 115) \| 162 \| (1, 110) \| 239 \| Single-stranded DNA-binding protein \| 2IHF\_A Single-stranded DNA-binding protein Single-stranded DNA binding protein (SSB); pdb70 \| 2FXQ\_A \| 99.7 \| 4.1e-21 \| 3.3e-25 \| 141.8 \| 109 \| (1, 114) \| 162 \| (1, 109) \| 264 \| Single-strand binding protein \| 2FXQ\_A Single-strand binding protein 5 strand beta-sheet extended loops; pdb70 \| 3K81\_C \| 99.7 \| 4.5e-21 \| 3.6e-25 \| 132.9 \| 117 \| (1, 119) \| 162 \| (18, 137) \| 164 \| MP18 RNA editing complex protein \| 3K81\_C MP18 RNA editing complex protein KREPA6, VHH, Single domain antibody; pdb70 \| 4APV\_A \| 99.7 \| 4.9e-21 \| 4e-25 \| 124.5 \| 105 \| (4, 117) \| 162 \| (1, 105) \| 112 \| PRIMOSOMAL REPLICATION PROTEIN N \| 4APV\_A PRIMOSOMAL REPLICATION PROTEIN N REPLICATION, PRIB PRIMOSOME SSDNA BINDING; pdb70 \| 1V1Q\_B \| 99.7 \| 5.3e-21 \| 4.2e-25 \| 128.4 \| 110 \| (1, 119) \| 162 \| (15, 124) \| 134 \| PRIMOSOMAL REPLICATION PROTEIN N \| 1V1Q\_B PRIMOSOMAL REPLICATION PROTEIN N PRIMOSOME, DNA REPLICATION, DNA BINDING HET: CYS; pdb70 \| 2IHF\_A \| 99.6 \| 6.7e-21 \| 5.5e-25 \| 138.7 \| 103 \| (2, 112) \| 162 \| (125, 227) \| 239 \| Single-stranded DNA-binding protein \| 2IHF\_A Single-stranded DNA-binding protein Single-stranded DNA binding protein (SSB); pdb70 \| 1SE8\_A \| 99.6 \| 7.7e-21 \| 6.2e-25 \| 143.0 \| 110 \| (1, 114) \| 162 \| (1, 111) \| 301 \| Single-strand binding protein \| 1SE8\_A Single-strand binding protein Single-strand binding protein, DNA BINDING; pdb70 \| 3UDG\_C \| 99.6 \| 7.7e-21 \| 6.2e-25 \| 143.0 \| 110 \| (1, 114) \| 162 \| (1, 111) \| 301 \| Single-stranded DNA-binding protein/DNA complex \| 3UDG\_C Single-stranded DNA-binding protein/DNA complex SSB, OB fold, beta-barrel, single-stranded HET: TMP; pdb70 \| 1WOC\_C \| 99.6 \| 8.7e-21 \| 7.1e-25 \| 121.0 \| 102 \| (5, 115) \| 162 \| (1, 102) \| 103 \| Primosomal replication protein n \| 1WOC\_C Primosomal replication protein n Oligonucleotide Binding fold, DNA BINDING; pdb70 \| 3K7U\_C \| 99.6 \| 1.5e-20 \| 1.2e-24 \| 128.1 \| 111 \| (1, 117) \| 162 \| (2, 119) \| 148 \| PROTEIN \| 3K7U\_C PROTEIN RNA-editing, OB-fold, RNA-editing proteins, kinetoplastids; pdb70 \| 3STB\_D \| 99.6 \| 2e-20 \| 1.6e-24 \| 126.9 \| 114 \| (3, 118) \| 162 \| (1, 117) \| 145 \| single domain antibody VHH, KREPA3 \| 3STB\_D single domain antibody VHH, KREPA3 RNA EDITING, Editosome, Nanobody, Single; pdb70 \| 4DNI\_A \| 99.6 \| 4.6e-20 \| 3.7e-24 \| 136.1 \| 102 \| (1, 104) \| 162 \| (142, 245) \| 257 \| Fusion protein of RNA-editing complex \| 4DNI\_A Fusion protein of RNA-editing complex KREPA3, KREPA6, Editosome, Protein/RNA binding; pdb70 \| 3STB\_C \| 99.6 \| 6.7e-20 \| 5.3e-24 \| 124.9 \| 97 \| (1, 103) \| 162 \| (40, 137) \| 148 \| single domain antibody VHH, KREPA3 \| 3STB\_C single domain antibody VHH, KREPA3 RNA EDITING, Editosome, Nanobody, Single; pdb70 \| 4GS3\_A \| 99.6 \| 9.4e-20 \| 7.7e-24 \| 117.2 \| 98 \| (2, 113) \| 162 \| (8, 106) \| 107 \| Single-stranded DNA-binding protein \| 4GS3\_A Single-stranded DNA-binding protein primosome, oligonucleotide/oligosaccharide binding domain, DNA; pdb70 \| 4DK6\_C \| 99.6 \| 1.3e-19 \| 1e-23 \| 116.3 \| 91 \| (3, 103) \| 162 \| (1, 93) \| 105 \| single domain antibody VHH, RNA-editing \| 4DK6\_C single domain antibody VHH, RNA-editing KREPA1, VHH, Single domain antibody; pdb70 \| 4DKA\_D \| 99.6 \| 1.3e-19 \| 1e-23 \| 116.3 \| 91 \| (3, 103) \| 162 \| (1, 93) \| 105 \| single domain antibody VHH, RNA-editing \| 4DKA\_D single domain antibody VHH, RNA-editing KREPA1, VHH, Single domain antibody; pdb70 \| 3KLW\_B \| 99.6 \| 1.6e-19 \| 1.3e-23 \| 114.1 \| 97 \| (5, 113) \| 162 \| (1, 97) \| 98 \| Primosomal replication protein n \| 3KLW\_B Primosomal replication protein n STRUCTURAL GENOMICS, PSI-2, PROTEIN STRUCTURE HET: MSE; pdb70 \| 4FDB\_A \| 99.6 \| 2.5e-19 \| 2.1e-23 \| 114.3 \| 85 \| (4, 91) \| 162 \| (1, 85) \| 101 \| Probable primosomal replication protein n \| 4FDB\_A Probable primosomal replication protein n Structural Genomics, PSI-Biology, Protein Structure; pdb70 \| 3FHW\_A \| 99.5 \| 3e-19 \| 2.4e-23 \| 116.5 \| 103 \| (5, 119) \| 162 \| (1, 103) \| 115 \| Primosomal replication protein n \| 3FHW\_A Primosomal replication protein n priB BpR162 X-RAY NESG, Structural HET: MSE; pdb70 \| 3KLW\_A \| 99.5 \| 9.4e-19 \| 7.7e-23 \| 110.6 \| 97 \| (5, 113) \| 162 \| (1, 97) \| 98 \| Primosomal replication protein n \| 3KLW\_A Primosomal replication protein n STRUCTURAL GENOMICS, PSI-2, PROTEIN STRUCTURE; pdb70 \| 2HQL\_B \| 99.5 \| 1.7e-18 \| 1.3e-22 \| 112.1 \| 93 \| (3, 111) \| 162 \| (6, 98) \| 110 \| Hypothetical protein MG376 homolog \| 2HQL\_B Hypothetical protein MG376 homolog Structural genomics, conserved hypothetical protein; pdb70 \| 2HQL\_D \| 99.5 \| 1.7e-18 \| 1.3e-22 \| 112.1 \| 93 \| (3, 111) \| 162 \| (6, 98) \| 110 \| Hypothetical protein MG376 homolog \| 2HQL\_D Hypothetical protein MG376 homolog Structural genomics, conserved hypothetical protein; pdb70 \| 4DNI\_A \| 99.3 \| 2e-16 \| 1.6e-20 \| 116.3 \| 97 \| (1, 103) \| 162 \| (26, 123) \| 257 \| Fusion protein of RNA-editing complex \| 4DNI\_A Fusion protein of RNA-editing complex KREPA3, KREPA6, Editosome, Protein/RNA binding |
| Top keywords  (threshold 1.00e-03 (evalue)) | **Single\_stranded, DNA\_binding, DNA, binding, Single\_strand, Fragment, Single, replication, complex, Primosomal** |
| Output files | ../../similar\_sequences/35\_FANPEZAQ\_CDS\_0035\_merged.svg ../../similar\_sequences/35\_FANPEZAQ\_CDS\_0035\_pdb70.a3m ../../similar\_sequences/35\_FANPEZAQ\_CDS\_0035\_pdb70.hhr ../../similar\_sequences/35\_FANPEZAQ\_CDS\_0035\_uniclust.a3m ../../similar\_sequences/35\_FANPEZAQ\_CDS\_0035\_uniclust.hhr |

#### Structure prediction (AlphaFold)^2^

|  |  |
| --- | --- |
| Stats | xml version="1.0" encoding="utf-8" standalone="no"?       2024-09-02T21:09:34.582616 image/svg+xml   Matplotlib v3.7.2, https://matplotlib.org/ |
| Predicted structure | **NGL Viewer Controls:**  - Center: *Left-Click* - Rotate: *Left-Click + Drag* - Translate: *Right-Click + Drag* - Zoom: *Shift + Left-Click + Drag* |
| Output files | ../../predicted\_structures/35\_FANPEZAQ\_CDS\_0035/features.pkl ../../predicted\_structures/35\_FANPEZAQ\_CDS\_0035/ranked\_0.pdb ../../predicted\_structures/35\_FANPEZAQ\_CDS\_0035/ranked\_0\_plots.svg ../../predicted\_structures/35\_FANPEZAQ\_CDS\_0035/result\_model\_1\_ptm\_pred\_0.pkl |

#### Structure similarity search results (Foldseek)^3^

|  |  |
| --- | --- |
| Structure databases searched | Pdb, Afdb-proteome, Afdb-uniprot50 |
| Results, scheme(s)  (Top layers only, threshold 1.00e-02 (evalue)) | xml version="1.0" encoding="utf-8" standalone="no"?       2024-09-02T21:11:11.159462 image/svg+xml   Matplotlib v3.7.2, https://matplotlib.org/ |
| Results, table  (threshold 1.00e-02 (evalue)) | db \| id \| prob \| evalue \| bits \| fident \| alnlen \| mismatch \| gapopen \| qstart \| qend \| tstart \| tend \| name \| description; pdb \| 7F2N\_A \| 1.0 \| 5.8e-16 \| 725 \| 0.625 \| 112 \| 42 \| 0 \| 2 \| 113 \| 1 \| 112 \| Single-stranded DNA-binding protein \| Single-stranded DNA-binding protein; pdb \| 4MZ9\_C \| 1.0 \| 8.448e-16 \| 712 \| 0.631 \| 114 \| 42 \| 0 \| 1 \| 114 \| 1 \| 114 \| Single-stranded DNA-binding protein \| Single-stranded DNA-binding protein; pdb \| 3PGZ\_B \| 1.0 \| 2.906e-15 \| 670 \| 0.631 \| 114 \| 40 \| 2 \| 2 \| 113 \| 1 \| 114 \| Single-stranded DNA-binding protein \| Single-stranded DNA-binding protein; pdb \| 3TQY\_C \| 1.0 \| 4.232e-15 \| 664 \| 0.618 \| 110 \| 39 \| 1 \| 4 \| 113 \| 2 \| 108 \| Single-stranded DNA-binding protein \| Single-stranded DNA-binding protein; pdb \| 7F2N\_C \| 1.0 \| 9.994e-15 \| 662 \| 0.603 \| 111 \| 40 \| 2 \| 2 \| 112 \| 1 \| 107 \| Single-stranded DNA-binding protein \| Single-stranded DNA-binding protein; pdb \| 1EYG\_D \| 1.0 \| 1.804e-14 \| 639 \| 0.616 \| 112 \| 40 \| 1 \| 1 \| 112 \| 1 \| 109 \| SINGLE-STRAND DNA-BINDING PROTEIN \| SINGLE-STRAND DNA-BINDING PROTEIN; pdb \| 5YUO\_C \| 1.0 \| 7.693e-14 \| 627 \| 0.6 \| 110 \| 38 \| 1 \| 4 \| 113 \| 2 \| 105 \| Single-stranded DNA-binding protein \| Single-stranded DNA-binding protein; pdb \| 1QVC\_D \| 1.0 \| 7.291e-14 \| 619 \| 0.571 \| 112 \| 47 \| 1 \| 1 \| 112 \| 1 \| 111 \| SINGLE STRANDED DNA BINDING PROTEIN MONOMER \| SINGLE STRANDED DNA BINDING PROTEIN MONOMER; pdb \| 6JDG\_D \| 1.0 \| 5.574e-14 \| 613 \| 0.6 \| 110 \| 37 \| 2 \| 4 \| 113 \| 2 \| 104 \| Single-stranded DNA-binding protein \| Single-stranded DNA-binding protein; pdb \| 6JDG\_A \| 1.0 \| 1.466e-13 \| 611 \| 0.572 \| 110 \| 37 \| 1 \| 4 \| 113 \| 2 \| 101 \| Single-stranded DNA-binding protein \| Single-stranded DNA-binding protein; pdb \| 7F2N\_D \| 1.0 \| 1.917e-13 \| 611 \| 0.571 \| 112 \| 39 \| 3 \| 2 \| 113 \| 1 \| 103 \| Single-stranded DNA-binding protein \| Single-stranded DNA-binding protein; pdb \| 5YUN\_B \| 1.0 \| 1.466e-13 \| 609 \| 0.576 \| 111 \| 39 \| 1 \| 4 \| 114 \| 1 \| 103 \| Single-stranded DNA-binding protein \| Single-stranded DNA-binding protein; pdb \| 5YUN\_A \| 1.0 \| 2.252e-13 \| 602 \| 0.576 \| 111 \| 40 \| 1 \| 4 \| 114 \| 1 \| 104 \| Single-stranded DNA-binding protein \| Single-stranded DNA-binding protein; pdb \| 7F2N\_B \| 1.0 \| 1.12e-13 \| 600 \| 0.543 \| 116 \| 43 \| 2 \| 2 \| 117 \| 1 \| 106 \| Single-stranded DNA-binding protein \| Single-stranded DNA-binding protein; pdb \| 5YUO\_D \| 1.0 \| 2.646e-13 \| 597 \| 0.581 \| 110 \| 37 \| 2 \| 4 \| 113 \| 2 \| 102 \| Single-stranded DNA-binding protein \| Single-stranded DNA-binding protein; pdb \| 1EYG\_B \| 1.0 \| 2.792e-13 \| 596 \| 0.587 \| 114 \| 36 \| 3 \| 1 \| 113 \| 1 \| 104 \| SINGLE-STRAND DNA-BINDING PROTEIN \| SINGLE-STRAND DNA-BINDING PROTEIN; pdb \| 7F25\_B \| 1.0 \| 2.252e-13 \| 596 \| 0.575 \| 113 \| 40 \| 3 \| 2 \| 113 \| 1 \| 106 \| Single-stranded DNA-binding protein 1 \| Single-stranded DNA-binding protein 1; pdb \| 7VUM\_A \| 1.0 \| 4.066e-13 \| 594 \| 0.572 \| 110 \| 37 \| 1 \| 4 \| 113 \| 2 \| 101 \| Single-stranded DNA-binding protein \| Single-stranded DNA-binding protein; pdb \| 5YUN\_D \| 1.0 \| 5.922e-13 \| 594 \| 0.585 \| 111 \| 37 \| 1 \| 2 \| 112 \| 1 \| 102 \| Single-stranded DNA-binding protein \| Single-stranded DNA-binding protein; pdb \| 3PGZ\_A \| 1.0 \| 1.546e-13 \| 594 \| 0.614 \| 114 \| 37 \| 3 \| 2 \| 113 \| 1 \| 109 \| Single-stranded DNA-binding protein \| Single-stranded DNA-binding protein; pdb \| 6IRQ\_C \| 1.0 \| 7.341e-13 \| 575 \| 0.581 \| 110 \| 39 \| 2 \| 4 \| 113 \| 2 \| 104 \| Single-stranded DNA-binding protein \| Single-stranded DNA-binding protein; pdb \| 5YUO\_B \| 1.0 \| 9.101e-13 \| 568 \| 0.545 \| 110 \| 39 \| 2 \| 4 \| 113 \| 2 \| 100 \| Single-stranded DNA-binding protein \| Single-stranded DNA-binding protein; pdb \| 5YUO\_A \| 1.0 \| 1.069e-12 \| 563 \| 0.558 \| 111 \| 38 \| 2 \| 4 \| 114 \| 1 \| 100 \| Single-stranded DNA-binding protein \| Single-stranded DNA-binding protein; pdb \| 1SRU\_D \| 1.0 \| 1.734e-12 \| 559 \| 0.584 \| 113 \| 34 \| 4 \| 1 \| 112 \| 1 \| 101 \| Single-strand binding protein \| Single-strand binding protein; pdb \| 1EQQ\_B \| 1.0 \| 5.612e-13 \| 558 \| 0.555 \| 117 \| 51 \| 1 \| 1 \| 117 \| 1 \| 116 \| SINGLE STRANDED DNA BINDING PROTEIN \| SINGLE STRANDED DNA BINDING PROTEIN; pdb \| 3TQY\_D \| 1.0 \| 1.399e-12 \| 548 \| 0.545 \| 110 \| 37 \| 3 \| 4 \| 113 \| 2 \| 98 \| Single-stranded DNA-binding protein \| Single-stranded DNA-binding protein; pdb \| 1SRU\_A \| 1.0 \| 4.559e-12 \| 543 \| 0.584 \| 113 \| 33 \| 4 \| 1 \| 112 \| 1 \| 100 \| Single-strand binding protein \| Single-strand binding protein; pdb \| 1SRU\_B \| 1.0 \| 3.13e-12 \| 541 \| 0.553 \| 112 \| 35 \| 3 \| 1 \| 112 \| 1 \| 97 \| Single-strand binding protein \| Single-strand binding protein; pdb \| 3ULP\_C \| 1.0 \| 2.967e-12 \| 539 \| 0.4 \| 115 \| 67 \| 2 \| 1 \| 113 \| 2 \| 116 \| Single-strand binding protein \| Single-strand binding protein; pdb \| 3ULP\_A \| 1.0 \| 3.13e-12 \| 534 \| 0.4 \| 115 \| 65 \| 3 \| 1 \| 113 \| 2 \| 114 \| Single-strand binding protein \| Single-strand binding protein; pdb \| 6JDG\_B \| 1.0 \| 7.006e-12 \| 532 \| 0.554 \| 110 \| 36 \| 3 \| 4 \| 113 \| 2 \| 98 \| Single-stranded DNA-binding protein \| Single-stranded DNA-binding protein; pdb \| 1SRU\_C \| 1.0 \| 9.67e-12 \| 529 \| 0.553 \| 112 \| 34 \| 3 \| 1 \| 112 \| 1 \| 96 \| Single-strand binding protein \| Single-strand binding protein; pdb \| 7VUM\_C \| 1.0 \| 1.408e-11 \| 524 \| 0.545 \| 110 \| 36 \| 3 \| 4 \| 113 \| 1 \| 96 \| Single-stranded DNA-binding protein \| Single-stranded DNA-binding protein; pdb \| 5ODN\_C \| 1.0 \| 1.265e-11 \| 524 \| 0.446 \| 112 \| 50 \| 2 \| 1 \| 112 \| 3 \| 102 \| Single-stranded DNA-binding protein \| Single-stranded DNA-binding protein; pdb \| 7VUM\_B \| 1.0 \| 1.944e-11 \| 522 \| 0.559 \| 109 \| 34 \| 2 \| 4 \| 112 \| 1 \| 95 \| Single-stranded DNA-binding protein \| Single-stranded DNA-binding protein; pdb \| 5ODN\_H \| 1.0 \| 1.265e-11 \| 512 \| 0.428 \| 112 \| 50 \| 3 \| 1 \| 112 \| 2 \| 99 \| Single-stranded DNA-binding protein \| Single-stranded DNA-binding protein; pdb \| 5ODN\_F \| 1.0 \| 6.292e-12 \| 504 \| 0.442 \| 113 \| 55 \| 3 \| 1 \| 113 \| 1 \| 105 \| Single-stranded DNA-binding protein \| Single-stranded DNA-binding protein; pdb \| 1KAW\_A \| 1.0 \| 9.164e-12 \| 503 \| 0.55 \| 109 \| 35 \| 4 \| 4 \| 112 \| 2 \| 96 \| SINGLE-STRANDED DNA BINDING PROTEIN \| SINGLE-STRANDED DNA BINDING PROTEIN; pdb \| 5ODN\_G \| 1.0 \| 1.335e-11 \| 503 \| 0.446 \| 112 \| 52 \| 4 \| 1 \| 112 \| 1 \| 102 \| Single-stranded DNA-binding protein \| Single-stranded DNA-binding protein; pdb \| 5ODN\_E \| 1.0 \| 4.095e-12 \| 503 \| 0.472 \| 110 \| 52 \| 3 \| 1 \| 110 \| 1 \| 104 \| Single-stranded DNA-binding protein \| Single-stranded DNA-binding protein; pdb \| 3ULP\_D \| 1.0 \| 1.568e-11 \| 503 \| 0.391 \| 115 \| 65 \| 4 \| 1 \| 113 \| 2 \| 113 \| Single-strand binding protein \| Single-strand binding protein; pdb \| 3ULP\_B \| 1.0 \| 1.408e-11 \| 502 \| 0.4 \| 115 \| 65 \| 4 \| 1 \| 113 \| 2 \| 114 \| Single-strand binding protein \| Single-strand binding protein; pdb \| 5ODN\_D \| 1.0 \| 9.67e-12 \| 498 \| 0.437 \| 112 \| 50 \| 3 \| 1 \| 112 \| 8 \| 106 \| Single-stranded DNA-binding protein \| Single-stranded DNA-binding protein; pdb \| 5ODN\_B \| 1.0 \| 4.321e-12 \| 497 \| 0.463 \| 110 \| 54 \| 3 \| 1 \| 110 \| 1 \| 105 \| Single-stranded DNA-binding protein \| Single-stranded DNA-binding protein; pdb \| 2VW9\_B \| 1.0 \| 2.41e-11 \| 476 \| 0.373 \| 107 \| 65 \| 2 \| 4 \| 110 \| 1 \| 105 \| SINGLE-STRANDED DNA BINDING PROTEIN \| SINGLE-STRANDED DNA BINDING PROTEIN; pdb \| 5ODP\_A \| 1.0 \| 4.623e-10 \| 450 \| 0.405 \| 111 \| 52 \| 3 \| 4 \| 114 \| 2 \| 98 \| Single-stranded DNA-binding protein \| Single-stranded DNA-binding protein; pdb \| 7F5Y\_B \| 1.0 \| 2.164e-11 \| 446 \| 0.313 \| 118 \| 78 \| 3 \| 1 \| 117 \| 1 \| 116 \| Single-stranded DNA-binding protein \| Single-stranded DNA-binding protein; pdb \| 7YM1\_B \| 1.0 \| 1.274e-10 \| 436 \| 0.336 \| 107 \| 65 \| 3 \| 4 \| 110 \| 1 \| 101 \| Single-stranded DNA-binding protein \| Single-stranded DNA-binding protein; pdb \| 1X3E\_B \| 1.0 \| 1.274e-10 \| 419 \| 0.284 \| 116 \| 78 \| 3 \| 6 \| 117 \| 4 \| 118 \| Single-strand binding protein \| Single-strand binding protein; pdb \| 5ODP\_G \| 1.0 \| 3.174e-10 \| 415 \| 0.401 \| 107 \| 50 \| 4 \| 2 \| 108 \| 1 \| 93 \| Single-stranded DNA-binding protein \| Single-stranded DNA-binding protein; pdb \| 3LGJ\_B \| 1.0 \| 1.152e-09 \| 409 \| 0.482 \| 114 \| 40 \| 5 \| 2 \| 113 \| 1 \| 97 \| Single-stranded DNA-binding protein \| Single-stranded DNA-binding protein; pdb \| 6BHW\_G \| 1.0 \| 8.346e-10 \| 409 \| 0.333 \| 105 \| 63 \| 3 \| 4 \| 108 \| 1 \| 98 \| Single-stranded DNA-binding protein A \| Single-stranded DNA-binding protein A; pdb \| 6RUP\_A \| 1.0 \| 1.353e-09 \| 408 \| 0.35 \| 114 \| 69 \| 3 \| 1 \| 113 \| 2 \| 111 \| Single-stranded DNA-binding protein, mitochondrial \| Single-stranded DNA-binding protein, mitochondrial; pdb \| 3LGJ\_A \| 1.0 \| 1.971e-09 \| 405 \| 0.46 \| 115 \| 40 \| 5 \| 2 \| 114 \| 1 \| 95 \| Single-stranded DNA-binding protein \| Single-stranded DNA-binding protein; pdb \| 7F5Z\_B \| 1.0 \| 3.008e-10 \| 401 \| 0.305 \| 118 \| 77 \| 4 \| 1 \| 117 \| 1 \| 114 \| Single-stranded DNA-binding protein \| Single-stranded DNA-binding protein; pdb \| 7D8J\_A \| 1.0 \| 6.047e-10 \| 393 \| 0.338 \| 118 \| 72 \| 3 \| 4 \| 121 \| 1 \| 112 \| Single-stranded DNA-binding protein \| Single-stranded DNA-binding protein; pdb \| 1S3O\_A \| 1.0 \| 2.443e-09 \| 391 \| 0.333 \| 114 \| 68 \| 3 \| 1 \| 113 \| 1 \| 107 \| Single-stranded DNA-binding protein, mitochondrial \| Single-stranded DNA-binding protein, mitochondrial; pdb \| 5YYU\_B \| 1.0 \| 2.194e-09 \| 387 \| 0.361 \| 105 \| 63 \| 3 \| 4 \| 108 \| 1 \| 101 \| Single-stranded DNA-binding protein \| Single-stranded DNA-binding protein; pdb \| 7F5Z\_A \| 1.0 \| 7.909e-10 \| 386 \| 0.305 \| 118 \| 75 \| 4 \| 1 \| 117 \| 1 \| 112 \| Single-stranded DNA-binding protein \| Single-stranded DNA-binding protein; pdb \| 6BHX\_C \| 1.0 \| 3.196e-09 \| 385 \| 0.3 \| 110 \| 67 \| 2 \| 1 \| 110 \| 1 \| 100 \| Single-stranded DNA-binding protein A \| Single-stranded DNA-binding protein A; pdb \| 3A5U\_A \| 1.0 \| 5.147e-10 \| 383 \| 0.305 \| 118 \| 78 \| 4 \| 1 \| 117 \| 1 \| 115 \| Single-stranded DNA-binding protein \| Single-stranded DNA-binding protein; pdb \| 7DEP\_A \| 1.0 \| 6.089e-09 \| 376 \| 0.33 \| 109 \| 63 \| 3 \| 4 \| 112 \| 1 \| 99 \| Single-stranded DNA-binding protein \| Single-stranded DNA-binding protein; pdb \| 6BHW\_E \| 1.0 \| 1.042e-08 \| 374 \| 0.308 \| 107 \| 65 \| 2 \| 4 \| 110 \| 2 \| 99 \| Single-stranded DNA-binding protein A \| Single-stranded DNA-binding protein A; pdb \| 1SE8\_A \| 1.0 \| 8.867e-09 \| 373 \| 0.336 \| 110 \| 62 \| 5 \| 5 \| 114 \| 115 \| 213 \| Single-strand binding protein \| Single-strand binding protein; pdb \| 6RUP\_B \| 1.0 \| 9.873e-09 \| 371 \| 0.333 \| 114 \| 63 \| 3 \| 1 \| 113 \| 8 \| 109 \| Single-stranded DNA-binding protein, mitochondrial \| Single-stranded DNA-binding protein, mitochondrial; pdb \| 1X3F\_B \| 1.0 \| 3.558e-09 \| 370 \| 0.282 \| 117 \| 72 \| 4 \| 6 \| 118 \| 4 \| 112 \| Single-strand binding protein \| Single-strand binding protein; pdb \| 2IHF\_A \| 1.0 \| 6.425e-09 \| 370 \| 0.286 \| 115 \| 77 \| 3 \| 1 \| 115 \| 1 \| 110 \| Single-stranded DNA-binding protein \| Single-stranded DNA-binding protein; pdb \| 6BHX\_D \| 1.0 \| 1.042e-08 \| 369 \| 0.299 \| 107 \| 64 \| 3 \| 4 \| 110 \| 1 \| 96 \| Single-stranded DNA-binding protein A \| Single-stranded DNA-binding protein A; pdb \| 6BHW\_B \| 1.0 \| 6.425e-09 \| 369 \| 0.327 \| 107 \| 63 \| 3 \| 4 \| 110 \| 2 \| 99 \| Single-stranded DNA-binding protein A \| Single-stranded DNA-binding protein A; pdb \| 3VDY\_A \| 1.0 \| 6.089e-09 \| 368 \| 0.289 \| 107 \| 68 \| 4 \| 4 \| 110 \| 2 \| 100 \| Single-stranded DNA-binding protein ssbB \| Single-stranded DNA-binding protein ssbB; pdb \| 6BHW\_F \| 1.0 \| 9.873e-09 \| 365 \| 0.295 \| 105 \| 63 \| 3 \| 4 \| 108 \| 2 \| 95 \| Single-stranded DNA-binding protein A \| Single-stranded DNA-binding protein A; pdb \| 6BHW\_C \| 1.0 \| 1.099e-08 \| 363 \| 0.29 \| 110 \| 68 \| 3 \| 1 \| 110 \| 1 \| 100 \| Single-stranded DNA-binding protein A \| Single-stranded DNA-binding protein A; pdb \| 5XGT\_A \| 1.0 \| 2.871e-09 \| 362 \| 0.342 \| 105 \| 65 \| 3 \| 4 \| 108 \| 1 \| 101 \| Single-stranded DNA-binding protein \| Single-stranded DNA-binding protein; pdb \| 1X3G\_A \| 1.0 \| 2.871e-09 \| 361 \| 0.26 \| 115 \| 76 \| 4 \| 7 \| 118 \| 4 \| 112 \| Single-strand binding protein \| Single-strand binding protein; pdb \| 3VDY\_B \| 1.0 \| 3.218e-08 \| 360 \| 0.27 \| 111 \| 67 \| 5 \| 4 \| 114 \| 1 \| 97 \| Single-stranded DNA-binding protein ssbB \| Single-stranded DNA-binding protein ssbB; pdb \| 7DEP\_B \| 1.0 \| 2.871e-09 \| 360 \| 0.361 \| 105 \| 59 \| 5 \| 4 \| 108 \| 1 \| 97 \| Single-stranded DNA-binding protein \| Single-stranded DNA-binding protein; pdb \| 3ULL\_A \| 1.0 \| 1.224e-08 \| 360 \| 0.289 \| 114 \| 72 \| 4 \| 1 \| 113 \| 1 \| 106 \| DNA BINDING PROTEIN \| DNA BINDING PROTEIN; pdb \| 3AFQ\_B \| 1.0 \| 7.964e-09 \| 358 \| 0.247 \| 121 \| 76 \| 5 \| 3 \| 117 \| 1 \| 112 \| Single-stranded DNA-binding protein \| Single-stranded DNA-binding protein; pdb \| 1X3G\_B \| 1.0 \| 9.357e-09 \| 357 \| 0.271 \| 118 \| 74 \| 4 \| 6 \| 117 \| 3 \| 114 \| Single-strand binding protein \| Single-strand binding protein; pdb \| 3EIV\_C \| 1.0 \| 4.655e-09 \| 355 \| 0.277 \| 108 \| 67 \| 4 \| 2 \| 108 \| 1 \| 98 \| Single-stranded DNA-binding protein 2 \| Single-stranded DNA-binding protein 2; pdb \| 6BHW\_A \| 1.0 \| 2.21e-08 \| 353 \| 0.277 \| 108 \| 66 \| 3 \| 1 \| 108 \| 1 \| 96 \| Single-stranded DNA-binding protein A \| Single-stranded DNA-binding protein A; pdb \| 1UE5\_B \| 1.0 \| 6.089e-09 \| 353 \| 0.271 \| 118 \| 68 \| 6 \| 6 \| 117 \| 3 \| 108 \| Single-strand binding protein \| Single-strand binding protein; pdb \| 5YYU\_C \| 1.0 \| 7.153e-09 \| 352 \| 0.323 \| 105 \| 63 \| 4 \| 4 \| 108 \| 1 \| 97 \| Single-stranded DNA-binding protein \| Single-stranded DNA-binding protein; pdb \| 3EIV\_B \| 1.0 \| 3.558e-09 \| 352 \| 0.282 \| 117 \| 74 \| 4 \| 1 \| 117 \| 1 \| 107 \| Single-stranded DNA-binding protein 2 \| Single-stranded DNA-binding protein 2; pdb \| 7D8J\_B \| 1.0 \| 8.867e-09 \| 351 \| 0.355 \| 107 \| 58 \| 5 \| 4 \| 110 \| 1 \| 96 \| Single-stranded DNA-binding protein \| Single-stranded DNA-binding protein; pdb \| 1UE1\_B \| 1.0 \| 5.468e-09 \| 342 \| 0.277 \| 119 \| 77 \| 6 \| 4 \| 117 \| 1 \| 115 \| Single-strand binding protein \| Single-strand binding protein; pdb \| 1UE1\_A \| 1.0 \| 4.411e-09 \| 341 \| 0.263 \| 114 \| 76 \| 3 \| 7 \| 117 \| 4 \| 112 \| Single-strand binding protein \| Single-strand binding protein; pdb \| 6BHW\_D \| 1.0 \| 2.21e-08 \| 338 \| 0.299 \| 107 \| 61 \| 4 \| 4 \| 110 \| 2 \| 94 \| Single-stranded DNA-binding protein A \| Single-stranded DNA-binding protein A; pdb \| 3A5U\_B \| 1.0 \| 8.867e-09 \| 338 \| 0.275 \| 120 \| 73 \| 6 \| 1 \| 117 \| 1 \| 109 \| Single-stranded DNA-binding protein \| Single-stranded DNA-binding protein; pdb \| 6BHX\_B \| 1.0 \| 4.21e-08 \| 336 \| 0.29 \| 110 \| 66 \| 4 \| 1 \| 110 \| 1 \| 98 \| Single-stranded DNA-binding protein A \| Single-stranded DNA-binding protein A; pdb \| 2CWA\_A \| 1.0 \| 8.929e-08 \| 335 \| 0.33 \| 112 \| 61 \| 4 \| 3 \| 114 \| 109 \| 206 \| Single-strand binding protein \| Single-strand binding protein; pdb \| 2DUD\_B \| 1.0 \| 9.422e-08 \| 331 \| 0.281 \| 110 \| 59 \| 6 \| 4 \| 112 \| 2 \| 92 \| Single-stranded DNA-binding protein \| Single-stranded DNA-binding protein; pdb \| 3AFP\_B \| 1.0 \| 2.094e-08 \| 329 \| 0.27 \| 122 \| 75 \| 7 \| 2 \| 117 \| 1 \| 114 \| Single-stranded DNA-binding protein \| Single-stranded DNA-binding protein; pdb \| 7YM1\_A \| 1.0 \| 2.739e-08 \| 324 \| 0.323 \| 105 \| 62 \| 3 \| 4 \| 108 \| 1 \| 96 \| Single-stranded DNA-binding protein \| Single-stranded DNA-binding protein; pdb \| 7F5Y\_A \| 1.0 \| 1.042e-08 \| 324 \| 0.25 \| 120 \| 74 \| 6 \| 1 \| 117 \| 3 \| 109 \| Single-stranded DNA-binding protein \| Single-stranded DNA-binding protein; pdb \| 3ULL\_B \| 1.0 \| 2.225e-07 \| 322 \| 0.286 \| 115 \| 69 \| 4 \| 1 \| 114 \| 1 \| 103 \| DNA BINDING PROTEIN \| DNA BINDING PROTEIN; pdb \| 1UE5\_A \| 1.0 \| 8.404e-09 \| 321 \| 0.252 \| 115 \| 79 \| 4 \| 6 \| 117 \| 4 \| 114 \| Single-strand binding protein \| Single-strand binding protein; pdb \| 3UDG\_B \| 1.0 \| 9.422e-08 \| 319 \| 0.336 \| 110 \| 58 \| 5 \| 5 \| 114 \| 120 \| 214 \| Single-stranded DNA-binding protein \| Single-stranded DNA-binding protein; pdb \| 6BHW\_H \| 1.0 \| 1.612e-07 \| 318 \| 0.296 \| 108 \| 66 \| 4 \| 4 \| 110 \| 2 \| 100 \| Single-stranded DNA-binding protein A \| Single-stranded DNA-binding protein A; pdb \| 2FXQ\_A \| 1.0 \| 2.89e-08 \| 318 \| 0.29 \| 117 \| 65 \| 6 \| 1 \| 114 \| 96 \| 197 \| Single-strand binding protein \| Single-strand binding protein; pdb \| 6BHX\_A \| 1.0 \| 9.942e-08 \| 316 \| 0.29 \| 110 \| 64 \| 4 \| 1 \| 110 \| 1 \| 96 \| Single-stranded DNA-binding protein A \| Single-stranded DNA-binding protein A; pdb \| 2DUD\_A \| 1.0 \| 1.998e-07 \| 315 \| 0.296 \| 108 \| 61 \| 5 \| 4 \| 110 \| 2 \| 95 \| Single-stranded DNA-binding protein \| Single-stranded DNA-binding protein; pdb \| 1X3E\_A \| 1.0 \| 1.689e-08 \| 314 \| 0.234 \| 115 \| 77 \| 5 \| 6 \| 117 \| 3 \| 109 \| Single-strand binding protein \| Single-strand binding protein; pdb \| 3AFP\_A \| 1.0 \| 1.881e-08 \| 311 \| 0.252 \| 111 \| 74 \| 4 \| 7 \| 117 \| 4 \| 105 \| Single-stranded DNA-binding protein \| Single-stranded DNA-binding protein; pdb \| 5YYU\_D \| 1.0 \| 4.473e-07 \| 308 \| 0.309 \| 110 \| 54 \| 5 \| 4 \| 113 \| 1 \| 88 \| Single-stranded DNA-binding protein \| Single-stranded DNA-binding protein; pdb \| 3UDG\_A \| 1.0 \| 3.99e-08 \| 306 \| 0.336 \| 107 \| 64 \| 4 \| 5 \| 110 \| 3 \| 103 \| Single-stranded DNA-binding protein \| Single-stranded DNA-binding protein; pdb \| 3UDG\_C \| 1.0 \| 6.826e-08 \| 306 \| 0.308 \| 107 \| 67 \| 3 \| 5 \| 110 \| 3 \| 103 \| Single-stranded DNA-binding protein \| Single-stranded DNA-binding protein; pdb \| 3AFQ\_C \| 1.0 \| 1.049e-07 \| 302 \| 0.213 \| 117 \| 71 \| 6 \| 6 \| 117 \| 4 \| 104 \| Single-stranded DNA-binding protein \| Single-stranded DNA-binding protein; pdb \| 1UE6\_C \| 1.0 \| 1.448e-07 \| 302 \| 0.267 \| 112 \| 69 \| 3 \| 6 \| 117 \| 3 \| 101 \| Single-strand binding protein \| Single-strand binding protein; pdb \| 2IHE\_A \| 1.0 \| 5.81e-08 \| 295 \| 0.299 \| 107 \| 70 \| 3 \| 5 \| 110 \| 4 \| 106 \| Single-stranded DNA-binding protein \| Single-stranded DNA-binding protein; pdb \| 4DNI\_A \| 1.0 \| 8.521e-07 \| 291 \| 0.208 \| 120 \| 87 \| 6 \| 1 \| 114 \| 139 \| 256 \| Fusion protein of RNA-editing complex proteins MP42 and MP18 \| Fusion protein of RNA-editing complex proteins MP42 and MP18; pdb \| 1UE6\_D \| 1.0 \| 5.545e-07 \| 286 \| 0.23 \| 117 \| 70 \| 5 \| 6 \| 117 \| 3 \| 104 \| Single-strand binding protein \| Single-strand binding protein; pdb \| 1X3F\_A \| 1.0 \| 1.049e-07 \| 286 \| 0.263 \| 114 \| 70 \| 4 \| 7 \| 117 \| 4 \| 106 \| Single-strand binding protein \| Single-strand binding protein; pdb \| 3AFQ\_D \| 1.0 \| 2.758e-07 \| 280 \| 0.217 \| 115 \| 75 \| 6 \| 6 \| 117 \| 4 \| 106 \| Single-stranded DNA-binding protein \| Single-stranded DNA-binding protein; pdb \| 3AFQ\_A \| 1.0 \| 3.807e-07 \| 276 \| 0.228 \| 118 \| 74 \| 5 \| 3 \| 117 \| 1 \| 104 \| Single-stranded DNA-binding protein \| Single-stranded DNA-binding protein; pdb \| 1WOC\_D \| 1.0 \| 3.608e-07 \| 272 \| 0.134 \| 104 \| 83 \| 4 \| 5 \| 108 \| 1 \| 97 \| Primosomal replication protein n \| Primosomal replication protein n; pdb \| 4APV\_A \| 1.0 \| 3.608e-07 \| 271 \| 0.16 \| 106 \| 79 \| 4 \| 5 \| 110 \| 1 \| 96 \| PRIMOSOMAL REPLICATION PROTEIN N \| PRIMOSOMAL REPLICATION PROTEIN N; pdb \| 3K80\_D \| 1.0 \| 2.364e-06 \| 270 \| 0.203 \| 113 \| 74 \| 7 \| 4 \| 114 \| 2 \| 100 \| MP18 RNA editing complex protein \| MP18 RNA editing complex protein; pdb \| 3K81\_D \| 1.0 \| 1.907e-06 \| 269 \| 0.203 \| 113 \| 80 \| 7 \| 4 \| 113 \| 1 \| 106 \| MP18 RNA editing complex protein \| MP18 RNA editing complex protein; pdb \| 3EIV\_A \| 1.0 \| 7.653e-07 \| 268 \| 0.23 \| 113 \| 68 \| 5 \| 5 \| 117 \| 1 \| 94 \| Single-stranded DNA-binding protein 2 \| Single-stranded DNA-binding protein 2; pdb \| 3K80\_C \| 1.0 \| 2.123e-06 \| 266 \| 0.203 \| 113 \| 77 \| 7 \| 4 \| 114 \| 1 \| 102 \| MP18 RNA editing complex protein \| MP18 RNA editing complex protein; pdb \| 1UE6\_B \| 1.0 \| 4.239e-07 \| 265 \| 0.226 \| 115 \| 69 \| 6 \| 6 \| 117 \| 3 \| 100 \| Single-strand binding protein \| Single-strand binding protein; pdb \| 1UE7\_C \| 1.0 \| 1.623e-06 \| 263 \| 0.25 \| 120 \| 64 \| 6 \| 2 \| 117 \| 1 \| 98 \| Single-strand binding protein \| Single-strand binding protein; pdb \| 1UE6\_A \| 1.0 \| 3.071e-07 \| 263 \| 0.243 \| 115 \| 72 \| 5 \| 6 \| 117 \| 4 \| 106 \| Single-strand binding protein \| Single-strand binding protein; pdb \| 2PNH\_A \| 1.0 \| 8.521e-07 \| 262 \| 0.121 \| 107 \| 83 \| 4 \| 4 \| 110 \| 1 \| 96 \| Primosomal replication protein n \| Primosomal replication protein n; pdb \| 1UE7\_D \| 1.0 \| 2.364e-06 \| 258 \| 0.226 \| 115 \| 61 \| 6 \| 7 \| 116 \| 4 \| 95 \| Single-strand binding protein \| Single-strand binding protein; pdb \| 4DAM\_B \| 1.0 \| 6.874e-07 \| 258 \| 0.232 \| 116 \| 76 \| 5 \| 7 \| 121 \| 5 \| 108 \| Single-stranded DNA-binding protein 1 \| Single-stranded DNA-binding protein 1; pdb \| 1Z9F\_A \| 1.0 \| 3.834e-06 \| 256 \| 0.3 \| 110 \| 55 \| 6 \| 1 \| 110 \| 1 \| 88 \| Single-strand binding protein \| Single-strand binding protein; pdb \| 1UE7\_B \| 1.0 \| 2.241e-06 \| 256 \| 0.232 \| 116 \| 59 \| 6 \| 7 \| 117 \| 4 \| 94 \| Single-strand binding protein \| Single-strand binding protein; pdb \| 5WQV\_A \| 1.0 \| 3.263e-06 \| 255 \| 0.111 \| 108 \| 82 \| 4 \| 5 \| 112 \| 1 \| 94 \| Primosomal replication protein N \| Primosomal replication protein N; pdb \| 1TXY\_A \| 1.0 \| 2.012e-06 \| 249 \| 0.121 \| 107 \| 79 \| 4 \| 4 \| 110 \| 1 \| 92 \| Primosomal replication protein n \| Primosomal replication protein n; pdb \| 2CCZ\_B \| 1.0 \| 1.115e-06 \| 246 \| 0.128 \| 101 \| 84 \| 2 \| 1 \| 101 \| 2 \| 98 \| PRIMOSOMAL REPLICATION PROTEIN N \| PRIMOSOMAL REPLICATION PROTEIN N; pdb \| 2PNH\_B \| 1.0 \| 1.807e-06 \| 243 \| 0.121 \| 107 \| 81 \| 4 \| 4 \| 110 \| 1 \| 94 \| Primosomal replication protein n \| Primosomal replication protein n; pdb \| 3K81\_C \| 1.0 \| 9.054e-06 \| 241 \| 0.201 \| 114 \| 78 \| 7 \| 4 \| 114 \| 2 \| 105 \| MP18 RNA editing complex protein \| MP18 RNA editing complex protein; pdb \| 3STB\_D \| 1.0 \| 9.054e-06 \| 240 \| 0.21 \| 114 \| 80 \| 8 \| 4 \| 114 \| 2 \| 108 \| MP18 RNA editing complex protein \| MP18 RNA editing complex protein; pdb \| 3KLW\_A \| 1.0 \| 1.391e-05 \| 239 \| 0.136 \| 110 \| 83 \| 5 \| 5 \| 114 \| 1 \| 98 \| Primosomal replication protein n \| Primosomal replication protein n; pdb \| 3STB\_C \| 1.0 \| 1.008e-05 \| 236 \| 0.173 \| 115 \| 86 \| 7 \| 1 \| 112 \| 3 \| 111 \| RNA-editing complex protein MP42 \| RNA-editing complex protein MP42; pdb \| 3EIV\_D \| 1.0 \| 2.123e-06 \| 232 \| 0.216 \| 111 \| 68 \| 6 \| 7 \| 117 \| 5 \| 96 \| Single-stranded DNA-binding protein 2 \| Single-stranded DNA-binding protein 2; pdb \| 1V1Q\_A \| 1.0 \| 1.807e-06 \| 232 \| 0.127 \| 102 \| 81 \| 4 \| 2 \| 102 \| 3 \| 97 \| PRIMOSOMAL REPLICATION PROTEIN N \| PRIMOSOMAL REPLICATION PROTEIN N; pdb \| 1UE7\_A \| 1.0 \| 5.583e-06 \| 231 \| 0.226 \| 115 \| 70 \| 5 \| 6 \| 117 \| 3 \| 101 \| Single-strand binding protein \| Single-strand binding protein; pdb \| 6CQM\_D \| 1.0 \| 6.921e-06 \| 229 \| 0.207 \| 111 \| 74 \| 5 \| 4 \| 113 \| 2 \| 99 \| Single-stranded DNA-binding protein RIM1, mitochondrial \| Single-stranded DNA-binding protein RIM1, mitochondrial; pdb \| 4DAM\_H \| 1.0 \| 5.255e-07 \| 229 \| 0.232 \| 116 \| 76 \| 5 \| 7 \| 121 \| 5 \| 108 \| Single-stranded DNA-binding protein 1 \| Single-stranded DNA-binding protein 1; pdb \| 3K8A\_A \| 1.0 \| 7.303e-06 \| 228 \| 0.151 \| 112 \| 82 \| 5 \| 3 \| 114 \| 2 \| 100 \| Putative primosomal replication protein \| Putative primosomal replication protein; pdb \| 3KOJ\_A \| 1.0 \| 4.535e-05 \| 221 \| 0.163 \| 110 \| 68 \| 6 \| 1 \| 109 \| 1 \| 87 \| uncharacterized protein ycf41 \| uncharacterized protein ycf41; pdb \| 6CQO\_D \| 1.0 \| 1.391e-05 \| 213 \| 0.207 \| 111 \| 74 \| 6 \| 4 \| 113 \| 2 \| 99 \| Single-stranded DNA-binding protein RIM1, mitochondrial \| Single-stranded DNA-binding protein RIM1, mitochondrial; pdb \| 5WQV\_B \| 1.0 \| 1.008e-05 \| 211 \| 0.122 \| 98 \| 73 \| 3 \| 5 \| 102 \| 1 \| 85 \| Primosomal replication protein N \| Primosomal replication protein N; pdb \| 6CQM\_F \| 1.0 \| 7.706e-06 \| 209 \| 0.216 \| 111 \| 70 \| 6 \| 4 \| 113 \| 2 \| 96 \| Single-stranded DNA-binding protein RIM1, mitochondrial \| Single-stranded DNA-binding protein RIM1, mitochondrial; pdb \| 3K7U\_C \| 1.0 \| 7.354e-05 \| 207 \| 0.191 \| 115 \| 74 \| 7 \| 4 \| 116 \| 1 \| 98 \| MP18 RNA editing complex protein \| MP18 RNA editing complex protein; pdb \| 3EN2\_A \| 1.0 \| 6.97e-05 \| 206 \| 0.119 \| 109 \| 78 \| 6 \| 4 \| 112 \| 1 \| 91 \| Probable primosomal replication protein n \| Probable primosomal replication protein n; pdb \| 3KOJ\_B \| 1.0 \| 8.188e-05 \| 205 \| 0.163 \| 110 \| 69 \| 6 \| 1 \| 109 \| 1 \| 88 \| uncharacterized protein ycf41 \| uncharacterized protein ycf41; pdb \| 4DAM\_E \| 1.0 \| 2.495e-06 \| 205 \| 0.243 \| 123 \| 78 \| 6 \| 4 \| 122 \| 1 \| 112 \| Single-stranded DNA-binding protein 1 \| Single-stranded DNA-binding protein 1; pdb \| 6CQM\_A \| 1.0 \| 1.635e-05 \| 204 \| 0.214 \| 112 \| 70 \| 6 \| 4 \| 114 \| 2 \| 96 \| Single-stranded DNA-binding protein RIM1, mitochondrial \| Single-stranded DNA-binding protein RIM1, mitochondrial; pdb \| 6CQO\_F \| 1.0 \| 1.184e-05 \| 203 \| 0.216 \| 111 \| 69 \| 7 \| 4 \| 113 \| 2 \| 95 \| Single-stranded DNA-binding protein RIM1, mitochondrial \| Single-stranded DNA-binding protein RIM1, mitochondrial; pdb \| 4FDB\_A \| 1.0 \| 6.26e-05 \| 203 \| 0.146 \| 109 \| 76 \| 7 \| 4 \| 111 \| 1 \| 93 \| Probable primosomal replication protein n \| Probable primosomal replication protein n; pdb \| 8FAK\_A \| 1.0 \| 3.114e-05 \| 196 \| 0.12 \| 108 \| 84 \| 4 \| 5 \| 112 \| 1 \| 97 \| Primosomal replication protein N \| Primosomal replication protein N; pdb \| 3FHW\_B \| 1.0 \| 3.86e-05 \| 189 \| 0.169 \| 112 \| 81 \| 6 \| 5 \| 116 \| 1 \| 100 \| Primosomal replication protein n \| Primosomal replication protein n; pdb \| 6CQM\_B \| 1.0 \| 2.951e-05 \| 188 \| 0.196 \| 112 \| 69 \| 6 \| 4 \| 114 \| 1 \| 92 \| Single-stranded DNA-binding protein RIM1, mitochondrial \| Single-stranded DNA-binding protein RIM1, mitochondrial; pdb \| 4DAM\_L \| 1.0 \| 2.651e-05 \| 183 \| 0.237 \| 97 \| 59 \| 6 \| 7 \| 101 \| 5 \| 88 \| Single-stranded DNA-binding protein 1 \| Single-stranded DNA-binding protein 1; pdb \| 3KLW\_B \| 1.0 \| 8.64e-05 \| 182 \| 0.16 \| 106 \| 73 \| 7 \| 5 \| 110 \| 1 \| 90 \| Primosomal replication protein n \| Primosomal replication protein n; pdb \| 6CQM\_C \| 1.0 \| 9.117e-05 \| 182 \| 0.205 \| 112 \| 72 \| 7 \| 4 \| 114 \| 1 \| 96 \| Single-stranded DNA-binding protein RIM1, mitochondrial \| Single-stranded DNA-binding protein RIM1, mitochondrial; pdb \| 4DAM\_J \| 1.0 \| 1.549e-05 \| 181 \| 0.228 \| 118 \| 74 \| 6 \| 7 \| 122 \| 5 \| 107 \| Single-stranded DNA-binding protein 1 \| Single-stranded DNA-binding protein 1; pdb \| 6CQO\_H \| 1.0 \| 9.117e-05 \| 176 \| 0.196 \| 112 \| 66 \| 7 \| 4 \| 113 \| 2 \| 91 \| Single-stranded DNA-binding protein RIM1, mitochondrial \| Single-stranded DNA-binding protein RIM1, mitochondrial; pdb \| 6CQK\_A \| 1.0 \| 0.0001478 \| 175 \| 0.2 \| 110 \| 71 \| 5 \| 4 \| 113 \| 1 \| 93 \| SsDNA-binding protein essential for mitochondrial genome maintenance \| SsDNA-binding protein essential for mitochondrial genome maintenance; pdb \| 1TXY\_B \| 1.0 \| 8.64e-05 \| 174 \| 0.135 \| 81 \| 62 \| 3 \| 4 \| 84 \| 1 \| 73 \| Primosomal replication protein n \| Primosomal replication protein n; pdb \| 6CQK\_B \| 1.0 \| 0.0001478 \| 166 \| 0.214 \| 112 \| 67 \| 7 \| 4 \| 113 \| 1 \| 93 \| SsDNA-binding protein essential for mitochondrial genome maintenance \| SsDNA-binding protein essential for mitochondrial genome maintenance; pdb \| 6CQO\_E \| 1.0 \| 0.0001071 \| 166 \| 0.203 \| 113 \| 68 \| 7 \| 4 \| 114 \| 2 \| 94 \| Single-stranded DNA-binding protein RIM1, mitochondrial \| Single-stranded DNA-binding protein RIM1, mitochondrial; pdb \| 4DAM\_C \| 1.0 \| 5.622e-05 \| 166 \| 0.264 \| 102 \| 56 \| 6 \| 4 \| 101 \| 1 \| 87 \| Single-stranded DNA-binding protein 1 \| Single-stranded DNA-binding protein 1; pdb \| 5GQO\_B \| 1.0 \| 0.0001478 \| 165 \| 0.171 \| 99 \| 65 \| 5 \| 6 \| 101 \| 3 \| 87 \| Single-stranded DNA-binding protein \| Single-stranded DNA-binding protein; pdb \| 3F2B\_A \| 1.0 \| 0.0002397 \| 161 \| 0.189 \| 116 \| 63 \| 9 \| 6 \| 117 \| 12 \| 100 \| DNA-directed DNA polymerase III alpha chain \| DNA-directed DNA polymerase III alpha chain; pdb \| 4GS3\_A \| 1.0 \| 0.0004328 \| 156 \| 0.171 \| 105 \| 68 \| 6 \| 6 \| 110 \| 3 \| 88 \| Single-stranded DNA-binding protein \| Single-stranded DNA-binding protein; pdb \| 4DAM\_G \| 1.0 \| 5.05e-05 \| 155 \| 0.267 \| 101 \| 56 \| 6 \| 4 \| 101 \| 1 \| 86 \| Single-stranded DNA-binding protein 1 \| Single-stranded DNA-binding protein 1; pdb \| 3F2D\_A \| 1.0 \| 0.0004567 \| 151 \| 0.198 \| 116 \| 62 \| 10 \| 6 \| 117 \| 12 \| 100 \| GEOBACILLUS KAUSTOPHILUS DNA POLC \| GEOBACILLUS KAUSTOPHILUS DNA POLC; pdb \| 2HQL\_E \| 1.0 \| 5.932e-05 \| 147 \| 0.134 \| 119 \| 75 \| 10 \| 4 \| 117 \| 1 \| 96 \| Hypothetical protein MG376 homolog \| Hypothetical protein MG376 homolog; pdb \| 5GQO\_A \| 1.0 \| 0.002288 \| 141 \| 0.158 \| 120 \| 72 \| 5 \| 5 \| 124 \| 1 \| 91 \| Single-stranded DNA-binding protein \| Single-stranded DNA-binding protein; pdb \| 4DK3\_C \| 1.0 \| 0.006347 \| 139 \| 0.15 \| 106 \| 66 \| 5 \| 7 \| 112 \| 2 \| 83 \| RNA-editing complex protein MP81 \| RNA-editing complex protein MP81; pdb \| 2HQL\_C \| 1.0 \| 0.0002972 \| 129 \| 0.161 \| 118 \| 67 \| 8 \| 4 \| 117 \| 1 \| 90 \| Hypothetical protein MG376 homolog \| Hypothetical protein MG376 homolog; pdb \| 2HQL\_F \| 1.0 \| 0.001845 \| 122 \| 0.144 \| 118 \| 67 \| 10 \| 4 \| 117 \| 1 \| 88 \| Hypothetical protein MG376 homolog \| Hypothetical protein MG376 homolog; pdb \| 8C5Z\_A \| 1.0 \| 0.001489 \| 119 \| 0.166 \| 108 \| 65 \| 6 \| 8 \| 111 \| 73 \| 159 \| Replication factor A \| Replication factor A; pdb \| 8OEL\_A \| 1.0 \| 0.006697 \| 113 \| 0.155 \| 109 \| 65 \| 7 \| 8 \| 111 \| 17 \| 103 \| Replication factor A \| Replication factor A; pdb \| 8AAJ\_A \| 1.0 \| 0.006347 \| 112 \| 0.155 \| 109 \| 65 \| 7 \| 8 \| 111 \| 77 \| 163 \| Replication factor A \| Replication factor A; pdb \| 8OEJ\_D \| 1.0 \| 0.006697 \| 109 \| 0.146 \| 116 \| 71 \| 8 \| 8 \| 117 \| 17 \| 110 \| Replication factor A \| Replication factor A; pdb \| 4YWK\_B \| 0.999 \| 0.002688 \| 98 \| 0.117 \| 102 \| 71 \| 5 \| 1 \| 101 \| 106 \| 189 \| Cell division control protein 21 \| Cell division control protein 21; pdb \| 6RAX\_4 \| 0.984 \| 0.00413 \| 80 \| 0.123 \| 162 \| 81 \| 9 \| 2 \| 110 \| 124 \| 277 \| DNA replication licensing factor MCM4 \| DNA replication licensing factor MCM4; afdb-proteome \| AF-A0A0H3GL04-F1-MODEL\_V4 \| 1.0 \| 8.933e-21 \| 770 \| 0.583 \| 173 \| 61 \| 2 \| 1 \| 162 \| 2 \| 174 \| Single-stranded DNA-binding protein \| Single-stranded DNA-binding protein; afdb-proteome \| AF-P0AGE0-F1-MODEL\_V4 \| 1.0 \| 2.349e-20 \| 757 \| 0.584 \| 178 \| 58 \| 3 \| 1 \| 162 \| 1 \| 178 \| Single-stranded DNA-binding protein \| Single-stranded DNA-binding protein; afdb-proteome \| AF-P0A2F6-F1-MODEL\_V4 \| 1.0 \| 3.421e-20 \| 750 \| 0.596 \| 176 \| 57 \| 5 \| 1 \| 162 \| 1 \| 176 \| Single-stranded DNA-binding protein 1 \| Single-stranded DNA-binding protein 1; afdb-proteome \| AF-Q327X2-F1-MODEL\_V4 \| 1.0 \| 2.478e-20 \| 739 \| 0.587 \| 177 \| 58 \| 3 \| 1 \| 162 \| 2 \| 178 \| Single-stranded DNA-binding protein \| Single-stranded DNA-binding protein; afdb-proteome \| AF-A0A0H3GZP4-F1-MODEL\_V4 \| 1.0 \| 9.058e-19 \| 723 \| 0.491 \| 171 \| 77 \| 2 \| 1 \| 161 \| 2 \| 172 \| Single-stranded DNA-binding protein \| Single-stranded DNA-binding protein; afdb-proteome \| AF-P40947-F1-MODEL\_V4 \| 1.0 \| 5.017e-19 \| 701 \| 0.535 \| 168 \| 69 \| 4 \| 1 \| 162 \| 1 \| 165 \| Single-stranded DNA-binding protein \| Single-stranded DNA-binding protein; afdb-proteome \| AF-Q93GP7-F1-MODEL\_V4 \| 1.0 \| 1.185e-18 \| 695 \| 0.508 \| 177 \| 66 \| 4 \| 1 \| 162 \| 2 \| 172 \| Single-stranded DNA-binding protein 2 \| Single-stranded DNA-binding protein 2; afdb-proteome \| AF-P44409-F1-MODEL\_V4 \| 1.0 \| 9.624e-18 \| 628 \| 0.537 \| 173 \| 64 \| 7 \| 1 \| 162 \| 1 \| 168 \| Single-stranded DNA-binding protein \| Single-stranded DNA-binding protein; afdb-proteome \| AF-Q5F7Y0-F1-MODEL\_V4 \| 1.0 \| 1.03e-15 \| 569 \| 0.406 \| 177 \| 83 \| 4 \| 4 \| 162 \| 2 \| 174 \| Single-stranded DNA-binding protein \| Single-stranded DNA-binding protein; afdb-proteome \| AF-O25841-F1-MODEL\_V4 \| 1.0 \| 5.823e-13 \| 513 \| 0.309 \| 152 \| 101 \| 2 \| 4 \| 155 \| 1 \| 148 \| Single-stranded DNA-binding protein \| Single-stranded DNA-binding protein; afdb-proteome \| AF-Q8I415-F1-MODEL\_V4 \| 1.0 \| 1.786e-13 \| 485 \| 0.317 \| 173 \| 104 \| 5 \| 4 \| 162 \| 81 \| 253 \| Single-stranded DNA-binding protein \| Single-stranded DNA-binding protein; afdb-proteome \| AF-P34496-F1-MODEL\_V4 \| 1.0 \| 9.574e-11 \| 462 \| 0.336 \| 116 \| 71 \| 4 \| 2 \| 115 \| 51 \| 162 \| Single-stranded DNA-binding protein, mitochondrial \| Single-stranded DNA-binding protein, mitochondrial; afdb-proteome \| AF-O69302-F1-MODEL\_V4 \| 1.0 \| 6.483e-13 \| 460 \| 0.308 \| 159 \| 98 \| 4 \| 4 \| 162 \| 1 \| 147 \| Single-stranded DNA-binding protein \| Single-stranded DNA-binding protein; afdb-proteome \| AF-B8A5I7-F1-MODEL\_V4 \| 1.0 \| 1.066e-10 \| 458 \| 0.344 \| 122 \| 71 \| 3 \| 1 \| 116 \| 25 \| 143 \| Single-stranded DNA-binding protein 1 \| Single-stranded DNA-binding protein 1; afdb-proteome \| AF-Q04837-F1-MODEL\_V4 \| 1.0 \| 1.125e-10 \| 451 \| 0.333 \| 120 \| 71 \| 3 \| 4 \| 117 \| 29 \| 145 \| Single-stranded DNA-binding protein, mitochondrial \| Single-stranded DNA-binding protein, mitochondrial; afdb-proteome \| AF-Q9CYR0-F1-MODEL\_V4 \| 1.0 \| 1.187e-10 \| 444 \| 0.333 \| 123 \| 74 \| 3 \| 1 \| 117 \| 26 \| 146 \| Single-stranded DNA-binding protein, mitochondrial \| Single-stranded DNA-binding protein, mitochondrial; afdb-proteome \| AF-A0A0K0DUQ6-F1-MODEL\_V4 \| 1.0 \| 7.37e-10 \| 442 \| 0.23 \| 117 \| 86 \| 2 \| 2 \| 118 \| 65 \| 177 \| Uncharacterized protein \| Uncharacterized protein; afdb-proteome \| AF-P54622-F1-MODEL\_V4 \| 1.0 \| 3.475e-10 \| 440 \| 0.296 \| 118 \| 77 \| 4 \| 1 \| 117 \| 34 \| 146 \| Single-stranded DNA-binding protein, mitochondrial \| Single-stranded DNA-binding protein, mitochondrial; afdb-proteome \| AF-P28042-F1-MODEL\_V4 \| 1.0 \| 1.01e-10 \| 440 \| 0.325 \| 129 \| 77 \| 4 \| 1 \| 122 \| 26 \| 151 \| Single-stranded DNA-binding protein, mitochondrial \| Single-stranded DNA-binding protein, mitochondrial; afdb-proteome \| AF-Q8DNH4-F1-MODEL\_V4 \| 1.0 \| 1.729e-10 \| 437 \| 0.33 \| 112 \| 68 \| 5 \| 4 \| 115 \| 1 \| 105 \| Single-stranded DNA-binding protein \| Single-stranded DNA-binding protein; afdb-proteome \| AF-Q2G112-F1-MODEL\_V4 \| 1.0 \| 3.248e-12 \| 419 \| 0.343 \| 157 \| 94 \| 6 \| 4 \| 159 \| 1 \| 149 \| Single-stranded DNA-binding protein \| Single-stranded DNA-binding protein; afdb-proteome \| AF-A0A133CLV7-F1-MODEL\_V4 \| 1.0 \| 3.248e-12 \| 414 \| 0.306 \| 160 \| 97 \| 7 \| 4 \| 159 \| 1 \| 150 \| Single-stranded DNA-binding protein \| Single-stranded DNA-binding protein; afdb-proteome \| AF-Q84J78-F1-MODEL\_V4 \| 1.0 \| 1.195e-09 \| 410 \| 0.266 \| 120 \| 80 \| 5 \| 4 \| 117 \| 70 \| 187 \| Single-stranded DNA-binding protein, mitochondrial \| Single-stranded DNA-binding protein, mitochondrial; afdb-proteome \| AF-P9WGD5-F1-MODEL\_V4 \| 1.0 \| 2.483e-12 \| 408 \| 0.288 \| 170 \| 107 \| 6 \| 1 \| 162 \| 1 \| 164 \| Single-stranded DNA-binding protein \| Single-stranded DNA-binding protein; afdb-proteome \| AF-X8FMG9-F1-MODEL\_V4 \| 1.0 \| 6.888e-12 \| 408 \| 0.254 \| 169 \| 117 \| 4 \| 1 \| 162 \| 1 \| 167 \| Single-stranded DNA-binding protein \| Single-stranded DNA-binding protein; afdb-proteome \| AF-I1LB73-F1-MODEL\_V4 \| 1.0 \| 1.482e-09 \| 402 \| 0.273 \| 117 \| 77 \| 5 \| 3 \| 113 \| 85 \| 199 \| Uncharacterized protein \| Uncharacterized protein; afdb-proteome \| AF-K0FER8-F1-MODEL\_V4 \| 1.0 \| 8.539e-12 \| 398 \| 0.294 \| 163 \| 107 \| 5 \| 1 \| 159 \| 1 \| 159 \| Single-stranded DNA-binding protein \| Single-stranded DNA-binding protein; afdb-proteome \| AF-P46390-F1-MODEL\_V4 \| 1.0 \| 6.888e-12 \| 396 \| 0.264 \| 170 \| 115 \| 5 \| 1 \| 162 \| 1 \| 168 \| Single-stranded DNA-binding protein \| Single-stranded DNA-binding protein; afdb-proteome \| AF-Q6L4W0-F1-MODEL\_V4 \| 1.0 \| 2.675e-09 \| 388 \| 0.237 \| 122 \| 87 \| 4 \| 4 \| 120 \| 75 \| 195 \| Os05g0509700 protein \| Os05g0509700 protein; afdb-proteome \| AF-Q9LII1-F1-MODEL\_V4 \| 1.0 \| 1.837e-09 \| 387 \| 0.244 \| 127 \| 86 \| 4 \| 1 \| 117 \| 76 \| 202 \| Nucleic acid-binding, OB-fold-like protein \| Nucleic acid-binding, OB-fold-like protein; afdb-proteome \| AF-P66855-F1-MODEL\_V4 \| 1.0 \| 6.574e-11 \| 385 \| 0.277 \| 162 \| 108 \| 6 \| 4 \| 162 \| 1 \| 156 \| Single-stranded DNA-binding protein \| Single-stranded DNA-binding protein; afdb-proteome \| AF-K7N5B5-F1-MODEL\_V4 \| 1.0 \| 4.11e-09 \| 381 \| 0.246 \| 130 \| 90 \| 4 \| 4 \| 129 \| 85 \| 210 \| Uncharacterized protein \| Uncharacterized protein; afdb-proteome \| AF-C4IYS0-F1-MODEL\_V4 \| 1.0 \| 3.896e-09 \| 380 \| 0.244 \| 127 \| 92 \| 4 \| 3 \| 125 \| 74 \| 200 \| Uncharacterized protein \| Uncharacterized protein; afdb-proteome \| AF-K0F0D9-F1-MODEL\_V4 \| 1.0 \| 3.099e-11 \| 379 \| 0.236 \| 165 \| 114 \| 6 \| 4 \| 162 \| 2 \| 160 \| Single-stranded DNA-binding protein \| Single-stranded DNA-binding protein; afdb-proteome \| AF-A0A132ZIN5-F1-MODEL\_V4 \| 1.0 \| 5.303e-11 \| 370 \| 0.272 \| 169 \| 109 \| 4 \| 4 \| 162 \| 1 \| 165 \| Single-stranded DNA-binding protein \| Single-stranded DNA-binding protein; afdb-proteome \| AF-Q2FWF7-F1-MODEL\_V4 \| 1.0 \| 8.263e-09 \| 369 \| 0.286 \| 115 \| 74 \| 5 \| 4 \| 118 \| 1 \| 107 \| Single-stranded DNA-binding protein \| Single-stranded DNA-binding protein; afdb-proteome \| AF-B4FRF4-F1-MODEL\_V4 \| 1.0 \| 5.096e-09 \| 365 \| 0.253 \| 126 \| 82 \| 4 \| 1 \| 115 \| 85 \| 209 \| Nucleic acid-binding OB-fold-like protein \| Nucleic acid-binding OB-fold-like protein; afdb-proteome \| AF-Q5VP75-F1-MODEL\_V4 \| 1.0 \| 1.492e-08 \| 359 \| 0.28 \| 125 \| 80 \| 3 \| 1 \| 115 \| 85 \| 209 \| Os01g0642900 protein \| Os01g0642900 protein; afdb-proteome \| AF-J9FJ05-F1-MODEL\_V4 \| 1.0 \| 3.164e-08 \| 355 \| 0.173 \| 115 \| 89 \| 3 \| 2 \| 116 \| 67 \| 175 \| Single-strand binding protein \| Single-strand binding protein; afdb-proteome \| AF-I1K195-F1-MODEL\_V4 \| 1.0 \| 8.263e-09 \| 353 \| 0.244 \| 135 \| 92 \| 4 \| 1 \| 125 \| 65 \| 199 \| Uncharacterized protein \| Uncharacterized protein; afdb-proteome \| AF-A0A0N4U7K8-F1-MODEL\_V4 \| 1.0 \| 9.64e-10 \| 343 \| 0.192 \| 171 \| 101 \| 4 \| 1 \| 136 \| 53 \| 221 \| Uncharacterized protein \| Uncharacterized protein; afdb-proteome \| AF-A0A0K0JYI2-F1-MODEL\_V4 \| 1.0 \| 1.024e-08 \| 336 \| 0.187 \| 139 \| 102 \| 4 \| 2 \| 136 \| 67 \| 198 \| Bm9218 \| Bm9218; afdb-proteome \| AF-K0F5S9-F1-MODEL\_V4 \| 1.0 \| 9.64e-10 \| 329 \| 0.193 \| 155 \| 105 \| 5 \| 10 \| 145 \| 2 \| 155 \| Single-stranded DNA-binding protein \| Single-stranded DNA-binding protein; afdb-proteome \| AF-A0A044UXX8-F1-MODEL\_V4 \| 1.0 \| 1.34e-08 \| 321 \| 0.137 \| 153 \| 121 \| 4 \| 2 \| 154 \| 67 \| 208 \| Uncharacterized protein \| Uncharacterized protein; afdb-proteome \| AF-Q0JDX1-F1-MODEL\_V4 \| 1.0 \| 2.862e-07 \| 317 \| 0.271 \| 114 \| 73 \| 6 \| 8 \| 114 \| 1 \| 111 \| Os04g0363700 protein \| Os04g0363700 protein; afdb-proteome \| AF-Q4CR03-F1-MODEL\_V4 \| 1.0 \| 7.885e-08 \| 305 \| 0.18 \| 144 \| 113 \| 5 \| 2 \| 142 \| 19 \| 160 \| RNA editing complex protein MP18, putative \| RNA editing complex protein MP18, putative; afdb-proteome \| AF-Q75GK2-F1-MODEL\_V4 \| 1.0 \| 2.693e-08 \| 298 \| 0.229 \| 157 \| 103 \| 4 \| 1 \| 157 \| 74 \| 212 \| Os03g0633900 protein \| Os03g0633900 protein; afdb-proteome \| AF-Q38B90-F1-MODEL\_V4 \| 1.0 \| 1.212e-07 \| 297 \| 0.195 \| 148 \| 113 \| 6 \| 2 \| 145 \| 19 \| 164 \| MP18 RNA editing complex protein \| MP18 RNA editing complex protein; afdb-proteome \| AF-Q4CZ11-F1-MODEL\_V4 \| 1.0 \| 1.031e-07 \| 296 \| 0.164 \| 146 \| 113 \| 7 \| 2 \| 141 \| 19 \| 161 \| RNA editing complex protein MP18, putative \| RNA editing complex protein MP18, putative; afdb-proteome \| AF-A4ICR4-F1-MODEL\_V4 \| 1.0 \| 8.779e-08 \| 296 \| 0.172 \| 151 \| 114 \| 6 \| 1 \| 142 \| 72 \| 220 \| MP18\_RNA\_editing\_complex\_protein\_-\_putative \| MP18\_RNA\_editing\_complex\_protein\_-\_putative; afdb-proteome \| AF-A0A0R0I0B6-F1-MODEL\_V4 \| 1.0 \| 8.32e-08 \| 295 \| 0.213 \| 145 \| 96 \| 4 \| 1 \| 145 \| 122 \| 248 \| Uncharacterized protein \| Uncharacterized protein; afdb-proteome \| AF-Q8ZK82-F1-MODEL\_V4 \| 1.0 \| 1.031e-07 \| 290 \| 0.131 \| 114 \| 89 \| 5 \| 4 \| 117 \| 1 \| 104 \| Primosomal replication protein N \| Primosomal replication protein N; afdb-proteome \| AF-C0NDW9-F1-MODEL\_V4 \| 1.0 \| 5.414e-08 \| 289 \| 0.273 \| 117 \| 74 \| 5 \| 2 \| 115 \| 27 \| 135 \| Single-stranded DNA-binding protein \| Single-stranded DNA-binding protein; afdb-proteome \| AF-I1JF51-F1-MODEL\_V4 \| 1.0 \| 1.031e-07 \| 289 \| 0.213 \| 145 \| 90 \| 5 \| 1 \| 141 \| 66 \| 190 \| Uncharacterized protein \| Uncharacterized protein; afdb-proteome \| AF-Q328J8-F1-MODEL\_V4 \| 1.0 \| 1.031e-07 \| 286 \| 0.131 \| 114 \| 89 \| 5 \| 4 \| 117 \| 1 \| 104 \| Primosomal replication protein N \| Primosomal replication protein N; afdb-proteome \| AF-A0A1C1D2M3-F1-MODEL\_V4 \| 1.0 \| 3.923e-08 \| 284 \| 0.225 \| 124 \| 83 \| 6 \| 1 \| 120 \| 27 \| 141 \| Single-stranded DNA-binding protein RIM1, mitochondrial \| Single-stranded DNA-binding protein RIM1, mitochondrial; afdb-proteome \| AF-C4J9S9-F1-MODEL\_V4 \| 1.0 \| 7.473e-08 \| 281 \| 0.207 \| 159 \| 112 \| 4 \| 1 \| 159 \| 71 \| 215 \| Protein OSB2 chloroplastic \| Protein OSB2 chloroplastic; afdb-proteome \| AF-U7PR10-F1-MODEL\_V4 \| 1.0 \| 1.502e-07 \| 280 \| 0.26 \| 119 \| 77 \| 4 \| 2 \| 117 \| 28 \| 138 \| Uncharacterized protein \| Uncharacterized protein; afdb-proteome \| AF-A0A0D2GAT6-F1-MODEL\_V4 \| 1.0 \| 5.131e-08 \| 279 \| 0.216 \| 125 \| 85 \| 6 \| 1 \| 121 \| 27 \| 142 \| Unplaced genomic scaffold supercont1.7, whole genome shotgun sequence \| Unplaced genomic scaffold supercont1.7, whole genome shotgun sequence; afdb-proteome \| AF-Q4DKA2-F1-MODEL\_V4 \| 1.0 \| 3.186e-07 \| 278 \| 0.181 \| 138 \| 89 \| 6 \| 4 \| 120 \| 13 \| 147 \| Uncharacterized protein \| Uncharacterized protein; afdb-proteome \| AF-P07013-F1-MODEL\_V4 \| 1.0 \| 2.712e-07 \| 276 \| 0.131 \| 114 \| 89 \| 5 \| 4 \| 117 \| 1 \| 104 \| Primosomal replication protein N \| Primosomal replication protein N; afdb-proteome \| AF-Q4D8F9-F1-MODEL\_V4 \| 1.0 \| 4.398e-07 \| 270 \| 0.169 \| 136 \| 89 \| 6 \| 6 \| 120 \| 38 \| 170 \| Uncharacterized protein \| Uncharacterized protein; afdb-proteome \| AF-A0A175VYV7-F1-MODEL\_V4 \| 1.0 \| 8.779e-08 \| 267 \| 0.255 \| 129 \| 82 \| 6 \| 1 \| 123 \| 25 \| 145 \| Single-stranded DNA-binding protein \| Single-stranded DNA-binding protein; afdb-proteome \| AF-K0ES78-F1-MODEL\_V4 \| 1.0 \| 4.863e-08 \| 267 \| 0.193 \| 165 \| 120 \| 5 \| 4 \| 156 \| 1 \| 164 \| Single-strand DNA binding protein (Modular protein) \| Single-strand DNA binding protein (Modular protein); afdb-proteome \| AF-O53205-F1-MODEL\_V4 \| 1.0 \| 8.32e-08 \| 259 \| 0.183 \| 158 \| 119 \| 6 \| 10 \| 162 \| 1 \| 153 \| Uncharacterized protein \| Uncharacterized protein; afdb-proteome \| AF-Q57YG2-F1-MODEL\_V4 \| 1.0 \| 9.328e-07 \| 252 \| 0.185 \| 140 \| 83 \| 7 \| 4 \| 115 \| 13 \| 149 \| Uncharacterized protein Tb927.8.680, mitochondrial \| Uncharacterized protein Tb927.8.680, mitochondrial; afdb-proteome \| AF-P44748-F1-MODEL\_V4 \| 1.0 \| 5.167e-07 \| 251 \| 0.157 \| 114 \| 85 \| 4 \| 1 \| 110 \| 1 \| 107 \| Primosomal replication protein N \| Primosomal replication protein N; afdb-proteome \| AF-X8F2P4-F1-MODEL\_V4 \| 1.0 \| 2.436e-07 \| 249 \| 0.14 \| 171 \| 123 \| 7 \| 6 \| 155 \| 4 \| 171 \| Single-stranded DNA-binding protein \| Single-stranded DNA-binding protein; afdb-proteome \| AF-A0A0K0JYI1-F1-MODEL\_V4 \| 1.0 \| 1.875e-06 \| 249 \| 0.181 \| 127 \| 90 \| 5 \| 2 \| 127 \| 55 \| 168 \| Bm9217 \| Bm9217; afdb-proteome \| AF-Q10GV8-F1-MODEL\_V4 \| 1.0 \| 6.805e-06 \| 243 \| 0.145 \| 103 \| 83 \| 3 \| 2 \| 102 \| 52 \| 151 \| Os03g0611700 protein \| Os03g0611700 protein; afdb-proteome \| AF-A0A3Q0KRK1-F1-MODEL\_V4 \| 1.0 \| 9.328e-07 \| 237 \| 0.17 \| 129 \| 96 \| 7 \| 2 \| 127 \| 33 \| 153 \| Uncharacterized protein \| Uncharacterized protein; afdb-proteome \| AF-Q5F924-F1-MODEL\_V4 \| 1.0 \| 7.181e-06 \| 236 \| 0.169 \| 112 \| 80 \| 6 \| 3 \| 114 \| 2 \| 100 \| Primosomal replication protein N \| Primosomal replication protein N; afdb-proteome \| AF-P32445-F1-MODEL\_V4 \| 1.0 \| 3.572e-06 \| 230 \| 0.208 \| 120 \| 85 \| 4 \| 2 \| 120 \| 16 \| 126 \| Single-stranded DNA-binding protein RIM1, mitochondrial \| Single-stranded DNA-binding protein RIM1, mitochondrial; afdb-proteome \| AF-A4ICD3-F1-MODEL\_V4 \| 1.0 \| 8.902e-06 \| 229 \| 0.184 \| 114 \| 78 \| 6 \| 2 \| 113 \| 526 \| 626 \| RNA editing comple protein MP63 \| RNA editing comple protein MP63; afdb-proteome \| AF-K7M5J7-F1-MODEL\_V4 \| 1.0 \| 4.197e-06 \| 223 \| 0.151 \| 112 \| 82 \| 5 \| 6 \| 115 \| 70 \| 170 \| Uncharacterized protein \| Uncharacterized protein; afdb-proteome \| AF-Q38AE3-F1-MODEL\_V4 \| 1.0 \| 2.47e-05 \| 223 \| 0.166 \| 114 \| 80 \| 7 \| 2 \| 113 \| 487 \| 587 \| RNA editing complex protein MP63 \| RNA editing complex protein MP63; afdb-proteome \| AF-A0A3Q0KDA2-F1-MODEL\_V4 \| 1.0 \| 1.979e-06 \| 221 \| 0.103 \| 135 \| 101 \| 4 \| 1 \| 117 \| 49 \| 181 \| Uncharacterized protein \| Uncharacterized protein; afdb-proteome \| AF-Q8GWJ4-F1-MODEL\_V4 \| 1.0 \| 1.359e-06 \| 215 \| 0.202 \| 148 \| 89 \| 9 \| 6 \| 148 \| 82 \| 205 \| Protein OSB3, chloroplastic/mitochondrial \| Protein OSB3, chloroplastic/mitochondrial; afdb-proteome \| AF-A0A0R0GAG9-F1-MODEL\_V4 \| 1.0 \| 4.931e-06 \| 212 \| 0.168 \| 113 \| 83 \| 4 \| 6 \| 116 \| 76 \| 179 \| Uncharacterized protein \| Uncharacterized protein; afdb-proteome \| AF-A0A1D6KXT6-F1-MODEL\_V4 \| 1.0 \| 2.341e-05 \| 206 \| 0.159 \| 119 \| 89 \| 4 \| 2 \| 117 \| 58 \| 168 \| Protein OSB1 mitochondrial \| Protein OSB1 mitochondrial; afdb-proteome \| AF-Q4E153-F1-MODEL\_V4 \| 1.0 \| 3.796e-05 \| 205 \| 0.131 \| 114 \| 84 \| 5 \| 2 \| 113 \| 490 \| 590 \| RNA editing complex protein MP63, putative \| RNA editing complex protein MP63, putative; afdb-proteome \| AF-Q8GXH3-F1-MODEL\_V4 \| 1.0 \| 9.911e-06 \| 203 \| 0.141 \| 156 \| 113 \| 4 \| 1 \| 156 \| 94 \| 228 \| Protein OSB2, chloroplastic \| Protein OSB2, chloroplastic; afdb-proteome \| AF-O14087-F1-MODEL\_V4 \| 1.0 \| 6.112e-06 \| 202 \| 0.177 \| 124 \| 90 \| 5 \| 7 \| 129 \| 27 \| 139 \| Single-stranded DNA-binding protein rim1, mitochondrial \| Single-stranded DNA-binding protein rim1, mitochondrial; afdb-proteome \| AF-C6T7T9-F1-MODEL\_V4 \| 1.0 \| 1.523e-05 \| 190 \| 0.197 \| 152 \| 98 \| 7 \| 6 \| 150 \| 37 \| 171 \| Uncharacterized protein \| Uncharacterized protein; afdb-proteome \| AF-Q5JKX9-F1-MODEL\_V4 \| 1.0 \| 1.296e-05 \| 188 \| 0.165 \| 157 \| 105 \| 7 \| 6 \| 156 \| 83 \| 219 \| Os01g0949060 protein \| Os01g0949060 protein; afdb-proteome \| AF-Q586L9-F1-MODEL\_V4 \| 1.0 \| 4.459e-05 \| 188 \| 0.161 \| 142 \| 85 \| 9 \| 2 \| 113 \| 625 \| 762 \| RNA-editing complex protein MP81 \| RNA-editing complex protein MP81; afdb-proteome \| AF-Q9FYJ2-F1-MODEL\_V4 \| 1.0 \| 1.104e-05 \| 184 \| 0.184 \| 163 \| 99 \| 7 \| 6 \| 151 \| 73 \| 218 \| Protein OSB4, chloroplastic \| Protein OSB4, chloroplastic; afdb-proteome \| AF-Q9SX99-F1-MODEL\_V4 \| 1.0 \| 0.0001378 \| 182 \| 0.094 \| 137 \| 109 \| 5 \| 1 \| 135 \| 51 \| 174 \| Protein OSB1, mitochondrial \| Protein OSB1, mitochondrial; afdb-proteome \| AF-A4HRK3-F1-MODEL\_V4 \| 1.0 \| 3.409e-05 \| 168 \| 0.128 \| 202 \| 130 \| 10 \| 2 \| 162 \| 242 \| 438 \| Hypothetical\_protein\_-\_conserved \| Hypothetical\_protein\_-\_conserved; afdb-proteome \| AF-A0A1D6G4L8-F1-MODEL\_V4 \| 1.0 \| 6.155e-05 \| 163 \| 0.122 \| 188 \| 120 \| 6 \| 6 \| 162 \| 94 \| 267 \| Protein OSB2 chloroplastic \| Protein OSB2 chloroplastic; afdb-proteome \| AF-Q9I5K8-F1-MODEL\_V4 \| 1.0 \| 0.0008554 \| 112 \| 0.142 \| 154 \| 116 \| 7 \| 1 \| 149 \| 2 \| 144 \| Helix destabilizing protein of bacteriophage Pf1 \| Helix destabilizing protein of bacteriophage Pf1; afdb-proteome \| AF-A0A1C1CYU6-F1-MODEL\_V4 \| 1.0 \| 0.00959 \| 108 \| 0.109 \| 119 \| 89 \| 4 \| 1 \| 116 \| 176 \| 280 \| Replication protein A subunit \| Replication protein A subunit; afdb-proteome \| AF-A0A0R0FI03-F1-MODEL\_V4 \| 0.997 \| 0.007332 \| 91 \| 0.121 \| 156 \| 109 \| 11 \| 3 \| 157 \| 258 \| 386 \| Uncharacterized protein \| Uncharacterized protein; afdb-proteome \| AF-A0A044UXX9-F1-MODEL\_V4 \| 0.991 \| 0.002504 \| 84 \| 0.155 \| 135 \| 86 \| 6 \| 12 \| 137 \| 65 \| 180 \| Uncharacterized protein \| Uncharacterized protein; afdb-uniprot50 \| AF-A0A8A7QY70-F1-MODEL\_V4 \| 1.0 \| 4.882e-17 \| 813 \| 0.643 \| 115 \| 41 \| 0 \| 1 \| 115 \| 2 \| 116 \| Single-stranded DNA-binding protein \| Single-stranded DNA-binding protein; afdb-uniprot50 \| AF-A0A5T2WJW7-F1-MODEL\_V4 \| 1.0 \| 1.318e-20 \| 813 \| 0.613 \| 163 \| 60 \| 2 \| 1 \| 162 \| 2 \| 162 \| Single-stranded DNA-binding protein \| Single-stranded DNA-binding protein; afdb-uniprot50 \| AF-Q8D254-F1-MODEL\_V4 \| 1.0 \| 2.181e-17 \| 810 \| 0.652 \| 118 \| 40 \| 1 \| 1 \| 117 \| 1 \| 118 \| Single-stranded DNA-binding protein \| Single-stranded DNA-binding protein; afdb-uniprot50 \| AF-A0A5T5YDS6-F1-MODEL\_V4 \| 1.0 \| 5.928e-20 \| 791 \| 0.586 \| 162 \| 65 \| 1 \| 1 \| 162 \| 2 \| 161 \| Single-stranded DNA-binding protein \| Single-stranded DNA-binding protein; afdb-uniprot50 \| AF-Q9RHF4-F1-MODEL\_V4 \| 1.0 \| 5.618e-20 \| 788 \| 0.601 \| 163 \| 62 \| 2 \| 1 \| 162 \| 2 \| 162 \| Single-stranded DNA-binding protein 2 \| Single-stranded DNA-binding protein 2; afdb-uniprot50 \| AF-A0A8B4JY02-F1-MODEL\_V4 \| 1.0 \| 9.11e-20 \| 787 \| 0.607 \| 163 \| 59 \| 3 \| 1 \| 162 \| 20 \| 178 \| Single-strand binding protein \| Single-strand binding protein; afdb-uniprot50 \| AF-A0A7X5QH72-F1-MODEL\_V4 \| 1.0 \| 8.633e-20 \| 784 \| 0.617 \| 162 \| 59 \| 2 \| 1 \| 162 \| 2 \| 160 \| Single-stranded DNA-binding protein \| Single-stranded DNA-binding protein; afdb-uniprot50 \| AF-A0A2C6DMN1-F1-MODEL\_V4 \| 1.0 \| 5.928e-20 \| 783 \| 0.603 \| 169 \| 60 \| 2 \| 1 \| 162 \| 2 \| 170 \| Single-stranded DNA-binding protein \| Single-stranded DNA-binding protein; afdb-uniprot50 \| AF-A0A759M963-F1-MODEL\_V4 \| 1.0 \| 2.949e-20 \| 782 \| 0.621 \| 164 \| 58 \| 3 \| 1 \| 162 \| 1 \| 162 \| Single-stranded DNA-binding protein \| Single-stranded DNA-binding protein; afdb-uniprot50 \| AF-V5Z4Q9-F1-MODEL\_V4 \| 1.0 \| 5.618e-20 \| 782 \| 0.573 \| 164 \| 67 \| 2 \| 1 \| 162 \| 2 \| 164 \| Single-stranded DNA-binding protein \| Single-stranded DNA-binding protein; afdb-uniprot50 \| AF-P28046-F1-MODEL\_V4 \| 1.0 \| 3.464e-20 \| 778 \| 0.612 \| 173 \| 56 \| 2 \| 1 \| 162 \| 2 \| 174 \| Single-stranded DNA-binding protein \| Single-stranded DNA-binding protein; afdb-uniprot50 \| AF-A0A8A9F1R6-F1-MODEL\_V4 \| 1.0 \| 1.014e-19 \| 777 \| 0.573 \| 164 \| 68 \| 2 \| 1 \| 162 \| 2 \| 165 \| Single-stranded DNA-binding protein \| Single-stranded DNA-binding protein; afdb-uniprot50 \| AF-A0A410X808-F1-MODEL\_V4 \| 1.0 \| 2.51e-20 \| 772 \| 0.622 \| 175 \| 53 \| 4 \| 1 \| 162 \| 2 \| 176 \| Single-stranded DNA-binding protein \| Single-stranded DNA-binding protein; afdb-uniprot50 \| AF-A0A6I7D2R0-F1-MODEL\_V4 \| 1.0 \| 4.815e-19 \| 770 \| 0.588 \| 163 \| 62 \| 3 \| 1 \| 162 \| 2 \| 160 \| Single-stranded DNA-binding protein \| Single-stranded DNA-binding protein; afdb-uniprot50 \| AF-A0A068YZG5-F1-MODEL\_V4 \| 1.0 \| 4.532e-20 \| 768 \| 0.581 \| 172 \| 62 \| 2 \| 1 \| 162 \| 2 \| 173 \| Single-stranded DNA-binding protein \| Single-stranded DNA-binding protein; afdb-uniprot50 \| AF-C0ARM9-F1-MODEL\_V4 \| 1.0 \| 1.919e-20 \| 767 \| 0.622 \| 172 \| 54 \| 3 \| 2 \| 162 \| 39 \| 210 \| Single-stranded DNA-binding protein \| Single-stranded DNA-binding protein; afdb-uniprot50 \| AF-E9CNB8-F1-MODEL\_V4 \| 1.0 \| 2.395e-19 \| 765 \| 0.576 \| 163 \| 66 \| 2 \| 1 \| 162 \| 2 \| 162 \| Single-stranded DNA-binding protein \| Single-stranded DNA-binding protein; afdb-uniprot50 \| AF-A0A763N3M4-F1-MODEL\_V4 \| 1.0 \| 6.094e-16 \| 764 \| 0.601 \| 113 \| 44 \| 1 \| 1 \| 113 \| 2 \| 113 \| Single-stranded DNA-binding protein \| Single-stranded DNA-binding protein; afdb-uniprot50 \| AF-A0A542C9S8-F1-MODEL\_V4 \| 1.0 \| 1.4e-19 \| 764 \| 0.586 \| 162 \| 65 \| 1 \| 1 \| 162 \| 2 \| 161 \| Single-stranded DNA-binding protein \| Single-stranded DNA-binding protein; afdb-uniprot50 \| AF-A0A085GLM7-F1-MODEL\_V4 \| 1.0 \| 2.025e-20 \| 764 \| 0.598 \| 177 \| 56 \| 3 \| 1 \| 162 \| 2 \| 178 \| Single-stranded DNA-binding protein \| Single-stranded DNA-binding protein; afdb-uniprot50 \| AF-A0A250B243-F1-MODEL\_V4 \| 1.0 \| 6.964e-20 \| 764 \| 0.589 \| 178 \| 57 \| 2 \| 1 \| 162 \| 2 \| 179 \| Single-stranded DNA-binding protein \| Single-stranded DNA-binding protein; afdb-uniprot50 \| AF-A0A3N6SJW5-F1-MODEL\_V4 \| 1.0 \| 5.657e-19 \| 763 \| 0.561 \| 162 \| 71 \| 0 \| 1 \| 162 \| 2 \| 163 \| Single-stranded DNA-binding protein \| Single-stranded DNA-binding protein; afdb-uniprot50 \| AF-A0A5V0FD94-F1-MODEL\_V4 \| 1.0 \| 2.039e-19 \| 762 \| 0.611 \| 162 \| 61 \| 1 \| 1 \| 162 \| 2 \| 161 \| Single-stranded DNA-binding protein \| Single-stranded DNA-binding protein; afdb-uniprot50 \| AF-A0A1W6B523-F1-MODEL\_V4 \| 1.0 \| 5.046e-20 \| 762 \| 0.607 \| 176 \| 55 \| 3 \| 1 \| 162 \| 2 \| 177 \| Single-stranded DNA-binding protein \| Single-stranded DNA-binding protein; afdb-uniprot50 \| AF-A0A090NBG8-F1-MODEL\_V4 \| 1.0 \| 6.6e-20 \| 760 \| 0.584 \| 178 \| 58 \| 3 \| 1 \| 162 \| 22 \| 199 \| Single-stranded DNA-binding protein \| Single-stranded DNA-binding protein; afdb-uniprot50 \| AF-A0A068RC77-F1-MODEL\_V4 \| 1.0 \| 8.633e-20 \| 759 \| 0.581 \| 172 \| 62 \| 2 \| 1 \| 162 \| 2 \| 173 \| Single-stranded DNA-binding protein \| Single-stranded DNA-binding protein; afdb-uniprot50 \| AF-A0A2W5S122-F1-MODEL\_V4 \| 1.0 \| 2.395e-19 \| 759 \| 0.521 \| 182 \| 66 \| 2 \| 1 \| 162 \| 2 \| 182 \| Single-stranded DNA-binding protein \| Single-stranded DNA-binding protein; afdb-uniprot50 \| AF-A0A2P8VNA9-F1-MODEL\_V4 \| 1.0 \| 2.649e-20 \| 758 \| 0.596 \| 181 \| 54 \| 3 \| 1 \| 162 \| 2 \| 182 \| Single-stranded DNA-binding protein \| Single-stranded DNA-binding protein; afdb-uniprot50 \| AF-A0A3A4CYF8-F1-MODEL\_V4 \| 1.0 \| 1.645e-19 \| 757 \| 0.586 \| 162 \| 64 \| 1 \| 1 \| 162 \| 2 \| 160 \| Single-stranded DNA-binding protein \| Single-stranded DNA-binding protein; afdb-uniprot50 \| AF-A0A6B1YL18-F1-MODEL\_V4 \| 1.0 \| 1.043e-15 \| 756 \| 0.548 \| 113 \| 51 \| 0 \| 1 \| 113 \| 1 \| 113 \| Single-stranded DNA-binding protein \| Single-stranded DNA-binding protein; afdb-uniprot50 \| AF-A0A6D2GFG8-F1-MODEL\_V4 \| 1.0 \| 8.182e-20 \| 756 \| 0.585 \| 176 \| 59 \| 4 \| 1 \| 162 \| 2 \| 177 \| Single-stranded DNA-binding protein \| Single-stranded DNA-binding protein; afdb-uniprot50 \| AF-T0NVG8-F1-MODEL\_V4 \| 1.0 \| 3.681e-19 \| 755 \| 0.581 \| 165 \| 65 \| 2 \| 1 \| 162 \| 2 \| 165 \| Single-stranded DNA-binding protein \| Single-stranded DNA-binding protein; afdb-uniprot50 \| AF-A0A7W3BKJ0-F1-MODEL\_V4 \| 1.0 \| 2.528e-19 \| 755 \| 0.545 \| 165 \| 69 \| 2 \| 1 \| 162 \| 2 \| 163 \| Single-stranded DNA-binding protein \| Single-stranded DNA-binding protein; afdb-uniprot50 \| AF-A0A850L2I0-F1-MODEL\_V4 \| 1.0 \| 5.081e-19 \| 754 \| 0.564 \| 163 \| 60 \| 3 \| 1 \| 162 \| 1 \| 153 \| Single-stranded DNA-binding protein \| Single-stranded DNA-binding protein; afdb-uniprot50 \| AF-A0A657IF21-F1-MODEL\_V4 \| 1.0 \| 4.815e-19 \| 754 \| 0.549 \| 162 \| 71 \| 1 \| 1 \| 162 \| 2 \| 161 \| Single-stranded DNA-binding protein \| Single-stranded DNA-binding protein; afdb-uniprot50 \| AF-A0A637XTZ4-F1-MODEL\_V4 \| 1.0 \| 1.257e-19 \| 754 \| 0.591 \| 174 \| 59 \| 4 \| 1 \| 162 \| 2 \| 175 \| Single-stranded DNA-binding protein \| Single-stranded DNA-binding protein; afdb-uniprot50 \| AF-C6CKS0-F1-MODEL\_V4 \| 1.0 \| 6.6e-20 \| 754 \| 0.582 \| 182 \| 56 \| 5 \| 1 \| 162 \| 1 \| 182 \| Single-stranded DNA-binding protein \| Single-stranded DNA-binding protein; afdb-uniprot50 \| AF-D4I2A3-F1-MODEL\_V4 \| 1.0 \| 5.618e-20 \| 752 \| 0.587 \| 182 \| 55 \| 4 \| 1 \| 162 \| 2 \| 183 \| Single-stranded DNA-binding protein \| Single-stranded DNA-binding protein; afdb-uniprot50 \| AF-A0A376WPU8-F1-MODEL\_V4 \| 1.0 \| 2.528e-19 \| 751 \| 0.6 \| 165 \| 60 \| 3 \| 1 \| 162 \| 1 \| 162 \| Single-stranded DNA-binding protein \| Single-stranded DNA-binding protein; afdb-uniprot50 \| AF-A0A2J0PNB2-F1-MODEL\_V4 \| 1.0 \| 1.4e-19 \| 751 \| 0.577 \| 168 \| 61 \| 3 \| 1 \| 162 \| 1 \| 164 \| Single-stranded DNA-binding protein \| Single-stranded DNA-binding protein; afdb-uniprot50 \| AF-A0A1Y0LD14-F1-MODEL\_V4 \| 1.0 \| 1.07e-19 \| 751 \| 0.576 \| 177 \| 60 \| 3 \| 1 \| 162 \| 2 \| 178 \| Single-stranded DNA-binding protein \| Single-stranded DNA-binding protein; afdb-uniprot50 \| AF-A0A2Z5T340-F1-MODEL\_V4 \| 1.0 \| 3.966e-16 \| 750 \| 0.561 \| 121 \| 52 \| 1 \| 1 \| 120 \| 2 \| 122 \| Single-stranded DNA-binding protein \| Single-stranded DNA-binding protein; afdb-uniprot50 \| AF-A0A4Q4A5A0-F1-MODEL\_V4 \| 1.0 \| 1.831e-19 \| 750 \| 0.584 \| 171 \| 55 \| 4 \| 1 \| 162 \| 2 \| 165 \| Single-stranded DNA-binding protein \| Single-stranded DNA-binding protein; afdb-uniprot50 \| AF-A0A348DL17-F1-MODEL\_V4 \| 1.0 \| 6.6e-20 \| 750 \| 0.576 \| 177 \| 60 \| 3 \| 1 \| 162 \| 2 \| 178 \| Single-stranded DNA-binding protein \| Single-stranded DNA-binding protein; afdb-uniprot50 \| AF-A0A0A2VXI7-F1-MODEL\_V4 \| 1.0 \| 5.618e-20 \| 749 \| 0.605 \| 175 \| 56 \| 2 \| 1 \| 162 \| 2 \| 176 \| Single-stranded DNA-binding protein \| Single-stranded DNA-binding protein; afdb-uniprot50 \| AF-A0A419B249-F1-MODEL\_V4 \| 1.0 \| 1.831e-19 \| 748 \| 0.612 \| 165 \| 61 \| 2 \| 1 \| 162 \| 2 \| 166 \| Single-stranded DNA-binding protein \| Single-stranded DNA-binding protein; afdb-uniprot50 \| AF-A0A5X6EUH9-F1-MODEL\_V4 \| 1.0 \| 1.327e-19 \| 748 \| 0.596 \| 166 \| 63 \| 2 \| 1 \| 162 \| 2 \| 167 \| Single-stranded DNA-binding protein \| Single-stranded DNA-binding protein; afdb-uniprot50 \| AF-A0A710F8C2-F1-MODEL\_V4 \| 1.0 \| 6.646e-19 \| 747 \| 0.543 \| 162 \| 72 \| 1 \| 1 \| 162 \| 2 \| 161 \| Single-stranded DNA-binding protein \| Single-stranded DNA-binding protein; afdb-uniprot50 \| AF-A0A5J6VYW7-F1-MODEL\_V4 \| 1.0 \| 3.133e-19 \| 746 \| 0.556 \| 178 \| 63 \| 3 \| 1 \| 162 \| 2 \| 179 \| Single-stranded DNA-binding protein \| Single-stranded DNA-binding protein; afdb-uniprot50 \| AF-A0A7W2V7Q4-F1-MODEL\_V4 \| 1.0 \| 6.964e-20 \| 745 \| 0.571 \| 175 \| 62 \| 2 \| 1 \| 162 \| 2 \| 176 \| Single-stranded DNA-binding protein \| Single-stranded DNA-binding protein; afdb-uniprot50 \| AF-A0A826P733-F1-MODEL\_V4 \| 1.0 \| 1.735e-19 \| 744 \| 0.6 \| 165 \| 60 \| 3 \| 1 \| 162 \| 1 \| 162 \| Single-stranded DNA-binding protein \| Single-stranded DNA-binding protein; afdb-uniprot50 \| AF-A0A1W6C1A7-F1-MODEL\_V4 \| 1.0 \| 3.681e-19 \| 743 \| 0.558 \| 170 \| 64 \| 3 \| 1 \| 162 \| 2 \| 168 \| Single-stranded DNA-binding protein \| Single-stranded DNA-binding protein; afdb-uniprot50 \| AF-A0A2V2SUM0-F1-MODEL\_V4 \| 1.0 \| 4.385e-17 \| 742 \| 0.551 \| 136 \| 59 \| 1 \| 1 \| 136 \| 1 \| 134 \| Single-stranded DNA-binding protein \| Single-stranded DNA-binding protein; afdb-uniprot50 \| AF-A0A4Q9D8C0-F1-MODEL\_V4 \| 1.0 \| 2.667e-19 \| 742 \| 0.613 \| 168 \| 55 \| 3 \| 1 \| 162 \| 1 \| 164 \| Single-stranded DNA-binding protein \| Single-stranded DNA-binding protein; afdb-uniprot50 \| AF-A0A0B6X7H2-F1-MODEL\_V4 \| 1.0 \| 8.693e-19 \| 742 \| 0.508 \| 169 \| 76 \| 2 \| 1 \| 162 \| 1 \| 169 \| Single-stranded DNA-binding protein \| Single-stranded DNA-binding protein; afdb-uniprot50 \| AF-A0A5U5MJG5-F1-MODEL\_V4 \| 1.0 \| 7.862e-18 \| 741 \| 0.565 \| 145 \| 58 \| 2 \| 1 \| 140 \| 2 \| 146 \| Single-stranded DNA-binding protein \| Single-stranded DNA-binding protein; afdb-uniprot50 \| AF-A0A5X2B0T4-F1-MODEL\_V4 \| 1.0 \| 6.298e-19 \| 741 \| 0.537 \| 162 \| 73 \| 1 \| 1 \| 162 \| 2 \| 161 \| Single-stranded DNA-binding protein \| Single-stranded DNA-binding protein; afdb-uniprot50 \| AF-A0A4P9VSW3-F1-MODEL\_V4 \| 1.0 \| 7.4e-19 \| 741 \| 0.559 \| 159 \| 69 \| 1 \| 4 \| 162 \| 3 \| 160 \| Single-stranded DNA-binding protein \| Single-stranded DNA-binding protein; afdb-uniprot50 \| AF-A0A3M4A4U1-F1-MODEL\_V4 \| 1.0 \| 2.096e-15 \| 740 \| 0.607 \| 112 \| 43 \| 1 \| 1 \| 112 \| 2 \| 112 \| Single-stranded DNA-binding protein \| Single-stranded DNA-binding protein; afdb-uniprot50 \| AF-A0A703UWP0-F1-MODEL\_V4 \| 1.0 \| 1.668e-17 \| 740 \| 0.566 \| 143 \| 53 \| 2 \| 1 \| 142 \| 2 \| 136 \| Single-stranded DNA-binding protein \| Single-stranded DNA-binding protein; afdb-uniprot50 \| AF-A0A0J9B2X7-F1-MODEL\_V4 \| 1.0 \| 5.081e-19 \| 739 \| 0.539 \| 165 \| 66 \| 2 \| 1 \| 162 \| 2 \| 159 \| Single-stranded DNA-binding protein \| Single-stranded DNA-binding protein; afdb-uniprot50 \| AF-A0A7S4LY53-F1-MODEL\_V4 \| 1.0 \| 6.964e-20 \| 739 \| 0.589 \| 178 \| 57 \| 4 \| 1 \| 162 \| 2 \| 179 \| Single-stranded DNA-binding protein \| Single-stranded DNA-binding protein; afdb-uniprot50 \| AF-A0A369WG84-F1-MODEL\_V4 \| 1.0 \| 3.489e-19 \| 739 \| 0.534 \| 187 \| 62 \| 3 \| 1 \| 162 \| 1 \| 187 \| Single-stranded DNA-binding protein \| Single-stranded DNA-binding protein; afdb-uniprot50 \| AF-A0A411WM66-F1-MODEL\_V4 \| 1.0 \| 2.723e-16 \| 737 \| 0.496 \| 129 \| 60 \| 1 \| 1 \| 129 \| 2 \| 125 \| Single-stranded DNA-binding protein \| Single-stranded DNA-binding protein; afdb-uniprot50 \| AF-A0A6G6PMY1-F1-MODEL\_V4 \| 1.0 \| 2.395e-19 \| 735 \| 0.603 \| 169 \| 60 \| 3 \| 1 \| 162 \| 1 \| 169 \| Single-stranded DNA-binding protein \| Single-stranded DNA-binding protein; afdb-uniprot50 \| AF-A0A2P5IIN3-F1-MODEL\_V4 \| 1.0 \| 3.133e-19 \| 734 \| 0.595 \| 168 \| 62 \| 3 \| 1 \| 162 \| 2 \| 169 \| Single-stranded DNA-binding protein \| Single-stranded DNA-binding protein; afdb-uniprot50 \| AF-A0A811DAC9-F1-MODEL\_V4 \| 1.0 \| 2.395e-19 \| 734 \| 0.572 \| 180 \| 59 \| 4 \| 1 \| 162 \| 1 \| 180 \| Single-stranded DNA-binding protein 1 \| Single-stranded DNA-binding protein 1; afdb-uniprot50 \| AF-C0Q532-F1-MODEL\_V4 \| 1.0 \| 2.814e-19 \| 734 \| 0.596 \| 176 \| 57 \| 5 \| 1 \| 162 \| 42 \| 217 \| Single-stranded DNA-binding protein \| Single-stranded DNA-binding protein; afdb-uniprot50 \| AF-A0A6N8F438-F1-MODEL\_V4 \| 1.0 \| 1.973e-16 \| 732 \| 0.573 \| 129 \| 53 \| 1 \| 1 \| 129 \| 1 \| 127 \| Single-stranded DNA-binding protein \| Single-stranded DNA-binding protein; afdb-uniprot50 \| AF-A0A1P8KIA2-F1-MODEL\_V4 \| 1.0 \| 1.487e-18 \| 732 \| 0.549 \| 162 \| 63 \| 3 \| 1 \| 162 \| 2 \| 153 \| Single-stranded DNA-binding protein \| Single-stranded DNA-binding protein; afdb-uniprot50 \| AF-Q9KUW2-F1-MODEL\_V4 \| 1.0 \| 5.969e-19 \| 732 \| 0.564 \| 179 \| 59 \| 5 \| 1 \| 162 \| 1 \| 177 \| Single-stranded DNA-binding protein \| Single-stranded DNA-binding protein; afdb-uniprot50 \| AF-A0A611K7F6-F1-MODEL\_V4 \| 1.0 \| 2.814e-19 \| 732 \| 0.568 \| 174 \| 60 \| 4 \| 1 \| 162 \| 1 \| 171 \| Single-stranded DNA-binding protein \| Single-stranded DNA-binding protein; afdb-uniprot50 \| AF-A0A2S6IZE8-F1-MODEL\_V4 \| 1.0 \| 7.972e-16 \| 731 \| 0.605 \| 119 \| 44 \| 2 \| 1 \| 117 \| 1 \| 118 \| Single-stranded DNA-binding protein \| Single-stranded DNA-binding protein; afdb-uniprot50 \| AF-A0A7G2IKD2-F1-MODEL\_V4 \| 1.0 \| 1.217e-16 \| 731 \| 0.601 \| 128 \| 42 \| 2 \| 1 \| 127 \| 15 \| 134 \| Single-stranded DNA-binding protein \| Single-stranded DNA-binding protein; afdb-uniprot50 \| AF-A0A419ARW4-F1-MODEL\_V4 \| 1.0 \| 1.07e-19 \| 730 \| 0.586 \| 184 \| 54 \| 3 \| 1 \| 162 \| 2 \| 185 \| Single-stranded DNA-binding protein \| Single-stranded DNA-binding protein; afdb-uniprot50 \| AF-A0A3D8ULM4-F1-MODEL\_V4 \| 1.0 \| 5.116e-18 \| 729 \| 0.524 \| 162 \| 69 \| 2 \| 1 \| 162 \| 2 \| 155 \| Single-stranded DNA-binding protein \| Single-stranded DNA-binding protein; afdb-uniprot50 \| AF-A0A629K0S7-F1-MODEL\_V4 \| 1.0 \| 1.2e-18 \| 728 \| 0.526 \| 167 \| 67 \| 3 \| 1 \| 155 \| 1 \| 167 \| Single-stranded DNA-binding protein \| Single-stranded DNA-binding protein; afdb-uniprot50 \| AF-V0AMD6-F1-MODEL\_V4 \| 1.0 \| 1.419e-17 \| 726 \| 0.575 \| 146 \| 52 \| 3 \| 1 \| 144 \| 1 \| 138 \| Single-stranded DNA-binding protein \| Single-stranded DNA-binding protein; afdb-uniprot50 \| AF-A0A2Z3HU03-F1-MODEL\_V4 \| 1.0 \| 4.595e-18 \| 726 \| 0.524 \| 164 \| 65 \| 2 \| 1 \| 162 \| 1 \| 153 \| Single-stranded DNA-binding protein \| Single-stranded DNA-binding protein; afdb-uniprot50 \| AF-A0A7H8I9U7-F1-MODEL\_V4 \| 1.0 \| 8.239e-19 \| 726 \| 0.577 \| 168 \| 65 \| 3 \| 1 \| 162 \| 1 \| 168 \| Single-stranded DNA-binding protein \| Single-stranded DNA-binding protein; afdb-uniprot50 \| AF-A0A630JIS0-F1-MODEL\_V4 \| 1.0 \| 6.011e-18 \| 725 \| 0.573 \| 150 \| 52 \| 4 \| 1 \| 138 \| 1 \| 150 \| Single-stranded DNA-binding protein \| Single-stranded DNA-binding protein; afdb-uniprot50 \| AF-A0A0C4WV47-F1-MODEL\_V4 \| 1.0 \| 3.707e-18 \| 725 \| 0.554 \| 164 \| 65 \| 3 \| 1 \| 162 \| 1 \| 158 \| Single-stranded DNA-binding protein \| Single-stranded DNA-binding protein; afdb-uniprot50 \| AF-A0A0F5ARE6-F1-MODEL\_V4 \| 1.0 \| 1.946e-18 \| 725 \| 0.524 \| 181 \| 66 \| 3 \| 1 \| 162 \| 2 \| 181 \| Single-stranded DNA-binding protein \| Single-stranded DNA-binding protein; afdb-uniprot50 \| AF-A0A2U8I3I1-F1-MODEL\_V4 \| 1.0 \| 2.166e-18 \| 725 \| 0.518 \| 164 \| 77 \| 1 \| 1 \| 162 \| 2 \| 165 \| Single-stranded DNA-binding protein \| Single-stranded DNA-binding protein; afdb-uniprot50 \| AF-A0A2D5T2D5-F1-MODEL\_V4 \| 1.0 \| 1.078e-18 \| 725 \| 0.55 \| 167 \| 70 \| 1 \| 1 \| 162 \| 1 \| 167 \| Single-stranded DNA-binding protein \| Single-stranded DNA-binding protein; afdb-uniprot50 \| AF-A0A139DMR2-F1-MODEL\_V4 \| 1.0 \| 5.474e-16 \| 724 \| 0.552 \| 123 \| 55 \| 0 \| 1 \| 123 \| 1 \| 123 \| Single-stranded DNA-binding protein \| Single-stranded DNA-binding protein; afdb-uniprot50 \| AF-G5SJV1-F1-MODEL\_V4 \| 1.0 \| 1.208e-17 \| 724 \| 0.564 \| 147 \| 55 \| 2 \| 1 \| 146 \| 2 \| 140 \| Single-stranded DNA-binding protein \| Single-stranded DNA-binding protein; afdb-uniprot50 \| AF-D3RW92-F1-MODEL\_V4 \| 1.0 \| 4.127e-18 \| 724 \| 0.518 \| 162 \| 70 \| 2 \| 1 \| 162 \| 1 \| 154 \| Single-stranded DNA-binding protein \| Single-stranded DNA-binding protein; afdb-uniprot50 \| AF-A0A2R4NNX4-F1-MODEL\_V4 \| 1.0 \| 2.286e-18 \| 723 \| 0.537 \| 162 \| 66 \| 2 \| 1 \| 162 \| 1 \| 153 \| Single-stranded DNA-binding protein \| Single-stranded DNA-binding protein; afdb-uniprot50 \| AF-A0A0H3GZP4-F1-MODEL\_V4 \| 1.0 \| 3.329e-18 \| 723 \| 0.491 \| 171 \| 77 \| 2 \| 1 \| 161 \| 2 \| 172 \| Single-stranded DNA-binding protein \| Single-stranded DNA-binding protein; afdb-uniprot50 \| AF-A0A2Z3HV94-F1-MODEL\_V4 \| 1.0 \| 1.656e-18 \| 723 \| 0.519 \| 181 \| 67 \| 4 \| 1 \| 162 \| 2 \| 181 \| Single-stranded DNA-binding protein \| Single-stranded DNA-binding protein; afdb-uniprot50 \| AF-A0A3D2B3S2-F1-MODEL\_V4 \| 1.0 \| 8.296e-18 \| 722 \| 0.567 \| 148 \| 55 \| 2 \| 1 \| 139 \| 2 \| 149 \| Single-stranded DNA-binding protein \| Single-stranded DNA-binding protein; afdb-uniprot50 \| AF-A0A7Y2N5T2-F1-MODEL\_V4 \| 1.0 \| 4.127e-18 \| 722 \| 0.546 \| 163 \| 70 \| 2 \| 1 \| 162 \| 1 \| 160 \| Single-stranded DNA-binding protein \| Single-stranded DNA-binding protein; afdb-uniprot50 \| AF-A0A4R7JHL5-F1-MODEL\_V4 \| 1.0 \| 1.487e-18 \| 721 \| 0.55 \| 169 \| 68 \| 3 \| 1 \| 162 \| 1 \| 168 \| Single-stranded DNA-binding protein \| Single-stranded DNA-binding protein; afdb-uniprot50 \| AF-A0A1B9LCJ4-F1-MODEL\_V4 \| 1.0 \| 8.239e-19 \| 721 \| 0.531 \| 173 \| 70 \| 4 \| 1 \| 162 \| 1 \| 173 \| Single-stranded DNA-binding protein \| Single-stranded DNA-binding protein; afdb-uniprot50 \| AF-A0A3B1E364-F1-MODEL\_V4 \| 1.0 \| 7.808e-19 \| 721 \| 0.529 \| 172 \| 71 \| 3 \| 1 \| 162 \| 2 \| 173 \| Single-stranded DNA-binding protein \| Single-stranded DNA-binding protein; afdb-uniprot50 \| AF-A0A4D7J836-F1-MODEL\_V4 \| 1.0 \| 1.857e-17 \| 720 \| 0.493 \| 162 \| 69 \| 1 \| 1 \| 162 \| 2 \| 150 \| Single-stranded DNA-binding protein \| Single-stranded DNA-binding protein; afdb-uniprot50 \| AF-A0A5K7YKA1-F1-MODEL\_V4 \| 1.0 \| 1.487e-18 \| 720 \| 0.544 \| 167 \| 67 \| 4 \| 1 \| 162 \| 1 \| 163 \| Single-stranded DNA-binding protein \| Single-stranded DNA-binding protein; afdb-uniprot50 \| AF-A0A2E9T352-F1-MODEL\_V4 \| 1.0 \| 1.2e-18 \| 720 \| 0.495 \| 200 \| 63 \| 4 \| 1 \| 162 \| 1 \| 200 \| Single-stranded DNA-binding protein \| Single-stranded DNA-binding protein; afdb-uniprot50 \| AF-A0A3S4HKI4-F1-MODEL\_V4 \| 1.0 \| 5.081e-19 \| 720 \| 0.558 \| 179 \| 62 \| 4 \| 1 \| 162 \| 1 \| 179 \| Single-stranded DNA-binding protein \| Single-stranded DNA-binding protein; afdb-uniprot50 \| AF-A3JEM4-F1-MODEL\_V4 \| 1.0 \| 1.419e-17 \| 719 \| 0.541 \| 155 \| 70 \| 1 \| 1 \| 155 \| 29 \| 182 \| Single-stranded DNA-binding protein \| Single-stranded DNA-binding protein; afdb-uniprot50 \| AF-A0A703J8T7-F1-MODEL\_V4 \| 1.0 \| 3.221e-15 \| 718 \| 0.647 \| 105 \| 37 \| 0 \| 1 \| 105 \| 2 \| 106 \| Single-stranded DNA-binding protein \| Single-stranded DNA-binding protein; afdb-uniprot50 \| AF-A0A0H4A0G0-F1-MODEL\_V4 \| 1.0 \| 2.873e-16 \| 718 \| 0.576 \| 130 \| 51 \| 2 \| 1 \| 129 \| 2 \| 128 \| Single-stranded DNA-binding protein \| Single-stranded DNA-binding protein; afdb-uniprot50 \| AF-A0A7U4TA53-F1-MODEL\_V4 \| 1.0 \| 5.361e-19 \| 718 \| 0.543 \| 184 \| 62 \| 4 \| 1 \| 162 \| 2 \| 185 \| Single-stranded DNA-binding protein \| Single-stranded DNA-binding protein; afdb-uniprot50 \| AF-A0A4V3X5D4-F1-MODEL\_V4 \| 1.0 \| 7.061e-18 \| 717 \| 0.536 \| 151 \| 68 \| 2 \| 1 \| 149 \| 1 \| 151 \| Single-stranded DNA-binding protein \| Single-stranded DNA-binding protein; afdb-uniprot50 \| AF-A0A709YGK6-F1-MODEL\_V4 \| 1.0 \| 3.329e-18 \| 717 \| 0.57 \| 156 \| 56 \| 3 \| 1 \| 145 \| 2 \| 157 \| Single-stranded DNA-binding protein \| Single-stranded DNA-binding protein; afdb-uniprot50 \| AF-A0A316FAR0-F1-MODEL\_V4 \| 1.0 \| 9.173e-19 \| 717 \| 0.553 \| 177 \| 61 \| 5 \| 1 \| 162 \| 2 \| 175 \| Single-stranded DNA-binding protein \| Single-stranded DNA-binding protein; afdb-uniprot50 \| AF-A0A5J6WRZ9-F1-MODEL\_V4 \| 1.0 \| 8.693e-19 \| 717 \| 0.573 \| 169 \| 61 \| 3 \| 1 \| 162 \| 2 \| 166 \| Single-stranded DNA-binding protein \| Single-stranded DNA-binding protein; afdb-uniprot50 \| AF-A0A3M1FWU7-F1-MODEL\_V4 \| 1.0 \| 2.053e-18 \| 716 \| 0.514 \| 169 \| 74 \| 2 \| 1 \| 162 \| 1 \| 168 \| Single-stranded DNA-binding protein \| Single-stranded DNA-binding protein; afdb-uniprot50 \| AF-A0A5V5DT66-F1-MODEL\_V4 \| 1.0 \| 3.993e-15 \| 715 \| 0.66 \| 103 \| 35 \| 0 \| 1 \| 103 \| 2 \| 104 \| Single-stranded DNA-binding protein \| Single-stranded DNA-binding protein; afdb-uniprot50 \| AF-A0A3G2N7B1-F1-MODEL\_V4 \| 1.0 \| 3.911e-18 \| 715 \| 0.539 \| 163 \| 63 \| 5 \| 1 \| 162 \| 1 \| 152 \| Single-stranded DNA-binding protein \| Single-stranded DNA-binding protein; afdb-uniprot50 \| AF-A0A1V0HPB3-F1-MODEL\_V4 \| 1.0 \| 2.286e-18 \| 715 \| 0.524 \| 166 \| 69 \| 3 \| 1 \| 162 \| 1 \| 160 \| Single-stranded DNA-binding protein \| Single-stranded DNA-binding protein; afdb-uniprot50 \| AF-A0A1X3M0N9-F1-MODEL\_V4 \| 1.0 \| 8.296e-18 \| 715 \| 0.519 \| 154 \| 71 \| 2 \| 1 \| 151 \| 14 \| 167 \| Single-stranded DNA-binding protein \| Single-stranded DNA-binding protein; afdb-uniprot50 \| AF-A0A379CU75-F1-MODEL\_V4 \| 1.0 \| 2.053e-18 \| 715 \| 0.548 \| 164 \| 67 \| 1 \| 1 \| 157 \| 2 \| 165 \| Single-stranded DNA-binding protein \| Single-stranded DNA-binding protein; afdb-uniprot50 \| AF-A0A3B8TV62-F1-MODEL\_V4 \| 1.0 \| 4.627e-17 \| 714 \| 0.568 \| 146 \| 60 \| 2 \| 1 \| 146 \| 1 \| 143 \| Single-stranded DNA-binding protein \| Single-stranded DNA-binding protein; afdb-uniprot50 \| AF-A0A2P5T016-F1-MODEL\_V4 \| 1.0 \| 2.545e-18 \| 714 \| 0.511 \| 170 \| 75 \| 3 \| 1 \| 162 \| 1 \| 170 \| Single-stranded DNA-binding protein \| Single-stranded DNA-binding protein; afdb-uniprot50 \| AF-A0A659R992-F1-MODEL\_V4 \| 1.0 \| 7.451e-18 \| 713 \| 0.552 \| 152 \| 59 \| 2 \| 1 \| 151 \| 2 \| 145 \| Single-stranded DNA-binding protein \| Single-stranded DNA-binding protein; afdb-uniprot50 \| AF-U7R096-F1-MODEL\_V4 \| 1.0 \| 8.754e-18 \| 713 \| 0.527 \| 163 \| 66 \| 3 \| 1 \| 162 \| 2 \| 154 \| Single-stranded DNA-binding protein \| Single-stranded DNA-binding protein; afdb-uniprot50 \| AF-A0A7Y8YBZ3-F1-MODEL\_V4 \| 1.0 \| 4.127e-18 \| 713 \| 0.549 \| 162 \| 65 \| 1 \| 1 \| 162 \| 1 \| 154 \| Single-stranded DNA-binding protein \| Single-stranded DNA-binding protein; afdb-uniprot50 \| AF-A0A707B4Y7-F1-MODEL\_V4 \| 1.0 \| 2.834e-18 \| 713 \| 0.524 \| 162 \| 68 \| 2 \| 1 \| 161 \| 2 \| 155 \| Single-stranded DNA-binding protein \| Single-stranded DNA-binding protein; afdb-uniprot50 \| AF-A0A610WMJ1-F1-MODEL\_V4 \| 1.0 \| 4.355e-18 \| 712 \| 0.521 \| 161 \| 68 \| 2 \| 1 \| 160 \| 2 \| 154 \| Single-stranded DNA-binding protein \| Single-stranded DNA-binding protein; afdb-uniprot50 \| AF-A0A6G2B591-F1-MODEL\_V4 \| 1.0 \| 5.398e-18 \| 712 \| 0.56 \| 164 \| 62 \| 4 \| 1 \| 162 \| 2 \| 157 \| Single-stranded DNA-binding protein \| Single-stranded DNA-binding protein; afdb-uniprot50 \| AF-A0A6G9HLE5-F1-MODEL\_V4 \| 1.0 \| 3.011e-17 \| 711 \| 0.537 \| 147 \| 64 \| 2 \| 1 \| 144 \| 2 \| 147 \| Single-stranded DNA-binding protein \| Single-stranded DNA-binding protein; afdb-uniprot50 \| AF-A0A6G7CNE3-F1-MODEL\_V4 \| 1.0 \| 7.451e-18 \| 711 \| 0.527 \| 163 \| 55 \| 3 \| 1 \| 162 \| 2 \| 143 \| Single-stranded DNA-binding protein \| Single-stranded DNA-binding protein; afdb-uniprot50 \| AF-A0A6C8WF48-F1-MODEL\_V4 \| 1.0 \| 3.329e-18 \| 711 \| 0.563 \| 158 \| 58 \| 3 \| 1 \| 147 \| 2 \| 159 \| Single-stranded DNA-binding protein \| Single-stranded DNA-binding protein; afdb-uniprot50 \| AF-A0A750BRM1-F1-MODEL\_V4 \| 1.0 \| 3.707e-18 \| 711 \| 0.481 \| 162 \| 81 \| 1 \| 1 \| 162 \| 2 \| 160 \| Single-stranded DNA-binding protein \| Single-stranded DNA-binding protein; afdb-uniprot50 \| AF-A0A370S7B0-F1-MODEL\_V4 \| 1.0 \| 1.844e-18 \| 711 \| 0.51 \| 186 \| 66 \| 4 \| 1 \| 162 \| 1 \| 185 \| Single-stranded DNA-binding protein \| Single-stranded DNA-binding protein; afdb-uniprot50 \| AF-A0A418YH34-F1-MODEL\_V4 \| 1.0 \| 1.487e-18 \| 710 \| 0.57 \| 163 \| 61 \| 4 \| 1 \| 162 \| 2 \| 156 \| Single-stranded DNA-binding protein \| Single-stranded DNA-binding protein; afdb-uniprot50 \| AF-E0WRV0-F1-MODEL\_V4 \| 1.0 \| 1.487e-18 \| 709 \| 0.5 \| 176 \| 74 \| 3 \| 1 \| 162 \| 2 \| 177 \| Single-stranded DNA-binding protein \| Single-stranded DNA-binding protein; afdb-uniprot50 \| AF-A0A383D6A9-F1-MODEL\_V4 \| 1.0 \| 1.987e-15 \| 708 \| 0.553 \| 112 \| 50 \| 0 \| 3 \| 114 \| 2 \| 113 \| Uncharacterized protein \| Uncharacterized protein; afdb-uniprot50 \| AF-A0A5U8SVX5-F1-MODEL\_V4 \| 1.0 \| 5.116e-18 \| 708 \| 0.635 \| 151 \| 49 \| 3 \| 1 \| 145 \| 1 \| 151 \| Single-stranded DNA-binding protein \| Single-stranded DNA-binding protein; afdb-uniprot50 \| AF-P28044-F1-MODEL\_V4 \| 1.0 \| 3.707e-18 \| 707 \| 0.502 \| 175 \| 73 \| 4 \| 1 \| 162 \| 2 \| 175 \| Plasmid-derived single-stranded DNA-binding protein \| Plasmid-derived single-stranded DNA-binding protein; afdb-uniprot50 \| AF-A0A2S5JEG1-F1-MODEL\_V4 \| 1.0 \| 3.376e-16 \| 706 \| 0.576 \| 130 \| 53 \| 2 \| 1 \| 129 \| 1 \| 129 \| Single-stranded DNA-binding protein \| Single-stranded DNA-binding protein; afdb-uniprot50 \| AF-A0A0H3ZW72-F1-MODEL\_V4 \| 1.0 \| 3.352e-17 \| 706 \| 0.533 \| 163 \| 54 \| 5 \| 1 \| 162 \| 2 \| 143 \| Single-stranded DNA-binding protein \| Single-stranded DNA-binding protein; afdb-uniprot50 \| AF-A0A2V4ZPU8-F1-MODEL\_V4 \| 1.0 \| 5.116e-18 \| 706 \| 0.527 \| 165 \| 74 \| 2 \| 1 \| 162 \| 1 \| 164 \| Single-stranded DNA-binding protein \| Single-stranded DNA-binding protein; afdb-uniprot50 \| AF-A0A2D0IKG4-F1-MODEL\_V4 \| 1.0 \| 6.692e-18 \| 706 \| 0.488 \| 170 \| 79 \| 3 \| 1 \| 162 \| 2 \| 171 \| Single-stranded DNA-binding protein \| Single-stranded DNA-binding protein; afdb-uniprot50 \| AF-P18022-F1-MODEL\_V4 \| 1.0 \| 4.127e-18 \| 706 \| 0.482 \| 176 \| 76 \| 4 \| 1 \| 162 \| 1 \| 175 \| Plasmid-derived single-stranded DNA-binding protein \| Plasmid-derived single-stranded DNA-binding protein; afdb-uniprot50 \| AF-A0A383C086-F1-MODEL\_V4 \| 1.0 \| 2.599e-15 \| 705 \| 0.616 \| 112 \| 41 \| 2 \| 1 \| 111 \| 1 \| 111 \| Uncharacterized protein \| Uncharacterized protein; afdb-uniprot50 \| AF-A0A369F188-F1-MODEL\_V4 \| 1.0 \| 1.429e-16 \| 705 \| 0.559 \| 143 \| 55 \| 3 \| 1 \| 143 \| 2 \| 136 \| Single-stranded DNA-binding protein \| Single-stranded DNA-binding protein; afdb-uniprot50 \| AF-A0A7W3G190-F1-MODEL\_V4 \| 1.0 \| 9.237e-18 \| 705 \| 0.53 \| 162 \| 64 \| 3 \| 1 \| 162 \| 2 \| 151 \| Single-stranded DNA-binding protein \| Single-stranded DNA-binding protein; afdb-uniprot50 \| AF-A0A379X3G7-F1-MODEL\_V4 \| 1.0 \| 1.208e-17 \| 705 \| 0.478 \| 163 \| 82 \| 2 \| 1 \| 162 \| 2 \| 162 \| Single-stranded DNA-binding protein \| Single-stranded DNA-binding protein; afdb-uniprot50 \| AF-A0A1I4SDU3-F1-MODEL\_V4 \| 1.0 \| 4.127e-18 \| 704 \| 0.533 \| 163 \| 75 \| 1 \| 1 \| 162 \| 14 \| 176 \| Single-stranded DNA-binding protein \| Single-stranded DNA-binding protein; afdb-uniprot50 \| AF-T1CAF7-F1-MODEL\_V4 \| 1.0 \| 7.608e-15 \| 702 \| 0.539 \| 113 \| 49 \| 1 \| 1 \| 113 \| 1 \| 110 \| Single-stranded DNA-binding protein \| Single-stranded DNA-binding protein; afdb-uniprot50 \| AF-A0A0F6TUU1-F1-MODEL\_V4 \| 1.0 \| 4.916e-16 \| 702 \| 0.492 \| 130 \| 66 \| 0 \| 1 \| 130 \| 2 \| 131 \| Single-stranded DNA-binding protein \| Single-stranded DNA-binding protein; afdb-uniprot50 \| AF-A0A2X2BQR0-F1-MODEL\_V4 \| 1.0 \| 1.208e-17 \| 702 \| 0.559 \| 161 \| 47 \| 3 \| 2 \| 162 \| 3 \| 139 \| Single-stranded DNA-binding protein \| Single-stranded DNA-binding protein; afdb-uniprot50 \| AF-A0A357KLJ1-F1-MODEL\_V4 \| 1.0 \| 4.355e-18 \| 701 \| 0.53 \| 164 \| 68 \| 3 \| 1 \| 162 \| 1 \| 157 \| Single-stranded DNA-binding protein \| Single-stranded DNA-binding protein; afdb-uniprot50 \| AF-A0A850RHP0-F1-MODEL\_V4 \| 1.0 \| 1.208e-17 \| 701 \| 0.478 \| 163 \| 81 \| 2 \| 1 \| 162 \| 1 \| 160 \| Single-stranded DNA-binding protein \| Single-stranded DNA-binding protein; afdb-uniprot50 \| AF-A0A5R9R0W8-F1-MODEL\_V4 \| 1.0 \| 1.208e-17 \| 701 \| 0.527 \| 163 \| 75 \| 2 \| 1 \| 162 \| 1 \| 162 \| Single-stranded DNA-binding protein \| Single-stranded DNA-binding protein; afdb-uniprot50 \| AF-A0A1T1H873-F1-MODEL\_V4 \| 1.0 \| 8.296e-18 \| 701 \| 0.494 \| 168 \| 79 \| 2 \| 1 \| 162 \| 1 \| 168 \| Single-stranded DNA-binding protein \| Single-stranded DNA-binding protein; afdb-uniprot50 \| AF-A0A7M2QME3-F1-MODEL\_V4 \| 1.0 \| 7.4e-19 \| 700 \| 0.568 \| 169 \| 60 \| 3 \| 1 \| 162 \| 2 \| 164 \| Single-stranded DNA-binding protein \| Single-stranded DNA-binding protein; afdb-uniprot50 \| AF-A0A7Z9CPI0-F1-MODEL\_V4 \| 1.0 \| 8.754e-18 \| 700 \| 0.469 \| 162 \| 83 \| 1 \| 1 \| 162 \| 2 \| 160 \| Single-stranded DNA-binding protein \| Single-stranded DNA-binding protein; afdb-uniprot50 \| AF-G5QBI0-F1-MODEL\_V4 \| 1.0 \| 3.155e-18 \| 700 \| 0.525 \| 175 \| 70 \| 4 \| 1 \| 162 \| 2 \| 176 \| Single-stranded DNA-binding protein \| Single-stranded DNA-binding protein; afdb-uniprot50 \| AF-A0A485A5Z5-F1-MODEL\_V4 \| 1.0 \| 6.692e-18 \| 699 \| 0.525 \| 160 \| 72 \| 2 \| 1 \| 156 \| 2 \| 161 \| Single-stranded DNA-binding protein \| Single-stranded DNA-binding protein; afdb-uniprot50 \| AF-A0A706BN23-F1-MODEL\_V4 \| 1.0 \| 7.061e-18 \| 698 \| 0.521 \| 161 \| 68 \| 2 \| 1 \| 160 \| 2 \| 154 \| Single-stranded DNA-binding protein \| Single-stranded DNA-binding protein; afdb-uniprot50 \| AF-A0A1B3B8T4-F1-MODEL\_V4 \| 1.0 \| 8.296e-18 \| 698 \| 0.533 \| 165 \| 68 \| 3 \| 1 \| 162 \| 2 \| 160 \| Single-stranded DNA-binding protein \| Single-stranded DNA-binding protein; afdb-uniprot50 \| AF-A0A1G5PGY1-F1-MODEL\_V4 \| 1.0 \| 3.329e-18 \| 698 \| 0.486 \| 187 \| 70 \| 4 \| 1 \| 162 \| 1 \| 186 \| Single-stranded DNA-binding protein \| Single-stranded DNA-binding protein; afdb-uniprot50 \| AF-A0A2X2V858-F1-MODEL\_V4 \| 1.0 \| 4.849e-18 \| 697 \| 0.53 \| 162 \| 72 \| 1 \| 1 \| 162 \| 2 \| 159 \| Single-stranded DNA-binding protein \| Single-stranded DNA-binding protein; afdb-uniprot50 \| AF-A0A2N3J8Y0-F1-MODEL\_V4 \| 1.0 \| 5.398e-18 \| 697 \| 0.551 \| 165 \| 67 \| 4 \| 1 \| 162 \| 1 \| 161 \| Single-stranded DNA-binding protein \| Single-stranded DNA-binding protein; afdb-uniprot50 \| AF-A0A7V7RBU2-F1-MODEL\_V4 \| 1.0 \| 3.155e-18 \| 696 \| 0.555 \| 162 \| 60 \| 3 \| 1 \| 162 \| 1 \| 150 \| Single-stranded DNA-binding protein \| Single-stranded DNA-binding protein; afdb-uniprot50 \| AF-A0A5R8ZPZ3-F1-MODEL\_V4 \| 1.0 \| 9.746e-18 \| 695 \| 0.496 \| 165 \| 69 \| 3 \| 1 \| 162 \| 1 \| 154 \| Single-stranded DNA-binding protein \| Single-stranded DNA-binding protein; afdb-uniprot50 \| AF-A0A3Q8D6N7-F1-MODEL\_V4 \| 1.0 \| 6.342e-18 \| 695 \| 0.461 \| 171 \| 81 \| 2 \| 1 \| 162 \| 2 \| 170 \| Single-stranded DNA-binding protein \| Single-stranded DNA-binding protein; afdb-uniprot50 \| AF-Q93GP7-F1-MODEL\_V4 \| 1.0 \| 4.355e-18 \| 695 \| 0.508 \| 177 \| 66 \| 4 \| 1 \| 162 \| 2 \| 172 \| Single-stranded DNA-binding protein 2 \| Single-stranded DNA-binding protein 2; afdb-uniprot50 \| AF-A0A659LI46-F1-MODEL\_V4 \| 1.0 \| 1.275e-17 \| 695 \| 0.481 \| 162 \| 77 \| 1 \| 1 \| 162 \| 2 \| 156 \| Single-stranded DNA-binding protein \| Single-stranded DNA-binding protein; afdb-uniprot50 \| AF-A0A5P9CCU7-F1-MODEL\_V4 \| 1.0 \| 1.085e-17 \| 694 \| 0.53 \| 164 \| 74 \| 3 \| 1 \| 162 \| 1 \| 163 \| Single-stranded DNA-binding protein \| Single-stranded DNA-binding protein; afdb-uniprot50 \| AF-U9Z7F7-F1-MODEL\_V4 \| 1.0 \| 7.451e-18 \| 694 \| 0.511 \| 174 \| 71 \| 3 \| 2 \| 162 \| 44 \| 216 \| Single-stranded DNA-binding protein \| Single-stranded DNA-binding protein; afdb-uniprot50 \| AF-A0A2E1TLI2-F1-MODEL\_V4 \| 1.0 \| 7.21e-15 \| 693 \| 0.581 \| 117 \| 45 \| 2 \| 1 \| 117 \| 1 \| 113 \| Single-stranded DNA-binding protein \| Single-stranded DNA-binding protein; afdb-uniprot50 \| AF-Q8K933-F1-MODEL\_V4 \| 1.0 \| 7.451e-18 \| 693 \| 0.518 \| 164 \| 75 \| 3 \| 1 \| 162 \| 1 \| 162 \| Single-stranded DNA-binding protein \| Single-stranded DNA-binding protein; afdb-uniprot50 \| AF-A0A379WHK7-F1-MODEL\_V4 \| 1.0 \| 1.76e-17 \| 693 \| 0.478 \| 161 \| 83 \| 1 \| 1 \| 160 \| 2 \| 162 \| Single-stranded DNA-binding protein \| Single-stranded DNA-binding protein; afdb-uniprot50 \| AF-A0A5Q0DJH1-F1-MODEL\_V4 \| 1.0 \| 7.608e-15 \| 692 \| 0.551 \| 116 \| 48 \| 2 \| 1 \| 116 \| 1 \| 112 \| Single-stranded DNA-binding protein \| Single-stranded DNA-binding protein; afdb-uniprot50 \| AF-Q8L2D1-F1-MODEL\_V4 \| 1.0 \| 1.58e-17 \| 692 \| 0.509 \| 157 \| 71 \| 3 \| 1 \| 154 \| 1 \| 154 \| Single-stranded DNA-binding protein \| Single-stranded DNA-binding protein; afdb-uniprot50 \| AF-A0A6G7VFM5-F1-MODEL\_V4 \| 1.0 \| 1.145e-17 \| 692 \| 0.512 \| 162 \| 74 \| 3 \| 1 \| 162 \| 2 \| 158 \| Single-stranded DNA-binding protein \| Single-stranded DNA-binding protein; afdb-uniprot50 \| AF-A0A375H7Q9-F1-MODEL\_V4 \| 1.0 \| 4.849e-18 \| 692 \| 0.568 \| 174 \| 56 \| 3 \| 1 \| 162 \| 1 \| 167 \| Single-stranded DNA-binding protein \| Single-stranded DNA-binding protein; afdb-uniprot50 \| AF-A0A1C4A5X5-F1-MODEL\_V4 \| 1.0 \| 4.156e-17 \| 691 \| 0.518 \| 164 \| 58 \| 5 \| 1 \| 162 \| 1 \| 145 \| Single-stranded DNA-binding protein \| Single-stranded DNA-binding protein; afdb-uniprot50 \| AF-A0A3G2HPE7-F1-MODEL\_V4 \| 1.0 \| 1.498e-17 \| 691 \| 0.506 \| 162 \| 78 \| 2 \| 1 \| 162 \| 1 \| 160 \| Single-stranded DNA-binding protein \| Single-stranded DNA-binding protein; afdb-uniprot50 \| AF-A0A661FAJ4-F1-MODEL\_V4 \| 1.0 \| 6.739e-17 \| 690 \| 0.515 \| 159 \| 59 \| 3 \| 4 \| 162 \| 1 \| 141 \| Single-stranded DNA-binding protein \| Single-stranded DNA-binding protein; afdb-uniprot50 \| AF-A0A485D3V8-F1-MODEL\_V4 \| 1.0 \| 4.882e-17 \| 690 \| 0.487 \| 154 \| 76 \| 1 \| 1 \| 154 \| 2 \| 152 \| Single-stranded DNA-binding protein \| Single-stranded DNA-binding protein; afdb-uniprot50 \| AF-A0A4Y3HW35-F1-MODEL\_V4 \| 1.0 \| 1.419e-17 \| 690 \| 0.509 \| 163 \| 72 \| 3 \| 1 \| 162 \| 2 \| 157 \| Single-stranded DNA-binding protein \| Single-stranded DNA-binding protein; afdb-uniprot50 \| AF-A0A7X9ENL8-F1-MODEL\_V4 \| 1.0 \| 3.707e-18 \| 690 \| 0.588 \| 158 \| 50 \| 3 \| 1 \| 143 \| 1 \| 158 \| Single-stranded DNA-binding protein \| Single-stranded DNA-binding protein; afdb-uniprot50 \| AF-A0A6B9GFR9-F1-MODEL\_V4 \| 1.0 \| 2.181e-17 \| 690 \| 0.469 \| 164 \| 84 \| 2 \| 1 \| 162 \| 2 \| 164 \| Single-stranded DNA-binding protein \| Single-stranded DNA-binding protein; afdb-uniprot50 \| AF-A0A1V4T3M5-F1-MODEL\_V4 \| 1.0 \| 1.208e-17 \| 690 \| 0.496 \| 163 \| 79 \| 2 \| 1 \| 162 \| 2 \| 162 \| Single-stranded DNA-binding protein \| Single-stranded DNA-binding protein; afdb-uniprot50 \| AF-A0A349ECJ1-F1-MODEL\_V4 \| 1.0 \| 5.512e-15 \| 689 \| 0.603 \| 116 \| 42 \| 2 \| 3 \| 117 \| 2 \| 114 \| Single-stranded DNA-binding protein \| Single-stranded DNA-binding protein; afdb-uniprot50 \| AF-A0A5C8BHE1-F1-MODEL\_V4 \| 1.0 \| 7.21e-15 \| 689 \| 0.533 \| 118 \| 51 \| 2 \| 1 \| 117 \| 1 \| 115 \| Single-stranded DNA-binding protein \| Single-stranded DNA-binding protein; afdb-uniprot50 \| AF-A0A5U2F1N9-F1-MODEL\_V4 \| 1.0 \| 3.177e-17 \| 689 \| 0.466 \| 163 \| 77 \| 2 \| 1 \| 162 \| 2 \| 155 \| Single-stranded DNA-binding protein \| Single-stranded DNA-binding protein; afdb-uniprot50 \| AF-A0A3M5NM76-F1-MODEL\_V4 \| 1.0 \| 1.217e-16 \| 688 \| 0.527 \| 146 \| 62 \| 3 \| 1 \| 146 \| 2 \| 140 \| Single-stranded DNA-binding protein \| Single-stranded DNA-binding protein; afdb-uniprot50 \| AF-A0A1I4SIB2-F1-MODEL\_V4 \| 1.0 \| 2.181e-17 \| 688 \| 0.527 \| 163 \| 72 \| 3 \| 1 \| 162 \| 1 \| 159 \| Single-stranded DNA-binding protein \| Single-stranded DNA-binding protein; afdb-uniprot50 \| AF-A0A0S2SIW1-F1-MODEL\_V4 \| 1.0 \| 1.844e-18 \| 688 \| 0.526 \| 186 \| 64 \| 5 \| 1 \| 162 \| 1 \| 186 \| Single-stranded DNA-binding protein \| Single-stranded DNA-binding protein; afdb-uniprot50 \| AF-C8CGN8-F1-MODEL\_V4 \| 1.0 \| 3.177e-17 \| 688 \| 0.487 \| 162 \| 82 \| 1 \| 1 \| 162 \| 2 \| 162 \| Single-stranded DNA-binding protein \| Single-stranded DNA-binding protein; afdb-uniprot50 \| AF-A0A6G9I4I2-F1-MODEL\_V4 \| 1.0 \| 3.352e-17 \| 688 \| 0.478 \| 161 \| 83 \| 1 \| 1 \| 161 \| 2 \| 161 \| Single-stranded DNA-binding protein \| Single-stranded DNA-binding protein; afdb-uniprot50 \| AF-A0A4V5J5Y1-F1-MODEL\_V4 \| 1.0 \| 5.187e-16 \| 687 \| 0.557 \| 140 \| 54 \| 3 \| 1 \| 140 \| 2 \| 133 \| Single-stranded DNA-binding protein \| Single-stranded DNA-binding protein; afdb-uniprot50 \| AF-A0A2P0QEJ1-F1-MODEL\_V4 \| 1.0 \| 2.853e-17 \| 687 \| 0.506 \| 162 \| 75 \| 2 \| 1 \| 162 \| 1 \| 157 \| Single-stranded DNA-binding protein \| Single-stranded DNA-binding protein; afdb-uniprot50 \| AF-A0A1S1HM16-F1-MODEL\_V4 \| 1.0 \| 5.398e-18 \| 686 \| 0.536 \| 164 \| 63 \| 5 \| 1 \| 162 \| 1 \| 153 \| Single-stranded DNA-binding protein \| Single-stranded DNA-binding protein; afdb-uniprot50 \| AF-A0A177YDV3-F1-MODEL\_V4 \| 1.0 \| 1.345e-17 \| 685 \| 0.527 \| 165 \| 72 \| 4 \| 1 \| 162 \| 1 \| 162 \| Single-stranded DNA-binding protein \| Single-stranded DNA-binding protein; afdb-uniprot50 \| AF-A0A6B1HYZ8-F1-MODEL\_V4 \| 1.0 \| 8.296e-18 \| 685 \| 0.517 \| 174 \| 70 \| 4 \| 1 \| 162 \| 2 \| 173 \| Single-stranded DNA-binding protein \| Single-stranded DNA-binding protein; afdb-uniprot50 \| AF-A0A3T9QRY1-F1-MODEL\_V4 \| 1.0 \| 9.746e-18 \| 685 \| 0.562 \| 169 \| 67 \| 5 \| 1 \| 162 \| 2 \| 170 \| Single-stranded DNA-binding protein \| Single-stranded DNA-binding protein; afdb-uniprot50 \| AF-A0A377BE37-F1-MODEL\_V4 \| 1.0 \| 1.028e-17 \| 684 \| 0.544 \| 169 \| 67 \| 4 \| 1 \| 162 \| 1 \| 166 \| Single-stranded DNA-binding protein \| Single-stranded DNA-binding protein; afdb-uniprot50 \| AF-A0A2K8U2W3-F1-MODEL\_V4 \| 1.0 \| 1.145e-17 \| 684 \| 0.494 \| 168 \| 76 \| 4 \| 1 \| 162 \| 30 \| 194 \| Single-stranded DNA-binding protein \| Single-stranded DNA-binding protein; afdb-uniprot50 \| AF-A0A377TIF1-F1-MODEL\_V4 \| 1.0 \| 6.692e-18 \| 684 \| 0.518 \| 162 \| 67 \| 3 \| 1 \| 162 \| 2 \| 152 \| Single-stranded DNA-binding protein \| Single-stranded DNA-binding protein; afdb-uniprot50 \| AF-A0A3C0J9N0-F1-MODEL\_V4 \| 1.0 \| 1.896e-14 \| 683 \| 0.525 \| 116 \| 52 \| 1 \| 1 \| 116 \| 1 \| 113 \| Single-stranded DNA-binding protein \| Single-stranded DNA-binding protein; afdb-uniprot50 \| AF-A0A7I6PS02-F1-MODEL\_V4 \| 1.0 \| 7.451e-18 \| 683 \| 0.541 \| 168 \| 64 \| 5 \| 1 \| 162 \| 2 \| 162 \| Single-stranded DNA-binding protein \| Single-stranded DNA-binding protein; afdb-uniprot50 \| AF-U3U1Y0-F1-MODEL\_V4 \| 1.0 \| 3.911e-18 \| 683 \| 0.537 \| 173 \| 67 \| 4 \| 1 \| 162 \| 1 \| 171 \| Single-stranded DNA-binding protein \| Single-stranded DNA-binding protein; afdb-uniprot50 \| AF-A0A7H4M7I9-F1-MODEL\_V4 \| 1.0 \| 1.374e-14 \| 681 \| 0.57 \| 107 \| 45 \| 1 \| 1 \| 106 \| 1 \| 107 \| Single-stranded DNA-binding protein \| Single-stranded DNA-binding protein; afdb-uniprot50 \| AF-A0A827CVI6-F1-MODEL\_V4 \| 1.0 \| 6.431e-16 \| 681 \| 0.424 \| 146 \| 83 \| 1 \| 1 \| 145 \| 2 \| 147 \| Single-stranded DNA-binding protein \| Single-stranded DNA-binding protein; afdb-uniprot50 \| AF-A0A850F6I9-F1-MODEL\_V4 \| 1.0 \| 9.746e-18 \| 681 \| 0.517 \| 174 \| 72 \| 4 \| 1 \| 162 \| 1 \| 174 \| Single-stranded DNA-binding protein \| Single-stranded DNA-binding protein; afdb-uniprot50 \| AF-A0A128F9E4-F1-MODEL\_V4 \| 1.0 \| 5.696e-18 \| 681 \| 0.528 \| 178 \| 65 \| 4 \| 1 \| 162 \| 2 \| 176 \| Single-stranded DNA-binding protein \| Single-stranded DNA-binding protein; afdb-uniprot50 \| AF-U1SCZ3-F1-MODEL\_V4 \| 1.0 \| 5.398e-18 \| 681 \| 0.542 \| 166 \| 68 \| 4 \| 1 \| 162 \| 25 \| 186 \| Single-stranded DNA-binding protein \| Single-stranded DNA-binding protein; afdb-uniprot50 \| AF-A0A447LUG1-F1-MODEL\_V4 \| 1.0 \| 8.754e-18 \| 680 \| 0.509 \| 163 \| 71 \| 2 \| 1 \| 162 \| 2 \| 156 \| Single-stranded DNA-binding protein \| Single-stranded DNA-binding protein; afdb-uniprot50 \| AF-A0A451D2Q2-F1-MODEL\_V4 \| 1.0 \| 2.302e-17 \| 680 \| 0.476 \| 170 \| 81 \| 1 \| 1 \| 162 \| 2 \| 171 \| Single-stranded DNA-binding protein \| Single-stranded DNA-binding protein; afdb-uniprot50 \| AF-Q8L2A6-F1-MODEL\_V4 \| 1.0 \| 3.329e-18 \| 680 \| 0.553 \| 179 \| 63 \| 5 \| 1 \| 162 \| 1 \| 179 \| Single-stranded DNA-binding protein \| Single-stranded DNA-binding protein; afdb-uniprot50 \| AF-A0A6F8PXD7-F1-MODEL\_V4 \| 1.0 \| 2.704e-17 \| 680 \| 0.468 \| 160 \| 84 \| 1 \| 4 \| 162 \| 3 \| 162 \| Single-stranded DNA-binding protein \| Single-stranded DNA-binding protein; afdb-uniprot50 \| AF-A0A0G4E5Q2-F1-MODEL\_V4 \| 1.0 \| 2.853e-17 \| 680 \| 0.533 \| 165 \| 73 \| 2 \| 1 \| 162 \| 61 \| 224 \| Single-stranded DNA-binding protein \| Single-stranded DNA-binding protein; afdb-uniprot50 \| AF-A7TUJ4-F1-MODEL\_V4 \| 1.0 \| 5.816e-15 \| 679 \| 0.631 \| 114 \| 36 \| 3 \| 1 \| 113 \| 1 \| 109 \| Single-stranded DNA-binding protein \| Single-stranded DNA-binding protein; afdb-uniprot50 \| AF-A0A4R1NRS4-F1-MODEL\_V4 \| 1.0 \| 7.111e-17 \| 679 \| 0.506 \| 162 \| 71 \| 4 \| 1 \| 162 \| 1 \| 153 \| Single-stranded DNA-binding protein \| Single-stranded DNA-binding protein; afdb-uniprot50 \| AF-A0A5Y2QF66-F1-MODEL\_V4 \| 1.0 \| 5.398e-18 \| 679 \| 0.565 \| 168 \| 54 \| 4 \| 5 \| 162 \| 1 \| 159 \| Single-stranded DNA-binding protein \| Single-stranded DNA-binding protein; afdb-uniprot50 \| AF-A0A5N7ZQN8-F1-MODEL\_V4 \| 1.0 \| 5.512e-15 \| 678 \| 0.552 \| 123 \| 50 \| 3 \| 1 \| 122 \| 1 \| 119 \| Single-stranded DNA-binding protein \| Single-stranded DNA-binding protein; afdb-uniprot50 \| AF-A0A3B8PC41-F1-MODEL\_V4 \| 1.0 \| 2.723e-16 \| 678 \| 0.542 \| 142 \| 60 \| 2 \| 1 \| 142 \| 1 \| 137 \| Single-stranded DNA-binding protein \| Single-stranded DNA-binding protein; afdb-uniprot50 \| AF-A0A2N1WWE7-F1-MODEL\_V4 \| 1.0 \| 9.301e-17 \| 678 \| 0.503 \| 151 \| 74 \| 1 \| 1 \| 151 \| 1 \| 150 \| Single-stranded DNA-binding protein \| Single-stranded DNA-binding protein; afdb-uniprot50 \| AF-X2JM87-F1-MODEL\_V4 \| 1.0 \| 1.959e-17 \| 678 \| 0.458 \| 170 \| 81 \| 2 \| 2 \| 162 \| 156 \| 323 \| Single-stranded DNA-binding protein \| Single-stranded DNA-binding protein; afdb-uniprot50 \| AF-Q1XGN9-F1-MODEL\_V4 \| 1.0 \| 9.814e-17 \| 677 \| 0.532 \| 154 \| 67 \| 2 \| 1 \| 154 \| 1 \| 149 \| Single-stranded DNA-binding protein \| Single-stranded DNA-binding protein; afdb-uniprot50 \| AF-A0A2D5QH01-F1-MODEL\_V4 \| 1.0 \| 3.938e-17 \| 677 \| 0.521 \| 165 \| 72 \| 3 \| 1 \| 162 \| 1 \| 161 \| Single-stranded DNA-binding protein \| Single-stranded DNA-binding protein; afdb-uniprot50 \| AF-A0A7C9GLE3-F1-MODEL\_V4 \| 1.0 \| 4.627e-17 \| 677 \| 0.472 \| 161 \| 83 \| 2 \| 1 \| 160 \| 1 \| 160 \| Single-stranded DNA-binding protein \| Single-stranded DNA-binding protein; afdb-uniprot50 \| AF-A0A2N7Y370-F1-MODEL\_V4 \| 1.0 \| 1.959e-17 \| 675 \| 0.539 \| 165 \| 65 \| 4 \| 1 \| 162 \| 1 \| 157 \| Single-stranded DNA-binding protein \| Single-stranded DNA-binding protein; afdb-uniprot50 \| AF-A0A427EMB1-F1-MODEL\_V4 \| 1.0 \| 2.853e-17 \| 675 \| 0.503 \| 165 \| 74 \| 3 \| 1 \| 162 \| 1 \| 160 \| Single-stranded DNA-binding protein \| Single-stranded DNA-binding protein; afdb-uniprot50 \| AF-A0A1H4ZXY2-F1-MODEL\_V4 \| 1.0 \| 4.882e-17 \| 675 \| 0.512 \| 162 \| 77 \| 2 \| 1 \| 162 \| 1 \| 160 \| Single-stranded DNA-binding protein \| Single-stranded DNA-binding protein; afdb-uniprot50 \| AF-A0A377WAK0-F1-MODEL\_V4 \| 1.0 \| 8.754e-18 \| 675 \| 0.497 \| 167 \| 71 \| 3 \| 1 \| 162 \| 1 \| 159 \| Single-stranded DNA-binding protein \| Single-stranded DNA-binding protein; afdb-uniprot50 \| AF-A0A5K1SDM4-F1-MODEL\_V4 \| 1.0 \| 6.052e-17 \| 674 \| 0.515 \| 159 \| 64 \| 3 \| 4 \| 162 \| 3 \| 148 \| Single-stranded DNA-binding protein \| Single-stranded DNA-binding protein; afdb-uniprot50 \| AF-A0A437JJK4-F1-MODEL\_V4 \| 1.0 \| 6.386e-17 \| 674 \| 0.545 \| 154 \| 65 \| 2 \| 1 \| 154 \| 1 \| 149 \| Single-stranded DNA-binding protein \| Single-stranded DNA-binding protein; afdb-uniprot50 \| AF-A0A346E026-F1-MODEL\_V4 \| 1.0 \| 9.814e-17 \| 673 \| 0.478 \| 163 \| 71 \| 3 \| 1 \| 162 \| 1 \| 150 \| Single-stranded DNA-binding protein \| Single-stranded DNA-binding protein; afdb-uniprot50 \| AF-A0A378N1F6-F1-MODEL\_V4 \| 1.0 \| 1.797e-14 \| 672 \| 0.598 \| 102 \| 40 \| 1 \| 1 \| 102 \| 1 \| 101 \| Single-stranded DNA-binding protein \| Single-stranded DNA-binding protein; afdb-uniprot50 \| AF-A0A5E5QEC3-F1-MODEL\_V4 \| 1.0 \| 9.951e-15 \| 672 \| 0.508 \| 120 \| 56 \| 3 \| 1 \| 118 \| 1 \| 119 \| Single-stranded DNA-binding protein \| Single-stranded DNA-binding protein; afdb-uniprot50 \| AF-A0A7G7WKN4-F1-MODEL\_V4 \| 1.0 \| 9.366e-16 \| 672 \| 0.609 \| 133 \| 50 \| 2 \| 1 \| 132 \| 1 \| 132 \| Single-stranded DNA-binding protein \| Single-stranded DNA-binding protein; afdb-uniprot50 \| AF-A0A7Z9WS61-F1-MODEL\_V4 \| 1.0 \| 1.76e-17 \| 672 \| 0.484 \| 165 \| 80 \| 2 \| 1 \| 162 \| 1 \| 163 \| Single-stranded DNA-binding protein \| Single-stranded DNA-binding protein; afdb-uniprot50 \| AF-A0A3Q9MM08-F1-MODEL\_V4 \| 1.0 \| 1.58e-17 \| 672 \| 0.473 \| 169 \| 82 \| 1 \| 1 \| 162 \| 2 \| 170 \| Single-stranded DNA-binding protein \| Single-stranded DNA-binding protein; afdb-uniprot50 \| AF-A0A844P9B2-F1-MODEL\_V4 \| 1.0 \| 4.021e-14 \| 671 \| 0.447 \| 114 \| 62 \| 1 \| 1 \| 113 \| 2 \| 115 \| Single-stranded DNA-binding protein \| Single-stranded DNA-binding protein; afdb-uniprot50 \| AF-A0A2D8P5X7-F1-MODEL\_V4 \| 1.0 \| 2.853e-17 \| 671 \| 0.487 \| 160 \| 77 \| 2 \| 4 \| 162 \| 2 \| 157 \| Single-stranded DNA-binding protein \| Single-stranded DNA-binding protein; afdb-uniprot50 \| AF-A0A4D6Y9S2-F1-MODEL\_V4 \| 1.0 \| 3.177e-17 \| 671 \| 0.45 \| 171 \| 79 \| 2 \| 1 \| 162 \| 1 \| 165 \| Single-stranded DNA-binding protein \| Single-stranded DNA-binding protein; afdb-uniprot50 \| AF-A0A6G7A8E4-F1-MODEL\_V4 \| 1.0 \| 1.275e-17 \| 670 \| 0.538 \| 169 \| 70 \| 4 \| 1 \| 162 \| 1 \| 168 \| Single-stranded DNA-binding protein \| Single-stranded DNA-binding protein; afdb-uniprot50 \| AF-A0A839HF94-F1-MODEL\_V4 \| 1.0 \| 4.627e-17 \| 670 \| 0.443 \| 160 \| 89 \| 0 \| 1 \| 160 \| 2 \| 161 \| Single-stranded DNA-binding protein \| Single-stranded DNA-binding protein; afdb-uniprot50 \| AF-A0A2V1GX48-F1-MODEL\_V4 \| 1.0 \| 2.704e-17 \| 670 \| 0.49 \| 163 \| 78 \| 3 \| 1 \| 162 \| 2 \| 160 \| Single-stranded DNA-binding protein \| Single-stranded DNA-binding protein; afdb-uniprot50 \| AF-A0A220SXH4-F1-MODEL\_V4 \| 1.0 \| 1.58e-17 \| 670 \| 0.483 \| 180 \| 69 \| 4 \| 1 \| 162 \| 2 \| 175 \| Single-stranded DNA-binding protein \| Single-stranded DNA-binding protein; afdb-uniprot50 \| AF-A0A6S6RS03-F1-MODEL\_V4 \| 1.0 \| 2.227e-14 \| 669 \| 0.548 \| 113 \| 48 \| 1 \| 1 \| 113 \| 1 \| 110 \| Single-stranded DNA-binding protein \| Single-stranded DNA-binding protein; afdb-uniprot50 \| AF-A0A6I7PW49-F1-MODEL\_V4 \| 1.0 \| 1.87e-16 \| 669 \| 0.435 \| 163 \| 75 \| 3 \| 1 \| 162 \| 2 \| 148 \| Single-stranded DNA-binding protein \| Single-stranded DNA-binding protein; afdb-uniprot50 \| AF-Q0E842-F1-MODEL\_V4 \| 1.0 \| 1.345e-17 \| 669 \| 0.537 \| 162 \| 62 \| 5 \| 1 \| 162 \| 15 \| 163 \| Single-stranded DNA-binding protein \| Single-stranded DNA-binding protein; afdb-uniprot50 \| AF-A0A023RQ42-F1-MODEL\_V4 \| 1.0 \| 1.498e-17 \| 669 \| 0.5 \| 190 \| 67 \| 5 \| 1 \| 162 \| 2 \| 191 \| Single-stranded DNA-binding protein \| Single-stranded DNA-binding protein; afdb-uniprot50 \| AF-A0A6I4WDE5-F1-MODEL\_V4 \| 1.0 \| 1.419e-17 \| 669 \| 0.53 \| 179 \| 62 \| 6 \| 1 \| 162 \| 1 \| 174 \| Single-stranded DNA-binding protein \| Single-stranded DNA-binding protein; afdb-uniprot50 \| AF-A0A3S9XDD4-F1-MODEL\_V4 \| 1.0 \| 2.096e-15 \| 668 \| 0.53 \| 130 \| 55 \| 2 \| 1 \| 130 \| 2 \| 125 \| Single-stranded DNA-binding protein \| Single-stranded DNA-binding protein; afdb-uniprot50 \| AF-A0A495VCG3-F1-MODEL\_V4 \| 1.0 \| 6.386e-17 \| 668 \| 0.469 \| 162 \| 78 \| 2 \| 1 \| 162 \| 1 \| 154 \| Single-stranded DNA-binding protein \| Single-stranded DNA-binding protein; afdb-uniprot50 \| AF-A0A1Z9NM89-F1-MODEL\_V4 \| 1.0 \| 3.032e-16 \| 667 \| 0.484 \| 163 \| 56 \| 4 \| 1 \| 162 \| 1 \| 136 \| Single-stranded DNA-binding protein \| Single-stranded DNA-binding protein; afdb-uniprot50 \| AF-A0A4V1B5F1-F1-MODEL\_V4 \| 1.0 \| 2.082e-16 \| 666 \| 0.515 \| 161 \| 62 \| 4 \| 3 \| 162 \| 2 \| 147 \| Single-stranded DNA-binding protein \| Single-stranded DNA-binding protein; afdb-uniprot50 \| AF-A0A443XDC5-F1-MODEL\_V4 \| 1.0 \| 5.736e-17 \| 666 \| 0.488 \| 172 \| 76 \| 4 \| 1 \| 162 \| 2 \| 171 \| Single-stranded DNA-binding protein \| Single-stranded DNA-binding protein; afdb-uniprot50 \| AF-R1F8G1-F1-MODEL\_V4 \| 1.0 \| 1.76e-17 \| 666 \| 0.489 \| 194 \| 67 \| 4 \| 1 \| 162 \| 2 \| 195 \| Single-stranded DNA-binding protein \| Single-stranded DNA-binding protein; afdb-uniprot50 \| AF-N6YXK7-F1-MODEL\_V4 \| 1.0 \| 5.776e-16 \| 665 \| 0.541 \| 146 \| 57 \| 4 \| 1 \| 146 \| 1 \| 136 \| Single-stranded DNA-binding protein \| Single-stranded DNA-binding protein; afdb-uniprot50 \| AF-A0A1F0UKE2-F1-MODEL\_V4 \| 1.0 \| 2.181e-17 \| 664 \| 0.57 \| 163 \| 61 \| 6 \| 1 \| 162 \| 1 \| 155 \| Single-stranded DNA-binding protein \| Single-stranded DNA-binding protein; afdb-uniprot50 \| AF-A0A5J5FSE3-F1-MODEL\_V4 \| 1.0 \| 1.519e-15 \| 664 \| 0.474 \| 135 \| 64 \| 1 \| 4 \| 138 \| 219 \| 346 \| Single-stranded DNA-binding protein \| Single-stranded DNA-binding protein; afdb-uniprot50 \| AF-A0A2E1N8X2-F1-MODEL\_V4 \| 1.0 \| 6.739e-17 \| 663 \| 0.524 \| 164 \| 62 \| 5 \| 1 \| 162 \| 1 \| 150 \| Single-stranded DNA-binding protein \| Single-stranded DNA-binding protein; afdb-uniprot50 \| AF-A0A2D5RVG7-F1-MODEL\_V4 \| 1.0 \| 4.882e-17 \| 663 \| 0.531 \| 160 \| 68 \| 3 \| 1 \| 160 \| 1 \| 153 \| Single-stranded DNA-binding protein \| Single-stranded DNA-binding protein; afdb-uniprot50 \| AF-A0A3G9IRR3-F1-MODEL\_V4 \| 1.0 \| 1.085e-17 \| 663 \| 0.492 \| 189 \| 69 \| 4 \| 1 \| 162 \| 35 \| 223 \| Single-stranded DNA-binding protein \| Single-stranded DNA-binding protein; afdb-uniprot50 \| AF-A0A7D5LTB4-F1-MODEL\_V4 \| 1.0 \| 1.161e-15 \| 662 \| 0.478 \| 142 \| 73 \| 1 \| 1 \| 142 \| 1 \| 141 \| Single-stranded DNA-binding protein \| Single-stranded DNA-binding protein; afdb-uniprot50 \| AF-A0A4R3W6J2-F1-MODEL\_V4 \| 1.0 \| 2.563e-17 \| 662 \| 0.533 \| 163 \| 61 \| 5 \| 1 \| 162 \| 1 \| 149 \| Single-stranded DNA-binding protein \| Single-stranded DNA-binding protein; afdb-uniprot50 \| AF-Q88QK5-F1-MODEL\_V4 \| 1.0 \| 2.302e-17 \| 662 \| 0.489 \| 182 \| 72 \| 4 \| 1 \| 162 \| 1 \| 181 \| Single-stranded DNA-binding protein \| Single-stranded DNA-binding protein; afdb-uniprot50 \| AF-U4SJQ3-F1-MODEL\_V4 \| 1.0 \| 6.386e-17 \| 661 \| 0.533 \| 163 \| 59 \| 5 \| 1 \| 162 \| 1 \| 147 \| Single-stranded DNA-binding protein \| Single-stranded DNA-binding protein; afdb-uniprot50 \| AF-A0A521H8Q0-F1-MODEL\_V4 \| 1.0 \| 1.093e-16 \| 661 \| 0.466 \| 163 \| 80 \| 4 \| 1 \| 162 \| 1 \| 157 \| Single-stranded DNA-binding protein \| Single-stranded DNA-binding protein; afdb-uniprot50 \| AF-D4HQH8-F1-MODEL\_V4 \| 1.0 \| 2.181e-17 \| 661 \| 0.494 \| 170 \| 76 \| 1 \| 2 \| 161 \| 43 \| 212 \| Single-stranded DNA-binding protein \| Single-stranded DNA-binding protein; afdb-uniprot50 \| AF-A0A7H9QSM4-F1-MODEL\_V4 \| 1.0 \| 2.893e-15 \| 660 \| 0.477 \| 136 \| 68 \| 2 \| 1 \| 135 \| 1 \| 134 \| Single-stranded DNA-binding protein \| Single-stranded DNA-binding protein; afdb-uniprot50 \| AF-A0A0E1CS86-F1-MODEL\_V4 \| 1.0 \| 3.732e-17 \| 660 \| 0.472 \| 180 \| 74 \| 4 \| 2 \| 162 \| 87 \| 264 \| Single-stranded DNA-binding protein \| Single-stranded DNA-binding protein; afdb-uniprot50 \| AF-A0A3S0XBA3-F1-MODEL\_V4 \| 1.0 \| 1.093e-16 \| 659 \| 0.524 \| 162 \| 64 \| 3 \| 1 \| 162 \| 1 \| 149 \| Single-stranded DNA-binding protein \| Single-stranded DNA-binding protein; afdb-uniprot50 \| AF-A0A2N1DPM0-F1-MODEL\_V4 \| 1.0 \| 2.853e-17 \| 659 \| 0.488 \| 168 \| 77 \| 3 \| 1 \| 162 \| 2 \| 166 \| Single-stranded DNA-binding protein \| Single-stranded DNA-binding protein; afdb-uniprot50 \| AF-A0A0B8Q6J5-F1-MODEL\_V4 \| 1.0 \| 2.318e-16 \| 658 \| 0.503 \| 147 \| 64 \| 2 \| 1 \| 146 \| 2 \| 140 \| Single-stranded DNA-binding protein \| Single-stranded DNA-binding protein; afdb-uniprot50 \| AF-A0A7Y0HDL1-F1-MODEL\_V4 \| 1.0 \| 1.275e-17 \| 658 \| 0.486 \| 181 \| 73 \| 5 \| 1 \| 162 \| 1 \| 180 \| Single-stranded DNA-binding protein \| Single-stranded DNA-binding protein; afdb-uniprot50 \| AF-A0A521VAV2-F1-MODEL\_V4 \| 1.0 \| 4.724e-14 \| 657 \| 0.558 \| 102 \| 44 \| 1 \| 1 \| 102 \| 1 \| 101 \| Single-stranded DNA-binding protein \| Single-stranded DNA-binding protein; afdb-uniprot50 \| AF-A0A5C2HL42-F1-MODEL\_V4 \| 1.0 \| 1.153e-16 \| 657 \| 0.457 \| 164 \| 84 \| 3 \| 1 \| 162 \| 1 \| 161 \| Single-stranded DNA-binding protein \| Single-stranded DNA-binding protein; afdb-uniprot50 \| AF-A0A379VN86-F1-MODEL\_V4 \| 1.0 \| 6.739e-17 \| 657 \| 0.506 \| 164 \| 67 \| 4 \| 1 \| 162 \| 1 \| 152 \| Single-stranded DNA-binding protein \| Single-stranded DNA-binding protein; afdb-uniprot50 \| AF-Q0I5U0-F1-MODEL\_V4 \| 1.0 \| 8.815e-17 \| 656 \| 0.527 \| 163 \| 62 \| 5 \| 1 \| 162 \| 1 \| 149 \| Single-stranded DNA-binding protein \| Single-stranded DNA-binding protein; afdb-uniprot50 \| AF-A0A6I4RC67-F1-MODEL\_V4 \| 1.0 \| 1.508e-16 \| 656 \| 0.49 \| 163 \| 72 \| 5 \| 1 \| 162 \| 1 \| 153 \| Single-stranded DNA-binding protein \| Single-stranded DNA-binding protein; afdb-uniprot50 \| AF-A0A1A5XLL0-F1-MODEL\_V4 \| 1.0 \| 2.446e-16 \| 656 \| 0.423 \| 196 \| 75 \| 4 \| 1 \| 162 \| 1 \| 192 \| Single-stranded DNA-binding protein \| Single-stranded DNA-binding protein; afdb-uniprot50 \| AF-A0A5J6Q657-F1-MODEL\_V4 \| 1.0 \| 7.16e-16 \| 655 \| 0.401 \| 162 \| 84 \| 2 \| 1 \| 162 \| 1 \| 149 \| Single-stranded DNA-binding protein \| Single-stranded DNA-binding protein; afdb-uniprot50 \| AF-A0A841HPC4-F1-MODEL\_V4 \| 1.0 \| 1.498e-17 \| 655 \| 0.518 \| 166 \| 64 \| 5 \| 1 \| 162 \| 1 \| 154 \| Single-strand DNA-binding protein \| Single-strand DNA-binding protein; afdb-uniprot50 \| AF-E1QMP1-F1-MODEL\_V4 \| 1.0 \| 6.386e-17 \| 655 \| 0.49 \| 161 \| 74 \| 3 \| 2 \| 162 \| 80 \| 232 \| Single-stranded DNA-binding protein \| Single-stranded DNA-binding protein; afdb-uniprot50 \| AF-A0A2T6GBE3-F1-MODEL\_V4 \| 1.0 \| 9.814e-17 \| 654 \| 0.439 \| 166 \| 88 \| 2 \| 1 \| 162 \| 1 \| 165 \| Single-stranded DNA-binding protein \| Single-stranded DNA-binding protein; afdb-uniprot50 \| AF-A0A432WBF7-F1-MODEL\_V4 \| 1.0 \| 6.386e-17 \| 654 \| 0.478 \| 165 \| 80 \| 3 \| 1 \| 162 \| 2 \| 163 \| Single-stranded DNA-binding protein \| Single-stranded DNA-binding protein; afdb-uniprot50 \| AF-A0A0E2KWP8-F1-MODEL\_V4 \| 1.0 \| 5.736e-17 \| 654 \| 0.493 \| 162 \| 74 \| 2 \| 1 \| 162 \| 2 \| 155 \| Single-stranded DNA-binding protein \| Single-stranded DNA-binding protein; afdb-uniprot50 \| AF-D9Z566-F1-MODEL\_V4 \| 1.0 \| 9.814e-17 \| 654 \| 0.494 \| 174 \| 74 \| 3 \| 2 \| 162 \| 38 \| 210 \| Single-stranded DNA-binding protein \| Single-stranded DNA-binding protein; afdb-uniprot50 \| AF-A0A376YLQ2-F1-MODEL\_V4 \| 1.0 \| 4.882e-17 \| 654 \| 0.478 \| 186 \| 65 \| 5 \| 1 \| 162 \| 41 \| 218 \| Single-stranded DNA-binding protein \| Single-stranded DNA-binding protein; afdb-uniprot50 \| AF-A0A382MFE2-F1-MODEL\_V4 \| 1.0 \| 3.612e-14 \| 653 \| 0.516 \| 122 \| 55 \| 2 \| 1 \| 122 \| 1 \| 118 \| Uncharacterized protein \| Uncharacterized protein; afdb-uniprot50 \| AF-A0A4V6Y5D9-F1-MODEL\_V4 \| 1.0 \| 7.21e-15 \| 653 \| 0.488 \| 127 \| 62 \| 2 \| 1 \| 124 \| 2 \| 128 \| Single-stranded DNA-binding protein \| Single-stranded DNA-binding protein; afdb-uniprot50 \| AF-A0A427E7X0-F1-MODEL\_V4 \| 1.0 \| 3.938e-17 \| 653 \| 0.517 \| 168 \| 72 \| 5 \| 1 \| 162 \| 1 \| 165 \| Single-stranded DNA-binding protein \| Single-stranded DNA-binding protein; afdb-uniprot50 \| AF-A0A6S6WKZ6-F1-MODEL\_V4 \| 1.0 \| 1.58e-17 \| 653 \| 0.497 \| 187 \| 66 \| 4 \| 1 \| 162 \| 2 \| 185 \| Single-stranded DNA-binding protein \| Single-stranded DNA-binding protein; afdb-uniprot50 \| AF-A0A6B0YI34-F1-MODEL\_V4 \| 1.0 \| 1.519e-15 \| 652 \| 0.521 \| 140 \| 58 \| 4 \| 1 \| 138 \| 1 \| 133 \| Single-stranded DNA-binding protein \| Single-stranded DNA-binding protein; afdb-uniprot50 \| AF-A0A1Z8N4J9-F1-MODEL\_V4 \| 1.0 \| 5.856e-14 \| 652 \| 0.456 \| 116 \| 60 \| 1 \| 1 \| 116 \| 1 \| 113 \| Single-stranded DNA-binding protein \| Single-stranded DNA-binding protein; afdb-uniprot50 \| AF-A0A6N8S0Z9-F1-MODEL\_V4 \| 1.0 \| 2.873e-16 \| 652 \| 0.509 \| 157 \| 69 \| 4 \| 1 \| 156 \| 1 \| 150 \| Single-stranded DNA-binding protein \| Single-stranded DNA-binding protein; afdb-uniprot50 \| AF-A0A0R2U7J0-F1-MODEL\_V4 \| 1.0 \| 8.47e-15 \| 651 \| 0.536 \| 138 \| 52 \| 2 \| 1 \| 138 \| 1 \| 126 \| Single-stranded DNA-binding protein \| Single-stranded DNA-binding protein; afdb-uniprot50 \| AF-A0A450WC46-F1-MODEL\_V4 \| 1.0 \| 1.093e-16 \| 651 \| 0.515 \| 165 \| 62 \| 4 \| 1 \| 162 \| 1 \| 150 \| Single-stranded DNA-binding protein \| Single-stranded DNA-binding protein; afdb-uniprot50 \| AF-A0A1E7Q8I5-F1-MODEL\_V4 \| 1.0 \| 9.814e-17 \| 651 \| 0.515 \| 157 \| 70 \| 2 \| 1 \| 157 \| 2 \| 152 \| Single-stranded DNA-binding protein \| Single-stranded DNA-binding protein; afdb-uniprot50 \| AF-A0A853I684-F1-MODEL\_V4 \| 1.0 \| 7.917e-17 \| 651 \| 0.512 \| 162 \| 74 \| 4 \| 1 \| 162 \| 1 \| 157 \| Single-stranded DNA-binding protein \| Single-stranded DNA-binding protein; afdb-uniprot50 \| AF-A0A4R6P349-F1-MODEL\_V4 \| 1.0 \| 4.95e-15 \| 650 \| 0.542 \| 118 \| 52 \| 2 \| 1 \| 117 \| 1 \| 117 \| Single-stranded DNA-binding protein \| Single-stranded DNA-binding protein; afdb-uniprot50 \| AF-A0A3A5L220-F1-MODEL\_V4 \| 1.0 \| 3.537e-17 \| 650 \| 0.537 \| 162 \| 71 \| 3 \| 1 \| 162 \| 1 \| 158 \| Single-stranded DNA-binding protein \| Single-stranded DNA-binding protein; afdb-uniprot50 \| AF-A0A6M8T6X9-F1-MODEL\_V4 \| 1.0 \| 9.301e-17 \| 650 \| 0.542 \| 164 \| 58 \| 5 \| 1 \| 162 \| 1 \| 149 \| Single-stranded DNA-binding protein \| Single-stranded DNA-binding protein; afdb-uniprot50 \| AF-A0A2G2KZ72-F1-MODEL\_V4 \| 1.0 \| 1.153e-16 \| 650 \| 0.461 \| 182 \| 78 \| 4 \| 1 \| 162 \| 2 \| 183 \| Single-stranded DNA-binding protein \| Single-stranded DNA-binding protein; afdb-uniprot50 \| AF-S6GE41-F1-MODEL\_V4 \| 1.0 \| 7.111e-17 \| 650 \| 0.512 \| 162 \| 74 \| 2 \| 1 \| 162 \| 1 \| 157 \| Single-stranded DNA-binding protein \| Single-stranded DNA-binding protein; afdb-uniprot50 \| AF-A0A1V3JSR9-F1-MODEL\_V4 \| 1.0 \| 1.987e-15 \| 649 \| 0.529 \| 136 \| 58 \| 3 \| 1 \| 135 \| 1 \| 131 \| Single-stranded DNA-binding protein \| Single-stranded DNA-binding protein; afdb-uniprot50 \| AF-A0A635JC17-F1-MODEL\_V4 \| 1.0 \| 1.691e-15 \| 649 \| 0.425 \| 162 \| 68 \| 3 \| 1 \| 162 \| 1 \| 137 \| Single-stranded DNA-binding protein \| Single-stranded DNA-binding protein; afdb-uniprot50 \| AF-A0A6N7E2V4-F1-MODEL\_V4 \| 1.0 \| 3.537e-17 \| 649 \| 0.507 \| 189 \| 61 \| 6 \| 4 \| 162 \| 3 \| 189 \| Single-stranded DNA-binding protein \| Single-stranded DNA-binding protein; afdb-uniprot50 \| AF-A0A3M0A0X9-F1-MODEL\_V4 \| 1.0 \| 9.301e-17 \| 649 \| 0.481 \| 164 \| 80 \| 2 \| 1 \| 162 \| 1 \| 161 \| Single-stranded DNA-binding protein \| Single-stranded DNA-binding protein; afdb-uniprot50 \| AF-H9ACF8-F1-MODEL\_V4 \| 1.0 \| 1.05e-14 \| 648 \| 0.564 \| 117 \| 44 \| 1 \| 1 \| 117 \| 2 \| 111 \| Single-stranded DNA-binding protein \| Single-stranded DNA-binding protein; afdb-uniprot50 \| AF-A0A1C3H441-F1-MODEL\_V4 \| 1.0 \| 1.508e-16 \| 648 \| 0.537 \| 162 \| 56 \| 7 \| 1 \| 162 \| 1 \| 143 \| Single-stranded DNA-binding protein \| Single-stranded DNA-binding protein; afdb-uniprot50 \| AF-G2FAV3-F1-MODEL\_V4 \| 1.0 \| 8.815e-17 \| 648 \| 0.5 \| 162 \| 67 \| 4 \| 1 \| 162 \| 1 \| 148 \| Single-stranded DNA-binding protein \| Single-stranded DNA-binding protein; afdb-uniprot50 \| AF-A0A7Y2XRP8-F1-MODEL\_V4 \| 1.0 \| 8.815e-17 \| 648 \| 0.515 \| 165 \| 70 \| 5 \| 1 \| 162 \| 1 \| 158 \| Single-stranded DNA-binding protein \| Single-stranded DNA-binding protein; afdb-uniprot50 \| AF-A0A1A9Z0W8-F1-MODEL\_V4 \| 1.0 \| 3.732e-17 \| 648 \| 0.464 \| 170 \| 83 \| 4 \| 1 \| 162 \| 1 \| 170 \| Uncharacterized protein \| Uncharacterized protein; afdb-uniprot50 \| AF-A0A023RSX4-F1-MODEL\_V4 \| 1.0 \| 4.627e-17 \| 648 \| 0.5 \| 172 \| 74 \| 5 \| 2 \| 162 \| 24 \| 194 \| Single-stranded DNA-binding protein \| Single-stranded DNA-binding protein; afdb-uniprot50 \| AF-A0A849HYH5-F1-MODEL\_V4 \| 1.0 \| 4.021e-14 \| 647 \| 0.504 \| 117 \| 52 \| 3 \| 1 \| 115 \| 1 \| 113 \| Single-stranded DNA-binding protein \| Single-stranded DNA-binding protein; afdb-uniprot50 \| AF-A0A5N0TC18-F1-MODEL\_V4 \| 1.0 \| 2.446e-16 \| 647 \| 0.503 \| 163 \| 72 \| 4 \| 1 \| 162 \| 1 \| 155 \| Single-stranded DNA-binding protein \| Single-stranded DNA-binding protein; afdb-uniprot50 \| AF-A0A4R3Y4S6-F1-MODEL\_V4 \| 1.0 \| 3.221e-15 \| 646 \| 0.541 \| 133 \| 54 \| 3 \| 1 \| 133 \| 1 \| 126 \| Single-stranded DNA-binding protein \| Single-stranded DNA-binding protein; afdb-uniprot50 \| AF-W1J8L6-F1-MODEL\_V4 \| 1.0 \| 1.036e-16 \| 646 \| 0.452 \| 170 \| 81 \| 4 \| 1 \| 162 \| 1 \| 166 \| Single-stranded DNA-binding protein \| Single-stranded DNA-binding protein; afdb-uniprot50 \| AF-A0A7C1ZZ21-F1-MODEL\_V4 \| 1.0 \| 4.243e-14 \| 645 \| 0.564 \| 108 \| 46 \| 1 \| 1 \| 108 \| 1 \| 107 \| Single-stranded DNA-binding protein \| Single-stranded DNA-binding protein; afdb-uniprot50 \| AF-A0A7D5QRG0-F1-MODEL\_V4 \| 1.0 \| 1.036e-16 \| 645 \| 0.459 \| 172 \| 80 \| 4 \| 1 \| 162 \| 1 \| 169 \| Single-stranded DNA-binding protein \| Single-stranded DNA-binding protein; afdb-uniprot50 \| AF-P59927-F1-MODEL\_V4 \| 1.0 \| 2.302e-17 \| 645 \| 0.526 \| 173 \| 67 \| 5 \| 1 \| 162 \| 1 \| 169 \| Single-stranded DNA-binding protein \| Single-stranded DNA-binding protein; afdb-uniprot50 \| AF-N6YC46-F1-MODEL\_V4 \| 1.0 \| 6.739e-17 \| 644 \| 0.538 \| 167 \| 65 \| 5 \| 1 \| 162 \| 1 \| 160 \| Single-stranded DNA-binding protein \| Single-stranded DNA-binding protein; afdb-uniprot50 \| AF-A0A455TMM6-F1-MODEL\_V4 \| 1.0 \| 6.052e-17 \| 644 \| 0.5 \| 170 \| 75 \| 1 \| 2 \| 161 \| 108 \| 277 \| Single-stranded DNA-binding protein \| Single-stranded DNA-binding protein; afdb-uniprot50 \| AF-A0A0M3G1E7-F1-MODEL\_V4 \| 1.0 \| 1.87e-16 \| 643 \| 0.503 \| 163 \| 63 \| 4 \| 1 \| 162 \| 1 \| 146 \| Single-stranded DNA-binding protein \| Single-stranded DNA-binding protein; afdb-uniprot50 \| AF-A0A524PZ59-F1-MODEL\_V4 \| 1.0 \| 2.873e-16 \| 643 \| 0.5 \| 162 \| 70 \| 4 \| 1 \| 162 \| 1 \| 151 \| Single-stranded DNA-binding protein \| Single-stranded DNA-binding protein; afdb-uniprot50 \| AF-A0A1H2DVA7-F1-MODEL\_V4 \| 1.0 \| 2.318e-16 \| 643 \| 0.451 \| 164 \| 86 \| 4 \| 1 \| 162 \| 1 \| 162 \| Single-stranded DNA-binding protein \| Single-stranded DNA-binding protein; afdb-uniprot50 \| AF-A0A0P7E436-F1-MODEL\_V4 \| 1.0 \| 3.938e-17 \| 643 \| 0.473 \| 188 \| 66 \| 5 \| 1 \| 162 \| 1 \| 181 \| Single-stranded DNA-binding protein \| Single-stranded DNA-binding protein; afdb-uniprot50 \| AF-S7U9Q9-F1-MODEL\_V4 \| 1.0 \| 1.772e-16 \| 642 \| 0.473 \| 167 \| 81 \| 3 \| 1 \| 162 \| 1 \| 165 \| Single-stranded DNA-binding protein \| Single-stranded DNA-binding protein; afdb-uniprot50 \| AF-A0A2D2CLI4-F1-MODEL\_V4 \| 1.0 \| 3.938e-17 \| 642 \| 0.492 \| 189 \| 69 \| 4 \| 1 \| 162 \| 67 \| 255 \| Single-stranded DNA-binding protein \| Single-stranded DNA-binding protein; afdb-uniprot50 \| AF-A0A7V5VC36-F1-MODEL\_V4 \| 1.0 \| 3.376e-16 \| 641 \| 0.509 \| 163 \| 65 \| 4 \| 1 \| 162 \| 2 \| 150 \| Single-stranded DNA-binding protein \| Single-stranded DNA-binding protein; afdb-uniprot50 \| AF-A0A3M3QJJ1-F1-MODEL\_V4 \| 1.0 \| 1.857e-17 \| 641 \| 0.486 \| 187 \| 70 \| 5 \| 1 \| 162 \| 7 \| 192 \| Single-stranded DNA-binding protein \| Single-stranded DNA-binding protein; afdb-uniprot50 \| AF-A0A2N0WZ92-F1-MODEL\_V4 \| 1.0 \| 1.284e-16 \| 640 \| 0.49 \| 165 \| 75 \| 5 \| 1 \| 162 \| 2 \| 160 \| Single-stranded DNA-binding protein \| Single-stranded DNA-binding protein; afdb-uniprot50 \| AF-A0A833GI75-F1-MODEL\_V4 \| 1.0 \| 2.599e-15 \| 639 \| 0.463 \| 151 \| 70 \| 3 \| 1 \| 151 \| 1 \| 140 \| Single-stranded DNA-binding protein \| Single-stranded DNA-binding protein; afdb-uniprot50 \| AF-A0A1D8KD04-F1-MODEL\_V4 \| 1.0 \| 4.659e-16 \| 639 \| 0.468 \| 160 \| 75 \| 3 \| 3 \| 162 \| 2 \| 151 \| Single-stranded DNA-binding protein \| Single-stranded DNA-binding protein; afdb-uniprot50 \| AF-A0A0N0IGT7-F1-MODEL\_V4 \| 1.0 \| 7.16e-16 \| 638 \| 0.454 \| 154 \| 71 \| 3 \| 1 \| 154 \| 1 \| 141 \| Single-stranded DNA-binding protein \| Single-stranded DNA-binding protein; afdb-uniprot50 \| AF-A0A523MI06-F1-MODEL\_V4 \| 1.0 \| 3.199e-16 \| 638 \| 0.496 \| 163 \| 68 \| 5 \| 1 \| 162 \| 1 \| 150 \| Single-stranded DNA-binding protein \| Single-stranded DNA-binding protein; afdb-uniprot50 \| AF-A0A482ZYE6-F1-MODEL\_V4 \| 1.0 \| 1.429e-16 \| 638 \| 0.527 \| 163 \| 68 \| 4 \| 1 \| 162 \| 1 \| 155 \| Single-stranded DNA-binding protein \| Single-stranded DNA-binding protein; afdb-uniprot50 \| AF-A0A7C8ECW2-F1-MODEL\_V4 \| 1.0 \| 1.374e-14 \| 638 \| 0.472 \| 125 \| 60 \| 2 \| 3 \| 127 \| 27 \| 145 \| Single-stranded DNA-binding protein \| Single-stranded DNA-binding protein; afdb-uniprot50 \| AF-A0A375IS87-F1-MODEL\_V4 \| 1.0 \| 1.093e-16 \| 638 \| 0.491 \| 175 \| 72 \| 4 \| 1 \| 162 \| 1 \| 171 \| Single-stranded DNA-binding protein \| Single-stranded DNA-binding protein; afdb-uniprot50 \| AF-A0A1S2DF59-F1-MODEL\_V4 \| 1.0 \| 6.386e-17 \| 638 \| 0.488 \| 176 \| 73 \| 4 \| 1 \| 162 \| 1 \| 173 \| Single-stranded DNA-binding protein \| Single-stranded DNA-binding protein; afdb-uniprot50 \| AF-A0A7C4XSU7-F1-MODEL\_V4 \| 1.0 \| 1.108e-14 \| 637 \| 0.589 \| 117 \| 45 \| 3 \| 1 \| 117 \| 1 \| 114 \| Single-stranded DNA-binding protein \| Single-stranded DNA-binding protein; afdb-uniprot50 \| AF-A0A2X4DRV2-F1-MODEL\_V4 \| 1.0 \| 2.873e-16 \| 637 \| 0.533 \| 163 \| 53 \| 5 \| 1 \| 162 \| 1 \| 141 \| Single-stranded DNA-binding protein \| Single-stranded DNA-binding protein; afdb-uniprot50 \| AF-A0A1V3K7V4-F1-MODEL\_V4 \| 1.0 \| 1.603e-15 \| 637 \| 0.492 \| 138 \| 63 \| 4 \| 1 \| 137 \| 1 \| 132 \| Single-stranded DNA-binding protein \| Single-stranded DNA-binding protein; afdb-uniprot50 \| AF-A0A1J5S6P7-F1-MODEL\_V4 \| 1.0 \| 2.723e-16 \| 637 \| 0.512 \| 162 \| 70 \| 5 \| 1 \| 162 \| 1 \| 153 \| Single-stranded DNA-binding protein \| Single-stranded DNA-binding protein; afdb-uniprot50 \| AF-A0A0F6L5J4-F1-MODEL\_V4 \| 1.0 \| 1.177e-13 \| 636 \| 0.526 \| 112 \| 49 \| 2 \| 4 \| 115 \| 1 \| 108 \| Single-stranded DNA-binding protein \| Single-stranded DNA-binding protein; afdb-uniprot50 \| AF-A0A832N208-F1-MODEL\_V4 \| 1.0 \| 1.217e-16 \| 636 \| 0.533 \| 165 \| 56 \| 7 \| 1 \| 162 \| 1 \| 147 \| Single-stranded DNA-binding protein \| Single-stranded DNA-binding protein; afdb-uniprot50 \| AF-A0A2H0DHP9-F1-MODEL\_V4 \| 1.0 \| 2.563e-17 \| 636 \| 0.511 \| 180 \| 66 \| 5 \| 1 \| 162 \| 1 \| 176 \| Single-stranded DNA-binding protein \| Single-stranded DNA-binding protein; afdb-uniprot50 \| AF-A0A7C8AUH4-F1-MODEL\_V4 \| 1.0 \| 8.876e-16 \| 635 \| 0.521 \| 163 \| 57 \| 5 \| 1 \| 162 \| 1 \| 143 \| Single-stranded DNA-binding protein \| Single-stranded DNA-binding protein; afdb-uniprot50 \| AF-A0A556UN06-F1-MODEL\_V4 \| 1.0 \| 7.972e-16 \| 635 \| 0.5 \| 162 \| 62 \| 4 \| 1 \| 162 \| 1 \| 143 \| Single-stranded DNA-binding protein \| Single-stranded DNA-binding protein; afdb-uniprot50 \| AF-A0A5E4RMM5-F1-MODEL\_V4 \| 1.0 \| 1.429e-16 \| 635 \| 0.488 \| 170 \| 73 \| 4 \| 1 \| 162 \| 1 \| 164 \| Single-stranded DNA-binding protein \| Single-stranded DNA-binding protein; afdb-uniprot50 \| AF-A0A165FGM2-F1-MODEL\_V4 \| 1.0 \| 2.082e-16 \| 635 \| 0.471 \| 174 \| 74 \| 5 \| 1 \| 162 \| 1 \| 168 \| Single-stranded DNA-binding protein \| Single-stranded DNA-binding protein; afdb-uniprot50 \| AF-E1VGN3-F1-MODEL\_V4 \| 1.0 \| 7.917e-17 \| 635 \| 0.466 \| 180 \| 75 \| 4 \| 1 \| 162 \| 2 \| 178 \| Single-stranded DNA-binding protein \| Single-stranded DNA-binding protein; afdb-uniprot50 \| AF-A0A832XC38-F1-MODEL\_V4 \| 1.0 \| 6.052e-17 \| 635 \| 0.429 \| 198 \| 72 \| 6 \| 1 \| 162 \| 1 \| 193 \| Single-stranded DNA-binding protein \| Single-stranded DNA-binding protein; afdb-uniprot50 \| AF-A0A1F4BVL2-F1-MODEL\_V4 \| 1.0 \| 1.987e-15 \| 634 \| 0.545 \| 132 \| 58 \| 2 \| 1 \| 132 \| 1 \| 130 \| Single-stranded DNA-binding protein \| Single-stranded DNA-binding protein; afdb-uniprot50 \| AF-A0A0Q8U118-F1-MODEL\_V4 \| 1.0 \| 7.503e-17 \| 634 \| 0.526 \| 167 \| 65 \| 5 \| 1 \| 162 \| 1 \| 158 \| Single-stranded DNA-binding protein \| Single-stranded DNA-binding protein; afdb-uniprot50 \| AF-A0A3T0K3E2-F1-MODEL\_V4 \| 1.0 \| 1.284e-16 \| 634 \| 0.473 \| 169 \| 75 \| 4 \| 1 \| 162 \| 1 \| 162 \| Single-stranded DNA-binding protein \| Single-stranded DNA-binding protein; afdb-uniprot50 \| AF-A0A3Q8WB38-F1-MODEL\_V4 \| 1.0 \| 1.153e-16 \| 634 \| 0.487 \| 162 \| 76 \| 2 \| 1 \| 162 \| 1 \| 155 \| Single-stranded DNA-binding protein \| Single-stranded DNA-binding protein; afdb-uniprot50 \| AF-A0A651I5E3-F1-MODEL\_V4 \| 1.0 \| 7.16e-16 \| 633 \| 0.455 \| 156 \| 78 \| 3 \| 1 \| 156 \| 14 \| 162 \| Single-stranded DNA-binding protein \| Single-stranded DNA-binding protein; afdb-uniprot50 \| AF-A0A353PC78-F1-MODEL\_V4 \| 1.0 \| 8.354e-17 \| 633 \| 0.477 \| 178 \| 74 \| 4 \| 1 \| 162 \| 2 \| 176 \| Single-stranded DNA-binding protein \| Single-stranded DNA-binding protein; afdb-uniprot50 \| AF-A0A379ANG6-F1-MODEL\_V4 \| 1.0 \| 4.156e-17 \| 633 \| 0.508 \| 183 \| 64 \| 5 \| 1 \| 162 \| 1 \| 178 \| Single-stranded DNA-binding protein \| Single-stranded DNA-binding protein; afdb-uniprot50 \| AF-A0A143WNF0-F1-MODEL\_V4 \| 1.0 \| 5.856e-14 \| 632 \| 0.527 \| 110 \| 51 \| 1 \| 1 \| 110 \| 1 \| 109 \| Single-stranded DNA-binding protein \| Single-stranded DNA-binding protein; afdb-uniprot50 \| AF-A0A0R2PAL9-F1-MODEL\_V4 \| 1.0 \| 3.562e-16 \| 632 \| 0.503 \| 163 \| 62 \| 5 \| 1 \| 162 \| 1 \| 145 \| Single-stranded DNA-binding protein \| Single-stranded DNA-binding protein; afdb-uniprot50 \| AF-A4VHR0-F1-MODEL\_V4 \| 1.0 \| 8.815e-17 \| 632 \| 0.518 \| 164 \| 68 \| 5 \| 5 \| 162 \| 1 \| 159 \| Single-stranded DNA-binding protein \| Single-stranded DNA-binding protein; afdb-uniprot50 \| AF-A0A7J6YTU1-F1-MODEL\_V4 \| 1.0 \| 2.742e-15 \| 632 \| 0.506 \| 144 \| 65 \| 4 \| 1 \| 142 \| 1 \| 140 \| BON domain-containing protein \| BON domain-containing protein; afdb-uniprot50 \| AF-B3PK66-F1-MODEL\_V4 \| 1.0 \| 1.093e-16 \| 631 \| 0.472 \| 199 \| 65 \| 5 \| 1 \| 162 \| 40 \| 235 \| Single-stranded DNA-binding protein \| Single-stranded DNA-binding protein; afdb-uniprot50 \| AF-W7Q024-F1-MODEL\_V4 \| 1.0 \| 4.659e-16 \| 630 \| 0.487 \| 162 \| 69 \| 3 \| 1 \| 162 \| 1 \| 148 \| Single-stranded DNA-binding protein \| Single-stranded DNA-binding protein; afdb-uniprot50 \| AF-G2JBS1-F1-MODEL\_V4 \| 1.0 \| 3.199e-16 \| 630 \| 0.521 \| 165 \| 65 \| 5 \| 1 \| 162 \| 1 \| 154 \| Single-stranded DNA-binding protein \| Single-stranded DNA-binding protein; afdb-uniprot50 \| AF-G2DCM5-F1-MODEL\_V4 \| 1.0 \| 6.739e-17 \| 630 \| 0.5 \| 162 \| 67 \| 3 \| 1 \| 162 \| 9 \| 156 \| Single-stranded DNA-binding protein \| Single-stranded DNA-binding protein; afdb-uniprot50 \| AF-A0A0J5P3F7-F1-MODEL\_V4 \| 1.0 \| 3.352e-17 \| 630 \| 0.524 \| 185 \| 57 \| 7 \| 1 \| 162 \| 1 \| 177 \| Single-stranded DNA-binding protein \| Single-stranded DNA-binding protein; afdb-uniprot50 \| AF-A0A375HE91-F1-MODEL\_V4 \| 1.0 \| 4.446e-15 \| 630 \| 0.517 \| 139 \| 54 \| 3 \| 1 \| 139 \| 1 \| 126 \| Single-stranded DNA-binding protein \| Single-stranded DNA-binding protein; afdb-uniprot50 \| AF-A0A3N2D590-F1-MODEL\_V4 \| 1.0 \| 2.082e-16 \| 630 \| 0.461 \| 169 \| 81 \| 2 \| 1 \| 162 \| 2 \| 167 \| Single-stranded DNA-binding protein \| Single-stranded DNA-binding protein; afdb-uniprot50 \| AF-A0A4Q3R8Z3-F1-MODEL\_V4 \| 1.0 \| 3.053e-15 \| 629 \| 0.5 \| 140 \| 61 \| 4 \| 1 \| 137 \| 1 \| 134 \| Single-stranded DNA-binding protein \| Single-stranded DNA-binding protein; afdb-uniprot50 \| AF-A0A1A7Q9A2-F1-MODEL\_V4 \| 1.0 \| 7.111e-17 \| 629 \| 0.52 \| 175 \| 66 \| 5 \| 1 \| 162 \| 1 \| 170 \| Single-stranded DNA-binding protein \| Single-stranded DNA-binding protein; afdb-uniprot50 \| AF-A0A2D6NS79-F1-MODEL\_V4 \| 1.0 \| 2.082e-16 \| 629 \| 0.5 \| 158 \| 69 \| 5 \| 1 \| 155 \| 1 \| 151 \| Single-stranded DNA-binding protein \| Single-stranded DNA-binding protein; afdb-uniprot50 \| AF-A0A849P2X6-F1-MODEL\_V4 \| 1.0 \| 3.399e-15 \| 628 \| 0.472 \| 161 \| 59 \| 3 \| 1 \| 161 \| 1 \| 135 \| Single-stranded DNA-binding protein \| Single-stranded DNA-binding protein; afdb-uniprot50 \| AF-A0A2E7MNJ4-F1-MODEL\_V4 \| 1.0 \| 1.1e-15 \| 628 \| 0.472 \| 161 \| 68 \| 3 \| 2 \| 162 \| 3 \| 146 \| Single-stranded DNA-binding protein \| Single-stranded DNA-binding protein; afdb-uniprot50 \| AF-A0A1F7FD92-F1-MODEL\_V4 \| 1.0 \| 3.562e-16 \| 628 \| 0.432 \| 162 \| 88 \| 3 \| 1 \| 162 \| 1 \| 158 \| Single-stranded DNA-binding protein \| Single-stranded DNA-binding protein; afdb-uniprot50 \| AF-A0A0E3Y7R2-F1-MODEL\_V4 \| 1.0 \| 2.723e-16 \| 628 \| 0.436 \| 174 \| 74 \| 3 \| 1 \| 162 \| 2 \| 163 \| Single-stranded DNA-binding protein \| Single-stranded DNA-binding protein; afdb-uniprot50 \| AF-A0A661EDJ0-F1-MODEL\_V4 \| 1.0 \| 6.094e-16 \| 628 \| 0.43 \| 165 \| 88 \| 2 \| 1 \| 162 \| 1 \| 162 \| Single-stranded DNA-binding protein \| Single-stranded DNA-binding protein; afdb-uniprot50 \| AF-A0A380N429-F1-MODEL\_V4 \| 1.0 \| 5.187e-16 \| 628 \| 0.457 \| 164 \| 83 \| 3 \| 1 \| 162 \| 2 \| 161 \| Single-stranded DNA-binding protein \| Single-stranded DNA-binding protein; afdb-uniprot50 \| AF-A0A154R4W2-F1-MODEL\_V4 \| 1.0 \| 4.185e-16 \| 628 \| 0.439 \| 166 \| 86 \| 2 \| 1 \| 162 \| 2 \| 164 \| Single-stranded DNA-binding protein \| Single-stranded DNA-binding protein; afdb-uniprot50 \| AF-A0A561JT25-F1-MODEL\_V4 \| 1.0 \| 1.439e-15 \| 627 \| 0.493 \| 162 \| 66 \| 3 \| 1 \| 162 \| 1 \| 146 \| Single-stranded DNA-binding protein \| Single-stranded DNA-binding protein; afdb-uniprot50 \| AF-A0A4U0YG63-F1-MODEL\_V4 \| 1.0 \| 7.16e-16 \| 627 \| 0.529 \| 155 \| 64 \| 4 \| 1 \| 154 \| 2 \| 148 \| Single-stranded DNA-binding protein \| Single-stranded DNA-binding protein; afdb-uniprot50 \| AF-A0A381ECU1-F1-MODEL\_V4 \| 1.0 \| 1.87e-16 \| 627 \| 0.45 \| 171 \| 79 \| 3 \| 4 \| 162 \| 3 \| 170 \| Single-stranded DNA-binding protein \| Single-stranded DNA-binding protein; afdb-uniprot50 \| AF-A0A3D1EPD5-F1-MODEL\_V4 \| 1.0 \| 1.311e-13 \| 626 \| 0.555 \| 108 \| 46 \| 2 \| 1 \| 108 \| 1 \| 106 \| Single-stranded DNA-binding protein \| Single-stranded DNA-binding protein; afdb-uniprot50 \| AF-V8G2X8-F1-MODEL\_V4 \| 1.0 \| 1.05e-14 \| 626 \| 0.511 \| 129 \| 55 \| 3 \| 1 \| 129 \| 1 \| 121 \| Single-stranded DNA-binding protein \| Single-stranded DNA-binding protein; afdb-uniprot50 \| AF-A0A7X7BUF0-F1-MODEL\_V4 \| 1.0 \| 1.603e-15 \| 626 \| 0.503 \| 163 \| 65 \| 4 \| 1 \| 162 \| 2 \| 149 \| Single-stranded DNA-binding protein \| Single-stranded DNA-binding protein; afdb-uniprot50 \| AF-A0A5Q0EJT3-F1-MODEL\_V4 \| 1.0 \| 3.199e-16 \| 626 \| 0.506 \| 166 \| 64 \| 4 \| 2 \| 162 \| 3 \| 155 \| Single-stranded DNA-binding protein \| Single-stranded DNA-binding protein; afdb-uniprot50 \| AF-A0A857TRT2-F1-MODEL\_V4 \| 1.0 \| 3.966e-16 \| 626 \| 0.442 \| 165 \| 85 \| 4 \| 1 \| 162 \| 2 \| 162 \| Single-stranded DNA-binding protein \| Single-stranded DNA-binding protein; afdb-uniprot50 \| AF-A0A2E9KR01-F1-MODEL\_V4 \| 1.0 \| 1.1e-15 \| 625 \| 0.466 \| 163 \| 68 \| 3 \| 1 \| 162 \| 1 \| 145 \| Single-stranded DNA-binding protein \| Single-stranded DNA-binding protein; afdb-uniprot50 \| AF-A0A1F9Y6H0-F1-MODEL\_V4 \| 1.0 \| 1.364e-15 \| 625 \| 0.401 \| 162 \| 91 \| 3 \| 1 \| 162 \| 1 \| 156 \| Single-stranded DNA-binding protein \| Single-stranded DNA-binding protein; afdb-uniprot50 \| AF-A0A1B4G7F0-F1-MODEL\_V4 \| 1.0 \| 9.301e-17 \| 625 \| 0.471 \| 174 \| 76 \| 5 \| 1 \| 162 \| 1 \| 170 \| Single-stranded DNA-binding protein \| Single-stranded DNA-binding protein; afdb-uniprot50 \| AF-G4MIH5-F1-MODEL\_V4 \| 1.0 \| 3.562e-16 \| 625 \| 0.443 \| 176 \| 81 \| 3 \| 1 \| 162 \| 28 \| 200 \| Single-stranded DNA-binding protein \| Single-stranded DNA-binding protein; afdb-uniprot50 \| AF-Q9PHE7-F1-MODEL\_V4 \| 1.0 \| 6.475e-15 \| 624 \| 0.432 \| 162 \| 66 \| 3 \| 1 \| 162 \| 1 \| 136 \| Single-stranded DNA-binding protein 1 \| Single-stranded DNA-binding protein 1; afdb-uniprot50 \| AF-A0A261RM57-F1-MODEL\_V4 \| 1.0 \| 3.785e-15 \| 624 \| 0.447 \| 163 \| 63 \| 5 \| 1 \| 162 \| 1 \| 137 \| Single-stranded DNA-binding protein \| Single-stranded DNA-binding protein; afdb-uniprot50 \| AF-G9ZJG1-F1-MODEL\_V4 \| 1.0 \| 3.785e-15 \| 624 \| 0.489 \| 143 \| 66 \| 2 \| 2 \| 140 \| 17 \| 156 \| Single-stranded DNA-binding protein \| Single-stranded DNA-binding protein; afdb-uniprot50 \| AF-A0A516V4K4-F1-MODEL\_V4 \| 1.0 \| 4.416e-16 \| 624 \| 0.439 \| 164 \| 86 \| 3 \| 1 \| 162 \| 1 \| 160 \| Single-stranded DNA-binding protein \| Single-stranded DNA-binding protein; afdb-uniprot50 \| AF-A0A451CMM4-F1-MODEL\_V4 \| 1.0 \| 3.199e-16 \| 624 \| 0.437 \| 160 \| 84 \| 2 \| 1 \| 160 \| 1 \| 154 \| Single-stranded DNA-binding protein \| Single-stranded DNA-binding protein; afdb-uniprot50 \| AF-A0A348HBX9-F1-MODEL\_V4 \| 1.0 \| 1.679e-16 \| 623 \| 0.482 \| 168 \| 76 \| 4 \| 1 \| 162 \| 26 \| 188 \| Single-stranded DNA-binding protein \| Single-stranded DNA-binding protein; afdb-uniprot50 \| AF-A0A522K774-F1-MODEL\_V4 \| 1.0 \| 3.266e-13 \| 622 \| 0.512 \| 117 \| 52 \| 3 \| 1 \| 116 \| 1 \| 113 \| Single-stranded DNA-binding protein \| Single-stranded DNA-binding protein; afdb-uniprot50 \| AF-A6EM42-F1-MODEL\_V4 \| 1.0 \| 9.496e-14 \| 622 \| 0.434 \| 122 \| 66 \| 1 \| 1 \| 122 \| 1 \| 119 \| Single-stranded DNA-binding protein \| Single-stranded DNA-binding protein; afdb-uniprot50 \| AF-A0A2G2QFS4-F1-MODEL\_V4 \| 1.0 \| 5.187e-16 \| 622 \| 0.471 \| 159 \| 75 \| 3 \| 4 \| 162 \| 2 \| 151 \| Single-stranded DNA-binding protein \| Single-stranded DNA-binding protein; afdb-uniprot50 \| AF-A0A1M5YJ27-F1-MODEL\_V4 \| 1.0 \| 1.987e-15 \| 622 \| 0.453 \| 163 \| 77 \| 6 \| 1 \| 162 \| 1 \| 152 \| Single-stranded DNA-binding protein \| Single-stranded DNA-binding protein; afdb-uniprot50 \| AF-A0A2M7SYI0-F1-MODEL\_V4 \| 1.0 \| 3.966e-16 \| 622 \| 0.469 \| 164 \| 76 \| 4 \| 1 \| 162 \| 1 \| 155 \| Single-stranded DNA-binding protein \| Single-stranded DNA-binding protein; afdb-uniprot50 \| AF-A0A3D4KFN2-F1-MODEL\_V4 \| 1.0 \| 8.412e-16 \| 622 \| 0.49 \| 159 \| 77 \| 2 \| 4 \| 162 \| 3 \| 157 \| Single-stranded DNA-binding protein \| Single-stranded DNA-binding protein; afdb-uniprot50 \| AF-A0A431KLH7-F1-MODEL\_V4 \| 1.0 \| 3.562e-16 \| 622 \| 0.475 \| 166 \| 78 \| 5 \| 1 \| 162 \| 1 \| 161 \| Single-stranded DNA-binding protein \| Single-stranded DNA-binding protein; afdb-uniprot50 \| AF-A0A380MSC4-F1-MODEL\_V4 \| 1.0 \| 5.776e-16 \| 621 \| 0.503 \| 163 \| 64 \| 5 \| 1 \| 162 \| 1 \| 147 \| Single-stranded DNA-binding protein \| Single-stranded DNA-binding protein; afdb-uniprot50 \| AF-A0A1E4ZSC0-F1-MODEL\_V4 \| 1.0 \| 4.416e-16 \| 621 \| 0.469 \| 162 \| 75 \| 5 \| 4 \| 162 \| 3 \| 156 \| Single-stranded DNA-binding protein \| Single-stranded DNA-binding protein; afdb-uniprot50 \| AF-A0A6H9SJ48-F1-MODEL\_V4 \| 1.0 \| 7.16e-16 \| 621 \| 0.484 \| 157 \| 74 \| 4 \| 1 \| 154 \| 1 \| 153 \| Single-stranded DNA-binding protein \| Single-stranded DNA-binding protein; afdb-uniprot50 \| AF-A0A2J9QVZ0-F1-MODEL\_V4 \| 1.0 \| 3.095e-13 \| 619 \| 0.457 \| 118 \| 61 \| 3 \| 1 \| 117 \| 1 \| 116 \| Single-stranded DNA-binding protein \| Single-stranded DNA-binding protein; afdb-uniprot50 \| AF-A0A7C4FIT9-F1-MODEL\_V4 \| 1.0 \| 1.529e-14 \| 619 \| 0.472 \| 127 \| 61 \| 3 \| 1 \| 127 \| 1 \| 121 \| Single-stranded DNA-binding protein \| Single-stranded DNA-binding protein; afdb-uniprot50 \| AF-A0A6L3NBL8-F1-MODEL\_V4 \| 1.0 \| 2.096e-15 \| 619 \| 0.487 \| 154 \| 68 \| 4 \| 1 \| 154 \| 1 \| 143 \| Single-stranded DNA-binding protein \| Single-stranded DNA-binding protein; afdb-uniprot50 \| AF-A0A1F9LDU6-F1-MODEL\_V4 \| 1.0 \| 3.447e-13 \| 618 \| 0.485 \| 107 \| 52 \| 2 \| 8 \| 114 \| 1 \| 104 \| Single-stranded DNA-binding protein \| Single-stranded DNA-binding protein; afdb-uniprot50 \| AF-A0A3D4GG74-F1-MODEL\_V4 \| 1.0 \| 1.459e-13 \| 618 \| 0.462 \| 121 \| 62 \| 2 \| 1 \| 121 \| 2 \| 119 \| Single-stranded DNA-binding protein \| Single-stranded DNA-binding protein; afdb-uniprot50 \| AF-A0A2D7RFL5-F1-MODEL\_V4 \| 1.0 \| 2.599e-15 \| 618 \| 0.493 \| 162 \| 65 \| 3 \| 1 \| 162 \| 1 \| 145 \| Single-stranded DNA-binding protein \| Single-stranded DNA-binding protein; afdb-uniprot50 \| AF-A0A4R7QAL6-F1-MODEL\_V4 \| 1.0 \| 1.909e-13 \| 618 \| 0.46 \| 115 \| 59 \| 1 \| 4 \| 118 \| 1 \| 112 \| Single-stranded DNA-binding protein \| Single-stranded DNA-binding protein; afdb-uniprot50 \| AF-S7UTG7-F1-MODEL\_V4 \| 1.0 \| 3.032e-16 \| 618 \| 0.446 \| 177 \| 83 \| 5 \| 1 \| 162 \| 1 \| 177 \| Single-stranded DNA-binding protein \| Single-stranded DNA-binding protein; afdb-uniprot50 \| AF-A0A7V8JJS2-F1-MODEL\_V4 \| 1.0 \| 1.987e-15 \| 617 \| 0.421 \| 173 \| 86 \| 3 \| 1 \| 162 \| 1 \| 170 \| Single-stranded DNA-binding protein \| Single-stranded DNA-binding protein; afdb-uniprot50 \| AF-A0A2G6T2T9-F1-MODEL\_V4 \| 1.0 \| 3.032e-16 \| 617 \| 0.514 \| 169 \| 69 \| 4 \| 1 \| 162 \| 1 \| 163 \| Single-stranded DNA-binding protein \| Single-stranded DNA-binding protein; afdb-uniprot50 \| AF-A0A3M8FH74-F1-MODEL\_V4 \| 1.0 \| 1.973e-16 \| 617 \| 0.444 \| 162 \| 85 \| 3 \| 1 \| 162 \| 1 \| 157 \| Single-stranded DNA-binding protein \| Single-stranded DNA-binding protein; afdb-uniprot50 \| AF-A0A369ZFS8-F1-MODEL\_V4 \| 1.0 \| 8.938e-15 \| 616 \| 0.531 \| 126 \| 53 \| 3 \| 4 \| 129 \| 2 \| 121 \| Single-stranded DNA-binding protein \| Single-stranded DNA-binding protein; afdb-uniprot50 \| AF-A0A1B8QYT8-F1-MODEL\_V4 \| 1.0 \| 7.972e-16 \| 616 \| 0.518 \| 162 \| 59 \| 6 \| 1 \| 162 \| 1 \| 143 \| Single-stranded DNA-binding protein \| Single-stranded DNA-binding protein; afdb-uniprot50 \| AF-H0Q1C2-F1-MODEL\_V4 \| 1.0 \| 1.439e-15 \| 616 \| 0.487 \| 162 \| 72 \| 4 \| 1 \| 162 \| 1 \| 151 \| Single-stranded DNA-binding protein \| Single-stranded DNA-binding protein; afdb-uniprot50 \| AF-A0A315BKE1-F1-MODEL\_V4 \| 1.0 \| 3.758e-16 \| 616 \| 0.509 \| 161 \| 72 \| 4 \| 4 \| 162 \| 1 \| 156 \| Single-stranded DNA-binding protein \| Single-stranded DNA-binding protein; afdb-uniprot50 \| AF-A0A2N1I3V0-F1-MODEL\_V4 \| 1.0 \| 2.197e-16 \| 616 \| 0.48 \| 179 \| 72 \| 6 \| 1 \| 162 \| 2 \| 176 \| Single-stranded DNA-binding protein \| Single-stranded DNA-binding protein; afdb-uniprot50 \| AF-A0A523Q0M1-F1-MODEL\_V4 \| 1.0 \| 8.139e-13 \| 615 \| 0.41 \| 117 \| 66 \| 1 \| 1 \| 117 \| 1 \| 114 \| Single-stranded DNA-binding protein \| Single-stranded DNA-binding protein; afdb-uniprot50 \| AF-A0A2A4T3P5-F1-MODEL\_V4 \| 1.0 \| 3.447e-13 \| 615 \| 0.435 \| 117 \| 63 \| 1 \| 1 \| 117 \| 1 \| 114 \| Single-stranded DNA-binding protein \| Single-stranded DNA-binding protein; afdb-uniprot50 \| AF-A0A418WRR6-F1-MODEL\_V4 \| 1.0 \| 5.187e-16 \| 615 \| 0.511 \| 174 \| 72 \| 5 \| 1 \| 162 \| 22 \| 194 \| Single-stranded DNA-binding protein \| Single-stranded DNA-binding protein; afdb-uniprot50 \| AF-A0A318L053-F1-MODEL\_V4 \| 1.0 \| 3.562e-16 \| 615 \| 0.49 \| 163 \| 76 \| 3 \| 1 \| 162 \| 1 \| 157 \| Single-stranded DNA-binding protein \| Single-stranded DNA-binding protein; afdb-uniprot50 \| AF-A4JUL0-F1-MODEL\_V4 \| 1.0 \| 4.916e-16 \| 615 \| 0.398 \| 188 \| 84 \| 3 \| 1 \| 162 \| 10 \| 194 \| Single-stranded DNA-binding protein \| Single-stranded DNA-binding protein; afdb-uniprot50 \| AF-A0A6N4TA35-F1-MODEL\_V4 \| 1.0 \| 1.883e-15 \| 614 \| 0.407 \| 162 \| 79 \| 4 \| 1 \| 162 \| 1 \| 145 \| Single-stranded DNA-binding protein \| Single-stranded DNA-binding protein; afdb-uniprot50 \| AF-A0A7W0C683-F1-MODEL\_V4 \| 1.0 \| 3.399e-15 \| 614 \| 0.456 \| 162 \| 73 \| 4 \| 1 \| 162 \| 1 \| 147 \| Single-stranded DNA-binding protein \| Single-stranded DNA-binding protein; afdb-uniprot50 \| AF-A0A6N7CZF7-F1-MODEL\_V4 \| 1.0 \| 4.416e-16 \| 614 \| 0.487 \| 162 \| 65 \| 4 \| 1 \| 162 \| 1 \| 144 \| Single-stranded DNA-binding protein \| Single-stranded DNA-binding protein; afdb-uniprot50 \| AF-A0A496NF27-F1-MODEL\_V4 \| 1.0 \| 2.581e-16 \| 614 \| 0.493 \| 160 \| 71 \| 4 \| 1 \| 160 \| 1 \| 150 \| Single-stranded DNA-binding protein \| Single-stranded DNA-binding protein; afdb-uniprot50 \| AF-A0A3A5ANI7-F1-MODEL\_V4 \| 1.0 \| 1.116e-13 \| 613 \| 0.453 \| 119 \| 62 \| 2 \| 4 \| 122 \| 1 \| 116 \| Single-stranded DNA-binding protein \| Single-stranded DNA-binding protein; afdb-uniprot50 \| AF-A0A369ZBV2-F1-MODEL\_V4 \| 1.0 \| 2.334e-15 \| 613 \| 0.484 \| 163 \| 58 \| 6 \| 1 \| 162 \| 1 \| 138 \| Single-stranded DNA-binding protein \| Single-stranded DNA-binding protein; afdb-uniprot50 \| AF-A0A2G9C242-F1-MODEL\_V4 \| 1.0 \| 1.691e-15 \| 613 \| 0.452 \| 157 \| 76 \| 4 \| 1 \| 157 \| 1 \| 147 \| Single-stranded DNA-binding protein \| Single-stranded DNA-binding protein; afdb-uniprot50 \| AF-A0A424WBH3-F1-MODEL\_V4 \| 1.0 \| 1.1e-15 \| 613 \| 0.453 \| 163 \| 75 \| 4 \| 1 \| 162 \| 1 \| 150 \| Single-stranded DNA-binding protein \| Single-stranded DNA-binding protein; afdb-uniprot50 \| AF-A0A6N8BTZ9-F1-MODEL\_V4 \| 1.0 \| 7.16e-16 \| 613 \| 0.402 \| 164 \| 89 \| 4 \| 1 \| 162 \| 1 \| 157 \| Single-stranded DNA-binding protein \| Single-stranded DNA-binding protein; afdb-uniprot50 \| AF-A0A109WX80-F1-MODEL\_V4 \| 1.0 \| 5.474e-16 \| 613 \| 0.435 \| 179 \| 77 \| 4 \| 1 \| 162 \| 1 \| 172 \| Single-stranded DNA-binding protein \| Single-stranded DNA-binding protein; afdb-uniprot50 \| AF-A0A252ER09-F1-MODEL\_V4 \| 1.0 \| 1.161e-15 \| 613 \| 0.431 \| 167 \| 87 \| 2 \| 3 \| 162 \| 2 \| 167 \| Single-stranded DNA-binding protein \| Single-stranded DNA-binding protein; afdb-uniprot50 \| AF-G0AGI2-F1-MODEL\_V4 \| 1.0 \| 4.416e-16 \| 613 \| 0.476 \| 170 \| 74 \| 4 \| 2 \| 162 \| 27 \| 190 \| Single-stranded DNA-binding protein \| Single-stranded DNA-binding protein; afdb-uniprot50 \| AF-A0A2N5LXF7-F1-MODEL\_V4 \| 1.0 \| 4.916e-16 \| 613 \| 0.438 \| 162 \| 88 \| 2 \| 1 \| 162 \| 1 \| 159 \| Single-stranded DNA-binding protein \| Single-stranded DNA-binding protein; afdb-uniprot50 \| AF-E6V3D0-F1-MODEL\_V4 \| 1.0 \| 3.562e-16 \| 613 \| 0.432 \| 192 \| 75 \| 3 \| 2 \| 162 \| 42 \| 230 \| Single-stranded DNA-binding protein \| Single-stranded DNA-binding protein; afdb-uniprot50 \| AF-V2UYE3-F1-MODEL\_V4 \| 1.0 \| 1.169e-14 \| 612 \| 0.5 \| 136 \| 60 \| 3 \| 4 \| 138 \| 3 \| 131 \| Single-stranded DNA-binding protein \| Single-stranded DNA-binding protein; afdb-uniprot50 \| AF-I2E0G5-F1-MODEL\_V4 \| 1.0 \| 8.027e-15 \| 612 \| 0.46 \| 139 \| 72 \| 1 \| 5 \| 143 \| 1 \| 136 \| Single-stranded DNA-binding protein \| Single-stranded DNA-binding protein; afdb-uniprot50 \| AF-A0A543Q277-F1-MODEL\_V4 \| 1.0 \| 1.093e-16 \| 612 \| 0.505 \| 170 \| 70 \| 7 \| 1 \| 162 \| 1 \| 164 \| Single-stranded DNA-binding protein \| Single-stranded DNA-binding protein; afdb-uniprot50 \| AF-A0A2T5MJC2-F1-MODEL\_V4 \| 1.0 \| 3.562e-16 \| 612 \| 0.502 \| 175 \| 68 \| 6 \| 1 \| 162 \| 1 \| 169 \| Single-stranded DNA-binding protein \| Single-stranded DNA-binding protein; afdb-uniprot50 \| AF-A0A7T4USW3-F1-MODEL\_V4 \| 1.0 \| 1.87e-16 \| 612 \| 0.46 \| 191 \| 69 \| 7 \| 1 \| 162 \| 1 \| 186 \| Single-stranded DNA-binding protein \| Single-stranded DNA-binding protein; afdb-uniprot50 \| AF-A0A661CUE1-F1-MODEL\_V4 \| 1.0 \| 1.364e-15 \| 611 \| 0.477 \| 159 \| 70 \| 5 \| 4 \| 162 \| 2 \| 147 \| Single-stranded DNA-binding protein \| Single-stranded DNA-binding protein; afdb-uniprot50 \| AF-A0A524BBB9-F1-MODEL\_V4 \| 1.0 \| 9.883e-16 \| 611 \| 0.484 \| 165 \| 71 \| 5 \| 1 \| 162 \| 1 \| 154 \| Single-stranded DNA-binding protein \| Single-stranded DNA-binding protein; afdb-uniprot50 \| AF-A0A259DP20-F1-MODEL\_V4 \| 1.0 \| 2.893e-15 \| 610 \| 0.427 \| 159 \| 74 \| 4 \| 4 \| 162 \| 2 \| 143 \| Single-stranded DNA-binding protein \| Single-stranded DNA-binding protein; afdb-uniprot50 \| AF-A0A450Z8W8-F1-MODEL\_V4 \| 1.0 \| 6.094e-16 \| 610 \| 0.463 \| 166 \| 78 \| 4 \| 1 \| 162 \| 1 \| 159 \| Single-stranded DNA-binding protein \| Single-stranded DNA-binding protein; afdb-uniprot50 \| AF-A0A7V8I1N6-F1-MODEL\_V4 \| 1.0 \| 4.185e-16 \| 610 \| 0.443 \| 185 \| 74 \| 7 \| 1 \| 162 \| 1 \| 179 \| Single-stranded DNA-binding protein \| Single-stranded DNA-binding protein; afdb-uniprot50 \| AF-A0A654KG85-F1-MODEL\_V4 \| 1.0 \| 2.934e-13 \| 609 \| 0.487 \| 121 \| 58 \| 2 \| 1 \| 121 \| 1 \| 117 \| Single-stranded DNA-binding protein \| Single-stranded DNA-binding protein; afdb-uniprot50 \| AF-A0A496W424-F1-MODEL\_V4 \| 1.0 \| 1.284e-16 \| 609 \| 0.487 \| 164 \| 76 \| 6 \| 1 \| 162 \| 1 \| 158 \| Single-stranded DNA-binding protein \| Single-stranded DNA-binding protein; afdb-uniprot50 \| AF-A0A157KWT1-F1-MODEL\_V4 \| 1.0 \| 1.896e-14 \| 608 \| 0.42 \| 157 \| 70 \| 5 \| 1 \| 156 \| 1 \| 137 \| Single-stranded DNA-binding protein \| Single-stranded DNA-binding protein; afdb-uniprot50 \| AF-A0A3D0DYT0-F1-MODEL\_V4 \| 1.0 \| 3.785e-15 \| 608 \| 0.457 \| 153 \| 70 \| 4 \| 1 \| 153 \| 1 \| 140 \| Single-stranded DNA-binding protein \| Single-stranded DNA-binding protein; afdb-uniprot50 \| AF-V4N8N1-F1-MODEL\_V4 \| 1.0 \| 1.797e-14 \| 608 \| 0.47 \| 136 \| 64 \| 3 \| 1 \| 136 \| 1 \| 128 \| Single-stranded DNA-binding protein \| Single-stranded DNA-binding protein; afdb-uniprot50 \| AF-A0A8B4QQV6-F1-MODEL\_V4 \| 1.0 \| 5.897e-13 \| 608 \| 0.407 \| 113 \| 64 \| 1 \| 1 \| 113 \| 1 \| 110 \| Helix-destabilizing protein \| Helix-destabilizing protein; afdb-uniprot50 \| AF-A0A6S6RZC4-F1-MODEL\_V4 \| 1.0 \| 2.082e-16 \| 608 \| 0.419 \| 162 \| 93 \| 1 \| 1 \| 162 \| 1 \| 161 \| Single-stranded DNA-binding protein \| Single-stranded DNA-binding protein; afdb-uniprot50 \| AF-A0A0F9C7M3-F1-MODEL\_V4 \| 1.0 \| 1.293e-15 \| 608 \| 0.397 \| 156 \| 85 \| 3 \| 1 \| 156 \| 1 \| 147 \| Uncharacterized protein \| Uncharacterized protein; afdb-uniprot50 \| AF-A0A1H6F8Q9-F1-MODEL\_V4 \| 1.0 \| 2.48e-14 \| 607 \| 0.456 \| 160 \| 62 \| 3 \| 3 \| 162 \| 2 \| 136 \| Single-stranded DNA-binding protein \| Single-stranded DNA-binding protein; afdb-uniprot50 \| AF-A0A2E1QNC3-F1-MODEL\_V4 \| 1.0 \| 1.108e-14 \| 607 \| 0.432 \| 164 \| 66 \| 6 \| 1 \| 162 \| 1 \| 139 \| Single-stranded DNA-binding protein \| Single-stranded DNA-binding protein; afdb-uniprot50 \| AF-G2Z404-F1-MODEL\_V4 \| 1.0 \| 2.934e-13 \| 607 \| 0.408 \| 120 \| 68 \| 1 \| 1 \| 120 \| 1 \| 117 \| Single-stranded DNA-binding protein \| Single-stranded DNA-binding protein; afdb-uniprot50 \| AF-A0A1V3KBP0-F1-MODEL\_V4 \| 1.0 \| 1.161e-15 \| 607 \| 0.49 \| 163 \| 63 \| 6 \| 1 \| 162 \| 1 \| 144 \| Single-stranded DNA-binding protein \| Single-stranded DNA-binding protein; afdb-uniprot50 \| AF-A0A383D837-F1-MODEL\_V4 \| 1.0 \| 4.446e-15 \| 607 \| 0.429 \| 163 \| 74 \| 4 \| 1 \| 162 \| 2 \| 146 \| Uncharacterized protein \| Uncharacterized protein; afdb-uniprot50 \| AF-A0A101GB98-F1-MODEL\_V4 \| 1.0 \| 2.096e-15 \| 607 \| 0.432 \| 162 \| 85 \| 2 \| 1 \| 162 \| 2 \| 156 \| Single-stranded DNA-binding protein \| Single-stranded DNA-binding protein; afdb-uniprot50 \| AF-A0A1D9CYH9-F1-MODEL\_V4 \| 1.0 \| 1.883e-15 \| 607 \| 0.481 \| 162 \| 76 \| 2 \| 4 \| 160 \| 3 \| 161 \| Single-stranded DNA-binding protein \| Single-stranded DNA-binding protein; afdb-uniprot50 \| AF-A0A151FIX3-F1-MODEL\_V4 \| 1.0 \| 7.972e-16 \| 606 \| 0.457 \| 177 \| 76 \| 6 \| 1 \| 162 \| 1 \| 172 \| Single-stranded DNA-binding protein \| Single-stranded DNA-binding protein; afdb-uniprot50 \| AF-A0A0S8BBA7-F1-MODEL\_V4 \| 1.0 \| 9.366e-16 \| 605 \| 0.46 \| 165 \| 71 \| 7 \| 1 \| 162 \| 1 \| 150 \| Single-stranded DNA-binding protein \| Single-stranded DNA-binding protein; afdb-uniprot50 \| AF-A0A345P7X1-F1-MODEL\_V4 \| 1.0 \| 7.16e-16 \| 605 \| 0.459 \| 159 \| 83 \| 2 \| 4 \| 162 \| 3 \| 158 \| Single-stranded DNA-binding protein \| Single-stranded DNA-binding protein; afdb-uniprot50 \| AF-A0A5P2H9B6-F1-MODEL\_V4 \| 1.0 \| 9.301e-17 \| 605 \| 0.491 \| 175 \| 74 \| 4 \| 1 \| 162 \| 1 \| 173 \| Single-stranded DNA-binding protein \| Single-stranded DNA-binding protein; afdb-uniprot50 \| AF-A0A2D9I903-F1-MODEL\_V4 \| 1.0 \| 3.447e-13 \| 604 \| 0.458 \| 120 \| 62 \| 1 \| 1 \| 120 \| 1 \| 117 \| Single-stranded DNA-binding protein \| Single-stranded DNA-binding protein; afdb-uniprot50 \| AF-A0A2P5L9K4-F1-MODEL\_V4 \| 1.0 \| 7.21e-15 \| 604 \| 0.427 \| 159 \| 72 \| 3 \| 4 \| 162 \| 1 \| 140 \| Single-stranded DNA-binding protein \| Single-stranded DNA-binding protein; afdb-uniprot50 \| AF-A0A5N3PH37-F1-MODEL\_V4 \| 1.0 \| 1.161e-15 \| 604 \| 0.485 \| 173 \| 77 \| 5 \| 1 \| 162 \| 1 \| 172 \| Single-stranded DNA-binding protein \| Single-stranded DNA-binding protein; afdb-uniprot50 \| AF-A0A1R4EI52-F1-MODEL\_V4 \| 1.0 \| 6.094e-16 \| 604 \| 0.512 \| 162 \| 73 \| 2 \| 4 \| 162 \| 3 \| 161 \| Single-stranded DNA-binding protein \| Single-stranded DNA-binding protein; afdb-uniprot50 \| AF-A0A7W3VKR4-F1-MODEL\_V4 \| 1.0 \| 8.589e-13 \| 603 \| 0.371 \| 113 \| 68 \| 1 \| 1 \| 113 \| 1 \| 110 \| Single-stranded DNA-binding protein \| Single-stranded DNA-binding protein; afdb-uniprot50 \| AF-A0A011Q4A3-F1-MODEL\_V4 \| 1.0 \| 3.587e-15 \| 603 \| 0.45 \| 151 \| 78 \| 2 \| 4 \| 154 \| 3 \| 148 \| Single-stranded DNA-binding protein \| Single-stranded DNA-binding protein; afdb-uniprot50 \| AF-A0A378T3Z6-F1-MODEL\_V4 \| 1.0 \| 2.446e-16 \| 603 \| 0.5 \| 166 \| 73 \| 5 \| 3 \| 162 \| 2 \| 163 \| Single-stranded DNA-binding protein \| Single-stranded DNA-binding protein; afdb-uniprot50 \| AF-A0A3D0SLK3-F1-MODEL\_V4 \| 1.0 \| 4.185e-16 \| 603 \| 0.467 \| 171 \| 77 \| 6 \| 1 \| 162 \| 1 \| 166 \| Single-stranded DNA-binding protein \| Single-stranded DNA-binding protein; afdb-uniprot50 \| AF-A0A0Q9PKG3-F1-MODEL\_V4 \| 1.0 \| 1.093e-16 \| 602 \| 0.466 \| 178 \| 78 \| 4 \| 1 \| 162 \| 1 \| 177 \| Single-stranded DNA-binding protein \| Single-stranded DNA-binding protein; afdb-uniprot50 \| AF-A0A2D3R7S5-F1-MODEL\_V4 \| 1.0 \| 3.993e-15 \| 602 \| 0.452 \| 157 \| 78 \| 3 \| 1 \| 157 \| 1 \| 149 \| Single-stranded DNA-binding protein \| Single-stranded DNA-binding protein; afdb-uniprot50 \| AF-K0MA94-F1-MODEL\_V4 \| 1.0 \| 8.412e-16 \| 602 \| 0.456 \| 173 \| 75 \| 4 \| 2 \| 162 \| 22 \| 187 \| Single-stranded DNA-binding protein \| Single-stranded DNA-binding protein; afdb-uniprot50 \| AF-A0A2V8KTP6-F1-MODEL\_V4 \| 1.0 \| 4.692e-15 \| 601 \| 0.423 \| 163 \| 80 \| 4 \| 1 \| 162 \| 1 \| 150 \| Single-stranded DNA-binding protein \| Single-stranded DNA-binding protein; afdb-uniprot50 \| AF-A0A522RZG9-F1-MODEL\_V4 \| 1.0 \| 1.161e-15 \| 601 \| 0.403 \| 186 \| 81 \| 4 \| 1 \| 162 \| 1 \| 180 \| Single-stranded DNA-binding protein \| Single-stranded DNA-binding protein; afdb-uniprot50 \| AF-A0A523HF25-F1-MODEL\_V4 \| 1.0 \| 9.062e-13 \| 600 \| 0.431 \| 116 \| 61 \| 2 \| 3 \| 117 \| 2 \| 113 \| Single-stranded DNA-binding protein \| Single-stranded DNA-binding protein; afdb-uniprot50 \| AF-A0A411SZB9-F1-MODEL\_V4 \| 1.0 \| 4.446e-15 \| 600 \| 0.447 \| 163 \| 77 \| 4 \| 1 \| 162 \| 1 \| 151 \| Single-stranded DNA-binding protein \| Single-stranded DNA-binding protein; afdb-uniprot50 \| AF-A0A238H9F9-F1-MODEL\_V4 \| 1.0 \| 2.318e-16 \| 600 \| 0.45 \| 173 \| 81 \| 4 \| 2 \| 162 \| 24 \| 194 \| Single-stranded DNA-binding protein \| Single-stranded DNA-binding protein; afdb-uniprot50 \| AF-A0A2S5CMA6-F1-MODEL\_V4 \| 1.0 \| 1.161e-15 \| 600 \| 0.45 \| 173 \| 78 \| 3 \| 2 \| 162 \| 46 \| 213 \| Single-stranded DNA-binding protein \| Single-stranded DNA-binding protein; afdb-uniprot50 \| AF-A0A4R6QQT1-F1-MODEL\_V4 \| 1.0 \| 1.043e-15 \| 600 \| 0.425 \| 188 \| 75 \| 4 \| 1 \| 162 \| 35 \| 215 \| Single-stranded DNA-binding protein \| Single-stranded DNA-binding protein; afdb-uniprot50 \| AF-A0A4R6DSH9-F1-MODEL\_V4 \| 1.0 \| 9.496e-14 \| 599 \| 0.457 \| 129 \| 63 \| 3 \| 1 \| 129 \| 1 \| 122 \| Single-stranded DNA-binding protein \| Single-stranded DNA-binding protein; afdb-uniprot50 \| AF-A0A1V3L736-F1-MODEL\_V4 \| 1.0 \| 9.366e-16 \| 599 \| 0.476 \| 170 \| 64 \| 6 \| 1 \| 162 \| 1 \| 153 \| Single-stranded DNA-binding protein \| Single-stranded DNA-binding protein; afdb-uniprot50 \| AF-A0A0G3ID38-F1-MODEL\_V4 \| 1.0 \| 7.555e-16 \| 599 \| 0.45 \| 171 \| 79 \| 5 \| 1 \| 162 \| 1 \| 165 \| Single-stranded DNA-binding protein \| Single-stranded DNA-binding protein; afdb-uniprot50 \| AF-A0A3N9EZF1-F1-MODEL\_V4 \| 1.0 \| 8.412e-16 \| 599 \| 0.441 \| 170 \| 83 \| 4 \| 1 \| 162 \| 1 \| 166 \| Single-stranded DNA-binding protein \| Single-stranded DNA-binding protein; afdb-uniprot50 \| AF-A0A833GPH7-F1-MODEL\_V4 \| 1.0 \| 1.87e-16 \| 599 \| 0.5 \| 166 \| 74 \| 5 \| 1 \| 162 \| 1 \| 161 \| Single-stranded DNA-binding protein \| Single-stranded DNA-binding protein; afdb-uniprot50 \| AF-S7TLS7-F1-MODEL\_V4 \| 1.0 \| 2.197e-16 \| 599 \| 0.479 \| 167 \| 76 \| 5 \| 1 \| 162 \| 1 \| 161 \| Single-stranded DNA-binding protein \| Single-stranded DNA-binding protein; afdb-uniprot50 \| AF-A0A5S4SIA5-F1-MODEL\_V4 \| 1.0 \| 2.35e-14 \| 598 \| 0.493 \| 146 \| 60 \| 4 \| 1 \| 145 \| 1 \| 133 \| Single-stranded DNA-binding protein \| Single-stranded DNA-binding protein; afdb-uniprot50 \| AF-A0A2E5IAI4-F1-MODEL\_V4 \| 1.0 \| 5.224e-15 \| 598 \| 0.417 \| 163 \| 79 \| 5 \| 1 \| 162 \| 1 \| 148 \| Single-stranded DNA-binding protein \| Single-stranded DNA-binding protein; afdb-uniprot50 \| AF-A0A2D9CGL3-F1-MODEL\_V4 \| 1.0 \| 2.873e-16 \| 598 \| 0.5 \| 162 \| 74 \| 3 \| 1 \| 159 \| 1 \| 158 \| Single-stranded DNA-binding protein \| Single-stranded DNA-binding protein; afdb-uniprot50 \| AF-A0A2W5QBP7-F1-MODEL\_V4 \| 1.0 \| 1.32e-12 \| 598 \| 0.347 \| 118 \| 76 \| 1 \| 1 \| 117 \| 157 \| 274 \| Single-stranded DNA-binding protein \| Single-stranded DNA-binding protein; afdb-uniprot50 \| AF-A0A0A1H6Y8-F1-MODEL\_V4 \| 1.0 \| 9.951e-15 \| 597 \| 0.447 \| 161 \| 65 \| 5 \| 3 \| 162 \| 2 \| 139 \| Single-stranded DNA-binding protein \| Single-stranded DNA-binding protein; afdb-uniprot50 \| AF-A0A5C7NPC6-F1-MODEL\_V4 \| 1.0 \| 8.027e-15 \| 597 \| 0.425 \| 160 \| 71 \| 4 \| 3 \| 162 \| 2 \| 140 \| Single-stranded DNA-binding protein \| Single-stranded DNA-binding protein; afdb-uniprot50 \| AF-A0A4Q7B5T0-F1-MODEL\_V4 \| 1.0 \| 4.049e-13 \| 596 \| 0.532 \| 107 \| 49 \| 1 \| 4 \| 110 \| 3 \| 108 \| Single-stranded DNA-binding protein \| Single-stranded DNA-binding protein; afdb-uniprot50 \| AF-A0A2N2Y9Q4-F1-MODEL\_V4 \| 1.0 \| 9.062e-13 \| 596 \| 0.435 \| 117 \| 63 \| 1 \| 1 \| 117 \| 1 \| 114 \| Single-stranded DNA-binding protein \| Single-stranded DNA-binding protein; afdb-uniprot50 \| AF-A0A318INE9-F1-MODEL\_V4 \| 1.0 \| 3.587e-15 \| 596 \| 0.445 \| 164 \| 68 \| 6 \| 1 \| 162 \| 1 \| 143 \| Single-stranded DNA-binding protein \| Single-stranded DNA-binding protein; afdb-uniprot50 \| AF-A0A6S6M805-F1-MODEL\_V4 \| 1.0 \| 1.169e-14 \| 596 \| 0.415 \| 159 \| 89 \| 2 \| 4 \| 162 \| 1 \| 155 \| Single-stranded DNA-binding protein \| Single-stranded DNA-binding protein; afdb-uniprot50 \| AF-A0A4Q4WRP0-F1-MODEL\_V4 \| 1.0 \| 4.477e-14 \| 596 \| 0.468 \| 126 \| 65 \| 2 \| 2 \| 127 \| 112 \| 235 \| Uncharacterized protein \| Uncharacterized protein; afdb-uniprot50 \| AF-A0A3D3HIB1-F1-MODEL\_V4 \| 1.0 \| 5.296e-13 \| 595 \| 0.367 \| 117 \| 71 \| 1 \| 1 \| 117 \| 1 \| 114 \| Single-stranded DNA-binding protein \| Single-stranded DNA-binding protein; afdb-uniprot50 \| AF-A0A1X7C379-F1-MODEL\_V4 \| 1.0 \| 7.661e-14 \| 595 \| 0.507 \| 130 \| 59 \| 3 \| 1 \| 130 \| 1 \| 125 \| Single-stranded DNA-binding protein \| Single-stranded DNA-binding protein; afdb-uniprot50 \| AF-A0A653A015-F1-MODEL\_V4 \| 1.0 \| 3.587e-15 \| 595 \| 0.463 \| 164 \| 62 \| 7 \| 1 \| 162 \| 1 \| 140 \| Single-stranded DNA-binding protein \| Single-stranded DNA-binding protein; afdb-uniprot50 \| AF-A0A2N0WIC8-F1-MODEL\_V4 \| 1.0 \| 4.416e-16 \| 595 \| 0.49 \| 165 \| 72 \| 4 \| 4 \| 160 \| 3 \| 163 \| Single-stranded DNA-binding protein \| Single-stranded DNA-binding protein; afdb-uniprot50 \| AF-A0A1Z4C5J3-F1-MODEL\_V4 \| 1.0 \| 5.776e-16 \| 595 \| 0.467 \| 169 \| 77 \| 3 \| 4 \| 162 \| 1 \| 166 \| Single-stranded DNA-binding protein \| Single-stranded DNA-binding protein; afdb-uniprot50 \| AF-A0A660N4U4-F1-MODEL\_V4 \| 1.0 \| 6.833e-15 \| 594 \| 0.443 \| 142 \| 74 \| 3 \| 4 \| 143 \| 2 \| 140 \| Single-stranded DNA-binding protein \| Single-stranded DNA-binding protein; afdb-uniprot50 \| AF-A0A536S686-F1-MODEL\_V4 \| 1.0 \| 6.566e-13 \| 594 \| 0.442 \| 113 \| 60 \| 1 \| 2 \| 114 \| 36 \| 145 \| Single-stranded DNA-binding protein \| Single-stranded DNA-binding protein; afdb-uniprot50 \| AF-A0A5J6PZX2-F1-MODEL\_V4 \| 1.0 \| 4.659e-16 \| 594 \| 0.444 \| 171 \| 81 \| 4 \| 3 \| 162 \| 2 \| 169 \| Single-stranded DNA-binding protein \| Single-stranded DNA-binding protein; afdb-uniprot50 \| AF-G4E3E4-F1-MODEL\_V4 \| 1.0 \| 6.786e-16 \| 594 \| 0.423 \| 170 \| 86 \| 4 \| 1 \| 162 \| 1 \| 166 \| Single-stranded DNA-binding protein \| Single-stranded DNA-binding protein; afdb-uniprot50 \| AF-A0A1V4GVS6-F1-MODEL\_V4 \| 1.0 \| 6.786e-16 \| 593 \| 0.494 \| 170 \| 72 \| 5 \| 1 \| 160 \| 1 \| 166 \| Single-stranded DNA-binding protein \| Single-stranded DNA-binding protein; afdb-uniprot50 \| AF-A0A2D5RWR9-F1-MODEL\_V4 \| 1.0 \| 5.474e-16 \| 592 \| 0.469 \| 162 \| 79 \| 3 \| 1 \| 158 \| 1 \| 159 \| Single-stranded DNA-binding protein \| Single-stranded DNA-binding protein; afdb-uniprot50 \| AF-A0A1V1X1A7-F1-MODEL\_V4 \| 1.0 \| 1.529e-14 \| 591 \| 0.421 \| 159 \| 73 \| 4 \| 4 \| 162 \| 1 \| 140 \| Single-stranded DNA-binding protein \| Single-stranded DNA-binding protein; afdb-uniprot50 \| AF-A0A645DBS0-F1-MODEL\_V4 \| 1.0 \| 4.446e-15 \| 591 \| 0.395 \| 162 \| 88 \| 3 \| 1 \| 162 \| 2 \| 153 \| Single-stranded DNA-binding protein \| Single-stranded DNA-binding protein; afdb-uniprot50 \| AF-F5T0V4-F1-MODEL\_V4 \| 1.0 \| 5.224e-15 \| 590 \| 0.479 \| 148 \| 70 \| 3 \| 4 \| 148 \| 2 \| 145 \| Single-stranded DNA-binding protein \| Single-stranded DNA-binding protein; afdb-uniprot50 \| AF-A0A6B1AVA2-F1-MODEL\_V4 \| 1.0 \| 1.784e-15 \| 590 \| 0.447 \| 161 \| 81 \| 4 \| 1 \| 160 \| 1 \| 154 \| Single-stranded DNA-binding protein \| Single-stranded DNA-binding protein; afdb-uniprot50 \| AF-A0A7W8HGC3-F1-MODEL\_V4 \| 1.0 \| 1.987e-15 \| 590 \| 0.437 \| 167 \| 84 \| 5 \| 1 \| 162 \| 1 \| 162 \| Single-stranded DNA-binding protein \| Single-stranded DNA-binding protein; afdb-uniprot50 \| AF-A0A7X4KLH9-F1-MODEL\_V4 \| 1.0 \| 1.519e-15 \| 589 \| 0.43 \| 179 \| 72 \| 4 \| 1 \| 162 \| 1 \| 166 \| Single-stranded DNA-binding protein \| Single-stranded DNA-binding protein; afdb-uniprot50 \| AF-A0A7V2HAB0-F1-MODEL\_V4 \| 1.0 \| 1.393e-12 \| 588 \| 0.376 \| 117 \| 70 \| 1 \| 4 \| 120 \| 2 \| 115 \| Single-stranded DNA-binding protein \| Single-stranded DNA-binding protein; afdb-uniprot50 \| AF-A0A3N5PE79-F1-MODEL\_V4 \| 1.0 \| 4.021e-14 \| 588 \| 0.477 \| 134 \| 61 \| 4 \| 1 \| 129 \| 1 \| 130 \| Single-stranded DNA-binding protein \| Single-stranded DNA-binding protein; afdb-uniprot50 \| AF-A0A1D2RY17-F1-MODEL\_V4 \| 1.0 \| 1.169e-14 \| 588 \| 0.413 \| 162 \| 78 \| 3 \| 1 \| 162 \| 1 \| 145 \| Single-stranded DNA-binding protein \| Single-stranded DNA-binding protein; afdb-uniprot50 \| AF-A1VVC7-F1-MODEL\_V4 \| 1.0 \| 3.758e-16 \| 588 \| 0.479 \| 167 \| 80 \| 3 \| 1 \| 162 \| 1 \| 165 \| Single-stranded DNA-binding protein \| Single-stranded DNA-binding protein; afdb-uniprot50 \| AF-E8LIW7-F1-MODEL\_V4 \| 1.0 \| 1.896e-14 \| 588 \| 0.4 \| 155 \| 89 \| 2 \| 1 \| 154 \| 1 \| 152 \| Single-stranded DNA-binding protein \| Single-stranded DNA-binding protein; afdb-uniprot50 \| AF-A0A7Y6YA81-F1-MODEL\_V4 \| 1.0 \| 4.243e-14 \| 587 \| 0.456 \| 162 \| 61 \| 4 \| 1 \| 162 \| 1 \| 135 \| Single-stranded DNA-binding protein \| Single-stranded DNA-binding protein; afdb-uniprot50 \| AF-A0A6A0IKH7-F1-MODEL\_V4 \| 1.0 \| 4.95e-15 \| 587 \| 0.429 \| 163 \| 69 \| 6 \| 1 \| 162 \| 1 \| 140 \| Single-stranded DNA-binding protein \| Single-stranded DNA-binding protein; afdb-uniprot50 \| AF-A0A3A5L1K9-F1-MODEL\_V4 \| 1.0 \| 2.096e-15 \| 587 \| 0.456 \| 171 \| 79 \| 6 \| 1 \| 162 \| 1 \| 166 \| Single-stranded DNA-binding protein \| Single-stranded DNA-binding protein; afdb-uniprot50 \| AF-A0A547EAB1-F1-MODEL\_V4 \| 1.0 \| 2.212e-15 \| 587 \| 0.443 \| 158 \| 71 \| 4 \| 1 \| 157 \| 1 \| 142 \| Single-stranded DNA-binding protein \| Single-stranded DNA-binding protein; afdb-uniprot50 \| AF-A0A523H4Y8-F1-MODEL\_V4 \| 1.0 \| 3.266e-13 \| 586 \| 0.428 \| 133 \| 71 \| 2 \| 1 \| 133 \| 1 \| 128 \| Single-stranded DNA-binding protein \| Single-stranded DNA-binding protein; afdb-uniprot50 \| AF-A0A1N7DQ25-F1-MODEL\_V4 \| 1.0 \| 1.883e-15 \| 586 \| 0.417 \| 163 \| 85 \| 3 \| 4 \| 162 \| 2 \| 158 \| Single-stranded DNA-binding protein \| Single-stranded DNA-binding protein; afdb-uniprot50 \| AF-V4P5D9-F1-MODEL\_V4 \| 1.0 \| 4.446e-15 \| 586 \| 0.53 \| 164 \| 71 \| 6 \| 1 \| 162 \| 1 \| 160 \| Single-stranded DNA-binding protein \| Single-stranded DNA-binding protein; afdb-uniprot50 \| AF-A0A179BNQ8-F1-MODEL\_V4 \| 1.0 \| 6.786e-16 \| 586 \| 0.491 \| 169 \| 73 \| 6 \| 1 \| 162 \| 1 \| 163 \| Single-stranded DNA-binding protein \| Single-stranded DNA-binding protein; afdb-uniprot50 \| AF-A0A7V0XHS0-F1-MODEL\_V4 \| 1.0 \| 1.987e-15 \| 586 \| 0.414 \| 169 \| 87 \| 4 \| 2 \| 162 \| 3 \| 167 \| Single-stranded DNA-binding protein \| Single-stranded DNA-binding protein; afdb-uniprot50 \| AF-V8G3X1-F1-MODEL\_V4 \| 1.0 \| 5.776e-16 \| 586 \| 0.456 \| 162 \| 78 \| 5 \| 1 \| 162 \| 1 \| 152 \| Single-stranded DNA-binding protein \| Single-stranded DNA-binding protein; afdb-uniprot50 \| AF-A0A1S9ZKL9-F1-MODEL\_V4 \| 1.0 \| 4.659e-16 \| 586 \| 0.493 \| 160 \| 72 \| 4 \| 4 \| 162 \| 2 \| 153 \| Single-stranded DNA-binding protein \| Single-stranded DNA-binding protein; afdb-uniprot50 \| AF-A0A3C0DYM7-F1-MODEL\_V4 \| 1.0 \| 6.88e-14 \| 585 \| 0.465 \| 131 \| 65 \| 3 \| 1 \| 131 \| 1 \| 126 \| Single-stranded DNA-binding protein \| Single-stranded DNA-binding protein; afdb-uniprot50 \| AF-A0A519KFE2-F1-MODEL\_V4 \| 1.0 \| 2.954e-12 \| 585 \| 0.424 \| 113 \| 62 \| 1 \| 1 \| 113 \| 1 \| 110 \| Single-stranded DNA-binding protein \| Single-stranded DNA-binding protein; afdb-uniprot50 \| AF-A0A7Y3IWH6-F1-MODEL\_V4 \| 1.0 \| 7.16e-16 \| 585 \| 0.47 \| 185 \| 62 \| 7 \| 4 \| 162 \| 3 \| 177 \| Single-stranded DNA-binding protein \| Single-stranded DNA-binding protein; afdb-uniprot50 \| AF-A0A1G2ATN0-F1-MODEL\_V4 \| 1.0 \| 3.095e-13 \| 584 \| 0.401 \| 117 \| 67 \| 2 \| 4 \| 120 \| 2 \| 115 \| Single-stranded DNA-binding protein \| Single-stranded DNA-binding protein; afdb-uniprot50 \| AF-A0A2S5QA54-F1-MODEL\_V4 \| 1.0 \| 4.692e-15 \| 584 \| 0.425 \| 162 \| 77 \| 5 \| 4 \| 162 \| 2 \| 150 \| Single-stranded DNA-binding protein \| Single-stranded DNA-binding protein; afdb-uniprot50 \| AF-A0A6L7MV45-F1-MODEL\_V4 \| 1.0 \| 9.883e-16 \| 584 \| 0.482 \| 168 \| 73 \| 5 \| 1 \| 162 \| 1 \| 160 \| Single-stranded DNA-binding protein \| Single-stranded DNA-binding protein; afdb-uniprot50 \| AF-A0A0T6W6Z2-F1-MODEL\_V4 \| 1.0 \| 5.512e-15 \| 584 \| 0.438 \| 162 \| 86 \| 2 \| 4 \| 162 \| 3 \| 162 \| Single-stranded DNA-binding protein \| Single-stranded DNA-binding protein; afdb-uniprot50 \| AF-A0A1G2BSC3-F1-MODEL\_V4 \| 1.0 \| 6.18e-14 \| 584 \| 0.402 \| 139 \| 76 \| 3 \| 1 \| 136 \| 38 \| 172 \| Single-stranded DNA-binding protein \| Single-stranded DNA-binding protein; afdb-uniprot50 \| AF-A0A7U3ZFE1-F1-MODEL\_V4 \| 1.0 \| 5.816e-15 \| 584 \| 0.404 \| 163 \| 91 \| 4 \| 2 \| 162 \| 15 \| 173 \| Single-stranded DNA-binding protein \| Single-stranded DNA-binding protein; afdb-uniprot50 \| AF-A0A6M8T455-F1-MODEL\_V4 \| 1.0 \| 5.856e-14 \| 583 \| 0.496 \| 125 \| 59 \| 2 \| 3 \| 127 \| 2 \| 122 \| Single-stranded DNA-binding protein \| Single-stranded DNA-binding protein; afdb-uniprot50 \| AF-A0A3M6R1S4-F1-MODEL\_V4 \| 1.0 \| 6.833e-15 \| 583 \| 0.481 \| 162 \| 60 \| 5 \| 1 \| 162 \| 1 \| 138 \| Single-stranded DNA-binding protein \| Single-stranded DNA-binding protein; afdb-uniprot50 \| AF-A0A7C1KV95-F1-MODEL\_V4 \| 1.0 \| 2.599e-15 \| 583 \| 0.366 \| 161 \| 97 \| 3 \| 4 \| 162 \| 2 \| 159 \| Single-stranded DNA-binding protein \| Single-stranded DNA-binding protein; afdb-uniprot50 \| AF-A0A6L4B9L0-F1-MODEL\_V4 \| 1.0 \| 9.366e-16 \| 583 \| 0.463 \| 192 \| 69 \| 5 \| 1 \| 162 \| 1 \| 188 \| Single-stranded DNA-binding protein \| Single-stranded DNA-binding protein; afdb-uniprot50 \| AF-A0A2W4U1S0-F1-MODEL\_V4 \| 1.0 \| 6.137e-15 \| 583 \| 0.45 \| 160 \| 81 \| 4 \| 4 \| 162 \| 1 \| 154 \| Single-stranded DNA-binding protein \| Single-stranded DNA-binding protein; afdb-uniprot50 \| AF-A0A1P8MPZ6-F1-MODEL\_V4 \| 1.0 \| 4.049e-13 \| 582 \| 0.42 \| 119 \| 62 \| 3 \| 1 \| 112 \| 2 \| 120 \| Single-stranded DNA-binding protein \| Single-stranded DNA-binding protein; afdb-uniprot50 \| AF-A0A661T5P7-F1-MODEL\_V4 \| 1.0 \| 8.47e-15 \| 582 \| 0.429 \| 163 \| 69 \| 6 \| 1 \| 162 \| 1 \| 140 \| Single-stranded DNA-binding protein \| Single-stranded DNA-binding protein; afdb-uniprot50 \| AF-A0A0M4SYI8-F1-MODEL\_V4 \| 1.0 \| 2.212e-15 \| 582 \| 0.452 \| 159 \| 79 \| 2 \| 4 \| 162 \| 3 \| 153 \| Single-stranded DNA-binding protein \| Single-stranded DNA-binding protein; afdb-uniprot50 \| AF-A0A011NB88-F1-MODEL\_V4 \| 1.0 \| 1.614e-14 \| 581 \| 0.425 \| 162 \| 74 \| 4 \| 5 \| 162 \| 1 \| 147 \| Single-stranded DNA-binding protein \| Single-stranded DNA-binding protein; afdb-uniprot50 \| AF-A0A1F3CA35-F1-MODEL\_V4 \| 1.0 \| 1.293e-15 \| 581 \| 0.425 \| 160 \| 82 \| 4 \| 4 \| 162 \| 1 \| 151 \| Single-stranded DNA-binding protein \| Single-stranded DNA-binding protein; afdb-uniprot50 \| AF-A0A6I2H1I0-F1-MODEL\_V4 \| 1.0 \| 2.318e-16 \| 581 \| 0.468 \| 173 \| 76 \| 6 \| 1 \| 162 \| 1 \| 168 \| Single-stranded DNA-binding protein \| Single-stranded DNA-binding protein; afdb-uniprot50 \| AF-A0A2E7E9P7-F1-MODEL\_V4 \| 1.0 \| 1.691e-15 \| 581 \| 0.442 \| 174 \| 81 \| 5 \| 1 \| 162 \| 1 \| 170 \| Single-stranded DNA-binding protein \| Single-stranded DNA-binding protein; afdb-uniprot50 \| AF-A0A4V1J3T3-F1-MODEL\_V4 \| 1.0 \| 6.52e-14 \| 580 \| 0.472 \| 129 \| 65 \| 3 \| 1 \| 129 \| 1 \| 126 \| Single-stranded DNA-binding protein \| Single-stranded DNA-binding protein; afdb-uniprot50 \| AF-A0A2G6LHZ8-F1-MODEL\_V4 \| 1.0 \| 8.412e-16 \| 580 \| 0.471 \| 178 \| 70 \| 5 \| 4 \| 162 \| 3 \| 175 \| Single-stranded DNA-binding protein \| Single-stranded DNA-binding protein; afdb-uniprot50 \| AF-A0A3C0C8R5-F1-MODEL\_V4 \| 1.0 \| 4.95e-15 \| 580 \| 0.386 \| 168 \| 94 \| 2 \| 1 \| 162 \| 1 \| 165 \| Single-stranded DNA-binding protein \| Single-stranded DNA-binding protein; afdb-uniprot50 \| AF-A0A349MDR0-F1-MODEL\_V4 \| 1.0 \| 6.52e-14 \| 579 \| 0.466 \| 135 \| 70 \| 2 \| 1 \| 135 \| 1 \| 133 \| Single-stranded DNA-binding protein \| Single-stranded DNA-binding protein; afdb-uniprot50 \| AF-A0A2X1LM92-F1-MODEL\_V4 \| 1.0 \| 2.35e-14 \| 579 \| 0.471 \| 142 \| 65 \| 3 \| 1 \| 142 \| 2 \| 133 \| Single-stranded DNA-binding protein \| Single-stranded DNA-binding protein; afdb-uniprot50 \| AF-A0A661IS78-F1-MODEL\_V4 \| 1.0 \| 8.47e-15 \| 579 \| 0.408 \| 159 \| 72 \| 6 \| 4 \| 162 \| 3 \| 139 \| Single-stranded DNA-binding protein \| Single-stranded DNA-binding protein; afdb-uniprot50 \| AF-A0A845GCZ9-F1-MODEL\_V4 \| 1.0 \| 6.137e-15 \| 579 \| 0.418 \| 160 \| 88 \| 4 \| 1 \| 160 \| 1 \| 155 \| Single-stranded DNA-binding protein \| Single-stranded DNA-binding protein; afdb-uniprot50 \| AF-A0A2D2LXU4-F1-MODEL\_V4 \| 1.0 \| 2.463e-15 \| 579 \| 0.456 \| 160 \| 78 \| 3 \| 4 \| 162 \| 3 \| 154 \| Single-stranded DNA-binding protein \| Single-stranded DNA-binding protein; afdb-uniprot50 \| AF-A0A535GRY2-F1-MODEL\_V4 \| 1.0 \| 9.562e-13 \| 578 \| 0.401 \| 117 \| 67 \| 1 \| 4 \| 120 \| 1 \| 114 \| Single-stranded DNA-binding protein \| Single-stranded DNA-binding protein; afdb-uniprot50 \| AF-A0A0M9TYI8-F1-MODEL\_V4 \| 1.0 \| 6.88e-14 \| 578 \| 0.427 \| 159 \| 68 \| 3 \| 4 \| 162 \| 1 \| 136 \| Single-stranded DNA-binding protein \| Single-stranded DNA-binding protein; afdb-uniprot50 \| AF-A0A383C8L9-F1-MODEL\_V4 \| 1.0 \| 9.431e-15 \| 578 \| 0.457 \| 164 \| 69 \| 7 \| 1 \| 162 \| 1 \| 146 \| Uncharacterized protein \| Uncharacterized protein; afdb-uniprot50 \| AF-A0A448TTN8-F1-MODEL\_V4 \| 1.0 \| 7.21e-15 \| 578 \| 0.472 \| 163 \| 69 \| 6 \| 1 \| 162 \| 1 \| 147 \| Single-stranded DNA-binding protein \| Single-stranded DNA-binding protein; afdb-uniprot50 \| AF-A0LIN0-F1-MODEL\_V4 \| 1.0 \| 1.519e-15 \| 578 \| 0.433 \| 173 \| 79 \| 6 \| 1 \| 162 \| 20 \| 184 \| Single-stranded DNA-binding protein \| Single-stranded DNA-binding protein; afdb-uniprot50 \| AF-A0A401FYS3-F1-MODEL\_V4 \| 1.0 \| 2e-14 \| 578 \| 0.515 \| 132 \| 51 \| 3 \| 1 \| 132 \| 1 \| 119 \| Single-stranded DNA-binding protein \| Single-stranded DNA-binding protein; afdb-uniprot50 \| AF-Q5I719-F1-MODEL\_V4 \| 1.0 \| 2e-14 \| 577 \| 0.415 \| 154 \| 79 \| 4 \| 1 \| 151 \| 1 \| 146 \| Single-stranded DNA-binding protein \| Single-stranded DNA-binding protein; afdb-uniprot50 \| AF-A0A2E8ZYW6-F1-MODEL\_V4 \| 1.0 \| 3.811e-14 \| 576 \| 0.383 \| 159 \| 78 \| 3 \| 4 \| 162 \| 2 \| 140 \| Single-stranded DNA-binding protein \| Single-stranded DNA-binding protein; afdb-uniprot50 \| AF-A0A654E3W4-F1-MODEL\_V4 \| 1.0 \| 5.512e-15 \| 576 \| 0.46 \| 163 \| 75 \| 6 \| 1 \| 162 \| 1 \| 151 \| Single-stranded DNA-binding protein \| Single-stranded DNA-binding protein; afdb-uniprot50 \| AF-A0A7X8GXX0-F1-MODEL\_V4 \| 1.0 \| 2.78e-13 \| 576 \| 0.448 \| 125 \| 63 \| 3 \| 1 \| 125 \| 1 \| 119 \| Single-stranded DNA-binding protein \| Single-stranded DNA-binding protein; afdb-uniprot50 \| AF-A0A2A2T4T3-F1-MODEL\_V4 \| 1.0 \| 1.449e-14 \| 574 \| 0.472 \| 161 \| 62 \| 6 \| 3 \| 162 \| 2 \| 140 \| Single-stranded DNA-binding protein \| Single-stranded DNA-binding protein; afdb-uniprot50 \| AF-B7J8U2-F1-MODEL\_V4 \| 1.0 \| 1.691e-15 \| 574 \| 0.466 \| 163 \| 73 \| 6 \| 1 \| 162 \| 1 \| 150 \| Single-stranded DNA-binding protein \| Single-stranded DNA-binding protein; afdb-uniprot50 \| AF-A0A448CRE5-F1-MODEL\_V4 \| 1.0 \| 7.21e-15 \| 574 \| 0.418 \| 160 \| 85 \| 3 \| 1 \| 157 \| 1 \| 155 \| Single-stranded DNA-binding protein \| Single-stranded DNA-binding protein; afdb-uniprot50 \| AF-A0A1A9X6E7-F1-MODEL\_V4 \| 1.0 \| 1.47e-12 \| 574 \| 0.637 \| 102 \| 37 \| 0 \| 16 \| 117 \| 433 \| 534 \| Replicative DNA helicase \| Replicative DNA helicase; afdb-uniprot50 \| AF-A0A812ILX3-F1-MODEL\_V4 \| 1.0 \| 2.742e-15 \| 573 \| 0.451 \| 177 \| 71 \| 6 \| 1 \| 162 \| 1 \| 166 \| Ssb protein \| Ssb protein; afdb-uniprot50 \| AF-A0A2U3KAX6-F1-MODEL\_V4 \| 1.0 \| 1.883e-15 \| 573 \| 0.419 \| 174 \| 83 \| 6 \| 1 \| 162 \| 1 \| 168 \| Single-stranded DNA-binding protein \| Single-stranded DNA-binding protein; afdb-uniprot50 \| AF-H6REC4-F1-MODEL\_V4 \| 1.0 \| 4.508e-13 \| 572 \| 0.4 \| 135 \| 73 \| 3 \| 1 \| 135 \| 1 \| 127 \| Single-stranded DNA-binding protein \| Single-stranded DNA-binding protein; afdb-uniprot50 \| AF-A0A424NW40-F1-MODEL\_V4 \| 1.0 \| 2.78e-13 \| 572 \| 0.351 \| 162 \| 76 \| 3 \| 1 \| 162 \| 1 \| 133 \| Single-stranded DNA-binding protein \| Single-stranded DNA-binding protein; afdb-uniprot50 \| AF-H0A2S4-F1-MODEL\_V4 \| 1.0 \| 5.816e-15 \| 572 \| 0.469 \| 164 \| 81 \| 5 \| 1 \| 162 \| 1 \| 160 \| Single-stranded DNA-binding protein \| Single-stranded DNA-binding protein; afdb-uniprot50 \| AF-A0A521PQ87-F1-MODEL\_V4 \| 1.0 \| 2.48e-14 \| 572 \| 0.383 \| 146 \| 86 \| 2 \| 2 \| 145 \| 16 \| 159 \| Single-stranded DNA-binding protein \| Single-stranded DNA-binding protein; afdb-uniprot50 \| AF-A0A2T2UL83-F1-MODEL\_V4 \| 1.0 \| 3.221e-15 \| 572 \| 0.388 \| 175 \| 89 \| 7 \| 1 \| 162 \| 1 \| 170 \| Single-stranded DNA-binding protein \| Single-stranded DNA-binding protein; afdb-uniprot50 \| AF-Q8Y2B4-F1-MODEL\_V4 \| 1.0 \| 3.587e-15 \| 572 \| 0.42 \| 183 \| 80 \| 7 \| 1 \| 162 \| 1 \| 178 \| Single-stranded DNA-binding protein \| Single-stranded DNA-binding protein; afdb-uniprot50 \| AF-A0A1W1H8U1-F1-MODEL\_V4 \| 1.0 \| 1.614e-14 \| 572 \| 0.408 \| 159 \| 86 \| 4 \| 1 \| 156 \| 1 \| 154 \| Single-stranded DNA-binding protein \| Single-stranded DNA-binding protein; afdb-uniprot50 \| AF-A0A662Z639-F1-MODEL\_V4 \| 1.0 \| 8.938e-15 \| 572 \| 0.406 \| 172 \| 89 \| 3 \| 1 \| 160 \| 1 \| 171 \| Single-stranded DNA-binding protein \| Single-stranded DNA-binding protein; afdb-uniprot50 \| AF-A0A831ZMD0-F1-MODEL\_V4 \| 1.0 \| 1.822e-12 \| 571 \| 0.436 \| 110 \| 60 \| 2 \| 1 \| 110 \| 1 \| 108 \| Single-stranded DNA-binding protein \| Single-stranded DNA-binding protein; afdb-uniprot50 \| AF-A0A496A9V7-F1-MODEL\_V4 \| 1.0 \| 1.449e-14 \| 571 \| 0.364 \| 162 \| 90 \| 4 \| 1 \| 162 \| 1 \| 149 \| Single-stranded DNA-binding protein \| Single-stranded DNA-binding protein; afdb-uniprot50 \| AF-A0A419IWL4-F1-MODEL\_V4 \| 1.0 \| 6.137e-15 \| 571 \| 0.396 \| 164 \| 93 \| 3 \| 1 \| 162 \| 2 \| 161 \| Single-stranded DNA-binding protein \| Single-stranded DNA-binding protein; afdb-uniprot50 \| AF-F0EYB2-F1-MODEL\_V4 \| 1.0 \| 7.714e-13 \| 570 \| 0.407 \| 113 \| 66 \| 1 \| 4 \| 116 \| 3 \| 114 \| Single-stranded DNA-binding protein \| Single-stranded DNA-binding protein; afdb-uniprot50 \| AF-A0A7V4A6M8-F1-MODEL\_V4 \| 1.0 \| 2.111e-14 \| 570 \| 0.408 \| 159 \| 71 \| 5 \| 4 \| 162 \| 3 \| 138 \| Single-stranded DNA-binding protein \| Single-stranded DNA-binding protein; afdb-uniprot50 \| AF-A0A0A2YFY3-F1-MODEL\_V4 \| 1.0 \| 4.95e-15 \| 570 \| 0.505 \| 170 \| 59 \| 7 \| 1 \| 162 \| 1 \| 153 \| Single-stranded DNA-binding protein \| Single-stranded DNA-binding protein; afdb-uniprot50 \| AF-F9GVF4-F1-MODEL\_V4 \| 1.0 \| 8.412e-16 \| 570 \| 0.469 \| 164 \| 75 \| 5 \| 1 \| 162 \| 1 \| 154 \| Single-stranded DNA-binding protein \| Single-stranded DNA-binding protein; afdb-uniprot50 \| AF-A0A0A6S9K9-F1-MODEL\_V4 \| 1.0 \| 1.691e-15 \| 570 \| 0.41 \| 178 \| 87 \| 5 \| 1 \| 162 \| 1 \| 176 \| Single-stranded DNA-binding protein \| Single-stranded DNA-binding protein; afdb-uniprot50 \| AF-A0A356VXD0-F1-MODEL\_V4 \| 1.0 \| 3.864e-12 \| 569 \| 0.656 \| 96 \| 32 \| 1 \| 4 \| 98 \| 3 \| 98 \| Single-stranded DNA-binding protein \| Single-stranded DNA-binding protein; afdb-uniprot50 \| AF-A0A7R9AGQ2-F1-MODEL\_V4 \| 1.0 \| 4.049e-13 \| 569 \| 0.466 \| 120 \| 63 \| 1 \| 3 \| 122 \| 2 \| 120 \| Hypothetical protein \| Hypothetical protein; afdb-uniprot50 \| AF-A0A2S4N7Z1-F1-MODEL\_V4 \| 1.0 \| 2.028e-12 \| 569 \| 0.391 \| 120 \| 70 \| 1 \| 1 \| 120 \| 1 \| 117 \| Single-stranded DNA-binding protein \| Single-stranded DNA-binding protein; afdb-uniprot50 \| AF-A0A831YQ24-F1-MODEL\_V4 \| 1.0 \| 4.692e-15 \| 569 \| 0.396 \| 174 \| 89 \| 4 \| 1 \| 162 \| 1 \| 170 \| Single-stranded DNA-binding protein \| Single-stranded DNA-binding protein; afdb-uniprot50 \| AF-A0A335FS28-F1-MODEL\_V4 \| 1.0 \| 4.049e-13 \| 568 \| 0.486 \| 115 \| 57 \| 2 \| 4 \| 118 \| 3 \| 115 \| Single-stranded DNA-binding protein \| Single-stranded DNA-binding protein; afdb-uniprot50 \| AF-A0A2V8QGU9-F1-MODEL\_V4 \| 1.0 \| 2.028e-12 \| 568 \| 0.439 \| 107 \| 59 \| 1 \| 4 \| 110 \| 2 \| 107 \| Single-stranded DNA-binding protein \| Single-stranded DNA-binding protein; afdb-uniprot50 \| AF-A0A2C6UNF7-F1-MODEL\_V4 \| 1.0 \| 1.123e-12 \| 568 \| 0.378 \| 132 \| 74 \| 2 \| 1 \| 132 \| 1 \| 124 \| Single-stranded DNA-binding protein \| Single-stranded DNA-binding protein; afdb-uniprot50 \| AF-A0A2W7INV0-F1-MODEL\_V4 \| 1.0 \| 4.214e-15 \| 568 \| 0.465 \| 172 \| 75 \| 7 \| 1 \| 162 \| 1 \| 165 \| Single-stranded DNA-binding protein \| Single-stranded DNA-binding protein; afdb-uniprot50 \| AF-A0A2N9WUQ3-F1-MODEL\_V4 \| 1.0 \| 3.47e-12 \| 567 \| 0.418 \| 117 \| 64 \| 2 \| 1 \| 117 \| 1 \| 113 \| Single-stranded DNA-binding protein \| Single-stranded DNA-binding protein; afdb-uniprot50 \| AF-A0A2D4XQA4-F1-MODEL\_V4 \| 1.0 \| 5.296e-13 \| 567 \| 0.408 \| 125 \| 66 \| 2 \| 5 \| 129 \| 1 \| 117 \| Single-stranded DNA-binding protein \| Single-stranded DNA-binding protein; afdb-uniprot50 \| AF-A0A535DEE2-F1-MODEL\_V4 \| 1.0 \| 4.273e-13 \| 567 \| 0.458 \| 120 \| 61 \| 3 \| 1 \| 120 \| 1 \| 116 \| Single-stranded DNA-binding protein \| Single-stranded DNA-binding protein; afdb-uniprot50 \| AF-A0A2E1JJ68-F1-MODEL\_V4 \| 1.0 \| 7.21e-15 \| 567 \| 0.453 \| 163 \| 71 \| 4 \| 1 \| 162 \| 1 \| 146 \| Single-stranded DNA-binding protein \| Single-stranded DNA-binding protein; afdb-uniprot50 \| AF-A0A7S8EHJ2-F1-MODEL\_V4 \| 1.0 \| 3.074e-14 \| 567 \| 0.414 \| 164 \| 83 \| 6 \| 1 \| 162 \| 1 \| 153 \| Single-stranded DNA-binding protein \| Single-stranded DNA-binding protein; afdb-uniprot50 \| AF-A0A2G4H6Y0-F1-MODEL\_V4 \| 1.0 \| 4.692e-15 \| 567 \| 0.425 \| 167 \| 79 \| 6 \| 4 \| 162 \| 3 \| 160 \| Single-stranded DNA-binding protein \| Single-stranded DNA-binding protein; afdb-uniprot50 \| AF-A1WP86-F1-MODEL\_V4 \| 1.0 \| 7.972e-16 \| 567 \| 0.491 \| 169 \| 77 \| 4 \| 1 \| 162 \| 1 \| 167 \| Single-stranded DNA-binding protein \| Single-stranded DNA-binding protein; afdb-uniprot50 \| AF-A0A7V4GHT9-F1-MODEL\_V4 \| 1.0 \| 3.587e-15 \| 567 \| 0.384 \| 177 \| 88 \| 6 \| 1 \| 162 \| 1 \| 171 \| Single-stranded DNA-binding protein \| Single-stranded DNA-binding protein; afdb-uniprot50 \| AF-A0A2R3QD86-F1-MODEL\_V4 \| 1.0 \| 1.883e-15 \| 567 \| 0.475 \| 162 \| 73 \| 3 \| 1 \| 162 \| 1 \| 150 \| Single-stranded DNA-binding protein \| Single-stranded DNA-binding protein; afdb-uniprot50 \| AF-A0A1D9HCR6-F1-MODEL\_V4 \| 1.0 \| 4.214e-15 \| 567 \| 0.434 \| 198 \| 72 \| 7 \| 1 \| 162 \| 1 \| 194 \| Single-stranded DNA-binding protein \| Single-stranded DNA-binding protein; afdb-uniprot50 \| AF-A0A1W9TJB4-F1-MODEL\_V4 \| 1.0 \| 4.95e-15 \| 566 \| 0.462 \| 162 \| 67 \| 6 \| 1 \| 162 \| 1 \| 142 \| Single-stranded DNA-binding protein \| Single-stranded DNA-binding protein; afdb-uniprot50 \| AF-A0A3D2TZ42-F1-MODEL\_V4 \| 1.0 \| 7.608e-15 \| 566 \| 0.386 \| 163 \| 88 \| 5 \| 1 \| 162 \| 1 \| 152 \| Single-stranded DNA-binding protein \| Single-stranded DNA-binding protein; afdb-uniprot50 \| AF-A0A3M4E9N4-F1-MODEL\_V4 \| 1.0 \| 4.54e-12 \| 566 \| 0.642 \| 98 \| 34 \| 1 \| 1 \| 98 \| 67 \| 163 \| Single-stranded DNA-binding protein \| Single-stranded DNA-binding protein; afdb-uniprot50 \| AF-A0A157QP91-F1-MODEL\_V4 \| 1.0 \| 4.95e-15 \| 566 \| 0.453 \| 183 \| 75 \| 6 \| 1 \| 162 \| 1 \| 179 \| Single-stranded DNA-binding protein \| Single-stranded DNA-binding protein; afdb-uniprot50 \| AF-A0A7H9WP57-F1-MODEL\_V4 \| 1.0 \| 4.757e-13 \| 565 \| 0.39 \| 128 \| 72 \| 3 \| 4 \| 131 \| 2 \| 123 \| Single-stranded DNA-binding protein \| Single-stranded DNA-binding protein; afdb-uniprot50 \| AF-A0A328B6D0-F1-MODEL\_V4 \| 1.0 \| 7.661e-14 \| 565 \| 0.414 \| 140 \| 79 \| 2 \| 4 \| 142 \| 1 \| 138 \| Single-stranded DNA-binding protein \| Single-stranded DNA-binding protein; afdb-uniprot50 \| AF-A0A328BGG8-F1-MODEL\_V4 \| 1.0 \| 5.55e-14 \| 565 \| 0.386 \| 150 \| 81 \| 4 \| 4 \| 150 \| 1 \| 142 \| Single-stranded DNA-binding protein \| Single-stranded DNA-binding protein; afdb-uniprot50 \| AF-A0A6J5FFK9-F1-MODEL\_V4 \| 1.0 \| 4.95e-15 \| 565 \| 0.447 \| 163 \| 67 \| 4 \| 4 \| 162 \| 2 \| 145 \| Single-stranded DNA-binding protein \| Single-stranded DNA-binding protein; afdb-uniprot50 \| AF-A0A2E4DF34-F1-MODEL\_V4 \| 1.0 \| 5.512e-15 \| 564 \| 0.431 \| 160 \| 76 \| 5 \| 4 \| 162 \| 2 \| 147 \| Single-stranded DNA-binding protein \| Single-stranded DNA-binding protein; afdb-uniprot50 \| AF-A7ILF7-F1-MODEL\_V4 \| 1.0 \| 2.48e-14 \| 564 \| 0.451 \| 166 \| 79 \| 6 \| 1 \| 162 \| 1 \| 158 \| Single-stranded DNA-binding protein \| Single-stranded DNA-binding protein; afdb-uniprot50 \| AF-A0A7V6WJS5-F1-MODEL\_V4 \| 1.0 \| 6.137e-15 \| 564 \| 0.413 \| 167 \| 82 \| 3 \| 4 \| 162 \| 2 \| 160 \| Single-stranded DNA-binding protein \| Single-stranded DNA-binding protein; afdb-uniprot50 \| AF-G4QAH2-F1-MODEL\_V4 \| 1.0 \| 7.608e-15 \| 564 \| 0.384 \| 169 \| 91 \| 4 \| 4 \| 162 \| 2 \| 167 \| Single-stranded DNA-binding protein \| Single-stranded DNA-binding protein; afdb-uniprot50 \| AF-A0A1F6IFU5-F1-MODEL\_V4 \| 1.0 \| 6.475e-15 \| 564 \| 0.358 \| 162 \| 97 \| 3 \| 1 \| 162 \| 1 \| 155 \| Single-stranded DNA-binding protein \| Single-stranded DNA-binding protein; afdb-uniprot50 \| AF-Q1N2F3-F1-MODEL\_V4 \| 1.0 \| 9.951e-15 \| 564 \| 0.46 \| 163 \| 79 \| 5 \| 1 \| 162 \| 1 \| 155 \| Single-stranded DNA-binding protein \| Single-stranded DNA-binding protein; afdb-uniprot50 \| AF-A0A370U549-F1-MODEL\_V4 \| 1.0 \| 1.302e-14 \| 564 \| 0.408 \| 164 \| 85 \| 4 \| 1 \| 160 \| 1 \| 156 \| Single-stranded DNA-binding protein \| Single-stranded DNA-binding protein; afdb-uniprot50 \| AF-A0A2E1MM44-F1-MODEL\_V4 \| 1.0 \| 2.913e-14 \| 563 \| 0.39 \| 164 \| 82 \| 6 \| 1 \| 162 \| 1 \| 148 \| Single-stranded DNA-binding protein \| Single-stranded DNA-binding protein; afdb-uniprot50 \| AF-A0A383HK71-F1-MODEL\_V4 \| 1.0 \| 1.703e-14 \| 563 \| 0.431 \| 183 \| 77 \| 4 \| 6 \| 162 \| 7 \| 188 \| Single-stranded DNA-binding protein \| Single-stranded DNA-binding protein; afdb-uniprot50 \| AF-R5ELR6-F1-MODEL\_V4 \| 1.0 \| 8.47e-15 \| 563 \| 0.381 \| 173 \| 93 \| 4 \| 1 \| 162 \| 1 \| 170 \| Single-stranded DNA-binding protein \| Single-stranded DNA-binding protein; afdb-uniprot50 \| AF-A0A257GV23-F1-MODEL\_V4 \| 1.0 \| 5.333e-12 \| 562 \| 0.427 \| 117 \| 64 \| 1 \| 1 \| 117 \| 1 \| 114 \| Single-stranded DNA-binding protein \| Single-stranded DNA-binding protein; afdb-uniprot50 \| AF-A0A2S0N5P2-F1-MODEL\_V4 \| 1.0 \| 5.55e-14 \| 562 \| 0.424 \| 153 \| 74 \| 2 \| 4 \| 156 \| 3 \| 141 \| Single-stranded DNA-binding protein \| Single-stranded DNA-binding protein; afdb-uniprot50 \| AF-A0A2E2SU00-F1-MODEL\_V4 \| 1.0 \| 4.477e-14 \| 562 \| 0.368 \| 163 \| 84 \| 4 \| 1 \| 162 \| 1 \| 145 \| Single-stranded DNA-binding protein \| Single-stranded DNA-binding protein; afdb-uniprot50 \| AF-A0A2G6HT26-F1-MODEL\_V4 \| 1.0 \| 4.214e-15 \| 562 \| 0.406 \| 172 \| 83 \| 6 \| 1 \| 162 \| 1 \| 163 \| Single-stranded DNA-binding protein \| Single-stranded DNA-binding protein; afdb-uniprot50 \| AF-A0A6N9BWC5-F1-MODEL\_V4 \| 1.0 \| 2.893e-15 \| 562 \| 0.434 \| 168 \| 88 \| 2 \| 1 \| 162 \| 1 \| 167 \| Single-stranded DNA-binding protein \| Single-stranded DNA-binding protein; afdb-uniprot50 \| AF-A0A7R8ZXC5-F1-MODEL\_V4 \| 1.0 \| 2.463e-15 \| 562 \| 0.421 \| 171 \| 80 \| 4 \| 5 \| 159 \| 1 \| 168 \| Hypothetical protein \| Hypothetical protein; afdb-uniprot50 \| AF-A0A7X9IAE8-F1-MODEL\_V4 \| 1.0 \| 1.065e-12 \| 561 \| 0.346 \| 127 \| 78 \| 2 \| 1 \| 127 \| 1 \| 122 \| Single-stranded DNA-binding protein \| Single-stranded DNA-binding protein; afdb-uniprot50 \| AF-A0A1B1Y8T8-F1-MODEL\_V4 \| 1.0 \| 2.243e-13 \| 561 \| 0.388 \| 162 \| 72 \| 5 \| 1 \| 162 \| 1 \| 135 \| Single-stranded DNA-binding protein \| Single-stranded DNA-binding protein; afdb-uniprot50 \| AF-A0A1E4K5Q6-F1-MODEL\_V4 \| 1.0 \| 6.137e-15 \| 561 \| 0.463 \| 179 \| 75 \| 5 \| 1 \| 162 \| 1 \| 175 \| Single-stranded DNA-binding protein \| Single-stranded DNA-binding protein; afdb-uniprot50 \| AF-L2F912-F1-MODEL\_V4 \| 1.0 \| 6.833e-15 \| 561 \| 0.407 \| 167 \| 88 \| 3 \| 4 \| 162 \| 3 \| 166 \| Single-stranded DNA-binding protein \| Single-stranded DNA-binding protein; afdb-uniprot50 \| AF-A0A535N5L0-F1-MODEL\_V4 \| 1.0 \| 3.864e-12 \| 561 \| 0.4 \| 115 \| 66 \| 1 \| 3 \| 117 \| 199 \| 310 \| Single-stranded DNA-binding protein \| Single-stranded DNA-binding protein; afdb-uniprot50 \| AF-A0A534RKS9-F1-MODEL\_V4 \| 1.0 \| 4.302e-12 \| 560 \| 0.401 \| 117 \| 67 \| 1 \| 1 \| 117 \| 115 \| 228 \| Single-stranded DNA-binding protein \| Single-stranded DNA-binding protein; afdb-uniprot50 \| AF-F0IH13-F1-MODEL\_V4 \| 1.0 \| 1.185e-12 \| 558 \| 0.418 \| 117 \| 66 \| 2 \| 1 \| 117 \| 13 \| 127 \| Single-stranded DNA-binding protein \| Single-stranded DNA-binding protein; afdb-uniprot50 \| AF-A0A4Q5VAN1-F1-MODEL\_V4 \| 1.0 \| 6.88e-14 \| 558 \| 0.459 \| 135 \| 65 \| 2 \| 1 \| 135 \| 1 \| 127 \| Single-stranded DNA-binding protein \| Single-stranded DNA-binding protein; afdb-uniprot50 \| AF-A0A1A7QB56-F1-MODEL\_V4 \| 1.0 \| 5.26e-14 \| 558 \| 0.468 \| 143 \| 70 \| 3 \| 1 \| 143 \| 1 \| 137 \| Single-stranded DNA-binding protein \| Single-stranded DNA-binding protein; afdb-uniprot50 \| AF-A0A2E9VE14-F1-MODEL\_V4 \| 1.0 \| 2.617e-14 \| 558 \| 0.444 \| 162 \| 70 \| 4 \| 1 \| 162 \| 2 \| 143 \| Single-stranded DNA-binding protein \| Single-stranded DNA-binding protein; afdb-uniprot50 \| AF-A0A1F8TSK2-F1-MODEL\_V4 \| 1.0 \| 1.529e-14 \| 558 \| 0.516 \| 151 \| 68 \| 2 \| 12 \| 162 \| 2 \| 147 \| Single-stranded DNA-binding protein \| Single-stranded DNA-binding protein; afdb-uniprot50 \| AF-E1YGH0-F1-MODEL\_V4 \| 1.0 \| 3.399e-15 \| 558 \| 0.425 \| 160 \| 77 \| 4 \| 3 \| 162 \| 21 \| 165 \| Single-stranded DNA-binding protein \| Single-stranded DNA-binding protein; afdb-uniprot50 \| AF-A0A7Z9G7J1-F1-MODEL\_V4 \| 1.0 \| 6.137e-15 \| 558 \| 0.4 \| 160 \| 89 \| 5 \| 4 \| 162 \| 2 \| 155 \| Single-stranded DNA-binding protein \| Single-stranded DNA-binding protein; afdb-uniprot50 \| AF-A0A661TMS4-F1-MODEL\_V4 \| 1.0 \| 6.137e-15 \| 557 \| 0.416 \| 168 \| 85 \| 6 \| 4 \| 162 \| 1 \| 164 \| Single-stranded DNA-binding protein \| Single-stranded DNA-binding protein; afdb-uniprot50 \| AF-A0A7X8D708-F1-MODEL\_V4 \| 1.0 \| 1.225e-15 \| 557 \| 0.47 \| 172 \| 79 \| 4 \| 1 \| 162 \| 1 \| 170 \| Single-stranded DNA-binding protein \| Single-stranded DNA-binding protein; afdb-uniprot50 \| AF-A0A3A1YSF5-F1-MODEL\_V4 \| 1.0 \| 1.043e-15 \| 557 \| 0.424 \| 179 \| 84 \| 5 \| 1 \| 162 \| 1 \| 177 \| Single-stranded DNA-binding protein \| Single-stranded DNA-binding protein; afdb-uniprot50 \| AF-A0A1W9L078-F1-MODEL\_V4 \| 1.0 \| 1.374e-14 \| 556 \| 0.425 \| 160 \| 80 \| 4 \| 3 \| 162 \| 2 \| 149 \| Single-stranded DNA-binding protein \| Single-stranded DNA-binding protein; afdb-uniprot50 \| AF-A0A662BHB8-F1-MODEL\_V4 \| 1.0 \| 6.928e-13 \| 555 \| 0.419 \| 131 \| 69 \| 4 \| 1 \| 131 \| 1 \| 124 \| Single-stranded DNA-binding protein \| Single-stranded DNA-binding protein; afdb-uniprot50 \| AF-A0A7K1X431-F1-MODEL\_V4 \| 1.0 \| 1.726e-12 \| 555 \| 0.406 \| 128 \| 68 \| 3 \| 1 \| 128 \| 1 \| 120 \| Single-stranded DNA-binding protein \| Single-stranded DNA-binding protein; afdb-uniprot50 \| AF-A0A1G2BK10-F1-MODEL\_V4 \| 1.0 \| 7.714e-13 \| 555 \| 0.429 \| 128 \| 65 \| 3 \| 4 \| 130 \| 2 \| 122 \| Single-stranded DNA-binding protein \| Single-stranded DNA-binding protein; afdb-uniprot50 \| AF-A0A2B8A0F0-F1-MODEL\_V4 \| 1.0 \| 2.014e-13 \| 555 \| 0.441 \| 145 \| 71 \| 5 \| 1 \| 144 \| 1 \| 136 \| Single-stranded DNA-binding protein \| Single-stranded DNA-binding protein; afdb-uniprot50 \| AF-G8R207-F1-MODEL\_V4 \| 1.0 \| 3.423e-14 \| 555 \| 0.45 \| 162 \| 73 \| 3 \| 1 \| 162 \| 1 \| 146 \| Single-stranded DNA-binding protein \| Single-stranded DNA-binding protein; afdb-uniprot50 \| AF-A0A5C8ILK4-F1-MODEL\_V4 \| 1.0 \| 3.612e-14 \| 555 \| 0.388 \| 162 \| 84 \| 5 \| 1 \| 162 \| 1 \| 147 \| Single-stranded DNA-binding protein \| Single-stranded DNA-binding protein; afdb-uniprot50 \| AF-A0A2H0L162-F1-MODEL\_V4 \| 1.0 \| 3.423e-14 \| 555 \| 0.422 \| 154 \| 82 \| 4 \| 1 \| 154 \| 1 \| 147 \| Single-stranded DNA-binding protein \| Single-stranded DNA-binding protein; afdb-uniprot50 \| AF-A0A3D1BFD1-F1-MODEL\_V4 \| 1.0 \| 3.221e-15 \| 555 \| 0.436 \| 165 \| 85 \| 5 \| 1 \| 162 \| 1 \| 160 \| Single-stranded DNA-binding protein \| Single-stranded DNA-binding protein; afdb-uniprot50 \| AF-A0A1F7BR67-F1-MODEL\_V4 \| 1.0 \| 1.529e-14 \| 555 \| 0.418 \| 160 \| 84 \| 3 \| 1 \| 160 \| 17 \| 167 \| Single-stranded DNA-binding protein \| Single-stranded DNA-binding protein; afdb-uniprot50 \| AF-A0A5B5VMW8-F1-MODEL\_V4 \| 1.0 \| 9.562e-13 \| 554 \| 0.436 \| 133 \| 69 \| 2 \| 4 \| 136 \| 1 \| 127 \| Single-stranded DNA-binding protein \| Single-stranded DNA-binding protein; afdb-uniprot50 \| AF-A0A1Z8MPJ0-F1-MODEL\_V4 \| 1.0 \| 1.909e-13 \| 554 \| 0.327 \| 159 \| 82 \| 3 \| 4 \| 162 \| 2 \| 135 \| Single-stranded DNA-binding protein \| Single-stranded DNA-binding protein; afdb-uniprot50 \| AF-A0A1F9RAQ1-F1-MODEL\_V4 \| 1.0 \| 4.446e-15 \| 554 \| 0.419 \| 162 \| 86 \| 5 \| 1 \| 162 \| 1 \| 154 \| Single-stranded DNA-binding protein \| Single-stranded DNA-binding protein; afdb-uniprot50 \| AF-A0A1F8B8P6-F1-MODEL\_V4 \| 1.0 \| 1.374e-14 \| 554 \| 0.37 \| 162 \| 96 \| 3 \| 1 \| 162 \| 1 \| 156 \| Single-stranded DNA-binding protein \| Single-stranded DNA-binding protein; afdb-uniprot50 \| AF-A0A3B9MCP7-F1-MODEL\_V4 \| 1.0 \| 1.636e-12 \| 553 \| 0.423 \| 118 \| 61 \| 3 \| 1 \| 114 \| 14 \| 128 \| Single-stranded DNA-binding protein \| Single-stranded DNA-binding protein; afdb-uniprot50 \| AF-A0A378XHW0-F1-MODEL\_V4 \| 1.0 \| 4.79e-12 \| 553 \| 0.347 \| 118 \| 72 \| 2 \| 1 \| 117 \| 1 \| 114 \| Single-stranded DNA-binding protein \| Single-stranded DNA-binding protein; afdb-uniprot50 \| AF-A0A5C7JQL9-F1-MODEL\_V4 \| 1.0 \| 3.811e-14 \| 553 \| 0.385 \| 174 \| 73 \| 6 \| 1 \| 162 \| 1 \| 152 \| Single-stranded DNA-binding protein \| Single-stranded DNA-binding protein; afdb-uniprot50 \| AF-A0A193KJZ2-F1-MODEL\_V4 \| 1.0 \| 2.35e-14 \| 553 \| 0.441 \| 163 \| 84 \| 5 \| 3 \| 162 \| 2 \| 160 \| Single-stranded DNA-binding protein \| Single-stranded DNA-binding protein; afdb-uniprot50 \| AF-A0A352LLW6-F1-MODEL\_V4 \| 1.0 \| 1.302e-14 \| 553 \| 0.442 \| 165 \| 81 \| 7 \| 1 \| 162 \| 1 \| 157 \| Single-stranded DNA-binding protein \| Single-stranded DNA-binding protein; afdb-uniprot50 \| AF-A0A0S8G146-F1-MODEL\_V4 \| 1.0 \| 1.715e-13 \| 552 \| 0.427 \| 159 \| 68 \| 4 \| 4 \| 162 \| 3 \| 138 \| Single-stranded DNA-binding protein \| Single-stranded DNA-binding protein; afdb-uniprot50 \| AF-A0A1G4UBM1-F1-MODEL\_V4 \| 1.0 \| 2.366e-13 \| 552 \| 0.424 \| 139 \| 72 \| 4 \| 1 \| 137 \| 1 \| 133 \| Single-stranded DNA-binding protein \| Single-stranded DNA-binding protein; afdb-uniprot50 \| AF-A0A7X3VRR7-F1-MODEL\_V4 \| 1.0 \| 2.497e-13 \| 552 \| 0.321 \| 146 \| 93 \| 2 \| 5 \| 150 \| 1 \| 140 \| Single-stranded DNA-binding protein \| Single-stranded DNA-binding protein; afdb-uniprot50 \| AF-A0A2H0BUB7-F1-MODEL\_V4 \| 1.0 \| 1.374e-14 \| 552 \| 0.385 \| 166 \| 94 \| 4 \| 1 \| 162 \| 2 \| 163 \| Single-stranded DNA-binding protein \| Single-stranded DNA-binding protein; afdb-uniprot50 \| AF-A0A4Y6J6J4-F1-MODEL\_V4 \| 1.0 \| 3.074e-14 \| 552 \| 0.341 \| 170 \| 96 \| 4 \| 1 \| 160 \| 2 \| 165 \| Single-stranded DNA-binding protein \| Single-stranded DNA-binding protein; afdb-uniprot50 \| AF-A0A0G1V161-F1-MODEL\_V4 \| 1.0 \| 4.508e-13 \| 551 \| 0.388 \| 126 \| 74 \| 2 \| 4 \| 127 \| 2 \| 126 \| Single-stranded DNA-binding protein \| Single-stranded DNA-binding protein; afdb-uniprot50 \| AF-A0A143XJT9-F1-MODEL\_V4 \| 1.0 \| 1.703e-14 \| 551 \| 0.392 \| 191 \| 81 \| 4 \| 4 \| 162 \| 1 \| 188 \| Single-stranded DNA-binding protein \| Single-stranded DNA-binding protein; afdb-uniprot50 \| AF-A0A517VMC9-F1-MODEL\_V4 \| 1.0 \| 3.244e-14 \| 550 \| 0.377 \| 167 \| 84 \| 7 \| 1 \| 162 \| 1 \| 152 \| Single-stranded DNA-binding protein \| Single-stranded DNA-binding protein; afdb-uniprot50 \| AF-A0A1Y1T5Q8-F1-MODEL\_V4 \| 1.0 \| 1.449e-14 \| 550 \| 0.325 \| 175 \| 102 \| 3 \| 1 \| 162 \| 1 \| 172 \| Single-stranded DNA-binding protein \| Single-stranded DNA-binding protein; afdb-uniprot50 \| AF-A0A1F6RWV3-F1-MODEL\_V4 \| 1.0 \| 2.334e-15 \| 550 \| 0.427 \| 166 \| 82 \| 4 \| 4 \| 162 \| 2 \| 161 \| Single-stranded DNA-binding protein \| Single-stranded DNA-binding protein; afdb-uniprot50 \| AF-A0A1W9NY84-F1-MODEL\_V4 \| 1.0 \| 5.333e-12 \| 550 \| 0.394 \| 119 \| 68 \| 2 \| 1 \| 118 \| 2 \| 117 \| Single-stranded DNA-binding protein \| Single-stranded DNA-binding protein; afdb-uniprot50 \| AF-A0A4P5V4F4-F1-MODEL\_V4 \| 1.0 \| 2.617e-14 \| 550 \| 0.373 \| 166 \| 95 \| 4 \| 1 \| 162 \| 1 \| 161 \| Single-stranded DNA-binding protein \| Single-stranded DNA-binding protein; afdb-uniprot50 \| AF-A0A2E9H291-F1-MODEL\_V4 \| 1.0 \| 6.52e-14 \| 550 \| 0.417 \| 163 \| 83 \| 4 \| 1 \| 155 \| 1 \| 159 \| Single-stranded DNA-binding protein \| Single-stranded DNA-binding protein; afdb-uniprot50 \| AF-A0A2H0SKG1-F1-MODEL\_V4 \| 1.0 \| 7.26e-14 \| 550 \| 0.351 \| 165 \| 101 \| 2 \| 1 \| 162 \| 2 \| 163 \| Single-stranded DNA-binding protein \| Single-stranded DNA-binding protein; afdb-uniprot50 \| AF-A0A554VRH5-F1-MODEL\_V4 \| 1.0 \| 1.726e-12 \| 549 \| 0.424 \| 125 \| 66 \| 3 \| 4 \| 128 \| 1 \| 119 \| Single-stranded DNA-binding protein \| Single-stranded DNA-binding protein; afdb-uniprot50 \| AF-A0A258BZL2-F1-MODEL\_V4 \| 1.0 \| 3.244e-14 \| 549 \| 0.409 \| 161 \| 78 \| 3 \| 3 \| 162 \| 2 \| 146 \| Single-stranded DNA-binding protein \| Single-stranded DNA-binding protein; afdb-uniprot50 \| AF-W7W8R5-F1-MODEL\_V4 \| 1.0 \| 2.334e-15 \| 549 \| 0.437 \| 167 \| 87 \| 4 \| 1 \| 162 \| 1 \| 165 \| Single-stranded DNA-binding protein \| Single-stranded DNA-binding protein; afdb-uniprot50 \| AF-A0A563D9X7-F1-MODEL\_V4 \| 1.0 \| 3.47e-12 \| 548 \| 0.401 \| 127 \| 68 \| 3 \| 1 \| 127 \| 1 \| 119 \| Single-stranded DNA-binding protein \| Single-stranded DNA-binding protein; afdb-uniprot50 \| AF-A0A2D6FJM0-F1-MODEL\_V4 \| 1.0 \| 8.529e-14 \| 548 \| 0.429 \| 163 \| 76 \| 5 \| 1 \| 162 \| 2 \| 148 \| Single-stranded DNA-binding protein \| Single-stranded DNA-binding protein; afdb-uniprot50 \| AF-A0A257H2V1-F1-MODEL\_V4 \| 1.0 \| 9.431e-15 \| 548 \| 0.404 \| 178 \| 78 \| 5 \| 1 \| 162 \| 1 \| 166 \| Single-stranded DNA-binding protein \| Single-stranded DNA-binding protein; afdb-uniprot50 \| AF-A0A5M4B7C0-F1-MODEL\_V4 \| 1.0 \| 1.636e-12 \| 547 \| 0.353 \| 133 \| 80 \| 2 \| 1 \| 130 \| 1 \| 130 \| Single-stranded DNA-binding protein \| Single-stranded DNA-binding protein; afdb-uniprot50 \| AF-R6A491-F1-MODEL\_V4 \| 1.0 \| 3.423e-14 \| 546 \| 0.422 \| 180 \| 81 \| 7 \| 1 \| 162 \| 1 \| 175 \| Single-stranded DNA-binding protein \| Single-stranded DNA-binding protein; afdb-uniprot50 \| AF-R5I4N3-F1-MODEL\_V4 \| 1.0 \| 6.137e-15 \| 546 \| 0.386 \| 194 \| 83 \| 4 \| 4 \| 162 \| 2 \| 194 \| Single-stranded DNA-binding protein \| Single-stranded DNA-binding protein; afdb-uniprot50 \| AF-A0A7C4TH37-F1-MODEL\_V4 \| 1.0 \| 5.897e-13 \| 545 \| 0.355 \| 135 \| 81 \| 3 \| 1 \| 134 \| 5 \| 134 \| Single-stranded DNA-binding protein \| Single-stranded DNA-binding protein; afdb-uniprot50 \| AF-A0A3N5P7G3-F1-MODEL\_V4 \| 1.0 \| 4.477e-14 \| 545 \| 0.419 \| 162 \| 72 \| 5 \| 1 \| 162 \| 1 \| 140 \| Single-stranded DNA-binding protein \| Single-stranded DNA-binding protein; afdb-uniprot50 \| AF-A0A1W9QM69-F1-MODEL\_V4 \| 1.0 \| 3.074e-14 \| 545 \| 0.364 \| 159 \| 83 \| 4 \| 4 \| 162 \| 2 \| 142 \| Single-stranded DNA-binding protein \| Single-stranded DNA-binding protein; afdb-uniprot50 \| AF-A0A6L7TFT4-F1-MODEL\_V4 \| 1.0 \| 3.423e-14 \| 545 \| 0.374 \| 171 \| 90 \| 4 \| 1 \| 162 \| 1 \| 163 \| Single-stranded DNA-binding protein \| Single-stranded DNA-binding protein; afdb-uniprot50 \| AF-A0A2N7RSM6-F1-MODEL\_V4 \| 1.0 \| 5.816e-15 \| 545 \| 0.391 \| 171 \| 90 \| 4 \| 1 \| 162 \| 1 \| 166 \| Single-stranded DNA-binding protein \| Single-stranded DNA-binding protein; afdb-uniprot50 \| AF-A0A535RBC2-F1-MODEL\_V4 \| 1.0 \| 4.077e-12 \| 544 \| 0.396 \| 111 \| 64 \| 2 \| 4 \| 114 \| 1 \| 108 \| Single-stranded DNA-binding protein \| Single-stranded DNA-binding protein; afdb-uniprot50 \| AF-A0A7Z9YCJ8-F1-MODEL\_V4 \| 1.0 \| 2.497e-13 \| 544 \| 0.36 \| 133 \| 83 \| 2 \| 1 \| 133 \| 1 \| 131 \| Single-stranded DNA-binding protein \| Single-stranded DNA-binding protein; afdb-uniprot50 \| AF-A0A351KW12-F1-MODEL\_V4 \| 1.0 \| 1.715e-13 \| 544 \| 0.381 \| 160 \| 78 \| 5 \| 4 \| 162 \| 3 \| 142 \| Single-stranded DNA-binding protein \| Single-stranded DNA-binding protein; afdb-uniprot50 \| AF-A0A3M0YZZ3-F1-MODEL\_V4 \| 1.0 \| 1.26e-11 \| 544 \| 0.393 \| 117 \| 68 \| 1 \| 1 \| 117 \| 3 \| 116 \| Single-stranded DNA-binding protein \| Single-stranded DNA-binding protein; afdb-uniprot50 \| AF-A0A3D4IIJ5-F1-MODEL\_V4 \| 1.0 \| 8.938e-15 \| 544 \| 0.402 \| 174 \| 83 \| 6 \| 1 \| 162 \| 2 \| 166 \| Single-stranded DNA-binding protein \| Single-stranded DNA-binding protein; afdb-uniprot50 \| AF-A0A2W6YVK2-F1-MODEL\_V4 \| 1.0 \| 1.169e-14 \| 544 \| 0.454 \| 176 \| 80 \| 6 \| 1 \| 162 \| 1 \| 174 \| Single-stranded DNA-binding protein \| Single-stranded DNA-binding protein; afdb-uniprot50 \| AF-A0A0F9SXG5-F1-MODEL\_V4 \| 1.0 \| 1.459e-13 \| 543 \| 0.401 \| 162 \| 78 \| 6 \| 1 \| 162 \| 1 \| 143 \| Uncharacterized protein \| Uncharacterized protein; afdb-uniprot50 \| AF-A0A077DE37-F1-MODEL\_V4 \| 1.0 \| 3.423e-14 \| 543 \| 0.406 \| 165 \| 86 \| 5 \| 3 \| 162 \| 2 \| 159 \| Single-stranded DNA-binding protein \| Single-stranded DNA-binding protein; afdb-uniprot50 \| AF-A0A1F7VHB8-F1-MODEL\_V4 \| 1.0 \| 1.374e-14 \| 543 \| 0.413 \| 167 \| 86 \| 5 \| 4 \| 162 \| 3 \| 165 \| Single-stranded DNA-binding protein \| Single-stranded DNA-binding protein; afdb-uniprot50 \| AF-A0A350CXW3-F1-MODEL\_V4 \| 1.0 \| 1.459e-13 \| 543 \| 0.387 \| 155 \| 88 \| 3 \| 1 \| 155 \| 20 \| 167 \| Single-stranded DNA-binding protein \| Single-stranded DNA-binding protein; afdb-uniprot50 \| AF-A0A4R3I482-F1-MODEL\_V4 \| 1.0 \| 2.334e-15 \| 543 \| 0.449 \| 187 \| 77 \| 5 \| 1 \| 162 \| 2 \| 187 \| Single-stranded DNA-binding protein \| Single-stranded DNA-binding protein; afdb-uniprot50 \| AF-A0A2M7DG48-F1-MODEL\_V4 \| 1.0 \| 2.243e-13 \| 542 \| 0.327 \| 162 \| 97 \| 2 \| 1 \| 162 \| 1 \| 150 \| Single-stranded DNA-binding protein \| Single-stranded DNA-binding protein; afdb-uniprot50 \| AF-A0A2M8EIY2-F1-MODEL\_V4 \| 1.0 \| 4.477e-14 \| 542 \| 0.367 \| 155 \| 92 \| 3 \| 1 \| 155 \| 1 \| 149 \| Single-stranded DNA-binding protein \| Single-stranded DNA-binding protein; afdb-uniprot50 \| AF-A0A1F1C449-F1-MODEL\_V4 \| 1.0 \| 7.714e-13 \| 540 \| 0.341 \| 117 \| 71 \| 2 \| 4 \| 120 \| 3 \| 113 \| Single-stranded DNA-binding protein \| Single-stranded DNA-binding protein; afdb-uniprot50 \| AF-R5E1D3-F1-MODEL\_V4 \| 1.0 \| 6.18e-14 \| 540 \| 0.389 \| 167 \| 88 \| 5 \| 1 \| 161 \| 1 \| 159 \| Single-stranded DNA-binding protein \| Single-stranded DNA-binding protein; afdb-uniprot50 \| AF-A0A4P6FUU1-F1-MODEL\_V4 \| 1.0 \| 4.985e-14 \| 540 \| 0.355 \| 163 \| 98 \| 6 \| 4 \| 162 \| 2 \| 161 \| Single-stranded DNA-binding protein \| Single-stranded DNA-binding protein; afdb-uniprot50 \| AF-S5RPZ6-F1-MODEL\_V4 \| 1.0 \| 6.137e-15 \| 540 \| 0.379 \| 182 \| 82 \| 4 \| 1 \| 162 \| 1 \| 171 \| Single-stranded DNA-binding protein \| Single-stranded DNA-binding protein; afdb-uniprot50 \| AF-A0A2H1YK25-F1-MODEL\_V4 \| 1.0 \| 6.266e-12 \| 539 \| 0.377 \| 127 \| 75 \| 2 \| 1 \| 127 \| 1 \| 123 \| Single-stranded DNA-binding protein \| Single-stranded DNA-binding protein; afdb-uniprot50 \| AF-A0A517YVP6-F1-MODEL\_V4 \| 1.0 \| 1.002e-13 \| 539 \| 0.368 \| 160 \| 89 \| 4 \| 4 \| 162 \| 3 \| 151 \| Single-stranded DNA-binding protein \| Single-stranded DNA-binding protein; afdb-uniprot50 \| AF-A0A7C4ZBJ7-F1-MODEL\_V4 \| 1.0 \| 2.617e-14 \| 539 \| 0.358 \| 159 \| 94 \| 3 \| 4 \| 162 \| 3 \| 153 \| Single-stranded DNA-binding protein \| Single-stranded DNA-binding protein; afdb-uniprot50 \| AF-A0A124FXW4-F1-MODEL\_V4 \| 1.0 \| 3.423e-14 \| 539 \| 0.337 \| 169 \| 96 \| 5 \| 1 \| 162 \| 2 \| 161 \| Single-stranded DNA-binding protein \| Single-stranded DNA-binding protein; afdb-uniprot50 \| AF-A0A1G1CF15-F1-MODEL\_V4 \| 1.0 \| 2.48e-14 \| 539 \| 0.366 \| 169 \| 96 \| 4 \| 1 \| 162 \| 2 \| 166 \| Single-stranded DNA-binding protein \| Single-stranded DNA-binding protein; afdb-uniprot50 \| AF-A0A4V0XBV8-F1-MODEL\_V4 \| 1.0 \| 9e-14 \| 539 \| 0.364 \| 162 \| 96 \| 3 \| 1 \| 161 \| 1 \| 156 \| Single-stranded DNA-binding protein \| Single-stranded DNA-binding protein; afdb-uniprot50 \| AF-A0A662ZJD1-F1-MODEL\_V4 \| 1.0 \| 4.021e-14 \| 539 \| 0.386 \| 163 \| 92 \| 4 \| 1 \| 159 \| 2 \| 160 \| Single-stranded DNA-binding protein \| Single-stranded DNA-binding protein; afdb-uniprot50 \| AF-X0YPY6-F1-MODEL\_V4 \| 1.0 \| 2.366e-13 \| 538 \| 0.363 \| 146 \| 90 \| 1 \| 5 \| 150 \| 1 \| 143 \| Uncharacterized protein \| Uncharacterized protein; afdb-uniprot50 \| AF-A0A660MIK8-F1-MODEL\_V4 \| 1.0 \| 2.8e-12 \| 537 \| 0.343 \| 128 \| 81 \| 1 \| 1 \| 128 \| 1 \| 125 \| Single-stranded DNA-binding protein \| Single-stranded DNA-binding protein; afdb-uniprot50 \| AF-A0A2D6STV9-F1-MODEL\_V4 \| 1.0 \| 6.88e-14 \| 537 \| 0.367 \| 166 \| 96 \| 6 \| 1 \| 162 \| 1 \| 161 \| Single-stranded DNA-binding protein \| Single-stranded DNA-binding protein; afdb-uniprot50 \| AF-A0A6H9LAI1-F1-MODEL\_V4 \| 1.0 \| 8.027e-15 \| 537 \| 0.4 \| 170 \| 90 \| 6 \| 1 \| 162 \| 1 \| 166 \| Single-stranded DNA-binding protein \| Single-stranded DNA-binding protein; afdb-uniprot50 \| AF-A0A0B7IME9-F1-MODEL\_V4 \| 1.0 \| 3.47e-12 \| 536 \| 0.375 \| 128 \| 74 \| 2 \| 2 \| 129 \| 3 \| 124 \| Single-stranded DNA-binding protein \| Single-stranded DNA-binding protein; afdb-uniprot50 \| AF-U2C4X8-F1-MODEL\_V4 \| 1.0 \| 2.635e-13 \| 536 \| 0.381 \| 139 \| 81 \| 2 \| 1 \| 139 \| 1 \| 134 \| Single-stranded DNA-binding protein \| Single-stranded DNA-binding protein; afdb-uniprot50 \| AF-A0A1F5BEK8-F1-MODEL\_V4 \| 1.0 \| 7.661e-14 \| 536 \| 0.376 \| 162 \| 88 \| 5 \| 2 \| 162 \| 4 \| 153 \| Single-stranded DNA-binding protein \| Single-stranded DNA-binding protein; afdb-uniprot50 \| AF-T1D461-F1-MODEL\_V4 \| 1.0 \| 1.529e-14 \| 535 \| 0.445 \| 164 \| 83 \| 4 \| 4 \| 162 \| 3 \| 163 \| Single-strand DNA-binding protein \| Single-strand DNA-binding protein; afdb-uniprot50 \| AF-A0A1Y1Q967-F1-MODEL\_V4 \| 1.0 \| 7.608e-15 \| 535 \| 0.396 \| 174 \| 91 \| 4 \| 1 \| 162 \| 1 \| 172 \| Single-stranded DNA-binding protein \| Single-stranded DNA-binding protein; afdb-uniprot50 \| AF-A0A2A4U0H7-F1-MODEL\_V4 \| 1.0 \| 2.48e-14 \| 535 \| 0.367 \| 185 \| 93 \| 2 \| 1 \| 162 \| 2 \| 185 \| Single-stranded DNA-binding protein \| Single-stranded DNA-binding protein; afdb-uniprot50 \| AF-A0A4Q7G2J8-F1-MODEL\_V4 \| 1.0 \| 1.311e-13 \| 535 \| 0.389 \| 167 \| 88 \| 6 \| 1 \| 162 \| 26 \| 183 \| Single-stranded DNA-binding protein \| Single-stranded DNA-binding protein; afdb-uniprot50 \| AF-A0A315EK95-F1-MODEL\_V4 \| 1.0 \| 3.637e-13 \| 534 \| 0.404 \| 146 \| 73 \| 3 \| 1 \| 146 \| 1 \| 132 \| Single-stranded DNA-binding protein \| Single-stranded DNA-binding protein; afdb-uniprot50 \| AF-A0A2M7CI87-F1-MODEL\_V4 \| 1.0 \| 3.074e-14 \| 534 \| 0.349 \| 163 \| 99 \| 3 \| 4 \| 162 \| 3 \| 162 \| Single-stranded DNA-binding protein \| Single-stranded DNA-binding protein; afdb-uniprot50 \| AF-A0A7L4ZN54-F1-MODEL\_V4 \| 1.0 \| 5.26e-14 \| 534 \| 0.329 \| 176 \| 101 \| 2 \| 1 \| 162 \| 1 \| 173 \| Single-stranded DNA-binding protein \| Single-stranded DNA-binding protein; afdb-uniprot50 \| AF-A0A7X6U5F3-F1-MODEL\_V4 \| 1.0 \| 4.823e-11 \| 533 \| 0.366 \| 101 \| 64 \| 0 \| 1 \| 101 \| 1 \| 101 \| Single-stranded DNA-binding protein \| Single-stranded DNA-binding protein; afdb-uniprot50 \| AF-A0A644Y581-F1-MODEL\_V4 \| 1.0 \| 4.077e-12 \| 533 \| 0.393 \| 127 \| 67 \| 3 \| 4 \| 129 \| 2 \| 119 \| Single-stranded DNA-binding protein \| Single-stranded DNA-binding protein; afdb-uniprot50 \| AF-A0A0B8QEM1-F1-MODEL\_V4 \| 1.0 \| 7.26e-14 \| 533 \| 0.457 \| 153 \| 75 \| 4 \| 12 \| 162 \| 2 \| 148 \| Single-stranded DNA-binding protein \| Single-stranded DNA-binding protein; afdb-uniprot50 \| AF-A0A2D8XDB3-F1-MODEL\_V4 \| 1.0 \| 1.116e-13 \| 533 \| 0.347 \| 161 \| 92 \| 5 \| 4 \| 162 \| 1 \| 150 \| Single-stranded DNA-binding protein \| Single-stranded DNA-binding protein; afdb-uniprot50 \| AF-A0A2E6JYQ6-F1-MODEL\_V4 \| 1.0 \| 3.244e-14 \| 533 \| 0.397 \| 161 \| 78 \| 4 \| 3 \| 162 \| 2 \| 144 \| Single-stranded DNA-binding protein \| Single-stranded DNA-binding protein; afdb-uniprot50 \| AF-A0A6L7VM16-F1-MODEL\_V4 \| 1.0 \| 1.449e-14 \| 533 \| 0.436 \| 181 \| 76 \| 9 \| 1 \| 162 \| 1 \| 174 \| Single-stranded DNA-binding protein \| Single-stranded DNA-binding protein; afdb-uniprot50 \| AF-A0A848FDV8-F1-MODEL\_V4 \| 1.0 \| 2.761e-14 \| 533 \| 0.371 \| 191 \| 87 \| 5 \| 1 \| 162 \| 1 \| 187 \| Single-stranded DNA-binding protein \| Single-stranded DNA-binding protein; afdb-uniprot50 \| AF-A0A365P1J8-F1-MODEL\_V4 \| 1.0 \| 4.79e-12 \| 532 \| 0.392 \| 130 \| 76 \| 1 \| 1 \| 130 \| 1 \| 127 \| Single-stranded DNA-binding protein \| Single-stranded DNA-binding protein; afdb-uniprot50 \| AF-A0A255ZVS7-F1-MODEL\_V4 \| 1.0 \| 1.636e-12 \| 532 \| 0.377 \| 135 \| 76 \| 2 \| 2 \| 136 \| 3 \| 129 \| Single-stranded DNA-binding protein \| Single-stranded DNA-binding protein; afdb-uniprot50 \| AF-A0A1F2Q8N9-F1-MODEL\_V4 \| 1.0 \| 2.14e-12 \| 532 \| 0.338 \| 139 \| 83 \| 3 \| 4 \| 138 \| 2 \| 135 \| Single-stranded DNA-binding protein \| Single-stranded DNA-binding protein; afdb-uniprot50 \| AF-A0A7Y4RCW9-F1-MODEL\_V4 \| 1.0 \| 9.496e-14 \| 532 \| 0.335 \| 164 \| 96 \| 5 \| 1 \| 162 \| 1 \| 153 \| Single-stranded DNA-binding protein \| Single-stranded DNA-binding protein; afdb-uniprot50 \| AF-A0A7T9BXX1-F1-MODEL\_V4 \| 1.0 \| 9.562e-13 \| 532 \| 0.328 \| 143 \| 90 \| 3 \| 1 \| 141 \| 1 \| 139 \| Single-stranded DNA-binding protein \| Single-stranded DNA-binding protein; afdb-uniprot50 \| AF-A0A081BZQ6-F1-MODEL\_V4 \| 1.0 \| 3.074e-14 \| 531 \| 0.376 \| 162 \| 97 \| 2 \| 1 \| 162 \| 2 \| 159 \| Single-stranded DNA-binding protein \| Single-stranded DNA-binding protein; afdb-uniprot50 \| AF-A0A662ZPE4-F1-MODEL\_V4 \| 1.0 \| 1.311e-13 \| 531 \| 0.351 \| 162 \| 98 \| 2 \| 1 \| 162 \| 1 \| 155 \| Single-stranded DNA-binding protein \| Single-stranded DNA-binding protein; afdb-uniprot50 \| AF-A0A1H9CG65-F1-MODEL\_V4 \| 1.0 \| 2.258e-12 \| 530 \| 0.345 \| 136 \| 85 \| 2 \| 1 \| 136 \| 1 \| 132 \| Single-stranded DNA-binding protein \| Single-stranded DNA-binding protein; afdb-uniprot50 \| AF-A0A0G0IUI9-F1-MODEL\_V4 \| 1.0 \| 1.809e-13 \| 530 \| 0.408 \| 159 \| 84 \| 3 \| 4 \| 162 \| 2 \| 150 \| Single-stranded DNA-binding protein \| Single-stranded DNA-binding protein; afdb-uniprot50 \| AF-K1ZMT3-F1-MODEL\_V4 \| 1.0 \| 8.529e-14 \| 530 \| 0.355 \| 163 \| 97 \| 4 \| 3 \| 162 \| 2 \| 159 \| Single-stranded DNA-binding protein \| Single-stranded DNA-binding protein; afdb-uniprot50 \| AF-A0A139KGR1-F1-MODEL\_V4 \| 1.0 \| 2.913e-14 \| 530 \| 0.395 \| 167 \| 91 \| 6 \| 1 \| 162 \| 1 \| 162 \| Single-stranded DNA-binding protein \| Single-stranded DNA-binding protein; afdb-uniprot50 \| AF-A0A4Q7V7A6-F1-MODEL\_V4 \| 1.0 \| 1.131e-11 \| 528 \| 0.394 \| 119 \| 68 \| 2 \| 4 \| 122 \| 1 \| 115 \| Single-stranded DNA-binding protein \| Single-stranded DNA-binding protein; afdb-uniprot50 \| AF-A0A847B064-F1-MODEL\_V4 \| 1.0 \| 4.508e-13 \| 527 \| 0.327 \| 162 \| 95 \| 4 \| 1 \| 162 \| 1 \| 148 \| Single-stranded DNA-binding protein \| Single-stranded DNA-binding protein; afdb-uniprot50 \| AF-A0A2H0VIG2-F1-MODEL\_V4 \| 1.0 \| 1.529e-14 \| 527 \| 0.359 \| 181 \| 93 \| 5 \| 1 \| 162 \| 2 \| 178 \| Single-stranded DNA-binding protein \| Single-stranded DNA-binding protein; afdb-uniprot50 \| AF-A6DBN5-F1-MODEL\_V4 \| 1.0 \| 1.057e-13 \| 526 \| 0.356 \| 160 \| 89 \| 5 \| 4 \| 162 \| 1 \| 147 \| Single-stranded DNA-binding protein \| Single-stranded DNA-binding protein; afdb-uniprot50 \| AF-A0A2H0YD16-F1-MODEL\_V4 \| 1.0 \| 3.423e-14 \| 526 \| 0.422 \| 161 \| 83 \| 4 \| 4 \| 162 \| 2 \| 154 \| Single-stranded DNA-binding protein \| Single-stranded DNA-binding protein; afdb-uniprot50 \| AF-A0A1V5SBT0-F1-MODEL\_V4 \| 1.0 \| 4.985e-14 \| 526 \| 0.391 \| 161 \| 88 \| 5 \| 4 \| 162 \| 3 \| 155 \| Single-stranded DNA-binding protein \| Single-stranded DNA-binding protein; afdb-uniprot50 \| AF-A0A660S5G9-F1-MODEL\_V4 \| 1.0 \| 1.402e-11 \| 524 \| 0.302 \| 119 \| 76 \| 3 \| 1 \| 115 \| 1 \| 116 \| Single-stranded DNA-binding protein \| Single-stranded DNA-binding protein; afdb-uniprot50 \| AF-A0A1G2R9I5-F1-MODEL\_V4 \| 1.0 \| 4.021e-14 \| 524 \| 0.408 \| 159 \| 88 \| 3 \| 4 \| 162 \| 2 \| 154 \| Single-stranded DNA-binding protein \| Single-stranded DNA-binding protein; afdb-uniprot50 \| AF-A0A2M6WFV0-F1-MODEL\_V4 \| 1.0 \| 3.074e-14 \| 524 \| 0.391 \| 184 \| 81 \| 7 \| 4 \| 162 \| 2 \| 179 \| Single-stranded DNA-binding protein \| Single-stranded DNA-binding protein; afdb-uniprot50 \| AF-A0A2S9Y8A6-F1-MODEL\_V4 \| 1.0 \| 1.169e-14 \| 524 \| 0.391 \| 166 \| 95 \| 3 \| 1 \| 162 \| 1 \| 164 \| Single-stranded DNA-binding protein \| Single-stranded DNA-binding protein; afdb-uniprot50 \| AF-A0A536IEW5-F1-MODEL\_V4 \| 1.0 \| 4.508e-13 \| 523 \| 0.413 \| 138 \| 75 \| 4 \| 1 \| 136 \| 1 \| 134 \| Single-stranded DNA-binding protein \| Single-stranded DNA-binding protein; afdb-uniprot50 \| AF-A0A257K0X4-F1-MODEL\_V4 \| 1.0 \| 5.02e-13 \| 523 \| 0.337 \| 154 \| 89 \| 5 \| 1 \| 154 \| 1 \| 141 \| Single-stranded DNA-binding protein \| Single-stranded DNA-binding protein; afdb-uniprot50 \| AF-A0A7V3EK99-F1-MODEL\_V4 \| 1.0 \| 5.26e-14 \| 523 \| 0.371 \| 167 \| 92 \| 6 \| 1 \| 162 \| 1 \| 159 \| Single-stranded DNA-binding protein \| Single-stranded DNA-binding protein; afdb-uniprot50 \| AF-A0A4P6X7E8-F1-MODEL\_V4 \| 1.0 \| 1.551e-12 \| 522 \| 0.372 \| 137 \| 81 \| 3 \| 1 \| 137 \| 1 \| 132 \| Single-stranded DNA-binding protein \| Single-stranded DNA-binding protein; afdb-uniprot50 \| AF-A0A7T9FFD8-F1-MODEL\_V4 \| 1.0 \| 1.116e-13 \| 522 \| 0.378 \| 164 \| 82 \| 7 \| 1 \| 162 \| 1 \| 146 \| Single-stranded DNA-binding protein \| Single-stranded DNA-binding protein; afdb-uniprot50 \| AF-A0A7Z9KRQ6-F1-MODEL\_V4 \| 1.0 \| 3.891e-11 \| 521 \| 0.272 \| 110 \| 77 \| 1 \| 4 \| 113 \| 3 \| 109 \| Single-stranded DNA-binding protein \| Single-stranded DNA-binding protein; afdb-uniprot50 \| AF-K1WZW4-F1-MODEL\_V4 \| 1.0 \| 1.48e-11 \| 520 \| 0.317 \| 129 \| 78 \| 4 \| 1 \| 128 \| 1 \| 120 \| Single-stranded DNA-binding protein \| Single-stranded DNA-binding protein; afdb-uniprot50 \| AF-A0A4U0GGN8-F1-MODEL\_V4 \| 1.0 \| 1.922e-12 \| 520 \| 0.337 \| 145 \| 82 \| 4 \| 1 \| 145 \| 2 \| 132 \| Single-stranded DNA-binding protein \| Single-stranded DNA-binding protein; afdb-uniprot50 \| AF-A0A2D5BSE2-F1-MODEL\_V4 \| 1.0 \| 4.243e-14 \| 520 \| 0.392 \| 163 \| 90 \| 6 \| 1 \| 162 \| 1 \| 155 \| Single-stranded DNA-binding protein \| Single-stranded DNA-binding protein; afdb-uniprot50 \| AF-A0A6N9D2R3-F1-MODEL\_V4 \| 1.0 \| 1.625e-13 \| 520 \| 0.397 \| 171 \| 88 \| 6 \| 1 \| 162 \| 1 \| 165 \| Single-stranded DNA-binding protein \| Single-stranded DNA-binding protein; afdb-uniprot50 \| AF-R4YT41-F1-MODEL\_V4 \| 1.0 \| 3.244e-14 \| 520 \| 0.419 \| 181 \| 84 \| 5 \| 1 \| 162 \| 1 \| 179 \| Single-stranded DNA-binding protein \| Single-stranded DNA-binding protein; afdb-uniprot50 \| AF-A0A1E8VZ74-F1-MODEL\_V4 \| 1.0 \| 2.8e-12 \| 519 \| 0.404 \| 126 \| 63 \| 2 \| 4 \| 129 \| 2 \| 115 \| Single-stranded DNA-binding protein \| Single-stranded DNA-binding protein; afdb-uniprot50 \| AF-A0A1V4QTT8-F1-MODEL\_V4 \| 1.0 \| 1.909e-13 \| 519 \| 0.381 \| 160 \| 91 \| 3 \| 1 \| 160 \| 1 \| 152 \| Single-stranded DNA-binding protein \| Single-stranded DNA-binding protein; afdb-uniprot50 \| AF-A0A2D6Q1P2-F1-MODEL\_V4 \| 1.0 \| 3.811e-14 \| 519 \| 0.357 \| 176 \| 97 \| 4 \| 1 \| 162 \| 1 \| 174 \| Single-stranded DNA-binding protein \| Single-stranded DNA-binding protein; afdb-uniprot50 \| AF-A0A1F3X205-F1-MODEL\_V4 \| 1.0 \| 4.302e-12 \| 518 \| 0.384 \| 125 \| 67 \| 3 \| 4 \| 128 \| 3 \| 117 \| Single-stranded DNA-binding protein \| Single-stranded DNA-binding protein; afdb-uniprot50 \| AF-A0A7Y4XJK4-F1-MODEL\_V4 \| 1.0 \| 1.116e-13 \| 518 \| 0.397 \| 166 \| 88 \| 4 \| 3 \| 162 \| 2 \| 161 \| Single-stranded DNA-binding protein \| Single-stranded DNA-binding protein; afdb-uniprot50 \| AF-A0A827P8J5-F1-MODEL\_V4 \| 1.0 \| 6.18e-14 \| 517 \| 0.611 \| 144 \| 51 \| 2 \| 21 \| 162 \| 1 \| 141 \| Single-stranded DNA-binding protein \| Single-stranded DNA-binding protein; afdb-uniprot50 \| AF-A0A3B9CYY7-F1-MODEL\_V4 \| 1.0 \| 4.54e-12 \| 516 \| 0.365 \| 134 \| 78 \| 4 \| 1 \| 132 \| 1 \| 129 \| Single-stranded DNA-binding protein \| Single-stranded DNA-binding protein; afdb-uniprot50 \| AF-A0A7X7RFW5-F1-MODEL\_V4 \| 1.0 \| 3.095e-13 \| 516 \| 0.31 \| 164 \| 101 \| 5 \| 1 \| 162 \| 1 \| 154 \| Single-stranded DNA-binding protein \| Single-stranded DNA-binding protein; afdb-uniprot50 \| AF-A0A3L7QDN6-F1-MODEL\_V4 \| 1.0 \| 4.724e-14 \| 516 \| 0.329 \| 173 \| 104 \| 4 \| 1 \| 162 \| 1 \| 172 \| Single-stranded DNA-binding protein \| Single-stranded DNA-binding protein; afdb-uniprot50 \| AF-A9AYW2-F1-MODEL\_V4 \| 1.0 \| 2.042e-11 \| 515 \| 0.375 \| 120 \| 69 \| 4 \| 1 \| 118 \| 3 \| 118 \| Single-stranded DNA-binding protein \| Single-stranded DNA-binding protein; afdb-uniprot50 \| AF-A0A660NA14-F1-MODEL\_V4 \| 1.0 \| 1.47e-12 \| 515 \| 0.308 \| 146 \| 91 \| 4 \| 4 \| 142 \| 3 \| 145 \| Single-stranded DNA-binding protein \| Single-stranded DNA-binding protein; afdb-uniprot50 \| AF-A0A7X6AFC1-F1-MODEL\_V4 \| 1.0 \| 2.366e-13 \| 515 \| 0.333 \| 168 \| 100 \| 5 \| 1 \| 161 \| 3 \| 165 \| Single-stranded DNA-binding protein \| Single-stranded DNA-binding protein; afdb-uniprot50 \| AF-A0A7C0ZN74-F1-MODEL\_V4 \| 1.0 \| 4.021e-14 \| 514 \| 0.353 \| 167 \| 99 \| 2 \| 4 \| 162 \| 2 \| 167 \| Single-stranded DNA-binding protein \| Single-stranded DNA-binding protein; afdb-uniprot50 \| AF-A0A432GXJ5-F1-MODEL\_V4 \| 1.0 \| 3.095e-13 \| 514 \| 0.3 \| 163 \| 105 \| 4 \| 1 \| 162 \| 75 \| 229 \| Single-stranded DNA-binding protein \| Single-stranded DNA-binding protein; afdb-uniprot50 \| AF-A0A317HH03-F1-MODEL\_V4 \| 1.0 \| 2.383e-12 \| 513 \| 0.387 \| 129 \| 77 \| 2 \| 1 \| 128 \| 8 \| 135 \| Single-stranded DNA-binding protein \| Single-stranded DNA-binding protein; afdb-uniprot50 \| AF-A0A2V2EFG8-F1-MODEL\_V4 \| 1.0 \| 6.566e-13 \| 513 \| 0.358 \| 162 \| 86 \| 5 \| 4 \| 162 \| 2 \| 148 \| Single-stranded DNA-binding protein \| Single-stranded DNA-binding protein; afdb-uniprot50 \| AF-A0A2E8VMN4-F1-MODEL\_V4 \| 1.0 \| 4.757e-13 \| 513 \| 0.333 \| 153 \| 92 \| 4 \| 1 \| 148 \| 6 \| 153 \| Single-stranded DNA-binding protein \| Single-stranded DNA-binding protein; afdb-uniprot50 \| AF-A0A7X8BKP9-F1-MODEL\_V4 \| 1.0 \| 2.028e-12 \| 512 \| 0.342 \| 146 \| 83 \| 3 \| 1 \| 146 \| 5 \| 137 \| Single-stranded DNA-binding protein \| Single-stranded DNA-binding protein; afdb-uniprot50 \| AF-A0A2M6ZHX2-F1-MODEL\_V4 \| 1.0 \| 6.222e-13 \| 512 \| 0.318 \| 157 \| 98 \| 4 \| 1 \| 156 \| 1 \| 149 \| Single-stranded DNA-binding protein \| Single-stranded DNA-binding protein; afdb-uniprot50 \| AF-F4T715-F1-MODEL\_V4 \| 1.0 \| 3.811e-14 \| 512 \| 0.506 \| 158 \| 67 \| 2 \| 16 \| 162 \| 1 \| 158 \| Single-stranded DNA-binding protein \| Single-stranded DNA-binding protein; afdb-uniprot50 \| AF-A0A3E4Z701-F1-MODEL\_V4 \| 1.0 \| 9e-14 \| 512 \| 0.379 \| 166 \| 94 \| 4 \| 4 \| 162 \| 1 \| 164 \| Single-stranded DNA-binding protein \| Single-stranded DNA-binding protein; afdb-uniprot50 \| AF-A0A1T0AXC1-F1-MODEL\_V4 \| 1.0 \| 6.657e-11 \| 511 \| 0.35 \| 117 \| 73 \| 1 \| 1 \| 117 \| 1 \| 114 \| Single-stranded DNA-binding protein \| Single-stranded DNA-binding protein; afdb-uniprot50 \| AF-A0A2E6Z2B5-F1-MODEL\_V4 \| 1.0 \| 7.714e-13 \| 511 \| 0.308 \| 159 \| 96 \| 3 \| 4 \| 162 \| 1 \| 145 \| Single-stranded DNA-binding protein \| Single-stranded DNA-binding protein; afdb-uniprot50 \| AF-A0A7C5KF93-F1-MODEL\_V4 \| 1.0 \| 1.177e-13 \| 511 \| 0.327 \| 168 \| 102 \| 4 \| 1 \| 162 \| 1 \| 163 \| Single-stranded DNA-binding protein \| Single-stranded DNA-binding protein; afdb-uniprot50 \| AF-A0A7C3URN1-F1-MODEL\_V4 \| 1.0 \| 1.26e-11 \| 510 \| 0.391 \| 115 \| 69 \| 1 \| 3 \| 117 \| 7 \| 120 \| Single-stranded DNA-binding protein \| Single-stranded DNA-binding protein; afdb-uniprot50 \| AF-A0A7W0J0Y0-F1-MODEL\_V4 \| 1.0 \| 1.123e-12 \| 510 \| 0.324 \| 154 \| 93 \| 6 \| 1 \| 154 \| 1 \| 143 \| Single-stranded DNA-binding protein \| Single-stranded DNA-binding protein; afdb-uniprot50 \| AF-A0A1L5KLY0-F1-MODEL\_V4 \| 1.0 \| 2.227e-14 \| 510 \| 0.584 \| 159 \| 51 \| 2 \| 19 \| 162 \| 1 \| 159 \| Single-stranded DNA-binding protein \| Single-stranded DNA-binding protein; afdb-uniprot50 \| AF-A0A1X4XY99-F1-MODEL\_V4 \| 1.0 \| 7.767e-12 \| 509 \| 0.325 \| 126 \| 81 \| 2 \| 4 \| 129 \| 2 \| 123 \| Single-stranded DNA-binding protein \| Single-stranded DNA-binding protein; afdb-uniprot50 \| AF-A0A1G0AMD8-F1-MODEL\_V4 \| 1.0 \| 5.054e-12 \| 509 \| 0.314 \| 143 \| 91 \| 2 \| 1 \| 143 \| 1 \| 136 \| Single-stranded DNA-binding protein \| Single-stranded DNA-binding protein; afdb-uniprot50 \| AF-A0A661XBR9-F1-MODEL\_V4 \| 1.0 \| 3.289e-12 \| 509 \| 0.343 \| 128 \| 78 \| 3 \| 1 \| 128 \| 1 \| 122 \| Single-stranded DNA-binding protein \| Single-stranded DNA-binding protein; afdb-uniprot50 \| AF-A0A7C5UPK2-F1-MODEL\_V4 \| 1.0 \| 1.715e-13 \| 509 \| 0.364 \| 159 \| 90 \| 3 \| 1 \| 156 \| 2 \| 152 \| Single-stranded DNA-binding protein \| Single-stranded DNA-binding protein; afdb-uniprot50 \| AF-A0A2E3HXA7-F1-MODEL\_V4 \| 1.0 \| 8.708e-11 \| 508 \| 0.343 \| 99 \| 65 \| 0 \| 4 \| 102 \| 3 \| 101 \| Single-stranded DNA-binding protein \| Single-stranded DNA-binding protein; afdb-uniprot50 \| AF-A0A250FUA3-F1-MODEL\_V4 \| 1.0 \| 7.31e-13 \| 508 \| 0.333 \| 156 \| 94 \| 3 \| 1 \| 156 \| 1 \| 146 \| Single-stranded DNA-binding protein \| Single-stranded DNA-binding protein; afdb-uniprot50 \| AF-A0A379MNE0-F1-MODEL\_V4 \| 1.0 \| 1.54e-13 \| 508 \| 0.4 \| 160 \| 88 \| 4 \| 4 \| 162 \| 1 \| 153 \| Single-stranded DNA-binding protein \| Single-stranded DNA-binding protein; afdb-uniprot50 \| AF-A0A1D3K7U2-F1-MODEL\_V4 \| 1.0 \| 4.508e-13 \| 508 \| 0.335 \| 173 \| 102 \| 6 \| 1 \| 162 \| 2 \| 172 \| Single-stranded DNA-binding protein \| Single-stranded DNA-binding protein; afdb-uniprot50 \| AF-A0A2E5LYI8-F1-MODEL\_V4 \| 1.0 \| 1.54e-13 \| 508 \| 0.356 \| 160 \| 95 \| 3 \| 3 \| 162 \| 2 \| 153 \| Multifunctional fusion protein \| Multifunctional fusion protein; afdb-uniprot50 \| AF-A0A0P6Y529-F1-MODEL\_V4 \| 1.0 \| 3.312e-11 \| 507 \| 0.336 \| 119 \| 75 \| 2 \| 1 \| 118 \| 3 \| 118 \| Single-stranded DNA-binding protein \| Single-stranded DNA-binding protein; afdb-uniprot50 \| AF-A0A7C4NQI9-F1-MODEL\_V4 \| 1.0 \| 1.561e-11 \| 506 \| 0.318 \| 116 \| 76 \| 2 \| 3 \| 116 \| 2 \| 116 \| Single-stranded DNA-binding protein \| Single-stranded DNA-binding protein; afdb-uniprot50 \| AF-A0A2S1QWE8-F1-MODEL\_V4 \| 1.0 \| 1.251e-12 \| 506 \| 0.349 \| 166 \| 85 \| 5 \| 1 \| 162 \| 1 \| 147 \| Single-stranded DNA-binding protein \| Single-stranded DNA-binding protein; afdb-uniprot50 \| AF-A0A653HVB8-F1-MODEL\_V4 \| 1.0 \| 6.222e-13 \| 506 \| 0.339 \| 159 \| 97 \| 4 \| 4 \| 162 \| 1 \| 151 \| Single-stranded DNA-binding protein \| Single-stranded DNA-binding protein; afdb-uniprot50 \| AF-A0A2T3ITI4-F1-MODEL\_V4 \| 1.0 \| 2.78e-13 \| 506 \| 0.331 \| 163 \| 101 \| 4 \| 4 \| 158 \| 1 \| 163 \| Single-stranded DNA-binding protein \| Single-stranded DNA-binding protein; afdb-uniprot50 \| AF-A0A0G0BK79-F1-MODEL\_V4 \| 1.0 \| 4.985e-14 \| 506 \| 0.329 \| 170 \| 105 \| 3 \| 1 \| 162 \| 2 \| 170 \| Single-stranded DNA-binding protein \| Single-stranded DNA-binding protein; afdb-uniprot50 \| AF-A0A3M1FQV6-F1-MODEL\_V4 \| 1.0 \| 2.383e-12 \| 505 \| 0.47 \| 136 \| 67 \| 2 \| 10 \| 145 \| 1 \| 131 \| Single-stranded DNA-binding protein \| Single-stranded DNA-binding protein; afdb-uniprot50 \| AF-A0A257LTW8-F1-MODEL\_V4 \| 1.0 \| 6.976e-12 \| 505 \| 0.338 \| 136 \| 86 \| 2 \| 1 \| 136 \| 5 \| 136 \| Single-stranded DNA-binding protein \| Single-stranded DNA-binding protein; afdb-uniprot50 \| AF-A0A7X3UCI6-F1-MODEL\_V4 \| 1.0 \| 1.715e-13 \| 505 \| 0.393 \| 160 \| 76 \| 4 \| 3 \| 162 \| 2 \| 140 \| Single-stranded DNA-binding protein \| Single-stranded DNA-binding protein; afdb-uniprot50 \| AF-A0A1J5HHA6-F1-MODEL\_V4 \| 1.0 \| 2.243e-13 \| 505 \| 0.312 \| 160 \| 101 \| 5 \| 4 \| 162 \| 3 \| 154 \| Single-stranded DNA-binding protein \| Single-stranded DNA-binding protein; afdb-uniprot50 \| AF-A0A7C7W444-F1-MODEL\_V4 \| 1.0 \| 3.139e-11 \| 505 \| 0.304 \| 115 \| 79 \| 1 \| 3 \| 117 \| 2 \| 115 \| Single-stranded DNA-binding protein \| Single-stranded DNA-binding protein; afdb-uniprot50 \| AF-A0A5C5X2Q6-F1-MODEL\_V4 \| 1.0 \| 1.32e-12 \| 504 \| 0.344 \| 145 \| 87 \| 5 \| 1 \| 144 \| 1 \| 138 \| Single-stranded DNA-binding protein \| Single-stranded DNA-binding protein; afdb-uniprot50 \| AF-A0A2N3G5B3-F1-MODEL\_V4 \| 1.0 \| 4.508e-13 \| 504 \| 0.339 \| 153 \| 94 \| 5 \| 1 \| 152 \| 1 \| 147 \| Single-stranded DNA-binding protein \| Single-stranded DNA-binding protein; afdb-uniprot50 \| AF-A0A661ED08-F1-MODEL\_V4 \| 1.0 \| 6.88e-14 \| 504 \| 0.382 \| 170 \| 89 \| 6 \| 4 \| 162 \| 3 \| 167 \| Single-stranded DNA-binding protein \| Single-stranded DNA-binding protein; afdb-uniprot50 \| AF-A0A2E7GB44-F1-MODEL\_V4 \| 1.0 \| 2.399e-11 \| 503 \| 0.375 \| 112 \| 69 \| 1 \| 4 \| 115 \| 3 \| 113 \| Single-stranded DNA-binding protein \| Single-stranded DNA-binding protein; afdb-uniprot50 \| AF-A0A060R893-F1-MODEL\_V4 \| 1.0 \| 2.8e-12 \| 503 \| 0.34 \| 141 \| 88 \| 3 \| 4 \| 143 \| 1 \| 137 \| Single-stranded DNA-binding protein \| Single-stranded DNA-binding protein; afdb-uniprot50 \| AF-A0A7W1TQ49-F1-MODEL\_V4 \| 1.0 \| 7.025e-11 \| 502 \| 0.356 \| 115 \| 71 \| 1 \| 1 \| 115 \| 4 \| 115 \| Single-stranded DNA-binding protein \| Single-stranded DNA-binding protein; afdb-uniprot50 \| AF-A0A6M8W133-F1-MODEL\_V4 \| 1.0 \| 9.062e-13 \| 502 \| 0.5 \| 154 \| 57 \| 7 \| 10 \| 162 \| 1 \| 135 \| Single-stranded DNA-binding protein \| Single-stranded DNA-binding protein; afdb-uniprot50 \| AF-A0A538CVG0-F1-MODEL\_V4 \| 1.0 \| 2.78e-13 \| 502 \| 0.329 \| 173 \| 91 \| 3 \| 2 \| 162 \| 80 \| 239 \| Single-stranded DNA-binding protein \| Single-stranded DNA-binding protein; afdb-uniprot50 \| AF-W0PFC5-F1-MODEL\_V4 \| 1.0 \| 3.495e-11 \| 501 \| 0.279 \| 118 \| 83 \| 2 \| 1 \| 117 \| 1 \| 117 \| Single-strand binding protein \| Single-strand binding protein; afdb-uniprot50 \| AF-A0A838E1P8-F1-MODEL\_V4 \| 1.0 \| 1.551e-12 \| 501 \| 0.301 \| 169 \| 86 \| 6 \| 1 \| 162 \| 1 \| 144 \| Single-stranded DNA-binding protein \| Single-stranded DNA-binding protein; afdb-uniprot50 \| AF-A0A7K2CRM9-F1-MODEL\_V4 \| 1.0 \| 3.837e-13 \| 501 \| 0.308 \| 162 \| 108 \| 2 \| 1 \| 162 \| 1 \| 158 \| Single-stranded DNA-binding protein \| Single-stranded DNA-binding protein; afdb-uniprot50 \| AF-A0A3M0Z921-F1-MODEL\_V4 \| 1.0 \| 3.266e-13 \| 500 \| 0.37 \| 178 \| 91 \| 6 \| 1 \| 162 \| 1 \| 173 \| Single-stranded DNA-binding protein \| Single-stranded DNA-binding protein; afdb-uniprot50 \| AF-A0A2H0DH32-F1-MODEL\_V4 \| 1.0 \| 5.628e-12 \| 499 \| 0.307 \| 140 \| 88 \| 3 \| 1 \| 136 \| 1 \| 135 \| Single-stranded DNA-binding protein \| Single-stranded DNA-binding protein; afdb-uniprot50 \| AF-I4B8K2-F1-MODEL\_V4 \| 1.0 \| 2.653e-12 \| 499 \| 0.296 \| 162 \| 99 \| 2 \| 1 \| 162 \| 1 \| 147 \| Single-stranded DNA-binding protein \| Single-stranded DNA-binding protein; afdb-uniprot50 \| AF-A0A1S9ZTE5-F1-MODEL\_V4 \| 1.0 \| 1.551e-12 \| 499 \| 0.293 \| 167 \| 91 \| 4 \| 4 \| 162 \| 2 \| 149 \| Single-stranded DNA-binding protein \| Single-stranded DNA-binding protein; afdb-uniprot50 \| AF-A0A2M7TEI6-F1-MODEL\_V4 \| 1.0 \| 9.062e-13 \| 499 \| 0.337 \| 163 \| 95 \| 5 \| 1 \| 162 \| 2 \| 152 \| Single-stranded DNA-binding protein \| Single-stranded DNA-binding protein; afdb-uniprot50 \| AF-A0A1H2FT79-F1-MODEL\_V4 \| 1.0 \| 1.809e-13 \| 499 \| 0.337 \| 172 \| 100 \| 6 \| 1 \| 162 \| 1 \| 168 \| Single-stranded DNA-binding protein \| Single-stranded DNA-binding protein; afdb-uniprot50 \| AF-A0A2N1TAT9-F1-MODEL\_V4 \| 1.0 \| 4.477e-14 \| 498 \| 0.353 \| 167 \| 98 \| 4 \| 4 \| 162 \| 3 \| 167 \| Single-stranded DNA-binding protein \| Single-stranded DNA-binding protein; afdb-uniprot50 \| AF-A0A836SQ28-F1-MODEL\_V4 \| 1.0 \| 5.054e-12 \| 497 \| 0.313 \| 137 \| 87 \| 3 \| 3 \| 135 \| 7 \| 140 \| Single-stranded DNA-binding protein \| Single-stranded DNA-binding protein; afdb-uniprot50 \| AF-A0A7V3VUI2-F1-MODEL\_V4 \| 1.0 \| 1.636e-12 \| 497 \| 0.299 \| 137 \| 91 \| 2 \| 3 \| 135 \| 7 \| 142 \| Single-stranded DNA-binding protein \| Single-stranded DNA-binding protein; afdb-uniprot50 \| AF-A0A7V3V0E3-F1-MODEL\_V4 \| 1.0 \| 6.611e-12 \| 497 \| 0.325 \| 126 \| 80 \| 2 \| 4 \| 129 \| 26 \| 146 \| Single-stranded DNA-binding protein \| Single-stranded DNA-binding protein; afdb-uniprot50 \| AF-A0A090LE13-F1-MODEL\_V4 \| 1.0 \| 1.023e-10 \| 497 \| 0.245 \| 118 \| 86 \| 1 \| 1 \| 118 \| 62 \| 176 \| Primosome PriB/single-strand DNA-binding family and Single-strand DNA-binding family and Nucleic acid-binding, OB-fold domain-containing protein \| Primosome PriB/single-strand DNA-binding family and Single-strand DNA-binding family and Nucleic acid-binding, OB-fold domain-containing protein; afdb-uniprot50 \| AF-A0A1Y4GFY9-F1-MODEL\_V4 \| 1.0 \| 6.222e-13 \| 496 \| 0.292 \| 157 \| 96 \| 4 \| 1 \| 157 \| 1 \| 142 \| Single-stranded DNA-binding protein \| Single-stranded DNA-binding protein; afdb-uniprot50 \| AF-A0A4Q7V6Z3-F1-MODEL\_V4 \| 1.0 \| 7.31e-13 \| 495 \| 0.356 \| 146 \| 83 \| 4 \| 1 \| 136 \| 1 \| 145 \| Single-stranded DNA-binding protein \| Single-stranded DNA-binding protein; afdb-uniprot50 \| AF-A0A523V6H0-F1-MODEL\_V4 \| 1.0 \| 1.822e-12 \| 495 \| 0.282 \| 163 \| 100 \| 5 \| 1 \| 162 \| 1 \| 147 \| Single-stranded DNA-binding protein \| Single-stranded DNA-binding protein; afdb-uniprot50 \| AF-A0A2D5WAR7-F1-MODEL\_V4 \| 1.0 \| 8.589e-13 \| 495 \| 0.312 \| 163 \| 101 \| 5 \| 1 \| 162 \| 1 \| 153 \| Single-stranded DNA-binding protein \| Single-stranded DNA-binding protein; afdb-uniprot50 \| AF-A0A381E7Q0-F1-MODEL\_V4 \| 1.0 \| 2.934e-13 \| 495 \| 0.309 \| 168 \| 105 \| 6 \| 1 \| 161 \| 2 \| 165 \| Single-stranded DNA-binding protein \| Single-stranded DNA-binding protein; afdb-uniprot50 \| AF-A0A7C3YLT4-F1-MODEL\_V4 \| 1.0 \| 5.589e-13 \| 494 \| 0.339 \| 162 \| 88 \| 5 \| 1 \| 162 \| 1 \| 143 \| Single-stranded DNA-binding protein \| Single-stranded DNA-binding protein; afdb-uniprot50 \| AF-A0A3M2CSC7-F1-MODEL\_V4 \| 1.0 \| 8.589e-13 \| 494 \| 0.304 \| 161 \| 106 \| 3 \| 4 \| 162 \| 3 \| 159 \| Single-stranded DNA-binding protein \| Single-stranded DNA-binding protein; afdb-uniprot50 \| AF-A0A1V6GP20-F1-MODEL\_V4 \| 1.0 \| 9.562e-13 \| 494 \| 0.345 \| 165 \| 99 \| 6 \| 1 \| 162 \| 1 \| 159 \| Single-stranded DNA-binding protein \| Single-stranded DNA-binding protein; afdb-uniprot50 \| AF-A0A2M8NDZ2-F1-MODEL\_V4 \| 1.0 \| 1.185e-12 \| 494 \| 0.297 \| 168 \| 107 \| 6 \| 1 \| 162 \| 2 \| 164 \| Single-stranded DNA-binding protein \| Single-stranded DNA-binding protein; afdb-uniprot50 \| AF-A0A7C2DEI9-F1-MODEL\_V4 \| 1.0 \| 9.062e-13 \| 493 \| 0.359 \| 164 \| 94 \| 5 \| 1 \| 162 \| 1 \| 155 \| Single-stranded DNA-binding protein \| Single-stranded DNA-binding protein; afdb-uniprot50 \| AF-A0A010YT66-F1-MODEL\_V4 \| 1.0 \| 1.123e-12 \| 493 \| 0.288 \| 163 \| 102 \| 4 \| 1 \| 162 \| 1 \| 150 \| Single-stranded DNA-binding protein \| Single-stranded DNA-binding protein; afdb-uniprot50 \| AF-A0A7M3PPS8-F1-MODEL\_V4 \| 1.0 \| 7.26e-14 \| 493 \| 0.457 \| 175 \| 77 \| 4 \| 1 \| 162 \| 26 \| 195 \| Single-stranded DNA-binding protein \| Single-stranded DNA-binding protein; afdb-uniprot50 \| AF-A0A523RKR1-F1-MODEL\_V4 \| 1.0 \| 4.508e-13 \| 493 \| 0.274 \| 208 \| 93 \| 7 \| 1 \| 162 \| 1 \| 196 \| Single-stranded DNA-binding protein \| Single-stranded DNA-binding protein; afdb-uniprot50 \| AF-A0A557PGW7-F1-MODEL\_V4 \| 1.0 \| 2.819e-11 \| 492 \| 0.267 \| 157 \| 87 \| 3 \| 4 \| 159 \| 3 \| 132 \| Single-stranded DNA-binding protein \| Single-stranded DNA-binding protein; afdb-uniprot50 \| AF-A0A7W1XXL3-F1-MODEL\_V4 \| 1.0 \| 1.393e-12 \| 492 \| 0.333 \| 159 \| 93 \| 5 \| 4 \| 162 \| 2 \| 147 \| Single-stranded DNA-binding protein \| Single-stranded DNA-binding protein; afdb-uniprot50 \| AF-A0A1Y4DPV1-F1-MODEL\_V4 \| 1.0 \| 2.653e-12 \| 491 \| 0.314 \| 159 \| 105 \| 2 \| 4 \| 162 \| 2 \| 156 \| Single-stranded DNA-binding protein \| Single-stranded DNA-binding protein; afdb-uniprot50 \| AF-A0A3M0WEQ2-F1-MODEL\_V4 \| 1.0 \| 7.767e-12 \| 490 \| 0.301 \| 159 \| 90 \| 3 \| 4 \| 162 \| 1 \| 138 \| Single-stranded DNA-binding protein \| Single-stranded DNA-binding protein; afdb-uniprot50 \| AF-A0A6L3YVY1-F1-MODEL\_V4 \| 1.0 \| 1.251e-12 \| 490 \| 0.364 \| 170 \| 96 \| 7 \| 1 \| 162 \| 1 \| 166 \| Single-stranded DNA-binding protein \| Single-stranded DNA-binding protein; afdb-uniprot50 \| AF-G2LEN2-F1-MODEL\_V4 \| 1.0 \| 1.185e-12 \| 489 \| 0.345 \| 159 \| 87 \| 3 \| 4 \| 162 \| 2 \| 143 \| Single-stranded DNA-binding protein \| Single-stranded DNA-binding protein; afdb-uniprot50 \| AF-A0A2N6QFG8-F1-MODEL\_V4 \| 1.0 \| 1.393e-12 \| 489 \| 0.317 \| 151 \| 95 \| 3 \| 1 \| 149 \| 1 \| 145 \| Single-stranded DNA-binding protein \| Single-stranded DNA-binding protein; afdb-uniprot50 \| AF-A0A1F9XKK2-F1-MODEL\_V4 \| 1.0 \| 2.042e-11 \| 488 \| 0.333 \| 123 \| 79 \| 2 \| 1 \| 123 \| 4 \| 123 \| Single-stranded DNA-binding protein \| Single-stranded DNA-binding protein; afdb-uniprot50 \| AF-A0A7Y2SQU7-F1-MODEL\_V4 \| 1.0 \| 1.48e-11 \| 488 \| 0.287 \| 132 \| 88 \| 3 \| 1 \| 129 \| 4 \| 132 \| Single-stranded DNA-binding protein \| Single-stranded DNA-binding protein; afdb-uniprot50 \| AF-A0A1H4XXA5-F1-MODEL\_V4 \| 1.0 \| 3.289e-12 \| 488 \| 0.32 \| 159 \| 85 \| 5 \| 4 \| 162 \| 1 \| 136 \| Single-stranded DNA-binding protein \| Single-stranded DNA-binding protein; afdb-uniprot50 \| AF-A0A1Y3UKN9-F1-MODEL\_V4 \| 1.0 \| 8.648e-12 \| 487 \| 0.3 \| 160 \| 81 \| 4 \| 4 \| 162 \| 2 \| 131 \| Single-stranded DNA-binding protein \| Single-stranded DNA-binding protein; afdb-uniprot50 \| AF-A0A7X6ICS6-F1-MODEL\_V4 \| 1.0 \| 2.042e-11 \| 487 \| 0.27 \| 159 \| 87 \| 4 \| 4 \| 162 \| 2 \| 131 \| Single-stranded DNA-binding protein \| Single-stranded DNA-binding protein; afdb-uniprot50 \| AF-A0A2E0L7G6-F1-MODEL\_V4 \| 1.0 \| 4.54e-12 \| 487 \| 0.308 \| 159 \| 85 \| 5 \| 4 \| 162 \| 2 \| 135 \| Single-stranded DNA-binding protein \| Single-stranded DNA-binding protein; afdb-uniprot50 \| AF-S3XEZ1-F1-MODEL\_V4 \| 1.0 \| 4.757e-13 \| 487 \| 0.358 \| 159 \| 91 \| 4 \| 4 \| 162 \| 1 \| 148 \| Single-stranded DNA-binding protein \| Single-stranded DNA-binding protein; afdb-uniprot50 \| AF-A0A354CZB3-F1-MODEL\_V4 \| 1.0 \| 2.954e-12 \| 486 \| 0.337 \| 154 \| 92 \| 4 \| 1 \| 154 \| 1 \| 144 \| Single-stranded DNA-binding protein \| Single-stranded DNA-binding protein; afdb-uniprot50 \| AF-A0A2E8DUA8-F1-MODEL\_V4 \| 1.0 \| 1.551e-12 \| 486 \| 0.246 \| 162 \| 100 \| 3 \| 1 \| 162 \| 16 \| 155 \| Single-stranded DNA-binding protein \| Single-stranded DNA-binding protein; afdb-uniprot50 \| AF-A0A073IMY4-F1-MODEL\_V4 \| 1.0 \| 8.589e-13 \| 486 \| 0.303 \| 181 \| 97 \| 6 \| 1 \| 162 \| 1 \| 171 \| Single-stranded DNA-binding protein \| Single-stranded DNA-binding protein; afdb-uniprot50 \| AF-A0A2M8NSD8-F1-MODEL\_V4 \| 1.0 \| 8.648e-12 \| 485 \| 0.385 \| 114 \| 68 \| 2 \| 1 \| 114 \| 1 \| 112 \| Single-stranded DNA-binding protein \| Single-stranded DNA-binding protein; afdb-uniprot50 \| AF-A0A257B1D5-F1-MODEL\_V4 \| 1.0 \| 4.302e-12 \| 485 \| 0.337 \| 151 \| 86 \| 2 \| 4 \| 154 \| 2 \| 138 \| Single-stranded DNA-binding protein \| Single-stranded DNA-binding protein; afdb-uniprot50 \| AF-A0A1F9VFL1-F1-MODEL\_V4 \| 1.0 \| 5.333e-12 \| 485 \| 0.341 \| 161 \| 89 \| 4 \| 3 \| 162 \| 2 \| 146 \| Single-stranded DNA-binding protein \| Single-stranded DNA-binding protein; afdb-uniprot50 \| AF-A0A2D6HC40-F1-MODEL\_V4 \| 1.0 \| 2.78e-13 \| 485 \| 0.319 \| 166 \| 102 \| 3 \| 3 \| 162 \| 2 \| 162 \| Single-stranded DNA-binding protein \| Single-stranded DNA-binding protein; afdb-uniprot50 \| AF-A0A495LT57-F1-MODEL\_V4 \| 1.0 \| 1.123e-12 \| 485 \| 0.333 \| 162 \| 92 \| 4 \| 1 \| 162 \| 21 \| 166 \| Single-stranded DNA-binding protein \| Single-stranded DNA-binding protein; afdb-uniprot50 \| AF-A0A351JNY7-F1-MODEL\_V4 \| 1.0 \| 3.662e-12 \| 484 \| 0.28 \| 146 \| 98 \| 2 \| 3 \| 148 \| 2 \| 140 \| Single-stranded DNA-binding protein \| Single-stranded DNA-binding protein; afdb-uniprot50 \| AF-A0A1F2X262-F1-MODEL\_V4 \| 1.0 \| 1.822e-12 \| 484 \| 0.287 \| 167 \| 97 \| 6 \| 1 \| 162 \| 1 \| 150 \| Single-stranded DNA-binding protein \| Single-stranded DNA-binding protein; afdb-uniprot50 \| AF-A0A7X8AGS9-F1-MODEL\_V4 \| 1.0 \| 4.79e-12 \| 483 \| 0.339 \| 159 \| 89 \| 4 \| 4 \| 162 \| 1 \| 143 \| Single-stranded DNA-binding protein \| Single-stranded DNA-binding protein; afdb-uniprot50 \| AF-A0A7C3RHW0-F1-MODEL\_V4 \| 1.0 \| 9.062e-13 \| 482 \| 0.341 \| 155 \| 95 \| 4 \| 1 \| 154 \| 3 \| 151 \| Single-stranded DNA-binding protein \| Single-stranded DNA-binding protein; afdb-uniprot50 \| AF-W5YUV6-F1-MODEL\_V4 \| 1.0 \| 6.222e-13 \| 482 \| 0.447 \| 161 \| 80 \| 3 \| 4 \| 162 \| 1 \| 154 \| Single-stranded DNA-binding protein \| Single-stranded DNA-binding protein; afdb-uniprot50 \| AF-A0A0K1F1N5-F1-MODEL\_V4 \| 1.0 \| 8.589e-13 \| 482 \| 0.331 \| 160 \| 97 \| 3 \| 4 \| 162 \| 2 \| 152 \| Single-stranded DNA-binding protein \| Single-stranded DNA-binding protein; afdb-uniprot50 \| AF-A0A845H8A1-F1-MODEL\_V4 \| 1.0 \| 1.123e-12 \| 481 \| 0.453 \| 152 \| 72 \| 4 \| 12 \| 162 \| 95 \| 236 \| Single-stranded DNA-binding protein \| Single-stranded DNA-binding protein; afdb-uniprot50 \| AF-A0A2A5A532-F1-MODEL\_V4 \| 1.0 \| 7.714e-13 \| 480 \| 0.293 \| 160 \| 101 \| 3 \| 4 \| 162 \| 3 \| 151 \| Single-stranded DNA-binding protein \| Single-stranded DNA-binding protein; afdb-uniprot50 \| AF-H0E9H1-F1-MODEL\_V4 \| 1.0 \| 4.508e-13 \| 480 \| 0.327 \| 162 \| 99 \| 4 \| 4 \| 162 \| 2 \| 156 \| Single-stranded DNA-binding protein \| Single-stranded DNA-binding protein; afdb-uniprot50 \| AF-A0A0S1SMJ0-F1-MODEL\_V4 \| 1.0 \| 1.123e-12 \| 480 \| 0.375 \| 141 \| 79 \| 2 \| 5 \| 137 \| 128 \| 267 \| Single-stranded DNA-binding protein \| Single-stranded DNA-binding protein; afdb-uniprot50 \| AF-A0A437UU05-F1-MODEL\_V4 \| 1.0 \| 6.222e-13 \| 479 \| 0.349 \| 169 \| 97 \| 5 \| 4 \| 162 \| 2 \| 167 \| Single-stranded DNA-binding protein \| Single-stranded DNA-binding protein; afdb-uniprot50 \| AF-A0A0F5PBD5-F1-MODEL\_V4 \| 1.0 \| 1.47e-12 \| 476 \| 0.355 \| 163 \| 97 \| 4 \| 1 \| 162 \| 1 \| 156 \| Single-stranded DNA-binding protein \| Single-stranded DNA-binding protein; afdb-uniprot50 \| AF-A0A1S9ZZ07-F1-MODEL\_V4 \| 1.0 \| 1.625e-13 \| 476 \| 0.383 \| 167 \| 89 \| 6 \| 4 \| 162 \| 1 \| 161 \| Single-stranded DNA-binding protein \| Single-stranded DNA-binding protein; afdb-uniprot50 \| AF-A0A3D0VSK9-F1-MODEL\_V4 \| 1.0 \| 2.839e-10 \| 475 \| 0.336 \| 113 \| 72 \| 2 \| 2 \| 114 \| 34 \| 143 \| Single-stranded DNA-binding protein \| Single-stranded DNA-binding protein; afdb-uniprot50 \| AF-A0A0G0L6C8-F1-MODEL\_V4 \| 1.0 \| 4.049e-13 \| 475 \| 0.344 \| 183 \| 86 \| 8 \| 4 \| 162 \| 3 \| 175 \| Single-stranded DNA-binding protein \| Single-stranded DNA-binding protein; afdb-uniprot50 \| AF-A0A1F3IVI4-F1-MODEL\_V4 \| 1.0 \| 3.918e-10 \| 474 \| 0.345 \| 110 \| 69 \| 2 \| 5 \| 114 \| 1 \| 107 \| Single-stranded DNA-binding protein \| Single-stranded DNA-binding protein; afdb-uniprot50 \| AF-A0A348YKY9-F1-MODEL\_V4 \| 1.0 \| 6.566e-13 \| 474 \| 0.359 \| 164 \| 93 \| 6 \| 4 \| 162 \| 3 \| 159 \| Single-stranded DNA-binding protein \| Single-stranded DNA-binding protein; afdb-uniprot50 \| AF-A0A1B0C486-F1-MODEL\_V4 \| 1.0 \| 4.273e-13 \| 474 \| 0.5 \| 150 \| 68 \| 2 \| 17 \| 161 \| 336 \| 483 \| Uncharacterized protein \| Uncharacterized protein; afdb-uniprot50 \| AF-A0A0H3CMN1-F1-MODEL\_V4 \| 1.0 \| 3.662e-12 \| 473 \| 0.535 \| 142 \| 52 \| 3 \| 21 \| 162 \| 2 \| 129 \| Single-stranded DNA-binding protein \| Single-stranded DNA-binding protein; afdb-uniprot50 \| AF-J0WGW8-F1-MODEL\_V4 \| 1.0 \| 4.54e-12 \| 471 \| 0.331 \| 154 \| 89 \| 4 \| 1 \| 154 \| 24 \| 163 \| Single-stranded DNA-binding protein \| Single-stranded DNA-binding protein; afdb-uniprot50 \| AF-A0A7S4G7Z5-F1-MODEL\_V4 \| 1.0 \| 2.258e-12 \| 471 \| 0.269 \| 167 \| 112 \| 4 \| 2 \| 162 \| 187 \| 349 \| Hypothetical protein \| Hypothetical protein; afdb-uniprot50 \| AF-A0A1F9TW11-F1-MODEL\_V4 \| 1.0 \| 5.938e-12 \| 470 \| 0.321 \| 168 \| 82 \| 7 \| 1 \| 162 \| 1 \| 142 \| Single-stranded DNA-binding protein \| Single-stranded DNA-binding protein; afdb-uniprot50 \| AF-A0A6I2VGK0-F1-MODEL\_V4 \| 1.0 \| 1.009e-12 \| 470 \| 0.333 \| 165 \| 94 \| 5 \| 2 \| 162 \| 19 \| 171 \| Single-stranded DNA-binding protein \| Single-stranded DNA-binding protein; afdb-uniprot50 \| AF-A0A1M3GUQ3-F1-MODEL\_V4 \| 1.0 \| 3.289e-12 \| 470 \| 0.308 \| 162 \| 106 \| 5 \| 4 \| 162 \| 1 \| 159 \| Single-stranded DNA-binding protein \| Single-stranded DNA-binding protein; afdb-uniprot50 \| AF-A0A5B8U1B7-F1-MODEL\_V4 \| 1.0 \| 4.757e-13 \| 467 \| 0.339 \| 171 \| 91 \| 4 \| 2 \| 162 \| 45 \| 203 \| Single-stranded DNA-binding protein \| Single-stranded DNA-binding protein; afdb-uniprot50 \| AF-A0A7J7BRE6-F1-MODEL\_V4 \| 1.0 \| 5.856e-14 \| 467 \| 0.443 \| 187 \| 69 \| 3 \| 10 \| 162 \| 543 \| 728 \| MFS domain-containing protein \| MFS domain-containing protein; afdb-uniprot50 \| AF-W0FMQ2-F1-MODEL\_V4 \| 1.0 \| 1.726e-12 \| 466 \| 0.352 \| 159 \| 94 \| 4 \| 4 \| 162 \| 2 \| 151 \| Single-stranded DNA-binding protein \| Single-stranded DNA-binding protein; afdb-uniprot50 \| AF-A0A354WRT5-F1-MODEL\_V4 \| 1.0 \| 3.837e-13 \| 466 \| 0.312 \| 176 \| 93 \| 5 \| 5 \| 162 \| 1 \| 166 \| Single-stranded DNA-binding protein \| Single-stranded DNA-binding protein; afdb-uniprot50 \| AF-A0A7V4E1N2-F1-MODEL\_V4 \| 1.0 \| 1.936e-11 \| 463 \| 0.302 \| 149 \| 90 \| 4 \| 1 \| 144 \| 1 \| 140 \| Single-stranded DNA-binding protein \| Single-stranded DNA-binding protein; afdb-uniprot50 \| AF-A0A1H1L2W3-F1-MODEL\_V4 \| 1.0 \| 1.26e-11 \| 463 \| 0.327 \| 159 \| 88 \| 5 \| 4 \| 162 \| 2 \| 141 \| Single-stranded DNA-binding protein \| Single-stranded DNA-binding protein; afdb-uniprot50 \| AF-A0A7T5RMB9-F1-MODEL\_V4 \| 1.0 \| 1.922e-12 \| 463 \| 0.339 \| 162 \| 90 \| 5 \| 1 \| 162 \| 5 \| 149 \| Single-stranded DNA-binding protein \| Single-stranded DNA-binding protein; afdb-uniprot50 \| AF-A0A7V2YYU3-F1-MODEL\_V4 \| 1.0 \| 3.289e-12 \| 463 \| 0.29 \| 165 \| 107 \| 4 \| 4 \| 162 \| 3 \| 163 \| Single-stranded DNA-binding protein \| Single-stranded DNA-binding protein; afdb-uniprot50 \| AF-A0A4T9TD94-F1-MODEL\_V4 \| 1.0 \| 4.302e-12 \| 463 \| 0.301 \| 159 \| 104 \| 3 \| 4 \| 162 \| 2 \| 153 \| Single-stranded DNA-binding protein \| Single-stranded DNA-binding protein; afdb-uniprot50 \| AF-A0A0F9VUL0-F1-MODEL\_V4 \| 1.0 \| 1.251e-12 \| 461 \| 0.279 \| 168 \| 107 \| 6 \| 1 \| 162 \| 18 \| 177 \| Uncharacterized protein \| Uncharacterized protein; afdb-uniprot50 \| AF-A0A210VRK1-F1-MODEL\_V4 \| 1.0 \| 1.726e-12 \| 460 \| 0.349 \| 166 \| 98 \| 6 \| 1 \| 160 \| 1 \| 162 \| Single-stranded DNA-binding protein \| Single-stranded DNA-binding protein; afdb-uniprot50 \| AF-A0A2E3VXK3-F1-MODEL\_V4 \| 1.0 \| 5.706e-10 \| 459 \| 0.327 \| 113 \| 73 \| 1 \| 4 \| 116 \| 2 \| 111 \| Single-stranded DNA-binding protein \| Single-stranded DNA-binding protein; afdb-uniprot50 \| AF-A0A1M2URQ5-F1-MODEL\_V4 \| 1.0 \| 1.48e-11 \| 458 \| 0.372 \| 161 \| 83 \| 4 \| 4 \| 162 \| 1 \| 145 \| Single-stranded DNA-binding protein \| Single-stranded DNA-binding protein; afdb-uniprot50 \| AF-A0A537YUJ5-F1-MODEL\_V4 \| 1.0 \| 1.123e-12 \| 458 \| 0.295 \| 159 \| 104 \| 4 \| 6 \| 162 \| 4 \| 156 \| Single-stranded DNA-binding protein \| Single-stranded DNA-binding protein; afdb-uniprot50 \| AF-A0A2H0S6Z3-F1-MODEL\_V4 \| 1.0 \| 8.139e-13 \| 458 \| 0.352 \| 156 \| 99 \| 2 \| 5 \| 160 \| 223 \| 376 \| Single-stranded DNA-binding protein \| Single-stranded DNA-binding protein; afdb-uniprot50 \| AF-A0A0P6YDB4-F1-MODEL\_V4 \| 1.0 \| 5.125e-10 \| 456 \| 0.3 \| 110 \| 75 \| 2 \| 1 \| 109 \| 1 \| 109 \| Single-stranded DNA-binding protein \| Single-stranded DNA-binding protein; afdb-uniprot50 \| AF-A0A7W2H3Y3-F1-MODEL\_V4 \| 1.0 \| 4.54e-12 \| 456 \| 0.283 \| 162 \| 105 \| 5 \| 1 \| 162 \| 1 \| 151 \| Single-stranded DNA-binding protein \| Single-stranded DNA-binding protein; afdb-uniprot50 \| AF-A0A1C5MNJ8-F1-MODEL\_V4 \| 1.0 \| 1.194e-11 \| 456 \| 0.283 \| 159 \| 107 \| 3 \| 4 \| 162 \| 2 \| 153 \| Single-stranded DNA-binding protein \| Single-stranded DNA-binding protein; afdb-uniprot50 \| AF-A0A7V2CKI6-F1-MODEL\_V4 \| 1.0 \| 1.32e-12 \| 456 \| 0.284 \| 172 \| 109 \| 6 \| 1 \| 162 \| 1 \| 168 \| Single-stranded DNA-binding protein \| Single-stranded DNA-binding protein; afdb-uniprot50 \| AF-A0A2N2XER4-F1-MODEL\_V4 \| 1.0 \| 5.37e-11 \| 454 \| 0.33 \| 139 \| 84 \| 2 \| 10 \| 142 \| 2 \| 137 \| Single-stranded DNA-binding protein \| Single-stranded DNA-binding protein; afdb-uniprot50 \| AF-A0A639KX21-F1-MODEL\_V4 \| 1.0 \| 2.69e-10 \| 453 \| 0.353 \| 113 \| 70 \| 3 \| 4 \| 115 \| 3 \| 113 \| Single-stranded DNA-binding protein \| Single-stranded DNA-binding protein; afdb-uniprot50 \| AF-A0A533RM11-F1-MODEL\_V4 \| 1.0 \| 3.687e-11 \| 453 \| 0.308 \| 159 \| 96 \| 4 \| 4 \| 162 \| 2 \| 146 \| Single-stranded DNA-binding protein \| Single-stranded DNA-binding protein; afdb-uniprot50 \| AF-A0A1G1CN45-F1-MODEL\_V4 \| 1.0 \| 2.954e-12 \| 453 \| 0.367 \| 158 \| 92 \| 3 \| 10 \| 162 \| 2 \| 156 \| Single-stranded DNA-binding protein \| Single-stranded DNA-binding protein; afdb-uniprot50 \| AF-A0A7X7NPE2-F1-MODEL\_V4 \| 1.0 \| 1.338e-10 \| 450 \| 0.313 \| 115 \| 76 \| 3 \| 4 \| 117 \| 2 \| 114 \| Single-stranded DNA-binding protein \| Single-stranded DNA-binding protein; afdb-uniprot50 \| AF-A0A2N4Z5P7-F1-MODEL\_V4 \| 1.0 \| 5.667e-11 \| 450 \| 0.25 \| 156 \| 115 \| 2 \| 1 \| 155 \| 2 \| 156 \| Single-stranded DNA-binding protein \| Single-stranded DNA-binding protein; afdb-uniprot50 \| AF-C4F8E6-F1-MODEL\_V4 \| 1.0 \| 1.268e-10 \| 449 \| 0.31 \| 161 \| 73 \| 5 \| 4 \| 162 \| 2 \| 126 \| Single-stranded DNA-binding protein \| Single-stranded DNA-binding protein; afdb-uniprot50 \| AF-A0A1G6KJG1-F1-MODEL\_V4 \| 1.0 \| 3.139e-11 \| 449 \| 0.341 \| 161 \| 87 \| 6 \| 4 \| 162 \| 2 \| 145 \| Single-stranded DNA-binding protein \| Single-stranded DNA-binding protein; afdb-uniprot50 \| AF-A0A7Y3KW57-F1-MODEL\_V4 \| 1.0 \| 1.072e-11 \| 449 \| 0.245 \| 163 \| 101 \| 4 \| 4 \| 162 \| 2 \| 146 \| Single-stranded DNA-binding protein \| Single-stranded DNA-binding protein; afdb-uniprot50 \| AF-A0A6M0LP34-F1-MODEL\_V4 \| 1.0 \| 1.131e-11 \| 449 \| 0.259 \| 162 \| 106 \| 6 \| 4 \| 160 \| 3 \| 155 \| Single-stranded DNA-binding protein \| Single-stranded DNA-binding protein; afdb-uniprot50 \| AF-A0A2D2LX13-F1-MODEL\_V4 \| 1.0 \| 4.54e-12 \| 449 \| 0.349 \| 163 \| 95 \| 4 \| 4 \| 162 \| 1 \| 156 \| Single-stranded DNA-binding protein \| Single-stranded DNA-binding protein; afdb-uniprot50 \| AF-A0A7V5N906-F1-MODEL\_V4 \| 1.0 \| 8.648e-12 \| 449 \| 0.275 \| 185 \| 102 \| 5 \| 4 \| 162 \| 3 \| 181 \| Single-stranded DNA-binding protein \| Single-stranded DNA-binding protein; afdb-uniprot50 \| AF-A0A349D2W1-F1-MODEL\_V4 \| 1.0 \| 1.922e-12 \| 448 \| 0.343 \| 172 \| 95 \| 6 \| 2 \| 162 \| 459 \| 623 \| Single-stranded DNA-binding protein \| Single-stranded DNA-binding protein; afdb-uniprot50 \| AF-A0A7X7YU40-F1-MODEL\_V4 \| 1.0 \| 3.139e-11 \| 446 \| 0.254 \| 165 \| 97 \| 5 \| 1 \| 162 \| 1 \| 142 \| Single-stranded DNA-binding protein \| Single-stranded DNA-binding protein; afdb-uniprot50 \| AF-L1NBJ9-F1-MODEL\_V4 \| 1.0 \| 2.057e-10 \| 446 \| 0.176 \| 136 \| 109 \| 1 \| 1 \| 136 \| 4 \| 136 \| Single-stranded DNA-binding protein \| Single-stranded DNA-binding protein; afdb-uniprot50 \| AF-A0A7C3PY12-F1-MODEL\_V4 \| 1.0 \| 4.54e-12 \| 446 \| 0.251 \| 175 \| 113 \| 5 \| 3 \| 162 \| 2 \| 173 \| Single-stranded DNA-binding protein \| Single-stranded DNA-binding protein; afdb-uniprot50 \| AF-A0A136NW82-F1-MODEL\_V4 \| 1.0 \| 6.657e-11 \| 445 \| 0.401 \| 122 \| 69 \| 4 \| 10 \| 130 \| 2 \| 120 \| Single-stranded DNA-binding protein \| Single-stranded DNA-binding protein; afdb-uniprot50 \| AF-T1A6S3-F1-MODEL\_V4 \| 1.0 \| 1.834e-11 \| 445 \| 0.308 \| 136 \| 87 \| 3 \| 3 \| 132 \| 2 \| 136 \| Single-strand DNA-binding protein \| Single-strand DNA-binding protein; afdb-uniprot50 \| AF-A0A7C1ENX2-F1-MODEL\_V4 \| 1.0 \| 7.361e-12 \| 445 \| 0.325 \| 132 \| 84 \| 3 \| 1 \| 129 \| 14 \| 143 \| Single-stranded DNA-binding protein \| Single-stranded DNA-binding protein; afdb-uniprot50 \| AF-A0A6B1E277-F1-MODEL\_V4 \| 1.0 \| 2.497e-13 \| 445 \| 0.318 \| 185 \| 99 \| 4 \| 4 \| 162 \| 2 \| 185 \| Single-stranded DNA-binding protein \| Single-stranded DNA-binding protein; afdb-uniprot50 \| AF-A0A6I2YPP7-F1-MODEL\_V4 \| 1.0 \| 1.338e-10 \| 444 \| 0.283 \| 120 \| 79 \| 2 \| 4 \| 117 \| 1 \| 119 \| Single-stranded DNA-binding protein \| Single-stranded DNA-binding protein; afdb-uniprot50 \| AF-A0A6L9IS86-F1-MODEL\_V4 \| 1.0 \| 1.936e-11 \| 444 \| 0.279 \| 129 \| 86 \| 3 \| 1 \| 129 \| 1 \| 122 \| Single-stranded DNA-binding protein \| Single-stranded DNA-binding protein; afdb-uniprot50 \| AF-A0A5C7U259-F1-MODEL\_V4 \| 1.0 \| 1.131e-11 \| 444 \| 0.288 \| 163 \| 103 \| 6 \| 1 \| 161 \| 23 \| 174 \| Single-stranded DNA-binding protein \| Single-stranded DNA-binding protein; afdb-uniprot50 \| AF-A0A415JIS1-F1-MODEL\_V4 \| 1.0 \| 1.147e-09 \| 443 \| 0.263 \| 114 \| 80 \| 2 \| 4 \| 117 \| 2 \| 111 \| Single-stranded DNA-binding protein \| Single-stranded DNA-binding protein; afdb-uniprot50 \| AF-A0A3M1DGA7-F1-MODEL\_V4 \| 1.0 \| 1.268e-10 \| 443 \| 0.22 \| 159 \| 101 \| 5 \| 5 \| 162 \| 1 \| 137 \| Single-stranded DNA-binding protein \| Single-stranded DNA-binding protein; afdb-uniprot50 \| AF-A0A0K0DUQ6-F1-MODEL\_V4 \| 1.0 \| 2.709e-09 \| 442 \| 0.23 \| 117 \| 86 \| 2 \| 2 \| 118 \| 65 \| 177 \| Uncharacterized protein \| Uncharacterized protein; afdb-uniprot50 \| AF-A0A1E3LMY7-F1-MODEL\_V4 \| 1.0 \| 5.706e-10 \| 441 \| 0.235 \| 136 \| 92 \| 2 \| 1 \| 136 \| 1 \| 124 \| Single-stranded DNA-binding protein \| Single-stranded DNA-binding protein; afdb-uniprot50 \| AF-I3UGJ2-F1-MODEL\_V4 \| 1.0 \| 3.139e-11 \| 441 \| 0.236 \| 152 \| 111 \| 3 \| 1 \| 151 \| 1 \| 148 \| Single-stranded DNA-binding protein \| Single-stranded DNA-binding protein; afdb-uniprot50 \| AF-A0A640Y9W9-F1-MODEL\_V4 \| 1.0 \| 7.767e-12 \| 441 \| 0.333 \| 162 \| 102 \| 4 \| 2 \| 162 \| 59 \| 215 \| Single-stranded DNA-binding protein \| Single-stranded DNA-binding protein; afdb-uniprot50 \| AF-A0A6J5JVU9-F1-MODEL\_V4 \| 1.0 \| 1.572e-10 \| 440 \| 0.37 \| 127 \| 72 \| 3 \| 1 \| 127 \| 1 \| 119 \| Single-stranded DNA-binding protein \| Single-stranded DNA-binding protein; afdb-uniprot50 \| AF-A0A7C3DLR0-F1-MODEL\_V4 \| 1.0 \| 4.302e-12 \| 440 \| 0.349 \| 166 \| 79 \| 7 \| 4 \| 162 \| 1 \| 144 \| Single-stranded DNA-binding protein \| Single-stranded DNA-binding protein; afdb-uniprot50 \| AF-J5MVP4-F1-MODEL\_V4 \| 1.0 \| 1.834e-11 \| 440 \| 0.447 \| 134 \| 66 \| 4 \| 1 \| 129 \| 1 \| 131 \| Single-stranded DNA-binding protein \| Single-stranded DNA-binding protein; afdb-uniprot50 \| AF-A0A0F9Z6S5-F1-MODEL\_V4 \| 1.0 \| 2.274e-11 \| 438 \| 0.333 \| 159 \| 91 \| 6 \| 4 \| 160 \| 3 \| 148 \| Single-stranded DNA-binding protein \| Single-stranded DNA-binding protein; afdb-uniprot50 \| AF-A0A532UZN4-F1-MODEL\_V4 \| 1.0 \| 2.274e-11 \| 437 \| 0.303 \| 165 \| 105 \| 6 \| 1 \| 160 \| 1 \| 160 \| Single-stranded DNA-binding protein \| Single-stranded DNA-binding protein; afdb-uniprot50 \| AF-A0A220QRB1-F1-MODEL\_V4 \| 1.0 \| 3.662e-12 \| 436 \| 0.367 \| 174 \| 89 \| 3 \| 2 \| 155 \| 156 \| 328 \| Single-stranded DNA-binding protein \| Single-stranded DNA-binding protein; afdb-uniprot50 \| AF-A0A2W6BUT5-F1-MODEL\_V4 \| 1.0 \| 5.333e-12 \| 435 \| 0.317 \| 173 \| 84 \| 6 \| 4 \| 162 \| 2 \| 154 \| Single-stranded DNA-binding protein \| Single-stranded DNA-binding protein; afdb-uniprot50 \| AF-S5DJE5-F1-MODEL\_V4 \| 1.0 \| 1.202e-10 \| 434 \| 0.267 \| 131 \| 87 \| 4 \| 4 \| 129 \| 2 \| 128 \| Single-stranded DNA-binding protein \| Single-stranded DNA-binding protein; afdb-uniprot50 \| AF-A0A2H0WY68-F1-MODEL\_V4 \| 1.0 \| 3.864e-12 \| 433 \| 0.343 \| 163 \| 94 \| 5 \| 10 \| 162 \| 2 \| 161 \| Single-stranded DNA-binding protein \| Single-stranded DNA-binding protein; afdb-uniprot50 \| AF-A0A0D8HFS3-F1-MODEL\_V4 \| 1.0 \| 7.767e-12 \| 431 \| 0.282 \| 163 \| 113 \| 2 \| 3 \| 162 \| 2 \| 163 \| Single-stranded DNA-binding protein \| Single-stranded DNA-binding protein; afdb-uniprot50 \| AF-A0A133ZE68-F1-MODEL\_V4 \| 1.0 \| 2.042e-11 \| 430 \| 0.253 \| 162 \| 113 \| 4 \| 4 \| 160 \| 2 \| 160 \| Single-stranded DNA-binding protein \| Single-stranded DNA-binding protein; afdb-uniprot50 \| AF-A0A7V1U542-F1-MODEL\_V4 \| 1.0 \| 4.332e-11 \| 427 \| 0.285 \| 161 \| 95 \| 7 \| 1 \| 157 \| 1 \| 145 \| Single-stranded DNA-binding protein \| Single-stranded DNA-binding protein; afdb-uniprot50 \| AF-A0A6A8M953-F1-MODEL\_V4 \| 1.0 \| 2.042e-11 \| 427 \| 0.327 \| 162 \| 99 \| 5 \| 5 \| 162 \| 1 \| 156 \| Single-stranded DNA-binding protein \| Single-stranded DNA-binding protein; afdb-uniprot50 \| AF-A0A6I1MPE9-F1-MODEL\_V4 \| 1.0 \| 1.03e-09 \| 427 \| 0.296 \| 118 \| 81 \| 2 \| 1 \| 117 \| 1 \| 117 \| Single-stranded DNA-binding protein \| Single-stranded DNA-binding protein; afdb-uniprot50 \| AF-A0A2V2G4L0-F1-MODEL\_V4 \| 1.0 \| 7.025e-11 \| 426 \| 0.265 \| 158 \| 105 \| 3 \| 5 \| 162 \| 1 \| 147 \| Single-stranded DNA-binding protein \| Single-stranded DNA-binding protein; afdb-uniprot50 \| AF-A0A3E2TPK5-F1-MODEL\_V4 \| 1.0 \| 4.362e-10 \| 425 \| 0.271 \| 162 \| 92 \| 5 \| 1 \| 162 \| 1 \| 136 \| Single-stranded DNA-binding protein \| Single-stranded DNA-binding protein; afdb-uniprot50 \| AF-A0A4U5MFD9-F1-MODEL\_V4 \| 1.0 \| 1.583e-09 \| 425 \| 0.28 \| 125 \| 84 \| 4 \| 2 \| 123 \| 56 \| 177 \| Uncharacterized protein \| Uncharacterized protein; afdb-uniprot50 \| AF-A0A4S2EXC4-F1-MODEL\_V4 \| 1.0 \| 1.072e-11 \| 425 \| 0.284 \| 186 \| 105 \| 5 \| 4 \| 162 \| 2 \| 186 \| Single-stranded DNA-binding protein \| Single-stranded DNA-binding protein; afdb-uniprot50 \| AF-A0A3A1YFT4-F1-MODEL\_V4 \| 1.0 \| 9.125e-12 \| 425 \| 0.284 \| 165 \| 110 \| 5 \| 1 \| 162 \| 1 \| 160 \| Single-stranded DNA-binding protein \| Single-stranded DNA-binding protein; afdb-uniprot50 \| AF-A0A1W6BYF7-F1-MODEL\_V4 \| 1.0 \| 4.332e-11 \| 424 \| 0.22 \| 154 \| 111 \| 4 \| 5 \| 154 \| 1 \| 149 \| Single-stranded DNA-binding protein \| Single-stranded DNA-binding protein; afdb-uniprot50 \| AF-A0A1Y3SED9-F1-MODEL\_V4 \| 1.0 \| 4.332e-11 \| 423 \| 0.228 \| 162 \| 110 \| 4 \| 1 \| 162 \| 29 \| 175 \| Single-stranded DNA-binding protein \| Single-stranded DNA-binding protein; afdb-uniprot50 \| AF-A0A356LBF3-F1-MODEL\_V4 \| 1.0 \| 1.023e-10 \| 422 \| 0.239 \| 146 \| 104 \| 4 \| 3 \| 142 \| 35 \| 179 \| Uncharacterized protein \| Uncharacterized protein; afdb-uniprot50 \| AF-A0A1L8N1R4-F1-MODEL\_V4 \| 1.0 \| 5.125e-10 \| 418 \| 0.205 \| 146 \| 110 \| 2 \| 3 \| 148 \| 2 \| 141 \| Single-stranded DNA-binding protein \| Single-stranded DNA-binding protein; afdb-uniprot50 \| AF-A0A5U6CUN8-F1-MODEL\_V4 \| 1.0 \| 3.312e-11 \| 417 \| 0.561 \| 139 \| 51 \| 2 \| 24 \| 162 \| 1 \| 129 \| Single-stranded DNA-binding protein \| Single-stranded DNA-binding protein; afdb-uniprot50 \| AF-A0A4S2ER89-F1-MODEL\_V4 \| 1.0 \| 1.139e-10 \| 417 \| 0.258 \| 147 \| 107 \| 2 \| 1 \| 146 \| 6 \| 151 \| Single-stranded DNA-binding protein \| Single-stranded DNA-binding protein; afdb-uniprot50 \| AF-A0A1F2YCG8-F1-MODEL\_V4 \| 1.0 \| 4.106e-11 \| 414 \| 0.281 \| 160 \| 93 \| 6 \| 1 \| 155 \| 1 \| 143 \| Single-stranded DNA-binding protein \| Single-stranded DNA-binding protein; afdb-uniprot50 \| AF-A0A2N3QHM9-F1-MODEL\_V4 \| 1.0 \| 8.196e-12 \| 414 \| 0.325 \| 160 \| 104 \| 4 \| 4 \| 162 \| 2 \| 158 \| Single-stranded DNA-binding protein \| Single-stranded DNA-binding protein; afdb-uniprot50 \| AF-A0A364VE05-F1-MODEL\_V4 \| 1.0 \| 3.139e-11 \| 414 \| 0.306 \| 150 \| 94 \| 5 \| 1 \| 144 \| 19 \| 164 \| Single-stranded DNA-binding protein \| Single-stranded DNA-binding protein; afdb-uniprot50 \| AF-A0A3C1CGQ0-F1-MODEL\_V4 \| 1.0 \| 1.08e-10 \| 413 \| 0.229 \| 144 \| 105 \| 2 \| 4 \| 142 \| 1 \| 143 \| Single-stranded DNA-binding protein \| Single-stranded DNA-binding protein; afdb-uniprot50 \| AF-A0A6G9HQU0-F1-MODEL\_V4 \| 1.0 \| 4.508e-13 \| 413 \| 0.448 \| 165 \| 80 \| 3 \| 1 \| 154 \| 1 \| 165 \| Single-stranded DNA-binding protein \| Single-stranded DNA-binding protein; afdb-uniprot50 \| AF-U2KLY9-F1-MODEL\_V4 \| 1.0 \| 4.106e-11 \| 412 \| 0.227 \| 158 \| 115 \| 3 \| 3 \| 160 \| 2 \| 152 \| Single-stranded DNA-binding protein \| Single-stranded DNA-binding protein; afdb-uniprot50 \| AF-A0A523Z5E7-F1-MODEL\_V4 \| 1.0 \| 5.979e-11 \| 410 \| 0.227 \| 158 \| 111 \| 4 \| 5 \| 162 \| 9 \| 155 \| Single-stranded DNA-binding protein \| Single-stranded DNA-binding protein; afdb-uniprot50 \| AF-A0A7C4ZDT3-F1-MODEL\_V4 \| 1.0 \| 1.268e-10 \| 409 \| 0.309 \| 152 \| 101 \| 2 \| 4 \| 155 \| 3 \| 150 \| Single-stranded DNA-binding protein \| Single-stranded DNA-binding protein; afdb-uniprot50 \| AF-A0A2H0Y1Q2-F1-MODEL\_V4 \| 1.0 \| 1.03e-09 \| 408 \| 0.259 \| 135 \| 95 \| 3 \| 1 \| 135 \| 21 \| 150 \| Single-stranded DNA-binding protein \| Single-stranded DNA-binding protein; afdb-uniprot50 \| AF-A0A5N9VXX7-F1-MODEL\_V4 \| 1.0 \| 1.572e-10 \| 407 \| 0.576 \| 125 \| 48 \| 2 \| 29 \| 151 \| 3 \| 124 \| Single-stranded DNA-binding protein \| Single-stranded DNA-binding protein; afdb-uniprot50 \| AF-A0A848J9W6-F1-MODEL\_V4 \| 1.0 \| 1.936e-11 \| 407 \| 0.273 \| 161 \| 101 \| 6 \| 4 \| 154 \| 2 \| 156 \| Single-stranded DNA-binding protein \| Single-stranded DNA-binding protein; afdb-uniprot50 \| AF-A0A1W9S8T2-F1-MODEL\_V4 \| 1.0 \| 2.672e-11 \| 407 \| 0.298 \| 184 \| 102 \| 6 \| 1 \| 162 \| 4 \| 182 \| Single-stranded DNA-binding protein \| Single-stranded DNA-binding protein; afdb-uniprot50 \| AF-A0A7V7XET6-F1-MODEL\_V4 \| 1.0 \| 3.495e-11 \| 405 \| 0.281 \| 160 \| 100 \| 5 \| 4 \| 162 \| 2 \| 147 \| Single-stranded DNA-binding protein \| Single-stranded DNA-binding protein; afdb-uniprot50 \| AF-A0A556R9X6-F1-MODEL\_V4 \| 1.0 \| 5.09e-11 \| 404 \| 0.289 \| 152 \| 100 \| 5 \| 6 \| 154 \| 3 \| 149 \| Single-stranded DNA-binding protein \| Single-stranded DNA-binding protein; afdb-uniprot50 \| AF-A0A7K3XWR7-F1-MODEL\_V4 \| 1.0 \| 1.847e-10 \| 403 \| 0.222 \| 162 \| 115 \| 4 \| 4 \| 162 \| 2 \| 155 \| Single-stranded DNA-binding protein \| Single-stranded DNA-binding protein; afdb-uniprot50 \| AF-A0A1Y3S4L9-F1-MODEL\_V4 \| 1.0 \| 1.202e-10 \| 402 \| 0.238 \| 159 \| 102 \| 5 \| 2 \| 160 \| 76 \| 215 \| Single-stranded DNA-binding protein \| Single-stranded DNA-binding protein; afdb-uniprot50 \| AF-X0UIT5-F1-MODEL\_V4 \| 1.0 \| 1.949e-10 \| 399 \| 0.269 \| 156 \| 98 \| 4 \| 8 \| 162 \| 1 \| 141 \| Uncharacterized protein \| Uncharacterized protein; afdb-uniprot50 \| AF-A0A509MKJ1-F1-MODEL\_V4 \| 1.0 \| 1.936e-11 \| 398 \| 0.31 \| 164 \| 93 \| 7 \| 7 \| 162 \| 4 \| 155 \| Single-stranded DNA-binding protein \| Single-stranded DNA-binding protein; afdb-uniprot50 \| AF-A0A7V9J1J1-F1-MODEL\_V4 \| 1.0 \| 3.312e-11 \| 398 \| 0.286 \| 171 \| 110 \| 6 \| 1 \| 162 \| 1 \| 168 \| Single-stranded DNA-binding protein \| Single-stranded DNA-binding protein; afdb-uniprot50 \| AF-V8QYI1-F1-MODEL\_V4 \| 1.0 \| 8.769e-10 \| 397 \| 0.263 \| 133 \| 92 \| 4 \| 1 \| 130 \| 1 \| 130 \| Uncharacterized protein \| Uncharacterized protein; afdb-uniprot50 \| AF-A0A2D7YMR2-F1-MODEL\_V4 \| 1.0 \| 1.422e-09 \| 397 \| 0.263 \| 133 \| 92 \| 4 \| 1 \| 131 \| 44 \| 172 \| Single-stranded DNA-binding protein \| Single-stranded DNA-binding protein; afdb-uniprot50 \| AF-A0A447TPT0-F1-MODEL\_V4 \| 1.0 \| 4.106e-11 \| 394 \| 0.545 \| 143 \| 52 \| 4 \| 29 \| 160 \| 5 \| 145 \| Single-stranded DNA-binding protein \| Single-stranded DNA-binding protein; afdb-uniprot50 \| AF-A0A5C7LUW6-F1-MODEL\_V4 \| 1.0 \| 7.821e-11 \| 394 \| 0.308 \| 159 \| 99 \| 5 \| 1 \| 152 \| 38 \| 192 \| Single-stranded DNA-binding protein \| Single-stranded DNA-binding protein; afdb-uniprot50 \| AF-A0A7U0YZF2-F1-MODEL\_V4 \| 1.0 \| 1.268e-10 \| 390 \| 0.279 \| 161 \| 101 \| 6 \| 7 \| 158 \| 5 \| 159 \| Single-stranded DNA-binding protein \| Single-stranded DNA-binding protein; afdb-uniprot50 \| AF-G5H4S1-F1-MODEL\_V4 \| 1.0 \| 6.353e-10 \| 389 \| 0.287 \| 160 \| 100 \| 6 \| 5 \| 162 \| 1 \| 148 \| Single-stranded DNA-binding protein \| Single-stranded DNA-binding protein; afdb-uniprot50 \| AF-A0A6L6LUW6-F1-MODEL\_V4 \| 1.0 \| 1.49e-10 \| 389 \| 0.26 \| 161 \| 107 \| 6 \| 3 \| 162 \| 104 \| 253 \| Single-stranded DNA-binding protein \| Single-stranded DNA-binding protein; afdb-uniprot50 \| AF-A0A415C335-F1-MODEL\_V4 \| 1.0 \| 2.399e-11 \| 388 \| 0.296 \| 162 \| 102 \| 8 \| 1 \| 155 \| 1 \| 157 \| Single-stranded DNA-binding protein \| Single-stranded DNA-binding protein; afdb-uniprot50 \| AF-A0A0F9XHB2-F1-MODEL\_V4 \| 1.0 \| 2.071e-09 \| 387 \| 0.212 \| 160 \| 102 \| 4 \| 4 \| 162 \| 2 \| 138 \| Uncharacterized protein \| Uncharacterized protein; afdb-uniprot50 \| AF-A0A6G2BJE3-F1-MODEL\_V4 \| 1.0 \| 2.399e-11 \| 387 \| 0.245 \| 163 \| 110 \| 4 \| 1 \| 154 \| 1 \| 159 \| Single-stranded DNA-binding protein \| Single-stranded DNA-binding protein; afdb-uniprot50 \| AF-A0A6N9CAY7-F1-MODEL\_V4 \| 1.0 \| 1.422e-09 \| 386 \| 0.331 \| 145 \| 85 \| 3 \| 18 \| 162 \| 1 \| 133 \| Single-stranded DNA-binding protein \| Single-stranded DNA-binding protein; afdb-uniprot50 \| AF-A0A1S2GGX7-F1-MODEL\_V4 \| 1.0 \| 2.819e-11 \| 384 \| 0.293 \| 160 \| 106 \| 5 \| 5 \| 160 \| 1 \| 157 \| Single-stranded DNA-binding protein \| Single-stranded DNA-binding protein; afdb-uniprot50 \| AF-A0A2I2AAY6-F1-MODEL\_V4 \| 1.0 \| 7.464e-10 \| 381 \| 0.192 \| 161 \| 123 \| 3 \| 4 \| 162 \| 3 \| 158 \| Single-stranded DNA-binding protein \| Single-stranded DNA-binding protein; afdb-uniprot50 \| AF-A0A3B9YJQ0-F1-MODEL\_V4 \| 1.0 \| 1.963e-09 \| 380 \| 0.26 \| 146 \| 94 \| 5 \| 1 \| 142 \| 4 \| 139 \| Single-stranded DNA-binding protein \| Single-stranded DNA-binding protein; afdb-uniprot50 \| AF-G9YJ09-F1-MODEL\_V4 \| 1.0 \| 1.147e-09 \| 378 \| 0.309 \| 152 \| 86 \| 4 \| 12 \| 162 \| 2 \| 135 \| Single-stranded DNA-binding protein \| Single-stranded DNA-binding protein; afdb-uniprot50 \| AF-A0A3G6ISV9-F1-MODEL\_V4 \| 1.0 \| 5.125e-10 \| 378 \| 0.256 \| 152 \| 103 \| 5 \| 5 \| 154 \| 1 \| 144 \| Single-stranded DNA-binding protein \| Single-stranded DNA-binding protein; afdb-uniprot50 \| AF-A0A2I1Y967-F1-MODEL\_V4 \| 1.0 \| 1.49e-10 \| 378 \| 0.169 \| 165 \| 126 \| 6 \| 4 \| 162 \| 1 \| 160 \| Single-stranded DNA-binding protein \| Single-stranded DNA-binding protein; afdb-uniprot50 \| AF-A0A7V9BRX3-F1-MODEL\_V4 \| 1.0 \| 3.945e-09 \| 377 \| 0.253 \| 130 \| 87 \| 6 \| 4 \| 130 \| 3 \| 125 \| Single-stranded DNA-binding protein \| Single-stranded DNA-binding protein; afdb-uniprot50 \| AF-A0A1Y2MLA3-F1-MODEL\_V4 \| 1.0 \| 1.751e-10 \| 377 \| 0.277 \| 155 \| 99 \| 7 \| 4 \| 154 \| 2 \| 147 \| Single-stranded DNA-binding protein \| Single-stranded DNA-binding protein; afdb-uniprot50 \| AF-A0A2D0IR08-F1-MODEL\_V4 \| 1.0 \| 2.839e-10 \| 376 \| 0.493 \| 144 \| 58 \| 2 \| 30 \| 162 \| 2 \| 141 \| Single-stranded DNA-binding protein \| Single-stranded DNA-binding protein; afdb-uniprot50 \| AF-A0A2N6T258-F1-MODEL\_V4 \| 1.0 \| 5.706e-10 \| 372 \| 0.253 \| 142 \| 100 \| 4 \| 4 \| 144 \| 3 \| 139 \| Single-stranded DNA-binding protein \| Single-stranded DNA-binding protein; afdb-uniprot50 \| AF-A0A2N6V0F4-F1-MODEL\_V4 \| 1.0 \| 9.696e-11 \| 372 \| 0.251 \| 183 \| 113 \| 8 \| 1 \| 162 \| 1 \| 180 \| Single-stranded DNA-binding protein \| Single-stranded DNA-binding protein; afdb-uniprot50 \| AF-A0A7V6GLJ9-F1-MODEL\_V4 \| 1.0 \| 1.49e-10 \| 371 \| 0.267 \| 168 \| 95 \| 7 \| 4 \| 162 \| 2 \| 150 \| Single-stranded DNA-binding protein \| Single-stranded DNA-binding protein; afdb-uniprot50 \| AF-A0A7Y0Y5A9-F1-MODEL\_V4 \| 1.0 \| 1.751e-10 \| 370 \| 0.288 \| 163 \| 106 \| 6 \| 4 \| 162 \| 2 \| 158 \| Single-stranded DNA-binding protein \| Single-stranded DNA-binding protein; afdb-uniprot50 \| AF-A0A3N2P4R5-F1-MODEL\_V4 \| 1.0 \| 1.572e-10 \| 370 \| 0.239 \| 171 \| 115 \| 7 \| 4 \| 162 \| 2 \| 169 \| Single-stranded DNA-binding protein \| Single-stranded DNA-binding protein; afdb-uniprot50 \| AF-A0A1T4P1L0-F1-MODEL\_V4 \| 1.0 \| 3.945e-09 \| 369 \| 0.33 \| 127 \| 76 \| 5 \| 5 \| 129 \| 1 \| 120 \| Single-stranded DNA-binding protein \| Single-stranded DNA-binding protein; afdb-uniprot50 \| AF-A0A6J5CGD1-F1-MODEL\_V4 \| 1.0 \| 5.125e-10 \| 369 \| 0.276 \| 163 \| 106 \| 5 \| 4 \| 162 \| 3 \| 157 \| Single-stranded DNA-binding protein \| Single-stranded DNA-binding protein; afdb-uniprot50 \| AF-A0A417WSE5-F1-MODEL\_V4 \| 1.0 \| 2.69e-10 \| 366 \| 0.283 \| 159 \| 108 \| 4 \| 5 \| 161 \| 1 \| 155 \| Single-stranded DNA-binding protein \| Single-stranded DNA-binding protein; afdb-uniprot50 \| AF-S3A3R1-F1-MODEL\_V4 \| 1.0 \| 1.087e-09 \| 365 \| 0.306 \| 137 \| 89 \| 5 \| 5 \| 139 \| 1 \| 133 \| Single-stranded DNA-binding protein \| Single-stranded DNA-binding protein; afdb-uniprot50 \| AF-A0A2N6TVJ7-F1-MODEL\_V4 \| 1.0 \| 1.5e-09 \| 365 \| 0.261 \| 149 \| 102 \| 5 \| 1 \| 146 \| 53 \| 196 \| Uncharacterized protein \| Uncharacterized protein; afdb-uniprot50 \| AF-A0A1X3AD52-F1-MODEL\_V4 \| 1.0 \| 7.074e-10 \| 364 \| 0.269 \| 156 \| 103 \| 5 \| 1 \| 152 \| 1 \| 149 \| Single-stranded DNA-binding protein \| Single-stranded DNA-binding protein; afdb-uniprot50 \| AF-A0A6L9SUC9-F1-MODEL\_V4 \| 1.0 \| 1.087e-09 \| 361 \| 0.243 \| 164 \| 114 \| 4 \| 1 \| 158 \| 1 \| 160 \| Single-stranded DNA-binding protein \| Single-stranded DNA-binding protein; afdb-uniprot50 \| AF-A0A1F6QQY9-F1-MODEL\_V4 \| 1.0 \| 7.986e-08 \| 360 \| 0.196 \| 112 \| 87 \| 2 \| 4 \| 115 \| 2 \| 110 \| Uncharacterized protein \| Uncharacterized protein; afdb-uniprot50 \| AF-A0A366INY2-F1-MODEL\_V4 \| 1.0 \| 3.713e-10 \| 360 \| 0.264 \| 155 \| 105 \| 6 \| 5 \| 154 \| 1 \| 151 \| Single-stranded DNA-binding protein \| Single-stranded DNA-binding protein; afdb-uniprot50 \| AF-A0A4Y6QZ46-F1-MODEL\_V4 \| 1.0 \| 5.408e-10 \| 359 \| 0.246 \| 162 \| 112 \| 5 \| 5 \| 162 \| 1 \| 156 \| Single-stranded DNA-binding protein \| Single-stranded DNA-binding protein; afdb-uniprot50 \| AF-A0A2S9Z3F5-F1-MODEL\_V4 \| 1.0 \| 1.087e-09 \| 358 \| 0.246 \| 166 \| 114 \| 5 \| 1 \| 162 \| 66 \| 224 \| Single-stranded DNA-binding protein \| Single-stranded DNA-binding protein; afdb-uniprot50 \| AF-A0A7T0PDH1-F1-MODEL\_V4 \| 1.0 \| 1.202e-10 \| 357 \| 0.291 \| 161 \| 106 \| 5 \| 6 \| 161 \| 3 \| 160 \| Single-stranded DNA-binding protein \| Single-stranded DNA-binding protein; afdb-uniprot50 \| AF-Q2MLT4-F1-MODEL\_V4 \| 1.0 \| 2.549e-10 \| 357 \| 0.248 \| 173 \| 116 \| 6 \| 1 \| 162 \| 1 \| 170 \| Single-stranded DNA-binding protein \| Single-stranded DNA-binding protein; afdb-uniprot50 \| AF-A0A1I2DXE7-F1-MODEL\_V4 \| 1.0 \| 2.433e-09 \| 357 \| 0.233 \| 133 \| 89 \| 6 \| 5 \| 129 \| 1 \| 128 \| Single-stranded DNA-binding protein \| Single-stranded DNA-binding protein; afdb-uniprot50 \| AF-B8DTM4-F1-MODEL\_V4 \| 1.0 \| 4.134e-10 \| 357 \| 0.244 \| 168 \| 117 \| 5 \| 1 \| 162 \| 1 \| 164 \| Single-stranded DNA-binding protein \| Single-stranded DNA-binding protein; afdb-uniprot50 \| AF-A0A1V3G0K3-F1-MODEL\_V4 \| 1.0 \| 3.161e-10 \| 356 \| 0.235 \| 153 \| 110 \| 6 \| 5 \| 154 \| 1 \| 149 \| Single-stranded DNA-binding protein \| Single-stranded DNA-binding protein; afdb-uniprot50 \| AF-A0A2W4PIC5-F1-MODEL\_V4 \| 1.0 \| 8.31e-10 \| 355 \| 0.238 \| 155 \| 102 \| 6 \| 7 \| 154 \| 4 \| 149 \| Single-stranded DNA-binding protein \| Single-stranded DNA-binding protein; afdb-uniprot50 \| AF-A0A4R7L5A8-F1-MODEL\_V4 \| 1.0 \| 2.071e-09 \| 354 \| 0.254 \| 161 \| 104 \| 7 \| 4 \| 162 \| 2 \| 148 \| Single-stranded DNA-binding protein \| Single-stranded DNA-binding protein; afdb-uniprot50 \| AF-A0A1E8CUC1-F1-MODEL\_V4 \| 1.0 \| 5.483e-08 \| 354 \| 0.31 \| 103 \| 68 \| 3 \| 7 \| 108 \| 5 \| 105 \| Single-stranded DNA-binding protein \| Single-stranded DNA-binding protein; afdb-uniprot50 \| AF-A0A6N7WM32-F1-MODEL\_V4 \| 1.0 \| 2.306e-09 \| 353 \| 0.272 \| 136 \| 93 \| 4 \| 5 \| 137 \| 1 \| 133 \| Single-stranded DNA-binding protein \| Single-stranded DNA-binding protein; afdb-uniprot50 \| AF-A0A0A1DMG2-F1-MODEL\_V4 \| 1.0 \| 2.071e-09 \| 352 \| 0.236 \| 152 \| 103 \| 5 \| 4 \| 154 \| 1 \| 140 \| Single-stranded DNA-binding protein \| Single-stranded DNA-binding protein; afdb-uniprot50 \| AF-A0A2J8GAC6-F1-MODEL\_V4 \| 1.0 \| 6.021e-10 \| 352 \| 0.236 \| 169 \| 118 \| 7 \| 1 \| 162 \| 39 \| 203 \| Single-stranded DNA-binding protein \| Single-stranded DNA-binding protein; afdb-uniprot50 \| AF-A0A2V4NL91-F1-MODEL\_V4 \| 1.0 \| 4.857e-10 \| 351 \| 0.254 \| 161 \| 109 \| 5 \| 2 \| 154 \| 28 \| 185 \| Single-stranded DNA-binding protein \| Single-stranded DNA-binding protein; afdb-uniprot50 \| AF-A0A2A3F440-F1-MODEL\_V4 \| 1.0 \| 1.348e-09 \| 350 \| 0.2 \| 165 \| 122 \| 6 \| 4 \| 162 \| 1 \| 161 \| Uncharacterized protein \| Uncharacterized protein; afdb-uniprot50 \| AF-A0A0G7ZN84-F1-MODEL\_V4 \| 1.0 \| 4.891e-09 \| 348 \| 0.225 \| 164 \| 115 \| 2 \| 4 \| 158 \| 1 \| 161 \| Single-stranded DNA-binding protein \| Single-stranded DNA-binding protein; afdb-uniprot50 \| AF-A0A212UIC7-F1-MODEL\_V4 \| 1.0 \| 4.362e-10 \| 347 \| 0.262 \| 164 \| 105 \| 7 \| 4 \| 154 \| 2 \| 162 \| Single-stranded DNA-binding protein \| Single-stranded DNA-binding protein; afdb-uniprot50 \| AF-A0A1S1D7V1-F1-MODEL\_V4 \| 1.0 \| 2.549e-10 \| 346 \| 0.239 \| 163 \| 115 \| 5 \| 6 \| 162 \| 3 \| 162 \| Single-stranded DNA-binding protein \| Single-stranded DNA-binding protein; afdb-uniprot50 \| AF-A0A0N1D9D7-F1-MODEL\_V4 \| 1.0 \| 1.671e-09 \| 345 \| 0.236 \| 161 \| 107 \| 6 \| 4 \| 162 \| 2 \| 148 \| Single-stranded DNA-binding protein \| Single-stranded DNA-binding protein; afdb-uniprot50 \| AF-M1UST2-F1-MODEL\_V4 \| 1.0 \| 1.422e-09 \| 345 \| 0.236 \| 152 \| 106 \| 4 \| 1 \| 144 \| 1 \| 150 \| Single-stranded DNA-binding protein \| Single-stranded DNA-binding protein; afdb-uniprot50 \| AF-A0A4Z1E4B9-F1-MODEL\_V4 \| 1.0 \| 5.125e-10 \| 345 \| 0.215 \| 167 \| 121 \| 5 \| 4 \| 162 \| 1 \| 165 \| Single-stranded DNA-binding protein \| Single-stranded DNA-binding protein; afdb-uniprot50 \| AF-A0A7W3IVP1-F1-MODEL\_V4 \| 1.0 \| 4.603e-10 \| 344 \| 0.245 \| 167 \| 114 \| 5 \| 6 \| 162 \| 3 \| 167 \| Single-stranded DNA-binding protein \| Single-stranded DNA-binding protein; afdb-uniprot50 \| AF-C6R1H3-F1-MODEL\_V4 \| 1.0 \| 3.161e-10 \| 342 \| 0.236 \| 165 \| 116 \| 5 \| 6 \| 162 \| 3 \| 165 \| Single-stranded DNA-binding protein \| Single-stranded DNA-binding protein; afdb-uniprot50 \| AF-A0A1F1ZNZ2-F1-MODEL\_V4 \| 1.0 \| 2.416e-10 \| 341 \| 0.275 \| 160 \| 104 \| 6 \| 5 \| 155 \| 1 \| 157 \| Single-stranded DNA-binding protein \| Single-stranded DNA-binding protein; afdb-uniprot50 \| AF-A0A1G3ZU57-F1-MODEL\_V4 \| 1.0 \| 1.5e-09 \| 341 \| 0.232 \| 176 \| 117 \| 6 \| 2 \| 162 \| 3 \| 175 \| Uncharacterized protein \| Uncharacterized protein; afdb-uniprot50 \| AF-A0A0K8Q012-F1-MODEL\_V4 \| 1.0 \| 9.253e-10 \| 340 \| 0.253 \| 166 \| 109 \| 6 \| 3 \| 156 \| 2 \| 164 \| Single-stranded DNA-binding protein \| Single-stranded DNA-binding protein; afdb-uniprot50 \| AF-A0A657BBA4-F1-MODEL\_V4 \| 1.0 \| 3.335e-10 \| 339 \| 0.482 \| 170 \| 55 \| 5 \| 23 \| 162 \| 2 \| 168 \| Single-stranded DNA-binding protein \| Single-stranded DNA-binding protein; afdb-uniprot50 \| AF-A0A6H9WNQ8-F1-MODEL\_V4 \| 1.0 \| 7.074e-10 \| 339 \| 0.206 \| 174 \| 123 \| 6 \| 1 \| 162 \| 50 \| 220 \| Single-stranded DNA-binding protein \| Single-stranded DNA-binding protein; afdb-uniprot50 \| AF-L1ME36-F1-MODEL\_V4 \| 1.0 \| 2.085e-08 \| 338 \| 0.203 \| 133 \| 99 \| 4 \| 2 \| 129 \| 5 \| 135 \| Single-strand binding family protein \| Single-strand binding family protein; afdb-uniprot50 \| AF-A0A3G6J8M0-F1-MODEL\_V4 \| 1.0 \| 5.445e-09 \| 337 \| 0.226 \| 137 \| 89 \| 5 \| 7 \| 128 \| 5 \| 139 \| Single-stranded DNA-binding protein \| Single-stranded DNA-binding protein; afdb-uniprot50 \| AF-A0A1D9FGE8-F1-MODEL\_V4 \| 1.0 \| 6.75e-09 \| 335 \| 0.208 \| 149 \| 107 \| 5 \| 8 \| 151 \| 1 \| 143 \| Single-stranded DNA-binding protein \| Single-stranded DNA-binding protein; afdb-uniprot50 \| AF-A0A449A2H2-F1-MODEL\_V4 \| 1.0 \| 1.357e-08 \| 335 \| 0.16 \| 162 \| 129 \| 3 \| 5 \| 162 \| 1 \| 159 \| Single-stranded DNA-binding protein \| Single-stranded DNA-binding protein; afdb-uniprot50 \| AF-A0A7Z0VU71-F1-MODEL\_V4 \| 1.0 \| 1.422e-09 \| 333 \| 0.209 \| 167 \| 117 \| 5 \| 1 \| 162 \| 1 \| 157 \| Uncharacterized protein \| Uncharacterized protein; afdb-uniprot50 \| AF-A0A3D5W1P2-F1-MODEL\_V4 \| 1.0 \| 1.348e-09 \| 333 \| 0.211 \| 161 \| 115 \| 6 \| 4 \| 154 \| 2 \| 160 \| Uncharacterized protein \| Uncharacterized protein; afdb-uniprot50 \| AF-A0A0M2LZ75-F1-MODEL\_V4 \| 1.0 \| 2.216e-07 \| 332 \| 0.187 \| 112 \| 85 \| 5 \| 4 \| 114 \| 1 \| 107 \| Uncharacterized protein \| Uncharacterized protein; afdb-uniprot50 \| AF-A0A6I7R8T6-F1-MODEL\_V4 \| 1.0 \| 3.568e-08 \| 331 \| 0.23 \| 130 \| 92 \| 5 \| 1 \| 129 \| 1 \| 123 \| Single-stranded DNA-binding protein \| Single-stranded DNA-binding protein; afdb-uniprot50 \| AF-A0A2G2FL91-F1-MODEL\_V4 \| 1.0 \| 1.095e-08 \| 329 \| 0.208 \| 139 \| 105 \| 3 \| 3 \| 138 \| 25 \| 161 \| Single-stranded DNA-binding protein \| Single-stranded DNA-binding protein; afdb-uniprot50 \| AF-A0A7M1SYA3-F1-MODEL\_V4 \| 1.0 \| 3.543e-09 \| 329 \| 0.189 \| 190 \| 123 \| 7 \| 1 \| 162 \| 1 \| 187 \| Single-stranded DNA-binding protein \| Single-stranded DNA-binding protein; afdb-uniprot50 \| AF-A0A7C6GI43-F1-MODEL\_V4 \| 1.0 \| 2.185e-09 \| 328 \| 0.229 \| 183 \| 118 \| 7 \| 1 \| 162 \| 36 \| 216 \| Single-stranded DNA-binding protein \| Single-stranded DNA-binding protein; afdb-uniprot50 \| AF-W6JX67-F1-MODEL\_V4 \| 1.0 \| 2.878e-08 \| 327 \| 0.269 \| 126 \| 88 \| 4 \| 4 \| 127 \| 1 \| 124 \| Single-stranded DNA-binding protein \| Single-stranded DNA-binding protein; afdb-uniprot50 \| AF-A0A2I1I393-F1-MODEL\_V4 \| 1.0 \| 4.635e-09 \| 327 \| 0.188 \| 175 \| 127 \| 5 \| 1 \| 162 \| 7 \| 179 \| Single-stranded DNA-binding protein \| Single-stranded DNA-binding protein; afdb-uniprot50 \| AF-A0A239QU82-F1-MODEL\_V4 \| 1.0 \| 3.358e-09 \| 326 \| 0.19 \| 168 \| 124 \| 7 \| 4 \| 162 \| 2 \| 166 \| Single-stranded DNA-binding protein \| Single-stranded DNA-binding protein; afdb-uniprot50 \| AF-A0A560W8C9-F1-MODEL\_V4 \| 1.0 \| 1.976e-08 \| 325 \| 0.224 \| 125 \| 87 \| 5 \| 4 \| 120 \| 1 \| 123 \| Single-stranded DNA-binding protein \| Single-stranded DNA-binding protein; afdb-uniprot50 \| AF-A0A2U1ZT47-F1-MODEL\_V4 \| 1.0 \| 3.016e-09 \| 325 \| 0.22 \| 163 \| 119 \| 5 \| 1 \| 162 \| 1 \| 156 \| Single-stranded DNA-binding protein \| Single-stranded DNA-binding protein; afdb-uniprot50 \| AF-A0A5B8ARK9-F1-MODEL\_V4 \| 1.0 \| 2.306e-09 \| 324 \| 0.202 \| 163 \| 112 \| 6 \| 3 \| 157 \| 2 \| 154 \| Single-stranded DNA-binding protein \| Single-stranded DNA-binding protein; afdb-uniprot50 \| AF-A0A5N5RH10-F1-MODEL\_V4 \| 1.0 \| 3.543e-09 \| 324 \| 0.226 \| 168 \| 118 \| 5 \| 1 \| 159 \| 1 \| 165 \| Single-stranded DNA-binding protein \| Single-stranded DNA-binding protein; afdb-uniprot50 \| AF-A0A5C7PTQ0-F1-MODEL\_V4 \| 1.0 \| 2.567e-09 \| 323 \| 0.252 \| 170 \| 106 \| 9 \| 1 \| 162 \| 1 \| 157 \| Single-stranded DNA-binding protein \| Single-stranded DNA-binding protein; afdb-uniprot50 \| AF-A0A1F0QJF3-F1-MODEL\_V4 \| 1.0 \| 3.205e-08 \| 322 \| 0.203 \| 123 \| 89 \| 4 \| 1 \| 120 \| 1 \| 117 \| Single-stranded DNA-binding protein \| Single-stranded DNA-binding protein; afdb-uniprot50 \| AF-A0A498C414-F1-MODEL\_V4 \| 1.0 \| 2.433e-09 \| 321 \| 0.247 \| 174 \| 112 \| 7 \| 5 \| 162 \| 1 \| 171 \| Single-stranded DNA-binding protein \| Single-stranded DNA-binding protein; afdb-uniprot50 \| AF-A0A7K1J6Z0-F1-MODEL\_V4 \| 1.0 \| 3.739e-09 \| 321 \| 0.229 \| 196 \| 115 \| 8 \| 1 \| 162 \| 1 \| 194 \| Single-stranded DNA-binding protein \| Single-stranded DNA-binding protein; afdb-uniprot50 \| AF-A0A4Q5B4F8-F1-MODEL\_V4 \| 1.0 \| 3.543e-09 \| 321 \| 0.231 \| 164 \| 114 \| 6 \| 1 \| 154 \| 1 \| 162 \| Single-strand binding protein \| Single-strand binding protein; afdb-uniprot50 \| AF-N6WBX6-F1-MODEL\_V4 \| 1.0 \| 3.358e-09 \| 321 \| 0.207 \| 169 \| 121 \| 7 \| 2 \| 162 \| 7 \| 170 \| Single-stranded DNA-binding protein \| Single-stranded DNA-binding protein; afdb-uniprot50 \| AF-A0A6N9YPB8-F1-MODEL\_V4 \| 1.0 \| 5.746e-09 \| 320 \| 0.235 \| 174 \| 115 \| 9 \| 1 \| 161 \| 1 \| 169 \| Single-stranded DNA-binding protein \| Single-stranded DNA-binding protein; afdb-uniprot50 \| AF-A0A0A7I9E0-F1-MODEL\_V4 \| 1.0 \| 1.037e-08 \| 318 \| 0.224 \| 169 \| 108 \| 7 \| 4 \| 154 \| 2 \| 165 \| Uncharacterized protein \| Uncharacterized protein; afdb-uniprot50 \| AF-A0A6G7Z2E4-F1-MODEL\_V4 \| 1.0 \| 3.358e-09 \| 318 \| 0.203 \| 167 \| 118 \| 6 \| 5 \| 162 \| 1 \| 161 \| Single-stranded DNA-binding protein \| Single-stranded DNA-binding protein; afdb-uniprot50 \| AF-A0A6A9V298-F1-MODEL\_V4 \| 1.0 \| 1.886e-07 \| 318 \| 0.213 \| 122 \| 86 \| 3 \| 2 \| 122 \| 56 \| 168 \| Single-stranded DNA-binding protein \| Single-stranded DNA-binding protein; afdb-uniprot50 \| AF-A0A2W0CY63-F1-MODEL\_V4 \| 1.0 \| 3.543e-09 \| 317 \| 0.204 \| 161 \| 118 \| 5 \| 4 \| 162 \| 2 \| 154 \| Single-stranded DNA-binding protein \| Single-stranded DNA-binding protein; afdb-uniprot50 \| AF-A0A3N4YNY9-F1-MODEL\_V4 \| 1.0 \| 1.422e-09 \| 317 \| 0.244 \| 176 \| 113 \| 8 \| 1 \| 162 \| 1 \| 170 \| Single-stranded DNA-binding protein \| Single-stranded DNA-binding protein; afdb-uniprot50 \| AF-A0A6I1P9S7-F1-MODEL\_V4 \| 1.0 \| 1.095e-08 \| 316 \| 0.182 \| 159 \| 125 \| 5 \| 1 \| 156 \| 1 \| 157 \| Single-stranded DNA-binding protein \| Single-stranded DNA-binding protein; afdb-uniprot50 \| AF-A0A7T0PDA0-F1-MODEL\_V4 \| 1.0 \| 7.123e-09 \| 315 \| 0.196 \| 168 \| 120 \| 7 \| 4 \| 162 \| 1 \| 162 \| Single-stranded DNA-binding protein \| Single-stranded DNA-binding protein; afdb-uniprot50 \| AF-A0A7T0LMC8-F1-MODEL\_V4 \| 1.0 \| 3.016e-09 \| 315 \| 0.215 \| 167 \| 117 \| 7 \| 4 \| 160 \| 2 \| 164 \| Single-stranded DNA-binding protein \| Single-stranded DNA-binding protein; afdb-uniprot50 \| AF-A0A6B8VYJ9-F1-MODEL\_V4 \| 1.0 \| 4.393e-09 \| 314 \| 0.211 \| 189 \| 115 \| 7 \| 6 \| 162 \| 4 \| 190 \| Single-stranded DNA-binding protein \| Single-stranded DNA-binding protein; afdb-uniprot50 \| AF-A0A087AGL6-F1-MODEL\_V4 \| 1.0 \| 9.317e-09 \| 314 \| 0.223 \| 170 \| 120 \| 7 \| 1 \| 162 \| 1 \| 166 \| Single-stranded DNA-binding protein \| Single-stranded DNA-binding protein; afdb-uniprot50 \| AF-A0A7K0K8X1-F1-MODEL\_V4 \| 1.0 \| 4.891e-09 \| 314 \| 0.182 \| 175 \| 126 \| 5 \| 4 \| 162 \| 2 \| 175 \| Single-stranded DNA-binding protein \| Single-stranded DNA-binding protein; afdb-uniprot50 \| AF-A0A1G0GPX9-F1-MODEL\_V4 \| 1.0 \| 2.747e-07 \| 313 \| 0.183 \| 120 \| 95 \| 1 \| 1 \| 120 \| 4 \| 120 \| Uncharacterized protein \| Uncharacterized protein; afdb-uniprot50 \| AF-A0A540R783-F1-MODEL\_V4 \| 1.0 \| 8.83e-09 \| 312 \| 0.254 \| 169 \| 110 \| 8 \| 5 \| 162 \| 1 \| 164 \| Single-stranded DNA-binding protein \| Single-stranded DNA-binding protein; afdb-uniprot50 \| AF-A0A1H5DK09-F1-MODEL\_V4 \| 1.0 \| 7.931e-09 \| 312 \| 0.198 \| 171 \| 122 \| 6 \| 1 \| 162 \| 1 \| 165 \| Single-stranded DNA-binding protein \| Single-stranded DNA-binding protein; afdb-uniprot50 \| AF-A0A852W0M8-F1-MODEL\_V4 \| 1.0 \| 5.786e-08 \| 312 \| 0.184 \| 125 \| 96 \| 4 \| 7 \| 127 \| 5 \| 127 \| Single-strand DNA-binding protein \| Single-strand DNA-binding protein; afdb-uniprot50 \| AF-D1NUV4-F1-MODEL\_V4 \| 1.0 \| 1.873e-08 \| 310 \| 0.222 \| 162 \| 118 \| 4 \| 1 \| 156 \| 1 \| 160 \| Single-strand binding family protein \| Single-strand binding family protein; afdb-uniprot50 \| AF-A0A6N9Z2N4-F1-MODEL\_V4 \| 1.0 \| 9.831e-09 \| 310 \| 0.21 \| 166 \| 119 \| 7 \| 1 \| 159 \| 1 \| 161 \| Single-stranded DNA-binding protein \| Single-stranded DNA-binding protein; afdb-uniprot50 \| AF-A0A2S8ZS37-F1-MODEL\_V4 \| 1.0 \| 4.635e-09 \| 308 \| 0.248 \| 165 \| 113 \| 7 \| 6 \| 162 \| 3 \| 164 \| Single-stranded DNA-binding protein \| Single-stranded DNA-binding protein; afdb-uniprot50 \| AF-A0A2N3QFT5-F1-MODEL\_V4 \| 1.0 \| 9.831e-09 \| 308 \| 0.225 \| 200 \| 115 \| 6 \| 1 \| 162 \| 1 \| 198 \| Single-stranded DNA-binding protein \| Single-stranded DNA-binding protein; afdb-uniprot50 \| AF-A0A1H4WAE0-F1-MODEL\_V4 \| 1.0 \| 7.931e-09 \| 307 \| 0.22 \| 159 \| 117 \| 5 \| 5 \| 162 \| 1 \| 153 \| Single-stranded DNA-binding protein \| Single-stranded DNA-binding protein; afdb-uniprot50 \| AF-A0A3D5VZ01-F1-MODEL\_V4 \| 1.0 \| 1.976e-08 \| 307 \| 0.181 \| 171 \| 125 \| 7 \| 1 \| 162 \| 1 \| 165 \| Single-stranded DNA-binding protein \| Single-stranded DNA-binding protein; afdb-uniprot50 \| AF-A0A3A4KSQ4-F1-MODEL\_V4 \| 1.0 \| 2.45e-08 \| 306 \| 0.217 \| 161 \| 105 \| 7 \| 1 \| 160 \| 6 \| 146 \| Single-stranded DNA-binding protein \| Single-stranded DNA-binding protein; afdb-uniprot50 \| AF-A0A1I6G343-F1-MODEL\_V4 \| 1.0 \| 1.286e-08 \| 306 \| 0.224 \| 156 \| 114 \| 5 \| 6 \| 156 \| 3 \| 156 \| Single-stranded DNA-binding protein \| Single-stranded DNA-binding protein; afdb-uniprot50 \| AF-A0A3L6ZP49-F1-MODEL\_V4 \| 1.0 \| 7.931e-09 \| 306 \| 0.188 \| 159 \| 116 \| 6 \| 4 \| 157 \| 3 \| 153 \| Single-stranded DNA-binding protein \| Single-stranded DNA-binding protein; afdb-uniprot50 \| AF-K6WX02-F1-MODEL\_V4 \| 1.0 \| 3.016e-09 \| 304 \| 0.218 \| 169 \| 115 \| 8 \| 4 \| 160 \| 1 \| 164 \| Single-stranded DNA-binding protein \| Single-stranded DNA-binding protein; afdb-uniprot50 \| AF-A0A1R4FAL8-F1-MODEL\_V4 \| 1.0 \| 5.161e-09 \| 304 \| 0.215 \| 181 \| 114 \| 9 \| 1 \| 162 \| 1 \| 172 \| Uncharacterized protein \| Uncharacterized protein; afdb-uniprot50 \| AF-A0A387BUL9-F1-MODEL\_V4 \| 1.0 \| 6.442e-08 \| 303 \| 0.223 \| 134 \| 98 \| 5 \| 1 \| 131 \| 1 \| 131 \| Single-stranded DNA-binding protein \| Single-stranded DNA-binding protein; afdb-uniprot50 \| AF-A0A1F6RE83-F1-MODEL\_V4 \| 1.0 \| 1.682e-08 \| 303 \| 0.214 \| 168 \| 121 \| 4 \| 4 \| 162 \| 2 \| 167 \| Single-stranded DNA-binding protein \| Single-stranded DNA-binding protein; afdb-uniprot50 \| AF-A0A7X8M9H6-F1-MODEL\_V4 \| 1.0 \| 8.426e-08 \| 303 \| 0.219 \| 141 \| 104 \| 5 \| 1 \| 138 \| 30 \| 167 \| Single-stranded DNA-binding protein \| Single-stranded DNA-binding protein; afdb-uniprot50 \| AF-A0A5C8I7Q5-F1-MODEL\_V4 \| 1.0 \| 8.368e-09 \| 303 \| 0.244 \| 172 \| 112 \| 7 \| 6 \| 162 \| 3 \| 171 \| Single-stranded DNA-binding protein \| Single-stranded DNA-binding protein; afdb-uniprot50 \| AF-A0A2A9D112-F1-MODEL\_V4 \| 1.0 \| 3.037e-08 \| 303 \| 0.203 \| 177 \| 115 \| 9 \| 1 \| 160 \| 17 \| 184 \| Single-strand DNA-binding protein \| Single-strand DNA-binding protein; afdb-uniprot50 \| AF-A0A6P0HKI8-F1-MODEL\_V4 \| 1.0 \| 1.682e-08 \| 302 \| 0.196 \| 173 \| 119 \| 6 \| 4 \| 162 \| 2 \| 168 \| Single-stranded DNA-binding protein \| Single-stranded DNA-binding protein; afdb-uniprot50 \| AF-A0A6I3JGH7-F1-MODEL\_V4 \| 1.0 \| 1.521e-07 \| 301 \| 0.155 \| 129 \| 101 \| 5 \| 1 \| 124 \| 1 \| 126 \| Single-stranded DNA-binding protein \| Single-stranded DNA-binding protein; afdb-uniprot50 \| AF-B1VGP3-F1-MODEL\_V4 \| 1.0 \| 4.192e-08 \| 301 \| 0.195 \| 169 \| 126 \| 6 \| 1 \| 161 \| 49 \| 215 \| Single-stranded DNA-binding protein \| Single-stranded DNA-binding protein; afdb-uniprot50 \| AF-A0A176QDX5-F1-MODEL\_V4 \| 1.0 \| 3.945e-09 \| 300 \| 0.235 \| 170 \| 112 \| 8 \| 4 \| 157 \| 1 \| 168 \| Single-stranded DNA-binding protein \| Single-stranded DNA-binding protein; afdb-uniprot50 \| AF-A0A7X7K7I0-F1-MODEL\_V4 \| 1.0 \| 1.775e-08 \| 299 \| 0.173 \| 161 \| 106 \| 5 \| 1 \| 136 \| 1 \| 159 \| Single-stranded DNA-binding protein \| Single-stranded DNA-binding protein; afdb-uniprot50 \| AF-A0A087BDX7-F1-MODEL\_V4 \| 1.0 \| 1.976e-08 \| 299 \| 0.205 \| 170 \| 123 \| 5 \| 1 \| 160 \| 1 \| 168 \| Single-stranded DNA-binding protein \| Single-stranded DNA-binding protein; afdb-uniprot50 \| AF-A0A430FWQ9-F1-MODEL\_V4 \| 1.0 \| 6.75e-09 \| 299 \| 0.225 \| 182 \| 117 \| 7 \| 1 \| 162 \| 1 \| 178 \| Single-stranded DNA-binding protein \| Single-stranded DNA-binding protein; afdb-uniprot50 \| AF-A0A2N6UPV5-F1-MODEL\_V4 \| 1.0 \| 1.286e-08 \| 298 \| 0.19 \| 163 \| 119 \| 6 \| 2 \| 154 \| 29 \| 188 \| Single-stranded DNA-binding protein \| Single-stranded DNA-binding protein; afdb-uniprot50 \| AF-A0A4Q9KLV9-F1-MODEL\_V4 \| 1.0 \| 2.728e-08 \| 297 \| 0.19 \| 173 \| 123 \| 8 \| 5 \| 162 \| 1 \| 171 \| Single-stranded DNA-binding protein \| Single-stranded DNA-binding protein; afdb-uniprot50 \| AF-A0A3G5T3X2-F1-MODEL\_V4 \| 1.0 \| 5.196e-08 \| 296 \| 0.174 \| 166 \| 125 \| 7 \| 1 \| 162 \| 1 \| 158 \| Single-stranded DNA-binding protein \| Single-stranded DNA-binding protein; afdb-uniprot50 \| AF-A0A6L4WZV7-F1-MODEL\_V4 \| 1.0 \| 2.085e-08 \| 296 \| 0.211 \| 170 \| 123 \| 7 \| 1 \| 162 \| 1 \| 167 \| Single-stranded DNA-binding protein \| Single-stranded DNA-binding protein; afdb-uniprot50 \| AF-A0A4Y8UKJ6-F1-MODEL\_V4 \| 1.0 \| 5.233e-07 \| 294 \| 0.21 \| 119 \| 86 \| 4 \| 3 \| 117 \| 2 \| 116 \| Uncharacterized protein \| Uncharacterized protein; afdb-uniprot50 \| AF-A0A1R4HUB9-F1-MODEL\_V4 \| 1.0 \| 8.891e-08 \| 294 \| 0.179 \| 156 \| 117 \| 6 \| 1 \| 151 \| 1 \| 150 \| Single-strand binding protein/Primosomal replication protein n \| Single-strand binding protein/Primosomal replication protein n; afdb-uniprot50 \| AF-A0A1X0E5M8-F1-MODEL\_V4 \| 1.0 \| 1.511e-08 \| 294 \| 0.201 \| 169 \| 117 \| 7 \| 6 \| 162 \| 4 \| 166 \| Single-stranded DNA-binding protein \| Single-stranded DNA-binding protein; afdb-uniprot50 \| AF-A0A849PFU8-F1-MODEL\_V4 \| 1.0 \| 4.925e-08 \| 294 \| 0.155 \| 180 \| 128 \| 8 \| 1 \| 162 \| 1 \| 174 \| Single-stranded DNA-binding protein \| Single-stranded DNA-binding protein; afdb-uniprot50 \| AF-A0A2P8EFA0-F1-MODEL\_V4 \| 1.0 \| 2.2e-08 \| 294 \| 0.223 \| 170 \| 117 \| 7 \| 4 \| 162 \| 1 \| 166 \| Single-stranded DNA-binding protein \| Single-stranded DNA-binding protein; afdb-uniprot50 \| AF-C0W1B7-F1-MODEL\_V4 \| 1.0 \| 1.511e-08 \| 294 \| 0.207 \| 178 \| 121 \| 8 \| 1 \| 162 \| 34 \| 207 \| Single-stranded DNA-binding protein \| Single-stranded DNA-binding protein; afdb-uniprot50 \| AF-A0A6I5N066-F1-MODEL\_V4 \| 1.0 \| 3.381e-08 \| 292 \| 0.226 \| 168 \| 119 \| 7 \| 1 \| 162 \| 1 \| 163 \| Single-stranded DNA-binding protein \| Single-stranded DNA-binding protein; afdb-uniprot50 \| AF-A0A2W5B054-F1-MODEL\_V4 \| 1.0 \| 7.172e-08 \| 291 \| 0.207 \| 169 \| 124 \| 6 \| 1 \| 161 \| 15 \| 181 \| Single-stranded DNA-binding protein \| Single-stranded DNA-binding protein; afdb-uniprot50 \| AF-X7ZHF1-F1-MODEL\_V4 \| 1.0 \| 3.765e-08 \| 291 \| 0.209 \| 162 \| 119 \| 7 \| 4 \| 162 \| 1 \| 156 \| Single-stranded DNA-binding protein \| Single-stranded DNA-binding protein; afdb-uniprot50 \| AF-A0A852XD35-F1-MODEL\_V4 \| 1.0 \| 1.594e-08 \| 290 \| 0.218 \| 169 \| 117 \| 6 \| 7 \| 162 \| 5 \| 171 \| Single-strand DNA-binding protein \| Single-strand DNA-binding protein; afdb-uniprot50 \| AF-A0A1A3DF40-F1-MODEL\_V4 \| 1.0 \| 2.878e-08 \| 288 \| 0.198 \| 166 \| 121 \| 7 \| 6 \| 162 \| 4 \| 166 \| Single-stranded DNA-binding protein \| Single-stranded DNA-binding protein; afdb-uniprot50 \| AF-A0A6N2TFT1-F1-MODEL\_V4 \| 1.0 \| 3.381e-08 \| 288 \| 0.169 \| 159 \| 120 \| 6 \| 7 \| 157 \| 6 \| 160 \| Single-stranded DNA-binding protein \| Single-stranded DNA-binding protein; afdb-uniprot50 \| AF-A0A1X3PBD3-F1-MODEL\_V4 \| 1.0 \| 3.973e-08 \| 287 \| 0.182 \| 164 \| 123 \| 8 \| 1 \| 162 \| 6 \| 160 \| Uncharacterized protein \| Uncharacterized protein; afdb-uniprot50 \| AF-A0A1X0DBK5-F1-MODEL\_V4 \| 1.0 \| 4.192e-08 \| 287 \| 0.204 \| 166 \| 117 \| 7 \| 6 \| 162 \| 4 \| 163 \| Single-stranded DNA-binding protein \| Single-stranded DNA-binding protein; afdb-uniprot50 \| AF-A0A087D7I9-F1-MODEL\_V4 \| 1.0 \| 2.585e-08 \| 287 \| 0.217 \| 170 \| 121 \| 7 \| 1 \| 162 \| 1 \| 166 \| Single-stranded DNA-binding protein \| Single-stranded DNA-binding protein; afdb-uniprot50 \| AF-A0A1H1ZTF3-F1-MODEL\_V4 \| 1.0 \| 2.085e-08 \| 286 \| 0.214 \| 177 \| 106 \| 10 \| 4 \| 154 \| 1 \| 170 \| Single-stranded DNA-binding protein \| Single-stranded DNA-binding protein; afdb-uniprot50 \| AF-A0A1Q4TRE2-F1-MODEL\_V4 \| 1.0 \| 2.585e-08 \| 285 \| 0.203 \| 172 \| 116 \| 7 \| 6 \| 162 \| 4 \| 169 \| Single-stranded DNA-binding protein \| Single-stranded DNA-binding protein; afdb-uniprot50 \| AF-A0A2S9YZ45-F1-MODEL\_V4 \| 1.0 \| 2.45e-08 \| 285 \| 0.209 \| 181 \| 119 \| 7 \| 4 \| 162 \| 2 \| 180 \| Uncharacterized protein \| Uncharacterized protein; afdb-uniprot50 \| AF-X8AMK4-F1-MODEL\_V4 \| 1.0 \| 8.891e-08 \| 284 \| 0.225 \| 151 \| 104 \| 6 \| 6 \| 151 \| 4 \| 146 \| Single-stranded DNA-binding protein \| Single-stranded DNA-binding protein; afdb-uniprot50 \| AF-A0A4R4ZQF2-F1-MODEL\_V4 \| 1.0 \| 6.797e-08 \| 284 \| 0.195 \| 164 \| 108 \| 5 \| 4 \| 162 \| 2 \| 146 \| Single-stranded DNA-binding protein \| Single-stranded DNA-binding protein; afdb-uniprot50 \| AF-A0A100XCE1-F1-MODEL\_V4 \| 1.0 \| 5.483e-08 \| 282 \| 0.191 \| 162 \| 121 \| 5 \| 6 \| 160 \| 4 \| 162 \| Single-stranded DNA-binding protein \| Single-stranded DNA-binding protein; afdb-uniprot50 \| AF-A0A8A6ALU5-F1-MODEL\_V4 \| 1.0 \| 4.925e-08 \| 281 \| 0.18 \| 166 \| 123 \| 7 \| 4 \| 162 \| 2 \| 161 \| Single-stranded DNA-binding protein \| Single-stranded DNA-binding protein; afdb-uniprot50 \| AF-A0A1W9ZUU9-F1-MODEL\_V4 \| 1.0 \| 4.925e-08 \| 280 \| 0.195 \| 164 \| 122 \| 7 \| 6 \| 162 \| 4 \| 164 \| Uncharacterized protein \| Uncharacterized protein; afdb-uniprot50 \| AF-A0A430FPW2-F1-MODEL\_V4 \| 1.0 \| 1.227e-07 \| 280 \| 0.232 \| 172 \| 118 \| 6 \| 1 \| 162 \| 2 \| 169 \| Single-strand binding protein \| Single-strand binding protein; afdb-uniprot50 \| AF-A0A3R6LGZ0-F1-MODEL\_V4 \| 1.0 \| 1.873e-08 \| 278 \| 0.194 \| 149 \| 105 \| 5 \| 2 \| 148 \| 16 \| 151 \| Single-stranded DNA-binding protein \| Single-stranded DNA-binding protein; afdb-uniprot50 \| AF-A0A1X0IMP9-F1-MODEL\_V4 \| 1.0 \| 9.9e-08 \| 278 \| 0.185 \| 156 \| 116 \| 6 \| 6 \| 156 \| 4 \| 153 \| Single-stranded DNA-binding protein \| Single-stranded DNA-binding protein; afdb-uniprot50 \| AF-A0A4U2YR72-F1-MODEL\_V4 \| 1.0 \| 1.694e-07 \| 278 \| 0.18 \| 166 \| 123 \| 8 \| 1 \| 160 \| 1 \| 159 \| Single-stranded DNA-binding protein \| Single-stranded DNA-binding protein; afdb-uniprot50 \| AF-A0A1X1TEZ9-F1-MODEL\_V4 \| 1.0 \| 1.366e-07 \| 277 \| 0.171 \| 157 \| 121 \| 6 \| 4 \| 154 \| 1 \| 154 \| Single-stranded DNA-binding protein \| Single-stranded DNA-binding protein; afdb-uniprot50 \| AF-A0A3A3Z1T3-F1-MODEL\_V4 \| 1.0 \| 4.192e-08 \| 277 \| 0.235 \| 153 \| 105 \| 3 \| 1 \| 151 \| 1 \| 143 \| Single-stranded DNA-binding protein \| Single-stranded DNA-binding protein; afdb-uniprot50 \| AF-A0A7Z0J5J7-F1-MODEL\_V4 \| 1.0 \| 5.196e-08 \| 277 \| 0.202 \| 163 \| 118 \| 7 \| 6 \| 162 \| 3 \| 159 \| Single-strand DNA-binding protein \| Single-strand DNA-binding protein; afdb-uniprot50 \| AF-C5BXK5-F1-MODEL\_V4 \| 1.0 \| 7.568e-08 \| 276 \| 0.18 \| 166 \| 125 \| 6 \| 4 \| 162 \| 2 \| 163 \| Single-stranded DNA-binding protein \| Single-stranded DNA-binding protein; afdb-uniprot50 \| AF-A0A7G6Z5V3-F1-MODEL\_V4 \| 1.0 \| 7.986e-08 \| 276 \| 0.164 \| 158 \| 119 \| 7 \| 4 \| 157 \| 3 \| 151 \| Single-stranded DNA-binding protein \| Single-stranded DNA-binding protein; afdb-uniprot50 \| AF-A0A101AME2-F1-MODEL\_V4 \| 1.0 \| 1.227e-07 \| 274 \| 0.175 \| 160 \| 122 \| 6 \| 6 \| 162 \| 4 \| 156 \| Single-stranded DNA-binding protein \| Single-stranded DNA-binding protein; afdb-uniprot50 \| AF-A0A3P1SET8-F1-MODEL\_V4 \| 1.0 \| 4.667e-08 \| 272 \| 0.17 \| 170 \| 124 \| 8 \| 1 \| 162 \| 1 \| 161 \| Single-stranded DNA-binding protein \| Single-stranded DNA-binding protein; afdb-uniprot50 \| AF-A0A0R3FMM0-F1-MODEL\_V4 \| 1.0 \| 1.163e-07 \| 271 \| 0.183 \| 158 \| 117 \| 7 \| 4 \| 155 \| 1 \| 152 \| Single-stranded DNA-binding protein \| Single-stranded DNA-binding protein; afdb-uniprot50 \| AF-A0A3N4ZQ97-F1-MODEL\_V4 \| 1.0 \| 1.045e-07 \| 270 \| 0.172 \| 185 \| 123 \| 6 \| 3 \| 162 \| 2 \| 181 \| Single-stranded DNA-binding protein \| Single-stranded DNA-binding protein; afdb-uniprot50 \| AF-A0A1Y0C1S9-F1-MODEL\_V4 \| 1.0 \| 9.9e-08 \| 269 \| 0.218 \| 160 \| 112 \| 5 \| 2 \| 160 \| 22 \| 169 \| Single-stranded DNA-binding protein \| Single-stranded DNA-binding protein; afdb-uniprot50 \| AF-A0A7S8MY85-F1-MODEL\_V4 \| 1.0 \| 9.382e-08 \| 268 \| 0.173 \| 161 \| 122 \| 7 \| 1 \| 154 \| 1 \| 157 \| Single-stranded DNA-binding protein \| Single-stranded DNA-binding protein; afdb-uniprot50 \| AF-A0A1Q4A4B6-F1-MODEL\_V4 \| 1.0 \| 1.787e-07 \| 266 \| 0.133 \| 165 \| 132 \| 6 \| 3 \| 162 \| 2 \| 160 \| Single-stranded DNA-binding protein \| Single-stranded DNA-binding protein; afdb-uniprot50 \| AF-A0A061LX28-F1-MODEL\_V4 \| 1.0 \| 2.1e-07 \| 266 \| 0.18 \| 166 \| 127 \| 6 \| 1 \| 160 \| 1 \| 163 \| Single-stranded DNA-binding protein \| Single-stranded DNA-binding protein; afdb-uniprot50 \| AF-A0A7T2TH27-F1-MODEL\_V4 \| 1.0 \| 3.791e-07 \| 265 \| 0.165 \| 157 \| 126 \| 5 \| 1 \| 156 \| 1 \| 153 \| Single-stranded DNA-binding protein \| Single-stranded DNA-binding protein; afdb-uniprot50 \| AF-A0A086ZVX5-F1-MODEL\_V4 \| 1.0 \| 6.105e-08 \| 265 \| 0.217 \| 179 \| 116 \| 9 \| 1 \| 159 \| 1 \| 175 \| Single-strand binding protein \| Single-strand binding protein; afdb-uniprot50 \| AF-A0A1E9HDT3-F1-MODEL\_V4 \| 1.0 \| 1.102e-07 \| 263 \| 0.215 \| 167 \| 117 \| 8 \| 1 \| 162 \| 5 \| 162 \| Uncharacterized protein \| Uncharacterized protein; afdb-uniprot50 \| AF-A0A4Q7NSB1-F1-MODEL\_V4 \| 1.0 \| 1.99e-07 \| 262 \| 0.168 \| 166 \| 120 \| 7 \| 4 \| 154 \| 1 \| 163 \| Single-stranded DNA-binding protein \| Single-stranded DNA-binding protein; afdb-uniprot50 \| AF-A0A7Y9YDV5-F1-MODEL\_V4 \| 1.0 \| 2.467e-07 \| 260 \| 0.14 \| 164 \| 129 \| 7 \| 1 \| 156 \| 1 \| 160 \| Single-stranded DNA-binding protein \| Single-stranded DNA-binding protein; afdb-uniprot50 \| AF-A0A178LXY2-F1-MODEL\_V4 \| 1.0 \| 1.99e-07 \| 258 \| 0.175 \| 160 \| 117 \| 5 \| 6 \| 160 \| 4 \| 153 \| Single-stranded DNA-binding protein \| Single-stranded DNA-binding protein; afdb-uniprot50 \| AF-A0A4P6MVX0-F1-MODEL\_V4 \| 1.0 \| 1.605e-07 \| 258 \| 0.171 \| 181 \| 116 \| 8 \| 5 \| 154 \| 1 \| 178 \| Single-stranded DNA-binding protein \| Single-stranded DNA-binding protein; afdb-uniprot50 \| AF-A0A2S0KF83-F1-MODEL\_V4 \| 1.0 \| 2.603e-07 \| 257 \| 0.191 \| 157 \| 115 \| 7 \| 7 \| 154 \| 5 \| 158 \| Single-stranded DNA-binding protein \| Single-stranded DNA-binding protein; afdb-uniprot50 \| AF-A0A7Z0EC79-F1-MODEL\_V4 \| 1.0 \| 6.845e-07 \| 254 \| 0.135 \| 155 \| 123 \| 7 \| 1 \| 150 \| 54 \| 202 \| Single-stranded DNA-binding protein \| Single-stranded DNA-binding protein; afdb-uniprot50 \| AF-A0A6N4W4B9-F1-MODEL\_V4 \| 1.0 \| 4.959e-07 \| 248 \| 0.155 \| 161 \| 122 \| 6 \| 7 \| 156 \| 5 \| 162 \| Single-stranded DNA-binding protein \| Single-stranded DNA-binding protein; afdb-uniprot50 \| AF-A0A2D9NY87-F1-MODEL\_V4 \| 1.0 \| 2.603e-07 \| 245 \| 0.173 \| 167 \| 125 \| 6 \| 1 \| 162 \| 14 \| 172 \| Uncharacterized protein \| Uncharacterized protein |
| Top keywords  (threshold 1.00e-02 (evalue)) | **DNA\_binding, Single\_stranded, Single\_strand, binding, replication, mitochondrial, Primosomal, n, A, complex** |
| Output files | ../../similar\_structures/35\_FANPEZAQ\_CDS\_0035\_afdb-proteome\_foldseek.tsv ../../similar\_structures/35\_FANPEZAQ\_CDS\_0035\_afdb-uniprot50\_foldseek.tsv ../../similar\_structures/35\_FANPEZAQ\_CDS\_0035\_merged.svg ../../similar\_structures/35\_FANPEZAQ\_CDS\_0035\_pdb\_foldseek.tsv |

  
  
  

Return to summary | Go to previous | Go to next

  


---

**Sequence/structure alignments coloring**  
Each object in the alignment figures is colored according to its E-value following this color coding:

1e-100
10
