## Supplementary material for "Completing the BASEL phage collection to unlock hidden diversity for systematic exploration of phage-host interactions": Data S2: 36.html

FANPEZAQ\_CDS\_0036


Return to summary | Go to previous | Go to next

|  |  |
| --- | --- |
| FANPEZAQ\_CDS\_0036  Page creation date: 02 Sep 2024, 12:00  Project folder: n/a  Input sequences file: Escherichia\_virus\_HeidiAbel.gb | dksa c4\_type domain\_containing trar transcriptional regulator zinc finger dnak rna conjugal transfer polymerase\_binding transcription factor fragment suppressor molecular chaperone dna\_binding general stress 16o duf2116 zn\_ribbon putative phage polymerase plasmid c\_4 type hypothetical dna\_directed alpha structural zn |

### Sequence information

|  |  |
| --- | --- |
| Name | FANPEZAQ\_CDS\_0036  36\_FANPEZAQ\_CDS\_0036 (pipeline id) |
| Imported annotations |  |
| Protein sequence | MTTNSRTIAALFAVSRMVTTTKSARLHLVNSGCTMADDIDRACEREEQERAAMIAAHLQK APAVKAVARGYCLNCGEDFPEGSKKIYCDADCANDHAAHLKRK |
| Number of residues | 103 |
| Molecular weight (Da) | 11242.75 |
| Output files | ../../query\_sequences/36\_FANPEZAQ\_CDS\_0036.fasta |

### Putative domain architecture and protein family

#### Search results (HHblits)^1^

|  |  |
| --- | --- |
| Domain family databases searched | Pfam, Ncbi-cd, Cath, Phrogs |
| Results, scheme(s)  (Top layers only; threshold 1.00e-03 (evalue)) | xml version="1.0" encoding="utf-8" standalone="no"?       2024-09-02T21:08:20.209596 image/svg+xml   Matplotlib v3.7.2, https://matplotlib.org/ |
| Results, table  (E-value ≤ 1.00e-03 (evalue)) | db \| id \| prob \| evalue \| pvalue \| score \| cols \| query \| query\_len \| template \| template\_len \| name \| description; phrogs \| 54 \| 99.7 \| 1.4e-21 \| 2e-25 \| 120.1 \| 62 \| (35, 98) \| 103 \| (1, 62) \| 63 \| DksA-like zinc-finger protein \| DksA-like zinc-finger protein; Category: other; MF490239\_p53 |
| Top keywords  (threshold 1.00e-03 (evalue)) | **DksA\_like, zinc\_finger, other, MF490239\_p53** |
| Output files | ../../domain\_architecture/36\_FANPEZAQ\_CDS\_0036\_cath.hhr ../../domain\_architecture/36\_FANPEZAQ\_CDS\_0036\_merged.svg ../../domain\_architecture/36\_FANPEZAQ\_CDS\_0036\_ncbi-cd.hhr ../../domain\_architecture/36\_FANPEZAQ\_CDS\_0036\_pfam.hhr ../../domain\_architecture/36\_FANPEZAQ\_CDS\_0036\_phrogs.hhr |

### Identical protein sequences/structures

#### Search results

|  |  |
| --- | --- |
| Protein sequence databases searched | Pdb, Swissprot, Refseq |
| Identical proteins found | -- |
| Top keywords | -- |
| Output files | -- |

### Similar protein sequences/structures

#### Sequence similarity search results (HHblits)^1^

|  |  |
| --- | --- |
| Sequence databases searched | Uniclust, Pdb70 |
| Results, scheme(s)  (Top layers only, threshold 1.00e-03 (evalue)) | xml version="1.0" encoding="utf-8" standalone="no"?       2024-09-02T21:08:45.027118 image/svg+xml   Matplotlib v3.7.2, https://matplotlib.org/ |
| Results, table(s)  (threshold 1.00e-03 (evalue)) | db \| id \| prob \| evalue \| pvalue \| score \| cols \| query \| query\_len \| template \| template\_len \| name \| description; uniclust \| UniRef100\_A0A061JN07 \| 99.5 \| 1.1e-17 \| 2.5e-23 \| 113.6 \| 90 \| (6, 101) \| 103 \| (6, 100) \| 109 \| DksA C4-type domain-containing protein \| DksA C4-type domain-containing protein; uniclust \| UniRef100\_A0A031IV80 \| 99.5 \| 1.4e-17 \| 3.5e-23 \| 108.5 \| 67 \| (33, 101) \| 103 \| (1, 71) \| 82 \| Prophage PSPPH06, DksA/TraR family C4-type zinc finger protein \| Prophage PSPPH06, DksA/TraR family C4-type zinc finger protein; uniclust \| UniRef100\_A0A084Z0P9 \| 99.5 \| 2.7e-17 \| 6.6e-23 \| 110.6 \| 72 \| (29, 102) \| 103 \| (2, 78) \| 98 \| DksA C4-type domain-containing protein \| DksA C4-type domain-containing protein; uniclust \| UniRef100\_A0A022PBT0 \| 99.5 \| 3.5e-17 \| 8.4e-23 \| 106.2 \| 67 \| (32, 102) \| 103 \| (1, 72) \| 79 \| Transcriptional regulator, TraR/DksA family \| Transcriptional regulator, TraR/DksA family; uniclust \| UniRef100\_A0A059ZYI4 \| 99.5 \| 1.8e-16 \| 4.3e-22 \| 108.5 \| 71 \| (28, 101) \| 103 \| (11, 87) \| 108 \| DksA C4-type domain-containing protein \| DksA C4-type domain-containing protein; uniclust \| UniRef100\_A0A009YU33 \| 99.4 \| 2.6e-16 \| 6.4e-22 \| 106.0 \| 71 \| (29, 102) \| 103 \| (2, 77) \| 98 \| Prokaryotic dksA/traR C4-type zinc finger family protein \| Prokaryotic dksA/traR C4-type zinc finger family protein; uniclust \| UniRef100\_A0A011MGP2 \| 99.4 \| 4.7e-16 \| 1.1e-21 \| 107.7 \| 68 \| (28, 99) \| 103 \| (8, 80) \| 117 \| Molecular chaperone DnaK \| Molecular chaperone DnaK; uniclust \| UniRef100\_A0A0F2KS34 \| 99.4 \| 8.5e-16 \| 2e-21 \| 102.0 \| 68 \| (30, 100) \| 103 \| (7, 79) \| 89 \| DksA C4-type domain-containing protein \| DksA C4-type domain-containing protein; uniclust \| UniRef100\_A0A0A8RLH0 \| 99.4 \| 1.1e-15 \| 2.6e-21 \| 100.0 \| 66 \| (32, 102) \| 103 \| (6, 76) \| 83 \| Uncharacterized protein \| Uncharacterized protein; uniclust \| UniRef100\_A0A086CZD8 \| 99.4 \| 2.2e-15 \| 5.1e-21 \| 100.1 \| 74 \| (27, 102) \| 103 \| (5, 83) \| 90 \| DksA C4-type domain-containing protein \| DksA C4-type domain-containing protein; uniclust \| UniRef100\_A0A023WVJ6 \| 99.3 \| 2.6e-15 \| 6e-21 \| 106.3 \| 71 \| (27, 101) \| 103 \| (42, 117) \| 136 \| Conjugal transfer protein TraR \| Conjugal transfer protein TraR; uniclust \| UniRef100\_A0A0H3PCF4 \| 99.3 \| 3.9e-15 \| 8.2e-21 \| 96.9 \| 69 \| (32, 102) \| 103 \| (2, 75) \| 83 \| DksA C4-type domain-containing protein \| DksA C4-type domain-containing protein; uniclust \| UniRef100\_A0A0C5DZD0 \| 99.3 \| 9.9e-15 \| 2.3e-20 \| 93.9 \| 65 \| (35, 102) \| 103 \| (1, 70) \| 74 \| DksA C4-type domain-containing protein \| DksA C4-type domain-containing protein; uniclust \| UniRef100\_A0A090IG19 \| 99.3 \| 1.4e-14 \| 3.3e-20 \| 94.9 \| 65 \| (35, 101) \| 103 \| (1, 74) \| 82 \| Uncharacterized phage protein \| Uncharacterized phage protein; uniclust \| UniRef100\_A0A011NC41 \| 99.2 \| 2.9e-14 \| 6.3e-20 \| 95.6 \| 71 \| (30, 101) \| 103 \| (7, 85) \| 98 \| DksA-like zinc finger domain containing protein \| DksA-like zinc finger domain containing protein; uniclust \| UniRef100\_A0A0C1EKQ1 \| 99.2 \| 3.3e-14 \| 6.5e-20 \| 94.4 \| 70 \| (27, 99) \| 103 \| (12, 86) \| 97 \| DksA C4-type domain-containing protein \| DksA C4-type domain-containing protein; uniclust \| UniRef100\_A0A2A5G8T1 \| 99.2 \| 3.8e-14 \| 8.9e-20 \| 92.8 \| 67 \| (35, 102) \| 103 \| (1, 75) \| 81 \| Conjugal transfer protein TraR \| Conjugal transfer protein TraR; uniclust \| UniRef100\_A0A140D8Z6 \| 99.2 \| 4.7e-14 \| 1.1e-19 \| 96.1 \| 69 \| (32, 102) \| 103 \| (1, 74) \| 104 \| Conjugal transfer protein TraR \| Conjugal transfer protein TraR; uniclust \| UniRef100\_A0A0N0E3Y8 \| 99.2 \| 6e-14 \| 1.3e-19 \| 90.2 \| 68 \| (34, 102) \| 103 \| (1, 73) \| 75 \| Transcriptional regulator, TraR/DksA family \| Transcriptional regulator, TraR/DksA family; uniclust \| UniRef100\_A0A0Q4N459 \| 99.2 \| 8.9e-14 \| 2.1e-19 \| 93.5 \| 68 \| (31, 102) \| 103 \| (5, 72) \| 95 \| DUF2116 family Zn-ribbon domain-containing protein \| DUF2116 family Zn-ribbon domain-containing protein; uniclust \| UniRef100\_A0A0A2X9Z0 \| 99.2 \| 1.1e-13 \| 2.4e-19 \| 91.9 \| 70 \| (31, 101) \| 103 \| (4, 80) \| 88 \| DksA C4-type domain-containing protein \| DksA C4-type domain-containing protein; uniclust \| UniRef100\_A0A3M1K5W8 \| 99.1 \| 2.2e-13 \| 4.9e-19 \| 90.1 \| 69 \| (32, 101) \| 103 \| (3, 80) \| 87 \| TraR/DksA family transcriptional regulator \| TraR/DksA family transcriptional regulator; uniclust \| UniRef100\_A0A060H8P7 \| 99.1 \| 3.7e-13 \| 8.5e-19 \| 107.5 \| 75 \| (27, 102) \| 103 \| (233, 320) \| 329 \| RNA polymerase-binding transcription factor DksA \| RNA polymerase-binding transcription factor DksA; uniclust \| UniRef100\_A0A317Q359 \| 99.1 \| 5e-13 \| 1e-18 \| 85.7 \| 64 \| (35, 102) \| 103 \| (3, 71) \| 75 \| Phage/conjugal plasmid C-4 type zinc finger TraR family protein (Fragment) \| Phage/conjugal plasmid C-4 type zinc finger TraR family protein (Fragment); uniclust \| UniRef100\_A0A010RU16 \| 99.1 \| 6.4e-13 \| 1.4e-18 \| 85.7 \| 64 \| (33, 101) \| 103 \| (1, 69) \| 74 \| Conjugal transfer protein TraR \| Conjugal transfer protein TraR; uniclust \| UniRef100\_A0A193QIP3 \| 99.1 \| 6.3e-13 \| 1.5e-18 \| 90.0 \| 69 \| (28, 102) \| 103 \| (9, 77) \| 97 \| Phage protein \| Phage protein; uniclust \| UniRef100\_A0A1W1XTU0 \| 99.1 \| 7.9e-13 \| 1.8e-18 \| 86.7 \| 63 \| (33, 101) \| 103 \| (5, 67) \| 81 \| DUF2116 family Zn-ribbon domain-containing protein \| DUF2116 family Zn-ribbon domain-containing protein; uniclust \| UniRef100\_A0A177QP03 \| 99.1 \| 9e-13 \| 2e-18 \| 87.1 \| 69 \| (30, 101) \| 103 \| (4, 75) \| 84 \| DksA C4-type domain-containing protein \| DksA C4-type domain-containing protein; uniclust \| UniRef100\_A0A034SXG4 \| 99.0 \| 1e-12 \| 2.4e-18 \| 89.6 \| 75 \| (15, 102) \| 103 \| (2, 76) \| 101 \| DUF2116 family Zn-ribbon domain-containing protein \| DUF2116 family Zn-ribbon domain-containing protein; uniclust \| UniRef100\_A0A062GXM3 \| 99.0 \| 1.1e-12 \| 2.4e-18 \| 86.8 \| 66 \| (32, 102) \| 103 \| (1, 70) \| 87 \| Prokaryotic dksA/traR C4-type zinc finger family protein \| Prokaryotic dksA/traR C4-type zinc finger family protein; uniclust \| UniRef100\_A0A0Q8QW33 \| 99.0 \| 1.5e-12 \| 3.4e-18 \| 86.9 \| 67 \| (31, 102) \| 103 \| (7, 73) \| 89 \| DUF2116 family Zn-ribbon domain-containing protein \| DUF2116 family Zn-ribbon domain-containing protein; uniclust \| UniRef100\_A0A1U9ZAF7 \| 99.0 \| 2.2e-12 \| 4.4e-18 \| 81.2 \| 63 \| (35, 101) \| 103 \| (1, 63) \| 67 \| Uncharacterized protein \| Uncharacterized protein; uniclust \| UniRef100\_A0A0J7J789 \| 99.0 \| 2.1e-12 \| 4.5e-18 \| 86.4 \| 60 \| (38, 101) \| 103 \| (21, 85) \| 92 \| Transcriptional regulator, TraR/DksA family \| Transcriptional regulator, TraR/DksA family; uniclust \| UniRef100\_A0A091AIU5 \| 99.0 \| 1.8e-12 \| 4.6e-18 \| 96.1 \| 65 \| (35, 100) \| 103 \| (76, 153) \| 174 \| DksA C4-type domain-containing protein \| DksA C4-type domain-containing protein; uniclust \| UniRef100\_A0A0H3A990 \| 99.0 \| 2.5e-12 \| 5e-18 \| 84.4 \| 68 \| (32, 100) \| 103 \| (3, 79) \| 87 \| Transcriptional regulator, TraR/DksA family \| Transcriptional regulator, TraR/DksA family; uniclust \| UniRef100\_A0A1G3M3M1 \| 99.0 \| 4e-12 \| 8e-18 \| 80.8 \| 62 \| (36, 99) \| 103 \| (1, 67) \| 72 \| DksA C4-type domain-containing protein (Fragment) \| DksA C4-type domain-containing protein (Fragment); uniclust \| UniRef100\_A0A031GDH3 \| 99.0 \| 3.7e-12 \| 9.6e-18 \| 94.2 \| 68 \| (32, 100) \| 103 \| (65, 145) \| 169 \| C4-type zinc finger protein, DksA/TraR family \| C4-type zinc finger protein, DksA/TraR family; uniclust \| UniRef100\_A0A1F2RT93 \| 99.0 \| 3.9e-12 \| 9.7e-18 \| 93.9 \| 69 \| (31, 100) \| 103 \| (61, 142) \| 168 \| DksA C4-type domain-containing protein \| DksA C4-type domain-containing protein; uniclust \| UniRef100\_A0A0B0VDY3 \| 98.9 \| 5.4e-12 \| 1.1e-17 \| 84.7 \| 70 \| (31, 102) \| 103 \| (7, 81) \| 96 \| Conjugal transfer protein TraR \| Conjugal transfer protein TraR; uniclust \| UniRef100\_A0A0H3JK08 \| 98.9 \| 6.6e-12 \| 1.5e-17 \| 84.4 \| 64 \| (33, 100) \| 103 \| (9, 78) \| 91 \| RNA polymerase-binding transcription factor \| RNA polymerase-binding transcription factor; uniclust \| UniRef100\_A0A1B9JT42 \| 98.9 \| 8.2e-12 \| 1.7e-17 \| 80.4 \| 64 \| (35, 100) \| 103 \| (1, 69) \| 74 \| DksA C4-type domain-containing protein \| DksA C4-type domain-containing protein; uniclust \| UniRef100\_A0A0A1VI61 \| 98.9 \| 7.6e-12 \| 2e-17 \| 92.0 \| 67 \| (34, 101) \| 103 \| (59, 138) \| 161 \| DnaK suppressor protein \| DnaK suppressor protein; uniclust \| UniRef100\_A0A076FE71 \| 98.9 \| 7.9e-12 \| 2e-17 \| 93.6 \| 70 \| (32, 102) \| 103 \| (81, 163) \| 184 \| DnaK suppressor protein \| DnaK suppressor protein; uniclust \| UniRef100\_A0A1G2UZM8 \| 98.9 \| 9.3e-12 \| 2.2e-17 \| 90.4 \| 71 \| (28, 99) \| 103 \| (48, 131) \| 149 \| DksA C4-type domain-containing protein \| DksA C4-type domain-containing protein; uniclust \| UniRef100\_A0A0C2UA16 \| 98.9 \| 1.1e-11 \| 2.4e-17 \| 80.9 \| 65 \| (35, 102) \| 103 \| (1, 70) \| 79 \| DksA C4-type domain-containing protein \| DksA C4-type domain-containing protein; uniclust \| UniRef100\_A0A059XN07 \| 98.9 \| 1e-11 \| 2.8e-17 \| 94.5 \| 65 \| (35, 100) \| 103 \| (74, 151) \| 208 \| Conjugal transfer protein TraR \| Conjugal transfer protein TraR; uniclust \| UniRef100\_A0A1G8NB44 \| 98.9 \| 1.3e-11 \| 2.8e-17 \| 81.1 \| 67 \| (32, 102) \| 103 \| (4, 77) \| 81 \| DksA/traR C4-type zinc finger \| DksA/traR C4-type zinc finger; uniclust \| UniRef100\_A0A1T4LZW9 \| 98.9 \| 1.5e-11 \| 3.3e-17 \| 94.3 \| 70 \| (31, 101) \| 103 \| (120, 202) \| 226 \| Transcriptional regulator, TraR/DksA family \| Transcriptional regulator, TraR/DksA family; uniclust \| UniRef100\_A0A2E4HLP3 \| 98.9 \| 1.6e-11 \| 3.4e-17 \| 84.1 \| 72 \| (30, 102) \| 103 \| (10, 94) \| 104 \| DksA C4-type domain-containing protein \| DksA C4-type domain-containing protein; uniclust \| UniRef100\_A0A0P0MAF7 \| 98.9 \| 1.6e-11 \| 3.5e-17 \| 101.7 \| 72 \| (29, 101) \| 103 \| (350, 434) \| 444 \| RNA polymerase-binding transcription factor DksA \| RNA polymerase-binding transcription factor DksA; uniclust \| UniRef100\_A0A0K2G9N8 \| 98.8 \| 1.9e-11 \| 4.6e-17 \| 94.1 \| 69 \| (31, 100) \| 103 \| (129, 210) \| 228 \| DnaK suppressor protein (Modular protein) \| DnaK suppressor protein (Modular protein); uniclust \| UniRef100\_A0A1I3XGG3 \| 98.8 \| 3.2e-11 \| 6.8e-17 \| 87.3 \| 77 \| (23, 100) \| 103 \| (50, 139) \| 155 \| Transcriptional regulator, TraR/DksA family \| Transcriptional regulator, TraR/DksA family; uniclust \| UniRef100\_A0A2H4TYU1 \| 98.8 \| 3.7e-11 \| 7.2e-17 \| 77.0 \| 65 \| (34, 102) \| 103 \| (1, 70) \| 76 \| Uncharacterized protein \| Uncharacterized protein; uniclust \| UniRef100\_A0A0G0NEV6 \| 98.8 \| 4.1e-11 \| 8.6e-17 \| 85.9 \| 68 \| (35, 103) \| 103 \| (62, 142) \| 144 \| Transcriptional regulator, TraR/DksA family \| Transcriptional regulator, TraR/DksA family; uniclust \| UniRef100\_A0A031GZT8 \| 98.8 \| 3.9e-11 \| 8.6e-17 \| 81.9 \| 68 \| (30, 101) \| 103 \| (9, 76) \| 101 \| DUF2116 family Zn-ribbon domain-containing protein \| DUF2116 family Zn-ribbon domain-containing protein; uniclust \| UniRef100\_A0A1F5P3U3 \| 98.8 \| 5.4e-11 \| 1.1e-16 \| 85.0 \| 67 \| (34, 101) \| 103 \| (40, 119) \| 139 \| DksA C4-type domain-containing protein \| DksA C4-type domain-containing protein; uniclust \| UniRef100\_A0A1F2RZ99 \| 98.8 \| 5e-11 \| 1.1e-16 \| 89.2 \| 64 \| (33, 97) \| 103 \| (81, 157) \| 183 \| DksA C4-type domain-containing protein \| DksA C4-type domain-containing protein; uniclust \| UniRef100\_A0A109RVY0 \| 98.8 \| 5.6e-11 \| 1.2e-16 \| 77.4 \| 66 \| (35, 102) \| 103 \| (1, 66) \| 78 \| Uncharacterized protein \| Uncharacterized protein; uniclust \| UniRef100\_A0A017TF53 \| 98.8 \| 4.8e-11 \| 1.2e-16 \| 90.8 \| 67 \| (34, 101) \| 103 \| (92, 171) \| 202 \| C4-type zinc finger protein, DksA/TraR family \| C4-type zinc finger protein, DksA/TraR family; uniclust \| UniRef100\_A0A072T172 \| 98.7 \| 5.8e-11 \| 1.5e-16 \| 87.6 \| 67 \| (33, 100) \| 103 \| (61, 140) \| 161 \| TraR/DksA family transcriptional regulator \| TraR/DksA family transcriptional regulator; uniclust \| UniRef100\_A0A1F9QKH6 \| 98.7 \| 6.5e-11 \| 1.5e-16 \| 88.5 \| 71 \| (29, 100) \| 103 \| (91, 174) \| 179 \| DksA C4-type domain-containing protein \| DksA C4-type domain-containing protein; uniclust \| UniRef100\_A0A023D0P2 \| 98.7 \| 7.6e-11 \| 1.6e-16 \| 88.8 \| 73 \| (28, 101) \| 103 \| (112, 197) \| 204 \| RNA polymerase-binding transcription factor DksA \| RNA polymerase-binding transcription factor DksA; uniclust \| UniRef100\_A0A099KCV7 \| 98.7 \| 7.7e-11 \| 1.7e-16 \| 84.8 \| 68 \| (32, 100) \| 103 \| (51, 131) \| 143 \| Transcriptional regulator, TraR/DksA family \| Transcriptional regulator, TraR/DksA family; uniclust \| UniRef100\_A0A021XD72 \| 98.7 \| 6.4e-11 \| 1.7e-16 \| 90.5 \| 63 \| (38, 101) \| 103 \| (102, 177) \| 209 \| DnaK suppressor protein \| DnaK suppressor protein; uniclust \| UniRef100\_A0A1F8GT99 \| 98.7 \| 7.4e-11 \| 1.8e-16 \| 86.6 \| 70 \| (31, 101) \| 103 \| (51, 133) \| 156 \| DksA C4-type domain-containing protein \| DksA C4-type domain-containing protein; uniclust \| UniRef100\_A0A0M9UC17 \| 98.7 \| 8.3e-11 \| 2e-16 \| 87.3 \| 70 \| (32, 102) \| 103 \| (59, 141) \| 169 \| DksA C4-type domain-containing protein \| DksA C4-type domain-containing protein; uniclust \| UniRef100\_A0A1F3SZI2 \| 98.7 \| 1.1e-10 \| 2.1e-16 \| 82.0 \| 65 \| (36, 101) \| 103 \| (43, 120) \| 132 \| DksA C4-type domain-containing protein \| DksA C4-type domain-containing protein; uniclust \| UniRef100\_A0A124FE87 \| 98.7 \| 9.6e-11 \| 2.2e-16 \| 86.8 \| 68 \| (33, 101) \| 103 \| (64, 144) \| 170 \| Transcriptional regulator, TraR/DksA family \| Transcriptional regulator, TraR/DksA family; uniclust \| UniRef100\_A0A0H3KWF3 \| 98.7 \| 1.3e-10 \| 2.4e-16 \| 79.3 \| 92 \| (5, 101) \| 103 \| (6, 105) \| 121 \| DksA C4-type domain-containing protein \| DksA C4-type domain-containing protein; uniclust \| UniRef100\_A0A011SK02 \| 98.7 \| 9.5e-11 \| 2.5e-16 \| 87.3 \| 64 \| (36, 100) \| 103 \| (78, 154) \| 173 \| Molecular chaperone DnaK \| Molecular chaperone DnaK; uniclust \| UniRef100\_A0A0D6QI25 \| 98.7 \| 1.1e-10 \| 2.5e-16 \| 85.4 \| 65 \| (32, 97) \| 103 \| (49, 126) \| 157 \| RNA polymerase-binding transcription factor DksA \| RNA polymerase-binding transcription factor DksA; uniclust \| UniRef100\_A0A085GNS1 \| 98.7 \| 1.4e-10 \| 2.9e-16 \| 87.2 \| 75 \| (27, 102) \| 103 \| (93, 180) \| 198 \| RNA polymerase-binding transcription factor DksA \| RNA polymerase-binding transcription factor DksA; uniclust \| UniRef100\_A0A010SR92 \| 98.7 \| 1.6e-10 \| 3.3e-16 \| 90.9 \| 75 \| (27, 102) \| 103 \| (190, 277) \| 286 \| RNA polymerase-binding transcription factor DksA \| RNA polymerase-binding transcription factor DksA; uniclust \| UniRef100\_A0A1B1EDL0 \| 98.7 \| 1.9e-10 \| 3.9e-16 \| 74.7 \| 66 \| (35, 101) \| 103 \| (1, 74) \| 78 \| DksA C4-type domain-containing protein \| DksA C4-type domain-containing protein; uniclust \| UniRef100\_A0A106QCK5 \| 98.7 \| 2e-10 \| 4.2e-16 \| 80.0 \| 73 \| (29, 102) \| 103 \| (22, 99) \| 115 \| DksA C4-type domain-containing protein \| DksA C4-type domain-containing protein; uniclust \| UniRef100\_A0A1F9ALS5 \| 98.7 \| 2e-10 \| 4.2e-16 \| 82.6 \| 69 \| (31, 100) \| 103 \| (58, 139) \| 145 \| DksA C4-type domain-containing protein \| DksA C4-type domain-containing protein; uniclust \| UniRef100\_A0A1V4XBY8 \| 98.7 \| 2.3e-10 \| 4.7e-16 \| 75.6 \| 65 \| (36, 101) \| 103 \| (1, 84) \| 86 \| RNA polymerase-binding transcription factor DksA \| RNA polymerase-binding transcription factor DksA; uniclust \| UniRef100\_A0A1G1JYH2 \| 98.6 \| 2.2e-10 \| 5e-16 \| 87.9 \| 65 \| (36, 101) \| 103 \| (135, 212) \| 218 \| DksA C4-type domain-containing protein \| DksA C4-type domain-containing protein; uniclust \| UniRef100\_A0A013XQT7 \| 98.6 \| 2.3e-10 \| 5.8e-16 \| 84.7 \| 66 \| (35, 101) \| 103 \| (74, 152) \| 163 \| DksA C4-type domain-containing protein \| DksA C4-type domain-containing protein; uniclust \| UniRef100\_A0A1J4Z2A6 \| 98.6 \| 2.5e-10 \| 6e-16 \| 82.9 \| 66 \| (35, 101) \| 103 \| (58, 136) \| 144 \| DksA C4-type domain-containing protein \| DksA C4-type domain-containing protein; uniclust \| UniRef100\_A0A523N3V7 \| 98.6 \| 3e-10 \| 6e-16 \| 78.0 \| 65 \| (34, 99) \| 103 \| (29, 106) \| 109 \| DksA C4-type domain-containing protein \| DksA C4-type domain-containing protein; uniclust \| UniRef100\_A0A011U6K4 \| 98.6 \| 2.9e-10 \| 6.5e-16 \| 88.9 \| 67 \| (34, 101) \| 103 \| (125, 204) \| 260 \| RNA polymerase-binding transcription factor DksA \| RNA polymerase-binding transcription factor DksA; uniclust \| UniRef100\_A0A656H975 \| 98.6 \| 3.7e-10 \| 7.3e-16 \| 74.3 \| 67 \| (35, 102) \| 103 \| (1, 73) \| 85 \| C4-type zinc finger DksA/TraR family protein \| C4-type zinc finger DksA/TraR family protein; uniclust \| UniRef100\_A0A0S7Z4A0 \| 98.6 \| 3.5e-10 \| 7.5e-16 \| 85.2 \| 70 \| (32, 102) \| 103 \| (98, 180) \| 198 \| DksA C4-type domain-containing protein \| DksA C4-type domain-containing protein; uniclust \| UniRef100\_A0A074M994 \| 98.6 \| 3.3e-10 \| 7.8e-16 \| 80.8 \| 64 \| (37, 101) \| 103 \| (43, 119) \| 126 \| DksA C4-type domain-containing protein \| DksA C4-type domain-containing protein; uniclust \| UniRef100\_A0A1F3BDN4 \| 98.6 \| 3.5e-10 \| 7.9e-16 \| 88.2 \| 67 \| (34, 101) \| 103 \| (145, 224) \| 246 \| DksA C4-type domain-containing protein \| DksA C4-type domain-containing protein; uniclust \| UniRef100\_A0A0U2WWH0 \| 98.6 \| 4.4e-10 \| 9e-16 \| 72.3 \| 63 \| (35, 102) \| 103 \| (1, 67) \| 72 \| DksA C4-type domain-containing protein \| DksA C4-type domain-containing protein; uniclust \| UniRef100\_A0A1W9HQR9 \| 98.6 \| 4.3e-10 \| 9.3e-16 \| 82.8 \| 66 \| (35, 101) \| 103 \| (42, 120) \| 164 \| DksA C4-type domain-containing protein \| DksA C4-type domain-containing protein; uniclust \| UniRef100\_A0A1J5DNA3 \| 98.6 \| 4.6e-10 \| 9.9e-16 \| 84.0 \| 71 \| (30, 101) \| 103 \| (42, 126) \| 187 \| DksA C4-type domain-containing protein \| DksA C4-type domain-containing protein; uniclust \| UniRef100\_A0A0F9CJ95 \| 98.6 \| 3.8e-10 \| 1e-15 \| 84.9 \| 65 \| (36, 101) \| 103 \| (86, 163) \| 183 \| DksA C4-type domain-containing protein \| DksA C4-type domain-containing protein; uniclust \| UniRef100\_A0A011NRS6 \| 98.6 \| 4.7e-10 \| 1e-15 \| 85.9 \| 74 \| (28, 102) \| 103 \| (108, 194) \| 222 \| RNA polymerase-binding transcription factor DksA \| RNA polymerase-binding transcription factor DksA; uniclust \| UniRef100\_A0A2M8KEH6 \| 98.6 \| 4.6e-10 \| 1e-15 \| 74.0 \| 38 \| (63, 101) \| 103 \| (33, 75) \| 79 \| DksA C4-type domain-containing protein (Fragment) \| DksA C4-type domain-containing protein (Fragment); uniclust \| UniRef100\_A0A0A6UQS0 \| 98.6 \| 5.1e-10 \| 1.1e-15 \| 82.6 \| 65 \| (36, 101) \| 103 \| (90, 167) \| 170 \| Conjugal transfer protein TraR \| Conjugal transfer protein TraR; uniclust \| UniRef100\_A0A1F2QVT2 \| 98.6 \| 4.9e-10 \| 1.2e-15 \| 83.1 \| 69 \| (32, 101) \| 103 \| (59, 140) \| 164 \| DksA C4-type domain-containing protein \| DksA C4-type domain-containing protein; uniclust \| UniRef100\_A0A0B0EK80 \| 98.6 \| 5.4e-10 \| 1.3e-15 \| 83.7 \| 70 \| (30, 100) \| 103 \| (46, 128) \| 176 \| DNA-binding protein \| DNA-binding protein; uniclust \| UniRef100\_A0A0S8CIM9 \| 98.6 \| 6e-10 \| 1.4e-15 \| 87.1 \| 72 \| (30, 102) \| 103 \| (90, 175) \| 249 \| DksA C4-type domain-containing protein \| DksA C4-type domain-containing protein; uniclust \| UniRef100\_A0A1Q7CPT7 \| 98.6 \| 6.7e-10 \| 1.4e-15 \| 75.5 \| 63 \| (38, 101) \| 103 \| (19, 94) \| 100 \| DksA C4-type domain-containing protein \| DksA C4-type domain-containing protein; uniclust \| UniRef100\_A0A1H9LGI8 \| 98.6 \| 6.8e-10 \| 1.4e-15 \| 71.5 \| 64 \| (33, 103) \| 103 \| (1, 66) \| 71 \| Phage/conjugal plasmid C-4 type zinc finger protein, TraR family \| Phage/conjugal plasmid C-4 type zinc finger protein, TraR family; uniclust \| UniRef100\_A0A2E1M7Z0 \| 98.5 \| 7.4e-10 \| 1.5e-15 \| 88.1 \| 69 \| (30, 99) \| 103 \| (232, 313) \| 318 \| DksA C4-type domain-containing protein \| DksA C4-type domain-containing protein; uniclust \| UniRef100\_A0A059KKD9 \| 98.5 \| 7e-10 \| 1.6e-15 \| 81.8 \| 62 \| (39, 101) \| 103 \| (73, 147) \| 160 \| Putative transcriptional regulator, TraR/DksA family \| Putative transcriptional regulator, TraR/DksA family; uniclust \| UniRef100\_A0A0B0EI80 \| 98.5 \| 7.9e-10 \| 1.7e-15 \| 79.7 \| 68 \| (32, 100) \| 103 \| (52, 132) \| 142 \| Transcriptional regulator \| Transcriptional regulator; uniclust \| UniRef100\_A0A0A2ZRM8 \| 98.5 \| 8.4e-10 \| 1.7e-15 \| 70.8 \| 63 \| (36, 101) \| 103 \| (1, 63) \| 70 \| DUF2116 family Zn-ribbon domain-containing protein \| DUF2116 family Zn-ribbon domain-containing protein; uniclust \| UniRef100\_A0A0Q1A6I1 \| 98.5 \| 7.6e-10 \| 1.8e-15 \| 82.1 \| 66 \| (36, 102) \| 103 \| (81, 159) \| 165 \| DksA C4-type domain-containing protein \| DksA C4-type domain-containing protein; uniclust \| UniRef100\_A0A0B5BBV4 \| 98.5 \| 7.8e-10 \| 1.8e-15 \| 82.1 \| 63 \| (37, 100) \| 103 \| (79, 154) \| 166 \| Molecular chaperone DnaK \| Molecular chaperone DnaK; uniclust \| UniRef100\_A0A1F9CTD9 \| 98.5 \| 9.5e-10 \| 1.9e-15 \| 79.1 \| 66 \| (35, 101) \| 103 \| (48, 126) \| 148 \| DksA C4-type domain-containing protein \| DksA C4-type domain-containing protein; uniclust \| UniRef100\_A0A0L8AA28 \| 98.5 \| 9.4e-10 \| 2e-15 \| 74.0 \| 73 \| (30, 102) \| 103 \| (8, 83) \| 91 \| Conjugal transfer protein TraR \| Conjugal transfer protein TraR; uniclust \| UniRef100\_A0A017HA46 \| 98.5 \| 9.2e-10 \| 2e-15 \| 78.6 \| 57 \| (40, 99) \| 103 \| (37, 98) \| 131 \| Putative Zinc-finger \| Putative Zinc-finger; uniclust \| UniRef100\_A0A1G2SET6 \| 98.5 \| 9.4e-10 \| 2e-15 \| 76.0 \| 40 \| (62, 102) \| 103 \| (56, 100) \| 106 \| DksA C4-type domain-containing protein \| DksA C4-type domain-containing protein; uniclust \| UniRef100\_A0A161WT96 \| 98.5 \| 1.1e-09 \| 2.1e-15 \| 71.2 \| 65 \| (35, 103) \| 103 \| (1, 66) \| 77 \| DksA C4-type domain-containing protein \| DksA C4-type domain-containing protein; uniclust \| UniRef100\_A0A2D8MXE8 \| 98.5 \| 1.1e-09 \| 2.1e-15 \| 82.9 \| 68 \| (34, 102) \| 103 \| (124, 204) \| 214 \| DksA C4-type domain-containing protein \| DksA C4-type domain-containing protein; uniclust \| UniRef100\_A0A094X2I6 \| 98.5 \| 1e-09 \| 2.4e-15 \| 82.0 \| 65 \| (34, 99) \| 103 \| (56, 133) \| 175 \| Putative dnaK suppressor \| Putative dnaK suppressor; uniclust \| UniRef100\_A0A0F9Z1F6 \| 98.5 \| 1.1e-09 \| 2.6e-15 \| 82.6 \| 65 \| (34, 99) \| 103 \| (70, 147) \| 186 \| Transcriptional regulator, TraR/DksA family \| Transcriptional regulator, TraR/DksA family; uniclust \| UniRef100\_A0A0A8R9Y5 \| 98.5 \| 1.5e-09 \| 2.8e-15 \| 76.6 \| 65 \| (31, 99) \| 103 \| (79, 148) \| 148 \| DksA C4-type domain-containing protein \| DksA C4-type domain-containing protein; uniclust \| UniRef100\_A0A0X8JHH5 \| 98.5 \| 1.7e-09 \| 3.6e-15 \| 79.9 \| 67 \| (31, 98) \| 103 \| (46, 125) \| 170 \| Transcriptional regulator \| Transcriptional regulator; uniclust \| UniRef100\_A0A0G0JDQ4 \| 98.5 \| 1.6e-09 \| 3.7e-15 \| 76.5 \| 66 \| (33, 99) \| 103 \| (35, 113) \| 118 \| Transcriptional regulator, TraR/DksA family \| Transcriptional regulator, TraR/DksA family; uniclust \| UniRef100\_A0A1F3SGG8 \| 98.5 \| 1.7e-09 \| 3.7e-15 \| 81.2 \| 69 \| (31, 100) \| 103 \| (69, 150) \| 180 \| DksA C4-type domain-containing protein \| DksA C4-type domain-containing protein; uniclust \| UniRef100\_A0A060NLE4 \| 98.4 \| 1.7e-09 \| 4.2e-15 \| 81.0 \| 61 \| (40, 101) \| 103 \| (87, 160) \| 173 \| DnaK suppressor protein \| DnaK suppressor protein; uniclust \| UniRef100\_A0A7C4ETC6 \| 98.4 \| 2.2e-09 \| 4.3e-15 \| 81.6 \| 66 \| (35, 101) \| 103 \| (137, 215) \| 242 \| TraR/DksA family transcriptional regulator \| TraR/DksA family transcriptional regulator; uniclust \| UniRef100\_A0A2G6EIW2 \| 98.4 \| 2.4e-09 \| 4.6e-15 \| 70.6 \| 71 \| (30, 101) \| 103 \| (10, 86) \| 87 \| Conjugal transfer protein TraR \| Conjugal transfer protein TraR; uniclust \| UniRef100\_A0A521SK58 \| 98.4 \| 2.4e-09 \| 5e-15 \| 75.2 \| 65 \| (35, 100) \| 103 \| (32, 109) \| 120 \| Transcriptional regulator, TraR/DksA family protein \| Transcriptional regulator, TraR/DksA family protein; uniclust \| UniRef100\_A0A2A3LML0 \| 98.4 \| 2.4e-09 \| 5.1e-15 \| 70.3 \| 63 \| (35, 100) \| 103 \| (1, 68) \| 78 \| DksA C4-type domain-containing protein \| DksA C4-type domain-containing protein; uniclust \| UniRef100\_UPI001CB70961 \| 98.4 \| 2.7e-09 \| 5.2e-15 \| 73.2 \| 71 \| (30, 101) \| 103 \| (33, 108) \| 112 \| TraR/DksA family transcriptional regulator \| TraR/DksA family transcriptional regulator; uniclust \| UniRef100\_A0A2E2AXQ7 \| 98.4 \| 2.5e-09 \| 5.2e-15 \| 76.8 \| 67 \| (32, 99) \| 103 \| (48, 127) \| 138 \| DksA C4-type domain-containing protein \| DksA C4-type domain-containing protein; uniclust \| UniRef100\_A0A1G3ZX42 \| 98.4 \| 2.3e-09 \| 5.4e-15 \| 86.3 \| 64 \| (36, 100) \| 103 \| (189, 265) \| 306 \| DksA C4-type domain-containing protein \| DksA C4-type domain-containing protein; uniclust \| UniRef100\_A0A068VTU4 \| 98.4 \| 2.9e-09 \| 6e-15 \| 82.7 \| 65 \| (36, 101) \| 103 \| (161, 238) \| 244 \| Transcriptional regulators, TraR/DksA family \| Transcriptional regulators, TraR/DksA family; uniclust \| UniRef100\_A0A0F2J1Q2 \| 98.4 \| 3e-09 \| 6.4e-15 \| 81.8 \| 68 \| (32, 100) \| 103 \| (126, 206) \| 220 \| RNA polymerase-binding, DksA \| RNA polymerase-binding, DksA; uniclust \| UniRef100\_A0A024YPC6 \| 98.4 \| 2.8e-09 \| 6.5e-15 \| 89.9 \| 65 \| (37, 102) \| 103 \| (402, 479) \| 481 \| DNA-binding protein \| DNA-binding protein; uniclust \| UniRef100\_A0A2A5B6B7 \| 98.4 \| 2.8e-09 \| 6.6e-15 \| 78.3 \| 69 \| (32, 101) \| 103 \| (64, 145) \| 150 \| DksA C4-type domain-containing protein \| DksA C4-type domain-containing protein; uniclust \| UniRef100\_A0A059IV93 \| 98.4 \| 3.1e-09 \| 7.3e-15 \| 81.2 \| 68 \| (33, 101) \| 103 \| (94, 174) \| 198 \| Putative dnaK suppressor protein \| Putative dnaK suppressor protein; uniclust \| UniRef100\_A0A1B4G123 \| 98.4 \| 4e-09 \| 8.5e-15 \| 68.5 \| 63 \| (35, 101) \| 103 \| (1, 64) \| 73 \| Conjugal transfer protein TraR \| Conjugal transfer protein TraR; uniclust \| UniRef100\_A0A5T2HSX0 \| 98.4 \| 4.6e-09 \| 8.7e-15 \| 67.2 \| 64 \| (32, 99) \| 103 \| (2, 70) \| 74 \| TraR/DksA family transcriptional regulator \| TraR/DksA family transcriptional regulator; uniclust \| UniRef100\_A0A2P5N6U8 \| 98.3 \| 5.3e-09 \| 1.1e-14 \| 70.6 \| 67 \| (35, 102) \| 103 \| (13, 84) \| 92 \| DksA C4-type domain-containing protein \| DksA C4-type domain-containing protein; uniclust \| UniRef100\_A0A0S3AHW3 \| 98.3 \| 5.2e-09 \| 1.1e-14 \| 76.1 \| 37 \| (63, 100) \| 103 \| (95, 136) \| 144 \| TraR/DksA family transcriptional regulator \| TraR/DksA family transcriptional regulator; uniclust \| UniRef100\_A0A081UWW5 \| 98.3 \| 4.7e-09 \| 1.2e-14 \| 79.8 \| 60 \| (40, 100) \| 103 \| (98, 170) \| 189 \| Molecular chaperone DnaK \| Molecular chaperone DnaK; uniclust \| UniRef100\_A0A0F2NQD6 \| 98.3 \| 5.3e-09 \| 1.2e-14 \| 76.7 \| 65 \| (36, 101) \| 103 \| (56, 133) \| 149 \| DksA C4-type domain-containing protein \| DksA C4-type domain-containing protein; uniclust \| UniRef100\_A0A0F9FJ65 \| 98.3 \| 5.7e-09 \| 1.3e-14 \| 81.0 \| 67 \| (33, 100) \| 103 \| (113, 192) \| 231 \| DksA C4-type domain-containing protein (Fragment) \| DksA C4-type domain-containing protein (Fragment); uniclust \| UniRef100\_A0A0B8WAV0 \| 98.3 \| 5.4e-09 \| 1.3e-14 \| 78.0 \| 66 \| (36, 102) \| 103 \| (62, 140) \| 168 \| Molecular chaperone DnaK \| Molecular chaperone DnaK; uniclust \| UniRef100\_A0A0A0GV12 \| 98.3 \| 6.9e-09 \| 1.3e-14 \| 68.9 \| 63 \| (35, 101) \| 103 \| (2, 69) \| 89 \| Protein traR \| Protein traR; uniclust \| UniRef100\_A0A171D6Y6 \| 98.3 \| 5.9e-09 \| 1.4e-14 \| 76.1 \| 62 \| (39, 101) \| 103 \| (61, 135) \| 143 \| Conjugal transfer protein TraR \| Conjugal transfer protein TraR; uniclust \| UniRef100\_A0A061NPW7 \| 98.3 \| 6.6e-09 \| 1.5e-14 \| 74.8 \| 37 \| (62, 99) \| 103 \| (88, 129) \| 131 \| DnaK suppressor protein \| DnaK suppressor protein; uniclust \| UniRef100\_A0A0N8QAU1 \| 98.3 \| 9.3e-09 \| 1.7e-14 \| 70.0 \| 65 \| (32, 99) \| 103 \| (40, 109) \| 110 \| TraR/DksA family transcriptional regulator \| TraR/DksA family transcriptional regulator; uniclust \| UniRef100\_A0A0R2XJZ4 \| 98.3 \| 8.5e-09 \| 1.8e-14 \| 84.9 \| 65 \| (36, 101) \| 103 \| (279, 356) \| 399 \| DksA C4-type domain-containing protein \| DksA C4-type domain-containing protein; uniclust \| UniRef100\_A0A1E4Q6M0 \| 98.3 \| 9.2e-09 \| 1.8e-14 \| 66.4 \| 61 \| (35, 101) \| 103 \| (1, 66) \| 73 \| DksA C4-type domain-containing protein \| DksA C4-type domain-containing protein; uniclust \| UniRef100\_A0A0D0WTJ9 \| 98.3 \| 8.1e-09 \| 1.9e-14 \| 75.3 \| 38 \| (62, 100) \| 103 \| (89, 131) \| 141 \| Conjugal transfer protein TraR \| Conjugal transfer protein TraR; uniclust \| UniRef100\_A0A3J9VMX4 \| 98.3 \| 1e-08 \| 2e-14 \| 66.2 \| 64 \| (36, 100) \| 103 \| (1, 70) \| 75 \| TraR/DksA family transcriptional regulator \| TraR/DksA family transcriptional regulator; uniclust \| UniRef100\_A0A0T5ZLC1 \| 98.3 \| 8.8e-09 \| 2e-14 \| 78.6 \| 68 \| (32, 100) \| 103 \| (105, 185) \| 197 \| TraR/DksA family transcriptional regulator, DnaK suppressor protein \| TraR/DksA family transcriptional regulator, DnaK suppressor protein; uniclust \| UniRef100\_A0A2D5I4F6 \| 98.3 \| 1e-08 \| 2.1e-14 \| 77.5 \| 74 \| (27, 101) \| 103 \| (117, 203) \| 207 \| RNA polymerase-binding transcription factor DksA \| RNA polymerase-binding transcription factor DksA; uniclust \| UniRef100\_UPI00218380DB \| 98.3 \| 1.1e-08 \| 2.1e-14 \| 70.4 \| 70 \| (27, 100) \| 103 \| (43, 117) \| 122 \| TraR/DksA family transcriptional regulator \| TraR/DksA family transcriptional regulator; uniclust \| UniRef100\_A0A106BVN3 \| 98.3 \| 1.1e-08 \| 2.2e-14 \| 66.4 \| 68 \| (34, 102) \| 103 \| (1, 68) \| 73 \| DUF2116 family Zn-ribbon domain-containing protein \| DUF2116 family Zn-ribbon domain-containing protein; uniclust \| UniRef100\_A0A0Q8PNH8 \| 98.3 \| 9.2e-09 \| 2.2e-14 \| 78.5 \| 64 \| (36, 100) \| 103 \| (110, 186) \| 194 \| DksA C4-type domain-containing protein \| DksA C4-type domain-containing protein; uniclust \| UniRef100\_A0A1M4XPA5 \| 98.3 \| 1.2e-08 \| 2.2e-14 \| 71.8 \| 64 \| (36, 100) \| 103 \| (26, 102) \| 135 \| Transcriptional regulator, TraR/DksA family \| Transcriptional regulator, TraR/DksA family; uniclust \| UniRef100\_A0A1H6CVK7 \| 98.3 \| 1.1e-08 \| 2.4e-14 \| 71.7 \| 35 \| (63, 98) \| 103 \| (74, 113) \| 116 \| DksA/traR C4-type zinc finger \| DksA/traR C4-type zinc finger; uniclust \| UniRef100\_A0A1Z9D4L2 \| 98.3 \| 1.1e-08 \| 2.4e-14 \| 80.9 \| 62 \| (37, 99) \| 103 \| (206, 280) \| 282 \| DksA C4-type domain-containing protein \| DksA C4-type domain-containing protein; uniclust \| UniRef100\_A0A0G3V2G2 \| 98.3 \| 1.1e-08 \| 2.5e-14 \| 81.5 \| 60 \| (39, 99) \| 103 \| (212, 284) \| 286 \| DnaK suppressor protein \| DnaK suppressor protein; uniclust \| UniRef100\_A0A0B6WW65 \| 98.3 \| 1.1e-08 \| 2.5e-14 \| 78.1 \| 37 \| (63, 100) \| 103 \| (134, 175) \| 206 \| Transcriptional regulator, TraR/DksA family \| Transcriptional regulator, TraR/DksA family; uniclust \| UniRef100\_A0A1Z8V918 \| 98.3 \| 1.1e-08 \| 2.5e-14 \| 73.5 \| 38 \| (63, 101) \| 103 \| (84, 126) \| 133 \| DksA C4-type domain-containing protein \| DksA C4-type domain-containing protein; uniclust \| UniRef100\_A0A1W9UTQ3 \| 98.3 \| 1.1e-08 \| 2.5e-14 \| 77.3 \| 59 \| (36, 95) \| 103 \| (105, 176) \| 185 \| DksA C4-type domain-containing protein \| DksA C4-type domain-containing protein; uniclust \| UniRef100\_A0A1F8N3L7 \| 98.3 \| 1.2e-08 \| 2.5e-14 \| 74.8 \| 68 \| (34, 102) \| 103 \| (67, 147) \| 149 \| DksA C4-type domain-containing protein \| DksA C4-type domain-containing protein; uniclust \| UniRef100\_A0A1G3XWY7 \| 98.3 \| 1.3e-08 \| 2.6e-14 \| 76.7 \| 37 \| (63, 100) \| 103 \| (138, 179) \| 190 \| DksA C4-type domain-containing protein \| DksA C4-type domain-containing protein; uniclust \| UniRef100\_A0A096H0I5 \| 98.2 \| 1.1e-08 \| 2.7e-14 \| 75.8 \| 38 \| (63, 101) \| 103 \| (102, 144) \| 155 \| Molecular chaperone DnaK \| Molecular chaperone DnaK; uniclust \| UniRef100\_A0A072N576 \| 98.2 \| 1.2e-08 \| 2.7e-14 \| 75.2 \| 65 \| (36, 101) \| 103 \| (56, 133) \| 151 \| Transcriptional regulator, TraR/DksA family \| Transcriptional regulator, TraR/DksA family; uniclust \| UniRef100\_A0A2W6C1C1 \| 98.2 \| 1.2e-08 \| 2.8e-14 \| 76.3 \| 69 \| (31, 100) \| 103 \| (52, 133) \| 172 \| DksA C4-type domain-containing protein \| DksA C4-type domain-containing protein; uniclust \| UniRef100\_A0A1F4QCC0 \| 98.2 \| 1.3e-08 \| 2.8e-14 \| 75.5 \| 67 \| (33, 100) \| 103 \| (55, 134) \| 165 \| DksA C4-type domain-containing protein \| DksA C4-type domain-containing protein; uniclust \| UniRef100\_A0A1F9TZQ5 \| 98.2 \| 1.4e-08 \| 2.9e-14 \| 74.9 \| 72 \| (27, 99) \| 103 \| (77, 161) \| 164 \| DksA C4-type domain-containing protein \| DksA C4-type domain-containing protein; uniclust \| UniRef100\_A0A1V8P4S1 \| 98.2 \| 1.5e-08 \| 3e-14 \| 68.8 \| 68 \| (31, 99) \| 103 \| (18, 91) \| 97 \| DksA C4-type domain-containing protein \| DksA C4-type domain-containing protein; uniclust \| UniRef100\_A0A0Q7CUW1 \| 98.2 \| 1.4e-08 \| 3e-14 \| 72.1 \| 38 \| (63, 101) \| 103 \| (76, 118) \| 123 \| DksA C4-type domain-containing protein \| DksA C4-type domain-containing protein; uniclust \| UniRef100\_A0A5Q0TJ53 \| 98.2 \| 1.4e-08 \| 3e-14 \| 65.0 \| 35 \| (67, 102) \| 103 \| (22, 61) \| 66 \| Transcriptional regulator \| Transcriptional regulator; uniclust \| UniRef100\_A0A497AXA6 \| 98.2 \| 1.6e-08 \| 3.2e-14 \| 68.3 \| 62 \| (40, 102) \| 103 \| (11, 85) \| 94 \| DksA C4-type domain-containing protein \| DksA C4-type domain-containing protein; uniclust \| UniRef100\_A0A1V6JBW9 \| 98.2 \| 1.4e-08 \| 3.2e-14 \| 82.3 \| 63 \| (39, 102) \| 103 \| (217, 292) \| 319 \| General stress protein 16O \| General stress protein 16O; uniclust \| UniRef100\_A0A1F8N7W5 \| 98.2 \| 1.6e-08 \| 3.3e-14 \| 74.9 \| 64 \| (36, 100) \| 103 \| (54, 130) \| 163 \| DksA C4-type domain-containing protein \| DksA C4-type domain-containing protein; uniclust \| UniRef100\_A0A522BF86 \| 98.2 \| 1.6e-08 \| 3.4e-14 \| 64.2 \| 36 \| (64, 100) \| 103 \| (18, 58) \| 64 \| DNA-binding protein (Fragment) \| DNA-binding protein (Fragment); uniclust \| UniRef100\_A0A021VUD3 \| 98.2 \| 1.6e-08 \| 3.6e-14 \| 77.2 \| 64 \| (36, 100) \| 103 \| (126, 202) \| 203 \| DNA-binding protein \| DNA-binding protein; uniclust \| UniRef100\_A0A0H1AEC4 \| 98.2 \| 1.7e-08 \| 3.7e-14 \| 77.3 \| 65 \| (36, 101) \| 103 \| (125, 202) \| 208 \| DNA-binding protein \| DNA-binding protein; uniclust \| UniRef100\_A0A2M9QWY4 \| 98.2 \| 1.8e-08 \| 3.7e-14 \| 66.4 \| 64 \| (35, 99) \| 103 \| (1, 71) \| 78 \| DksA C4-type domain-containing protein \| DksA C4-type domain-containing protein; uniclust \| UniRef100\_A0A327X3U0 \| 98.2 \| 1.9e-08 \| 3.8e-14 \| 64.3 \| 61 \| (35, 101) \| 103 \| (1, 61) \| 69 \| Uncharacterized protein \| Uncharacterized protein; uniclust \| UniRef100\_A0A022LFJ2 \| 98.2 \| 1.6e-08 \| 3.9e-14 \| 77.6 \| 39 \| (62, 101) \| 103 \| (139, 182) \| 200 \| Molecular chaperone DnaK \| Molecular chaperone DnaK; uniclust \| UniRef100\_A0A014MPS1 \| 98.2 \| 1.8e-08 \| 4.5e-14 \| 75.0 \| 39 \| (62, 101) \| 103 \| (101, 144) \| 160 \| Molecular chaperone DnaK \| Molecular chaperone DnaK; uniclust \| UniRef100\_A0A089YWC5 \| 98.2 \| 2e-08 \| 4.5e-14 \| 86.2 \| 62 \| (37, 99) \| 103 \| (494, 568) \| 569 \| DksA C4-type domain-containing protein \| DksA C4-type domain-containing protein; uniclust \| UniRef100\_A0A139TQP8 \| 98.2 \| 2.1e-08 \| 4.6e-14 \| 81.6 \| 65 \| (36, 101) \| 103 \| (233, 310) \| 349 \| C4-type zinc finger protein, DksA/TraR family \| C4-type zinc finger protein, DksA/TraR family; uniclust \| UniRef100\_A0A099TCY4 \| 98.2 \| 1.7e-08 \| 4.6e-14 \| 74.3 \| 62 \| (39, 101) \| 103 \| (61, 135) \| 150 \| Dimethylmenaquinone methyltransferase \| Dimethylmenaquinone methyltransferase; uniclust \| UniRef100\_A0A1M6SXT8 \| 98.2 \| 2.3e-08 \| 4.8e-14 \| 72.8 \| 64 \| (35, 99) \| 103 \| (44, 120) \| 151 \| Transcriptional regulator, TraR/DksA family \| Transcriptional regulator, TraR/DksA family; uniclust \| UniRef100\_A0A011U294 \| 98.2 \| 2.1e-08 \| 4.8e-14 \| 75.4 \| 38 \| (63, 101) \| 103 \| (118, 160) \| 172 \| Conjugal transfer protein TraR \| Conjugal transfer protein TraR; uniclust \| UniRef100\_A0A1J4SK56 \| 98.2 \| 2.5e-08 \| 4.9e-14 \| 70.2 \| 66 \| (33, 99) \| 103 \| (39, 117) \| 123 \| DksA C4-type domain-containing protein \| DksA C4-type domain-containing protein; uniclust \| UniRef100\_UPI00226C8043 \| 98.2 \| 2.8e-08 \| 5.1e-14 \| 68.3 \| 64 \| (36, 102) \| 103 \| (48, 116) \| 118 \| TraR/DksA C4-type zinc finger protein \| TraR/DksA C4-type zinc finger protein; uniclust \| UniRef100\_A0A3A0CR10 \| 98.2 \| 2.6e-08 \| 5.2e-14 \| 77.4 \| 78 \| (23, 101) \| 103 \| (153, 243) \| 260 \| DksA C4-type domain-containing protein \| DksA C4-type domain-containing protein; uniclust \| UniRef100\_A0A015MT88 \| 98.2 \| 2.1e-08 \| 5.2e-14 \| 82.2 \| 60 \| (37, 97) \| 103 \| (99, 171) \| 336 \| Molecular chaperone DnaK \| Molecular chaperone DnaK; uniclust \| UniRef100\_A0A0K1JR25 \| 98.2 \| 2.6e-08 \| 5.4e-14 \| 81.3 \| 63 \| (37, 100) \| 103 \| (283, 358) \| 359 \| DksA C4-type domain-containing protein \| DksA C4-type domain-containing protein; uniclust \| UniRef100\_A0A378VW00 \| 98.2 \| 3e-08 \| 5.5e-14 \| 68.9 \| 60 \| (35, 97) \| 103 \| (1, 65) \| 127 \| Phage associated protein \| Phage associated protein; uniclust \| UniRef100\_A0A2E2EEP1 \| 98.2 \| 2.9e-08 \| 5.6e-14 \| 72.8 \| 65 \| (34, 99) \| 103 \| (45, 122) \| 168 \| DksA C4-type domain-containing protein \| DksA C4-type domain-containing protein; uniclust \| UniRef100\_UPI0009DCB1FE \| 98.2 \| 3.2e-08 \| 5.9e-14 \| 66.5 \| 65 \| (34, 99) \| 103 \| (29, 101) \| 102 \| TraR/DksA C4-type zinc finger protein \| TraR/DksA C4-type zinc finger protein; uniclust \| UniRef100\_A0A2V7YIT4 \| 98.2 \| 3e-08 \| 6.1e-14 \| 67.8 \| 38 \| (62, 100) \| 103 \| (51, 93) \| 100 \| DksA C4-type domain-containing protein \| DksA C4-type domain-containing protein; uniclust \| UniRef100\_A0A024QBA7 \| 98.1 \| 2.8e-08 \| 6.4e-14 \| 73.0 \| 56 \| (38, 94) \| 103 \| (63, 131) \| 146 \| General stress protein 16O \| General stress protein 16O; uniclust \| UniRef100\_A0A126SXD5 \| 98.1 \| 3.4e-08 \| 6.9e-14 \| 75.6 \| 74 \| (28, 102) \| 103 \| (129, 215) \| 218 \| RNA polymerase-binding transcription factor DksA \| RNA polymerase-binding transcription factor DksA; uniclust \| UniRef100\_A0A6J5L1I0 \| 98.1 \| 3.9e-08 \| 7.1e-14 \| 61.0 \| 57 \| (35, 94) \| 103 \| (1, 62) \| 63 \| DksA-like zinc finger domain containing protein \| DksA-like zinc finger domain containing protein; uniclust \| UniRef100\_A0A359KPZ9 \| 98.1 \| 3.5e-08 \| 7.2e-14 \| 71.4 \| 37 \| (63, 100) \| 103 \| (87, 128) \| 142 \| DksA C4-type domain-containing protein \| DksA C4-type domain-containing protein; uniclust \| UniRef100\_A0A031FUW9 \| 98.1 \| 3.4e-08 \| 7.3e-14 \| 72.1 \| 37 \| (63, 100) \| 103 \| (93, 134) \| 145 \| DnaK suppressor protein \| DnaK suppressor protein; uniclust \| UniRef100\_A0A099KYM7 \| 98.1 \| 3.1e-08 \| 7.5e-14 \| 72.4 \| 66 \| (35, 101) \| 103 \| (57, 135) \| 140 \| Transcriptional regulator, TraR/DksA family \| Transcriptional regulator, TraR/DksA family; uniclust \| UniRef100\_A0A1F3T813 \| 98.1 \| 3.5e-08 \| 7.5e-14 \| 74.8 \| 67 \| (33, 100) \| 103 \| (56, 135) \| 189 \| DksA C4-type domain-containing protein \| DksA C4-type domain-containing protein; uniclust \| UniRef100\_A0A2A4PPZ5 \| 98.1 \| 3.9e-08 \| 7.7e-14 \| 63.8 \| 61 \| (35, 101) \| 103 \| (1, 66) \| 74 \| DksA C4-type domain-containing protein \| DksA C4-type domain-containing protein; uniclust \| UniRef100\_A0A085FRZ0 \| 98.1 \| 3.4e-08 \| 7.8e-14 \| 72.5 \| 66 \| (35, 101) \| 103 \| (51, 129) \| 145 \| C4-type zinc finger protein, DksA/TraR family \| C4-type zinc finger protein, DksA/TraR family; uniclust \| UniRef100\_A0A084SHE3 \| 98.1 \| 3.4e-08 \| 7.9e-14 \| 71.4 \| 63 \| (36, 99) \| 103 \| (52, 127) \| 131 \| DksA C4-type domain-containing protein \| DksA C4-type domain-containing protein; uniclust \| UniRef100\_A0A1I9LJW4 \| 98.1 \| 4.1e-08 \| 8.2e-14 \| 59.8 \| 44 \| (35, 81) \| 103 \| (1, 44) \| 52 \| TraR/DksA family transcriptional regulator \| TraR/DksA family transcriptional regulator; uniclust \| UniRef100\_A0A7C5UR01 \| 98.1 \| 4.3e-08 \| 8.3e-14 \| 65.7 \| 60 \| (38, 98) \| 103 \| (20, 92) \| 93 \| TraR/DksA family transcriptional regulator \| TraR/DksA family transcriptional regulator; uniclust \| UniRef100\_A0A0D8L7R9 \| 98.1 \| 4.2e-08 \| 8.3e-14 \| 63.8 \| 63 \| (35, 102) \| 103 \| (1, 63) \| 74 \| DUF2116 family Zn-ribbon domain-containing protein \| DUF2116 family Zn-ribbon domain-containing protein; uniclust \| UniRef100\_A0A940WD45 \| 98.1 \| 4.7e-08 \| 8.6e-14 \| 65.5 \| 65 \| (36, 101) \| 103 \| (2, 79) \| 99 \| Molecular chaperone DnaK \| Molecular chaperone DnaK; uniclust \| UniRef100\_A0A2A4WBF5 \| 98.1 \| 4.6e-08 \| 9.1e-14 \| 62.1 \| 38 \| (63, 101) \| 103 \| (19, 61) \| 65 \| DksA C4-type domain-containing protein \| DksA C4-type domain-containing protein; uniclust \| UniRef100\_A0A3A0EJC5 \| 98.1 \| 4.7e-08 \| 9.3e-14 \| 73.5 \| 70 \| (30, 100) \| 103 \| (105, 187) \| 194 \| DksA C4-type domain-containing protein \| DksA C4-type domain-containing protein; uniclust \| UniRef100\_UPI0004B56B7F \| 98.1 \| 5.2e-08 \| 9.5e-14 \| 58.2 \| 36 \| (63, 99) \| 103 \| (7, 47) \| 51 \| TraR/DksA family transcriptional regulator \| TraR/DksA family transcriptional regulator; uniclust \| UniRef100\_A0A031IJ02 \| 98.1 \| 4e-08 \| 9.5e-14 \| 77.8 \| 62 \| (37, 99) \| 103 \| (59, 133) \| 252 \| Conjugal transfer protein TraR \| Conjugal transfer protein TraR; uniclust \| UniRef100\_A0A8T5Z470 \| 98.1 \| 5.2e-08 \| 9.6e-14 \| 61.2 \| 62 \| (35, 98) \| 103 \| (1, 67) \| 67 \| Conjugal transfer protein TraR \| Conjugal transfer protein TraR; uniclust \| UniRef100\_A0A063BFQ5 \| 98.1 \| 4.4e-08 \| 9.8e-14 \| 73.4 \| 38 \| (62, 100) \| 103 \| (119, 161) \| 169 \| Transcriptional regulator, TraR/DksA family \| Transcriptional regulator, TraR/DksA family; uniclust \| UniRef100\_A0A059ZVK0 \| 98.1 \| 4.5e-08 \| 9.8e-14 \| 69.9 \| 71 \| (30, 101) \| 103 \| (5, 84) \| 124 \| DksA C4-type domain-containing protein \| DksA C4-type domain-containing protein; uniclust \| UniRef100\_A0A7C5QEY6 \| 98.1 \| 4.9e-08 \| 1e-13 \| 68.8 \| 69 \| (33, 102) \| 103 \| (35, 116) \| 120 \| Conjugal transfer protein TraR \| Conjugal transfer protein TraR; uniclust \| UniRef100\_A0A1G1F137 \| 98.1 \| 4.8e-08 \| 1e-13 \| 77.4 \| 66 \| (33, 99) \| 103 \| (179, 257) \| 265 \| DksA C4-type domain-containing protein \| DksA C4-type domain-containing protein; uniclust \| UniRef100\_A0A1Q6YPU8 \| 98.1 \| 4.7e-08 \| 1.1e-13 \| 73.1 \| 63 \| (37, 100) \| 103 \| (74, 149) \| 164 \| DksA C4-type domain-containing protein \| DksA C4-type domain-containing protein; uniclust \| UniRef100\_A0A1B1LRN6 \| 98.1 \| 4.9e-08 \| 1.1e-13 \| 71.1 \| 71 \| (30, 101) \| 103 \| (40, 125) \| 140 \| DksA C4-type domain-containing protein \| DksA C4-type domain-containing protein; uniclust \| UniRef100\_A0A0D5NN62 \| 98.1 \| 4.9e-08 \| 1.1e-13 \| 76.4 \| 57 \| (37, 94) \| 103 \| (52, 121) \| 230 \| Conjugal transfer protein TraR \| Conjugal transfer protein TraR; uniclust \| UniRef100\_A0A024P9L5 \| 98.1 \| 5.2e-08 \| 1.2e-13 \| 76.0 \| 63 \| (37, 100) \| 103 \| (65, 140) \| 229 \| General stress protein 16O \| General stress protein 16O; uniclust \| UniRef100\_A0A936IL68 \| 98.1 \| 6.1e-08 \| 1.2e-13 \| 68.8 \| 61 \| (39, 100) \| 103 \| (48, 121) \| 132 \| TraR/DksA family transcriptional regulator \| TraR/DksA family transcriptional regulator; uniclust \| UniRef100\_A0A095UJM2 \| 98.1 \| 5.2e-08 \| 1.2e-13 \| 70.4 \| 40 \| (62, 102) \| 103 \| (84, 128) \| 131 \| DnaK suppressor protein \| DnaK suppressor protein; uniclust \| UniRef100\_A0A3C0KQS3 \| 98.1 \| 6.3e-08 \| 1.2e-13 \| 63.9 \| 65 \| (36, 101) \| 103 \| (5, 75) \| 81 \| DksA C4-type domain-containing protein \| DksA C4-type domain-containing protein; uniclust \| UniRef100\_A0A0M7FD39 \| 98.1 \| 5.8e-08 \| 1.3e-13 \| 64.5 \| 69 \| (31, 101) \| 103 \| (6, 74) \| 79 \| Conjugal transfer protein TraR \| Conjugal transfer protein TraR; uniclust \| UniRef100\_A0A011NM41 \| 98.1 \| 5.7e-08 \| 1.3e-13 \| 76.8 \| 70 \| (31, 101) \| 103 \| (151, 233) \| 250 \| RNA polymerase-binding transcription factor DksA \| RNA polymerase-binding transcription factor DksA; uniclust \| UniRef100\_A0A0K2SKL3 \| 98.1 \| 6.2e-08 \| 1.4e-13 \| 77.9 \| 65 \| (37, 102) \| 103 \| (70, 147) \| 296 \| Transcriptional regulator \| Transcriptional regulator; uniclust \| UniRef100\_A0A063KX61 \| 98.1 \| 6.8e-08 \| 1.4e-13 \| 72.9 \| 74 \| (27, 101) \| 103 \| (94, 180) \| 197 \| RNA polymerase-binding transcription factor DksA \| RNA polymerase-binding transcription factor DksA; uniclust \| UniRef100\_A0A094JEJ5 \| 98.0 \| 6.9e-08 \| 1.6e-13 \| 70.6 \| 39 \| (63, 102) \| 103 \| (94, 137) \| 141 \| DksA C4-type domain-containing protein \| DksA C4-type domain-containing protein; uniclust \| UniRef100\_A0A1G0XCN3 \| 98.0 \| 7.5e-08 \| 1.6e-13 \| 71.9 \| 38 \| (63, 101) \| 103 \| (113, 155) \| 169 \| DksA C4-type domain-containing protein \| DksA C4-type domain-containing protein; uniclust \| UniRef100\_A0A084SZT3 \| 98.0 \| 7.7e-08 \| 1.7e-13 \| 71.6 \| 34 \| (65, 99) \| 103 \| (110, 148) \| 163 \| Conjugal transfer protein TraR \| Conjugal transfer protein TraR; uniclust \| UniRef100\_A0A354P9W1 \| 98.0 \| 9.2e-08 \| 1.7e-13 \| 65.1 \| 68 \| (33, 101) \| 103 \| (10, 90) \| 108 \| DksA C4-type domain-containing protein (Fragment) \| DksA C4-type domain-containing protein (Fragment); uniclust \| UniRef100\_A0A1I5WIC7 \| 98.0 \| 8e-08 \| 1.7e-13 \| 64.9 \| 66 \| (35, 101) \| 103 \| (4, 82) \| 88 \| DnaK suppressor protein \| DnaK suppressor protein; uniclust \| UniRef100\_A0A014N519 \| 98.0 \| 8e-08 \| 1.7e-13 \| 77.3 \| 62 \| (37, 99) \| 103 \| (229, 303) \| 304 \| DNA-binding protein \| DNA-binding protein; uniclust \| UniRef100\_A0A1E7ZE78 \| 98.0 \| 8.2e-08 \| 1.7e-13 \| 63.4 \| 66 \| (36, 102) \| 103 \| (1, 75) \| 77 \| DksA C4-type domain-containing protein \| DksA C4-type domain-containing protein; uniclust \| UniRef100\_A0A4P6WX91 \| 98.0 \| 9.5e-08 \| 1.8e-13 \| 64.2 \| 61 \| (37, 101) \| 103 \| (31, 96) \| 100 \| TraR/DksA family transcriptional regulator \| TraR/DksA family transcriptional regulator; uniclust \| UniRef100\_A0A5M8P5R7 \| 98.0 \| 9.5e-08 \| 1.8e-13 \| 56.9 \| 37 \| (64, 101) \| 103 \| (7, 48) \| 49 \| DksA C4-type domain-containing protein (Fragment) \| DksA C4-type domain-containing protein (Fragment); uniclust \| UniRef100\_A0A250L6W6 \| 98.0 \| 8.6e-08 \| 1.8e-13 \| 72.0 \| 36 \| (63, 99) \| 103 \| (132, 172) \| 181 \| DksA C4-type domain-containing protein \| DksA C4-type domain-containing protein; uniclust \| UniRef100\_A0A136KHD1 \| 98.0 \| 8.1e-08 \| 1.8e-13 \| 68.7 \| 62 \| (36, 98) \| 103 \| (48, 122) \| 123 \| General stress protein 16O \| General stress protein 16O; uniclust \| UniRef100\_A0A3M0YYI6 \| 98.0 \| 8.6e-08 \| 1.8e-13 \| 70.0 \| 36 \| (63, 99) \| 103 \| (86, 126) \| 146 \| TraR/DksA family transcriptional regulator \| TraR/DksA family transcriptional regulator; uniclust \| UniRef100\_UPI000A104C83 \| 98.0 \| 8.9e-08 \| 1.8e-13 \| 61.2 \| 34 \| (67, 101) \| 103 \| (26, 64) \| 65 \| TraR/DksA C4-type zinc finger protein \| TraR/DksA C4-type zinc finger protein; uniclust \| UniRef100\_A0A1G2BEF3 \| 98.0 \| 8e-08 \| 1.9e-13 \| 72.0 \| 37 \| (62, 99) \| 103 \| (110, 151) \| 164 \| DksA C4-type domain-containing protein \| DksA C4-type domain-containing protein; uniclust \| UniRef100\_A0A081FU55 \| 98.0 \| 1e-07 \| 1.9e-13 \| 63.8 \| 39 \| (63, 102) \| 103 \| (43, 86) \| 91 \| DksA C4-type domain-containing protein \| DksA C4-type domain-containing protein; uniclust \| UniRef100\_A0A358PEC4 \| 98.0 \| 9.2e-08 \| 2e-13 \| 67.0 \| 59 \| (36, 95) \| 103 \| (37, 108) \| 110 \| Conjugal transfer protein TraR \| Conjugal transfer protein TraR; uniclust \| UniRef100\_UPI000681A24E \| 98.0 \| 1e-07 \| 2e-13 \| 62.1 \| 65 \| (36, 101) \| 103 \| (1, 70) \| 77 \| TraR/DksA C4-type zinc finger protein \| TraR/DksA C4-type zinc finger protein; uniclust \| UniRef100\_A0A521HPH1 \| 98.0 \| 1e-07 \| 2.1e-13 \| 62.6 \| 32 \| (67, 99) \| 103 \| (21, 57) \| 75 \| DksA C4-type domain-containing protein (Fragment) \| DksA C4-type domain-containing protein (Fragment); uniclust \| UniRef100\_A0A3N5HTP4 \| 98.0 \| 1e-07 \| 2.2e-13 \| 62.6 \| 38 \| (63, 101) \| 103 \| (29, 71) \| 75 \| DksA C4-type domain-containing protein (Fragment) \| DksA C4-type domain-containing protein (Fragment); uniclust \| UniRef100\_A0A6A7Y5C6 \| 98.0 \| 1.2e-07 \| 2.2e-13 \| 65.7 \| 66 \| (35, 102) \| 103 \| (50, 120) \| 122 \| DksA C4-type domain-containing protein \| DksA C4-type domain-containing protein; uniclust \| UniRef100\_A0A1E7J167 \| 98.0 \| 1e-07 \| 2.2e-13 \| 76.5 \| 37 \| (63, 100) \| 103 \| (86, 127) \| 282 \| DksA C4-type domain-containing protein \| DksA C4-type domain-containing protein; uniclust \| UniRef100\_A0A1F4UBE2 \| 98.0 \| 1.2e-07 \| 2.3e-13 \| 61.5 \| 60 \| (39, 99) \| 103 \| (2, 74) \| 75 \| DksA C4-type domain-containing protein \| DksA C4-type domain-containing protein; uniclust \| UniRef100\_A0A0Q5ZHN8 \| 98.0 \| 1.1e-07 \| 2.4e-13 \| 69.4 \| 63 \| (33, 99) \| 103 \| (63, 130) \| 139 \| DksA C4-type domain-containing protein \| DksA C4-type domain-containing protein; uniclust \| UniRef100\_A0A0F7GCS4 \| 98.0 \| 1.1e-07 \| 2.4e-13 \| 70.3 \| 62 \| (39, 101) \| 103 \| (64, 138) \| 149 \| Putative DNA-binding protein \| Putative DNA-binding protein; uniclust \| UniRef100\_A0A2S5R2M0 \| 98.0 \| 1.2e-07 \| 2.5e-13 \| 63.3 \| 34 \| (67, 101) \| 103 \| (37, 75) \| 83 \| DksA C4-type domain-containing protein \| DksA C4-type domain-containing protein; uniclust \| UniRef100\_A0A2T5CG19 \| 98.0 \| 1.4e-07 \| 2.5e-13 \| 68.0 \| 64 \| (35, 99) \| 103 \| (67, 143) \| 152 \| DksA C4-type domain-containing protein \| DksA C4-type domain-containing protein; uniclust \| UniRef100\_A0A0M4DJJ5 \| 98.0 \| 1.2e-07 \| 2.6e-13 \| 82.0 \| 62 \| (37, 99) \| 103 \| (512, 586) \| 587 \| DNA-binding protein \| DNA-binding protein; uniclust \| UniRef100\_A0A9E5MJY6 \| 98.0 \| 1.4e-07 \| 2.7e-13 \| 60.8 \| 63 \| (36, 99) \| 103 \| (1, 69) \| 73 \| Conjugal transfer protein TraR \| Conjugal transfer protein TraR; uniclust \| UniRef100\_UPI001C11592D \| 98.0 \| 1.5e-07 \| 2.8e-13 \| 63.1 \| 68 \| (30, 101) \| 103 \| (12, 84) \| 98 \| TraR/DksA C4-type zinc finger protein \| TraR/DksA C4-type zinc finger protein; uniclust \| UniRef100\_A0A2W4MHC5 \| 98.0 \| 1.3e-07 \| 2.8e-13 \| 68.4 \| 38 \| (63, 101) \| 103 \| (84, 126) \| 129 \| DksA C4-type domain-containing protein \| DksA C4-type domain-containing protein; uniclust \| UniRef100\_A0A3N7CE60 \| 98.0 \| 1.5e-07 \| 2.9e-13 \| 62.5 \| 65 \| (35, 100) \| 103 \| (1, 80) \| 88 \| DksA C4-type domain-containing protein \| DksA C4-type domain-containing protein; uniclust \| UniRef100\_A0A349E9S1 \| 98.0 \| 1.4e-07 \| 2.9e-13 \| 66.1 \| 70 \| (31, 101) \| 103 \| (29, 111) \| 112 \| DksA C4-type domain-containing protein \| DksA C4-type domain-containing protein; uniclust \| UniRef100\_A0A2V6NZ32 \| 98.0 \| 1.4e-07 \| 3e-13 \| 72.0 \| 63 \| (34, 97) \| 103 \| (48, 123) \| 194 \| DksA C4-type domain-containing protein \| DksA C4-type domain-containing protein; uniclust \| UniRef100\_A0A2G6DF11 \| 98.0 \| 1.6e-07 \| 3e-13 \| 63.2 \| 68 \| (35, 103) \| 103 \| (1, 81) \| 100 \| DksA C4-type domain-containing protein \| DksA C4-type domain-containing protein; uniclust \| UniRef100\_A0A136NY37 \| 98.0 \| 1.4e-07 \| 3e-13 \| 67.2 \| 63 \| (36, 99) \| 103 \| (43, 118) \| 121 \| TraR/DksA family transcriptional regulator \| TraR/DksA family transcriptional regulator; uniclust \| UniRef100\_A0A023WNM1 \| 98.0 \| 1.2e-07 \| 3.1e-13 \| 72.1 \| 38 \| (62, 100) \| 103 \| (123, 165) \| 181 \| Conjugal transfer protein TraR \| Conjugal transfer protein TraR; uniclust \| UniRef100\_A0A2G4KCU6 \| 97.9 \| 1.7e-07 \| 3.1e-13 \| 65.9 \| 69 \| (30, 100) \| 103 \| (54, 127) \| 133 \| DksA C4-type domain-containing protein \| DksA C4-type domain-containing protein; uniclust \| UniRef100\_A0A6F8XJD3 \| 97.9 \| 1.6e-07 \| 3.2e-13 \| 75.6 \| 62 \| (37, 99) \| 103 \| (247, 321) \| 321 \| DksA C4-type domain-containing protein \| DksA C4-type domain-containing protein; uniclust \| UniRef100\_A0A1N7MQG6 \| 97.9 \| 1.7e-07 \| 3.5e-13 \| 61.8 \| 51 \| (46, 99) \| 103 \| (15, 70) \| 78 \| Transcriptional regulator, TraR/DksA family \| Transcriptional regulator, TraR/DksA family; uniclust \| UniRef100\_A0A5Q0EGG8 \| 97.9 \| 1.9e-07 \| 3.6e-13 \| 59.6 \| 37 \| (63, 100) \| 103 \| (20, 61) \| 68 \| TraR/DksA C4-type zinc finger protein \| TraR/DksA C4-type zinc finger protein; uniclust \| UniRef100\_A0A2D5B804 \| 97.9 \| 1.7e-07 \| 3.6e-13 \| 71.5 \| 70 \| (31, 101) \| 103 \| (110, 192) \| 194 \| DksA C4-type domain-containing protein \| DksA C4-type domain-containing protein; uniclust \| UniRef100\_A0A1Q6VM36 \| 97.9 \| 1.9e-07 \| 3.7e-13 \| 67.5 \| 70 \| (31, 101) \| 103 \| (65, 147) \| 150 \| DksA C4-type domain-containing protein (Fragment) \| DksA C4-type domain-containing protein (Fragment); uniclust \| UniRef100\_A0A3M1Q0B7 \| 97.9 \| 2e-07 \| 3.8e-13 \| 69.9 \| 64 \| (35, 99) \| 103 \| (57, 133) \| 198 \| TraR/DksA family transcriptional regulator \| TraR/DksA family transcriptional regulator; uniclust \| UniRef100\_UPI001D046260 \| 97.9 \| 2.1e-07 \| 3.9e-13 \| 62.5 \| 66 \| (31, 100) \| 103 \| (11, 81) \| 98 \| TraR/DksA C4-type zinc finger protein \| TraR/DksA C4-type zinc finger protein; uniclust \| UniRef100\_A0A2S3QGI0 \| 97.9 \| 1.9e-07 \| 4e-13 \| 72.9 \| 61 \| (37, 98) \| 103 \| (50, 123) \| 235 \| Conjugal transfer protein TraR \| Conjugal transfer protein TraR; uniclust \| UniRef100\_A0A101FKR2 \| 97.9 \| 1.8e-07 \| 4e-13 \| 69.9 \| 63 \| (37, 100) \| 103 \| (68, 145) \| 163 \| DNA binding protein, DksA/TraR family \| DNA binding protein, DksA/TraR family; uniclust \| UniRef100\_A0A3D3CQ74 \| 97.9 \| 2.2e-07 \| 4.2e-13 \| 58.0 \| 37 \| (63, 100) \| 103 \| (10, 51) \| 60 \| Molecular chaperone DnaK (Fragment) \| Molecular chaperone DnaK (Fragment); uniclust \| UniRef100\_A0A7C6PEY8 \| 97.9 \| 2.4e-07 \| 4.4e-13 \| 71.1 \| 68 \| (31, 99) \| 103 \| (178, 250) \| 257 \| TraR/DksA family transcriptional regulator \| TraR/DksA family transcriptional regulator; uniclust \| UniRef100\_A0A2E2XIR7 \| 97.9 \| 2.1e-07 \| 4.7e-13 \| 67.4 \| 65 \| (34, 99) \| 103 \| (47, 124) \| 129 \| Dimethylmenaquinone methyltransferase \| Dimethylmenaquinone methyltransferase; uniclust \| UniRef100\_A0A254TH52 \| 97.9 \| 2.3e-07 \| 4.8e-13 \| 66.9 \| 37 \| (63, 100) \| 103 \| (85, 126) \| 134 \| DksA C4-type domain-containing protein \| DksA C4-type domain-containing protein; uniclust \| UniRef100\_S9TQT2 \| 97.9 \| 2.5e-07 \| 4.8e-13 \| 60.4 \| 65 \| (35, 102) \| 103 \| (1, 70) \| 77 \| Conjugal transfer protein TraR \| Conjugal transfer protein TraR; uniclust \| UniRef100\_A0A7X5VPI8 \| 97.9 \| 2.5e-07 \| 5e-13 \| 73.1 \| 69 \| (30, 99) \| 103 \| (142, 223) \| 279 \| DksA C4-type domain-containing protein \| DksA C4-type domain-containing protein; uniclust \| UniRef100\_A0A096BHG0 \| 97.9 \| 2.2e-07 \| 5e-13 \| 74.2 \| 62 \| (37, 99) \| 103 \| (74, 148) \| 259 \| Molecular chaperone DnaK \| Molecular chaperone DnaK; uniclust \| UniRef100\_A0A2N9MYM1 \| 97.9 \| 2.8e-07 \| 5.1e-13 \| 55.9 \| 35 \| (64, 99) \| 103 \| (9, 48) \| 54 \| DksA C4-type domain-containing protein \| DksA C4-type domain-containing protein; uniclust \| UniRef100\_A0A258HF81 \| 97.9 \| 2.8e-07 \| 5.2e-13 \| 61.7 \| 60 \| (39, 99) \| 103 \| (16, 88) \| 96 \| DksA C4-type domain-containing protein \| DksA C4-type domain-containing protein; uniclust \| UniRef100\_A0A2E3VVZ9 \| 97.9 \| 2.5e-07 \| 5.2e-13 \| 68.7 \| 66 \| (34, 100) \| 103 \| (78, 156) \| 159 \| Conjugal transfer protein TraR \| Conjugal transfer protein TraR; uniclust \| UniRef100\_A0A3A1P1K3 \| 97.9 \| 2.7e-07 \| 5.3e-13 \| 57.1 \| 32 \| (68, 100) \| 103 \| (15, 51) \| 55 \| DksA C4-type domain-containing protein \| DksA C4-type domain-containing protein; uniclust \| UniRef100\_A0A1G6Q4B8 \| 97.9 \| 2.5e-07 \| 5.4e-13 \| 65.4 \| 66 \| (30, 96) \| 103 \| (30, 108) \| 113 \| DksA/traR C4-type zinc finger \| DksA/traR C4-type zinc finger; uniclust \| UniRef100\_A0A0P7RX32 \| 97.9 \| 2.6e-07 \| 5.4e-13 \| 61.8 \| 66 \| (32, 101) \| 103 \| (6, 76) \| 83 \| TraR/DksA family transcriptional regulator \| TraR/DksA family transcriptional regulator; uniclust \| UniRef100\_A0A2V8LIK5 \| 97.9 \| 2.7e-07 \| 5.4e-13 \| 60.3 \| 37 \| (63, 100) \| 103 \| (22, 63) \| 72 \| DksA C4-type domain-containing protein \| DksA C4-type domain-containing protein; uniclust \| UniRef100\_A0A2W4MDT4 \| 97.9 \| 2.9e-07 \| 5.5e-13 \| 66.7 \| 62 \| (37, 99) \| 103 \| (51, 125) \| 150 \| Conjugal transfer protein TraR (Fragment) \| Conjugal transfer protein TraR (Fragment); uniclust \| UniRef100\_A0A524JED7 \| 97.9 \| 2.8e-07 \| 5.5e-13 \| 64.6 \| 55 \| (39, 97) \| 103 \| (57, 116) \| 118 \| TraR/DksA family transcriptional regulator \| TraR/DksA family transcriptional regulator; uniclust \| UniRef100\_A0A1F2RVJ7 \| 97.9 \| 2.8e-07 \| 5.6e-13 \| 66.6 \| 38 \| (62, 100) \| 103 \| (78, 120) \| 136 \| DksA C4-type domain-containing protein \| DksA C4-type domain-containing protein; uniclust \| UniRef100\_A0A259NJV5 \| 97.9 \| 2.5e-07 \| 5.7e-13 \| 66.1 \| 36 \| (63, 99) \| 103 \| (77, 117) \| 119 \| DksA C4-type domain-containing protein \| DksA C4-type domain-containing protein; uniclust \| UniRef100\_A0A519BE40 \| 97.9 \| 3e-07 \| 5.8e-13 \| 66.9 \| 70 \| (32, 102) \| 103 \| (50, 132) \| 159 \| TraR/DksA family transcriptional regulator \| TraR/DksA family transcriptional regulator; uniclust \| UniRef100\_A0A2E9I554 \| 97.9 \| 2.8e-07 \| 5.9e-13 \| 61.6 \| 34 \| (63, 97) \| 103 \| (42, 80) \| 82 \| DksA C4-type domain-containing protein \| DksA C4-type domain-containing protein; uniclust \| UniRef100\_UPI0009BB9C06 \| 97.9 \| 3.1e-07 \| 5.9e-13 \| 57.3 \| 35 \| (67, 102) \| 103 \| (13, 52) \| 60 \| TraR/DksA C4-type zinc finger protein \| TraR/DksA C4-type zinc finger protein; uniclust \| UniRef100\_A0A521I243 \| 97.9 \| 2.9e-07 \| 5.9e-13 \| 66.4 \| 66 \| (34, 100) \| 103 \| (41, 119) \| 132 \| TraR/DksA family transcriptional regulator \| TraR/DksA family transcriptional regulator; uniclust \| UniRef100\_A0A7C3RPK9 \| 97.9 \| 2.9e-07 \| 5.9e-13 \| 70.0 \| 63 \| (32, 95) \| 103 \| (43, 118) \| 195 \| TraR/DksA family transcriptional regulator \| TraR/DksA family transcriptional regulator; uniclust \| UniRef100\_A0A1D9H525 \| 97.9 \| 2.9e-07 \| 6.3e-13 \| 71.3 \| 64 \| (36, 100) \| 103 \| (78, 154) \| 210 \| DksA C4-type domain-containing protein \| DksA C4-type domain-containing protein; uniclust \| UniRef100\_A0A2D6ISF3 \| 97.9 \| 3.4e-07 \| 6.4e-13 \| 64.0 \| 67 \| (34, 101) \| 103 \| (44, 115) \| 119 \| DksA C4-type domain-containing protein \| DksA C4-type domain-containing protein; uniclust \| UniRef100\_A0A379YCN4 \| 97.8 \| 3.5e-07 \| 6.5e-13 \| 61.4 \| 61 \| (35, 99) \| 103 \| (1, 66) \| 97 \| DnaK suppressor protein \| DnaK suppressor protein; uniclust \| UniRef100\_A0A060NKP5 \| 97.8 \| 2.9e-07 \| 6.6e-13 \| 78.1 \| 70 \| (32, 102) \| 103 \| (361, 443) \| 448 \| RNA polymerase-binding transcription factor DksA \| RNA polymerase-binding transcription factor DksA; uniclust \| UniRef100\_A0A2D8UDS4 \| 97.8 \| 3.3e-07 \| 6.8e-13 \| 69.9 \| 64 \| (36, 100) \| 103 \| (79, 159) \| 196 \| DksA C4-type domain-containing protein \| DksA C4-type domain-containing protein; uniclust \| UniRef100\_A0A3D3KCQ9 \| 97.8 \| 3.7e-07 \| 6.8e-13 \| 60.1 \| 65 \| (29, 99) \| 103 \| (16, 85) \| 86 \| Molecular chaperone DnaK \| Molecular chaperone DnaK; uniclust \| UniRef100\_A0A2V7CK04 \| 97.8 \| 3.8e-07 \| 7.1e-13 \| 67.2 \| 64 \| (35, 99) \| 103 \| (96, 172) \| 174 \| DksA C4-type domain-containing protein \| DksA C4-type domain-containing protein; uniclust \| UniRef100\_A0A1Q7D0Z5 \| 97.8 \| 3.8e-07 \| 7.1e-13 \| 66.4 \| 66 \| (35, 101) \| 103 \| (76, 154) \| 157 \| DksA C4-type domain-containing protein \| DksA C4-type domain-containing protein; uniclust \| UniRef100\_A0A014QDA9 \| 97.8 \| 3.5e-07 \| 7.4e-13 \| 74.4 \| 70 \| (31, 101) \| 103 \| (244, 326) \| 331 \| RNA polymerase-binding transcription factor DksA \| RNA polymerase-binding transcription factor DksA; uniclust \| UniRef100\_A0A098S5Y4 \| 97.8 \| 3e-07 \| 7.6e-13 \| 68.3 \| 38 \| (63, 101) \| 103 \| (103, 145) \| 151 \| DksA C4-type domain-containing protein \| DksA C4-type domain-containing protein; uniclust \| UniRef100\_A0A9D1R109 \| 97.8 \| 4e-07 \| 7.6e-13 \| 65.8 \| 66 \| (34, 100) \| 103 \| (32, 110) \| 149 \| TraR/DksA family transcriptional regulator \| TraR/DksA family transcriptional regulator; uniclust \| UniRef100\_A0A6N6N474 \| 97.8 \| 4.3e-07 \| 7.8e-13 \| 66.6 \| 67 \| (31, 98) \| 103 \| (91, 170) \| 176 \| TraR/DksA family transcriptional regulator \| TraR/DksA family transcriptional regulator; uniclust \| UniRef100\_A0A368U9L5 \| 97.8 \| 4.3e-07 \| 8e-13 \| 60.1 \| 70 \| (31, 103) \| 103 \| (11, 85) \| 89 \| TraR/DksA family transcriptional regulator \| TraR/DksA family transcriptional regulator; uniclust \| UniRef100\_A0A520BYY6 \| 97.8 \| 4.2e-07 \| 8.1e-13 \| 62.7 \| 36 \| (63, 99) \| 103 \| (54, 94) \| 105 \| TraR/DksA family transcriptional regulator \| TraR/DksA family transcriptional regulator; uniclust \| UniRef100\_A0A4Q7DA94 \| 97.8 \| 4.1e-07 \| 8.2e-13 \| 57.8 \| 37 \| (63, 100) \| 103 \| (21, 62) \| 63 \| RNA polymerase-binding transcription factor DksA \| RNA polymerase-binding transcription factor DksA; uniclust \| UniRef100\_A0A0S8E9J3 \| 97.8 \| 4.2e-07 \| 8.4e-13 \| 68.7 \| 62 \| (37, 99) \| 103 \| (91, 165) \| 187 \| DksA C4-type domain-containing protein \| DksA C4-type domain-containing protein; uniclust \| UniRef100\_A0A537JIW5 \| 97.8 \| 4.7e-07 \| 8.6e-13 \| 60.0 \| 64 \| (36, 100) \| 103 \| (9, 85) \| 89 \| DksA C4-type domain-containing protein \| DksA C4-type domain-containing protein; uniclust \| UniRef100\_A0A2I8QNI8 \| 97.8 \| 4.8e-07 \| 8.8e-13 \| 62.0 \| 66 \| (32, 101) \| 103 \| (34, 104) \| 110 \| DksA C4-type domain-containing protein \| DksA C4-type domain-containing protein; uniclust \| UniRef100\_A0A061A4Y2 \| 97.8 \| 4e-07 \| 9.1e-13 \| 67.6 \| 36 \| (63, 99) \| 103 \| (104, 144) \| 152 \| DNA-binding protein \| DNA-binding protein; uniclust \| UniRef100\_A0A931US23 \| 97.8 \| 5e-07 \| 9.2e-13 \| 69.2 \| 69 \| (31, 100) \| 103 \| (36, 117) \| 249 \| TraR/DksA family transcriptional regulator \| TraR/DksA family transcriptional regulator; uniclust \| UniRef100\_A0A0A6UJY1 \| 97.8 \| 4.7e-07 \| 9.4e-13 \| 68.2 \| 63 \| (37, 100) \| 103 \| (107, 182) \| 184 \| Conjugal transfer protein TraR \| Conjugal transfer protein TraR; uniclust \| UniRef100\_A0A017SZY0 \| 97.8 \| 4.4e-07 \| 9.8e-13 \| 69.4 \| 69 \| (33, 102) \| 103 \| (94, 175) \| 188 \| C4-type zinc finger protein, DksA/TraR family \| C4-type zinc finger protein, DksA/TraR family; uniclust \| UniRef100\_A0A059V4A5 \| 97.8 \| 5e-07 \| 1e-12 \| 61.0 \| 65 \| (35, 102) \| 103 \| (14, 83) \| 89 \| DksA C4-type domain-containing protein \| DksA C4-type domain-containing protein; uniclust \| UniRef100\_I2NPZ6 \| 97.8 \| 5.5e-07 \| 1e-12 \| 66.8 \| 86 \| (15, 101) \| 103 \| (90, 188) \| 192 \| RNA polymerase-binding transcription factor DksA \| RNA polymerase-binding transcription factor DksA; uniclust \| UniRef100\_A0A1Q7DI09 \| 97.8 \| 5.3e-07 \| 1e-12 \| 59.4 \| 35 \| (66, 101) \| 103 \| (37, 76) \| 78 \| DksA C4-type domain-containing protein \| DksA C4-type domain-containing protein; uniclust \| UniRef100\_A0A0P0FQ45 \| 97.8 \| 5.2e-07 \| 1.1e-12 \| 71.9 \| 65 \| (36, 101) \| 103 \| (172, 249) \| 282 \| General stress protein 16O \| General stress protein 16O; uniclust \| UniRef100\_A0A147EFJ8 \| 97.8 \| 4.9e-07 \| 1.1e-12 \| 60.4 \| 36 \| (63, 99) \| 103 \| (36, 76) \| 79 \| DksA C4-type domain-containing protein \| DksA C4-type domain-containing protein; uniclust \| UniRef100\_A0A7Y8L3J6 \| 97.8 \| 5.9e-07 \| 1.1e-12 \| 54.4 \| 34 \| (64, 98) \| 103 \| (4, 42) \| 53 \| TraR/DksA C4-type zinc finger protein \| TraR/DksA C4-type zinc finger protein; uniclust \| UniRef100\_A0A1M6JC87 \| 97.8 \| 5.8e-07 \| 1.2e-12 \| 68.0 \| 60 \| (38, 98) \| 103 \| (61, 133) \| 189 \| Transcriptional regulator, TraR/DksA family \| Transcriptional regulator, TraR/DksA family; uniclust \| UniRef100\_A0A3C0SA11 \| 97.8 \| 6.1e-07 \| 1.2e-12 \| 52.0 \| 33 \| (66, 99) \| 103 \| (2, 39) \| 40 \| DksA C4-type domain-containing protein (Fragment) \| DksA C4-type domain-containing protein (Fragment); uniclust \| UniRef100\_A0A0J5GIB6 \| 97.8 \| 6.4e-07 \| 1.2e-12 \| 56.8 \| 31 \| (68, 99) \| 103 \| (31, 66) \| 68 \| DksA C4-type domain-containing protein \| DksA C4-type domain-containing protein; uniclust \| UniRef100\_K2K8W1 \| 97.8 \| 6.4e-07 \| 1.2e-12 \| 56.9 \| 63 \| (35, 101) \| 103 \| (1, 64) \| 69 \| Uncharacterized protein \| Uncharacterized protein; uniclust \| UniRef100\_A0A091ATI3 \| 97.8 \| 6.4e-07 \| 1.2e-12 \| 58.0 \| 63 \| (35, 102) \| 103 \| (1, 66) \| 73 \| DksA C4-type domain-containing protein \| DksA C4-type domain-containing protein; uniclust \| UniRef100\_A0A0G4E598 \| 97.8 \| 5.7e-07 \| 1.2e-12 \| 63.6 \| 73 \| (28, 101) \| 103 \| (15, 100) \| 112 \| C4-type zinc finger protein, DksA/TraR family \| C4-type zinc finger protein, DksA/TraR family; uniclust \| UniRef100\_A0A2V9DB68 \| 97.8 \| 6.3e-07 \| 1.2e-12 \| 59.6 \| 36 \| (63, 99) \| 103 \| (34, 74) \| 82 \| DksA C4-type domain-containing protein \| DksA C4-type domain-containing protein; uniclust \| UniRef100\_A0A149QS87 \| 97.8 \| 6.7e-07 \| 1.3e-12 \| 58.6 \| 63 \| (36, 99) \| 103 \| (6, 76) \| 78 \| DksA C4-type domain-containing protein \| DksA C4-type domain-containing protein; uniclust \| UniRef100\_A0A0S7ZBX2 \| 97.8 \| 6e-07 \| 1.3e-12 \| 68.1 \| 62 \| (38, 100) \| 103 \| (82, 158) \| 180 \| DksA C4-type domain-containing protein \| DksA C4-type domain-containing protein; uniclust \| UniRef100\_A0A1Y3QGI6 \| 97.7 \| 6.8e-07 \| 1.3e-12 \| 67.5 \| 61 \| (37, 98) \| 103 \| (49, 122) \| 189 \| DksA C4-type domain-containing protein \| DksA C4-type domain-containing protein; uniclust \| UniRef100\_A0A949QK11 \| 97.7 \| 7e-07 \| 1.3e-12 \| 54.4 \| 38 \| (63, 101) \| 103 \| (6, 48) \| 52 \| TraR/DksA C4-type zinc finger protein \| TraR/DksA C4-type zinc finger protein; uniclust \| UniRef100\_A0A3L7RXF9 \| 97.7 \| 7.1e-07 \| 1.4e-12 \| 62.3 \| 37 \| (63, 100) \| 103 \| (41, 82) \| 114 \| TraR/DksA family transcriptional regulator (Fragment) \| TraR/DksA family transcriptional regulator (Fragment); uniclust \| UniRef100\_UPI000376C2D9 \| 97.7 \| 7.4e-07 \| 1.4e-12 \| 58.0 \| 65 \| (35, 100) \| 103 \| (1, 72) \| 74 \| hypothetical protein \| hypothetical protein; uniclust \| UniRef100\_A0A0K6H4T7 \| 97.7 \| 7.7e-07 \| 1.5e-12 \| 59.6 \| 35 \| (63, 98) \| 103 \| (39, 78) \| 84 \| Transcriptional regulator, TraR/DksA family \| Transcriptional regulator, TraR/DksA family; uniclust \| UniRef100\_UPI001D0B57A9 \| 97.7 \| 8.4e-07 \| 1.6e-12 \| 54.5 \| 35 \| (66, 101) \| 103 \| (11, 50) \| 57 \| TraR/DksA family transcriptional regulator \| TraR/DksA family transcriptional regulator; uniclust \| UniRef100\_A0A1H1X644 \| 97.7 \| 8.1e-07 \| 1.6e-12 \| 71.3 \| 62 \| (37, 99) \| 103 \| (233, 307) \| 307 \| RNA polymerase-binding protein DksA \| RNA polymerase-binding protein DksA; uniclust \| UniRef100\_A0A447JDU8 \| 97.7 \| 8.8e-07 \| 1.6e-12 \| 62.1 \| 63 \| (35, 101) \| 103 \| (1, 68) \| 127 \| site-specific DNA-methyltransferase (adenine-specific) \| site-specific DNA-methyltransferase (adenine-specific); uniclust \| UniRef100\_A0A2V9X0D4 \| 97.7 \| 8.6e-07 \| 1.6e-12 \| 53.9 \| 34 \| (64, 98) \| 103 \| (11, 49) \| 51 \| DksA C4-type domain-containing protein \| DksA C4-type domain-containing protein; uniclust \| UniRef100\_A0A0K8QN02 \| 97.7 \| 8.6e-07 \| 1.6e-12 \| 59.3 \| 65 \| (35, 100) \| 103 \| (1, 76) \| 87 \| C4-type zinc finger DksA/TraR family protein \| C4-type zinc finger DksA/TraR family protein; uniclust \| UniRef100\_A0A7C2IE66 \| 97.7 \| 9.5e-07 \| 1.7e-12 \| 60.9 \| 39 \| (63, 102) \| 103 \| (54, 97) \| 113 \| DksA C4-type domain-containing protein \| DksA C4-type domain-containing protein; uniclust \| UniRef100\_A0A375H4V9 \| 97.7 \| 8e-07 \| 1.7e-12 \| 58.5 \| 67 \| (35, 102) \| 103 \| (1, 70) \| 72 \| Uncharacterized protein \| Uncharacterized protein; uniclust \| UniRef100\_A0A090ARX1 \| 97.7 \| 8.7e-07 \| 1.8e-12 \| 59.0 \| 60 \| (36, 101) \| 103 \| (1, 61) \| 80 \| Conserved uncharacterized protein \| Conserved uncharacterized protein; uniclust \| UniRef100\_A0A0A6CYW5 \| 97.7 \| 8.7e-07 \| 1.8e-12 \| 58.4 \| 60 \| (35, 99) \| 103 \| (1, 60) \| 73 \| Conjugal transfer protein TraR \| Conjugal transfer protein TraR; uniclust \| UniRef100\_A0A7Y5VRL4 \| 97.7 \| 1e-06 \| 1.9e-12 \| 63.9 \| 70 \| (30, 100) \| 103 \| (63, 145) \| 160 \| TraR/DksA family transcriptional regulator \| TraR/DksA family transcriptional regulator; uniclust \| UniRef100\_A0A177HY54 \| 97.7 \| 9.4e-07 \| 1.9e-12 \| 77.6 \| 61 \| (38, 99) \| 103 \| (686, 759) \| 760 \| RNA polymerase-binding transcription factor DksA \| RNA polymerase-binding transcription factor DksA; uniclust \| UniRef100\_A0A142XHA1 \| 97.7 \| 9e-07 \| 1.9e-12 \| 64.0 \| 66 \| (33, 99) \| 103 \| (48, 126) \| 129 \| General stress protein 16O \| General stress protein 16O; uniclust \| UniRef100\_A0A7Y6CRV9 \| 97.7 \| 1e-06 \| 2e-12 \| 66.5 \| 62 \| (36, 98) \| 103 \| (122, 196) \| 197 \| TraR/DksA family transcriptional regulator \| TraR/DksA family transcriptional regulator; uniclust \| UniRef100\_A0A7V2X9Q0 \| 97.7 \| 1.1e-06 \| 2.1e-12 \| 63.3 \| 67 \| (32, 99) \| 103 \| (56, 135) \| 152 \| TraR/DksA family transcriptional regulator \| TraR/DksA family transcriptional regulator; uniclust \| UniRef100\_A0A0H2KR58 \| 97.7 \| 9.8e-07 \| 2.1e-12 \| 65.7 \| 36 \| (63, 99) \| 103 \| (113, 153) \| 158 \| DksA C4-type domain-containing protein \| DksA C4-type domain-containing protein; uniclust \| UniRef100\_A0A9E2N3C8 \| 97.7 \| 1.2e-06 \| 2.1e-12 \| 63.3 \| 38 \| (64, 102) \| 103 \| (89, 131) \| 153 \| TraR/DksA family transcriptional regulator \| TraR/DksA family transcriptional regulator; uniclust \| UniRef100\_A0A9D6TUP0 \| 97.7 \| 1.2e-06 \| 2.1e-12 \| 58.3 \| 39 \| (63, 102) \| 103 \| (33, 76) \| 89 \| TraR/DksA C4-type zinc finger protein \| TraR/DksA C4-type zinc finger protein; uniclust \| UniRef100\_A0A3D1TEN9 \| 97.7 \| 1e-06 \| 2.1e-12 \| 60.9 \| 38 \| (63, 101) \| 103 \| (47, 89) \| 96 \| DksA C4-type domain-containing protein (Fragment) \| DksA C4-type domain-containing protein (Fragment); uniclust \| UniRef100\_A0A929HNH4 \| 97.7 \| 1.2e-06 \| 2.2e-12 \| 67.6 \| 70 \| (31, 101) \| 103 \| (152, 234) \| 260 \| TraR/DksA family transcriptional regulator \| TraR/DksA family transcriptional regulator; uniclust \| UniRef100\_A0A146G6K6 \| 97.7 \| 1.2e-06 \| 2.2e-12 \| 67.4 \| 65 \| (36, 101) \| 103 \| (105, 182) \| 218 \| RNA polymerase-binding transcription factor DksA \| RNA polymerase-binding transcription factor DksA; uniclust \| UniRef100\_A0A536HB94 \| 97.7 \| 1.2e-06 \| 2.3e-12 \| 56.1 \| 36 \| (65, 101) \| 103 \| (29, 69) \| 72 \| DksA C4-type domain-containing protein \| DksA C4-type domain-containing protein; uniclust \| UniRef100\_A0A1G8LDJ8 \| 97.7 \| 1.1e-06 \| 2.3e-12 \| 68.8 \| 60 \| (37, 97) \| 103 \| (75, 147) \| 219 \| RNA polymerase-binding protein DksA \| RNA polymerase-binding protein DksA; uniclust \| UniRef100\_A0A7Y2DU12 \| 97.7 \| 1.2e-06 \| 2.4e-12 \| 63.2 \| 62 \| (37, 99) \| 103 \| (48, 122) \| 131 \| DksA C4-type domain-containing protein \| DksA C4-type domain-containing protein; uniclust \| UniRef100\_A0A4Q3NJL8 \| 97.7 \| 1.2e-06 \| 2.4e-12 \| 66.4 \| 36 \| (63, 99) \| 103 \| (136, 176) \| 193 \| TraR/DksA family transcriptional regulator \| TraR/DksA family transcriptional regulator; uniclust \| UniRef100\_A0A6B0XPH2 \| 97.7 \| 1.2e-06 \| 2.4e-12 \| 72.8 \| 68 \| (31, 99) \| 103 \| (377, 457) \| 463 \| TraR/DksA family transcriptional regulator \| TraR/DksA family transcriptional regulator; uniclust \| UniRef100\_A0A7C5ZAZ5 \| 97.7 \| 1.2e-06 \| 2.4e-12 \| 71.3 \| 68 \| (31, 99) \| 103 \| (293, 373) \| 375 \| DksA C4-type domain-containing protein \| DksA C4-type domain-containing protein; uniclust \| UniRef100\_A0A1V4XFM2 \| 97.7 \| 1.3e-06 \| 2.4e-12 \| 64.2 \| 65 \| (35, 100) \| 103 \| (58, 135) \| 159 \| RNA polymerase-binding transcription factor DksA \| RNA polymerase-binding transcription factor DksA; uniclust \| UniRef100\_A0A1G3QAA7 \| 97.7 \| 1.2e-06 \| 2.5e-12 \| 65.0 \| 66 \| (34, 100) \| 103 \| (58, 136) \| 158 \| DksA C4-type domain-containing protein \| DksA C4-type domain-containing protein; uniclust \| UniRef100\_UPI001FE60161 \| 97.7 \| 1.4e-06 \| 2.5e-12 \| 59.0 \| 67 \| (31, 101) \| 103 \| (20, 91) \| 99 \| TraR/DksA family transcriptional regulator \| TraR/DksA family transcriptional regulator; uniclust \| UniRef100\_A0A889IPF0 \| 97.6 \| 1.3e-06 \| 2.6e-12 \| 58.4 \| 59 \| (40, 99) \| 103 \| (9, 74) \| 87 \| DksA C4-type domain-containing protein \| DksA C4-type domain-containing protein; uniclust \| UniRef100\_A0A081K6D0 \| 97.6 \| 1.2e-06 \| 2.6e-12 \| 64.6 \| 38 \| (63, 101) \| 103 \| (97, 139) \| 141 \| Molecular chaperone DnaK \| Molecular chaperone DnaK; uniclust \| UniRef100\_A0A0F7JX09 \| 97.6 \| 1.5e-06 \| 2.8e-12 \| 59.2 \| 64 \| (36, 100) \| 103 \| (1, 72) \| 96 \| DksA C4-type domain-containing protein \| DksA C4-type domain-containing protein; uniclust \| UniRef100\_UPI001F35BB70 \| 97.6 \| 1.6e-06 \| 2.9e-12 \| 53.3 \| 31 \| (68, 99) \| 103 \| (15, 50) \| 56 \| TraR/DksA family transcriptional regulator \| TraR/DksA family transcriptional regulator; uniclust \| UniRef100\_A0A336N326 \| 97.6 \| 1.5e-06 \| 2.9e-12 \| 56.5 \| 66 \| (35, 101) \| 103 \| (1, 72) \| 73 \| DksA C4-type domain-containing protein \| DksA C4-type domain-containing protein; uniclust \| UniRef100\_A0A0K3R167 \| 97.6 \| 1.6e-06 \| 2.9e-12 \| 59.5 \| 70 \| (30, 101) \| 103 \| (10, 84) \| 103 \| Conjugal transfer protein TraR \| Conjugal transfer protein TraR; uniclust \| UniRef100\_A0A7V3BK14 \| 97.6 \| 1.5e-06 \| 2.9e-12 \| 63.8 \| 67 \| (32, 99) \| 103 \| (74, 153) \| 159 \| TraR/DksA family transcriptional regulator \| TraR/DksA family transcriptional regulator; uniclust \| UniRef100\_A0A846Q3K0 \| 97.6 \| 1.6e-06 \| 3e-12 \| 57.1 \| 37 \| (63, 100) \| 103 \| (33, 74) \| 77 \| DksA C4-type domain-containing protein (Fragment) \| DksA C4-type domain-containing protein (Fragment); uniclust \| UniRef100\_A0A8H9EJZ3 \| 97.6 \| 1.6e-06 \| 3.1e-12 \| 57.1 \| 35 \| (67, 102) \| 103 \| (33, 72) \| 76 \| TraR/DksA family transcriptional regulator \| TraR/DksA family transcriptional regulator; uniclust \| UniRef100\_A0A3B9NYN9 \| 97.6 \| 1.7e-06 \| 3.2e-12 \| 53.3 \| 34 \| (68, 102) \| 103 \| (8, 48) \| 54 \| DksA C4-type domain-containing protein \| DksA C4-type domain-containing protein; uniclust \| UniRef100\_UPI0002E76F17 \| 97.6 \| 1.7e-06 \| 3.3e-12 \| 55.8 \| 62 \| (34, 102) \| 103 \| (2, 63) \| 70 \| hypothetical protein \| hypothetical protein; uniclust \| UniRef100\_UPI001D0D1C56 \| 97.6 \| 1.7e-06 \| 3.3e-12 \| 52.8 \| 35 \| (65, 100) \| 103 \| (12, 50) \| 51 \| TraR/DksA C4-type zinc finger protein \| TraR/DksA C4-type zinc finger protein; uniclust \| UniRef100\_A0A1V4S0C0 \| 97.6 \| 1.8e-06 \| 3.4e-12 \| 66.0 \| 68 \| (31, 99) \| 103 \| (38, 119) \| 238 \| DksA C4-type domain-containing protein \| DksA C4-type domain-containing protein; uniclust \| UniRef100\_B9BF14 \| 97.6 \| 1.8e-06 \| 3.6e-12 \| 67.9 \| 64 \| (36, 100) \| 103 \| (167, 243) \| 252 \| Transcriptional regulator, TraR/DksA family \| Transcriptional regulator, TraR/DksA family; uniclust \| UniRef100\_A0A0G0PVY8 \| 97.6 \| 1.7e-06 \| 3.7e-12 \| 66.7 \| 63 \| (37, 100) \| 103 \| (100, 175) \| 200 \| DnaK suppressor protein (Modular protein) \| DnaK suppressor protein (Modular protein); uniclust \| UniRef100\_A0A3A4A6E0 \| 97.6 \| 1.9e-06 \| 3.7e-12 \| 54.7 \| 35 \| (63, 98) \| 103 \| (20, 59) \| 61 \| DksA C4-type domain-containing protein \| DksA C4-type domain-containing protein; uniclust \| UniRef100\_A0A2D9HEW6 \| 97.6 \| 1.8e-06 \| 3.8e-12 \| 64.5 \| 35 \| (64, 99) \| 103 \| (116, 155) \| 157 \| DksA C4-type domain-containing protein \| DksA C4-type domain-containing protein; uniclust \| UniRef100\_M4SM17 \| 97.6 \| 2e-06 \| 3.8e-12 \| 59.4 \| 69 \| (28, 99) \| 103 \| (14, 82) \| 109 \| Uncharacterized protein \| Uncharacterized protein; uniclust \| UniRef100\_A0A2E4X1U5 \| 97.6 \| 1.8e-06 \| 3.8e-12 \| 63.9 \| 64 \| (36, 100) \| 103 \| (46, 122) \| 148 \| RNA polymerase-binding protein DksA \| RNA polymerase-binding protein DksA; uniclust \| UniRef100\_A0A1F3SKA7 \| 97.6 \| 1.9e-06 \| 3.9e-12 \| 62.8 \| 67 \| (32, 99) \| 103 \| (39, 118) \| 141 \| DksA C4-type domain-containing protein \| DksA C4-type domain-containing protein; uniclust \| UniRef100\_A0A1V5UVG4 \| 97.6 \| 2e-06 \| 3.9e-12 \| 67.7 \| 62 \| (37, 99) \| 103 \| (175, 249) \| 257 \| General stress protein 16O \| General stress protein 16O; uniclust \| UniRef100\_A0A953US15 \| 97.6 \| 2.2e-06 \| 4e-12 \| 62.6 \| 74 \| (24, 98) \| 103 \| (24, 110) \| 165 \| TraR/DksA family transcriptional regulator \| TraR/DksA family transcriptional regulator; uniclust \| UniRef100\_A0A7C4I3K4 \| 97.6 \| 2.1e-06 \| 4e-12 \| 52.2 \| 38 \| (63, 101) \| 103 \| (3, 45) \| 49 \| Transcriptional regulator, TraR/DksA family protein (Fragment) \| Transcriptional regulator, TraR/DksA family protein (Fragment); uniclust \| UniRef100\_UPI001FEB62E6 \| 97.6 \| 2.2e-06 \| 4.1e-12 \| 51.9 \| 33 \| (67, 100) \| 103 \| (4, 41) \| 52 \| TraR/DksA C4-type zinc finger protein \| TraR/DksA C4-type zinc finger protein; uniclust \| UniRef100\_A0A5Q0M9K3 \| 97.6 \| 2.2e-06 \| 4.1e-12 \| 62.6 \| 63 \| (36, 99) \| 103 \| (67, 142) \| 153 \| TraR/DksA family transcriptional regulator \| TraR/DksA family transcriptional regulator; uniclust \| UniRef100\_A0A258C2E0 \| 97.6 \| 2.2e-06 \| 4.1e-12 \| 52.6 \| 33 \| (67, 100) \| 103 \| (14, 51) \| 52 \| DksA C4-type domain-containing protein \| DksA C4-type domain-containing protein; uniclust \| UniRef100\_A0A0J0V0N8 \| 97.6 \| 2e-06 \| 4.2e-12 \| 67.6 \| 61 \| (39, 100) \| 103 \| (150, 223) \| 225 \| Transcriptional regulator, TraR/DksA family \| Transcriptional regulator, TraR/DksA family; uniclust \| UniRef100\_A0A0G0L1H4 \| 97.6 \| 1.8e-06 \| 4.3e-12 \| 64.6 \| 62 \| (36, 98) \| 103 \| (72, 146) \| 153 \| DksA C4-type domain-containing protein \| DksA C4-type domain-containing protein; uniclust \| UniRef100\_A0A345GTX4 \| 97.6 \| 2.3e-06 \| 4.4e-12 \| 56.0 \| 63 \| (35, 100) \| 103 \| (4, 66) \| 74 \| DUF2116 family Zn-ribbon domain-containing protein \| DUF2116 family Zn-ribbon domain-containing protein; uniclust \| UniRef100\_A0A9C8GJI6 \| 97.6 \| 2.4e-06 \| 4.4e-12 \| 61.6 \| 36 \| (63, 99) \| 103 \| (110, 150) \| 151 \| Molecular chaperone DnaK \| Molecular chaperone DnaK; uniclust \| UniRef100\_A0A944V385 \| 97.6 \| 2.4e-06 \| 4.5e-12 \| 53.9 \| 40 \| (63, 103) \| 103 \| (20, 64) \| 65 \| DksA C4-type domain-containing protein \| DksA C4-type domain-containing protein; uniclust \| UniRef100\_A0A250KX83 \| 97.6 \| 2.3e-06 \| 4.5e-12 \| 58.7 \| 67 \| (34, 101) \| 103 \| (2, 82) \| 96 \| Transcriptional regulator, TraR/DksA family \| Transcriptional regulator, TraR/DksA family; uniclust \| UniRef100\_A0A3E0P096 \| 97.6 \| 2.2e-06 \| 4.5e-12 \| 65.3 \| 66 \| (33, 99) \| 103 \| (105, 183) \| 189 \| TraR/DksA family transcriptional regulator \| TraR/DksA family transcriptional regulator; uniclust \| UniRef100\_A0A0J7LZ52 \| 97.6 \| 2.2e-06 \| 4.7e-12 \| 56.3 \| 51 \| (42, 95) \| 103 \| (11, 66) \| 70 \| DksA C4-type domain-containing protein \| DksA C4-type domain-containing protein; uniclust \| UniRef100\_A0A010ZWH0 \| 97.5 \| 2.1e-06 \| 4.9e-12 \| 65.8 \| 35 \| (63, 98) \| 103 \| (123, 162) \| 182 \| DnaK suppressor protein \| DnaK suppressor protein; uniclust \| UniRef100\_A0A2E3W236 \| 97.5 \| 2.8e-06 \| 5.2e-12 \| 63.8 \| 65 \| (36, 101) \| 103 \| (81, 158) \| 202 \| DksA C4-type domain-containing protein \| DksA C4-type domain-containing protein; uniclust \| UniRef100\_A0A2V6NWY6 \| 97.5 \| 2.8e-06 \| 5.2e-12 \| 54.7 \| 39 \| (63, 102) \| 103 \| (15, 58) \| 73 \| DksA C4-type domain-containing protein \| DksA C4-type domain-containing protein; uniclust \| UniRef100\_A0A142WZJ9 \| 97.5 \| 2.3e-06 \| 5.2e-12 \| 68.9 \| 64 \| (34, 98) \| 103 \| (49, 125) \| 264 \| General stress protein 16O \| General stress protein 16O; uniclust \| UniRef100\_A0A086P6H6 \| 97.5 \| 2.3e-06 \| 5.2e-12 \| 65.3 \| 35 \| (62, 97) \| 103 \| (96, 135) \| 177 \| DnaK suppressor protein \| DnaK suppressor protein; uniclust \| UniRef100\_A0A5Q2RKE7 \| 97.5 \| 2.6e-06 \| 5.2e-12 \| 60.7 \| 35 \| (63, 98) \| 103 \| (83, 122) \| 123 \| DksA C4-type domain-containing protein \| DksA C4-type domain-containing protein; uniclust \| UniRef100\_A0A965KYD4 \| 97.5 \| 2.9e-06 \| 5.2e-12 \| 61.5 \| 65 \| (35, 100) \| 103 \| (80, 149) \| 154 \| DksA C4-type domain-containing protein \| DksA C4-type domain-containing protein; uniclust \| UniRef100\_A0A3D1BL39 \| 97.5 \| 2.7e-06 \| 5.3e-12 \| 57.1 \| 36 \| (63, 99) \| 103 \| (30, 70) \| 86 \| Transcriptional regulator, TraR/DksA family protein (Fragment) \| Transcriptional regulator, TraR/DksA family protein (Fragment); uniclust \| UniRef100\_A0A1F3X404 \| 97.5 \| 2.9e-06 \| 5.6e-12 \| 55.4 \| 37 \| (63, 100) \| 103 \| (26, 67) \| 75 \| DksA C4-type domain-containing protein \| DksA C4-type domain-containing protein; uniclust \| UniRef100\_A0A3D8HS68 \| 97.5 \| 3.3e-06 \| 6e-12 \| 61.0 \| 66 \| (35, 101) \| 103 \| (52, 130) \| 150 \| RNA polymerase-binding protein DksA \| RNA polymerase-binding protein DksA; uniclust \| UniRef100\_A0A0G0G8U5 \| 97.5 \| 2.7e-06 \| 6.1e-12 \| 63.8 \| 59 \| (38, 97) \| 103 \| (80, 151) \| 155 \| Transcriptional regulator, TraR/DksA family \| Transcriptional regulator, TraR/DksA family; uniclust \| UniRef100\_A0A0G0CFG9 \| 97.5 \| 2.6e-06 \| 6.2e-12 \| 62.8 \| 64 \| (33, 97) \| 103 \| (57, 133) \| 138 \| Transcriptional regulator, TraR/DksA family \| Transcriptional regulator, TraR/DksA family; uniclust \| UniRef100\_UPI0003651E65 \| 97.5 \| 3.1e-06 \| 6.2e-12 \| 56.4 \| 63 \| (39, 102) \| 103 \| (8, 75) \| 78 \| hypothetical protein \| hypothetical protein; uniclust \| UniRef100\_A0A2R4VRC7 \| 97.5 \| 3.3e-06 \| 6.3e-12 \| 52.3 \| 33 \| (68, 101) \| 103 \| (11, 48) \| 54 \| DksA C4-type domain-containing protein \| DksA C4-type domain-containing protein; uniclust \| UniRef100\_A0A0F3Q901 \| 97.5 \| 3.1e-06 \| 6.4e-12 \| 62.9 \| 71 \| (30, 101) \| 103 \| (65, 148) \| 153 \| Putative dnaK suppressor-like protein \| Putative dnaK suppressor-like protein; uniclust \| UniRef100\_A0A350NQT1 \| 97.5 \| 3.4e-06 \| 7e-12 \| 60.9 \| 37 \| (63, 100) \| 103 \| (27, 68) \| 130 \| DksA C4-type domain-containing protein \| DksA C4-type domain-containing protein; uniclust \| UniRef100\_G5LKE7 \| 97.5 \| 3.6e-06 \| 7.1e-12 \| 52.9 \| 34 \| (67, 101) \| 103 \| (13, 51) \| 58 \| Putative Zinc-finger containing protein (Fragment) \| Putative Zinc-finger containing protein (Fragment); uniclust \| UniRef100\_A0A1H9NNW1 \| 97.5 \| 3.5e-06 \| 7.3e-12 \| 61.9 \| 58 \| (37, 95) \| 103 \| (45, 115) \| 140 \| Transcriptional regulator, TraR/DksA family \| Transcriptional regulator, TraR/DksA family; uniclust \| UniRef100\_A0A518LQ83 \| 97.5 \| 3.4e-06 \| 7.3e-12 \| 64.8 \| 64 \| (36, 100) \| 103 \| (109, 185) \| 189 \| General stress protein 16O \| General stress protein 16O; uniclust \| UniRef100\_A0A1E7HCG8 \| 97.5 \| 3.9e-06 \| 7.5e-12 \| 64.1 \| 37 \| (64, 101) \| 103 \| (75, 116) \| 203 \| DksA C4-type domain-containing protein \| DksA C4-type domain-containing protein; uniclust \| UniRef100\_A0A0K1NCL4 \| 97.5 \| 3.8e-06 \| 7.6e-12 \| 62.1 \| 38 \| (63, 101) \| 103 \| (104, 146) \| 152 \| DksA C4-type domain-containing protein \| DksA C4-type domain-containing protein; uniclust \| UniRef100\_A0A1Q7U7Y8 \| 97.5 \| 4.2e-06 \| 7.7e-12 \| 60.6 \| 71 \| (30, 101) \| 103 \| (24, 107) \| 153 \| DksA C4-type domain-containing protein \| DksA C4-type domain-containing protein; uniclust \| UniRef100\_A0A0W8E1D5 \| 97.5 \| 3.7e-06 \| 8e-12 \| 63.6 \| 61 \| (38, 99) \| 103 \| (48, 121) \| 167 \| Dnak suppressor protein \| Dnak suppressor protein; uniclust \| UniRef100\_A0A1Q7SQ33 \| 97.5 \| 4.1e-06 \| 8.4e-12 \| 63.7 \| 38 \| (63, 101) \| 103 \| (104, 146) \| 181 \| DksA C4-type domain-containing protein \| DksA C4-type domain-containing protein; uniclust \| UniRef100\_A0A2E3P7D0 \| 97.5 \| 4.4e-06 \| 8.5e-12 \| 53.4 \| 54 \| (34, 93) \| 103 \| (2, 60) \| 65 \| Molecular chaperone DnaK \| Molecular chaperone DnaK; uniclust \| UniRef100\_A0A223PH24 \| 97.5 \| 4.2e-06 \| 8.7e-12 \| 59.2 \| 37 \| (63, 100) \| 103 \| (68, 109) \| 111 \| Molecular chaperone DnaK \| Molecular chaperone DnaK; uniclust \| UniRef100\_A0A1V1W784 \| 97.4 \| 4.4e-06 \| 9e-12 \| 59.9 \| 35 \| (63, 98) \| 103 \| (74, 113) \| 122 \| C4-type zinc finger protein, DksA/TraR family \| C4-type zinc finger protein, DksA/TraR family; uniclust \| UniRef100\_A0A7Y8LK13 \| 97.4 \| 4.9e-06 \| 9.4e-12 \| 53.4 \| 35 \| (64, 99) \| 103 \| (14, 53) \| 67 \| TraR/DksA C4-type zinc finger protein \| TraR/DksA C4-type zinc finger protein; uniclust \| UniRef100\_A0A0D8FU18 \| 97.4 \| 4.6e-06 \| 9.6e-12 \| 65.5 \| 54 \| (44, 98) \| 103 \| (148, 214) \| 232 \| General stress protein 16O \| General stress protein 16O; uniclust \| UniRef100\_A0A0F7HEF4 \| 97.4 \| 4.9e-06 \| 9.6e-12 \| 58.8 \| 52 \| (46, 100) \| 103 \| (15, 71) \| 117 \| DksA C4-type domain-containing protein \| DksA C4-type domain-containing protein; uniclust \| UniRef100\_A0A3N5HZ15 \| 97.4 \| 5.3e-06 \| 9.8e-12 \| 60.3 \| 63 \| (36, 99) \| 103 \| (79, 154) \| 157 \| TraR/DksA family transcriptional regulator \| TraR/DksA family transcriptional regulator; uniclust \| UniRef100\_A0A925I2M1 \| 97.4 \| 5.1e-06 \| 9.8e-12 \| 57.4 \| 38 \| (63, 101) \| 103 \| (54, 96) \| 103 \| TraR/DksA family transcriptional regulator \| TraR/DksA family transcriptional regulator; uniclust \| UniRef100\_A0A1G1XR92 \| 97.4 \| 5.2e-06 \| 1e-11 \| 60.2 \| 39 \| (63, 102) \| 103 \| (82, 125) \| 140 \| DksA C4-type domain-containing protein \| DksA C4-type domain-containing protein; uniclust \| UniRef100\_A0A2H0RQV3 \| 97.4 \| 5.6e-06 \| 1e-11 \| 57.1 \| 36 \| (63, 99) \| 103 \| (50, 90) \| 109 \| DksA C4-type domain-containing protein \| DksA C4-type domain-containing protein; uniclust \| UniRef100\_A0A0S4KQC6 \| 97.4 \| 5e-06 \| 1.1e-11 \| 63.1 \| 69 \| (31, 100) \| 103 \| (84, 165) \| 176 \| DksA C4-type domain-containing protein \| DksA C4-type domain-containing protein; uniclust \| UniRef100\_A0A0G0A276 \| 97.4 \| 5.2e-06 \| 1.1e-11 \| 63.7 \| 38 \| (63, 101) \| 103 \| (123, 165) \| 187 \| DNA-binding protein \| DNA-binding protein; uniclust \| UniRef100\_A0A0G1FXN7 \| 97.4 \| 4.6e-06 \| 1.1e-11 \| 61.4 \| 36 \| (62, 98) \| 103 \| (91, 131) \| 135 \| DksA C4-type domain-containing protein \| DksA C4-type domain-containing protein; uniclust \| UniRef100\_A0A7C3SEN0 \| 97.4 \| 5.3e-06 \| 1.1e-11 \| 55.1 \| 35 \| (63, 98) \| 103 \| (35, 74) \| 75 \| DksA C4-type domain-containing protein \| DksA C4-type domain-containing protein; uniclust \| UniRef100\_A0A067LRQ7 \| 97.4 \| 5.1e-06 \| 1.1e-11 \| 61.9 \| 36 \| (63, 99) \| 103 \| (103, 143) \| 148 \| Suppressor protein DnaK \| Suppressor protein DnaK; uniclust \| UniRef100\_UPI0019149785 \| 97.4 \| 6.3e-06 \| 1.2e-11 \| 53.1 \| 63 \| (35, 101) \| 103 \| (1, 64) \| 72 \| hypothetical protein \| hypothetical protein; uniclust \| UniRef100\_A0A1V3RQS2 \| 97.4 \| 5.8e-06 \| 1.2e-11 \| 56.6 \| 64 \| (33, 97) \| 103 \| (11, 82) \| 92 \| DksA C4-type domain-containing protein \| DksA C4-type domain-containing protein; uniclust \| UniRef100\_A0A1G2XD98 \| 97.4 \| 5.6e-06 \| 1.2e-11 \| 64.9 \| 67 \| (32, 99) \| 103 \| (82, 161) \| 222 \| DksA C4-type domain-containing protein \| DksA C4-type domain-containing protein; uniclust \| UniRef100\_UPI0009FAB1EC \| 97.4 \| 6.4e-06 \| 1.2e-11 \| 61.1 \| 59 \| (37, 96) \| 103 \| (49, 120) \| 181 \| TraR/DksA C4-type zinc finger protein \| TraR/DksA C4-type zinc finger protein; uniclust \| UniRef100\_UPI001F4393EB \| 97.4 \| 6.5e-06 \| 1.2e-11 \| 52.9 \| 32 \| (67, 99) \| 103 \| (30, 66) \| 71 \| TraR/DksA C4-type zinc finger protein \| TraR/DksA C4-type zinc finger protein; uniclust \| UniRef100\_A0A1U9NPL5 \| 97.4 \| 5.5e-06 \| 1.2e-11 \| 67.2 \| 65 \| (34, 99) \| 103 \| (141, 218) \| 284 \| General stress protein 16O \| General stress protein 16O; uniclust \| UniRef100\_A0A411YA84 \| 97.4 \| 6.6e-06 \| 1.2e-11 \| 60.9 \| 62 \| (34, 96) \| 103 \| (99, 173) \| 177 \| TraR/DksA family transcriptional regulator \| TraR/DksA family transcriptional regulator; uniclust \| UniRef100\_UPI002362D03C \| 97.4 \| 6.6e-06 \| 1.2e-11 \| 50.2 \| 33 \| (68, 101) \| 103 \| (11, 48) \| 53 \| TraR/DksA C4-type zinc finger protein \| TraR/DksA C4-type zinc finger protein; uniclust \| UniRef100\_A0A0S4V0E8 \| 97.4 \| 6.7e-06 \| 1.2e-11 \| 54.4 \| 36 \| (63, 99) \| 103 \| (42, 82) \| 84 \| DnaK suppressor protein \| DnaK suppressor protein; uniclust \| UniRef100\_A0A356T9N2 \| 97.4 \| 6.2e-06 \| 1.3e-11 \| 67.8 \| 71 \| (30, 101) \| 103 \| (251, 334) \| 346 \| RNA polymerase-binding protein DksA \| RNA polymerase-binding protein DksA; uniclust \| UniRef100\_A0A7V5HPI1 \| 97.4 \| 6.3e-06 \| 1.3e-11 \| 51.6 \| 34 \| (63, 97) \| 103 \| (14, 52) \| 55 \| TraR/DksA family transcriptional regulator \| TraR/DksA family transcriptional regulator; uniclust \| UniRef100\_A0A377TLM7 \| 97.4 \| 7.1e-06 \| 1.3e-11 \| 61.6 \| 74 \| (27, 101) \| 103 \| (42, 128) \| 197 \| RNA polymerase-binding transcription factor DksA \| RNA polymerase-binding transcription factor DksA; uniclust \| UniRef100\_A0A7C3B7W3 \| 97.4 \| 6.8e-06 \| 1.3e-11 \| 47.9 \| 31 \| (64, 95) \| 103 \| (3, 38) \| 40 \| TraR/DksA family transcriptional regulator \| TraR/DksA family transcriptional regulator; uniclust \| UniRef100\_A0A3N5XHU1 \| 97.4 \| 7.2e-06 \| 1.3e-11 \| 56.6 \| 35 \| (63, 98) \| 103 \| (60, 99) \| 109 \| DksA C4-type domain-containing protein (Fragment) \| DksA C4-type domain-containing protein (Fragment); uniclust \| UniRef100\_A0A1Z3LXK0 \| 97.4 \| 6.7e-06 \| 1.3e-11 \| 58.3 \| 53 \| (44, 99) \| 103 \| (43, 100) \| 117 \| DksA C4-type domain-containing protein \| DksA C4-type domain-containing protein; uniclust \| UniRef100\_A0A2X1UWH1 \| 97.4 \| 6.9e-06 \| 1.3e-11 \| 55.0 \| 67 \| (33, 100) \| 103 \| (3, 77) \| 83 \| Phage/conjugal plasmid C-4 type zinc finger protein, TraR family \| Phage/conjugal plasmid C-4 type zinc finger protein, TraR family; uniclust \| UniRef100\_A0A2V8PR07 \| 97.4 \| 7.5e-06 \| 1.4e-11 \| 55.2 \| 62 \| (36, 98) \| 103 \| (9, 83) \| 94 \| DksA C4-type domain-containing protein \| DksA C4-type domain-containing protein; uniclust \| UniRef100\_A0A951CX43 \| 97.4 \| 7.7e-06 \| 1.4e-11 \| 52.5 \| 36 \| (63, 99) \| 103 \| (16, 56) \| 70 \| TraR/DksA C4-type zinc finger protein \| TraR/DksA C4-type zinc finger protein; uniclust \| UniRef100\_A0A7V2MWM1 \| 97.4 \| 7.9e-06 \| 1.5e-11 \| 59.2 \| 38 \| (63, 101) \| 103 \| (100, 142) \| 152 \| TraR/DksA family transcriptional regulator \| TraR/DksA family transcriptional regulator; uniclust \| UniRef100\_A0A1G3ZPZ8 \| 97.4 \| 7.2e-06 \| 1.5e-11 \| 69.7 \| 65 \| (36, 101) \| 103 \| (352, 429) \| 477 \| DksA C4-type domain-containing protein \| DksA C4-type domain-containing protein; uniclust \| UniRef100\_A0A0J6YR85 \| 97.4 \| 7.3e-06 \| 1.5e-11 \| 64.3 \| 68 \| (31, 99) \| 103 \| (149, 229) \| 231 \| DksA C4-type domain-containing protein \| DksA C4-type domain-containing protein; uniclust \| UniRef100\_A0A7C5J814 \| 97.4 \| 8.1e-06 \| 1.5e-11 \| 54.5 \| 64 \| (36, 100) \| 103 \| (1, 82) \| 88 \| DksA C4-type domain-containing protein \| DksA C4-type domain-containing protein; uniclust \| UniRef100\_A0A2V8ELD6 \| 97.4 \| 7.6e-06 \| 1.5e-11 \| 63.1 \| 37 \| (63, 100) \| 103 \| (144, 185) \| 209 \| DksA C4-type domain-containing protein \| DksA C4-type domain-containing protein; uniclust \| UniRef100\_A0A540V863 \| 97.4 \| 7.8e-06 \| 1.5e-11 \| 52.7 \| 62 \| (36, 102) \| 103 \| (1, 62) \| 67 \| DUF2116 family Zn-ribbon domain-containing protein \| DUF2116 family Zn-ribbon domain-containing protein; uniclust \| UniRef100\_A0A2V7UG18 \| 97.4 \| 8.2e-06 \| 1.5e-11 \| 58.9 \| 63 \| (36, 99) \| 103 \| (71, 146) \| 148 \| DksA C4-type domain-containing protein \| DksA C4-type domain-containing protein; uniclust \| UniRef100\_A0A0C1W9X3 \| 97.4 \| 7.4e-06 \| 1.5e-11 \| 60.4 \| 67 \| (35, 102) \| 103 \| (51, 130) \| 142 \| DksA C4-type domain-containing protein \| DksA C4-type domain-containing protein; uniclust \| UniRef100\_A0A2N8GFW6 \| 97.3 \| 8.7e-06 \| 1.6e-11 \| 48.6 \| 46 \| (35, 81) \| 103 \| (1, 47) \| 47 \| Conjugal transfer protein TraR (Fragment) \| Conjugal transfer protein TraR (Fragment); uniclust \| UniRef100\_UPI0022357C3F \| 97.3 \| 8.9e-06 \| 1.6e-11 \| 53.2 \| 35 \| (63, 98) \| 103 \| (7, 46) \| 78 \| TraR/DksA family transcriptional regulator \| TraR/DksA family transcriptional regulator; uniclust \| UniRef100\_A0A940R1G3 \| 97.3 \| 9e-06 \| 1.6e-11 \| 59.7 \| 65 \| (35, 100) \| 103 \| (39, 116) \| 165 \| C4-type zinc finger protein, DksA/TraR family \| C4-type zinc finger protein, DksA/TraR family; uniclust \| UniRef100\_A0A2E4ZEP8 \| 97.3 \| 8.8e-06 \| 1.7e-11 \| 56.7 \| 35 \| (63, 98) \| 103 \| (65, 104) \| 109 \| DksA C4-type domain-containing protein \| DksA C4-type domain-containing protein; uniclust \| UniRef100\_A0A2D6N8B5 \| 97.3 \| 9.1e-06 \| 1.8e-11 \| 58.2 \| 36 \| (63, 99) \| 103 \| (76, 116) \| 125 \| Conjugal transfer protein TraR \| Conjugal transfer protein TraR; uniclust \| UniRef100\_A0A836SUD0 \| 97.3 \| 9.5e-06 \| 1.8e-11 \| 53.0 \| 36 \| (64, 100) \| 103 \| (10, 50) \| 73 \| DksA C4-type domain-containing protein \| DksA C4-type domain-containing protein; uniclust \| UniRef100\_A0A1F9A4Z3 \| 97.3 \| 9.1e-06 \| 1.8e-11 \| 57.9 \| 71 \| (31, 102) \| 103 \| (35, 118) \| 119 \| DksA C4-type domain-containing protein \| DksA C4-type domain-containing protein; uniclust \| UniRef100\_A0A2T2V596 \| 97.3 \| 9.1e-06 \| 1.8e-11 \| 56.8 \| 64 \| (37, 101) \| 103 \| (25, 101) \| 104 \| DksA C4-type domain-containing protein \| DksA C4-type domain-containing protein; uniclust \| UniRef100\_A0A4R2CHW8 \| 97.3 \| 8.9e-06 \| 1.8e-11 \| 60.7 \| 38 \| (63, 101) \| 103 \| (112, 154) \| 155 \| TraR/DksA family transcriptional regulator \| TraR/DksA family transcriptional regulator; uniclust \| UniRef100\_A0A0G0VR09 \| 97.3 \| 7.9e-06 \| 1.8e-11 \| 68.6 \| 35 \| (63, 98) \| 103 \| (310, 349) \| 365 \| DNA-binding protein \| DNA-binding protein; uniclust \| UniRef100\_A0A497A8C1 \| 97.3 \| 9.7e-06 \| 1.9e-11 \| 54.4 \| 36 \| (63, 99) \| 103 \| (36, 76) \| 84 \| Molecular chaperone DnaK (Fragment) \| Molecular chaperone DnaK (Fragment); uniclust \| UniRef100\_A0A2E7BH60 \| 97.3 \| 1e-05 \| 1.9e-11 \| 58.2 \| 68 \| (31, 99) \| 103 \| (39, 119) \| 143 \| DksA C4-type domain-containing protein \| DksA C4-type domain-containing protein; uniclust \| UniRef100\_A0A7Z9JQQ5 \| 97.3 \| 9.9e-06 \| 1.9e-11 \| 52.8 \| 35 \| (66, 101) \| 103 \| (31, 70) \| 71 \| RNA polymerase-binding protein DksA \| RNA polymerase-binding protein DksA; uniclust \| UniRef100\_A0A1H7ZBE4 \| 97.3 \| 9.8e-06 \| 1.9e-11 \| 52.6 \| 63 \| (35, 99) \| 103 \| (1, 66) \| 67 \| Transcriptional regulator, TraR/DksA family \| Transcriptional regulator, TraR/DksA family; uniclust \| UniRef100\_A0A0S2HZL1 \| 97.3 \| 8.8e-06 \| 1.9e-11 \| 60.1 \| 38 \| (63, 101) \| 103 \| (94, 136) \| 138 \| RNA polymerase-binding protein DksA \| RNA polymerase-binding protein DksA; uniclust \| UniRef100\_A0A934NEW0 \| 97.3 \| 1.1e-05 \| 2e-11 \| 51.0 \| 32 \| (63, 95) \| 103 \| (23, 59) \| 63 \| TraR/DksA C4-type zinc finger protein \| TraR/DksA C4-type zinc finger protein; uniclust \| UniRef100\_A0A1Z9AE00 \| 97.3 \| 9.9e-06 \| 2e-11 \| 64.7 \| 65 \| (36, 101) \| 103 \| (158, 235) \| 264 \| DksA C4-type domain-containing protein (Fragment) \| DksA C4-type domain-containing protein (Fragment); uniclust \| UniRef100\_A0A317E8Z2 \| 97.3 \| 1.1e-05 \| 2.1e-11 \| 51.5 \| 60 \| (36, 99) \| 103 \| (1, 67) \| 68 \| Conjugal transfer protein TraR \| Conjugal transfer protein TraR; uniclust \| UniRef100\_A0A177RA40 \| 97.3 \| 1.2e-05 \| 2.1e-11 \| 58.8 \| 62 \| (36, 98) \| 103 \| (44, 118) \| 158 \| DksA C4-type domain-containing protein (Fragment) \| DksA C4-type domain-containing protein (Fragment); uniclust \| UniRef100\_A0A0B8P4P1 \| 97.3 \| 1.1e-05 \| 2.2e-11 \| 51.4 \| 39 \| (63, 102) \| 103 \| (17, 60) \| 63 \| C4-type zinc finger protein \| C4-type zinc finger protein; uniclust \| UniRef100\_A0A084SYV1 \| 97.3 \| 1.1e-05 \| 2.2e-11 \| 58.7 \| 37 \| (63, 100) \| 103 \| (80, 121) \| 133 \| DksA \| DksA; uniclust \| UniRef100\_A0A1G1E393 \| 97.3 \| 9.8e-06 \| 2.2e-11 \| 66.0 \| 33 \| (63, 96) \| 103 \| (89, 126) \| 281 \| DksA C4-type domain-containing protein \| DksA C4-type domain-containing protein; uniclust \| UniRef100\_A0A496XS46 \| 97.3 \| 1.2e-05 \| 2.2e-11 \| 56.2 \| 69 \| (29, 98) \| 103 \| (32, 113) \| 114 \| DksA C4-type domain-containing protein \| DksA C4-type domain-containing protein; uniclust \| UniRef100\_A0A4R4LN85 \| 97.3 \| 1.2e-05 \| 2.2e-11 \| 59.2 \| 32 \| (63, 95) \| 103 \| (130, 166) \| 168 \| Molecular chaperone DnaK \| Molecular chaperone DnaK; uniclust \| UniRef100\_A0A2D5E8I5 \| 97.3 \| 1e-05 \| 2.3e-11 \| 68.1 \| 64 \| (36, 100) \| 103 \| (286, 362) \| 375 \| DksA C4-type domain-containing protein \| DksA C4-type domain-containing protein; uniclust \| UniRef100\_A0A0U2YW82 \| 97.3 \| 1.2e-05 \| 2.3e-11 \| 54.5 \| 34 \| (63, 97) \| 103 \| (57, 95) \| 97 \| C4\_traR\_proteo: phage/conjugal plasmid C-4 type zinc finger protein, TraR \| C4\_traR\_proteo: phage/conjugal plasmid C-4 type zinc finger protein, TraR; uniclust \| UniRef100\_A0A1A9UK84 \| 97.3 \| 1.3e-05 \| 2.4e-11 \| 56.8 \| 65 \| (27, 99) \| 103 \| (54, 118) \| 128 \| Ribosomal protein L9 domain-containing protein \| Ribosomal protein L9 domain-containing protein; uniclust \| UniRef100\_A0A257H7G2 \| 97.3 \| 1.3e-05 \| 2.4e-11 \| 46.6 \| 33 \| (66, 99) \| 103 \| (3, 40) \| 41 \| DksA C4-type domain-containing protein \| DksA C4-type domain-containing protein; uniclust \| UniRef100\_A0A357XRK3 \| 97.3 \| 1.3e-05 \| 2.4e-11 \| 64.6 \| 64 \| (35, 99) \| 103 \| (267, 343) \| 356 \| DksA C4-type domain-containing protein \| DksA C4-type domain-containing protein; uniclust \| UniRef100\_A0A081NL60 \| 97.3 \| 1.3e-05 \| 2.5e-11 \| 52.8 \| 65 \| (36, 101) \| 103 \| (1, 71) \| 75 \| DksA C4-type domain-containing protein \| DksA C4-type domain-containing protein; uniclust \| UniRef100\_A0A7Y6Z4L2 \| 97.3 \| 1.4e-05 \| 2.5e-11 \| 57.3 \| 65 \| (35, 101) \| 103 \| (62, 134) \| 137 \| TraR/DksA C4-type zinc finger protein \| TraR/DksA C4-type zinc finger protein; uniclust \| UniRef100\_A0A257XAG9 \| 97.3 \| 1.2e-05 \| 2.6e-11 \| 59.5 \| 65 \| (35, 100) \| 103 \| (56, 133) \| 144 \| DksA C4-type domain-containing protein \| DksA C4-type domain-containing protein; uniclust \| UniRef100\_Q31HW7 \| 97.3 \| 1.4e-05 \| 2.7e-11 \| 55.4 \| 68 \| (31, 102) \| 103 \| (42, 110) \| 111 \| Uncharacterized protein \| Uncharacterized protein; uniclust \| UniRef100\_A0A845GB54 \| 97.2 \| 1.5e-05 \| 2.8e-11 \| 53.4 \| 37 \| (63, 100) \| 103 \| (47, 88) \| 89 \| TraR/DksA family transcriptional regulator \| TraR/DksA family transcriptional regulator; uniclust \| UniRef100\_A0A968KN48 \| 97.2 \| 1.5e-05 \| 2.8e-11 \| 49.2 \| 35 \| (66, 101) \| 103 \| (4, 43) \| 56 \| YteA family sporulation protein \| YteA family sporulation protein; uniclust \| UniRef100\_A0A535JRC0 \| 97.2 \| 1.5e-05 \| 2.8e-11 \| 54.8 \| 33 \| (65, 98) \| 103 \| (67, 104) \| 105 \| DksA C4-type domain-containing protein \| DksA C4-type domain-containing protein; pdb70 \| 5W1S\_M \| 99.4 \| 7.3e-18 \| 7.1e-22 \| 106.1 \| 66 \| (35, 102) \| 103 \| (1, 71) \| 79 \| DNA-directed RNA polymerase subunit alpha \| 5W1S\_M DNA-directed RNA polymerase subunit alpha RNA polymerase, TraR, TRANSFERASE; pdb70 \| 6PST\_N \| 99.4 \| 2e-17 \| 2e-21 \| 101.4 \| 65 \| (36, 102) \| 103 \| (1, 70) \| 72 \| DNA-directed RNA polymerase subunit alpha \| 6PST\_N DNA-directed RNA polymerase subunit alpha TRANSCRIPTION, transcription-dna complex HET: 1N7; pdb70 \| 6PSV\_N \| 99.4 \| 2e-17 \| 2e-21 \| 101.4 \| 65 \| (36, 102) \| 103 \| (1, 70) \| 72 \| DNA-directed RNA polymerase subunit alpha \| 6PSV\_N DNA-directed RNA polymerase subunit alpha TRANSCRIPTION, transcription-dna complex HET: 1N7; pdb70 \| 4IJJ\_B \| 99.2 \| 1e-15 \| 1e-19 \| 104.1 \| 51 \| (48, 102) \| 103 \| (77, 132) \| 136 \| Putative C4-type zinc finger protein \| 4IJJ\_B Putative C4-type zinc finger protein DksA fold, transcription factor, RNA HET: SO4; pdb70 \| 6PTG\_B \| 99.2 \| 1.7e-15 \| 1.7e-19 \| 102.7 \| 50 \| (48, 101) \| 103 \| (75, 129) \| 132 \| DnaK Suppressor \| 6PTG\_B DnaK Suppressor SSGCID, Structural Genomics, Seattle Structural; pdb70 \| 1TJL\_B \| 99.2 \| 2.5e-15 \| 2.4e-19 \| 104.1 \| 51 \| (48, 102) \| 103 \| (93, 148) \| 151 \| DnaK suppressor protein \| 1TJL\_B DnaK suppressor protein DksA, transcription factor, RNA polymerase HET: ZN; pdb70 \| 5VSW\_M \| 99.2 \| 2.5e-15 \| 2.4e-19 \| 104.1 \| 51 \| (48, 102) \| 103 \| (93, 148) \| 151 \| DNA-directed RNA polymerase subunit alpha \| 5VSW\_M DNA-directed RNA polymerase subunit alpha RNA polymerase, DksA, ppGpp, transferase-dna HET: G4P; pdb70 \| 2KQ9\_A \| 99.1 \| 1.5e-14 \| 1.5e-18 \| 95.2 \| 44 \| (48, 95) \| 103 \| (63, 111) \| 112 \| DnaK suppressor protein \| 2KQ9\_A DnaK suppressor protein Zinc binding protein, Structural Genomics HET: ZN; pdb70 \| 2KGO\_A \| 99.0 \| 3.6e-14 \| 3.3e-18 \| 94.7 \| 60 \| (40, 102) \| 103 \| (29, 93) \| 108 \| Uncharacterized protein ybiI \| 2KGO\_A Uncharacterized protein ybiI Zn finger, partially disordered, structural HET: ZN |
| Top keywords  (threshold 1.00e-03 (evalue)) | **DksA, C4\_type, domain\_containing, TraR, transcriptional, regulator, zinc, finger, DnaK, RNA** |
| Output files | ../../similar\_sequences/36\_FANPEZAQ\_CDS\_0036\_merged.svg ../../similar\_sequences/36\_FANPEZAQ\_CDS\_0036\_pdb70.a3m ../../similar\_sequences/36\_FANPEZAQ\_CDS\_0036\_pdb70.hhr ../../similar\_sequences/36\_FANPEZAQ\_CDS\_0036\_uniclust.a3m ../../similar\_sequences/36\_FANPEZAQ\_CDS\_0036\_uniclust.hhr |

#### Structure prediction (AlphaFold)^2^

|  |  |
| --- | --- |
| Stats | xml version="1.0" encoding="utf-8" standalone="no"?       2024-09-02T21:09:35.392415 image/svg+xml   Matplotlib v3.7.2, https://matplotlib.org/ |
| Predicted structure | **NGL Viewer Controls:**  - Center: *Left-Click* - Rotate: *Left-Click + Drag* - Translate: *Right-Click + Drag* - Zoom: *Shift + Left-Click + Drag* |
| Output files | ../../predicted\_structures/36\_FANPEZAQ\_CDS\_0036/features.pkl ../../predicted\_structures/36\_FANPEZAQ\_CDS\_0036/ranked\_0.pdb ../../predicted\_structures/36\_FANPEZAQ\_CDS\_0036/ranked\_0\_plots.svg ../../predicted\_structures/36\_FANPEZAQ\_CDS\_0036/result\_model\_1\_ptm\_pred\_0.pkl |

#### Structure similarity search results (Foldseek)^3^

|  |  |
| --- | --- |
| Structure databases searched | Pdb, Afdb-proteome, Afdb-uniprot50 |
| Results, scheme(s)  (Top layers only, threshold 1.00e-02 (evalue)) | xml version="1.0" encoding="utf-8" standalone="no"?       2024-09-02T21:11:11.331828 image/svg+xml   Matplotlib v3.7.2, https://matplotlib.org/ |
| Results, table  (threshold 1.00e-02 (evalue)) | db \| id \| prob \| evalue \| bits \| fident \| alnlen \| mismatch \| gapopen \| qstart \| qend \| tstart \| tend \| name \| description; afdb-uniprot50 \| AF-Q31HW7-F1-MODEL\_V4 \| 1.0 \| 0.002348 \| 108 \| 0.281 \| 103 \| 63 \| 5 \| 1 \| 103 \| 18 \| 109 \| Uncharacterized protein \| Uncharacterized protein |
| Top keywords  (threshold 1.00e-02 (evalue)) | -- |
| Output files | ../../similar\_structures/36\_FANPEZAQ\_CDS\_0036\_afdb-proteome\_foldseek.tsv ../../similar\_structures/36\_FANPEZAQ\_CDS\_0036\_afdb-uniprot50\_foldseek.tsv ../../similar\_structures/36\_FANPEZAQ\_CDS\_0036\_merged.svg ../../similar\_structures/36\_FANPEZAQ\_CDS\_0036\_pdb\_foldseek.tsv |

  
  
  

Return to summary | Go to previous | Go to next

  


---

**Sequence/structure alignments coloring**  
Each object in the alignment figures is colored according to its E-value following this color coding:

1e-100
10
