## Supplementary material for "Completing the BASEL phage collection to unlock hidden diversity for systematic exploration of phage-host interactions": Data S2: 38.html

FANPEZAQ\_CDS\_0038


Return to summary | Go to previous | Go to next

|  |  |
| --- | --- |
| FANPEZAQ\_CDS\_0038  Page creation date: 02 Sep 2024, 12:00  Project folder: n/a  Input sequences file: Escherichia\_virus\_HeidiAbel.gb | domain\_containing helix\_turn\_helix dna\_binding regulator hth\_17 transcriptional excisionase dna binding alpa merr phage a putative tp and regulatory ap yes cp gp multidrug\_efflux transporter engineered transcription complex escherichia hth coli organism\_taxid arch excision expressed in expression\_system\_taxid alpha integration gene packaging mainly orthogonal bundle b terminase activator hth\_type d xis \_3' c |

### Sequence information

|  |  |
| --- | --- |
| Name | FANPEZAQ\_CDS\_0038  38\_FANPEZAQ\_CDS\_0038 (pipeline id) |
| Imported annotations |  |
| Protein sequence | MAQPAQKQSTYLTPQELVLRWNGAVTTGTLANWRNKGKGPAYTKFGSRVRYSITSVEAYE AKHMIGANDNEEC |
| Number of residues | 73 |
| Molecular weight (Da) | 8136.05 |
| Output files | ../../query\_sequences/38\_FANPEZAQ\_CDS\_0038.fasta |

### Putative domain architecture and protein family

#### Search results (HHblits)^1^

|  |  |
| --- | --- |
| Domain family databases searched | Pfam, Ncbi-cd, Cath, Phrogs |
| Results, scheme(s)  (Top layers only; threshold 1.00e-03 (evalue)) | xml version="1.0" encoding="utf-8" standalone="no"?       2024-09-02T21:08:20.579766 image/svg+xml   Matplotlib v3.7.2, https://matplotlib.org/ |
| Results, table  (E-value ≤ 1.00e-03 (evalue)) | db \| id \| prob \| evalue \| pvalue \| score \| cols \| query \| query\_len \| template \| template\_len \| name \| description; pfam \| PF09035 \| 97.8 \| 4.7e-09 \| 7.7e-13 \| 41.5 \| 56 \| (6, 63) \| 73 \| (10, 66) \| 69 \| Tn916-Xis \| Excisionase from transposon Tn916; pfam \| PF06806 \| 97.8 \| 7.2e-09 \| 1.2e-12 \| 41.3 \| 57 \| (5, 63) \| 73 \| (3, 69) \| 70 \| DUF1233 \| Putative excisionase (DUF1233); pfam \| PF10743 \| 97.7 \| 1e-08 \| 1.7e-12 \| 42.7 \| 62 \| (1, 64) \| 73 \| (1, 70) \| 87 \| Phage\_Cox \| Regulatory phage protein cox; pfam \| PF04936 \| 97.4 \| 7e-08 \| 1.2e-11 \| 45.9 \| 55 \| (8, 64) \| 73 \| (11, 66) \| 186 \| DUF658 \| Protein of unknown function (DUF658); pfam \| PF04645 \| 97.2 \| 1.8e-07 \| 2.9e-11 \| 44.0 \| 51 \| (8, 63) \| 73 \| (16, 70) \| 181 \| DUF603 \| Protein of unknown function, DUF603; pfam \| PF07618 \| 97.1 \| 3.5e-07 \| 5.7e-11 \| 35.0 \| 47 \| (13, 62) \| 73 \| (1, 55) \| 57 \| DUF1580 \| Protein of unknown function (DUF1580); pfam \| PF04936 \| 96.9 \| 1.1e-06 \| 1.8e-10 \| 41.6 \| 39 \| (12, 52) \| 73 \| (89, 127) \| 186 \| DUF658 \| Protein of unknown function (DUF658); pfam \| PF12964 \| 96.9 \| 1.2e-06 \| 2e-10 \| 36.3 \| 53 \| (8, 62) \| 73 \| (42, 95) \| 96 \| DUF3853 \| Protein of unknown function (DUF3853); pfam \| PF12728 \| 96.8 \| 1.5e-06 \| 2.4e-10 \| 30.8 \| 47 \| (12, 62) \| 73 \| (2, 49) \| 51 \| HTH\_17 \| Helix-turn-helix domain; pfam \| PF07278 \| 96.8 \| 1.5e-06 \| 2.5e-10 \| 39.1 \| 50 \| (12, 64) \| 73 \| (2, 51) \| 147 \| DUF1441 \| Protein of unknown function (DUF1441); pfam \| PF05930 \| 96.8 \| 1.8e-06 \| 3e-10 \| 30.5 \| 48 \| (10, 59) \| 73 \| (2, 51) \| 52 \| Phage\_AlpA \| Prophage CP4-57 regulatory protein (AlpA); pfam \| PF07037 \| 96.7 \| 2.5e-06 \| 4.1e-10 \| 36.4 \| 50 \| (12, 63) \| 73 \| (1, 55) \| 113 \| DUF1323 \| Putative transcription regulator (DUF1323); pfam \| PF13411 \| 96.6 \| 3.7e-06 \| 6e-10 \| 31.9 \| 48 \| (12, 61) \| 73 \| (1, 49) \| 70 \| MerR\_1 \| MerR HTH family regulatory protein; pfam \| PF20038 \| 96.5 \| 4.3e-06 \| 7.1e-10 \| 31.2 \| 54 \| (8, 63) \| 73 \| (4, 64) \| 66 \| HTH\_59 \| Helix-turn-helix domain; pfam \| PF07471 \| 96.4 \| 6.6e-06 \| 1.1e-09 \| 37.1 \| 52 \| (11, 64) \| 73 \| (2, 54) \| 162 \| Phage\_Nu1 \| Phage DNA packaging protein Nu1; pfam \| PF20063 \| 96.4 \| 7.9e-06 \| 1.3e-09 \| 28.2 \| 48 \| (10, 61) \| 73 \| (2, 49) \| 51 \| DUF6462 \| Family of unknown function (DUF6462); pfam \| PF13171 \| 96.3 \| 9.6e-06 \| 1.6e-09 \| 38.3 \| 44 \| (11, 56) \| 73 \| (1, 51) \| 199 \| DUF4004 \| Protein of unknown function (DUF4004); pfam \| PF07825 \| 96.2 \| 1.6e-05 \| 2.6e-09 \| 31.3 \| 44 \| (11, 56) \| 73 \| (2, 48) \| 72 \| Exc \| Excisionase-like protein; pfam \| PF13591 \| 95.3 \| 0.00015 \| 2.4e-08 \| 28.3 \| 46 \| (12, 59) \| 73 \| (1, 47) \| 86 \| MerR\_2 \| MerR HTH family regulatory protein; pfam \| PF11242 \| 94.5 \| 0.00046 \| 7.7e-08 \| 27.0 \| 29 \| (7, 37) \| 73 \| (10, 38) \| 64 \| DUF2774 \| Protein of unknown function (DUF2774); pfam \| PF17443 \| 94.2 \| 0.0007 \| 1.1e-07 \| 25.3 \| 32 \| (6, 39) \| 73 \| (3, 34) \| 62 \| pXO2-72 \| Uncharacterized protein pXO2-72; pfam \| PF00376 \| 94.0 \| 0.00083 \| 1.4e-07 \| 20.7 \| 26 \| (14, 41) \| 73 \| (2, 27) \| 38 \| MerR \| MerR family regulatory protein; pfam \| PF08876 \| 93.8 \| 0.001 \| 1.7e-07 \| 28.5 \| 32 \| (10, 43) \| 73 \| (23, 54) \| 109 \| DUF1836 \| Domain of unknown function (DUF1836); cath \| 4lhfA00 \| 98.2 \| 2.7e-10 \| 3.6e-14 \| 55.0 \| 61 \| (1, 64) \| 73 \| (1, 61) \| 91 \| Regulatory protein cox \| CATHCODE: 6.10.200.10 NAME: Regulatory protein cox. Chain: a. Engineered: yes SOURCE: Enterobacteria phage p2. Organism\_taxid: 10679. Gene: cox. Expressed in: escherichia coli. Expression\_system\_taxid: 469008. CLASS: Special, ARCH: Helix non-globular, TOPOL: Multidrug-efflux Transporter Regulator; Chain: A; Domain 2, HOMOL: Regulatory phage protein Cox; cath \| 2kvvA00 \| 98.2 \| 4.7e-10 \| 6.3e-14 \| 53.4 \| 60 \| (3, 65) \| 73 \| (2, 72) \| 78 \| Putative excisionase \| CATHCODE: 1.10.1660.60 NAME: Putative excisionase. Chain: a. Engineered: yes SOURCE: Klebsiella pneumoniae. Organism\_taxid: 272620. Strain: atcc 700721 / mgh 78578. Gene: kpn78578\_13480, kpn\_01377. Expressed in: escherichia coli. Expression\_system\_taxid: 469008. CLASS: Mainly Alpha, ARCH: Orthogonal Bundle, TOPOL: Multidrug-efflux Transporter Regulator; Chain: A; Domain 2, HOMOL: Putative excisionased domain DUF1233; cath \| 1y6uA01 \| 97.4 \| 1.1e-07 \| 1.5e-11 \| 38.5 \| 47 \| (10, 59) \| 73 \| (1, 49) \| 49 \| Excisionase from transposon tn916 \| CATHCODE: 3.90.105.50 NAME: Excisionase from transposon tn916. Chain: a. Synonym: xis. Engineered:yes SOURCE: Enterococcus faecalis. Organism\_taxid: 1351. Gene: xis. Expressed in:escherichia coli. Expression\_system\_taxid: 562. CLASS: Alpha Beta, ARCH: Alpha-Beta Complex, TOPOL: Molybdopterin biosynthesis moea protein, domain 2, HOMOL: Molybdopterin biosynthesis moea protein, domain 2; cath \| 1z4hA01 \| 97.3 \| 1.6e-07 \| 2.2e-11 \| 39.0 \| 52 \| (11, 65) \| 73 \| (2, 54) \| 58 \| Tor inhibition protein \| CATHCODE: 1.10.238.160 NAME: Tor inhibition protein. Chain: a. Synonym: tori. Engineered: yes SOURCE: Escherichia coli. Organism\_taxid: 562. Expressed in: escherichia colibl21(de3). Expression\_system\_taxid: 469008. CLASS: Mainly Alpha, ARCH: Orthogonal Bundle, TOPOL: Recoverin; domain 1, HOMOL: Recoverin; domain 1; cath \| 1j9iA00 \| 96.0 \| 2.9e-05 \| 4.1e-09 \| 32.6 \| 52 \| (11, 66) \| 73 \| (2, 56) \| 68 \| Terminase small subunit \| CATHCODE: 1.10.10.10 NAME: Terminase small subunit. Chain: a, b. Fragment: DNA binding domain, residues 1-68. Synonym: DNA packaging protein nu1. Gpnu1 dbd. Engineered: yes SOURCE: Enterobacteria phage lambda. Organism\_taxid: 10710. Gene: nu1. Expressed in: escherichia coli. Expression\_system\_taxid: 562. Expression\_system\_vector\_type: plasmid CLASS: Mainly Alpha, ARCH: Orthogonal Bundle, TOPOL: Arc Repressor Mutant, subunit A, HOMOL: Winged helix-like DNA-binding domain superfamily/Winged helix DNA-binding domain; cath \| 1pm6A00 \| 95.9 \| 4.3e-05 \| 5.9e-09 \| 33.4 \| 43 \| (11, 56) \| 73 \| (2, 48) \| 72 \| Excisionase \| CATHCODE: 1.10.1660.20 NAME: Excisionase. Chain: a. Engineered: yes. Mutation: yes SOURCE: Enterobacteria phage hk022. Organism\_taxid: 10742. Gene: xis. Expressed in: escherichia coli. Expression\_system\_taxid: 562. CLASS: Mainly Alpha, ARCH: Orthogonal Bundle, TOPOL: Multidrug-efflux Transporter Regulator; Chain: A; Domain 2, HOMOL: Multidrug-efflux Transporter Regulator; Chain: A; Domain 2; cath \| 3gpvA00 \| 95.6 \| 7.9e-05 \| 1.1e-08 \| 35.6 \| 55 \| (5, 62) \| 73 \| (10, 66) \| 128 \| Transcriptional regulator, merr family \| CATHCODE: 1.10.1660.10 NAME: Transcriptional regulator, merr family. Chain: a, b. Engineered: yes SOURCE: Bacillus thuringiensis serovar konkukian. Organism\_taxid: 180856. Gene: bt9727\_1615. Expressed in: escherichia coli. Expression\_system\_taxid: 562. CLASS: Mainly Alpha, ARCH: Orthogonal Bundle, TOPOL: Multidrug-efflux Transporter Regulator; Chain: A; Domain 2, HOMOL: Multidrug-efflux Transporter Regulator; Chain: A; Domain 2; cath \| 4r4eB00 \| 95.5 \| 0.00011 \| 1.6e-08 \| 31.7 \| 56 \| (2, 60) \| 73 \| (2, 58) \| 84 \| Hth-type transcriptional regulator glnr \| CATHCODE: 1.10.1660.10 NAME: Hth-type transcriptional regulator glnr. Chain: b, a. Engineered: yes.Dna (5'-d(\*ap\*tp\*tp\*cp\*tp\*gp\*ap\*cp\*a)-3'). Chain: d. Engineered: yes.Other\_details: cognate DNA. Dna (5'-d(\*tp\*gp\*tp\*cp\*ap\*gp\*tp\*a)-3'). Chain: e. Engineered: yes. Other\_details: cognate DNA (complement to 2) SOURCE: Bacillus subtilis subsp. Subtilis. Organism\_taxid: 224308. Strain: 168. Gene: glnr, bsu17450. Expressed in: escherichia coli. Expression\_system\_taxid: 562. Yes. Organism\_scientific: synthetic DNA. Organism\_taxid: 32630. CLASS: Mainly Alpha, ARCH: Orthogonal Bundle, TOPOL: Multidrug-efflux Transporter Regulator; Chain: A; Domain 2, HOMOL: Multidrug-efflux Transporter Regulator; Chain: A; Domain 2; cath \| 1r8eA02 \| 95.3 \| 0.00017 \| 2.4e-08 \| 29.9 \| 51 \| (9, 62) \| 73 \| (3, 56) \| 75 \| 5'-d(\*gp\*ap\*cp\*cp\*cp\*tp\*cp\*cp\*cp\*cp\*tp\*tp\*ap\*gp\*gp\*gp\*gp\*ap \*gp\*gp\*gp\*tp\*c)-3' \| CATHCODE: 1.10.1660.10 NAME: 5'-d(\*gp\*ap\*cp\*cp\*cp\*tp\*cp\*cp\*cp\*cp\*tp\*tp\*ap\*gp\*gp\*gp\*gp\*ap \*gp\*gp\*gp\*tp\*c)-3'. Chain: b. Engineered: yes. Multidrug-efflux transporter regulator. Chain: a. Engineered: yes SOURCE: Yes. CLASS: Mainly Alpha, ARCH: Orthogonal Bundle, TOPOL: Multidrug-efflux Transporter Regulator; Chain: A; Domain 2, HOMOL: Multidrug-efflux Transporter Regulator; Chain: A; Domain 2; cath \| 5i41B00 \| 95.1 \| 0.00024 \| 3.4e-08 \| 28.6 \| 50 \| (10, 62) \| 73 \| (2, 52) \| 69 \| Chromosome-anchoring protein raca \| CATHCODE: 1.10.1660.10 NAME: Chromosome-anchoring protein raca. Chain: b. Engineered: yes SOURCE: Bacillus subtilis. Organism\_taxid: 224308. Strain: 168. Gene: raca, ywkc, bsu37030. Expressed in: escherichia coli. Expression\_system\_taxid: 562 CLASS: Mainly Alpha, ARCH: Orthogonal Bundle, TOPOL: Multidrug-efflux Transporter Regulator; Chain: A; Domain 2, HOMOL: Multidrug-efflux Transporter Regulator; Chain: A; Domain 2; cath \| 2zhgA00 \| 95.0 \| 0.00029 \| 3.9e-08 \| 34.0 \| 55 \| (3, 60) \| 73 \| (3, 58) \| 135 \| Redox-sensitive transcriptional activator soxr \| CATHCODE: 1.10.1660.10 NAME: Redox-sensitive transcriptional activator soxr. Chain: a. Engineered: yes. Dna (5'- d(\*dgp\*dcp\*dcp\*dtp\*dcp\*dap\*dap\*dgp\*dtp\*dtp\*dap\*dap\*dcp\*dtp\*dtp\*dgp\*da p\*dgp\*dgp\*dc)-3'). Chain: b. Engineered: yes SOURCE: Escherichia coli. Organism\_taxid: 83333. Strain: k12. Gene: soxr. Expressed in: escherichia coli. Expression\_system\_taxid: 562. Expression\_system\_vector\_type: plasmid. CLASS: Mainly Alpha, ARCH: Orthogonal Bundle, TOPOL: Multidrug-efflux Transporter Regulator; Chain: A; Domain 2, HOMOL: Multidrug-efflux Transporter Regulator; Chain: A; Domain 2; cath \| 3hh0A01 \| 94.8 \| 0.00036 \| 5e-08 \| 28.9 \| 49 \| (10, 61) \| 73 \| (3, 53) \| 74 \| Transcriptional regulator, merr family \| CATHCODE: 1.10.1660.10 NAME: Transcriptional regulator, merr family. Chain: a, b, c, d. Engineered:yes SOURCE: Bacillus cereus atcc 14579. Organism\_taxid: 226900. Atcc: 14579. Gene:bc\_0953. Expressed in: escherichia coli. Expression\_system\_taxid: 562. CLASS: Mainly Alpha, ARCH: Orthogonal Bundle, TOPOL: Multidrug-efflux Transporter Regulator; Chain: A; Domain 2, HOMOL: Multidrug-efflux Transporter Regulator; Chain: A; Domain 2; cath \| 2jmlA00 \| 94.8 \| 0.00037 \| 5.1e-08 \| 29.7 \| 52 \| (9, 62) \| 73 \| (3, 56) \| 81 \| Dna binding domain/transcriptional regulator \| CATHCODE: 1.10.1660.10 NAME: Dna binding domain/transcriptional regulator. Chain: a. Fragment: n-terminal domain, residues 1-78. Engineered: yes SOURCE: Myxococcus xanthus. Organism\_taxid: 246197. Strain: dk 1622. Expressed in: escherichia coli bl21(de3). Expression\_system\_taxid: 469008. CLASS: Mainly Alpha, ARCH: Orthogonal Bundle, TOPOL: Multidrug-efflux Transporter Regulator; Chain: A; Domain 2, HOMOL: Multidrug-efflux Transporter Regulator; Chain: A; Domain 2; cath \| 5d8cA00 \| 94.7 \| 0.00043 \| 5.9e-08 \| 33.3 \| 49 \| (10, 61) \| 73 \| (3, 53) \| 137 \| Merr family regulator protein \| CATHCODE: 1.10.1660.10 NAME: Merr family regulator protein. Chain: a, b. Engineered: yes. Dna (5'- d(\*cp\*tp\*tp\*ap\*gp\*ap\*gp\*tp\*tp\*cp\*ap\*cp\*tp\*cp\*tp\*ap\*ap\*g)-3'). Chain: c. Engineered: yes. Other\_details: promoter sequence for adhc-estd operon and nmlr gene.. Dna (5'- d(\*cp\*tp\*tp\*ap\*gp\*ap\*gp\*tp\*gp\*ap\*ap\*cp\*tp\*cp\*tp\*ap\*ap\*g)-3'). Chain: d. Engineered: yes SOURCE: Haemophilus influenzae (strain atcc 51907 / dsm 11121 / kw20 / rd). Organism\_taxid: 71421. Strain: atcc 51907 / dsm 11121 / kw20 / rd. Gene:hi\_0186. Expressed in: escherichia coli. Expression\_system\_taxid: 469008. CLASS: Mainly Alpha, ARCH: Orthogonal Bundle, TOPOL: Multidrug-efflux Transporter Regulator; Chain: A; Domain 2, HOMOL: Multidrug-efflux Transporter Regulator; Chain: A; Domain 2; cath \| 3vw4A02 \| 94.1 \| 0.00096 \| 1.3e-07 \| 24.4 \| 27 \| (9, 37) \| 73 \| (5, 31) \| 33 \| Rep \| CATHCODE: 1.10.1220.10 NAME: Rep. Chain: a, b. Fragment: DNA-bindig domain. Synonym: replication initiator cole2-rep. Engineered: yes. Dna (5'- d(p\*ap\*ap\*tp\*gp\*ap\*gp\*ap\*cp\*cp\*ap\*gp\*ap\*tp\*ap\*ap\*gp\*cp\*cp\*tp\*tp\*ap\*tp \*c)-3'). Chain: c, e. Engineered: yes. Dna (5'- d(p\*gp\*ap\*tp\*ap\*ap\*gp\*gp\*cp\*tp\*tp\*ap\*tp\*cp\*tp\*gp\*gp\*tp\*cp\*tp\*cp\*ap\*tp \*t)-3'). Chain: d, f. Engineered: yes SOURCE: Escherichia coli. Organism\_taxid: 562. Gene: rep. Expressed in: escherichia coli. Expression\_system\_taxid: 562. CLASS: Mainly Alpha, ARCH: Orthogonal Bundle, TOPOL: Arc Repressor Mutant, HOMOL: Met repressor-like; phrogs \| 66 \| 99.1 \| 7e-15 \| 8.7e-19 \| 73.9 \| 67 \| (2, 70) \| 73 \| (8, 77) \| 81 \| excisionase and transcriptional regulator \| excisionase and transcriptional regulator; Category: integration and excision; NC\_023692\_p35; phrogs \| 6401 \| 98.6 \| 8.6e-12 \| 9.9e-16 \| 60.1 \| 62 \| (3, 66) \| 73 \| (3, 64) \| 66 \| DNA binding protein \| DNA binding protein; Category: DNA, RNA and nucleotide metabolism; MG757155\_p60; phrogs \| 10045 \| 98.6 \| 1.2e-11 \| 1.4e-15 \| 58.5 \| 54 \| (5, 61) \| 73 \| (3, 57) \| 62 \| HTH DNA binding protein \| HTH DNA binding protein; Category: DNA, RNA and nucleotide metabolism; KU963258\_p53; phrogs \| 15534 \| 98.4 \| 4e-11 \| 4.6e-15 \| 60.6 \| 58 \| (7, 68) \| 73 \| (32, 89) \| 93 \| HTH DNA binding protein \| HTH DNA binding protein; Category: DNA, RNA and nucleotide metabolism; NC\_028930\_p36; phrogs \| 7338 \| 98.2 \| 3e-10 \| 3.5e-14 \| 58.0 \| 60 \| (7, 69) \| 73 \| (34, 93) \| 97 \| NA \| NA; Category: unknown function; p137519 VI\_04448; phrogs \| 2907 \| 98.2 \| 3e-10 \| 3.5e-14 \| 59.0 \| 59 \| (5, 66) \| 73 \| (3, 65) \| 103 \| excisionase \| excisionase; Category: integration and excision; p256170 VI\_09967; phrogs \| 668 \| 98.1 \| 5.9e-10 \| 7.3e-14 \| 56.1 \| 53 \| (6, 63) \| 73 \| (31, 84) \| 85 \| HTH DNA binding protein \| HTH DNA binding protein; Category: DNA, RNA and nucleotide metabolism; p164379 VI\_04615; phrogs \| 2790 \| 98.1 \| 8.2e-10 \| 9.3e-14 \| 55.8 \| 63 \| (6, 70) \| 73 \| (20, 83) \| 91 \| excisionase and transcriptional regulator \| excisionase and transcriptional regulator; Category: integration and excision; p43040 VI\_12007; phrogs \| 2474 \| 97.9 \| 5.3e-09 \| 6.1e-13 \| 51.1 \| 55 \| (3, 59) \| 73 \| (7, 62) \| 74 \| MerR-like transcriptional regulator \| MerR-like transcriptional regulator; Category: transcription regulation; NC\_026585\_p102; phrogs \| 57 \| 97.8 \| 5.5e-09 \| 6.8e-13 \| 58.4 \| 57 \| (9, 67) \| 73 \| (2, 60) \| 173 \| terminase small subunit \| terminase small subunit; Category: head and packaging; p277333 VI\_02593; phrogs \| 6620 \| 97.8 \| 7e-09 \| 8.1e-13 \| 57.6 \| 52 \| (9, 62) \| 73 \| (3, 55) \| 170 \| transposase \| transposase; Category: integration and excision; p336225 VI\_06314; phrogs \| 4691 \| 97.8 \| 9.2e-09 \| 1e-12 \| 49.5 \| 54 \| (6, 61) \| 73 \| (13, 66) \| 70 \| excisionase \| excisionase; Category: integration and excision; p211020 VI\_00698; phrogs \| 4745 \| 97.7 \| 1.2e-08 \| 1.4e-12 \| 48.7 \| 54 \| (9, 64) \| 73 \| (2, 63) \| 65 \| DNA binding protein \| DNA binding protein; Category: DNA, RNA and nucleotide metabolism; KR093652\_p84; phrogs \| 25262 \| 97.7 \| 1.9e-08 \| 2.1e-12 \| 50.5 \| 60 \| (6, 68) \| 73 \| (21, 80) \| 96 \| NA \| NA; Category: unknown function; p39263 VI\_01014; phrogs \| 9834 \| 97.5 \| 5.2e-08 \| 5.9e-12 \| 46.5 \| 58 \| (8, 68) \| 73 \| (3, 62) \| 67 \| excisionase and transcriptional regulator \| excisionase and transcriptional regulator; Category: integration and excision; p40253 VI\_09193; phrogs \| 11671 \| 97.5 \| 6.7e-08 \| 7.7e-12 \| 51.1 \| 57 \| (4, 63) \| 73 \| (12, 69) \| 122 \| NA \| NA; Category: unknown function; p36461 VI\_11179; phrogs \| 2655 \| 97.5 \| 6.9e-08 \| 8.1e-12 \| 46.9 \| 56 \| (9, 66) \| 73 \| (3, 58) \| 70 \| excisionase \| excisionase; Category: integration and excision; p346156 VI\_04054; phrogs \| 575 \| 97.4 \| 1.1e-07 \| 1.3e-11 \| 45.9 \| 33 \| (7, 41) \| 73 \| (10, 42) \| 67 \| Cox-like excisionase and repressor \| Cox-like excisionase and repressor; Category: integration and excision; p264496 VI\_02100; phrogs \| 1969 \| 97.2 \| 2.6e-07 \| 2.9e-11 \| 47.4 \| 52 \| (7, 61) \| 73 \| (50, 101) \| 103 \| NA \| NA; Category: unknown function; NC\_023689\_p106; phrogs \| 1864 \| 97.2 \| 3.4e-07 \| 3.9e-11 \| 44.0 \| 51 \| (9, 63) \| 73 \| (18, 68) \| 71 \| NA \| NA; Category: unknown function; p434791 VI\_03009; phrogs \| 7735 \| 97.1 \| 4.6e-07 \| 5.2e-11 \| 43.8 \| 54 \| (6, 62) \| 73 \| (12, 66) \| 74 \| excisionase \| excisionase; Category: integration and excision; p109736 VI\_12389; phrogs \| 2620 \| 97.0 \| 1e-06 \| 1.1e-10 \| 48.5 \| 60 \| (6, 67) \| 73 \| (74, 139) \| 187 \| excisionase and transcriptional regulator \| excisionase and transcriptional regulator; Category: integration and excision; p31239 VI\_02649; phrogs \| 7506 \| 96.8 \| 2.4e-06 \| 2.7e-10 \| 41.6 \| 56 \| (1, 59) \| 73 \| (4, 59) \| 85 \| recombination directionality factor \| recombination directionality factor; Category: integration and excision; NC\_012788\_p34; phrogs \| 1729 \| 96.7 \| 3.2e-06 \| 3.7e-10 \| 45.5 \| 53 \| (9, 63) \| 73 \| (95, 148) \| 152 \| NA \| NA; Category: unknown function; NC\_016435\_p45; phrogs \| 5866 \| 96.6 \| 4.6e-06 \| 5.1e-10 \| 41.1 \| 57 \| (8, 66) \| 73 \| (36, 93) \| 96 \| NA \| NA; Category: unknown function; JX409894\_p1; phrogs \| 6355 \| 96.3 \| 1.4e-05 \| 1.6e-09 \| 39.7 \| 57 \| (8, 66) \| 73 \| (16, 78) \| 89 \| NA \| NA; Category: unknown function; NC\_031098\_p35; phrogs \| 2528 \| 96.2 \| 1.9e-05 \| 2.2e-09 \| 39.0 \| 61 \| (7, 69) \| 73 \| (24, 84) \| 89 \| NA \| NA; Category: unknown function; p416682 VI\_10147; phrogs \| 2538 \| 96.2 \| 2.3e-05 \| 2.8e-09 \| 40.8 \| 30 \| (9, 40) \| 73 \| (14, 43) \| 113 \| terminase small subunit \| terminase small subunit; Category: head and packaging; p133796 VI\_05051; phrogs \| 151 \| 96.0 \| 3.7e-05 \| 4.6e-09 \| 38.2 \| 28 \| (9, 38) \| 73 \| (20, 47) \| 91 \| transposase \| transposase; Category: integration and excision; p268658 VI\_04551; phrogs \| 10458 \| 96.0 \| 4.2e-05 \| 4.8e-09 \| 38.0 \| 50 \| (9, 61) \| 73 \| (39, 90) \| 93 \| NA \| NA; Category: unknown function; p298856 VI\_05329; phrogs \| 7676 \| 95.9 \| 4.4e-05 \| 5e-09 \| 36.4 \| 31 \| (6, 38) \| 73 \| (11, 41) \| 75 \| NA \| NA; Category: unknown function; NC\_023688\_p227; phrogs \| 24579 \| 95.9 \| 5e-05 \| 5.7e-09 \| 39.0 \| 34 \| (5, 40) \| 73 \| (25, 58) \| 126 \| terminase small subunit \| terminase small subunit; Category: head and packaging; KP836356\_p30; phrogs \| 28156 \| 95.8 \| 6.2e-05 \| 6.9e-09 \| 39.2 \| 53 \| (6, 61) \| 73 \| (4, 57) \| 156 \| NA \| NA; Category: unknown function; JF937093\_p45; phrogs \| 20793 \| 95.8 \| 7.4e-05 \| 8.3e-09 \| 36.1 \| 52 \| (9, 62) \| 73 \| (5, 57) \| 86 \| NA \| NA; Category: unknown function; NC\_022918\_p114; phrogs \| 629 \| 95.7 \| 8.4e-05 \| 9.8e-09 \| 36.8 \| 45 \| (11, 56) \| 73 \| (3, 50) \| 86 \| excisionase \| excisionase; Category: integration and excision; p127807 VI\_03659; phrogs \| 22784 \| 95.7 \| 9.3e-05 \| 1e-08 \| 34.9 \| 54 \| (9, 68) \| 73 \| (15, 68) \| 76 \| NA \| NA; Category: unknown function; p30848 VI\_02556; phrogs \| 17173 \| 95.7 \| 9.5e-05 \| 1.1e-08 \| 35.8 \| 55 \| (9, 67) \| 73 \| (41, 96) \| 98 \| NA \| NA; Category: unknown function; p170583 VI\_00608; phrogs \| 7950 \| 95.6 \| 0.00011 \| 1.3e-08 \| 33.2 \| 27 \| (8, 36) \| 73 \| (10, 36) \| 53 \| transcriptional repressor \| transcriptional repressor; Category: transcription regulation; p141193 VI\_04895; phrogs \| 2951 \| 95.5 \| 0.00012 \| 1.3e-08 \| 35.7 \| 50 \| (8, 60) \| 73 \| (23, 75) \| 90 \| NA \| NA; Category: unknown function; p434966 VI\_08277; phrogs \| 19880 \| 95.5 \| 0.00015 \| 1.6e-08 \| 38.4 \| 47 \| (9, 57) \| 73 \| (84, 130) \| 151 \| transcriptional regulator \| transcriptional regulator; Category: transcription regulation; MG720309\_p38; phrogs \| 33310 \| 95.3 \| 0.00021 \| 2.4e-08 \| 37.4 \| 49 \| (9, 61) \| 73 \| (99, 147) \| 151 \| NA \| NA; Category: unknown function; p240132 VI\_01320; phrogs \| 11967 \| 95.2 \| 0.00025 \| 2.8e-08 \| 34.3 \| 46 \| (8, 56) \| 73 \| (14, 62) \| 87 \| NA \| NA; Category: unknown function; KX898400\_p53; phrogs \| 534 \| 95.1 \| 0.00026 \| 3e-08 \| 34.2 \| 27 \| (9, 37) \| 73 \| (23, 49) \| 87 \| DNA binding protein \| DNA binding protein; Category: DNA, RNA and nucleotide metabolism; p132807 VI\_04689; phrogs \| 9099 \| 94.9 \| 0.00037 \| 4.1e-08 \| 34.8 \| 59 \| (6, 66) \| 73 \| (52, 111) \| 115 \| NA \| NA; Category: unknown function; p72437 VI\_12339; phrogs \| 35 \| 94.9 \| 0.00036 \| 4.4e-08 \| 32.8 \| 27 \| (12, 40) \| 73 \| (17, 43) \| 75 \| transcriptional repressor \| transcriptional repressor; Category: transcription regulation; p323300 VI\_10199; phrogs \| 4494 \| 94.9 \| 0.00041 \| 4.7e-08 \| 34.1 \| 31 \| (8, 40) \| 73 \| (14, 44) \| 88 \| NA \| NA; Category: unknown function; p409661 VI\_06907; phrogs \| 9848 \| 94.7 \| 0.00053 \| 6e-08 \| 33.8 \| 49 \| (11, 62) \| 73 \| (32, 81) \| 89 \| NA \| NA; Category: unknown function; KC595516\_p66; phrogs \| 12527 \| 94.7 \| 0.00057 \| 6.3e-08 \| 34.4 \| 53 \| (10, 65) \| 73 \| (28, 80) \| 130 \| excisionase \| excisionase; Category: integration and excision; p35305 VI\_03424; phrogs \| 17819 \| 94.4 \| 0.00081 \| 9.2e-08 \| 30.9 \| 30 \| (7, 38) \| 73 \| (18, 47) \| 64 \| NA \| NA; Category: unknown function; p411564 VI\_01840; phrogs \| 8013 \| 94.3 \| 0.00087 \| 9.9e-08 \| 38.4 \| 50 \| (8, 60) \| 73 \| (195, 244) \| 259 \| NA \| NA; Category: unknown function; p437881 VI\_04144; phrogs \| 5751 \| 94.3 \| 0.00092 \| 1.1e-07 \| 36.1 \| 27 \| (10, 38) \| 73 \| (23, 49) \| 164 \| terminase large subunit \| terminase large subunit; Category: head and packaging; p273217 VI\_06917; phrogs \| 309 \| 94.2 \| 0.00092 \| 1.1e-07 \| 35.9 \| 26 \| (10, 37) \| 73 \| (22, 47) \| 153 \| terminase small subunit \| terminase small subunit; Category: head and packaging; p301521 VI\_00938; phrogs \| 3218 \| 94.2 \| 0.00099 \| 1.1e-07 \| 35.2 \| 56 \| (10, 67) \| 73 \| (32, 93) \| 152 \| terminase small subunit \| terminase small subunit; Category: head and packaging; p174882 VI\_05927 |
| Top keywords  (threshold 1.00e-03 (evalue)) | **A, tp, Regulator, and, ap, DNA, yes, cp, gp, Engineered** |
| Output files | ../../domain\_architecture/38\_FANPEZAQ\_CDS\_0038\_cath.hhr ../../domain\_architecture/38\_FANPEZAQ\_CDS\_0038\_merged.svg ../../domain\_architecture/38\_FANPEZAQ\_CDS\_0038\_ncbi-cd.hhr ../../domain\_architecture/38\_FANPEZAQ\_CDS\_0038\_pfam.hhr ../../domain\_architecture/38\_FANPEZAQ\_CDS\_0038\_phrogs.hhr |

### Identical protein sequences/structures

#### Search results

|  |  |
| --- | --- |
| Protein sequence databases searched | Pdb, Swissprot, Refseq |
| Identical proteins found | -- |
| Top keywords | -- |
| Output files | -- |

### Similar protein sequences/structures

#### Sequence similarity search results (HHblits)^1^

|  |  |
| --- | --- |
| Sequence databases searched | Uniclust, Pdb70 |
| Results, scheme(s)  (Top layers only, threshold 1.00e-03 (evalue)) | xml version="1.0" encoding="utf-8" standalone="no"?       2024-09-02T21:08:45.791884 image/svg+xml   Matplotlib v3.7.2, https://matplotlib.org/ |
| Results, table(s)  (threshold 1.00e-03 (evalue)) | db \| id \| prob \| evalue \| pvalue \| score \| cols \| query \| query\_len \| template \| template\_len \| name \| description; uniclust \| UniRef100\_A0A0N7LYK3 \| 99.8 \| 4.9e-24 \| 9.7e-30 \| 110.8 \| 68 \| (1, 70) \| 73 \| (8, 75) \| 82 \| Helix-turn-helix domain protein \| Helix-turn-helix domain protein; uniclust \| UniRef100\_A0A0H5Q954 \| 99.8 \| 2.3e-23 \| 4.6e-29 \| 110.0 \| 66 \| (2, 69) \| 73 \| (5, 70) \| 89 \| Helix-turn-helix domain-containing protein \| Helix-turn-helix domain-containing protein; uniclust \| UniRef100\_A0A0F2N5I2 \| 99.8 \| 3.4e-22 \| 6.9e-28 \| 106.0 \| 63 \| (4, 68) \| 73 \| (17, 79) \| 90 \| Helix-turn-helix domain-containing protein \| Helix-turn-helix domain-containing protein; uniclust \| UniRef100\_A0A0B5I6H3 \| 99.8 \| 6.1e-22 \| 1.2e-27 \| 100.7 \| 67 \| (2, 70) \| 73 \| (1, 67) \| 72 \| Helix-turn-helix domain-containing protein \| Helix-turn-helix domain-containing protein; uniclust \| UniRef100\_A0A1F8XFD9 \| 99.8 \| 8.2e-22 \| 1.6e-27 \| 103.9 \| 61 \| (6, 68) \| 73 \| (14, 74) \| 88 \| Helix-turn-helix domain-containing protein \| Helix-turn-helix domain-containing protein; uniclust \| UniRef100\_A0A0M4QCQ6 \| 99.8 \| 8.9e-22 \| 1.7e-27 \| 105.1 \| 63 \| (4, 68) \| 73 \| (29, 91) \| 99 \| Excisionase \| Excisionase; uniclust \| UniRef100\_A0A059G6X6 \| 99.7 \| 1.7e-21 \| 3.3e-27 \| 102.5 \| 67 \| (1, 69) \| 73 \| (15, 81) \| 88 \| Helix-turn-helix domain-containing protein \| Helix-turn-helix domain-containing protein; uniclust \| UniRef100\_A0A0Q8DCD6 \| 99.7 \| 2.1e-21 \| 4.1e-27 \| 98.5 \| 65 \| (2, 68) \| 73 \| (1, 65) \| 72 \| Helix-turn-helix domain-containing protein \| Helix-turn-helix domain-containing protein; uniclust \| UniRef100\_A0A072MSX1 \| 99.7 \| 2.1e-21 \| 4.3e-27 \| 103.1 \| 62 \| (6, 69) \| 73 \| (12, 77) \| 89 \| Helix-turn-helix domain-containing protein \| Helix-turn-helix domain-containing protein; uniclust \| UniRef100\_A0A059G7C8 \| 99.7 \| 2.2e-21 \| 4.5e-27 \| 104.1 \| 63 \| (6, 70) \| 73 \| (11, 73) \| 95 \| Helix-turn-helix domain-containing protein \| Helix-turn-helix domain-containing protein; uniclust \| UniRef100\_A0A087AQ78 \| 99.7 \| 2.4e-21 \| 4.9e-27 \| 106.2 \| 64 \| (4, 69) \| 73 \| (32, 95) \| 110 \| Helix-turn-helix domain-containing protein \| Helix-turn-helix domain-containing protein; uniclust \| UniRef100\_A0A099FD25 \| 99.7 \| 2.8e-21 \| 5.6e-27 \| 105.0 \| 66 \| (2, 69) \| 73 \| (11, 76) \| 104 \| Helix-turn-helix domain-containing protein \| Helix-turn-helix domain-containing protein; uniclust \| UniRef100\_A0A0C9PRY0 \| 99.7 \| 2.9e-21 \| 5.9e-27 \| 102.3 \| 63 \| (2, 67) \| 73 \| (7, 69) \| 89 \| Phosphotransferase system mannitol/fructose-specific IIA domain \| Phosphotransferase system mannitol/fructose-specific IIA domain; uniclust \| UniRef100\_A0A0F5PY42 \| 99.7 \| 3.3e-21 \| 6.7e-27 \| 105.4 \| 64 \| (5, 70) \| 73 \| (21, 84) \| 106 \| Transcriptional regulator, AlpA family \| Transcriptional regulator, AlpA family; uniclust \| UniRef100\_A0A0J5S0Y2 \| 99.7 \| 3.4e-21 \| 6.9e-27 \| 100.8 \| 65 \| (2, 68) \| 73 \| (8, 72) \| 82 \| Helix-turn-helix domain-containing protein \| Helix-turn-helix domain-containing protein; uniclust \| UniRef100\_A0A0F9Z934 \| 99.7 \| 5.9e-21 \| 1.2e-26 \| 99.4 \| 64 \| (4, 70) \| 73 \| (10, 73) \| 80 \| Helix-turn-helix domain-containing protein \| Helix-turn-helix domain-containing protein; uniclust \| UniRef100\_A0A031GGX7 \| 99.7 \| 6.2e-21 \| 1.2e-26 \| 99.1 \| 62 \| (5, 68) \| 73 \| (10, 71) \| 80 \| Helix-turn-helix domain protein \| Helix-turn-helix domain protein; uniclust \| UniRef100\_A0A0K3BH05 \| 99.7 \| 8.9e-21 \| 1.8e-26 \| 99.2 \| 65 \| (3, 69) \| 73 \| (14, 78) \| 84 \| Helix-turn-helix domain-containing protein \| Helix-turn-helix domain-containing protein; uniclust \| UniRef100\_A0A0H1R5M7 \| 99.7 \| 9.5e-21 \| 1.9e-26 \| 99.9 \| 65 \| (3, 69) \| 73 \| (11, 75) \| 89 \| Helix-turn-helix domain-containing protein \| Helix-turn-helix domain-containing protein; uniclust \| UniRef100\_A0A0Q8Y888 \| 99.7 \| 1.1e-20 \| 2.1e-26 \| 99.7 \| 59 \| (6, 66) \| 73 \| (15, 73) \| 89 \| Helix-turn-helix domain-containing protein \| Helix-turn-helix domain-containing protein; uniclust \| UniRef100\_A0A013WJ46 \| 99.7 \| 1.1e-20 \| 2.2e-26 \| 101.8 \| 68 \| (1, 70) \| 73 \| (12, 80) \| 101 \| Helix-turn-helix domain-containing protein \| Helix-turn-helix domain-containing protein; uniclust \| UniRef100\_A0A1E7YNR3 \| 99.7 \| 1.5e-20 \| 3e-26 \| 99.6 \| 63 \| (2, 67) \| 73 \| (12, 74) \| 93 \| DNA-binding protein \| DNA-binding protein; uniclust \| UniRef100\_A0A0S7XKK0 \| 99.7 \| 1.6e-20 \| 3.1e-26 \| 97.5 \| 64 \| (2, 68) \| 73 \| (1, 64) \| 85 \| Helix-turn-helix domain-containing protein \| Helix-turn-helix domain-containing protein; uniclust \| UniRef100\_A0A0H1R6E4 \| 99.7 \| 1.8e-20 \| 3.5e-26 \| 99.1 \| 62 \| (7, 70) \| 73 \| (15, 76) \| 91 \| Helix-turn-helix domain-containing protein \| Helix-turn-helix domain-containing protein; uniclust \| UniRef100\_A0A0W8GA41 \| 99.7 \| 1.8e-20 \| 3.5e-26 \| 99.2 \| 66 \| (3, 70) \| 73 \| (12, 77) \| 91 \| Helix-turn-helix domain-containing protein \| Helix-turn-helix domain-containing protein; uniclust \| UniRef100\_A0A0S8E603 \| 99.7 \| 1.8e-20 \| 3.7e-26 \| 99.0 \| 61 \| (3, 66) \| 73 \| (9, 69) \| 88 \| Helix-turn-helix domain-containing protein \| Helix-turn-helix domain-containing protein; uniclust \| UniRef100\_A0A7X6PMZ9 \| 99.7 \| 2.2e-20 \| 4.2e-26 \| 94.8 \| 65 \| (2, 68) \| 73 \| (5, 70) \| 73 \| Helix-turn-helix domain-containing protein \| Helix-turn-helix domain-containing protein; uniclust \| UniRef100\_A0A1F1ED30 \| 99.7 \| 2.3e-20 \| 4.6e-26 \| 97.2 \| 66 \| (2, 69) \| 73 \| (10, 75) \| 80 \| Helix-turn-helix domain-containing protein \| Helix-turn-helix domain-containing protein; uniclust \| UniRef100\_A0A0D8HFS2 \| 99.7 \| 2.8e-20 \| 5.6e-26 \| 99.7 \| 64 \| (2, 68) \| 73 \| (13, 76) \| 95 \| Helix-turn-helix domain protein \| Helix-turn-helix domain protein; uniclust \| UniRef100\_A0A059G1X5 \| 99.7 \| 2.8e-20 \| 5.7e-26 \| 97.9 \| 68 \| (1, 70) \| 73 \| (5, 73) \| 84 \| Helix-turn-helix domain-containing protein \| Helix-turn-helix domain-containing protein; uniclust \| UniRef100\_A0A0D6JKC0 \| 99.7 \| 3.1e-20 \| 6.1e-26 \| 98.4 \| 60 \| (9, 70) \| 73 \| (24, 84) \| 91 \| Helix-turn-helix domain-containing protein \| Helix-turn-helix domain-containing protein; uniclust \| UniRef100\_A0A126NYS1 \| 99.7 \| 3.2e-20 \| 6.5e-26 \| 97.8 \| 67 \| (2, 70) \| 73 \| (7, 74) \| 85 \| Helix-turn-helix domain-containing protein \| Helix-turn-helix domain-containing protein; uniclust \| UniRef100\_A0A077AYW7 \| 99.7 \| 3.4e-20 \| 6.6e-26 \| 95.4 \| 65 \| (3, 69) \| 73 \| (6, 70) \| 76 \| Helix-turn-helix domain-containing protein \| Helix-turn-helix domain-containing protein; uniclust \| UniRef100\_A0A0K6HIE1 \| 99.7 \| 3.4e-20 \| 6.9e-26 \| 101.9 \| 59 \| (5, 65) \| 73 \| (24, 82) \| 112 \| Helix-turn-helix domain \| Helix-turn-helix domain; uniclust \| UniRef100\_A0A094ZE53 \| 99.7 \| 4.2e-20 \| 8.3e-26 \| 97.4 \| 63 \| (6, 70) \| 73 \| (11, 74) \| 87 \| AlpA family phage regulatory protein \| AlpA family phage regulatory protein; uniclust \| UniRef100\_A0A062WXZ1 \| 99.7 \| 4.2e-20 \| 8.3e-26 \| 101.0 \| 62 \| (6, 69) \| 73 \| (31, 92) \| 110 \| Transcriptional regulator, AlpA family \| Transcriptional regulator, AlpA family; uniclust \| UniRef100\_A0A0A0BTB2 \| 99.7 \| 4.2e-20 \| 8.4e-26 \| 97.4 \| 57 \| (9, 67) \| 73 \| (18, 74) \| 85 \| Excisionase \| Excisionase; uniclust \| UniRef100\_A0A059DTE2 \| 99.7 \| 5.1e-20 \| 1.1e-25 \| 100.5 \| 61 \| (3, 66) \| 73 \| (14, 74) \| 103 \| Helix-turn-helix domain-containing protein \| Helix-turn-helix domain-containing protein; uniclust \| UniRef100\_A0A022MTG3 \| 99.7 \| 5.2e-20 \| 1.1e-25 \| 97.5 \| 57 \| (10, 68) \| 73 \| (18, 74) \| 86 \| DNA-binding protein \| DNA-binding protein; uniclust \| UniRef100\_A0A0J6SUB7 \| 99.7 \| 5.6e-20 \| 1.1e-25 \| 99.1 \| 61 \| (8, 70) \| 73 \| (28, 88) \| 99 \| Helix-turn-helix domain-containing protein \| Helix-turn-helix domain-containing protein; uniclust \| UniRef100\_A0A076JIU9 \| 99.7 \| 6.3e-20 \| 1.2e-25 \| 98.9 \| 60 \| (9, 70) \| 73 \| (26, 85) \| 102 \| Putative transcriptional regulator \| Putative transcriptional regulator; uniclust \| UniRef100\_A0A0Q0HYW1 \| 99.7 \| 6.3e-20 \| 1.3e-25 \| 96.4 \| 64 \| (3, 68) \| 73 \| (5, 69) \| 85 \| Helix-turn-helix domain protein \| Helix-turn-helix domain protein; uniclust \| UniRef100\_A0A0B5ERS0 \| 99.7 \| 6.6e-20 \| 1.3e-25 \| 98.2 \| 59 \| (8, 68) \| 73 \| (31, 89) \| 98 \| DNA-binding protein \| DNA-binding protein; uniclust \| UniRef100\_A0A0G3GU35 \| 99.7 \| 6.9e-20 \| 1.3e-25 \| 92.4 \| 65 \| (3, 69) \| 73 \| (2, 66) \| 70 \| Transcriptional regulator, AlpA family \| Transcriptional regulator, AlpA family; uniclust \| UniRef100\_A0A0K2GIG6 \| 99.7 \| 7.8e-20 \| 1.5e-25 \| 98.7 \| 63 \| (3, 68) \| 73 \| (12, 74) \| 102 \| Helix-turn-helix domain-containing protein \| Helix-turn-helix domain-containing protein; uniclust \| UniRef100\_A0A099JLS6 \| 99.7 \| 8.7e-20 \| 1.7e-25 \| 97.3 \| 60 \| (8, 69) \| 73 \| (17, 77) \| 92 \| Prophage regulatory protein \| Prophage regulatory protein; uniclust \| UniRef100\_A0A0A6UQH0 \| 99.7 \| 9.6e-20 \| 1.9e-25 \| 93.5 \| 66 \| (2, 69) \| 73 \| (10, 75) \| 79 \| Excisionase \| Excisionase; uniclust \| UniRef100\_A0A081HUQ7 \| 99.7 \| 1.4e-19 \| 2.8e-25 \| 95.6 \| 62 \| (4, 67) \| 73 \| (12, 73) \| 88 \| Helix-turn-helix domain-containing protein \| Helix-turn-helix domain-containing protein; uniclust \| UniRef100\_A0A094WFN5 \| 99.7 \| 1.4e-19 \| 2.8e-25 \| 98.3 \| 64 \| (5, 70) \| 73 \| (18, 81) \| 106 \| DNA-binding protein \| DNA-binding protein; uniclust \| UniRef100\_A0A1L7AKC8 \| 99.7 \| 1.4e-19 \| 2.8e-25 \| 98.0 \| 61 \| (4, 66) \| 73 \| (21, 82) \| 102 \| Helix-turn-helix domain-containing protein \| Helix-turn-helix domain-containing protein; uniclust \| UniRef100\_A0A1F5RZJ7 \| 99.7 \| 1.4e-19 \| 2.9e-25 \| 95.5 \| 65 \| (1, 68) \| 73 \| (5, 69) \| 87 \| Helix-turn-helix domain-containing protein \| Helix-turn-helix domain-containing protein; uniclust \| UniRef100\_A0A3A5AND8 \| 99.7 \| 1.6e-19 \| 3.1e-25 \| 92.9 \| 65 \| (2, 69) \| 73 \| (2, 66) \| 76 \| DNA-binding protein \| DNA-binding protein; uniclust \| UniRef100\_A0A1M4LEU1 \| 99.7 \| 1.7e-19 \| 3.3e-25 \| 94.6 \| 62 \| (6, 69) \| 73 \| (14, 75) \| 86 \| Helix-turn-helix domain-containing protein \| Helix-turn-helix domain-containing protein; uniclust \| UniRef100\_A0A0L6JWJ1 \| 99.7 \| 1.7e-19 \| 3.3e-25 \| 94.8 \| 65 \| (1, 68) \| 73 \| (6, 70) \| 85 \| DNA binding domain protein, excisionase family \| DNA binding domain protein, excisionase family; uniclust \| UniRef100\_A0A062VY92 \| 99.7 \| 2e-19 \| 3.8e-25 \| 92.5 \| 59 \| (9, 69) \| 73 \| (15, 73) \| 79 \| Helix-turn-helix domain-containing protein \| Helix-turn-helix domain-containing protein; uniclust \| UniRef100\_A0A0F9NKP1 \| 99.7 \| 2e-19 \| 4e-25 \| 92.9 \| 64 \| (2, 68) \| 73 \| (3, 66) \| 75 \| Helix-turn-helix domain-containing protein \| Helix-turn-helix domain-containing protein; uniclust \| UniRef100\_A0A0B9AT96 \| 99.7 \| 2.2e-19 \| 4.3e-25 \| 91.3 \| 65 \| (1, 68) \| 73 \| (1, 65) \| 69 \| Helix-turn-helix domain-containing protein \| Helix-turn-helix domain-containing protein; uniclust \| UniRef100\_A0A090MQQ0 \| 99.6 \| 2.3e-19 \| 4.6e-25 \| 95.2 \| 66 \| (3, 70) \| 73 \| (9, 74) \| 89 \| Helix-turn-helix domain protein \| Helix-turn-helix domain protein; uniclust \| UniRef100\_A0A1A9CVS8 \| 99.6 \| 2.4e-19 \| 4.6e-25 \| 93.8 \| 66 \| (1, 68) \| 73 \| (9, 74) \| 85 \| Helix-turn-helix domain-containing protein \| Helix-turn-helix domain-containing protein; uniclust \| UniRef100\_A0A0A7I9V8 \| 99.6 \| 2.4e-19 \| 4.8e-25 \| 96.5 \| 65 \| (3, 69) \| 73 \| (14, 78) \| 98 \| Helix-turn-helix domain-containing protein \| Helix-turn-helix domain-containing protein; uniclust \| UniRef100\_A0A0K0N7A4 \| 99.6 \| 2.6e-19 \| 5.3e-25 \| 95.6 \| 61 \| (7, 69) \| 73 \| (26, 86) \| 94 \| Helix-turn-helix domain-containing protein \| Helix-turn-helix domain-containing protein; uniclust \| UniRef100\_A0A059E4I3 \| 99.6 \| 2.9e-19 \| 5.7e-25 \| 95.8 \| 65 \| (4, 70) \| 73 \| (21, 85) \| 98 \| Helix-turn-helix domain-containing protein \| Helix-turn-helix domain-containing protein; uniclust \| UniRef100\_A0A0A1CWK4 \| 99.6 \| 2.9e-19 \| 5.7e-25 \| 91.0 \| 64 \| (1, 66) \| 73 \| (1, 64) \| 72 \| Helix-turn-helix domain-containing protein \| Helix-turn-helix domain-containing protein; uniclust \| UniRef100\_A0A066PHX5 \| 99.6 \| 3e-19 \| 5.9e-25 \| 96.8 \| 66 \| (3, 70) \| 73 \| (18, 83) \| 107 \| Helix-turn-helix domain-containing protein \| Helix-turn-helix domain-containing protein; uniclust \| UniRef100\_A0A089LM03 \| 99.6 \| 3.3e-19 \| 6.6e-25 \| 92.1 \| 63 \| (3, 68) \| 73 \| (1, 63) \| 76 \| Excisionase \| Excisionase; uniclust \| UniRef100\_A0A1S9CZ14 \| 99.6 \| 3.5e-19 \| 6.8e-25 \| 89.9 \| 63 \| (5, 69) \| 73 \| (4, 66) \| 70 \| Helix-turn-helix domain-containing protein \| Helix-turn-helix domain-containing protein; uniclust \| UniRef100\_A0A0M0BEZ1 \| 99.6 \| 3.5e-19 \| 6.9e-25 \| 93.2 \| 60 \| (1, 63) \| 73 \| (1, 60) \| 82 \| Helix-turn-helix domain-containing protein \| Helix-turn-helix domain-containing protein; uniclust \| UniRef100\_A0A068F4K0 \| 99.6 \| 3.4e-19 \| 6.9e-25 \| 93.0 \| 65 \| (1, 67) \| 73 \| (4, 69) \| 79 \| Excise \| Excise; uniclust \| UniRef100\_A0A1U7GLT6 \| 99.6 \| 3.6e-19 \| 7.1e-25 \| 92.4 \| 62 \| (5, 68) \| 73 \| (8, 69) \| 80 \| Helix-turn-helix domain-containing protein \| Helix-turn-helix domain-containing protein; uniclust \| UniRef100\_A0A0M2HKS0 \| 99.6 \| 3.9e-19 \| 7.5e-25 \| 96.4 \| 66 \| (2, 69) \| 73 \| (7, 72) \| 107 \| Helix-turn-helix domain protein \| Helix-turn-helix domain protein; uniclust \| UniRef100\_A0A077M4I0 \| 99.6 \| 4e-19 \| 8e-25 \| 97.0 \| 57 \| (9, 67) \| 73 \| (25, 81) \| 107 \| Helix-turn-helix domain-containing protein \| Helix-turn-helix domain-containing protein; uniclust \| UniRef100\_A0A061LXH0 \| 99.6 \| 4.3e-19 \| 8.6e-25 \| 94.6 \| 61 \| (8, 70) \| 73 \| (22, 82) \| 91 \| Helix-turn-helix domain-containing protein \| Helix-turn-helix domain-containing protein; uniclust \| UniRef100\_A0A1F6T9P4 \| 99.6 \| 4.4e-19 \| 8.6e-25 \| 94.3 \| 61 \| (8, 70) \| 73 \| (17, 77) \| 91 \| Helix-turn-helix domain-containing protein \| Helix-turn-helix domain-containing protein; uniclust \| UniRef100\_A0A087BZX1 \| 99.6 \| 4.7e-19 \| 9.3e-25 \| 92.0 \| 59 \| (9, 69) \| 73 \| (16, 74) \| 80 \| DNA binding domain, excisionase family \| DNA binding domain, excisionase family; uniclust \| UniRef100\_A0A068Z4E0 \| 99.6 \| 5.5e-19 \| 1.1e-24 \| 97.1 \| 61 \| (7, 69) \| 73 \| (33, 94) \| 108 \| Helix-turn-helix domain-containing protein \| Helix-turn-helix domain-containing protein; uniclust \| UniRef100\_A0A133L1R7 \| 99.6 \| 5.7e-19 \| 1.1e-24 \| 92.7 \| 62 \| (6, 69) \| 73 \| (14, 76) \| 83 \| Transcriptional regulator \| Transcriptional regulator; uniclust \| UniRef100\_A0A0H1RFW3 \| 99.6 \| 6e-19 \| 1.2e-24 \| 92.2 \| 62 \| (6, 69) \| 73 \| (14, 75) \| 80 \| Helix-turn-helix domain-containing protein \| Helix-turn-helix domain-containing protein; uniclust \| UniRef100\_A0A1F4N4R1 \| 99.6 \| 6.3e-19 \| 1.3e-24 \| 92.2 \| 63 \| (3, 68) \| 73 \| (1, 63) \| 80 \| Helix-turn-helix domain-containing protein \| Helix-turn-helix domain-containing protein; uniclust \| UniRef100\_A0A094S3N1 \| 99.6 \| 6.4e-19 \| 1.3e-24 \| 93.4 \| 66 \| (2, 69) \| 73 \| (8, 73) \| 87 \| AlpA family transcriptional regulator \| AlpA family transcriptional regulator; uniclust \| UniRef100\_A0A061LYD6 \| 99.6 \| 6.9e-19 \| 1.4e-24 \| 95.5 \| 64 \| (4, 69) \| 73 \| (29, 92) \| 106 \| Ethanolamine utilization protein EutA \| Ethanolamine utilization protein EutA; uniclust \| UniRef100\_A0A0B2Y500 \| 99.6 \| 7.1e-19 \| 1.4e-24 \| 92.2 \| 68 \| (1, 70) \| 73 \| (4, 71) \| 85 \| DNA-binding protein \| DNA-binding protein; uniclust \| UniRef100\_A0A0K8PHS6 \| 99.6 \| 7.4e-19 \| 1.5e-24 \| 96.1 \| 58 \| (8, 67) \| 73 \| (28, 86) \| 109 \| Helix-turn-helix domain-containing protein \| Helix-turn-helix domain-containing protein; uniclust \| UniRef100\_A0A0S4LPK6 \| 99.6 \| 8.7e-19 \| 1.7e-24 \| 94.4 \| 62 \| (3, 67) \| 73 \| (17, 78) \| 101 \| Putative DNA binding domain, excisionase family \| Putative DNA binding domain, excisionase family; uniclust \| UniRef100\_A0A011UV86 \| 99.6 \| 8.9e-19 \| 1.8e-24 \| 97.1 \| 61 \| (7, 69) \| 73 \| (27, 87) \| 116 \| Transcriptional regulator \| Transcriptional regulator; uniclust \| UniRef100\_A0A069E1R7 \| 99.6 \| 1e-18 \| 2e-24 \| 92.0 \| 65 \| (4, 70) \| 73 \| (14, 78) \| 82 \| Helix-turn-helix domain-containing protein \| Helix-turn-helix domain-containing protein; uniclust \| UniRef100\_A0A1Y0BBP6 \| 99.6 \| 1.1e-18 \| 2.1e-24 \| 90.2 \| 61 \| (5, 68) \| 73 \| (2, 62) \| 75 \| B42 \| B42; uniclust \| UniRef100\_A0A1A3ELP5 \| 99.6 \| 1.2e-18 \| 2.3e-24 \| 90.9 \| 65 \| (4, 70) \| 73 \| (11, 75) \| 82 \| Helix-turn-helix domain-containing protein \| Helix-turn-helix domain-containing protein; uniclust \| UniRef100\_A0A090REP0 \| 99.6 \| 1.2e-18 \| 2.4e-24 \| 89.2 \| 63 \| (5, 69) \| 73 \| (2, 65) \| 72 \| Transcriptional regulator \| Transcriptional regulator; uniclust \| UniRef100\_A0A099KUC7 \| 99.6 \| 1.2e-18 \| 2.4e-24 \| 92.6 \| 65 \| (1, 67) \| 73 \| (2, 67) \| 89 \| Helix-turn-helix domain-containing protein \| Helix-turn-helix domain-containing protein; uniclust \| UniRef100\_A0A0L1KD87 \| 99.6 \| 1.3e-18 \| 2.6e-24 \| 92.3 \| 62 \| (6, 69) \| 73 \| (18, 79) \| 93 \| Helix-turn-helix domain-containing protein \| Helix-turn-helix domain-containing protein; uniclust \| UniRef100\_A0A0S8JD43 \| 99.6 \| 1.4e-18 \| 2.8e-24 \| 91.4 \| 56 \| (2, 60) \| 73 \| (6, 61) \| 83 \| Helix-turn-helix domain-containing protein \| Helix-turn-helix domain-containing protein; uniclust \| UniRef100\_A0A1G9V0Q2 \| 99.6 \| 1.5e-18 \| 2.9e-24 \| 91.2 \| 64 \| (4, 69) \| 73 \| (8, 71) \| 82 \| Helix-turn-helix domain-containing protein \| Helix-turn-helix domain-containing protein; uniclust \| UniRef100\_A0A444PYM9 \| 99.6 \| 1.5e-18 \| 2.9e-24 \| 89.1 \| 63 \| (2, 66) \| 73 \| (1, 63) \| 72 \| DNA-binding protein \| DNA-binding protein; uniclust \| UniRef100\_A0A014NQA7 \| 99.6 \| 1.6e-18 \| 3.1e-24 \| 91.1 \| 57 \| (8, 66) \| 73 \| (18, 74) \| 89 \| Helix-turn-helix domain-containing protein \| Helix-turn-helix domain-containing protein; uniclust \| UniRef100\_A0A0N8HY41 \| 99.6 \| 1.7e-18 \| 3.4e-24 \| 90.4 \| 60 \| (8, 69) \| 73 \| (18, 77) \| 80 \| Helix-turn-helix domain-containing protein \| Helix-turn-helix domain-containing protein; uniclust \| UniRef100\_A0A0Q2LK60 \| 99.6 \| 1.8e-18 \| 3.4e-24 \| 88.6 \| 64 \| (5, 70) \| 73 \| (3, 66) \| 74 \| Helix-turn-helix domain-containing protein \| Helix-turn-helix domain-containing protein; uniclust \| UniRef100\_A0A083XX69 \| 99.6 \| 1.8e-18 \| 3.5e-24 \| 91.5 \| 61 \| (7, 69) \| 73 \| (12, 72) \| 88 \| Helix-turn-helix domain-containing protein \| Helix-turn-helix domain-containing protein; uniclust \| UniRef100\_A0A176YVV7 \| 99.6 \| 1.9e-18 \| 3.7e-24 \| 89.1 \| 64 \| (4, 69) \| 73 \| (6, 69) \| 77 \| HTH merR-type domain-containing protein \| HTH merR-type domain-containing protein; uniclust \| UniRef100\_A0A1F9V6F5 \| 99.6 \| 1.9e-18 \| 3.7e-24 \| 93.0 \| 60 \| (6, 68) \| 73 \| (19, 78) \| 98 \| Helix-turn-helix domain-containing protein \| Helix-turn-helix domain-containing protein; uniclust \| UniRef100\_A0A2T3IMR8 \| 99.6 \| 2.1e-18 \| 4.1e-24 \| 89.9 \| 65 \| (2, 68) \| 73 \| (9, 74) \| 80 \| AlpA family phage regulatory protein \| AlpA family phage regulatory protein; uniclust \| UniRef100\_A0A059DSQ7 \| 99.6 \| 2.3e-18 \| 4.5e-24 \| 92.1 \| 62 \| (6, 69) \| 73 \| (17, 78) \| 98 \| Helix-turn-helix domain-containing protein \| Helix-turn-helix domain-containing protein; uniclust \| UniRef100\_A0A022LRD1 \| 99.6 \| 2.3e-18 \| 4.5e-24 \| 92.4 \| 63 \| (4, 68) \| 73 \| (26, 88) \| 97 \| Helix-turn-helix domain-containing protein \| Helix-turn-helix domain-containing protein; uniclust \| UniRef100\_A0A0S7XS04 \| 99.6 \| 2.4e-18 \| 4.8e-24 \| 88.3 \| 65 \| (3, 70) \| 73 \| (2, 66) \| 73 \| Helix-turn-helix domain-containing protein \| Helix-turn-helix domain-containing protein; uniclust \| UniRef100\_A0A0A8B4D0 \| 99.6 \| 2.6e-18 \| 5e-24 \| 94.9 \| 60 \| (7, 68) \| 73 \| (35, 95) \| 120 \| Transcriptional regulator \| Transcriptional regulator; uniclust \| UniRef100\_A0A0R0LNK5 \| 99.6 \| 2.6e-18 \| 5.1e-24 \| 90.8 \| 67 \| (1, 69) \| 73 \| (1, 67) \| 87 \| Helix-turn-helix domain-containing protein \| Helix-turn-helix domain-containing protein; uniclust \| UniRef100\_A0A086PDB8 \| 99.6 \| 2.6e-18 \| 5.2e-24 \| 92.7 \| 58 \| (8, 67) \| 73 \| (27, 84) \| 99 \| Helix-turn-helix domain-containing protein \| Helix-turn-helix domain-containing protein; uniclust \| UniRef100\_A0A099KYA7 \| 99.6 \| 2.6e-18 \| 5.3e-24 \| 91.2 \| 65 \| (2, 68) \| 73 \| (12, 77) \| 86 \| Phage transcriptional regulator, AlpA \| Phage transcriptional regulator, AlpA; uniclust \| UniRef100\_A0A1H6A7S1 \| 99.6 \| 3.5e-18 \| 6.8e-24 \| 88.6 \| 59 \| (9, 69) \| 73 \| (16, 74) \| 82 \| Helix-turn-helix domain-containing protein \| Helix-turn-helix domain-containing protein; uniclust \| UniRef100\_A0A078BLR7 \| 99.6 \| 3.7e-18 \| 7.3e-24 \| 90.5 \| 63 \| (5, 69) \| 73 \| (22, 84) \| 88 \| Putative Bacteriophage related protein \| Putative Bacteriophage related protein; uniclust \| UniRef100\_A0A069S043 \| 99.6 \| 4.4e-18 \| 8.7e-24 \| 89.2 \| 65 \| (2, 69) \| 73 \| (16, 80) \| 83 \| MerR HTH regulatory family protein \| MerR HTH regulatory family protein; uniclust \| UniRef100\_A0A0F9NA78 \| 99.6 \| 4.4e-18 \| 8.8e-24 \| 88.9 \| 65 \| (1, 68) \| 73 \| (2, 66) \| 79 \| Helix-turn-helix domain-containing protein \| Helix-turn-helix domain-containing protein; uniclust \| UniRef100\_A0A081FVL7 \| 99.6 \| 4.5e-18 \| 8.8e-24 \| 91.3 \| 62 \| (3, 67) \| 73 \| (17, 78) \| 96 \| Putative phage protein \| Putative phage protein; uniclust \| UniRef100\_A0A0B5FUY6 \| 99.6 \| 5.1e-18 \| 9.8e-24 \| 89.5 \| 60 \| (3, 65) \| 73 \| (7, 66) \| 89 \| DNA-binding protein \| DNA-binding protein; uniclust \| UniRef100\_A0A011TBA6 \| 99.6 \| 5.2e-18 \| 1e-23 \| 91.7 \| 65 \| (3, 69) \| 73 \| (13, 77) \| 102 \| Helix-turn-helix domain-containing protein \| Helix-turn-helix domain-containing protein; uniclust \| UniRef100\_A0A357ZGF1 \| 99.6 \| 5.3e-18 \| 1e-23 \| 87.2 \| 62 \| (7, 70) \| 73 \| (2, 64) \| 76 \| Helix-turn-helix domain-containing protein \| Helix-turn-helix domain-containing protein; uniclust \| UniRef100\_A0A132H9Y5 \| 99.6 \| 5.8e-18 \| 1.1e-23 \| 91.7 \| 62 \| (3, 67) \| 73 \| (22, 83) \| 110 \| Helix-turn-helix domain-containing protein \| Helix-turn-helix domain-containing protein; uniclust \| UniRef100\_A0A1G7WEL0 \| 99.6 \| 6.1e-18 \| 1.2e-23 \| 85.7 \| 61 \| (4, 66) \| 73 \| (6, 67) \| 69 \| Helix-turn-helix domain-containing protein \| Helix-turn-helix domain-containing protein; uniclust \| UniRef100\_A0A0R3MKB8 \| 99.6 \| 6.2e-18 \| 1.2e-23 \| 90.2 \| 60 \| (9, 70) \| 73 \| (16, 75) \| 94 \| Helix-turn-helix domain-containing protein \| Helix-turn-helix domain-containing protein; uniclust \| UniRef100\_A0A0F5MUR4 \| 99.6 \| 6.2e-18 \| 1.2e-23 \| 90.3 \| 60 \| (6, 67) \| 73 \| (21, 80) \| 93 \| DNA-binding protein \| DNA-binding protein; uniclust \| UniRef100\_A0A0J1FJT4 \| 99.6 \| 6.2e-18 \| 1.2e-23 \| 85.0 \| 56 \| (9, 68) \| 73 \| (3, 58) \| 64 \| Helix-turn-helix domain protein \| Helix-turn-helix domain protein; uniclust \| UniRef100\_A0A0C9PVN7 \| 99.6 \| 6.2e-18 \| 1.3e-23 \| 88.0 \| 66 \| (1, 69) \| 73 \| (3, 68) \| 77 \| Phosphotransferase system mannitol/fructose-specific IIA domain \| Phosphotransferase system mannitol/fructose-specific IIA domain; uniclust \| UniRef100\_A0A1X0ZRW0 \| 99.6 \| 6.8e-18 \| 1.3e-23 \| 84.2 \| 60 \| (8, 69) \| 73 \| (3, 63) \| 68 \| Helix-turn-helix domain-containing protein \| Helix-turn-helix domain-containing protein; uniclust \| UniRef100\_A0A0B3S532 \| 99.6 \| 6.7e-18 \| 1.3e-23 \| 91.6 \| 62 \| (7, 70) \| 73 \| (34, 96) \| 102 \| Phage regulator-like protein \| Phage regulator-like protein; uniclust \| UniRef100\_A0A0J6YTM7 \| 99.6 \| 7e-18 \| 1.4e-23 \| 87.2 \| 65 \| (4, 70) \| 73 \| (1, 66) \| 77 \| Helix-turn-helix domain protein \| Helix-turn-helix domain protein; uniclust \| UniRef100\_A0A075NWM2 \| 99.6 \| 7.1e-18 \| 1.4e-23 \| 93.8 \| 62 \| (5, 68) \| 73 \| (28, 95) \| 123 \| Helix-turn-helix domain-containing protein \| Helix-turn-helix domain-containing protein; uniclust \| UniRef100\_A0A259S763 \| 99.6 \| 7.3e-18 \| 1.4e-23 \| 87.5 \| 63 \| (2, 67) \| 73 \| (6, 68) \| 79 \| Helix-turn-helix domain-containing protein \| Helix-turn-helix domain-containing protein; uniclust \| UniRef100\_A0A380TD81 \| 99.6 \| 7.2e-18 \| 1.4e-23 \| 92.5 \| 60 \| (10, 71) \| 73 \| (34, 94) \| 106 \| Transcriptional regulator \| Transcriptional regulator; uniclust \| UniRef100\_A0A060UPH2 \| 99.6 \| 7.4e-18 \| 1.5e-23 \| 91.6 \| 60 \| (9, 70) \| 73 \| (36, 96) \| 103 \| Helix-turn-helix domain-containing protein \| Helix-turn-helix domain-containing protein; uniclust \| UniRef100\_A0A0H3LYR5 \| 99.6 \| 7.6e-18 \| 1.5e-23 \| 87.7 \| 59 \| (8, 68) \| 73 \| (18, 76) \| 80 \| Helix-turn-helix domain-containing protein \| Helix-turn-helix domain-containing protein; uniclust \| UniRef100\_A0A087F9Y2 \| 99.6 \| 7.6e-18 \| 1.5e-23 \| 90.6 \| 61 \| (7, 69) \| 73 \| (26, 87) \| 95 \| Helix-turn-helix domain-containing protein \| Helix-turn-helix domain-containing protein; uniclust \| UniRef100\_A0A0F7N9N1 \| 99.6 \| 8e-18 \| 1.5e-23 \| 90.9 \| 58 \| (8, 67) \| 73 \| (25, 82) \| 104 \| Helix-turn-helix domain-containing protein \| Helix-turn-helix domain-containing protein; uniclust \| UniRef100\_A0A1E4NHF0 \| 99.6 \| 7.9e-18 \| 1.6e-23 \| 89.6 \| 61 \| (7, 69) \| 73 \| (26, 86) \| 89 \| Helix-turn-helix domain-containing protein \| Helix-turn-helix domain-containing protein; uniclust \| UniRef100\_A0A0S7X7T0 \| 99.6 \| 8.1e-18 \| 1.6e-23 \| 87.4 \| 62 \| (4, 68) \| 73 \| (6, 67) \| 78 \| Helix-turn-helix domain-containing protein \| Helix-turn-helix domain-containing protein; uniclust \| UniRef100\_A0A010YHV7 \| 99.6 \| 8.2e-18 \| 1.6e-23 \| 89.5 \| 63 \| (5, 69) \| 73 \| (24, 87) \| 92 \| DNA-binding protein, excisionase family \| DNA-binding protein, excisionase family; uniclust \| UniRef100\_A0A0M9EFS7 \| 99.6 \| 8.8e-18 \| 1.7e-23 \| 89.1 \| 61 \| (8, 70) \| 73 \| (19, 80) \| 93 \| Helix-turn-helix domain protein \| Helix-turn-helix domain protein; uniclust \| UniRef100\_A0A2A6RG20 \| 99.6 \| 8.7e-18 \| 1.7e-23 \| 90.2 \| 62 \| (4, 68) \| 73 \| (10, 71) \| 98 \| Helix-turn-helix domain-containing protein \| Helix-turn-helix domain-containing protein; uniclust \| UniRef100\_A0A327KP88 \| 99.6 \| 8.9e-18 \| 1.7e-23 \| 87.8 \| 62 \| (6, 69) \| 73 \| (1, 62) \| 86 \| Helix-turn-helix domain-containing protein \| Helix-turn-helix domain-containing protein; uniclust \| UniRef100\_A0A2H9N9A0 \| 99.6 \| 8.8e-18 \| 1.7e-23 \| 88.2 \| 62 \| (6, 69) \| 73 \| (13, 78) \| 85 \| Helix-turn-helix domain-containing protein \| Helix-turn-helix domain-containing protein; uniclust \| UniRef100\_A0A097IEB9 \| 99.6 \| 9.6e-18 \| 1.9e-23 \| 88.7 \| 65 \| (4, 70) \| 73 \| (14, 78) \| 87 \| MerR family transcriptional regulator \| MerR family transcriptional regulator; uniclust \| UniRef100\_A0A0G1M4Q2 \| 99.6 \| 1e-17 \| 2e-23 \| 85.2 \| 64 \| (4, 70) \| 73 \| (1, 64) \| 67 \| Excisionase/Xis, DNA-binding protein \| Excisionase/Xis, DNA-binding protein; uniclust \| UniRef100\_A0A0Q5ZYK8 \| 99.6 \| 1.1e-17 \| 2.1e-23 \| 87.7 \| 60 \| (9, 70) \| 73 \| (18, 77) \| 84 \| Helix-turn-helix domain-containing protein \| Helix-turn-helix domain-containing protein; uniclust \| UniRef100\_A0A061STS5 \| 99.5 \| 1.1e-17 \| 2.3e-23 \| 87.9 \| 62 \| (7, 70) \| 73 \| (12, 73) \| 82 \| AlpA family transcriptional regulator \| AlpA family transcriptional regulator; uniclust \| UniRef100\_A0A2S5MS32 \| 99.5 \| 1.2e-17 \| 2.4e-23 \| 86.4 \| 63 \| (5, 69) \| 73 \| (13, 76) \| 79 \| DNA-binding protein \| DNA-binding protein; uniclust \| UniRef100\_A0A0F6TRE5 \| 99.5 \| 1.2e-17 \| 2.4e-23 \| 85.9 \| 62 \| (5, 68) \| 73 \| (10, 72) \| 75 \| Helix-turn-helix domain-containing protein \| Helix-turn-helix domain-containing protein; uniclust \| UniRef100\_A0A1X1MNM6 \| 99.5 \| 1.3e-17 \| 2.4e-23 \| 84.1 \| 57 \| (8, 66) \| 73 \| (15, 71) \| 74 \| Helix-turn-helix domain-containing protein \| Helix-turn-helix domain-containing protein; uniclust \| UniRef100\_A0A1H8Y5U8 \| 99.5 \| 1.2e-17 \| 2.4e-23 \| 86.8 \| 65 \| (4, 70) \| 73 \| (6, 70) \| 83 \| Helix-turn-helix domain-containing protein \| Helix-turn-helix domain-containing protein; uniclust \| UniRef100\_A0A2D5PMF7 \| 99.5 \| 1.3e-17 \| 2.5e-23 \| 86.8 \| 63 \| (3, 67) \| 73 \| (4, 66) \| 79 \| Terminase \| Terminase; uniclust \| UniRef100\_A0A080N3X4 \| 99.5 \| 1.4e-17 \| 2.7e-23 \| 86.1 \| 61 \| (8, 70) \| 73 \| (16, 76) \| 79 \| DNA-binding protein \| DNA-binding protein; uniclust \| UniRef100\_A0A062VFU0 \| 99.5 \| 1.4e-17 \| 2.8e-23 \| 91.0 \| 62 \| (3, 67) \| 73 \| (14, 75) \| 106 \| DNA-binding protein \| DNA-binding protein; uniclust \| UniRef100\_A0A0F4QZF8 \| 99.5 \| 1.4e-17 \| 2.8e-23 \| 86.7 \| 63 \| (6, 70) \| 73 \| (11, 74) \| 81 \| Helix-turn-helix domain-containing protein \| Helix-turn-helix domain-containing protein; uniclust \| UniRef100\_A0A0F2RWG3 \| 99.5 \| 1.5e-17 \| 2.9e-23 \| 86.7 \| 62 \| (5, 69) \| 73 \| (13, 74) \| 78 \| Helix-turn-helix domain-containing protein \| Helix-turn-helix domain-containing protein; uniclust \| UniRef100\_A0A075MKU4 \| 99.5 \| 1.5e-17 \| 3e-23 \| 87.0 \| 64 \| (4, 69) \| 73 \| (2, 65) \| 84 \| Helix-turn-helix domain-containing protein \| Helix-turn-helix domain-containing protein; uniclust \| UniRef100\_A0A2S8SD78 \| 99.5 \| 1.5e-17 \| 3e-23 \| 86.9 \| 65 \| (3, 69) \| 73 \| (7, 72) \| 81 \| Helix-turn-helix protein \| Helix-turn-helix protein; uniclust \| UniRef100\_A0A0Q0XEN5 \| 99.5 \| 1.6e-17 \| 3.1e-23 \| 90.1 \| 65 \| (4, 70) \| 73 \| (24, 88) \| 105 \| Transcriptional regulator \| Transcriptional regulator; uniclust \| UniRef100\_A0A061M0H8 \| 99.5 \| 1.7e-17 \| 3.2e-23 \| 89.7 \| 57 \| (8, 66) \| 73 \| (31, 87) \| 102 \| Excisionase \| Excisionase; uniclust \| UniRef100\_A0A1J4YZ76 \| 99.5 \| 1.6e-17 \| 3.3e-23 \| 89.4 \| 62 \| (3, 67) \| 73 \| (9, 70) \| 96 \| Helix-turn-helix domain-containing protein \| Helix-turn-helix domain-containing protein; uniclust \| UniRef100\_A0A071MIW0 \| 99.5 \| 1.6e-17 \| 3.3e-23 \| 88.9 \| 61 \| (6, 68) \| 73 \| (15, 76) \| 92 \| Helix-turn-helix domain-containing protein \| Helix-turn-helix domain-containing protein; uniclust \| UniRef100\_A0A081N1X2 \| 99.5 \| 1.8e-17 \| 3.5e-23 \| 87.2 \| 66 \| (2, 69) \| 73 \| (13, 79) \| 85 \| AlpA family transcriptional regulator \| AlpA family transcriptional regulator; uniclust \| UniRef100\_A0A0N8K6P2 \| 99.5 \| 1.8e-17 \| 3.5e-23 \| 84.8 \| 59 \| (6, 67) \| 73 \| (1, 59) \| 69 \| DNA binding domain, excisionase family \| DNA binding domain, excisionase family; uniclust \| UniRef100\_A0A0S7XYB7 \| 99.5 \| 1.8e-17 \| 3.6e-23 \| 87.6 \| 61 \| (6, 68) \| 73 \| (21, 82) \| 89 \| Helix-turn-helix domain-containing protein \| Helix-turn-helix domain-containing protein; uniclust \| UniRef100\_A0A098RX04 \| 99.5 \| 1.8e-17 \| 3.6e-23 \| 88.4 \| 63 \| (4, 68) \| 73 \| (18, 81) \| 92 \| AlpA family phage regulatory protein \| AlpA family phage regulatory protein; uniclust \| UniRef100\_A0A0U3NKV4 \| 99.5 \| 1.9e-17 \| 3.7e-23 \| 88.0 \| 62 \| (7, 70) \| 73 \| (17, 78) \| 91 \| Helix-turn-helix domain-containing protein \| Helix-turn-helix domain-containing protein; uniclust \| UniRef100\_A0A074TFL5 \| 99.5 \| 2e-17 \| 3.9e-23 \| 90.4 \| 59 \| (10, 70) \| 73 \| (33, 91) \| 113 \| Helix-turn-helix domain-containing protein \| Helix-turn-helix domain-containing protein; uniclust \| UniRef100\_A0A096CY27 \| 99.5 \| 2.1e-17 \| 4e-23 \| 87.2 \| 61 \| (4, 67) \| 73 \| (20, 80) \| 95 \| AlpA family transcriptional regulator \| AlpA family transcriptional regulator; uniclust \| UniRef100\_A0A0T5PC48 \| 99.5 \| 2.2e-17 \| 4.3e-23 \| 87.2 \| 63 \| (3, 67) \| 73 \| (1, 64) \| 88 \| Putative transcriptional regulator \| Putative transcriptional regulator; uniclust \| UniRef100\_A0A0I9TFY0 \| 99.5 \| 2.3e-17 \| 4.4e-23 \| 82.8 \| 62 \| (6, 69) \| 73 \| (2, 63) \| 67 \| DNA-binding protein \| DNA-binding protein; uniclust \| UniRef100\_A0A142XZZ0 \| 99.5 \| 2.3e-17 \| 4.5e-23 \| 89.0 \| 57 \| (6, 64) \| 73 \| (28, 84) \| 100 \| Prophage CP4-57 regulatory protein (AlpA) \| Prophage CP4-57 regulatory protein (AlpA); uniclust \| UniRef100\_A0A0F9IQP4 \| 99.5 \| 2.3e-17 \| 4.5e-23 \| 86.0 \| 62 \| (4, 68) \| 73 \| (12, 73) \| 79 \| Helix-turn-helix domain-containing protein \| Helix-turn-helix domain-containing protein; uniclust \| UniRef100\_A0A1H1FCL2 \| 99.5 \| 2.3e-17 \| 4.6e-23 \| 88.5 \| 63 \| (5, 69) \| 73 \| (11, 74) \| 91 \| Helix-turn-helix domain-containing protein \| Helix-turn-helix domain-containing protein; uniclust \| UniRef100\_A0A024EAZ7 \| 99.5 \| 2.4e-17 \| 4.6e-23 \| 89.6 \| 62 \| (3, 67) \| 73 \| (15, 76) \| 106 \| Phage transcriptional regulator, AlpA \| Phage transcriptional regulator, AlpA; uniclust \| UniRef100\_A0A0F3IQQ3 \| 99.5 \| 2.3e-17 \| 4.7e-23 \| 88.3 \| 66 \| (1, 68) \| 73 \| (6, 72) \| 90 \| Transcriptional regulator \| Transcriptional regulator; uniclust \| UniRef100\_A0A024P892 \| 99.5 \| 2.4e-17 \| 4.8e-23 \| 89.6 \| 61 \| (5, 68) \| 73 \| (18, 78) \| 98 \| DNA binding domain, excisionase family \| DNA binding domain, excisionase family; uniclust \| UniRef100\_A0A1F7JH96 \| 99.5 \| 2.5e-17 \| 4.9e-23 \| 82.1 \| 60 \| (8, 70) \| 73 \| (1, 60) \| 63 \| Helix-turn-helix domain-containing protein \| Helix-turn-helix domain-containing protein; uniclust \| UniRef100\_A0A011Q608 \| 99.5 \| 2.6e-17 \| 5.1e-23 \| 87.9 \| 60 \| (9, 70) \| 73 \| (27, 87) \| 92 \| Helix-turn-helix domain protein \| Helix-turn-helix domain protein; uniclust \| UniRef100\_A0A151AMP3 \| 99.5 \| 2.5e-17 \| 5.1e-23 \| 88.0 \| 62 \| (3, 67) \| 73 \| (12, 73) \| 90 \| Helix-turn-helix domain protein \| Helix-turn-helix domain protein; uniclust \| UniRef100\_A0A0D1MDJ0 \| 99.5 \| 2.9e-17 \| 5.6e-23 \| 86.6 \| 62 \| (3, 66) \| 73 \| (6, 67) \| 90 \| Helix-turn-helix domain-containing protein \| Helix-turn-helix domain-containing protein; uniclust \| UniRef100\_A0A0K8JIR0 \| 99.5 \| 2.9e-17 \| 5.6e-23 \| 86.0 \| 64 \| (1, 67) \| 73 \| (1, 64) \| 82 \| Helix-turn-helix domain-containing protein \| Helix-turn-helix domain-containing protein; uniclust \| UniRef100\_A0A087BB61 \| 99.5 \| 2.8e-17 \| 5.6e-23 \| 88.6 \| 63 \| (6, 70) \| 73 \| (24, 86) \| 97 \| Putative phage excisionase \| Putative phage excisionase; uniclust \| UniRef100\_A0A060NMA2 \| 99.5 \| 2.9e-17 \| 5.7e-23 \| 86.0 \| 66 \| (2, 69) \| 73 \| (10, 75) \| 85 \| Helix-turn-helix domain-containing protein \| Helix-turn-helix domain-containing protein; uniclust \| UniRef100\_A0A2U2MS54 \| 99.5 \| 3.1e-17 \| 6.1e-23 \| 85.9 \| 64 \| (5, 70) \| 73 \| (11, 74) \| 81 \| Excisionase \| Excisionase; uniclust \| UniRef100\_A0A0Q6FGU9 \| 99.5 \| 3.4e-17 \| 6.6e-23 \| 87.9 \| 58 \| (7, 66) \| 73 \| (24, 81) \| 96 \| DNA-binding protein \| DNA-binding protein; uniclust \| UniRef100\_A0A078BM66 \| 99.5 \| 3.3e-17 \| 6.6e-23 \| 85.3 \| 63 \| (1, 65) \| 73 \| (5, 67) \| 78 \| Putative Phage transcriptional regulator, AlpA \| Putative Phage transcriptional regulator, AlpA; uniclust \| UniRef100\_A0A353H9N9 \| 99.5 \| 3.3e-17 \| 6.6e-23 \| 86.7 \| 59 \| (5, 66) \| 73 \| (6, 64) \| 87 \| DNA-binding protein \| DNA-binding protein; uniclust \| UniRef100\_A0A0N1B023 \| 99.5 \| 3.3e-17 \| 6.7e-23 \| 86.0 \| 64 \| (4, 69) \| 73 \| (9, 72) \| 80 \| AlpA family transcriptional regulator \| AlpA family transcriptional regulator; uniclust \| UniRef100\_A0A068YKC6 \| 99.5 \| 3.5e-17 \| 6.9e-23 \| 84.9 \| 66 \| (1, 68) \| 73 \| (1, 68) \| 78 \| Helix-turn-helix domain-containing protein \| Helix-turn-helix domain-containing protein; uniclust \| UniRef100\_A0A090MSV2 \| 99.5 \| 3.8e-17 \| 7.3e-23 \| 88.0 \| 65 \| (4, 68) \| 73 \| (13, 77) \| 102 \| Helix-turn-helix domain-containing protein \| Helix-turn-helix domain-containing protein; uniclust \| UniRef100\_A0A1V5E809 \| 99.5 \| 3.8e-17 \| 7.3e-23 \| 87.3 \| 63 \| (2, 67) \| 73 \| (11, 73) \| 97 \| Helix-turn-helix domain protein \| Helix-turn-helix domain protein; uniclust \| UniRef100\_A0A2E3AS91 \| 99.5 \| 3.8e-17 \| 7.4e-23 \| 85.1 \| 68 \| (2, 71) \| 73 \| (8, 77) \| 79 \| DNA-binding protein \| DNA-binding protein; uniclust \| UniRef100\_A0A0F8ZPA5 \| 99.5 \| 3.9e-17 \| 7.6e-23 \| 83.3 \| 60 \| (6, 68) \| 73 \| (8, 67) \| 71 \| Helix-turn-helix domain-containing protein (Fragment) \| Helix-turn-helix domain-containing protein (Fragment); uniclust \| UniRef100\_A0A066UD20 \| 99.5 \| 4e-17 \| 7.9e-23 \| 88.0 \| 62 \| (6, 69) \| 73 \| (18, 80) \| 97 \| Prophage CP4-57 regulatory protein AlpA \| Prophage CP4-57 regulatory protein AlpA; uniclust \| UniRef100\_A0A0U2WJB7 \| 99.5 \| 4.1e-17 \| 8e-23 \| 85.7 \| 62 \| (5, 69) \| 73 \| (9, 70) \| 87 \| Excisionase family DNA-binding domain-containing protein \| Excisionase family DNA-binding domain-containing protein; uniclust \| UniRef100\_A0A086PAC3 \| 99.5 \| 4.2e-17 \| 8.1e-23 \| 84.5 \| 64 \| (1, 66) \| 73 \| (1, 66) \| 79 \| Phage transcriptional regulator AlpA \| Phage transcriptional regulator AlpA; uniclust \| UniRef100\_A0A1C0A4S8 \| 99.5 \| 4.2e-17 \| 8.3e-23 \| 85.2 \| 61 \| (5, 68) \| 73 \| (14, 74) \| 79 \| DNA-binding protein \| DNA-binding protein; uniclust \| UniRef100\_A0A1G3X112 \| 99.5 \| 4.2e-17 \| 8.4e-23 \| 86.0 \| 64 \| (2, 68) \| 73 \| (9, 72) \| 84 \| Helix-turn-helix domain-containing protein \| Helix-turn-helix domain-containing protein; uniclust \| UniRef100\_A0A1G5P793 \| 99.5 \| 4.6e-17 \| 9e-23 \| 85.6 \| 63 \| (4, 68) \| 73 \| (6, 69) \| 86 \| Transcriptional regulator, AlpA family \| Transcriptional regulator, AlpA family; uniclust \| UniRef100\_A0A086MZX1 \| 99.5 \| 4.6e-17 \| 9e-23 \| 87.1 \| 56 \| (10, 67) \| 73 \| (35, 90) \| 96 \| Excisionase \| Excisionase; uniclust \| UniRef100\_A0A1I4TIB0 \| 99.5 \| 4.6e-17 \| 9.1e-23 \| 88.0 \| 60 \| (10, 71) \| 73 \| (30, 89) \| 99 \| Helix-turn-helix domain-containing protein \| Helix-turn-helix domain-containing protein; uniclust \| UniRef100\_A0A931WHN7 \| 99.5 \| 4.9e-17 \| 9.3e-23 \| 83.5 \| 63 \| (3, 68) \| 73 \| (3, 65) \| 80 \| Helix-turn-helix domain-containing protein \| Helix-turn-helix domain-containing protein; uniclust \| UniRef100\_A0A2W4KKW8 \| 99.5 \| 5.3e-17 \| 1e-22 \| 85.3 \| 58 \| (9, 69) \| 73 \| (12, 69) \| 84 \| Excisionase \| Excisionase; uniclust \| UniRef100\_A0A1G8RV34 \| 99.5 \| 5.4e-17 \| 1.1e-22 \| 84.4 \| 61 \| (3, 66) \| 73 \| (2, 62) \| 80 \| DNA binding domain-containing protein, excisionase family \| DNA binding domain-containing protein, excisionase family; uniclust \| UniRef100\_A0A0C2YUN8 \| 99.5 \| 5.6e-17 \| 1.1e-22 \| 83.8 \| 65 \| (1, 67) \| 73 \| (1, 65) \| 79 \| Helix-turn-helix domain-containing protein \| Helix-turn-helix domain-containing protein; uniclust \| UniRef100\_A0A2N9JAS9 \| 99.5 \| 5.5e-17 \| 1.1e-22 \| 83.8 \| 61 \| (6, 69) \| 73 \| (8, 68) \| 75 \| DNA binding domain-containing protein, excisionase family \| DNA binding domain-containing protein, excisionase family; uniclust \| UniRef100\_A0A099GKU9 \| 99.5 \| 5.8e-17 \| 1.1e-22 \| 86.3 \| 59 \| (9, 69) \| 73 \| (22, 80) \| 91 \| Helix-turn-helix domain-containing protein \| Helix-turn-helix domain-containing protein; uniclust \| UniRef100\_A0A0P6XBV6 \| 99.5 \| 6.1e-17 \| 1.2e-22 \| 86.9 \| 60 \| (4, 66) \| 73 \| (17, 76) \| 95 \| Helix-turn-helix domain-containing protein \| Helix-turn-helix domain-containing protein; uniclust \| UniRef100\_A0A0F5ESX0 \| 99.5 \| 6.1e-17 \| 1.2e-22 \| 87.3 \| 62 \| (5, 68) \| 73 \| (16, 78) \| 93 \| AlpA family phage regulatory protein \| AlpA family phage regulatory protein; uniclust \| UniRef100\_A0A164B9G5 \| 99.5 \| 6.5e-17 \| 1.3e-22 \| 86.0 \| 61 \| (7, 69) \| 73 \| (11, 71) \| 94 \| Helix-turn-helix domain-containing protein \| Helix-turn-helix domain-containing protein; uniclust \| UniRef100\_A0A2E7GW86 \| 99.5 \| 6.5e-17 \| 1.3e-22 \| 83.0 \| 60 \| (7, 68) \| 73 \| (6, 65) \| 75 \| DNA-binding protein \| DNA-binding protein; uniclust \| UniRef100\_A0A1F8ZYZ8 \| 99.5 \| 6.6e-17 \| 1.3e-22 \| 85.9 \| 60 \| (4, 66) \| 73 \| (8, 67) \| 89 \| Helix-turn-helix domain-containing protein \| Helix-turn-helix domain-containing protein; uniclust \| UniRef100\_A0A0F7HAF8 \| 99.5 \| 6.6e-17 \| 1.3e-22 \| 84.4 \| 59 \| (7, 67) \| 73 \| (10, 68) \| 78 \| Response regulator inhibitor for tor operon \| Response regulator inhibitor for tor operon; uniclust \| UniRef100\_A0A2A3YGA8 \| 99.5 \| 7.1e-17 \| 1.3e-22 \| 81.4 \| 60 \| (6, 67) \| 73 \| (11, 70) \| 72 \| Helix-turn-helix domain-containing protein \| Helix-turn-helix domain-containing protein; uniclust \| UniRef100\_A0A1H6SP13 \| 99.5 \| 7e-17 \| 1.4e-22 \| 85.9 \| 62 \| (6, 69) \| 73 \| (25, 87) \| 92 \| Transcriptional regulator, AlpA family \| Transcriptional regulator, AlpA family; uniclust \| UniRef100\_A0A0Q9SUY6 \| 99.5 \| 6.9e-17 \| 1.4e-22 \| 85.1 \| 60 \| (7, 68) \| 73 \| (23, 82) \| 84 \| Excisionase \| Excisionase; uniclust \| UniRef100\_A0A0F9MAA1 \| 99.5 \| 7e-17 \| 1.4e-22 \| 84.9 \| 58 \| (7, 67) \| 73 \| (2, 59) \| 85 \| Helix-turn-helix domain-containing protein \| Helix-turn-helix domain-containing protein; uniclust \| UniRef100\_A0A2S5CX75 \| 99.5 \| 8.2e-17 \| 1.6e-22 \| 82.6 \| 60 \| (3, 65) \| 73 \| (3, 62) \| 73 \| Helix-turn-helix domain-containing protein \| Helix-turn-helix domain-containing protein; uniclust \| UniRef100\_A0A0R3E754 \| 99.5 \| 8.6e-17 \| 1.7e-22 \| 84.3 \| 61 \| (6, 69) \| 73 \| (18, 78) \| 88 \| Helix-turn-helix domain-containing protein \| Helix-turn-helix domain-containing protein; uniclust \| UniRef100\_A0A2T6CQ99 \| 99.5 \| 8.5e-17 \| 1.7e-22 \| 83.3 \| 64 \| (3, 68) \| 73 \| (12, 75) \| 78 \| Helix-turn-helix domain-containing protein \| Helix-turn-helix domain-containing protein; uniclust \| UniRef100\_A0A1A3DQH9 \| 99.5 \| 8.6e-17 \| 1.7e-22 \| 82.6 \| 64 \| (3, 68) \| 73 \| (6, 69) \| 75 \| Helix-turn-helix domain-containing protein \| Helix-turn-helix domain-containing protein; uniclust \| UniRef100\_A0A255R1K7 \| 99.5 \| 8.6e-17 \| 1.7e-22 \| 86.2 \| 55 \| (8, 64) \| 73 \| (27, 81) \| 94 \| Helix-turn-helix domain-containing protein \| Helix-turn-helix domain-containing protein; uniclust \| UniRef100\_A0A1F0QJQ0 \| 99.5 \| 8.8e-17 \| 1.7e-22 \| 83.2 \| 57 \| (11, 69) \| 73 \| (18, 74) \| 79 \| Helix-turn-helix domain-containing protein \| Helix-turn-helix domain-containing protein; uniclust \| UniRef100\_A0A0C9PAJ7 \| 99.5 \| 8.7e-17 \| 1.7e-22 \| 84.8 \| 58 \| (9, 69) \| 73 \| (22, 79) \| 85 \| Helix-turn-helix domain-containing protein \| Helix-turn-helix domain-containing protein; uniclust \| UniRef100\_A0A1G9DMJ4 \| 99.5 \| 8.8e-17 \| 1.7e-22 \| 83.4 \| 61 \| (7, 70) \| 73 \| (12, 72) \| 76 \| DNA binding domain-containing protein, excisionase family \| DNA binding domain-containing protein, excisionase family; uniclust \| UniRef100\_A0A0G0M802 \| 99.5 \| 9.4e-17 \| 1.8e-22 \| 83.2 \| 62 \| (4, 68) \| 73 \| (4, 66) \| 77 \| Binding domain protein, excisionase family protein \| Binding domain protein, excisionase family protein; uniclust \| UniRef100\_A0A017HNG0 \| 99.5 \| 9.5e-17 \| 1.8e-22 \| 86.6 \| 61 \| (6, 68) \| 73 \| (20, 81) \| 101 \| Phage transcriptional regulator, AlpA \| Phage transcriptional regulator, AlpA; uniclust \| UniRef100\_A0A2H0DID7 \| 99.5 \| 1e-16 \| 1.9e-22 \| 81.4 \| 62 \| (3, 67) \| 73 \| (1, 63) \| 70 \| Helix-turn-helix domain-containing protein \| Helix-turn-helix domain-containing protein; uniclust \| UniRef100\_A0A2V9PDW9 \| 99.5 \| 1e-16 \| 1.9e-22 \| 83.5 \| 64 \| (4, 68) \| 73 \| (11, 74) \| 82 \| DNA-binding protein \| DNA-binding protein; uniclust \| UniRef100\_A0A0B8Q8F9 \| 99.5 \| 1e-16 \| 2e-22 \| 88.1 \| 65 \| (3, 69) \| 73 \| (24, 89) \| 108 \| Transcriptional regulator \| Transcriptional regulator; uniclust \| UniRef100\_A0A2N1ZXV4 \| 99.5 \| 1.1e-16 \| 2.1e-22 \| 83.6 \| 65 \| (1, 67) \| 73 \| (1, 66) \| 80 \| Transcriptional regulator \| Transcriptional regulator; uniclust \| UniRef100\_A0A1F2R0F2 \| 99.5 \| 1e-16 \| 2.1e-22 \| 84.8 \| 57 \| (8, 67) \| 73 \| (11, 67) \| 82 \| Helix-turn-helix domain-containing protein \| Helix-turn-helix domain-containing protein; uniclust \| UniRef100\_A0A1X1PI46 \| 99.5 \| 1.1e-16 \| 2.1e-22 \| 82.3 \| 61 \| (6, 68) \| 73 \| (7, 67) \| 75 \| Helix-turn-helix domain-containing protein \| Helix-turn-helix domain-containing protein; uniclust \| UniRef100\_A0A2V9WFU4 \| 99.5 \| 1.1e-16 \| 2.2e-22 \| 84.5 \| 66 \| (1, 68) \| 73 \| (1, 67) \| 86 \| Helix-turn-helix domain-containing protein \| Helix-turn-helix domain-containing protein; uniclust \| UniRef100\_A0A017HTQ8 \| 99.5 \| 1.1e-16 \| 2.2e-22 \| 86.3 \| 61 \| (6, 68) \| 73 \| (21, 82) \| 94 \| DNA-binding protein, putative \| DNA-binding protein, putative; uniclust \| UniRef100\_A0A0B4XAG0 \| 99.5 \| 1.1e-16 \| 2.2e-22 \| 87.2 \| 60 \| (6, 67) \| 73 \| (24, 83) \| 108 \| Helix-turn-helix domain-containing protein \| Helix-turn-helix domain-containing protein; uniclust \| UniRef100\_A0A011QGD0 \| 99.5 \| 1.2e-16 \| 2.3e-22 \| 87.0 \| 60 \| (5, 67) \| 73 \| (30, 89) \| 114 \| DNA binding domain, excisionase family \| DNA binding domain, excisionase family; uniclust \| UniRef100\_A0A109IMN0 \| 99.5 \| 1.2e-16 \| 2.3e-22 \| 81.5 \| 60 \| (7, 68) \| 73 \| (11, 70) \| 78 \| Transcriptional regulator, AlpA family \| Transcriptional regulator, AlpA family; uniclust \| UniRef100\_A0A0P6X5W5 \| 99.5 \| 1.2e-16 \| 2.3e-22 \| 85.5 \| 60 \| (4, 66) \| 73 \| (9, 68) \| 95 \| DNA-binding protein \| DNA-binding protein; uniclust \| UniRef100\_A0A077M1T7 \| 99.5 \| 1.2e-16 \| 2.3e-22 \| 84.3 \| 63 \| (4, 68) \| 73 \| (18, 80) \| 86 \| Helix-turn-helix domain-containing protein \| Helix-turn-helix domain-containing protein; uniclust \| UniRef100\_A0A0F7PLT3 \| 99.5 \| 1.2e-16 \| 2.4e-22 \| 85.7 \| 66 \| (1, 68) \| 73 \| (3, 73) \| 96 \| Helix-turn-helix domain \| Helix-turn-helix domain; uniclust \| UniRef100\_A0A0F5Q0G5 \| 99.5 \| 1.2e-16 \| 2.4e-22 \| 84.2 \| 64 \| (3, 69) \| 73 \| (1, 64) \| 84 \| DNA binding domain-containing protein, excisionase family \| DNA binding domain-containing protein, excisionase family; uniclust \| UniRef100\_A0A136K2Y7 \| 99.5 \| 1.2e-16 \| 2.4e-22 \| 82.2 \| 64 \| (2, 68) \| 73 \| (1, 64) \| 74 \| Helix-turn-helix domain protein \| Helix-turn-helix domain protein; uniclust \| UniRef100\_A0A022LBF6 \| 99.5 \| 1.3e-16 \| 2.5e-22 \| 83.4 \| 61 \| (7, 68) \| 73 \| (16, 76) \| 84 \| Helix-turn-helix domain-containing protein \| Helix-turn-helix domain-containing protein; uniclust \| UniRef100\_A0A0K8MEP3 \| 99.5 \| 1.3e-16 \| 2.5e-22 \| 84.8 \| 61 \| (7, 69) \| 73 \| (24, 85) \| 93 \| Helix-turn-helix domain protein \| Helix-turn-helix domain protein; uniclust \| UniRef100\_A0A258IUU4 \| 99.5 \| 1.4e-16 \| 2.7e-22 \| 83.3 \| 67 \| (2, 70) \| 73 \| (8, 74) \| 82 \| DNA-binding protein \| DNA-binding protein; uniclust \| UniRef100\_A0A496RWF2 \| 99.5 \| 1.4e-16 \| 2.7e-22 \| 81.2 \| 62 \| (5, 68) \| 73 \| (3, 64) \| 74 \| Helix-turn-helix domain-containing protein \| Helix-turn-helix domain-containing protein; uniclust \| UniRef100\_A0A077M848 \| 99.5 \| 1.5e-16 \| 2.9e-22 \| 86.6 \| 62 \| (2, 66) \| 73 \| (12, 73) \| 110 \| Phage transcriptional regulator, AlpA \| Phage transcriptional regulator, AlpA; uniclust \| UniRef100\_A0A0T6APZ4 \| 99.5 \| 1.5e-16 \| 2.9e-22 \| 85.8 \| 62 \| (5, 69) \| 73 \| (25, 86) \| 94 \| Excisionase \| Excisionase; uniclust \| UniRef100\_A0A066UG88 \| 99.5 \| 1.5e-16 \| 3e-22 \| 85.3 \| 59 \| (9, 69) \| 73 \| (32, 91) \| 95 \| Excisionase \| Excisionase; uniclust \| UniRef100\_A0A0H4J0C0 \| 99.5 \| 1.6e-16 \| 3.1e-22 \| 87.1 \| 61 \| (6, 68) \| 73 \| (21, 82) \| 106 \| AlpA family transcriptional regulator \| AlpA family transcriptional regulator; uniclust \| UniRef100\_A0A4R9BK06 \| 99.5 \| 1.6e-16 \| 3.1e-22 \| 86.2 \| 60 \| (10, 71) \| 73 \| (38, 97) \| 101 \| DNA-binding protein \| DNA-binding protein; uniclust \| UniRef100\_A0A127K3W8 \| 99.5 \| 1.6e-16 \| 3.2e-22 \| 88.0 \| 61 \| (7, 69) \| 73 \| (45, 105) \| 109 \| Helix-turn-helix domain-containing protein \| Helix-turn-helix domain-containing protein; uniclust \| UniRef100\_A0A066ZRH0 \| 99.5 \| 1.7e-16 \| 3.3e-22 \| 84.5 \| 65 \| (3, 69) \| 73 \| (7, 72) \| 87 \| AlpA family transcriptional regulator \| AlpA family transcriptional regulator; uniclust \| UniRef100\_A0A0G3G4R0 \| 99.5 \| 1.7e-16 \| 3.3e-22 \| 83.3 \| 64 \| (3, 68) \| 73 \| (9, 73) \| 82 \| AlpA family transcriptional regulator \| AlpA family transcriptional regulator; uniclust \| UniRef100\_A0A1V6CFG2 \| 99.5 \| 1.8e-16 \| 3.4e-22 \| 79.9 \| 58 \| (6, 66) \| 73 \| (2, 59) \| 73 \| Helix-turn-helix domain protein \| Helix-turn-helix domain protein; uniclust \| UniRef100\_A0A0F8YFR4 \| 99.5 \| 1.8e-16 \| 3.5e-22 \| 82.8 \| 62 \| (3, 67) \| 73 \| (10, 71) \| 84 \| HTH merR-type domain-containing protein \| HTH merR-type domain-containing protein; uniclust \| UniRef100\_A0A167I6B1 \| 99.5 \| 1.9e-16 \| 3.8e-22 \| 86.0 \| 59 \| (8, 68) \| 73 \| (25, 83) \| 98 \| Helix-turn-helix domain-containing protein \| Helix-turn-helix domain-containing protein; uniclust \| UniRef100\_A0A086Y3N3 \| 99.5 \| 2e-16 \| 3.9e-22 \| 84.1 \| 64 \| (2, 67) \| 73 \| (14, 79) \| 86 \| AlpA family transcriptional regulator \| AlpA family transcriptional regulator; uniclust \| UniRef100\_A0A0Q7G9F4 \| 99.5 \| 2.1e-16 \| 4e-22 \| 78.2 \| 60 \| (6, 67) \| 73 \| (1, 60) \| 62 \| Helix-turn-helix domain-containing protein \| Helix-turn-helix domain-containing protein; uniclust \| UniRef100\_A0A098B1F2 \| 99.5 \| 2.1e-16 \| 4e-22 \| 80.8 \| 64 \| (2, 68) \| 73 \| (6, 69) \| 78 \| DNA binding domain, putative \| DNA binding domain, putative; uniclust \| UniRef100\_A0A136KDH9 \| 99.5 \| 2e-16 \| 4e-22 \| 85.7 \| 58 \| (6, 66) \| 73 \| (28, 85) \| 99 \| Helix-turn-helix domain protein \| Helix-turn-helix domain protein; uniclust \| UniRef100\_A0A261TDA7 \| 99.5 \| 2.1e-16 \| 4.1e-22 \| 86.8 \| 58 \| (5, 64) \| 73 \| (7, 65) \| 110 \| Helix-turn-helix domain-containing protein \| Helix-turn-helix domain-containing protein; uniclust \| UniRef100\_A0A1I5LC99 \| 99.5 \| 2e-16 \| 4.1e-22 \| 84.3 \| 64 \| (3, 69) \| 73 \| (16, 79) \| 88 \| DNA binding domain-containing protein, excisionase family \| DNA binding domain-containing protein, excisionase family; uniclust \| UniRef100\_A0A1Q7SV79 \| 99.5 \| 2e-16 \| 4.1e-22 \| 82.3 \| 64 \| (2, 68) \| 73 \| (2, 66) \| 76 \| Helix-turn-helix domain-containing protein \| Helix-turn-helix domain-containing protein; uniclust \| UniRef100\_A0A0G0RVY6 \| 99.5 \| 2.2e-16 \| 4.3e-22 \| 83.2 \| 62 \| (3, 67) \| 73 \| (7, 68) \| 86 \| Binding domain protein, excisionase family protein \| Binding domain protein, excisionase family protein; uniclust \| UniRef100\_A0A2N2NCM9 \| 99.5 \| 2.2e-16 \| 4.3e-22 \| 82.5 \| 59 \| (7, 68) \| 73 \| (13, 71) \| 79 \| DNA-binding protein \| DNA-binding protein; uniclust \| UniRef100\_A0A086ZPG8 \| 99.5 \| 2.3e-16 \| 4.4e-22 \| 85.0 \| 60 \| (4, 66) \| 73 \| (12, 71) \| 101 \| DNA binding domain-containing protein, excisionase family \| DNA binding domain-containing protein, excisionase family; uniclust \| UniRef100\_A0A085ADN8 \| 99.4 \| 2.3e-16 \| 4.5e-22 \| 82.4 \| 64 \| (3, 68) \| 73 \| (3, 66) \| 79 \| AlpA family phage regulatory protein \| AlpA family phage regulatory protein; uniclust \| UniRef100\_A0A031JQZ3 \| 99.4 \| 2.3e-16 \| 4.5e-22 \| 87.0 \| 59 \| (9, 69) \| 73 \| (36, 94) \| 107 \| Helix-turn-helix domain-containing protein \| Helix-turn-helix domain-containing protein; uniclust \| UniRef100\_A0A010YJ93 \| 99.4 \| 2.3e-16 \| 4.6e-22 \| 83.7 \| 61 \| (5, 67) \| 73 \| (13, 73) \| 88 \| DNA-binding protein, excisionase family \| DNA-binding protein, excisionase family; uniclust \| UniRef100\_A0A0A3IXC5 \| 99.4 \| 2.4e-16 \| 4.8e-22 \| 83.7 \| 62 \| (4, 68) \| 73 \| (12, 73) \| 89 \| Excisionase \| Excisionase; uniclust \| UniRef100\_A0A086MTU0 \| 99.4 \| 2.4e-16 \| 4.8e-22 \| 84.4 \| 60 \| (7, 68) \| 73 \| (29, 89) \| 92 \| Excisionase \| Excisionase; uniclust \| UniRef100\_A0A0W0SED0 \| 99.4 \| 2.5e-16 \| 4.9e-22 \| 82.4 \| 64 \| (4, 69) \| 73 \| (1, 64) \| 84 \| DNA-binding protein \| DNA-binding protein; uniclust \| UniRef100\_A0A059XS14 \| 99.4 \| 2.5e-16 \| 5e-22 \| 85.1 \| 64 \| (2, 67) \| 73 \| (11, 75) \| 95 \| AlpA family transcriptional regulator \| AlpA family transcriptional regulator; uniclust \| UniRef100\_A0A1J5D8F8 \| 99.4 \| 2.5e-16 \| 5.1e-22 \| 82.4 \| 63 \| (2, 67) \| 73 \| (1, 63) \| 79 \| Helix-turn-helix domain-containing protein \| Helix-turn-helix domain-containing protein; uniclust \| UniRef100\_A0A0F2RJT8 \| 99.4 \| 2.7e-16 \| 5.3e-22 \| 84.2 \| 63 \| (6, 70) \| 73 \| (23, 86) \| 100 \| Helix-turn-helix domain-containing protein \| Helix-turn-helix domain-containing protein; uniclust \| UniRef100\_A0A3M0XFV3 \| 99.4 \| 2.8e-16 \| 5.3e-22 \| 83.1 \| 59 \| (5, 66) \| 73 \| (4, 62) \| 93 \| DNA-binding protein \| DNA-binding protein; uniclust \| UniRef100\_A0A1V4VR13 \| 99.4 \| 2.8e-16 \| 5.4e-22 \| 81.1 \| 65 \| (2, 69) \| 73 \| (1, 65) \| 75 \| Helix-turn-helix domain protein \| Helix-turn-helix domain protein; uniclust \| UniRef100\_A0A1N7PI06 \| 99.4 \| 2.8e-16 \| 5.5e-22 \| 84.9 \| 62 \| (2, 65) \| 73 \| (16, 78) \| 101 \| Prophage regulatory protein \| Prophage regulatory protein; uniclust \| UniRef100\_A0A0F2R716 \| 99.4 \| 2.8e-16 \| 5.6e-22 \| 83.1 \| 64 \| (7, 72) \| 73 \| (18, 81) \| 82 \| Helix-turn-helix domain-containing protein \| Helix-turn-helix domain-containing protein; uniclust \| UniRef100\_A0A1G2ZI97 \| 99.4 \| 2.9e-16 \| 5.6e-22 \| 83.7 \| 53 \| (8, 63) \| 73 \| (22, 74) \| 91 \| Helix-turn-helix domain-containing protein \| Helix-turn-helix domain-containing protein; uniclust \| UniRef100\_A0A1F1E662 \| 99.4 \| 2.9e-16 \| 5.7e-22 \| 83.2 \| 59 \| (9, 69) \| 73 \| (19, 80) \| 88 \| Helix-turn-helix domain-containing protein \| Helix-turn-helix domain-containing protein; uniclust \| UniRef100\_A0A0G3V2N5 \| 99.4 \| 2.9e-16 \| 5.7e-22 \| 84.9 \| 58 \| (8, 68) \| 73 \| (22, 79) \| 97 \| Transcriptional regulator, AlpA family \| Transcriptional regulator, AlpA family; uniclust \| UniRef100\_A0A089IG40 \| 99.4 \| 3.2e-16 \| 6.3e-22 \| 79.7 \| 65 \| (1, 69) \| 73 \| (1, 65) \| 69 \| Helix-turn-helix domain-containing protein \| Helix-turn-helix domain-containing protein; uniclust \| UniRef100\_A0A1F3TX82 \| 99.4 \| 3.3e-16 \| 6.3e-22 \| 81.8 \| 61 \| (4, 67) \| 73 \| (11, 71) \| 84 \| Excisionase \| Excisionase; uniclust \| UniRef100\_A0A191WES9 \| 99.4 \| 3.4e-16 \| 6.7e-22 \| 83.6 \| 58 \| (5, 65) \| 73 \| (11, 68) \| 90 \| Helix-turn-helix domain-containing protein \| Helix-turn-helix domain-containing protein; uniclust \| UniRef100\_A0A0C1RLB9 \| 99.4 \| 3.4e-16 \| 6.8e-22 \| 82.4 \| 58 \| (8, 68) \| 73 \| (19, 76) \| 84 \| Helix-turn-helix domain-containing protein \| Helix-turn-helix domain-containing protein; uniclust \| UniRef100\_A0A0C1X3E4 \| 99.4 \| 3.6e-16 \| 7e-22 \| 82.9 \| 65 \| (4, 70) \| 73 \| (16, 84) \| 93 \| DNA-binding protein \| DNA-binding protein; uniclust \| UniRef100\_A0A1G1K0V6 \| 99.4 \| 3.6e-16 \| 7e-22 \| 82.9 \| 57 \| (6, 65) \| 73 \| (9, 65) \| 93 \| Helix-turn-helix domain-containing protein \| Helix-turn-helix domain-containing protein; uniclust \| UniRef100\_A0A067A1U4 \| 99.4 \| 3.5e-16 \| 7e-22 \| 86.4 \| 65 \| (2, 68) \| 73 \| (28, 93) \| 109 \| AlpA family transcriptional regulator \| AlpA family transcriptional regulator; uniclust \| UniRef100\_A0A2E7A109 \| 99.4 \| 3.8e-16 \| 7.2e-22 \| 80.6 \| 57 \| (7, 65) \| 73 \| (22, 80) \| 82 \| Helix-turn-helix domain-containing protein \| Helix-turn-helix domain-containing protein; uniclust \| UniRef100\_A0A060UYJ0 \| 99.4 \| 3.7e-16 \| 7.2e-22 \| 83.1 \| 63 \| (4, 68) \| 73 \| (15, 78) \| 87 \| AlpA family transcriptional regulator \| AlpA family transcriptional regulator; uniclust \| UniRef100\_A0A060ZVZ6 \| 99.4 \| 4e-16 \| 7.7e-22 \| 86.2 \| 60 \| (8, 69) \| 73 \| (53, 112) \| 120 \| Helix-turn-helix domain-containing protein \| Helix-turn-helix domain-containing protein; uniclust \| UniRef100\_A0A1F8XWQ5 \| 99.4 \| 3.9e-16 \| 7.8e-22 \| 88.8 \| 52 \| (10, 64) \| 73 \| (19, 70) \| 132 \| Helix-turn-helix domain-containing protein \| Helix-turn-helix domain-containing protein; uniclust \| UniRef100\_A0A1U9USI9 \| 99.4 \| 4e-16 \| 7.9e-22 \| 80.6 \| 62 \| (3, 66) \| 73 \| (9, 70) \| 76 \| DNA-binding protein \| DNA-binding protein; uniclust \| UniRef100\_A0A6P0HA59 \| 99.4 \| 4.2e-16 \| 7.9e-22 \| 81.7 \| 60 \| (7, 68) \| 73 \| (11, 70) \| 93 \| Helix-turn-helix domain-containing protein \| Helix-turn-helix domain-containing protein; uniclust \| UniRef100\_A0A1Y0UY49 \| 99.4 \| 4.2e-16 \| 8.3e-22 \| 79.9 \| 66 \| (1, 68) \| 73 \| (1, 68) \| 72 \| Helix-turn-helix domain-containing protein \| Helix-turn-helix domain-containing protein; uniclust \| UniRef100\_A0A3A5M606 \| 99.4 \| 4.4e-16 \| 8.4e-22 \| 80.8 \| 57 \| (8, 66) \| 73 \| (20, 76) \| 80 \| DNA-binding protein \| DNA-binding protein; uniclust \| UniRef100\_A0A1V5FMB9 \| 99.4 \| 4.4e-16 \| 8.5e-22 \| 79.9 \| 64 \| (1, 66) \| 73 \| (1, 65) \| 77 \| Helix-turn-helix domain protein \| Helix-turn-helix domain protein; uniclust \| UniRef100\_A0A1V5QDI4 \| 99.4 \| 4.2e-16 \| 8.5e-22 \| 81.2 \| 64 \| (3, 69) \| 73 \| (4, 67) \| 75 \| Helix-turn-helix domain protein \| Helix-turn-helix domain protein; uniclust \| UniRef100\_A0A095XXS8 \| 99.4 \| 4.4e-16 \| 8.7e-22 \| 81.4 \| 64 \| (4, 68) \| 73 \| (8, 71) \| 78 \| HTH merR-type domain-containing protein \| HTH merR-type domain-containing protein; uniclust \| UniRef100\_A0A5B0D528 \| 99.4 \| 4.5e-16 \| 8.8e-22 \| 77.9 \| 59 \| (5, 65) \| 73 \| (2, 60) \| 63 \| Helix-turn-helix domain-containing protein \| Helix-turn-helix domain-containing protein; uniclust \| UniRef100\_A0A1I3XCD0 \| 99.4 \| 4.5e-16 \| 8.9e-22 \| 83.4 \| 54 \| (9, 64) \| 73 \| (20, 73) \| 92 \| Helix-turn-helix domain-containing protein \| Helix-turn-helix domain-containing protein; uniclust \| UniRef100\_A0A1W9XLA3 \| 99.4 \| 4.5e-16 \| 8.9e-22 \| 85.1 \| 63 \| (6, 70) \| 73 \| (31, 94) \| 103 \| Helix-turn-helix domain-containing protein \| Helix-turn-helix domain-containing protein; uniclust \| UniRef100\_A0A0Q8V938 \| 99.4 \| 4.7e-16 \| 8.9e-22 \| 84.0 \| 58 \| (9, 68) \| 73 \| (42, 99) \| 108 \| Helix-turn-helix domain-containing protein \| Helix-turn-helix domain-containing protein; uniclust \| UniRef100\_A0A0F4ICJ3 \| 99.4 \| 4.8e-16 \| 9.2e-22 \| 82.8 \| 60 \| (9, 70) \| 73 \| (38, 97) \| 100 \| Helix-turn-helix domain-containing protein \| Helix-turn-helix domain-containing protein; uniclust \| UniRef100\_A0A011U9H2 \| 99.4 \| 4.9e-16 \| 9.5e-22 \| 85.3 \| 60 \| (8, 69) \| 73 \| (21, 80) \| 112 \| Transcriptional regulator \| Transcriptional regulator; uniclust \| UniRef100\_A0A0C1R836 \| 99.4 \| 5e-16 \| 9.7e-22 \| 85.7 \| 59 \| (7, 68) \| 73 \| (28, 86) \| 120 \| Helix-turn-helix domain-containing protein \| Helix-turn-helix domain-containing protein; uniclust \| UniRef100\_A0A0D6ZBS2 \| 99.4 \| 5.1e-16 \| 9.8e-22 \| 79.7 \| 60 \| (6, 68) \| 73 \| (7, 66) \| 75 \| Helix-turn-helix domain-containing protein \| Helix-turn-helix domain-containing protein; uniclust \| UniRef100\_A0A096BCL5 \| 99.4 \| 5.1e-16 \| 1e-21 \| 82.5 \| 59 \| (7, 68) \| 73 \| (18, 76) \| 89 \| Helix-turn-helix domain-containing protein \| Helix-turn-helix domain-containing protein; uniclust \| UniRef100\_A0A2E3YEM9 \| 99.4 \| 5.3e-16 \| 1e-21 \| 82.4 \| 59 \| (5, 66) \| 73 \| (6, 64) \| 96 \| Helix-turn-helix domain-containing protein \| Helix-turn-helix domain-containing protein; uniclust \| UniRef100\_A0A1H3K8W7 \| 99.4 \| 5.4e-16 \| 1e-21 \| 79.0 \| 59 \| (7, 67) \| 73 \| (9, 68) \| 70 \| Transcriptional regulator, AlpA family \| Transcriptional regulator, AlpA family; uniclust \| UniRef100\_A0A0G1DFI7 \| 99.4 \| 5.4e-16 \| 1.1e-21 \| 83.8 \| 62 \| (3, 67) \| 73 \| (16, 82) \| 95 \| Helix-turn-helix domain-containing protein \| Helix-turn-helix domain-containing protein; uniclust \| UniRef100\_A0A953LVT0 \| 99.4 \| 5.7e-16 \| 1.1e-21 \| 83.7 \| 63 \| (5, 69) \| 73 \| (32, 94) \| 103 \| Helix-turn-helix domain-containing protein \| Helix-turn-helix domain-containing protein; uniclust \| UniRef100\_A0A136KUN2 \| 99.4 \| 6.2e-16 \| 1.2e-21 \| 96.0 \| 65 \| (1, 67) \| 73 \| (18, 82) \| 272 \| MerR family transcriptional regulator \| MerR family transcriptional regulator; uniclust \| UniRef100\_A0A0J1BH83 \| 99.4 \| 6.2e-16 \| 1.2e-21 \| 82.8 \| 55 \| (9, 65) \| 73 \| (21, 75) \| 88 \| Helix-turn-helix domain-containing protein \| Helix-turn-helix domain-containing protein; uniclust \| UniRef100\_A0A0F7GG26 \| 99.4 \| 6.4e-16 \| 1.3e-21 \| 95.5 \| 61 \| (1, 64) \| 73 \| (38, 98) \| 256 \| Nitrogen regulation protein NR(I) \| Nitrogen regulation protein NR(I); uniclust \| UniRef100\_A0A1V1PRD8 \| 99.4 \| 7e-16 \| 1.4e-21 \| 82.9 \| 61 \| (7, 69) \| 73 \| (26, 86) \| 97 \| Helix-turn-helix domain-containing protein \| Helix-turn-helix domain-containing protein; uniclust \| UniRef100\_A0A023WP26 \| 99.4 \| 6.6e-16 \| 1.4e-21 \| 84.1 \| 63 \| (5, 69) \| 73 \| (19, 82) \| 96 \| Transcriptional regulator \| Transcriptional regulator; uniclust \| UniRef100\_A0A062UK24 \| 99.4 \| 7.1e-16 \| 1.4e-21 \| 81.9 \| 60 \| (8, 69) \| 73 \| (16, 75) \| 90 \| Helix-turn-helix domain-containing protein \| Helix-turn-helix domain-containing protein; uniclust \| UniRef100\_A0A7L8RVP0 \| 99.4 \| 7.5e-16 \| 1.4e-21 \| 81.6 \| 62 \| (6, 69) \| 73 \| (27, 88) \| 95 \| Helix-turn-helix domain-containing protein \| Helix-turn-helix domain-containing protein; uniclust \| UniRef100\_A0A2E6LLT9 \| 99.4 \| 7.6e-16 \| 1.4e-21 \| 75.8 \| 55 \| (10, 66) \| 73 \| (2, 57) \| 60 \| AlpA family transcriptional regulator \| AlpA family transcriptional regulator; uniclust \| UniRef100\_A0A0A0B9Q1 \| 99.4 \| 7.6e-16 \| 1.5e-21 \| 80.2 \| 59 \| (4, 64) \| 73 \| (19, 78) \| 81 \| Excisionase \| Excisionase; uniclust \| UniRef100\_A0A0M0T6P6 \| 99.4 \| 7.7e-16 \| 1.5e-21 \| 81.1 \| 63 \| (4, 68) \| 73 \| (9, 71) \| 85 \| Uncharacterized protein \| Uncharacterized protein; uniclust \| UniRef100\_A0A1Q6XGV3 \| 99.4 \| 8.2e-16 \| 1.6e-21 \| 80.8 \| 64 \| (3, 68) \| 73 \| (17, 80) \| 87 \| Helix-turn-helix domain-containing protein \| Helix-turn-helix domain-containing protein; uniclust \| UniRef100\_A0A0G0MGN7 \| 99.4 \| 8.4e-16 \| 1.6e-21 \| 79.9 \| 61 \| (3, 67) \| 73 \| (8, 68) \| 77 \| Excise: DNA binding domain, excisionase fami \| Excise: DNA binding domain, excisionase fami; uniclust \| UniRef100\_A0A1F6Q9F2 \| 99.4 \| 8.7e-16 \| 1.7e-21 \| 79.5 \| 58 \| (8, 68) \| 73 \| (20, 77) \| 81 \| Helix-turn-helix domain-containing protein \| Helix-turn-helix domain-containing protein; uniclust \| UniRef100\_A0A0Q5WB56 \| 99.4 \| 8.5e-16 \| 1.7e-21 \| 81.8 \| 57 \| (9, 68) \| 73 \| (23, 79) \| 88 \| Helix-turn-helix domain-containing protein \| Helix-turn-helix domain-containing protein; uniclust \| UniRef100\_A0A542ZXW8 \| 99.4 \| 8.9e-16 \| 1.7e-21 \| 77.0 \| 61 \| (6, 68) \| 73 \| (2, 63) \| 66 \| Helix-turn-helix protein \| Helix-turn-helix protein; uniclust \| UniRef100\_A0A0Q9QUK2 \| 99.4 \| 9.4e-16 \| 1.8e-21 \| 80.3 \| 57 \| (9, 67) \| 73 \| (18, 74) \| 85 \| Helix-turn-helix domain-containing protein \| Helix-turn-helix domain-containing protein; uniclust \| UniRef100\_A0A1Q6JKS2 \| 99.4 \| 9.9e-16 \| 1.9e-21 \| 78.1 \| 61 \| (5, 67) \| 73 \| (9, 70) \| 75 \| Helix-turn-helix domain-containing protein \| Helix-turn-helix domain-containing protein; uniclust \| UniRef100\_A0A3B7DB92 \| 99.4 \| 1e-15 \| 1.9e-21 \| 77.1 \| 58 \| (8, 67) \| 73 \| (11, 68) \| 71 \| DNA-binding protein \| DNA-binding protein; uniclust \| UniRef100\_A0A1C5D3R9 \| 99.4 \| 1e-15 \| 2e-21 \| 79.6 \| 59 \| (5, 66) \| 73 \| (13, 71) \| 80 \| DNA binding domain-containing protein, excisionase family \| DNA binding domain-containing protein, excisionase family; uniclust \| UniRef100\_A0A068Z9N5 \| 99.4 \| 1e-15 \| 2e-21 \| 81.7 \| 64 \| (3, 68) \| 73 \| (6, 70) \| 87 \| AlpA family transcriptional regulator \| AlpA family transcriptional regulator; uniclust \| UniRef100\_A0A1A9BV43 \| 99.4 \| 1.1e-15 \| 2.1e-21 \| 77.5 \| 60 \| (5, 66) \| 73 \| (2, 62) \| 67 \| Helix-turn-helix domain-containing protein \| Helix-turn-helix domain-containing protein; uniclust \| UniRef100\_A0A2H1STW7 \| 99.4 \| 1.1e-15 \| 2.1e-21 \| 80.6 \| 62 \| (4, 68) \| 73 \| (29, 90) \| 97 \| Excisionase (Modular protein) \| Excisionase (Modular protein); uniclust \| UniRef100\_A0A0G0QHG5 \| 99.4 \| 1.1e-15 \| 2.1e-21 \| 79.8 \| 61 \| (5, 68) \| 73 \| (6, 70) \| 78 \| Binding domain protein, excisionase family protein \| Binding domain protein, excisionase family protein; uniclust \| UniRef100\_A0A0B3SSZ9 \| 99.4 \| 1.1e-15 \| 2.2e-21 \| 79.5 \| 59 \| (6, 66) \| 73 \| (21, 79) \| 82 \| Transcriptional repressor, MerR family protein \| Transcriptional repressor, MerR family protein; uniclust \| UniRef100\_A0A6I5VMA1 \| 99.4 \| 1.2e-15 \| 2.2e-21 \| 81.0 \| 60 \| (8, 69) \| 73 \| (25, 85) \| 93 \| Helix-turn-helix domain-containing protein \| Helix-turn-helix domain-containing protein; uniclust \| UniRef100\_A0A1E3YDB6 \| 99.4 \| 1.2e-15 \| 2.3e-21 \| 91.7 \| 63 \| (4, 69) \| 73 \| (28, 90) \| 229 \| Response regulatory domain-containing protein \| Response regulatory domain-containing protein; uniclust \| UniRef100\_A0A1H8HT30 \| 99.4 \| 1.2e-15 \| 2.3e-21 \| 80.1 \| 62 \| (1, 64) \| 73 \| (1, 63) \| 81 \| Transcriptional regulator, AlpA family \| Transcriptional regulator, AlpA family; uniclust \| UniRef100\_A0A1H6INB4 \| 99.4 \| 1.2e-15 \| 2.4e-21 \| 81.9 \| 61 \| (5, 68) \| 73 \| (18, 78) \| 90 \| DNA binding domain-containing protein, excisionase family \| DNA binding domain-containing protein, excisionase family; uniclust \| UniRef100\_A0A0F8ZBU4 \| 99.4 \| 1.3e-15 \| 2.5e-21 \| 78.3 \| 58 \| (7, 67) \| 73 \| (1, 58) \| 76 \| Helix-turn-helix domain-containing protein \| Helix-turn-helix domain-containing protein; uniclust \| UniRef100\_A0A1J1EMG5 \| 99.4 \| 1.3e-15 \| 2.5e-21 \| 78.3 \| 57 \| (7, 66) \| 73 \| (5, 61) \| 71 \| DNA binding domain, excisionase family \| DNA binding domain, excisionase family; uniclust \| UniRef100\_A0A353GQK3 \| 99.4 \| 1.3e-15 \| 2.5e-21 \| 77.4 \| 62 \| (6, 69) \| 73 \| (1, 65) \| 70 \| Helix-turn-helix domain-containing protein \| Helix-turn-helix domain-containing protein; uniclust \| UniRef100\_A0A5N8YNY4 \| 99.4 \| 1.3e-15 \| 2.5e-21 \| 81.8 \| 59 \| (7, 67) \| 73 \| (23, 81) \| 98 \| Helix-turn-helix domain-containing protein \| Helix-turn-helix domain-containing protein; uniclust \| UniRef100\_A0A1M3GUP1 \| 99.4 \| 1.4e-15 \| 2.6e-21 \| 80.3 \| 59 \| (6, 66) \| 73 \| (2, 60) \| 88 \| Helix-turn-helix domain-containing protein \| Helix-turn-helix domain-containing protein; uniclust \| UniRef100\_A0A1C2G2R7 \| 99.4 \| 1.3e-15 \| 2.6e-21 \| 81.6 \| 63 \| (2, 66) \| 73 \| (5, 68) \| 89 \| Helix-turn-helix domain-containing protein \| Helix-turn-helix domain-containing protein; uniclust \| UniRef100\_A0A023D901 \| 99.4 \| 1.5e-15 \| 2.8e-21 \| 81.1 \| 60 \| (8, 69) \| 73 \| (13, 72) \| 98 \| Helix-turn-helix domain-containing protein \| Helix-turn-helix domain-containing protein; uniclust \| UniRef100\_A0A061YB04 \| 99.4 \| 1.4e-15 \| 2.9e-21 \| 81.6 \| 65 \| (3, 69) \| 73 \| (9, 75) \| 91 \| AlpA family phage regulatory protein \| AlpA family phage regulatory protein; uniclust \| UniRef100\_A0A1I0GNV9 \| 99.4 \| 1.5e-15 \| 2.9e-21 \| 79.6 \| 55 \| (8, 64) \| 73 \| (15, 69) \| 88 \| Helix-turn-helix domain-containing protein \| Helix-turn-helix domain-containing protein; uniclust \| UniRef100\_A0A4Q2YA13 \| 99.4 \| 1.5e-15 \| 2.9e-21 \| 79.7 \| 59 \| (8, 68) \| 73 \| (20, 78) \| 83 \| DNA-binding protein (Fragment) \| DNA-binding protein (Fragment); uniclust \| UniRef100\_A0A1M6UHM6 \| 99.4 \| 1.5e-15 \| 2.9e-21 \| 81.6 \| 62 \| (6, 69) \| 73 \| (24, 85) \| 98 \| Helix-turn-helix domain-containing protein \| Helix-turn-helix domain-containing protein; uniclust \| UniRef100\_A0A0C1C693 \| 99.4 \| 1.5e-15 \| 2.9e-21 \| 79.9 \| 63 \| (4, 68) \| 73 \| (5, 68) \| 79 \| Uncharacterized protein ORF88 \| Uncharacterized protein ORF88; uniclust \| UniRef100\_A0A099F695 \| 99.4 \| 1.5e-15 \| 3e-21 \| 81.6 \| 61 \| (7, 69) \| 73 \| (18, 78) \| 97 \| Helix-turn-helix domain-containing protein \| Helix-turn-helix domain-containing protein; uniclust \| UniRef100\_A0A0K1EPI1 \| 99.4 \| 1.6e-15 \| 3e-21 \| 81.0 \| 62 \| (4, 68) \| 73 \| (12, 73) \| 98 \| DNA-binding protein \| DNA-binding protein; uniclust \| UniRef100\_A0A2D8NLB9 \| 99.4 \| 1.6e-15 \| 3.2e-21 \| 79.1 \| 60 \| (7, 68) \| 73 \| (12, 71) \| 82 \| Excisionase \| Excisionase; uniclust \| UniRef100\_A0A1H3INP0 \| 99.4 \| 1.7e-15 \| 3.2e-21 \| 77.3 \| 62 \| (6, 69) \| 73 \| (2, 63) \| 74 \| Helix-turn-helix domain-containing protein \| Helix-turn-helix domain-containing protein; uniclust \| UniRef100\_A0A2H9U2J9 \| 99.4 \| 1.7e-15 \| 3.2e-21 \| 75.0 \| 59 \| (8, 68) \| 73 \| (1, 59) \| 63 \| AlpA family transcriptional regulator \| AlpA family transcriptional regulator; uniclust \| UniRef100\_A0A085DXN1 \| 99.4 \| 1.6e-15 \| 3.2e-21 \| 83.5 \| 63 \| (4, 68) \| 73 \| (17, 80) \| 106 \| AlpA family transcriptional regulator \| AlpA family transcriptional regulator; uniclust \| UniRef100\_A0A0F2RL48 \| 99.4 \| 1.7e-15 \| 3.3e-21 \| 83.5 \| 59 \| (9, 69) \| 73 \| (39, 99) \| 109 \| Helix-turn-helix domain-containing protein \| Helix-turn-helix domain-containing protein; uniclust \| UniRef100\_A0A0B2F8R1 \| 99.4 \| 1.7e-15 \| 3.3e-21 \| 79.8 \| 60 \| (5, 67) \| 73 \| (12, 71) \| 83 \| Helix-turn-helix domain-containing protein \| Helix-turn-helix domain-containing protein; uniclust \| UniRef100\_A0A1H1ZMM5 \| 99.4 \| 1.8e-15 \| 3.3e-21 \| 78.6 \| 60 \| (8, 69) \| 73 \| (23, 82) \| 85 \| DNA binding domain-containing protein, excisionase family \| DNA binding domain-containing protein, excisionase family; uniclust \| UniRef100\_A0A7W8EEW1 \| 99.4 \| 1.8e-15 \| 3.4e-21 \| 78.1 \| 62 \| (6, 69) \| 73 \| (14, 75) \| 78 \| Putative DNA-binding transcriptional regulator AlpA \| Putative DNA-binding transcriptional regulator AlpA; uniclust \| UniRef100\_A0A1A0KLC7 \| 99.4 \| 1.8e-15 \| 3.4e-21 \| 79.6 \| 58 \| (9, 68) \| 73 \| (21, 78) \| 87 \| Helix-turn-helix domain-containing protein \| Helix-turn-helix domain-containing protein; uniclust \| UniRef100\_A0A0M9CB32 \| 99.4 \| 1.8e-15 \| 3.6e-21 \| 82.0 \| 60 \| (6, 67) \| 73 \| (19, 79) \| 100 \| Helix-turn-helix domain-containing protein \| Helix-turn-helix domain-containing protein; uniclust \| UniRef100\_A0A011NTG0 \| 99.4 \| 1.9e-15 \| 3.6e-21 \| 81.9 \| 59 \| (4, 65) \| 73 \| (25, 83) \| 109 \| DNA binding domain, excisionase family \| DNA binding domain, excisionase family; uniclust \| UniRef100\_A0A2A4I9V0 \| 99.4 \| 1.9e-15 \| 3.7e-21 \| 78.7 \| 62 \| (7, 70) \| 73 \| (17, 78) \| 83 \| DNA-binding protein \| DNA-binding protein; uniclust \| UniRef100\_A0A087DCF9 \| 99.4 \| 1.9e-15 \| 3.7e-21 \| 78.4 \| 61 \| (5, 67) \| 73 \| (9, 70) \| 78 \| Helix-turn-helix domain-containing protein \| Helix-turn-helix domain-containing protein; uniclust \| UniRef100\_A0A1F9ABL8 \| 99.4 \| 1.9e-15 \| 3.7e-21 \| 78.8 \| 58 \| (7, 67) \| 73 \| (19, 76) \| 86 \| Helix-turn-helix domain-containing protein \| Helix-turn-helix domain-containing protein; uniclust \| UniRef100\_A0A1C6IJV6 \| 99.4 \| 1.9e-15 \| 3.7e-21 \| 77.3 \| 63 \| (1, 66) \| 73 \| (1, 64) \| 69 \| DNA binding domain, excisionase family \| DNA binding domain, excisionase family; uniclust \| UniRef100\_A0A1V5YRC9 \| 99.4 \| 2e-15 \| 3.9e-21 \| 78.5 \| 58 \| (5, 64) \| 73 \| (7, 64) \| 80 \| Helix-turn-helix domain protein \| Helix-turn-helix domain protein; uniclust \| UniRef100\_A0A0K9ETV7 \| 99.4 \| 2e-15 \| 3.9e-21 \| 76.2 \| 61 \| (5, 68) \| 73 \| (2, 62) \| 66 \| Helix-turn-helix domain-containing protein \| Helix-turn-helix domain-containing protein; uniclust \| UniRef100\_A0A011UIH5 \| 99.4 \| 2e-15 \| 4.1e-21 \| 78.8 \| 63 \| (2, 67) \| 73 \| (6, 68) \| 77 \| Excisionase \| Excisionase; uniclust \| UniRef100\_A0A291ILY0 \| 99.4 \| 2e-15 \| 4.1e-21 \| 81.1 \| 60 \| (8, 69) \| 73 \| (21, 81) \| 91 \| Helix-turn-helix domain-containing protein \| Helix-turn-helix domain-containing protein; uniclust \| UniRef100\_A0A081HYW1 \| 99.4 \| 2.1e-15 \| 4.1e-21 \| 78.9 \| 63 \| (2, 66) \| 73 \| (9, 71) \| 81 \| Helix-turn-helix domain-containing protein \| Helix-turn-helix domain-containing protein; uniclust \| UniRef100\_A0A229VYW8 \| 99.4 \| 2.1e-15 \| 4.2e-21 \| 82.9 \| 61 \| (8, 70) \| 73 \| (28, 88) \| 111 \| Phage transcriptional regulator AlpA \| Phage transcriptional regulator AlpA; uniclust \| UniRef100\_A0A0D6EU63 \| 99.4 \| 2.1e-15 \| 4.2e-21 \| 81.0 \| 57 \| (10, 68) \| 73 \| (22, 80) \| 92 \| Phage transcriptional regulator, AlpA \| Phage transcriptional regulator, AlpA; uniclust \| UniRef100\_A0A2S8GGC5 \| 99.4 \| 2.2e-15 \| 4.2e-21 \| 78.0 \| 57 \| (7, 65) \| 73 \| (9, 65) \| 76 \| Excisionase \| Excisionase; uniclust \| UniRef100\_A0A1F9CTC9 \| 99.4 \| 2.2e-15 \| 4.3e-21 \| 78.0 \| 58 \| (3, 63) \| 73 \| (1, 58) \| 77 \| Helix-turn-helix domain-containing protein \| Helix-turn-helix domain-containing protein; uniclust \| UniRef100\_A0A1D2R863 \| 99.4 \| 2.2e-15 \| 4.3e-21 \| 78.6 \| 59 \| (8, 68) \| 73 \| (11, 70) \| 84 \| Helix-turn-helix domain-containing protein \| Helix-turn-helix domain-containing protein; uniclust \| UniRef100\_A0A0K8QST4 \| 99.4 \| 2.2e-15 \| 4.3e-21 \| 79.2 \| 58 \| (5, 65) \| 73 \| (6, 63) \| 83 \| Transcriptional regulator, AlpA family \| Transcriptional regulator, AlpA family; uniclust \| UniRef100\_A0A522NUE7 \| 99.4 \| 2.2e-15 \| 4.4e-21 \| 76.7 \| 54 \| (9, 65) \| 73 \| (10, 64) \| 67 \| DNA-binding protein \| DNA-binding protein; uniclust \| UniRef100\_A0A135IGU9 \| 99.4 \| 2.3e-15 \| 4.5e-21 \| 75.2 \| 56 \| (12, 69) \| 73 \| (2, 57) \| 66 \| Helix-turn-helix domain-containing protein \| Helix-turn-helix domain-containing protein; uniclust \| UniRef100\_A0A6N4ULT1 \| 99.4 \| 2.3e-15 \| 4.5e-21 \| 77.3 \| 63 \| (4, 68) \| 73 \| (2, 66) \| 74 \| Helix-turn-helix domain-containing protein \| Helix-turn-helix domain-containing protein; uniclust \| UniRef100\_A0A024H216 \| 99.4 \| 2.3e-15 \| 4.5e-21 \| 80.9 \| 59 \| (9, 70) \| 73 \| (26, 84) \| 94 \| Uncharacterized domain protein \| Uncharacterized domain protein; uniclust \| UniRef100\_A0A0F2JEK6 \| 99.4 \| 2.3e-15 \| 4.6e-21 \| 77.5 \| 60 \| (6, 68) \| 73 \| (2, 63) \| 72 \| Helix-turn-helix domain-containing protein \| Helix-turn-helix domain-containing protein; uniclust \| UniRef100\_A0A1I4YVV8 \| 99.4 \| 2.4e-15 \| 4.7e-21 \| 82.1 \| 64 \| (4, 69) \| 73 \| (20, 83) \| 102 \| DNA binding domain-containing protein, excisionase family \| DNA binding domain-containing protein, excisionase family; uniclust \| UniRef100\_A0A0K2ZQU6 \| 99.4 \| 2.4e-15 \| 4.7e-21 \| 81.7 \| 62 \| (5, 68) \| 73 \| (22, 85) \| 99 \| Uncharacterized protein \| Uncharacterized protein; uniclust \| UniRef100\_A0A961Y0H3 \| 99.4 \| 2.5e-15 \| 4.7e-21 \| 78.7 \| 67 \| (1, 69) \| 73 \| (1, 71) \| 88 \| Helix-turn-helix domain-containing protein \| Helix-turn-helix domain-containing protein; uniclust \| UniRef100\_A0A0K2SNP2 \| 99.4 \| 2.5e-15 \| 4.8e-21 \| 82.7 \| 59 \| (1, 62) \| 73 \| (4, 62) \| 110 \| Molybdenum-pterin binding protein \| Molybdenum-pterin binding protein; uniclust \| UniRef100\_A0A1H8TNU4 \| 99.4 \| 2.6e-15 \| 4.8e-21 \| 77.7 \| 59 \| (8, 68) \| 73 \| (20, 78) \| 83 \| Transcriptional regulator, AlpA family \| Transcriptional regulator, AlpA family; uniclust \| UniRef100\_A0A1F2WGB9 \| 99.4 \| 2.5e-15 \| 4.8e-21 \| 81.2 \| 60 \| (6, 68) \| 73 \| (19, 78) \| 103 \| Helix-turn-helix domain-containing protein (Fragment) \| Helix-turn-helix domain-containing protein (Fragment); uniclust \| UniRef100\_A0A0S7WQI0 \| 99.4 \| 2.5e-15 \| 4.9e-21 \| 80.7 \| 57 \| (7, 66) \| 73 \| (23, 79) \| 94 \| Helix-turn-helix domain-containing protein \| Helix-turn-helix domain-containing protein; uniclust \| UniRef100\_A0A3E0Q7Z1 \| 99.4 \| 2.6e-15 \| 5e-21 \| 78.0 \| 64 \| (1, 67) \| 73 \| (1, 64) \| 78 \| DNA-binding protein \| DNA-binding protein; uniclust \| UniRef100\_A0A0C2VMS3 \| 99.4 \| 2.6e-15 \| 5.1e-21 \| 77.1 \| 61 \| (6, 69) \| 73 \| (2, 62) \| 72 \| Helix-turn-helix domain-containing protein \| Helix-turn-helix domain-containing protein; uniclust \| UniRef100\_A0A089N5B4 \| 99.4 \| 2.7e-15 \| 5.2e-21 \| 79.4 \| 62 \| (5, 69) \| 73 \| (14, 75) \| 88 \| Helix-turn-helix domain-containing protein \| Helix-turn-helix domain-containing protein; uniclust \| UniRef100\_A0A2C8F565 \| 99.4 \| 2.8e-15 \| 5.3e-21 \| 79.6 \| 59 \| (7, 67) \| 73 \| (30, 88) \| 92 \| Helix-turn-helix domain-containing protein \| Helix-turn-helix domain-containing protein; uniclust \| UniRef100\_A0A068NIB7 \| 99.4 \| 2.8e-15 \| 5.4e-21 \| 82.5 \| 59 \| (6, 67) \| 73 \| (32, 90) \| 109 \| Excisionase \| Excisionase; uniclust \| UniRef100\_A0A0M6XPV5 \| 99.4 \| 2.9e-15 \| 5.5e-21 \| 74.5 \| 54 \| (7, 62) \| 73 \| (10, 63) \| 66 \| DNA binding domain, excisionase family \| DNA binding domain, excisionase family; uniclust \| UniRef100\_A0A0G3BTD9 \| 99.4 \| 3e-15 \| 5.8e-21 \| 77.2 \| 64 \| (5, 70) \| 73 \| (4, 67) \| 78 \| Helix-turn-helix domain-containing protein \| Helix-turn-helix domain-containing protein; uniclust \| UniRef100\_A0A085K4B9 \| 99.4 \| 3e-15 \| 5.8e-21 \| 83.4 \| 61 \| (7, 69) \| 73 \| (24, 84) \| 121 \| DNA-binding protein \| DNA-binding protein; uniclust \| UniRef100\_A0A0C1AYG1 \| 99.4 \| 3e-15 \| 5.8e-21 \| 81.0 \| 59 \| (10, 70) \| 73 \| (37, 96) \| 100 \| Helix-turn-helix domain-containing protein \| Helix-turn-helix domain-containing protein; uniclust \| UniRef100\_A0A326TZW7 \| 99.3 \| 3.2e-15 \| 6.2e-21 \| 81.6 \| 59 \| (6, 67) \| 73 \| (14, 72) \| 104 \| PTS system nitrogen regulatory IIA component \| PTS system nitrogen regulatory IIA component; uniclust \| UniRef100\_A0A7T0C1C5 \| 99.3 \| 3.2e-15 \| 6.2e-21 \| 79.7 \| 60 \| (6, 68) \| 73 \| (1, 60) \| 96 \| Helix-turn-helix domain-containing protein \| Helix-turn-helix domain-containing protein; uniclust \| UniRef100\_A0A1F5N9I8 \| 99.3 \| 3.2e-15 \| 6.2e-21 \| 79.3 \| 62 \| (3, 67) \| 73 \| (16, 77) \| 88 \| Helix-turn-helix domain-containing protein \| Helix-turn-helix domain-containing protein; uniclust \| UniRef100\_A0A0A2WTF0 \| 99.3 \| 3.3e-15 \| 6.3e-21 \| 77.9 \| 60 \| (3, 65) \| 73 \| (7, 66) \| 80 \| Helix-turn-helix domain-containing protein \| Helix-turn-helix domain-containing protein; uniclust \| UniRef100\_A0A0A5HUG6 \| 99.3 \| 3.2e-15 \| 6.4e-21 \| 80.8 \| 62 \| (6, 69) \| 73 \| (21, 83) \| 95 \| AlpA family transcriptional regulator \| AlpA family transcriptional regulator; uniclust \| UniRef100\_L0J4B7 \| 99.3 \| 3.4e-15 \| 6.5e-21 \| 80.2 \| 64 \| (2, 67) \| 73 \| (1, 64) \| 103 \| Helix-turn-helix domain-containing protein \| Helix-turn-helix domain-containing protein; uniclust \| UniRef100\_A0A2E2QBI3 \| 99.3 \| 3.4e-15 \| 6.7e-21 \| 78.7 \| 61 \| (3, 65) \| 73 \| (7, 72) \| 86 \| DNA-binding protein \| DNA-binding protein; uniclust \| UniRef100\_A0A0G0BKN9 \| 99.3 \| 3.5e-15 \| 6.8e-21 \| 78.7 \| 60 \| (5, 67) \| 73 \| (17, 78) \| 84 \| Transcriptional regulator, MerR family \| Transcriptional regulator, MerR family; uniclust \| UniRef100\_A0A0D5M2K3 \| 99.3 \| 3.4e-15 \| 6.8e-21 \| 80.5 \| 56 \| (8, 65) \| 73 \| (20, 76) \| 92 \| Phage transcriptional regulator, AlpA \| Phage transcriptional regulator, AlpA; uniclust \| UniRef100\_A0A0A2XY55 \| 99.3 \| 3.5e-15 \| 6.9e-21 \| 80.3 \| 61 \| (6, 68) \| 73 \| (23, 84) \| 94 \| AlpA family phage regulatory protein \| AlpA family phage regulatory protein; uniclust \| UniRef100\_A0A3D3UV23 \| 99.3 \| 3.6e-15 \| 7e-21 \| 79.4 \| 60 \| (2, 63) \| 73 \| (7, 66) \| 90 \| Helix-turn-helix domain-containing protein \| Helix-turn-helix domain-containing protein; uniclust \| UniRef100\_A0A1A0M401 \| 99.3 \| 3.7e-15 \| 7.1e-21 \| 78.8 \| 65 \| (5, 70) \| 73 \| (9, 73) \| 89 \| Helix-turn-helix domain-containing protein \| Helix-turn-helix domain-containing protein; uniclust \| UniRef100\_A0A143B7R0 \| 99.3 \| 3.7e-15 \| 7.1e-21 \| 80.3 \| 60 \| (5, 67) \| 73 \| (30, 89) \| 99 \| DNA-binding protein \| DNA-binding protein; uniclust \| UniRef100\_A0A485B3H2 \| 99.3 \| 3.8e-15 \| 7.1e-21 \| 74.2 \| 56 \| (8, 66) \| 73 \| (2, 57) \| 69 \| Predicted transcriptional regulator \| Predicted transcriptional regulator; uniclust \| UniRef100\_A0A0Q7JLQ8 \| 99.3 \| 3.7e-15 \| 7.1e-21 \| 76.0 \| 65 \| (1, 67) \| 73 \| (1, 66) \| 70 \| Helix-turn-helix domain-containing protein \| Helix-turn-helix domain-containing protein; uniclust \| UniRef100\_A0A644TYX9 \| 99.3 \| 3.7e-15 \| 7.4e-21 \| 80.8 \| 59 \| (9, 70) \| 73 \| (38, 96) \| 98 \| Helix-turn-helix domain-containing protein \| Helix-turn-helix domain-containing protein; uniclust \| UniRef100\_A0A0K8QVH0 \| 99.3 \| 3.9e-15 \| 7.5e-21 \| 80.2 \| 58 \| (5, 65) \| 73 \| (18, 75) \| 106 \| Protein containing DNA binding domain, excisionase family \| Protein containing DNA binding domain, excisionase family; uniclust \| UniRef100\_A0A1G8QBS7 \| 99.3 \| 4e-15 \| 7.7e-21 \| 77.9 \| 62 \| (6, 69) \| 73 \| (13, 74) \| 83 \| Transcriptional regulator, AlpA family \| Transcriptional regulator, AlpA family; uniclust \| UniRef100\_A0A2E8B578 \| 99.3 \| 4.2e-15 \| 7.8e-21 \| 79.4 \| 58 \| (7, 67) \| 73 \| (38, 95) \| 104 \| DNA-binding protein \| DNA-binding protein; uniclust \| UniRef100\_A0A0F8YEC9 \| 99.3 \| 4e-15 \| 8e-21 \| 79.4 \| 58 \| (6, 65) \| 73 \| (14, 71) \| 88 \| Helix-turn-helix domain-containing protein \| Helix-turn-helix domain-containing protein; uniclust \| UniRef100\_A0A0B0DF89 \| 99.3 \| 4.1e-15 \| 8e-21 \| 77.4 \| 65 \| (3, 69) \| 73 \| (11, 75) \| 79 \| Helix-turn-helix domain-containing protein \| Helix-turn-helix domain-containing protein; uniclust \| UniRef100\_A0A177QRW9 \| 99.3 \| 4.2e-15 \| 8.2e-21 \| 83.8 \| 59 \| (6, 67) \| 73 \| (21, 79) \| 131 \| Helix-turn-helix domain-containing protein \| Helix-turn-helix domain-containing protein; uniclust \| UniRef100\_A0A0T6A0K1 \| 99.3 \| 4.2e-15 \| 8.3e-21 \| 76.4 \| 60 \| (4, 66) \| 73 \| (3, 62) \| 72 \| Helix-turn-helix domain-containing protein \| Helix-turn-helix domain-containing protein; uniclust \| UniRef100\_A0A1X1MGL1 \| 99.3 \| 4.6e-15 \| 8.8e-21 \| 77.0 \| 59 \| (5, 65) \| 73 \| (3, 61) \| 84 \| Helix-turn-helix domain-containing protein \| Helix-turn-helix domain-containing protein; uniclust \| UniRef100\_A0A059E7V9 \| 99.3 \| 4.6e-15 \| 8.9e-21 \| 82.0 \| 61 \| (7, 69) \| 73 \| (42, 102) \| 118 \| DNA-binding protein \| DNA-binding protein; uniclust \| UniRef100\_A0A109LZ39 \| 99.3 \| 4.6e-15 \| 8.9e-21 \| 75.9 \| 57 \| (7, 65) \| 73 \| (13, 70) \| 73 \| Helix-turn-helix domain-containing protein \| Helix-turn-helix domain-containing protein; uniclust \| UniRef100\_A0A022GBV9 \| 99.3 \| 4.6e-15 \| 8.9e-21 \| 82.1 \| 61 \| (8, 70) \| 73 \| (38, 98) \| 117 \| Transcriptional regulator \| Transcriptional regulator; uniclust \| UniRef100\_A0A087L5X3 \| 99.3 \| 4.5e-15 \| 8.9e-21 \| 79.8 \| 64 \| (3, 68) \| 73 \| (8, 73) \| 92 \| AlpA family transcriptional regulator \| AlpA family transcriptional regulator; uniclust \| UniRef100\_A0A0M7B8D3 \| 99.3 \| 4.7e-15 \| 9e-21 \| 81.3 \| 59 \| (7, 67) \| 73 \| (2, 60) \| 115 \| Helix-turn-helix domain protein \| Helix-turn-helix domain protein; uniclust \| UniRef100\_A0A094PUR9 \| 99.3 \| 4.7e-15 \| 9e-21 \| 78.0 \| 61 \| (5, 67) \| 73 \| (13, 73) \| 88 \| Helix-turn-helix domain-containing protein \| Helix-turn-helix domain-containing protein; uniclust \| UniRef100\_A0A1N6NC24 \| 99.3 \| 4.6e-15 \| 9e-21 \| 77.7 \| 59 \| (8, 68) \| 73 \| (19, 77) \| 83 \| Helix-turn-helix domain-containing protein \| Helix-turn-helix domain-containing protein; uniclust \| UniRef100\_A0A078LJ41 \| 99.3 \| 4.7e-15 \| 9.4e-21 \| 80.4 \| 66 \| (1, 68) \| 73 \| (5, 71) \| 96 \| Transcriptional regulator \| Transcriptional regulator; uniclust \| UniRef100\_A0A366HET7 \| 99.3 \| 5e-15 \| 9.5e-21 \| 78.3 \| 60 \| (8, 69) \| 73 \| (18, 77) \| 92 \| Helix-turn-helix protein \| Helix-turn-helix protein; uniclust \| UniRef100\_A0A1B4S5B3 \| 99.3 \| 5e-15 \| 9.7e-21 \| 85.1 \| 65 \| (2, 68) \| 73 \| (14, 78) \| 146 \| Helix-turn-helix domain-containing protein \| Helix-turn-helix domain-containing protein; uniclust \| UniRef100\_A0A1H1RBV9 \| 99.3 \| 4.9e-15 \| 9.7e-21 \| 77.5 \| 55 \| (7, 64) \| 73 \| (14, 68) \| 79 \| DNA binding domain-containing protein, excisionase family \| DNA binding domain-containing protein, excisionase family; uniclust \| UniRef100\_A0A2G2K1G3 \| 99.3 \| 5e-15 \| 9.8e-21 \| 80.2 \| 60 \| (6, 68) \| 73 \| (18, 77) \| 97 \| DNA-binding protein \| DNA-binding protein; uniclust \| UniRef100\_A0A290ZH97 \| 99.3 \| 5.2e-15 \| 9.9e-21 \| 76.1 \| 67 \| (1, 69) \| 73 \| (1, 68) \| 79 \| Excisionase \| Excisionase; uniclust \| UniRef100\_A0A382E9C4 \| 99.3 \| 5.4e-15 \| 1.1e-20 \| 78.2 \| 64 \| (5, 70) \| 73 \| (14, 77) \| 87 \| Helix-turn-helix domain-containing protein \| Helix-turn-helix domain-containing protein; uniclust \| UniRef100\_A0A2G2GU36 \| 99.3 \| 5.5e-15 \| 1.1e-20 \| 78.1 \| 62 \| (7, 70) \| 73 \| (13, 79) \| 86 \| Helix-turn-helix domain-containing protein \| Helix-turn-helix domain-containing protein; uniclust \| UniRef100\_A0A9E1W7L4 \| 99.3 \| 5.7e-15 \| 1.1e-20 \| 75.9 \| 61 \| (8, 70) \| 73 \| (14, 74) \| 79 \| Helix-turn-helix domain-containing protein \| Helix-turn-helix domain-containing protein; uniclust \| UniRef100\_A0A011V604 \| 99.3 \| 5.5e-15 \| 1.1e-20 \| 79.2 \| 61 \| (6, 68) \| 73 \| (19, 79) \| 93 \| AlpA family transcriptional regulator \| AlpA family transcriptional regulator; uniclust \| UniRef100\_A0A1H3REZ3 \| 99.3 \| 5.6e-15 \| 1.1e-20 \| 74.2 \| 59 \| (5, 66) \| 73 \| (4, 62) \| 66 \| Putative molybdopterin biosynthesis protein \| Putative molybdopterin biosynthesis protein; uniclust \| UniRef100\_A0A4D7DJZ9 \| 99.3 \| 5.8e-15 \| 1.1e-20 \| 73.8 \| 61 \| (5, 67) \| 73 \| (3, 63) \| 66 \| Helix-turn-helix domain-containing protein \| Helix-turn-helix domain-containing protein; uniclust \| UniRef100\_A0A4U1KY23 \| 99.3 \| 5.9e-15 \| 1.1e-20 \| 73.2 \| 57 \| (12, 70) \| 73 \| (1, 58) \| 64 \| Helix-turn-helix domain-containing protein \| Helix-turn-helix domain-containing protein; uniclust \| UniRef100\_A0A1A9SZU8 \| 99.3 \| 5.8e-15 \| 1.1e-20 \| 77.5 \| 59 \| (8, 68) \| 73 \| (2, 61) \| 81 \| Helix-turn-helix domain-containing protein \| Helix-turn-helix domain-containing protein; uniclust \| UniRef100\_A0A0K2GF18 \| 99.3 \| 5.9e-15 \| 1.1e-20 \| 77.2 \| 58 \| (7, 67) \| 73 \| (11, 68) \| 81 \| Helix-turn-helix domain-containing protein \| Helix-turn-helix domain-containing protein; uniclust \| UniRef100\_A0A087MDQ3 \| 99.3 \| 6e-15 \| 1.2e-20 \| 77.2 \| 64 \| (2, 67) \| 73 \| (13, 76) \| 81 \| Helix-turn-helix domain-containing protein \| Helix-turn-helix domain-containing protein; uniclust \| UniRef100\_A0A9D8HGG9 \| 99.3 \| 6.3e-15 \| 1.2e-20 \| 78.9 \| 62 \| (5, 68) \| 73 \| (5, 66) \| 106 \| Helix-turn-helix domain-containing protein \| Helix-turn-helix domain-containing protein; uniclust \| UniRef100\_A0A399Y389 \| 99.3 \| 6.1e-15 \| 1.2e-20 \| 76.9 \| 59 \| (7, 68) \| 73 \| (1, 59) \| 80 \| AlpA family transcriptional regulator \| AlpA family transcriptional regulator; uniclust \| UniRef100\_A0A097ELU5 \| 99.3 \| 6.1e-15 \| 1.2e-20 \| 78.9 \| 58 \| (8, 67) \| 73 \| (19, 76) \| 93 \| AlpA family transcriptional regulator \| AlpA family transcriptional regulator; uniclust \| UniRef100\_A0A084EIT9 \| 99.3 \| 6.4e-15 \| 1.2e-20 \| 77.2 \| 63 \| (6, 68) \| 73 \| (20, 82) \| 95 \| Helix-turn-helix domain-containing protein \| Helix-turn-helix domain-containing protein; uniclust \| UniRef100\_A0A356W9G0 \| 99.3 \| 6.2e-15 \| 1.2e-20 \| 77.9 \| 60 \| (8, 69) \| 73 \| (23, 83) \| 89 \| Helix-turn-helix domain-containing protein \| Helix-turn-helix domain-containing protein; uniclust \| UniRef100\_A0A1M5BQS2 \| 99.3 \| 6.4e-15 \| 1.2e-20 \| 80.1 \| 62 \| (5, 68) \| 73 \| (22, 87) \| 109 \| Helix-turn-helix domain-containing protein \| Helix-turn-helix domain-containing protein; uniclust \| UniRef100\_A0A0K2B6L2 \| 99.3 \| 6.4e-15 \| 1.2e-20 \| 78.8 \| 64 \| (3, 68) \| 73 \| (14, 78) \| 95 \| PCQ3\_68 \| PCQ3\_68; uniclust \| UniRef100\_A0A7W7ZCS8 \| 99.3 \| 6.5e-15 \| 1.2e-20 \| 77.8 \| 56 \| (9, 66) \| 73 \| (16, 71) \| 90 \| Putative DNA-binding transcriptional regulator AlpA \| Putative DNA-binding transcriptional regulator AlpA; uniclust \| UniRef100\_A0A1Q5VLA1 \| 99.3 \| 6.5e-15 \| 1.3e-20 \| 78.2 \| 64 \| (3, 68) \| 73 \| (8, 72) \| 86 \| AlpA family transcriptional regulator \| AlpA family transcriptional regulator; uniclust \| UniRef100\_A0A017HLU1 \| 99.3 \| 6.5e-15 \| 1.3e-20 \| 78.6 \| 58 \| (9, 68) \| 73 \| (23, 83) \| 89 \| Helix-turn-helix domain-containing protein \| Helix-turn-helix domain-containing protein; uniclust \| UniRef100\_A0A0C1GY24 \| 99.3 \| 6.5e-15 \| 1.3e-20 \| 78.1 \| 64 \| (3, 68) \| 73 \| (6, 70) \| 84 \| DNA-binding protein \| DNA-binding protein; uniclust \| UniRef100\_A0A1H1H733 \| 99.3 \| 6.8e-15 \| 1.3e-20 \| 76.0 \| 59 \| (7, 67) \| 73 \| (10, 68) \| 78 \| Helix-turn-helix domain-containing protein \| Helix-turn-helix domain-containing protein; uniclust \| UniRef100\_A0A0P7CAH9 \| 99.3 \| 6.9e-15 \| 1.4e-20 \| 78.4 \| 62 \| (4, 68) \| 73 \| (6, 67) \| 88 \| Helix-turn-helix domain-containing protein \| Helix-turn-helix domain-containing protein; uniclust \| UniRef100\_A0A0K1JJ15 \| 99.3 \| 6.8e-15 \| 1.4e-20 \| 80.0 \| 62 \| (5, 68) \| 73 \| (16, 77) \| 97 \| Helix-turn-helix domain-containing protein \| Helix-turn-helix domain-containing protein; uniclust \| UniRef100\_A0A011QUG7 \| 99.3 \| 7e-15 \| 1.4e-20 \| 77.4 \| 61 \| (7, 69) \| 73 \| (15, 75) \| 85 \| Helix-turn-helix domain protein \| Helix-turn-helix domain protein; uniclust \| UniRef100\_A0A0D4DU24 \| 99.3 \| 7.2e-15 \| 1.4e-20 \| 80.4 \| 58 \| (9, 68) \| 73 \| (21, 78) \| 115 \| DNA-binding protein \| DNA-binding protein; uniclust \| UniRef100\_A0A2H0LJP8 \| 99.3 \| 7.2e-15 \| 1.4e-20 \| 76.8 \| 56 \| (8, 66) \| 73 \| (6, 61) \| 84 \| Helix-turn-helix domain-containing protein \| Helix-turn-helix domain-containing protein; uniclust \| UniRef100\_A0A1A7QHD2 \| 99.3 \| 7.2e-15 \| 1.4e-20 \| 75.8 \| 57 \| (4, 62) \| 73 \| (10, 66) \| 76 \| Helix-turn-helix domain-containing protein \| Helix-turn-helix domain-containing protein; uniclust \| UniRef100\_A0A2N3QTQ8 \| 99.3 \| 7.5e-15 \| 1.4e-20 \| 75.3 \| 57 \| (8, 66) \| 73 \| (14, 71) \| 77 \| Phage transcriptional regulator, AlpA \| Phage transcriptional regulator, AlpA; uniclust \| UniRef100\_A0A1M6DCH5 \| 99.3 \| 7.6e-15 \| 1.4e-20 \| 74.7 \| 62 \| (6, 69) \| 73 \| (8, 69) \| 74 \| Helix-turn-helix domain-containing protein \| Helix-turn-helix domain-containing protein; uniclust \| UniRef100\_A0A381XKI2 \| 99.3 \| 7.5e-15 \| 1.5e-20 \| 74.0 \| 62 \| (2, 66) \| 73 \| (2, 63) \| 66 \| Helix-turn-helix domain-containing protein \| Helix-turn-helix domain-containing protein; uniclust \| UniRef100\_A0A1V4VK00 \| 99.3 \| 7.5e-15 \| 1.5e-20 \| 74.9 \| 58 \| (6, 66) \| 73 \| (5, 62) \| 71 \| Helix-turn-helix domain protein \| Helix-turn-helix domain protein; uniclust \| UniRef100\_A0A060ZRS2 \| 99.3 \| 7.6e-15 \| 1.5e-20 \| 77.3 \| 61 \| (6, 69) \| 73 \| (16, 76) \| 81 \| DNA binding domain protein, excisionase family \| DNA binding domain protein, excisionase family; uniclust \| UniRef100\_A0A0Q8DF19 \| 99.3 \| 7.7e-15 \| 1.5e-20 \| 79.3 \| 56 \| (9, 66) \| 73 \| (30, 85) \| 98 \| Helix-turn-helix domain-containing protein \| Helix-turn-helix domain-containing protein; uniclust \| UniRef100\_A0A079YS49 \| 99.3 \| 7.7e-15 \| 1.5e-20 \| 80.9 \| 56 \| (7, 65) \| 73 \| (46, 101) \| 104 \| DNA-binding protein \| DNA-binding protein; uniclust \| UniRef100\_A0A095UZ92 \| 99.3 \| 7.7e-15 \| 1.6e-20 \| 82.4 \| 58 \| (7, 67) \| 73 \| (52, 109) \| 115 \| HTH merR-type domain-containing protein \| HTH merR-type domain-containing protein; uniclust \| UniRef100\_A0A1G9J2G5 \| 99.3 \| 8.3e-15 \| 1.6e-20 \| 73.3 \| 58 \| (6, 65) \| 73 \| (2, 59) \| 65 \| Helix-turn-helix domain-containing protein \| Helix-turn-helix domain-containing protein; uniclust \| UniRef100\_A0A1H8Y2E5 \| 99.3 \| 8.7e-15 \| 1.6e-20 \| 77.6 \| 56 \| (9, 66) \| 73 \| (41, 96) \| 98 \| DNA binding domain-containing protein, excisionase family \| DNA binding domain-containing protein, excisionase family; uniclust \| UniRef100\_A0A0P7YQL3 \| 99.3 \| 8.4e-15 \| 1.7e-20 \| 80.4 \| 60 \| (9, 70) \| 73 \| (42, 102) \| 105 \| Helix-turn-helix domain-containing protein \| Helix-turn-helix domain-containing protein; uniclust \| UniRef100\_A0A8J2ZB08 \| 99.3 \| 8.8e-15 \| 1.7e-20 \| 74.3 \| 67 \| (1, 69) \| 73 \| (1, 67) \| 77 \| Helix-turn-helix domain-containing protein \| Helix-turn-helix domain-containing protein; uniclust \| UniRef100\_A0A1F9Y074 \| 99.3 \| 9e-15 \| 1.7e-20 \| 76.1 \| 62 \| (6, 69) \| 73 \| (13, 76) \| 81 \| Helix-turn-helix domain-containing protein \| Helix-turn-helix domain-containing protein; uniclust \| UniRef100\_A0A149UPN1 \| 99.3 \| 9e-15 \| 1.7e-20 \| 76.6 \| 62 \| (7, 70) \| 73 \| (15, 77) \| 81 \| AlpA family phage regulatory protein \| AlpA family phage regulatory protein; uniclust \| UniRef100\_A0A1H0TUD0 \| 99.3 \| 9.6e-15 \| 1.9e-20 \| 75.1 \| 64 \| (4, 69) \| 73 \| (5, 69) \| 73 \| Transcriptional regulator, AlpA family \| Transcriptional regulator, AlpA family; uniclust \| UniRef100\_A0A2W4KKL3 \| 99.3 \| 9.4e-15 \| 1.9e-20 \| 77.0 \| 59 \| (6, 67) \| 73 \| (8, 66) \| 81 \| DNA-binding protein \| DNA-binding protein; uniclust \| UniRef100\_A0A371RLY0 \| 99.3 \| 1e-14 \| 1.9e-20 \| 75.2 \| 61 \| (6, 68) \| 73 \| (15, 76) \| 78 \| DNA-binding protein \| DNA-binding protein; uniclust \| UniRef100\_A0A098U8C9 \| 99.3 \| 1e-14 \| 2e-20 \| 78.2 \| 66 \| (2, 69) \| 73 \| (14, 80) \| 91 \| AlpA family transcriptional regulator \| AlpA family transcriptional regulator; uniclust \| UniRef100\_A0A839A0Y5 \| 99.3 \| 1e-14 \| 2e-20 \| 73.7 \| 58 \| (7, 66) \| 73 \| (1, 58) \| 69 \| Helix-turn-helix domain-containing protein \| Helix-turn-helix domain-containing protein; uniclust \| UniRef100\_A0A2D6GE75 \| 99.3 \| 1e-14 \| 2e-20 \| 76.6 \| 64 \| (2, 67) \| 73 \| (7, 70) \| 79 \| DNA-binding protein \| DNA-binding protein; uniclust \| UniRef100\_A0A0W1L165 \| 99.3 \| 1e-14 \| 2e-20 \| 78.5 \| 57 \| (6, 64) \| 73 \| (21, 78) \| 92 \| AlpA family transcriptional regulator \| AlpA family transcriptional regulator; uniclust \| UniRef100\_A0A2N1W432 \| 99.3 \| 1e-14 \| 2e-20 \| 76.3 \| 58 \| (7, 67) \| 73 \| (1, 58) \| 83 \| DNA-binding protein \| DNA-binding protein; uniclust \| UniRef100\_A0A139D2J3 \| 99.3 \| 1e-14 \| 2e-20 \| 73.9 \| 57 \| (9, 68) \| 73 \| (2, 58) \| 65 \| Helix-turn-helix domain-containing protein \| Helix-turn-helix domain-containing protein; uniclust \| UniRef100\_A0A1I0F1X2 \| 99.3 \| 1.1e-14 \| 2.1e-20 \| 72.3 \| 57 \| (6, 65) \| 73 \| (2, 58) \| 62 \| DNA binding domain-containing protein, excisionase family \| DNA binding domain-containing protein, excisionase family; uniclust \| UniRef100\_A0A1E7HLJ3 \| 99.3 \| 1.1e-14 \| 2.1e-20 \| 78.5 \| 61 \| (6, 68) \| 73 \| (27, 91) \| 95 \| Helix-turn-helix domain-containing protein \| Helix-turn-helix domain-containing protein; uniclust \| UniRef100\_A0A1Q5RFL0 \| 99.3 \| 1.1e-14 \| 2.1e-20 \| 80.1 \| 59 \| (9, 69) \| 73 \| (27, 85) \| 112 \| Helix-turn-helix domain-containing protein \| Helix-turn-helix domain-containing protein; uniclust \| UniRef100\_A0A5C5RVC0 \| 99.3 \| 1.2e-14 \| 2.2e-20 \| 77.5 \| 59 \| (8, 68) \| 73 \| (37, 95) \| 98 \| Helix-turn-helix domain-containing protein \| Helix-turn-helix domain-containing protein; uniclust \| UniRef100\_A0A1P8XFB5 \| 99.3 \| 1.1e-14 \| 2.2e-20 \| 76.3 \| 58 \| (10, 69) \| 73 \| (11, 69) \| 83 \| DNA-binding protein \| DNA-binding protein; uniclust \| UniRef100\_A0A085DXK9 \| 99.3 \| 1.1e-14 \| 2.2e-20 \| 81.1 \| 62 \| (5, 68) \| 73 \| (28, 90) \| 113 \| AlpA family transcriptional regulator \| AlpA family transcriptional regulator; uniclust \| UniRef100\_A0A011NBN1 \| 99.3 \| 1.1e-14 \| 2.3e-20 \| 81.5 \| 61 \| (6, 68) \| 73 \| (25, 86) \| 112 \| DNA-binding protein \| DNA-binding protein; uniclust \| UniRef100\_A0A0K9EV25 \| 99.3 \| 1.2e-14 \| 2.3e-20 \| 76.6 \| 63 \| (2, 66) \| 73 \| (1, 63) \| 84 \| Helix-turn-helix domain-containing protein \| Helix-turn-helix domain-containing protein; uniclust \| UniRef100\_A0A085AAA5 \| 99.3 \| 1.1e-14 \| 2.3e-20 \| 79.1 \| 64 \| (3, 68) \| 73 \| (16, 80) \| 97 \| DNA-binding protein \| DNA-binding protein; uniclust \| UniRef100\_A0A074TCW2 \| 99.3 \| 1.2e-14 \| 2.3e-20 \| 77.9 \| 64 \| (5, 70) \| 73 \| (14, 79) \| 89 \| Prophage CP4-57 regulatory \| Prophage CP4-57 regulatory; uniclust \| UniRef100\_A0A142WZF1 \| 99.3 \| 1.2e-14 \| 2.3e-20 \| 78.0 \| 61 \| (4, 66) \| 73 \| (10, 71) \| 94 \| MerR family regulatory protein \| MerR family regulatory protein; uniclust \| UniRef100\_A0A954DPY9 \| 99.3 \| 1.2e-14 \| 2.3e-20 \| 74.5 \| 62 \| (5, 68) \| 73 \| (2, 63) \| 81 \| Helix-turn-helix domain-containing protein \| Helix-turn-helix domain-containing protein; pdb70 \| 1Z4H\_A \| 98.6 \| 9.9e-12 \| 7.2e-16 \| 53.1 \| 62 \| (3, 66) \| 73 \| (2, 63) \| 66 \| Tor inhibition protein \| 1Z4H\_A Tor inhibition protein winged helix, reverse turn, PROTEIN; pdb70 \| 6AMA\_Y \| 98.5 \| 3.7e-11 \| 2.7e-15 \| 51.4 \| 64 \| (2, 67) \| 73 \| (5, 68) \| 71 \| DNA binding domain-containing protein, excisionase \| 6AMA\_Y DNA binding domain-containing protein, excisionase BldC, S. coelicolor, developmental switch; pdb70 \| 6AMA\_A \| 98.5 \| 4.4e-11 \| 3.2e-15 \| 51.3 \| 62 \| (4, 67) \| 73 \| (7, 68) \| 71 \| DNA binding domain-containing protein, excisionase \| 6AMA\_A DNA binding domain-containing protein, excisionase BldC, S. coelicolor, developmental switch; pdb70 \| 6AMK\_A \| 98.4 \| 5.9e-11 \| 4.3e-15 \| 51.1 \| 60 \| (6, 67) \| 73 \| (10, 69) \| 72 \| DNA binding domain-containing protein, excisionase \| 6AMK\_A DNA binding domain-containing protein, excisionase BldC, Streptomyces, MerR-like, DNA BINDING HET: MSE; pdb70 \| 6HLK\_A \| 98.4 \| 1.2e-10 \| 8.8e-15 \| 57.0 \| 58 \| (6, 66) \| 73 \| (2, 60) \| 153 \| Redirecting phage packaging protein C \| 6HLK\_A Redirecting phage packaging protein C Redirecting packaging protein, DNA Binding HET: MSE; pdb70 \| 6HN7\_A \| 98.4 \| 1.2e-10 \| 8.8e-15 \| 57.0 \| 58 \| (6, 66) \| 73 \| (2, 60) \| 153 \| Redirecting phage packaging protein C \| 6HN7\_A Redirecting phage packaging protein C Redirecting packaging protein, DNA Binding; pdb70 \| 2ZHG\_A \| 98.3 \| 2.9e-10 \| 2.1e-14 \| 56.4 \| 57 \| (1, 59) \| 73 \| (1, 57) \| 154 \| Redox-sensitive transcriptional activator soxR/DNA Complex \| 2ZHG\_A Redox-sensitive transcriptional activator soxR/DNA Complex oxidative stress, MerR family, Activator; pdb70 \| 2ZHH\_A \| 98.3 \| 2.9e-10 \| 2.1e-14 \| 56.4 \| 57 \| (1, 59) \| 73 \| (1, 57) \| 154 \| Redox-sensitive transcriptional activator soxR \| 2ZHH\_A Redox-sensitive transcriptional activator soxR oxidative stress, MerR family, Activator; pdb70 \| 1Y6U\_A \| 98.2 \| 3.8e-10 \| 2.8e-14 \| 48.6 \| 56 \| (6, 64) \| 73 \| (11, 68) \| 70 \| Excisionase from transposon Tn916 \| 1Y6U\_A Excisionase from transposon Tn916 DNA architectural protein, Tyrosine recombinase; pdb70 \| 4J2N\_B \| 98.1 \| 9.9e-10 \| 7.2e-14 \| 44.7 \| 54 \| (8, 63) \| 73 \| (3, 56) \| 58 \| Gp37 \| 4J2N\_B Gp37 Winged-Helix, Doman Swap, Filament, VIRAL HET: SO4; pdb70 \| 4J2N\_C \| 98.1 \| 9.9e-10 \| 7.2e-14 \| 44.7 \| 54 \| (8, 63) \| 73 \| (3, 56) \| 58 \| Gp37 \| 4J2N\_C Gp37 Winged-Helix, Doman Swap, Filament, VIRAL HET: SO4; pdb70 \| 3GPV\_B \| 98.1 \| 1.1e-09 \| 7.9e-14 \| 53.7 \| 58 \| (2, 61) \| 73 \| (7, 65) \| 148 \| Transcriptional regulator, MerR family \| 3GPV\_B Transcriptional regulator, MerR family transcriptional regulator, MerR family, Protein; pdb70 \| 4LHF\_A \| 98.1 \| 1.9e-09 \| 1.4e-13 \| 49.3 \| 57 \| (5, 63) \| 73 \| (5, 69) \| 91 \| Regulatory protein cox \| 4LHF\_A Regulatory protein cox helix-turn-helix, DNA binding, VIRAL PROTEIN; pdb70 \| 4R4E\_A \| 98.0 \| 2.2e-09 \| 1.6e-13 \| 47.2 \| 57 \| (3, 61) \| 73 \| (3, 59) \| 84 \| HTH-type transcriptional regulator GlnR \| 4R4E\_A HTH-type transcriptional regulator GlnR TnrA/GlnR family, B. subtilis, nitrogen HET: CXS; pdb70 \| 5C8E\_G \| 98.0 \| 2.3e-09 \| 1.6e-13 \| 57.8 \| 59 \| (2, 62) \| 73 \| (17, 76) \| 305 \| Light-dependent transcriptional regulator CarH/DNA Complex \| 5C8E\_G Light-dependent transcriptional regulator CarH/DNA Complex Transcription factor, light sensor, adenosylcobalamin-binding HET: B12, 5AD; pdb70 \| 5C8F\_A \| 98.0 \| 2.3e-09 \| 1.6e-13 \| 57.8 \| 59 \| (2, 62) \| 73 \| (17, 76) \| 305 \| Light-dependent transcriptional regulator CarH \| 5C8F\_A Light-dependent transcriptional regulator CarH Transcription factor, light sensor, adenosylcobalamin-binding HET: B12; pdb70 \| 6HN7\_B \| 98.0 \| 3.2e-09 \| 2.3e-13 \| 49.2 \| 56 \| (8, 66) \| 73 \| (2, 59) \| 101 \| Redirecting phage packaging protein C \| 6HN7\_B Redirecting phage packaging protein C Redirecting packaging protein, DNA Binding; pdb70 \| 4R24\_B \| 98.0 \| 3.3e-09 \| 2.4e-13 \| 46.7 \| 54 \| (5, 60) \| 73 \| (2, 55) \| 85 \| HTH-type transcriptional regulator TnrA \| 4R24\_B HTH-type transcriptional regulator TnrA TnrA, GS, B. subtilis, GlnK; pdb70 \| 3UCS\_B \| 97.9 \| 5.3e-09 \| 3.9e-13 \| 48.1 \| 54 \| (6, 61) \| 73 \| (3, 56) \| 102 \| Chaperone-modulator protein CbpM, Curved DNA-binding \| 3UCS\_B Chaperone-modulator protein CbpM, Curved DNA-binding protein-protein complex, Structural Genomics, Montreal-Kingston; pdb70 \| 5CRL\_B \| 97.9 \| 6.2e-09 \| 4.4e-13 \| 50.1 \| 54 \| (6, 61) \| 73 \| (2, 56) \| 134 \| Mercuric resistance operon regulatory protein \| 5CRL\_B Mercuric resistance operon regulatory protein transcription activator, MerR, mercury, P.; pdb70 \| 1J9I\_A \| 97.8 \| 1.2e-08 \| 8.9e-13 \| 43.2 \| 54 \| (11, 67) \| 73 \| (2, 57) \| 68 \| TERMINASE SMALL SUBUNIT \| 1J9I\_A TERMINASE SMALL SUBUNIT DNA BINDING DOMAIN, HOMODIMER, VIRAL; pdb70 \| 4R22\_B \| 97.8 \| 1.7e-08 \| 1.2e-12 \| 43.8 \| 54 \| (6, 61) \| 73 \| (2, 55) \| 82 \| HTH-type transcriptional regulator TnrA \| 4R22\_B HTH-type transcriptional regulator TnrA New family of transcription regulators; pdb70 \| 5XQL\_A \| 97.7 \| 2.3e-08 \| 1.6e-12 \| 52.8 \| 57 \| (2, 60) \| 73 \| (12, 70) \| 291 \| Multidrug-efflux transporter 1 regulator \| 5XQL\_A Multidrug-efflux transporter 1 regulator Pseudomonas aeruginosa, transcriptional regulator, TRANSCRIPTION HET: C2E; pdb70 \| 5GPE\_C \| 97.7 \| 2.4e-08 \| 1.7e-12 \| 47.6 \| 49 \| (11, 61) \| 73 \| (1, 50) \| 129 \| Transcriptional regulator, MerR-family \| 5GPE\_C Transcriptional regulator, MerR-family PbrR691, MerR family, Transcription activator; pdb70 \| 5GPE\_D \| 97.7 \| 2.4e-08 \| 1.7e-12 \| 47.6 \| 49 \| (11, 61) \| 73 \| (1, 50) \| 129 \| Transcriptional regulator, MerR-family \| 5GPE\_D Transcriptional regulator, MerR-family PbrR691, MerR family, Transcription activator; pdb70 \| 4UA1\_A \| 97.7 \| 2.5e-08 \| 1.8e-12 \| 47.8 \| 50 \| (11, 62) \| 73 \| (2, 52) \| 132 \| Transcriptional regulator MerR \| 4UA1\_A Transcriptional regulator MerR Metalloregulatory protein, DNA BINDING PROTEIN; pdb70 \| 4UA1\_B \| 97.7 \| 2.5e-08 \| 1.8e-12 \| 47.8 \| 50 \| (11, 62) \| 73 \| (2, 52) \| 132 \| Transcriptional regulator MerR \| 4UA1\_B Transcriptional regulator MerR Metalloregulatory protein, DNA BINDING PROTEIN; pdb70 \| 4UA2\_C \| 97.7 \| 2.5e-08 \| 1.8e-12 \| 47.8 \| 50 \| (11, 62) \| 73 \| (2, 52) \| 132 \| Transcriptional regulator MerR \| 4UA2\_C Transcriptional regulator MerR Metalloregulatory protein, DNA BINDING PROTEIN HET: MSE; pdb70 \| 3GP4\_A \| 97.7 \| 2.8e-08 \| 2e-12 \| 48.2 \| 49 \| (11, 61) \| 73 \| (2, 51) \| 142 \| Transcriptional regulator, MerR family \| 3GP4\_A Transcriptional regulator, MerR family structural genomics, MerR, transcriptional regulator HET: GOL; pdb70 \| 3GP4\_B \| 97.7 \| 2.8e-08 \| 2e-12 \| 48.2 \| 49 \| (11, 61) \| 73 \| (2, 51) \| 142 \| Transcriptional regulator, MerR family \| 3GP4\_B Transcriptional regulator, MerR family structural genomics, MerR, transcriptional regulator HET: GOL; pdb70 \| 3HH0\_A \| 97.7 \| 2.9e-08 \| 2.1e-12 \| 48.6 \| 50 \| (9, 60) \| 73 \| (2, 52) \| 146 \| Transcriptional regulator, MerR family \| 3HH0\_A Transcriptional regulator, MerR family Transcriptional regulator, MerR, Protein Structure; pdb70 \| 3HH0\_B \| 97.7 \| 2.9e-08 \| 2.1e-12 \| 48.6 \| 50 \| (9, 60) \| 73 \| (2, 52) \| 146 \| Transcriptional regulator, MerR family \| 3HH0\_B Transcriptional regulator, MerR family Transcriptional regulator, MerR, Protein Structure HET: MSE; pdb70 \| 6JGV\_A \| 97.7 \| 3.2e-08 \| 2.3e-12 \| 48.4 \| 48 \| (12, 61) \| 73 \| (1, 49) \| 147 \| CadR \| 6JGV\_A CadR CadR, MerR family, cadmium regulator; pdb70 \| 6JGX\_A \| 97.7 \| 3.2e-08 \| 2.3e-12 \| 48.4 \| 48 \| (12, 61) \| 73 \| (1, 49) \| 147 \| CadR/DNA \| 6JGX\_A CadR/DNA CadR, MerR family, cadmium regulator; pdb70 \| 6JNI\_A \| 97.7 \| 3.2e-08 \| 2.3e-12 \| 48.4 \| 48 \| (12, 61) \| 73 \| (1, 49) \| 147 \| CadR/DNA \| 6JNI\_A CadR/DNA CadR, MerR family, cadmium regulator HET: ZN; pdb70 \| 3QAO\_A \| 97.7 \| 3.5e-08 \| 2.5e-12 \| 51.6 \| 49 \| (10, 60) \| 73 \| (2, 51) \| 249 \| MerR-like transcriptional regulator \| 3QAO\_A MerR-like transcriptional regulator structural genomics, The Center for HET: GOL; pdb70 \| 5E01\_A \| 97.6 \| 4e-08 \| 2.9e-12 \| 46.7 \| 50 \| (9, 60) \| 73 \| (2, 52) \| 128 \| Uncharacterized HTH-type transcriptional regulator HI\_0186/DNA \| 5E01\_A Uncharacterized HTH-type transcriptional regulator HI\_0186/DNA Transcription factor, MerR, thiol-based genetic; pdb70 \| 5E01\_B \| 97.6 \| 4e-08 \| 2.9e-12 \| 46.7 \| 50 \| (9, 60) \| 73 \| (2, 52) \| 128 \| Uncharacterized HTH-type transcriptional regulator HI\_0186/DNA \| 5E01\_B Uncharacterized HTH-type transcriptional regulator HI\_0186/DNA Transcription factor, MerR, thiol-based genetic; pdb70 \| 1R8D\_A \| 97.6 \| 4.1e-08 \| 2.9e-12 \| 45.2 \| 50 \| (11, 62) \| 73 \| (2, 52) \| 109 \| transcription activator MtaN/DNA Complex \| 1R8D\_A transcription activator MtaN/DNA Complex protein-DNA complex, Transcription-DNA COMPLEX HET: SO4; pdb70 \| 1Q06\_B \| 97.6 \| 4.2e-08 \| 3e-12 \| 47.4 \| 48 \| (12, 61) \| 73 \| (1, 49) \| 135 \| Transcriptional regulator cueR \| 1Q06\_B Transcriptional regulator cueR MerR family transcriptional regulator, copper; pdb70 \| 1Q07\_A \| 97.6 \| 4.2e-08 \| 3e-12 \| 47.4 \| 48 \| (12, 61) \| 73 \| (1, 49) \| 135 \| Transcriptional regulator cueR \| 1Q07\_A Transcriptional regulator cueR MerR family transcriptional regulator, copper; pdb70 \| 4WLW\_A \| 97.6 \| 4.2e-08 \| 3e-12 \| 47.4 \| 48 \| (12, 61) \| 73 \| (1, 49) \| 135 \| TRANSCRIPTIONAL REGULATOR CUER/DNA Complex \| 4WLW\_A TRANSCRIPTIONAL REGULATOR CUER/DNA Complex PROTEIN-DNA COMPLEX, MERR-FAMILY TRANSCRIPTION REGULATOR HET: MSE; pdb70 \| 6JGF\_A \| 97.6 \| 4.6e-08 \| 3.4e-12 \| 46.4 \| 48 \| (12, 61) \| 73 \| (1, 49) \| 126 \| CadR \| 6JGF\_A CadR CadR, MerR family, cadmium regulator HET: MSE; pdb70 \| 1PM6\_A \| 97.6 \| 5e-08 \| 3.6e-12 \| 42.6 \| 47 \| (11, 57) \| 73 \| (2, 49) \| 72 \| Excisionase \| 1PM6\_A Excisionase ANTIPARALLEL BETA-SHEET, WINGED-HELIX, CIS-TRANS-TRANS TRIPROLINE; pdb70 \| 5D8C\_A \| 97.5 \| 7.1e-08 \| 5.1e-12 \| 46.4 \| 50 \| (9, 60) \| 73 \| (2, 52) \| 137 \| Uncharacterized HTH-type transcriptional regulator HI\_0186/DNA \| 5D8C\_A Uncharacterized HTH-type transcriptional regulator HI\_0186/DNA Transcription factor, MerR, thiol-based genetic; pdb70 \| 3Q2Y\_A \| 97.5 \| 7.9e-08 \| 5.7e-12 \| 50.6 \| 51 \| (8, 60) \| 73 \| (2, 54) \| 284 \| Multidrug-efflux transporter 1 regulator/DNA complex \| 3Q2Y\_A Multidrug-efflux transporter 1 regulator/DNA complex Protein DNA complex, transcription regulator HET: GOL, ET; pdb70 \| 2OG0\_B \| 97.5 \| 8.7e-08 \| 6.3e-12 \| 38.8 \| 46 \| (11, 56) \| 73 \| (2, 48) \| 52 \| Excisionase/DNA Complex \| 2OG0\_B Excisionase/DNA Complex PROTEIN-DNA COMPLEX, DNA ARCHITECTURAL PROTEIN; pdb70 \| 1R8E\_A \| 97.5 \| 9e-08 \| 6.5e-12 \| 49.9 \| 51 \| (8, 60) \| 73 \| (2, 54) \| 278 \| multidrug-efflux transporter regulator/DNA Complex \| 1R8E\_A multidrug-efflux transporter regulator/DNA Complex PROTEIN-DNA COMPLEX, MERR-FAMILY TRANSCRIPTION ACTIVATOR HET: P4P, GOL, IMD; pdb70 \| 2JML\_A \| 97.5 \| 1e-07 \| 7.3e-12 \| 41.2 \| 53 \| (8, 62) \| 73 \| (2, 56) \| 81 \| DNA BINDING DOMAIN/TRANSCRIPTIONAL REGULATOR \| 2JML\_A DNA BINDING DOMAIN/TRANSCRIPTIONAL REGULATOR anti-repressor, MerR, carotenogenesis, TRANSCRIPTION; pdb70 \| 3WHP\_A \| 97.5 \| 1.1e-07 \| 8e-12 \| 50.6 \| 57 \| (5, 63) \| 73 \| (5, 62) \| 290 \| Probable transcriptional regulator \| 3WHP\_A Probable transcriptional regulator B12-binding domain, Rossmann fold, Four HET: B12; pdb70 \| 2VZ4\_A \| 97.5 \| 1.2e-07 \| 8.3e-12 \| 43.6 \| 49 \| (11, 61) \| 73 \| (1, 50) \| 108 \| HTH-TYPE TRANSCRIPTIONAL ACTIVATOR TIPA \| 2VZ4\_A HTH-TYPE TRANSCRIPTIONAL ACTIVATOR TIPA TRANSCRIPTION, RESISTANCE, ANTIBIOTIC, DNA-BINDING, STREPTOMYCES; pdb70 \| 2DG6\_A \| 97.5 \| 1.2e-07 \| 8.7e-12 \| 49.4 \| 49 \| (12, 62) \| 73 \| (1, 50) \| 222 \| putative transcriptional regulator \| 2DG6\_A putative transcriptional regulator Winged-helix motif, MerR family, GENE; pdb70 \| 1RH6\_B \| 97.4 \| 2e-07 \| 1.4e-11 \| 38.2 \| 47 \| (11, 57) \| 73 \| (2, 49) \| 55 \| Excisionase/DNA Complex \| 1RH6\_B Excisionase/DNA Complex Protein-DNA complex, DNA architectural protein; pdb70 \| 6P0U\_E \| 97.4 \| 2e-07 \| 1.4e-11 \| 38.2 \| 47 \| (11, 57) \| 73 \| (2, 49) \| 55 \| DNA-binding protein Fis, Excisionase/DNA Complex \| 6P0U\_E DNA-binding protein Fis, Excisionase/DNA Complex Protein-DNA ternary complex, DNA shape; pdb70 \| 5I41\_B \| 97.2 \| 4.5e-07 \| 3.3e-11 \| 37.3 \| 48 \| (10, 59) \| 73 \| (2, 49) \| 69 \| Chromosome-anchoring protein RacA \| 5I41\_B Chromosome-anchoring protein RacA RacA, axial filament, sporulation, B.; pdb70 \| 5I44\_A \| 97.2 \| 4.5e-07 \| 3.3e-11 \| 37.3 \| 48 \| (10, 59) \| 73 \| (2, 49) \| 69 \| Chromosome-anchoring protein RacA/DNA Complex \| 5I44\_A Chromosome-anchoring protein RacA/DNA Complex RacA, B. subtilis, axial filament; pdb70 \| 5XBT\_B \| 97.2 \| 5.4e-07 \| 3.9e-11 \| 46.7 \| 50 \| (10, 61) \| 73 \| (2, 53) \| 272 \| Probable transcriptional regulator \| 5XBT\_B Probable transcriptional regulator c-di-GMP, Receptor, HTH domain, DNA HET: GOL, PEG, IMD, C2E; pdb70 \| 2KFS\_A \| 97.1 \| 7.7e-07 \| 5.6e-11 \| 43.6 \| 48 \| (8, 58) \| 73 \| (28, 75) \| 148 \| Conserved hypothetical regulatory protein \| 2KFS\_A Conserved hypothetical regulatory protein wHTH, DNA binding, Phosphorylation, DNA-binding; pdb70 \| 5YDC\_B \| 96.9 \| 2.5e-06 \| 1.8e-10 \| 41.4 \| 49 \| (5, 59) \| 73 \| (16, 64) \| 142 \| Uncharacterized HTH-type transcriptional regulator Rv1828 \| 5YDC\_B Uncharacterized HTH-type transcriptional regulator Rv1828 DNA binding, HTH motif, promoter; pdb70 \| 5YDD\_B \| 96.9 \| 2.5e-06 \| 1.8e-10 \| 41.4 \| 49 \| (5, 59) \| 73 \| (16, 64) \| 142 \| Uncharacterized HTH-type transcriptional regulator Rv1828 \| 5YDD\_B Uncharacterized HTH-type transcriptional regulator Rv1828 DNA binding, HTH motif, promoter HET: 8TU; pdb70 \| 5AF3\_B \| 96.6 \| 7.4e-06 \| 5.3e-10 \| 42.9 \| 52 \| (6, 59) \| 73 \| (12, 78) \| 241 \| VAPBC49 \| 5AF3\_B VAPBC49 MYCOBACTERIUM TUBERCULOSIS, DNA BINDING, TA; pdb70 \| 6LND\_K \| 96.4 \| 1.5e-05 \| 1.1e-09 \| 44.1 \| 62 \| (8, 71) \| 73 \| (321, 389) \| 396 \| transposition protein TniQ \| 6LND\_K transposition protein TniQ Type I-F CRISPR-Cas system: Csy HET: MSE; pdb70 \| 6PIG\_J \| 96.3 \| 2.4e-05 \| 1.7e-09 \| 42.9 \| 60 \| (9, 70) \| 73 \| (296, 362) \| 369 \| cas7 type I-F CRISPR-associated protein \| 6PIG\_J cas7 type I-F CRISPR-associated protein CRISPR/Cas, Cascade, RNA BINDING PROTEIN; pdb70 \| 6V9P\_A \| 96.3 \| 2.6e-05 \| 1.9e-09 \| 43.1 \| 60 \| (9, 70) \| 73 \| (321, 387) \| 395 \| TniQ family protein \| 6V9P\_A TniQ family protein Type I-F CRISPR-Cas system, transposition |
| Top keywords  (threshold 1.00e-03 (evalue)) | **Helix\_turn\_helix, domain\_containing, regulator, transcriptional, DNA, excisionase, DNA\_binding, AlpA, binding, MerR** |
| Output files | ../../similar\_sequences/38\_FANPEZAQ\_CDS\_0038\_merged.svg ../../similar\_sequences/38\_FANPEZAQ\_CDS\_0038\_pdb70.a3m ../../similar\_sequences/38\_FANPEZAQ\_CDS\_0038\_pdb70.hhr ../../similar\_sequences/38\_FANPEZAQ\_CDS\_0038\_uniclust.a3m ../../similar\_sequences/38\_FANPEZAQ\_CDS\_0038\_uniclust.hhr |

#### Structure prediction (AlphaFold)^2^

|  |  |
| --- | --- |
| Stats | xml version="1.0" encoding="utf-8" standalone="no"?       2024-09-02T21:09:37.075729 image/svg+xml   Matplotlib v3.7.2, https://matplotlib.org/ |
| Predicted structure | **NGL Viewer Controls:**  - Center: *Left-Click* - Rotate: *Left-Click + Drag* - Translate: *Right-Click + Drag* - Zoom: *Shift + Left-Click + Drag* |
| Output files | ../../predicted\_structures/38\_FANPEZAQ\_CDS\_0038/features.pkl ../../predicted\_structures/38\_FANPEZAQ\_CDS\_0038/ranked\_0.pdb ../../predicted\_structures/38\_FANPEZAQ\_CDS\_0038/ranked\_0\_plots.svg ../../predicted\_structures/38\_FANPEZAQ\_CDS\_0038/result\_model\_1\_ptm\_pred\_0.pkl |

#### Structure similarity search results (Foldseek)^3^

|  |  |
| --- | --- |
| Structure databases searched | Pdb, Afdb-proteome, Afdb-uniprot50 |
| Results, scheme(s)  (Top layers only, threshold 1.00e-02 (evalue)) | xml version="1.0" encoding="utf-8" standalone="no"?       2024-09-02T21:11:13.160571 image/svg+xml   Matplotlib v3.7.2, https://matplotlib.org/ |
| Results, table  (threshold 1.00e-02 (evalue)) | db \| id \| prob \| evalue \| bits \| fident \| alnlen \| mismatch \| gapopen \| qstart \| qend \| tstart \| tend \| name \| description; pdb \| 8DGL\_B \| 1.0 \| 0.008615 \| 128 \| 0.272 \| 55 \| 38 \| 1 \| 9 \| 63 \| 16 \| 68 \| Recombination Directionality Factor RdfS \| Recombination Directionality Factor RdfS; afdb-proteome \| AF-O53399-F1-MODEL\_V4 \| 1.0 \| 0.002637 \| 147 \| 0.262 \| 61 \| 43 \| 1 \| 5 \| 65 \| 189 \| 247 \| HTH\_17 domain-containing protein \| HTH\_17 domain-containing protein; afdb-uniprot50 \| AF-A0A2Z3IG45-F1-MODEL\_V4 \| 1.0 \| 5.132e-07 \| 269 \| 0.569 \| 72 \| 27 \| 1 \| 1 \| 72 \| 1 \| 68 \| Uncharacterized protein \| Uncharacterized protein; afdb-uniprot50 \| AF-A0A158S0W0-F1-MODEL\_V4 \| 1.0 \| 1.744e-05 \| 254 \| 0.547 \| 53 \| 24 \| 0 \| 11 \| 63 \| 6 \| 58 \| Uncharacterized protein \| Uncharacterized protein; afdb-uniprot50 \| AF-A0A375H718-F1-MODEL\_V4 \| 1.0 \| 6.265e-07 \| 250 \| 0.534 \| 73 \| 32 \| 1 \| 1 \| 73 \| 1 \| 71 \| Uncharacterized protein \| Uncharacterized protein; afdb-uniprot50 \| AF-A0A5E7ZHS3-F1-MODEL\_V4 \| 1.0 \| 2.276e-05 \| 235 \| 0.39 \| 64 \| 38 \| 1 \| 1 \| 64 \| 1 \| 63 \| DNA-binding protein \| DNA-binding protein; afdb-uniprot50 \| AF-A0A0S4UXE0-F1-MODEL\_V4 \| 1.0 \| 2.433e-05 \| 230 \| 0.365 \| 63 \| 40 \| 0 \| 11 \| 73 \| 5 \| 67 \| HTH\_17 domain-containing protein \| HTH\_17 domain-containing protein; afdb-uniprot50 \| AF-A0A1Q4EAJ3-F1-MODEL\_V4 \| 1.0 \| 9.203e-05 \| 228 \| 0.396 \| 58 \| 35 \| 0 \| 7 \| 64 \| 2 \| 59 \| DNA-binding protein \| DNA-binding protein; afdb-uniprot50 \| AF-A0A2T4HLL8-F1-MODEL\_V4 \| 1.0 \| 0.000179 \| 220 \| 0.368 \| 57 \| 36 \| 0 \| 7 \| 63 \| 2 \| 58 \| DNA-binding protein \| DNA-binding protein; afdb-uniprot50 \| AF-A0A1C3NJQ6-F1-MODEL\_V4 \| 1.0 \| 1.527e-05 \| 215 \| 0.452 \| 73 \| 35 \| 1 \| 1 \| 73 \| 18 \| 85 \| Uncharacterized protein \| Uncharacterized protein; afdb-uniprot50 \| AF-H4F9I4-F1-MODEL\_V4 \| 1.0 \| 1.864e-05 \| 215 \| 0.319 \| 72 \| 49 \| 0 \| 1 \| 72 \| 46 \| 117 \| Uncharacterized protein \| Uncharacterized protein; afdb-uniprot50 \| AF-A0A841M6V1-F1-MODEL\_V4 \| 1.0 \| 5.057e-05 \| 214 \| 0.333 \| 69 \| 45 \| 1 \| 1 \| 69 \| 29 \| 96 \| Uncharacterized protein \| Uncharacterized protein; afdb-uniprot50 \| AF-A0A5S4YCH6-F1-MODEL\_V4 \| 1.0 \| 9.203e-05 \| 211 \| 0.298 \| 67 \| 47 \| 0 \| 7 \| 73 \| 2 \| 68 \| Helix-turn-helix domain-containing protein \| Helix-turn-helix domain-containing protein; afdb-uniprot50 \| AF-A0A227JR88-F1-MODEL\_V4 \| 1.0 \| 0.0002852 \| 210 \| 0.537 \| 54 \| 25 \| 0 \| 10 \| 63 \| 6 \| 59 \| HTH\_17 domain-containing protein \| HTH\_17 domain-containing protein; afdb-uniprot50 \| AF-A0A1V1PTW5-F1-MODEL\_V4 \| 1.0 \| 7.538e-05 \| 210 \| 0.365 \| 63 \| 40 \| 0 \| 1 \| 63 \| 3 \| 65 \| Uncharacterized protein \| Uncharacterized protein; afdb-uniprot50 \| AF-A0A2Z5UBR4-F1-MODEL\_V4 \| 1.0 \| 0.0003721 \| 209 \| 0.333 \| 57 \| 38 \| 0 \| 7 \| 63 \| 2 \| 58 \| Transcriptional regulator MerR family \| Transcriptional regulator MerR family; afdb-uniprot50 \| AF-A0A145VQA6-F1-MODEL\_V4 \| 1.0 \| 6.174e-05 \| 208 \| 0.461 \| 65 \| 35 \| 0 \| 6 \| 70 \| 2 \| 66 \| Uncharacterized protein \| Uncharacterized protein; afdb-uniprot50 \| AF-A0A356RFX9-F1-MODEL\_V4 \| 1.0 \| 0.0001466 \| 208 \| 0.344 \| 58 \| 36 \| 2 \| 7 \| 63 \| 1 \| 57 \| HTH\_17 domain-containing protein \| HTH\_17 domain-containing protein; afdb-uniprot50 \| AF-A0A177IQV5-F1-MODEL\_V4 \| 1.0 \| 0.0001124 \| 207 \| 0.38 \| 63 \| 34 \| 1 \| 1 \| 63 \| 1 \| 58 \| HTH\_17 domain-containing protein \| HTH\_17 domain-containing protein; afdb-uniprot50 \| AF-A0A6G4QV14-F1-MODEL\_V4 \| 1.0 \| 0.0001466 \| 205 \| 0.307 \| 65 \| 45 \| 0 \| 7 \| 71 \| 2 \| 66 \| Helix-turn-helix domain-containing protein \| Helix-turn-helix domain-containing protein; afdb-uniprot50 \| AF-A0A504ERG4-F1-MODEL\_V4 \| 1.0 \| 9.836e-05 \| 203 \| 0.295 \| 71 \| 50 \| 0 \| 2 \| 72 \| 12 \| 82 \| Helix-turn-helix domain-containing protein \| Helix-turn-helix domain-containing protein; afdb-uniprot50 \| AF-A0A2X0SMX0-F1-MODEL\_V4 \| 1.0 \| 0.0001372 \| 202 \| 0.38 \| 63 \| 37 \| 1 \| 1 \| 63 \| 17 \| 77 \| Uncharacterized protein \| Uncharacterized protein; afdb-uniprot50 \| AF-A0A6L5FK90-F1-MODEL\_V4 \| 1.0 \| 0.000179 \| 201 \| 0.363 \| 66 \| 42 \| 0 \| 7 \| 72 \| 2 \| 67 \| DNA-binding protein \| DNA-binding protein; afdb-uniprot50 \| AF-A0A1X7FPK0-F1-MODEL\_V4 \| 1.0 \| 0.0004543 \| 200 \| 0.363 \| 55 \| 35 \| 0 \| 7 \| 61 \| 2 \| 56 \| Uncharacterized protein \| Uncharacterized protein; afdb-uniprot50 \| AF-A0A1M6DCH5-F1-MODEL\_V4 \| 1.0 \| 0.0001051 \| 200 \| 0.349 \| 63 \| 37 \| 1 \| 1 \| 63 \| 1 \| 59 \| Helix-turn-helix domain-containing protein \| Helix-turn-helix domain-containing protein; afdb-uniprot50 \| AF-A0A1D2SFS9-F1-MODEL\_V4 \| 1.0 \| 3.875e-05 \| 200 \| 0.333 \| 72 \| 44 \| 2 \| 1 \| 72 \| 1 \| 68 \| Uncharacterized protein \| Uncharacterized protein; afdb-uniprot50 \| AF-A0A0W8FLL0-F1-MODEL\_V4 \| 1.0 \| 0.0001372 \| 199 \| 0.333 \| 66 \| 43 \| 1 \| 7 \| 72 \| 3 \| 67 \| Uncharacterized protein \| Uncharacterized protein; afdb-uniprot50 \| AF-A0A2S0XML1-F1-MODEL\_V4 \| 1.0 \| 0.0001124 \| 198 \| 0.367 \| 68 \| 38 \| 1 \| 1 \| 63 \| 23 \| 90 \| Uncharacterized protein \| Uncharacterized protein; afdb-uniprot50 \| AF-T0GF19-F1-MODEL\_V4 \| 1.0 \| 0.000179 \| 198 \| 0.292 \| 65 \| 46 \| 0 \| 9 \| 73 \| 4 \| 68 \| Uncharacterized protein \| Uncharacterized protein; afdb-uniprot50 \| AF-A0A0H3J1R4-F1-MODEL\_V4 \| 1.0 \| 0.0001124 \| 197 \| 0.333 \| 69 \| 41 \| 1 \| 1 \| 69 \| 1 \| 64 \| HTH transcriptional regulator \| HTH transcriptional regulator; afdb-uniprot50 \| AF-A0A3B0N048-F1-MODEL\_V4 \| 1.0 \| 0.0002336 \| 196 \| 0.393 \| 61 \| 35 \| 1 \| 7 \| 67 \| 1 \| 59 \| HTH\_17 domain-containing protein \| HTH\_17 domain-containing protein; afdb-uniprot50 \| AF-A0A222X839-F1-MODEL\_V4 \| 1.0 \| 0.0001567 \| 196 \| 0.409 \| 61 \| 34 \| 1 \| 11 \| 71 \| 5 \| 63 \| Uncharacterized protein \| Uncharacterized protein; afdb-uniprot50 \| AF-A0A4P8HF99-F1-MODEL\_V4 \| 1.0 \| 0.0002668 \| 194 \| 0.333 \| 66 \| 42 \| 2 \| 1 \| 66 \| 1 \| 64 \| Helix-turn-helix domain-containing protein \| Helix-turn-helix domain-containing protein; afdb-uniprot50 \| AF-A0A4Q2SXB6-F1-MODEL\_V4 \| 1.0 \| 0.0002852 \| 194 \| 0.328 \| 67 \| 45 \| 0 \| 1 \| 67 \| 24 \| 90 \| DNA-binding protein \| DNA-binding protein; afdb-uniprot50 \| AF-A0A8B3L4A2-F1-MODEL\_V4 \| 1.0 \| 0.0003482 \| 193 \| 0.38 \| 63 \| 34 \| 1 \| 1 \| 63 \| 1 \| 58 \| DNA-binding protein \| DNA-binding protein; afdb-uniprot50 \| AF-K2KUD9-F1-MODEL\_V4 \| 1.0 \| 0.0001372 \| 192 \| 0.396 \| 63 \| 36 \| 1 \| 11 \| 73 \| 5 \| 65 \| Uncharacterized protein \| Uncharacterized protein; afdb-uniprot50 \| AF-A0A4Q3WF84-F1-MODEL\_V4 \| 1.0 \| 0.0002496 \| 192 \| 0.353 \| 65 \| 40 \| 1 \| 2 \| 66 \| 7 \| 69 \| DNA-binding protein \| DNA-binding protein; afdb-uniprot50 \| AF-A0A7Y3WDS6-F1-MODEL\_V4 \| 1.0 \| 0.001153 \| 192 \| 0.381 \| 55 \| 34 \| 0 \| 9 \| 63 \| 4 \| 58 \| Helix-turn-helix domain-containing protein \| Helix-turn-helix domain-containing protein; afdb-uniprot50 \| AF-A0A8A6KER8-F1-MODEL\_V4 \| 1.0 \| 0.0001675 \| 191 \| 0.417 \| 67 \| 34 \| 1 \| 12 \| 73 \| 1 \| 67 \| DNA-binding protein \| DNA-binding protein; afdb-uniprot50 \| AF-A0A7Z8LR69-F1-MODEL\_V4 \| 1.0 \| 0.0002668 \| 190 \| 0.409 \| 61 \| 34 \| 1 \| 11 \| 71 \| 5 \| 63 \| Helix-turn-helix protein \| Helix-turn-helix protein; afdb-uniprot50 \| AF-A0A099F695-F1-MODEL\_V4 \| 1.0 \| 0.0003048 \| 189 \| 0.328 \| 67 \| 43 \| 1 \| 7 \| 73 \| 1 \| 65 \| HTH\_17 domain-containing protein \| HTH\_17 domain-containing protein; afdb-uniprot50 \| AF-A0A525JL77-F1-MODEL\_V4 \| 1.0 \| 0.0003977 \| 189 \| 0.349 \| 63 \| 39 \| 1 \| 1 \| 63 \| 1 \| 61 \| DNA-binding protein \| DNA-binding protein; afdb-uniprot50 \| AF-A0A3A5K4U6-F1-MODEL\_V4 \| 1.0 \| 0.0001372 \| 189 \| 0.323 \| 68 \| 41 \| 1 \| 11 \| 73 \| 6 \| 73 \| DNA-binding protein \| DNA-binding protein; afdb-uniprot50 \| AF-A0A2G5QHB5-F1-MODEL\_V4 \| 1.0 \| 0.0004251 \| 189 \| 0.311 \| 61 \| 42 \| 0 \| 11 \| 71 \| 4 \| 64 \| HTH\_17 domain-containing protein \| HTH\_17 domain-containing protein; afdb-uniprot50 \| AF-A0A4R1R9W0-F1-MODEL\_V4 \| 1.0 \| 0.0008268 \| 187 \| 0.379 \| 58 \| 35 \| 1 \| 7 \| 63 \| 1 \| 58 \| Uncharacterized protein \| Uncharacterized protein; afdb-uniprot50 \| AF-A0A1N6NC24-F1-MODEL\_V4 \| 1.0 \| 9.836e-05 \| 187 \| 0.378 \| 74 \| 40 \| 3 \| 1 \| 73 \| 6 \| 74 \| Uncharacterized protein \| Uncharacterized protein; afdb-uniprot50 \| AF-R0EA79-F1-MODEL\_V4 \| 1.0 \| 0.0003977 \| 187 \| 0.333 \| 66 \| 42 \| 1 \| 1 \| 66 \| 16 \| 79 \| Uncharacterized protein \| Uncharacterized protein; afdb-uniprot50 \| AF-A0A2N3AXX5-F1-MODEL\_V4 \| 1.0 \| 0.0002185 \| 187 \| 0.409 \| 61 \| 34 \| 1 \| 11 \| 71 \| 5 \| 63 \| HTH\_17 domain-containing protein \| HTH\_17 domain-containing protein; afdb-uniprot50 \| AF-A0A1M3H0S1-F1-MODEL\_V4 \| 1.0 \| 0.000519 \| 186 \| 0.444 \| 54 \| 28 \| 1 \| 9 \| 62 \| 2 \| 53 \| Uncharacterized protein \| Uncharacterized protein; afdb-uniprot50 \| AF-A0A7J5CN46-F1-MODEL\_V4 \| 1.0 \| 0.0002496 \| 185 \| 0.323 \| 65 \| 42 \| 1 \| 2 \| 66 \| 5 \| 67 \| Helix-turn-helix domain-containing protein \| Helix-turn-helix domain-containing protein; afdb-uniprot50 \| AF-A0A1M3GUP1-F1-MODEL\_V4 \| 1.0 \| 0.0006772 \| 185 \| 0.444 \| 54 \| 28 \| 1 \| 10 \| 63 \| 4 \| 55 \| HTH\_17 domain-containing protein \| HTH\_17 domain-containing protein; afdb-uniprot50 \| AF-A0A7Y4Y2M7-F1-MODEL\_V4 \| 1.0 \| 0.0001675 \| 185 \| 0.301 \| 73 \| 48 \| 2 \| 1 \| 73 \| 6 \| 75 \| DNA-binding protein \| DNA-binding protein; afdb-uniprot50 \| AF-A0A2E2ZU57-F1-MODEL\_V4 \| 1.0 \| 0.0005928 \| 184 \| 0.333 \| 63 \| 37 \| 1 \| 1 \| 63 \| 1 \| 58 \| DNA-binding protein \| DNA-binding protein; afdb-uniprot50 \| AF-A0A1H8ZIM8-F1-MODEL\_V4 \| 1.0 \| 0.0008836 \| 184 \| 0.389 \| 59 \| 34 \| 1 \| 7 \| 65 \| 3 \| 59 \| Uncharacterized protein \| Uncharacterized protein; afdb-uniprot50 \| AF-A0A2E8PB88-F1-MODEL\_V4 \| 1.0 \| 0.0001466 \| 184 \| 0.367 \| 68 \| 40 \| 2 \| 2 \| 68 \| 8 \| 73 \| HTH\_17 domain-containing protein \| HTH\_17 domain-containing protein; afdb-uniprot50 \| AF-A0A373FQ25-F1-MODEL\_V4 \| 1.0 \| 0.0004856 \| 183 \| 0.349 \| 63 \| 39 \| 1 \| 11 \| 73 \| 2 \| 62 \| DNA-binding protein \| DNA-binding protein; afdb-uniprot50 \| AF-A0A4Q3YNY5-F1-MODEL\_V4 \| 1.0 \| 0.000519 \| 183 \| 0.359 \| 64 \| 39 \| 1 \| 10 \| 73 \| 4 \| 65 \| DNA-binding protein \| DNA-binding protein; afdb-uniprot50 \| AF-A0A840STV4-F1-MODEL\_V4 \| 1.0 \| 0.0004543 \| 183 \| 0.403 \| 62 \| 35 \| 1 \| 10 \| 71 \| 4 \| 63 \| Putative site-specific integrase-resolvase \| Putative site-specific integrase-resolvase; afdb-uniprot50 \| AF-A0A4R6RSK0-F1-MODEL\_V4 \| 1.0 \| 0.0001124 \| 183 \| 0.287 \| 80 \| 50 \| 2 \| 1 \| 73 \| 3 \| 82 \| Helix-turn-helix protein \| Helix-turn-helix protein; afdb-uniprot50 \| AF-A0A554SP46-F1-MODEL\_V4 \| 1.0 \| 0.0008836 \| 182 \| 0.363 \| 55 \| 33 \| 1 \| 9 \| 63 \| 2 \| 54 \| Helix-turn-helix domain-containing protein \| Helix-turn-helix domain-containing protein; afdb-uniprot50 \| AF-A0A5A7Y4Z2-F1-MODEL\_V4 \| 1.0 \| 0.0008836 \| 182 \| 0.403 \| 57 \| 32 \| 1 \| 11 \| 67 \| 8 \| 62 \| DNA-binding protein \| DNA-binding protein; afdb-uniprot50 \| AF-A0A442F8A9-F1-MODEL\_V4 \| 1.0 \| 0.0003257 \| 182 \| 0.315 \| 73 \| 45 \| 1 \| 1 \| 73 \| 1 \| 68 \| DNA-binding protein \| DNA-binding protein; afdb-uniprot50 \| AF-A0A2E8L6W0-F1-MODEL\_V4 \| 1.0 \| 0.0004543 \| 182 \| 0.384 \| 65 \| 38 \| 1 \| 9 \| 73 \| 14 \| 76 \| HTH\_17 domain-containing protein \| HTH\_17 domain-containing protein; afdb-uniprot50 \| AF-A0A495BRN6-F1-MODEL\_V4 \| 1.0 \| 0.001009 \| 182 \| 0.392 \| 51 \| 29 \| 1 \| 11 \| 61 \| 5 \| 53 \| Uncharacterized protein \| Uncharacterized protein; afdb-uniprot50 \| AF-A0A1L5BSX1-F1-MODEL\_V4 \| 1.0 \| 0.0005547 \| 181 \| 0.362 \| 58 \| 35 \| 1 \| 11 \| 68 \| 4 \| 59 \| DNA-binding protein \| DNA-binding protein; afdb-uniprot50 \| AF-A0A1Y1R0V8-F1-MODEL\_V4 \| 1.0 \| 0.0003257 \| 181 \| 0.428 \| 63 \| 34 \| 1 \| 11 \| 73 \| 5 \| 65 \| HTH\_17 domain-containing protein \| HTH\_17 domain-containing protein; afdb-uniprot50 \| AF-A0A4R6UD26-F1-MODEL\_V4 \| 1.0 \| 0.0003721 \| 181 \| 0.315 \| 73 \| 48 \| 1 \| 1 \| 73 \| 16 \| 86 \| Uncharacterized protein \| Uncharacterized protein; afdb-uniprot50 \| AF-A0A5C7Y246-F1-MODEL\_V4 \| 1.0 \| 0.0005928 \| 180 \| 0.36 \| 61 \| 37 \| 1 \| 9 \| 69 \| 3 \| 61 \| DNA-binding protein \| DNA-binding protein; afdb-uniprot50 \| AF-A0A1M3GVS8-F1-MODEL\_V4 \| 1.0 \| 0.0003257 \| 180 \| 0.388 \| 67 \| 38 \| 2 \| 7 \| 73 \| 1 \| 64 \| HTH\_17 domain-containing protein \| HTH\_17 domain-containing protein; afdb-uniprot50 \| AF-A0A0G3IYW9-F1-MODEL\_V4 \| 1.0 \| 0.0002185 \| 180 \| 0.328 \| 73 \| 44 \| 2 \| 1 \| 73 \| 1 \| 68 \| Uncharacterized protein \| Uncharacterized protein; afdb-uniprot50 \| AF-A0A2A2GKF9-F1-MODEL\_V4 \| 1.0 \| 0.0003257 \| 180 \| 0.264 \| 68 \| 48 \| 1 \| 2 \| 69 \| 6 \| 71 \| Terminase \| Terminase; afdb-uniprot50 \| AF-A0A6V8MMJ1-F1-MODEL\_V4 \| 1.0 \| 0.0002336 \| 180 \| 0.285 \| 70 \| 48 \| 1 \| 1 \| 70 \| 7 \| 74 \| HTH\_17 domain-containing protein \| HTH\_17 domain-containing protein; afdb-uniprot50 \| AF-A0A3M6RS62-F1-MODEL\_V4 \| 1.0 \| 0.0003977 \| 180 \| 0.328 \| 64 \| 39 \| 2 \| 1 \| 62 \| 14 \| 75 \| DNA-binding protein \| DNA-binding protein; afdb-uniprot50 \| AF-A0A239NMV0-F1-MODEL\_V4 \| 1.0 \| 0.0002045 \| 180 \| 0.25 \| 80 \| 53 \| 1 \| 1 \| 73 \| 3 \| 82 \| HTH\_17 domain-containing protein \| HTH\_17 domain-containing protein; afdb-uniprot50 \| AF-A0A7Y1SMM4-F1-MODEL\_V4 \| 1.0 \| 0.0004251 \| 180 \| 0.319 \| 72 \| 46 \| 2 \| 1 \| 72 \| 1 \| 69 \| Helix-turn-helix domain-containing protein \| Helix-turn-helix domain-containing protein; afdb-uniprot50 \| AF-A0A347ULU5-F1-MODEL\_V4 \| 1.0 \| 0.0008268 \| 179 \| 0.393 \| 61 \| 35 \| 1 \| 11 \| 71 \| 5 \| 63 \| DNA-binding protein \| DNA-binding protein; afdb-uniprot50 \| AF-A0A539DI59-F1-MODEL\_V4 \| 1.0 \| 0.0009444 \| 179 \| 0.317 \| 63 \| 43 \| 0 \| 1 \| 63 \| 7 \| 69 \| Uncharacterized protein \| Uncharacterized protein; afdb-uniprot50 \| AF-A0A3D1ZSG9-F1-MODEL\_V4 \| 1.0 \| 0.0005547 \| 178 \| 0.383 \| 60 \| 35 \| 1 \| 11 \| 70 \| 5 \| 62 \| HTH\_17 domain-containing protein \| HTH\_17 domain-containing protein; afdb-uniprot50 \| AF-A0A7J9VS78-F1-MODEL\_V4 \| 1.0 \| 0.0003257 \| 178 \| 0.357 \| 70 \| 42 \| 2 \| 4 \| 73 \| 2 \| 68 \| DNA-binding protein \| DNA-binding protein; afdb-uniprot50 \| AF-A0A4P5VMN8-F1-MODEL\_V4 \| 1.0 \| 0.0004251 \| 178 \| 0.358 \| 67 \| 41 \| 1 \| 7 \| 73 \| 2 \| 66 \| Uncharacterized protein \| Uncharacterized protein; afdb-uniprot50 \| AF-B1YV21-F1-MODEL\_V4 \| 1.0 \| 0.0003482 \| 177 \| 0.342 \| 70 \| 40 \| 2 \| 7 \| 72 \| 2 \| 69 \| HTH\_17 domain-containing protein \| HTH\_17 domain-containing protein; afdb-uniprot50 \| AF-A0A653J2V4-F1-MODEL\_V4 \| 1.0 \| 0.0002852 \| 177 \| 0.318 \| 66 \| 41 \| 2 \| 2 \| 65 \| 3 \| 66 \| DNA-binding protein \| DNA-binding protein; afdb-uniprot50 \| AF-A0A3A4KJX6-F1-MODEL\_V4 \| 1.0 \| 0.0006772 \| 177 \| 0.365 \| 63 \| 39 \| 1 \| 1 \| 63 \| 6 \| 67 \| DNA-binding protein \| DNA-binding protein; afdb-uniprot50 \| AF-A0A807ZGS9-F1-MODEL\_V4 \| 1.0 \| 0.001505 \| 177 \| 0.327 \| 58 \| 37 \| 1 \| 8 \| 65 \| 43 \| 98 \| DNA-binding protein \| DNA-binding protein; afdb-uniprot50 \| AF-E5Y5F8-F1-MODEL\_V4 \| 1.0 \| 0.0004543 \| 177 \| 0.277 \| 72 \| 50 \| 1 \| 1 \| 72 \| 33 \| 102 \| Uncharacterized protein \| Uncharacterized protein; afdb-uniprot50 \| AF-A0A833D1M0-F1-MODEL\_V4 \| 1.0 \| 0.0005928 \| 176 \| 0.349 \| 63 \| 39 \| 1 \| 11 \| 73 \| 2 \| 62 \| Helix-turn-helix domain-containing protein \| Helix-turn-helix domain-containing protein; afdb-uniprot50 \| AF-A0A2N6B1S7-F1-MODEL\_V4 \| 1.0 \| 0.0006336 \| 176 \| 0.333 \| 63 \| 40 \| 1 \| 11 \| 73 \| 5 \| 65 \| HTH\_17 domain-containing protein \| HTH\_17 domain-containing protein; afdb-uniprot50 \| AF-A0A4V1RMP8-F1-MODEL\_V4 \| 1.0 \| 0.0009444 \| 176 \| 0.322 \| 59 \| 38 \| 1 \| 4 \| 62 \| 3 \| 59 \| DNA-binding protein \| DNA-binding protein; afdb-uniprot50 \| AF-A0A383AGQ8-F1-MODEL\_V4 \| 1.0 \| 0.0005547 \| 176 \| 0.235 \| 68 \| 50 \| 1 \| 4 \| 71 \| 1 \| 66 \| Uncharacterized protein \| Uncharacterized protein; afdb-uniprot50 \| AF-A0A433X345-F1-MODEL\_V4 \| 1.0 \| 0.0007736 \| 176 \| 0.318 \| 66 \| 43 \| 1 \| 8 \| 73 \| 4 \| 67 \| DNA-binding protein \| DNA-binding protein; afdb-uniprot50 \| AF-A0A840XSZ3-F1-MODEL\_V4 \| 1.0 \| 0.0006772 \| 176 \| 0.365 \| 63 \| 38 \| 1 \| 11 \| 73 \| 5 \| 65 \| Putative site-specific integrase-resolvase \| Putative site-specific integrase-resolvase; afdb-uniprot50 \| AF-A0A353PQN1-F1-MODEL\_V4 \| 1.0 \| 0.0003482 \| 176 \| 0.349 \| 63 \| 37 \| 2 \| 11 \| 73 \| 5 \| 63 \| HTH\_17 domain-containing protein \| HTH\_17 domain-containing protein; afdb-uniprot50 \| AF-A0A2E7K219-F1-MODEL\_V4 \| 1.0 \| 0.0004543 \| 176 \| 0.343 \| 67 \| 42 \| 1 \| 7 \| 73 \| 2 \| 66 \| HTH\_17 domain-containing protein \| HTH\_17 domain-containing protein; afdb-uniprot50 \| AF-A0A7L9SIQ9-F1-MODEL\_V4 \| 1.0 \| 0.0006336 \| 175 \| 0.269 \| 63 \| 44 \| 1 \| 11 \| 73 \| 5 \| 65 \| Helix-turn-helix domain-containing protein \| Helix-turn-helix domain-containing protein; afdb-uniprot50 \| AF-A0A1M3IL58-F1-MODEL\_V4 \| 1.0 \| 0.001009 \| 175 \| 0.407 \| 54 \| 30 \| 1 \| 11 \| 64 \| 5 \| 56 \| Uncharacterized protein \| Uncharacterized protein; afdb-uniprot50 \| AF-A0A2E3P4L9-F1-MODEL\_V4 \| 1.0 \| 0.0005928 \| 175 \| 0.419 \| 62 \| 33 \| 2 \| 11 \| 72 \| 5 \| 63 \| Helix-turn-helix domain-containing protein \| Helix-turn-helix domain-containing protein; afdb-uniprot50 \| AF-A0A3D9U6K0-F1-MODEL\_V4 \| 1.0 \| 0.0002852 \| 175 \| 0.266 \| 75 \| 52 \| 2 \| 1 \| 73 \| 4 \| 77 \| Uncharacterized protein \| Uncharacterized protein; afdb-uniprot50 \| AF-A0A329YBV5-F1-MODEL\_V4 \| 1.0 \| 0.0008836 \| 175 \| 0.306 \| 62 \| 43 \| 0 \| 11 \| 72 \| 6 \| 67 \| DNA-binding protein \| DNA-binding protein; afdb-uniprot50 \| AF-A0A6C1KV97-F1-MODEL\_V4 \| 1.0 \| 0.0003721 \| 175 \| 0.315 \| 73 \| 45 \| 1 \| 1 \| 73 \| 1 \| 68 \| Helix-turn-helix domain-containing protein \| Helix-turn-helix domain-containing protein; afdb-uniprot50 \| AF-A0A509Y4U3-F1-MODEL\_V4 \| 1.0 \| 0.0003048 \| 175 \| 0.367 \| 68 \| 41 \| 1 \| 2 \| 69 \| 17 \| 82 \| Helix-turn-helix domain-containing protein \| Helix-turn-helix domain-containing protein; afdb-uniprot50 \| AF-A0A315ECV1-F1-MODEL\_V4 \| 1.0 \| 0.0001913 \| 175 \| 0.306 \| 75 \| 46 \| 2 \| 1 \| 71 \| 15 \| 87 \| HTH\_17 domain-containing protein \| HTH\_17 domain-containing protein; afdb-uniprot50 \| AF-A0A6P0HA59-F1-MODEL\_V4 \| 1.0 \| 0.0004251 \| 175 \| 0.328 \| 64 \| 41 \| 1 \| 2 \| 65 \| 6 \| 67 \| Helix-turn-helix domain-containing protein \| Helix-turn-helix domain-containing protein; afdb-uniprot50 \| AF-L7L686-F1-MODEL\_V4 \| 1.0 \| 0.001009 \| 174 \| 0.333 \| 57 \| 36 \| 1 \| 7 \| 63 \| 3 \| 57 \| Uncharacterized protein \| Uncharacterized protein; afdb-uniprot50 \| AF-A0A5C7MH74-F1-MODEL\_V4 \| 1.0 \| 0.0003048 \| 174 \| 0.393 \| 66 \| 38 \| 1 \| 4 \| 69 \| 1 \| 64 \| DNA-binding protein \| DNA-binding protein; afdb-uniprot50 \| AF-A0A2G2CSM9-F1-MODEL\_V4 \| 1.0 \| 0.0005547 \| 174 \| 0.235 \| 68 \| 50 \| 1 \| 1 \| 68 \| 1 \| 66 \| DNA-binding protein \| DNA-binding protein; afdb-uniprot50 \| AF-A0A2G1YBU3-F1-MODEL\_V4 \| 1.0 \| 0.001153 \| 174 \| 0.317 \| 63 \| 41 \| 1 \| 4 \| 66 \| 2 \| 62 \| DNA-binding protein \| DNA-binding protein; afdb-uniprot50 \| AF-A0A7G6WVR5-F1-MODEL\_V4 \| 1.0 \| 0.0002852 \| 174 \| 0.263 \| 72 \| 51 \| 1 \| 1 \| 72 \| 4 \| 73 \| Helix-turn-helix domain-containing protein \| Helix-turn-helix domain-containing protein; afdb-uniprot50 \| AF-A0A7V7KBR4-F1-MODEL\_V4 \| 1.0 \| 0.0001567 \| 174 \| 0.378 \| 74 \| 42 \| 3 \| 1 \| 73 \| 3 \| 73 \| Helix-turn-helix domain-containing protein \| Helix-turn-helix domain-containing protein; afdb-uniprot50 \| AF-A0A1F1WCH5-F1-MODEL\_V4 \| 1.0 \| 0.0003977 \| 174 \| 0.301 \| 73 \| 48 \| 2 \| 1 \| 73 \| 9 \| 78 \| HTH\_17 domain-containing protein \| HTH\_17 domain-containing protein; afdb-uniprot50 \| AF-A0A1M6C0J0-F1-MODEL\_V4 \| 1.0 \| 0.0005928 \| 173 \| 0.313 \| 67 \| 44 \| 1 \| 7 \| 73 \| 2 \| 66 \| DNA binding domain-containing protein, excisionase family \| DNA binding domain-containing protein, excisionase family; afdb-uniprot50 \| AF-A0A5B8IWQ0-F1-MODEL\_V4 \| 1.0 \| 0.0005928 \| 173 \| 0.343 \| 64 \| 39 \| 2 \| 1 \| 63 \| 3 \| 64 \| Helix-turn-helix domain-containing protein \| Helix-turn-helix domain-containing protein; afdb-uniprot50 \| AF-A0A7V3JQH2-F1-MODEL\_V4 \| 1.0 \| 0.0004856 \| 173 \| 0.236 \| 72 \| 53 \| 1 \| 1 \| 72 \| 3 \| 72 \| DNA-binding protein \| DNA-binding protein; afdb-uniprot50 \| AF-A0A4R8ZYI3-F1-MODEL\_V4 \| 1.0 \| 0.001009 \| 173 \| 0.41 \| 56 \| 30 \| 2 \| 11 \| 66 \| 9 \| 61 \| DNA-binding protein \| DNA-binding protein; afdb-uniprot50 \| AF-A0A1I7L0T8-F1-MODEL\_V4 \| 1.0 \| 0.001079 \| 173 \| 0.303 \| 66 \| 44 \| 1 \| 8 \| 73 \| 23 \| 86 \| Uncharacterized protein \| Uncharacterized protein; afdb-uniprot50 \| AF-A0A7W1M9X1-F1-MODEL\_V4 \| 1.0 \| 0.0001913 \| 173 \| 0.391 \| 74 \| 38 \| 3 \| 1 \| 70 \| 1 \| 71 \| Helix-turn-helix domain-containing protein \| Helix-turn-helix domain-containing protein; afdb-uniprot50 \| AF-A0A521H311-F1-MODEL\_V4 \| 1.0 \| 0.0008836 \| 172 \| 0.327 \| 61 \| 39 \| 1 \| 11 \| 71 \| 6 \| 64 \| DNA-binding protein \| DNA-binding protein; afdb-uniprot50 \| AF-A0A2D8GJA3-F1-MODEL\_V4 \| 1.0 \| 0.0008268 \| 171 \| 0.306 \| 62 \| 41 \| 1 \| 11 \| 72 \| 4 \| 63 \| DNA-binding protein \| DNA-binding protein; afdb-uniprot50 \| AF-A0A317L7L9-F1-MODEL\_V4 \| 1.0 \| 0.0004543 \| 171 \| 0.378 \| 66 \| 36 \| 2 \| 11 \| 73 \| 5 \| 68 \| Helix-turn-helix domain-containing protein \| Helix-turn-helix domain-containing protein; afdb-uniprot50 \| AF-A0A4S3M5G0-F1-MODEL\_V4 \| 1.0 \| 0.0006336 \| 171 \| 0.313 \| 67 \| 41 \| 2 \| 7 \| 73 \| 2 \| 63 \| DNA-binding protein \| DNA-binding protein; afdb-uniprot50 \| AF-A0A2W7C0P0-F1-MODEL\_V4 \| 1.0 \| 0.004081 \| 171 \| 0.377 \| 53 \| 33 \| 0 \| 11 \| 63 \| 3 \| 55 \| DNA-binding protein \| DNA-binding protein; afdb-uniprot50 \| AF-A0A1S2PGN3-F1-MODEL\_V4 \| 1.0 \| 0.001009 \| 171 \| 0.315 \| 57 \| 38 \| 1 \| 9 \| 65 \| 21 \| 76 \| DNA-binding protein \| DNA-binding protein; afdb-uniprot50 \| AF-A0A0Q7BDA8-F1-MODEL\_V4 \| 1.0 \| 0.0004543 \| 171 \| 0.328 \| 67 \| 43 \| 1 \| 1 \| 67 \| 19 \| 83 \| HTH merR-type domain-containing protein \| HTH merR-type domain-containing protein; afdb-uniprot50 \| AF-A0A4Y8PHP0-F1-MODEL\_V4 \| 1.0 \| 0.0004251 \| 171 \| 0.333 \| 69 \| 43 \| 2 \| 2 \| 69 \| 19 \| 85 \| DNA-binding protein \| DNA-binding protein; afdb-uniprot50 \| AF-A0A2W7BSX5-F1-MODEL\_V4 \| 1.0 \| 0.001153 \| 171 \| 0.313 \| 67 \| 41 \| 1 \| 1 \| 67 \| 1 \| 62 \| Uncharacterized protein \| Uncharacterized protein; afdb-uniprot50 \| AF-A0A2H5VV35-F1-MODEL\_V4 \| 1.0 \| 0.001009 \| 170 \| 0.322 \| 62 \| 38 \| 2 \| 11 \| 72 \| 4 \| 61 \| HTH\_17 domain-containing protein \| HTH\_17 domain-containing protein; afdb-uniprot50 \| AF-G1V6J5-F1-MODEL\_V4 \| 1.0 \| 0.001153 \| 170 \| 0.303 \| 66 \| 42 \| 2 \| 1 \| 66 \| 1 \| 62 \| HTH\_17 domain-containing protein \| HTH\_17 domain-containing protein; afdb-uniprot50 \| AF-A0A495A877-F1-MODEL\_V4 \| 1.0 \| 0.0005547 \| 170 \| 0.285 \| 70 \| 48 \| 1 \| 4 \| 73 \| 1 \| 68 \| DNA-binding protein \| DNA-binding protein; afdb-uniprot50 \| AF-A0A6N7ZTL0-F1-MODEL\_V4 \| 1.0 \| 0.002243 \| 170 \| 0.285 \| 56 \| 38 \| 1 \| 8 \| 63 \| 9 \| 62 \| DNA-binding protein \| DNA-binding protein; afdb-uniprot50 \| AF-A0A5B7QT10-F1-MODEL\_V4 \| 1.0 \| 0.0007736 \| 170 \| 0.333 \| 60 \| 38 \| 1 \| 3 \| 62 \| 6 \| 63 \| Helix-turn-helix domain-containing protein \| Helix-turn-helix domain-containing protein; afdb-uniprot50 \| AF-A3WRZ7-F1-MODEL\_V4 \| 1.0 \| 0.0004251 \| 170 \| 0.337 \| 83 \| 40 \| 2 \| 1 \| 73 \| 1 \| 78 \| HTH\_17 domain-containing protein \| HTH\_17 domain-containing protein; afdb-uniprot50 \| AF-M3AAY3-F1-MODEL\_V4 \| 1.0 \| 0.0006772 \| 170 \| 0.365 \| 63 \| 38 \| 1 \| 1 \| 63 \| 11 \| 71 \| HTH\_17 domain-containing protein \| HTH\_17 domain-containing protein; afdb-uniprot50 \| AF-A3X0W1-F1-MODEL\_V4 \| 1.0 \| 0.001009 \| 170 \| 0.344 \| 61 \| 38 \| 1 \| 11 \| 71 \| 5 \| 63 \| Uncharacterized protein \| Uncharacterized protein; afdb-uniprot50 \| AF-A0A7I7SIE3-F1-MODEL\_V4 \| 1.0 \| 0.0006772 \| 170 \| 0.291 \| 72 \| 49 \| 1 \| 1 \| 72 \| 10 \| 79 \| HTH\_17 domain-containing protein \| HTH\_17 domain-containing protein; afdb-uniprot50 \| AF-A0A2D5PMF7-F1-MODEL\_V4 \| 1.0 \| 0.0005547 \| 169 \| 0.369 \| 65 \| 39 \| 1 \| 4 \| 68 \| 1 \| 63 \| Terminase \| Terminase; afdb-uniprot50 \| AF-A0A562QVB6-F1-MODEL\_V4 \| 1.0 \| 0.001317 \| 169 \| 0.298 \| 67 \| 45 \| 1 \| 7 \| 73 \| 1 \| 65 \| Excisionase family DNA binding protein \| Excisionase family DNA binding protein; afdb-uniprot50 \| AF-A0A5B8KXC5-F1-MODEL\_V4 \| 1.0 \| 0.0008836 \| 169 \| 0.396 \| 63 \| 36 \| 1 \| 11 \| 73 \| 5 \| 65 \| Helix-turn-helix domain-containing protein \| Helix-turn-helix domain-containing protein; afdb-uniprot50 \| AF-H8YVZ3-F1-MODEL\_V4 \| 1.0 \| 0.0007736 \| 169 \| 0.278 \| 61 \| 41 \| 2 \| 4 \| 63 \| 3 \| 61 \| HTH\_17 domain-containing protein \| HTH\_17 domain-containing protein; afdb-uniprot50 \| AF-A0A2S5P8W1-F1-MODEL\_V4 \| 1.0 \| 0.0004251 \| 169 \| 0.283 \| 74 \| 50 \| 2 \| 1 \| 73 \| 5 \| 76 \| HTH\_17 domain-containing protein \| HTH\_17 domain-containing protein; afdb-uniprot50 \| AF-A0A1X1T3U5-F1-MODEL\_V4 \| 1.0 \| 0.001009 \| 169 \| 0.311 \| 61 \| 40 \| 1 \| 2 \| 62 \| 15 \| 73 \| Uncharacterized protein \| Uncharacterized protein; afdb-uniprot50 \| AF-A0A7U7GDB5-F1-MODEL\_V4 \| 1.0 \| 0.002397 \| 169 \| 0.301 \| 53 \| 35 \| 1 \| 11 \| 63 \| 4 \| 54 \| HTH\_17 domain-containing protein \| HTH\_17 domain-containing protein; afdb-uniprot50 \| AF-A0A6N1BNC1-F1-MODEL\_V4 \| 1.0 \| 0.0003257 \| 169 \| 0.311 \| 77 \| 46 \| 2 \| 1 \| 72 \| 26 \| 100 \| Helix-turn-helix domain-containing protein \| Helix-turn-helix domain-containing protein; afdb-uniprot50 \| AF-A0A0J0YNG3-F1-MODEL\_V4 \| 1.0 \| 0.001317 \| 168 \| 0.349 \| 63 \| 38 \| 2 \| 7 \| 69 \| 3 \| 62 \| Transcriptional regulator, AlpA family \| Transcriptional regulator, AlpA family; afdb-uniprot50 \| AF-A0A7M2X5E3-F1-MODEL\_V4 \| 1.0 \| 0.0007238 \| 168 \| 0.353 \| 65 \| 38 \| 2 \| 11 \| 73 \| 5 \| 67 \| Helix-turn-helix domain-containing protein \| Helix-turn-helix domain-containing protein; afdb-uniprot50 \| AF-A0A7S7SJL7-F1-MODEL\_V4 \| 1.0 \| 0.0007736 \| 168 \| 0.301 \| 73 \| 47 \| 2 \| 1 \| 73 \| 1 \| 69 \| Helix-turn-helix domain-containing protein \| Helix-turn-helix domain-containing protein; afdb-uniprot50 \| AF-A0A2D6TB11-F1-MODEL\_V4 \| 1.0 \| 0.0009444 \| 168 \| 0.349 \| 63 \| 39 \| 1 \| 11 \| 73 \| 5 \| 65 \| HTH\_17 domain-containing protein \| HTH\_17 domain-containing protein; afdb-uniprot50 \| AF-A0A543I5H3-F1-MODEL\_V4 \| 1.0 \| 0.0006336 \| 168 \| 0.291 \| 72 \| 49 \| 1 \| 2 \| 73 \| 5 \| 74 \| Helix-turn-helix protein \| Helix-turn-helix protein; afdb-uniprot50 \| AF-A0A7W1M9Q7-F1-MODEL\_V4 \| 1.0 \| 0.0007736 \| 168 \| 0.358 \| 67 \| 38 \| 2 \| 7 \| 73 \| 1 \| 62 \| Helix-turn-helix domain-containing protein \| Helix-turn-helix domain-containing protein; afdb-uniprot50 \| AF-A0A3P1T187-F1-MODEL\_V4 \| 1.0 \| 0.001079 \| 168 \| 0.333 \| 63 \| 40 \| 1 \| 11 \| 73 \| 5 \| 65 \| DNA-binding protein \| DNA-binding protein; afdb-uniprot50 \| AF-A0A7V2T8E4-F1-MODEL\_V4 \| 1.0 \| 0.0002668 \| 168 \| 0.257 \| 70 \| 49 \| 2 \| 4 \| 73 \| 2 \| 68 \| DNA-binding protein \| DNA-binding protein; afdb-uniprot50 \| AF-A0A7K2THJ8-F1-MODEL\_V4 \| 1.0 \| 0.0009444 \| 168 \| 0.291 \| 72 \| 49 \| 1 \| 1 \| 72 \| 3 \| 72 \| DNA-binding protein \| DNA-binding protein; afdb-uniprot50 \| AF-A0A2N8LFG3-F1-MODEL\_V4 \| 1.0 \| 0.002562 \| 167 \| 0.333 \| 54 \| 34 \| 1 \| 10 \| 63 \| 3 \| 54 \| DNA-binding protein \| DNA-binding protein; afdb-uniprot50 \| AF-A0A2I1V145-F1-MODEL\_V4 \| 1.0 \| 0.001009 \| 167 \| 0.278 \| 61 \| 41 \| 2 \| 7 \| 66 \| 2 \| 60 \| DNA-binding protein \| DNA-binding protein; afdb-uniprot50 \| AF-A0A5C7L536-F1-MODEL\_V4 \| 1.0 \| 0.0006772 \| 167 \| 0.26 \| 73 \| 49 \| 2 \| 1 \| 73 \| 1 \| 68 \| DNA-binding protein \| DNA-binding protein; afdb-uniprot50 \| AF-A0A5P9H8R0-F1-MODEL\_V4 \| 1.0 \| 0.0005928 \| 167 \| 0.205 \| 73 \| 56 \| 1 \| 1 \| 73 \| 1 \| 71 \| Helix-turn-helix domain protein \| Helix-turn-helix domain protein; afdb-uniprot50 \| AF-A0A5B2Z9L9-F1-MODEL\_V4 \| 1.0 \| 0.0003482 \| 167 \| 0.308 \| 68 \| 44 \| 2 \| 7 \| 73 \| 1 \| 66 \| Helix-turn-helix domain-containing protein \| Helix-turn-helix domain-containing protein; afdb-uniprot50 \| AF-A0A850RC49-F1-MODEL\_V4 \| 1.0 \| 0.0007238 \| 167 \| 0.271 \| 70 \| 49 \| 1 \| 1 \| 70 \| 4 \| 71 \| Helix-turn-helix domain-containing protein \| Helix-turn-helix domain-containing protein; afdb-uniprot50 \| AF-A0A2N3VFX6-F1-MODEL\_V4 \| 1.0 \| 0.001009 \| 167 \| 0.343 \| 64 \| 38 \| 2 \| 1 \| 62 \| 5 \| 66 \| Helix-turn-helix protein \| Helix-turn-helix protein; afdb-uniprot50 \| AF-A0A1Q3UQ13-F1-MODEL\_V4 \| 1.0 \| 0.0006772 \| 167 \| 0.256 \| 78 \| 47 \| 1 \| 7 \| 73 \| 2 \| 79 \| DNA-binding protein \| DNA-binding protein; afdb-uniprot50 \| AF-A0A2H6HB99-F1-MODEL\_V4 \| 1.0 \| 0.001009 \| 167 \| 0.241 \| 62 \| 45 \| 1 \| 11 \| 72 \| 7 \| 66 \| Helix-turn-helix domain protein \| Helix-turn-helix domain protein; afdb-uniprot50 \| AF-A0A7W3P5A7-F1-MODEL\_V4 \| 1.0 \| 0.002927 \| 166 \| 0.32 \| 53 \| 34 \| 1 \| 11 \| 63 \| 6 \| 56 \| Excisionase family DNA binding protein \| Excisionase family DNA binding protein; afdb-uniprot50 \| AF-A0A1A0UL62-F1-MODEL\_V4 \| 1.0 \| 0.001317 \| 166 \| 0.343 \| 64 \| 40 \| 1 \| 10 \| 73 \| 4 \| 65 \| HTH\_17 domain-containing protein \| HTH\_17 domain-containing protein; afdb-uniprot50 \| AF-A0A6G9XNP3-F1-MODEL\_V4 \| 1.0 \| 0.001719 \| 166 \| 0.25 \| 64 \| 46 \| 1 \| 9 \| 72 \| 5 \| 66 \| Helix-turn-helix domain-containing protein \| Helix-turn-helix domain-containing protein; afdb-uniprot50 \| AF-A0A167I6B1-F1-MODEL\_V4 \| 1.0 \| 0.002098 \| 166 \| 0.292 \| 65 \| 44 \| 1 \| 9 \| 73 \| 6 \| 68 \| HTH\_17 domain-containing protein \| HTH\_17 domain-containing protein; afdb-uniprot50 \| AF-A0A3A4K0Q0-F1-MODEL\_V4 \| 1.0 \| 0.001837 \| 166 \| 0.246 \| 65 \| 47 \| 1 \| 9 \| 73 \| 6 \| 68 \| DNA-binding protein \| DNA-binding protein; afdb-uniprot50 \| AF-A0A060NMA2-F1-MODEL\_V4 \| 1.0 \| 0.0003482 \| 166 \| 0.328 \| 73 \| 44 \| 2 \| 4 \| 73 \| 1 \| 71 \| Uncharacterized protein \| Uncharacterized protein; afdb-uniprot50 \| AF-A0A1G7RCE4-F1-MODEL\_V4 \| 1.0 \| 0.000519 \| 166 \| 0.25 \| 76 \| 50 \| 2 \| 1 \| 71 \| 2 \| 75 \| Helix-turn-helix domain-containing protein \| Helix-turn-helix domain-containing protein; afdb-uniprot50 \| AF-A0A2Z6GBU1-F1-MODEL\_V4 \| 1.0 \| 0.001079 \| 166 \| 0.306 \| 62 \| 41 \| 1 \| 11 \| 72 \| 5 \| 64 \| Uncharacterized protein \| Uncharacterized protein; afdb-uniprot50 \| AF-A0A1G6T1N9-F1-MODEL\_V4 \| 1.0 \| 0.0008268 \| 166 \| 0.287 \| 73 \| 50 \| 1 \| 1 \| 73 \| 11 \| 81 \| Helix-turn-helix domain-containing protein \| Helix-turn-helix domain-containing protein; afdb-uniprot50 \| AF-A0A5M8SNF4-F1-MODEL\_V4 \| 1.0 \| 0.0004856 \| 166 \| 0.275 \| 80 \| 49 \| 3 \| 1 \| 73 \| 13 \| 90 \| Helix-turn-helix domain-containing protein \| Helix-turn-helix domain-containing protein; afdb-uniprot50 \| AF-A0A1E4MWD2-F1-MODEL\_V4 \| 1.0 \| 0.003343 \| 165 \| 0.321 \| 56 \| 36 \| 1 \| 11 \| 66 \| 2 \| 55 \| HTH\_17 domain-containing protein \| HTH\_17 domain-containing protein; afdb-uniprot50 \| AF-A0A0H2KQU9-F1-MODEL\_V4 \| 1.0 \| 0.002397 \| 165 \| 0.288 \| 52 \| 35 \| 1 \| 11 \| 62 \| 4 \| 53 \| HTH\_17 domain-containing protein \| HTH\_17 domain-containing protein; afdb-uniprot50 \| AF-A0A5C7YER4-F1-MODEL\_V4 \| 1.0 \| 0.000519 \| 165 \| 0.315 \| 73 \| 44 \| 4 \| 1 \| 73 \| 1 \| 67 \| DNA-binding protein \| DNA-binding protein; afdb-uniprot50 \| AF-A0A2U1SGD4-F1-MODEL\_V4 \| 1.0 \| 0.0007736 \| 165 \| 0.315 \| 73 \| 48 \| 1 \| 1 \| 73 \| 2 \| 72 \| Excisionase \| Excisionase; afdb-uniprot50 \| AF-A0A7R7GJG0-F1-MODEL\_V4 \| 1.0 \| 0.000519 \| 165 \| 0.365 \| 63 \| 34 \| 2 \| 11 \| 73 \| 5 \| 61 \| Uncharacterized protein \| Uncharacterized protein; afdb-uniprot50 \| AF-A0A7Y9R7W6-F1-MODEL\_V4 \| 1.0 \| 0.0003977 \| 165 \| 0.328 \| 67 \| 39 \| 2 \| 1 \| 63 \| 3 \| 67 \| Uncharacterized protein \| Uncharacterized protein; afdb-uniprot50 \| AF-A0A2E3RBG2-F1-MODEL\_V4 \| 1.0 \| 0.006084 \| 165 \| 0.346 \| 52 \| 34 \| 0 \| 11 \| 62 \| 4 \| 55 \| DNA-binding protein \| DNA-binding protein; afdb-uniprot50 \| AF-A0A2X4RW04-F1-MODEL\_V4 \| 1.0 \| 0.001079 \| 165 \| 0.285 \| 63 \| 43 \| 1 \| 1 \| 63 \| 1 \| 61 \| Regulatory protein MerR \| Regulatory protein MerR; afdb-uniprot50 \| AF-A0A1F9M8Y5-F1-MODEL\_V4 \| 1.0 \| 0.001153 \| 165 \| 0.285 \| 63 \| 43 \| 1 \| 1 \| 63 \| 6 \| 66 \| HTH\_17 domain-containing protein \| HTH\_17 domain-containing protein; afdb-uniprot50 \| AF-A0A5J6ERQ1-F1-MODEL\_V4 \| 1.0 \| 0.0007238 \| 165 \| 0.287 \| 66 \| 46 \| 1 \| 2 \| 67 \| 13 \| 77 \| DNA-binding protein \| DNA-binding protein; afdb-uniprot50 \| AF-A0A1E3Y8P5-F1-MODEL\_V4 \| 1.0 \| 0.0002336 \| 165 \| 0.217 \| 78 \| 54 \| 3 \| 1 \| 73 \| 1 \| 76 \| HTH\_17 domain-containing protein \| HTH\_17 domain-containing protein; afdb-uniprot50 \| AF-A0A4P9SC28-F1-MODEL\_V4 \| 1.0 \| 0.0005547 \| 165 \| 0.26 \| 73 \| 51 \| 2 \| 1 \| 72 \| 11 \| 81 \| Helix-turn-helix domain-containing protein \| Helix-turn-helix domain-containing protein; afdb-uniprot50 \| AF-A0A1M3GW08-F1-MODEL\_V4 \| 1.0 \| 0.000519 \| 165 \| 0.31 \| 74 \| 47 \| 2 \| 1 \| 72 \| 2 \| 73 \| Uncharacterized protein \| Uncharacterized protein; afdb-uniprot50 \| AF-A0A7I7T0P9-F1-MODEL\_V4 \| 1.0 \| 0.0003482 \| 165 \| 0.283 \| 74 \| 49 \| 2 \| 1 \| 72 \| 48 \| 119 \| Uncharacterized protein \| Uncharacterized protein; afdb-uniprot50 \| AF-A0A0M9VKJ4-F1-MODEL\_V4 \| 1.0 \| 0.0009444 \| 164 \| 0.412 \| 63 \| 35 \| 1 \| 1 \| 63 \| 1 \| 61 \| HTH\_17 domain-containing protein \| HTH\_17 domain-containing protein; afdb-uniprot50 \| AF-A0A3T0DJV1-F1-MODEL\_V4 \| 1.0 \| 0.001837 \| 164 \| 0.266 \| 60 \| 41 \| 2 \| 7 \| 65 \| 2 \| 59 \| Excisionase \| Excisionase; afdb-uniprot50 \| AF-A0A522CQA7-F1-MODEL\_V4 \| 1.0 \| 0.002562 \| 164 \| 0.271 \| 59 \| 41 \| 1 \| 11 \| 69 \| 5 \| 61 \| DNA-binding protein \| DNA-binding protein; afdb-uniprot50 \| AF-A0A291RRP8-F1-MODEL\_V4 \| 1.0 \| 0.001317 \| 164 \| 0.272 \| 66 \| 45 \| 2 \| 1 \| 65 \| 1 \| 64 \| DNA-binding protein \| DNA-binding protein; afdb-uniprot50 \| AF-A0A6I1KHJ9-F1-MODEL\_V4 \| 1.0 \| 0.0008268 \| 164 \| 0.314 \| 70 \| 46 \| 1 \| 4 \| 73 \| 1 \| 68 \| HTH\_17 domain-containing protein \| HTH\_17 domain-containing protein; afdb-uniprot50 \| AF-A0A1J5V6Q7-F1-MODEL\_V4 \| 1.0 \| 0.0008268 \| 164 \| 0.313 \| 67 \| 43 \| 2 \| 1 \| 66 \| 1 \| 65 \| HTH\_17 domain-containing protein \| HTH\_17 domain-containing protein; afdb-uniprot50 \| AF-A0A7Y6T9Q4-F1-MODEL\_V4 \| 1.0 \| 0.001317 \| 164 \| 0.276 \| 65 \| 43 \| 2 \| 1 \| 63 \| 1 \| 63 \| Helix-turn-helix domain-containing protein \| Helix-turn-helix domain-containing protein; afdb-uniprot50 \| AF-A0A0N7LYK3-F1-MODEL\_V4 \| 1.0 \| 0.004081 \| 164 \| 0.309 \| 55 \| 36 \| 1 \| 9 \| 63 \| 5 \| 57 \| Helix-turn-helix domain protein \| Helix-turn-helix domain protein; afdb-uniprot50 \| AF-F8AAE3-F1-MODEL\_V4 \| 1.0 \| 0.0007736 \| 164 \| 0.333 \| 66 \| 42 \| 2 \| 7 \| 72 \| 13 \| 76 \| HTH\_17 domain-containing protein \| HTH\_17 domain-containing protein; afdb-uniprot50 \| AF-A0A0P7XPZ0-F1-MODEL\_V4 \| 1.0 \| 0.001009 \| 164 \| 0.276 \| 65 \| 45 \| 1 \| 2 \| 66 \| 13 \| 75 \| Putative site-specific integrase-resolvase \| Putative site-specific integrase-resolvase; afdb-uniprot50 \| AF-A0A1M3H9S1-F1-MODEL\_V4 \| 1.0 \| 0.001009 \| 164 \| 0.349 \| 63 \| 36 \| 2 \| 11 \| 73 \| 5 \| 62 \| Uncharacterized protein \| Uncharacterized protein; afdb-uniprot50 \| AF-A0A853UBA9-F1-MODEL\_V4 \| 1.0 \| 0.0002496 \| 164 \| 0.276 \| 76 \| 51 \| 2 \| 1 \| 72 \| 2 \| 77 \| Uncharacterized protein \| Uncharacterized protein; afdb-uniprot50 \| AF-A0A843S3B5-F1-MODEL\_V4 \| 1.0 \| 0.0005547 \| 164 \| 0.253 \| 75 \| 52 \| 2 \| 1 \| 73 \| 1 \| 73 \| Helix-turn-helix domain-containing protein \| Helix-turn-helix domain-containing protein; afdb-uniprot50 \| AF-A0A318WSP9-F1-MODEL\_V4 \| 1.0 \| 0.0006772 \| 164 \| 0.21 \| 76 \| 54 \| 2 \| 1 \| 72 \| 26 \| 99 \| AlpA family transcriptional regulator \| AlpA family transcriptional regulator; afdb-uniprot50 \| AF-K2EJ62-F1-MODEL\_V4 \| 1.0 \| 0.0003721 \| 164 \| 0.315 \| 73 \| 44 \| 3 \| 1 \| 73 \| 58 \| 124 \| Uncharacterized protein \| Uncharacterized protein; afdb-uniprot50 \| AF-D8F5B5-F1-MODEL\_V4 \| 1.0 \| 0.0001913 \| 164 \| 0.297 \| 84 \| 46 \| 4 \| 1 \| 73 \| 20 \| 101 \| Transcriptional regulator, MerR family \| Transcriptional regulator, MerR family; afdb-uniprot50 \| AF-A0A1Y0CB63-F1-MODEL\_V4 \| 1.0 \| 0.001608 \| 163 \| 0.372 \| 59 \| 34 \| 2 \| 11 \| 69 \| 6 \| 61 \| DNA-binding protein \| DNA-binding protein; afdb-uniprot50 \| AF-A0A7V0J3F2-F1-MODEL\_V4 \| 1.0 \| 0.003128 \| 163 \| 0.272 \| 55 \| 38 \| 1 \| 11 \| 65 \| 7 \| 59 \| DNA-binding protein \| DNA-binding protein; afdb-uniprot50 \| AF-A0A387BVQ2-F1-MODEL\_V4 \| 1.0 \| 0.001719 \| 163 \| 0.295 \| 61 \| 41 \| 1 \| 3 \| 63 \| 2 \| 60 \| DNA-binding protein \| DNA-binding protein; afdb-uniprot50 \| AF-A0A1X0H7C2-F1-MODEL\_V4 \| 1.0 \| 0.0005547 \| 163 \| 0.289 \| 69 \| 45 \| 2 \| 1 \| 67 \| 2 \| 68 \| DNA-binding protein \| DNA-binding protein; afdb-uniprot50 \| AF-A0A347Q4W5-F1-MODEL\_V4 \| 1.0 \| 0.004662 \| 163 \| 0.327 \| 55 \| 35 \| 1 \| 9 \| 63 \| 7 \| 59 \| HTH\_17 domain-containing protein \| HTH\_17 domain-containing protein; afdb-uniprot50 \| AF-A0A7Y9KJU0-F1-MODEL\_V4 \| 1.0 \| 0.0005928 \| 163 \| 0.32 \| 78 \| 47 \| 2 \| 1 \| 73 \| 1 \| 77 \| Putative DNA-binding transcriptional regulator AlpA \| Putative DNA-binding transcriptional regulator AlpA; afdb-uniprot50 \| AF-A0A849C2U3-F1-MODEL\_V4 \| 1.0 \| 0.002397 \| 163 \| 0.265 \| 64 \| 45 \| 1 \| 10 \| 73 \| 7 \| 68 \| Helix-turn-helix domain-containing protein \| Helix-turn-helix domain-containing protein; afdb-uniprot50 \| AF-A0A7Y6EM70-F1-MODEL\_V4 \| 1.0 \| 0.0004856 \| 163 \| 0.319 \| 72 \| 47 \| 1 \| 1 \| 72 \| 14 \| 83 \| Helix-turn-helix domain-containing protein \| Helix-turn-helix domain-containing protein; afdb-uniprot50 \| AF-A0A316MH91-F1-MODEL\_V4 \| 1.0 \| 0.0005547 \| 163 \| 0.256 \| 78 \| 51 \| 2 \| 1 \| 73 \| 7 \| 82 \| DNA-binding protein \| DNA-binding protein; afdb-uniprot50 \| AF-L0J4B7-F1-MODEL\_V4 \| 1.0 \| 0.001009 \| 163 \| 0.333 \| 63 \| 40 \| 1 \| 11 \| 73 \| 9 \| 69 \| Uncharacterized protein \| Uncharacterized protein; afdb-uniprot50 \| AF-A0A3N6F8T7-F1-MODEL\_V4 \| 1.0 \| 0.002098 \| 163 \| 0.215 \| 65 \| 49 \| 1 \| 2 \| 66 \| 3 \| 65 \| Helix-turn-helix domain protein \| Helix-turn-helix domain protein; afdb-uniprot50 \| AF-A0A6L5XLJ8-F1-MODEL\_V4 \| 1.0 \| 0.001317 \| 162 \| 0.36 \| 61 \| 37 \| 1 \| 12 \| 72 \| 2 \| 60 \| DNA-binding protein \| DNA-binding protein; afdb-uniprot50 \| AF-A0A7C5CBG1-F1-MODEL\_V4 \| 1.0 \| 0.001408 \| 162 \| 0.344 \| 61 \| 38 \| 1 \| 12 \| 72 \| 1 \| 59 \| DNA-binding protein \| DNA-binding protein; afdb-uniprot50 \| AF-A0A372MC99-F1-MODEL\_V4 \| 1.0 \| 0.002738 \| 162 \| 0.322 \| 59 \| 38 \| 1 \| 9 \| 67 \| 7 \| 63 \| DNA-binding protein \| DNA-binding protein; afdb-uniprot50 \| AF-A0A4Q5Y382-F1-MODEL\_V4 \| 1.0 \| 0.001408 \| 162 \| 0.366 \| 60 \| 36 \| 1 \| 12 \| 71 \| 1 \| 58 \| DNA-binding protein \| DNA-binding protein; afdb-uniprot50 \| AF-A0A0F2QGE5-F1-MODEL\_V4 \| 1.0 \| 0.001505 \| 162 \| 0.26 \| 73 \| 52 \| 1 \| 1 \| 73 \| 1 \| 71 \| HTH\_17 domain-containing protein \| HTH\_17 domain-containing protein; afdb-uniprot50 \| AF-A0A853IUT5-F1-MODEL\_V4 \| 1.0 \| 0.0006772 \| 162 \| 0.305 \| 72 \| 47 \| 2 \| 1 \| 72 \| 1 \| 69 \| Helix-turn-helix domain-containing protein \| Helix-turn-helix domain-containing protein; afdb-uniprot50 \| AF-A0A0S7CWQ4-F1-MODEL\_V4 \| 1.0 \| 0.001079 \| 162 \| 0.292 \| 65 \| 44 \| 1 \| 1 \| 65 \| 6 \| 68 \| HTH\_17 domain-containing protein \| HTH\_17 domain-containing protein; afdb-uniprot50 \| AF-A0A321LRP0-F1-MODEL\_V4 \| 1.0 \| 0.0008268 \| 162 \| 0.3 \| 70 \| 47 \| 1 \| 4 \| 73 \| 2 \| 69 \| Excisionase \| Excisionase; afdb-uniprot50 \| AF-A0A7X6SV09-F1-MODEL\_V4 \| 1.0 \| 0.0006336 \| 162 \| 0.226 \| 75 \| 54 \| 3 \| 1 \| 73 \| 2 \| 74 \| Helix-turn-helix domain-containing protein \| Helix-turn-helix domain-containing protein; afdb-uniprot50 \| AF-A0A840A4S4-F1-MODEL\_V4 \| 1.0 \| 0.0008268 \| 162 \| 0.348 \| 66 \| 38 \| 2 \| 11 \| 73 \| 5 \| 68 \| Putative site-specific integrase-resolvase \| Putative site-specific integrase-resolvase; afdb-uniprot50 \| AF-A0A7K1XH10-F1-MODEL\_V4 \| 1.0 \| 0.0009444 \| 162 \| 0.313 \| 67 \| 45 \| 1 \| 1 \| 67 \| 12 \| 77 \| Helix-turn-helix domain-containing protein \| Helix-turn-helix domain-containing protein; afdb-uniprot50 \| AF-A0A259S496-F1-MODEL\_V4 \| 1.0 \| 0.0003257 \| 162 \| 0.297 \| 74 \| 49 \| 2 \| 1 \| 73 \| 1 \| 72 \| HTH\_17 domain-containing protein \| HTH\_17 domain-containing protein; afdb-uniprot50 \| AF-A0A6V8MPT2-F1-MODEL\_V4 \| 1.0 \| 0.0008836 \| 162 \| 0.232 \| 73 \| 54 \| 1 \| 1 \| 73 \| 7 \| 77 \| HTH\_17 domain-containing protein \| HTH\_17 domain-containing protein; afdb-uniprot50 \| AF-A0A7W1M9N3-F1-MODEL\_V4 \| 1.0 \| 0.001408 \| 162 \| 0.301 \| 63 \| 42 \| 1 \| 11 \| 73 \| 5 \| 65 \| DNA-binding protein \| DNA-binding protein; afdb-uniprot50 \| AF-Q82S53-F1-MODEL\_V4 \| 1.0 \| 0.0005547 \| 162 \| 0.277 \| 72 \| 49 \| 2 \| 1 \| 71 \| 15 \| 84 \| Uncharacterized protein \| Uncharacterized protein; afdb-uniprot50 \| AF-A0A4Q7G7Y9-F1-MODEL\_V4 \| 1.0 \| 0.001009 \| 162 \| 0.267 \| 71 \| 47 \| 1 \| 1 \| 71 \| 1 \| 66 \| DNA-binding protein \| DNA-binding protein; afdb-uniprot50 \| AF-A0A315EIU2-F1-MODEL\_V4 \| 1.0 \| 0.0009444 \| 162 \| 0.348 \| 66 \| 38 \| 3 \| 1 \| 63 \| 15 \| 78 \| Uncharacterized protein \| Uncharacterized protein; afdb-uniprot50 \| AF-A0A842HLN5-F1-MODEL\_V4 \| 1.0 \| 0.001505 \| 161 \| 0.301 \| 63 \| 42 \| 1 \| 11 \| 73 \| 5 \| 65 \| DNA-binding protein \| DNA-binding protein; afdb-uniprot50 \| AF-A0A0Q4BJD0-F1-MODEL\_V4 \| 1.0 \| 0.002562 \| 161 \| 0.365 \| 63 \| 38 \| 1 \| 11 \| 73 \| 5 \| 65 \| HTH\_17 domain-containing protein \| HTH\_17 domain-containing protein; afdb-uniprot50 \| AF-A0A4R2IPL2-F1-MODEL\_V4 \| 1.0 \| 0.001408 \| 161 \| 0.283 \| 67 \| 46 \| 1 \| 1 \| 67 \| 2 \| 66 \| Excisionase family DNA binding protein \| Excisionase family DNA binding protein; afdb-uniprot50 \| AF-A0A1B8SCM6-F1-MODEL\_V4 \| 1.0 \| 0.001153 \| 161 \| 0.285 \| 63 \| 41 \| 2 \| 11 \| 73 \| 10 \| 68 \| HTH\_17 domain-containing protein \| HTH\_17 domain-containing protein; afdb-uniprot50 \| AF-A0A7J9W6U6-F1-MODEL\_V4 \| 1.0 \| 0.003819 \| 161 \| 0.246 \| 65 \| 49 \| 0 \| 1 \| 65 \| 2 \| 66 \| Helix-turn-helix domain-containing protein \| Helix-turn-helix domain-containing protein; afdb-uniprot50 \| AF-A0A2V8UL43-F1-MODEL\_V4 \| 1.0 \| 0.004081 \| 161 \| 0.333 \| 51 \| 32 \| 1 \| 11 \| 61 \| 4 \| 52 \| DNA-binding protein \| DNA-binding protein; afdb-uniprot50 \| AF-A0A2H5F3C0-F1-MODEL\_V4 \| 1.0 \| 0.001317 \| 161 \| 0.191 \| 73 \| 57 \| 1 \| 1 \| 73 \| 1 \| 71 \| AlpA family transcriptional regulator \| AlpA family transcriptional regulator; afdb-uniprot50 \| AF-A0A2N9N2Z5-F1-MODEL\_V4 \| 1.0 \| 0.001232 \| 161 \| 0.301 \| 63 \| 41 \| 2 \| 11 \| 73 \| 4 \| 63 \| HTH\_17 domain-containing protein \| HTH\_17 domain-containing protein; afdb-uniprot50 \| AF-A0A315DB40-F1-MODEL\_V4 \| 1.0 \| 0.0009444 \| 161 \| 0.307 \| 65 \| 40 \| 2 \| 11 \| 72 \| 5 \| 67 \| Uncharacterized protein \| Uncharacterized protein; afdb-uniprot50 \| AF-A0A7T1I020-F1-MODEL\_V4 \| 1.0 \| 0.001232 \| 161 \| 0.272 \| 66 \| 45 \| 2 \| 1 \| 65 \| 10 \| 73 \| Helix-turn-helix domain-containing protein \| Helix-turn-helix domain-containing protein; afdb-uniprot50 \| AF-A0A0N1H048-F1-MODEL\_V4 \| 1.0 \| 0.0008836 \| 161 \| 0.298 \| 67 \| 46 \| 1 \| 2 \| 68 \| 13 \| 78 \| HTH\_17 domain-containing protein \| HTH\_17 domain-containing protein; afdb-uniprot50 \| AF-A0A0Q8Q4S6-F1-MODEL\_V4 \| 1.0 \| 0.0005928 \| 161 \| 0.308 \| 68 \| 45 \| 2 \| 1 \| 67 \| 10 \| 76 \| HTH\_17 domain-containing protein \| HTH\_17 domain-containing protein; afdb-uniprot50 \| AF-A3XET4-F1-MODEL\_V4 \| 1.0 \| 0.0008268 \| 161 \| 0.291 \| 72 \| 48 \| 2 \| 2 \| 73 \| 10 \| 78 \| Uncharacterized protein \| Uncharacterized protein; afdb-uniprot50 \| AF-A0A2A4Z6F6-F1-MODEL\_V4 \| 1.0 \| 0.001079 \| 161 \| 0.26 \| 73 \| 49 \| 2 \| 1 \| 73 \| 1 \| 68 \| DNA-binding protein \| DNA-binding protein; afdb-uniprot50 \| AF-A0A2X0SBJ5-F1-MODEL\_V4 \| 1.0 \| 0.000519 \| 161 \| 0.287 \| 73 \| 49 \| 2 \| 1 \| 73 \| 16 \| 85 \| Uncharacterized protein \| Uncharacterized protein; afdb-uniprot50 \| AF-W0RB88-F1-MODEL\_V4 \| 1.0 \| 0.001232 \| 161 \| 0.253 \| 71 \| 51 \| 1 \| 1 \| 71 \| 12 \| 80 \| HTH\_17 domain-containing protein \| HTH\_17 domain-containing protein; afdb-uniprot50 \| AF-A0A2V1NPH5-F1-MODEL\_V4 \| 1.0 \| 0.001608 \| 161 \| 0.311 \| 61 \| 41 \| 1 \| 9 \| 69 \| 30 \| 89 \| DNA-binding protein \| DNA-binding protein; afdb-uniprot50 \| AF-A0A4S2B6Y8-F1-MODEL\_V4 \| 1.0 \| 0.0009444 \| 161 \| 0.343 \| 64 \| 38 \| 2 \| 2 \| 63 \| 10 \| 71 \| DNA-binding protein \| DNA-binding protein; afdb-uniprot50 \| AF-A0A437M628-F1-MODEL\_V4 \| 1.0 \| 0.001009 \| 161 \| 0.243 \| 74 \| 53 \| 2 \| 1 \| 73 \| 16 \| 87 \| DNA-binding protein \| DNA-binding protein; afdb-uniprot50 \| AF-S9ZET1-F1-MODEL\_V4 \| 1.0 \| 0.0006772 \| 161 \| 0.267 \| 71 \| 50 \| 1 \| 1 \| 71 \| 7 \| 75 \| Uncharacterized protein \| Uncharacterized protein; afdb-uniprot50 \| AF-A0A7Z0JE43-F1-MODEL\_V4 \| 1.0 \| 0.002397 \| 161 \| 0.264 \| 68 \| 48 \| 1 \| 2 \| 69 \| 36 \| 101 \| HTH\_17 domain-containing protein \| HTH\_17 domain-containing protein; afdb-uniprot50 \| AF-A0A1A2QLL9-F1-MODEL\_V4 \| 1.0 \| 0.001232 \| 160 \| 0.294 \| 68 \| 44 \| 2 \| 1 \| 68 \| 1 \| 64 \| HTH\_17 domain-containing protein \| HTH\_17 domain-containing protein; afdb-uniprot50 \| AF-A0A4P8SWN0-F1-MODEL\_V4 \| 1.0 \| 0.001963 \| 160 \| 0.3 \| 60 \| 40 \| 1 \| 4 \| 63 \| 1 \| 58 \| Uncharacterized protein \| Uncharacterized protein; afdb-uniprot50 \| AF-A0A3E0NRW3-F1-MODEL\_V4 \| 1.0 \| 0.001837 \| 160 \| 0.26 \| 69 \| 49 \| 1 \| 4 \| 72 \| 1 \| 67 \| DNA-binding protein \| DNA-binding protein; afdb-uniprot50 \| AF-A0A7C6R8I4-F1-MODEL\_V4 \| 1.0 \| 0.001153 \| 160 \| 0.342 \| 73 \| 43 \| 3 \| 1 \| 73 \| 1 \| 68 \| Helix-turn-helix domain-containing protein \| Helix-turn-helix domain-containing protein; afdb-uniprot50 \| AF-A0A850RHE4-F1-MODEL\_V4 \| 1.0 \| 0.001505 \| 160 \| 0.333 \| 63 \| 40 \| 1 \| 1 \| 63 \| 2 \| 62 \| Helix-turn-helix domain-containing protein \| Helix-turn-helix domain-containing protein; afdb-uniprot50 \| AF-A0A2T7TRJ2-F1-MODEL\_V4 \| 1.0 \| 0.001505 \| 160 \| 0.328 \| 67 \| 40 \| 2 \| 7 \| 73 \| 2 \| 63 \| HTH\_17 domain-containing protein \| HTH\_17 domain-containing protein; afdb-uniprot50 \| AF-A0A7C2GW26-F1-MODEL\_V4 \| 1.0 \| 0.001505 \| 160 \| 0.287 \| 73 \| 48 \| 2 \| 1 \| 73 \| 1 \| 69 \| Helix-turn-helix domain-containing protein \| Helix-turn-helix domain-containing protein; afdb-uniprot50 \| AF-A0A6I3ZMZ5-F1-MODEL\_V4 \| 1.0 \| 0.001317 \| 160 \| 0.273 \| 73 \| 50 \| 2 \| 1 \| 73 \| 1 \| 70 \| Helix-turn-helix domain-containing protein \| Helix-turn-helix domain-containing protein; afdb-uniprot50 \| AF-A0A4Q8QL74-F1-MODEL\_V4 \| 1.0 \| 0.001408 \| 160 \| 0.27 \| 74 \| 50 \| 2 \| 1 \| 72 \| 1 \| 72 \| DNA-binding protein \| DNA-binding protein; afdb-uniprot50 \| AF-A0A1P8NCH9-F1-MODEL\_V4 \| 1.0 \| 0.001837 \| 160 \| 0.263 \| 72 \| 51 \| 1 \| 1 \| 72 \| 7 \| 76 \| HTH\_17 domain-containing protein \| HTH\_17 domain-containing protein; afdb-uniprot50 \| AF-J0MNL2-F1-MODEL\_V4 \| 1.0 \| 0.003343 \| 160 \| 0.253 \| 63 \| 45 \| 1 \| 11 \| 73 \| 19 \| 79 \| Uncharacterized protein \| Uncharacterized protein; afdb-uniprot50 \| AF-A0A1H1ZMM5-F1-MODEL\_V4 \| 1.0 \| 0.001232 \| 160 \| 0.263 \| 72 \| 51 \| 1 \| 1 \| 72 \| 18 \| 87 \| DNA binding domain-containing protein, excisionase family \| DNA binding domain-containing protein, excisionase family; afdb-uniprot50 \| AF-K2DX72-F1-MODEL\_V4 \| 1.0 \| 0.002098 \| 160 \| 0.36 \| 61 \| 37 \| 1 \| 3 \| 63 \| 2 \| 60 \| Uncharacterized protein \| Uncharacterized protein; afdb-uniprot50 \| AF-A0A437M765-F1-MODEL\_V4 \| 1.0 \| 0.002738 \| 160 \| 0.238 \| 67 \| 49 \| 1 \| 1 \| 67 \| 19 \| 83 \| Uncharacterized protein \| Uncharacterized protein; afdb-uniprot50 \| AF-A0A2V9QRN0-F1-MODEL\_V4 \| 1.0 \| 0.001232 \| 160 \| 0.273 \| 73 \| 51 \| 1 \| 1 \| 73 \| 23 \| 93 \| HTH\_17 domain-containing protein \| HTH\_17 domain-containing protein; afdb-uniprot50 \| AF-A0A238VQI5-F1-MODEL\_V4 \| 1.0 \| 0.002738 \| 159 \| 0.263 \| 72 \| 45 \| 2 \| 1 \| 72 \| 1 \| 64 \| Helix-turn-helix domain-containing protein \| Helix-turn-helix domain-containing protein; afdb-uniprot50 \| AF-A0A7V9MYE0-F1-MODEL\_V4 \| 1.0 \| 0.002738 \| 159 \| 0.257 \| 66 \| 47 \| 1 \| 7 \| 72 \| 1 \| 64 \| Helix-turn-helix domain-containing protein \| Helix-turn-helix domain-containing protein; afdb-uniprot50 \| AF-A0A1I1BCB6-F1-MODEL\_V4 \| 1.0 \| 0.001719 \| 159 \| 0.268 \| 67 \| 46 \| 2 \| 1 \| 66 \| 2 \| 66 \| Transcriptional regulator, AlpA family \| Transcriptional regulator, AlpA family; afdb-uniprot50 \| AF-A0A1A2GHT1-F1-MODEL\_V4 \| 1.0 \| 0.004081 \| 159 \| 0.295 \| 61 \| 41 \| 1 \| 8 \| 68 \| 9 \| 67 \| HTH\_17 domain-containing protein \| HTH\_17 domain-containing protein; afdb-uniprot50 \| AF-A0A4P9S7Y4-F1-MODEL\_V4 \| 1.0 \| 0.001317 \| 159 \| 0.363 \| 66 \| 39 \| 2 \| 7 \| 71 \| 2 \| 65 \| Helix-turn-helix domain-containing protein \| Helix-turn-helix domain-containing protein; afdb-uniprot50 \| AF-I2ACQ1-F1-MODEL\_V4 \| 1.0 \| 0.001719 \| 159 \| 0.285 \| 63 \| 42 \| 2 \| 11 \| 73 \| 4 \| 63 \| HTH\_17 domain-containing protein \| HTH\_17 domain-containing protein; afdb-uniprot50 \| AF-A0A7Y2W1R5-F1-MODEL\_V4 \| 1.0 \| 0.001963 \| 159 \| 0.306 \| 62 \| 41 \| 1 \| 1 \| 62 \| 1 \| 60 \| Helix-turn-helix domain-containing protein \| Helix-turn-helix domain-containing protein; afdb-uniprot50 \| AF-A0A7L9BRT5-F1-MODEL\_V4 \| 1.0 \| 0.001232 \| 159 \| 0.328 \| 70 \| 45 \| 1 \| 4 \| 73 \| 1 \| 68 \| Helix-turn-helix domain-containing protein \| Helix-turn-helix domain-containing protein; afdb-uniprot50 \| AF-A0A7M3MAM1-F1-MODEL\_V4 \| 1.0 \| 0.004983 \| 159 \| 0.24 \| 54 \| 41 \| 0 \| 10 \| 63 \| 17 \| 70 \| Uncharacterized protein \| Uncharacterized protein; afdb-uniprot50 \| AF-A0A2G1QH89-F1-MODEL\_V4 \| 1.0 \| 0.0004543 \| 159 \| 0.269 \| 78 \| 50 \| 3 \| 1 \| 73 \| 1 \| 76 \| Transcriptional regulator \| Transcriptional regulator; afdb-uniprot50 \| AF-A0A4Q7FM96-F1-MODEL\_V4 \| 1.0 \| 0.003343 \| 159 \| 0.311 \| 61 \| 40 \| 1 \| 5 \| 65 \| 13 \| 71 \| Transcriptional regulator \| Transcriptional regulator; afdb-uniprot50 \| AF-A0A6N9BF07-F1-MODEL\_V4 \| 1.0 \| 0.0008268 \| 159 \| 0.266 \| 75 \| 51 \| 2 \| 1 \| 73 \| 1 \| 73 \| Helix-turn-helix domain-containing protein \| Helix-turn-helix domain-containing protein; afdb-uniprot50 \| AF-A0A5N0E6B6-F1-MODEL\_V4 \| 1.0 \| 0.004662 \| 159 \| 0.28 \| 57 \| 39 \| 1 \| 9 \| 65 \| 5 \| 59 \| Helix-turn-helix domain-containing protein \| Helix-turn-helix domain-containing protein; afdb-uniprot50 \| AF-A0A2U9NH93-F1-MODEL\_V4 \| 1.0 \| 0.0009444 \| 159 \| 0.292 \| 65 \| 42 \| 2 \| 1 \| 63 \| 9 \| 71 \| DNA-binding protein \| DNA-binding protein; afdb-uniprot50 \| AF-A0A2E1IAE1-F1-MODEL\_V4 \| 1.0 \| 0.0005928 \| 158 \| 0.306 \| 62 \| 41 \| 1 \| 4 \| 65 \| 2 \| 61 \| DNA-binding protein \| DNA-binding protein; afdb-uniprot50 \| AF-A0A366EM39-F1-MODEL\_V4 \| 1.0 \| 0.001719 \| 158 \| 0.349 \| 63 \| 39 \| 1 \| 11 \| 73 \| 7 \| 67 \| Helix-turn-helix protein \| Helix-turn-helix protein; afdb-uniprot50 \| AF-A0A1H1LN62-F1-MODEL\_V4 \| 1.0 \| 0.001608 \| 158 \| 0.257 \| 66 \| 47 \| 1 \| 2 \| 67 \| 4 \| 67 \| Helix-turn-helix domain-containing protein \| Helix-turn-helix domain-containing protein; afdb-uniprot50 \| AF-A0A2K8UDT2-F1-MODEL\_V4 \| 1.0 \| 0.0003721 \| 158 \| 0.301 \| 73 \| 43 \| 3 \| 1 \| 73 \| 1 \| 65 \| HTH\_17 domain-containing protein \| HTH\_17 domain-containing protein; afdb-uniprot50 \| AF-T0HS16-F1-MODEL\_V4 \| 1.0 \| 0.001232 \| 158 \| 0.208 \| 72 \| 55 \| 1 \| 2 \| 73 \| 6 \| 75 \| Uncharacterized protein \| Uncharacterized protein; afdb-uniprot50 \| AF-A0A6P0GKB9-F1-MODEL\_V4 \| 1.0 \| 0.002243 \| 158 \| 0.219 \| 73 \| 55 \| 1 \| 1 \| 73 \| 4 \| 74 \| Helix-turn-helix domain-containing protein \| Helix-turn-helix domain-containing protein; afdb-uniprot50 \| AF-A0A286EZU0-F1-MODEL\_V4 \| 1.0 \| 0.001608 \| 158 \| 0.318 \| 66 \| 43 \| 1 \| 2 \| 67 \| 13 \| 76 \| Helix-turn-helix domain-containing protein \| Helix-turn-helix domain-containing protein; afdb-uniprot50 \| AF-A0A7U3LS04-F1-MODEL\_V4 \| 1.0 \| 0.001837 \| 158 \| 0.287 \| 66 \| 43 \| 2 \| 10 \| 73 \| 4 \| 67 \| Uncharacterized protein \| Uncharacterized protein; afdb-uniprot50 \| AF-A0A562C335-F1-MODEL\_V4 \| 1.0 \| 0.0008268 \| 158 \| 0.26 \| 73 \| 49 \| 2 \| 1 \| 73 \| 11 \| 78 \| AlpA family transcriptional regulator \| AlpA family transcriptional regulator; afdb-uniprot50 \| AF-A0A4T0UKW9-F1-MODEL\_V4 \| 1.0 \| 0.0004251 \| 158 \| 0.309 \| 71 \| 44 \| 3 \| 1 \| 67 \| 8 \| 77 \| Helix-turn-helix domain-containing protein \| Helix-turn-helix domain-containing protein; afdb-uniprot50 \| AF-A0A1M3IN55-F1-MODEL\_V4 \| 1.0 \| 0.001837 \| 158 \| 0.323 \| 65 \| 42 \| 1 \| 7 \| 71 \| 1 \| 63 \| HTH\_17 domain-containing protein \| HTH\_17 domain-containing protein; afdb-uniprot50 \| AF-A0A1G7WPJ9-F1-MODEL\_V4 \| 1.0 \| 0.001505 \| 158 \| 0.26 \| 73 \| 51 \| 2 \| 1 \| 73 \| 13 \| 82 \| DNA binding domain-containing protein, excisionase family \| DNA binding domain-containing protein, excisionase family; afdb-uniprot50 \| AF-A0A512T4M0-F1-MODEL\_V4 \| 1.0 \| 0.001408 \| 158 \| 0.26 \| 69 \| 49 \| 1 \| 4 \| 72 \| 1 \| 67 \| HTH\_17 domain-containing protein \| HTH\_17 domain-containing protein; afdb-uniprot50 \| AF-A0A7Z0EDB2-F1-MODEL\_V4 \| 1.0 \| 0.001963 \| 158 \| 0.338 \| 62 \| 39 \| 1 \| 1 \| 62 \| 42 \| 101 \| Putative DNA-binding transcriptional regulator AlpA \| Putative DNA-binding transcriptional regulator AlpA; afdb-uniprot50 \| AF-A0A7Y9SJU6-F1-MODEL\_V4 \| 1.0 \| 0.002738 \| 157 \| 0.338 \| 62 \| 39 \| 1 \| 1 \| 62 \| 1 \| 60 \| Excisionase family DNA binding protein \| Excisionase family DNA binding protein; afdb-uniprot50 \| AF-A0A1B8SBD0-F1-MODEL\_V4 \| 1.0 \| 0.002243 \| 157 \| 0.285 \| 70 \| 44 \| 2 \| 4 \| 73 \| 1 \| 64 \| Excisionase \| Excisionase; afdb-uniprot50 \| AF-A0A1V3NRC7-F1-MODEL\_V4 \| 1.0 \| 0.002243 \| 157 \| 0.206 \| 63 \| 48 \| 1 \| 11 \| 73 \| 6 \| 66 \| DNA-binding protein \| DNA-binding protein; afdb-uniprot50 \| AF-Q5Z3A8-F1-MODEL\_V4 \| 1.0 \| 0.002243 \| 157 \| 0.317 \| 63 \| 40 \| 2 \| 1 \| 63 \| 1 \| 60 \| HTH\_17 domain-containing protein \| HTH\_17 domain-containing protein; afdb-uniprot50 \| AF-A0A7G6U3R5-F1-MODEL\_V4 \| 1.0 \| 0.002562 \| 157 \| 0.222 \| 63 \| 47 \| 1 \| 1 \| 63 \| 3 \| 63 \| Helix-turn-helix domain-containing protein \| Helix-turn-helix domain-containing protein; afdb-uniprot50 \| AF-A0A5N6S6V5-F1-MODEL\_V4 \| 1.0 \| 0.0006336 \| 157 \| 0.287 \| 73 \| 49 \| 2 \| 1 \| 73 \| 2 \| 71 \| DNA-binding protein \| DNA-binding protein; afdb-uniprot50 \| AF-A0A4R5PE23-F1-MODEL\_V4 \| 1.0 \| 0.0009444 \| 157 \| 0.301 \| 73 \| 47 \| 3 \| 1 \| 72 \| 1 \| 70 \| DNA-binding protein \| DNA-binding protein; afdb-uniprot50 \| AF-A0A2A3D3G3-F1-MODEL\_V4 \| 1.0 \| 0.001232 \| 157 \| 0.297 \| 74 \| 49 \| 2 \| 1 \| 73 \| 1 \| 72 \| Uncharacterized protein \| Uncharacterized protein; afdb-uniprot50 \| AF-A0A4Y3UN51-F1-MODEL\_V4 \| 1.0 \| 0.0005547 \| 157 \| 0.306 \| 75 \| 48 \| 2 \| 1 \| 73 \| 1 \| 73 \| Uncharacterized protein \| Uncharacterized protein; afdb-uniprot50 \| AF-A0A1I6Z5G2-F1-MODEL\_V4 \| 1.0 \| 0.001317 \| 157 \| 0.323 \| 68 \| 41 \| 1 \| 11 \| 73 \| 6 \| 73 \| Uncharacterized protein \| Uncharacterized protein; afdb-uniprot50 \| AF-A0A3S0GSK5-F1-MODEL\_V4 \| 1.0 \| 0.002562 \| 157 \| 0.264 \| 68 \| 48 \| 1 \| 5 \| 72 \| 11 \| 76 \| DNA-binding protein \| DNA-binding protein; afdb-uniprot50 \| AF-A0A7C3Q0U4-F1-MODEL\_V4 \| 1.0 \| 0.0007238 \| 157 \| 0.25 \| 76 \| 51 \| 2 \| 2 \| 73 \| 10 \| 83 \| DNA-binding protein \| DNA-binding protein; afdb-uniprot50 \| AF-A0A1V4XWG2-F1-MODEL\_V4 \| 1.0 \| 0.002098 \| 157 \| 0.328 \| 64 \| 40 \| 2 \| 2 \| 62 \| 17 \| 80 \| Uncharacterized protein \| Uncharacterized protein; afdb-uniprot50 \| AF-A0A7D5MY55-F1-MODEL\_V4 \| 1.0 \| 0.001963 \| 157 \| 0.303 \| 66 \| 43 \| 2 \| 9 \| 73 \| 2 \| 65 \| Helix-turn-helix domain-containing protein \| Helix-turn-helix domain-containing protein; afdb-uniprot50 \| AF-A0A315BJ48-F1-MODEL\_V4 \| 1.0 \| 0.0003721 \| 157 \| 0.301 \| 83 \| 46 \| 4 \| 1 \| 73 \| 40 \| 120 \| Uncharacterized protein \| Uncharacterized protein; afdb-uniprot50 \| AF-A0A1H3G4F0-F1-MODEL\_V4 \| 1.0 \| 0.003128 \| 156 \| 0.245 \| 61 \| 44 \| 1 \| 7 \| 67 \| 1 \| 59 \| Transcriptional regulator, AlpA family \| Transcriptional regulator, AlpA family; afdb-uniprot50 \| AF-A0A1I6Y1L1-F1-MODEL\_V4 \| 1.0 \| 0.001009 \| 156 \| 0.338 \| 62 \| 37 \| 2 \| 11 \| 72 \| 6 \| 63 \| Transcriptional regulator, AlpA family \| Transcriptional regulator, AlpA family; afdb-uniprot50 \| AF-S3AQA5-F1-MODEL\_V4 \| 1.0 \| 0.002927 \| 156 \| 0.253 \| 63 \| 45 \| 1 \| 1 \| 63 \| 3 \| 63 \| Excisionase family DNA binding domain-containing protein \| Excisionase family DNA binding domain-containing protein; afdb-uniprot50 \| AF-A0A380MTT9-F1-MODEL\_V4 \| 1.0 \| 0.002243 \| 156 \| 0.292 \| 65 \| 42 \| 2 \| 1 \| 65 \| 2 \| 62 \| Excisionase or Xis \| Excisionase or Xis; afdb-uniprot50 \| AF-A0A1Q8JCF0-F1-MODEL\_V4 \| 1.0 \| 0.0009444 \| 156 \| 0.268 \| 67 \| 46 \| 2 \| 1 \| 66 \| 1 \| 65 \| HTH\_17 domain-containing protein \| HTH\_17 domain-containing protein; afdb-uniprot50 \| AF-A0A7Y0DBL9-F1-MODEL\_V4 \| 1.0 \| 0.001408 \| 156 \| 0.303 \| 66 \| 42 \| 2 \| 1 \| 64 \| 1 \| 64 \| Helix-turn-helix domain-containing protein \| Helix-turn-helix domain-containing protein; afdb-uniprot50 \| AF-A0A150HHP1-F1-MODEL\_V4 \| 1.0 \| 0.001837 \| 156 \| 0.253 \| 63 \| 44 \| 2 \| 11 \| 72 \| 7 \| 67 \| Helix-turn-helix domain protein \| Helix-turn-helix domain protein; afdb-uniprot50 \| AF-A0A494RQT9-F1-MODEL\_V4 \| 1.0 \| 0.001408 \| 156 \| 0.367 \| 68 \| 40 \| 2 \| 1 \| 67 \| 1 \| 66 \| DNA-binding protein \| DNA-binding protein; afdb-uniprot50 \| AF-A0A2A4V5K8-F1-MODEL\_V4 \| 1.0 \| 0.001408 \| 156 \| 0.253 \| 71 \| 48 \| 3 \| 4 \| 73 \| 2 \| 68 \| HTH\_17 domain-containing protein \| HTH\_17 domain-containing protein; afdb-uniprot50 \| AF-A0A3S4FDS9-F1-MODEL\_V4 \| 1.0 \| 0.002243 \| 156 \| 0.313 \| 67 \| 43 \| 2 \| 7 \| 72 \| 2 \| 66 \| HTH\_17 domain-containing protein \| HTH\_17 domain-containing protein; afdb-uniprot50 \| AF-A0A5C7LIH8-F1-MODEL\_V4 \| 1.0 \| 0.001232 \| 156 \| 0.232 \| 73 \| 51 \| 3 \| 1 \| 73 \| 1 \| 68 \| DNA-binding protein \| DNA-binding protein; afdb-uniprot50 \| AF-A0A6L5G2Y0-F1-MODEL\_V4 \| 1.0 \| 0.003819 \| 156 \| 0.3 \| 60 \| 39 \| 2 \| 7 \| 65 \| 2 \| 59 \| Helix-turn-helix domain-containing protein \| Helix-turn-helix domain-containing protein; afdb-uniprot50 \| AF-A0A1A0KLC7-F1-MODEL\_V4 \| 1.0 \| 0.004662 \| 156 \| 0.222 \| 63 \| 47 \| 1 \| 11 \| 73 \| 7 \| 67 \| HTH\_17 domain-containing protein \| HTH\_17 domain-containing protein; afdb-uniprot50 \| AF-A0A4U2Z9B1-F1-MODEL\_V4 \| 1.0 \| 0.002562 \| 156 \| 0.238 \| 63 \| 46 \| 1 \| 1 \| 63 \| 5 \| 65 \| Helix-turn-helix domain-containing protein \| Helix-turn-helix domain-containing protein; afdb-uniprot50 \| AF-A0A4Y8QXZ4-F1-MODEL\_V4 \| 1.0 \| 0.001505 \| 156 \| 0.26 \| 69 \| 48 \| 2 \| 1 \| 68 \| 2 \| 68 \| DNA-binding protein \| DNA-binding protein; afdb-uniprot50 \| AF-A0A2W7IHD9-F1-MODEL\_V4 \| 1.0 \| 0.0008836 \| 156 \| 0.315 \| 76 \| 43 \| 2 \| 1 \| 69 \| 1 \| 74 \| Helix-turn-helix protein \| Helix-turn-helix protein; afdb-uniprot50 \| AF-A0A5B8RKC5-F1-MODEL\_V4 \| 1.0 \| 0.0007736 \| 156 \| 0.26 \| 73 \| 50 \| 2 \| 3 \| 73 \| 6 \| 76 \| HTH\_17 domain-containing protein \| HTH\_17 domain-containing protein; afdb-uniprot50 \| AF-A0A4Q2SX89-F1-MODEL\_V4 \| 1.0 \| 0.001408 \| 156 \| 0.257 \| 70 \| 47 \| 2 \| 2 \| 67 \| 5 \| 73 \| DNA-binding protein \| DNA-binding protein; afdb-uniprot50 \| AF-A0A7G7BH93-F1-MODEL\_V4 \| 1.0 \| 0.004362 \| 156 \| 0.305 \| 59 \| 39 \| 1 \| 9 \| 67 \| 21 \| 77 \| Helix-turn-helix domain-containing protein \| Helix-turn-helix domain-containing protein; afdb-uniprot50 \| AF-A0A6M3XPC7-F1-MODEL\_V4 \| 1.0 \| 0.0009444 \| 156 \| 0.273 \| 73 \| 49 \| 3 \| 1 \| 73 \| 8 \| 76 \| Putative DNA binding, helix-turn-helix domain containing protein \| Putative DNA binding, helix-turn-helix domain containing protein; afdb-uniprot50 \| AF-A0A1T4VUB8-F1-MODEL\_V4 \| 1.0 \| 0.004983 \| 156 \| 0.245 \| 61 \| 44 \| 1 \| 2 \| 62 \| 19 \| 77 \| Helix-turn-helix domain-containing protein \| Helix-turn-helix domain-containing protein; afdb-uniprot50 \| AF-A0A4R9QI88-F1-MODEL\_V4 \| 1.0 \| 0.002098 \| 156 \| 0.275 \| 69 \| 48 \| 1 \| 5 \| 73 \| 11 \| 77 \| DNA-binding protein \| DNA-binding protein; afdb-uniprot50 \| AF-A0A285VVA6-F1-MODEL\_V4 \| 1.0 \| 0.001963 \| 156 \| 0.253 \| 67 \| 47 \| 2 \| 1 \| 67 \| 28 \| 91 \| DNA binding domain-containing protein, excisionase family \| DNA binding domain-containing protein, excisionase family; afdb-uniprot50 \| AF-A0A7X7D5U4-F1-MODEL\_V4 \| 1.0 \| 0.001079 \| 156 \| 0.293 \| 75 \| 49 \| 2 \| 1 \| 73 \| 3 \| 75 \| Helix-turn-helix domain-containing protein \| Helix-turn-helix domain-containing protein; afdb-uniprot50 \| AF-A0A7X4A2Z2-F1-MODEL\_V4 \| 1.0 \| 0.0008268 \| 156 \| 0.256 \| 74 \| 52 \| 2 \| 1 \| 73 \| 34 \| 105 \| Helix-turn-helix domain-containing protein \| Helix-turn-helix domain-containing protein; afdb-uniprot50 \| AF-A0A6N6VFE6-F1-MODEL\_V4 \| 1.0 \| 0.003343 \| 155 \| 0.285 \| 63 \| 43 \| 1 \| 11 \| 73 \| 4 \| 64 \| Helix-turn-helix domain-containing protein \| Helix-turn-helix domain-containing protein; afdb-uniprot50 \| AF-A0A412KDG0-F1-MODEL\_V4 \| 1.0 \| 0.002098 \| 155 \| 0.333 \| 63 \| 38 \| 2 \| 12 \| 72 \| 2 \| 62 \| DNA-binding protein \| DNA-binding protein; afdb-uniprot50 \| AF-A0A087BZX1-F1-MODEL\_V4 \| 1.0 \| 0.002397 \| 155 \| 0.318 \| 66 \| 43 \| 1 \| 7 \| 72 \| 2 \| 65 \| DNA binding domain, excisionase family \| DNA binding domain, excisionase family; afdb-uniprot50 \| AF-A0A2U2MS54-F1-MODEL\_V4 \| 1.0 \| 0.003573 \| 155 \| 0.328 \| 64 \| 40 \| 3 \| 10 \| 73 \| 6 \| 66 \| Excisionase \| Excisionase; afdb-uniprot50 \| AF-A0A1F1ZUF8-F1-MODEL\_V4 \| 1.0 \| 0.002397 \| 155 \| 0.246 \| 65 \| 46 \| 2 \| 10 \| 73 \| 4 \| 66 \| HTH\_17 domain-containing protein \| HTH\_17 domain-containing protein; afdb-uniprot50 \| AF-F5XSJ0-F1-MODEL\_V4 \| 1.0 \| 0.001719 \| 155 \| 0.242 \| 70 \| 50 \| 2 \| 4 \| 72 \| 2 \| 69 \| HTH\_17 domain-containing protein \| HTH\_17 domain-containing protein; afdb-uniprot50 \| AF-A0A4R6RE54-F1-MODEL\_V4 \| 1.0 \| 0.002927 \| 155 \| 0.262 \| 61 \| 43 \| 1 \| 11 \| 71 \| 5 \| 63 \| Helix-turn-helix protein \| Helix-turn-helix protein; afdb-uniprot50 \| AF-A0A2G1YXW8-F1-MODEL\_V4 \| 1.0 \| 0.001408 \| 155 \| 0.318 \| 69 \| 41 \| 2 \| 9 \| 73 \| 2 \| 68 \| DNA-binding protein \| DNA-binding protein; afdb-uniprot50 \| AF-A0A2V9L6Q6-F1-MODEL\_V4 \| 1.0 \| 0.001963 \| 155 \| 0.285 \| 70 \| 48 \| 1 \| 4 \| 73 \| 2 \| 69 \| HTH\_17 domain-containing protein \| HTH\_17 domain-containing protein; afdb-uniprot50 \| AF-A0A0T1UKU9-F1-MODEL\_V4 \| 1.0 \| 0.001719 \| 155 \| 0.257 \| 66 \| 48 \| 1 \| 2 \| 67 \| 7 \| 71 \| DNA-binding protein \| DNA-binding protein; afdb-uniprot50 \| AF-A0A526W0E5-F1-MODEL\_V4 \| 1.0 \| 0.001079 \| 155 \| 0.342 \| 73 \| 45 \| 2 \| 1 \| 73 \| 1 \| 70 \| Helix-turn-helix domain-containing protein \| Helix-turn-helix domain-containing protein; afdb-uniprot50 \| AF-A0A6N0YFG5-F1-MODEL\_V4 \| 1.0 \| 0.003343 \| 155 \| 0.344 \| 58 \| 36 \| 1 \| 6 \| 63 \| 12 \| 67 \| DNA-binding protein \| DNA-binding protein; afdb-uniprot50 \| AF-A0A4P6KDJ9-F1-MODEL\_V4 \| 1.0 \| 0.001153 \| 155 \| 0.205 \| 73 \| 55 \| 2 \| 1 \| 73 \| 4 \| 73 \| DNA-binding protein \| DNA-binding protein; afdb-uniprot50 \| AF-A0A3G9G7L5-F1-MODEL\_V4 \| 1.0 \| 0.004081 \| 155 \| 0.258 \| 62 \| 42 \| 2 \| 9 \| 70 \| 15 \| 72 \| HTH\_17 domain-containing protein \| HTH\_17 domain-containing protein; afdb-uniprot50 \| AF-A0A6A0QUD0-F1-MODEL\_V4 \| 1.0 \| 0.003128 \| 155 \| 0.25 \| 64 \| 46 \| 1 \| 2 \| 65 \| 15 \| 76 \| DNA-binding protein \| DNA-binding protein; afdb-uniprot50 \| AF-A0A7Y0HTB1-F1-MODEL\_V4 \| 1.0 \| 0.003573 \| 155 \| 0.261 \| 65 \| 45 \| 1 \| 9 \| 73 \| 20 \| 81 \| Uncharacterized protein \| Uncharacterized protein; afdb-uniprot50 \| AF-H6RNM7-F1-MODEL\_V4 \| 1.0 \| 0.002738 \| 155 \| 0.235 \| 68 \| 50 \| 1 \| 2 \| 69 \| 18 \| 83 \| Transcriptional regulator, MerR family \| Transcriptional regulator, MerR family; afdb-uniprot50 \| AF-B7J9L7-F1-MODEL\_V4 \| 1.0 \| 0.001079 \| 155 \| 0.291 \| 72 \| 49 \| 1 \| 2 \| 73 \| 35 \| 104 \| Uncharacterized protein \| Uncharacterized protein; afdb-uniprot50 \| AF-A0A2G4YWK8-F1-MODEL\_V4 \| 1.0 \| 0.003128 \| 155 \| 0.208 \| 67 \| 51 \| 1 \| 5 \| 71 \| 21 \| 85 \| HTH\_17 domain-containing protein \| HTH\_17 domain-containing protein; afdb-uniprot50 \| AF-A0A3A5AUL8-F1-MODEL\_V4 \| 1.0 \| 0.003343 \| 154 \| 0.253 \| 63 \| 46 \| 1 \| 11 \| 73 \| 3 \| 64 \| Uncharacterized protein \| Uncharacterized protein; afdb-uniprot50 \| AF-A0A1X1TZH7-F1-MODEL\_V4 \| 1.0 \| 0.002397 \| 154 \| 0.318 \| 66 \| 41 \| 2 \| 1 \| 66 \| 1 \| 62 \| Uncharacterized protein \| Uncharacterized protein; afdb-uniprot50 \| AF-A0A100VZK0-F1-MODEL\_V4 \| 1.0 \| 0.001963 \| 154 \| 0.301 \| 73 \| 46 \| 2 \| 1 \| 73 \| 1 \| 68 \| HTH\_17 domain-containing protein \| HTH\_17 domain-containing protein; afdb-uniprot50 \| AF-A0A2D6BJI6-F1-MODEL\_V4 \| 1.0 \| 0.001505 \| 154 \| 0.285 \| 63 \| 39 \| 2 \| 11 \| 73 \| 4 \| 60 \| HTH\_17 domain-containing protein \| HTH\_17 domain-containing protein; afdb-uniprot50 \| AF-A0A517SMF5-F1-MODEL\_V4 \| 1.0 \| 0.0007736 \| 154 \| 0.243 \| 74 \| 53 \| 2 \| 1 \| 73 \| 1 \| 72 \| Helix-turn-helix domain protein \| Helix-turn-helix domain protein; afdb-uniprot50 \| AF-A0A833FY91-F1-MODEL\_V4 \| 1.0 \| 0.002098 \| 154 \| 0.277 \| 72 \| 46 \| 2 \| 1 \| 72 \| 1 \| 66 \| Helix-turn-helix domain-containing protein \| Helix-turn-helix domain-containing protein; afdb-uniprot50 \| AF-A0A849DSH8-F1-MODEL\_V4 \| 1.0 \| 0.0007736 \| 154 \| 0.232 \| 73 \| 54 \| 2 \| 1 \| 73 \| 4 \| 74 \| Helix-turn-helix domain-containing protein \| Helix-turn-helix domain-containing protein; afdb-uniprot50 \| AF-A0A7X5U2M3-F1-MODEL\_V4 \| 1.0 \| 0.0006336 \| 154 \| 0.31 \| 74 \| 43 \| 3 \| 2 \| 73 \| 10 \| 77 \| Excisionase family DNA binding protein \| Excisionase family DNA binding protein; afdb-uniprot50 \| AF-A0A4Q2KLM2-F1-MODEL\_V4 \| 1.0 \| 0.002397 \| 154 \| 0.311 \| 77 \| 45 \| 3 \| 1 \| 72 \| 1 \| 74 \| DNA-binding protein \| DNA-binding protein; afdb-uniprot50 \| AF-A0A1G6EPZ3-F1-MODEL\_V4 \| 1.0 \| 0.0008268 \| 154 \| 0.258 \| 85 \| 49 \| 3 \| 1 \| 73 \| 1 \| 83 \| Uncharacterized protein \| Uncharacterized protein; afdb-uniprot50 \| AF-A0A024H216-F1-MODEL\_V4 \| 1.0 \| 0.002562 \| 154 \| 0.265 \| 64 \| 45 \| 1 \| 10 \| 73 \| 23 \| 84 \| Uncharacterized domain protein \| Uncharacterized domain protein; afdb-uniprot50 \| AF-A0A4R4DRZ7-F1-MODEL\_V4 \| 1.0 \| 0.001505 \| 154 \| 0.3 \| 70 \| 47 \| 1 \| 2 \| 71 \| 10 \| 77 \| DNA-binding protein \| DNA-binding protein; afdb-uniprot50 \| AF-A0A2N3GRQ9-F1-MODEL\_V4 \| 1.0 \| 0.002243 \| 154 \| 0.26 \| 69 \| 48 \| 2 \| 5 \| 73 \| 5 \| 70 \| HTH\_17 domain-containing protein \| HTH\_17 domain-containing protein; afdb-uniprot50 \| AF-A0A2A9DZ10-F1-MODEL\_V4 \| 1.0 \| 0.001079 \| 154 \| 0.191 \| 73 \| 56 \| 2 \| 1 \| 73 \| 28 \| 97 \| Excisionase family DNA binding protein \| Excisionase family DNA binding protein; afdb-uniprot50 \| AF-A0A2L2Q153-F1-MODEL\_V4 \| 1.0 \| 0.001837 \| 154 \| 0.257 \| 66 \| 46 \| 2 \| 1 \| 65 \| 31 \| 94 \| DNA-binding protein \| DNA-binding protein; afdb-uniprot50 \| AF-A0A2A4SJ14-F1-MODEL\_V4 \| 1.0 \| 0.001317 \| 154 \| 0.194 \| 77 \| 55 \| 2 \| 1 \| 72 \| 17 \| 91 \| HTH\_17 domain-containing protein \| HTH\_17 domain-containing protein; afdb-uniprot50 \| AF-A0A2D6EPB2-F1-MODEL\_V4 \| 1.0 \| 0.004662 \| 154 \| 0.258 \| 62 \| 44 \| 1 \| 2 \| 63 \| 40 \| 99 \| HTH\_17 domain-containing protein \| HTH\_17 domain-containing protein; afdb-uniprot50 \| AF-A0A7W9JAP3-F1-MODEL\_V4 \| 1.0 \| 0.002098 \| 154 \| 0.289 \| 76 \| 49 \| 2 \| 1 \| 73 \| 1 \| 74 \| Uncharacterized protein \| Uncharacterized protein; afdb-uniprot50 \| AF-A0A4R4WWB4-F1-MODEL\_V4 \| 1.0 \| 0.005692 \| 153 \| 0.327 \| 55 \| 35 \| 2 \| 11 \| 65 \| 8 \| 60 \| DNA-binding protein \| DNA-binding protein; afdb-uniprot50 \| AF-A0A0B2AB94-F1-MODEL\_V4 \| 1.0 \| 0.003343 \| 153 \| 0.317 \| 63 \| 41 \| 2 \| 12 \| 73 \| 1 \| 62 \| Uncharacterized protein \| Uncharacterized protein; afdb-uniprot50 \| AF-A0A7S7TXP9-F1-MODEL\_V4 \| 1.0 \| 0.004983 \| 153 \| 0.285 \| 63 \| 42 \| 2 \| 11 \| 73 \| 4 \| 63 \| DNA-binding protein \| DNA-binding protein; afdb-uniprot50 \| AF-A3VLE0-F1-MODEL\_V4 \| 1.0 \| 0.002243 \| 153 \| 0.242 \| 70 \| 48 \| 3 \| 4 \| 73 \| 2 \| 66 \| HTH\_17 domain-containing protein \| HTH\_17 domain-containing protein; afdb-uniprot50 \| AF-A0A1X1RN46-F1-MODEL\_V4 \| 1.0 \| 0.001009 \| 153 \| 0.314 \| 70 \| 45 \| 2 \| 4 \| 72 \| 1 \| 68 \| HTH merR-type domain-containing protein \| HTH merR-type domain-containing protein; afdb-uniprot50 \| AF-A0A7X6MSW7-F1-MODEL\_V4 \| 1.0 \| 0.005692 \| 153 \| 0.254 \| 59 \| 42 \| 1 \| 4 \| 62 \| 2 \| 58 \| Helix-turn-helix domain-containing protein \| Helix-turn-helix domain-containing protein; afdb-uniprot50 \| AF-A0A6I0ENH1-F1-MODEL\_V4 \| 1.0 \| 0.003343 \| 153 \| 0.228 \| 70 \| 52 \| 1 \| 4 \| 73 \| 2 \| 69 \| Helix-turn-helix domain-containing protein \| Helix-turn-helix domain-containing protein; afdb-uniprot50 \| AF-A0A1Q9W072-F1-MODEL\_V4 \| 1.0 \| 0.002738 \| 153 \| 0.189 \| 74 \| 55 \| 3 \| 1 \| 73 \| 1 \| 70 \| HTH\_17 domain-containing protein \| HTH\_17 domain-containing protein; afdb-uniprot50 \| AF-A0A1H3DCW2-F1-MODEL\_V4 \| 1.0 \| 0.0007736 \| 153 \| 0.356 \| 73 \| 41 \| 3 \| 1 \| 73 \| 1 \| 67 \| Helix-turn-helix domain-containing protein \| Helix-turn-helix domain-containing protein; afdb-uniprot50 \| AF-H5USK6-F1-MODEL\_V4 \| 1.0 \| 0.002397 \| 153 \| 0.222 \| 63 \| 46 \| 2 \| 1 \| 63 \| 12 \| 71 \| HTH\_17 domain-containing protein \| HTH\_17 domain-containing protein; afdb-uniprot50 \| AF-A0A0N8HY41-F1-MODEL\_V4 \| 1.0 \| 0.001408 \| 153 \| 0.279 \| 68 \| 46 \| 2 \| 1 \| 67 \| 8 \| 73 \| Uncharacterized protein \| Uncharacterized protein; afdb-uniprot50 \| AF-A0A2W7C802-F1-MODEL\_V4 \| 1.0 \| 0.0005928 \| 153 \| 0.283 \| 74 \| 49 \| 2 \| 1 \| 72 \| 4 \| 75 \| Uncharacterized protein \| Uncharacterized protein; afdb-uniprot50 \| AF-A0A355T8Q2-F1-MODEL\_V4 \| 1.0 \| 0.001317 \| 153 \| 0.323 \| 71 \| 42 \| 2 \| 7 \| 73 \| 2 \| 70 \| DNA-binding protein \| DNA-binding protein; afdb-uniprot50 \| AF-A0A366ZRD9-F1-MODEL\_V4 \| 1.0 \| 0.002397 \| 153 \| 0.235 \| 68 \| 49 \| 2 \| 1 \| 67 \| 6 \| 71 \| DNA-binding protein \| DNA-binding protein; afdb-uniprot50 \| AF-A0A0Q8WXY6-F1-MODEL\_V4 \| 1.0 \| 0.002243 \| 153 \| 0.24 \| 79 \| 52 \| 2 \| 1 \| 73 \| 1 \| 77 \| HTH\_17 domain-containing protein \| HTH\_17 domain-containing protein; afdb-uniprot50 \| AF-A0A2A3I5W9-F1-MODEL\_V4 \| 1.0 \| 0.003343 \| 153 \| 0.287 \| 66 \| 45 \| 1 \| 2 \| 67 \| 17 \| 80 \| Excisionase family DNA binding protein \| Excisionase family DNA binding protein; afdb-uniprot50 \| AF-A0A2N8K0S6-F1-MODEL\_V4 \| 1.0 \| 0.001153 \| 153 \| 0.266 \| 75 \| 51 \| 3 \| 1 \| 73 \| 8 \| 80 \| Excisionase \| Excisionase; afdb-uniprot50 \| AF-A0A521H308-F1-MODEL\_V4 \| 1.0 \| 0.001963 \| 153 \| 0.3 \| 70 \| 47 \| 1 \| 4 \| 73 \| 2 \| 69 \| DNA-binding protein \| DNA-binding protein; afdb-uniprot50 \| AF-J1HLR3-F1-MODEL\_V4 \| 1.0 \| 0.002397 \| 153 \| 0.291 \| 72 \| 49 \| 1 \| 2 \| 73 \| 12 \| 81 \| DNA-binding helix-turn-helix protein \| DNA-binding helix-turn-helix protein; afdb-uniprot50 \| AF-A0A0Q8Q995-F1-MODEL\_V4 \| 1.0 \| 0.007428 \| 152 \| 0.214 \| 56 \| 42 \| 1 \| 11 \| 66 \| 4 \| 57 \| HTH\_17 domain-containing protein \| HTH\_17 domain-containing protein; afdb-uniprot50 \| AF-A0A1A2P8X3-F1-MODEL\_V4 \| 1.0 \| 0.002397 \| 152 \| 0.276 \| 65 \| 44 \| 2 \| 10 \| 73 \| 3 \| 65 \| Uncharacterized protein \| Uncharacterized protein; afdb-uniprot50 \| AF-A0A2T4JGE1-F1-MODEL\_V4 \| 1.0 \| 0.001837 \| 152 \| 0.328 \| 67 \| 38 \| 2 \| 12 \| 73 \| 1 \| 65 \| Uncharacterized protein \| Uncharacterized protein; afdb-uniprot50 \| AF-A0A355T7V6-F1-MODEL\_V4 \| 1.0 \| 0.002098 \| 152 \| 0.375 \| 64 \| 38 \| 1 \| 7 \| 70 \| 2 \| 63 \| DNA-binding protein \| DNA-binding protein; afdb-uniprot50 \| AF-A0A5R8PA16-F1-MODEL\_V4 \| 1.0 \| 0.002243 \| 152 \| 0.26 \| 73 \| 50 \| 3 \| 1 \| 72 \| 1 \| 70 \| Helix-turn-helix domain-containing protein \| Helix-turn-helix domain-containing protein; afdb-uniprot50 \| AF-A0A4Y4C815-F1-MODEL\_V4 \| 1.0 \| 0.0009444 \| 152 \| 0.297 \| 74 \| 45 \| 3 \| 1 \| 73 \| 2 \| 69 \| HTH\_17 domain-containing protein \| HTH\_17 domain-containing protein; afdb-uniprot50 \| AF-A0A0C2YAS8-F1-MODEL\_V4 \| 1.0 \| 0.001608 \| 152 \| 0.301 \| 73 \| 47 \| 2 \| 1 \| 73 \| 1 \| 69 \| Uncharacterized protein \| Uncharacterized protein; afdb-uniprot50 \| AF-A0A161VAV6-F1-MODEL\_V4 \| 1.0 \| 0.001963 \| 152 \| 0.205 \| 73 \| 56 \| 1 \| 1 \| 73 \| 2 \| 72 \| Helix-turn-helix domain protein \| Helix-turn-helix domain protein; afdb-uniprot50 \| AF-A0A7W8EEW1-F1-MODEL\_V4 \| 1.0 \| 0.002243 \| 152 \| 0.243 \| 74 \| 52 \| 2 \| 1 \| 72 \| 3 \| 74 \| Putative DNA-binding transcriptional regulator AlpA \| Putative DNA-binding transcriptional regulator AlpA; afdb-uniprot50 \| AF-A0A0M9CB32-F1-MODEL\_V4 \| 1.0 \| 0.003819 \| 152 \| 0.268 \| 67 \| 47 \| 2 \| 7 \| 73 \| 9 \| 73 \| HTH\_17 domain-containing protein \| HTH\_17 domain-containing protein; afdb-uniprot50 \| AF-A0A542NQW3-F1-MODEL\_V4 \| 1.0 \| 0.004362 \| 152 \| 0.265 \| 64 \| 44 \| 2 \| 1 \| 63 \| 6 \| 67 \| Helix-turn-helix protein \| Helix-turn-helix protein; afdb-uniprot50 \| AF-A0A7W0NGD1-F1-MODEL\_V4 \| 1.0 \| 0.002397 \| 152 \| 0.281 \| 64 \| 43 \| 2 \| 1 \| 63 \| 250 \| 311 \| Helix-turn-helix domain-containing protein \| Helix-turn-helix domain-containing protein; afdb-uniprot50 \| AF-A0A0Q9KLL4-F1-MODEL\_V4 \| 1.0 \| 0.002098 \| 151 \| 0.272 \| 66 \| 42 \| 3 \| 1 \| 65 \| 1 \| 61 \| HTH\_17 domain-containing protein \| HTH\_17 domain-containing protein; afdb-uniprot50 \| AF-A0A6G7PU95-F1-MODEL\_V4 \| 1.0 \| 0.002927 \| 151 \| 0.283 \| 67 \| 44 \| 2 \| 7 \| 73 \| 1 \| 63 \| Helix-turn-helix domain-containing protein \| Helix-turn-helix domain-containing protein; afdb-uniprot50 \| AF-A0A7X6CXT2-F1-MODEL\_V4 \| 1.0 \| 0.003343 \| 151 \| 0.308 \| 68 \| 43 \| 2 \| 1 \| 68 \| 1 \| 64 \| Helix-turn-helix domain-containing protein \| Helix-turn-helix domain-containing protein; afdb-uniprot50 \| AF-A0A059G3P2-F1-MODEL\_V4 \| 1.0 \| 0.004362 \| 151 \| 0.268 \| 67 \| 46 \| 2 \| 7 \| 73 \| 1 \| 64 \| HTH\_17 domain-containing protein \| HTH\_17 domain-containing protein; afdb-uniprot50 \| AF-A0A661MGY5-F1-MODEL\_V4 \| 1.0 \| 0.003128 \| 151 \| 0.301 \| 63 \| 42 \| 1 \| 11 \| 73 \| 4 \| 64 \| DNA-binding protein \| DNA-binding protein; afdb-uniprot50 \| AF-A0A1A2D498-F1-MODEL\_V4 \| 1.0 \| 0.003819 \| 151 \| 0.261 \| 65 \| 44 \| 2 \| 11 \| 73 \| 3 \| 65 \| Transcriptional regulator \| Transcriptional regulator; afdb-uniprot50 \| AF-A0A1I4RPU8-F1-MODEL\_V4 \| 1.0 \| 0.004983 \| 151 \| 0.166 \| 66 \| 51 \| 2 \| 6 \| 69 \| 2 \| 65 \| Uncharacterized protein \| Uncharacterized protein; afdb-uniprot50 \| AF-A0A2N6F548-F1-MODEL\_V4 \| 1.0 \| 0.003343 \| 151 \| 0.23 \| 65 \| 48 \| 1 \| 1 \| 65 \| 2 \| 64 \| HTH\_17 domain-containing protein \| HTH\_17 domain-containing protein; afdb-uniprot50 \| AF-A0A258BKJ5-F1-MODEL\_V4 \| 1.0 \| 0.004081 \| 151 \| 0.285 \| 63 \| 43 \| 2 \| 11 \| 73 \| 4 \| 64 \| Uncharacterized protein \| Uncharacterized protein; afdb-uniprot50 \| AF-A0A380MLV7-F1-MODEL\_V4 \| 1.0 \| 0.002738 \| 151 \| 0.287 \| 66 \| 44 \| 2 \| 1 \| 65 \| 2 \| 65 \| Excisionase or Xis \| Excisionase or Xis; afdb-uniprot50 \| AF-A0A0N7M3V3-F1-MODEL\_V4 \| 1.0 \| 0.001153 \| 151 \| 0.357 \| 70 \| 42 \| 2 \| 4 \| 72 \| 2 \| 69 \| Helix-turn-helix domain protein \| Helix-turn-helix domain protein; afdb-uniprot50 \| AF-A0A7C8G298-F1-MODEL\_V4 \| 1.0 \| 0.003343 \| 151 \| 0.287 \| 73 \| 49 \| 2 \| 1 \| 73 \| 2 \| 71 \| Helix-turn-helix domain-containing protein \| Helix-turn-helix domain-containing protein; afdb-uniprot50 \| AF-A0A2G2M150-F1-MODEL\_V4 \| 1.0 \| 0.008485 \| 151 \| 0.254 \| 55 \| 39 \| 1 \| 10 \| 64 \| 4 \| 56 \| HTH\_17 domain-containing protein \| HTH\_17 domain-containing protein; afdb-uniprot50 \| AF-A0A8A9YMU8-F1-MODEL\_V4 \| 1.0 \| 0.001079 \| 151 \| 0.205 \| 73 \| 55 \| 2 \| 2 \| 73 \| 6 \| 76 \| Helix-turn-helix domain-containing protein \| Helix-turn-helix domain-containing protein; afdb-uniprot50 \| AF-A0A245ZQK2-F1-MODEL\_V4 \| 1.0 \| 0.001232 \| 151 \| 0.243 \| 78 \| 52 \| 3 \| 1 \| 73 \| 1 \| 76 \| Helix-turn-helix domain protein \| Helix-turn-helix domain protein; afdb-uniprot50 \| AF-A0A1R1IPT8-F1-MODEL\_V4 \| 1.0 \| 0.003819 \| 151 \| 0.23 \| 65 \| 48 \| 1 \| 9 \| 73 \| 20 \| 82 \| DNA-binding protein \| DNA-binding protein; afdb-uniprot50 \| AF-A0A429YW83-F1-MODEL\_V4 \| 1.0 \| 0.006084 \| 151 \| 0.301 \| 63 \| 39 \| 1 \| 1 \| 63 \| 1 \| 58 \| DNA-binding protein \| DNA-binding protein; afdb-uniprot50 \| AF-A0A7W1M9M0-F1-MODEL\_V4 \| 1.0 \| 0.001837 \| 151 \| 0.308 \| 68 \| 44 \| 2 \| 6 \| 73 \| 2 \| 66 \| DNA-binding protein \| DNA-binding protein; afdb-uniprot50 \| AF-A0A6N9RRQ5-F1-MODEL\_V4 \| 1.0 \| 0.003573 \| 151 \| 0.219 \| 73 \| 55 \| 1 \| 1 \| 73 \| 9 \| 79 \| Helix-turn-helix domain-containing protein \| Helix-turn-helix domain-containing protein; afdb-uniprot50 \| AF-A0A6G4WQN4-F1-MODEL\_V4 \| 1.0 \| 0.0006772 \| 151 \| 0.246 \| 77 \| 52 \| 2 \| 1 \| 73 \| 14 \| 88 \| Helix-turn-helix domain-containing protein \| Helix-turn-helix domain-containing protein; afdb-uniprot50 \| AF-A0A2G6Y1E4-F1-MODEL\_V4 \| 1.0 \| 0.003573 \| 151 \| 0.253 \| 67 \| 46 \| 2 \| 1 \| 65 \| 21 \| 85 \| Excisionase family DNA binding protein \| Excisionase family DNA binding protein; afdb-uniprot50 \| AF-A0A2A3HXH1-F1-MODEL\_V4 \| 1.0 \| 0.002562 \| 151 \| 0.294 \| 68 \| 45 \| 2 \| 1 \| 67 \| 24 \| 89 \| Helix-turn-helix protein \| Helix-turn-helix protein; afdb-uniprot50 \| AF-A0A386UE03-F1-MODEL\_V4 \| 1.0 \| 0.001232 \| 151 \| 0.333 \| 63 \| 35 \| 2 \| 11 \| 73 \| 110 \| 165 \| HTH\_17 domain-containing protein \| HTH\_17 domain-containing protein; afdb-uniprot50 \| AF-A0A423UVZ0-F1-MODEL\_V4 \| 1.0 \| 0.003343 \| 151 \| 0.323 \| 65 \| 40 \| 2 \| 1 \| 65 \| 174 \| 234 \| DNA-binding protein \| DNA-binding protein; afdb-uniprot50 \| AF-A0A2U9S8C5-F1-MODEL\_V4 \| 1.0 \| 0.002397 \| 150 \| 0.365 \| 63 \| 36 \| 2 \| 11 \| 73 \| 4 \| 62 \| DNA-binding protein \| DNA-binding protein; afdb-uniprot50 \| AF-A0A2E7B4B3-F1-MODEL\_V4 \| 1.0 \| 0.005692 \| 150 \| 0.312 \| 64 \| 40 \| 2 \| 10 \| 73 \| 6 \| 65 \| DNA-binding protein \| DNA-binding protein; afdb-uniprot50 \| AF-A0A1I1SH13-F1-MODEL\_V4 \| 1.0 \| 0.003343 \| 150 \| 0.232 \| 73 \| 49 \| 3 \| 1 \| 73 \| 1 \| 66 \| DNA binding domain-containing protein, excisionase family \| DNA binding domain-containing protein, excisionase family; afdb-uniprot50 \| AF-A0A2D8ZUW2-F1-MODEL\_V4 \| 1.0 \| 0.001153 \| 150 \| 0.328 \| 73 \| 41 \| 3 \| 1 \| 73 \| 1 \| 65 \| DNA-binding protein \| DNA-binding protein; afdb-uniprot50 \| AF-A0A6G2N519-F1-MODEL\_V4 \| 1.0 \| 0.0009444 \| 150 \| 0.253 \| 67 \| 48 \| 2 \| 1 \| 66 \| 2 \| 67 \| Helix-turn-helix domain-containing protein \| Helix-turn-helix domain-containing protein; afdb-uniprot50 \| AF-A0A1X1RFI8-F1-MODEL\_V4 \| 1.0 \| 0.002098 \| 150 \| 0.205 \| 73 \| 53 \| 2 \| 1 \| 73 \| 1 \| 68 \| HTH\_17 domain-containing protein \| HTH\_17 domain-containing protein; afdb-uniprot50 \| AF-A0A3N4Z623-F1-MODEL\_V4 \| 1.0 \| 0.002397 \| 150 \| 0.257 \| 70 \| 50 \| 1 \| 4 \| 73 \| 2 \| 69 \| Excisionase family DNA binding protein \| Excisionase family DNA binding protein; afdb-uniprot50 \| AF-A0A846W105-F1-MODEL\_V4 \| 1.0 \| 0.004662 \| 150 \| 0.228 \| 70 \| 52 \| 1 \| 4 \| 73 \| 3 \| 70 \| Helix-turn-helix domain-containing protein \| Helix-turn-helix domain-containing protein; afdb-uniprot50 \| AF-A0A809IK81-F1-MODEL\_V4 \| 1.0 \| 0.002562 \| 150 \| 0.285 \| 63 \| 43 \| 1 \| 1 \| 63 \| 7 \| 67 \| DNA-binding protein \| DNA-binding protein; afdb-uniprot50 \| AF-A0A2E5BQF7-F1-MODEL\_V4 \| 1.0 \| 0.002397 \| 150 \| 0.271 \| 70 \| 48 \| 2 \| 2 \| 70 \| 10 \| 77 \| HTH\_17 domain-containing protein \| HTH\_17 domain-containing protein; afdb-uniprot50 \| AF-A0A1C6LZ92-F1-MODEL\_V4 \| 1.0 \| 0.001963 \| 150 \| 0.216 \| 74 \| 52 \| 3 \| 1 \| 73 \| 10 \| 78 \| Helix-turn-helix domain-containing protein \| Helix-turn-helix domain-containing protein; afdb-uniprot50 \| AF-A0A1G9CHM9-F1-MODEL\_V4 \| 1.0 \| 0.001963 \| 150 \| 0.243 \| 78 \| 54 \| 2 \| 1 \| 73 \| 1 \| 78 \| Uncharacterized protein \| Uncharacterized protein; afdb-uniprot50 \| AF-A0A1G6EMA2-F1-MODEL\_V4 \| 1.0 \| 0.0006772 \| 150 \| 0.36 \| 75 \| 43 \| 3 \| 1 \| 72 \| 2 \| 74 \| Uncharacterized protein \| Uncharacterized protein; afdb-uniprot50 \| AF-A0A2D8NLB9-F1-MODEL\_V4 \| 1.0 \| 0.002562 \| 150 \| 0.282 \| 78 \| 47 \| 3 \| 1 \| 73 \| 1 \| 74 \| Excisionase \| Excisionase; afdb-uniprot50 \| AF-A0A4Y9QPR4-F1-MODEL\_V4 \| 1.0 \| 0.002738 \| 150 \| 0.25 \| 76 \| 52 \| 3 \| 1 \| 73 \| 2 \| 75 \| DNA-binding protein \| DNA-binding protein; afdb-uniprot50 \| AF-A0A559VPY9-F1-MODEL\_V4 \| 1.0 \| 0.001505 \| 150 \| 0.287 \| 73 \| 47 \| 2 \| 1 \| 73 \| 16 \| 83 \| Excisionase family DNA binding protein \| Excisionase family DNA binding protein; afdb-uniprot50 \| AF-A0A7W0L409-F1-MODEL\_V4 \| 1.0 \| 0.003819 \| 150 \| 0.232 \| 73 \| 54 \| 1 \| 1 \| 73 \| 16 \| 86 \| Helix-turn-helix domain-containing protein \| Helix-turn-helix domain-containing protein; afdb-uniprot50 \| AF-A0A2E2QLK1-F1-MODEL\_V4 \| 1.0 \| 0.002738 \| 150 \| 0.228 \| 70 \| 51 \| 2 \| 1 \| 70 \| 18 \| 84 \| HTH\_17 domain-containing protein \| HTH\_17 domain-containing protein; afdb-uniprot50 \| AF-A8LA07-F1-MODEL\_V4 \| 1.0 \| 0.001963 \| 150 \| 0.242 \| 66 \| 47 \| 3 \| 1 \| 65 \| 25 \| 88 \| Excision promoter, Xis \| Excision promoter, Xis; afdb-uniprot50 \| AF-A0A0P1F1V2-F1-MODEL\_V4 \| 1.0 \| 0.004983 \| 150 \| 0.281 \| 71 \| 49 \| 1 \| 3 \| 73 \| 14 \| 82 \| Helix-turn-helix domain protein \| Helix-turn-helix domain protein; afdb-uniprot50 \| AF-A0A2W1TE65-F1-MODEL\_V4 \| 1.0 \| 0.006084 \| 149 \| 0.295 \| 61 \| 41 \| 1 \| 12 \| 72 \| 1 \| 59 \| HTH\_17 domain-containing protein \| HTH\_17 domain-containing protein; afdb-uniprot50 \| AF-A0A2W4LWT2-F1-MODEL\_V4 \| 1.0 \| 0.008485 \| 149 \| 0.298 \| 57 \| 38 \| 1 \| 11 \| 67 \| 4 \| 58 \| DNA-binding protein \| DNA-binding protein; afdb-uniprot50 \| AF-A0A7C5TS90-F1-MODEL\_V4 \| 1.0 \| 0.004362 \| 149 \| 0.253 \| 67 \| 48 \| 1 \| 6 \| 72 \| 2 \| 66 \| DNA-binding protein \| DNA-binding protein; afdb-uniprot50 \| AF-A0A7W7GUH3-F1-MODEL\_V4 \| 1.0 \| 0.004983 \| 149 \| 0.218 \| 64 \| 48 \| 1 \| 1 \| 64 \| 3 \| 64 \| Excisionase family DNA binding protein \| Excisionase family DNA binding protein; afdb-uniprot50 \| AF-A0A2M8B589-F1-MODEL\_V4 \| 1.0 \| 0.009692 \| 149 \| 0.213 \| 61 \| 46 \| 1 \| 9 \| 69 \| 5 \| 63 \| Transcriptional regulator \| Transcriptional regulator; afdb-uniprot50 \| AF-A0A7G2SRX6-F1-MODEL\_V4 \| 1.0 \| 0.003128 \| 149 \| 0.246 \| 73 \| 53 \| 1 \| 1 \| 73 \| 1 \| 71 \| HTH\_17 domain-containing protein \| HTH\_17 domain-containing protein; afdb-uniprot50 \| AF-A0A2P8E071-F1-MODEL\_V4 \| 1.0 \| 0.002927 \| 149 \| 0.25 \| 68 \| 49 \| 1 \| 1 \| 68 \| 7 \| 72 \| Excisionase family DNA binding protein \| Excisionase family DNA binding protein; afdb-uniprot50 \| AF-A0A2N3GRP0-F1-MODEL\_V4 \| 1.0 \| 0.003343 \| 149 \| 0.212 \| 66 \| 49 \| 2 \| 1 \| 65 \| 9 \| 72 \| HTH\_17 domain-containing protein \| HTH\_17 domain-containing protein; afdb-uniprot50 \| AF-A0A7W0KKL6-F1-MODEL\_V4 \| 1.0 \| 0.002243 \| 149 \| 0.226 \| 75 \| 54 \| 2 \| 1 \| 73 \| 3 \| 75 \| Helix-turn-helix domain-containing protein \| Helix-turn-helix domain-containing protein; afdb-uniprot50 \| AF-A0A3P3VTI3-F1-MODEL\_V4 \| 1.0 \| 0.001963 \| 149 \| 0.219 \| 73 \| 54 \| 2 \| 2 \| 73 \| 5 \| 75 \| DNA-binding protein \| DNA-binding protein; afdb-uniprot50 \| AF-A0A399UEI2-F1-MODEL\_V4 \| 1.0 \| 0.003819 \| 149 \| 0.232 \| 73 \| 54 \| 1 \| 1 \| 73 \| 7 \| 77 \| DNA-binding protein \| DNA-binding protein; afdb-uniprot50 \| AF-A0A7X5F027-F1-MODEL\_V4 \| 1.0 \| 0.004662 \| 149 \| 0.228 \| 70 \| 52 \| 1 \| 1 \| 70 \| 3 \| 70 \| Helix-turn-helix domain-containing protein \| Helix-turn-helix domain-containing protein; afdb-uniprot50 \| AF-A0A0B5ERS0-F1-MODEL\_V4 \| 1.0 \| 0.001837 \| 149 \| 0.275 \| 69 \| 45 \| 2 \| 1 \| 65 \| 8 \| 75 \| DNA-binding protein \| DNA-binding protein; afdb-uniprot50 \| AF-F6F0P7-F1-MODEL\_V4 \| 1.0 \| 0.005692 \| 149 \| 0.215 \| 65 \| 49 \| 1 \| 9 \| 73 \| 16 \| 78 \| Uncharacterized protein \| Uncharacterized protein; afdb-uniprot50 \| AF-B6JJF5-F1-MODEL\_V4 \| 1.0 \| 0.002738 \| 149 \| 0.307 \| 65 \| 41 \| 2 \| 1 \| 63 \| 11 \| 73 \| HTH\_17 domain-containing protein \| HTH\_17 domain-containing protein; afdb-uniprot50 \| AF-A0A2W5ETA6-F1-MODEL\_V4 \| 1.0 \| 0.002397 \| 149 \| 0.328 \| 73 \| 46 \| 2 \| 1 \| 73 \| 13 \| 82 \| DNA-binding protein \| DNA-binding protein; afdb-uniprot50 \| AF-A0A1G6DPZ1-F1-MODEL\_V4 \| 1.0 \| 0.003819 \| 149 \| 0.333 \| 63 \| 40 \| 1 \| 11 \| 73 \| 14 \| 74 \| Uncharacterized protein \| Uncharacterized protein; afdb-uniprot50 \| AF-A0A1M3Q4B2-F1-MODEL\_V4 \| 1.0 \| 0.005692 \| 149 \| 0.241 \| 62 \| 44 \| 2 \| 1 \| 62 \| 6 \| 64 \| DNA-binding protein \| DNA-binding protein; afdb-uniprot50 \| AF-A0A316AUK9-F1-MODEL\_V4 \| 1.0 \| 0.006084 \| 149 \| 0.253 \| 63 \| 45 \| 1 \| 1 \| 63 \| 9 \| 69 \| Helix-turn-helix protein \| Helix-turn-helix protein; afdb-uniprot50 \| AF-A0A7W3J5K9-F1-MODEL\_V4 \| 1.0 \| 0.002098 \| 149 \| 0.232 \| 73 \| 54 \| 1 \| 1 \| 73 \| 17 \| 87 \| Excisionase family DNA binding protein \| Excisionase family DNA binding protein; afdb-uniprot50 \| AF-A0A435X2Z5-F1-MODEL\_V4 \| 1.0 \| 0.002243 \| 149 \| 0.253 \| 75 \| 51 \| 2 \| 2 \| 73 \| 32 \| 104 \| DNA-binding protein \| DNA-binding protein; afdb-uniprot50 \| AF-A0A5A7WVK6-F1-MODEL\_V4 \| 1.0 \| 0.004983 \| 148 \| 0.205 \| 68 \| 52 \| 1 \| 6 \| 73 \| 2 \| 67 \| DNA-binding protein \| DNA-binding protein; afdb-uniprot50 \| AF-A0A371P3H7-F1-MODEL\_V4 \| 1.0 \| 0.004662 \| 148 \| 0.235 \| 68 \| 49 \| 2 \| 1 \| 68 \| 1 \| 65 \| DNA-binding protein \| DNA-binding protein; afdb-uniprot50 \| AF-A0A2H6H355-F1-MODEL\_V4 \| 1.0 \| 0.003573 \| 148 \| 0.279 \| 68 \| 47 \| 1 \| 6 \| 73 \| 2 \| 67 \| Helix-turn-helix domain protein \| Helix-turn-helix domain protein; afdb-uniprot50 \| AF-A0A349HMZ8-F1-MODEL\_V4 \| 1.0 \| 0.001719 \| 148 \| 0.318 \| 66 \| 42 \| 2 \| 9 \| 73 \| 2 \| 65 \| DNA-binding protein \| DNA-binding protein; afdb-uniprot50 \| AF-A0A4Q7ZEX0-F1-MODEL\_V4 \| 1.0 \| 0.001079 \| 148 \| 0.25 \| 72 \| 48 \| 3 \| 1 \| 72 \| 4 \| 69 \| AlpA family transcriptional regulator \| AlpA family transcriptional regulator; afdb-uniprot50 \| AF-A0A7U3UU80-F1-MODEL\_V4 \| 1.0 \| 0.001608 \| 148 \| 0.279 \| 68 \| 44 \| 2 \| 1 \| 65 \| 1 \| 66 \| Uncharacterized protein \| Uncharacterized protein; afdb-uniprot50 \| AF-A0A7T2WNU1-F1-MODEL\_V4 \| 1.0 \| 0.002927 \| 148 \| 0.273 \| 73 \| 47 \| 3 \| 1 \| 70 \| 1 \| 70 \| Helix-turn-helix domain-containing protein \| Helix-turn-helix domain-containing protein; afdb-uniprot50 \| AF-A0A512JRF1-F1-MODEL\_V4 \| 1.0 \| 0.002243 \| 148 \| 0.239 \| 71 \| 52 \| 1 \| 2 \| 72 \| 3 \| 71 \| HTH\_17 domain-containing protein \| HTH\_17 domain-containing protein; afdb-uniprot50 \| AF-A0A090MQQ0-F1-MODEL\_V4 \| 1.0 \| 0.004662 \| 148 \| 0.285 \| 63 \| 43 \| 1 \| 11 \| 73 \| 5 \| 65 \| Helix-turn-helix domain protein \| Helix-turn-helix domain protein; afdb-uniprot50 \| AF-A0A229SBU2-F1-MODEL\_V4 \| 1.0 \| 0.002397 \| 148 \| 0.267 \| 71 \| 49 \| 2 \| 4 \| 73 \| 3 \| 71 \| HTH\_17 domain-containing protein \| HTH\_17 domain-containing protein; afdb-uniprot50 \| AF-A0A4Q8TE58-F1-MODEL\_V4 \| 1.0 \| 0.003128 \| 148 \| 0.276 \| 65 \| 45 \| 1 \| 1 \| 65 \| 6 \| 68 \| DNA-binding protein \| DNA-binding protein; afdb-uniprot50 \| AF-U5VTE3-F1-MODEL\_V4 \| 1.0 \| 0.003819 \| 148 \| 0.242 \| 66 \| 48 \| 1 \| 2 \| 67 \| 7 \| 70 \| Excisionase/Xis, DNA-binding protein \| Excisionase/Xis, DNA-binding protein; afdb-uniprot50 \| AF-A0A1A2CNF7-F1-MODEL\_V4 \| 1.0 \| 0.002927 \| 148 \| 0.246 \| 73 \| 53 \| 1 \| 1 \| 73 \| 2 \| 72 \| HTH\_17 domain-containing protein \| HTH\_17 domain-containing protein; afdb-uniprot50 \| AF-A0A238ZEI8-F1-MODEL\_V4 \| 1.0 \| 0.005326 \| 148 \| 0.265 \| 64 \| 45 \| 1 \| 10 \| 73 \| 15 \| 76 \| Uncharacterized protein \| Uncharacterized protein; afdb-uniprot50 \| AF-A0A7C9M524-F1-MODEL\_V4 \| 1.0 \| 0.002562 \| 148 \| 0.328 \| 73 \| 47 \| 2 \| 1 \| 73 \| 1 \| 71 \| Helix-turn-helix domain-containing protein \| Helix-turn-helix domain-containing protein; afdb-uniprot50 \| AF-A0A059GC80-F1-MODEL\_V4 \| 1.0 \| 0.004662 \| 148 \| 0.232 \| 73 \| 54 \| 1 \| 1 \| 73 \| 8 \| 78 \| HTH\_17 domain-containing protein \| HTH\_17 domain-containing protein; afdb-uniprot50 \| AF-A0A349CXT0-F1-MODEL\_V4 \| 1.0 \| 0.001009 \| 148 \| 0.27 \| 74 \| 49 \| 3 \| 1 \| 73 \| 1 \| 70 \| HTH\_17 domain-containing protein \| HTH\_17 domain-containing protein; afdb-uniprot50 \| AF-A0A2A4ZEI9-F1-MODEL\_V4 \| 1.0 \| 0.002098 \| 148 \| 0.256 \| 74 \| 50 \| 2 \| 1 \| 71 \| 13 \| 84 \| Uncharacterized protein \| Uncharacterized protein; afdb-uniprot50 \| AF-E8RUQ5-F1-MODEL\_V4 \| 1.0 \| 0.004662 \| 148 \| 0.26 \| 69 \| 47 \| 2 \| 5 \| 73 \| 30 \| 94 \| HTH\_17 domain-containing protein \| HTH\_17 domain-containing protein; afdb-uniprot50 \| AF-A0A6N8XP61-F1-MODEL\_V4 \| 1.0 \| 0.002738 \| 148 \| 0.24 \| 75 \| 53 \| 2 \| 1 \| 73 \| 26 \| 98 \| Helix-turn-helix domain-containing protein \| Helix-turn-helix domain-containing protein; afdb-uniprot50 \| AF-A0A2N5C742-F1-MODEL\_V4 \| 1.0 \| 0.001608 \| 148 \| 0.287 \| 73 \| 47 \| 2 \| 1 \| 70 \| 13 \| 83 \| DNA-binding protein \| DNA-binding protein; afdb-uniprot50 \| AF-A0A445MWH5-F1-MODEL\_V4 \| 1.0 \| 0.003573 \| 147 \| 0.294 \| 68 \| 46 \| 1 \| 6 \| 73 \| 2 \| 67 \| HTH\_17 domain-containing protein \| HTH\_17 domain-containing protein; afdb-uniprot50 \| AF-A0A520X4P5-F1-MODEL\_V4 \| 1.0 \| 0.003573 \| 147 \| 0.246 \| 73 \| 49 \| 3 \| 1 \| 73 \| 1 \| 67 \| DNA-binding protein \| DNA-binding protein; afdb-uniprot50 \| AF-A0A7X0FN29-F1-MODEL\_V4 \| 1.0 \| 0.003573 \| 147 \| 0.271 \| 70 \| 48 \| 3 \| 4 \| 73 \| 2 \| 68 \| Uncharacterized protein \| Uncharacterized protein; afdb-uniprot50 \| AF-A0A4Z0KEZ5-F1-MODEL\_V4 \| 1.0 \| 0.002397 \| 147 \| 0.217 \| 69 \| 51 \| 2 \| 1 \| 69 \| 3 \| 68 \| DNA-binding protein \| DNA-binding protein; afdb-uniprot50 \| AF-A0A243RM94-F1-MODEL\_V4 \| 1.0 \| 0.001963 \| 147 \| 0.25 \| 72 \| 52 \| 1 \| 1 \| 72 \| 1 \| 70 \| DNA-binding protein \| DNA-binding protein; afdb-uniprot50 \| AF-A0A2A9K2C8-F1-MODEL\_V4 \| 1.0 \| 0.002098 \| 147 \| 0.253 \| 71 \| 49 \| 2 \| 1 \| 69 \| 1 \| 69 \| AlpA family transcriptional regulator \| AlpA family transcriptional regulator; afdb-uniprot50 \| AF-A0A7J5BMM3-F1-MODEL\_V4 \| 1.0 \| 0.002243 \| 147 \| 0.238 \| 67 \| 47 \| 2 \| 1 \| 65 \| 2 \| 66 \| Helix-turn-helix domain-containing protein \| Helix-turn-helix domain-containing protein; afdb-uniprot50 \| AF-A0A840EWM2-F1-MODEL\_V4 \| 1.0 \| 0.003819 \| 147 \| 0.219 \| 73 \| 53 \| 3 \| 1 \| 73 \| 1 \| 69 \| Uncharacterized protein \| Uncharacterized protein; afdb-uniprot50 \| AF-A0A7Y7M2Z0-F1-MODEL\_V4 \| 1.0 \| 0.003343 \| 147 \| 0.223 \| 67 \| 48 \| 2 \| 1 \| 65 \| 6 \| 70 \| Helix-turn-helix domain-containing protein \| Helix-turn-helix domain-containing protein; afdb-uniprot50 \| AF-A0A2K8Y4P9-F1-MODEL\_V4 \| 1.0 \| 0.005692 \| 147 \| 0.322 \| 59 \| 38 \| 1 \| 4 \| 62 \| 1 \| 57 \| Excisionase \| Excisionase; afdb-uniprot50 \| AF-A0A2D6SS17-F1-MODEL\_V4 \| 1.0 \| 0.003128 \| 147 \| 0.257 \| 70 \| 50 \| 1 \| 4 \| 73 \| 1 \| 68 \| HTH\_17 domain-containing protein \| HTH\_17 domain-containing protein; afdb-uniprot50 \| AF-A0A4Y9NPZ2-F1-MODEL\_V4 \| 1.0 \| 0.004662 \| 147 \| 0.222 \| 63 \| 47 \| 1 \| 11 \| 73 \| 13 \| 73 \| DNA-binding protein \| DNA-binding protein; afdb-uniprot50 \| AF-A0A2V5JGJ3-F1-MODEL\_V4 \| 1.0 \| 0.003343 \| 147 \| 0.27 \| 74 \| 51 \| 2 \| 1 \| 73 \| 2 \| 73 \| DNA-binding protein \| DNA-binding protein; afdb-uniprot50 \| AF-A0A522NW80-F1-MODEL\_V4 \| 1.0 \| 0.004983 \| 147 \| 0.25 \| 72 \| 52 \| 1 \| 2 \| 73 \| 8 \| 77 \| DNA-binding protein \| DNA-binding protein; afdb-uniprot50 \| AF-A0A0D7CQX3-F1-MODEL\_V4 \| 1.0 \| 0.001317 \| 147 \| 0.295 \| 71 \| 42 \| 2 \| 2 \| 65 \| 7 \| 76 \| DNA-binding protein \| DNA-binding protein; afdb-uniprot50 \| AF-A0A397LT87-F1-MODEL\_V4 \| 1.0 \| 0.002243 \| 147 \| 0.21 \| 76 \| 55 \| 2 \| 1 \| 73 \| 9 \| 82 \| AlpA family transcriptional regulator \| AlpA family transcriptional regulator; afdb-uniprot50 \| AF-G2JA45-F1-MODEL\_V4 \| 1.0 \| 0.001963 \| 147 \| 0.315 \| 73 \| 45 \| 3 \| 1 \| 73 \| 1 \| 68 \| Uncharacterized protein \| Uncharacterized protein; afdb-uniprot50 \| AF-A0A286GLX8-F1-MODEL\_V4 \| 1.0 \| 0.003343 \| 147 \| 0.208 \| 72 \| 55 \| 1 \| 1 \| 72 \| 16 \| 85 \| Helix-turn-helix domain-containing protein \| Helix-turn-helix domain-containing protein; afdb-uniprot50 \| AF-A0A1V9K8R2-F1-MODEL\_V4 \| 1.0 \| 0.005692 \| 147 \| 0.271 \| 59 \| 41 \| 1 \| 9 \| 67 \| 30 \| 86 \| DNA-binding protein \| DNA-binding protein; afdb-uniprot50 \| AF-A0A7Z0AIF4-F1-MODEL\_V4 \| 1.0 \| 0.001408 \| 147 \| 0.191 \| 73 \| 56 \| 2 \| 2 \| 73 \| 20 \| 90 \| Excisionase family DNA binding protein \| Excisionase family DNA binding protein; afdb-uniprot50 \| AF-Q5P854-F1-MODEL\_V4 \| 1.0 \| 0.001153 \| 147 \| 0.32 \| 75 \| 46 \| 3 \| 1 \| 72 \| 17 \| 89 \| HTH\_17 domain-containing protein \| HTH\_17 domain-containing protein; afdb-uniprot50 \| AF-A0A1G7ELU0-F1-MODEL\_V4 \| 1.0 \| 0.005692 \| 147 \| 0.38 \| 63 \| 36 \| 2 \| 11 \| 73 \| 41 \| 100 \| Helix-turn-helix domain-containing protein \| Helix-turn-helix domain-containing protein; afdb-uniprot50 \| AF-A0A2W6S3C3-F1-MODEL\_V4 \| 1.0 \| 0.001408 \| 147 \| 0.273 \| 73 \| 50 \| 2 \| 1 \| 72 \| 19 \| 89 \| DNA-binding protein \| DNA-binding protein; afdb-uniprot50 \| AF-M7A7X6-F1-MODEL\_V4 \| 1.0 \| 0.002927 \| 146 \| 0.312 \| 64 \| 39 \| 3 \| 1 \| 62 \| 1 \| 61 \| HTH\_17 domain-containing protein \| HTH\_17 domain-containing protein; afdb-uniprot50 \| AF-A0A2K9DLX8-F1-MODEL\_V4 \| 1.0 \| 0.008485 \| 146 \| 0.312 \| 64 \| 41 \| 2 \| 1 \| 63 \| 2 \| 63 \| DNA-binding protein \| DNA-binding protein; afdb-uniprot50 \| AF-A0A348G1J1-F1-MODEL\_V4 \| 1.0 \| 0.001408 \| 146 \| 0.231 \| 69 \| 46 \| 2 \| 4 \| 72 \| 2 \| 63 \| HTH\_17 domain-containing protein \| HTH\_17 domain-containing protein; afdb-uniprot50 \| AF-A0A2S5X2A3-F1-MODEL\_V4 \| 1.0 \| 0.002927 \| 146 \| 0.271 \| 70 \| 45 \| 2 \| 1 \| 66 \| 1 \| 68 \| HTH\_17 domain-containing protein \| HTH\_17 domain-containing protein; afdb-uniprot50 \| AF-A0A2U2AVN2-F1-MODEL\_V4 \| 1.0 \| 0.003819 \| 146 \| 0.295 \| 61 \| 39 \| 2 \| 2 \| 62 \| 9 \| 65 \| Excisionase \| Excisionase; afdb-uniprot50 \| AF-A0A0B5HSA4-F1-MODEL\_V4 \| 1.0 \| 0.001963 \| 146 \| 0.257 \| 70 \| 47 \| 2 \| 1 \| 67 \| 3 \| 70 \| DNA-binding protein \| DNA-binding protein; afdb-uniprot50 \| AF-A0A344WL72-F1-MODEL\_V4 \| 1.0 \| 0.002927 \| 146 \| 0.322 \| 62 \| 40 \| 2 \| 1 \| 62 \| 1 \| 60 \| Uncharacterized protein \| Uncharacterized protein; afdb-uniprot50 \| AF-A0A6I7Y267-F1-MODEL\_V4 \| 1.0 \| 0.007428 \| 146 \| 0.265 \| 64 \| 45 \| 1 \| 1 \| 64 \| 8 \| 69 \| DNA-binding protein \| DNA-binding protein; afdb-uniprot50 \| AF-A0A3A0G615-F1-MODEL\_V4 \| 1.0 \| 0.002927 \| 146 \| 0.246 \| 73 \| 51 \| 2 \| 1 \| 73 \| 2 \| 70 \| HTH\_17 domain-containing protein \| HTH\_17 domain-containing protein; afdb-uniprot50 \| AF-A0A841CXJ6-F1-MODEL\_V4 \| 1.0 \| 0.001719 \| 146 \| 0.246 \| 77 \| 48 \| 2 \| 1 \| 69 \| 1 \| 75 \| Uncharacterized protein \| Uncharacterized protein; afdb-uniprot50 \| AF-A0A6A7LAR2-F1-MODEL\_V4 \| 1.0 \| 0.003819 \| 146 \| 0.23 \| 65 \| 46 \| 2 \| 1 \| 63 \| 6 \| 68 \| Helix-turn-helix domain-containing protein \| Helix-turn-helix domain-containing protein; afdb-uniprot50 \| AF-B0URF6-F1-MODEL\_V4 \| 1.0 \| 0.003128 \| 146 \| 0.242 \| 70 \| 51 \| 1 \| 1 \| 70 \| 2 \| 69 \| HTH\_17 domain-containing protein \| HTH\_17 domain-containing protein; afdb-uniprot50 \| AF-A0A1H1VI02-F1-MODEL\_V4 \| 1.0 \| 0.001837 \| 146 \| 0.243 \| 74 \| 53 \| 2 \| 1 \| 73 \| 8 \| 79 \| DNA binding domain-containing protein, excisionase family \| DNA binding domain-containing protein, excisionase family; afdb-uniprot50 \| AF-A0A3R8VZY2-F1-MODEL\_V4 \| 1.0 \| 0.001837 \| 146 \| 0.242 \| 70 \| 50 \| 2 \| 2 \| 70 \| 16 \| 83 \| DNA-binding protein \| DNA-binding protein; afdb-uniprot50 \| AF-A0A846XD13-F1-MODEL\_V4 \| 1.0 \| 0.002098 \| 146 \| 0.275 \| 69 \| 46 \| 3 \| 2 \| 68 \| 21 \| 87 \| Helix-turn-helix domain-containing protein \| Helix-turn-helix domain-containing protein; afdb-uniprot50 \| AF-A0A1C3RET6-F1-MODEL\_V4 \| 1.0 \| 0.002927 \| 146 \| 0.246 \| 73 \| 53 \| 1 \| 1 \| 73 \| 9 \| 79 \| Transcriptional regulator (Modular protein) \| Transcriptional regulator (Modular protein); afdb-uniprot50 \| AF-A0A2A2D798-F1-MODEL\_V4 \| 1.0 \| 0.003573 \| 146 \| 0.227 \| 79 \| 53 \| 2 \| 1 \| 73 \| 1 \| 77 \| Uncharacterized protein \| Uncharacterized protein; afdb-uniprot50 \| AF-A0A6L7X0Z1-F1-MODEL\_V4 \| 1.0 \| 0.001608 \| 146 \| 0.246 \| 73 \| 52 \| 2 \| 1 \| 72 \| 45 \| 115 \| Helix-turn-helix domain-containing protein \| Helix-turn-helix domain-containing protein; afdb-uniprot50 \| AF-A0A1W2FSX5-F1-MODEL\_V4 \| 1.0 \| 0.004983 \| 145 \| 0.25 \| 64 \| 45 \| 2 \| 1 \| 63 \| 2 \| 63 \| Helix-turn-helix domain-containing protein \| Helix-turn-helix domain-containing protein; afdb-uniprot50 \| AF-A0A5B8C8T0-F1-MODEL\_V4 \| 1.0 \| 0.003573 \| 145 \| 0.277 \| 72 \| 46 \| 3 \| 1 \| 72 \| 1 \| 66 \| Helix-turn-helix domain-containing protein \| Helix-turn-helix domain-containing protein; afdb-uniprot50 \| AF-A0A1V0KSN6-F1-MODEL\_V4 \| 1.0 \| 0.004081 \| 145 \| 0.268 \| 67 \| 45 \| 3 \| 7 \| 72 \| 2 \| 65 \| Excisionase \| Excisionase; afdb-uniprot50 \| AF-A0A6L9SHQ0-F1-MODEL\_V4 \| 1.0 \| 0.007939 \| 145 \| 0.283 \| 67 \| 46 \| 1 \| 1 \| 67 \| 2 \| 66 \| Helix-turn-helix domain-containing protein \| Helix-turn-helix domain-containing protein; afdb-uniprot50 \| AF-A0A7Y0CGF4-F1-MODEL\_V4 \| 1.0 \| 0.001153 \| 145 \| 0.287 \| 73 \| 45 \| 3 \| 1 \| 73 \| 3 \| 68 \| Helix-turn-helix domain-containing protein \| Helix-turn-helix domain-containing protein; afdb-uniprot50 \| AF-A0A2W4I437-F1-MODEL\_V4 \| 1.0 \| 0.004983 \| 145 \| 0.257 \| 66 \| 46 \| 2 \| 1 \| 65 \| 3 \| 66 \| DNA-binding protein \| DNA-binding protein; afdb-uniprot50 \| AF-A0A2W2MKA0-F1-MODEL\_V4 \| 1.0 \| 0.001963 \| 145 \| 0.246 \| 73 \| 52 \| 2 \| 1 \| 73 \| 2 \| 71 \| HTH\_17 domain-containing protein \| HTH\_17 domain-containing protein; afdb-uniprot50 \| AF-A0A1P8X724-F1-MODEL\_V4 \| 1.0 \| 0.004081 \| 145 \| 0.232 \| 73 \| 52 \| 2 \| 1 \| 73 \| 2 \| 70 \| DNA-binding protein \| DNA-binding protein; afdb-uniprot50 \| AF-A0A7W5FGQ6-F1-MODEL\_V4 \| 1.0 \| 0.003128 \| 145 \| 0.279 \| 68 \| 46 \| 2 \| 1 \| 67 \| 2 \| 67 \| Putative DNA-binding transcriptional regulator AlpA \| Putative DNA-binding transcriptional regulator AlpA; afdb-uniprot50 \| AF-A0A7Y9GK84-F1-MODEL\_V4 \| 1.0 \| 0.003128 \| 145 \| 0.219 \| 73 \| 53 \| 2 \| 3 \| 73 \| 2 \| 72 \| Putative DNA-binding transcriptional regulator AlpA \| Putative DNA-binding transcriptional regulator AlpA; afdb-uniprot50 \| AF-A0A645HPJ7-F1-MODEL\_V4 \| 1.0 \| 0.002738 \| 145 \| 0.219 \| 73 \| 55 \| 1 \| 1 \| 73 \| 1 \| 71 \| HTH\_17 domain-containing protein \| HTH\_17 domain-containing protein; afdb-uniprot50 \| AF-A0A515KRJ5-F1-MODEL\_V4 \| 1.0 \| 0.003128 \| 145 \| 0.226 \| 75 \| 54 \| 2 \| 1 \| 73 \| 1 \| 73 \| Helix-turn-helix domain-containing protein \| Helix-turn-helix domain-containing protein; afdb-uniprot50 \| AF-A0A2U3QJS9-F1-MODEL\_V4 \| 1.0 \| 0.003819 \| 145 \| 0.26 \| 69 \| 45 \| 2 \| 9 \| 73 \| 2 \| 68 \| Uncharacterized protein \| Uncharacterized protein; afdb-uniprot50 \| AF-A0A511AET4-F1-MODEL\_V4 \| 1.0 \| 0.004662 \| 145 \| 0.26 \| 73 \| 53 \| 1 \| 1 \| 73 \| 6 \| 77 \| Uncharacterized protein \| Uncharacterized protein; afdb-uniprot50 \| AF-A0A140GVH6-F1-MODEL\_V4 \| 1.0 \| 0.002927 \| 145 \| 0.177 \| 79 \| 57 \| 3 \| 1 \| 73 \| 1 \| 77 \| Phage transcriptional regulator AlpA \| Phage transcriptional regulator AlpA; afdb-uniprot50 \| AF-A0A7X0VA92-F1-MODEL\_V4 \| 1.0 \| 0.002562 \| 145 \| 0.276 \| 76 \| 44 \| 2 \| 1 \| 67 \| 1 \| 74 \| Helix-turn-helix domain-containing protein \| Helix-turn-helix domain-containing protein; afdb-uniprot50 \| AF-A0A7X3S6D7-F1-MODEL\_V4 \| 1.0 \| 0.004662 \| 145 \| 0.281 \| 71 \| 49 \| 1 \| 2 \| 72 \| 14 \| 82 \| Helix-turn-helix domain-containing protein \| Helix-turn-helix domain-containing protein; afdb-uniprot50 \| AF-A0A1R1SIM1-F1-MODEL\_V4 \| 1.0 \| 0.007939 \| 145 \| 0.28 \| 57 \| 39 \| 1 \| 9 \| 65 \| 35 \| 89 \| DNA-binding protein \| DNA-binding protein; afdb-uniprot50 \| AF-A0A2P2BX08-F1-MODEL\_V4 \| 1.0 \| 0.002243 \| 145 \| 0.305 \| 72 \| 45 \| 3 \| 2 \| 71 \| 4 \| 72 \| Transcriptional repressor, dcmR family (Modular protein) \| Transcriptional repressor, dcmR family (Modular protein); afdb-uniprot50 \| AF-A0A4R8WX82-F1-MODEL\_V4 \| 1.0 \| 0.005692 \| 145 \| 0.322 \| 62 \| 38 \| 2 \| 2 \| 61 \| 40 \| 99 \| Helix-turn-helix domain-containing protein \| Helix-turn-helix domain-containing protein; afdb-uniprot50 \| AF-A0A2D9H295-F1-MODEL\_V4 \| 1.0 \| 0.0008268 \| 145 \| 0.276 \| 76 \| 46 \| 4 \| 1 \| 73 \| 44 \| 113 \| HTH\_17 domain-containing protein \| HTH\_17 domain-containing protein; afdb-uniprot50 \| AF-D6KAR1-F1-MODEL\_V4 \| 1.0 \| 0.003573 \| 145 \| 0.214 \| 70 \| 48 \| 2 \| 1 \| 65 \| 44 \| 111 \| Excisionase family DNA-binding domain-containing protein \| Excisionase family DNA-binding domain-containing protein; afdb-uniprot50 \| AF-A0A1Q7VQ48-F1-MODEL\_V4 \| 1.0 \| 0.005326 \| 144 \| 0.238 \| 63 \| 45 \| 2 \| 1 \| 63 \| 1 \| 60 \| HTH\_17 domain-containing protein \| HTH\_17 domain-containing protein; afdb-uniprot50 \| AF-A0A562V6Z3-F1-MODEL\_V4 \| 1.0 \| 0.003819 \| 144 \| 0.25 \| 68 \| 46 \| 3 \| 1 \| 68 \| 1 \| 63 \| AlpA family transcriptional regulator \| AlpA family transcriptional regulator; afdb-uniprot50 \| AF-A0A2N0TJM6-F1-MODEL\_V4 \| 1.0 \| 0.005326 \| 144 \| 0.257 \| 66 \| 45 \| 2 \| 9 \| 73 \| 6 \| 68 \| Uncharacterized protein \| Uncharacterized protein; afdb-uniprot50 \| AF-A0A7Y4L6Z8-F1-MODEL\_V4 \| 1.0 \| 0.004662 \| 144 \| 0.185 \| 70 \| 54 \| 2 \| 1 \| 70 \| 1 \| 67 \| Helix-turn-helix domain-containing protein \| Helix-turn-helix domain-containing protein; afdb-uniprot50 \| AF-A0A7I9Y7H1-F1-MODEL\_V4 \| 1.0 \| 0.005326 \| 144 \| 0.314 \| 70 \| 45 \| 2 \| 1 \| 69 \| 1 \| 68 \| HTH\_17 domain-containing protein \| HTH\_17 domain-containing protein; afdb-uniprot50 \| AF-A0A3M2M828-F1-MODEL\_V4 \| 1.0 \| 0.004081 \| 144 \| 0.222 \| 72 \| 52 \| 2 \| 1 \| 72 \| 3 \| 70 \| Helix-turn-helix domain-containing protein \| Helix-turn-helix domain-containing protein; afdb-uniprot50 \| AF-A0A1M5QRE5-F1-MODEL\_V4 \| 1.0 \| 0.004983 \| 144 \| 0.257 \| 70 \| 50 \| 1 \| 4 \| 73 \| 2 \| 69 \| Uncharacterized protein \| Uncharacterized protein; afdb-uniprot50 \| AF-A0A445MRB1-F1-MODEL\_V4 \| 1.0 \| 0.002927 \| 144 \| 0.307 \| 65 \| 41 \| 2 \| 1 \| 63 \| 1 \| 63 \| HTH\_17 domain-containing protein \| HTH\_17 domain-containing protein; afdb-uniprot50 \| AF-A0A077MB24-F1-MODEL\_V4 \| 1.0 \| 0.005326 \| 144 \| 0.238 \| 67 \| 49 \| 1 \| 1 \| 67 \| 6 \| 70 \| HTH\_17 domain-containing protein \| HTH\_17 domain-containing protein; afdb-uniprot50 \| AF-B8EI03-F1-MODEL\_V4 \| 1.0 \| 0.001837 \| 144 \| 0.256 \| 74 \| 52 \| 2 \| 1 \| 73 \| 2 \| 73 \| Phage transcriptional regulator, AlpA \| Phage transcriptional regulator, AlpA; afdb-uniprot50 \| AF-A0A0F3ILT4-F1-MODEL\_V4 \| 1.0 \| 0.004983 \| 144 \| 0.258 \| 62 \| 43 \| 2 \| 2 \| 62 \| 9 \| 68 \| Uncharacterized protein \| Uncharacterized protein; afdb-uniprot50 \| AF-I9WCR2-F1-MODEL\_V4 \| 1.0 \| 0.006502 \| 144 \| 0.202 \| 69 \| 53 \| 1 \| 5 \| 73 \| 13 \| 79 \| HTH\_17 domain-containing protein \| HTH\_17 domain-containing protein; afdb-uniprot50 \| AF-A0A3G7HH05-F1-MODEL\_V4 \| 1.0 \| 0.001963 \| 144 \| 0.256 \| 74 \| 52 \| 2 \| 1 \| 73 \| 9 \| 80 \| HTH\_17 domain-containing protein \| HTH\_17 domain-containing protein; afdb-uniprot50 \| AF-A0A225SLN1-F1-MODEL\_V4 \| 1.0 \| 0.004983 \| 144 \| 0.303 \| 66 \| 44 \| 1 \| 8 \| 73 \| 13 \| 76 \| Uncharacterized protein \| Uncharacterized protein; afdb-uniprot50 \| AF-A0A1H3I1Q4-F1-MODEL\_V4 \| 1.0 \| 0.001963 \| 144 \| 0.243 \| 74 \| 53 \| 2 \| 1 \| 73 \| 8 \| 79 \| DNA binding domain-containing protein, excisionase family \| DNA binding domain-containing protein, excisionase family; afdb-uniprot50 \| AF-A0A2N0FF21-F1-MODEL\_V4 \| 1.0 \| 0.002243 \| 144 \| 0.253 \| 71 \| 47 \| 2 \| 1 \| 66 \| 12 \| 81 \| Helix-turn-helix protein \| Helix-turn-helix protein; afdb-uniprot50 \| AF-A0A2T3GGH2-F1-MODEL\_V4 \| 1.0 \| 0.002562 \| 144 \| 0.241 \| 87 \| 49 \| 3 \| 1 \| 73 \| 2 \| 85 \| DNA-binding protein \| DNA-binding protein; afdb-uniprot50 \| AF-A0A844B6Q9-F1-MODEL\_V4 \| 1.0 \| 0.005326 \| 144 \| 0.205 \| 73 \| 55 \| 2 \| 1 \| 73 \| 13 \| 82 \| Helix-turn-helix domain-containing protein \| Helix-turn-helix domain-containing protein; afdb-uniprot50 \| AF-A0A5C8ULD5-F1-MODEL\_V4 \| 1.0 \| 0.004662 \| 144 \| 0.253 \| 71 \| 51 \| 1 \| 1 \| 71 \| 17 \| 85 \| DNA-binding protein \| DNA-binding protein; afdb-uniprot50 \| AF-A0A4Y3VET6-F1-MODEL\_V4 \| 1.0 \| 0.004081 \| 144 \| 0.208 \| 72 \| 52 \| 2 \| 2 \| 73 \| 24 \| 90 \| HTH\_17 domain-containing protein \| HTH\_17 domain-containing protein; afdb-uniprot50 \| AF-A0A846WWH9-F1-MODEL\_V4 \| 1.0 \| 0.002927 \| 144 \| 0.314 \| 70 \| 44 \| 2 \| 2 \| 69 \| 24 \| 91 \| Helix-turn-helix domain-containing protein \| Helix-turn-helix domain-containing protein; afdb-uniprot50 \| AF-A0A8B5TVS5-F1-MODEL\_V4 \| 1.0 \| 0.003573 \| 144 \| 0.253 \| 75 \| 52 \| 2 \| 1 \| 73 \| 24 \| 96 \| Helix-turn-helix domain-containing protein \| Helix-turn-helix domain-containing protein; afdb-uniprot50 \| AF-A0A7V8K6X7-F1-MODEL\_V4 \| 1.0 \| 0.008485 \| 144 \| 0.184 \| 65 \| 51 \| 1 \| 9 \| 73 \| 29 \| 91 \| Uncharacterized protein \| Uncharacterized protein; afdb-uniprot50 \| AF-A0A503YCB6-F1-MODEL\_V4 \| 1.0 \| 0.001963 \| 144 \| 0.253 \| 75 \| 52 \| 2 \| 1 \| 73 \| 26 \| 98 \| Helix-turn-helix domain-containing protein \| Helix-turn-helix domain-containing protein; afdb-uniprot50 \| AF-A0A2Z6AYS9-F1-MODEL\_V4 \| 1.0 \| 0.004362 \| 144 \| 0.267 \| 71 \| 49 \| 2 \| 3 \| 73 \| 40 \| 107 \| HTH\_17 domain-containing protein \| HTH\_17 domain-containing protein; afdb-uniprot50 \| AF-D2S8Y0-F1-MODEL\_V4 \| 1.0 \| 0.002243 \| 144 \| 0.233 \| 77 \| 53 \| 3 \| 1 \| 73 \| 1 \| 75 \| Putative transcriptional regulator, MerR family \| Putative transcriptional regulator, MerR family; afdb-uniprot50 \| AF-A0A6I6EB25-F1-MODEL\_V4 \| 1.0 \| 0.007939 \| 143 \| 0.269 \| 63 \| 44 \| 1 \| 1 \| 63 \| 1 \| 61 \| DNA-binding protein \| DNA-binding protein; afdb-uniprot50 \| AF-A0A7W1TAK0-F1-MODEL\_V4 \| 1.0 \| 0.004081 \| 143 \| 0.317 \| 63 \| 41 \| 2 \| 1 \| 63 \| 1 \| 61 \| Helix-turn-helix domain-containing protein \| Helix-turn-helix domain-containing protein; afdb-uniprot50 \| AF-A0A0F9SYS0-F1-MODEL\_V4 \| 1.0 \| 0.002927 \| 143 \| 0.25 \| 72 \| 48 \| 3 \| 1 \| 69 \| 1 \| 69 \| Uncharacterized protein \| Uncharacterized protein; afdb-uniprot50 \| AF-A0A1C4T028-F1-MODEL\_V4 \| 1.0 \| 0.006502 \| 143 \| 0.261 \| 65 \| 46 \| 1 \| 1 \| 65 \| 6 \| 68 \| DNA binding domain-containing protein, excisionase family \| DNA binding domain-containing protein, excisionase family; afdb-uniprot50 \| AF-A0A382E9C4-F1-MODEL\_V4 \| 1.0 \| 0.002243 \| 143 \| 0.297 \| 74 \| 48 \| 2 \| 1 \| 72 \| 1 \| 72 \| Uncharacterized protein \| Uncharacterized protein; afdb-uniprot50 \| AF-A0A1P8XFC1-F1-MODEL\_V4 \| 1.0 \| 0.001153 \| 143 \| 0.333 \| 75 \| 41 \| 3 \| 1 \| 73 \| 2 \| 69 \| Uncharacterized protein \| Uncharacterized protein; afdb-uniprot50 \| AF-A0A7W1GPM6-F1-MODEL\_V4 \| 1.0 \| 0.005692 \| 143 \| 0.285 \| 63 \| 42 \| 2 \| 11 \| 73 \| 4 \| 63 \| Helix-turn-helix domain-containing protein \| Helix-turn-helix domain-containing protein; afdb-uniprot50 \| AF-A0A127ENA5-F1-MODEL\_V4 \| 1.0 \| 0.003573 \| 143 \| 0.256 \| 74 \| 52 \| 2 \| 1 \| 73 \| 1 \| 72 \| Excisionase family DNA binding domain-containing protein \| Excisionase family DNA binding domain-containing protein; afdb-uniprot50 \| AF-A0A1C5D9B9-F1-MODEL\_V4 \| 1.0 \| 0.003343 \| 143 \| 0.246 \| 73 \| 47 \| 2 \| 1 \| 67 \| 1 \| 71 \| Helix-turn-helix domain-containing protein \| Helix-turn-helix domain-containing protein; afdb-uniprot50 \| AF-A0A5H2Y9R4-F1-MODEL\_V4 \| 1.0 \| 0.004081 \| 143 \| 0.232 \| 73 \| 54 \| 1 \| 1 \| 73 \| 1 \| 71 \| HTH\_17 domain-containing protein \| HTH\_17 domain-containing protein; afdb-uniprot50 \| AF-W6KAN3-F1-MODEL\_V4 \| 1.0 \| 0.002562 \| 143 \| 0.302 \| 76 \| 48 \| 2 \| 1 \| 73 \| 1 \| 74 \| Putative DNA binding domain protein, excisionase family \| Putative DNA binding domain protein, excisionase family; afdb-uniprot50 \| AF-A0A512JRE3-F1-MODEL\_V4 \| 1.0 \| 0.002738 \| 143 \| 0.306 \| 75 \| 48 \| 2 \| 1 \| 73 \| 1 \| 73 \| HTH\_17 domain-containing protein \| HTH\_17 domain-containing protein; afdb-uniprot50 \| AF-A0A4Z1CN04-F1-MODEL\_V4 \| 1.0 \| 0.002562 \| 143 \| 0.246 \| 73 \| 51 \| 3 \| 1 \| 72 \| 7 \| 76 \| DNA-binding protein \| DNA-binding protein; afdb-uniprot50 \| AF-A0A6A8W264-F1-MODEL\_V4 \| 1.0 \| 0.00695 \| 143 \| 0.217 \| 69 \| 51 \| 2 \| 1 \| 69 \| 11 \| 76 \| AlpA family phage regulatory protein \| AlpA family phage regulatory protein; afdb-uniprot50 \| AF-A0A6L7XK63-F1-MODEL\_V4 \| 1.0 \| 0.003343 \| 143 \| 0.285 \| 77 \| 49 \| 2 \| 1 \| 73 \| 1 \| 75 \| Helix-turn-helix domain-containing protein \| Helix-turn-helix domain-containing protein; afdb-uniprot50 \| AF-A0A839UPM0-F1-MODEL\_V4 \| 1.0 \| 0.001719 \| 143 \| 0.222 \| 72 \| 53 \| 2 \| 3 \| 73 \| 12 \| 81 \| Putative DNA-binding transcriptional regulator AlpA \| Putative DNA-binding transcriptional regulator AlpA; afdb-uniprot50 \| AF-A0A7I7VI49-F1-MODEL\_V4 \| 1.0 \| 0.006084 \| 143 \| 0.307 \| 65 \| 42 \| 2 \| 2 \| 65 \| 10 \| 72 \| HTH\_17 domain-containing protein \| HTH\_17 domain-containing protein; afdb-uniprot50 \| AF-A0A0S3F4D4-F1-MODEL\_V4 \| 1.0 \| 0.003573 \| 143 \| 0.208 \| 67 \| 49 \| 2 \| 9 \| 73 \| 17 \| 81 \| Uncharacterized protein \| Uncharacterized protein; afdb-uniprot50 \| AF-A0A2N3BAH4-F1-MODEL\_V4 \| 1.0 \| 0.003573 \| 143 \| 0.222 \| 81 \| 53 \| 2 \| 1 \| 73 \| 1 \| 79 \| Transcriptional regulator \| Transcriptional regulator; afdb-uniprot50 \| AF-A0A062VUZ0-F1-MODEL\_V4 \| 1.0 \| 0.002098 \| 143 \| 0.229 \| 74 \| 54 \| 2 \| 1 \| 73 \| 7 \| 78 \| HTH\_17 domain-containing protein \| HTH\_17 domain-containing protein; afdb-uniprot50 \| AF-A0A2A2ZNT2-F1-MODEL\_V4 \| 1.0 \| 0.003128 \| 143 \| 0.275 \| 69 \| 42 \| 3 \| 1 \| 63 \| 13 \| 79 \| DNA-binding protein \| DNA-binding protein; afdb-uniprot50 \| AF-A0A420WJK2-F1-MODEL\_V4 \| 1.0 \| 0.004362 \| 143 \| 0.232 \| 73 \| 52 \| 2 \| 1 \| 71 \| 10 \| 80 \| Helix-turn-helix protein \| Helix-turn-helix protein; afdb-uniprot50 \| AF-A0A367A4J1-F1-MODEL\_V4 \| 1.0 \| 0.006502 \| 143 \| 0.264 \| 68 \| 48 \| 1 \| 1 \| 68 \| 30 \| 95 \| HTH\_17 domain-containing protein \| HTH\_17 domain-containing protein; afdb-uniprot50 \| AF-A0A1H8Y2E5-F1-MODEL\_V4 \| 1.0 \| 0.001608 \| 143 \| 0.253 \| 75 \| 44 \| 2 \| 1 \| 65 \| 24 \| 96 \| DNA binding domain-containing protein, excisionase family \| DNA binding domain-containing protein, excisionase family; afdb-uniprot50 \| AF-A0A841NDL1-F1-MODEL\_V4 \| 1.0 \| 0.006502 \| 143 \| 0.261 \| 65 \| 44 \| 2 \| 2 \| 64 \| 39 \| 101 \| Excisionase family DNA binding protein \| Excisionase family DNA binding protein; afdb-uniprot50 \| AF-A0A329LI23-F1-MODEL\_V4 \| 1.0 \| 0.004362 \| 143 \| 0.238 \| 67 \| 49 \| 1 \| 1 \| 67 \| 61 \| 125 \| HTH\_17 domain-containing protein \| HTH\_17 domain-containing protein; afdb-uniprot50 \| AF-A0A2W4JHC9-F1-MODEL\_V4 \| 1.0 \| 0.006084 \| 143 \| 0.202 \| 74 \| 56 \| 2 \| 1 \| 73 \| 3 \| 74 \| HTH\_17 domain-containing protein \| HTH\_17 domain-containing protein; afdb-uniprot50 \| AF-A0A2K4LH34-F1-MODEL\_V4 \| 1.0 \| 0.006084 \| 142 \| 0.269 \| 63 \| 44 \| 1 \| 11 \| 73 \| 3 \| 63 \| DNA-binding protein \| DNA-binding protein; afdb-uniprot50 \| AF-A0A2H9N9A0-F1-MODEL\_V4 \| 1.0 \| 0.007939 \| 142 \| 0.285 \| 63 \| 42 \| 2 \| 11 \| 72 \| 4 \| 64 \| HTH\_17 domain-containing protein \| HTH\_17 domain-containing protein; afdb-uniprot50 \| AF-A0A126UVT8-F1-MODEL\_V4 \| 1.0 \| 0.005326 \| 142 \| 0.236 \| 72 \| 51 \| 3 \| 1 \| 72 \| 1 \| 68 \| HTH\_17 domain-containing protein \| HTH\_17 domain-containing protein; afdb-uniprot50 \| AF-A0A449CYP3-F1-MODEL\_V4 \| 1.0 \| 0.004081 \| 142 \| 0.231 \| 69 \| 49 \| 2 \| 1 \| 67 \| 1 \| 67 \| Uncharacterized protein \| Uncharacterized protein; afdb-uniprot50 \| AF-A0A7Y0UIH7-F1-MODEL\_V4 \| 1.0 \| 0.006502 \| 142 \| 0.242 \| 66 \| 47 \| 2 \| 7 \| 71 \| 1 \| 64 \| Helix-turn-helix domain-containing protein \| Helix-turn-helix domain-containing protein; afdb-uniprot50 \| AF-A0A7X5VIB6-F1-MODEL\_V4 \| 1.0 \| 0.004081 \| 142 \| 0.26 \| 73 \| 49 \| 2 \| 1 \| 73 \| 1 \| 68 \| Excisionase family DNA binding protein \| Excisionase family DNA binding protein; afdb-uniprot50 \| AF-A0A7Z9V025-F1-MODEL\_V4 \| 1.0 \| 0.004983 \| 142 \| 0.274 \| 62 \| 43 \| 1 \| 2 \| 63 \| 4 \| 63 \| DNA-binding protein \| DNA-binding protein; afdb-uniprot50 \| AF-A0A1B2A925-F1-MODEL\_V4 \| 1.0 \| 0.009692 \| 142 \| 0.234 \| 64 \| 46 \| 2 \| 10 \| 73 \| 8 \| 68 \| Helix-turn-helix domain protein \| Helix-turn-helix domain protein; afdb-uniprot50 \| AF-A0A4Z1CGY7-F1-MODEL\_V4 \| 1.0 \| 0.003573 \| 142 \| 0.279 \| 68 \| 43 \| 3 \| 1 \| 64 \| 1 \| 66 \| DNA-binding protein \| DNA-binding protein; afdb-uniprot50 \| AF-A0A6N6T802-F1-MODEL\_V4 \| 1.0 \| 0.001608 \| 142 \| 0.31 \| 74 \| 48 \| 2 \| 1 \| 73 \| 1 \| 72 \| Helix-turn-helix domain-containing protein \| Helix-turn-helix domain-containing protein; afdb-uniprot50 \| AF-A0A382R345-F1-MODEL\_V4 \| 1.0 \| 0.004983 \| 142 \| 0.296 \| 64 \| 42 \| 2 \| 1 \| 63 \| 2 \| 63 \| HTH\_17 domain-containing protein \| HTH\_17 domain-containing protein; afdb-uniprot50 \| AF-A0A1W6NWZ4-F1-MODEL\_V4 \| 1.0 \| 0.006084 \| 142 \| 0.186 \| 75 \| 57 \| 2 \| 1 \| 73 \| 3 \| 75 \| Phenylglyoxylate:acceptor oxidoreductase \| Phenylglyoxylate:acceptor oxidoreductase; afdb-uniprot50 \| AF-A0A4S5ESI4-F1-MODEL\_V4 \| 1.0 \| 0.004662 \| 142 \| 0.218 \| 64 \| 47 \| 2 \| 1 \| 63 \| 13 \| 74 \| Helix-turn-helix domain-containing protein \| Helix-turn-helix domain-containing protein; afdb-uniprot50 \| AF-A0A4V4RAK6-F1-MODEL\_V4 \| 1.0 \| 0.008485 \| 142 \| 0.246 \| 65 \| 44 \| 1 \| 2 \| 66 \| 20 \| 79 \| Uncharacterized protein \| Uncharacterized protein; afdb-uniprot50 \| AF-A0A3N5ZHA9-F1-MODEL\_V4 \| 1.0 \| 0.003128 \| 142 \| 0.24 \| 75 \| 52 \| 2 \| 2 \| 73 \| 8 \| 80 \| DNA-binding protein \| DNA-binding protein; afdb-uniprot50 \| AF-A0A7G8YVT3-F1-MODEL\_V4 \| 1.0 \| 0.001963 \| 142 \| 0.202 \| 79 \| 55 \| 3 \| 1 \| 73 \| 2 \| 78 \| Helix-turn-helix domain-containing protein \| Helix-turn-helix domain-containing protein; afdb-uniprot50 \| AF-A0A7W1WKP4-F1-MODEL\_V4 \| 1.0 \| 0.002098 \| 142 \| 0.181 \| 77 \| 56 \| 3 \| 1 \| 72 \| 5 \| 79 \| Helix-turn-helix domain-containing protein \| Helix-turn-helix domain-containing protein; afdb-uniprot50 \| AF-H5Y6A6-F1-MODEL\_V4 \| 1.0 \| 0.006084 \| 142 \| 0.154 \| 71 \| 57 \| 2 \| 1 \| 71 \| 15 \| 82 \| Uncharacterized protein \| Uncharacterized protein; afdb-uniprot50 \| AF-A0A512NPT7-F1-MODEL\_V4 \| 1.0 \| 0.001719 \| 142 \| 0.256 \| 74 \| 48 \| 3 \| 1 \| 72 \| 17 \| 85 \| HTH\_17 domain-containing protein \| HTH\_17 domain-containing protein; afdb-uniprot50 \| AF-A0A2V9IUE2-F1-MODEL\_V4 \| 1.0 \| 0.007939 \| 142 \| 0.222 \| 63 \| 47 \| 1 \| 11 \| 73 \| 3 \| 63 \| HTH\_17 domain-containing protein \| HTH\_17 domain-containing protein; afdb-uniprot50 \| AF-A0A2S8CG40-F1-MODEL\_V4 \| 1.0 \| 0.004362 \| 142 \| 0.191 \| 68 \| 52 \| 2 \| 2 \| 68 \| 35 \| 100 \| HTH\_17 domain-containing protein \| HTH\_17 domain-containing protein; afdb-uniprot50 \| AF-A0A328N1Y7-F1-MODEL\_V4 \| 1.0 \| 0.001719 \| 142 \| 0.281 \| 71 \| 45 \| 3 \| 2 \| 72 \| 43 \| 107 \| HTH\_17 domain-containing protein \| HTH\_17 domain-containing protein; afdb-uniprot50 \| AF-A0A7Y9ZK00-F1-MODEL\_V4 \| 1.0 \| 0.009692 \| 142 \| 0.2 \| 65 \| 50 \| 1 \| 1 \| 65 \| 45 \| 107 \| Putative DNA-binding transcriptional regulator AlpA \| Putative DNA-binding transcriptional regulator AlpA; afdb-uniprot50 \| AF-A0A327JJ67-F1-MODEL\_V4 \| 1.0 \| 0.003573 \| 142 \| 0.277 \| 72 \| 50 \| 1 \| 2 \| 73 \| 43 \| 112 \| HTH\_17 domain-containing protein \| HTH\_17 domain-containing protein; afdb-uniprot50 \| AF-A0A2E8YX83-F1-MODEL\_V4 \| 1.0 \| 0.001719 \| 142 \| 0.263 \| 76 \| 51 \| 3 \| 1 \| 73 \| 44 \| 117 \| HTH\_17 domain-containing protein \| HTH\_17 domain-containing protein; afdb-uniprot50 \| AF-A0A2G7CQS8-F1-MODEL\_V4 \| 1.0 \| 0.003819 \| 142 \| 0.228 \| 70 \| 49 \| 2 \| 1 \| 67 \| 62 \| 129 \| Helix-turn-helix protein \| Helix-turn-helix protein; afdb-uniprot50 \| AF-A0A5B0V902-F1-MODEL\_V4 \| 1.0 \| 0.009692 \| 141 \| 0.222 \| 63 \| 47 \| 1 \| 11 \| 73 \| 4 \| 64 \| Helix-turn-helix domain-containing protein \| Helix-turn-helix domain-containing protein; afdb-uniprot50 \| AF-A0A3N1AY40-F1-MODEL\_V4 \| 1.0 \| 0.007428 \| 141 \| 0.184 \| 65 \| 51 \| 1 \| 1 \| 65 \| 3 \| 65 \| Helix-turn-helix protein \| Helix-turn-helix protein; afdb-uniprot50 \| AF-A0A1H8K0G4-F1-MODEL\_V4 \| 1.0 \| 0.003573 \| 141 \| 0.214 \| 70 \| 50 \| 2 \| 4 \| 73 \| 1 \| 65 \| DNA binding domain, excisionase family \| DNA binding domain, excisionase family; afdb-uniprot50 \| AF-A0A1X0XIA3-F1-MODEL\_V4 \| 1.0 \| 0.003128 \| 141 \| 0.298 \| 67 \| 42 \| 2 \| 10 \| 73 \| 3 \| 67 \| Uncharacterized protein \| Uncharacterized protein; afdb-uniprot50 \| AF-A0A2W6CXR4-F1-MODEL\_V4 \| 1.0 \| 0.004662 \| 141 \| 0.25 \| 72 \| 45 \| 3 \| 1 \| 66 \| 1 \| 69 \| HTH\_17 domain-containing protein \| HTH\_17 domain-containing protein; afdb-uniprot50 \| AF-A0A7W0SNE2-F1-MODEL\_V4 \| 1.0 \| 0.006502 \| 141 \| 0.2 \| 65 \| 50 \| 1 \| 1 \| 65 \| 2 \| 64 \| Helix-turn-helix domain-containing protein \| Helix-turn-helix domain-containing protein; afdb-uniprot50 \| AF-A0A2R4WUM3-F1-MODEL\_V4 \| 1.0 \| 0.001232 \| 141 \| 0.294 \| 78 \| 45 \| 3 \| 1 \| 73 \| 10 \| 82 \| DNA-binding protein \| DNA-binding protein; afdb-uniprot50 \| AF-A0A386UJQ9-F1-MODEL\_V4 \| 1.0 \| 0.006502 \| 141 \| 0.171 \| 76 \| 58 \| 2 \| 1 \| 73 \| 6 \| 79 \| DNA-binding protein \| DNA-binding protein; afdb-uniprot50 \| AF-A0A7Y0B853-F1-MODEL\_V4 \| 1.0 \| 0.002098 \| 140 \| 0.333 \| 69 \| 38 \| 3 \| 1 \| 69 \| 1 \| 61 \| Helix-turn-helix domain-containing protein \| Helix-turn-helix domain-containing protein; afdb-uniprot50 \| AF-A0A1Q8KIY1-F1-MODEL\_V4 \| 1.0 \| 0.007939 \| 140 \| 0.276 \| 65 \| 42 \| 2 \| 3 \| 67 \| 2 \| 61 \| DNA binding domain, excisionase/Xis \| DNA binding domain, excisionase/Xis; afdb-uniprot50 \| AF-A0A6G4X2R8-F1-MODEL\_V4 \| 1.0 \| 0.009068 \| 140 \| 0.268 \| 67 \| 45 \| 2 \| 1 \| 67 \| 5 \| 67 \| Helix-turn-helix domain-containing protein \| Helix-turn-helix domain-containing protein; afdb-uniprot50 \| AF-K2AU10-F1-MODEL\_V4 \| 1.0 \| 0.006502 \| 140 \| 0.292 \| 65 \| 43 \| 1 \| 11 \| 72 \| 4 \| 68 \| Uncharacterized protein \| Uncharacterized protein; afdb-uniprot50 \| AF-A0A543P9T2-F1-MODEL\_V4 \| 1.0 \| 0.003819 \| 140 \| 0.289 \| 69 \| 46 \| 2 \| 1 \| 68 \| 2 \| 68 \| Excisionase family DNA binding protein \| Excisionase family DNA binding protein; afdb-uniprot50 \| AF-A0A2M8W6L7-F1-MODEL\_V4 \| 1.0 \| 0.009692 \| 140 \| 0.222 \| 63 \| 47 \| 1 \| 1 \| 63 \| 7 \| 67 \| AlpA family transcriptional regulator \| AlpA family transcriptional regulator; afdb-uniprot50 \| AF-A0A0R3E397-F1-MODEL\_V4 \| 1.0 \| 0.004662 \| 140 \| 0.273 \| 73 \| 49 \| 2 \| 3 \| 73 \| 5 \| 75 \| Uncharacterized protein \| Uncharacterized protein; afdb-uniprot50 \| AF-A0A1X1MGL1-F1-MODEL\_V4 \| 1.0 \| 0.005692 \| 140 \| 0.205 \| 73 \| 54 \| 2 \| 1 \| 73 \| 1 \| 69 \| Uncharacterized protein \| Uncharacterized protein; afdb-uniprot50 \| AF-A0A2N1I4M7-F1-MODEL\_V4 \| 1.0 \| 0.004983 \| 140 \| 0.268 \| 67 \| 45 \| 2 \| 2 \| 66 \| 9 \| 73 \| Uncharacterized protein \| Uncharacterized protein; afdb-uniprot50 \| AF-A0A516Q5F3-F1-MODEL\_V4 \| 1.0 \| 0.005326 \| 140 \| 0.202 \| 69 \| 51 \| 2 \| 1 \| 67 \| 1 \| 67 \| Helix-turn-helix domain-containing protein \| Helix-turn-helix domain-containing protein; afdb-uniprot50 \| AF-A0A7J0BY77-F1-MODEL\_V4 \| 1.0 \| 0.001837 \| 140 \| 0.293 \| 75 \| 49 \| 3 \| 1 \| 73 \| 9 \| 81 \| HTH\_17 domain-containing protein \| HTH\_17 domain-containing protein; afdb-uniprot50 \| AF-A0A530LMQ4-F1-MODEL\_V4 \| 1.0 \| 0.00695 \| 140 \| 0.246 \| 69 \| 50 \| 1 \| 5 \| 73 \| 11 \| 77 \| Helix-turn-helix domain-containing protein \| Helix-turn-helix domain-containing protein; afdb-uniprot50 \| AF-A0A1E4PMA0-F1-MODEL\_V4 \| 1.0 \| 0.003819 \| 140 \| 0.279 \| 68 \| 42 \| 2 \| 1 \| 63 \| 4 \| 69 \| HTH\_17 domain-containing protein \| HTH\_17 domain-containing protein; afdb-uniprot50 \| AF-A0A0V2FH72-F1-MODEL\_V4 \| 1.0 \| 0.001963 \| 140 \| 0.337 \| 74 \| 43 \| 4 \| 1 \| 73 \| 1 \| 69 \| Transcriptional regulator \| Transcriptional regulator; afdb-uniprot50 \| AF-A5EBE1-F1-MODEL\_V4 \| 1.0 \| 0.002927 \| 140 \| 0.26 \| 73 \| 49 \| 3 \| 1 \| 73 \| 3 \| 70 \| Uncharacterized protein \| Uncharacterized protein; afdb-uniprot50 \| AF-A0A7K0CZS0-F1-MODEL\_V4 \| 1.0 \| 0.005326 \| 140 \| 0.207 \| 77 \| 55 \| 3 \| 1 \| 73 \| 31 \| 105 \| HTH\_17 domain-containing protein \| HTH\_17 domain-containing protein; afdb-uniprot50 \| AF-A0A653Z374-F1-MODEL\_V4 \| 1.0 \| 0.006084 \| 140 \| 0.285 \| 63 \| 42 \| 2 \| 1 \| 62 \| 33 \| 93 \| HTH\_17 domain-containing protein \| HTH\_17 domain-containing protein; afdb-uniprot50 \| AF-A0A087KGM0-F1-MODEL\_V4 \| 1.0 \| 0.009068 \| 139 \| 0.225 \| 62 \| 46 \| 1 \| 6 \| 67 \| 2 \| 61 \| MerR family transcriptional regulator \| MerR family transcriptional regulator; afdb-uniprot50 \| AF-A0A1Y4CKH7-F1-MODEL\_V4 \| 1.0 \| 0.006502 \| 139 \| 0.268 \| 67 \| 45 \| 2 \| 1 \| 65 \| 2 \| 66 \| HTH\_17 domain-containing protein \| HTH\_17 domain-containing protein; afdb-uniprot50 \| AF-A0A7W8FGC6-F1-MODEL\_V4 \| 1.0 \| 0.00695 \| 139 \| 0.278 \| 61 \| 41 \| 2 \| 4 \| 63 \| 2 \| 60 \| Uncharacterized protein \| Uncharacterized protein; afdb-uniprot50 \| AF-A0A0J6XLH8-F1-MODEL\_V4 \| 1.0 \| 0.006084 \| 139 \| 0.268 \| 67 \| 46 \| 2 \| 1 \| 67 \| 1 \| 64 \| DNA-binding protein \| DNA-binding protein; afdb-uniprot50 \| AF-A0A0U3FN39-F1-MODEL\_V4 \| 1.0 \| 0.005326 \| 139 \| 0.236 \| 72 \| 49 \| 2 \| 1 \| 72 \| 5 \| 70 \| HTH\_17 domain-containing protein \| HTH\_17 domain-containing protein; afdb-uniprot50 \| AF-A0A238Y6F1-F1-MODEL\_V4 \| 1.0 \| 0.008485 \| 139 \| 0.268 \| 67 \| 47 \| 1 \| 1 \| 67 \| 2 \| 66 \| DNA binding domain-containing protein, excisionase family \| DNA binding domain-containing protein, excisionase family; afdb-uniprot50 \| AF-A0A7Y7M0V9-F1-MODEL\_V4 \| 1.0 \| 0.005326 \| 139 \| 0.253 \| 63 \| 44 \| 2 \| 1 \| 62 \| 6 \| 66 \| Helix-turn-helix domain-containing protein \| Helix-turn-helix domain-containing protein; afdb-uniprot50 \| AF-A0A1H0BM39-F1-MODEL\_V4 \| 1.0 \| 0.003819 \| 139 \| 0.289 \| 69 \| 45 \| 3 \| 1 \| 67 \| 1 \| 67 \| DNA binding domain-containing protein, excisionase family \| DNA binding domain-containing protein, excisionase family; afdb-uniprot50 \| AF-A0A4R8S6P8-F1-MODEL\_V4 \| 1.0 \| 0.003819 \| 139 \| 0.226 \| 75 \| 53 \| 3 \| 1 \| 73 \| 1 \| 72 \| Helix-turn-helix domain protein \| Helix-turn-helix domain protein; afdb-uniprot50 \| AF-A0A1M5SRH2-F1-MODEL\_V4 \| 1.0 \| 0.005326 \| 139 \| 0.24 \| 75 \| 53 \| 2 \| 1 \| 73 \| 1 \| 73 \| Helix-turn-helix domain-containing protein \| Helix-turn-helix domain-containing protein; afdb-uniprot50 \| AF-A0A4Q2RK28-F1-MODEL\_V4 \| 1.0 \| 0.008485 \| 139 \| 0.223 \| 67 \| 50 \| 1 \| 1 \| 67 \| 5 \| 69 \| DNA-binding protein \| DNA-binding protein; afdb-uniprot50 \| AF-A0A200HEW4-F1-MODEL\_V4 \| 1.0 \| 0.004662 \| 139 \| 0.219 \| 73 \| 54 \| 2 \| 1 \| 73 \| 4 \| 73 \| HTH\_17 domain-containing protein \| HTH\_17 domain-containing protein; afdb-uniprot50 \| AF-A0A2W6C6E5-F1-MODEL\_V4 \| 1.0 \| 0.004662 \| 139 \| 0.303 \| 79 \| 46 \| 2 \| 1 \| 72 \| 1 \| 77 \| DNA-binding protein \| DNA-binding protein; afdb-uniprot50 \| AF-A0A1I1ZX04-F1-MODEL\_V4 \| 1.0 \| 0.003819 \| 139 \| 0.191 \| 73 \| 52 \| 2 \| 1 \| 73 \| 2 \| 67 \| DNA binding domain-containing protein, excisionase family \| DNA binding domain-containing protein, excisionase family; afdb-uniprot50 \| AF-A0A1H9I537-F1-MODEL\_V4 \| 1.0 \| 0.003573 \| 139 \| 0.228 \| 70 \| 50 \| 3 \| 1 \| 69 \| 10 \| 76 \| Helix-turn-helix domain-containing protein \| Helix-turn-helix domain-containing protein; afdb-uniprot50 \| AF-A0A7Z0J817-F1-MODEL\_V4 \| 1.0 \| 0.003819 \| 139 \| 0.26 \| 73 \| 50 \| 3 \| 1 \| 73 \| 11 \| 79 \| Putative DNA-binding transcriptional regulator AlpA \| Putative DNA-binding transcriptional regulator AlpA; afdb-uniprot50 \| AF-A0A430F7K9-F1-MODEL\_V4 \| 1.0 \| 0.005692 \| 139 \| 0.211 \| 71 \| 51 \| 3 \| 4 \| 71 \| 2 \| 70 \| HTH\_17 domain-containing protein \| HTH\_17 domain-containing protein; afdb-uniprot50 \| AF-A0A1I0Z939-F1-MODEL\_V4 \| 1.0 \| 0.003573 \| 139 \| 0.25 \| 68 \| 45 \| 3 \| 2 \| 66 \| 20 \| 84 \| Helix-turn-helix domain-containing protein \| Helix-turn-helix domain-containing protein; afdb-uniprot50 \| AF-A0A4U7MSR7-F1-MODEL\_V4 \| 1.0 \| 0.007939 \| 139 \| 0.205 \| 73 \| 55 \| 2 \| 2 \| 73 \| 13 \| 83 \| Helix-turn-helix domain-containing protein \| Helix-turn-helix domain-containing protein; afdb-uniprot50 \| AF-A0A5M9ZFW5-F1-MODEL\_V4 \| 1.0 \| 0.006502 \| 139 \| 0.206 \| 63 \| 48 \| 1 \| 1 \| 63 \| 25 \| 85 \| DNA-binding protein \| DNA-binding protein; afdb-uniprot50 \| AF-A0A4R1K3A1-F1-MODEL\_V4 \| 1.0 \| 0.005692 \| 139 \| 0.238 \| 67 \| 47 \| 2 \| 6 \| 72 \| 42 \| 104 \| Uncharacterized protein \| Uncharacterized protein; afdb-uniprot50 \| AF-A0A1I1T9U6-F1-MODEL\_V4 \| 1.0 \| 0.009692 \| 139 \| 0.257 \| 66 \| 46 \| 2 \| 2 \| 67 \| 41 \| 103 \| Helix-turn-helix domain-containing protein \| Helix-turn-helix domain-containing protein; afdb-uniprot50 \| AF-A0A544Y1F3-F1-MODEL\_V4 \| 1.0 \| 0.00695 \| 138 \| 0.253 \| 63 \| 43 \| 3 \| 2 \| 62 \| 4 \| 64 \| Helix-turn-helix domain-containing protein \| Helix-turn-helix domain-containing protein; afdb-uniprot50 \| AF-A0A7W3U6U0-F1-MODEL\_V4 \| 1.0 \| 0.006502 \| 138 \| 0.278 \| 61 \| 41 \| 2 \| 4 \| 63 \| 1 \| 59 \| Helix-turn-helix domain-containing protein \| Helix-turn-helix domain-containing protein; afdb-uniprot50 \| AF-A0A4Y3WPK9-F1-MODEL\_V4 \| 1.0 \| 0.005692 \| 138 \| 0.238 \| 67 \| 45 \| 2 \| 11 \| 73 \| 5 \| 69 \| HTH\_17 domain-containing protein \| HTH\_17 domain-containing protein; afdb-uniprot50 \| AF-A0A081HUQ7-F1-MODEL\_V4 \| 1.0 \| 0.005692 \| 138 \| 0.235 \| 68 \| 45 \| 2 \| 11 \| 73 \| 4 \| 69 \| HTH\_17 domain-containing protein \| HTH\_17 domain-containing protein; afdb-uniprot50 \| AF-A0A2S5U1P4-F1-MODEL\_V4 \| 1.0 \| 0.001505 \| 138 \| 0.243 \| 74 \| 50 \| 3 \| 1 \| 73 \| 1 \| 69 \| Excisionase \| Excisionase; afdb-uniprot50 \| AF-H6RJB0-F1-MODEL\_V4 \| 1.0 \| 0.003819 \| 138 \| 0.26 \| 69 \| 47 \| 3 \| 1 \| 67 \| 1 \| 67 \| Transcriptional regulator, MerR family \| Transcriptional regulator, MerR family; afdb-uniprot50 \| AF-A0A0Q5J0X5-F1-MODEL\_V4 \| 1.0 \| 0.004362 \| 138 \| 0.24 \| 75 \| 50 \| 3 \| 1 \| 73 \| 6 \| 75 \| HTH\_17 domain-containing protein \| HTH\_17 domain-containing protein; afdb-uniprot50 \| AF-A0A7I7MBG1-F1-MODEL\_V4 \| 1.0 \| 0.009692 \| 138 \| 0.223 \| 67 \| 51 \| 1 \| 1 \| 67 \| 8 \| 73 \| HTH\_17 domain-containing protein \| HTH\_17 domain-containing protein; afdb-uniprot50 \| AF-A0A6D1I828-F1-MODEL\_V4 \| 1.0 \| 0.004983 \| 138 \| 0.222 \| 72 \| 54 \| 2 \| 2 \| 72 \| 6 \| 76 \| DNA-binding protein \| DNA-binding protein; afdb-uniprot50 \| AF-A0A238JFU8-F1-MODEL\_V4 \| 1.0 \| 0.007939 \| 138 \| 0.218 \| 64 \| 47 \| 2 \| 1 \| 63 \| 12 \| 73 \| Uncharacterized protein \| Uncharacterized protein; afdb-uniprot50 \| AF-A0A7Y9MMD5-F1-MODEL\_V4 \| 1.0 \| 0.002098 \| 138 \| 0.278 \| 79 \| 49 \| 3 \| 1 \| 73 \| 1 \| 77 \| Putative DNA-binding transcriptional regulator AlpA \| Putative DNA-binding transcriptional regulator AlpA; afdb-uniprot50 \| AF-E9USC6-F1-MODEL\_V4 \| 1.0 \| 0.003573 \| 138 \| 0.246 \| 73 \| 51 \| 3 \| 1 \| 72 \| 12 \| 81 \| Conserved domain protein \| Conserved domain protein; afdb-uniprot50 \| AF-A0A1V0KL81-F1-MODEL\_V4 \| 1.0 \| 0.004983 \| 138 \| 0.253 \| 75 \| 52 \| 2 \| 1 \| 73 \| 5 \| 77 \| HTH\_17 domain-containing protein \| HTH\_17 domain-containing protein; afdb-uniprot50 \| AF-A0A7Y9J849-F1-MODEL\_V4 \| 1.0 \| 0.002738 \| 138 \| 0.194 \| 72 \| 52 \| 2 \| 2 \| 73 \| 18 \| 83 \| Putative DNA-binding transcriptional regulator AlpA \| Putative DNA-binding transcriptional regulator AlpA; afdb-uniprot50 \| AF-A0A0F2QH64-F1-MODEL\_V4 \| 1.0 \| 0.007428 \| 138 \| 0.27 \| 74 \| 48 \| 3 \| 2 \| 73 \| 21 \| 90 \| HTH\_17 domain-containing protein \| HTH\_17 domain-containing protein; afdb-uniprot50 \| AF-A0A2N5CW05-F1-MODEL\_V4 \| 1.0 \| 0.005326 \| 138 \| 0.26 \| 73 \| 50 \| 4 \| 1 \| 73 \| 11 \| 79 \| DNA-binding protein \| DNA-binding protein; afdb-uniprot50 \| AF-A0A285KSA3-F1-MODEL\_V4 \| 1.0 \| 0.002243 \| 138 \| 0.266 \| 75 \| 50 \| 2 \| 1 \| 72 \| 24 \| 96 \| DNA binding domain-containing protein, excisionase family \| DNA binding domain-containing protein, excisionase family; afdb-uniprot50 \| AF-A0A850Q143-F1-MODEL\_V4 \| 1.0 \| 0.004362 \| 137 \| 0.26 \| 73 \| 46 \| 3 \| 1 \| 73 \| 1 \| 65 \| Helix-turn-helix domain-containing protein \| Helix-turn-helix domain-containing protein; afdb-uniprot50 \| AF-A0A1G7GIV7-F1-MODEL\_V4 \| 1.0 \| 0.009692 \| 137 \| 0.211 \| 71 \| 53 \| 2 \| 1 \| 71 \| 1 \| 68 \| Uncharacterized protein \| Uncharacterized protein; afdb-uniprot50 \| AF-A0A7T8MJK6-F1-MODEL\_V4 \| 1.0 \| 0.005326 \| 137 \| 0.256 \| 74 \| 52 \| 2 \| 1 \| 73 \| 1 \| 72 \| Helix-turn-helix domain-containing protein \| Helix-turn-helix domain-containing protein; afdb-uniprot50 \| AF-A0A101VLE6-F1-MODEL\_V4 \| 1.0 \| 0.006502 \| 137 \| 0.219 \| 73 \| 53 \| 2 \| 1 \| 73 \| 1 \| 69 \| HTH\_17 domain-containing protein \| HTH\_17 domain-containing protein; afdb-uniprot50 \| AF-A0A1H0JZA4-F1-MODEL\_V4 \| 1.0 \| 0.005692 \| 137 \| 0.219 \| 73 \| 52 \| 3 \| 1 \| 73 \| 6 \| 73 \| DNA binding domain-containing protein, excisionase family \| DNA binding domain-containing protein, excisionase family; afdb-uniprot50 \| AF-A0A543C838-F1-MODEL\_V4 \| 1.0 \| 0.004983 \| 137 \| 0.246 \| 73 \| 49 \| 3 \| 1 \| 73 \| 2 \| 68 \| Helix-turn-helix protein \| Helix-turn-helix protein; afdb-uniprot50 \| AF-A0A3G2N4L3-F1-MODEL\_V4 \| 1.0 \| 0.003128 \| 137 \| 0.266 \| 75 \| 51 \| 3 \| 1 \| 73 \| 1 \| 73 \| DNA-binding protein \| DNA-binding protein; afdb-uniprot50 \| AF-A0A101AJT1-F1-MODEL\_V4 \| 1.0 \| 0.008485 \| 137 \| 0.215 \| 65 \| 47 \| 2 \| 1 \| 63 \| 8 \| 70 \| HTH\_17 domain-containing protein \| HTH\_17 domain-containing protein; afdb-uniprot50 \| AF-A0A846MX55-F1-MODEL\_V4 \| 1.0 \| 0.004662 \| 137 \| 0.207 \| 77 \| 45 \| 2 \| 1 \| 63 \| 1 \| 75 \| Uncharacterized protein \| Uncharacterized protein; afdb-uniprot50 \| AF-A0A6N7ZMY0-F1-MODEL\_V4 \| 1.0 \| 0.004983 \| 137 \| 0.25 \| 76 \| 50 \| 2 \| 2 \| 72 \| 5 \| 78 \| Helix-turn-helix domain-containing protein \| Helix-turn-helix domain-containing protein; afdb-uniprot50 \| AF-A0A0N1NIG7-F1-MODEL\_V4 \| 1.0 \| 0.004983 \| 137 \| 0.278 \| 79 \| 43 \| 2 \| 1 \| 67 \| 1 \| 77 \| HTH\_17 domain-containing protein \| HTH\_17 domain-containing protein; afdb-uniprot50 \| AF-A0A2S9F4K8-F1-MODEL\_V4 \| 1.0 \| 0.007939 \| 137 \| 0.268 \| 67 \| 45 \| 3 \| 1 \| 66 \| 16 \| 79 \| DNA-binding protein \| DNA-binding protein; afdb-uniprot50 \| AF-A0A7Z0BSY9-F1-MODEL\_V4 \| 1.0 \| 0.007939 \| 137 \| 0.226 \| 75 \| 54 \| 2 \| 1 \| 73 \| 10 \| 82 \| Putative DNA-binding transcriptional regulator AlpA \| Putative DNA-binding transcriptional regulator AlpA; afdb-uniprot50 \| AF-A0A1G7SMF3-F1-MODEL\_V4 \| 1.0 \| 0.009068 \| 137 \| 0.178 \| 73 \| 57 \| 2 \| 1 \| 72 \| 9 \| 79 \| Uncharacterized protein \| Uncharacterized protein; afdb-uniprot50 \| AF-A0A7W7ZCS8-F1-MODEL\_V4 \| 1.0 \| 0.006084 \| 137 \| 0.232 \| 73 \| 53 \| 3 \| 1 \| 72 \| 4 \| 74 \| Putative DNA-binding transcriptional regulator AlpA \| Putative DNA-binding transcriptional regulator AlpA; afdb-uniprot50 \| AF-A0A2P2C867-F1-MODEL\_V4 \| 1.0 \| 0.003343 \| 137 \| 0.185 \| 81 \| 55 \| 3 \| 1 \| 73 \| 4 \| 81 \| Phage transcriptional regulator, AlpA \| Phage transcriptional regulator, AlpA; afdb-uniprot50 \| AF-A0A0Q4WSX8-F1-MODEL\_V4 \| 1.0 \| 0.006502 \| 137 \| 0.24 \| 79 \| 52 \| 2 \| 3 \| 73 \| 17 \| 95 \| Uncharacterized protein \| Uncharacterized protein; afdb-uniprot50 \| AF-A0A148N521-F1-MODEL\_V4 \| 1.0 \| 0.004662 \| 136 \| 0.279 \| 68 \| 45 \| 3 \| 7 \| 73 \| 2 \| 66 \| Excisionase \| Excisionase; afdb-uniprot50 \| AF-A0A158A0S3-F1-MODEL\_V4 \| 1.0 \| 0.009068 \| 136 \| 0.23 \| 65 \| 48 \| 1 \| 9 \| 73 \| 2 \| 64 \| Uncharacterized protein \| Uncharacterized protein; afdb-uniprot50 \| AF-A0A2P6WDG2-F1-MODEL\_V4 \| 1.0 \| 0.00695 \| 136 \| 0.194 \| 67 \| 51 \| 2 \| 7 \| 73 \| 3 \| 66 \| HTH\_17 domain-containing protein \| HTH\_17 domain-containing protein; afdb-uniprot50 \| AF-A0A2G2QSP1-F1-MODEL\_V4 \| 1.0 \| 0.008485 \| 136 \| 0.257 \| 66 \| 46 \| 2 \| 11 \| 73 \| 3 \| 68 \| Excisionase \| Excisionase; afdb-uniprot50 \| AF-A0A428YP86-F1-MODEL\_V4 \| 1.0 \| 0.005326 \| 136 \| 0.267 \| 71 \| 48 \| 2 \| 1 \| 69 \| 1 \| 69 \| HTH\_17 domain-containing protein \| HTH\_17 domain-containing protein; afdb-uniprot50 \| AF-A0A3S3TAN4-F1-MODEL\_V4 \| 1.0 \| 0.009692 \| 136 \| 0.274 \| 62 \| 42 \| 2 \| 2 \| 62 \| 10 \| 69 \| DNA-binding protein \| DNA-binding protein; afdb-uniprot50 \| AF-A0A1J0S147-F1-MODEL\_V4 \| 1.0 \| 0.003573 \| 136 \| 0.352 \| 71 \| 41 \| 3 \| 1 \| 70 \| 1 \| 67 \| DNA-binding protein \| DNA-binding protein; afdb-uniprot50 \| AF-A0A165EL06-F1-MODEL\_V4 \| 1.0 \| 0.003573 \| 136 \| 0.25 \| 76 \| 51 \| 4 \| 1 \| 73 \| 1 \| 73 \| HTH\_17 domain-containing protein \| HTH\_17 domain-containing protein; afdb-uniprot50 \| AF-A0A327KQE5-F1-MODEL\_V4 \| 1.0 \| 0.003128 \| 136 \| 0.24 \| 79 \| 52 \| 3 \| 1 \| 73 \| 1 \| 77 \| HTH\_17 domain-containing protein \| HTH\_17 domain-containing protein; afdb-uniprot50 \| AF-A0A841L8L4-F1-MODEL\_V4 \| 1.0 \| 0.001719 \| 136 \| 0.24 \| 75 \| 53 \| 3 \| 1 \| 73 \| 6 \| 78 \| Uncharacterized protein \| Uncharacterized protein; afdb-uniprot50 \| AF-A0A0L0LTJ3-F1-MODEL\_V4 \| 1.0 \| 0.004362 \| 136 \| 0.223 \| 85 \| 51 \| 3 \| 1 \| 73 \| 1 \| 82 \| Uncharacterized protein \| Uncharacterized protein; afdb-uniprot50 \| AF-A0A1Q5AXY7-F1-MODEL\_V4 \| 1.0 \| 0.007428 \| 136 \| 0.208 \| 67 \| 50 \| 2 \| 1 \| 67 \| 15 \| 78 \| DNA-binding protein \| DNA-binding protein; afdb-uniprot50 \| AF-A0A2P6WDT5-F1-MODEL\_V4 \| 1.0 \| 0.004362 \| 136 \| 0.272 \| 77 \| 49 \| 3 \| 1 \| 73 \| 8 \| 81 \| DNA-binding protein \| DNA-binding protein; afdb-uniprot50 \| AF-A0A229VYW8-F1-MODEL\_V4 \| 1.0 \| 0.002243 \| 136 \| 0.252 \| 87 \| 48 \| 4 \| 1 \| 72 \| 1 \| 85 \| Phage transcriptional regulator AlpA \| Phage transcriptional regulator AlpA; afdb-uniprot50 \| AF-A0A259MIW6-F1-MODEL\_V4 \| 1.0 \| 0.004362 \| 136 \| 0.253 \| 71 \| 49 \| 1 \| 1 \| 67 \| 29 \| 99 \| Uncharacterized protein \| Uncharacterized protein; afdb-uniprot50 \| AF-A0A2T6M167-F1-MODEL\_V4 \| 1.0 \| 0.005692 \| 136 \| 0.298 \| 67 \| 43 \| 2 \| 2 \| 66 \| 89 \| 153 \| Uncharacterized protein \| Uncharacterized protein; afdb-uniprot50 \| AF-A0A6S7F7W2-F1-MODEL\_V4 \| 1.0 \| 0.002562 \| 136 \| 0.253 \| 75 \| 52 \| 3 \| 1 \| 73 \| 63 \| 135 \| HTH\_17 domain-containing protein \| HTH\_17 domain-containing protein; afdb-uniprot50 \| AF-A0A7V9RN05-F1-MODEL\_V4 \| 1.0 \| 0.007939 \| 135 \| 0.235 \| 68 \| 47 \| 2 \| 6 \| 73 \| 2 \| 64 \| Helix-turn-helix domain-containing protein \| Helix-turn-helix domain-containing protein; afdb-uniprot50 \| AF-A0A7V1HES1-F1-MODEL\_V4 \| 1.0 \| 0.003819 \| 135 \| 0.239 \| 71 \| 51 \| 2 \| 4 \| 73 \| 1 \| 69 \| DNA-binding protein \| DNA-binding protein; afdb-uniprot50 \| AF-A0A418KYP8-F1-MODEL\_V4 \| 1.0 \| 0.008485 \| 135 \| 0.257 \| 66 \| 44 \| 2 \| 1 \| 63 \| 5 \| 68 \| Uncharacterized protein \| Uncharacterized protein; afdb-uniprot50 \| AF-A0A066PHX5-F1-MODEL\_V4 \| 1.0 \| 0.004362 \| 135 \| 0.225 \| 71 \| 52 \| 2 \| 4 \| 73 \| 1 \| 69 \| Helix-turn-helix domain-containing protein \| Helix-turn-helix domain-containing protein; afdb-uniprot50 \| AF-A0A2J0TSF3-F1-MODEL\_V4 \| 1.0 \| 0.004983 \| 135 \| 0.26 \| 73 \| 48 \| 3 \| 1 \| 73 \| 8 \| 74 \| HTH\_17 domain-containing protein \| HTH\_17 domain-containing protein; afdb-uniprot50 \| AF-A0A498B7D3-F1-MODEL\_V4 \| 1.0 \| 0.005326 \| 135 \| 0.298 \| 67 \| 44 \| 2 \| 2 \| 67 \| 12 \| 76 \| Excisionase family DNA binding protein \| Excisionase family DNA binding protein; afdb-uniprot50 \| AF-A0A7V9PX81-F1-MODEL\_V4 \| 1.0 \| 0.009692 \| 135 \| 0.253 \| 79 \| 51 \| 2 \| 1 \| 73 \| 4 \| 80 \| Helix-turn-helix domain-containing protein \| Helix-turn-helix domain-containing protein; afdb-uniprot50 \| AF-A0A2E2MPT9-F1-MODEL\_V4 \| 1.0 \| 0.004983 \| 135 \| 0.243 \| 74 \| 50 \| 4 \| 1 \| 72 \| 6 \| 75 \| DNA-binding protein \| DNA-binding protein; afdb-uniprot50 \| AF-A0A3S0LBJ7-F1-MODEL\_V4 \| 1.0 \| 0.002927 \| 135 \| 0.263 \| 76 \| 50 \| 3 \| 2 \| 73 \| 17 \| 90 \| DNA-binding protein \| DNA-binding protein; afdb-uniprot50 \| AF-A0A844ZUN2-F1-MODEL\_V4 \| 1.0 \| 0.00695 \| 135 \| 0.273 \| 73 \| 51 \| 1 \| 1 \| 73 \| 24 \| 94 \| Uncharacterized protein \| Uncharacterized protein; afdb-uniprot50 \| AF-A0A6G6Z347-F1-MODEL\_V4 \| 1.0 \| 0.005692 \| 135 \| 0.202 \| 74 \| 56 \| 3 \| 1 \| 73 \| 3 \| 74 \| Helix-turn-helix domain-containing protein \| Helix-turn-helix domain-containing protein; afdb-uniprot50 \| AF-A0A6H9WTP2-F1-MODEL\_V4 \| 1.0 \| 0.008485 \| 135 \| 0.268 \| 67 \| 47 \| 1 \| 1 \| 67 \| 40 \| 104 \| Helix-turn-helix domain-containing protein \| Helix-turn-helix domain-containing protein; afdb-uniprot50 \| AF-A0A7W6IQJ9-F1-MODEL\_V4 \| 1.0 \| 0.00695 \| 135 \| 0.246 \| 69 \| 46 \| 2 \| 1 \| 69 \| 1 \| 63 \| Putative DNA-binding transcriptional regulator AlpA \| Putative DNA-binding transcriptional regulator AlpA; afdb-uniprot50 \| AF-A0A2N2KGC4-F1-MODEL\_V4 \| 1.0 \| 0.004983 \| 135 \| 0.236 \| 76 \| 51 \| 2 \| 2 \| 72 \| 59 \| 132 \| Uncharacterized protein \| Uncharacterized protein; afdb-uniprot50 \| AF-A0A401Z9H7-F1-MODEL\_V4 \| 1.0 \| 0.005692 \| 134 \| 0.2 \| 75 \| 51 \| 3 \| 1 \| 73 \| 1 \| 68 \| HTH\_17 domain-containing protein \| HTH\_17 domain-containing protein; afdb-uniprot50 \| AF-A0A1H9VIH3-F1-MODEL\_V4 \| 1.0 \| 0.009068 \| 134 \| 0.26 \| 69 \| 49 \| 2 \| 1 \| 69 \| 1 \| 67 \| Prophage CP4-57 regulatory protein (AlpA) \| Prophage CP4-57 regulatory protein (AlpA); afdb-uniprot50 \| AF-A0A512DG42-F1-MODEL\_V4 \| 1.0 \| 0.007428 \| 134 \| 0.231 \| 69 \| 48 \| 3 \| 1 \| 67 \| 1 \| 66 \| HTH\_17 domain-containing protein \| HTH\_17 domain-containing protein; afdb-uniprot50 \| AF-A0A0M2HKS0-F1-MODEL\_V4 \| 1.0 \| 0.005326 \| 134 \| 0.26 \| 73 \| 51 \| 2 \| 1 \| 71 \| 3 \| 74 \| Uncharacterized protein \| Uncharacterized protein; afdb-uniprot50 \| AF-A0A525HJG6-F1-MODEL\_V4 \| 1.0 \| 0.004662 \| 134 \| 0.287 \| 73 \| 48 \| 3 \| 3 \| 73 \| 2 \| 72 \| DNA-binding protein \| DNA-binding protein; afdb-uniprot50 \| AF-A0A1A2DCW1-F1-MODEL\_V4 \| 1.0 \| 0.005326 \| 134 \| 0.226 \| 75 \| 51 \| 3 \| 1 \| 70 \| 1 \| 73 \| Uncharacterized protein \| Uncharacterized protein; afdb-uniprot50 \| AF-A0A1N7FDQ6-F1-MODEL\_V4 \| 1.0 \| 0.004983 \| 134 \| 0.231 \| 69 \| 45 \| 2 \| 1 \| 63 \| 3 \| 69 \| Uncharacterized protein \| Uncharacterized protein; afdb-uniprot50 \| AF-A0A1F2UUZ3-F1-MODEL\_V4 \| 1.0 \| 0.004983 \| 134 \| 0.282 \| 78 \| 47 \| 3 \| 1 \| 73 \| 1 \| 74 \| HTH\_17 domain-containing protein \| HTH\_17 domain-containing protein; afdb-uniprot50 \| AF-A0A1S6ERV9-F1-MODEL\_V4 \| 1.0 \| 0.008485 \| 134 \| 0.219 \| 73 \| 53 \| 2 \| 2 \| 72 \| 8 \| 78 \| HTH\_17 domain-containing protein \| HTH\_17 domain-containing protein; afdb-uniprot50 \| AF-A0A8A3LV53-F1-MODEL\_V4 \| 1.0 \| 0.005326 \| 134 \| 0.21 \| 76 \| 57 \| 2 \| 1 \| 73 \| 5 \| 80 \| Helix-turn-helix domain-containing protein \| Helix-turn-helix domain-containing protein; afdb-uniprot50 \| AF-A0A6B8MDK0-F1-MODEL\_V4 \| 1.0 \| 0.003573 \| 134 \| 0.27 \| 74 \| 50 \| 3 \| 1 \| 72 \| 10 \| 81 \| Helix-turn-helix domain-containing protein \| Helix-turn-helix domain-containing protein; afdb-uniprot50 \| AF-A0A1W6ZN77-F1-MODEL\_V4 \| 1.0 \| 0.004983 \| 134 \| 0.205 \| 78 \| 54 \| 2 \| 2 \| 73 \| 4 \| 79 \| HTH\_17 domain-containing protein \| HTH\_17 domain-containing protein; afdb-uniprot50 \| AF-A0A382CIH3-F1-MODEL\_V4 \| 1.0 \| 0.004662 \| 134 \| 0.213 \| 75 \| 54 \| 3 \| 1 \| 73 \| 4 \| 75 \| HTH\_17 domain-containing protein \| HTH\_17 domain-containing protein; afdb-uniprot50 \| AF-A0A7Y7M2F2-F1-MODEL\_V4 \| 1.0 \| 0.004662 \| 134 \| 0.223 \| 76 \| 54 \| 3 \| 1 \| 73 \| 6 \| 79 \| Helix-turn-helix domain-containing protein \| Helix-turn-helix domain-containing protein; afdb-uniprot50 \| AF-H6RRY8-F1-MODEL\_V4 \| 1.0 \| 0.00695 \| 134 \| 0.303 \| 66 \| 43 \| 2 \| 1 \| 65 \| 17 \| 80 \| Transcriptional regulator, MerR family \| Transcriptional regulator, MerR family; afdb-uniprot50 \| AF-A0A7I7UXQ3-F1-MODEL\_V4 \| 1.0 \| 0.006502 \| 134 \| 0.205 \| 73 \| 55 \| 3 \| 1 \| 73 \| 21 \| 90 \| HTH\_17 domain-containing protein \| HTH\_17 domain-containing protein; afdb-uniprot50 \| AF-A0A2P7ACK4-F1-MODEL\_V4 \| 1.0 \| 0.005326 \| 133 \| 0.202 \| 79 \| 53 \| 3 \| 1 \| 73 \| 1 \| 75 \| DNA-binding protein \| DNA-binding protein; afdb-uniprot50 \| AF-A0A6V8MGW9-F1-MODEL\_V4 \| 1.0 \| 0.006084 \| 133 \| 0.26 \| 73 \| 51 \| 2 \| 1 \| 73 \| 7 \| 76 \| HTH\_17 domain-containing protein \| HTH\_17 domain-containing protein; afdb-uniprot50 \| AF-A0A354W361-F1-MODEL\_V4 \| 1.0 \| 0.007428 \| 133 \| 0.212 \| 66 \| 47 \| 3 \| 1 \| 63 \| 2 \| 65 \| DNA-binding protein \| DNA-binding protein; afdb-uniprot50 \| AF-A0A178HUC9-F1-MODEL\_V4 \| 1.0 \| 0.005326 \| 133 \| 0.285 \| 77 \| 47 \| 3 \| 1 \| 73 \| 8 \| 80 \| HTH\_17 domain-containing protein \| HTH\_17 domain-containing protein; afdb-uniprot50 \| AF-A0A3A0FTT9-F1-MODEL\_V4 \| 1.0 \| 0.006502 \| 133 \| 0.236 \| 72 \| 49 \| 2 \| 2 \| 73 \| 16 \| 81 \| DNA-binding protein \| DNA-binding protein; afdb-uniprot50 \| AF-A0A6L9W7M6-F1-MODEL\_V4 \| 1.0 \| 0.009692 \| 133 \| 0.205 \| 73 \| 56 \| 1 \| 1 \| 73 \| 2 \| 72 \| Helix-turn-helix domain-containing protein \| Helix-turn-helix domain-containing protein; afdb-uniprot50 \| AF-A0A7Y7LZW5-F1-MODEL\_V4 \| 1.0 \| 0.005692 \| 133 \| 0.205 \| 73 \| 53 \| 2 \| 1 \| 73 \| 7 \| 74 \| Helix-turn-helix domain-containing protein \| Helix-turn-helix domain-containing protein; afdb-uniprot50 \| AF-A0A6V8MQ78-F1-MODEL\_V4 \| 1.0 \| 0.004362 \| 133 \| 0.246 \| 77 \| 50 \| 3 \| 2 \| 72 \| 12 \| 86 \| HTH\_17 domain-containing protein \| HTH\_17 domain-containing protein; afdb-uniprot50 \| AF-A0A4R0Z385-F1-MODEL\_V4 \| 1.0 \| 0.009068 \| 133 \| 0.25 \| 72 \| 52 \| 1 \| 2 \| 73 \| 29 \| 98 \| DNA-binding protein \| DNA-binding protein; afdb-uniprot50 \| AF-A0A3M0HKQ5-F1-MODEL\_V4 \| 1.0 \| 0.009068 \| 133 \| 0.223 \| 67 \| 47 \| 2 \| 1 \| 67 \| 46 \| 107 \| AlpA family phage regulatory protein \| AlpA family phage regulatory protein; afdb-uniprot50 \| AF-A0A0U5BBK8-F1-MODEL\_V4 \| 1.0 \| 0.005692 \| 133 \| 0.216 \| 74 \| 53 \| 3 \| 2 \| 72 \| 41 \| 112 \| Phage transcriptional regulator, AlpA \| Phage transcriptional regulator, AlpA; afdb-uniprot50 \| AF-A0A7Y4MFA2-F1-MODEL\_V4 \| 1.0 \| 0.008485 \| 133 \| 0.238 \| 67 \| 48 \| 2 \| 2 \| 67 \| 5 \| 69 \| Helix-turn-helix domain-containing protein \| Helix-turn-helix domain-containing protein; afdb-uniprot50 \| AF-A0A1Q8KWK1-F1-MODEL\_V4 \| 1.0 \| 0.003573 \| 132 \| 0.275 \| 69 \| 42 \| 3 \| 1 \| 69 \| 1 \| 61 \| HTH\_17 domain-containing protein \| HTH\_17 domain-containing protein; afdb-uniprot50 \| AF-A0A7Y6NP98-F1-MODEL\_V4 \| 1.0 \| 0.007428 \| 132 \| 0.246 \| 73 \| 51 \| 3 \| 1 \| 72 \| 1 \| 70 \| Helix-turn-helix domain-containing protein \| Helix-turn-helix domain-containing protein; afdb-uniprot50 \| AF-A0A7Y0VX78-F1-MODEL\_V4 \| 1.0 \| 0.006084 \| 132 \| 0.216 \| 74 \| 52 \| 3 \| 1 \| 73 \| 3 \| 71 \| Helix-turn-helix domain-containing protein \| Helix-turn-helix domain-containing protein; afdb-uniprot50 \| AF-A0A2V5IYM0-F1-MODEL\_V4 \| 1.0 \| 0.009692 \| 132 \| 0.205 \| 68 \| 52 \| 1 \| 1 \| 68 \| 8 \| 73 \| Excisionase \| Excisionase; afdb-uniprot50 \| AF-A0A7G9L3X4-F1-MODEL\_V4 \| 1.0 \| 0.009692 \| 132 \| 0.2 \| 65 \| 46 \| 2 \| 9 \| 73 \| 15 \| 73 \| Helix-turn-helix domain-containing protein \| Helix-turn-helix domain-containing protein; afdb-uniprot50 \| AF-A0A2T0TPB9-F1-MODEL\_V4 \| 1.0 \| 0.00695 \| 132 \| 0.192 \| 78 \| 55 \| 2 \| 1 \| 72 \| 2 \| 77 \| Excisionase family DNA binding protein \| Excisionase family DNA binding protein; afdb-uniprot50 \| AF-A0A1W1D8J7-F1-MODEL\_V4 \| 1.0 \| 0.003573 \| 132 \| 0.293 \| 75 \| 47 \| 3 \| 1 \| 73 \| 5 \| 75 \| HTH\_17 domain-containing protein \| HTH\_17 domain-containing protein; afdb-uniprot50 \| AF-A0A7L5Z0X3-F1-MODEL\_V4 \| 1.0 \| 0.007939 \| 132 \| 0.232 \| 73 \| 53 \| 2 \| 2 \| 73 \| 5 \| 75 \| Helix-turn-helix domain-containing protein \| Helix-turn-helix domain-containing protein; afdb-uniprot50 \| AF-A0A2V8IKF4-F1-MODEL\_V4 \| 1.0 \| 0.002562 \| 132 \| 0.209 \| 86 \| 53 \| 3 \| 1 \| 73 \| 38 \| 121 \| HTH\_17 domain-containing protein \| HTH\_17 domain-containing protein; afdb-uniprot50 \| AF-A0A2E5FA21-F1-MODEL\_V4 \| 1.0 \| 0.006084 \| 131 \| 0.246 \| 73 \| 51 \| 3 \| 1 \| 72 \| 7 \| 76 \| Uncharacterized protein \| Uncharacterized protein; afdb-uniprot50 \| AF-A0A365H3H9-F1-MODEL\_V4 \| 1.0 \| 0.006502 \| 131 \| 0.291 \| 72 \| 48 \| 2 \| 1 \| 72 \| 10 \| 78 \| DNA-binding protein \| DNA-binding protein; afdb-uniprot50 \| AF-A0A1T4K0K2-F1-MODEL\_V4 \| 1.0 \| 0.004362 \| 131 \| 0.259 \| 77 \| 51 \| 3 \| 1 \| 73 \| 7 \| 81 \| Transcriptional regulator, AlpA family \| Transcriptional regulator, AlpA family; afdb-uniprot50 \| AF-A0A7Z0GLK2-F1-MODEL\_V4 \| 1.0 \| 0.00695 \| 131 \| 0.285 \| 77 \| 49 \| 5 \| 1 \| 73 \| 9 \| 83 \| Uncharacterized protein \| Uncharacterized protein; afdb-uniprot50 \| AF-A0A3S4WXW0-F1-MODEL\_V4 \| 1.0 \| 0.003819 \| 131 \| 0.294 \| 78 \| 48 \| 3 \| 1 \| 73 \| 1 \| 76 \| Predicted transcriptional regulator \| Predicted transcriptional regulator; afdb-uniprot50 \| AF-A0A090MHP8-F1-MODEL\_V4 \| 1.0 \| 0.009068 \| 131 \| 0.18 \| 72 \| 57 \| 1 \| 2 \| 73 \| 27 \| 96 \| Uncultured bacterium genome assembly Metasoil\_fosmids\_resub \| Uncultured bacterium genome assembly Metasoil\_fosmids\_resub; afdb-uniprot50 \| AF-A0A7Z7RKN0-F1-MODEL\_V4 \| 1.0 \| 0.009692 \| 131 \| 0.235 \| 68 \| 46 \| 2 \| 6 \| 73 \| 2 \| 63 \| Predicted transcriptional regulator \| Predicted transcriptional regulator; afdb-uniprot50 \| AF-A0A239JAJ2-F1-MODEL\_V4 \| 1.0 \| 0.007428 \| 130 \| 0.246 \| 69 \| 44 \| 2 \| 11 \| 73 \| 5 \| 71 \| Uncharacterized protein \| Uncharacterized protein; afdb-uniprot50 \| AF-A0A5C4W3D3-F1-MODEL\_V4 \| 1.0 \| 0.009068 \| 130 \| 0.25 \| 64 \| 44 \| 2 \| 1 \| 62 \| 4 \| 65 \| Helix-turn-helix domain-containing protein \| Helix-turn-helix domain-containing protein; afdb-uniprot50 \| AF-A0A7V3NCW0-F1-MODEL\_V4 \| 1.0 \| 0.004983 \| 130 \| 0.227 \| 79 \| 53 \| 3 \| 1 \| 73 \| 51 \| 127 \| DNA-binding protein \| DNA-binding protein; afdb-uniprot50 \| AF-A0A4Q2S2U5-F1-MODEL\_V4 \| 1.0 \| 0.006502 \| 130 \| 0.208 \| 67 \| 45 \| 2 \| 2 \| 62 \| 192 \| 256 \| Helix-turn-helix domain-containing protein \| Helix-turn-helix domain-containing protein; afdb-uniprot50 \| AF-A0A1M6YKR5-F1-MODEL\_V4 \| 1.0 \| 0.008485 \| 129 \| 0.338 \| 68 \| 42 \| 3 \| 7 \| 73 \| 2 \| 67 \| Transcriptional regulator, AlpA family \| Transcriptional regulator, AlpA family; afdb-uniprot50 \| AF-A0A355EX70-F1-MODEL\_V4 \| 1.0 \| 0.006084 \| 129 \| 0.297 \| 74 \| 48 \| 3 \| 1 \| 72 \| 1 \| 72 \| Uncharacterized protein \| Uncharacterized protein; afdb-uniprot50 \| AF-A0A557SM49-F1-MODEL\_V4 \| 1.0 \| 0.009068 \| 129 \| 0.229 \| 74 \| 53 \| 3 \| 1 \| 73 \| 2 \| 72 \| Helix-turn-helix domain-containing protein \| Helix-turn-helix domain-containing protein; afdb-uniprot50 \| AF-A0A0Q7K039-F1-MODEL\_V4 \| 1.0 \| 0.007428 \| 129 \| 0.205 \| 78 \| 53 \| 3 \| 1 \| 72 \| 2 \| 76 \| HTH\_17 domain-containing protein \| HTH\_17 domain-containing protein; afdb-uniprot50 \| AF-I3ZJK5-F1-MODEL\_V4 \| 1.0 \| 0.00695 \| 129 \| 0.297 \| 74 \| 48 \| 2 \| 2 \| 73 \| 7 \| 78 \| HTH\_17 domain-containing protein \| HTH\_17 domain-containing protein; afdb-uniprot50 \| AF-A0A522CQL4-F1-MODEL\_V4 \| 1.0 \| 0.007939 \| 129 \| 0.181 \| 77 \| 59 \| 2 \| 1 \| 73 \| 3 \| 79 \| DNA-binding protein \| DNA-binding protein; afdb-uniprot50 \| AF-A0A3D9YWA9-F1-MODEL\_V4 \| 1.0 \| 0.009692 \| 129 \| 0.253 \| 75 \| 52 \| 3 \| 1 \| 73 \| 13 \| 85 \| Uncharacterized protein \| Uncharacterized protein; afdb-uniprot50 \| AF-A0A3S8T1N0-F1-MODEL\_V4 \| 1.0 \| 0.007428 \| 128 \| 0.253 \| 71 \| 50 \| 2 \| 4 \| 73 \| 3 \| 71 \| DNA-binding protein \| DNA-binding protein; afdb-uniprot50 \| AF-A0A840ARV2-F1-MODEL\_V4 \| 1.0 \| 0.006084 \| 128 \| 0.232 \| 73 \| 52 \| 2 \| 1 \| 73 \| 4 \| 72 \| Excisionase family DNA binding protein \| Excisionase family DNA binding protein; afdb-uniprot50 \| AF-A0A382YY28-F1-MODEL\_V4 \| 1.0 \| 0.004362 \| 128 \| 0.23 \| 78 \| 54 \| 3 \| 1 \| 73 \| 1 \| 77 \| Uncharacterized protein \| Uncharacterized protein; afdb-uniprot50 \| AF-A0A4R0IXT6-F1-MODEL\_V4 \| 1.0 \| 0.008485 \| 128 \| 0.228 \| 70 \| 50 \| 2 \| 4 \| 73 \| 2 \| 67 \| DNA-binding protein \| DNA-binding protein; afdb-uniprot50 \| AF-A0A1M7TPV6-F1-MODEL\_V4 \| 1.0 \| 0.00695 \| 128 \| 0.236 \| 72 \| 52 \| 2 \| 2 \| 72 \| 9 \| 78 \| Uncharacterized protein \| Uncharacterized protein; afdb-uniprot50 \| AF-F6EZQ7-F1-MODEL\_V4 \| 1.0 \| 0.009068 \| 128 \| 0.256 \| 74 \| 50 \| 3 \| 1 \| 73 \| 7 \| 76 \| Uncharacterized protein \| Uncharacterized protein; afdb-uniprot50 \| AF-T0ZSB8-F1-MODEL\_V4 \| 1.0 \| 0.002243 \| 128 \| 0.272 \| 77 \| 46 \| 4 \| 4 \| 72 \| 3 \| 77 \| Uncharacterized protein \| Uncharacterized protein; afdb-uniprot50 \| AF-A0A0K8MEP3-F1-MODEL\_V4 \| 1.0 \| 0.007939 \| 127 \| 0.213 \| 75 \| 54 \| 3 \| 1 \| 72 \| 1 \| 73 \| Helix-turn-helix domain protein \| Helix-turn-helix domain protein; afdb-uniprot50 \| AF-A0A3S0AX38-F1-MODEL\_V4 \| 1.0 \| 0.008485 \| 127 \| 0.298 \| 77 \| 48 \| 2 \| 1 \| 73 \| 6 \| 80 \| DNA-binding protein \| DNA-binding protein; afdb-uniprot50 \| AF-A0A2N2KEI7-F1-MODEL\_V4 \| 1.0 \| 0.009068 \| 127 \| 0.227 \| 79 \| 53 \| 3 \| 1 \| 73 \| 4 \| 80 \| DNA-binding protein \| DNA-binding protein; afdb-uniprot50 \| AF-A0A8A7CSH7-F1-MODEL\_V4 \| 1.0 \| 0.009068 \| 127 \| 0.31 \| 74 \| 46 \| 2 \| 1 \| 70 \| 1 \| 73 \| Uncharacterized protein \| Uncharacterized protein; afdb-uniprot50 \| AF-A0A1M3AAN7-F1-MODEL\_V4 \| 1.0 \| 0.005692 \| 127 \| 0.28 \| 82 \| 47 \| 4 \| 2 \| 73 \| 5 \| 84 \| Uncharacterized protein \| Uncharacterized protein; afdb-uniprot50 \| AF-A0A1H3NPD0-F1-MODEL\_V4 \| 1.0 \| 0.008485 \| 127 \| 0.246 \| 73 \| 51 \| 3 \| 1 \| 71 \| 1 \| 71 \| DNA binding domain-containing protein, excisionase family \| DNA binding domain-containing protein, excisionase family; afdb-uniprot50 \| AF-A0A2W4XEV9-F1-MODEL\_V4 \| 1.0 \| 0.008485 \| 127 \| 0.2 \| 75 \| 57 \| 1 \| 1 \| 72 \| 5 \| 79 \| Uncharacterized protein \| Uncharacterized protein; afdb-uniprot50 \| AF-F4G8U6-F1-MODEL\_V4 \| 1.0 \| 0.007939 \| 127 \| 0.246 \| 69 \| 44 \| 2 \| 11 \| 73 \| 5 \| 71 \| Uncharacterized protein \| Uncharacterized protein; afdb-uniprot50 \| AF-A0A3N0GII4-F1-MODEL\_V4 \| 1.0 \| 0.009692 \| 126 \| 0.21 \| 76 \| 55 \| 3 \| 1 \| 73 \| 1 \| 74 \| DNA-binding protein \| DNA-binding protein; afdb-uniprot50 \| AF-A0A1V3R129-F1-MODEL\_V4 \| 1.0 \| 0.007428 \| 126 \| 0.26 \| 73 \| 48 \| 3 \| 1 \| 73 \| 3 \| 69 \| HTH\_17 domain-containing protein \| HTH\_17 domain-containing protein; afdb-uniprot50 \| AF-A0A5S4WPV7-F1-MODEL\_V4 \| 1.0 \| 0.009692 \| 126 \| 0.259 \| 77 \| 51 \| 3 \| 1 \| 72 \| 1 \| 76 \| Uncharacterized protein \| Uncharacterized protein; afdb-uniprot50 \| AF-A0A1M3DSQ0-F1-MODEL\_V4 \| 1.0 \| 0.007939 \| 126 \| 0.235 \| 89 \| 49 \| 3 \| 1 \| 72 \| 3 \| 89 \| HTH\_17 domain-containing protein \| HTH\_17 domain-containing protein; afdb-uniprot50 \| AF-A0A7H4GTR2-F1-MODEL\_V4 \| 1.0 \| 0.008485 \| 126 \| 0.226 \| 75 \| 51 \| 4 \| 1 \| 73 \| 33 \| 102 \| Helix-turn-helix domain-containing protein \| Helix-turn-helix domain-containing protein; afdb-uniprot50 \| AF-A0A2V8WJS3-F1-MODEL\_V4 \| 1.0 \| 0.006502 \| 126 \| 0.25 \| 76 \| 51 \| 3 \| 1 \| 72 \| 12 \| 85 \| HTH\_17 domain-containing protein \| HTH\_17 domain-containing protein; afdb-uniprot50 \| AF-A0A1I7DBH4-F1-MODEL\_V4 \| 1.0 \| 0.009692 \| 125 \| 0.243 \| 74 \| 52 \| 3 \| 1 \| 72 \| 1 \| 72 \| Transcriptional regulator, AlpA family \| Transcriptional regulator, AlpA family; afdb-uniprot50 \| AF-A0A516NU13-F1-MODEL\_V4 \| 1.0 \| 0.007428 \| 125 \| 0.277 \| 72 \| 44 \| 3 \| 1 \| 72 \| 1 \| 64 \| DNA-binding protein \| DNA-binding protein; afdb-uniprot50 \| AF-A0A2W6VUQ2-F1-MODEL\_V4 \| 1.0 \| 0.009692 \| 125 \| 0.283 \| 74 \| 50 \| 2 \| 1 \| 73 \| 3 \| 74 \| HTH\_17 domain-containing protein \| HTH\_17 domain-containing protein; afdb-uniprot50 \| AF-A0A3S0D818-F1-MODEL\_V4 \| 1.0 \| 0.006502 \| 125 \| 0.236 \| 76 \| 53 \| 3 \| 1 \| 73 \| 5 \| 78 \| DNA-binding protein \| DNA-binding protein; afdb-uniprot50 \| AF-A0A5D4XRR2-F1-MODEL\_V4 \| 1.0 \| 0.00695 \| 125 \| 0.215 \| 79 \| 54 \| 3 \| 1 \| 73 \| 1 \| 77 \| Helix-turn-helix domain-containing protein \| Helix-turn-helix domain-containing protein; afdb-uniprot50 \| AF-A0A2E7UH17-F1-MODEL\_V4 \| 1.0 \| 0.007428 \| 125 \| 0.283 \| 81 \| 48 \| 3 \| 1 \| 73 \| 1 \| 79 \| DNA-binding protein \| DNA-binding protein; afdb-uniprot50 \| AF-A0A3F3GUT7-F1-MODEL\_V4 \| 1.0 \| 0.009692 \| 125 \| 0.213 \| 75 \| 55 \| 3 \| 1 \| 73 \| 2 \| 74 \| Excisionase DNA binding domain-containing protein \| Excisionase DNA binding domain-containing protein; afdb-uniprot50 \| AF-A0A7C9KIR3-F1-MODEL\_V4 \| 1.0 \| 0.007939 \| 124 \| 0.246 \| 77 \| 50 \| 3 \| 1 \| 73 \| 6 \| 78 \| Helix-turn-helix domain-containing protein \| Helix-turn-helix domain-containing protein; afdb-uniprot50 \| AF-A0A2V2R565-F1-MODEL\_V4 \| 1.0 \| 0.009692 \| 124 \| 0.256 \| 82 \| 50 \| 4 \| 1 \| 73 \| 2 \| 81 \| Uncharacterized protein \| Uncharacterized protein; afdb-uniprot50 \| AF-E2S8B6-F1-MODEL\_V4 \| 1.0 \| 0.009692 \| 124 \| 0.226 \| 75 \| 54 \| 2 \| 1 \| 73 \| 18 \| 90 \| HTH\_17 domain-containing protein \| HTH\_17 domain-containing protein; afdb-uniprot50 \| AF-A0A2T5JVL9-F1-MODEL\_V4 \| 1.0 \| 0.009692 \| 123 \| 0.243 \| 74 \| 53 \| 2 \| 1 \| 73 \| 8 \| 79 \| Helix-turn-helix protein \| Helix-turn-helix protein; afdb-uniprot50 \| AF-A0A833LJH9-F1-MODEL\_V4 \| 1.0 \| 0.007939 \| 122 \| 0.232 \| 73 \| 53 \| 2 \| 1 \| 73 \| 6 \| 75 \| Helix-turn-helix domain-containing protein \| Helix-turn-helix domain-containing protein; afdb-uniprot50 \| AF-A0A2E7ISN4-F1-MODEL\_V4 \| 1.0 \| 0.009692 \| 121 \| 0.214 \| 70 \| 51 \| 3 \| 1 \| 68 \| 1 \| 68 \| Transcriptional regulator \| Transcriptional regulator; afdb-uniprot50 \| AF-A0A4R6WVG7-F1-MODEL\_V4 \| 1.0 \| 0.004662 \| 121 \| 0.217 \| 78 \| 54 \| 3 \| 1 \| 73 \| 1 \| 76 \| Uncharacterized protein \| Uncharacterized protein; afdb-uniprot50 \| AF-A0A4P5WYP2-F1-MODEL\_V4 \| 1.0 \| 0.009692 \| 119 \| 0.23 \| 78 \| 51 \| 5 \| 1 \| 73 \| 1 \| 74 \| HTH\_17 domain-containing protein \| HTH\_17 domain-containing protein; afdb-uniprot50 \| AF-F4CSF2-F1-MODEL\_V4 \| 1.0 \| 0.007428 \| 118 \| 0.277 \| 72 \| 41 \| 3 \| 1 \| 68 \| 22 \| 86 \| Prophage CP4-57 regulatory \| Prophage CP4-57 regulatory; afdb-uniprot50 \| AF-A0A1Q4X2L2-F1-MODEL\_V4 \| 1.0 \| 0.003819 \| 118 \| 0.259 \| 77 \| 51 \| 3 \| 1 \| 73 \| 4 \| 78 \| HTH merR-type domain-containing protein \| HTH merR-type domain-containing protein; afdb-uniprot50 \| AF-A0A087CGI1-F1-MODEL\_V4 \| 1.0 \| 0.008485 \| 117 \| 0.25 \| 72 \| 47 \| 3 \| 1 \| 72 \| 3 \| 67 \| Phage transcriptional regulator AlpA \| Phage transcriptional regulator AlpA |
| Top keywords  (threshold 1.00e-02 (evalue)) | **domain\_containing, DNA\_binding, Helix\_turn\_helix, HTH\_17, Excisionase, transcriptional, regulator, DNA, binding, AlpA** |
| Output files | ../../similar\_structures/38\_FANPEZAQ\_CDS\_0038\_afdb-proteome\_foldseek.tsv ../../similar\_structures/38\_FANPEZAQ\_CDS\_0038\_afdb-uniprot50\_foldseek.tsv ../../similar\_structures/38\_FANPEZAQ\_CDS\_0038\_merged.svg ../../similar\_structures/38\_FANPEZAQ\_CDS\_0038\_pdb\_foldseek.tsv |

  
  
  

Return to summary | Go to previous | Go to next

  


---

**Sequence/structure alignments coloring**  
Each object in the alignment figures is colored according to its E-value following this color coding:

1e-100
10
