## Supplementary material for "Completing the BASEL phage collection to unlock hidden diversity for systematic exploration of phage-host interactions": Data S2: 39.html

FANPEZAQ\_CDS\_0039


Return to summary | Go to previous | Go to next

|  |  |
| --- | --- |
| FANPEZAQ\_CDS\_0039  Page creation date: 02 Sep 2024, 12:00  Project folder: n/a  Input sequences file: Escherichia\_virus\_HeidiAbel.gb | kaic domain\_containing aaa atpase dna circadian clock reca kinase repair recombination fragment and atp helicase rad51 profile rada synthase in beta homologous recombinase homolog atp\_binding complex putative nucleotide a signal the replicative alpha containing serine radb adp mg threonine rna metabolism c non\_specific recognition p\_loop triphosphate e primase particle fold |

### Sequence information

|  |  |
| --- | --- |
| Name | FANPEZAQ\_CDS\_0039  39\_FANPEZAQ\_CDS\_0039 (pipeline id) |
| Imported annotations |  |
| Protein sequence | MAVINIRKAEREGARLVIGLAGISGSGKTLSALMLAYGLANGDGTKVGFLDTENRRGSLY ASDDTYDKIQESLGLKERPDAFWIGDLEPPFSPQRYIDAIREFEQAGVEVLVIDSISHEW EGTGGCEEIATLANPMKPQWNRAKAEHKRFMNALLQSNMHIICCIRAREKVKLVKVNGKT EYEPQGIMPVTEKNVMFEMTASLMMWDSGSAQQVMKCPEELRGILGRETGYISAQDGLAL RQWVQGGNKLDPKVEAYRNKLRGVTEQGEQYTQECWDKTPPAIRKKLGDEFKQSLLQAAR AYDEQRANANGDAQDVDDLNSQVMGTGAQ |
| Number of residues | 329 |
| Molecular weight (Da) | 36565.06 |
| Output files | ../../query\_sequences/39\_FANPEZAQ\_CDS\_0039.fasta |

### Putative domain architecture and protein family

#### Search results (HHblits)^1^

|  |  |
| --- | --- |
| Domain family databases searched | Pfam, Ncbi-cd, Cath, Phrogs |
| Results, scheme(s)  (Top layers only; threshold 1.00e-03 (evalue)) | xml version="1.0" encoding="utf-8" standalone="no"?       2024-09-02T21:08:20.907143 image/svg+xml   Matplotlib v3.7.2, https://matplotlib.org/ |
| Results, table  (E-value ≤ 1.00e-03 (evalue)) | db \| id \| prob \| evalue \| pvalue \| score \| cols \| query \| query\_len \| template \| template\_len \| name \| description; pfam \| PF08423 \| 96.2 \| 9.4e-06 \| 2e-09 \| 65.4 \| 146 \| (13, 170) \| 329 \| (35, 189) \| 255 \| Rad51 \| Rad51; pfam \| PF06745 \| 96.1 \| 1.5e-05 \| 3.3e-09 \| 61.5 \| 135 \| (13, 169) \| 329 \| (17, 166) \| 232 \| ATPase \| KaiC; pfam \| PF00154 \| 95.4 \| 8.1e-05 \| 1.7e-08 \| 60.7 \| 141 \| (14, 171) \| 329 \| (51, 192) \| 261 \| RecA \| recA bacterial DNA recombination protein; pfam \| PF09807 \| 94.6 \| 0.00031 \| 6.7e-08 \| 55.7 \| 133 \| (14, 169) \| 329 \| (17, 182) \| 248 \| ELP6 \| Elongation complex protein 6; ncbi-cd \| cd01124 \| 96.6 \| 2.5e-06 \| 6.6e-10 \| 62.9 \| 42 \| (15, 57) \| 329 \| (19, 60) \| 222 \| KaiC-like \| cd01124 KaiC-like; Circadian Clock Protein KaiC. KaiC is a circadian clock protein, most studied in cyanobacteria.; ncbi-cd \| cd19475 \| 96.6 \| 2.7e-06 \| 6.9e-10 \| 66.4 \| 102 \| (14, 123) \| 329 \| (18, 128) \| 220 \| FlaH \| cd19475 FlaH; flagellar accessory protein FlaH. Flagellar accessory protein FlaH is part of the motor of the archaellum membrane-anchored archaeal motility structure, together with FlaX and FlaI.; ncbi-cd \| cd19484 \| 96.3 \| 6.5e-06 \| 1.7e-09 \| 61.8 \| 101 \| (14, 122) \| 329 \| (19, 125) \| 218 \| KaiC\_C \| cd19484 KaiC\_C; C-terminal domain of Circadian Clock Protein KaiC. KaiC is a circadian clock protein, most studied in cyanobacteria.; ncbi-cd \| cd19495 \| 96.3 \| 7.1e-06 \| 1.9e-09 \| 62.4 \| 42 \| (17, 59) \| 329 \| (2, 44) \| 228 \| Elp6 \| cd19495 Elp6; Elongator subcomplex subunit Elp6. Elongator is a highly conserved multiprotein complex involved in RNA polymerase II-mediated transcriptional elongation and many other processes, including cytoskeleton organization, exocytosis, and tRNA modification.; ncbi-cd \| cd19488 \| 96.2 \| 7.8e-06 \| 2.1e-09 \| 61.0 \| 42 \| (14, 56) \| 329 \| (18, 59) \| 225 \| KaiC-like\_N \| cd19488 KaiC-like\_N; N-terminal domain of KaiC family protein; uncharacterized subfamily. KaiC is a circadian clock protein, most studied in cyanobacteria.; ncbi-cd \| cd01394 \| 96.2 \| 8.6e-06 \| 2.3e-09 \| 60.1 \| 39 \| (15, 54) \| 329 \| (19, 57) \| 216 \| archRadB \| cd01394 archRadB; archaeal RadB. The archaeal protein RadB shares similarity RadA, the archaeal functional homologue to the bacterial RecA.; ncbi-cd \| cd19486 \| 96.2 \| 9e-06 \| 2.4e-09 \| 60.2 \| 40 \| (15, 55) \| 329 \| (19, 58) \| 230 \| KaiC\_arch \| cd19486 KaiC\_arch; KaiC family protein; uncharacterized subfamily similar to Pyrococcus horikoshii PH0284.; ncbi-cd \| cd01393 \| 96.1 \| 1.1e-05 \| 2.9e-09 \| 56.6 \| 37 \| (17, 54) \| 329 \| (3, 39) \| 185 \| RecA-like \| cd01393 RecA-like; RecA family. RecA is a bacterial enzyme which has roles in homologous recombination, DNA repair, and the induction of the SOS response.; ncbi-cd \| cd19487 \| 96.1 \| 1.2e-05 \| 3.3e-09 \| 59.1 \| 41 \| (15, 56) \| 329 \| (19, 59) \| 219 \| KaiC-like\_C \| cd19487 KaiC-like\_C; C-terminal domain of KaiC family protein; uncharacterized subfamily. KaiC is a circadian clock protein, most studied in cyanobacteria.; ncbi-cd \| cd19485 \| 95.9 \| 2e-05 \| 5.3e-09 \| 59.5 \| 44 \| (15, 58) \| 329 \| (19, 62) \| 226 \| KaiC-N \| cd19485 KaiC-N; N-terminal domain of Circadian Clock Protein Kaic. KaiC is a circadian clock protein, most studied in cyanobacteria.; ncbi-cd \| cd19513 \| 95.8 \| 2.9e-05 \| 7.4e-09 \| 58.8 \| 41 \| (14, 55) \| 329 \| (18, 64) \| 235 \| Rad51 \| cd19513 Rad51; RAD51D recombinase. RAD51 recombinase plays an essential role in DNA repair by homologous recombination (HR).; ncbi-cd \| cd19493 \| 95.8 \| 2.9e-05 \| 7.7e-09 \| 57.0 \| 39 \| (15, 54) \| 329 \| (11, 55) \| 222 \| Rad51B \| cd19493 Rad51B; RAD51B recombinase. RAD51B recombinase, a RAD51 paralog, plays an important role in DNA repair by homologous recombination (HR).; ncbi-cd \| cd19492 \| 95.8 \| 3e-05 \| 8e-09 \| 55.3 \| 39 \| (16, 54) \| 329 \| (2, 45) \| 172 \| Rad51C \| cd19492 Rad51C; RAD51C recombinase. RAD51C recombinase, a RAD51 paralog, plays an important role in DNA repair by homologous recombination (HR).; ncbi-cd \| cd19515 \| 95.7 \| 3.3e-05 \| 8.4e-09 \| 59.1 \| 41 \| (14, 55) \| 329 \| (18, 64) \| 233 \| archRadA \| cd19515 archRadA; archaeal recombinase Rad51/RadA. This group includes the archaeal protein RadA which is a homolog of Rad51.; ncbi-cd \| cd01123 \| 95.4 \| 6.4e-05 \| 1.7e-08 \| 56.1 \| 41 \| (15, 55) \| 329 \| (19, 64) \| 234 \| Rad51\_DMC1\_archRadA \| cd01123 Rad51\_DMC1\_archRadA; recombinase Rad51, DMC1, and archaeal RadA. This group of recombinases includes the eukaryotic proteins RAD51, RAD55/57 and the meiosis-specific protein DMC1, and the archaeal protein RadA.; ncbi-cd \| cd19514 \| 95.4 \| 6.9e-05 \| 1.8e-08 \| 56.9 \| 41 \| (15, 55) \| 329 \| (19, 64) \| 236 \| DMC1 \| cd19514 DMC1; homologous-pairing protein DMC1. DMC1 has a central role in homologous recombination in meiosis.; ncbi-cd \| cd19483 \| 95.2 \| 9.2e-05 \| 2.5e-08 \| 53.8 \| 37 \| (18, 55) \| 329 \| (1, 38) \| 231 \| RecA-like\_Gp4D\_helicase \| cd19483 RecA-like\_Gp4D\_helicase; RecA-like domain of Escherichia coli bacteriophage T7 Gp4D helicase. This family includes the RecA-like domain of the Gp4D fragment of the Gene4 helicase-primase (Gp4) from bacteriophage T7.; ncbi-cd \| cd01132 \| 95.2 \| 0.00011 \| 2.9e-08 \| 58.6 \| 44 \| (15, 59) \| 329 \| (69, 112) \| 274 \| F1-ATPase\_alpha\_CD \| cd01132 F1-ATPase\_alpha\_CD; F1 ATP synthase alpha subunit, central domain. The F-ATPase is found in bacterial plasma membranes, mitochondrial inner membranes and in chloroplast thylakoid membranes.; ncbi-cd \| cd01121 \| 94.7 \| 0.00023 \| 5.7e-08 \| 56.0 \| 40 \| (14, 54) \| 329 \| (81, 120) \| 268 \| RadA\_SMS\_N \| cd01121 RadA\_SMS\_N; bacterial RadA DNA repair protein. Sms or bacterial RadA is a DNA repair protein that plays a role in recombination and recombinational repair of DNA damaged by UV radiation, X-rays, and chemical agent and is responsible for the stabilization or processing of branched DNA molecules.; ncbi-cd \| cd01125 \| 94.6 \| 0.00028 \| 7.2e-08 \| 53.6 \| 40 \| (16, 55) \| 329 \| (2, 50) \| 238 \| RepA\_RSF1010\_like \| cd01125 RepA\_RSF1010\_like; Hexameric Replicative Helicase RepA of plasmid RSF1010 and related proteins. This family includes the homo-hexameric replicative helicase RepA encoded by plasmid RSF1010.; ncbi-cd \| cd17875 \| 94.4 \| 0.00033 \| 8.8e-08 \| 49.9 \| 37 \| (17, 54) \| 329 \| (2, 38) \| 193 \| SRP54\_G \| cd17875 SRP54\_G; GTPase domain of the signal recognition 54 kDa subunit. The signal recognition particle (SRP) mediates the transport to or across the plasma membrane in bacteria and the endoplasmic reticulum in eukaryotes.; ncbi-cd \| cd00544 \| 94.2 \| 0.00042 \| 1.1e-07 \| 49.6 \| 33 \| (18, 54) \| 329 \| (1, 33) \| 166 \| CobU \| cd00544 CobU; Adenosylcobinamide kinase / adenosylcobinamide phosphate guanyltransferase (CobU).; ncbi-cd \| cd18539 \| 93.4 \| 0.00098 \| 2.6e-07 \| 47.5 \| 37 \| (17, 54) \| 329 \| (2, 38) \| 193 \| SRP\_G \| cd18539 SRP\_G; GTPase domain of signal recognition particle protein. The signal recognition particle (SRP) mediates the transport to or across the plasma membrane in bacteria and the endoplasmic reticulum in eukaryotes.; cath \| 1g19A01 \| 95.7 \| 5.4e-05 \| 1e-08 \| 59.1 \| 98 \| (12, 124) \| 329 \| (30, 128) \| 242 \| Reca protein \| CATHCODE: 3.40.50.300 NAME: Reca protein. Chain: a. Synonym: recombination protein reca. Ec: 3.1.-.- SOURCE: Mycobacterium tuberculosis. Organism\_taxid: 1773 CLASS: Alpha Beta, ARCH: 3-Layer(aba) Sandwich, TOPOL: Rossmann fold, HOMOL: P-loop containing nucleotide triphosphate hydrolases; cath \| 1szpB02 \| 95.6 \| 5.3e-05 \| 1.1e-08 \| 57.9 \| 105 \| (12, 122) \| 329 \| (24, 137) \| 249 \| Dna repair protein rad51 \| CATHCODE: 3.40.50.300 NAME: Dna repair protein rad51. Chain: a, b, c, d, e, f. Fragment: del(1-79). Engineered: yes SOURCE: Saccharomyces cerevisiae. Baker's yeast. Organism\_taxid: 4932. Gene: rad51, yer095w. Expressed in: escherichia coli. Expression\_system\_taxid: 562. CLASS: Alpha Beta, ARCH: 3-Layer(aba) Sandwich, TOPOL: Rossmann fold, HOMOL: P-loop containing nucleotide triphosphate hydrolases; cath \| 4ydsA00 \| 95.6 \| 6.6e-05 \| 1.3e-08 \| 56.9 \| 104 \| (12, 123) \| 329 \| (17, 131) \| 228 \| Flagella-related protein h \| CATHCODE: 3.40.50.300 NAME: Flagella-related protein h. Chain: a. Engineered: yes SOURCE: Sulfolobus acidocaldarius. Organism\_taxid: 330779. Strain: atcc 33909 / dsm 639 / jcm 8929 / nbrc 15157 / ncimb 11770. Gene: saci\_1174. Expressed in: escherichia coli. Expression\_system\_taxid: 511693. CLASS: Alpha Beta, ARCH: 3-Layer(aba) Sandwich, TOPOL: Rossmann fold, HOMOL: P-loop containing nucleotide triphosphate hydrolases; cath \| 4wiaA00 \| 95.5 \| 6.8e-05 \| 1.4e-08 \| 56.8 \| 105 \| (12, 124) \| 329 \| (24, 137) \| 233 \| Putative flagella-related protein h \| CATHCODE: 3.40.50.300 NAME: Putative flagella-related protein h. Chain: a, b, c. Engineered: yes SOURCE: Methanocaldococcus jannaschii dsm 2661. Organism\_taxid: 243232. Gene: flah, mj0899. Expressed in: escherichia coli. Expression\_system\_taxid: 562. CLASS: Alpha Beta, ARCH: 3-Layer(aba) Sandwich, TOPOL: Rossmann fold, HOMOL: P-loop containing nucleotide triphosphate hydrolases; cath \| 3ewaA02 \| 95.4 \| 9.8e-05 \| 1.9e-08 \| 57.3 \| 105 \| (12, 122) \| 329 \| (29, 153) \| 256 \| Dna repair and recombination protein rada \| CATHCODE: 3.40.50.300 NAME: Dna repair and recombination protein rada. Chain: a. Engineered: yes. Mutation: yes SOURCE: Methanococcus maripaludis. Organism\_taxid: 39152. Gene: mmp1222, rada.Expressed in: escherichia coli. Expression\_system\_taxid: 562. CLASS: Alpha Beta, ARCH: 3-Layer(aba) Sandwich, TOPOL: Rossmann fold, HOMOL: P-loop containing nucleotide triphosphate hydrolases; cath \| 1tf7A02 \| 95.4 \| 9.7e-05 \| 1.9e-08 \| 56.8 \| 104 \| (12, 123) \| 329 \| (29, 138) \| 248 \| Kaic \| CATHCODE: 3.40.50.300 NAME: Kaic. Chain: a, b, c, d, e, f. Engineered: yes SOURCE: Synechococcus sp.. Organism\_taxid: 1131. Gene: pcc7942. Expressed in:escherichia coli bl21(de3). Expression\_system\_taxid: 469008. CLASS: Alpha Beta, ARCH: 3-Layer(aba) Sandwich, TOPOL: Rossmann fold, HOMOL: P-loop containing nucleotide triphosphate hydrolases; cath \| 1tf7A01 \| 95.3 \| 0.00011 \| 2.1e-08 \| 56.2 \| 102 \| (13, 122) \| 329 \| (37, 153) \| 249 \| Kaic \| CATHCODE: 3.40.50.300 NAME: Kaic. Chain: a, b, c, d, e, f. Engineered: yes SOURCE: Synechococcus sp.. Organism\_taxid: 1131. Gene: pcc7942. Expressed in:escherichia coli bl21(de3). Expression\_system\_taxid: 469008. CLASS: Alpha Beta, ARCH: 3-Layer(aba) Sandwich, TOPOL: Rossmann fold, HOMOL: P-loop containing nucleotide triphosphate hydrolases; cath \| 3io5A00 \| 95.3 \| 0.00013 \| 2.4e-08 \| 61.0 \| 99 \| (12, 123) \| 329 \| (26, 127) \| 315 \| Recombination and repair protein \| CATHCODE: 3.40.50.300 NAME: Recombination and repair protein. Chain: a, b. Fragment: unp residues 30-358. Engineered: yes SOURCE: Enterobacteria phage t4. Bacteriophage t4. Organism\_taxid: 10665. Gene: fdsa, uvsx. Expressed in: escherichia coli. Expression\_system\_taxid: 562. CLASS: Alpha Beta, ARCH: 3-Layer(aba) Sandwich, TOPOL: Rossmann fold, HOMOL: P-loop containing nucleotide triphosphate hydrolases; cath \| 2dr3A00 \| 95.1 \| 0.00016 \| 3.3e-08 \| 54.3 \| 45 \| (12, 57) \| 329 \| (20, 64) \| 247 \| Upf0273 protein ph0284 \| CATHCODE: 3.40.50.300 NAME: Upf0273 protein ph0284. Chain: a, b, c, d, e, f. Engineered: yes SOURCE: Pyrococcus horikoshii. Organism\_taxid: 70601. Strain: ot3. Gene: ph0284. Expressed in: escherichia coli. Expression\_system\_taxid: 562. CLASS: Alpha Beta, ARCH: 3-Layer(aba) Sandwich, TOPOL: Rossmann fold, HOMOL: P-loop containing nucleotide triphosphate hydrolases; cath \| 2cvhA01 \| 94.5 \| 0.00039 \| 7.5e-08 \| 52.6 \| 103 \| (12, 123) \| 329 \| (11, 115) \| 214 \| Dna repair and recombination protein radb \| CATHCODE: 3.40.50.300 NAME: Dna repair and recombination protein radb. Chain: a, b. Engineered: yes SOURCE: Thermococcus kodakarensis. Organism\_taxid: 311400. Gene: pk-rec. Expressed in: escherichia coli. Expression\_system\_taxid: 562. CLASS: Alpha Beta, ARCH: 3-Layer(aba) Sandwich, TOPOL: Rossmann fold, HOMOL: P-loop containing nucleotide triphosphate hydrolases; cath \| 2w0mA00 \| 93.8 \| 0.00083 \| 1.7e-07 \| 50.5 \| 45 \| (12, 57) \| 329 \| (20, 64) \| 235 \| Sso2452 \| CATHCODE: 3.40.50.300 NAME: Sso2452. Chain: a. Fragment: residues 1-235. Engineered: yes SOURCE: Sulfolobus solfataricus p2. Organism\_taxid: 273057. Expressed in: escherichia coli bl21(de3). Expression\_system\_taxid: 469008. CLASS: Alpha Beta, ARCH: 3-Layer(aba) Sandwich, TOPOL: Rossmann fold, HOMOL: P-loop containing nucleotide triphosphate hydrolases; phrogs \| 1366 \| 99.9 \| 1.2e-29 \| 1.4e-33 \| 226.2 \| 281 \| (3, 302) \| 329 \| (1, 297) \| 312 \| NA \| NA; Category: unknown function; p165524 VI\_04513; phrogs \| 243 \| 99.5 \| 1.2e-18 \| 1.4e-22 \| 151.9 \| 194 \| (7, 225) \| 329 \| (29, 247) \| 296 \| DnaB-like replicative helicase \| DnaB-like replicative helicase; Category: DNA, RNA and nucleotide metabolism; NC\_028904\_p36; phrogs \| 239 \| 99.5 \| 3.4e-18 \| 4.3e-22 \| 164.0 \| 192 \| (10, 221) \| 329 \| (303, 526) \| 543 \| DNA primase/helicase \| DNA primase/helicase; Category: DNA, RNA and nucleotide metabolism; NC\_031029\_p115; phrogs \| 19 \| 99.4 \| 2.2e-17 \| 2.8e-21 \| 155.1 \| 193 \| (10, 224) \| 329 \| (193, 416) \| 441 \| DnaB-like replicative helicase \| DnaB-like replicative helicase; Category: DNA, RNA and nucleotide metabolism; p414569 VI\_10123; phrogs \| 97 \| 99.2 \| 1e-15 \| 1.3e-19 \| 141.5 \| 183 \| (9, 209) \| 329 \| (56, 249) \| 421 \| UvsX-like recombinase \| UvsX-like recombinase; Category: other; NC\_023573\_p129; phrogs \| 124 \| 99.1 \| 6.1e-15 \| 7.6e-19 \| 133.7 \| 171 \| (8, 210) \| 329 \| (11, 201) \| 348 \| Sak4-like ssDNA annealing protein \| Sak4-like ssDNA annealing protein; Category: DNA, RNA and nucleotide metabolism; KY554773\_p62; phrogs \| 2597 \| 99.0 \| 5.1e-14 \| 5.9e-18 \| 130.9 \| 187 \| (11, 222) \| 329 \| (367, 565) \| 759 \| DNA primase \| DNA primase; Category: DNA, RNA and nucleotide metabolism; p160743 VI\_04926; phrogs \| 3123 \| 98.9 \| 2e-13 \| 2.4e-17 \| 125.9 \| 180 \| (10, 211) \| 329 \| (341, 541) \| 641 \| NA \| NA; Category: unknown function; NC\_021063\_p56; phrogs \| 8337 \| 98.7 \| 3.5e-12 \| 4.1e-16 \| 107.2 \| 170 \| (4, 199) \| 329 \| (39, 218) \| 307 \| DNA repair protein \| DNA repair protein; Category: DNA, RNA and nucleotide metabolism; NC\_028762\_p33; phrogs \| 3757 \| 98.5 \| 1.2e-11 \| 1.5e-15 \| 109.0 \| 172 \| (10, 249) \| 329 \| (127, 299) \| 379 \| NA \| NA; Category: unknown function; p40711 VI\_00882; phrogs \| 6891 \| 98.5 \| 1.2e-11 \| 1.5e-15 \| 100.8 \| 117 \| (12, 168) \| 329 \| (31, 150) \| 205 \| ATPase \| ATPase; Category: other; p126525 VI\_01034; phrogs \| 906 \| 98.4 \| 8.3e-11 \| 9.7e-15 \| 110.2 \| 172 \| (12, 209) \| 329 \| (387, 585) \| 741 \| DNA polymerase/primase \| DNA polymerase/primase; Category: DNA, RNA and nucleotide metabolism; p30025 VI\_00108; phrogs \| 6357 \| 98.3 \| 1.1e-10 \| 1.2e-14 \| 97.2 \| 179 \| (5, 224) \| 329 \| (21, 213) \| 320 \| NA \| NA; Category: unknown function; p178901 VI\_06273; phrogs \| 11530 \| 98.2 \| 4.6e-10 \| 5.1e-14 \| 89.3 \| 180 \| (26, 225) \| 329 \| (1, 213) \| 259 \| NA \| NA; Category: unknown function; p393811 VI\_07291; phrogs \| 1284 \| 98.1 \| 7.7e-10 \| 8.8e-14 \| 91.3 \| 174 \| (11, 211) \| 329 \| (32, 221) \| 298 \| NA \| NA; Category: unknown function; FJ848885\_p44; phrogs \| 26214 \| 98.1 \| 8.8e-10 \| 9.8e-14 \| 90.6 \| 177 \| (3, 224) \| 329 \| (43, 226) \| 420 \| NA \| NA; Category: unknown function; p118985 VI\_00400; phrogs \| 2850 \| 98.1 \| 8.9e-10 \| 1e-13 \| 95.4 \| 169 \| (9, 211) \| 329 \| (156, 341) \| 466 \| NA \| NA; Category: unknown function; p280403 VI\_02056; phrogs \| 2872 \| 98.1 \| 8.7e-10 \| 1e-13 \| 88.9 \| 159 \| (15, 210) \| 329 \| (2, 163) \| 212 \| Sak4-like ssDNA annealing protein \| Sak4-like ssDNA annealing protein; Category: DNA, RNA and nucleotide metabolism; NC\_024365\_p27; phrogs \| 6012 \| 98.0 \| 2.1e-09 \| 2.4e-13 \| 94.2 \| 169 \| (11, 211) \| 329 \| (133, 318) \| 433 \| DNA helicase \| DNA helicase; Category: DNA, RNA and nucleotide metabolism; JF937101\_p149; phrogs \| 296 \| 97.9 \| 3.1e-09 \| 3.9e-13 \| 93.6 \| 115 \| (13, 172) \| 329 \| (91, 208) \| 312 \| DNA transposition protein \| DNA transposition protein; Category: integration and excision; p199288 VI\_08191; phrogs \| 13004 \| 97.6 \| 3.2e-08 \| 3.6e-12 \| 84.0 \| 185 \| (8, 210) \| 329 \| (258, 460) \| 485 \| NA \| NA; Category: unknown function; p26785 VI\_00069; phrogs \| 3228 \| 97.6 \| 4.1e-08 \| 4.7e-12 \| 89.6 \| 187 \| (12, 227) \| 329 \| (348, 549) \| 728 \| replicative helicase-primase \| replicative helicase-primase; Category: DNA, RNA and nucleotide metabolism; NC\_023608\_p43; phrogs \| 8097 \| 97.3 \| 1.5e-07 \| 1.7e-11 \| 82.9 \| 179 \| (9, 210) \| 329 \| (270, 460) \| 561 \| exonuclease \| exonuclease; Category: DNA, RNA and nucleotide metabolism; p356704 VI\_06144; phrogs \| 5482 \| 97.3 \| 1.5e-07 \| 1.7e-11 \| 76.5 \| 145 \| (44, 211) \| 329 \| (9, 163) \| 249 \| NA \| NA; Category: unknown function; p303059 VI\_08688; phrogs \| 18375 \| 97.1 \| 4.5e-07 \| 5.1e-11 \| 78.7 \| 182 \| (15, 211) \| 329 \| (329, 526) \| 598 \| DNA primase/helicase \| DNA primase/helicase; Category: DNA, RNA and nucleotide metabolism; p356848 VI\_04074; phrogs \| 25693 \| 97.0 \| 7e-07 \| 7.9e-11 \| 74.3 \| 144 \| (12, 172) \| 329 \| (47, 214) \| 405 \| NA \| NA; Category: unknown function; p114671 VI\_04236; phrogs \| 293 \| 97.0 \| 7.1e-07 \| 8.7e-11 \| 75.4 \| 48 \| (105, 171) \| 329 \| (151, 199) \| 253 \| ABC transporter \| ABC transporter; Category: moron, auxiliary metabolic gene and host takeover; p21813 VI\_04201; phrogs \| 9172 \| 97.0 \| 1e-06 \| 1.2e-10 \| 78.2 \| 120 \| (14, 172) \| 329 \| (97, 221) \| 518 \| NA \| NA; Category: unknown function; KX607102\_p48; phrogs \| 11143 \| 96.9 \| 1.5e-06 \| 1.7e-10 \| 73.4 \| 101 \| (14, 124) \| 329 \| (47, 156) \| 382 \| NA \| NA; Category: unknown function; p430754 VI\_01265; phrogs \| 1979 \| 96.9 \| 1.6e-06 \| 1.9e-10 \| 75.9 \| 115 \| (13, 172) \| 329 \| (120, 242) \| 384 \| NA \| NA; Category: unknown function; p439672 VI\_12223; phrogs \| 2666 \| 96.7 \| 3.4e-06 \| 4.1e-10 \| 68.8 \| 139 \| (12, 172) \| 329 \| (4, 153) \| 212 \| terminase large subunit \| terminase large subunit; Category: head and packaging; p394408 VI\_00688; phrogs \| 2392 \| 96.6 \| 5.6e-06 \| 6.7e-10 \| 79.1 \| 28 \| (12, 39) \| 329 \| (508, 535) \| 726 \| ABC transporter \| ABC transporter; Category: moron, auxiliary metabolic gene and host takeover; p58017 VI\_07050; phrogs \| 30773 \| 96.5 \| 8e-06 \| 9e-10 \| 70.6 \| 114 \| (17, 172) \| 329 \| (182, 297) \| 647 \| NA \| NA; Category: unknown function; p438992 VI\_07698; phrogs \| 5846 \| 96.4 \| 8.7e-06 \| 1e-09 \| 69.4 \| 25 \| (15, 39) \| 329 \| (36, 60) \| 306 \| clamp loader of DNA polymerase \| clamp loader of DNA polymerase; Category: DNA, RNA and nucleotide metabolism; KY606587\_p99; phrogs \| 10401 \| 96.3 \| 1.6e-05 \| 1.8e-09 \| 68.9 \| 78 \| (14, 124) \| 329 \| (198, 275) \| 385 \| ATPase \| ATPase; Category: other; MG189906\_p41; phrogs \| 36804 \| 96.0 \| 4e-05 \| 4.4e-09 \| 54.7 \| 84 \| (19, 122) \| 329 \| (2, 85) \| 162 \| NA \| NA; Category: unknown function; p120336 VI\_12480; phrogs \| 7485 \| 95.9 \| 4.3e-05 \| 5e-09 \| 66.5 \| 45 \| (9, 57) \| 329 \| (213, 257) \| 408 \| NA \| NA; Category: unknown function; NC\_019406\_p235; phrogs \| 3277 \| 95.9 \| 5e-05 \| 5.8e-09 \| 59.6 \| 27 \| (14, 40) \| 329 \| (65, 91) \| 194 \| DnaC-like helicase loader \| DnaC-like helicase loader; Category: DNA, RNA and nucleotide metabolism; p17234 VI\_01038; phrogs \| 164 \| 95.7 \| 7.6e-05 \| 9.6e-09 \| 62.5 \| 43 \| (16, 59) \| 329 \| (4, 46) \| 258 \| ParA-like partition protein \| ParA-like partition protein; Category: DNA, RNA and nucleotide metabolism; p190960 VI\_07687; phrogs \| 27930 \| 95.6 \| 0.00011 \| 1.2e-08 \| 61.3 \| 195 \| (18, 243) \| 329 \| (252, 472) \| 506 \| NA \| NA; Category: unknown function; NC\_021327\_p98; phrogs \| 50 \| 94.9 \| 0.00033 \| 4.2e-08 \| 58.7 \| 40 \| (14, 54) \| 329 \| (109, 148) \| 271 \| DnaC-like helicase loader \| DnaC-like helicase loader; Category: DNA, RNA and nucleotide metabolism; NC\_025421\_p36; phrogs \| 8638 \| 94.8 \| 0.00043 \| 4.9e-08 \| 60.7 \| 118 \| (10, 170) \| 329 \| (308, 425) \| 566 \| NA \| NA; Category: unknown function; NC\_004685\_p96; phrogs \| 7998 \| 94.8 \| 0.00046 \| 5.3e-08 \| 58.5 \| 28 \| (13, 40) \| 329 \| (38, 65) \| 358 \| NA \| NA; Category: unknown function; NC\_008695\_p16; phrogs \| 2071 \| 94.7 \| 0.00048 \| 5.9e-08 \| 51.4 \| 22 \| (16, 37) \| 329 \| (3, 24) \| 134 \| ATPase \| ATPase; Category: other; KX552041\_p115; phrogs \| 27475 \| 94.6 \| 0.00057 \| 6.4e-08 \| 57.9 \| 42 \| (13, 55) \| 329 \| (481, 522) \| 570 \| NA \| NA; Category: unknown function; p89643 VI\_08714; phrogs \| 5484 \| 94.5 \| 0.00071 \| 8.3e-08 \| 54.2 \| 24 \| (16, 39) \| 329 \| (23, 46) \| 231 \| DnaA-like replication initiation protein \| DnaA-like replication initiation protein; Category: DNA, RNA and nucleotide metabolism; NC\_005361\_p6; phrogs \| 325 \| 94.3 \| 0.00076 \| 9.5e-08 \| 59.8 \| 43 \| (14, 57) \| 329 \| (31, 73) \| 405 \| Dda-like helicase \| Dda-like helicase; Category: DNA, RNA and nucleotide metabolism; MF360957\_p200 |
| Top keywords  (threshold 1.00e-03 (evalue)) | **and, DNA, in, nucleotide, the, a, RNA, metabolism, KaiC, is** |
| Output files | ../../domain\_architecture/39\_FANPEZAQ\_CDS\_0039\_cath.hhr ../../domain\_architecture/39\_FANPEZAQ\_CDS\_0039\_merged.svg ../../domain\_architecture/39\_FANPEZAQ\_CDS\_0039\_ncbi-cd.hhr ../../domain\_architecture/39\_FANPEZAQ\_CDS\_0039\_pfam.hhr ../../domain\_architecture/39\_FANPEZAQ\_CDS\_0039\_phrogs.hhr |

### Identical protein sequences/structures

#### Search results

|  |  |
| --- | --- |
| Protein sequence databases searched | Pdb, Swissprot, Refseq |
| Identical proteins found | -- |
| Top keywords | -- |
| Output files | -- |

### Similar protein sequences/structures

#### Sequence similarity search results (HHblits)^1^

|  |  |
| --- | --- |
| Sequence databases searched | Uniclust, Pdb70 |
| Results, scheme(s)  (Top layers only, threshold 1.00e-03 (evalue)) | xml version="1.0" encoding="utf-8" standalone="no"?       2024-09-02T21:08:46.455137 image/svg+xml   Matplotlib v3.7.2, https://matplotlib.org/ |
| Results, table(s)  (threshold 1.00e-03 (evalue)) | db \| id \| prob \| evalue \| pvalue \| score \| cols \| query \| query\_len \| template \| template\_len \| name \| description; uniclust \| UniRef100\_A0A013U8Q2 \| 100.0 \| 2.4e-57 \| 5.1e-63 \| 417.9 \| 257 \| (1, 280) \| 329 \| (26, 288) \| 375 \| AAA domain protein \| AAA domain protein; uniclust \| UniRef100\_A0A077KDR8 \| 100.0 \| 1.1e-54 \| 2.4e-60 \| 403.4 \| 266 \| (1, 289) \| 329 \| (52, 324) \| 400 \| AAA family ATPase \| AAA family ATPase; uniclust \| UniRef100\_A0A088FQU0 \| 100.0 \| 4.1e-54 \| 8.5e-60 \| 398.7 \| 319 \| (1, 328) \| 329 \| (38, 359) \| 429 \| Replicative DNA helicase \| Replicative DNA helicase; uniclust \| UniRef100\_A0A1H3YZW9 \| 100.0 \| 8.8e-52 \| 1.8e-57 \| 364.5 \| 225 \| (1, 249) \| 329 \| (17, 243) \| 306 \| Signal recognition particle subunit FFH/SRP54 (Srp54) \| Signal recognition particle subunit FFH/SRP54 (Srp54); uniclust \| UniRef100\_A0A1I4QA63 \| 100.0 \| 2.4e-51 \| 4.7e-57 \| 366.1 \| 227 \| (1, 251) \| 329 \| (1, 234) \| 353 \| AAA domain-containing protein \| AAA domain-containing protein; uniclust \| UniRef100\_A0A0D6TKS4 \| 100.0 \| 4.1e-51 \| 7.9e-57 \| 355.0 \| 260 \| (4, 289) \| 329 \| (20, 282) \| 312 \| ATPase AAA \| ATPase AAA; uniclust \| UniRef100\_A0A0U1KY64 \| 100.0 \| 1.8e-49 \| 3.6e-55 \| 360.0 \| 227 \| (1, 250) \| 329 \| (1, 233) \| 382 \| Phage protein \| Phage protein; uniclust \| UniRef100\_A0A482RRG4 \| 100.0 \| 1.2e-48 \| 2.5e-54 \| 359.3 \| 229 \| (1, 250) \| 329 \| (37, 268) \| 378 \| AAA family ATPase (Fragment) \| AAA family ATPase (Fragment); uniclust \| UniRef100\_A0A2S3QKN9 \| 100.0 \| 3.1e-48 \| 6.5e-54 \| 349.7 \| 227 \| (1, 251) \| 329 \| (6, 238) \| 326 \| AAA family ATPase \| AAA family ATPase; uniclust \| UniRef100\_A0A086MRB5 \| 100.0 \| 1.4e-46 \| 3.1e-52 \| 353.4 \| 251 \| (1, 278) \| 329 \| (56, 314) \| 433 \| Uncharacterized protein \| Uncharacterized protein; uniclust \| UniRef100\_A0A1G8G4H4 \| 100.0 \| 4.6e-46 \| 9.3e-52 \| 317.1 \| 228 \| (3, 250) \| 329 \| (1, 230) \| 242 \| AAA+ ATPase domain-containing protein \| AAA+ ATPase domain-containing protein; uniclust \| UniRef100\_A0A380WK58 \| 100.0 \| 2.7e-45 \| 5.5e-51 \| 343.1 \| 290 \| (1, 308) \| 329 \| (37, 332) \| 514 \| AAA+ ATPase domain-containing protein \| AAA+ ATPase domain-containing protein; uniclust \| UniRef100\_A0A1V5Z756 \| 100.0 \| 1.9e-44 \| 3.8e-50 \| 320.8 \| 223 \| (3, 250) \| 329 \| (14, 241) \| 352 \| AAA+ ATPase domain-containing protein \| AAA+ ATPase domain-containing protein; uniclust \| UniRef100\_A0A142X8B7 \| 100.0 \| 2.3e-44 \| 4.8e-50 \| 328.5 \| 228 \| (2, 250) \| 329 \| (41, 275) \| 384 \| Signal recognition particle 54 kDa protein \| Signal recognition particle 54 kDa protein; uniclust \| UniRef100\_A0A1C5G6R7 \| 100.0 \| 2.5e-44 \| 5.1e-50 \| 326.9 \| 269 \| (1, 279) \| 329 \| (5, 285) \| 378 \| RecA-superfamily ATPase, KaiC/GvpD/RAD55 family \| RecA-superfamily ATPase, KaiC/GvpD/RAD55 family; uniclust \| UniRef100\_A0A060M5P6 \| 100.0 \| 3.8e-44 \| 7.8e-50 \| 324.9 \| 237 \| (1, 251) \| 329 \| (2, 249) \| 342 \| AAA+ ATPase domain-containing protein \| AAA+ ATPase domain-containing protein; uniclust \| UniRef100\_A0A3E0KAZ1 \| 100.0 \| 6.5e-44 \| 1.3e-49 \| 318.7 \| 227 \| (1, 250) \| 329 \| (1, 236) \| 388 \| ATP-binding protein \| ATP-binding protein; uniclust \| UniRef100\_A0A1V5X060 \| 100.0 \| 8.3e-44 \| 1.7e-49 \| 326.6 \| 223 \| (3, 250) \| 329 \| (30, 257) \| 396 \| AAA+ ATPase domain-containing protein \| AAA+ ATPase domain-containing protein; uniclust \| UniRef100\_A0A0F8ZUD5 \| 100.0 \| 2.5e-43 \| 5.3e-49 \| 328.3 \| 288 \| (1, 307) \| 329 \| (44, 370) \| 407 \| AAA+ ATPase domain-containing protein (Fragment) \| AAA+ ATPase domain-containing protein (Fragment); uniclust \| UniRef100\_A0A1J5PJ53 \| 100.0 \| 4.2e-43 \| 8.2e-49 \| 302.8 \| 260 \| (4, 289) \| 329 \| (1, 262) \| 294 \| Uncharacterized protein \| Uncharacterized protein; uniclust \| UniRef100\_A0A256B8D6 \| 100.0 \| 5.9e-43 \| 1.2e-48 \| 327.3 \| 226 \| (1, 250) \| 329 \| (43, 276) \| 446 \| AAA+ ATPase domain-containing protein \| AAA+ ATPase domain-containing protein; uniclust \| UniRef100\_A0A101TWA6 \| 100.0 \| 2.9e-41 \| 5.6e-47 \| 305.2 \| 232 \| (3, 250) \| 329 \| (49, 288) \| 404 \| AAA family ATPase \| AAA family ATPase; uniclust \| UniRef100\_A0A060N9D0 \| 100.0 \| 3.6e-40 \| 7.6e-46 \| 301.8 \| 231 \| (1, 250) \| 329 \| (27, 265) \| 353 \| Uncharacterized protein \| Uncharacterized protein; uniclust \| UniRef100\_A0A944V0U4 \| 100.0 \| 6.2e-40 \| 1.2e-45 \| 290.4 \| 222 \| (4, 250) \| 329 \| (18, 247) \| 358 \| AAA family ATPase \| AAA family ATPase; uniclust \| UniRef100\_A0A2T2WVC8 \| 100.0 \| 4.7e-39 \| 9.2e-45 \| 273.2 \| 220 \| (3, 245) \| 329 \| (1, 225) \| 239 \| AAA+ ATPase domain-containing protein \| AAA+ ATPase domain-containing protein; uniclust \| UniRef100\_A0A4D8QWP6 \| 100.0 \| 1.7e-38 \| 3.2e-44 \| 286.7 \| 290 \| (1, 307) \| 329 \| (3, 298) \| 393 \| AAA domain-containing protein \| AAA domain-containing protein; uniclust \| UniRef100\_A0A061SUU8 \| 100.0 \| 1e-37 \| 2e-43 \| 288.0 \| 289 \| (4, 307) \| 329 \| (31, 351) \| 436 \| AAA domain-containing protein \| AAA domain-containing protein; uniclust \| UniRef100\_A0A143C2N2 \| 100.0 \| 8.6e-37 \| 1.7e-42 \| 285.3 \| 232 \| (1, 251) \| 329 \| (86, 323) \| 526 \| AAA+ ATPase domain-containing protein \| AAA+ ATPase domain-containing protein; uniclust \| UniRef100\_A0A494TFH0 \| 100.0 \| 1e-36 \| 2.1e-42 \| 277.9 \| 288 \| (1, 307) \| 329 \| (12, 343) \| 358 \| AAA+ ATPase domain-containing protein \| AAA+ ATPase domain-containing protein; uniclust \| UniRef100\_A0A141GF08 \| 100.0 \| 1.5e-36 \| 2.7e-42 \| 273.9 \| 287 \| (4, 308) \| 329 \| (40, 332) \| 482 \| Recombinational DNA repair protein RecT \| Recombinational DNA repair protein RecT; uniclust \| UniRef100\_A0A1G0HWF8 \| 100.0 \| 7.7e-36 \| 1.5e-41 \| 269.4 \| 231 \| (1, 250) \| 329 \| (1, 253) \| 363 \| AAA+ ATPase domain-containing protein \| AAA+ ATPase domain-containing protein; uniclust \| UniRef100\_A0A832BEQ0 \| 100.0 \| 7e-35 \| 1.3e-40 \| 255.6 \| 230 \| (1, 251) \| 329 \| (1, 232) \| 406 \| AAA+ ATPase domain-containing protein \| AAA+ ATPase domain-containing protein; uniclust \| UniRef100\_A0A401YHK5 \| 100.0 \| 1.6e-34 \| 3.1e-40 \| 263.2 \| 255 \| (1, 278) \| 329 \| (1, 263) \| 516 \| AAA+ ATPase domain-containing protein \| AAA+ ATPase domain-containing protein; uniclust \| UniRef100\_A0A661JH92 \| 100.0 \| 3e-34 \| 5.8e-40 \| 231.6 \| 169 \| (3, 191) \| 329 \| (1, 173) \| 173 \| AAA+ ATPase domain-containing protein (Fragment) \| AAA+ ATPase domain-containing protein (Fragment); uniclust \| UniRef100\_A0A0F9SZA2 \| 100.0 \| 1.2e-33 \| 2.2e-39 \| 247.8 \| 249 \| (4, 279) \| 329 \| (1, 265) \| 402 \| AAA+ ATPase domain-containing protein \| AAA+ ATPase domain-containing protein; uniclust \| UniRef100\_A0A0J1FG05 \| 100.0 \| 8e-33 \| 1.5e-38 \| 242.2 \| 232 \| (4, 251) \| 329 \| (2, 239) \| 393 \| AAA+ ATPase domain-containing protein \| AAA+ ATPase domain-containing protein; uniclust \| UniRef100\_A9B384 \| 100.0 \| 1.1e-32 \| 2.1e-38 \| 235.3 \| 251 \| (5, 279) \| 329 \| (1, 271) \| 316 \| AAA ATPase \| AAA ATPase; uniclust \| UniRef100\_A0A0F8WAX0 \| 99.9 \| 2.6e-32 \| 5.1e-38 \| 232.2 \| 229 \| (1, 250) \| 329 \| (1, 245) \| 251 \| AAA+ ATPase domain-containing protein (Fragment) \| AAA+ ATPase domain-containing protein (Fragment); uniclust \| UniRef100\_A0A2H9UQR6 \| 99.9 \| 4e-32 \| 7.5e-38 \| 236.7 \| 226 \| (1, 250) \| 329 \| (1, 232) \| 358 \| AAA family ATPase \| AAA family ATPase; uniclust \| UniRef100\_A0A6J5N212 \| 99.9 \| 4.9e-32 \| 9e-38 \| 232.1 \| 213 \| (17, 250) \| 329 \| (2, 223) \| 322 \| AAA domain containing protein \| AAA domain containing protein; uniclust \| UniRef100\_A0A0F9T6J3 \| 99.9 \| 2.4e-31 \| 4.5e-37 \| 232.6 \| 221 \| (4, 248) \| 329 \| (2, 230) \| 385 \| AAA+ ATPase domain-containing protein \| AAA+ ATPase domain-containing protein; uniclust \| UniRef100\_A0A1M4ZD27 \| 99.9 \| 1.8e-30 \| 3.3e-36 \| 235.4 \| 231 \| (4, 248) \| 329 \| (1, 238) \| 544 \| AAA+ ATPase domain-containing protein \| AAA+ ATPase domain-containing protein; uniclust \| UniRef100\_A0A1T4YH29 \| 99.9 \| 2.9e-30 \| 5.4e-36 \| 212.2 \| 192 \| (4, 222) \| 329 \| (1, 198) \| 207 \| AAA domain-containing protein (Fragment) \| AAA domain-containing protein (Fragment); uniclust \| UniRef100\_A0A955XKB8 \| 99.9 \| 6.1e-30 \| 1.1e-35 \| 213.6 \| 207 \| (21, 248) \| 329 \| (1, 221) \| 258 \| AAA family ATPase \| AAA family ATPase; uniclust \| UniRef100\_A0A1Y2P4Y3 \| 99.9 \| 2.7e-29 \| 5e-35 \| 224.8 \| 222 \| (5, 248) \| 329 \| (50, 281) \| 423 \| AAA+ ATPase domain-containing protein \| AAA+ ATPase domain-containing protein; uniclust \| UniRef100\_A0A0F9WQ23 \| 99.9 \| 3e-29 \| 5.6e-35 \| 226.9 \| 231 \| (5, 249) \| 329 \| (1, 242) \| 464 \| AAA+ ATPase domain-containing protein \| AAA+ ATPase domain-containing protein; uniclust \| UniRef100\_UPI001902FDC2 \| 99.9 \| 4.1e-29 \| 7.6e-35 \| 219.2 \| 220 \| (4, 250) \| 329 \| (8, 233) \| 388 \| ATP-binding protein \| ATP-binding protein; uniclust \| UniRef100\_UPI002147EB62 \| 99.9 \| 4.2e-29 \| 7.8e-35 \| 225.9 \| 234 \| (1, 250) \| 329 \| (1, 239) \| 522 \| ATP-binding protein \| ATP-binding protein; uniclust \| UniRef100\_A0A4Q3F7I8 \| 99.9 \| 4.3e-29 \| 8.1e-35 \| 207.7 \| 184 \| (91, 281) \| 329 \| (8, 193) \| 227 \| AAA family ATPase (Fragment) \| AAA family ATPase (Fragment); uniclust \| UniRef100\_A0A6M3M0Y2 \| 99.9 \| 9.1e-28 \| 1.7e-33 \| 210.2 \| 221 \| (4, 249) \| 329 \| (1, 227) \| 376 \| Putative ATPase domain containing protein \| Putative ATPase domain containing protein; uniclust \| UniRef100\_A0A847YW20 \| 99.9 \| 9.3e-28 \| 1.7e-33 \| 209.6 \| 216 \| (3, 248) \| 329 \| (1, 224) \| 367 \| ATP-binding protein \| ATP-binding protein; uniclust \| UniRef100\_A0A7K1B1R5 \| 99.9 \| 1.2e-27 \| 2.1e-33 \| 215.9 \| 226 \| (4, 250) \| 329 \| (3, 234) \| 499 \| AAA family ATPase \| AAA family ATPase; uniclust \| UniRef100\_A0A2I0CWX1 \| 99.9 \| 1.2e-27 \| 2.2e-33 \| 192.2 \| 177 \| (52, 248) \| 329 \| (3, 186) \| 191 \| AAA family ATPase \| AAA family ATPase; uniclust \| UniRef100\_A0A4Q3VBQ7 \| 99.9 \| 1.3e-27 \| 2.4e-33 \| 215.8 \| 226 \| (4, 249) \| 329 \| (3, 233) \| 507 \| AAA family ATPase \| AAA family ATPase; uniclust \| UniRef100\_A0A1H5LQB0 \| 99.9 \| 1.9e-27 \| 3.5e-33 \| 205.6 \| 229 \| (4, 251) \| 329 \| (82, 318) \| 335 \| AAA domain-containing protein \| AAA domain-containing protein; uniclust \| UniRef100\_A0A3R6YIJ8 \| 99.9 \| 6.5e-27 \| 1.3e-32 \| 197.6 \| 182 \| (1, 193) \| 329 \| (1, 198) \| 212 \| AAA+ ATPase domain-containing protein \| AAA+ ATPase domain-containing protein; uniclust \| UniRef100\_A0A7X2BBX7 \| 99.9 \| 9.6e-27 \| 1.8e-32 \| 192.2 \| 197 \| (3, 208) \| 329 \| (1, 208) \| 211 \| AAA family ATPase (Fragment) \| AAA family ATPase (Fragment); uniclust \| UniRef100\_A0A3M1BKJ4 \| 99.9 \| 1.2e-26 \| 2.2e-32 \| 193.6 \| 233 \| (3, 249) \| 329 \| (1, 239) \| 250 \| AAA+ ATPase domain-containing protein \| AAA+ ATPase domain-containing protein; uniclust \| UniRef100\_A0A0F9GWA1 \| 99.9 \| 5e-26 \| 1.1e-31 \| 211.3 \| 201 \| (2, 239) \| 329 \| (29, 247) \| 347 \| AAA+ ATPase domain-containing protein \| AAA+ ATPase domain-containing protein; uniclust \| UniRef100\_A0A0Q6CAF9 \| 99.9 \| 1.3e-25 \| 2.4e-31 \| 199.9 \| 230 \| (1, 248) \| 329 \| (1, 252) \| 376 \| AAA+ ATPase domain-containing protein \| AAA+ ATPase domain-containing protein; uniclust \| UniRef100\_A0A9D8XCB7 \| 99.9 \| 3.6e-25 \| 6.7e-31 \| 191.2 \| 221 \| (12, 250) \| 329 \| (8, 239) \| 325 \| AAA family ATPase \| AAA family ATPase; uniclust \| UniRef100\_A0A945NPG0 \| 99.9 \| 4.3e-25 \| 7.8e-31 \| 178.3 \| 183 \| (109, 299) \| 329 \| (1, 185) \| 194 \| AAA family ATPase \| AAA family ATPase; uniclust \| UniRef100\_A0A6J5T713 \| 99.8 \| 3.6e-24 \| 6.6e-30 \| 189.6 \| 225 \| (4, 248) \| 329 \| (8, 249) \| 398 \| AAA domain containing protein \| AAA domain containing protein; uniclust \| UniRef100\_A0A0F9EYI3 \| 99.8 \| 5.1e-24 \| 9.3e-30 \| 178.5 \| 190 \| (45, 248) \| 329 \| (26, 222) \| 251 \| AAA domain-containing protein \| AAA domain-containing protein; uniclust \| UniRef100\_A0A4Q3WWG6 \| 99.8 \| 3e-23 \| 5.8e-29 \| 162.7 \| 121 \| (4, 143) \| 329 \| (6, 132) \| 132 \| ATP-binding protein (Fragment) \| ATP-binding protein (Fragment); uniclust \| UniRef100\_A0A0F9G2Y5 \| 99.8 \| 4e-23 \| 8.1e-29 \| 182.8 \| 209 \| (1, 247) \| 329 \| (33, 253) \| 266 \| AAA domain-containing protein (Fragment) \| AAA domain-containing protein (Fragment); uniclust \| UniRef100\_UPI00187D605D \| 99.8 \| 4.5e-23 \| 8.3e-29 \| 186.7 \| 281 \| (4, 307) \| 329 \| (3, 301) \| 488 \| AAA family ATPase \| AAA family ATPase; uniclust \| UniRef100\_A0A1Y3GD00 \| 99.8 \| 7.7e-23 \| 1.6e-28 \| 180.6 \| 180 \| (12, 219) \| 329 \| (20, 200) \| 227 \| DNA repair and recombination protein RadB \| DNA repair and recombination protein RadB; uniclust \| UniRef100\_A0A3N5F2N2 \| 99.8 \| 9e-23 \| 1.7e-28 \| 182.6 \| 267 \| (1, 282) \| 329 \| (1, 300) \| 330 \| AAA+ ATPase domain-containing protein \| AAA+ ATPase domain-containing protein; uniclust \| UniRef100\_A0A4Q3TR60 \| 99.8 \| 1e-22 \| 1.9e-28 \| 177.9 \| 187 \| (45, 248) \| 329 \| (6, 197) \| 347 \| AAA family ATPase (Fragment) \| AAA family ATPase (Fragment); uniclust \| UniRef100\_UPI0011A87DE2 \| 99.8 \| 1.6e-22 \| 3e-28 \| 169.9 \| 167 \| (80, 250) \| 329 \| (13, 181) \| 245 \| AAA family ATPase \| AAA family ATPase; uniclust \| UniRef100\_A0A075H6A9 \| 99.8 \| 1.5e-22 \| 3.3e-28 \| 190.6 \| 180 \| (13, 219) \| 329 \| (81, 264) \| 323 \| DNA repair and recombination protein RadB \| DNA repair and recombination protein RadB; uniclust \| UniRef100\_A0A933YJK3 \| 99.8 \| 1.9e-22 \| 3.5e-28 \| 171.6 \| 165 \| (80, 248) \| 329 \| (7, 177) \| 278 \| AAA family ATPase \| AAA family ATPase; uniclust \| UniRef100\_UPI001CD76CA4 \| 99.8 \| 2.2e-22 \| 4e-28 \| 177.9 \| 224 \| (4, 250) \| 329 \| (3, 231) \| 383 \| AAA family ATPase \| AAA family ATPase; uniclust \| UniRef100\_A0A0D8J3Z1 \| 99.8 \| 2.4e-22 \| 5e-28 \| 187.8 \| 228 \| (3, 245) \| 329 \| (1, 279) \| 394 \| Uncharacterized protein \| Uncharacterized protein; uniclust \| UniRef100\_A0A062V191 \| 99.8 \| 2.5e-22 \| 5.5e-28 \| 188.5 \| 182 \| (13, 222) \| 329 \| (79, 264) \| 326 \| DNA repair and recombination protein RadB \| DNA repair and recombination protein RadB; uniclust \| UniRef100\_UPI00207722C5 \| 99.8 \| 3.1e-22 \| 5.8e-28 \| 166.6 \| 187 \| (88, 280) \| 329 \| (6, 199) \| 234 \| AAA family ATPase \| AAA family ATPase; uniclust \| UniRef100\_A0A522DM60 \| 99.8 \| 3.2e-22 \| 5.9e-28 \| 171.8 \| 185 \| (45, 250) \| 329 \| (18, 210) \| 299 \| AAA family ATPase (Fragment) \| AAA family ATPase (Fragment); uniclust \| UniRef100\_A0A957C2B4 \| 99.8 \| 4e-22 \| 7.4e-28 \| 181.5 \| 189 \| (6, 210) \| 329 \| (17, 212) \| 507 \| AAA family ATPase \| AAA family ATPase; uniclust \| UniRef100\_A0A1V5ITA8 \| 99.8 \| 4.7e-22 \| 8.7e-28 \| 169.9 \| 150 \| (97, 250) \| 329 \| (3, 160) \| 286 \| Uncharacterized protein \| Uncharacterized protein; uniclust \| UniRef100\_A0A0E1T1I5 \| 99.8 \| 1e-21 \| 2.2e-27 \| 187.5 \| 226 \| (4, 247) \| 329 \| (24, 265) \| 413 \| Putative bacteriophage protein \| Putative bacteriophage protein; uniclust \| UniRef100\_A0A0U4INR8 \| 99.8 \| 1.2e-21 \| 2.2e-27 \| 172.4 \| 250 \| (10, 278) \| 329 \| (11, 267) \| 365 \| ATPase \| ATPase; uniclust \| UniRef100\_A0A073KND6 \| 99.8 \| 1.1e-21 \| 2.3e-27 \| 182.0 \| 219 \| (1, 248) \| 329 \| (13, 252) \| 340 \| Uncharacterized protein \| Uncharacterized protein; uniclust \| UniRef100\_UPI001F0E2617 \| 99.7 \| 1.5e-21 \| 2.8e-27 \| 160.1 \| 161 \| (81, 248) \| 329 \| (3, 165) \| 209 \| AAA family ATPase \| AAA family ATPase; uniclust \| UniRef100\_UPI001C64201A \| 99.7 \| 2e-21 \| 3.6e-27 \| 159.3 \| 148 \| (137, 289) \| 329 \| (43, 192) \| 207 \| AAA family ATPase \| AAA family ATPase; uniclust \| UniRef100\_A0A7C1KN99 \| 99.7 \| 3.4e-21 \| 6.3e-27 \| 170.4 \| 232 \| (3, 250) \| 329 \| (22, 260) \| 345 \| AAA family ATPase \| AAA family ATPase; uniclust \| UniRef100\_UPI0019EE306B \| 99.7 \| 5.6e-21 \| 1e-26 \| 162.2 \| 225 \| (97, 325) \| 329 \| (3, 242) \| 267 \| ATP-binding protein \| ATP-binding protein; uniclust \| UniRef100\_A0A832QGK7 \| 99.7 \| 6.8e-21 \| 1.3e-26 \| 170.4 \| 270 \| (4, 282) \| 329 \| (8, 296) \| 322 \| AAA family ATPase \| AAA family ATPase; uniclust \| UniRef100\_UPI00225E2A99 \| 99.7 \| 7.9e-21 \| 1.5e-26 \| 158.0 \| 193 \| (4, 218) \| 329 \| (25, 226) \| 228 \| AAA family ATPase \| AAA family ATPase; uniclust \| UniRef100\_A0A2N5JRX4 \| 99.7 \| 1.4e-20 \| 2.6e-26 \| 161.3 \| 227 \| (3, 248) \| 329 \| (1, 232) \| 288 \| AAA+ ATPase domain-containing protein \| AAA+ ATPase domain-containing protein; uniclust \| UniRef100\_A0A151DXG7 \| 99.7 \| 1.6e-20 \| 3.2e-26 \| 163.4 \| 181 \| (13, 224) \| 329 \| (26, 208) \| 226 \| DNA repair and recombination protein RadB \| DNA repair and recombination protein RadB; uniclust \| UniRef100\_A0A3S0CCS2 \| 99.7 \| 3.2e-20 \| 5.8e-26 \| 165.0 \| 226 \| (9, 248) \| 329 \| (10, 250) \| 390 \| AAA family ATPase \| AAA family ATPase; uniclust \| UniRef100\_UPI0007C66D04 \| 99.7 \| 3.2e-20 \| 5.9e-26 \| 155.5 \| 169 \| (103, 278) \| 329 \| (9, 182) \| 239 \| AAA family ATPase \| AAA family ATPase; uniclust \| UniRef100\_A0A073JQI6 \| 99.7 \| 3.7e-20 \| 7.5e-26 \| 169.2 \| 213 \| (4, 247) \| 329 \| (7, 238) \| 332 \| AAA family ATPase \| AAA family ATPase; uniclust \| UniRef100\_UPI000E5BAA8A \| 99.7 \| 1.2e-19 \| 2.2e-25 \| 158.9 \| 224 \| (4, 249) \| 329 \| (5, 250) \| 337 \| AAA family ATPase \| AAA family ATPase; uniclust \| UniRef100\_A0A424XXY3 \| 99.7 \| 1.4e-19 \| 2.6e-25 \| 154.3 \| 133 \| (113, 248) \| 329 \| (1, 148) \| 252 \| AAA family ATPase (Fragment) \| AAA family ATPase (Fragment); uniclust \| UniRef100\_A0A1F5UBV1 \| 99.7 \| 1.5e-19 \| 2.9e-25 \| 160.9 \| 209 \| (4, 244) \| 329 \| (2, 231) \| 326 \| AAA domain-containing protein \| AAA domain-containing protein; uniclust \| UniRef100\_A0A519UE97 \| 99.7 \| 1.5e-19 \| 2.9e-25 \| 140.1 \| 111 \| (4, 131) \| 329 \| (1, 111) \| 121 \| AAA family ATPase (Fragment) \| AAA family ATPase (Fragment); uniclust \| UniRef100\_UPI0014873B00 \| 99.7 \| 2.3e-19 \| 4.3e-25 \| 160.1 \| 223 \| (4, 251) \| 329 \| (4, 235) \| 398 \| AAA family ATPase \| AAA family ATPase; uniclust \| UniRef100\_A0A0M0BRB8 \| 99.6 \| 2.7e-19 \| 5.9e-25 \| 162.0 \| 170 \| (13, 209) \| 329 \| (37, 210) \| 248 \| DNA repair and recombination protein RadB \| DNA repair and recombination protein RadB; uniclust \| UniRef100\_A0A1F6X2V9 \| 99.6 \| 6.6e-19 \| 1.4e-24 \| 156.9 \| 163 \| (15, 208) \| 329 \| (22, 188) \| 222 \| DNA repair and recombination protein RadB \| DNA repair and recombination protein RadB; uniclust \| UniRef100\_A0A0F3GVT5 \| 99.6 \| 1.4e-18 \| 2.7e-24 \| 157.1 \| 189 \| (5, 227) \| 329 \| (51, 254) \| 351 \| Uncharacterized protein \| Uncharacterized protein; uniclust \| UniRef100\_UPI0009E9984F \| 99.6 \| 1.7e-18 \| 3.1e-24 \| 145.9 \| 149 \| (123, 278) \| 329 \| (27, 183) \| 216 \| AAA family ATPase \| AAA family ATPase; uniclust \| UniRef100\_UPI00051C4C85 \| 99.6 \| 2.6e-18 \| 4.8e-24 \| 145.6 \| 136 \| (108, 250) \| 329 \| (1, 144) \| 253 \| AAA family ATPase \| AAA family ATPase; uniclust \| UniRef100\_A0A0F9FK63 \| 99.6 \| 5.4e-18 \| 1.1e-23 \| 156.7 \| 183 \| (9, 227) \| 329 \| (22, 209) \| 313 \| AAA domain-containing protein \| AAA domain-containing protein; uniclust \| UniRef100\_UPI0022FF9F80 \| 99.5 \| 1.4e-17 \| 2.6e-23 \| 143.4 \| 170 \| (80, 250) \| 329 \| (5, 196) \| 284 \| hypothetical protein \| hypothetical protein; uniclust \| UniRef100\_UPI001BAED1AF \| 99.5 \| 1.6e-17 \| 2.9e-23 \| 126.9 \| 114 \| (50, 182) \| 329 \| (1, 115) \| 116 \| AAA family ATPase \| AAA family ATPase; uniclust \| UniRef100\_A0A375H3M6 \| 99.5 \| 1.8e-17 \| 3.2e-23 \| 134.7 \| 177 \| (152, 328) \| 329 \| (2, 183) \| 184 \| Uncharacterized protein \| Uncharacterized protein; uniclust \| UniRef100\_A0A1R1CCR1 \| 99.5 \| 2.3e-17 \| 4.5e-23 \| 150.8 \| 210 \| (4, 244) \| 329 \| (1, 231) \| 324 \| Uncharacterized protein \| Uncharacterized protein; uniclust \| UniRef100\_A0A0Q5C8F9 \| 99.5 \| 2.9e-17 \| 6e-23 \| 155.4 \| 187 \| (1, 211) \| 329 \| (27, 233) \| 385 \| AAA+ ATPase domain-containing protein \| AAA+ ATPase domain-containing protein; uniclust \| UniRef100\_A0A4Q3X6T5 \| 99.5 \| 3.3e-17 \| 6.4e-23 \| 122.2 \| 96 \| (4, 120) \| 329 \| (1, 96) \| 96 \| AAA family ATPase (Fragment) \| AAA family ATPase (Fragment); uniclust \| UniRef100\_A0A6A6K1N9 \| 99.5 \| 5.7e-17 \| 1e-22 \| 148.7 \| 151 \| (9, 174) \| 329 \| (328, 481) \| 490 \| Putative exodeoxyribonuclease 8 PDDEXK-like domain-containing protein \| Putative exodeoxyribonuclease 8 PDDEXK-like domain-containing protein; uniclust \| UniRef100\_A0A024YQ28 \| 99.5 \| 7.3e-17 \| 1.5e-22 \| 155.7 \| 186 \| (5, 213) \| 329 \| (41, 258) \| 445 \| Uncharacterized protein \| Uncharacterized protein; uniclust \| UniRef100\_A0A4Q3H202 \| 99.5 \| 1e-16 \| 1.9e-22 \| 144.4 \| 235 \| (11, 279) \| 329 \| (37, 272) \| 316 \| AAA family ATPase \| AAA family ATPase; uniclust \| UniRef100\_UPI000F843E5B \| 99.5 \| 1.1e-16 \| 2.1e-22 \| 131.6 \| 108 \| (176, 289) \| 329 \| (9, 124) \| 156 \| hypothetical protein \| hypothetical protein; uniclust \| UniRef100\_A0A7W3XXB5 \| 99.5 \| 1.2e-16 \| 2.3e-22 \| 150.8 \| 229 \| (4, 249) \| 329 \| (5, 245) \| 572 \| AAA family ATPase \| AAA family ATPase; uniclust \| UniRef100\_A0A5C7LUL3 \| 99.5 \| 1.4e-16 \| 2.6e-22 \| 148.9 \| 206 \| (4, 227) \| 329 \| (5, 230) \| 416 \| AAA family ATPase \| AAA family ATPase; uniclust \| UniRef100\_UPI00196AFCA5 \| 99.5 \| 2e-16 \| 3.7e-22 \| 129.3 \| 112 \| (137, 251) \| 329 \| (1, 114) \| 175 \| hypothetical protein \| hypothetical protein; uniclust \| UniRef100\_UPI0021F7B6B2 \| 99.4 \| 3.9e-16 \| 7.3e-22 \| 119.3 \| 104 \| (3, 122) \| 329 \| (1, 104) \| 112 \| ATP-binding protein \| ATP-binding protein; uniclust \| UniRef100\_UPI0006D7CC93 \| 99.4 \| 5.6e-16 \| 1e-21 \| 132.4 \| 177 \| (48, 251) \| 329 \| (3, 185) \| 258 \| hypothetical protein \| hypothetical protein; uniclust \| UniRef100\_UPI001EF63525 \| 99.4 \| 6.4e-16 \| 1.2e-21 \| 137.4 \| 232 \| (6, 248) \| 329 \| (8, 266) \| 356 \| AAA family ATPase \| AAA family ATPase; uniclust \| UniRef100\_A0A2D6RLK7 \| 99.4 \| 7.6e-16 \| 1.5e-21 \| 131.0 \| 153 \| (15, 208) \| 329 \| (9, 162) \| 205 \| RecA family profile 1 domain-containing protein \| RecA family profile 1 domain-containing protein; uniclust \| UniRef100\_A0A5C7M288 \| 99.4 \| 8.4e-16 \| 1.6e-21 \| 134.0 \| 189 \| (4, 210) \| 329 \| (16, 217) \| 230 \| AAA family ATPase (Fragment) \| AAA family ATPase (Fragment); uniclust \| UniRef100\_A0A081CB03 \| 99.4 \| 8.9e-16 \| 1.9e-21 \| 145.1 \| 184 \| (11, 226) \| 329 \| (61, 265) \| 336 \| Gp70 protein \| Gp70 protein; uniclust \| UniRef100\_A0A0S8I6U9 \| 99.4 \| 1.2e-15 \| 2.7e-21 \| 143.2 \| 166 \| (12, 211) \| 329 \| (72, 243) \| 305 \| KaiC domain-containing protein \| KaiC domain-containing protein; uniclust \| UniRef100\_A0A967WJV0 \| 99.4 \| 1.7e-15 \| 3.1e-21 \| 122.0 \| 115 \| (1, 131) \| 329 \| (1, 115) \| 168 \| AAA family ATPase \| AAA family ATPase; uniclust \| UniRef100\_A0A0G0FNL2 \| 99.4 \| 1.9e-15 \| 3.8e-21 \| 138.9 \| 172 \| (12, 208) \| 329 \| (99, 271) \| 306 \| Protein RecA (Fragment) \| Protein RecA (Fragment); uniclust \| UniRef100\_A0A011NUY9 \| 99.4 \| 1.9e-15 \| 4e-21 \| 148.6 \| 172 \| (12, 208) \| 329 \| (130, 302) \| 481 \| Protein RecA \| Protein RecA; uniclust \| UniRef100\_A0A1I2ZH41 \| 99.4 \| 2.1e-15 \| 4.1e-21 \| 133.7 \| 187 \| (13, 223) \| 329 \| (47, 235) \| 263 \| DNA repair and recombination protein RadB \| DNA repair and recombination protein RadB; uniclust \| UniRef100\_A0A955WSM5 \| 99.4 \| 3.2e-15 \| 5.9e-21 \| 113.0 \| 103 \| (107, 209) \| 329 \| (2, 106) \| 113 \| AAA family ATPase (Fragment) \| AAA family ATPase (Fragment); uniclust \| UniRef100\_A0A087RS38 \| 99.3 \| 2.8e-15 \| 6.2e-21 \| 143.5 \| 162 \| (12, 208) \| 329 \| (125, 301) \| 353 \| Circadian clock protein kinase KaiC \| Circadian clock protein kinase KaiC; uniclust \| UniRef100\_A0A1C6RH26 \| 99.3 \| 3.8e-15 \| 7.6e-21 \| 142.8 \| 183 \| (8, 213) \| 329 \| (14, 224) \| 441 \| AAA domain-containing protein \| AAA domain-containing protein; uniclust \| UniRef100\_A0A0Q8MKG9 \| 99.3 \| 4e-15 \| 7.7e-21 \| 137.1 \| 171 \| (12, 207) \| 329 \| (69, 240) \| 364 \| Protein RecA \| Protein RecA; uniclust \| UniRef100\_A0A1D8MRM1 \| 99.3 \| 3.9e-15 \| 8.4e-21 \| 136.8 \| 162 \| (12, 209) \| 329 \| (48, 212) \| 257 \| KaiC domain-containing protein \| KaiC domain-containing protein; uniclust \| UniRef100\_A0A9D7RLT2 \| 99.3 \| 5.9e-15 \| 1.1e-20 \| 121.3 \| 117 \| (24, 160) \| 329 \| (22, 145) \| 191 \| AAA family ATPase \| AAA family ATPase; uniclust \| UniRef100\_A0A256Z8Y2 \| 99.3 \| 5.5e-15 \| 1.2e-20 \| 133.6 \| 160 \| (17, 213) \| 329 \| (37, 199) \| 231 \| AAA+ ATPase domain-containing protein \| AAA+ ATPase domain-containing protein; uniclust \| UniRef100\_A0A1Q9N6T9 \| 99.3 \| 6.4e-15 \| 1.3e-20 \| 132.4 \| 180 \| (12, 212) \| 329 \| (38, 222) \| 249 \| DNA repair and recombination protein RadB \| DNA repair and recombination protein RadB; uniclust \| UniRef100\_A0A0B3ADI5 \| 99.3 \| 6.8e-15 \| 1.5e-20 \| 139.7 \| 161 \| (12, 209) \| 329 \| (83, 255) \| 336 \| Circadian clock protein KaiC \| Circadian clock protein KaiC; uniclust \| UniRef100\_A0A010TBU8 \| 99.3 \| 7.2e-15 \| 1.5e-20 \| 146.4 \| 173 \| (12, 209) \| 329 \| (178, 351) \| 558 \| Protein RecA \| Protein RecA; uniclust \| UniRef100\_A0A062VA97 \| 99.3 \| 8.5e-15 \| 1.9e-20 \| 146.2 \| 161 \| (12, 209) \| 329 \| (176, 354) \| 474 \| RecA-superfamily ATPase possibly involved in signal transduction \| RecA-superfamily ATPase possibly involved in signal transduction; uniclust \| UniRef100\_A0A2R6MJT5 \| 99.3 \| 9.6e-15 \| 1.9e-20 \| 123.0 \| 135 \| (13, 167) \| 329 \| (29, 168) \| 170 \| DNA repair protein RadB (Fragment) \| DNA repair protein RadB (Fragment); uniclust \| UniRef100\_A0A077HSX7 \| 99.3 \| 9.8e-15 \| 1.9e-20 \| 138.0 \| 252 \| (6, 281) \| 329 \| (32, 307) \| 429 \| Uncharacterized protein \| Uncharacterized protein; uniclust \| UniRef100\_A0A0F2LA32 \| 99.3 \| 1.1e-14 \| 2.3e-20 \| 132.3 \| 160 \| (13, 208) \| 329 \| (43, 208) \| 267 \| Recombination protein RecA (Fragment) \| Recombination protein RecA (Fragment); uniclust \| UniRef100\_A0A256YLX7 \| 99.3 \| 1.1e-14 \| 2.3e-20 \| 129.4 \| 184 \| (13, 234) \| 329 \| (18, 201) \| 211 \| AAA family ATPase \| AAA family ATPase; uniclust \| UniRef100\_A0A2V2U9P3 \| 99.3 \| 1.1e-14 \| 2.3e-20 \| 132.6 \| 160 \| (12, 207) \| 329 \| (25, 190) \| 246 \| KaiC domain-containing protein \| KaiC domain-containing protein; uniclust \| UniRef100\_A0A0B3B298 \| 99.3 \| 1.4e-14 \| 3.1e-20 \| 138.7 \| 164 \| (12, 211) \| 329 \| (95, 267) \| 340 \| KaiC domain-containing protein \| KaiC domain-containing protein; uniclust \| UniRef100\_A0A3N5F8L2 \| 99.3 \| 2.1e-14 \| 3.9e-20 \| 125.1 \| 188 \| (11, 233) \| 329 \| (46, 248) \| 291 \| ATP-binding protein \| ATP-binding protein; uniclust \| UniRef100\_A0A0F9UF35 \| 99.3 \| 2.2e-14 \| 4e-20 \| 132.6 \| 232 \| (4, 245) \| 329 \| (5, 253) \| 490 \| AAA+ ATPase domain-containing protein \| AAA+ ATPase domain-containing protein; uniclust \| UniRef100\_UPI000507A9AD \| 99.3 \| 2.6e-14 \| 4.9e-20 \| 128.8 \| 221 \| (4, 244) \| 329 \| (2, 240) \| 389 \| AAA family ATPase \| AAA family ATPase; uniclust \| UniRef100\_A0A4Q6ASF8 \| 99.2 \| 3.7e-14 \| 7.1e-20 \| 125.0 \| 231 \| (12, 279) \| 329 \| (9, 241) \| 274 \| Uncharacterized protein \| Uncharacterized protein; uniclust \| UniRef100\_A0A420ZB44 \| 99.2 \| 4.7e-14 \| 9.4e-20 \| 129.1 \| 179 \| (14, 213) \| 329 \| (65, 252) \| 311 \| RecA family profile 1 domain-containing protein \| RecA family profile 1 domain-containing protein; uniclust \| UniRef100\_A0A256ZN40 \| 99.2 \| 6.3e-14 \| 1.2e-19 \| 123.1 \| 176 \| (14, 219) \| 329 \| (32, 210) \| 244 \| RecA family profile 1 domain-containing protein \| RecA family profile 1 domain-containing protein; uniclust \| UniRef100\_A0A0U3SJ19 \| 99.2 \| 6.1e-14 \| 1.3e-19 \| 134.1 \| 180 \| (13, 227) \| 329 \| (49, 236) \| 362 \| Circadian clock protein KaiC \| Circadian clock protein KaiC; uniclust \| UniRef100\_A0A256YSC6 \| 99.2 \| 6.7e-14 \| 1.4e-19 \| 118.1 \| 130 \| (14, 160) \| 329 \| (32, 165) \| 166 \| DNA repair protein RadB (Fragment) \| DNA repair protein RadB (Fragment); uniclust \| UniRef100\_A0A0F5I4I8 \| 99.2 \| 6.3e-14 \| 1.4e-19 \| 124.5 \| 137 \| (12, 167) \| 329 \| (37, 179) \| 192 \| Circadian clock protein KaiC \| Circadian clock protein KaiC; uniclust \| UniRef100\_A0A084SEJ6 \| 99.2 \| 6.7e-14 \| 1.4e-19 \| 137.3 \| 172 \| (12, 208) \| 329 \| (145, 317) \| 487 \| Protein RecA \| Protein RecA; uniclust \| UniRef100\_A0A062V187 \| 99.2 \| 6.9e-14 \| 1.6e-19 \| 136.0 \| 179 \| (12, 226) \| 329 \| (102, 303) \| 353 \| RecA-superfamily ATPase possibly involved in signal transduction \| RecA-superfamily ATPase possibly involved in signal transduction; uniclust \| UniRef100\_A0A075MT78 \| 99.2 \| 8.8e-14 \| 1.8e-19 \| 131.1 \| 162 \| (14, 208) \| 329 \| (91, 258) \| 336 \| RecA-superfamily ATPase possibly involved in signal transduction \| RecA-superfamily ATPase possibly involved in signal transduction; uniclust \| UniRef100\_A0A023NGB4 \| 99.2 \| 1.1e-13 \| 2.3e-19 \| 126.1 \| 162 \| (13, 207) \| 329 \| (19, 184) \| 267 \| AAA+ ATPase domain-containing protein \| AAA+ ATPase domain-containing protein; uniclust \| UniRef100\_A0A0F9D4W2 \| 99.2 \| 1.2e-13 \| 2.4e-19 \| 131.4 \| 188 \| (12, 234) \| 329 \| (75, 278) \| 395 \| KaiC domain-containing protein (Fragment) \| KaiC domain-containing protein (Fragment); uniclust \| UniRef100\_A0A142XSV4 \| 99.2 \| 1.2e-13 \| 2.4e-19 \| 130.4 \| 172 \| (12, 208) \| 329 \| (102, 274) \| 414 \| Protein RecA \| Protein RecA; uniclust \| UniRef100\_A0A132MHS1 \| 99.2 \| 1.3e-13 \| 2.6e-19 \| 133.8 \| 244 \| (9, 278) \| 329 \| (19, 306) \| 477 \| Uncharacterized protein \| Uncharacterized protein; uniclust \| UniRef100\_A0A221NLH3 \| 99.2 \| 1.4e-13 \| 2.8e-19 \| 132.3 \| 215 \| (4, 245) \| 329 \| (11, 245) \| 406 \| Uncharacterized protein \| Uncharacterized protein; uniclust \| UniRef100\_A0A151ENH4 \| 99.2 \| 1.3e-13 \| 2.8e-19 \| 130.2 \| 178 \| (13, 226) \| 329 \| (57, 250) \| 297 \| KaiC domain-containing protein \| KaiC domain-containing protein; uniclust \| UniRef100\_A0A534KY56 \| 99.2 \| 1.7e-13 \| 3e-19 \| 128.2 \| 170 \| (13, 211) \| 329 \| (336, 510) \| 539 \| DNA repair and recombination protein RadB \| DNA repair and recombination protein RadB; uniclust \| UniRef100\_A0A0P0N116 \| 99.2 \| 1.5e-13 \| 3.1e-19 \| 122.0 \| 157 \| (15, 213) \| 329 \| (32, 188) \| 220 \| RecA/RadA recombinase \| RecA/RadA recombinase; uniclust \| UniRef100\_A0A075LS49 \| 99.1 \| 1.9e-13 \| 3.9e-19 \| 126.3 \| 169 \| (13, 227) \| 329 \| (57, 246) \| 277 \| KaiC domain-containing protein \| KaiC domain-containing protein; uniclust \| UniRef100\_A0A062V7X4 \| 99.1 \| 1.8e-13 \| 4.1e-19 \| 142.3 \| 195 \| (13, 242) \| 329 \| (346, 547) \| 685 \| non-specific serine/threonine protein kinase \| non-specific serine/threonine protein kinase; uniclust \| UniRef100\_A0A0S8KLX2 \| 99.1 \| 2.1e-13 \| 4.3e-19 \| 125.8 \| 155 \| (14, 207) \| 329 \| (48, 212) \| 289 \| KaiC domain-containing protein \| KaiC domain-containing protein; uniclust \| UniRef100\_A0A059LQV8 \| 99.1 \| 2.2e-13 \| 4.5e-19 \| 134.9 \| 173 \| (11, 208) \| 329 \| (189, 362) \| 512 \| DNA repair DNA-dependent ATPase RecA \| DNA repair DNA-dependent ATPase RecA; uniclust \| UniRef100\_A0A2T2WUP4 \| 99.1 \| 2.6e-13 \| 4.8e-19 \| 112.1 \| 141 \| (133, 278) \| 329 \| (6, 151) \| 191 \| AAA family ATPase (Fragment) \| AAA family ATPase (Fragment); uniclust \| UniRef100\_A0A1F2P412 \| 99.1 \| 2.5e-13 \| 5.4e-19 \| 131.3 \| 162 \| (12, 208) \| 329 \| (95, 270) \| 358 \| ATPase \| ATPase; uniclust \| UniRef100\_A0A023ZVC7 \| 99.1 \| 2.9e-13 \| 5.8e-19 \| 129.2 \| 172 \| (12, 208) \| 329 \| (94, 266) \| 438 \| Protein RecA (Fragment) \| Protein RecA (Fragment); uniclust \| UniRef100\_A0A017HR86 \| 99.1 \| 2.9e-13 \| 5.9e-19 \| 131.1 \| 172 \| (12, 208) \| 329 \| (144, 316) \| 492 \| Protein RecA \| Protein RecA; uniclust \| UniRef100\_A0A090VYE0 \| 99.1 \| 3.1e-13 \| 6.4e-19 \| 127.0 \| 167 \| (13, 223) \| 329 \| (89, 258) \| 323 \| DNA repair protein RadA \| DNA repair protein RadA; uniclust \| UniRef100\_A0A4V1U163 \| 99.1 \| 3.6e-13 \| 6.9e-19 \| 119.6 \| 190 \| (1, 213) \| 329 \| (1, 218) \| 272 \| ATP-binding protein (Fragment) \| ATP-binding protein (Fragment); uniclust \| UniRef100\_A0A0F9TSC2 \| 99.1 \| 3.5e-13 \| 7.5e-19 \| 132.2 \| 173 \| (12, 209) \| 329 \| (85, 258) \| 427 \| RecA family profile 2 domain-containing protein \| RecA family profile 2 domain-containing protein; uniclust \| UniRef100\_A0A097QT34 \| 99.1 \| 3.6e-13 \| 8e-19 \| 136.7 \| 132 \| (13, 166) \| 329 \| (286, 421) \| 526 \| non-specific serine/threonine protein kinase \| non-specific serine/threonine protein kinase; uniclust \| UniRef100\_A0A062V6P2 \| 99.1 \| 3.6e-13 \| 8e-19 \| 128.8 \| 134 \| (12, 167) \| 329 \| (84, 233) \| 321 \| RecA-superfamily ATPase possibly involved in signal transduction \| RecA-superfamily ATPase possibly involved in signal transduction; uniclust \| UniRef100\_A0A6J5R7H7 \| 99.1 \| 4.4e-13 \| 8e-19 \| 121.0 \| 239 \| (2, 247) \| 329 \| (102, 359) \| 378 \| AAA domain containing protein \| AAA domain containing protein; uniclust \| UniRef100\_A0A133VEK1 \| 99.1 \| 3.7e-13 \| 8.1e-19 \| 123.8 \| 187 \| (13, 234) \| 329 \| (27, 222) \| 248 \| KaiC domain-containing protein \| KaiC domain-containing protein; uniclust \| UniRef100\_A0A101HXY9 \| 99.1 \| 3.9e-13 \| 8.2e-19 \| 128.0 \| 169 \| (12, 223) \| 329 \| (96, 266) \| 350 \| DNA repair protein radA (Fragment) \| DNA repair protein radA (Fragment); uniclust \| UniRef100\_A0A099T178 \| 99.1 \| 4.4e-13 \| 8.7e-19 \| 125.2 \| 156 \| (12, 209) \| 329 \| (91, 248) \| 360 \| Circadian clock protein KaiC \| Circadian clock protein KaiC; uniclust \| UniRef100\_A0A009YIA0 \| 99.1 \| 4.2e-13 \| 8.9e-19 \| 139.0 \| 168 \| (13, 221) \| 329 \| (243, 411) \| 721 \| DNA repair protein RadA \| DNA repair protein RadA; uniclust \| UniRef100\_A0A2H5XG24 \| 99.1 \| 4.3e-13 \| 9e-19 \| 133.1 \| 155 \| (13, 208) \| 329 \| (33, 197) \| 503 \| non-specific serine/threonine protein kinase \| non-specific serine/threonine protein kinase; uniclust \| UniRef100\_A0A550GPC2 \| 99.1 \| 4.8e-13 \| 9.2e-19 \| 113.0 \| 144 \| (13, 172) \| 329 \| (30, 177) \| 185 \| RecA family profile 1 domain-containing protein (Fragment) \| RecA family profile 1 domain-containing protein (Fragment); uniclust \| UniRef100\_A0A0F9LPC4 \| 99.1 \| 4.5e-13 \| 9.3e-19 \| 127.9 \| 175 \| (14, 212) \| 329 \| (109, 296) \| 372 \| DNA repair and recombination protein RadA (Fragment) \| DNA repair and recombination protein RadA (Fragment); uniclust \| UniRef100\_A0A0N7JCY7 \| 99.1 \| 4.3e-13 \| 9.6e-19 \| 135.4 \| 163 \| (12, 208) \| 329 \| (35, 202) \| 491 \| KaiC domain-containing protein \| KaiC domain-containing protein; uniclust \| UniRef100\_A0A0M8NHN2 \| 99.1 \| 5e-13 \| 1e-18 \| 127.5 \| 154 \| (14, 210) \| 329 \| (109, 264) \| 376 \| DNA repair protein RadA (Fragment) \| DNA repair protein RadA (Fragment); uniclust \| UniRef100\_A0A1Y4D5T3 \| 99.1 \| 5.3e-13 \| 1.1e-18 \| 125.1 \| 177 \| (4, 208) \| 329 \| (18, 212) \| 291 \| AAA+ ATPase domain-containing protein \| AAA+ ATPase domain-containing protein; uniclust \| UniRef100\_A0A062XPL1 \| 99.1 \| 6.3e-13 \| 1.3e-18 \| 121.2 \| 177 \| (4, 208) \| 329 \| (8, 193) \| 262 \| NTP-binding protein \| NTP-binding protein; uniclust \| UniRef100\_A0A0C1NQ08 \| 99.1 \| 6.4e-13 \| 1.4e-18 \| 118.2 \| 139 \| (12, 167) \| 329 \| (40, 188) \| 205 \| KaiC domain-containing protein (Fragment) \| KaiC domain-containing protein (Fragment); uniclust \| UniRef100\_A0A2W4NC12 \| 99.1 \| 7.8e-13 \| 1.4e-18 \| 103.4 \| 108 \| (4, 128) \| 329 \| (23, 130) \| 132 \| AAA family ATPase (Fragment) \| AAA family ATPase (Fragment); uniclust \| UniRef100\_A0A0G3M7Z9 \| 99.1 \| 6.7e-13 \| 1.5e-18 \| 131.0 \| 165 \| (13, 209) \| 329 \| (93, 273) \| 421 \| AAA family ATPase \| AAA family ATPase; uniclust \| UniRef100\_A0A011RPQ0 \| 99.1 \| 7.5e-13 \| 1.6e-18 \| 136.7 \| 168 \| (13, 221) \| 329 \| (206, 374) \| 652 \| DNA repair protein RadA \| DNA repair protein RadA; uniclust \| UniRef100\_A0A017RU46 \| 99.1 \| 8.2e-13 \| 1.6e-18 \| 128.3 \| 173 \| (12, 209) \| 329 \| (133, 306) \| 495 \| Protein RecA \| Protein RecA; uniclust \| UniRef100\_A0A0C2V8R7 \| 99.1 \| 7.7e-13 \| 1.6e-18 \| 131.9 \| 159 \| (12, 210) \| 329 \| (34, 202) \| 491 \| Circadian clock protein KaiC \| Circadian clock protein KaiC; uniclust \| UniRef100\_A0A075LY79 \| 99.1 \| 8.4e-13 \| 1.8e-18 \| 123.1 \| 180 \| (12, 226) \| 329 \| (38, 237) \| 292 \| ATPase \| ATPase; uniclust \| UniRef100\_A0A1Y4TW55 \| 99.1 \| 9.8e-13 \| 1.9e-18 \| 97.7 \| 84 \| (3, 102) \| 329 \| (1, 84) \| 85 \| AAA family ATPase (Fragment) \| AAA family ATPase (Fragment); uniclust \| UniRef100\_A0A2K2V568 \| 99.0 \| 1e-12 \| 2.1e-18 \| 124.3 \| 179 \| (13, 213) \| 329 \| (113, 301) \| 333 \| DNA repair and recombination protein RadA \| DNA repair and recombination protein RadA; uniclust \| UniRef100\_A0A059T7T3 \| 99.0 \| 1.1e-12 \| 2.4e-18 \| 124.7 \| 179 \| (1, 209) \| 329 \| (33, 221) \| 312 \| AAA\_24 domain-containing protein \| AAA\_24 domain-containing protein; uniclust \| UniRef100\_A0A067XSV1 \| 99.0 \| 1.3e-12 \| 2.4e-18 \| 120.7 \| 172 \| (12, 208) \| 329 \| (59, 231) \| 379 \| Protein RecA (Fragment) \| Protein RecA (Fragment); uniclust \| UniRef100\_A0A0L0LD37 \| 99.0 \| 1.4e-12 \| 2.5e-18 \| 114.9 \| 197 \| (13, 227) \| 329 \| (29, 240) \| 301 \| Uncharacterized protein (Fragment) \| Uncharacterized protein (Fragment); uniclust \| UniRef100\_A0A257AQ51 \| 99.0 \| 1.3e-12 \| 2.7e-18 \| 120.0 \| 171 \| (13, 212) \| 329 \| (54, 228) \| 266 \| RecA family profile 1 domain-containing protein \| RecA family profile 1 domain-containing protein; uniclust \| UniRef100\_A0A0E4FH58 \| 99.0 \| 1.4e-12 \| 2.7e-18 \| 116.9 \| 145 \| (12, 173) \| 329 \| (64, 209) \| 298 \| Protein RecA (Fragment) \| Protein RecA (Fragment); uniclust \| UniRef100\_A0A0E0TB11 \| 99.0 \| 1.4e-12 \| 2.8e-18 \| 114.0 \| 140 \| (12, 168) \| 329 \| (79, 219) \| 222 \| Protein RecA \| Protein RecA; uniclust \| UniRef100\_A0A1F5S0T5 \| 99.0 \| 1.4e-12 \| 3e-18 \| 129.5 \| 181 \| (12, 227) \| 329 \| (284, 484) \| 514 \| KaiC domain-containing protein \| KaiC domain-containing protein; uniclust \| UniRef100\_A0A3A3FWN4 \| 99.0 \| 1.6e-12 \| 3e-18 \| 95.8 \| 87 \| (20, 124) \| 329 \| (1, 87) \| 93 \| AAA domain-containing protein \| AAA domain-containing protein; uniclust \| UniRef100\_A0A059KPY5 \| 99.0 \| 1.5e-12 \| 3.2e-18 \| 130.4 \| 167 \| (14, 223) \| 329 \| (158, 327) \| 534 \| DNA repair protein RadA \| DNA repair protein RadA; uniclust \| UniRef100\_A0A448ZC34 \| 99.0 \| 1.6e-12 \| 3.2e-18 \| 126.4 \| 171 \| (12, 207) \| 329 \| (166, 337) \| 481 \| RecA family profile 1 domain-containing protein \| RecA family profile 1 domain-containing protein; uniclust \| UniRef100\_A0A023H9M0 \| 99.0 \| 1.9e-12 \| 3.6e-18 \| 116.3 \| 144 \| (12, 172) \| 329 \| (86, 230) \| 280 \| Protein RecA (Fragment) \| Protein RecA (Fragment); uniclust \| UniRef100\_A0A662JTN0 \| 99.0 \| 1.9e-12 \| 3.8e-18 \| 116.6 \| 178 \| (12, 209) \| 329 \| (43, 229) \| 267 \| KaiC domain-containing protein (Fragment) \| KaiC domain-containing protein (Fragment); uniclust \| UniRef100\_A0A0W0VSK8 \| 99.0 \| 1.9e-12 \| 3.9e-18 \| 120.3 \| 181 \| (13, 227) \| 329 \| (39, 232) \| 310 \| DNA integration/recombination/inversion protein \| DNA integration/recombination/inversion protein; uniclust \| UniRef100\_A0A013VEU8 \| 99.0 \| 1.9e-12 \| 4.2e-18 \| 135.3 \| 198 \| (13, 243) \| 329 \| (379, 586) \| 728 \| non-specific serine/threonine protein kinase \| non-specific serine/threonine protein kinase; uniclust \| UniRef100\_A0A5R2ATA9 \| 99.0 \| 2.4e-12 \| 4.4e-18 \| 115.6 \| 195 \| (1, 213) \| 329 \| (1, 200) \| 351 \| Uncharacterized protein \| Uncharacterized protein; uniclust \| UniRef100\_A0A061SUX7 \| 99.0 \| 2.3e-12 \| 4.4e-18 \| 125.1 \| 172 \| (12, 208) \| 329 \| (151, 323) \| 588 \| Protein RecA \| Protein RecA; uniclust \| UniRef100\_A0A016QUY6 \| 99.0 \| 2.3e-12 \| 4.9e-18 \| 130.8 \| 168 \| (14, 222) \| 329 \| (143, 311) \| 581 \| DNA repair protein RadA \| DNA repair protein RadA; uniclust \| UniRef100\_A0A101DJW8 \| 99.0 \| 2.4e-12 \| 5e-18 \| 124.6 \| 189 \| (14, 234) \| 329 \| (148, 343) \| 380 \| Putative circadian clock protein, KaiC \| Putative circadian clock protein, KaiC; uniclust \| UniRef100\_A0A1U9WRA5 \| 99.0 \| 2.4e-12 \| 5e-18 \| 124.6 \| 173 \| (11, 208) \| 329 \| (60, 241) \| 396 \| RecA \| RecA; uniclust \| UniRef100\_A0A0B3ADK1 \| 99.0 \| 2.4e-12 \| 5.1e-18 \| 124.0 \| 195 \| (12, 243) \| 329 \| (123, 342) \| 346 \| Circadian clock protein, KaiC \| Circadian clock protein, KaiC; uniclust \| UniRef100\_A0A2D6KXY2 \| 99.0 \| 2.5e-12 \| 5.2e-18 \| 119.4 \| 179 \| (12, 223) \| 329 \| (60, 243) \| 287 \| KaiC domain-containing protein \| KaiC domain-containing protein; uniclust \| UniRef100\_A0A011T6A1 \| 99.0 \| 2.7e-12 \| 5.7e-18 \| 127.0 \| 157 \| (14, 210) \| 329 \| (102, 258) \| 499 \| DNA repair protein RadA \| DNA repair protein RadA; uniclust \| UniRef100\_A0A256XEU0 \| 99.0 \| 2.8e-12 \| 5.8e-18 \| 118.7 \| 176 \| (12, 208) \| 329 \| (55, 243) \| 290 \| KaiC domain-containing protein \| KaiC domain-containing protein; uniclust \| UniRef100\_A0A0F8YVJ1 \| 99.0 \| 3.2e-12 \| 5.8e-18 \| 100.7 \| 111 \| (4, 131) \| 329 \| (1, 111) \| 136 \| AAA domain-containing protein (Fragment) \| AAA domain-containing protein (Fragment); uniclust \| UniRef100\_A0A0N0D162 \| 99.0 \| 3.2e-12 \| 6.5e-18 \| 128.0 \| 200 \| (13, 247) \| 329 \| (280, 486) \| 595 \| KaiC 1 (Fragment) \| KaiC 1 (Fragment); uniclust \| UniRef100\_A0A2R7Y1F6 \| 99.0 \| 3.1e-12 \| 7e-18 \| 130.4 \| 161 \| (13, 208) \| 329 \| (59, 221) \| 516 \| KaiC domain-containing protein \| KaiC domain-containing protein; uniclust \| UniRef100\_A0A1H8MGT9 \| 99.0 \| 3.5e-12 \| 7.3e-18 \| 126.3 \| 183 \| (13, 227) \| 329 \| (267, 453) \| 500 \| Circadian clock protein KaiC \| Circadian clock protein KaiC; uniclust \| UniRef100\_A0A1D2RFQ8 \| 99.0 \| 3.5e-12 \| 7.4e-18 \| 115.1 \| 138 \| (13, 167) \| 329 \| (35, 183) \| 223 \| KaiC domain-containing protein \| KaiC domain-containing protein; uniclust \| UniRef100\_A0A075FP22 \| 99.0 \| 3.6e-12 \| 7.8e-18 \| 121.5 \| 138 \| (12, 167) \| 329 \| (74, 224) \| 315 \| KaiC domain-containing protein \| KaiC domain-containing protein; uniclust \| UniRef100\_A0A0Q4BEX0 \| 99.0 \| 3.9e-12 \| 8.1e-18 \| 117.6 \| 138 \| (12, 167) \| 329 \| (51, 202) \| 280 \| Circadian clock protein KaiC \| Circadian clock protein KaiC; uniclust \| UniRef100\_A0A151E288 \| 99.0 \| 4.7e-12 \| 8.6e-18 \| 112.1 \| 191 \| (4, 227) \| 329 \| (2, 202) \| 310 \| Uncharacterized protein \| Uncharacterized protein; uniclust \| UniRef100\_UPI001CCF95CF \| 99.0 \| 4.8e-12 \| 8.7e-18 \| 106.3 \| 118 \| (152, 278) \| 329 \| (31, 152) \| 207 \| hypothetical protein \| hypothetical protein; uniclust \| UniRef100\_A0A077LYA0 \| 99.0 \| 4.3e-12 \| 8.8e-18 \| 131.6 \| 145 \| (12, 173) \| 329 \| (92, 237) \| 820 \| Protein RecA \| Protein RecA; uniclust \| UniRef100\_A0A5P6NYW8 \| 99.0 \| 4.7e-12 \| 8.9e-18 \| 120.0 \| 204 \| (6, 227) \| 329 \| (103, 325) \| 501 \| AAA family ATPase \| AAA family ATPase; uniclust \| UniRef100\_A0A7C5ZM63 \| 99.0 \| 5e-12 \| 9.1e-18 \| 119.6 \| 112 \| (135, 250) \| 329 \| (7, 121) \| 573 \| IS481 family transposase \| IS481 family transposase; uniclust \| UniRef100\_A0A0B7MPK7 \| 99.0 \| 4.5e-12 \| 9.3e-18 \| 127.9 \| 137 \| (12, 167) \| 329 \| (274, 413) \| 571 \| non-specific serine/threonine protein kinase \| non-specific serine/threonine protein kinase; uniclust \| UniRef100\_G2JBT2 \| 99.0 \| 5.2e-12 \| 9.6e-18 \| 102.1 \| 83 \| (165, 250) \| 329 \| (1, 84) \| 160 \| Uncharacterized protein \| Uncharacterized protein; uniclust \| UniRef100\_A0A089NFY9 \| 98.9 \| 5.2e-12 \| 9.9e-18 \| 116.6 \| 173 \| (11, 208) \| 329 \| (69, 242) \| 361 \| Protein RecA (Fragment) \| Protein RecA (Fragment); uniclust \| UniRef100\_A0A011NVD5 \| 98.9 \| 4.8e-12 \| 1e-17 \| 131.6 \| 136 \| (13, 167) \| 329 \| (395, 532) \| 715 \| non-specific serine/threonine protein kinase \| non-specific serine/threonine protein kinase; uniclust \| UniRef100\_A0A031LJV2 \| 98.9 \| 4.9e-12 \| 1.1e-17 \| 128.2 \| 173 \| (14, 211) \| 329 \| (233, 420) \| 535 \| DNA repair and recombination protein RadA \| DNA repair and recombination protein RadA; uniclust \| UniRef100\_UPI0022E10B1A \| 98.9 \| 6.2e-12 \| 1.1e-17 \| 103.9 \| 113 \| (135, 251) \| 329 \| (9, 122) \| 185 \| hypothetical protein \| hypothetical protein; uniclust \| UniRef100\_A0A2D6SQ13 \| 98.9 \| 6.3e-12 \| 1.2e-17 \| 111.5 \| 175 \| (15, 212) \| 329 \| (99, 276) \| 314 \| DNA repair and recombination protein RadB \| DNA repair and recombination protein RadB; uniclust \| UniRef100\_A0A068RTV8 \| 98.9 \| 5.8e-12 \| 1.2e-17 \| 125.0 \| 175 \| (11, 213) \| 329 \| (93, 278) \| 483 \| Dna repair protein rad51 homolog 3 \| Dna repair protein rad51 homolog 3; uniclust \| UniRef100\_A0A139CHR2 \| 98.9 \| 5.9e-12 \| 1.2e-17 \| 114.0 \| 192 \| (14, 234) \| 329 \| (23, 224) \| 241 \| Circadian clock protein KaiC \| Circadian clock protein KaiC; uniclust \| UniRef100\_A0A075G1W7 \| 98.9 \| 6.8e-12 \| 1.4e-17 \| 115.8 \| 143 \| (12, 168) \| 329 \| (69, 211) \| 274 \| RecA/RadA recombinase-like protein (RAD51) \| RecA/RadA recombinase-like protein (RAD51); uniclust \| UniRef100\_A0A062UYH3 \| 98.9 \| 7.5e-12 \| 1.6e-17 \| 128.2 \| 200 \| (13, 247) \| 329 \| (267, 475) \| 570 \| non-specific serine/threonine protein kinase \| non-specific serine/threonine protein kinase; uniclust \| UniRef100\_A0A059X0Y6 \| 98.9 \| 8.2e-12 \| 1.7e-17 \| 124.0 \| 166 \| (14, 222) \| 329 \| (197, 365) \| 562 \| AAA domain protein \| AAA domain protein; uniclust \| UniRef100\_A0A3S1ZGV7 \| 98.9 \| 8.8e-12 \| 1.8e-17 \| 117.5 \| 182 \| (12, 227) \| 329 \| (103, 291) \| 344 \| Serine/threonine protein kinase (Fragment) \| Serine/threonine protein kinase (Fragment); uniclust \| UniRef100\_A0A0B8WUF2 \| 98.9 \| 8.8e-12 \| 1.8e-17 \| 124.4 \| 166 \| (13, 221) \| 329 \| (93, 261) \| 577 \| DNA repair protein RadA \| DNA repair protein RadA; uniclust \| UniRef100\_A0A1W9U784 \| 98.9 \| 9.4e-12 \| 1.9e-17 \| 111.2 \| 168 \| (4, 206) \| 329 \| (9, 181) \| 234 \| Uncharacterized protein \| Uncharacterized protein; uniclust \| UniRef100\_A0A0F8Y5Y5 \| 98.9 \| 9.5e-12 \| 2e-17 \| 113.2 \| 175 \| (12, 210) \| 329 \| (18, 203) \| 237 \| DNA repair and recombination protein RadB \| DNA repair and recombination protein RadB; uniclust \| UniRef100\_A0A010S4V8 \| 98.9 \| 9.4e-12 \| 2e-17 \| 125.0 \| 175 \| (13, 209) \| 329 \| (190, 378) \| 502 \| DNA repair protein RAD51 homolog \| DNA repair protein RAD51 homolog; uniclust \| UniRef100\_A0A0F8W083 \| 98.9 \| 1.1e-11 \| 2.2e-17 \| 118.1 \| 173 \| (12, 209) \| 329 \| (57, 230) \| 389 \| RecA family profile 2 domain-containing protein (Fragment) \| RecA family profile 2 domain-containing protein (Fragment); uniclust \| UniRef100\_A0A1G0WQ90 \| 98.9 \| 1e-11 \| 2.2e-17 \| 125.1 \| 162 \| (11, 210) \| 329 \| (79, 250) \| 500 \| AAA+ ATPase domain-containing protein \| AAA+ ATPase domain-containing protein; uniclust \| UniRef100\_A0A0F7PFT0 \| 98.9 \| 1.1e-11 \| 2.2e-17 \| 120.0 \| 161 \| (12, 207) \| 329 \| (220, 386) \| 432 \| Circadian clock protein, KaiC \| Circadian clock protein, KaiC; uniclust \| UniRef100\_A0A8T7MHD9 \| 98.9 \| 1.4e-11 \| 2.6e-17 \| 112.6 \| 202 \| (12, 245) \| 329 \| (16, 233) \| 400 \| AAA family ATPase \| AAA family ATPase; uniclust \| UniRef100\_A0A0K8Q481 \| 98.9 \| 1.3e-11 \| 2.6e-17 \| 122.4 \| 171 \| (12, 207) \| 329 \| (132, 303) \| 609 \| Multifunctional fusion protein \| Multifunctional fusion protein; uniclust \| UniRef100\_A0A0F9LI83 \| 98.9 \| 1.4e-11 \| 2.6e-17 \| 112.3 \| 236 \| (4, 248) \| 329 \| (3, 252) \| 394 \| AAA+ ATPase domain-containing protein \| AAA+ ATPase domain-containing protein; uniclust \| UniRef100\_A0A017N5Y4 \| 98.9 \| 1.3e-11 \| 2.7e-17 \| 119.1 \| 172 \| (1, 210) \| 329 \| (62, 243) \| 419 \| AAA domain protein \| AAA domain protein; uniclust \| UniRef100\_A0A258HM16 \| 98.9 \| 1.4e-11 \| 2.7e-17 \| 106.2 \| 109 \| (12, 127) \| 329 \| (31, 142) \| 196 \| KaiC domain-containing protein \| KaiC domain-containing protein; uniclust \| UniRef100\_A0A0D6JWN7 \| 98.9 \| 1.4e-11 \| 2.8e-17 \| 120.3 \| 161 \| (12, 207) \| 329 \| (261, 427) \| 473 \| Circadian clock protein kinase KaiC \| Circadian clock protein kinase KaiC; uniclust \| UniRef100\_A0A0Q7FJN5 \| 98.9 \| 1.4e-11 \| 2.9e-17 \| 118.2 \| 200 \| (13, 248) \| 329 \| (60, 268) \| 385 \| KaiC domain-containing protein \| KaiC domain-containing protein; uniclust \| UniRef100\_A0A0E3ZFC2 \| 98.9 \| 1.5e-11 \| 3e-17 \| 120.7 \| 136 \| (13, 167) \| 329 \| (287, 424) \| 510 \| non-specific serine/threonine protein kinase \| non-specific serine/threonine protein kinase; uniclust \| UniRef100\_A0A2J8A2H5 \| 98.9 \| 1.6e-11 \| 3.1e-17 \| 113.5 \| 144 \| (12, 172) \| 329 \| (126, 270) \| 346 \| Protein RecA \| Protein RecA; uniclust \| UniRef100\_A0A023X4H3 \| 98.9 \| 1.7e-11 \| 3.6e-17 \| 122.7 \| 166 \| (13, 221) \| 329 \| (155, 323) \| 552 \| DNA repair protein RadA \| DNA repair protein RadA; uniclust \| UniRef100\_A0A0G0QNB5 \| 98.9 \| 1.8e-11 \| 3.7e-17 \| 120.7 \| 164 \| (14, 221) \| 329 \| (94, 260) \| 449 \| DNA repair protein RadA \| DNA repair protein RadA; uniclust \| UniRef100\_A0A369T1U4 \| 98.9 \| 1.9e-11 \| 3.8e-17 \| 113.0 \| 138 \| (13, 167) \| 329 \| (101, 250) \| 314 \| DNA repair and recombination protein RadA \| DNA repair and recombination protein RadA; uniclust \| UniRef100\_A0A176VER6 \| 98.9 \| 1.9e-11 \| 3.9e-17 \| 119.8 \| 172 \| (11, 207) \| 329 \| (181, 353) \| 513 \| Uncharacterized protein \| Uncharacterized protein; uniclust \| UniRef100\_A0A2D6P3X9 \| 98.9 \| 1.9e-11 \| 3.9e-17 \| 115.3 \| 159 \| (14, 211) \| 329 \| (125, 293) \| 333 \| KaiC domain-containing protein \| KaiC domain-containing protein; uniclust \| UniRef100\_A0A0F9FGH6 \| 98.8 \| 2.1e-11 \| 4.1e-17 \| 106.3 \| 144 \| (12, 173) \| 329 \| (59, 204) \| 218 \| RecA family profile 1 domain-containing protein (Fragment) \| RecA family profile 1 domain-containing protein (Fragment); uniclust \| UniRef100\_A0A2G9PAU5 \| 98.8 \| 2e-11 \| 4.4e-17 \| 112.9 \| 164 \| (12, 212) \| 329 \| (32, 213) \| 251 \| KaiC-like domain-containing protein \| KaiC-like domain-containing protein; uniclust \| UniRef100\_A0A023Y4B6 \| 98.8 \| 2.3e-11 \| 4.4e-17 \| 113.9 \| 172 \| (12, 207) \| 329 \| (130, 303) \| 438 \| Protein RecA \| Protein RecA; uniclust \| UniRef100\_A0A3N5EQ31 \| 98.8 \| 2.3e-11 \| 4.7e-17 \| 110.0 \| 136 \| (7, 168) \| 329 \| (10, 148) \| 252 \| ATP-binding protein \| ATP-binding protein; uniclust \| UniRef100\_A0A3A1Y8F9 \| 98.8 \| 2.7e-11 \| 5.2e-17 \| 110.7 \| 190 \| (5, 208) \| 329 \| (6, 203) \| 325 \| AAA domain-containing protein \| AAA domain-containing protein; uniclust \| UniRef100\_A0A3M1F472 \| 98.8 \| 2.8e-11 \| 5.2e-17 \| 110.0 \| 167 \| (15, 210) \| 329 \| (178, 347) \| 379 \| RecA family profile 1 domain-containing protein \| RecA family profile 1 domain-containing protein; uniclust \| UniRef100\_A0A0B3AV81 \| 98.8 \| 2.8e-11 \| 5.4e-17 \| 114.9 \| 166 \| (12, 207) \| 329 \| (183, 364) \| 416 \| Circadian clock protein KaiC \| Circadian clock protein KaiC; uniclust \| UniRef100\_A0A662VFD8 \| 98.8 \| 2.7e-11 \| 5.5e-17 \| 115.1 \| 162 \| (13, 208) \| 329 \| (32, 195) \| 368 \| KaiC domain-containing protein (Fragment) \| KaiC domain-containing protein (Fragment); uniclust \| UniRef100\_A0A0E3SP83 \| 98.8 \| 2.7e-11 \| 5.6e-17 \| 105.7 \| 154 \| (43, 223) \| 329 \| (3, 159) \| 184 \| DNA repair and recombination protein RadB \| DNA repair and recombination protein RadB; uniclust \| UniRef100\_A0A023NM81 \| 98.8 \| 3e-11 \| 5.7e-17 \| 104.2 \| 145 \| (12, 173) \| 329 \| (20, 165) \| 208 \| Protein RecA (Fragment) \| Protein RecA (Fragment); uniclust \| UniRef100\_A0A011NVD5 \| 98.8 \| 2.9e-11 \| 6.2e-17 \| 126.1 \| 163 \| (12, 208) \| 329 \| (152, 326) \| 715 \| non-specific serine/threonine protein kinase \| non-specific serine/threonine protein kinase; uniclust \| UniRef100\_A0A0F9CU89 \| 98.8 \| 3.2e-11 \| 6.5e-17 \| 110.6 \| 207 \| (5, 227) \| 329 \| (5, 239) \| 282 \| AAA domain-containing protein \| AAA domain-containing protein; uniclust \| UniRef100\_A0A382IL75 \| 98.8 \| 3.4e-11 \| 6.6e-17 \| 113.2 \| 167 \| (13, 222) \| 329 \| (68, 237) \| 369 \| RecA family profile 1 domain-containing protein (Fragment) \| RecA family profile 1 domain-containing protein (Fragment); uniclust \| UniRef100\_A0A2R6D0K4 \| 98.8 \| 3.5e-11 \| 7e-17 \| 107.0 \| 137 \| (13, 169) \| 329 \| (45, 192) \| 229 \| KaiC-like domain-containing protein \| KaiC-like domain-containing protein; uniclust \| UniRef100\_A0A0E3T6D1 \| 98.8 \| 3.9e-11 \| 7.7e-17 \| 112.2 \| 176 \| (7, 211) \| 329 \| (18, 207) \| 337 \| ATP-binding protein \| ATP-binding protein; uniclust \| UniRef100\_A0A1H6L1N6 \| 98.8 \| 2.7e-11 \| 7.7e-17 \| 134.1 \| 169 \| (14, 225) \| 329 \| (88, 259) \| 846 \| DNA repair protein RadA \| DNA repair protein RadA; uniclust \| UniRef100\_A0A088F6N8 \| 98.8 \| 3.9e-11 \| 7.7e-17 \| 111.3 \| 187 \| (4, 227) \| 329 \| (44, 251) \| 300 \| ATPase \| ATPase; uniclust \| UniRef100\_A0A2H9PL68 \| 98.8 \| 4.2e-11 \| 8e-17 \| 112.4 \| 157 \| (14, 210) \| 329 \| (177, 343) \| 386 \| KaiC domain-containing protein \| KaiC domain-containing protein; uniclust \| UniRef100\_A0A328S0V5 \| 98.8 \| 4.2e-11 \| 8.7e-17 \| 113.9 \| 169 \| (15, 206) \| 329 \| (116, 298) \| 338 \| DNA repair and recombination protein RadA (Fragment) \| DNA repair and recombination protein RadA (Fragment); uniclust \| UniRef100\_A0A7C5P7L2 \| 98.8 \| 4.9e-11 \| 9.5e-17 \| 104.9 \| 138 \| (12, 173) \| 329 \| (25, 164) \| 223 \| Circadian clock KaiC-like protein \| Circadian clock KaiC-like protein; uniclust \| UniRef100\_A0A0E3LEL1 \| 98.8 \| 4.9e-11 \| 1e-16 \| 100.8 \| 104 \| (12, 119) \| 329 \| (42, 147) \| 151 \| Circadian clock protein KaiC \| Circadian clock protein KaiC; uniclust \| UniRef100\_A0A0G2ZHU4 \| 98.8 \| 5.3e-11 \| 1.1e-16 \| 123.6 \| 137 \| (12, 167) \| 329 \| (313, 452) \| 785 \| Circadian clock protein KaiC \| Circadian clock protein KaiC; uniclust \| UniRef100\_A0A0A1VGZ2 \| 98.8 \| 5.3e-11 \| 1.1e-16 \| 114.1 \| 186 \| (12, 234) \| 329 \| (45, 240) \| 343 \| RecA-superfamily ATPases implicated in signal transduction \| RecA-superfamily ATPases implicated in signal transduction; uniclust \| UniRef100\_A0A497KWF6 \| 98.8 \| 5.8e-11 \| 1.1e-16 \| 104.5 \| 135 \| (12, 166) \| 329 \| (49, 195) \| 228 \| KaiC domain-containing protein (Fragment) \| KaiC domain-containing protein (Fragment); uniclust \| UniRef100\_A0A1C8HS01 \| 98.8 \| 5.8e-11 \| 1.2e-16 \| 112.2 \| 181 \| (12, 212) \| 329 \| (16, 233) \| 321 \| Putative ATPase \| Putative ATPase; uniclust \| UniRef100\_A0A1G9ZZ45 \| 98.8 \| 6.1e-11 \| 1.2e-16 \| 111.4 \| 161 \| (12, 207) \| 329 \| (154, 320) \| 366 \| KaiC domain protein, AF\_0351 family \| KaiC domain protein, AF\_0351 family; uniclust \| UniRef100\_A0A067DDE5 \| 98.8 \| 6.4e-11 \| 1.3e-16 \| 110.3 \| 137 \| (12, 168) \| 329 \| (115, 255) \| 307 \| RecA family profile 1 domain-containing protein (Fragment) \| RecA family profile 1 domain-containing protein (Fragment); uniclust \| UniRef100\_A0A0J1D9A8 \| 98.8 \| 6.2e-11 \| 1.3e-16 \| 121.5 \| 161 \| (12, 207) \| 329 \| (321, 485) \| 612 \| non-specific serine/threonine protein kinase \| non-specific serine/threonine protein kinase; uniclust \| UniRef100\_A0A938TZZ1 \| 98.8 \| 7.1e-11 \| 1.3e-16 \| 110.4 \| 186 \| (4, 221) \| 329 \| (82, 282) \| 419 \| Uncharacterized protein \| Uncharacterized protein; uniclust \| UniRef100\_A0A0A0RM32 \| 98.8 \| 7e-11 \| 1.4e-16 \| 112.6 \| 197 \| (13, 241) \| 329 \| (49, 250) \| 371 \| RecA-like DNA recombinase \| RecA-like DNA recombinase; uniclust \| UniRef100\_A0A497QXX0 \| 98.7 \| 7.5e-11 \| 1.5e-16 \| 106.1 \| 171 \| (13, 212) \| 329 \| (44, 217) \| 262 \| RecA family profile 1 domain-containing protein \| RecA family profile 1 domain-containing protein; uniclust \| UniRef100\_A0A7J4EQR4 \| 98.7 \| 8.4e-11 \| 1.6e-16 \| 102.6 \| 209 \| (10, 244) \| 329 \| (28, 249) \| 265 \| AAA family ATPase \| AAA family ATPase; uniclust \| UniRef100\_A0A062V0T1 \| 98.7 \| 7.6e-11 \| 1.6e-16 \| 113.5 \| 173 \| (13, 223) \| 329 \| (77, 259) \| 352 \| RecA-superfamily ATPase possibly involved in signal transduction \| RecA-superfamily ATPase possibly involved in signal transduction; uniclust \| UniRef100\_A0A0W0VNN5 \| 98.7 \| 8.5e-11 \| 1.7e-16 \| 112.5 \| 195 \| (13, 242) \| 329 \| (44, 245) \| 357 \| DNA integration/recombination/inversion protein \| DNA integration/recombination/inversion protein; uniclust \| UniRef100\_A0A534KBF9 \| 98.7 \| 9.7e-11 \| 1.8e-16 \| 103.9 \| 99 \| (13, 122) \| 329 \| (29, 132) \| 302 \| RecA family profile 1 domain-containing protein \| RecA family profile 1 domain-containing protein; uniclust \| UniRef100\_UPI000408E576 \| 98.7 \| 9.7e-11 \| 1.8e-16 \| 110.1 \| 205 \| (4, 226) \| 329 \| (1, 228) \| 504 \| AAA family ATPase \| AAA family ATPase; uniclust \| UniRef100\_A0A034WVC2 \| 98.7 \| 8.5e-11 \| 1.8e-16 \| 114.2 \| 209 \| (12, 243) \| 329 \| (99, 334) \| 380 \| DNA repair protein RAD51-like protein 3 \| DNA repair protein RAD51-like protein 3; uniclust \| UniRef100\_A0A2M8NG75 \| 98.7 \| 9.1e-11 \| 1.8e-16 \| 105.2 \| 153 \| (17, 210) \| 329 \| (1, 153) \| 248 \| DNA repair protein RadA (Fragment) \| DNA repair protein RadA (Fragment); uniclust \| UniRef100\_A0A7J4N8G8 \| 98.7 \| 9.9e-11 \| 1.8e-16 \| 103.6 \| 166 \| (13, 208) \| 329 \| (84, 254) \| 295 \| DNA repair and recombination protein RadB \| DNA repair and recombination protein RadB; uniclust \| UniRef100\_A0A059DH68 \| 98.7 \| 9.4e-11 \| 1.8e-16 \| 113.6 \| 173 \| (12, 208) \| 329 \| (156, 331) \| 479 \| RecA family profile 2 domain-containing protein \| RecA family profile 2 domain-containing protein; uniclust \| UniRef100\_A0A2X2K0Q1 \| 98.7 \| 9.4e-11 \| 1.9e-16 \| 107.2 \| 155 \| (13, 211) \| 329 \| (47, 203) \| 260 \| DNA repair protein RadA \| DNA repair protein RadA; uniclust \| UniRef100\_A0A6H1ZJT6 \| 98.7 \| 9.3e-11 \| 2e-16 \| 106.8 \| 172 \| (15, 208) \| 329 \| (8, 186) \| 229 \| Putative ATPase domain containing protein \| Putative ATPase domain containing protein; uniclust \| UniRef100\_A0A0F9GA24 \| 98.7 \| 9.3e-11 \| 2e-16 \| 115.1 \| 171 \| (12, 207) \| 329 \| (60, 246) \| 395 \| AAA+ ATPase domain-containing protein (Fragment) \| AAA+ ATPase domain-containing protein (Fragment); uniclust \| UniRef100\_A0A259P4E8 \| 98.7 \| 9.6e-11 \| 2e-16 \| 117.2 \| 134 \| (13, 167) \| 329 \| (37, 177) \| 489 \| Protein kinase (Fragment) \| Protein kinase (Fragment); uniclust \| UniRef100\_A0A0K1E4U1 \| 98.7 \| 1e-10 \| 2.1e-16 \| 113.6 \| 141 \| (16, 208) \| 329 \| (9, 152) \| 389 \| RecA-superfamily ATPase implicated in signal transduction \| RecA-superfamily ATPase implicated in signal transduction; uniclust \| UniRef100\_A0A0P9DUN6 \| 98.7 \| 1.1e-10 \| 2.3e-16 \| 100.7 \| 138 \| (13, 167) \| 329 \| (26, 166) \| 185 \| Circadian clock protein KaiC \| Circadian clock protein KaiC; uniclust \| UniRef100\_A0A2V7UA84 \| 98.7 \| 1.2e-10 \| 2.4e-16 \| 108.8 \| 143 \| (12, 173) \| 329 \| (35, 189) \| 342 \| Circadian clock protein KaiC (Fragment) \| Circadian clock protein KaiC (Fragment); uniclust \| UniRef100\_U2YUY2 \| 98.7 \| 1.4e-10 \| 2.5e-16 \| 106.9 \| 184 \| (13, 220) \| 329 \| (202, 387) \| 414 \| DNA repair and recombination protein RadB \| DNA repair and recombination protein RadB; uniclust \| UniRef100\_A0A151BLE4 \| 98.7 \| 1.2e-10 \| 2.5e-16 \| 117.2 \| 164 \| (12, 208) \| 329 \| (104, 271) \| 558 \| KaiC domain-containing protein \| KaiC domain-containing protein; uniclust \| UniRef100\_A0A1J8P3X0 \| 98.7 \| 1.3e-10 \| 2.5e-16 \| 108.0 \| 172 \| (12, 208) \| 329 \| (61, 233) \| 355 \| Protein RecA \| Protein RecA; uniclust \| UniRef100\_A0A062XNQ7 \| 98.7 \| 1.2e-10 \| 2.6e-16 \| 118.3 \| 162 \| (13, 207) \| 329 \| (319, 482) \| 541 \| non-specific serine/threonine protein kinase \| non-specific serine/threonine protein kinase; uniclust \| UniRef100\_A0A0F9NK85 \| 98.7 \| 1.4e-10 \| 2.8e-16 \| 104.4 \| 167 \| (12, 207) \| 329 \| (24, 196) \| 252 \| Phage nucleotide-binding protein \| Phage nucleotide-binding protein; uniclust \| UniRef100\_A0A0C9PX14 \| 98.7 \| 1.4e-10 \| 2.8e-16 \| 113.1 \| 190 \| (13, 219) \| 329 \| (111, 318) \| 391 \| RadA protein \| RadA protein; uniclust \| UniRef100\_A0A031LLC8 \| 98.7 \| 1.4e-10 \| 2.9e-16 \| 102.9 \| 145 \| (13, 208) \| 329 \| (26, 170) \| 196 \| ATPase AAA \| ATPase AAA; uniclust \| UniRef100\_A0A062UYH3 \| 98.7 \| 1.3e-10 \| 2.9e-16 \| 119.6 \| 155 \| (13, 208) \| 329 \| (39, 198) \| 570 \| non-specific serine/threonine protein kinase \| non-specific serine/threonine protein kinase; uniclust \| UniRef100\_A0A6J5NUX5 \| 98.7 \| 1.5e-10 \| 2.9e-16 \| 111.2 \| 171 \| (12, 208) \| 329 \| (56, 235) \| 408 \| RecA RecA/RadA recombinase \| RecA RecA/RadA recombinase; uniclust \| UniRef100\_A0A017TDB8 \| 98.7 \| 1.3e-10 \| 3e-16 \| 122.8 \| 191 \| (12, 234) \| 329 \| (348, 546) \| 653 \| non-specific serine/threonine protein kinase \| non-specific serine/threonine protein kinase; uniclust \| UniRef100\_A0A0M0BPC7 \| 98.7 \| 1.4e-10 \| 3e-16 \| 118.1 \| 164 \| (12, 208) \| 329 \| (48, 215) \| 527 \| KaiC domain-containing protein \| KaiC domain-containing protein; uniclust \| UniRef100\_A0A0S8EHQ0 \| 98.7 \| 1.5e-10 \| 3.1e-16 \| 105.5 \| 163 \| (12, 211) \| 329 \| (38, 205) \| 248 \| KaiC-like domain-containing protein \| KaiC-like domain-containing protein; uniclust \| UniRef100\_A0A014N6S2 \| 98.7 \| 1.6e-10 \| 3.2e-16 \| 116.1 \| 182 \| (7, 210) \| 329 \| (16, 224) \| 553 \| Uncharacterized protein \| Uncharacterized protein; uniclust \| UniRef100\_A0A0P6Y9P5 \| 98.7 \| 1.7e-10 \| 3.3e-16 \| 108.5 \| 178 \| (5, 211) \| 329 \| (1, 192) \| 350 \| AAA+ ATPase domain-containing protein \| AAA+ ATPase domain-containing protein; uniclust \| UniRef100\_A0A0F9HTH3 \| 98.7 \| 1.7e-10 \| 3.3e-16 \| 111.8 \| 167 \| (13, 206) \| 329 \| (116, 284) \| 439 \| RecA family profile 2 domain-containing protein \| RecA family profile 2 domain-containing protein; uniclust \| UniRef100\_A0A0B5FRR7 \| 98.7 \| 1.6e-10 \| 3.3e-16 \| 109.1 \| 156 \| (15, 207) \| 329 \| (47, 212) \| 295 \| ATP-binding protein \| ATP-binding protein; uniclust \| UniRef100\_A0A0S6W1H8 \| 98.7 \| 1.7e-10 \| 3.4e-16 \| 115.4 \| 161 \| (15, 207) \| 329 \| (266, 428) \| 552 \| Circadian clock protein KaiC \| Circadian clock protein KaiC; uniclust \| UniRef100\_A0A7C2V9T1 \| 98.7 \| 1.8e-10 \| 3.4e-16 \| 94.4 \| 102 \| (15, 122) \| 329 \| (22, 126) \| 142 \| Recombinase RecA (Fragment) \| Recombinase RecA (Fragment); uniclust \| UniRef100\_A0A2G6VT70 \| 98.7 \| 1.7e-10 \| 3.6e-16 \| 112.9 \| 164 \| (16, 208) \| 329 \| (122, 303) \| 418 \| AAA domain-containing protein \| AAA domain-containing protein; uniclust \| UniRef100\_A0A0F9RAF4 \| 98.7 \| 1.8e-10 \| 3.7e-16 \| 110.2 \| 174 \| (12, 208) \| 329 \| (70, 244) \| 372 \| RecA family profile 2 domain-containing protein \| RecA family profile 2 domain-containing protein; uniclust \| UniRef100\_A0A3B9JM63 \| 98.7 \| 2e-10 \| 3.8e-16 \| 102.5 \| 168 \| (10, 211) \| 329 \| (9, 184) \| 285 \| ATP-binding protein \| ATP-binding protein; uniclust \| UniRef100\_A0A257A9J6 \| 98.7 \| 2e-10 \| 3.9e-16 \| 105.8 \| 148 \| (12, 169) \| 329 \| (43, 193) \| 281 \| RecA family profile 1 domain-containing protein \| RecA family profile 1 domain-containing protein; uniclust \| UniRef100\_A0A0F0HMR4 \| 98.7 \| 1.8e-10 \| 4e-16 \| 114.4 \| 169 \| (12, 209) \| 329 \| (63, 233) \| 406 \| Protein RecA \| Protein RecA; uniclust \| UniRef100\_A0A0B5HAI0 \| 98.7 \| 2.1e-10 \| 4.1e-16 \| 98.6 \| 96 \| (12, 122) \| 329 \| (77, 173) \| 178 \| Protein RecA (Fragment) \| Protein RecA (Fragment); uniclust \| UniRef100\_A0A0P9H5V8 \| 98.7 \| 2.4e-10 \| 4.5e-16 \| 111.0 \| 169 \| (12, 208) \| 329 \| (70, 242) \| 518 \| Protein RecA \| Protein RecA; uniclust \| UniRef100\_A0A060RFG9 \| 98.7 \| 2.2e-10 \| 4.6e-16 \| 105.3 \| 170 \| (1, 207) \| 329 \| (5, 184) \| 240 \| RecA \| RecA; uniclust \| UniRef100\_A0A4Q3IJ75 \| 98.6 \| 2.6e-10 \| 4.8e-16 \| 92.7 \| 83 \| (165, 251) \| 329 \| (1, 85) \| 158 \| Uncharacterized protein \| Uncharacterized protein; uniclust \| UniRef100\_A0A1F5S7A6 \| 98.6 \| 2.5e-10 \| 4.9e-16 \| 111.6 \| 157 \| (13, 209) \| 329 \| (91, 248) \| 468 \| DNA repair protein RadA \| DNA repair protein RadA; uniclust \| UniRef100\_A0A838VNY1 \| 98.6 \| 2.7e-10 \| 5.1e-16 \| 101.6 \| 110 \| (133, 246) \| 329 \| (10, 124) \| 266 \| Uncharacterized protein \| Uncharacterized protein; uniclust \| UniRef100\_A0A497T6C2 \| 98.6 \| 2.6e-10 \| 5.1e-16 \| 95.9 \| 100 \| (12, 118) \| 329 \| (27, 128) \| 151 \| KaiC domain-containing protein (Fragment) \| KaiC domain-containing protein (Fragment); uniclust \| UniRef100\_A0A022QKB8 \| 98.6 \| 2.7e-10 \| 5.2e-16 \| 112.7 \| 173 \| (12, 207) \| 329 \| (236, 409) \| 607 \| RecA family profile 2 domain-containing protein \| RecA family profile 2 domain-containing protein; uniclust \| UniRef100\_A0A0B7MPK7 \| 98.6 \| 2.5e-10 \| 5.3e-16 \| 116.1 \| 156 \| (13, 208) \| 329 \| (37, 202) \| 571 \| non-specific serine/threonine protein kinase \| non-specific serine/threonine protein kinase; uniclust \| UniRef100\_A0A5R9EA46 \| 98.6 \| 3.1e-10 \| 5.6e-16 \| 106.9 \| 244 \| (15, 282) \| 329 \| (164, 443) \| 501 \| AAA family ATPase \| AAA family ATPase; uniclust \| UniRef100\_A0A5A8DKM2 \| 98.6 \| 3e-10 \| 5.6e-16 \| 108.8 \| 172 \| (12, 208) \| 329 \| (122, 294) \| 504 \| RecA family profile 2 domain-containing protein \| RecA family profile 2 domain-containing protein; uniclust \| UniRef100\_A0A8S5MWZ3 \| 98.6 \| 3e-10 \| 5.8e-16 \| 102.7 \| 210 \| (12, 248) \| 329 \| (41, 269) \| 280 \| AAA domain protein \| AAA domain protein; uniclust \| UniRef100\_A0A4R6LFL0 \| 98.6 \| 2.9e-10 \| 5.9e-16 \| 106.6 \| 167 \| (15, 227) \| 329 \| (34, 220) \| 317 \| KaiC/GvpD/RAD55 family RecA-like ATPase \| KaiC/GvpD/RAD55 family RecA-like ATPase; uniclust \| UniRef100\_A0A022PSD5 \| 98.6 \| 3.3e-10 \| 6.5e-16 \| 112.4 \| 172 \| (12, 208) \| 329 \| (205, 380) \| 542 \| RecA family profile 1 domain-containing protein \| RecA family profile 1 domain-containing protein; uniclust \| UniRef100\_A0A4Q6DJA5 \| 98.6 \| 3.7e-10 \| 6.8e-16 \| 97.0 \| 191 \| (31, 247) \| 329 \| (26, 223) \| 229 \| Uncharacterized protein \| Uncharacterized protein; uniclust \| UniRef100\_A0A256Z3M3 \| 98.6 \| 3.5e-10 \| 6.8e-16 \| 101.1 \| 162 \| (13, 208) \| 329 \| (23, 186) \| 243 \| KaiC domain-containing protein (Fragment) \| KaiC domain-containing protein (Fragment); uniclust \| UniRef100\_A0A0F9FQA9 \| 98.6 \| 3.2e-10 \| 6.8e-16 \| 109.3 \| 174 \| (12, 208) \| 329 \| (55, 232) \| 350 \| RecA family profile 1 domain-containing protein (Fragment) \| RecA family profile 1 domain-containing protein (Fragment); uniclust \| UniRef100\_A0A1X3I5A2 \| 98.6 \| 3.7e-10 \| 6.9e-16 \| 92.7 \| 141 \| (4, 164) \| 329 \| (2, 156) \| 161 \| Putative bacteriophage protein \| Putative bacteriophage protein; uniclust \| UniRef100\_A0A135VKT4 \| 98.6 \| 3.4e-10 \| 7e-16 \| 110.0 \| 179 \| (13, 210) \| 329 \| (101, 287) \| 396 \| DNA repair and recombination protein RadA (Fragment) \| DNA repair and recombination protein RadA (Fragment); uniclust \| UniRef100\_A0A024G263 \| 98.6 \| 3.6e-10 \| 7.1e-16 \| 112.9 \| 172 \| (13, 223) \| 329 \| (161, 346) \| 607 \| RecA family profile 1 domain-containing protein \| RecA family profile 1 domain-containing protein; uniclust \| UniRef100\_A0A497GIR7 \| 98.6 \| 3.9e-10 \| 7.4e-16 \| 101.2 \| 136 \| (13, 166) \| 329 \| (49, 202) \| 272 \| Recombinase RecA (Fragment) \| Recombinase RecA (Fragment); uniclust \| UniRef100\_A0A0K0N6N5 \| 98.6 \| 3.7e-10 \| 7.6e-16 \| 105.3 \| 209 \| (14, 244) \| 329 \| (33, 252) \| 291 \| RecA-like DNA recombinase \| RecA-like DNA recombinase; uniclust \| UniRef100\_A0A2W4IIX8 \| 98.6 \| 4.2e-10 \| 7.8e-16 \| 100.6 \| 201 \| (12, 248) \| 329 \| (9, 219) \| 313 \| AAA family ATPase \| AAA family ATPase; uniclust \| UniRef100\_A0A0D6E0K0 \| 98.6 \| 3.7e-10 \| 7.9e-16 \| 111.4 \| 128 \| (11, 170) \| 329 \| (34, 170) \| 390 \| Putative phage nucleotide-binding protein \| Putative phage nucleotide-binding protein; uniclust \| UniRef100\_A0A497JUW8 \| 98.6 \| 3.9e-10 \| 8e-16 \| 109.0 \| 139 \| (12, 167) \| 329 \| (29, 171) \| 397 \| KaiC domain-containing protein (Fragment) \| KaiC domain-containing protein (Fragment); uniclust \| UniRef100\_A0A7M6UVN9 \| 98.6 \| 4.9e-10 \| 9.9e-16 \| 105.9 \| 185 \| (13, 219) \| 329 \| (93, 296) \| 320 \| RecA family profile 1 domain-containing protein \| RecA family profile 1 domain-containing protein; uniclust \| UniRef100\_A0A015L9R8 \| 98.6 \| 5e-10 \| 1e-15 \| 110.1 \| 180 \| (15, 216) \| 329 \| (202, 395) \| 480 \| Dmc1p \| Dmc1p; uniclust \| UniRef100\_D8U914 \| 98.6 \| 5.5e-10 \| 1e-15 \| 106.7 \| 173 \| (12, 210) \| 329 \| (275, 449) \| 576 \| Uncharacterized protein \| Uncharacterized protein; uniclust \| UniRef100\_A0A010ZS25 \| 98.6 \| 5e-10 \| 1e-15 \| 112.1 \| 208 \| (14, 246) \| 329 \| (133, 346) \| 512 \| RecA/RadA recombinase \| RecA/RadA recombinase; uniclust \| UniRef100\_A0A059AII9 \| 98.6 \| 5e-10 \| 1e-15 \| 117.3 \| 159 \| (13, 211) \| 329 \| (314, 476) \| 817 \| RecA family profile 1 domain-containing protein \| RecA family profile 1 domain-containing protein; uniclust \| UniRef100\_A0A0H5AZJ9 \| 98.6 \| 5.7e-10 \| 1.1e-15 \| 111.5 \| 182 \| (13, 227) \| 329 \| (314, 500) \| 545 \| non-specific serine/threonine protein kinase \| non-specific serine/threonine protein kinase; uniclust \| UniRef100\_A0A074TZL4 \| 98.6 \| 5.5e-10 \| 1.2e-15 \| 114.2 \| 180 \| (13, 223) \| 329 \| (112, 307) \| 518 \| AAA+ ATPase domain-containing protein \| AAA+ ATPase domain-containing protein; uniclust \| UniRef100\_A0A024VLC0 \| 98.6 \| 6.7e-10 \| 1.3e-15 \| 105.1 \| 176 \| (13, 210) \| 329 \| (143, 332) \| 390 \| Meiotic recombinase Dmc1 \| Meiotic recombinase Dmc1; uniclust \| UniRef100\_A0A2D6N6W6 \| 98.6 \| 7.1e-10 \| 1.4e-15 \| 101.9 \| 166 \| (13, 221) \| 329 \| (83, 251) \| 302 \| DNA repair protein RadA (Fragment) \| DNA repair protein RadA (Fragment); uniclust \| UniRef100\_A0A0Q4BB21 \| 98.6 \| 6.5e-10 \| 1.4e-15 \| 106.8 \| 163 \| (12, 210) \| 329 \| (88, 262) \| 314 \| KaiC-like domain-containing protein \| KaiC-like domain-containing protein; uniclust \| UniRef100\_A0A151EFE0 \| 98.6 \| 7.4e-10 \| 1.4e-15 \| 102.6 \| 163 \| (11, 208) \| 329 \| (127, 308) \| 353 \| KaiC domain-containing protein \| KaiC domain-containing protein; uniclust \| UniRef100\_A0A063Z806 \| 98.5 \| 6.9e-10 \| 1.5e-15 \| 105.6 \| 163 \| (12, 211) \| 329 \| (90, 260) \| 306 \| Flagellar accessory protein FlaH \| Flagellar accessory protein FlaH; uniclust \| UniRef100\_A0A1V4YX98 \| 98.5 \| 7.3e-10 \| 1.5e-15 \| 109.9 \| 139 \| (12, 167) \| 329 \| (33, 189) \| 491 \| Circadian clock protein KaiC \| Circadian clock protein KaiC; uniclust \| UniRef100\_A0A024US34 \| 98.5 \| 6.9e-10 \| 1.5e-15 \| 110.4 \| 149 \| (13, 172) \| 329 \| (119, 282) \| 414 \| RecA family profile 1 domain-containing protein \| RecA family profile 1 domain-containing protein; uniclust \| UniRef100\_UPI000B11D97B \| 98.5 \| 8.6e-10 \| 1.6e-15 \| 77.6 \| 64 \| (3, 66) \| 329 \| (1, 64) \| 68 \| AAA family ATPase \| AAA family ATPase; uniclust \| UniRef100\_A0A182F8F8 \| 98.5 \| 8.1e-10 \| 1.6e-15 \| 101.8 \| 160 \| (13, 182) \| 329 \| (55, 223) \| 288 \| RecA family profile 1 domain-containing protein \| RecA family profile 1 domain-containing protein; uniclust \| UniRef100\_A0A059D4I4 \| 98.5 \| 7.7e-10 \| 1.7e-15 \| 114.2 \| 150 \| (12, 172) \| 329 \| (199, 363) \| 578 \| RecA family profile 1 domain-containing protein \| RecA family profile 1 domain-containing protein; uniclust \| UniRef100\_A0A022QJU5 \| 98.5 \| 9e-10 \| 1.7e-15 \| 107.9 \| 147 \| (11, 174) \| 329 \| (193, 340) \| 588 \| RecA family profile 2 domain-containing protein \| RecA family profile 2 domain-containing protein; uniclust \| UniRef100\_A0A0F9W5I4 \| 98.5 \| 8.4e-10 \| 1.8e-15 \| 110.0 \| 184 \| (13, 234) \| 329 \| (115, 306) \| 433 \| AAA+ ATPase domain-containing protein \| AAA+ ATPase domain-containing protein; uniclust \| UniRef100\_A0A0F9CDA0 \| 98.5 \| 9.1e-10 \| 1.8e-15 \| 100.8 \| 177 \| (13, 210) \| 329 \| (10, 204) \| 259 \| AAA+ ATPase domain-containing protein \| AAA+ ATPase domain-containing protein; uniclust \| UniRef100\_A0A1H1AH51 \| 98.5 \| 9.2e-10 \| 1.9e-15 \| 110.1 \| 179 \| (13, 227) \| 329 \| (271, 454) \| 500 \| non-specific serine/threonine protein kinase \| non-specific serine/threonine protein kinase; uniclust \| UniRef100\_A0A0G1VH57 \| 98.5 \| 1e-09 \| 2e-15 \| 107.7 \| 167 \| (14, 221) \| 329 \| (91, 260) \| 490 \| Repair protein radA protein \| Repair protein radA protein; uniclust \| UniRef100\_A0A022QH90 \| 98.5 \| 1.1e-09 \| 2.1e-15 \| 109.1 \| 175 \| (12, 208) \| 329 \| (256, 431) \| 648 \| RecA family profile 2 domain-containing protein (Fragment) \| RecA family profile 2 domain-containing protein (Fragment); uniclust \| UniRef100\_A0A2H5XG24 \| 98.5 \| 1e-09 \| 2.1e-15 \| 110.6 \| 134 \| (14, 167) \| 329 \| (270, 407) \| 503 \| non-specific serine/threonine protein kinase \| non-specific serine/threonine protein kinase; uniclust \| UniRef100\_A0A2D7G538 \| 98.5 \| 1.1e-09 \| 2.2e-15 \| 94.2 \| 96 \| (12, 122) \| 329 \| (70, 166) \| 176 \| Protein RecA (Fragment) \| Protein RecA (Fragment); uniclust \| UniRef100\_A0A496QMI5 \| 98.5 \| 1.2e-09 \| 2.2e-15 \| 103.2 \| 183 \| (12, 227) \| 329 \| (47, 241) \| 376 \| KaiC 1 (Fragment) \| KaiC 1 (Fragment); uniclust \| UniRef100\_A0A022QWM4 \| 98.5 \| 1.1e-09 \| 2.3e-15 \| 110.5 \| 181 \| (12, 210) \| 329 \| (157, 361) \| 468 \| RecA family profile 1 domain-containing protein \| RecA family profile 1 domain-containing protein; uniclust \| UniRef100\_A0A176VLD6 \| 98.5 \| 1.2e-09 \| 2.3e-15 \| 105.4 \| 175 \| (12, 208) \| 329 \| (167, 343) \| 480 \| RecA family profile 2 domain-containing protein \| RecA family profile 2 domain-containing protein; uniclust \| UniRef100\_UPI001BF05A99 \| 98.5 \| 1.3e-09 \| 2.3e-15 \| 85.5 \| 87 \| (188, 278) \| 329 \| (2, 90) \| 123 \| hypothetical protein \| hypothetical protein; uniclust \| UniRef100\_A0A2T2RW71 \| 98.5 \| 1.3e-09 \| 2.4e-15 \| 96.6 \| 142 \| (12, 172) \| 329 \| (36, 189) \| 255 \| Circadian clock protein KaiC (Fragment) \| Circadian clock protein KaiC (Fragment); uniclust \| UniRef100\_A0A1Q7Z991 \| 98.5 \| 1.1e-09 \| 2.4e-15 \| 107.1 \| 190 \| (12, 234) \| 329 \| (125, 322) \| 371 \| KaiC domain-containing protein \| KaiC domain-containing protein; uniclust \| UniRef100\_A0A117M3S6 \| 98.5 \| 1.3e-09 \| 2.5e-15 \| 98.0 \| 131 \| (14, 167) \| 329 \| (44, 182) \| 234 \| RecA-superfamily ATPase (Fragment) \| RecA-superfamily ATPase (Fragment); uniclust \| UniRef100\_A0A2S3QK73 \| 98.5 \| 1.4e-09 \| 2.6e-15 \| 88.9 \| 91 \| (156, 250) \| 329 \| (2, 94) \| 160 \| Uncharacterized protein \| Uncharacterized protein; uniclust \| UniRef100\_A0A1F9QK12 \| 98.5 \| 1.4e-09 \| 2.7e-15 \| 98.5 \| 156 \| (14, 211) \| 329 \| (37, 194) \| 246 \| RecA family profile 1 domain-containing protein \| RecA family profile 1 domain-containing protein; uniclust \| UniRef100\_A0A1T5JGQ2 \| 98.5 \| 1.3e-09 \| 2.7e-15 \| 110.5 \| 191 \| (14, 227) \| 329 \| (184, 376) \| 574 \| AAA domain-containing protein \| AAA domain-containing protein; uniclust \| UniRef100\_A0A328EIC8 \| 98.5 \| 1.5e-09 \| 2.8e-15 \| 101.5 \| 198 \| (11, 247) \| 329 \| (9, 213) \| 455 \| AAA+ ATPase domain-containing protein \| AAA+ ATPase domain-containing protein; uniclust \| UniRef100\_A0A077WH29 \| 98.5 \| 1.3e-09 \| 2.8e-15 \| 107.9 \| 153 \| (13, 172) \| 329 \| (127, 291) \| 405 \| RecA family profile 1 domain-containing protein \| RecA family profile 1 domain-containing protein; uniclust \| UniRef100\_A0A0E0D408 \| 98.5 \| 1.5e-09 \| 2.9e-15 \| 104.7 \| 146 \| (11, 173) \| 329 \| (127, 273) \| 428 \| RecA family profile 1 domain-containing protein \| RecA family profile 1 domain-containing protein; uniclust \| UniRef100\_A0A256ZNM9 \| 98.5 \| 1.5e-09 \| 2.9e-15 \| 95.2 \| 136 \| (12, 167) \| 329 \| (45, 199) \| 210 \| KaiC domain-containing protein (Fragment) \| KaiC domain-containing protein (Fragment); uniclust \| UniRef100\_A0A7J7B7P2 \| 98.5 \| 1.6e-09 \| 3e-15 \| 99.9 \| 172 \| (12, 208) \| 329 \| (165, 337) \| 398 \| Protein RecA \| Protein RecA; uniclust \| UniRef100\_A0A0D0THD4 \| 98.5 \| 1.5e-09 \| 3e-15 \| 103.3 \| 176 \| (15, 212) \| 329 \| (129, 319) \| 357 \| Meiotic recombinase Dmc1 \| Meiotic recombinase Dmc1; uniclust \| UniRef100\_A0A097F767 \| 98.5 \| 1.6e-09 \| 3e-15 \| 92.4 \| 96 \| (12, 122) \| 329 \| (57, 153) \| 178 \| Protein RecA (Fragment) \| Protein RecA (Fragment); uniclust \| UniRef100\_A0A2U2HNZ2 \| 98.5 \| 1.5e-09 \| 3.1e-15 \| 104.0 \| 184 \| (12, 227) \| 329 \| (94, 284) \| 341 \| Protein kinase (Fragment) \| Protein kinase (Fragment); uniclust \| UniRef100\_UPI001F57B665 \| 98.5 \| 1.7e-09 \| 3.1e-15 \| 100.6 \| 269 \| (7, 299) \| 329 \| (22, 324) \| 430 \| hypothetical protein \| hypothetical protein; uniclust \| UniRef100\_A0A0D3DWG8 \| 98.5 \| 1.8e-09 \| 3.4e-15 \| 110.4 \| 175 \| (12, 208) \| 329 \| (581, 756) \| 905 \| RecA family profile 2 domain-containing protein \| RecA family profile 2 domain-containing protein; uniclust \| UniRef100\_A0A087F9L5 \| 98.5 \| 1.8e-09 \| 3.6e-15 \| 92.9 \| 96 \| (12, 122) \| 329 \| (73, 169) \| 174 \| Protein RecA \| Protein RecA; uniclust \| UniRef100\_A0A096CMX0 \| 98.5 \| 1.7e-09 \| 3.6e-15 \| 107.3 \| 132 \| (1, 168) \| 329 \| (11, 155) \| 402 \| Phage nucleotide-binding protein \| Phage nucleotide-binding protein; uniclust \| UniRef100\_A0A7S3XE34 \| 98.5 \| 1.9e-09 \| 3.6e-15 \| 101.1 \| 175 \| (12, 208) \| 329 \| (169, 344) \| 465 \| Uncharacterized protein \| Uncharacterized protein; uniclust \| UniRef100\_A0A0B8QMU1 \| 98.5 \| 1.9e-09 \| 3.6e-15 \| 98.5 \| 167 \| (2, 197) \| 329 \| (18, 191) \| 323 \| Predicted ATP-dependent serine protease \| Predicted ATP-dependent serine protease; uniclust \| UniRef100\_A0A194PGT0 \| 98.5 \| 1.8e-09 \| 3.6e-15 \| 105.6 \| 144 \| (13, 172) \| 329 \| (114, 262) \| 402 \| DNA repair protein XRCC3 \| DNA repair protein XRCC3; uniclust \| UniRef100\_A0A1Q3SF85 \| 98.5 \| 1.9e-09 \| 3.8e-15 \| 102.2 \| 205 \| (12, 248) \| 329 \| (22, 247) \| 364 \| AAA+ ATPase domain-containing protein \| AAA+ ATPase domain-containing protein; uniclust \| UniRef100\_A0A351QB39 \| 98.4 \| 2.2e-09 \| 4.1e-15 \| 101.1 \| 171 \| (12, 207) \| 329 \| (65, 236) \| 436 \| Protein RecA (Fragment) \| Protein RecA (Fragment); uniclust \| UniRef100\_A0A140L9N3 \| 98.4 \| 2.3e-09 \| 4.2e-15 \| 86.7 \| 136 \| (12, 166) \| 329 \| (6, 144) \| 148 \| Circadian clock protein kinase KaiC \| Circadian clock protein kinase KaiC; uniclust \| UniRef100\_A0A932C5E2 \| 98.4 \| 2.3e-09 \| 4.2e-15 \| 103.6 \| 172 \| (12, 209) \| 329 \| (72, 245) \| 630 \| Protein RecA \| Protein RecA; uniclust \| UniRef100\_A0A497RG95 \| 98.4 \| 2.2e-09 \| 4.2e-15 \| 91.3 \| 105 \| (12, 122) \| 329 \| (21, 144) \| 170 \| KaiC domain-containing protein (Fragment) \| KaiC domain-containing protein (Fragment); uniclust \| UniRef100\_A0A0T7BLU2 \| 98.4 \| 2.2e-09 \| 4.4e-15 \| 110.4 \| 105 \| (12, 122) \| 329 \| (262, 370) \| 731 \| Recombinase RecA \| Recombinase RecA; uniclust \| UniRef100\_A0A0F9CML5 \| 98.4 \| 2.2e-09 \| 4.5e-15 \| 103.8 \| 173 \| (12, 209) \| 329 \| (68, 247) \| 364 \| RecA family profile 1 domain-containing protein \| RecA family profile 1 domain-containing protein; uniclust \| UniRef100\_A0A0U4B4X9 \| 98.4 \| 2.3e-09 \| 4.5e-15 \| 102.5 \| 212 \| (14, 248) \| 329 \| (69, 315) \| 371 \| RecA-like DNA recombinase \| RecA-like DNA recombinase; uniclust \| UniRef100\_A0A929RAQ8 \| 98.4 \| 2.2e-09 \| 4.5e-15 \| 112.1 \| 193 \| (13, 227) \| 329 \| (398, 590) \| 742 \| Bifunctional DNA primase/polymerase \| Bifunctional DNA primase/polymerase; uniclust \| UniRef100\_A0A022L896 \| 98.4 \| 2.3e-09 \| 4.6e-15 \| 110.6 \| 188 \| (14, 226) \| 329 \| (409, 598) \| 798 \| DNA primase/polymerase bifunctional N-terminal domain-containing protein \| DNA primase/polymerase bifunctional N-terminal domain-containing protein; uniclust \| UniRef100\_A0A2V8NE51 \| 98.4 \| 2.3e-09 \| 4.7e-15 \| 102.0 \| 185 \| (13, 234) \| 329 \| (48, 242) \| 317 \| Recombinase RecA (Fragment) \| Recombinase RecA (Fragment); uniclust \| UniRef100\_A0A1F2T428 \| 98.4 \| 2.2e-09 \| 4.8e-15 \| 116.4 \| 204 \| (12, 246) \| 329 \| (342, 579) \| 902 \| AAA+ ATPase domain-containing protein \| AAA+ ATPase domain-containing protein; uniclust \| UniRef100\_A0A0M3K0W3 \| 98.4 \| 2.3e-09 \| 4.8e-15 \| 102.7 \| 175 \| (13, 205) \| 329 \| (78, 264) \| 338 \| Hus1-like protein (inferred by orthology to a C. elegans protein) \| Hus1-like protein (inferred by orthology to a C. elegans protein); uniclust \| UniRef100\_A0A174TWV9 \| 98.4 \| 2.4e-09 \| 4.9e-15 \| 102.9 \| 129 \| (12, 172) \| 329 \| (12, 150) \| 383 \| ATP-binding protein \| ATP-binding protein; uniclust \| UniRef100\_A0A024FWZ3 \| 98.4 \| 2.4e-09 \| 5e-15 \| 110.2 \| 181 \| (12, 211) \| 329 \| (226, 432) \| 582 \| RecA family profile 1 domain-containing protein \| RecA family profile 1 domain-containing protein; uniclust \| UniRef100\_A0A2D6X309 \| 98.4 \| 2.6e-09 \| 5.3e-15 \| 97.5 \| 161 \| (11, 206) \| 329 \| (5, 169) \| 237 \| UDP-N-acetylglucosamine kinase \| UDP-N-acetylglucosamine kinase; uniclust \| UniRef100\_A0A0Q8QVK5 \| 98.4 \| 2.8e-09 \| 5.4e-15 \| 96.3 \| 177 \| (18, 227) \| 329 \| (2, 183) \| 245 \| KaiC-like domain-containing protein \| KaiC-like domain-containing protein; uniclust \| UniRef100\_A0A1F9U338 \| 98.4 \| 2.8e-09 \| 5.6e-15 \| 104.0 \| 138 \| (13, 169) \| 329 \| (211, 350) \| 450 \| KaiC domain-containing protein \| KaiC domain-containing protein; uniclust \| UniRef100\_UPI001FD21D5B \| 98.4 \| 3e-09 \| 5.8e-15 \| 89.3 \| 104 \| (152, 260) \| 329 \| (3, 108) \| 167 \| hypothetical protein \| hypothetical protein; uniclust \| UniRef100\_A0A3S0KB12 \| 98.4 \| 3.1e-09 \| 6e-15 \| 102.4 \| 134 \| (13, 167) \| 329 \| (95, 236) \| 405 \| AAA family ATPase \| AAA family ATPase; uniclust \| UniRef100\_A0A3M1RHG9 \| 98.4 \| 3.3e-09 \| 6.2e-15 \| 99.3 \| 135 \| (13, 167) \| 329 \| (170, 322) \| 399 \| EVE domain-containing protein \| EVE domain-containing protein; uniclust \| UniRef100\_A0A973P365 \| 98.4 \| 3.6e-09 \| 6.8e-15 \| 103.0 \| 176 \| (10, 208) \| 329 \| (25, 216) \| 535 \| AAA family ATPase \| AAA family ATPase; uniclust \| UniRef100\_A0A015SWH5 \| 98.4 \| 3.2e-09 \| 7.1e-15 \| 101.5 \| 91 \| (14, 118) \| 329 \| (75, 166) \| 285 \| ATP-binding protein \| ATP-binding protein; uniclust \| UniRef100\_A0A257A7A1 \| 98.4 \| 3.8e-09 \| 7.4e-15 \| 103.4 \| 152 \| (14, 211) \| 329 \| (263, 418) \| 492 \| KaiA-binding protein \| KaiA-binding protein; uniclust \| UniRef100\_A0A0B5HJS6 \| 98.4 \| 3.9e-09 \| 7.4e-15 \| 100.2 \| 158 \| (14, 207) \| 329 \| (170, 334) \| 383 \| KaiC domain-containing protein \| KaiC domain-containing protein; uniclust \| UniRef100\_A0A1F2P869 \| 98.4 \| 3.9e-09 \| 7.5e-15 \| 96.3 \| 160 \| (12, 206) \| 329 \| (57, 222) \| 278 \| Circadian clock protein KaiC \| Circadian clock protein KaiC; uniclust \| UniRef100\_A0A3N5LY41 \| 98.4 \| 3.9e-09 \| 7.7e-15 \| 95.6 \| 162 \| (13, 209) \| 329 \| (23, 187) \| 241 \| KaiC 1 (Fragment) \| KaiC 1 (Fragment); uniclust \| UniRef100\_A0A060WM17 \| 98.4 \| 3.9e-09 \| 7.8e-15 \| 106.0 \| 152 \| (12, 171) \| 329 \| (170, 335) \| 547 \| RecA family profile 1 domain-containing protein \| RecA family profile 1 domain-containing protein; uniclust \| UniRef100\_A0A101IMN0 \| 98.4 \| 3.7e-09 \| 7.8e-15 \| 96.4 \| 94 \| (13, 122) \| 329 \| (97, 190) \| 218 \| DNA repair protein radA \| DNA repair protein radA; uniclust \| UniRef100\_A0A2C9W427 \| 98.4 \| 4.4e-09 \| 8.2e-15 \| 91.0 \| 145 \| (13, 174) \| 329 \| (29, 174) \| 222 \| RecA family profile 1 domain-containing protein \| RecA family profile 1 domain-containing protein; uniclust \| UniRef100\_A0A0I9SDJ7 \| 98.4 \| 3.9e-09 \| 8.3e-15 \| 104.0 \| 168 \| (13, 209) \| 329 \| (77, 257) \| 404 \| AAA family ATPase \| AAA family ATPase; uniclust \| UniRef100\_A0A067EHN3 \| 98.4 \| 4.4e-09 \| 8.3e-15 \| 100.7 \| 146 \| (11, 173) \| 329 \| (170, 316) \| 480 \| Uncharacterized protein \| Uncharacterized protein; uniclust \| UniRef100\_A0A6P6H533 \| 98.4 \| 4.6e-09 \| 8.5e-15 \| 90.7 \| 168 \| (17, 209) \| 329 \| (2, 170) \| 230 \| Uncharacterized protein LOC112851458 \| Uncharacterized protein LOC112851458; uniclust \| UniRef100\_A0A061ASR8 \| 98.4 \| 4.5e-09 \| 8.6e-15 \| 101.5 \| 174 \| (13, 208) \| 329 \| (250, 437) \| 475 \| DNA repair protein RAD51 homolog \| DNA repair protein RAD51 homolog; uniclust \| UniRef100\_A0A2M4ASR0 \| 98.4 \| 4.6e-09 \| 9e-15 \| 94.2 \| 146 \| (13, 167) \| 329 \| (39, 203) \| 234 \| Putative meiotic recombination protein dmc1 (Fragment) \| Putative meiotic recombination protein dmc1 (Fragment); uniclust \| UniRef100\_UPI001FE17A8B \| 98.4 \| 4.9e-09 \| 9e-15 \| 76.9 \| 81 \| (4, 102) \| 329 \| (2, 82) \| 83 \| AAA family ATPase \| AAA family ATPase; uniclust \| UniRef100\_A0A660ZY62 \| 98.4 \| 4.5e-09 \| 9e-15 \| 96.3 \| 190 \| (16, 223) \| 329 \| (11, 238) \| 263 \| AAA family ATPase \| AAA family ATPase; uniclust \| UniRef100\_A0A497FRI7 \| 98.4 \| 5.1e-09 \| 9.6e-15 \| 93.1 \| 202 \| (14, 244) \| 329 \| (24, 243) \| 252 \| Uncharacterized protein \| Uncharacterized protein; uniclust \| UniRef100\_A0A497G818 \| 98.4 \| 5.1e-09 \| 9.6e-15 \| 98.5 \| 129 \| (12, 167) \| 329 \| (209, 339) \| 399 \| Uncharacterized protein (Fragment) \| Uncharacterized protein (Fragment); uniclust \| UniRef100\_A0A259P4E8 \| 98.4 \| 4.6e-09 \| 9.7e-15 \| 106.0 \| 136 \| (13, 168) \| 329 \| (273, 412) \| 489 \| Protein kinase (Fragment) \| Protein kinase (Fragment); uniclust \| UniRef100\_A0A0F9ILE9 \| 98.4 \| 4.8e-09 \| 9.7e-15 \| 102.4 \| 176 \| (10, 209) \| 329 \| (89, 271) \| 413 \| RecA family profile 2 domain-containing protein \| RecA family profile 2 domain-containing protein; uniclust \| UniRef100\_A0A1G6XCP8 \| 98.4 \| 4.6e-09 \| 9.7e-15 \| 108.1 \| 157 \| (12, 208) \| 329 \| (53, 239) \| 587 \| RecA-superfamily ATPase, KaiC/GvpD/RAD55 family \| RecA-superfamily ATPase, KaiC/GvpD/RAD55 family; uniclust \| UniRef100\_A0A075MRR8 \| 98.4 \| 4.9e-09 \| 1e-14 \| 96.6 \| 140 \| (13, 171) \| 329 \| (38, 179) \| 251 \| RecA/RadA recombinase \| RecA/RadA recombinase; uniclust \| UniRef100\_A0A0J1FYZ4 \| 98.4 \| 4.6e-09 \| 1e-14 \| 105.9 \| 133 \| (12, 167) \| 329 \| (133, 280) \| 449 \| AAA+ ATPase domain-containing protein \| AAA+ ATPase domain-containing protein; uniclust \| UniRef100\_A0A0F2LDE9 \| 98.4 \| 5e-09 \| 1e-14 \| 102.7 \| 93 \| (17, 122) \| 329 \| (23, 116) \| 410 \| AAA+ ATPase domain-containing protein \| AAA+ ATPase domain-containing protein; uniclust \| UniRef100\_A0A165HV77 \| 98.4 \| 5.1e-09 \| 1e-14 \| 98.8 \| 188 \| (7, 208) \| 329 \| (26, 224) \| 307 \| AAA+ ATPase domain-containing protein \| AAA+ ATPase domain-containing protein; uniclust \| UniRef100\_A0A1V5WB44 \| 98.4 \| 5e-09 \| 1e-14 \| 105.3 \| 140 \| (12, 168) \| 329 \| (103, 253) \| 498 \| Uncharacterized protein \| Uncharacterized protein; uniclust \| UniRef100\_A0A2N1IGH6 \| 98.4 \| 5.2e-09 \| 1e-14 \| 99.7 \| 130 \| (15, 167) \| 329 \| (42, 190) \| 357 \| AAA+ ATPase domain-containing protein \| AAA+ ATPase domain-containing protein; uniclust \| UniRef100\_A0A0Q3RAV7 \| 98.4 \| 5.5e-09 \| 1e-14 \| 100.8 \| 144 \| (12, 172) \| 329 \| (197, 340) \| 475 \| RecA family profile 2 domain-containing protein \| RecA family profile 2 domain-containing protein; uniclust \| UniRef100\_A0A0K8Q9N1 \| 98.4 \| 5.3e-09 \| 1.1e-14 \| 103.2 \| 161 \| (15, 212) \| 329 \| (86, 247) \| 487 \| DNA repair protein RadA homolog \| DNA repair protein RadA homolog; uniclust \| UniRef100\_A0A3M1F514 \| 98.4 \| 5.4e-09 \| 1.1e-14 \| 102.9 \| 137 \| (12, 167) \| 329 \| (39, 181) \| 494 \| KaiC domain-containing protein (Fragment) \| KaiC domain-containing protein (Fragment); uniclust \| UniRef100\_A0A535E7X6 \| 98.3 \| 5.4e-09 \| 1.1e-14 \| 102.8 \| 156 \| (13, 206) \| 329 \| (257, 416) \| 465 \| KaiC domain-containing protein \| KaiC domain-containing protein; uniclust \| UniRef100\_A0A535HST7 \| 98.3 \| 6e-09 \| 1.1e-14 \| 98.2 \| 145 \| (12, 173) \| 329 \| (179, 324) \| 475 \| Protein RecA \| Protein RecA; uniclust \| UniRef100\_A0A061GLK6 \| 98.3 \| 6e-09 \| 1.1e-14 \| 102.1 \| 172 \| (12, 207) \| 329 \| (261, 435) \| 577 \| RecA DNA recombination family protein \| RecA DNA recombination family protein; uniclust \| UniRef100\_A0A967YFN2 \| 98.3 \| 6.2e-09 \| 1.2e-14 \| 88.8 \| 152 \| (13, 207) \| 329 \| (38, 189) \| 190 \| AAA family ATPase (Fragment) \| AAA family ATPase (Fragment); uniclust \| UniRef100\_A0A6A5KQJ0 \| 98.3 \| 6.5e-09 \| 1.2e-14 \| 101.9 \| 171 \| (12, 208) \| 329 \| (441, 613) \| 716 \| Recombinase A \| Recombinase A; uniclust \| UniRef100\_A0A165ZYR3 \| 98.3 \| 6e-09 \| 1.2e-14 \| 92.4 \| 134 \| (4, 168) \| 329 \| (14, 149) \| 203 \| Uncharacterized protein \| Uncharacterized protein; uniclust \| UniRef100\_A0A1V5WES9 \| 98.3 \| 5.8e-09 \| 1.2e-14 \| 101.6 \| 172 \| (11, 206) \| 329 \| (52, 232) \| 381 \| Protein RecA \| Protein RecA; uniclust \| UniRef100\_A0A819BHB5 \| 98.3 \| 6.6e-09 \| 1.2e-14 \| 93.7 \| 145 \| (12, 173) \| 329 \| (58, 203) \| 319 \| Protein RecA \| Protein RecA; uniclust \| UniRef100\_A0A2T8IIP5 \| 98.3 \| 6.5e-09 \| 1.2e-14 \| 95.4 \| 145 \| (12, 173) \| 329 \| (133, 277) \| 311 \| RecA family profile 1 domain-containing protein \| RecA family profile 1 domain-containing protein; uniclust \| UniRef100\_A0A062V3E0 \| 98.3 \| 5.7e-09 \| 1.3e-14 \| 100.7 \| 137 \| (12, 167) \| 329 \| (74, 215) \| 298 \| Putative ATPase involved in flagella biogenesis \| Putative ATPase involved in flagella biogenesis; uniclust \| UniRef100\_A0A8T6WHX9 \| 98.3 \| 6.8e-09 \| 1.3e-14 \| 85.2 \| 128 \| (12, 167) \| 329 \| (12, 149) \| 161 \| AAA family ATPase (Fragment) \| AAA family ATPase (Fragment); uniclust \| UniRef100\_A0A023NLX7 \| 98.3 \| 6.6e-09 \| 1.3e-14 \| 86.2 \| 96 \| (12, 122) \| 329 \| (30, 126) \| 150 \| Protein RecA (Fragment) \| Protein RecA (Fragment); uniclust \| UniRef100\_A0A1Y1I963 \| 98.3 \| 6.9e-09 \| 1.3e-14 \| 98.6 \| 144 \| (12, 173) \| 329 \| (190, 335) \| 514 \| Putative DNA repair and recombination protein RecA \| Putative DNA repair and recombination protein RecA; uniclust \| UniRef100\_A0A6G1C7K2 \| 98.3 \| 6.9e-09 \| 1.3e-14 \| 93.4 \| 146 \| (12, 174) \| 329 \| (87, 233) \| 279 \| RecA family profile 1 domain-containing protein (Fragment) \| RecA family profile 1 domain-containing protein (Fragment); uniclust \| UniRef100\_A0A088GGJ1 \| 98.3 \| 6.7e-09 \| 1.3e-14 \| 92.2 \| 157 \| (11, 208) \| 329 \| (14, 176) \| 226 \| Phage nucleotide-binding protein \| Phage nucleotide-binding protein; uniclust \| UniRef100\_A0A7X8AP94 \| 98.3 \| 6.8e-09 \| 1.3e-14 \| 101.6 \| 193 \| (14, 227) \| 329 \| (112, 304) \| 503 \| AAA family ATPase \| AAA family ATPase; uniclust \| UniRef100\_A0A0F2L3G7 \| 98.3 \| 6.8e-09 \| 1.3e-14 \| 97.8 \| 165 \| (12, 209) \| 329 \| (64, 271) \| 335 \| Recombination protein RecA \| Recombination protein RecA; uniclust \| UniRef100\_A0A059VKV7 \| 98.3 \| 6.4e-09 \| 1.3e-14 \| 101.6 \| 131 \| (13, 171) \| 329 \| (37, 182) \| 365 \| AAA-ATPase \| AAA-ATPase; uniclust \| UniRef100\_A0A2E0FRW8 \| 98.3 \| 6.6e-09 \| 1.4e-14 \| 101.5 \| 122 \| (15, 169) \| 329 \| (205, 326) \| 400 \| Flagellar biosynthesis protein FlhF \| Flagellar biosynthesis protein FlhF; uniclust \| UniRef100\_A0A3M1V3N6 \| 98.3 \| 7.4e-09 \| 1.4e-14 \| 93.7 \| 177 \| (13, 216) \| 329 \| (118, 305) \| 327 \| AAA family ATPase \| AAA family ATPase; uniclust \| UniRef100\_A0A0K1E313 \| 98.3 \| 6.8e-09 \| 1.4e-14 \| 99.8 \| 92 \| (15, 122) \| 329 \| (26, 118) \| 365 \| RecA-superfamily ATPase implicated in signal transduction \| RecA-superfamily ATPase implicated in signal transduction; uniclust \| UniRef100\_A0A017SP12 \| 98.3 \| 6.8e-09 \| 1.4e-14 \| 101.2 \| 158 \| (12, 172) \| 329 \| (68, 243) \| 404 \| p-loop containing nucleoside triphosphate hydrolase protein \| p-loop containing nucleoside triphosphate hydrolase protein; uniclust \| UniRef100\_A0A1G8FKD8 \| 98.3 \| 7.5e-09 \| 1.4e-14 \| 70.7 \| 55 \| (4, 62) \| 329 \| (1, 55) \| 57 \| AAA domain-containing protein (Fragment) \| AAA domain-containing protein (Fragment); uniclust \| UniRef100\_A0A117SN30 \| 98.3 \| 7.4e-09 \| 1.4e-14 \| 90.8 \| 121 \| (14, 167) \| 329 \| (9, 132) \| 220 \| AAA+ ATPase domain-containing protein \| AAA+ ATPase domain-containing protein; uniclust \| UniRef100\_A0A0Q4KZI2 \| 98.3 \| 6.8e-09 \| 1.5e-14 \| 103.5 \| 135 \| (13, 167) \| 329 \| (80, 235) \| 405 \| AAA+ ATPase domain-containing protein \| AAA+ ATPase domain-containing protein; uniclust \| UniRef100\_A0A6A5LD20 \| 98.3 \| 8e-09 \| 1.5e-14 \| 101.6 \| 171 \| (12, 207) \| 329 \| (444, 615) \| 738 \| RecA family profile 1 domain-containing protein \| RecA family profile 1 domain-containing protein; uniclust \| UniRef100\_A0A059WSK7 \| 98.3 \| 7.5e-09 \| 1.5e-14 \| 103.6 \| 166 \| (14, 222) \| 329 \| (68, 236) \| 473 \| Subunit ChlI of Mg-chelatase (Fragment) \| Subunit ChlI of Mg-chelatase (Fragment); uniclust \| UniRef100\_A0A060HIS1 \| 98.3 \| 7.4e-09 \| 1.6e-14 \| 98.0 \| 136 \| (12, 167) \| 329 \| (61, 205) \| 275 \| Flagella protein FlaH \| Flagella protein FlaH; uniclust \| UniRef100\_A0A059AIS9 \| 98.3 \| 8.1e-09 \| 1.6e-14 \| 101.4 \| 159 \| (13, 211) \| 329 \| (267, 429) \| 460 \| RecA family profile 1 domain-containing protein \| RecA family profile 1 domain-containing protein; uniclust \| UniRef100\_A0A0P0N535 \| 98.3 \| 7.5e-09 \| 1.6e-14 \| 99.8 \| 163 \| (12, 210) \| 329 \| (52, 237) \| 308 \| KaiC domain-containing protein \| KaiC domain-containing protein; uniclust \| UniRef100\_A0A2N1PDZ8 \| 98.3 \| 8.5e-09 \| 1.6e-14 \| 86.7 \| 96 \| (12, 122) \| 329 \| (57, 153) \| 159 \| Protein RecA (Fragment) \| Protein RecA (Fragment); uniclust \| UniRef100\_A0A0F7L738 \| 98.3 \| 7.8e-09 \| 1.7e-14 \| 108.9 \| 175 \| (13, 218) \| 329 \| (324, 514) \| 694 \| Putative AAA ATPase \| Putative AAA ATPase; uniclust \| UniRef100\_A0A0S6W5U3 \| 98.3 \| 8.5e-09 \| 1.7e-14 \| 89.6 \| 136 \| (12, 166) \| 329 \| (33, 181) \| 185 \| Circadian clock protein KaiC \| Circadian clock protein KaiC; uniclust \| UniRef100\_A0A2D6VTU5 \| 98.3 \| 7.7e-09 \| 1.7e-14 \| 110.1 \| 137 \| (13, 167) \| 329 \| (383, 534) \| 730 \| DNA primase/polymerase bifunctional N-terminal domain-containing protein \| DNA primase/polymerase bifunctional N-terminal domain-containing protein; uniclust \| UniRef100\_A0A089ZBC3 \| 98.3 \| 8.4e-09 \| 1.7e-14 \| 95.4 \| 162 \| (12, 209) \| 329 \| (33, 216) \| 250 \| ATPase \| ATPase; pdb70 \| 2ZUC\_B \| 97.2 \| 2.5e-07 \| 2.9e-11 \| 81.3 \| 108 \| (10, 122) \| 329 \| (102, 218) \| 324 \| DNA repair and recombination protein \| 2ZUC\_B DNA repair and recombination protein ARCHAEA, FILAMENT, LEFT-HANDED, DNA BINDING; pdb70 \| 2BKE\_A \| 97.2 \| 2.6e-07 \| 3e-11 \| 81.3 \| 108 \| (10, 122) \| 329 \| (102, 218) \| 324 \| DNA REPAIR AND RECOMBINATION PROTEIN \| 2BKE\_A DNA REPAIR AND RECOMBINATION PROTEIN DNA-BINDING PROTEIN, HOMOLOGOUS RECOMBINATION, DNA; pdb70 \| 2Z43\_A \| 97.2 \| 2.6e-07 \| 3e-11 \| 81.3 \| 108 \| (10, 122) \| 329 \| (102, 218) \| 324 \| DNA repair and recombination protein \| 2Z43\_A DNA repair and recombination protein ARCHAEA, FILAMENT, DNA BINDING, RECOMBINATION; pdb70 \| 4WIA\_B \| 97.2 \| 2.7e-07 \| 3.4e-11 \| 74.9 \| 108 \| (10, 125) \| 329 \| (22, 138) \| 233 \| Putative flagella-related protein H \| 4WIA\_B Putative flagella-related protein H archaea, flagella, ATP-binding protein HET: SO4, MSE; pdb70 \| 4DC9\_F \| 97.2 \| 3.7e-07 \| 4.3e-11 \| 77.1 \| 108 \| (10, 122) \| 329 \| (37, 163) \| 266 \| DNA repair and recombination protein \| 4DC9\_F DNA repair and recombination protein Hexamer, RadA, Recombinase, Homologous Recombination HET: NO3; pdb70 \| 4QKQ\_I \| 97.2 \| 3.7e-07 \| 4.3e-11 \| 77.1 \| 108 \| (10, 122) \| 329 \| (37, 163) \| 266 \| DNA repair and recombination protein \| 4QKQ\_I DNA repair and recombination protein RadA, Rad51, DMC1, RecA, ATPase HET: NO3, 35N; pdb70 \| 3HR8\_A \| 97.0 \| 7e-07 \| 7.9e-11 \| 80.3 \| 145 \| (10, 171) \| 329 \| (56, 201) \| 356 \| Protein recA \| 3HR8\_A Protein recA Alpha and beta proteins (a/b; pdb70 \| 5JZC\_F \| 97.0 \| 9.9e-07 \| 1.1e-10 \| 77.7 \| 109 \| (10, 123) \| 329 \| (115, 231) \| 339 \| DNA repair protein RAD51 homolog \| 5JZC\_F DNA repair protein RAD51 homolog cryoEM DNA repair recombinase, CELL; pdb70 \| 5NWL\_K \| 97.0 \| 9.9e-07 \| 1.1e-10 \| 77.7 \| 109 \| (10, 123) \| 329 \| (115, 231) \| 339 \| DNA repair protein RAD51 homolog \| 5NWL\_K DNA repair protein RAD51 homolog ATPase, DNA-strand exchange, Homologous Recombination HET: ATP, MG; pdb70 \| 2G88\_A \| 97.0 \| 1.1e-06 \| 1.3e-10 \| 78.6 \| 145 \| (10, 171) \| 329 \| (56, 201) \| 349 \| Protein recA \| 2G88\_A Protein recA RECOMBINATION, DNA-REPAIR HET: DTP, CIT; pdb70 \| 1XU4\_A \| 96.9 \| 1.1e-06 \| 1.3e-10 \| 76.1 \| 109 \| (10, 123) \| 329 \| (93, 220) \| 322 \| ATPASE IN COMPLEX WITH AMP-PNP \| 1XU4\_A ATPASE IN COMPLEX WITH AMP-PNP ATPASE, PROTEIN-ATP COMPLEX, CO-FACTORS, POTASSIUM-DEPENDENCE HET: ANP; pdb70 \| 1SZP\_D \| 96.9 \| 1.3e-06 \| 1.6e-10 \| 75.3 \| 110 \| (10, 124) \| 329 \| (94, 211) \| 321 \| DNA repair protein RAD51 \| 1SZP\_D DNA repair protein RAD51 homologous recombination, asymmetry, Rad51 filament HET: SO4; pdb70 \| 3LDA\_A \| 96.9 \| 1.5e-06 \| 1.6e-10 \| 80.6 \| 109 \| (10, 123) \| 329 \| (173, 289) \| 400 \| DNA repair protein RAD51 (E.C.3.1.22.4) \| 3LDA\_A DNA repair protein RAD51 (E.C.3.1.22.4) DNA BINDING PROTEIN, ATP-binding, DNA; pdb70 \| 1SZP\_A \| 96.9 \| 1.3e-06 \| 1.6e-10 \| 75.3 \| 109 \| (10, 123) \| 329 \| (94, 210) \| 321 \| DNA repair protein RAD51 \| 1SZP\_A DNA repair protein RAD51 homologous recombination, asymmetry, Rad51 filament HET: SO4; pdb70 \| 4PPF\_A \| 96.9 \| 1.7e-06 \| 1.9e-10 \| 77.4 \| 99 \| (10, 123) \| 329 \| (55, 154) \| 350 \| Protein RecA, 1st part, 2nd \| 4PPF\_A Protein RecA, 1st part, 2nd HOMOLOGOUS RECOMBINATION, DNA REPAIR, ATPASE HET: FLC, EDO; pdb70 \| 4PSV\_A \| 96.9 \| 1.7e-06 \| 1.9e-10 \| 77.4 \| 99 \| (10, 123) \| 329 \| (55, 154) \| 350 \| Protein RecA, 1st part, 2nd \| 4PSV\_A Protein RecA, 1st part, 2nd HOMOLOGOUS RECOMBINATION, DNA REPAIR, ATPASE; pdb70 \| 2I1Q\_A \| 96.8 \| 1.8e-06 \| 2.2e-10 \| 74.6 \| 109 \| (10, 123) \| 329 \| (93, 220) \| 322 \| DNA repair and recombination protein \| 2I1Q\_A DNA repair and recombination protein ATPASE, RECOMBINASE, ATP COMPLEX, Calcium HET: ANP; pdb70 \| 1V5W\_B \| 96.8 \| 2.2e-06 \| 2.6e-10 \| 75.8 \| 110 \| (10, 124) \| 329 \| (117, 236) \| 343 \| Meiotic recombination protein DMC1/LIM15 homolog \| 1V5W\_B Meiotic recombination protein DMC1/LIM15 homolog DNA-binding protein, ring protein, octamer; pdb70 \| 1XP8\_A \| 96.8 \| 2.4e-06 \| 2.7e-10 \| 77.0 \| 99 \| (10, 123) \| 329 \| (69, 168) \| 366 \| RecA protein \| 1XP8\_A RecA protein Recombination, Radioresistance, DNA-repair, ATPase, DNA-binding HET: AGS; pdb70 \| 5JRJ\_A \| 96.8 \| 2.5e-06 \| 2.8e-10 \| 76.0 \| 99 \| (10, 123) \| 329 \| (61, 160) \| 351 \| Protein RecA \| 5JRJ\_A Protein RecA DNA-binding protein, ATP-dependent DNA protein HET: ATP, ADP; pdb70 \| 2ZTS\_B \| 96.8 \| 2.3e-06 \| 2.9e-10 \| 69.0 \| 45 \| (10, 54) \| 329 \| (25, 69) \| 251 \| Putative uncharacterized protein PH0186 \| 2ZTS\_B Putative uncharacterized protein PH0186 KaiC like protein, ATP-binding, Nucleotide-binding HET: ADP, MSE; pdb70 \| 2ZTS\_C \| 96.8 \| 2.3e-06 \| 2.9e-10 \| 69.0 \| 45 \| (10, 54) \| 329 \| (25, 69) \| 251 \| Putative uncharacterized protein PH0186 \| 2ZTS\_C Putative uncharacterized protein PH0186 KaiC like protein, ATP-binding, Nucleotide-binding HET: ADP; pdb70 \| 2ZJB\_A \| 96.7 \| 2.8e-06 \| 3.2e-10 \| 75.0 \| 109 \| (10, 123) \| 329 \| (117, 235) \| 343 \| Meiotic recombination protein DMC1/LIM15 homolog \| 2ZJB\_A Meiotic recombination protein DMC1/LIM15 homolog DNA-BINDING PROTEIN, RING PROTEIN, OCTAMER; pdb70 \| 1PZN\_A \| 96.7 \| 2.9e-06 \| 3.3e-10 \| 75.1 \| 108 \| (10, 122) \| 329 \| (126, 246) \| 349 \| DNA repair and recombination protein \| 1PZN\_A DNA repair and recombination protein HEPTAMERIC RING HEPTAMER HET: IMD, MPD, GOL, SO4; pdb70 \| 1PZN\_G \| 96.7 \| 2.9e-06 \| 3.3e-10 \| 75.1 \| 108 \| (10, 122) \| 329 \| (126, 246) \| 349 \| DNA repair and recombination protein \| 1PZN\_G DNA repair and recombination protein HEPTAMERIC RING HEPTAMER HET: GOL, SO4, MPD, MSE, IMD; pdb70 \| 6R3P\_A \| 96.7 \| 2.9e-06 \| 3.5e-10 \| 69.9 \| 45 \| (10, 54) \| 329 \| (35, 84) \| 261 \| Meiotic recombination protein DMC1/LIM15 homolog \| 6R3P\_A Meiotic recombination protein DMC1/LIM15 homolog Recombinase, Homologous recombination, Strand invasion HET: PG4, 1PE, EDO, PEG, P6G; pdb70 \| 6R3P\_C \| 96.7 \| 3.3e-06 \| 4e-10 \| 69.6 \| 45 \| (10, 54) \| 329 \| (35, 84) \| 261 \| Meiotic recombination protein DMC1/LIM15 homolog \| 6R3P\_C Meiotic recombination protein DMC1/LIM15 homolog Recombinase, Homologous recombination, Strand invasion HET: PG4, P6G, 1PE, PEG, EDO; pdb70 \| 4TWZ\_A \| 96.7 \| 3.6e-06 \| 4.1e-10 \| 74.8 \| 99 \| (10, 123) \| 329 \| (54, 153) \| 352 \| PROTEIN RECA \| 4TWZ\_A PROTEIN RECA HOMOLOGOUS RECOMBINATION, DNA BINDING, RECOMBINATION; pdb70 \| 1N0W\_A \| 96.6 \| 4e-06 \| 5.1e-10 \| 67.0 \| 109 \| (10, 123) \| 329 \| (19, 135) \| 243 \| DNA repair protein RAD51 homolog \| 1N0W\_A DNA repair protein RAD51 homolog DNA repair, homologous recombination, breast HET: EDO, MSE; pdb70 \| 4YDS\_A \| 96.6 \| 5e-06 \| 6.3e-10 \| 66.2 \| 45 \| (10, 55) \| 329 \| (15, 59) \| 228 \| Flagella-related protein H \| 4YDS\_A Flagella-related protein H RecA superfamily ATPase, HYDROLASE HET: ATP; pdb70 \| 5JWR\_A \| 96.5 \| 6e-06 \| 7.7e-10 \| 65.5 \| 45 \| (10, 54) \| 329 \| (27, 71) \| 247 \| Circadian clock protein kinase KaiC \| 5JWR\_A Circadian clock protein kinase KaiC Transcription Regulator, foldswitch HET: PO4; pdb70 \| 5JEC\_B \| 96.2 \| 1.7e-05 \| 2.1e-09 \| 62.3 \| 44 \| (11, 54) \| 329 \| (21, 69) \| 231 \| DNA repair and recombination protein \| 5JEC\_B DNA repair and recombination protein DNA repair, fragment based drug HET: SO4; pdb70 \| 4IJM\_A \| 96.2 \| 1.8e-05 \| 2.2e-09 \| 72.0 \| 106 \| (10, 123) \| 329 \| (263, 374) \| 512 \| Circadian clock protein kinase KaiC \| 4IJM\_A Circadian clock protein kinase KaiC ATP binding, biological rhythms, auto-kinase HET: TPO, PO4, ATP; pdb70 \| 2W0M\_A \| 96.2 \| 1.8e-05 \| 2.3e-09 \| 61.5 \| 44 \| (10, 54) \| 329 \| (18, 61) \| 235 \| SSO2452 \| 2W0M\_A SSO2452 SSO2452, RECA, SULFOLOBUS SOLFATARICUS P2; pdb70 \| 2DR3\_D \| 96.2 \| 1.8e-05 \| 2.3e-09 \| 62.5 \| 43 \| (11, 54) \| 329 \| (19, 61) \| 247 \| UPF0273 protein PH0284 \| 2DR3\_D UPF0273 protein PH0284 RecA superfamily ATPase, Hexamer, Structural HET: ADP; pdb70 \| 2CVH\_B \| 96.2 \| 2e-05 \| 2.5e-09 \| 61.1 \| 40 \| (11, 54) \| 329 \| (16, 55) \| 220 \| DNA repair and recombination protein \| 2CVH\_B DNA repair and recombination protein RadB, filament formation, homologous recombination; pdb70 \| 3IO5\_A \| 96.1 \| 2.6e-05 \| 2.9e-09 \| 69.5 \| 46 \| (10, 56) \| 329 \| (24, 70) \| 333 \| Recombination and repair protein \| 3IO5\_A Recombination and repair protein storage dimer, inactive conformation, RecA HET: PO4; pdb70 \| 3IO5\_B \| 96.1 \| 2.6e-05 \| 2.9e-09 \| 69.5 \| 46 \| (10, 56) \| 329 \| (24, 70) \| 333 \| Recombination and repair protein \| 3IO5\_B Recombination and repair protein storage dimer, inactive conformation, RecA HET: PO4; pdb70 \| 4TL6\_C \| 96.1 \| 2.4e-05 \| 3.1e-09 \| 61.8 \| 44 \| (11, 54) \| 329 \| (35, 78) \| 253 \| Circadian clock protein kinase KaiC \| 4TL6\_C Circadian clock protein kinase KaiC Serine/threonine-protein kinase, TRANSFERASE HET: ANP; pdb70 \| 4TL7\_B \| 96.1 \| 2.4e-05 \| 3.1e-09 \| 61.8 \| 44 \| (11, 54) \| 329 \| (35, 78) \| 253 \| Circadian clock protein kinase KaiC \| 4TL7\_B Circadian clock protein kinase KaiC Serine/threonine-protein kinase, TRANSFERASE HET: SO4, ATP; pdb70 \| 4IJM\_A \| 96.0 \| 3.3e-05 \| 3.9e-09 \| 70.2 \| 46 \| (10, 55) \| 329 \| (21, 66) \| 512 \| Circadian clock protein kinase KaiC \| 4IJM\_A Circadian clock protein kinase KaiC ATP binding, biological rhythms, auto-kinase HET: TPO, PO4, ATP; pdb70 \| 5J4L\_A \| 96.0 \| 3.6e-05 \| 4.6e-09 \| 59.7 \| 45 \| (10, 54) \| 329 \| (20, 69) \| 231 \| DNA repair and recombination protein \| 5J4L\_A DNA repair and recombination protein DNA repair, fragment based drug; pdb70 \| 3CMV\_H \| 96.0 \| 4e-05 \| 4.7e-09 \| 79.0 \| 97 \| (11, 122) \| 329 \| (30, 127) \| 1357 \| PROTEIN recA \| 3CMV\_H PROTEIN recA homologous recombination, recombination HET: ANP, MG; pdb70 \| 6CQP\_A \| 95.9 \| 5.1e-05 \| 6e-09 \| 64.8 \| 42 \| (12, 54) \| 329 \| (96, 137) \| 303 \| Signal recognition particle receptor FtsY \| 6CQP\_A Signal recognition particle receptor FtsY cotranslational delivery SRP, PROTEIN TRANSPORT HET: GOL; pdb70 \| 6N9B\_B \| 95.9 \| 5.1e-05 \| 6e-09 \| 64.8 \| 42 \| (12, 54) \| 329 \| (96, 137) \| 303 \| Signal recognition particle receptor FtsY \| 6N9B\_B Signal recognition particle receptor FtsY FtsY, SRP, Signal recognition particle HET: ACT; pdb70 \| 1Q57\_G \| 95.8 \| 5.5e-05 \| 6.5e-09 \| 69.1 \| 45 \| (11, 55) \| 329 \| (238, 282) \| 503 \| DNA primase/helicase (E.C.2.7.7.-) \| 1Q57\_G DNA primase/helicase (E.C.2.7.7.-) primase, helicase, dNTPase, DNA replication; pdb70 \| 1U9I\_F \| 95.8 \| 5.9e-05 \| 7.1e-09 \| 68.5 \| 44 \| (11, 54) \| 329 \| (35, 78) \| 519 \| KaiC \| 1U9I\_F KaiC homohexamer, circadian, kaia, kaib, kaic HET: SEP, ATP, TPO; pdb70 \| 3CMV\_A \| 95.8 \| 6.5e-05 \| 7.6e-09 \| 77.4 \| 97 \| (11, 122) \| 329 \| (30, 127) \| 1357 \| PROTEIN recA \| 3CMV\_A PROTEIN recA homologous recombination, recombination HET: ANP, MG; pdb70 \| 3CMV\_A \| 95.8 \| 6.9e-05 \| 8e-09 \| 77.2 \| 97 \| (11, 122) \| 329 \| (1078, 1175) \| 1357 \| PROTEIN recA \| 3CMV\_A PROTEIN recA homologous recombination, recombination HET: ANP, MG; pdb70 \| 3CMV\_H \| 95.8 \| 6.9e-05 \| 8.1e-09 \| 77.1 \| 97 \| (11, 122) \| 329 \| (1078, 1175) \| 1357 \| PROTEIN recA \| 3CMV\_H PROTEIN recA homologous recombination, recombination HET: ANP, MG; pdb70 \| 3CMT\_D \| 95.7 \| 7.4e-05 \| 8.6e-09 \| 79.0 \| 97 \| (11, 122) \| 329 \| (1078, 1175) \| 1706 \| PROTEIN recA/DNA Complex \| 3CMT\_D PROTEIN recA/DNA Complex homologous recombination, ATP-binding, Cytoplasm, DNA HET: ADP, MG, ALF; pdb70 \| 3CMW\_C \| 95.7 \| 7.9e-05 \| 9.2e-09 \| 78.8 \| 97 \| (11, 122) \| 329 \| (1078, 1175) \| 1706 \| PROTEIN recA/DNA Complex \| 3CMW\_C PROTEIN recA/DNA Complex homologous recombination, recombination-DNA COMPLEX HET: ADP, MG, ALF; pdb70 \| 3CMU\_A \| 95.7 \| 8.2e-05 \| 9.3e-09 \| 80.5 \| 97 \| (11, 122) \| 329 \| (379, 476) \| 2050 \| PROTEIN recA/DNA Complex \| 3CMU\_A PROTEIN recA/DNA Complex homologous recombination, recombination-DNA COMPLEX HET: ADP, ALF; pdb70 \| 3CMW\_C \| 95.7 \| 8.6e-05 \| 1e-08 \| 78.5 \| 98 \| (10, 122) \| 329 \| (727, 825) \| 1706 \| PROTEIN recA/DNA Complex \| 3CMW\_C PROTEIN recA/DNA Complex homologous recombination, recombination-DNA COMPLEX HET: ADP, MG, ALF; pdb70 \| 3CMT\_D \| 95.6 \| 9.2e-05 \| 1.1e-08 \| 78.3 \| 97 \| (11, 122) \| 329 \| (1427, 1524) \| 1706 \| PROTEIN recA/DNA Complex \| 3CMT\_D PROTEIN recA/DNA Complex homologous recombination, ATP-binding, Cytoplasm, DNA HET: ADP, MG, ALF; pdb70 \| 3CMV\_A \| 95.6 \| 0.00011 \| 1.2e-08 \| 75.7 \| 98 \| (10, 122) \| 329 \| (378, 476) \| 1357 \| PROTEIN recA \| 3CMV\_A PROTEIN recA homologous recombination, recombination HET: ANP, MG; pdb70 \| 5LKM\_A \| 95.6 \| 0.00011 \| 1.3e-08 \| 66.1 \| 42 \| (11, 54) \| 329 \| (84, 125) \| 452 \| DNA repair protein RadA \| 5LKM\_A DNA repair protein RadA Helicase, recombination, DNA-binding protein, Lon-protease HET: TYD; pdb70 \| 3CMV\_H \| 95.5 \| 0.00012 \| 1.4e-08 \| 75.3 \| 98 \| (10, 122) \| 329 \| (727, 825) \| 1357 \| PROTEIN recA \| 3CMV\_H PROTEIN recA homologous recombination, recombination HET: ANP, MG; pdb70 \| 5IKN\_E \| 95.5 \| 0.00012 \| 1.4e-08 \| 65.7 \| 45 \| (11, 55) \| 329 \| (238, 282) \| 486 \| DNA-directed DNA polymerase (E.C.2.7.7.7,3.1.11.-), DNA \| 5IKN\_E DNA-directed DNA polymerase (E.C.2.7.7.7,3.1.11.-), DNA Replisome, TRANSFERASE; pdb70 \| 5IKN\_H \| 95.5 \| 0.00012 \| 1.4e-08 \| 65.7 \| 45 \| (11, 55) \| 329 \| (238, 282) \| 486 \| DNA-directed DNA polymerase (E.C.2.7.7.7,3.1.11.-), DNA \| 5IKN\_H DNA-directed DNA polymerase (E.C.2.7.7.7,3.1.11.-), DNA Replisome, TRANSFERASE; pdb70 \| 3CMW\_C \| 95.5 \| 0.00012 \| 1.4e-08 \| 77.2 \| 97 \| (11, 122) \| 329 \| (1427, 1524) \| 1706 \| PROTEIN recA/DNA Complex \| 3CMW\_C PROTEIN recA/DNA Complex homologous recombination, recombination-DNA COMPLEX HET: ADP, MG, ALF; pdb70 \| 3CMV\_A \| 95.5 \| 0.00013 \| 1.5e-08 \| 75.0 \| 97 \| (11, 122) \| 329 \| (728, 825) \| 1357 \| PROTEIN recA \| 3CMV\_A PROTEIN recA homologous recombination, recombination HET: ANP, MG; pdb70 \| 3CMT\_D \| 95.5 \| 0.00014 \| 1.6e-08 \| 76.9 \| 98 \| (11, 123) \| 329 \| (30, 128) \| 1706 \| PROTEIN recA/DNA Complex \| 3CMT\_D PROTEIN recA/DNA Complex homologous recombination, ATP-binding, Cytoplasm, DNA HET: ADP, MG, ALF; pdb70 \| 3CMT\_D \| 95.5 \| 0.00014 \| 1.6e-08 \| 76.9 \| 98 \| (10, 122) \| 329 \| (727, 825) \| 1706 \| PROTEIN recA/DNA Complex \| 3CMT\_D PROTEIN recA/DNA Complex homologous recombination, ATP-binding, Cytoplasm, DNA HET: ADP, MG, ALF; pdb70 \| 3CMW\_C \| 95.5 \| 0.00014 \| 1.6e-08 \| 76.8 \| 98 \| (11, 123) \| 329 \| (30, 128) \| 1706 \| PROTEIN recA/DNA Complex \| 3CMW\_C PROTEIN recA/DNA Complex homologous recombination, recombination-DNA COMPLEX HET: ADP, MG, ALF; pdb70 \| 3CMV\_H \| 95.4 \| 0.00014 \| 1.7e-08 \| 74.7 \| 97 \| (11, 122) \| 329 \| (379, 476) \| 1357 \| PROTEIN recA \| 3CMV\_H PROTEIN recA homologous recombination, recombination HET: ANP, MG; pdb70 \| 3CMU\_A \| 95.4 \| 0.00015 \| 1.7e-08 \| 78.4 \| 97 \| (11, 122) \| 329 \| (728, 825) \| 2050 \| PROTEIN recA/DNA Complex \| 3CMU\_A PROTEIN recA/DNA Complex homologous recombination, recombination-DNA COMPLEX HET: ADP, ALF; pdb70 \| 1NLF\_B \| 95.4 \| 0.00015 \| 1.8e-08 \| 59.9 \| 45 \| (11, 55) \| 329 \| (26, 79) \| 279 \| Regulatory protein repA \| 1NLF\_B Regulatory protein repA replicative DNA helicase structural changes HET: SO4; pdb70 \| 1OLO\_B \| 95.4 \| 0.00015 \| 1.8e-08 \| 59.9 \| 45 \| (11, 55) \| 329 \| (26, 79) \| 279 \| REGULATORY PROTEIN REPA \| 1OLO\_B REGULATORY PROTEIN REPA DNA HELICASE, ATPASE, MOTOR PROTEIN HET: SO4; pdb70 \| 3CMU\_A \| 95.3 \| 0.00018 \| 2e-08 \| 77.8 \| 97 \| (11, 122) \| 329 \| (1423, 1520) \| 2050 \| PROTEIN recA/DNA Complex \| 3CMU\_A PROTEIN recA/DNA Complex homologous recombination, recombination-DNA COMPLEX HET: ADP, ALF; pdb70 \| 1CR2\_A \| 95.3 \| 0.00018 \| 2.2e-08 \| 58.6 \| 45 \| (10, 54) \| 329 \| (30, 74) \| 296 \| DNA PRIMASE/HELICASE (E.C.2.7.7.-) \| 1CR2\_A DNA PRIMASE/HELICASE (E.C.2.7.7.-) RECA-TYPE PROTEIN FOLD, TRANSFERASE HET: SO4, DTP; pdb70 \| 1CR4\_A \| 95.3 \| 0.00018 \| 2.2e-08 \| 58.6 \| 45 \| (10, 54) \| 329 \| (30, 74) \| 296 \| DNA PRIMASE/HELICASE (E.C.2.7.7.-) \| 1CR4\_A DNA PRIMASE/HELICASE (E.C.2.7.7.-) RECA-TYPE PROTEIN FOLD, TRANSFERASE HET: SO4, TYD; pdb70 \| 1U9I\_F \| 95.3 \| 0.00019 \| 2.3e-08 \| 64.9 \| 106 \| (10, 123) \| 329 \| (276, 387) \| 519 \| KaiC \| 1U9I\_F KaiC homohexamer, circadian, kaia, kaib, kaic HET: SEP, ATP, TPO; pdb70 \| 3ZN8\_D \| 95.2 \| 0.00021 \| 2.5e-08 \| 60.6 \| 41 \| (12, 53) \| 329 \| (90, 130) \| 295 \| SIGNAL RECOGNITION PARTICLE PROTEIN (E.C.3.6.5.4) \| 3ZN8\_D SIGNAL RECOGNITION PARTICLE PROTEIN (E.C.3.6.5.4) PROTEIN TRANSPORT, HYDROLASE; pdb70 \| 1E0K\_B \| 95.1 \| 0.00023 \| 3e-08 \| 57.3 \| 44 \| (11, 54) \| 329 \| (41, 84) \| 289 \| DNA HELICASE \| 1E0K\_B DNA HELICASE HELICASE, ATPASE, DNA REPLICATION; pdb70 \| 1KMH\_A \| 95.1 \| 0.00029 \| 3.1e-08 \| 66.9 \| 43 \| (11, 54) \| 329 \| (159, 201) \| 507 \| ATPase alpha subunit (E.C.3.6.3.14)/ATPase beta \| 1KMH\_A ATPase alpha subunit (E.C.3.6.3.14)/ATPase beta Protein-Inhibitor Complex, HYDROLASE HET: TTX; pdb70 \| 6FKF\_E \| 95.1 \| 0.00029 \| 3.1e-08 \| 66.9 \| 43 \| (11, 54) \| 329 \| (159, 201) \| 507 \| ATP synthase subunit alpha, chloroplastic \| 6FKF\_E ATP synthase subunit alpha, chloroplastic ATP synthase, membrane protein complex HET: ADP, ATP; pdb70 \| 6FKH\_A \| 95.1 \| 0.00029 \| 3.1e-08 \| 66.9 \| 43 \| (11, 54) \| 329 \| (159, 201) \| 507 \| ATP synthase subunit a, chloroplastic \| 6FKH\_A ATP synthase subunit a, chloroplastic ATP synthase, membrane protein complex HET: ATP, ADP; pdb70 \| 2R9V\_A \| 95.0 \| 0.00034 \| 3.6e-08 \| 66.5 \| 42 \| (11, 53) \| 329 \| (171, 212) \| 515 \| ATP synthase subunit alpha (E.C.3.6.3.14) \| 2R9V\_A ATP synthase subunit alpha (E.C.3.6.3.14) TM1612, ATP synthase subunit alpha HET: PG4, ATP; pdb70 \| 3CMU\_A \| 95.0 \| 0.00033 \| 3.7e-08 \| 75.6 \| 98 \| (11, 123) \| 329 \| (30, 128) \| 2050 \| PROTEIN recA/DNA Complex \| 3CMU\_A PROTEIN recA/DNA Complex homologous recombination, recombination-DNA COMPLEX HET: ADP, ALF; pdb70 \| 5CDF\_A \| 94.9 \| 0.00043 \| 4.5e-08 \| 65.7 \| 43 \| (11, 53) \| 329 \| (158, 208) \| 511 \| ATP synthase subunit alpha (E.C.3.6.3.14) \| 5CDF\_A ATP synthase subunit alpha (E.C.3.6.3.14) hydrolase HET: GOL; pdb70 \| 3OE7\_B \| 94.8 \| 0.00049 \| 5.1e-08 \| 65.2 \| 43 \| (11, 53) \| 329 \| (160, 209) \| 510 \| ATP synthase subunit alpha (E.C.3.6.3.14) \| 3OE7\_B ATP synthase subunit alpha (E.C.3.6.3.14) ATP SYNTHASE, ATP PHOSPHATASE, F1F0 HET: ANP, MG; pdb70 \| 6Q45\_A \| 94.8 \| 0.00051 \| 5.4e-08 \| 64.9 \| 43 \| (11, 54) \| 329 \| (158, 200) \| 500 \| ATP synthase subunit alpha (E.C.3.6.3.14) \| 6Q45\_A ATP synthase subunit alpha (E.C.3.6.3.14) Complex, Fusobacterium, HYDROLASE, ATPase HET: MG, ADP, ATP; pdb70 \| 4B2Q\_C \| 94.8 \| 0.00052 \| 5.5e-08 \| 64.4 \| 43 \| (11, 53) \| 329 \| (135, 184) \| 485 \| ATP SYNTHASE SUBUNIT ALPHA, MITOCHONDRIAL \| 4B2Q\_C ATP SYNTHASE SUBUNIT ALPHA, MITOCHONDRIAL HYDROLASE, SUBTOMOGRAM AVERAGE HET: ADP, ATP, MG; pdb70 \| 5NCO\_l \| 94.7 \| 0.00048 \| 5.8e-08 \| 56.0 \| 41 \| (12, 53) \| 329 \| (67, 107) \| 271 \| 50S ribosomal protein L2, 50S \| 5NCO\_l 50S ribosomal protein L2, 50S Ribosome, SRP, Sec translocon, SRP HET: GDP, ALF; pdb70 \| 3BH0\_A \| 94.5 \| 0.0007 \| 8.2e-08 \| 57.6 \| 44 \| (11, 55) \| 329 \| (64, 107) \| 315 \| B.subtilis phage SPP1 DNA sequence \| 3BH0\_A B.subtilis phage SPP1 DNA sequence Helicase, ATPase, REPLICATION; pdb70 \| 1LS1\_A \| 94.4 \| 0.00072 \| 8.5e-08 \| 57.0 \| 41 \| (13, 54) \| 329 \| (96, 136) \| 295 \| SIGNAL RECOGNITION PARTICLE PROTEIN \| 1LS1\_A SIGNAL RECOGNITION PARTICLE PROTEIN Ffh, SRP54, SRP, GTPase, ultrahigh; pdb70 \| 6N2Z\_A \| 94.4 \| 0.00081 \| 8.5e-08 \| 63.6 \| 43 \| (11, 54) \| 329 \| (158, 200) \| 502 \| Bacterial ATP synthase \| 6N2Z\_A Bacterial ATP synthase Bacterial ATP synthase, HYDROLASE HET: ADP, ATP; pdb70 \| 6N7I\_D \| 94.4 \| 0.00076 \| 9.1e-08 \| 61.5 \| 44 \| (11, 54) \| 329 \| (301, 344) \| 566 \| DNA primase/helicase/DNA Complex \| 6N7I\_D DNA primase/helicase/DNA Complex helicase, ATPase, hexamer, DNA replication HET: TTP; pdb70 \| 2JDI\_A \| 94.4 \| 0.0009 \| 9.5e-08 \| 63.4 \| 44 \| (11, 54) \| 329 \| (158, 208) \| 510 \| ATP SYNTHASE SUBUNIT ALPHA HEART \| 2JDI\_A ATP SYNTHASE SUBUNIT ALPHA HEART ATP PHOSPHORYLASE, HYDROLASE, ATP SYNTHESIS HET: ANP; pdb70 \| 4C7O\_D \| 94.3 \| 0.00082 \| 1e-07 \| 54.8 \| 41 \| (12, 53) \| 329 \| (71, 111) \| 278 \| SIGNAL RECOGNITION PARTICLE PROTEIN, SIGNAL \| 4C7O\_D SIGNAL RECOGNITION PARTICLE PROTEIN, SIGNAL NUCLEAR PROTEIN-RNA COMPLEX, NUCLEAR PROTEIN HET: ALF, GDP; pdb70 \| 6N9X\_B \| 94.3 \| 0.00088 \| 1.1e-07 \| 61.0 \| 45 \| (10, 54) \| 329 \| (300, 344) \| 566 \| DNA-directed DNA polymerase, TrxA/DNA Complex \| 6N9X\_B DNA-directed DNA polymerase, TrxA/DNA Complex DNA polymerase, primase, helicase, DNA HET: DOC, TTP; pdb70 \| 6J5K\_BA \| 94.2 \| 0.001 \| 1.1e-07 \| 62.9 \| 44 \| (11, 54) \| 329 \| (157, 207) \| 509 \| ATP synthase F1 subunit alpha \| 6J5K\_BA ATP synthase F1 subunit alpha membrane protein HET: ADP, ATP, MG; pdb70 \| 1JPN\_A \| 94.2 \| 0.00099 \| 1.2e-07 \| 55.9 \| 41 \| (13, 54) \| 329 \| (96, 136) \| 296 \| SIGNAL RECOGNITION PARTICLE PROTEIN \| 1JPN\_A SIGNAL RECOGNITION PARTICLE PROTEIN Ffh, SRP, GMPPNP, Signal Recognition HET: ACY, GNP |
| Top keywords  (threshold 1.00e-03 (evalue)) | **ATPase, domain\_containing, AAA, RecA, DNA, recombination, Fragment, KaiC, repair, profile** |
| Output files | ../../similar\_sequences/39\_FANPEZAQ\_CDS\_0039\_merged.svg ../../similar\_sequences/39\_FANPEZAQ\_CDS\_0039\_pdb70.a3m ../../similar\_sequences/39\_FANPEZAQ\_CDS\_0039\_pdb70.hhr ../../similar\_sequences/39\_FANPEZAQ\_CDS\_0039\_uniclust.a3m ../../similar\_sequences/39\_FANPEZAQ\_CDS\_0039\_uniclust.hhr |

#### Structure prediction (AlphaFold)^2^

|  |  |
| --- | --- |
| Stats | xml version="1.0" encoding="utf-8" standalone="no"?       2024-09-02T21:09:38.105150 image/svg+xml   Matplotlib v3.7.2, https://matplotlib.org/ |
| Predicted structure | **NGL Viewer Controls:**  - Center: *Left-Click* - Rotate: *Left-Click + Drag* - Translate: *Right-Click + Drag* - Zoom: *Shift + Left-Click + Drag* |
| Output files | ../../predicted\_structures/39\_FANPEZAQ\_CDS\_0039/features.pkl ../../predicted\_structures/39\_FANPEZAQ\_CDS\_0039/ranked\_0.pdb ../../predicted\_structures/39\_FANPEZAQ\_CDS\_0039/ranked\_0\_plots.svg ../../predicted\_structures/39\_FANPEZAQ\_CDS\_0039/result\_model\_1\_ptm\_pred\_0.pkl |

#### Structure similarity search results (Foldseek)^3^

|  |  |
| --- | --- |
| Structure databases searched | Pdb, Afdb-proteome, Afdb-uniprot50 |
| Results, scheme(s)  (Top layers only, threshold 1.00e-02 (evalue)) | xml version="1.0" encoding="utf-8" standalone="no"?       2024-09-02T21:11:13.957999 image/svg+xml   Matplotlib v3.7.2, https://matplotlib.org/ |
| Results, table  (threshold 1.00e-02 (evalue)) | db \| id \| prob \| evalue \| bits \| fident \| alnlen \| mismatch \| gapopen \| qstart \| qend \| tstart \| tend \| name \| description; pdb \| 4TL7\_A \| 1.0 \| 3.47e-06 \| 167 \| 0.14 \| 227 \| 143 \| 9 \| 16 \| 233 \| 25 \| 208 \| Circadian clock protein kinase KaiC \| Circadian clock protein kinase KaiC; pdb \| 8FAZ\_D \| 1.0 \| 4.1e-06 \| 154 \| 0.131 \| 243 \| 142 \| 18 \| 12 \| 231 \| 98 \| 294 \| DNA repair protein RAD51 homolog 4 \| DNA repair protein RAD51 homolog 4; pdb \| 4TL7\_D \| 1.0 \| 1.948e-05 \| 150 \| 0.14 \| 228 \| 143 \| 10 \| 16 \| 233 \| 24 \| 208 \| Circadian clock protein kinase KaiC \| Circadian clock protein kinase KaiC; pdb \| 4TLB\_F \| 1.0 \| 1.648e-05 \| 148 \| 0.135 \| 228 \| 139 \| 13 \| 16 \| 233 \| 24 \| 203 \| Circadian clock protein kinase KaiC \| Circadian clock protein kinase KaiC; pdb \| 8OUZ\_C \| 1.0 \| 1.559e-05 \| 145 \| 0.127 \| 243 \| 142 \| 17 \| 16 \| 231 \| 100 \| 299 \| DNA repair protein RAD51 homolog 4 \| DNA repair protein RAD51 homolog 4; pdb \| 8GBJ\_D \| 1.0 \| 2.627e-06 \| 144 \| 0.151 \| 244 \| 134 \| 18 \| 16 \| 231 \| 100 \| 298 \| DNA repair protein RAD51 homolog 4 \| DNA repair protein RAD51 homolog 4; pdb \| 7V3X\_G \| 1.0 \| 2.177e-05 \| 144 \| 0.146 \| 218 \| 130 \| 12 \| 16 \| 219 \| 24 \| 199 \| Circadian clock protein kinase KaiC \| Circadian clock protein kinase KaiC; pdb \| 5YZ8\_A \| 1.0 \| 3.592e-05 \| 142 \| 0.144 \| 215 \| 135 \| 12 \| 16 \| 219 \| 24 \| 200 \| Circadian Clock Protein Kinase KaiC \| Circadian Clock Protein Kinase KaiC; pdb \| 4TLB\_D \| 1.0 \| 2.302e-05 \| 140 \| 0.132 \| 256 \| 149 \| 13 \| 5 \| 233 \| 1 \| 210 \| Circadian clock protein kinase KaiC \| Circadian clock protein kinase KaiC; pdb \| 7DY2\_D \| 1.0 \| 4.745e-05 \| 138 \| 0.136 \| 212 \| 137 \| 10 \| 16 \| 219 \| 23 \| 196 \| Circadian clock protein kinase KaiC \| Circadian clock protein kinase KaiC; pdb \| 4TL8\_B \| 1.0 \| 3.04e-05 \| 137 \| 0.142 \| 246 \| 146 \| 10 \| 1 \| 233 \| 6 \| 199 \| Circadian clock protein kinase KaiC \| Circadian clock protein kinase KaiC; pdb \| 4TLA\_B \| 1.0 \| 3.398e-05 \| 136 \| 0.131 \| 228 \| 136 \| 12 \| 16 \| 233 \| 25 \| 200 \| Circadian clock protein kinase KaiC \| Circadian clock protein kinase KaiC; pdb \| 4TLC\_B \| 1.0 \| 5.016e-05 \| 136 \| 0.139 \| 229 \| 140 \| 13 \| 16 \| 233 \| 26 \| 208 \| Circadian clock protein kinase KaiC \| Circadian clock protein kinase KaiC; pdb \| 4TL8\_F \| 1.0 \| 4.745e-05 \| 135 \| 0.14 \| 227 \| 133 \| 10 \| 16 \| 233 \| 24 \| 197 \| Circadian clock protein kinase KaiC \| Circadian clock protein kinase KaiC; pdb \| 7DY1\_C \| 1.0 \| 4.488e-05 \| 135 \| 0.154 \| 233 \| 134 \| 13 \| 12 \| 233 \| 22 \| 202 \| Circadian clock protein kinase KaiC \| Circadian clock protein kinase KaiC; pdb \| 7DY2\_B \| 1.0 \| 9.988e-06 \| 135 \| 0.157 \| 229 \| 132 \| 13 \| 16 \| 233 \| 23 \| 201 \| Circadian clock protein kinase KaiC \| Circadian clock protein kinase KaiC; pdb \| 4TLB\_A \| 1.0 \| 4.745e-05 \| 134 \| 0.142 \| 253 \| 150 \| 16 \| 1 \| 233 \| 2 \| 207 \| Circadian clock protein kinase KaiC \| Circadian clock protein kinase KaiC; pdb \| 4TLB\_C \| 1.0 \| 6.267e-05 \| 134 \| 0.122 \| 228 \| 144 \| 11 \| 16 \| 233 \| 26 \| 207 \| Circadian clock protein kinase KaiC \| Circadian clock protein kinase KaiC; pdb \| 5YZ8\_C \| 1.0 \| 0.0001292 \| 132 \| 0.14 \| 214 \| 138 \| 12 \| 16 \| 219 \| 24 \| 201 \| Circadian Clock Protein Kinase KaiC \| Circadian Clock Protein Kinase KaiC; pdb \| 7V3X\_A \| 1.0 \| 4.488e-05 \| 132 \| 0.145 \| 233 \| 132 \| 14 \| 16 \| 233 \| 23 \| 203 \| Circadian clock protein kinase KaiC \| Circadian clock protein kinase KaiC; pdb \| 4TLD\_F \| 1.0 \| 9.252e-05 \| 130 \| 0.14 \| 227 \| 135 \| 10 \| 16 \| 233 \| 24 \| 199 \| Circadian clock protein kinase KaiC \| Circadian clock protein kinase KaiC; pdb \| 4TLB\_B \| 1.0 \| 3.592e-05 \| 130 \| 0.152 \| 230 \| 133 \| 11 \| 16 \| 233 \| 26 \| 205 \| Circadian clock protein kinase KaiC \| Circadian clock protein kinase KaiC; pdb \| 4TLA\_C \| 1.0 \| 5.928e-05 \| 130 \| 0.148 \| 229 \| 140 \| 14 \| 16 \| 233 \| 27 \| 211 \| Circadian clock protein kinase KaiC \| Circadian clock protein kinase KaiC; pdb \| 7DXQ\_E \| 1.0 \| 4.488e-05 \| 130 \| 0.152 \| 216 \| 131 \| 11 \| 16 \| 219 \| 24 \| 199 \| Circadian clock protein kinase KaiC \| Circadian clock protein kinase KaiC; pdb \| 4TLE\_B \| 1.0 \| 9.252e-05 \| 129 \| 0.14 \| 228 \| 133 \| 11 \| 16 \| 233 \| 26 \| 200 \| Circadian clock protein kinase KaiC \| Circadian clock protein kinase KaiC; pdb \| 3HR8\_A \| 1.0 \| 2.433e-05 \| 129 \| 0.189 \| 201 \| 121 \| 12 \| 12 \| 204 \| 57 \| 223 \| Protein recA \| Protein recA; pdb \| 4TLE\_C \| 1.0 \| 7.005e-05 \| 128 \| 0.147 \| 231 \| 131 \| 14 \| 16 \| 233 \| 28 \| 205 \| Circadian clock protein kinase KaiC \| Circadian clock protein kinase KaiC; pdb \| 2ZR0\_A \| 1.0 \| 5.607e-05 \| 128 \| 0.208 \| 197 \| 112 \| 12 \| 12 \| 204 \| 56 \| 212 \| Protein recA \| Protein recA; pdb \| 4TL9\_F \| 1.0 \| 0.0001093 \| 127 \| 0.135 \| 228 \| 135 \| 12 \| 16 \| 233 \| 23 \| 198 \| Circadian clock protein kinase KaiC \| Circadian clock protein kinase KaiC; pdb \| 4TLD\_D \| 1.0 \| 5.303e-05 \| 127 \| 0.134 \| 231 \| 133 \| 13 \| 16 \| 233 \| 28 \| 204 \| Circadian clock protein kinase KaiC \| Circadian clock protein kinase KaiC; pdb \| 4TLC\_D \| 1.0 \| 6.267e-05 \| 127 \| 0.139 \| 229 \| 137 \| 13 \| 16 \| 233 \| 28 \| 207 \| Circadian clock protein kinase KaiC \| Circadian clock protein kinase KaiC; pdb \| 7DY2\_E \| 1.0 \| 0.0001907 \| 127 \| 0.148 \| 215 \| 135 \| 11 \| 16 \| 219 \| 23 \| 200 \| Circadian clock protein kinase KaiC \| Circadian clock protein kinase KaiC; pdb \| 4TL6\_C \| 1.0 \| 7.829e-05 \| 126 \| 0.138 \| 252 \| 147 \| 13 \| 1 \| 233 \| 1 \| 201 \| Circadian clock protein kinase KaiC \| Circadian clock protein kinase KaiC; pdb \| 7DY2\_A \| 1.0 \| 8.751e-05 \| 126 \| 0.122 \| 229 \| 145 \| 12 \| 16 \| 233 \| 248 \| 431 \| Circadian clock protein kinase KaiC \| Circadian clock protein kinase KaiC; pdb \| 7V3X\_B \| 1.0 \| 7.406e-05 \| 126 \| 0.144 \| 228 \| 138 \| 14 \| 16 \| 233 \| 24 \| 204 \| Circadian clock protein kinase KaiC \| Circadian clock protein kinase KaiC; pdb \| 4TL8\_C \| 1.0 \| 0.0001034 \| 125 \| 0.13 \| 230 \| 136 \| 11 \| 16 \| 233 \| 26 \| 203 \| Circadian clock protein kinase KaiC \| Circadian clock protein kinase KaiC; pdb \| 7V3X\_H \| 1.0 \| 6.267e-05 \| 125 \| 0.152 \| 229 \| 136 \| 15 \| 16 \| 233 \| 24 \| 205 \| Circadian clock protein kinase KaiC \| Circadian clock protein kinase KaiC; pdb \| 7DXQ\_A \| 1.0 \| 7.829e-05 \| 124 \| 0.142 \| 232 \| 133 \| 13 \| 16 \| 233 \| 23 \| 202 \| Circadian clock protein kinase KaiC \| Circadian clock protein kinase KaiC; pdb \| 5YZ8\_B \| 1.0 \| 0.0004912 \| 123 \| 0.135 \| 214 \| 139 \| 12 \| 16 \| 219 \| 24 \| 201 \| Circadian Clock Protein Kinase KaiC \| Circadian Clock Protein Kinase KaiC; pdb \| 4TLC\_F \| 1.0 \| 8.277e-05 \| 122 \| 0.139 \| 229 \| 136 \| 14 \| 16 \| 233 \| 24 \| 202 \| Circadian clock protein kinase KaiC \| Circadian clock protein kinase KaiC; pdb \| 4TLE\_D \| 1.0 \| 0.0001292 \| 122 \| 0.148 \| 229 \| 129 \| 14 \| 16 \| 233 \| 28 \| 201 \| Circadian clock protein kinase KaiC \| Circadian clock protein kinase KaiC; pdb \| 4TL9\_D \| 1.0 \| 0.0001222 \| 122 \| 0.134 \| 246 \| 148 \| 12 \| 1 \| 233 \| 9 \| 202 \| Circadian clock protein kinase KaiC \| Circadian clock protein kinase KaiC; pdb \| 4TLC\_C \| 1.0 \| 0.0001156 \| 122 \| 0.131 \| 228 \| 144 \| 12 \| 16 \| 233 \| 26 \| 209 \| Circadian clock protein kinase KaiC \| Circadian clock protein kinase KaiC; pdb \| 7V3X\_I \| 1.0 \| 7.406e-05 \| 121 \| 0.133 \| 254 \| 152 \| 15 \| 1 \| 233 \| 1 \| 207 \| Circadian clock protein kinase KaiC \| Circadian clock protein kinase KaiC; pdb \| 4IJM\_D \| 1.0 \| 8.277e-05 \| 121 \| 0.135 \| 244 \| 134 \| 15 \| 16 \| 233 \| 27 \| 219 \| Circadian clock protein kinase KaiC \| Circadian clock protein kinase KaiC; pdb \| 4O0M\_C \| 1.0 \| 0.0002132 \| 121 \| 0.157 \| 241 \| 138 \| 15 \| 12 \| 233 \| 29 \| 223 \| Circadian clock protein kinase KaiC \| Circadian clock protein kinase KaiC; pdb \| 3DVL\_F \| 1.0 \| 0.0002016 \| 121 \| 0.143 \| 237 \| 140 \| 14 \| 16 \| 233 \| 27 \| 219 \| Circadian clock protein kinase kaiC \| Circadian clock protein kinase kaiC; pdb \| 4TL8\_E \| 1.0 \| 7.406e-05 \| 120 \| 0.151 \| 231 \| 127 \| 12 \| 16 \| 233 \| 25 \| 199 \| Circadian clock protein kinase KaiC \| Circadian clock protein kinase KaiC; pdb \| 7DYK\_A \| 1.0 \| 0.0002519 \| 120 \| 0.13 \| 199 \| 120 \| 9 \| 16 \| 186 \| 242 \| 415 \| Circadian clock protein kinase KaiC \| Circadian clock protein kinase KaiC; pdb \| 7DYJ\_B \| 1.0 \| 0.0001907 \| 120 \| 0.14 \| 214 \| 134 \| 12 \| 16 \| 219 \| 23 \| 196 \| Circadian clock protein kinase KaiC \| Circadian clock protein kinase KaiC; pdb \| 7V3X\_C \| 1.0 \| 8.277e-05 \| 120 \| 0.13 \| 253 \| 154 \| 14 \| 1 \| 233 \| 1 \| 207 \| Circadian clock protein kinase KaiC \| Circadian clock protein kinase KaiC; pdb \| 7DY1\_D \| 1.0 \| 0.0001034 \| 119 \| 0.14 \| 235 \| 133 \| 12 \| 12 \| 233 \| 22 \| 200 \| Circadian clock protein kinase KaiC \| Circadian clock protein kinase KaiC; pdb \| 7DY2\_H \| 1.0 \| 7.829e-05 \| 119 \| 0.144 \| 236 \| 125 \| 13 \| 16 \| 233 \| 23 \| 199 \| Circadian clock protein kinase KaiC \| Circadian clock protein kinase KaiC; pdb \| 4TLE\_F \| 1.0 \| 0.0001034 \| 118 \| 0.15 \| 232 \| 125 \| 11 \| 16 \| 233 \| 24 \| 197 \| Circadian clock protein kinase KaiC \| Circadian clock protein kinase KaiC; pdb \| 5YZ8\_E \| 1.0 \| 0.0004646 \| 118 \| 0.135 \| 214 \| 136 \| 11 \| 16 \| 219 \| 23 \| 197 \| Circadian Clock Protein Kinase KaiC \| Circadian Clock Protein Kinase KaiC; pdb \| 7V3X\_J \| 1.0 \| 0.0002016 \| 118 \| 0.135 \| 222 \| 130 \| 14 \| 16 \| 219 \| 23 \| 200 \| Circadian clock protein kinase KaiC \| Circadian clock protein kinase KaiC; pdb \| 4TL9\_C \| 1.0 \| 0.0002816 \| 117 \| 0.125 \| 231 \| 137 \| 13 \| 16 \| 233 \| 27 \| 205 \| Circadian clock protein kinase KaiC \| Circadian clock protein kinase KaiC; pdb \| 4TL6\_B \| 1.0 \| 6.626e-05 \| 117 \| 0.147 \| 230 \| 138 \| 10 \| 16 \| 233 \| 24 \| 207 \| Circadian clock protein kinase KaiC \| Circadian clock protein kinase KaiC; pdb \| 7V3X\_R \| 1.0 \| 0.0003147 \| 117 \| 0.135 \| 229 \| 141 \| 13 \| 16 \| 233 \| 23 \| 205 \| Circadian clock protein kinase KaiC \| Circadian clock protein kinase KaiC; pdb \| 7V3X\_M \| 1.0 \| 0.0003327 \| 117 \| 0.13 \| 214 \| 137 \| 11 \| 16 \| 217 \| 248 \| 424 \| Circadian clock protein kinase KaiC \| Circadian clock protein kinase KaiC; pdb \| 7V3X\_L \| 1.0 \| 0.0001093 \| 117 \| 0.144 \| 235 \| 138 \| 15 \| 12 \| 233 \| 24 \| 208 \| Circadian clock protein kinase KaiC \| Circadian clock protein kinase KaiC; pdb \| 2ZTS\_B \| 1.0 \| 9.781e-05 \| 116 \| 0.16 \| 224 \| 122 \| 18 \| 12 \| 219 \| 22 \| 195 \| Putative uncharacterized protein PH0186 \| Putative uncharacterized protein PH0186; pdb \| 4TLD\_B \| 1.0 \| 0.0002816 \| 116 \| 0.131 \| 228 \| 137 \| 12 \| 16 \| 233 \| 25 \| 201 \| Circadian clock protein kinase KaiC \| Circadian clock protein kinase KaiC; pdb \| 4TL6\_A \| 1.0 \| 0.0003518 \| 116 \| 0.144 \| 228 \| 134 \| 11 \| 16 \| 233 \| 26 \| 202 \| Circadian clock protein kinase KaiC \| Circadian clock protein kinase KaiC; pdb \| 3K0C\_D \| 1.0 \| 0.0001614 \| 116 \| 0.13 \| 237 \| 143 \| 13 \| 16 \| 233 \| 27 \| 219 \| Circadian clock protein kinase KaiC \| Circadian clock protein kinase KaiC; pdb \| 7DYI\_A \| 1.0 \| 0.0001907 \| 115 \| 0.134 \| 231 \| 133 \| 12 \| 16 \| 233 \| 24 \| 200 \| Circadian clock protein kinase KaiC \| Circadian clock protein kinase KaiC; pdb \| 7DXQ\_D \| 1.0 \| 0.0003518 \| 115 \| 0.131 \| 236 \| 135 \| 15 \| 16 \| 233 \| 248 \| 431 \| Circadian clock protein kinase KaiC \| Circadian clock protein kinase KaiC; pdb \| 7DXQ\_B \| 1.0 \| 0.0001366 \| 115 \| 0.137 \| 233 \| 135 \| 14 \| 16 \| 233 \| 27 \| 208 \| Circadian clock protein kinase KaiC \| Circadian clock protein kinase KaiC; pdb \| 4TLA\_A \| 1.0 \| 0.0002132 \| 114 \| 0.141 \| 234 \| 127 \| 14 \| 16 \| 233 \| 25 \| 200 \| Circadian clock protein kinase KaiC \| Circadian clock protein kinase KaiC; pdb \| 7X1Y\_D \| 1.0 \| 5.607e-05 \| 114 \| 0.131 \| 228 \| 130 \| 13 \| 16 \| 216 \| 269 \| 455 \| Circadian clock oscillator protein KaiC \| Circadian clock oscillator protein KaiC; pdb \| 7V3X\_D \| 1.0 \| 0.0002016 \| 114 \| 0.135 \| 237 \| 132 \| 13 \| 16 \| 233 \| 24 \| 206 \| Circadian clock protein kinase KaiC \| Circadian clock protein kinase KaiC; pdb \| 4TLD\_C \| 1.0 \| 0.0004395 \| 113 \| 0.126 \| 230 \| 137 \| 12 \| 16 \| 233 \| 27 \| 204 \| Circadian clock protein kinase KaiC \| Circadian clock protein kinase KaiC; pdb \| 7DYE\_A \| 1.0 \| 0.0002016 \| 113 \| 0.13 \| 230 \| 134 \| 12 \| 16 \| 233 \| 24 \| 199 \| Circadian clock protein kinase KaiC \| Circadian clock protein kinase KaiC; pdb \| 7V3X\_X \| 1.0 \| 0.001414 \| 113 \| 0.136 \| 213 \| 141 \| 11 \| 16 \| 219 \| 246 \| 424 \| Circadian clock protein kinase KaiC \| Circadian clock protein kinase KaiC; pdb \| 7DY2\_I \| 1.0 \| 0.0001804 \| 113 \| 0.15 \| 239 \| 132 \| 17 \| 12 \| 233 \| 21 \| 205 \| Circadian clock protein kinase KaiC \| Circadian clock protein kinase KaiC; pdb \| 7V3X\_F \| 1.0 \| 0.0001222 \| 113 \| 0.147 \| 231 \| 134 \| 14 \| 16 \| 233 \| 26 \| 206 \| Circadian clock protein kinase KaiC \| Circadian clock protein kinase KaiC; pdb \| 3K0F\_D \| 1.0 \| 0.0001907 \| 113 \| 0.145 \| 241 \| 135 \| 12 \| 16 \| 233 \| 27 \| 219 \| Circadian clock protein kinase KaiC \| Circadian clock protein kinase KaiC; pdb \| 7DY2\_G \| 1.0 \| 0.0003932 \| 112 \| 0.119 \| 243 \| 142 \| 13 \| 16 \| 235 \| 244 \| 437 \| Circadian clock protein kinase KaiC \| Circadian clock protein kinase KaiC; pdb \| 7S65\_E \| 1.0 \| 0.0002977 \| 112 \| 0.152 \| 236 \| 139 \| 16 \| 16 \| 233 \| 26 \| 218 \| Circadian clock protein kinase KaiC \| Circadian clock protein kinase KaiC; pdb \| 7DXQ\_F \| 1.0 \| 0.0002663 \| 112 \| 0.141 \| 233 \| 134 \| 16 \| 16 \| 233 \| 27 \| 208 \| Circadian clock protein kinase KaiC \| Circadian clock protein kinase KaiC; pdb \| 5YZ8\_F \| 1.0 \| 0.001197 \| 111 \| 0.14 \| 213 \| 134 \| 11 \| 16 \| 219 \| 24 \| 196 \| Circadian Clock Protein Kinase KaiC \| Circadian Clock Protein Kinase KaiC; pdb \| 6X61\_I \| 1.0 \| 0.001265 \| 111 \| 0.153 \| 222 \| 136 \| 11 \| 12 \| 216 \| 20 \| 206 \| Circadian clock protein kinase KaiC \| Circadian clock protein kinase KaiC; pdb \| 3S1A\_A \| 1.0 \| 9.781e-05 \| 111 \| 0.144 \| 242 \| 134 \| 14 \| 16 \| 233 \| 27 \| 219 \| Circadian clock protein kinase kaiC \| Circadian clock protein kinase kaiC; pdb \| 2ZTS\_C \| 1.0 \| 0.0002254 \| 110 \| 0.172 \| 226 \| 127 \| 15 \| 12 \| 219 \| 22 \| 205 \| Putative uncharacterized protein PH0186 \| Putative uncharacterized protein PH0186; pdb \| 7DYJ\_A \| 1.0 \| 0.0004646 \| 110 \| 0.131 \| 197 \| 116 \| 9 \| 15 \| 182 \| 242 \| 412 \| Circadian clock protein kinase KaiC \| Circadian clock protein kinase KaiC; pdb \| 8FWI\_A \| 1.0 \| 6.4e-06 \| 110 \| 0.158 \| 335 \| 203 \| 21 \| 16 \| 326 \| 265 \| 544 \| Circadian clock protein KaiC \| Circadian clock protein KaiC; pdb \| 4TLD\_A \| 1.0 \| 0.0005193 \| 109 \| 0.123 \| 227 \| 139 \| 10 \| 16 \| 233 \| 26 \| 201 \| Circadian clock protein kinase KaiC \| Circadian clock protein kinase KaiC; pdb \| 7DY2\_F \| 1.0 \| 0.0003719 \| 109 \| 0.127 \| 243 \| 140 \| 15 \| 16 \| 235 \| 252 \| 445 \| Circadian clock protein kinase KaiC \| Circadian clock protein kinase KaiC; pdb \| 7V3X\_E \| 1.0 \| 0.0002816 \| 109 \| 0.143 \| 237 \| 135 \| 16 \| 12 \| 233 \| 22 \| 205 \| Circadian clock protein kinase KaiC \| Circadian clock protein kinase KaiC; pdb \| 8DB3\_B \| 1.0 \| 4.015e-05 \| 108 \| 0.147 \| 238 \| 122 \| 17 \| 12 \| 216 \| 250 \| 439 \| Circadian clock protein KaiC \| Circadian clock protein KaiC; pdb \| 7V3X\_W \| 1.0 \| 0.0003147 \| 108 \| 0.12 \| 200 \| 123 \| 14 \| 16 \| 204 \| 248 \| 405 \| Circadian clock protein kinase KaiC \| Circadian clock protein kinase KaiC; pdb \| 7V3X\_V \| 1.0 \| 0.0005805 \| 107 \| 0.142 \| 232 \| 137 \| 15 \| 16 \| 233 \| 222 \| 405 \| Circadian clock protein kinase KaiC \| Circadian clock protein kinase KaiC; pdb \| 7DY1\_B \| 1.0 \| 0.0004912 \| 107 \| 0.144 \| 221 \| 130 \| 15 \| 16 \| 219 \| 237 \| 415 \| Circadian clock protein kinase KaiC \| Circadian clock protein kinase KaiC; pdb \| 7V3X\_Q \| 1.0 \| 0.0002383 \| 107 \| 0.137 \| 232 \| 133 \| 14 \| 16 \| 233 \| 23 \| 201 \| Circadian clock protein kinase KaiC \| Circadian clock protein kinase KaiC; pdb \| 3JZM\_B \| 1.0 \| 0.0003147 \| 107 \| 0.137 \| 240 \| 138 \| 13 \| 16 \| 233 \| 27 \| 219 \| Circadian clock protein kinase kaiC \| Circadian clock protein kinase kaiC; pdb \| 7WDC\_B \| 1.0 \| 0.001013 \| 106 \| 0.136 \| 213 \| 136 \| 11 \| 16 \| 219 \| 23 \| 196 \| Circadian clock protein kinase KaiC \| Circadian clock protein kinase KaiC; pdb \| 3K0A\_D \| 1.0 \| 0.0006137 \| 106 \| 0.141 \| 240 \| 137 \| 12 \| 16 \| 233 \| 27 \| 219 \| Circadian clock protein kinase KaiC \| Circadian clock protein kinase KaiC; pdb \| 4TLB\_E \| 1.0 \| 0.0002519 \| 105 \| 0.14 \| 235 \| 136 \| 15 \| 16 \| 233 \| 26 \| 211 \| Circadian clock protein kinase KaiC \| Circadian clock protein kinase KaiC; pdb \| 1CR1\_A \| 1.0 \| 0.0002254 \| 105 \| 0.136 \| 250 \| 132 \| 15 \| 10 \| 230 \| 32 \| 226 \| DNA PRIMASE/HELICASE \| DNA PRIMASE/HELICASE; pdb \| 7WDC\_A \| 1.0 \| 0.0003719 \| 105 \| 0.137 \| 218 \| 130 \| 12 \| 16 \| 219 \| 23 \| 196 \| Circadian clock protein kinase KaiC \| Circadian clock protein kinase KaiC; pdb \| 4TLA\_F \| 1.0 \| 0.0009578 \| 103 \| 0.139 \| 236 \| 133 \| 14 \| 16 \| 233 \| 24 \| 207 \| Circadian clock protein kinase KaiC \| Circadian clock protein kinase KaiC; pdb \| 3K09\_B \| 1.0 \| 0.0005193 \| 103 \| 0.145 \| 241 \| 135 \| 14 \| 16 \| 233 \| 27 \| 219 \| Circadian clock protein kinase kaiC \| Circadian clock protein kinase kaiC; pdb \| 4TLA\_E \| 1.0 \| 0.000857 \| 102 \| 0.135 \| 244 \| 134 \| 15 \| 16 \| 233 \| 25 \| 217 \| Circadian clock protein kinase KaiC \| Circadian clock protein kinase KaiC; pdb \| 2Q6T\_D \| 1.0 \| 0.0006859 \| 102 \| 0.145 \| 241 \| 139 \| 19 \| 8 \| 219 \| 180 \| 382 \| DnaB replication fork helicase \| DnaB replication fork helicase; pdb \| 7DYK\_B \| 1.0 \| 0.0004646 \| 102 \| 0.13 \| 238 \| 138 \| 14 \| 16 \| 219 \| 244 \| 446 \| Circadian clock protein kinase KaiC \| Circadian clock protein kinase KaiC; pdb \| 4TL9\_E \| 1.0 \| 0.0009578 \| 101 \| 0.143 \| 230 \| 132 \| 15 \| 16 \| 233 \| 25 \| 201 \| Circadian clock protein kinase KaiC \| Circadian clock protein kinase KaiC; pdb \| 4ESV\_C \| 1.0 \| 0.0003719 \| 101 \| 0.12 \| 224 \| 141 \| 17 \| 9 \| 204 \| 193 \| 388 \| Replicative helicase \| Replicative helicase; pdb \| 7V3X\_N \| 1.0 \| 0.002608 \| 101 \| 0.132 \| 219 \| 135 \| 12 \| 16 \| 219 \| 239 \| 417 \| Circadian clock protein kinase KaiC \| Circadian clock protein kinase KaiC; pdb \| 7DY2\_C \| 1.0 \| 0.001581 \| 101 \| 0.135 \| 222 \| 127 \| 14 \| 16 \| 217 \| 246 \| 422 \| Circadian clock protein kinase KaiC \| Circadian clock protein kinase KaiC; pdb \| 7DXQ\_C \| 1.0 \| 0.0006488 \| 101 \| 0.147 \| 231 \| 134 \| 13 \| 16 \| 233 \| 27 \| 207 \| Circadian clock protein kinase KaiC \| Circadian clock protein kinase KaiC; pdb \| 8FWJ\_A \| 1.0 \| 1.056e-05 \| 101 \| 0.157 \| 355 \| 194 \| 24 \| 12 \| 329 \| 263 \| 549 \| Circadian clock protein KaiC \| Circadian clock protein KaiC; pdb \| 5JWO\_A \| 1.0 \| 0.0002977 \| 100 \| 0.163 \| 214 \| 114 \| 17 \| 12 \| 189 \| 21 \| 205 \| Circadian clock protein kinase KaiC \| Circadian clock protein kinase KaiC; pdb \| 4TLE\_E \| 1.0 \| 0.0005805 \| 100 \| 0.139 \| 229 \| 131 \| 16 \| 16 \| 233 \| 26 \| 199 \| Circadian clock protein kinase KaiC \| Circadian clock protein kinase KaiC; pdb \| 7V3X\_O \| 1.0 \| 0.001013 \| 100 \| 0.139 \| 230 \| 132 \| 13 \| 16 \| 233 \| 25 \| 200 \| Circadian clock protein kinase KaiC \| Circadian clock protein kinase KaiC; pdb \| 8H9E\_D \| 1.0 \| 4.245e-05 \| 100 \| 0.143 \| 314 \| 182 \| 27 \| 16 \| 298 \| 143 \| 400 \| ATP synthase subunit beta, mitochondrial \| ATP synthase subunit beta, mitochondrial; pdb \| 3K0E\_C \| 1.0 \| 0.0006488 \| 100 \| 0.145 \| 241 \| 135 \| 13 \| 16 \| 233 \| 27 \| 219 \| Circadian clock protein kinase KaiC \| Circadian clock protein kinase KaiC; pdb \| 7DYI\_B \| 0.999 \| 0.0003147 \| 99 \| 0.131 \| 243 \| 134 \| 15 \| 16 \| 220 \| 241 \| 444 \| Circadian clock protein kinase KaiC \| Circadian clock protein kinase KaiC; pdb \| 7DY2\_L \| 0.999 \| 0.0006859 \| 99 \| 0.146 \| 232 \| 129 \| 15 \| 16 \| 233 \| 23 \| 199 \| Circadian clock protein kinase KaiC \| Circadian clock protein kinase KaiC; pdb \| 1CR2\_A \| 0.999 \| 0.0004157 \| 98 \| 0.153 \| 254 \| 117 \| 19 \| 10 \| 230 \| 32 \| 220 \| DNA PRIMASE/HELICASE \| DNA PRIMASE/HELICASE; pdb \| 7DY1\_A \| 0.999 \| 0.0006137 \| 98 \| 0.14 \| 249 \| 130 \| 18 \| 16 \| 235 \| 239 \| 432 \| Circadian clock protein kinase KaiC \| Circadian clock protein kinase KaiC; pdb \| 4ESV\_A \| 0.999 \| 0.0006488 \| 97 \| 0.13 \| 223 \| 140 \| 18 \| 9 \| 204 \| 177 \| 372 \| Replicative helicase \| Replicative helicase; pdb \| 4ESV\_B \| 0.999 \| 0.0006859 \| 97 \| 0.114 \| 218 \| 138 \| 15 \| 11 \| 204 \| 194 \| 380 \| Replicative helicase \| Replicative helicase; pdb \| 8H9I\_D \| 0.999 \| 9.252e-05 \| 97 \| 0.151 \| 316 \| 177 \| 25 \| 16 \| 298 \| 143 \| 400 \| ATP synthase subunit beta, mitochondrial \| ATP synthase subunit beta, mitochondrial; pdb \| 7S65\_C \| 0.999 \| 0.001975 \| 97 \| 0.141 \| 241 \| 135 \| 16 \| 16 \| 235 \| 268 \| 457 \| Circadian clock protein kinase KaiC \| Circadian clock protein kinase KaiC; pdb \| 7DY2\_J \| 0.999 \| 0.0003719 \| 97 \| 0.133 \| 239 \| 126 \| 14 \| 16 \| 214 \| 249 \| 446 \| Circadian clock protein kinase KaiC \| Circadian clock protein kinase KaiC; pdb \| 4ESV\_E \| 0.999 \| 0.0007667 \| 96 \| 0.109 \| 228 \| 139 \| 18 \| 9 \| 204 \| 177 \| 372 \| Replicative helicase \| Replicative helicase; pdb \| 4ESV\_D \| 0.999 \| 0.0007252 \| 96 \| 0.13 \| 223 \| 140 \| 18 \| 9 \| 204 \| 190 \| 385 \| Replicative helicase \| Replicative helicase; pdb \| 3OEH\_D \| 0.999 \| 6.267e-05 \| 96 \| 0.144 \| 319 \| 181 \| 21 \| 12 \| 298 \| 144 \| 402 \| ATP synthase subunit beta \| ATP synthase subunit beta; pdb \| 1XP8\_A \| 0.998 \| 0.0003147 \| 95 \| 0.136 \| 285 \| 143 \| 19 \| 12 \| 246 \| 56 \| 287 \| RecA protein \| RecA protein; pdb \| 5DN6\_F \| 0.998 \| 0.0002016 \| 95 \| 0.159 \| 320 \| 177 \| 24 \| 12 \| 298 \| 140 \| 400 \| ATP synthase subunit beta \| ATP synthase subunit beta; pdb \| 5FL7\_D \| 0.998 \| 0.0002132 \| 95 \| 0.134 \| 304 \| 169 \| 21 \| 16 \| 281 \| 146 \| 393 \| ATP SYNTHASE SUBUNIT BETA \| ATP SYNTHASE SUBUNIT BETA; pdb \| 8DB3\_A \| 0.998 \| 0.0002519 \| 93 \| 0.154 \| 227 \| 121 \| 14 \| 16 \| 216 \| 253 \| 434 \| Circadian clock protein KaiC \| Circadian clock protein KaiC; pdb \| 7TJX\_E \| 0.998 \| 5.928e-05 \| 93 \| 0.151 \| 311 \| 182 \| 21 \| 12 \| 298 \| 141 \| 393 \| ATP synthase subunit beta \| ATP synthase subunit beta; pdb \| 8DBA\_G \| 0.998 \| 3.798e-05 \| 93 \| 0.147 \| 332 \| 177 \| 21 \| 16 \| 324 \| 265 \| 513 \| Circadian clock protein KaiC \| Circadian clock protein KaiC; pdb \| 4TLD\_E \| 0.997 \| 0.001265 \| 91 \| 0.141 \| 234 \| 130 \| 17 \| 16 \| 233 \| 24 \| 202 \| Circadian clock protein kinase KaiC \| Circadian clock protein kinase KaiC; pdb \| 4TLC\_A \| 0.997 \| 0.0005491 \| 91 \| 0.133 \| 262 \| 146 \| 19 \| 1 \| 233 \| 2 \| 211 \| Circadian clock protein kinase KaiC \| Circadian clock protein kinase KaiC; pdb \| 2Q6T\_B \| 0.997 \| 0.002088 \| 91 \| 0.156 \| 223 \| 121 \| 15 \| 12 \| 204 \| 183 \| 368 \| DnaB replication fork helicase \| DnaB replication fork helicase; pdb \| 4ESV\_J \| 0.996 \| 0.001495 \| 90 \| 0.122 \| 229 \| 142 \| 16 \| 11 \| 209 \| 176 \| 375 \| Replicative helicase \| Replicative helicase; pdb \| 4ESV\_H \| 0.996 \| 0.001868 \| 90 \| 0.13 \| 223 \| 137 \| 17 \| 11 \| 204 \| 181 \| 375 \| Replicative helicase \| Replicative helicase; pdb \| 4ESV\_I \| 0.996 \| 0.001071 \| 89 \| 0.132 \| 226 \| 133 \| 18 \| 11 \| 204 \| 178 \| 372 \| Replicative helicase \| Replicative helicase; pdb \| 2Q6T\_C \| 0.995 \| 0.001868 \| 88 \| 0.156 \| 224 \| 121 \| 19 \| 8 \| 204 \| 178 \| 360 \| DnaB replication fork helicase \| DnaB replication fork helicase; pdb \| 8DBA\_F \| 0.995 \| 0.0001156 \| 88 \| 0.152 \| 334 \| 175 \| 18 \| 16 \| 324 \| 246 \| 496 \| Circadian clock protein KaiC \| Circadian clock protein KaiC; pdb \| 8DBA\_J \| 0.995 \| 0.0001366 \| 88 \| 0.158 \| 329 \| 178 \| 18 \| 16 \| 324 \| 251 \| 500 \| Circadian clock protein KaiC \| Circadian clock protein KaiC; pdb \| 2ZRF\_A \| 0.994 \| 5.928e-05 \| 87 \| 0.162 \| 295 \| 153 \| 16 \| 16 \| 260 \| 58 \| 308 \| Protein recA \| Protein recA; pdb \| 4ESV\_L \| 0.994 \| 0.001414 \| 87 \| 0.109 \| 228 \| 136 \| 15 \| 11 \| 204 \| 179 \| 373 \| Replicative helicase \| Replicative helicase; pdb \| 8DB3\_C \| 0.994 \| 0.0004157 \| 87 \| 0.143 \| 230 \| 131 \| 16 \| 12 \| 216 \| 258 \| 446 \| Circadian clock protein KaiC \| Circadian clock protein KaiC; pdb \| 8DBA\_B \| 0.993 \| 0.0001614 \| 86 \| 0.174 \| 327 \| 180 \| 22 \| 16 \| 315 \| 241 \| 504 \| Circadian clock protein KaiC \| Circadian clock protein KaiC; pdb \| 8DBA\_K \| 0.993 \| 0.0002016 \| 86 \| 0.143 \| 335 \| 180 \| 17 \| 16 \| 324 \| 249 \| 502 \| Circadian clock protein KaiC \| Circadian clock protein KaiC; pdb \| 4YDS\_A \| 0.992 \| 0.0002816 \| 85 \| 0.137 \| 232 \| 141 \| 15 \| 15 \| 216 \| 20 \| 222 \| Flagella-related protein H \| Flagella-related protein H; pdb \| 3IO5\_B \| 0.992 \| 0.002757 \| 85 \| 0.134 \| 216 \| 113 \| 13 \| 15 \| 217 \| 23 \| 177 \| Recombination and repair protein \| Recombination and repair protein; pdb \| 6N7S\_B \| 0.992 \| 0.002915 \| 85 \| 0.145 \| 248 \| 135 \| 15 \| 10 \| 217 \| 35 \| 245 \| DNA primase/helicase \| DNA primase/helicase; pdb \| 4ESV\_K \| 0.992 \| 0.001975 \| 85 \| 0.112 \| 231 \| 132 \| 14 \| 11 \| 204 \| 181 \| 375 \| Replicative helicase \| Replicative helicase; pdb \| 6YO0\_F1 \| 0.992 \| 0.0004157 \| 85 \| 0.155 \| 316 \| 182 \| 25 \| 12 \| 298 \| 140 \| 399 \| ATP synthase subunit beta \| ATP synthase subunit beta; pdb \| 2ZRD\_A \| 0.991 \| 0.0001804 \| 84 \| 0.168 \| 291 \| 143 \| 18 \| 12 \| 246 \| 56 \| 303 \| Protein recA \| Protein recA; pdb \| 8H9T\_D \| 0.991 \| 0.0007252 \| 84 \| 0.159 \| 289 \| 153 \| 26 \| 16 \| 261 \| 143 \| 384 \| ATP synthase subunit beta, mitochondrial \| ATP synthase subunit beta, mitochondrial; pdb \| 3FKS\_D \| 0.991 \| 0.0007252 \| 84 \| 0.133 \| 308 \| 167 \| 25 \| 12 \| 279 \| 144 \| 391 \| ATP synthase subunit beta, mitochondrial \| ATP synthase subunit beta, mitochondrial; pdb \| 6FKH\_B \| 0.991 \| 0.0006137 \| 84 \| 0.14 \| 313 \| 185 \| 25 \| 16 \| 298 \| 150 \| 408 \| ATP synthase subunit b, chloroplastic \| ATP synthase subunit b, chloroplastic; pdb \| 4PQR\_A \| 0.99 \| 0.0001093 \| 83 \| 0.14 \| 292 \| 151 \| 19 \| 11 \| 246 \| 54 \| 301 \| Protein RecA, 1st part, 2nd part \| Protein RecA, 1st part, 2nd part; pdb \| 2ZRE\_A \| 0.99 \| 0.0001222 \| 83 \| 0.16 \| 292 \| 142 \| 18 \| 12 \| 246 \| 56 \| 301 \| Protein recA \| Protein recA; pdb \| 8DBA\_I \| 0.99 \| 0.0001907 \| 83 \| 0.148 \| 337 \| 173 \| 24 \| 16 \| 324 \| 240 \| 490 \| Circadian clock protein KaiC \| Circadian clock protein KaiC; pdb \| 8DBA\_H \| 0.99 \| 0.0003719 \| 83 \| 0.152 \| 340 \| 172 \| 21 \| 16 \| 324 \| 251 \| 505 \| Circadian clock protein KaiC \| Circadian clock protein KaiC; pdb \| 5JWR\_C \| 0.988 \| 0.001197 \| 82 \| 0.178 \| 224 \| 102 \| 16 \| 12 \| 189 \| 22 \| 209 \| Circadian clock protein kinase KaiC \| Circadian clock protein kinase KaiC; pdb \| 2R6C\_B \| 0.988 \| 0.001414 \| 82 \| 0.123 \| 267 \| 146 \| 15 \| 10 \| 250 \| 154 \| 358 \| Replicative helicase \| Replicative helicase; pdb \| 4ESV\_G \| 0.988 \| 0.002333 \| 82 \| 0.124 \| 233 \| 127 \| 17 \| 11 \| 204 \| 181 \| 375 \| Replicative helicase \| Replicative helicase; pdb \| 2F43\_B \| 0.988 \| 0.001767 \| 82 \| 0.124 \| 282 \| 169 \| 20 \| 16 \| 265 \| 150 \| 385 \| ATP synthase beta chain, mitochondrial \| ATP synthase beta chain, mitochondrial; pdb \| 8DBA\_A \| 0.988 \| 0.0003518 \| 82 \| 0.146 \| 335 \| 174 \| 21 \| 16 \| 324 \| 255 \| 503 \| Circadian clock protein KaiC \| Circadian clock protein KaiC; pdb \| 8DBA\_C \| 0.986 \| 0.0001366 \| 81 \| 0.153 \| 338 \| 183 \| 23 \| 12 \| 318 \| 242 \| 507 \| Circadian clock protein KaiC \| Circadian clock protein KaiC; pdb \| 8DBA\_L \| 0.981 \| 0.001071 \| 79 \| 0.141 \| 332 \| 181 \| 18 \| 16 \| 324 \| 249 \| 499 \| Circadian clock protein KaiC \| Circadian clock protein KaiC; pdb \| 3OFN\_V \| 0.978 \| 0.00385 \| 78 \| 0.148 \| 282 \| 161 \| 25 \| 6 \| 246 \| 116 \| 359 \| ATP synthase subunit beta \| ATP synthase subunit beta; pdb \| 6RD4\_X \| 0.975 \| 0.001265 \| 77 \| 0.15 \| 319 \| 172 \| 23 \| 18 \| 298 \| 147 \| 404 \| ATP synthase subunit beta \| ATP synthase subunit beta; pdb \| 4KFR\_C \| 0.971 \| 0.00481 \| 76 \| 0.142 \| 238 \| 130 \| 14 \| 15 \| 245 \| 4 \| 174 \| Genome packaging NTPase B204 \| Genome packaging NTPase B204; pdb \| 8H9S\_E \| 0.971 \| 0.002088 \| 76 \| 0.142 \| 315 \| 179 \| 22 \| 12 \| 298 \| 140 \| 391 \| ATP synthase subunit beta, mitochondrial \| ATP synthase subunit beta, mitochondrial; pdb \| 1NBM\_E \| 0.971 \| 0.0004395 \| 76 \| 0.159 \| 326 \| 168 \| 27 \| 12 \| 298 \| 140 \| 398 \| F1-ATPASE \| F1-ATPASE; pdb \| 8DBA\_D \| 0.967 \| 0.001071 \| 75 \| 0.15 \| 340 \| 165 \| 22 \| 16 \| 324 \| 262 \| 508 \| Circadian clock protein KaiC \| Circadian clock protein KaiC; pdb \| 1MAB\_B \| 0.956 \| 0.004071 \| 73 \| 0.142 \| 280 \| 172 \| 19 \| 12 \| 265 \| 148 \| 385 \| PROTEIN (F1-ATPASE BETA CHAIN) \| PROTEIN (F1-ATPASE BETA CHAIN); pdb \| 2R6A\_B \| 0.941 \| 0.005684 \| 71 \| 0.129 \| 217 \| 116 \| 14 \| 10 \| 181 \| 184 \| 372 \| Replicative helicase \| Replicative helicase; pdb \| 4KFT\_A \| 0.933 \| 0.005086 \| 70 \| 0.15 \| 239 \| 120 \| 17 \| 15 \| 241 \| 4 \| 171 \| Genome packaging NTPase B204 \| Genome packaging NTPase B204; pdb \| 4NMN\_B \| 0.912 \| 0.006354 \| 68 \| 0.156 \| 230 \| 116 \| 16 \| 10 \| 204 \| 182 \| 368 \| Replicative DNA helicase \| Replicative DNA helicase; pdb \| 4ZC0\_A \| 0.912 \| 0.006354 \| 68 \| 0.119 \| 235 \| 125 \| 20 \| 10 \| 204 \| 180 \| 372 \| Replicative DNA helicase \| Replicative DNA helicase; pdb \| 4KFR\_A \| 0.9 \| 0.005377 \| 67 \| 0.137 \| 248 \| 124 \| 19 \| 15 \| 245 \| 4 \| 178 \| Genome packaging NTPase B204 \| Genome packaging NTPase B204; pdb \| 4KFU\_A \| 0.9 \| 0.006354 \| 67 \| 0.145 \| 247 \| 126 \| 19 \| 15 \| 245 \| 3 \| 180 \| Genome packaging NTPase B204 \| Genome packaging NTPase B204; pdb \| 4R2H\_A \| 0.9 \| 0.006717 \| 67 \| 0.122 \| 253 \| 133 \| 21 \| 11 \| 246 \| 2 \| 182 \| STIV B204 ATPase \| STIV B204 ATPase; pdb \| 4KFT\_B \| 0.872 \| 0.005086 \| 65 \| 0.146 \| 252 \| 112 \| 22 \| 15 \| 245 \| 4 \| 173 \| Genome packaging NTPase B204 \| Genome packaging NTPase B204; pdb \| 6TE0\_D \| 0.872 \| 0.006717 \| 65 \| 0.141 \| 319 \| 182 \| 26 \| 12 \| 298 \| 145 \| 403 \| ATP synthase subunit beta \| ATP synthase subunit beta; pdb \| 5CDF\_E \| 0.663 \| 0.00481 \| 56 \| 0.147 \| 333 \| 156 \| 24 \| 12 \| 298 \| 140 \| 390 \| ATP synthase subunit beta \| ATP synthase subunit beta; afdb-proteome \| AF-A0A158Q445-F1-MODEL\_V4 \| 1.0 \| 6.327e-07 \| 189 \| 0.129 \| 224 \| 140 \| 15 \| 12 \| 217 \| 44 \| 230 \| RECA\_2 domain-containing protein \| RECA\_2 domain-containing protein; afdb-proteome \| AF-A0A0J9XTJ7-F1-MODEL\_V4 \| 1.0 \| 1.234e-06 \| 182 \| 0.122 \| 221 \| 138 \| 14 \| 12 \| 214 \| 44 \| 226 \| BMA-RFS-1 \| BMA-RFS-1; afdb-proteome \| AF-A0A0N4UFL0-F1-MODEL\_V4 \| 1.0 \| 1.167e-06 \| 172 \| 0.155 \| 232 \| 142 \| 19 \| 12 \| 219 \| 127 \| 328 \| DNA repair protein RAD51 homolog \| DNA repair protein RAD51 homolog; afdb-proteome \| AF-A0A0P0VCR4-F1-MODEL\_V4 \| 1.0 \| 9.875e-07 \| 166 \| 0.171 \| 221 \| 141 \| 14 \| 12 \| 217 \| 136 \| 329 \| Os01g0945001 protein \| Os01g0945001 protein; afdb-proteome \| AF-Q6YU07-F1-MODEL\_V4 \| 1.0 \| 6.925e-06 \| 163 \| 0.161 \| 236 \| 136 \| 16 \| 16 \| 219 \| 62 \| 267 \| Os02g0562100 protein \| Os02g0562100 protein; afdb-proteome \| AF-A0A2K6W9N3-F1-MODEL\_V4 \| 1.0 \| 5.543e-06 \| 163 \| 0.159 \| 232 \| 141 \| 19 \| 12 \| 219 \| 134 \| 335 \| DNA repair protein RAD51 homolog \| DNA repair protein RAD51 homolog; afdb-proteome \| AF-A0A044UQU3-F1-MODEL\_V4 \| 1.0 \| 6.195e-06 \| 161 \| 0.113 \| 238 \| 145 \| 15 \| 16 \| 219 \| 38 \| 243 \| RECA\_2 domain-containing protein \| RECA\_2 domain-containing protein; afdb-proteome \| AF-Q4CYE3-F1-MODEL\_V4 \| 1.0 \| 1.143e-05 \| 154 \| 0.137 \| 232 \| 146 \| 17 \| 12 \| 219 \| 150 \| 351 \| DNA repair protein RAD51 homolog \| DNA repair protein RAD51 homolog; afdb-proteome \| AF-A0A1D6HAQ9-F1-MODEL\_V4 \| 1.0 \| 8.183e-06 \| 153 \| 0.142 \| 231 \| 136 \| 14 \| 16 \| 214 \| 58 \| 258 \| DNA repair protein XRCC3-like protein \| DNA repair protein XRCC3-like protein; afdb-proteome \| AF-B4FBZ5-F1-MODEL\_V4 \| 1.0 \| 3.177e-06 \| 153 \| 0.142 \| 238 \| 137 \| 19 \| 12 \| 219 \| 120 \| 320 \| DNA repair protein RAD51 homolog \| DNA repair protein RAD51 homolog; afdb-proteome \| AF-I1LYB2-F1-MODEL\_V4 \| 1.0 \| 3.359e-06 \| 153 \| 0.158 \| 234 \| 138 \| 17 \| 12 \| 219 \| 124 \| 324 \| DNA repair protein RAD51 homolog \| DNA repair protein RAD51 homolog; afdb-proteome \| AF-Q32CM9-F1-MODEL\_V4 \| 1.0 \| 1.022e-05 \| 152 \| 0.172 \| 209 \| 133 \| 14 \| 12 \| 213 \| 59 \| 234 \| Protein RecA \| Protein RecA; afdb-proteome \| AF-O75771-F1-MODEL\_V4 \| 1.0 \| 6.195e-06 \| 151 \| 0.123 \| 268 \| 151 \| 19 \| 12 \| 245 \| 99 \| 316 \| DNA repair protein RAD51 homolog 4 \| DNA repair protein RAD51 homolog 4; afdb-proteome \| AF-O55230-F1-MODEL\_V4 \| 1.0 \| 6.925e-06 \| 151 \| 0.124 \| 265 \| 145 \| 20 \| 12 \| 237 \| 99 \| 315 \| DNA repair protein RAD51 homolog 4 \| DNA repair protein RAD51 homolog 4; afdb-proteome \| AF-Q8IIS8-F1-MODEL\_V4 \| 1.0 \| 5.543e-06 \| 151 \| 0.149 \| 234 \| 140 \| 18 \| 12 \| 219 \| 129 \| 329 \| DNA repair protein RAD51 homolog \| DNA repair protein RAD51 homolog; afdb-proteome \| AF-Q95Q25-F1-MODEL\_V4 \| 1.0 \| 1.208e-05 \| 151 \| 0.122 \| 236 \| 145 \| 17 \| 12 \| 219 \| 173 \| 374 \| DNA repair protein RAD51 homolog \| DNA repair protein RAD51 homolog; afdb-proteome \| AF-F1R474-F1-MODEL\_V4 \| 1.0 \| 9.146e-06 \| 150 \| 0.1 \| 258 \| 158 \| 16 \| 12 \| 238 \| 99 \| 313 \| DNA repair protein \| DNA repair protein; afdb-proteome \| AF-P0A452-F1-MODEL\_V4 \| 1.0 \| 1.783e-05 \| 150 \| 0.176 \| 198 \| 127 \| 11 \| 12 \| 204 \| 71 \| 237 \| Protein RecA \| Protein RecA; afdb-proteome \| AF-Q9XED7-F1-MODEL\_V4 \| 1.0 \| 5.543e-06 \| 149 \| 0.142 \| 238 \| 137 \| 19 \| 12 \| 219 \| 120 \| 320 \| DNA repair protein RAD51 homolog B \| DNA repair protein RAD51 homolog B; afdb-proteome \| AF-A0A077YZT8-F1-MODEL\_V4 \| 1.0 \| 8.651e-06 \| 149 \| 0.132 \| 241 \| 137 \| 17 \| 12 \| 219 \| 140 \| 341 \| DNA repair protein RAD51 homolog \| DNA repair protein RAD51 homolog; afdb-proteome \| AF-Q2FZ09-F1-MODEL\_V4 \| 1.0 \| 1.596e-05 \| 148 \| 0.18 \| 210 \| 136 \| 12 \| 12 \| 216 \| 32 \| 210 \| Protein RecA \| Protein RecA; afdb-proteome \| AF-I1MS33-F1-MODEL\_V4 \| 1.0 \| 6.925e-06 \| 148 \| 0.16 \| 237 \| 134 \| 18 \| 12 \| 219 \| 123 \| 323 \| DNA repair protein RAD51 homolog \| DNA repair protein RAD51 homolog; afdb-proteome \| AF-Q67EU8-F1-MODEL\_V4 \| 1.0 \| 6.925e-06 \| 147 \| 0.141 \| 234 \| 142 \| 17 \| 12 \| 219 \| 120 \| 320 \| DNA repair protein RAD51 homolog A \| DNA repair protein RAD51 homolog A; afdb-proteome \| AF-Q0P7V6-F1-MODEL\_V4 \| 1.0 \| 1.783e-05 \| 147 \| 0.184 \| 195 \| 130 \| 13 \| 12 \| 204 \| 57 \| 224 \| Protein RecA \| Protein RecA; afdb-proteome \| AF-Q384K0-F1-MODEL\_V4 \| 1.0 \| 2.108e-05 \| 147 \| 0.135 \| 229 \| 144 \| 15 \| 16 \| 219 \| 154 \| 353 \| DNA repair protein RAD51 homolog \| DNA repair protein RAD51 homolog; afdb-proteome \| AF-Q8SZ30-F1-MODEL\_V4 \| 1.0 \| 3.005e-06 \| 146 \| 0.138 \| 238 \| 139 \| 14 \| 16 \| 219 \| 65 \| 270 \| RE19845p \| RE19845p; afdb-proteome \| AF-Q8SBB9-F1-MODEL\_V4 \| 1.0 \| 4.691e-06 \| 146 \| 0.137 \| 240 \| 136 \| 20 \| 12 \| 219 \| 119 \| 319 \| DNA repair protein RAD51 homolog \| DNA repair protein RAD51 homolog; afdb-proteome \| AF-A0A132YZI1-F1-MODEL\_V4 \| 1.0 \| 2.356e-05 \| 146 \| 0.176 \| 198 \| 127 \| 12 \| 12 \| 204 \| 57 \| 223 \| Protein RecA \| Protein RecA; afdb-proteome \| AF-A4I3C9-F1-MODEL\_V4 \| 1.0 \| 2.108e-05 \| 145 \| 0.149 \| 234 \| 141 \| 20 \| 12 \| 219 \| 155 \| 356 \| DNA repair protein RAD51 homolog \| DNA repair protein RAD51 homolog; afdb-proteome \| AF-Q2FX33-F1-MODEL\_V4 \| 1.0 \| 1.509e-05 \| 142 \| 0.12 \| 225 \| 143 \| 12 \| 11 \| 214 \| 24 \| 214 \| Conserved hypothetical phage protein \| Conserved hypothetical phage protein; afdb-proteome \| AF-Q8RY99-F1-MODEL\_V4 \| 1.0 \| 2.275e-06 \| 140 \| 0.177 \| 293 \| 148 \| 20 \| 12 \| 244 \| 115 \| 374 \| DNA repair protein recA homolog 2, mitochondrial \| DNA repair protein recA homolog 2, mitochondrial; afdb-proteome \| AF-Q53QJ3-F1-MODEL\_V4 \| 1.0 \| 9.146e-06 \| 140 \| 0.21 \| 200 \| 121 \| 11 \| 12 \| 204 \| 121 \| 290 \| Os11g0302700 protein \| Os11g0302700 protein; afdb-proteome \| AF-B5DFH5-F1-MODEL\_V4 \| 1.0 \| 2.633e-05 \| 135 \| 0.112 \| 266 \| 149 \| 19 \| 12 \| 238 \| 99 \| 316 \| DNA repair protein \| DNA repair protein; afdb-proteome \| AF-Q9UUL2-F1-MODEL\_V4 \| 1.0 \| 5.134e-05 \| 134 \| 0.146 \| 253 \| 155 \| 16 \| 16 \| 218 \| 94 \| 335 \| DNA repair protein rhp57 \| DNA repair protein rhp57; afdb-proteome \| AF-A0A0H3GS46-F1-MODEL\_V4 \| 1.0 \| 1.022e-05 \| 130 \| 0.155 \| 329 \| 182 \| 24 \| 12 \| 299 \| 59 \| 332 \| Protein RecA \| Protein RecA; afdb-proteome \| AF-A0A0R0EV99-F1-MODEL\_V4 \| 1.0 \| 1.925e-06 \| 130 \| 0.165 \| 326 \| 161 \| 23 \| 12 \| 261 \| 113 \| 403 \| Uncharacterized protein \| Uncharacterized protein; afdb-proteome \| AF-Q555F1-F1-MODEL\_V4 \| 1.0 \| 1.143e-05 \| 128 \| 0.117 \| 315 \| 165 \| 19 \| 16 \| 244 \| 105 \| 392 \| RECA\_2 domain-containing protein \| RECA\_2 domain-containing protein; afdb-proteome \| AF-P65977-F1-MODEL\_V4 \| 1.0 \| 1.886e-05 \| 127 \| 0.151 \| 329 \| 183 \| 22 \| 12 \| 299 \| 59 \| 332 \| Protein RecA \| Protein RecA; afdb-proteome \| AF-Q5N6Y9-F1-MODEL\_V4 \| 1.0 \| 6.414e-05 \| 127 \| 0.185 \| 227 \| 136 \| 12 \| 12 \| 219 \| 118 \| 314 \| Os01g0901200 protein \| Os01g0901200 protein; afdb-proteome \| AF-Q2FWQ9-F1-MODEL\_V4 \| 1.0 \| 9.469e-05 \| 125 \| 0.144 \| 228 \| 136 \| 16 \| 11 \| 214 \| 24 \| 216 \| Conserved hypothetical phage protein \| Conserved hypothetical phage protein; afdb-proteome \| AF-K7MBY5-F1-MODEL\_V4 \| 1.0 \| 5.543e-06 \| 125 \| 0.144 \| 368 \| 206 \| 23 \| 12 \| 329 \| 116 \| 424 \| Uncharacterized protein \| Uncharacterized protein; afdb-proteome \| AF-I1L1J2-F1-MODEL\_V4 \| 1.0 \| 7.321e-06 \| 125 \| 0.136 \| 365 \| 208 \| 22 \| 12 \| 329 \| 116 \| 420 \| Uncharacterized protein \| Uncharacterized protein; afdb-proteome \| AF-A0A0R0FHZ4-F1-MODEL\_V4 \| 1.0 \| 9.67e-06 \| 125 \| 0.162 \| 356 \| 196 \| 23 \| 12 \| 329 \| 136 \| 427 \| Uncharacterized protein \| Uncharacterized protein; afdb-proteome \| AF-Q9ZUP2-F1-MODEL\_V4 \| 1.0 \| 1.022e-05 \| 121 \| 0.148 \| 351 \| 197 \| 22 \| 12 \| 319 \| 111 \| 402 \| DNA repair protein recA homolog 3, mitochondrial \| DNA repair protein recA homolog 3, mitochondrial; afdb-proteome \| AF-A0A0N4UIX0-F1-MODEL\_V4 \| 1.0 \| 8.472e-05 \| 120 \| 0.118 \| 236 \| 136 \| 15 \| 12 \| 218 \| 46 \| 238 \| Checkpoint protein \| Checkpoint protein; afdb-proteome \| AF-A0A0K0DTA7-F1-MODEL\_V4 \| 1.0 \| 5.134e-05 \| 119 \| 0.155 \| 219 \| 123 \| 16 \| 16 \| 204 \| 37 \| 223 \| Rad51 domain-containing protein \| Rad51 domain-containing protein; afdb-proteome \| AF-U7Q6E5-F1-MODEL\_V4 \| 1.0 \| 0.0002307 \| 115 \| 0.148 \| 236 \| 143 \| 17 \| 1 \| 204 \| 111 \| 320 \| RECA\_2 domain-containing protein \| RECA\_2 domain-containing protein; afdb-proteome \| AF-Q58754-F1-MODEL\_V4 \| 1.0 \| 0.0001652 \| 113 \| 0.117 \| 238 \| 138 \| 14 \| 10 \| 219 \| 21 \| 214 \| UPF0273 protein MJ1359 \| UPF0273 protein MJ1359; afdb-proteome \| AF-Q386Q5-F1-MODEL\_V4 \| 1.0 \| 0.0001652 \| 105 \| 0.111 \| 305 \| 145 \| 15 \| 16 \| 204 \| 125 \| 419 \| Recombinase Rad51, putative \| Recombinase Rad51, putative; afdb-proteome \| AF-F4J2X9-F1-MODEL\_V4 \| 1.0 \| 0.001096 \| 101 \| 0.119 \| 234 \| 149 \| 17 \| 3 \| 216 \| 5 \| 201 \| tRNA dimethylallyltransferase \| tRNA dimethylallyltransferase; afdb-proteome \| AF-A0A133CMJ6-F1-MODEL\_V4 \| 1.0 \| 0.0004255 \| 101 \| 0.128 \| 225 \| 136 \| 16 \| 10 \| 204 \| 200 \| 394 \| Replicative DNA helicase \| Replicative DNA helicase; afdb-proteome \| AF-P35901-F1-MODEL\_V4 \| 1.0 \| 0.0001846 \| 100 \| 0.185 \| 221 \| 130 \| 15 \| 12 \| 214 \| 59 \| 247 \| Protein RecA \| Protein RecA; afdb-proteome \| AF-A0A175VRV5-F1-MODEL\_V4 \| 0.999 \| 0.002388 \| 98 \| 0.206 \| 179 \| 109 \| 10 \| 12 \| 170 \| 145 \| 310 \| DNA repair protein rhp57 \| DNA repair protein rhp57; afdb-proteome \| AF-A4I499-F1-MODEL\_V4 \| 0.998 \| 0.0009273 \| 93 \| 0.099 \| 353 \| 154 \| 18 \| 12 \| 219 \| 298 \| 631 \| Rad51/recA\_bacterial\_DNA\_recombination\_protein /KaiC\_-\_putative \| Rad51/recA\_bacterial\_DNA\_recombination\_protein /KaiC\_-\_putative; afdb-proteome \| AF-Q58309-F1-MODEL\_V4 \| 0.996 \| 0.002984 \| 90 \| 0.15 \| 206 \| 116 \| 13 \| 12 \| 186 \| 26 \| 203 \| Putative flagella-related protein H \| Putative flagella-related protein H; afdb-proteome \| AF-P10719-F1-MODEL\_V4 \| 0.995 \| 0.0004498 \| 88 \| 0.151 \| 317 \| 182 \| 22 \| 12 \| 298 \| 198 \| 457 \| ATP synthase subunit beta, mitochondrial \| ATP synthase subunit beta, mitochondrial; afdb-proteome \| AF-A0A077Z2B0-F1-MODEL\_V4 \| 0.994 \| 0.005503 \| 87 \| 0.146 \| 239 \| 144 \| 15 \| 11 \| 234 \| 265 \| 458 \| Alpha:beta hydrolase fold protein \| Alpha:beta hydrolase fold protein; afdb-proteome \| AF-Q8T4C4-F1-MODEL\_V4 \| 0.993 \| 0.0007847 \| 86 \| 0.156 \| 314 \| 184 \| 21 \| 12 \| 298 \| 242 \| 501 \| ATP synthase subunit beta \| ATP synthase subunit beta; afdb-proteome \| AF-O25916-F1-MODEL\_V4 \| 0.992 \| 0.0006281 \| 85 \| 0.118 \| 303 \| 159 \| 17 \| 10 \| 278 \| 193 \| 421 \| Replicative DNA helicase \| Replicative DNA helicase; afdb-proteome \| AF-Q2G1S4-F1-MODEL\_V4 \| 0.991 \| 0.0007422 \| 84 \| 0.136 \| 235 \| 126 \| 15 \| 10 \| 204 \| 200 \| 397 \| Replicative DNA helicase \| Replicative DNA helicase; afdb-proteome \| AF-A8WGC6-F1-MODEL\_V4 \| 0.991 \| 0.0007847 \| 84 \| 0.145 \| 316 \| 185 \| 23 \| 12 \| 298 \| 187 \| 446 \| ATP synthase subunit beta \| ATP synthase subunit beta; afdb-proteome \| AF-A0A381MM20-F1-MODEL\_V4 \| 0.991 \| 0.001447 \| 84 \| 0.148 \| 390 \| 212 \| 26 \| 12 \| 324 \| 178 \| 524 \| ATP synthase subunit beta \| ATP synthase subunit beta; afdb-proteome \| AF-P00827-F1-MODEL\_V4 \| 0.986 \| 0.0005315 \| 81 \| 0.157 \| 323 \| 178 \| 28 \| 10 \| 298 \| 162 \| 424 \| ATP synthase subunit beta, chloroplastic \| ATP synthase subunit beta, chloroplastic; afdb-proteome \| AF-P19366-F1-MODEL\_V4 \| 0.978 \| 0.00171 \| 78 \| 0.161 \| 303 \| 170 \| 24 \| 16 \| 284 \| 166 \| 418 \| ATP synthase subunit beta, chloroplastic \| ATP synthase subunit beta, chloroplastic; afdb-proteome \| AF-Q2PMV0-F1-MODEL\_V4 \| 0.978 \| 0.0005028 \| 78 \| 0.159 \| 319 \| 162 \| 25 \| 10 \| 284 \| 162 \| 418 \| ATP synthase subunit beta, chloroplastic \| ATP synthase subunit beta, chloroplastic; afdb-proteome \| AF-A0A3Q0KEM2-F1-MODEL\_V4 \| 0.978 \| 0.001808 \| 78 \| 0.134 \| 319 \| 185 \| 22 \| 12 \| 298 \| 187 \| 446 \| ATP synthase subunit beta \| ATP synthase subunit beta; afdb-proteome \| AF-Q05825-F1-MODEL\_V4 \| 0.967 \| 0.001912 \| 75 \| 0.151 \| 324 \| 174 \| 26 \| 12 \| 298 \| 175 \| 434 \| ATP synthase subunit beta, mitochondrial \| ATP synthase subunit beta, mitochondrial; afdb-proteome \| AF-A0A0P0Y832-F1-MODEL\_V4 \| 0.961 \| 0.002388 \| 74 \| 0.161 \| 316 \| 165 \| 27 \| 10 \| 284 \| 84 \| 340 \| ATP synthase subunit beta \| ATP synthase subunit beta; afdb-proteome \| AF-A0A1D8PI62-F1-MODEL\_V4 \| 0.949 \| 0.002822 \| 72 \| 0.122 \| 221 \| 122 \| 13 \| 12 \| 179 \| 102 \| 303 \| Putative DNA-dependent ATPase \| Putative DNA-dependent ATPase; afdb-proteome \| AF-Q5Z9S8-F1-MODEL\_V4 \| 0.9 \| 0.004405 \| 67 \| 0.132 \| 346 \| 165 \| 18 \| 16 \| 300 \| 933 \| 1204 \| ABC transporter G family member 42 \| ABC transporter G family member 42; afdb-proteome \| AF-Q55GB1-F1-MODEL\_V4 \| 0.817 \| 0.003526 \| 62 \| 0.121 \| 346 \| 175 \| 21 \| 10 \| 299 \| 869 \| 1141 \| ABC transporter G family member 15 \| ABC transporter G family member 15; afdb-uniprot50 \| AF-A0A8A6KEC5-F1-MODEL\_V4 \| 1.0 \| 1.139e-41 \| 1647 \| 0.595 \| 331 \| 115 \| 6 \| 1 \| 326 \| 1 \| 317 \| AAA family ATPase \| AAA family ATPase; afdb-uniprot50 \| AF-A0A0E9N151-F1-MODEL\_V4 \| 1.0 \| 6.391e-33 \| 1260 \| 0.424 \| 332 \| 167 \| 8 \| 1 \| 329 \| 1 \| 311 \| Uncharacterized protein \| Uncharacterized protein; afdb-uniprot50 \| AF-A0A5C1A6A2-F1-MODEL\_V4 \| 1.0 \| 7.498e-30 \| 1031 \| 0.414 \| 311 \| 168 \| 8 \| 1 \| 309 \| 1 \| 299 \| Uncharacterized protein \| Uncharacterized protein; afdb-uniprot50 \| AF-A0A7Y7GN48-F1-MODEL\_V4 \| 1.0 \| 6.433e-28 \| 1015 \| 0.341 \| 337 \| 193 \| 8 \| 4 \| 329 \| 1 \| 319 \| AAA family ATPase \| AAA family ATPase; afdb-uniprot50 \| AF-A0A4Y8QD62-F1-MODEL\_V4 \| 1.0 \| 5.331e-27 \| 979 \| 0.35 \| 340 \| 187 \| 12 \| 4 \| 329 \| 1 \| 320 \| Uncharacterized protein \| Uncharacterized protein; afdb-uniprot50 \| AF-M1NXB7-F1-MODEL\_V4 \| 1.0 \| 2.266e-26 \| 978 \| 0.355 \| 307 \| 173 \| 8 \| 4 \| 304 \| 1 \| 288 \| Putative replicative DNA helicase \| Putative replicative DNA helicase; afdb-uniprot50 \| AF-A0A4D8QWP6-F1-MODEL\_V4 \| 1.0 \| 4.67e-26 \| 904 \| 0.329 \| 343 \| 195 \| 14 \| 1 \| 329 \| 1 \| 322 \| Uncharacterized protein \| Uncharacterized protein; afdb-uniprot50 \| AF-H8FP22-F1-MODEL\_V4 \| 1.0 \| 2.93e-25 \| 889 \| 0.335 \| 340 \| 195 \| 12 \| 1 \| 329 \| 3 \| 322 \| Uncharacterized protein \| Uncharacterized protein; afdb-uniprot50 \| AF-A0A3E1NQ56-F1-MODEL\_V4 \| 1.0 \| 7.39e-24 \| 787 \| 0.288 \| 312 \| 182 \| 11 \| 1 \| 304 \| 1 \| 280 \| AAA family ATPase \| AAA family ATPase; afdb-uniprot50 \| AF-A0A2U8QUP8-F1-MODEL\_V4 \| 1.0 \| 1.441e-23 \| 784 \| 0.287 \| 327 \| 188 \| 9 \| 4 \| 329 \| 1 \| 283 \| AAA family ATPase \| AAA family ATPase; afdb-uniprot50 \| AF-A0A1M4VWU6-F1-MODEL\_V4 \| 1.0 \| 3.321e-23 \| 776 \| 0.29 \| 303 \| 186 \| 9 \| 4 \| 303 \| 1 \| 277 \| AAA domain-containing protein \| AAA domain-containing protein; afdb-uniprot50 \| AF-A0A1H5JJL2-F1-MODEL\_V4 \| 1.0 \| 1.289e-23 \| 772 \| 0.352 \| 321 \| 162 \| 10 \| 4 \| 299 \| 3 \| 302 \| AAA domain-containing protein \| AAA domain-containing protein; afdb-uniprot50 \| AF-A0A4V2ADJ8-F1-MODEL\_V4 \| 1.0 \| 6.124e-23 \| 768 \| 0.291 \| 312 \| 187 \| 10 \| 4 \| 308 \| 1 \| 285 \| AAA family ATPase \| AAA family ATPase; afdb-uniprot50 \| AF-A0A1H5LQB0-F1-MODEL\_V4 \| 1.0 \| 1.025e-20 \| 765 \| 0.393 \| 254 \| 124 \| 9 \| 4 \| 248 \| 82 \| 314 \| AAA domain-containing protein \| AAA domain-containing protein; afdb-uniprot50 \| AF-G2KMZ3-F1-MODEL\_V4 \| 1.0 \| 4.541e-22 \| 765 \| 0.295 \| 311 \| 186 \| 12 \| 4 \| 307 \| 8 \| 292 \| Uncharacterized protein \| Uncharacterized protein; afdb-uniprot50 \| AF-A0A1G8G4H4-F1-MODEL\_V4 \| 1.0 \| 3.118e-20 \| 763 \| 0.352 \| 241 \| 132 \| 6 \| 4 \| 241 \| 1 \| 220 \| AAA domain-containing protein \| AAA domain-containing protein; afdb-uniprot50 \| AF-A0A444WBT6-F1-MODEL\_V4 \| 1.0 \| 1.971e-22 \| 756 \| 0.284 \| 309 \| 192 \| 9 \| 4 \| 311 \| 1 \| 281 \| AAA ATPase \| AAA ATPase; afdb-uniprot50 \| AF-A0A832BEQ0-F1-MODEL\_V4 \| 1.0 \| 8.089e-23 \| 746 \| 0.295 \| 308 \| 182 \| 9 \| 1 \| 299 \| 1 \| 282 \| Uncharacterized protein \| Uncharacterized protein; afdb-uniprot50 \| AF-A0A362XEI9-F1-MODEL\_V4 \| 1.0 \| 1.577e-22 \| 742 \| 0.295 \| 298 \| 180 \| 9 \| 4 \| 298 \| 34 \| 304 \| Signal recognition particle subunit FFH/SRP54 (Srp54) \| Signal recognition particle subunit FFH/SRP54 (Srp54); afdb-uniprot50 \| AF-E2ZIG4-F1-MODEL\_V4 \| 1.0 \| 1.353e-20 \| 740 \| 0.316 \| 256 \| 151 \| 8 \| 4 \| 251 \| 2 \| 241 \| Uncharacterized protein \| Uncharacterized protein; afdb-uniprot50 \| AF-A0A4Q3D406-F1-MODEL\_V4 \| 1.0 \| 7.493e-22 \| 731 \| 0.257 \| 307 \| 204 \| 8 \| 4 \| 309 \| 1 \| 284 \| AAA family ATPase \| AAA family ATPase; afdb-uniprot50 \| AF-A0A7C1KN99-F1-MODEL\_V4 \| 1.0 \| 1.253e-19 \| 731 \| 0.322 \| 248 \| 145 \| 9 \| 4 \| 244 \| 1 \| 232 \| AAA family ATPase \| AAA family ATPase; afdb-uniprot50 \| AF-A0A4Q4B7Y5-F1-MODEL\_V4 \| 1.0 \| 7.087e-22 \| 723 \| 0.274 \| 313 \| 200 \| 8 \| 4 \| 315 \| 1 \| 287 \| AAA family ATPase \| AAA family ATPase; afdb-uniprot50 \| AF-A0A1I4QA63-F1-MODEL\_V4 \| 1.0 \| 5.145e-20 \| 720 \| 0.328 \| 253 \| 141 \| 10 \| 6 \| 253 \| 4 \| 232 \| AAA domain-containing protein \| AAA domain-containing protein; afdb-uniprot50 \| AF-A0A2T7U531-F1-MODEL\_V4 \| 1.0 \| 2.127e-23 \| 720 \| 0.309 \| 352 \| 170 \| 12 \| 4 \| 302 \| 41 \| 372 \| Uncharacterized protein \| Uncharacterized protein; afdb-uniprot50 \| AF-A0A841K3V8-F1-MODEL\_V4 \| 1.0 \| 2.81e-23 \| 716 \| 0.312 \| 349 \| 172 \| 12 \| 4 \| 305 \| 3 \| 330 \| Uncharacterized protein \| Uncharacterized protein; afdb-uniprot50 \| AF-A0A447CPV3-F1-MODEL\_V4 \| 1.0 \| 5.006e-24 \| 714 \| 0.317 \| 372 \| 175 \| 13 \| 1 \| 306 \| 1 \| 359 \| Uncharacterized protein \| Uncharacterized protein; afdb-uniprot50 \| AF-A0A1D9LC38-F1-MODEL\_V4 \| 1.0 \| 3.252e-22 \| 710 \| 0.252 \| 313 \| 202 \| 8 \| 4 \| 308 \| 1 \| 289 \| Uncharacterized protein \| Uncharacterized protein; afdb-uniprot50 \| AF-A0A1U7MFH0-F1-MODEL\_V4 \| 1.0 \| 5.751e-20 \| 705 \| 0.281 \| 249 \| 151 \| 7 \| 4 \| 246 \| 1 \| 227 \| AAA domain-containing protein \| AAA domain-containing protein; afdb-uniprot50 \| AF-A0A0J1FG05-F1-MODEL\_V4 \| 1.0 \| 8.031e-20 \| 705 \| 0.307 \| 254 \| 154 \| 7 \| 4 \| 251 \| 2 \| 239 \| Uncharacterized protein \| Uncharacterized protein; afdb-uniprot50 \| AF-A0A0Q5Z437-F1-MODEL\_V4 \| 1.0 \| 9.042e-23 \| 698 \| 0.293 \| 348 \| 190 \| 9 \| 4 \| 306 \| 3 \| 339 \| AAA domain-containing protein \| AAA domain-containing protein; afdb-uniprot50 \| AF-A0A1V5QK79-F1-MODEL\_V4 \| 1.0 \| 1.046e-21 \| 696 \| 0.286 \| 318 \| 190 \| 11 \| 4 \| 310 \| 1 \| 292 \| AAA domain-containing protein \| AAA domain-containing protein; afdb-uniprot50 \| AF-A0A3N5PH66-F1-MODEL\_V4 \| 1.0 \| 6.08e-20 \| 694 \| 0.313 \| 265 \| 146 \| 10 \| 1 \| 249 \| 1 \| 245 \| Uncharacterized protein \| Uncharacterized protein; afdb-uniprot50 \| AF-A0A3G6REK7-F1-MODEL\_V4 \| 1.0 \| 4.206e-21 \| 689 \| 0.275 \| 312 \| 180 \| 9 \| 4 \| 298 \| 1 \| 283 \| AAA family ATPase \| AAA family ATPase; afdb-uniprot50 \| AF-A0A7V1BX95-F1-MODEL\_V4 \| 1.0 \| 7.493e-22 \| 680 \| 0.262 \| 339 \| 200 \| 13 \| 4 \| 309 \| 23 \| 344 \| Uncharacterized protein \| Uncharacterized protein; afdb-uniprot50 \| AF-A0A1V3JIB2-F1-MODEL\_V4 \| 1.0 \| 1.236e-21 \| 676 \| 0.266 \| 323 \| 190 \| 10 \| 4 \| 306 \| 3 \| 298 \| AAA family ATPase \| AAA family ATPase; afdb-uniprot50 \| AF-A0A847VJL2-F1-MODEL\_V4 \| 1.0 \| 3.95e-18 \| 672 \| 0.319 \| 244 \| 137 \| 8 \| 4 \| 241 \| 1 \| 221 \| ATP-binding protein \| ATP-binding protein; afdb-uniprot50 \| AF-A0A2T5YD05-F1-MODEL\_V4 \| 1.0 \| 5.366e-22 \| 672 \| 0.302 \| 324 \| 179 \| 11 \| 4 \| 314 \| 1 \| 290 \| Signal recognition particle subunit FFH/SRP54 (Srp54) \| Signal recognition particle subunit FFH/SRP54 (Srp54); afdb-uniprot50 \| AF-A0A0G3CIG4-F1-MODEL\_V4 \| 1.0 \| 8.201e-21 \| 671 \| 0.267 \| 310 \| 193 \| 10 \| 4 \| 304 \| 1 \| 285 \| AAA domain-containing protein \| AAA domain-containing protein; afdb-uniprot50 \| AF-A0A242NEU9-F1-MODEL\_V4 \| 1.0 \| 5.555e-21 \| 671 \| 0.278 \| 320 \| 185 \| 14 \| 4 \| 309 \| 2 \| 289 \| AAA domain-containing protein \| AAA domain-containing protein; afdb-uniprot50 \| AF-A0A7C8LT02-F1-MODEL\_V4 \| 1.0 \| 1.93e-21 \| 667 \| 0.233 \| 330 \| 215 \| 12 \| 4 \| 314 \| 1 \| 311 \| AAA family ATPase \| AAA family ATPase; afdb-uniprot50 \| AF-A0A0F8WAX0-F1-MODEL\_V4 \| 1.0 \| 4.508e-19 \| 666 \| 0.324 \| 271 \| 142 \| 11 \| 1 \| 253 \| 1 \| 248 \| AAA domain-containing protein \| AAA domain-containing protein; afdb-uniprot50 \| AF-A0A512JIX0-F1-MODEL\_V4 \| 1.0 \| 6.341e-22 \| 661 \| 0.26 \| 403 \| 194 \| 16 \| 1 \| 313 \| 1 \| 389 \| Uncharacterized protein \| Uncharacterized protein; afdb-uniprot50 \| AF-A0A1M4ZD27-F1-MODEL\_V4 \| 1.0 \| 4.033e-19 \| 661 \| 0.292 \| 256 \| 158 \| 9 \| 4 \| 251 \| 1 \| 241 \| AAA domain-containing protein \| AAA domain-containing protein; afdb-uniprot50 \| AF-R5T4I4-F1-MODEL\_V4 \| 1.0 \| 2.584e-19 \| 660 \| 0.281 \| 259 \| 153 \| 9 \| 1 \| 251 \| 1 \| 234 \| AAA domain-containing protein \| AAA domain-containing protein; afdb-uniprot50 \| AF-N9H8P7-F1-MODEL\_V4 \| 1.0 \| 8.375e-22 \| 660 \| 0.26 \| 334 \| 210 \| 11 \| 1 \| 324 \| 1 \| 307 \| AAA domain-containing protein \| AAA domain-containing protein; afdb-uniprot50 \| AF-A0A843H9K7-F1-MODEL\_V4 \| 1.0 \| 3.608e-19 \| 659 \| 0.297 \| 259 \| 155 \| 10 \| 4 \| 251 \| 3 \| 245 \| AAA family ATPase \| AAA family ATPase; afdb-uniprot50 \| AF-A0A4Q3Y5G8-F1-MODEL\_V4 \| 1.0 \| 2.62e-17 \| 656 \| 0.306 \| 248 \| 146 \| 7 \| 4 \| 247 \| 1 \| 226 \| AAA family ATPase \| AAA family ATPase; afdb-uniprot50 \| AF-A0A5M9TR59-F1-MODEL\_V4 \| 1.0 \| 3.184e-21 \| 656 \| 0.293 \| 303 \| 187 \| 10 \| 4 \| 295 \| 3 \| 289 \| AAA family ATPase \| AAA family ATPase; afdb-uniprot50 \| AF-A0A7V9QIV5-F1-MODEL\_V4 \| 1.0 \| 2.28e-21 \| 656 \| 0.299 \| 344 \| 170 \| 12 \| 4 \| 298 \| 3 \| 324 \| AAA family ATPase \| AAA family ATPase; afdb-uniprot50 \| AF-A0A1Q6RAL6-F1-MODEL\_V4 \| 1.0 \| 3.815e-19 \| 654 \| 0.332 \| 256 \| 139 \| 8 \| 1 \| 249 \| 1 \| 231 \| Uncharacterized protein \| Uncharacterized protein; afdb-uniprot50 \| AF-A0A2W4MPK8-F1-MODEL\_V4 \| 1.0 \| 2.675e-18 \| 653 \| 0.407 \| 211 \| 102 \| 5 \| 4 \| 209 \| 3 \| 195 \| AAA domain-containing protein \| AAA domain-containing protein; afdb-uniprot50 \| AF-A0A1Z4C5W6-F1-MODEL\_V4 \| 1.0 \| 3.118e-20 \| 653 \| 0.288 \| 309 \| 177 \| 11 \| 5 \| 307 \| 1 \| 272 \| AAA family ATPase \| AAA family ATPase; afdb-uniprot50 \| AF-A0A8B5WUU9-F1-MODEL\_V4 \| 1.0 \| 3.843e-22 \| 652 \| 0.285 \| 368 \| 187 \| 14 \| 1 \| 310 \| 1 \| 350 \| Uncharacterized protein \| Uncharacterized protein; afdb-uniprot50 \| AF-A0A0P0CW41-F1-MODEL\_V4 \| 1.0 \| 2.04e-21 \| 650 \| 0.274 \| 332 \| 201 \| 10 \| 4 \| 329 \| 1 \| 298 \| AAA domain-containing protein \| AAA domain-containing protein; afdb-uniprot50 \| AF-A4J1U0-F1-MODEL\_V4 \| 1.0 \| 1.956e-19 \| 648 \| 0.314 \| 261 \| 147 \| 10 \| 1 \| 256 \| 1 \| 234 \| Uncharacterized protein \| Uncharacterized protein; afdb-uniprot50 \| AF-A0A5S4ZQJ3-F1-MODEL\_V4 \| 1.0 \| 4.033e-19 \| 646 \| 0.319 \| 257 \| 144 \| 9 \| 1 \| 252 \| 1 \| 231 \| AAA domain-containing protein \| AAA domain-containing protein; afdb-uniprot50 \| AF-A0A554XCB7-F1-MODEL\_V4 \| 1.0 \| 8.67e-21 \| 645 \| 0.306 \| 313 \| 177 \| 11 \| 4 \| 307 \| 1 \| 282 \| AAA domain protein \| AAA domain protein; afdb-uniprot50 \| AF-A0A843HQ30-F1-MODEL\_V4 \| 1.0 \| 7.864e-19 \| 644 \| 0.31 \| 258 \| 153 \| 10 \| 4 \| 251 \| 3 \| 245 \| AAA family ATPase \| AAA family ATPase; afdb-uniprot50 \| AF-A0A7H1NU03-F1-MODEL\_V4 \| 1.0 \| 6.703e-22 \| 644 \| 0.331 \| 347 \| 156 \| 15 \| 1 \| 299 \| 1 \| 319 \| AAA domain protein \| AAA domain protein; afdb-uniprot50 \| AF-V2TRU2-F1-MODEL\_V4 \| 1.0 \| 1.998e-20 \| 643 \| 0.283 \| 318 \| 187 \| 10 \| 1 \| 308 \| 1 \| 287 \| AAA domain-containing protein \| AAA domain-containing protein; afdb-uniprot50 \| AF-A0A241VL01-F1-MODEL\_V4 \| 1.0 \| 4.97e-21 \| 643 \| 0.26 \| 330 \| 206 \| 12 \| 1 \| 321 \| 1 \| 301 \| AAA family ATPase \| AAA family ATPase; afdb-uniprot50 \| AF-M1P9I7-F1-MODEL\_V4 \| 1.0 \| 1.145e-20 \| 638 \| 0.25 \| 316 \| 199 \| 9 \| 6 \| 305 \| 4 \| 297 \| AAA domain-containing protein \| AAA domain-containing protein; afdb-uniprot50 \| AF-D6SV02-F1-MODEL\_V4 \| 1.0 \| 1.691e-20 \| 636 \| 0.301 \| 338 \| 178 \| 18 \| 5 \| 323 \| 1 \| 299 \| AAA ATPase \| AAA ATPase; afdb-uniprot50 \| AF-A0A1W7ACF2-F1-MODEL\_V4 \| 1.0 \| 1.061e-19 \| 635 \| 0.256 \| 320 \| 193 \| 11 \| 4 \| 299 \| 1 \| 299 \| AAA domain-containing protein \| AAA domain-containing protein; afdb-uniprot50 \| AF-A0A850RF01-F1-MODEL\_V4 \| 1.0 \| 3.012e-21 \| 635 \| 0.284 \| 323 \| 191 \| 9 \| 5 \| 312 \| 1 \| 298 \| ATP-binding protein \| ATP-binding protein; afdb-uniprot50 \| AF-A0A0F9IND8-F1-MODEL\_V4 \| 1.0 \| 7.284e-18 \| 634 \| 0.319 \| 260 \| 142 \| 10 \| 4 \| 244 \| 3 \| 246 \| Uncharacterized protein \| Uncharacterized protein; afdb-uniprot50 \| AF-A0A1G6IL95-F1-MODEL\_V4 \| 1.0 \| 4.867e-20 \| 634 \| 0.259 \| 320 \| 192 \| 13 \| 4 \| 298 \| 2 \| 301 \| AAA domain-containing protein \| AAA domain-containing protein; afdb-uniprot50 \| AF-C3X1R4-F1-MODEL\_V4 \| 1.0 \| 2.04e-21 \| 633 \| 0.28 \| 328 \| 185 \| 14 \| 4 \| 311 \| 3 \| 299 \| AAA domain-containing protein \| AAA domain-containing protein; afdb-uniprot50 \| AF-B5ZFT3-F1-MODEL\_V4 \| 1.0 \| 1.106e-21 \| 633 \| 0.325 \| 353 \| 180 \| 18 \| 4 \| 313 \| 3 \| 340 \| Uncharacterized protein \| Uncharacterized protein; afdb-uniprot50 \| AF-A0A3D6BFW0-F1-MODEL\_V4 \| 1.0 \| 6.428e-20 \| 631 \| 0.264 \| 329 \| 193 \| 12 \| 4 \| 329 \| 1 \| 283 \| AAA family ATPase \| AAA family ATPase; afdb-uniprot50 \| AF-A0A2W4BX75-F1-MODEL\_V4 \| 1.0 \| 2.496e-20 \| 631 \| 0.268 \| 335 \| 194 \| 12 \| 1 \| 309 \| 1 \| 310 \| Uncharacterized protein \| Uncharacterized protein; afdb-uniprot50 \| AF-B0VT02-F1-MODEL\_V4 \| 1.0 \| 4.701e-21 \| 631 \| 0.249 \| 333 \| 208 \| 13 \| 4 \| 324 \| 3 \| 305 \| Putative bacteriophage protein \| Putative bacteriophage protein; afdb-uniprot50 \| AF-G4QCR6-F1-MODEL\_V4 \| 1.0 \| 2.949e-20 \| 630 \| 0.283 \| 318 \| 176 \| 11 \| 4 \| 299 \| 1 \| 288 \| Phage protein \| Phage protein; afdb-uniprot50 \| AF-A0A6J4I179-F1-MODEL\_V4 \| 1.0 \| 7.284e-18 \| 627 \| 0.295 \| 247 \| 148 \| 7 \| 4 \| 245 \| 1 \| 226 \| Phage protein (ACLAME 621) \| Phage protein (ACLAME 621); afdb-uniprot50 \| AF-A0A3D6AQ56-F1-MODEL\_V4 \| 1.0 \| 3.559e-21 \| 626 \| 0.282 \| 347 \| 192 \| 16 \| 1 \| 328 \| 1 \| 309 \| AAA domain-containing protein \| AAA domain-containing protein; afdb-uniprot50 \| AF-A0A098AXD5-F1-MODEL\_V4 \| 1.0 \| 9.49e-20 \| 625 \| 0.3 \| 320 \| 181 \| 11 \| 1 \| 308 \| 1 \| 289 \| AAA ATPase \| AAA ATPase; afdb-uniprot50 \| AF-A0A2B3NNN2-F1-MODEL\_V4 \| 1.0 \| 2.77e-17 \| 622 \| 0.284 \| 260 \| 148 \| 10 \| 4 \| 245 \| 2 \| 241 \| Uncharacterized protein \| Uncharacterized protein; afdb-uniprot50 \| AF-A0A519PNY2-F1-MODEL\_V4 \| 1.0 \| 1.253e-19 \| 622 \| 0.26 \| 303 \| 184 \| 14 \| 4 \| 299 \| 1 \| 270 \| AAA family ATPase \| AAA family ATPase; afdb-uniprot50 \| AF-A0A7X7C3C3-F1-MODEL\_V4 \| 1.0 \| 1.053e-16 \| 620 \| 0.287 \| 254 \| 146 \| 8 \| 4 \| 251 \| 1 \| 225 \| AAA family ATPase \| AAA family ATPase; afdb-uniprot50 \| AF-A0A2T0SYP3-F1-MODEL\_V4 \| 1.0 \| 2.888e-19 \| 619 \| 0.246 \| 308 \| 194 \| 17 \| 4 \| 299 \| 1 \| 282 \| AAA domain-containing protein \| AAA domain-containing protein; afdb-uniprot50 \| AF-A0A0U5IPQ9-F1-MODEL\_V4 \| 1.0 \| 1.787e-20 \| 618 \| 0.263 \| 326 \| 199 \| 10 \| 5 \| 314 \| 1 \| 301 \| AAA ATPase \| AAA ATPase; afdb-uniprot50 \| AF-A0A4R3HQG4-F1-MODEL\_V4 \| 1.0 \| 3.366e-21 \| 615 \| 0.294 \| 333 \| 188 \| 16 \| 1 \| 316 \| 36 \| 338 \| AAA domain-containing protein \| AAA domain-containing protein; afdb-uniprot50 \| AF-A0A136Q9Y5-F1-MODEL\_V4 \| 1.0 \| 1.621e-18 \| 612 \| 0.317 \| 280 \| 156 \| 11 \| 5 \| 276 \| 1 \| 253 \| ATPase AAA \| ATPase AAA; afdb-uniprot50 \| AF-G2JBT6-F1-MODEL\_V4 \| 1.0 \| 3.485e-20 \| 612 \| 0.292 \| 321 \| 182 \| 11 \| 4 \| 310 \| 1 \| 290 \| AAA domain-containing protein \| AAA domain-containing protein; afdb-uniprot50 \| AF-A0A1Y4GR65-F1-MODEL\_V4 \| 1.0 \| 2.949e-20 \| 612 \| 0.283 \| 325 \| 180 \| 12 \| 4 \| 299 \| 3 \| 303 \| AAA domain-containing protein \| AAA domain-containing protein; afdb-uniprot50 \| AF-A0A0F8ZUD5-F1-MODEL\_V4 \| 1.0 \| 8.491e-20 \| 610 \| 0.299 \| 311 \| 172 \| 13 \| 11 \| 300 \| 2 \| 287 \| AAA domain-containing protein \| AAA domain-containing protein; afdb-uniprot50 \| AF-A0A519PTH0-F1-MODEL\_V4 \| 1.0 \| 2.888e-19 \| 610 \| 0.291 \| 295 \| 175 \| 11 \| 4 \| 294 \| 1 \| 265 \| AAA family ATPase \| AAA family ATPase; afdb-uniprot50 \| AF-A0A3M1BKJ4-F1-MODEL\_V4 \| 1.0 \| 4.323e-17 \| 609 \| 0.271 \| 254 \| 155 \| 11 \| 4 \| 246 \| 2 \| 236 \| AAA domain-containing protein \| AAA domain-containing protein; afdb-uniprot50 \| AF-A0A1V5IYB1-F1-MODEL\_V4 \| 1.0 \| 3.342e-18 \| 609 \| 0.25 \| 287 \| 184 \| 9 \| 2 \| 280 \| 1 \| 264 \| Uncharacterized protein \| Uncharacterized protein; afdb-uniprot50 \| AF-A0A6N6ZWE1-F1-MODEL\_V4 \| 1.0 \| 7.185e-20 \| 604 \| 0.285 \| 319 \| 181 \| 17 \| 4 \| 302 \| 9 \| 300 \| Uncharacterized protein \| Uncharacterized protein; afdb-uniprot50 \| AF-A0A512AWV3-F1-MODEL\_V4 \| 1.0 \| 8.031e-20 \| 603 \| 0.271 \| 335 \| 201 \| 13 \| 4 \| 329 \| 1 \| 301 \| Uncharacterized protein \| Uncharacterized protein; afdb-uniprot50 \| AF-A0A5M6IBX0-F1-MODEL\_V4 \| 1.0 \| 1.28e-20 \| 597 \| 0.271 \| 365 \| 177 \| 15 \| 4 \| 307 \| 3 \| 339 \| AAA family ATPase \| AAA family ATPase; afdb-uniprot50 \| AF-A0A3C1G7Q9-F1-MODEL\_V4 \| 1.0 \| 1.121e-19 \| 596 \| 0.282 \| 340 \| 185 \| 15 \| 1 \| 309 \| 1 \| 312 \| Uncharacterized protein \| Uncharacterized protein; afdb-uniprot50 \| AF-A0A4Q2ZQB1-F1-MODEL\_V4 \| 1.0 \| 1.916e-18 \| 594 \| 0.27 \| 303 \| 182 \| 11 \| 1 \| 303 \| 1 \| 264 \| AAA domain-containing protein \| AAA domain-containing protein; afdb-uniprot50 \| AF-A0A3C0TWV1-F1-MODEL\_V4 \| 1.0 \| 3.297e-20 \| 593 \| 0.291 \| 367 \| 185 \| 13 \| 4 \| 316 \| 3 \| 348 \| Uncharacterized protein \| Uncharacterized protein; afdb-uniprot50 \| AF-A0A3E0KAZ1-F1-MODEL\_V4 \| 1.0 \| 2.928e-17 \| 591 \| 0.292 \| 256 \| 150 \| 8 \| 4 \| 250 \| 2 \| 235 \| ATP-binding protein \| ATP-binding protein; afdb-uniprot50 \| AF-A0A327WUV5-F1-MODEL\_V4 \| 1.0 \| 3.608e-19 \| 589 \| 0.28 \| 303 \| 173 \| 12 \| 4 \| 297 \| 1 \| 267 \| AAA domain-containing protein \| AAA domain-containing protein; afdb-uniprot50 \| AF-A0A1G8RUQ5-F1-MODEL\_V4 \| 1.0 \| 2.584e-19 \| 588 \| 0.234 \| 328 \| 207 \| 13 \| 4 \| 311 \| 1 \| 304 \| Uncharacterized protein \| Uncharacterized protein; afdb-uniprot50 \| AF-A0A4U6DCX4-F1-MODEL\_V4 \| 1.0 \| 2.828e-18 \| 585 \| 0.266 \| 300 \| 172 \| 12 \| 19 \| 310 \| 2 \| 261 \| AAA family ATPase \| AAA family ATPase; afdb-uniprot50 \| AF-A0A3N5VX40-F1-MODEL\_V4 \| 1.0 \| 5.401e-17 \| 583 \| 0.295 \| 257 \| 151 \| 9 \| 4 \| 252 \| 3 \| 237 \| AAA domain-containing protein \| AAA domain-containing protein; afdb-uniprot50 \| AF-X0UF94-F1-MODEL\_V4 \| 1.0 \| 3.95e-18 \| 581 \| 0.273 \| 304 \| 178 \| 13 \| 4 \| 297 \| 1 \| 271 \| AAA domain-containing protein \| AAA domain-containing protein; afdb-uniprot50 \| AF-A0A2I1K2F4-F1-MODEL\_V4 \| 1.0 \| 4.508e-19 \| 581 \| 0.236 \| 326 \| 197 \| 11 \| 4 \| 303 \| 1 \| 300 \| AAA domain-containing protein \| AAA domain-containing protein; afdb-uniprot50 \| AF-A0A5C7QJD8-F1-MODEL\_V4 \| 1.0 \| 1.161e-18 \| 581 \| 0.252 \| 329 \| 210 \| 9 \| 1 \| 306 \| 1 \| 316 \| Uncharacterized protein \| Uncharacterized protein; afdb-uniprot50 \| AF-A0A7Y5WDD0-F1-MODEL\_V4 \| 1.0 \| 5.632e-19 \| 581 \| 0.25 \| 335 \| 184 \| 13 \| 4 \| 298 \| 1 \| 308 \| AAA family ATPase \| AAA family ATPase; afdb-uniprot50 \| AF-A0A7K1B1R5-F1-MODEL\_V4 \| 1.0 \| 3.161e-18 \| 580 \| 0.237 \| 312 \| 180 \| 15 \| 4 \| 295 \| 3 \| 276 \| AAA family ATPase \| AAA family ATPase; afdb-uniprot50 \| AF-A0A7Y7NCN0-F1-MODEL\_V4 \| 1.0 \| 6.428e-20 \| 578 \| 0.271 \| 321 \| 187 \| 13 \| 4 \| 299 \| 1 \| 299 \| AAA family ATPase \| AAA family ATPase; afdb-uniprot50 \| AF-A0A3R7B9V5-F1-MODEL\_V4 \| 1.0 \| 1.837e-16 \| 574 \| 0.27 \| 251 \| 157 \| 7 \| 5 \| 250 \| 1 \| 230 \| AAA family ATPase \| AAA family ATPase; afdb-uniprot50 \| AF-A0A482RRG4-F1-MODEL\_V4 \| 1.0 \| 1.534e-18 \| 572 \| 0.24 \| 299 \| 188 \| 12 \| 4 \| 298 \| 40 \| 303 \| Uncharacterized protein \| Uncharacterized protein; afdb-uniprot50 \| AF-A0A060N9D0-F1-MODEL\_V4 \| 1.0 \| 2.312e-19 \| 571 \| 0.225 \| 359 \| 186 \| 15 \| 4 \| 299 \| 1 \| 330 \| Uncharacterized protein \| Uncharacterized protein; afdb-uniprot50 \| AF-A0A6V8M265-F1-MODEL\_V4 \| 1.0 \| 5.751e-20 \| 571 \| 0.247 \| 359 \| 194 \| 14 \| 1 \| 298 \| 1 \| 344 \| Uncharacterized protein \| Uncharacterized protein; afdb-uniprot50 \| AF-A0A2T2WVC8-F1-MODEL\_V4 \| 1.0 \| 1.61e-15 \| 570 \| 0.254 \| 251 \| 153 \| 9 \| 4 \| 246 \| 2 \| 226 \| AAA domain-containing protein \| AAA domain-containing protein; afdb-uniprot50 \| AF-A0A2D9FRJ3-F1-MODEL\_V4 \| 1.0 \| 2.171e-16 \| 570 \| 0.269 \| 249 \| 152 \| 9 \| 1 \| 244 \| 12 \| 235 \| AAA family ATPase \| AAA family ATPase; afdb-uniprot50 \| AF-A0A7X9JFT3-F1-MODEL\_V4 \| 1.0 \| 5.327e-19 \| 570 \| 0.237 \| 379 \| 197 \| 14 \| 4 \| 329 \| 2 \| 341 \| ATP-binding protein \| ATP-binding protein; afdb-uniprot50 \| AF-A0A4Q6ABA1-F1-MODEL\_V4 \| 1.0 \| 3.228e-19 \| 569 \| 0.229 \| 340 \| 193 \| 15 \| 4 \| 305 \| 1 \| 309 \| ATP-binding protein \| ATP-binding protein; afdb-uniprot50 \| AF-A9B384-F1-MODEL\_V4 \| 1.0 \| 2.565e-16 \| 565 \| 0.262 \| 251 \| 157 \| 9 \| 5 \| 250 \| 1 \| 228 \| AAA ATPase \| AAA ATPase; afdb-uniprot50 \| AF-L7U581-F1-MODEL\_V4 \| 1.0 \| 5.515e-18 \| 562 \| 0.247 \| 327 \| 199 \| 18 \| 5 \| 316 \| 3 \| 297 \| Uncharacterized protein \| Uncharacterized protein; afdb-uniprot50 \| AF-A0A1V5UIG9-F1-MODEL\_V4 \| 1.0 \| 1.644e-16 \| 562 \| 0.262 \| 251 \| 153 \| 8 \| 4 \| 247 \| 1 \| 226 \| AAA domain-containing protein \| AAA domain-containing protein; afdb-uniprot50 \| AF-A0A225E1D1-F1-MODEL\_V4 \| 1.0 \| 3.95e-18 \| 559 \| 0.269 \| 334 \| 191 \| 13 \| 1 \| 308 \| 1 \| 307 \| Uncharacterized protein \| Uncharacterized protein; afdb-uniprot50 \| AF-A0A847H377-F1-MODEL\_V4 \| 1.0 \| 2.928e-17 \| 558 \| 0.235 \| 306 \| 201 \| 8 \| 1 \| 299 \| 1 \| 280 \| AAA family ATPase \| AAA family ATPase; afdb-uniprot50 \| AF-A0A5C7LG64-F1-MODEL\_V4 \| 1.0 \| 2.99e-18 \| 558 \| 0.268 \| 332 \| 189 \| 15 \| 4 \| 310 \| 2 \| 304 \| ATP-binding protein \| ATP-binding protein; afdb-uniprot50 \| AF-A0A7V9ZER8-F1-MODEL\_V4 \| 1.0 \| 1.227e-18 \| 554 \| 0.227 \| 334 \| 197 \| 13 \| 4 \| 307 \| 2 \| 304 \| AAA family ATPase \| AAA family ATPase; afdb-uniprot50 \| AF-A0A3D9B4I0-F1-MODEL\_V4 \| 1.0 \| 1.161e-18 \| 554 \| 0.248 \| 334 \| 181 \| 13 \| 4 \| 296 \| 1 \| 305 \| AAA family ATPase \| AAA family ATPase; afdb-uniprot50 \| AF-A0A0Q6CAF9-F1-MODEL\_V4 \| 1.0 \| 1.098e-18 \| 554 \| 0.255 \| 387 \| 194 \| 16 \| 4 \| 328 \| 3 \| 357 \| Uncharacterized protein \| Uncharacterized protein; afdb-uniprot50 \| AF-A0A5C0VJB0-F1-MODEL\_V4 \| 1.0 \| 1.644e-16 \| 552 \| 0.234 \| 307 \| 191 \| 13 \| 4 \| 304 \| 1 \| 269 \| AAA family ATPase \| AAA family ATPase; afdb-uniprot50 \| AF-A0A2S8SVY0-F1-MODEL\_V4 \| 1.0 \| 1.837e-16 \| 552 \| 0.255 \| 258 \| 151 \| 12 \| 1 \| 246 \| 1 \| 229 \| AAA domain-containing protein \| AAA domain-containing protein; afdb-uniprot50 \| AF-D4IQQ6-F1-MODEL\_V4 \| 1.0 \| 1.61e-15 \| 551 \| 0.289 \| 235 \| 133 \| 8 \| 18 \| 246 \| 3 \| 209 \| Uncharacterized protein \| Uncharacterized protein; afdb-uniprot50 \| AF-A0A1G8EH60-F1-MODEL\_V4 \| 1.0 \| 7.542e-17 \| 550 \| 0.238 \| 281 \| 183 \| 11 \| 4 \| 277 \| 3 \| 259 \| AAA domain-containing protein \| AAA domain-containing protein; afdb-uniprot50 \| AF-A0A4R3DYZ1-F1-MODEL\_V4 \| 1.0 \| 3.736e-18 \| 549 \| 0.254 \| 342 \| 191 \| 14 \| 4 \| 303 \| 3 \| 322 \| Uncharacterized protein \| Uncharacterized protein; afdb-uniprot50 \| AF-A0A410K460-F1-MODEL\_V4 \| 1.0 \| 3.161e-18 \| 548 \| 0.229 \| 327 \| 194 \| 11 \| 4 \| 298 \| 2 \| 302 \| Uncharacterized protein \| Uncharacterized protein; afdb-uniprot50 \| AF-A0A2D1KV30-F1-MODEL\_V4 \| 1.0 \| 6.517e-18 \| 548 \| 0.194 \| 324 \| 201 \| 11 \| 4 \| 295 \| 1 \| 296 \| Uncharacterized protein \| Uncharacterized protein; afdb-uniprot50 \| AF-A0A3N1GT73-F1-MODEL\_V4 \| 1.0 \| 2.53e-18 \| 548 \| 0.228 \| 319 \| 202 \| 13 \| 6 \| 299 \| 7 \| 306 \| KaiC/GvpD/RAD55 family RecA-like ATPase \| KaiC/GvpD/RAD55 family RecA-like ATPase; afdb-uniprot50 \| AF-A0A3N2NXN1-F1-MODEL\_V4 \| 1.0 \| 1.812e-18 \| 547 \| 0.215 \| 357 \| 231 \| 18 \| 1 \| 328 \| 5 \| 341 \| Uncharacterized protein \| Uncharacterized protein; afdb-uniprot50 \| AF-A0A142XCB2-F1-MODEL\_V4 \| 1.0 \| 2.675e-18 \| 546 \| 0.298 \| 332 \| 175 \| 13 \| 4 \| 302 \| 8 \| 314 \| Uncharacterized protein \| Uncharacterized protein; afdb-uniprot50 \| AF-A0A3D5KDT0-F1-MODEL\_V4 \| 1.0 \| 2.025e-18 \| 545 \| 0.229 \| 348 \| 202 \| 14 \| 1 \| 311 \| 1 \| 319 \| Uncharacterized protein \| Uncharacterized protein; afdb-uniprot50 \| AF-A0A512J936-F1-MODEL\_V4 \| 1.0 \| 3.95e-18 \| 545 \| 0.237 \| 350 \| 200 \| 13 \| 4 \| 308 \| 7 \| 334 \| Uncharacterized protein \| Uncharacterized protein; afdb-uniprot50 \| AF-A0A7Y4U332-F1-MODEL\_V4 \| 1.0 \| 1.09e-15 \| 544 \| 0.274 \| 255 \| 149 \| 8 \| 4 \| 246 \| 1 \| 231 \| AAA family ATPase \| AAA family ATPase; afdb-uniprot50 \| AF-A0A661JH92-F1-MODEL\_V4 \| 1.0 \| 1.01e-14 \| 543 \| 0.359 \| 192 \| 99 \| 4 \| 4 \| 191 \| 2 \| 173 \| AAA domain-containing protein \| AAA domain-containing protein; afdb-uniprot50 \| AF-A0A0F9T6J3-F1-MODEL\_V4 \| 1.0 \| 1.61e-15 \| 540 \| 0.26 \| 250 \| 154 \| 9 \| 6 \| 248 \| 4 \| 229 \| AAA domain-containing protein \| AAA domain-containing protein; afdb-uniprot50 \| AF-A0A2N3UTJ3-F1-MODEL\_V4 \| 1.0 \| 1.244e-16 \| 538 \| 0.233 \| 291 \| 183 \| 13 \| 1 \| 277 \| 1 \| 265 \| AAA domain-containing protein \| AAA domain-containing protein; afdb-uniprot50 \| AF-A0A1G0HWF8-F1-MODEL\_V4 \| 1.0 \| 2.828e-18 \| 535 \| 0.259 \| 355 \| 193 \| 12 \| 1 \| 307 \| 1 \| 333 \| Uncharacterized protein \| Uncharacterized protein; afdb-uniprot50 \| AF-A0A6P1WJL1-F1-MODEL\_V4 \| 1.0 \| 4.934e-18 \| 535 \| 0.289 \| 384 \| 187 \| 18 \| 1 \| 325 \| 1 \| 357 \| AAA family ATPase \| AAA family ATPase; afdb-uniprot50 \| AF-I3IJD0-F1-MODEL\_V4 \| 1.0 \| 3.342e-18 \| 534 \| 0.222 \| 346 \| 206 \| 12 \| 6 \| 313 \| 20 \| 340 \| ATPase \| ATPase; afdb-uniprot50 \| AF-A0A7W6JAD3-F1-MODEL\_V4 \| 1.0 \| 2.675e-18 \| 534 \| 0.265 \| 380 \| 198 \| 18 \| 4 \| 311 \| 3 \| 373 \| Uncharacterized protein \| Uncharacterized protein; afdb-uniprot50 \| AF-A0A1V6GAY5-F1-MODEL\_V4 \| 1.0 \| 6.89e-18 \| 533 \| 0.21 \| 337 \| 212 \| 11 \| 5 \| 309 \| 1 \| 315 \| AAA domain-containing protein \| AAA domain-containing protein; afdb-uniprot50 \| AF-A0A4Q3VBQ7-F1-MODEL\_V4 \| 1.0 \| 1.031e-15 \| 532 \| 0.248 \| 254 \| 164 \| 9 \| 4 \| 251 \| 3 \| 235 \| Uncharacterized protein \| Uncharacterized protein; afdb-uniprot50 \| AF-A0A7X9GKV1-F1-MODEL\_V4 \| 1.0 \| 1.316e-16 \| 531 \| 0.283 \| 293 \| 167 \| 9 \| 6 \| 282 \| 4 \| 269 \| AAA family ATPase \| AAA family ATPase; afdb-uniprot50 \| AF-A0A7C6ZLP1-F1-MODEL\_V4 \| 1.0 \| 2.171e-16 \| 530 \| 0.247 \| 255 \| 158 \| 9 \| 5 \| 251 \| 1 \| 229 \| AAA family ATPase \| AAA family ATPase; afdb-uniprot50 \| AF-A0A2Z3HGM3-F1-MODEL\_V4 \| 1.0 \| 2.77e-17 \| 530 \| 0.217 \| 312 \| 221 \| 7 \| 4 \| 307 \| 2 \| 298 \| Uncharacterized protein \| Uncharacterized protein; afdb-uniprot50 \| AF-A0A535F1B2-F1-MODEL\_V4 \| 1.0 \| 3.342e-18 \| 529 \| 0.233 \| 342 \| 207 \| 15 \| 4 \| 316 \| 2 \| 317 \| Uncharacterized protein \| Uncharacterized protein; afdb-uniprot50 \| AF-A0A4V6M1G0-F1-MODEL\_V4 \| 1.0 \| 3.032e-16 \| 529 \| 0.229 \| 283 \| 180 \| 10 \| 4 \| 267 \| 3 \| 266 \| AAA ATPase \| AAA ATPase; afdb-uniprot50 \| AF-A0A2T2XCV9-F1-MODEL\_V4 \| 1.0 \| 1.876e-17 \| 526 \| 0.246 \| 308 \| 191 \| 10 \| 6 \| 298 \| 11 \| 292 \| AAA family ATPase \| AAA family ATPase; afdb-uniprot50 \| AF-A0A6G2PR85-F1-MODEL\_V4 \| 1.0 \| 3.582e-16 \| 524 \| 0.228 \| 258 \| 162 \| 11 \| 4 \| 251 \| 23 \| 253 \| AAA family ATPase \| AAA family ATPase; afdb-uniprot50 \| AF-A0A7J4N567-F1-MODEL\_V4 \| 1.0 \| 2.928e-17 \| 523 \| 0.232 \| 322 \| 190 \| 16 \| 4 \| 299 \| 1 \| 291 \| AAA family ATPase \| AAA family ATPase; afdb-uniprot50 \| AF-A0A0F9WQ23-F1-MODEL\_V4 \| 1.0 \| 4.383e-15 \| 523 \| 0.25 \| 252 \| 166 \| 8 \| 5 \| 252 \| 1 \| 233 \| AAA domain-containing protein \| AAA domain-containing protein; afdb-uniprot50 \| AF-A0A6A6K1N9-F1-MODEL\_V4 \| 1.0 \| 1.193e-14 \| 523 \| 0.522 \| 174 \| 61 \| 5 \| 4 \| 172 \| 323 \| 479 \| DUF3799 domain-containing protein \| DUF3799 domain-containing protein; afdb-uniprot50 \| AF-A0A651G1T1-F1-MODEL\_V4 \| 1.0 \| 7.284e-18 \| 520 \| 0.268 \| 361 \| 182 \| 14 \| 5 \| 308 \| 1 \| 336 \| AAA family ATPase \| AAA family ATPase; afdb-uniprot50 \| AF-A0A1L8CRN1-F1-MODEL\_V4 \| 1.0 \| 6.89e-18 \| 516 \| 0.244 \| 343 \| 188 \| 12 \| 4 \| 308 \| 1 \| 310 \| AAA domain-containing protein \| AAA domain-containing protein; afdb-uniprot50 \| AF-A0A2T0KP83-F1-MODEL\_V4 \| 1.0 \| 1.271e-17 \| 515 \| 0.208 \| 345 \| 211 \| 16 \| 7 \| 328 \| 1 \| 306 \| AAA domain-containing protein \| AAA domain-containing protein; afdb-uniprot50 \| AF-A0A142XQT8-F1-MODEL\_V4 \| 1.0 \| 3.658e-17 \| 512 \| 0.208 \| 321 \| 212 \| 13 \| 1 \| 299 \| 1 \| 301 \| Signal recognition particle protein Srp54 \| Signal recognition particle protein Srp54; afdb-uniprot50 \| AF-A0A7X2N516-F1-MODEL\_V4 \| 1.0 \| 1.316e-16 \| 510 \| 0.214 \| 331 \| 197 \| 11 \| 4 \| 298 \| 3 \| 306 \| AAA family ATPase \| AAA family ATPase; afdb-uniprot50 \| AF-F5YIK8-F1-MODEL\_V4 \| 1.0 \| 2.011e-15 \| 508 \| 0.295 \| 257 \| 147 \| 10 \| 4 \| 251 \| 1 \| 232 \| AAA domain-containing protein \| AAA domain-containing protein; afdb-uniprot50 \| AF-A0A3N5GT73-F1-MODEL\_V4 \| 1.0 \| 8.142e-18 \| 505 \| 0.232 \| 331 \| 195 \| 14 \| 4 \| 303 \| 21 \| 323 \| ATP-binding protein \| ATP-binding protein; afdb-uniprot50 \| AF-A0A1L6L855-F1-MODEL\_V4 \| 1.0 \| 1.679e-17 \| 505 \| 0.24 \| 333 \| 207 \| 13 \| 1 \| 309 \| 14 \| 324 \| AAA domain-containing protein \| AAA domain-containing protein; afdb-uniprot50 \| AF-A0A350BEK3-F1-MODEL\_V4 \| 1.0 \| 9.621e-18 \| 502 \| 0.246 \| 361 \| 190 \| 17 \| 4 \| 310 \| 1 \| 333 \| ATP-binding protein \| ATP-binding protein; afdb-uniprot50 \| AF-A0A1V9K0D4-F1-MODEL\_V4 \| 1.0 \| 2.126e-15 \| 499 \| 0.227 \| 255 \| 160 \| 12 \| 4 \| 248 \| 22 \| 249 \| AAA domain-containing protein \| AAA domain-containing protein; afdb-uniprot50 \| AF-A0A544U8C9-F1-MODEL\_V4 \| 1.0 \| 9.961e-17 \| 499 \| 0.251 \| 310 \| 198 \| 13 \| 4 \| 296 \| 1 \| 293 \| Uncharacterized protein \| Uncharacterized protein; afdb-uniprot50 \| AF-A0A0Q4XY38-F1-MODEL\_V4 \| 1.0 \| 1.555e-16 \| 499 \| 0.252 \| 333 \| 190 \| 15 \| 1 \| 303 \| 1 \| 304 \| Uncharacterized protein \| Uncharacterized protein; afdb-uniprot50 \| AF-A0A3N4SMN9-F1-MODEL\_V4 \| 1.0 \| 2.217e-17 \| 498 \| 0.252 \| 348 \| 194 \| 16 \| 4 \| 318 \| 22 \| 336 \| AAA domain-containing protein \| AAA domain-containing protein; afdb-uniprot50 \| AF-A0A1L6LWE2-F1-MODEL\_V4 \| 1.0 \| 1.837e-16 \| 498 \| 0.214 \| 326 \| 211 \| 12 \| 4 \| 299 \| 8 \| 318 \| Uncharacterized protein \| Uncharacterized protein; afdb-uniprot50 \| AF-A0A0F9SZA2-F1-MODEL\_V4 \| 1.0 \| 3.096e-17 \| 498 \| 0.255 \| 317 \| 182 \| 16 \| 4 \| 298 \| 1 \| 285 \| AAA domain-containing protein \| AAA domain-containing protein; afdb-uniprot50 \| AF-T0HD80-F1-MODEL\_V4 \| 1.0 \| 7.133e-17 \| 496 \| 0.237 \| 354 \| 180 \| 14 \| 11 \| 302 \| 2 \| 327 \| AAA domain-containing protein \| AAA domain-containing protein; afdb-uniprot50 \| AF-A0A1V0UC96-F1-MODEL\_V4 \| 1.0 \| 8.255e-16 \| 496 \| 0.244 \| 278 \| 173 \| 14 \| 4 \| 267 \| 28 \| 282 \| Uncharacterized protein \| Uncharacterized protein; afdb-uniprot50 \| AF-A0A2N5JRX4-F1-MODEL\_V4 \| 1.0 \| 3.709e-15 \| 493 \| 0.244 \| 303 \| 203 \| 10 \| 4 \| 299 \| 2 \| 285 \| AAA domain-containing protein \| AAA domain-containing protein; afdb-uniprot50 \| AF-A0A560BNE4-F1-MODEL\_V4 \| 1.0 \| 2.097e-17 \| 491 \| 0.282 \| 347 \| 167 \| 19 \| 1 \| 303 \| 1 \| 309 \| AAA domain-containing protein \| AAA domain-containing protein; afdb-uniprot50 \| AF-A0A842HGV6-F1-MODEL\_V4 \| 1.0 \| 1.053e-16 \| 488 \| 0.231 \| 345 \| 201 \| 13 \| 5 \| 311 \| 1 \| 319 \| ATP-binding protein \| ATP-binding protein; afdb-uniprot50 \| AF-A0A7Y0Q481-F1-MODEL\_V4 \| 1.0 \| 1.576e-14 \| 487 \| 0.306 \| 225 \| 124 \| 8 \| 1 \| 217 \| 1 \| 201 \| ATP-binding protein \| ATP-binding protein; afdb-uniprot50 \| AF-A0A5C7PD19-F1-MODEL\_V4 \| 1.0 \| 1.837e-16 \| 486 \| 0.224 \| 366 \| 197 \| 15 \| 1 \| 307 \| 2 \| 339 \| Uncharacterized protein \| Uncharacterized protein; afdb-uniprot50 \| AF-A0A1V5Z756-F1-MODEL\_V4 \| 1.0 \| 7.542e-17 \| 480 \| 0.258 \| 352 \| 176 \| 14 \| 6 \| 298 \| 4 \| 329 \| AAA domain-containing protein \| AAA domain-containing protein; afdb-uniprot50 \| AF-A0A133ZYS2-F1-MODEL\_V4 \| 1.0 \| 3.205e-16 \| 479 \| 0.245 \| 310 \| 199 \| 10 \| 4 \| 298 \| 6 \| 295 \| Uncharacterized protein \| Uncharacterized protein; afdb-uniprot50 \| AF-A0A7X7JYV6-F1-MODEL\_V4 \| 1.0 \| 8.912e-17 \| 477 \| 0.252 \| 336 \| 199 \| 18 \| 4 \| 309 \| 5 \| 318 \| AAA family ATPase \| AAA family ATPase; afdb-uniprot50 \| AF-A0A353M2R8-F1-MODEL\_V4 \| 1.0 \| 1.177e-16 \| 476 \| 0.222 \| 355 \| 201 \| 14 \| 5 \| 309 \| 1 \| 330 \| AAA family ATPase \| AAA family ATPase; afdb-uniprot50 \| AF-A0A6M3M0Y2-F1-MODEL\_V4 \| 1.0 \| 5.401e-17 \| 469 \| 0.238 \| 344 \| 193 \| 13 \| 4 \| 312 \| 1 \| 310 \| Putative ATPase domain containing protein \| Putative ATPase domain containing protein; afdb-uniprot50 \| AF-A0A7K2YW75-F1-MODEL\_V4 \| 1.0 \| 4.004e-16 \| 469 \| 0.224 \| 339 \| 199 \| 18 \| 4 \| 314 \| 3 \| 305 \| AAA family ATPase \| AAA family ATPase; afdb-uniprot50 \| AF-A0A2K8ZC48-F1-MODEL\_V4 \| 1.0 \| 6.12e-15 \| 465 \| 0.244 \| 286 \| 169 \| 12 \| 21 \| 298 \| 2 \| 248 \| AAA family ATPase \| AAA family ATPase; afdb-uniprot50 \| AF-A0A1V5X060-F1-MODEL\_V4 \| 1.0 \| 1.09e-15 \| 464 \| 0.222 \| 315 \| 196 \| 13 \| 6 \| 298 \| 21 \| 308 \| AAA domain-containing protein \| AAA domain-containing protein; afdb-uniprot50 \| AF-A0A7J5GPD8-F1-MODEL\_V4 \| 1.0 \| 1.689e-12 \| 463 \| 0.334 \| 203 \| 104 \| 6 \| 4 \| 200 \| 2 \| 179 \| AAA family ATPase \| AAA family ATPase; afdb-uniprot50 \| AF-A0A353K9R3-F1-MODEL\_V4 \| 1.0 \| 1.09e-15 \| 462 \| 0.224 \| 339 \| 204 \| 17 \| 1 \| 314 \| 58 \| 362 \| Uncharacterized protein \| Uncharacterized protein; afdb-uniprot50 \| AF-A0A652LKG1-F1-MODEL\_V4 \| 1.0 \| 5.002e-16 \| 461 \| 0.231 \| 328 \| 192 \| 18 \| 1 \| 300 \| 1 \| 296 \| Uncharacterized protein \| Uncharacterized protein; afdb-uniprot50 \| AF-A0A2E5S350-F1-MODEL\_V4 \| 1.0 \| 2.376e-15 \| 460 \| 0.224 \| 330 \| 202 \| 13 \| 4 \| 315 \| 2 \| 295 \| AAA family ATPase \| AAA family ATPase; afdb-uniprot50 \| AF-A0A0F9EYI3-F1-MODEL\_V4 \| 1.0 \| 3.182e-13 \| 455 \| 0.221 \| 230 \| 152 \| 6 \| 31 \| 246 \| 4 \| 220 \| Uncharacterized protein \| Uncharacterized protein; afdb-uniprot50 \| AF-A0A4Q3H9L2-F1-MODEL\_V4 \| 1.0 \| 9.035e-15 \| 452 \| 0.187 \| 315 \| 211 \| 12 \| 14 \| 325 \| 8 \| 280 \| Uncharacterized protein \| Uncharacterized protein; afdb-uniprot50 \| AF-L8MVA5-F1-MODEL\_V4 \| 1.0 \| 3.632e-14 \| 451 \| 0.224 \| 250 \| 153 \| 11 \| 4 \| 245 \| 3 \| 219 \| AAA ATPase \| AAA ATPase; afdb-uniprot50 \| AF-A0A1C5FLC7-F1-MODEL\_V4 \| 1.0 \| 6.12e-15 \| 447 \| 0.192 \| 338 \| 218 \| 13 \| 1 \| 315 \| 21 \| 326 \| AAA domain-containing protein \| AAA domain-containing protein; afdb-uniprot50 \| AF-A0A554L1H4-F1-MODEL\_V4 \| 1.0 \| 4.797e-14 \| 446 \| 0.323 \| 247 \| 122 \| 8 \| 4 \| 250 \| 1 \| 202 \| AAA domain-containing protein \| AAA domain-containing protein; afdb-uniprot50 \| AF-A0A1B8H5H1-F1-MODEL\_V4 \| 1.0 \| 1.01e-14 \| 441 \| 0.228 \| 293 \| 182 \| 13 \| 40 \| 316 \| 3 \| 267 \| Uncharacterized protein \| Uncharacterized protein; afdb-uniprot50 \| AF-A0A2Z5X544-F1-MODEL\_V4 \| 1.0 \| 8.369e-14 \| 441 \| 0.232 \| 250 \| 151 \| 11 \| 4 \| 245 \| 3 \| 219 \| AAA ATPase \| AAA ATPase; afdb-uniprot50 \| AF-A0A4Q5Y1S6-F1-MODEL\_V4 \| 1.0 \| 2.637e-12 \| 440 \| 0.189 \| 237 \| 161 \| 9 \| 11 \| 246 \| 38 \| 244 \| Uncharacterized protein \| Uncharacterized protein; afdb-uniprot50 \| AF-A0A644T0C1-F1-MODEL\_V4 \| 1.0 \| 2.248e-15 \| 440 \| 0.226 \| 322 \| 194 \| 15 \| 4 \| 302 \| 1 \| 290 \| AAA domain-containing protein \| AAA domain-containing protein; afdb-uniprot50 \| AF-A0A1Z4RCR8-F1-MODEL\_V4 \| 1.0 \| 2.295e-16 \| 437 \| 0.215 \| 358 \| 207 \| 16 \| 4 \| 325 \| 1 \| 320 \| ERF family protein \| ERF family protein; afdb-uniprot50 \| AF-A0A6L9HL74-F1-MODEL\_V4 \| 1.0 \| 5.789e-15 \| 433 \| 0.2 \| 335 \| 197 \| 13 \| 4 \| 300 \| 5 \| 306 \| Uncharacterized protein \| Uncharacterized protein; afdb-uniprot50 \| AF-A0A375H3M6-F1-MODEL\_V4 \| 1.0 \| 4.443e-13 \| 432 \| 0.433 \| 187 \| 93 \| 3 \| 152 \| 329 \| 2 \| 184 \| Uncharacterized protein \| Uncharacterized protein; afdb-uniprot50 \| AF-D3DCQ5-F1-MODEL\_V4 \| 1.0 \| 1.41e-14 \| 428 \| 0.206 \| 325 \| 208 \| 15 \| 4 \| 302 \| 6 \| 306 \| Uncharacterized protein \| Uncharacterized protein; afdb-uniprot50 \| AF-A0A4Q3H202-F1-MODEL\_V4 \| 1.0 \| 1.46e-13 \| 421 \| 0.183 \| 306 \| 205 \| 16 \| 11 \| 313 \| 4 \| 267 \| Uncharacterized protein \| Uncharacterized protein; afdb-uniprot50 \| AF-A0A2P2GMU8-F1-MODEL\_V4 \| 1.0 \| 1.491e-14 \| 412 \| 0.221 \| 334 \| 183 \| 17 \| 18 \| 317 \| 2 \| 292 \| Uncharacterized protein \| Uncharacterized protein; afdb-uniprot50 \| AF-A0A1Y4U0G7-F1-MODEL\_V4 \| 1.0 \| 1.48e-11 \| 399 \| 0.289 \| 207 \| 113 \| 8 \| 43 \| 243 \| 2 \| 180 \| AAA family ATPase \| AAA family ATPase; afdb-uniprot50 \| AF-H0A058-F1-MODEL\_V4 \| 1.0 \| 6.699e-14 \| 381 \| 0.205 \| 389 \| 196 \| 19 \| 1 \| 312 \| 1 \| 353 \| AAA domain-containing protein \| AAA domain-containing protein; afdb-uniprot50 \| AF-A0A520A7W8-F1-MODEL\_V4 \| 1.0 \| 5.323e-11 \| 380 \| 0.205 \| 209 \| 136 \| 10 \| 14 \| 219 \| 9 \| 190 \| Uncharacterized protein \| Uncharacterized protein; afdb-uniprot50 \| AF-A0A840V466-F1-MODEL\_V4 \| 1.0 \| 7.332e-13 \| 375 \| 0.253 \| 264 \| 162 \| 9 \| 51 \| 297 \| 1 \| 246 \| Uncharacterized protein \| Uncharacterized protein; afdb-uniprot50 \| AF-R8E138-F1-MODEL\_V4 \| 1.0 \| 1.532e-10 \| 359 \| 0.292 \| 198 \| 108 \| 7 \| 4 \| 185 \| 2 \| 183 \| Uncharacterized protein \| Uncharacterized protein; afdb-uniprot50 \| AF-A0A2I0CWX1-F1-MODEL\_V4 \| 1.0 \| 5.827e-10 \| 351 \| 0.246 \| 203 \| 124 \| 7 \| 52 \| 246 \| 3 \| 184 \| Uncharacterized protein \| Uncharacterized protein; afdb-uniprot50 \| AF-A0A1V5T3V6-F1-MODEL\_V4 \| 1.0 \| 6.424e-12 \| 341 \| 0.195 \| 373 \| 217 \| 15 \| 6 \| 317 \| 7 \| 357 \| Uncharacterized protein \| Uncharacterized protein; afdb-uniprot50 \| AF-A0A522DM60-F1-MODEL\_V4 \| 1.0 \| 6.076e-12 \| 339 \| 0.203 \| 315 \| 197 \| 17 \| 27 \| 320 \| 3 \| 284 \| Uncharacterized protein \| Uncharacterized protein; afdb-uniprot50 \| AF-A0A658Z6H7-F1-MODEL\_V4 \| 1.0 \| 3.812e-11 \| 339 \| 0.253 \| 272 \| 159 \| 16 \| 1 \| 250 \| 1 \| 250 \| Putative bacteriophage protein \| Putative bacteriophage protein; afdb-uniprot50 \| AF-A0A6M3LFY1-F1-MODEL\_V4 \| 1.0 \| 6.935e-13 \| 337 \| 0.234 \| 350 \| 189 \| 19 \| 6 \| 303 \| 5 \| 327 \| Putative ATPase domain containing protein \| Putative ATPase domain containing protein; afdb-uniprot50 \| AF-A0A832HSN5-F1-MODEL\_V4 \| 1.0 \| 1.003e-11 \| 331 \| 0.188 \| 361 \| 212 \| 21 \| 4 \| 299 \| 14 \| 358 \| Uncharacterized protein \| Uncharacterized protein; afdb-uniprot50 \| AF-A0A2H9V4N0-F1-MODEL\_V4 \| 1.0 \| 3.893e-12 \| 324 \| 0.209 \| 329 \| 186 \| 20 \| 6 \| 286 \| 67 \| 369 \| AAA domain-containing protein \| AAA domain-containing protein; afdb-uniprot50 \| AF-A0A6L7X1V5-F1-MODEL\_V4 \| 1.0 \| 4.116e-12 \| 323 \| 0.22 \| 358 \| 201 \| 23 \| 1 \| 305 \| 1 \| 333 \| AAA family ATPase \| AAA family ATPase; afdb-uniprot50 \| AF-A0A2M8C2I9-F1-MODEL\_V4 \| 1.0 \| 5.436e-12 \| 322 \| 0.181 \| 385 \| 215 \| 24 \| 6 \| 324 \| 10 \| 360 \| Uncharacterized protein \| Uncharacterized protein; afdb-uniprot50 \| AF-A0A5R2ATA9-F1-MODEL\_V4 \| 1.0 \| 5.747e-12 \| 321 \| 0.157 \| 369 \| 230 \| 11 \| 4 \| 329 \| 3 \| 333 \| Uncharacterized protein \| Uncharacterized protein; afdb-uniprot50 \| AF-A0A1F5UT66-F1-MODEL\_V4 \| 1.0 \| 3.182e-13 \| 307 \| 0.147 \| 353 \| 214 \| 18 \| 4 \| 306 \| 2 \| 317 \| Uncharacterized protein \| Uncharacterized protein; afdb-uniprot50 \| AF-A0A1F5UBV1-F1-MODEL\_V4 \| 1.0 \| 2.693e-13 \| 302 \| 0.164 \| 359 \| 213 \| 19 \| 4 \| 311 \| 2 \| 324 \| Uncharacterized protein \| Uncharacterized protein; afdb-uniprot50 \| AF-A0A6L5MSV1-F1-MODEL\_V4 \| 1.0 \| 1.608e-07 \| 301 \| 0.266 \| 120 \| 87 \| 1 \| 75 \| 193 \| 7 \| 126 \| AAA family ATPase \| AAA family ATPase; afdb-uniprot50 \| AF-A0A6F9WZH7-F1-MODEL\_V4 \| 1.0 \| 5.035e-11 \| 300 \| 0.203 \| 379 \| 205 \| 21 \| 1 \| 309 \| 1 \| 352 \| Bacteriophage protein \| Bacteriophage protein; afdb-uniprot50 \| AF-A0A3A1YKB4-F1-MODEL\_V4 \| 1.0 \| 4.173e-10 \| 295 \| 0.172 \| 261 \| 174 \| 11 \| 6 \| 250 \| 7 \| 241 \| Uncharacterized protein \| Uncharacterized protein; afdb-uniprot50 \| AF-A0A4Q3TR60-F1-MODEL\_V4 \| 1.0 \| 5.628e-11 \| 290 \| 0.194 \| 329 \| 182 \| 14 \| 46 \| 329 \| 7 \| 297 \| Uncharacterized protein \| Uncharacterized protein; afdb-uniprot50 \| AF-A0A2H0NBL3-F1-MODEL\_V4 \| 1.0 \| 1.297e-10 \| 286 \| 0.165 \| 362 \| 211 \| 22 \| 2 \| 298 \| 3 \| 338 \| Uncharacterized protein \| Uncharacterized protein; afdb-uniprot50 \| AF-A0A522DTH7-F1-MODEL\_V4 \| 1.0 \| 1.16e-10 \| 272 \| 0.177 \| 377 \| 205 \| 18 \| 4 \| 299 \| 7 \| 359 \| Uncharacterized protein \| Uncharacterized protein; afdb-uniprot50 \| AF-A0A3R6YIJ8-F1-MODEL\_V4 \| 1.0 \| 2.009e-07 \| 269 \| 0.259 \| 177 \| 107 \| 6 \| 4 \| 166 \| 1 \| 167 \| Uncharacterized protein \| Uncharacterized protein; afdb-uniprot50 \| AF-A0A430AW13-F1-MODEL\_V4 \| 1.0 \| 1.587e-09 \| 265 \| 0.174 \| 316 \| 175 \| 19 \| 1 \| 267 \| 1 \| 279 \| Uncharacterized protein \| Uncharacterized protein; afdb-uniprot50 \| AF-A0A844U1W6-F1-MODEL\_V4 \| 1.0 \| 3.785e-08 \| 254 \| 0.228 \| 219 \| 143 \| 9 \| 89 \| 298 \| 32 \| 233 \| AAA family ATPase \| AAA family ATPase; afdb-uniprot50 \| AF-A0A4R0EH56-F1-MODEL\_V4 \| 1.0 \| 4.829e-09 \| 249 \| 0.191 \| 266 \| 189 \| 9 \| 75 \| 324 \| 8 \| 263 \| AAA family ATPase \| AAA family ATPase; afdb-uniprot50 \| AF-A0A149SNW0-F1-MODEL\_V4 \| 1.0 \| 2.967e-07 \| 244 \| 0.428 \| 126 \| 71 \| 1 \| 196 \| 320 \| 1 \| 126 \| Uncharacterized protein \| Uncharacterized protein; afdb-uniprot50 \| AF-A0A7X2BBX7-F1-MODEL\_V4 \| 1.0 \| 1.968e-06 \| 241 \| 0.364 \| 162 \| 80 \| 6 \| 4 \| 153 \| 2 \| 152 \| Uncharacterized protein \| Uncharacterized protein; afdb-uniprot50 \| AF-A0A3D6BEG5-F1-MODEL\_V4 \| 1.0 \| 1.643e-08 \| 241 \| 0.138 \| 369 \| 234 \| 18 \| 7 \| 314 \| 8 \| 353 \| Uncharacterized protein \| Uncharacterized protein; afdb-uniprot50 \| AF-A0A521Z886-F1-MODEL\_V4 \| 1.0 \| 1.941e-08 \| 236 \| 0.187 \| 256 \| 150 \| 15 \| 4 \| 228 \| 53 \| 281 \| Uncharacterized protein \| Uncharacterized protein; afdb-uniprot50 \| AF-A0A4Q3F7I8-F1-MODEL\_V4 \| 1.0 \| 9.22e-08 \| 232 \| 0.242 \| 210 \| 142 \| 7 \| 107 \| 311 \| 4 \| 201 \| AAA family ATPase \| AAA family ATPase; afdb-uniprot50 \| AF-A0A7V1ZNT5-F1-MODEL\_V4 \| 1.0 \| 1.47e-08 \| 231 \| 0.15 \| 259 \| 168 \| 15 \| 4 \| 245 \| 20 \| 243 \| Uncharacterized protein \| Uncharacterized protein; afdb-uniprot50 \| AF-A0A1J5PJ53-F1-MODEL\_V4 \| 1.0 \| 8.54e-07 \| 230 \| 0.195 \| 194 \| 141 \| 4 \| 108 \| 299 \| 1 \| 181 \| Uncharacterized protein \| Uncharacterized protein; afdb-uniprot50 \| AF-A0A7Y5QKL6-F1-MODEL\_V4 \| 1.0 \| 1.554e-08 \| 230 \| 0.142 \| 267 \| 172 \| 16 \| 1 \| 245 \| 1 \| 232 \| AAA family ATPase \| AAA family ATPase; afdb-uniprot50 \| AF-A0A1V5X4J5-F1-MODEL\_V4 \| 1.0 \| 4.23e-08 \| 228 \| 0.145 \| 220 \| 142 \| 11 \| 4 \| 204 \| 1 \| 193 \| Uncharacterized protein \| Uncharacterized protein; afdb-uniprot50 \| AF-A0A132MHS1-F1-MODEL\_V4 \| 1.0 \| 2.926e-09 \| 216 \| 0.148 \| 385 \| 215 \| 22 \| 1 \| 310 \| 1 \| 347 \| Uncharacterized protein \| Uncharacterized protein; afdb-uniprot50 \| AF-A0A7V5SU14-F1-MODEL\_V4 \| 1.0 \| 2.564e-08 \| 213 \| 0.128 \| 303 \| 186 \| 23 \| 11 \| 268 \| 56 \| 325 \| Uncharacterized protein \| Uncharacterized protein; afdb-uniprot50 \| AF-A0A0F8ZTA4-F1-MODEL\_V4 \| 1.0 \| 5.472e-07 \| 204 \| 0.139 \| 223 \| 154 \| 8 \| 15 \| 207 \| 6 \| 220 \| Uncharacterized protein \| Uncharacterized protein; afdb-uniprot50 \| AF-A0A1Y4D5T3-F1-MODEL\_V4 \| 1.0 \| 8.721e-08 \| 197 \| 0.156 \| 294 \| 155 \| 19 \| 1 \| 244 \| 1 \| 251 \| AAA domain-containing protein \| AAA domain-containing protein; afdb-uniprot50 \| AF-A0A7X6VZ73-F1-MODEL\_V4 \| 1.0 \| 8.54e-07 \| 195 \| 0.164 \| 225 \| 132 \| 14 \| 15 \| 214 \| 14 \| 207 \| AAA family ATPase \| AAA family ATPase; afdb-uniprot50 \| AF-A0A0P6Y9P5-F1-MODEL\_V4 \| 1.0 \| 6.603e-08 \| 194 \| 0.163 \| 319 \| 195 \| 20 \| 15 \| 298 \| 11 \| 292 \| Uncharacterized protein \| Uncharacterized protein; afdb-uniprot50 \| AF-A0A2A7EE04-F1-MODEL\_V4 \| 1.0 \| 0.0001429 \| 193 \| 0.312 \| 112 \| 68 \| 2 \| 52 \| 163 \| 1 \| 103 \| Uncharacterized protein \| Uncharacterized protein; afdb-uniprot50 \| AF-A0A2E5HJ52-F1-MODEL\_V4 \| 1.0 \| 8.54e-07 \| 186 \| 0.127 \| 275 \| 164 \| 24 \| 1 \| 246 \| 1 \| 228 \| AAA domain-containing protein \| AAA domain-containing protein; afdb-uniprot50 \| AF-A0A0F8VWM4-F1-MODEL\_V4 \| 1.0 \| 8.54e-07 \| 184 \| 0.123 \| 235 \| 141 \| 10 \| 15 \| 204 \| 10 \| 224 \| Uncharacterized protein \| Uncharacterized protein; afdb-uniprot50 \| AF-A0A420Z9U9-F1-MODEL\_V4 \| 1.0 \| 3.247e-06 \| 182 \| 0.109 \| 283 \| 177 \| 15 \| 7 \| 241 \| 40 \| 295 \| Uncharacterized protein \| Uncharacterized protein; afdb-uniprot50 \| AF-A0A4P9Y712-F1-MODEL\_V4 \| 1.0 \| 4.289e-06 \| 180 \| 0.152 \| 236 \| 141 \| 16 \| 16 \| 231 \| 43 \| 239 \| P-loop containing nucleoside triphosphate hydrolase protein \| P-loop containing nucleoside triphosphate hydrolase protein; afdb-uniprot50 \| AF-A0A374DWF3-F1-MODEL\_V4 \| 1.0 \| 1.49e-06 \| 178 \| 0.142 \| 239 \| 143 \| 11 \| 1 \| 204 \| 1 \| 212 \| Uncharacterized protein \| Uncharacterized protein; afdb-uniprot50 \| AF-A0A7S0RNJ2-F1-MODEL\_V4 \| 1.0 \| 8.842e-06 \| 177 \| 0.135 \| 221 \| 145 \| 12 \| 16 \| 216 \| 102 \| 296 \| Hypothetical protein \| Hypothetical protein; afdb-uniprot50 \| AF-A0A5E4LTZ5-F1-MODEL\_V4 \| 1.0 \| 8.363e-06 \| 175 \| 0.198 \| 217 \| 128 \| 10 \| 15 \| 219 \| 36 \| 218 \| Circadian clock protein kinase KaiC \| Circadian clock protein kinase KaiC; afdb-uniprot50 \| AF-A0A2N3LFU5-F1-MODEL\_V4 \| 1.0 \| 3.386e-08 \| 174 \| 0.142 \| 408 \| 214 \| 24 \| 1 \| 300 \| 1 \| 380 \| Uncharacterized protein \| Uncharacterized protein; afdb-uniprot50 \| AF-A0A417LQP8-F1-MODEL\_V4 \| 1.0 \| 1.861e-06 \| 172 \| 0.145 \| 240 \| 141 \| 11 \| 1 \| 204 \| 1 \| 212 \| Uncharacterized protein \| Uncharacterized protein; afdb-uniprot50 \| AF-A0A5E4LUF9-F1-MODEL\_V4 \| 1.0 \| 1.927e-05 \| 169 \| 0.128 \| 218 \| 143 \| 11 \| 12 \| 217 \| 48 \| 230 \| Circadian clock protein kinase KaiC \| Circadian clock protein kinase KaiC; afdb-uniprot50 \| AF-A0A7V1G7H0-F1-MODEL\_V4 \| 1.0 \| 4.2e-05 \| 166 \| 0.14 \| 220 \| 129 \| 11 \| 15 \| 215 \| 3 \| 181 \| Adenosylcobinamide kinase \| Adenosylcobinamide kinase; afdb-uniprot50 \| AF-A0A0F9END0-F1-MODEL\_V4 \| 1.0 \| 7.077e-06 \| 165 \| 0.101 \| 247 \| 142 \| 12 \| 14 \| 207 \| 1 \| 220 \| Uncharacterized protein \| Uncharacterized protein; afdb-uniprot50 \| AF-A0A535BIN7-F1-MODEL\_V4 \| 1.0 \| 4.694e-05 \| 163 \| 0.15 \| 220 \| 143 \| 11 \| 9 \| 219 \| 13 \| 197 \| Recombinase RecA \| Recombinase RecA; afdb-uniprot50 \| AF-A0A0F9W451-F1-MODEL\_V4 \| 1.0 \| 5.358e-06 \| 163 \| 0.131 \| 221 \| 136 \| 14 \| 6 \| 204 \| 24 \| 210 \| Uncharacterized protein \| Uncharacterized protein; afdb-uniprot50 \| AF-A0A497IGW5-F1-MODEL\_V4 \| 1.0 \| 5.865e-05 \| 161 \| 0.127 \| 244 \| 155 \| 13 \| 20 \| 229 \| 2 \| 221 \| Uncharacterized protein \| Uncharacterized protein; afdb-uniprot50 \| AF-M0PFA0-F1-MODEL\_V4 \| 1.0 \| 2.691e-05 \| 161 \| 0.155 \| 218 \| 137 \| 11 \| 15 \| 219 \| 214 \| 397 \| Circadian clock protein, KaiC \| Circadian clock protein, KaiC; afdb-uniprot50 \| AF-A0A7C2AU95-F1-MODEL\_V4 \| 1.0 \| 2.154e-05 \| 161 \| 0.175 \| 217 \| 134 \| 12 \| 12 \| 219 \| 1 \| 181 \| PAS domain S-box protein \| PAS domain S-box protein; afdb-uniprot50 \| AF-A4BYK4-F1-MODEL\_V4 \| 1.0 \| 6.2e-05 \| 160 \| 0.173 \| 207 \| 143 \| 7 \| 108 \| 307 \| 1 \| 186 \| Uncharacterized protein \| Uncharacterized protein; afdb-uniprot50 \| AF-A0A256JKU2-F1-MODEL\_V4 \| 1.0 \| 4.44e-05 \| 159 \| 0.159 \| 219 \| 137 \| 9 \| 14 \| 219 \| 33 \| 217 \| KaiC domain-containing protein \| KaiC domain-containing protein; afdb-uniprot50 \| AF-A0A5E4LUA0-F1-MODEL\_V4 \| 1.0 \| 1.542e-05 \| 159 \| 0.158 \| 234 \| 144 \| 16 \| 12 \| 219 \| 25 \| 231 \| Circadian clock protein kinase KaiC \| Circadian clock protein kinase KaiC; afdb-uniprot50 \| AF-A0A534TRW8-F1-MODEL\_V4 \| 1.0 \| 1.724e-05 \| 159 \| 0.18 \| 227 \| 135 \| 12 \| 16 \| 233 \| 266 \| 450 \| Recombinase RecA \| Recombinase RecA; afdb-uniprot50 \| AF-A0A1X4G9F5-F1-MODEL\_V4 \| 1.0 \| 5.865e-05 \| 157 \| 0.159 \| 219 \| 137 \| 10 \| 14 \| 219 \| 61 \| 245 \| KaiC domain-containing protein \| KaiC domain-containing protein; afdb-uniprot50 \| AF-A0A2E7DLC9-F1-MODEL\_V4 \| 1.0 \| 3.362e-05 \| 156 \| 0.147 \| 223 \| 139 \| 11 \| 14 \| 215 \| 17 \| 209 \| Uncharacterized protein \| Uncharacterized protein; afdb-uniprot50 \| AF-A0A7C3LMJ7-F1-MODEL\_V4 \| 1.0 \| 5.665e-06 \| 155 \| 0.161 \| 217 \| 130 \| 12 \| 11 \| 214 \| 5 \| 182 \| Uncharacterized protein \| Uncharacterized protein; afdb-uniprot50 \| AF-A0A0F9TKH8-F1-MODEL\_V4 \| 1.0 \| 1.861e-06 \| 155 \| 0.114 \| 261 \| 145 \| 15 \| 16 \| 214 \| 1 \| 237 \| Uncharacterized protein \| Uncharacterized protein; afdb-uniprot50 \| AF-A0A5E4LSB6-F1-MODEL\_V4 \| 1.0 \| 2.691e-05 \| 155 \| 0.152 \| 229 \| 132 \| 15 \| 12 \| 219 \| 23 \| 210 \| Circadian clock protein kinase KaiC \| Circadian clock protein kinase KaiC; afdb-uniprot50 \| AF-A0A5E4HZV1-F1-MODEL\_V4 \| 1.0 \| 3.18e-05 \| 155 \| 0.138 \| 217 \| 139 \| 12 \| 16 \| 219 \| 28 \| 209 \| DNA repair and recombination protein RadB \| DNA repair and recombination protein RadB; afdb-uniprot50 \| AF-A0A843BZ86-F1-MODEL\_V4 \| 1.0 \| 4.694e-05 \| 155 \| 0.172 \| 220 \| 133 \| 11 \| 12 \| 219 \| 40 \| 222 \| AAA family ATPase \| AAA family ATPase; afdb-uniprot50 \| AF-K1XPL3-F1-MODEL\_V4 \| 1.0 \| 1.305e-05 \| 154 \| 0.133 \| 254 \| 136 \| 18 \| 15 \| 214 \| 16 \| 239 \| Uncharacterized protein \| Uncharacterized protein; afdb-uniprot50 \| AF-A0A658JC52-F1-MODEL\_V4 \| 1.0 \| 1.105e-05 \| 152 \| 0.157 \| 216 \| 132 \| 14 \| 4 \| 198 \| 8 \| 194 \| Uncharacterized protein \| Uncharacterized protein; afdb-uniprot50 \| AF-A0A0F9BRG4-F1-MODEL\_V4 \| 1.0 \| 5.665e-06 \| 152 \| 0.102 \| 301 \| 173 \| 17 \| 14 \| 246 \| 1 \| 272 \| Uncharacterized protein \| Uncharacterized protein; afdb-uniprot50 \| AF-A0A662PXI2-F1-MODEL\_V4 \| 1.0 \| 4.694e-05 \| 152 \| 0.187 \| 219 \| 125 \| 14 \| 16 \| 219 \| 21 \| 201 \| KaiC domain-containing protein \| KaiC domain-containing protein; afdb-uniprot50 \| AF-A0A7V5YVJ5-F1-MODEL\_V4 \| 1.0 \| 0.000223 \| 152 \| 0.131 \| 213 \| 143 \| 9 \| 16 \| 219 \| 238 \| 417 \| Circadian clock protein KaiC \| Circadian clock protein KaiC; afdb-uniprot50 \| AF-A0A2T9XZ21-F1-MODEL\_V4 \| 1.0 \| 4.534e-06 \| 151 \| 0.15 \| 239 \| 139 \| 17 \| 10 \| 204 \| 481 \| 699 \| alpha-1,2-Mannosidase \| alpha-1,2-Mannosidase; afdb-uniprot50 \| AF-A0A2D6SN79-F1-MODEL\_V4 \| 1.0 \| 3.758e-05 \| 150 \| 0.142 \| 217 \| 141 \| 11 \| 16 \| 219 \| 25 \| 209 \| KaiC domain-containing protein \| KaiC domain-containing protein; afdb-uniprot50 \| AF-A0A6M3KQ16-F1-MODEL\_V4 \| 1.0 \| 4.794e-06 \| 150 \| 0.134 \| 215 \| 132 \| 10 \| 9 \| 204 \| 40 \| 219 \| Putative ATPase domain containing protein \| Putative ATPase domain containing protein; afdb-uniprot50 \| AF-A0A419KUX3-F1-MODEL\_V4 \| 1.0 \| 1.823e-05 \| 149 \| 0.161 \| 235 \| 132 \| 15 \| 12 \| 231 \| 32 \| 216 \| KaiC domain-containing protein \| KaiC domain-containing protein; afdb-uniprot50 \| AF-A0A1H6WCT0-F1-MODEL\_V4 \| 1.0 \| 9.678e-05 \| 149 \| 0.165 \| 218 \| 135 \| 11 \| 15 \| 219 \| 331 \| 514 \| KaiC domain protein, AF\_0351 family \| KaiC domain protein, AF\_0351 family; afdb-uniprot50 \| AF-A0A7C5RLF6-F1-MODEL\_V4 \| 1.0 \| 4.694e-05 \| 148 \| 0.147 \| 224 \| 134 \| 15 \| 12 \| 219 \| 22 \| 204 \| KaiC domain-containing protein \| KaiC domain-containing protein; afdb-uniprot50 \| AF-A0A497IAX5-F1-MODEL\_V4 \| 1.0 \| 4.44e-05 \| 148 \| 0.147 \| 223 \| 137 \| 12 \| 12 \| 219 \| 28 \| 212 \| KaiC domain-containing protein \| KaiC domain-containing protein; afdb-uniprot50 \| AF-A0A2R4X0W3-F1-MODEL\_V4 \| 1.0 \| 5.547e-05 \| 148 \| 0.177 \| 220 \| 127 \| 13 \| 16 \| 219 \| 94 \| 275 \| KaiC domain-containing protein \| KaiC domain-containing protein; afdb-uniprot50 \| AF-A0A897MUN6-F1-MODEL\_V4 \| 1.0 \| 4.2e-05 \| 148 \| 0.176 \| 227 \| 124 \| 15 \| 14 \| 219 \| 203 \| 387 \| RecA-superfamily ATPase implicated in signal transduction \| RecA-superfamily ATPase implicated in signal transduction; afdb-uniprot50 \| AF-A0A1D2QW89-F1-MODEL\_V4 \| 1.0 \| 9.678e-05 \| 148 \| 0.168 \| 220 \| 129 \| 10 \| 16 \| 220 \| 265 \| 445 \| Uncharacterized protein \| Uncharacterized protein; afdb-uniprot50 \| AF-A0A7C3ZCY4-F1-MODEL\_V4 \| 1.0 \| 4.694e-05 \| 147 \| 0.156 \| 217 \| 135 \| 12 \| 16 \| 219 \| 73 \| 254 \| KaiC domain-containing protein \| KaiC domain-containing protein; afdb-uniprot50 \| AF-A0A5N5UE56-F1-MODEL\_V4 \| 1.0 \| 6.555e-05 \| 147 \| 0.18 \| 222 \| 128 \| 12 \| 14 \| 219 \| 177 \| 360 \| KaiC domain-containing protein \| KaiC domain-containing protein; afdb-uniprot50 \| AF-A0A842X9A4-F1-MODEL\_V4 \| 1.0 \| 5.865e-05 \| 146 \| 0.18 \| 200 \| 115 \| 13 \| 16 \| 204 \| 25 \| 186 \| Uncharacterized protein \| Uncharacterized protein; afdb-uniprot50 \| AF-A0A7C6EJA8-F1-MODEL\_V4 \| 1.0 \| 5.547e-05 \| 146 \| 0.114 \| 218 \| 140 \| 11 \| 16 \| 219 \| 28 \| 206 \| KaiC domain-containing protein \| KaiC domain-containing protein; afdb-uniprot50 \| AF-A0A238V873-F1-MODEL\_V4 \| 1.0 \| 6.2e-05 \| 146 \| 0.175 \| 228 \| 121 \| 14 \| 15 \| 219 \| 254 \| 437 \| KaiC domain protein, AF\_0351 family \| KaiC domain protein, AF\_0351 family; afdb-uniprot50 \| AF-A0A662Q2C8-F1-MODEL\_V4 \| 1.0 \| 2.407e-05 \| 146 \| 0.198 \| 227 \| 115 \| 18 \| 12 \| 217 \| 23 \| 203 \| KaiC domain-containing protein \| KaiC domain-containing protein; afdb-uniprot50 \| AF-A0A832YZT3-F1-MODEL\_V4 \| 1.0 \| 0.0001351 \| 146 \| 0.158 \| 196 \| 124 \| 9 \| 15 \| 204 \| 24 \| 184 \| Uncharacterized protein \| Uncharacterized protein; afdb-uniprot50 \| AF-A0A846PET3-F1-MODEL\_V4 \| 1.0 \| 0.0001429 \| 145 \| 0.167 \| 221 \| 132 \| 12 \| 12 \| 219 \| 30 \| 211 \| AAA family ATPase \| AAA family ATPase; afdb-uniprot50 \| AF-A0A7J2ZGV2-F1-MODEL\_V4 \| 1.0 \| 3.973e-05 \| 145 \| 0.186 \| 231 \| 131 \| 13 \| 16 \| 233 \| 45 \| 231 \| KaiC domain-containing protein \| KaiC domain-containing protein; afdb-uniprot50 \| AF-A0A842SU21-F1-MODEL\_V4 \| 1.0 \| 9.154e-05 \| 145 \| 0.152 \| 223 \| 139 \| 11 \| 12 \| 219 \| 24 \| 211 \| Uncharacterized protein \| Uncharacterized protein; afdb-uniprot50 \| AF-A0A4U7CWT0-F1-MODEL\_V4 \| 1.0 \| 8.658e-05 \| 145 \| 0.182 \| 225 \| 123 \| 15 \| 15 \| 219 \| 101 \| 284 \| KaiC domain-containing protein \| KaiC domain-containing protein; afdb-uniprot50 \| AF-A0A6C0UJT1-F1-MODEL\_V4 \| 1.0 \| 9.154e-05 \| 145 \| 0.174 \| 223 \| 133 \| 13 \| 10 \| 219 \| 210 \| 394 \| KaiC domain-containing protein \| KaiC domain-containing protein; afdb-uniprot50 \| AF-A0A662Q6K3-F1-MODEL\_V4 \| 1.0 \| 8.19e-05 \| 144 \| 0.15 \| 232 \| 128 \| 14 \| 9 \| 219 \| 18 \| 201 \| KaiC domain-containing protein \| KaiC domain-containing protein; afdb-uniprot50 \| AF-A0A497IXB1-F1-MODEL\_V4 \| 1.0 \| 6.555e-05 \| 144 \| 0.164 \| 231 \| 119 \| 13 \| 15 \| 219 \| 22 \| 204 \| KaiC domain-containing protein \| KaiC domain-containing protein; afdb-uniprot50 \| AF-A0A497KE38-F1-MODEL\_V4 \| 1.0 \| 0.0001429 \| 144 \| 0.151 \| 204 \| 120 \| 13 \| 12 \| 204 \| 23 \| 184 \| Circadian clock KaiC-like protein \| Circadian clock KaiC-like protein; afdb-uniprot50 \| AF-A0A2E6QQJ8-F1-MODEL\_V4 \| 1.0 \| 0.0001278 \| 144 \| 0.17 \| 217 \| 130 \| 13 \| 16 \| 219 \| 230 \| 409 \| KaiC domain-containing protein \| KaiC domain-containing protein; afdb-uniprot50 \| AF-A0A1G3MAD4-F1-MODEL\_V4 \| 1.0 \| 6.555e-05 \| 144 \| 0.177 \| 209 \| 126 \| 12 \| 12 \| 204 \| 105 \| 283 \| RECA\_3 domain-containing protein \| RECA\_3 domain-containing protein; afdb-uniprot50 \| AF-A0A2E1XF51-F1-MODEL\_V4 \| 1.0 \| 0.000151 \| 144 \| 0.146 \| 218 \| 137 \| 12 \| 11 \| 217 \| 260 \| 439 \| Serine/threonine protein kinase \| Serine/threonine protein kinase; afdb-uniprot50 \| AF-A0A5E4IP08-F1-MODEL\_V4 \| 1.0 \| 0.0001144 \| 143 \| 0.156 \| 224 \| 134 \| 11 \| 12 \| 219 \| 66 \| 250 \| DNA repair and recombination protein RadB \| DNA repair and recombination protein RadB; afdb-uniprot50 \| AF-A0A1C2JI50-F1-MODEL\_V4 \| 1.0 \| 3.837e-06 \| 143 \| 0.134 \| 245 \| 137 \| 15 \| 4 \| 215 \| 7 \| 209 \| Uncharacterized protein \| Uncharacterized protein; afdb-uniprot50 \| AF-A0A7C4HE07-F1-MODEL\_V4 \| 1.0 \| 0.0001995 \| 143 \| 0.131 \| 220 \| 140 \| 11 \| 12 \| 219 \| 21 \| 201 \| KaiC domain-containing protein \| KaiC domain-containing protein; afdb-uniprot50 \| AF-A0A497FEI0-F1-MODEL\_V4 \| 1.0 \| 6.555e-05 \| 143 \| 0.171 \| 227 \| 124 \| 15 \| 9 \| 219 \| 17 \| 195 \| Uncharacterized protein \| Uncharacterized protein; afdb-uniprot50 \| AF-A0A497KSX2-F1-MODEL\_V4 \| 1.0 \| 0.000151 \| 143 \| 0.149 \| 214 \| 132 \| 13 \| 15 \| 217 \| 8 \| 182 \| KaiC domain-containing protein \| KaiC domain-containing protein; afdb-uniprot50 \| AF-A0A7C7QD37-F1-MODEL\_V4 \| 1.0 \| 8.19e-05 \| 142 \| 0.157 \| 228 \| 121 \| 16 \| 16 \| 219 \| 26 \| 206 \| KaiC domain-containing protein \| KaiC domain-containing protein; afdb-uniprot50 \| AF-A0A2H9L029-F1-MODEL\_V4 \| 1.0 \| 2.691e-05 \| 142 \| 0.16 \| 237 \| 130 \| 14 \| 16 \| 233 \| 25 \| 211 \| KaiC domain-containing protein \| KaiC domain-containing protein; afdb-uniprot50 \| AF-A0A7C1VTC4-F1-MODEL\_V4 \| 1.0 \| 5.865e-05 \| 142 \| 0.186 \| 204 \| 130 \| 11 \| 12 \| 204 \| 61 \| 239 \| Uncharacterized protein \| Uncharacterized protein; afdb-uniprot50 \| AF-A0A151Z912-F1-MODEL\_V4 \| 1.0 \| 0.000223 \| 141 \| 0.108 \| 249 \| 154 \| 15 \| 12 \| 217 \| 106 \| 329 \| Putative DNA repair protein \| Putative DNA repair protein; afdb-uniprot50 \| AF-A0A842U289-F1-MODEL\_V4 \| 1.0 \| 3.758e-05 \| 140 \| 0.129 \| 217 \| 130 \| 14 \| 13 \| 204 \| 17 \| 199 \| Uncharacterized protein \| Uncharacterized protein; afdb-uniprot50 \| AF-A0A662SDZ3-F1-MODEL\_V4 \| 1.0 \| 0.0003292 \| 140 \| 0.149 \| 214 \| 132 \| 10 \| 16 \| 219 \| 26 \| 199 \| KaiC domain-containing protein \| KaiC domain-containing protein; afdb-uniprot50 \| AF-A0A7J4SSQ8-F1-MODEL\_V4 \| 1.0 \| 6.555e-05 \| 140 \| 0.126 \| 245 \| 153 \| 12 \| 1 \| 219 \| 1 \| 210 \| KaiC domain-containing protein \| KaiC domain-containing protein; afdb-uniprot50 \| AF-A0A5E4LAZ7-F1-MODEL\_V4 \| 1.0 \| 0.0001209 \| 140 \| 0.188 \| 217 \| 129 \| 13 \| 16 \| 219 \| 25 \| 207 \| Circadian clock protein kinase KaiC \| Circadian clock protein kinase KaiC; afdb-uniprot50 \| AF-A0A534MTG6-F1-MODEL\_V4 \| 1.0 \| 1.823e-05 \| 140 \| 0.154 \| 278 \| 143 \| 17 \| 1 \| 235 \| 8 \| 236 \| KaiC domain-containing protein \| KaiC domain-containing protein; afdb-uniprot50 \| AF-A0A357C3H2-F1-MODEL\_V4 \| 1.0 \| 1.459e-05 \| 140 \| 0.144 \| 243 \| 146 \| 15 \| 5 \| 214 \| 1 \| 214 \| Uncharacterized protein \| Uncharacterized protein; afdb-uniprot50 \| AF-A0A354WT19-F1-MODEL\_V4 \| 1.0 \| 1.128e-06 \| 140 \| 0.123 \| 380 \| 205 \| 21 \| 4 \| 296 \| 2 \| 340 \| Uncharacterized protein \| Uncharacterized protein; afdb-uniprot50 \| AF-A0A7C5XQS7-F1-MODEL\_V4 \| 1.0 \| 8.658e-05 \| 140 \| 0.179 \| 217 \| 131 \| 14 \| 12 \| 217 \| 22 \| 202 \| KaiC domain-containing protein \| KaiC domain-containing protein; afdb-uniprot50 \| AF-A0A133UDP6-F1-MODEL\_V4 \| 1.0 \| 0.0001209 \| 139 \| 0.144 \| 222 \| 131 \| 14 \| 15 \| 219 \| 23 \| 202 \| KaiC domain-containing protein \| KaiC domain-containing protein; afdb-uniprot50 \| AF-A0A7C3V282-F1-MODEL\_V4 \| 1.0 \| 0.0003292 \| 139 \| 0.146 \| 218 \| 134 \| 10 \| 16 \| 219 \| 29 \| 208 \| KaiC domain-containing protein \| KaiC domain-containing protein; afdb-uniprot50 \| AF-A0A1M4ZBV9-F1-MODEL\_V4 \| 1.0 \| 4.694e-05 \| 139 \| 0.127 \| 251 \| 155 \| 15 \| 1 \| 231 \| 36 \| 242 \| AAA domain-containing protein \| AAA domain-containing protein; afdb-uniprot50 \| AF-A0A7Y6UET4-F1-MODEL\_V4 \| 1.0 \| 5.247e-05 \| 139 \| 0.144 \| 229 \| 142 \| 13 \| 16 \| 233 \| 263 \| 448 \| AAA family ATPase \| AAA family ATPase; afdb-uniprot50 \| AF-A0A5E4LUJ3-F1-MODEL\_V4 \| 1.0 \| 0.0002109 \| 138 \| 0.155 \| 225 \| 132 \| 13 \| 12 \| 219 \| 21 \| 204 \| DNA repair and recombination protein RadB \| DNA repair and recombination protein RadB; afdb-uniprot50 \| AF-A0A3N5NHE5-F1-MODEL\_V4 \| 1.0 \| 6.2e-05 \| 138 \| 0.171 \| 210 \| 115 \| 13 \| 12 \| 191 \| 24 \| 204 \| Protein kinase \| Protein kinase; afdb-uniprot50 \| AF-A0A662U9F7-F1-MODEL\_V4 \| 1.0 \| 0.0001082 \| 138 \| 0.151 \| 225 \| 134 \| 14 \| 12 \| 219 \| 25 \| 209 \| Uncharacterized protein \| Uncharacterized protein; afdb-uniprot50 \| AF-A0A7W1H037-F1-MODEL\_V4 \| 1.0 \| 0.000348 \| 137 \| 0.178 \| 218 \| 129 \| 13 \| 16 \| 219 \| 7 \| 188 \| Protein kinase \| Protein kinase; afdb-uniprot50 \| AF-A0A4Q3GU50-F1-MODEL\_V4 \| 1.0 \| 0.0001278 \| 137 \| 0.16 \| 218 \| 139 \| 12 \| 11 \| 219 \| 13 \| 195 \| Circadian clock protein KaiC \| Circadian clock protein KaiC; afdb-uniprot50 \| AF-A0A660V9I0-F1-MODEL\_V4 \| 1.0 \| 5.247e-05 \| 137 \| 0.17 \| 229 \| 122 \| 15 \| 12 \| 219 \| 186 \| 367 \| Uncharacterized protein \| Uncharacterized protein; afdb-uniprot50 \| AF-A0A1Q7P4E9-F1-MODEL\_V4 \| 1.0 \| 0.0001785 \| 137 \| 0.182 \| 225 \| 118 \| 15 \| 16 \| 219 \| 24 \| 203 \| KaiC domain-containing protein \| KaiC domain-containing protein; afdb-uniprot50 \| AF-A0A6J4UUK0-F1-MODEL\_V4 \| 1.0 \| 3.973e-05 \| 137 \| 0.15 \| 232 \| 128 \| 15 \| 12 \| 219 \| 288 \| 474 \| Uncharacterized protein \| Uncharacterized protein; afdb-uniprot50 \| AF-A0A3D4ZSM3-F1-MODEL\_V4 \| 1.0 \| 8.19e-05 \| 137 \| 0.157 \| 222 \| 127 \| 12 \| 16 \| 219 \| 259 \| 438 \| Uncharacterized protein \| Uncharacterized protein; afdb-uniprot50 \| AF-A0A7C4EZ54-F1-MODEL\_V4 \| 1.0 \| 0.0001429 \| 136 \| 0.146 \| 226 \| 131 \| 15 \| 12 \| 219 \| 21 \| 202 \| KaiC domain-containing protein \| KaiC domain-containing protein; afdb-uniprot50 \| AF-A0A353NQU0-F1-MODEL\_V4 \| 1.0 \| 0.0001995 \| 136 \| 0.153 \| 234 \| 137 \| 13 \| 12 \| 233 \| 21 \| 205 \| ATPase \| ATPase; afdb-uniprot50 \| AF-A0A180F7Q6-F1-MODEL\_V4 \| 1.0 \| 0.0001597 \| 135 \| 0.161 \| 229 \| 143 \| 14 \| 3 \| 219 \| 43 \| 234 \| Uncharacterized protein \| Uncharacterized protein; afdb-uniprot50 \| AF-A0A848LTY7-F1-MODEL\_V4 \| 1.0 \| 0.0001429 \| 135 \| 0.15 \| 239 \| 141 \| 13 \| 16 \| 235 \| 267 \| 462 \| AAA family ATPase \| AAA family ATPase; afdb-uniprot50 \| AF-A0A101DYX4-F1-MODEL\_V4 \| 1.0 \| 8.658e-05 \| 134 \| 0.168 \| 226 \| 134 \| 14 \| 11 \| 219 \| 14 \| 202 \| KaiC domain-containing protein \| KaiC domain-containing protein; afdb-uniprot50 \| AF-A0A345E6S3-F1-MODEL\_V4 \| 1.0 \| 0.0001082 \| 134 \| 0.167 \| 227 \| 124 \| 14 \| 15 \| 219 \| 151 \| 334 \| KaiC domain-containing protein \| KaiC domain-containing protein; afdb-uniprot50 \| AF-A0A7J2HFW6-F1-MODEL\_V4 \| 1.0 \| 0.000151 \| 134 \| 0.183 \| 218 \| 127 \| 14 \| 16 \| 219 \| 21 \| 201 \| KaiC domain-containing protein \| KaiC domain-containing protein; afdb-uniprot50 \| AF-A0A7J3XLY6-F1-MODEL\_V4 \| 1.0 \| 0.0001688 \| 134 \| 0.139 \| 223 \| 135 \| 14 \| 12 \| 219 \| 159 \| 339 \| KaiC domain-containing protein \| KaiC domain-containing protein; afdb-uniprot50 \| AF-A0A7C5UV01-F1-MODEL\_V4 \| 1.0 \| 0.0001887 \| 133 \| 0.159 \| 244 \| 138 \| 16 \| 16 \| 237 \| 22 \| 220 \| KaiC domain-containing protein \| KaiC domain-containing protein; afdb-uniprot50 \| AF-A0A1F5DE45-F1-MODEL\_V4 \| 1.0 \| 0.0002492 \| 133 \| 0.151 \| 218 \| 134 \| 10 \| 16 \| 219 \| 24 \| 204 \| KaiC domain-containing protein \| KaiC domain-containing protein; afdb-uniprot50 \| AF-A0A3C1QFQ6-F1-MODEL\_V4 \| 1.0 \| 6.93e-05 \| 133 \| 0.143 \| 293 \| 168 \| 18 \| 15 \| 244 \| 20 \| 292 \| Uncharacterized protein \| Uncharacterized protein; afdb-uniprot50 \| AF-A0A7W0YP95-F1-MODEL\_V4 \| 1.0 \| 1.105e-05 \| 133 \| 0.168 \| 321 \| 178 \| 21 \| 9 \| 287 \| 14 \| 287 \| AAA family ATPase \| AAA family ATPase; afdb-uniprot50 \| AF-A0A5D5ANE0-F1-MODEL\_V4 \| 1.0 \| 0.0002786 \| 133 \| 0.168 \| 220 \| 128 \| 10 \| 15 \| 217 \| 152 \| 333 \| KaiC domain-containing protein \| KaiC domain-containing protein; afdb-uniprot50 \| AF-A0A1Y2E1Y0-F1-MODEL\_V4 \| 1.0 \| 0.0001023 \| 133 \| 0.151 \| 238 \| 140 \| 16 \| 12 \| 217 \| 72 \| 279 \| p-loop containing nucleoside triphosphate hydrolase protein \| p-loop containing nucleoside triphosphate hydrolase protein; afdb-uniprot50 \| AF-A0A838ULP5-F1-MODEL\_V4 \| 1.0 \| 0.0007175 \| 133 \| 0.179 \| 217 \| 131 \| 15 \| 16 \| 217 \| 279 \| 463 \| AAA family ATPase \| AAA family ATPase; afdb-uniprot50 \| AF-A0A533RVL5-F1-MODEL\_V4 \| 1.0 \| 0.000223 \| 132 \| 0.15 \| 199 \| 119 \| 12 \| 16 \| 204 \| 24 \| 182 \| KaiC domain-containing protein \| KaiC domain-containing protein; afdb-uniprot50 \| AF-A0A7C3A7E9-F1-MODEL\_V4 \| 1.0 \| 0.000151 \| 132 \| 0.161 \| 235 \| 119 \| 14 \| 12 \| 219 \| 21 \| 204 \| KaiC domain-containing protein \| KaiC domain-containing protein; afdb-uniprot50 \| AF-A0A2R6IYK1-F1-MODEL\_V4 \| 1.0 \| 0.0002945 \| 132 \| 0.18 \| 221 \| 125 \| 14 \| 14 \| 217 \| 34 \| 215 \| KaiC domain-containing protein \| KaiC domain-containing protein; afdb-uniprot50 \| AF-A0A4Q3K7Q1-F1-MODEL\_V4 \| 1.0 \| 0.0005432 \| 132 \| 0.17 \| 217 \| 131 \| 16 \| 12 \| 217 \| 34 \| 212 \| Circadian clock protein KaiC \| Circadian clock protein KaiC; afdb-uniprot50 \| AF-A0A7J3MCA1-F1-MODEL\_V4 \| 1.0 \| 0.0002945 \| 132 \| 0.169 \| 230 \| 124 \| 18 \| 11 \| 219 \| 18 \| 201 \| KaiC domain-containing protein \| KaiC domain-containing protein; afdb-uniprot50 \| AF-A0A349YSR0-F1-MODEL\_V4 \| 1.0 \| 1.542e-05 \| 132 \| 0.12 \| 416 \| 220 \| 21 \| 1 \| 300 \| 1 \| 386 \| Uncharacterized protein \| Uncharacterized protein; afdb-uniprot50 \| AF-A0A662Q5Z7-F1-MODEL\_V4 \| 1.0 \| 0.0001082 \| 132 \| 0.147 \| 223 \| 127 \| 14 \| 12 \| 217 \| 24 \| 200 \| KaiC domain-containing protein \| KaiC domain-containing protein; afdb-uniprot50 \| AF-A0A419JWC1-F1-MODEL\_V4 \| 1.0 \| 0.000368 \| 132 \| 0.174 \| 218 \| 129 \| 13 \| 16 \| 219 \| 121 \| 301 \| KaiC domain-containing protein \| KaiC domain-containing protein; afdb-uniprot50 \| AF-O29797-F1-MODEL\_V4 \| 1.0 \| 0.0001278 \| 132 \| 0.17 \| 223 \| 132 \| 14 \| 11 \| 219 \| 19 \| 202 \| Uncharacterized protein \| Uncharacterized protein; afdb-uniprot50 \| AF-A0A662IS64-F1-MODEL\_V4 \| 1.0 \| 0.000151 \| 132 \| 0.137 \| 225 \| 139 \| 14 \| 11 \| 219 \| 12 \| 197 \| KaiC domain-containing protein \| KaiC domain-containing protein; afdb-uniprot50 \| AF-A0A534VV70-F1-MODEL\_V4 \| 1.0 \| 0.000486 \| 132 \| 0.119 \| 218 \| 135 \| 12 \| 18 \| 219 \| 2 \| 178 \| Uncharacterized protein \| Uncharacterized protein; afdb-uniprot50 \| AF-A0A7J3BH97-F1-MODEL\_V4 \| 1.0 \| 0.0001688 \| 132 \| 0.157 \| 229 \| 123 \| 15 \| 10 \| 216 \| 19 \| 199 \| KaiC domain-containing protein \| KaiC domain-containing protein; afdb-uniprot50 \| AF-A0A497P1I2-F1-MODEL\_V4 \| 1.0 \| 0.0001887 \| 132 \| 0.191 \| 225 \| 116 \| 14 \| 16 \| 219 \| 21 \| 200 \| KaiC domain-containing protein \| KaiC domain-containing protein; afdb-uniprot50 \| AF-A0A7C5TP83-F1-MODEL\_V4 \| 1.0 \| 0.0001887 \| 132 \| 0.148 \| 222 \| 136 \| 13 \| 12 \| 219 \| 39 \| 221 \| KaiC domain-containing protein \| KaiC domain-containing protein; afdb-uniprot50 \| AF-A0A7J3LXP7-F1-MODEL\_V4 \| 1.0 \| 0.0001887 \| 132 \| 0.156 \| 223 \| 127 \| 13 \| 12 \| 216 \| 105 \| 284 \| KaiC domain-containing protein \| KaiC domain-containing protein; afdb-uniprot50 \| AF-A0A7C3R271-F1-MODEL\_V4 \| 1.0 \| 0.0002635 \| 131 \| 0.143 \| 230 \| 139 \| 12 \| 16 \| 233 \| 24 \| 207 \| KaiC domain-containing protein \| KaiC domain-containing protein; afdb-uniprot50 \| AF-A0A7J2Z1S8-F1-MODEL\_V4 \| 1.0 \| 0.0007586 \| 131 \| 0.111 \| 215 \| 148 \| 9 \| 16 \| 219 \| 36 \| 218 \| KaiC domain-containing protein \| KaiC domain-containing protein; afdb-uniprot50 \| AF-A0A842UGP0-F1-MODEL\_V4 \| 1.0 \| 0.000223 \| 131 \| 0.119 \| 252 \| 147 \| 15 \| 12 \| 233 \| 30 \| 236 \| Uncharacterized protein \| Uncharacterized protein; afdb-uniprot50 \| AF-A0A5E4HXF3-F1-MODEL\_V4 \| 1.0 \| 0.0001995 \| 131 \| 0.149 \| 221 \| 132 \| 14 \| 16 \| 219 \| 27 \| 208 \| Circadian clock protein kinase KaiC \| Circadian clock protein kinase KaiC; afdb-uniprot50 \| AF-A0A838RIV3-F1-MODEL\_V4 \| 1.0 \| 2.407e-05 \| 131 \| 0.175 \| 325 \| 177 \| 23 \| 12 \| 297 \| 83 \| 355 \| AAA family ATPase \| AAA family ATPase; afdb-uniprot50 \| AF-A0A2G4DS72-F1-MODEL\_V4 \| 1.0 \| 0.0001429 \| 131 \| 0.151 \| 258 \| 139 \| 14 \| 16 \| 251 \| 201 \| 400 \| Serine/threonine protein kinase \| Serine/threonine protein kinase; afdb-uniprot50 \| AF-A0A7X8FBR8-F1-MODEL\_V4 \| 1.0 \| 9.154e-05 \| 131 \| 0.146 \| 225 \| 132 \| 13 \| 11 \| 219 \| 20 \| 200 \| AAA family ATPase \| AAA family ATPase; afdb-uniprot50 \| AF-F5Y0R5-F1-MODEL\_V4 \| 1.0 \| 9.154e-05 \| 131 \| 0.17 \| 235 \| 132 \| 14 \| 12 \| 233 \| 264 \| 448 \| KaiC domain-containing protein \| KaiC domain-containing protein; afdb-uniprot50 \| AF-A0A497L1L9-F1-MODEL\_V4 \| 1.0 \| 0.000389 \| 131 \| 0.142 \| 225 \| 133 \| 15 \| 12 \| 219 \| 23 \| 204 \| KaiC domain-containing protein \| KaiC domain-containing protein; afdb-uniprot50 \| AF-A0A497EX83-F1-MODEL\_V4 \| 1.0 \| 0.0002109 \| 130 \| 0.16 \| 218 \| 127 \| 13 \| 16 \| 217 \| 24 \| 201 \| KaiC domain-containing protein \| KaiC domain-containing protein; afdb-uniprot50 \| AF-I7ZAC5-F1-MODEL\_V4 \| 1.0 \| 0.0002109 \| 130 \| 0.15 \| 226 \| 134 \| 15 \| 8 \| 217 \| 3 \| 186 \| KaiC domain-containing protein \| KaiC domain-containing protein; afdb-uniprot50 \| AF-A0A7J2T6C6-F1-MODEL\_V4 \| 1.0 \| 0.0004597 \| 130 \| 0.115 \| 216 \| 145 \| 11 \| 16 \| 219 \| 25 \| 206 \| KaiC domain-containing protein \| KaiC domain-containing protein; afdb-uniprot50 \| AF-A0A2D6XGW7-F1-MODEL\_V4 \| 1.0 \| 2.691e-05 \| 130 \| 0.117 \| 315 \| 177 \| 20 \| 1 \| 246 \| 6 \| 288 \| Uncharacterized protein \| Uncharacterized protein; afdb-uniprot50 \| AF-A0A497JUW8-F1-MODEL\_V4 \| 1.0 \| 0.000348 \| 130 \| 0.149 \| 227 \| 131 \| 15 \| 12 \| 219 \| 23 \| 206 \| KaiC domain-containing protein \| KaiC domain-containing protein; afdb-uniprot50 \| AF-A0A847TZG1-F1-MODEL\_V4 \| 1.0 \| 0.0002786 \| 130 \| 0.177 \| 225 \| 128 \| 15 \| 12 \| 219 \| 199 \| 383 \| KaiC domain-containing protein \| KaiC domain-containing protein; afdb-uniprot50 \| AF-A0A415N119-F1-MODEL\_V4 \| 1.0 \| 5.989e-06 \| 130 \| 0.118 \| 353 \| 177 \| 17 \| 14 \| 280 \| 1 \| 305 \| Uncharacterized protein \| Uncharacterized protein; afdb-uniprot50 \| AF-A0A838IMW4-F1-MODEL\_V4 \| 1.0 \| 0.0001023 \| 130 \| 0.159 \| 220 \| 136 \| 14 \| 12 \| 219 \| 255 \| 437 \| Uncharacterized protein \| Uncharacterized protein; afdb-uniprot50 \| AF-A0A7V5Z082-F1-MODEL\_V4 \| 1.0 \| 0.0002635 \| 130 \| 0.157 \| 229 \| 116 \| 15 \| 16 \| 219 \| 261 \| 437 \| ATPase domain-containing protein \| ATPase domain-containing protein; afdb-uniprot50 \| AF-A0A497LB97-F1-MODEL\_V4 \| 1.0 \| 0.0002358 \| 130 \| 0.169 \| 218 \| 130 \| 11 \| 16 \| 219 \| 24 \| 204 \| KaiC domain-containing protein \| KaiC domain-containing protein; afdb-uniprot50 \| AF-A0A7C5QII8-F1-MODEL\_V4 \| 1.0 \| 6.2e-05 \| 130 \| 0.144 \| 256 \| 138 \| 16 \| 1 \| 219 \| 234 \| 445 \| Circadian clock protein KaiC \| Circadian clock protein KaiC; afdb-uniprot50 \| AF-A0A3D0N2Q1-F1-MODEL\_V4 \| 1.0 \| 0.0002492 \| 130 \| 0.152 \| 230 \| 138 \| 13 \| 16 \| 233 \| 262 \| 446 \| Uncharacterized protein \| Uncharacterized protein; afdb-uniprot50 \| AF-A0A1Q7ZS30-F1-MODEL\_V4 \| 1.0 \| 0.0001785 \| 129 \| 0.173 \| 225 \| 133 \| 13 \| 9 \| 219 \| 20 \| 205 \| KaiC domain-containing protein \| KaiC domain-containing protein; afdb-uniprot50 \| AF-A0A7T9DJA6-F1-MODEL\_V4 \| 1.0 \| 0.0002945 \| 129 \| 0.136 \| 227 \| 132 \| 15 \| 16 \| 219 \| 40 \| 225 \| KaiC domain-containing protein \| KaiC domain-containing protein; afdb-uniprot50 \| AF-A0A3R7AIS7-F1-MODEL\_V4 \| 1.0 \| 0.0004113 \| 129 \| 0.144 \| 228 \| 132 \| 13 \| 10 \| 219 \| 23 \| 205 \| Uncharacterized protein \| Uncharacterized protein; afdb-uniprot50 \| AF-A0A7J2U3Z7-F1-MODEL\_V4 \| 1.0 \| 0.000486 \| 129 \| 0.125 \| 223 \| 136 \| 11 \| 15 \| 219 \| 20 \| 201 \| KaiC domain-containing protein \| KaiC domain-containing protein; afdb-uniprot50 \| AF-A0A419KUI7-F1-MODEL\_V4 \| 1.0 \| 6.93e-05 \| 129 \| 0.158 \| 227 \| 124 \| 19 \| 12 \| 217 \| 23 \| 203 \| KaiC domain-containing protein \| KaiC domain-containing protein; afdb-uniprot50 \| AF-A0A2V7X6T5-F1-MODEL\_V4 \| 1.0 \| 0.0001351 \| 129 \| 0.131 \| 259 \| 146 \| 16 \| 4 \| 233 \| 242 \| 450 \| Uncharacterized protein \| Uncharacterized protein; afdb-uniprot50 \| AF-A0A6B0SKH5-F1-MODEL\_V4 \| 1.0 \| 0.000389 \| 128 \| 0.157 \| 242 \| 127 \| 13 \| 5 \| 219 \| 1 \| 192 \| KaiC domain-containing protein \| KaiC domain-containing protein; afdb-uniprot50 \| AF-Q3IR97-F1-MODEL\_V4 \| 1.0 \| 0.0001144 \| 128 \| 0.166 \| 234 \| 121 \| 15 \| 10 \| 219 \| 84 \| 267 \| KaiC domain protein \| KaiC domain protein; afdb-uniprot50 \| AF-A0A109C679-F1-MODEL\_V4 \| 1.0 \| 0.0002358 \| 128 \| 0.135 \| 222 \| 137 \| 15 \| 12 \| 219 \| 49 \| 229 \| Circadian clock protein KaiC \| Circadian clock protein KaiC; afdb-uniprot50 \| AF-A0A422QQ32-F1-MODEL\_V4 \| 1.0 \| 0.0001785 \| 128 \| 0.151 \| 225 \| 130 \| 16 \| 12 \| 219 \| 128 \| 308 \| KaiC domain-containing protein \| KaiC domain-containing protein; afdb-uniprot50 \| AF-A0A832G139-F1-MODEL\_V4 \| 1.0 \| 0.0005432 \| 128 \| 0.144 \| 215 \| 134 \| 12 \| 16 \| 216 \| 36 \| 214 \| Uncharacterized protein \| Uncharacterized protein; afdb-uniprot50 \| AF-E4RMH3-F1-MODEL\_V4 \| 1.0 \| 3.758e-05 \| 128 \| 0.143 \| 251 \| 156 \| 13 \| 12 \| 232 \| 258 \| 479 \| Non-specific serine/threonine protein kinase \| Non-specific serine/threonine protein kinase; afdb-uniprot50 \| AF-A0A534NIL1-F1-MODEL\_V4 \| 1.0 \| 0.0003114 \| 128 \| 0.118 \| 254 \| 146 \| 15 \| 1 \| 219 \| 189 \| 399 \| Uncharacterized protein \| Uncharacterized protein; afdb-uniprot50 \| AF-A0A349M2U1-F1-MODEL\_V4 \| 1.0 \| 0.0004113 \| 127 \| 0.166 \| 234 \| 133 \| 16 \| 16 \| 233 \| 46 \| 233 \| Circadian clock protein KaiC \| Circadian clock protein KaiC; afdb-uniprot50 \| AF-A0A5C3M5K1-F1-MODEL\_V4 \| 1.0 \| 0.001653 \| 127 \| 0.146 \| 232 \| 144 \| 14 \| 15 \| 219 \| 101 \| 305 \| P-loop containing nucleoside triphosphate hydrolase protein \| P-loop containing nucleoside triphosphate hydrolase protein; afdb-uniprot50 \| AF-A0A7C2L107-F1-MODEL\_V4 \| 1.0 \| 0.0004597 \| 127 \| 0.126 \| 222 \| 139 \| 12 \| 12 \| 219 \| 26 \| 206 \| KaiC domain-containing protein \| KaiC domain-containing protein; afdb-uniprot50 \| AF-A0A660VS27-F1-MODEL\_V4 \| 1.0 \| 9.154e-05 \| 127 \| 0.155 \| 231 \| 123 \| 16 \| 12 \| 219 \| 252 \| 433 \| Uncharacterized protein \| Uncharacterized protein; afdb-uniprot50 \| AF-A0A2V8PAG4-F1-MODEL\_V4 \| 1.0 \| 0.0001429 \| 127 \| 0.118 \| 254 \| 147 \| 14 \| 1 \| 219 \| 303 \| 514 \| Uncharacterized protein \| Uncharacterized protein; afdb-uniprot50 \| AF-A0A0F9J633-F1-MODEL\_V4 \| 1.0 \| 0.0002786 \| 126 \| 0.144 \| 305 \| 162 \| 19 \| 4 \| 245 \| 7 \| 275 \| Uncharacterized protein \| Uncharacterized protein; afdb-uniprot50 \| AF-A0A7C4DZ70-F1-MODEL\_V4 \| 1.0 \| 0.0004348 \| 126 \| 0.137 \| 226 \| 128 \| 14 \| 12 \| 217 \| 23 \| 201 \| KaiC domain-containing protein \| KaiC domain-containing protein; afdb-uniprot50 \| AF-I3R747-F1-MODEL\_V4 \| 1.0 \| 0.000389 \| 126 \| 0.178 \| 224 \| 125 \| 15 \| 15 \| 219 \| 202 \| 385 \| Circadian clock protein KaiC \| Circadian clock protein KaiC; afdb-uniprot50 \| AF-A0A7J2M1T3-F1-MODEL\_V4 \| 1.0 \| 0.0001785 \| 126 \| 0.146 \| 225 \| 133 \| 14 \| 12 \| 219 \| 30 \| 212 \| Uncharacterized protein \| Uncharacterized protein; afdb-uniprot50 \| AF-A0A497LWX5-F1-MODEL\_V4 \| 1.0 \| 0.0001278 \| 126 \| 0.156 \| 224 \| 126 \| 16 \| 16 \| 219 \| 24 \| 204 \| KaiC domain-containing protein \| KaiC domain-containing protein; afdb-uniprot50 \| AF-A0A497NIN7-F1-MODEL\_V4 \| 1.0 \| 0.000368 \| 126 \| 0.162 \| 227 \| 128 \| 18 \| 12 \| 219 \| 23 \| 206 \| KaiC domain-containing protein \| KaiC domain-containing protein; afdb-uniprot50 \| AF-A0A7W0A236-F1-MODEL\_V4 \| 1.0 \| 0.0006419 \| 126 \| 0.181 \| 198 \| 121 \| 13 \| 16 \| 204 \| 297 \| 462 \| AAA family ATPase \| AAA family ATPase; afdb-uniprot50 \| AF-A0A0C5BY25-F1-MODEL\_V4 \| 1.0 \| 0.000389 \| 125 \| 0.185 \| 226 \| 120 \| 15 \| 12 \| 219 \| 23 \| 202 \| KaiC domain-containing protein \| KaiC domain-containing protein; afdb-uniprot50 \| AF-A0A497NB40-F1-MODEL\_V4 \| 1.0 \| 0.0006072 \| 125 \| 0.169 \| 224 \| 129 \| 15 \| 12 \| 219 \| 22 \| 204 \| KaiC domain-containing protein \| KaiC domain-containing protein; afdb-uniprot50 \| AF-A0A662UES6-F1-MODEL\_V4 \| 1.0 \| 0.0007586 \| 125 \| 0.156 \| 223 \| 134 \| 17 \| 11 \| 216 \| 30 \| 215 \| KaiC domain-containing protein \| KaiC domain-containing protein; afdb-uniprot50 \| AF-A0A1F5YWC1-F1-MODEL\_V4 \| 1.0 \| 0.0002945 \| 125 \| 0.131 \| 235 \| 138 \| 13 \| 16 \| 233 \| 23 \| 208 \| Uncharacterized protein \| Uncharacterized protein; afdb-uniprot50 \| AF-A0A4D8QES0-F1-MODEL\_V4 \| 1.0 \| 0.0008964 \| 124 \| 0.139 \| 229 \| 135 \| 15 \| 9 \| 219 \| 7 \| 191 \| KaiC domain-containing protein \| KaiC domain-containing protein; afdb-uniprot50 \| AF-A0A365TER5-F1-MODEL\_V4 \| 1.0 \| 0.0001351 \| 124 \| 0.189 \| 232 \| 114 \| 15 \| 15 \| 219 \| 124 \| 308 \| KaiC domain-containing protein \| KaiC domain-containing protein; afdb-uniprot50 \| AF-A0A1B2I3A7-F1-MODEL\_V4 \| 1.0 \| 2.545e-05 \| 124 \| 0.128 \| 374 \| 205 \| 26 \| 12 \| 329 \| 61 \| 369 \| Protein RecA \| Protein RecA; afdb-uniprot50 \| AF-A0A7J2QYE4-F1-MODEL\_V4 \| 1.0 \| 0.0006072 \| 124 \| 0.157 \| 210 \| 133 \| 13 \| 12 \| 204 \| 59 \| 241 \| RECA\_3 domain-containing protein \| RECA\_3 domain-containing protein; afdb-uniprot50 \| AF-A0A662Q9S3-F1-MODEL\_V4 \| 1.0 \| 0.0005743 \| 124 \| 0.154 \| 220 \| 129 \| 14 \| 15 \| 217 \| 150 \| 329 \| KaiC domain-containing protein \| KaiC domain-containing protein; afdb-uniprot50 \| AF-A0A537GIY2-F1-MODEL\_V4 \| 1.0 \| 0.000802 \| 123 \| 0.115 \| 243 \| 151 \| 13 \| 1 \| 216 \| 16 \| 221 \| KaiC domain-containing protein \| KaiC domain-containing protein; afdb-uniprot50 \| AF-A0A2U0RYE1-F1-MODEL\_V4 \| 1.0 \| 0.000368 \| 123 \| 0.13 \| 214 \| 146 \| 11 \| 16 \| 219 \| 25 \| 208 \| KaiC domain-containing protein \| KaiC domain-containing protein; afdb-uniprot50 \| AF-A0A3B9KJ87-F1-MODEL\_V4 \| 1.0 \| 0.001059 \| 123 \| 0.139 \| 229 \| 145 \| 10 \| 16 \| 233 \| 122 \| 309 \| KaiC domain-containing protein \| KaiC domain-containing protein; afdb-uniprot50 \| AF-A0A662S5U5-F1-MODEL\_V4 \| 1.0 \| 4.694e-05 \| 123 \| 0.151 \| 231 \| 123 \| 12 \| 16 \| 219 \| 258 \| 442 \| Uncharacterized protein \| Uncharacterized protein; afdb-uniprot50 \| AF-A0A498G4R8-F1-MODEL\_V4 \| 1.0 \| 0.000348 \| 123 \| 0.161 \| 247 \| 133 \| 16 \| 15 \| 235 \| 246 \| 444 \| KaiC domain-containing protein \| KaiC domain-containing protein; afdb-uniprot50 \| AF-A0A7C5P3J9-F1-MODEL\_V4 \| 1.0 \| 0.0004348 \| 123 \| 0.159 \| 219 \| 133 \| 12 \| 12 \| 217 \| 28 \| 208 \| KaiC domain-containing protein \| KaiC domain-containing protein; afdb-uniprot50 \| AF-A0A6I2H1L9-F1-MODEL\_V4 \| 1.0 \| 0.000368 \| 123 \| 0.135 \| 236 \| 139 \| 13 \| 16 \| 233 \| 79 \| 267 \| AAA family ATPase \| AAA family ATPase; afdb-uniprot50 \| AF-A0A7C7I8H2-F1-MODEL\_V4 \| 1.0 \| 0.001564 \| 122 \| 0.161 \| 161 \| 101 \| 7 \| 16 \| 166 \| 28 \| 164 \| ATPase domain-containing protein \| ATPase domain-containing protein; afdb-uniprot50 \| AF-A0A7C4DT93-F1-MODEL\_V4 \| 1.0 \| 7.327e-05 \| 122 \| 0.18 \| 255 \| 125 \| 18 \| 12 \| 237 \| 21 \| 220 \| KaiC domain-containing protein \| KaiC domain-containing protein; afdb-uniprot50 \| AF-A0A537G127-F1-MODEL\_V4 \| 1.0 \| 0.0006072 \| 122 \| 0.159 \| 226 \| 135 \| 12 \| 9 \| 219 \| 20 \| 205 \| KaiC domain-containing protein \| KaiC domain-containing protein; afdb-uniprot50 \| AF-A0A7C4W5N6-F1-MODEL\_V4 \| 1.0 \| 0.0005743 \| 122 \| 0.136 \| 227 \| 132 \| 13 \| 12 \| 219 \| 23 \| 204 \| KaiC domain-containing protein \| KaiC domain-containing protein; afdb-uniprot50 \| AF-A0A3M1F514-F1-MODEL\_V4 \| 1.0 \| 0.0002358 \| 122 \| 0.161 \| 247 \| 129 \| 16 \| 16 \| 235 \| 43 \| 238 \| Uncharacterized protein \| Uncharacterized protein; afdb-uniprot50 \| AF-A0A7C3TCM6-F1-MODEL\_V4 \| 1.0 \| 0.0001995 \| 122 \| 0.128 \| 233 \| 128 \| 15 \| 12 \| 219 \| 23 \| 205 \| KaiC domain-containing protein \| KaiC domain-containing protein; afdb-uniprot50 \| AF-A0A537F4V6-F1-MODEL\_V4 \| 1.0 \| 0.0004597 \| 122 \| 0.159 \| 226 \| 129 \| 13 \| 12 \| 219 \| 33 \| 215 \| Circadian clock KaiC-like protein \| Circadian clock KaiC-like protein; afdb-uniprot50 \| AF-A0A7C5UBI0-F1-MODEL\_V4 \| 1.0 \| 9.154e-05 \| 122 \| 0.144 \| 318 \| 182 \| 20 \| 16 \| 298 \| 266 \| 528 \| Uncharacterized protein \| Uncharacterized protein; afdb-uniprot50 \| AF-A0A3D1TYK0-F1-MODEL\_V4 \| 1.0 \| 0.000368 \| 121 \| 0.146 \| 232 \| 136 \| 13 \| 9 \| 219 \| 1 \| 191 \| Circadian clock protein KaiC \| Circadian clock protein KaiC; afdb-uniprot50 \| AF-A0A7C5P6R4-F1-MODEL\_V4 \| 1.0 \| 0.0001023 \| 121 \| 0.174 \| 223 \| 130 \| 16 \| 11 \| 219 \| 19 \| 201 \| Circadian clock KaiC-like protein \| Circadian clock KaiC-like protein; afdb-uniprot50 \| AF-A0A2N1WRM6-F1-MODEL\_V4 \| 1.0 \| 0.0001597 \| 121 \| 0.164 \| 268 \| 138 \| 16 \| 4 \| 234 \| 252 \| 470 \| Circadian clock protein KaiC \| Circadian clock protein KaiC; afdb-uniprot50 \| AF-A0A7J4B2L6-F1-MODEL\_V4 \| 1.0 \| 0.0001597 \| 120 \| 0.135 \| 206 \| 122 \| 10 \| 16 \| 193 \| 28 \| 205 \| KaiC domain-containing protein \| KaiC domain-containing protein; afdb-uniprot50 \| AF-A0A7J4B336-F1-MODEL\_V4 \| 1.0 \| 0.0005743 \| 120 \| 0.158 \| 234 \| 134 \| 16 \| 16 \| 233 \| 25 \| 211 \| KaiC domain-containing protein \| KaiC domain-containing protein; afdb-uniprot50 \| AF-A0A1Q8BEF9-F1-MODEL\_V4 \| 1.0 \| 0.001748 \| 120 \| 0.165 \| 223 \| 132 \| 13 \| 12 \| 219 \| 12 \| 195 \| KaiC domain-containing protein \| KaiC domain-containing protein; afdb-uniprot50 \| AF-A0A497NEQ1-F1-MODEL\_V4 \| 1.0 \| 0.000802 \| 120 \| 0.168 \| 219 \| 129 \| 13 \| 16 \| 219 \| 25 \| 205 \| KaiC domain-containing protein \| KaiC domain-containing protein; afdb-uniprot50 \| AF-A0A662ILG2-F1-MODEL\_V4 \| 1.0 \| 0.0006072 \| 120 \| 0.158 \| 239 \| 127 \| 15 \| 16 \| 233 \| 28 \| 213 \| Uncharacterized protein \| Uncharacterized protein; afdb-uniprot50 \| AF-A0A497IS76-F1-MODEL\_V4 \| 1.0 \| 0.0005743 \| 119 \| 0.122 \| 228 \| 131 \| 14 \| 16 \| 219 \| 10 \| 192 \| KaiC domain-containing protein \| KaiC domain-containing protein; afdb-uniprot50 \| AF-A0A662KAI4-F1-MODEL\_V4 \| 1.0 \| 0.000802 \| 119 \| 0.161 \| 217 \| 133 \| 13 \| 16 \| 219 \| 32 \| 212 \| KaiC domain-containing protein \| KaiC domain-containing protein; afdb-uniprot50 \| AF-A0A1I1D661-F1-MODEL\_V4 \| 1.0 \| 0.0001351 \| 119 \| 0.177 \| 242 \| 119 \| 16 \| 15 \| 231 \| 253 \| 439 \| KaiC domain protein, AF\_0351 family \| KaiC domain protein, AF\_0351 family; afdb-uniprot50 \| AF-A0A7J2ZH85-F1-MODEL\_V4 \| 1.0 \| 0.000389 \| 119 \| 0.171 \| 222 \| 131 \| 13 \| 12 \| 219 \| 22 \| 204 \| KaiC domain-containing protein \| KaiC domain-containing protein; afdb-uniprot50 \| AF-A0A7C4H9Y4-F1-MODEL\_V4 \| 1.0 \| 0.001002 \| 119 \| 0.13 \| 222 \| 138 \| 13 \| 12 \| 219 \| 26 \| 206 \| KaiC domain-containing protein \| KaiC domain-containing protein; afdb-uniprot50 \| AF-A0A7J3I9A3-F1-MODEL\_V4 \| 1.0 \| 0.00112 \| 119 \| 0.172 \| 203 \| 122 \| 14 \| 12 \| 206 \| 21 \| 185 \| KaiC domain-containing protein \| KaiC domain-containing protein; afdb-uniprot50 \| AF-A0A2R7Y1F6-F1-MODEL\_V4 \| 1.0 \| 0.002184 \| 119 \| 0.155 \| 200 \| 118 \| 12 \| 16 \| 204 \| 48 \| 207 \| KaiC domain-containing protein \| KaiC domain-containing protein; afdb-uniprot50 \| AF-A0A6J4JV05-F1-MODEL\_V4 \| 1.0 \| 0.0009477 \| 119 \| 0.172 \| 215 \| 133 \| 13 \| 16 \| 219 \| 273 \| 453 \| Circadian clock protein KaiC \| Circadian clock protein KaiC; afdb-uniprot50 \| AF-A0A6I2GST4-F1-MODEL\_V4 \| 1.0 \| 0.000368 \| 119 \| 0.167 \| 251 \| 138 \| 16 \| 1 \| 219 \| 249 \| 460 \| AAA family ATPase \| AAA family ATPase; afdb-uniprot50 \| AF-A0A1G6QEK8-F1-MODEL\_V4 \| 1.0 \| 3.554e-05 \| 119 \| 0.126 \| 349 \| 193 \| 22 \| 16 \| 316 \| 270 \| 554 \| Circadian clock protein KaiC \| Circadian clock protein KaiC; afdb-uniprot50 \| AF-A0A7J5W1L4-F1-MODEL\_V4 \| 1.0 \| 0.0001429 \| 118 \| 0.122 \| 244 \| 152 \| 14 \| 16 \| 227 \| 278 \| 491 \| Putative circadian clock protein \| Putative circadian clock protein; afdb-uniprot50 \| AF-A0A7T5UKT8-F1-MODEL\_V4 \| 1.0 \| 4.694e-05 \| 118 \| 0.164 \| 340 \| 176 \| 23 \| 12 \| 299 \| 273 \| 556 \| Circadian clock protein KaiC \| Circadian clock protein KaiC; afdb-uniprot50 \| AF-A0A3D6C4X9-F1-MODEL\_V4 \| 1.0 \| 0.0005743 \| 117 \| 0.143 \| 244 \| 126 \| 17 \| 16 \| 231 \| 38 \| 226 \| Uncharacterized protein \| Uncharacterized protein; afdb-uniprot50 \| AF-A0A7J3A042-F1-MODEL\_V4 \| 1.0 \| 0.001653 \| 116 \| 0.135 \| 199 \| 134 \| 8 \| 12 \| 207 \| 23 \| 186 \| KaiC domain-containing protein \| KaiC domain-containing protein; afdb-uniprot50 \| AF-A0A7C5Y1U1-F1-MODEL\_V4 \| 1.0 \| 0.002309 \| 116 \| 0.154 \| 227 \| 141 \| 12 \| 16 \| 233 \| 24 \| 208 \| KaiC domain-containing protein \| KaiC domain-containing protein; afdb-uniprot50 \| AF-A0A7C4B8K6-F1-MODEL\_V4 \| 1.0 \| 0.001399 \| 116 \| 0.136 \| 220 \| 136 \| 14 \| 12 \| 218 \| 30 \| 208 \| Uncharacterized protein \| Uncharacterized protein; afdb-uniprot50 \| AF-A0A662URN2-F1-MODEL\_V4 \| 1.0 \| 0.0006072 \| 116 \| 0.08 \| 223 \| 145 \| 15 \| 12 \| 219 \| 23 \| 200 \| KaiC domain-containing protein \| KaiC domain-containing protein; afdb-uniprot50 \| AF-A0A832KH48-F1-MODEL\_V4 \| 1.0 \| 0.0006787 \| 116 \| 0.144 \| 236 \| 121 \| 15 \| 12 \| 219 \| 22 \| 204 \| Uncharacterized protein \| Uncharacterized protein; afdb-uniprot50 \| AF-A0A7J3F1A2-F1-MODEL\_V4 \| 1.0 \| 0.001479 \| 116 \| 0.13 \| 222 \| 136 \| 14 \| 15 \| 219 \| 28 \| 209 \| KaiC domain-containing protein \| KaiC domain-containing protein; afdb-uniprot50 \| AF-A0A2N5KIB9-F1-MODEL\_V4 \| 1.0 \| 0.0004348 \| 115 \| 0.168 \| 208 \| 117 \| 12 \| 16 \| 193 \| 4 \| 185 \| KaiC domain-containing protein \| KaiC domain-containing protein; afdb-uniprot50 \| AF-A0A5J4KQT9-F1-MODEL\_V4 \| 1.0 \| 0.0004113 \| 115 \| 0.13 \| 237 \| 138 \| 14 \| 16 \| 233 \| 33 \| 220 \| KaiC domain-containing protein \| KaiC domain-containing protein; afdb-uniprot50 \| AF-A0A550HA37-F1-MODEL\_V4 \| 1.0 \| 0.002184 \| 115 \| 0.127 \| 228 \| 146 \| 14 \| 12 \| 219 \| 62 \| 256 \| Rad51 domain-containing protein \| Rad51 domain-containing protein; afdb-uniprot50 \| AF-A0A6J4MRF5-F1-MODEL\_V4 \| 1.0 \| 0.001002 \| 115 \| 0.14 \| 271 \| 148 \| 16 \| 4 \| 235 \| 247 \| 471 \| KaiC domain-containing protein \| KaiC domain-containing protein; afdb-uniprot50 \| AF-A0A4P7BXA5-F1-MODEL\_V4 \| 1.0 \| 0.002065 \| 115 \| 0.14 \| 220 \| 129 \| 15 \| 16 \| 217 \| 273 \| 450 \| Circadian clock protein KaiC \| Circadian clock protein KaiC; afdb-uniprot50 \| AF-A0A7C4DEG0-F1-MODEL\_V4 \| 1.0 \| 0.0005743 \| 114 \| 0.16 \| 231 \| 123 \| 17 \| 12 \| 219 \| 22 \| 204 \| KaiC domain-containing protein \| KaiC domain-containing protein; afdb-uniprot50 \| AF-A0A662HB48-F1-MODEL\_V4 \| 1.0 \| 0.001323 \| 114 \| 0.11 \| 227 \| 138 \| 16 \| 12 \| 219 \| 37 \| 218 \| Uncharacterized protein \| Uncharacterized protein; afdb-uniprot50 \| AF-A0A497JZ60-F1-MODEL\_V4 \| 1.0 \| 8.19e-05 \| 114 \| 0.14 \| 334 \| 177 \| 23 \| 16 \| 299 \| 25 \| 298 \| KaiC domain-containing protein \| KaiC domain-containing protein; afdb-uniprot50 \| AF-A0A1H8MGT9-F1-MODEL\_V4 \| 1.0 \| 0.0004113 \| 114 \| 0.153 \| 235 \| 122 \| 16 \| 11 \| 219 \| 181 \| 364 \| Circadian clock protein KaiC \| Circadian clock protein KaiC; afdb-uniprot50 \| AF-A0A2V6WLC1-F1-MODEL\_V4 \| 1.0 \| 0.002184 \| 114 \| 0.121 \| 197 \| 135 \| 9 \| 16 \| 207 \| 267 \| 430 \| Uncharacterized protein \| Uncharacterized protein; afdb-uniprot50 \| AF-A0A2W5QUD6-F1-MODEL\_V4 \| 1.0 \| 0.0001023 \| 114 \| 0.133 \| 359 \| 204 \| 23 \| 12 \| 329 \| 270 \| 562 \| Uncharacterized protein \| Uncharacterized protein; afdb-uniprot50 \| AF-A0A7C1M4D4-F1-MODEL\_V4 \| 1.0 \| 0.0002945 \| 113 \| 0.17 \| 229 \| 132 \| 15 \| 7 \| 219 \| 15 \| 201 \| Circadian clock KaiC-like protein \| Circadian clock KaiC-like protein; afdb-uniprot50 \| AF-A0A564WRY8-F1-MODEL\_V4 \| 1.0 \| 3.554e-05 \| 113 \| 0.118 \| 363 \| 188 \| 15 \| 14 \| 301 \| 1 \| 306 \| Uncharacterized protein \| Uncharacterized protein; afdb-uniprot50 \| AF-A0A346S328-F1-MODEL\_V4 \| 1.0 \| 0.0002492 \| 112 \| 0.141 \| 247 \| 155 \| 15 \| 16 \| 231 \| 9 \| 229 \| KaiC domain-containing protein \| KaiC domain-containing protein; afdb-uniprot50 \| AF-A0A843CIY9-F1-MODEL\_V4 \| 1.0 \| 0.0007175 \| 112 \| 0.146 \| 267 \| 138 \| 19 \| 12 \| 252 \| 36 \| 238 \| AAA family ATPase \| AAA family ATPase; afdb-uniprot50 \| AF-A0A1F5UAY4-F1-MODEL\_V4 \| 1.0 \| 0.0009477 \| 112 \| 0.152 \| 223 \| 135 \| 13 \| 12 \| 219 \| 25 \| 208 \| KaiC domain-containing protein \| KaiC domain-containing protein; afdb-uniprot50 \| AF-A0A7C4NE86-F1-MODEL\_V4 \| 1.0 \| 0.000802 \| 112 \| 0.156 \| 223 \| 133 \| 15 \| 12 \| 219 \| 26 \| 208 \| KaiC domain-containing protein \| KaiC domain-containing protein; afdb-uniprot50 \| AF-A0A537FDB0-F1-MODEL\_V4 \| 1.0 \| 0.0009477 \| 112 \| 0.15 \| 239 \| 133 \| 16 \| 12 \| 233 \| 23 \| 208 \| Uncharacterized protein \| Uncharacterized protein; afdb-uniprot50 \| AF-A0A352NFF9-F1-MODEL\_V4 \| 1.0 \| 0.0001995 \| 110 \| 0.189 \| 237 \| 125 \| 17 \| 12 \| 216 \| 256 \| 457 \| ATPase \| ATPase; afdb-uniprot50 \| AF-A0A2R7Y6U6-F1-MODEL\_V4 \| 1.0 \| 0.001252 \| 110 \| 0.087 \| 216 \| 135 \| 13 \| 13 \| 189 \| 24 \| 216 \| KaiC domain-containing protein \| KaiC domain-containing protein; afdb-uniprot50 \| AF-A0A661S0K0-F1-MODEL\_V4 \| 1.0 \| 9.678e-05 \| 110 \| 0.125 \| 239 \| 132 \| 14 \| 16 \| 216 \| 330 \| 529 \| KaiC 1 \| KaiC 1; afdb-uniprot50 \| AF-A0A355UR33-F1-MODEL\_V4 \| 1.0 \| 1.927e-05 \| 109 \| 0.141 \| 361 \| 202 \| 22 \| 12 \| 329 \| 145 \| 440 \| KaiC domain-containing protein \| KaiC domain-containing protein; afdb-uniprot50 \| AF-A0A7J3QBJ0-F1-MODEL\_V4 \| 1.0 \| 0.0008964 \| 109 \| 0.154 \| 207 \| 127 \| 17 \| 7 \| 204 \| 18 \| 185 \| KaiC domain-containing protein \| KaiC domain-containing protein; afdb-uniprot50 \| AF-A0A0A6UQ55-F1-MODEL\_V4 \| 1.0 \| 0.000802 \| 109 \| 0.121 \| 247 \| 144 \| 16 \| 5 \| 219 \| 234 \| 439 \| Circadian clock protein KaiC \| Circadian clock protein KaiC; afdb-uniprot50 \| AF-A0A849T9L3-F1-MODEL\_V4 \| 1.0 \| 0.002884 \| 108 \| 0.133 \| 225 \| 135 \| 13 \| 12 \| 219 \| 8 \| 189 \| Uncharacterized protein \| Uncharacterized protein; afdb-uniprot50 \| AF-A0A7J3A6W1-F1-MODEL\_V4 \| 1.0 \| 0.0004113 \| 108 \| 0.167 \| 227 \| 125 \| 18 \| 12 \| 219 \| 22 \| 203 \| KaiC domain-containing protein \| KaiC domain-containing protein; afdb-uniprot50 \| AF-B0ED98-F1-MODEL\_V4 \| 1.0 \| 0.0007586 \| 108 \| 0.126 \| 230 \| 139 \| 18 \| 12 \| 218 \| 104 \| 294 \| Uncharacterized protein \| Uncharacterized protein; afdb-uniprot50 \| AF-A2BKH9-F1-MODEL\_V4 \| 1.0 \| 0.002065 \| 108 \| 0.146 \| 218 \| 135 \| 11 \| 16 \| 219 \| 32 \| 212 \| Putative RecA ATPase \| Putative RecA ATPase; afdb-uniprot50 \| AF-A0A534PG62-F1-MODEL\_V4 \| 1.0 \| 0.001954 \| 108 \| 0.187 \| 224 \| 125 \| 14 \| 12 \| 219 \| 269 \| 451 \| Recombinase RecA \| Recombinase RecA; afdb-uniprot50 \| AF-A0A1B1UC61-F1-MODEL\_V4 \| 1.0 \| 0.004759 \| 107 \| 0.139 \| 179 \| 128 \| 9 \| 12 \| 185 \| 38 \| 195 \| ATPase domain-containing protein \| ATPase domain-containing protein; afdb-uniprot50 \| AF-A0A0U3SJ19-F1-MODEL\_V4 \| 1.0 \| 0.000368 \| 107 \| 0.168 \| 232 \| 125 \| 18 \| 9 \| 219 \| 17 \| 201 \| Circadian clock protein KaiC \| Circadian clock protein KaiC; afdb-uniprot50 \| AF-A0A524KT06-F1-MODEL\_V4 \| 1.0 \| 0.004759 \| 107 \| 0.138 \| 216 \| 137 \| 14 \| 16 \| 219 \| 90 \| 268 \| KaiC domain-containing protein \| KaiC domain-containing protein; afdb-uniprot50 \| AF-A0A6A8FI11-F1-MODEL\_V4 \| 1.0 \| 0.0006419 \| 107 \| 0.129 \| 232 \| 145 \| 14 \| 16 \| 220 \| 260 \| 461 \| AAA family ATPase \| AAA family ATPase; afdb-uniprot50 \| AF-A0A4Q3NXW5-F1-MODEL\_V4 \| 1.0 \| 0.002728 \| 106 \| 0.162 \| 227 \| 126 \| 16 \| 12 \| 218 \| 18 \| 200 \| AAA domain-containing protein \| AAA domain-containing protein; afdb-uniprot50 \| AF-A0A7V8XVW8-F1-MODEL\_V4 \| 1.0 \| 0.001748 \| 106 \| 0.128 \| 233 \| 133 \| 15 \| 10 \| 219 \| 83 \| 268 \| KaiC domain-containing protein \| KaiC domain-containing protein; afdb-uniprot50 \| AF-A0A2E2DQY8-F1-MODEL\_V4 \| 1.0 \| 0.0007175 \| 106 \| 0.13 \| 230 \| 119 \| 14 \| 16 \| 193 \| 42 \| 242 \| ATPase domain-containing protein \| ATPase domain-containing protein; afdb-uniprot50 \| AF-A0A2J6XK50-F1-MODEL\_V4 \| 1.0 \| 7.327e-05 \| 106 \| 0.18 \| 233 \| 111 \| 15 \| 16 \| 194 \| 25 \| 231 \| KaiC domain-containing protein \| KaiC domain-containing protein; afdb-uniprot50 \| AF-A0A7J3BDE4-F1-MODEL\_V4 \| 1.0 \| 0.001184 \| 106 \| 0.14 \| 235 \| 123 \| 16 \| 12 \| 219 \| 51 \| 233 \| KaiC domain-containing protein \| KaiC domain-containing protein; afdb-uniprot50 \| AF-A0A7W1AFM2-F1-MODEL\_V4 \| 1.0 \| 0.0002635 \| 106 \| 0.121 \| 347 \| 196 \| 25 \| 1 \| 287 \| 204 \| 501 \| AAA family ATPase \| AAA family ATPase; afdb-uniprot50 \| AF-A0A4Y7WTN4-F1-MODEL\_V4 \| 1.0 \| 0.0002109 \| 105 \| 0.132 \| 265 \| 155 \| 17 \| 16 \| 243 \| 10 \| 236 \| KaiC domain-containing protein \| KaiC domain-containing protein; afdb-uniprot50 \| AF-A0A523ZRI4-F1-MODEL\_V4 \| 1.0 \| 0.000802 \| 105 \| 0.146 \| 226 \| 131 \| 11 \| 16 \| 198 \| 100 \| 306 \| KaiC domain-containing protein \| KaiC domain-containing protein; afdb-uniprot50 \| AF-A0A7J3B871-F1-MODEL\_V4 \| 1.0 \| 0.0002635 \| 104 \| 0.148 \| 236 \| 117 \| 14 \| 12 \| 193 \| 21 \| 226 \| KaiC domain-containing protein \| KaiC domain-containing protein; afdb-uniprot50 \| AF-A0A1Q7MI04-F1-MODEL\_V4 \| 1.0 \| 0.001848 \| 104 \| 0.142 \| 266 \| 140 \| 19 \| 12 \| 252 \| 35 \| 237 \| KaiC domain-containing protein \| KaiC domain-containing protein; afdb-uniprot50 \| AF-A0A537H318-F1-MODEL\_V4 \| 1.0 \| 0.001323 \| 104 \| 0.154 \| 214 \| 118 \| 13 \| 12 \| 193 \| 20 \| 202 \| ATPase \| ATPase; afdb-uniprot50 \| AF-A0A1J4UF65-F1-MODEL\_V4 \| 1.0 \| 0.001399 \| 104 \| 0.132 \| 249 \| 147 \| 14 \| 1 \| 217 \| 29 \| 240 \| KaiC domain-containing protein \| KaiC domain-containing protein; afdb-uniprot50 \| AF-B3PNA4-F1-MODEL\_V4 \| 1.0 \| 0.001479 \| 104 \| 0.149 \| 214 \| 122 \| 16 \| 16 \| 204 \| 42 \| 220 \| Bacteriophage MAV1 replication protein RepB \| Bacteriophage MAV1 replication protein RepB; afdb-uniprot50 \| AF-A0A085WJA9-F1-MODEL\_V4 \| 1.0 \| 0.0006072 \| 104 \| 0.138 \| 238 \| 146 \| 13 \| 10 \| 219 \| 231 \| 437 \| KaiC domain-containing protein \| KaiC domain-containing protein; afdb-uniprot50 \| AF-A0A662UBR9-F1-MODEL\_V4 \| 1.0 \| 0.00381 \| 104 \| 0.117 \| 230 \| 134 \| 15 \| 12 \| 219 \| 35 \| 217 \| KaiC domain-containing protein \| KaiC domain-containing protein; afdb-uniprot50 \| AF-A0A211YR69-F1-MODEL\_V4 \| 1.0 \| 0.0006787 \| 104 \| 0.144 \| 270 \| 149 \| 13 \| 4 \| 224 \| 13 \| 249 \| KaiC domain-containing protein \| KaiC domain-containing protein; afdb-uniprot50 \| AF-A0A7W1PRM7-F1-MODEL\_V4 \| 1.0 \| 0.002065 \| 104 \| 0.113 \| 238 \| 141 \| 12 \| 4 \| 214 \| 275 \| 469 \| AAA family ATPase \| AAA family ATPase; afdb-uniprot50 \| AF-A0A7C4DZK8-F1-MODEL\_V4 \| 1.0 \| 0.0003114 \| 103 \| 0.15 \| 239 \| 132 \| 15 \| 16 \| 219 \| 26 \| 228 \| KaiC domain-containing protein \| KaiC domain-containing protein; afdb-uniprot50 \| AF-A0A1Q8BK86-F1-MODEL\_V4 \| 1.0 \| 0.004028 \| 103 \| 0.134 \| 230 \| 131 \| 16 \| 12 \| 219 \| 167 \| 350 \| KaiC domain-containing protein \| KaiC domain-containing protein; afdb-uniprot50 \| AF-A0A832KI38-F1-MODEL\_V4 \| 1.0 \| 0.0006419 \| 103 \| 0.154 \| 227 \| 108 \| 14 \| 12 \| 189 \| 20 \| 211 \| Uncharacterized protein \| Uncharacterized protein; afdb-uniprot50 \| AF-A0A662WCX8-F1-MODEL\_V4 \| 1.0 \| 0.001479 \| 102 \| 0.165 \| 200 \| 115 \| 12 \| 12 \| 180 \| 22 \| 200 \| ATPase \| ATPase; afdb-uniprot50 \| AF-A0A534Y3E6-F1-MODEL\_V4 \| 1.0 \| 0.0008964 \| 102 \| 0.154 \| 253 \| 130 \| 19 \| 12 \| 235 \| 204 \| 401 \| Recombinase RecA \| Recombinase RecA; afdb-uniprot50 \| AF-A0A7C2Q0A6-F1-MODEL\_V4 \| 1.0 \| 0.001564 \| 102 \| 0.152 \| 210 \| 114 \| 12 \| 16 \| 189 \| 24 \| 205 \| KaiC domain-containing protein \| KaiC domain-containing protein; afdb-uniprot50 \| AF-A0A0M9E807-F1-MODEL\_V4 \| 1.0 \| 8.19e-05 \| 102 \| 0.13 \| 361 \| 194 \| 20 \| 16 \| 328 \| 281 \| 569 \| KaiC 1 \| KaiC 1; afdb-uniprot50 \| AF-A0A4Q3H1A7-F1-MODEL\_V4 \| 1.0 \| 0.002184 \| 102 \| 0.125 \| 240 \| 133 \| 12 \| 13 \| 198 \| 283 \| 499 \| AAA domain-containing protein \| AAA domain-containing protein; afdb-uniprot50 \| AF-A0A497HTL3-F1-MODEL\_V4 \| 1.0 \| 0.001748 \| 101 \| 0.153 \| 196 \| 117 \| 10 \| 12 \| 182 \| 61 \| 232 \| AAA family ATPase \| AAA family ATPase; afdb-uniprot50 \| AF-I7JA90-F1-MODEL\_V4 \| 1.0 \| 0.001479 \| 101 \| 0.138 \| 216 \| 137 \| 14 \| 16 \| 215 \| 35 \| 217 \| p-loop containing nucleoside triphosphate hydrolase, putative \| p-loop containing nucleoside triphosphate hydrolase, putative; afdb-uniprot50 \| AF-A0A256Z1J6-F1-MODEL\_V4 \| 1.0 \| 0.000802 \| 101 \| 0.147 \| 244 \| 145 \| 15 \| 12 \| 224 \| 25 \| 236 \| Uncharacterized protein \| Uncharacterized protein; afdb-uniprot50 \| AF-A0A662VFD8-F1-MODEL\_V4 \| 1.0 \| 0.0006072 \| 100 \| 0.15 \| 246 \| 127 \| 17 \| 16 \| 219 \| 27 \| 232 \| KaiC domain-containing protein \| KaiC domain-containing protein; afdb-uniprot50 \| AF-A0A4Q5ZS73-F1-MODEL\_V4 \| 1.0 \| 0.002184 \| 100 \| 0.153 \| 222 \| 119 \| 13 \| 16 \| 189 \| 150 \| 350 \| Circadian clock protein KaiC \| Circadian clock protein KaiC; afdb-uniprot50 \| AF-A0A058Z4P2-F1-MODEL\_V4 \| 1.0 \| 0.002309 \| 100 \| 0.142 \| 288 \| 171 \| 18 \| 11 \| 251 \| 163 \| 421 \| RECA\_2 domain-containing protein \| RECA\_2 domain-containing protein; afdb-uniprot50 \| AF-A0A7W0Y1W2-F1-MODEL\_V4 \| 0.999 \| 0.00381 \| 99 \| 0.122 \| 221 \| 129 \| 11 \| 18 \| 193 \| 2 \| 202 \| KaiC domain-containing protein \| KaiC domain-containing protein; afdb-uniprot50 \| AF-A0A160VU18-F1-MODEL\_V4 \| 0.999 \| 0.0007175 \| 99 \| 0.17 \| 235 \| 117 \| 18 \| 11 \| 219 \| 20 \| 202 \| Circadian clock KaiC-like protein \| Circadian clock KaiC-like protein; afdb-uniprot50 \| AF-A0A3S0DQB7-F1-MODEL\_V4 \| 0.999 \| 0.001323 \| 99 \| 0.112 \| 445 \| 210 \| 25 \| 15 \| 311 \| 16 \| 423 \| Uncharacterized protein \| Uncharacterized protein; afdb-uniprot50 \| AF-A0A662MQT2-F1-MODEL\_V4 \| 0.999 \| 0.0006072 \| 98 \| 0.145 \| 233 \| 125 \| 17 \| 11 \| 219 \| 19 \| 201 \| Circadian clock KaiC-like protein \| Circadian clock KaiC-like protein; afdb-uniprot50 \| AF-A0A6N7PSS2-F1-MODEL\_V4 \| 0.999 \| 0.001184 \| 98 \| 0.162 \| 253 \| 129 \| 18 \| 10 \| 235 \| 258 \| 454 \| Uncharacterized protein \| Uncharacterized protein; afdb-uniprot50 \| AF-A0A534Z7Q3-F1-MODEL\_V4 \| 0.999 \| 0.001252 \| 98 \| 0.148 \| 235 \| 122 \| 16 \| 12 \| 219 \| 269 \| 452 \| Uncharacterized protein \| Uncharacterized protein; afdb-uniprot50 \| AF-A0A7C7RJU4-F1-MODEL\_V4 \| 0.999 \| 0.0009477 \| 97 \| 0.163 \| 318 \| 168 \| 23 \| 16 \| 299 \| 25 \| 278 \| Uncharacterized protein \| Uncharacterized protein; afdb-uniprot50 \| AF-A0A1G1G598-F1-MODEL\_V4 \| 0.999 \| 0.000151 \| 97 \| 0.127 \| 360 \| 216 \| 26 \| 8 \| 329 \| 4 \| 303 \| KaiC domain-containing protein \| KaiC domain-containing protein; afdb-uniprot50 \| AF-A0A3N0BQY0-F1-MODEL\_V4 \| 0.999 \| 0.0003114 \| 97 \| 0.138 \| 333 \| 167 \| 23 \| 12 \| 287 \| 72 \| 341 \| LuxR family transcriptional regulator \| LuxR family transcriptional regulator; afdb-uniprot50 \| AF-A0A662RV12-F1-MODEL\_V4 \| 0.999 \| 0.003603 \| 97 \| 0.142 \| 224 \| 125 \| 17 \| 16 \| 217 \| 40 \| 218 \| Uncharacterized protein \| Uncharacterized protein; afdb-uniprot50 \| AF-A0A7C4CKA9-F1-MODEL\_V4 \| 0.999 \| 0.00928 \| 97 \| 0.135 \| 228 \| 127 \| 14 \| 12 \| 217 \| 27 \| 206 \| KaiC domain-containing protein \| KaiC domain-containing protein; afdb-uniprot50 \| AF-A0A842V294-F1-MODEL\_V4 \| 0.999 \| 0.00112 \| 97 \| 0.149 \| 247 \| 126 \| 19 \| 12 \| 233 \| 263 \| 450 \| AAA family ATPase \| AAA family ATPase; afdb-uniprot50 \| AF-A0A7C4HA07-F1-MODEL\_V4 \| 0.998 \| 0.004502 \| 94 \| 0.17 \| 241 \| 120 \| 12 \| 1 \| 219 \| 1 \| 183 \| KaiC domain-containing protein \| KaiC domain-containing protein; afdb-uniprot50 \| AF-A0A0A8UMG1-F1-MODEL\_V4 \| 0.998 \| 0.000348 \| 94 \| 0.151 \| 324 \| 183 \| 21 \| 16 \| 306 \| 259 \| 523 \| Uncharacterized protein \| Uncharacterized protein; afdb-uniprot50 \| AF-A0A6J4UVX8-F1-MODEL\_V4 \| 0.998 \| 0.001184 \| 93 \| 0.133 \| 254 \| 129 \| 13 \| 4 \| 192 \| 231 \| 458 \| Circadian clock protein KaiC \| Circadian clock protein KaiC; afdb-uniprot50 \| AF-A0A6J4UDQ5-F1-MODEL\_V4 \| 0.997 \| 0.004028 \| 92 \| 0.145 \| 240 \| 140 \| 14 \| 14 \| 220 \| 264 \| 471 \| Uncharacterized protein \| Uncharacterized protein; afdb-uniprot50 \| AF-A0A3A6QRG0-F1-MODEL\_V4 \| 0.997 \| 0.007026 \| 91 \| 0.149 \| 227 \| 132 \| 11 \| 16 \| 219 \| 33 \| 221 \| ATPase \| ATPase; afdb-uniprot50 \| AF-A0A7C3A453-F1-MODEL\_V4 \| 0.997 \| 0.0005138 \| 91 \| 0.151 \| 278 \| 138 \| 19 \| 9 \| 231 \| 23 \| 257 \| Circadian clock KaiC-like protein \| Circadian clock KaiC-like protein; afdb-uniprot50 \| AF-A0A5E4K8W4-F1-MODEL\_V4 \| 0.997 \| 0.0006787 \| 91 \| 0.133 \| 263 \| 142 \| 16 \| 16 \| 227 \| 146 \| 373 \| Circadian clock protein kinase KaiC \| Circadian clock protein kinase KaiC; afdb-uniprot50 \| AF-A0A7C4H7G7-F1-MODEL\_V4 \| 0.997 \| 0.001564 \| 91 \| 0.161 \| 235 \| 113 \| 17 \| 10 \| 194 \| 248 \| 448 \| Uncharacterized protein \| Uncharacterized protein; afdb-uniprot50 \| AF-A0A838ND27-F1-MODEL\_V4 \| 0.997 \| 0.0005432 \| 91 \| 0.117 \| 356 \| 215 \| 20 \| 12 \| 329 \| 275 \| 569 \| Circadian clock protein KaiC \| Circadian clock protein KaiC; afdb-uniprot50 \| AF-A0A7C3LFH4-F1-MODEL\_V4 \| 0.996 \| 0.000368 \| 90 \| 0.114 \| 359 \| 214 \| 22 \| 12 \| 329 \| 115 \| 410 \| Circadian clock protein KaiC \| Circadian clock protein KaiC; afdb-uniprot50 \| AF-A0A7C4CR27-F1-MODEL\_V4 \| 0.996 \| 0.0004348 \| 89 \| 0.12 \| 275 \| 147 \| 16 \| 16 \| 231 \| 24 \| 262 \| KaiC domain-containing protein \| KaiC domain-containing protein; afdb-uniprot50 \| AF-A0A1I0BGA0-F1-MODEL\_V4 \| 0.995 \| 0.0006419 \| 88 \| 0.106 \| 320 \| 186 \| 22 \| 31 \| 298 \| 393 \| 664 \| Homing endonuclease \| Homing endonuclease; afdb-uniprot50 \| AF-A0A662RV64-F1-MODEL\_V4 \| 0.994 \| 0.002309 \| 87 \| 0.136 \| 213 \| 124 \| 13 \| 11 \| 189 \| 33 \| 219 \| KaiC domain-containing protein \| KaiC domain-containing protein; afdb-uniprot50 \| AF-A0A838UR59-F1-MODEL\_V4 \| 0.993 \| 0.006646 \| 86 \| 0.117 \| 246 \| 132 \| 15 \| 12 \| 197 \| 281 \| 501 \| AAA family ATPase \| AAA family ATPase; afdb-uniprot50 \| AF-A0A7J2JS41-F1-MODEL\_V4 \| 0.992 \| 0.001252 \| 85 \| 0.136 \| 314 \| 179 \| 18 \| 1 \| 251 \| 25 \| 309 \| AAA domain-containing protein \| AAA domain-containing protein; afdb-uniprot50 \| AF-A0A4V2B6G6-F1-MODEL\_V4 \| 0.991 \| 0.00532 \| 84 \| 0.107 \| 232 \| 118 \| 12 \| 28 \| 193 \| 2 \| 210 \| Protein kinase \| Protein kinase; afdb-uniprot50 \| AF-A0A3D1T9M1-F1-MODEL\_V4 \| 0.991 \| 0.004028 \| 84 \| 0.127 \| 235 \| 124 \| 12 \| 16 \| 195 \| 71 \| 279 \| ATPase domain-containing protein \| ATPase domain-containing protein; afdb-uniprot50 \| AF-L0AA79-F1-MODEL\_V4 \| 0.988 \| 0.0001023 \| 82 \| 0.145 \| 296 \| 146 \| 22 \| 12 \| 233 \| 23 \| 285 \| RecA-superfamily ATPase possibly involved in signal transduction \| RecA-superfamily ATPase possibly involved in signal transduction; afdb-uniprot50 \| AF-A0A133VMH7-F1-MODEL\_V4 \| 0.956 \| 0.005946 \| 73 \| 0.142 \| 224 \| 124 \| 13 \| 12 \| 193 \| 57 \| 254 \| ATPase domain-containing protein \| ATPase domain-containing protein; afdb-uniprot50 \| AF-A0A2N5K4H2-F1-MODEL\_V4 \| 0.956 \| 0.005946 \| 73 \| 0.145 \| 240 \| 121 \| 16 \| 12 \| 195 \| 265 \| 476 \| Uncharacterized protein \| Uncharacterized protein |
| Top keywords  (threshold 1.00e-02 (evalue)) | **KaiC, domain\_containing, Circadian, clock, AAA, kinase, ATPase, DNA, repair, beta** |
| Output files | ../../similar\_structures/39\_FANPEZAQ\_CDS\_0039\_afdb-proteome\_foldseek.tsv ../../similar\_structures/39\_FANPEZAQ\_CDS\_0039\_afdb-uniprot50\_foldseek.tsv ../../similar\_structures/39\_FANPEZAQ\_CDS\_0039\_merged.svg ../../similar\_structures/39\_FANPEZAQ\_CDS\_0039\_pdb\_foldseek.tsv |

  
  
  

Return to summary | Go to previous | Go to next

  


---

**Sequence/structure alignments coloring**  
Each object in the alignment figures is colored according to its E-value following this color coding:

1e-100
10
