## Supplementary material for "Completing the BASEL phage collection to unlock hidden diversity for systematic exploration of phage-host interactions": Data S2: 41.html

FANPEZAQ\_CDS\_0041


Return to summary | Go to previous | Go to next

|  |  |
| --- | --- |
| FANPEZAQ\_CDS\_0041  Page creation date: 02 Sep 2024, 12:00  Project folder: n/a  Input sequences file: Escherichia\_virus\_HeidiAbel.gb | domain\_containing duf3799 exodeoxyribonuclease putative pddexk\_like e d pd\_ xk dna helicase nuclease\_like viii exonuclease pddexk\_1 nuclease fragment breaking\_rejoining 3'\_5' endonuclease\_like a superfamily atp\_dependent rece \_ cas4 crispr\_associated complex csa1 recb hypothetical phage type c containing i\_a yes lambda uvrd sap beta anp rep mitochondrial genome maintenance engineered escherichia coli gp60 |

### Sequence information

|  |  |
| --- | --- |
| Name | FANPEZAQ\_CDS\_0041  41\_FANPEZAQ\_CDS\_0041 (pipeline id) |
| Imported annotations |  |
| Protein sequence | MKPGVYEGIPNAEYHGGPGISKSGLDLVHRSPMHYHAVVTAANDRTPTAAQELGTAAHAL ILEPDVFADTYCLALRRSDVPDAIDDREVLVEMVNKLNEGRLAKLPTSGNKNELIARIME FTGWTGETSELEAMKGAELKAIIEKANESRTGLLPTSGSRHDLAYLLRSNGVKVTLWSDV LAEWTENNPGRMVLTPETWDQLHAMAAAVHAHPAAGSLLTSCPGEAEKSVYWIDATTGVL CRCRPDWWRDDNVIVDLKTTEDASPEGFAKSIANWRYDVQAAYYLDGVQQATGKRPKAFV FIAVEKKPPYGVGVYVLDSESVDLGRAQYQHDLRVYAECVRTGVWPGYGDKIQTISLPGW HANKNQHLLGAV |
| Number of residues | 372 |
| Molecular weight (Da) | 40833.65 |
| Output files | ../../query\_sequences/41\_FANPEZAQ\_CDS\_0041.fasta |

### Putative domain architecture and protein family

#### Search results (HHblits)^1^

|  |  |
| --- | --- |
| Domain family databases searched | Pfam, Ncbi-cd, Cath, Phrogs |
| Results, scheme(s)  (Top layers only; threshold 1.00e-03 (evalue)) | xml version="1.0" encoding="utf-8" standalone="no"?       2024-09-02T21:08:21.329101 image/svg+xml   Matplotlib v3.7.2, https://matplotlib.org/ |
| Results, table  (E-value ≤ 1.00e-03 (evalue)) | db \| id \| prob \| evalue \| pvalue \| score \| cols \| query \| query\_len \| template \| template\_len \| name \| description; pfam \| PF12684 \| 97.5 \| 2.2e-08 \| 5.4e-12 \| 79.8 \| 122 \| (225, 348) \| 372 \| (95, 225) \| 235 \| DUF3799 \| PDDEXK-like domain of unknown function (DUF3799); pfam \| PF10926 \| 96.8 \| 1.1e-06 \| 2.5e-10 \| 78.1 \| 91 \| (238, 339) \| 372 \| (121, 218) \| 406 \| DUF2800 \| Protein of unknown function (DUF2800); pfam \| PF06023 \| 96.5 \| 3e-06 \| 7.1e-10 \| 68.9 \| 105 \| (226, 348) \| 372 \| (157, 266) \| 281 \| Csa1 \| CRISPR-associated exonuclease Csa1; pfam \| PF12705 \| 95.5 \| 6.7e-05 \| 1.6e-08 \| 59.4 \| 63 \| (238, 304) \| 372 \| (136, 202) \| 272 \| PDDEXK\_1 \| PD-(D/E)XK nuclease superfamily; pfam \| PF01930 \| 93.7 \| 0.00077 \| 1.8e-07 \| 48.9 \| 86 \| (243, 347) \| 372 \| (57, 144) \| 162 \| Cas\_Cas4 \| Domain of unknown function DUF83; cath \| 3h4rA00 \| 99.7 \| 1.3e-21 \| 2.1e-25 \| 170.9 \| 225 \| (8, 365) \| 372 \| (1, 229) \| 231 \| Exodeoxyribonuclease 8 \| CATHCODE: 3.90.320.10 NAME: Exodeoxyribonuclease 8. Chain: a. Fragment: c-terminal domain: unp residues 606-866. Synonym: exodeoxyribonuclease viii, exo viii. Engineered: yes. Mutation: yes SOURCE: Escherichia coli. Organism\_taxid: 83333. Strain: k-12. Gene: b1350, jw1344, rece. Expressed in: escherichia coli. Expression\_system\_taxid: 562. CLASS: Alpha Beta, ARCH: Alpha-Beta Complex, TOPOL: Lambda Exonuclease; Chain A, HOMOL: Lambda Exonuclease; Chain A; cath \| 3l0aA00 \| 99.2 \| 5.9e-16 \| 9.4e-20 \| 137.0 \| 221 \| (7, 348) \| 372 \| (2, 240) \| 266 \| Putative exonuclease \| CATHCODE: 3.90.320.10 NAME: Putative exonuclease. Chain: a. Engineered: yes SOURCE: Eubacterium rectale. Organism\_taxid: 515619. Strain: atcc 33656 / vpi 0990. Gene: eubrec\_2131, rer070207002219. Expressed in: escherichia coli. Expression\_system\_taxid: 562. CLASS: Alpha Beta, ARCH: Alpha-Beta Complex, TOPOL: Lambda Exonuclease; Chain A, HOMOL: Lambda Exonuclease; Chain A; cath \| 3u4qA06 \| 97.6 \| 2.5e-08 \| 4e-12 \| 85.8 \| 58 \| (238, 296) \| 372 \| (163, 228) \| 244 \| Atp-dependent helicase/nuclease subunit a \| CATHCODE: 3.90.320.10 NAME: Atp-dependent helicase/nuclease subunit a. Chain: a. Synonym: atp-dependent helicase/nuclease adda. Ec: 3.1.-.-, 3.6.4.12. Engineered: yes. Mutation: yes. Atp-dependent helicase/deoxyribonuclease subunit b. Chain: b. Synonym: atp-dependent helicase/nuclease addb. Ec: 3.1.-.-, 3.6.4.12. Engineered: yes. Mutation: yes. Dna (27-mer). Chain: x. Engineered: yes SOURCE: Bacillus subtilis. Organism\_taxid: 1423. Gene: adda, bsu10630. Expressed in: escherichia coli. Expression\_system\_taxid: 562. Bacillus subtilis. Organism\_taxid: 1423. Gene: addb, bsu10620. Expressed in: escherichia coli. Expression\_system\_taxid: 562. CLASS: Alpha Beta, ARCH: Alpha-Beta Complex, TOPOL: Lambda Exonuclease; Chain A, HOMOL: Lambda Exonuclease; Chain A; cath \| 4r5qA00 \| 96.6 \| 4.3e-06 \| 6.9e-10 \| 68.8 \| 48 \| (16, 63) \| 372 \| (52, 99) \| 216 \| Crispr-associated exonuclease, cas4 family \| CATHCODE: 3.90.320.10 NAME: Crispr-associated exonuclease, cas4 family. Chain: a. Engineered: yes SOURCE: Pyrobaculum calidifontis jcm 11548. Organism\_taxid: 410359. Strain: jcm 11548 / va1. Gene: pcal\_0546. Expressed in: escherichia coli. Expression\_system\_taxid: 469008. CLASS: Alpha Beta, ARCH: Alpha-Beta Complex, TOPOL: Lambda Exonuclease; Chain A, HOMOL: Lambda Exonuclease; Chain A; cath \| 4ic1A00 \| 96.2 \| 1.4e-05 \| 2.3e-09 \| 63.7 \| 49 \| (237, 291) \| 372 \| (94, 146) \| 206 \| Uncharacterized protein \| CATHCODE: 3.90.320.10 NAME: Uncharacterized protein. Chain: a, b, c, d, f, g, h, i, j, k. Engineered: yes SOURCE: Sulfolobus solfataricus. Organism\_taxid: 273057. Strain: atcc 35092 / dsm 1617 / jcm 11322 / p2. Gene: sso0001. Expressed in: escherichia coli. Expression\_system\_taxid: 469008. CLASS: Alpha Beta, ARCH: Alpha-Beta Complex, TOPOL: Lambda Exonuclease; Chain A, HOMOL: Lambda Exonuclease; Chain A; phrogs \| 412 \| 100.0 \| 1.1e-63 \| 1.4e-67 \| 473.5 \| 248 \| (2, 353) \| 372 \| (4, 265) \| 267 \| exonuclease VIII \| exonuclease VIII; Category: DNA, RNA and nucleotide metabolism; p123465 VI\_04287; phrogs \| 28222 \| 99.9 \| 1.5e-29 \| 1.6e-33 \| 250.5 \| 276 \| (4, 348) \| 372 \| (639, 924) \| 941 \| exonuclease \| exonuclease; Category: DNA, RNA and nucleotide metabolism; NC\_020850\_p8; phrogs \| 1227 \| 99.9 \| 6.3e-27 \| 8.1e-31 \| 218.9 \| 150 \| (195, 347) \| 372 \| (77, 246) \| 268 \| exonuclease \| exonuclease; Category: DNA, RNA and nucleotide metabolism; KX456208\_p10 |
| Top keywords  (threshold 1.00e-03 (evalue)) | **Exonuclease, A, yes, Lambda, Engineered, escherichia, coli, Organism\_taxid, Gene, Expressed** |
| Output files | ../../domain\_architecture/41\_FANPEZAQ\_CDS\_0041\_cath.hhr ../../domain\_architecture/41\_FANPEZAQ\_CDS\_0041\_merged.svg ../../domain\_architecture/41\_FANPEZAQ\_CDS\_0041\_ncbi-cd.hhr ../../domain\_architecture/41\_FANPEZAQ\_CDS\_0041\_pfam.hhr ../../domain\_architecture/41\_FANPEZAQ\_CDS\_0041\_phrogs.hhr |

### Identical protein sequences/structures

#### Search results

|  |  |
| --- | --- |
| Protein sequence databases searched | Pdb, Swissprot, Refseq |
| Identical proteins found | -- |
| Top keywords | -- |
| Output files | -- |

### Similar protein sequences/structures

#### Sequence similarity search results (HHblits)^1^

|  |  |
| --- | --- |
| Sequence databases searched | Uniclust, Pdb70 |
| Results, scheme(s)  (Top layers only, threshold 1.00e-03 (evalue)) | xml version="1.0" encoding="utf-8" standalone="no"?       2024-09-02T21:08:47.281446 image/svg+xml   Matplotlib v3.7.2, https://matplotlib.org/ |
| Results, table(s)  (threshold 1.00e-03 (evalue)) | db \| id \| prob \| evalue \| pvalue \| score \| cols \| query \| query\_len \| template \| template\_len \| name \| description; uniclust \| UniRef100\_A0A088FVF0 \| 100.0 \| 3.3e-70 \| 6.9e-76 \| 514.8 \| 357 \| (1, 371) \| 372 \| (95, 456) \| 476 \| Putative exodeoxyribonuclease 8 PDDEXK-like domain-containing protein \| Putative exodeoxyribonuclease 8 PDDEXK-like domain-containing protein; uniclust \| UniRef100\_A0A0F9QDF5 \| 100.0 \| 1.4e-63 \| 3.1e-69 \| 467.4 \| 271 \| (2, 368) \| 372 \| (73, 344) \| 358 \| Putative exodeoxyribonuclease 8 PDDEXK-like domain-containing protein \| Putative exodeoxyribonuclease 8 PDDEXK-like domain-containing protein; uniclust \| UniRef100\_A0A0F6SEZ4 \| 100.0 \| 3.8e-63 \| 8.2e-69 \| 454.4 \| 263 \| (6, 365) \| 372 \| (34, 296) \| 318 \| Exodeoxyribonuclease VIII \| Exodeoxyribonuclease VIII; uniclust \| UniRef100\_A0A0F5JU14 \| 100.0 \| 7.1e-63 \| 1.5e-68 \| 442.3 \| 286 \| (2, 370) \| 372 \| (23, 314) \| 325 \| Exonuclease VIII \| Exonuclease VIII; uniclust \| UniRef100\_A0A0F9TR59 \| 100.0 \| 4.9e-60 \| 1.1e-65 \| 435.5 \| 279 \| (1, 368) \| 372 \| (25, 304) \| 321 \| Putative exodeoxyribonuclease 8 PDDEXK-like domain-containing protein \| Putative exodeoxyribonuclease 8 PDDEXK-like domain-containing protein; uniclust \| UniRef100\_A0A0K6IY38 \| 100.0 \| 1.8e-59 \| 3.8e-65 \| 424.5 \| 273 \| (2, 369) \| 372 \| (61, 335) \| 343 \| Putative exodeoxyribonuclease 8 PDDEXK-like domain-containing protein \| Putative exodeoxyribonuclease 8 PDDEXK-like domain-containing protein; uniclust \| UniRef100\_A0A023W6M4 \| 100.0 \| 9.6e-59 \| 2e-64 \| 432.0 \| 269 \| (1, 370) \| 372 \| (84, 358) \| 384 \| RecB-like exonuclease/helicase \| RecB-like exonuclease/helicase; uniclust \| UniRef100\_A0A009G867 \| 100.0 \| 2.4e-58 \| 4.9e-64 \| 426.0 \| 266 \| (1, 368) \| 372 \| (87, 359) \| 378 \| Putative exodeoxyribonuclease 8 PDDEXK-like domain-containing protein \| Putative exodeoxyribonuclease 8 PDDEXK-like domain-containing protein; uniclust \| UniRef100\_A0A011UJY5 \| 100.0 \| 1.4e-56 \| 2.9e-62 \| 430.1 \| 266 \| (2, 370) \| 372 \| (148, 419) \| 504 \| Putative exodeoxyribonuclease 8 PDDEXK-like domain-containing protein \| Putative exodeoxyribonuclease 8 PDDEXK-like domain-containing protein; uniclust \| UniRef100\_A0A3D2A6G1 \| 100.0 \| 3.4e-54 \| 6.5e-60 \| 367.5 \| 265 \| (2, 361) \| 372 \| (3, 275) \| 275 \| Putative exodeoxyribonuclease 8 PDDEXK-like domain-containing protein \| Putative exodeoxyribonuclease 8 PDDEXK-like domain-containing protein; uniclust \| UniRef100\_A0A011VDN1 \| 100.0 \| 1.2e-53 \| 2.6e-59 \| 395.3 \| 280 \| (2, 370) \| 372 \| (47, 339) \| 361 \| Putative exodeoxyribonuclease 8 PDDEXK-like domain-containing protein \| Putative exodeoxyribonuclease 8 PDDEXK-like domain-containing protein; uniclust \| UniRef100\_A0A0F9EGN9 \| 100.0 \| 1.5e-53 \| 3e-59 \| 379.5 \| 263 \| (1, 364) \| 372 \| (8, 272) \| 295 \| Putative exodeoxyribonuclease 8 PDDEXK-like domain-containing protein (Fragment) \| Putative exodeoxyribonuclease 8 PDDEXK-like domain-containing protein (Fragment); uniclust \| UniRef100\_A0A1K1LD38 \| 100.0 \| 2.5e-51 \| 5.1e-57 \| 365.4 \| 270 \| (1, 368) \| 372 \| (9, 283) \| 312 \| Exodeoxyribonuclease VIII \| Exodeoxyribonuclease VIII; uniclust \| UniRef100\_A0A318CJ92 \| 100.0 \| 3.9e-51 \| 7.7e-57 \| 361.4 \| 265 \| (1, 367) \| 372 \| (48, 312) \| 317 \| Putative exodeoxyribonuclease 8 PDDEXK-like domain-containing protein \| Putative exodeoxyribonuclease 8 PDDEXK-like domain-containing protein; uniclust \| UniRef100\_A0A011MMW1 \| 100.0 \| 7.7e-51 \| 1.4e-56 \| 345.0 \| 265 \| (3, 367) \| 372 \| (6, 272) \| 296 \| Exodeoxyribonuclease 8 \| Exodeoxyribonuclease 8; uniclust \| UniRef100\_UPI0018DF9E2C \| 100.0 \| 3.4e-50 \| 6.3e-56 \| 351.4 \| 369 \| (2, 371) \| 372 \| (12, 454) \| 459 \| PD-(D/E)XK nuclease-like domain-containing protein \| PD-(D/E)XK nuclease-like domain-containing protein; uniclust \| UniRef100\_A0A023ZU70 \| 100.0 \| 4.8e-48 \| 1e-53 \| 355.6 \| 302 \| (7, 370) \| 372 \| (50, 359) \| 365 \| Putative exodeoxyribonuclease VIII \| Putative exodeoxyribonuclease VIII; uniclust \| UniRef100\_A0A2U9P084 \| 100.0 \| 2.4e-47 \| 4.6e-53 \| 351.3 \| 268 \| (2, 369) \| 372 \| (214, 491) \| 499 \| Putative exodeoxyribonuclease 8 PDDEXK-like domain-containing protein \| Putative exodeoxyribonuclease 8 PDDEXK-like domain-containing protein; uniclust \| UniRef100\_A0A0F6B083 \| 100.0 \| 2.8e-47 \| 5.6e-53 \| 376.5 \| 273 \| (1, 372) \| 372 \| (610, 886) \| 889 \| Exodeoxyribonuclease 8 \| Exodeoxyribonuclease 8; uniclust \| UniRef100\_A0A1Z7ZMG2 \| 100.0 \| 4.8e-47 \| 9.4e-53 \| 330.7 \| 261 \| (2, 370) \| 372 \| (21, 285) \| 296 \| Putative exodeoxyribonuclease 8 PDDEXK-like domain-containing protein \| Putative exodeoxyribonuclease 8 PDDEXK-like domain-containing protein; uniclust \| UniRef100\_A0A1E4WVT7 \| 100.0 \| 8e-47 \| 1.5e-52 \| 344.3 \| 357 \| (2, 368) \| 372 \| (100, 522) \| 562 \| Putative exodeoxyribonuclease 8 PDDEXK-like domain-containing protein \| Putative exodeoxyribonuclease 8 PDDEXK-like domain-containing protein; uniclust \| UniRef100\_A0A076LUB6 \| 100.0 \| 1.1e-46 \| 2.3e-52 \| 385.2 \| 354 \| (2, 369) \| 372 \| (689, 1110) \| 1223 \| Exodeoxyribonuclease VIII \| Exodeoxyribonuclease VIII; uniclust \| UniRef100\_A0A2W1SGM9 \| 100.0 \| 4.7e-46 \| 9e-52 \| 337.8 \| 271 \| (2, 368) \| 372 \| (166, 442) \| 444 \| Putative exodeoxyribonuclease 8 PDDEXK-like domain-containing protein \| Putative exodeoxyribonuclease 8 PDDEXK-like domain-containing protein; uniclust \| UniRef100\_A0A6J5P3Q0 \| 100.0 \| 1.7e-45 \| 3.5e-51 \| 328.7 \| 257 \| (9, 365) \| 372 \| (33, 290) \| 296 \| Exodeoxyribonuclease 8, PDDEXK-like domain containing protein \| Exodeoxyribonuclease 8, PDDEXK-like domain containing protein; uniclust \| UniRef100\_A0A061SV84 \| 100.0 \| 2e-45 \| 4.1e-51 \| 343.5 \| 264 \| (1, 368) \| 372 \| (91, 367) \| 404 \| Putative exodeoxyribonuclease 8 PDDEXK-like domain-containing protein \| Putative exodeoxyribonuclease 8 PDDEXK-like domain-containing protein; uniclust \| UniRef100\_A0A0A0Q3N4 \| 100.0 \| 3.5e-45 \| 6.7e-51 \| 319.9 \| 278 \| (3, 371) \| 372 \| (78, 360) \| 368 \| PDDEXK-like domain \| PDDEXK-like domain; uniclust \| UniRef100\_A0A0D8L788 \| 100.0 \| 8.2e-45 \| 1.6e-50 \| 335.4 \| 352 \| (2, 368) \| 372 \| (150, 569) \| 577 \| DNA breaking-rejoining protein \| DNA breaking-rejoining protein; uniclust \| UniRef100\_A0A0F9G9B6 \| 100.0 \| 8.3e-45 \| 1.7e-50 \| 329.1 \| 265 \| (2, 368) \| 372 \| (15, 295) \| 330 \| Putative exodeoxyribonuclease 8 PDDEXK-like domain-containing protein \| Putative exodeoxyribonuclease 8 PDDEXK-like domain-containing protein; uniclust \| UniRef100\_A0A0J6S4G7 \| 100.0 \| 3.7e-44 \| 7.7e-50 \| 330.6 \| 295 \| (1, 365) \| 372 \| (37, 368) \| 373 \| Putative exodeoxyribonuclease 8 PDDEXK-like domain-containing protein \| Putative exodeoxyribonuclease 8 PDDEXK-like domain-containing protein; uniclust \| UniRef100\_A0A523QSH7 \| 100.0 \| 3.1e-43 \| 5.9e-49 \| 300.7 \| 264 \| (1, 364) \| 372 \| (21, 292) \| 297 \| Putative exodeoxyribonuclease 8 PDDEXK-like domain-containing protein \| Putative exodeoxyribonuclease 8 PDDEXK-like domain-containing protein; uniclust \| UniRef100\_A0A0F9RJV3 \| 100.0 \| 8.3e-43 \| 1.6e-48 \| 299.9 \| 266 \| (3, 365) \| 372 \| (7, 273) \| 291 \| Putative exodeoxyribonuclease 8 PDDEXK-like domain-containing protein \| Putative exodeoxyribonuclease 8 PDDEXK-like domain-containing protein; uniclust \| UniRef100\_A0A0F7L8T5 \| 100.0 \| 1.2e-42 \| 2.4e-48 \| 309.4 \| 234 \| (9, 349) \| 372 \| (39, 274) \| 293 \| Putative exodeoxyribonuclease 8 PDDEXK-like domain-containing protein \| Putative exodeoxyribonuclease 8 PDDEXK-like domain-containing protein; uniclust \| UniRef100\_A0A7X5Z2D5 \| 100.0 \| 2.8e-42 \| 5.5e-48 \| 312.4 \| 266 \| (2, 369) \| 372 \| (124, 394) \| 407 \| Putative exodeoxyribonuclease 8 PDDEXK-like domain-containing protein \| Putative exodeoxyribonuclease 8 PDDEXK-like domain-containing protein; uniclust \| UniRef100\_A0A501Y166 \| 100.0 \| 3.5e-42 \| 6.8e-48 \| 316.2 \| 336 \| (1, 357) \| 372 \| (86, 430) \| 441 \| Putative exodeoxyribonuclease 8 PDDEXK-like domain-containing protein \| Putative exodeoxyribonuclease 8 PDDEXK-like domain-containing protein; uniclust \| UniRef100\_A0A4R4A4G8 \| 100.0 \| 6.9e-42 \| 1.3e-47 \| 285.8 \| 255 \| (3, 359) \| 372 \| (12, 270) \| 273 \| Exodeoxyribonuclease VIII \| Exodeoxyribonuclease VIII; uniclust \| UniRef100\_A0A2D7NC52 \| 100.0 \| 8e-42 \| 1.6e-47 \| 297.1 \| 260 \| (3, 367) \| 372 \| (16, 277) \| 289 \| Putative exodeoxyribonuclease 8 PDDEXK-like domain-containing protein \| Putative exodeoxyribonuclease 8 PDDEXK-like domain-containing protein; uniclust \| UniRef100\_A0A192A857 \| 100.0 \| 1.2e-41 \| 2.5e-47 \| 308.8 \| 260 \| (5, 366) \| 372 \| (110, 375) \| 392 \| Putative exodeoxyribonuclease 8 PDDEXK-like domain-containing protein \| Putative exodeoxyribonuclease 8 PDDEXK-like domain-containing protein; uniclust \| UniRef100\_A0A254QH28 \| 100.0 \| 2.3e-41 \| 4.5e-47 \| 282.1 \| 206 \| (1, 306) \| 372 \| (6, 211) \| 211 \| Putative exodeoxyribonuclease 8 PDDEXK-like domain-containing protein (Fragment) \| Putative exodeoxyribonuclease 8 PDDEXK-like domain-containing protein (Fragment); uniclust \| UniRef100\_A0A661RLU8 \| 100.0 \| 7.1e-41 \| 1.4e-46 \| 289.6 \| 272 \| (1, 366) \| 372 \| (15, 296) \| 306 \| Putative exodeoxyribonuclease 8 PDDEXK-like domain-containing protein \| Putative exodeoxyribonuclease 8 PDDEXK-like domain-containing protein; uniclust \| UniRef100\_A0A1X0YIJ6 \| 100.0 \| 1.2e-40 \| 2.3e-46 \| 300.8 \| 353 \| (2, 368) \| 372 \| (49, 469) \| 481 \| Exonuclease VIII (Fragment) \| Exonuclease VIII (Fragment); uniclust \| UniRef100\_A0A378FZ40 \| 100.0 \| 1.7e-40 \| 3.3e-46 \| 322.1 \| 359 \| (2, 369) \| 372 \| (579, 1000) \| 1005 \| Putative exodeoxyribonuclease VIII \| Putative exodeoxyribonuclease VIII; uniclust \| UniRef100\_A0A2I7RCC9 \| 100.0 \| 2.1e-40 \| 4.1e-46 \| 282.9 \| 208 \| (1, 306) \| 372 \| (36, 247) \| 253 \| Uncharacterized protein \| Uncharacterized protein; uniclust \| UniRef100\_A0A139BNH5 \| 100.0 \| 4.1e-40 \| 8.1e-46 \| 274.2 \| 187 \| (4, 288) \| 372 \| (15, 201) \| 202 \| Putative exodeoxyribonuclease 8 PDDEXK-like domain-containing protein (Fragment) \| Putative exodeoxyribonuclease 8 PDDEXK-like domain-containing protein (Fragment); uniclust \| UniRef100\_A0A090EAA3 \| 100.0 \| 5e-40 \| 1e-45 \| 303.5 \| 307 \| (1, 365) \| 372 \| (63, 407) \| 415 \| Putative exodeoxyribonuclease 8 PDDEXK-like domain-containing protein \| Putative exodeoxyribonuclease 8 PDDEXK-like domain-containing protein; uniclust \| UniRef100\_A0A142XW82 \| 100.0 \| 1.8e-39 \| 3.4e-45 \| 276.2 \| 258 \| (10, 363) \| 372 \| (5, 268) \| 285 \| Exodeoxyribonuclease 8 \| Exodeoxyribonuclease 8; uniclust \| UniRef100\_UPI0005E1294F \| 100.0 \| 2e-39 \| 3.6e-45 \| 279.6 \| 261 \| (3, 366) \| 372 \| (94, 361) \| 384 \| PD-(D/E)XK nuclease-like domain-containing protein \| PD-(D/E)XK nuclease-like domain-containing protein; uniclust \| UniRef100\_A0A0F6B562 \| 100.0 \| 3e-39 \| 5.7e-45 \| 314.3 \| 353 \| (2, 368) \| 372 \| (558, 978) \| 987 \| Exodeoxyribonuclease VIII-like protein \| Exodeoxyribonuclease VIII-like protein; uniclust \| UniRef100\_E6QW59 \| 100.0 \| 3.8e-39 \| 6.9e-45 \| 281.2 \| 354 \| (5, 365) \| 372 \| (4, 424) \| 427 \| Putative exodeoxyribonuclease 8 PDDEXK-like domain-containing protein \| Putative exodeoxyribonuclease 8 PDDEXK-like domain-containing protein; uniclust \| UniRef100\_A0A071LVK8 \| 100.0 \| 7.3e-39 \| 1.4e-44 \| 301.8 \| 353 \| (2, 368) \| 372 \| (324, 744) \| 769 \| Exodeoxyribonuclease VIII \| Exodeoxyribonuclease VIII; uniclust \| UniRef100\_A0A370CF63 \| 100.0 \| 1e-38 \| 1.9e-44 \| 285.5 \| 263 \| (1, 362) \| 372 \| (11, 275) \| 541 \| Putative exodeoxyribonuclease 8 PDDEXK-like domain-containing protein \| Putative exodeoxyribonuclease 8 PDDEXK-like domain-containing protein; uniclust \| UniRef100\_A0A076JDP6 \| 100.0 \| 9.3e-39 \| 1.9e-44 \| 285.6 \| 261 \| (6, 371) \| 372 \| (17, 296) \| 299 \| Phage-related exonuclease, PD-(D/E)XK superfamily \| Phage-related exonuclease, PD-(D/E)XK superfamily; uniclust \| UniRef100\_A0A0S8J9S9 \| 100.0 \| 9.6e-39 \| 1.9e-44 \| 287.6 \| 257 \| (6, 359) \| 372 \| (69, 328) \| 340 \| Putative exodeoxyribonuclease 8 PDDEXK-like domain-containing protein \| Putative exodeoxyribonuclease 8 PDDEXK-like domain-containing protein; uniclust \| UniRef100\_A0A448S1F5 \| 100.0 \| 1.7e-38 \| 3.2e-44 \| 278.8 \| 267 \| (1, 368) \| 372 \| (1, 273) \| 451 \| Exodeoxyribonuclease 8 \| Exodeoxyribonuclease 8; uniclust \| UniRef100\_A0A505CFK2 \| 100.0 \| 3.8e-38 \| 7e-44 \| 288.1 \| 265 \| (1, 364) \| 372 \| (414, 682) \| 685 \| Exodeoxyribonuclease 8 \| Exodeoxyribonuclease 8; uniclust \| UniRef100\_A0A1B3AZZ2 \| 100.0 \| 4.9e-38 \| 9.1e-44 \| 277.9 \| 259 \| (3, 370) \| 372 \| (168, 429) \| 440 \| Exonuclease \| Exonuclease; uniclust \| UniRef100\_A0A2D8XAW4 \| 100.0 \| 6.3e-38 \| 1.2e-43 \| 268.5 \| 261 \| (3, 364) \| 372 \| (13, 283) \| 309 \| Putative exodeoxyribonuclease 8 PDDEXK-like domain-containing protein \| Putative exodeoxyribonuclease 8 PDDEXK-like domain-containing protein; uniclust \| UniRef100\_A0A077ND06 \| 100.0 \| 1.6e-37 \| 3.1e-43 \| 264.0 \| 261 \| (1, 364) \| 372 \| (4, 268) \| 276 \| Phage-related exonuclease \| Phage-related exonuclease; uniclust \| UniRef100\_A0A376LPM8 \| 100.0 \| 2e-37 \| 3.8e-43 \| 298.1 \| 264 \| (1, 363) \| 372 \| (505, 772) \| 836 \| Exonuclease VIII \| Exonuclease VIII; uniclust \| UniRef100\_A0A1F9UXV9 \| 100.0 \| 2.8e-37 \| 5.6e-43 \| 269.3 \| 253 \| (8, 369) \| 372 \| (9, 265) \| 273 \| Uncharacterized protein \| Uncharacterized protein; uniclust \| UniRef100\_A0A1Z9H6K7 \| 100.0 \| 3e-37 \| 5.9e-43 \| 266.5 \| 239 \| (4, 345) \| 372 \| (36, 276) \| 280 \| Putative exodeoxyribonuclease 8 PDDEXK-like domain-containing protein \| Putative exodeoxyribonuclease 8 PDDEXK-like domain-containing protein; uniclust \| UniRef100\_A0A0P0IK53 \| 100.0 \| 6.1e-37 \| 1.2e-42 \| 268.2 \| 307 \| (7, 369) \| 372 \| (19, 342) \| 349 \| Exonuclease \| Exonuclease; uniclust \| UniRef100\_A0A3B8UEV6 \| 100.0 \| 6.7e-37 \| 1.2e-42 \| 266.4 \| 273 \| (2, 370) \| 372 \| (111, 396) \| 412 \| Putative exodeoxyribonuclease 8 PDDEXK-like domain-containing protein \| Putative exodeoxyribonuclease 8 PDDEXK-like domain-containing protein; uniclust \| UniRef100\_Q11ZS8 \| 100.0 \| 9.6e-37 \| 1.8e-42 \| 266.7 \| 253 \| (8, 360) \| 372 \| (82, 340) \| 358 \| Putative exodeoxyribonuclease 8 PDDEXK-like domain-containing protein \| Putative exodeoxyribonuclease 8 PDDEXK-like domain-containing protein; uniclust \| UniRef100\_A0A379GM57 \| 100.0 \| 1.7e-36 \| 3.1e-42 \| 276.0 \| 352 \| (2, 368) \| 372 \| (150, 570) \| 575 \| Exodeoxyribonuclease 8 \| Exodeoxyribonuclease 8; uniclust \| UniRef100\_A0A0J9D3R0 \| 100.0 \| 2.7e-36 \| 5.1e-42 \| 268.8 \| 275 \| (2, 369) \| 372 \| (121, 432) \| 442 \| Putative exodeoxyribonuclease 8 PDDEXK-like domain-containing protein \| Putative exodeoxyribonuclease 8 PDDEXK-like domain-containing protein; uniclust \| UniRef100\_UPI001C312083 \| 100.0 \| 5.7e-36 \| 1e-41 \| 256.7 \| 262 \| (2, 362) \| 372 \| (91, 358) \| 359 \| PD-(D/E)XK nuclease-like domain-containing protein \| PD-(D/E)XK nuclease-like domain-containing protein; uniclust \| UniRef100\_A0A2X7FM56 \| 100.0 \| 5.8e-36 \| 1.1e-41 \| 281.6 \| 355 \| (2, 368) \| 372 \| (353, 773) \| 776 \| Putative phage exodeoxyribonuclease \| Putative phage exodeoxyribonuclease; uniclust \| UniRef100\_A0A0F9KSW2 \| 100.0 \| 5.7e-36 \| 1.1e-41 \| 257.3 \| 260 \| (8, 365) \| 372 \| (10, 272) \| 277 \| Putative exodeoxyribonuclease 8 PDDEXK-like domain-containing protein \| Putative exodeoxyribonuclease 8 PDDEXK-like domain-containing protein; uniclust \| UniRef100\_UPI001961519C \| 100.0 \| 1.5e-35 \| 2.8e-41 \| 253.2 \| 261 \| (8, 369) \| 372 \| (78, 347) \| 349 \| PD-(D/E)XK nuclease-like domain-containing protein \| PD-(D/E)XK nuclease-like domain-containing protein; uniclust \| UniRef100\_A0A3N0CLS5 \| 100.0 \| 1.8e-35 \| 3.5e-41 \| 256.6 \| 263 \| (2, 362) \| 372 \| (12, 280) \| 294 \| Putative exodeoxyribonuclease 8 PDDEXK-like domain-containing protein \| Putative exodeoxyribonuclease 8 PDDEXK-like domain-containing protein; uniclust \| UniRef100\_A0A2R5F5H8 \| 100.0 \| 1.9e-35 \| 3.6e-41 \| 254.4 \| 256 \| (7, 362) \| 372 \| (31, 289) \| 292 \| Coproporphyrinogen III oxidase \| Coproporphyrinogen III oxidase; uniclust \| UniRef100\_H8FP23 \| 100.0 \| 2.4e-35 \| 4.4e-41 \| 254.4 \| 267 \| (2, 365) \| 372 \| (64, 350) \| 378 \| Putative exodeoxyribonuclease 8 PDDEXK-like domain-containing protein \| Putative exodeoxyribonuclease 8 PDDEXK-like domain-containing protein; uniclust \| UniRef100\_A0A1V5GYE4 \| 100.0 \| 2.4e-35 \| 4.7e-41 \| 257.5 \| 259 \| (1, 365) \| 372 \| (9, 272) \| 283 \| Exodeoxyribonuclease 8 \| Exodeoxyribonuclease 8; uniclust \| UniRef100\_UPI001C47B09F \| 100.0 \| 3.7e-35 \| 6.8e-41 \| 254.1 \| 258 \| (7, 362) \| 372 \| (100, 378) \| 389 \| PD-(D/E)XK nuclease-like domain-containing protein \| PD-(D/E)XK nuclease-like domain-containing protein; uniclust \| UniRef100\_A0A6J7VNU5 \| 100.0 \| 4.8e-35 \| 9.2e-41 \| 251.4 \| 261 \| (3, 365) \| 372 \| (9, 270) \| 276 \| Exodeoxyribonuclease 8, PDDEXK-like domain containing protein \| Exodeoxyribonuclease 8, PDDEXK-like domain containing protein; uniclust \| UniRef100\_A0A0F8VZZ2 \| 100.0 \| 4.9e-35 \| 9.4e-41 \| 241.7 \| 178 \| (187, 367) \| 372 \| (33, 214) \| 215 \| Putative exodeoxyribonuclease 8 PDDEXK-like domain-containing protein (Fragment) \| Putative exodeoxyribonuclease 8 PDDEXK-like domain-containing protein (Fragment); uniclust \| UniRef100\_A0A2G2G2Z2 \| 100.0 \| 7.8e-35 \| 1.5e-40 \| 253.2 \| 249 \| (2, 348) \| 372 \| (5, 264) \| 289 \| Putative exodeoxyribonuclease 8 PDDEXK-like domain-containing protein \| Putative exodeoxyribonuclease 8 PDDEXK-like domain-containing protein; uniclust \| UniRef100\_A0A437M7Z1 \| 100.0 \| 9.6e-35 \| 1.8e-40 \| 259.8 \| 304 \| (2, 363) \| 372 \| (45, 384) \| 387 \| Exodeoxyribonuclease 8 PDDEXK-like domain-containing protein \| Exodeoxyribonuclease 8 PDDEXK-like domain-containing protein; uniclust \| UniRef100\_A0A377Q5C7 \| 100.0 \| 1.3e-34 \| 2.3e-40 \| 245.1 \| 288 \| (2, 353) \| 372 \| (10, 301) \| 319 \| Exodeoxyribonuclease 8 \| Exodeoxyribonuclease 8; uniclust \| UniRef100\_A0A086A3E8 \| 100.0 \| 1.5e-34 \| 3.1e-40 \| 268.8 \| 308 \| (3, 365) \| 372 \| (83, 405) \| 416 \| Uncharacterized protein \| Uncharacterized protein; uniclust \| UniRef100\_A0A2I5T572 \| 100.0 \| 6e-34 \| 1.2e-39 \| 281.6 \| 359 \| (2, 369) \| 372 \| (552, 973) \| 980 \| Exodeoxyribonuclease VIII \| Exodeoxyribonuclease VIII; uniclust \| UniRef100\_A0A7K0GP64 \| 100.0 \| 6.7e-34 \| 1.4e-39 \| 231.7 \| 142 \| (225, 366) \| 372 \| (9, 151) \| 156 \| Exodeoxyribonuclease VIII \| Exodeoxyribonuclease VIII; uniclust \| UniRef100\_UPI001438A939 \| 100.0 \| 7.4e-34 \| 1.4e-39 \| 244.3 \| 260 \| (2, 365) \| 372 \| (49, 312) \| 318 \| PD-(D/E)XK nuclease-like domain-containing protein \| PD-(D/E)XK nuclease-like domain-containing protein; uniclust \| UniRef100\_UPI0009ACE8C2 \| 100.0 \| 9e-34 \| 1.7e-39 \| 235.8 \| 211 \| (1, 306) \| 372 \| (41, 252) \| 260 \| PD-(D/E)XK nuclease-like domain-containing protein \| PD-(D/E)XK nuclease-like domain-containing protein; uniclust \| UniRef100\_A0A433FYR1 \| 100.0 \| 9.2e-34 \| 1.7e-39 \| 245.3 \| 281 \| (4, 369) \| 372 \| (93, 379) \| 384 \| Putative exodeoxyribonuclease 8 PDDEXK-like domain-containing protein \| Putative exodeoxyribonuclease 8 PDDEXK-like domain-containing protein; uniclust \| UniRef100\_A0A7Y6TYS2 \| 100.0 \| 1.6e-33 \| 2.9e-39 \| 243.2 \| 260 \| (4, 361) \| 372 \| (112, 374) \| 374 \| PD-(D/E)XK nuclease-like domain-containing protein \| PD-(D/E)XK nuclease-like domain-containing protein; uniclust \| UniRef100\_A0A2D7BYZ1 \| 100.0 \| 1.6e-33 \| 3e-39 \| 240.9 \| 248 \| (2, 356) \| 372 \| (30, 279) \| 284 \| Putative exodeoxyribonuclease 8 PDDEXK-like domain-containing protein \| Putative exodeoxyribonuclease 8 PDDEXK-like domain-containing protein; uniclust \| UniRef100\_A0A2S6AMK5 \| 100.0 \| 2.1e-33 \| 3.8e-39 \| 240.8 \| 257 \| (2, 361) \| 372 \| (81, 347) \| 353 \| RecE \| RecE; uniclust \| UniRef100\_A0A759M5V4 \| 100.0 \| 3.3e-33 \| 6.2e-39 \| 248.9 \| 309 \| (3, 368) \| 372 \| (140, 461) \| 474 \| Putative exodeoxyribonuclease 8 PDDEXK-like domain-containing protein \| Putative exodeoxyribonuclease 8 PDDEXK-like domain-containing protein; uniclust \| UniRef100\_A0A3S0UCX3 \| 100.0 \| 1.5e-32 \| 2.8e-38 \| 235.1 \| 234 \| (9, 349) \| 372 \| (1, 238) \| 272 \| Putative exodeoxyribonuclease 8 PDDEXK-like domain-containing protein \| Putative exodeoxyribonuclease 8 PDDEXK-like domain-containing protein; uniclust \| UniRef100\_A0A0K8Q1C6 \| 100.0 \| 1.5e-32 \| 3.1e-38 \| 226.3 \| 137 \| (229, 366) \| 372 \| (1, 138) \| 159 \| Exodeoxyribonuclease 8 \| Exodeoxyribonuclease 8; uniclust \| UniRef100\_A0A6P1CUJ9 \| 99.9 \| 3.3e-32 \| 6.1e-38 \| 240.9 \| 257 \| (2, 363) \| 372 \| (6, 269) \| 464 \| Putative exodeoxyribonuclease 8 PDDEXK-like domain-containing protein \| Putative exodeoxyribonuclease 8 PDDEXK-like domain-containing protein; uniclust \| UniRef100\_A0A5Q4ZYH7 \| 99.9 \| 3.1e-32 \| 6.1e-38 \| 266.1 \| 369 \| (2, 371) \| 372 \| (57, 623) \| 683 \| Uncharacterized protein \| Uncharacterized protein; uniclust \| UniRef100\_A0A6H1ZNC6 \| 99.9 \| 3.6e-32 \| 6.8e-38 \| 235.8 \| 287 \| (2, 368) \| 372 \| (29, 324) \| 325 \| Putative exodeoxyribonuclease 8 PDDEXK-like domain-containing protein \| Putative exodeoxyribonuclease 8 PDDEXK-like domain-containing protein; uniclust \| UniRef100\_A0A936Z580 \| 99.9 \| 4.8e-32 \| 8.7e-38 \| 228.7 \| 264 \| (5, 365) \| 372 \| (8, 301) \| 307 \| PD-(D/E)XK nuclease-like domain-containing protein \| PD-(D/E)XK nuclease-like domain-containing protein; uniclust \| UniRef100\_A0A196PBF4 \| 99.9 \| 5.1e-32 \| 1e-37 \| 242.1 \| 282 \| (2, 365) \| 372 \| (20, 315) \| 326 \| Putative exodeoxyribonuclease 8 PDDEXK-like domain-containing protein \| Putative exodeoxyribonuclease 8 PDDEXK-like domain-containing protein; uniclust \| UniRef100\_A0A9C7CW69 \| 99.9 \| 8.7e-32 \| 1.6e-37 \| 225.0 \| 261 \| (6, 362) \| 372 \| (17, 283) \| 284 \| Exodeoxyribonuclease 8 \| Exodeoxyribonuclease 8; uniclust \| UniRef100\_A0A0S8DSQ9 \| 99.9 \| 1.1e-31 \| 2e-37 \| 223.1 \| 262 \| (2, 364) \| 372 \| (3, 268) \| 272 \| Putative exodeoxyribonuclease 8 PDDEXK-like domain-containing protein \| Putative exodeoxyribonuclease 8 PDDEXK-like domain-containing protein; uniclust \| UniRef100\_UPI0006FF5E4D \| 99.9 \| 1.8e-31 \| 3.4e-37 \| 230.2 \| 179 \| (186, 368) \| 372 \| (46, 231) \| 256 \| PD-(D/E)XK nuclease-like domain-containing protein \| PD-(D/E)XK nuclease-like domain-containing protein; uniclust \| UniRef100\_A0A146GG86 \| 99.9 \| 2.1e-31 \| 3.8e-37 \| 224.2 \| 269 \| (1, 368) \| 372 \| (15, 293) \| 298 \| Putative exodeoxyribonuclease 8 PDDEXK-like domain-containing protein \| Putative exodeoxyribonuclease 8 PDDEXK-like domain-containing protein; uniclust \| UniRef100\_A0A7C5SKK8 \| 99.9 \| 4.2e-31 \| 7.9e-37 \| 213.7 \| 162 \| (207, 370) \| 372 \| (5, 169) \| 182 \| Putative exodeoxyribonuclease 8 PDDEXK-like domain-containing protein \| Putative exodeoxyribonuclease 8 PDDEXK-like domain-containing protein; uniclust \| UniRef100\_UPI001C485F61 \| 99.9 \| 8.2e-31 \| 1.5e-36 \| 225.9 \| 280 \| (2, 288) \| 372 \| (24, 362) \| 362 \| PD-(D/E)XK nuclease-like domain-containing protein \| PD-(D/E)XK nuclease-like domain-containing protein; uniclust \| UniRef100\_I6APA3 \| 99.9 \| 9e-31 \| 1.7e-36 \| 223.8 \| 285 \| (4, 371) \| 372 \| (13, 317) \| 320 \| Putative exodeoxyribonuclease 8 PDDEXK-like domain-containing protein \| Putative exodeoxyribonuclease 8 PDDEXK-like domain-containing protein; uniclust \| UniRef100\_A0A965K260 \| 99.9 \| 1.2e-30 \| 2.2e-36 \| 221.4 \| 279 \| (2, 363) \| 372 \| (10, 314) \| 317 \| Putative exodeoxyribonuclease 8 PDDEXK-like domain-containing protein \| Putative exodeoxyribonuclease 8 PDDEXK-like domain-containing protein; uniclust \| UniRef100\_UPI00190851CF \| 99.9 \| 1.9e-30 \| 3.4e-36 \| 220.0 \| 292 \| (2, 363) \| 372 \| (14, 314) \| 314 \| PD-(D/E)XK nuclease-like domain-containing protein \| PD-(D/E)XK nuclease-like domain-containing protein; uniclust \| UniRef100\_A0A0C1ZBA7 \| 99.9 \| 3.3e-30 \| 6.5e-36 \| 237.0 \| 308 \| (2, 366) \| 372 \| (62, 381) \| 385 \| Putative exodeoxyribonuclease 8 PDDEXK-like domain-containing protein \| Putative exodeoxyribonuclease 8 PDDEXK-like domain-containing protein; uniclust \| UniRef100\_A0A2E7BLG2 \| 99.9 \| 3.8e-30 \| 6.9e-36 \| 217.6 \| 257 \| (7, 367) \| 372 \| (42, 304) \| 307 \| Putative exodeoxyribonuclease 8 PDDEXK-like domain-containing protein \| Putative exodeoxyribonuclease 8 PDDEXK-like domain-containing protein; uniclust \| UniRef100\_A0A0K1H0Q4 \| 99.9 \| 5.9e-30 \| 1.1e-35 \| 241.8 \| 263 \| (5, 368) \| 372 \| (423, 692) \| 700 \| Exodeoxyribonuclease VIII \| Exodeoxyribonuclease VIII; uniclust \| UniRef100\_UPI0015E1F6CA \| 99.9 \| 8e-30 \| 1.7e-35 \| 219.2 \| 172 \| (3, 265) \| 372 \| (7, 179) \| 202 \| PD-(D/E)XK nuclease-like domain-containing protein \| PD-(D/E)XK nuclease-like domain-containing protein; uniclust \| UniRef100\_A0A2E5C063 \| 99.9 \| 9e-30 \| 1.7e-35 \| 211.3 \| 231 \| (3, 336) \| 372 \| (7, 239) \| 243 \| Putative exodeoxyribonuclease 8 PDDEXK-like domain-containing protein \| Putative exodeoxyribonuclease 8 PDDEXK-like domain-containing protein; uniclust \| UniRef100\_A0A177W5I7 \| 99.9 \| 1e-29 \| 1.9e-35 \| 218.3 \| 300 \| (4, 365) \| 372 \| (15, 320) \| 324 \| Exonuclease VIII \| Exonuclease VIII; uniclust \| UniRef100\_A0A0F8ZM74 \| 99.9 \| 1.3e-29 \| 2.4e-35 \| 208.2 \| 180 \| (184, 368) \| 372 \| (26, 205) \| 215 \| Putative exodeoxyribonuclease 8 PDDEXK-like domain-containing protein (Fragment) \| Putative exodeoxyribonuclease 8 PDDEXK-like domain-containing protein (Fragment); uniclust \| UniRef100\_UPI001E48B583 \| 99.9 \| 1.4e-29 \| 2.5e-35 \| 200.3 \| 146 \| (225, 370) \| 372 \| (7, 162) \| 167 \| PD-(D/E)XK nuclease-like domain-containing protein \| PD-(D/E)XK nuclease-like domain-containing protein; uniclust \| UniRef100\_A0A2A2HMK2 \| 99.9 \| 1.8e-29 \| 3.5e-35 \| 232.7 \| 262 \| (5, 370) \| 372 \| (254, 522) \| 531 \| Putative exodeoxyribonuclease 8 PDDEXK-like domain-containing protein \| Putative exodeoxyribonuclease 8 PDDEXK-like domain-containing protein; uniclust \| UniRef100\_A0A3M2CPY5 \| 99.9 \| 2.2e-29 \| 4e-35 \| 212.8 \| 267 \| (3, 362) \| 372 \| (16, 284) \| 303 \| Putative exodeoxyribonuclease 8 PDDEXK-like domain-containing protein \| Putative exodeoxyribonuclease 8 PDDEXK-like domain-containing protein; uniclust \| UniRef100\_UPI0021473028 \| 99.9 \| 2.5e-29 \| 4.6e-35 \| 205.0 \| 175 \| (188, 367) \| 372 \| (56, 230) \| 232 \| PD-(D/E)XK nuclease-like domain-containing protein \| PD-(D/E)XK nuclease-like domain-containing protein; uniclust \| UniRef100\_UPI00226FB7C9 \| 99.9 \| 3.3e-29 \| 6.1e-35 \| 221.4 \| 268 \| (7, 370) \| 372 \| (156, 440) \| 445 \| PD-(D/E)XK nuclease-like domain-containing protein \| PD-(D/E)XK nuclease-like domain-containing protein; uniclust \| UniRef100\_A0A0B8T7Q5 \| 99.9 \| 4.8e-29 \| 9.2e-35 \| 217.8 \| 243 \| (19, 361) \| 372 \| (37, 286) \| 288 \| Exonuclease VIII, 5'-3' specific dsDNA exonuclease \| Exonuclease VIII, 5'-3' specific dsDNA exonuclease; uniclust \| UniRef100\_A0A6H2A032 \| 99.9 \| 4.9e-29 \| 9.3e-35 \| 211.5 \| 249 \| (9, 363) \| 372 \| (2, 252) \| 267 \| Putative exodeoxyribonuclease 8 PDDEXK-like domain-containing protein \| Putative exodeoxyribonuclease 8 PDDEXK-like domain-containing protein; uniclust \| UniRef100\_A0A3A1WJX1 \| 99.9 \| 7e-29 \| 1.3e-34 \| 217.9 \| 272 \| (2, 369) \| 372 \| (120, 408) \| 418 \| Putative exodeoxyribonuclease 8 PDDEXK-like domain-containing protein \| Putative exodeoxyribonuclease 8 PDDEXK-like domain-containing protein; uniclust \| UniRef100\_A0A966V4M2 \| 99.9 \| 8.8e-29 \| 1.6e-34 \| 210.6 \| 259 \| (9, 363) \| 372 \| (1, 263) \| 279 \| Putative exodeoxyribonuclease 8 PDDEXK-like domain-containing protein \| Putative exodeoxyribonuclease 8 PDDEXK-like domain-containing protein; uniclust \| UniRef100\_UPI0005A261FB \| 99.9 \| 9.2e-29 \| 1.7e-34 \| 203.6 \| 215 \| (2, 315) \| 372 \| (16, 241) \| 248 \| PD-(D/E)XK nuclease-like domain-containing protein \| PD-(D/E)XK nuclease-like domain-containing protein; uniclust \| UniRef100\_A0A1B1IP21 \| 99.9 \| 1e-28 \| 2.1e-34 \| 223.1 \| 211 \| (11, 340) \| 372 \| (34, 247) \| 269 \| Putative exodeoxyribonuclease 8 PDDEXK-like domain-containing protein \| Putative exodeoxyribonuclease 8 PDDEXK-like domain-containing protein; uniclust \| UniRef100\_A0A076HWB5 \| 99.9 \| 1.1e-28 \| 2.3e-34 \| 223.6 \| 141 \| (195, 341) \| 372 \| (97, 244) \| 283 \| Putative exodeoxyribonuclease 8 PDDEXK-like domain-containing protein \| Putative exodeoxyribonuclease 8 PDDEXK-like domain-containing protein; uniclust \| UniRef100\_A0A2D9QDN0 \| 99.9 \| 1.4e-28 \| 2.6e-34 \| 202.5 \| 238 \| (19, 362) \| 372 \| (7, 248) \| 248 \| Putative exodeoxyribonuclease 8 PDDEXK-like domain-containing protein \| Putative exodeoxyribonuclease 8 PDDEXK-like domain-containing protein; uniclust \| UniRef100\_A0A2Z3HVS0 \| 99.9 \| 1.5e-28 \| 2.8e-34 \| 199.4 \| 168 \| (189, 364) \| 372 \| (4, 175) \| 180 \| Putative exodeoxyribonuclease 8 PDDEXK-like domain-containing protein \| Putative exodeoxyribonuclease 8 PDDEXK-like domain-containing protein; uniclust \| UniRef100\_A0A354CHA7 \| 99.9 \| 2.2e-28 \| 4.2e-34 \| 218.0 \| 270 \| (7, 369) \| 372 \| (53, 333) \| 343 \| Putative exodeoxyribonuclease 8 PDDEXK-like domain-containing protein \| Putative exodeoxyribonuclease 8 PDDEXK-like domain-containing protein; uniclust \| UniRef100\_A0A497PDQ5 \| 99.9 \| 3.3e-28 \| 6.1e-34 \| 208.0 \| 257 \| (3, 362) \| 372 \| (9, 265) \| 330 \| Putative exodeoxyribonuclease 8 PDDEXK-like domain-containing protein \| Putative exodeoxyribonuclease 8 PDDEXK-like domain-containing protein; uniclust \| UniRef100\_A0A8G2BHZ2 \| 99.9 \| 4.2e-28 \| 7.7e-34 \| 206.4 \| 273 \| (1, 370) \| 372 \| (3, 296) \| 318 \| Putative exodeoxyribonuclease 8 PDDEXK-like domain-containing protein \| Putative exodeoxyribonuclease 8 PDDEXK-like domain-containing protein; uniclust \| UniRef100\_A0A6J5MSZ1 \| 99.9 \| 5.3e-28 \| 9.7e-34 \| 205.7 \| 273 \| (8, 364) \| 372 \| (30, 309) \| 316 \| Exodeoxyribonuclease 8, PDDEXK-like domain containing protein \| Exodeoxyribonuclease 8, PDDEXK-like domain containing protein; uniclust \| UniRef100\_A0A7C1PFD3 \| 99.9 \| 6.2e-28 \| 1.2e-33 \| 213.2 \| 255 \| (8, 351) \| 372 \| (18, 291) \| 344 \| Putative exodeoxyribonuclease 8 PDDEXK-like domain-containing protein \| Putative exodeoxyribonuclease 8 PDDEXK-like domain-containing protein; uniclust \| UniRef100\_A0A7T5UDE4 \| 99.9 \| 6.8e-28 \| 1.3e-33 \| 205.2 \| 249 \| (7, 353) \| 372 \| (57, 312) \| 319 \| PD-(D/E)XK nuclease-like domain-containing protein \| PD-(D/E)XK nuclease-like domain-containing protein; uniclust \| UniRef100\_A0A0F9TVJ5 \| 99.9 \| 7e-28 \| 1.3e-33 \| 200.9 \| 251 \| (2, 353) \| 372 \| (15, 266) \| 270 \| Putative exodeoxyribonuclease 8 PDDEXK-like domain-containing protein \| Putative exodeoxyribonuclease 8 PDDEXK-like domain-containing protein; uniclust \| UniRef100\_A0A2L0VPK0 \| 99.9 \| 1.2e-27 \| 2.2e-33 \| 207.6 \| 263 \| (2, 356) \| 372 \| (31, 316) \| 344 \| Putative exodeoxyribonuclease 8 PDDEXK-like domain-containing protein \| Putative exodeoxyribonuclease 8 PDDEXK-like domain-containing protein; uniclust \| UniRef100\_A0A843HS31 \| 99.9 \| 1.4e-27 \| 2.6e-33 \| 207.1 \| 237 \| (8, 341) \| 372 \| (4, 249) \| 289 \| PD-(D/E)XK nuclease-like domain-containing protein \| PD-(D/E)XK nuclease-like domain-containing protein; uniclust \| UniRef100\_A0A1G3FSC3 \| 99.9 \| 2e-27 \| 3.8e-33 \| 202.7 \| 267 \| (2, 364) \| 372 \| (9, 297) \| 300 \| Putative exodeoxyribonuclease 8 PDDEXK-like domain-containing protein \| Putative exodeoxyribonuclease 8 PDDEXK-like domain-containing protein; uniclust \| UniRef100\_A0A0F9FLX8 \| 99.9 \| 2.5e-27 \| 4.5e-33 \| 197.8 \| 260 \| (1, 363) \| 372 \| (1, 267) \| 271 \| Putative exodeoxyribonuclease 8 PDDEXK-like domain-containing protein \| Putative exodeoxyribonuclease 8 PDDEXK-like domain-containing protein; uniclust \| UniRef100\_UPI0015869DB5 \| 99.9 \| 2.6e-27 \| 4.7e-33 \| 194.4 \| 182 \| (175, 362) \| 372 \| (55, 237) \| 239 \| PD-(D/E)XK nuclease-like domain-containing protein \| PD-(D/E)XK nuclease-like domain-containing protein; uniclust \| UniRef100\_A0A2D7PF36 \| 99.9 \| 2.6e-27 \| 4.9e-33 \| 204.2 \| 235 \| (6, 347) \| 372 \| (22, 274) \| 294 \| Putative exodeoxyribonuclease 8 PDDEXK-like domain-containing protein \| Putative exodeoxyribonuclease 8 PDDEXK-like domain-containing protein; uniclust \| UniRef100\_A0A3M1GEU5 \| 99.9 \| 3.7e-27 \| 6.9e-33 \| 195.4 \| 214 \| (4, 310) \| 372 \| (24, 243) \| 257 \| Putative exodeoxyribonuclease 8 PDDEXK-like domain-containing protein (Fragment) \| Putative exodeoxyribonuclease 8 PDDEXK-like domain-containing protein (Fragment); uniclust \| UniRef100\_A0A3R8K6A3 \| 99.9 \| 3.6e-27 \| 6.9e-33 \| 203.1 \| 265 \| (3, 369) \| 372 \| (11, 293) \| 296 \| Putative exodeoxyribonuclease 8 PDDEXK-like domain-containing protein \| Putative exodeoxyribonuclease 8 PDDEXK-like domain-containing protein; uniclust \| UniRef100\_A0A2D7GWI6 \| 99.9 \| 5.1e-27 \| 9.3e-33 \| 205.3 \| 321 \| (2, 350) \| 372 \| (35, 362) \| 376 \| SAP domain-containing protein \| SAP domain-containing protein; uniclust \| UniRef100\_A0A2E5F7Y4 \| 99.9 \| 9.7e-27 \| 1.8e-32 \| 198.2 \| 271 \| (3, 364) \| 372 \| (24, 308) \| 316 \| Putative exodeoxyribonuclease 8 PDDEXK-like domain-containing protein \| Putative exodeoxyribonuclease 8 PDDEXK-like domain-containing protein; uniclust \| UniRef100\_A0A6H1ZNA7 \| 99.9 \| 1.3e-26 \| 2.4e-32 \| 195.4 \| 271 \| (2, 363) \| 372 \| (9, 287) \| 291 \| Putative exodeoxyribonuclease 8 PDDEXK-like domain-containing protein \| Putative exodeoxyribonuclease 8 PDDEXK-like domain-containing protein; uniclust \| UniRef100\_A0A524RVX3 \| 99.9 \| 1.8e-26 \| 3.3e-32 \| 204.2 \| 296 \| (6, 371) \| 372 \| (107, 430) \| 437 \| Putative exodeoxyribonuclease 8 PDDEXK-like domain-containing protein \| Putative exodeoxyribonuclease 8 PDDEXK-like domain-containing protein; uniclust \| UniRef100\_A0A2N2TMM3 \| 99.9 \| 2.8e-26 \| 5.1e-32 \| 198.6 \| 258 \| (6, 365) \| 372 \| (89, 352) \| 360 \| Putative exodeoxyribonuclease 8 PDDEXK-like domain-containing protein \| Putative exodeoxyribonuclease 8 PDDEXK-like domain-containing protein; uniclust \| UniRef100\_A0A3A0DGM8 \| 99.9 \| 2.8e-26 \| 5.3e-32 \| 209.5 \| 249 \| (8, 353) \| 372 \| (233, 485) \| 490 \| Putative exodeoxyribonuclease 8 PDDEXK-like domain-containing protein \| Putative exodeoxyribonuclease 8 PDDEXK-like domain-containing protein; uniclust \| UniRef100\_UPI001CF244C9 \| 99.9 \| 3.5e-26 \| 6.4e-32 \| 187.1 \| 187 \| (181, 369) \| 372 \| (39, 229) \| 232 \| PD-(D/E)XK nuclease-like domain-containing protein \| PD-(D/E)XK nuclease-like domain-containing protein; uniclust \| UniRef100\_UPI001675557B \| 99.9 \| 5.8e-26 \| 1.1e-31 \| 199.9 \| 266 \| (1, 363) \| 372 \| (92, 369) \| 415 \| PD-(D/E)XK nuclease-like domain-containing protein \| PD-(D/E)XK nuclease-like domain-containing protein; uniclust \| UniRef100\_A0A496WM50 \| 99.9 \| 8.2e-26 \| 1.5e-31 \| 190.0 \| 264 \| (1, 353) \| 372 \| (1, 264) \| 283 \| Putative exodeoxyribonuclease 8 PDDEXK-like domain-containing protein \| Putative exodeoxyribonuclease 8 PDDEXK-like domain-containing protein; uniclust \| UniRef100\_UPI001FE141F0 \| 99.9 \| 8.9e-26 \| 1.6e-31 \| 186.6 \| 178 \| (187, 367) \| 372 \| (52, 233) \| 249 \| PD-(D/E)XK nuclease-like domain-containing protein \| PD-(D/E)XK nuclease-like domain-containing protein; uniclust \| UniRef100\_A0A966W4D5 \| 99.9 \| 9.8e-26 \| 1.8e-31 \| 183.1 \| 176 \| (186, 364) \| 372 \| (33, 211) \| 219 \| Putative exodeoxyribonuclease 8 PDDEXK-like domain-containing protein \| Putative exodeoxyribonuclease 8 PDDEXK-like domain-containing protein; uniclust \| UniRef100\_A0A2E2D771 \| 99.9 \| 9.6e-26 \| 1.9e-31 \| 202.0 \| 282 \| (17, 364) \| 372 \| (40, 326) \| 336 \| Putative exodeoxyribonuclease 8 PDDEXK-like domain-containing protein \| Putative exodeoxyribonuclease 8 PDDEXK-like domain-containing protein; uniclust \| UniRef100\_A0A1H5JK49 \| 99.9 \| 1e-25 \| 1.9e-31 \| 194.5 \| 263 \| (2, 366) \| 372 \| (5, 288) \| 306 \| Putative exodeoxyribonuclease 8 PDDEXK-like domain-containing protein \| Putative exodeoxyribonuclease 8 PDDEXK-like domain-containing protein; uniclust \| UniRef100\_A0A518EYZ7 \| 99.9 \| 1.3e-25 \| 2.4e-31 \| 190.8 \| 267 \| (2, 366) \| 372 \| (11, 285) \| 306 \| Exodeoxyribonuclease 8 \| Exodeoxyribonuclease 8; uniclust \| UniRef100\_A0A1H6WGJ5 \| 99.9 \| 1.4e-25 \| 2.6e-31 \| 199.9 \| 252 \| (2, 351) \| 372 \| (32, 294) \| 462 \| Putative exodeoxyribonuclease 8 PDDEXK-like domain-containing protein \| Putative exodeoxyribonuclease 8 PDDEXK-like domain-containing protein; uniclust \| UniRef100\_A0A2E8RC03 \| 99.9 \| 1.6e-25 \| 2.9e-31 \| 190.8 \| 252 \| (7, 349) \| 372 \| (5, 263) \| 277 \| Putative exodeoxyribonuclease 8 PDDEXK-like domain-containing protein \| Putative exodeoxyribonuclease 8 PDDEXK-like domain-containing protein; uniclust \| UniRef100\_A0A0F9N5B6 \| 99.9 \| 1.9e-25 \| 3.8e-31 \| 199.7 \| 271 \| (7, 367) \| 372 \| (14, 320) \| 327 \| Putative exodeoxyribonuclease 8 PDDEXK-like domain-containing protein \| Putative exodeoxyribonuclease 8 PDDEXK-like domain-containing protein; uniclust \| UniRef100\_A0A022N7N2 \| 99.9 \| 1.9e-25 \| 4.1e-31 \| 211.0 \| 230 \| (7, 348) \| 372 \| (60, 302) \| 368 \| Putative exodeoxyribonuclease 8 PDDEXK-like domain-containing protein \| Putative exodeoxyribonuclease 8 PDDEXK-like domain-containing protein; uniclust \| UniRef100\_UPI001BB88D46 \| 99.9 \| 2.9e-25 \| 5.4e-31 \| 193.0 \| 323 \| (2, 351) \| 372 \| (21, 344) \| 369 \| PD-(D/E)XK nuclease-like domain-containing protein \| PD-(D/E)XK nuclease-like domain-containing protein; uniclust \| UniRef100\_A0A2D7FJB4 \| 99.9 \| 4.5e-25 \| 8.2e-31 \| 185.9 \| 256 \| (5, 359) \| 372 \| (18, 277) \| 285 \| Putative exodeoxyribonuclease 8 PDDEXK-like domain-containing protein \| Putative exodeoxyribonuclease 8 PDDEXK-like domain-containing protein; uniclust \| UniRef100\_A0A433CKI3 \| 99.9 \| 4.3e-25 \| 8.4e-31 \| 194.0 \| 228 \| (8, 343) \| 372 \| (24, 256) \| 277 \| Putative exodeoxyribonuclease 8 PDDEXK-like domain-containing protein \| Putative exodeoxyribonuclease 8 PDDEXK-like domain-containing protein; uniclust \| UniRef100\_A0A0K1LMW8 \| 99.9 \| 4.7e-25 \| 8.6e-31 \| 196.0 \| 230 \| (2, 291) \| 372 \| (31, 269) \| 447 \| Nucleic acid-binding protein \| Nucleic acid-binding protein; uniclust \| UniRef100\_A0A5M6B4U3 \| 99.9 \| 4.8e-25 \| 9e-31 \| 192.0 \| 268 \| (10, 364) \| 372 \| (21, 307) \| 314 \| Putative exodeoxyribonuclease 8 PDDEXK-like domain-containing protein \| Putative exodeoxyribonuclease 8 PDDEXK-like domain-containing protein; uniclust \| UniRef100\_A0A516LLF0 \| 99.9 \| 5.2e-25 \| 9.6e-31 \| 184.6 \| 265 \| (2, 364) \| 372 \| (5, 271) \| 274 \| Putative exodeoxyribonuclease 8 \| Putative exodeoxyribonuclease 8; uniclust \| UniRef100\_A0A4Q7G2P0 \| 99.8 \| 7.4e-25 \| 1.4e-30 \| 195.0 \| 301 \| (2, 362) \| 372 \| (114, 449) \| 453 \| Putative exodeoxyribonuclease 8 PDDEXK-like domain-containing protein \| Putative exodeoxyribonuclease 8 PDDEXK-like domain-containing protein; uniclust \| UniRef100\_A0A7X6FQD1 \| 99.8 \| 1.5e-24 \| 2.8e-30 \| 183.7 \| 235 \| (1, 291) \| 372 \| (16, 264) \| 296 \| Putative exodeoxyribonuclease 8 PDDEXK-like domain-containing protein \| Putative exodeoxyribonuclease 8 PDDEXK-like domain-containing protein; uniclust \| UniRef100\_A0A385DTB7 \| 99.8 \| 2.7e-24 \| 5.6e-30 \| 201.6 \| 294 \| (7, 347) \| 372 \| (23, 340) \| 365 \| Exonuclease, PD-(D/E)XK superfamily \| Exonuclease, PD-(D/E)XK superfamily; uniclust \| UniRef100\_A0A2D6XBJ2 \| 99.8 \| 3.1e-24 \| 5.6e-30 \| 179.7 \| 262 \| (7, 364) \| 372 \| (3, 267) \| 269 \| Putative exodeoxyribonuclease 8 PDDEXK-like domain-containing protein \| Putative exodeoxyribonuclease 8 PDDEXK-like domain-containing protein; uniclust \| UniRef100\_A0A0F9U718 \| 99.8 \| 3.1e-24 \| 5.6e-30 \| 179.5 \| 258 \| (2, 356) \| 372 \| (4, 262) \| 267 \| Putative exodeoxyribonuclease 8 PDDEXK-like domain-containing protein \| Putative exodeoxyribonuclease 8 PDDEXK-like domain-containing protein; uniclust \| UniRef100\_A0A0J6BHA2 \| 99.8 \| 4.2e-24 \| 8e-30 \| 183.6 \| 248 \| (7, 350) \| 372 \| (2, 256) \| 282 \| Putative exodeoxyribonuclease 8 PDDEXK-like domain-containing protein \| Putative exodeoxyribonuclease 8 PDDEXK-like domain-containing protein; uniclust \| UniRef100\_UPI001E36E344 \| 99.8 \| 4.6e-24 \| 8.5e-30 \| 183.5 \| 267 \| (81, 368) \| 372 \| (15, 293) \| 332 \| PD-(D/E)XK nuclease-like domain-containing protein \| PD-(D/E)XK nuclease-like domain-containing protein; uniclust \| UniRef100\_A0A1Q6ECM9 \| 99.8 \| 5.8e-24 \| 1.1e-29 \| 188.4 \| 278 \| (7, 365) \| 372 \| (37, 346) \| 356 \| Putative exodeoxyribonuclease 8 PDDEXK-like domain-containing protein \| Putative exodeoxyribonuclease 8 PDDEXK-like domain-containing protein; uniclust \| UniRef100\_A0A2M9PH52 \| 99.8 \| 6.4e-24 \| 1.2e-29 \| 189.1 \| 267 \| (4, 367) \| 372 \| (9, 290) \| 450 \| Putative exodeoxyribonuclease 8 PDDEXK-like domain-containing protein \| Putative exodeoxyribonuclease 8 PDDEXK-like domain-containing protein; uniclust \| UniRef100\_A0A1P8VVD3 \| 99.8 \| 7.3e-24 \| 1.4e-29 \| 194.4 \| 232 \| (2, 291) \| 372 \| (172, 413) \| 550 \| Nucleic acid-binding protein \| Nucleic acid-binding protein; uniclust \| UniRef100\_A0A1F6GLK6 \| 99.8 \| 9.7e-24 \| 1.8e-29 \| 183.5 \| 317 \| (2, 364) \| 372 \| (17, 354) \| 363 \| Putative exodeoxyribonuclease 8 PDDEXK-like domain-containing protein \| Putative exodeoxyribonuclease 8 PDDEXK-like domain-containing protein; uniclust \| UniRef100\_A0A7M1RQF0 \| 99.8 \| 9.7e-24 \| 1.9e-29 \| 191.6 \| 291 \| (8, 347) \| 372 \| (13, 324) \| 347 \| Exonuclease \| Exonuclease; uniclust \| UniRef100\_A0A5C7PQS1 \| 99.8 \| 1.1e-23 \| 2e-29 \| 176.6 \| 256 \| (9, 362) \| 372 \| (1, 269) \| 271 \| Putative exodeoxyribonuclease 8 PDDEXK-like domain-containing protein \| Putative exodeoxyribonuclease 8 PDDEXK-like domain-containing protein; uniclust \| UniRef100\_UPI001BC86F65 \| 99.8 \| 1.1e-23 \| 2.1e-29 \| 176.3 \| 154 \| (1, 246) \| 372 \| (115, 268) \| 268 \| PD-(D/E)XK nuclease-like domain-containing protein \| PD-(D/E)XK nuclease-like domain-containing protein; uniclust \| UniRef100\_A0A5C7P550 \| 99.8 \| 1.3e-23 \| 2.4e-29 \| 180.3 \| 230 \| (19, 355) \| 372 \| (71, 301) \| 324 \| Putative exodeoxyribonuclease 8 PDDEXK-like domain-containing protein \| Putative exodeoxyribonuclease 8 PDDEXK-like domain-containing protein; uniclust \| UniRef100\_UPI0007C549EE \| 99.8 \| 1.5e-23 \| 2.8e-29 \| 181.6 \| 257 \| (5, 365) \| 372 \| (80, 344) \| 350 \| PD-(D/E)XK nuclease-like domain-containing protein \| PD-(D/E)XK nuclease-like domain-containing protein; uniclust \| UniRef100\_A0A1G3LP96 \| 99.8 \| 2e-23 \| 3.8e-29 \| 181.4 \| 251 \| (6, 362) \| 372 \| (4, 264) \| 266 \| Putative exodeoxyribonuclease 8 PDDEXK-like domain-containing protein \| Putative exodeoxyribonuclease 8 PDDEXK-like domain-containing protein; uniclust \| UniRef100\_A0A2A2GZ94 \| 99.8 \| 2.6e-23 \| 4.8e-29 \| 174.3 \| 167 \| (191, 370) \| 372 \| (66, 238) \| 247 \| Putative exodeoxyribonuclease 8 PDDEXK-like domain-containing protein \| Putative exodeoxyribonuclease 8 PDDEXK-like domain-containing protein; uniclust \| UniRef100\_A0A150AJT1 \| 99.8 \| 2.6e-23 \| 5e-29 \| 180.2 \| 137 \| (195, 337) \| 372 \| (93, 231) \| 250 \| Putative exodeoxyribonuclease 8 PDDEXK-like domain-containing protein \| Putative exodeoxyribonuclease 8 PDDEXK-like domain-containing protein; uniclust \| UniRef100\_A0A966PZB7 \| 99.8 \| 2.7e-23 \| 5e-29 \| 174.5 \| 248 \| (19, 367) \| 372 \| (17, 268) \| 273 \| Putative exodeoxyribonuclease 8 PDDEXK-like domain-containing protein \| Putative exodeoxyribonuclease 8 PDDEXK-like domain-containing protein; uniclust \| UniRef100\_A0A2D5TTD4 \| 99.8 \| 2.9e-23 \| 5.4e-29 \| 186.2 \| 229 \| (1, 283) \| 372 \| (7, 244) \| 391 \| Exodeoxyribonuclease 8 PDDEXK-like domain-containing protein \| Exodeoxyribonuclease 8 PDDEXK-like domain-containing protein; uniclust \| UniRef100\_A0A3B8U8D5 \| 99.8 \| 3.1e-23 \| 5.7e-29 \| 177.8 \| 269 \| (19, 349) \| 372 \| (16, 287) \| 320 \| SAP domain-containing protein \| SAP domain-containing protein; uniclust \| UniRef100\_A0A357XXN4 \| 99.8 \| 3.6e-23 \| 6.7e-29 \| 175.7 \| 247 \| (5, 348) \| 372 \| (8, 257) \| 267 \| Putative exodeoxyribonuclease 8 PDDEXK-like domain-containing protein \| Putative exodeoxyribonuclease 8 PDDEXK-like domain-containing protein; uniclust \| UniRef100\_M3JBV3 \| 99.8 \| 3.9e-23 \| 7.3e-29 \| 174.5 \| 224 \| (8, 365) \| 372 \| (24, 249) \| 253 \| Bacteriophage protein \| Bacteriophage protein; uniclust \| UniRef100\_A0A6J5S1P5 \| 99.8 \| 4.6e-23 \| 8.4e-29 \| 181.7 \| 238 \| (6, 347) \| 372 \| (5, 246) \| 405 \| Exodeoxyribonuclease 8, PDDEXK-like domain containing protein \| Exodeoxyribonuclease 8, PDDEXK-like domain containing protein; uniclust \| UniRef100\_A0A2E7BUE9 \| 99.8 \| 4.6e-23 \| 8.4e-29 \| 177.1 \| 263 \| (6, 362) \| 372 \| (36, 313) \| 325 \| Putative exodeoxyribonuclease 8 PDDEXK-like domain-containing protein \| Putative exodeoxyribonuclease 8 PDDEXK-like domain-containing protein; uniclust \| UniRef100\_A0A350PI72 \| 99.8 \| 5e-23 \| 9.1e-29 \| 168.1 \| 178 \| (10, 290) \| 372 \| (38, 216) \| 221 \| Putative exodeoxyribonuclease 8 PDDEXK-like domain-containing protein (Fragment) \| Putative exodeoxyribonuclease 8 PDDEXK-like domain-containing protein (Fragment); uniclust \| UniRef100\_A0A0F9ABF3 \| 99.8 \| 5.5e-23 \| 1e-28 \| 173.1 \| 179 \| (10, 291) \| 372 \| (97, 275) \| 277 \| Putative exodeoxyribonuclease 8 PDDEXK-like domain-containing protein (Fragment) \| Putative exodeoxyribonuclease 8 PDDEXK-like domain-containing protein (Fragment); uniclust \| UniRef100\_UPI000AEFFF0F \| 99.8 \| 5.5e-23 \| 1e-28 \| 168.9 \| 179 \| (187, 369) \| 372 \| (14, 196) \| 205 \| PD-(D/E)XK nuclease-like domain-containing protein \| PD-(D/E)XK nuclease-like domain-containing protein; uniclust \| UniRef100\_A0A8T3QJZ4 \| 99.8 \| 6.3e-23 \| 1.2e-28 \| 179.6 \| 263 \| (5, 367) \| 372 \| (35, 312) \| 320 \| PD-(D/E)XK nuclease-like domain-containing protein \| PD-(D/E)XK nuclease-like domain-containing protein; uniclust \| UniRef100\_UPI0013CB2100 \| 99.8 \| 8.3e-23 \| 1.5e-28 \| 172.7 \| 211 \| (154, 368) \| 372 \| (4, 224) \| 263 \| PD-(D/E)XK nuclease-like domain-containing protein \| PD-(D/E)XK nuclease-like domain-containing protein; uniclust \| UniRef100\_A0A3C0EHM3 \| 99.8 \| 8.1e-23 \| 1.6e-28 \| 166.9 \| 163 \| (186, 355) \| 372 \| (12, 174) \| 177 \| Putative exodeoxyribonuclease 8 PDDEXK-like domain-containing protein \| Putative exodeoxyribonuclease 8 PDDEXK-like domain-containing protein; uniclust \| UniRef100\_A0A2D6PI96 \| 99.8 \| 7.9e-23 \| 1.6e-28 \| 188.8 \| 287 \| (11, 347) \| 372 \| (8, 323) \| 355 \| PD-(D/E)XK endonuclease-like domain-containing protein \| PD-(D/E)XK endonuclease-like domain-containing protein; uniclust \| UniRef100\_A0A661I8B6 \| 99.8 \| 1.4e-22 \| 2.6e-28 \| 171.5 \| 242 \| (3, 348) \| 372 \| (23, 271) \| 286 \| Putative exodeoxyribonuclease 8 PDDEXK-like domain-containing protein \| Putative exodeoxyribonuclease 8 PDDEXK-like domain-containing protein; uniclust \| UniRef100\_A0A395XFE4 \| 99.8 \| 1.5e-22 \| 2.8e-28 \| 178.3 \| 261 \| (5, 366) \| 372 \| (95, 396) \| 400 \| Nuclease \| Nuclease; uniclust \| UniRef100\_A0A516LE23 \| 99.8 \| 1.7e-22 \| 3e-28 \| 170.1 \| 242 \| (5, 343) \| 372 \| (9, 252) \| 274 \| Putative exodeoxyribonuclease 8 \| Putative exodeoxyribonuclease 8; uniclust \| UniRef100\_A0A966VAZ2 \| 99.8 \| 2e-22 \| 3.6e-28 \| 172.0 \| 259 \| (5, 359) \| 372 \| (35, 298) \| 305 \| Putative exodeoxyribonuclease 8 PDDEXK-like domain-containing protein \| Putative exodeoxyribonuclease 8 PDDEXK-like domain-containing protein; uniclust \| UniRef100\_A0A7X8XER7 \| 99.8 \| 2.2e-22 \| 4.1e-28 \| 168.3 \| 198 \| (2, 295) \| 372 \| (20, 220) \| 235 \| Putative exodeoxyribonuclease 8 PDDEXK-like domain-containing protein (Fragment) \| Putative exodeoxyribonuclease 8 PDDEXK-like domain-containing protein (Fragment); uniclust \| UniRef100\_A0A6J5LX80 \| 99.8 \| 3.2e-22 \| 6e-28 \| 169.6 \| 246 \| (3, 350) \| 372 \| (5, 255) \| 268 \| Exodeoxyribonuclease 8, PDDEXK-like domain containing protein \| Exodeoxyribonuclease 8, PDDEXK-like domain containing protein; uniclust \| UniRef100\_A0A968CQ55 \| 99.8 \| 3.6e-22 \| 6.6e-28 \| 165.5 \| 227 \| (2, 354) \| 372 \| (5, 243) \| 244 \| Putative exodeoxyribonuclease 8 PDDEXK-like domain-containing protein \| Putative exodeoxyribonuclease 8 PDDEXK-like domain-containing protein; uniclust \| UniRef100\_A0A6I0ESC0 \| 99.8 \| 3.7e-22 \| 6.8e-28 \| 176.6 \| 245 \| (10, 351) \| 372 \| (158, 407) \| 414 \| Putative exodeoxyribonuclease 8 PDDEXK-like domain-containing protein \| Putative exodeoxyribonuclease 8 PDDEXK-like domain-containing protein; uniclust \| UniRef100\_A0A2D7YTB8 \| 99.8 \| 4.1e-22 \| 7.6e-28 \| 168.2 \| 258 \| (4, 364) \| 372 \| (7, 278) \| 280 \| Putative exodeoxyribonuclease 8 PDDEXK-like domain-containing protein \| Putative exodeoxyribonuclease 8 PDDEXK-like domain-containing protein; uniclust \| UniRef100\_A0A5C5X348 \| 99.8 \| 5.6e-22 \| 1e-27 \| 167.6 \| 257 \| (5, 365) \| 372 \| (12, 273) \| 282 \| Putative exodeoxyribonuclease 8 PDDEXK-like domain-containing protein \| Putative exodeoxyribonuclease 8 PDDEXK-like domain-containing protein; uniclust \| UniRef100\_A0A2E8IY15 \| 99.8 \| 6.6e-22 \| 1.2e-27 \| 166.4 \| 177 \| (187, 365) \| 372 \| (93, 270) \| 272 \| Putative exodeoxyribonuclease 8 PDDEXK-like domain-containing protein \| Putative exodeoxyribonuclease 8 PDDEXK-like domain-containing protein; uniclust \| UniRef100\_A0A935XAD7 \| 99.8 \| 7.9e-22 \| 1.4e-27 \| 164.8 \| 146 \| (188, 338) \| 372 \| (93, 240) \| 258 \| PD-(D/E)XK nuclease-like domain-containing protein \| PD-(D/E)XK nuclease-like domain-containing protein; uniclust \| UniRef100\_A0A485A5W0 \| 99.8 \| 8.6e-22 \| 1.6e-27 \| 176.9 \| 245 \| (2, 249) \| 372 \| (163, 460) \| 470 \| Exodeoxyribonuclease 8 \| Exodeoxyribonuclease 8; uniclust \| UniRef100\_A0A2E4HLS4 \| 99.8 \| 1.1e-21 \| 2.1e-27 \| 178.2 \| 282 \| (11, 335) \| 372 \| (20, 319) \| 343 \| PD-(D/E)XK endonuclease-like domain-containing protein \| PD-(D/E)XK endonuclease-like domain-containing protein; uniclust \| UniRef100\_A0A0G9KAG8 \| 99.7 \| 1.9e-21 \| 3.6e-27 \| 167.7 \| 235 \| (2, 341) \| 372 \| (20, 262) \| 284 \| Putative exodeoxyribonuclease 8 PDDEXK-like domain-containing protein \| Putative exodeoxyribonuclease 8 PDDEXK-like domain-containing protein; uniclust \| UniRef100\_A0A6B9SPH0 \| 99.7 \| 2.7e-21 \| 5e-27 \| 161.0 \| 225 \| (7, 291) \| 372 \| (9, 241) \| 249 \| Putative exonuclease \| Putative exonuclease; uniclust \| UniRef100\_UPI000C7CBD65 \| 99.7 \| 2.9e-21 \| 5.3e-27 \| 190.0 \| 334 \| (6, 348) \| 372 \| (868, 1312) \| 1341 \| hypothetical protein \| hypothetical protein; uniclust \| UniRef100\_A0A5C7PJN5 \| 99.7 \| 3.1e-21 \| 5.8e-27 \| 163.0 \| 262 \| (7, 362) \| 372 \| (6, 274) \| 278 \| Putative exodeoxyribonuclease 8 PDDEXK-like domain-containing protein \| Putative exodeoxyribonuclease 8 PDDEXK-like domain-containing protein; uniclust \| UniRef100\_A0A966RAL2 \| 99.7 \| 3.3e-21 \| 6.1e-27 \| 164.2 \| 269 \| (1, 370) \| 372 \| (2, 293) \| 297 \| Putative exodeoxyribonuclease 8 PDDEXK-like domain-containing protein \| Putative exodeoxyribonuclease 8 PDDEXK-like domain-containing protein; uniclust \| UniRef100\_A0A948KE74 \| 99.7 \| 3.8e-21 \| 7e-27 \| 160.8 \| 175 \| (187, 368) \| 372 \| (57, 236) \| 257 \| PD-(D/E)XK nuclease-like domain-containing protein \| PD-(D/E)XK nuclease-like domain-containing protein; uniclust \| UniRef100\_A0A076HWD4 \| 99.7 \| 3.7e-21 \| 7e-27 \| 169.2 \| 126 \| (205, 338) \| 372 \| (139, 268) \| 309 \| Putative exodeoxyribonuclease 8 PDDEXK-like domain-containing protein \| Putative exodeoxyribonuclease 8 PDDEXK-like domain-containing protein; uniclust \| UniRef100\_A0A970WNN9 \| 99.7 \| 4.7e-21 \| 8.7e-27 \| 151.6 \| 160 \| (205, 366) \| 372 \| (1, 165) \| 175 \| Putative exodeoxyribonuclease 8 PDDEXK-like domain-containing protein \| Putative exodeoxyribonuclease 8 PDDEXK-like domain-containing protein; uniclust \| UniRef100\_A0A956M5I9 \| 99.7 \| 5.4e-21 \| 1e-26 \| 161.9 \| 247 \| (2, 350) \| 372 \| (9, 275) \| 283 \| PD-(D/E)XK nuclease-like domain-containing protein (Fragment) \| PD-(D/E)XK nuclease-like domain-containing protein (Fragment); uniclust \| UniRef100\_A0A5T5VGN4 \| 99.7 \| 7.1e-21 \| 1.3e-26 \| 148.4 \| 137 \| (1, 233) \| 372 \| (1, 140) \| 140 \| Exodeoxyribonuclease 8 (Fragment) \| Exodeoxyribonuclease 8 (Fragment); uniclust \| UniRef100\_A0A0Q5QGX0 \| 99.7 \| 6.9e-21 \| 1.3e-26 \| 172.6 \| 286 \| (19, 367) \| 372 \| (46, 334) \| 350 \| Putative exodeoxyribonuclease 8 PDDEXK-like domain-containing protein \| Putative exodeoxyribonuclease 8 PDDEXK-like domain-containing protein; uniclust \| UniRef100\_A0A2D8XKX9 \| 99.7 \| 7.9e-21 \| 1.5e-26 \| 165.1 \| 242 \| (17, 345) \| 372 \| (24, 267) \| 278 \| Putative exodeoxyribonuclease 8 PDDEXK-like domain-containing protein \| Putative exodeoxyribonuclease 8 PDDEXK-like domain-containing protein; uniclust \| UniRef100\_A0A097ENH1 \| 99.7 \| 9.2e-21 \| 1.7e-26 \| 159.1 \| 246 \| (13, 364) \| 372 \| (2, 255) \| 263 \| Putative exodeoxyribonuclease 8 PDDEXK-like domain-containing protein \| Putative exodeoxyribonuclease 8 PDDEXK-like domain-containing protein; uniclust \| UniRef100\_UPI0005F02F46 \| 99.7 \| 9.6e-21 \| 1.8e-26 \| 185.4 \| 330 \| (6, 349) \| 372 \| (540, 946) \| 988 \| hypothetical protein \| hypothetical protein; uniclust \| UniRef100\_A0A965MNM5 \| 99.7 \| 9.8e-21 \| 1.9e-26 \| 166.6 \| 265 \| (6, 369) \| 372 \| (29, 302) \| 304 \| Putative exodeoxyribonuclease 8 PDDEXK-like domain-containing protein \| Putative exodeoxyribonuclease 8 PDDEXK-like domain-containing protein; uniclust \| UniRef100\_A0A0F3IMN9 \| 99.7 \| 2.6e-20 \| 4.7e-26 \| 159.3 \| 235 \| (8, 343) \| 372 \| (47, 287) \| 301 \| Putative exodeoxyribonuclease 8 PDDEXK-like domain-containing protein \| Putative exodeoxyribonuclease 8 PDDEXK-like domain-containing protein; uniclust \| UniRef100\_A0A936DK79 \| 99.7 \| 2.6e-20 \| 4.8e-26 \| 158.4 \| 267 \| (10, 364) \| 372 \| (2, 273) \| 289 \| PD-(D/E)XK nuclease-like domain-containing protein \| PD-(D/E)XK nuclease-like domain-containing protein; uniclust \| UniRef100\_A0A3M1NIT7 \| 99.7 \| 3.4e-20 \| 6.2e-26 \| 156.9 \| 258 \| (8, 366) \| 372 \| (4, 264) \| 277 \| Putative exodeoxyribonuclease 8 PDDEXK-like domain-containing protein \| Putative exodeoxyribonuclease 8 PDDEXK-like domain-containing protein; uniclust \| UniRef100\_A0A0F9AK76 \| 99.7 \| 3.5e-20 \| 6.5e-26 \| 154.8 \| 176 \| (184, 364) \| 372 \| (67, 247) \| 252 \| Putative exodeoxyribonuclease 8 PDDEXK-like domain-containing protein (Fragment) \| Putative exodeoxyribonuclease 8 PDDEXK-like domain-containing protein (Fragment); uniclust \| UniRef100\_A0A7Y6MP93 \| 99.7 \| 3.6e-20 \| 6.7e-26 \| 177.9 \| 283 \| (7, 348) \| 372 \| (641, 935) \| 970 \| PD-(D/E)XK nuclease-like domain-containing protein \| PD-(D/E)XK nuclease-like domain-containing protein; uniclust \| UniRef100\_A0A517YVS6 \| 99.7 \| 3.9e-20 \| 7.1e-26 \| 158.0 \| 260 \| (3, 363) \| 372 \| (23, 294) \| 298 \| Exodeoxyribonuclease 8 \| Exodeoxyribonuclease 8; uniclust \| UniRef100\_A0A4V0Z515 \| 99.7 \| 4e-20 \| 7.4e-26 \| 148.1 \| 171 \| (191, 362) \| 372 \| (4, 183) \| 188 \| Putative exodeoxyribonuclease 8 PDDEXK-like domain-containing protein \| Putative exodeoxyribonuclease 8 PDDEXK-like domain-containing protein; uniclust \| UniRef100\_A0A962Q8Y1 \| 99.7 \| 4.4e-20 \| 8.1e-26 \| 156.6 \| 260 \| (6, 365) \| 372 \| (4, 273) \| 282 \| PD-(D/E)XK nuclease-like domain-containing protein \| PD-(D/E)XK nuclease-like domain-containing protein; uniclust \| UniRef100\_A0A6J5PGR2 \| 99.7 \| 5.3e-20 \| 9.8e-26 \| 155.9 \| 263 \| (3, 368) \| 372 \| (9, 276) \| 279 \| Exodeoxyribonuclease 8, PDDEXK-like domain containing protein \| Exodeoxyribonuclease 8, PDDEXK-like domain containing protein; uniclust \| UniRef100\_A0A485AXQ3 \| 99.7 \| 5.9e-20 \| 1.1e-25 \| 142.1 \| 78 \| (1, 78) \| 372 \| (4, 81) \| 136 \| Exodeoxyribonuclease 8 \| Exodeoxyribonuclease 8; uniclust \| UniRef100\_A0A3S9D4G6 \| 99.7 \| 5.9e-20 \| 1.1e-25 \| 149.4 \| 148 \| (3, 261) \| 372 \| (21, 177) \| 185 \| Uncharacterized protein \| Uncharacterized protein; uniclust \| UniRef100\_A0A955HL80 \| 99.7 \| 5.9e-20 \| 1.1e-25 \| 159.9 \| 230 \| (18, 348) \| 372 \| (26, 262) \| 277 \| PD-(D/E)XK nuclease-like domain-containing protein \| PD-(D/E)XK nuclease-like domain-containing protein; uniclust \| UniRef100\_A0A1E4ZGR8 \| 99.7 \| 6.4e-20 \| 1.2e-25 \| 156.9 \| 269 \| (2, 364) \| 372 \| (7, 296) \| 301 \| Putative exodeoxyribonuclease 8 PDDEXK-like domain-containing protein \| Putative exodeoxyribonuclease 8 PDDEXK-like domain-containing protein; uniclust \| UniRef100\_A0A511B8A3 \| 99.7 \| 7.2e-20 \| 1.4e-25 \| 147.1 \| 147 \| (215, 367) \| 372 \| (7, 156) \| 168 \| Putative exodeoxyribonuclease 8 PDDEXK-like domain-containing protein \| Putative exodeoxyribonuclease 8 PDDEXK-like domain-containing protein; uniclust \| UniRef100\_A0A914CHP8 \| 99.7 \| 7.7e-20 \| 1.4e-25 \| 184.4 \| 215 \| (3, 313) \| 372 \| (6, 224) \| 2182 \| Putative exodeoxyribonuclease 8 PDDEXK-like domain-containing protein \| Putative exodeoxyribonuclease 8 PDDEXK-like domain-containing protein; uniclust \| UniRef100\_A0A0P0DG67 \| 99.7 \| 8e-20 \| 1.5e-25 \| 161.4 \| 278 \| (1, 365) \| 372 \| (6, 324) \| 337 \| Putative exodeoxyribonuclease 8 PDDEXK-like domain-containing protein \| Putative exodeoxyribonuclease 8 PDDEXK-like domain-containing protein; uniclust \| UniRef100\_A0A5C7PU20 \| 99.7 \| 8.2e-20 \| 1.5e-25 \| 154.7 \| 254 \| (13, 361) \| 372 \| (19, 277) \| 278 \| Putative exodeoxyribonuclease 8 PDDEXK-like domain-containing protein \| Putative exodeoxyribonuclease 8 PDDEXK-like domain-containing protein; uniclust \| UniRef100\_Q6LJU7 \| 99.7 \| 1e-19 \| 1.8e-25 \| 177.1 \| 335 \| (1, 349) \| 372 \| (661, 1116) \| 1148 \| Uncharacterized protein \| Uncharacterized protein; uniclust \| UniRef100\_A0A447XY29 \| 99.7 \| 1.1e-19 \| 2.1e-25 \| 156.2 \| 210 \| (153, 368) \| 372 \| (31, 250) \| 254 \| Putative phage exodeoxyribonuclease \| Putative phage exodeoxyribonuclease; uniclust \| UniRef100\_A0A976KWG8 \| 99.7 \| 1.3e-19 \| 2.4e-25 \| 155.8 \| 249 \| (6, 338) \| 372 \| (9, 266) \| 274 \| Putative exodeoxyribonuclease 8 PDDEXK-like domain-containing protein \| Putative exodeoxyribonuclease 8 PDDEXK-like domain-containing protein; uniclust \| UniRef100\_A0A2E1MZJ8 \| 99.7 \| 1.4e-19 \| 2.7e-25 \| 156.4 \| 287 \| (9, 347) \| 372 \| (1, 316) \| 326 \| Putative exodeoxyribonuclease 8 PDDEXK-like domain-containing protein \| Putative exodeoxyribonuclease 8 PDDEXK-like domain-containing protein; uniclust \| UniRef100\_UPI0014383DDD \| 99.7 \| 1.8e-19 \| 3.4e-25 \| 152.0 \| 186 \| (5, 286) \| 372 \| (75, 264) \| 269 \| PD-(D/E)XK nuclease-like domain-containing protein \| PD-(D/E)XK nuclease-like domain-containing protein; uniclust \| UniRef100\_A0A1I5SZ79 \| 99.7 \| 2.2e-19 \| 4e-25 \| 153.0 \| 271 \| (2, 362) \| 372 \| (4, 288) \| 289 \| Putative exodeoxyribonuclease 8 PDDEXK-like domain-containing protein \| Putative exodeoxyribonuclease 8 PDDEXK-like domain-containing protein; uniclust \| UniRef100\_A0A7V6NHJ0 \| 99.6 \| 2.6e-19 \| 5e-25 \| 153.9 \| 197 \| (6, 315) \| 372 \| (8, 217) \| 220 \| Putative exodeoxyribonuclease 8 PDDEXK-like domain-containing protein \| Putative exodeoxyribonuclease 8 PDDEXK-like domain-containing protein; uniclust \| UniRef100\_A0A963CJ35 \| 99.6 \| 2.8e-19 \| 5.2e-25 \| 140.8 \| 135 \| (2, 233) \| 372 \| (23, 158) \| 166 \| PD-(D/E)XK nuclease-like domain-containing protein (Fragment) \| PD-(D/E)XK nuclease-like domain-containing protein (Fragment); uniclust \| UniRef100\_A0A843GNI8 \| 99.6 \| 3.2e-19 \| 5.8e-25 \| 150.6 \| 252 \| (7, 361) \| 372 \| (15, 268) \| 269 \| PD-(D/E)XK nuclease-like domain-containing protein \| PD-(D/E)XK nuclease-like domain-containing protein; uniclust \| UniRef100\_A0A0M4SKY7 \| 99.6 \| 3.1e-19 \| 5.9e-25 \| 155.6 \| 233 \| (6, 344) \| 372 \| (26, 261) \| 294 \| Putative exodeoxyribonuclease 8 PDDEXK-like domain-containing protein \| Putative exodeoxyribonuclease 8 PDDEXK-like domain-containing protein; uniclust \| UniRef100\_UPI00158604F7 \| 99.6 \| 3.3e-19 \| 6.4e-25 \| 133.8 \| 97 \| (268, 365) \| 372 \| (2, 99) \| 104 \| PD-(D/E)XK nuclease-like domain-containing protein \| PD-(D/E)XK nuclease-like domain-containing protein; uniclust \| UniRef100\_A0A3D1XDL4 \| 99.6 \| 3.7e-19 \| 6.8e-25 \| 145.3 \| 161 \| (187, 348) \| 372 \| (31, 193) \| 211 \| Putative exodeoxyribonuclease 8 PDDEXK-like domain-containing protein (Fragment) \| Putative exodeoxyribonuclease 8 PDDEXK-like domain-containing protein (Fragment); uniclust \| UniRef100\_A0A0F9I7X6 \| 99.6 \| 4.3e-19 \| 7.9e-25 \| 152.7 \| 262 \| (4, 362) \| 372 \| (18, 284) \| 311 \| Putative exodeoxyribonuclease 8 PDDEXK-like domain-containing protein \| Putative exodeoxyribonuclease 8 PDDEXK-like domain-containing protein; uniclust \| UniRef100\_A0A2I7RBQ8 \| 99.6 \| 4.6e-19 \| 8.4e-25 \| 167.3 \| 279 \| (5, 349) \| 372 \| (445, 732) \| 750 \| Coil containing protein \| Coil containing protein; uniclust \| UniRef100\_A0A3C0F448 \| 99.6 \| 4.5e-19 \| 8.5e-25 \| 147.3 \| 167 \| (11, 288) \| 372 \| (18, 187) \| 194 \| Putative exodeoxyribonuclease 8 PDDEXK-like domain-containing protein \| Putative exodeoxyribonuclease 8 PDDEXK-like domain-containing protein; uniclust \| UniRef100\_UPI0005B4C7AB \| 99.6 \| 5e-19 \| 9.2e-25 \| 157.2 \| 144 \| (1, 236) \| 372 \| (244, 387) \| 406 \| PD-(D/E)XK nuclease-like domain-containing protein \| PD-(D/E)XK nuclease-like domain-containing protein; uniclust \| UniRef100\_A0A967T6U3 \| 99.6 \| 6.1e-19 \| 1.1e-24 \| 140.8 \| 165 \| (197, 369) \| 372 \| (2, 179) \| 181 \| Putative exodeoxyribonuclease 8 PDDEXK-like domain-containing protein \| Putative exodeoxyribonuclease 8 PDDEXK-like domain-containing protein; uniclust \| UniRef100\_A0A516LFV1 \| 99.6 \| 8.6e-19 \| 1.6e-24 \| 152.6 \| 295 \| (18, 347) \| 372 \| (4, 310) \| 340 \| Putative exodeoxyribonuclease 8 \| Putative exodeoxyribonuclease 8; uniclust \| UniRef100\_A0A482N4I2 \| 99.6 \| 3e-18 \| 5.5e-24 \| 135.5 \| 160 \| (206, 367) \| 372 \| (2, 167) \| 168 \| Putative exodeoxyribonuclease 8 PDDEXK-like domain-containing protein \| Putative exodeoxyribonuclease 8 PDDEXK-like domain-containing protein; uniclust \| UniRef100\_R1HTJ0 \| 99.6 \| 3.9e-18 \| 7.1e-24 \| 131.9 \| 128 \| (242, 370) \| 372 \| (1, 134) \| 146 \| Gp60 protein (Fragment) \| Gp60 protein (Fragment); uniclust \| UniRef100\_A0A2H5BFV1 \| 99.6 \| 4e-18 \| 7.3e-24 \| 165.0 \| 275 \| (5, 348) \| 372 \| (635, 919) \| 936 \| Exodeoxyribonuclease type VIII \| Exodeoxyribonuclease type VIII; uniclust \| UniRef100\_A0A098RLH1 \| 99.6 \| 4.2e-18 \| 7.7e-24 \| 139.4 \| 189 \| (2, 291) \| 372 \| (8, 204) \| 211 \| Putative exodeoxyribonuclease 8 PDDEXK-like domain-containing protein (Fragment) \| Putative exodeoxyribonuclease 8 PDDEXK-like domain-containing protein (Fragment); uniclust \| UniRef100\_A0A827N2G5 \| 99.6 \| 4.9e-18 \| 8.9e-24 \| 148.1 \| 138 \| (1, 232) \| 372 \| (1, 140) \| 339 \| Adenine methylase \| Adenine methylase; uniclust \| UniRef100\_A0A6A6K1N9 \| 99.6 \| 7e-18 \| 1.3e-23 \| 153.3 \| 124 \| (225, 350) \| 372 \| (200, 323) \| 490 \| Putative exodeoxyribonuclease 8 PDDEXK-like domain-containing protein \| Putative exodeoxyribonuclease 8 PDDEXK-like domain-containing protein; uniclust \| UniRef100\_A0A351GFU4 \| 99.6 \| 8.3e-18 \| 1.6e-23 \| 149.8 \| 293 \| (10, 346) \| 372 \| (2, 322) \| 337 \| Putative exodeoxyribonuclease 8 PDDEXK-like domain-containing protein \| Putative exodeoxyribonuclease 8 PDDEXK-like domain-containing protein; uniclust \| UniRef100\_UPI00189A7229 \| 99.6 \| 8.7e-18 \| 1.6e-23 \| 136.6 \| 170 \| (193, 363) \| 372 \| (9, 181) \| 190 \| PD-(D/E)XK nuclease-like domain-containing protein \| PD-(D/E)XK nuclease-like domain-containing protein; uniclust \| UniRef100\_UPI000C27CD9A \| 99.6 \| 1e-17 \| 1.9e-23 \| 131.5 \| 145 \| (201, 348) \| 372 \| (2, 148) \| 159 \| PD-(D/E)XK nuclease-like domain-containing protein \| PD-(D/E)XK nuclease-like domain-containing protein; uniclust \| UniRef100\_A0A5P8D2N7 \| 99.5 \| 1.2e-17 \| 2.3e-23 \| 151.3 \| 179 \| (187, 368) \| 372 \| (210, 394) \| 396 \| Exonuclease \| Exonuclease; uniclust \| UniRef100\_A0A345LVI2 \| 99.5 \| 1.3e-17 \| 2.3e-23 \| 131.8 \| 148 \| (1, 250) \| 372 \| (1, 148) \| 155 \| PD-(D/E)XK nuclease-like domain-containing protein \| PD-(D/E)XK nuclease-like domain-containing protein; uniclust \| UniRef100\_A0A150JKQ2 \| 99.5 \| 1.3e-17 \| 2.5e-23 \| 153.0 \| 274 \| (8, 330) \| 372 \| (22, 313) \| 382 \| Uncharacterized protein \| Uncharacterized protein; uniclust \| UniRef100\_A0A1T4X5K6 \| 99.5 \| 1.4e-17 \| 2.6e-23 \| 147.3 \| 270 \| (7, 317) \| 372 \| (23, 309) \| 380 \| Putative exodeoxyribonuclease 8 PDDEXK-like domain-containing protein \| Putative exodeoxyribonuclease 8 PDDEXK-like domain-containing protein; uniclust \| UniRef100\_A0A3D2IK29 \| 99.5 \| 1.5e-17 \| 2.7e-23 \| 134.4 \| 165 \| (188, 357) \| 372 \| (14, 187) \| 192 \| Putative exodeoxyribonuclease 8 PDDEXK-like domain-containing protein (Fragment) \| Putative exodeoxyribonuclease 8 PDDEXK-like domain-containing protein (Fragment); uniclust \| UniRef100\_UPI001BAEF683 \| 99.5 \| 2.1e-17 \| 3.8e-23 \| 135.0 \| 167 \| (190, 359) \| 372 \| (32, 202) \| 206 \| PD-(D/E)XK nuclease-like domain-containing protein \| PD-(D/E)XK nuclease-like domain-containing protein; uniclust \| UniRef100\_UPI00102DF95B \| 99.5 \| 2.8e-17 \| 5.1e-23 \| 142.3 \| 237 \| (2, 341) \| 372 \| (20, 272) \| 315 \| PD-(D/E)XK nuclease-like domain-containing protein \| PD-(D/E)XK nuclease-like domain-containing protein; uniclust \| UniRef100\_UPI001EE8661C \| 99.5 \| 2.8e-17 \| 5.2e-23 \| 140.0 \| 241 \| (19, 347) \| 372 \| (22, 263) \| 278 \| PD-(D/E)XK nuclease-like domain-containing protein \| PD-(D/E)XK nuclease-like domain-containing protein; uniclust \| UniRef100\_A0A4U9HTL9 \| 99.5 \| 5.2e-17 \| 9.8e-23 \| 160.2 \| 227 \| (2, 231) \| 372 \| (699, 978) \| 980 \| Exodeoxyribonuclease 8 \| Exodeoxyribonuclease 8; uniclust \| UniRef100\_A0A0E3B9I3 \| 99.5 \| 6.5e-17 \| 1.2e-22 \| 134.4 \| 176 \| (184, 363) \| 372 \| (31, 218) \| 230 \| Putative exodeoxyribonuclease 8 PDDEXK-like domain-containing protein \| Putative exodeoxyribonuclease 8 PDDEXK-like domain-containing protein; uniclust \| UniRef100\_A0A4Q3BZQ0 \| 99.5 \| 8.5e-17 \| 1.6e-22 \| 139.9 \| 207 \| (2, 312) \| 372 \| (45, 266) \| 276 \| PD-(D/E)XK endonuclease-like domain-containing protein \| PD-(D/E)XK endonuclease-like domain-containing protein; uniclust \| UniRef100\_A0A160CAK3 \| 99.5 \| 9.8e-17 \| 1.8e-22 \| 135.5 \| 181 \| (185, 367) \| 372 \| (45, 256) \| 258 \| Exonuclease \| Exonuclease; uniclust \| UniRef100\_A0A6S6PGI9 \| 99.5 \| 1e-16 \| 1.8e-22 \| 127.2 \| 142 \| (1, 235) \| 372 \| (4, 151) \| 168 \| Putative exodeoxyribonuclease 8 PDDEXK-like domain-containing protein \| Putative exodeoxyribonuclease 8 PDDEXK-like domain-containing protein; uniclust \| UniRef100\_A0A955A2S3 \| 99.5 \| 1.2e-16 \| 2.2e-22 \| 141.5 \| 303 \| (2, 364) \| 372 \| (27, 361) \| 378 \| PD-(D/E)XK nuclease-like domain-containing protein \| PD-(D/E)XK nuclease-like domain-containing protein; uniclust \| UniRef100\_A0A5C7PD05 \| 99.5 \| 1.4e-16 \| 2.5e-22 \| 139.4 \| 260 \| (1, 365) \| 372 \| (51, 324) \| 339 \| Putative exodeoxyribonuclease 8 PDDEXK-like domain-containing protein \| Putative exodeoxyribonuclease 8 PDDEXK-like domain-containing protein; uniclust \| UniRef100\_A0A5C8B335 \| 99.5 \| 1.7e-16 \| 3.1e-22 \| 134.6 \| 251 \| (9, 366) \| 372 \| (1, 254) \| 265 \| Putative exodeoxyribonuclease 8 PDDEXK-like domain-containing protein \| Putative exodeoxyribonuclease 8 PDDEXK-like domain-containing protein; uniclust \| UniRef100\_A0A0K8Q145 \| 99.5 \| 1.7e-16 \| 3.1e-22 \| 126.2 \| 138 \| (2, 242) \| 372 \| (4, 142) \| 170 \| Exodeoxyribonuclease 8 \| Exodeoxyribonuclease 8; uniclust \| UniRef100\_UPI001E46E591 \| 99.5 \| 1.8e-16 \| 3.4e-22 \| 132.2 \| 167 \| (45, 262) \| 372 \| (14, 185) \| 235 \| PD-(D/E)XK nuclease-like domain-containing protein \| PD-(D/E)XK nuclease-like domain-containing protein; uniclust \| UniRef100\_UPI001FEF4C2F \| 99.5 \| 2e-16 \| 3.6e-22 \| 129.7 \| 168 \| (192, 362) \| 372 \| (24, 196) \| 207 \| PD-(D/E)XK nuclease-like domain-containing protein \| PD-(D/E)XK nuclease-like domain-containing protein; uniclust \| UniRef100\_A0A965PNL8 \| 99.5 \| 2.1e-16 \| 3.9e-22 \| 127.0 \| 148 \| (215, 364) \| 372 \| (4, 162) \| 182 \| Putative exodeoxyribonuclease 8 PDDEXK-like domain-containing protein \| Putative exodeoxyribonuclease 8 PDDEXK-like domain-containing protein; uniclust \| UniRef100\_A0A218KSW6 \| 99.4 \| 2.5e-16 \| 4.7e-22 \| 136.3 \| 130 \| (209, 341) \| 372 \| (102, 233) \| 261 \| Putative exonuclease \| Putative exonuclease; uniclust \| UniRef100\_UPI0018805882 \| 99.4 \| 2.8e-16 \| 5.1e-22 \| 121.0 \| 78 \| (1, 78) \| 372 \| (40, 117) \| 138 \| PD-(D/E)XK nuclease-like domain-containing protein \| PD-(D/E)XK nuclease-like domain-containing protein; uniclust \| UniRef100\_UPI001354BAA8 \| 99.4 \| 2.9e-16 \| 5.4e-22 \| 122.8 \| 101 \| (186, 289) \| 372 \| (34, 135) \| 152 \| PD-(D/E)XK nuclease-like domain-containing protein \| PD-(D/E)XK nuclease-like domain-containing protein; uniclust \| UniRef100\_A0A379GH20 \| 99.4 \| 3.1e-16 \| 5.7e-22 \| 135.9 \| 131 \| (2, 163) \| 372 \| (149, 280) \| 312 \| Exodeoxyribonuclease 8 \| Exodeoxyribonuclease 8; uniclust \| UniRef100\_A0A3U7CCZ6 \| 99.4 \| 3.1e-16 \| 5.7e-22 \| 128.7 \| 78 \| (1, 78) \| 372 \| (78, 155) \| 208 \| Exodeoxyribonuclease 8 (Fragment) \| Exodeoxyribonuclease 8 (Fragment); uniclust \| UniRef100\_A0A6I1JKT6 \| 99.4 \| 3.3e-16 \| 6.1e-22 \| 130.4 \| 165 \| (186, 366) \| 372 \| (45, 213) \| 231 \| Phage protein \| Phage protein; uniclust \| UniRef100\_G5S9W1 \| 99.4 \| 3.8e-16 \| 7e-22 \| 131.5 \| 78 \| (1, 78) \| 372 \| (113, 190) \| 243 \| Exodeoxyribonuclease 8 \| Exodeoxyribonuclease 8; uniclust \| UniRef100\_UPI0021E8A40C \| 99.4 \| 3.9e-16 \| 7.1e-22 \| 139.1 \| 169 \| (190, 359) \| 372 \| (192, 383) \| 391 \| PD-(D/E)XK nuclease-like domain-containing protein \| PD-(D/E)XK nuclease-like domain-containing protein; uniclust \| UniRef100\_A0A2E3D7E6 \| 99.4 \| 4.1e-16 \| 7.6e-22 \| 132.9 \| 242 \| (7, 358) \| 372 \| (3, 248) \| 273 \| Uncharacterized protein \| Uncharacterized protein; uniclust \| UniRef100\_A0A3B9ZLY2 \| 99.4 \| 4.6e-16 \| 8.5e-22 \| 138.1 \| 297 \| (8, 350) \| 372 \| (15, 336) \| 378 \| Putative exodeoxyribonuclease 8 PDDEXK-like domain-containing protein \| Putative exodeoxyribonuclease 8 PDDEXK-like domain-containing protein; uniclust \| UniRef100\_A0A3S4K5S1 \| 99.4 \| 4.7e-16 \| 8.7e-22 \| 126.9 \| 130 \| (2, 162) \| 372 \| (7, 137) \| 179 \| Putative phage exodeoxyribonuclease \| Putative phage exodeoxyribonuclease; uniclust \| UniRef100\_UPI0004091708 \| 99.4 \| 5.1e-16 \| 9.3e-22 \| 128.5 \| 130 \| (1, 139) \| 372 \| (78, 213) \| 220 \| SAP domain-containing protein \| SAP domain-containing protein; uniclust \| UniRef100\_A0A946HKP6 \| 99.4 \| 6.2e-16 \| 1.1e-21 \| 123.6 \| 161 \| (199, 364) \| 372 \| (4, 169) \| 174 \| Putative exodeoxyribonuclease 8 PDDEXK-like domain-containing protein \| Putative exodeoxyribonuclease 8 PDDEXK-like domain-containing protein; uniclust \| UniRef100\_A0A359MF53 \| 99.4 \| 6.9e-16 \| 1.3e-21 \| 127.9 \| 145 \| (187, 343) \| 372 \| (36, 182) \| 198 \| Putative exodeoxyribonuclease 8 PDDEXK-like domain-containing protein (Fragment) \| Putative exodeoxyribonuclease 8 PDDEXK-like domain-containing protein (Fragment); uniclust \| UniRef100\_A0A942R8U6 \| 99.4 \| 7.1e-16 \| 1.3e-21 \| 122.4 \| 127 \| (1, 136) \| 372 \| (14, 141) \| 166 \| Uncharacterized protein (Fragment) \| Uncharacterized protein (Fragment); uniclust \| UniRef100\_M7RB49 \| 99.4 \| 8.2e-16 \| 1.5e-21 \| 124.4 \| 78 \| (1, 78) \| 372 \| (96, 173) \| 188 \| Uncharacterized protein (Fragment) \| Uncharacterized protein (Fragment); uniclust \| UniRef100\_UPI001CC2243F \| 99.4 \| 9.9e-16 \| 1.9e-21 \| 123.0 \| 125 \| (205, 337) \| 372 \| (1, 129) \| 158 \| PD-(D/E)XK nuclease-like domain-containing protein \| PD-(D/E)XK nuclease-like domain-containing protein; uniclust \| UniRef100\_A0A1L3I482 \| 99.4 \| 1.6e-15 \| 3e-21 \| 116.4 \| 103 \| (205, 307) \| 372 \| (1, 110) \| 134 \| Exodeoxyribonuclease 8 \| Exodeoxyribonuclease 8; uniclust \| UniRef100\_A0A853IND3 \| 99.4 \| 1.6e-15 \| 3e-21 \| 125.1 \| 119 \| (187, 307) \| 372 \| (48, 166) \| 189 \| PD-(D/E)XK nuclease-like domain-containing protein (Fragment) \| PD-(D/E)XK nuclease-like domain-containing protein (Fragment); uniclust \| UniRef100\_A0A2A6DEY2 \| 99.4 \| 1.6e-15 \| 3e-21 \| 136.5 \| 78 \| (1, 78) \| 372 \| (293, 370) \| 423 \| Exodeoxyribonuclease 8 \| Exodeoxyribonuclease 8; uniclust \| UniRef100\_A0A139TR47 \| 99.4 \| 1.8e-15 \| 3.7e-21 \| 140.9 \| 145 \| (198, 349) \| 372 \| (142, 286) \| 314 \| PD-(D/E)XK endonuclease-like domain-containing protein \| PD-(D/E)XK endonuclease-like domain-containing protein; uniclust \| UniRef100\_A0A377K6Z1 \| 99.4 \| 2.1e-15 \| 3.9e-21 \| 119.2 \| 72 \| (2, 73) \| 372 \| (46, 118) \| 146 \| Putative phage exodeoxyribonuclease \| Putative phage exodeoxyribonuclease; uniclust \| UniRef100\_A0A1I7N9A7 \| 99.4 \| 2e-15 \| 4e-21 \| 153.7 \| 149 \| (188, 338) \| 372 \| (512, 674) \| 717 \| DNA 3'-5' helicase \| DNA 3'-5' helicase; uniclust \| UniRef100\_UPI001CECFBF9 \| 99.4 \| 2.2e-15 \| 4e-21 \| 141.2 \| 161 \| (1, 169) \| 372 \| (415, 602) \| 626 \| DNA breaking-rejoining protein \| DNA breaking-rejoining protein; uniclust \| UniRef100\_K1U9P6 \| 99.4 \| 2.5e-15 \| 4.5e-21 \| 141.0 \| 291 \| (13, 347) \| 372 \| (307, 614) \| 628 \| Protein containing Metallophosphoesterase domain protein \| Protein containing Metallophosphoesterase domain protein; uniclust \| UniRef100\_UPI001A94C8F2 \| 99.4 \| 2.5e-15 \| 4.7e-21 \| 131.8 \| 78 \| (1, 78) \| 372 \| (213, 290) \| 309 \| PD-(D/E)XK nuclease-like domain-containing protein \| PD-(D/E)XK nuclease-like domain-containing protein; uniclust \| UniRef100\_A0A021VU21 \| 99.4 \| 2.6e-15 \| 5.3e-21 \| 149.0 \| 116 \| (225, 350) \| 372 \| (272, 391) \| 503 \| Exodeoxyribonuclease V subunit beta \| Exodeoxyribonuclease V subunit beta; uniclust \| UniRef100\_A0A0R1M3A8 \| 99.3 \| 3.1e-15 \| 6e-21 \| 129.2 \| 148 \| (195, 347) \| 372 \| (19, 175) \| 203 \| Putative exodeoxyribonuclease 8 PDDEXK-like domain-containing protein \| Putative exodeoxyribonuclease 8 PDDEXK-like domain-containing protein; uniclust \| UniRef100\_UPI00210133BA \| 99.3 \| 4.6e-15 \| 8.4e-21 \| 115.3 \| 127 \| (187, 315) \| 372 \| (4, 139) \| 143 \| PD-(D/E)XK nuclease-like domain-containing protein \| PD-(D/E)XK nuclease-like domain-containing protein; uniclust \| UniRef100\_A0A524P3G8 \| 99.3 \| 5e-15 \| 9.2e-21 \| 117.9 \| 115 \| (235, 351) \| 372 \| (6, 122) \| 167 \| Putative exodeoxyribonuclease 8 PDDEXK-like domain-containing protein (Fragment) \| Putative exodeoxyribonuclease 8 PDDEXK-like domain-containing protein (Fragment); uniclust \| UniRef100\_X1D8E3 \| 99.3 \| 5.4e-15 \| 9.9e-21 \| 100.9 \| 65 \| (2, 66) \| 372 \| (4, 70) \| 70 \| Uncharacterized protein (Fragment) \| Uncharacterized protein (Fragment); uniclust \| UniRef100\_A0A0R2JVU7 \| 99.3 \| 5.1e-15 \| 1e-20 \| 126.2 \| 148 \| (195, 348) \| 372 \| (11, 166) \| 191 \| Putative exodeoxyribonuclease 8 PDDEXK-like domain-containing protein \| Putative exodeoxyribonuclease 8 PDDEXK-like domain-containing protein; uniclust \| UniRef100\_UPI000E170CDF \| 99.3 \| 5.9e-15 \| 1.1e-20 \| 108.4 \| 74 \| (5, 78) \| 372 \| (13, 87) \| 103 \| PD-(D/E)XK nuclease-like domain-containing protein \| PD-(D/E)XK nuclease-like domain-containing protein; uniclust \| UniRef100\_A0A7X7QZ66 \| 99.3 \| 6.1e-15 \| 1.1e-20 \| 106.5 \| 89 \| (274, 362) \| 372 \| (3, 91) \| 94 \| Putative exodeoxyribonuclease 8 PDDEXK-like domain-containing protein \| Putative exodeoxyribonuclease 8 PDDEXK-like domain-containing protein; uniclust \| UniRef100\_A0A238X2A8 \| 99.3 \| 6.8e-15 \| 1.2e-20 \| 126.5 \| 140 \| (199, 346) \| 372 \| (71, 215) \| 285 \| Putative exodeoxyribonuclease 8 PDDEXK-like domain-containing protein \| Putative exodeoxyribonuclease 8 PDDEXK-like domain-containing protein; uniclust \| UniRef100\_A0A098LCC7 \| 99.3 \| 6.6e-15 \| 1.4e-20 \| 160.5 \| 149 \| (188, 338) \| 372 \| (891, 1053) \| 1107 \| DNA 3'-5' helicase \| DNA 3'-5' helicase; uniclust \| UniRef100\_A0A0S7WJM6 \| 99.3 \| 6.9e-15 \| 1.4e-20 \| 139.4 \| 212 \| (19, 349) \| 372 \| (54, 272) \| 348 \| PD-(D/E)XK endonuclease-like domain-containing protein \| PD-(D/E)XK endonuclease-like domain-containing protein; uniclust \| UniRef100\_A0A2D6B2Q9 \| 99.3 \| 8.7e-15 \| 1.6e-20 \| 126.1 \| 252 \| (3, 348) \| 372 \| (2, 277) \| 288 \| Putative exodeoxyribonuclease 8 PDDEXK-like domain-containing protein \| Putative exodeoxyribonuclease 8 PDDEXK-like domain-containing protein; uniclust \| UniRef100\_A0A3D2IFL8 \| 99.3 \| 1e-14 \| 1.9e-20 \| 109.3 \| 71 \| (187, 260) \| 372 \| (44, 114) \| 115 \| Putative exodeoxyribonuclease 8 PDDEXK-like domain-containing protein (Fragment) \| Putative exodeoxyribonuclease 8 PDDEXK-like domain-containing protein (Fragment); uniclust \| UniRef100\_A0A0V8BEL4 \| 99.3 \| 1e-14 \| 1.9e-20 \| 128.7 \| 227 \| (5, 348) \| 372 \| (22, 270) \| 293 \| Prophage Lp1 protein 19 \| Prophage Lp1 protein 19; uniclust \| UniRef100\_UPI0022659236 \| 99.3 \| 1.4e-14 \| 2.6e-20 \| 118.8 \| 175 \| (188, 363) \| 372 \| (12, 193) \| 201 \| PD-(D/E)XK nuclease-like domain-containing protein \| PD-(D/E)XK nuclease-like domain-containing protein; uniclust \| UniRef100\_A0A8S0I2N3 \| 99.3 \| 1.5e-14 \| 2.8e-20 \| 106.2 \| 82 \| (225, 308) \| 372 \| (14, 96) \| 102 \| Putative exodeoxyribonuclease 8 PDDEXK-like domain-containing protein \| Putative exodeoxyribonuclease 8 PDDEXK-like domain-containing protein; uniclust \| UniRef100\_A0A1M5H3U8 \| 99.3 \| 1.5e-14 \| 2.9e-20 \| 142.6 \| 149 \| (188, 338) \| 372 \| (457, 619) \| 658 \| PD-(D/E)XK nuclease superfamily protein \| PD-(D/E)XK nuclease superfamily protein; uniclust \| UniRef100\_A0A447JGP7 \| 99.3 \| 1.7e-14 \| 3.2e-20 \| 105.3 \| 93 \| (194, 289) \| 372 \| (2, 95) \| 99 \| Exonuclease VIII \| Exonuclease VIII; uniclust \| UniRef100\_A0A368JHD6 \| 99.3 \| 1.7e-14 \| 3.3e-20 \| 120.9 \| 133 \| (201, 340) \| 372 \| (59, 194) \| 215 \| Putative exodeoxyribonuclease 8 PDDEXK-like domain-containing protein \| Putative exodeoxyribonuclease 8 PDDEXK-like domain-containing protein; uniclust \| UniRef100\_A0A2A4RDJ8 \| 99.3 \| 2e-14 \| 4.1e-20 \| 153.9 \| 224 \| (19, 349) \| 372 \| (880, 1114) \| 1152 \| DNA 3'-5' helicase \| DNA 3'-5' helicase; uniclust \| UniRef100\_A0A081PKL5 \| 99.3 \| 2.1e-14 \| 4.3e-20 \| 128.9 \| 140 \| (191, 337) \| 372 \| (76, 223) \| 249 \| Uncharacterized protein \| Uncharacterized protein; uniclust \| UniRef100\_A0A3D8IYQ2 \| 99.3 \| 2.4e-14 \| 4.3e-20 \| 121.9 \| 217 \| (9, 335) \| 372 \| (1, 226) \| 260 \| Putative exodeoxyribonuclease 8 PDDEXK-like domain-containing protein \| Putative exodeoxyribonuclease 8 PDDEXK-like domain-containing protein; uniclust \| UniRef100\_A0A1F9B6U3 \| 99.3 \| 2.3e-14 \| 4.9e-20 \| 137.7 \| 140 \| (199, 349) \| 372 \| (173, 321) \| 362 \| PD-(D/E)XK endonuclease-like domain-containing protein (Fragment) \| PD-(D/E)XK endonuclease-like domain-containing protein (Fragment); uniclust \| UniRef100\_A0A0C6EI61 \| 99.3 \| 2.5e-14 \| 5e-20 \| 140.9 \| 114 \| (225, 352) \| 372 \| (200, 317) \| 491 \| PD-(D/E)XK endonuclease-like domain-containing protein \| PD-(D/E)XK endonuclease-like domain-containing protein; uniclust \| UniRef100\_A0A2X3JIG8 \| 99.3 \| 2.7e-14 \| 5e-20 \| 121.1 \| 152 \| (2, 160) \| 372 \| (67, 244) \| 253 \| Putative phage exodeoxyribonuclease \| Putative phage exodeoxyribonuclease; uniclust \| UniRef100\_A0A350BS60 \| 99.2 \| 3e-14 \| 5.6e-20 \| 121.0 \| 163 \| (184, 348) \| 372 \| (57, 247) \| 256 \| Putative exodeoxyribonuclease 8 PDDEXK-like domain-containing protein \| Putative exodeoxyribonuclease 8 PDDEXK-like domain-containing protein; uniclust \| UniRef100\_UPI001B805189 \| 99.2 \| 3.1e-14 \| 5.6e-20 \| 100.7 \| 80 \| (205, 285) \| 372 \| (1, 83) \| 83 \| PD-(D/E)XK nuclease-like domain-containing protein \| PD-(D/E)XK nuclease-like domain-containing protein; uniclust \| UniRef100\_A0A1V5NKN1 \| 99.2 \| 3.1e-14 \| 6.3e-20 \| 142.0 \| 232 \| (8, 339) \| 372 \| (244, 493) \| 538 \| ATP-dependent helicase/deoxyribonuclease subunit B \| ATP-dependent helicase/deoxyribonuclease subunit B; uniclust \| UniRef100\_A0A379CLP5 \| 99.2 \| 3.6e-14 \| 6.5e-20 \| 120.4 \| 158 \| (2, 165) \| 372 \| (24, 205) \| 253 \| Exodeoxyribonuclease 8 \| Exodeoxyribonuclease 8; uniclust \| UniRef100\_M4QBX2 \| 99.2 \| 4.2e-14 \| 7.7e-20 \| 125.6 \| 176 \| (31, 260) \| 372 \| (3, 182) \| 361 \| Putative exodeoxyribonuclease 8 PDDEXK-like domain-containing protein \| Putative exodeoxyribonuclease 8 PDDEXK-like domain-containing protein; uniclust \| UniRef100\_A0A962Z9F4 \| 99.2 \| 4.4e-14 \| 8e-20 \| 116.4 \| 134 \| (191, 326) \| 372 \| (65, 205) \| 205 \| PD-(D/E)XK nuclease family protein (Fragment) \| PD-(D/E)XK nuclease family protein (Fragment); uniclust \| UniRef100\_A0A7C1JUS1 \| 99.2 \| 4.6e-14 \| 8.4e-20 \| 117.2 \| 138 \| (205, 349) \| 372 \| (1, 138) \| 216 \| Putative exodeoxyribonuclease 8 PDDEXK-like domain-containing protein \| Putative exodeoxyribonuclease 8 PDDEXK-like domain-containing protein; uniclust \| UniRef100\_A0A098SBQ5 \| 99.2 \| 4.1e-14 \| 9e-20 \| 156.2 \| 217 \| (18, 338) \| 372 \| (824, 1056) \| 1105 \| DNA 3'-5' helicase \| DNA 3'-5' helicase; uniclust \| UniRef100\_A0A059XC20 \| 99.2 \| 5.5e-14 \| 1.1e-19 \| 134.2 \| 110 \| (225, 350) \| 372 \| (199, 311) \| 344 \| PD-(D/E)XK nuclease superfamily (Fragment) \| PD-(D/E)XK nuclease superfamily (Fragment); uniclust \| UniRef100\_A0A0G0LEM3 \| 99.2 \| 6.2e-14 \| 1.3e-19 \| 136.1 \| 231 \| (14, 364) \| 372 \| (58, 303) \| 429 \| PD-(D/E)XK endonuclease-like domain-containing protein \| PD-(D/E)XK endonuclease-like domain-containing protein; uniclust \| UniRef100\_A0A1T5CLE9 \| 99.2 \| 7.4e-14 \| 1.4e-19 \| 115.7 \| 171 \| (20, 306) \| 372 \| (1, 179) \| 182 \| Putative exodeoxyribonuclease 8 PDDEXK-like domain-containing protein \| Putative exodeoxyribonuclease 8 PDDEXK-like domain-containing protein; uniclust \| UniRef100\_X7ZHD3 \| 99.2 \| 7.8e-14 \| 1.4e-19 \| 109.5 \| 94 \| (253, 346) \| 372 \| (9, 102) \| 149 \| Gp60 domain protein \| Gp60 domain protein; uniclust \| UniRef100\_A0A6J5Q956 \| 99.2 \| 8.4e-14 \| 1.5e-19 \| 115.9 \| 163 \| (184, 350) \| 372 \| (15, 190) \| 219 \| Exodeoxyribonuclease 8, PDDEXK-like domain containing protein \| Exodeoxyribonuclease 8, PDDEXK-like domain containing protein; uniclust \| UniRef100\_A0A6M3ITI9 \| 99.2 \| 8.4e-14 \| 1.5e-19 \| 122.0 \| 259 \| (8, 348) \| 372 \| (5, 290) \| 321 \| Putative exodeoxyribonuclease 8 PDDEXK-like domain-containing protein \| Putative exodeoxyribonuclease 8 PDDEXK-like domain-containing protein; uniclust \| UniRef100\_A0A949QL58 \| 99.2 \| 8.6e-14 \| 1.6e-19 \| 104.4 \| 109 \| (254, 362) \| 372 \| (2, 113) \| 114 \| PD-(D/E)XK nuclease-like domain-containing protein \| PD-(D/E)XK nuclease-like domain-containing protein; uniclust \| UniRef100\_UPI0015868581 \| 99.2 \| 8.6e-14 \| 1.6e-19 \| 98.5 \| 66 \| (1, 67) \| 372 \| (18, 83) \| 83 \| PD-(D/E)XK nuclease-like domain-containing protein \| PD-(D/E)XK nuclease-like domain-containing protein; uniclust \| UniRef100\_UPI00215A86D0 \| 99.2 \| 9.2e-14 \| 1.7e-19 \| 99.5 \| 75 \| (229, 306) \| 372 \| (1, 76) \| 88 \| PD-(D/E)XK nuclease-like domain-containing protein \| PD-(D/E)XK nuclease-like domain-containing protein; uniclust \| UniRef100\_A0A376LMM8 \| 99.2 \| 1.1e-13 \| 2.1e-19 \| 133.4 \| 140 \| (1, 232) \| 372 \| (613, 752) \| 752 \| Exonuclease VIII \| Exonuclease VIII; uniclust \| UniRef100\_A0A958WPB8 \| 99.2 \| 1.2e-13 \| 2.2e-19 \| 119.0 \| 137 \| (194, 339) \| 372 \| (134, 276) \| 280 \| PD-(D/E)XK nuclease-like domain-containing protein \| PD-(D/E)XK nuclease-like domain-containing protein; uniclust \| UniRef100\_UPI001FEDDF68 \| 99.2 \| 1.2e-13 \| 2.3e-19 \| 107.6 \| 117 \| (192, 310) \| 372 \| (24, 142) \| 142 \| PD-(D/E)XK nuclease-like domain-containing protein \| PD-(D/E)XK nuclease-like domain-containing protein; uniclust \| UniRef100\_A0A5C7JQK3 \| 99.2 \| 1.3e-13 \| 2.4e-19 \| 120.3 \| 249 \| (15, 328) \| 372 \| (11, 265) \| 310 \| Putative exodeoxyribonuclease 8 PDDEXK-like domain-containing protein \| Putative exodeoxyribonuclease 8 PDDEXK-like domain-containing protein; uniclust \| UniRef100\_UPI000DFC6B13 \| 99.2 \| 1.3e-13 \| 2.5e-19 \| 113.9 \| 132 \| (2, 142) \| 372 \| (37, 174) \| 207 \| SAP domain-containing protein \| SAP domain-containing protein; uniclust \| UniRef100\_A0A1V1PGN8 \| 99.2 \| 1.6e-13 \| 3.1e-19 \| 122.0 \| 255 \| (7, 338) \| 372 \| (2, 277) \| 300 \| Exodeoxyribonuclease 8 PDDEXK-like domain-containing protein \| Exodeoxyribonuclease 8 PDDEXK-like domain-containing protein; uniclust \| UniRef100\_A0A0F9EWN7 \| 99.2 \| 1.5e-13 \| 3.2e-19 \| 129.6 \| 137 \| (200, 352) \| 372 \| (133, 271) \| 296 \| PD-(D/E)XK endonuclease-like domain-containing protein (Fragment) \| PD-(D/E)XK endonuclease-like domain-containing protein (Fragment); uniclust \| UniRef100\_A0A2E9HYF9 \| 99.2 \| 1.8e-13 \| 3.3e-19 \| 121.6 \| 177 \| (184, 365) \| 372 \| (159, 346) \| 357 \| Putative exodeoxyribonuclease 8 PDDEXK-like domain-containing protein \| Putative exodeoxyribonuclease 8 PDDEXK-like domain-containing protein; uniclust \| UniRef100\_A0A0S7ZE36 \| 99.2 \| 1.7e-13 \| 3.4e-19 \| 135.0 \| 109 \| (225, 349) \| 372 \| (323, 434) \| 473 \| DNA 3'-5' helicase (Fragment) \| DNA 3'-5' helicase (Fragment); uniclust \| UniRef100\_A0A8S5NP51 \| 99.2 \| 1.9e-13 \| 3.5e-19 \| 106.6 \| 107 \| (255, 362) \| 372 \| (33, 140) \| 142 \| Exodeoxyribonuclease 8 \| Exodeoxyribonuclease 8; uniclust \| UniRef100\_UPI001FDECBD7 \| 99.2 \| 2e-13 \| 3.6e-19 \| 109.1 \| 118 \| (245, 362) \| 372 \| (1, 127) \| 165 \| PD-(D/E)XK nuclease-like domain-containing protein \| PD-(D/E)XK nuclease-like domain-containing protein; uniclust \| UniRef100\_A0A0G0NCS9 \| 99.1 \| 2.9e-13 \| 5.8e-19 \| 130.6 \| 231 \| (9, 348) \| 372 \| (143, 391) \| 414 \| DNA 3'-5' helicase (Fragment) \| DNA 3'-5' helicase (Fragment); uniclust \| UniRef100\_A0A0G1A9X7 \| 99.1 \| 3.6e-13 \| 7.4e-19 \| 141.2 \| 218 \| (18, 338) \| 372 \| (497, 731) \| 777 \| DNA 3'-5' helicase \| DNA 3'-5' helicase; uniclust \| UniRef100\_A0A447U5S1 \| 99.1 \| 4.2e-13 \| 7.8e-19 \| 113.4 \| 201 \| (84, 308) \| 372 \| (22, 231) \| 239 \| Exodeoxyribonuclease VIII \| Exodeoxyribonuclease VIII; uniclust \| UniRef100\_A0A139BNJ2 \| 99.1 \| 4.2e-13 \| 8.2e-19 \| 99.9 \| 79 \| (291, 370) \| 372 \| (4, 82) \| 86 \| Putative exodeoxyribonuclease 8 PDDEXK-like domain-containing protein \| Putative exodeoxyribonuclease 8 PDDEXK-like domain-containing protein; uniclust \| UniRef100\_UPI001E519A63 \| 99.1 \| 4.6e-13 \| 8.5e-19 \| 109.4 \| 150 \| (190, 347) \| 372 \| (6, 161) \| 189 \| PD-(D/E)XK nuclease-like domain-containing protein \| PD-(D/E)XK nuclease-like domain-containing protein; uniclust \| UniRef100\_A0A843FLM9 \| 99.1 \| 5.1e-13 \| 9.9e-19 \| 124.5 \| 286 \| (7, 343) \| 372 \| (23, 346) \| 361 \| PD-(D/E)XK nuclease-like domain-containing protein \| PD-(D/E)XK nuclease-like domain-containing protein; uniclust \| UniRef100\_A0A0D5A873 \| 99.1 \| 5.2e-13 \| 1e-18 \| 125.7 \| 105 \| (236, 349) \| 372 \| (213, 321) \| 391 \| RecB family exonuclease \| RecB family exonuclease; uniclust \| UniRef100\_UPI001F4ADC0F \| 99.1 \| 5.4e-13 \| 1e-18 \| 114.4 \| 149 \| (195, 348) \| 372 \| (49, 205) \| 229 \| PD-(D/E)XK nuclease-like domain-containing protein \| PD-(D/E)XK nuclease-like domain-containing protein; uniclust \| UniRef100\_B6YS92 \| 99.1 \| 5.9e-13 \| 1.1e-18 \| 119.3 \| 226 \| (8, 292) \| 372 \| (15, 250) \| 327 \| Putative exodeoxyribonuclease 8 PDDEXK-like domain-containing protein \| Putative exodeoxyribonuclease 8 PDDEXK-like domain-containing protein; uniclust \| UniRef100\_A0A160VHU9 \| 99.1 \| 5.8e-13 \| 1.2e-18 \| 124.0 \| 120 \| (225, 362) \| 372 \| (190, 324) \| 327 \| RecB family exonuclease \| RecB family exonuclease; uniclust \| UniRef100\_A0A8E0IFE8 \| 99.1 \| 6.9e-13 \| 1.3e-18 \| 123.8 \| 147 \| (195, 347) \| 372 \| (370, 524) \| 549 \| Putative exodeoxyribonuclease 8 PDDEXK-like domain-containing protein \| Putative exodeoxyribonuclease 8 PDDEXK-like domain-containing protein; uniclust \| UniRef100\_UPI0018CF9690 \| 99.1 \| 7.6e-13 \| 1.4e-18 \| 103.9 \| 83 \| (2, 84) \| 372 \| (54, 140) \| 146 \| hypothetical protein \| hypothetical protein; uniclust \| UniRef100\_A0A3C0G174 \| 99.1 \| 8.2e-13 \| 1.5e-18 \| 99.3 \| 93 \| (253, 347) \| 372 \| (2, 94) \| 113 \| Putative exodeoxyribonuclease 8 PDDEXK-like domain-containing protein \| Putative exodeoxyribonuclease 8 PDDEXK-like domain-containing protein; uniclust \| UniRef100\_UPI0009718900 \| 99.1 \| 8.2e-13 \| 1.5e-18 \| 107.1 \| 163 \| (201, 370) \| 372 \| (3, 170) \| 179 \| PD-(D/E)XK nuclease-like domain-containing protein \| PD-(D/E)XK nuclease-like domain-containing protein; uniclust \| UniRef100\_A0A024YLP4 \| 99.1 \| 9.1e-13 \| 1.7e-18 \| 123.8 \| 105 \| (236, 349) \| 372 \| (253, 361) \| 430 \| Recombinase RecB \| Recombinase RecB; uniclust \| UniRef100\_A0A4P0Y455 \| 99.1 \| 9.4e-13 \| 1.8e-18 \| 113.1 \| 99 \| (2, 102) \| 372 \| (130, 229) \| 235 \| Putative exodeoxyribonuclease VIII \| Putative exodeoxyribonuclease VIII; uniclust \| UniRef100\_A0A177Q060 \| 99.0 \| 1e-12 \| 2.1e-18 \| 123.0 \| 153 \| (200, 364) \| 372 \| (125, 289) \| 300 \| PD-(D/E)XK endonuclease-like domain-containing protein \| PD-(D/E)XK endonuclease-like domain-containing protein; uniclust \| UniRef100\_A0A066U5F6 \| 99.0 \| 1e-12 \| 2.2e-18 \| 128.5 \| 126 \| (199, 339) \| 372 \| (142, 270) \| 367 \| Recombinase RecB \| Recombinase RecB; uniclust \| UniRef100\_A0A060HEP5 \| 99.0 \| 1.1e-12 \| 2.2e-18 \| 141.9 \| 138 \| (196, 349) \| 372 \| (926, 1067) \| 1094 \| DNA 3'-5' helicase \| DNA 3'-5' helicase; uniclust \| UniRef100\_A0A0G0B930 \| 99.0 \| 1.1e-12 \| 2.3e-18 \| 129.6 \| 110 \| (225, 350) \| 372 \| (357, 470) \| 489 \| DNA 3'-5' helicase (Fragment) \| DNA 3'-5' helicase (Fragment); uniclust \| UniRef100\_A0A257LV29 \| 99.0 \| 1.3e-12 \| 2.7e-18 \| 119.6 \| 113 \| (225, 349) \| 372 \| (112, 227) \| 258 \| PD-(D/E)XK endonuclease-like domain-containing protein (Fragment) \| PD-(D/E)XK endonuclease-like domain-containing protein (Fragment); uniclust \| UniRef100\_A0A4R3AZ53 \| 99.0 \| 1.7e-12 \| 3.2e-18 \| 124.4 \| 141 \| (196, 349) \| 372 \| (340, 484) \| 528 \| RecB family exonuclease \| RecB family exonuclease; uniclust \| UniRef100\_A0A0F9QNT3 \| 99.0 \| 1.6e-12 \| 3.3e-18 \| 142.8 \| 110 \| (225, 350) \| 372 \| (932, 1045) \| 1088 \| DNA 3'-5' helicase \| DNA 3'-5' helicase; uniclust \| UniRef100\_A0A0F9YKX6 \| 99.0 \| 1.8e-12 \| 3.6e-18 \| 123.5 \| 134 \| (199, 347) \| 372 \| (140, 278) \| 340 \| PD-(D/E)XK endonuclease-like domain-containing protein \| PD-(D/E)XK endonuclease-like domain-containing protein; uniclust \| UniRef100\_A0A0G1WC88 \| 99.0 \| 1.8e-12 \| 3.7e-18 \| 123.3 \| 110 \| (225, 349) \| 372 \| (190, 303) \| 357 \| UvrD/REP helicase (Fragment) \| UvrD/REP helicase (Fragment); uniclust \| UniRef100\_A0A6M3LWC3 \| 99.0 \| 2.3e-12 \| 4.1e-18 \| 113.4 \| 280 \| (19, 346) \| 372 \| (3, 296) \| 317 \| Putative exodeoxyribonuclease 8 PDDEXK-like domain-containing protein \| Putative exodeoxyribonuclease 8 PDDEXK-like domain-containing protein; uniclust \| UniRef100\_UPI000E016246 \| 99.0 \| 2.3e-12 \| 4.3e-18 \| 109.7 \| 134 \| (2, 231) \| 372 \| (19, 162) \| 248 \| hypothetical protein \| hypothetical protein; uniclust \| UniRef100\_A0A5V7WNI2 \| 99.0 \| 2.2e-12 \| 4.4e-18 \| 128.6 \| 101 \| (2, 104) \| 372 \| (329, 430) \| 550 \| Exodeoxyribonuclease (Fragment) \| Exodeoxyribonuclease (Fragment); uniclust \| UniRef100\_A0A327J2B3 \| 99.0 \| 2.3e-12 \| 4.5e-18 \| 136.4 \| 110 \| (225, 338) \| 372 \| (893, 1006) \| 1032 \| DNA 3'-5' helicase \| DNA 3'-5' helicase; uniclust \| UniRef100\_A0A0F9YNH9 \| 99.0 \| 2.3e-12 \| 4.7e-18 \| 139.7 \| 225 \| (9, 337) \| 372 \| (814, 1056) \| 1108 \| DNA 3'-5' helicase \| DNA 3'-5' helicase; uniclust \| UniRef100\_A0A117IC06 \| 99.0 \| 2.5e-12 \| 4.7e-18 \| 93.3 \| 78 \| (292, 369) \| 372 \| (5, 82) \| 84 \| RecE (Fragment) \| RecE (Fragment); uniclust \| UniRef100\_A0A0A0B9Q3 \| 99.0 \| 2.5e-12 \| 4.8e-18 \| 123.7 \| 116 \| (225, 350) \| 372 \| (185, 304) \| 477 \| Recombinase RecB \| Recombinase RecB; uniclust \| UniRef100\_A0A9D7FBP7 \| 99.0 \| 2.7e-12 \| 4.9e-18 \| 96.1 \| 70 \| (7, 77) \| 372 \| (25, 95) \| 109 \| PD-(D/E)XK nuclease-like domain-containing protein \| PD-(D/E)XK nuclease-like domain-containing protein; uniclust \| UniRef100\_A0A963IY46 \| 99.0 \| 2.7e-12 \| 5e-18 \| 98.4 \| 106 \| (253, 360) \| 372 \| (2, 108) \| 125 \| Uncharacterized protein \| Uncharacterized protein; uniclust \| UniRef100\_A0A965VWR8 \| 99.0 \| 2.9e-12 \| 5.4e-18 \| 111.0 \| 239 \| (11, 290) \| 372 \| (16, 271) \| 282 \| PD-(D/E)XK endonuclease-like domain-containing protein (Fragment) \| PD-(D/E)XK endonuclease-like domain-containing protein (Fragment); uniclust \| UniRef100\_UPI001E64B484 \| 99.0 \| 3e-12 \| 5.4e-18 \| 102.9 \| 96 \| (2, 100) \| 372 \| (24, 119) \| 166 \| hypothetical protein \| hypothetical protein; uniclust \| UniRef100\_A0A2E7DJB3 \| 99.0 \| 2.8e-12 \| 5.5e-18 \| 117.7 \| 121 \| (235, 363) \| 372 \| (174, 304) \| 306 \| PD-(D/E)XK endonuclease-like domain-containing protein \| PD-(D/E)XK endonuclease-like domain-containing protein; uniclust \| UniRef100\_A0A4Q3PZX2 \| 99.0 \| 3.2e-12 \| 6e-18 \| 105.2 \| 114 \| (225, 338) \| 372 \| (36, 161) \| 176 \| Putative exodeoxyribonuclease 8 PDDEXK-like domain-containing protein (Fragment) \| Putative exodeoxyribonuclease 8 PDDEXK-like domain-containing protein (Fragment); uniclust \| UniRef100\_A0A349YBV5 \| 99.0 \| 3.5e-12 \| 6.5e-18 \| 109.6 \| 138 \| (195, 337) \| 372 \| (92, 231) \| 242 \| Putative exodeoxyribonuclease 8 PDDEXK-like domain-containing protein \| Putative exodeoxyribonuclease 8 PDDEXK-like domain-containing protein; uniclust \| UniRef100\_A0A1I0PE57 \| 99.0 \| 3.5e-12 \| 6.7e-18 \| 120.9 \| 149 \| (188, 338) \| 372 \| (273, 438) \| 468 \| PD-(D/E)XK nuclease superfamily protein \| PD-(D/E)XK nuclease superfamily protein; uniclust \| UniRef100\_A0A081S3H6 \| 99.0 \| 3.5e-12 \| 7.1e-18 \| 131.1 \| 112 \| (225, 349) \| 372 \| (585, 700) \| 722 \| DNA 3'-5' helicase (Fragment) \| DNA 3'-5' helicase (Fragment); uniclust \| UniRef100\_UPI000EC7A980 \| 99.0 \| 4.1e-12 \| 7.5e-18 \| 88.1 \| 64 \| (225, 288) \| 372 \| (5, 72) \| 72 \| PD-(D/E)XK nuclease-like domain-containing protein \| PD-(D/E)XK nuclease-like domain-containing protein; uniclust \| UniRef100\_A0A5C7PXG6 \| 99.0 \| 4.3e-12 \| 8e-18 \| 94.7 \| 70 \| (3, 74) \| 372 \| (30, 100) \| 107 \| Putative exodeoxyribonuclease 8 PDDEXK-like domain-containing protein (Fragment) \| Putative exodeoxyribonuclease 8 PDDEXK-like domain-containing protein (Fragment); uniclust \| UniRef100\_A0A0G0K286 \| 99.0 \| 4.5e-12 \| 8.9e-18 \| 124.3 \| 230 \| (11, 347) \| 372 \| (239, 490) \| 510 \| DNA 3'-5' helicase \| DNA 3'-5' helicase; uniclust \| UniRef100\_A0A968V6E1 \| 99.0 \| 5.1e-12 \| 9.4e-18 \| 111.8 \| 262 \| (7, 349) \| 372 \| (3, 293) \| 329 \| Uncharacterized protein \| Uncharacterized protein; uniclust \| UniRef100\_A0A6M3J0N3 \| 98.9 \| 5.4e-12 \| 9.9e-18 \| 104.5 \| 166 \| (2, 262) \| 372 \| (8, 193) \| 202 \| Uncharacterized protein (Fragment) \| Uncharacterized protein (Fragment); uniclust \| UniRef100\_A0A0G0EV96 \| 98.9 \| 4.9e-12 \| 1e-17 \| 132.5 \| 109 \| (225, 349) \| 372 \| (606, 718) \| 741 \| DNA 3'-5' helicase \| DNA 3'-5' helicase; uniclust \| UniRef100\_UPI002165B33C \| 98.9 \| 5.8e-12 \| 1.1e-17 \| 108.8 \| 146 \| (216, 363) \| 372 \| (104, 261) \| 272 \| PD-(D/E)XK nuclease-like domain-containing protein \| PD-(D/E)XK nuclease-like domain-containing protein; uniclust \| UniRef100\_UPI000B1595CC \| 98.9 \| 5.9e-12 \| 1.1e-17 \| 111.9 \| 211 \| (4, 318) \| 372 \| (37, 262) \| 341 \| PD-(D/E)XK nuclease-like domain-containing protein \| PD-(D/E)XK nuclease-like domain-containing protein; uniclust \| UniRef100\_A0A0F9D9H6 \| 98.9 \| 5.6e-12 \| 1.2e-17 \| 120.5 \| 148 \| (190, 351) \| 372 \| (132, 281) \| 324 \| PD-(D/E)XK endonuclease-like domain-containing protein \| PD-(D/E)XK endonuclease-like domain-containing protein; uniclust \| UniRef100\_A0A1M7D7C6 \| 98.9 \| 6.5e-12 \| 1.2e-17 \| 109.0 \| 138 \| (192, 337) \| 372 \| (129, 273) \| 282 \| Putative exodeoxyribonuclease 8 PDDEXK-like domain-containing protein \| Putative exodeoxyribonuclease 8 PDDEXK-like domain-containing protein; uniclust \| UniRef100\_A0A1F2WZJ7 \| 98.9 \| 6.2e-12 \| 1.2e-17 \| 117.0 \| 162 \| (187, 364) \| 372 \| (174, 347) \| 353 \| PD-(D/E)XK endonuclease-like domain-containing protein \| PD-(D/E)XK endonuclease-like domain-containing protein; uniclust \| UniRef100\_A0A485BQR4 \| 98.9 \| 6.7e-12 \| 1.2e-17 \| 96.7 \| 90 \| (2, 93) \| 372 \| (36, 126) \| 128 \| Exodeoxyribonuclease 8 \| Exodeoxyribonuclease 8; uniclust \| UniRef100\_A0A0F9EXF6 \| 98.9 \| 6.7e-12 \| 1.3e-17 \| 106.4 \| 166 \| (2, 269) \| 372 \| (15, 199) \| 211 \| Exodeoxyribonuclease 8 PDDEXK-like domain-containing protein (Fragment) \| Exodeoxyribonuclease 8 PDDEXK-like domain-containing protein (Fragment); uniclust \| UniRef100\_A0A920Q8L2 \| 98.9 \| 6.8e-12 \| 1.3e-17 \| 108.9 \| 126 \| (8, 233) \| 372 \| (138, 263) \| 283 \| Putative exodeoxyribonuclease 8 PDDEXK-like domain-containing protein \| Putative exodeoxyribonuclease 8 PDDEXK-like domain-containing protein; uniclust \| UniRef100\_UPI0022DE3438 \| 98.9 \| 6.9e-12 \| 1.3e-17 \| 104.9 \| 133 \| (199, 339) \| 372 \| (47, 186) \| 215 \| PD-(D/E)XK nuclease-like domain-containing protein \| PD-(D/E)XK nuclease-like domain-containing protein; uniclust \| UniRef100\_A0A095Y853 \| 98.9 \| 7.2e-12 \| 1.5e-17 \| 135.1 \| 137 \| (195, 350) \| 372 \| (906, 1047) \| 1071 \| DNA 3'-5' helicase \| DNA 3'-5' helicase; uniclust \| UniRef100\_A0A136KRS2 \| 98.9 \| 8.1e-12 \| 1.5e-17 \| 113.6 \| 148 \| (191, 339) \| 372 \| (128, 291) \| 316 \| ATP-dependent helicase/deoxyribonuclease subunit B \| ATP-dependent helicase/deoxyribonuclease subunit B; uniclust \| UniRef100\_A0A0G3WJV3 \| 98.9 \| 8.4e-12 \| 1.7e-17 \| 117.7 \| 132 \| (199, 348) \| 372 \| (141, 276) \| 309 \| PD-(D/E)XK endonuclease-like domain-containing protein \| PD-(D/E)XK endonuclease-like domain-containing protein; uniclust \| UniRef100\_A0A0F9YF59 \| 98.9 \| 8.5e-12 \| 1.7e-17 \| 124.1 \| 127 \| (225, 362) \| 372 \| (176, 316) \| 482 \| PD-(D/E)XK endonuclease-like domain-containing protein \| PD-(D/E)XK endonuclease-like domain-containing protein; uniclust \| UniRef100\_A0A379YD88 \| 98.9 \| 9.3e-12 \| 1.7e-17 \| 101.7 \| 153 \| (210, 368) \| 372 \| (2, 164) \| 168 \| Exonuclease VIII \| Exonuclease VIII; uniclust \| UniRef100\_A0A094PIS3 \| 98.9 \| 9.1e-12 \| 1.8e-17 \| 116.1 \| 106 \| (236, 350) \| 372 \| (161, 268) \| 329 \| PD-(D/E)XK endonuclease-like domain-containing protein \| PD-(D/E)XK endonuclease-like domain-containing protein; uniclust \| UniRef100\_UPI0020C55E59 \| 98.9 \| 1e-11 \| 1.9e-17 \| 94.1 \| 90 \| (187, 279) \| 372 \| (23, 113) \| 113 \| PD-(D/E)XK nuclease-like domain-containing protein \| PD-(D/E)XK nuclease-like domain-containing protein; uniclust \| UniRef100\_A0A1J4YR24 \| 98.9 \| 1.1e-11 \| 2.1e-17 \| 85.0 \| 62 \| (5, 68) \| 372 \| (5, 66) \| 68 \| Uncharacterized protein \| Uncharacterized protein; uniclust \| UniRef100\_A0A0G0RCS7 \| 98.9 \| 1.1e-11 \| 2.2e-17 \| 123.0 \| 112 \| (225, 348) \| 372 \| (387, 501) \| 531 \| DNA 3'-5' helicase \| DNA 3'-5' helicase; uniclust \| UniRef100\_A0A1G0YSF3 \| 98.9 \| 1.3e-11 \| 2.7e-17 \| 117.1 \| 229 \| (8, 349) \| 372 \| (6, 249) \| 300 \| PD-(D/E)XK endonuclease-like domain-containing protein \| PD-(D/E)XK endonuclease-like domain-containing protein; uniclust \| UniRef100\_A0A5S9M3W7 \| 98.9 \| 1.6e-11 \| 3.1e-17 \| 101.2 \| 148 \| (2, 259) \| 372 \| (2, 155) \| 155 \| Putative exodeoxyribonuclease 8 PDDEXK-like domain-containing protein \| Putative exodeoxyribonuclease 8 PDDEXK-like domain-containing protein; uniclust \| UniRef100\_UPI0017820ABF \| 98.9 \| 1.9e-11 \| 3.5e-17 \| 106.1 \| 176 \| (183, 361) \| 372 \| (63, 274) \| 277 \| PD-(D/E)XK nuclease-like domain-containing protein \| PD-(D/E)XK nuclease-like domain-containing protein; uniclust \| UniRef100\_A0A024JZZ9 \| 98.9 \| 1.9e-11 \| 3.5e-17 \| 113.3 \| 147 \| (194, 349) \| 372 \| (124, 277) \| 356 \| Recombinase B \| Recombinase B; uniclust \| UniRef100\_A0A5C5X7Z0 \| 98.8 \| 2.2e-11 \| 4.1e-17 \| 108.2 \| 241 \| (8, 351) \| 372 \| (13, 323) \| 333 \| Putative exodeoxyribonuclease 8 PDDEXK-like domain-containing protein \| Putative exodeoxyribonuclease 8 PDDEXK-like domain-containing protein; uniclust \| UniRef100\_A0A350BX12 \| 98.8 \| 2.2e-11 \| 4.3e-17 \| 111.0 \| 148 \| (189, 338) \| 372 \| (53, 214) \| 260 \| DNA helicase UvrD (Fragment) \| DNA helicase UvrD (Fragment); uniclust \| UniRef100\_X0YC96 \| 98.8 \| 2.4e-11 \| 4.4e-17 \| 96.0 \| 74 \| (2, 78) \| 372 \| (6, 80) \| 146 \| Uncharacterized protein (Fragment) \| Uncharacterized protein (Fragment); uniclust \| UniRef100\_A0A1G6MCA1 \| 98.8 \| 2.4e-11 \| 4.9e-17 \| 130.7 \| 141 \| (197, 350) \| 372 \| (961, 1108) \| 1131 \| DNA 3'-5' helicase \| DNA 3'-5' helicase; uniclust \| UniRef100\_A0A068NW26 \| 98.8 \| 2.6e-11 \| 5.2e-17 \| 114.6 \| 213 \| (17, 347) \| 372 \| (45, 268) \| 323 \| RecB family exonuclease \| RecB family exonuclease; uniclust \| UniRef100\_A0A377E4A8 \| 98.8 \| 3.1e-11 \| 5.7e-17 \| 97.2 \| 64 \| (1, 64) \| 372 \| (94, 157) \| 165 \| Exonuclease VIII \| Exonuclease VIII; uniclust \| UniRef100\_A0A962VA62 \| 98.8 \| 3.2e-11 \| 5.9e-17 \| 89.6 \| 69 \| (4, 77) \| 372 \| (22, 90) \| 102 \| Uncharacterized protein (Fragment) \| Uncharacterized protein (Fragment); uniclust \| UniRef100\_UPI000F6DF811 \| 98.8 \| 3.4e-11 \| 6.2e-17 \| 90.9 \| 70 \| (1, 75) \| 372 \| (1, 70) \| 111 \| hypothetical protein \| hypothetical protein; uniclust \| UniRef100\_A0A2U3ERY8 \| 98.8 \| 3.8e-11 \| 6.9e-17 \| 109.5 \| 166 \| (188, 363) \| 372 \| (223, 394) \| 413 \| Exodeoxyribonuclease 8 PDDEXK-like domain-containing protein \| Exodeoxyribonuclease 8 PDDEXK-like domain-containing protein; uniclust \| UniRef100\_A0A1G3BLT7 \| 98.8 \| 3.5e-11 \| 6.9e-17 \| 129.0 \| 145 \| (191, 348) \| 372 \| (963, 1112) \| 1169 \| DNA 3'-5' helicase (Fragment) \| DNA 3'-5' helicase (Fragment); uniclust \| UniRef100\_A0A0S8E471 \| 98.8 \| 3.6e-11 \| 7.3e-17 \| 129.3 \| 112 \| (225, 350) \| 372 \| (879, 994) \| 1014 \| DNA 3'-5' helicase \| DNA 3'-5' helicase; uniclust \| UniRef100\_UPI0020CB8203 \| 98.8 \| 4.4e-11 \| 8.1e-17 \| 99.6 \| 189 \| (34, 290) \| 372 \| (3, 193) \| 204 \| PD-(D/E)XK nuclease-like domain-containing protein \| PD-(D/E)XK nuclease-like domain-containing protein; uniclust \| UniRef100\_UPI0011161D0A \| 98.8 \| 4.5e-11 \| 8.2e-17 \| 88.7 \| 77 \| (2, 78) \| 372 \| (15, 92) \| 101 \| hypothetical protein \| hypothetical protein; uniclust \| UniRef100\_UPI0007C58163 \| 98.8 \| 4.5e-11 \| 8.3e-17 \| 94.2 \| 95 \| (188, 291) \| 372 \| (10, 107) \| 143 \| PD-(D/E)XK nuclease-like domain-containing protein \| PD-(D/E)XK nuclease-like domain-containing protein; uniclust \| UniRef100\_A0A2D9CKI8 \| 98.8 \| 4.6e-11 \| 8.5e-17 \| 108.6 \| 138 \| (198, 335) \| 372 \| (186, 348) \| 399 \| Putative exodeoxyribonuclease 8 PDDEXK-like domain-containing protein \| Putative exodeoxyribonuclease 8 PDDEXK-like domain-containing protein; uniclust \| UniRef100\_UPI0020A48B88 \| 98.8 \| 5.3e-11 \| 9.8e-17 \| 102.2 \| 95 \| (231, 326) \| 372 \| (132, 230) \| 254 \| PD-(D/E)XK nuclease-like domain-containing protein \| PD-(D/E)XK nuclease-like domain-containing protein; uniclust \| UniRef100\_A0A1V1FV31 \| 98.8 \| 5.3e-11 \| 9.8e-17 \| 105.1 \| 152 \| (188, 342) \| 372 \| (120, 290) \| 314 \| Exonucleaase VIII \| Exonucleaase VIII; uniclust \| UniRef100\_A0A0F9G4G2 \| 98.8 \| 5.5e-11 \| 1e-16 \| 89.9 \| 91 \| (272, 363) \| 372 \| (4, 96) \| 112 \| Putative exodeoxyribonuclease 8 PDDEXK-like domain-containing protein (Fragment) \| Putative exodeoxyribonuclease 8 PDDEXK-like domain-containing protein (Fragment); uniclust \| UniRef100\_A0A3D2SYI2 \| 98.8 \| 5.8e-11 \| 1.1e-16 \| 101.9 \| 70 \| (236, 307) \| 372 \| (147, 216) \| 252 \| Putative exodeoxyribonuclease 8 PDDEXK-like domain-containing protein \| Putative exodeoxyribonuclease 8 PDDEXK-like domain-containing protein; uniclust \| UniRef100\_A0A3N5V3F8 \| 98.8 \| 6e-11 \| 1.1e-16 \| 95.2 \| 74 \| (2, 77) \| 372 \| (28, 102) \| 160 \| Uncharacterized protein (Fragment) \| Uncharacterized protein (Fragment); uniclust \| UniRef100\_A0A1F2QKG7 \| 98.8 \| 5.5e-11 \| 1.1e-16 \| 113.5 \| 101 \| (236, 346) \| 372 \| (158, 264) \| 320 \| PD-(D/E)XK endonuclease-like domain-containing protein \| PD-(D/E)XK endonuclease-like domain-containing protein; uniclust \| UniRef100\_A0A1M3BII6 \| 98.8 \| 5.5e-11 \| 1.1e-16 \| 133.0 \| 145 \| (193, 348) \| 372 \| (1236, 1383) \| 1448 \| DNA 3'-5' helicase (Fragment) \| DNA 3'-5' helicase (Fragment); uniclust \| UniRef100\_A0A1F9WEL2 \| 98.8 \| 6.9e-11 \| 1.3e-16 \| 111.8 \| 103 \| (198, 312) \| 372 \| (105, 211) \| 399 \| PD-(D/E)XK endonuclease-like domain-containing protein (Fragment) \| PD-(D/E)XK endonuclease-like domain-containing protein (Fragment); uniclust \| UniRef100\_A0A377TQV8 \| 98.8 \| 7.6e-11 \| 1.4e-16 \| 105.4 \| 100 \| (2, 103) \| 372 \| (239, 339) \| 344 \| Putative exodeoxyribonuclease VIII \| Putative exodeoxyribonuclease VIII; uniclust \| UniRef100\_UPI001F44166F \| 98.7 \| 8e-11 \| 1.5e-16 \| 92.9 \| 117 \| (253, 370) \| 372 \| (12, 140) \| 143 \| PD-(D/E)XK nuclease-like domain-containing protein \| PD-(D/E)XK nuclease-like domain-containing protein; uniclust \| UniRef100\_A0A1Z8ZBW6 \| 98.7 \| 7.7e-11 \| 1.6e-16 \| 128.0 \| 121 \| (225, 362) \| 372 \| (881, 1015) \| 1018 \| DNA 3'-5' helicase \| DNA 3'-5' helicase; uniclust \| UniRef100\_UPI00208F1201 \| 98.7 \| 8.9e-11 \| 1.6e-16 \| 85.6 \| 64 \| (1, 69) \| 372 \| (1, 64) \| 91 \| hypothetical protein \| hypothetical protein; uniclust \| UniRef100\_A0A0G2BII5 \| 98.7 \| 8.3e-11 \| 1.6e-16 \| 108.7 \| 101 \| (225, 337) \| 372 \| (143, 246) \| 285 \| UvrD/REP helicase (Fragment) \| UvrD/REP helicase (Fragment); uniclust \| UniRef100\_A0A3N0GKK1 \| 98.7 \| 8.7e-11 \| 1.7e-16 \| 110.1 \| 103 \| (235, 348) \| 372 \| (159, 264) \| 332 \| PD-(D/E)XK endonuclease-like domain-containing protein \| PD-(D/E)XK endonuclease-like domain-containing protein; uniclust \| UniRef100\_UPI000B0585E9 \| 98.7 \| 1.1e-10 \| 2e-16 \| 96.0 \| 73 \| (188, 262) \| 372 \| (101, 175) \| 185 \| PD-(D/E)XK nuclease-like domain-containing protein \| PD-(D/E)XK nuclease-like domain-containing protein; uniclust \| UniRef100\_A0A1Q7DWU8 \| 98.7 \| 1e-10 \| 2.1e-16 \| 119.7 \| 110 \| (225, 350) \| 372 \| (606, 719) \| 738 \| DNA 3'-5' helicase \| DNA 3'-5' helicase; uniclust \| UniRef100\_A0A0G1VV66 \| 98.7 \| 1.1e-10 \| 2.1e-16 \| 121.3 \| 146 \| (190, 337) \| 372 \| (617, 778) \| 814 \| DNA 3'-5' helicase (Fragment) \| DNA 3'-5' helicase (Fragment); uniclust \| UniRef100\_A0A0P6X7G0 \| 98.7 \| 1.3e-10 \| 2.6e-16 \| 108.0 \| 104 \| (236, 350) \| 372 \| (155, 262) \| 288 \| PD-(D/E)XK endonuclease-like domain-containing protein \| PD-(D/E)XK endonuclease-like domain-containing protein; uniclust \| UniRef100\_UPI00214D1424 \| 98.7 \| 1.4e-10 \| 2.6e-16 \| 86.1 \| 69 \| (298, 366) \| 372 \| (2, 70) \| 100 \| PD-(D/E)XK nuclease-like domain-containing protein \| PD-(D/E)XK nuclease-like domain-containing protein; uniclust \| UniRef100\_UPI000DFB18C6 \| 98.7 \| 1.5e-10 \| 2.7e-16 \| 88.4 \| 105 \| (263, 368) \| 372 \| (3, 111) \| 117 \| PD-(D/E)XK nuclease-like domain-containing protein \| PD-(D/E)XK nuclease-like domain-containing protein; uniclust \| UniRef100\_A0A537TML3 \| 98.7 \| 1.5e-10 \| 2.8e-16 \| 116.8 \| 146 \| (193, 351) \| 372 \| (508, 658) \| 692 \| DNA 3'-5' helicase (Fragment) \| DNA 3'-5' helicase (Fragment); uniclust \| UniRef100\_A0A0F9KKT2 \| 98.7 \| 1.4e-10 \| 3e-16 \| 107.8 \| 117 \| (237, 366) \| 372 \| (134, 260) \| 265 \| PD-(D/E)XK endonuclease-like domain-containing protein \| PD-(D/E)XK endonuclease-like domain-containing protein; uniclust \| UniRef100\_UPI001E3ED308 \| 98.7 \| 1.7e-10 \| 3.1e-16 \| 95.0 \| 164 \| (7, 230) \| 372 \| (12, 185) \| 185 \| hypothetical protein \| hypothetical protein; uniclust \| UniRef100\_A0A2V9BQB0 \| 98.7 \| 1.7e-10 \| 3.3e-16 \| 119.3 \| 110 \| (225, 350) \| 372 \| (632, 745) \| 778 \| DNA 3'-5' helicase \| DNA 3'-5' helicase; uniclust \| UniRef100\_A0A1Q7DKN2 \| 98.7 \| 1.7e-10 \| 3.4e-16 \| 122.4 \| 110 \| (225, 350) \| 372 \| (855, 968) \| 1013 \| DNA 3'-5' helicase \| DNA 3'-5' helicase; uniclust \| UniRef100\_A0A0C1UKG6 \| 98.7 \| 1.6e-10 \| 3.4e-16 \| 130.1 \| 157 \| (188, 350) \| 372 \| (859, 1034) \| 1119 \| PD-(D/E)XK endonuclease-like domain-containing protein \| PD-(D/E)XK endonuclease-like domain-containing protein; uniclust \| UniRef100\_A0A973F465 \| 98.7 \| 1.9e-10 \| 3.5e-16 \| 100.2 \| 228 \| (9, 339) \| 372 \| (4, 241) \| 277 \| Putative exodeoxyribonuclease 8 PDDEXK-like domain-containing protein \| Putative exodeoxyribonuclease 8 PDDEXK-like domain-containing protein; uniclust \| UniRef100\_UPI0020BD75F5 \| 98.7 \| 1.9e-10 \| 3.5e-16 \| 96.9 \| 146 \| (195, 346) \| 372 \| (32, 193) \| 217 \| PD-(D/E)XK nuclease-like domain-containing protein \| PD-(D/E)XK nuclease-like domain-containing protein; uniclust \| UniRef100\_A0A1V5YSQ9 \| 98.7 \| 1.8e-10 \| 3.8e-16 \| 127.5 \| 130 \| (196, 337) \| 372 \| (1038, 1167) \| 1194 \| DNA 3'-5' helicase \| DNA 3'-5' helicase; uniclust \| UniRef100\_A0A0B3AHC3 \| 98.7 \| 2e-10 \| 4e-16 \| 111.8 \| 105 \| (225, 349) \| 372 \| (193, 300) \| 375 \| PD-(D/E)XK endonuclease-like domain-containing protein \| PD-(D/E)XK endonuclease-like domain-containing protein; uniclust \| UniRef100\_A0A101HUX6 \| 98.7 \| 2e-10 \| 4.1e-16 \| 121.5 \| 112 \| (226, 349) \| 372 \| (626, 740) \| 766 \| DNA 3'-5' helicase (Fragment) \| DNA 3'-5' helicase (Fragment); uniclust \| UniRef100\_UPI0007C48233 \| 98.7 \| 2.3e-10 \| 4.2e-16 \| 104.6 \| 134 \| (5, 231) \| 372 \| (254, 389) \| 392 \| hypothetical protein \| hypothetical protein; uniclust \| UniRef100\_A0A1G1TB27 \| 98.7 \| 2.1e-10 \| 4.2e-16 \| 123.7 \| 146 \| (191, 338) \| 372 \| (865, 1040) \| 1092 \| DNA 3'-5' helicase \| DNA 3'-5' helicase; uniclust \| UniRef100\_A0A074LL59 \| 98.7 \| 2e-10 \| 4.5e-16 \| 130.0 \| 146 \| (196, 349) \| 372 \| (1035, 1188) \| 1220 \| DNA 3'-5' helicase \| DNA 3'-5' helicase; uniclust \| UniRef100\_A0A3Q8YTG7 \| 98.7 \| 2.5e-10 \| 4.7e-16 \| 89.9 \| 106 \| (253, 359) \| 372 \| (14, 124) \| 140 \| Putative exodeoxyribonuclease 8 PDDEXK-like domain-containing protein \| Putative exodeoxyribonuclease 8 PDDEXK-like domain-containing protein; uniclust \| UniRef100\_A0A0G0MTT2 \| 98.6 \| 2.4e-10 \| 4.9e-16 \| 108.8 \| 135 \| (197, 350) \| 372 \| (118, 264) \| 302 \| RecB family exonuclease \| RecB family exonuclease; uniclust \| UniRef100\_A0A1F2X2I5 \| 98.6 \| 2.5e-10 \| 5.3e-16 \| 126.8 \| 159 \| (188, 350) \| 372 \| (888, 1062) \| 1142 \| PD-(D/E)XK endonuclease-like domain-containing protein \| PD-(D/E)XK endonuclease-like domain-containing protein; uniclust \| UniRef100\_A0A2W4MB63 \| 98.6 \| 2.8e-10 \| 5.4e-16 \| 112.3 \| 158 \| (190, 364) \| 372 \| (390, 558) \| 563 \| DNA 3'-5' helicase \| DNA 3'-5' helicase; uniclust \| UniRef100\_A0A199YTG5 \| 98.6 \| 2.9e-10 \| 5.6e-16 \| 120.0 \| 140 \| (195, 338) \| 372 \| (827, 994) \| 1330 \| DNA 3'-5' helicase \| DNA 3'-5' helicase; uniclust \| UniRef100\_A0A0S7XKG7 \| 98.6 \| 2.8e-10 \| 6.1e-16 \| 126.2 \| 141 \| (198, 344) \| 372 \| (917, 1061) \| 1088 \| DNA 3'-5' helicase (Fragment) \| DNA 3'-5' helicase (Fragment); uniclust \| UniRef100\_UPI001E3DABD6 \| 98.6 \| 3.7e-10 \| 6.8e-16 \| 84.6 \| 86 \| (252, 338) \| 372 \| (7, 94) \| 104 \| PD-(D/E)XK nuclease-like domain-containing protein \| PD-(D/E)XK nuclease-like domain-containing protein; uniclust \| UniRef100\_A0A2E4L226 \| 98.6 \| 3.6e-10 \| 6.8e-16 \| 114.3 \| 121 \| (225, 361) \| 372 \| (654, 791) \| 793 \| DNA 3'-5' helicase \| DNA 3'-5' helicase; uniclust \| UniRef100\_A0A0G1JH46 \| 98.6 \| 3.4e-10 \| 6.9e-16 \| 124.8 \| 104 \| (225, 338) \| 372 \| (1253, 1360) \| 1396 \| DNA 3'-5' helicase \| DNA 3'-5' helicase; uniclust \| UniRef100\_A0A6N8NPZ1 \| 98.6 \| 4.2e-10 \| 7.7e-16 \| 93.6 \| 112 \| (136, 247) \| 372 \| (47, 189) \| 190 \| Exodeoxyribonuclease VIII (Fragment) \| Exodeoxyribonuclease VIII (Fragment); uniclust \| UniRef100\_A0A0B3ADA3 \| 98.6 \| 4.1e-10 \| 8.4e-16 \| 110.0 \| 113 \| (226, 350) \| 372 \| (170, 288) \| 374 \| PD-(D/E)XK endonuclease-like domain-containing protein \| PD-(D/E)XK endonuclease-like domain-containing protein; uniclust \| UniRef100\_UPI000AE98C0E \| 98.6 \| 4.4e-10 \| 8.4e-16 \| 94.6 \| 87 \| (196, 287) \| 372 \| (71, 165) \| 175 \| PD-(D/E)XK nuclease-like domain-containing protein \| PD-(D/E)XK nuclease-like domain-containing protein; pdb70 \| 3H4R\_A \| 99.2 \| 2.6e-15 \| 3.1e-19 \| 131.8 \| 133 \| (225, 364) \| 372 \| (126, 262) \| 265 \| Exodeoxyribonuclease 8 (E.C.3.1.11.-) \| 3H4R\_A Exodeoxyribonuclease 8 (E.C.3.1.11.-) Exonuclease, Recombination, Hydrolase, Nuclease; pdb70 \| 5ZYU\_A \| 97.7 \| 1.5e-08 \| 1.7e-12 \| 89.6 \| 116 \| (224, 348) \| 372 \| (130, 250) \| 254 \| Mitochondrial genome maintenance exonuclease 1/DNA \| 5ZYU\_A Mitochondrial genome maintenance exonuclease 1/DNA huamnMGME1, DNA complex, DNA exonuclease; pdb70 \| 5ZYW\_A \| 97.7 \| 1.5e-08 \| 1.7e-12 \| 89.6 \| 116 \| (224, 348) \| 372 \| (130, 250) \| 254 \| Mitochondrial genome maintenance exonuclease 1 \| 5ZYW\_A Mitochondrial genome maintenance exonuclease 1 huamnMGME1, DNA complex, DNA exonuclease HET: TLA; pdb70 \| 5ZYT\_D \| 97.6 \| 3.2e-08 \| 3.4e-12 \| 92.3 \| 115 \| (224, 347) \| 372 \| (200, 319) \| 324 \| Mitochondrial genome maintenance exonuclease 1/DNA \| 5ZYT\_D Mitochondrial genome maintenance exonuclease 1/DNA human MGME1, DNA complex, DNA; pdb70 \| 3L0A\_A \| 97.4 \| 1.2e-07 \| 1.4e-11 \| 83.5 \| 119 \| (225, 349) \| 372 \| (114, 240) \| 266 \| Putative exonuclease \| 3L0A\_A Putative exonuclease RER070207002219, Putative exonuclease, Structural Genomics HET: PE4; pdb70 \| 6PPJ\_A \| 96.9 \| 1.2e-06 \| 1.5e-10 \| 88.3 \| 103 \| (237, 349) \| 372 \| (911, 1023) \| 1045 \| UvrD/REP helicase (E.C.3.6.4.12), ATP-dependent DNA \| 6PPJ\_A UvrD/REP helicase (E.C.3.6.4.12), ATP-dependent DNA DNA BINDING PROTEIN HET: ANP; pdb70 \| 6PPR\_A \| 96.9 \| 1.2e-06 \| 1.5e-10 \| 88.3 \| 103 \| (237, 349) \| 372 \| (911, 1023) \| 1045 \| UvrD/REP helicase, ATP-dependent DNA helicase \| 6PPR\_A UvrD/REP helicase, ATP-dependent DNA helicase DNA, DNA BINDING PROTEIN, DNA HET: ANP; pdb70 \| 6PPU\_A \| 96.9 \| 1.3e-06 \| 1.6e-10 \| 85.3 \| 103 \| (237, 349) \| 372 \| (590, 702) \| 724 \| UvrD/REP helicase, ATP-dependent DNA helicase \| 6PPU\_A UvrD/REP helicase, ATP-dependent DNA helicase DNA, DNA BINDING PROTEIN, DNA; pdb70 \| 6PPR\_B \| 96.6 \| 4e-06 \| 4.9e-10 \| 85.4 \| 104 \| (223, 341) \| 372 \| (979, 1088) \| 1095 \| UvrD/REP helicase, ATP-dependent DNA helicase \| 6PPR\_B UvrD/REP helicase, ATP-dependent DNA helicase DNA, DNA BINDING PROTEIN, DNA HET: ANP; pdb70 \| 6PPJ\_B \| 96.6 \| 3.9e-06 \| 4.9e-10 \| 85.1 \| 77 \| (224, 307) \| 372 \| (980, 1062) \| 1095 \| UvrD/REP helicase (E.C.3.6.4.12), ATP-dependent DNA \| 6PPJ\_B UvrD/REP helicase (E.C.3.6.4.12), ATP-dependent DNA DNA BINDING PROTEIN HET: ANP; pdb70 \| 3U4Q\_B \| 96.3 \| 1.5e-05 \| 1.9e-09 \| 81.3 \| 50 \| (238, 291) \| 372 \| (936, 992) \| 1166 \| ATP-dependent helicase/nuclease subunit A (E.C.3.1.-.- \| 3U4Q\_B ATP-dependent helicase/nuclease subunit A (E.C.3.1.-.- Helicase, nuclease, Double strand DNA; pdb70 \| 4CEI\_B \| 96.3 \| 1.5e-05 \| 1.9e-09 \| 81.3 \| 50 \| (238, 291) \| 372 \| (936, 992) \| 1166 \| ATP-DEPENDENT HELICASE/NUCLEASE SUBUNIT A (E.C.3.1.-.- \| 4CEI\_B ATP-DEPENDENT HELICASE/NUCLEASE SUBUNIT A (E.C.3.1.-.- HYDROLASE-DNA COMPLEX, HELICASE-NUCLEASE, BACTERIAL PROTEINS HET: ANP; pdb70 \| 4CEJ\_B \| 96.3 \| 1.5e-05 \| 1.9e-09 \| 81.3 \| 50 \| (238, 291) \| 372 \| (936, 992) \| 1166 \| ATP-DEPENDENT HELICASE/NUCLEASE SUBUNIT A (E.C.3.1.-.- \| 4CEJ\_B ATP-DEPENDENT HELICASE/NUCLEASE SUBUNIT A (E.C.3.1.-.- HYDROLASE-DNA COMPLEX, HELICASE-NUCLEASE, DNA BREAKS HET: ANP; pdb70 \| 4IC1\_D \| 95.9 \| 4.3e-05 \| 5.2e-09 \| 62.7 \| 58 \| (238, 307) \| 372 \| (95, 156) \| 206 \| Uncharacterized protein \| 4IC1\_D Uncharacterized protein Cas4, CRISPR, MCSG, Exonuclease, PSI-Biology HET: MN, SF4; pdb70 \| 4IC1\_I \| 95.9 \| 4.3e-05 \| 5.2e-09 \| 62.7 \| 58 \| (238, 307) \| 372 \| (95, 156) \| 206 \| Uncharacterized protein \| 4IC1\_I Uncharacterized protein Cas4, CRISPR, MCSG, Exonuclease, PSI-Biology HET: SF4, MN, MSE; pdb70 \| 3U4Q\_A \| 95.8 \| 6.8e-05 \| 8.1e-09 \| 77.9 \| 83 \| (222, 306) \| 372 \| (1124, 1226) \| 1232 \| ATP-dependent helicase/nuclease subunit A (E.C.3.1.-.- \| 3U4Q\_A ATP-dependent helicase/nuclease subunit A (E.C.3.1.-.- Helicase, nuclease, Double strand DNA; pdb70 \| 4CEJ\_A \| 95.8 \| 6.8e-05 \| 8.1e-09 \| 77.9 \| 83 \| (222, 306) \| 372 \| (1124, 1226) \| 1232 \| ATP-DEPENDENT HELICASE/NUCLEASE SUBUNIT A (E.C.3.1.-.- \| 4CEJ\_A ATP-DEPENDENT HELICASE/NUCLEASE SUBUNIT A (E.C.3.1.-.- HYDROLASE-DNA COMPLEX, HELICASE-NUCLEASE, DNA BREAKS HET: ANP; pdb70 \| 4CEI\_A \| 95.8 \| 7e-05 \| 8.3e-09 \| 77.8 \| 83 \| (222, 306) \| 372 \| (1124, 1226) \| 1232 \| ATP-DEPENDENT HELICASE/NUCLEASE SUBUNIT A (E.C.3.1.-.- \| 4CEI\_A ATP-DEPENDENT HELICASE/NUCLEASE SUBUNIT A (E.C.3.1.-.- HYDROLASE-DNA COMPLEX, HELICASE-NUCLEASE, BACTERIAL PROTEINS HET: ANP; pdb70 \| 5EAN\_A \| 95.6 \| 0.0001 \| 1.2e-08 \| 76.4 \| 71 \| (223, 307) \| 372 \| (262, 343) \| 1059 \| DNA replication ATP-dependent helicase/nuclease DNA2/DNA \| 5EAN\_A DNA replication ATP-dependent helicase/nuclease DNA2/DNA DNA binding protein, Hydrolase-DNA complex HET: ADP; pdb70 \| 1W36\_B \| 95.2 \| 0.00021 \| 2.5e-08 \| 74.0 \| 54 \| (238, 293) \| 372 \| (1059, 1123) \| 1180 \| EXODEOXYRIBONUCLEASE V BETA CHAIN (E.C.3.1.11.5) \| 1W36\_B EXODEOXYRIBONUCLEASE V BETA CHAIN (E.C.3.1.11.5) RECOMBINATION, HELICASE, NUCLEASE, HYDROLASE, DNA |
| Top keywords  (threshold 1.00e-03 (evalue)) | **domain\_containing, exodeoxyribonuclease, Putative, PDDEXK\_like, E, PD\_, D, XK, nuclease\_like, DNA** |
| Output files | ../../similar\_sequences/41\_FANPEZAQ\_CDS\_0041\_merged.svg ../../similar\_sequences/41\_FANPEZAQ\_CDS\_0041\_pdb70.a3m ../../similar\_sequences/41\_FANPEZAQ\_CDS\_0041\_pdb70.hhr ../../similar\_sequences/41\_FANPEZAQ\_CDS\_0041\_uniclust.a3m ../../similar\_sequences/41\_FANPEZAQ\_CDS\_0041\_uniclust.hhr |

#### Structure prediction (AlphaFold)^2^

|  |  |
| --- | --- |
| Stats | xml version="1.0" encoding="utf-8" standalone="no"?       2024-09-02T21:09:40.359380 image/svg+xml   Matplotlib v3.7.2, https://matplotlib.org/ |
| Predicted structure | **NGL Viewer Controls:**  - Center: *Left-Click* - Rotate: *Left-Click + Drag* - Translate: *Right-Click + Drag* - Zoom: *Shift + Left-Click + Drag* |
| Output files | ../../predicted\_structures/41\_FANPEZAQ\_CDS\_0041/features.pkl ../../predicted\_structures/41\_FANPEZAQ\_CDS\_0041/ranked\_0.pdb ../../predicted\_structures/41\_FANPEZAQ\_CDS\_0041/ranked\_0\_plots.svg ../../predicted\_structures/41\_FANPEZAQ\_CDS\_0041/result\_model\_1\_ptm\_pred\_0.pkl |

#### Structure similarity search results (Foldseek)^3^

|  |  |
| --- | --- |
| Structure databases searched | Pdb, Afdb-proteome, Afdb-uniprot50 |
| Results, scheme(s)  (Top layers only, threshold 1.00e-02 (evalue)) | xml version="1.0" encoding="utf-8" standalone="no"?       2024-09-02T21:11:17.549784 image/svg+xml   Matplotlib v3.7.2, https://matplotlib.org/ |
| Results, table  (threshold 1.00e-02 (evalue)) | db \| id \| prob \| evalue \| bits \| fident \| alnlen \| mismatch \| gapopen \| qstart \| qend \| tstart \| tend \| name \| description; pdb \| 3H4R\_A \| 1.0 \| 9.159e-15 \| 534 \| 0.236 \| 347 \| 128 \| 8 \| 18 \| 360 \| 1 \| 214 \| Exodeoxyribonuclease 8 \| Exodeoxyribonuclease 8; pdb \| 3L0A\_A \| 1.0 \| 1.235e-07 \| 191 \| 0.161 \| 241 \| 154 \| 13 \| 142 \| 356 \| 40 \| 258 \| Putative exonuclease \| Putative exonuclease; pdb \| 5ZYU\_B \| 0.998 \| 0.001179 \| 93 \| 0.14 \| 264 \| 163 \| 17 \| 98 \| 344 \| 7 \| 223 \| Mitochondrial genome maintenance exonuclease 1 \| Mitochondrial genome maintenance exonuclease 1; pdb \| 5ZYT\_A \| 0.997 \| 0.001054 \| 92 \| 0.123 \| 252 \| 164 \| 16 \| 97 \| 342 \| 19 \| 219 \| Mitochondrial genome maintenance exonuclease 1 \| Mitochondrial genome maintenance exonuclease 1; pdb \| 4IC1\_C \| 0.991 \| 0.008437 \| 84 \| 0.133 \| 142 \| 99 \| 8 \| 199 \| 331 \| 51 \| 177 \| Uncharacterized protein \| Uncharacterized protein; pdb \| 3SZ5\_A \| 0.933 \| 0.004809 \| 70 \| 0.125 \| 200 \| 132 \| 12 \| 155 \| 343 \| 22 \| 189 \| Exonuclease \| Exonuclease; pdb \| 4CEH\_A \| 0.837 \| 0.004809 \| 63 \| 0.123 \| 323 \| 149 \| 19 \| 6 \| 315 \| 953 \| 1154 \| ATP-DEPENDENT HELICASE/NUCLEASE SUBUNIT A \| ATP-DEPENDENT HELICASE/NUCLEASE SUBUNIT A; afdb-proteome \| AF-Q8XF26-F1-MODEL\_V4 \| 1.0 \| 1.427e-24 \| 759 \| 0.222 \| 436 \| 265 \| 12 \| 1 \| 372 \| 536 \| 961 \| Gifsy-1 prophage protein \| Gifsy-1 prophage protein; afdb-proteome \| AF-P15032-F1-MODEL\_V4 \| 1.0 \| 5.85e-22 \| 668 \| 0.274 \| 368 \| 164 \| 9 \| 1 \| 364 \| 595 \| 863 \| Exodeoxyribonuclease 8 \| Exodeoxyribonuclease 8; afdb-proteome \| AF-Q7AQ72-F1-MODEL\_V4 \| 1.0 \| 7.175e-08 \| 194 \| 0.119 \| 351 \| 202 \| 15 \| 7 \| 351 \| 4 \| 253 \| PDDEXK\_1 domain-containing protein \| PDDEXK\_1 domain-containing protein; afdb-proteome \| AF-O33254-F1-MODEL\_V4 \| 1.0 \| 1.764e-07 \| 188 \| 0.141 \| 353 \| 180 \| 17 \| 9 \| 351 \| 9 \| 248 \| PDDEXK\_1 domain-containing protein \| PDDEXK\_1 domain-containing protein; afdb-proteome \| AF-A0A077ZMK2-F1-MODEL\_V4 \| 1.0 \| 3.29e-05 \| 127 \| 0.181 \| 248 \| 164 \| 13 \| 101 \| 322 \| 878 \| 1112 \| DUF4494 and SNF2 N domain containing protein \| DUF4494 and SNF2 N domain containing protein; afdb-proteome \| AF-K0F1Q7-F1-MODEL\_V4 \| 1.0 \| 2.484e-05 \| 114 \| 0.129 \| 286 \| 185 \| 14 \| 86 \| 364 \| 2 \| 230 \| Putative RecB family exonuclease \| Putative RecB family exonuclease; afdb-proteome \| AF-A0A3Q0KJI8-F1-MODEL\_V4 \| 0.999 \| 0.000437 \| 99 \| 0.154 \| 253 \| 159 \| 20 \| 108 \| 342 \| 41 \| 256 \| Mitochondrial genome maintenance exonuclease 1 \| Mitochondrial genome maintenance exonuclease 1; afdb-proteome \| AF-Q55EN1-F1-MODEL\_V4 \| 0.999 \| 6.834e-05 \| 98 \| 0.127 \| 360 \| 228 \| 20 \| 11 \| 360 \| 72 \| 355 \| PDDEXK\_1 domain-containing protein \| PDDEXK\_1 domain-containing protein; afdb-proteome \| AF-A9QY30-F1-MODEL\_V4 \| 0.998 \| 0.0005472 \| 95 \| 0.136 \| 292 \| 168 \| 19 \| 99 \| 352 \| 39 \| 284 \| Mitochondrial genome maintenance exonuclease 1 \| Mitochondrial genome maintenance exonuclease 1; afdb-proteome \| AF-Q5PPI6-F1-MODEL\_V4 \| 0.975 \| 0.002955 \| 77 \| 0.135 \| 281 \| 154 \| 18 \| 102 \| 353 \| 118 \| 338 \| Mitochondrial genome maintenance exonuclease 1 \| Mitochondrial genome maintenance exonuclease 1; afdb-uniprot50 \| AF-A0A158E8Q1-F1-MODEL\_V4 \| 1.0 \| 8.188e-48 \| 1795 \| 0.577 \| 372 \| 149 \| 5 \| 1 \| 369 \| 1 \| 367 \| Exonuclease VIII \| Exonuclease VIII; afdb-uniprot50 \| AF-A0A8A6KNI9-F1-MODEL\_V4 \| 1.0 \| 2.488e-30 \| 1192 \| 0.36 \| 374 \| 136 \| 6 \| 1 \| 366 \| 4 \| 282 \| PD-(D/E)XK nuclease-like domain-containing protein \| PD-(D/E)XK nuclease-like domain-containing protein; afdb-uniprot50 \| AF-A0A1V1UMV8-F1-MODEL\_V4 \| 1.0 \| 1.496e-31 \| 1136 \| 0.355 \| 385 \| 179 \| 7 \| 1 \| 372 \| 7 \| 335 \| Exodeoxyribonuclease 8 \| Exodeoxyribonuclease 8; afdb-uniprot50 \| AF-A0A4R2KU57-F1-MODEL\_V4 \| 1.0 \| 5.141e-32 \| 1115 \| 0.388 \| 381 \| 165 \| 7 \| 1 \| 371 \| 7 \| 329 \| PDDEXK-like uncharacterized protein DUF3799 \| PDDEXK-like uncharacterized protein DUF3799; afdb-uniprot50 \| AF-A0A2M7C2P3-F1-MODEL\_V4 \| 1.0 \| 8.184e-26 \| 994 \| 0.313 \| 361 \| 147 \| 7 \| 5 \| 363 \| 4 \| 265 \| Exodeoxyribonuclease VIII \| Exodeoxyribonuclease VIII; afdb-uniprot50 \| AF-R1H4T8-F1-MODEL\_V4 \| 1.0 \| 2.112e-28 \| 993 \| 0.291 \| 436 \| 229 \| 11 \| 2 \| 369 \| 100 \| 523 \| DUF3799 domain-containing protein \| DUF3799 domain-containing protein; afdb-uniprot50 \| AF-A0A7D5NWS8-F1-MODEL\_V4 \| 1.0 \| 3.131e-28 \| 992 \| 0.264 \| 435 \| 244 \| 10 \| 1 \| 369 \| 21 \| 445 \| PD-(D/E)XK nuclease-like domain-containing protein \| PD-(D/E)XK nuclease-like domain-containing protein; afdb-uniprot50 \| AF-A0A1D2X562-F1-MODEL\_V4 \| 1.0 \| 1.691e-27 \| 973 \| 0.297 \| 363 \| 193 \| 10 \| 1 \| 360 \| 1 \| 304 \| DUF3799 domain-containing protein \| DUF3799 domain-containing protein; afdb-uniprot50 \| AF-A0A349EF37-F1-MODEL\_V4 \| 1.0 \| 2.252e-25 \| 954 \| 0.327 \| 363 \| 142 \| 6 \| 4 \| 364 \| 7 \| 269 \| Exodeoxyribonuclease VIII \| Exodeoxyribonuclease VIII; afdb-uniprot50 \| AF-D0XHM1-F1-MODEL\_V4 \| 1.0 \| 2.826e-24 \| 946 \| 0.308 \| 357 \| 144 \| 7 \| 8 \| 363 \| 2 \| 256 \| DUF3799 domain-containing protein \| DUF3799 domain-containing protein; afdb-uniprot50 \| AF-A0A1A7Q996-F1-MODEL\_V4 \| 1.0 \| 7.332e-25 \| 945 \| 0.355 \| 352 \| 125 \| 6 \| 9 \| 360 \| 1 \| 250 \| Exodeoxyribonuclease VIII \| Exodeoxyribonuclease VIII; afdb-uniprot50 \| AF-A0A1H0UVZ5-F1-MODEL\_V4 \| 1.0 \| 5.233e-25 \| 920 \| 0.327 \| 372 \| 148 \| 10 \| 1 \| 366 \| 4 \| 279 \| Exodeoxyribonuclease VIII \| Exodeoxyribonuclease VIII; afdb-uniprot50 \| AF-A0A6J4XAC9-F1-MODEL\_V4 \| 1.0 \| 1.611e-24 \| 917 \| 0.297 \| 360 \| 156 \| 6 \| 1 \| 360 \| 5 \| 267 \| DUF3799 domain-containing protein \| DUF3799 domain-containing protein; afdb-uniprot50 \| AF-A0A4Q6DHW1-F1-MODEL\_V4 \| 1.0 \| 6.569e-24 \| 914 \| 0.234 \| 371 \| 184 \| 6 \| 1 \| 370 \| 1 \| 272 \| DUF3799 domain-containing protein \| DUF3799 domain-containing protein; afdb-uniprot50 \| AF-A0A227JR50-F1-MODEL\_V4 \| 1.0 \| 4.653e-27 \| 906 \| 0.341 \| 372 \| 158 \| 12 \| 5 \| 370 \| 46 \| 336 \| DUF3799 domain-containing protein \| DUF3799 domain-containing protein; afdb-uniprot50 \| AF-A0A4D8QPW1-F1-MODEL\_V4 \| 1.0 \| 3.163e-24 \| 903 \| 0.287 \| 362 \| 154 \| 9 \| 1 \| 361 \| 9 \| 267 \| DUF3799 domain-containing protein \| DUF3799 domain-containing protein; afdb-uniprot50 \| AF-A0A0K1QBP3-F1-MODEL\_V4 \| 1.0 \| 1.44e-24 \| 903 \| 0.308 \| 363 \| 152 \| 10 \| 1 \| 360 \| 19 \| 285 \| Exodeoxyribonuclease VIII \| Exodeoxyribonuclease VIII; afdb-uniprot50 \| AF-A0A800EYB9-F1-MODEL\_V4 \| 1.0 \| 3.971e-23 \| 897 \| 0.252 \| 372 \| 178 \| 7 \| 1 \| 369 \| 3 \| 277 \| DUF3799 domain-containing protein \| DUF3799 domain-containing protein; afdb-uniprot50 \| AF-A0A6M3KVF3-F1-MODEL\_V4 \| 1.0 \| 2.022e-23 \| 896 \| 0.281 \| 373 \| 167 \| 10 \| 1 \| 371 \| 6 \| 279 \| DUF3799 domain-containing protein \| DUF3799 domain-containing protein; afdb-uniprot50 \| AF-A0A158DVS7-F1-MODEL\_V4 \| 1.0 \| 1.523e-24 \| 891 \| 0.267 \| 366 \| 164 \| 10 \| 2 \| 361 \| 8 \| 275 \| Exonuclease VIII \| Exonuclease VIII; afdb-uniprot50 \| AF-A0A2M7J5F5-F1-MODEL\_V4 \| 1.0 \| 1.607e-25 \| 889 \| 0.313 \| 361 \| 147 \| 7 \| 5 \| 363 \| 65 \| 326 \| Exodeoxyribonuclease VIII \| Exodeoxyribonuclease VIII; afdb-uniprot50 \| AF-A0A3B7N4J6-F1-MODEL\_V4 \| 1.0 \| 6.194e-25 \| 887 \| 0.301 \| 371 \| 156 \| 11 \| 1 \| 367 \| 1 \| 272 \| Exodeoxyribonuclease VIII \| Exodeoxyribonuclease VIII; afdb-uniprot50 \| AF-A0A7W8M817-F1-MODEL\_V4 \| 1.0 \| 4.959e-24 \| 887 \| 0.276 \| 373 \| 165 \| 9 \| 1 \| 367 \| 9 \| 282 \| Exodeoxyribonuclease VIII \| Exodeoxyribonuclease VIII; afdb-uniprot50 \| AF-A0A7C5FTZ9-F1-MODEL\_V4 \| 1.0 \| 2.394e-23 \| 883 \| 0.298 \| 362 \| 155 \| 8 \| 5 \| 366 \| 2 \| 264 \| Exodeoxyribonuclease VIII \| Exodeoxyribonuclease VIII; afdb-uniprot50 \| AF-C5A7V4-F1-MODEL\_V4 \| 1.0 \| 4.2e-23 \| 883 \| 0.282 \| 364 \| 152 \| 7 \| 5 \| 360 \| 2 \| 264 \| Exonuclease VIII, 5'-3' specific dsDNA exonuclease \| Exonuclease VIII, 5'-3' specific dsDNA exonuclease; afdb-uniprot50 \| AF-A0A5C5DDI0-F1-MODEL\_V4 \| 1.0 \| 8.679e-25 \| 874 \| 0.239 \| 389 \| 265 \| 10 \| 1 \| 372 \| 118 \| 492 \| DUF3799 domain-containing protein \| DUF3799 domain-containing protein; afdb-uniprot50 \| AF-A0A838BVQ7-F1-MODEL\_V4 \| 1.0 \| 4.179e-25 \| 872 \| 0.328 \| 362 \| 138 \| 8 \| 1 \| 360 \| 1 \| 259 \| PD-(D/E)XK nuclease-like domain-containing protein \| PD-(D/E)XK nuclease-like domain-containing protein; afdb-uniprot50 \| AF-A0A2G2QFT7-F1-MODEL\_V4 \| 1.0 \| 5.246e-24 \| 871 \| 0.291 \| 371 \| 161 \| 6 \| 1 \| 371 \| 14 \| 282 \| DUF3799 domain-containing protein \| DUF3799 domain-containing protein; afdb-uniprot50 \| AF-A0A838LRU1-F1-MODEL\_V4 \| 1.0 \| 7.737e-26 \| 869 \| 0.317 \| 369 \| 183 \| 15 \| 2 \| 363 \| 5 \| 311 \| Uncharacterized protein \| Uncharacterized protein; afdb-uniprot50 \| AF-D0Z7N0-F1-MODEL\_V4 \| 1.0 \| 4.168e-26 \| 869 \| 0.224 \| 436 \| 260 \| 11 \| 1 \| 370 \| 401 \| 824 \| Gifsy-1 prophage protein \| Gifsy-1 prophage protein; afdb-uniprot50 \| AF-R7GUS0-F1-MODEL\_V4 \| 1.0 \| 2.532e-23 \| 867 \| 0.317 \| 356 \| 146 \| 10 \| 9 \| 364 \| 1 \| 259 \| DUF3799 domain-containing protein \| DUF3799 domain-containing protein; afdb-uniprot50 \| AF-A0A2D7CSN2-F1-MODEL\_V4 \| 1.0 \| 9.205e-24 \| 865 \| 0.327 \| 373 \| 143 \| 8 \| 4 \| 372 \| 18 \| 286 \| Exodeoxyribonuclease VIII \| Exodeoxyribonuclease VIII; afdb-uniprot50 \| AF-A0A6M3JML4-F1-MODEL\_V4 \| 1.0 \| 1.293e-22 \| 864 \| 0.267 \| 367 \| 162 \| 9 \| 1 \| 364 \| 2 \| 264 \| DUF3799 domain-containing protein \| DUF3799 domain-containing protein; afdb-uniprot50 \| AF-A0A2E1R914-F1-MODEL\_V4 \| 1.0 \| 3.548e-23 \| 864 \| 0.304 \| 364 \| 149 \| 9 \| 2 \| 360 \| 8 \| 272 \| Exodeoxyribonuclease VIII \| Exodeoxyribonuclease VIII; afdb-uniprot50 \| AF-A0A762GQR2-F1-MODEL\_V4 \| 1.0 \| 7.314e-26 \| 862 \| 0.249 \| 433 \| 250 \| 15 \| 1 \| 369 \| 687 \| 1108 \| Exodeoxyribonuclease \| Exodeoxyribonuclease; afdb-uniprot50 \| AF-A0A149U469-F1-MODEL\_V4 \| 1.0 \| 2.819e-25 \| 857 \| 0.316 \| 373 \| 177 \| 16 \| 1 \| 364 \| 10 \| 313 \| DUF3799 domain-containing protein \| DUF3799 domain-containing protein; afdb-uniprot50 \| AF-A0A612C5H5-F1-MODEL\_V4 \| 1.0 \| 1.519e-25 \| 853 \| 0.236 \| 439 \| 254 \| 13 \| 1 \| 372 \| 546 \| 970 \| Exodeoxyribonuclease \| Exodeoxyribonuclease; afdb-uniprot50 \| AF-A0A5J0RYS5-F1-MODEL\_V4 \| 1.0 \| 2.382e-25 \| 848 \| 0.23 \| 439 \| 257 \| 13 \| 1 \| 372 \| 260 \| 684 \| Exodeoxyribonuclease \| Exodeoxyribonuclease; afdb-uniprot50 \| AF-A0A0F9KCK9-F1-MODEL\_V4 \| 1.0 \| 1.527e-23 \| 847 \| 0.254 \| 366 \| 171 \| 10 \| 1 \| 361 \| 1 \| 269 \| DUF3799 domain-containing protein \| DUF3799 domain-containing protein; afdb-uniprot50 \| AF-A0A0F9QDF5-F1-MODEL\_V4 \| 1.0 \| 2.841e-22 \| 845 \| 0.279 \| 354 \| 150 \| 10 \| 9 \| 361 \| 1 \| 250 \| DUF3799 domain-containing protein \| DUF3799 domain-containing protein; afdb-uniprot50 \| AF-A0A1F9B5U7-F1-MODEL\_V4 \| 1.0 \| 1.092e-22 \| 845 \| 0.268 \| 365 \| 162 \| 8 \| 1 \| 361 \| 1 \| 264 \| DUF3799 domain-containing protein \| DUF3799 domain-containing protein; afdb-uniprot50 \| AF-B7UGT8-F1-MODEL\_V4 \| 1.0 \| 1.615e-23 \| 841 \| 0.285 \| 371 \| 162 \| 10 \| 1 \| 367 \| 1 \| 272 \| Predicted exonuclease VIII/RecE-like protein \| Predicted exonuclease VIII/RecE-like protein; afdb-uniprot50 \| AF-A0A2G2PPI1-F1-MODEL\_V4 \| 1.0 \| 1.09e-23 \| 841 \| 0.273 \| 373 \| 188 \| 9 \| 1 \| 370 \| 1 \| 293 \| DUF3799 domain-containing protein \| DUF3799 domain-containing protein; afdb-uniprot50 \| AF-A0A2G2KSA4-F1-MODEL\_V4 \| 1.0 \| 3.548e-23 \| 834 \| 0.288 \| 367 \| 161 \| 7 \| 5 \| 371 \| 20 \| 286 \| DUF3799 domain-containing protein \| DUF3799 domain-containing protein; afdb-uniprot50 \| AF-A0A5J2KB08-F1-MODEL\_V4 \| 1.0 \| 2.834e-23 \| 831 \| 0.285 \| 371 \| 162 \| 9 \| 1 \| 367 \| 1 \| 272 \| Exodeoxyribonuclease VIII \| Exodeoxyribonuclease VIII; afdb-uniprot50 \| AF-A0A1Y6CQ16-F1-MODEL\_V4 \| 1.0 \| 6.967e-23 \| 831 \| 0.231 \| 372 \| 183 \| 7 \| 1 \| 371 \| 22 \| 291 \| Exodeoxyribonuclease VIII \| Exodeoxyribonuclease VIII; afdb-uniprot50 \| AF-A0A6H0ITG2-F1-MODEL\_V4 \| 1.0 \| 4.7e-23 \| 830 \| 0.302 \| 360 \| 150 \| 8 \| 4 \| 361 \| 14 \| 274 \| DUF3799 domain-containing protein \| DUF3799 domain-containing protein; afdb-uniprot50 \| AF-A0A1W9QRF2-F1-MODEL\_V4 \| 1.0 \| 1.225e-21 \| 829 \| 0.248 \| 366 \| 171 \| 8 \| 5 \| 367 \| 2 \| 266 \| DUF3799 domain-containing protein \| DUF3799 domain-containing protein; afdb-uniprot50 \| AF-A0A4P5VJR3-F1-MODEL\_V4 \| 1.0 \| 9.228e-23 \| 827 \| 0.299 \| 364 \| 150 \| 10 \| 3 \| 360 \| 2 \| 266 \| DUF3799 domain-containing protein \| DUF3799 domain-containing protein; afdb-uniprot50 \| AF-A0A1G2ZKS6-F1-MODEL\_V4 \| 1.0 \| 2.4e-22 \| 827 \| 0.255 \| 364 \| 166 \| 10 \| 1 \| 360 \| 4 \| 266 \| DUF3799 domain-containing protein \| DUF3799 domain-containing protein; afdb-uniprot50 \| AF-A0A348HFN7-F1-MODEL\_V4 \| 1.0 \| 6.586e-23 \| 827 \| 0.283 \| 370 \| 153 \| 10 \| 1 \| 362 \| 6 \| 271 \| Dehydrogenases \| Dehydrogenases; afdb-uniprot50 \| AF-A0A7U1K146-F1-MODEL\_V4 \| 1.0 \| 1.364e-23 \| 825 \| 0.281 \| 376 \| 157 \| 14 \| 1 \| 367 \| 1 \| 272 \| PD-(D/E)XK nuclease-like domain-containing protein \| PD-(D/E)XK nuclease-like domain-containing protein; afdb-uniprot50 \| AF-A0A1J8PMA9-F1-MODEL\_V4 \| 1.0 \| 9.762e-23 \| 825 \| 0.28 \| 363 \| 160 \| 12 \| 1 \| 361 \| 11 \| 274 \| DUF3799 domain-containing protein \| DUF3799 domain-containing protein; afdb-uniprot50 \| AF-A0A6G6JMC2-F1-MODEL\_V4 \| 1.0 \| 2.519e-25 \| 825 \| 0.238 \| 436 \| 250 \| 13 \| 1 \| 368 \| 480 \| 901 \| Exodeoxyribonuclease \| Exodeoxyribonuclease; afdb-uniprot50 \| AF-E6QW59-F1-MODEL\_V4 \| 1.0 \| 3.512e-27 \| 824 \| 0.293 \| 447 \| 204 \| 16 \| 3 \| 363 \| 2 \| 422 \| Uncharacterized protein \| Uncharacterized protein; afdb-uniprot50 \| AF-A0A4V2X429-F1-MODEL\_V4 \| 1.0 \| 6.586e-23 \| 823 \| 0.264 \| 374 \| 163 \| 12 \| 1 \| 365 \| 1 \| 271 \| Exodeoxyribonuclease VIII \| Exodeoxyribonuclease VIII; afdb-uniprot50 \| AF-A0A1H3GZQ2-F1-MODEL\_V4 \| 1.0 \| 5.885e-23 \| 823 \| 0.267 \| 367 \| 181 \| 10 \| 1 \| 361 \| 3 \| 287 \| DUF3799 domain-containing protein \| DUF3799 domain-containing protein; afdb-uniprot50 \| AF-A0A6M3LLV5-F1-MODEL\_V4 \| 1.0 \| 4.984e-22 \| 820 \| 0.27 \| 377 \| 162 \| 10 \| 1 \| 369 \| 1 \| 272 \| DUF3799 domain-containing protein \| DUF3799 domain-containing protein; afdb-uniprot50 \| AF-A0A7T8DU22-F1-MODEL\_V4 \| 1.0 \| 1.027e-24 \| 820 \| 0.236 \| 435 \| 253 \| 13 \| 1 \| 369 \| 139 \| 560 \| PD-(D/E)XK nuclease-like domain-containing protein \| PD-(D/E)XK nuclease-like domain-containing protein; afdb-uniprot50 \| AF-A0A7G8SH51-F1-MODEL\_V4 \| 1.0 \| 1.907e-24 \| 818 \| 0.216 \| 438 \| 259 \| 14 \| 1 \| 369 \| 498 \| 920 \| PD-(D/E)XK nuclease-like domain-containing protein \| PD-(D/E)XK nuclease-like domain-containing protein; afdb-uniprot50 \| AF-A0A5E4PG85-F1-MODEL\_V4 \| 1.0 \| 3.354e-23 \| 817 \| 0.287 \| 362 \| 165 \| 9 \| 1 \| 360 \| 3 \| 273 \| Exodeoxyribonuclease 8 \| Exodeoxyribonuclease 8; afdb-uniprot50 \| AF-A0A4P7C0W9-F1-MODEL\_V4 \| 1.0 \| 1.158e-21 \| 813 \| 0.297 \| 343 \| 139 \| 8 \| 1 \| 342 \| 30 \| 271 \| Exodeoxyribonuclease VIII \| Exodeoxyribonuclease VIII; afdb-uniprot50 \| AF-A0A2D7YR06-F1-MODEL\_V4 \| 1.0 \| 1.451e-21 \| 809 \| 0.298 \| 355 \| 137 \| 8 \| 9 \| 361 \| 1 \| 245 \| DUF3799 domain-containing protein \| DUF3799 domain-containing protein; afdb-uniprot50 \| AF-A0A6M3IX22-F1-MODEL\_V4 \| 1.0 \| 1.156e-22 \| 808 \| 0.239 \| 372 \| 184 \| 10 \| 1 \| 370 \| 4 \| 278 \| DUF3799 domain-containing protein \| DUF3799 domain-containing protein; afdb-uniprot50 \| AF-A0A663BKX6-F1-MODEL\_V4 \| 1.0 \| 7.332e-25 \| 807 \| 0.225 \| 443 \| 255 \| 14 \| 1 \| 372 \| 493 \| 918 \| Exodeoxyribonuclease 8 \| Exodeoxyribonuclease 8; afdb-uniprot50 \| AF-A0A4Q6DHU3-F1-MODEL\_V4 \| 1.0 \| 3.179e-22 \| 806 \| 0.206 \| 377 \| 194 \| 9 \| 1 \| 371 \| 5 \| 282 \| DUF3799 domain-containing protein \| DUF3799 domain-containing protein; afdb-uniprot50 \| AF-A0A149SNY1-F1-MODEL\_V4 \| 1.0 \| 8.746e-22 \| 805 \| 0.254 \| 369 \| 161 \| 11 \| 9 \| 368 \| 1 \| 264 \| DUF3799 domain-containing protein \| DUF3799 domain-containing protein; afdb-uniprot50 \| AF-A0A1C6FHM2-F1-MODEL\_V4 \| 1.0 \| 4.984e-22 \| 804 \| 0.28 \| 364 \| 156 \| 8 \| 7 \| 364 \| 3 \| 266 \| Exodeoxyribonuclease 8 \| Exodeoxyribonuclease 8; afdb-uniprot50 \| AF-A0A656QBR2-F1-MODEL\_V4 \| 1.0 \| 1.035e-21 \| 804 \| 0.248 \| 358 \| 171 \| 7 \| 3 \| 360 \| 10 \| 269 \| DUF3799 domain-containing protein \| DUF3799 domain-containing protein; afdb-uniprot50 \| AF-A0A5C9BVQ4-F1-MODEL\_V4 \| 1.0 \| 6.241e-22 \| 803 \| 0.279 \| 365 \| 156 \| 9 \| 4 \| 362 \| 31 \| 294 \| DUF3799 domain-containing protein \| DUF3799 domain-containing protein; afdb-uniprot50 \| AF-A0A2H9TC24-F1-MODEL\_V4 \| 1.0 \| 3.548e-23 \| 803 \| 0.252 \| 372 \| 213 \| 12 \| 1 \| 368 \| 15 \| 325 \| Exodeoxyribonuclease 8 \| Exodeoxyribonuclease 8; afdb-uniprot50 \| AF-A0A5M7LBW2-F1-MODEL\_V4 \| 1.0 \| 2.257e-24 \| 802 \| 0.231 \| 436 \| 253 \| 13 \| 1 \| 368 \| 529 \| 950 \| Exodeoxyribonuclease \| Exodeoxyribonuclease; afdb-uniprot50 \| AF-A0A7S9S4Y9-F1-MODEL\_V4 \| 1.0 \| 1.451e-21 \| 801 \| 0.28 \| 364 \| 154 \| 10 \| 8 \| 363 \| 1 \| 264 \| Exodeoxyribonuclease VIII \| Exodeoxyribonuclease VIII; afdb-uniprot50 \| AF-A0A1H2DRY6-F1-MODEL\_V4 \| 1.0 \| 1.033e-22 \| 801 \| 0.274 \| 361 \| 178 \| 9 \| 1 \| 361 \| 1 \| 277 \| Exodeoxyribonuclease VIII \| Exodeoxyribonuclease VIII; afdb-uniprot50 \| AF-A0A2N3KY65-F1-MODEL\_V4 \| 1.0 \| 9.228e-23 \| 801 \| 0.228 \| 380 \| 191 \| 10 \| 1 \| 370 \| 3 \| 290 \| DUF3799 domain-containing protein \| DUF3799 domain-containing protein; afdb-uniprot50 \| AF-A0A767X0T4-F1-MODEL\_V4 \| 1.0 \| 6.931e-25 \| 801 \| 0.227 \| 440 \| 258 \| 12 \| 1 \| 372 \| 1 \| 426 \| DNA breaking-rejoining protein \| DNA breaking-rejoining protein; afdb-uniprot50 \| AF-A0A2Z3HXT5-F1-MODEL\_V4 \| 1.0 \| 4.984e-22 \| 800 \| 0.273 \| 365 \| 162 \| 10 \| 1 \| 361 \| 1 \| 266 \| Exodeoxyribonuclease VIII \| Exodeoxyribonuclease VIII; afdb-uniprot50 \| AF-A0A5M6ITC2-F1-MODEL\_V4 \| 1.0 \| 5.9e-22 \| 800 \| 0.286 \| 360 \| 154 \| 7 \| 1 \| 358 \| 3 \| 261 \| DUF3799 domain-containing protein \| DUF3799 domain-containing protein; afdb-uniprot50 \| AF-A0A742LIF7-F1-MODEL\_V4 \| 1.0 \| 8.679e-25 \| 798 \| 0.207 \| 438 \| 270 \| 11 \| 1 \| 372 \| 462 \| 888 \| DNA breaking-rejoining protein \| DNA breaking-rejoining protein; afdb-uniprot50 \| AF-A0A2D6JHQ6-F1-MODEL\_V4 \| 1.0 \| 2.545e-21 \| 797 \| 0.243 \| 361 \| 165 \| 9 \| 9 \| 363 \| 5 \| 263 \| Exodeoxyribonuclease VIII \| Exodeoxyribonuclease VIII; afdb-uniprot50 \| AF-A0A7Z1AHF7-F1-MODEL\_V4 \| 1.0 \| 2.145e-22 \| 796 \| 0.278 \| 373 \| 165 \| 11 \| 1 \| 370 \| 11 \| 282 \| DUF3799 domain-containing protein \| DUF3799 domain-containing protein; afdb-uniprot50 \| AF-A0A741YHL3-F1-MODEL\_V4 \| 1.0 \| 6.552e-25 \| 796 \| 0.226 \| 438 \| 261 \| 12 \| 1 \| 372 \| 358 \| 783 \| Exodeoxyribonuclease \| Exodeoxyribonuclease; afdb-uniprot50 \| AF-Q9AKZ1-F1-MODEL\_V4 \| 1.0 \| 5.578e-22 \| 795 \| 0.226 \| 371 \| 185 \| 10 \| 1 \| 369 \| 6 \| 276 \| DUF3799 domain-containing protein \| DUF3799 domain-containing protein; afdb-uniprot50 \| AF-A0A212KLV9-F1-MODEL\_V4 \| 1.0 \| 3.971e-23 \| 795 \| 0.236 \| 381 \| 184 \| 11 \| 1 \| 371 \| 5 \| 288 \| DUF3799 domain-containing protein \| DUF3799 domain-containing protein; afdb-uniprot50 \| AF-A0A1E4HKX6-F1-MODEL\_V4 \| 1.0 \| 7.815e-22 \| 794 \| 0.28 \| 342 \| 149 \| 9 \| 23 \| 363 \| 13 \| 258 \| DUF3799 domain-containing protein \| DUF3799 domain-containing protein; afdb-uniprot50 \| AF-A0A1V2RAV6-F1-MODEL\_V4 \| 1.0 \| 3.557e-22 \| 793 \| 0.266 \| 379 \| 166 \| 12 \| 1 \| 370 \| 6 \| 281 \| Exodeoxyribonuclease 8 \| Exodeoxyribonuclease 8; afdb-uniprot50 \| AF-A0A2E2DCA2-F1-MODEL\_V4 \| 1.0 \| 1.921e-21 \| 793 \| 0.257 \| 365 \| 170 \| 8 \| 5 \| 369 \| 19 \| 282 \| DUF3799 domain-containing protein \| DUF3799 domain-containing protein; afdb-uniprot50 \| AF-A0A250BLY8-F1-MODEL\_V4 \| 1.0 \| 1.216e-24 \| 793 \| 0.231 \| 437 \| 256 \| 13 \| 1 \| 370 \| 511 \| 934 \| Exodeoxyribonuclease \| Exodeoxyribonuclease; afdb-uniprot50 \| AF-A0A6H0K5E3-F1-MODEL\_V4 \| 1.0 \| 2.012e-25 \| 793 \| 0.26 \| 438 \| 245 \| 14 \| 1 \| 372 \| 570 \| 994 \| Exodeoxyribonuclease \| Exodeoxyribonuclease; afdb-uniprot50 \| AF-A0A3S0A9S4-F1-MODEL\_V4 \| 1.0 \| 1.713e-22 \| 792 \| 0.274 \| 379 \| 160 \| 11 \| 1 \| 366 \| 6 \| 282 \| DUF3799 domain-containing protein \| DUF3799 domain-containing protein; afdb-uniprot50 \| AF-A0A346DXG2-F1-MODEL\_V4 \| 1.0 \| 6.586e-23 \| 792 \| 0.241 \| 385 \| 178 \| 14 \| 1 \| 371 \| 4 \| 288 \| DUF3799 domain-containing protein \| DUF3799 domain-containing protein; afdb-uniprot50 \| AF-A0A5X6IFM5-F1-MODEL\_V4 \| 1.0 \| 2.819e-25 \| 791 \| 0.224 \| 436 \| 264 \| 12 \| 1 \| 372 \| 54 \| 479 \| Exodeoxyribonuclease \| Exodeoxyribonuclease; afdb-uniprot50 \| AF-A0A6P1K602-F1-MODEL\_V4 \| 1.0 \| 7.332e-25 \| 790 \| 0.217 \| 437 \| 260 \| 15 \| 1 \| 369 \| 600 \| 1022 \| Exodeoxyribonuclease VIII \| Exodeoxyribonuclease VIII; afdb-uniprot50 \| AF-A0A3D0NKI8-F1-MODEL\_V4 \| 1.0 \| 1.153e-23 \| 789 \| 0.277 \| 367 \| 192 \| 10 \| 1 \| 359 \| 9 \| 310 \| DUF3799 domain-containing protein \| DUF3799 domain-containing protein; afdb-uniprot50 \| AF-G5S9R5-F1-MODEL\_V4 \| 1.0 \| 3.337e-25 \| 789 \| 0.221 \| 438 \| 263 \| 13 \| 1 \| 372 \| 177 \| 602 \| Exodeoxyribonuclease 8 \| Exodeoxyribonuclease 8; afdb-uniprot50 \| AF-A0A1Y2SD06-F1-MODEL\_V4 \| 1.0 \| 7.388e-22 \| 788 \| 0.266 \| 368 \| 167 \| 9 \| 1 \| 364 \| 1 \| 269 \| Exodeoxyribonuclease VIII \| Exodeoxyribonuclease VIII; afdb-uniprot50 \| AF-A0A5W2M3A0-F1-MODEL\_V4 \| 1.0 \| 4.421e-25 \| 787 \| 0.224 \| 436 \| 264 \| 12 \| 1 \| 372 \| 143 \| 568 \| DNA breaking-rejoining protein \| DNA breaking-rejoining protein; afdb-uniprot50 \| AF-A0A0F9LUR6-F1-MODEL\_V4 \| 1.0 \| 4.2e-23 \| 786 \| 0.271 \| 364 \| 177 \| 9 \| 1 \| 364 \| 25 \| 300 \| DUF3799 domain-containing protein \| DUF3799 domain-containing protein; afdb-uniprot50 \| AF-A0A767X2R0-F1-MODEL\_V4 \| 1.0 \| 5.233e-25 \| 786 \| 0.224 \| 436 \| 264 \| 12 \| 1 \| 372 \| 99 \| 524 \| Exodeoxyribonuclease \| Exodeoxyribonuclease; afdb-uniprot50 \| AF-A0A5V9PFN1-F1-MODEL\_V4 \| 1.0 \| 4.947e-25 \| 785 \| 0.224 \| 436 \| 264 \| 12 \| 1 \| 372 \| 474 \| 899 \| Exodeoxyribonuclease \| Exodeoxyribonuclease; afdb-uniprot50 \| AF-A0A5C6BNI7-F1-MODEL\_V4 \| 1.0 \| 2.139e-23 \| 784 \| 0.254 \| 377 \| 183 \| 11 \| 2 \| 366 \| 20 \| 310 \| Exodeoxyribonuclease 8 \| Exodeoxyribonuclease 8; afdb-uniprot50 \| AF-A0A754MTK6-F1-MODEL\_V4 \| 1.0 \| 3.53e-25 \| 784 \| 0.228 \| 438 \| 260 \| 13 \| 1 \| 372 \| 142 \| 567 \| Exodeoxyribonuclease \| Exodeoxyribonuclease; afdb-uniprot50 \| AF-A0A515CTF2-F1-MODEL\_V4 \| 1.0 \| 9.712e-25 \| 784 \| 0.215 \| 437 \| 261 \| 13 \| 1 \| 369 \| 612 \| 1034 \| DUF3799 domain-containing protein \| DUF3799 domain-containing protein; afdb-uniprot50 \| AF-A0A0T9L9Z5-F1-MODEL\_V4 \| 1.0 \| 9.712e-25 \| 783 \| 0.228 \| 438 \| 254 \| 14 \| 1 \| 369 \| 159 \| 581 \| Exodeoxyribonuclease 8 \| Exodeoxyribonuclease 8; afdb-uniprot50 \| AF-A0A5V3EY84-F1-MODEL\_V4 \| 1.0 \| 3.53e-25 \| 782 \| 0.228 \| 438 \| 260 \| 13 \| 1 \| 372 \| 347 \| 772 \| Exodeoxyribonuclease \| Exodeoxyribonuclease; afdb-uniprot50 \| AF-A0A4P8C6B6-F1-MODEL\_V4 \| 1.0 \| 1.907e-24 \| 782 \| 0.224 \| 437 \| 257 \| 15 \| 1 \| 369 \| 618 \| 1040 \| Exodeoxyribonuclease VIII \| Exodeoxyribonuclease VIII; afdb-uniprot50 \| AF-A0A5Z8QTS3-F1-MODEL\_V4 \| 1.0 \| 6.552e-25 \| 781 \| 0.224 \| 436 \| 264 \| 12 \| 1 \| 372 \| 142 \| 567 \| Exodeoxyribonuclease \| Exodeoxyribonuclease; afdb-uniprot50 \| AF-A0A5V8U3D0-F1-MODEL\_V4 \| 1.0 \| 1.087e-24 \| 781 \| 0.209 \| 439 \| 268 \| 13 \| 1 \| 372 \| 246 \| 672 \| DNA breaking-rejoining protein \| DNA breaking-rejoining protein; afdb-uniprot50 \| AF-A0A2S4QN94-F1-MODEL\_V4 \| 1.0 \| 2.145e-22 \| 781 \| 0.218 \| 385 \| 231 \| 10 \| 1 \| 372 \| 749 \| 1076 \| Exodeoxyribonuclease \| Exodeoxyribonuclease; afdb-uniprot50 \| AF-A0A2E0Z4I9-F1-MODEL\_V4 \| 1.0 \| 5.273e-22 \| 780 \| 0.258 \| 367 \| 173 \| 10 \| 3 \| 364 \| 8 \| 280 \| DUF3799 domain-containing protein \| DUF3799 domain-containing protein; afdb-uniprot50 \| AF-A0A826KR83-F1-MODEL\_V4 \| 1.0 \| 2.99e-24 \| 780 \| 0.226 \| 437 \| 256 \| 15 \| 1 \| 369 \| 107 \| 529 \| Exonuclease VIII \| Exonuclease VIII; afdb-uniprot50 \| AF-A0A726YLN1-F1-MODEL\_V4 \| 1.0 \| 4.421e-25 \| 778 \| 0.225 \| 434 \| 266 \| 14 \| 1 \| 372 \| 546 \| 971 \| Exodeoxyribonuclease \| Exodeoxyribonuclease; afdb-uniprot50 \| AF-W1I0H3-F1-MODEL\_V4 \| 1.0 \| 5.233e-25 \| 777 \| 0.226 \| 438 \| 261 \| 12 \| 1 \| 372 \| 115 \| 540 \| Exodeoxyribonuclease VIII \| Exodeoxyribonuclease VIII; afdb-uniprot50 \| AF-A0A378FZ40-F1-MODEL\_V4 \| 1.0 \| 3.961e-24 \| 777 \| 0.22 \| 440 \| 261 \| 12 \| 1 \| 372 \| 195 \| 620 \| Putative exodeoxyribonuclease VIII \| Putative exodeoxyribonuclease VIII; afdb-uniprot50 \| AF-A0A723W028-F1-MODEL\_V4 \| 1.0 \| 5.535e-25 \| 777 \| 0.223 \| 434 \| 267 \| 14 \| 1 \| 372 \| 550 \| 975 \| Exodeoxyribonuclease \| Exodeoxyribonuclease; afdb-uniprot50 \| AF-A0A839L3J4-F1-MODEL\_V4 \| 1.0 \| 6.586e-23 \| 776 \| 0.272 \| 370 \| 166 \| 9 \| 1 \| 366 \| 36 \| 306 \| PD-(D/E)XK nuclease-like domain-containing protein \| PD-(D/E)XK nuclease-like domain-containing protein; afdb-uniprot50 \| AF-A0A808IEU1-F1-MODEL\_V4 \| 1.0 \| 9.181e-25 \| 775 \| 0.22 \| 439 \| 262 \| 14 \| 1 \| 372 \| 481 \| 906 \| Exodeoxyribonuclease VIII \| Exodeoxyribonuclease VIII; afdb-uniprot50 \| AF-A0A759BTV9-F1-MODEL\_V4 \| 1.0 \| 1.286e-24 \| 774 \| 0.222 \| 436 \| 265 \| 12 \| 1 \| 372 \| 129 \| 554 \| Exodeoxyribonuclease \| Exodeoxyribonuclease; afdb-uniprot50 \| AF-A0A5W6KM85-F1-MODEL\_V4 \| 1.0 \| 5.535e-25 \| 773 \| 0.21 \| 438 \| 269 \| 15 \| 1 \| 372 \| 166 \| 592 \| DNA breaking-rejoining protein \| DNA breaking-rejoining protein; afdb-uniprot50 \| AF-A0A1G4AE34-F1-MODEL\_V4 \| 1.0 \| 9.275e-21 \| 772 \| 0.249 \| 365 \| 167 \| 10 \| 1 \| 361 \| 1 \| 262 \| DUF3799 domain-containing protein \| DUF3799 domain-containing protein; afdb-uniprot50 \| AF-A0A7Z9CR49-F1-MODEL\_V4 \| 1.0 \| 1.027e-24 \| 772 \| 0.221 \| 438 \| 263 \| 12 \| 1 \| 372 \| 100 \| 525 \| Putative exodeoxyribonuclease VIII \| Putative exodeoxyribonuclease VIII; afdb-uniprot50 \| AF-A0A839HGV9-F1-MODEL\_V4 \| 1.0 \| 2.274e-21 \| 771 \| 0.26 \| 357 \| 163 \| 11 \| 5 \| 360 \| 11 \| 267 \| PD-(D/E)XK nuclease-like domain-containing protein \| PD-(D/E)XK nuclease-like domain-containing protein; afdb-uniprot50 \| AF-A0A764YUZ2-F1-MODEL\_V4 \| 1.0 \| 1.704e-24 \| 770 \| 0.222 \| 436 \| 265 \| 12 \| 1 \| 372 \| 271 \| 696 \| Exodeoxyribonuclease \| Exodeoxyribonuclease; afdb-uniprot50 \| AF-A0A7I6ZDW9-F1-MODEL\_V4 \| 1.0 \| 2.841e-22 \| 770 \| 0.23 \| 382 \| 224 \| 10 \| 1 \| 369 \| 506 \| 830 \| Uncharacterized protein \| Uncharacterized protein; afdb-uniprot50 \| AF-A0A0F9TR59-F1-MODEL\_V4 \| 1.0 \| 1.296e-21 \| 769 \| 0.246 \| 369 \| 186 \| 10 \| 1 \| 367 \| 11 \| 289 \| DUF3799 domain-containing protein \| DUF3799 domain-containing protein; afdb-uniprot50 \| AF-A0A609CCL8-F1-MODEL\_V4 \| 1.0 \| 6.931e-25 \| 768 \| 0.212 \| 438 \| 268 \| 15 \| 1 \| 372 \| 487 \| 913 \| Exodeoxyribonuclease \| Exodeoxyribonuclease; afdb-uniprot50 \| AF-A0A430DP05-F1-MODEL\_V4 \| 1.0 \| 4.984e-22 \| 767 \| 0.241 \| 397 \| 182 \| 10 \| 2 \| 371 \| 7 \| 311 \| DUF3799 domain-containing protein \| DUF3799 domain-containing protein; afdb-uniprot50 \| AF-A0A3D0NII1-F1-MODEL\_V4 \| 1.0 \| 3.548e-23 \| 765 \| 0.311 \| 360 \| 145 \| 11 \| 5 \| 360 \| 51 \| 311 \| DUF3799 domain-containing protein \| DUF3799 domain-containing protein; afdb-uniprot50 \| AF-A0A221DNX1-F1-MODEL\_V4 \| 1.0 \| 9.712e-25 \| 765 \| 0.216 \| 435 \| 262 \| 13 \| 1 \| 369 \| 456 \| 877 \| DUF3799 domain-containing protein \| DUF3799 domain-containing protein; afdb-uniprot50 \| AF-A0A5V7BL52-F1-MODEL\_V4 \| 1.0 \| 5.564e-23 \| 764 \| 0.261 \| 375 \| 164 \| 11 \| 1 \| 366 \| 34 \| 304 \| Exonuclease VIII \| Exonuclease VIII; afdb-uniprot50 \| AF-A0A7X4MGN6-F1-MODEL\_V4 \| 1.0 \| 4.19e-24 \| 764 \| 0.218 \| 440 \| 262 \| 14 \| 1 \| 372 \| 166 \| 591 \| DUF3799 domain-containing protein \| DUF3799 domain-containing protein; afdb-uniprot50 \| AF-A0A6Y1QS66-F1-MODEL\_V4 \| 1.0 \| 1.216e-24 \| 764 \| 0.222 \| 440 \| 260 \| 14 \| 1 \| 372 \| 546 \| 971 \| Exodeoxyribonuclease \| Exodeoxyribonuclease; afdb-uniprot50 \| AF-A0A377Q5C7-F1-MODEL\_V4 \| 1.0 \| 1.619e-22 \| 763 \| 0.252 \| 368 \| 203 \| 14 \| 1 \| 363 \| 9 \| 309 \| Exodeoxyribonuclease 8 \| Exodeoxyribonuclease 8; afdb-uniprot50 \| AF-A0A0P8LGK8-F1-MODEL\_V4 \| 1.0 \| 3.539e-24 \| 762 \| 0.217 \| 436 \| 267 \| 12 \| 1 \| 372 \| 656 \| 1081 \| Exodeoxyribonuclease \| Exodeoxyribonuclease; afdb-uniprot50 \| AF-A0A0M8PNS2-F1-MODEL\_V4 \| 1.0 \| 2.15e-21 \| 760 \| 0.282 \| 375 \| 160 \| 11 \| 2 \| 369 \| 7 \| 279 \| RecE \| RecE; afdb-uniprot50 \| AF-A0A4R0ER10-F1-MODEL\_V4 \| 1.0 \| 1.161e-20 \| 760 \| 0.208 \| 375 \| 195 \| 8 \| 2 \| 368 \| 9 \| 289 \| DUF3799 domain-containing protein \| DUF3799 domain-containing protein; afdb-uniprot50 \| AF-Q8XF26-F1-MODEL\_V4 \| 1.0 \| 5.246e-24 \| 759 \| 0.222 \| 436 \| 265 \| 12 \| 1 \| 372 \| 536 \| 961 \| Gifsy-1 prophage protein \| Gifsy-1 prophage protein; afdb-uniprot50 \| AF-A0A523QSH7-F1-MODEL\_V4 \| 1.0 \| 2.274e-21 \| 758 \| 0.239 \| 371 \| 171 \| 10 \| 1 \| 363 \| 21 \| 288 \| DUF3799 domain-containing protein \| DUF3799 domain-containing protein; afdb-uniprot50 \| AF-A0A7V5Q1K7-F1-MODEL\_V4 \| 1.0 \| 2.406e-21 \| 758 \| 0.234 \| 367 \| 194 \| 12 \| 1 \| 367 \| 5 \| 284 \| DUF3799 domain-containing protein \| DUF3799 domain-containing protein; afdb-uniprot50 \| AF-A0A2I8NWI0-F1-MODEL\_V4 \| 1.0 \| 5.246e-24 \| 758 \| 0.222 \| 436 \| 265 \| 11 \| 1 \| 372 \| 571 \| 996 \| Exodeoxyribonuclease VIII \| Exodeoxyribonuclease VIII; afdb-uniprot50 \| AF-A0A318FG45-F1-MODEL\_V4 \| 1.0 \| 2.526e-24 \| 758 \| 0.227 \| 440 \| 258 \| 13 \| 1 \| 372 \| 594 \| 1019 \| Exodeoxyribonuclease VIII \| Exodeoxyribonuclease VIII; afdb-uniprot50 \| AF-A0A6C7DAJ4-F1-MODEL\_V4 \| 1.0 \| 1.44e-24 \| 757 \| 0.224 \| 437 \| 263 \| 14 \| 1 \| 372 \| 542 \| 967 \| Exodeoxyribonuclease \| Exodeoxyribonuclease; afdb-uniprot50 \| AF-A0A420S4T2-F1-MODEL\_V4 \| 1.0 \| 1.721e-20 \| 756 \| 0.241 \| 373 \| 175 \| 10 \| 8 \| 372 \| 10 \| 282 \| DUF3799 domain-containing protein \| DUF3799 domain-containing protein; afdb-uniprot50 \| AF-D6PKU1-F1-MODEL\_V4 \| 1.0 \| 3.575e-20 \| 755 \| 0.249 \| 365 \| 163 \| 10 \| 1 \| 360 \| 1 \| 259 \| Exonuclease VIII 5 > 3 specific dsDNA exonuclease \| Exonuclease VIII 5 > 3 specific dsDNA exonuclease; afdb-uniprot50 \| AF-A0A6N7BWK4-F1-MODEL\_V4 \| 1.0 \| 4.466e-21 \| 755 \| 0.254 \| 370 \| 171 \| 9 \| 1 \| 360 \| 1 \| 275 \| DUF3799 domain-containing protein \| DUF3799 domain-containing protein; afdb-uniprot50 \| AF-A0A1I3YJV0-F1-MODEL\_V4 \| 1.0 \| 4.466e-21 \| 755 \| 0.246 \| 373 \| 175 \| 11 \| 1 \| 371 \| 5 \| 273 \| DUF3799 domain-containing protein \| DUF3799 domain-containing protein; afdb-uniprot50 \| AF-A0A564TYU2-F1-MODEL\_V4 \| 1.0 \| 6.21e-24 \| 752 \| 0.221 \| 433 \| 263 \| 13 \| 1 \| 369 \| 203 \| 625 \| Exodeoxyribonuclease 8 \| Exodeoxyribonuclease 8; afdb-uniprot50 \| AF-A0A5E9SPN3-F1-MODEL\_V4 \| 1.0 \| 1.527e-23 \| 752 \| 0.217 \| 436 \| 259 \| 15 \| 1 \| 368 \| 427 \| 848 \| Exonuclease VIII \| Exonuclease VIII; afdb-uniprot50 \| AF-A0A2D9CGT1-F1-MODEL\_V4 \| 1.0 \| 1.161e-20 \| 750 \| 0.262 \| 358 \| 158 \| 10 \| 5 \| 360 \| 3 \| 256 \| DUF3799 domain-containing protein \| DUF3799 domain-containing protein; afdb-uniprot50 \| AF-A0A0P0EB10-F1-MODEL\_V4 \| 1.0 \| 1.821e-20 \| 750 \| 0.252 \| 372 \| 174 \| 9 \| 3 \| 371 \| 2 \| 272 \| DUF3799 domain-containing protein \| DUF3799 domain-containing protein; afdb-uniprot50 \| AF-A0A756I411-F1-MODEL\_V4 \| 1.0 \| 2.388e-24 \| 749 \| 0.223 \| 438 \| 262 \| 12 \| 1 \| 372 \| 54 \| 479 \| DNA breaking-rejoining protein \| DNA breaking-rejoining protein; afdb-uniprot50 \| AF-A0A2S9I8G0-F1-MODEL\_V4 \| 1.0 \| 9.712e-25 \| 749 \| 0.232 \| 439 \| 255 \| 14 \| 1 \| 371 \| 430 \| 854 \| Exodeoxyribonuclease VIII \| Exodeoxyribonuclease VIII; afdb-uniprot50 \| AF-A0A754B5G1-F1-MODEL\_V4 \| 1.0 \| 3.013e-21 \| 749 \| 0.233 \| 385 \| 225 \| 10 \| 1 \| 372 \| 569 \| 896 \| Exodeoxyribonuclease VIII \| Exodeoxyribonuclease VIII; afdb-uniprot50 \| AF-A0A480BQL0-F1-MODEL\_V4 \| 1.0 \| 2.552e-20 \| 747 \| 0.229 \| 357 \| 173 \| 6 \| 9 \| 364 \| 1 \| 256 \| DUF3799 domain-containing protein \| DUF3799 domain-containing protein; afdb-uniprot50 \| AF-A0A0Q9C5I5-F1-MODEL\_V4 \| 1.0 \| 7.002e-21 \| 746 \| 0.27 \| 374 \| 166 \| 11 \| 1 \| 372 \| 2 \| 270 \| DUF3799 domain-containing protein \| DUF3799 domain-containing protein; afdb-uniprot50 \| AF-A0A7T8PWF9-F1-MODEL\_V4 \| 1.0 \| 3.991e-21 \| 746 \| 0.265 \| 343 \| 170 \| 9 \| 20 \| 361 \| 2 \| 263 \| PD-(D/E)XK nuclease-like domain-containing protein \| PD-(D/E)XK nuclease-like domain-containing protein; afdb-uniprot50 \| AF-A0A2L1VE72-F1-MODEL\_V4 \| 1.0 \| 7.407e-21 \| 746 \| 0.238 \| 378 \| 180 \| 11 \| 1 \| 368 \| 2 \| 281 \| Nuclease \| Nuclease; afdb-uniprot50 \| AF-A0A090V4Q6-F1-MODEL\_V4 \| 1.0 \| 4.432e-24 \| 746 \| 0.231 \| 436 \| 261 \| 12 \| 1 \| 372 \| 814 \| 1239 \| Putative exodeoxyribonuclease VIII \| Putative exodeoxyribonuclease VIII; afdb-uniprot50 \| AF-A0A0F9EGN9-F1-MODEL\_V4 \| 1.0 \| 2.848e-21 \| 745 \| 0.258 \| 363 \| 165 \| 9 \| 1 \| 360 \| 4 \| 265 \| DUF3799 domain-containing protein \| DUF3799 domain-containing protein; afdb-uniprot50 \| AF-A0A3C1SW29-F1-MODEL\_V4 \| 1.0 \| 9.786e-22 \| 745 \| 0.28 \| 363 \| 153 \| 10 \| 1 \| 363 \| 44 \| 298 \| DUF3799 domain-containing protein \| DUF3799 domain-containing protein; afdb-uniprot50 \| AF-A0A7W3KNY4-F1-MODEL\_V4 \| 1.0 \| 4.466e-21 \| 744 \| 0.286 \| 366 \| 154 \| 10 \| 1 \| 360 \| 11 \| 275 \| Exodeoxyribonuclease VIII \| Exodeoxyribonuclease VIII; afdb-uniprot50 \| AF-A0A376RV81-F1-MODEL\_V4 \| 1.0 \| 6.967e-23 \| 744 \| 0.219 \| 438 \| 258 \| 15 \| 1 \| 369 \| 136 \| 558 \| Putative phage exodeoxyribonuclease \| Putative phage exodeoxyribonuclease; afdb-uniprot50 \| AF-A0A6M3KS37-F1-MODEL\_V4 \| 1.0 \| 4.232e-20 \| 743 \| 0.26 \| 361 \| 163 \| 10 \| 3 \| 360 \| 2 \| 261 \| DUF3799 domain-containing protein \| DUF3799 domain-containing protein; afdb-uniprot50 \| AF-A0A7W3F3C1-F1-MODEL\_V4 \| 1.0 \| 1.619e-22 \| 743 \| 0.227 \| 382 \| 225 \| 11 \| 1 \| 369 \| 664 \| 988 \| PD-(D/E)XK nuclease-like domain-containing protein \| PD-(D/E)XK nuclease-like domain-containing protein; afdb-uniprot50 \| AF-A0A6B1U9G5-F1-MODEL\_V4 \| 1.0 \| 2.257e-24 \| 743 \| 0.217 \| 437 \| 266 \| 15 \| 1 \| 372 \| 680 \| 1105 \| Exodeoxyribonuclease \| Exodeoxyribonuclease; afdb-uniprot50 \| AF-A0A0Q5C9J8-F1-MODEL\_V4 \| 1.0 \| 2.156e-20 \| 742 \| 0.24 \| 370 \| 183 \| 9 \| 1 \| 369 \| 1 \| 273 \| DUF3799 domain-containing protein \| DUF3799 domain-containing protein; afdb-uniprot50 \| AF-A0A7W3A5W8-F1-MODEL\_V4 \| 1.0 \| 2.545e-21 \| 742 \| 0.214 \| 382 \| 228 \| 10 \| 1 \| 368 \| 535 \| 858 \| PD-(D/E)XK nuclease-like domain-containing protein \| PD-(D/E)XK nuclease-like domain-containing protein; afdb-uniprot50 \| AF-A0A5W0BDJ2-F1-MODEL\_V4 \| 1.0 \| 1.03e-23 \| 742 \| 0.24 \| 440 \| 252 \| 15 \| 1 \| 371 \| 561 \| 987 \| Exonuclease VIII \| Exonuclease VIII; afdb-uniprot50 \| AF-A0A448S1F5-F1-MODEL\_V4 \| 1.0 \| 9.205e-24 \| 741 \| 0.282 \| 375 \| 166 \| 9 \| 1 \| 371 \| 1 \| 276 \| Exodeoxyribonuclease 8 \| Exodeoxyribonuclease 8; afdb-uniprot50 \| AF-A0A3S2UWS8-F1-MODEL\_V4 \| 1.0 \| 3.005e-22 \| 740 \| 0.284 \| 394 \| 148 \| 12 \| 2 \| 361 \| 24 \| 317 \| Uncharacterized protein \| Uncharacterized protein; afdb-uniprot50 \| AF-A0A7Y7QAL0-F1-MODEL\_V4 \| 1.0 \| 2.263e-23 \| 740 \| 0.264 \| 367 \| 207 \| 7 \| 1 \| 359 \| 97 \| 408 \| PD-(D/E)XK nuclease-like domain-containing protein \| PD-(D/E)XK nuclease-like domain-containing protein; afdb-uniprot50 \| AF-A0A4Q3L0R6-F1-MODEL\_V4 \| 1.0 \| 7.835e-21 \| 739 \| 0.239 \| 364 \| 192 \| 11 \| 6 \| 369 \| 1 \| 279 \| DUF3799 domain-containing protein \| DUF3799 domain-containing protein; afdb-uniprot50 \| AF-A0A2S4QV97-F1-MODEL\_V4 \| 1.0 \| 2.139e-23 \| 737 \| 0.217 \| 436 \| 267 \| 11 \| 1 \| 372 \| 658 \| 1083 \| Exodeoxyribonuclease \| Exodeoxyribonuclease; afdb-uniprot50 \| AF-A0A0F9TVJ5-F1-MODEL\_V4 \| 1.0 \| 6.636e-20 \| 736 \| 0.215 \| 358 \| 179 \| 7 \| 1 \| 357 \| 14 \| 270 \| DUF3799 domain-containing protein \| DUF3799 domain-containing protein; afdb-uniprot50 \| AF-A0A7Y9J8Y8-F1-MODEL\_V4 \| 1.0 \| 1.717e-21 \| 735 \| 0.256 \| 374 \| 171 \| 12 \| 1 \| 372 \| 33 \| 301 \| DUF3799 domain-containing protein \| DUF3799 domain-containing protein; afdb-uniprot50 \| AF-A0A5U5MI79-F1-MODEL\_V4 \| 1.0 \| 4.688e-24 \| 734 \| 0.229 \| 418 \| 248 \| 12 \| 1 \| 354 \| 142 \| 549 \| Exodeoxyribonuclease \| Exodeoxyribonuclease; afdb-uniprot50 \| AF-A0A3L9YEK8-F1-MODEL\_V4 \| 1.0 \| 8.768e-21 \| 733 \| 0.247 \| 371 \| 171 \| 9 \| 8 \| 368 \| 36 \| 308 \| PDDEXK-like uncharacterized protein DUF3799 \| PDDEXK-like uncharacterized protein DUF3799; afdb-uniprot50 \| AF-A0A542ZT56-F1-MODEL\_V4 \| 1.0 \| 2.699e-20 \| 732 \| 0.222 \| 377 \| 179 \| 13 \| 1 \| 369 \| 4 \| 274 \| PDDEXK-like uncharacterized protein DUF3799 \| PDDEXK-like uncharacterized protein DUF3799; afdb-uniprot50 \| AF-A0A3M1FF26-F1-MODEL\_V4 \| 1.0 \| 3.195e-20 \| 730 \| 0.25 \| 363 \| 159 \| 12 \| 1 \| 361 \| 1 \| 252 \| DUF3799 domain-containing protein \| DUF3799 domain-containing protein; afdb-uniprot50 \| AF-A0A1E4IEB0-F1-MODEL\_V4 \| 1.0 \| 7.002e-21 \| 730 \| 0.249 \| 361 \| 184 \| 10 \| 3 \| 360 \| 7 \| 283 \| DUF3799 domain-containing protein \| DUF3799 domain-containing protein; afdb-uniprot50 \| AF-A0A1X0DYE3-F1-MODEL\_V4 \| 1.0 \| 3.991e-21 \| 729 \| 0.257 \| 376 \| 171 \| 14 \| 1 \| 365 \| 4 \| 282 \| DUF3799 domain-containing protein \| DUF3799 domain-containing protein; afdb-uniprot50 \| AF-M3VCA4-F1-MODEL\_V4 \| 1.0 \| 9.275e-21 \| 728 \| 0.281 \| 362 \| 154 \| 13 \| 1 \| 360 \| 6 \| 263 \| DUF3799 domain-containing protein \| DUF3799 domain-containing protein; afdb-uniprot50 \| AF-A0A828EP49-F1-MODEL\_V4 \| 1.0 \| 3.354e-23 \| 727 \| 0.229 \| 436 \| 256 \| 16 \| 1 \| 369 \| 258 \| 680 \| Exonuclease VIII \| Exonuclease VIII; afdb-uniprot50 \| AF-B8FNJ4-F1-MODEL\_V4 \| 1.0 \| 5.93e-20 \| 726 \| 0.212 \| 357 \| 177 \| 7 \| 5 \| 360 \| 4 \| 257 \| Putative bacteriophage protein \| Putative bacteriophage protein; afdb-uniprot50 \| AF-A0A2W1PQ23-F1-MODEL\_V4 \| 1.0 \| 1.375e-20 \| 726 \| 0.252 \| 372 \| 177 \| 13 \| 1 \| 367 \| 1 \| 276 \| RecE \| RecE; afdb-uniprot50 \| AF-A0A100Y6B0-F1-MODEL\_V4 \| 1.0 \| 7.407e-21 \| 724 \| 0.279 \| 368 \| 157 \| 12 \| 9 \| 368 \| 1 \| 268 \| DUF3799 domain-containing protein \| DUF3799 domain-containing protein; afdb-uniprot50 \| AF-A0A2J6NWE3-F1-MODEL\_V4 \| 1.0 \| 8.768e-21 \| 724 \| 0.256 \| 382 \| 187 \| 14 \| 1 \| 371 \| 2 \| 297 \| DUF3799 domain-containing protein \| DUF3799 domain-containing protein; afdb-uniprot50 \| AF-A0A6H1ZNA7-F1-MODEL\_V4 \| 1.0 \| 1.038e-20 \| 723 \| 0.221 \| 366 \| 186 \| 11 \| 3 \| 360 \| 10 \| 284 \| DUF3799 domain-containing protein \| DUF3799 domain-containing protein; afdb-uniprot50 \| AF-A0A2E3N1L6-F1-MODEL\_V4 \| 1.0 \| 1.826e-19 \| 722 \| 0.245 \| 366 \| 168 \| 10 \| 9 \| 361 \| 3 \| 273 \| DUF3799 domain-containing protein \| DUF3799 domain-containing protein; afdb-uniprot50 \| AF-A0A6M3JA96-F1-MODEL\_V4 \| 1.0 \| 5.915e-21 \| 721 \| 0.247 \| 371 \| 199 \| 10 \| 1 \| 368 \| 3 \| 296 \| DUF3799 domain-containing protein \| DUF3799 domain-containing protein; afdb-uniprot50 \| AF-A0A0F9U718-F1-MODEL\_V4 \| 1.0 \| 1.931e-19 \| 720 \| 0.202 \| 360 \| 189 \| 7 \| 1 \| 359 \| 3 \| 265 \| DUF3799 domain-containing protein \| DUF3799 domain-containing protein; afdb-uniprot50 \| AF-A0A6M3Y0Y1-F1-MODEL\_V4 \| 1.0 \| 2.863e-19 \| 720 \| 0.237 \| 367 \| 174 \| 8 \| 1 \| 360 \| 2 \| 269 \| DUF3799 domain-containing protein \| DUF3799 domain-containing protein; afdb-uniprot50 \| AF-A0A653VM09-F1-MODEL\_V4 \| 1.0 \| 2.848e-21 \| 719 \| 0.232 \| 370 \| 184 \| 11 \| 5 \| 370 \| 7 \| 280 \| DUF3799 domain-containing protein \| DUF3799 domain-containing protein; afdb-uniprot50 \| AF-A0A7W6CSA3-F1-MODEL\_V4 \| 1.0 \| 6.257e-21 \| 719 \| 0.244 \| 385 \| 181 \| 10 \| 2 \| 368 \| 6 \| 298 \| DUF3799 domain-containing protein \| DUF3799 domain-containing protein; afdb-uniprot50 \| AF-A0A654C1H5-F1-MODEL\_V4 \| 1.0 \| 1.807e-23 \| 719 \| 0.221 \| 437 \| 258 \| 14 \| 1 \| 369 \| 603 \| 1025 \| DUF3799 domain-containing protein \| DUF3799 domain-containing protein; afdb-uniprot50 \| AF-A0A2T3BX21-F1-MODEL\_V4 \| 1.0 \| 1.101e-19 \| 718 \| 0.246 \| 365 \| 167 \| 14 \| 1 \| 364 \| 3 \| 260 \| DUF3799 domain-containing protein \| DUF3799 domain-containing protein; afdb-uniprot50 \| AF-A0A4P6GRI8-F1-MODEL\_V4 \| 1.0 \| 2.4e-22 \| 718 \| 0.261 \| 363 \| 181 \| 8 \| 1 \| 360 \| 60 \| 338 \| DUF3799 domain-containing protein \| DUF3799 domain-containing protein; afdb-uniprot50 \| AF-D4LDT1-F1-MODEL\_V4 \| 1.0 \| 1.378e-19 \| 717 \| 0.243 \| 362 \| 163 \| 9 \| 9 \| 361 \| 1 \| 260 \| DUF3799 domain-containing protein \| DUF3799 domain-containing protein; afdb-uniprot50 \| AF-A0A5B9W8V5-F1-MODEL\_V4 \| 1.0 \| 8.79e-20 \| 717 \| 0.247 \| 367 \| 172 \| 9 \| 5 \| 369 \| 2 \| 266 \| Exodeoxyribonuclease 8 \| Exodeoxyribonuclease 8; afdb-uniprot50 \| AF-A0A370CF63-F1-MODEL\_V4 \| 1.0 \| 4.7e-23 \| 717 \| 0.253 \| 386 \| 174 \| 13 \| 1 \| 371 \| 11 \| 297 \| DUF3799 domain-containing protein \| DUF3799 domain-containing protein; afdb-uniprot50 \| AF-A0A6B3LMA0-F1-MODEL\_V4 \| 1.0 \| 4.477e-20 \| 716 \| 0.254 \| 365 \| 169 \| 14 \| 5 \| 360 \| 14 \| 284 \| DUF3799 domain-containing protein \| DUF3799 domain-containing protein; afdb-uniprot50 \| AF-A0A5C8A0A4-F1-MODEL\_V4 \| 1.0 \| 3.773e-21 \| 716 \| 0.245 \| 374 \| 175 \| 10 \| 1 \| 370 \| 45 \| 315 \| DUF3799 domain-containing protein \| DUF3799 domain-containing protein; afdb-uniprot50 \| AF-A0A1Z9X1Y0-F1-MODEL\_V4 \| 1.0 \| 5.273e-22 \| 714 \| 0.214 \| 382 \| 218 \| 8 \| 1 \| 358 \| 39 \| 362 \| DUF3799 domain-containing protein \| DUF3799 domain-containing protein; afdb-uniprot50 \| AF-A0A2E8DZF3-F1-MODEL\_V4 \| 1.0 \| 3.38e-20 \| 712 \| 0.271 \| 364 \| 147 \| 11 \| 7 \| 360 \| 10 \| 265 \| DUF3799 domain-containing protein \| DUF3799 domain-containing protein; afdb-uniprot50 \| AF-A0A759M5V4-F1-MODEL\_V4 \| 1.0 \| 3.763e-22 \| 712 \| 0.237 \| 383 \| 226 \| 12 \| 1 \| 372 \| 116 \| 443 \| DUF3799 domain-containing protein \| DUF3799 domain-containing protein; afdb-uniprot50 \| AF-A0A0F9K1Z8-F1-MODEL\_V4 \| 1.0 \| 4.736e-20 \| 711 \| 0.23 \| 368 \| 182 \| 9 \| 1 \| 361 \| 22 \| 295 \| DUF3799 domain-containing protein \| DUF3799 domain-containing protein; afdb-uniprot50 \| AF-A0A2D4TU41-F1-MODEL\_V4 \| 1.0 \| 9.228e-23 \| 711 \| 0.238 \| 382 \| 248 \| 12 \| 1 \| 358 \| 41 \| 403 \| Uncharacterized protein \| Uncharacterized protein; afdb-uniprot50 \| AF-B6XEX5-F1-MODEL\_V4 \| 1.0 \| 4.984e-22 \| 709 \| 0.252 \| 384 \| 219 \| 13 \| 1 \| 372 \| 148 \| 475 \| DUF3799 domain-containing protein \| DUF3799 domain-containing protein; afdb-uniprot50 \| AF-A0A7Z1T1G3-F1-MODEL\_V4 \| 1.0 \| 2.027e-22 \| 708 \| 0.283 \| 371 \| 163 \| 9 \| 1 \| 367 \| 567 \| 838 \| Exonuclease VIII \| Exonuclease VIII; afdb-uniprot50 \| AF-A0A0Q6ECD3-F1-MODEL\_V4 \| 1.0 \| 1.375e-20 \| 707 \| 0.232 \| 375 \| 188 \| 11 \| 1 \| 371 \| 4 \| 282 \| DUF3799 domain-containing protein \| DUF3799 domain-containing protein; afdb-uniprot50 \| AF-A0A722GKL2-F1-MODEL\_V4 \| 1.0 \| 7.37e-23 \| 707 \| 0.211 \| 420 \| 253 \| 11 \| 19 \| 372 \| 2 \| 409 \| DNA breaking-rejoining protein \| DNA breaking-rejoining protein; afdb-uniprot50 \| AF-A0A1C6GFL6-F1-MODEL\_V4 \| 1.0 \| 3.028e-19 \| 706 \| 0.242 \| 363 \| 169 \| 10 \| 9 \| 364 \| 1 \| 264 \| Exodeoxyribonuclease 8 \| Exodeoxyribonuclease 8; afdb-uniprot50 \| AF-E6WYE1-F1-MODEL\_V4 \| 1.0 \| 5.93e-20 \| 701 \| 0.273 \| 355 \| 154 \| 10 \| 9 \| 361 \| 1 \| 253 \| DUF3799 domain-containing protein \| DUF3799 domain-containing protein; afdb-uniprot50 \| AF-C7R228-F1-MODEL\_V4 \| 1.0 \| 1.931e-19 \| 701 \| 0.248 \| 382 \| 170 \| 12 \| 1 \| 369 \| 3 \| 280 \| DUF3799 domain-containing protein \| DUF3799 domain-containing protein; afdb-uniprot50 \| AF-A0A845GHL0-F1-MODEL\_V4 \| 1.0 \| 1.232e-19 \| 699 \| 0.236 \| 363 \| 172 \| 12 \| 2 \| 360 \| 16 \| 277 \| Uncharacterized protein \| Uncharacterized protein; afdb-uniprot50 \| AF-A0A0S8DSQ9-F1-MODEL\_V4 \| 1.0 \| 6.653e-19 \| 698 \| 0.205 \| 365 \| 185 \| 7 \| 1 \| 361 \| 2 \| 265 \| DUF3799 domain-containing protein \| DUF3799 domain-containing protein; afdb-uniprot50 \| AF-A0A0C1AUR5-F1-MODEL\_V4 \| 1.0 \| 2.706e-19 \| 697 \| 0.237 \| 379 \| 174 \| 14 \| 1 \| 370 \| 4 \| 276 \| DUF3799 domain-containing protein \| DUF3799 domain-containing protein; afdb-uniprot50 \| AF-A0A7S9HDT4-F1-MODEL\_V4 \| 1.0 \| 4.011e-19 \| 697 \| 0.241 \| 377 \| 177 \| 12 \| 1 \| 372 \| 20 \| 292 \| PD-(D/E)XK nuclease-like domain-containing protein \| PD-(D/E)XK nuclease-like domain-containing protein; afdb-uniprot50 \| AF-A0A1V5Z6N5-F1-MODEL\_V4 \| 1.0 \| 1.627e-20 \| 696 \| 0.276 \| 383 \| 159 \| 14 \| 1 \| 370 \| 10 \| 287 \| Exodeoxyribonuclease 8 \| Exodeoxyribonuclease 8; afdb-uniprot50 \| AF-A0A742LDT8-F1-MODEL\_V4 \| 1.0 \| 2.145e-22 \| 696 \| 0.265 \| 369 \| 159 \| 13 \| 1 \| 360 \| 196 \| 461 \| DUF3799 domain-containing protein \| DUF3799 domain-containing protein; afdb-uniprot50 \| AF-A0A6G7XGW4-F1-MODEL\_V4 \| 1.0 \| 7.875e-19 \| 695 \| 0.231 \| 372 \| 181 \| 11 \| 1 \| 370 \| 3 \| 271 \| DUF3799 domain-containing protein \| DUF3799 domain-containing protein; afdb-uniprot50 \| AF-A0A366DC85-F1-MODEL\_V4 \| 1.0 \| 8.309e-20 \| 695 \| 0.238 \| 378 \| 175 \| 15 \| 1 \| 368 \| 1 \| 275 \| PDDEXK-like uncharacterized protein DUF3799 \| PDDEXK-like uncharacterized protein DUF3799; afdb-uniprot50 \| AF-A0A0J8U4U2-F1-MODEL\_V4 \| 1.0 \| 1.458e-19 \| 694 \| 0.243 \| 374 \| 178 \| 12 \| 2 \| 371 \| 7 \| 279 \| DUF3799 domain-containing protein \| DUF3799 domain-containing protein; afdb-uniprot50 \| AF-A0A3D3U8P6-F1-MODEL\_V4 \| 1.0 \| 4.748e-19 \| 693 \| 0.235 \| 361 \| 164 \| 8 \| 9 \| 358 \| 1 \| 260 \| DUF3799 domain-containing protein \| DUF3799 domain-containing protein; afdb-uniprot50 \| AF-A0A606Y4Y5-F1-MODEL\_V4 \| 1.0 \| 1.531e-22 \| 693 \| 0.27 \| 370 \| 167 \| 10 \| 1 \| 366 \| 541 \| 811 \| Exonuclease VIII \| Exonuclease VIII; afdb-uniprot50 \| AF-A0A0F9FLX8-F1-MODEL\_V4 \| 1.0 \| 7.445e-19 \| 690 \| 0.198 \| 373 \| 189 \| 10 \| 1 \| 366 \| 1 \| 270 \| DUF3799 domain-containing protein \| DUF3799 domain-containing protein; afdb-uniprot50 \| AF-A0A6Y5LE31-F1-MODEL\_V4 \| 1.0 \| 3.179e-22 \| 690 \| 0.262 \| 369 \| 160 \| 13 \| 1 \| 360 \| 501 \| 766 \| DUF1391 domain-containing protein \| DUF1391 domain-containing protein; afdb-uniprot50 \| AF-A0A7V9ZES2-F1-MODEL\_V4 \| 1.0 \| 1.636e-18 \| 689 \| 0.174 \| 361 \| 195 \| 7 \| 1 \| 360 \| 7 \| 265 \| PD-(D/E)XK nuclease-like domain-containing protein \| PD-(D/E)XK nuclease-like domain-containing protein; afdb-uniprot50 \| AF-A0A2E7ZS37-F1-MODEL\_V4 \| 1.0 \| 1.717e-21 \| 689 \| 0.266 \| 372 \| 188 \| 12 \| 5 \| 364 \| 10 \| 308 \| DUF3799 domain-containing protein \| DUF3799 domain-containing protein; afdb-uniprot50 \| AF-A0A2I1I759-F1-MODEL\_V4 \| 1.0 \| 2.286e-19 \| 688 \| 0.223 \| 372 \| 172 \| 13 \| 1 \| 363 \| 6 \| 269 \| DUF3799 domain-containing protein \| DUF3799 domain-containing protein; afdb-uniprot50 \| AF-A0A5C7NLS0-F1-MODEL\_V4 \| 1.0 \| 5.01e-20 \| 688 \| 0.242 \| 383 \| 183 \| 14 \| 1 \| 371 \| 11 \| 298 \| DUF3799 domain-containing protein \| DUF3799 domain-containing protein; afdb-uniprot50 \| AF-A0A2E5HBV2-F1-MODEL\_V4 \| 1.0 \| 5.62e-19 \| 686 \| 0.261 \| 367 \| 165 \| 10 \| 1 \| 364 \| 2 \| 265 \| DUF3799 domain-containing protein \| DUF3799 domain-containing protein; afdb-uniprot50 \| AF-A0A0W7W9N2-F1-MODEL\_V4 \| 1.0 \| 5.01e-20 \| 685 \| 0.228 \| 380 \| 189 \| 10 \| 2 \| 371 \| 7 \| 292 \| DUF3799 domain-containing protein \| DUF3799 domain-containing protein; afdb-uniprot50 \| AF-A0A6M3JG38-F1-MODEL\_V4 \| 1.0 \| 1.382e-18 \| 683 \| 0.243 \| 365 \| 170 \| 12 \| 1 \| 360 \| 4 \| 267 \| DUF3799 domain-containing protein \| DUF3799 domain-containing protein; afdb-uniprot50 \| AF-A0A6I4WMS0-F1-MODEL\_V4 \| 1.0 \| 1.615e-23 \| 683 \| 0.226 \| 398 \| 252 \| 9 \| 3 \| 358 \| 34 \| 417 \| PD-(D/E)XK nuclease-like domain-containing protein \| PD-(D/E)XK nuclease-like domain-containing protein; afdb-uniprot50 \| AF-A0A0K6IY38-F1-MODEL\_V4 \| 1.0 \| 5.62e-19 \| 682 \| 0.299 \| 321 \| 120 \| 9 \| 52 \| 363 \| 2 \| 226 \| DUF3799 domain-containing protein \| DUF3799 domain-containing protein; afdb-uniprot50 \| AF-A0A8B5X8J1-F1-MODEL\_V4 \| 1.0 \| 6.653e-19 \| 682 \| 0.224 \| 356 \| 175 \| 6 \| 8 \| 359 \| 1 \| 259 \| Uncharacterized protein \| Uncharacterized protein; afdb-uniprot50 \| AF-A0A1Q5A5E3-F1-MODEL\_V4 \| 1.0 \| 1.826e-19 \| 682 \| 0.237 \| 375 \| 179 \| 10 \| 1 \| 368 \| 16 \| 290 \| DUF3799 domain-containing protein \| DUF3799 domain-containing protein; afdb-uniprot50 \| AF-A0A5X3VD53-F1-MODEL\_V4 \| 1.0 \| 1.447e-22 \| 682 \| 0.216 \| 424 \| 250 \| 12 \| 17 \| 372 \| 1 \| 410 \| Exodeoxyribonuclease \| Exodeoxyribonuclease; afdb-uniprot50 \| AF-A0A418GY69-F1-MODEL\_V4 \| 1.0 \| 4.454e-22 \| 682 \| 0.267 \| 370 \| 168 \| 9 \| 1 \| 366 \| 277 \| 547 \| Exodeoxyribonuclease 8 \| Exodeoxyribonuclease 8; afdb-uniprot50 \| AF-A0A7U5YQG6-F1-MODEL\_V4 \| 1.0 \| 1.619e-22 \| 682 \| 0.262 \| 369 \| 160 \| 11 \| 1 \| 360 \| 567 \| 832 \| Exonuclease VIII \| Exonuclease VIII; afdb-uniprot50 \| AF-A0A494WBY5-F1-MODEL\_V4 \| 1.0 \| 7.02e-20 \| 681 \| 0.244 \| 384 \| 183 \| 10 \| 2 \| 371 \| 17 \| 307 \| DUF3799 domain-containing protein \| DUF3799 domain-containing protein; afdb-uniprot50 \| AF-A0A2D6F114-F1-MODEL\_V4 \| 1.0 \| 1.038e-20 \| 681 \| 0.251 \| 366 \| 201 \| 13 \| 5 \| 360 \| 8 \| 310 \| DUF3799 domain-containing protein \| DUF3799 domain-containing protein; afdb-uniprot50 \| AF-A0A827ZGB3-F1-MODEL\_V4 \| 1.0 \| 1.917e-22 \| 681 \| 0.265 \| 373 \| 165 \| 11 \| 1 \| 366 \| 501 \| 771 \| Exodeoxyribonuclease VIII \| Exodeoxyribonuclease VIII; afdb-uniprot50 \| AF-A0A7U0UP56-F1-MODEL\_V4 \| 1.0 \| 2.418e-19 \| 680 \| 0.231 \| 371 \| 173 \| 14 \| 2 \| 367 \| 5 \| 268 \| PD-(D/E)XK nuclease-like domain-containing protein \| PD-(D/E)XK nuclease-like domain-containing protein; afdb-uniprot50 \| AF-A0A7C5WUZ4-F1-MODEL\_V4 \| 1.0 \| 3.584e-19 \| 680 \| 0.254 \| 365 \| 161 \| 12 \| 6 \| 364 \| 11 \| 270 \| DUF3799 domain-containing protein \| DUF3799 domain-containing protein; afdb-uniprot50 \| AF-A0A5C1APE4-F1-MODEL\_V4 \| 1.0 \| 1.3e-20 \| 679 \| 0.209 \| 411 \| 218 \| 15 \| 1 \| 367 \| 9 \| 356 \| Exodeoxyribonuclease 8 \| Exodeoxyribonuclease 8; afdb-uniprot50 \| AF-A0A1K1LD38-F1-MODEL\_V4 \| 1.0 \| 2.706e-19 \| 678 \| 0.232 \| 375 \| 178 \| 14 \| 1 \| 366 \| 1 \| 274 \| Exodeoxyribonuclease VIII \| Exodeoxyribonuclease VIII; afdb-uniprot50 \| AF-A0A730CAU1-F1-MODEL\_V4 \| 1.0 \| 1.534e-21 \| 678 \| 0.205 \| 418 \| 254 \| 11 \| 21 \| 372 \| 1 \| 406 \| Exodeoxyribonuclease \| Exodeoxyribonuclease; afdb-uniprot50 \| AF-A0A6M3J3Y5-F1-MODEL\_V4 \| 1.0 \| 1.041e-19 \| 677 \| 0.238 \| 360 \| 169 \| 10 \| 7 \| 361 \| 3 \| 262 \| DUF3799 domain-containing protein \| DUF3799 domain-containing protein; afdb-uniprot50 \| AF-A0A829DNE5-F1-MODEL\_V4 \| 1.0 \| 3.005e-22 \| 677 \| 0.265 \| 373 \| 165 \| 11 \| 1 \| 366 \| 616 \| 886 \| Uncharacterized protein \| Uncharacterized protein; afdb-uniprot50 \| AF-A0A731HUQ6-F1-MODEL\_V4 \| 1.0 \| 2.027e-22 \| 676 \| 0.214 \| 424 \| 251 \| 12 \| 17 \| 372 \| 1 \| 410 \| Exodeoxyribonuclease \| Exodeoxyribonuclease; afdb-uniprot50 \| AF-A0A2A7UMV7-F1-MODEL\_V4 \| 1.0 \| 5.62e-19 \| 675 \| 0.22 \| 376 \| 181 \| 12 \| 1 \| 370 \| 6 \| 275 \| RecE \| RecE; afdb-uniprot50 \| AF-A0A0A1DF13-F1-MODEL\_V4 \| 1.0 \| 1.232e-19 \| 675 \| 0.25 \| 376 \| 178 \| 14 \| 1 \| 372 \| 18 \| 293 \| Phage exonuclease \| Phage exonuclease; afdb-uniprot50 \| AF-A0A849A8M1-F1-MODEL\_V4 \| 1.0 \| 1.458e-19 \| 673 \| 0.272 \| 370 \| 169 \| 10 \| 1 \| 365 \| 1 \| 275 \| Uncharacterized protein \| Uncharacterized protein; afdb-uniprot50 \| AF-A0A2N3VGV8-F1-MODEL\_V4 \| 1.0 \| 3.584e-19 \| 673 \| 0.238 \| 382 \| 175 \| 13 \| 2 \| 368 \| 6 \| 286 \| PDDEXK-like uncharacterized protein DUF3799 \| PDDEXK-like uncharacterized protein DUF3799; afdb-uniprot50 \| AF-A0A6X8AL85-F1-MODEL\_V4 \| 1.0 \| 6.984e-22 \| 673 \| 0.264 \| 374 \| 164 \| 11 \| 1 \| 366 \| 237 \| 507 \| Exodeoxyribonuclease 8 \| Exodeoxyribonuclease 8; afdb-uniprot50 \| AF-A0A7H9CND5-F1-MODEL\_V4 \| 1.0 \| 4.748e-19 \| 672 \| 0.225 \| 360 \| 169 \| 11 \| 9 \| 358 \| 1 \| 260 \| Putative exonuclease VIII \| Putative exonuclease VIII; afdb-uniprot50 \| AF-A0A2D9C7V6-F1-MODEL\_V4 \| 1.0 \| 5.326e-18 \| 671 \| 0.222 \| 359 \| 172 \| 12 \| 4 \| 360 \| 2 \| 255 \| DUF3799 domain-containing protein \| DUF3799 domain-containing protein; afdb-uniprot50 \| AF-A0A2D5F9Z8-F1-MODEL\_V4 \| 1.0 \| 2.699e-20 \| 671 \| 0.253 \| 355 \| 169 \| 9 \| 5 \| 357 \| 2 \| 262 \| DUF3799 domain-containing protein \| DUF3799 domain-containing protein; afdb-uniprot50 \| AF-A0A730GR09-F1-MODEL\_V4 \| 1.0 \| 8.746e-22 \| 671 \| 0.21 \| 418 \| 252 \| 11 \| 21 \| 372 \| 1 \| 406 \| DNA breaking-rejoining protein \| DNA breaking-rejoining protein; afdb-uniprot50 \| AF-A0A378Y5Z6-F1-MODEL\_V4 \| 1.0 \| 8.79e-20 \| 670 \| 0.249 \| 373 \| 170 \| 13 \| 1 \| 367 \| 1 \| 269 \| Exodeoxyribonuclease 8 \| Exodeoxyribonuclease 8; afdb-uniprot50 \| AF-A0A3M9MM27-F1-MODEL\_V4 \| 1.0 \| 3.028e-19 \| 669 \| 0.255 \| 368 \| 169 \| 15 \| 2 \| 360 \| 10 \| 281 \| DUF3799 domain-containing protein \| DUF3799 domain-containing protein; afdb-uniprot50 \| AF-A0A3A5MMF7-F1-MODEL\_V4 \| 1.0 \| 1.041e-19 \| 669 \| 0.237 \| 371 \| 183 \| 11 \| 2 \| 367 \| 36 \| 311 \| DUF3799 domain-containing protein \| DUF3799 domain-containing protein; afdb-uniprot50 \| AF-A0A638M5J8-F1-MODEL\_V4 \| 1.0 \| 3.005e-22 \| 669 \| 0.264 \| 375 \| 163 \| 12 \| 1 \| 366 \| 314 \| 584 \| Exodeoxyribonuclease 8 \| Exodeoxyribonuclease 8; afdb-uniprot50 \| AF-A0A854GB88-F1-MODEL\_V4 \| 1.0 \| 7.855e-20 \| 668 \| 0.229 \| 370 \| 174 \| 12 \| 8 \| 370 \| 34 \| 299 \| Uncharacterized protein \| Uncharacterized protein; afdb-uniprot50 \| AF-P15032-F1-MODEL\_V4 \| 1.0 \| 2.15e-21 \| 668 \| 0.274 \| 368 \| 164 \| 9 \| 1 \| 364 \| 595 \| 863 \| Exodeoxyribonuclease 8 \| Exodeoxyribonuclease 8; afdb-uniprot50 \| AF-A0A0N0A2F3-F1-MODEL\_V4 \| 1.0 \| 4.477e-20 \| 667 \| 0.257 \| 381 \| 169 \| 14 \| 1 \| 369 \| 23 \| 301 \| DUF3799 domain-containing protein \| DUF3799 domain-containing protein; afdb-uniprot50 \| AF-A0A3L7AHV8-F1-MODEL\_V4 \| 1.0 \| 8.309e-20 \| 667 \| 0.224 \| 383 \| 187 \| 12 \| 2 \| 372 \| 18 \| 302 \| DUF3799 domain-containing protein \| DUF3799 domain-containing protein; afdb-uniprot50 \| AF-A0A7Z0J579-F1-MODEL\_V4 \| 1.0 \| 1.726e-19 \| 666 \| 0.263 \| 376 \| 171 \| 13 \| 1 \| 368 \| 5 \| 282 \| DUF3799 domain-containing protein \| DUF3799 domain-containing protein; afdb-uniprot50 \| AF-Q6LJU7-F1-MODEL\_V4 \| 1.0 \| 1.296e-21 \| 666 \| 0.185 \| 501 \| 266 \| 17 \| 1 \| 372 \| 661 \| 1148 \| DUF3799 domain-containing protein \| DUF3799 domain-containing protein; afdb-uniprot50 \| AF-A0A709X174-F1-MODEL\_V4 \| 1.0 \| 6.619e-21 \| 665 \| 0.206 \| 411 \| 248 \| 11 \| 28 \| 372 \| 2 \| 400 \| DUF3799 domain-containing protein \| DUF3799 domain-containing protein; afdb-uniprot50 \| AF-A0A2U9P084-F1-MODEL\_V4 \| 1.0 \| 1.296e-21 \| 665 \| 0.273 \| 377 \| 164 \| 11 \| 2 \| 368 \| 163 \| 439 \| DUF3799 domain-containing protein \| DUF3799 domain-containing protein; afdb-uniprot50 \| AF-A0A1X3IAK9-F1-MODEL\_V4 \| 1.0 \| 9.251e-22 \| 665 \| 0.268 \| 373 \| 164 \| 11 \| 1 \| 366 \| 319 \| 589 \| Exodeoxyribonuclease 8 (Exodeoxyribonuclease VIII) (EXOVIII) \| Exodeoxyribonuclease 8 (Exodeoxyribonuclease VIII) (EXOVIII); afdb-uniprot50 \| AF-A0A536UDP7-F1-MODEL\_V4 \| 1.0 \| 5.62e-19 \| 664 \| 0.23 \| 373 \| 164 \| 11 \| 20 \| 370 \| 1 \| 272 \| DUF3799 domain-containing protein \| DUF3799 domain-containing protein; afdb-uniprot50 \| AF-A0A1Z9Q5Q2-F1-MODEL\_V4 \| 1.0 \| 1.306e-18 \| 663 \| 0.199 \| 366 \| 186 \| 10 \| 1 \| 361 \| 5 \| 268 \| DUF3799 domain-containing protein \| DUF3799 domain-containing protein; afdb-uniprot50 \| AF-A0A1G3M6I0-F1-MODEL\_V4 \| 1.0 \| 1.826e-19 \| 663 \| 0.254 \| 370 \| 181 \| 13 \| 1 \| 360 \| 22 \| 306 \| DUF3799 domain-containing protein \| DUF3799 domain-containing protein; afdb-uniprot50 \| AF-A0A735S3T4-F1-MODEL\_V4 \| 1.0 \| 2.692e-21 \| 663 \| 0.272 \| 370 \| 166 \| 9 \| 1 \| 366 \| 121 \| 391 \| Exonuclease VIII \| Exonuclease VIII; afdb-uniprot50 \| AF-A0A2W6R8J9-F1-MODEL\_V4 \| 1.0 \| 2.558e-19 \| 662 \| 0.235 \| 373 \| 174 \| 13 \| 2 \| 371 \| 11 \| 275 \| RecE \| RecE; afdb-uniprot50 \| AF-A0A2W1QND0-F1-MODEL\_V4 \| 1.0 \| 1.164e-19 \| 661 \| 0.262 \| 373 \| 176 \| 11 \| 1 \| 370 \| 24 \| 300 \| DUF3799 domain-containing protein \| DUF3799 domain-containing protein; afdb-uniprot50 \| AF-A0A1M5I3B0-F1-MODEL\_V4 \| 1.0 \| 2.043e-19 \| 661 \| 0.24 \| 366 \| 171 \| 9 \| 1 \| 360 \| 53 \| 317 \| Exodeoxyribonuclease VIII \| Exodeoxyribonuclease VIII; afdb-uniprot50 \| AF-A0A7Z7VL18-F1-MODEL\_V4 \| 1.0 \| 2.841e-22 \| 661 \| 0.259 \| 374 \| 232 \| 11 \| 1 \| 359 \| 77 \| 420 \| SAP domain-containing protein \| SAP domain-containing protein; afdb-uniprot50 \| AF-A0A0L6CK16-F1-MODEL\_V4 \| 1.0 \| 7.445e-19 \| 660 \| 0.241 \| 368 \| 166 \| 14 \| 5 \| 362 \| 9 \| 273 \| DUF3799 domain-containing protein \| DUF3799 domain-containing protein; afdb-uniprot50 \| AF-A0A4Q8AK80-F1-MODEL\_V4 \| 1.0 \| 1.103e-18 \| 659 \| 0.236 \| 363 \| 179 \| 11 \| 1 \| 360 \| 2 \| 269 \| PDDEXK-like uncharacterized protein DUF3799 \| PDDEXK-like uncharacterized protein DUF3799; afdb-uniprot50 \| AF-A0A759U666-F1-MODEL\_V4 \| 1.0 \| 5.592e-21 \| 659 \| 0.203 \| 417 \| 254 \| 11 \| 22 \| 372 \| 1 \| 405 \| Exodeoxyribonuclease \| Exodeoxyribonuclease; afdb-uniprot50 \| AF-A0A5T3EKQ1-F1-MODEL\_V4 \| 1.0 \| 3.557e-22 \| 659 \| 0.231 \| 411 \| 234 \| 14 \| 1 \| 343 \| 550 \| 946 \| Exodeoxyribonuclease \| Exodeoxyribonuclease; afdb-uniprot50 \| AF-L7VX35-F1-MODEL\_V4 \| 1.0 \| 4.243e-19 \| 658 \| 0.211 \| 355 \| 176 \| 9 \| 9 \| 361 \| 1 \| 253 \| Exodeoxyribonuclease VIII \| Exodeoxyribonuclease VIII; afdb-uniprot50 \| AF-A0A1H1XHG2-F1-MODEL\_V4 \| 1.0 \| 1.232e-19 \| 658 \| 0.231 \| 372 \| 187 \| 14 \| 1 \| 368 \| 14 \| 290 \| DUF3799 domain-containing protein \| DUF3799 domain-containing protein; afdb-uniprot50 \| AF-A0A4V2NXX9-F1-MODEL\_V4 \| 1.0 \| 1.926e-20 \| 658 \| 0.269 \| 371 \| 165 \| 11 \| 2 \| 370 \| 61 \| 327 \| DUF3799 domain-containing protein \| DUF3799 domain-containing protein; afdb-uniprot50 \| AF-A0A191WF22-F1-MODEL\_V4 \| 1.0 \| 5.035e-18 \| 657 \| 0.237 \| 370 \| 174 \| 14 \| 1 \| 369 \| 4 \| 266 \| DUF3799 domain-containing protein \| DUF3799 domain-containing protein; afdb-uniprot50 \| AF-A0A367HZT2-F1-MODEL\_V4 \| 1.0 \| 4.748e-19 \| 657 \| 0.247 \| 372 \| 167 \| 12 \| 9 \| 368 \| 1 \| 271 \| DUF3799 domain-containing protein \| DUF3799 domain-containing protein; afdb-uniprot50 \| AF-A0A2G8BG73-F1-MODEL\_V4 \| 1.0 \| 2.424e-18 \| 656 \| 0.247 \| 367 \| 167 \| 13 \| 1 \| 361 \| 4 \| 267 \| DUF3799 domain-containing protein \| DUF3799 domain-containing protein; afdb-uniprot50 \| AF-A0A7D7WG90-F1-MODEL\_V4 \| 1.0 \| 4.243e-19 \| 656 \| 0.252 \| 364 \| 171 \| 11 \| 4 \| 366 \| 2 \| 265 \| DUF3799 domain-containing protein \| DUF3799 domain-containing protein; afdb-uniprot50 \| AF-A0A497R8U6-F1-MODEL\_V4 \| 1.0 \| 2.043e-19 \| 655 \| 0.242 \| 371 \| 170 \| 12 \| 1 \| 365 \| 45 \| 310 \| DUF3799 domain-containing protein \| DUF3799 domain-containing protein; afdb-uniprot50 \| AF-A0A7W4YZB4-F1-MODEL\_V4 \| 1.0 \| 6.653e-19 \| 654 \| 0.224 \| 379 \| 187 \| 9 \| 2 \| 370 \| 4 \| 285 \| DUF3799 domain-containing protein \| DUF3799 domain-containing protein; afdb-uniprot50 \| AF-A0A4Y3NBY8-F1-MODEL\_V4 \| 1.0 \| 1.164e-19 \| 654 \| 0.252 \| 376 \| 175 \| 14 \| 2 \| 372 \| 53 \| 327 \| DUF3799 domain-containing protein \| DUF3799 domain-containing protein; afdb-uniprot50 \| AF-A0A634SIY9-F1-MODEL\_V4 \| 1.0 \| 8.288e-21 \| 654 \| 0.27 \| 374 \| 162 \| 12 \| 1 \| 366 \| 85 \| 355 \| Exonuclease VIII \| Exonuclease VIII; afdb-uniprot50 \| AF-A0A1T4USH8-F1-MODEL\_V4 \| 1.0 \| 5.286e-21 \| 653 \| 0.242 \| 375 \| 209 \| 12 \| 1 \| 361 \| 44 \| 357 \| Exodeoxyribonuclease 8 \| Exodeoxyribonuclease 8; afdb-uniprot50 \| AF-A0A7S7FKE8-F1-MODEL\_V4 \| 1.0 \| 4.243e-19 \| 652 \| 0.255 \| 376 \| 160 \| 15 \| 1 \| 360 \| 5 \| 276 \| PD-(D/E)XK nuclease-like domain-containing protein \| PD-(D/E)XK nuclease-like domain-containing protein; afdb-uniprot50 \| AF-A0A0M2HG29-F1-MODEL\_V4 \| 1.0 \| 1.926e-20 \| 652 \| 0.258 \| 372 \| 175 \| 13 \| 4 \| 371 \| 37 \| 311 \| Exodeoxyribonuclease 8 \| Exodeoxyribonuclease 8; afdb-uniprot50 \| AF-A0A142XVC8-F1-MODEL\_V4 \| 1.0 \| 7.875e-19 \| 651 \| 0.216 \| 360 \| 178 \| 12 \| 10 \| 363 \| 3 \| 264 \| Exodeoxyribonuclease 8 \| Exodeoxyribonuclease 8; afdb-uniprot50 \| AF-A0A843H5C9-F1-MODEL\_V4 \| 1.0 \| 6.653e-19 \| 651 \| 0.239 \| 355 \| 198 \| 12 \| 1 \| 351 \| 1 \| 287 \| PD-(D/E)XK nuclease-like domain-containing protein \| PD-(D/E)XK nuclease-like domain-containing protein; afdb-uniprot50 \| AF-A0A6I5FGL7-F1-MODEL\_V4 \| 1.0 \| 1.167e-18 \| 650 \| 0.228 \| 377 \| 182 \| 12 \| 1 \| 368 \| 13 \| 289 \| DUF3799 domain-containing protein \| DUF3799 domain-containing protein; afdb-uniprot50 \| AF-A0A0F9G9B6-F1-MODEL\_V4 \| 1.0 \| 3.388e-19 \| 650 \| 0.211 \| 388 \| 192 \| 14 \| 1 \| 372 \| 5 \| 294 \| DUF3799 domain-containing protein \| DUF3799 domain-containing protein; afdb-uniprot50 \| AF-A0A2W5NKZ8-F1-MODEL\_V4 \| 1.0 \| 1.926e-20 \| 650 \| 0.249 \| 373 \| 188 \| 9 \| 5 \| 371 \| 15 \| 301 \| DUF3799 domain-containing protein \| DUF3799 domain-containing protein; afdb-uniprot50 \| AF-H8FP23-F1-MODEL\_V4 \| 1.0 \| 9.811e-21 \| 650 \| 0.248 \| 391 \| 177 \| 13 \| 1 \| 371 \| 63 \| 356 \| DUF3799 domain-containing protein \| DUF3799 domain-containing protein; afdb-uniprot50 \| AF-A0A7Y5UCT3-F1-MODEL\_V4 \| 1.0 \| 1.462e-18 \| 649 \| 0.243 \| 365 \| 167 \| 13 \| 1 \| 361 \| 3 \| 262 \| DUF3799 domain-containing protein \| DUF3799 domain-containing protein; afdb-uniprot50 \| AF-A0A661DE24-F1-MODEL\_V4 \| 1.0 \| 1.31e-17 \| 649 \| 0.224 \| 361 \| 172 \| 8 \| 5 \| 360 \| 2 \| 259 \| DUF3799 domain-containing protein \| DUF3799 domain-containing protein; afdb-uniprot50 \| AF-A0A2E9H4D8-F1-MODEL\_V4 \| 1.0 \| 2.848e-21 \| 649 \| 0.238 \| 382 \| 213 \| 14 \| 1 \| 359 \| 47 \| 373 \| SAP domain-containing protein \| SAP domain-containing protein; afdb-uniprot50 \| AF-A0A524KUU5-F1-MODEL\_V4 \| 1.0 \| 2.877e-17 \| 648 \| 0.242 \| 342 \| 153 \| 9 \| 1 \| 336 \| 1 \| 242 \| DUF3799 domain-containing protein \| DUF3799 domain-containing protein; afdb-uniprot50 \| AF-A0A848KZI4-F1-MODEL\_V4 \| 1.0 \| 1.936e-18 \| 648 \| 0.236 \| 380 \| 171 \| 14 \| 1 \| 369 \| 12 \| 283 \| Uncharacterized protein \| Uncharacterized protein; afdb-uniprot50 \| AF-A0A5C7Q151-F1-MODEL\_V4 \| 1.0 \| 5.313e-19 \| 648 \| 0.246 \| 369 \| 196 \| 12 \| 2 \| 366 \| 7 \| 297 \| DUF3799 domain-containing protein \| DUF3799 domain-containing protein; afdb-uniprot50 \| AF-A0A447PEW2-F1-MODEL\_V4 \| 1.0 \| 1.816e-21 \| 648 \| 0.265 \| 376 \| 162 \| 12 \| 1 \| 367 \| 391 \| 661 \| Exonuclease VIII \| Exonuclease VIII; afdb-uniprot50 \| AF-A0A354UC05-F1-MODEL\_V4 \| 1.0 \| 4.021e-18 \| 647 \| 0.212 \| 353 \| 180 \| 8 \| 9 \| 357 \| 13 \| 271 \| DUF3799 domain-containing protein \| DUF3799 domain-containing protein; afdb-uniprot50 \| AF-A0A5C2FIG8-F1-MODEL\_V4 \| 1.0 \| 2.713e-18 \| 647 \| 0.219 \| 374 \| 179 \| 15 \| 4 \| 369 \| 3 \| 271 \| DUF3799 domain-containing protein \| DUF3799 domain-containing protein; afdb-uniprot50 \| AF-A0A5C7QK33-F1-MODEL\_V4 \| 1.0 \| 1.826e-19 \| 646 \| 0.232 \| 375 \| 193 \| 12 \| 1 \| 364 \| 6 \| 296 \| DUF3799 domain-containing protein \| DUF3799 domain-containing protein; afdb-uniprot50 \| AF-A0A7X0NGZ1-F1-MODEL\_V4 \| 1.0 \| 5.606e-20 \| 646 \| 0.209 \| 397 \| 216 \| 8 \| 1 \| 357 \| 34 \| 372 \| Exodeoxyribonuclease VIII \| Exodeoxyribonuclease VIII; afdb-uniprot50 \| AF-A0A164JW40-F1-MODEL\_V4 \| 1.0 \| 5.945e-19 \| 645 \| 0.238 \| 369 \| 169 \| 12 \| 9 \| 368 \| 1 \| 266 \| DUF3799 domain-containing protein \| DUF3799 domain-containing protein; afdb-uniprot50 \| AF-A0A7Y5JXA5-F1-MODEL\_V4 \| 1.0 \| 8.331e-19 \| 645 \| 0.237 \| 362 \| 173 \| 13 \| 1 \| 359 \| 5 \| 266 \| RecE \| RecE; afdb-uniprot50 \| AF-A0A0J9D3R0-F1-MODEL\_V4 \| 1.0 \| 2.552e-20 \| 645 \| 0.226 \| 406 \| 186 \| 11 \| 2 \| 371 \| 118 \| 431 \| DUF3799 domain-containing protein \| DUF3799 domain-containing protein; afdb-uniprot50 \| AF-A0A7X9ENY2-F1-MODEL\_V4 \| 1.0 \| 3.397e-18 \| 644 \| 0.241 \| 365 \| 165 \| 13 \| 3 \| 359 \| 2 \| 262 \| DUF3799 domain-containing protein \| DUF3799 domain-containing protein; afdb-uniprot50 \| AF-A0A1S2WIB5-F1-MODEL\_V4 \| 1.0 \| 5.035e-18 \| 644 \| 0.229 \| 374 \| 181 \| 12 \| 2 \| 369 \| 8 \| 280 \| RecE \| RecE; afdb-uniprot50 \| AF-A0A7X7ZLD9-F1-MODEL\_V4 \| 1.0 \| 2.167e-18 \| 644 \| 0.24 \| 362 \| 172 \| 8 \| 1 \| 361 \| 11 \| 270 \| DUF3799 domain-containing protein \| DUF3799 domain-containing protein; afdb-uniprot50 \| AF-A0A022LJE0-F1-MODEL\_V4 \| 1.0 \| 4.748e-19 \| 644 \| 0.231 \| 384 \| 193 \| 15 \| 1 \| 372 \| 3 \| 296 \| DUF3799 domain-containing protein \| DUF3799 domain-containing protein; afdb-uniprot50 \| AF-A0A2D7NC52-F1-MODEL\_V4 \| 1.0 \| 1.17e-17 \| 643 \| 0.203 \| 358 \| 177 \| 10 \| 3 \| 357 \| 5 \| 257 \| DUF3799 domain-containing protein \| DUF3799 domain-containing protein; afdb-uniprot50 \| AF-A0A1N2I403-F1-MODEL\_V4 \| 1.0 \| 4.243e-19 \| 643 \| 0.219 \| 369 \| 185 \| 15 \| 1 \| 360 \| 5 \| 279 \| RecE \| RecE; afdb-uniprot50 \| AF-A0A753B4R6-F1-MODEL\_V4 \| 1.0 \| 2.545e-21 \| 643 \| 0.268 \| 376 \| 161 \| 12 \| 1 \| 367 \| 621 \| 891 \| Exodeoxyribonuclease VIII \| Exodeoxyribonuclease VIII; afdb-uniprot50 \| AF-A0A0F6SEZ4-F1-MODEL\_V4 \| 1.0 \| 9.346e-18 \| 642 \| 0.197 \| 354 \| 186 \| 8 \| 9 \| 360 \| 1 \| 258 \| Exodeoxyribonuclease VIII \| Exodeoxyribonuclease VIII; afdb-uniprot50 \| AF-A0A2T3IFT8-F1-MODEL\_V4 \| 1.0 \| 8.768e-21 \| 640 \| 0.242 \| 375 \| 209 \| 12 \| 1 \| 361 \| 89 \| 402 \| DUF3799 domain-containing protein \| DUF3799 domain-containing protein; afdb-uniprot50 \| AF-D2U156-F1-MODEL\_V4 \| 1.0 \| 8.352e-18 \| 639 \| 0.193 \| 366 \| 190 \| 9 \| 1 \| 363 \| 1 \| 264 \| Exodeoxyribonuclease 8 \| Exodeoxyribonuclease 8; afdb-uniprot50 \| AF-N6YA75-F1-MODEL\_V4 \| 1.0 \| 5.313e-19 \| 639 \| 0.243 \| 358 \| 159 \| 9 \| 8 \| 360 \| 43 \| 293 \| DUF3799 domain-containing protein \| DUF3799 domain-containing protein; afdb-uniprot50 \| AF-A0A2D8W508-F1-MODEL\_V4 \| 1.0 \| 1.73e-18 \| 639 \| 0.225 \| 381 \| 186 \| 11 \| 2 \| 372 \| 15 \| 296 \| DUF3799 domain-containing protein \| DUF3799 domain-containing protein; afdb-uniprot50 \| AF-A0A646KLD2-F1-MODEL\_V4 \| 1.0 \| 1.101e-19 \| 639 \| 0.259 \| 381 \| 166 \| 14 \| 1 \| 368 \| 45 \| 322 \| DUF3799 domain-containing protein \| DUF3799 domain-containing protein; afdb-uniprot50 \| AF-A0A7Y6TYS2-F1-MODEL\_V4 \| 1.0 \| 5.01e-20 \| 639 \| 0.23 \| 365 \| 176 \| 9 \| 1 \| 360 \| 109 \| 373 \| PD-(D/E)XK nuclease-like domain-containing protein \| PD-(D/E)XK nuclease-like domain-containing protein; afdb-uniprot50 \| AF-A0A7Y5TGE9-F1-MODEL\_V4 \| 1.0 \| 3.036e-18 \| 638 \| 0.223 \| 363 \| 171 \| 13 \| 1 \| 357 \| 3 \| 260 \| DUF3799 domain-containing protein \| DUF3799 domain-containing protein; afdb-uniprot50 \| AF-A0A518EYZ7-F1-MODEL\_V4 \| 1.0 \| 1.382e-18 \| 637 \| 0.184 \| 373 \| 198 \| 12 \| 2 \| 366 \| 11 \| 285 \| Exodeoxyribonuclease 8 \| Exodeoxyribonuclease 8; afdb-uniprot50 \| AF-A0A560CFR0-F1-MODEL\_V4 \| 1.0 \| 2.424e-18 \| 637 \| 0.232 \| 378 \| 180 \| 11 \| 1 \| 371 \| 7 \| 281 \| PDDEXK-like uncharacterized protein DUF3799 \| PDDEXK-like uncharacterized protein DUF3799; afdb-uniprot50 \| AF-A0A2D7BYZ1-F1-MODEL\_V4 \| 1.0 \| 2.048e-18 \| 636 \| 0.218 \| 361 \| 171 \| 10 \| 1 \| 357 \| 16 \| 269 \| DUF3799 domain-containing protein \| DUF3799 domain-containing protein; afdb-uniprot50 \| AF-A0A3M8FHL9-F1-MODEL\_V4 \| 1.0 \| 5.606e-20 \| 636 \| 0.231 \| 371 \| 214 \| 15 \| 5 \| 368 \| 13 \| 319 \| DUF3799 domain-containing protein \| DUF3799 domain-containing protein; afdb-uniprot50 \| AF-A0A2W5UA26-F1-MODEL\_V4 \| 1.0 \| 2.706e-19 \| 636 \| 0.192 \| 384 \| 242 \| 12 \| 1 \| 372 \| 21 \| 348 \| DUF3799 domain-containing protein \| DUF3799 domain-containing protein; afdb-uniprot50 \| AF-A0A8B2U0W5-F1-MODEL\_V4 \| 1.0 \| 9.251e-22 \| 635 \| 0.241 \| 397 \| 247 \| 13 \| 3 \| 358 \| 99 \| 482 \| Uncharacterized protein \| Uncharacterized protein; afdb-uniprot50 \| AF-A0A2W6WBB9-F1-MODEL\_V4 \| 1.0 \| 2.855e-20 \| 634 \| 0.246 \| 385 \| 184 \| 11 \| 1 \| 371 \| 78 \| 370 \| DUF3799 domain-containing protein \| DUF3799 domain-containing protein; afdb-uniprot50 \| AF-Q6LHC4-F1-MODEL\_V4 \| 1.0 \| 7.895e-18 \| 633 \| 0.377 \| 196 \| 111 \| 3 \| 178 \| 370 \| 26 \| 213 \| DUF3799 domain-containing protein \| DUF3799 domain-containing protein; afdb-uniprot50 \| AF-A0A7W4B1R6-F1-MODEL\_V4 \| 1.0 \| 3.773e-21 \| 633 \| 0.262 \| 381 \| 231 \| 11 \| 1 \| 357 \| 34 \| 388 \| PD-(D/E)XK nuclease-like domain-containing protein \| PD-(D/E)XK nuclease-like domain-containing protein; afdb-uniprot50 \| AF-A0A100WID5-F1-MODEL\_V4 \| 1.0 \| 7.463e-18 \| 632 \| 0.237 \| 375 \| 174 \| 17 \| 2 \| 366 \| 7 \| 279 \| DUF3799 domain-containing protein \| DUF3799 domain-containing protein; afdb-uniprot50 \| AF-A0A6M3LIW3-F1-MODEL\_V4 \| 1.0 \| 1.83e-18 \| 632 \| 0.22 \| 381 \| 188 \| 13 \| 1 \| 370 \| 3 \| 285 \| DUF3799 domain-containing protein \| DUF3799 domain-containing protein; afdb-uniprot50 \| AF-A0A3S9D344-F1-MODEL\_V4 \| 1.0 \| 1.382e-18 \| 632 \| 0.212 \| 382 \| 200 \| 11 \| 1 \| 371 \| 15 \| 306 \| DUF3799 domain-containing protein \| DUF3799 domain-containing protein; afdb-uniprot50 \| AF-A0A2X1UAB7-F1-MODEL\_V4 \| 1.0 \| 5.313e-19 \| 632 \| 0.225 \| 377 \| 182 \| 13 \| 1 \| 370 \| 57 \| 330 \| Gp60 protein \| Gp60 protein; afdb-uniprot50 \| AF-A0A1G4X2S6-F1-MODEL\_V4 \| 1.0 \| 7.426e-20 \| 632 \| 0.257 \| 376 \| 170 \| 13 \| 2 \| 371 \| 73 \| 345 \| PD-(D/E)XK nuclease superfamily protein \| PD-(D/E)XK nuclease superfamily protein; afdb-uniprot50 \| AF-A0A6G7XI10-F1-MODEL\_V4 \| 1.0 \| 7.895e-18 \| 630 \| 0.232 \| 375 \| 187 \| 11 \| 2 \| 371 \| 7 \| 285 \| DUF3799 domain-containing protein \| DUF3799 domain-containing protein; afdb-uniprot50 \| AF-A0A5R8NB53-F1-MODEL\_V4 \| 1.0 \| 9.322e-19 \| 629 \| 0.223 \| 367 \| 173 \| 13 \| 1 \| 360 \| 5 \| 266 \| DUF3799 domain-containing protein \| DUF3799 domain-containing protein; afdb-uniprot50 \| AF-A0A3N5PIH7-F1-MODEL\_V4 \| 1.0 \| 5.945e-19 \| 629 \| 0.222 \| 391 \| 183 \| 17 \| 1 \| 372 \| 6 \| 294 \| DUF3799 domain-containing protein \| DUF3799 domain-containing protein; afdb-uniprot50 \| AF-A0A7Y9DWL9-F1-MODEL\_V4 \| 1.0 \| 5.3e-20 \| 629 \| 0.217 \| 400 \| 182 \| 17 \| 1 \| 371 \| 1 \| 298 \| DUF3799 domain-containing protein \| DUF3799 domain-containing protein; afdb-uniprot50 \| AF-N1MBF6-F1-MODEL\_V4 \| 1.0 \| 3.036e-18 \| 628 \| 0.4 \| 205 \| 114 \| 4 \| 172 \| 369 \| 19 \| 221 \| DUF3799 domain-containing protein \| DUF3799 domain-containing protein; afdb-uniprot50 \| AF-A0A7W7XTG8-F1-MODEL\_V4 \| 1.0 \| 2.87e-18 \| 628 \| 0.248 \| 370 \| 168 \| 13 \| 1 \| 364 \| 11 \| 276 \| DUF3799 domain-containing protein \| DUF3799 domain-containing protein; afdb-uniprot50 \| AF-A0A142XW82-F1-MODEL\_V4 \| 1.0 \| 7.915e-17 \| 628 \| 0.206 \| 368 \| 189 \| 11 \| 10 \| 371 \| 5 \| 275 \| Exodeoxyribonuclease 8 \| Exodeoxyribonuclease 8; afdb-uniprot50 \| AF-A0A0F9IPA0-F1-MODEL\_V4 \| 1.0 \| 5.035e-18 \| 628 \| 0.198 \| 377 \| 195 \| 10 \| 1 \| 367 \| 10 \| 289 \| DUF3799 domain-containing protein \| DUF3799 domain-containing protein; afdb-uniprot50 \| AF-A0A7J5DV78-F1-MODEL\_V4 \| 1.0 \| 1.103e-18 \| 628 \| 0.255 \| 379 \| 180 \| 12 \| 1 \| 369 \| 7 \| 293 \| RecE \| RecE; afdb-uniprot50 \| AF-A0A2D8XAW4-F1-MODEL\_V4 \| 1.0 \| 4.243e-19 \| 628 \| 0.269 \| 368 \| 152 \| 12 \| 11 \| 365 \| 7 \| 270 \| DUF3799 domain-containing protein \| DUF3799 domain-containing protein; afdb-uniprot50 \| AF-W7WYY6-F1-MODEL\_V4 \| 1.0 \| 3.594e-18 \| 628 \| 0.185 \| 372 \| 188 \| 12 \| 1 \| 360 \| 13 \| 281 \| Exodeoxyribonuclease 8 \| Exodeoxyribonuclease 8; afdb-uniprot50 \| AF-A0A4D8R4H4-F1-MODEL\_V4 \| 1.0 \| 1.726e-19 \| 628 \| 0.217 \| 390 \| 194 \| 15 \| 1 \| 371 \| 7 \| 304 \| DUF3799 domain-containing protein \| DUF3799 domain-containing protein; afdb-uniprot50 \| AF-A0A7G8J5Q1-F1-MODEL\_V4 \| 1.0 \| 1.826e-19 \| 626 \| 0.25 \| 371 \| 192 \| 12 \| 2 \| 360 \| 10 \| 306 \| DUF3799 domain-containing protein \| DUF3799 domain-containing protein; afdb-uniprot50 \| AF-A0A291RDB1-F1-MODEL\_V4 \| 1.0 \| 2.706e-19 \| 625 \| 0.246 \| 373 \| 174 \| 11 \| 3 \| 370 \| 46 \| 316 \| RecE \| RecE; afdb-uniprot50 \| AF-A0A269PKN2-F1-MODEL\_V4 \| 1.0 \| 3.792e-19 \| 625 \| 0.198 \| 387 \| 223 \| 12 \| 1 \| 360 \| 1 \| 327 \| DUF3799 domain-containing protein \| DUF3799 domain-containing protein; afdb-uniprot50 \| AF-A0A6G2NMW0-F1-MODEL\_V4 \| 1.0 \| 3.38e-20 \| 625 \| 0.27 \| 377 \| 165 \| 14 \| 1 \| 368 \| 160 \| 435 \| DUF3799 domain-containing protein \| DUF3799 domain-containing protein; afdb-uniprot50 \| AF-A0A069T421-F1-MODEL\_V4 \| 1.0 \| 1.64e-17 \| 624 \| 0.213 \| 366 \| 177 \| 15 \| 1 \| 361 \| 3 \| 262 \| DUF3799 domain-containing protein \| DUF3799 domain-containing protein; afdb-uniprot50 \| AF-A0A2A9CPI8-F1-MODEL\_V4 \| 1.0 \| 3.02e-20 \| 623 \| 0.264 \| 390 \| 161 \| 18 \| 2 \| 372 \| 17 \| 299 \| PDDEXK-like uncharacterized protein DUF3799 \| PDDEXK-like uncharacterized protein DUF3799; afdb-uniprot50 \| AF-A0A5C8H0D8-F1-MODEL\_V4 \| 1.0 \| 7.426e-20 \| 623 \| 0.265 \| 377 \| 168 \| 12 \| 1 \| 368 \| 231 \| 507 \| DUF3799 domain-containing protein \| DUF3799 domain-containing protein; afdb-uniprot50 \| AF-A0A852R0H0-F1-MODEL\_V4 \| 1.0 \| 7.463e-18 \| 622 \| 0.248 \| 354 \| 163 \| 12 \| 1 \| 347 \| 5 \| 262 \| Uncharacterized protein \| Uncharacterized protein; afdb-uniprot50 \| AF-A0A3D6C748-F1-MODEL\_V4 \| 1.0 \| 2.424e-18 \| 622 \| 0.211 \| 369 \| 176 \| 13 \| 3 \| 361 \| 7 \| 270 \| Exonuclease VIII \| Exonuclease VIII; afdb-uniprot50 \| AF-A0A178MMD6-F1-MODEL\_V4 \| 1.0 \| 3.594e-18 \| 619 \| 0.223 \| 362 \| 171 \| 12 \| 18 \| 369 \| 12 \| 273 \| DUF3799 domain-containing protein \| DUF3799 domain-containing protein; afdb-uniprot50 \| AF-W5TN31-F1-MODEL\_V4 \| 1.0 \| 9.861e-19 \| 619 \| 0.228 \| 368 \| 172 \| 12 \| 1 \| 360 \| 36 \| 299 \| DUF3799 domain-containing protein \| DUF3799 domain-containing protein; afdb-uniprot50 \| AF-A0A1F8JB47-F1-MODEL\_V4 \| 1.0 \| 7.875e-19 \| 618 \| 0.22 \| 381 \| 179 \| 16 \| 1 \| 367 \| 3 \| 279 \| DUF3799 domain-containing protein \| DUF3799 domain-containing protein; afdb-uniprot50 \| AF-A0A3M2CPY5-F1-MODEL\_V4 \| 1.0 \| 1.735e-17 \| 618 \| 0.194 \| 370 \| 203 \| 9 \| 1 \| 368 \| 14 \| 290 \| DUF3799 domain-containing protein \| DUF3799 domain-containing protein; afdb-uniprot50 \| AF-A0A635X7L7-F1-MODEL\_V4 \| 1.0 \| 5.286e-21 \| 618 \| 0.208 \| 412 \| 248 \| 11 \| 27 \| 372 \| 3 \| 402 \| DNA breaking-rejoining protein \| DNA breaking-rejoining protein; afdb-uniprot50 \| AF-A0A7U6J3P3-F1-MODEL\_V4 \| 1.0 \| 5.93e-20 \| 618 \| 0.146 \| 545 \| 286 \| 19 \| 1 \| 371 \| 56 \| 595 \| DUF3799 domain-containing protein \| DUF3799 domain-containing protein; afdb-uniprot50 \| AF-A0A7X5U491-F1-MODEL\_V4 \| 1.0 \| 8.352e-18 \| 617 \| 0.22 \| 372 \| 192 \| 12 \| 9 \| 372 \| 1 \| 282 \| DUF3799 domain-containing protein \| DUF3799 domain-containing protein; afdb-uniprot50 \| AF-A0A5B8R4A6-F1-MODEL\_V4 \| 1.0 \| 1.926e-20 \| 617 \| 0.241 \| 372 \| 244 \| 13 \| 1 \| 359 \| 64 \| 410 \| DUF3799 domain-containing protein \| DUF3799 domain-containing protein; afdb-uniprot50 \| AF-A0A0F9N017-F1-MODEL\_V4 \| 1.0 \| 9.346e-18 \| 615 \| 0.218 \| 379 \| 187 \| 17 \| 2 \| 369 \| 5 \| 285 \| DUF3799 domain-containing protein \| DUF3799 domain-containing protein; afdb-uniprot50 \| AF-A0A7U2VEE6-F1-MODEL\_V4 \| 1.0 \| 1.926e-20 \| 615 \| 0.242 \| 380 \| 183 \| 12 \| 1 \| 369 \| 196 \| 481 \| DUF3799 domain-containing protein \| DUF3799 domain-containing protein; afdb-uniprot50 \| AF-A0A7L5Z1X1-F1-MODEL\_V4 \| 1.0 \| 3.036e-18 \| 614 \| 0.251 \| 362 \| 168 \| 14 \| 2 \| 361 \| 5 \| 265 \| PD-(D/E)XK nuclease-like domain-containing protein \| PD-(D/E)XK nuclease-like domain-containing protein; afdb-uniprot50 \| AF-A0A6M6JHN5-F1-MODEL\_V4 \| 1.0 \| 3.801e-18 \| 614 \| 0.214 \| 391 \| 191 \| 14 \| 1 \| 371 \| 7 \| 301 \| DUF3799 domain-containing protein \| DUF3799 domain-containing protein; afdb-uniprot50 \| AF-A0A5M8H4X0-F1-MODEL\_V4 \| 1.0 \| 3.792e-19 \| 614 \| 0.251 \| 378 \| 170 \| 13 \| 1 \| 367 \| 57 \| 332 \| DUF3799 domain-containing protein \| DUF3799 domain-containing protein; afdb-uniprot50 \| AF-A0A2D7FJB4-F1-MODEL\_V4 \| 1.0 \| 4.265e-17 \| 612 \| 0.165 \| 368 \| 208 \| 10 \| 1 \| 366 \| 14 \| 284 \| DUF3799 domain-containing protein \| DUF3799 domain-containing protein; afdb-uniprot50 \| AF-A0A0P0YX84-F1-MODEL\_V4 \| 1.0 \| 3.801e-18 \| 612 \| 0.221 \| 384 \| 190 \| 13 \| 2 \| 371 \| 16 \| 304 \| DUF3799 domain-containing protein \| DUF3799 domain-containing protein; afdb-uniprot50 \| AF-A0A7M1R1P4-F1-MODEL\_V4 \| 1.0 \| 2.713e-18 \| 611 \| 0.218 \| 380 \| 187 \| 15 \| 1 \| 371 \| 5 \| 283 \| PD-(D/E)XK nuclease-like domain-containing protein \| PD-(D/E)XK nuclease-like domain-containing protein; afdb-uniprot50 \| AF-A0A6H2A4F4-F1-MODEL\_V4 \| 1.0 \| 1.936e-18 \| 611 \| 0.217 \| 372 \| 191 \| 14 \| 2 \| 360 \| 8 \| 292 \| DUF3799 domain-containing protein \| DUF3799 domain-containing protein; afdb-uniprot50 \| AF-A0A2E7BLG2-F1-MODEL\_V4 \| 1.0 \| 1.106e-17 \| 611 \| 0.23 \| 373 \| 173 \| 12 \| 5 \| 369 \| 40 \| 306 \| DUF3799 domain-containing protein \| DUF3799 domain-containing protein; afdb-uniprot50 \| AF-A0A4Q6UDT3-F1-MODEL\_V4 \| 1.0 \| 1.462e-18 \| 611 \| 0.264 \| 375 \| 166 \| 13 \| 3 \| 368 \| 49 \| 322 \| DUF3799 domain-containing protein \| DUF3799 domain-containing protein; afdb-uniprot50 \| AF-A0A7K1LBD0-F1-MODEL\_V4 \| 1.0 \| 5.023e-19 \| 611 \| 0.242 \| 379 \| 175 \| 12 \| 1 \| 369 \| 54 \| 330 \| DUF3799 domain-containing protein \| DUF3799 domain-containing protein; afdb-uniprot50 \| AF-A0A7W5AMD2-F1-MODEL\_V4 \| 1.0 \| 1.462e-18 \| 610 \| 0.237 \| 383 \| 179 \| 17 \| 1 \| 369 \| 11 \| 294 \| DUF3799 domain-containing protein \| DUF3799 domain-containing protein; afdb-uniprot50 \| AF-A0A3D1IV21-F1-MODEL\_V4 \| 1.0 \| 1.167e-18 \| 610 \| 0.238 \| 381 \| 184 \| 14 \| 2 \| 369 \| 7 \| 294 \| DUF3799 domain-containing protein \| DUF3799 domain-containing protein; afdb-uniprot50 \| AF-A0A3B9WYA0-F1-MODEL\_V4 \| 1.0 \| 1.644e-16 \| 609 \| 0.193 \| 357 \| 179 \| 9 \| 2 \| 357 \| 4 \| 252 \| DUF3799 domain-containing protein \| DUF3799 domain-containing protein; afdb-uniprot50 \| AF-A0A1V5V1L8-F1-MODEL\_V4 \| 1.0 \| 4.511e-17 \| 607 \| 0.233 \| 359 \| 172 \| 13 \| 5 \| 357 \| 6 \| 267 \| Exonuclease VIII \| Exonuclease VIII; afdb-uniprot50 \| AF-A0A4Q3G6R8-F1-MODEL\_V4 \| 1.0 \| 4.265e-17 \| 607 \| 0.22 \| 385 \| 184 \| 14 \| 1 \| 372 \| 5 \| 286 \| DUF3799 domain-containing protein \| DUF3799 domain-containing protein; afdb-uniprot50 \| AF-A0A3C0M8N0-F1-MODEL\_V4 \| 1.0 \| 3.036e-18 \| 607 \| 0.214 \| 382 \| 186 \| 10 \| 2 \| 371 \| 14 \| 293 \| DUF3799 domain-containing protein \| DUF3799 domain-containing protein; afdb-uniprot50 \| AF-A0A7G8A5U5-F1-MODEL\_V4 \| 1.0 \| 1.631e-19 \| 607 \| 0.24 \| 375 \| 187 \| 15 \| 3 \| 371 \| 73 \| 355 \| DUF3799 domain-containing protein \| DUF3799 domain-containing protein; afdb-uniprot50 \| AF-A0A2E7BUE9-F1-MODEL\_V4 \| 1.0 \| 1.306e-18 \| 605 \| 0.215 \| 376 \| 186 \| 12 \| 5 \| 365 \| 35 \| 316 \| DUF3799 domain-containing protein \| DUF3799 domain-containing protein; afdb-uniprot50 \| AF-A0A0F9KSW2-F1-MODEL\_V4 \| 1.0 \| 2.298e-17 \| 604 \| 0.21 \| 371 \| 190 \| 11 \| 1 \| 366 \| 1 \| 273 \| DUF3799 domain-containing protein \| DUF3799 domain-containing protein; afdb-uniprot50 \| AF-A0A231HCK3-F1-MODEL\_V4 \| 1.0 \| 2.292e-18 \| 604 \| 0.23 \| 368 \| 175 \| 14 \| 1 \| 361 \| 30 \| 296 \| Exodeoxyribonuclease 8 \| Exodeoxyribonuclease 8; afdb-uniprot50 \| AF-A0A1M5MMI5-F1-MODEL\_V4 \| 1.0 \| 3.397e-18 \| 604 \| 0.222 \| 382 \| 190 \| 12 \| 2 \| 371 \| 14 \| 300 \| DUF3799 domain-containing protein \| DUF3799 domain-containing protein; afdb-uniprot50 \| AF-A0A6L5VLS5-F1-MODEL\_V4 \| 1.0 \| 7.895e-18 \| 603 \| 0.247 \| 340 \| 153 \| 9 \| 31 \| 366 \| 4 \| 244 \| DUF3799 domain-containing protein \| DUF3799 domain-containing protein; afdb-uniprot50 \| AF-A0A3B8U8D5-F1-MODEL\_V4 \| 1.0 \| 2.161e-19 \| 603 \| 0.208 \| 365 \| 221 \| 12 \| 5 \| 364 \| 3 \| 304 \| SAP domain-containing protein \| SAP domain-containing protein; afdb-uniprot50 \| AF-A0A350P1N0-F1-MODEL\_V4 \| 1.0 \| 1.941e-17 \| 600 \| 0.238 \| 369 \| 162 \| 13 \| 5 \| 361 \| 4 \| 265 \| DUF3799 domain-containing protein \| DUF3799 domain-containing protein; afdb-uniprot50 \| AF-N6UG32-F1-MODEL\_V4 \| 1.0 \| 2.713e-18 \| 600 \| 0.244 \| 377 \| 177 \| 8 \| 1 \| 366 \| 42 \| 321 \| DUF3799 domain-containing protein \| DUF3799 domain-containing protein; afdb-uniprot50 \| AF-A0A4P6HS96-F1-MODEL\_V4 \| 1.0 \| 2.713e-18 \| 600 \| 0.231 \| 376 \| 189 \| 10 \| 1 \| 366 \| 38 \| 323 \| DUF3799 domain-containing protein \| DUF3799 domain-containing protein; afdb-uniprot50 \| AF-A0A7Y0U3A1-F1-MODEL\_V4 \| 1.0 \| 2.431e-17 \| 599 \| 0.221 \| 370 \| 178 \| 13 \| 1 \| 361 \| 8 \| 276 \| DUF3799 domain-containing protein \| DUF3799 domain-containing protein; afdb-uniprot50 \| AF-A0A705UGF7-F1-MODEL\_V4 \| 1.0 \| 2.855e-20 \| 599 \| 0.209 \| 406 \| 243 \| 11 \| 33 \| 372 \| 2 \| 395 \| DNA breaking-rejoining protein \| DNA breaking-rejoining protein; afdb-uniprot50 \| AF-A0A3V4IUN9-F1-MODEL\_V4 \| 1.0 \| 4.997e-21 \| 599 \| 0.217 \| 410 \| 243 \| 14 \| 25 \| 368 \| 2 \| 399 \| DUF3799 domain-containing protein \| DUF3799 domain-containing protein; afdb-uniprot50 \| AF-A0A3D6BSW6-F1-MODEL\_V4 \| 1.0 \| 7.073e-17 \| 598 \| 0.206 \| 358 \| 170 \| 10 \| 3 \| 348 \| 10 \| 265 \| DUF3799 domain-containing protein \| DUF3799 domain-containing protein; afdb-uniprot50 \| AF-A0A0F9RJV3-F1-MODEL\_V4 \| 1.0 \| 7.073e-17 \| 598 \| 0.192 \| 363 \| 184 \| 8 \| 1 \| 361 \| 5 \| 260 \| DUF3799 domain-containing protein \| DUF3799 domain-containing protein; afdb-uniprot50 \| AF-A0A524RVX3-F1-MODEL\_V4 \| 1.0 \| 2.558e-19 \| 598 \| 0.228 \| 399 \| 210 \| 15 \| 1 \| 371 \| 102 \| 430 \| DUF3799 domain-containing protein \| DUF3799 domain-containing protein; afdb-uniprot50 \| AF-A0A2T6LPT7-F1-MODEL\_V4 \| 1.0 \| 3.792e-19 \| 597 \| 0.25 \| 387 \| 178 \| 14 \| 2 \| 371 \| 60 \| 351 \| PDDEXK-like uncharacterized protein DUF3799 \| PDDEXK-like uncharacterized protein DUF3799; afdb-uniprot50 \| AF-A0A5C7P0Q3-F1-MODEL\_V4 \| 1.0 \| 7.915e-17 \| 596 \| 0.204 \| 376 \| 175 \| 13 \| 1 \| 358 \| 2 \| 271 \| DUF3799 domain-containing protein \| DUF3799 domain-containing protein; afdb-uniprot50 \| AF-A0A2E5F7Y4-F1-MODEL\_V4 \| 1.0 \| 2.87e-18 \| 596 \| 0.231 \| 385 \| 188 \| 17 \| 1 \| 370 \| 22 \| 313 \| DUF3799 domain-containing protein \| DUF3799 domain-containing protein; afdb-uniprot50 \| AF-A0A7K3LW35-F1-MODEL\_V4 \| 1.0 \| 9.322e-19 \| 596 \| 0.223 \| 371 \| 180 \| 12 \| 2 \| 371 \| 58 \| 321 \| DUF3799 domain-containing protein \| DUF3799 domain-containing protein; afdb-uniprot50 \| AF-A0A100W6Y0-F1-MODEL\_V4 \| 1.0 \| 3.028e-19 \| 595 \| 0.22 \| 363 \| 191 \| 13 \| 1 \| 360 \| 53 \| 326 \| Gp60 \| Gp60; afdb-uniprot50 \| AF-A0A757V7J2-F1-MODEL\_V4 \| 1.0 \| 2.699e-20 \| 595 \| 0.204 \| 410 \| 250 \| 13 \| 27 \| 372 \| 2 \| 399 \| DNA breaking-rejoining protein \| DNA breaking-rejoining protein; afdb-uniprot50 \| AF-A0A7Y8WZA1-F1-MODEL\_V4 \| 1.0 \| 4.031e-17 \| 593 \| 0.226 \| 371 \| 174 \| 14 \| 1 \| 364 \| 6 \| 270 \| PD-(D/E)XK nuclease-like domain-containing protein \| PD-(D/E)XK nuclease-like domain-containing protein; afdb-uniprot50 \| AF-A0A496WM50-F1-MODEL\_V4 \| 1.0 \| 1.469e-16 \| 593 \| 0.205 \| 370 \| 203 \| 11 \| 1 \| 370 \| 1 \| 279 \| DUF3799 domain-containing protein \| DUF3799 domain-containing protein; afdb-uniprot50 \| AF-A0A7Y6MP93-F1-MODEL\_V4 \| 1.0 \| 5.62e-19 \| 593 \| 0.174 \| 384 \| 240 \| 12 \| 5 \| 369 \| 639 \| 964 \| PD-(D/E)XK nuclease-like domain-containing protein \| PD-(D/E)XK nuclease-like domain-containing protein; afdb-uniprot50 \| AF-A0A2D5TXR6-F1-MODEL\_V4 \| 1.0 \| 6.321e-17 \| 591 \| 0.219 \| 351 \| 161 \| 9 \| 1 \| 344 \| 12 \| 256 \| DUF3799 domain-containing protein \| DUF3799 domain-containing protein; afdb-uniprot50 \| AF-G2KMZ2-F1-MODEL\_V4 \| 1.0 \| 1.238e-17 \| 590 \| 0.203 \| 402 \| 220 \| 15 \| 1 \| 363 \| 20 \| 360 \| SAP domain-containing protein \| SAP domain-containing protein; afdb-uniprot50 \| AF-A0A6M3IKZ8-F1-MODEL\_V4 \| 1.0 \| 4.265e-17 \| 589 \| 0.228 \| 367 \| 174 \| 12 \| 2 \| 361 \| 9 \| 273 \| DUF3799 domain-containing protein \| DUF3799 domain-containing protein; afdb-uniprot50 \| AF-A0A5R8P622-F1-MODEL\_V4 \| 1.0 \| 6.289e-19 \| 589 \| 0.218 \| 375 \| 181 \| 16 \| 1 \| 368 \| 77 \| 346 \| DUF3799 domain-containing protein \| DUF3799 domain-containing protein; afdb-uniprot50 \| AF-A0A7K2J285-F1-MODEL\_V4 \| 1.0 \| 4.031e-17 \| 588 \| 0.371 \| 202 \| 116 \| 6 \| 177 \| 369 \| 61 \| 260 \| DUF3799 domain-containing protein \| DUF3799 domain-containing protein; afdb-uniprot50 \| AF-A0A6H1Z9N2-F1-MODEL\_V4 \| 1.0 \| 2.87e-18 \| 588 \| 0.229 \| 362 \| 210 \| 13 \| 9 \| 362 \| 1 \| 301 \| DUF3799 domain-containing protein \| DUF3799 domain-containing protein; afdb-uniprot50 \| AF-A0A1F6GLK6-F1-MODEL\_V4 \| 1.0 \| 1.378e-19 \| 587 \| 0.21 \| 408 \| 225 \| 17 \| 1 \| 372 \| 16 \| 362 \| DUF3799 domain-containing protein \| DUF3799 domain-containing protein; afdb-uniprot50 \| AF-A0A359JZU9-F1-MODEL\_V4 \| 1.0 \| 8.309e-20 \| 587 \| 0.218 \| 380 \| 230 \| 14 \| 3 \| 359 \| 35 \| 370 \| SAP domain-containing protein \| SAP domain-containing protein; afdb-uniprot50 \| AF-A0A2E1XRD1-F1-MODEL\_V4 \| 1.0 \| 3.811e-17 \| 586 \| 0.193 \| 377 \| 189 \| 13 \| 2 \| 367 \| 5 \| 277 \| DUF3799 domain-containing protein \| DUF3799 domain-containing protein; afdb-uniprot50 \| AF-A0A161XAN9-F1-MODEL\_V4 \| 1.0 \| 5.023e-19 \| 586 \| 0.235 \| 391 \| 190 \| 12 \| 2 \| 371 \| 13 \| 315 \| Exodeoxyribonuclease 8 \| Exodeoxyribonuclease 8; afdb-uniprot50 \| AF-A0A8B3WJL6-F1-MODEL\_V4 \| 1.0 \| 1.389e-16 \| 585 \| 0.2 \| 374 \| 190 \| 13 \| 1 \| 365 \| 5 \| 278 \| RecE \| RecE; afdb-uniprot50 \| AF-A0A853KWC9-F1-MODEL\_V4 \| 1.0 \| 1.936e-18 \| 585 \| 0.206 \| 397 \| 219 \| 14 \| 1 \| 361 \| 39 \| 375 \| Uncharacterized protein \| Uncharacterized protein; afdb-uniprot50 \| AF-A0A2W1SGM9-F1-MODEL\_V4 \| 1.0 \| 5.62e-19 \| 585 \| 0.247 \| 372 \| 180 \| 11 \| 1 \| 367 \| 110 \| 386 \| DUF3799 domain-containing protein \| DUF3799 domain-containing protein; afdb-uniprot50 \| AF-A0A5Y7B1G5-F1-MODEL\_V4 \| 1.0 \| 1.926e-20 \| 585 \| 0.21 \| 408 \| 244 \| 13 \| 31 \| 372 \| 1 \| 396 \| Exodeoxyribonuclease \| Exodeoxyribonuclease; afdb-uniprot50 \| AF-A0A661RLU8-F1-MODEL\_V4 \| 1.0 \| 9.886e-18 \| 584 \| 0.217 \| 381 \| 184 \| 13 \| 1 \| 366 \| 8 \| 289 \| DUF3799 domain-containing protein \| DUF3799 domain-containing protein; afdb-uniprot50 \| AF-A0A2D4XKL8-F1-MODEL\_V4 \| 1.0 \| 3.228e-16 \| 583 \| 0.2 \| 354 \| 169 \| 12 \| 1 \| 352 \| 1 \| 242 \| DUF3799 domain-containing protein \| DUF3799 domain-containing protein; afdb-uniprot50 \| AF-A0A7T4IB75-F1-MODEL\_V4 \| 1.0 \| 4.265e-17 \| 583 \| 0.232 \| 365 \| 174 \| 7 \| 12 \| 366 \| 2 \| 270 \| PD-(D/E)XK nuclease-like domain-containing protein \| PD-(D/E)XK nuclease-like domain-containing protein; afdb-uniprot50 \| AF-A0A762RJE4-F1-MODEL\_V4 \| 1.0 \| 4.001e-20 \| 582 \| 0.232 \| 391 \| 218 \| 12 \| 1 \| 323 \| 12 \| 388 \| Exodeoxyribonuclease \| Exodeoxyribonuclease; afdb-uniprot50 \| AF-A0A4Y8YA29-F1-MODEL\_V4 \| 1.0 \| 3.584e-19 \| 582 \| 0.263 \| 380 \| 166 \| 16 \| 1 \| 368 \| 235 \| 512 \| DUF3799 domain-containing protein \| DUF3799 domain-containing protein; afdb-uniprot50 \| AF-A0A146GG86-F1-MODEL\_V4 \| 1.0 \| 1.385e-17 \| 581 \| 0.194 \| 380 \| 197 \| 11 \| 1 \| 370 \| 15 \| 295 \| DUF3799 domain-containing protein \| DUF3799 domain-containing protein; afdb-uniprot50 \| AF-A0A2T0JIJ1-F1-MODEL\_V4 \| 1.0 \| 3.211e-18 \| 581 \| 0.225 \| 385 \| 183 \| 17 \| 1 \| 372 \| 17 \| 299 \| PDDEXK-like uncharacterized protein DUF3799 \| PDDEXK-like uncharacterized protein DUF3799; afdb-uniprot50 \| AF-I6APA3-F1-MODEL\_V4 \| 1.0 \| 2.571e-17 \| 581 \| 0.206 \| 387 \| 204 \| 16 \| 1 \| 366 \| 8 \| 312 \| DUF3799 domain-containing protein \| DUF3799 domain-containing protein; afdb-uniprot50 \| AF-A0A6P1IM96-F1-MODEL\_V4 \| 1.0 \| 2.571e-17 \| 580 \| 0.198 \| 363 \| 183 \| 11 \| 5 \| 361 \| 33 \| 293 \| DUF3799 domain-containing protein \| DUF3799 domain-containing protein; afdb-uniprot50 \| AF-A0A2E0GW99-F1-MODEL\_V4 \| 1.0 \| 1.469e-16 \| 578 \| 0.206 \| 382 \| 184 \| 15 \| 1 \| 368 \| 1 \| 277 \| DUF3799 domain-containing protein \| DUF3799 domain-containing protein; afdb-uniprot50 \| AF-A0A3B8UEV6-F1-MODEL\_V4 \| 1.0 \| 4.488e-19 \| 578 \| 0.238 \| 386 \| 181 \| 15 \| 1 \| 371 \| 110 \| 397 \| DUF3799 domain-containing protein \| DUF3799 domain-containing protein; afdb-uniprot50 \| AF-A0A0F9QMY9-F1-MODEL\_V4 \| 1.0 \| 1.109e-16 \| 577 \| 0.211 \| 364 \| 180 \| 13 \| 9 \| 360 \| 12 \| 280 \| DUF3799 domain-containing protein \| DUF3799 domain-containing protein; afdb-uniprot50 \| AF-A0A7X5Z2D5-F1-MODEL\_V4 \| 1.0 \| 8.812e-19 \| 576 \| 0.239 \| 376 \| 178 \| 11 \| 1 \| 371 \| 124 \| 396 \| DUF3799 domain-containing protein \| DUF3799 domain-containing protein; afdb-uniprot50 \| AF-Q11ZS8-F1-MODEL\_V4 \| 1.0 \| 2.172e-17 \| 575 \| 0.196 \| 381 \| 192 \| 9 \| 2 \| 368 \| 41 \| 321 \| DUF3799 domain-containing protein \| DUF3799 domain-containing protein; afdb-uniprot50 \| AF-A0A844B7Y8-F1-MODEL\_V4 \| 1.0 \| 1.167e-18 \| 574 \| 0.191 \| 382 \| 192 \| 12 \| 3 \| 367 \| 82 \| 363 \| Uncharacterized protein \| Uncharacterized protein; afdb-uniprot50 \| AF-A0A2D6XBJ2-F1-MODEL\_V4 \| 1.0 \| 8.88e-16 \| 573 \| 0.198 \| 362 \| 191 \| 13 \| 6 \| 364 \| 2 \| 267 \| DUF3799 domain-containing protein \| DUF3799 domain-containing protein; afdb-uniprot50 \| AF-A0A7X6NUU3-F1-MODEL\_V4 \| 1.0 \| 2.172e-17 \| 573 \| 0.255 \| 384 \| 169 \| 15 \| 1 \| 371 \| 13 \| 292 \| DUF3799 domain-containing protein \| DUF3799 domain-containing protein; afdb-uniprot50 \| AF-A0A2U1AQD0-F1-MODEL\_V4 \| 1.0 \| 5.991e-16 \| 572 \| 0.2 \| 359 \| 185 \| 12 \| 5 \| 358 \| 8 \| 269 \| PDDEXK-like uncharacterized protein DUF3799 \| PDDEXK-like uncharacterized protein DUF3799; afdb-uniprot50 \| AF-T0HIE6-F1-MODEL\_V4 \| 1.0 \| 8.373e-17 \| 572 \| 0.189 \| 386 \| 196 \| 15 \| 1 \| 371 \| 19 \| 302 \| DUF3799 domain-containing protein \| DUF3799 domain-containing protein; afdb-uniprot50 \| AF-A0A5V6MQJ5-F1-MODEL\_V4 \| 1.0 \| 1.631e-19 \| 571 \| 0.212 \| 404 \| 240 \| 11 \| 35 \| 372 \| 2 \| 393 \| Exodeoxyribonuclease \| Exodeoxyribonuclease; afdb-uniprot50 \| AF-J9SAI7-F1-MODEL\_V4 \| 1.0 \| 5.945e-19 \| 570 \| 0.259 \| 370 \| 167 \| 13 \| 1 \| 369 \| 192 \| 455 \| DUF3799 domain-containing protein \| DUF3799 domain-containing protein; afdb-uniprot50 \| AF-A0A559PZG8-F1-MODEL\_V4 \| 1.0 \| 1.17e-17 \| 569 \| 0.232 \| 387 \| 204 \| 14 \| 2 \| 372 \| 14 \| 323 \| PDDEXK-like uncharacterized protein DUF3799 \| PDDEXK-like uncharacterized protein DUF3799; afdb-uniprot50 \| AF-A0A2Z5YJC4-F1-MODEL\_V4 \| 1.0 \| 4.511e-17 \| 569 \| 0.216 \| 375 \| 187 \| 12 \| 2 \| 369 \| 55 \| 329 \| DUF3799 domain-containing protein \| DUF3799 domain-containing protein; afdb-uniprot50 \| AF-A0A433FYR1-F1-MODEL\_V4 \| 1.0 \| 9.861e-19 \| 569 \| 0.238 \| 381 \| 190 \| 15 \| 3 \| 372 \| 92 \| 383 \| DUF3799 domain-containing protein \| DUF3799 domain-containing protein; afdb-uniprot50 \| AF-A0A3N0V111-F1-MODEL\_V4 \| 1.0 \| 1.554e-16 \| 568 \| 0.203 \| 349 \| 174 \| 11 \| 1 \| 346 \| 13 \| 260 \| DUF3799 domain-containing protein \| DUF3799 domain-containing protein; afdb-uniprot50 \| AF-A0A7Z8LAH0-F1-MODEL\_V4 \| 1.0 \| 2.584e-15 \| 568 \| 0.198 \| 352 \| 177 \| 8 \| 9 \| 360 \| 8 \| 254 \| DUF3799 domain-containing protein \| DUF3799 domain-containing protein; afdb-uniprot50 \| AF-A0A2S6AMK5-F1-MODEL\_V4 \| 1.0 \| 1.462e-18 \| 568 \| 0.247 \| 368 \| 164 \| 13 \| 1 \| 358 \| 80 \| 344 \| RecE \| RecE; afdb-uniprot50 \| AF-A0A5R8NEL4-F1-MODEL\_V4 \| 1.0 \| 1.167e-18 \| 568 \| 0.205 \| 374 \| 187 \| 14 \| 1 \| 368 \| 163 \| 432 \| DUF3799 domain-containing protein \| DUF3799 domain-containing protein; afdb-uniprot50 \| AF-A0A6H2A032-F1-MODEL\_V4 \| 1.0 \| 7.501e-16 \| 567 \| 0.203 \| 358 \| 170 \| 13 \| 9 \| 361 \| 1 \| 248 \| DUF3799 domain-containing protein \| DUF3799 domain-containing protein; afdb-uniprot50 \| AF-A0A4Y9QDI6-F1-MODEL\_V4 \| 1.0 \| 1.554e-16 \| 566 \| 0.222 \| 342 \| 165 \| 9 \| 32 \| 366 \| 8 \| 255 \| DUF3799 domain-containing protein \| DUF3799 domain-containing protein; afdb-uniprot50 \| AF-A0A2E1MYG3-F1-MODEL\_V4 \| 1.0 \| 1.558e-15 \| 566 \| 0.204 \| 366 \| 176 \| 14 \| 1 \| 360 \| 1 \| 257 \| DUF3799 domain-containing protein \| DUF3799 domain-containing protein; afdb-uniprot50 \| AF-A0A5C5X348-F1-MODEL\_V4 \| 1.0 \| 2.584e-15 \| 566 \| 0.18 \| 376 \| 199 \| 9 \| 1 \| 371 \| 8 \| 279 \| DUF3799 domain-containing protein \| DUF3799 domain-containing protein; afdb-uniprot50 \| AF-A0A2N2TMM3-F1-MODEL\_V4 \| 1.0 \| 6.305e-18 \| 566 \| 0.197 \| 365 \| 183 \| 11 \| 5 \| 362 \| 88 \| 349 \| DUF3799 domain-containing protein \| DUF3799 domain-containing protein; afdb-uniprot50 \| AF-W0E2G2-F1-MODEL\_V4 \| 1.0 \| 1.558e-15 \| 565 \| 0.382 \| 188 \| 106 \| 6 \| 178 \| 361 \| 22 \| 203 \| Exonuclease VIII \| Exonuclease VIII; afdb-uniprot50 \| AF-A0A5W6TLU2-F1-MODEL\_V4 \| 1.0 \| 1.84e-16 \| 565 \| 0.247 \| 340 \| 153 \| 10 \| 31 \| 366 \| 1 \| 241 \| Exonuclease VIII \| Exonuclease VIII; afdb-uniprot50 \| AF-H0A060-F1-MODEL\_V4 \| 1.0 \| 5.649e-17 \| 565 \| 0.21 \| 389 \| 188 \| 16 \| 1 \| 371 \| 1 \| 288 \| DUF3799 domain-containing protein \| DUF3799 domain-containing protein; afdb-uniprot50 \| AF-A0A7W7MM07-F1-MODEL\_V4 \| 1.0 \| 1.64e-17 \| 563 \| 0.243 \| 399 \| 170 \| 17 \| 1 \| 372 \| 6 \| 299 \| DUF3799 domain-containing protein \| DUF3799 domain-containing protein; afdb-uniprot50 \| AF-A0A0F8VZZ2-F1-MODEL\_V4 \| 1.0 \| 3.83e-15 \| 562 \| 0.396 \| 179 \| 103 \| 4 \| 189 \| 364 \| 15 \| 191 \| DUF3799 domain-containing protein \| DUF3799 domain-containing protein; afdb-uniprot50 \| AF-A0A2S4KG23-F1-MODEL\_V4 \| 1.0 \| 3.603e-17 \| 562 \| 0.197 \| 369 \| 179 \| 12 \| 5 \| 360 \| 11 \| 275 \| DUF3799 domain-containing protein \| DUF3799 domain-containing protein; afdb-uniprot50 \| AF-Q21QE7-F1-MODEL\_V4 \| 1.0 \| 7.895e-18 \| 562 \| 0.213 \| 379 \| 178 \| 16 \| 5 \| 364 \| 53 \| 330 \| DUF3799 domain-containing protein \| DUF3799 domain-containing protein; afdb-uniprot50 \| AF-A0A3D4T735-F1-MODEL\_V4 \| 1.0 \| 3.821e-16 \| 561 \| 0.254 \| 318 \| 134 \| 8 \| 53 \| 364 \| 4 \| 224 \| DUF3799 domain-containing protein \| DUF3799 domain-containing protein; afdb-uniprot50 \| AF-A0A705Y2E6-F1-MODEL\_V4 \| 1.0 \| 4.477e-20 \| 559 \| 0.224 \| 387 \| 226 \| 12 \| 1 \| 323 \| 80 \| 456 \| Exodeoxyribonuclease \| Exodeoxyribonuclease; afdb-uniprot50 \| AF-A0A3C0TWV8-F1-MODEL\_V4 \| 1.0 \| 1.31e-17 \| 558 \| 0.203 \| 383 \| 189 \| 17 \| 1 \| 368 \| 24 \| 305 \| DUF3799 domain-containing protein \| DUF3799 domain-containing protein; afdb-uniprot50 \| AF-A0A759Y1M9-F1-MODEL\_V4 \| 1.0 \| 3.575e-20 \| 557 \| 0.226 \| 389 \| 223 \| 13 \| 1 \| 323 \| 80 \| 456 \| DNA breaking-rejoining protein \| DNA breaking-rejoining protein; afdb-uniprot50 \| AF-A0A509MJM3-F1-MODEL\_V4 \| 1.0 \| 1.946e-16 \| 556 \| 0.203 \| 369 \| 185 \| 11 \| 1 \| 360 \| 6 \| 274 \| RecE \| RecE; afdb-uniprot50 \| AF-A0A845UWG1-F1-MODEL\_V4 \| 1.0 \| 1.941e-17 \| 556 \| 0.227 \| 383 \| 186 \| 16 \| 1 \| 371 \| 5 \| 289 \| Uncharacterized protein \| Uncharacterized protein; afdb-uniprot50 \| AF-A0A316RHU8-F1-MODEL\_V4 \| 1.0 \| 4.011e-19 \| 555 \| 0.218 \| 380 \| 212 \| 13 \| 1 \| 364 \| 57 \| 367 \| DUF3799 domain-containing protein \| DUF3799 domain-containing protein; afdb-uniprot50 \| AF-A0A7W4CX72-F1-MODEL\_V4 \| 1.0 \| 8.857e-17 \| 554 \| 0.223 \| 380 \| 184 \| 17 \| 1 \| 370 \| 11 \| 289 \| PD-(D/E)XK nuclease-like domain-containing protein \| PD-(D/E)XK nuclease-like domain-containing protein; afdb-uniprot50 \| AF-A0A761RJR1-F1-MODEL\_V4 \| 1.0 \| 2.855e-20 \| 554 \| 0.22 \| 385 \| 230 \| 14 \| 1 \| 323 \| 191 \| 567 \| Exodeoxyribonuclease \| Exodeoxyribonuclease; afdb-uniprot50 \| AF-A0A316RSE2-F1-MODEL\_V4 \| 1.0 \| 1.244e-15 \| 553 \| 0.2 \| 359 \| 186 \| 12 \| 5 \| 358 \| 8 \| 270 \| DUF3799 domain-containing protein \| DUF3799 domain-containing protein; afdb-uniprot50 \| AF-A0A7C5FX24-F1-MODEL\_V4 \| 1.0 \| 3.821e-16 \| 553 \| 0.189 \| 354 \| 188 \| 11 \| 5 \| 356 \| 45 \| 301 \| DUF3799 domain-containing protein \| DUF3799 domain-containing protein; afdb-uniprot50 \| AF-A0A0F9I7X6-F1-MODEL\_V4 \| 1.0 \| 1.109e-16 \| 553 \| 0.185 \| 388 \| 199 \| 14 \| 4 \| 371 \| 17 \| 307 \| DUF3799 domain-containing protein \| DUF3799 domain-containing protein; afdb-uniprot50 \| AF-A0A3R8K6A3-F1-MODEL\_V4 \| 1.0 \| 3.414e-16 \| 552 \| 0.194 \| 391 \| 193 \| 15 \| 1 \| 372 \| 9 \| 296 \| DUF3799 domain-containing protein \| DUF3799 domain-containing protein; afdb-uniprot50 \| AF-A0A2E2S9B9-F1-MODEL\_V4 \| 1.0 \| 8.394e-16 \| 552 \| 0.196 \| 382 \| 185 \| 11 \| 1 \| 364 \| 7 \| 284 \| DUF3799 domain-containing protein \| DUF3799 domain-containing protein; afdb-uniprot50 \| AF-E0MRN4-F1-MODEL\_V4 \| 1.0 \| 3.821e-16 \| 552 \| 0.17 \| 388 \| 207 \| 11 \| 2 \| 371 \| 15 \| 305 \| DUF3799 domain-containing protein \| DUF3799 domain-containing protein; afdb-uniprot50 \| AF-A0A7J5DTI8-F1-MODEL\_V4 \| 1.0 \| 2.713e-18 \| 552 \| 0.246 \| 378 \| 183 \| 10 \| 1 \| 368 \| 84 \| 369 \| RecE \| RecE; afdb-uniprot50 \| AF-A0A2D9QDN0-F1-MODEL\_V4 \| 1.0 \| 4.796e-15 \| 551 \| 0.21 \| 346 \| 161 \| 10 \| 20 \| 360 \| 8 \| 246 \| DUF3799 domain-containing protein \| DUF3799 domain-containing protein; afdb-uniprot50 \| AF-A0A2V3UAZ7-F1-MODEL\_V4 \| 1.0 \| 5.354e-16 \| 551 \| 0.187 \| 399 \| 226 \| 14 \| 1 \| 362 \| 8 \| 345 \| PDDEXK-like uncharacterized protein DUF3799 \| PDDEXK-like uncharacterized protein DUF3799; afdb-uniprot50 \| AF-A0A7V9TIF0-F1-MODEL\_V4 \| 1.0 \| 1.739e-16 \| 550 \| 0.214 \| 341 \| 163 \| 10 \| 18 \| 358 \| 27 \| 262 \| PD-(D/E)XK nuclease-like domain-containing protein \| PD-(D/E)XK nuclease-like domain-containing protein; afdb-uniprot50 \| AF-A0A163TGH6-F1-MODEL\_V4 \| 1.0 \| 6.321e-17 \| 550 \| 0.217 \| 386 \| 196 \| 14 \| 2 \| 370 \| 14 \| 310 \| DUF3799 domain-containing protein \| DUF3799 domain-containing protein; afdb-uniprot50 \| AF-A0A2A7UDS5-F1-MODEL\_V4 \| 1.0 \| 5.635e-18 \| 550 \| 0.219 \| 365 \| 174 \| 15 \| 1 \| 358 \| 149 \| 409 \| DUF3799 domain-containing protein \| DUF3799 domain-containing protein; afdb-uniprot50 \| AF-A0A2D7VYX6-F1-MODEL\_V4 \| 1.0 \| 1.392e-15 \| 548 \| 0.213 \| 356 \| 171 \| 11 \| 14 \| 361 \| 3 \| 257 \| DUF3799 domain-containing protein \| DUF3799 domain-containing protein; afdb-uniprot50 \| AF-A0A2N3FUY9-F1-MODEL\_V4 \| 1.0 \| 1.55e-17 \| 548 \| 0.205 \| 380 \| 189 \| 13 \| 1 \| 368 \| 87 \| 365 \| DUF3799 domain-containing protein \| DUF3799 domain-containing protein; afdb-uniprot50 \| AF-A0A757U2L4-F1-MODEL\_V4 \| 1.0 \| 1.101e-19 \| 547 \| 0.226 \| 389 \| 223 \| 13 \| 1 \| 323 \| 119 \| 495 \| Exodeoxyribonuclease \| Exodeoxyribonuclease; afdb-uniprot50 \| AF-A0A5C7P550-F1-MODEL\_V4 \| 1.0 \| 4.772e-17 \| 545 \| 0.206 \| 348 \| 168 \| 8 \| 19 \| 365 \| 71 \| 311 \| DUF3799 domain-containing protein \| DUF3799 domain-containing protein; afdb-uniprot50 \| AF-A0A1E4ZGR8-F1-MODEL\_V4 \| 1.0 \| 1.946e-16 \| 544 \| 0.19 \| 388 \| 201 \| 13 \| 1 \| 368 \| 6 \| 300 \| DUF3799 domain-containing protein \| DUF3799 domain-containing protein; afdb-uniprot50 \| AF-A0A0Q6ZRJ8-F1-MODEL\_V4 \| 1.0 \| 2.178e-16 \| 544 \| 0.209 \| 386 \| 196 \| 13 \| 2 \| 371 \| 16 \| 308 \| DUF3799 domain-containing protein \| DUF3799 domain-containing protein; afdb-uniprot50 \| AF-A0A3A1WJX1-F1-MODEL\_V4 \| 1.0 \| 8.352e-18 \| 544 \| 0.196 \| 387 \| 198 \| 10 \| 2 \| 371 \| 120 \| 410 \| DUF3799 domain-containing protein \| DUF3799 domain-containing protein; afdb-uniprot50 \| AF-A0A2D6XD80-F1-MODEL\_V4 \| 1.0 \| 2.298e-17 \| 543 \| 0.211 \| 355 \| 184 \| 9 \| 8 \| 359 \| 4 \| 265 \| DUF3799 domain-containing protein \| DUF3799 domain-containing protein; afdb-uniprot50 \| AF-A0A7W4AYG5-F1-MODEL\_V4 \| 1.0 \| 1.936e-18 \| 543 \| 0.212 \| 372 \| 252 \| 14 \| 1 \| 357 \| 89 \| 434 \| PD-(D/E)XK nuclease-like domain-containing protein \| PD-(D/E)XK nuclease-like domain-containing protein; afdb-uniprot50 \| AF-A0A5Q4ZYH7-F1-MODEL\_V4 \| 1.0 \| 1.546e-18 \| 543 \| 0.145 \| 548 \| 282 \| 21 \| 8 \| 371 \| 30 \| 574 \| DUF3799 domain-containing protein \| DUF3799 domain-containing protein; afdb-uniprot50 \| AF-A0A5C7PJN5-F1-MODEL\_V4 \| 1.0 \| 6.704e-16 \| 542 \| 0.21 \| 366 \| 186 \| 13 \| 5 \| 362 \| 4 \| 274 \| DUF3799 domain-containing protein \| DUF3799 domain-containing protein; afdb-uniprot50 \| AF-A0A1I5SZ79-F1-MODEL\_V4 \| 1.0 \| 5.991e-16 \| 540 \| 0.172 \| 376 \| 203 \| 15 \| 1 \| 360 \| 3 \| 286 \| DUF3799 domain-containing protein \| DUF3799 domain-containing protein; afdb-uniprot50 \| AF-A0A6L3A789-F1-MODEL\_V4 \| 1.0 \| 3.612e-16 \| 536 \| 0.198 \| 363 \| 184 \| 13 \| 8 \| 358 \| 40 \| 307 \| DUF3799 domain-containing protein \| DUF3799 domain-containing protein; afdb-uniprot50 \| AF-A0A562ZN70-F1-MODEL\_V4 \| 1.0 \| 2.172e-17 \| 536 \| 0.201 \| 363 \| 180 \| 13 \| 5 \| 360 \| 104 \| 363 \| DUF3799 domain-containing protein \| DUF3799 domain-containing protein; afdb-uniprot50 \| AF-A0A833IUC5-F1-MODEL\_V4 \| 1.0 \| 1.473e-15 \| 535 \| 0.333 \| 225 \| 138 \| 6 \| 154 \| 369 \| 4 \| 225 \| Uncharacterized protein \| Uncharacterized protein; afdb-uniprot50 \| AF-A0A846T0U2-F1-MODEL\_V4 \| 1.0 \| 7.935e-16 \| 534 \| 0.177 \| 366 \| 192 \| 9 \| 9 \| 371 \| 1 \| 260 \| Uncharacterized protein \| Uncharacterized protein; afdb-uniprot50 \| AF-A0A356J474-F1-MODEL\_V4 \| 1.0 \| 1.051e-15 \| 533 \| 0.213 \| 360 \| 181 \| 12 \| 9 \| 362 \| 30 \| 293 \| DUF3799 domain-containing protein \| DUF3799 domain-containing protein; afdb-uniprot50 \| AF-A0A846LX61-F1-MODEL\_V4 \| 1.0 \| 5.991e-16 \| 533 \| 0.195 \| 379 \| 191 \| 15 \| 1 \| 371 \| 30 \| 302 \| Uncharacterized protein \| Uncharacterized protein; afdb-uniprot50 \| AF-A0A7C1PFD3-F1-MODEL\_V4 \| 1.0 \| 3.228e-16 \| 532 \| 0.181 \| 358 \| 185 \| 14 \| 20 \| 360 \| 1 \| 267 \| DUF3799 domain-containing protein \| DUF3799 domain-containing protein; afdb-uniprot50 \| AF-A0A0G1R8M5-F1-MODEL\_V4 \| 1.0 \| 1.956e-14 \| 531 \| 0.341 \| 193 \| 122 \| 4 \| 178 \| 370 \| 20 \| 207 \| GP60 protein \| GP60 protein; afdb-uniprot50 \| AF-A0A2E5C063-F1-MODEL\_V4 \| 1.0 \| 3.83e-15 \| 531 \| 0.21 \| 342 \| 161 \| 10 \| 1 \| 338 \| 5 \| 241 \| DUF3799 domain-containing protein \| DUF3799 domain-containing protein; afdb-uniprot50 \| AF-A0A429MIW0-F1-MODEL\_V4 \| 1.0 \| 2.892e-15 \| 530 \| 0.293 \| 201 \| 127 \| 5 \| 178 \| 368 \| 22 \| 217 \| Nuclease \| Nuclease; afdb-uniprot50 \| AF-A0A7Y6AGS1-F1-MODEL\_V4 \| 1.0 \| 1.648e-15 \| 530 \| 0.302 \| 218 \| 127 \| 3 \| 157 \| 368 \| 37 \| 235 \| RecE \| RecE; afdb-uniprot50 \| AF-A0A355H9R2-F1-MODEL\_V4 \| 1.0 \| 1.844e-15 \| 529 \| 0.21 \| 342 \| 165 \| 10 \| 1 \| 340 \| 5 \| 243 \| DUF3799 domain-containing protein \| DUF3799 domain-containing protein; afdb-uniprot50 \| AF-A0A6M3J5F8-F1-MODEL\_V4 \| 1.0 \| 2.727e-16 \| 529 \| 0.209 \| 387 \| 192 \| 18 \| 7 \| 367 \| 3 \| 301 \| DUF3799 domain-containing protein \| DUF3799 domain-containing protein; afdb-uniprot50 \| AF-A0A7V1BWR4-F1-MODEL\_V4 \| 1.0 \| 1.739e-16 \| 529 \| 0.184 \| 385 \| 193 \| 15 \| 9 \| 361 \| 12 \| 307 \| DUF3799 domain-containing protein \| DUF3799 domain-containing protein; afdb-uniprot50 \| AF-A0A752A1T4-F1-MODEL\_V4 \| 1.0 \| 3.811e-17 \| 529 \| 0.199 \| 407 \| 219 \| 15 \| 5 \| 367 \| 22 \| 365 \| Exodeoxyribonuclease VIII \| Exodeoxyribonuclease VIII; afdb-uniprot50 \| AF-A0A318CJ92-F1-MODEL\_V4 \| 1.0 \| 8.415e-15 \| 528 \| 0.239 \| 359 \| 151 \| 12 \| 1 \| 357 \| 5 \| 243 \| DUF3799 domain-containing protein \| DUF3799 domain-containing protein; afdb-uniprot50 \| AF-A0A7D8UL27-F1-MODEL\_V4 \| 1.0 \| 4.042e-16 \| 528 \| 0.201 \| 357 \| 181 \| 13 \| 8 \| 358 \| 60 \| 318 \| DUF3799 domain-containing protein \| DUF3799 domain-containing protein; afdb-uniprot50 \| AF-A0A3B9NQ75-F1-MODEL\_V4 \| 1.0 \| 2.178e-16 \| 527 \| 0.192 \| 380 \| 189 \| 13 \| 8 \| 371 \| 11 \| 288 \| DUF3799 domain-containing protein \| DUF3799 domain-containing protein; afdb-uniprot50 \| AF-A0A7K1SKS8-F1-MODEL\_V4 \| 1.0 \| 1.306e-18 \| 527 \| 0.204 \| 382 \| 223 \| 14 \| 1 \| 361 \| 55 \| 376 \| DUF3799 domain-containing protein \| DUF3799 domain-containing protein; afdb-uniprot50 \| AF-A0A497PDQ5-F1-MODEL\_V4 \| 1.0 \| 1.469e-16 \| 526 \| 0.171 \| 367 \| 196 \| 13 \| 1 \| 363 \| 4 \| 266 \| DUF3799 domain-containing protein \| DUF3799 domain-containing protein; afdb-uniprot50 \| AF-A0A2E8IY15-F1-MODEL\_V4 \| 1.0 \| 3.236e-15 \| 524 \| 0.179 \| 367 \| 185 \| 13 \| 5 \| 363 \| 10 \| 268 \| DUF3799 domain-containing protein \| DUF3799 domain-containing protein; afdb-uniprot50 \| AF-A0A7R7HDE4-F1-MODEL\_V4 \| 1.0 \| 4.275e-16 \| 524 \| 0.19 \| 398 \| 228 \| 15 \| 1 \| 361 \| 7 \| 347 \| Uncharacterized protein \| Uncharacterized protein; afdb-uniprot50 \| AF-A0A3E5H9L3-F1-MODEL\_V4 \| 1.0 \| 9.346e-18 \| 524 \| 0.175 \| 381 \| 229 \| 13 \| 1 \| 361 \| 53 \| 368 \| DUF3799 domain-containing protein \| DUF3799 domain-containing protein; afdb-uniprot50 \| AF-A0A4U8SHN3-F1-MODEL\_V4 \| 1.0 \| 1.316e-15 \| 523 \| 0.198 \| 362 \| 168 \| 14 \| 9 \| 363 \| 4 \| 250 \| DUF3799 domain-containing protein \| DUF3799 domain-containing protein; afdb-uniprot50 \| AF-A0A6P1CUJ9-F1-MODEL\_V4 \| 1.0 \| 5.96e-18 \| 523 \| 0.217 \| 368 \| 176 \| 13 \| 1 \| 361 \| 5 \| 267 \| DUF3799 domain-containing protein \| DUF3799 domain-containing protein; afdb-uniprot50 \| AF-A0A501WTJ5-F1-MODEL\_V4 \| 1.0 \| 1.389e-16 \| 522 \| 0.209 \| 387 \| 182 \| 17 \| 1 \| 368 \| 4 \| 285 \| DUF3799 domain-containing protein \| DUF3799 domain-containing protein; afdb-uniprot50 \| AF-A0A0Q6C361-F1-MODEL\_V4 \| 1.0 \| 8.88e-16 \| 522 \| 0.174 \| 390 \| 216 \| 13 \| 2 \| 371 \| 14 \| 317 \| DUF3799 domain-containing protein \| DUF3799 domain-containing protein; afdb-uniprot50 \| AF-A0A2D7EEB4-F1-MODEL\_V4 \| 1.0 \| 1.051e-15 \| 522 \| 0.17 \| 386 \| 224 \| 16 \| 1 \| 371 \| 13 \| 317 \| DUF3799 domain-containing protein \| DUF3799 domain-containing protein; afdb-uniprot50 \| AF-A0A521STB6-F1-MODEL\_V4 \| 1.0 \| 3.236e-15 \| 521 \| 0.18 \| 354 \| 180 \| 10 \| 9 \| 359 \| 1 \| 247 \| DUF3799 domain-containing protein \| DUF3799 domain-containing protein; afdb-uniprot50 \| AF-A0A6M3JLX9-F1-MODEL\_V4 \| 1.0 \| 7.091e-16 \| 521 \| 0.191 \| 386 \| 194 \| 13 \| 3 \| 366 \| 7 \| 296 \| DUF3799 domain-containing protein \| DUF3799 domain-containing protein; afdb-uniprot50 \| AF-A0A5M6INY5-F1-MODEL\_V4 \| 1.0 \| 3.051e-16 \| 520 \| 0.181 \| 386 \| 195 \| 17 \| 2 \| 369 \| 7 \| 289 \| DUF3799 domain-containing protein \| DUF3799 domain-containing protein; afdb-uniprot50 \| AF-A0A0Q6PBB0-F1-MODEL\_V4 \| 1.0 \| 2.178e-16 \| 518 \| 0.177 \| 400 \| 210 \| 16 \| 1 \| 371 \| 21 \| 330 \| DUF3799 domain-containing protein \| DUF3799 domain-containing protein; afdb-uniprot50 \| AF-A0A354GE15-F1-MODEL\_V4 \| 1.0 \| 4.052e-15 \| 517 \| 0.193 \| 372 \| 175 \| 15 \| 3 \| 360 \| 2 \| 262 \| DUF3799 domain-containing protein \| DUF3799 domain-containing protein; afdb-uniprot50 \| AF-A0A2W6TFB9-F1-MODEL\_V4 \| 1.0 \| 9.911e-17 \| 517 \| 0.206 \| 388 \| 206 \| 15 \| 2 \| 371 \| 45 \| 348 \| DUF3799 domain-containing protein \| DUF3799 domain-containing protein; afdb-uniprot50 \| AF-A0A261QAX9-F1-MODEL\_V4 \| 1.0 \| 4.021e-18 \| 517 \| 0.212 \| 381 \| 220 \| 14 \| 1 \| 361 \| 62 \| 382 \| DUF3799 domain-containing protein \| DUF3799 domain-containing protein; afdb-uniprot50 \| AF-A0A7K0K0E5-F1-MODEL\_V4 \| 1.0 \| 2.183e-15 \| 516 \| 0.247 \| 371 \| 164 \| 18 \| 1 \| 360 \| 16 \| 282 \| DUF3799 domain-containing protein \| DUF3799 domain-containing protein; afdb-uniprot50 \| AF-F4SH48-F1-MODEL\_V4 \| 1.0 \| 7.875e-19 \| 516 \| 0.204 \| 382 \| 231 \| 11 \| 54 \| 372 \| 2 \| 373 \| Exodeoxyribonuclease VIII \| Exodeoxyribonuclease VIII; afdb-uniprot50 \| AF-A0A8B5WVL2-F1-MODEL\_V4 \| 1.0 \| 4.772e-17 \| 515 \| 0.225 \| 408 \| 173 \| 20 \| 1 \| 371 \| 7 \| 308 \| Uncharacterized protein \| Uncharacterized protein; afdb-uniprot50 \| AF-A0A447CPW6-F1-MODEL\_V4 \| 1.0 \| 5.663e-16 \| 514 \| 0.169 \| 383 \| 202 \| 15 \| 1 \| 367 \| 5 \| 287 \| DUF3799 domain-containing protein \| DUF3799 domain-containing protein; afdb-uniprot50 \| AF-A0A850CSY3-F1-MODEL\_V4 \| 1.0 \| 2.315e-14 \| 511 \| 0.319 \| 188 \| 118 \| 6 \| 179 \| 361 \| 24 \| 206 \| Uncharacterized protein \| Uncharacterized protein; afdb-uniprot50 \| AF-A0A844H0G1-F1-MODEL\_V4 \| 1.0 \| 6.337e-16 \| 510 \| 0.209 \| 386 \| 192 \| 18 \| 1 \| 371 \| 5 \| 292 \| Uncharacterized protein \| Uncharacterized protein; afdb-uniprot50 \| AF-A0A6M3XHK2-F1-MODEL\_V4 \| 1.0 \| 4.275e-16 \| 509 \| 0.202 \| 381 \| 203 \| 15 \| 1 \| 366 \| 28 \| 322 \| DUF3799 domain-containing protein \| DUF3799 domain-containing protein; afdb-uniprot50 \| AF-A0A3R6CYF7-F1-MODEL\_V4 \| 1.0 \| 6.305e-18 \| 509 \| 0.17 \| 392 \| 230 \| 14 \| 1 \| 361 \| 63 \| 390 \| DUF3799 domain-containing protein \| DUF3799 domain-containing protein; afdb-uniprot50 \| AF-A0A843GNI8-F1-MODEL\_V4 \| 1.0 \| 2.183e-15 \| 508 \| 0.143 \| 363 \| 205 \| 14 \| 1 \| 361 \| 10 \| 268 \| PD-(D/E)XK nuclease-like domain-containing protein \| PD-(D/E)XK nuclease-like domain-containing protein; afdb-uniprot50 \| AF-A0A5X8YNM2-F1-MODEL\_V4 \| 1.0 \| 6.006e-15 \| 507 \| 0.247 \| 315 \| 133 \| 11 \| 57 \| 367 \| 2 \| 216 \| Exonuclease VIII \| Exonuclease VIII; afdb-uniprot50 \| AF-A0A5M6IB30-F1-MODEL\_V4 \| 1.0 \| 3.414e-16 \| 507 \| 0.206 \| 382 \| 188 \| 13 \| 2 \| 367 \| 18 \| 300 \| DUF3799 domain-containing protein \| DUF3799 domain-containing protein; afdb-uniprot50 \| AF-A0A369QT25-F1-MODEL\_V4 \| 1.0 \| 5.991e-16 \| 507 \| 0.226 \| 384 \| 186 \| 17 \| 1 \| 371 \| 47 \| 332 \| DUF3799 domain-containing protein \| DUF3799 domain-containing protein; afdb-uniprot50 \| AF-A0A2D7PF36-F1-MODEL\_V4 \| 1.0 \| 4.534e-15 \| 506 \| 0.207 \| 376 \| 168 \| 15 \| 5 \| 359 \| 20 \| 286 \| DUF3799 domain-containing protein \| DUF3799 domain-containing protein; afdb-uniprot50 \| AF-A0A744K618-F1-MODEL\_V4 \| 1.0 \| 9.417e-15 \| 505 \| 0.242 \| 313 \| 134 \| 9 \| 58 \| 366 \| 2 \| 215 \| Exonuclease VIII \| Exonuclease VIII; afdb-uniprot50 \| AF-A0A7C7TML4-F1-MODEL\_V4 \| 1.0 \| 6.738e-14 \| 505 \| 0.161 \| 341 \| 166 \| 10 \| 1 \| 338 \| 8 \| 231 \| DUF3799 domain-containing protein \| DUF3799 domain-containing protein; afdb-uniprot50 \| AF-A0A1G0HWF1-F1-MODEL\_V4 \| 1.0 \| 1.84e-16 \| 505 \| 0.209 \| 372 \| 190 \| 17 \| 12 \| 368 \| 2 \| 284 \| DUF3799 domain-containing protein \| DUF3799 domain-containing protein; afdb-uniprot50 \| AF-A0A317H245-F1-MODEL\_V4 \| 1.0 \| 1.051e-15 \| 505 \| 0.213 \| 393 \| 196 \| 16 \| 2 \| 371 \| 18 \| 320 \| DUF3799 domain-containing protein \| DUF3799 domain-containing protein; afdb-uniprot50 \| AF-A0A7W6J8M0-F1-MODEL\_V4 \| 1.0 \| 3.414e-16 \| 504 \| 0.197 \| 395 \| 190 \| 16 \| 1 \| 367 \| 11 \| 306 \| DUF3799 domain-containing protein \| DUF3799 domain-containing protein; afdb-uniprot50 \| AF-A0A8B3HGI6-F1-MODEL\_V4 \| 1.0 \| 8.437e-14 \| 503 \| 0.337 \| 181 \| 108 \| 6 \| 190 \| 366 \| 1 \| 173 \| Uncharacterized protein \| Uncharacterized protein; afdb-uniprot50 \| AF-A0A1Z7ZMG2-F1-MODEL\_V4 \| 1.0 \| 1.247e-14 \| 502 \| 0.194 \| 376 \| 188 \| 13 \| 1 \| 371 \| 13 \| 278 \| DUF3799 domain-containing protein \| DUF3799 domain-containing protein; afdb-uniprot50 \| AF-A0A0S9D1Q2-F1-MODEL\_V4 \| 1.0 \| 1.247e-14 \| 501 \| 0.279 \| 326 \| 129 \| 12 \| 43 \| 364 \| 2 \| 225 \| DUF3799 domain-containing protein \| DUF3799 domain-containing protein; afdb-uniprot50 \| AF-A0A3A0DGM8-F1-MODEL\_V4 \| 1.0 \| 1.051e-15 \| 500 \| 0.205 \| 355 \| 181 \| 12 \| 8 \| 358 \| 233 \| 490 \| Uncharacterized protein \| Uncharacterized protein; afdb-uniprot50 \| AF-S2W335-F1-MODEL\_V4 \| 1.0 \| 5.074e-15 \| 498 \| 0.191 \| 361 \| 192 \| 13 \| 2 \| 361 \| 3 \| 264 \| DUF3799 domain-containing protein \| DUF3799 domain-containing protein; afdb-uniprot50 \| AF-A0A6N6ZVV7-F1-MODEL\_V4 \| 1.0 \| 1.316e-15 \| 498 \| 0.192 \| 395 \| 193 \| 15 \| 2 \| 369 \| 4 \| 299 \| DUF3799 domain-containing protein \| DUF3799 domain-containing protein; afdb-uniprot50 \| AF-A0A3N0CLS5-F1-MODEL\_V4 \| 1.0 \| 3.83e-15 \| 497 \| 0.201 \| 368 \| 190 \| 15 \| 2 \| 363 \| 12 \| 281 \| DUF3799 domain-containing protein \| DUF3799 domain-containing protein; afdb-uniprot50 \| AF-A0A7U2KNN2-F1-MODEL\_V4 \| 1.0 \| 1.179e-14 \| 497 \| 0.177 \| 384 \| 222 \| 12 \| 1 \| 371 \| 19 \| 321 \| PD-(D/E)XK nuclease-like domain-containing protein \| PD-(D/E)XK nuclease-like domain-containing protein; afdb-uniprot50 \| AF-A0A2D7EZG3-F1-MODEL\_V4 \| 1.0 \| 2.315e-14 \| 496 \| 0.236 \| 355 \| 159 \| 13 \| 5 \| 354 \| 6 \| 253 \| DUF3799 domain-containing protein \| DUF3799 domain-containing protein; afdb-uniprot50 \| AF-A0A1H5JK49-F1-MODEL\_V4 \| 1.0 \| 3.236e-15 \| 496 \| 0.204 \| 387 \| 189 \| 18 \| 1 \| 371 \| 4 \| 287 \| DUF3799 domain-containing protein \| DUF3799 domain-containing protein; afdb-uniprot50 \| AF-A0A7V4U9Y0-F1-MODEL\_V4 \| 1.0 \| 1.844e-15 \| 496 \| 0.202 \| 356 \| 182 \| 12 \| 8 \| 358 \| 78 \| 336 \| DUF3799 domain-containing protein \| DUF3799 domain-containing protein; afdb-uniprot50 \| AF-A0A373FBR7-F1-MODEL\_V4 \| 1.0 \| 4.052e-15 \| 495 \| 0.162 \| 375 \| 187 \| 13 \| 5 \| 360 \| 7 \| 273 \| DUF3799 domain-containing protein \| DUF3799 domain-containing protein; afdb-uniprot50 \| AF-A0A5C0AA35-F1-MODEL\_V4 \| 1.0 \| 2.309e-15 \| 495 \| 0.179 \| 385 \| 195 \| 17 \| 1 \| 369 \| 6 \| 285 \| DUF3799 domain-containing protein \| DUF3799 domain-containing protein; afdb-uniprot50 \| AF-A0A7Y0EXA6-F1-MODEL\_V4 \| 1.0 \| 4.796e-15 \| 493 \| 0.201 \| 383 \| 182 \| 15 \| 5 \| 371 \| 4 \| 278 \| DUF3799 domain-containing protein \| DUF3799 domain-containing protein; afdb-uniprot50 \| AF-A0A4Q8QN88-F1-MODEL\_V4 \| 1.0 \| 2.069e-14 \| 493 \| 0.189 \| 385 \| 201 \| 17 \| 1 \| 369 \| 12 \| 301 \| DUF3799 domain-containing protein \| DUF3799 domain-containing protein; afdb-uniprot50 \| AF-A0A509JFD6-F1-MODEL\_V4 \| 1.0 \| 2.178e-16 \| 492 \| 0.166 \| 390 \| 205 \| 11 \| 1 \| 366 \| 1 \| 294 \| Uncharacterized protein \| Uncharacterized protein; afdb-uniprot50 \| AF-A0A5C7PU20-F1-MODEL\_V4 \| 1.0 \| 1.32e-14 \| 491 \| 0.188 \| 361 \| 193 \| 13 \| 6 \| 361 \| 12 \| 277 \| DUF3799 domain-containing protein \| DUF3799 domain-containing protein; afdb-uniprot50 \| AF-A0A2D7YTB8-F1-MODEL\_V4 \| 1.0 \| 1.752e-13 \| 491 \| 0.176 \| 375 \| 192 \| 15 \| 1 \| 361 \| 4 \| 275 \| DUF3799 domain-containing protein \| DUF3799 domain-containing protein; afdb-uniprot50 \| AF-A0A097ENH1-F1-MODEL\_V4 \| 1.0 \| 4.297e-14 \| 488 \| 0.196 \| 366 \| 178 \| 15 \| 14 \| 370 \| 3 \| 261 \| DUF3799 domain-containing protein \| DUF3799 domain-containing protein; afdb-uniprot50 \| AF-A0A7C1JNN6-F1-MODEL\_V4 \| 1.0 \| 1.854e-13 \| 487 \| 0.192 \| 333 \| 170 \| 11 \| 25 \| 356 \| 2 \| 236 \| DUF3799 domain-containing protein \| DUF3799 domain-containing protein; afdb-uniprot50 \| AF-Q4LBT7-F1-MODEL\_V4 \| 1.0 \| 4.275e-16 \| 487 \| 0.24 \| 375 \| 174 \| 14 \| 5 \| 371 \| 292 \| 563 \| RecE protein \| RecE protein; afdb-uniprot50 \| AF-A0A3S0D0M8-F1-MODEL\_V4 \| 1.0 \| 1.056e-13 \| 482 \| 0.149 \| 402 \| 201 \| 14 \| 1 \| 369 \| 6 \| 299 \| Uncharacterized protein \| Uncharacterized protein; afdb-uniprot50 \| AF-A0A5B8CGE1-F1-MODEL\_V4 \| 1.0 \| 1.562e-14 \| 482 \| 0.201 \| 382 \| 192 \| 15 \| 1 \| 369 \| 38 \| 319 \| DUF3799 domain-containing protein \| DUF3799 domain-containing protein; afdb-uniprot50 \| AF-A0A077NJB7-F1-MODEL\_V4 \| 1.0 \| 1.059e-12 \| 480 \| 0.264 \| 174 \| 118 \| 4 \| 194 \| 361 \| 10 \| 179 \| Phage-related exonuclease \| Phage-related exonuclease; afdb-uniprot50 \| AF-A0A318MLD0-F1-MODEL\_V4 \| 1.0 \| 2.069e-14 \| 480 \| 0.189 \| 386 \| 186 \| 15 \| 1 \| 369 \| 1 \| 276 \| DUF3799 domain-containing protein \| DUF3799 domain-containing protein; afdb-uniprot50 \| AF-A0A6L9STJ4-F1-MODEL\_V4 \| 1.0 \| 3.83e-15 \| 479 \| 0.161 \| 378 \| 188 \| 15 \| 1 \| 361 \| 1 \| 266 \| DUF3799 domain-containing protein \| DUF3799 domain-containing protein; afdb-uniprot50 \| AF-A0A3M1NIT7-F1-MODEL\_V4 \| 1.0 \| 1.251e-13 \| 478 \| 0.172 \| 359 \| 194 \| 11 \| 5 \| 361 \| 2 \| 259 \| DUF3799 domain-containing protein \| DUF3799 domain-containing protein; afdb-uniprot50 \| AF-A0A437M7Z1-F1-MODEL\_V4 \| 1.0 \| 6.353e-15 \| 478 \| 0.162 \| 413 \| 256 \| 14 \| 1 \| 370 \| 44 \| 409 \| Uncharacterized protein \| Uncharacterized protein; afdb-uniprot50 \| AF-A0A3G2V3T1-F1-MODEL\_V4 \| 1.0 \| 1.389e-16 \| 476 \| 0.188 \| 408 \| 219 \| 22 \| 1 \| 361 \| 15 \| 357 \| DUF3799 domain-containing protein \| DUF3799 domain-containing protein; afdb-uniprot50 \| AF-A0A517YVS6-F1-MODEL\_V4 \| 1.0 \| 2.069e-14 \| 475 \| 0.17 \| 382 \| 192 \| 13 \| 1 \| 364 \| 21 \| 295 \| Exodeoxyribonuclease 8 \| Exodeoxyribonuclease 8; afdb-uniprot50 \| AF-A0A8A5F1F3-F1-MODEL\_V4 \| 1.0 \| 4.052e-15 \| 475 \| 0.196 \| 387 \| 199 \| 18 \| 1 \| 364 \| 10 \| 307 \| PD-(D/E)XK nuclease-like domain-containing protein \| PD-(D/E)XK nuclease-like domain-containing protein; afdb-uniprot50 \| AF-A0A843HS31-F1-MODEL\_V4 \| 1.0 \| 7.127e-14 \| 474 \| 0.156 \| 377 \| 200 \| 14 \| 5 \| 366 \| 2 \| 275 \| PD-(D/E)XK nuclease-like domain-containing protein \| PD-(D/E)XK nuclease-like domain-containing protein; afdb-uniprot50 \| AF-A0A5Y1VD64-F1-MODEL\_V4 \| 1.0 \| 1.12e-12 \| 473 \| 0.339 \| 171 \| 101 \| 6 \| 200 \| 366 \| 2 \| 164 \| Exonuclease VIII \| Exonuclease VIII; afdb-uniprot50 \| AF-A0A2A5D9T2-F1-MODEL\_V4 \| 1.0 \| 2.747e-13 \| 473 \| 0.182 \| 368 \| 187 \| 16 \| 1 \| 358 \| 10 \| 273 \| DUF3799 domain-containing protein \| DUF3799 domain-containing protein; afdb-uniprot50 \| AF-A0A358N4E2-F1-MODEL\_V4 \| 1.0 \| 2.189e-14 \| 473 \| 0.181 \| 363 \| 191 \| 14 \| 10 \| 360 \| 3 \| 271 \| DUF3799 domain-containing protein \| DUF3799 domain-containing protein; afdb-uniprot50 \| AF-A0A1V5IXR4-F1-MODEL\_V4 \| 1.0 \| 8.925e-14 \| 473 \| 0.193 \| 372 \| 181 \| 14 \| 6 \| 355 \| 21 \| 295 \| Exonuclease VIII \| Exonuclease VIII; afdb-uniprot50 \| AF-A0A6I0ESC0-F1-MODEL\_V4 \| 1.0 \| 2.584e-15 \| 473 \| 0.213 \| 356 \| 178 \| 12 \| 8 \| 358 \| 156 \| 414 \| DUF3799 domain-containing protein \| DUF3799 domain-containing protein; afdb-uniprot50 \| AF-R5BQV7-F1-MODEL\_V4 \| 1.0 \| 3.228e-16 \| 472 \| 0.167 \| 381 \| 235 \| 19 \| 6 \| 355 \| 2 \| 331 \| DUF3799 domain-containing protein \| DUF3799 domain-containing protein; afdb-uniprot50 \| AF-A0A1H4NL54-F1-MODEL\_V4 \| 1.0 \| 2.892e-15 \| 471 \| 0.16 \| 411 \| 235 \| 16 \| 1 \| 360 \| 16 \| 367 \| DUF3799 domain-containing protein \| DUF3799 domain-containing protein; afdb-uniprot50 \| AF-A0A5N9VVT2-F1-MODEL\_V4 \| 1.0 \| 1.484e-12 \| 469 \| 0.349 \| 169 \| 98 \| 6 \| 194 \| 358 \| 1 \| 161 \| DUF3799 domain-containing protein \| DUF3799 domain-containing protein; afdb-uniprot50 \| AF-A0A0Q6CAS4-F1-MODEL\_V4 \| 1.0 \| 3.059e-15 \| 469 \| 0.178 \| 387 \| 195 \| 19 \| 1 \| 367 \| 14 \| 297 \| DUF3799 domain-containing protein \| DUF3799 domain-containing protein; afdb-uniprot50 \| AF-A0A0K1H0Q4-F1-MODEL\_V4 \| 1.0 \| 3.228e-16 \| 469 \| 0.206 \| 377 \| 185 \| 13 \| 5 \| 371 \| 416 \| 688 \| Exodeoxyribonuclease VIII \| Exodeoxyribonuclease VIII; afdb-uniprot50 \| AF-A0A5V0VKT4-F1-MODEL\_V4 \| 1.0 \| 2.315e-14 \| 468 \| 0.232 \| 293 \| 191 \| 7 \| 90 \| 372 \| 2 \| 270 \| DUF3799 domain-containing protein \| DUF3799 domain-containing protein; afdb-uniprot50 \| AF-A0A5C7PQS1-F1-MODEL\_V4 \| 1.0 \| 4.808e-14 \| 468 \| 0.184 \| 364 \| 188 \| 14 \| 9 \| 360 \| 1 \| 267 \| DUF3799 domain-containing protein \| DUF3799 domain-containing protein; afdb-uniprot50 \| AF-A0A1F9UXV9-F1-MODEL\_V4 \| 1.0 \| 6.369e-14 \| 468 \| 0.157 \| 363 \| 176 \| 11 \| 17 \| 359 \| 8 \| 260 \| DUF3799 domain-containing protein \| DUF3799 domain-containing protein; afdb-uniprot50 \| AF-A0A381SAS3-F1-MODEL\_V4 \| 1.0 \| 1.326e-12 \| 465 \| 0.21 \| 290 \| 132 \| 8 \| 52 \| 340 \| 2 \| 195 \| DUF3799 domain-containing protein \| DUF3799 domain-containing protein; afdb-uniprot50 \| AF-A0A2U1TII1-F1-MODEL\_V4 \| 1.0 \| 8.902e-15 \| 465 \| 0.222 \| 288 \| 180 \| 7 \| 98 \| 370 \| 2 \| 260 \| Exodeoxyribonuclease VIII \| Exodeoxyribonuclease VIII; afdb-uniprot50 \| AF-A0A0Q4WZ62-F1-MODEL\_V4 \| 1.0 \| 6.721e-15 \| 465 \| 0.181 \| 386 \| 193 \| 20 \| 2 \| 367 \| 13 \| 295 \| DUF3799 domain-containing protein \| DUF3799 domain-containing protein; afdb-uniprot50 \| AF-A0A7R6V9M0-F1-MODEL\_V4 \| 1.0 \| 1.323e-13 \| 463 \| 0.172 \| 354 \| 185 \| 13 \| 9 \| 358 \| 1 \| 250 \| Exodeoxyribonuclease VIII \| Exodeoxyribonuclease VIII; afdb-uniprot50 \| AF-A0A4P7L2A5-F1-MODEL\_V4 \| 1.0 \| 3.236e-15 \| 463 \| 0.213 \| 374 \| 183 \| 13 \| 5 \| 371 \| 254 \| 523 \| Exodeoxyribonuclease 8 \| Exodeoxyribonuclease 8; afdb-uniprot50 \| AF-A0A0Q8VDF8-F1-MODEL\_V4 \| 1.0 \| 4.083e-12 \| 462 \| 0.445 \| 146 \| 75 \| 2 \| 229 \| 370 \| 2 \| 145 \| DUF3799 domain-containing protein \| DUF3799 domain-containing protein; afdb-uniprot50 \| AF-A0A6P1PXM4-F1-MODEL\_V4 \| 1.0 \| 6.353e-15 \| 462 \| 0.232 \| 378 \| 164 \| 15 \| 1 \| 354 \| 1 \| 276 \| Exodeoxyribonuclease 8 \| Exodeoxyribonuclease 8; afdb-uniprot50 \| AF-A0A8B3TCB0-F1-MODEL\_V4 \| 1.0 \| 1.477e-14 \| 461 \| 0.157 \| 393 \| 207 \| 14 \| 1 \| 371 \| 19 \| 309 \| Uncharacterized protein \| Uncharacterized protein; afdb-uniprot50 \| AF-A0A0F9N5B6-F1-MODEL\_V4 \| 1.0 \| 8.902e-15 \| 459 \| 0.183 \| 403 \| 203 \| 19 \| 3 \| 369 \| 4 \| 316 \| DUF3799 domain-containing protein \| DUF3799 domain-containing protein; afdb-uniprot50 \| AF-A0A841AAK3-F1-MODEL\_V4 \| 1.0 \| 4.796e-15 \| 459 \| 0.203 \| 388 \| 186 \| 21 \| 2 \| 367 \| 13 \| 299 \| Uncharacterized protein \| Uncharacterized protein; afdb-uniprot50 \| AF-A0A165XES0-F1-MODEL\_V4 \| 1.0 \| 1.396e-14 \| 459 \| 0.149 \| 401 \| 243 \| 17 \| 1 \| 363 \| 10 \| 350 \| Exonuclease VIII \| Exonuclease VIII; afdb-uniprot50 \| AF-A0A2Z5YI50-F1-MODEL\_V4 \| 1.0 \| 9.987e-14 \| 458 \| 0.206 \| 368 \| 188 \| 16 \| 1 \| 361 \| 5 \| 275 \| DUF3799 domain-containing protein \| DUF3799 domain-containing protein; afdb-uniprot50 \| AF-A0A0F3IMN9-F1-MODEL\_V4 \| 1.0 \| 1.656e-13 \| 457 \| 0.181 \| 353 \| 181 \| 11 \| 9 \| 356 \| 48 \| 297 \| DUF3799 domain-containing protein \| DUF3799 domain-containing protein; afdb-uniprot50 \| AF-A0A2Z3HVS0-F1-MODEL\_V4 \| 1.0 \| 3.449e-12 \| 455 \| 0.341 \| 164 \| 97 \| 5 \| 204 \| 361 \| 2 \| 160 \| DUF3799 domain-containing protein \| DUF3799 domain-containing protein; afdb-uniprot50 \| AF-A0A354CHA7-F1-MODEL\_V4 \| 1.0 \| 8.415e-15 \| 453 \| 0.203 \| 363 \| 189 \| 12 \| 7 \| 360 \| 53 \| 324 \| DUF3799 domain-containing protein \| DUF3799 domain-containing protein; afdb-uniprot50 \| AF-A0A832HXX1-F1-MODEL\_V4 \| 1.0 \| 1.051e-15 \| 453 \| 0.184 \| 402 \| 204 \| 19 \| 1 \| 371 \| 10 \| 318 \| Uncharacterized protein \| Uncharacterized protein; afdb-uniprot50 \| AF-A0A2E8RC03-F1-MODEL\_V4 \| 1.0 \| 1.182e-13 \| 452 \| 0.207 \| 367 \| 179 \| 14 \| 8 \| 363 \| 5 \| 270 \| DUF3799 domain-containing protein \| DUF3799 domain-containing protein; afdb-uniprot50 \| AF-X0VAD8-F1-MODEL\_V4 \| 1.0 \| 1.406e-11 \| 451 \| 0.417 \| 134 \| 77 \| 1 \| 232 \| 364 \| 2 \| 135 \| DUF3799 domain-containing protein \| DUF3799 domain-containing protein; afdb-uniprot50 \| AF-A0A2A6NRI3-F1-MODEL\_V4 \| 1.0 \| 1.396e-14 \| 448 \| 0.164 \| 407 \| 235 \| 16 \| 1 \| 361 \| 64 \| 411 \| DUF3799 domain-containing protein \| DUF3799 domain-containing protein; afdb-uniprot50 \| AF-A0A0F9AK76-F1-MODEL\_V4 \| 1.0 \| 1.966e-12 \| 447 \| 0.287 \| 188 \| 124 \| 7 \| 181 \| 363 \| 64 \| 246 \| DUF3799 domain-containing protein \| DUF3799 domain-containing protein; afdb-uniprot50 \| AF-A0A5C7PD05-F1-MODEL\_V4 \| 1.0 \| 1.652e-14 \| 444 \| 0.162 \| 381 \| 200 \| 15 \| 1 \| 367 \| 51 \| 326 \| DUF3799 domain-containing protein \| DUF3799 domain-containing protein; afdb-uniprot50 \| AF-A0A244CWF3-F1-MODEL\_V4 \| 1.0 \| 8.88e-16 \| 444 \| 0.156 \| 409 \| 242 \| 16 \| 1 \| 372 \| 54 \| 396 \| DUF3799 domain-containing protein \| DUF3799 domain-containing protein; afdb-uniprot50 \| AF-A0A2D6B2Q9-F1-MODEL\_V4 \| 1.0 \| 9.987e-14 \| 443 \| 0.185 \| 382 \| 191 \| 15 \| 1 \| 356 \| 1 \| 288 \| DUF3799 domain-containing protein \| DUF3799 domain-containing protein; afdb-uniprot50 \| AF-A0A3D1XDL4-F1-MODEL\_V4 \| 1.0 \| 2.194e-13 \| 442 \| 0.242 \| 210 \| 136 \| 5 \| 155 \| 361 \| 17 \| 206 \| DUF3799 domain-containing protein \| DUF3799 domain-containing protein; afdb-uniprot50 \| AF-A0A7V9TKI5-F1-MODEL\_V4 \| 1.0 \| 3.44e-13 \| 440 \| 0.173 \| 381 \| 197 \| 16 \| 9 \| 371 \| 1 \| 281 \| Uncharacterized protein \| Uncharacterized protein; afdb-uniprot50 \| AF-A0A2E9HYF9-F1-MODEL\_V4 \| 1.0 \| 1.112e-15 \| 440 \| 0.172 \| 388 \| 212 \| 17 \| 2 \| 371 \| 56 \| 352 \| DUF3799 domain-containing protein \| DUF3799 domain-containing protein; afdb-uniprot50 \| AF-A0A4P7L155-F1-MODEL\_V4 \| 1.0 \| 1.656e-13 \| 436 \| 0.212 \| 325 \| 146 \| 12 \| 53 \| 371 \| 2 \| 222 \| Exodeoxyribonuclease 8 \| Exodeoxyribonuclease 8; afdb-uniprot50 \| AF-A0A709TYJ6-F1-MODEL\_V4 \| 1.0 \| 5.354e-16 \| 434 \| 0.223 \| 335 \| 205 \| 8 \| 83 \| 372 \| 4 \| 328 \| DNA breaking-rejoining protein \| DNA breaking-rejoining protein; afdb-uniprot50 \| AF-A0A626E748-F1-MODEL\_V4 \| 1.0 \| 6.337e-16 \| 434 \| 0.223 \| 335 \| 205 \| 8 \| 83 \| 372 \| 7 \| 331 \| DNA breaking-rejoining protein \| DNA breaking-rejoining protein; afdb-uniprot50 \| AF-M3JBV3-F1-MODEL\_V4 \| 1.0 \| 2.604e-12 \| 433 \| 0.164 \| 359 \| 166 \| 9 \| 9 \| 366 \| 25 \| 250 \| Bacteriophage protein \| Bacteriophage protein; afdb-uniprot50 \| AF-A0A825MRR2-F1-MODEL\_V4 \| 1.0 \| 2.906e-13 \| 432 \| 0.193 \| 331 \| 157 \| 12 \| 40 \| 361 \| 5 \| 234 \| Uncharacterized protein \| Uncharacterized protein; afdb-uniprot50 \| AF-A0A348FZE1-F1-MODEL\_V4 \| 1.0 \| 3.432e-14 \| 432 \| 0.196 \| 391 \| 190 \| 19 \| 2 \| 371 \| 15 \| 302 \| DUF3799 domain-containing protein \| DUF3799 domain-containing protein; afdb-uniprot50 \| AF-A0A705SAZ7-F1-MODEL\_V4 \| 1.0 \| 3.821e-16 \| 432 \| 0.223 \| 335 \| 205 \| 8 \| 83 \| 372 \| 5 \| 329 \| DNA breaking-rejoining protein \| DNA breaking-rejoining protein; afdb-uniprot50 \| AF-A0A741MSY8-F1-MODEL\_V4 \| 1.0 \| 3.821e-16 \| 431 \| 0.223 \| 335 \| 205 \| 9 \| 83 \| 372 \| 3 \| 327 \| DNA breaking-rejoining protein \| DNA breaking-rejoining protein; afdb-uniprot50 \| AF-A0A3N2N6Q1-F1-MODEL\_V4 \| 1.0 \| 3.423e-15 \| 430 \| 0.181 \| 392 \| 214 \| 19 \| 5 \| 367 \| 35 \| 348 \| DUF3799 domain-containing protein \| DUF3799 domain-containing protein; afdb-uniprot50 \| AF-A0A5U6R0D4-F1-MODEL\_V4 \| 1.0 \| 3.821e-16 \| 429 \| 0.22 \| 336 \| 205 \| 9 \| 83 \| 372 \| 4 \| 328 \| Exodeoxyribonuclease \| Exodeoxyribonuclease; afdb-uniprot50 \| AF-A0A5X2NJW9-F1-MODEL\_V4 \| 1.0 \| 6.337e-16 \| 429 \| 0.223 \| 335 \| 205 \| 8 \| 83 \| 372 \| 5 \| 329 \| DNA breaking-rejoining protein \| DNA breaking-rejoining protein; afdb-uniprot50 \| AF-A0A706H7T9-F1-MODEL\_V4 \| 1.0 \| 5.991e-16 \| 428 \| 0.222 \| 337 \| 203 \| 9 \| 83 \| 372 \| 6 \| 330 \| DNA breaking-rejoining protein \| DNA breaking-rejoining protein; afdb-uniprot50 \| AF-A0A722TRM3-F1-MODEL\_V4 \| 1.0 \| 5.991e-16 \| 427 \| 0.223 \| 335 \| 205 \| 9 \| 83 \| 372 \| 3 \| 327 \| DNA breaking-rejoining protein \| DNA breaking-rejoining protein; afdb-uniprot50 \| AF-A0A710PYI5-F1-MODEL\_V4 \| 1.0 \| 5.991e-16 \| 426 \| 0.223 \| 335 \| 205 \| 8 \| 83 \| 372 \| 5 \| 329 \| DNA breaking-rejoining protein \| DNA breaking-rejoining protein; afdb-uniprot50 \| AF-A0A5C7M3T7-F1-MODEL\_V4 \| 1.0 \| 5.394e-13 \| 425 \| 0.186 \| 359 \| 188 \| 17 \| 9 \| 362 \| 1 \| 260 \| DUF3799 domain-containing protein \| DUF3799 domain-containing protein; afdb-uniprot50 \| AF-A0A704YVF3-F1-MODEL\_V4 \| 1.0 \| 1.473e-15 \| 425 \| 0.22 \| 335 \| 206 \| 8 \| 83 \| 372 \| 7 \| 331 \| DNA breaking-rejoining protein \| DNA breaking-rejoining protein; afdb-uniprot50 \| AF-A0A710ZUB8-F1-MODEL\_V4 \| 1.0 \| 5.061e-16 \| 424 \| 0.22 \| 336 \| 205 \| 9 \| 83 \| 372 \| 4 \| 328 \| Exodeoxyribonuclease \| Exodeoxyribonuclease; afdb-uniprot50 \| AF-A0A705GFT1-F1-MODEL\_V4 \| 1.0 \| 1.176e-15 \| 424 \| 0.22 \| 336 \| 205 \| 9 \| 83 \| 372 \| 7 \| 331 \| Exodeoxyribonuclease \| Exodeoxyribonuclease; afdb-uniprot50 \| AF-A0A705K906-F1-MODEL\_V4 \| 1.0 \| 6.337e-16 \| 424 \| 0.223 \| 335 \| 205 \| 8 \| 83 \| 372 \| 11 \| 335 \| DNA breaking-rejoining protein \| DNA breaking-rejoining protein; afdb-uniprot50 \| AF-A0A6W0EZW0-F1-MODEL\_V4 \| 1.0 \| 1.316e-15 \| 422 \| 0.22 \| 335 \| 206 \| 9 \| 83 \| 372 \| 3 \| 327 \| Exodeoxyribonuclease \| Exodeoxyribonuclease; afdb-uniprot50 \| AF-A0A5X0DDY5-F1-MODEL\_V4 \| 1.0 \| 5.991e-16 \| 422 \| 0.22 \| 336 \| 205 \| 9 \| 83 \| 372 \| 7 \| 331 \| Exodeoxyribonuclease \| Exodeoxyribonuclease; afdb-uniprot50 \| AF-A0A7K0GP64-F1-MODEL\_V4 \| 1.0 \| 1.333e-10 \| 421 \| 0.52 \| 121 \| 58 \| 0 \| 241 \| 361 \| 1 \| 121 \| Exodeoxyribonuclease VIII \| Exodeoxyribonuclease VIII; afdb-uniprot50 \| AF-A0A661I8B6-F1-MODEL\_V4 \| 1.0 \| 3.261e-12 \| 421 \| 0.179 \| 357 \| 182 \| 13 \| 9 \| 358 \| 29 \| 281 \| DUF3799 domain-containing protein \| DUF3799 domain-containing protein; afdb-uniprot50 \| AF-A0A3T5P7X6-F1-MODEL\_V4 \| 1.0 \| 8.88e-16 \| 421 \| 0.223 \| 335 \| 205 \| 8 \| 83 \| 372 \| 7 \| 331 \| DNA breaking-rejoining protein \| DNA breaking-rejoining protein; afdb-uniprot50 \| AF-A0A2U1SSX7-F1-MODEL\_V4 \| 1.0 \| 1.118e-13 \| 421 \| 0.195 \| 405 \| 203 \| 17 \| 1 \| 371 \| 14 \| 329 \| DUF3799 domain-containing protein \| DUF3799 domain-containing protein; afdb-uniprot50 \| AF-A0A5W4VQX4-F1-MODEL\_V4 \| 1.0 \| 7.091e-16 \| 421 \| 0.219 \| 337 \| 204 \| 9 \| 83 \| 372 \| 18 \| 342 \| DNA breaking-rejoining protein \| DNA breaking-rejoining protein; afdb-uniprot50 \| AF-B2KDD5-F1-MODEL\_V4 \| 1.0 \| 2.597e-13 \| 420 \| 0.144 \| 387 \| 206 \| 15 \| 6 \| 367 \| 1 \| 287 \| DUF3799 domain-containing protein \| DUF3799 domain-containing protein; afdb-uniprot50 \| AF-A0A721STD1-F1-MODEL\_V4 \| 1.0 \| 1.244e-15 \| 420 \| 0.223 \| 335 \| 205 \| 8 \| 83 \| 372 \| 6 \| 330 \| DNA breaking-rejoining protein \| DNA breaking-rejoining protein; afdb-uniprot50 \| AF-A0A630QVP1-F1-MODEL\_V4 \| 1.0 \| 4.319e-12 \| 419 \| 0.261 \| 226 \| 151 \| 5 \| 157 \| 372 \| 1 \| 220 \| DUF3799 domain-containing protein \| DUF3799 domain-containing protein; afdb-uniprot50 \| AF-A0A5C8B335-F1-MODEL\_V4 \| 1.0 \| 1.966e-12 \| 419 \| 0.147 \| 379 \| 190 \| 16 \| 9 \| 370 \| 1 \| 263 \| DUF3799 domain-containing protein \| DUF3799 domain-containing protein; afdb-uniprot50 \| AF-A0A722NJL0-F1-MODEL\_V4 \| 1.0 \| 1.051e-15 \| 419 \| 0.22 \| 335 \| 206 \| 9 \| 83 \| 372 \| 3 \| 327 \| Exodeoxyribonuclease \| Exodeoxyribonuclease; afdb-uniprot50 \| AF-A0A3B9ZLY2-F1-MODEL\_V4 \| 1.0 \| 5.663e-16 \| 419 \| 0.193 \| 387 \| 241 \| 21 \| 7 \| 368 \| 13 \| 353 \| DUF3799 domain-containing protein \| DUF3799 domain-containing protein; afdb-uniprot50 \| AF-A0A3U7DT70-F1-MODEL\_V4 \| 1.0 \| 2.085e-11 \| 418 \| 0.354 \| 155 \| 89 \| 5 \| 216 \| 366 \| 2 \| 149 \| Exonuclease VIII \| Exonuclease VIII; afdb-uniprot50 \| AF-A0A2E1MZJ8-F1-MODEL\_V4 \| 1.0 \| 2.899e-14 \| 418 \| 0.183 \| 382 \| 220 \| 18 \| 9 \| 354 \| 1 \| 326 \| DUF3799 domain-containing protein \| DUF3799 domain-containing protein; afdb-uniprot50 \| AF-A0A709BXU1-F1-MODEL\_V4 \| 1.0 \| 2.443e-15 \| 418 \| 0.22 \| 336 \| 205 \| 9 \| 83 \| 372 \| 3 \| 327 \| Exodeoxyribonuclease \| Exodeoxyribonuclease; afdb-uniprot50 \| AF-A0A722CY88-F1-MODEL\_V4 \| 1.0 \| 8.394e-16 \| 417 \| 0.22 \| 336 \| 205 \| 9 \| 83 \| 372 \| 5 \| 329 \| Exodeoxyribonuclease \| Exodeoxyribonuclease; afdb-uniprot50 \| AF-A0A6Y0XSK0-F1-MODEL\_V4 \| 1.0 \| 8.394e-16 \| 417 \| 0.22 \| 336 \| 205 \| 9 \| 83 \| 372 \| 5 \| 329 \| Exodeoxyribonuclease \| Exodeoxyribonuclease; afdb-uniprot50 \| AF-A0A087B4Z0-F1-MODEL\_V4 \| 1.0 \| 4.557e-13 \| 416 \| 0.173 \| 381 \| 188 \| 20 \| 5 \| 368 \| 2 \| 272 \| Phage protein \| Phage protein; afdb-uniprot50 \| AF-A0A6L6J4B9-F1-MODEL\_V4 \| 1.0 \| 2.194e-13 \| 416 \| 0.19 \| 373 \| 176 \| 18 \| 1 \| 359 \| 8 \| 268 \| DUF3799 domain-containing protein \| DUF3799 domain-containing protein; afdb-uniprot50 \| AF-X8AH60-F1-MODEL\_V4 \| 1.0 \| 5.422e-11 \| 413 \| 0.345 \| 168 \| 102 \| 6 \| 207 \| 369 \| 3 \| 167 \| DUF3799 domain-containing protein \| DUF3799 domain-containing protein; afdb-uniprot50 \| AF-A0A705X8M6-F1-MODEL\_V4 \| 1.0 \| 7.935e-16 \| 413 \| 0.215 \| 338 \| 204 \| 10 \| 83 \| 372 \| 11 \| 335 \| Exodeoxyribonuclease \| Exodeoxyribonuclease; afdb-uniprot50 \| AF-A0A2E2D771-F1-MODEL\_V4 \| 1.0 \| 1.956e-14 \| 409 \| 0.175 \| 353 \| 214 \| 16 \| 19 \| 359 \| 24 \| 311 \| DUF3799 domain-containing protein \| DUF3799 domain-containing protein; afdb-uniprot50 \| AF-A0A3S5DAA5-F1-MODEL\_V4 \| 1.0 \| 3.658e-11 \| 407 \| 0.299 \| 147 \| 100 \| 2 \| 221 \| 364 \| 10 \| 156 \| Exodeoxyribonuclease 8 \| Exodeoxyribonuclease 8; afdb-uniprot50 \| AF-A0A6N8TQN8-F1-MODEL\_V4 \| 1.0 \| 3.84e-14 \| 407 \| 0.126 \| 331 \| 225 \| 12 \| 15 \| 343 \| 43 \| 311 \| Uncharacterized protein \| Uncharacterized protein; afdb-uniprot50 \| AF-A0A5W8UZN8-F1-MODEL\_V4 \| 1.0 \| 7.109e-15 \| 404 \| 0.23 \| 321 \| 192 \| 9 \| 97 \| 372 \| 2 \| 312 \| Exodeoxyribonuclease \| Exodeoxyribonuclease; afdb-uniprot50 \| AF-A0A2E3D7E6-F1-MODEL\_V4 \| 1.0 \| 2.327e-12 \| 402 \| 0.174 \| 373 \| 191 \| 14 \| 8 \| 372 \| 4 \| 267 \| DUF3799 domain-containing protein \| DUF3799 domain-containing protein; afdb-uniprot50 \| AF-A0A1M6GAE2-F1-MODEL\_V4 \| 1.0 \| 5.381e-14 \| 397 \| 0.163 \| 355 \| 228 \| 13 \| 19 \| 370 \| 49 \| 337 \| DUF3799 domain-containing protein \| DUF3799 domain-containing protein; afdb-uniprot50 \| AF-A0A1H6WGJ5-F1-MODEL\_V4 \| 1.0 \| 3.84e-14 \| 396 \| 0.243 \| 373 \| 167 \| 18 \| 1 \| 359 \| 31 \| 302 \| DUF3799 domain-containing protein \| DUF3799 domain-containing protein; afdb-uniprot50 \| AF-A0A5W2I5P1-F1-MODEL\_V4 \| 1.0 \| 3.63e-14 \| 395 \| 0.23 \| 321 \| 192 \| 9 \| 97 \| 372 \| 2 \| 312 \| Exodeoxyribonuclease \| Exodeoxyribonuclease; afdb-uniprot50 \| AF-A0A495SNW1-F1-MODEL\_V4 \| 1.0 \| 4.297e-14 \| 391 \| 0.151 \| 364 \| 233 \| 15 \| 6 \| 360 \| 22 \| 318 \| PDDEXK-like uncharacterized protein DUF3799 \| PDDEXK-like uncharacterized protein DUF3799; afdb-uniprot50 \| AF-A0A318TK26-F1-MODEL\_V4 \| 1.0 \| 1.059e-12 \| 390 \| 0.185 \| 400 \| 220 \| 23 \| 3 \| 361 \| 10 \| 344 \| PDDEXK-like uncharacterized protein DUF3799 \| PDDEXK-like uncharacterized protein DUF3799; afdb-uniprot50 \| AF-A0A4Q2UH62-F1-MODEL\_V4 \| 1.0 \| 1.188e-11 \| 388 \| 0.189 \| 364 \| 144 \| 10 \| 9 \| 359 \| 3 \| 228 \| DUF3799 domain-containing protein \| DUF3799 domain-containing protein; afdb-uniprot50 \| AF-A0A5V8R8Y3-F1-MODEL\_V4 \| 1.0 \| 3.63e-14 \| 388 \| 0.231 \| 315 \| 187 \| 9 \| 103 \| 372 \| 3 \| 307 \| DNA breaking-rejoining protein \| DNA breaking-rejoining protein; afdb-uniprot50 \| AF-A0A369XIG4-F1-MODEL\_V4 \| 1.0 \| 8.057e-10 \| 386 \| 0.453 \| 128 \| 66 \| 3 \| 238 \| 363 \| 2 \| 127 \| DUF3799 domain-containing protein \| DUF3799 domain-containing protein; afdb-uniprot50 \| AF-A0A3Z7BBV9-F1-MODEL\_V4 \| 1.0 \| 8.925e-14 \| 386 \| 0.231 \| 315 \| 187 \| 9 \| 103 \| 372 \| 1 \| 305 \| Exodeoxyribonuclease \| Exodeoxyribonuclease; afdb-uniprot50 \| AF-A0A553GCW7-F1-MODEL\_V4 \| 1.0 \| 2.205e-11 \| 385 \| 0.161 \| 341 \| 150 \| 8 \| 1 \| 338 \| 2 \| 209 \| DUF3799 domain-containing protein \| DUF3799 domain-containing protein; afdb-uniprot50 \| AF-A0A379GHA6-F1-MODEL\_V4 \| 1.0 \| 1.484e-12 \| 385 \| 0.242 \| 264 \| 166 \| 9 \| 105 \| 358 \| 5 \| 244 \| Phage-related exonuclease \| Phage-related exonuclease; afdb-uniprot50 \| AF-A0A6M3ITI9-F1-MODEL\_V4 \| 1.0 \| 1.001e-12 \| 385 \| 0.182 \| 390 \| 203 \| 15 \| 5 \| 358 \| 2 \| 311 \| DUF3799 domain-containing protein \| DUF3799 domain-containing protein; afdb-uniprot50 \| AF-A0A7U4E4Q7-F1-MODEL\_V4 \| 1.0 \| 7.597e-11 \| 384 \| 0.179 \| 334 \| 139 \| 10 \| 7 \| 338 \| 9 \| 209 \| DUF3799 domain-containing protein \| DUF3799 domain-containing protein; afdb-uniprot50 \| AF-A0A662AYQ9-F1-MODEL\_V4 \| 1.0 \| 4.569e-12 \| 384 \| 0.153 \| 391 \| 194 \| 20 \| 1 \| 356 \| 1 \| 289 \| DUF3799 domain-containing protein \| DUF3799 domain-containing protein; afdb-uniprot50 \| AF-A0A2K8Z643-F1-MODEL\_V4 \| 1.0 \| 2.211e-10 \| 379 \| 0.175 \| 331 \| 131 \| 12 \| 14 \| 338 \| 4 \| 198 \| DUF3799 domain-containing protein \| DUF3799 domain-containing protein; afdb-uniprot50 \| AF-A0A2N7BEU4-F1-MODEL\_V4 \| 1.0 \| 1.578e-10 \| 378 \| 0.182 \| 334 \| 137 \| 9 \| 7 \| 340 \| 3 \| 200 \| DUF3799 domain-containing protein \| DUF3799 domain-containing protein; afdb-uniprot50 \| AF-A0A2D8XKX9-F1-MODEL\_V4 \| 1.0 \| 9.489e-12 \| 378 \| 0.15 \| 340 \| 192 \| 13 \| 19 \| 355 \| 16 \| 261 \| DUF3799 domain-containing protein \| DUF3799 domain-containing protein; afdb-uniprot50 \| AF-A0A857VCF5-F1-MODEL\_V4 \| 1.0 \| 1.863e-11 \| 376 \| 0.162 \| 358 \| 186 \| 16 \| 9 \| 359 \| 1 \| 251 \| Uncharacterized protein \| Uncharacterized protein; afdb-uniprot50 \| AF-A0A0E3B9I3-F1-MODEL\_V4 \| 1.0 \| 1.574e-11 \| 375 \| 0.244 \| 196 \| 132 \| 7 \| 177 \| 360 \| 24 \| 215 \| DUF3799 domain-containing protein \| DUF3799 domain-containing protein; afdb-uniprot50 \| AF-A0A433CKI3-F1-MODEL\_V4 \| 1.0 \| 4.58e-11 \| 375 \| 0.159 \| 327 \| 160 \| 12 \| 17 \| 341 \| 3 \| 216 \| DUF3799 domain-containing protein \| DUF3799 domain-containing protein; afdb-uniprot50 \| AF-A0A0F8ZM74-F1-MODEL\_V4 \| 1.0 \| 2.211e-10 \| 374 \| 0.302 \| 192 \| 123 \| 8 \| 183 \| 372 \| 2 \| 184 \| DUF3799 domain-containing protein \| DUF3799 domain-containing protein; afdb-uniprot50 \| AF-A0A721W8G7-F1-MODEL\_V4 \| 1.0 \| 6.082e-10 \| 373 \| 0.259 \| 189 \| 124 \| 5 \| 194 \| 372 \| 3 \| 185 \| DUF3799 domain-containing protein \| DUF3799 domain-containing protein; afdb-uniprot50 \| AF-A0A7J5TSP0-F1-MODEL\_V4 \| 1.0 \| 3.869e-11 \| 373 \| 0.183 \| 371 \| 150 \| 10 \| 6 \| 361 \| 1 \| 233 \| DUF3799 domain-containing protein \| DUF3799 domain-containing protein; afdb-uniprot50 \| AF-A0A2D9CGG8-F1-MODEL\_V4 \| 1.0 \| 1.123e-11 \| 373 \| 0.15 \| 338 \| 195 \| 13 \| 19 \| 356 \| 15 \| 260 \| DUF3799 domain-containing protein \| DUF3799 domain-containing protein; afdb-uniprot50 \| AF-A0A1B9YEN4-F1-MODEL\_V4 \| 1.0 \| 7.145e-13 \| 373 \| 0.151 \| 370 \| 233 \| 13 \| 6 \| 360 \| 12 \| 315 \| DUF3799 domain-containing protein \| DUF3799 domain-containing protein; afdb-uniprot50 \| AF-A0A7X9RXT8-F1-MODEL\_V4 \| 1.0 \| 4.845e-11 \| 372 \| 0.173 \| 345 \| 147 \| 7 \| 1 \| 341 \| 27 \| 237 \| DUF3799 domain-containing protein \| DUF3799 domain-containing protein; afdb-uniprot50 \| AF-A0A7C2PXI9-F1-MODEL\_V4 \| 1.0 \| 1.26e-10 \| 370 \| 0.278 \| 187 \| 125 \| 6 \| 174 \| 358 \| 105 \| 283 \| DUF3799 domain-containing protein \| DUF3799 domain-containing protein; afdb-uniprot50 \| AF-A0A7Z9YY02-F1-MODEL\_V4 \| 1.0 \| 3.475e-09 \| 369 \| 0.49 \| 108 \| 55 \| 0 \| 254 \| 361 \| 1 \| 108 \| Exodeoxyribonuclease VIII \| Exodeoxyribonuclease VIII; afdb-uniprot50 \| AF-A0A352REG3-F1-MODEL\_V4 \| 1.0 \| 2.217e-09 \| 368 \| 0.294 \| 139 \| 93 \| 4 \| 203 \| 340 \| 2 \| 136 \| DUF3799 domain-containing protein \| DUF3799 domain-containing protein; afdb-uniprot50 \| AF-A0A519U1A8-F1-MODEL\_V4 \| 1.0 \| 1.976e-10 \| 367 \| 0.159 \| 377 \| 157 \| 10 \| 1 \| 371 \| 10 \| 232 \| DUF3799 domain-containing protein \| DUF3799 domain-containing protein; afdb-uniprot50 \| AF-I8ANZ2-F1-MODEL\_V4 \| 1.0 \| 1.574e-11 \| 367 \| 0.17 \| 386 \| 197 \| 12 \| 6 \| 372 \| 1 \| 282 \| Phage protein \| Phage protein; afdb-uniprot50 \| AF-A0A4V0Z515-F1-MODEL\_V4 \| 1.0 \| 6.082e-10 \| 366 \| 0.287 \| 181 \| 119 \| 4 \| 189 \| 360 \| 2 \| 181 \| DUF3799 domain-containing protein \| DUF3799 domain-containing protein; afdb-uniprot50 \| AF-A0A6X8JIS0-F1-MODEL\_V4 \| 1.0 \| 1.182e-13 \| 364 \| 0.228 \| 306 \| 174 \| 7 \| 1 \| 250 \| 457 \| 756 \| Exodeoxyribonuclease \| Exodeoxyribonuclease; afdb-uniprot50 \| AF-A0A6Y3UI71-F1-MODEL\_V4 \| 1.0 \| 5.087e-14 \| 363 \| 0.219 \| 328 \| 201 \| 10 \| 1 \| 273 \| 219 \| 546 \| Exodeoxyribonuclease \| Exodeoxyribonuclease; afdb-uniprot50 \| AF-A0A3D2IK29-F1-MODEL\_V4 \| 1.0 \| 1.766e-10 \| 360 \| 0.29 \| 196 \| 119 \| 10 \| 177 \| 359 \| 1 \| 189 \| DUF3799 domain-containing protein \| DUF3799 domain-containing protein; afdb-uniprot50 \| AF-A0A485BQ07-F1-MODEL\_V4 \| 1.0 \| 2.455e-13 \| 359 \| 0.227 \| 312 \| 188 \| 10 \| 100 \| 368 \| 28 \| 329 \| Exonuclease VIII \| Exonuclease VIII; afdb-uniprot50 \| AF-X0ST36-F1-MODEL\_V4 \| 1.0 \| 2.095e-09 \| 357 \| 0.357 \| 137 \| 87 \| 1 \| 234 \| 369 \| 5 \| 141 \| DUF3799 domain-containing protein \| DUF3799 domain-containing protein; afdb-uniprot50 \| AF-A0A1W6ECB7-F1-MODEL\_V4 \| 1.0 \| 3.098e-10 \| 356 \| 0.158 \| 359 \| 156 \| 11 \| 8 \| 358 \| 2 \| 222 \| DUF3799 domain-containing protein \| DUF3799 domain-containing protein; afdb-uniprot50 \| AF-A0A238X2A8-F1-MODEL\_V4 \| 1.0 \| 2.468e-11 \| 355 \| 0.196 \| 357 \| 140 \| 12 \| 1 \| 349 \| 1 \| 218 \| DUF3799 domain-containing protein \| DUF3799 domain-containing protein; afdb-uniprot50 \| AF-A0A705E0Z0-F1-MODEL\_V4 \| 1.0 \| 1.182e-13 \| 354 \| 0.216 \| 300 \| 185 \| 8 \| 1 \| 250 \| 63 \| 362 \| Exodeoxyribonuclease \| Exodeoxyribonuclease; afdb-uniprot50 \| AF-A0A758PAR8-F1-MODEL\_V4 \| 1.0 \| 2.074e-13 \| 350 \| 0.208 \| 321 \| 177 \| 10 \| 68 \| 323 \| 2 \| 310 \| DNA breaking-rejoining protein \| DNA breaking-rejoining protein; afdb-uniprot50 \| AF-A0A7Y0FMZ2-F1-MODEL\_V4 \| 1.0 \| 2.211e-10 \| 347 \| 0.17 \| 375 \| 161 \| 14 \| 1 \| 358 \| 1 \| 242 \| DUF3799 domain-containing protein \| DUF3799 domain-containing protein; afdb-uniprot50 \| AF-A0A7W4DRQ1-F1-MODEL\_V4 \| 1.0 \| 3.667e-10 \| 343 \| 0.153 \| 359 \| 182 \| 15 \| 9 \| 355 \| 1 \| 249 \| PD-(D/E)XK nuclease-like domain-containing protein \| PD-(D/E)XK nuclease-like domain-containing protein; afdb-uniprot50 \| AF-A0A1G2Z6Y7-F1-MODEL\_V4 \| 1.0 \| 2.474e-10 \| 341 \| 0.22 \| 200 \| 143 \| 7 \| 164 \| 359 \| 25 \| 215 \| DUF3799 domain-containing protein \| DUF3799 domain-containing protein; afdb-uniprot50 \| AF-A0A447XY29-F1-MODEL\_V4 \| 1.0 \| 4.093e-11 \| 340 \| 0.214 \| 280 \| 170 \| 9 \| 105 \| 372 \| 1 \| 242 \| Putative phage exodeoxyribonuclease \| Putative phage exodeoxyribonuclease; afdb-uniprot50 \| AF-A0A1B1G3H1-F1-MODEL\_V4 \| 1.0 \| 1.067e-09 \| 340 \| 0.158 \| 341 \| 149 \| 8 \| 2 \| 338 \| 25 \| 231 \| DUF3799 domain-containing protein \| DUF3799 domain-containing protein; afdb-uniprot50 \| AF-A0A5V5KL18-F1-MODEL\_V4 \| 1.0 \| 2.597e-13 \| 340 \| 0.222 \| 328 \| 200 \| 10 \| 1 \| 273 \| 39 \| 366 \| Exodeoxyribonuclease \| Exodeoxyribonuclease; afdb-uniprot50 \| AF-A0A4S3LPF3-F1-MODEL\_V4 \| 1.0 \| 1.254e-12 \| 339 \| 0.214 \| 308 \| 176 \| 8 \| 1 \| 250 \| 485 \| 784 \| Uncharacterized protein \| Uncharacterized protein; afdb-uniprot50 \| AF-A0A4Q2UMS3-F1-MODEL\_V4 \| 1.0 \| 6.082e-10 \| 338 \| 0.184 \| 347 \| 140 \| 11 \| 14 \| 355 \| 3 \| 211 \| DUF3799 domain-containing protein \| DUF3799 domain-containing protein; afdb-uniprot50 \| AF-A0A1G3PFN4-F1-MODEL\_V4 \| 1.0 \| 1.333e-10 \| 337 \| 0.146 \| 369 \| 187 \| 18 \| 5 \| 359 \| 4 \| 258 \| DUF3799 domain-containing protein \| DUF3799 domain-containing protein; afdb-uniprot50 \| AF-A0A485A5W0-F1-MODEL\_V4 \| 1.0 \| 1.757e-12 \| 337 \| 0.214 \| 308 \| 176 \| 9 \| 1 \| 250 \| 162 \| 461 \| Exodeoxyribonuclease 8 \| Exodeoxyribonuclease 8; afdb-uniprot50 \| AF-A0A5C5X7Z0-F1-MODEL\_V4 \| 1.0 \| 3.869e-11 \| 336 \| 0.154 \| 415 \| 191 \| 18 \| 6 \| 356 \| 11 \| 329 \| DUF3799 domain-containing protein \| DUF3799 domain-containing protein; afdb-uniprot50 \| AF-A0A2U3ERY8-F1-MODEL\_V4 \| 1.0 \| 3.269e-11 \| 335 \| 0.159 \| 376 \| 176 \| 12 \| 5 \| 369 \| 154 \| 400 \| Uncharacterized protein \| Uncharacterized protein; afdb-uniprot50 \| AF-A0A5X5VH27-F1-MODEL\_V4 \| 1.0 \| 1.326e-12 \| 334 \| 0.223 \| 304 \| 178 \| 7 \| 1 \| 250 \| 634 \| 933 \| DNA breaking-rejoining protein \| DNA breaking-rejoining protein; afdb-uniprot50 \| AF-A0A7H1NU02-F1-MODEL\_V4 \| 1.0 \| 1.197e-08 \| 333 \| 0.261 \| 149 \| 103 \| 6 \| 224 \| 369 \| 8 \| 152 \| DUF3799 domain-containing protein \| DUF3799 domain-containing protein; afdb-uniprot50 \| AF-A0A2E4HLS4-F1-MODEL\_V4 \| 1.0 \| 4.319e-12 \| 332 \| 0.135 \| 361 \| 241 \| 17 \| 8 \| 345 \| 2 \| 314 \| PDDEXK\_1 domain-containing protein \| PDDEXK\_1 domain-containing protein; afdb-uniprot50 \| AF-A0A6Y4WD11-F1-MODEL\_V4 \| 1.0 \| 1.059e-12 \| 330 \| 0.218 \| 297 \| 182 \| 8 \| 1 \| 247 \| 110 \| 406 \| Exodeoxyribonuclease \| Exodeoxyribonuclease; afdb-uniprot50 \| AF-A0A2D8BQ94-F1-MODEL\_V4 \| 1.0 \| 2.217e-09 \| 319 \| 0.13 \| 353 \| 180 \| 14 \| 11 \| 357 \| 10 \| 241 \| DUF3799 domain-containing protein \| DUF3799 domain-containing protein; afdb-uniprot50 \| AF-A0A060RHE4-F1-MODEL\_V4 \| 1.0 \| 2.624e-09 \| 318 \| 0.133 \| 381 \| 194 \| 15 \| 9 \| 363 \| 1 \| 271 \| DUF3799 domain-containing protein \| DUF3799 domain-containing protein; afdb-uniprot50 \| AF-A0A447MUU1-F1-MODEL\_V4 \| 1.0 \| 2.747e-13 \| 318 \| 0.179 \| 340 \| 213 \| 11 \| 33 \| 311 \| 9 \| 343 \| Exodeoxyribonuclease VIII \| Exodeoxyribonuclease VIII; afdb-uniprot50 \| AF-A0A4Q2UGG0-F1-MODEL\_V4 \| 1.0 \| 1.126e-10 \| 314 \| 0.248 \| 205 \| 134 \| 8 \| 167 \| 358 \| 19 \| 216 \| DUF3799 domain-containing protein \| DUF3799 domain-containing protein; afdb-uniprot50 \| AF-A0A0K8Q1C6-F1-MODEL\_V4 \| 1.0 \| 2.222e-08 \| 309 \| 0.431 \| 116 \| 64 \| 2 \| 256 \| 371 \| 2 \| 115 \| Exodeoxyribonuclease 8 \| Exodeoxyribonuclease 8; afdb-uniprot50 \| AF-A0A6G4MZF1-F1-MODEL\_V4 \| 1.0 \| 2.48e-09 \| 306 \| 0.125 \| 391 \| 200 \| 15 \| 6 \| 370 \| 1 \| 275 \| Nuclease \| Nuclease; afdb-uniprot50 \| AF-A0A3M1KL91-F1-MODEL\_V4 \| 1.0 \| 5.164e-08 \| 302 \| 0.288 \| 142 \| 93 \| 5 \| 217 \| 356 \| 4 \| 139 \| DUF3799 domain-containing protein \| DUF3799 domain-containing protein; afdb-uniprot50 \| AF-A0A7C1JUS1-F1-MODEL\_V4 \| 1.0 \| 9.585e-08 \| 302 \| 0.267 \| 153 \| 105 \| 4 \| 207 \| 359 \| 3 \| 148 \| DUF3799 domain-containing protein \| DUF3799 domain-containing protein; afdb-uniprot50 \| AF-A0A5Y2EHS0-F1-MODEL\_V4 \| 1.0 \| 2.929e-10 \| 302 \| 0.254 \| 283 \| 106 \| 8 \| 1 \| 277 \| 49 \| 232 \| Exonuclease VIII \| Exonuclease VIII; afdb-uniprot50 \| AF-A0A511B8A3-F1-MODEL\_V4 \| 1.0 \| 2.351e-08 \| 297 \| 0.26 \| 161 \| 105 \| 7 \| 222 \| 372 \| 12 \| 168 \| DUF3799 domain-containing protein \| DUF3799 domain-containing protein; afdb-uniprot50 \| AF-A0A5M6A0E3-F1-MODEL\_V4 \| 1.0 \| 1.263e-09 \| 297 \| 0.218 \| 215 \| 133 \| 10 \| 177 \| 364 \| 21 \| 227 \| DUF3799 domain-containing protein \| DUF3799 domain-containing protein; afdb-uniprot50 \| AF-A0A5X5NRN7-F1-MODEL\_V4 \| 1.0 \| 6.806e-10 \| 295 \| 0.244 \| 229 \| 139 \| 7 \| 108 \| 326 \| 2 \| 206 \| DUF3799 domain-containing protein \| DUF3799 domain-containing protein; afdb-uniprot50 \| AF-A0A377TT21-F1-MODEL\_V4 \| 1.0 \| 1.971e-11 \| 295 \| 0.22 \| 286 \| 185 \| 8 \| 84 \| 359 \| 15 \| 272 \| Putative exodeoxyribonuclease VIII \| Putative exodeoxyribonuclease VIII; afdb-uniprot50 \| AF-A0A350BS60-F1-MODEL\_V4 \| 1.0 \| 3.098e-10 \| 291 \| 0.218 \| 275 \| 159 \| 13 \| 114 \| 354 \| 4 \| 256 \| DUF3799 domain-containing protein \| DUF3799 domain-containing protein; afdb-uniprot50 \| AF-A0A349YBV5-F1-MODEL\_V4 \| 1.0 \| 2.351e-08 \| 288 \| 0.226 \| 159 \| 116 \| 5 \| 187 \| 343 \| 84 \| 237 \| DUF3799 domain-containing protein \| DUF3799 domain-containing protein; afdb-uniprot50 \| AF-A0A3R7HPJ5-F1-MODEL\_V4 \| 1.0 \| 2.211e-10 \| 286 \| 0.176 \| 375 \| 200 \| 21 \| 9 \| 358 \| 1 \| 291 \| DUF3799 domain-containing protein \| DUF3799 domain-containing protein; afdb-uniprot50 \| AF-A0A2S8GSL0-F1-MODEL\_V4 \| 1.0 \| 3.475e-09 \| 286 \| 0.129 \| 354 \| 181 \| 14 \| 6 \| 346 \| 29 \| 268 \| Uncharacterized protein \| Uncharacterized protein; afdb-uniprot50 \| AF-A0A7C5SKK8-F1-MODEL\_V4 \| 1.0 \| 3.493e-07 \| 285 \| 0.315 \| 130 \| 88 \| 1 \| 232 \| 360 \| 1 \| 130 \| DUF3799 domain-containing protein \| DUF3799 domain-containing protein; afdb-uniprot50 \| AF-A0A352NZ68-F1-MODEL\_V4 \| 1.0 \| 1.986e-08 \| 279 \| 0.123 \| 381 \| 185 \| 16 \| 8 \| 361 \| 7 \| 265 \| DUF3799 domain-containing protein \| DUF3799 domain-containing protein; afdb-uniprot50 \| AF-A0A1L3I482-F1-MODEL\_V4 \| 1.0 \| 2.487e-08 \| 274 \| 0.451 \| 135 \| 60 \| 2 \| 207 \| 334 \| 3 \| 130 \| Exodeoxyribonuclease 8 \| Exodeoxyribonuclease 8; afdb-uniprot50 \| AF-A0A706A9B5-F1-MODEL\_V4 \| 1.0 \| 7.2e-10 \| 271 \| 0.227 \| 251 \| 160 \| 7 \| 83 \| 323 \| 12 \| 238 \| Exodeoxyribonuclease \| Exodeoxyribonuclease; afdb-uniprot50 \| AF-A0A1S8PH38-F1-MODEL\_V4 \| 1.0 \| 5.463e-08 \| 270 \| 0.137 \| 370 \| 179 \| 17 \| 8 \| 357 \| 2 \| 251 \| PD-(D/E)XK nuclease superfamily protein \| PD-(D/E)XK nuclease superfamily protein; afdb-uniprot50 \| AF-Z9JM14-F1-MODEL\_V4 \| 1.0 \| 1.273e-06 \| 265 \| 0.394 \| 109 \| 64 \| 2 \| 255 \| 363 \| 8 \| 114 \| DUF3799 domain-containing protein \| DUF3799 domain-containing protein; afdb-uniprot50 \| AF-A0A069CWL0-F1-MODEL\_V4 \| 1.0 \| 1.503e-07 \| 264 \| 0.113 \| 360 \| 184 \| 12 \| 6 \| 344 \| 1 \| 246 \| Phage-related protein \| Phage-related protein; afdb-uniprot50 \| AF-A0A1V1PGN8-F1-MODEL\_V4 \| 1.0 \| 5.151e-09 \| 261 \| 0.158 \| 379 \| 211 \| 19 \| 6 \| 358 \| 1 \| 297 \| Uncharacterized protein \| Uncharacterized protein; afdb-uniprot50 \| AF-A0A853IND3-F1-MODEL\_V4 \| 1.0 \| 1.014e-07 \| 260 \| 0.251 \| 159 \| 110 \| 5 \| 155 \| 307 \| 11 \| 166 \| PD-(D/E)XK nuclease-like domain-containing protein \| PD-(D/E)XK nuclease-like domain-containing protein; afdb-uniprot50 \| AF-A0A8A3QEV9-F1-MODEL\_V4 \| 1.0 \| 8.566e-08 \| 260 \| 0.122 \| 360 \| 173 \| 16 \| 20 \| 357 \| 1 \| 239 \| PD-(D/E)XK nuclease-like domain-containing protein \| PD-(D/E)XK nuclease-like domain-containing protein; afdb-uniprot50 \| AF-A0A6M3IM01-F1-MODEL\_V4 \| 1.0 \| 1.981e-09 \| 260 \| 0.133 \| 352 \| 183 \| 15 \| 6 \| 348 \| 4 \| 242 \| Putative PD-(D/E)XK nuclease superfamily protein \| Putative PD-(D/E)XK nuclease superfamily protein; afdb-uniprot50 \| AF-A0A524P3G8-F1-MODEL\_V4 \| 1.0 \| 1.779e-07 \| 258 \| 0.29 \| 148 \| 97 \| 4 \| 230 \| 371 \| 1 \| 146 \| DUF3799 domain-containing protein \| DUF3799 domain-containing protein; afdb-uniprot50 \| AF-A0A6G9Q4G4-F1-MODEL\_V4 \| 1.0 \| 3.484e-08 \| 257 \| 0.143 \| 384 \| 178 \| 18 \| 7 \| 358 \| 3 \| 267 \| DUF3799 domain-containing protein \| DUF3799 domain-containing protein; afdb-uniprot50 \| AF-A0A0R1R1I5-F1-MODEL\_V4 \| 1.0 \| 1.503e-07 \| 254 \| 0.113 \| 360 \| 188 \| 14 \| 20 \| 361 \| 1 \| 247 \| DUF3799 domain-containing protein \| DUF3799 domain-containing protein; afdb-uniprot50 \| AF-A0A2N5WAQ2-F1-MODEL\_V4 \| 1.0 \| 2.493e-07 \| 252 \| 0.121 \| 369 \| 176 \| 12 \| 18 \| 356 \| 3 \| 253 \| PD-(D/E)XK nuclease superfamily protein \| PD-(D/E)XK nuclease superfamily protein; afdb-uniprot50 \| AF-A0A512PLA9-F1-MODEL\_V4 \| 1.0 \| 3.114e-08 \| 252 \| 0.124 \| 379 \| 188 \| 18 \| 7 \| 359 \| 41 \| 301 \| HTH cro/C1-type domain-containing protein \| HTH cro/C1-type domain-containing protein; afdb-uniprot50 \| AF-F5VG10-F1-MODEL\_V4 \| 1.0 \| 3.106e-09 \| 250 \| 0.14 \| 398 \| 219 \| 21 \| 11 \| 360 \| 4 \| 326 \| DUF3799 domain-containing protein \| DUF3799 domain-containing protein; afdb-uniprot50 \| AF-A0A3Q8YTG7-F1-MODEL\_V4 \| 1.0 \| 2.958e-06 \| 246 \| 0.306 \| 111 \| 73 \| 2 \| 253 \| 359 \| 14 \| 124 \| DUF3799 domain-containing protein \| DUF3799 domain-containing protein; afdb-uniprot50 \| AF-X0TZX0-F1-MODEL\_V4 \| 1.0 \| 1.686e-06 \| 246 \| 0.176 \| 249 \| 110 \| 4 \| 1 \| 249 \| 6 \| 159 \| DUF3799 domain-containing protein \| DUF3799 domain-containing protein; afdb-uniprot50 \| AF-A0A3C0G174-F1-MODEL\_V4 \| 1.0 \| 1.273e-06 \| 243 \| 0.339 \| 106 \| 68 \| 2 \| 253 \| 358 \| 2 \| 105 \| DUF3799 domain-containing protein \| DUF3799 domain-containing protein; afdb-uniprot50 \| AF-R1HTJ0-F1-MODEL\_V4 \| 1.0 \| 7.674e-07 \| 243 \| 0.274 \| 142 \| 89 \| 3 \| 231 \| 371 \| 7 \| 135 \| Gp60 protein \| Gp60 protein; afdb-uniprot50 \| AF-A0A1E3KRK4-F1-MODEL\_V4 \| 1.0 \| 6.858e-07 \| 240 \| 0.119 \| 360 \| 178 \| 15 \| 20 \| 357 \| 1 \| 243 \| DUF3799 domain-containing protein \| DUF3799 domain-containing protein; afdb-uniprot50 \| AF-A0A842HJB6-F1-MODEL\_V4 \| 1.0 \| 1.877e-08 \| 240 \| 0.117 \| 367 \| 203 \| 17 \| 1 \| 350 \| 2 \| 264 \| PD-(D/E)XK nuclease family protein \| PD-(D/E)XK nuclease family protein; afdb-uniprot50 \| AF-A0A379YD88-F1-MODEL\_V4 \| 1.0 \| 3.502e-06 \| 239 \| 0.225 \| 151 \| 101 \| 5 \| 229 \| 369 \| 2 \| 146 \| Exonuclease VIII \| Exonuclease VIII; afdb-uniprot50 \| AF-A0A6I1JKT6-F1-MODEL\_V4 \| 1.0 \| 1.266e-08 \| 239 \| 0.278 \| 201 \| 117 \| 9 \| 178 \| 371 \| 39 \| 218 \| Phage protein \| Phage protein; afdb-uniprot50 \| AF-A0A7C6QXB3-F1-MODEL\_V4 \| 1.0 \| 4.124e-08 \| 237 \| 0.123 \| 340 \| 177 \| 12 \| 16 \| 347 \| 3 \| 229 \| PD-(D/E)XK nuclease family protein \| PD-(D/E)XK nuclease family protein; afdb-uniprot50 \| AF-A0A7X6AC15-F1-MODEL\_V4 \| 1.0 \| 9.585e-08 \| 236 \| 0.128 \| 343 \| 183 \| 15 \| 15 \| 350 \| 5 \| 238 \| PDDEXK\_1 domain-containing protein \| PDDEXK\_1 domain-containing protein; afdb-uniprot50 \| AF-A0A3Z0FCG3-F1-MODEL\_V4 \| 1.0 \| 1.067e-09 \| 234 \| 0.215 \| 279 \| 157 \| 8 \| 53 \| 273 \| 5 \| 279 \| Exodeoxyribonuclease \| Exodeoxyribonuclease; afdb-uniprot50 \| AF-A0A645IKQ3-F1-MODEL\_V4 \| 1.0 \| 6.876e-06 \| 232 \| 0.324 \| 114 \| 71 \| 5 \| 246 \| 357 \| 2 \| 111 \| DUF3799 domain-containing protein \| DUF3799 domain-containing protein; afdb-uniprot50 \| AF-A0A8A7W4S7-F1-MODEL\_V4 \| 1.0 \| 1.27e-07 \| 232 \| 0.134 \| 364 \| 195 \| 16 \| 17 \| 371 \| 2 \| 254 \| PD-(D/E)XK nuclease family protein \| PD-(D/E)XK nuclease family protein; afdb-uniprot50 \| AF-X0YVD2-F1-MODEL\_V4 \| 1.0 \| 8.077e-09 \| 230 \| 0.175 \| 280 \| 171 \| 16 \| 90 \| 354 \| 3 \| 237 \| DUF3799 domain-containing protein \| DUF3799 domain-containing protein; afdb-uniprot50 \| AF-A0A139TR47-F1-MODEL\_V4 \| 1.0 \| 8.098e-08 \| 230 \| 0.131 \| 374 \| 211 \| 12 \| 1 \| 371 \| 1 \| 263 \| PDDEXK\_1 domain-containing protein \| PDDEXK\_1 domain-containing protein; afdb-uniprot50 \| AF-A0A6B2LYD2-F1-MODEL\_V4 \| 1.0 \| 7.655e-08 \| 230 \| 0.117 \| 332 \| 186 \| 13 \| 17 \| 343 \| 19 \| 248 \| PD-(D/E)XK nuclease family protein \| PD-(D/E)XK nuclease family protein; afdb-uniprot50 \| AF-X0SG33-F1-MODEL\_V4 \| 1.0 \| 2.624e-09 \| 228 \| 0.164 \| 352 \| 200 \| 18 \| 43 \| 368 \| 33 \| 316 \| Uncharacterized protein \| Uncharacterized protein; afdb-uniprot50 \| AF-A0A1F9X0L3-F1-MODEL\_V4 \| 1.0 \| 1.343e-07 \| 223 \| 0.12 \| 340 \| 183 \| 11 \| 18 \| 350 \| 4 \| 234 \| PDDEXK\_1 domain-containing protein \| PDDEXK\_1 domain-containing protein; afdb-uniprot50 \| AF-A0A1M3CJF3-F1-MODEL\_V4 \| 1.0 \| 2.493e-07 \| 221 \| 0.125 \| 343 \| 180 \| 16 \| 18 \| 348 \| 10 \| 244 \| PDDEXK\_1 domain-containing protein \| PDDEXK\_1 domain-containing protein; afdb-uniprot50 \| AF-A0A661TMP3-F1-MODEL\_V4 \| 1.0 \| 2.943e-08 \| 221 \| 0.109 \| 348 \| 191 \| 13 \| 18 \| 351 \| 11 \| 253 \| PDDEXK\_1 domain-containing protein \| PDDEXK\_1 domain-containing protein; afdb-uniprot50 \| AF-A0A2T6D714-F1-MODEL\_V4 \| 1.0 \| 1.2e-07 \| 220 \| 0.133 \| 351 \| 176 \| 16 \| 14 \| 348 \| 6 \| 244 \| PD-(D/E)XK nuclease family protein \| PD-(D/E)XK nuclease family protein; afdb-uniprot50 \| AF-A0A517QQP5-F1-MODEL\_V4 \| 1.0 \| 1.503e-07 \| 219 \| 0.125 \| 360 \| 203 \| 15 \| 1 \| 350 \| 3 \| 260 \| PD-(D/E)XK nuclease superfamily protein \| PD-(D/E)XK nuclease superfamily protein; afdb-uniprot50 \| AF-A0A1M6GGK6-F1-MODEL\_V4 \| 1.0 \| 6.841e-08 \| 218 \| 0.144 \| 380 \| 202 \| 18 \| 1 \| 372 \| 4 \| 268 \| Putative RecB family exonuclease \| Putative RecB family exonuclease; afdb-uniprot50 \| AF-A0A831P8V5-F1-MODEL\_V4 \| 1.0 \| 2.351e-08 \| 218 \| 0.135 \| 361 \| 190 \| 12 \| 1 \| 350 \| 8 \| 257 \| PD-(D/E)XK nuclease family protein \| PD-(D/E)XK nuclease family protein; afdb-uniprot50 \| AF-A0A1V9R6K6-F1-MODEL\_V4 \| 1.0 \| 4.374e-07 \| 218 \| 0.121 \| 386 \| 192 \| 17 \| 11 \| 367 \| 8 \| 275 \| DUF3799 domain-containing protein \| DUF3799 domain-containing protein; afdb-uniprot50 \| AF-A0A722BMP5-F1-MODEL\_V4 \| 1.0 \| 2.369e-05 \| 217 \| 0.26 \| 150 \| 95 \| 5 \| 233 \| 372 \| 1 \| 144 \| DNA breaking-rejoining protein \| DNA breaking-rejoining protein; afdb-uniprot50 \| AF-A0A0R2B3T1-F1-MODEL\_V4 \| 1.0 \| 2.357e-07 \| 217 \| 0.162 \| 222 \| 142 \| 11 \| 157 \| 357 \| 18 \| 216 \| DUF3799 domain-containing protein \| DUF3799 domain-containing protein; afdb-uniprot50 \| AF-A0A2H0X8D0-F1-MODEL\_V4 \| 1.0 \| 1.779e-07 \| 217 \| 0.102 \| 333 \| 189 \| 13 \| 19 \| 348 \| 11 \| 236 \| PDDEXK\_1 domain-containing protein \| PDDEXK\_1 domain-containing protein; afdb-uniprot50 \| AF-A0A3N9MVW9-F1-MODEL\_V4 \| 1.0 \| 9.061e-08 \| 217 \| 0.115 \| 345 \| 185 \| 14 \| 17 \| 350 \| 2 \| 237 \| PD-(D/E)XK nuclease family protein \| PD-(D/E)XK nuclease family protein; afdb-uniprot50 \| AF-A0A5V0PBZ2-F1-MODEL\_V4 \| 1.0 \| 9.585e-08 \| 216 \| 0.195 \| 235 \| 128 \| 5 \| 71 \| 250 \| 2 \| 230 \| Uncharacterized protein \| Uncharacterized protein; afdb-uniprot50 \| AF-X0YAJ3-F1-MODEL\_V4 \| 1.0 \| 3.909e-07 \| 215 \| 0.129 \| 341 \| 173 \| 17 \| 19 \| 343 \| 2 \| 234 \| PDDEXK\_1 domain-containing protein \| PDDEXK\_1 domain-containing protein; afdb-uniprot50 \| AF-A0A6A7VXV8-F1-MODEL\_V4 \| 1.0 \| 2.797e-06 \| 214 \| 0.137 \| 348 \| 164 \| 13 \| 6 \| 340 \| 1 \| 225 \| Uncharacterized protein \| Uncharacterized protein; afdb-uniprot50 \| AF-A0A2A2QKQ5-F1-MODEL\_V4 \| 1.0 \| 1.882e-07 \| 214 \| 0.132 \| 378 \| 210 \| 16 \| 1 \| 372 \| 9 \| 274 \| PDDEXK\_1 domain-containing protein \| PDDEXK\_1 domain-containing protein; afdb-uniprot50 \| AF-A0A7X7QZ66-F1-MODEL\_V4 \| 1.0 \| 2.966e-05 \| 213 \| 0.348 \| 89 \| 58 \| 0 \| 273 \| 361 \| 2 \| 90 \| DUF3799 domain-containing protein \| DUF3799 domain-containing protein; afdb-uniprot50 \| AF-A0A5W5XL97-F1-MODEL\_V4 \| 1.0 \| 2.369e-05 \| 211 \| 0.253 \| 146 \| 93 \| 5 \| 237 \| 372 \| 3 \| 142 \| DNA breaking-rejoining protein \| DNA breaking-rejoining protein; afdb-uniprot50 \| AF-A0A0F9F1T3-F1-MODEL\_V4 \| 1.0 \| 4.65e-05 \| 211 \| 0.228 \| 127 \| 90 \| 4 \| 222 \| 343 \| 4 \| 127 \| DUF3799 domain-containing protein \| DUF3799 domain-containing protein; afdb-uniprot50 \| AF-A0A496LXW8-F1-MODEL\_V4 \| 1.0 \| 7.674e-07 \| 211 \| 0.118 \| 380 \| 191 \| 19 \| 16 \| 372 \| 3 \| 261 \| DUF3799 domain-containing protein \| DUF3799 domain-containing protein; afdb-uniprot50 \| AF-A0A3C0NN37-F1-MODEL\_V4 \| 1.0 \| 8.118e-07 \| 210 \| 0.131 \| 379 \| 173 \| 17 \| 6 \| 361 \| 1 \| 246 \| DUF3799 domain-containing protein \| DUF3799 domain-containing protein; afdb-uniprot50 \| AF-A0A1M5WQQ9-F1-MODEL\_V4 \| 1.0 \| 2.637e-07 \| 206 \| 0.139 \| 374 \| 211 \| 20 \| 19 \| 357 \| 2 \| 299 \| Uncharacterized protein \| Uncharacterized protein; afdb-uniprot50 \| AF-A0A6H2A5D7-F1-MODEL\_V4 \| 1.0 \| 1.073e-07 \| 204 \| 0.121 \| 353 \| 194 \| 16 \| 6 \| 348 \| 1 \| 247 \| Putative PD-(D/E)XK nuclease superfamily protein \| Putative PD-(D/E)XK nuclease superfamily protein; afdb-uniprot50 \| AF-A0A2E0QSB0-F1-MODEL\_V4 \| 1.0 \| 3.493e-07 \| 203 \| 0.143 \| 349 \| 188 \| 15 \| 8 \| 348 \| 20 \| 265 \| PDDEXK\_1 domain-containing protein \| PDDEXK\_1 domain-containing protein; afdb-uniprot50 \| AF-H6SML8-F1-MODEL\_V4 \| 1.0 \| 0.0002006 \| 202 \| 0.289 \| 69 \| 49 \| 0 \| 250 \| 318 \| 10 \| 78 \| DUF3799 domain-containing protein \| DUF3799 domain-containing protein; afdb-uniprot50 \| AF-A0A512M8Q3-F1-MODEL\_V4 \| 1.0 \| 1.273e-06 \| 201 \| 0.117 \| 340 \| 186 \| 15 \| 18 \| 350 \| 24 \| 256 \| PDDEXK\_1 domain-containing protein \| PDDEXK\_1 domain-containing protein; afdb-uniprot50 \| AF-A0A7C3TTB3-F1-MODEL\_V4 \| 1.0 \| 0.0001209 \| 200 \| 0.257 \| 105 \| 76 \| 1 \| 254 \| 358 \| 14 \| 116 \| DUF3799 domain-containing protein \| DUF3799 domain-containing protein; afdb-uniprot50 \| AF-A0A7L6WNN6-F1-MODEL\_V4 \| 1.0 \| 4.894e-07 \| 200 \| 0.148 \| 250 \| 171 \| 13 \| 137 \| 360 \| 17 \| 250 \| PD-(D/E)XK nuclease-like domain-containing protein \| PD-(D/E)XK nuclease-like domain-containing protein; afdb-uniprot50 \| AF-A0A1Z8QGD4-F1-MODEL\_V4 \| 1.0 \| 8.588e-07 \| 200 \| 0.101 \| 345 \| 183 \| 13 \| 18 \| 347 \| 5 \| 237 \| PDDEXK\_1 domain-containing protein \| PDDEXK\_1 domain-containing protein; afdb-uniprot50 \| AF-A0A851GBJ6-F1-MODEL\_V4 \| 1.0 \| 1.27e-07 \| 198 \| 0.158 \| 373 \| 183 \| 23 \| 7 \| 352 \| 20 \| 288 \| PD-(D/E)XK nuclease family protein \| PD-(D/E)XK nuclease family protein; afdb-uniprot50 \| AF-A0A556QJB4-F1-MODEL\_V4 \| 1.0 \| 1.347e-06 \| 197 \| 0.149 \| 362 \| 189 \| 16 \| 1 \| 348 \| 7 \| 263 \| PD-(D/E)XK nuclease family protein \| PD-(D/E)XK nuclease family protein; afdb-uniprot50 \| AF-A0A0F9HJD9-F1-MODEL\_V4 \| 1.0 \| 1.014e-07 \| 196 \| 0.142 \| 344 \| 186 \| 15 \| 19 \| 348 \| 14 \| 262 \| PDDEXK\_1 domain-containing protein \| PDDEXK\_1 domain-containing protein; afdb-uniprot50 \| AF-A0A758AID2-F1-MODEL\_V4 \| 1.0 \| 8.159e-05 \| 195 \| 0.236 \| 127 \| 84 \| 4 \| 253 \| 372 \| 20 \| 140 \| DNA breaking-rejoining protein \| DNA breaking-rejoining protein; afdb-uniprot50 \| AF-A0A6A6K1N9-F1-MODEL\_V4 \| 1.0 \| 1.678e-08 \| 195 \| 0.179 \| 318 \| 152 \| 9 \| 43 \| 359 \| 123 \| 332 \| DUF3799 domain-containing protein \| DUF3799 domain-containing protein; afdb-uniprot50 \| AF-A0A0R2JVU7-F1-MODEL\_V4 \| 1.0 \| 5.491e-06 \| 194 \| 0.185 \| 183 \| 118 \| 10 \| 197 \| 357 \| 3 \| 176 \| DUF3799 domain-containing protein \| DUF3799 domain-containing protein; afdb-uniprot50 \| AF-A0A3A4UKW7-F1-MODEL\_V4 \| 1.0 \| 5.794e-07 \| 193 \| 0.131 \| 373 \| 187 \| 20 \| 17 \| 359 \| 4 \| 269 \| PD-(D/E)XK nuclease family protein \| PD-(D/E)XK nuclease family protein; afdb-uniprot50 \| AF-A0A3A4NCG6-F1-MODEL\_V4 \| 1.0 \| 2.493e-07 \| 193 \| 0.112 \| 355 \| 206 \| 12 \| 1 \| 351 \| 15 \| 264 \| PD-(D/E)XK nuclease family protein \| PD-(D/E)XK nuclease family protein; afdb-uniprot50 \| AF-A0A3R8KJC4-F1-MODEL\_V4 \| 1.0 \| 8.118e-07 \| 193 \| 0.127 \| 352 \| 198 \| 15 \| 7 \| 351 \| 47 \| 296 \| PD-(D/E)XK nuclease family protein \| PD-(D/E)XK nuclease family protein; afdb-uniprot50 \| AF-A0A5T2DQK0-F1-MODEL\_V4 \| 1.0 \| 0.0001022 \| 190 \| 0.226 \| 128 \| 86 \| 4 \| 252 \| 372 \| 6 \| 127 \| DNA breaking-rejoining protein \| DNA breaking-rejoining protein; afdb-uniprot50 \| AF-A0A1B1UKD2-F1-MODEL\_V4 \| 1.0 \| 0.000818 \| 188 \| 0.252 \| 91 \| 67 \| 1 \| 252 \| 342 \| 6 \| 95 \| Uncharacterized protein \| Uncharacterized protein; afdb-uniprot50 \| AF-A0A6C2U3G2-F1-MODEL\_V4 \| 1.0 \| 3.302e-07 \| 187 \| 0.127 \| 353 \| 183 \| 18 \| 12 \| 348 \| 6 \| 249 \| PDDEXK\_1 domain-containing protein \| PDDEXK\_1 domain-containing protein; afdb-uniprot50 \| AF-A0A5Y3WNM2-F1-MODEL\_V4 \| 1.0 \| 0.0001209 \| 186 \| 0.234 \| 128 \| 85 \| 4 \| 252 \| 372 \| 3 \| 124 \| DNA breaking-rejoining protein \| DNA breaking-rejoining protein; afdb-uniprot50 \| AF-A0A4R1S4K0-F1-MODEL\_V4 \| 1.0 \| 1.996e-06 \| 186 \| 0.128 \| 350 \| 181 \| 15 \| 6 \| 347 \| 17 \| 250 \| RecB family exonuclease \| RecB family exonuclease; afdb-uniprot50 \| AF-A0A5W2H069-F1-MODEL\_V4 \| 1.0 \| 9.061e-08 \| 185 \| 0.199 \| 261 \| 151 \| 7 \| 14 \| 220 \| 2 \| 258 \| Uncharacterized protein \| Uncharacterized protein; afdb-uniprot50 \| AF-A0A5Y0QKR5-F1-MODEL\_V4 \| 1.0 \| 2.637e-07 \| 185 \| 0.19 \| 257 \| 150 \| 7 \| 1 \| 203 \| 21 \| 273 \| Exodeoxyribonuclease \| Exodeoxyribonuclease; afdb-uniprot50 \| AF-A0A610YZ68-F1-MODEL\_V4 \| 1.0 \| 9.659e-05 \| 184 \| 0.228 \| 127 \| 85 \| 4 \| 253 \| 372 \| 18 \| 138 \| DUF3799 domain-containing protein \| DUF3799 domain-containing protein; afdb-uniprot50 \| AF-A0A5P0ZGJ0-F1-MODEL\_V4 \| 1.0 \| 7.236e-08 \| 184 \| 0.15 \| 346 \| 177 \| 20 \| 43 \| 357 \| 4 \| 263 \| DUF3799 domain-containing protein \| DUF3799 domain-containing protein; afdb-uniprot50 \| AF-A0A410SCK6-F1-MODEL\_V4 \| 1.0 \| 6.5e-06 \| 180 \| 0.168 \| 196 \| 142 \| 5 \| 178 \| 369 \| 6 \| 184 \| PD-(D/E)XK nuclease family protein \| PD-(D/E)XK nuclease family protein; afdb-uniprot50 \| AF-A0A3A4A5A1-F1-MODEL\_V4 \| 1.0 \| 1.594e-06 \| 176 \| 0.134 \| 358 \| 203 \| 16 \| 1 \| 350 \| 1 \| 259 \| PD-(D/E)XK nuclease family protein \| PD-(D/E)XK nuclease family protein; afdb-uniprot50 \| AF-A0A853QFX1-F1-MODEL\_V4 \| 1.0 \| 2.363e-06 \| 175 \| 0.152 \| 275 \| 149 \| 14 \| 116 \| 356 \| 2 \| 226 \| Uncharacterized protein \| Uncharacterized protein; afdb-uniprot50 \| AF-A0A6M3IPF6-F1-MODEL\_V4 \| 1.0 \| 6.483e-07 \| 174 \| 0.109 \| 346 \| 182 \| 15 \| 8 \| 342 \| 1 \| 231 \| Putative PD-(D/E)XK nuclease superfamily protein \| Putative PD-(D/E)XK nuclease superfamily protein; afdb-uniprot50 \| AF-A0A5S9M6A0-F1-MODEL\_V4 \| 1.0 \| 0.0001279 \| 172 \| 0.187 \| 144 \| 99 \| 8 \| 222 \| 353 \| 4 \| 141 \| DUF3799 domain-containing protein \| DUF3799 domain-containing protein; afdb-uniprot50 \| AF-A0A7J9WA11-F1-MODEL\_V4 \| 1.0 \| 4.135e-07 \| 170 \| 0.122 \| 342 \| 185 \| 17 \| 18 \| 351 \| 164 \| 398 \| PDDEXK\_1 domain-containing protein \| PDDEXK\_1 domain-containing protein; afdb-uniprot50 \| AF-A0A1V5IYR0-F1-MODEL\_V4 \| 1.0 \| 6.144e-06 \| 169 \| 0.109 \| 338 \| 187 \| 16 \| 18 \| 347 \| 5 \| 236 \| PD-(D/E)XK nuclease superfamily protein \| PD-(D/E)XK nuclease superfamily protein; afdb-uniprot50 \| AF-A0A6M3LHK4-F1-MODEL\_V4 \| 1.0 \| 1.276e-05 \| 168 \| 0.142 \| 218 \| 135 \| 13 \| 177 \| 357 \| 13 \| 215 \| DUF3799 domain-containing protein \| DUF3799 domain-containing protein; afdb-uniprot50 \| AF-A0A1I4S4P9-F1-MODEL\_V4 \| 1.0 \| 9.084e-07 \| 168 \| 0.101 \| 345 \| 191 \| 17 \| 7 \| 343 \| 158 \| 391 \| DNA helicase \| DNA helicase; afdb-uniprot50 \| AF-X7ZHD3-F1-MODEL\_V4 \| 1.0 \| 0.0003939 \| 167 \| 0.258 \| 124 \| 75 \| 1 \| 253 \| 359 \| 9 \| 132 \| Gp60 domain protein \| Gp60 domain protein; afdb-uniprot50 \| AF-A0A2N2ISB4-F1-MODEL\_V4 \| 1.0 \| 4.907e-06 \| 166 \| 0.121 \| 361 \| 179 \| 17 \| 1 \| 341 \| 1 \| 243 \| PDDEXK\_1 domain-containing protein \| PDDEXK\_1 domain-containing protein; afdb-uniprot50 \| AF-A0A3N5JX30-F1-MODEL\_V4 \| 1.0 \| 0.0002811 \| 165 \| 0.19 \| 147 \| 103 \| 6 \| 196 \| 338 \| 2 \| 136 \| PDDEXK\_1 domain-containing protein \| PDDEXK\_1 domain-containing protein; afdb-uniprot50 \| AF-A0A4Q3AU83-F1-MODEL\_V4 \| 1.0 \| 6.16e-05 \| 163 \| 0.15 \| 179 \| 136 \| 10 \| 178 \| 349 \| 6 \| 175 \| PD-(D/E)XK nuclease family protein \| PD-(D/E)XK nuclease family protein; afdb-uniprot50 \| AF-A0A7V3L4A6-F1-MODEL\_V4 \| 1.0 \| 1.887e-06 \| 163 \| 0.121 \| 361 \| 205 \| 19 \| 19 \| 350 \| 10 \| 287 \| PDDEXK\_1 domain-containing protein \| PDDEXK\_1 domain-containing protein; afdb-uniprot50 \| AF-A0A7C3FI44-F1-MODEL\_V4 \| 1.0 \| 1.511e-05 \| 162 \| 0.141 \| 226 \| 134 \| 9 \| 105 \| 319 \| 5 \| 181 \| PDDEXK\_1 domain-containing protein \| PDDEXK\_1 domain-containing protein; afdb-uniprot50 \| AF-A0LJW0-F1-MODEL\_V4 \| 1.0 \| 3.511e-05 \| 161 \| 0.138 \| 159 \| 108 \| 9 \| 223 \| 372 \| 1 \| 139 \| PDDEXK\_1 domain-containing protein \| PDDEXK\_1 domain-containing protein; afdb-uniprot50 \| AF-A0A842HHQ9-F1-MODEL\_V4 \| 1.0 \| 8.159e-05 \| 160 \| 0.17 \| 182 \| 124 \| 12 \| 182 \| 348 \| 5 \| 174 \| PD-(D/E)XK nuclease family protein \| PD-(D/E)XK nuclease family protein; afdb-uniprot50 \| AF-A0A0F9T4J1-F1-MODEL\_V4 \| 1.0 \| 9.634e-06 \| 160 \| 0.111 \| 270 \| 177 \| 11 \| 111 \| 370 \| 8 \| 224 \| Uncharacterized protein \| Uncharacterized protein; afdb-uniprot50 \| AF-A0A3C0ZKX1-F1-MODEL\_V4 \| 1.0 \| 2.111e-06 \| 160 \| 0.11 \| 336 \| 183 \| 16 \| 16 \| 343 \| 360 \| 587 \| DNA helicase \| DNA helicase; afdb-uniprot50 \| AF-A0A6M3INL5-F1-MODEL\_V4 \| 1.0 \| 1.69e-05 \| 159 \| 0.106 \| 244 \| 169 \| 8 \| 111 \| 349 \| 8 \| 207 \| Putative PD-(D/E)XK nuclease superfamily protein \| Putative PD-(D/E)XK nuclease superfamily protein; afdb-uniprot50 \| AF-A0A6J7E3Q5-F1-MODEL\_V4 \| 1.0 \| 3.129e-06 \| 158 \| 0.112 \| 373 \| 222 \| 20 \| 2 \| 350 \| 10 \| 297 \| Unannotated protein \| Unannotated protein; afdb-uniprot50 \| AF-A0A5C7RJB2-F1-MODEL\_V4 \| 1.0 \| 1.347e-06 \| 157 \| 0.084 \| 343 \| 200 \| 15 \| 8 \| 343 \| 68 \| 303 \| DNA helicase \| DNA helicase; afdb-uniprot50 \| AF-A0A6G2KJL8-F1-MODEL\_V4 \| 1.0 \| 7.292e-05 \| 157 \| 0.174 \| 183 \| 117 \| 8 \| 159 \| 337 \| 954 \| 1106 \| DNA helicase \| DNA helicase; afdb-uniprot50 \| AF-A0A0F9G4G2-F1-MODEL\_V4 \| 1.0 \| 0.0008653 \| 155 \| 0.329 \| 97 \| 56 \| 4 \| 270 \| 361 \| 2 \| 94 \| DUF3799 domain-containing protein \| DUF3799 domain-containing protein; afdb-uniprot50 \| AF-A0A7X9KIB3-F1-MODEL\_V4 \| 1.0 \| 0.0001695 \| 155 \| 0.167 \| 137 \| 102 \| 7 \| 201 \| 333 \| 7 \| 135 \| PDDEXK\_1 domain-containing protein \| PDDEXK\_1 domain-containing protein; afdb-uniprot50 \| AF-A0A6M3IV43-F1-MODEL\_V4 \| 1.0 \| 1.276e-05 \| 154 \| 0.143 \| 265 \| 165 \| 12 \| 111 \| 368 \| 9 \| 218 \| Putative PD-(D/E)XK nuclease superfamily protein \| Putative PD-(D/E)XK nuclease superfamily protein; afdb-uniprot50 \| AF-A0A3N5PX14-F1-MODEL\_V4 \| 1.0 \| 2.499e-06 \| 154 \| 0.113 \| 352 \| 175 \| 17 \| 17 \| 346 \| 337 \| 573 \| DNA helicase \| DNA helicase; afdb-uniprot50 \| AF-A0A7W0FIM0-F1-MODEL\_V4 \| 1.0 \| 5.491e-06 \| 150 \| 0.11 \| 335 \| 180 \| 16 \| 16 \| 338 \| 568 \| 796 \| DNA helicase \| DNA helicase; afdb-uniprot50 \| AF-A0A6L2R4U5-F1-MODEL\_V4 \| 1.0 \| 6.5e-06 \| 150 \| 0.093 \| 353 \| 201 \| 18 \| 7 \| 351 \| 702 \| 943 \| DNA helicase \| DNA helicase; afdb-uniprot50 \| AF-R6ZK78-F1-MODEL\_V4 \| 1.0 \| 8.139e-06 \| 149 \| 0.118 \| 355 \| 190 \| 12 \| 6 \| 350 \| 80 \| 321 \| PDDEXK\_1 domain-containing protein \| PDDEXK\_1 domain-containing protein; afdb-uniprot50 \| AF-A0A7W9ZL67-F1-MODEL\_V4 \| 1.0 \| 6.5e-06 \| 149 \| 0.098 \| 376 \| 213 \| 18 \| 6 \| 370 \| 115 \| 375 \| DNA helicase \| DNA helicase; afdb-uniprot50 \| AF-A0A2H5VL30-F1-MODEL\_V4 \| 1.0 \| 4.135e-07 \| 149 \| 0.158 \| 353 \| 200 \| 20 \| 7 \| 350 \| 940 \| 1204 \| DNA helicase \| DNA helicase; afdb-uniprot50 \| AF-A0A537ZYU4-F1-MODEL\_V4 \| 1.0 \| 4.894e-07 \| 148 \| 0.128 \| 350 \| 217 \| 16 \| 6 \| 354 \| 871 \| 1133 \| DNA helicase \| DNA helicase; afdb-uniprot50 \| AF-A0A0R1TRL8-F1-MODEL\_V4 \| 1.0 \| 0.0004932 \| 147 \| 0.14 \| 164 \| 114 \| 9 \| 225 \| 368 \| 2 \| 158 \| DUF3799 domain-containing protein \| DUF3799 domain-containing protein; afdb-uniprot50 \| AF-A0A706XGY7-F1-MODEL\_V4 \| 1.0 \| 4.639e-06 \| 147 \| 0.225 \| 204 \| 117 \| 4 \| 83 \| 250 \| 6 \| 204 \| Uncharacterized protein \| Uncharacterized protein; afdb-uniprot50 \| AF-A0A7W1BLJ0-F1-MODEL\_V4 \| 1.0 \| 1.594e-06 \| 147 \| 0.129 \| 348 \| 193 \| 21 \| 7 \| 338 \| 637 \| 890 \| DNA helicase \| DNA helicase; afdb-uniprot50 \| AF-A0A6G0A6A2-F1-MODEL\_V4 \| 1.0 \| 0.0001279 \| 147 \| 0.213 \| 183 \| 102 \| 9 \| 162 \| 336 \| 991 \| 1139 \| DNA helicase \| DNA helicase; afdb-uniprot50 \| AF-A0A2H0LI12-F1-MODEL\_V4 \| 1.0 \| 0.0002512 \| 146 \| 0.156 \| 147 \| 107 \| 5 \| 208 \| 351 \| 2 \| 134 \| PDDEXK\_1 domain-containing protein \| PDDEXK\_1 domain-containing protein; afdb-uniprot50 \| AF-A0A5Y3U2L9-F1-MODEL\_V4 \| 1.0 \| 6.144e-06 \| 146 \| 0.226 \| 203 \| 118 \| 3 \| 83 \| 250 \| 5 \| 203 \| DNA breaking-rejoining protein \| DNA breaking-rejoining protein; afdb-uniprot50 \| AF-A0A2E7SPB8-F1-MODEL\_V4 \| 1.0 \| 1.69e-05 \| 146 \| 0.106 \| 367 \| 198 \| 20 \| 10 \| 347 \| 14 \| 279 \| PDDEXK\_1 domain-containing protein \| PDDEXK\_1 domain-containing protein; afdb-uniprot50 \| AF-A0A843GQU3-F1-MODEL\_V4 \| 1.0 \| 0.003153 \| 145 \| 0.19 \| 100 \| 76 \| 4 \| 254 \| 351 \| 1 \| 97 \| PD-(D/E)XK nuclease-like domain-containing protein \| PD-(D/E)XK nuclease-like domain-containing protein; afdb-uniprot50 \| AF-A0A3D5RXV7-F1-MODEL\_V4 \| 1.0 \| 0.0003327 \| 145 \| 0.174 \| 149 \| 102 \| 9 \| 201 \| 344 \| 7 \| 139 \| PDDEXK\_1 domain-containing protein \| PDDEXK\_1 domain-containing protein; afdb-uniprot50 \| AF-A0A1V5U7X3-F1-MODEL\_V4 \| 1.0 \| 0.0001353 \| 145 \| 0.137 \| 204 \| 139 \| 9 \| 154 \| 342 \| 1 \| 182 \| PDDEXK\_1 domain-containing protein \| PDDEXK\_1 domain-containing protein; afdb-uniprot50 \| AF-A0A6Y1UZP9-F1-MODEL\_V4 \| 1.0 \| 3.919e-06 \| 145 \| 0.226 \| 203 \| 118 \| 4 \| 83 \| 250 \| 3 \| 201 \| DNA breaking-rejoining protein \| DNA breaking-rejoining protein; afdb-uniprot50 \| AF-A0A709G0R9-F1-MODEL\_V4 \| 1.0 \| 6.144e-06 \| 145 \| 0.226 \| 203 \| 118 \| 3 \| 83 \| 250 \| 6 \| 204 \| DNA breaking-rejoining protein \| DNA breaking-rejoining protein; afdb-uniprot50 \| AF-U2QAP5-F1-MODEL\_V4 \| 1.0 \| 1.35e-05 \| 145 \| 0.074 \| 334 \| 197 \| 15 \| 16 \| 343 \| 109 \| 336 \| DNA helicase \| DNA helicase; afdb-uniprot50 \| AF-A0A708WCW9-F1-MODEL\_V4 \| 1.0 \| 6.876e-06 \| 144 \| 0.226 \| 203 \| 118 \| 3 \| 83 \| 250 \| 4 \| 202 \| DNA breaking-rejoining protein \| DNA breaking-rejoining protein; afdb-uniprot50 \| AF-A0A726Q5Q9-F1-MODEL\_V4 \| 1.0 \| 6.876e-06 \| 144 \| 0.226 \| 203 \| 118 \| 3 \| 83 \| 250 \| 4 \| 202 \| DNA breaking-rejoining protein \| DNA breaking-rejoining protein; afdb-uniprot50 \| AF-A0A702ARL0-F1-MODEL\_V4 \| 1.0 \| 4.385e-06 \| 144 \| 0.225 \| 204 \| 117 \| 4 \| 83 \| 250 \| 5 \| 203 \| Uncharacterized protein \| Uncharacterized protein; afdb-uniprot50 \| AF-A0A3M1HHZ9-F1-MODEL\_V4 \| 1.0 \| 8.61e-06 \| 144 \| 0.103 \| 338 \| 195 \| 15 \| 17 \| 346 \| 94 \| 331 \| PD-(D/E)XK nuclease family protein \| PD-(D/E)XK nuclease family protein; afdb-uniprot50 \| AF-A0A3A2J0B7-F1-MODEL\_V4 \| 1.0 \| 1.598e-05 \| 144 \| 0.117 \| 356 \| 197 \| 16 \| 6 \| 353 \| 414 \| 660 \| DNA helicase \| DNA helicase; afdb-uniprot50 \| AF-A0A7X8LCA3-F1-MODEL\_V4 \| 1.0 \| 1.019e-05 \| 144 \| 0.095 \| 344 \| 195 \| 17 \| 16 \| 351 \| 502 \| 737 \| DNA helicase \| DNA helicase; afdb-uniprot50 \| AF-D8FG64-F1-MODEL\_V4 \| 1.0 \| 3.31e-06 \| 143 \| 0.122 \| 336 \| 188 \| 18 \| 16 \| 335 \| 8 \| 252 \| PDDEXK\_1 domain-containing protein \| PDDEXK\_1 domain-containing protein; afdb-uniprot50 \| AF-A0A359LS55-F1-MODEL\_V4 \| 1.0 \| 9.107e-06 \| 143 \| 0.119 \| 352 \| 184 \| 15 \| 6 \| 342 \| 579 \| 819 \| PDDEXK\_1 domain-containing protein \| PDDEXK\_1 domain-containing protein; afdb-uniprot50 \| AF-A0A327X879-F1-MODEL\_V4 \| 1.0 \| 3.929e-05 \| 143 \| 0.113 \| 345 \| 191 \| 17 \| 18 \| 352 \| 707 \| 946 \| DNA helicase \| DNA helicase; afdb-uniprot50 \| AF-A0A722ZM29-F1-MODEL\_V4 \| 1.0 \| 6.876e-06 \| 142 \| 0.226 \| 203 \| 118 \| 4 \| 83 \| 250 \| 2 \| 200 \| DNA breaking-rejoining protein \| DNA breaking-rejoining protein; afdb-uniprot50 \| AF-A0A607LSL1-F1-MODEL\_V4 \| 1.0 \| 8.61e-06 \| 142 \| 0.221 \| 203 \| 119 \| 4 \| 83 \| 250 \| 3 \| 201 \| Uncharacterized protein \| Uncharacterized protein; afdb-uniprot50 \| AF-A0A614G6N4-F1-MODEL\_V4 \| 1.0 \| 8.61e-06 \| 142 \| 0.225 \| 204 \| 117 \| 4 \| 83 \| 250 \| 4 \| 202 \| Uncharacterized protein \| Uncharacterized protein; afdb-uniprot50 \| AF-A0A611B465-F1-MODEL\_V4 \| 1.0 \| 9.107e-06 \| 142 \| 0.226 \| 203 \| 118 \| 3 \| 83 \| 250 \| 7 \| 205 \| DNA breaking-rejoining protein \| DNA breaking-rejoining protein; afdb-uniprot50 \| AF-A0A1E5IMV9-F1-MODEL\_V4 \| 1.0 \| 9.634e-06 \| 142 \| 0.125 \| 344 \| 204 \| 16 \| 6 \| 341 \| 31 \| 285 \| PDDEXK\_1 domain-containing protein \| PDDEXK\_1 domain-containing protein; afdb-uniprot50 \| AF-A0A538KGZ7-F1-MODEL\_V4 \| 1.0 \| 4.385e-06 \| 142 \| 0.116 \| 359 \| 181 \| 14 \| 6 \| 341 \| 912 \| 1157 \| DNA helicase \| DNA helicase; afdb-uniprot50 \| AF-A0A708ZUJ8-F1-MODEL\_V4 \| 1.0 \| 9.634e-06 \| 141 \| 0.226 \| 203 \| 118 \| 4 \| 83 \| 250 \| 3 \| 201 \| DNA breaking-rejoining protein \| DNA breaking-rejoining protein; afdb-uniprot50 \| AF-A0A704U7J8-F1-MODEL\_V4 \| 1.0 \| 4.907e-06 \| 141 \| 0.226 \| 203 \| 118 \| 3 \| 83 \| 250 \| 7 \| 205 \| DNA breaking-rejoining protein \| DNA breaking-rejoining protein; afdb-uniprot50 \| AF-A0A704ZNC4-F1-MODEL\_V4 \| 1.0 \| 9.634e-06 \| 141 \| 0.225 \| 204 \| 117 \| 4 \| 83 \| 250 \| 7 \| 205 \| Uncharacterized protein \| Uncharacterized protein; afdb-uniprot50 \| AF-A0A317GV55-F1-MODEL\_V4 \| 1.0 \| 7.694e-06 \| 141 \| 0.116 \| 360 \| 212 \| 21 \| 9 \| 343 \| 1 \| 279 \| PDDEXK\_1 domain-containing protein \| PDDEXK\_1 domain-containing protein; afdb-uniprot50 \| AF-A0A3Z3JMT0-F1-MODEL\_V4 \| 1.0 \| 4.639e-06 \| 140 \| 0.225 \| 204 \| 117 \| 4 \| 83 \| 250 \| 7 \| 205 \| Uncharacterized protein \| Uncharacterized protein; afdb-uniprot50 \| AF-I4EFM0-F1-MODEL\_V4 \| 1.0 \| 5.505e-05 \| 140 \| 0.167 \| 173 \| 122 \| 10 \| 193 \| 353 \| 32 \| 194 \| PDDEXK\_1 domain-containing protein \| PDDEXK\_1 domain-containing protein; afdb-uniprot50 \| AF-R7JEN0-F1-MODEL\_V4 \| 1.0 \| 3.137e-05 \| 140 \| 0.119 \| 351 \| 188 \| 16 \| 11 \| 351 \| 336 \| 575 \| DNA helicase \| DNA helicase; afdb-uniprot50 \| AF-A0A523W971-F1-MODEL\_V4 \| 1.0 \| 3.319e-05 \| 140 \| 0.128 \| 359 \| 187 \| 19 \| 7 \| 350 \| 659 \| 906 \| Dna2/Cas4 domain-containing protein \| Dna2/Cas4 domain-containing protein; afdb-uniprot50 \| AF-A0A625UR15-F1-MODEL\_V4 \| 1.0 \| 1.14e-05 \| 139 \| 0.225 \| 204 \| 117 \| 4 \| 83 \| 250 \| 7 \| 205 \| Uncharacterized protein \| Uncharacterized protein; afdb-uniprot50 \| AF-A0A0B5GGD9-F1-MODEL\_V4 \| 1.0 \| 2.001e-05 \| 139 \| 0.074 \| 336 \| 195 \| 18 \| 16 \| 343 \| 206 \| 433 \| DNA helicase \| DNA helicase; afdb-uniprot50 \| AF-A0A254S109-F1-MODEL\_V4 \| 1.0 \| 1.14e-05 \| 138 \| 0.111 \| 342 \| 187 \| 17 \| 7 \| 338 \| 712 \| 946 \| DNA helicase \| DNA helicase; afdb-uniprot50 \| AF-A0A538LVJ5-F1-MODEL\_V4 \| 1.0 \| 6.5e-06 \| 138 \| 0.151 \| 330 \| 175 \| 14 \| 17 \| 338 \| 854 \| 1086 \| DNA helicase \| DNA helicase; afdb-uniprot50 \| AF-A0A2H0RW27-F1-MODEL\_V4 \| 1.0 \| 0.0001695 \| 137 \| 0.131 \| 343 \| 173 \| 17 \| 17 \| 342 \| 2 \| 236 \| PDDEXK\_1 domain-containing protein \| PDDEXK\_1 domain-containing protein; afdb-uniprot50 \| AF-A0A7K0RIH0-F1-MODEL\_V4 \| 1.0 \| 9.107e-06 \| 137 \| 0.145 \| 357 \| 182 \| 17 \| 6 \| 347 \| 212 \| 460 \| Uncharacterized protein \| Uncharacterized protein; afdb-uniprot50 \| AF-A0A1I6KGK5-F1-MODEL\_V4 \| 1.0 \| 4.65e-05 \| 137 \| 0.084 \| 330 \| 191 \| 14 \| 18 \| 342 \| 705 \| 928 \| DNA helicase \| DNA helicase; afdb-uniprot50 \| AF-A0A1E5J1P3-F1-MODEL\_V4 \| 1.0 \| 4.65e-05 \| 137 \| 0.099 \| 333 \| 185 \| 16 \| 18 \| 343 \| 702 \| 926 \| DNA helicase \| DNA helicase; afdb-uniprot50 \| AF-A0A2W5Y867-F1-MODEL\_V4 \| 1.0 \| 1.35e-05 \| 137 \| 0.142 \| 364 \| 167 \| 16 \| 9 \| 338 \| 902 \| 1154 \| DNA helicase \| DNA helicase; afdb-uniprot50 \| AF-A0A3D0QKZ3-F1-MODEL\_V4 \| 1.0 \| 1.428e-05 \| 136 \| 0.106 \| 338 \| 206 \| 15 \| 6 \| 334 \| 1 \| 251 \| PDDEXK\_1 domain-containing protein \| PDDEXK\_1 domain-containing protein; afdb-uniprot50 \| AF-A0A7S7SGN9-F1-MODEL\_V4 \| 1.0 \| 9.634e-06 \| 136 \| 0.091 \| 382 \| 204 \| 16 \| 9 \| 371 \| 609 \| 866 \| PD-(D/E)XK nuclease family protein \| PD-(D/E)XK nuclease family protein; afdb-uniprot50 \| AF-R5VS20-F1-MODEL\_V4 \| 1.0 \| 7.292e-05 \| 135 \| 0.075 \| 343 \| 200 \| 16 \| 18 \| 352 \| 710 \| 943 \| DNA helicase \| DNA helicase; afdb-uniprot50 \| AF-A0A538TBY8-F1-MODEL\_V4 \| 1.0 \| 0.001699 \| 134 \| 0.206 \| 126 \| 85 \| 6 \| 220 \| 341 \| 5 \| 119 \| PDDEXK\_1 domain-containing protein \| PDDEXK\_1 domain-containing protein; afdb-uniprot50 \| AF-B1GYT7-F1-MODEL\_V4 \| 1.0 \| 1.511e-05 \| 134 \| 0.09 \| 342 \| 209 \| 16 \| 6 \| 334 \| 1 \| 253 \| PD-(D/E)XK nuclease family protein \| PD-(D/E)XK nuclease family protein; afdb-uniprot50 \| AF-A0A7V4SV88-F1-MODEL\_V4 \| 1.0 \| 9.107e-06 \| 134 \| 0.136 \| 345 \| 216 \| 16 \| 17 \| 350 \| 5 \| 278 \| PD-(D/E)XK nuclease family protein \| PD-(D/E)XK nuclease family protein; afdb-uniprot50 \| AF-A0A1G9NLF5-F1-MODEL\_V4 \| 1.0 \| 1.276e-05 \| 134 \| 0.126 \| 378 \| 188 \| 19 \| 20 \| 358 \| 3 \| 277 \| PD-(D/E)XK nuclease superfamily protein \| PD-(D/E)XK nuclease superfamily protein; afdb-uniprot50 \| AF-A0A2G0E7V9-F1-MODEL\_V4 \| 1.0 \| 4.65e-05 \| 134 \| 0.077 \| 335 \| 194 \| 17 \| 18 \| 345 \| 360 \| 586 \| DNA helicase \| DNA helicase; afdb-uniprot50 \| AF-A0A5F0K2G5-F1-MODEL\_V4 \| 1.0 \| 2.001e-05 \| 134 \| 0.098 \| 344 \| 189 \| 16 \| 8 \| 342 \| 665 \| 896 \| DNA helicase \| DNA helicase; afdb-uniprot50 \| AF-R7LXF4-F1-MODEL\_V4 \| 1.0 \| 1.276e-05 \| 134 \| 0.129 \| 370 \| 197 \| 19 \| 11 \| 371 \| 762 \| 1015 \| DNA helicase \| DNA helicase; afdb-uniprot50 \| AF-A0A383DLI3-F1-MODEL\_V4 \| 1.0 \| 0.0008653 \| 133 \| 0.179 \| 167 \| 108 \| 9 \| 182 \| 343 \| 3 \| 145 \| PDDEXK\_1 domain-containing protein \| PDDEXK\_1 domain-containing protein; afdb-uniprot50 \| AF-K4ZR42-F1-MODEL\_V4 \| 1.0 \| 0.001084 \| 133 \| 0.175 \| 148 \| 96 \| 8 \| 238 \| 363 \| 7 \| 150 \| DUF3799 domain-containing protein \| DUF3799 domain-containing protein; afdb-uniprot50 \| AF-A0A5C7U316-F1-MODEL\_V4 \| 1.0 \| 7.292e-05 \| 133 \| 0.1 \| 377 \| 199 \| 20 \| 18 \| 370 \| 3 \| 263 \| PD-(D/E)XK nuclease family protein \| PD-(D/E)XK nuclease family protein; afdb-uniprot50 \| AF-A0A0S4SUN1-F1-MODEL\_V4 \| 1.0 \| 7.713e-05 \| 133 \| 0.117 \| 410 \| 182 \| 15 \| 9 \| 344 \| 6 \| 309 \| PD-(D/E)XK nuclease superfamily \| PD-(D/E)XK nuclease superfamily; afdb-uniprot50 \| AF-A0A2E6AAK1-F1-MODEL\_V4 \| 1.0 \| 0.001797 \| 132 \| 0.137 \| 153 \| 102 \| 10 \| 202 \| 344 \| 98 \| 230 \| Uncharacterized protein \| Uncharacterized protein; afdb-uniprot50 \| AF-A0A1W1IJA3-F1-MODEL\_V4 \| 1.0 \| 5.505e-05 \| 132 \| 0.093 \| 332 \| 188 \| 15 \| 18 \| 343 \| 705 \| 929 \| DNA helicase \| DNA helicase; afdb-uniprot50 \| AF-A0A7Y1VN02-F1-MODEL\_V4 \| 1.0 \| 8.159e-05 \| 131 \| 0.158 \| 246 \| 152 \| 13 \| 115 \| 351 \| 18 \| 217 \| Uncharacterized protein \| Uncharacterized protein; afdb-uniprot50 \| AF-A0A011A8K6-F1-MODEL\_V4 \| 1.0 \| 6.16e-05 \| 131 \| 0.149 \| 207 \| 134 \| 12 \| 161 \| 343 \| 44 \| 232 \| PDDEXK\_1 domain-containing protein \| PDDEXK\_1 domain-containing protein; afdb-uniprot50 \| AF-A0A7C6NF67-F1-MODEL\_V4 \| 1.0 \| 5.505e-05 \| 131 \| 0.12 \| 332 \| 159 \| 13 \| 16 \| 338 \| 6 \| 213 \| PD-(D/E)XK nuclease family protein \| PD-(D/E)XK nuclease family protein; afdb-uniprot50 \| AF-A0A7C1SR97-F1-MODEL\_V4 \| 1.0 \| 2.117e-05 \| 131 \| 0.093 \| 375 \| 226 \| 21 \| 19 \| 372 \| 14 \| 295 \| PD-(D/E)XK nuclease family protein \| PD-(D/E)XK nuclease family protein; afdb-uniprot50 \| AF-A0A609LX09-F1-MODEL\_V4 \| 1.0 \| 0.001213 \| 130 \| 0.25 \| 148 \| 93 \| 2 \| 103 \| 250 \| 3 \| 132 \| Uncharacterized protein \| Uncharacterized protein; afdb-uniprot50 \| AF-A0A5X8LAM5-F1-MODEL\_V4 \| 1.0 \| 0.0004407 \| 130 \| 0.231 \| 190 \| 123 \| 4 \| 88 \| 272 \| 6 \| 177 \| Uncharacterized protein \| Uncharacterized protein; afdb-uniprot50 \| AF-A0A6N8NPZ1-F1-MODEL\_V4 \| 1.0 \| 3.929e-05 \| 130 \| 0.221 \| 194 \| 108 \| 4 \| 91 \| 247 \| 2 \| 189 \| Exodeoxyribonuclease VIII \| Exodeoxyribonuclease VIII; afdb-uniprot50 \| AF-A0A0S8KUQ7-F1-MODEL\_V4 \| 1.0 \| 0.0001432 \| 130 \| 0.111 \| 207 \| 143 \| 11 \| 157 \| 343 \| 34 \| 219 \| Uncharacterized protein \| Uncharacterized protein; afdb-uniprot50 \| AF-A0A6G0AGT9-F1-MODEL\_V4 \| 1.0 \| 2.966e-05 \| 130 \| 0.133 \| 337 \| 202 \| 20 \| 17 \| 347 \| 398 \| 650 \| DNA helicase \| DNA helicase; afdb-uniprot50 \| AF-A0A2M7MLQ6-F1-MODEL\_V4 \| 1.0 \| 9.634e-06 \| 130 \| 0.107 \| 343 \| 209 \| 16 \| 18 \| 347 \| 920 \| 1178 \| DNA helicase \| DNA helicase; afdb-uniprot50 \| AF-A0A3C0F7Z2-F1-MODEL\_V4 \| 1.0 \| 0.0004932 \| 129 \| 0.127 \| 204 \| 157 \| 8 \| 178 \| 370 \| 24 \| 217 \| Uncharacterized protein \| Uncharacterized protein; afdb-uniprot50 \| AF-A0A832GQ63-F1-MODEL\_V4 \| 1.0 \| 9.634e-06 \| 129 \| 0.129 \| 354 \| 195 \| 19 \| 16 \| 351 \| 44 \| 302 \| Type I-A CRISPR-associated protein Cas4/Csa1 \| Type I-A CRISPR-associated protein Cas4/Csa1; afdb-uniprot50 \| AF-A0A0F8WAD0-F1-MODEL\_V4 \| 1.0 \| 0.004177 \| 128 \| 0.304 \| 92 \| 59 \| 1 \| 284 \| 370 \| 2 \| 93 \| Uncharacterized protein \| Uncharacterized protein; afdb-uniprot50 \| AF-A0A382D5K0-F1-MODEL\_V4 \| 1.0 \| 0.0002657 \| 128 \| 0.176 \| 187 \| 106 \| 12 \| 164 \| 343 \| 34 \| 179 \| PDDEXK\_1 domain-containing protein \| PDDEXK\_1 domain-containing protein; afdb-uniprot50 \| AF-K1UR72-F1-MODEL\_V4 \| 1.0 \| 0.0002122 \| 128 \| 0.106 \| 254 \| 169 \| 12 \| 114 \| 364 \| 5 \| 203 \| UvrD/REP helicase \| UvrD/REP helicase; afdb-uniprot50 \| AF-A0A2M7X4F5-F1-MODEL\_V4 \| 1.0 \| 2.804e-05 \| 127 \| 0.12 \| 348 \| 188 \| 18 \| 7 \| 342 \| 749 \| 990 \| DNA helicase \| DNA helicase; afdb-uniprot50 \| AF-A0A660LK76-F1-MODEL\_V4 \| 1.0 \| 3.319e-05 \| 127 \| 0.126 \| 341 \| 182 \| 16 \| 17 \| 347 \| 878 \| 1112 \| DNA helicase \| DNA helicase; afdb-uniprot50 \| AF-A0A077ZMK2-F1-MODEL\_V4 \| 1.0 \| 0.0001209 \| 127 \| 0.181 \| 248 \| 164 \| 13 \| 101 \| 322 \| 878 \| 1112 \| DUF4494 and SNF2 N domain containing protein \| DUF4494 and SNF2 N domain containing protein; afdb-uniprot50 \| AF-A0A7C4ZXI5-F1-MODEL\_V4 \| 1.0 \| 2.111e-06 \| 126 \| 0.131 \| 364 \| 196 \| 24 \| 22 \| 370 \| 59 \| 317 \| Type I-A CRISPR-associated protein Cas4/Csa1 \| Type I-A CRISPR-associated protein Cas4/Csa1; afdb-uniprot50 \| AF-A0A0F9SQ52-F1-MODEL\_V4 \| 1.0 \| 4.396e-05 \| 126 \| 0.122 \| 384 \| 193 \| 18 \| 9 \| 370 \| 945 \| 1206 \| DNA helicase \| DNA helicase; afdb-uniprot50 \| AF-A0A6S6Y1F3-F1-MODEL\_V4 \| 1.0 \| 0.000818 \| 125 \| 0.307 \| 127 \| 71 \| 4 \| 254 \| 372 \| 43 \| 160 \| DUF3799 domain-containing protein \| DUF3799 domain-containing protein; afdb-uniprot50 \| AF-A0A3B9YZQ0-F1-MODEL\_V4 \| 1.0 \| 3.929e-05 \| 125 \| 0.132 \| 256 \| 154 \| 15 \| 99 \| 333 \| 423 \| 631 \| DNA helicase \| DNA helicase; afdb-uniprot50 \| AF-A0A6J6ESD5-F1-MODEL\_V4 \| 1.0 \| 7.713e-05 \| 125 \| 0.121 \| 353 \| 167 \| 16 \| 18 \| 351 \| 775 \| 1003 \| DNA helicase \| DNA helicase; afdb-uniprot50 \| AF-A0A7V9VQS8-F1-MODEL\_V4 \| 1.0 \| 0.001084 \| 125 \| 0.167 \| 143 \| 112 \| 4 \| 196 \| 335 \| 958 \| 1096 \| DNA helicase \| DNA helicase; afdb-uniprot50 \| AF-A0A2D6SUJ8-F1-MODEL\_V4 \| 1.0 \| 2.369e-05 \| 124 \| 0.122 \| 367 \| 177 \| 19 \| 6 \| 343 \| 287 \| 537 \| DNA helicase \| DNA helicase; afdb-uniprot50 \| AF-A0A7W1NG24-F1-MODEL\_V4 \| 1.0 \| 0.001357 \| 124 \| 0.167 \| 143 \| 112 \| 4 \| 196 \| 335 \| 758 \| 896 \| DNA helicase \| DNA helicase; afdb-uniprot50 \| AF-A0A506XSD7-F1-MODEL\_V4 \| 1.0 \| 1.35e-05 \| 124 \| 0.121 \| 380 \| 229 \| 22 \| 3 \| 365 \| 683 \| 974 \| PD-(D/E)XK nuclease family protein \| PD-(D/E)XK nuclease family protein; afdb-uniprot50 \| AF-A0A7Y2X9K5-F1-MODEL\_V4 \| 1.0 \| 2.369e-05 \| 124 \| 0.085 \| 397 \| 224 \| 18 \| 6 \| 358 \| 903 \| 1204 \| DNA helicase \| DNA helicase; afdb-uniprot50 \| AF-A0A7X7UZ96-F1-MODEL\_V4 \| 1.0 \| 2.966e-05 \| 124 \| 0.099 \| 352 \| 230 \| 17 \| 6 \| 347 \| 925 \| 1199 \| DNA helicase \| DNA helicase; afdb-uniprot50 \| AF-A0A5U2XD06-F1-MODEL\_V4 \| 1.0 \| 0.004674 \| 123 \| 0.281 \| 110 \| 79 \| 0 \| 141 \| 250 \| 2 \| 111 \| Uncharacterized protein \| Uncharacterized protein; afdb-uniprot50 \| AF-F3KJA7-F1-MODEL\_V4 \| 1.0 \| 0.0006533 \| 123 \| 0.128 \| 164 \| 122 \| 9 \| 182 \| 343 \| 35 \| 179 \| PDDEXK\_1 domain-containing protein \| PDDEXK\_1 domain-containing protein; afdb-uniprot50 \| AF-A0A1Z9PHH6-F1-MODEL\_V4 \| 1.0 \| 0.0002973 \| 122 \| 0.123 \| 227 \| 132 \| 11 \| 178 \| 343 \| 38 \| 258 \| Uncharacterized protein \| Uncharacterized protein; afdb-uniprot50 \| AF-A0A350P152-F1-MODEL\_V4 \| 1.0 \| 0.0005838 \| 122 \| 0.149 \| 174 \| 122 \| 9 \| 217 \| 371 \| 76 \| 242 \| Uncharacterized protein \| Uncharacterized protein; afdb-uniprot50 \| AF-A0A1W9VNN7-F1-MODEL\_V4 \| 1.0 \| 6.893e-05 \| 122 \| 0.105 \| 332 \| 204 \| 18 \| 16 \| 338 \| 11 \| 258 \| PDDEXK\_1 domain-containing protein \| PDDEXK\_1 domain-containing protein; afdb-uniprot50 \| AF-A0A7W0AR45-F1-MODEL\_V4 \| 1.0 \| 3.511e-05 \| 122 \| 0.127 \| 352 \| 193 \| 16 \| 7 \| 347 \| 565 \| 813 \| DNA helicase \| DNA helicase; afdb-uniprot50 \| AF-A0A350UHQ6-F1-MODEL\_V4 \| 1.0 \| 4.919e-05 \| 122 \| 0.111 \| 360 \| 199 \| 21 \| 6 \| 349 \| 583 \| 837 \| PDDEXK\_1 domain-containing protein \| PDDEXK\_1 domain-containing protein; afdb-uniprot50 \| AF-A0A3D3R7C6-F1-MODEL\_V4 \| 1.0 \| 7.292e-05 \| 122 \| 0.122 \| 270 \| 175 \| 11 \| 87 \| 343 \| 948 \| 1168 \| DNA helicase \| DNA helicase; afdb-uniprot50 \| AF-A0A3M2E2F5-F1-MODEL\_V4 \| 1.0 \| 0.000731 \| 121 \| 0.131 \| 183 \| 137 \| 10 \| 196 \| 367 \| 6 \| 177 \| Double-strand break repair protein AddB \| Double-strand break repair protein AddB; afdb-uniprot50 \| AF-A0A6N8BXU9-F1-MODEL\_V4 \| 1.0 \| 0.001699 \| 121 \| 0.1 \| 149 \| 113 \| 7 \| 200 \| 338 \| 80 \| 217 \| PD-(D/E)XK nuclease family protein \| PD-(D/E)XK nuclease family protein; afdb-uniprot50 \| AF-A0A1Q4FIU6-F1-MODEL\_V4 \| 1.0 \| 0.0002006 \| 121 \| 0.2 \| 200 \| 129 \| 13 \| 167 \| 346 \| 31 \| 219 \| PDDEXK\_1 domain-containing protein \| PDDEXK\_1 domain-containing protein; afdb-uniprot50 \| AF-A0A497SU83-F1-MODEL\_V4 \| 1.0 \| 2.506e-05 \| 121 \| 0.112 \| 348 \| 208 \| 18 \| 18 \| 353 \| 41 \| 299 \| Type I-A CRISPR-associated protein Cas4/Csa1 \| Type I-A CRISPR-associated protein Cas4/Csa1; afdb-uniprot50 \| AF-A0A7V7XH75-F1-MODEL\_V4 \| 1.0 \| 0.0001022 \| 121 \| 0.114 \| 350 \| 205 \| 17 \| 19 \| 353 \| 787 \| 1046 \| DNA helicase \| DNA helicase; afdb-uniprot50 \| AF-A0A3E0QRG2-F1-MODEL\_V4 \| 1.0 \| 0.001283 \| 120 \| 0.14 \| 206 \| 129 \| 10 \| 179 \| 343 \| 14 \| 212 \| Uncharacterized protein \| Uncharacterized protein; afdb-uniprot50 \| AF-A0A662HTM3-F1-MODEL\_V4 \| 1.0 \| 0.0001209 \| 120 \| 0.108 \| 342 \| 191 \| 17 \| 19 \| 353 \| 40 \| 274 \| Type I-A CRISPR-associated protein Cas4/Csa1 \| Type I-A CRISPR-associated protein Cas4/Csa1; afdb-uniprot50 \| AF-A0A7C4GM66-F1-MODEL\_V4 \| 1.0 \| 8.631e-05 \| 120 \| 0.12 \| 349 \| 201 \| 18 \| 14 \| 351 \| 42 \| 295 \| Type I-A CRISPR-associated protein Cas4/Csa1 \| Type I-A CRISPR-associated protein Cas4/Csa1; afdb-uniprot50 \| AF-A0A497Q0N2-F1-MODEL\_V4 \| 1.0 \| 4.919e-05 \| 120 \| 0.091 \| 337 \| 199 \| 14 \| 18 \| 336 \| 45 \| 292 \| Type I-A CRISPR-associated protein Cas4/Csa1 \| Type I-A CRISPR-associated protein Cas4/Csa1; afdb-uniprot50 \| AF-A0A7W1M4Q3-F1-MODEL\_V4 \| 1.0 \| 4.156e-05 \| 120 \| 0.123 \| 348 \| 179 \| 14 \| 7 \| 339 \| 250 \| 486 \| DNA helicase \| DNA helicase; afdb-uniprot50 \| AF-A0A7C3DWA4-F1-MODEL\_V4 \| 1.0 \| 2.65e-05 \| 120 \| 0.101 \| 366 \| 207 \| 17 \| 11 \| 368 \| 390 \| 641 \| PD-(D/E)XK nuclease family protein \| PD-(D/E)XK nuclease family protein; afdb-uniprot50 \| AF-A0A3L7WDI4-F1-MODEL\_V4 \| 1.0 \| 0.0001353 \| 120 \| 0.129 \| 254 \| 155 \| 13 \| 99 \| 334 \| 1033 \| 1238 \| DNA helicase \| DNA helicase; afdb-uniprot50 \| AF-A0A136PHC7-F1-MODEL\_V4 \| 1.0 \| 0.001146 \| 119 \| 0.12 \| 166 \| 130 \| 8 \| 203 \| 364 \| 2 \| 155 \| Exonuclease V subunit beta \| Exonuclease V subunit beta; afdb-uniprot50 \| AF-A0A7V9FD96-F1-MODEL\_V4 \| 1.0 \| 8.631e-05 \| 119 \| 0.122 \| 383 \| 173 \| 18 \| 18 \| 369 \| 7 \| 257 \| PD-(D/E)XK nuclease family protein \| PD-(D/E)XK nuclease family protein; afdb-uniprot50 \| AF-A0A2P5LHJ5-F1-MODEL\_V4 \| 1.0 \| 0.0002375 \| 119 \| 0.111 \| 394 \| 205 \| 22 \| 1 \| 368 \| 4 \| 278 \| Double-strand break repair protein AddB \| Double-strand break repair protein AddB; afdb-uniprot50 \| AF-A0A1F8NK16-F1-MODEL\_V4 \| 1.0 \| 3.714e-05 \| 119 \| 0.124 \| 353 \| 200 \| 17 \| 1 \| 338 \| 1 \| 259 \| PDDEXK\_1 domain-containing protein \| PDDEXK\_1 domain-containing protein; afdb-uniprot50 \| AF-A0A0F9C255-F1-MODEL\_V4 \| 1.0 \| 5.204e-05 \| 119 \| 0.117 \| 390 \| 221 \| 23 \| 6 \| 371 \| 9 \| 299 \| PDDEXK\_1 domain-containing protein \| PDDEXK\_1 domain-containing protein; afdb-uniprot50 \| AF-A0A3B8MFY8-F1-MODEL\_V4 \| 1.0 \| 8.631e-05 \| 119 \| 0.131 \| 289 \| 168 \| 15 \| 76 \| 337 \| 595 \| 827 \| DNA helicase \| DNA helicase; afdb-uniprot50 \| AF-A0A357CMJ9-F1-MODEL\_V4 \| 1.0 \| 6.16e-05 \| 119 \| 0.126 \| 372 \| 183 \| 17 \| 20 \| 372 \| 895 \| 1143 \| DNA helicase \| DNA helicase; afdb-uniprot50 \| AF-X0RWR8-F1-MODEL\_V4 \| 1.0 \| 0.0002006 \| 118 \| 0.14 \| 227 \| 140 \| 13 \| 157 \| 364 \| 7 \| 197 \| PDDEXK\_1 domain-containing protein \| PDDEXK\_1 domain-containing protein; afdb-uniprot50 \| AF-E3HAC0-F1-MODEL\_V4 \| 1.0 \| 0.0005217 \| 118 \| 0.103 \| 376 \| 180 \| 18 \| 9 \| 369 \| 1 \| 234 \| PDDEXK\_1 domain-containing protein \| PDDEXK\_1 domain-containing protein; afdb-uniprot50 \| AF-A0A1B8SLC9-F1-MODEL\_V4 \| 1.0 \| 0.0001896 \| 118 \| 0.155 \| 186 \| 133 \| 11 \| 180 \| 354 \| 67 \| 239 \| Uncharacterized protein \| Uncharacterized protein; afdb-uniprot50 \| AF-A0A7C5JZ99-F1-MODEL\_V4 \| 1.0 \| 0.0004662 \| 118 \| 0.1 \| 340 \| 178 \| 15 \| 19 \| 350 \| 8 \| 227 \| PD-(D/E)XK nuclease family protein \| PD-(D/E)XK nuclease family protein; afdb-uniprot50 \| AF-A0A1W9UX36-F1-MODEL\_V4 \| 1.0 \| 0.001901 \| 118 \| 0.176 \| 159 \| 109 \| 9 \| 200 \| 346 \| 94 \| 242 \| PDDEXK\_1 domain-containing protein \| PDDEXK\_1 domain-containing protein; afdb-uniprot50 \| AF-A0A843DE36-F1-MODEL\_V4 \| 1.0 \| 0.0001793 \| 118 \| 0.107 \| 354 \| 198 \| 14 \| 6 \| 349 \| 728 \| 973 \| ATP-dependent helicase \| ATP-dependent helicase; afdb-uniprot50 \| AF-A0A662I0G3-F1-MODEL\_V4 \| 1.0 \| 9.131e-05 \| 117 \| 0.128 \| 351 \| 203 \| 19 \| 17 \| 355 \| 44 \| 303 \| Type I-A CRISPR-associated protein Cas4/Csa1 \| Type I-A CRISPR-associated protein Cas4/Csa1; afdb-uniprot50 \| AF-A0A7V9KBP8-F1-MODEL\_V4 \| 1.0 \| 0.0001081 \| 117 \| 0.13 \| 361 \| 182 \| 20 \| 10 \| 349 \| 461 \| 710 \| DNA helicase \| DNA helicase; afdb-uniprot50 \| AF-A0A1M3KGY8-F1-MODEL\_V4 \| 1.0 \| 0.0003939 \| 116 \| 0.091 \| 337 \| 184 \| 15 \| 19 \| 350 \| 8 \| 227 \| PDDEXK\_1 domain-containing protein \| PDDEXK\_1 domain-containing protein; afdb-uniprot50 \| AF-A0A431HDA1-F1-MODEL\_V4 \| 1.0 \| 0.0001022 \| 116 \| 0.09 \| 387 \| 215 \| 19 \| 1 \| 372 \| 626 \| 890 \| PDDEXK\_1 domain-containing protein \| PDDEXK\_1 domain-containing protein; afdb-uniprot50 \| AF-A0A351EJF5-F1-MODEL\_V4 \| 1.0 \| 3.929e-05 \| 116 \| 0.136 \| 322 \| 181 \| 17 \| 45 \| 346 \| 687 \| 931 \| DNA helicase \| DNA helicase; afdb-uniprot50 \| AF-A0A6M8UY77-F1-MODEL\_V4 \| 1.0 \| 4.156e-05 \| 116 \| 0.123 \| 355 \| 205 \| 20 \| 6 \| 347 \| 907 \| 1168 \| DNA helicase \| DNA helicase; afdb-uniprot50 \| AF-A0A7V4J4G6-F1-MODEL\_V4 \| 1.0 \| 0.0008653 \| 115 \| 0.158 \| 196 \| 124 \| 12 \| 196 \| 370 \| 60 \| 235 \| PDDEXK\_1 domain-containing protein \| PDDEXK\_1 domain-containing protein; afdb-uniprot50 \| AF-A0A7V3LX21-F1-MODEL\_V4 \| 1.0 \| 0.0001793 \| 115 \| 0.138 \| 239 \| 173 \| 11 \| 111 \| 338 \| 44 \| 260 \| PDDEXK\_1 domain-containing protein \| PDDEXK\_1 domain-containing protein; afdb-uniprot50 \| AF-A0A3M1GPK9-F1-MODEL\_V4 \| 1.0 \| 5.823e-05 \| 115 \| 0.117 \| 332 \| 174 \| 16 \| 20 \| 337 \| 938 \| 1164 \| DNA helicase \| DNA helicase; afdb-uniprot50 \| AF-A0A4Q6FWM5-F1-MODEL\_V4 \| 1.0 \| 0.002011 \| 114 \| 0.202 \| 153 \| 103 \| 9 \| 205 \| 343 \| 28 \| 175 \| ATP-dependent DNA helicase \| ATP-dependent DNA helicase; afdb-uniprot50 \| AF-A0A846AL31-F1-MODEL\_V4 \| 1.0 \| 0.002011 \| 114 \| 0.1 \| 368 \| 187 \| 17 \| 16 \| 369 \| 2 \| 239 \| PD-(D/E)XK nuclease family protein \| PD-(D/E)XK nuclease family protein; afdb-uniprot50 \| AF-A0A0L8EQ63-F1-MODEL\_V4 \| 1.0 \| 0.0002512 \| 114 \| 0.088 \| 326 \| 203 \| 15 \| 43 \| 354 \| 10 \| 255 \| PDDEXK\_1 domain-containing protein \| PDDEXK\_1 domain-containing protein; afdb-uniprot50 \| AF-A0A0F8WPB7-F1-MODEL\_V4 \| 1.0 \| 0.0002122 \| 114 \| 0.075 \| 369 \| 195 \| 19 \| 9 \| 347 \| 1 \| 253 \| Uncharacterized protein \| Uncharacterized protein; afdb-uniprot50 \| AF-E0STX3-F1-MODEL\_V4 \| 1.0 \| 0.0001209 \| 114 \| 0.106 \| 356 \| 177 \| 15 \| 6 \| 356 \| 70 \| 289 \| CRISPR-associated protein, Csa1 family \| CRISPR-associated protein, Csa1 family; afdb-uniprot50 \| AF-A0A2D5TWL9-F1-MODEL\_V4 \| 1.0 \| 0.0002512 \| 113 \| 0.141 \| 233 \| 139 \| 10 \| 111 \| 338 \| 51 \| 227 \| PDDEXK\_1 domain-containing protein \| PDDEXK\_1 domain-containing protein; afdb-uniprot50 \| AF-A0A841KPM6-F1-MODEL\_V4 \| 1.0 \| 0.0009154 \| 113 \| 0.106 \| 365 \| 175 \| 18 \| 19 \| 354 \| 10 \| 252 \| CRISPR/Cas system-associated exonuclease Cas4 (RecB family) \| CRISPR/Cas system-associated exonuclease Cas4 (RecB family); afdb-uniprot50 \| AF-A0A7C0UD41-F1-MODEL\_V4 \| 1.0 \| 0.0001209 \| 113 \| 0.13 \| 368 \| 211 \| 22 \| 1 \| 351 \| 5 \| 280 \| Type I-A CRISPR-associated protein Cas4/Csa1 \| Type I-A CRISPR-associated protein Cas4/Csa1; afdb-uniprot50 \| AF-A0A7Y2DTL4-F1-MODEL\_V4 \| 1.0 \| 0.0002245 \| 113 \| 0.107 \| 345 \| 187 \| 17 \| 19 \| 338 \| 12 \| 260 \| PDDEXK\_1 domain-containing protein \| PDDEXK\_1 domain-containing protein; afdb-uniprot50 \| AF-A0A850I7L6-F1-MODEL\_V4 \| 1.0 \| 0.0003145 \| 113 \| 0.131 \| 205 \| 139 \| 12 \| 180 \| 371 \| 81 \| 259 \| Uncharacterized protein \| Uncharacterized protein; afdb-uniprot50 \| AF-R5VWW7-F1-MODEL\_V4 \| 1.0 \| 0.0006533 \| 113 \| 0.093 \| 330 \| 189 \| 15 \| 16 \| 338 \| 510 \| 736 \| DNA helicase \| DNA helicase; afdb-uniprot50 \| AF-A0A554JBM9-F1-MODEL\_V4 \| 1.0 \| 0.0002006 \| 113 \| 0.11 \| 380 \| 211 \| 23 \| 12 \| 369 \| 758 \| 1032 \| DNA helicase \| DNA helicase; afdb-uniprot50 \| AF-A0A2D4YUG9-F1-MODEL\_V4 \| 1.0 \| 0.0001896 \| 112 \| 0.119 \| 251 \| 159 \| 16 \| 104 \| 344 \| 26 \| 224 \| Exonuclease \| Exonuclease; afdb-uniprot50 \| AF-A0A4P5V7J0-F1-MODEL\_V4 \| 1.0 \| 3.714e-05 \| 112 \| 0.125 \| 310 \| 174 \| 19 \| 63 \| 333 \| 240 \| 491 \| DNA helicase \| DNA helicase; afdb-uniprot50 \| AF-A0A7W2B9I7-F1-MODEL\_V4 \| 1.0 \| 0.0001143 \| 112 \| 0.109 \| 384 \| 207 \| 22 \| 1 \| 366 \| 665 \| 931 \| PD-(D/E)XK nuclease family protein \| PD-(D/E)XK nuclease family protein; afdb-uniprot50 \| AF-A0A497EM85-F1-MODEL\_V4 \| 1.0 \| 0.0006176 \| 111 \| 0.111 \| 341 \| 188 \| 16 \| 19 \| 354 \| 65 \| 295 \| Type I-A CRISPR-associated protein Cas4/Csa1 \| Type I-A CRISPR-associated protein Cas4/Csa1; afdb-uniprot50 \| AF-A0A2E5G5K7-F1-MODEL\_V4 \| 1.0 \| 0.0002811 \| 111 \| 0.123 \| 371 \| 205 \| 20 \| 17 \| 370 \| 677 \| 944 \| PDDEXK\_1 domain-containing protein \| PDDEXK\_1 domain-containing protein; afdb-uniprot50 \| AF-A0A5E4HM91-F1-MODEL\_V4 \| 1.0 \| 0.002381 \| 110 \| 0.161 \| 198 \| 121 \| 12 \| 164 \| 353 \| 65 \| 225 \| Uncharacterized protein \| Uncharacterized protein; afdb-uniprot50 \| AF-A0A496U7R8-F1-MODEL\_V4 \| 1.0 \| 0.0006911 \| 109 \| 0.101 \| 353 \| 202 \| 17 \| 17 \| 363 \| 627 \| 870 \| PDDEXK\_1 domain-containing protein \| PDDEXK\_1 domain-containing protein; afdb-uniprot50 \| AF-A0A7C5FEZ8-F1-MODEL\_V4 \| 1.0 \| 0.007329 \| 108 \| 0.096 \| 176 \| 123 \| 8 \| 220 \| 372 \| 7 \| 169 \| PDDEXK\_1 domain-containing protein \| PDDEXK\_1 domain-containing protein; afdb-uniprot50 \| AF-A0A1E5IMZ6-F1-MODEL\_V4 \| 1.0 \| 0.0004662 \| 108 \| 0.104 \| 307 \| 189 \| 16 \| 81 \| 368 \| 14 \| 253 \| PDDEXK\_1 domain-containing protein \| PDDEXK\_1 domain-containing protein; afdb-uniprot50 \| AF-A0A7C4A432-F1-MODEL\_V4 \| 1.0 \| 0.0001602 \| 108 \| 0.11 \| 345 \| 205 \| 21 \| 17 \| 341 \| 6 \| 268 \| PDDEXK\_1 domain-containing protein \| PDDEXK\_1 domain-containing protein; afdb-uniprot50 \| AF-A0A2N8HCS8-F1-MODEL\_V4 \| 1.0 \| 0.005853 \| 107 \| 0.188 \| 106 \| 72 \| 4 \| 249 \| 348 \| 32 \| 129 \| PDDEXK\_1 domain-containing protein \| PDDEXK\_1 domain-containing protein; afdb-uniprot50 \| AF-A0A2E9QA28-F1-MODEL\_V4 \| 1.0 \| 0.0002122 \| 107 \| 0.117 \| 282 \| 166 \| 19 \| 86 \| 346 \| 49 \| 268 \| Uncharacterized protein \| Uncharacterized protein; afdb-uniprot50 \| AF-A0A3M2AIE2-F1-MODEL\_V4 \| 1.0 \| 0.0001793 \| 107 \| 0.137 \| 393 \| 194 \| 23 \| 6 \| 366 \| 486 \| 765 \| PD-(D/E)XK nuclease family protein \| PD-(D/E)XK nuclease family protein; afdb-uniprot50 \| AF-A0A1U9K8J1-F1-MODEL\_V4 \| 1.0 \| 3.137e-05 \| 107 \| 0.127 \| 393 \| 217 \| 20 \| 1 \| 351 \| 875 \| 1183 \| DNA helicase \| DNA helicase; afdb-uniprot50 \| AF-A0A2M7TJF4-F1-MODEL\_V4 \| 1.0 \| 0.001518 \| 106 \| 0.1 \| 319 \| 167 \| 17 \| 31 \| 341 \| 3 \| 209 \| PDDEXK\_1 domain-containing protein \| PDDEXK\_1 domain-containing protein; afdb-uniprot50 \| AF-A0A2M6ZV13-F1-MODEL\_V4 \| 1.0 \| 0.0006533 \| 106 \| 0.1 \| 300 \| 188 \| 14 \| 87 \| 370 \| 11 \| 244 \| PDDEXK\_1 domain-containing protein \| PDDEXK\_1 domain-containing protein; afdb-uniprot50 \| AF-A0A7T8V446-F1-MODEL\_V4 \| 1.0 \| 0.002011 \| 106 \| 0.075 \| 330 \| 173 \| 13 \| 19 \| 338 \| 7 \| 214 \| PD-(D/E)XK nuclease family protein \| PD-(D/E)XK nuclease family protein; afdb-uniprot50 \| AF-A0A7C7QVR4-F1-MODEL\_V4 \| 1.0 \| 0.0002657 \| 106 \| 0.125 \| 366 \| 190 \| 22 \| 18 \| 366 \| 39 \| 291 \| PDDEXK\_1 domain-containing protein \| PDDEXK\_1 domain-containing protein; afdb-uniprot50 \| AF-A0A3L7YM13-F1-MODEL\_V4 \| 1.0 \| 0.0001793 \| 106 \| 0.117 \| 247 \| 151 \| 16 \| 111 \| 352 \| 870 \| 1054 \| DNA helicase \| DNA helicase; afdb-uniprot50 \| AF-A0A382UHU8-F1-MODEL\_V4 \| 1.0 \| 0.001797 \| 105 \| 0.142 \| 140 \| 99 \| 7 \| 225 \| 355 \| 2 \| 129 \| PDDEXK\_1 domain-containing protein \| PDDEXK\_1 domain-containing protein; afdb-uniprot50 \| AF-A0A2E5ASM7-F1-MODEL\_V4 \| 1.0 \| 0.0007733 \| 105 \| 0.15 \| 226 \| 136 \| 15 \| 137 \| 346 \| 44 \| 229 \| PDDEXK\_1 domain-containing protein \| PDDEXK\_1 domain-containing protein; afdb-uniprot50 \| AF-A0A523LAM6-F1-MODEL\_V4 \| 1.0 \| 0.0004662 \| 105 \| 0.128 \| 280 \| 133 \| 13 \| 44 \| 316 \| 63 \| 238 \| PDDEXK\_1 domain-containing protein \| PDDEXK\_1 domain-containing protein; afdb-uniprot50 \| AF-A0A160T011-F1-MODEL\_V4 \| 1.0 \| 0.0002122 \| 105 \| 0.149 \| 287 \| 165 \| 12 \| 83 \| 350 \| 6 \| 232 \| Uncharacterized protein \| Uncharacterized protein; afdb-uniprot50 \| AF-A0A1V4IB94-F1-MODEL\_V4 \| 1.0 \| 0.004674 \| 104 \| 0.095 \| 199 \| 147 \| 8 \| 183 \| 370 \| 62 \| 238 \| PD-(D/E)XK nuclease superfamily protein \| PD-(D/E)XK nuclease superfamily protein; afdb-uniprot50 \| AF-A0A7X9J9S1-F1-MODEL\_V4 \| 1.0 \| 0.000731 \| 104 \| 0.11 \| 325 \| 163 \| 15 \| 19 \| 338 \| 8 \| 211 \| PD-(D/E)XK nuclease family protein \| PD-(D/E)XK nuclease family protein; afdb-uniprot50 \| AF-A0A3B9QJZ0-F1-MODEL\_V4 \| 1.0 \| 0.0003145 \| 104 \| 0.133 \| 285 \| 165 \| 16 \| 82 \| 346 \| 285 \| 507 \| DNA helicase \| DNA helicase; afdb-uniprot50 \| AF-A0A7V7XI94-F1-MODEL\_V4 \| 1.0 \| 0.001797 \| 103 \| 0.093 \| 342 \| 169 \| 17 \| 16 \| 346 \| 16 \| 227 \| PD-(D/E)XK nuclease family protein \| PD-(D/E)XK nuclease family protein; afdb-uniprot50 \| AF-A0A2E0ZUF9-F1-MODEL\_V4 \| 1.0 \| 0.0001896 \| 103 \| 0.147 \| 338 \| 178 \| 16 \| 6 \| 332 \| 232 \| 470 \| DNA helicase \| DNA helicase; afdb-uniprot50 \| AF-A0A1Z8S8D1-F1-MODEL\_V4 \| 1.0 \| 0.001283 \| 102 \| 0.142 \| 245 \| 138 \| 14 \| 134 \| 352 \| 1 \| 199 \| PDDEXK\_1 domain-containing protein \| PDDEXK\_1 domain-containing protein; afdb-uniprot50 \| AF-A0A2D9L0E1-F1-MODEL\_V4 \| 1.0 \| 0.0006176 \| 102 \| 0.136 \| 234 \| 153 \| 14 \| 168 \| 372 \| 52 \| 265 \| PDDEXK\_1 domain-containing protein \| PDDEXK\_1 domain-containing protein; afdb-uniprot50 \| AF-A0A381XZY0-F1-MODEL\_V4 \| 1.0 \| 0.001901 \| 101 \| 0.162 \| 209 \| 119 \| 10 \| 197 \| 369 \| 15 \| 203 \| PDDEXK\_1 domain-containing protein \| PDDEXK\_1 domain-containing protein; afdb-uniprot50 \| AF-A0A0F9FF23-F1-MODEL\_V4 \| 1.0 \| 0.006549 \| 101 \| 0.13 \| 192 \| 122 \| 14 \| 179 \| 359 \| 58 \| 215 \| Uncharacterized protein \| Uncharacterized protein; afdb-uniprot50 \| AF-H0E406-F1-MODEL\_V4 \| 1.0 \| 0.0001353 \| 101 \| 0.14 \| 300 \| 159 \| 16 \| 82 \| 338 \| 691 \| 934 \| DNA helicase \| DNA helicase; afdb-uniprot50 \| AF-A0A2P5NMI1-F1-MODEL\_V4 \| 1.0 \| 0.001901 \| 101 \| 0.111 \| 384 \| 224 \| 17 \| 1 \| 371 \| 744 \| 1023 \| Double-strand break repair protein AddB \| Double-strand break repair protein AddB; afdb-uniprot50 \| AF-A0A292YCD1-F1-MODEL\_V4 \| 1.0 \| 0.0002122 \| 101 \| 0.142 \| 379 \| 177 \| 21 \| 7 \| 350 \| 952 \| 1217 \| DNA helicase \| DNA helicase; afdb-uniprot50 \| AF-A0A705R219-F1-MODEL\_V4 \| 1.0 \| 0.004419 \| 100 \| 0.201 \| 174 \| 98 \| 4 \| 83 \| 220 \| 7 \| 175 \| Uncharacterized protein \| Uncharacterized protein; afdb-uniprot50 \| AF-A0A7W1EQU2-F1-MODEL\_V4 \| 0.999 \| 0.0004166 \| 99 \| 0.123 \| 299 \| 167 \| 20 \| 84 \| 356 \| 38 \| 267 \| Uncharacterized protein \| Uncharacterized protein; afdb-uniprot50 \| AF-A0A840DK55-F1-MODEL\_V4 \| 0.999 \| 0.0004662 \| 99 \| 0.135 \| 400 \| 189 \| 24 \| 9 \| 368 \| 663 \| 945 \| RecB family exonuclease \| RecB family exonuclease; afdb-uniprot50 \| AF-A0A832FQV0-F1-MODEL\_V4 \| 0.999 \| 0.001283 \| 98 \| 0.169 \| 206 \| 119 \| 13 \| 161 \| 362 \| 14 \| 171 \| DUF911 domain-containing protein \| DUF911 domain-containing protein; afdb-uniprot50 \| AF-A0A537W8D4-F1-MODEL\_V4 \| 0.999 \| 0.002127 \| 98 \| 0.122 \| 204 \| 140 \| 12 \| 154 \| 351 \| 8 \| 178 \| PDDEXK\_1 domain-containing protein \| PDDEXK\_1 domain-containing protein; afdb-uniprot50 \| AF-A0A162LB28-F1-MODEL\_V4 \| 0.999 \| 0.003336 \| 98 \| 0.077 \| 322 \| 168 \| 16 \| 19 \| 334 \| 5 \| 203 \| PD-(D/E)XK nuclease superfamily protein \| PD-(D/E)XK nuclease superfamily protein; afdb-uniprot50 \| AF-A0A1U9NQE7-F1-MODEL\_V4 \| 0.999 \| 0.004419 \| 98 \| 0.125 \| 207 \| 131 \| 10 \| 197 \| 369 \| 44 \| 234 \| PD-(D/E)XK nuclease superfamily protein \| PD-(D/E)XK nuclease superfamily protein; afdb-uniprot50 \| AF-A0A350REJ8-F1-MODEL\_V4 \| 0.999 \| 0.0004407 \| 98 \| 0.16 \| 331 \| 194 \| 17 \| 32 \| 334 \| 73 \| 347 \| PDDEXK\_1 domain-containing protein \| PDDEXK\_1 domain-containing protein; afdb-uniprot50 \| AF-A0A4P5SGY1-F1-MODEL\_V4 \| 0.999 \| 0.001084 \| 98 \| 0.13 \| 284 \| 197 \| 14 \| 104 \| 372 \| 670 \| 918 \| PDDEXK\_1 domain-containing protein \| PDDEXK\_1 domain-containing protein; afdb-uniprot50 \| AF-U6B6L1-F1-MODEL\_V4 \| 0.999 \| 0.0006911 \| 98 \| 0.093 \| 376 \| 215 \| 22 \| 15 \| 368 \| 758 \| 1029 \| Inactivated superfamily I helicase \| Inactivated superfamily I helicase; afdb-uniprot50 \| AF-A0A2S5MP45-F1-MODEL\_V4 \| 0.999 \| 0.00523 \| 97 \| 0.14 \| 185 \| 132 \| 11 \| 199 \| 369 \| 3 \| 174 \| Double-strand break repair protein AddB \| Double-strand break repair protein AddB; afdb-uniprot50 \| AF-A0A662HJV4-F1-MODEL\_V4 \| 0.999 \| 0.003153 \| 97 \| 0.109 \| 339 \| 177 \| 18 \| 19 \| 351 \| 47 \| 266 \| Type I-A CRISPR-associated protein Cas4/Csa1 \| Type I-A CRISPR-associated protein Cas4/Csa1; afdb-uniprot50 \| AF-A0A1W9S0B8-F1-MODEL\_V4 \| 0.999 \| 0.0002245 \| 97 \| 0.16 \| 286 \| 166 \| 13 \| 38 \| 319 \| 557 \| 772 \| DNA helicase \| DNA helicase; afdb-uniprot50 \| AF-A0A1Q3MWE1-F1-MODEL\_V4 \| 0.999 \| 0.003153 \| 97 \| 0.126 \| 348 \| 179 \| 21 \| 17 \| 351 \| 624 \| 859 \| PDDEXK\_1 domain-containing protein \| PDDEXK\_1 domain-containing protein; afdb-uniprot50 \| AF-A0A7V4APT1-F1-MODEL\_V4 \| 0.999 \| 0.0004407 \| 96 \| 0.099 \| 352 \| 215 \| 20 \| 6 \| 335 \| 1 \| 272 \| PD-(D/E)XK nuclease family protein \| PD-(D/E)XK nuclease family protein; afdb-uniprot50 \| AF-D9PF80-F1-MODEL\_V4 \| 0.998 \| 0.0005519 \| 95 \| 0.16 \| 237 \| 128 \| 15 \| 100 \| 316 \| 52 \| 237 \| PDDEXK\_1 domain-containing protein \| PDDEXK\_1 domain-containing protein; afdb-uniprot50 \| AF-A0A6N8W5D8-F1-MODEL\_V4 \| 0.998 \| 0.0004932 \| 95 \| 0.121 \| 355 \| 162 \| 17 \| 18 \| 354 \| 358 \| 580 \| DNA helicase \| DNA helicase; afdb-uniprot50 \| AF-A0A661U7B8-F1-MODEL\_V4 \| 0.998 \| 0.001146 \| 95 \| 0.144 \| 242 \| 145 \| 10 \| 116 \| 319 \| 862 \| 1079 \| DNA helicase \| DNA helicase; afdb-uniprot50 \| AF-A0A3M1Y1D5-F1-MODEL\_V4 \| 0.998 \| 0.0004166 \| 94 \| 0.109 \| 293 \| 184 \| 16 \| 66 \| 334 \| 219 \| 458 \| DNA helicase \| DNA helicase; afdb-uniprot50 \| AF-A0A356N865-F1-MODEL\_V4 \| 0.998 \| 0.002381 \| 93 \| 0.125 \| 352 \| 175 \| 22 \| 19 \| 351 \| 798 \| 1035 \| DNA helicase \| DNA helicase; afdb-uniprot50 \| AF-A0A2M6YXS7-F1-MODEL\_V4 \| 0.997 \| 0.001901 \| 92 \| 0.128 \| 367 \| 173 \| 15 \| 17 \| 349 \| 956 \| 1209 \| DNA helicase \| DNA helicase; afdb-uniprot50 \| AF-A0A7C1B943-F1-MODEL\_V4 \| 0.997 \| 0.0005217 \| 91 \| 0.168 \| 285 \| 155 \| 14 \| 43 \| 319 \| 691 \| 901 \| DNA helicase \| DNA helicase; afdb-uniprot50 \| AF-A0A6I1QTZ0-F1-MODEL\_V4 \| 0.997 \| 0.001901 \| 91 \| 0.138 \| 368 \| 199 \| 21 \| 12 \| 353 \| 724 \| 999 \| UvrD-like helicase C-terminal domain-containing protein \| UvrD-like helicase C-terminal domain-containing protein; afdb-uniprot50 \| AF-A0A0F8YJ77-F1-MODEL\_V4 \| 0.996 \| 0.008201 \| 90 \| 0.152 \| 190 \| 137 \| 9 \| 194 \| 369 \| 10 \| 189 \| Uncharacterized protein \| Uncharacterized protein; afdb-uniprot50 \| AF-A0A2E5NP53-F1-MODEL\_V4 \| 0.996 \| 0.002818 \| 90 \| 0.098 \| 283 \| 144 \| 12 \| 43 \| 318 \| 927 \| 1105 \| DNA helicase \| DNA helicase; afdb-uniprot50 \| AF-A0A5J4EAI4-F1-MODEL\_V4 \| 0.995 \| 0.002011 \| 88 \| 0.096 \| 291 \| 143 \| 16 \| 43 \| 318 \| 332 \| 517 \| DNA helicase \| DNA helicase; afdb-uniprot50 \| AF-A0A2T4UF80-F1-MODEL\_V4 \| 0.995 \| 0.0006533 \| 88 \| 0.127 \| 415 \| 192 \| 23 \| 6 \| 359 \| 905 \| 1210 \| DNA helicase \| DNA helicase; afdb-uniprot50 \| AF-A0A7C5FTP4-F1-MODEL\_V4 \| 0.994 \| 0.005853 \| 87 \| 0.135 \| 229 \| 144 \| 13 \| 182 \| 372 \| 11 \| 223 \| PDDEXK\_1 domain-containing protein \| PDDEXK\_1 domain-containing protein; afdb-uniprot50 \| AF-A0A256Z4F0-F1-MODEL\_V4 \| 0.993 \| 0.001213 \| 86 \| 0.107 \| 280 \| 180 \| 15 \| 104 \| 351 \| 240 \| 481 \| Cas\_Cas4 domain-containing protein \| Cas\_Cas4 domain-containing protein; afdb-uniprot50 \| AF-A0A1F8V201-F1-MODEL\_V4 \| 0.991 \| 0.004944 \| 84 \| 0.116 \| 223 \| 151 \| 15 \| 109 \| 318 \| 910 \| 1099 \| DNA helicase \| DNA helicase; afdb-uniprot50 \| AF-A0A662G4I6-F1-MODEL\_V4 \| 0.99 \| 0.0003939 \| 83 \| 0.125 \| 288 \| 174 \| 14 \| 88 \| 364 \| 2 \| 222 \| Type I-A CRISPR-associated protein Cas4/Csa1 \| Type I-A CRISPR-associated protein Cas4/Csa1; afdb-uniprot50 \| AF-A0A7C5H0W5-F1-MODEL\_V4 \| 0.99 \| 0.0009683 \| 83 \| 0.121 \| 347 \| 230 \| 18 \| 23 \| 351 \| 13 \| 302 \| Dna2/Cas4 domain-containing protein \| Dna2/Cas4 domain-containing protein; afdb-uniprot50 \| AF-A0A0S8BQY4-F1-MODEL\_V4 \| 0.99 \| 0.001797 \| 83 \| 0.122 \| 237 \| 130 \| 14 \| 87 \| 316 \| 270 \| 435 \| DNA helicase \| DNA helicase; afdb-uniprot50 \| AF-A0A7C1YNJ4-F1-MODEL\_V4 \| 0.99 \| 0.003949 \| 83 \| 0.101 \| 247 \| 146 \| 15 \| 82 \| 319 \| 963 \| 1142 \| DNA helicase \| DNA helicase; afdb-uniprot50 \| AF-A0A3C1LBA3-F1-MODEL\_V4 \| 0.988 \| 0.001606 \| 82 \| 0.145 \| 337 \| 183 \| 20 \| 35 \| 333 \| 353 \| 622 \| DNA helicase \| DNA helicase; afdb-uniprot50 \| AF-A0A2N6N5L1-F1-MODEL\_V4 \| 0.988 \| 0.006928 \| 82 \| 0.104 \| 286 \| 140 \| 14 \| 45 \| 318 \| 596 \| 777 \| DNA helicase \| DNA helicase; afdb-uniprot50 \| AF-A0A849E0V5-F1-MODEL\_V4 \| 0.986 \| 0.004674 \| 81 \| 0.095 \| 240 \| 142 \| 15 \| 87 \| 319 \| 223 \| 394 \| Uncharacterized protein \| Uncharacterized protein; afdb-uniprot50 \| AF-Q83C98-F1-MODEL\_V4 \| 0.986 \| 0.006928 \| 81 \| 0.091 \| 284 \| 147 \| 13 \| 43 \| 319 \| 923 \| 1102 \| DNA helicase \| DNA helicase; afdb-uniprot50 \| AF-A0A0F9E004-F1-MODEL\_V4 \| 0.984 \| 0.003949 \| 80 \| 0.112 \| 213 \| 130 \| 10 \| 159 \| 343 \| 13 \| 194 \| Uncharacterized protein \| Uncharacterized protein; afdb-uniprot50 \| AF-A0A812L764-F1-MODEL\_V4 \| 0.984 \| 0.001024 \| 80 \| 0.15 \| 378 \| 222 \| 22 \| 42 \| 359 \| 27 \| 365 \| SWA2 protein \| SWA2 protein; afdb-uniprot50 \| AF-A0A3C1TRC6-F1-MODEL\_V4 \| 0.984 \| 0.004944 \| 80 \| 0.126 \| 252 \| 130 \| 15 \| 82 \| 316 \| 584 \| 762 \| DNA helicase \| DNA helicase; afdb-uniprot50 \| AF-A0A1G1K328-F1-MODEL\_V4 \| 0.984 \| 0.004674 \| 80 \| 0.128 \| 357 \| 199 \| 19 \| 7 \| 338 \| 860 \| 1129 \| DNA helicase \| DNA helicase; afdb-uniprot50 \| AF-A0A1F8RNI5-F1-MODEL\_V4 \| 0.981 \| 0.005533 \| 79 \| 0.128 \| 280 \| 147 \| 15 \| 43 \| 318 \| 838 \| 1024 \| DNA helicase \| DNA helicase; afdb-uniprot50 \| AF-A0A6B1AB89-F1-MODEL\_V4 \| 0.981 \| 0.001283 \| 79 \| 0.155 \| 329 \| 179 \| 21 \| 44 \| 351 \| 847 \| 1097 \| DNA helicase \| DNA helicase; afdb-uniprot50 \| AF-A0A0F9TAF0-F1-MODEL\_V4 \| 0.978 \| 0.007329 \| 78 \| 0.134 \| 276 \| 157 \| 14 \| 105 \| 359 \| 17 \| 231 \| YqaJ domain-containing protein \| YqaJ domain-containing protein; afdb-uniprot50 \| AF-A0A521I6S1-F1-MODEL\_V4 \| 0.978 \| 0.0009683 \| 78 \| 0.135 \| 287 \| 189 \| 14 \| 43 \| 318 \| 181 \| 419 \| DNA helicase \| DNA helicase; afdb-uniprot50 \| AF-A0A5C7V023-F1-MODEL\_V4 \| 0.975 \| 0.007329 \| 77 \| 0.113 \| 281 \| 142 \| 15 \| 43 \| 315 \| 486 \| 667 \| DNA helicase \| DNA helicase; afdb-uniprot50 \| AF-A0A812ZGY8-F1-MODEL\_V4 \| 0.975 \| 0.00225 \| 77 \| 0.123 \| 308 \| 166 \| 18 \| 100 \| 372 \| 280 \| 518 \| Hypothetical protein \| Hypothetical protein; afdb-uniprot50 \| AF-A0A6M3JLX4-F1-MODEL\_V4 \| 0.971 \| 0.003733 \| 76 \| 0.118 \| 185 \| 136 \| 11 \| 170 \| 343 \| 34 \| 202 \| Putative exonuclease \| Putative exonuclease; afdb-uniprot50 \| AF-A0A2D9JQL8-F1-MODEL\_V4 \| 0.971 \| 0.008675 \| 76 \| 0.114 \| 287 \| 144 \| 16 \| 42 \| 319 \| 951 \| 1136 \| DNA helicase \| DNA helicase; afdb-uniprot50 \| AF-A0A1W9K967-F1-MODEL\_V4 \| 0.933 \| 0.009708 \| 70 \| 0.105 \| 304 \| 182 \| 16 \| 35 \| 316 \| 585 \| 820 \| DNA helicase \| DNA helicase; afdb-uniprot50 \| AF-A0A812XXC2-F1-MODEL\_V4 \| 0.933 \| 0.003733 \| 70 \| 0.129 \| 317 \| 191 \| 23 \| 81 \| 359 \| 15 \| 284 \| Hypothetical protein \| Hypothetical protein; afdb-uniprot50 \| AF-A0A2P4PVM3-F1-MODEL\_V4 \| 0.772 \| 0.00523 \| 60 \| 0.142 \| 338 \| 202 \| 21 \| 75 \| 370 \| 1 \| 292 \| Uncharacterized protein \| Uncharacterized protein |
| Top keywords  (threshold 1.00e-02 (evalue)) | **domain\_containing, DUF3799, Exodeoxyribonuclease, DNA, helicase, PD\_, D, E, XK, VIII** |
| Output files | ../../similar\_structures/41\_FANPEZAQ\_CDS\_0041\_afdb-proteome\_foldseek.tsv ../../similar\_structures/41\_FANPEZAQ\_CDS\_0041\_afdb-uniprot50\_foldseek.tsv ../../similar\_structures/41\_FANPEZAQ\_CDS\_0041\_merged.svg ../../similar\_structures/41\_FANPEZAQ\_CDS\_0041\_pdb\_foldseek.tsv |

  
  
  

Return to summary | Go to previous | Go to next

  


---

**Sequence/structure alignments coloring**  
Each object in the alignment figures is colored according to its E-value following this color coding:

1e-100
10
