## Supplementary material for "Completing the BASEL phage collection to unlock hidden diversity for systematic exploration of phage-host interactions": Data S2: 42.html

FANPEZAQ\_CDS\_0042


Return to summary | Go to previous | Go to next

|  |  |
| --- | --- |
| FANPEZAQ\_CDS\_0042  Page creation date: 02 Sep 2024, 12:00  Project folder: n/a  Input sequences file: Escherichia\_virus\_HeidiAbel.gb | transcriptional domain\_containing regulator repressor hth cro c1\_type peptidase helix\_turn\_helix s24 phage putative c peptidase\_s24 and lexa peptisase transcription s24\_like dna hth\_type signal ci contains s26a s26b xre domains dna\_binding complex with regulation s26c structural binding |

### Sequence information

|  |  |
| --- | --- |
| Name | FANPEZAQ\_CDS\_0042  42\_FANPEZAQ\_CDS\_0042 (pipeline id) |
| Imported annotations |  |
| Protein sequence | MPNRIRERRQAAGMTLQDVAEKLGTTAVTVSRWEREPQRVTLPILDRLADAIGCRKEELL SATAVTANALFFNDGVMSTMSAFYGLPAESLAVVKVVTDSMEPTLLNGDSCVIDKSISWI DNAGIYAIAMNGEARMVRCQRRIDGNVRVLCDNELYKFDEVCTDDDLEVIGKVIGLNRKI |
| Number of residues | 180 |
| Molecular weight (Da) | 19913.70 |
| Output files | ../../query\_sequences/42\_FANPEZAQ\_CDS\_0042.fasta |

### Putative domain architecture and protein family

#### Search results (HHblits)^1^

|  |  |
| --- | --- |
| Domain family databases searched | Pfam, Ncbi-cd, Cath, Phrogs |
| Results, scheme(s)  (Top layers only; threshold 1.00e-03 (evalue)) | xml version="1.0" encoding="utf-8" standalone="no"?       2024-09-02T21:08:21.681344 image/svg+xml   Matplotlib v3.7.2, https://matplotlib.org/ |
| Results, table  (E-value ≤ 1.00e-03 (evalue)) | db \| id \| prob \| evalue \| pvalue \| score \| cols \| query \| query\_len \| template \| template\_len \| name \| description; pfam \| PF00717 \| 98.2 \| 3e-10 \| 4.8e-14 \| 64.0 \| 84 \| (89, 174) \| 180 \| (32, 116) \| 116 \| Peptidase\_S24 \| Peptidase S24-like; pfam \| PF08667 \| 98.1 \| 6.6e-10 \| 1.1e-13 \| 68.0 \| 121 \| (4, 140) \| 180 \| (9, 137) \| 174 \| BetR \| BetR domain; pfam \| PF16452 \| 97.4 \| 8.5e-08 \| 1.4e-11 \| 52.6 \| 66 \| (105, 175) \| 180 \| (36, 101) \| 101 \| Phage\_CI\_C \| Bacteriophage CI repressor C-terminal domain; ncbi-cd \| cd06529 \| 97.0 \| 4.8e-07 \| 9.1e-11 \| 49.3 \| 79 \| (94, 174) \| 180 \| (3, 81) \| 81 \| S24\_LexA-like \| cd06529 S24\_LexA-like; Peptidase S24 LexA-like proteins are involved in the SOS response leading to the repair of single-stranded DNA within the bacterial cell.; ncbi-cd \| cd06462 \| 96.2 \| 1.1e-05 \| 2.1e-09 \| 43.9 \| 64 \| (94, 157) \| 180 \| (3, 68) \| 84 \| Peptidase\_S24\_S26 \| cd06462 Peptidase\_S24\_S26; The S24, S26 LexA/signal peptidase superfamily contains LexA-related and type I signal peptidase families.; ncbi-cd \| cd06530 \| 94.9 \| 0.00023 \| 4.3e-08 \| 38.4 \| 55 \| (94, 153) \| 180 \| (3, 63) \| 85 \| S26\_SPase\_I \| cd06530 S26\_SPase\_I; The S26 Type I signal peptidase (SPase; LepB; leader peptidase B; leader peptidase I.; cath \| 1jheA00 \| 98.5 \| 2.7e-11 \| 3.2e-15 \| 70.0 \| 89 \| (88, 179) \| 180 \| (40, 131) \| 135 \| Lexa repressor \| CATHCODE: 2.10.109.10 NAME: Lexa repressor. Chain: a, b. Fragment: c-terminus, residues 68-202. Engineered: yes. Mutation: yes SOURCE: Escherichia coli. Organism\_taxid: 562. Gene: lexa. Expressed in: escherichia coli. Expression\_system\_taxid: 562 CLASS: Mainly Beta, ARCH: Ribbon, TOPOL: Umud Fragment, subunit A, HOMOL: Umud Fragment, subunit A; cath \| 1ay9A00 \| 98.4 \| 7.2e-11 \| 8.5e-15 \| 65.6 \| 90 \| (86, 179) \| 180 \| (15, 106) \| 108 \| Umud protein \| CATHCODE: 2.10.109.10 NAME: Umud protein. Chain: a, b. SOURCE: Escherichia coli. Organism\_taxid: 562 CLASS: Mainly Beta, ARCH: Ribbon, TOPOL: Umud Fragment, subunit A, HOMOL: Umud Fragment, subunit A; cath \| 3k2zA02 \| 98.4 \| 9e-11 \| 1.1e-14 \| 65.9 \| 87 \| (90, 179) \| 180 \| (31, 118) \| 119 \| Lexa repressor \| CATHCODE: 2.10.109.10 NAME: Lexa repressor. Chain: a, b. Engineered: yes. Mutation: yes SOURCE: Thermotoga maritima. Organism\_taxid: 2336. Gene: lexa, tm\_1082. Expressed in: escherichia coli. Expression\_system\_taxid: 562. CLASS: Mainly Beta, ARCH: Ribbon, TOPOL: Umud Fragment, subunit A, HOMOL: Umud Fragment, subunit A; cath \| 1f39A00 \| 98.2 \| 3.7e-10 \| 4.3e-14 \| 61.7 \| 86 \| (89, 178) \| 180 \| (3, 95) \| 101 \| Repressor protein ci \| CATHCODE: 2.10.109.10 NAME: Repressor protein ci. Chain: a, b. Fragment: c-terminal domain. Engineered: yes SOURCE: Enterobacteria phage lambda. Organism\_taxid: 10710. Expressed in: escherichia coli. Expression\_system\_taxid: 562. Expression\_system\_vector\_type: plasmid. CLASS: Mainly Beta, ARCH: Ribbon, TOPOL: Umud Fragment, subunit A, HOMOL: Umud Fragment, subunit A; cath \| 1b12C01 \| 97.9 \| 5.4e-09 \| 6.4e-13 \| 60.7 \| 89 \| (90, 179) \| 180 \| (5, 118) \| 137 \| Signal peptidase i \| CATHCODE: 2.10.109.10 NAME: Signal peptidase i. Chain: a, b, c, d. Fragment: catalytic domain. Synonym: spase i, leader peptidase i. Engineered: yes SOURCE: Escherichia coli. Organism\_taxid: 469008. Strain: bl21(de3). Cellular\_location: periplasm. Gene: lepb. Expressed in: escherichia coli bl21(de3). Expression\_system\_taxid: 469008. CLASS: Mainly Beta, ARCH: Ribbon, TOPOL: Umud Fragment, subunit A, HOMOL: Umud Fragment, subunit A; cath \| 2fjrA02 \| 97.2 \| 3.4e-07 \| 4.1e-11 \| 50.8 \| 67 \| (106, 179) \| 180 \| (44, 111) \| 113 \| Repressor protein ci \| CATHCODE: 2.10.109.10 NAME: Repressor protein ci. Chain: a, b. Engineered: yes. Mutation: yes SOURCE: Enterobacteria phage 186. Organism\_taxid: 29252. Gene: ci. Expressed in: escherichia coli. Expression\_system\_taxid: 562 CLASS: Mainly Beta, ARCH: Ribbon, TOPOL: Umud Fragment, subunit A, HOMOL: Umud Fragment, subunit A; cath \| 4k8wA00 \| 94.7 \| 0.00054 \| 6.4e-08 \| 37.7 \| 44 \| (98, 141) \| 180 \| (2, 45) \| 129 \| Lepa \| CATHCODE: 2.10.109.10 NAME: Lepa. Chain: a. Fragment: unp residues 45-173. Engineered: yes SOURCE: Streptococcus pyogenes. Organism\_taxid: 1314. Strain: 90/306s. Gene: lepa, sipa. Expressed in: escherichia coli. Expression\_system\_taxid: 469008. CLASS: Mainly Beta, ARCH: Ribbon, TOPOL: Umud Fragment, subunit A, HOMOL: Umud Fragment, subunit A; phrogs \| 4 \| 99.7 \| 4.3e-23 \| 5.2e-27 \| 148.6 \| 160 \| (14, 179) \| 180 \| (22, 208) \| 209 \| transcriptional repressor \| transcriptional repressor; Category: transcription regulation; p259758 VI\_06270; phrogs \| 811 \| 99.7 \| 2.2e-21 \| 2.5e-25 \| 131.1 \| 92 \| (85, 179) \| 180 \| (68, 161) \| 165 \| UmuD-like protein \| UmuD-like protein; Category: DNA, RNA and nucleotide metabolism; p6911 VI\_09232; phrogs \| 6201 \| 99.7 \| 2.6e-21 \| 3e-25 \| 125.7 \| 92 \| (85, 178) \| 180 \| (38, 132) \| 139 \| CI-like repressor \| CI-like repressor; Category: transcription regulation; p89843 VI\_01594; phrogs \| 10470 \| 99.0 \| 6.3e-14 \| 7e-18 \| 79.2 \| 73 \| (103, 179) \| 180 \| (4, 77) \| 79 \| UmuD-like protein \| UmuD-like protein; Category: DNA, RNA and nucleotide metabolism; p434438 VI\_09939; phrogs \| 2328 \| 98.7 \| 1.9e-12 \| 2.2e-16 \| 83.0 \| 63 \| (1, 64) \| 180 \| (19, 82) \| 123 \| transcriptional repressor \| transcriptional repressor; Category: transcription regulation; MG099936\_p34; phrogs \| 8558 \| 98.7 \| 3.8e-12 \| 4.3e-16 \| 74.6 \| 73 \| (102, 179) \| 180 \| (8, 82) \| 87 \| NA \| NA; Category: unknown function; p316397 VI\_07324; phrogs \| 1373 \| 98.5 \| 1.6e-11 \| 1.9e-15 \| 75.6 \| 57 \| (2, 59) \| 180 \| (35, 92) \| 95 \| transcriptional regulator \| transcriptional regulator; Category: transcription regulation; NC\_005857\_p48; phrogs \| 2967 \| 98.4 \| 7.4e-11 \| 8.6e-15 \| 79.5 \| 62 \| (1, 63) \| 180 \| (7, 69) \| 167 \| transcriptional regulator \| transcriptional regulator; Category: transcription regulation; p123578 VI\_00485; phrogs \| 7241 \| 98.4 \| 8.2e-11 \| 9.3e-15 \| 73.6 \| 57 \| (5, 62) \| 180 \| (66, 123) \| 128 \| transcriptional regulator \| transcriptional regulator; Category: transcription regulation; p63620 VI\_05511; phrogs \| 4583 \| 98.2 \| 4.8e-10 \| 5.5e-14 \| 65.3 \| 62 \| (2, 64) \| 180 \| (12, 74) \| 77 \| transcriptional repressor \| transcriptional repressor; Category: transcription regulation; NC\_031243\_p35; phrogs \| 1381 \| 98.0 \| 1.7e-09 \| 1.9e-13 \| 66.4 \| 60 \| (3, 65) \| 180 \| (11, 72) \| 99 \| plasmid antitoxin with HTH domain \| plasmid antitoxin with HTH domain; Category: moron, auxiliary metabolic gene and host takeover; p79485 VI\_11488; phrogs \| 4836 \| 98.0 \| 1.8e-09 \| 2e-13 \| 73.0 \| 65 \| (2, 67) \| 180 \| (84, 149) \| 233 \| CII-like regulator \| CII-like regulator; Category: transcription regulation; p13677 VI\_07483; phrogs \| 2439 \| 97.8 \| 6.2e-09 \| 7.2e-13 \| 78.7 \| 61 \| (2, 63) \| 180 \| (9, 70) \| 385 \| transcriptional regulator \| transcriptional regulator; Category: transcription regulation; p298790 VI\_12393; phrogs \| 147 \| 97.8 \| 8.4e-09 \| 9.7e-13 \| 64.3 \| 67 \| (1, 68) \| 180 \| (13, 82) \| 112 \| transcriptional regulator \| transcriptional regulator; Category: transcription regulation; p315289 VI\_07968; phrogs \| 2104 \| 97.8 \| 1e-08 \| 1.2e-12 \| 61.5 \| 39 \| (3, 42) \| 180 \| (33, 71) \| 92 \| NA \| NA; Category: unknown function; p72070 VI\_03178; phrogs \| 1468 \| 97.6 \| 2.5e-08 \| 2.9e-12 \| 56.5 \| 56 \| (2, 58) \| 180 \| (12, 67) \| 69 \| transcriptional repressor \| transcriptional repressor; Category: transcription regulation; p307668 VI\_06058; phrogs \| 4779 \| 97.6 \| 3.3e-08 \| 3.8e-12 \| 57.6 \| 35 \| (2, 36) \| 180 \| (3, 37) \| 78 \| transcriptional regulator \| transcriptional regulator; Category: transcription regulation; p225361 VI\_06249; phrogs \| 5823 \| 97.6 \| 3.8e-08 \| 4.4e-12 \| 74.0 \| 63 \| (1, 64) \| 180 \| (9, 74) \| 410 \| transcriptional regulator \| transcriptional regulator; Category: transcription regulation; NC\_028952\_p26; phrogs \| 2339 \| 97.5 \| 5.4e-08 \| 6.3e-12 \| 58.5 \| 53 \| (4, 56) \| 180 \| (32, 85) \| 93 \| HTH DNA binding protein \| HTH DNA binding protein; Category: DNA, RNA and nucleotide metabolism; p425338 VI\_07226; phrogs \| 6831 \| 97.4 \| 7.8e-08 \| 8.9e-12 \| 53.3 \| 43 \| (8, 50) \| 180 \| (10, 54) \| 66 \| transcriptional regulator \| transcriptional regulator; Category: transcription regulation; p431926 VI\_12195; phrogs \| 769 \| 97.4 \| 8e-08 \| 9.2e-12 \| 57.4 \| 63 \| (3, 67) \| 180 \| (4, 67) \| 90 \| transcriptional repressor \| transcriptional repressor; Category: transcription regulation; p166433 VI\_07600; phrogs \| 1337 \| 97.4 \| 1.1e-07 \| 1.2e-11 \| 52.8 \| 52 \| (2, 53) \| 180 \| (4, 57) \| 61 \| DNA binding protein \| DNA binding protein; Category: DNA, RNA and nucleotide metabolism; p177899 VI\_00920; phrogs \| 638 \| 97.4 \| 1.1e-07 \| 1.3e-11 \| 62.5 \| 62 \| (3, 65) \| 180 \| (5, 68) \| 148 \| transcriptional regulator \| transcriptional regulator; Category: transcription regulation; p438490 VI\_02648; phrogs \| 34939 \| 97.2 \| 3.3e-07 \| 3.7e-11 \| 61.7 \| 66 \| (3, 68) \| 180 \| (12, 78) \| 276 \| NA \| NA; Category: unknown function; p54066 VI\_01167; phrogs \| 3037 \| 97.0 \| 8.4e-07 \| 9.6e-11 \| 60.8 \| 54 \| (13, 67) \| 180 \| (29, 83) \| 215 \| transcriptional repressor \| transcriptional repressor; Category: transcription regulation; p321370 VI\_07179; phrogs \| 13326 \| 97.0 \| 8.6e-07 \| 9.6e-11 \| 55.8 \| 64 \| (3, 67) \| 180 \| (5, 71) \| 161 \| NA \| NA; Category: unknown function; p201792 VI\_05516; phrogs \| 4441 \| 97.0 \| 1e-06 \| 1.2e-10 \| 53.9 \| 52 \| (11, 63) \| 180 \| (14, 67) \| 109 \| HTH DNA binding protein \| HTH DNA binding protein; Category: DNA, RNA and nucleotide metabolism; p152087 VI\_05530; phrogs \| 9103 \| 96.9 \| 1.2e-06 \| 1.3e-10 \| 50.0 \| 60 \| (5, 65) \| 180 \| (13, 74) \| 78 \| transcriptional repressor \| transcriptional repressor; Category: transcription regulation; KY624611\_p11; phrogs \| 3642 \| 96.9 \| 1.3e-06 \| 1.5e-10 \| 51.5 \| 51 \| (4, 55) \| 180 \| (9, 60) \| 90 \| HTH DNA binding protein \| HTH DNA binding protein; Category: DNA, RNA and nucleotide metabolism; p184909 VI\_05451; phrogs \| 4739 \| 96.9 \| 1.4e-06 \| 1.6e-10 \| 57.0 \| 64 \| (3, 67) \| 180 \| (71, 135) \| 216 \| NA \| NA; Category: unknown function; p402726 VI\_08646; phrogs \| 35 \| 96.8 \| 1.7e-06 \| 2.1e-10 \| 49.9 \| 57 \| (4, 62) \| 180 \| (8, 67) \| 75 \| transcriptional repressor \| transcriptional repressor; Category: transcription regulation; p323300 VI\_10199; phrogs \| 3557 \| 96.8 \| 1.9e-06 \| 2.1e-10 \| 53.9 \| 62 \| (2, 65) \| 180 \| (15, 77) \| 148 \| transcriptional regulator \| transcriptional regulator; Category: transcription regulation; p437537 VI\_09606; phrogs \| 7950 \| 96.8 \| 2.6e-06 \| 2.9e-10 \| 45.0 \| 39 \| (2, 41) \| 180 \| (2, 40) \| 53 \| transcriptional repressor \| transcriptional repressor; Category: transcription regulation; p141193 VI\_04895; phrogs \| 18342 \| 96.7 \| 2.9e-06 \| 3.2e-10 \| 51.7 \| 59 \| (5, 63) \| 180 \| (23, 83) \| 123 \| transcriptional repressor \| transcriptional repressor; Category: transcription regulation; p158151 VI\_11365; phrogs \| 1702 \| 96.7 \| 2.9e-06 \| 3.3e-10 \| 51.6 \| 63 \| (2, 65) \| 180 \| (30, 96) \| 112 \| transcriptional repressor \| transcriptional repressor; Category: transcription regulation; NC\_022052\_p39; phrogs \| 6463 \| 96.7 \| 3.8e-06 \| 4.3e-10 \| 47.4 \| 57 \| (5, 62) \| 180 \| (3, 66) \| 77 \| HTH DNA binding protein \| HTH DNA binding protein; Category: DNA, RNA and nucleotide metabolism; NC\_004683\_p47; phrogs \| 5736 \| 96.6 \| 4.8e-06 \| 5.3e-10 \| 51.8 \| 60 \| (5, 65) \| 180 \| (11, 71) \| 151 \| transcriptional regulator \| transcriptional regulator; Category: transcription regulation; p339615 VI\_06353; phrogs \| 7126 \| 96.6 \| 4.9e-06 \| 5.6e-10 \| 51.6 \| 52 \| (13, 64) \| 180 \| (31, 83) \| 123 \| HTH DNA binding protein \| HTH DNA binding protein; Category: DNA, RNA and nucleotide metabolism; NC\_031107\_p106; phrogs \| 925 \| 96.6 \| 5.3e-06 \| 6.1e-10 \| 47.7 \| 56 \| (2, 58) \| 180 \| (15, 71) \| 76 \| HTH DNA binding protein \| HTH DNA binding protein; Category: DNA, RNA and nucleotide metabolism; p18585 VI\_03080; phrogs \| 1994 \| 96.5 \| 7.1e-06 \| 7.9e-10 \| 52.5 \| 61 \| (3, 64) \| 180 \| (2, 63) \| 170 \| DNA binding protein \| DNA binding protein; Category: DNA, RNA and nucleotide metabolism; NC\_031023\_p45; phrogs \| 18104 \| 96.4 \| 1.3e-05 \| 1.4e-09 \| 42.0 \| 48 \| (3, 53) \| 180 \| (5, 53) \| 56 \| transcriptional regulator \| transcriptional regulator; Category: transcription regulation; NC\_025471\_p63; phrogs \| 3949 \| 96.3 \| 1.4e-05 \| 1.6e-09 \| 51.5 \| 58 \| (5, 63) \| 180 \| (5, 64) \| 163 \| transcriptional repressor \| transcriptional repressor; Category: transcription regulation; JQ300538\_p45; phrogs \| 6223 \| 96.3 \| 1.7e-05 \| 1.9e-09 \| 44.9 \| 33 \| (10, 42) \| 180 \| (14, 46) \| 77 \| transcriptional repressor \| transcriptional repressor; Category: transcription regulation; p192116 VI\_09192; phrogs \| 955 \| 96.1 \| 3.2e-05 \| 3.6e-09 \| 50.2 \| 65 \| (1, 65) \| 180 \| (56, 123) \| 174 \| HTH DNA binding protein \| HTH DNA binding protein; Category: DNA, RNA and nucleotide metabolism; JN699010\_p46; phrogs \| 4739 \| 96.1 \| 3.3e-05 \| 3.7e-09 \| 50.3 \| 64 \| (1, 65) \| 180 \| (149, 213) \| 216 \| NA \| NA; Category: unknown function; p402726 VI\_08646; phrogs \| 7396 \| 96.0 \| 3.5e-05 \| 4e-09 \| 52.1 \| 66 \| (1, 67) \| 180 \| (13, 81) \| 233 \| NA \| NA; Category: unknown function; p13562 VI\_08594; phrogs \| 4268 \| 96.0 \| 3.7e-05 \| 4.2e-09 \| 45.4 \| 50 \| (4, 56) \| 180 \| (39, 89) \| 94 \| transcriptional repressor \| transcriptional repressor; Category: transcription regulation; p79532 VI\_11488; phrogs \| 8 \| 96.0 \| 3.8e-05 \| 4.5e-09 \| 46.1 \| 41 \| (14, 55) \| 180 \| (1, 42) \| 95 \| transcriptional regulator \| transcriptional regulator; Category: transcription regulation; p309030 VI\_07697; phrogs \| 18318 \| 95.9 \| 5.1e-05 \| 5.8e-09 \| 41.8 \| 54 \| (4, 57) \| 180 \| (12, 66) \| 76 \| HTH DNA binding protein \| HTH DNA binding protein; Category: DNA, RNA and nucleotide metabolism; KU574722\_p511; phrogs \| 38027 \| 95.9 \| 5.3e-05 \| 5.9e-09 \| 44.7 \| 58 \| (4, 62) \| 180 \| (61, 119) \| 132 \| transcriptional repressor \| transcriptional repressor; Category: transcription regulation; NC\_030931\_p18; phrogs \| 18068 \| 95.9 \| 5.4e-05 \| 6e-09 \| 46.1 \| 60 \| (5, 67) \| 180 \| (6, 66) \| 139 \| NA \| NA; Category: unknown function; KR093642\_p39; phrogs \| 14076 \| 95.8 \| 6.5e-05 \| 7.3e-09 \| 42.4 \| 34 \| (3, 36) \| 180 \| (17, 55) \| 81 \| transcriptional regulator \| transcriptional regulator; Category: transcription regulation; p109717 VI\_00077; phrogs \| 1529 \| 95.8 \| 6.5e-05 \| 7.4e-09 \| 41.3 \| 38 \| (2, 41) \| 180 \| (2, 39) \| 65 \| NA \| NA; Category: unknown function; p439749 VI\_04529; phrogs \| 32768 \| 95.8 \| 7.3e-05 \| 8.1e-09 \| 53.0 \| 62 \| (1, 62) \| 180 \| (10, 72) \| 446 \| NA \| NA; Category: unknown function; KT151957\_p71; phrogs \| 5301 \| 95.6 \| 9.9e-05 \| 1.1e-08 \| 45.8 \| 52 \| (10, 62) \| 180 \| (21, 73) \| 143 \| transcriptional repressor \| transcriptional repressor; Category: transcription regulation; p166315 VI\_04146; phrogs \| 38278 \| 95.6 \| 0.00011 \| 1.2e-08 \| 46.3 \| 53 \| (1, 55) \| 180 \| (18, 71) \| 180 \| NA \| NA; Category: unknown function; p17072 VI\_01301; phrogs \| 296 \| 95.6 \| 0.00011 \| 1.3e-08 \| 53.5 \| 35 \| (2, 36) \| 180 \| (7, 41) \| 312 \| DNA transposition protein \| DNA transposition protein; Category: integration and excision; p199288 VI\_08191; phrogs \| 517 \| 95.5 \| 0.00013 \| 1.5e-08 \| 43.9 \| 31 \| (5, 36) \| 180 \| (73, 103) \| 107 \| HTH DNA binding protein \| HTH DNA binding protein; Category: DNA, RNA and nucleotide metabolism; MF668278\_p45; phrogs \| 6962 \| 95.5 \| 0.00014 \| 1.6e-08 \| 38.4 \| 33 \| (4, 36) \| 180 \| (5, 37) \| 59 \| NA \| NA; Category: unknown function; p75562 VI\_02384; phrogs \| 725 \| 95.4 \| 0.00016 \| 1.8e-08 \| 43.7 \| 41 \| (12, 55) \| 180 \| (28, 68) \| 106 \| CII-like regulator \| CII-like regulator; Category: transcription regulation; p72240 VI\_06449; phrogs \| 3 \| 95.3 \| 0.00018 \| 2.2e-08 \| 41.3 \| 50 \| (2, 54) \| 180 \| (7, 59) \| 80 \| transcriptional repressor \| transcriptional repressor; Category: transcription regulation; p29899 VI\_00998; phrogs \| 4031 \| 95.2 \| 0.00022 \| 2.5e-08 \| 54.5 \| 55 \| (3, 58) \| 180 \| (518, 574) \| 622 \| NA \| NA; Category: unknown function; p115970 VI\_05096; phrogs \| 6499 \| 95.1 \| 0.00026 \| 3e-08 \| 45.7 \| 51 \| (5, 56) \| 180 \| (13, 70) \| 157 \| transcriptional regulator \| transcriptional regulator; Category: transcription regulation; p14253 VI\_01278; phrogs \| 4151 \| 95.1 \| 0.00028 \| 3.2e-08 \| 46.2 \| 27 \| (10, 36) \| 180 \| (17, 43) \| 168 \| DNA binding protein \| DNA binding protein; Category: DNA, RNA and nucleotide metabolism; p284557 VI\_05952; phrogs \| 34341 \| 95.0 \| 0.00032 \| 3.6e-08 \| 40.6 \| 34 \| (3, 36) \| 180 \| (74, 107) \| 122 \| transcriptional regulator \| transcriptional regulator; Category: transcription regulation; MF360957\_p141; phrogs \| 24325 \| 95.0 \| 0.00033 \| 3.6e-08 \| 41.2 \| 61 \| (2, 62) \| 180 \| (54, 115) \| 133 \| transcriptional regulator \| transcriptional regulator; Category: transcription regulation; p318984 VI\_12108; phrogs \| 2538 \| 94.8 \| 0.00043 \| 5e-08 \| 42.5 \| 28 \| (9, 36) \| 180 \| (12, 39) \| 113 \| terminase small subunit \| terminase small subunit; Category: head and packaging; p133796 VI\_05051; phrogs \| 17822 \| 94.8 \| 0.00047 \| 5.3e-08 \| 46.3 \| 63 \| (3, 67) \| 180 \| (10, 73) \| 267 \| NA \| NA; Category: unknown function; p205501 VI\_03162; phrogs \| 15562 \| 94.8 \| 0.00048 \| 5.5e-08 \| 44.4 \| 49 \| (94, 145) \| 180 \| (31, 80) \| 185 \| NA \| NA; Category: unknown function; p342664 VI\_05383; phrogs \| 1124 \| 94.7 \| 0.00051 \| 5.9e-08 \| 45.7 \| 39 \| (12, 52) \| 180 \| (16, 55) \| 182 \| transcriptional regulator \| transcriptional regulator; Category: transcription regulation; p275302 VI\_05197; phrogs \| 409 \| 94.7 \| 0.00053 \| 6.2e-08 \| 38.6 \| 28 \| (7, 36) \| 180 \| (5, 32) \| 72 \| transcriptional regulator \| transcriptional regulator; Category: transcription regulation; p88852 VI\_07592; phrogs \| 19579 \| 94.6 \| 0.00059 \| 6.6e-08 \| 44.7 \| 51 \| (12, 63) \| 180 \| (14, 65) \| 225 \| transcriptional regulator \| transcriptional regulator; Category: transcription regulation; p68909 VI\_04167; phrogs \| 7676 \| 94.4 \| 0.0008 \| 9e-08 \| 36.9 \| 26 \| (11, 36) \| 180 \| (14, 39) \| 75 \| NA \| NA; Category: unknown function; NC\_023688\_p227 |
| Top keywords  (threshold 1.00e-03 (evalue)) | **transcription, regulation, transcriptional, DNA, repressor, regulator, A, and, Fragment, Umud** |
| Output files | ../../domain\_architecture/42\_FANPEZAQ\_CDS\_0042\_cath.hhr ../../domain\_architecture/42\_FANPEZAQ\_CDS\_0042\_merged.svg ../../domain\_architecture/42\_FANPEZAQ\_CDS\_0042\_ncbi-cd.hhr ../../domain\_architecture/42\_FANPEZAQ\_CDS\_0042\_pfam.hhr ../../domain\_architecture/42\_FANPEZAQ\_CDS\_0042\_phrogs.hhr |

### Identical protein sequences/structures

#### Search results

|  |  |
| --- | --- |
| Protein sequence databases searched | Pdb, Swissprot, Refseq |
| Identical proteins found | -- |
| Top keywords | -- |
| Output files | -- |

### Similar protein sequences/structures

#### Sequence similarity search results (HHblits)^1^

|  |  |
| --- | --- |
| Sequence databases searched | Uniclust, Pdb70 |
| Results, scheme(s)  (Top layers only, threshold 1.00e-03 (evalue)) | xml version="1.0" encoding="utf-8" standalone="no"?       2024-09-02T21:08:48.043791 image/svg+xml   Matplotlib v3.7.2, https://matplotlib.org/ |
| Results, table(s)  (threshold 1.00e-03 (evalue)) | db \| id \| prob \| evalue \| pvalue \| score \| cols \| query \| query\_len \| template \| template\_len \| name \| description; uniclust \| UniRef100\_A0A078LJD5 \| 100.0 \| 3.8e-37 \| 7.1e-43 \| 225.0 \| 179 \| (1, 180) \| 180 \| (78, 299) \| 306 \| Phage repressor protein \| Phage repressor protein; uniclust \| UniRef100\_A0A0C1R042 \| 100.0 \| 2.3e-35 \| 4.3e-41 \| 203.2 \| 177 \| (1, 180) \| 180 \| (27, 225) \| 228 \| Cro/Cl family transcriptional regulator \| Cro/Cl family transcriptional regulator; uniclust \| UniRef100\_A0A0F9H182 \| 100.0 \| 5.1e-35 \| 9.5e-41 \| 208.8 \| 179 \| (1, 180) \| 180 \| (37, 251) \| 253 \| HTH cro/C1-type domain-containing protein \| HTH cro/C1-type domain-containing protein; uniclust \| UniRef100\_A0A521VUF4 \| 100.0 \| 4.6e-34 \| 8.6e-40 \| 194.6 \| 178 \| (1, 180) \| 180 \| (24, 234) \| 236 \| Peptidase S24 (Fragment) \| Peptidase S24 (Fragment); uniclust \| UniRef100\_A0A1M7T7T0 \| 100.0 \| 5.7e-34 \| 1.1e-39 \| 196.7 \| 178 \| (1, 179) \| 180 \| (30, 265) \| 267 \| Phage repressor protein C, contains Cro/C1-type HTH and peptisase s24 domains \| Phage repressor protein C, contains Cro/C1-type HTH and peptisase s24 domains; uniclust \| UniRef100\_A0A078BMD6 \| 100.0 \| 7.9e-34 \| 1.5e-39 \| 205.2 \| 177 \| (1, 180) \| 180 \| (42, 256) \| 280 \| Putative phage repressor \| Putative phage repressor; uniclust \| UniRef100\_A0A2G2CXL5 \| 100.0 \| 1.1e-33 \| 2.1e-39 \| 197.9 \| 178 \| (1, 180) \| 180 \| (33, 250) \| 253 \| HTH cro/C1-type domain-containing protein \| HTH cro/C1-type domain-containing protein; uniclust \| UniRef100\_A0A075WVE9 \| 100.0 \| 1.8e-33 \| 3.4e-39 \| 196.5 \| 178 \| (1, 179) \| 180 \| (37, 241) \| 244 \| HTH cro/C1-type domain-containing protein \| HTH cro/C1-type domain-containing protein; uniclust \| UniRef100\_A0A1W0D753 \| 100.0 \| 3.1e-33 \| 5.7e-39 \| 191.2 \| 178 \| (1, 179) \| 180 \| (29, 248) \| 258 \| HTH cro/C1-type domain-containing protein \| HTH cro/C1-type domain-containing protein; uniclust \| UniRef100\_A0A139KEG7 \| 100.0 \| 3.5e-33 \| 6.4e-39 \| 188.4 \| 176 \| (2, 177) \| 180 \| (11, 222) \| 227 \| Peptidase S24-like protein \| Peptidase S24-like protein; uniclust \| UniRef100\_A0A1V4GWD0 \| 100.0 \| 7.8e-33 \| 1.5e-38 \| 194.8 \| 177 \| (1, 179) \| 180 \| (23, 230) \| 241 \| HTH cro/C1-type domain-containing protein \| HTH cro/C1-type domain-containing protein; uniclust \| UniRef100\_A0A066TBR7 \| 100.0 \| 9.6e-33 \| 1.8e-38 \| 197.0 \| 178 \| (1, 180) \| 180 \| (68, 299) \| 315 \| Putative transcriptional regulator \| Putative transcriptional regulator; uniclust \| UniRef100\_A0A099K7Y6 \| 100.0 \| 1e-32 \| 1.9e-38 \| 204.6 \| 177 \| (1, 180) \| 180 \| (85, 301) \| 327 \| Phage C1 repressor \| Phage C1 repressor; uniclust \| UniRef100\_A0A0C1W593 \| 100.0 \| 1.3e-32 \| 2.4e-38 \| 190.8 \| 176 \| (1, 177) \| 180 \| (16, 218) \| 233 \| Phage repressor protein \| Phage repressor protein; uniclust \| UniRef100\_A0A022N9X9 \| 100.0 \| 2.3e-32 \| 4.2e-38 \| 196.3 \| 178 \| (1, 180) \| 180 \| (40, 271) \| 275 \| Peptidase S24 \| Peptidase S24; uniclust \| UniRef100\_A0A075CI58 \| 99.9 \| 3.7e-32 \| 6.9e-38 \| 192.3 \| 165 \| (15, 180) \| 180 \| (55, 256) \| 275 \| C repressor \| C repressor; uniclust \| UniRef100\_A0A1H9B3Z2 \| 99.9 \| 7.6e-32 \| 1.4e-37 \| 184.9 \| 178 \| (1, 180) \| 180 \| (20, 228) \| 230 \| Helix-turn-helix \| Helix-turn-helix; uniclust \| UniRef100\_A0A0S4SC53 \| 99.9 \| 8.1e-32 \| 1.5e-37 \| 190.9 \| 178 \| (1, 179) \| 180 \| (10, 233) \| 253 \| DNA-binding protein \| DNA-binding protein; uniclust \| UniRef100\_A0A0J5GKK0 \| 99.9 \| 8.7e-32 \| 1.6e-37 \| 189.0 \| 178 \| (1, 178) \| 180 \| (29, 260) \| 263 \| HTH cro/C1-type domain-containing protein \| HTH cro/C1-type domain-containing protein; uniclust \| UniRef100\_A0A0J7Y5V7 \| 99.9 \| 1.1e-31 \| 2.1e-37 \| 188.1 \| 177 \| (2, 180) \| 180 \| (37, 252) \| 255 \| HTH cro/C1-type domain-containing protein \| HTH cro/C1-type domain-containing protein; uniclust \| UniRef100\_A0A011TVH5 \| 99.9 \| 1.3e-31 \| 2.5e-37 \| 187.6 \| 179 \| (1, 179) \| 180 \| (40, 259) \| 264 \| Transcriptional regulator \| Transcriptional regulator; uniclust \| UniRef100\_A0A086WQD3 \| 99.9 \| 1.4e-31 \| 2.6e-37 \| 188.4 \| 177 \| (2, 179) \| 180 \| (26, 245) \| 254 \| Prophage transcriptional regulator \| Prophage transcriptional regulator; uniclust \| UniRef100\_A0A013SC04 \| 99.9 \| 1.7e-31 \| 3.2e-37 \| 194.6 \| 178 \| (1, 178) \| 180 \| (49, 275) \| 309 \| Peptidase S24-like family protein \| Peptidase S24-like family protein; uniclust \| UniRef100\_A0A060H7F8 \| 99.9 \| 3.1e-31 \| 5.7e-37 \| 190.7 \| 173 \| (1, 176) \| 180 \| (42, 269) \| 281 \| Peptidase S24 \| Peptidase S24; uniclust \| UniRef100\_A0A0G3CSZ6 \| 99.9 \| 3.6e-31 \| 6.7e-37 \| 182.3 \| 176 \| (1, 179) \| 180 \| (31, 243) \| 246 \| HTH cro/C1-type domain-containing protein \| HTH cro/C1-type domain-containing protein; uniclust \| UniRef100\_A0A090IB00 \| 99.9 \| 4.4e-31 \| 8.1e-37 \| 187.4 \| 178 \| (1, 179) \| 180 \| (36, 251) \| 254 \| Uncharacterized phage protein, peptidase S24 family \| Uncharacterized phage protein, peptidase S24 family; uniclust \| UniRef100\_A0A081NQ11 \| 99.9 \| 4.8e-31 \| 8.9e-37 \| 186.7 \| 177 \| (1, 179) \| 180 \| (55, 257) \| 264 \| Helix-turn-helix domain-containing protein \| Helix-turn-helix domain-containing protein; uniclust \| UniRef100\_A0A095YT12 \| 99.9 \| 5.2e-31 \| 9.7e-37 \| 185.9 \| 176 \| (1, 179) \| 180 \| (23, 226) \| 236 \| HTH cro/C1-type domain-containing protein \| HTH cro/C1-type domain-containing protein; uniclust \| UniRef100\_A0A0B1TYP4 \| 99.9 \| 6.3e-31 \| 1.2e-36 \| 188.3 \| 176 \| (3, 180) \| 180 \| (48, 270) \| 284 \| Putative transcriptional regulator \| Putative transcriptional regulator; uniclust \| UniRef100\_A0A011VKK5 \| 99.9 \| 7.3e-31 \| 1.3e-36 \| 189.8 \| 178 \| (2, 180) \| 180 \| (69, 294) \| 295 \| Transcriptional regulator \| Transcriptional regulator; uniclust \| UniRef100\_A0A098ELK1 \| 99.9 \| 8.1e-31 \| 1.5e-36 \| 188.5 \| 178 \| (1, 179) \| 180 \| (42, 245) \| 255 \| LexA repressor \| LexA repressor; uniclust \| UniRef100\_A0A0K6IU92 \| 99.9 \| 9.8e-31 \| 1.8e-36 \| 188.8 \| 177 \| (2, 179) \| 180 \| (23, 313) \| 316 \| Peptidase S24-like/Helix-turn-helix \| Peptidase S24-like/Helix-turn-helix; uniclust \| UniRef100\_A0A074V8P9 \| 99.9 \| 1e-30 \| 1.9e-36 \| 193.1 \| 177 \| (1, 179) \| 180 \| (54, 275) \| 304 \| Putative transcriptional regulator \| Putative transcriptional regulator; uniclust \| UniRef100\_A0A0A2XT17 \| 99.9 \| 1.1e-30 \| 2e-36 \| 183.7 \| 174 \| (1, 176) \| 180 \| (22, 234) \| 245 \| Transcriptional regulator \| Transcriptional regulator; uniclust \| UniRef100\_A0A1Y6CX45 \| 99.9 \| 1.1e-30 \| 2e-36 \| 180.5 \| 176 \| (1, 180) \| 180 \| (26, 240) \| 242 \| Phage repressor protein C, contains Cro/C1-type HTH and peptisase s24 domains \| Phage repressor protein C, contains Cro/C1-type HTH and peptisase s24 domains; uniclust \| UniRef100\_A0A096ALY3 \| 99.9 \| 1.4e-30 \| 2.6e-36 \| 184.2 \| 177 \| (1, 180) \| 180 \| (34, 239) \| 248 \| HTH cro/C1-type domain-containing protein \| HTH cro/C1-type domain-containing protein; uniclust \| UniRef100\_A0A094YT33 \| 99.9 \| 2e-30 \| 3.8e-36 \| 181.5 \| 178 \| (2, 180) \| 180 \| (28, 253) \| 254 \| Putative transcriptional regulator \| Putative transcriptional regulator; uniclust \| UniRef100\_A0A101ETK8 \| 99.9 \| 2.2e-30 \| 4e-36 \| 173.5 \| 176 \| (1, 178) \| 180 \| (11, 213) \| 230 \| Putative prophage repressor \| Putative prophage repressor; uniclust \| UniRef100\_A0A132DZT8 \| 99.9 \| 2.1e-30 \| 4e-36 \| 185.2 \| 175 \| (5, 180) \| 180 \| (47, 261) \| 276 \| Uncharacterized protein \| Uncharacterized protein; uniclust \| UniRef100\_A0A1C5ZA08 \| 99.9 \| 2.2e-30 \| 4.1e-36 \| 176.4 \| 178 \| (1, 180) \| 180 \| (12, 219) \| 222 \| LexA repressor \| LexA repressor; uniclust \| UniRef100\_A0A0H2MAD8 \| 99.9 \| 2.7e-30 \| 5e-36 \| 181.9 \| 178 \| (1, 180) \| 180 \| (37, 253) \| 255 \| Putative HTH-type transcriptional regulator \| Putative HTH-type transcriptional regulator; uniclust \| UniRef100\_A0A084EVF3 \| 99.9 \| 2.9e-30 \| 5.4e-36 \| 182.7 \| 178 \| (2, 180) \| 180 \| (90, 312) \| 313 \| Putative HTH-type transcriptional regulator \| Putative HTH-type transcriptional regulator; uniclust \| UniRef100\_A0A142BB31 \| 99.9 \| 3.6e-30 \| 6.6e-36 \| 174.1 \| 175 \| (1, 179) \| 180 \| (12, 200) \| 203 \| S24 family peptidase \| S24 family peptidase; uniclust \| UniRef100\_A0A0Q2XP48 \| 99.9 \| 3.7e-30 \| 6.9e-36 \| 180.4 \| 174 \| (3, 180) \| 180 \| (35, 244) \| 246 \| HTH cro/C1-type domain-containing protein \| HTH cro/C1-type domain-containing protein; uniclust \| UniRef100\_A0A1C3RIU3 \| 99.9 \| 4e-30 \| 7.3e-36 \| 175.5 \| 177 \| (1, 180) \| 180 \| (22, 235) \| 237 \| Putative Transcriptional regulator \| Putative Transcriptional regulator; uniclust \| UniRef100\_A0A0A2XGJ6 \| 99.9 \| 4.3e-30 \| 8e-36 \| 178.9 \| 177 \| (1, 178) \| 180 \| (23, 241) \| 247 \| Transcriptional regulator \| Transcriptional regulator; uniclust \| UniRef100\_A0A011VZ09 \| 99.9 \| 5.1e-30 \| 9.4e-36 \| 185.3 \| 178 \| (1, 180) \| 180 \| (56, 264) \| 272 \| LexA family transcriptional regulator \| LexA family transcriptional regulator; uniclust \| UniRef100\_A0A173YIP3 \| 99.9 \| 5.4e-30 \| 9.9e-36 \| 176.0 \| 177 \| (1, 180) \| 180 \| (19, 227) \| 230 \| LexA repressor \| LexA repressor; uniclust \| UniRef100\_A0A0Q7I1K8 \| 99.9 \| 5.9e-30 \| 1.1e-35 \| 179.1 \| 177 \| (4, 180) \| 180 \| (44, 261) \| 262 \| HTH cro/C1-type domain-containing protein \| HTH cro/C1-type domain-containing protein; uniclust \| UniRef100\_A0A074VC69 \| 99.9 \| 6.1e-30 \| 1.1e-35 \| 183.0 \| 175 \| (1, 176) \| 180 \| (25, 243) \| 280 \| Putative transcriptional regulator \| Putative transcriptional regulator; uniclust \| UniRef100\_A0A086PBI9 \| 99.9 \| 6.9e-30 \| 1.3e-35 \| 185.1 \| 177 \| (2, 180) \| 180 \| (83, 301) \| 302 \| Putative phage repressor \| Putative phage repressor; uniclust \| UniRef100\_A0A248LHK2 \| 99.9 \| 7.4e-30 \| 1.4e-35 \| 172.7 \| 178 \| (2, 180) \| 180 \| (37, 260) \| 261 \| Repressor \| Repressor; uniclust \| UniRef100\_A0A089HUS8 \| 99.9 \| 9.4e-30 \| 1.7e-35 \| 182.6 \| 175 \| (1, 179) \| 180 \| (61, 259) \| 289 \| HTH cro/C1-type domain-containing protein \| HTH cro/C1-type domain-containing protein; uniclust \| UniRef100\_A0A0D8J1P7 \| 99.9 \| 9.5e-30 \| 1.7e-35 \| 171.1 \| 174 \| (1, 177) \| 180 \| (12, 213) \| 219 \| Helix-turn-helix domain-containing protein \| Helix-turn-helix domain-containing protein; uniclust \| UniRef100\_A0A193GZ18 \| 99.9 \| 1e-29 \| 1.8e-35 \| 165.0 \| 179 \| (2, 180) \| 180 \| (11, 189) \| 193 \| Putative DNA binding helix-turn helix protein \| Putative DNA binding helix-turn helix protein; uniclust \| UniRef100\_A0A017H4C1 \| 99.9 \| 1e-29 \| 1.9e-35 \| 181.7 \| 177 \| (1, 179) \| 180 \| (22, 231) \| 271 \| Peptidase S24 \| Peptidase S24; uniclust \| UniRef100\_A0A2E3AY80 \| 99.9 \| 1.2e-29 \| 2.2e-35 \| 169.9 \| 174 \| (2, 176) \| 180 \| (7, 216) \| 222 \| Peptidase S24 \| Peptidase S24; uniclust \| UniRef100\_A0A099GKA4 \| 99.9 \| 1.2e-29 \| 2.2e-35 \| 179.6 \| 178 \| (1, 179) \| 180 \| (46, 269) \| 288 \| Peptidase S24/S26A/S26B/S26C domain-containing protein \| Peptidase S24/S26A/S26B/S26C domain-containing protein; uniclust \| UniRef100\_A0A1G0SGA6 \| 99.9 \| 1.3e-29 \| 2.4e-35 \| 169.7 \| 177 \| (1, 180) \| 180 \| (15, 226) \| 227 \| HTH cro/C1-type domain-containing protein \| HTH cro/C1-type domain-containing protein; uniclust \| UniRef100\_A0A0Q1A481 \| 99.9 \| 1.6e-29 \| 2.9e-35 \| 173.5 \| 180 \| (1, 180) \| 180 \| (7, 221) \| 226 \| Uncharacterized protein \| Uncharacterized protein; uniclust \| UniRef100\_A0A352NLU9 \| 99.9 \| 3.5e-29 \| 6.5e-35 \| 172.2 \| 176 \| (1, 179) \| 180 \| (59, 264) \| 265 \| LexA family transcriptional regulator \| LexA family transcriptional regulator; uniclust \| UniRef100\_A0A073IPP6 \| 99.9 \| 3.6e-29 \| 6.6e-35 \| 175.6 \| 176 \| (1, 178) \| 180 \| (22, 244) \| 252 \| HTH cro/C1-type domain-containing protein \| HTH cro/C1-type domain-containing protein; uniclust \| UniRef100\_A0A059L2Y0 \| 99.9 \| 5e-29 \| 9.3e-35 \| 184.3 \| 175 \| (2, 177) \| 180 \| (90, 320) \| 370 \| XRE family transcriptional regulator \| XRE family transcriptional regulator; uniclust \| UniRef100\_A0A1G6D4T4 \| 99.9 \| 5.4e-29 \| 1e-34 \| 179.9 \| 172 \| (1, 178) \| 180 \| (75, 309) \| 353 \| Peptidase S24-like \| Peptidase S24-like; uniclust \| UniRef100\_A0A080NM98 \| 99.9 \| 5.5e-29 \| 1e-34 \| 173.7 \| 176 \| (3, 180) \| 180 \| (53, 262) \| 265 \| LexA family transcriptional regulator \| LexA family transcriptional regulator; uniclust \| UniRef100\_A0A062XFL7 \| 99.9 \| 5.7e-29 \| 1e-34 \| 178.3 \| 176 \| (1, 178) \| 180 \| (37, 267) \| 275 \| HTH cro/C1-type domain-containing protein \| HTH cro/C1-type domain-containing protein; uniclust \| UniRef100\_A0A0Q8U4Q3 \| 99.9 \| 6.1e-29 \| 1.1e-34 \| 180.4 \| 176 \| (2, 179) \| 180 \| (33, 248) \| 309 \| Phage repressor protein \| Phage repressor protein; uniclust \| UniRef100\_A0A009Z9C0 \| 99.9 \| 7.8e-29 \| 1.4e-34 \| 180.8 \| 176 \| (1, 176) \| 180 \| (55, 287) \| 308 \| Repressor protein CI \| Repressor protein CI; uniclust \| UniRef100\_A0A062H0E0 \| 99.9 \| 9.4e-29 \| 1.7e-34 \| 167.0 \| 178 \| (1, 179) \| 180 \| (23, 235) \| 238 \| Helix-turn-helix family protein \| Helix-turn-helix family protein; uniclust \| UniRef100\_A0A143B3W3 \| 99.9 \| 1e-28 \| 1.9e-34 \| 176.4 \| 171 \| (10, 180) \| 180 \| (58, 273) \| 276 \| LexA-related transcriptional regulator \| LexA-related transcriptional regulator; uniclust \| UniRef100\_A0A3C1VZE1 \| 99.9 \| 1.4e-28 \| 2.6e-34 \| 168.7 \| 174 \| (1, 175) \| 180 \| (10, 225) \| 233 \| HTH cro/C1-type domain-containing protein \| HTH cro/C1-type domain-containing protein; uniclust \| UniRef100\_A0A009FWP1 \| 99.9 \| 1.5e-28 \| 2.7e-34 \| 172.2 \| 176 \| (1, 177) \| 180 \| (16, 240) \| 245 \| Helix-turn-helix family protein \| Helix-turn-helix family protein; uniclust \| UniRef100\_A0A009GET6 \| 99.9 \| 1.5e-28 \| 2.7e-34 \| 171.0 \| 174 \| (2, 176) \| 180 \| (43, 271) \| 278 \| Peptidase S24-like family protein \| Peptidase S24-like family protein; uniclust \| UniRef100\_A0A060HCU5 \| 99.9 \| 1.6e-28 \| 3e-34 \| 175.5 \| 176 \| (1, 178) \| 180 \| (48, 264) \| 271 \| HTH cro/C1-type domain-containing protein \| HTH cro/C1-type domain-containing protein; uniclust \| UniRef100\_A0A023D7Y1 \| 99.9 \| 2e-28 \| 3.6e-34 \| 172.6 \| 175 \| (2, 178) \| 180 \| (31, 251) \| 256 \| Transcriptional regulator phage repressor \| Transcriptional regulator phage repressor; uniclust \| UniRef100\_A0A0K1F1X2 \| 99.9 \| 2.2e-28 \| 4.1e-34 \| 170.3 \| 175 \| (1, 177) \| 180 \| (23, 231) \| 240 \| HTH cro/C1-type domain-containing protein \| HTH cro/C1-type domain-containing protein; uniclust \| UniRef100\_A0A023KYN1 \| 99.9 \| 2.3e-28 \| 4.3e-34 \| 178.6 \| 175 \| (1, 177) \| 180 \| (65, 282) \| 314 \| Helix-turn-helix transcriptional regulator \| Helix-turn-helix transcriptional regulator; uniclust \| UniRef100\_A0A124G6Q3 \| 99.9 \| 2.5e-28 \| 4.5e-34 \| 167.5 \| 178 \| (1, 179) \| 180 \| (9, 243) \| 245 \| XRE family transcriptional regulator \| XRE family transcriptional regulator; uniclust \| UniRef100\_A0A068A299 \| 99.9 \| 2.7e-28 \| 5e-34 \| 176.4 \| 177 \| (1, 179) \| 180 \| (34, 271) \| 275 \| Repressor \| Repressor; uniclust \| UniRef100\_A0A013WWL1 \| 99.9 \| 3e-28 \| 5.5e-34 \| 175.5 \| 177 \| (3, 180) \| 180 \| (53, 262) \| 275 \| Peptidase S24 \| Peptidase S24; uniclust \| UniRef100\_A0A1F8XTB5 \| 99.9 \| 3e-28 \| 5.6e-34 \| 170.2 \| 178 \| (1, 180) \| 180 \| (46, 262) \| 263 \| HTH cro/C1-type domain-containing protein \| HTH cro/C1-type domain-containing protein; uniclust \| UniRef100\_A0A1Y5FD13 \| 99.9 \| 4.3e-28 \| 8e-34 \| 163.6 \| 174 \| (4, 180) \| 180 \| (14, 217) \| 218 \| Peptidase S24/S26A/S26B/S26C domain-containing protein \| Peptidase S24/S26A/S26B/S26C domain-containing protein; uniclust \| UniRef100\_A0A0F2J2G4 \| 99.9 \| 4.3e-28 \| 8e-34 \| 170.3 \| 175 \| (1, 178) \| 180 \| (44, 244) \| 276 \| Repressor protein \| Repressor protein; uniclust \| UniRef100\_A0A0G4C027 \| 99.9 \| 4.5e-28 \| 8.3e-34 \| 172.1 \| 177 \| (1, 178) \| 180 \| (18, 249) \| 255 \| Peptidase S24-like protein \| Peptidase S24-like protein; uniclust \| UniRef100\_A0A0B5QLH2 \| 99.9 \| 5.4e-28 \| 9.9e-34 \| 169.1 \| 177 \| (1, 179) \| 180 \| (21, 236) \| 241 \| HTH cro/C1-type domain-containing protein \| HTH cro/C1-type domain-containing protein; uniclust \| UniRef100\_A0A073INQ6 \| 99.9 \| 6.6e-28 \| 1.2e-33 \| 162.9 \| 176 \| (1, 179) \| 180 \| (5, 228) \| 238 \| HTH cro/C1-type domain-containing protein \| HTH cro/C1-type domain-containing protein; uniclust \| UniRef100\_A0A0R1Q5L8 \| 99.9 \| 7.1e-28 \| 1.3e-33 \| 163.9 \| 174 \| (2, 177) \| 180 \| (14, 211) \| 218 \| Peptidase s24-like protein \| Peptidase s24-like protein; uniclust \| UniRef100\_A0A011VZ04 \| 99.9 \| 7.1e-28 \| 1.3e-33 \| 170.9 \| 178 \| (1, 179) \| 180 \| (14, 227) \| 234 \| DNA-binding protein \| DNA-binding protein; uniclust \| UniRef100\_A0A011PI42 \| 99.9 \| 8.1e-28 \| 1.5e-33 \| 175.2 \| 173 \| (2, 176) \| 180 \| (86, 309) \| 320 \| Pyocin repressor protein \| Pyocin repressor protein; uniclust \| UniRef100\_A0A1V5P064 \| 99.9 \| 8.3e-28 \| 1.5e-33 \| 169.0 \| 179 \| (1, 179) \| 180 \| (39, 248) \| 265 \| LexA repressor \| LexA repressor; uniclust \| UniRef100\_A0A085ERU0 \| 99.9 \| 8.9e-28 \| 1.7e-33 \| 171.9 \| 177 \| (2, 179) \| 180 \| (36, 253) \| 261 \| Putative transcriptional regulator \| Putative transcriptional regulator; uniclust \| UniRef100\_A0A134A0B4 \| 99.9 \| 9.9e-28 \| 1.8e-33 \| 172.0 \| 176 \| (1, 179) \| 180 \| (17, 260) \| 267 \| Peptidase S24-like protein \| Peptidase S24-like protein; uniclust \| UniRef100\_A0A0M9GHV2 \| 99.9 \| 1.5e-27 \| 2.8e-33 \| 159.9 \| 176 \| (2, 179) \| 180 \| (27, 237) \| 239 \| Putative transcriptional regulator \| Putative transcriptional regulator; uniclust \| UniRef100\_A0A1R1MJE6 \| 99.9 \| 1.6e-27 \| 3e-33 \| 162.4 \| 177 \| (1, 179) \| 180 \| (21, 213) \| 216 \| HTH cro/C1-type domain-containing protein \| HTH cro/C1-type domain-containing protein; uniclust \| UniRef100\_A0A1F9FXD4 \| 99.9 \| 1.8e-27 \| 3.4e-33 \| 162.8 \| 176 \| (1, 179) \| 180 \| (26, 230) \| 232 \| HTH cro/C1-type domain-containing protein \| HTH cro/C1-type domain-containing protein; uniclust \| UniRef100\_A0A074VB08 \| 99.9 \| 2.2e-27 \| 4.1e-33 \| 161.2 \| 176 \| (1, 178) \| 180 \| (24, 241) \| 245 \| HTH cro/C1-type domain-containing protein \| HTH cro/C1-type domain-containing protein; uniclust \| UniRef100\_A0A084J761 \| 99.9 \| 2.2e-27 \| 4.1e-33 \| 174.8 \| 175 \| (1, 177) \| 180 \| (52, 283) \| 335 \| HTH cro/C1-type domain-containing protein \| HTH cro/C1-type domain-containing protein; uniclust \| UniRef100\_A0A7C2V4D0 \| 99.9 \| 2.3e-27 \| 4.2e-33 \| 159.4 \| 174 \| (1, 177) \| 180 \| (19, 215) \| 224 \| Helix-turn-helix domain-containing protein \| Helix-turn-helix domain-containing protein; uniclust \| UniRef100\_A0A081NDK8 \| 99.9 \| 3.1e-27 \| 5.7e-33 \| 158.6 \| 166 \| (14, 180) \| 180 \| (33, 231) \| 232 \| Repressor \| Repressor; uniclust \| UniRef100\_A0A031IUW7 \| 99.9 \| 3.7e-27 \| 6.9e-33 \| 172.0 \| 173 \| (2, 176) \| 180 \| (78, 308) \| 321 \| Putative phage repressor \| Putative phage repressor; uniclust \| UniRef100\_A0A926HWW3 \| 99.9 \| 3.8e-27 \| 7e-33 \| 153.6 \| 176 \| (2, 180) \| 180 \| (3, 185) \| 186 \| Helix-turn-helix domain-containing protein \| Helix-turn-helix domain-containing protein; uniclust \| UniRef100\_A0A0Q8TCT1 \| 99.9 \| 4.6e-27 \| 8.5e-33 \| 163.0 \| 175 \| (5, 180) \| 180 \| (38, 241) \| 242 \| Peptidase S24/S26A/S26B/S26C domain-containing protein \| Peptidase S24/S26A/S26B/S26C domain-containing protein; uniclust \| UniRef100\_A0A072NUH5 \| 99.9 \| 5e-27 \| 9.3e-33 \| 169.4 \| 175 \| (1, 178) \| 180 \| (15, 280) \| 298 \| SOS response transcriptional repressor, RecA-mediated autopeptidase \| SOS response transcriptional repressor, RecA-mediated autopeptidase; uniclust \| UniRef100\_A0A075SG92 \| 99.9 \| 5.4e-27 \| 1e-32 \| 173.0 \| 177 \| (1, 179) \| 180 \| (76, 319) \| 337 \| Repressor \| Repressor; uniclust \| UniRef100\_A0A0R1VT03 \| 99.9 \| 5.9e-27 \| 1.1e-32 \| 162.0 \| 173 \| (1, 175) \| 180 \| (21, 252) \| 253 \| Phage repressor like XRE family transcriptional regulator \| Phage repressor like XRE family transcriptional regulator; uniclust \| UniRef100\_A0A060H2Y3 \| 99.9 \| 6.3e-27 \| 1.2e-32 \| 168.8 \| 174 \| (1, 175) \| 180 \| (82, 314) \| 358 \| Peptidase S24, S26A and S26B \| Peptidase S24, S26A and S26B; uniclust \| UniRef100\_A0A0R1LCU9 \| 99.9 \| 6.4e-27 \| 1.2e-32 \| 163.8 \| 172 \| (1, 175) \| 180 \| (37, 269) \| 286 \| Peptidase S24-like protein \| Peptidase S24-like protein; uniclust \| UniRef100\_A0A1B3JHZ4 \| 99.9 \| 6.7e-27 \| 1.2e-32 \| 159.8 \| 175 \| (1, 176) \| 180 \| (14, 238) \| 247 \| Phage repressor \| Phage repressor; uniclust \| UniRef100\_A0A1C6GZH6 \| 99.9 \| 6.8e-27 \| 1.3e-32 \| 166.7 \| 168 \| (1, 176) \| 180 \| (22, 249) \| 306 \| LexA repressor \| LexA repressor; uniclust \| UniRef100\_A0A0J6S859 \| 99.9 \| 7.4e-27 \| 1.4e-32 \| 166.6 \| 171 \| (1, 174) \| 180 \| (31, 275) \| 283 \| HTH cro/C1-type domain-containing protein \| HTH cro/C1-type domain-containing protein; uniclust \| UniRef100\_A0A1W1Y2G5 \| 99.9 \| 8.4e-27 \| 1.6e-32 \| 165.7 \| 177 \| (1, 179) \| 180 \| (23, 247) \| 265 \| Phage repressor protein C, contains Cro/C1-type HTH and peptisase s24 domains \| Phage repressor protein C, contains Cro/C1-type HTH and peptisase s24 domains; uniclust \| UniRef100\_A0A212QLE3 \| 99.9 \| 8.8e-27 \| 1.6e-32 \| 163.4 \| 178 \| (1, 179) \| 180 \| (38, 276) \| 277 \| Phage repressor protein C, contains Cro/C1-type HTH and peptisase s24 domains \| Phage repressor protein C, contains Cro/C1-type HTH and peptisase s24 domains; uniclust \| UniRef100\_A0A024L3W0 \| 99.9 \| 9.1e-27 \| 1.7e-32 \| 171.1 \| 175 \| (1, 179) \| 180 \| (69, 281) \| 312 \| Helix-turn-helix domain-containing protein \| Helix-turn-helix domain-containing protein; uniclust \| UniRef100\_A0A015NCW0 \| 99.9 \| 9.8e-27 \| 1.8e-32 \| 171.3 \| 177 \| (2, 178) \| 180 \| (62, 292) \| 297 \| HTH cro/C1-type domain-containing protein \| HTH cro/C1-type domain-containing protein; uniclust \| UniRef100\_A0A1G7D0E1 \| 99.9 \| 1.3e-26 \| 2.4e-32 \| 154.8 \| 164 \| (16, 179) \| 180 \| (35, 224) \| 226 \| Peptidase S24-like \| Peptidase S24-like; uniclust \| UniRef100\_A0A968IL67 \| 99.9 \| 1.7e-26 \| 3.1e-32 \| 151.1 \| 175 \| (3, 180) \| 180 \| (28, 235) \| 237 \| Helix-turn-helix domain-containing protein \| Helix-turn-helix domain-containing protein; uniclust \| UniRef100\_A0A009PV59 \| 99.9 \| 1.7e-26 \| 3.2e-32 \| 162.4 \| 178 \| (1, 179) \| 180 \| (47, 256) \| 271 \| Helix-turn-helix family protein \| Helix-turn-helix family protein; uniclust \| UniRef100\_A0A0D0RL49 \| 99.9 \| 1.9e-26 \| 3.6e-32 \| 162.7 \| 175 \| (1, 177) \| 180 \| (19, 242) \| 251 \| CI-like repressor, phage associated \| CI-like repressor, phage associated; uniclust \| UniRef100\_A0A1V6DM68 \| 99.9 \| 2.4e-26 \| 4.4e-32 \| 159.0 \| 180 \| (1, 180) \| 180 \| (2, 241) \| 244 \| LexA repressor \| LexA repressor; uniclust \| UniRef100\_A0A099I651 \| 99.9 \| 2.6e-26 \| 4.9e-32 \| 169.4 \| 170 \| (1, 176) \| 180 \| (53, 295) \| 312 \| HTH cro/C1-type domain-containing protein \| HTH cro/C1-type domain-containing protein; uniclust \| UniRef100\_A0A0Q4FWG9 \| 99.9 \| 2.7e-26 \| 5e-32 \| 161.9 \| 177 \| (3, 180) \| 180 \| (57, 272) \| 272 \| HTH cro/C1-type domain-containing protein \| HTH cro/C1-type domain-containing protein; uniclust \| UniRef100\_A0A1G3BMS2 \| 99.9 \| 2.7e-26 \| 5e-32 \| 158.4 \| 176 \| (1, 176) \| 180 \| (60, 259) \| 265 \| HTH cro/C1-type domain-containing protein \| HTH cro/C1-type domain-containing protein; uniclust \| UniRef100\_A0A1G3L741 \| 99.9 \| 2.9e-26 \| 5.3e-32 \| 161.9 \| 175 \| (1, 177) \| 180 \| (10, 244) \| 249 \| HTH cro/C1-type domain-containing protein \| HTH cro/C1-type domain-containing protein; uniclust \| UniRef100\_A0A090V113 \| 99.9 \| 3.1e-26 \| 5.7e-32 \| 163.8 \| 175 \| (1, 179) \| 180 \| (31, 239) \| 253 \| LexA repressor \| LexA repressor; uniclust \| UniRef100\_UPI000FD88B00 \| 99.9 \| 3.3e-26 \| 6.1e-32 \| 155.2 \| 174 \| (2, 177) \| 180 \| (19, 237) \| 241 \| S24 family peptidase \| S24 family peptidase; uniclust \| UniRef100\_A0A2S5N385 \| 99.9 \| 4.4e-26 \| 8.2e-32 \| 161.2 \| 177 \| (1, 179) \| 180 \| (66, 291) \| 292 \| HTH cro/C1-type domain-containing protein \| HTH cro/C1-type domain-containing protein; uniclust \| UniRef100\_A0A0U3VIS1 \| 99.9 \| 4.6e-26 \| 8.5e-32 \| 156.4 \| 176 \| (1, 178) \| 180 \| (9, 211) \| 226 \| LexA family transcriptional regulator \| LexA family transcriptional regulator; uniclust \| UniRef100\_A0A0S4R308 \| 99.9 \| 5.3e-26 \| 9.8e-32 \| 154.4 \| 174 \| (2, 177) \| 180 \| (10, 222) \| 230 \| DNA-binding protein \| DNA-binding protein; uniclust \| UniRef100\_A0A023CWD3 \| 99.9 \| 5.4e-26 \| 1e-31 \| 153.2 \| 176 \| (2, 179) \| 180 \| (18, 215) \| 225 \| Xre family DNA-binding protein \| Xre family DNA-binding protein; uniclust \| UniRef100\_A0A0Q2Y0V6 \| 99.9 \| 5.8e-26 \| 1.1e-31 \| 157.2 \| 165 \| (14, 179) \| 180 \| (1, 216) \| 236 \| Peptidase S24/S26A/S26B/S26C domain-containing protein \| Peptidase S24/S26A/S26B/S26C domain-containing protein; uniclust \| UniRef100\_A0A068RDG1 \| 99.9 \| 5.8e-26 \| 1.1e-31 \| 156.8 \| 174 \| (1, 176) \| 180 \| (23, 222) \| 232 \| Helix-turn-helix/Peptidase S24-like domain-containing protein \| Helix-turn-helix/Peptidase S24-like domain-containing protein; uniclust \| UniRef100\_A0A0A5HTQ8 \| 99.9 \| 6.4e-26 \| 1.2e-31 \| 157.4 \| 162 \| (15, 180) \| 180 \| (60, 243) \| 250 \| Transcriptional regulator \| Transcriptional regulator; uniclust \| UniRef100\_A0A024E9Q4 \| 99.9 \| 7.9e-26 \| 1.5e-31 \| 164.5 \| 176 \| (3, 178) \| 180 \| (47, 277) \| 284 \| Phage repressor \| Phage repressor; uniclust \| UniRef100\_A0A0Q3EP03 \| 99.9 \| 8.8e-26 \| 1.6e-31 \| 162.3 \| 175 \| (2, 176) \| 180 \| (52, 265) \| 276 \| Helix-turn-helix domain-containing protein \| Helix-turn-helix domain-containing protein; uniclust \| UniRef100\_A0A377PS33 \| 99.9 \| 9e-26 \| 1.7e-31 \| 153.2 \| 175 \| (1, 180) \| 180 \| (4, 247) \| 249 \| LexA repressor \| LexA repressor; uniclust \| UniRef100\_A0A369T885 \| 99.9 \| 9.5e-26 \| 1.7e-31 \| 153.6 \| 176 \| (1, 180) \| 180 \| (38, 255) \| 257 \| XRE family transcriptional regulator \| XRE family transcriptional regulator; uniclust \| UniRef100\_A0A0C9MTW4 \| 99.9 \| 1.1e-25 \| 2e-31 \| 158.3 \| 174 \| (2, 179) \| 180 \| (36, 244) \| 251 \| DNA, contig: SP630 \| DNA, contig: SP630; uniclust \| UniRef100\_A0A098RK60 \| 99.9 \| 1.1e-25 \| 2e-31 \| 160.0 \| 176 \| (2, 178) \| 180 \| (23, 277) \| 279 \| Repressor \| Repressor; uniclust \| UniRef100\_A0A081MYG4 \| 99.9 \| 1.1e-25 \| 2.1e-31 \| 158.0 \| 178 \| (1, 179) \| 180 \| (15, 219) \| 225 \| HTH cro/C1-type domain-containing protein \| HTH cro/C1-type domain-containing protein; uniclust \| UniRef100\_A0A1F4HU16 \| 99.9 \| 1.2e-25 \| 2.2e-31 \| 158.4 \| 175 \| (1, 177) \| 180 \| (63, 281) \| 286 \| Repressor \| Repressor; uniclust \| UniRef100\_A0A0V8LX08 \| 99.9 \| 1.2e-25 \| 2.2e-31 \| 154.7 \| 172 \| (1, 180) \| 180 \| (13, 216) \| 228 \| Transcriptional regulator \| Transcriptional regulator; uniclust \| UniRef100\_A0A0K1HH28 \| 99.9 \| 1.2e-25 \| 2.3e-31 \| 155.0 \| 174 \| (2, 175) \| 180 \| (8, 237) \| 256 \| Putative transcriptional regulator, XRE family (Peptidase S24 LexA-like domain) \| Putative transcriptional regulator, XRE family (Peptidase S24 LexA-like domain); uniclust \| UniRef100\_A0A0C2YQB8 \| 99.9 \| 1.3e-25 \| 2.3e-31 \| 161.2 \| 177 \| (3, 180) \| 180 \| (54, 273) \| 278 \| Peptidase \| Peptidase; uniclust \| UniRef100\_A0A0C3RMF8 \| 99.9 \| 1.3e-25 \| 2.4e-31 \| 151.3 \| 101 \| (79, 180) \| 180 \| (70, 172) \| 175 \| Peptidase S24/S26A/S26B/S26C domain-containing protein \| Peptidase S24/S26A/S26B/S26C domain-containing protein; uniclust \| UniRef100\_A0A0D6H1B7 \| 99.9 \| 1.3e-25 \| 2.4e-31 \| 158.5 \| 177 \| (1, 177) \| 180 \| (21, 233) \| 243 \| LexA family transcriptional regulator \| LexA family transcriptional regulator; uniclust \| UniRef100\_A0A090HXE6 \| 99.9 \| 1.3e-25 \| 2.5e-31 \| 158.1 \| 176 \| (1, 176) \| 180 \| (23, 235) \| 242 \| LexA protein \| LexA protein; uniclust \| UniRef100\_A0A937YQE5 \| 99.9 \| 1.5e-25 \| 2.7e-31 \| 147.1 \| 178 \| (2, 180) \| 180 \| (35, 246) \| 246 \| Helix-turn-helix transcriptional regulator \| Helix-turn-helix transcriptional regulator; uniclust \| UniRef100\_A0A1A9RLP6 \| 99.9 \| 1.5e-25 \| 2.8e-31 \| 156.4 \| 175 \| (2, 177) \| 180 \| (35, 253) \| 264 \| Peptidase S24/S26A/S26B/S26C domain-containing protein \| Peptidase S24/S26A/S26B/S26C domain-containing protein; uniclust \| UniRef100\_A0A060RFY0 \| 99.9 \| 1.6e-25 \| 3e-31 \| 162.5 \| 177 \| (1, 179) \| 180 \| (29, 262) \| 272 \| Pleiotropic regulator of exopolysaccharide synthe sis, competence and biofilm formation Ftr, XRE family \| Pleiotropic regulator of exopolysaccharide synthe sis, competence and biofilm formation Ftr, XRE family; uniclust \| UniRef100\_A0A1Q6T3X1 \| 99.9 \| 2.1e-25 \| 3.9e-31 \| 153.2 \| 175 \| (4, 179) \| 180 \| (10, 222) \| 226 \| Peptidase S24/S26A/S26B/S26C domain-containing protein \| Peptidase S24/S26A/S26B/S26C domain-containing protein; uniclust \| UniRef100\_A0A061QGB9 \| 99.9 \| 2.2e-25 \| 4e-31 \| 160.0 \| 176 \| (2, 179) \| 180 \| (63, 252) \| 265 \| LexA repressor \| LexA repressor; uniclust \| UniRef100\_A0A212IUA3 \| 99.9 \| 2.2e-25 \| 4.1e-31 \| 146.0 \| 175 \| (1, 179) \| 180 \| (3, 217) \| 218 \| Putative Phage repressor \| Putative Phage repressor; uniclust \| UniRef100\_A0A0U4BEI9 \| 99.9 \| 2.3e-25 \| 4.3e-31 \| 159.2 \| 175 \| (1, 177) \| 180 \| (33, 256) \| 266 \| HTH cro/C1-type domain-containing protein \| HTH cro/C1-type domain-containing protein; uniclust \| UniRef100\_A0A066SSH7 \| 99.9 \| 2.3e-25 \| 4.3e-31 \| 157.7 \| 175 \| (1, 179) \| 180 \| (37, 248) \| 254 \| Helix-turn-helix domain-containing protein \| Helix-turn-helix domain-containing protein; uniclust \| UniRef100\_A0A0B1TSB9 \| 99.9 \| 2.9e-25 \| 5.4e-31 \| 156.1 \| 177 \| (2, 180) \| 180 \| (35, 281) \| 282 \| Putative transcriptional regulator \| Putative transcriptional regulator; uniclust \| UniRef100\_A0A1I1VLL5 \| 99.9 \| 3e-25 \| 5.5e-31 \| 152.4 \| 177 \| (2, 179) \| 180 \| (21, 252) \| 256 \| Phage repressor protein C, contains Cro/C1-type HTH and peptisase s24 domains \| Phage repressor protein C, contains Cro/C1-type HTH and peptisase s24 domains; uniclust \| UniRef100\_A0A077PXM9 \| 99.9 \| 3e-25 \| 5.5e-31 \| 157.3 \| 172 \| (2, 176) \| 180 \| (89, 319) \| 324 \| Repressor protein CI (Modular protein) \| Repressor protein CI (Modular protein); uniclust \| UniRef100\_A0A0F8XX58 \| 99.9 \| 3.7e-25 \| 6.8e-31 \| 157.5 \| 177 \| (2, 178) \| 180 \| (31, 267) \| 277 \| HTH cro/C1-type domain-containing protein (Fragment) \| HTH cro/C1-type domain-containing protein (Fragment); uniclust \| UniRef100\_A0A1C5L5I0 \| 99.9 \| 3.9e-25 \| 7.3e-31 \| 150.8 \| 177 \| (1, 178) \| 180 \| (12, 212) \| 219 \| LexA repressor \| LexA repressor; uniclust \| UniRef100\_A0A0J1F8E5 \| 99.9 \| 4e-25 \| 7.3e-31 \| 158.2 \| 176 \| (1, 179) \| 180 \| (21, 226) \| 291 \| HTH-type transcriptional regulator Xre \| HTH-type transcriptional regulator Xre; uniclust \| UniRef100\_A0A149SNV7 \| 99.9 \| 4.9e-25 \| 9e-31 \| 152.7 \| 172 \| (2, 175) \| 180 \| (18, 233) \| 238 \| HTH cro/C1-type domain-containing protein \| HTH cro/C1-type domain-containing protein; uniclust \| UniRef100\_A0A352MTY7 \| 99.9 \| 4.9e-25 \| 9.1e-31 \| 147.7 \| 175 \| (1, 177) \| 180 \| (5, 193) \| 195 \| HTH cro/C1-type domain-containing protein \| HTH cro/C1-type domain-containing protein; uniclust \| UniRef100\_A0A061QF65 \| 99.9 \| 5e-25 \| 9.1e-31 \| 150.3 \| 169 \| (12, 180) \| 180 \| (23, 226) \| 227 \| Repressor protein CI \| Repressor protein CI; uniclust \| UniRef100\_A0A1V5PB98 \| 99.9 \| 5.2e-25 \| 9.7e-31 \| 148.8 \| 172 \| (3, 177) \| 180 \| (23, 226) \| 230 \| LexA repressor \| LexA repressor; uniclust \| UniRef100\_A0A5C1MF53 \| 99.8 \| 5.5e-25 \| 1e-30 \| 150.4 \| 175 \| (2, 180) \| 180 \| (45, 263) \| 263 \| Helix-turn-helix transcriptional regulator \| Helix-turn-helix transcriptional regulator; uniclust \| UniRef100\_A0A0Y7KSX8 \| 99.8 \| 5.6e-25 \| 1e-30 \| 148.9 \| 179 \| (1, 180) \| 180 \| (30, 256) \| 256 \| Helix-turn-helix domain-containing protein \| Helix-turn-helix domain-containing protein; uniclust \| UniRef100\_A0A0K6I8B4 \| 99.8 \| 6.3e-25 \| 1.2e-30 \| 155.1 \| 173 \| (1, 175) \| 180 \| (62, 282) \| 290 \| Peptidase S24-like \| Peptidase S24-like; uniclust \| UniRef100\_A0A419V0E4 \| 99.8 \| 6.4e-25 \| 1.2e-30 \| 146.3 \| 175 \| (1, 177) \| 180 \| (7, 206) \| 210 \| SOS-response transcriptional repressor LexA \| SOS-response transcriptional repressor LexA; uniclust \| UniRef100\_A0A1M5MNV0 \| 99.8 \| 7.7e-25 \| 1.4e-30 \| 153.5 \| 173 \| (2, 175) \| 180 \| (29, 258) \| 266 \| Peptidase S24-like \| Peptidase S24-like; uniclust \| UniRef100\_A0A133N482 \| 99.8 \| 8.6e-25 \| 1.6e-30 \| 154.2 \| 176 \| (1, 178) \| 180 \| (88, 282) \| 286 \| Repressor LexA \| Repressor LexA; uniclust \| UniRef100\_A0A014MMR0 \| 99.8 \| 1.1e-24 \| 2e-30 \| 147.5 \| 178 \| (2, 179) \| 180 \| (4, 216) \| 219 \| Peptidase S24/S26A/S26B/S26C domain-containing protein \| Peptidase S24/S26A/S26B/S26C domain-containing protein; uniclust \| UniRef100\_A0A099WLJ4 \| 99.8 \| 1.1e-24 \| 2.1e-30 \| 154.6 \| 177 \| (1, 177) \| 180 \| (60, 271) \| 284 \| HTH cro/C1-type domain-containing protein \| HTH cro/C1-type domain-containing protein; uniclust \| UniRef100\_A0A0V8LXT6 \| 99.8 \| 1.2e-24 \| 2.2e-30 \| 148.7 \| 175 \| (1, 180) \| 180 \| (16, 231) \| 233 \| HTH cro/C1-type domain-containing protein \| HTH cro/C1-type domain-containing protein; uniclust \| UniRef100\_A0A023D3P8 \| 99.8 \| 1.2e-24 \| 2.2e-30 \| 147.1 \| 174 \| (2, 178) \| 180 \| (36, 234) \| 238 \| Phage repressor \| Phage repressor; uniclust \| UniRef100\_A0A071M1S3 \| 99.8 \| 1.6e-24 \| 2.9e-30 \| 151.0 \| 175 \| (1, 179) \| 180 \| (30, 239) \| 241 \| XRE family transcriptional regulator \| XRE family transcriptional regulator; uniclust \| UniRef100\_A0A091C634 \| 99.8 \| 1.6e-24 \| 2.9e-30 \| 148.3 \| 171 \| (2, 174) \| 180 \| (8, 231) \| 232 \| Phage repressor \| Phage repressor; uniclust \| UniRef100\_A0A1B1PWH0 \| 99.8 \| 2.1e-24 \| 3.9e-30 \| 142.6 \| 177 \| (2, 178) \| 180 \| (5, 227) \| 229 \| HTH cro/C1-type domain-containing protein \| HTH cro/C1-type domain-containing protein; uniclust \| UniRef100\_A0A098F7G8 \| 99.8 \| 2.2e-24 \| 4e-30 \| 157.6 \| 175 \| (1, 177) \| 180 \| (14, 322) \| 340 \| SOS-response transcriptional repressor \| SOS-response transcriptional repressor; uniclust \| UniRef100\_A0A015NHE4 \| 99.8 \| 2.3e-24 \| 4.3e-30 \| 153.3 \| 177 \| (1, 177) \| 180 \| (24, 242) \| 256 \| HTH cro/C1-type domain-containing protein \| HTH cro/C1-type domain-containing protein; uniclust \| UniRef100\_F6DTI3 \| 99.8 \| 3.3e-24 \| 6.1e-30 \| 150.4 \| 174 \| (1, 177) \| 180 \| (8, 277) \| 294 \| LexA repressor \| LexA repressor; uniclust \| UniRef100\_A0A2C9D6H6 \| 99.8 \| 3.6e-24 \| 6.6e-30 \| 142.6 \| 176 \| (2, 179) \| 180 \| (8, 207) \| 209 \| Pyocin repressor protein \| Pyocin repressor protein; uniclust \| UniRef100\_A0A087MCA7 \| 99.8 \| 3.7e-24 \| 6.8e-30 \| 152.1 \| 165 \| (13, 179) \| 180 \| (53, 246) \| 288 \| Phage repressor protein \| Phage repressor protein; uniclust \| UniRef100\_A0A523J2I9 \| 99.8 \| 4.2e-24 \| 7.7e-30 \| 149.3 \| 176 \| (3, 179) \| 180 \| (35, 245) \| 246 \| Helix-turn-helix domain-containing protein \| Helix-turn-helix domain-containing protein; uniclust \| UniRef100\_A0A084UBN0 \| 99.8 \| 4.2e-24 \| 7.8e-30 \| 155.5 \| 173 \| (2, 179) \| 180 \| (44, 295) \| 297 \| Putative transcriptional regulator \| Putative transcriptional regulator; uniclust \| UniRef100\_A0A1V5GIA7 \| 99.8 \| 5e-24 \| 9.1e-30 \| 153.7 \| 172 \| (1, 178) \| 180 \| (65, 296) \| 330 \| HTH-type transcriptional regulator ImmR \| HTH-type transcriptional regulator ImmR; uniclust \| UniRef100\_A0A143Z9V0 \| 99.8 \| 5.3e-24 \| 9.7e-30 \| 144.9 \| 177 \| (1, 179) \| 180 \| (9, 256) \| 261 \| Phage repressor protein C, contains Cro/C1-type HTH and peptisase s24 domains \| Phage repressor protein C, contains Cro/C1-type HTH and peptisase s24 domains; uniclust \| UniRef100\_A0A353R1K5 \| 99.8 \| 5.3e-24 \| 9.7e-30 \| 144.4 \| 175 \| (1, 180) \| 180 \| (27, 230) \| 231 \| HTH cro/C1-type domain-containing protein \| HTH cro/C1-type domain-containing protein; uniclust \| UniRef100\_A0A3A4K5N6 \| 99.8 \| 5.5e-24 \| 1e-29 \| 141.3 \| 175 \| (1, 178) \| 180 \| (10, 203) \| 205 \| LexA family transcriptional regulator \| LexA family transcriptional regulator; uniclust \| UniRef100\_A0A1H4AEP1 \| 99.8 \| 6.5e-24 \| 1.2e-29 \| 140.4 \| 175 \| (1, 180) \| 180 \| (5, 193) \| 195 \| SOS-response transcriptional repressor LexA (RecA-mediated autopeptidase) \| SOS-response transcriptional repressor LexA (RecA-mediated autopeptidase); uniclust \| UniRef100\_A0A2U0TA89 \| 99.8 \| 6.7e-24 \| 1.2e-29 \| 141.1 \| 174 \| (3, 179) \| 180 \| (28, 215) \| 225 \| Helix-turn-helix protein \| Helix-turn-helix protein; uniclust \| UniRef100\_A0A2U3L084 \| 99.8 \| 7.8e-24 \| 1.4e-29 \| 144.9 \| 180 \| (1, 180) \| 180 \| (16, 246) \| 247 \| Putative phage repressor \| Putative phage repressor; uniclust \| UniRef100\_A0A0K2GBJ9 \| 99.8 \| 1.5e-23 \| 2.7e-29 \| 149.1 \| 176 \| (2, 180) \| 180 \| (43, 231) \| 256 \| LexA repressor \| LexA repressor; uniclust \| UniRef100\_A0A0S4W647 \| 99.8 \| 1.5e-23 \| 2.8e-29 \| 145.8 \| 172 \| (3, 176) \| 180 \| (12, 254) \| 260 \| Peptidase S24/S26A/S26B/S26C domain-containing protein \| Peptidase S24/S26A/S26B/S26C domain-containing protein; uniclust \| UniRef100\_A0A101VVJ0 \| 99.8 \| 1.6e-23 \| 3e-29 \| 144.4 \| 176 \| (1, 179) \| 180 \| (18, 221) \| 245 \| Peptidase S24/S26A/S26B/S26C domain-containing protein \| Peptidase S24/S26A/S26B/S26C domain-containing protein; uniclust \| UniRef100\_A0A1B7XER7 \| 99.8 \| 1.7e-23 \| 3.1e-29 \| 147.9 \| 164 \| (15, 180) \| 180 \| (37, 262) \| 265 \| Transcriptional regulator \| Transcriptional regulator; uniclust \| UniRef100\_A0A098G5U5 \| 99.8 \| 1.9e-23 \| 3.5e-29 \| 148.2 \| 178 \| (1, 179) \| 180 \| (47, 262) \| 274 \| HTH cro/C1-type domain-containing protein \| HTH cro/C1-type domain-containing protein; uniclust \| UniRef100\_A0A0Q5M1T3 \| 99.8 \| 1.9e-23 \| 3.6e-29 \| 142.6 \| 179 \| (1, 180) \| 180 \| (22, 248) \| 248 \| Peptidase S24 \| Peptidase S24; uniclust \| UniRef100\_A0A523U297 \| 99.8 \| 1.9e-23 \| 3.6e-29 \| 144.5 \| 175 \| (1, 177) \| 180 \| (23, 244) \| 257 \| Helix-turn-helix domain-containing protein \| Helix-turn-helix domain-containing protein; uniclust \| UniRef100\_A0A072CT56 \| 99.8 \| 1.9e-23 \| 3.6e-29 \| 154.4 \| 177 \| (1, 179) \| 180 \| (61, 276) \| 319 \| Repressor \| Repressor; uniclust \| UniRef100\_A0A084INR4 \| 99.8 \| 1.9e-23 \| 3.6e-29 \| 149.1 \| 171 \| (2, 178) \| 180 \| (41, 245) \| 250 \| Repressor protein c2 \| Repressor protein c2; uniclust \| UniRef100\_A0A1Q6U9K6 \| 99.8 \| 2e-23 \| 3.7e-29 \| 138.2 \| 176 \| (3, 178) \| 180 \| (7, 222) \| 225 \| HTH cro/C1-type domain-containing protein \| HTH cro/C1-type domain-containing protein; uniclust \| UniRef100\_A0A3D4YWI1 \| 99.8 \| 2.2e-23 \| 4.1e-29 \| 141.5 \| 172 \| (1, 174) \| 180 \| (7, 213) \| 233 \| HTH cro/C1-type domain-containing protein \| HTH cro/C1-type domain-containing protein; uniclust \| UniRef100\_A0A098EGV2 \| 99.8 \| 2.2e-23 \| 4.1e-29 \| 139.4 \| 178 \| (2, 180) \| 180 \| (12, 213) \| 214 \| DNA-binding protein \| DNA-binding protein; uniclust \| UniRef100\_A0A1B8QAD9 \| 99.8 \| 2.3e-23 \| 4.2e-29 \| 143.1 \| 178 \| (1, 179) \| 180 \| (16, 224) \| 230 \| HTH cro/C1-type domain-containing protein \| HTH cro/C1-type domain-containing protein; uniclust \| UniRef100\_A0A1G5MEX5 \| 99.8 \| 2.5e-23 \| 4.6e-29 \| 144.1 \| 177 \| (1, 179) \| 180 \| (39, 274) \| 275 \| Phage repressor protein C, contains Cro/C1-type HTH and peptisase s24 domains \| Phage repressor protein C, contains Cro/C1-type HTH and peptisase s24 domains; uniclust \| UniRef100\_A0A023BWP2 \| 99.8 \| 2.5e-23 \| 4.7e-29 \| 152.5 \| 164 \| (2, 166) \| 180 \| (38, 241) \| 302 \| DNA-binding protein \| DNA-binding protein; uniclust \| UniRef100\_A0A133XV83 \| 99.8 \| 2.6e-23 \| 4.8e-29 \| 143.4 \| 175 \| (1, 177) \| 180 \| (29, 234) \| 250 \| DNA-binding helix-turn-helix protein \| DNA-binding helix-turn-helix protein; uniclust \| UniRef100\_A0A1F1DBG0 \| 99.8 \| 2.7e-23 \| 5e-29 \| 143.2 \| 176 \| (2, 179) \| 180 \| (23, 263) \| 273 \| Repressor \| Repressor; uniclust \| UniRef100\_A0A014B1G1 \| 99.8 \| 2.7e-23 \| 5e-29 \| 147.4 \| 179 \| (1, 179) \| 180 \| (13, 234) \| 243 \| Helix-turn-helix family protein \| Helix-turn-helix family protein; uniclust \| UniRef100\_A0A2S7JR70 \| 99.8 \| 2.9e-23 \| 5.4e-29 \| 146.1 \| 178 \| (2, 180) \| 180 \| (25, 299) \| 299 \| Peptidase S24/S26A/S26B/S26C domain-containing protein \| Peptidase S24/S26A/S26B/S26C domain-containing protein; uniclust \| UniRef100\_A0A2X0V8H6 \| 99.8 \| 3e-23 \| 5.6e-29 \| 134.5 \| 177 \| (1, 180) \| 180 \| (15, 202) \| 203 \| LexA repressor \| LexA repressor; uniclust \| UniRef100\_A0A0E3BVD5 \| 99.8 \| 4.3e-23 \| 7.9e-29 \| 142.2 \| 175 \| (2, 177) \| 180 \| (10, 231) \| 236 \| Repressor \| Repressor; uniclust \| UniRef100\_A0A2N7Q1N3 \| 99.8 \| 4.4e-23 \| 8.1e-29 \| 136.9 \| 175 \| (1, 176) \| 180 \| (16, 210) \| 220 \| Phage repressor protein \| Phage repressor protein; uniclust \| UniRef100\_A0A118HZG0 \| 99.8 \| 4.6e-23 \| 8.5e-29 \| 149.3 \| 159 \| (20, 180) \| 180 \| (172, 377) \| 377 \| HTH cro/C1-type domain-containing protein \| HTH cro/C1-type domain-containing protein; uniclust \| UniRef100\_A0A077LIM9 \| 99.8 \| 4.7e-23 \| 8.8e-29 \| 147.2 \| 173 \| (2, 177) \| 180 \| (59, 271) \| 277 \| Repressor protein cI \| Repressor protein cI; uniclust \| UniRef100\_A0A0D0JL38 \| 99.8 \| 6.4e-23 \| 1.2e-28 \| 142.6 \| 172 \| (2, 175) \| 180 \| (42, 254) \| 264 \| Phage-related repressor protein \| Phage-related repressor protein; uniclust \| UniRef100\_A0A2D3WLJ9 \| 99.8 \| 6.7e-23 \| 1.2e-28 \| 142.4 \| 175 \| (1, 176) \| 180 \| (28, 252) \| 262 \| HTH cro/C1-type domain-containing protein \| HTH cro/C1-type domain-containing protein; uniclust \| UniRef100\_A0A066TCP8 \| 99.8 \| 6.9e-23 \| 1.3e-28 \| 140.2 \| 168 \| (12, 180) \| 180 \| (39, 237) \| 238 \| HTH cro/C1-type domain-containing protein \| HTH cro/C1-type domain-containing protein; uniclust \| UniRef100\_A0A0H4A1L4 \| 99.8 \| 7.1e-23 \| 1.3e-28 \| 137.4 \| 173 \| (1, 178) \| 180 \| (6, 202) \| 211 \| Phage repressor \| Phage repressor; uniclust \| UniRef100\_A0A059V0W4 \| 99.8 \| 8.2e-23 \| 1.5e-28 \| 149.6 \| 175 \| (1, 177) \| 180 \| (60, 329) \| 334 \| Repressor \| Repressor; uniclust \| UniRef100\_A0A0Q5CIB1 \| 99.8 \| 8.5e-23 \| 1.6e-28 \| 144.5 \| 169 \| (5, 176) \| 180 \| (48, 280) \| 289 \| Peptidase S24/S26A/S26B/S26C domain-containing protein \| Peptidase S24/S26A/S26B/S26C domain-containing protein; uniclust \| UniRef100\_A0A162BGE3 \| 99.8 \| 1e-22 \| 1.9e-28 \| 137.7 \| 172 \| (1, 178) \| 180 \| (3, 214) \| 225 \| Repressor \| Repressor; uniclust \| UniRef100\_A0A1G8JBR8 \| 99.8 \| 1.1e-22 \| 2e-28 \| 147.3 \| 175 \| (2, 176) \| 180 \| (101, 335) \| 347 \| Phage repressor protein C, contains Cro/C1-type HTH and peptisase s24 domains \| Phage repressor protein C, contains Cro/C1-type HTH and peptisase s24 domains; uniclust \| UniRef100\_A0A0M2DVU2 \| 99.8 \| 1.1e-22 \| 2e-28 \| 141.7 \| 176 \| (4, 179) \| 180 \| (17, 250) \| 254 \| Peptidase S24/S26A/S26B/S26C domain-containing protein \| Peptidase S24/S26A/S26B/S26C domain-containing protein; uniclust \| UniRef100\_A0A172WXK4 \| 99.8 \| 1.3e-22 \| 2.5e-28 \| 140.6 \| 173 \| (2, 176) \| 180 \| (23, 252) \| 260 \| XRE family transcriptional regulator \| XRE family transcriptional regulator; uniclust \| UniRef100\_A0A1L8R4K9 \| 99.8 \| 1.4e-22 \| 2.6e-28 \| 138.7 \| 174 \| (1, 176) \| 180 \| (10, 240) \| 244 \| Multidrug transporter \| Multidrug transporter; uniclust \| UniRef100\_A0A2M9FSW7 \| 99.8 \| 1.7e-22 \| 3.1e-28 \| 139.2 \| 169 \| (1, 174) \| 180 \| (13, 230) \| 231 \| HTH cro/C1-type domain-containing protein \| HTH cro/C1-type domain-containing protein; uniclust \| UniRef100\_A0A085JF28 \| 99.8 \| 1.8e-22 \| 3.3e-28 \| 143.8 \| 175 \| (1, 178) \| 180 \| (47, 255) \| 266 \| Phage repressor protein C2 \| Phage repressor protein C2; uniclust \| UniRef100\_A0A2A2HM07 \| 99.8 \| 2e-22 \| 3.6e-28 \| 138.0 \| 175 \| (1, 177) \| 180 \| (17, 240) \| 248 \| Phage repressor protein \| Phage repressor protein; uniclust \| UniRef100\_A0A0T9T2W8 \| 99.8 \| 2e-22 \| 3.6e-28 \| 138.2 \| 173 \| (2, 178) \| 180 \| (29, 219) \| 236 \| Uncharacterized HTH-type transcriptional regulator HI\_1476 \| Uncharacterized HTH-type transcriptional regulator HI\_1476; uniclust \| UniRef100\_A0A191HUA0 \| 99.8 \| 2e-22 \| 3.7e-28 \| 136.3 \| 178 \| (1, 180) \| 180 \| (35, 236) \| 237 \| HTH cro/C1-type domain-containing protein \| HTH cro/C1-type domain-containing protein; uniclust \| UniRef100\_A0A0N8P2N3 \| 99.8 \| 2.3e-22 \| 4.2e-28 \| 121.7 \| 92 \| (89, 180) \| 180 \| (4, 97) \| 101 \| Peptidase S24/S26A/S26B/S26C domain-containing protein (Fragment) \| Peptidase S24/S26A/S26B/S26C domain-containing protein (Fragment); uniclust \| UniRef100\_A0A7X5Y7A7 \| 99.8 \| 2.4e-22 \| 4.4e-28 \| 136.3 \| 177 \| (2, 180) \| 180 \| (89, 306) \| 306 \| Phage repressor protein C with HTH and peptisase S24 domain \| Phage repressor protein C with HTH and peptisase S24 domain; uniclust \| UniRef100\_UPI002147CCE2 \| 99.8 \| 2.8e-22 \| 5.1e-28 \| 136.2 \| 175 \| (2, 176) \| 180 \| (70, 295) \| 309 \| S24 family peptidase \| S24 family peptidase; uniclust \| UniRef100\_A0A1E4G2P2 \| 99.8 \| 2.8e-22 \| 5.1e-28 \| 140.2 \| 174 \| (1, 178) \| 180 \| (34, 236) \| 245 \| HTH cro/C1-type domain-containing protein \| HTH cro/C1-type domain-containing protein; uniclust \| UniRef100\_A0A1R1LYN7 \| 99.8 \| 2.9e-22 \| 5.3e-28 \| 133.8 \| 174 \| (1, 176) \| 180 \| (3, 221) \| 227 \| HTH cro/C1-type domain-containing protein \| HTH cro/C1-type domain-containing protein; uniclust \| UniRef100\_A0A1C2DEK8 \| 99.8 \| 3.3e-22 \| 6.1e-28 \| 135.3 \| 98 \| (83, 180) \| 180 \| (86, 185) \| 196 \| Peptidase S24/S26A/S26B/S26C domain-containing protein \| Peptidase S24/S26A/S26B/S26C domain-containing protein; uniclust \| UniRef100\_A0A2A2R811 \| 99.8 \| 3.4e-22 \| 6.3e-28 \| 140.0 \| 176 \| (1, 180) \| 180 \| (26, 241) \| 262 \| Peptidase S24/S26A/S26B/S26C domain-containing protein \| Peptidase S24/S26A/S26B/S26C domain-containing protein; uniclust \| UniRef100\_A0A1D9GLV6 \| 99.8 \| 3.5e-22 \| 6.4e-28 \| 147.1 \| 174 \| (2, 176) \| 180 \| (65, 357) \| 369 \| Peptidase S24/S26A/S26B/S26C domain-containing protein \| Peptidase S24/S26A/S26B/S26C domain-containing protein; uniclust \| UniRef100\_A0A4Q2J692 \| 99.8 \| 3.5e-22 \| 6.5e-28 \| 135.0 \| 174 \| (1, 178) \| 180 \| (4, 245) \| 256 \| LexA family transcriptional regulator \| LexA family transcriptional regulator; uniclust \| UniRef100\_A0A0W8E3K7 \| 99.8 \| 3.6e-22 \| 6.6e-28 \| 147.5 \| 176 \| (1, 179) \| 180 \| (29, 274) \| 333 \| Sos-response repressor and protease lexa \| Sos-response repressor and protease lexa; uniclust \| UniRef100\_A0A1H0VW04 \| 99.8 \| 3.8e-22 \| 6.9e-28 \| 137.9 \| 172 \| (4, 178) \| 180 \| (20, 235) \| 240 \| SOS-response transcriptional repressor LexA (RecA-mediated autopeptidase) \| SOS-response transcriptional repressor LexA (RecA-mediated autopeptidase); uniclust \| UniRef100\_A0A0K4HVA9 \| 99.8 \| 4e-22 \| 7.4e-28 \| 140.5 \| 174 \| (2, 176) \| 180 \| (48, 244) \| 264 \| Helix-turn-helix domain-containing protein \| Helix-turn-helix domain-containing protein; uniclust \| UniRef100\_A0A1H6C042 \| 99.8 \| 4.2e-22 \| 7.8e-28 \| 143.4 \| 176 \| (2, 178) \| 180 \| (42, 282) \| 309 \| Helix-turn-helix \| Helix-turn-helix; uniclust \| UniRef100\_A0A1D2UGZ7 \| 99.8 \| 5e-22 \| 9.2e-28 \| 143.3 \| 174 \| (1, 175) \| 180 \| (101, 327) \| 333 \| HTH cro/C1-type domain-containing protein \| HTH cro/C1-type domain-containing protein; uniclust \| UniRef100\_A0A023WV23 \| 99.8 \| 5.8e-22 \| 1.1e-27 \| 143.6 \| 173 \| (1, 175) \| 180 \| (43, 247) \| 260 \| Cro/Cl family transcriptional regulator \| Cro/Cl family transcriptional regulator; uniclust \| UniRef100\_A0A013SVQ8 \| 99.8 \| 6e-22 \| 1.1e-27 \| 148.7 \| 174 \| (1, 178) \| 180 \| (88, 300) \| 332 \| Helix-turn-helix family protein \| Helix-turn-helix family protein; uniclust \| UniRef100\_A0A2W4N736 \| 99.8 \| 6.1e-22 \| 1.1e-27 \| 138.0 \| 174 \| (1, 176) \| 180 \| (68, 277) \| 340 \| HTH cro/C1-type domain-containing protein \| HTH cro/C1-type domain-containing protein; uniclust \| UniRef100\_A0A196NVN3 \| 99.8 \| 7.3e-22 \| 1.4e-27 \| 132.9 \| 176 \| (4, 180) \| 180 \| (20, 235) \| 236 \| Peptidase S24/S26A/S26B/S26C domain-containing protein \| Peptidase S24/S26A/S26B/S26C domain-containing protein; uniclust \| UniRef100\_A0A1R0YXM3 \| 99.8 \| 8.1e-22 \| 1.5e-27 \| 129.2 \| 174 \| (1, 179) \| 180 \| (4, 192) \| 198 \| HTH cro/C1-type domain-containing protein \| HTH cro/C1-type domain-containing protein; uniclust \| UniRef100\_A0A292QUL5 \| 99.8 \| 8.9e-22 \| 1.6e-27 \| 133.4 \| 163 \| (12, 179) \| 180 \| (17, 201) \| 210 \| Peptidase S24/S26A/S26B/S26C domain-containing protein \| Peptidase S24/S26A/S26B/S26C domain-containing protein; uniclust \| UniRef100\_A0A0E2BPP9 \| 99.8 \| 8.9e-22 \| 1.6e-27 \| 138.2 \| 174 \| (1, 176) \| 180 \| (38, 252) \| 259 \| SOS-response repressor/protease \| SOS-response repressor/protease; uniclust \| UniRef100\_A0A077QL17 \| 99.8 \| 9.8e-22 \| 1.8e-27 \| 140.3 \| 173 \| (1, 177) \| 180 \| (14, 216) \| 281 \| Helix-turn-helix/peptidase S24-like domain protein (Modular protein) \| Helix-turn-helix/peptidase S24-like domain protein (Modular protein); uniclust \| UniRef100\_A0A268THE3 \| 99.8 \| 1.1e-21 \| 2.1e-27 \| 135.3 \| 88 \| (87, 177) \| 180 \| (175, 262) \| 267 \| HTH cro/C1-type domain-containing protein \| HTH cro/C1-type domain-containing protein; uniclust \| UniRef100\_A0A0Q5GW81 \| 99.7 \| 1.4e-21 \| 2.5e-27 \| 127.8 \| 97 \| (84, 180) \| 180 \| (62, 160) \| 161 \| Peptidase S24/S26A/S26B/S26C domain-containing protein \| Peptidase S24/S26A/S26B/S26C domain-containing protein; uniclust \| UniRef100\_A0A0A8GVX7 \| 99.7 \| 1.4e-21 \| 2.6e-27 \| 134.9 \| 159 \| (4, 177) \| 180 \| (30, 225) \| 232 \| Peptidase S24 LexA-like protein \| Peptidase S24 LexA-like protein; uniclust \| UniRef100\_A0A084A6X1 \| 99.7 \| 1.4e-21 \| 2.6e-27 \| 137.8 \| 176 \| (1, 177) \| 180 \| (13, 254) \| 297 \| Putative transcriptional regulator \| Putative transcriptional regulator; uniclust \| UniRef100\_A0A0U3FB36 \| 99.7 \| 1.5e-21 \| 2.7e-27 \| 132.0 \| 178 \| (1, 179) \| 180 \| (12, 219) \| 227 \| HTH/peptidase S24 family domain-containing protein \| HTH/peptidase S24 family domain-containing protein; uniclust \| UniRef100\_A0A1G3FIK6 \| 99.7 \| 1.5e-21 \| 2.7e-27 \| 131.1 \| 178 \| (1, 179) \| 180 \| (16, 242) \| 243 \| Transcriptional regulator \| Transcriptional regulator; uniclust \| UniRef100\_A0A1F9A3V9 \| 99.7 \| 1.5e-21 \| 2.8e-27 \| 126.2 \| 93 \| (87, 180) \| 180 \| (62, 156) \| 158 \| Peptidase S24/S26A/S26B/S26C domain-containing protein \| Peptidase S24/S26A/S26B/S26C domain-containing protein; uniclust \| UniRef100\_A0A2E9H220 \| 99.7 \| 1.6e-21 \| 2.9e-27 \| 136.4 \| 174 \| (1, 175) \| 180 \| (39, 260) \| 275 \| HTH cro/C1-type domain-containing protein \| HTH cro/C1-type domain-containing protein; uniclust \| UniRef100\_A0A2I0GID1 \| 99.7 \| 1.8e-21 \| 3.3e-27 \| 139.3 \| 173 \| (1, 174) \| 180 \| (11, 288) \| 298 \| HTH cro/C1-type domain-containing protein \| HTH cro/C1-type domain-containing protein; uniclust \| UniRef100\_A0A4U0PYZ7 \| 99.7 \| 1.9e-21 \| 3.5e-27 \| 136.3 \| 176 \| (2, 178) \| 180 \| (96, 319) \| 322 \| Helix-turn-helix transcriptional regulator \| Helix-turn-helix transcriptional regulator; uniclust \| UniRef100\_A0A1Q6I010 \| 99.7 \| 2e-21 \| 3.6e-27 \| 137.6 \| 165 \| (2, 168) \| 180 \| (54, 258) \| 274 \| HTH cro/C1-type domain-containing protein \| HTH cro/C1-type domain-containing protein; uniclust \| UniRef100\_B6AS86 \| 99.7 \| 2e-21 \| 3.6e-27 \| 128.7 \| 176 \| (2, 179) \| 180 \| (7, 228) \| 229 \| Putative phage repressor \| Putative phage repressor; uniclust \| UniRef100\_A0A1Y2K654 \| 99.7 \| 2e-21 \| 3.6e-27 \| 122.6 \| 96 \| (84, 180) \| 180 \| (30, 127) \| 129 \| Putative Cro/CI family transcriptional regulator \| Putative Cro/CI family transcriptional regulator; uniclust \| UniRef100\_A0A016QME0 \| 99.7 \| 2e-21 \| 3.7e-27 \| 134.9 \| 171 \| (2, 176) \| 180 \| (30, 243) \| 251 \| LexA repressor \| LexA repressor; uniclust \| UniRef100\_A0A640WCV9 \| 99.7 \| 2.1e-21 \| 3.9e-27 \| 129.2 \| 173 \| (2, 174) \| 180 \| (5, 234) \| 242 \| Helix-turn-helix domain-containing protein \| Helix-turn-helix domain-containing protein; uniclust \| UniRef100\_UPI0009857A7F \| 99.7 \| 2.4e-21 \| 4.4e-27 \| 129.7 \| 168 \| (1, 176) \| 180 \| (10, 219) \| 228 \| XRE family transcriptional regulator \| XRE family transcriptional regulator; uniclust \| UniRef100\_A0A071LTP5 \| 99.7 \| 2.4e-21 \| 4.5e-27 \| 140.9 \| 100 \| (80, 180) \| 180 \| (171, 272) \| 279 \| HTH Mu-type domain-containing protein \| HTH Mu-type domain-containing protein; uniclust \| UniRef100\_A0A1E7YT01 \| 99.7 \| 2.8e-21 \| 5.1e-27 \| 137.7 \| 171 \| (5, 176) \| 180 \| (70, 292) \| 305 \| Peptidase S24/S26A/S26B/S26C domain-containing protein \| Peptidase S24/S26A/S26B/S26C domain-containing protein; uniclust \| UniRef100\_A0A356M362 \| 99.7 \| 2.8e-21 \| 5.1e-27 \| 131.7 \| 177 \| (1, 179) \| 180 \| (12, 222) \| 229 \| HTH cro/C1-type domain-containing protein \| HTH cro/C1-type domain-containing protein; uniclust \| UniRef100\_A0A081RAB9 \| 99.7 \| 2.9e-21 \| 5.3e-27 \| 120.2 \| 99 \| (80, 180) \| 180 \| (23, 122) \| 123 \| Putative phage repressor \| Putative phage repressor; uniclust \| UniRef100\_A0A519BE94 \| 99.7 \| 2.9e-21 \| 5.4e-27 \| 130.3 \| 178 \| (1, 180) \| 180 \| (14, 234) \| 234 \| Helix-turn-helix domain-containing protein \| Helix-turn-helix domain-containing protein; uniclust \| UniRef100\_A0A946HKU8 \| 99.7 \| 3.7e-21 \| 6.8e-27 \| 122.5 \| 176 \| (4, 180) \| 180 \| (2, 189) \| 190 \| Helix-turn-helix domain-containing protein \| Helix-turn-helix domain-containing protein; uniclust \| UniRef100\_A0A011NA96 \| 99.7 \| 3.8e-21 \| 7e-27 \| 131.5 \| 156 \| (19, 176) \| 180 \| (1, 200) \| 210 \| Transcriptional regulator \| Transcriptional regulator; uniclust \| UniRef100\_UPI001432A832 \| 99.7 \| 4e-21 \| 7.4e-27 \| 124.7 \| 180 \| (1, 180) \| 180 \| (3, 207) \| 207 \| helix-turn-helix domain-containing protein \| helix-turn-helix domain-containing protein; uniclust \| UniRef100\_A0A971PC19 \| 99.7 \| 4.2e-21 \| 7.7e-27 \| 131.8 \| 173 \| (1, 176) \| 180 \| (3, 253) \| 255 \| LexA family transcriptional regulator \| LexA family transcriptional regulator; uniclust \| UniRef100\_UPI0004923115 \| 99.7 \| 4.8e-21 \| 8.9e-27 \| 130.7 \| 177 \| (1, 179) \| 180 \| (10, 249) \| 250 \| helix-turn-helix domain-containing protein \| helix-turn-helix domain-containing protein; uniclust \| UniRef100\_A0A0K2XET6 \| 99.7 \| 5e-21 \| 9.3e-27 \| 131.2 \| 175 \| (3, 178) \| 180 \| (38, 242) \| 246 \| Transcriptional regulator \| Transcriptional regulator; uniclust \| UniRef100\_A0A9D8S324 \| 99.7 \| 5.7e-21 \| 1.1e-26 \| 123.9 \| 174 \| (2, 176) \| 180 \| (2, 185) \| 192 \| LexA family transcriptional regulator \| LexA family transcriptional regulator; uniclust \| UniRef100\_A0A4Q2X6Y6 \| 99.7 \| 5.7e-21 \| 1.1e-26 \| 130.6 \| 175 \| (1, 178) \| 180 \| (12, 220) \| 221 \| LexA family transcriptional regulator \| LexA family transcriptional regulator; uniclust \| UniRef100\_A0A0H0MNU5 \| 99.7 \| 5.7e-21 \| 1.1e-26 \| 131.8 \| 171 \| (2, 176) \| 180 \| (27, 232) \| 246 \| Helix-turn-helix domain-containing protein \| Helix-turn-helix domain-containing protein; uniclust \| UniRef100\_A0A158HTA6 \| 99.7 \| 6.2e-21 \| 1.1e-26 \| 130.0 \| 178 \| (1, 179) \| 180 \| (8, 245) \| 249 \| LexA repressor \| LexA repressor; uniclust \| UniRef100\_A0A239PSQ2 \| 99.7 \| 6.2e-21 \| 1.1e-26 \| 134.6 \| 176 \| (1, 179) \| 180 \| (59, 262) \| 265 \| Transcriptional regulator, contains XRE-family HTH domain \| Transcriptional regulator, contains XRE-family HTH domain; uniclust \| UniRef100\_A0A136P7M0 \| 99.7 \| 7.2e-21 \| 1.3e-26 \| 132.0 \| 179 \| (1, 180) \| 180 \| (15, 221) \| 222 \| LexA repressor \| LexA repressor; uniclust \| UniRef100\_A0A0J6YUX8 \| 99.7 \| 7.3e-21 \| 1.3e-26 \| 136.2 \| 177 \| (1, 179) \| 180 \| (95, 313) \| 314 \| HTH cro/C1-type domain-containing protein \| HTH cro/C1-type domain-containing protein; uniclust \| UniRef100\_A0A068QRI4 \| 99.7 \| 9.3e-21 \| 1.7e-26 \| 137.6 \| 173 \| (1, 177) \| 180 \| (49, 261) \| 270 \| Repressor protein C2 \| Repressor protein C2; uniclust \| UniRef100\_A0A258GFY3 \| 99.7 \| 9.9e-21 \| 1.8e-26 \| 128.8 \| 176 \| (2, 177) \| 180 \| (6, 243) \| 246 \| HTH cro/C1-type domain-containing protein \| HTH cro/C1-type domain-containing protein; uniclust \| UniRef100\_A0A367LYV8 \| 99.7 \| 1e-20 \| 1.8e-26 \| 114.4 \| 96 \| (84, 180) \| 180 \| (15, 113) \| 114 \| S24 family peptidase (Fragment) \| S24 family peptidase (Fragment); uniclust \| UniRef100\_A0A174MQZ3 \| 99.7 \| 1.1e-20 \| 2.1e-26 \| 129.4 \| 175 \| (1, 177) \| 180 \| (7, 215) \| 226 \| Antitoxin PezA \| Antitoxin PezA; uniclust \| UniRef100\_A0A2P1S2C8 \| 99.7 \| 1.2e-20 \| 2.2e-26 \| 130.6 \| 177 \| (1, 179) \| 180 \| (7, 220) \| 259 \| HTH cro/C1-type domain-containing protein \| HTH cro/C1-type domain-containing protein; uniclust \| UniRef100\_A0A0F2RP23 \| 99.7 \| 1.2e-20 \| 2.2e-26 \| 124.3 \| 102 \| (78, 180) \| 180 \| (58, 160) \| 161 \| Peptidase S24/S26A/S26B/S26C domain-containing protein \| Peptidase S24/S26A/S26B/S26C domain-containing protein; uniclust \| UniRef100\_A0A258AUA7 \| 99.7 \| 1.2e-20 \| 2.2e-26 \| 137.8 \| 175 \| (3, 179) \| 180 \| (90, 307) \| 313 \| HTH cro/C1-type domain-containing protein \| HTH cro/C1-type domain-containing protein; uniclust \| UniRef100\_UPI0013D78761 \| 99.7 \| 1.2e-20 \| 2.3e-26 \| 134.7 \| 178 \| (1, 180) \| 180 \| (4, 323) \| 324 \| S24 family peptidase \| S24 family peptidase; uniclust \| UniRef100\_A0A3G9GCF8 \| 99.7 \| 1.3e-20 \| 2.4e-26 \| 117.3 \| 100 \| (80, 180) \| 180 \| (36, 138) \| 138 \| Probable transcription regulator \| Probable transcription regulator; uniclust \| UniRef100\_A0A0N0MC17 \| 99.7 \| 1.5e-20 \| 2.8e-26 \| 126.0 \| 98 \| (82, 180) \| 180 \| (80, 180) \| 180 \| Peptidase S24/S26A/S26B/S26C domain-containing protein \| Peptidase S24/S26A/S26B/S26C domain-containing protein; uniclust \| UniRef100\_A0A193FV67 \| 99.7 \| 1.5e-20 \| 2.8e-26 \| 140.5 \| 174 \| (3, 176) \| 180 \| (139, 384) \| 392 \| HTH cro/C1-type domain-containing protein \| HTH cro/C1-type domain-containing protein; uniclust \| UniRef100\_A0A2P5MV31 \| 99.7 \| 1.7e-20 \| 3e-26 \| 131.1 \| 177 \| (1, 179) \| 180 \| (17, 322) \| 323 \| HTH cro/C1-type domain-containing protein \| HTH cro/C1-type domain-containing protein; uniclust \| UniRef100\_A0A1B2I1E8 \| 99.7 \| 1.7e-20 \| 3e-26 \| 131.0 \| 176 \| (1, 179) \| 180 \| (32, 258) \| 265 \| HTH cro/C1-type domain-containing protein \| HTH cro/C1-type domain-containing protein; uniclust \| UniRef100\_A0A0B1U779 \| 99.7 \| 1.8e-20 \| 3.3e-26 \| 134.8 \| 177 \| (3, 180) \| 180 \| (48, 269) \| 273 \| Putative DNA-binding protein \| Putative DNA-binding protein; uniclust \| UniRef100\_A0A8S5PML4 \| 99.7 \| 1.9e-20 \| 3.5e-26 \| 134.1 \| 174 \| (2, 176) \| 180 \| (129, 345) \| 355 \| Helix-turn-helix domain protein \| Helix-turn-helix domain protein; uniclust \| UniRef100\_A0A010SKQ2 \| 99.7 \| 2.1e-20 \| 3.9e-26 \| 137.2 \| 174 \| (1, 178) \| 180 \| (151, 361) \| 368 \| Sulfurtransferase \| Sulfurtransferase; uniclust \| UniRef100\_A0A971C5L3 \| 99.7 \| 2.1e-20 \| 3.9e-26 \| 128.7 \| 174 \| (1, 176) \| 180 \| (10, 286) \| 291 \| Helix-turn-helix domain-containing protein \| Helix-turn-helix domain-containing protein; uniclust \| UniRef100\_A0A6B1CMI7 \| 99.7 \| 2.1e-20 \| 3.9e-26 \| 131.9 \| 173 \| (4, 180) \| 180 \| (27, 261) \| 278 \| Helix-turn-helix transcriptional regulator \| Helix-turn-helix transcriptional regulator; uniclust \| UniRef100\_A0A069PBS4 \| 99.7 \| 2.2e-20 \| 4.1e-26 \| 137.5 \| 169 \| (2, 174) \| 180 \| (62, 279) \| 290 \| Phage repressor protein \| Phage repressor protein; uniclust \| UniRef100\_A0A087NEV6 \| 99.7 \| 2.2e-20 \| 4.1e-26 \| 114.0 \| 93 \| (87, 180) \| 180 \| (13, 108) \| 109 \| Peptidase S24/S26A/S26B/S26C domain-containing protein \| Peptidase S24/S26A/S26B/S26C domain-containing protein; uniclust \| UniRef100\_A0A292SMX3 \| 99.7 \| 2.3e-20 \| 4.3e-26 \| 125.5 \| 166 \| (12, 179) \| 180 \| (27, 212) \| 222 \| Peptidase S24/S26A/S26B/S26C domain-containing protein \| Peptidase S24/S26A/S26B/S26C domain-containing protein; uniclust \| UniRef100\_A0A1E7ZE28 \| 99.7 \| 2.5e-20 \| 4.6e-26 \| 129.4 \| 175 \| (1, 177) \| 180 \| (13, 202) \| 206 \| Peptidase S24/S26A/S26B/S26C domain-containing protein \| Peptidase S24/S26A/S26B/S26C domain-containing protein; uniclust \| UniRef100\_A0A1C5MIN5 \| 99.7 \| 2.9e-20 \| 5.4e-26 \| 123.9 \| 167 \| (1, 179) \| 180 \| (7, 178) \| 190 \| Bifunctional HTH-domain containing protein/aminotransferase \| Bifunctional HTH-domain containing protein/aminotransferase; uniclust \| UniRef100\_A0A9E5G3I3 \| 99.7 \| 2.9e-20 \| 5.4e-26 \| 126.7 \| 179 \| (1, 180) \| 180 \| (17, 246) \| 251 \| LexA family transcriptional regulator \| LexA family transcriptional regulator; uniclust \| UniRef100\_A4U201 \| 99.7 \| 3.1e-20 \| 5.7e-26 \| 121.2 \| 178 \| (2, 180) \| 180 \| (6, 222) \| 222 \| Transcriptional regulator \| Transcriptional regulator; uniclust \| UniRef100\_A0A3S0D8L6 \| 99.7 \| 3.7e-20 \| 6.8e-26 \| 125.9 \| 166 \| (12, 177) \| 180 \| (25, 219) \| 224 \| LexA family transcriptional regulator \| LexA family transcriptional regulator; uniclust \| UniRef100\_A0A0E2NFI1 \| 99.7 \| 3.8e-20 \| 7.1e-26 \| 135.3 \| 173 \| (2, 176) \| 180 \| (46, 267) \| 274 \| Repressor \| Repressor; uniclust \| UniRef100\_A0A2D5TJQ4 \| 99.7 \| 4e-20 \| 7.3e-26 \| 124.7 \| 172 \| (1, 176) \| 180 \| (12, 217) \| 223 \| HTH cro/C1-type domain-containing protein \| HTH cro/C1-type domain-containing protein; uniclust \| UniRef100\_A0A1C3H508 \| 99.7 \| 4.4e-20 \| 8.1e-26 \| 126.6 \| 91 \| (86, 178) \| 180 \| (191, 281) \| 284 \| Phage repressor \| Phage repressor; uniclust \| UniRef100\_A0A1F9LJ24 \| 99.7 \| 4.4e-20 \| 8.2e-26 \| 130.8 \| 177 \| (1, 177) \| 180 \| (36, 274) \| 283 \| HTH cro/C1-type domain-containing protein \| HTH cro/C1-type domain-containing protein; uniclust \| UniRef100\_A0A1H5VJF6 \| 99.7 \| 4.6e-20 \| 8.5e-26 \| 127.8 \| 174 \| (2, 179) \| 180 \| (5, 274) \| 290 \| Peptidase S24-like \| Peptidase S24-like; uniclust \| UniRef100\_A0A1G3PJ90 \| 99.7 \| 5.2e-20 \| 9.6e-26 \| 122.0 \| 175 \| (2, 178) \| 180 \| (4, 216) \| 219 \| HTH cro/C1-type domain-containing protein \| HTH cro/C1-type domain-containing protein; uniclust \| UniRef100\_A0A7R6VQ36 \| 99.7 \| 5.3e-20 \| 9.8e-26 \| 123.8 \| 177 \| (1, 177) \| 180 \| (4, 229) \| 239 \| Phage repressor \| Phage repressor; uniclust \| UniRef100\_A0A0T2MLF1 \| 99.7 \| 5.7e-20 \| 1e-25 \| 134.4 \| 178 \| (1, 178) \| 180 \| (68, 280) \| 339 \| Phage repressor \| Phage repressor; uniclust \| UniRef100\_A0A3P1Y7B9 \| 99.7 \| 6.6e-20 \| 1.2e-25 \| 122.6 \| 174 \| (1, 175) \| 180 \| (13, 226) \| 237 \| LexA family transcriptional regulator \| LexA family transcriptional regulator; uniclust \| UniRef100\_A0A1I2BPM5 \| 99.7 \| 6.8e-20 \| 1.3e-25 \| 131.6 \| 175 \| (1, 176) \| 180 \| (20, 249) \| 299 \| Phage repressor protein C, contains Cro/C1-type HTH and peptisase s24 domains \| Phage repressor protein C, contains Cro/C1-type HTH and peptisase s24 domains; uniclust \| UniRef100\_A0A010RVE5 \| 99.7 \| 7.5e-20 \| 1.4e-25 \| 134.3 \| 174 \| (2, 178) \| 180 \| (85, 294) \| 339 \| Cro/Cl family transcriptional regulator \| Cro/Cl family transcriptional regulator; uniclust \| UniRef100\_A0A7C6H9D8 \| 99.7 \| 7.7e-20 \| 1.4e-25 \| 122.7 \| 174 \| (4, 180) \| 180 \| (3, 214) \| 215 \| Helix-turn-helix transcriptional regulator \| Helix-turn-helix transcriptional regulator; uniclust \| UniRef100\_A0A009PCU9 \| 99.7 \| 8.2e-20 \| 1.5e-25 \| 130.7 \| 177 \| (2, 179) \| 180 \| (50, 261) \| 281 \| Peptidase S24-like family protein \| Peptidase S24-like family protein; uniclust \| UniRef100\_A0A2Z4PPI4 \| 99.7 \| 8.4e-20 \| 1.6e-25 \| 109.6 \| 90 \| (90, 180) \| 180 \| (8, 101) \| 102 \| Peptidase S24/S26A/S26B/S26C domain-containing protein \| Peptidase S24/S26A/S26B/S26C domain-containing protein; uniclust \| UniRef100\_A0A2P8KL91 \| 99.7 \| 8.5e-20 \| 1.6e-25 \| 126.9 \| 170 \| (3, 177) \| 180 \| (35, 249) \| 254 \| Uncharacterized protein \| Uncharacterized protein; uniclust \| UniRef100\_A0A1I1N4A7 \| 99.7 \| 1.1e-19 \| 1.9e-25 \| 122.0 \| 178 \| (1, 179) \| 180 \| (17, 232) \| 235 \| Phage repressor protein C, contains Cro/C1-type HTH and peptisase s24 domains \| Phage repressor protein C, contains Cro/C1-type HTH and peptisase s24 domains; uniclust \| UniRef100\_A0A1B2Z1M4 \| 99.7 \| 1.1e-19 \| 2e-25 \| 128.9 \| 176 \| (1, 176) \| 180 \| (43, 248) \| 291 \| HTH cro/C1-type domain-containing protein \| HTH cro/C1-type domain-containing protein; uniclust \| UniRef100\_A0A1Q3MNN2 \| 99.7 \| 1.1e-19 \| 2.1e-25 \| 124.3 \| 170 \| (2, 178) \| 180 \| (14, 202) \| 222 \| Peptidase S24/S26A/S26B/S26C domain-containing protein \| Peptidase S24/S26A/S26B/S26C domain-containing protein; uniclust \| UniRef100\_A0A1U9LFW6 \| 99.7 \| 1.2e-19 \| 2.2e-25 \| 117.3 \| 100 \| (80, 180) \| 180 \| (61, 163) \| 163 \| Peptidase S24/S26A/S26B/S26C domain-containing protein \| Peptidase S24/S26A/S26B/S26C domain-containing protein; uniclust \| UniRef100\_A0A847BTK1 \| 99.7 \| 1.3e-19 \| 2.3e-25 \| 106.4 \| 83 \| (95, 178) \| 180 \| (2, 84) \| 88 \| S24 family peptidase \| S24 family peptidase; uniclust \| UniRef100\_A0A081J3X7 \| 99.7 \| 1.3e-19 \| 2.3e-25 \| 128.4 \| 168 \| (11, 179) \| 180 \| (66, 282) \| 285 \| Transcriptional regulator \| Transcriptional regulator; uniclust \| UniRef100\_A0A942GU57 \| 99.7 \| 1.3e-19 \| 2.3e-25 \| 124.8 \| 174 \| (2, 178) \| 180 \| (23, 233) \| 262 \| Helix-turn-helix domain-containing protein \| Helix-turn-helix domain-containing protein; uniclust \| UniRef100\_A0A0D6I1W4 \| 99.7 \| 1.3e-19 \| 2.4e-25 \| 110.6 \| 93 \| (87, 180) \| 180 \| (13, 108) \| 110 \| Repressor LexA \| Repressor LexA; uniclust \| UniRef100\_A0A173T592 \| 99.7 \| 1.3e-19 \| 2.4e-25 \| 128.4 \| 174 \| (1, 175) \| 180 \| (36, 281) \| 284 \| Peptidase S24-like \| Peptidase S24-like; uniclust \| UniRef100\_A0A1V5IGK5 \| 99.7 \| 1.5e-19 \| 2.8e-25 \| 126.5 \| 172 \| (1, 174) \| 180 \| (18, 248) \| 259 \| LexA repressor \| LexA repressor; uniclust \| UniRef100\_A0A0W0R9T4 \| 99.7 \| 1.6e-19 \| 3e-25 \| 124.5 \| 173 \| (3, 177) \| 180 \| (10, 209) \| 215 \| HTH-type transcriptional regulator \| HTH-type transcriptional regulator; uniclust \| UniRef100\_A0A0A1AFM4 \| 99.7 \| 1.6e-19 \| 3e-25 \| 120.8 \| 177 \| (1, 178) \| 180 \| (12, 216) \| 218 \| Helix-turn-helix domain-containing protein \| Helix-turn-helix domain-containing protein; uniclust \| UniRef100\_A0A1F7T9S1 \| 99.7 \| 1.6e-19 \| 3e-25 \| 125.1 \| 177 \| (2, 179) \| 180 \| (26, 245) \| 282 \| HTH cro/C1-type domain-containing protein \| HTH cro/C1-type domain-containing protein; uniclust \| UniRef100\_A0A0U2IWK9 \| 99.7 \| 1.7e-19 \| 3e-25 \| 115.2 \| 98 \| (82, 180) \| 180 \| (34, 134) \| 134 \| Putative peptidase S24 \| Putative peptidase S24; uniclust \| UniRef100\_A0A1C6E8P6 \| 99.7 \| 1.8e-19 \| 3.4e-25 \| 128.0 \| 169 \| (2, 177) \| 180 \| (8, 325) \| 329 \| Peptidase S24-like \| Peptidase S24-like; uniclust \| UniRef100\_A0A0R6PHW4 \| 99.7 \| 1.9e-19 \| 3.6e-25 \| 128.2 \| 175 \| (3, 179) \| 180 \| (40, 258) \| 278 \| HTH cro/C1-type domain-containing protein \| HTH cro/C1-type domain-containing protein; uniclust \| UniRef100\_A0A220S122 \| 99.7 \| 2e-19 \| 3.7e-25 \| 122.1 \| 173 \| (1, 174) \| 180 \| (19, 233) \| 243 \| HTH cro/C1-type domain-containing protein \| HTH cro/C1-type domain-containing protein; uniclust \| UniRef100\_A0A1I5LCX6 \| 99.7 \| 2.1e-19 \| 3.8e-25 \| 131.5 \| 176 \| (1, 179) \| 180 \| (126, 369) \| 374 \| Peptidase S24-like \| Peptidase S24-like; uniclust \| UniRef100\_A0A432UMV4 \| 99.7 \| 2.1e-19 \| 3.9e-25 \| 125.2 \| 164 \| (13, 177) \| 180 \| (50, 275) \| 281 \| Peptidase S24/S26A/S26B/S26C domain-containing protein \| Peptidase S24/S26A/S26B/S26C domain-containing protein; uniclust \| UniRef100\_A0A848G1C2 \| 99.7 \| 2.2e-19 \| 4e-25 \| 119.7 \| 176 \| (1, 178) \| 180 \| (2, 222) \| 224 \| LexA family transcriptional regulator \| LexA family transcriptional regulator; uniclust \| UniRef100\_A0A963CV82 \| 99.7 \| 2.2e-19 \| 4e-25 \| 116.1 \| 176 \| (1, 178) \| 180 \| (4, 203) \| 206 \| LexA family transcriptional regulator \| LexA family transcriptional regulator; uniclust \| UniRef100\_A0A8T3RZ60 \| 99.7 \| 2.2e-19 \| 4.1e-25 \| 123.2 \| 164 \| (15, 179) \| 180 \| (27, 280) \| 282 \| LexA family transcriptional regulator \| LexA family transcriptional regulator; uniclust \| UniRef100\_A0A2E3MGU9 \| 99.6 \| 2.7e-19 \| 5e-25 \| 119.3 \| 176 \| (2, 178) \| 180 \| (9, 222) \| 226 \| HTH cro/C1-type domain-containing protein \| HTH cro/C1-type domain-containing protein; uniclust \| UniRef100\_A0A0B0SD21 \| 99.6 \| 3e-19 \| 5.5e-25 \| 121.9 \| 173 \| (3, 179) \| 180 \| (13, 220) \| 231 \| Peptidase S24 \| Peptidase S24; uniclust \| UniRef100\_A0A0A1H208 \| 99.6 \| 3e-19 \| 5.6e-25 \| 122.6 \| 174 \| (1, 176) \| 180 \| (14, 228) \| 243 \| Putative phage repressor \| Putative phage repressor; uniclust \| UniRef100\_A0A085FSU8 \| 99.6 \| 3.1e-19 \| 5.7e-25 \| 129.2 \| 171 \| (1, 174) \| 180 \| (45, 262) \| 271 \| Putative phage repressor \| Putative phage repressor; uniclust \| UniRef100\_A0A3D5VC65 \| 99.6 \| 3.2e-19 \| 5.9e-25 \| 122.0 \| 178 \| (3, 180) \| 180 \| (25, 262) \| 262 \| HTH cro/C1-type domain-containing protein \| HTH cro/C1-type domain-containing protein; uniclust \| UniRef100\_A0A011UQX8 \| 99.6 \| 3.4e-19 \| 6.3e-25 \| 132.3 \| 177 \| (2, 179) \| 180 \| (96, 315) \| 329 \| Peptidase S24 \| Peptidase S24; uniclust \| UniRef100\_A0A1M7HEK4 \| 99.6 \| 3.6e-19 \| 6.6e-25 \| 124.6 \| 175 \| (2, 179) \| 180 \| (13, 259) \| 271 \| Peptidase S24-like \| Peptidase S24-like; uniclust \| UniRef100\_A0A0K9N6J2 \| 99.6 \| 3.9e-19 \| 7.3e-25 \| 127.0 \| 169 \| (1, 177) \| 180 \| (12, 255) \| 319 \| Putative repressor \| Putative repressor; uniclust \| UniRef100\_A0A0Y7I845 \| 99.6 \| 4.3e-19 \| 7.9e-25 \| 121.8 \| 165 \| (9, 174) \| 180 \| (28, 243) \| 253 \| Peptidase S24-like protein \| Peptidase S24-like protein; uniclust \| UniRef100\_A0A095FKS9 \| 99.6 \| 4.3e-19 \| 7.9e-25 \| 120.5 \| 176 \| (1, 179) \| 180 \| (5, 229) \| 232 \| Transcriptional regulator \| Transcriptional regulator; uniclust \| UniRef100\_G0EP36 \| 99.6 \| 5.2e-19 \| 9.7e-25 \| 121.3 \| 174 \| (1, 177) \| 180 \| (19, 254) \| 262 \| Putative phage repressor \| Putative phage repressor; uniclust \| UniRef100\_A0A9D1SLB7 \| 99.6 \| 5.4e-19 \| 1e-24 \| 119.5 \| 176 \| (1, 180) \| 180 \| (8, 209) \| 211 \| LexA family transcriptional regulator \| LexA family transcriptional regulator; uniclust \| UniRef100\_A0A0S6X5H7 \| 99.6 \| 6.2e-19 \| 1.1e-24 \| 106.3 \| 95 \| (83, 179) \| 180 \| (20, 114) \| 115 \| Peptidase S24/S26A/S26B/S26C domain-containing protein \| Peptidase S24/S26A/S26B/S26C domain-containing protein; uniclust \| UniRef100\_A0A6S7CQU9 \| 99.6 \| 7.2e-19 \| 1.3e-24 \| 123.2 \| 164 \| (16, 180) \| 180 \| (114, 340) \| 340 \| Peptidase S24/S26A/S26B/S26C domain-containing protein \| Peptidase S24/S26A/S26B/S26C domain-containing protein; uniclust \| UniRef100\_A0A0C6FN11 \| 99.6 \| 7.1e-19 \| 1.3e-24 \| 128.1 \| 171 \| (1, 174) \| 180 \| (41, 253) \| 261 \| Putative phage repressor \| Putative phage repressor; uniclust \| UniRef100\_A0A1F3L8R9 \| 99.6 \| 7.6e-19 \| 1.4e-24 \| 121.2 \| 158 \| (1, 159) \| 180 \| (10, 210) \| 238 \| HTH cro/C1-type domain-containing protein \| HTH cro/C1-type domain-containing protein; uniclust \| UniRef100\_A0A1F4NGE6 \| 99.6 \| 7.9e-19 \| 1.5e-24 \| 122.9 \| 172 \| (3, 174) \| 180 \| (35, 256) \| 264 \| HTH cro/C1-type domain-containing protein \| HTH cro/C1-type domain-containing protein; uniclust \| UniRef100\_A0A1U9JWC3 \| 99.6 \| 8.1e-19 \| 1.5e-24 \| 123.5 \| 170 \| (1, 174) \| 180 \| (49, 265) \| 271 \| Putative DNA-binding protein \| Putative DNA-binding protein; uniclust \| UniRef100\_A0A1F5AT30 \| 99.6 \| 9.3e-19 \| 1.7e-24 \| 120.3 \| 178 \| (2, 179) \| 180 \| (7, 240) \| 251 \| HTH cro/C1-type domain-containing protein \| HTH cro/C1-type domain-containing protein; uniclust \| UniRef100\_UPI0009838998 \| 99.6 \| 9.6e-19 \| 1.8e-24 \| 116.9 \| 169 \| (3, 179) \| 180 \| (13, 214) \| 218 \| S24 family peptidase \| S24 family peptidase; uniclust \| UniRef100\_A0A352SWL9 \| 99.6 \| 1.1e-18 \| 2e-24 \| 119.0 \| 172 \| (1, 175) \| 180 \| (5, 248) \| 259 \| HTH cro/C1-type domain-containing protein \| HTH cro/C1-type domain-containing protein; uniclust \| UniRef100\_A0A1J5NH46 \| 99.6 \| 1.1e-18 \| 2e-24 \| 110.4 \| 92 \| (87, 179) \| 180 \| (45, 140) \| 142 \| Putative HTH-type transcriptional regulator \| Putative HTH-type transcriptional regulator; uniclust \| UniRef100\_A0A1A7C1A4 \| 99.6 \| 1.2e-18 \| 2.2e-24 \| 123.9 \| 159 \| (15, 176) \| 180 \| (64, 257) \| 266 \| Peptidase S24-like protein \| Peptidase S24-like protein; uniclust \| UniRef100\_A0A0F9HCG5 \| 99.6 \| 1.3e-18 \| 2.4e-24 \| 124.9 \| 177 \| (2, 178) \| 180 \| (67, 278) \| 284 \| HTH cro/C1-type domain-containing protein \| HTH cro/C1-type domain-containing protein; uniclust \| UniRef100\_A0A1X0W151 \| 99.6 \| 1.7e-18 \| 3.2e-24 \| 122.7 \| 172 \| (3, 177) \| 180 \| (24, 227) \| 247 \| HTH cro/C1-type domain-containing protein \| HTH cro/C1-type domain-containing protein; uniclust \| UniRef100\_A0A5B9DX04 \| 99.6 \| 1.8e-18 \| 3.3e-24 \| 118.6 \| 175 \| (2, 177) \| 180 \| (34, 251) \| 262 \| Helix-turn-helix transcriptional regulator \| Helix-turn-helix transcriptional regulator; uniclust \| UniRef100\_A0A060I420 \| 99.6 \| 1.8e-18 \| 3.3e-24 \| 126.7 \| 174 \| (2, 177) \| 180 \| (79, 279) \| 292 \| Peptidase S24/S26 family protein \| Peptidase S24/S26 family protein; uniclust \| UniRef100\_A0A077N0G1 \| 99.6 \| 1.8e-18 \| 3.3e-24 \| 125.2 \| 161 \| (1, 162) \| 180 \| (44, 235) \| 295 \| Helix-turn-helix/peptidase S24-like domain protein \| Helix-turn-helix/peptidase S24-like domain protein; uniclust \| UniRef100\_A0A329BE56 \| 99.6 \| 2e-18 \| 3.6e-24 \| 121.1 \| 170 \| (1, 174) \| 180 \| (102, 306) \| 316 \| SOS-response transcriptional repressor LexA \| SOS-response transcriptional repressor LexA; uniclust \| UniRef100\_A0A2A4NKI2 \| 99.6 \| 2.1e-18 \| 3.8e-24 \| 125.3 \| 164 \| (2, 166) \| 180 \| (52, 273) \| 339 \| Peptidase S24/S26A/S26B/S26C domain-containing protein \| Peptidase S24/S26A/S26B/S26C domain-containing protein; uniclust \| UniRef100\_A0A085JGV9 \| 99.6 \| 2.2e-18 \| 4.1e-24 \| 122.6 \| 170 \| (2, 176) \| 180 \| (38, 239) \| 307 \| Phage repressor \| Phage repressor; uniclust \| UniRef100\_A0A068SNY7 \| 99.6 \| 2.3e-18 \| 4.2e-24 \| 122.2 \| 163 \| (16, 178) \| 180 \| (90, 298) \| 300 \| Prophage MuSo2, transcriptional regulator, Cro/CI family \| Prophage MuSo2, transcriptional regulator, Cro/CI family; uniclust \| UniRef100\_A0A1Y5TZI9 \| 99.6 \| 2.3e-18 \| 4.2e-24 \| 111.4 \| 100 \| (80, 180) \| 180 \| (61, 163) \| 164 \| Peptidase S24-like protein \| Peptidase S24-like protein; uniclust \| UniRef100\_A0A098CTP1 \| 99.6 \| 2.4e-18 \| 4.4e-24 \| 120.3 \| 173 \| (1, 176) \| 180 \| (23, 266) \| 277 \| Transcriptional repressor DicA \| Transcriptional repressor DicA; uniclust \| UniRef100\_A0A258L743 \| 99.6 \| 2.5e-18 \| 4.6e-24 \| 119.8 \| 172 \| (2, 177) \| 180 \| (26, 209) \| 221 \| HTH cro/C1-type domain-containing protein \| HTH cro/C1-type domain-containing protein; uniclust \| UniRef100\_A0A2N1S0V3 \| 99.6 \| 2.5e-18 \| 4.6e-24 \| 117.3 \| 172 \| (2, 175) \| 180 \| (7, 227) \| 236 \| HTH cro/C1-type domain-containing protein \| HTH cro/C1-type domain-containing protein; uniclust \| UniRef100\_A0A959R3J2 \| 99.6 \| 2.7e-18 \| 4.9e-24 \| 114.2 \| 177 \| (1, 179) \| 180 \| (4, 215) \| 218 \| Helix-turn-helix domain-containing protein \| Helix-turn-helix domain-containing protein; uniclust \| UniRef100\_A0A0Q6W6Y0 \| 99.6 \| 2.7e-18 \| 4.9e-24 \| 123.5 \| 160 \| (17, 177) \| 180 \| (71, 251) \| 267 \| Peptidase S24/S26A/S26B/S26C domain-containing protein \| Peptidase S24/S26A/S26B/S26C domain-containing protein; uniclust \| UniRef100\_A0A1I5T8W6 \| 99.6 \| 2.8e-18 \| 5.2e-24 \| 120.4 \| 156 \| (2, 157) \| 180 \| (25, 213) \| 237 \| Helix-turn-helix \| Helix-turn-helix; uniclust \| UniRef100\_UPI0004669D61 \| 99.6 \| 3e-18 \| 5.4e-24 \| 111.5 \| 130 \| (1, 132) \| 180 \| (4, 167) \| 171 \| S24 family peptidase \| S24 family peptidase; uniclust \| UniRef100\_A0A1F7T224 \| 99.6 \| 3.1e-18 \| 5.6e-24 \| 117.6 \| 177 \| (2, 179) \| 180 \| (39, 265) \| 267 \| HTH cro/C1-type domain-containing protein \| HTH cro/C1-type domain-containing protein; uniclust \| UniRef100\_A0A940P805 \| 99.6 \| 3.3e-18 \| 6e-24 \| 111.8 \| 143 \| (1, 144) \| 180 \| (4, 172) \| 176 \| Helix-turn-helix transcriptional regulator (Fragment) \| Helix-turn-helix transcriptional regulator (Fragment); uniclust \| UniRef100\_A0A126NZ97 \| 99.6 \| 3.5e-18 \| 6.4e-24 \| 118.5 \| 177 \| (1, 178) \| 180 \| (12, 228) \| 252 \| HTH cro/C1-type domain-containing protein \| HTH cro/C1-type domain-containing protein; uniclust \| UniRef100\_A0A3C0EH31 \| 99.6 \| 3.6e-18 \| 6.7e-24 \| 115.3 \| 165 \| (2, 167) \| 180 \| (11, 208) \| 221 \| HTH cro/C1-type domain-containing protein \| HTH cro/C1-type domain-containing protein; uniclust \| UniRef100\_A0A1C3K7V4 \| 99.6 \| 3.7e-18 \| 6.8e-24 \| 124.9 \| 178 \| (1, 178) \| 180 \| (15, 343) \| 357 \| Putative phage repressor protein \| Putative phage repressor protein; uniclust \| UniRef100\_A0A223PBQ1 \| 99.6 \| 3.7e-18 \| 6.8e-24 \| 120.0 \| 163 \| (13, 177) \| 180 \| (24, 233) \| 241 \| Transcriptional regulator \| Transcriptional regulator; uniclust \| UniRef100\_A0A0C1MWF1 \| 99.6 \| 3.7e-18 \| 6.8e-24 \| 105.7 \| 91 \| (87, 179) \| 180 \| (8, 99) \| 110 \| Peptidase S24/S26A/S26B/S26C domain-containing protein \| Peptidase S24/S26A/S26B/S26C domain-containing protein; uniclust \| UniRef100\_A0A2E4Q3Q2 \| 99.6 \| 3.9e-18 \| 7.2e-24 \| 116.5 \| 174 \| (1, 176) \| 180 \| (69, 298) \| 302 \| XRE family transcriptional regulator \| XRE family transcriptional regulator; uniclust \| UniRef100\_A0A0S4SRP9 \| 99.6 \| 4.2e-18 \| 7.7e-24 \| 119.8 \| 174 \| (2, 177) \| 180 \| (7, 233) \| 254 \| Signal peptidase I \| Signal peptidase I; uniclust \| UniRef100\_A0A352PSP8 \| 99.6 \| 5e-18 \| 9.2e-24 \| 112.9 \| 100 \| (80, 179) \| 180 \| (66, 169) \| 171 \| Peptidase S24/S26A/S26B/S26C domain-containing protein \| Peptidase S24/S26A/S26B/S26C domain-containing protein; uniclust \| UniRef100\_A0A1M7RMF5 \| 99.6 \| 5.2e-18 \| 9.6e-24 \| 114.8 \| 92 \| (87, 180) \| 180 \| (81, 173) \| 177 \| SOS regulatory protein LexA \| SOS regulatory protein LexA; uniclust \| UniRef100\_A0A0S2EQH0 \| 99.6 \| 5.6e-18 \| 1e-23 \| 120.5 \| 173 \| (2, 177) \| 180 \| (64, 256) \| 268 \| HTH cro/C1-type domain-containing protein \| HTH cro/C1-type domain-containing protein; uniclust \| UniRef100\_A0A011Q9M4 \| 99.6 \| 6.7e-18 \| 1.2e-23 \| 115.2 \| 174 \| (2, 178) \| 180 \| (21, 217) \| 222 \| Putative HTH-type transcriptional regulator \| Putative HTH-type transcriptional regulator; uniclust \| UniRef100\_UPI001403D558 \| 99.6 \| 6.9e-18 \| 1.3e-23 \| 120.4 \| 176 \| (1, 177) \| 180 \| (130, 390) \| 394 \| helix-turn-helix transcriptional regulator \| helix-turn-helix transcriptional regulator; uniclust \| UniRef100\_A0A2P9HMT6 \| 99.6 \| 6.9e-18 \| 1.3e-23 \| 117.1 \| 173 \| (3, 176) \| 180 \| (8, 227) \| 231 \| HTH cro/C1-type domain-containing protein \| HTH cro/C1-type domain-containing protein; uniclust \| UniRef100\_A0A2V2G387 \| 99.6 \| 7e-18 \| 1.3e-23 \| 120.8 \| 167 \| (12, 180) \| 180 \| (54, 251) \| 251 \| HTH cro/C1-type domain-containing protein \| HTH cro/C1-type domain-containing protein; uniclust \| UniRef100\_A0A084A767 \| 99.6 \| 7.8e-18 \| 1.4e-23 \| 117.6 \| 174 \| (2, 177) \| 180 \| (20, 292) \| 295 \| Phage repressor \| Phage repressor; uniclust \| UniRef100\_A0A927Y316 \| 99.6 \| 7.8e-18 \| 1.4e-23 \| 120.1 \| 176 \| (1, 178) \| 180 \| (3, 307) \| 309 \| HTH cro/C1-type domain-containing protein \| HTH cro/C1-type domain-containing protein; uniclust \| UniRef100\_A0A0U3G7P0 \| 99.6 \| 7.9e-18 \| 1.5e-23 \| 111.7 \| 174 \| (3, 178) \| 180 \| (2, 195) \| 198 \| Helix-turn-helix domain-containing protein \| Helix-turn-helix domain-containing protein; uniclust \| UniRef100\_A0A845JXB3 \| 99.6 \| 8e-18 \| 1.5e-23 \| 111.2 \| 100 \| (80, 179) \| 180 \| (129, 228) \| 229 \| Transcriptional regulator \| Transcriptional regulator; uniclust \| UniRef100\_A0A1Q6G157 \| 99.6 \| 8.3e-18 \| 1.5e-23 \| 117.7 \| 175 \| (3, 179) \| 180 \| (25, 233) \| 237 \| Peptidase S24/S26A/S26B/S26C domain-containing protein \| Peptidase S24/S26A/S26B/S26C domain-containing protein; uniclust \| UniRef100\_A0A5S3XEB8 \| 99.6 \| 8.4e-18 \| 1.6e-23 \| 114.3 \| 176 \| (2, 178) \| 180 \| (25, 211) \| 217 \| HTH cro/C1-type domain-containing protein \| HTH cro/C1-type domain-containing protein; uniclust \| UniRef100\_A0A8I2F569 \| 99.6 \| 9e-18 \| 1.7e-23 \| 114.9 \| 174 \| (1, 175) \| 180 \| (17, 216) \| 229 \| Helix-turn-helix domain-containing protein \| Helix-turn-helix domain-containing protein; uniclust \| UniRef100\_A0A1I7BNY7 \| 99.6 \| 1e-17 \| 1.9e-23 \| 115.1 \| 93 \| (85, 177) \| 180 \| (124, 219) \| 223 \| Peptidase S24-like \| Peptidase S24-like; uniclust \| UniRef100\_A0A0S2SLA0 \| 99.6 \| 1.1e-17 \| 2e-23 \| 110.0 \| 100 \| (80, 179) \| 180 \| (53, 156) \| 158 \| Peptidase S24/S26A/S26B/S26C domain-containing protein \| Peptidase S24/S26A/S26B/S26C domain-containing protein; uniclust \| UniRef100\_A0A172U149 \| 99.6 \| 1.1e-17 \| 2e-23 \| 118.1 \| 165 \| (3, 167) \| 180 \| (21, 235) \| 260 \| HTH cro/C1-type domain-containing protein \| HTH cro/C1-type domain-containing protein; uniclust \| UniRef100\_A0A1F9G5D9 \| 99.5 \| 1.2e-17 \| 2.3e-23 \| 118.9 \| 174 \| (2, 176) \| 180 \| (31, 258) \| 276 \| HTH cro/C1-type domain-containing protein \| HTH cro/C1-type domain-containing protein; uniclust \| UniRef100\_A0A2P7TC32 \| 99.5 \| 1.3e-17 \| 2.4e-23 \| 111.7 \| 175 \| (1, 178) \| 180 \| (10, 222) \| 224 \| Helix-turn-helix domain-containing protein \| Helix-turn-helix domain-containing protein; uniclust \| UniRef100\_A0A0C2R644 \| 99.5 \| 1.3e-17 \| 2.4e-23 \| 122.6 \| 87 \| (88, 177) \| 180 \| (234, 322) \| 330 \| HTH cro/C1-type domain-containing protein \| HTH cro/C1-type domain-containing protein; uniclust \| UniRef100\_A0A1I6T486 \| 99.5 \| 1.4e-17 \| 2.6e-23 \| 117.8 \| 173 \| (3, 178) \| 180 \| (5, 236) \| 242 \| Peptidase S24-like \| Peptidase S24-like; uniclust \| UniRef100\_A0A381KSX4 \| 99.5 \| 1.5e-17 \| 2.8e-23 \| 112.8 \| 179 \| (1, 179) \| 180 \| (15, 215) \| 216 \| Helix-turn-helix \| Helix-turn-helix; uniclust \| UniRef100\_A0A1F3SQC4 \| 99.5 \| 1.5e-17 \| 2.8e-23 \| 109.6 \| 94 \| (85, 180) \| 180 \| (55, 148) \| 152 \| Peptidase S24/S26A/S26B/S26C domain-containing protein \| Peptidase S24/S26A/S26B/S26C domain-containing protein; uniclust \| UniRef100\_A0A031G478 \| 99.5 \| 1.6e-17 \| 2.9e-23 \| 118.3 \| 177 \| (3, 179) \| 180 \| (46, 251) \| 254 \| Peptidase s24-like domain protein \| Peptidase s24-like domain protein; uniclust \| UniRef100\_A0A136LGU8 \| 99.5 \| 1.6e-17 \| 3e-23 \| 122.4 \| 151 \| (15, 166) \| 180 \| (36, 232) \| 305 \| Transcriptional regulator \| Transcriptional regulator; uniclust \| UniRef100\_A0A066ZT37 \| 99.5 \| 1.7e-17 \| 3.1e-23 \| 120.0 \| 99 \| (80, 180) \| 180 \| (140, 239) \| 258 \| DNA polymerase V \| DNA polymerase V; uniclust \| UniRef100\_A0A085EU53 \| 99.5 \| 1.7e-17 \| 3.2e-23 \| 120.8 \| 179 \| (1, 180) \| 180 \| (39, 263) \| 278 \| Helix-turn-helix protein \| Helix-turn-helix protein; uniclust \| UniRef100\_A0A0E4A064 \| 99.5 \| 1.8e-17 \| 3.3e-23 \| 118.1 \| 175 \| (2, 177) \| 180 \| (38, 254) \| 261 \| Repressor \| Repressor; uniclust \| UniRef100\_A0A554XCA7 \| 99.5 \| 1.9e-17 \| 3.5e-23 \| 116.5 \| 88 \| (89, 176) \| 180 \| (227, 316) \| 325 \| Peptidase S24-like protein \| Peptidase S24-like protein; uniclust \| UniRef100\_A0A031INB5 \| 99.5 \| 1.9e-17 \| 3.5e-23 \| 120.6 \| 165 \| (11, 177) \| 180 \| (48, 302) \| 307 \| Prophage PSPPH03, Cro/CI family transcriptional regulator \| Prophage PSPPH03, Cro/CI family transcriptional regulator; uniclust \| UniRef100\_A0A1Q3QX90 \| 99.5 \| 1.9e-17 \| 3.6e-23 \| 115.8 \| 175 \| (2, 177) \| 180 \| (49, 268) \| 276 \| HTH cro/C1-type domain-containing protein \| HTH cro/C1-type domain-containing protein; uniclust \| UniRef100\_A0A1Q3VUE3 \| 99.5 \| 2e-17 \| 3.6e-23 \| 116.1 \| 160 \| (16, 175) \| 180 \| (47, 231) \| 241 \| HTH cro/C1-type domain-containing protein \| HTH cro/C1-type domain-containing protein; uniclust \| UniRef100\_A0A6I4YRF4 \| 99.5 \| 2e-17 \| 3.7e-23 \| 118.2 \| 160 \| (15, 175) \| 180 \| (151, 358) \| 370 \| Helix-turn-helix domain-containing protein \| Helix-turn-helix domain-containing protein; uniclust \| UniRef100\_A0A0B6CXM0 \| 99.5 \| 2.1e-17 \| 3.8e-23 \| 113.5 \| 173 \| (4, 177) \| 180 \| (11, 216) \| 228 \| Helix-turn-helix family protein \| Helix-turn-helix family protein; uniclust \| UniRef100\_A0A0K8JF70 \| 99.5 \| 2.4e-17 \| 4.4e-23 \| 114.8 \| 175 \| (1, 177) \| 180 \| (10, 295) \| 298 \| HTH cro/C1-type domain-containing protein \| HTH cro/C1-type domain-containing protein; uniclust \| UniRef100\_A0A940CVJ9 \| 99.5 \| 2.4e-17 \| 4.4e-23 \| 111.5 \| 175 \| (1, 177) \| 180 \| (3, 229) \| 244 \| LexA family transcriptional regulator \| LexA family transcriptional regulator; uniclust \| UniRef100\_A0A315CMB6 \| 99.5 \| 2.4e-17 \| 4.5e-23 \| 110.6 \| 174 \| (1, 178) \| 180 \| (5, 201) \| 206 \| HTH cro/C1-type domain-containing protein \| HTH cro/C1-type domain-containing protein; uniclust \| UniRef100\_A0A380LI00 \| 99.5 \| 2.6e-17 \| 4.7e-23 \| 109.7 \| 169 \| (9, 179) \| 180 \| (15, 198) \| 201 \| CI-like repressor, S. pneumoniae bacteriophage EJ-1 \| CI-like repressor, S. pneumoniae bacteriophage EJ-1; uniclust \| UniRef100\_A0A844PAU3 \| 99.5 \| 2.7e-17 \| 5e-23 \| 109.7 \| 171 \| (6, 180) \| 180 \| (28, 214) \| 214 \| Transcriptional regulator \| Transcriptional regulator; uniclust \| UniRef100\_A0A7W2GRI4 \| 99.5 \| 3e-17 \| 5.5e-23 \| 114.1 \| 162 \| (16, 178) \| 180 \| (56, 265) \| 272 \| Helix-turn-helix domain-containing protein \| Helix-turn-helix domain-containing protein; uniclust \| UniRef100\_A0A0A8HUB8 \| 99.5 \| 3e-17 \| 5.6e-23 \| 113.6 \| 171 \| (4, 175) \| 180 \| (18, 225) \| 243 \| Peptidase S24 LexA-like protein \| Peptidase S24 LexA-like protein; uniclust \| UniRef100\_A0A8J8BJY7 \| 99.5 \| 3e-17 \| 5.6e-23 \| 112.4 \| 173 \| (4, 179) \| 180 \| (9, 211) \| 245 \| LexA family transcriptional regulator \| LexA family transcriptional regulator; uniclust \| UniRef100\_A0A101VSQ7 \| 99.5 \| 3.1e-17 \| 5.7e-23 \| 114.7 \| 172 \| (1, 176) \| 180 \| (11, 213) \| 228 \| HTH cro/C1-type domain-containing protein \| HTH cro/C1-type domain-containing protein; uniclust \| UniRef100\_A0A2D5Y4P2 \| 99.5 \| 3.2e-17 \| 6e-23 \| 112.3 \| 164 \| (13, 177) \| 180 \| (15, 197) \| 200 \| Peptidase S24/S26A/S26B/S26C domain-containing protein \| Peptidase S24/S26A/S26B/S26C domain-containing protein; uniclust \| UniRef100\_A0A2S0UMT0 \| 99.5 \| 3.3e-17 \| 6.1e-23 \| 95.5 \| 79 \| (101, 179) \| 180 \| (1, 82) \| 85 \| Peptidase S24/S26A/S26B/S26C domain-containing protein \| Peptidase S24/S26A/S26B/S26C domain-containing protein; uniclust \| UniRef100\_E5BEW3 \| 99.5 \| 3.6e-17 \| 6.6e-23 \| 116.0 \| 173 \| (4, 179) \| 180 \| (9, 310) \| 345 \| Putative phage head-tail adaptor \| Putative phage head-tail adaptor; uniclust \| UniRef100\_A0A2A4XVH8 \| 99.5 \| 3.7e-17 \| 6.9e-23 \| 113.6 \| 171 \| (1, 174) \| 180 \| (4, 223) \| 231 \| HTH cro/C1-type domain-containing protein \| HTH cro/C1-type domain-containing protein; uniclust \| UniRef100\_A0A0K2Y4A7 \| 99.5 \| 4.6e-17 \| 8.5e-23 \| 119.7 \| 176 \| (1, 177) \| 180 \| (10, 317) \| 339 \| LexA-family transcriptional regulator \| LexA-family transcriptional regulator; uniclust \| UniRef100\_A0A2E9YV30 \| 99.5 \| 4.7e-17 \| 8.6e-23 \| 112.1 \| 176 \| (1, 176) \| 180 \| (10, 222) \| 230 \| HTH cro/C1-type domain-containing protein \| HTH cro/C1-type domain-containing protein; uniclust \| UniRef100\_A0A0U5N406 \| 99.5 \| 4.8e-17 \| 8.9e-23 \| 117.6 \| 156 \| (2, 157) \| 180 \| (61, 281) \| 303 \| Putative Repressor protein C \| Putative Repressor protein C; uniclust \| UniRef100\_A0A011VG60 \| 99.5 \| 5.2e-17 \| 9.6e-23 \| 117.5 \| 168 \| (3, 177) \| 180 \| (46, 234) \| 281 \| Peptidase S24 \| Peptidase S24; uniclust \| UniRef100\_A0A1A0FJR0 \| 99.5 \| 5.5e-17 \| 1e-22 \| 107.3 \| 94 \| (84, 177) \| 180 \| (82, 178) \| 181 \| Peptidase S24/S26A/S26B/S26C domain-containing protein \| Peptidase S24/S26A/S26B/S26C domain-containing protein; uniclust \| UniRef100\_A0A2A4N3F3 \| 99.5 \| 5.6e-17 \| 1e-22 \| 114.9 \| 176 \| (2, 179) \| 180 \| (27, 281) \| 293 \| Peptidase S24/S26A/S26B/S26C domain-containing protein \| Peptidase S24/S26A/S26B/S26C domain-containing protein; uniclust \| UniRef100\_A0A0A1VGJ5 \| 99.5 \| 5.6e-17 \| 1e-22 \| 106.4 \| 99 \| (80, 180) \| 180 \| (44, 143) \| 153 \| SOS-response transcriptional repressor \| SOS-response transcriptional repressor; uniclust \| UniRef100\_A0A017RV89 \| 99.5 \| 5.6e-17 \| 1e-22 \| 117.1 \| 176 \| (1, 178) \| 180 \| (25, 243) \| 268 \| HTH cro/C1-type domain-containing protein \| HTH cro/C1-type domain-containing protein; uniclust \| UniRef100\_A0A0N0ZPS4 \| 99.5 \| 5.9e-17 \| 1.1e-22 \| 114.6 \| 176 \| (2, 179) \| 180 \| (29, 241) \| 259 \| HTH cro/C1-type domain-containing protein \| HTH cro/C1-type domain-containing protein; uniclust \| UniRef100\_A0A1F3XYF0 \| 99.5 \| 6.4e-17 \| 1.2e-22 \| 113.5 \| 177 \| (1, 177) \| 180 \| (20, 252) \| 258 \| HTH cro/C1-type domain-containing protein \| HTH cro/C1-type domain-containing protein; uniclust \| UniRef100\_UPI0018DFAD8D \| 99.5 \| 6.5e-17 \| 1.2e-22 \| 101.6 \| 159 \| (2, 177) \| 180 \| (3, 162) \| 164 \| LexA family transcriptional regulator \| LexA family transcriptional regulator; uniclust \| UniRef100\_A0A1A5RXV5 \| 99.5 \| 6.7e-17 \| 1.2e-22 \| 112.7 \| 172 \| (2, 174) \| 180 \| (76, 266) \| 272 \| HTH cro/C1-type domain-containing protein \| HTH cro/C1-type domain-containing protein; uniclust \| UniRef100\_A0A0T2QE85 \| 99.5 \| 6.9e-17 \| 1.3e-22 \| 112.3 \| 172 \| (2, 179) \| 180 \| (31, 224) \| 235 \| HTH cro/C1-type domain-containing protein \| HTH cro/C1-type domain-containing protein; uniclust \| UniRef100\_A0A7W1G0U4 \| 99.5 \| 7.5e-17 \| 1.4e-22 \| 108.9 \| 160 \| (6, 166) \| 180 \| (13, 200) \| 238 \| LexA family transcriptional regulator \| LexA family transcriptional regulator; uniclust \| UniRef100\_A0A085EXQ7 \| 99.5 \| 8.2e-17 \| 1.5e-22 \| 118.5 \| 171 \| (2, 175) \| 180 \| (52, 269) \| 279 \| Putative phage repressor \| Putative phage repressor; uniclust \| UniRef100\_A0A4Q3HVL5 \| 99.5 \| 8.3e-17 \| 1.5e-22 \| 97.5 \| 99 \| (80, 180) \| 180 \| (10, 109) \| 111 \| Error-prone repair protein UmuD (Fragment) \| Error-prone repair protein UmuD (Fragment); uniclust \| UniRef100\_A0A133MBD3 \| 99.5 \| 8.3e-17 \| 1.5e-22 \| 120.6 \| 175 \| (1, 178) \| 180 \| (25, 365) \| 375 \| DNA-binding helix-turn-helix protein \| DNA-binding helix-turn-helix protein; uniclust \| UniRef100\_A0A7Z9U9M0 \| 99.5 \| 8.4e-17 \| 1.6e-22 \| 107.3 \| 95 \| (85, 180) \| 180 \| (76, 175) \| 175 \| Peptidase S24/S26A/S26B/S26C domain-containing protein \| Peptidase S24/S26A/S26B/S26C domain-containing protein; uniclust \| UniRef100\_A0A7V3HNI8 \| 99.5 \| 8.6e-17 \| 1.6e-22 \| 108.2 \| 174 \| (1, 177) \| 180 \| (19, 257) \| 262 \| Helix-turn-helix domain-containing protein \| Helix-turn-helix domain-containing protein; uniclust \| UniRef100\_A0A090CZE3 \| 99.5 \| 8.8e-17 \| 1.6e-22 \| 110.4 \| 99 \| (80, 180) \| 180 \| (96, 195) \| 204 \| Protein UmuD \| Protein UmuD; uniclust \| UniRef100\_A0A124BRC6 \| 99.5 \| 9.7e-17 \| 1.8e-22 \| 112.8 \| 172 \| (2, 177) \| 180 \| (24, 223) \| 233 \| Helix-turn-helix domain protein \| Helix-turn-helix domain protein; uniclust \| UniRef100\_A0A1B7JZC6 \| 99.5 \| 1e-16 \| 1.9e-22 \| 106.4 \| 91 \| (85, 176) \| 180 \| (68, 160) \| 169 \| Phage repressor \| Phage repressor; uniclust \| UniRef100\_A0A2N9XWK5 \| 99.5 \| 1.1e-16 \| 1.9e-22 \| 106.9 \| 91 \| (85, 176) \| 180 \| (76, 167) \| 174 \| Peptidase S24/S26A/S26B/S26C domain-containing protein \| Peptidase S24/S26A/S26B/S26C domain-containing protein; uniclust \| UniRef100\_A0A143XNR7 \| 99.5 \| 1.1e-16 \| 2.1e-22 \| 114.8 \| 173 \| (2, 178) \| 180 \| (20, 234) \| 270 \| Peptidase S24/S26A/S26B/S26C domain-containing protein \| Peptidase S24/S26A/S26B/S26C domain-containing protein; uniclust \| UniRef100\_A0A1C3EE82 \| 99.5 \| 1.2e-16 \| 2.2e-22 \| 110.2 \| 175 \| (2, 176) \| 180 \| (89, 312) \| 318 \| HTH cro/C1-type domain-containing protein \| HTH cro/C1-type domain-containing protein; uniclust \| UniRef100\_A0A1C2K2G3 \| 99.5 \| 1.3e-16 \| 2.3e-22 \| 110.9 \| 174 \| (1, 178) \| 180 \| (15, 221) \| 229 \| HTH cro/C1-type domain-containing protein \| HTH cro/C1-type domain-containing protein; uniclust \| UniRef100\_A0A494X9D3 \| 99.5 \| 1.3e-16 \| 2.4e-22 \| 109.0 \| 177 \| (1, 177) \| 180 \| (13, 249) \| 254 \| LexA family transcriptional regulator \| LexA family transcriptional regulator; uniclust \| UniRef100\_A0A351J0K7 \| 99.5 \| 1.3e-16 \| 2.4e-22 \| 107.4 \| 171 \| (2, 174) \| 180 \| (4, 237) \| 238 \| HTH cro/C1-type domain-containing protein \| HTH cro/C1-type domain-containing protein; uniclust \| UniRef100\_A0A060BDL7 \| 99.5 \| 1.4e-16 \| 2.6e-22 \| 112.4 \| 175 \| (1, 179) \| 180 \| (17, 207) \| 245 \| XRE family transcriptional regulator \| XRE family transcriptional regulator; uniclust \| UniRef100\_A0A1R1BA90 \| 99.5 \| 1.4e-16 \| 2.6e-22 \| 117.5 \| 87 \| (88, 177) \| 180 \| (279, 367) \| 370 \| HTH cro/C1-type domain-containing protein \| HTH cro/C1-type domain-containing protein; uniclust \| UniRef100\_A0A123TEK7 \| 99.5 \| 1.4e-16 \| 2.6e-22 \| 107.2 \| 91 \| (87, 179) \| 180 \| (76, 166) \| 174 \| Putative phage repressor \| Putative phage repressor; uniclust \| UniRef100\_A0A0U5F099 \| 99.5 \| 1.5e-16 \| 2.8e-22 \| 115.1 \| 89 \| (87, 176) \| 180 \| (204, 296) \| 300 \| Putative phage repressor \| Putative phage repressor; uniclust \| UniRef100\_A0A0G0TYJ6 \| 99.5 \| 1.5e-16 \| 2.8e-22 \| 110.7 \| 175 \| (1, 179) \| 180 \| (11, 247) \| 248 \| LexA repressor \| LexA repressor; uniclust \| UniRef100\_UPI001F49D1FF \| 99.5 \| 1.6e-16 \| 2.9e-22 \| 91.1 \| 86 \| (95, 180) \| 180 \| (3, 90) \| 90 \| S24 family peptidase \| S24 family peptidase; uniclust \| UniRef100\_A0A098GCJ7 \| 99.5 \| 1.6e-16 \| 2.9e-22 \| 116.8 \| 175 \| (2, 178) \| 180 \| (41, 244) \| 276 \| Putative repressor protein from phage origin \| Putative repressor protein from phage origin; uniclust \| UniRef100\_UPI0013CFBFF4 \| 99.5 \| 1.6e-16 \| 2.9e-22 \| 105.6 \| 177 \| (1, 179) \| 180 \| (18, 210) \| 211 \| LexA family transcriptional regulator \| LexA family transcriptional regulator; uniclust \| UniRef100\_A0A031HR16 \| 99.5 \| 1.6e-16 \| 2.9e-22 \| 109.5 \| 155 \| (25, 180) \| 180 \| (62, 242) \| 249 \| Peptidase S24/S26A/S26B/S26C domain-containing protein \| Peptidase S24/S26A/S26B/S26C domain-containing protein; uniclust \| UniRef100\_A0A6L7WV89 \| 99.5 \| 1.6e-16 \| 3e-22 \| 109.9 \| 177 \| (1, 180) \| 180 \| (29, 288) \| 288 \| LexA family transcriptional regulator \| LexA family transcriptional regulator; uniclust \| UniRef100\_A0A179BUG2 \| 99.5 \| 1.7e-16 \| 3.1e-22 \| 115.2 \| 173 \| (1, 176) \| 180 \| (32, 250) \| 257 \| HTH cro/C1-type domain-containing protein \| HTH cro/C1-type domain-containing protein; uniclust \| UniRef100\_UPI001E65DB02 \| 99.5 \| 1.8e-16 \| 3.2e-22 \| 105.7 \| 173 \| (1, 175) \| 180 \| (7, 228) \| 235 \| LexA family transcriptional regulator \| LexA family transcriptional regulator; uniclust \| UniRef100\_A0A1H0WB76 \| 99.5 \| 1.8e-16 \| 3.4e-22 \| 98.5 \| 90 \| (88, 177) \| 180 \| (20, 114) \| 122 \| Peptidase S24-like \| Peptidase S24-like; uniclust \| UniRef100\_A0A437MC40 \| 99.5 \| 1.8e-16 \| 3.4e-22 \| 113.2 \| 154 \| (2, 155) \| 180 \| (49, 254) \| 279 \| Helix-turn-helix transcriptional regulator \| Helix-turn-helix transcriptional regulator; uniclust \| UniRef100\_A0A0Q4YH48 \| 99.5 \| 1.9e-16 \| 3.5e-22 \| 109.3 \| 159 \| (17, 175) \| 180 \| (57, 228) \| 238 \| Peptidase S24 \| Peptidase S24; uniclust \| UniRef100\_A0A0L6J7F1 \| 99.5 \| 1.9e-16 \| 3.6e-22 \| 115.4 \| 169 \| (2, 177) \| 180 \| (61, 243) \| 266 \| Xre family transcriptional regulator \| Xre family transcriptional regulator; uniclust \| UniRef100\_A0A1I4Z2S5 \| 99.5 \| 2e-16 \| 3.6e-22 \| 99.1 \| 98 \| (80, 179) \| 180 \| (28, 126) \| 132 \| DNA polymerase V \| DNA polymerase V; uniclust \| UniRef100\_A0A3C1EDA2 \| 99.5 \| 2e-16 \| 3.7e-22 \| 103.0 \| 173 \| (1, 180) \| 180 \| (3, 179) \| 180 \| HTH cro/C1-type domain-containing protein \| HTH cro/C1-type domain-containing protein; uniclust \| UniRef100\_A0A191HVY2 \| 99.5 \| 2.1e-16 \| 3.8e-22 \| 99.6 \| 92 \| (87, 180) \| 180 \| (19, 115) \| 127 \| Peptidase S24/S26A/S26B/S26C domain-containing protein \| Peptidase S24/S26A/S26B/S26C domain-containing protein; uniclust \| UniRef100\_A0A4R7DWV3 \| 99.5 \| 2.2e-16 \| 4e-22 \| 110.5 \| 173 \| (1, 177) \| 180 \| (16, 286) \| 292 \| Helix-turn-helix protein \| Helix-turn-helix protein; uniclust \| UniRef100\_A0A0S4S9Z8 \| 99.5 \| 2.2e-16 \| 4.1e-22 \| 107.8 \| 163 \| (13, 180) \| 180 \| (21, 210) \| 213 \| Phage repressor protein \| Phage repressor protein; uniclust \| UniRef100\_A0A0H3ZMC9 \| 99.5 \| 2.4e-16 \| 4.4e-22 \| 112.7 \| 96 \| (81, 176) \| 180 \| (141, 240) \| 280 \| HTH Mu-type domain-containing protein \| HTH Mu-type domain-containing protein; uniclust \| UniRef100\_A0A0R6PGT9 \| 99.5 \| 2.4e-16 \| 4.5e-22 \| 106.0 \| 173 \| (6, 180) \| 180 \| (13, 196) \| 197 \| Peptidase S24/S26A/S26B/S26C domain-containing protein \| Peptidase S24/S26A/S26B/S26C domain-containing protein; uniclust \| UniRef100\_A0A8X8TGA2 \| 99.4 \| 2.5e-16 \| 4.7e-22 \| 109.9 \| 171 \| (2, 178) \| 180 \| (7, 203) \| 220 \| Helix-turn-helix transcriptional regulator \| Helix-turn-helix transcriptional regulator; uniclust \| UniRef100\_A0A6L7U9Q8 \| 99.4 \| 2.6e-16 \| 4.8e-22 \| 106.1 \| 93 \| (85, 179) \| 180 \| (146, 238) \| 239 \| LexA family transcriptional regulator \| LexA family transcriptional regulator; uniclust \| UniRef100\_UPI000688D37D \| 99.4 \| 2.7e-16 \| 4.9e-22 \| 101.7 \| 84 \| (90, 176) \| 180 \| (67, 151) \| 158 \| S24/S26 family peptidase \| S24/S26 family peptidase; pdb70 \| 3BDN\_A \| 99.7 \| 2.1e-22 \| 1.4e-26 \| 132.0 \| 173 \| (3, 179) \| 180 \| (16, 231) \| 236 \| PROTEIN/DNA Complex \| 3BDN\_A PROTEIN/DNA Complex lambda, repressor, allostery, cooperativity, DNA; pdb70 \| 2FJR\_A \| 99.6 \| 1.1e-19 \| 7.3e-24 \| 115.0 \| 167 \| (2, 179) \| 180 \| (8, 187) \| 189 \| Repressor protein CI \| 2FJR\_A Repressor protein CI genetic switch, regulation, repressor, cooperativity; pdb70 \| 1JHC\_A \| 99.0 \| 3.9e-14 \| 2.6e-18 \| 84.6 \| 88 \| (89, 179) \| 180 \| (41, 131) \| 137 \| LEXA REPRESSOR (E.C.3.4.21.88) \| 1JHC\_A LEXA REPRESSOR (E.C.3.4.21.88) LexA SOS repressor, HYDROLASE; pdb70 \| 6A2Q\_A \| 99.0 \| 6.1e-14 \| 4.1e-18 \| 81.6 \| 86 \| (89, 179) \| 180 \| (31, 117) \| 118 \| LexA repressor (E.C.3.4.21.88) \| 6A2Q\_A LexA repressor (E.C.3.4.21.88) Mycobacterium tuberculosis, LexA, SOS response HET: CME; pdb70 \| 6A2S\_C \| 99.0 \| 7.3e-14 \| 4.9e-18 \| 80.3 \| 86 \| (89, 179) \| 180 \| (24, 110) \| 111 \| LexA repressor (E.C.3.4.21.88) \| 6A2S\_C LexA repressor (E.C.3.4.21.88) Mycobacterium tuberculosis, LexA, SOS response HET: CME, PEG, P6G; pdb70 \| 1F39\_A \| 99.0 \| 8e-14 \| 5.4e-18 \| 78.8 \| 87 \| (89, 179) \| 180 \| (3, 96) \| 101 \| REPRESSOR PROTEIN CI \| 1F39\_A REPRESSOR PROTEIN CI COOPERATIVE OPERATOR BINDING, RECA-MEDIATED SELF-CLEAVAGE; pdb70 \| 4Y42\_I \| 99.0 \| 1.3e-13 \| 8.5e-18 \| 84.2 \| 126 \| (3, 138) \| 180 \| (16, 155) \| 156 \| Cyanase (E.C.4.2.1.104) \| 4Y42\_I Cyanase (E.C.4.2.1.104) CynS, cyanase, lyase HET: GOL; pdb70 \| 6B6M\_A \| 99.0 \| 1.3e-13 \| 8.5e-18 \| 84.2 \| 126 \| (3, 138) \| 180 \| (16, 155) \| 156 \| Cyanate hydratase (E.C.4.2.1.104) \| 6B6M\_A Cyanate hydratase (E.C.4.2.1.104) Cyanase, LYASE; pdb70 \| 3K2Z\_B \| 98.9 \| 1.8e-13 \| 1.2e-17 \| 86.4 \| 88 \| (89, 179) \| 180 \| (107, 195) \| 196 \| LexA repressor (E.C.3.4.21.88) \| 3K2Z\_B LexA repressor (E.C.3.4.21.88) winged helix-turn-helix, repressor, LexA, SOS; pdb70 \| 1AY9\_B \| 98.9 \| 1.9e-13 \| 1.3e-17 \| 78.2 \| 87 \| (89, 179) \| 180 \| (18, 106) \| 108 \| UMUD PROTEIN \| 1AY9\_B UMUD PROTEIN MUTAGENESIS PROTEIN, DNA REPAIR, HYDROLASE; pdb70 \| 1UMU\_B \| 98.9 \| 4.4e-13 \| 2.9e-17 \| 77.7 \| 87 \| (89, 179) \| 180 \| (25, 113) \| 116 \| UMUD' \| 1UMU\_B UMUD' INDUCED MUTAGENESIS, SOS MUTAGENESIS, DNA HET: MSE; pdb70 \| 1JHF\_A \| 98.8 \| 6.4e-13 \| 4.3e-17 \| 84.3 \| 159 \| (12, 179) \| 180 \| (24, 198) \| 202 \| LEXA REPRESSOR (E.C.3.4.21.88) \| 1JHF\_A LEXA REPRESSOR (E.C.3.4.21.88) LexA SOS repressor, HYDROLASE; pdb70 \| 2HNF\_A \| 98.8 \| 1.2e-12 \| 8.1e-17 \| 77.6 \| 84 \| (89, 176) \| 180 \| (42, 132) \| 133 \| Phage lambda repressor protein CI \| 2HNF\_A Phage lambda repressor protein CI VIRAL PROTEIN; pdb70 \| 4N31\_B \| 98.5 \| 3.8e-11 \| 2.6e-15 \| 73.5 \| 60 \| (91, 151) \| 180 \| (29, 89) \| 163 \| SipA \| 4N31\_B SipA Streptococcus pyogenes, pilus polymerisation, Signal HET: PO4, PTY; pdb70 \| 4N31\_A \| 98.5 \| 4.8e-11 \| 3.2e-15 \| 73.0 \| 60 \| (91, 151) \| 180 \| (29, 89) \| 163 \| SipA \| 4N31\_A SipA Streptococcus pyogenes, pilus polymerisation, Signal HET: PO4, PTY; pdb70 \| 4ME8\_A \| 98.2 \| 7.1e-10 \| 4.8e-14 \| 66.8 \| 57 \| (94, 151) \| 180 \| (2, 60) \| 151 \| Signal peptidase I (E.C.3.4.21.89) \| 4ME8\_A Signal peptidase I (E.C.3.4.21.89) PF00717 family, Peptidase\_S24, Structural Genomics; pdb70 \| 4NV4\_A \| 98.1 \| 2e-09 \| 1.3e-13 \| 66.5 \| 59 \| (91, 150) \| 180 \| (20, 83) \| 176 \| Signal peptidase I (E.C.3.4.21.89) \| 4NV4\_A Signal peptidase I (E.C.3.4.21.89) Structural Genomics, NIAID, National Institute HET: PEG, PG4; pdb70 \| 4NV4\_B \| 98.0 \| 2.2e-09 \| 1.5e-13 \| 66.3 \| 59 \| (91, 150) \| 180 \| (20, 83) \| 176 \| Signal peptidase I (E.C.3.4.21.89) \| 4NV4\_B Signal peptidase I (E.C.3.4.21.89) Structural Genomics, NIAID, National Institute HET: PG4, PEG; pdb70 \| 1B12\_D \| 98.0 \| 2.5e-09 \| 1.7e-13 \| 69.6 \| 60 \| (92, 152) \| 180 \| (7, 85) \| 248 \| SIGNAL PEPTIDASE I (E.C.3.9.21.89) \| 1B12\_D SIGNAL PEPTIDASE I (E.C.3.9.21.89) SERINE PROTEINASE, SERINE-DEPENDANT HYDROLASE, SIGNAL HET: 1PN; pdb70 \| 4K8W\_A \| 98.0 \| 3.1e-09 \| 2.1e-13 \| 62.1 \| 55 \| (97, 152) \| 180 \| (1, 56) \| 129 \| LepA \| 4K8W\_A LepA Predominately beta-strand, S26A signal peptidase; pdb70 \| 2FKD\_F \| 98.0 \| 3.3e-09 \| 2.2e-13 \| 60.3 \| 69 \| (105, 179) \| 180 \| (40, 108) \| 110 \| Repressor protein CI \| 2FKD\_F Repressor protein CI genetic switch, regulation, cooperativity, repressor; pdb70 \| 6B88\_B \| 97.9 \| 7.5e-09 \| 5.1e-13 \| 67.2 \| 59 \| (92, 151) \| 180 \| (6, 83) \| 247 \| Signal peptidase I (E.C.3.4.21.89), ILE-THR-LEU-LEU \| 6B88\_B Signal peptidase I (E.C.3.4.21.89), ILE-THR-LEU-LEU Signal peptidase, SBDD, antibiotic, HYDROLASE HET: 1PE, CZD; pdb70 \| 2K9Q\_B \| 97.7 \| 2.2e-08 \| 1.4e-12 \| 52.9 \| 62 \| (2, 63) \| 180 \| (4, 65) \| 77 \| uncharacterized protein \| 2K9Q\_B uncharacterized protein all helix, helix-turn-helix, Plasmid, Structural; pdb70 \| 4WVI\_A \| 97.7 \| 2.2e-08 \| 1.5e-12 \| 71.9 \| 58 \| (93, 151) \| 180 \| (380, 442) \| 533 \| Maltose binding protein - \| 4WVI\_A Maltose binding protein - SpsB Type-I signal peptidase, Peptide HET: MAL; pdb70 \| 5J2Y\_A \| 97.7 \| 2.3e-08 \| 1.5e-12 \| 53.3 \| 61 \| (2, 62) \| 180 \| (13, 74) \| 80 \| Regulatory protein/DNA Complex \| 5J2Y\_A Regulatory protein/DNA Complex Quorum-sensing repressor, gene regulation, RsaL-DNA HET: MSE; pdb70 \| 5J2Y\_B \| 97.7 \| 2.3e-08 \| 1.5e-12 \| 53.3 \| 61 \| (2, 62) \| 180 \| (13, 74) \| 80 \| Regulatory protein/DNA Complex \| 5J2Y\_B Regulatory protein/DNA Complex Quorum-sensing repressor, gene regulation, RsaL-DNA HET: MSE; pdb70 \| 2KPJ\_A \| 97.7 \| 3.5e-08 \| 2.4e-12 \| 54.2 \| 61 \| (3, 63) \| 180 \| (12, 72) \| 94 \| SOS-response transcriptional repressor, LexA \| 2KPJ\_A SOS-response transcriptional repressor, LexA NESG, GFT, Structural Genomics, PSI-2; pdb70 \| 1LLI\_B \| 97.6 \| 3.8e-08 \| 2.6e-12 \| 53.9 \| 53 \| (10, 62) \| 180 \| (27, 79) \| 92 \| LAMBDA REPRESSOR (TRIPLE MUTANT)/DNA COMPLEX \| 1LLI\_B LAMBDA REPRESSOR (TRIPLE MUTANT)/DNA COMPLEX PROTEIN-DNA COMPLEX, DOUBLE HELIX, TRANSCRIPTION-DNA; pdb70 \| 3KZ3\_B \| 97.6 \| 7e-08 \| 4.7e-12 \| 51.1 \| 60 \| (4, 63) \| 180 \| (16, 75) \| 80 \| Repressor protein CI \| 3KZ3\_B Repressor protein CI Five helix bundle, DNA-binding, Repressor; pdb70 \| 4I6U\_A \| 97.6 \| 7.1e-08 \| 4.8e-12 \| 51.3 \| 61 \| (2, 62) \| 180 \| (13, 73) \| 82 \| Regulatory protein \| 4I6U\_A Regulatory protein Restriction-modification, helix-turn-helix, transcriptional regulato, DNA HET: GOL, PEG, ACT; pdb70 \| 4I6R\_B \| 97.5 \| 9.9e-08 \| 6.6e-12 \| 50.7 \| 62 \| (2, 63) \| 180 \| (13, 74) \| 82 \| Regulatory protein \| 4I6R\_B Regulatory protein Restriction-modification, helix-turn-helix, transcriptional regulator, DNA HET: GOL, SO4; pdb70 \| 6B9R\_C \| 97.4 \| 1.4e-07 \| 9.3e-12 \| 66.5 \| 63 \| (1, 63) \| 180 \| (17, 79) \| 450 \| Methylphosphonate synthase \| 6B9R\_C Methylphosphonate synthase Phosphonate, Hydroxymethylphosphonate, Iron, OXIDOREDUCTASE HET: 2HE, GOL; pdb70 \| 6B9R\_D \| 97.4 \| 1.4e-07 \| 9.3e-12 \| 66.5 \| 63 \| (1, 63) \| 180 \| (17, 79) \| 450 \| Methylphosphonate synthase \| 6B9R\_D Methylphosphonate synthase Phosphonate, Hydroxymethylphosphonate, Iron, OXIDOREDUCTASE HET: GOL, 2HE; pdb70 \| 1Y9Q\_A \| 97.4 \| 1.4e-07 \| 9.7e-12 \| 58.6 \| 62 \| (2, 63) \| 180 \| (13, 74) \| 192 \| transcriptional regulator, HTH\_3 family \| 1Y9Q\_A transcriptional regulator, HTH\_3 family HTH\_3 family, Transcriptional Regulaator, Strucutral; pdb70 \| 2EF8\_A \| 97.3 \| 2.7e-07 \| 1.8e-11 \| 49.3 \| 62 \| (2, 63) \| 180 \| (12, 77) \| 84 \| Putative transcription factor \| 2EF8\_A Putative transcription factor helix-turn-helix, DNA binding protein, TRANSCRIPTION; pdb70 \| 3B7H\_A \| 97.3 \| 2.8e-07 \| 1.9e-11 \| 48.2 \| 61 \| (3, 63) \| 180 \| (10, 71) \| 78 \| Prophage Lp1 protein 11 \| 3B7H\_A Prophage Lp1 protein 11 prophage Lp1 protein 11, structural; pdb70 \| 2P5T\_E \| 97.3 \| 2.9e-07 \| 1.9e-11 \| 55.5 \| 62 \| (2, 63) \| 180 \| (3, 64) \| 158 \| fragment of PezA helix-turn-helix motif \| 2P5T\_E fragment of PezA helix-turn-helix motif postsegregational killing system, phosphoryltransferase, helix-turn-helix; pdb70 \| 2WIU\_B \| 97.3 \| 3e-07 \| 2e-11 \| 49.4 \| 61 \| (3, 63) \| 180 \| (15, 75) \| 88 \| PROTEIN HIPA (E.C.2.7.11.1), HTH-TYPE TRANSCRIPTIONAL \| 2WIU\_B PROTEIN HIPA (E.C.2.7.11.1), HTH-TYPE TRANSCRIPTIONAL TRANSFERASE TRANSCRIPTION COMPLEX, SERINE KINASE HET: CL; pdb70 \| 5D4Z\_M \| 97.3 \| 3.2e-07 \| 2.2e-11 \| 51.5 \| 61 \| (2, 62) \| 180 \| (6, 68) \| 107 \| Repressor \| 5D4Z\_M Repressor Repressor, DNA BINDING PROTEIN; pdb70 \| 2XI8\_B \| 97.3 \| 3.3e-07 \| 2.2e-11 \| 45.9 \| 61 \| (2, 62) \| 180 \| (3, 63) \| 66 \| PUTATIVE TRANSCRIPTION REGULATOR \| 2XI8\_B PUTATIVE TRANSCRIPTION REGULATOR TRANSCRIPTION, HTH DNA-BINDING MOTIF HET: GOL; pdb70 \| 4MCT\_C \| 97.3 \| 3.5e-07 \| 2.4e-11 \| 52.8 \| 62 \| (2, 63) \| 180 \| (10, 71) \| 125 \| Antidote protein, Killer protein \| 4MCT\_C Antidote protein, Killer protein bacterial toxins, biofilms, cell metabolism; pdb70 \| 3OMT\_A \| 97.3 \| 3.9e-07 \| 2.6e-11 \| 46.7 \| 60 \| (3, 62) \| 180 \| (11, 70) \| 73 \| uncharacterized protein \| 3OMT\_A uncharacterized protein structural genomics, PSI-2, Protein Structure; pdb70 \| 1Y7Y\_A \| 97.3 \| 4.1e-07 \| 2.7e-11 \| 46.8 \| 58 \| (3, 60) \| 180 \| (16, 73) \| 74 \| C.AhdI \| 1Y7Y\_A C.AhdI HELIX-TURN-HELIX, DNA-BINDING PROTEIN, TRANSCRIPTIONAL REGULATOR; pdb70 \| 3OMT\_B \| 97.3 \| 4.1e-07 \| 2.7e-11 \| 46.7 \| 60 \| (3, 62) \| 180 \| (11, 70) \| 73 \| uncharacterized protein \| 3OMT\_B uncharacterized protein structural genomics, PSI-2, Protein Structure; pdb70 \| 6B9T\_D \| 97.2 \| 4.2e-07 \| 2.8e-11 \| 64.1 \| 64 \| (1, 64) \| 180 \| (16, 79) \| 457 \| Methylphosphonate synthase (E.C.1.13.11.73) \| 6B9T\_D Methylphosphonate synthase (E.C.1.13.11.73) Phosphonate, Methylphosphonate, Iron, OXIDOREDUCTASE HET: 2HE, FMT; pdb70 \| 6B9T\_C \| 97.2 \| 4.5e-07 \| 3e-11 \| 64.0 \| 64 \| (1, 64) \| 180 \| (16, 79) \| 457 \| Methylphosphonate synthase (E.C.1.13.11.73) \| 6B9T\_C Methylphosphonate synthase (E.C.1.13.11.73) Phosphonate, Methylphosphonate, Iron, OXIDOREDUCTASE HET: FMT, 2HE; pdb70 \| 3F6W\_C \| 97.2 \| 4.6e-07 \| 3.1e-11 \| 48.0 \| 61 \| (3, 63) \| 180 \| (17, 77) \| 83 \| XRE-family like protein \| 3F6W\_C XRE-family like protein helix-turn-helix DNA binding protein HET: BTB, MSE; pdb70 \| 5UK3\_G \| 97.2 \| 4.8e-07 \| 3.2e-11 \| 55.5 \| 62 \| (2, 63) \| 180 \| (21, 82) \| 176 \| Uncharacterized protein \| 5UK3\_G Uncharacterized protein Tetranychus urticae, cyanase, agricultural pest; pdb70 \| 5UK3\_I \| 97.2 \| 4.8e-07 \| 3.2e-11 \| 55.5 \| 62 \| (2, 63) \| 180 \| (21, 82) \| 176 \| Uncharacterized protein \| 5UK3\_I Uncharacterized protein Tetranychus urticae, cyanase, agricultural pest; pdb70 \| 1ADR\_A \| 97.2 \| 5.9e-07 \| 3.9e-11 \| 46.6 \| 61 \| (3, 63) \| 180 \| (8, 68) \| 76 \| P22 C2 REPRESSOR (AMINO-TERMINAL DNA-BINDING \| 1ADR\_A P22 C2 REPRESSOR (AMINO-TERMINAL DNA-BINDING TRANSCRIPTION REGULATION; pdb70 \| 6RNZ\_B \| 97.2 \| 6.5e-07 \| 4.3e-11 \| 44.7 \| 59 \| (3, 61) \| 180 \| (5, 63) \| 66 \| HTH-type transcriptional regulator DdrOC \| 6RNZ\_B HTH-type transcriptional regulator DdrOC ----, DNA BINDING PROTEIN HET: GOL; pdb70 \| 3JXC\_L \| 97.2 \| 7e-07 \| 4.6e-11 \| 44.8 \| 59 \| (3, 61) \| 180 \| (7, 65) \| 67 \| Repressor protein C2/DNA Complex, THALLIUM \| 3JXC\_L Repressor protein C2/DNA Complex, THALLIUM protein-DNA complex, DNA-binding, Repressor, Transcription; pdb70 \| 4RYK\_A \| 97.2 \| 7e-07 \| 4.7e-11 \| 59.0 \| 62 \| (1, 63) \| 180 \| (10, 71) \| 306 \| Lmo0325 protein \| 4RYK\_A Lmo0325 protein Structural Genomics, PSI-Biology, Midwest Center HET: TLA, SUC; pdb70 \| 6JQ1\_B \| 97.2 \| 7e-07 \| 4.7e-11 \| 51.8 \| 62 \| (2, 63) \| 180 \| (7, 68) \| 132 \| Transcriptional regulator, XRE family \| 6JQ1\_B Transcriptional regulator, XRE family Transcription factor, Xre, HTH, Dimerization; pdb70 \| 6QER\_C \| 97.1 \| 7.6e-07 \| 5e-11 \| 58.3 \| 61 \| (1, 61) \| 180 \| (7, 68) \| 310 \| Transcriptional regulator \| 6QER\_C Transcriptional regulator Quorum sensing, Competence, TPR domain; pdb70 \| 5JUF\_A \| 97.1 \| 8.2e-07 \| 5.4e-11 \| 58.2 \| 61 \| (1, 61) \| 180 \| (7, 68) \| 310 \| Transcriptional regulator \| 5JUF\_A Transcriptional regulator RNPP, Transcriptional activator, TPR, HTH HET: SO4; pdb70 \| 3JXB\_C \| 97.1 \| 8.7e-07 \| 5.7e-11 \| 44.4 \| 60 \| (2, 61) \| 180 \| (6, 65) \| 67 \| Repressor protein C2/DNA Complex \| 3JXB\_C Repressor protein C2/DNA Complex protein-DNA complex, DNA-binding, Repressor, Transcription; pdb70 \| 2R1J\_R \| 97.1 \| 8.9e-07 \| 5.9e-11 \| 44.5 \| 59 \| (3, 61) \| 180 \| (8, 66) \| 68 \| Repressor protein C2/DNA Complex \| 2R1J\_R Repressor protein C2/DNA Complex Protein-DNA complex, Helix-turn-helix, DNA-binding, Repressor; pdb70 \| 2XIU\_A \| 97.1 \| 9.3e-07 \| 6.1e-11 \| 44.0 \| 61 \| (2, 62) \| 180 \| (3, 63) \| 66 \| CYLR2 \| 2XIU\_A CYLR2 DNA BINDING PROTEIN, HTH-DNA BINDING HET: MTN; pdb70 \| 6HU8\_A \| 97.1 \| 1e-06 \| 6.6e-11 \| 58.0 \| 62 \| (1, 62) \| 180 \| (7, 69) \| 310 \| Uncharacterized protein \| 6HU8\_A Uncharacterized protein RNPP family TPR domain HTH; pdb70 \| 6H49\_A \| 97.1 \| 9.9e-07 \| 6.7e-11 \| 53.0 \| 62 \| (2, 63) \| 180 \| (16, 77) \| 157 \| Orf20 \| 6H49\_A Orf20 SaPI, Repressor, STRUCTURAL PROTEIN HET: SO4; pdb70 \| 6H4C\_D \| 97.1 \| 9.9e-07 \| 6.7e-11 \| 53.0 \| 62 \| (2, 63) \| 180 \| (16, 77) \| 157 \| dUTPase, Orf20 \| 6H4C\_D dUTPase, Orf20 SaPI, Repressor, Complex, STRUCTURAL PROTEIN HET: PEG; pdb70 \| 5W4M\_B \| 97.1 \| 1.1e-06 \| 7.1e-11 \| 57.0 \| 61 \| (1, 62) \| 180 \| (5, 65) \| 280 \| Transcriptional regulator \| 5W4M\_B Transcriptional regulator DNA BINDING, PHEROMONE BINDING, REPEAT HET: GOL, DMS; pdb70 \| 4Z5H\_A \| 97.1 \| 1.1e-06 \| 7.5e-11 \| 44.8 \| 58 \| (3, 60) \| 180 \| (13, 70) \| 72 \| E.coli transcription repressor, HipB subunit \| 4Z5H\_A E.coli transcription repressor, HipB subunit HTH motif, Transcription repressor, HipA; pdb70 \| 2B5A\_C \| 97.1 \| 1.2e-06 \| 7.7e-11 \| 45.4 \| 60 \| (3, 62) \| 180 \| (13, 72) \| 77 \| C.BclI \| 2B5A\_C C.BclI HELIX-TURN-HELIX motif, GENE REGULATION; pdb70 \| 3BS3\_A \| 97.0 \| 1.3e-06 \| 9e-11 \| 44.9 \| 60 \| (3, 62) \| 180 \| (13, 72) \| 76 \| Putative DNA-binding protein \| 3BS3\_A Putative DNA-binding protein DNA-binding, XRE-family, Structural Genomics, PSI-2; pdb70 \| 5WOQ\_B \| 97.0 \| 1.4e-06 \| 9.4e-11 \| 49.8 \| 62 \| (2, 63) \| 180 \| (18, 79) \| 120 \| Transcriptional regulator ClgR \| 5WOQ\_B Transcriptional regulator ClgR NIAID, structural genomics, transcription factor; pdb70 \| 5WOQ\_C \| 97.0 \| 1.4e-06 \| 9.4e-11 \| 49.8 \| 62 \| (2, 63) \| 180 \| (18, 79) \| 120 \| Transcriptional regulator ClgR \| 5WOQ\_C Transcriptional regulator ClgR NIAID, structural genomics, transcription factor; pdb70 \| 4YG7\_C \| 97.0 \| 1.4e-06 \| 9.4e-11 \| 44.4 \| 57 \| (3, 59) \| 180 \| (12, 68) \| 71 \| HipB, HipA/DNA Complex \| 4YG7\_C HipB, HipA/DNA Complex persistence, multidrug tolerance, autorepression, promoter; pdb70 \| 4Z59\_A \| 97.0 \| 1.4e-06 \| 9.4e-11 \| 44.4 \| 57 \| (3, 59) \| 180 \| (12, 68) \| 71 \| E.coli transcription repressor, HipB/DNA Complex \| 4Z59\_A E.coli transcription repressor, HipB/DNA Complex HTH motif, Transcription repressor, HipA; pdb70 \| 2GRM\_A \| 97.0 \| 1.5e-06 \| 9.9e-11 \| 57.9 \| 61 \| (1, 62) \| 180 \| (4, 64) \| 317 \| PrgX, peptide \| 2GRM\_A PrgX, peptide receptor, inhibitor, TRANSCRIPTION; pdb70 \| 5Y6A\_A \| 97.0 \| 1.5e-06 \| 1e-10 \| 51.8 \| 62 \| (2, 63) \| 180 \| (5, 67) \| 151 \| chain A and B \| 5Y6A\_A chain A and B Unknown function, VIRAL PROTEIN; pdb70 \| 2BNM\_B \| 97.0 \| 1.5e-06 \| 1e-10 \| 54.1 \| 62 \| (2, 63) \| 180 \| (12, 74) \| 198 \| EPOXIDASE \| 2BNM\_B EPOXIDASE OXIDOREDUCTASE, EPOXIDASE, CUPIN, HTH, CATION-DEPENDANT HET: SO4; pdb70 \| 4J1X\_C \| 97.0 \| 1.7e-06 \| 1.2e-10 \| 53.8 \| 62 \| (2, 63) \| 180 \| (11, 73) \| 197 \| Epoxidase \| 4J1X\_C Epoxidase keto product, Hydroxypropylphosphonic acid epoxidase HET: 1JJ; pdb70 \| 1B0N\_A \| 97.0 \| 1.8e-06 \| 1.2e-10 \| 48.4 \| 62 \| (2, 63) \| 180 \| (3, 65) \| 111 \| PROTEIN (SINR PROTEIN) \| 1B0N\_A PROTEIN (SINR PROTEIN) TRANSCRIPTION REGULATOR, ANTAGONIST, SPORULATION; pdb70 \| 3KXA\_A \| 97.0 \| 1.8e-06 \| 1.2e-10 \| 50.9 \| 61 \| (3, 63) \| 180 \| (71, 131) \| 141 \| Putative uncharacterized protein \| 3KXA\_A Putative uncharacterized protein NGO0477, Neisseria gonorrhoeae, New protein HET: CL, ASN, UNX, MSE; pdb70 \| 4YV9\_D \| 97.0 \| 1.8e-06 \| 1.2e-10 \| 56.1 \| 61 \| (2, 63) \| 180 \| (6, 66) \| 284 \| Transcriptional regulator, Cyclosporin A \| 4YV9\_D Transcriptional regulator, Cyclosporin A DNA binding, pheromone binding, repeat HET: DAL, SO4, MVA, SAR, MSE, ABA, BMT, MLE; pdb70 \| 5TN0\_A \| 97.0 \| 1.9e-06 \| 1.2e-10 \| 43.3 \| 62 \| (2, 63) \| 180 \| (3, 65) \| 69 \| HTH-type transcriptional regulator SinR \| 5TN0\_A HTH-type transcriptional regulator SinR biofilm formation, TRANSCRIPTION; pdb70 \| 2IUO\_B \| 96.9 \| 1.9e-06 \| 1.3e-10 \| 51.7 \| 61 \| (2, 62) \| 180 \| (15, 75) \| 156 \| CYANATE HYDRATASE (E.C.4.2.1.104) \| 2IUO\_B CYANATE HYDRATASE (E.C.4.2.1.104) LYASE, CYANATE DEGRADATION HET: CL, SO4, BR; pdb70 \| 1R63\_A \| 96.9 \| 2.1e-06 \| 1.4e-10 \| 42.3 \| 59 \| (2, 61) \| 180 \| (3, 61) \| 63 \| REPRESSOR PROTEIN FROM BACTERIOPHAGE 434 \| 1R63\_A REPRESSOR PROTEIN FROM BACTERIOPHAGE 434 GENE REGULATING PROTEIN, PHAGE 434; pdb70 \| 1DW9\_C \| 96.9 \| 2.1e-06 \| 1.4e-10 \| 51.5 \| 61 \| (2, 62) \| 180 \| (15, 75) \| 156 \| CYANATE LYASE (E.C.4.3.99.1) \| 1DW9\_C CYANATE LYASE (E.C.4.3.99.1) LYASE, CYANATE DEGRADATION, STRUCTURAL GENOMICS HET: CL, SO4, MSE; pdb70 \| 1DWK\_J \| 96.9 \| 2.1e-06 \| 1.4e-10 \| 51.5 \| 61 \| (2, 62) \| 180 \| (15, 75) \| 156 \| CYANATE HYDRATASE (E.C.4.2.1.104) \| 1DWK\_J CYANATE HYDRATASE (E.C.4.2.1.104) LYASE, CYANATE DEGRADATION, PSI, PROTEIN HET: OXL, SO4, MSE; pdb70 \| 3OP9\_A \| 96.9 \| 2.2e-06 \| 1.5e-10 \| 48.1 \| 61 \| (3, 63) \| 180 \| (12, 72) \| 114 \| Pli0006 protein \| 3OP9\_A Pli0006 protein Structural Genomics, PSI-2, Protein Structure; pdb70 \| 6F8H\_B \| 96.9 \| 2.4e-06 \| 1.6e-10 \| 47.6 \| 62 \| (2, 63) \| 180 \| (18, 80) \| 105 \| XRE family transcriptional regulator \| 6F8H\_B XRE family transcriptional regulator GraA, HigA, antitoxin; pdb70 \| 3MLF\_A \| 96.9 \| 2.5e-06 \| 1.7e-10 \| 48.0 \| 61 \| (3, 63) \| 180 \| (26, 86) \| 111 \| transcriptional regulator \| 3MLF\_A transcriptional regulator structural genomics, Helix-turn-helix XRE-family like; pdb70 \| 3MLF\_D \| 96.9 \| 2.5e-06 \| 1.7e-10 \| 48.0 \| 61 \| (3, 63) \| 180 \| (26, 86) \| 111 \| transcriptional regulator \| 3MLF\_D transcriptional regulator structural genomics, Helix-turn-helix XRE-family like HET: MSE; pdb70 \| 1RIO\_B \| 96.9 \| 2.6e-06 \| 1.8e-10 \| 46.6 \| 55 \| (9, 63) \| 180 \| (27, 81) \| 98 \| sigma factor SigA/Repressor protein CI/DNA \| 1RIO\_B sigma factor SigA/Repressor protein CI/DNA HELIX-TURN-HELIX, TRANSCRIPTION ACTIVATION, transcription-DNA COMPLEX HET: MSE; pdb70 \| 2ICT\_A \| 96.9 \| 2.7e-06 \| 1.8e-10 \| 46.2 \| 61 \| (3, 63) \| 180 \| (11, 71) \| 94 \| Putative HTH-type transcriptional regulator yddM \| 2ICT\_A Putative HTH-type transcriptional regulator yddM helix-turn-helix, Structural Genomics, PSI-2, Protein; pdb70 \| 1R69\_A \| 96.9 \| 2.7e-06 \| 1.8e-10 \| 42.8 \| 61 \| (2, 63) \| 180 \| (3, 63) \| 69 \| 434 REPRESSOR (AMINO-TERMINAL DOMAIN) (R1-69) \| 1R69\_A 434 REPRESSOR (AMINO-TERMINAL DOMAIN) (R1-69) GENE REGULATING PROTEIN; pdb70 \| 5D50\_A \| 96.9 \| 2.9e-06 \| 1.9e-10 \| 52.8 \| 63 \| (2, 64) \| 180 \| (10, 73) \| 199 \| Repressor, Anti-repressor protein \| 5D50\_A Repressor, Anti-repressor protein Repressor, Anti-repressor, complex, DNA BINDING; pdb70 \| 5D50\_B \| 96.9 \| 2.9e-06 \| 1.9e-10 \| 52.8 \| 63 \| (2, 64) \| 180 \| (10, 73) \| 199 \| Repressor, Anti-repressor protein \| 5D50\_B Repressor, Anti-repressor protein Repressor, Anti-repressor, complex, DNA BINDING; pdb70 \| 3IVP\_D \| 96.9 \| 2.9e-06 \| 2e-10 \| 48.6 \| 61 \| (3, 63) \| 180 \| (15, 75) \| 126 \| Putative transposon-related DNA-binding protein \| 3IVP\_D Putative transposon-related DNA-binding protein APC62618, transposon-related DNA-binding, Clostridium difficile HET: MSE, PG4; pdb70 \| 3FYA\_A \| 96.9 \| 2.9e-06 \| 2e-10 \| 46.5 \| 61 \| (3, 63) \| 180 \| (31, 91) \| 99 \| Regulatory protein \| 3FYA\_A Regulatory protein transcriptional regulator, helix-turn-helix, Restriction-modification, TRANSCRIPTION; pdb70 \| 2MEZ\_A \| 96.8 \| 3e-06 \| 2e-10 \| 47.2 \| 61 \| (3, 63) \| 180 \| (13, 73) \| 106 \| Multiprotein Bridging Factor (MBP-like) \| 2MEZ\_A Multiprotein Bridging Factor (MBP-like) Multiple Binding Factor MBF1, Helix-turn-Helix; pdb70 \| 2EWT\_A \| 96.8 \| 3.2e-06 \| 2.1e-10 \| 42.7 \| 57 \| (4, 60) \| 180 \| (12, 70) \| 71 \| putative DNA-binding protein \| 2EWT\_A putative DNA-binding protein the DNA-binding domain of BldD HET: SO4; pdb70 \| 4O8B\_A \| 96.8 \| 3.4e-06 \| 2.3e-10 \| 48.3 \| 62 \| (2, 63) \| 180 \| (3, 68) \| 122 \| Uncharacterized protein \| 4O8B\_A Uncharacterized protein HTH motif, transcriptional activator, promoter; pdb70 \| 1X57\_A \| 96.8 \| 3.5e-06 \| 2.3e-10 \| 45.2 \| 61 \| (3, 63) \| 180 \| (16, 76) \| 91 \| Endothelial differentiation-related factor 1 \| 1X57\_A Endothelial differentiation-related factor 1 EDF1, HMBF1alpha, helix-turn-helix, Structural Genomics; pdb70 \| 3O9X\_A \| 96.8 \| 4.1e-06 \| 2.7e-10 \| 48.4 \| 57 \| (3, 60) \| 180 \| (74, 130) \| 133 \| Uncharacterized HTH-type transcriptional regulator ygiT/DNA \| 3O9X\_A Uncharacterized HTH-type transcriptional regulator ygiT/DNA HTH-XRE DNA binding motif, Transcriptional HET: GOL; pdb70 \| 5FD4\_B \| 96.8 \| 4.1e-06 \| 2.7e-10 \| 55.8 \| 61 \| (2, 62) \| 180 \| (32, 93) \| 324 \| ComR \| 5FD4\_B ComR Streptococcus, Competence, Quorum sensing, ComR; pdb70 \| 3FYM\_A \| 96.8 \| 4.5e-06 \| 3e-10 \| 48.3 \| 62 \| (2, 63) \| 180 \| (5, 72) \| 130 \| Putative uncharacterized protein \| 3FYM\_A Putative uncharacterized protein HTH DNA binding, DNA BINDING; pdb70 \| 1ZUG\_A \| 96.7 \| 4.8e-06 \| 3.2e-10 \| 42.2 \| 60 \| (3, 63) \| 180 \| (6, 65) \| 71 \| PHAGE 434 CRO PROTEIN \| 1ZUG\_A PHAGE 434 CRO PROTEIN GENE REGULATING PROTEIN, TRANSCRIPTION REGULATION; pdb70 \| 2CRO\_A \| 96.7 \| 4.8e-06 \| 3.2e-10 \| 42.2 \| 60 \| (3, 63) \| 180 \| (6, 65) \| 71 \| 434 CRO PROTEIN \| 2CRO\_A 434 CRO PROTEIN GENE REGULATING PROTEIN; pdb70 \| 5Y69\_A \| 96.7 \| 5.4e-06 \| 3.6e-10 \| 49.2 \| 62 \| (2, 63) \| 180 \| (5, 67) \| 151 \| chain A \| 5Y69\_A chain A UNKNOWN FUCTION, VIRAL PROTEIN HET: MSE; pdb70 \| 3CEC\_A \| 96.7 \| 5.8e-06 \| 3.9e-10 \| 45.8 \| 61 \| (3, 63) \| 180 \| (21, 81) \| 104 \| Putative antidote protein of plasmid \| 3CEC\_A Putative antidote protein of plasmid Structural genomics, Joint Center for; pdb70 \| 1S4K\_B \| 96.7 \| 6.3e-06 \| 4.2e-10 \| 46.9 \| 59 \| (2, 60) \| 180 \| (5, 65) \| 120 \| putative cytoplasmic protein ydil \| 1S4K\_B putative cytoplasmic protein ydil structural genomics, MCSG, Salmonella typhimurium HET: MSE; pdb70 \| 3G5G\_G \| 96.7 \| 6.4e-06 \| 4.3e-10 \| 45.1 \| 60 \| (3, 62) \| 180 \| (31, 90) \| 99 \| Regulatory protein \| 3G5G\_G Regulatory protein transcriptional regulator, helix-turn-helix, Restriction-modification, TRANSCRIPTION; pdb70 \| 3G5G\_H \| 96.7 \| 6.4e-06 \| 4.3e-10 \| 45.1 \| 60 \| (3, 62) \| 180 \| (31, 90) \| 99 \| Regulatory protein \| 3G5G\_H Regulatory protein transcriptional regulator, helix-turn-helix, Restriction-modification, TRANSCRIPTION; pdb70 \| 1LMB\_4 \| 96.7 \| 6.6e-06 \| 4.4e-10 \| 44.0 \| 55 \| (9, 63) \| 180 \| (26, 80) \| 92 \| LAMBDA REPRESSOR/DNA COMPLEX \| 1LMB\_4 LAMBDA REPRESSOR/DNA COMPLEX PROTEIN-DNA COMPLEX, DOUBLE HELIX, TRANSCRIPTION-DNA; pdb70 \| 5IFG\_B \| 96.7 \| 6.5e-06 \| 4.4e-10 \| 48.2 \| 56 \| (3, 60) \| 180 \| (82, 137) \| 138 \| mRNA interferase HigB (E.C.3.1.-.-), Antitoxin \| 5IFG\_B mRNA interferase HigB (E.C.3.1.-.-), Antitoxin mRNA interferase, HYDROLASE-ANTITOXIN complex HET: MSE; pdb70 \| 6IRP\_A \| 96.7 \| 6.5e-06 \| 4.4e-10 \| 48.2 \| 56 \| (3, 60) \| 180 \| (82, 137) \| 138 \| Antitoxin HigA \| 6IRP\_A Antitoxin HigA Antoxin, ANTITOXIN HET: MSE; pdb70 \| 6JQ4\_A \| 96.7 \| 6.5e-06 \| 4.4e-10 \| 48.2 \| 56 \| (3, 60) \| 180 \| (82, 137) \| 138 \| Antitoxin HigA \| 6JQ4\_A Antitoxin HigA Helix-Turn-Helix Domain, ANTITOXIN; pdb70 \| 5V5T\_A \| 96.7 \| 6.8e-06 \| 4.6e-10 \| 54.6 \| 61 \| (2, 63) \| 180 \| (11, 71) \| 306 \| Conserved domain protein \| 5V5T\_A Conserved domain protein TRANSCRIPTION REGULATOR, TRANSCRIPTION HET: MSE; pdb70 \| 5V5T\_D \| 96.7 \| 6.8e-06 \| 4.6e-10 \| 54.6 \| 61 \| (2, 63) \| 180 \| (11, 71) \| 306 \| Conserved domain protein \| 5V5T\_D Conserved domain protein TRANSCRIPTION REGULATOR, TRANSCRIPTION HET: MSE; pdb70 \| 5V5U\_B \| 96.7 \| 6.8e-06 \| 4.6e-10 \| 54.6 \| 61 \| (2, 63) \| 180 \| (11, 71) \| 306 \| Conserved domain protein \| 5V5U\_B Conserved domain protein TRANSCRIPTION REGULATOR, TRANSCRIPTION; pdb70 \| 3ZHI\_A \| 96.6 \| 8e-06 \| 5.4e-10 \| 43.3 \| 60 \| (3, 62) \| 180 \| (8, 76) \| 83 \| CI \| 3ZHI\_A CI TRANSCRIPTION, TRANSCRIPTION REGULATION; pdb70 \| 3F51\_E \| 96.6 \| 8.4e-06 \| 5.6e-10 \| 46.0 \| 61 \| (3, 63) \| 180 \| (31, 91) \| 117 \| clp gene regulator (ClgR) \| 3F51\_E clp gene regulator (ClgR) gene regulator, helix-turn-helix, transcriptional activator HET: MPD; pdb70 \| 3F52\_E \| 96.6 \| 8.4e-06 \| 5.6e-10 \| 46.0 \| 61 \| (3, 63) \| 180 \| (31, 91) \| 117 \| clp gene regulator (ClgR) \| 3F52\_E clp gene regulator (ClgR) gene regulator, helix-turn-helix motif, transcriptional HET: GOL; pdb70 \| 4MCX\_C \| 96.6 \| 8.4e-06 \| 5.6e-10 \| 44.9 \| 61 \| (3, 63) \| 180 \| (11, 71) \| 104 \| Host inhibition of growth A \| 4MCX\_C Host inhibition of growth A bacterial toxins, biofilms, cell metabolism; pdb70 \| 3PXP\_A \| 96.6 \| 8.9e-06 \| 6e-10 \| 53.8 \| 62 \| (2, 63) \| 180 \| (8, 79) \| 292 \| Helix-turn-helix domain protein \| 3PXP\_A Helix-turn-helix domain protein DNA-BINDING, BASIC HELIX-LOOP-HELIX MOTIF, BHLH HET: MYR, EDO; pdb70 \| 3R1F\_N \| 96.6 \| 9.7e-06 \| 6.5e-10 \| 47.2 \| 60 \| (3, 62) \| 180 \| (10, 81) \| 135 \| ESX-1 secretion-associated regulator EspR \| 3R1F\_N ESX-1 secretion-associated regulator EspR helix-turn-helix, transcription factor, helix-turn-helix transcription HET: MSE; pdb70 \| 2L49\_A \| 96.6 \| 9.8e-06 \| 6.6e-10 \| 44.1 \| 56 \| (3, 58) \| 180 \| (7, 62) \| 99 \| C protein \| 2L49\_A C protein P2 bacteriophage, P2 C, direct; pdb70 \| 2XCJ\_B \| 96.6 \| 9.8e-06 \| 6.6e-10 \| 44.1 \| 56 \| (3, 58) \| 180 \| (7, 62) \| 99 \| C PROTEIN \| 2XCJ\_B C PROTEIN DIRECT REPEATS, REPRESSOR, HELIX-TURN-HELIX, VIRAL HET: GOL; pdb70 \| 3VK0\_A \| 96.6 \| 1e-05 \| 6.7e-10 \| 45.6 \| 62 \| (3, 64) \| 180 \| (24, 85) \| 114 \| Transcriptional regulator \| 3VK0\_A Transcriptional regulator HTH motif, XRE transcription factor; pdb70 \| 4MCX\_A \| 96.6 \| 1e-05 \| 6.7e-10 \| 44.5 \| 61 \| (3, 63) \| 180 \| (11, 71) \| 104 \| Host inhibition of growth A \| 4MCX\_A Host inhibition of growth A bacterial toxins, biofilms, cell metabolism; pdb70 \| 3TYS\_A \| 96.5 \| 1.1e-05 \| 7.1e-10 \| 43.2 \| 58 \| (4, 62) \| 180 \| (28, 85) \| 88 \| Predicted transcriptional regulator \| 3TYS\_A Predicted transcriptional regulator Structural Genomics, Center for Structural; pdb70 \| 2JVL\_A \| 96.5 \| 1.1e-05 \| 7.3e-10 \| 44.9 \| 61 \| (3, 63) \| 180 \| (37, 99) \| 107 \| TrMBF1 \| 2JVL\_A TrMBF1 MBF1, Coactivator, Trichoderma reesei, helix-turn-helix; pdb70 \| 6AF3\_D \| 96.5 \| 1.1e-05 \| 7.6e-10 \| 43.8 \| 59 \| (4, 62) \| 180 \| (35, 93) \| 97 \| HigB toxin, HigA antitoxin \| 6AF3\_D HigB toxin, HigA antitoxin Toxin-Antitoxin complex, TOXIN HET: MSE; pdb70 \| 6AF4\_B \| 96.5 \| 1.1e-05 \| 7.6e-10 \| 43.8 \| 59 \| (4, 62) \| 180 \| (35, 93) \| 97 \| HigB, HigA \| 6AF4\_B HigB, HigA Toxin-Antitoxin complex, TOXIN; pdb70 \| 3QYX\_B \| 96.5 \| 1.1e-05 \| 7.6e-10 \| 46.8 \| 61 \| (2, 62) \| 180 \| (7, 79) \| 133 \| ESX-1 secretion-associated regulator EspR/DNA complex \| 3QYX\_B ESX-1 secretion-associated regulator EspR/DNA complex N-terminal HTH motif, C-terminal dimerization; pdb70 \| 4NDW\_B \| 96.5 \| 1.2e-05 \| 8.5e-10 \| 47.9 \| 60 \| (3, 62) \| 180 \| (28, 99) \| 153 \| Nucleoid-associated protein EspR \| 4NDW\_B Nucleoid-associated protein EspR ALL HELICAL, HELIX-TURN-HELIX MOTIF, TRANSACTIONAL; pdb70 \| 6CF1\_B \| 96.5 \| 1.3e-05 \| 8.5e-10 \| 47.0 \| 61 \| (3, 63) \| 180 \| (45, 105) \| 138 \| Antitoxin HigA \| 6CF1\_B Antitoxin HigA Helix-turn-helix motif, transcriptional regulator, toxin-antitoxin; pdb70 \| 3PXP\_B \| 96.5 \| 1.3e-05 \| 9e-10 \| 52.9 \| 62 \| (2, 63) \| 180 \| (8, 79) \| 292 \| Helix-turn-helix domain protein \| 3PXP\_B Helix-turn-helix domain protein DNA-BINDING, BASIC HELIX-LOOP-HELIX MOTIF, BHLH HET: MSE, EDO, MYR; pdb70 \| 3FMY\_A \| 96.5 \| 1.4e-05 \| 9.3e-10 \| 40.5 \| 57 \| (3, 60) \| 180 \| (14, 70) \| 73 \| Uncharacterized HTH-type transcriptional regulator ygiT \| 3FMY\_A Uncharacterized HTH-type transcriptional regulator ygiT helix-turn-helix, DNA-binding, Transcription, Transcription regulation; pdb70 \| 2AW6\_B \| 96.5 \| 1.5e-05 \| 9.8e-10 \| 53.1 \| 60 \| (2, 62) \| 180 \| (5, 64) \| 317 \| PrgX \| 2AW6\_B PrgX Repressor, pheromone, DNA binding, regulatory; pdb70 \| 5A7L\_A \| 96.4 \| 1.6e-05 \| 1e-09 \| 41.3 \| 61 \| (3, 63) \| 180 \| (7, 76) \| 80 \| CI \| 5A7L\_A CI TRANSCRIPTION; pdb70 \| 3QWG\_A \| 96.4 \| 1.6e-05 \| 1.1e-09 \| 45.5 \| 62 \| (2, 63) \| 180 \| (7, 80) \| 123 \| ESX-1 secretion-associated regulator EspR \| 3QWG\_A ESX-1 secretion-associated regulator EspR N-terminal helix-turn-helix motif, Transcription factor; pdb70 \| 2KZ8\_A \| 96.4 \| 1.7e-05 \| 1.2e-09 \| 45.5 \| 58 \| (3, 61) \| 180 \| (72, 129) \| 131 \| Uncharacterized HTH-type transcriptional regulator ygiT \| 2KZ8\_A Uncharacterized HTH-type transcriptional regulator ygiT Zinc finger, Helix turn helix; pdb70 \| 2H8R\_B \| 96.4 \| 1.8e-05 \| 1.2e-09 \| 50.2 \| 61 \| (2, 62) \| 180 \| (33, 93) \| 221 \| Hepatocyte nuclear factor 1-beta/DNA Complex \| 2H8R\_B Hepatocyte nuclear factor 1-beta/DNA Complex trasncription factor, POU, homeo, protein-DNA; pdb70 \| 2EBY\_A \| 96.4 \| 2e-05 \| 1.4e-09 \| 44.0 \| 58 \| (6, 63) \| 180 \| (17, 74) \| 113 \| Putative HTH-type transcriptional regulator ybaQ \| 2EBY\_A Putative HTH-type transcriptional regulator ybaQ Hypothetical protein, JW0472, Structural Genomics HET: SO4; pdb70 \| 6HPC\_A \| 96.4 \| 2.2e-05 \| 1.4e-09 \| 45.2 \| 55 \| (4, 58) \| 180 \| (81, 135) \| 138 \| Antitoxin HicB \| 6HPC\_A Antitoxin HicB dna binding, helix-turn-helix, HTH, ANTITOXIN HET: MSE; pdb70 \| 1NEQ\_A \| 96.3 \| 2.3e-05 \| 1.5e-09 \| 39.8 \| 58 \| (3, 63) \| 180 \| (12, 69) \| 74 \| DNA-BINDING PROTEIN NER \| 1NEQ\_A DNA-BINDING PROTEIN NER DNA-BINDING PROTEIN; pdb70 \| 2AUW\_B \| 96.3 \| 2.3e-05 \| 1.5e-09 \| 47.5 \| 59 \| (3, 62) \| 180 \| (93, 151) \| 170 \| hypothetical protein NE0471 \| 2AUW\_B hypothetical protein NE0471 alpha-beta structure, Structural Genomics, PSI HET: GOL; pdb70 \| 3QQ6\_A \| 96.3 \| 2.4e-05 \| 1.6e-09 \| 40.4 \| 59 \| (4, 62) \| 180 \| (14, 73) \| 78 \| HTH-type transcriptional regulator sinR \| 3QQ6\_A HTH-type transcriptional regulator sinR Helix-Turn-Helix motif, biofilm, repressor, transcriptional; pdb70 \| 4ICH\_A \| 96.3 \| 2.5e-05 \| 1.7e-09 \| 51.7 \| 62 \| (3, 64) \| 180 \| (33, 94) \| 311 \| Transcriptional regulator \| 4ICH\_A Transcriptional regulator Structural Genomics, PSI-Biology, Midwest Center HET: MSE, B3P; pdb70 \| 6CHV\_B \| 96.3 \| 2.5e-05 \| 1.7e-09 \| 44.5 \| 60 \| (3, 62) \| 180 \| (28, 87) \| 121 \| Antitoxin HigA/DNA Complex \| 6CHV\_B Antitoxin HigA/DNA Complex Helix-turn-helix DNA binding protein, DNA-protein; pdb70 \| 4PU7\_B \| 96.3 \| 2.6e-05 \| 1.7e-09 \| 44.3 \| 59 \| (4, 63) \| 180 \| (53, 111) \| 118 \| Toxin-antitoxin system antidote transcriptional repressor \| 4PU7\_B Toxin-antitoxin system antidote transcriptional repressor Toxin Antitoxin System, TOXIN; pdb70 \| 3EUS\_A \| 96.3 \| 2.6e-05 \| 1.7e-09 \| 40.9 \| 59 \| (3, 61) \| 180 \| (17, 76) \| 86 \| DNA-binding protein \| 3EUS\_A DNA-binding protein DNA binding protein, STRUCTURAL GENOMICS; pdb70 \| 2A6C\_A \| 96.3 \| 2.7e-05 \| 1.8e-09 \| 40.8 \| 58 \| (4, 61) \| 180 \| (22, 80) \| 83 \| Helix-turn-helix motif \| 2A6C\_A Helix-turn-helix motif PUTATIVE TRANSCRIPTIONAL REGULATOR, STRUCTURAL GENOMICS HET: GOL, CIT, MSE; pdb70 \| 4PU4\_D \| 96.3 \| 2.8e-05 \| 1.9e-09 \| 44.1 \| 59 \| (4, 63) \| 180 \| (53, 111) \| 118 \| Toxin-antitoxin system toxin HipA family \| 4PU4\_D Toxin-antitoxin system toxin HipA family Toxin Antitoxin System, TOXIN-ANTITOXIN-DNA complex; pdb70 \| 4GQM\_A \| 96.3 \| 2.8e-05 \| 1.9e-09 \| 44.0 \| 62 \| (2, 63) \| 180 \| (15, 82) \| 118 \| CT009 \| 4GQM\_A CT009 helix-turn-helix, UNKNOWN FUNCTION; pdb70 \| 2WUS\_R \| 96.3 \| 2.9e-05 \| 2e-09 \| 43.5 \| 61 \| (3, 63) \| 180 \| (10, 76) \| 112 \| ROD SHAPE-DETERMINING PROTEIN MREB, PUTATIVE \| 2WUS\_R ROD SHAPE-DETERMINING PROTEIN MREB, PUTATIVE STRUCTURAL PROTEIN, CELL WALL MORPHOGENESIS; pdb70 \| 2WUS\_S \| 96.3 \| 2.9e-05 \| 2e-09 \| 43.5 \| 61 \| (3, 63) \| 180 \| (10, 76) \| 112 \| ROD SHAPE-DETERMINING PROTEIN MREB, PUTATIVE \| 2WUS\_S ROD SHAPE-DETERMINING PROTEIN MREB, PUTATIVE STRUCTURAL PROTEIN, CELL WALL MORPHOGENESIS; pdb70 \| 6U0I\_B \| 96.3 \| 3e-05 \| 2e-09 \| 45.5 \| 54 \| (4, 57) \| 180 \| (91, 144) \| 148 \| Antitoxin HicB \| 6U0I\_B Antitoxin HicB structural genomics, Center for Structural HET: MSE; pdb70 \| 6F8S\_C \| 96.2 \| 3e-05 \| 2e-09 \| 42.1 \| 60 \| (4, 63) \| 180 \| (14, 74) \| 99 \| XRE family transcriptional regulator, Putative \| 6F8S\_C XRE family transcriptional regulator, Putative GraA, GraT, HigA, HigB, GraTA HET: PEG, SO4; pdb70 \| 2O38\_A \| 96.2 \| 3.3e-05 \| 2.2e-09 \| 43.8 \| 60 \| (4, 63) \| 180 \| (44, 104) \| 120 \| Hypothetical protein \| 2O38\_A Hypothetical protein alpha-beta, helix-turn-helix, Structural Genomics, PSI-2 HET: ACY, MSE; pdb70 \| 2O38\_B \| 96.2 \| 3.3e-05 \| 2.2e-09 \| 43.8 \| 60 \| (4, 63) \| 180 \| (44, 104) \| 120 \| Hypothetical protein \| 2O38\_B Hypothetical protein alpha-beta, helix-turn-helix, Structural Genomics, PSI-2 HET: ACY; pdb70 \| 4YBA\_B \| 96.2 \| 3.6e-05 \| 2.4e-09 \| 41.8 \| 53 \| (3, 55) \| 180 \| (12, 64) \| 99 \| C.kpn2I controller protein \| 4YBA\_B C.kpn2I controller protein DNA-protein interactions, helix-turn-helix, restriction-modification, controller; pdb70 \| 6CHV\_D \| 96.2 \| 3.6e-05 \| 2.4e-09 \| 43.8 \| 60 \| (3, 62) \| 180 \| (28, 87) \| 121 \| Antitoxin HigA/DNA Complex \| 6CHV\_D Antitoxin HigA/DNA Complex Helix-turn-helix DNA binding protein, DNA-protein; pdb70 \| 4JCY\_A \| 96.2 \| 4e-05 \| 2.6e-09 \| 41.3 \| 62 \| (2, 63) \| 180 \| (3, 68) \| 98 \| Csp231I C protein \| 4JCY\_A Csp231I C protein helix-turn-helix, C controller protein, restriction-modification; pdb70 \| 4JQD\_E \| 96.2 \| 4.1e-05 \| 2.7e-09 \| 41.2 \| 62 \| (2, 63) \| 180 \| (3, 68) \| 98 \| Csp231I C protein \| 4JQD\_E Csp231I C protein helix-turn-helix, C controller protein, restriction-modification; pdb70 \| 2PPX\_A \| 96.1 \| 4.9e-05 \| 3.3e-09 \| 41.4 \| 57 \| (4, 61) \| 180 \| (34, 90) \| 99 \| Uncharacterized protein Atu1735 \| 2PPX\_A Uncharacterized protein Atu1735 Agrobacterium tumefaciens, HTH-motif, XRE-family, structural HET: GOL, SO4; pdb70 \| 2AWI\_F \| 96.1 \| 4.9e-05 \| 3.3e-09 \| 50.4 \| 60 \| (2, 62) \| 180 \| (5, 64) \| 317 \| PrgX \| 2AWI\_F PrgX Repressor, pheromone, DNA binding, regulatory HET: MSE; pdb70 \| 4GHJ\_A \| 96.1 \| 5e-05 \| 3.4e-09 \| 41.7 \| 57 \| (3, 61) \| 180 \| (39, 95) \| 101 \| Probable transcriptional regulator \| 4GHJ\_A Probable transcriptional regulator Structural Genomics, NIAID, National Institute HET: MSE; pdb70 \| 4GHJ\_B \| 96.1 \| 5e-05 \| 3.4e-09 \| 41.7 \| 57 \| (3, 61) \| 180 \| (39, 95) \| 101 \| Probable transcriptional regulator \| 4GHJ\_B Probable transcriptional regulator Structural Genomics, NIAID, National Institute; pdb70 \| 6B9R\_D \| 96.0 \| 5.7e-05 \| 3.8e-09 \| 53.0 \| 63 \| (2, 64) \| 180 \| (239, 301) \| 450 \| Methylphosphonate synthase \| 6B9R\_D Methylphosphonate synthase Phosphonate, Hydroxymethylphosphonate, Iron, OXIDOREDUCTASE HET: GOL, 2HE; pdb70 \| 6B9R\_C \| 96.0 \| 5.7e-05 \| 3.8e-09 \| 53.0 \| 63 \| (2, 64) \| 180 \| (239, 301) \| 450 \| Methylphosphonate synthase \| 6B9R\_C Methylphosphonate synthase Phosphonate, Hydroxymethylphosphonate, Iron, OXIDOREDUCTASE HET: 2HE, GOL; pdb70 \| 2HIN\_A \| 96.0 \| 5.7e-05 \| 3.9e-09 \| 38.5 \| 54 \| (5, 60) \| 180 \| (3, 58) \| 71 \| Repressor protein \| 2HIN\_A Repressor protein TRANSCRIPTION FACTOR, DIMER INTERFACE, HELIX-TURN-HELIX HET: SO4; pdb70 \| 2HIN\_B \| 96.0 \| 5.7e-05 \| 3.9e-09 \| 38.5 \| 54 \| (5, 60) \| 180 \| (3, 58) \| 71 \| Repressor protein \| 2HIN\_B Repressor protein TRANSCRIPTION FACTOR, DIMER INTERFACE, HELIX-TURN-HELIX HET: SO4; pdb70 \| 3BD1\_A \| 96.0 \| 5.8e-05 \| 3.9e-09 \| 39.1 \| 58 \| (4, 63) \| 180 \| (3, 62) \| 79 \| Cro protein \| 3BD1\_A Cro protein transcription factor, helix-turn-helix, prophage, structural HET: GOL, SO4; pdb70 \| 2OFY\_A \| 96.0 \| 6.2e-05 \| 4.1e-09 \| 39.5 \| 50 \| (13, 62) \| 180 \| (27, 77) \| 86 \| Putative XRE-family transcriptional regulator \| 2OFY\_A Putative XRE-family transcriptional regulator TRANSCRIPTION REGULATOR, XRE-FAMILY, STRUCTURAL GENOMICS; pdb70 \| 2OFY\_B \| 96.0 \| 6.3e-05 \| 4.2e-09 \| 39.4 \| 50 \| (13, 62) \| 180 \| (27, 77) \| 86 \| Putative XRE-family transcriptional regulator \| 2OFY\_B Putative XRE-family transcriptional regulator TRANSCRIPTION REGULATOR, XRE-FAMILY, STRUCTURAL GENOMICS HET: MSE; pdb70 \| 5Y24\_B \| 96.0 \| 6.6e-05 \| 4.4e-09 \| 51.4 \| 56 \| (1, 56) \| 180 \| (4, 62) \| 396 \| AimR transcriptional regulator, GLY-MET-PRO-ARG-GLY-ALA \| 5Y24\_B AimR transcriptional regulator, GLY-MET-PRO-ARG-GLY-ALA DNA binding protein, peptide binding HET: MSE; pdb70 \| 5D50\_A \| 96.0 \| 7.1e-05 \| 4.7e-09 \| 46.2 \| 60 \| (5, 64) \| 180 \| (101, 162) \| 199 \| Repressor, Anti-repressor protein \| 5D50\_A Repressor, Anti-repressor protein Repressor, Anti-repressor, complex, DNA BINDING; pdb70 \| 5D50\_B \| 96.0 \| 7.1e-05 \| 4.7e-09 \| 46.2 \| 60 \| (5, 64) \| 180 \| (101, 162) \| 199 \| Repressor, Anti-repressor protein \| 5D50\_B Repressor, Anti-repressor protein Repressor, Anti-repressor, complex, DNA BINDING; pdb70 \| 3TRB\_B \| 95.9 \| 7.5e-05 \| 5e-09 \| 41.0 \| 59 \| (5, 63) \| 180 \| (18, 77) \| 104 \| Virulence-associated protein I \| 3TRB\_B Virulence-associated protein I Mobile and extrachromosomal element functions HET: MSE; pdb70 \| 2OX6\_C \| 95.9 \| 7.4e-05 \| 5e-09 \| 45.0 \| 51 \| (3, 53) \| 180 \| (10, 60) \| 166 \| Hypothetical protein \| 2OX6\_C Hypothetical protein Shewanella oneidensis, Structural genomics, PSI-2 HET: MG; pdb70 \| 1IC8\_B \| 95.9 \| 8e-05 \| 5.4e-09 \| 46.0 \| 59 \| (2, 60) \| 180 \| (32, 90) \| 194 \| HEPATOCYTE NUCLEAR FACTOR 1-ALPHA/DNA Complex \| 1IC8\_B HEPATOCYTE NUCLEAR FACTOR 1-ALPHA/DNA Complex Transcription regulation DNA-binding; pdb70 \| 6FKG\_C \| 95.9 \| 8.2e-05 \| 5.5e-09 \| 41.5 \| 57 \| (3, 59) \| 180 \| (10, 69) \| 115 \| Uncharacterized protein Rv1989c, Uncharacterized protein \| 6FKG\_C Uncharacterized protein Rv1989c, Uncharacterized protein Toxin-Antitoxin system Phosphorylase NAD+-binding, TOXIN HET: GOL; pdb70 \| 2OX6\_A \| 95.9 \| 8.4e-05 \| 5.7e-09 \| 44.8 \| 51 \| (3, 53) \| 180 \| (10, 60) \| 166 \| Hypothetical protein \| 2OX6\_A Hypothetical protein Shewanella oneidensis, Structural genomics, PSI-2 HET: MG; pdb70 \| 5W8Y\_A \| 95.9 \| 8.5e-05 \| 5.7e-09 \| 37.7 \| 56 \| (4, 61) \| 180 \| (3, 60) \| 73 \| protein XPH1 \| 5W8Y\_A protein XPH1 helix-turn-helix, structural evolution, conformational switch; pdb70 \| 3VWB\_A \| 95.9 \| 8.4e-05 \| 5.8e-09 \| 43.6 \| 34 \| (3, 37) \| 180 \| (32, 65) \| 143 \| Virulence regulon transcriptional activator virB/DNA \| 3VWB\_A Virulence regulon transcriptional activator virB/DNA HTH DNA binding motif, dsDNA; pdb70 \| 3W2A\_A \| 95.9 \| 8.4e-05 \| 5.8e-09 \| 43.6 \| 34 \| (3, 37) \| 180 \| (32, 65) \| 143 \| Virulence regulon transcriptional activator VirB/DNA \| 3W2A\_A Virulence regulon transcriptional activator VirB/DNA ParS like DNA binding sites; pdb70 \| 4YAR\_A \| 95.9 \| 8.5e-05 \| 5.8e-09 \| 51.9 \| 61 \| (3, 63) \| 180 \| (6, 67) \| 443 \| 2-Hydroxyethyl-phosphonate Dioxygenase (HEPD) \| 4YAR\_A 2-Hydroxyethyl-phosphonate Dioxygenase (HEPD) Dioxygenase, OXIDOREDUCTASE HET: CD; pdb70 \| 5ZW6\_B \| 95.8 \| 0.0001 \| 6.7e-09 \| 50.3 \| 56 \| (1, 56) \| 180 \| (4, 62) \| 391 \| AimR transcriptional regulator, GLY-MET-PRO-ARG-GLY-ALA \| 5ZW6\_B AimR transcriptional regulator, GLY-MET-PRO-ARG-GLY-ALA dimer, phage SPbeta, TRANSCRIPTION REGULATOR; pdb70 \| 2LCV\_A \| 95.8 \| 0.00011 \| 7.2e-09 \| 36.1 \| 54 \| (7, 60) \| 180 \| (3, 59) \| 67 \| HTH-type transcriptional repressor CytR \| 2LCV\_A HTH-type transcriptional repressor CytR BACTERIAL GENE REPRESSOR, HELIX TURN; pdb70 \| 2MQK\_A \| 95.7 \| 0.00014 \| 9.1e-09 \| 35.6 \| 37 \| (3, 39) \| 180 \| (10, 46) \| 65 \| ATP-dependent target DNA activator B \| 2MQK\_A ATP-dependent target DNA activator B HYDROLASE; pdb70 \| 1NR3\_A \| 95.7 \| 0.00014 \| 9.2e-09 \| 41.3 \| 52 \| (11, 62) \| 180 \| (3, 56) \| 122 \| DNA-BINDING PROTEIN TFX \| 1NR3\_A DNA-BINDING PROTEIN TFX NORTHEAST STRUCTURAL GENOMICS CONSORTIUM, REDUCED-DIMENSIONALITY; pdb70 \| 1CJG\_A \| 95.7 \| 0.00014 \| 9.2e-09 \| 35.5 \| 46 \| (12, 57) \| 180 \| (2, 50) \| 62 \| LAC REPRESSOR HP62/DNA COMPLEX \| 1CJG\_A LAC REPRESSOR HP62/DNA COMPLEX TRANSCRIPTION REGULATION, LAC OPERON, LAC; pdb70 \| 4YAR\_A \| 95.7 \| 0.00015 \| 1.1e-08 \| 50.6 \| 62 \| (2, 63) \| 180 \| (232, 294) \| 443 \| 2-Hydroxyethyl-phosphonate Dioxygenase (HEPD) \| 4YAR\_A 2-Hydroxyethyl-phosphonate Dioxygenase (HEPD) Dioxygenase, OXIDOREDUCTASE HET: CD; pdb70 \| 2L1P\_A \| 95.6 \| 0.00017 \| 1.1e-08 \| 38.0 \| 42 \| (11, 54) \| 180 \| (30, 71) \| 83 \| DNA-binding protein SATB1 \| 2L1P\_A DNA-binding protein SATB1 PSI-Biology, NESG, Structural Genomics, Protein; pdb70 \| 3NZL\_A \| 95.6 \| 0.00017 \| 1.1e-08 \| 38.0 \| 42 \| (11, 54) \| 180 \| (30, 71) \| 83 \| DNA-binding protein SATB1 \| 3NZL\_A DNA-binding protein SATB1 alpha-helical domain, Structural Genomics, PSI-Biology; pdb70 \| 3ME5\_A \| 95.6 \| 0.00017 \| 1.2e-08 \| 51.0 \| 40 \| (2, 41) \| 180 \| (18, 63) \| 482 \| Cytosine-specific methyltransferase (E.C.2.1.1.37) \| 3ME5\_A Cytosine-specific methyltransferase (E.C.2.1.1.37) STRUCTURAL GENOMICS, PROTEIN STRUCTURE INITIATIVE; pdb70 \| 6B9T\_C \| 95.6 \| 0.00018 \| 1.2e-08 \| 50.5 \| 62 \| (2, 63) \| 180 \| (244, 305) \| 457 \| Methylphosphonate synthase (E.C.1.13.11.73) \| 6B9T\_C Methylphosphonate synthase (E.C.1.13.11.73) Phosphonate, Methylphosphonate, Iron, OXIDOREDUCTASE HET: FMT, 2HE; pdb70 \| 6B9T\_D \| 95.6 \| 0.00018 \| 1.2e-08 \| 50.4 \| 61 \| (3, 63) \| 180 \| (245, 305) \| 457 \| Methylphosphonate synthase (E.C.1.13.11.73) \| 6B9T\_D Methylphosphonate synthase (E.C.1.13.11.73) Phosphonate, Methylphosphonate, Iron, OXIDOREDUCTASE HET: 2HE, FMT; pdb70 \| 6JG8\_A \| 95.5 \| 0.00021 \| 1.4e-08 \| 48.8 \| 55 \| (2, 56) \| 180 \| (6, 63) \| 395 \| AimR transcriptional regulator/DNA Complex \| 6JG8\_A AimR transcriptional regulator/DNA Complex AimR, Apo, HTH, PEPTIDE BINDING; pdb70 \| 5J9I\_D \| 95.5 \| 0.00021 \| 1.4e-08 \| 39.1 \| 58 \| (3, 61) \| 180 \| (43, 100) \| 104 \| Antitoxin igA-2 \| 5J9I\_D Antitoxin igA-2 toxin-antitoxin system, antitoxin; pdb70 \| 2QFC\_A \| 95.4 \| 0.0003 \| 2e-08 \| 45.0 \| 58 \| (2, 60) \| 180 \| (7, 64) \| 293 \| PlcR protein, C-terminus pentapeptide from \| 2QFC\_A PlcR protein, C-terminus pentapeptide from TPR HTH, TRANSCRIPTION REGULATION HET: PEG; pdb70 \| 3U3W\_B \| 95.4 \| 0.0003 \| 2e-08 \| 45.0 \| 58 \| (2, 60) \| 180 \| (7, 64) \| 293 \| Transcriptional activator PlcR protein, C-terminus \| 3U3W\_B Transcriptional activator PlcR protein, C-terminus ternary complex, PlcR-PAPR7-DNA, HTH DNA-binding; pdb70 \| 3D1N\_I \| 95.3 \| 0.00035 \| 2.3e-08 \| 40.6 \| 50 \| (2, 53) \| 180 \| (10, 65) \| 151 \| POU domain, class 6, transcription \| 3D1N\_I POU domain, class 6, transcription Protein-DNA complex, Helix-turn-helix (HTH), DNA-binding HET: MSE; pdb70 \| 5ZVV\_B \| 95.2 \| 0.00037 \| 2.5e-08 \| 46.8 \| 55 \| (2, 56) \| 180 \| (3, 60) \| 382 \| AimR transcriptional regulator \| 5ZVV\_B AimR transcriptional regulator dimer, phage phi3T, TRANSCRIPTION REGULATOR HET: GOL, MSE; pdb70 \| 1RZS\_A \| 95.2 \| 0.00044 \| 2.9e-08 \| 33.0 \| 45 \| (5, 50) \| 180 \| (3, 47) \| 61 \| Regulatory protein cro \| 1RZS\_A Regulatory protein cro HELIX-TURN-HELIX, DNA-BINDING PROTEIN, STRUCTURAL EVOLUTION; pdb70 \| 5JAA\_A \| 95.1 \| 0.00044 \| 2.9e-08 \| 37.7 \| 57 \| (3, 60) \| 180 \| (42, 98) \| 103 \| Antitoxin igA-2, Toxin HigB-2 \| 5JAA\_A Antitoxin igA-2, Toxin HigB-2 toxin-antitoxin system, toxin HET: MSE; pdb70 \| 3D1N\_N \| 95.1 \| 0.00046 \| 3e-08 \| 40.1 \| 51 \| (2, 54) \| 180 \| (10, 66) \| 151 \| POU domain, class 6, transcription \| 3D1N\_N POU domain, class 6, transcription Protein-DNA complex, Helix-turn-helix (HTH), DNA-binding HET: MSE; pdb70 \| 1GT0\_C \| 94.6 \| 0.001 \| 7e-08 \| 39.0 \| 49 \| (2, 52) \| 180 \| (13, 67) \| 159 \| OCTAMER-BINDING TRANSCRIPTION FACTOR 1, TRANSCRIPTION \| 1GT0\_C OCTAMER-BINDING TRANSCRIPTION FACTOR 1, TRANSCRIPTION TRANSCRIPTION FACTOR, POU FACTORS, SOX |
| Top keywords  (threshold 1.00e-03 (evalue)) | **domain\_containing, repressor, transcriptional, regulator, HTH, cro, C1\_type, Peptidase, Helix\_turn\_helix, S24** |
| Output files | ../../similar\_sequences/42\_FANPEZAQ\_CDS\_0042\_merged.svg ../../similar\_sequences/42\_FANPEZAQ\_CDS\_0042\_pdb70.a3m ../../similar\_sequences/42\_FANPEZAQ\_CDS\_0042\_pdb70.hhr ../../similar\_sequences/42\_FANPEZAQ\_CDS\_0042\_uniclust.a3m ../../similar\_sequences/42\_FANPEZAQ\_CDS\_0042\_uniclust.hhr |

#### Structure prediction (AlphaFold)^2^

|  |  |
| --- | --- |
| Stats | xml version="1.0" encoding="utf-8" standalone="no"?       2024-09-02T21:09:41.312135 image/svg+xml   Matplotlib v3.7.2, https://matplotlib.org/ |
| Predicted structure | **NGL Viewer Controls:**  - Center: *Left-Click* - Rotate: *Left-Click + Drag* - Translate: *Right-Click + Drag* - Zoom: *Shift + Left-Click + Drag* |
| Output files | ../../predicted\_structures/42\_FANPEZAQ\_CDS\_0042/features.pkl ../../predicted\_structures/42\_FANPEZAQ\_CDS\_0042/ranked\_0.pdb ../../predicted\_structures/42\_FANPEZAQ\_CDS\_0042/ranked\_0\_plots.svg ../../predicted\_structures/42\_FANPEZAQ\_CDS\_0042/result\_model\_1\_ptm\_pred\_0.pkl |

#### Structure similarity search results (Foldseek)^3^

|  |  |
| --- | --- |
| Structure databases searched | Pdb, Afdb-proteome, Afdb-uniprot50 |
| Results, scheme(s)  (Top layers only, threshold 1.00e-02 (evalue)) | xml version="1.0" encoding="utf-8" standalone="no"?       2024-09-02T21:11:21.799040 image/svg+xml   Matplotlib v3.7.2, https://matplotlib.org/ |
| Results, table  (threshold 1.00e-02 (evalue)) | db \| id \| prob \| evalue \| bits \| fident \| alnlen \| mismatch \| gapopen \| qstart \| qend \| tstart \| tend \| name \| description; pdb \| 3K2Z\_A \| 1.0 \| 8.188e-06 \| 226 \| 0.252 \| 91 \| 64 \| 4 \| 91 \| 180 \| 105 \| 192 \| LexA repressor \| LexA repressor; pdb \| 6A2R\_A \| 1.0 \| 1.14e-06 \| 222 \| 0.3 \| 103 \| 61 \| 5 \| 84 \| 180 \| 2 \| 99 \| LexA repressor \| LexA repressor; pdb \| 6A2S\_A \| 1.0 \| 2.157e-06 \| 220 \| 0.297 \| 101 \| 60 \| 5 \| 86 \| 180 \| 5 \| 100 \| LexA repressor \| LexA repressor; pdb \| 3K2Z\_B \| 1.0 \| 5.782e-06 \| 217 \| 0.235 \| 102 \| 68 \| 5 \| 86 \| 180 \| 97 \| 195 \| LexA repressor \| LexA repressor; pdb \| 6A2T\_A \| 1.0 \| 1.229e-05 \| 194 \| 0.307 \| 91 \| 57 \| 4 \| 91 \| 180 \| 9 \| 94 \| LexA repressor \| LexA repressor; pdb \| 1UMU\_B \| 1.0 \| 6.232e-05 \| 189 \| 0.235 \| 102 \| 72 \| 5 \| 80 \| 179 \| 9 \| 106 \| UMUD' \| UMUD'; pdb \| 8GMT\_A \| 1.0 \| 7.859e-05 \| 184 \| 0.237 \| 101 \| 71 \| 5 \| 80 \| 178 \| 2 \| 98 \| DNA polymerase V subunit UmuD \| DNA polymerase V subunit UmuD; pdb \| 1JHH\_B \| 1.0 \| 8.825e-05 \| 182 \| 0.177 \| 90 \| 68 \| 5 \| 93 \| 179 \| 32 \| 118 \| LEXA REPRESSOR \| LEXA REPRESSOR; pdb \| 1JHE\_B \| 1.0 \| 0.000105 \| 180 \| 0.166 \| 90 \| 69 \| 5 \| 93 \| 179 \| 38 \| 124 \| LEXA REPRESSOR \| LEXA REPRESSOR; pdb \| 7ZRA\_B \| 1.0 \| 0.0001179 \| 179 \| 0.188 \| 90 \| 67 \| 5 \| 93 \| 179 \| 38 \| 124 \| LexA repressor \| LexA repressor; pdb \| 1JHF\_B \| 1.0 \| 0.0001179 \| 178 \| 0.188 \| 90 \| 67 \| 5 \| 93 \| 179 \| 24 \| 110 \| LEXA REPRESSOR \| LEXA REPRESSOR; pdb \| 8GMS\_B \| 1.0 \| 0.000105 \| 175 \| 0.188 \| 90 \| 67 \| 5 \| 93 \| 179 \| 27 \| 113 \| LexA repressor \| LexA repressor; pdb \| 1KCA\_A \| 1.0 \| 9.911e-05 \| 173 \| 0.197 \| 96 \| 66 \| 6 \| 89 \| 177 \| 3 \| 94 \| REPRESSOR PROTEIN CI \| REPRESSOR PROTEIN CI; pdb \| 2HO0\_A \| 1.0 \| 0.000177 \| 165 \| 0.217 \| 92 \| 65 \| 7 \| 89 \| 177 \| 41 \| 128 \| Repressor protein cI101-229DM-K192A \| Repressor protein cI101-229DM-K192A; pdb \| 1JHH\_A \| 1.0 \| 0.0002815 \| 165 \| 0.177 \| 90 \| 68 \| 5 \| 93 \| 179 \| 111 \| 197 \| LEXA REPRESSOR \| LEXA REPRESSOR; pdb \| 1F39\_B \| 1.0 \| 0.0001988 \| 164 \| 0.202 \| 94 \| 64 \| 6 \| 90 \| 176 \| 4 \| 93 \| REPRESSOR PROTEIN CI \| REPRESSOR PROTEIN CI; pdb \| 8GMS\_A \| 1.0 \| 0.0002983 \| 163 \| 0.188 \| 90 \| 67 \| 5 \| 93 \| 179 \| 37 \| 123 \| LexA repressor \| LexA repressor; pdb \| 8GMU\_A \| 1.0 \| 0.0001876 \| 159 \| 0.202 \| 94 \| 64 \| 6 \| 90 \| 176 \| 39 \| 128 \| Repressor protein cI \| Repressor protein cI; pdb \| 3K3R\_F \| 1.0 \| 0.0002815 \| 159 \| 0.191 \| 89 \| 66 \| 5 \| 93 \| 178 \| 86 \| 171 \| LexA repressor \| LexA repressor; pdb \| 2HNF\_A \| 1.0 \| 0.0002365 \| 158 \| 0.204 \| 93 \| 66 \| 7 \| 89 \| 177 \| 42 \| 130 \| Repressor protein cI101-229DM-K192A \| Repressor protein cI101-229DM-K192A; pdb \| 3OMT\_B \| 1.0 \| 0.004296 \| 157 \| 0.278 \| 61 \| 44 \| 0 \| 3 \| 63 \| 6 \| 66 \| uncharacterized protein \| uncharacterized protein; pdb \| 7JVT\_C \| 1.0 \| 4.777e-07 \| 154 \| 0.155 \| 193 \| 130 \| 6 \| 6 \| 180 \| 20 \| 197 \| Repressor protein CI \| Repressor protein CI; pdb \| 3BS3\_A \| 1.0 \| 0.006084 \| 153 \| 0.254 \| 59 \| 44 \| 0 \| 3 \| 61 \| 4 \| 62 \| Putative DNA-binding protein \| Putative DNA-binding protein; pdb \| 3JXC\_L \| 1.0 \| 0.003407 \| 149 \| 0.293 \| 58 \| 41 \| 0 \| 4 \| 61 \| 7 \| 64 \| Repressor protein C2 \| Repressor protein C2; pdb \| 1ADR\_A \| 1.0 \| 0.001132 \| 148 \| 0.264 \| 68 \| 50 \| 0 \| 4 \| 71 \| 9 \| 76 \| P22 C2 REPRESSOR \| P22 C2 REPRESSOR; pdb \| 3JSO\_A \| 1.0 \| 1.16e-05 \| 144 \| 0.164 \| 195 \| 122 \| 13 \| 5 \| 179 \| 14 \| 187 \| LexA repressor \| LexA repressor; pdb \| 3JSO\_B \| 1.0 \| 8.677e-06 \| 140 \| 0.168 \| 196 \| 118 \| 12 \| 5 \| 179 \| 14 \| 185 \| LexA repressor \| LexA repressor; pdb \| 2FKD\_B \| 1.0 \| 0.002406 \| 139 \| 0.202 \| 74 \| 53 \| 2 \| 107 \| 180 \| 42 \| 109 \| Repressor protein CI \| Repressor protein CI; pdb \| 3K3R\_E \| 1.0 \| 0.0012 \| 138 \| 0.179 \| 89 \| 67 \| 5 \| 93 \| 178 \| 87 \| 172 \| LexA repressor \| LexA repressor; pdb \| 1ADR\_A \| 1.0 \| 0.002863 \| 134 \| 0.264 \| 68 \| 50 \| 0 \| 4 \| 71 \| 9 \| 76 \| P22 C2 REPRESSOR \| P22 C2 REPRESSOR; pdb \| 1R63\_A \| 1.0 \| 0.007672 \| 132 \| 0.275 \| 58 \| 41 \| 1 \| 4 \| 61 \| 5 \| 61 \| REPRESSOR PROTEIN FROM BACTERIOPHAGE 434 \| REPRESSOR PROTEIN FROM BACTERIOPHAGE 434; pdb \| 1ADR\_A \| 1.0 \| 0.008616 \| 124 \| 0.264 \| 68 \| 50 \| 0 \| 4 \| 71 \| 9 \| 76 \| P22 C2 REPRESSOR \| P22 C2 REPRESSOR; pdb \| 3JSP\_B \| 1.0 \| 8.328e-05 \| 122 \| 0.166 \| 198 \| 119 \| 13 \| 5 \| 179 \| 14 \| 188 \| LexA repressor \| LexA repressor; pdb \| 4N31\_A \| 1.0 \| 0.0008976 \| 121 \| 0.118 \| 135 \| 72 \| 5 \| 90 \| 179 \| 7 \| 139 \| SipA \| SipA; pdb \| 1ADR\_A \| 1.0 \| 0.008616 \| 120 \| 0.264 \| 68 \| 50 \| 0 \| 4 \| 71 \| 9 \| 76 \| P22 C2 REPRESSOR \| P22 C2 REPRESSOR; pdb \| 3JSP\_A \| 1.0 \| 8.825e-05 \| 119 \| 0.172 \| 197 \| 124 \| 13 \| 5 \| 179 \| 14 \| 193 \| LexA repressor \| LexA repressor; pdb \| 6JQ1\_A \| 1.0 \| 0.004296 \| 112 \| 0.264 \| 87 \| 63 \| 1 \| 4 \| 89 \| 7 \| 93 \| Transcriptional regulator, XRE family \| Transcriptional regulator, XRE family; pdb \| 2FJR\_B \| 1.0 \| 4.153e-05 \| 112 \| 0.133 \| 202 \| 126 \| 10 \| 5 \| 180 \| 10 \| 188 \| Repressor protein CI \| Repressor protein CI; pdb \| 3BDN\_A \| 1.0 \| 0.000167 \| 112 \| 0.14 \| 235 \| 131 \| 10 \| 2 \| 180 \| 11 \| 230 \| Lambda Repressor \| Lambda Repressor; pdb \| 4N31\_B \| 1.0 \| 0.00227 \| 104 \| 0.133 \| 135 \| 68 \| 5 \| 91 \| 180 \| 3 \| 133 \| SipA \| SipA; pdb \| 1ZZ6\_A \| 0.999 \| 7.859e-05 \| 99 \| 0.134 \| 208 \| 110 \| 13 \| 5 \| 178 \| 10 \| 181 \| Hydroxypropylphosphonic Acid Epoxidase \| Hydroxypropylphosphonic Acid Epoxidase; pdb \| 1ZZ7\_B \| 0.999 \| 0.000167 \| 96 \| 0.134 \| 201 \| 112 \| 13 \| 5 \| 178 \| 9 \| 174 \| Hydroxyprophylphosphonic Acid Epoxidase \| Hydroxyprophylphosphonic Acid Epoxidase; pdb \| 3OP9\_A \| 0.998 \| 0.00813 \| 93 \| 0.15 \| 93 \| 71 \| 2 \| 1 \| 89 \| 1 \| 89 \| Pli0006 protein \| Pli0006 protein; pdb \| 1ZZ7\_A \| 0.998 \| 0.0002232 \| 93 \| 0.122 \| 187 \| 119 \| 9 \| 5 \| 169 \| 9 \| 172 \| Hydroxyprophylphosphonic Acid Epoxidase \| Hydroxyprophylphosphonic Acid Epoxidase; pdb \| 2BNO\_A \| 0.998 \| 0.0002506 \| 93 \| 0.128 \| 202 \| 112 \| 10 \| 5 \| 175 \| 12 \| 180 \| EPOXIDASE \| EPOXIDASE; pdb \| 7P2P\_A \| 0.997 \| 0.003407 \| 91 \| 0.184 \| 119 \| 81 \| 6 \| 76 \| 178 \| 17 \| 135 \| Signal peptidase complex catalytic subunit SEC11A \| Signal peptidase complex catalytic subunit SEC11A; pdb \| 1ZZ6\_B \| 0.997 \| 0.0002106 \| 91 \| 0.134 \| 193 \| 114 \| 11 \| 5 \| 169 \| 10 \| 177 \| Hydroxypropylphosphonic Acid Epoxidase \| Hydroxypropylphosphonic Acid Epoxidase; pdb \| 2BNN\_B \| 0.997 \| 0.0002106 \| 91 \| 0.129 \| 208 \| 114 \| 12 \| 5 \| 174 \| 11 \| 189 \| EPOXIDASE \| EPOXIDASE; pdb \| 1ZZ9\_C \| 0.997 \| 0.0002656 \| 91 \| 0.125 \| 200 \| 115 \| 11 \| 5 \| 169 \| 10 \| 184 \| Hydroxypropylphosphonic Acid Epoxidase \| Hydroxypropylphosphonic Acid Epoxidase; pdb \| 1ZZC\_A \| 0.996 \| 0.0004224 \| 90 \| 0.134 \| 193 \| 112 \| 11 \| 5 \| 173 \| 11 \| 172 \| hydroxypropylphosphonic acid epoxidase \| hydroxypropylphosphonic acid epoxidase; pdb \| 1ZZ9\_A \| 0.996 \| 0.0004224 \| 89 \| 0.133 \| 203 \| 112 \| 12 \| 5 \| 169 \| 10 \| 186 \| Hydroxypropylphosphonic Acid Epoxidase \| Hydroxypropylphosphonic Acid Epoxidase; pdb \| 1ZZ8\_C \| 0.995 \| 0.0003549 \| 88 \| 0.129 \| 208 \| 118 \| 13 \| 5 \| 174 \| 10 \| 192 \| Hydroxypropylphosphonic Acid Epoxidase \| Hydroxypropylphosphonic Acid Epoxidase; pdb \| 1ZZB\_A \| 0.994 \| 0.0005327 \| 87 \| 0.131 \| 197 \| 113 \| 11 \| 5 \| 176 \| 10 \| 173 \| Hydroxypropylphosphonic Acid Epoxidase \| Hydroxypropylphosphonic Acid Epoxidase; pdb \| 4J1X\_C \| 0.994 \| 0.0005982 \| 87 \| 0.121 \| 206 \| 112 \| 10 \| 5 \| 169 \| 12 \| 189 \| Epoxidase \| Epoxidase; pdb \| 6B9R\_C \| 0.949 \| 0.003407 \| 72 \| 0.153 \| 202 \| 107 \| 13 \| 5 \| 173 \| 238 \| 408 \| Hydroxyethylphosphonate dioxygenase \| Hydroxyethylphosphonate dioxygenase; pdb \| 3GBF\_A \| 0.933 \| 0.0005645 \| 70 \| 0.121 \| 214 \| 113 \| 15 \| 4 \| 173 \| 7 \| 189 \| PhpD \| PhpD; pdb \| 3RZZ\_A \| 0.923 \| 0.001132 \| 69 \| 0.105 \| 208 \| 117 \| 14 \| 4 \| 173 \| 7 \| 183 \| Hydroxyethylphoshphonate Dioxygenase (PhpD) \| Hydroxyethylphoshphonate Dioxygenase (PhpD); pdb \| 1Y9Q\_A \| 0.872 \| 0.006832 \| 65 \| 0.117 \| 187 \| 115 \| 12 \| 4 \| 169 \| 12 \| 169 \| transcriptional regulator, HTH\_3 family \| transcriptional regulator, HTH\_3 family; pdb \| 3G7D\_A \| 0.772 \| 0.00361 \| 60 \| 0.119 \| 210 \| 111 \| 14 \| 4 \| 173 \| 7 \| 182 \| PhpD \| PhpD; pdb \| 4YAR\_A \| 0.772 \| 0.004296 \| 60 \| 0.109 \| 220 \| 118 \| 14 \| 4 \| 173 \| 5 \| 196 \| 2-hydroxyethylphosphonate dioxygenase \| 2-hydroxyethylphosphonate dioxygenase; afdb-proteome \| AF-Q8ZQI4-F1-MODEL\_V4 \| 1.0 \| 4.604e-13 \| 350 \| 0.23 \| 221 \| 124 \| 5 \| 4 \| 180 \| 8 \| 226 \| Putative Fels-1 prophage transcriptional regulator \| Putative Fels-1 prophage transcriptional regulator; afdb-proteome \| AF-Q5F9C0-F1-MODEL\_V4 \| 1.0 \| 6.006e-11 \| 302 \| 0.2 \| 210 \| 132 \| 8 \| 5 \| 180 \| 29 \| 236 \| Cro/Cl family transcriptional regulator \| Cro/Cl family transcriptional regulator; afdb-proteome \| AF-Q9I551-F1-MODEL\_V4 \| 1.0 \| 1.915e-10 \| 283 \| 0.206 \| 233 \| 128 \| 5 \| 4 \| 180 \| 6 \| 237 \| Probable transcriptional regulator \| Probable transcriptional regulator; afdb-proteome \| AF-A0A0H3GP86-F1-MODEL\_V4 \| 1.0 \| 1.005e-07 \| 281 \| 0.298 \| 114 \| 74 \| 3 \| 70 \| 180 \| 7 \| 117 \| Regulatory protein CI bacteriophage origin \| Regulatory protein CI bacteriophage origin; afdb-proteome \| AF-P44207-F1-MODEL\_V4 \| 1.0 \| 2.874e-10 \| 272 \| 0.195 \| 230 \| 129 \| 8 \| 4 \| 180 \| 12 \| 238 \| Uncharacterized HTH-type transcriptional regulator HI\_1476 \| Uncharacterized HTH-type transcriptional regulator HI\_1476; afdb-proteome \| AF-Q5F7Q4-F1-MODEL\_V4 \| 1.0 \| 9.167e-10 \| 270 \| 0.147 \| 231 \| 142 \| 5 \| 4 \| 180 \| 7 \| 236 \| Repressor \| Repressor; afdb-proteome \| AF-Q5F6C9-F1-MODEL\_V4 \| 1.0 \| 2.03e-10 \| 263 \| 0.168 \| 225 \| 135 \| 7 \| 4 \| 176 \| 19 \| 243 \| Repressor \| Repressor; afdb-proteome \| AF-Q06553-F1-MODEL\_V4 \| 1.0 \| 1.458e-09 \| 256 \| 0.149 \| 234 \| 141 \| 7 \| 4 \| 180 \| 15 \| 247 \| HTH-type transcriptional regulator PrtR \| HTH-type transcriptional regulator PrtR; afdb-proteome \| AF-Q5F6W7-F1-MODEL\_V4 \| 1.0 \| 2.759e-09 \| 245 \| 0.177 \| 231 \| 136 \| 9 \| 1 \| 180 \| 1 \| 228 \| Repressor \| Repressor; afdb-proteome \| AF-P22493-F1-MODEL\_V4 \| 1.0 \| 1.237e-05 \| 216 \| 0.262 \| 103 \| 70 \| 5 \| 80 \| 180 \| 40 \| 138 \| Protein UmuD \| Protein UmuD; afdb-proteome \| AF-Q2FX22-F1-MODEL\_V4 \| 1.0 \| 6.977e-09 \| 211 \| 0.174 \| 229 \| 128 \| 9 \| 4 \| 174 \| 13 \| 238 \| Phage repressor protein, putative \| Phage repressor protein, putative; afdb-proteome \| AF-Q32H33-F1-MODEL\_V4 \| 1.0 \| 3.316e-05 \| 204 \| 0.233 \| 103 \| 73 \| 5 \| 80 \| 180 \| 40 \| 138 \| UmuD \| UmuD; afdb-proteome \| AF-Q32F74-F1-MODEL\_V4 \| 1.0 \| 4.211e-08 \| 204 \| 0.151 \| 218 \| 132 \| 10 \| 6 \| 179 \| 24 \| 232 \| Repressor protein CI \| Repressor protein CI; afdb-proteome \| AF-Q2FY89-F1-MODEL\_V4 \| 1.0 \| 2.648e-08 \| 200 \| 0.161 \| 229 \| 128 \| 9 \| 5 \| 174 \| 13 \| 236 \| Helix-turn-helix domain protein \| Helix-turn-helix domain protein; afdb-proteome \| AF-P0AG11-F1-MODEL\_V4 \| 1.0 \| 5.588e-05 \| 198 \| 0.233 \| 103 \| 73 \| 5 \| 80 \| 180 \| 40 \| 138 \| Protein UmuD \| Protein UmuD; afdb-proteome \| AF-P23831-F1-MODEL\_V4 \| 1.0 \| 4.976e-05 \| 189 \| 0.227 \| 101 \| 71 \| 4 \| 80 \| 179 \| 41 \| 135 \| Protein SamA \| Protein SamA; afdb-proteome \| AF-A0A0H3H1L6-F1-MODEL\_V4 \| 1.0 \| 0.0001334 \| 184 \| 0.178 \| 101 \| 78 \| 4 \| 80 \| 179 \| 42 \| 138 \| Protein impA \| Protein impA; afdb-proteome \| AF-P75974-F1-MODEL\_V4 \| 1.0 \| 5.401e-07 \| 181 \| 0.127 \| 211 \| 145 \| 11 \| 4 \| 179 \| 11 \| 217 \| Prophage repressor CohE \| Prophage repressor CohE; afdb-proteome \| AF-P0A273-F1-MODEL\_V4 \| 1.0 \| 0.0001782 \| 178 \| 0.177 \| 90 \| 68 \| 5 \| 93 \| 179 \| 112 \| 198 \| LexA repressor \| LexA repressor; afdb-proteome \| AF-A0A0H3GQ57-F1-MODEL\_V4 \| 1.0 \| 0.0002121 \| 176 \| 0.188 \| 90 \| 67 \| 5 \| 93 \| 179 \| 112 \| 198 \| LexA repressor \| LexA repressor; afdb-proteome \| AF-P0A7C2-F1-MODEL\_V4 \| 1.0 \| 0.0002524 \| 173 \| 0.188 \| 90 \| 67 \| 5 \| 93 \| 179 \| 112 \| 198 \| LexA repressor \| LexA repressor; afdb-proteome \| AF-Q327V0-F1-MODEL\_V4 \| 1.0 \| 0.0003182 \| 171 \| 0.188 \| 90 \| 67 \| 5 \| 93 \| 179 \| 112 \| 198 \| LexA repressor \| LexA repressor; afdb-proteome \| AF-A0A0H3GN19-F1-MODEL\_V4 \| 1.0 \| 0.0004013 \| 170 \| 0.203 \| 103 \| 76 \| 5 \| 80 \| 180 \| 40 \| 138 \| DNA polymerase V subunit UmuD \| DNA polymerase V subunit UmuD; afdb-proteome \| AF-P37452-F1-MODEL\_V4 \| 1.0 \| 0.0007167 \| 167 \| 0.188 \| 90 \| 67 \| 5 \| 93 \| 179 \| 118 \| 204 \| LexA repressor \| LexA repressor; afdb-proteome \| AF-A0A2S7MUS3-F1-MODEL\_V4 \| 1.0 \| 3.661e-06 \| 156 \| 0.186 \| 214 \| 112 \| 16 \| 5 \| 180 \| 19 \| 208 \| LexA repressor \| LexA repressor; afdb-proteome \| AF-Q9HWV1-F1-MODEL\_V4 \| 1.0 \| 0.002286 \| 153 \| 0.287 \| 66 \| 47 \| 0 \| 1 \| 66 \| 1 \| 66 \| Probable transcriptional regulator \| Probable transcriptional regulator; afdb-proteome \| AF-A0A0H3H1N6-F1-MODEL\_V4 \| 1.0 \| 9.646e-07 \| 142 \| 0.185 \| 199 \| 126 \| 11 \| 5 \| 179 \| 10 \| 196 \| CI repressor \| CI repressor; afdb-proteome \| AF-A0A133CLZ3-F1-MODEL\_V4 \| 1.0 \| 0.0009038 \| 140 \| 0.264 \| 87 \| 57 \| 1 \| 5 \| 84 \| 10 \| 96 \| Helix-turn-helix transcriptional regulator \| Helix-turn-helix transcriptional regulator; afdb-proteome \| AF-Q57720-F1-MODEL\_V4 \| 1.0 \| 0.009193 \| 139 \| 0.23 \| 65 \| 50 \| 0 \| 1 \| 65 \| 1 \| 65 \| Uncharacterized HTH-type transcriptional regulator MJ0272 \| Uncharacterized HTH-type transcriptional regulator MJ0272; afdb-proteome \| AF-Q2FYU1-F1-MODEL\_V4 \| 1.0 \| 3.514e-05 \| 138 \| 0.189 \| 216 \| 112 \| 15 \| 5 \| 180 \| 15 \| 207 \| LexA repressor \| LexA repressor; afdb-proteome \| AF-Q2G1V7-F1-MODEL\_V4 \| 1.0 \| 0.008186 \| 137 \| 0.212 \| 66 \| 52 \| 0 \| 1 \| 66 \| 1 \| 66 \| HTH cro/C1-type domain-containing protein \| HTH cro/C1-type domain-containing protein; afdb-proteome \| AF-A0A132ZE74-F1-MODEL\_V4 \| 1.0 \| 0.006491 \| 137 \| 0.205 \| 68 \| 54 \| 0 \| 1 \| 68 \| 1 \| 68 \| Helix-turn-helix transcriptional regulator \| Helix-turn-helix transcriptional regulator; afdb-proteome \| AF-Q4AAY3-F1-MODEL\_V4 \| 1.0 \| 8.244e-06 \| 125 \| 0.162 \| 209 \| 128 \| 11 \| 5 \| 180 \| 16 \| 210 \| Fels-2 prophage protein \| Fels-2 prophage protein; afdb-proteome \| AF-Q8ZNJ5-F1-MODEL\_V4 \| 1.0 \| 0.0007167 \| 121 \| 0.136 \| 125 \| 90 \| 4 \| 61 \| 179 \| 43 \| 155 \| Putative peptidase \| Putative peptidase; afdb-proteome \| AF-K7M8T7-F1-MODEL\_V4 \| 1.0 \| 0.0005683 \| 120 \| 0.196 \| 117 \| 74 \| 7 \| 76 \| 177 \| 9 \| 120 \| Signal peptidase I \| Signal peptidase I; afdb-proteome \| AF-Q2FX38-F1-MODEL\_V4 \| 1.0 \| 0.006491 \| 119 \| 0.246 \| 77 \| 54 \| 2 \| 5 \| 80 \| 11 \| 84 \| Helix-turn-helix DNA binding protein \| Helix-turn-helix DNA binding protein; afdb-proteome \| AF-I1KFJ2-F1-MODEL\_V4 \| 1.0 \| 0.0009038 \| 116 \| 0.196 \| 117 \| 74 \| 7 \| 76 \| 177 \| 36 \| 147 \| Signal peptidase I \| Signal peptidase I; afdb-proteome \| AF-Q0DJD8-F1-MODEL\_V4 \| 1.0 \| 0.0007167 \| 115 \| 0.185 \| 124 \| 77 \| 7 \| 74 \| 177 \| 28 \| 147 \| Signal peptidase I \| Signal peptidase I; afdb-proteome \| AF-P23873-F1-MODEL\_V4 \| 1.0 \| 0.009742 \| 113 \| 0.217 \| 69 \| 49 \| 1 \| 5 \| 73 \| 17 \| 80 \| Antitoxin HipB \| Antitoxin HipB; afdb-proteome \| AF-Q9R0P6-F1-MODEL\_V4 \| 1.0 \| 0.0001682 \| 113 \| 0.209 \| 153 \| 91 \| 8 \| 45 \| 177 \| 4 \| 146 \| Signal peptidase complex catalytic subunit SEC11A \| Signal peptidase complex catalytic subunit SEC11A; afdb-proteome \| AF-A0A0H3GXD8-F1-MODEL\_V4 \| 1.0 \| 0.009742 \| 112 \| 0.275 \| 69 \| 49 \| 1 \| 5 \| 73 \| 16 \| 83 \| DNA-binding transcriptional regulator HipB \| DNA-binding transcriptional regulator HipB; afdb-proteome \| AF-A0A044QWB3-F1-MODEL\_V4 \| 1.0 \| 0.001813 \| 111 \| 0.17 \| 117 \| 77 \| 7 \| 76 \| 177 \| 2 \| 113 \| Signal peptidase complex catalytic subunit SEC11 \| Signal peptidase complex catalytic subunit SEC11; afdb-proteome \| AF-Q5A869-F1-MODEL\_V4 \| 1.0 \| 0.002422 \| 111 \| 0.162 \| 117 \| 81 \| 6 \| 74 \| 177 \| 21 \| 133 \| Signal peptidase complex catalytic subunit SEC11 \| Signal peptidase complex catalytic subunit SEC11; afdb-proteome \| AF-P06966-F1-MODEL\_V4 \| 1.0 \| 0.008186 \| 110 \| 0.255 \| 90 \| 61 \| 1 \| 4 \| 87 \| 11 \| 100 \| HTH-type transcriptional regulator DicA \| HTH-type transcriptional regulator DicA; afdb-proteome \| AF-P15367-F1-MODEL\_V4 \| 1.0 \| 0.001921 \| 106 \| 0.158 \| 120 \| 79 \| 9 \| 74 \| 177 \| 21 \| 134 \| Signal peptidase complex catalytic subunit SEC11 \| Signal peptidase complex catalytic subunit SEC11; afdb-proteome \| AF-P42667-F1-MODEL\_V4 \| 1.0 \| 0.0001334 \| 106 \| 0.181 \| 160 \| 93 \| 12 \| 35 \| 177 \| 8 \| 146 \| Signal peptidase complex catalytic subunit SEC11A \| Signal peptidase complex catalytic subunit SEC11A; afdb-proteome \| AF-K0F7R0-F1-MODEL\_V4 \| 1.0 \| 0.007725 \| 105 \| 0.215 \| 93 \| 61 \| 3 \| 5 \| 85 \| 40 \| 132 \| Putative DNA-binding protein \| Putative DNA-binding protein; afdb-proteome \| AF-O97066-F1-MODEL\_V4 \| 1.0 \| 0.0001057 \| 104 \| 0.149 \| 167 \| 104 \| 12 \| 28 \| 177 \| 7 \| 152 \| Signal peptidase complex catalytic subunit SEC11 \| Signal peptidase complex catalytic subunit SEC11; afdb-proteome \| AF-A0A0D2F1V0-F1-MODEL\_V4 \| 1.0 \| 0.001921 \| 104 \| 0.161 \| 130 \| 79 \| 7 \| 74 \| 177 \| 30 \| 155 \| Signal peptidase complex catalytic subunit SEC11 \| Signal peptidase complex catalytic subunit SEC11; afdb-proteome \| AF-I1KAD4-F1-MODEL\_V4 \| 1.0 \| 0.001614 \| 104 \| 0.128 \| 171 \| 92 \| 8 \| 64 \| 180 \| 24 \| 191 \| Peptidase\_S26 domain-containing protein \| Peptidase\_S26 domain-containing protein; afdb-proteome \| AF-P67812-F1-MODEL\_V4 \| 1.0 \| 0.0002834 \| 103 \| 0.219 \| 155 \| 87 \| 11 \| 45 \| 177 \| 4 \| 146 \| Signal peptidase complex catalytic subunit SEC11A \| Signal peptidase complex catalytic subunit SEC11A; afdb-proteome \| AF-B8A1P4-F1-MODEL\_V4 \| 1.0 \| 0.0003787 \| 102 \| 0.159 \| 163 \| 103 \| 9 \| 30 \| 177 \| 4 \| 147 \| Signal peptidase I \| Signal peptidase I; afdb-proteome \| AF-Q9BY50-F1-MODEL\_V4 \| 1.0 \| 0.0002001 \| 102 \| 0.168 \| 160 \| 95 \| 12 \| 35 \| 177 \| 20 \| 158 \| Signal peptidase complex catalytic subunit SEC11C \| Signal peptidase complex catalytic subunit SEC11C; afdb-proteome \| AF-Q1LVK8-F1-MODEL\_V4 \| 1.0 \| 0.0001587 \| 100 \| 0.164 \| 164 \| 99 \| 12 \| 31 \| 177 \| 4 \| 146 \| Signal peptidase complex catalytic subunit SEC11 \| Signal peptidase complex catalytic subunit SEC11; afdb-proteome \| AF-Q9I6R3-F1-MODEL\_V4 \| 0.999 \| 0.0002247 \| 98 \| 0.136 \| 198 \| 113 \| 8 \| 1 \| 169 \| 1 \| 169 \| Probable transcriptional regulator \| Probable transcriptional regulator; afdb-proteome \| AF-A0A132P6Q0-F1-MODEL\_V4 \| 0.999 \| 0.00729 \| 97 \| 0.123 \| 113 \| 84 \| 7 \| 74 \| 177 \| 17 \| 123 \| Signal peptidase \| Signal peptidase; afdb-proteome \| AF-Q9N3D0-F1-MODEL\_V4 \| 0.999 \| 0.0009038 \| 97 \| 0.164 \| 158 \| 98 \| 9 \| 35 \| 177 \| 11 \| 149 \| Signal peptidase complex catalytic subunit SEC11 \| Signal peptidase complex catalytic subunit SEC11; afdb-proteome \| AF-B4FU77-F1-MODEL\_V4 \| 0.999 \| 0.001356 \| 97 \| 0.13 \| 168 \| 89 \| 7 \| 64 \| 177 \| 32 \| 196 \| Thylakoidal processing peptidase 1 chloroplastic \| Thylakoidal processing peptidase 1 chloroplastic; afdb-proteome \| AF-Q9D8V7-F1-MODEL\_V4 \| 0.999 \| 0.0003003 \| 96 \| 0.168 \| 160 \| 95 \| 12 \| 35 \| 177 \| 20 \| 158 \| Signal peptidase complex catalytic subunit SEC11C \| Signal peptidase complex catalytic subunit SEC11C; afdb-proteome \| AF-Q8ZP82-F1-MODEL\_V4 \| 0.999 \| 0.0002834 \| 96 \| 0.124 \| 193 \| 109 \| 8 \| 4 \| 170 \| 25 \| 183 \| Putative transcriptional regulator \| Putative transcriptional regulator; afdb-proteome \| AF-Q54RP1-F1-MODEL\_V4 \| 0.999 \| 0.006491 \| 96 \| 0.176 \| 125 \| 75 \| 6 \| 74 \| 173 \| 151 \| 272 \| Uncharacterized protein \| Uncharacterized protein; afdb-proteome \| AF-A0A077ZFD1-F1-MODEL\_V4 \| 0.999 \| 0.0002001 \| 96 \| 0.154 \| 220 \| 129 \| 14 \| 8 \| 177 \| 154 \| 366 \| Signal peptidase complex catalytic subunit SEC11 \| Signal peptidase complex catalytic subunit SEC11; afdb-proteome \| AF-Q5VN14-F1-MODEL\_V4 \| 0.998 \| 0.0009038 \| 95 \| 0.141 \| 163 \| 106 \| 9 \| 30 \| 177 \| 4 \| 147 \| Signal peptidase I \| Signal peptidase I; afdb-proteome \| AF-I1MSH9-F1-MODEL\_V4 \| 0.998 \| 0.0006023 \| 95 \| 0.166 \| 186 \| 110 \| 9 \| 7 \| 177 \| 3 \| 158 \| Signal peptidase I \| Signal peptidase I; afdb-proteome \| AF-Q86JD4-F1-MODEL\_V4 \| 0.998 \| 0.0008529 \| 94 \| 0.144 \| 166 \| 108 \| 9 \| 26 \| 177 \| 1 \| 146 \| Signal peptidase complex catalytic subunit sec11 \| Signal peptidase complex catalytic subunit sec11; afdb-proteome \| AF-Q9WTR7-F1-MODEL\_V4 \| 0.998 \| 0.0004253 \| 94 \| 0.168 \| 160 \| 95 \| 12 \| 35 \| 177 \| 20 \| 158 \| Signal peptidase complex catalytic subunit SEC11C \| Signal peptidase complex catalytic subunit SEC11C; afdb-proteome \| AF-K0EK64-F1-MODEL\_V4 \| 0.998 \| 0.0002524 \| 94 \| 0.13 \| 207 \| 118 \| 10 \| 4 \| 180 \| 16 \| 190 \| XRE family transcriptional regulator \| XRE family transcriptional regulator; afdb-proteome \| AF-A0A1D6QAA1-F1-MODEL\_V4 \| 0.998 \| 0.001208 \| 94 \| 0.135 \| 170 \| 103 \| 8 \| 28 \| 177 \| 2 \| 147 \| Signal peptidase I \| Signal peptidase I; afdb-proteome \| AF-B4F8H6-F1-MODEL\_V4 \| 0.998 \| 0.00114 \| 93 \| 0.165 \| 163 \| 102 \| 9 \| 30 \| 177 \| 4 \| 147 \| Signal peptidase I \| Signal peptidase I; afdb-proteome \| AF-Q9HVS2-F1-MODEL\_V4 \| 0.997 \| 0.0002001 \| 92 \| 0.165 \| 200 \| 110 \| 12 \| 4 \| 175 \| 9 \| 179 \| Probable transcriptional regulator \| Probable transcriptional regulator; afdb-proteome \| AF-A0A1C1CFT4-F1-MODEL\_V4 \| 0.997 \| 0.0008048 \| 92 \| 0.157 \| 178 \| 101 \| 10 \| 26 \| 177 \| 1 \| 155 \| Signal peptidase complex catalytic subunit SEC11 \| Signal peptidase complex catalytic subunit SEC11; afdb-proteome \| AF-K0EMM4-F1-MODEL\_V4 \| 0.997 \| 0.0003574 \| 92 \| 0.141 \| 191 \| 117 \| 7 \| 4 \| 175 \| 17 \| 179 \| XRE family transcriptional regulator \| XRE family transcriptional regulator; afdb-proteome \| AF-Q9SSR2-F1-MODEL\_V4 \| 0.996 \| 0.001614 \| 90 \| 0.165 \| 163 \| 102 \| 9 \| 30 \| 177 \| 4 \| 147 \| Signal peptidase I \| Signal peptidase I; afdb-proteome \| AF-Q4DIR9-F1-MODEL\_V4 \| 0.996 \| 0.0006023 \| 90 \| 0.176 \| 181 \| 102 \| 10 \| 27 \| 177 \| 10 \| 173 \| Signal peptidase I \| Signal peptidase I; afdb-proteome \| AF-A0A1D6L610-F1-MODEL\_V4 \| 0.996 \| 0.00729 \| 90 \| 0.141 \| 162 \| 74 \| 8 \| 72 \| 173 \| 278 \| 434 \| Peptidase/ serine-type peptidase \| Peptidase/ serine-type peptidase; afdb-proteome \| AF-Q4DSS4-F1-MODEL\_V4 \| 0.996 \| 0.0008048 \| 89 \| 0.197 \| 177 \| 103 \| 11 \| 27 \| 177 \| 10 \| 173 \| Signal peptidase I \| Signal peptidase I; afdb-proteome \| AF-Q9LW08-F1-MODEL\_V4 \| 0.995 \| 0.002286 \| 88 \| 0.153 \| 163 \| 104 \| 9 \| 30 \| 177 \| 4 \| 147 \| Signal peptidase I \| Signal peptidase I; afdb-proteome \| AF-A0A0H3H3B8-F1-MODEL\_V4 \| 0.995 \| 0.001208 \| 88 \| 0.144 \| 166 \| 85 \| 6 \| 4 \| 137 \| 25 \| 165 \| Helix-turn-helix motif-containing protein \| Helix-turn-helix motif-containing protein; afdb-proteome \| AF-A0A133CSK7-F1-MODEL\_V4 \| 0.994 \| 0.0008048 \| 87 \| 0.1 \| 199 \| 115 \| 9 \| 4 \| 173 \| 6 \| 169 \| Cro/Cl family transcriptional regulator \| Cro/Cl family transcriptional regulator; afdb-proteome \| AF-C0NKT8-F1-MODEL\_V4 \| 0.994 \| 0.0008048 \| 87 \| 0.172 \| 180 \| 95 \| 13 \| 26 \| 177 \| 1 \| 154 \| Signal peptidase complex catalytic subunit SEC11 \| Signal peptidase complex catalytic subunit SEC11; afdb-proteome \| AF-A0A0H3GK51-F1-MODEL\_V4 \| 0.994 \| 0.001921 \| 87 \| 0.144 \| 152 \| 96 \| 8 \| 5 \| 136 \| 30 \| 167 \| Putative helix-turn-helix regulatory protein \| Putative helix-turn-helix regulatory protein; afdb-proteome \| AF-Q57Z71-F1-MODEL\_V4 \| 0.994 \| 0.0009038 \| 87 \| 0.156 \| 192 \| 118 \| 11 \| 9 \| 177 \| 5 \| 175 \| Signal peptidase I \| Signal peptidase I; afdb-proteome \| AF-K0F3Z4-F1-MODEL\_V4 \| 0.993 \| 0.00114 \| 86 \| 0.133 \| 202 \| 100 \| 10 \| 4 \| 173 \| 20 \| 178 \| DNA binding regulatory protein \| DNA binding regulatory protein; afdb-proteome \| AF-Q9HUH8-F1-MODEL\_V4 \| 0.993 \| 0.0007595 \| 86 \| 0.142 \| 197 \| 108 \| 7 \| 4 \| 169 \| 23 \| 189 \| Probable transcriptional regulator \| Probable transcriptional regulator; afdb-proteome \| AF-Q7Y0D0-F1-MODEL\_V4 \| 0.993 \| 0.008186 \| 86 \| 0.153 \| 163 \| 73 \| 9 \| 72 \| 174 \| 300 \| 457 \| Os03g0765200 protein \| Os03g0765200 protein; afdb-proteome \| AF-A0A1D6GGF0-F1-MODEL\_V4 \| 0.993 \| 0.007725 \| 86 \| 0.148 \| 162 \| 75 \| 8 \| 72 \| 174 \| 293 \| 450 \| Thylakoidal processing peptidase 1 chloroplastic \| Thylakoidal processing peptidase 1 chloroplastic; afdb-proteome \| AF-P77626-F1-MODEL\_V4 \| 0.992 \| 0.002035 \| 85 \| 0.14 \| 150 \| 100 \| 5 \| 5 \| 140 \| 12 \| 146 \| HTH-type transcriptional regulator SutR \| HTH-type transcriptional regulator SutR; afdb-proteome \| AF-O04348-F1-MODEL\_V4 \| 0.992 \| 0.005781 \| 85 \| 0.16 \| 175 \| 82 \| 9 \| 57 \| 174 \| 147 \| 313 \| Thylakoidal processing peptidase 1, chloroplastic \| Thylakoidal processing peptidase 1, chloroplastic; afdb-proteome \| AF-I1JTG1-F1-MODEL\_V4 \| 0.992 \| 0.002567 \| 85 \| 0.133 \| 202 \| 91 \| 10 \| 26 \| 173 \| 162 \| 333 \| Peptidase\_S26 domain-containing protein \| Peptidase\_S26 domain-containing protein; afdb-proteome \| AF-X8FMT1-F1-MODEL\_V4 \| 0.99 \| 0.0008529 \| 83 \| 0.128 \| 202 \| 131 \| 12 \| 1 \| 179 \| 1 \| 180 \| Helix-turn-helix family protein \| Helix-turn-helix family protein; afdb-proteome \| AF-Q32FP6-F1-MODEL\_V4 \| 0.986 \| 0.00343 \| 81 \| 0.14 \| 150 \| 100 \| 5 \| 5 \| 140 \| 12 \| 146 \| HTH cro/C1-type domain-containing protein \| HTH cro/C1-type domain-containing protein; afdb-proteome \| AF-C1GU90-F1-MODEL\_V4 \| 0.986 \| 0.005148 \| 81 \| 0.155 \| 187 \| 100 \| 10 \| 26 \| 177 \| 1 \| 164 \| Signal peptidase complex catalytic subunit SEC11 \| Signal peptidase complex catalytic subunit SEC11; afdb-proteome \| AF-Q9I601-F1-MODEL\_V4 \| 0.984 \| 0.004584 \| 80 \| 0.162 \| 154 \| 91 \| 6 \| 5 \| 136 \| 15 \| 152 \| Probable transcriptional regulator \| Probable transcriptional regulator; afdb-proteome \| AF-Q9I1G3-F1-MODEL\_V4 \| 0.984 \| 0.003237 \| 80 \| 0.101 \| 158 \| 97 \| 7 \| 5 \| 136 \| 17 \| 155 \| Probable transcriptional regulator \| Probable transcriptional regulator; afdb-proteome \| AF-K0ES92-F1-MODEL\_V4 \| 0.975 \| 0.001523 \| 77 \| 0.163 \| 190 \| 109 \| 11 \| 5 \| 172 \| 12 \| 173 \| DNA-binding protein \| DNA-binding protein; afdb-proteome \| AF-Q9I3Y5-F1-MODEL\_V4 \| 0.975 \| 0.003055 \| 77 \| 0.147 \| 190 \| 104 \| 10 \| 5 \| 173 \| 49 \| 201 \| Probable transcriptional regulator \| Probable transcriptional regulator; afdb-proteome \| AF-A0A5K4EAL6-F1-MODEL\_V4 \| 0.971 \| 0.009193 \| 76 \| 0.165 \| 139 \| 79 \| 8 \| 68 \| 173 \| 7 \| 141 \| Mitochondrial signal peptidase (S26 family) \| Mitochondrial signal peptidase (S26 family); afdb-proteome \| AF-Q2G2A6-F1-MODEL\_V4 \| 0.967 \| 0.005455 \| 75 \| 0.125 \| 176 \| 111 \| 9 \| 4 \| 153 \| 6 \| 164 \| HTH cro/C1-type domain-containing protein \| HTH cro/C1-type domain-containing protein; afdb-proteome \| AF-Q9I2L3-F1-MODEL\_V4 \| 0.967 \| 0.003055 \| 75 \| 0.16 \| 200 \| 109 \| 11 \| 4 \| 171 \| 10 \| 182 \| Probable transcriptional regulator \| Probable transcriptional regulator; afdb-proteome \| AF-K0EYC2-F1-MODEL\_V4 \| 0.967 \| 0.002422 \| 75 \| 0.158 \| 202 \| 118 \| 10 \| 5 \| 179 \| 19 \| 195 \| Regulatory protein \| Regulatory protein; afdb-proteome \| AF-A0A0H3GYK3-F1-MODEL\_V4 \| 0.961 \| 0.002157 \| 74 \| 0.162 \| 197 \| 113 \| 11 \| 4 \| 173 \| 15 \| 186 \| Putative regulator \| Putative regulator; afdb-proteome \| AF-A0A0H3GP19-F1-MODEL\_V4 \| 0.956 \| 0.004082 \| 73 \| 0.135 \| 184 \| 113 \| 8 \| 5 \| 169 \| 11 \| 167 \| Putative regulator \| Putative regulator; afdb-proteome \| AF-Q8ZPC6-F1-MODEL\_V4 \| 0.949 \| 0.004584 \| 72 \| 0.121 \| 197 \| 121 \| 11 \| 5 \| 179 \| 12 \| 178 \| Putative oxidoreductase \| Putative oxidoreductase; afdb-proteome \| AF-Q32F57-F1-MODEL\_V4 \| 0.949 \| 0.005455 \| 72 \| 0.13 \| 192 \| 119 \| 10 \| 4 \| 176 \| 11 \| 173 \| HTH cro/C1-type domain-containing protein \| HTH cro/C1-type domain-containing protein; afdb-proteome \| AF-Q8H0W1-F1-MODEL\_V4 \| 0.933 \| 0.003237 \| 70 \| 0.14 \| 178 \| 86 \| 10 \| 61 \| 174 \| 94 \| 268 \| Chloroplast processing peptidase \| Chloroplast processing peptidase; afdb-proteome \| AF-Q8ZKS3-F1-MODEL\_V4 \| 0.9 \| 0.006879 \| 67 \| 0.11 \| 199 \| 104 \| 8 \| 5 \| 172 \| 18 \| 174 \| HTH cro/C1-type domain-containing protein \| HTH cro/C1-type domain-containing protein; afdb-proteome \| AF-A0A133CSL8-F1-MODEL\_V4 \| 0.872 \| 0.008675 \| 65 \| 0.116 \| 248 \| 115 \| 14 \| 4 \| 166 \| 11 \| 239 \| Helix-turn-helix domain-containing protein \| Helix-turn-helix domain-containing protein; afdb-uniprot50 \| AF-A0A139JVH4-F1-MODEL\_V4 \| 1.0 \| 3.747e-13 \| 391 \| 0.215 \| 190 \| 135 \| 5 \| 4 \| 180 \| 12 \| 200 \| DNA-binding helix-turn-helix protein \| DNA-binding helix-turn-helix protein; afdb-uniprot50 \| AF-A0A316C0P5-F1-MODEL\_V4 \| 1.0 \| 5.306e-13 \| 371 \| 0.2 \| 200 \| 137 \| 3 \| 1 \| 180 \| 1 \| 197 \| DNA-binding XRE family transcriptional regulator \| DNA-binding XRE family transcriptional regulator; afdb-uniprot50 \| AF-A0A1C7W9W0-F1-MODEL\_V4 \| 1.0 \| 5.622e-13 \| 366 \| 0.25 \| 216 \| 119 \| 6 \| 4 \| 180 \| 6 \| 217 \| HTH cro/C1-type domain-containing protein \| HTH cro/C1-type domain-containing protein; afdb-uniprot50 \| AF-A0A661FLS5-F1-MODEL\_V4 \| 1.0 \| 4.458e-13 \| 365 \| 0.207 \| 222 \| 129 \| 7 \| 4 \| 180 \| 24 \| 243 \| HTH cro/C1-type domain-containing protein \| HTH cro/C1-type domain-containing protein; afdb-uniprot50 \| AF-A0A2E3AY80-F1-MODEL\_V4 \| 1.0 \| 1.9e-12 \| 362 \| 0.276 \| 210 \| 117 \| 7 \| 4 \| 180 \| 9 \| 216 \| Peptidase S24 \| Peptidase S24; afdb-uniprot50 \| AF-A0A1S8YLB7-F1-MODEL\_V4 \| 1.0 \| 3.148e-13 \| 360 \| 0.243 \| 222 \| 121 \| 7 \| 4 \| 180 \| 6 \| 225 \| Transcriptional regulator \| Transcriptional regulator; afdb-uniprot50 \| AF-A0A1Q6SZL5-F1-MODEL\_V4 \| 1.0 \| 1.422e-12 \| 359 \| 0.255 \| 180 \| 122 \| 3 \| 1 \| 180 \| 1 \| 168 \| HTH cro/C1-type domain-containing protein \| HTH cro/C1-type domain-containing protein; afdb-uniprot50 \| AF-A0A7U4Q650-F1-MODEL\_V4 \| 1.0 \| 8.941e-13 \| 359 \| 0.226 \| 221 \| 125 \| 6 \| 4 \| 180 \| 6 \| 224 \| Helix-turn-helix transcriptional regulator \| Helix-turn-helix transcriptional regulator; afdb-uniprot50 \| AF-Q57QA9-F1-MODEL\_V4 \| 1.0 \| 3.97e-13 \| 359 \| 0.27 \| 222 \| 110 \| 7 \| 4 \| 177 \| 6 \| 223 \| Regulatory protein \| Regulatory protein; afdb-uniprot50 \| AF-A0A327JZK4-F1-MODEL\_V4 \| 1.0 \| 1.355e-08 \| 350 \| 0.349 \| 106 \| 65 \| 2 \| 78 \| 180 \| 49 \| 153 \| Peptidase\_S24 domain-containing protein \| Peptidase\_S24 domain-containing protein; afdb-uniprot50 \| AF-A0A7V8G8N6-F1-MODEL\_V4 \| 1.0 \| 3.022e-12 \| 348 \| 0.215 \| 213 \| 130 \| 6 \| 4 \| 180 \| 28 \| 239 \| Putative HTH-type transcriptional regulator \| Putative HTH-type transcriptional regulator; afdb-uniprot50 \| AF-A0A6F8VBR1-F1-MODEL\_V4 \| 1.0 \| 4.038e-12 \| 348 \| 0.213 \| 239 \| 124 \| 6 \| 4 \| 180 \| 10 \| 246 \| Repressor \| Repressor; afdb-uniprot50 \| AF-A0A1H0M9N5-F1-MODEL\_V4 \| 1.0 \| 2.691e-12 \| 348 \| 0.222 \| 216 \| 128 \| 6 \| 4 \| 180 \| 51 \| 265 \| Phage repressor protein C, contains Cro/C1-type HTH and peptisase s24 domains \| Phage repressor protein C, contains Cro/C1-type HTH and peptisase s24 domains; afdb-uniprot50 \| AF-A0LD08-F1-MODEL\_V4 \| 1.0 \| 4.279e-12 \| 346 \| 0.252 \| 214 \| 117 \| 7 \| 4 \| 180 \| 8 \| 215 \| Putative phage repressor \| Putative phage repressor; afdb-uniprot50 \| AF-A0A2A4XTD6-F1-MODEL\_V4 \| 1.0 \| 3.393e-12 \| 345 \| 0.226 \| 212 \| 127 \| 6 \| 4 \| 180 \| 13 \| 222 \| HTH cro/C1-type domain-containing protein \| HTH cro/C1-type domain-containing protein; afdb-uniprot50 \| AF-K2J5W2-F1-MODEL\_V4 \| 1.0 \| 6.757e-09 \| 344 \| 0.318 \| 113 \| 69 \| 3 \| 71 \| 180 \| 50 \| 157 \| Phage repressor \| Phage repressor; afdb-uniprot50 \| AF-K2JV89-F1-MODEL\_V4 \| 1.0 \| 1.597e-12 \| 343 \| 0.252 \| 206 \| 121 \| 6 \| 4 \| 180 \| 25 \| 226 \| Putative phage repressor \| Putative phage repressor; afdb-uniprot50 \| AF-A0A2X1U723-F1-MODEL\_V4 \| 1.0 \| 4.279e-12 \| 343 \| 0.21 \| 219 \| 131 \| 7 \| 2 \| 180 \| 230 \| 446 \| Putative regulatory protein \| Putative regulatory protein; afdb-uniprot50 \| AF-A0A1I4Q661-F1-MODEL\_V4 \| 1.0 \| 7.642e-12 \| 341 \| 0.211 \| 222 \| 125 \| 6 \| 4 \| 180 \| 8 \| 224 \| HTH cro/C1-type domain-containing protein \| HTH cro/C1-type domain-containing protein; afdb-uniprot50 \| AF-D4BUI1-F1-MODEL\_V4 \| 1.0 \| 7.212e-12 \| 341 \| 0.201 \| 223 \| 131 \| 4 \| 4 \| 180 \| 7 \| 228 \| DNA-binding helix-turn-helix protein \| DNA-binding helix-turn-helix protein; afdb-uniprot50 \| AF-A0A315BKI3-F1-MODEL\_V4 \| 1.0 \| 5.143e-08 \| 339 \| 0.294 \| 95 \| 66 \| 1 \| 87 \| 180 \| 80 \| 174 \| Peptidase\_S24 domain-containing protein \| Peptidase\_S24 domain-containing protein; afdb-uniprot50 \| AF-A0A495RIU6-F1-MODEL\_V4 \| 1.0 \| 3.202e-12 \| 339 \| 0.207 \| 212 \| 130 \| 7 \| 4 \| 180 \| 6 \| 214 \| Phage repressor protein C with HTH and peptisase S24 domain \| Phage repressor protein C with HTH and peptisase S24 domain; afdb-uniprot50 \| AF-A0A2D5PK86-F1-MODEL\_V4 \| 1.0 \| 3.022e-12 \| 339 \| 0.213 \| 225 \| 129 \| 5 \| 4 \| 180 \| 6 \| 230 \| HTH cro/C1-type domain-containing protein \| HTH cro/C1-type domain-containing protein; afdb-uniprot50 \| AF-U0ZRD1-F1-MODEL\_V4 \| 1.0 \| 5.719e-12 \| 337 \| 0.201 \| 208 \| 128 \| 5 \| 4 \| 178 \| 7 \| 209 \| HTH cro/C1-type domain-containing protein \| HTH cro/C1-type domain-containing protein; afdb-uniprot50 \| AF-A0A2Z4R558-F1-MODEL\_V4 \| 1.0 \| 6.422e-12 \| 337 \| 0.192 \| 213 \| 135 \| 5 \| 4 \| 180 \| 9 \| 220 \| HTH cro/C1-type domain-containing protein \| HTH cro/C1-type domain-containing protein; afdb-uniprot50 \| AF-A0A2R3QQH4-F1-MODEL\_V4 \| 1.0 \| 5.719e-12 \| 337 \| 0.206 \| 208 \| 131 \| 6 \| 4 \| 180 \| 28 \| 232 \| HTH cro/C1-type domain-containing protein \| HTH cro/C1-type domain-containing protein; afdb-uniprot50 \| AF-A0A844BRP4-F1-MODEL\_V4 \| 1.0 \| 7.642e-12 \| 335 \| 0.228 \| 197 \| 128 \| 4 \| 4 \| 180 \| 9 \| 201 \| Helix-turn-helix domain-containing protein \| Helix-turn-helix domain-containing protein; afdb-uniprot50 \| AF-A0A4U0PX94-F1-MODEL\_V4 \| 1.0 \| 8.582e-12 \| 335 \| 0.208 \| 230 \| 128 \| 4 \| 4 \| 180 \| 7 \| 235 \| Helix-turn-helix domain-containing protein \| Helix-turn-helix domain-containing protein; afdb-uniprot50 \| AF-A0A836WXJ3-F1-MODEL\_V4 \| 1.0 \| 1.288e-11 \| 334 \| 0.221 \| 226 \| 126 \| 5 \| 1 \| 180 \| 1 \| 222 \| Helix-turn-helix transcriptional regulator \| Helix-turn-helix transcriptional regulator; afdb-uniprot50 \| AF-A0A7T9LJB7-F1-MODEL\_V4 \| 1.0 \| 1.365e-11 \| 333 \| 0.207 \| 222 \| 131 \| 6 \| 2 \| 180 \| 5 \| 224 \| Helix-turn-helix transcriptional regulator \| Helix-turn-helix transcriptional regulator; afdb-uniprot50 \| AF-A0A2P5N6V1-F1-MODEL\_V4 \| 1.0 \| 4.279e-12 \| 333 \| 0.19 \| 242 \| 129 \| 6 \| 5 \| 180 \| 41 \| 281 \| HTH cro/C1-type domain-containing protein \| HTH cro/C1-type domain-containing protein; afdb-uniprot50 \| AF-A0A6N7BSU4-F1-MODEL\_V4 \| 1.0 \| 1.082e-11 \| 332 \| 0.218 \| 197 \| 127 \| 4 \| 4 \| 180 \| 8 \| 197 \| HTH cro/C1-type domain-containing protein \| HTH cro/C1-type domain-containing protein; afdb-uniprot50 \| AF-A0A6F8VD58-F1-MODEL\_V4 \| 1.0 \| 7.642e-12 \| 332 \| 0.235 \| 212 \| 125 \| 6 \| 5 \| 180 \| 7 \| 217 \| HTH cro/C1-type domain-containing protein \| HTH cro/C1-type domain-containing protein; afdb-uniprot50 \| AF-A0A2N9XPR0-F1-MODEL\_V4 \| 1.0 \| 5.45e-08 \| 331 \| 0.284 \| 95 \| 64 \| 2 \| 89 \| 180 \| 6 \| 99 \| Peptidase\_S24 domain-containing protein \| Peptidase\_S24 domain-containing protein; afdb-uniprot50 \| AF-A0A2N9XWM8-F1-MODEL\_V4 \| 1.0 \| 2.565e-08 \| 331 \| 0.273 \| 106 \| 73 \| 2 \| 78 \| 180 \| 7 \| 111 \| Peptidase\_S24 domain-containing protein \| Peptidase\_S24 domain-containing protein; afdb-uniprot50 \| AF-A0A6N7AJK2-F1-MODEL\_V4 \| 1.0 \| 1.824e-11 \| 331 \| 0.231 \| 207 \| 124 \| 7 \| 5 \| 180 \| 20 \| 222 \| Prophage MuMc02 S24 family peptidase \| Prophage MuMc02 S24 family peptidase; afdb-uniprot50 \| AF-A0A1G0CQ54-F1-MODEL\_V4 \| 1.0 \| 5.45e-08 \| 330 \| 0.345 \| 107 \| 66 \| 2 \| 77 \| 180 \| 69 \| 174 \| Peptidase\_S24 domain-containing protein \| Peptidase\_S24 domain-containing protein; afdb-uniprot50 \| AF-A0A139KEG7-F1-MODEL\_V4 \| 1.0 \| 1.365e-11 \| 330 \| 0.197 \| 213 \| 134 \| 5 \| 4 \| 180 \| 6 \| 217 \| Peptidase S24-like protein \| Peptidase S24-like protein; afdb-uniprot50 \| AF-A0A2U0YHI6-F1-MODEL\_V4 \| 1.0 \| 1.933e-11 \| 329 \| 0.248 \| 209 \| 122 \| 8 \| 4 \| 180 \| 20 \| 225 \| Phage repressor protein C with HTH and peptisase S24 domain \| Phage repressor protein C with HTH and peptisase S24 domain; afdb-uniprot50 \| AF-A0A1H7XMA1-F1-MODEL\_V4 \| 1.0 \| 5.092e-12 \| 329 \| 0.2 \| 220 \| 130 \| 8 \| 4 \| 180 \| 7 \| 223 \| Phage repressor protein C, contains Cro/C1-type HTH and peptisase s24 domains \| Phage repressor protein C, contains Cro/C1-type HTH and peptisase s24 domains; afdb-uniprot50 \| AF-A0A7Y5VHZ7-F1-MODEL\_V4 \| 1.0 \| 1.215e-11 \| 329 \| 0.188 \| 228 \| 132 \| 5 \| 4 \| 179 \| 7 \| 233 \| Helix-turn-helix domain-containing protein \| Helix-turn-helix domain-containing protein; afdb-uniprot50 \| AF-A0A2N9Y298-F1-MODEL\_V4 \| 1.0 \| 1.933e-11 \| 328 \| 0.197 \| 192 \| 137 \| 3 \| 4 \| 180 \| 6 \| 195 \| HTH cro/C1-type domain-containing protein \| HTH cro/C1-type domain-containing protein; afdb-uniprot50 \| AF-A0A6S5X474-F1-MODEL\_V4 \| 1.0 \| 3.876e-11 \| 328 \| 0.2 \| 209 \| 131 \| 4 \| 4 \| 180 \| 11 \| 215 \| HTH cro/C1-type domain-containing protein \| HTH cro/C1-type domain-containing protein; afdb-uniprot50 \| AF-A0A7Z6VI80-F1-MODEL\_V4 \| 1.0 \| 2.048e-11 \| 328 \| 0.2 \| 215 \| 130 \| 6 \| 4 \| 180 \| 8 \| 218 \| Helix-turn-helix transcriptional regulator \| Helix-turn-helix transcriptional regulator; afdb-uniprot50 \| AF-B9Z333-F1-MODEL\_V4 \| 1.0 \| 8.582e-12 \| 328 \| 0.2 \| 224 \| 130 \| 6 \| 4 \| 180 \| 6 \| 227 \| Putative phage repressor \| Putative phage repressor; afdb-uniprot50 \| AF-G8NRH6-F1-MODEL\_V4 \| 1.0 \| 8.582e-12 \| 328 \| 0.251 \| 235 \| 116 \| 9 \| 1 \| 176 \| 1 \| 234 \| Putative phage repressor \| Putative phage repressor; afdb-uniprot50 \| AF-A0A7W5EVC8-F1-MODEL\_V4 \| 1.0 \| 2.88e-08 \| 327 \| 0.304 \| 105 \| 68 \| 3 \| 79 \| 180 \| 36 \| 138 \| Phage repressor protein C with HTH and peptisase S24 domain \| Phage repressor protein C with HTH and peptisase S24 domain; afdb-uniprot50 \| AF-A0A1C4CK44-F1-MODEL\_V4 \| 1.0 \| 9.637e-12 \| 327 \| 0.226 \| 216 \| 123 \| 6 \| 5 \| 180 \| 9 \| 220 \| Peptidase S24-like \| Peptidase S24-like; afdb-uniprot50 \| AF-G2ZXI3-F1-MODEL\_V4 \| 1.0 \| 2.17e-11 \| 327 \| 0.209 \| 224 \| 125 \| 7 \| 4 \| 180 \| 7 \| 225 \| Putative phage repressor \| Putative phage repressor; afdb-uniprot50 \| AF-A0A3S2YW43-F1-MODEL\_V4 \| 1.0 \| 1.365e-11 \| 327 \| 0.205 \| 238 \| 127 \| 7 \| 4 \| 180 \| 22 \| 258 \| Peptidase\_S24 domain-containing protein \| Peptidase\_S24 domain-containing protein; afdb-uniprot50 \| AF-A0A1H5XPH2-F1-MODEL\_V4 \| 1.0 \| 5.092e-12 \| 327 \| 0.212 \| 231 \| 126 \| 8 \| 5 \| 180 \| 33 \| 262 \| Phage repressor protein C, contains Cro/C1-type HTH and peptisase s24 domains \| Phage repressor protein C, contains Cro/C1-type HTH and peptisase s24 domains; afdb-uniprot50 \| AF-D5RM48-F1-MODEL\_V4 \| 1.0 \| 1.082e-11 \| 327 \| 0.201 \| 228 \| 129 \| 5 \| 4 \| 180 \| 39 \| 264 \| Peptidase S24-like protein \| Peptidase S24-like protein; afdb-uniprot50 \| AF-X2JS37-F1-MODEL\_V4 \| 1.0 \| 3.876e-11 \| 327 \| 0.218 \| 220 \| 127 \| 7 \| 4 \| 180 \| 72 \| 289 \| HTH cro/C1-type domain-containing protein \| HTH cro/C1-type domain-containing protein; afdb-uniprot50 \| AF-A0A0Q5YYS1-F1-MODEL\_V4 \| 1.0 \| 9.637e-12 \| 326 \| 0.219 \| 210 \| 131 \| 7 \| 4 \| 180 \| 7 \| 216 \| HTH cro/C1-type domain-containing protein \| HTH cro/C1-type domain-containing protein; afdb-uniprot50 \| AF-F3BKE6-F1-MODEL\_V4 \| 1.0 \| 9.094e-12 \| 325 \| 0.255 \| 192 \| 124 \| 5 \| 4 \| 180 \| 19 \| 206 \| HTH cro/C1-type domain-containing protein \| HTH cro/C1-type domain-containing protein; afdb-uniprot50 \| AF-A0A1H2HQC4-F1-MODEL\_V4 \| 1.0 \| 5.775e-08 \| 325 \| 0.307 \| 104 \| 68 \| 2 \| 80 \| 180 \| 126 \| 228 \| Peptidase S24-like \| Peptidase S24-like; afdb-uniprot50 \| AF-A0A3M2H9H5-F1-MODEL\_V4 \| 1.0 \| 1.082e-11 \| 325 \| 0.202 \| 207 \| 134 \| 6 \| 4 \| 180 \| 25 \| 230 \| XRE family transcriptional regulator \| XRE family transcriptional regulator; afdb-uniprot50 \| AF-A0A1I3KW04-F1-MODEL\_V4 \| 1.0 \| 9.637e-12 \| 325 \| 0.196 \| 224 \| 133 \| 4 \| 4 \| 180 \| 7 \| 230 \| Peptidase S24-like \| Peptidase S24-like; afdb-uniprot50 \| AF-A0A348HFP4-F1-MODEL\_V4 \| 1.0 \| 1.824e-11 \| 325 \| 0.221 \| 221 \| 127 \| 7 \| 4 \| 180 \| 12 \| 231 \| Predicted transcriptional regulator \| Predicted transcriptional regulator; afdb-uniprot50 \| AF-A0A2I6S9H6-F1-MODEL\_V4 \| 1.0 \| 3.658e-11 \| 325 \| 0.189 \| 232 \| 131 \| 6 \| 4 \| 180 \| 6 \| 235 \| HTH cro/C1-type domain-containing protein \| HTH cro/C1-type domain-containing protein; afdb-uniprot50 \| AF-A0A2G5FSI2-F1-MODEL\_V4 \| 1.0 \| 1.288e-11 \| 325 \| 0.209 \| 215 \| 127 \| 8 \| 4 \| 180 \| 37 \| 246 \| C repressor \| C repressor; afdb-uniprot50 \| AF-A5EUX6-F1-MODEL\_V4 \| 1.0 \| 1.824e-11 \| 325 \| 0.188 \| 228 \| 131 \| 7 \| 5 \| 180 \| 80 \| 305 \| Regulatory protein RegA2 \| Regulatory protein RegA2; afdb-uniprot50 \| AF-A0A1Q6UEY7-F1-MODEL\_V4 \| 1.0 \| 6.922e-11 \| 324 \| 0.2 \| 215 \| 133 \| 7 \| 1 \| 180 \| 1 \| 211 \| HTH cro/C1-type domain-containing protein \| HTH cro/C1-type domain-containing protein; afdb-uniprot50 \| AF-A0A2D9F6P6-F1-MODEL\_V4 \| 1.0 \| 2.048e-11 \| 324 \| 0.222 \| 220 \| 126 \| 7 \| 2 \| 180 \| 8 \| 223 \| HTH cro/C1-type domain-containing protein \| HTH cro/C1-type domain-containing protein; afdb-uniprot50 \| AF-A0A1Y2S5X0-F1-MODEL\_V4 \| 1.0 \| 1.288e-11 \| 324 \| 0.19 \| 221 \| 129 \| 9 \| 4 \| 178 \| 9 \| 225 \| Transcriptional regulator \| Transcriptional regulator; afdb-uniprot50 \| AF-A0A7I6ZJ48-F1-MODEL\_V4 \| 1.0 \| 6.12e-08 \| 324 \| 0.245 \| 106 \| 76 \| 2 \| 78 \| 180 \| 133 \| 237 \| Uncharacterized protein \| Uncharacterized protein; afdb-uniprot50 \| AF-A0A7W5EKL1-F1-MODEL\_V4 \| 1.0 \| 2.437e-11 \| 324 \| 0.229 \| 231 \| 122 \| 6 \| 4 \| 179 \| 30 \| 259 \| Phage repressor protein C with HTH and peptisase S24 domain \| Phage repressor protein C with HTH and peptisase S24 domain; afdb-uniprot50 \| AF-A0A0K6IU92-F1-MODEL\_V4 \| 1.0 \| 1.533e-11 \| 324 \| 0.192 \| 265 \| 124 \| 5 \| 4 \| 180 \| 23 \| 285 \| Peptidase S24-like/Helix-turn-helix \| Peptidase S24-like/Helix-turn-helix; afdb-uniprot50 \| AF-J6DSP2-F1-MODEL\_V4 \| 1.0 \| 1.093e-07 \| 323 \| 0.304 \| 105 \| 69 \| 2 \| 79 \| 180 \| 36 \| 139 \| Prophage MuSo1, transcriptional regulator, Cro/CI family protein \| Prophage MuSo1, transcriptional regulator, Cro/CI family protein; afdb-uniprot50 \| AF-A0A0F2RP23-F1-MODEL\_V4 \| 1.0 \| 1.227e-07 \| 323 \| 0.361 \| 105 \| 64 \| 3 \| 78 \| 180 \| 40 \| 143 \| Peptidase\_S24 domain-containing protein \| Peptidase\_S24 domain-containing protein; afdb-uniprot50 \| AF-A0A1S6HW15-F1-MODEL\_V4 \| 1.0 \| 1.824e-11 \| 323 \| 0.2 \| 205 \| 134 \| 6 \| 4 \| 180 \| 16 \| 218 \| Helix-turn-helix domain \| Helix-turn-helix domain; afdb-uniprot50 \| AF-A0A378NHI5-F1-MODEL\_V4 \| 1.0 \| 1.365e-11 \| 323 \| 0.21 \| 214 \| 129 \| 8 \| 4 \| 180 \| 7 \| 217 \| Uncharacterized HTH-type transcriptional regulator HI\_1476 \| Uncharacterized HTH-type transcriptional regulator HI\_1476; afdb-uniprot50 \| AF-A0A3B7PQJ3-F1-MODEL\_V4 \| 1.0 \| 1.533e-11 \| 323 \| 0.19 \| 220 \| 131 \| 6 \| 4 \| 178 \| 6 \| 223 \| Helix-turn-helix transcriptional regulator \| Helix-turn-helix transcriptional regulator; afdb-uniprot50 \| AF-H8W5A1-F1-MODEL\_V4 \| 1.0 \| 5.45e-08 \| 322 \| 0.301 \| 106 \| 70 \| 2 \| 78 \| 180 \| 70 \| 174 \| Putative transcriptional regulator, Cro/CI family \| Putative transcriptional regulator, Cro/CI family; afdb-uniprot50 \| AF-H8L1T9-F1-MODEL\_V4 \| 1.0 \| 2.9e-11 \| 322 \| 0.228 \| 219 \| 126 \| 6 \| 4 \| 180 \| 8 \| 225 \| Putative transcriptional regulator \| Putative transcriptional regulator; afdb-uniprot50 \| AF-A0A443VFS7-F1-MODEL\_V4 \| 1.0 \| 2.17e-11 \| 322 \| 0.232 \| 219 \| 124 \| 8 \| 4 \| 180 \| 8 \| 224 \| HTH cro/C1-type domain-containing protein \| HTH cro/C1-type domain-containing protein; afdb-uniprot50 \| AF-A0A2D6GSD7-F1-MODEL\_V4 \| 1.0 \| 9.637e-12 \| 322 \| 0.235 \| 217 \| 125 \| 6 \| 4 \| 180 \| 18 \| 233 \| Phage repressor protein C \| Phage repressor protein C; afdb-uniprot50 \| AF-A0A367LYV8-F1-MODEL\_V4 \| 1.0 \| 1.031e-07 \| 321 \| 0.292 \| 99 \| 66 \| 2 \| 85 \| 180 \| 5 \| 102 \| S24 family peptidase \| S24 family peptidase; afdb-uniprot50 \| AF-A0A5B9EHR9-F1-MODEL\_V4 \| 1.0 \| 7.283e-08 \| 321 \| 0.257 \| 105 \| 74 \| 2 \| 78 \| 179 \| 51 \| 154 \| S24 family peptidase \| S24 family peptidase; afdb-uniprot50 \| AF-A0A1R1LYN7-F1-MODEL\_V4 \| 1.0 \| 1.446e-11 \| 321 \| 0.22 \| 222 \| 122 \| 7 \| 4 \| 180 \| 6 \| 221 \| HTH cro/C1-type domain-containing protein \| HTH cro/C1-type domain-containing protein; afdb-uniprot50 \| AF-A0A561H5B3-F1-MODEL\_V4 \| 1.0 \| 1.824e-11 \| 321 \| 0.233 \| 201 \| 129 \| 7 \| 4 \| 180 \| 92 \| 291 \| Phage repressor protein C with HTH and peptisase S24 domain \| Phage repressor protein C with HTH and peptisase S24 domain; afdb-uniprot50 \| AF-A0A192IPT4-F1-MODEL\_V4 \| 1.0 \| 3.234e-08 \| 320 \| 0.291 \| 120 \| 81 \| 2 \| 64 \| 180 \| 17 \| 135 \| Peptidase\_S24 domain-containing protein \| Peptidase\_S24 domain-containing protein; afdb-uniprot50 \| AF-A0A0Q5GW81-F1-MODEL\_V4 \| 1.0 \| 6.12e-08 \| 320 \| 0.263 \| 114 \| 77 \| 2 \| 70 \| 180 \| 28 \| 137 \| Peptidase\_S24 domain-containing protein \| Peptidase\_S24 domain-containing protein; afdb-uniprot50 \| AF-A0A317EGV6-F1-MODEL\_V4 \| 1.0 \| 1.721e-11 \| 320 \| 0.237 \| 223 \| 125 \| 7 \| 1 \| 180 \| 1 \| 221 \| HTH cro/C1-type domain-containing protein \| HTH cro/C1-type domain-containing protein; afdb-uniprot50 \| AF-A0A4R2GU03-F1-MODEL\_V4 \| 1.0 \| 1.624e-11 \| 320 \| 0.202 \| 217 \| 130 \| 5 \| 3 \| 180 \| 11 \| 223 \| Helix-turn-helix protein \| Helix-turn-helix protein; afdb-uniprot50 \| AF-G2IX61-F1-MODEL\_V4 \| 1.0 \| 1.933e-11 \| 320 \| 0.191 \| 219 \| 133 \| 4 \| 4 \| 178 \| 7 \| 225 \| Putative phage repressor \| Putative phage repressor; afdb-uniprot50 \| AF-C6XCG7-F1-MODEL\_V4 \| 1.0 \| 1.031e-07 \| 320 \| 0.35 \| 100 \| 61 \| 2 \| 85 \| 180 \| 131 \| 230 \| Putative phage repressor \| Putative phage repressor; afdb-uniprot50 \| AF-A0A3M1A2A7-F1-MODEL\_V4 \| 1.0 \| 1.64e-07 \| 319 \| 0.257 \| 105 \| 74 \| 2 \| 79 \| 180 \| 70 \| 173 \| Helix-turn-helix transcriptional regulator \| Helix-turn-helix transcriptional regulator; afdb-uniprot50 \| AF-A0A7C3DGY8-F1-MODEL\_V4 \| 1.0 \| 1.824e-11 \| 319 \| 0.205 \| 219 \| 130 \| 6 \| 4 \| 180 \| 10 \| 226 \| Helix-turn-helix transcriptional regulator \| Helix-turn-helix transcriptional regulator; afdb-uniprot50 \| AF-A0A439P541-F1-MODEL\_V4 \| 1.0 \| 7.335e-11 \| 319 \| 0.204 \| 205 \| 131 \| 6 \| 1 \| 180 \| 39 \| 236 \| Helix-turn-helix domain-containing protein \| Helix-turn-helix domain-containing protein; afdb-uniprot50 \| AF-A0A2W6V6P1-F1-MODEL\_V4 \| 1.0 \| 1.933e-11 \| 319 \| 0.237 \| 215 \| 124 \| 5 \| 4 \| 180 \| 30 \| 242 \| Helix-turn-helix transcriptional regulator \| Helix-turn-helix transcriptional regulator; afdb-uniprot50 \| AF-A0A1S1NYL5-F1-MODEL\_V4 \| 1.0 \| 1.933e-11 \| 319 \| 0.221 \| 239 \| 121 \| 5 \| 4 \| 178 \| 8 \| 245 \| Helix-turn-helix transcriptional regulator \| Helix-turn-helix transcriptional regulator; afdb-uniprot50 \| AF-A0A5B8R9J4-F1-MODEL\_V4 \| 1.0 \| 4.888e-11 \| 318 \| 0.247 \| 202 \| 125 \| 6 \| 4 \| 180 \| 28 \| 227 \| Putative HTH-type transcriptional regulator \| Putative HTH-type transcriptional regulator; afdb-uniprot50 \| AF-A0A5P1R9H6-F1-MODEL\_V4 \| 1.0 \| 2.737e-11 \| 318 \| 0.228 \| 223 \| 124 \| 8 \| 4 \| 180 \| 6 \| 226 \| Helix-turn-helix domain-containing protein \| Helix-turn-helix domain-containing protein; afdb-uniprot50 \| AF-A0A077F5X8-F1-MODEL\_V4 \| 1.0 \| 2.437e-11 \| 318 \| 0.167 \| 233 \| 138 \| 3 \| 4 \| 180 \| 7 \| 239 \| Phage transcriptional regulator \| Phage transcriptional regulator; afdb-uniprot50 \| AF-A0A1S6HSY5-F1-MODEL\_V4 \| 1.0 \| 1.365e-11 \| 318 \| 0.188 \| 228 \| 133 \| 5 \| 4 \| 180 \| 17 \| 243 \| Putative transcriptional regulator \| Putative transcriptional regulator; afdb-uniprot50 \| AF-A0A5C1E747-F1-MODEL\_V4 \| 1.0 \| 2.737e-11 \| 318 \| 0.2 \| 240 \| 129 \| 8 \| 4 \| 180 \| 19 \| 258 \| HTH cro/C1-type domain-containing protein \| HTH cro/C1-type domain-containing protein; afdb-uniprot50 \| AF-A0A2D2D3J4-F1-MODEL\_V4 \| 1.0 \| 2.3e-11 \| 317 \| 0.237 \| 215 \| 124 \| 6 \| 4 \| 180 \| 18 \| 230 \| Phage repressor protein \| Phage repressor protein; afdb-uniprot50 \| AF-P71159-F1-MODEL\_V4 \| 1.0 \| 4.612e-11 \| 317 \| 0.192 \| 228 \| 130 \| 5 \| 5 \| 180 \| 7 \| 232 \| Regulatory protein \| Regulatory protein; afdb-uniprot50 \| AF-A0A345Y6T3-F1-MODEL\_V4 \| 1.0 \| 2.583e-11 \| 317 \| 0.191 \| 230 \| 131 \| 6 \| 5 \| 180 \| 13 \| 241 \| Helix-turn-helix transcriptional regulator \| Helix-turn-helix transcriptional regulator; afdb-uniprot50 \| AF-A0A0C1Q293-F1-MODEL\_V4 \| 1.0 \| 2.437e-11 \| 317 \| 0.209 \| 220 \| 128 \| 7 \| 5 \| 180 \| 26 \| 243 \| Repressor \| Repressor; afdb-uniprot50 \| AF-A0A7Z8ZKU1-F1-MODEL\_V4 \| 1.0 \| 2.9e-11 \| 317 \| 0.186 \| 220 \| 134 \| 7 \| 4 \| 180 \| 32 \| 249 \| DNA-binding/peptidase S24 domain protein \| DNA-binding/peptidase S24 domain protein; afdb-uniprot50 \| AF-A0A545SG22-F1-MODEL\_V4 \| 1.0 \| 3.257e-11 \| 317 \| 0.224 \| 223 \| 127 \| 7 \| 1 \| 179 \| 37 \| 257 \| Helix-turn-helix transcriptional regulator \| Helix-turn-helix transcriptional regulator; afdb-uniprot50 \| AF-A0A1Q3UZ38-F1-MODEL\_V4 \| 1.0 \| 7.718e-08 \| 316 \| 0.292 \| 113 \| 72 \| 3 \| 71 \| 180 \| 3 \| 110 \| Peptidase\_S24 domain-containing protein \| Peptidase\_S24 domain-containing protein; afdb-uniprot50 \| AF-A0A2G6PXF8-F1-MODEL\_V4 \| 1.0 \| 3.658e-11 \| 316 \| 0.248 \| 209 \| 122 \| 8 \| 4 \| 180 \| 7 \| 212 \| HTH cro/C1-type domain-containing protein \| HTH cro/C1-type domain-containing protein; afdb-uniprot50 \| AF-F3QJB3-F1-MODEL\_V4 \| 1.0 \| 3.257e-11 \| 316 \| 0.203 \| 221 \| 129 \| 6 \| 4 \| 180 \| 7 \| 224 \| Peptidase S24-like protein \| Peptidase S24-like protein; afdb-uniprot50 \| AF-A0A4Q9D5B2-F1-MODEL\_V4 \| 1.0 \| 1.721e-11 \| 316 \| 0.217 \| 221 \| 128 \| 6 \| 4 \| 180 \| 6 \| 225 \| Helix-turn-helix transcriptional regulator \| Helix-turn-helix transcriptional regulator; afdb-uniprot50 \| AF-A0A7T0G1L7-F1-MODEL\_V4 \| 1.0 \| 3.658e-11 \| 316 \| 0.189 \| 216 \| 136 \| 5 \| 4 \| 180 \| 15 \| 230 \| Helix-turn-helix transcriptional regulator \| Helix-turn-helix transcriptional regulator; afdb-uniprot50 \| AF-A0A516SAZ0-F1-MODEL\_V4 \| 1.0 \| 4.612e-11 \| 316 \| 0.213 \| 234 \| 126 \| 7 \| 4 \| 180 \| 7 \| 239 \| Helix-turn-helix domain-containing protein \| Helix-turn-helix domain-containing protein; afdb-uniprot50 \| AF-A0A5C1MF53-F1-MODEL\_V4 \| 1.0 \| 4.612e-11 \| 316 \| 0.212 \| 216 \| 129 \| 8 \| 4 \| 179 \| 30 \| 244 \| Helix-turn-helix transcriptional regulator \| Helix-turn-helix transcriptional regulator; afdb-uniprot50 \| AF-A4N1A7-F1-MODEL\_V4 \| 1.0 \| 6.486e-08 \| 315 \| 0.322 \| 96 \| 61 \| 2 \| 88 \| 180 \| 93 \| 187 \| Putative transcriptional regulator \| Putative transcriptional regulator; afdb-uniprot50 \| AF-A0A562C8X5-F1-MODEL\_V4 \| 1.0 \| 2.737e-11 \| 315 \| 0.221 \| 212 \| 124 \| 7 \| 4 \| 180 \| 9 \| 214 \| Phage repressor protein C with HTH and peptisase S24 domain \| Phage repressor protein C with HTH and peptisase S24 domain; afdb-uniprot50 \| AF-A0A318MXH2-F1-MODEL\_V4 \| 1.0 \| 3.257e-11 \| 315 \| 0.195 \| 215 \| 130 \| 7 \| 5 \| 180 \| 9 \| 219 \| Transcriptional regulator \| Transcriptional regulator; afdb-uniprot50 \| AF-A0A7R6SVB1-F1-MODEL\_V4 \| 1.0 \| 1.039e-10 \| 315 \| 0.236 \| 220 \| 123 \| 7 \| 4 \| 180 \| 8 \| 225 \| Peptidase S24 \| Peptidase S24; afdb-uniprot50 \| AF-A0A1B9KJY8-F1-MODEL\_V4 \| 1.0 \| 3.876e-11 \| 315 \| 0.185 \| 226 \| 133 \| 6 \| 4 \| 180 \| 7 \| 230 \| HTH cro/C1-type domain-containing protein \| HTH cro/C1-type domain-containing protein; afdb-uniprot50 \| AF-A0A3R8T613-F1-MODEL\_V4 \| 1.0 \| 1.147e-11 \| 315 \| 0.202 \| 232 \| 133 \| 10 \| 1 \| 180 \| 1 \| 232 \| Helix-turn-helix transcriptional regulator \| Helix-turn-helix transcriptional regulator; afdb-uniprot50 \| AF-A0A1B3E902-F1-MODEL\_V4 \| 1.0 \| 3.451e-11 \| 315 \| 0.169 \| 224 \| 137 \| 5 \| 4 \| 179 \| 19 \| 241 \| Cro/Cl family transcriptional regulator \| Cro/Cl family transcriptional regulator; afdb-uniprot50 \| AF-A0A358RV08-F1-MODEL\_V4 \| 1.0 \| 4.322e-08 \| 314 \| 0.271 \| 114 \| 75 \| 3 \| 70 \| 180 \| 5 \| 113 \| Peptidase\_S24 domain-containing protein \| Peptidase\_S24 domain-containing protein; afdb-uniprot50 \| AF-A0A423DG37-F1-MODEL\_V4 \| 1.0 \| 1.166e-10 \| 314 \| 0.213 \| 206 \| 128 \| 5 \| 6 \| 180 \| 2 \| 204 \| HTH cro/C1-type domain-containing protein \| HTH cro/C1-type domain-containing protein; afdb-uniprot50 \| AF-A0A5E4YQN1-F1-MODEL\_V4 \| 1.0 \| 6.532e-11 \| 314 \| 0.2 \| 205 \| 136 \| 4 \| 4 \| 180 \| 19 \| 223 \| Putative HTH-type transcriptional regulator \| Putative HTH-type transcriptional regulator; afdb-uniprot50 \| AF-A0A6G6Q5W5-F1-MODEL\_V4 \| 1.0 \| 3.451e-11 \| 314 \| 0.177 \| 225 \| 136 \| 5 \| 4 \| 180 \| 6 \| 229 \| Helix-turn-helix transcriptional regulator \| Helix-turn-helix transcriptional regulator; afdb-uniprot50 \| AF-A0A3D5BE89-F1-MODEL\_V4 \| 1.0 \| 3.074e-11 \| 314 \| 0.2 \| 234 \| 130 \| 5 \| 4 \| 180 \| 6 \| 239 \| XRE family transcriptional regulator \| XRE family transcriptional regulator; afdb-uniprot50 \| AF-A0A2M8RY32-F1-MODEL\_V4 \| 1.0 \| 3.632e-08 \| 313 \| 0.232 \| 116 \| 82 \| 4 \| 70 \| 180 \| 9 \| 122 \| Peptidase\_S24 domain-containing protein \| Peptidase\_S24 domain-containing protein; afdb-uniprot50 \| AF-U7P2I5-F1-MODEL\_V4 \| 1.0 \| 1.461e-07 \| 313 \| 0.285 \| 105 \| 71 \| 2 \| 79 \| 180 \| 49 \| 152 \| Peptidase\_S24 domain-containing protein \| Peptidase\_S24 domain-containing protein; afdb-uniprot50 \| AF-A0A1L9CZL4-F1-MODEL\_V4 \| 1.0 \| 2.034e-08 \| 313 \| 0.251 \| 127 \| 84 \| 4 \| 60 \| 180 \| 75 \| 196 \| Peptidase\_S24 domain-containing protein \| Peptidase\_S24 domain-containing protein; afdb-uniprot50 \| AF-E6W350-F1-MODEL\_V4 \| 1.0 \| 3.257e-11 \| 313 \| 0.235 \| 212 \| 125 \| 6 \| 4 \| 180 \| 6 \| 215 \| Peptidase S24/S26A/S26B, conserved region \| Peptidase S24/S26A/S26B, conserved region; afdb-uniprot50 \| AF-A0A7Y6VAX6-F1-MODEL\_V4 \| 1.0 \| 1.824e-11 \| 313 \| 0.208 \| 225 \| 130 \| 8 \| 1 \| 180 \| 3 \| 224 \| Helix-turn-helix transcriptional regulator \| Helix-turn-helix transcriptional regulator; afdb-uniprot50 \| AF-A0A1C0A0X4-F1-MODEL\_V4 \| 1.0 \| 6.532e-11 \| 313 \| 0.172 \| 226 \| 136 \| 5 \| 4 \| 180 \| 7 \| 230 \| HTH cro/C1-type domain-containing protein \| HTH cro/C1-type domain-containing protein; afdb-uniprot50 \| AF-A0A2R3QPF1-F1-MODEL\_V4 \| 1.0 \| 3.451e-11 \| 313 \| 0.209 \| 205 \| 131 \| 6 \| 4 \| 180 \| 29 \| 230 \| C repressor \| C repressor; afdb-uniprot50 \| AF-A0A1L3JAI5-F1-MODEL\_V4 \| 1.0 \| 3.658e-11 \| 313 \| 0.192 \| 223 \| 131 \| 6 \| 6 \| 180 \| 11 \| 232 \| Peptidase\_S24 domain-containing protein \| Peptidase\_S24 domain-containing protein; afdb-uniprot50 \| AF-A0A2W6GVH0-F1-MODEL\_V4 \| 1.0 \| 4.612e-11 \| 313 \| 0.222 \| 225 \| 124 \| 7 \| 4 \| 179 \| 10 \| 232 \| Peptidase\_S24 domain-containing protein \| Peptidase\_S24 domain-containing protein; afdb-uniprot50 \| AF-A0A318DG58-F1-MODEL\_V4 \| 1.0 \| 3.257e-11 \| 313 \| 0.214 \| 233 \| 123 \| 6 \| 5 \| 178 \| 8 \| 239 \| Phage repressor protein C with HTH and peptisase S24 domain \| Phage repressor protein C with HTH and peptisase S24 domain; afdb-uniprot50 \| AF-A0A1C3H797-F1-MODEL\_V4 \| 1.0 \| 5.489e-11 \| 313 \| 0.218 \| 215 \| 129 \| 6 \| 4 \| 180 \| 28 \| 241 \| Putative transcriptional regulator \| Putative transcriptional regulator; afdb-uniprot50 \| AF-A0A060H678-F1-MODEL\_V4 \| 1.0 \| 5.18e-11 \| 313 \| 0.176 \| 243 \| 131 \| 7 \| 5 \| 179 \| 7 \| 248 \| Peptidase S24 \| Peptidase S24; afdb-uniprot50 \| AF-A0A7T8EGD1-F1-MODEL\_V4 \| 1.0 \| 1.031e-07 \| 312 \| 0.292 \| 106 \| 70 \| 3 \| 78 \| 180 \| 76 \| 179 \| S24 family peptidase \| S24 family peptidase; afdb-uniprot50 \| AF-A9L6K9-F1-MODEL\_V4 \| 1.0 \| 3.451e-11 \| 312 \| 0.197 \| 208 \| 134 \| 5 \| 4 \| 180 \| 7 \| 212 \| Putative phage repressor \| Putative phage repressor; afdb-uniprot50 \| AF-A0A7X6EK24-F1-MODEL\_V4 \| 1.0 \| 9.25e-11 \| 312 \| 0.195 \| 215 \| 130 \| 7 \| 4 \| 180 \| 11 \| 220 \| Helix-turn-helix domain-containing protein \| Helix-turn-helix domain-containing protein; afdb-uniprot50 \| AF-R5YMJ3-F1-MODEL\_V4 \| 1.0 \| 4.888e-11 \| 312 \| 0.164 \| 213 \| 139 \| 4 \| 4 \| 178 \| 10 \| 221 \| Putative transcriptional regulator repressor \| Putative transcriptional regulator repressor; afdb-uniprot50 \| AF-A0A249DYA3-F1-MODEL\_V4 \| 1.0 \| 3.451e-11 \| 312 \| 0.194 \| 226 \| 133 \| 4 \| 4 \| 180 \| 6 \| 231 \| XRE family transcriptional regulator \| XRE family transcriptional regulator; afdb-uniprot50 \| AF-A0A2W6WL33-F1-MODEL\_V4 \| 1.0 \| 6.532e-11 \| 312 \| 0.169 \| 213 \| 137 \| 6 \| 4 \| 180 \| 27 \| 235 \| HTH cro/C1-type domain-containing protein \| HTH cro/C1-type domain-containing protein; afdb-uniprot50 \| AF-A0A1I1Q734-F1-MODEL\_V4 \| 1.0 \| 4.612e-11 \| 312 \| 0.211 \| 227 \| 126 \| 7 \| 5 \| 180 \| 31 \| 255 \| Phage repressor protein C, contains Cro/C1-type HTH and peptisase s24 domains \| Phage repressor protein C, contains Cro/C1-type HTH and peptisase s24 domains; afdb-uniprot50 \| AF-A0A0F7KKY2-F1-MODEL\_V4 \| 1.0 \| 2.323e-07 \| 311 \| 0.317 \| 107 \| 69 \| 2 \| 77 \| 180 \| 35 \| 140 \| Peptidase\_S24 domain-containing protein \| Peptidase\_S24 domain-containing protein; afdb-uniprot50 \| AF-A0A855FN50-F1-MODEL\_V4 \| 1.0 \| 4.58e-08 \| 311 \| 0.254 \| 114 \| 78 \| 4 \| 70 \| 180 \| 83 \| 192 \| Uncharacterized protein \| Uncharacterized protein; afdb-uniprot50 \| AF-A0A7H1NQ78-F1-MODEL\_V4 \| 1.0 \| 3.658e-11 \| 311 \| 0.199 \| 221 \| 131 \| 5 \| 4 \| 180 \| 8 \| 226 \| HTH-type transcriptional regulator PrtR \| HTH-type transcriptional regulator PrtR; afdb-uniprot50 \| AF-A0A437QDQ5-F1-MODEL\_V4 \| 1.0 \| 4.612e-11 \| 311 \| 0.235 \| 225 \| 123 \| 7 \| 4 \| 180 \| 8 \| 231 \| Helix-turn-helix transcriptional regulator \| Helix-turn-helix transcriptional regulator; afdb-uniprot50 \| AF-A0A847S398-F1-MODEL\_V4 \| 1.0 \| 4.612e-11 \| 311 \| 0.243 \| 230 \| 119 \| 8 \| 4 \| 180 \| 7 \| 234 \| Helix-turn-helix transcriptional regulator \| Helix-turn-helix transcriptional regulator; afdb-uniprot50 \| AF-A0A0D5N774-F1-MODEL\_V4 \| 1.0 \| 1.471e-10 \| 310 \| 0.215 \| 200 \| 128 \| 8 \| 5 \| 180 \| 8 \| 202 \| Peptidase\_S24 domain-containing protein \| Peptidase\_S24 domain-containing protein; afdb-uniprot50 \| AF-A0A3D5BAV1-F1-MODEL\_V4 \| 1.0 \| 8.729e-11 \| 310 \| 0.193 \| 202 \| 134 \| 4 \| 5 \| 180 \| 10 \| 208 \| Peptidase\_S24 domain-containing protein \| Peptidase\_S24 domain-containing protein; afdb-uniprot50 \| AF-A0A252A0T3-F1-MODEL\_V4 \| 1.0 \| 4.888e-11 \| 310 \| 0.21 \| 204 \| 127 \| 7 \| 4 \| 180 \| 14 \| 210 \| Repressor \| Repressor; afdb-uniprot50 \| AF-A0A246HR45-F1-MODEL\_V4 \| 1.0 \| 2.048e-11 \| 310 \| 0.199 \| 221 \| 130 \| 9 \| 4 \| 180 \| 6 \| 223 \| Transcriptional regulator \| Transcriptional regulator; afdb-uniprot50 \| AF-A0A3D3SH38-F1-MODEL\_V4 \| 1.0 \| 5.18e-11 \| 310 \| 0.2 \| 219 \| 131 \| 7 \| 4 \| 180 \| 7 \| 223 \| HTH cro/C1-type domain-containing protein \| HTH cro/C1-type domain-containing protein; afdb-uniprot50 \| AF-A0A379CA04-F1-MODEL\_V4 \| 1.0 \| 4.612e-11 \| 310 \| 0.184 \| 217 \| 134 \| 8 \| 4 \| 180 \| 7 \| 220 \| Uncharacterized HTH-type transcriptional regulator HI\_1476 \| Uncharacterized HTH-type transcriptional regulator HI\_1476; afdb-uniprot50 \| AF-A0A1J0D5X4-F1-MODEL\_V4 \| 1.0 \| 6.922e-11 \| 310 \| 0.181 \| 220 \| 131 \| 7 \| 4 \| 179 \| 10 \| 224 \| HTH cro/C1-type domain-containing protein \| HTH cro/C1-type domain-containing protein; afdb-uniprot50 \| AF-A0A495RI07-F1-MODEL\_V4 \| 1.0 \| 8.237e-11 \| 310 \| 0.195 \| 225 \| 131 \| 4 \| 4 \| 180 \| 5 \| 227 \| Phage repressor protein C with HTH and peptisase S24 domain \| Phage repressor protein C with HTH and peptisase S24 domain; afdb-uniprot50 \| AF-A0A2E2KKV0-F1-MODEL\_V4 \| 1.0 \| 4.612e-11 \| 310 \| 0.251 \| 215 \| 118 \| 5 \| 4 \| 180 \| 19 \| 228 \| HTH cro/C1-type domain-containing protein \| HTH cro/C1-type domain-containing protein; afdb-uniprot50 \| AF-D0IHH2-F1-MODEL\_V4 \| 1.0 \| 3.451e-11 \| 310 \| 0.205 \| 229 \| 130 \| 5 \| 4 \| 180 \| 9 \| 237 \| Predicted transcriptional regulator \| Predicted transcriptional regulator; afdb-uniprot50 \| AF-A0A011NA96-F1-MODEL\_V4 \| 1.0 \| 9.733e-08 \| 309 \| 0.226 \| 106 \| 77 \| 3 \| 78 \| 180 \| 31 \| 134 \| Transcriptional regulator \| Transcriptional regulator; afdb-uniprot50 \| AF-A0A1H2PS65-F1-MODEL\_V4 \| 1.0 \| 1.039e-10 \| 309 \| 0.201 \| 218 \| 128 \| 8 \| 4 \| 180 \| 6 \| 218 \| Phage repressor protein C, contains Cro/C1-type HTH and peptisase s24 domains \| Phage repressor protein C, contains Cro/C1-type HTH and peptisase s24 domains; afdb-uniprot50 \| AF-A0A2J6JA73-F1-MODEL\_V4 \| 1.0 \| 6.164e-11 \| 309 \| 0.185 \| 221 \| 134 \| 6 \| 5 \| 180 \| 7 \| 226 \| Transcriptional regulator \| Transcriptional regulator; afdb-uniprot50 \| AF-A0A346JU79-F1-MODEL\_V4 \| 1.0 \| 3.451e-11 \| 309 \| 0.223 \| 224 \| 120 \| 9 \| 4 \| 177 \| 7 \| 226 \| Helix-turn-helix transcriptional regulator \| Helix-turn-helix transcriptional regulator; afdb-uniprot50 \| AF-A0A2U1TLL0-F1-MODEL\_V4 \| 1.0 \| 1.039e-10 \| 308 \| 0.206 \| 208 \| 132 \| 6 \| 4 \| 180 \| 12 \| 217 \| HTH cro/C1-type domain-containing protein \| HTH cro/C1-type domain-containing protein; afdb-uniprot50 \| AF-A0A522BFE5-F1-MODEL\_V4 \| 1.0 \| 1.855e-10 \| 308 \| 0.22 \| 213 \| 127 \| 6 \| 4 \| 180 \| 14 \| 223 \| Helix-turn-helix transcriptional regulator \| Helix-turn-helix transcriptional regulator; afdb-uniprot50 \| AF-A0A1G5CCK6-F1-MODEL\_V4 \| 1.0 \| 4.612e-11 \| 308 \| 0.235 \| 212 \| 127 \| 8 \| 2 \| 180 \| 17 \| 226 \| Phage repressor protein C, contains Cro/C1-type HTH and peptisase s24 domains \| Phage repressor protein C, contains Cro/C1-type HTH and peptisase s24 domains; afdb-uniprot50 \| AF-A0A222FGK0-F1-MODEL\_V4 \| 1.0 \| 7.335e-11 \| 308 \| 0.204 \| 215 \| 131 \| 6 \| 4 \| 180 \| 18 \| 230 \| Transcriptional regulator \| Transcriptional regulator; afdb-uniprot50 \| AF-A0A1N7DPA7-F1-MODEL\_V4 \| 1.0 \| 4.107e-11 \| 308 \| 0.193 \| 227 \| 129 \| 6 \| 4 \| 178 \| 6 \| 230 \| Phage repressor protein C, contains Cro/C1-type HTH and peptisase s24 domains \| Phage repressor protein C, contains Cro/C1-type HTH and peptisase s24 domains; afdb-uniprot50 \| AF-A0A3T0ZVY7-F1-MODEL\_V4 \| 1.0 \| 3.658e-11 \| 308 \| 0.201 \| 218 \| 130 \| 5 \| 4 \| 179 \| 31 \| 246 \| Transcriptional regulator \| Transcriptional regulator; afdb-uniprot50 \| AF-A0A149QP86-F1-MODEL\_V4 \| 1.0 \| 1.548e-07 \| 307 \| 0.301 \| 93 \| 61 \| 2 \| 88 \| 177 \| 46 \| 137 \| Repressor \| Repressor; afdb-uniprot50 \| AF-A0A546XR11-F1-MODEL\_V4 \| 1.0 \| 5.775e-08 \| 307 \| 0.26 \| 123 \| 80 \| 4 \| 64 \| 180 \| 48 \| 165 \| Helix-turn-helix transcriptional regulator \| Helix-turn-helix transcriptional regulator; afdb-uniprot50 \| AF-A0A2D9DW47-F1-MODEL\_V4 \| 1.0 \| 4.353e-11 \| 307 \| 0.188 \| 207 \| 134 \| 8 \| 5 \| 180 \| 10 \| 213 \| Peptidase\_S24 domain-containing protein \| Peptidase\_S24 domain-containing protein; afdb-uniprot50 \| AF-A0A7X5QU28-F1-MODEL\_V4 \| 1.0 \| 8.729e-11 \| 307 \| 0.177 \| 214 \| 138 \| 6 \| 4 \| 180 \| 4 \| 216 \| Helix-turn-helix transcriptional regulator \| Helix-turn-helix transcriptional regulator; afdb-uniprot50 \| AF-F3QJB4-F1-MODEL\_V4 \| 1.0 \| 1.236e-10 \| 307 \| 0.201 \| 213 \| 129 \| 6 \| 4 \| 178 \| 13 \| 222 \| Peptidase S24-like protein \| Peptidase S24-like protein; afdb-uniprot50 \| AF-A0A657K4X9-F1-MODEL\_V4 \| 1.0 \| 7.335e-11 \| 307 \| 0.193 \| 232 \| 129 \| 6 \| 4 \| 180 \| 7 \| 235 \| HTH cro/C1-type domain-containing protein \| HTH cro/C1-type domain-containing protein; afdb-uniprot50 \| AF-W6W8I6-F1-MODEL\_V4 \| 1.0 \| 1.166e-10 \| 307 \| 0.201 \| 238 \| 128 \| 6 \| 4 \| 180 \| 9 \| 245 \| Putative phage repressor \| Putative phage repressor; afdb-uniprot50 \| AF-A0A8B3TEF1-F1-MODEL\_V4 \| 1.0 \| 5.18e-11 \| 307 \| 0.18 \| 249 \| 134 \| 8 \| 1 \| 180 \| 1 \| 248 \| Helix-turn-helix domain-containing protein \| Helix-turn-helix domain-containing protein; afdb-uniprot50 \| AF-A0A7T5A214-F1-MODEL\_V4 \| 1.0 \| 4.612e-11 \| 307 \| 0.232 \| 219 \| 122 \| 7 \| 4 \| 180 \| 37 \| 251 \| LexA family transcriptional regulator \| LexA family transcriptional regulator; afdb-uniprot50 \| AF-A0A286C7A4-F1-MODEL\_V4 \| 1.0 \| 6.12e-08 \| 307 \| 0.318 \| 113 \| 69 \| 3 \| 71 \| 180 \| 154 \| 261 \| Mu DNA-binding domain-containing protein \| Mu DNA-binding domain-containing protein; afdb-uniprot50 \| AF-A0A743J265-F1-MODEL\_V4 \| 1.0 \| 4.853e-08 \| 306 \| 0.309 \| 113 \| 70 \| 4 \| 71 \| 180 \| 56 \| 163 \| Helix-turn-helix transcriptional regulator \| Helix-turn-helix transcriptional regulator; afdb-uniprot50 \| AF-A0A2P1VM30-F1-MODEL\_V4 \| 1.0 \| 1.227e-07 \| 306 \| 0.245 \| 106 \| 75 \| 3 \| 78 \| 180 \| 104 \| 207 \| Transcriptional regulator \| Transcriptional regulator; afdb-uniprot50 \| AF-A0A4R2GXK3-F1-MODEL\_V4 \| 1.0 \| 1.855e-10 \| 306 \| 0.212 \| 212 \| 127 \| 9 \| 4 \| 180 \| 18 \| 224 \| Phage repressor protein C with HTH and peptisase S24 domain \| Phage repressor protein C with HTH and peptisase S24 domain; afdb-uniprot50 \| AF-A0A366FEU9-F1-MODEL\_V4 \| 1.0 \| 2.583e-11 \| 306 \| 0.222 \| 220 \| 123 \| 8 \| 4 \| 180 \| 6 \| 220 \| Phage repressor protein C with HTH and peptisase S24 domain \| Phage repressor protein C with HTH and peptisase S24 domain; afdb-uniprot50 \| AF-A0A2J9DWQ2-F1-MODEL\_V4 \| 1.0 \| 7.335e-11 \| 306 \| 0.2 \| 220 \| 131 \| 6 \| 4 \| 180 \| 9 \| 226 \| Helix-turn-helix transcriptional regulator \| Helix-turn-helix transcriptional regulator; afdb-uniprot50 \| AF-A0A7X2GXD4-F1-MODEL\_V4 \| 1.0 \| 4.107e-11 \| 306 \| 0.194 \| 226 \| 134 \| 8 \| 1 \| 180 \| 1 \| 224 \| Helix-turn-helix domain-containing protein \| Helix-turn-helix domain-containing protein; afdb-uniprot50 \| AF-A0A852PQB2-F1-MODEL\_V4 \| 1.0 \| 3.451e-11 \| 306 \| 0.174 \| 223 \| 136 \| 7 \| 4 \| 180 \| 7 \| 227 \| Helix-turn-helix transcriptional regulator \| Helix-turn-helix transcriptional regulator; afdb-uniprot50 \| AF-A0A7W6G492-F1-MODEL\_V4 \| 1.0 \| 4.178e-10 \| 306 \| 0.172 \| 238 \| 129 \| 7 \| 5 \| 180 \| 7 \| 238 \| Phage repressor protein C with HTH and peptisase S24 domain \| Phage repressor protein C with HTH and peptisase S24 domain; afdb-uniprot50 \| AF-A0A495YMK1-F1-MODEL\_V4 \| 1.0 \| 1.166e-10 \| 306 \| 0.164 \| 237 \| 134 \| 7 \| 4 \| 180 \| 7 \| 239 \| Phage repressor protein \| Phage repressor protein; afdb-uniprot50 \| AF-A0A1Y2K654-F1-MODEL\_V4 \| 1.0 \| 7.718e-08 \| 305 \| 0.211 \| 118 \| 84 \| 4 \| 66 \| 180 \| 1 \| 112 \| Putative Cro/CI family transcriptional regulator \| Putative Cro/CI family transcriptional regulator; afdb-uniprot50 \| AF-A0A2W6YVV3-F1-MODEL\_V4 \| 1.0 \| 1.75e-10 \| 305 \| 0.198 \| 212 \| 133 \| 6 \| 4 \| 180 \| 4 \| 213 \| HTH cro/C1-type domain-containing protein \| HTH cro/C1-type domain-containing protein; afdb-uniprot50 \| AF-A0A1X0TH41-F1-MODEL\_V4 \| 1.0 \| 1.559e-10 \| 305 \| 0.17 \| 211 \| 139 \| 6 \| 4 \| 180 \| 7 \| 215 \| HTH cro/C1-type domain-containing protein \| HTH cro/C1-type domain-containing protein; afdb-uniprot50 \| AF-A0A4Q7BGU0-F1-MODEL\_V4 \| 1.0 \| 1.31e-10 \| 305 \| 0.213 \| 220 \| 123 \| 5 \| 4 \| 180 \| 6 \| 218 \| Peptidase S24 \| Peptidase S24; afdb-uniprot50 \| AF-A0A212KKI8-F1-MODEL\_V4 \| 1.0 \| 8.237e-11 \| 305 \| 0.22 \| 209 \| 129 \| 6 \| 5 \| 180 \| 14 \| 221 \| Putative Repressor protein CI \| Putative Repressor protein CI; afdb-uniprot50 \| AF-A0A6N9SY62-F1-MODEL\_V4 \| 1.0 \| 1.471e-10 \| 305 \| 0.213 \| 220 \| 129 \| 5 \| 4 \| 180 \| 12 \| 230 \| Helix-turn-helix domain-containing protein \| Helix-turn-helix domain-containing protein; afdb-uniprot50 \| AF-A0A7Y2QFS9-F1-MODEL\_V4 \| 1.0 \| 1.166e-10 \| 305 \| 0.171 \| 228 \| 132 \| 6 \| 4 \| 178 \| 8 \| 231 \| Helix-turn-helix transcriptional regulator \| Helix-turn-helix transcriptional regulator; afdb-uniprot50 \| AF-A0A516SD59-F1-MODEL\_V4 \| 1.0 \| 5.489e-11 \| 305 \| 0.231 \| 233 \| 122 \| 9 \| 4 \| 180 \| 6 \| 237 \| Helix-turn-helix transcriptional regulator \| Helix-turn-helix transcriptional regulator; afdb-uniprot50 \| AF-I7C2G6-F1-MODEL\_V4 \| 1.0 \| 2.737e-11 \| 305 \| 0.184 \| 260 \| 128 \| 7 \| 4 \| 180 \| 8 \| 266 \| Putative phage repressor \| Putative phage repressor; afdb-uniprot50 \| AF-C4GFS1-F1-MODEL\_V4 \| 1.0 \| 1.236e-10 \| 305 \| 0.161 \| 229 \| 138 \| 7 \| 4 \| 180 \| 125 \| 351 \| DNA-binding helix-turn-helix protein \| DNA-binding helix-turn-helix protein; afdb-uniprot50 \| AF-A0A554RHB1-F1-MODEL\_V4 \| 1.0 \| 1.952e-07 \| 304 \| 0.245 \| 106 \| 75 \| 3 \| 79 \| 180 \| 65 \| 169 \| Helix-turn-helix transcriptional regulator \| Helix-turn-helix transcriptional regulator; afdb-uniprot50 \| AF-A0A345DBG2-F1-MODEL\_V4 \| 1.0 \| 1.471e-10 \| 304 \| 0.195 \| 200 \| 135 \| 4 \| 5 \| 179 \| 7 \| 205 \| HTH-type transcriptional regulator PrtR \| HTH-type transcriptional regulator PrtR; afdb-uniprot50 \| AF-A0A2X1U757-F1-MODEL\_V4 \| 1.0 \| 8.729e-11 \| 304 \| 0.206 \| 213 \| 130 \| 9 \| 4 \| 180 \| 7 \| 216 \| Phage repressor protein, phage associated protein \| Phage repressor protein, phage associated protein; afdb-uniprot50 \| AF-V2QBR0-F1-MODEL\_V4 \| 1.0 \| 5.489e-11 \| 304 \| 0.2 \| 215 \| 130 \| 7 \| 6 \| 180 \| 12 \| 224 \| HTH cro/C1-type domain-containing protein \| HTH cro/C1-type domain-containing protein; afdb-uniprot50 \| AF-A0A376BT84-F1-MODEL\_V4 \| 1.0 \| 6.532e-11 \| 304 \| 0.233 \| 214 \| 125 \| 7 \| 4 \| 180 \| 16 \| 227 \| Uncharacterized HTH-type transcriptional regulator HI\_1476 \| Uncharacterized HTH-type transcriptional regulator HI\_1476; afdb-uniprot50 \| AF-A0A369YGT9-F1-MODEL\_V4 \| 1.0 \| 1.75e-10 \| 304 \| 0.185 \| 221 \| 133 \| 7 \| 4 \| 180 \| 7 \| 224 \| Helix-turn-helix transcriptional regulator \| Helix-turn-helix transcriptional regulator; afdb-uniprot50 \| AF-A0A369T885-F1-MODEL\_V4 \| 1.0 \| 9.802e-11 \| 304 \| 0.169 \| 218 \| 136 \| 8 \| 4 \| 180 \| 41 \| 254 \| XRE family transcriptional regulator \| XRE family transcriptional regulator; afdb-uniprot50 \| AF-A0A349XYC5-F1-MODEL\_V4 \| 1.0 \| 1.64e-07 \| 303 \| 0.271 \| 114 \| 75 \| 3 \| 70 \| 180 \| 9 \| 117 \| Helix-turn-helix transcriptional regulator \| Helix-turn-helix transcriptional regulator; afdb-uniprot50 \| AF-A0A828SCT0-F1-MODEL\_V4 \| 1.0 \| 1.548e-07 \| 303 \| 0.287 \| 108 \| 72 \| 3 \| 77 \| 180 \| 45 \| 151 \| Peptidase S24-like family protein \| Peptidase S24-like family protein; afdb-uniprot50 \| AF-U3U086-F1-MODEL\_V4 \| 1.0 \| 1.158e-07 \| 303 \| 0.28 \| 107 \| 71 \| 4 \| 78 \| 180 \| 60 \| 164 \| Repressor protein CI \| Repressor protein CI; afdb-uniprot50 \| AF-A0A718TEP2-F1-MODEL\_V4 \| 1.0 \| 6.12e-08 \| 303 \| 0.309 \| 113 \| 70 \| 4 \| 71 \| 180 \| 63 \| 170 \| Helix-turn-helix transcriptional regulator \| Helix-turn-helix transcriptional regulator; afdb-uniprot50 \| AF-A0A515D4V4-F1-MODEL\_V4 \| 1.0 \| 1.158e-07 \| 303 \| 0.303 \| 112 \| 71 \| 4 \| 72 \| 180 \| 80 \| 187 \| Helix-turn-helix transcriptional regulator \| Helix-turn-helix transcriptional regulator; afdb-uniprot50 \| AF-C0N8P1-F1-MODEL\_V4 \| 1.0 \| 2.764e-07 \| 303 \| 0.256 \| 109 \| 75 \| 2 \| 77 \| 180 \| 102 \| 209 \| Peptidase S24-like domain protein \| Peptidase S24-like domain protein; afdb-uniprot50 \| AF-A0A2L2BE93-F1-MODEL\_V4 \| 1.0 \| 9.802e-11 \| 303 \| 0.199 \| 216 \| 131 \| 4 \| 4 \| 178 \| 7 \| 221 \| HTH cro/C1-type domain-containing protein \| HTH cro/C1-type domain-containing protein; afdb-uniprot50 \| AF-A0A8B5S1H8-F1-MODEL\_V4 \| 1.0 \| 6.164e-11 \| 303 \| 0.177 \| 225 \| 136 \| 6 \| 4 \| 180 \| 6 \| 229 \| Helix-turn-helix transcriptional regulator \| Helix-turn-helix transcriptional regulator; afdb-uniprot50 \| AF-A0A0F4R425-F1-MODEL\_V4 \| 1.0 \| 5.489e-11 \| 303 \| 0.202 \| 237 \| 128 \| 6 \| 4 \| 180 \| 6 \| 241 \| HTH cro/C1-type domain-containing protein \| HTH cro/C1-type domain-containing protein; afdb-uniprot50 \| AF-A0A3Z8HGE4-F1-MODEL\_V4 \| 1.0 \| 2.929e-07 \| 302 \| 0.34 \| 100 \| 61 \| 3 \| 83 \| 179 \| 3 \| 100 \| Helix-turn-helix transcriptional regulator \| Helix-turn-helix transcriptional regulator; afdb-uniprot50 \| AF-A0A6L3ERI8-F1-MODEL\_V4 \| 1.0 \| 1.207e-08 \| 302 \| 0.236 \| 127 \| 87 \| 3 \| 60 \| 178 \| 1 \| 125 \| S24 family peptidase \| S24 family peptidase; afdb-uniprot50 \| AF-A0A160BZ99-F1-MODEL\_V4 \| 1.0 \| 1.842e-07 \| 302 \| 0.224 \| 107 \| 78 \| 3 \| 79 \| 180 \| 49 \| 155 \| Peptidase\_S24 domain-containing protein \| Peptidase\_S24 domain-containing protein; afdb-uniprot50 \| AF-A0A1C3EAH7-F1-MODEL\_V4 \| 1.0 \| 1.75e-10 \| 302 \| 0.242 \| 194 \| 127 \| 4 \| 4 \| 180 \| 18 \| 208 \| HTH cro/C1-type domain-containing protein \| HTH cro/C1-type domain-containing protein; afdb-uniprot50 \| AF-K9RVQ1-F1-MODEL\_V4 \| 1.0 \| 8.237e-11 \| 302 \| 0.21 \| 209 \| 130 \| 7 \| 3 \| 180 \| 11 \| 215 \| Putative transcriptional regulator \| Putative transcriptional regulator; afdb-uniprot50 \| AF-A0A2T1AI87-F1-MODEL\_V4 \| 1.0 \| 8.729e-11 \| 302 \| 0.221 \| 212 \| 129 \| 4 \| 4 \| 180 \| 9 \| 219 \| Helix-turn-helix protein \| Helix-turn-helix protein; afdb-uniprot50 \| AF-B4RKJ1-F1-MODEL\_V4 \| 1.0 \| 8.729e-11 \| 302 \| 0.219 \| 210 \| 128 \| 6 \| 5 \| 180 \| 11 \| 218 \| Putative lambda repressor protein cI, putative phage associated protein \| Putative lambda repressor protein cI, putative phage associated protein; afdb-uniprot50 \| AF-A0A1N6R0A9-F1-MODEL\_V4 \| 1.0 \| 1.652e-10 \| 302 \| 0.218 \| 220 \| 123 \| 6 \| 5 \| 180 \| 8 \| 222 \| Phage repressor protein C, contains Cro/C1-type HTH and peptisase s24 domains \| Phage repressor protein C, contains Cro/C1-type HTH and peptisase s24 domains; afdb-uniprot50 \| AF-A0A0Q4U5S7-F1-MODEL\_V4 \| 1.0 \| 1.166e-10 \| 302 \| 0.184 \| 211 \| 137 \| 5 \| 4 \| 180 \| 15 \| 224 \| HTH cro/C1-type domain-containing protein \| HTH cro/C1-type domain-containing protein; afdb-uniprot50 \| AF-A0A380TXN7-F1-MODEL\_V4 \| 1.0 \| 4.353e-11 \| 302 \| 0.189 \| 216 \| 134 \| 7 \| 4 \| 180 \| 12 \| 225 \| Putative phage repressor \| Putative phage repressor; afdb-uniprot50 \| AF-A0A1R3U4A3-F1-MODEL\_V4 \| 1.0 \| 4.107e-11 \| 302 \| 0.213 \| 225 \| 127 \| 7 \| 4 \| 180 \| 12 \| 234 \| Putative HTH-type transcriptional regulator \| Putative HTH-type transcriptional regulator; afdb-uniprot50 \| AF-A0A7G5DM43-F1-MODEL\_V4 \| 1.0 \| 6.532e-11 \| 302 \| 0.18 \| 227 \| 138 \| 6 \| 1 \| 180 \| 16 \| 241 \| Helix-turn-helix transcriptional regulator \| Helix-turn-helix transcriptional regulator; afdb-uniprot50 \| AF-U1I7S2-F1-MODEL\_V4 \| 1.0 \| 3.074e-11 \| 302 \| 0.171 \| 222 \| 135 \| 6 \| 6 \| 180 \| 26 \| 245 \| Transcriptional regulator \| Transcriptional regulator; afdb-uniprot50 \| AF-A0A154U565-F1-MODEL\_V4 \| 1.0 \| 2.608e-07 \| 301 \| 0.285 \| 105 \| 71 \| 2 \| 79 \| 180 \| 62 \| 165 \| Transcriptional regulator \| Transcriptional regulator; afdb-uniprot50 \| AF-Q0FYV8-F1-MODEL\_V4 \| 1.0 \| 2.608e-07 \| 301 \| 0.273 \| 106 \| 74 \| 1 \| 78 \| 180 \| 78 \| 183 \| Prophage MuSo1, transcriptional regulator, Cro/CI family protein \| Prophage MuSo1, transcriptional regulator, Cro/CI family protein; afdb-uniprot50 \| AF-A0A6G8C4E6-F1-MODEL\_V4 \| 1.0 \| 3.104e-07 \| 301 \| 0.237 \| 97 \| 71 \| 2 \| 87 \| 180 \| 95 \| 191 \| Helix-turn-helix transcriptional regulator \| Helix-turn-helix transcriptional regulator; afdb-uniprot50 \| AF-A0A212J1Z6-F1-MODEL\_V4 \| 1.0 \| 8.237e-11 \| 301 \| 0.213 \| 201 \| 128 \| 9 \| 5 \| 180 \| 7 \| 202 \| Putative Phage repressor \| Putative Phage repressor; afdb-uniprot50 \| AF-A0A7Y8SNM8-F1-MODEL\_V4 \| 1.0 \| 1.236e-10 \| 301 \| 0.199 \| 216 \| 123 \| 8 \| 6 \| 180 \| 2 \| 208 \| Helix-turn-helix transcriptional regulator \| Helix-turn-helix transcriptional regulator; afdb-uniprot50 \| AF-A0A1G0HXS8-F1-MODEL\_V4 \| 1.0 \| 1.652e-10 \| 301 \| 0.209 \| 210 \| 129 \| 6 \| 7 \| 180 \| 3 \| 211 \| HTH cro/C1-type domain-containing protein \| HTH cro/C1-type domain-containing protein; afdb-uniprot50 \| AF-A0A0M3G0U6-F1-MODEL\_V4 \| 1.0 \| 1.236e-10 \| 301 \| 0.211 \| 217 \| 130 \| 6 \| 4 \| 180 \| 7 \| 222 \| HTH cro/C1-type domain-containing protein \| HTH cro/C1-type domain-containing protein; afdb-uniprot50 \| AF-G9ZDL4-F1-MODEL\_V4 \| 1.0 \| 1.559e-10 \| 301 \| 0.209 \| 215 \| 129 \| 6 \| 4 \| 180 \| 22 \| 233 \| DNA-binding helix-turn-helix protein \| DNA-binding helix-turn-helix protein; afdb-uniprot50 \| AF-A0A1S7MQB7-F1-MODEL\_V4 \| 1.0 \| 1.388e-10 \| 301 \| 0.211 \| 218 \| 129 \| 7 \| 4 \| 180 \| 26 \| 241 \| Putative transcriptional regulator \| Putative transcriptional regulator; afdb-uniprot50 \| AF-A0A7V8F8X1-F1-MODEL\_V4 \| 1.0 \| 2.479e-10 \| 301 \| 0.191 \| 225 \| 132 \| 6 \| 5 \| 180 \| 28 \| 251 \| HTH cro/C1-type domain-containing protein \| HTH cro/C1-type domain-containing protein; afdb-uniprot50 \| AF-A0A077LIF9-F1-MODEL\_V4 \| 1.0 \| 4.888e-11 \| 301 \| 0.151 \| 271 \| 135 \| 6 \| 4 \| 180 \| 8 \| 277 \| Prophage PSPPH03, Cro/CI family transcriptional regulator \| Prophage PSPPH03, Cro/CI family transcriptional regulator; afdb-uniprot50 \| AF-A0A808X786-F1-MODEL\_V4 \| 1.0 \| 2.764e-07 \| 300 \| 0.257 \| 105 \| 73 \| 3 \| 79 \| 180 \| 62 \| 164 \| Helix-turn-helix transcriptional regulator \| Helix-turn-helix transcriptional regulator; afdb-uniprot50 \| AF-A0A380RBK7-F1-MODEL\_V4 \| 1.0 \| 3.914e-07 \| 300 \| 0.288 \| 104 \| 72 \| 2 \| 78 \| 180 \| 71 \| 173 \| Peptidase S24-like \| Peptidase S24-like; afdb-uniprot50 \| AF-A0A4V1P805-F1-MODEL\_V4 \| 1.0 \| 1.471e-10 \| 300 \| 0.219 \| 219 \| 130 \| 6 \| 1 \| 180 \| 1 \| 217 \| Peptidase\_S24 domain-containing protein \| Peptidase\_S24 domain-containing protein; afdb-uniprot50 \| AF-A0A5A7NJR8-F1-MODEL\_V4 \| 1.0 \| 2.339e-10 \| 300 \| 0.179 \| 223 \| 137 \| 8 \| 1 \| 180 \| 1 \| 220 \| Peptidase\_S24 domain-containing protein \| Peptidase\_S24 domain-containing protein; afdb-uniprot50 \| AF-A0A377QCZ4-F1-MODEL\_V4 \| 1.0 \| 8.729e-11 \| 300 \| 0.19 \| 215 \| 131 \| 8 \| 8 \| 180 \| 10 \| 223 \| Uncharacterized HTH-type transcriptional regulator HI\_1476 \| Uncharacterized HTH-type transcriptional regulator HI\_1476; afdb-uniprot50 \| AF-A0A256C1Y2-F1-MODEL\_V4 \| 1.0 \| 2.784e-10 \| 300 \| 0.156 \| 223 \| 137 \| 6 \| 4 \| 180 \| 6 \| 223 \| HTH cro/C1-type domain-containing protein \| HTH cro/C1-type domain-containing protein; afdb-uniprot50 \| AF-A0A256CUE4-F1-MODEL\_V4 \| 1.0 \| 2.083e-10 \| 300 \| 0.172 \| 232 \| 135 \| 5 \| 4 \| 180 \| 5 \| 234 \| HTH cro/C1-type domain-containing protein \| HTH cro/C1-type domain-containing protein; afdb-uniprot50 \| AF-A0A7W2R0E4-F1-MODEL\_V4 \| 1.0 \| 1.236e-10 \| 300 \| 0.187 \| 235 \| 130 \| 4 \| 4 \| 180 \| 7 \| 238 \| Helix-turn-helix transcriptional regulator \| Helix-turn-helix transcriptional regulator; afdb-uniprot50 \| AF-B7J7H3-F1-MODEL\_V4 \| 1.0 \| 1.039e-10 \| 300 \| 0.231 \| 229 \| 123 \| 5 \| 4 \| 180 \| 6 \| 233 \| Transcriptional regulator, putative \| Transcriptional regulator, putative; afdb-uniprot50 \| AF-A0A6B0DRA6-F1-MODEL\_V4 \| 1.0 \| 1.855e-10 \| 300 \| 0.221 \| 217 \| 126 \| 7 \| 4 \| 180 \| 49 \| 262 \| Helix-turn-helix domain-containing protein \| Helix-turn-helix domain-containing protein; afdb-uniprot50 \| AF-A0A376PCJ4-F1-MODEL\_V4 \| 1.0 \| 2.929e-07 \| 299 \| 0.252 \| 107 \| 74 \| 4 \| 78 \| 180 \| 46 \| 150 \| Phage repressor protein CI \| Phage repressor protein CI; afdb-uniprot50 \| AF-A0A4R1YUP2-F1-MODEL\_V4 \| 1.0 \| 1.952e-07 \| 299 \| 0.266 \| 105 \| 73 \| 2 \| 79 \| 180 \| 59 \| 162 \| Peptidase S24-like protein \| Peptidase S24-like protein; afdb-uniprot50 \| AF-Q9CNL1-F1-MODEL\_V4 \| 1.0 \| 2.461e-07 \| 299 \| 0.285 \| 105 \| 69 \| 4 \| 80 \| 180 \| 113 \| 215 \| HTH Mu-type domain-containing protein \| HTH Mu-type domain-containing protein; afdb-uniprot50 \| AF-A0A556UNU0-F1-MODEL\_V4 \| 1.0 \| 9.25e-11 \| 299 \| 0.229 \| 196 \| 112 \| 6 \| 19 \| 180 \| 28 \| 218 \| Helix-turn-helix transcriptional regulator \| Helix-turn-helix transcriptional regulator; afdb-uniprot50 \| AF-A0A3G3L9K2-F1-MODEL\_V4 \| 1.0 \| 1.559e-10 \| 299 \| 0.184 \| 222 \| 132 \| 6 \| 4 \| 178 \| 7 \| 226 \| Helix-turn-helix transcriptional regulator \| Helix-turn-helix transcriptional regulator; afdb-uniprot50 \| AF-A0A2V1H275-F1-MODEL\_V4 \| 1.0 \| 1.966e-10 \| 299 \| 0.196 \| 229 \| 131 \| 4 \| 4 \| 180 \| 6 \| 233 \| Phage repressor protein \| Phage repressor protein; afdb-uniprot50 \| AF-A0A2G2PID2-F1-MODEL\_V4 \| 1.0 \| 8.237e-11 \| 299 \| 0.203 \| 226 \| 130 \| 5 \| 5 \| 180 \| 8 \| 233 \| XRE family transcriptional regulator \| XRE family transcriptional regulator; afdb-uniprot50 \| AF-C4LBA5-F1-MODEL\_V4 \| 1.0 \| 1.039e-10 \| 299 \| 0.198 \| 227 \| 130 \| 6 \| 4 \| 180 \| 31 \| 255 \| Putative phage repressor \| Putative phage repressor; afdb-uniprot50 \| AF-A0A7X4W2Y3-F1-MODEL\_V4 \| 1.0 \| 8.237e-11 \| 299 \| 0.206 \| 242 \| 124 \| 8 \| 4 \| 180 \| 23 \| 261 \| Helix-turn-helix domain-containing protein \| Helix-turn-helix domain-containing protein; afdb-uniprot50 \| AF-A0A2W1MLG5-F1-MODEL\_V4 \| 1.0 \| 3.451e-11 \| 299 \| 0.19 \| 278 \| 123 \| 7 \| 4 \| 180 \| 6 \| 282 \| Transcriptional regulator \| Transcriptional regulator; afdb-uniprot50 \| AF-A0A7H5FDH5-F1-MODEL\_V4 \| 1.0 \| 2.608e-07 \| 298 \| 0.254 \| 106 \| 75 \| 2 \| 78 \| 180 \| 9 \| 113 \| Cro/Cl family transcriptional regulator \| Cro/Cl family transcriptional regulator; afdb-uniprot50 \| AF-A0A1N7AVQ2-F1-MODEL\_V4 \| 1.0 \| 2.764e-07 \| 298 \| 0.245 \| 102 \| 75 \| 2 \| 79 \| 180 \| 53 \| 152 \| Peptidase S24-like \| Peptidase S24-like; afdb-uniprot50 \| AF-A0A7S7LXX0-F1-MODEL\_V4 \| 1.0 \| 2.207e-10 \| 298 \| 0.187 \| 213 \| 134 \| 6 \| 5 \| 180 \| 8 \| 218 \| LexA family transcriptional regulator \| LexA family transcriptional regulator; afdb-uniprot50 \| AF-A0A031LXB0-F1-MODEL\_V4 \| 1.0 \| 1.855e-10 \| 298 \| 0.19 \| 221 \| 135 \| 5 \| 4 \| 180 \| 7 \| 227 \| Peptidase S24 \| Peptidase S24; afdb-uniprot50 \| AF-A0A380MRE3-F1-MODEL\_V4 \| 1.0 \| 1.855e-10 \| 298 \| 0.195 \| 230 \| 131 \| 6 \| 4 \| 180 \| 6 \| 234 \| Uncharacterized HTH-type transcriptional regulator HI\_1476 \| Uncharacterized HTH-type transcriptional regulator HI\_1476; afdb-uniprot50 \| AF-A0A625Y966-F1-MODEL\_V4 \| 1.0 \| 1.75e-10 \| 298 \| 0.2 \| 220 \| 131 \| 8 \| 4 \| 180 \| 13 \| 230 \| Helix-turn-helix transcriptional regulator \| Helix-turn-helix transcriptional regulator; afdb-uniprot50 \| AF-A0LBS2-F1-MODEL\_V4 \| 1.0 \| 2.784e-10 \| 298 \| 0.181 \| 232 \| 132 \| 8 \| 4 \| 180 \| 36 \| 264 \| Putative phage repressor \| Putative phage repressor; afdb-uniprot50 \| AF-Q82XF6-F1-MODEL\_V4 \| 1.0 \| 2.323e-07 \| 298 \| 0.261 \| 107 \| 75 \| 2 \| 78 \| 180 \| 157 \| 263 \| Peptidase\_S24 domain-containing protein \| Peptidase\_S24 domain-containing protein; afdb-uniprot50 \| AF-A0A514BU83-F1-MODEL\_V4 \| 1.0 \| 7.773e-11 \| 298 \| 0.173 \| 231 \| 136 \| 11 \| 4 \| 180 \| 38 \| 267 \| Helix-turn-helix transcriptional regulator \| Helix-turn-helix transcriptional regulator; afdb-uniprot50 \| AF-A0A5Y2E1G9-F1-MODEL\_V4 \| 1.0 \| 7.718e-08 \| 297 \| 0.31 \| 116 \| 73 \| 4 \| 68 \| 180 \| 5 \| 116 \| Helix-turn-helix transcriptional regulator \| Helix-turn-helix transcriptional regulator; afdb-uniprot50 \| AF-A0A556SZ41-F1-MODEL\_V4 \| 1.0 \| 1.031e-07 \| 297 \| 0.279 \| 111 \| 72 \| 4 \| 71 \| 178 \| 14 \| 119 \| Helix-turn-helix transcriptional regulator \| Helix-turn-helix transcriptional regulator; afdb-uniprot50 \| AF-A0A1W9QTR7-F1-MODEL\_V4 \| 1.0 \| 1.301e-07 \| 297 \| 0.3 \| 103 \| 71 \| 1 \| 78 \| 180 \| 51 \| 152 \| Phage repressor protein \| Phage repressor protein; afdb-uniprot50 \| AF-A3NLY9-F1-MODEL\_V4 \| 1.0 \| 1.31e-10 \| 297 \| 0.211 \| 217 \| 127 \| 8 \| 4 \| 179 \| 7 \| 220 \| Helix-turn-helix motif:Peptidase S24, S26A and S26B \| Helix-turn-helix motif:Peptidase S24, S26A and S26B; afdb-uniprot50 \| AF-E6W293-F1-MODEL\_V4 \| 1.0 \| 2.339e-10 \| 297 \| 0.2 \| 220 \| 130 \| 7 \| 5 \| 180 \| 7 \| 224 \| CI repressor \| CI repressor; afdb-uniprot50 \| AF-A0A0D5LT23-F1-MODEL\_V4 \| 1.0 \| 3.313e-10 \| 297 \| 0.187 \| 219 \| 129 \| 5 \| 5 \| 180 \| 12 \| 224 \| Peptidase\_S24 domain-containing protein \| Peptidase\_S24 domain-containing protein; afdb-uniprot50 \| AF-I3YGS2-F1-MODEL\_V4 \| 1.0 \| 1.166e-10 \| 297 \| 0.238 \| 218 \| 123 \| 6 \| 4 \| 180 \| 12 \| 227 \| Putative transcriptional regulator \| Putative transcriptional regulator; afdb-uniprot50 \| AF-A0A6G9IAJ9-F1-MODEL\_V4 \| 1.0 \| 2.479e-10 \| 297 \| 0.178 \| 207 \| 137 \| 7 \| 5 \| 180 \| 28 \| 232 \| Helix-turn-helix transcriptional regulator \| Helix-turn-helix transcriptional regulator; afdb-uniprot50 \| AF-A0A1V0B9I0-F1-MODEL\_V4 \| 1.0 \| 4.107e-11 \| 297 \| 0.19 \| 231 \| 132 \| 7 \| 4 \| 180 \| 14 \| 243 \| HTH cro/C1-type domain-containing protein \| HTH cro/C1-type domain-containing protein; afdb-uniprot50 \| AF-A0A3N9U3X2-F1-MODEL\_V4 \| 1.0 \| 1.039e-10 \| 297 \| 0.224 \| 227 \| 125 \| 8 \| 4 \| 180 \| 22 \| 247 \| Helix-turn-helix domain-containing protein \| Helix-turn-helix domain-containing protein; afdb-uniprot50 \| AF-A0A2M9G0Q7-F1-MODEL\_V4 \| 1.0 \| 1.236e-10 \| 297 \| 0.209 \| 243 \| 121 \| 7 \| 4 \| 180 \| 21 \| 258 \| HTH cro/C1-type domain-containing protein \| HTH cro/C1-type domain-containing protein; afdb-uniprot50 \| AF-A0A1B6W082-F1-MODEL\_V4 \| 1.0 \| 4.396e-07 \| 296 \| 0.254 \| 106 \| 75 \| 2 \| 78 \| 180 \| 22 \| 126 \| Peptidase\_S24 domain-containing protein \| Peptidase\_S24 domain-containing protein; afdb-uniprot50 \| AF-A0A560TXL0-F1-MODEL\_V4 \| 1.0 \| 8.179e-08 \| 296 \| 0.26 \| 115 \| 77 \| 3 \| 70 \| 180 \| 28 \| 138 \| Peptidase S24-like protein \| Peptidase S24-like protein; afdb-uniprot50 \| AF-A0A212KJY0-F1-MODEL\_V4 \| 1.0 \| 3.486e-07 \| 296 \| 0.273 \| 106 \| 72 \| 3 \| 78 \| 180 \| 49 \| 152 \| Phage repressor \| Phage repressor; afdb-uniprot50 \| AF-A0A1F9A3V9-F1-MODEL\_V4 \| 1.0 \| 4.396e-07 \| 296 \| 0.291 \| 103 \| 70 \| 2 \| 80 \| 180 \| 55 \| 156 \| Peptidase\_S24 domain-containing protein \| Peptidase\_S24 domain-containing protein; afdb-uniprot50 \| AF-N8ZST7-F1-MODEL\_V4 \| 1.0 \| 1.559e-10 \| 296 \| 0.188 \| 196 \| 139 \| 4 \| 4 \| 180 \| 7 \| 201 \| HTH cro/C1-type domain-containing protein \| HTH cro/C1-type domain-containing protein; afdb-uniprot50 \| AF-A0A377R622-F1-MODEL\_V4 \| 1.0 \| 1.039e-10 \| 296 \| 0.188 \| 217 \| 135 \| 6 \| 3 \| 179 \| 2 \| 217 \| Uncharacterized HTH-type transcriptional regulator HI\_1476 \| Uncharacterized HTH-type transcriptional regulator HI\_1476; afdb-uniprot50 \| AF-A0A839LDZ9-F1-MODEL\_V4 \| 1.0 \| 1.75e-10 \| 296 \| 0.217 \| 207 \| 130 \| 5 \| 4 \| 180 \| 18 \| 222 \| Helix-turn-helix domain-containing protein \| Helix-turn-helix domain-containing protein; afdb-uniprot50 \| AF-U5DI73-F1-MODEL\_V4 \| 1.0 \| 6.269e-10 \| 296 \| 0.202 \| 207 \| 132 \| 5 \| 4 \| 180 \| 18 \| 221 \| Putative transcriptional regulator \| Putative transcriptional regulator; afdb-uniprot50 \| AF-A0A7X4HWJ8-F1-MODEL\_V4 \| 1.0 \| 2.784e-10 \| 296 \| 0.179 \| 228 \| 134 \| 5 \| 4 \| 180 \| 7 \| 232 \| Helix-turn-helix domain-containing protein \| Helix-turn-helix domain-containing protein; afdb-uniprot50 \| AF-A0A7W6EDZ3-F1-MODEL\_V4 \| 1.0 \| 3.126e-10 \| 296 \| 0.224 \| 209 \| 126 \| 8 \| 4 \| 180 \| 30 \| 234 \| Phage repressor protein C with HTH and peptisase S24 domain \| Phage repressor protein C with HTH and peptisase S24 domain; afdb-uniprot50 \| AF-A0A1H5YB65-F1-MODEL\_V4 \| 1.0 \| 2.929e-07 \| 295 \| 0.261 \| 107 \| 74 \| 2 \| 79 \| 180 \| 29 \| 135 \| Peptidase S24-like \| Peptidase S24-like; afdb-uniprot50 \| AF-A0A1I7CAN9-F1-MODEL\_V4 \| 1.0 \| 2.068e-07 \| 295 \| 0.28 \| 114 \| 74 \| 3 \| 70 \| 180 \| 28 \| 136 \| Peptidase S24-like \| Peptidase S24-like; afdb-uniprot50 \| AF-A0A0D9LFD7-F1-MODEL\_V4 \| 1.0 \| 1.966e-10 \| 295 \| 0.207 \| 212 \| 131 \| 7 \| 4 \| 180 \| 7 \| 216 \| Peptidase S24-like protein \| Peptidase S24-like protein; afdb-uniprot50 \| AF-R1F2E5-F1-MODEL\_V4 \| 1.0 \| 1.31e-10 \| 295 \| 0.22 \| 218 \| 126 \| 7 \| 4 \| 179 \| 7 \| 222 \| Prophage MuSo1, transcriptional regulator, Cro/CI family protein \| Prophage MuSo1, transcriptional regulator, Cro/CI family protein; afdb-uniprot50 \| AF-A0A0Q6T7L7-F1-MODEL\_V4 \| 1.0 \| 1.471e-10 \| 295 \| 0.184 \| 222 \| 133 \| 7 \| 4 \| 179 \| 6 \| 225 \| HTH cro/C1-type domain-containing protein \| HTH cro/C1-type domain-containing protein; afdb-uniprot50 \| AF-A0A680P814-F1-MODEL\_V4 \| 1.0 \| 2.784e-10 \| 295 \| 0.207 \| 222 \| 121 \| 8 \| 4 \| 176 \| 6 \| 221 \| Helix-turn-helix transcriptional regulator \| Helix-turn-helix transcriptional regulator; afdb-uniprot50 \| AF-A0A7Y7MC10-F1-MODEL\_V4 \| 1.0 \| 2.339e-10 \| 295 \| 0.247 \| 226 \| 121 \| 6 \| 4 \| 180 \| 6 \| 231 \| Helix-turn-helix transcriptional regulator \| Helix-turn-helix transcriptional regulator; afdb-uniprot50 \| AF-A0A1Q3VUE3-F1-MODEL\_V4 \| 1.0 \| 7.335e-11 \| 295 \| 0.195 \| 210 \| 126 \| 7 \| 5 \| 177 \| 28 \| 231 \| HTH cro/C1-type domain-containing protein \| HTH cro/C1-type domain-containing protein; afdb-uniprot50 \| AF-A0A202B2B8-F1-MODEL\_V4 \| 1.0 \| 1.966e-10 \| 295 \| 0.214 \| 219 \| 126 \| 7 \| 4 \| 178 \| 20 \| 236 \| HTH cro/C1-type domain-containing protein \| HTH cro/C1-type domain-containing protein; afdb-uniprot50 \| AF-F9RZN7-F1-MODEL\_V4 \| 1.0 \| 2.784e-10 \| 295 \| 0.159 \| 264 \| 133 \| 6 \| 4 \| 180 \| 7 \| 268 \| Prophage MuSo1, Cro/CI family transcriptional regulator \| Prophage MuSo1, Cro/CI family transcriptional regulator; afdb-uniprot50 \| AF-A0A5P1R1X3-F1-MODEL\_V4 \| 1.0 \| 1.652e-10 \| 295 \| 0.197 \| 233 \| 128 \| 6 \| 4 \| 180 \| 53 \| 282 \| Helix-turn-helix domain-containing protein \| Helix-turn-helix domain-containing protein; afdb-uniprot50 \| AF-A0A1H9EDS2-F1-MODEL\_V4 \| 1.0 \| 2.207e-10 \| 295 \| 0.155 \| 296 \| 129 \| 6 \| 4 \| 180 \| 6 \| 299 \| Phage repressor protein C, contains Cro/C1-type HTH and peptisase s24 domains \| Phage repressor protein C, contains Cro/C1-type HTH and peptisase s24 domains; afdb-uniprot50 \| AF-A0A1V3RQU3-F1-MODEL\_V4 \| 1.0 \| 8.179e-08 \| 294 \| 0.274 \| 124 \| 82 \| 4 \| 60 \| 180 \| 2 \| 120 \| Peptidase\_S24 domain-containing protein \| Peptidase\_S24 domain-containing protein; afdb-uniprot50 \| AF-A0A257JTL0-F1-MODEL\_V4 \| 1.0 \| 9.342e-07 \| 294 \| 0.245 \| 106 \| 76 \| 2 \| 78 \| 180 \| 66 \| 170 \| Transcriptional regulator \| Transcriptional regulator; afdb-uniprot50 \| AF-A0A7L9WS46-F1-MODEL\_V4 \| 1.0 \| 3.289e-07 \| 294 \| 0.269 \| 104 \| 72 \| 2 \| 80 \| 180 \| 99 \| 201 \| Uncharacterized protein \| Uncharacterized protein; afdb-uniprot50 \| AF-A0A256C4D7-F1-MODEL\_V4 \| 1.0 \| 2.339e-10 \| 294 \| 0.186 \| 215 \| 138 \| 7 \| 1 \| 180 \| 1 \| 213 \| Transcriptional regulator \| Transcriptional regulator; afdb-uniprot50 \| AF-A0A2L0I076-F1-MODEL\_V4 \| 1.0 \| 2.627e-10 \| 294 \| 0.218 \| 211 \| 125 \| 7 \| 4 \| 179 \| 8 \| 213 \| Peptidase S24 \| Peptidase S24; afdb-uniprot50 \| AF-A0A832USA2-F1-MODEL\_V4 \| 1.0 \| 4.427e-10 \| 294 \| 0.165 \| 218 \| 136 \| 6 \| 4 \| 180 \| 9 \| 221 \| Helix-turn-helix transcriptional regulator \| Helix-turn-helix transcriptional regulator; afdb-uniprot50 \| AF-A0A1I0CD38-F1-MODEL\_V4 \| 1.0 \| 3.126e-10 \| 294 \| 0.193 \| 217 \| 133 \| 6 \| 4 \| 180 \| 9 \| 223 \| Phage repressor protein C, contains Cro/C1-type HTH and peptisase s24 domains \| Phage repressor protein C, contains Cro/C1-type HTH and peptisase s24 domains; afdb-uniprot50 \| AF-A0A2G6CHA8-F1-MODEL\_V4 \| 1.0 \| 2.339e-10 \| 294 \| 0.214 \| 219 \| 129 \| 5 \| 4 \| 180 \| 6 \| 223 \| Transcriptional regulator \| Transcriptional regulator; afdb-uniprot50 \| AF-A0A425B209-F1-MODEL\_V4 \| 1.0 \| 2.339e-10 \| 294 \| 0.2 \| 224 \| 128 \| 6 \| 5 \| 180 \| 3 \| 223 \| XRE family transcriptional regulator \| XRE family transcriptional regulator; afdb-uniprot50 \| AF-A0A1A7PW04-F1-MODEL\_V4 \| 1.0 \| 5.489e-11 \| 294 \| 0.173 \| 225 \| 136 \| 8 \| 4 \| 180 \| 7 \| 229 \| Repressor \| Repressor; afdb-uniprot50 \| AF-A0A2T5R3U0-F1-MODEL\_V4 \| 1.0 \| 5.489e-11 \| 294 \| 0.238 \| 226 \| 116 \| 9 \| 5 \| 180 \| 15 \| 234 \| Phage repressor protein C with HTH and peptisase S24 domain \| Phage repressor protein C with HTH and peptisase S24 domain; afdb-uniprot50 \| AF-A0A139SW86-F1-MODEL\_V4 \| 1.0 \| 2.207e-10 \| 294 \| 0.172 \| 232 \| 137 \| 5 \| 4 \| 180 \| 7 \| 238 \| HTH cro/C1-type domain-containing protein \| HTH cro/C1-type domain-containing protein; afdb-uniprot50 \| AF-A0A246GMR8-F1-MODEL\_V4 \| 1.0 \| 1.855e-10 \| 294 \| 0.193 \| 238 \| 128 \| 7 \| 4 \| 180 \| 8 \| 242 \| XRE family transcriptional regulator \| XRE family transcriptional regulator; afdb-uniprot50 \| AF-A0A5E9N793-F1-MODEL\_V4 \| 1.0 \| 1.75e-10 \| 294 \| 0.193 \| 248 \| 128 \| 6 \| 5 \| 180 \| 8 \| 255 \| Helix-turn-helix domain-containing protein \| Helix-turn-helix domain-containing protein; afdb-uniprot50 \| AF-A0A486QK96-F1-MODEL\_V4 \| 1.0 \| 1.093e-07 \| 293 \| 0.268 \| 123 \| 84 \| 3 \| 61 \| 180 \| 4 \| 123 \| Putative phage repressor protein CI \| Putative phage repressor protein CI; afdb-uniprot50 \| AF-A0A653B0F6-F1-MODEL\_V4 \| 1.0 \| 2.068e-07 \| 293 \| 0.2 \| 105 \| 80 \| 2 \| 79 \| 180 \| 49 \| 152 \| Phage repressor protein C, contains Cro/C1-type HTH and peptisase s24 domains \| Phage repressor protein C, contains Cro/C1-type HTH and peptisase s24 domains; afdb-uniprot50 \| AF-A0A5C4SB52-F1-MODEL\_V4 \| 1.0 \| 4.148e-07 \| 293 \| 0.276 \| 105 \| 72 \| 3 \| 79 \| 180 \| 56 \| 159 \| S24 family peptidase \| S24 family peptidase; afdb-uniprot50 \| AF-A0A2J4SU95-F1-MODEL\_V4 \| 1.0 \| 1.64e-07 \| 293 \| 0.283 \| 113 \| 73 \| 4 \| 71 \| 180 \| 68 \| 175 \| Phage repressor protein \| Phage repressor protein; afdb-uniprot50 \| AF-A0A5D3WFX3-F1-MODEL\_V4 \| 1.0 \| 2.339e-10 \| 293 \| 0.255 \| 215 \| 115 \| 10 \| 4 \| 180 \| 7 \| 214 \| Phage repressor protein C with HTH and peptisase S24 domain \| Phage repressor protein C with HTH and peptisase S24 domain; afdb-uniprot50 \| AF-A0A2M7FQV6-F1-MODEL\_V4 \| 1.0 \| 8.237e-11 \| 293 \| 0.199 \| 226 \| 128 \| 5 \| 1 \| 178 \| 3 \| 223 \| HTH cro/C1-type domain-containing protein \| HTH cro/C1-type domain-containing protein; afdb-uniprot50 \| AF-A0A1Y2Q9C5-F1-MODEL\_V4 \| 1.0 \| 1.652e-10 \| 293 \| 0.185 \| 216 \| 133 \| 5 \| 4 \| 178 \| 7 \| 220 \| HTH cro/C1-type domain-containing protein \| HTH cro/C1-type domain-containing protein; afdb-uniprot50 \| AF-E6W0K3-F1-MODEL\_V4 \| 1.0 \| 2.784e-10 \| 293 \| 0.208 \| 221 \| 130 \| 5 \| 4 \| 180 \| 5 \| 224 \| Helix-turn-helix domain protein \| Helix-turn-helix domain protein; afdb-uniprot50 \| AF-A0A1V3JA05-F1-MODEL\_V4 \| 1.0 \| 2.784e-10 \| 293 \| 0.178 \| 219 \| 135 \| 8 \| 4 \| 180 \| 7 \| 222 \| Transcriptional regulator \| Transcriptional regulator; afdb-uniprot50 \| AF-A0A357I973-F1-MODEL\_V4 \| 1.0 \| 1.75e-10 \| 293 \| 0.183 \| 218 \| 133 \| 6 \| 5 \| 178 \| 11 \| 227 \| HTH cro/C1-type domain-containing protein \| HTH cro/C1-type domain-containing protein; afdb-uniprot50 \| AF-A0A6P1WJJ3-F1-MODEL\_V4 \| 1.0 \| 6.269e-10 \| 293 \| 0.196 \| 224 \| 130 \| 6 \| 5 \| 180 \| 6 \| 227 \| Helix-turn-helix transcriptional regulator \| Helix-turn-helix transcriptional regulator; afdb-uniprot50 \| AF-A0A2G6CCY5-F1-MODEL\_V4 \| 1.0 \| 3.126e-10 \| 293 \| 0.218 \| 229 \| 122 \| 7 \| 4 \| 180 \| 10 \| 233 \| Transcriptional regulator \| Transcriptional regulator; afdb-uniprot50 \| AF-A0A1H1QPG0-F1-MODEL\_V4 \| 1.0 \| 1.559e-10 \| 293 \| 0.161 \| 223 \| 140 \| 6 \| 4 \| 180 \| 19 \| 240 \| Phage repressor protein C, contains Cro/C1-type HTH and peptisase s24 domains \| Phage repressor protein C, contains Cro/C1-type HTH and peptisase s24 domains; afdb-uniprot50 \| AF-A0A4R1MGX5-F1-MODEL\_V4 \| 1.0 \| 2.784e-10 \| 293 \| 0.185 \| 242 \| 129 \| 5 \| 4 \| 178 \| 7 \| 247 \| Phage repressor protein C with HTH and peptisase S24 domain \| Phage repressor protein C with HTH and peptisase S24 domain; afdb-uniprot50 \| AF-A0A5D3YA70-F1-MODEL\_V4 \| 1.0 \| 6.12e-08 \| 293 \| 0.276 \| 130 \| 78 \| 4 \| 63 \| 180 \| 123 \| 248 \| Phage repressor protein C with HTH and peptisase S24 domain \| Phage repressor protein C with HTH and peptisase S24 domain; afdb-uniprot50 \| AF-A0A1F0H5F4-F1-MODEL\_V4 \| 1.0 \| 3.876e-11 \| 293 \| 0.21 \| 209 \| 128 \| 7 \| 5 \| 176 \| 37 \| 245 \| HTH cro/C1-type domain-containing protein \| HTH cro/C1-type domain-containing protein; afdb-uniprot50 \| AF-A0A516WA19-F1-MODEL\_V4 \| 1.0 \| 1.388e-10 \| 293 \| 0.177 \| 225 \| 128 \| 6 \| 5 \| 179 \| 50 \| 267 \| Helix-turn-helix transcriptional regulator \| Helix-turn-helix transcriptional regulator; afdb-uniprot50 \| AF-A0A2C9D6H6-F1-MODEL\_V4 \| 1.0 \| 2.192e-07 \| 292 \| 0.272 \| 110 \| 73 \| 4 \| 77 \| 180 \| 75 \| 183 \| Pyocin repressor protein \| Pyocin repressor protein; afdb-uniprot50 \| AF-A0A5B8KTY6-F1-MODEL\_V4 \| 1.0 \| 5.583e-10 \| 292 \| 0.207 \| 217 \| 132 \| 6 \| 1 \| 180 \| 1 \| 214 \| Helix-turn-helix transcriptional regulator \| Helix-turn-helix transcriptional regulator; afdb-uniprot50 \| AF-B9JYW3-F1-MODEL\_V4 \| 1.0 \| 1.652e-10 \| 292 \| 0.187 \| 219 \| 139 \| 6 \| 1 \| 180 \| 1 \| 219 \| Transcriptional regulator \| Transcriptional regulator; afdb-uniprot50 \| AF-A0A1B8PUD1-F1-MODEL\_V4 \| 1.0 \| 1.559e-10 \| 292 \| 0.165 \| 218 \| 135 \| 6 \| 6 \| 178 \| 9 \| 224 \| HTH cro/C1-type domain-containing protein \| HTH cro/C1-type domain-containing protein; afdb-uniprot50 \| AF-A0A844RUM5-F1-MODEL\_V4 \| 1.0 \| 3.511e-10 \| 292 \| 0.221 \| 217 \| 126 \| 6 \| 4 \| 180 \| 12 \| 225 \| Helix-turn-helix domain-containing protein \| Helix-turn-helix domain-containing protein; afdb-uniprot50 \| AF-A0A2Z4RAG6-F1-MODEL\_V4 \| 1.0 \| 1.471e-10 \| 292 \| 0.191 \| 219 \| 133 \| 7 \| 4 \| 180 \| 9 \| 225 \| Peptidase\_S24 domain-containing protein \| Peptidase\_S24 domain-containing protein; afdb-uniprot50 \| AF-A0A6M4XFP4-F1-MODEL\_V4 \| 1.0 \| 2.95e-10 \| 292 \| 0.197 \| 223 \| 126 \| 7 \| 5 \| 180 \| 16 \| 232 \| Helix-turn-helix transcriptional regulator \| Helix-turn-helix transcriptional regulator; afdb-uniprot50 \| AF-A0A6V8N692-F1-MODEL\_V4 \| 1.0 \| 3.126e-10 \| 292 \| 0.222 \| 234 \| 119 \| 7 \| 5 \| 179 \| 3 \| 232 \| Transcriptional regulator \| Transcriptional regulator; afdb-uniprot50 \| AF-A0A653HZL2-F1-MODEL\_V4 \| 1.0 \| 8.878e-10 \| 292 \| 0.165 \| 223 \| 137 \| 5 \| 4 \| 178 \| 13 \| 234 \| HTH cro/C1-type domain-containing protein \| HTH cro/C1-type domain-containing protein; afdb-uniprot50 \| AF-A0A354IYV1-F1-MODEL\_V4 \| 1.0 \| 2.339e-10 \| 292 \| 0.168 \| 231 \| 137 \| 6 \| 4 \| 180 \| 7 \| 236 \| Peptidase \| Peptidase; afdb-uniprot50 \| AF-A1W671-F1-MODEL\_V4 \| 1.0 \| 2.437e-11 \| 292 \| 0.247 \| 202 \| 119 \| 7 \| 4 \| 180 \| 44 \| 237 \| Putative phage repressor \| Putative phage repressor; afdb-uniprot50 \| AF-A0A7W4FX54-F1-MODEL\_V4 \| 1.0 \| 2.083e-10 \| 292 \| 0.234 \| 230 \| 123 \| 5 \| 4 \| 180 \| 8 \| 237 \| Helix-turn-helix transcriptional regulator \| Helix-turn-helix transcriptional regulator; afdb-uniprot50 \| AF-A0A7Y7YNG6-F1-MODEL\_V4 \| 1.0 \| 4.178e-10 \| 292 \| 0.179 \| 234 \| 133 \| 8 \| 4 \| 180 \| 14 \| 245 \| Helix-turn-helix transcriptional regulator \| Helix-turn-helix transcriptional regulator; afdb-uniprot50 \| AF-A0A0J5QMV2-F1-MODEL\_V4 \| 1.0 \| 8.729e-11 \| 292 \| 0.206 \| 247 \| 124 \| 6 \| 4 \| 180 \| 7 \| 251 \| Putative transcriptional regulator \| Putative transcriptional regulator; afdb-uniprot50 \| AF-Q2RVM0-F1-MODEL\_V4 \| 1.0 \| 1.388e-10 \| 292 \| 0.21 \| 237 \| 127 \| 8 \| 2 \| 180 \| 31 \| 265 \| Phage repressor \| Phage repressor; afdb-uniprot50 \| AF-E1SGK7-F1-MODEL\_V4 \| 1.0 \| 2.929e-07 \| 291 \| 0.263 \| 114 \| 77 \| 4 \| 70 \| 180 \| 30 \| 139 \| 26 kDa repressor protein (Regulatory protein CI) \| 26 kDa repressor protein (Regulatory protein CI); afdb-uniprot50 \| AF-A0A1G7D0E1-F1-MODEL\_V4 \| 1.0 \| 2.339e-10 \| 291 \| 0.271 \| 199 \| 116 \| 6 \| 4 \| 180 \| 6 \| 197 \| Peptidase S24-like \| Peptidase S24-like; afdb-uniprot50 \| AF-A0A2D5S2B6-F1-MODEL\_V4 \| 1.0 \| 2.627e-10 \| 291 \| 0.204 \| 205 \| 131 \| 6 \| 4 \| 180 \| 12 \| 212 \| HTH cro/C1-type domain-containing protein \| HTH cro/C1-type domain-containing protein; afdb-uniprot50 \| AF-A0A829P4G5-F1-MODEL\_V4 \| 1.0 \| 7.46e-10 \| 291 \| 0.201 \| 208 \| 133 \| 3 \| 6 \| 180 \| 10 \| 217 \| Peptidase S24 \| Peptidase S24; afdb-uniprot50 \| AF-A0A1Q8ZSD3-F1-MODEL\_V4 \| 1.0 \| 1.855e-10 \| 291 \| 0.191 \| 219 \| 138 \| 6 \| 1 \| 180 \| 1 \| 219 \| Transcriptional regulator \| Transcriptional regulator; afdb-uniprot50 \| AF-A0A368ALS6-F1-MODEL\_V4 \| 1.0 \| 4.691e-10 \| 291 \| 0.192 \| 213 \| 132 \| 6 \| 4 \| 180 \| 11 \| 219 \| HTH cro/C1-type domain-containing protein \| HTH cro/C1-type domain-containing protein; afdb-uniprot50 \| AF-A0A329E845-F1-MODEL\_V4 \| 1.0 \| 3.72e-10 \| 291 \| 0.2 \| 215 \| 133 \| 4 \| 4 \| 180 \| 7 \| 220 \| Phage repressor protein C with HTH and peptisase S24 domain \| Phage repressor protein C with HTH and peptisase S24 domain; afdb-uniprot50 \| AF-A0A1Q5TGM7-F1-MODEL\_V4 \| 1.0 \| 2.95e-10 \| 291 \| 0.163 \| 220 \| 138 \| 8 \| 4 \| 180 \| 9 \| 225 \| Transcriptional regulator \| Transcriptional regulator; afdb-uniprot50 \| AF-A0A0C5S4V4-F1-MODEL\_V4 \| 1.0 \| 2.784e-10 \| 291 \| 0.184 \| 233 \| 131 \| 5 \| 4 \| 180 \| 7 \| 236 \| XRE family transcriptional regulator \| XRE family transcriptional regulator; afdb-uniprot50 \| AF-A0A2D5RVH7-F1-MODEL\_V4 \| 1.0 \| 2.339e-10 \| 291 \| 0.214 \| 233 \| 127 \| 5 \| 4 \| 180 \| 8 \| 240 \| XRE family transcriptional regulator \| XRE family transcriptional regulator; afdb-uniprot50 \| AF-A0A829TAF0-F1-MODEL\_V4 \| 1.0 \| 2.083e-10 \| 291 \| 0.185 \| 232 \| 134 \| 5 \| 4 \| 180 \| 14 \| 245 \| Predicted transcriptional regulator \| Predicted transcriptional regulator; afdb-uniprot50 \| AF-A0A7T1GX48-F1-MODEL\_V4 \| 1.0 \| 2.627e-10 \| 291 \| 0.192 \| 229 \| 132 \| 8 \| 4 \| 180 \| 18 \| 245 \| Helix-turn-helix domain-containing protein \| Helix-turn-helix domain-containing protein; afdb-uniprot50 \| AF-A0A1H6KDH3-F1-MODEL\_V4 \| 1.0 \| 4.396e-07 \| 291 \| 0.235 \| 106 \| 76 \| 3 \| 78 \| 180 \| 152 \| 255 \| Peptidase S24-like \| Peptidase S24-like; afdb-uniprot50 \| AF-A0A015NHE4-F1-MODEL\_V4 \| 1.0 \| 4.691e-10 \| 291 \| 0.176 \| 249 \| 132 \| 7 \| 4 \| 180 \| 7 \| 254 \| HTH cro/C1-type domain-containing protein \| HTH cro/C1-type domain-containing protein; afdb-uniprot50 \| AF-A0A0J1HD31-F1-MODEL\_V4 \| 1.0 \| 3.72e-10 \| 291 \| 0.17 \| 252 \| 131 \| 8 \| 4 \| 180 \| 26 \| 274 \| Peptidase\_S24 domain-containing protein \| Peptidase\_S24 domain-containing protein; afdb-uniprot50 \| AF-A0A1Y1IYP3-F1-MODEL\_V4 \| 1.0 \| 1.652e-10 \| 291 \| 0.193 \| 222 \| 129 \| 10 \| 2 \| 180 \| 106 \| 320 \| Prophage transcriptional regulator \| Prophage transcriptional regulator; afdb-uniprot50 \| AF-A0A377R2D0-F1-MODEL\_V4 \| 1.0 \| 1.461e-07 \| 290 \| 0.268 \| 123 \| 85 \| 3 \| 60 \| 179 \| 5 \| 125 \| Uncharacterized HTH-type transcriptional regulator HI\_1476 \| Uncharacterized HTH-type transcriptional regulator HI\_1476; afdb-uniprot50 \| AF-A0A2N7RSW1-F1-MODEL\_V4 \| 1.0 \| 4.936e-07 \| 290 \| 0.235 \| 102 \| 76 \| 2 \| 79 \| 180 \| 30 \| 129 \| LexA repressor \| LexA repressor; afdb-uniprot50 \| AF-A0A6B8QMJ2-F1-MODEL\_V4 \| 1.0 \| 6.597e-07 \| 290 \| 0.285 \| 105 \| 71 \| 2 \| 79 \| 180 \| 71 \| 174 \| S24 family peptidase \| S24 family peptidase; afdb-uniprot50 \| AF-A0A7W0C9Y1-F1-MODEL\_V4 \| 1.0 \| 1.668e-06 \| 290 \| 0.234 \| 94 \| 72 \| 0 \| 87 \| 180 \| 82 \| 175 \| Phage repressor protein C with HTH and peptisase S24 domain \| Phage repressor protein C with HTH and peptisase S24 domain; afdb-uniprot50 \| AF-A0A6C1MWJ7-F1-MODEL\_V4 \| 1.0 \| 5.583e-10 \| 290 \| 0.184 \| 211 \| 136 \| 6 \| 4 \| 180 \| 7 \| 215 \| LexA family transcriptional regulator \| LexA family transcriptional regulator; afdb-uniprot50 \| AF-A0A2N9XVJ2-F1-MODEL\_V4 \| 1.0 \| 3.313e-10 \| 290 \| 0.188 \| 218 \| 134 \| 5 \| 4 \| 180 \| 5 \| 220 \| HTH cro/C1-type domain-containing protein \| HTH cro/C1-type domain-containing protein; afdb-uniprot50 \| AF-A0A6F8QDC6-F1-MODEL\_V4 \| 1.0 \| 4.971e-10 \| 290 \| 0.165 \| 224 \| 138 \| 7 \| 6 \| 180 \| 2 \| 225 \| HTH cro/C1-type domain-containing protein \| HTH cro/C1-type domain-containing protein; afdb-uniprot50 \| AF-A0A6N9DJF1-F1-MODEL\_V4 \| 1.0 \| 7.46e-10 \| 290 \| 0.174 \| 212 \| 137 \| 6 \| 4 \| 180 \| 25 \| 233 \| Helix-turn-helix domain-containing protein \| Helix-turn-helix domain-containing protein; afdb-uniprot50 \| AF-A0A3A4NEQ1-F1-MODEL\_V4 \| 1.0 \| 2.784e-10 \| 290 \| 0.174 \| 223 \| 136 \| 5 \| 4 \| 180 \| 17 \| 237 \| Helix-turn-helix transcriptional regulator \| Helix-turn-helix transcriptional regulator; afdb-uniprot50 \| AF-A0A3M3NJT7-F1-MODEL\_V4 \| 1.0 \| 2.083e-10 \| 290 \| 0.188 \| 249 \| 128 \| 5 \| 5 \| 180 \| 8 \| 255 \| HTH cro/C1-type domain-containing protein \| HTH cro/C1-type domain-containing protein; afdb-uniprot50 \| AF-A0A8B5ZRH6-F1-MODEL\_V4 \| 1.0 \| 7.04e-10 \| 290 \| 0.157 \| 228 \| 140 \| 5 \| 4 \| 180 \| 30 \| 256 \| Helix-turn-helix transcriptional regulator \| Helix-turn-helix transcriptional regulator; afdb-uniprot50 \| AF-A0A6M4XRT2-F1-MODEL\_V4 \| 1.0 \| 1.952e-07 \| 289 \| 0.28 \| 114 \| 74 \| 3 \| 70 \| 180 \| 1 \| 109 \| S24 family peptidase \| S24 family peptidase; afdb-uniprot50 \| AF-A0A376TJ78-F1-MODEL\_V4 \| 1.0 \| 2.068e-07 \| 289 \| 0.283 \| 113 \| 74 \| 3 \| 71 \| 180 \| 48 \| 156 \| Regulatory protein CI from bacteriophage origin \| Regulatory protein CI from bacteriophage origin; afdb-uniprot50 \| AF-A0A1E3WK34-F1-MODEL\_V4 \| 1.0 \| 3.511e-10 \| 289 \| 0.202 \| 212 \| 131 \| 7 \| 4 \| 180 \| 10 \| 218 \| HTH cro/C1-type domain-containing protein \| HTH cro/C1-type domain-containing protein; afdb-uniprot50 \| AF-A0A127MKS3-F1-MODEL\_V4 \| 1.0 \| 4.971e-10 \| 289 \| 0.184 \| 217 \| 131 \| 7 \| 4 \| 180 \| 8 \| 218 \| Putative HTH-type transcriptional regulator \| Putative HTH-type transcriptional regulator; afdb-uniprot50 \| AF-A0A496L3E2-F1-MODEL\_V4 \| 1.0 \| 4.691e-10 \| 289 \| 0.21 \| 223 \| 129 \| 4 \| 4 \| 180 \| 7 \| 228 \| Helix-turn-helix transcriptional regulator \| Helix-turn-helix transcriptional regulator; afdb-uniprot50 \| AF-A0A261R781-F1-MODEL\_V4 \| 1.0 \| 3.72e-10 \| 289 \| 0.187 \| 224 \| 133 \| 6 \| 4 \| 179 \| 6 \| 228 \| HTH cro/C1-type domain-containing protein \| HTH cro/C1-type domain-containing protein; afdb-uniprot50 \| AF-A0A443J1K1-F1-MODEL\_V4 \| 1.0 \| 2.339e-10 \| 289 \| 0.186 \| 225 \| 134 \| 6 \| 4 \| 180 \| 10 \| 233 \| LexA family transcriptional regulator \| LexA family transcriptional regulator; afdb-uniprot50 \| AF-A0A198XTH8-F1-MODEL\_V4 \| 1.0 \| 3.313e-10 \| 289 \| 0.201 \| 228 \| 130 \| 7 \| 4 \| 180 \| 6 \| 232 \| Transcriptional regulator \| Transcriptional regulator; afdb-uniprot50 \| AF-A0A7X3VZG0-F1-MODEL\_V4 \| 1.0 \| 1.855e-10 \| 289 \| 0.214 \| 238 \| 117 \| 8 \| 5 \| 180 \| 10 \| 239 \| Helix-turn-helix transcriptional regulator \| Helix-turn-helix transcriptional regulator; afdb-uniprot50 \| AF-A0A3L0WWP8-F1-MODEL\_V4 \| 1.0 \| 1.855e-10 \| 289 \| 0.199 \| 241 \| 126 \| 5 \| 4 \| 180 \| 17 \| 254 \| LexA family transcriptional regulator \| LexA family transcriptional regulator; afdb-uniprot50 \| AF-A0A6I6QEU7-F1-MODEL\_V4 \| 1.0 \| 1.166e-10 \| 289 \| 0.18 \| 271 \| 127 \| 5 \| 4 \| 180 \| 6 \| 275 \| Helix-turn-helix domain-containing protein \| Helix-turn-helix domain-containing protein; afdb-uniprot50 \| AF-A0A5E9U8S2-F1-MODEL\_V4 \| 1.0 \| 4.971e-10 \| 289 \| 0.149 \| 267 \| 136 \| 4 \| 4 \| 180 \| 16 \| 281 \| Helix-turn-helix domain-containing protein \| Helix-turn-helix domain-containing protein; afdb-uniprot50 \| AF-A0A251ZV98-F1-MODEL\_V4 \| 1.0 \| 5.874e-07 \| 288 \| 0.311 \| 93 \| 60 \| 2 \| 87 \| 176 \| 5 \| 96 \| Repressor \| Repressor; afdb-uniprot50 \| AF-A0A0Q4PTP6-F1-MODEL\_V4 \| 1.0 \| 2.461e-07 \| 288 \| 0.22 \| 118 \| 85 \| 3 \| 65 \| 180 \| 2 \| 114 \| Peptidase\_S24 domain-containing protein \| Peptidase\_S24 domain-containing protein; afdb-uniprot50 \| AF-K1JKB5-F1-MODEL\_V4 \| 1.0 \| 1.227e-07 \| 288 \| 0.243 \| 123 \| 84 \| 4 \| 61 \| 180 \| 3 \| 119 \| Peptidase\_S24 domain-containing protein \| Peptidase\_S24 domain-containing protein; afdb-uniprot50 \| AF-A0A7H4P4T5-F1-MODEL\_V4 \| 1.0 \| 2.323e-07 \| 288 \| 0.283 \| 113 \| 73 \| 4 \| 71 \| 180 \| 33 \| 140 \| Phage repressor \| Phage repressor; afdb-uniprot50 \| AF-A0A1E4EGW2-F1-MODEL\_V4 \| 1.0 \| 6.225e-07 \| 288 \| 0.266 \| 105 \| 73 \| 2 \| 79 \| 180 \| 43 \| 146 \| Peptidase\_S24 domain-containing protein \| Peptidase\_S24 domain-containing protein; afdb-uniprot50 \| AF-A0A515EKI5-F1-MODEL\_V4 \| 1.0 \| 5.231e-07 \| 288 \| 0.219 \| 105 \| 78 \| 2 \| 79 \| 180 \| 50 \| 153 \| Helix-turn-helix transcriptional regulator \| Helix-turn-helix transcriptional regulator; afdb-uniprot50 \| AF-A0A5K1I4S0-F1-MODEL\_V4 \| 1.0 \| 4.658e-07 \| 288 \| 0.262 \| 103 \| 73 \| 1 \| 78 \| 177 \| 60 \| 162 \| HTH-type transcriptional regulator PrtR \| HTH-type transcriptional regulator PrtR; afdb-uniprot50 \| AF-A0A7H5FFX3-F1-MODEL\_V4 \| 1.0 \| 6.99e-07 \| 288 \| 0.235 \| 102 \| 78 \| 0 \| 79 \| 180 \| 79 \| 180 \| Transcriptional regulator \| Transcriptional regulator; afdb-uniprot50 \| AF-N9N570-F1-MODEL\_V4 \| 1.0 \| 3.511e-10 \| 288 \| 0.198 \| 191 \| 134 \| 3 \| 4 \| 180 \| 7 \| 192 \| HTH cro/C1-type domain-containing protein \| HTH cro/C1-type domain-containing protein; afdb-uniprot50 \| AF-A0A285TSU5-F1-MODEL\_V4 \| 1.0 \| 4.691e-10 \| 288 \| 0.171 \| 216 \| 137 \| 7 \| 5 \| 180 \| 7 \| 220 \| Phage repressor protein C, contains Cro/C1-type HTH and peptisase s24 domains \| Phage repressor protein C, contains Cro/C1-type HTH and peptisase s24 domains; afdb-uniprot50 \| AF-A0A1X0TJB0-F1-MODEL\_V4 \| 1.0 \| 1.388e-10 \| 288 \| 0.204 \| 220 \| 123 \| 5 \| 5 \| 180 \| 9 \| 220 \| HTH cro/C1-type domain-containing protein \| HTH cro/C1-type domain-containing protein; afdb-uniprot50 \| AF-A0A2N9XSA8-F1-MODEL\_V4 \| 1.0 \| 5.583e-10 \| 288 \| 0.14 \| 228 \| 142 \| 5 \| 6 \| 180 \| 3 \| 229 \| HTH cro/C1-type domain-containing protein \| HTH cro/C1-type domain-containing protein; afdb-uniprot50 \| AF-A0A376I3L8-F1-MODEL\_V4 \| 1.0 \| 4.691e-10 \| 288 \| 0.17 \| 211 \| 140 \| 6 \| 4 \| 180 \| 22 \| 231 \| DNA-binding/peptidase S24 domain protein \| DNA-binding/peptidase S24 domain protein; afdb-uniprot50 \| AF-A0A3B0MMP0-F1-MODEL\_V4 \| 1.0 \| 3.313e-10 \| 288 \| 0.149 \| 228 \| 142 \| 4 \| 4 \| 180 \| 20 \| 246 \| HTH-type transcriptional regulator PrtR \| HTH-type transcriptional regulator PrtR; afdb-uniprot50 \| AF-A0A5C8AJZ0-F1-MODEL\_V4 \| 1.0 \| 3.511e-10 \| 288 \| 0.194 \| 252 \| 124 \| 6 \| 4 \| 180 \| 7 \| 254 \| XRE family transcriptional regulator \| XRE family transcriptional regulator; afdb-uniprot50 \| AF-A0A6L4AN93-F1-MODEL\_V4 \| 1.0 \| 8.878e-10 \| 288 \| 0.18 \| 210 \| 136 \| 4 \| 4 \| 180 \| 68 \| 274 \| Helix-turn-helix transcriptional regulator \| Helix-turn-helix transcriptional regulator; afdb-uniprot50 \| AF-A0A510U6Q2-F1-MODEL\_V4 \| 1.0 \| 2.479e-10 \| 287 \| 0.213 \| 183 \| 135 \| 4 \| 5 \| 180 \| 15 \| 195 \| HTH cro/C1-type domain-containing protein \| HTH cro/C1-type domain-containing protein; afdb-uniprot50 \| AF-F9T596-F1-MODEL\_V4 \| 1.0 \| 3.511e-10 \| 287 \| 0.211 \| 208 \| 126 \| 4 \| 4 \| 180 \| 7 \| 207 \| Repressor \| Repressor; afdb-uniprot50 \| AF-A0A640W7B7-F1-MODEL\_V4 \| 1.0 \| 4.178e-10 \| 287 \| 0.166 \| 210 \| 136 \| 7 \| 4 \| 180 \| 11 \| 214 \| Helix-turn-helix domain-containing protein \| Helix-turn-helix domain-containing protein; afdb-uniprot50 \| AF-A0A099KPN4-F1-MODEL\_V4 \| 1.0 \| 2.479e-10 \| 287 \| 0.211 \| 227 \| 122 \| 9 \| 1 \| 180 \| 1 \| 217 \| Putative phage repressor \| Putative phage repressor; afdb-uniprot50 \| AF-A0A418WU97-F1-MODEL\_V4 \| 1.0 \| 3.126e-10 \| 287 \| 0.196 \| 214 \| 133 \| 4 \| 4 \| 180 \| 6 \| 217 \| XRE family transcriptional regulator \| XRE family transcriptional regulator; afdb-uniprot50 \| AF-A0A7V2F687-F1-MODEL\_V4 \| 1.0 \| 2.784e-10 \| 287 \| 0.218 \| 224 \| 125 \| 6 \| 4 \| 180 \| 8 \| 228 \| Helix-turn-helix transcriptional regulator \| Helix-turn-helix transcriptional regulator; afdb-uniprot50 \| AF-A0A2N2R9N5-F1-MODEL\_V4 \| 1.0 \| 4.612e-11 \| 287 \| 0.26 \| 196 \| 124 \| 7 \| 4 \| 180 \| 34 \| 227 \| Peptidase S24 \| Peptidase S24; afdb-uniprot50 \| AF-A0A7W2VIL3-F1-MODEL\_V4 \| 1.0 \| 8.378e-10 \| 287 \| 0.223 \| 219 \| 127 \| 6 \| 4 \| 180 \| 18 \| 235 \| Helix-turn-helix transcriptional regulator \| Helix-turn-helix transcriptional regulator; afdb-uniprot50 \| AF-A0A2D6RTW2-F1-MODEL\_V4 \| 1.0 \| 2.95e-10 \| 287 \| 0.175 \| 240 \| 134 \| 8 \| 5 \| 180 \| 4 \| 243 \| Phage repressor protein \| Phage repressor protein; afdb-uniprot50 \| AF-A0A1C3H9I3-F1-MODEL\_V4 \| 1.0 \| 5.916e-10 \| 287 \| 0.197 \| 228 \| 131 \| 4 \| 5 \| 180 \| 19 \| 246 \| HTH-type transcriptional regulator PrtR \| HTH-type transcriptional regulator PrtR; afdb-uniprot50 \| AF-A0A2E7Q619-F1-MODEL\_V4 \| 1.0 \| 8.378e-10 \| 287 \| 0.213 \| 230 \| 123 \| 9 \| 5 \| 180 \| 40 \| 265 \| HTH cro/C1-type domain-containing protein \| HTH cro/C1-type domain-containing protein; afdb-uniprot50 \| AF-A0A2I2MFF4-F1-MODEL\_V4 \| 1.0 \| 2.784e-10 \| 287 \| 0.193 \| 233 \| 130 \| 8 \| 4 \| 180 \| 106 \| 336 \| Helix-turn-helix protein \| Helix-turn-helix protein; afdb-uniprot50 \| AF-A0A377M491-F1-MODEL\_V4 \| 1.0 \| 3.694e-07 \| 286 \| 0.271 \| 114 \| 76 \| 4 \| 70 \| 180 \| 10 \| 119 \| Putative repressor protein \| Putative repressor protein; afdb-uniprot50 \| AF-A0A0Q4G0B2-F1-MODEL\_V4 \| 1.0 \| 6.99e-07 \| 286 \| 0.264 \| 106 \| 73 \| 2 \| 79 \| 179 \| 54 \| 159 \| Peptidase\_S24 domain-containing protein \| Peptidase\_S24 domain-containing protein; afdb-uniprot50 \| AF-A0A5C7FWN9-F1-MODEL\_V4 \| 1.0 \| 5.231e-07 \| 286 \| 0.266 \| 105 \| 72 \| 3 \| 79 \| 180 \| 64 \| 166 \| S24 family peptidase \| S24 family peptidase; afdb-uniprot50 \| AF-A0A1C3H412-F1-MODEL\_V4 \| 1.0 \| 4.148e-07 \| 286 \| 0.245 \| 106 \| 72 \| 3 \| 79 \| 178 \| 70 \| 173 \| Phage repressor \| Phage repressor; afdb-uniprot50 \| AF-A0A264VM17-F1-MODEL\_V4 \| 1.0 \| 3.486e-07 \| 286 \| 0.223 \| 112 \| 80 \| 4 \| 72 \| 180 \| 77 \| 184 \| Peptidase\_S24 domain-containing protein \| Peptidase\_S24 domain-containing protein; afdb-uniprot50 \| AF-A0A660NC27-F1-MODEL\_V4 \| 1.0 \| 5.583e-10 \| 286 \| 0.183 \| 218 \| 134 \| 7 \| 4 \| 180 \| 6 \| 220 \| S24 family peptidase \| S24 family peptidase; afdb-uniprot50 \| AF-Q72BW5-F1-MODEL\_V4 \| 1.0 \| 1.559e-10 \| 286 \| 0.205 \| 209 \| 127 \| 8 \| 5 \| 180 \| 22 \| 224 \| Transcriptional regulator, putative \| Transcriptional regulator, putative; afdb-uniprot50 \| AF-A0A220S1V6-F1-MODEL\_V4 \| 1.0 \| 2.479e-10 \| 286 \| 0.171 \| 228 \| 137 \| 6 \| 4 \| 180 \| 3 \| 229 \| Chromophore lyase \| Chromophore lyase; afdb-uniprot50 \| AF-A0A5E4V8H1-F1-MODEL\_V4 \| 1.0 \| 4.971e-10 \| 286 \| 0.181 \| 215 \| 128 \| 7 \| 4 \| 180 \| 31 \| 235 \| Putative HTH-type transcriptional regulator \| Putative HTH-type transcriptional regulator; afdb-uniprot50 \| AF-A0A7L5XVV2-F1-MODEL\_V4 \| 1.0 \| 4.691e-10 \| 286 \| 0.211 \| 213 \| 131 \| 5 \| 4 \| 180 \| 33 \| 244 \| Helix-turn-helix transcriptional regulator \| Helix-turn-helix transcriptional regulator; afdb-uniprot50 \| AF-A0A1V3NCJ2-F1-MODEL\_V4 \| 1.0 \| 1.257e-09 \| 286 \| 0.202 \| 222 \| 129 \| 6 \| 5 \| 180 \| 29 \| 248 \| Peptidase\_S24 domain-containing protein \| Peptidase\_S24 domain-containing protein; afdb-uniprot50 \| AF-A0A7T7Q6W5-F1-MODEL\_V4 \| 1.0 \| 3.72e-10 \| 286 \| 0.175 \| 239 \| 134 \| 6 \| 5 \| 180 \| 10 \| 248 \| Helix-turn-helix transcriptional regulator \| Helix-turn-helix transcriptional regulator; afdb-uniprot50 \| AF-A0A0P7CEB0-F1-MODEL\_V4 \| 1.0 \| 3.72e-10 \| 286 \| 0.194 \| 252 \| 124 \| 6 \| 4 \| 180 \| 7 \| 254 \| XRE family transcriptional regulator \| XRE family transcriptional regulator; afdb-uniprot50 \| AF-A0A7R7BN55-F1-MODEL\_V4 \| 1.0 \| 4.178e-10 \| 285 \| 0.198 \| 202 \| 136 \| 4 \| 1 \| 180 \| 1 \| 198 \| Uncharacterized protein \| Uncharacterized protein; afdb-uniprot50 \| AF-A0A1V4T976-F1-MODEL\_V4 \| 1.0 \| 6.269e-10 \| 285 \| 0.2 \| 215 \| 132 \| 6 \| 4 \| 180 \| 5 \| 217 \| Peptidase\_S24 domain-containing protein \| Peptidase\_S24 domain-containing protein; afdb-uniprot50 \| AF-A0A3S8YQ18-F1-MODEL\_V4 \| 1.0 \| 7.906e-10 \| 285 \| 0.223 \| 210 \| 125 \| 9 \| 4 \| 180 \| 13 \| 217 \| S24 family peptidase \| S24 family peptidase; afdb-uniprot50 \| AF-A0A6G8YM25-F1-MODEL\_V4 \| 1.0 \| 1.471e-10 \| 285 \| 0.202 \| 217 \| 127 \| 10 \| 5 \| 180 \| 11 \| 222 \| Peptidase\_S24 domain-containing protein \| Peptidase\_S24 domain-containing protein; afdb-uniprot50 \| AF-A0A7X2KYD5-F1-MODEL\_V4 \| 1.0 \| 1.323e-06 \| 285 \| 0.186 \| 102 \| 83 \| 0 \| 79 \| 180 \| 136 \| 237 \| DUF723 domain-containing protein \| DUF723 domain-containing protein; afdb-uniprot50 \| AF-A0A7D7NB06-F1-MODEL\_V4 \| 1.0 \| 3.126e-10 \| 285 \| 0.149 \| 234 \| 141 \| 5 \| 4 \| 180 \| 5 \| 237 \| Chromophore lyase \| Chromophore lyase; afdb-uniprot50 \| AF-A0A1B3JHZ4-F1-MODEL\_V4 \| 1.0 \| 2.784e-10 \| 285 \| 0.152 \| 242 \| 139 \| 9 \| 4 \| 180 \| 7 \| 247 \| Phage repressor \| Phage repressor; afdb-uniprot50 \| AF-A0A4Q0YMX5-F1-MODEL\_V4 \| 1.0 \| 4.427e-10 \| 285 \| 0.202 \| 227 \| 130 \| 7 \| 4 \| 180 \| 91 \| 316 \| Uncharacterized protein \| Uncharacterized protein; afdb-uniprot50 \| AF-A0A2N0WL24-F1-MODEL\_V4 \| 1.0 \| 1.64e-07 \| 284 \| 0.278 \| 104 \| 72 \| 3 \| 79 \| 180 \| 5 \| 107 \| Transcriptional regulator \| Transcriptional regulator; afdb-uniprot50 \| AF-A0A1Z4C411-F1-MODEL\_V4 \| 1.0 \| 1.093e-07 \| 284 \| 0.316 \| 117 \| 71 \| 4 \| 65 \| 177 \| 4 \| 115 \| Peptidase\_S24 domain-containing protein \| Peptidase\_S24 domain-containing protein; afdb-uniprot50 \| AF-W1ED85-F1-MODEL\_V4 \| 1.0 \| 3.289e-07 \| 284 \| 0.298 \| 114 \| 74 \| 3 \| 70 \| 180 \| 7 \| 117 \| COG2932: Predicted transcriptional regulator \| COG2932: Predicted transcriptional regulator; afdb-uniprot50 \| AF-X1P3M5-F1-MODEL\_V4 \| 1.0 \| 4.148e-07 \| 284 \| 0.247 \| 113 \| 79 \| 3 \| 71 \| 180 \| 27 \| 136 \| Peptidase\_S24 domain-containing protein \| Peptidase\_S24 domain-containing protein; afdb-uniprot50 \| AF-A0A0H3EGA9-F1-MODEL\_V4 \| 1.0 \| 3.313e-10 \| 284 \| 0.224 \| 209 \| 118 \| 7 \| 14 \| 180 \| 1 \| 207 \| Regulatory protein CI from bacteriophage origin \| Regulatory protein CI from bacteriophage origin; afdb-uniprot50 \| AF-U7H6U4-F1-MODEL\_V4 \| 1.0 \| 2.207e-10 \| 284 \| 0.228 \| 219 \| 121 \| 6 \| 6 \| 180 \| 3 \| 217 \| HTH cro/C1-type domain-containing protein \| HTH cro/C1-type domain-containing protein; afdb-uniprot50 \| AF-A0A7W7Y2E7-F1-MODEL\_V4 \| 1.0 \| 9.408e-10 \| 284 \| 0.208 \| 216 \| 128 \| 8 \| 5 \| 180 \| 7 \| 219 \| Phage repressor protein C with HTH and peptisase S24 domain \| Phage repressor protein C with HTH and peptisase S24 domain; afdb-uniprot50 \| AF-A0A2X4G6V1-F1-MODEL\_V4 \| 1.0 \| 5.874e-07 \| 284 \| 0.252 \| 107 \| 74 \| 6 \| 78 \| 180 \| 113 \| 217 \| Peptidase S24-like protein \| Peptidase S24-like protein; afdb-uniprot50 \| AF-A0A7W4B885-F1-MODEL\_V4 \| 1.0 \| 3.72e-10 \| 284 \| 0.201 \| 233 \| 130 \| 7 \| 1 \| 180 \| 1 \| 230 \| LexA family transcriptional regulator \| LexA family transcriptional regulator; afdb-uniprot50 \| AF-A0A0H2MCK7-F1-MODEL\_V4 \| 1.0 \| 4.178e-10 \| 284 \| 0.186 \| 215 \| 136 \| 6 \| 4 \| 180 \| 18 \| 231 \| Putative HTH-type transcriptional regulator \| Putative HTH-type transcriptional regulator; afdb-uniprot50 \| AF-A0A147GPB6-F1-MODEL\_V4 \| 1.0 \| 5.916e-10 \| 284 \| 0.168 \| 225 \| 139 \| 8 \| 4 \| 180 \| 8 \| 232 \| HTH cro/C1-type domain-containing protein \| HTH cro/C1-type domain-containing protein; afdb-uniprot50 \| AF-A0A4Q5YJ50-F1-MODEL\_V4 \| 1.0 \| 2.627e-10 \| 284 \| 0.216 \| 231 \| 127 \| 5 \| 4 \| 180 \| 6 \| 236 \| Helix-turn-helix transcriptional regulator \| Helix-turn-helix transcriptional regulator; afdb-uniprot50 \| AF-A4JD17-F1-MODEL\_V4 \| 1.0 \| 3.942e-10 \| 284 \| 0.222 \| 225 \| 126 \| 10 \| 4 \| 180 \| 15 \| 238 \| Putative phage repressor \| Putative phage repressor; afdb-uniprot50 \| AF-N9M239-F1-MODEL\_V4 \| 1.0 \| 4.691e-10 \| 284 \| 0.193 \| 233 \| 129 \| 6 \| 4 \| 178 \| 7 \| 238 \| HTH cro/C1-type domain-containing protein \| HTH cro/C1-type domain-containing protein; afdb-uniprot50 \| AF-A0A3S0MVQ0-F1-MODEL\_V4 \| 1.0 \| 4.971e-10 \| 284 \| 0.213 \| 229 \| 127 \| 6 \| 5 \| 180 \| 14 \| 242 \| XRE family transcriptional regulator \| XRE family transcriptional regulator; afdb-uniprot50 \| AF-A0A6P2FI33-F1-MODEL\_V4 \| 1.0 \| 4.178e-10 \| 284 \| 0.172 \| 232 \| 133 \| 9 \| 5 \| 179 \| 22 \| 251 \| Pyocin repressor protein \| Pyocin repressor protein; afdb-uniprot50 \| AF-A0A060H7F8-F1-MODEL\_V4 \| 1.0 \| 3.72e-10 \| 284 \| 0.205 \| 229 \| 125 \| 9 \| 5 \| 178 \| 29 \| 255 \| Peptidase S24 \| Peptidase S24; afdb-uniprot50 \| AF-A0A2S6NDQ8-F1-MODEL\_V4 \| 1.0 \| 5.916e-10 \| 284 \| 0.165 \| 224 \| 138 \| 5 \| 5 \| 180 \| 44 \| 266 \| HTH cro/C1-type domain-containing protein \| HTH cro/C1-type domain-containing protein; afdb-uniprot50 \| AF-A0A6M3M5L7-F1-MODEL\_V4 \| 1.0 \| 1.559e-10 \| 284 \| 0.166 \| 265 \| 132 \| 6 \| 4 \| 180 \| 6 \| 269 \| Putative peptidase \| Putative peptidase; afdb-uniprot50 \| AF-A0A1Y0G6S2-F1-MODEL\_V4 \| 1.0 \| 6.597e-07 \| 284 \| 0.31 \| 100 \| 65 \| 2 \| 78 \| 173 \| 201 \| 300 \| Peptidase\_S24 domain-containing protein \| Peptidase\_S24 domain-containing protein; afdb-uniprot50 \| AF-A0A6B1DL16-F1-MODEL\_V4 \| 1.0 \| 2.339e-10 \| 284 \| 0.204 \| 230 \| 130 \| 4 \| 4 \| 180 \| 163 \| 392 \| Helix-turn-helix domain-containing protein \| Helix-turn-helix domain-containing protein; afdb-uniprot50 \| AF-A0A429MKG1-F1-MODEL\_V4 \| 1.0 \| 5.231e-07 \| 283 \| 0.214 \| 112 \| 80 \| 3 \| 70 \| 178 \| 28 \| 134 \| S24 family peptidase \| S24 family peptidase; afdb-uniprot50 \| AF-A0A839VM87-F1-MODEL\_V4 \| 1.0 \| 5.874e-07 \| 283 \| 0.247 \| 105 \| 74 \| 3 \| 79 \| 180 \| 100 \| 202 \| Uncharacterized protein \| Uncharacterized protein; afdb-uniprot50 \| AF-A0A379LIF7-F1-MODEL\_V4 \| 1.0 \| 8.878e-10 \| 283 \| 0.159 \| 226 \| 135 \| 8 \| 5 \| 178 \| 10 \| 232 \| Uncharacterized HTH-type transcriptional regulator HI\_1476 \| Uncharacterized HTH-type transcriptional regulator HI\_1476; afdb-uniprot50 \| AF-A0A3D0LWT4-F1-MODEL\_V4 \| 1.0 \| 9.408e-10 \| 283 \| 0.195 \| 230 \| 130 \| 8 \| 4 \| 180 \| 6 \| 233 \| Peptidase\_S24 domain-containing protein \| Peptidase\_S24 domain-containing protein; afdb-uniprot50 \| AF-A0A7G2S2Y5-F1-MODEL\_V4 \| 1.0 \| 2.627e-10 \| 283 \| 0.189 \| 227 \| 133 \| 6 \| 5 \| 180 \| 10 \| 236 \| XRE family transcriptional regulator \| XRE family transcriptional regulator; afdb-uniprot50 \| AF-Q9I551-F1-MODEL\_V4 \| 1.0 \| 7.04e-10 \| 283 \| 0.206 \| 233 \| 128 \| 5 \| 4 \| 180 \| 6 \| 237 \| Probable transcriptional regulator \| Probable transcriptional regulator; afdb-uniprot50 \| AF-A0A443LNA1-F1-MODEL\_V4 \| 1.0 \| 5.268e-10 \| 283 \| 0.2 \| 225 \| 131 \| 7 \| 4 \| 180 \| 15 \| 238 \| LexA family transcriptional regulator \| LexA family transcriptional regulator; afdb-uniprot50 \| AF-A0A5D9DBC8-F1-MODEL\_V4 \| 1.0 \| 6.269e-10 \| 283 \| 0.195 \| 235 \| 125 \| 6 \| 4 \| 177 \| 68 \| 299 \| Helix-turn-helix transcriptional regulator \| Helix-turn-helix transcriptional regulator; afdb-uniprot50 \| AF-V9H5H0-F1-MODEL\_V4 \| 1.0 \| 1.057e-09 \| 282 \| 0.205 \| 200 \| 135 \| 8 \| 1 \| 180 \| 1 \| 196 \| Peptidase\_S24 domain-containing protein \| Peptidase\_S24 domain-containing protein; afdb-uniprot50 \| AF-A0A7T0BVD4-F1-MODEL\_V4 \| 1.0 \| 5.583e-10 \| 282 \| 0.19 \| 221 \| 132 \| 7 \| 5 \| 180 \| 3 \| 221 \| Helix-turn-helix transcriptional regulator \| Helix-turn-helix transcriptional regulator; afdb-uniprot50 \| AF-A0A3B0IWS9-F1-MODEL\_V4 \| 1.0 \| 4.178e-10 \| 282 \| 0.157 \| 228 \| 139 \| 5 \| 4 \| 180 \| 6 \| 231 \| HTH-type transcriptional regulator PrtR \| HTH-type transcriptional regulator PrtR; afdb-uniprot50 \| AF-A0A0N9VWL4-F1-MODEL\_V4 \| 1.0 \| 1.652e-10 \| 282 \| 0.173 \| 236 \| 133 \| 7 \| 6 \| 180 \| 6 \| 240 \| HTH cro/C1-type domain-containing protein \| HTH cro/C1-type domain-containing protein; afdb-uniprot50 \| AF-A0A6V8IAU7-F1-MODEL\_V4 \| 1.0 \| 2.95e-10 \| 282 \| 0.206 \| 232 \| 118 \| 8 \| 4 \| 180 \| 19 \| 239 \| Putative HTH-type transcriptional regulator \| Putative HTH-type transcriptional regulator; afdb-uniprot50 \| AF-A0A5C8WGV8-F1-MODEL\_V4 \| 1.0 \| 9.408e-10 \| 282 \| 0.193 \| 232 \| 126 \| 6 \| 4 \| 180 \| 20 \| 245 \| LexA family transcriptional regulator \| LexA family transcriptional regulator; afdb-uniprot50 \| AF-C0N2Z7-F1-MODEL\_V4 \| 1.0 \| 1.057e-09 \| 282 \| 0.181 \| 265 \| 127 \| 6 \| 4 \| 180 \| 10 \| 272 \| Peptidase S24-like domain protein \| Peptidase S24-like domain protein; afdb-uniprot50 \| AF-Q3JTQ6-F1-MODEL\_V4 \| 1.0 \| 4.178e-10 \| 282 \| 0.211 \| 208 \| 130 \| 6 \| 5 \| 180 \| 84 \| 289 \| Repressor protein, putative \| Repressor protein, putative; afdb-uniprot50 \| AF-A0A511FR38-F1-MODEL\_V4 \| 1.0 \| 3.72e-10 \| 281 \| 0.217 \| 198 \| 125 \| 7 \| 4 \| 175 \| 15 \| 208 \| Repressor \| Repressor; afdb-uniprot50 \| AF-A0A255ZI18-F1-MODEL\_V4 \| 1.0 \| 5.583e-10 \| 281 \| 0.179 \| 217 \| 129 \| 5 \| 5 \| 179 \| 7 \| 216 \| HTH cro/C1-type domain-containing protein \| HTH cro/C1-type domain-containing protein; afdb-uniprot50 \| AF-A0A0S4TX81-F1-MODEL\_V4 \| 1.0 \| 2.627e-10 \| 281 \| 0.215 \| 218 \| 127 \| 7 \| 5 \| 180 \| 6 \| 221 \| Peptidase\_S24 domain-containing protein \| Peptidase\_S24 domain-containing protein; afdb-uniprot50 \| AF-A0A7C4YFR3-F1-MODEL\_V4 \| 1.0 \| 1.559e-10 \| 281 \| 0.222 \| 225 \| 128 \| 9 \| 1 \| 180 \| 3 \| 225 \| Helix-turn-helix transcriptional regulator \| Helix-turn-helix transcriptional regulator; afdb-uniprot50 \| AF-A0A1H0LEQ9-F1-MODEL\_V4 \| 1.0 \| 4.178e-10 \| 281 \| 0.229 \| 227 \| 126 \| 7 \| 1 \| 180 \| 1 \| 225 \| Phage repressor protein C, contains Cro/C1-type HTH and peptisase s24 domains \| Phage repressor protein C, contains Cro/C1-type HTH and peptisase s24 domains; afdb-uniprot50 \| AF-A3JLF3-F1-MODEL\_V4 \| 1.0 \| 2.245e-09 \| 281 \| 0.163 \| 226 \| 139 \| 5 \| 4 \| 180 \| 13 \| 237 \| Probable transcriptional regulator \| Probable transcriptional regulator; afdb-uniprot50 \| AF-A0A1C3ELF7-F1-MODEL\_V4 \| 1.0 \| 5.916e-10 \| 281 \| 0.167 \| 233 \| 137 \| 6 \| 4 \| 180 \| 7 \| 238 \| XRE family transcriptional regulator \| XRE family transcriptional regulator; afdb-uniprot50 \| AF-A0A7Z0UXS2-F1-MODEL\_V4 \| 1.0 \| 7.46e-10 \| 281 \| 0.132 \| 242 \| 145 \| 5 \| 4 \| 180 \| 10 \| 251 \| Transcriptional regulator \| Transcriptional regulator; afdb-uniprot50 \| AF-A0A7U2YMF5-F1-MODEL\_V4 \| 1.0 \| 1.057e-09 \| 281 \| 0.18 \| 238 \| 132 \| 8 \| 5 \| 180 \| 19 \| 255 \| Peptidase\_S24 domain-containing protein \| Peptidase\_S24 domain-containing protein; afdb-uniprot50 \| AF-A0A554U561-F1-MODEL\_V4 \| 1.0 \| 1.057e-09 \| 281 \| 0.164 \| 255 \| 133 \| 8 \| 4 \| 180 \| 5 \| 257 \| LexA family transcriptional regulator \| LexA family transcriptional regulator; afdb-uniprot50 \| AF-A0A1Q9R2D7-F1-MODEL\_V4 \| 1.0 \| 3.511e-10 \| 281 \| 0.158 \| 240 \| 137 \| 5 \| 6 \| 180 \| 22 \| 261 \| HTH-type transcriptional regulator PrtR \| HTH-type transcriptional regulator PrtR; afdb-uniprot50 \| AF-B2U8I8-F1-MODEL\_V4 \| 1.0 \| 1.75e-10 \| 281 \| 0.186 \| 241 \| 130 \| 7 \| 4 \| 180 \| 62 \| 300 \| Putative phage repressor \| Putative phage repressor; afdb-uniprot50 \| AF-T5KQ06-F1-MODEL\_V4 \| 1.0 \| 2.764e-07 \| 280 \| 0.276 \| 112 \| 74 \| 6 \| 73 \| 180 \| 1 \| 109 \| Peptidase\_S24 domain-containing protein \| Peptidase\_S24 domain-containing protein; afdb-uniprot50 \| AF-A0A2U2BY69-F1-MODEL\_V4 \| 1.0 \| 6.225e-07 \| 280 \| 0.271 \| 114 \| 79 \| 2 \| 70 \| 180 \| 6 \| 118 \| Phage repressor protein \| Phage repressor protein; afdb-uniprot50 \| AF-A0A127QZI9-F1-MODEL\_V4 \| 1.0 \| 2.764e-07 \| 280 \| 0.258 \| 120 \| 80 \| 4 \| 64 \| 180 \| 16 \| 129 \| Peptidase S24-like family protein \| Peptidase S24-like family protein; afdb-uniprot50 \| AF-A0A6B2D5R2-F1-MODEL\_V4 \| 1.0 \| 1.12e-09 \| 280 \| 0.206 \| 199 \| 132 \| 6 \| 7 \| 180 \| 3 \| 200 \| HTH cro/C1-type domain-containing protein \| HTH cro/C1-type domain-containing protein; afdb-uniprot50 \| AF-A0A3S4Z798-F1-MODEL\_V4 \| 1.0 \| 1.585e-09 \| 280 \| 0.18 \| 211 \| 138 \| 4 \| 4 \| 180 \| 6 \| 215 \| Transcriptional regulator \| Transcriptional regulator; afdb-uniprot50 \| AF-A0A1Q8SUI1-F1-MODEL\_V4 \| 1.0 \| 6.644e-10 \| 280 \| 0.196 \| 219 \| 128 \| 7 \| 4 \| 180 \| 7 \| 219 \| HTH cro/C1-type domain-containing protein \| HTH cro/C1-type domain-containing protein; afdb-uniprot50 \| AF-A0A7S7P6L2-F1-MODEL\_V4 \| 1.0 \| 5.916e-10 \| 280 \| 0.2 \| 220 \| 133 \| 5 \| 1 \| 178 \| 1 \| 219 \| Helix-turn-helix transcriptional regulator \| Helix-turn-helix transcriptional regulator; afdb-uniprot50 \| AF-A0A3R9FTS3-F1-MODEL\_V4 \| 1.0 \| 9.97e-10 \| 280 \| 0.16 \| 230 \| 139 \| 5 \| 4 \| 180 \| 7 \| 235 \| XRE family transcriptional regulator \| XRE family transcriptional regulator; afdb-uniprot50 \| AF-A0A806J3Z0-F1-MODEL\_V4 \| 1.0 \| 4.971e-10 \| 280 \| 0.165 \| 230 \| 135 \| 9 \| 5 \| 180 \| 12 \| 238 \| Bacteriophage transcriptional regulator \| Bacteriophage transcriptional regulator; afdb-uniprot50 \| AF-A0A0C1K7W0-F1-MODEL\_V4 \| 1.0 \| 1.768e-06 \| 279 \| 0.217 \| 92 \| 71 \| 1 \| 89 \| 180 \| 12 \| 102 \| Repressor \| Repressor; afdb-uniprot50 \| AF-A0A6G6J700-F1-MODEL\_V4 \| 1.0 \| 1.402e-06 \| 279 \| 0.274 \| 102 \| 74 \| 0 \| 79 \| 180 \| 82 \| 183 \| Helix-turn-helix transcriptional regulator \| Helix-turn-helix transcriptional regulator; afdb-uniprot50 \| AF-A0A1I4RVT8-F1-MODEL\_V4 \| 1.0 \| 5.268e-10 \| 279 \| 0.211 \| 227 \| 127 \| 7 \| 4 \| 180 \| 6 \| 230 \| Peptidase S24-like \| Peptidase S24-like; afdb-uniprot50 \| AF-A0A4D7AWP4-F1-MODEL\_V4 \| 1.0 \| 9.97e-10 \| 279 \| 0.204 \| 220 \| 130 \| 7 \| 4 \| 180 \| 16 \| 233 \| Helix-turn-helix transcriptional regulator \| Helix-turn-helix transcriptional regulator; afdb-uniprot50 \| AF-A0A6G6JTQ1-F1-MODEL\_V4 \| 1.0 \| 4.971e-10 \| 279 \| 0.193 \| 227 \| 131 \| 9 \| 4 \| 180 \| 14 \| 238 \| Chromophore lyase \| Chromophore lyase; afdb-uniprot50 \| AF-A0A2T6F9B1-F1-MODEL\_V4 \| 1.0 \| 4.691e-10 \| 279 \| 0.196 \| 214 \| 132 \| 8 \| 5 \| 180 \| 34 \| 245 \| HTH cro/C1-type domain-containing protein \| HTH cro/C1-type domain-containing protein; afdb-uniprot50 \| AF-A0A3D4KIJ4-F1-MODEL\_V4 \| 1.0 \| 1.999e-09 \| 279 \| 0.152 \| 243 \| 133 \| 8 \| 4 \| 178 \| 14 \| 251 \| HTH cro/C1-type domain-containing protein \| HTH cro/C1-type domain-containing protein; afdb-uniprot50 \| AF-A0A2L0WYT4-F1-MODEL\_V4 \| 1.0 \| 6.644e-10 \| 279 \| 0.211 \| 232 \| 127 \| 5 \| 4 \| 180 \| 20 \| 250 \| HTH cro/C1-type domain-containing protein \| HTH cro/C1-type domain-containing protein; afdb-uniprot50 \| AF-A0A3E0WQD8-F1-MODEL\_V4 \| 1.0 \| 5.916e-10 \| 279 \| 0.224 \| 241 \| 121 \| 9 \| 5 \| 180 \| 15 \| 254 \| HTH cro/C1-type domain-containing protein \| HTH cro/C1-type domain-containing protein; afdb-uniprot50 \| AF-A0A2D5F736-F1-MODEL\_V4 \| 1.0 \| 4.427e-10 \| 279 \| 0.223 \| 224 \| 126 \| 7 \| 4 \| 180 \| 38 \| 260 \| Peptidase \| Peptidase; afdb-uniprot50 \| AF-A0A4S2Q5B1-F1-MODEL\_V4 \| 1.0 \| 7.04e-10 \| 279 \| 0.166 \| 222 \| 138 \| 8 \| 4 \| 180 \| 65 \| 284 \| XRE family transcriptional regulator \| XRE family transcriptional regulator; afdb-uniprot50 \| AF-A0A2J4NYR4-F1-MODEL\_V4 \| 1.0 \| 3.486e-07 \| 278 \| 0.272 \| 125 \| 83 \| 5 \| 60 \| 180 \| 1 \| 121 \| Phage repressor protein \| Phage repressor protein; afdb-uniprot50 \| AF-A0A1Y1QLB8-F1-MODEL\_V4 \| 1.0 \| 1.68e-09 \| 278 \| 0.183 \| 202 \| 138 \| 4 \| 5 \| 180 \| 10 \| 210 \| HTH cro/C1-type domain-containing protein \| HTH cro/C1-type domain-containing protein; afdb-uniprot50 \| AF-A0A2S5TZG8-F1-MODEL\_V4 \| 1.0 \| 6.644e-10 \| 278 \| 0.221 \| 212 \| 114 \| 6 \| 15 \| 180 \| 5 \| 211 \| Helix-turn-helix transcriptional regulator \| Helix-turn-helix transcriptional regulator; afdb-uniprot50 \| AF-A0A6G8AHU4-F1-MODEL\_V4 \| 1.0 \| 7.04e-10 \| 278 \| 0.195 \| 225 \| 134 \| 8 \| 1 \| 180 \| 1 \| 223 \| Helix-turn-helix transcriptional regulator \| Helix-turn-helix transcriptional regulator; afdb-uniprot50 \| AF-A0A2L0PMA1-F1-MODEL\_V4 \| 1.0 \| 6.269e-10 \| 278 \| 0.209 \| 210 \| 128 \| 9 \| 5 \| 177 \| 15 \| 223 \| Putative transcriptional repressor \| Putative transcriptional repressor; afdb-uniprot50 \| AF-A0A1N6NSV1-F1-MODEL\_V4 \| 1.0 \| 8.378e-10 \| 278 \| 0.188 \| 223 \| 126 \| 5 \| 4 \| 180 \| 20 \| 233 \| Phage repressor protein C, contains Cro/C1-type HTH and peptisase s24 domains \| Phage repressor protein C, contains Cro/C1-type HTH and peptisase s24 domains; afdb-uniprot50 \| AF-A0A3C0XA60-F1-MODEL\_V4 \| 1.0 \| 7.906e-10 \| 278 \| 0.194 \| 226 \| 131 \| 10 \| 4 \| 180 \| 8 \| 231 \| Peptidase\_S24 domain-containing protein \| Peptidase\_S24 domain-containing protein; afdb-uniprot50 \| AF-A0A1D9BCV5-F1-MODEL\_V4 \| 1.0 \| 4.691e-10 \| 278 \| 0.2 \| 224 \| 129 \| 8 \| 4 \| 178 \| 7 \| 229 \| Peptidase\_S24 domain-containing protein \| Peptidase\_S24 domain-containing protein; afdb-uniprot50 \| AF-A0A1N6XE22-F1-MODEL\_V4 \| 1.0 \| 1.966e-10 \| 278 \| 0.19 \| 231 \| 129 \| 8 \| 5 \| 180 \| 12 \| 239 \| Phage repressor protein C, contains Cro/C1-type HTH and peptisase s24 domains \| Phage repressor protein C, contains Cro/C1-type HTH and peptisase s24 domains; afdb-uniprot50 \| AF-D7N153-F1-MODEL\_V4 \| 1.0 \| 5.583e-10 \| 278 \| 0.183 \| 212 \| 135 \| 9 \| 5 \| 180 \| 36 \| 245 \| Peptidase\_S24 domain-containing protein \| Peptidase\_S24 domain-containing protein; afdb-uniprot50 \| AF-A0A1G9CRS8-F1-MODEL\_V4 \| 1.0 \| 9.185e-08 \| 278 \| 0.291 \| 127 \| 81 \| 4 \| 61 \| 180 \| 127 \| 251 \| Peptidase S24-like \| Peptidase S24-like; afdb-uniprot50 \| AF-A0A2N8C2G3-F1-MODEL\_V4 \| 1.0 \| 1.966e-10 \| 278 \| 0.177 \| 236 \| 132 \| 7 \| 6 \| 180 \| 22 \| 256 \| Helix-turn-helix domain-containing protein \| Helix-turn-helix domain-containing protein; afdb-uniprot50 \| AF-A0A2E8IH19-F1-MODEL\_V4 \| 1.0 \| 8.878e-10 \| 278 \| 0.184 \| 228 \| 134 \| 7 \| 4 \| 180 \| 37 \| 263 \| Transcriptional regulator \| Transcriptional regulator; afdb-uniprot50 \| AF-A0A1Y3KYL7-F1-MODEL\_V4 \| 1.0 \| 8.378e-10 \| 278 \| 0.193 \| 243 \| 129 \| 8 \| 4 \| 180 \| 44 \| 285 \| HTH cro/C1-type domain-containing protein \| HTH cro/C1-type domain-containing protein; afdb-uniprot50 \| AF-A0A7C2AYL5-F1-MODEL\_V4 \| 1.0 \| 1.75e-10 \| 278 \| 0.156 \| 274 \| 133 \| 6 \| 4 \| 180 \| 14 \| 286 \| LexA family transcriptional regulator \| LexA family transcriptional regulator; afdb-uniprot50 \| AF-A0A1N7DHM0-F1-MODEL\_V4 \| 1.0 \| 5.583e-10 \| 277 \| 0.195 \| 215 \| 129 \| 7 \| 4 \| 180 \| 7 \| 215 \| Phage repressor protein C, contains Cro/C1-type HTH and peptisase s24 domains \| Phage repressor protein C, contains Cro/C1-type HTH and peptisase s24 domains; afdb-uniprot50 \| AF-Q1JZ74-F1-MODEL\_V4 \| 1.0 \| 1.257e-09 \| 277 \| 0.183 \| 207 \| 136 \| 5 \| 4 \| 180 \| 27 \| 230 \| Putative phage repressor \| Putative phage repressor; afdb-uniprot50 \| AF-Q1H1H4-F1-MODEL\_V4 \| 1.0 \| 9.899e-07 \| 277 \| 0.309 \| 97 \| 64 \| 1 \| 87 \| 180 \| 140 \| 236 \| Putative phage repressor \| Putative phage repressor; afdb-uniprot50 \| AF-A0A2N2QYU2-F1-MODEL\_V4 \| 1.0 \| 1.31e-10 \| 277 \| 0.275 \| 196 \| 121 \| 8 \| 4 \| 180 \| 42 \| 235 \| Peptidase S24 \| Peptidase S24; afdb-uniprot50 \| AF-A0A561MAZ7-F1-MODEL\_V4 \| 1.0 \| 1.585e-09 \| 277 \| 0.164 \| 237 \| 130 \| 7 \| 5 \| 179 \| 7 \| 237 \| Phage repressor protein C with HTH and peptisase S24 domain \| Phage repressor protein C with HTH and peptisase S24 domain; afdb-uniprot50 \| AF-A0A6M8SNY5-F1-MODEL\_V4 \| 1.0 \| 5.268e-10 \| 277 \| 0.19 \| 236 \| 131 \| 10 \| 4 \| 180 \| 10 \| 244 \| Helix-turn-helix transcriptional regulator \| Helix-turn-helix transcriptional regulator; afdb-uniprot50 \| AF-A0A510XCN0-F1-MODEL\_V4 \| 1.0 \| 1.186e-09 \| 277 \| 0.222 \| 238 \| 124 \| 6 \| 4 \| 180 \| 9 \| 246 \| HTH cro/C1-type domain-containing protein \| HTH cro/C1-type domain-containing protein; afdb-uniprot50 \| AF-A0A843YNI7-F1-MODEL\_V4 \| 1.0 \| 2.784e-10 \| 277 \| 0.166 \| 240 \| 135 \| 7 \| 4 \| 180 \| 12 \| 249 \| Uncharacterized protein \| Uncharacterized protein; afdb-uniprot50 \| AF-A0A6P2YEJ5-F1-MODEL\_V4 \| 1.0 \| 1.496e-09 \| 277 \| 0.168 \| 232 \| 137 \| 5 \| 4 \| 179 \| 17 \| 248 \| Putative phage repressor \| Putative phage repressor; afdb-uniprot50 \| AF-A0A562RKT0-F1-MODEL\_V4 \| 1.0 \| 8.878e-10 \| 277 \| 0.175 \| 256 \| 130 \| 6 \| 4 \| 179 \| 4 \| 258 \| Peptidase S24-like protein \| Peptidase S24-like protein; afdb-uniprot50 \| AF-A0A285VR71-F1-MODEL\_V4 \| 1.0 \| 1.412e-09 \| 277 \| 0.213 \| 225 \| 127 \| 6 \| 4 \| 180 \| 49 \| 271 \| Phage repressor protein C, contains Cro/C1-type HTH and peptisase s24 domains \| Phage repressor protein C, contains Cro/C1-type HTH and peptisase s24 domains; afdb-uniprot50 \| AF-A0A846VQU4-F1-MODEL\_V4 \| 1.0 \| 4.148e-07 \| 276 \| 0.285 \| 119 \| 77 \| 4 \| 65 \| 180 \| 19 \| 132 \| S24 family peptidase \| S24 family peptidase; afdb-uniprot50 \| AF-H0KGR1-F1-MODEL\_V4 \| 1.0 \| 2.831e-09 \| 276 \| 0.159 \| 194 \| 138 \| 7 \| 5 \| 180 \| 10 \| 196 \| DNA-binding protein RDGA \| DNA-binding protein RDGA; afdb-uniprot50 \| AF-A0A848G1C2-F1-MODEL\_V4 \| 1.0 \| 2.119e-09 \| 276 \| 0.185 \| 226 \| 127 \| 8 \| 5 \| 180 \| 6 \| 224 \| LexA family transcriptional regulator \| LexA family transcriptional regulator; afdb-uniprot50 \| AF-A0A2G6EVX0-F1-MODEL\_V4 \| 1.0 \| 4.971e-10 \| 276 \| 0.198 \| 212 \| 128 \| 6 \| 4 \| 174 \| 12 \| 222 \| HTH cro/C1-type domain-containing protein \| HTH cro/C1-type domain-containing protein; afdb-uniprot50 \| AF-A0A1X0TFD1-F1-MODEL\_V4 \| 1.0 \| 6.644e-10 \| 276 \| 0.156 \| 223 \| 136 \| 8 \| 4 \| 178 \| 7 \| 225 \| HTH cro/C1-type domain-containing protein \| HTH cro/C1-type domain-containing protein; afdb-uniprot50 \| AF-B0C586-F1-MODEL\_V4 \| 1.0 \| 1.332e-09 \| 276 \| 0.168 \| 214 \| 138 \| 6 \| 4 \| 180 \| 17 \| 227 \| Peptidase, S24 family \| Peptidase, S24 family; afdb-uniprot50 \| AF-A0A1Z4BY84-F1-MODEL\_V4 \| 1.0 \| 1.057e-09 \| 276 \| 0.191 \| 224 \| 132 \| 7 \| 5 \| 180 \| 8 \| 230 \| XRE family transcriptional regulator \| XRE family transcriptional regulator; afdb-uniprot50 \| AF-A0A4R0EQW7-F1-MODEL\_V4 \| 1.0 \| 1.257e-09 \| 276 \| 0.158 \| 227 \| 141 \| 3 \| 4 \| 180 \| 6 \| 232 \| Helix-turn-helix transcriptional regulator \| Helix-turn-helix transcriptional regulator; afdb-uniprot50 \| AF-A0A7U9F4N3-F1-MODEL\_V4 \| 1.0 \| 9.97e-10 \| 276 \| 0.174 \| 229 \| 135 \| 4 \| 5 \| 180 \| 8 \| 235 \| Transcriptional regulator \| Transcriptional regulator; afdb-uniprot50 \| AF-A0A7D5D6C4-F1-MODEL\_V4 \| 1.0 \| 6.269e-10 \| 276 \| 0.194 \| 206 \| 135 \| 6 \| 4 \| 180 \| 34 \| 237 \| Helix-turn-helix domain-containing protein \| Helix-turn-helix domain-containing protein; afdb-uniprot50 \| AF-A0A0D6Q481-F1-MODEL\_V4 \| 1.0 \| 4.691e-10 \| 276 \| 0.206 \| 232 \| 125 \| 7 \| 4 \| 179 \| 22 \| 250 \| Transcriptional regulator phage repressor \| Transcriptional regulator phage repressor; afdb-uniprot50 \| AF-A0A1G8G7E8-F1-MODEL\_V4 \| 1.0 \| 1.585e-09 \| 276 \| 0.16 \| 250 \| 134 \| 7 \| 5 \| 180 \| 11 \| 258 \| Phage repressor protein C, contains Cro/C1-type HTH and peptisase s24 domains \| Phage repressor protein C, contains Cro/C1-type HTH and peptisase s24 domains; afdb-uniprot50 \| AF-A0A4U8YGI0-F1-MODEL\_V4 \| 1.0 \| 1.887e-09 \| 276 \| 0.205 \| 234 \| 128 \| 4 \| 4 \| 180 \| 35 \| 267 \| Peptidase s24/s26a/s26b/s26c \| Peptidase s24/s26a/s26b/s26c; afdb-uniprot50 \| AF-B8L9T0-F1-MODEL\_V4 \| 1.0 \| 5.916e-10 \| 276 \| 0.194 \| 226 \| 131 \| 9 \| 4 \| 180 \| 74 \| 297 \| Phage-related repressor protein \| Phage-related repressor protein; afdb-uniprot50 \| AF-A0A354J173-F1-MODEL\_V4 \| 1.0 \| 1.574e-06 \| 275 \| 0.2 \| 105 \| 79 \| 2 \| 79 \| 179 \| 25 \| 128 \| Helix-turn-helix transcriptional regulator \| Helix-turn-helix transcriptional regulator; afdb-uniprot50 \| AF-B9BQH7-F1-MODEL\_V4 \| 1.0 \| 7.408e-07 \| 275 \| 0.259 \| 108 \| 74 \| 4 \| 78 \| 180 \| 38 \| 144 \| Prophage transcriptional regulator \| Prophage transcriptional regulator; afdb-uniprot50 \| AF-A0A8B3VSP7-F1-MODEL\_V4 \| 1.0 \| 9.899e-07 \| 275 \| 0.23 \| 104 \| 74 \| 4 \| 79 \| 179 \| 51 \| 151 \| Putative phage repressor \| Putative phage repressor; afdb-uniprot50 \| AF-C8N745-F1-MODEL\_V4 \| 1.0 \| 6.644e-10 \| 275 \| 0.218 \| 211 \| 124 \| 9 \| 4 \| 180 \| 6 \| 209 \| Peptidase S24-like protein \| Peptidase S24-like protein; afdb-uniprot50 \| AF-A0A0Q3EYM7-F1-MODEL\_V4 \| 1.0 \| 9.408e-10 \| 275 \| 0.181 \| 215 \| 137 \| 5 \| 4 \| 180 \| 6 \| 219 \| Peptidase\_S24 domain-containing protein \| Peptidase\_S24 domain-containing protein; afdb-uniprot50 \| AF-A0A6G9RL62-F1-MODEL\_V4 \| 1.0 \| 4.178e-10 \| 275 \| 0.221 \| 217 \| 123 \| 12 \| 4 \| 180 \| 13 \| 223 \| Helix-turn-helix transcriptional regulator \| Helix-turn-helix transcriptional regulator; afdb-uniprot50 \| AF-A0A1X0TKP2-F1-MODEL\_V4 \| 1.0 \| 1.332e-09 \| 275 \| 0.169 \| 224 \| 131 \| 8 \| 4 \| 178 \| 7 \| 224 \| HTH cro/C1-type domain-containing protein \| HTH cro/C1-type domain-containing protein; afdb-uniprot50 \| AF-A0A3S5GHX0-F1-MODEL\_V4 \| 1.0 \| 1.471e-10 \| 275 \| 0.19 \| 210 \| 127 \| 7 \| 4 \| 172 \| 7 \| 214 \| Transcriptional regulator \| Transcriptional regulator; afdb-uniprot50 \| AF-A0A077NT47-F1-MODEL\_V4 \| 1.0 \| 5.916e-10 \| 275 \| 0.162 \| 221 \| 139 \| 9 \| 4 \| 180 \| 11 \| 229 \| 26 kDa repressor protein \| 26 kDa repressor protein; afdb-uniprot50 \| AF-A0A522KYU9-F1-MODEL\_V4 \| 1.0 \| 1.412e-09 \| 275 \| 0.145 \| 233 \| 140 \| 7 \| 4 \| 178 \| 6 \| 237 \| LexA family transcriptional regulator \| LexA family transcriptional regulator; afdb-uniprot50 \| AF-A0A846UWQ4-F1-MODEL\_V4 \| 1.0 \| 8.378e-10 \| 275 \| 0.184 \| 228 \| 127 \| 8 \| 5 \| 179 \| 14 \| 235 \| Phage repressor protein C with HTH and peptisase S24 domain \| Phage repressor protein C with HTH and peptisase S24 domain; afdb-uniprot50 \| AF-A0A432UMV4-F1-MODEL\_V4 \| 1.0 \| 4.971e-10 \| 275 \| 0.163 \| 233 \| 133 \| 7 \| 7 \| 178 \| 12 \| 243 \| Peptidase\_S24 domain-containing protein \| Peptidase\_S24 domain-containing protein; afdb-uniprot50 \| AF-A0A104MVH2-F1-MODEL\_V4 \| 1.0 \| 2.379e-09 \| 275 \| 0.178 \| 230 \| 133 \| 6 \| 4 \| 178 \| 20 \| 248 \| HTH cro/C1-type domain-containing protein \| HTH cro/C1-type domain-containing protein; afdb-uniprot50 \| AF-A0A364H0J3-F1-MODEL\_V4 \| 1.0 \| 1.68e-09 \| 275 \| 0.159 \| 232 \| 139 \| 5 \| 4 \| 179 \| 29 \| 260 \| Peptidase S24-like protein \| Peptidase S24-like protein; afdb-uniprot50 \| AF-A0A2E4Q3Q2-F1-MODEL\_V4 \| 1.0 \| 1.496e-09 \| 275 \| 0.215 \| 232 \| 126 \| 6 \| 4 \| 180 \| 72 \| 302 \| XRE family transcriptional regulator \| XRE family transcriptional regulator; afdb-uniprot50 \| AF-A0A1H6SGR5-F1-MODEL\_V4 \| 1.0 \| 1.75e-10 \| 275 \| 0.153 \| 273 \| 133 \| 8 \| 5 \| 180 \| 35 \| 306 \| Peptidase S24-like \| Peptidase S24-like; afdb-uniprot50 \| AF-A0A060H3F4-F1-MODEL\_V4 \| 1.0 \| 1.332e-09 \| 275 \| 0.169 \| 248 \| 132 \| 6 \| 4 \| 178 \| 69 \| 315 \| Uncharacterized protein \| Uncharacterized protein; afdb-uniprot50 \| AF-A0A7U4R302-F1-MODEL\_V4 \| 1.0 \| 5.543e-07 \| 274 \| 0.258 \| 112 \| 75 \| 5 \| 70 \| 179 \| 25 \| 130 \| Peptidase\_S24 domain-containing protein \| Peptidase\_S24 domain-containing protein; afdb-uniprot50 \| AF-A0A0S2SFV9-F1-MODEL\_V4 \| 1.0 \| 6.644e-10 \| 274 \| 0.175 \| 216 \| 131 \| 7 \| 10 \| 180 \| 2 \| 215 \| Transcriptional regulator \| Transcriptional regulator; afdb-uniprot50 \| AF-A0A4Q4AGJ7-F1-MODEL\_V4 \| 1.0 \| 7.46e-10 \| 274 \| 0.207 \| 217 \| 131 \| 7 \| 4 \| 180 \| 12 \| 227 \| Helix-turn-helix transcriptional regulator \| Helix-turn-helix transcriptional regulator; afdb-uniprot50 \| AF-A0A2U8I4J9-F1-MODEL\_V4 \| 1.0 \| 1.999e-09 \| 274 \| 0.154 \| 226 \| 138 \| 5 \| 4 \| 180 \| 8 \| 229 \| HTH cro/C1-type domain-containing protein \| HTH cro/C1-type domain-containing protein; afdb-uniprot50 \| AF-A0A352Z9M4-F1-MODEL\_V4 \| 1.0 \| 1.412e-09 \| 274 \| 0.199 \| 221 \| 130 \| 8 \| 4 \| 180 \| 13 \| 230 \| HTH cro/C1-type domain-containing protein \| HTH cro/C1-type domain-containing protein; afdb-uniprot50 \| AF-A0A2A7U2T6-F1-MODEL\_V4 \| 1.0 \| 1.585e-09 \| 274 \| 0.152 \| 230 \| 141 \| 5 \| 4 \| 180 \| 8 \| 236 \| Phage repressor protein C \| Phage repressor protein C; afdb-uniprot50 \| AF-A0A346MWA8-F1-MODEL\_V4 \| 1.0 \| 1.257e-09 \| 274 \| 0.207 \| 222 \| 124 \| 7 \| 6 \| 180 \| 29 \| 245 \| Helix-turn-helix domain-containing protein \| Helix-turn-helix domain-containing protein; afdb-uniprot50 \| AF-A0A069PBS4-F1-MODEL\_V4 \| 1.0 \| 6.644e-10 \| 274 \| 0.159 \| 232 \| 137 \| 7 \| 4 \| 180 \| 17 \| 245 \| Phage repressor protein \| Phage repressor protein; afdb-uniprot50 \| AF-A0A7T4WC91-F1-MODEL\_V4 \| 1.0 \| 5.583e-10 \| 274 \| 0.168 \| 238 \| 134 \| 8 \| 5 \| 180 \| 25 \| 260 \| S24 family peptidase \| S24 family peptidase; afdb-uniprot50 \| AF-A0A7Z0ACD2-F1-MODEL\_V4 \| 1.0 \| 6.644e-10 \| 274 \| 0.189 \| 243 \| 130 \| 8 \| 5 \| 180 \| 19 \| 261 \| Phage repressor protein C with HTH and peptisase S24 domain \| Phage repressor protein C with HTH and peptisase S24 domain; afdb-uniprot50 \| AF-A0A2N9XWK5-F1-MODEL\_V4 \| 1.0 \| 3.486e-07 \| 273 \| 0.192 \| 125 \| 94 \| 2 \| 62 \| 180 \| 27 \| 150 \| Peptidase\_S24 domain-containing protein \| Peptidase\_S24 domain-containing protein; afdb-uniprot50 \| AF-A0A806J765-F1-MODEL\_V4 \| 1.0 \| 7.85e-07 \| 273 \| 0.278 \| 104 \| 70 \| 3 \| 77 \| 176 \| 62 \| 164 \| LexA family repressor/S24 family protease \| LexA family repressor/S24 family protease; afdb-uniprot50 \| AF-A0A1I4GAU7-F1-MODEL\_V4 \| 1.0 \| 2.245e-09 \| 273 \| 0.176 \| 215 \| 133 \| 7 \| 4 \| 180 \| 11 \| 219 \| HTH cro/C1-type domain-containing protein \| HTH cro/C1-type domain-containing protein; afdb-uniprot50 \| AF-A0A261U5A9-F1-MODEL\_V4 \| 1.0 \| 1.12e-09 \| 273 \| 0.199 \| 221 \| 131 \| 6 \| 4 \| 180 \| 7 \| 225 \| HTH cro/C1-type domain-containing protein \| HTH cro/C1-type domain-containing protein; afdb-uniprot50 \| AF-A0A1X0TEC1-F1-MODEL\_V4 \| 1.0 \| 8.878e-10 \| 273 \| 0.161 \| 217 \| 140 \| 8 \| 4 \| 180 \| 11 \| 225 \| Peptidase\_S24 domain-containing protein \| Peptidase\_S24 domain-containing protein; afdb-uniprot50 \| AF-A0A425B2V4-F1-MODEL\_V4 \| 1.0 \| 1.057e-09 \| 273 \| 0.163 \| 232 \| 139 \| 5 \| 3 \| 180 \| 2 \| 232 \| Chromophore lyase \| Chromophore lyase; afdb-uniprot50 \| AF-C3X398-F1-MODEL\_V4 \| 1.0 \| 6.269e-10 \| 273 \| 0.16 \| 225 \| 139 \| 9 \| 4 \| 180 \| 7 \| 229 \| HTH cro/C1-type domain-containing protein \| HTH cro/C1-type domain-containing protein; afdb-uniprot50 \| AF-A0A3D5UZY1-F1-MODEL\_V4 \| 1.0 \| 1.412e-09 \| 273 \| 0.182 \| 235 \| 133 \| 6 \| 4 \| 180 \| 6 \| 239 \| HTH cro/C1-type domain-containing protein \| HTH cro/C1-type domain-containing protein; afdb-uniprot50 \| AF-A0A2S5QN69-F1-MODEL\_V4 \| 1.0 \| 4.691e-10 \| 273 \| 0.184 \| 233 \| 132 \| 9 \| 5 \| 180 \| 11 \| 242 \| Phage repressor protein \| Phage repressor protein; afdb-uniprot50 \| AF-A0A7Z1UV24-F1-MODEL\_V4 \| 1.0 \| 1.68e-09 \| 273 \| 0.171 \| 227 \| 136 \| 7 \| 5 \| 180 \| 27 \| 252 \| HTH cro/C1-type domain-containing protein \| HTH cro/C1-type domain-containing protein; afdb-uniprot50 \| AF-A0A1I1VLZ3-F1-MODEL\_V4 \| 1.0 \| 4.936e-07 \| 273 \| 0.336 \| 113 \| 67 \| 3 \| 71 \| 180 \| 161 \| 268 \| Peptidase S24-like \| Peptidase S24-like; afdb-uniprot50 \| AF-A0A4R7VY96-F1-MODEL\_V4 \| 1.0 \| 4.178e-10 \| 273 \| 0.151 \| 258 \| 134 \| 6 \| 6 \| 180 \| 22 \| 277 \| Phage repressor protein C with HTH and peptisase S24 domain \| Phage repressor protein C with HTH and peptisase S24 domain; afdb-uniprot50 \| AF-A0A1G8LL79-F1-MODEL\_V4 \| 1.0 \| 2.95e-10 \| 273 \| 0.148 \| 269 \| 135 \| 7 \| 5 \| 180 \| 22 \| 289 \| Phage repressor protein C, contains Cro/C1-type HTH and peptisase s24 domains \| Phage repressor protein C, contains Cro/C1-type HTH and peptisase s24 domains; afdb-uniprot50 \| AF-A0A2J0UCK8-F1-MODEL\_V4 \| 1.0 \| 4.738e-06 \| 272 \| 0.23 \| 91 \| 69 \| 1 \| 90 \| 180 \| 18 \| 107 \| Uncharacterized protein \| Uncharacterized protein; afdb-uniprot50 \| AF-A0A1F1UL81-F1-MODEL\_V4 \| 1.0 \| 5.874e-07 \| 272 \| 0.247 \| 109 \| 77 \| 3 \| 77 \| 180 \| 45 \| 153 \| Peptidase\_S24 domain-containing protein \| Peptidase\_S24 domain-containing protein; afdb-uniprot50 \| AF-A0A5E7AN99-F1-MODEL\_V4 \| 1.0 \| 3.914e-07 \| 272 \| 0.3 \| 110 \| 68 \| 4 \| 79 \| 180 \| 65 \| 173 \| Peptidase\_S24 domain-containing protein \| Peptidase\_S24 domain-containing protein; afdb-uniprot50 \| AF-A0A1B1A083-F1-MODEL\_V4 \| 1.0 \| 1.842e-07 \| 272 \| 0.253 \| 126 \| 83 \| 5 \| 61 \| 180 \| 54 \| 174 \| Transcriptional regulator \| Transcriptional regulator; afdb-uniprot50 \| AF-A0A661J126-F1-MODEL\_V4 \| 1.0 \| 1.496e-09 \| 272 \| 0.203 \| 211 \| 127 \| 6 \| 4 \| 180 \| 8 \| 211 \| Peptidase S24 \| Peptidase S24; afdb-uniprot50 \| AF-A0A0Q6LRH4-F1-MODEL\_V4 \| 1.0 \| 7.46e-10 \| 272 \| 0.185 \| 226 \| 128 \| 8 \| 5 \| 179 \| 7 \| 227 \| HTH cro/C1-type domain-containing protein \| HTH cro/C1-type domain-containing protein; afdb-uniprot50 \| AF-A0A1G0MXC6-F1-MODEL\_V4 \| 1.0 \| 6.269e-10 \| 272 \| 0.219 \| 219 \| 123 \| 8 \| 5 \| 180 \| 17 \| 230 \| Repressor \| Repressor; afdb-uniprot50 \| AF-A0A2A4V065-F1-MODEL\_V4 \| 1.0 \| 1.186e-09 \| 272 \| 0.163 \| 238 \| 131 \| 8 \| 4 \| 180 \| 9 \| 239 \| Peptidase S24 \| Peptidase S24; afdb-uniprot50 \| AF-A0A2J7TQM8-F1-MODEL\_V4 \| 1.0 \| 1.585e-09 \| 272 \| 0.185 \| 248 \| 130 \| 4 \| 5 \| 180 \| 8 \| 255 \| XRE family transcriptional regulator \| XRE family transcriptional regulator; afdb-uniprot50 \| AF-A0A2E1HCN9-F1-MODEL\_V4 \| 1.0 \| 2.672e-09 \| 272 \| 0.17 \| 241 \| 134 \| 6 \| 4 \| 180 \| 43 \| 281 \| HTH cro/C1-type domain-containing protein \| HTH cro/C1-type domain-containing protein; afdb-uniprot50 \| AF-A0A1D3JVS7-F1-MODEL\_V4 \| 1.0 \| 5.268e-10 \| 272 \| 0.121 \| 264 \| 141 \| 5 \| 7 \| 180 \| 24 \| 286 \| Propanediol utilization protein \| Propanediol utilization protein; afdb-uniprot50 \| AF-A0A6I6SSB5-F1-MODEL\_V4 \| 1.0 \| 1.186e-09 \| 272 \| 0.204 \| 235 \| 125 \| 6 \| 4 \| 177 \| 58 \| 291 \| Helix-turn-helix transcriptional regulator \| Helix-turn-helix transcriptional regulator; afdb-uniprot50 \| AF-A0A1B5D1A6-F1-MODEL\_V4 \| 1.0 \| 2.363e-06 \| 271 \| 0.254 \| 106 \| 75 \| 2 \| 78 \| 180 \| 66 \| 170 \| Putative HTH-type transcriptional regulator \| Putative HTH-type transcriptional regulator; afdb-uniprot50 \| AF-A0A173HMY6-F1-MODEL\_V4 \| 1.0 \| 7.85e-07 \| 271 \| 0.22 \| 109 \| 78 \| 3 \| 78 \| 180 \| 100 \| 207 \| CI repressor protein \| CI repressor protein; afdb-uniprot50 \| AF-A0A1C0A407-F1-MODEL\_V4 \| 1.0 \| 1.12e-09 \| 271 \| 0.184 \| 206 \| 127 \| 7 \| 14 \| 180 \| 6 \| 209 \| HTH cro/C1-type domain-containing protein \| HTH cro/C1-type domain-containing protein; afdb-uniprot50 \| AF-A0A4R4G357-F1-MODEL\_V4 \| 1.0 \| 2.521e-09 \| 271 \| 0.198 \| 196 \| 121 \| 5 \| 17 \| 180 \| 27 \| 218 \| Phage repressor protein \| Phage repressor protein; afdb-uniprot50 \| AF-A0A6I1ZNW8-F1-MODEL\_V4 \| 1.0 \| 2.245e-09 \| 271 \| 0.192 \| 213 \| 135 \| 4 \| 4 \| 180 \| 19 \| 230 \| Peptidase\_S24 domain-containing protein \| Peptidase\_S24 domain-containing protein; afdb-uniprot50 \| AF-A0A742DF20-F1-MODEL\_V4 \| 1.0 \| 3.18e-09 \| 271 \| 0.159 \| 226 \| 140 \| 6 \| 4 \| 180 \| 8 \| 232 \| Helix-turn-helix transcriptional regulator \| Helix-turn-helix transcriptional regulator; afdb-uniprot50 \| AF-A0A1S9ZSJ1-F1-MODEL\_V4 \| 1.0 \| 8.378e-10 \| 271 \| 0.167 \| 233 \| 136 \| 7 \| 4 \| 180 \| 7 \| 237 \| Uncharacterized HTH-type transcriptional regulator HI\_1476 \| Uncharacterized HTH-type transcriptional regulator HI\_1476; afdb-uniprot50 \| AF-A0A6I5AJN6-F1-MODEL\_V4 \| 1.0 \| 2.245e-09 \| 271 \| 0.213 \| 220 \| 124 \| 9 \| 4 \| 178 \| 13 \| 228 \| Helix-turn-helix domain-containing protein \| Helix-turn-helix domain-containing protein; afdb-uniprot50 \| AF-A0A4P0TEV4-F1-MODEL\_V4 \| 1.0 \| 1.402e-06 \| 270 \| 0.254 \| 106 \| 75 \| 1 \| 79 \| 180 \| 67 \| 172 \| Transcriptional regulator \| Transcriptional regulator; afdb-uniprot50 \| AF-A0A011TVH5-F1-MODEL\_V4 \| 1.0 \| 1.057e-09 \| 270 \| 0.206 \| 203 \| 125 \| 6 \| 14 \| 180 \| 1 \| 203 \| Transcriptional regulator \| Transcriptional regulator; afdb-uniprot50 \| AF-A0A7D0IFV9-F1-MODEL\_V4 \| 1.0 \| 1.186e-09 \| 270 \| 0.19 \| 226 \| 132 \| 9 \| 4 \| 180 \| 6 \| 229 \| XRE family transcriptional regulator \| XRE family transcriptional regulator; afdb-uniprot50 \| AF-A0A1I1ETS9-F1-MODEL\_V4 \| 1.0 \| 1.412e-09 \| 270 \| 0.208 \| 225 \| 128 \| 7 \| 4 \| 180 \| 6 \| 228 \| Phage repressor protein C, contains Cro/C1-type HTH and peptisase s24 domains \| Phage repressor protein C, contains Cro/C1-type HTH and peptisase s24 domains; afdb-uniprot50 \| AF-A0A1M5I2V5-F1-MODEL\_V4 \| 1.0 \| 7.906e-10 \| 270 \| 0.194 \| 221 \| 122 \| 5 \| 15 \| 180 \| 14 \| 233 \| Phage repressor protein C, contains Cro/C1-type HTH and peptisase s24 domains \| Phage repressor protein C, contains Cro/C1-type HTH and peptisase s24 domains; afdb-uniprot50 \| AF-A0A1N7UFW9-F1-MODEL\_V4 \| 1.0 \| 4.971e-10 \| 270 \| 0.132 \| 272 \| 140 \| 6 \| 4 \| 180 \| 14 \| 284 \| Propanediol utilization protein \| Propanediol utilization protein; afdb-uniprot50 \| AF-A0A822VGQ0-F1-MODEL\_V4 \| 1.0 \| 1.049e-06 \| 269 \| 0.247 \| 105 \| 74 \| 5 \| 77 \| 178 \| 55 \| 157 \| Peptidase S24-like protein \| Peptidase S24-like protein; afdb-uniprot50 \| AF-A0A1H3E809-F1-MODEL\_V4 \| 1.0 \| 1.887e-09 \| 269 \| 0.192 \| 213 \| 126 \| 8 \| 12 \| 180 \| 14 \| 224 \| Phage repressor protein C, contains Cro/C1-type HTH and peptisase s24 domains \| Phage repressor protein C, contains Cro/C1-type HTH and peptisase s24 domains; afdb-uniprot50 \| AF-A0A5B9DY71-F1-MODEL\_V4 \| 1.0 \| 9.408e-10 \| 269 \| 0.214 \| 224 \| 126 \| 9 \| 5 \| 180 \| 4 \| 225 \| Helix-turn-helix transcriptional regulator \| Helix-turn-helix transcriptional regulator; afdb-uniprot50 \| AF-A0A4R8FYR2-F1-MODEL\_V4 \| 1.0 \| 1.412e-09 \| 269 \| 0.199 \| 226 \| 131 \| 6 \| 4 \| 180 \| 8 \| 232 \| Phage repressor protein C with HTH and peptisase S24 domain \| Phage repressor protein C with HTH and peptisase S24 domain; afdb-uniprot50 \| AF-A0A1L2B3E0-F1-MODEL\_V4 \| 1.0 \| 1.496e-09 \| 269 \| 0.19 \| 247 \| 128 \| 6 \| 4 \| 178 \| 6 \| 252 \| Helix-turn-helix transcriptional regulator \| Helix-turn-helix transcriptional regulator; afdb-uniprot50 \| AF-A0A3M2VAP0-F1-MODEL\_V4 \| 1.0 \| 1.257e-09 \| 269 \| 0.178 \| 224 \| 136 \| 5 \| 5 \| 180 \| 31 \| 254 \| Putative prophage PSSB64-02, repressor protein \| Putative prophage PSSB64-02, repressor protein; afdb-uniprot50 \| AF-A0A2E1XTS7-F1-MODEL\_V4 \| 1.0 \| 2.831e-09 \| 269 \| 0.175 \| 211 \| 138 \| 6 \| 5 \| 180 \| 78 \| 287 \| HTH cro/C1-type domain-containing protein \| HTH cro/C1-type domain-containing protein; afdb-uniprot50 \| AF-A0A1I3KCB5-F1-MODEL\_V4 \| 1.0 \| 8.378e-10 \| 269 \| 0.139 \| 266 \| 138 \| 4 \| 5 \| 180 \| 22 \| 286 \| Helix-turn-helix \| Helix-turn-helix; afdb-uniprot50 \| AF-A0A379GG64-F1-MODEL\_V4 \| 1.0 \| 2.229e-06 \| 268 \| 0.218 \| 110 \| 82 \| 1 \| 70 \| 179 \| 1 \| 106 \| Phage repressor protein \| Phage repressor protein; afdb-uniprot50 \| AF-K9RXU0-F1-MODEL\_V4 \| 1.0 \| 4.396e-07 \| 268 \| 0.266 \| 120 \| 82 \| 3 \| 64 \| 180 \| 8 \| 124 \| Putative transcriptional regulator \| Putative transcriptional regulator; afdb-uniprot50 \| AF-A0A839A2N0-F1-MODEL\_V4 \| 1.0 \| 6.017e-09 \| 268 \| 0.177 \| 220 \| 134 \| 7 \| 4 \| 179 \| 7 \| 223 \| Helix-turn-helix domain-containing protein \| Helix-turn-helix domain-containing protein; afdb-uniprot50 \| AF-A0A0A0FDV9-F1-MODEL\_V4 \| 1.0 \| 4.503e-09 \| 268 \| 0.151 \| 225 \| 142 \| 7 \| 4 \| 180 \| 7 \| 230 \| Putative transcriptional regulator \| Putative transcriptional regulator; afdb-uniprot50 \| AF-A0A7U0N6L3-F1-MODEL\_V4 \| 1.0 \| 1.412e-09 \| 268 \| 0.188 \| 223 \| 130 \| 10 \| 4 \| 180 \| 13 \| 230 \| Helix-turn-helix transcriptional regulator \| Helix-turn-helix transcriptional regulator; afdb-uniprot50 \| AF-A0A1B8PKT6-F1-MODEL\_V4 \| 1.0 \| 2.521e-09 \| 268 \| 0.178 \| 230 \| 135 \| 5 \| 4 \| 180 \| 7 \| 235 \| HTH cro/C1-type domain-containing protein \| HTH cro/C1-type domain-containing protein; afdb-uniprot50 \| AF-A0A1B1EDQ4-F1-MODEL\_V4 \| 1.0 \| 2.521e-09 \| 268 \| 0.186 \| 225 \| 132 \| 8 \| 5 \| 180 \| 18 \| 240 \| HTH cro/C1-type domain-containing protein \| HTH cro/C1-type domain-containing protein; afdb-uniprot50 \| AF-A0A3N0V596-F1-MODEL\_V4 \| 1.0 \| 1.057e-09 \| 268 \| 0.182 \| 230 \| 134 \| 9 \| 4 \| 180 \| 16 \| 244 \| XRE family transcriptional regulator \| XRE family transcriptional regulator; afdb-uniprot50 \| AF-A0A348DDK2-F1-MODEL\_V4 \| 1.0 \| 9.408e-10 \| 268 \| 0.2 \| 224 \| 130 \| 7 \| 4 \| 180 \| 35 \| 256 \| Repressor \| Repressor; afdb-uniprot50 \| AF-A0A7Y7J3L6-F1-MODEL\_V4 \| 1.0 \| 2.379e-09 \| 268 \| 0.204 \| 230 \| 130 \| 6 \| 4 \| 180 \| 41 \| 270 \| Helix-turn-helix transcriptional regulator \| Helix-turn-helix transcriptional regulator; afdb-uniprot50 \| AF-A0A2S9AU56-F1-MODEL\_V4 \| 1.0 \| 1.496e-09 \| 268 \| 0.125 \| 278 \| 141 \| 7 \| 4 \| 180 \| 14 \| 290 \| Propanediol utilization protein \| Propanediol utilization protein; afdb-uniprot50 \| AF-A0A2D0KAP2-F1-MODEL\_V4 \| 1.0 \| 2.831e-09 \| 268 \| 0.157 \| 235 \| 139 \| 5 \| 4 \| 180 \| 97 \| 330 \| Repressor \| Repressor; afdb-uniprot50 \| AF-A0A2W6VYN7-F1-MODEL\_V4 \| 1.0 \| 1.668e-06 \| 267 \| 0.26 \| 100 \| 67 \| 4 \| 87 \| 180 \| 5 \| 103 \| Uncharacterized protein \| Uncharacterized protein; afdb-uniprot50 \| AF-A0A7G2K0K2-F1-MODEL\_V4 \| 1.0 \| 2.229e-06 \| 267 \| 0.252 \| 111 \| 79 \| 1 \| 70 \| 180 \| 1 \| 107 \| Prophage LambdaSo, transcriptional regulator, Cro/CI family \| Prophage LambdaSo, transcriptional regulator, Cro/CI family; afdb-uniprot50 \| AF-A0A4Q8L9Y8-F1-MODEL\_V4 \| 1.0 \| 5.32e-06 \| 267 \| 0.191 \| 94 \| 74 \| 2 \| 87 \| 180 \| 20 \| 111 \| Uncharacterized protein \| Uncharacterized protein; afdb-uniprot50 \| AF-A0A7W4SU23-F1-MODEL\_V4 \| 1.0 \| 1.496e-09 \| 267 \| 0.181 \| 198 \| 131 \| 7 \| 4 \| 180 \| 10 \| 197 \| Transcriptional regulator with XRE-family HTH domain \| Transcriptional regulator with XRE-family HTH domain; afdb-uniprot50 \| AF-A0A349EQG1-F1-MODEL\_V4 \| 1.0 \| 4.249e-09 \| 267 \| 0.209 \| 210 \| 130 \| 5 \| 4 \| 180 \| 6 \| 212 \| HTH cro/C1-type domain-containing protein \| HTH cro/C1-type domain-containing protein; afdb-uniprot50 \| AF-A0A7T7HH85-F1-MODEL\_V4 \| 1.0 \| 4.772e-09 \| 267 \| 0.18 \| 211 \| 138 \| 6 \| 4 \| 180 \| 15 \| 224 \| Helix-turn-helix domain-containing protein \| Helix-turn-helix domain-containing protein; afdb-uniprot50 \| AF-A0A4R1J3T7-F1-MODEL\_V4 \| 1.0 \| 4.249e-09 \| 267 \| 0.206 \| 228 \| 122 \| 8 \| 5 \| 178 \| 7 \| 229 \| Peptidase S24-like protein \| Peptidase S24-like protein; afdb-uniprot50 \| AF-A0A3N4NGB5-F1-MODEL\_V4 \| 1.0 \| 6.269e-10 \| 267 \| 0.208 \| 221 \| 125 \| 7 \| 4 \| 176 \| 7 \| 225 \| XRE family transcriptional regulator \| XRE family transcriptional regulator; afdb-uniprot50 \| AF-A0A2A2B7J3-F1-MODEL\_V4 \| 1.0 \| 5.678e-09 \| 267 \| 0.135 \| 228 \| 135 \| 6 \| 5 \| 178 \| 16 \| 235 \| Peptidase\_S24 domain-containing protein \| Peptidase\_S24 domain-containing protein; afdb-uniprot50 \| AF-A0A4U9HM21-F1-MODEL\_V4 \| 1.0 \| 2.653e-06 \| 266 \| 0.245 \| 106 \| 76 \| 1 \| 79 \| 180 \| 73 \| 178 \| Peptidase S24-like \| Peptidase S24-like; afdb-uniprot50 \| AF-A0A377ICP7-F1-MODEL\_V4 \| 1.0 \| 5.143e-08 \| 266 \| 0.248 \| 141 \| 90 \| 5 \| 48 \| 180 \| 48 \| 180 \| LexA family repressor/S24 family protease \| LexA family repressor/S24 family protease; afdb-uniprot50 \| AF-A0A7Y9BX19-F1-MODEL\_V4 \| 1.0 \| 6.757e-09 \| 266 \| 0.152 \| 230 \| 138 \| 6 \| 4 \| 180 \| 9 \| 234 \| Helix-turn-helix transcriptional regulator \| Helix-turn-helix transcriptional regulator; afdb-uniprot50 \| AF-A0A1Y5STX6-F1-MODEL\_V4 \| 1.0 \| 1.496e-09 \| 266 \| 0.163 \| 226 \| 138 \| 9 \| 4 \| 180 \| 14 \| 237 \| HTH-type transcriptional regulator PrtR \| HTH-type transcriptional regulator PrtR; afdb-uniprot50 \| AF-A0A7S9VBU4-F1-MODEL\_V4 \| 1.0 \| 3.257e-11 \| 266 \| 0.21 \| 238 \| 124 \| 8 \| 4 \| 179 \| 7 \| 242 \| Helix-turn-helix domain-containing protein \| Helix-turn-helix domain-containing protein; afdb-uniprot50 \| AF-A0A369TDZ1-F1-MODEL\_V4 \| 1.0 \| 5.056e-09 \| 266 \| 0.181 \| 242 \| 135 \| 6 \| 1 \| 180 \| 8 \| 248 \| LexA family transcriptional regulator \| LexA family transcriptional regulator; afdb-uniprot50 \| AF-A0A1W6K966-F1-MODEL\_V4 \| 1.0 \| 3.784e-09 \| 266 \| 0.192 \| 250 \| 128 \| 5 \| 4 \| 180 \| 6 \| 254 \| HTH-type transcriptional regulator PrtR \| HTH-type transcriptional regulator PrtR; afdb-uniprot50 \| AF-A0A2G2PKM8-F1-MODEL\_V4 \| 1.0 \| 3e-09 \| 266 \| 0.188 \| 239 \| 129 \| 6 \| 4 \| 180 \| 30 \| 265 \| Cro/Cl family transcriptional regulator \| Cro/Cl family transcriptional regulator; afdb-uniprot50 \| AF-A0A7G2K0R3-F1-MODEL\_V4 \| 1.0 \| 7.11e-06 \| 265 \| 0.271 \| 92 \| 67 \| 0 \| 89 \| 180 \| 5 \| 96 \| Prophage LambdaSo, transcriptional regulator, Cro/CI family \| Prophage LambdaSo, transcriptional regulator, Cro/CI family; afdb-uniprot50 \| AF-A0A0E1W9H6-F1-MODEL\_V4 \| 1.0 \| 1.68e-09 \| 265 \| 0.217 \| 207 \| 127 \| 6 \| 6 \| 180 \| 11 \| 214 \| Putative repressor protein \| Putative repressor protein; afdb-uniprot50 \| AF-A0A7M1S497-F1-MODEL\_V4 \| 1.0 \| 1.057e-09 \| 265 \| 0.182 \| 208 \| 136 \| 7 \| 5 \| 180 \| 12 \| 217 \| LexA family transcriptional regulator \| LexA family transcriptional regulator; afdb-uniprot50 \| AF-E7H635-F1-MODEL\_V4 \| 1.0 \| 8.878e-10 \| 265 \| 0.219 \| 214 \| 128 \| 7 \| 4 \| 180 \| 7 \| 218 \| HTH cro/C1-type domain-containing protein \| HTH cro/C1-type domain-containing protein; afdb-uniprot50 \| AF-A0A286U3Z1-F1-MODEL\_V4 \| 1.0 \| 2.379e-09 \| 265 \| 0.169 \| 212 \| 135 \| 5 \| 9 \| 180 \| 11 \| 221 \| Peptidase S24-like domain protein \| Peptidase S24-like domain protein; afdb-uniprot50 \| AF-X2H633-F1-MODEL\_V4 \| 1.0 \| 1.999e-09 \| 265 \| 0.151 \| 231 \| 143 \| 4 \| 3 \| 180 \| 2 \| 232 \| Putative transcriptional regulator \| Putative transcriptional regulator; afdb-uniprot50 \| AF-A0A212QJ24-F1-MODEL\_V4 \| 1.0 \| 2.672e-09 \| 265 \| 0.191 \| 225 \| 128 \| 7 \| 5 \| 180 \| 14 \| 233 \| Phage repressor protein C, contains Cro/C1-type HTH and peptisase s24 domains \| Phage repressor protein C, contains Cro/C1-type HTH and peptisase s24 domains; afdb-uniprot50 \| AF-A0A2S5KUI7-F1-MODEL\_V4 \| 1.0 \| 1.887e-09 \| 265 \| 0.22 \| 231 \| 125 \| 9 \| 4 \| 180 \| 10 \| 239 \| Peptidase\_S24 domain-containing protein \| Peptidase\_S24 domain-containing protein; afdb-uniprot50 \| AF-A0A2T5PGI5-F1-MODEL\_V4 \| 1.0 \| 2.379e-09 \| 265 \| 0.184 \| 239 \| 131 \| 5 \| 4 \| 180 \| 10 \| 246 \| Peptidase \| Peptidase; afdb-uniprot50 \| AF-A0A1Q9R370-F1-MODEL\_V4 \| 1.0 \| 1.412e-09 \| 265 \| 0.14 \| 277 \| 137 \| 9 \| 4 \| 180 \| 6 \| 281 \| HTH cro/C1-type domain-containing protein \| HTH cro/C1-type domain-containing protein; afdb-uniprot50 \| AF-A0A0W0N0W8-F1-MODEL\_V4 \| 1.0 \| 6.269e-10 \| 265 \| 0.14 \| 270 \| 138 \| 6 \| 4 \| 180 \| 14 \| 282 \| Propanediol utilization protein \| Propanediol utilization protein; afdb-uniprot50 \| AF-A0A379D1W1-F1-MODEL\_V4 \| 1.0 \| 3.914e-07 \| 264 \| 0.248 \| 125 \| 85 \| 5 \| 60 \| 180 \| 2 \| 121 \| Uncharacterized HTH-type transcriptional regulator HI\_1476 \| Uncharacterized HTH-type transcriptional regulator HI\_1476; afdb-uniprot50 \| AF-A0A5E7J7C6-F1-MODEL\_V4 \| 1.0 \| 6.99e-07 \| 264 \| 0.213 \| 122 \| 91 \| 3 \| 63 \| 180 \| 6 \| 126 \| Peptidase\_S24 domain-containing protein \| Peptidase\_S24 domain-containing protein; afdb-uniprot50 \| AF-A0A317DV49-F1-MODEL\_V4 \| 1.0 \| 2.521e-09 \| 264 \| 0.226 \| 199 \| 114 \| 6 \| 20 \| 180 \| 2 \| 198 \| HTH cro/C1-type domain-containing protein \| HTH cro/C1-type domain-containing protein; afdb-uniprot50 \| AF-A0A3A8EIV5-F1-MODEL\_V4 \| 1.0 \| 1.186e-09 \| 264 \| 0.231 \| 203 \| 123 \| 7 \| 4 \| 174 \| 6 \| 207 \| Peptidase S24 \| Peptidase S24; afdb-uniprot50 \| AF-A0A850JR98-F1-MODEL\_V4 \| 1.0 \| 3.369e-09 \| 264 \| 0.207 \| 212 \| 129 \| 7 \| 4 \| 180 \| 10 \| 217 \| Helix-turn-helix transcriptional regulator \| Helix-turn-helix transcriptional regulator; afdb-uniprot50 \| AF-J9Z8Z9-F1-MODEL\_V4 \| 1.0 \| 2.521e-09 \| 264 \| 0.216 \| 217 \| 122 \| 10 \| 5 \| 178 \| 8 \| 219 \| Repressor protein \| Repressor protein; afdb-uniprot50 \| AF-A0A2P5LAB6-F1-MODEL\_V4 \| 1.0 \| 2.521e-09 \| 264 \| 0.191 \| 219 \| 133 \| 7 \| 4 \| 180 \| 7 \| 223 \| HTH cro/C1-type domain-containing protein \| HTH cro/C1-type domain-containing protein; afdb-uniprot50 \| AF-A0A840M8M5-F1-MODEL\_V4 \| 1.0 \| 2.831e-09 \| 264 \| 0.188 \| 218 \| 136 \| 6 \| 4 \| 180 \| 15 \| 232 \| Phage repressor protein C with HTH and peptisase S24 domain \| Phage repressor protein C with HTH and peptisase S24 domain; afdb-uniprot50 \| AF-A0A0U2B2W6-F1-MODEL\_V4 \| 1.0 \| 1.257e-09 \| 264 \| 0.19 \| 236 \| 131 \| 8 \| 1 \| 180 \| 3 \| 234 \| Peptidase\_S24 domain-containing protein \| Peptidase\_S24 domain-containing protein; afdb-uniprot50 \| AF-A0A2N5ZG77-F1-MODEL\_V4 \| 1.0 \| 3.571e-09 \| 264 \| 0.173 \| 213 \| 138 \| 6 \| 4 \| 180 \| 36 \| 246 \| HTH cro/C1-type domain-containing protein \| HTH cro/C1-type domain-containing protein; afdb-uniprot50 \| AF-A0A0S4IE42-F1-MODEL\_V4 \| 1.0 \| 1.496e-09 \| 264 \| 0.146 \| 273 \| 135 \| 6 \| 5 \| 180 \| 9 \| 280 \| LexA repressor \| LexA repressor; afdb-uniprot50 \| AF-A0A0B5K5I8-F1-MODEL\_V4 \| 1.0 \| 1.12e-09 \| 264 \| 0.141 \| 275 \| 136 \| 7 \| 5 \| 180 \| 22 \| 295 \| Cro/Cl family transcriptional regulator \| Cro/Cl family transcriptional regulator; afdb-uniprot50 \| AF-A0A7C9KFV0-F1-MODEL\_V4 \| 1.0 \| 1.186e-09 \| 264 \| 0.157 \| 292 \| 126 \| 9 \| 4 \| 179 \| 27 \| 314 \| Helix-turn-helix domain-containing protein \| Helix-turn-helix domain-containing protein; afdb-uniprot50 \| AF-A0A1Q4D1Y9-F1-MODEL\_V4 \| 1.0 \| 8.319e-07 \| 263 \| 0.258 \| 116 \| 76 \| 4 \| 70 \| 180 \| 28 \| 138 \| Peptidase\_S24 domain-containing protein \| Peptidase\_S24 domain-containing protein; afdb-uniprot50 \| AF-W7PA95-F1-MODEL\_V4 \| 1.0 \| 7.535e-06 \| 263 \| 0.196 \| 102 \| 82 \| 0 \| 79 \| 180 \| 87 \| 188 \| HTH-type transcriptional regulator RdgA \| HTH-type transcriptional regulator RdgA; afdb-uniprot50 \| AF-A0A7V8JTY9-F1-MODEL\_V4 \| 1.0 \| 2.245e-09 \| 263 \| 0.212 \| 202 \| 122 \| 6 \| 5 \| 173 \| 15 \| 212 \| Putative HTH-type transcriptional regulator \| Putative HTH-type transcriptional regulator; afdb-uniprot50 \| AF-A0A2D4RVR8-F1-MODEL\_V4 \| 1.0 \| 7.588e-09 \| 263 \| 0.196 \| 214 \| 134 \| 5 \| 4 \| 180 \| 7 \| 219 \| HTH cro/C1-type domain-containing protein \| HTH cro/C1-type domain-containing protein; afdb-uniprot50 \| AF-A0A154WGA2-F1-MODEL\_V4 \| 1.0 \| 4.249e-09 \| 263 \| 0.161 \| 223 \| 140 \| 11 \| 1 \| 180 \| 4 \| 222 \| Peptidase S24 \| Peptidase S24; afdb-uniprot50 \| AF-A0A836MN97-F1-MODEL\_V4 \| 1.0 \| 8.878e-10 \| 263 \| 0.193 \| 238 \| 130 \| 10 \| 1 \| 180 \| 1 \| 234 \| Putative transcriptional regulator \| Putative transcriptional regulator; afdb-uniprot50 \| AF-Q5F6C9-F1-MODEL\_V4 \| 1.0 \| 7.46e-10 \| 263 \| 0.168 \| 225 \| 135 \| 7 \| 4 \| 176 \| 19 \| 243 \| Repressor \| Repressor; afdb-uniprot50 \| AF-A0A849P416-F1-MODEL\_V4 \| 1.0 \| 3.571e-09 \| 263 \| 0.203 \| 221 \| 127 \| 7 \| 5 \| 180 \| 38 \| 254 \| S24 family peptidase \| S24 family peptidase; afdb-uniprot50 \| AF-A0A845BW56-F1-MODEL\_V4 \| 1.0 \| 3.571e-09 \| 263 \| 0.201 \| 238 \| 129 \| 5 \| 4 \| 180 \| 27 \| 264 \| Helix-turn-helix domain-containing protein \| Helix-turn-helix domain-containing protein; afdb-uniprot50 \| AF-A0A5M8F983-F1-MODEL\_V4 \| 1.0 \| 4.427e-10 \| 263 \| 0.146 \| 266 \| 133 \| 7 \| 4 \| 176 \| 14 \| 278 \| Helix-turn-helix domain-containing protein \| Helix-turn-helix domain-containing protein; afdb-uniprot50 \| AF-A0A1H9EPH6-F1-MODEL\_V4 \| 1.0 \| 3.313e-10 \| 263 \| 0.161 \| 273 \| 129 \| 7 \| 7 \| 180 \| 24 \| 295 \| Phage repressor protein C, contains Cro/C1-type HTH and peptisase s24 domains \| Phage repressor protein C, contains Cro/C1-type HTH and peptisase s24 domains; afdb-uniprot50 \| AF-A0A239LDH9-F1-MODEL\_V4 \| 1.0 \| 2.379e-09 \| 263 \| 0.189 \| 232 \| 129 \| 7 \| 5 \| 179 \| 85 \| 314 \| Phage repressor protein C, contains Cro/C1-type HTH and peptisase s24 domains \| Phage repressor protein C, contains Cro/C1-type HTH and peptisase s24 domains; afdb-uniprot50 \| AF-Q4QKX1-F1-MODEL\_V4 \| 1.0 \| 2.672e-09 \| 263 \| 0.138 \| 246 \| 142 \| 6 \| 4 \| 180 \| 74 \| 318 \| Uncharacterized protein \| Uncharacterized protein; afdb-uniprot50 \| AF-A0A549SHD6-F1-MODEL\_V4 \| 1.0 \| 3.545e-06 \| 262 \| 0.204 \| 98 \| 71 \| 3 \| 89 \| 180 \| 13 \| 109 \| S24 family peptidase \| S24 family peptidase; afdb-uniprot50 \| AF-A0A6G9VXL7-F1-MODEL\_V4 \| 1.0 \| 2.229e-06 \| 262 \| 0.254 \| 106 \| 75 \| 2 \| 78 \| 180 \| 40 \| 144 \| S24 family peptidase \| S24 family peptidase; afdb-uniprot50 \| AF-A0A166UAP7-F1-MODEL\_V4 \| 1.0 \| 6.709e-06 \| 262 \| 0.288 \| 104 \| 72 \| 2 \| 78 \| 180 \| 70 \| 172 \| Peptidase\_S24 domain-containing protein \| Peptidase\_S24 domain-containing protein; afdb-uniprot50 \| AF-A0A3B9IP20-F1-MODEL\_V4 \| 1.0 \| 1.999e-09 \| 262 \| 0.198 \| 196 \| 121 \| 5 \| 20 \| 180 \| 2 \| 196 \| Transcriptional regulator \| Transcriptional regulator; afdb-uniprot50 \| AF-A0A554GDH4-F1-MODEL\_V4 \| 1.0 \| 1.768e-06 \| 262 \| 0.271 \| 107 \| 73 \| 1 \| 79 \| 180 \| 90 \| 196 \| Helix-turn-helix transcriptional regulator \| Helix-turn-helix transcriptional regulator; afdb-uniprot50 \| AF-A0A5K1S6Z7-F1-MODEL\_V4 \| 1.0 \| 1.985e-06 \| 262 \| 0.257 \| 105 \| 75 \| 1 \| 79 \| 180 \| 98 \| 202 \| Helix-turn-helix transcriptional regulator \| Helix-turn-helix transcriptional regulator; afdb-uniprot50 \| AF-A0A317EHX9-F1-MODEL\_V4 \| 1.0 \| 2.831e-09 \| 262 \| 0.19 \| 215 \| 134 \| 6 \| 4 \| 180 \| 7 \| 219 \| HTH cro/C1-type domain-containing protein \| HTH cro/C1-type domain-containing protein; afdb-uniprot50 \| AF-A0A7H8UUY2-F1-MODEL\_V4 \| 1.0 \| 2.521e-09 \| 262 \| 0.19 \| 215 \| 129 \| 8 \| 6 \| 180 \| 14 \| 223 \| Helix-turn-helix transcriptional regulator \| Helix-turn-helix transcriptional regulator; afdb-uniprot50 \| AF-A0A1I2E7J1-F1-MODEL\_V4 \| 1.0 \| 2.245e-09 \| 262 \| 0.172 \| 237 \| 135 \| 8 \| 1 \| 180 \| 1 \| 233 \| Peptidase S24-like \| Peptidase S24-like; afdb-uniprot50 \| AF-A0A515DE75-F1-MODEL\_V4 \| 1.0 \| 8.378e-10 \| 262 \| 0.248 \| 197 \| 126 \| 9 \| 4 \| 180 \| 52 \| 246 \| S24 family peptidase \| S24 family peptidase; afdb-uniprot50 \| AF-A0A7M3AWF2-F1-MODEL\_V4 \| 1.0 \| 2.672e-09 \| 262 \| 0.165 \| 241 \| 136 \| 7 \| 4 \| 180 \| 18 \| 257 \| LexA family transcriptional regulator \| LexA family transcriptional regulator; afdb-uniprot50 \| AF-A0A5E4YYK6-F1-MODEL\_V4 \| 1.0 \| 3.369e-09 \| 262 \| 0.138 \| 259 \| 140 \| 7 \| 4 \| 180 \| 19 \| 276 \| Putative phage repressor \| Putative phage repressor; afdb-uniprot50 \| AF-A0A266Q680-F1-MODEL\_V4 \| 1.0 \| 2.521e-09 \| 262 \| 0.191 \| 267 \| 121 \| 11 \| 4 \| 180 \| 19 \| 280 \| Peptidase\_S24 domain-containing protein \| Peptidase\_S24 domain-containing protein; afdb-uniprot50 \| AF-A0A6D0P928-F1-MODEL\_V4 \| 1.0 \| 1.112e-06 \| 261 \| 0.241 \| 116 \| 81 \| 4 \| 68 \| 180 \| 5 \| 116 \| Peptidase\_S24 domain-containing protein \| Peptidase\_S24 domain-containing protein; afdb-uniprot50 \| AF-A1VX46-F1-MODEL\_V4 \| 1.0 \| 7.161e-09 \| 261 \| 0.172 \| 203 \| 126 \| 5 \| 19 \| 180 \| 3 \| 204 \| Putative phage repressor \| Putative phage repressor; afdb-uniprot50 \| AF-A0A2S9TGE6-F1-MODEL\_V4 \| 1.0 \| 5.056e-09 \| 261 \| 0.183 \| 218 \| 134 \| 6 \| 5 \| 180 \| 7 \| 222 \| Phage repressor protein \| Phage repressor protein; afdb-uniprot50 \| AF-A0A2E5FYH6-F1-MODEL\_V4 \| 1.0 \| 8.041e-09 \| 261 \| 0.162 \| 240 \| 137 \| 6 \| 4 \| 180 \| 13 \| 251 \| HTH cro/C1-type domain-containing protein \| HTH cro/C1-type domain-containing protein; afdb-uniprot50 \| AF-A0A317EB58-F1-MODEL\_V4 \| 1.0 \| 2.672e-09 \| 261 \| 0.191 \| 225 \| 128 \| 10 \| 5 \| 180 \| 34 \| 253 \| Peptidase\_S24 domain-containing protein \| Peptidase\_S24 domain-containing protein; afdb-uniprot50 \| AF-A0A2S4J7K4-F1-MODEL\_V4 \| 1.0 \| 3e-09 \| 261 \| 0.176 \| 255 \| 131 \| 7 \| 4 \| 180 \| 6 \| 259 \| XRE family transcriptional regulator \| XRE family transcriptional regulator; afdb-uniprot50 \| AF-A0A2N0ZNY9-F1-MODEL\_V4 \| 1.0 \| 7.161e-09 \| 261 \| 0.128 \| 257 \| 143 \| 4 \| 4 \| 180 \| 12 \| 267 \| HTH cro/C1-type domain-containing protein \| HTH cro/C1-type domain-containing protein; afdb-uniprot50 \| AF-A0A3B8ZF36-F1-MODEL\_V4 \| 1.0 \| 1.412e-09 \| 261 \| 0.188 \| 239 \| 127 \| 8 \| 8 \| 180 \| 42 \| 279 \| Peptidase\_S24 domain-containing protein \| Peptidase\_S24 domain-containing protein; afdb-uniprot50 \| AF-A0A0U4VN79-F1-MODEL\_V4 \| 1.0 \| 2.672e-09 \| 261 \| 0.149 \| 274 \| 133 \| 7 \| 5 \| 180 \| 9 \| 280 \| Repressor \| Repressor; afdb-uniprot50 \| AF-A0A062GNF5-F1-MODEL\_V4 \| 1.0 \| 5.975e-06 \| 260 \| 0.247 \| 89 \| 63 \| 2 \| 93 \| 178 \| 3 \| 90 \| Peptidase S24-like family protein \| Peptidase S24-like family protein; afdb-uniprot50 \| AF-A0A7T8S055-F1-MODEL\_V4 \| 1.0 \| 3.289e-07 \| 260 \| 0.195 \| 128 \| 93 \| 5 \| 61 \| 180 \| 41 \| 166 \| Helix-turn-helix transcriptional regulator \| Helix-turn-helix transcriptional regulator; afdb-uniprot50 \| AF-A0A7U9HK71-F1-MODEL\_V4 \| 1.0 \| 2.521e-09 \| 260 \| 0.164 \| 219 \| 138 \| 9 \| 4 \| 180 \| 20 \| 235 \| Peptidase\_S24 domain-containing protein \| Peptidase\_S24 domain-containing protein; afdb-uniprot50 \| AF-A0A7X5UB38-F1-MODEL\_V4 \| 1.0 \| 2.831e-09 \| 260 \| 0.157 \| 229 \| 136 \| 8 \| 5 \| 180 \| 15 \| 239 \| Peptidase\_S24 domain-containing protein \| Peptidase\_S24 domain-containing protein; afdb-uniprot50 \| AF-A0A031LX11-F1-MODEL\_V4 \| 1.0 \| 3.571e-09 \| 260 \| 0.141 \| 241 \| 138 \| 7 \| 5 \| 178 \| 4 \| 242 \| Alkaline phosphatase \| Alkaline phosphatase; afdb-uniprot50 \| AF-A0A258UIF6-F1-MODEL\_V4 \| 1.0 \| 7.161e-09 \| 260 \| 0.211 \| 217 \| 129 \| 8 \| 5 \| 180 \| 36 \| 251 \| HTH cro/C1-type domain-containing protein \| HTH cro/C1-type domain-containing protein; afdb-uniprot50 \| AF-A0A6J4EE08-F1-MODEL\_V4 \| 1.0 \| 3.18e-09 \| 260 \| 0.163 \| 238 \| 138 \| 5 \| 4 \| 180 \| 16 \| 253 \| HTH-type transcriptional regulator PrtR \| HTH-type transcriptional regulator PrtR; afdb-uniprot50 \| AF-A0A0P9ZT73-F1-MODEL\_V4 \| 1.0 \| 1.585e-09 \| 260 \| 0.169 \| 266 \| 130 \| 7 \| 5 \| 180 \| 3 \| 267 \| Cro/Cl family transcriptional regulator \| Cro/Cl family transcriptional regulator; afdb-uniprot50 \| AF-A0A1L7NES5-F1-MODEL\_V4 \| 1.0 \| 2.504e-06 \| 259 \| 0.233 \| 107 \| 77 \| 1 \| 79 \| 180 \| 90 \| 196 \| Transcriptional regulator \| Transcriptional regulator; afdb-uniprot50 \| AF-A0A081MYG4-F1-MODEL\_V4 \| 1.0 \| 1.014e-08 \| 259 \| 0.195 \| 200 \| 124 \| 6 \| 13 \| 180 \| 4 \| 198 \| HTH cro/C1-type domain-containing protein \| HTH cro/C1-type domain-containing protein; afdb-uniprot50 \| AF-A0A419U229-F1-MODEL\_V4 \| 1.0 \| 1.257e-09 \| 259 \| 0.213 \| 215 \| 111 \| 9 \| 19 \| 178 \| 4 \| 215 \| Phage repressor protein C with HTH and peptisase S24 domain \| Phage repressor protein C with HTH and peptisase S24 domain; afdb-uniprot50 \| AF-A0A2G2Q498-F1-MODEL\_V4 \| 1.0 \| 2.245e-09 \| 259 \| 0.195 \| 225 \| 128 \| 8 \| 1 \| 180 \| 5 \| 221 \| Transcriptional regulator \| Transcriptional regulator; afdb-uniprot50 \| AF-A0A7Y1GCX0-F1-MODEL\_V4 \| 1.0 \| 2.104e-06 \| 259 \| 0.242 \| 107 \| 76 \| 1 \| 79 \| 180 \| 130 \| 236 \| Helix-turn-helix transcriptional regulator \| Helix-turn-helix transcriptional regulator; afdb-uniprot50 \| AF-A0A094SR02-F1-MODEL\_V4 \| 1.0 \| 3.784e-09 \| 259 \| 0.21 \| 228 \| 125 \| 11 \| 4 \| 180 \| 13 \| 236 \| Peptidase\_S24 domain-containing protein \| Peptidase\_S24 domain-containing protein; afdb-uniprot50 \| AF-A0A4Q3MRY5-F1-MODEL\_V4 \| 1.0 \| 3.369e-09 \| 259 \| 0.191 \| 245 \| 131 \| 9 \| 1 \| 180 \| 1 \| 243 \| LexA family transcriptional repressor \| LexA family transcriptional repressor; afdb-uniprot50 \| AF-A0A7H1NPH5-F1-MODEL\_V4 \| 1.0 \| 6.757e-09 \| 259 \| 0.162 \| 234 \| 136 \| 8 \| 4 \| 180 \| 28 \| 258 \| HTH-type transcriptional regulator PrtR \| HTH-type transcriptional regulator PrtR; afdb-uniprot50 \| AF-A0A1H8PP99-F1-MODEL\_V4 \| 1.0 \| 1.496e-09 \| 258 \| 0.2 \| 215 \| 119 \| 6 \| 14 \| 180 \| 1 \| 210 \| Helix-turn-helix \| Helix-turn-helix; afdb-uniprot50 \| AF-A0A1I5S1T1-F1-MODEL\_V4 \| 1.0 \| 4.503e-09 \| 258 \| 0.205 \| 224 \| 130 \| 9 \| 1 \| 180 \| 1 \| 220 \| Phage repressor protein C, contains Cro/C1-type HTH and peptisase s24 domains \| Phage repressor protein C, contains Cro/C1-type HTH and peptisase s24 domains; afdb-uniprot50 \| AF-A0A165VS19-F1-MODEL\_V4 \| 1.0 \| 2.504e-06 \| 258 \| 0.235 \| 106 \| 77 \| 1 \| 79 \| 180 \| 107 \| 212 \| Peptidase\_S24 domain-containing protein \| Peptidase\_S24 domain-containing protein; afdb-uniprot50 \| AF-A0A378PYP5-F1-MODEL\_V4 \| 1.0 \| 5.678e-09 \| 258 \| 0.163 \| 233 \| 139 \| 6 \| 1 \| 180 \| 3 \| 232 \| Uncharacterized HTH-type transcriptional regulator HI\_1476 \| Uncharacterized HTH-type transcriptional regulator HI\_1476; afdb-uniprot50 \| AF-A0A386DUQ7-F1-MODEL\_V4 \| 1.0 \| 9.03e-09 \| 258 \| 0.206 \| 213 \| 128 \| 4 \| 4 \| 176 \| 39 \| 250 \| Putative phage repressor \| Putative phage repressor; afdb-uniprot50 \| AF-A0A0P9KR81-F1-MODEL\_V4 \| 1.0 \| 2.831e-09 \| 258 \| 0.166 \| 258 \| 134 \| 9 \| 1 \| 180 \| 2 \| 256 \| HTH cro/C1-type domain-containing protein \| HTH cro/C1-type domain-containing protein; afdb-uniprot50 \| AF-A0A7D5QHE9-F1-MODEL\_V4 \| 1.0 \| 4.249e-09 \| 258 \| 0.148 \| 249 \| 138 \| 3 \| 6 \| 180 \| 22 \| 270 \| Helix-turn-helix transcriptional regulator \| Helix-turn-helix transcriptional regulator; afdb-uniprot50 \| AF-A0A8B6UYU7-F1-MODEL\_V4 \| 1.0 \| 1.057e-09 \| 258 \| 0.145 \| 274 \| 136 \| 8 \| 4 \| 180 \| 14 \| 286 \| Helix-turn-helix domain-containing protein \| Helix-turn-helix domain-containing protein; afdb-uniprot50 \| AF-A0A4U1KS30-F1-MODEL\_V4 \| 1.0 \| 1.78e-09 \| 258 \| 0.15 \| 285 \| 132 \| 7 \| 4 \| 180 \| 6 \| 288 \| LexA family transcriptional regulator \| LexA family transcriptional regulator; afdb-uniprot50 \| AF-A0A285Q3X1-F1-MODEL\_V4 \| 1.0 \| 4.249e-09 \| 258 \| 0.138 \| 289 \| 136 \| 5 \| 4 \| 180 \| 7 \| 294 \| Phage repressor protein C, contains Cro/C1-type HTH and peptisase s24 domains \| Phage repressor protein C, contains Cro/C1-type HTH and peptisase s24 domains; afdb-uniprot50 \| AF-A0A3D4Z7E1-F1-MODEL\_V4 \| 1.0 \| 4.738e-06 \| 257 \| 0.27 \| 96 \| 66 \| 4 \| 87 \| 180 \| 17 \| 110 \| Peptidase\_S24 domain-containing protein \| Peptidase\_S24 domain-containing protein; afdb-uniprot50 \| AF-A0A0R0C0L3-F1-MODEL\_V4 \| 1.0 \| 2.504e-06 \| 257 \| 0.257 \| 101 \| 66 \| 5 \| 87 \| 180 \| 22 \| 120 \| Uncharacterized protein \| Uncharacterized protein; afdb-uniprot50 \| AF-F4G9T8-F1-MODEL\_V4 \| 1.0 \| 4.691e-10 \| 257 \| 0.256 \| 187 \| 110 \| 7 \| 15 \| 180 \| 8 \| 186 \| Peptidase S24/S26A/S26B, conserved region \| Peptidase S24/S26A/S26B, conserved region; afdb-uniprot50 \| AF-A0A4P6X1R6-F1-MODEL\_V4 \| 1.0 \| 6.644e-10 \| 257 \| 0.251 \| 187 \| 111 \| 7 \| 15 \| 180 \| 8 \| 186 \| Peptidase S24-like protein \| Peptidase S24-like protein; afdb-uniprot50 \| AF-A0A1G7S5X8-F1-MODEL\_V4 \| 1.0 \| 3.784e-09 \| 257 \| 0.227 \| 202 \| 121 \| 7 \| 13 \| 180 \| 2 \| 202 \| Helix-turn-helix domain-containing protein \| Helix-turn-helix domain-containing protein; afdb-uniprot50 \| AF-A0A4R1V102-F1-MODEL\_V4 \| 1.0 \| 1.075e-08 \| 257 \| 0.168 \| 202 \| 141 \| 4 \| 5 \| 180 \| 36 \| 236 \| Phage repressor protein C with HTH and peptisase S24 domain \| Phage repressor protein C with HTH and peptisase S24 domain; afdb-uniprot50 \| AF-A0A1A9RFF4-F1-MODEL\_V4 \| 1.0 \| 6.377e-09 \| 257 \| 0.181 \| 231 \| 135 \| 5 \| 4 \| 180 \| 7 \| 237 \| HTH cro/C1-type domain-containing protein \| HTH cro/C1-type domain-containing protein; afdb-uniprot50 \| AF-A0A848FZX3-F1-MODEL\_V4 \| 1.0 \| 3.571e-09 \| 257 \| 0.188 \| 239 \| 131 \| 7 \| 4 \| 180 \| 7 \| 244 \| Helix-turn-helix transcriptional regulator \| Helix-turn-helix transcriptional regulator; afdb-uniprot50 \| AF-A0A2E8IKW7-F1-MODEL\_V4 \| 1.0 \| 4.322e-08 \| 257 \| 0.141 \| 254 \| 139 \| 5 \| 4 \| 180 \| 9 \| 260 \| XRE family transcriptional regulator \| XRE family transcriptional regulator; afdb-uniprot50 \| AF-A0A3M4BSU0-F1-MODEL\_V4 \| 1.0 \| 3e-09 \| 257 \| 0.173 \| 236 \| 131 \| 6 \| 6 \| 180 \| 35 \| 267 \| HTH cro/C1-type domain-containing protein \| HTH cro/C1-type domain-containing protein; afdb-uniprot50 \| AF-A0A2Z4RT63-F1-MODEL\_V4 \| 1.0 \| 2.521e-09 \| 257 \| 0.138 \| 267 \| 136 \| 7 \| 7 \| 180 \| 24 \| 289 \| Helix-turn-helix domain-containing protein \| Helix-turn-helix domain-containing protein; afdb-uniprot50 \| AF-A0A1E8PLB2-F1-MODEL\_V4 \| 1.0 \| 1.323e-06 \| 256 \| 0.228 \| 118 \| 83 \| 4 \| 60 \| 174 \| 2 \| 114 \| Peptidase\_S24 domain-containing protein \| Peptidase\_S24 domain-containing protein; afdb-uniprot50 \| AF-A0A2S9K153-F1-MODEL\_V4 \| 1.0 \| 1.697e-05 \| 256 \| 0.2 \| 100 \| 75 \| 2 \| 86 \| 180 \| 31 \| 130 \| Peptidase\_S24 domain-containing protein \| Peptidase\_S24 domain-containing protein; afdb-uniprot50 \| AF-A0A5C8BLB1-F1-MODEL\_V4 \| 1.0 \| 8.461e-06 \| 256 \| 0.21 \| 100 \| 77 \| 2 \| 78 \| 176 \| 51 \| 149 \| S24 family peptidase \| S24 family peptidase; afdb-uniprot50 \| AF-A0A0F4NKC0-F1-MODEL\_V4 \| 1.0 \| 3.571e-09 \| 256 \| 0.193 \| 207 \| 131 \| 7 \| 4 \| 179 \| 5 \| 206 \| HTH cro/C1-type domain-containing protein \| HTH cro/C1-type domain-containing protein; afdb-uniprot50 \| AF-R6P6E9-F1-MODEL\_V4 \| 1.0 \| 6.017e-09 \| 256 \| 0.186 \| 214 \| 127 \| 8 \| 5 \| 180 \| 7 \| 211 \| Phage transcriptional regulator \| Phage transcriptional regulator; afdb-uniprot50 \| AF-D4XHG0-F1-MODEL\_V4 \| 1.0 \| 6.017e-09 \| 256 \| 0.168 \| 220 \| 136 \| 9 \| 4 \| 180 \| 8 \| 223 \| DNA-binding helix-turn-helix protein \| DNA-binding helix-turn-helix protein; afdb-uniprot50 \| AF-A0A6N4T9B0-F1-MODEL\_V4 \| 1.0 \| 4.01e-09 \| 256 \| 0.168 \| 225 \| 136 \| 8 \| 5 \| 179 \| 7 \| 230 \| HTH cro/C1-type domain-containing protein \| HTH cro/C1-type domain-containing protein; afdb-uniprot50 \| AF-A0A318DXJ5-F1-MODEL\_V4 \| 1.0 \| 8.521e-09 \| 256 \| 0.159 \| 219 \| 139 \| 8 \| 6 \| 180 \| 14 \| 231 \| HTH cro/C1-type domain-containing protein \| HTH cro/C1-type domain-containing protein; afdb-uniprot50 \| AF-F2JZ54-F1-MODEL\_V4 \| 1.0 \| 1.436e-08 \| 256 \| 0.173 \| 236 \| 131 \| 6 \| 4 \| 180 \| 6 \| 236 \| Putative phage repressor \| Putative phage repressor; afdb-uniprot50 \| AF-A0A2N6F112-F1-MODEL\_V4 \| 1.0 \| 2.521e-09 \| 256 \| 0.216 \| 236 \| 122 \| 8 \| 6 \| 180 \| 14 \| 247 \| Peptidase\_S24 domain-containing protein \| Peptidase\_S24 domain-containing protein; afdb-uniprot50 \| AF-Q06553-F1-MODEL\_V4 \| 1.0 \| 5.358e-09 \| 256 \| 0.149 \| 234 \| 141 \| 7 \| 4 \| 180 \| 15 \| 247 \| HTH-type transcriptional regulator PrtR \| HTH-type transcriptional regulator PrtR; afdb-uniprot50 \| AF-A0A7Z2ZDG1-F1-MODEL\_V4 \| 1.0 \| 1.999e-09 \| 256 \| 0.186 \| 247 \| 129 \| 8 \| 5 \| 180 \| 10 \| 255 \| Cro/Cl family transcriptional regulator \| Cro/Cl family transcriptional regulator; afdb-uniprot50 \| AF-A0A3M3D8N0-F1-MODEL\_V4 \| 1.0 \| 5.678e-09 \| 256 \| 0.169 \| 236 \| 137 \| 6 \| 2 \| 180 \| 41 \| 274 \| Cro/CI family transcriptional regulator \| Cro/CI family transcriptional regulator; afdb-uniprot50 \| AF-A0A6S7B3R8-F1-MODEL\_V4 \| 1.0 \| 4.772e-09 \| 256 \| 0.166 \| 265 \| 131 \| 6 \| 4 \| 178 \| 16 \| 280 \| HTH-type transcriptional regulator PrtR \| HTH-type transcriptional regulator PrtR; afdb-uniprot50 \| AF-A0A3D3HTG8-F1-MODEL\_V4 \| 1.0 \| 6.757e-09 \| 256 \| 0.17 \| 229 \| 136 \| 7 \| 4 \| 179 \| 62 \| 289 \| Phage repressor protein \| Phage repressor protein; afdb-uniprot50 \| AF-A0A7X3HDH1-F1-MODEL\_V4 \| 1.0 \| 7.588e-09 \| 256 \| 0.165 \| 229 \| 137 \| 5 \| 5 \| 180 \| 163 \| 390 \| Helix-turn-helix domain-containing protein \| Helix-turn-helix domain-containing protein; afdb-uniprot50 \| AF-A0A6A7Z0J1-F1-MODEL\_V4 \| 1.0 \| 4.219e-06 \| 255 \| 0.245 \| 106 \| 76 \| 1 \| 79 \| 180 \| 65 \| 170 \| Peptidase\_S24 domain-containing protein \| Peptidase\_S24 domain-containing protein; afdb-uniprot50 \| AF-A0A346N4Q6-F1-MODEL\_V4 \| 1.0 \| 5.678e-09 \| 255 \| 0.181 \| 220 \| 130 \| 7 \| 4 \| 178 \| 6 \| 220 \| Helix-turn-helix transcriptional regulator \| Helix-turn-helix transcriptional regulator; afdb-uniprot50 \| AF-A0A1H2PQK0-F1-MODEL\_V4 \| 1.0 \| 3.18e-09 \| 255 \| 0.172 \| 226 \| 136 \| 8 \| 4 \| 180 \| 5 \| 228 \| Phage repressor protein C, contains Cro/C1-type HTH and peptisase s24 domains \| Phage repressor protein C, contains Cro/C1-type HTH and peptisase s24 domains; afdb-uniprot50 \| AF-A0A2L0S440-F1-MODEL\_V4 \| 1.0 \| 1.279e-08 \| 255 \| 0.163 \| 220 \| 143 \| 5 \| 1 \| 180 \| 10 \| 228 \| HTH cro/C1-type domain-containing protein \| HTH cro/C1-type domain-containing protein; afdb-uniprot50 \| AF-A0A4R3NNU1-F1-MODEL\_V4 \| 1.0 \| 1.811e-08 \| 255 \| 0.165 \| 217 \| 141 \| 5 \| 4 \| 180 \| 14 \| 230 \| Phage repressor protein C with HTH and peptisase S24 domain \| Phage repressor protein C with HTH and peptisase S24 domain; afdb-uniprot50 \| AF-A0A7Y1BXH5-F1-MODEL\_V4 \| 1.0 \| 8.521e-09 \| 255 \| 0.157 \| 242 \| 139 \| 6 \| 4 \| 180 \| 14 \| 255 \| LexA family transcriptional regulator \| LexA family transcriptional regulator; afdb-uniprot50 \| AF-W6W2S2-F1-MODEL\_V4 \| 1.0 \| 2.521e-09 \| 255 \| 0.143 \| 265 \| 135 \| 7 \| 7 \| 180 \| 24 \| 287 \| Putative phage repressor \| Putative phage repressor; afdb-uniprot50 \| AF-A0A4U0PYZ7-F1-MODEL\_V4 \| 1.0 \| 4.249e-09 \| 255 \| 0.171 \| 239 \| 135 \| 7 \| 5 \| 180 \| 88 \| 326 \| Helix-turn-helix transcriptional regulator \| Helix-turn-helix transcriptional regulator; afdb-uniprot50 \| AF-A0A2T5S3K6-F1-MODEL\_V4 \| 1.0 \| 1.112e-06 \| 254 \| 0.228 \| 118 \| 85 \| 3 \| 68 \| 180 \| 17 \| 133 \| Peptidase S24-like protein \| Peptidase S24-like protein; afdb-uniprot50 \| AF-A0A845JXB3-F1-MODEL\_V4 \| 1.0 \| 5.678e-09 \| 254 \| 0.183 \| 224 \| 133 \| 7 \| 5 \| 180 \| 8 \| 229 \| Transcriptional regulator \| Transcriptional regulator; afdb-uniprot50 \| AF-B5JYF6-F1-MODEL\_V4 \| 1.0 \| 1.139e-08 \| 254 \| 0.18 \| 216 \| 135 \| 7 \| 4 \| 180 \| 18 \| 230 \| Prophage MuMc02, peptidase, family S24 \| Prophage MuMc02, peptidase, family S24; afdb-uniprot50 \| AF-A0A370QQ69-F1-MODEL\_V4 \| 1.0 \| 5.056e-09 \| 254 \| 0.157 \| 228 \| 141 \| 6 \| 4 \| 180 \| 12 \| 239 \| Helix-turn-helix protein \| Helix-turn-helix protein; afdb-uniprot50 \| AF-A0A1G5MEX5-F1-MODEL\_V4 \| 1.0 \| 6.757e-09 \| 254 \| 0.19 \| 236 \| 129 \| 8 \| 4 \| 180 \| 10 \| 242 \| Phage repressor protein C, contains Cro/C1-type HTH and peptisase s24 domains \| Phage repressor protein C, contains Cro/C1-type HTH and peptisase s24 domains; afdb-uniprot50 \| AF-A0A3M5WRJ3-F1-MODEL\_V4 \| 1.0 \| 4.503e-09 \| 254 \| 0.166 \| 246 \| 134 \| 6 \| 6 \| 180 \| 99 \| 344 \| Putative transcriptional regulator \| Putative transcriptional regulator; afdb-uniprot50 \| AF-A0A096AWJ2-F1-MODEL\_V4 \| 1.0 \| 1.402e-06 \| 253 \| 0.237 \| 118 \| 82 \| 5 \| 70 \| 180 \| 1 \| 117 \| Peptidase\_S24 domain-containing protein \| Peptidase\_S24 domain-containing protein; afdb-uniprot50 \| AF-A0A8B3GBH4-F1-MODEL\_V4 \| 1.0 \| 3.757e-06 \| 253 \| 0.198 \| 106 \| 80 \| 1 \| 80 \| 180 \| 47 \| 152 \| Uncharacterized protein \| Uncharacterized protein; afdb-uniprot50 \| AF-A0A4V1KJS4-F1-MODEL\_V4 \| 1.0 \| 1.522e-08 \| 253 \| 0.236 \| 173 \| 96 \| 6 \| 43 \| 180 \| 5 \| 176 \| Helix-turn-helix transcriptional regulator \| Helix-turn-helix transcriptional regulator; afdb-uniprot50 \| AF-A0A1N7Q2M4-F1-MODEL\_V4 \| 1.0 \| 6.597e-07 \| 253 \| 0.236 \| 127 \| 86 \| 4 \| 61 \| 180 \| 75 \| 197 \| Peptidase S24-like \| Peptidase S24-like; afdb-uniprot50 \| AF-A0A173JAB6-F1-MODEL\_V4 \| 1.0 \| 4.738e-06 \| 253 \| 0.177 \| 107 \| 83 \| 1 \| 78 \| 179 \| 83 \| 189 \| Transcriptional regulator \| Transcriptional regulator; afdb-uniprot50 \| AF-A0A7X8V275-F1-MODEL\_V4 \| 1.0 \| 1.355e-08 \| 253 \| 0.219 \| 210 \| 123 \| 7 \| 4 \| 176 \| 6 \| 211 \| Helix-turn-helix domain-containing protein \| Helix-turn-helix domain-containing protein; afdb-uniprot50 \| AF-A0A5C7QA30-F1-MODEL\_V4 \| 1.0 \| 8.521e-09 \| 253 \| 0.184 \| 206 \| 133 \| 7 \| 5 \| 180 \| 18 \| 218 \| Helix-turn-helix transcriptional regulator \| Helix-turn-helix transcriptional regulator; afdb-uniprot50 \| AF-A0A449IEK0-F1-MODEL\_V4 \| 1.0 \| 6.377e-09 \| 253 \| 0.174 \| 229 \| 134 \| 6 \| 4 \| 180 \| 14 \| 239 \| Peptidase S24, S26A and S26B \| Peptidase S24, S26A and S26B; afdb-uniprot50 \| AF-A0A7Y4UN23-F1-MODEL\_V4 \| 1.0 \| 2.245e-09 \| 253 \| 0.242 \| 198 \| 121 \| 6 \| 4 \| 180 \| 78 \| 267 \| S24 family peptidase \| S24 family peptidase; afdb-uniprot50 \| AF-A0A0P7CW07-F1-MODEL\_V4 \| 1.0 \| 9.03e-09 \| 253 \| 0.163 \| 239 \| 137 \| 6 \| 5 \| 180 \| 29 \| 267 \| Transcriptional regulator \| Transcriptional regulator; afdb-uniprot50 \| AF-A0A4R6DX28-F1-MODEL\_V4 \| 1.0 \| 4.503e-09 \| 253 \| 0.153 \| 260 \| 134 \| 8 \| 4 \| 178 \| 19 \| 277 \| Peptidase S24-like protein \| Peptidase S24-like protein; afdb-uniprot50 \| AF-A0A1H2N8M0-F1-MODEL\_V4 \| 1.0 \| 2.672e-09 \| 253 \| 0.136 \| 264 \| 138 \| 5 \| 6 \| 180 \| 23 \| 285 \| Helix-turn-helix \| Helix-turn-helix; afdb-uniprot50 \| AF-A0A3S4MVN3-F1-MODEL\_V4 \| 1.0 \| 4.01e-09 \| 253 \| 0.209 \| 244 \| 124 \| 10 \| 4 \| 180 \| 66 \| 307 \| CI repressor protein \| CI repressor protein; afdb-uniprot50 \| AF-S6WAL0-F1-MODEL\_V4 \| 1.0 \| 2.104e-06 \| 252 \| 0.21 \| 114 \| 81 \| 2 \| 72 \| 180 \| 1 \| 110 \| Transcriptional regulator \| Transcriptional regulator; afdb-uniprot50 \| AF-A0A1X0T0I3-F1-MODEL\_V4 \| 1.0 \| 1.112e-06 \| 252 \| 0.225 \| 124 \| 88 \| 3 \| 61 \| 180 \| 13 \| 132 \| Phage repressor \| Phage repressor; afdb-uniprot50 \| AF-A0A5R9NXX0-F1-MODEL\_V4 \| 1.0 \| 8.521e-09 \| 252 \| 0.18 \| 183 \| 131 \| 7 \| 13 \| 180 \| 3 \| 181 \| XRE family transcriptional regulator \| XRE family transcriptional regulator; afdb-uniprot50 \| AF-A0A839ILY4-F1-MODEL\_V4 \| 1.0 \| 1.139e-08 \| 252 \| 0.184 \| 228 \| 133 \| 6 \| 4 \| 180 \| 11 \| 236 \| Helix-turn-helix transcriptional regulator \| Helix-turn-helix transcriptional regulator; afdb-uniprot50 \| AF-A0A2N9XC71-F1-MODEL\_V4 \| 1.0 \| 6.017e-09 \| 252 \| 0.167 \| 233 \| 137 \| 8 \| 4 \| 180 \| 8 \| 239 \| Chromophore lyase \| Chromophore lyase; afdb-uniprot50 \| AF-A0A2Z6EVF8-F1-MODEL\_V4 \| 1.0 \| 4.249e-09 \| 252 \| 0.178 \| 219 \| 132 \| 9 \| 4 \| 179 \| 28 \| 241 \| Peptidase S24, S26A and S26B with Helix-turn-helix motif \| Peptidase S24, S26A and S26B with Helix-turn-helix motif; afdb-uniprot50 \| AF-A0A1G4ZS22-F1-MODEL\_V4 \| 1.0 \| 1.279e-08 \| 252 \| 0.148 \| 229 \| 141 \| 4 \| 4 \| 180 \| 9 \| 235 \| Phage repressor protein C, contains Cro/C1-type HTH and peptisase s24 domains \| Phage repressor protein C, contains Cro/C1-type HTH and peptisase s24 domains; afdb-uniprot50 \| AF-A0A7W8HG37-F1-MODEL\_V4 \| 1.0 \| 2.672e-09 \| 252 \| 0.168 \| 249 \| 133 \| 8 \| 5 \| 180 \| 12 \| 259 \| Phage repressor protein C with HTH and peptisase S24 domain \| Phage repressor protein C with HTH and peptisase S24 domain; afdb-uniprot50 \| AF-W6WJM4-F1-MODEL\_V4 \| 1.0 \| 5.056e-09 \| 252 \| 0.165 \| 224 \| 139 \| 5 \| 4 \| 180 \| 37 \| 259 \| Putative phage repressor \| Putative phage repressor; afdb-uniprot50 \| AF-K2C6Z1-F1-MODEL\_V4 \| 1.0 \| 2.155e-08 \| 252 \| 0.173 \| 231 \| 134 \| 6 \| 4 \| 180 \| 31 \| 258 \| HTH cro/C1-type domain-containing protein \| HTH cro/C1-type domain-containing protein; afdb-uniprot50 \| AF-A0A0M7LNV7-F1-MODEL\_V4 \| 1.0 \| 5.678e-09 \| 252 \| 0.187 \| 219 \| 135 \| 7 \| 4 \| 180 \| 54 \| 271 \| DNA polymerase V subunit UmuD \| DNA polymerase V subunit UmuD; afdb-uniprot50 \| AF-A0A2S8Y6M7-F1-MODEL\_V4 \| 1.0 \| 1.78e-09 \| 252 \| 0.138 \| 267 \| 136 \| 7 \| 7 \| 180 \| 24 \| 289 \| Propanediol utilization protein \| Propanediol utilization protein; afdb-uniprot50 \| AF-A0A833N995-F1-MODEL\_V4 \| 1.0 \| 8.041e-09 \| 252 \| 0.146 \| 314 \| 134 \| 6 \| 1 \| 180 \| 61 \| 374 \| Phage cI repressor (ACLAME 5) \| Phage cI repressor (ACLAME 5); afdb-uniprot50 \| AF-A0A1K1W354-F1-MODEL\_V4 \| 1.0 \| 8.521e-09 \| 251 \| 0.23 \| 182 \| 93 \| 5 \| 41 \| 180 \| 3 \| 179 \| Phage repressor protein C, contains Cro/C1-type HTH and peptisase s24 domains \| Phage repressor protein C, contains Cro/C1-type HTH and peptisase s24 domains; afdb-uniprot50 \| AF-U1ZQ79-F1-MODEL\_V4 \| 1.0 \| 8.461e-06 \| 251 \| 0.25 \| 100 \| 68 \| 3 \| 87 \| 179 \| 91 \| 190 \| Peptidase\_S24 domain-containing protein \| Peptidase\_S24 domain-containing protein; afdb-uniprot50 \| AF-A0A7S9UK10-F1-MODEL\_V4 \| 1.0 \| 6.709e-06 \| 251 \| 0.223 \| 103 \| 78 \| 2 \| 79 \| 180 \| 98 \| 199 \| S24 family peptidase \| S24 family peptidase; afdb-uniprot50 \| AF-A0A0A7S0B9-F1-MODEL\_V4 \| 1.0 \| 8.521e-09 \| 251 \| 0.202 \| 222 \| 124 \| 9 \| 8 \| 180 \| 3 \| 220 \| Putative transcriptional regulator \| Putative transcriptional regulator; afdb-uniprot50 \| AF-A0A2E6QFD6-F1-MODEL\_V4 \| 1.0 \| 1.014e-08 \| 251 \| 0.186 \| 215 \| 131 \| 7 \| 4 \| 180 \| 16 \| 224 \| HTH cro/C1-type domain-containing protein \| HTH cro/C1-type domain-containing protein; afdb-uniprot50 \| AF-A0A171KSF2-F1-MODEL\_V4 \| 1.0 \| 5.056e-09 \| 251 \| 0.173 \| 219 \| 131 \| 9 \| 7 \| 180 \| 6 \| 219 \| HTH cro/C1-type domain-containing protein \| HTH cro/C1-type domain-containing protein; afdb-uniprot50 \| AF-A0A3R8XKJ5-F1-MODEL\_V4 \| 1.0 \| 6.377e-09 \| 251 \| 0.159 \| 232 \| 129 \| 6 \| 10 \| 180 \| 2 \| 228 \| XRE family transcriptional regulator \| XRE family transcriptional regulator; afdb-uniprot50 \| AF-A0A3S0YRJ1-F1-MODEL\_V4 \| 1.0 \| 8.041e-09 \| 251 \| 0.194 \| 226 \| 130 \| 8 \| 4 \| 180 \| 9 \| 231 \| LexA family transcriptional regulator \| LexA family transcriptional regulator; afdb-uniprot50 \| AF-A0A1G6NPZ1-F1-MODEL\_V4 \| 1.0 \| 6.377e-09 \| 251 \| 0.159 \| 245 \| 134 \| 6 \| 5 \| 179 \| 12 \| 254 \| Phage repressor protein C, contains Cro/C1-type HTH and peptisase s24 domains \| Phage repressor protein C, contains Cro/C1-type HTH and peptisase s24 domains; afdb-uniprot50 \| AF-A0A2G4E5F7-F1-MODEL\_V4 \| 1.0 \| 9.97e-10 \| 251 \| 0.174 \| 269 \| 128 \| 8 \| 5 \| 180 \| 3 \| 270 \| Cro/Cl family transcriptional regulator \| Cro/Cl family transcriptional regulator; afdb-uniprot50 \| AF-A0A8A8PHC1-F1-MODEL\_V4 \| 1.0 \| 1.919e-08 \| 251 \| 0.186 \| 263 \| 127 \| 7 \| 4 \| 180 \| 29 \| 290 \| Helix-turn-helix transcriptional regulator \| Helix-turn-helix transcriptional regulator; afdb-uniprot50 \| AF-A0A8B4I5H6-F1-MODEL\_V4 \| 1.0 \| 8.521e-09 \| 251 \| 0.168 \| 237 \| 135 \| 6 \| 4 \| 180 \| 92 \| 326 \| Putative phage repressor \| Putative phage repressor; afdb-uniprot50 \| AF-A0A2N9Y4B0-F1-MODEL\_V4 \| 1.0 \| 7.85e-07 \| 250 \| 0.184 \| 125 \| 93 \| 4 \| 57 \| 178 \| 8 \| 126 \| Peptidase\_S24 domain-containing protein \| Peptidase\_S24 domain-containing protein; afdb-uniprot50 \| AF-A0A0V8T744-F1-MODEL\_V4 \| 1.0 \| 4.219e-06 \| 250 \| 0.152 \| 105 \| 85 \| 2 \| 79 \| 180 \| 82 \| 185 \| Cro/Cl family transcriptional regulator \| Cro/Cl family transcriptional regulator; afdb-uniprot50 \| AF-C0EP73-F1-MODEL\_V4 \| 1.0 \| 4.772e-09 \| 250 \| 0.17 \| 211 \| 131 \| 7 \| 10 \| 180 \| 5 \| 211 \| Peptidase S24-like protein \| Peptidase S24-like protein; afdb-uniprot50 \| AF-A0A1G5V1P4-F1-MODEL\_V4 \| 1.0 \| 1.075e-08 \| 250 \| 0.181 \| 226 \| 137 \| 7 \| 1 \| 180 \| 2 \| 225 \| Phage repressor protein C, contains Cro/C1-type HTH and peptisase s24 domains \| Phage repressor protein C, contains Cro/C1-type HTH and peptisase s24 domains; afdb-uniprot50 \| AF-A0A1D2QRN9-F1-MODEL\_V4 \| 1.0 \| 4.772e-09 \| 250 \| 0.214 \| 224 \| 124 \| 8 \| 5 \| 179 \| 7 \| 227 \| Peptidase\_S24 domain-containing protein \| Peptidase\_S24 domain-containing protein; afdb-uniprot50 \| AF-A0A6N8AT90-F1-MODEL\_V4 \| 1.0 \| 4.01e-09 \| 250 \| 0.196 \| 229 \| 124 \| 10 \| 5 \| 180 \| 11 \| 232 \| Helix-turn-helix transcriptional regulator \| Helix-turn-helix transcriptional regulator; afdb-uniprot50 \| AF-A0A530R6J1-F1-MODEL\_V4 \| 1.0 \| 1.522e-08 \| 250 \| 0.185 \| 221 \| 133 \| 8 \| 5 \| 180 \| 26 \| 244 \| Helix-turn-helix domain-containing protein \| Helix-turn-helix domain-containing protein; afdb-uniprot50 \| AF-A0A1B8Q949-F1-MODEL\_V4 \| 1.0 \| 4.249e-09 \| 250 \| 0.146 \| 245 \| 136 \| 7 \| 5 \| 178 \| 5 \| 247 \| HTH cro/C1-type domain-containing protein \| HTH cro/C1-type domain-containing protein; afdb-uniprot50 \| AF-A0A7U3YNF5-F1-MODEL\_V4 \| 1.0 \| 1.207e-08 \| 250 \| 0.16 \| 243 \| 135 \| 6 \| 4 \| 180 \| 6 \| 245 \| Putative phage repressor \| Putative phage repressor; afdb-uniprot50 \| AF-A0A433C752-F1-MODEL\_V4 \| 1.0 \| 2.245e-09 \| 250 \| 0.188 \| 202 \| 137 \| 8 \| 4 \| 180 \| 47 \| 246 \| S24 family peptidase \| S24 family peptidase; afdb-uniprot50 \| AF-A0A2S7E1D1-F1-MODEL\_V4 \| 1.0 \| 4.01e-09 \| 250 \| 0.182 \| 241 \| 132 \| 5 \| 5 \| 180 \| 35 \| 275 \| HTH cro/C1-type domain-containing protein \| HTH cro/C1-type domain-containing protein; afdb-uniprot50 \| AF-A0A495IUG7-F1-MODEL\_V4 \| 1.0 \| 2.155e-08 \| 250 \| 0.139 \| 323 \| 132 \| 5 \| 4 \| 180 \| 22 \| 344 \| Phage repressor protein C with HTH and peptisase S24 domain \| Phage repressor protein C with HTH and peptisase S24 domain; afdb-uniprot50 \| AF-A0A837MMU5-F1-MODEL\_V4 \| 1.0 \| 5.975e-06 \| 249 \| 0.194 \| 108 \| 82 \| 1 \| 78 \| 180 \| 90 \| 197 \| Transcriptional regulator \| Transcriptional regulator; afdb-uniprot50 \| AF-A0A2D5S0G5-F1-MODEL\_V4 \| 1.0 \| 5.358e-09 \| 249 \| 0.199 \| 231 \| 128 \| 6 \| 5 \| 180 \| 9 \| 237 \| Transcriptional regulator \| Transcriptional regulator; afdb-uniprot50 \| AF-A0A6N8RY26-F1-MODEL\_V4 \| 1.0 \| 4.58e-08 \| 249 \| 0.17 \| 234 \| 136 \| 7 \| 4 \| 180 \| 18 \| 250 \| Helix-turn-helix domain-containing protein \| Helix-turn-helix domain-containing protein; afdb-uniprot50 \| AF-A0A7T8M381-F1-MODEL\_V4 \| 1.0 \| 1.049e-06 \| 248 \| 0.258 \| 120 \| 73 \| 4 \| 64 \| 180 \| 6 \| 112 \| S24 family peptidase \| S24 family peptidase; afdb-uniprot50 \| AF-A0A4S1MQ12-F1-MODEL\_V4 \| 1.0 \| 2.019e-05 \| 248 \| 0.184 \| 92 \| 73 \| 2 \| 89 \| 180 \| 27 \| 116 \| Uncharacterized protein \| Uncharacterized protein; afdb-uniprot50 \| AF-A0A4R7VYY6-F1-MODEL\_V4 \| 1.0 \| 3.981e-06 \| 248 \| 0.224 \| 116 \| 81 \| 2 \| 70 \| 180 \| 26 \| 137 \| Peptidase S24-like protein \| Peptidase S24-like protein; afdb-uniprot50 \| AF-A0A7G2JK01-F1-MODEL\_V4 \| 1.0 \| 1.355e-08 \| 248 \| 0.199 \| 211 \| 120 \| 9 \| 15 \| 180 \| 3 \| 209 \| Putative HTH-type transcriptional regulator \| Putative HTH-type transcriptional regulator; afdb-uniprot50 \| AF-A0A2X1U872-F1-MODEL\_V4 \| 1.0 \| 4.772e-09 \| 248 \| 0.173 \| 213 \| 133 \| 8 \| 5 \| 176 \| 11 \| 221 \| Phage repressor protein, phage associated protein \| Phage repressor protein, phage associated protein; afdb-uniprot50 \| AF-A0A3B7M1G8-F1-MODEL\_V4 \| 1.0 \| 1.075e-08 \| 248 \| 0.179 \| 223 \| 134 \| 5 \| 5 \| 180 \| 11 \| 231 \| Helix-turn-helix transcriptional regulator \| Helix-turn-helix transcriptional regulator; afdb-uniprot50 \| AF-N8XK06-F1-MODEL\_V4 \| 1.0 \| 1.014e-08 \| 248 \| 0.145 \| 227 \| 144 \| 5 \| 1 \| 180 \| 14 \| 237 \| HTH cro/C1-type domain-containing protein \| HTH cro/C1-type domain-containing protein; afdb-uniprot50 \| AF-A0A258NSS6-F1-MODEL\_V4 \| 1.0 \| 1.355e-08 \| 248 \| 0.222 \| 225 \| 129 \| 6 \| 2 \| 180 \| 58 \| 282 \| XRE family transcriptional regulator \| XRE family transcriptional regulator; afdb-uniprot50 \| AF-A0A844RU21-F1-MODEL\_V4 \| 1.0 \| 6.017e-09 \| 248 \| 0.211 \| 208 \| 121 \| 7 \| 4 \| 172 \| 49 \| 252 \| Helix-turn-helix domain-containing protein \| Helix-turn-helix domain-containing protein; afdb-uniprot50 \| AF-A0A2P5MV31-F1-MODEL\_V4 \| 1.0 \| 1.139e-08 \| 248 \| 0.158 \| 290 \| 129 \| 8 \| 4 \| 180 \| 6 \| 293 \| HTH cro/C1-type domain-containing protein \| HTH cro/C1-type domain-containing protein; afdb-uniprot50 \| AF-A0A178J3C6-F1-MODEL\_V4 \| 1.0 \| 4.503e-09 \| 247 \| 0.187 \| 197 \| 121 \| 8 \| 14 \| 180 \| 3 \| 190 \| HTH cro/C1-type domain-containing protein \| HTH cro/C1-type domain-containing protein; afdb-uniprot50 \| AF-A0A833L7F1-F1-MODEL\_V4 \| 1.0 \| 1.522e-08 \| 247 \| 0.184 \| 206 \| 133 \| 6 \| 4 \| 180 \| 9 \| 208 \| Uncharacterized protein \| Uncharacterized protein; afdb-uniprot50 \| AF-A0A1H8RCZ9-F1-MODEL\_V4 \| 1.0 \| 5.358e-09 \| 247 \| 0.175 \| 233 \| 139 \| 7 \| 1 \| 180 \| 2 \| 234 \| Peptidase S24-like \| Peptidase S24-like; afdb-uniprot50 \| AF-A0A7W6RC19-F1-MODEL\_V4 \| 1.0 \| 2.155e-08 \| 247 \| 0.186 \| 241 \| 121 \| 6 \| 14 \| 180 \| 1 \| 240 \| Phage repressor protein C with HTH and peptisase S24 domain \| Phage repressor protein C with HTH and peptisase S24 domain; afdb-uniprot50 \| AF-B2I5H5-F1-MODEL\_V4 \| 1.0 \| 6.377e-09 \| 247 \| 0.172 \| 243 \| 132 \| 9 \| 4 \| 178 \| 7 \| 248 \| Putative phage repressor \| Putative phage repressor; afdb-uniprot50 \| AF-A0A6G6IUW8-F1-MODEL\_V4 \| 1.0 \| 1.279e-08 \| 247 \| 0.159 \| 245 \| 137 \| 5 \| 5 \| 180 \| 10 \| 254 \| Helix-turn-helix domain-containing protein \| Helix-turn-helix domain-containing protein; afdb-uniprot50 \| AF-A0A2E6JYY0-F1-MODEL\_V4 \| 1.0 \| 1.355e-08 \| 247 \| 0.146 \| 226 \| 142 \| 4 \| 5 \| 180 \| 34 \| 258 \| HTH cro/C1-type domain-containing protein \| HTH cro/C1-type domain-containing protein; afdb-uniprot50 \| AF-A0A7Y4W621-F1-MODEL\_V4 \| 1.0 \| 5.358e-09 \| 247 \| 0.182 \| 257 \| 121 \| 10 \| 12 \| 180 \| 11 \| 266 \| Helix-turn-helix transcriptional regulator \| Helix-turn-helix transcriptional regulator; afdb-uniprot50 \| AF-A0A143ZKM8-F1-MODEL\_V4 \| 1.0 \| 1.014e-08 \| 246 \| 0.2 \| 204 \| 124 \| 5 \| 16 \| 180 \| 6 \| 209 \| Phage repressor protein C, contains Cro/C1-type HTH and peptisase s24 domains \| Phage repressor protein C, contains Cro/C1-type HTH and peptisase s24 domains; afdb-uniprot50 \| AF-A0A2D3WEV3-F1-MODEL\_V4 \| 1.0 \| 6.757e-09 \| 246 \| 0.187 \| 208 \| 133 \| 5 \| 5 \| 180 \| 15 \| 218 \| Phage repressor protein \| Phage repressor protein; afdb-uniprot50 \| AF-C3X7V2-F1-MODEL\_V4 \| 1.0 \| 1.613e-08 \| 246 \| 0.208 \| 216 \| 130 \| 6 \| 4 \| 180 \| 6 \| 219 \| Peptidase S24-like protein \| Peptidase S24-like protein; afdb-uniprot50 \| AF-A0A126Z9R7-F1-MODEL\_V4 \| 1.0 \| 5.056e-09 \| 246 \| 0.158 \| 221 \| 139 \| 4 \| 5 \| 180 \| 14 \| 232 \| Peptidase\_S24 domain-containing protein \| Peptidase\_S24 domain-containing protein; afdb-uniprot50 \| AF-A0A1U9JXL2-F1-MODEL\_V4 \| 1.0 \| 5.45e-08 \| 246 \| 0.156 \| 224 \| 142 \| 7 \| 4 \| 180 \| 7 \| 230 \| Peptidase\_S24 domain-containing protein \| Peptidase\_S24 domain-containing protein; afdb-uniprot50 \| AF-A0A832UN00-F1-MODEL\_V4 \| 1.0 \| 2.565e-08 \| 246 \| 0.2 \| 235 \| 124 \| 6 \| 5 \| 180 \| 10 \| 239 \| Uncharacterized protein \| Uncharacterized protein; afdb-uniprot50 \| AF-A0A068T867-F1-MODEL\_V4 \| 1.0 \| 2.42e-08 \| 245 \| 0.191 \| 204 \| 121 \| 6 \| 16 \| 179 \| 5 \| 204 \| Putative transcriptional regulator, repressor \| Putative transcriptional regulator, repressor; afdb-uniprot50 \| AF-A0A0Q9EUS2-F1-MODEL\_V4 \| 1.0 \| 1.522e-08 \| 245 \| 0.142 \| 231 \| 142 \| 8 \| 5 \| 179 \| 5 \| 235 \| HTH cro/C1-type domain-containing protein \| HTH cro/C1-type domain-containing protein; afdb-uniprot50 \| AF-A0A1H3KSL9-F1-MODEL\_V4 \| 1.0 \| 2.88e-08 \| 245 \| 0.137 \| 261 \| 137 \| 6 \| 4 \| 180 \| 8 \| 264 \| Peptidase S24-like \| Peptidase S24-like; afdb-uniprot50 \| AF-A0A2N2KCH0-F1-MODEL\_V4 \| 1.0 \| 8.041e-09 \| 245 \| 0.148 \| 263 \| 137 \| 5 \| 5 \| 180 \| 13 \| 275 \| Peptidase\_S24 domain-containing protein \| Peptidase\_S24 domain-containing protein; afdb-uniprot50 \| AF-A0A327JDI9-F1-MODEL\_V4 \| 1.0 \| 2.718e-08 \| 244 \| 0.139 \| 208 \| 140 \| 7 \| 4 \| 180 \| 5 \| 204 \| Peptidase\_S24 domain-containing protein \| Peptidase\_S24 domain-containing protein; afdb-uniprot50 \| AF-A0A4Z0BX65-F1-MODEL\_V4 \| 1.0 \| 1.613e-08 \| 244 \| 0.152 \| 210 \| 143 \| 3 \| 4 \| 180 \| 19 \| 226 \| Helix-turn-helix transcriptional regulator \| Helix-turn-helix transcriptional regulator; afdb-uniprot50 \| AF-A0A1H1UI98-F1-MODEL\_V4 \| 1.0 \| 2.88e-08 \| 244 \| 0.165 \| 242 \| 132 \| 7 \| 4 \| 180 \| 12 \| 248 \| Peptidase\_S24 domain-containing protein \| Peptidase\_S24 domain-containing protein; afdb-uniprot50 \| AF-J2XH41-F1-MODEL\_V4 \| 1.0 \| 1.709e-08 \| 244 \| 0.155 \| 232 \| 138 \| 4 \| 4 \| 179 \| 76 \| 305 \| Putative transcriptional regulator \| Putative transcriptional regulator; afdb-uniprot50 \| AF-A0A1M7LJF6-F1-MODEL\_V4 \| 1.0 \| 7.588e-09 \| 244 \| 0.129 \| 254 \| 131 \| 5 \| 16 \| 180 \| 116 \| 368 \| Peptidase S24-like \| Peptidase S24-like; afdb-uniprot50 \| AF-A0A0G9JXZ6-F1-MODEL\_V4 \| 1.0 \| 2.034e-08 \| 243 \| 0.197 \| 197 \| 121 \| 6 \| 19 \| 180 \| 3 \| 197 \| Phage repressor protein \| Phage repressor protein; afdb-uniprot50 \| AF-A0A1Q4CNZ7-F1-MODEL\_V4 \| 1.0 \| 2.034e-08 \| 243 \| 0.172 \| 214 \| 120 \| 8 \| 6 \| 180 \| 2 \| 197 \| HTH cro/C1-type domain-containing protein \| HTH cro/C1-type domain-containing protein; afdb-uniprot50 \| AF-A0A378CDN0-F1-MODEL\_V4 \| 1.0 \| 1.436e-08 \| 243 \| 0.233 \| 193 \| 109 \| 6 \| 26 \| 180 \| 4 \| 195 \| Putative phage repressor protein CI \| Putative phage repressor protein CI; afdb-uniprot50 \| AF-A0A1B9LJP2-F1-MODEL\_V4 \| 1.0 \| 3.052e-08 \| 243 \| 0.177 \| 220 \| 131 \| 5 \| 4 \| 180 \| 10 \| 222 \| Transcriptional regulator \| Transcriptional regulator; afdb-uniprot50 \| AF-A0A7U5AQA5-F1-MODEL\_V4 \| 1.0 \| 1.436e-08 \| 243 \| 0.181 \| 209 \| 136 \| 8 \| 5 \| 180 \| 28 \| 234 \| Peptidase S24-like family protein \| Peptidase S24-like family protein; afdb-uniprot50 \| AF-A0A1F0J3A6-F1-MODEL\_V4 \| 1.0 \| 1.279e-08 \| 243 \| 0.169 \| 230 \| 133 \| 7 \| 9 \| 180 \| 14 \| 243 \| Transcriptional regulator \| Transcriptional regulator; afdb-uniprot50 \| AF-A0A2R7RTB9-F1-MODEL\_V4 \| 1.0 \| 2.284e-08 \| 243 \| 0.15 \| 232 \| 139 \| 5 \| 4 \| 179 \| 9 \| 238 \| Transcriptional regulator \| Transcriptional regulator; afdb-uniprot50 \| AF-A0A855KF29-F1-MODEL\_V4 \| 1.0 \| 1.075e-08 \| 243 \| 0.18 \| 255 \| 130 \| 6 \| 4 \| 180 \| 6 \| 259 \| Phage repressor protein C \| Phage repressor protein C; afdb-uniprot50 \| AF-A0A015NCW0-F1-MODEL\_V4 \| 1.0 \| 7.161e-09 \| 242 \| 0.211 \| 232 \| 128 \| 10 \| 3 \| 180 \| 8 \| 238 \| HTH cro/C1-type domain-containing protein \| HTH cro/C1-type domain-containing protein; afdb-uniprot50 \| AF-A0A2V4FY31-F1-MODEL\_V4 \| 1.0 \| 3.052e-08 \| 242 \| 0.182 \| 235 \| 134 \| 5 \| 4 \| 180 \| 14 \| 248 \| Helix-turn-helix transcriptional regulator \| Helix-turn-helix transcriptional regulator; afdb-uniprot50 \| AF-A0A1S8DCK6-F1-MODEL\_V4 \| 1.0 \| 4.503e-09 \| 242 \| 0.207 \| 246 \| 124 \| 9 \| 5 \| 180 \| 3 \| 247 \| Peptidase\_S24 domain-containing protein \| Peptidase\_S24 domain-containing protein; afdb-uniprot50 \| AF-A0A6J4E657-F1-MODEL\_V4 \| 1.0 \| 7.588e-09 \| 242 \| 0.167 \| 245 \| 123 \| 7 \| 16 \| 180 \| 24 \| 267 \| HTH cro/C1-type domain-containing protein \| HTH cro/C1-type domain-containing protein; afdb-uniprot50 \| AF-A0A7X1TFZ1-F1-MODEL\_V4 \| 1.0 \| 8.041e-09 \| 242 \| 0.16 \| 281 \| 131 \| 11 \| 4 \| 180 \| 6 \| 285 \| Peptidase\_S24 domain-containing protein \| Peptidase\_S24 domain-containing protein; afdb-uniprot50 \| AF-A0A4Q7D2I1-F1-MODEL\_V4 \| 1.0 \| 3.632e-08 \| 242 \| 0.167 \| 215 \| 139 \| 6 \| 5 \| 180 \| 103 \| 316 \| Helix-turn-helix domain-containing protein \| Helix-turn-helix domain-containing protein; afdb-uniprot50 \| AF-A0A081B6G2-F1-MODEL\_V4 \| 1.0 \| 3.234e-08 \| 241 \| 0.202 \| 202 \| 121 \| 7 \| 16 \| 180 \| 7 \| 205 \| Putative phage repressor \| Putative phage repressor; afdb-uniprot50 \| AF-A0A388SDE2-F1-MODEL\_V4 \| 1.0 \| 1.436e-08 \| 241 \| 0.18 \| 194 \| 140 \| 4 \| 4 \| 180 \| 14 \| 205 \| HTH cro/C1-type domain-containing protein \| HTH cro/C1-type domain-containing protein; afdb-uniprot50 \| AF-A0A1C3GJA4-F1-MODEL\_V4 \| 1.0 \| 1.131e-05 \| 241 \| 0.205 \| 107 \| 80 \| 1 \| 79 \| 180 \| 83 \| 189 \| Pyocin repressor protein \| Pyocin repressor protein; afdb-uniprot50 \| AF-A0A1E7QYY0-F1-MODEL\_V4 \| 1.0 \| 9.569e-09 \| 241 \| 0.174 \| 241 \| 133 \| 7 \| 5 \| 180 \| 8 \| 247 \| Helix-turn-helix transcriptional regulator \| Helix-turn-helix transcriptional regulator; afdb-uniprot50 \| AF-A0A6S5P8N1-F1-MODEL\_V4 \| 1.0 \| 7.588e-09 \| 241 \| 0.156 \| 237 \| 138 \| 7 \| 5 \| 180 \| 15 \| 250 \| Transcriptional regulator \| Transcriptional regulator; afdb-uniprot50 \| AF-A0A7Y0VS83-F1-MODEL\_V4 \| 1.0 \| 2.284e-08 \| 241 \| 0.172 \| 232 \| 137 \| 6 \| 1 \| 179 \| 1 \| 230 \| Helix-turn-helix transcriptional regulator \| Helix-turn-helix transcriptional regulator; afdb-uniprot50 \| AF-A0A1F8VMR4-F1-MODEL\_V4 \| 1.0 \| 3.052e-08 \| 240 \| 0.211 \| 199 \| 120 \| 6 \| 13 \| 180 \| 4 \| 196 \| Peptidase\_S24 domain-containing protein \| Peptidase\_S24 domain-containing protein; afdb-uniprot50 \| AF-A5UES4-F1-MODEL\_V4 \| 1.0 \| 4.078e-08 \| 240 \| 0.179 \| 212 \| 125 \| 4 \| 18 \| 180 \| 2 \| 213 \| Predicted transcriptional regulator \| Predicted transcriptional regulator; afdb-uniprot50 \| AF-A0A1F0F535-F1-MODEL\_V4 \| 1.0 \| 1.919e-08 \| 240 \| 0.154 \| 233 \| 139 \| 7 \| 4 \| 180 \| 3 \| 233 \| Repressor \| Repressor; afdb-uniprot50 \| AF-A0A2T5HFF1-F1-MODEL\_V4 \| 1.0 \| 1.207e-08 \| 240 \| 0.206 \| 237 \| 127 \| 5 \| 5 \| 180 \| 3 \| 239 \| Phage repressor protein C with HTH and peptisase S24 domain \| Phage repressor protein C with HTH and peptisase S24 domain; afdb-uniprot50 \| AF-A0A809N3S2-F1-MODEL\_V4 \| 1.0 \| 1.014e-08 \| 240 \| 0.159 \| 232 \| 137 \| 10 \| 5 \| 180 \| 24 \| 253 \| Uncharacterized protein \| Uncharacterized protein; afdb-uniprot50 \| AF-A0A853GEI4-F1-MODEL\_V4 \| 1.0 \| 8.179e-08 \| 240 \| 0.151 \| 238 \| 141 \| 8 \| 4 \| 180 \| 18 \| 255 \| Helix-turn-helix transcriptional regulator \| Helix-turn-helix transcriptional regulator; afdb-uniprot50 \| AF-A0A246GY71-F1-MODEL\_V4 \| 1.0 \| 6.017e-09 \| 240 \| 0.132 \| 272 \| 139 \| 8 \| 5 \| 180 \| 3 \| 273 \| Phage repressor protein \| Phage repressor protein; afdb-uniprot50 \| AF-A0A1H8V9H3-F1-MODEL\_V4 \| 1.0 \| 1.279e-08 \| 240 \| 0.165 \| 247 \| 135 \| 8 \| 5 \| 180 \| 32 \| 278 \| Phage repressor protein C, contains Cro/C1-type HTH and peptisase s24 domains \| Phage repressor protein C, contains Cro/C1-type HTH and peptisase s24 domains; afdb-uniprot50 \| AF-A0A653E6P4-F1-MODEL\_V4 \| 1.0 \| 8.521e-09 \| 240 \| 0.155 \| 244 \| 138 \| 7 \| 5 \| 180 \| 94 \| 337 \| HTH-type transcriptional regulator PrtR (Modular protein) \| HTH-type transcriptional regulator PrtR (Modular protein); afdb-uniprot50 \| AF-A0A543HLB3-F1-MODEL\_V4 \| 1.0 \| 1.709e-08 \| 239 \| 0.168 \| 232 \| 134 \| 9 \| 4 \| 180 \| 7 \| 234 \| Phage repressor protein C with HTH and peptisase S24 domain \| Phage repressor protein C with HTH and peptisase S24 domain; afdb-uniprot50 \| AF-A0A6M3M6C4-F1-MODEL\_V4 \| 1.0 \| 2.284e-08 \| 239 \| 0.181 \| 237 \| 134 \| 6 \| 4 \| 180 \| 5 \| 241 \| Putative peptidase \| Putative peptidase; afdb-uniprot50 \| AF-E6QCB5-F1-MODEL\_V4 \| 1.0 \| 1.613e-08 \| 239 \| 0.182 \| 235 \| 128 \| 7 \| 4 \| 180 \| 14 \| 242 \| Putative 26 kDa repressor protein (Regulatory protein CI) \| Putative 26 kDa repressor protein (Regulatory protein CI); afdb-uniprot50 \| AF-A0A1M4WAA2-F1-MODEL\_V4 \| 1.0 \| 1.436e-08 \| 239 \| 0.181 \| 220 \| 124 \| 6 \| 16 \| 180 \| 25 \| 243 \| Peptidase S24-like \| Peptidase S24-like; afdb-uniprot50 \| AF-A0A329VHA2-F1-MODEL\_V4 \| 1.0 \| 1.709e-08 \| 239 \| 0.176 \| 232 \| 131 \| 8 \| 7 \| 180 \| 16 \| 245 \| Phage repressor protein \| Phage repressor protein; afdb-uniprot50 \| AF-A0A653E7Y4-F1-MODEL\_V4 \| 1.0 \| 1.198e-05 \| 238 \| 0.229 \| 109 \| 77 \| 2 \| 79 \| 180 \| 52 \| 160 \| HTH-type transcriptional regulator PrtR \| HTH-type transcriptional regulator PrtR; afdb-uniprot50 \| AF-A0A5A7N7S5-F1-MODEL\_V4 \| 1.0 \| 1.548e-07 \| 238 \| 0.189 \| 179 \| 102 \| 8 \| 41 \| 180 \| 2 \| 176 \| Peptidase\_S24 domain-containing protein \| Peptidase\_S24 domain-containing protein; afdb-uniprot50 \| AF-A0A6I1NN78-F1-MODEL\_V4 \| 1.0 \| 1.426e-05 \| 238 \| 0.214 \| 107 \| 79 \| 1 \| 79 \| 180 \| 94 \| 200 \| Transcriptional regulator \| Transcriptional regulator; afdb-uniprot50 \| AF-A0A359LPK0-F1-MODEL\_V4 \| 1.0 \| 3.234e-08 \| 238 \| 0.171 \| 204 \| 139 \| 7 \| 4 \| 180 \| 9 \| 209 \| Peptidase\_S24 domain-containing protein \| Peptidase\_S24 domain-containing protein; afdb-uniprot50 \| AF-A0A222FJU3-F1-MODEL\_V4 \| 1.0 \| 3.571e-09 \| 238 \| 0.194 \| 211 \| 132 \| 7 \| 6 \| 180 \| 9 \| 217 \| LexA family transcriptional repressor \| LexA family transcriptional repressor; afdb-uniprot50 \| AF-G4QCW3-F1-MODEL\_V4 \| 1.0 \| 2.565e-08 \| 238 \| 0.153 \| 222 \| 139 \| 8 \| 5 \| 180 \| 4 \| 222 \| Putative phage repressor \| Putative phage repressor; afdb-uniprot50 \| AF-A0A7U3BSK1-F1-MODEL\_V4 \| 1.0 \| 2.565e-08 \| 238 \| 0.186 \| 220 \| 136 \| 7 \| 4 \| 180 \| 19 \| 238 \| Transcriptional regulator \| Transcriptional regulator; afdb-uniprot50 \| AF-A0A7D5DA53-F1-MODEL\_V4 \| 1.0 \| 3.427e-08 \| 238 \| 0.134 \| 230 \| 144 \| 4 \| 4 \| 180 \| 7 \| 234 \| Helix-turn-helix transcriptional regulator \| Helix-turn-helix transcriptional regulator; afdb-uniprot50 \| AF-A0A7S9VER3-F1-MODEL\_V4 \| 1.0 \| 1.709e-08 \| 238 \| 0.173 \| 242 \| 131 \| 5 \| 5 \| 177 \| 4 \| 245 \| Helix-turn-helix transcriptional regulator \| Helix-turn-helix transcriptional regulator; afdb-uniprot50 \| AF-A0A2W5EWM8-F1-MODEL\_V4 \| 1.0 \| 1.522e-08 \| 238 \| 0.176 \| 250 \| 133 \| 8 \| 4 \| 180 \| 14 \| 263 \| XRE family transcriptional regulator \| XRE family transcriptional regulator; afdb-uniprot50 \| AF-A0A285N7N5-F1-MODEL\_V4 \| 1.0 \| 4.772e-09 \| 238 \| 0.172 \| 267 \| 132 \| 10 \| 1 \| 180 \| 1 \| 265 \| Phage repressor protein C, contains Cro/C1-type HTH and peptisase s24 domains \| Phage repressor protein C, contains Cro/C1-type HTH and peptisase s24 domains; afdb-uniprot50 \| AF-B9TGR6-F1-MODEL\_V4 \| 1.0 \| 1.139e-08 \| 238 \| 0.19 \| 221 \| 125 \| 8 \| 5 \| 174 \| 21 \| 238 \| HTH cro/C1-type domain-containing protein \| HTH cro/C1-type domain-containing protein; afdb-uniprot50 \| AF-A0A6I4YRF4-F1-MODEL\_V4 \| 1.0 \| 1.139e-08 \| 238 \| 0.207 \| 226 \| 127 \| 10 \| 4 \| 180 \| 133 \| 355 \| Helix-turn-helix domain-containing protein \| Helix-turn-helix domain-containing protein; afdb-uniprot50 \| AF-A0A2R8CKN3-F1-MODEL\_V4 \| 1.0 \| 1.139e-08 \| 237 \| 0.173 \| 207 \| 134 \| 9 \| 4 \| 180 \| 7 \| 206 \| HTH cro/C1-type domain-containing protein \| HTH cro/C1-type domain-containing protein; afdb-uniprot50 \| AF-A0A292SJV4-F1-MODEL\_V4 \| 1.0 \| 2.42e-08 \| 237 \| 0.186 \| 209 \| 127 \| 5 \| 4 \| 179 \| 7 \| 205 \| Helix-turn-helix domain-containing protein \| Helix-turn-helix domain-containing protein; afdb-uniprot50 \| AF-A0A345PBQ8-F1-MODEL\_V4 \| 1.0 \| 2.718e-08 \| 237 \| 0.144 \| 214 \| 137 \| 4 \| 13 \| 180 \| 5 \| 218 \| Helix-turn-helix transcriptional regulator \| Helix-turn-helix transcriptional regulator; afdb-uniprot50 \| AF-A0A427G804-F1-MODEL\_V4 \| 1.0 \| 2.718e-08 \| 237 \| 0.159 \| 232 \| 129 \| 6 \| 10 \| 180 \| 2 \| 228 \| XRE family transcriptional regulator \| XRE family transcriptional regulator; afdb-uniprot50 \| AF-A0A853SDR3-F1-MODEL\_V4 \| 1.0 \| 3.052e-08 \| 237 \| 0.161 \| 204 \| 140 \| 7 \| 4 \| 180 \| 29 \| 228 \| Uncharacterized protein \| Uncharacterized protein; afdb-uniprot50 \| AF-A0A241W2E3-F1-MODEL\_V4 \| 1.0 \| 1.014e-08 \| 237 \| 0.161 \| 223 \| 132 \| 9 \| 4 \| 180 \| 53 \| 266 \| HTH cro/C1-type domain-containing protein \| HTH cro/C1-type domain-containing protein; afdb-uniprot50 \| AF-A0A2S9API7-F1-MODEL\_V4 \| 1.0 \| 2.565e-08 \| 237 \| 0.126 \| 276 \| 141 \| 8 \| 4 \| 180 \| 6 \| 280 \| HTH cro/C1-type domain-containing protein \| HTH cro/C1-type domain-containing protein; afdb-uniprot50 \| AF-A0A5E6XVR5-F1-MODEL\_V4 \| 1.0 \| 2.42e-08 \| 237 \| 0.16 \| 230 \| 135 \| 5 \| 4 \| 177 \| 9 \| 236 \| HTH-type transcriptional regulator PrtR \| HTH-type transcriptional regulator PrtR; afdb-uniprot50 \| AF-A0A2W0ES33-F1-MODEL\_V4 \| 1.0 \| 9.03e-09 \| 237 \| 0.141 \| 275 \| 136 \| 8 \| 5 \| 180 \| 11 \| 284 \| Cro/Cl family transcriptional regulator \| Cro/Cl family transcriptional regulator; afdb-uniprot50 \| AF-A0A212IUA3-F1-MODEL\_V4 \| 1.0 \| 5.143e-08 \| 236 \| 0.206 \| 199 \| 111 \| 7 \| 24 \| 180 \| 3 \| 196 \| Putative Phage repressor \| Putative Phage repressor; afdb-uniprot50 \| AF-A0A4R1PLM3-F1-MODEL\_V4 \| 1.0 \| 8.521e-09 \| 236 \| 0.148 \| 216 \| 125 \| 7 \| 7 \| 180 \| 24 \| 222 \| Phage repressor protein C with HTH and peptisase S24 domain \| Phage repressor protein C with HTH and peptisase S24 domain; afdb-uniprot50 \| AF-A0A2Z4PXZ2-F1-MODEL\_V4 \| 1.0 \| 6.757e-09 \| 236 \| 0.183 \| 224 \| 124 \| 10 \| 14 \| 180 \| 4 \| 225 \| Cro/Cl family transcriptional regulator \| Cro/Cl family transcriptional regulator; afdb-uniprot50 \| AF-A0A080NQX9-F1-MODEL\_V4 \| 1.0 \| 6.486e-08 \| 236 \| 0.175 \| 217 \| 138 \| 7 \| 5 \| 180 \| 19 \| 235 \| Peptidase S24-like family protein \| Peptidase S24-like family protein; afdb-uniprot50 \| AF-A0A286BZZ3-F1-MODEL\_V4 \| 1.0 \| 6.873e-08 \| 236 \| 0.156 \| 236 \| 136 \| 6 \| 4 \| 180 \| 6 \| 237 \| Phage repressor protein C, contains Cro/C1-type HTH and peptisase s24 domains \| Phage repressor protein C, contains Cro/C1-type HTH and peptisase s24 domains; afdb-uniprot50 \| AF-A0A1I7KDD2-F1-MODEL\_V4 \| 1.0 \| 3.052e-08 \| 236 \| 0.14 \| 250 \| 138 \| 7 \| 5 \| 180 \| 5 \| 251 \| Peptidase S24-like \| Peptidase S24-like; afdb-uniprot50 \| AF-A0A3D2LZR0-F1-MODEL\_V4 \| 1.0 \| 1.873e-06 \| 235 \| 0.153 \| 124 \| 100 \| 2 \| 62 \| 180 \| 2 \| 125 \| Peptidase\_S24 domain-containing protein \| Peptidase\_S24 domain-containing protein; afdb-uniprot50 \| AF-A0A3G6YKD9-F1-MODEL\_V4 \| 1.0 \| 3.346e-06 \| 235 \| 0.195 \| 128 \| 90 \| 4 \| 61 \| 180 \| 4 \| 126 \| Helix-turn-helix transcriptional regulator \| Helix-turn-helix transcriptional regulator; afdb-uniprot50 \| AF-A0A837E4W0-F1-MODEL\_V4 \| 1.0 \| 1.574e-06 \| 235 \| 0.186 \| 129 \| 95 \| 3 \| 62 \| 180 \| 38 \| 166 \| Uncharacterized protein \| Uncharacterized protein; afdb-uniprot50 \| AF-A9H6C0-F1-MODEL\_V4 \| 1.0 \| 1.279e-08 \| 235 \| 0.188 \| 217 \| 112 \| 7 \| 16 \| 179 \| 5 \| 210 \| Putative peptidase \| Putative peptidase; afdb-uniprot50 \| AF-A0A072T9Z9-F1-MODEL\_V4 \| 1.0 \| 1.093e-07 \| 235 \| 0.159 \| 219 \| 131 \| 8 \| 14 \| 180 \| 1 \| 218 \| HTH cro/C1-type domain-containing protein \| HTH cro/C1-type domain-containing protein; afdb-uniprot50 \| AF-A0A5M9J6V1-F1-MODEL\_V4 \| 1.0 \| 1.613e-08 \| 235 \| 0.176 \| 227 \| 127 \| 7 \| 10 \| 180 \| 2 \| 224 \| HTH-type transcriptional regulator PrtR \| HTH-type transcriptional regulator PrtR; afdb-uniprot50 \| AF-A0A519E7W6-F1-MODEL\_V4 \| 1.0 \| 1.075e-08 \| 235 \| 0.192 \| 218 \| 133 \| 9 \| 4 \| 180 \| 55 \| 270 \| S24 family peptidase \| S24 family peptidase; afdb-uniprot50 \| AF-A0A3R9Z5S5-F1-MODEL\_V4 \| 1.0 \| 1.031e-07 \| 235 \| 0.179 \| 239 \| 132 \| 6 \| 4 \| 180 \| 53 \| 289 \| Helix-turn-helix transcriptional regulator \| Helix-turn-helix transcriptional regulator; afdb-uniprot50 \| AF-V8RAQ2-F1-MODEL\_V4 \| 1.0 \| 9.569e-09 \| 235 \| 0.121 \| 271 \| 140 \| 6 \| 7 \| 180 \| 24 \| 293 \| Propanediol utilization protein \| Propanediol utilization protein; afdb-uniprot50 \| AF-A0A157S832-F1-MODEL\_V4 \| 1.0 \| 7.984e-06 \| 234 \| 0.221 \| 122 \| 92 \| 3 \| 60 \| 180 \| 8 \| 127 \| Transcriptional regulator \| Transcriptional regulator; afdb-uniprot50 \| AF-A0A1E4G2P2-F1-MODEL\_V4 \| 1.0 \| 4.58e-08 \| 234 \| 0.161 \| 211 \| 137 \| 7 \| 4 \| 180 \| 26 \| 230 \| HTH cro/C1-type domain-containing protein \| HTH cro/C1-type domain-containing protein; afdb-uniprot50 \| AF-A0A2N3B9K3-F1-MODEL\_V4 \| 1.0 \| 2.88e-08 \| 234 \| 0.173 \| 231 \| 135 \| 6 \| 4 \| 180 \| 13 \| 241 \| HTH cro/C1-type domain-containing protein \| HTH cro/C1-type domain-containing protein; afdb-uniprot50 \| AF-A0A2E3KX44-F1-MODEL\_V4 \| 1.0 \| 2.155e-08 \| 234 \| 0.166 \| 252 \| 137 \| 8 \| 1 \| 180 \| 2 \| 252 \| Transcriptional regulator \| Transcriptional regulator; afdb-uniprot50 \| AF-J8RZK5-F1-MODEL\_V4 \| 1.0 \| 1.697e-05 \| 233 \| 0.166 \| 108 \| 85 \| 1 \| 78 \| 180 \| 25 \| 132 \| Transcriptional regulator \| Transcriptional regulator; afdb-uniprot50 \| AF-A0A7Y6QLG8-F1-MODEL\_V4 \| 1.0 \| 2.284e-08 \| 233 \| 0.18 \| 210 \| 132 \| 9 \| 4 \| 180 \| 29 \| 231 \| Peptidase\_S24 domain-containing protein \| Peptidase\_S24 domain-containing protein; afdb-uniprot50 \| AF-A0A542RP09-F1-MODEL\_V4 \| 1.0 \| 2.42e-08 \| 233 \| 0.166 \| 228 \| 136 \| 7 \| 6 \| 180 \| 11 \| 237 \| Phage repressor protein C with HTH and peptisase S24 domain \| Phage repressor protein C with HTH and peptisase S24 domain; afdb-uniprot50 \| AF-A0A109Q3B0-F1-MODEL\_V4 \| 1.0 \| 7.283e-08 \| 233 \| 0.163 \| 232 \| 140 \| 6 \| 1 \| 180 \| 1 \| 230 \| Peptidase\_S24 domain-containing protein \| Peptidase\_S24 domain-containing protein; afdb-uniprot50 \| AF-A0A1I1CFP1-F1-MODEL\_V4 \| 1.0 \| 5.45e-08 \| 233 \| 0.161 \| 229 \| 139 \| 8 \| 5 \| 180 \| 11 \| 239 \| Peptidase S24-like \| Peptidase S24-like; afdb-uniprot50 \| AF-A0A2D2DKA6-F1-MODEL\_V4 \| 1.0 \| 3.052e-08 \| 233 \| 0.131 \| 235 \| 142 \| 4 \| 7 \| 180 \| 26 \| 259 \| HTH cro/C1-type domain-containing protein \| HTH cro/C1-type domain-containing protein; afdb-uniprot50 \| AF-A0A367M7U5-F1-MODEL\_V4 \| 1.0 \| 2.155e-08 \| 233 \| 0.132 \| 271 \| 139 \| 8 \| 5 \| 180 \| 11 \| 280 \| Phage repressor protein \| Phage repressor protein; afdb-uniprot50 \| AF-A0A443K7M6-F1-MODEL\_V4 \| 1.0 \| 9.733e-08 \| 233 \| 0.149 \| 294 \| 130 \| 8 \| 4 \| 179 \| 7 \| 298 \| HTH cro/C1-type domain-containing protein \| HTH cro/C1-type domain-containing protein; afdb-uniprot50 \| AF-A0A1Z5YTF2-F1-MODEL\_V4 \| 1.0 \| 4.853e-08 \| 232 \| 0.166 \| 216 \| 120 \| 6 \| 20 \| 180 \| 2 \| 212 \| HTH cro/C1-type domain-containing protein \| HTH cro/C1-type domain-containing protein; afdb-uniprot50 \| AF-A0A1Q6U9K6-F1-MODEL\_V4 \| 1.0 \| 5.143e-08 \| 232 \| 0.171 \| 216 \| 134 \| 8 \| 6 \| 180 \| 9 \| 220 \| HTH cro/C1-type domain-containing protein \| HTH cro/C1-type domain-containing protein; afdb-uniprot50 \| AF-A0A5C7JG03-F1-MODEL\_V4 \| 1.0 \| 3.234e-08 \| 232 \| 0.206 \| 194 \| 132 \| 7 \| 4 \| 179 \| 45 \| 234 \| Peptidase\_S24 domain-containing protein \| Peptidase\_S24 domain-containing protein; afdb-uniprot50 \| AF-D5CU95-F1-MODEL\_V4 \| 1.0 \| 3.427e-08 \| 232 \| 0.226 \| 225 \| 114 \| 8 \| 4 \| 174 \| 15 \| 233 \| Putative phage repressor \| Putative phage repressor; afdb-uniprot50 \| AF-A0A6B0DM06-F1-MODEL\_V4 \| 1.0 \| 9.569e-09 \| 232 \| 0.235 \| 195 \| 111 \| 4 \| 4 \| 163 \| 12 \| 203 \| Helix-turn-helix domain-containing protein \| Helix-turn-helix domain-containing protein; afdb-uniprot50 \| AF-A0A5B8CV78-F1-MODEL\_V4 \| 1.0 \| 1.709e-08 \| 231 \| 0.198 \| 181 \| 106 \| 4 \| 33 \| 174 \| 4 \| 184 \| Helix-turn-helix transcriptional regulator \| Helix-turn-helix transcriptional regulator; afdb-uniprot50 \| AF-A0A1Y5SGE5-F1-MODEL\_V4 \| 1.0 \| 2.88e-08 \| 231 \| 0.219 \| 228 \| 111 \| 10 \| 14 \| 180 \| 1 \| 222 \| Putative HTH-type transcriptional regulator \| Putative HTH-type transcriptional regulator; afdb-uniprot50 \| AF-A0A6I1H3E1-F1-MODEL\_V4 \| 1.0 \| 7.283e-08 \| 231 \| 0.169 \| 206 \| 138 \| 7 \| 4 \| 180 \| 28 \| 229 \| Peptidase\_S24 domain-containing protein \| Peptidase\_S24 domain-containing protein; afdb-uniprot50 \| AF-A0A845GQS4-F1-MODEL\_V4 \| 1.0 \| 4.322e-08 \| 231 \| 0.175 \| 211 \| 130 \| 8 \| 4 \| 180 \| 30 \| 230 \| Uncharacterized protein \| Uncharacterized protein; afdb-uniprot50 \| AF-A0A2S7JR70-F1-MODEL\_V4 \| 1.0 \| 2.88e-08 \| 231 \| 0.191 \| 282 \| 121 \| 10 \| 4 \| 180 \| 5 \| 284 \| Uncharacterized protein \| Uncharacterized protein; afdb-uniprot50 \| AF-A0A7Y5EC04-F1-MODEL\_V4 \| 1.0 \| 2.14e-05 \| 230 \| 0.166 \| 108 \| 85 \| 1 \| 78 \| 180 \| 51 \| 158 \| Transcriptional regulator \| Transcriptional regulator; afdb-uniprot50 \| AF-A0A7U3GDM4-F1-MODEL\_V4 \| 1.0 \| 6.873e-08 \| 230 \| 0.125 \| 215 \| 150 \| 5 \| 4 \| 180 \| 8 \| 222 \| LexA family transcriptional regulator \| LexA family transcriptional regulator; afdb-uniprot50 \| AF-Q1IFI8-F1-MODEL\_V4 \| 1.0 \| 6.486e-08 \| 230 \| 0.141 \| 226 \| 140 \| 6 \| 4 \| 180 \| 7 \| 227 \| Putative transcriptional regulator \| Putative transcriptional regulator; afdb-uniprot50 \| AF-A0A1F7FIP4-F1-MODEL\_V4 \| 1.0 \| 4.078e-08 \| 230 \| 0.129 \| 232 \| 145 \| 7 \| 5 \| 180 \| 22 \| 252 \| HTH cro/C1-type domain-containing protein \| HTH cro/C1-type domain-containing protein; afdb-uniprot50 \| AF-A0A679GE06-F1-MODEL\_V4 \| 1.0 \| 2.155e-08 \| 230 \| 0.132 \| 271 \| 139 \| 7 \| 5 \| 180 \| 6 \| 275 \| Peptidase\_S24 domain-containing protein \| Peptidase\_S24 domain-containing protein; afdb-uniprot50 \| AF-A0A1F4NAY1-F1-MODEL\_V4 \| 1.0 \| 2.546e-05 \| 229 \| 0.221 \| 104 \| 77 \| 4 \| 79 \| 180 \| 78 \| 179 \| Peptidase\_S24 domain-containing protein \| Peptidase\_S24 domain-containing protein; afdb-uniprot50 \| AF-A0A0M7MMT4-F1-MODEL\_V4 \| 1.0 \| 1.798e-05 \| 229 \| 0.257 \| 97 \| 66 \| 2 \| 87 \| 180 \| 90 \| 183 \| DNA polymerase V subunit UmuD \| DNA polymerase V subunit UmuD; afdb-uniprot50 \| AF-A0A1P8END7-F1-MODEL\_V4 \| 1.0 \| 3.234e-08 \| 229 \| 0.168 \| 219 \| 139 \| 6 \| 4 \| 180 \| 11 \| 228 \| HTH cro/C1-type domain-containing protein \| HTH cro/C1-type domain-containing protein; afdb-uniprot50 \| AF-A0A3D0L6H5-F1-MODEL\_V4 \| 1.0 \| 7.283e-08 \| 229 \| 0.17 \| 217 \| 133 \| 7 \| 8 \| 180 \| 15 \| 228 \| Peptidase\_S24 domain-containing protein \| Peptidase\_S24 domain-containing protein; afdb-uniprot50 \| AF-A0A2X1WHQ7-F1-MODEL\_V4 \| 1.0 \| 2.42e-08 \| 229 \| 0.178 \| 235 \| 131 \| 7 \| 5 \| 180 \| 4 \| 235 \| Uncharacterized HTH-type transcriptional regulator HI\_1476 \| Uncharacterized HTH-type transcriptional regulator HI\_1476; afdb-uniprot50 \| AF-A0A3A5J7R3-F1-MODEL\_V4 \| 1.0 \| 6.873e-08 \| 229 \| 0.152 \| 229 \| 136 \| 6 \| 4 \| 180 \| 16 \| 238 \| LexA family transcriptional regulator \| LexA family transcriptional regulator; afdb-uniprot50 \| AF-A0A358AXE2-F1-MODEL\_V4 \| 1.0 \| 8.667e-08 \| 229 \| 0.16 \| 225 \| 141 \| 6 \| 4 \| 180 \| 19 \| 243 \| Transcriptional regulator \| Transcriptional regulator; afdb-uniprot50 \| AF-A0A074V4S2-F1-MODEL\_V4 \| 1.0 \| 9.185e-08 \| 229 \| 0.183 \| 240 \| 131 \| 8 \| 4 \| 180 \| 11 \| 248 \| Peptidase S24-like \| Peptidase S24-like; afdb-uniprot50 \| AF-A0A6L7X9K4-F1-MODEL\_V4 \| 1.0 \| 2.284e-08 \| 229 \| 0.122 \| 253 \| 139 \| 6 \| 5 \| 180 \| 4 \| 250 \| S24 family peptidase \| S24 family peptidase; afdb-uniprot50 \| AF-A0A6G6IVE1-F1-MODEL\_V4 \| 1.0 \| 4.58e-08 \| 229 \| 0.158 \| 234 \| 135 \| 6 \| 5 \| 177 \| 44 \| 276 \| Helix-turn-helix domain-containing protein \| Helix-turn-helix domain-containing protein; afdb-uniprot50 \| AF-A0A348SFI4-F1-MODEL\_V4 \| 1.0 \| 5.45e-08 \| 229 \| 0.142 \| 246 \| 139 \| 5 \| 5 \| 178 \| 34 \| 279 \| HTH cro/C1-type domain-containing protein \| HTH cro/C1-type domain-containing protein; afdb-uniprot50 \| AF-A0A7Z8D4G4-F1-MODEL\_V4 \| 1.0 \| 6.12e-08 \| 228 \| 0.159 \| 207 \| 121 \| 7 \| 26 \| 180 \| 1 \| 206 \| XRE family transcriptional regulator \| XRE family transcriptional regulator; afdb-uniprot50 \| AF-A0A4D7B2Y1-F1-MODEL\_V4 \| 1.0 \| 3.052e-08 \| 228 \| 0.2 \| 214 \| 130 \| 9 \| 4 \| 180 \| 15 \| 224 \| HTH cro/C1-type domain-containing protein \| HTH cro/C1-type domain-containing protein; afdb-uniprot50 \| AF-A0A7W7NEM4-F1-MODEL\_V4 \| 1.0 \| 4.078e-08 \| 228 \| 0.133 \| 263 \| 141 \| 8 \| 4 \| 180 \| 7 \| 268 \| Phage repressor protein C with HTH and peptisase S24 domain \| Phage repressor protein C with HTH and peptisase S24 domain; afdb-uniprot50 \| AF-A0A1Y0ER74-F1-MODEL\_V4 \| 1.0 \| 3.234e-08 \| 227 \| 0.186 \| 220 \| 128 \| 14 \| 4 \| 178 \| 7 \| 220 \| Peptidase\_S24 domain-containing protein \| Peptidase\_S24 domain-containing protein; afdb-uniprot50 \| AF-A0A1V5QRV5-F1-MODEL\_V4 \| 1.0 \| 4.58e-08 \| 227 \| 0.207 \| 231 \| 116 \| 9 \| 15 \| 180 \| 9 \| 237 \| HTH-type transcriptional regulator PrtR \| HTH-type transcriptional regulator PrtR; afdb-uniprot50 \| AF-A0A0Q5KKZ7-F1-MODEL\_V4 \| 1.0 \| 4.322e-08 \| 227 \| 0.137 \| 232 \| 144 \| 8 \| 4 \| 180 \| 13 \| 243 \| Peptidase\_S24 domain-containing protein \| Peptidase\_S24 domain-containing protein; afdb-uniprot50 \| AF-A0A220RCN3-F1-MODEL\_V4 \| 1.0 \| 7.718e-08 \| 227 \| 0.19 \| 241 \| 126 \| 7 \| 4 \| 177 \| 7 \| 245 \| Peptidase\_S24 domain-containing protein \| Peptidase\_S24 domain-containing protein; afdb-uniprot50 \| AF-A0A1E7YT01-F1-MODEL\_V4 \| 1.0 \| 4.58e-08 \| 227 \| 0.163 \| 238 \| 137 \| 6 \| 4 \| 180 \| 8 \| 244 \| Peptidase\_S24 domain-containing protein \| Peptidase\_S24 domain-containing protein; afdb-uniprot50 \| AF-A0A854CXB7-F1-MODEL\_V4 \| 1.0 \| 2.718e-08 \| 227 \| 0.175 \| 245 \| 130 \| 9 \| 2 \| 178 \| 5 \| 245 \| Transcriptional regulator \| Transcriptional regulator; afdb-uniprot50 \| AF-A0A1H5XW02-F1-MODEL\_V4 \| 1.0 \| 6.12e-08 \| 227 \| 0.159 \| 251 \| 135 \| 6 \| 6 \| 180 \| 12 \| 262 \| Helix-turn-helix \| Helix-turn-helix; afdb-uniprot50 \| AF-A0A239BRE0-F1-MODEL\_V4 \| 1.0 \| 2.034e-08 \| 227 \| 0.126 \| 261 \| 143 \| 8 \| 4 \| 180 \| 7 \| 266 \| Peptidase S24-like \| Peptidase S24-like; afdb-uniprot50 \| AF-A0A1N7HEF4-F1-MODEL\_V4 \| 1.0 \| 3.234e-08 \| 227 \| 0.137 \| 262 \| 140 \| 8 \| 4 \| 180 \| 22 \| 282 \| Phage repressor protein C, contains Cro/C1-type HTH and peptisase s24 domains \| Phage repressor protein C, contains Cro/C1-type HTH and peptisase s24 domains; afdb-uniprot50 \| AF-A0A1E7PYC2-F1-MODEL\_V4 \| 1.0 \| 7.283e-08 \| 227 \| 0.141 \| 269 \| 141 \| 6 \| 1 \| 180 \| 26 \| 293 \| HTH cro/C1-type domain-containing protein \| HTH cro/C1-type domain-containing protein; afdb-uniprot50 \| AF-A0A1X0ZUR4-F1-MODEL\_V4 \| 1.0 \| 2.88e-08 \| 226 \| 0.179 \| 223 \| 126 \| 6 \| 4 \| 179 \| 7 \| 219 \| Transcriptional regulator \| Transcriptional regulator; afdb-uniprot50 \| AF-A0A521EXD5-F1-MODEL\_V4 \| 1.0 \| 1.031e-07 \| 226 \| 0.157 \| 222 \| 141 \| 7 \| 4 \| 180 \| 10 \| 230 \| Peptidase S24-like \| Peptidase S24-like; afdb-uniprot50 \| AF-A0A7C7ACW4-F1-MODEL\_V4 \| 1.0 \| 1.811e-08 \| 226 \| 0.156 \| 230 \| 140 \| 5 \| 4 \| 180 \| 7 \| 235 \| Helix-turn-helix transcriptional regulator \| Helix-turn-helix transcriptional regulator; afdb-uniprot50 \| AF-A0A4R1XGS3-F1-MODEL\_V4 \| 1.0 \| 8.667e-08 \| 226 \| 0.181 \| 232 \| 132 \| 10 \| 5 \| 180 \| 7 \| 236 \| Phage repressor protein C with HTH and peptisase S24 domain \| Phage repressor protein C with HTH and peptisase S24 domain; afdb-uniprot50 \| AF-A0A379JE71-F1-MODEL\_V4 \| 1.0 \| 7.283e-08 \| 226 \| 0.164 \| 225 \| 130 \| 6 \| 8 \| 179 \| 2 \| 221 \| Putative regulatory protein \| Putative regulatory protein; afdb-uniprot50 \| AF-A0A2S2F875-F1-MODEL\_V4 \| 1.0 \| 9.733e-08 \| 226 \| 0.146 \| 218 \| 139 \| 5 \| 4 \| 180 \| 34 \| 245 \| Helix-turn-helix domain-containing protein \| Helix-turn-helix domain-containing protein; afdb-uniprot50 \| AF-A0A1F4NGE6-F1-MODEL\_V4 \| 1.0 \| 2.192e-07 \| 226 \| 0.131 \| 235 \| 145 \| 8 \| 4 \| 180 \| 17 \| 250 \| HTH cro/C1-type domain-containing protein \| HTH cro/C1-type domain-containing protein; afdb-uniprot50 \| AF-A0A7U9I271-F1-MODEL\_V4 \| 1.0 \| 4.322e-08 \| 226 \| 0.132 \| 265 \| 141 \| 8 \| 4 \| 180 \| 7 \| 270 \| Peptidase\_S24 domain-containing protein \| Peptidase\_S24 domain-containing protein; afdb-uniprot50 \| AF-A0A7W8WGP6-F1-MODEL\_V4 \| 1.0 \| 5.143e-08 \| 226 \| 0.16 \| 275 \| 114 \| 11 \| 16 \| 180 \| 8 \| 275 \| Phage repressor protein C with HTH and peptisase S24 domain \| Phage repressor protein C with HTH and peptisase S24 domain; afdb-uniprot50 \| AF-F2K1S3-F1-MODEL\_V4 \| 1.0 \| 6.486e-08 \| 225 \| 0.196 \| 204 \| 119 \| 4 \| 20 \| 179 \| 2 \| 204 \| Putative phage repressor \| Putative phage repressor; afdb-uniprot50 \| AF-A0A6S7DEF1-F1-MODEL\_V4 \| 1.0 \| 3.234e-08 \| 225 \| 0.187 \| 229 \| 130 \| 8 \| 5 \| 180 \| 7 \| 232 \| HTH cro/C1-type domain-containing protein \| HTH cro/C1-type domain-containing protein; afdb-uniprot50 \| AF-A0A7U5XY97-F1-MODEL\_V4 \| 1.0 \| 2.565e-08 \| 225 \| 0.214 \| 210 \| 123 \| 10 \| 4 \| 180 \| 37 \| 237 \| Peptidase\_S24 domain-containing protein \| Peptidase\_S24 domain-containing protein; afdb-uniprot50 \| AF-A0A3G7V6R7-F1-MODEL\_V4 \| 1.0 \| 8.667e-08 \| 225 \| 0.163 \| 239 \| 136 \| 5 \| 4 \| 180 \| 14 \| 250 \| Phage cI repressor \| Phage cI repressor; afdb-uniprot50 \| AF-A0A080NQD6-F1-MODEL\_V4 \| 1.0 \| 5.45e-08 \| 225 \| 0.153 \| 247 \| 137 \| 11 \| 4 \| 180 \| 8 \| 252 \| Peptidase S24-like family protein \| Peptidase S24-like family protein; afdb-uniprot50 \| AF-A0A6M9CND3-F1-MODEL\_V4 \| 1.0 \| 7.718e-08 \| 225 \| 0.148 \| 249 \| 137 \| 12 \| 5 \| 180 \| 13 \| 259 \| Peptidase\_S24 domain-containing protein \| Peptidase\_S24 domain-containing protein; afdb-uniprot50 \| AF-D8IV29-F1-MODEL\_V4 \| 1.0 \| 3.849e-08 \| 225 \| 0.157 \| 266 \| 132 \| 8 \| 4 \| 180 \| 6 \| 268 \| Peptidase S24 LexA-like protein \| Peptidase S24 LexA-like protein; afdb-uniprot50 \| AF-A0A6B1CR96-F1-MODEL\_V4 \| 1.0 \| 1.031e-07 \| 225 \| 0.146 \| 267 \| 134 \| 8 \| 5 \| 180 \| 7 \| 270 \| S24 family peptidase \| S24 family peptidase; afdb-uniprot50 \| AF-A0A1W2AAM9-F1-MODEL\_V4 \| 1.0 \| 1.919e-08 \| 225 \| 0.142 \| 274 \| 132 \| 8 \| 5 \| 180 \| 11 \| 279 \| Phage repressor protein C, contains Cro/C1-type HTH and peptisase s24 domains \| Phage repressor protein C, contains Cro/C1-type HTH and peptisase s24 domains; afdb-uniprot50 \| AF-A0A4R5UKI9-F1-MODEL\_V4 \| 1.0 \| 1.093e-07 \| 225 \| 0.16 \| 230 \| 139 \| 5 \| 4 \| 180 \| 79 \| 307 \| LexA family transcriptional regulator \| LexA family transcriptional regulator; afdb-uniprot50 \| AF-A0A7G2K0G5-F1-MODEL\_V4 \| 1.0 \| 7.232e-05 \| 224 \| 0.277 \| 83 \| 60 \| 0 \| 98 \| 180 \| 5 \| 87 \| Prophage LambdaSo, transcriptional regulator, Cro/CI family \| Prophage LambdaSo, transcriptional regulator, Cro/CI family; afdb-uniprot50 \| AF-A0A3G6YX37-F1-MODEL\_V4 \| 1.0 \| 6.12e-08 \| 224 \| 0.158 \| 195 \| 118 \| 4 \| 29 \| 178 \| 4 \| 197 \| LexA family transcriptional regulator \| LexA family transcriptional regulator; afdb-uniprot50 \| AF-A0A3B0MER7-F1-MODEL\_V4 \| 1.0 \| 9.185e-08 \| 224 \| 0.161 \| 211 \| 140 \| 7 \| 5 \| 179 \| 7 \| 216 \| HTH-type transcriptional regulator PrtR \| HTH-type transcriptional regulator PrtR; afdb-uniprot50 \| AF-A0A3P3E8B0-F1-MODEL\_V4 \| 1.0 \| 1.158e-07 \| 224 \| 0.179 \| 217 \| 136 \| 9 \| 4 \| 180 \| 29 \| 243 \| Helix-turn-helix transcriptional regulator \| Helix-turn-helix transcriptional regulator; afdb-uniprot50 \| AF-A0A5C7VD74-F1-MODEL\_V4 \| 1.0 \| 1.158e-07 \| 224 \| 0.173 \| 213 \| 134 \| 7 \| 5 \| 180 \| 38 \| 245 \| Helix-turn-helix transcriptional regulator \| Helix-turn-helix transcriptional regulator; afdb-uniprot50 \| AF-A0A2U0YK76-F1-MODEL\_V4 \| 1.0 \| 1.031e-07 \| 224 \| 0.155 \| 218 \| 141 \| 6 \| 5 \| 180 \| 37 \| 253 \| Phage repressor protein C with HTH and peptisase S24 domain \| Phage repressor protein C with HTH and peptisase S24 domain; afdb-uniprot50 \| AF-A0A1H9E7D8-F1-MODEL\_V4 \| 1.0 \| 4.078e-08 \| 224 \| 0.161 \| 248 \| 133 \| 12 \| 5 \| 179 \| 12 \| 257 \| Phage repressor protein C, contains Cro/C1-type HTH and peptisase s24 domains \| Phage repressor protein C, contains Cro/C1-type HTH and peptisase s24 domains; afdb-uniprot50 \| AF-A0A105T6I1-F1-MODEL\_V4 \| 1.0 \| 1.548e-07 \| 224 \| 0.161 \| 236 \| 137 \| 5 \| 4 \| 180 \| 33 \| 266 \| Pyocin repressor protein \| Pyocin repressor protein; afdb-uniprot50 \| AF-E6QGP3-F1-MODEL\_V4 \| 1.0 \| 9.733e-08 \| 224 \| 0.133 \| 233 \| 142 \| 8 \| 4 \| 180 \| 66 \| 294 \| Peptidase\_S24 domain-containing protein \| Peptidase\_S24 domain-containing protein; afdb-uniprot50 \| AF-A0A661II41-F1-MODEL\_V4 \| 1.0 \| 1.131e-05 \| 223 \| 0.166 \| 108 \| 86 \| 2 \| 74 \| 178 \| 19 \| 125 \| Peptidase\_S24 domain-containing protein \| Peptidase\_S24 domain-containing protein; afdb-uniprot50 \| AF-A0A7W9RYE0-F1-MODEL\_V4 \| 1.0 \| 4.078e-08 \| 223 \| 0.188 \| 207 \| 133 \| 11 \| 4 \| 179 \| 12 \| 214 \| Phage repressor protein C with HTH and peptisase S24 domain \| Phage repressor protein C with HTH and peptisase S24 domain; afdb-uniprot50 \| AF-A0A379J5I1-F1-MODEL\_V4 \| 1.0 \| 4.853e-08 \| 223 \| 0.169 \| 189 \| 121 \| 4 \| 27 \| 180 \| 37 \| 224 \| Prophage PSPPH03, Cro/CI family transcriptional \| Prophage PSPPH03, Cro/CI family transcriptional; afdb-uniprot50 \| AF-D9Y858-F1-MODEL\_V4 \| 1.0 \| 5.45e-08 \| 223 \| 0.137 \| 225 \| 137 \| 10 \| 5 \| 180 \| 12 \| 228 \| Uncharacterized protein \| Uncharacterized protein; afdb-uniprot50 \| AF-A0A512L9E3-F1-MODEL\_V4 \| 1.0 \| 3.632e-08 \| 223 \| 0.19 \| 205 \| 136 \| 10 \| 4 \| 180 \| 36 \| 238 \| Peptidase\_S24 domain-containing protein \| Peptidase\_S24 domain-containing protein; afdb-uniprot50 \| AF-A0A8B4GDE0-F1-MODEL\_V4 \| 1.0 \| 1.227e-07 \| 223 \| 0.171 \| 239 \| 134 \| 8 \| 5 \| 180 \| 11 \| 248 \| Peptidase S24-like \| Peptidase S24-like; afdb-uniprot50 \| AF-A0A2D9ESJ0-F1-MODEL\_V4 \| 1.0 \| 5.775e-08 \| 223 \| 0.172 \| 237 \| 133 \| 13 \| 5 \| 180 \| 11 \| 245 \| Peptidase\_S24 domain-containing protein \| Peptidase\_S24 domain-containing protein; afdb-uniprot50 \| AF-A0A086D105-F1-MODEL\_V4 \| 1.0 \| 9.185e-08 \| 223 \| 0.183 \| 256 \| 128 \| 10 \| 5 \| 180 \| 3 \| 257 \| Peptidase\_S24 domain-containing protein \| Peptidase\_S24 domain-containing protein; afdb-uniprot50 \| AF-A0A149V4K2-F1-MODEL\_V4 \| 1.0 \| 2.068e-07 \| 222 \| 0.213 \| 173 \| 109 \| 6 \| 29 \| 176 \| 2 \| 172 \| Repressor \| Repressor; afdb-uniprot50 \| AF-A0A7G2K117-F1-MODEL\_V4 \| 1.0 \| 4.936e-07 \| 222 \| 0.21 \| 157 \| 117 \| 3 \| 25 \| 180 \| 33 \| 183 \| Prophage LambdaSo, transcriptional regulator, Cro/CI family \| Prophage LambdaSo, transcriptional regulator, Cro/CI family; afdb-uniprot50 \| AF-A0A7W5JLM8-F1-MODEL\_V4 \| 1.0 \| 6.873e-08 \| 222 \| 0.2 \| 190 \| 109 \| 7 \| 32 \| 180 \| 3 \| 190 \| Phage repressor protein C with HTH and peptisase S24 domain \| Phage repressor protein C with HTH and peptisase S24 domain; afdb-uniprot50 \| AF-A0A5N7XEU1-F1-MODEL\_V4 \| 1.0 \| 2.192e-07 \| 222 \| 0.187 \| 229 \| 132 \| 8 \| 4 \| 180 \| 22 \| 248 \| Peptidase S24 \| Peptidase S24; afdb-uniprot50 \| AF-A0A385N4F7-F1-MODEL\_V4 \| 1.0 \| 3.849e-08 \| 222 \| 0.109 \| 219 \| 146 \| 8 \| 4 \| 180 \| 52 \| 263 \| LexA family transcriptional regulator \| LexA family transcriptional regulator; afdb-uniprot50 \| AF-A0A7V1K4M1-F1-MODEL\_V4 \| 1.0 \| 7.283e-08 \| 221 \| 0.189 \| 195 \| 121 \| 5 \| 18 \| 180 \| 1 \| 190 \| Phage repressor protein \| Phage repressor protein; afdb-uniprot50 \| AF-A0A8B3EZK2-F1-MODEL\_V4 \| 1.0 \| 1.227e-07 \| 221 \| 0.183 \| 202 \| 116 \| 4 \| 28 \| 180 \| 13 \| 214 \| Helix-turn-helix transcriptional regulator \| Helix-turn-helix transcriptional regulator; afdb-uniprot50 \| AF-A0A1H6XI94-F1-MODEL\_V4 \| 1.0 \| 1.709e-08 \| 221 \| 0.159 \| 226 \| 126 \| 8 \| 1 \| 180 \| 18 \| 225 \| Phage repressor protein C, contains Cro/C1-type HTH and peptisase s24 domains \| Phage repressor protein C, contains Cro/C1-type HTH and peptisase s24 domains; afdb-uniprot50 \| AF-A0A543BE40-F1-MODEL\_V4 \| 1.0 \| 2.461e-07 \| 221 \| 0.139 \| 222 \| 145 \| 8 \| 4 \| 180 \| 12 \| 232 \| Phage repressor protein C with HTH and peptisase S24 domain \| Phage repressor protein C with HTH and peptisase S24 domain; afdb-uniprot50 \| AF-A0A103DW42-F1-MODEL\_V4 \| 1.0 \| 9.733e-08 \| 221 \| 0.15 \| 233 \| 139 \| 9 \| 4 \| 179 \| 10 \| 240 \| Repressor \| Repressor; afdb-uniprot50 \| AF-A0A1E4LU30-F1-MODEL\_V4 \| 1.0 \| 6.486e-08 \| 220 \| 0.186 \| 231 \| 126 \| 11 \| 4 \| 179 \| 5 \| 228 \| Repressor \| Repressor; afdb-uniprot50 \| AF-A0A6L7U9Q8-F1-MODEL\_V4 \| 1.0 \| 4.322e-08 \| 220 \| 0.162 \| 240 \| 130 \| 8 \| 5 \| 180 \| 7 \| 239 \| LexA family transcriptional regulator \| LexA family transcriptional regulator; afdb-uniprot50 \| AF-A0A3D1CH01-F1-MODEL\_V4 \| 1.0 \| 1.093e-07 \| 219 \| 0.172 \| 226 \| 135 \| 8 \| 5 \| 180 \| 12 \| 235 \| Transcriptional regulator \| Transcriptional regulator; afdb-uniprot50 \| AF-A0A2C9WXH1-F1-MODEL\_V4 \| 1.0 \| 5.45e-08 \| 219 \| 0.161 \| 241 \| 137 \| 10 \| 4 \| 180 \| 8 \| 247 \| Peptidase\_S24 domain-containing protein \| Peptidase\_S24 domain-containing protein; afdb-uniprot50 \| AF-A0A5E7B5Z1-F1-MODEL\_V4 \| 1.0 \| 7.718e-08 \| 218 \| 0.147 \| 197 \| 126 \| 7 \| 17 \| 180 \| 41 \| 228 \| Peptidase\_S24 domain-containing protein \| Peptidase\_S24 domain-containing protein; afdb-uniprot50 \| AF-A0A1M3JND1-F1-MODEL\_V4 \| 1.0 \| 7.283e-08 \| 218 \| 0.215 \| 209 \| 127 \| 9 \| 4 \| 180 \| 21 \| 224 \| Peptidase\_S24 domain-containing protein \| Peptidase\_S24 domain-containing protein; afdb-uniprot50 \| AF-B4RJ87-F1-MODEL\_V4 \| 1.0 \| 1.158e-07 \| 218 \| 0.142 \| 231 \| 139 \| 5 \| 9 \| 180 \| 4 \| 234 \| Putative phage repressor protein, putative phage associated protein \| Putative phage repressor protein, putative phage associated protein; afdb-uniprot50 \| AF-A0A844BRX6-F1-MODEL\_V4 \| 1.0 \| 3.427e-08 \| 218 \| 0.208 \| 235 \| 118 \| 7 \| 4 \| 171 \| 14 \| 247 \| Transcriptional regulator \| Transcriptional regulator; afdb-uniprot50 \| AF-A0A3D3KGL4-F1-MODEL\_V4 \| 1.0 \| 5.775e-08 \| 218 \| 0.137 \| 291 \| 136 \| 7 \| 4 \| 180 \| 11 \| 300 \| HTH cro/C1-type domain-containing protein \| HTH cro/C1-type domain-containing protein; afdb-uniprot50 \| AF-A0A6M3XFS2-F1-MODEL\_V4 \| 1.0 \| 4.078e-08 \| 218 \| 0.161 \| 211 \| 124 \| 8 \| 7 \| 180 \| 118 \| 312 \| Putative peptidase \| Putative peptidase; afdb-uniprot50 \| AF-A0A3B9HQD5-F1-MODEL\_V4 \| 1.0 \| 1.378e-07 \| 218 \| 0.146 \| 232 \| 138 \| 7 \| 8 \| 180 \| 85 \| 315 \| HTH cro/C1-type domain-containing protein \| HTH cro/C1-type domain-containing protein; afdb-uniprot50 \| AF-A0A812QVM4-F1-MODEL\_V4 \| 1.0 \| 8.179e-08 \| 218 \| 0.157 \| 203 \| 127 \| 4 \| 16 \| 175 \| 5 \| 206 \| CI protein \| CI protein; afdb-uniprot50 \| AF-A0A0D6SZZ1-F1-MODEL\_V4 \| 1.0 \| 1.158e-07 \| 217 \| 0.178 \| 185 \| 116 \| 5 \| 31 \| 180 \| 4 \| 187 \| Peptidase\_S24 domain-containing protein \| Peptidase\_S24 domain-containing protein; afdb-uniprot50 \| AF-G1V581-F1-MODEL\_V4 \| 1.0 \| 1.842e-07 \| 217 \| 0.186 \| 236 \| 125 \| 9 \| 5 \| 180 \| 13 \| 241 \| HTH cro/C1-type domain-containing protein \| HTH cro/C1-type domain-containing protein; afdb-uniprot50 \| AF-A0A853YD01-F1-MODEL\_V4 \| 1.0 \| 1.461e-07 \| 217 \| 0.202 \| 247 \| 124 \| 8 \| 6 \| 180 \| 2 \| 247 \| Uncharacterized protein \| Uncharacterized protein; afdb-uniprot50 \| AF-A0A4U8YR75-F1-MODEL\_V4 \| 1.0 \| 2.929e-07 \| 217 \| 0.174 \| 252 \| 131 \| 7 \| 4 \| 180 \| 5 \| 254 \| Bacteriophage ci repressor \| Bacteriophage ci repressor; afdb-uniprot50 \| AF-A0A4Q7NCE9-F1-MODEL\_V4 \| 1.0 \| 5.45e-08 \| 217 \| 0.147 \| 264 \| 126 \| 7 \| 14 \| 180 \| 33 \| 294 \| Phage repressor protein C with HTH and peptisase S24 domain \| Phage repressor protein C with HTH and peptisase S24 domain; afdb-uniprot50 \| AF-A0A5C8BK61-F1-MODEL\_V4 \| 1.0 \| 2.88e-08 \| 216 \| 0.178 \| 207 \| 128 \| 7 \| 5 \| 176 \| 7 \| 206 \| Helix-turn-helix transcriptional regulator \| Helix-turn-helix transcriptional regulator; afdb-uniprot50 \| AF-A0A3B0RY47-F1-MODEL\_V4 \| 1.0 \| 6.486e-08 \| 216 \| 0.14 \| 221 \| 140 \| 8 \| 5 \| 180 \| 6 \| 221 \| Uncharacterized protein \| Uncharacterized protein; afdb-uniprot50 \| AF-A0A4V3FW68-F1-MODEL\_V4 \| 1.0 \| 1.842e-07 \| 216 \| 0.143 \| 230 \| 140 \| 5 \| 4 \| 180 \| 7 \| 232 \| Peptidase S24-like protein \| Peptidase S24-like protein; afdb-uniprot50 \| AF-A0A3F3I4Y1-F1-MODEL\_V4 \| 1.0 \| 1.378e-07 \| 216 \| 0.155 \| 238 \| 135 \| 6 \| 8 \| 180 \| 6 \| 242 \| Peptidase\_S24 domain-containing protein \| Peptidase\_S24 domain-containing protein; afdb-uniprot50 \| AF-A0A1V2K4Z0-F1-MODEL\_V4 \| 1.0 \| 9.569e-09 \| 216 \| 0.151 \| 245 \| 139 \| 8 \| 4 \| 180 \| 6 \| 249 \| HTH cro/C1-type domain-containing protein \| HTH cro/C1-type domain-containing protein; afdb-uniprot50 \| AF-A0A2E7GR68-F1-MODEL\_V4 \| 1.0 \| 1.227e-07 \| 215 \| 0.239 \| 192 \| 114 \| 9 \| 5 \| 180 \| 10 \| 185 \| Peptidase\_S24 domain-containing protein \| Peptidase\_S24 domain-containing protein; afdb-uniprot50 \| AF-A0A2X1VHT4-F1-MODEL\_V4 \| 1.0 \| 2.929e-07 \| 215 \| 0.199 \| 216 \| 130 \| 10 \| 5 \| 180 \| 16 \| 228 \| Uncharacterized HTH-type transcriptional regulator HI\_1476 \| Uncharacterized HTH-type transcriptional regulator HI\_1476; afdb-uniprot50 \| AF-A0A7V4MJH9-F1-MODEL\_V4 \| 1.0 \| 4.148e-07 \| 215 \| 0.183 \| 234 \| 127 \| 7 \| 5 \| 180 \| 8 \| 235 \| Helix-turn-helix transcriptional regulator \| Helix-turn-helix transcriptional regulator; afdb-uniprot50 \| AF-A0A410UEY1-F1-MODEL\_V4 \| 1.0 \| 6.873e-08 \| 215 \| 0.144 \| 228 \| 139 \| 10 \| 5 \| 180 \| 15 \| 238 \| Uncharacterized protein \| Uncharacterized protein; afdb-uniprot50 \| AF-A0A1H7KZP5-F1-MODEL\_V4 \| 1.0 \| 1.548e-07 \| 215 \| 0.198 \| 232 \| 128 \| 10 \| 6 \| 180 \| 17 \| 247 \| Peptidase S24-like \| Peptidase S24-like; afdb-uniprot50 \| AF-A0A4R3NQD2-F1-MODEL\_V4 \| 1.0 \| 2.764e-07 \| 214 \| 0.145 \| 199 \| 129 \| 5 \| 23 \| 180 \| 2 \| 200 \| Phage repressor protein C with HTH and peptisase S24 domain \| Phage repressor protein C with HTH and peptisase S24 domain; afdb-uniprot50 \| AF-A0A7Y4J6H1-F1-MODEL\_V4 \| 1.0 \| 1.27e-05 \| 213 \| 0.188 \| 122 \| 91 \| 2 \| 59 \| 180 \| 4 \| 117 \| Uncharacterized protein \| Uncharacterized protein; afdb-uniprot50 \| AF-A0A7U6R1V1-F1-MODEL\_V4 \| 1.0 \| 9.733e-08 \| 213 \| 0.208 \| 206 \| 107 \| 8 \| 25 \| 180 \| 1 \| 200 \| Transcriptional regulator \| Transcriptional regulator; afdb-uniprot50 \| AF-A0A0W0WMM9-F1-MODEL\_V4 \| 1.0 \| 1.227e-07 \| 213 \| 0.106 \| 207 \| 155 \| 7 \| 3 \| 180 \| 2 \| 207 \| HTH-type transcriptional regulator \| HTH-type transcriptional regulator; afdb-uniprot50 \| AF-A0A2S5QA13-F1-MODEL\_V4 \| 1.0 \| 1.093e-07 \| 213 \| 0.184 \| 211 \| 137 \| 7 \| 4 \| 180 \| 7 \| 216 \| HTH cro/C1-type domain-containing protein \| HTH cro/C1-type domain-containing protein; afdb-uniprot50 \| AF-A0A2E8YIR4-F1-MODEL\_V4 \| 1.0 \| 1.436e-08 \| 213 \| 0.18 \| 238 \| 121 \| 7 \| 1 \| 180 \| 1 \| 222 \| HTH cro/C1-type domain-containing protein \| HTH cro/C1-type domain-containing protein; afdb-uniprot50 \| AF-A0A132C1D9-F1-MODEL\_V4 \| 1.0 \| 1.64e-07 \| 213 \| 0.234 \| 192 \| 116 \| 10 \| 5 \| 180 \| 115 \| 291 \| Putative HTH-type transcriptional regulator \| Putative HTH-type transcriptional regulator; afdb-uniprot50 \| AF-A0A2G6DFY9-F1-MODEL\_V4 \| 1.0 \| 9.733e-08 \| 212 \| 0.141 \| 240 \| 135 \| 8 \| 6 \| 180 \| 4 \| 237 \| Peptidase S24 \| Peptidase S24; afdb-uniprot50 \| AF-A0A496JQM1-F1-MODEL\_V4 \| 1.0 \| 2.608e-07 \| 212 \| 0.141 \| 248 \| 138 \| 8 \| 4 \| 180 \| 13 \| 256 \| Peptidase\_S24 domain-containing protein \| Peptidase\_S24 domain-containing protein; afdb-uniprot50 \| AF-A0A0A8RA87-F1-MODEL\_V4 \| 1.0 \| 2.068e-07 \| 212 \| 0.152 \| 236 \| 141 \| 5 \| 4 \| 180 \| 33 \| 268 \| Repressor protein CI \| Repressor protein CI; afdb-uniprot50 \| AF-A0A1A9WSH4-F1-MODEL\_V4 \| 1.0 \| 2.764e-07 \| 212 \| 0.173 \| 230 \| 131 \| 9 \| 4 \| 175 \| 525 \| 753 \| HTH cro/C1-type domain-containing protein \| HTH cro/C1-type domain-containing protein; afdb-uniprot50 \| AF-A0A1A7Q739-F1-MODEL\_V4 \| 1.0 \| 9.185e-08 \| 211 \| 0.186 \| 225 \| 120 \| 8 \| 13 \| 180 \| 11 \| 229 \| HTH Mu-type domain-containing protein \| HTH Mu-type domain-containing protein; afdb-uniprot50 \| AF-A0A7U4SE54-F1-MODEL\_V4 \| 1.0 \| 2.608e-07 \| 210 \| 0.149 \| 227 \| 140 \| 10 \| 4 \| 180 \| 7 \| 230 \| Peptidase S24 \| Peptidase S24; afdb-uniprot50 \| AF-A0A1H7A0H9-F1-MODEL\_V4 \| 1.0 \| 8.667e-08 \| 209 \| 0.15 \| 186 \| 112 \| 7 \| 28 \| 180 \| 18 \| 190 \| Phage repressor protein C, contains Cro/C1-type HTH and peptisase s24 domains \| Phage repressor protein C, contains Cro/C1-type HTH and peptisase s24 domains; afdb-uniprot50 \| AF-A0A2A3MNA1-F1-MODEL\_V4 \| 1.0 \| 3.289e-07 \| 209 \| 0.172 \| 214 \| 118 \| 4 \| 24 \| 180 \| 2 \| 213 \| Peptidase \| Peptidase; afdb-uniprot50 \| AF-I2NH04-F1-MODEL\_V4 \| 1.0 \| 1.842e-07 \| 209 \| 0.188 \| 217 \| 126 \| 11 \| 12 \| 180 \| 4 \| 218 \| Peptidase S24-like protein \| Peptidase S24-like protein; afdb-uniprot50 \| AF-A0A431J112-F1-MODEL\_V4 \| 1.0 \| 4.396e-07 \| 209 \| 0.146 \| 211 \| 140 \| 10 \| 5 \| 179 \| 20 \| 226 \| S24 family peptidase \| S24 family peptidase; afdb-uniprot50 \| AF-A0A4P6QMF0-F1-MODEL\_V4 \| 1.0 \| 1.301e-07 \| 209 \| 0.142 \| 224 \| 126 \| 9 \| 16 \| 180 \| 14 \| 230 \| S24 family peptidase \| S24 family peptidase; afdb-uniprot50 \| AF-A0A439VN75-F1-MODEL\_V4 \| 1.0 \| 7.283e-08 \| 209 \| 0.142 \| 280 \| 134 \| 10 \| 5 \| 180 \| 26 \| 303 \| Helix-turn-helix domain-containing protein \| Helix-turn-helix domain-containing protein; afdb-uniprot50 \| AF-A0A1F6GLC5-F1-MODEL\_V4 \| 1.0 \| 2.608e-07 \| 209 \| 0.216 \| 203 \| 123 \| 9 \| 9 \| 180 \| 112 \| 309 \| Peptidase\_S24 domain-containing protein \| Peptidase\_S24 domain-containing protein; afdb-uniprot50 \| AF-A0A0M7MMN5-F1-MODEL\_V4 \| 1.0 \| 2.461e-07 \| 208 \| 0.175 \| 222 \| 133 \| 9 \| 4 \| 180 \| 13 \| 229 \| DNA polymerase V subunit UmuD \| DNA polymerase V subunit UmuD; afdb-uniprot50 \| AF-A0A7X3U948-F1-MODEL\_V4 \| 1.0 \| 1.378e-07 \| 208 \| 0.149 \| 255 \| 127 \| 8 \| 5 \| 180 \| 8 \| 251 \| S24 family peptidase \| S24 family peptidase; afdb-uniprot50 \| AF-A0A7X3H7P9-F1-MODEL\_V4 \| 1.0 \| 5.543e-07 \| 208 \| 0.116 \| 284 \| 142 \| 8 \| 5 \| 180 \| 9 \| 291 \| Helix-turn-helix domain-containing protein \| Helix-turn-helix domain-containing protein; afdb-uniprot50 \| AF-A0A554XCA7-F1-MODEL\_V4 \| 1.0 \| 2.323e-07 \| 208 \| 0.136 \| 308 \| 129 \| 9 \| 5 \| 176 \| 8 \| 314 \| Peptidase S24-like protein \| Peptidase S24-like protein; afdb-uniprot50 \| AF-A0A0Q5MIV8-F1-MODEL\_V4 \| 1.0 \| 9.12e-05 \| 207 \| 0.171 \| 105 \| 84 \| 3 \| 78 \| 180 \| 137 \| 240 \| Peptidase\_S24 domain-containing protein \| Peptidase\_S24 domain-containing protein; afdb-uniprot50 \| AF-A0A2N6IH07-F1-MODEL\_V4 \| 1.0 \| 1.738e-07 \| 207 \| 0.154 \| 227 \| 127 \| 7 \| 7 \| 180 \| 24 \| 238 \| HTH cro/C1-type domain-containing protein \| HTH cro/C1-type domain-containing protein; afdb-uniprot50 \| AF-A0A1G8AY88-F1-MODEL\_V4 \| 1.0 \| 2.068e-07 \| 207 \| 0.156 \| 243 \| 134 \| 11 \| 5 \| 180 \| 3 \| 241 \| Peptidase S24-like \| Peptidase S24-like; afdb-uniprot50 \| AF-A0A2H0SA59-F1-MODEL\_V4 \| 1.0 \| 0.0001369 \| 206 \| 0.204 \| 98 \| 71 \| 2 \| 90 \| 180 \| 110 \| 207 \| Peptidase\_S24 domain-containing protein \| Peptidase\_S24 domain-containing protein; afdb-uniprot50 \| AF-A0A0Q8TCT1-F1-MODEL\_V4 \| 1.0 \| 4.658e-07 \| 206 \| 0.144 \| 236 \| 141 \| 9 \| 5 \| 180 \| 4 \| 238 \| Peptidase\_S24 domain-containing protein \| Peptidase\_S24 domain-containing protein; afdb-uniprot50 \| AF-A0A2E9SLM5-F1-MODEL\_V4 \| 1.0 \| 7.85e-07 \| 205 \| 0.125 \| 287 \| 139 \| 5 \| 5 \| 180 \| 11 \| 296 \| Peptidase\_S24 domain-containing protein \| Peptidase\_S24 domain-containing protein; afdb-uniprot50 \| AF-C0AQU8-F1-MODEL\_V4 \| 1.0 \| 0.0002745 \| 204 \| 0.237 \| 80 \| 61 \| 0 \| 101 \| 180 \| 1 \| 80 \| Peptidase S24-like protein \| Peptidase S24-like protein; afdb-uniprot50 \| AF-A0A545B7V3-F1-MODEL\_V4 \| 1.0 \| 8.121e-05 \| 204 \| 0.137 \| 102 \| 85 \| 3 \| 80 \| 180 \| 43 \| 142 \| Peptidase S24 \| Peptidase S24; afdb-uniprot50 \| AF-A0A0Q4GGM9-F1-MODEL\_V4 \| 1.0 \| 4.396e-07 \| 203 \| 0.183 \| 207 \| 112 \| 9 \| 9 \| 179 \| 11 \| 196 \| Peptidase\_S24 domain-containing protein \| Peptidase\_S24 domain-containing protein; afdb-uniprot50 \| AF-A0A0W0R1A0-F1-MODEL\_V4 \| 1.0 \| 4.658e-07 \| 203 \| 0.118 \| 194 \| 147 \| 7 \| 5 \| 180 \| 7 \| 194 \| HTH-type transcriptional regulator \| HTH-type transcriptional regulator; afdb-uniprot50 \| AF-A0A1S7SGT8-F1-MODEL\_V4 \| 1.0 \| 9.899e-07 \| 202 \| 0.207 \| 178 \| 99 \| 6 \| 28 \| 180 \| 27 \| 187 \| Transcriptional regulator \| Transcriptional regulator; afdb-uniprot50 \| AF-A0A2A5YC16-F1-MODEL\_V4 \| 1.0 \| 2.323e-07 \| 202 \| 0.162 \| 209 \| 137 \| 7 \| 5 \| 180 \| 7 \| 210 \| Uncharacterized protein \| Uncharacterized protein; afdb-uniprot50 \| AF-A0A427PUZ0-F1-MODEL\_V4 \| 1.0 \| 5.543e-07 \| 202 \| 0.163 \| 226 \| 132 \| 6 \| 7 \| 176 \| 16 \| 240 \| S24 family peptidase \| S24 family peptidase; afdb-uniprot50 \| AF-A0A3N0V7P7-F1-MODEL\_V4 \| 1.0 \| 3.289e-07 \| 202 \| 0.146 \| 253 \| 137 \| 9 \| 5 \| 180 \| 14 \| 264 \| Peptidase\_S24 domain-containing protein \| Peptidase\_S24 domain-containing protein; afdb-uniprot50 \| AF-A0A346YJJ6-F1-MODEL\_V4 \| 1.0 \| 5.543e-07 \| 202 \| 0.129 \| 217 \| 141 \| 7 \| 4 \| 173 \| 44 \| 259 \| Peptidase\_S24 domain-containing protein \| Peptidase\_S24 domain-containing protein; afdb-uniprot50 \| AF-A0A4Y9RWB5-F1-MODEL\_V4 \| 1.0 \| 1.64e-07 \| 201 \| 0.18 \| 205 \| 137 \| 7 \| 4 \| 180 \| 23 \| 224 \| S24 family peptidase \| S24 family peptidase; afdb-uniprot50 \| AF-A0A1I1CVZ8-F1-MODEL\_V4 \| 1.0 \| 3.694e-07 \| 201 \| 0.152 \| 236 \| 139 \| 8 \| 5 \| 180 \| 3 \| 237 \| Phage repressor protein C, contains Cro/C1-type HTH and peptisase s24 domains \| Phage repressor protein C, contains Cro/C1-type HTH and peptisase s24 domains; afdb-uniprot50 \| AF-A0A410UEA8-F1-MODEL\_V4 \| 1.0 \| 1.952e-07 \| 201 \| 0.175 \| 234 \| 134 \| 9 \| 4 \| 180 \| 38 \| 269 \| Helix-turn-helix transcriptional regulator \| Helix-turn-helix transcriptional regulator; afdb-uniprot50 \| AF-A0A6S6N4B6-F1-MODEL\_V4 \| 1.0 \| 8.815e-07 \| 200 \| 0.155 \| 180 \| 112 \| 6 \| 41 \| 180 \| 5 \| 184 \| Peptidase\_S24 domain-containing protein \| Peptidase\_S24 domain-containing protein; afdb-uniprot50 \| AF-A0A1N6QCR8-F1-MODEL\_V4 \| 1.0 \| 1.402e-06 \| 200 \| 0.191 \| 193 \| 106 \| 6 \| 38 \| 180 \| 5 \| 197 \| Phage repressor protein C, contains Cro/C1-type HTH and peptisase s24 domains \| Phage repressor protein C, contains Cro/C1-type HTH and peptisase s24 domains; afdb-uniprot50 \| AF-A0A0G9HCL5-F1-MODEL\_V4 \| 1.0 \| 5.874e-07 \| 200 \| 0.165 \| 229 \| 135 \| 8 \| 4 \| 180 \| 15 \| 239 \| Uncharacterized protein \| Uncharacterized protein; afdb-uniprot50 \| AF-A0A6L8GJ02-F1-MODEL\_V4 \| 1.0 \| 6.331e-06 \| 200 \| 0.225 \| 155 \| 109 \| 3 \| 27 \| 180 \| 38 \| 182 \| Peptidase\_S24 domain-containing protein \| Peptidase\_S24 domain-containing protein; afdb-uniprot50 \| AF-A0A2W5FGP3-F1-MODEL\_V4 \| 1.0 \| 2.068e-07 \| 199 \| 0.183 \| 185 \| 133 \| 6 \| 5 \| 180 \| 6 \| 181 \| Peptidase\_S24 domain-containing protein \| Peptidase\_S24 domain-containing protein; afdb-uniprot50 \| AF-A0A3B0J7Y0-F1-MODEL\_V4 \| 1.0 \| 1.486e-06 \| 199 \| 0.18 \| 188 \| 105 \| 4 \| 41 \| 179 \| 3 \| 190 \| HTH-type transcriptional regulator PrtR \| HTH-type transcriptional regulator PrtR; afdb-uniprot50 \| AF-E3I329-F1-MODEL\_V4 \| 1.0 \| 4.853e-08 \| 199 \| 0.2 \| 190 \| 123 \| 9 \| 3 \| 174 \| 20 \| 198 \| Helix-turn-helix domain protein \| Helix-turn-helix domain protein; afdb-uniprot50 \| AF-A0A550ESR5-F1-MODEL\_V4 \| 1.0 \| 2.323e-07 \| 199 \| 0.15 \| 206 \| 126 \| 8 \| 20 \| 180 \| 68 \| 269 \| S24 family peptidase \| S24 family peptidase; afdb-uniprot50 \| AF-A0A2C9WTG2-F1-MODEL\_V4 \| 1.0 \| 5.874e-07 \| 198 \| 0.16 \| 212 \| 130 \| 9 \| 4 \| 180 \| 8 \| 206 \| Peptidase S24 \| Peptidase S24; afdb-uniprot50 \| AF-A0A1B4V7W1-F1-MODEL\_V4 \| 1.0 \| 1.842e-07 \| 198 \| 0.177 \| 281 \| 123 \| 10 \| 4 \| 180 \| 21 \| 297 \| Peptidase S24 \| Peptidase S24; afdb-uniprot50 \| AF-A0A6L8HJW3-F1-MODEL\_V4 \| 1.0 \| 4.658e-07 \| 197 \| 0.229 \| 183 \| 104 \| 7 \| 27 \| 180 \| 42 \| 216 \| S24 family peptidase \| S24 family peptidase; afdb-uniprot50 \| AF-A0A1B8TEV8-F1-MODEL\_V4 \| 1.0 \| 4.936e-07 \| 197 \| 0.16 \| 250 \| 119 \| 10 \| 20 \| 180 \| 25 \| 272 \| Peptidase\_S24 domain-containing protein \| Peptidase\_S24 domain-containing protein; afdb-uniprot50 \| AF-A0A349Y0P3-F1-MODEL\_V4 \| 1.0 \| 6.597e-07 \| 196 \| 0.186 \| 214 \| 130 \| 8 \| 5 \| 180 \| 8 \| 215 \| Transcriptional regulator \| Transcriptional regulator; afdb-uniprot50 \| AF-A0A3L7AJ75-F1-MODEL\_V4 \| 1.0 \| 4.396e-07 \| 196 \| 0.175 \| 205 \| 126 \| 7 \| 4 \| 180 \| 41 \| 230 \| HTH cro/C1-type domain-containing protein \| HTH cro/C1-type domain-containing protein; afdb-uniprot50 \| AF-A0A1I2A0G8-F1-MODEL\_V4 \| 1.0 \| 7.408e-07 \| 196 \| 0.15 \| 239 \| 129 \| 10 \| 4 \| 180 \| 34 \| 260 \| Phage repressor protein C, contains Cro/C1-type HTH and peptisase s24 domains \| Phage repressor protein C, contains Cro/C1-type HTH and peptisase s24 domains; afdb-uniprot50 \| AF-A0A5M6I649-F1-MODEL\_V4 \| 1.0 \| 0.0001219 \| 195 \| 0.235 \| 85 \| 63 \| 1 \| 1 \| 85 \| 1 \| 83 \| Helix-turn-helix transcriptional regulator \| Helix-turn-helix transcriptional regulator; afdb-uniprot50 \| AF-A0A1E7ZE28-F1-MODEL\_V4 \| 1.0 \| 4.936e-07 \| 195 \| 0.157 \| 197 \| 125 \| 7 \| 5 \| 180 \| 10 \| 186 \| Peptidase\_S24 domain-containing protein \| Peptidase\_S24 domain-containing protein; afdb-uniprot50 \| AF-X6BLE3-F1-MODEL\_V4 \| 1.0 \| 1.402e-06 \| 195 \| 0.189 \| 190 \| 109 \| 6 \| 35 \| 180 \| 2 \| 190 \| Peptidase\_S24 domain-containing protein \| Peptidase\_S24 domain-containing protein; afdb-uniprot50 \| AF-A0A143Z9V0-F1-MODEL\_V4 \| 1.0 \| 6.225e-07 \| 195 \| 0.175 \| 233 \| 127 \| 9 \| 10 \| 180 \| 3 \| 232 \| Phage repressor protein C, contains Cro/C1-type HTH and peptisase s24 domains \| Phage repressor protein C, contains Cro/C1-type HTH and peptisase s24 domains; afdb-uniprot50 \| AF-A0A5B9EMT8-F1-MODEL\_V4 \| 1.0 \| 6.99e-07 \| 194 \| 0.168 \| 220 \| 135 \| 8 \| 5 \| 180 \| 8 \| 223 \| S24 family peptidase \| S24 family peptidase; afdb-uniprot50 \| AF-A0A6P2FXV7-F1-MODEL\_V4 \| 1.0 \| 0.000259 \| 194 \| 0.215 \| 93 \| 71 \| 2 \| 85 \| 176 \| 139 \| 230 \| Uncharacterized protein \| Uncharacterized protein; afdb-uniprot50 \| AF-D5AQW8-F1-MODEL\_V4 \| 1.0 \| 1.486e-06 \| 193 \| 0.172 \| 203 \| 121 \| 6 \| 25 \| 180 \| 1 \| 203 \| Peptidase, S24 family \| Peptidase, S24 family; afdb-uniprot50 \| AF-A0A2D0AFF9-F1-MODEL\_V4 \| 1.0 \| 1.248e-06 \| 192 \| 0.155 \| 232 \| 132 \| 7 \| 13 \| 180 \| 5 \| 236 \| Transcriptional regulator \| Transcriptional regulator; afdb-uniprot50 \| AF-A0A7Y6JDF7-F1-MODEL\_V4 \| 1.0 \| 1.768e-06 \| 191 \| 0.169 \| 201 \| 116 \| 5 \| 30 \| 180 \| 2 \| 201 \| LexA family transcriptional regulator \| LexA family transcriptional regulator; afdb-uniprot50 \| AF-A0A3B1AL92-F1-MODEL\_V4 \| 1.0 \| 3.104e-07 \| 191 \| 0.185 \| 221 \| 131 \| 9 \| 4 \| 180 \| 7 \| 222 \| HTH cro/C1-type domain-containing protein \| HTH cro/C1-type domain-containing protein; afdb-uniprot50 \| AF-A0A3D4G9A8-F1-MODEL\_V4 \| 1.0 \| 6.225e-07 \| 191 \| 0.114 \| 228 \| 145 \| 10 \| 5 \| 180 \| 12 \| 234 \| Uncharacterized protein \| Uncharacterized protein; afdb-uniprot50 \| AF-S7UV51-F1-MODEL\_V4 \| 1.0 \| 1.486e-06 \| 191 \| 0.159 \| 251 \| 133 \| 12 \| 4 \| 180 \| 27 \| 273 \| Phage C1 repressor \| Phage C1 repressor; afdb-uniprot50 \| AF-A0A4R3I013-F1-MODEL\_V4 \| 1.0 \| 5.874e-07 \| 191 \| 0.182 \| 208 \| 125 \| 10 \| 7 \| 173 \| 73 \| 276 \| Phage repressor protein C with HTH and peptisase S24 domain \| Phage repressor protein C with HTH and peptisase S24 domain; afdb-uniprot50 \| AF-A0A7X0AWH3-F1-MODEL\_V4 \| 1.0 \| 0.0002176 \| 189 \| 0.31 \| 87 \| 57 \| 1 \| 2 \| 85 \| 50 \| 136 \| Transcriptional regulator with XRE-family HTH domain \| Transcriptional regulator with XRE-family HTH domain; afdb-uniprot50 \| AF-A0A444LGP6-F1-MODEL\_V4 \| 1.0 \| 5.975e-06 \| 187 \| 0.097 \| 226 \| 150 \| 4 \| 8 \| 180 \| 39 \| 263 \| S24 family peptidase \| S24 family peptidase; afdb-uniprot50 \| AF-A0A5N7S1F1-F1-MODEL\_V4 \| 1.0 \| 8.815e-07 \| 187 \| 0.131 \| 289 \| 137 \| 9 \| 5 \| 180 \| 4 \| 291 \| S24 family peptidase \| S24 family peptidase; afdb-uniprot50 \| AF-A0A2Z4RTP7-F1-MODEL\_V4 \| 1.0 \| 1.768e-06 \| 186 \| 0.179 \| 212 \| 111 \| 6 \| 28 \| 180 \| 15 \| 222 \| XRE family transcriptional regulator \| XRE family transcriptional regulator; afdb-uniprot50 \| AF-A0A352VC09-F1-MODEL\_V4 \| 1.0 \| 1.323e-06 \| 186 \| 0.171 \| 239 \| 127 \| 7 \| 4 \| 180 \| 6 \| 235 \| Peptidase\_S24 domain-containing protein \| Peptidase\_S24 domain-containing protein; afdb-uniprot50 \| AF-A0A1H3E1Y6-F1-MODEL\_V4 \| 1.0 \| 0.0001537 \| 185 \| 0.115 \| 121 \| 100 \| 5 \| 61 \| 180 \| 13 \| 127 \| Uncharacterized protein \| Uncharacterized protein; afdb-uniprot50 \| AF-A0A4Q2IVG0-F1-MODEL\_V4 \| 1.0 \| 7.85e-07 \| 185 \| 0.157 \| 260 \| 132 \| 8 \| 5 \| 180 \| 10 \| 266 \| Helix-turn-helix transcriptional regulator \| Helix-turn-helix transcriptional regulator; afdb-uniprot50 \| AF-A0A1B8P414-F1-MODEL\_V4 \| 1.0 \| 0.0002306 \| 184 \| 0.288 \| 90 \| 62 \| 1 \| 79 \| 166 \| 32 \| 121 \| HTH-type transcriptional regulator PrtR \| HTH-type transcriptional regulator PrtR; afdb-uniprot50 \| AF-A0A7X4ALL0-F1-MODEL\_V4 \| 1.0 \| 1.112e-06 \| 184 \| 0.195 \| 189 \| 119 \| 5 \| 20 \| 180 \| 14 \| 197 \| Helix-turn-helix transcriptional regulator \| Helix-turn-helix transcriptional regulator; afdb-uniprot50 \| AF-A0A0H4KUA3-F1-MODEL\_V4 \| 1.0 \| 1.323e-06 \| 184 \| 0.125 \| 207 \| 149 \| 6 \| 5 \| 180 \| 7 \| 212 \| Uncharacterized protein \| Uncharacterized protein; afdb-uniprot50 \| AF-A0A3D8V7V0-F1-MODEL\_V4 \| 1.0 \| 0.0001219 \| 183 \| 0.3 \| 90 \| 62 \| 1 \| 1 \| 89 \| 1 \| 90 \| XRE family transcriptional regulator \| XRE family transcriptional regulator; afdb-uniprot50 \| AF-A0A7W6RHR6-F1-MODEL\_V4 \| 1.0 \| 0.0003887 \| 183 \| 0.273 \| 84 \| 60 \| 1 \| 2 \| 84 \| 5 \| 88 \| Transcriptional regulator with XRE-family HTH domain \| Transcriptional regulator with XRE-family HTH domain; afdb-uniprot50 \| AF-A0A146H4F4-F1-MODEL\_V4 \| 1.0 \| 7.85e-07 \| 179 \| 0.17 \| 229 \| 129 \| 11 \| 4 \| 178 \| 19 \| 240 \| MASE1 domain-containing protein \| MASE1 domain-containing protein; afdb-uniprot50 \| AF-A0A845M0L5-F1-MODEL\_V4 \| 1.0 \| 1.548e-07 \| 178 \| 0.18 \| 216 \| 124 \| 10 \| 1 \| 180 \| 1 \| 199 \| Helix-turn-helix domain-containing protein \| Helix-turn-helix domain-containing protein; afdb-uniprot50 \| AF-A0A4P6L3Y5-F1-MODEL\_V4 \| 1.0 \| 6.331e-06 \| 177 \| 0.17 \| 176 \| 114 \| 6 \| 16 \| 180 \| 18 \| 172 \| Peptidase\_S24 domain-containing protein \| Peptidase\_S24 domain-containing protein; afdb-uniprot50 \| AF-A0A6L6YJW9-F1-MODEL\_V4 \| 1.0 \| 5.874e-07 \| 177 \| 0.143 \| 237 \| 128 \| 14 \| 5 \| 176 \| 5 \| 231 \| Uncharacterized protein \| Uncharacterized protein; afdb-uniprot50 \| AF-A0A2S1XL62-F1-MODEL\_V4 \| 1.0 \| 0.0003461 \| 175 \| 0.279 \| 86 \| 56 \| 2 \| 2 \| 87 \| 32 \| 111 \| HTH cro/C1-type domain-containing protein \| HTH cro/C1-type domain-containing protein; afdb-uniprot50 \| AF-A0A2V9FMP7-F1-MODEL\_V4 \| 1.0 \| 0.0003887 \| 171 \| 0.258 \| 85 \| 62 \| 1 \| 4 \| 87 \| 41 \| 125 \| HTH cro/C1-type domain-containing protein \| HTH cro/C1-type domain-containing protein; afdb-uniprot50 \| AF-A0A2V9A659-F1-MODEL\_V4 \| 1.0 \| 0.0004625 \| 169 \| 0.27 \| 85 \| 61 \| 1 \| 4 \| 87 \| 29 \| 113 \| HTH cro/C1-type domain-containing protein \| HTH cro/C1-type domain-containing protein; afdb-uniprot50 \| AF-A0A7C5S2C7-F1-MODEL\_V4 \| 1.0 \| 0.0007795 \| 168 \| 0.25 \| 88 \| 59 \| 2 \| 2 \| 85 \| 6 \| 90 \| XRE family transcriptional regulator \| XRE family transcriptional regulator; afdb-uniprot50 \| AF-A0A2D6LJL1-F1-MODEL\_V4 \| 1.0 \| 0.002634 \| 167 \| 0.172 \| 87 \| 70 \| 1 \| 91 \| 177 \| 2 \| 86 \| Peptidase\_S24 domain-containing protein \| Peptidase\_S24 domain-containing protein; afdb-uniprot50 \| AF-A0A0J8AY60-F1-MODEL\_V4 \| 1.0 \| 0.0003668 \| 167 \| 0.265 \| 94 \| 64 \| 2 \| 2 \| 90 \| 4 \| 97 \| Cro/Cl family transcriptional regulator \| Cro/Cl family transcriptional regulator; afdb-uniprot50 \| AF-A0A1C7HIQ4-F1-MODEL\_V4 \| 1.0 \| 1.131e-05 \| 167 \| 0.119 \| 242 \| 143 \| 12 \| 5 \| 180 \| 19 \| 256 \| Peptidase\_S24 domain-containing protein \| Peptidase\_S24 domain-containing protein; afdb-uniprot50 \| AF-F3C821-F1-MODEL\_V4 \| 1.0 \| 1.323e-06 \| 167 \| 0.138 \| 209 \| 127 \| 6 \| 4 \| 163 \| 7 \| 211 \| Cro/CI family transcriptional regulator \| Cro/CI family transcriptional regulator; afdb-uniprot50 \| AF-A0A353J872-F1-MODEL\_V4 \| 1.0 \| 0.0006941 \| 163 \| 0.197 \| 86 \| 66 \| 2 \| 1 \| 85 \| 1 \| 84 \| Transcriptional regulator \| Transcriptional regulator; afdb-uniprot50 \| AF-A0A1H6UM04-F1-MODEL\_V4 \| 1.0 \| 0.002214 \| 161 \| 0.223 \| 85 \| 63 \| 1 \| 1 \| 85 \| 2 \| 83 \| DNA-binding transcriptional regulator, XRE-family HTH domain \| DNA-binding transcriptional regulator, XRE-family HTH domain; afdb-uniprot50 \| AF-A0A009HGE2-F1-MODEL\_V4 \| 1.0 \| 2.546e-05 \| 161 \| 0.113 \| 203 \| 126 \| 9 \| 9 \| 180 \| 8 \| 187 \| HTH cro/C1-type domain-containing protein \| HTH cro/C1-type domain-containing protein; afdb-uniprot50 \| AF-A0A844NU01-F1-MODEL\_V4 \| 1.0 \| 2.019e-05 \| 161 \| 0.141 \| 191 \| 117 \| 7 \| 33 \| 180 \| 4 \| 190 \| S24 family peptidase \| S24 family peptidase; afdb-uniprot50 \| AF-A0A7I0GSB9-F1-MODEL\_V4 \| 1.0 \| 0.001314 \| 159 \| 0.188 \| 85 \| 69 \| 0 \| 1 \| 85 \| 1 \| 85 \| HTH cro/C1-type domain-containing protein \| HTH cro/C1-type domain-containing protein; afdb-uniprot50 \| AF-A0A0F9DHM8-F1-MODEL\_V4 \| 1.0 \| 0.002089 \| 155 \| 0.238 \| 84 \| 64 \| 0 \| 4 \| 87 \| 87 \| 170 \| HTH cro/C1-type domain-containing protein \| HTH cro/C1-type domain-containing protein; afdb-uniprot50 \| AF-A0A0G1HFB0-F1-MODEL\_V4 \| 1.0 \| 6.824e-05 \| 155 \| 0.113 \| 193 \| 116 \| 7 \| 40 \| 180 \| 14 \| 203 \| Peptidase\_S24 domain-containing protein \| Peptidase\_S24 domain-containing protein; afdb-uniprot50 \| AF-A0A511XD88-F1-MODEL\_V4 \| 1.0 \| 0.0001291 \| 154 \| 0.162 \| 123 \| 81 \| 3 \| 61 \| 177 \| 3 \| 109 \| Uncharacterized protein \| Uncharacterized protein; afdb-uniprot50 \| AF-A0A7V6M500-F1-MODEL\_V4 \| 1.0 \| 0.00186 \| 153 \| 0.173 \| 104 \| 80 \| 2 \| 74 \| 177 \| 15 \| 112 \| Signal peptidase I \| Signal peptidase I; afdb-uniprot50 \| AF-A0A1H4C6J2-F1-MODEL\_V4 \| 1.0 \| 0.001563 \| 152 \| 0.191 \| 89 \| 68 \| 1 \| 3 \| 87 \| 19 \| 107 \| DNA-binding transcriptional regulator, XRE-family HTH domain \| DNA-binding transcriptional regulator, XRE-family HTH domain; afdb-uniprot50 \| AF-A0A1N7RAL9-F1-MODEL\_V4 \| 1.0 \| 0.002634 \| 150 \| 0.216 \| 83 \| 65 \| 0 \| 3 \| 85 \| 7 \| 89 \| Cro/C1-type HTH DNA-binding domain-containing protein \| Cro/C1-type HTH DNA-binding domain-containing protein; afdb-uniprot50 \| AF-A0A371J4W6-F1-MODEL\_V4 \| 1.0 \| 0.002214 \| 148 \| 0.16 \| 87 \| 67 \| 2 \| 2 \| 84 \| 4 \| 88 \| Transcriptional regulator \| Transcriptional regulator; afdb-uniprot50 \| AF-A0A352CT79-F1-MODEL\_V4 \| 1.0 \| 0.00186 \| 147 \| 0.194 \| 103 \| 75 \| 4 \| 77 \| 177 \| 19 \| 115 \| Signal peptidase I \| Signal peptidase I; afdb-uniprot50 \| AF-A0A2P4V625-F1-MODEL\_V4 \| 1.0 \| 1.426e-05 \| 146 \| 0.132 \| 226 \| 140 \| 8 \| 4 \| 180 \| 60 \| 278 \| Peptidase\_S24 domain-containing protein \| Peptidase\_S24 domain-containing protein; afdb-uniprot50 \| AF-A0A1F2RWP3-F1-MODEL\_V4 \| 1.0 \| 0.004189 \| 145 \| 0.226 \| 84 \| 65 \| 0 \| 4 \| 87 \| 40 \| 123 \| HTH cro/C1-type domain-containing protein \| HTH cro/C1-type domain-containing protein; afdb-uniprot50 \| AF-A0A1I5C6A5-F1-MODEL\_V4 \| 1.0 \| 0.001656 \| 143 \| 0.209 \| 86 \| 64 \| 1 \| 4 \| 85 \| 5 \| 90 \| DNA-binding transcriptional regulator, XRE-family HTH domain \| DNA-binding transcriptional regulator, XRE-family HTH domain; afdb-uniprot50 \| AF-A0A1C5W6V3-F1-MODEL\_V4 \| 1.0 \| 0.002214 \| 143 \| 0.15 \| 106 \| 83 \| 4 \| 75 \| 177 \| 44 \| 145 \| Signal peptidase I W \| Signal peptidase I W; afdb-uniprot50 \| AF-A0A135PBJ9-F1-MODEL\_V4 \| 1.0 \| 0.002634 \| 141 \| 0.2 \| 85 \| 64 \| 2 \| 5 \| 87 \| 7 \| 89 \| DNA-binding helix-turn-helix protein \| DNA-binding helix-turn-helix protein; afdb-uniprot50 \| AF-A0A0P8AQ76-F1-MODEL\_V4 \| 1.0 \| 0.003135 \| 140 \| 0.255 \| 90 \| 59 \| 2 \| 4 \| 89 \| 30 \| 115 \| Helix-turn-helix domain/PilZ domain \| Helix-turn-helix domain/PilZ domain; afdb-uniprot50 \| AF-A0A417R8M7-F1-MODEL\_V4 \| 1.0 \| 0.001392 \| 140 \| 0.211 \| 118 \| 84 \| 5 \| 63 \| 177 \| 3 \| 114 \| Signal peptidase I \| Signal peptidase I; afdb-uniprot50 \| AF-A0A841M4G8-F1-MODEL\_V4 \| 1.0 \| 0.007481 \| 139 \| 0.178 \| 84 \| 69 \| 0 \| 2 \| 85 \| 21 \| 104 \| Transcriptional regulator with XRE-family HTH domain \| Transcriptional regulator with XRE-family HTH domain; afdb-uniprot50 \| AF-A0A417SP50-F1-MODEL\_V4 \| 1.0 \| 0.002958 \| 136 \| 0.157 \| 108 \| 80 \| 5 \| 74 \| 176 \| 10 \| 111 \| Signal peptidase I \| Signal peptidase I; afdb-uniprot50 \| AF-A0A564VS75-F1-MODEL\_V4 \| 1.0 \| 0.00706 \| 131 \| 0.15 \| 106 \| 83 \| 4 \| 75 \| 177 \| 53 \| 154 \| Signal peptidase I W \| Signal peptidase I W; afdb-uniprot50 \| AF-A0A533RLY2-F1-MODEL\_V4 \| 1.0 \| 0.001755 \| 130 \| 0.161 \| 118 \| 81 \| 8 \| 70 \| 176 \| 1 \| 111 \| Peptidase\_S26 domain-containing protein \| Peptidase\_S26 domain-containing protein; afdb-uniprot50 \| AF-D4LGL3-F1-MODEL\_V4 \| 1.0 \| 0.007928 \| 130 \| 0.15 \| 106 \| 83 \| 4 \| 75 \| 177 \| 14 \| 115 \| Signal peptidase I, archaeal type \| Signal peptidase I, archaeal type; afdb-uniprot50 \| AF-E1IGW0-F1-MODEL\_V4 \| 1.0 \| 0.001563 \| 130 \| 0.168 \| 119 \| 84 \| 5 \| 70 \| 177 \| 14 \| 128 \| Peptidase S26B, signal peptidase \| Peptidase S26B, signal peptidase; afdb-uniprot50 \| AF-A0A374IVV9-F1-MODEL\_V4 \| 1.0 \| 0.006287 \| 127 \| 0.144 \| 111 \| 86 \| 4 \| 72 \| 177 \| 9 \| 115 \| Signal peptidase I \| Signal peptidase I; afdb-uniprot50 \| AF-A0A4Y8SEM8-F1-MODEL\_V4 \| 1.0 \| 0.006287 \| 121 \| 0.154 \| 97 \| 74 \| 3 \| 3 \| 96 \| 8 \| 99 \| XRE family transcriptional regulator \| XRE family transcriptional regulator; afdb-uniprot50 \| AF-A0A7J3QAC2-F1-MODEL\_V4 \| 1.0 \| 0.0003887 \| 110 \| 0.165 \| 211 \| 128 \| 13 \| 6 \| 177 \| 157 \| 358 \| Signal peptidase I \| Signal peptidase I; afdb-uniprot50 \| AF-A0A5S4WPM9-F1-MODEL\_V4 \| 1.0 \| 0.0001829 \| 105 \| 0.171 \| 187 \| 119 \| 9 \| 4 \| 173 \| 36 \| 203 \| Helix-turn-helix domain-containing protein \| Helix-turn-helix domain-containing protein; afdb-uniprot50 \| AF-A0A7C3AT77-F1-MODEL\_V4 \| 1.0 \| 0.002958 \| 104 \| 0.165 \| 175 \| 116 \| 8 \| 6 \| 177 \| 557 \| 704 \| Signal peptidase I \| Signal peptidase I; afdb-uniprot50 \| AF-A0A7Y8ZJC6-F1-MODEL\_V4 \| 1.0 \| 0.001563 \| 102 \| 0.094 \| 170 \| 97 \| 7 \| 5 \| 166 \| 7 \| 127 \| Transcriptional regulator with XRE-family HTH domain \| Transcriptional regulator with XRE-family HTH domain |
| Top keywords  (threshold 1.00e-02 (evalue)) | **transcriptional, regulator, domain\_containing, HTH, repressor, cro, C1\_type, Helix\_turn\_helix, Peptidase, Phage** |
| Output files | ../../similar\_structures/42\_FANPEZAQ\_CDS\_0042\_afdb-proteome\_foldseek.tsv ../../similar\_structures/42\_FANPEZAQ\_CDS\_0042\_afdb-uniprot50\_foldseek.tsv ../../similar\_structures/42\_FANPEZAQ\_CDS\_0042\_merged.svg ../../similar\_structures/42\_FANPEZAQ\_CDS\_0042\_pdb\_foldseek.tsv |

  
  
  

Return to summary | Go to previous | Go to next

  


---

**Sequence/structure alignments coloring**  
Each object in the alignment figures is colored according to its E-value following this color coding:

1e-100
10
