## Supplementary material for "Completing the BASEL phage collection to unlock hidden diversity for systematic exploration of phage-host interactions": Data S2: 44.html

FANPEZAQ\_CDS\_0044


Return to summary | Go to previous | Go to next

|  |  |
| --- | --- |
| FANPEZAQ\_CDS\_0044  Page creation date: 02 Sep 2024, 12:00  Project folder: n/a  Input sequences file: Escherichia\_virus\_HeidiAbel.gb | helicase rna dna a atp\_dependent dead box and in the hydrolase i deah complex c yes nucleotide fragment dead\_box e containing engineered factor p\_loop triphosphate adp repair fold atpase alpha or beta rossmann arch 3\_layer aba sandwich dexh\_box also domain\_containing proteins organism\_taxid transcription expressed gene escherichia coli type |

### Sequence information

|  |  |
| --- | --- |
| Name | FANPEZAQ\_CDS\_0044  44\_FANPEZAQ\_CDS\_0044 (pipeline id) |
| Imported annotations | Escherichia\_virus\_HeidiAbel Bas97 |
| Protein sequence | MGVVLRDYQQDIIDKGRQAMRRHKRILLQAPTGAGKTALASFMAAQTSQKGQQVWFICHR AELVLQTSLTFRKFGIDHGFIAAGYPMNLRAGVQVCSIDTLKNRLHTLPAPKLAIMDEAH HCGAAGWARVNAWLFENGAYVIGLSATPKRLDGTGLDDHFDDIVLGPSVRWLMDNGHLSD YRMFCPDIPDMKGVRKQMGDFSKGETAEKMDKPKLTGNIISHWLKYAKGMKTVGFAVNIA HSRHLTQAFIDSGIRAAHLDGGTDKAERKRIIQDYANDNLDVIFNVSLFGEGFDLSAIAQ TDVTIDCLIDAQPTQSLSLVLQKDGRVLRPSKGKTAIILDHAGNALRHGFPDDEREWSLA GDDSGSKSSSAGGPPPPVICKGCFNAIRRPLPPKCPHCQKELQAEAKEIEVAEGELREMD ERAKESLRAKMRREEDACKDIGALTALAMKRGLPNPSGWAIAKMGGRRFKR |
| Number of residues | 471 |
| Molecular weight (Da) | 51873.00 |
| Output files | ../../query\_sequences/44\_FANPEZAQ\_CDS\_0044.fasta |

### Putative domain architecture and protein family

#### Search results (HHblits)^1^

|  |  |
| --- | --- |
| Domain family databases searched | Pfam, Ncbi-cd, Cath, Phrogs |
| Results, scheme(s)  (Top layers only; threshold 1.00e-03 (evalue)) | xml version="1.0" encoding="utf-8" standalone="no"?       2024-09-02T21:08:22.220930 image/svg+xml   Matplotlib v3.7.2, https://matplotlib.org/ |
| Results, table  (E-value ≤ 1.00e-03 (evalue)) | db \| id \| prob \| evalue \| pvalue \| score \| cols \| query \| query\_len \| template \| template\_len \| name \| description; pfam \| PF06862 \| 99.7 \| 1.9e-22 \| 3.5e-26 \| 201.9 \| 317 \| (6, 343) \| 471 \| (1, 469) \| 495 \| UTP25 \| Utp25, U3 small nucleolar RNA-associated SSU processome protein 25; pfam \| PF02399 \| 99.6 \| 1.3e-20 \| 2.3e-24 \| 201.0 \| 309 \| (7, 345) \| 471 \| (37, 393) \| 832 \| Herpes\_ori\_bp \| Origin of replication binding protein; pfam \| PF04851 \| 97.4 \| 7.5e-08 \| 1.4e-11 \| 73.8 \| 143 \| (3, 149) \| 471 \| (2, 150) \| 151 \| ResIII \| Type III restriction enzyme, res subunit; pfam \| PF00270 \| 97.2 \| 2.1e-07 \| 4e-11 \| 72.4 \| 138 \| (7, 149) \| 471 \| (2, 161) \| 167 \| DEAD \| DEAD/DEAH box helicase; pfam \| PF07652 \| 96.9 \| 1e-06 \| 2e-10 \| 67.2 \| 131 \| (22, 156) \| 471 \| (3, 144) \| 148 \| Flavi\_DEAD \| Flavivirus DEAD domain; pfam \| PF18766 \| 96.8 \| 1.3e-06 \| 2.4e-10 \| 73.5 \| 128 \| (23, 152) \| 471 \| (23, 174) \| 239 \| SWI2\_SNF2 \| SWI2/SNF2 ATPase; pfam \| PF02562 \| 96.8 \| 1.3e-06 \| 2.5e-10 \| 70.8 \| 60 \| (4, 67) \| 471 \| (4, 65) \| 205 \| PhoH \| PhoH-like protein; pfam \| PF00176 \| 96.7 \| 1.9e-06 \| 3.6e-10 \| 76.4 \| 143 \| (8, 153) \| 471 \| (1, 164) \| 318 \| SNF2-rel\_dom \| SNF2-related domain; pfam \| PF00271 \| 96.6 \| 2.5e-06 \| 4.7e-10 \| 60.5 \| 94 \| (229, 330) \| 471 \| (15, 109) \| 110 \| Helicase\_C \| Helicase conserved C-terminal domain; pfam \| PF16203 \| 96.6 \| 2.8e-06 \| 5.2e-10 \| 73.5 \| 92 \| (229, 332) \| 471 \| (61, 154) \| 256 \| ERCC3\_RAD25\_C \| ERCC3/RAD25/XPB C-terminal helicase; pfam \| PF12340 \| 96.3 \| 8.8e-06 \| 1.6e-09 \| 69.5 \| 71 \| (2, 74) \| 471 \| (20, 95) \| 229 \| DUF3638 \| Protein of unknown function (DUF3638); pfam \| PF13872 \| 96.3 \| 8.7e-06 \| 1.6e-09 \| 71.2 \| 71 \| (4, 74) \| 471 \| (35, 112) \| 306 \| AAA\_34 \| P-loop containing NTP hydrolase pore-1; pfam \| PF03354 \| 96.0 \| 2.2e-05 \| 4.1e-09 \| 62.1 \| 68 \| (7, 74) \| 471 \| (1, 78) \| 175 \| TerL\_ATPase \| Terminase large subunit, ATPase domain; pfam \| PF05127 \| 95.5 \| 7.3e-05 \| 1.4e-08 \| 57.3 \| 46 \| (27, 72) \| 471 \| (1, 48) \| 168 \| Helicase\_RecD \| Helicase; pfam \| PF05876 \| 95.5 \| 7.5e-05 \| 1.4e-08 \| 62.9 \| 69 \| (3, 73) \| 471 \| (9, 79) \| 247 \| GpA\_ATPase \| Phage terminase large subunit gpA, ATPase domain; pfam \| PF13871 \| 95.1 \| 0.00017 \| 3.2e-08 \| 62.5 \| 74 \| (255, 335) \| 471 \| (26, 118) \| 277 \| Helicase\_C\_4 \| C-terminal domain on Strawberry notch homologue; pfam \| PF13086 \| 95.1 \| 0.00017 \| 3.3e-08 \| 59.6 \| 65 \| (5, 73) \| 471 \| (2, 70) \| 273 \| AAA\_11 \| AAA domain; pfam \| PF07517 \| 95.1 \| 0.00019 \| 3.5e-08 \| 68.9 \| 115 \| (3, 123) \| 471 \| (133, 267) \| 548 \| SecA\_DEAD \| SecA DEAD-like domain; pfam \| PF13245 \| 95.0 \| 0.00019 \| 3.6e-08 \| 52.3 \| 26 \| (22, 47) \| 471 \| (10, 35) \| 141 \| AAA\_19 \| AAA domain; pfam \| PF04466 \| 94.9 \| 0.00022 \| 4.1e-08 \| 57.5 \| 126 \| (24, 151) \| 471 \| (2, 143) \| 202 \| Terminase\_3 \| Phage terminase large subunit; pfam \| PF14617 \| 94.5 \| 0.00043 \| 8e-08 \| 58.3 \| 101 \| (22, 122) \| 471 \| (76, 205) \| 246 \| CMS1 \| U3-containing 90S pre-ribosomal complex subunit; pfam \| PF00580 \| 93.8 \| 0.00091 \| 1.7e-07 \| 55.3 \| 63 \| (5, 73) \| 471 \| (1, 67) \| 245 \| UvrD-helicase \| UvrD/REP helicase N-terminal domain; ncbi-cd \| cd09696 \| 99.8 \| 1.5e-24 \| 3e-28 \| 232.4 \| 301 \| (2, 341) \| 471 \| (12, 389) \| 843 \| Cas3\_I \| cd09696 Cas3\_I; CRISPR/Cas system-associated protein Cas3; Distinct Cas3 family with HD domain fused to C-termus of Helicase domain.; ncbi-cd \| cd09710 \| 99.7 \| 1.4e-23 \| 2.9e-27 \| 196.1 \| 294 \| (8, 328) \| 471 \| (1, 353) \| 353 \| Cas3\_I-D \| cd09710 Cas3\_I-D; CRISPR/Cas system-associated protein Cas3; Distinct diverged subfamily of Cas3 helicase domain.; ncbi-cd \| cd09639 \| 99.7 \| 3.4e-23 \| 6.7e-27 \| 194.7 \| 283 \| (26, 345) \| 471 \| (2, 332) \| 353 \| Cas3\_I \| cd09639 Cas3\_I; CRISPR/Cas system-associated protein Cas3. CRISPR (Clustered Regularly Interspaced Short Palindromic Repeats) and associated Cas proteins comprise a system for heritable host defense by prokaryotic cells against phage and other foreign DNA.; ncbi-cd \| cd09673 \| 99.5 \| 3.5e-19 \| 6.8e-23 \| 194.0 \| 328 \| (4, 347) \| 471 \| (404, 896) \| 1106 \| Cas3\_Cas2\_I-F \| cd09673 Cas3\_Cas2\_I-F; CRISPR/Cas system-associated protein Cas3/Cas2. CRISPR (Clustered Regularly Interspaced Short Palindromic Repeats) and associated Cas proteins comprise a system for heritable host defense by prokaryotic cells against phage and other foreign DNA.; ncbi-cd \| cd18013 \| 98.8 \| 4.8e-13 \| 9.4e-17 \| 116.7 \| 144 \| (5, 152) \| 471 \| (1, 160) \| 218 \| DEXQc\_bact\_SNF2 \| cd18013 DEXQc\_bact\_SNF2; DEXQ-box helicase domain of bacterial SNF2 family proteins. Proteins belonging to the SNF2 family of DNA dependent ATPases are important members of the chromatin remodeling complexes that are implicated in epigenetic control of gene expression.; ncbi-cd \| cd18073 \| 98.6 \| 5.3e-12 \| 1e-15 \| 106.2 \| 147 \| (4, 154) \| 471 \| (2, 178) \| 202 \| DEXHc\_RIG-I\_DDX58 \| cd18073 DEXHc\_RIG-I\_DDX58; DEXH-box helicase domain of RIG-I. RIG-I (Retinoic acid-inducible gene I protein), also called DEAD box protein 58 (DDX58), is a pathogen-recognition receptor that recognizes viral 5'-triphosphates carrying double-stranded RNA.; ncbi-cd \| cd17951 \| 98.5 \| 9.3e-12 \| 1.9e-15 \| 104.1 \| 149 \| (4, 156) \| 471 \| (12, 195) \| 206 \| DEADc\_DDX41 \| cd17951 DEADc\_DDX41; DEAD-box helicase domain of DEAD box protein 41. DDX41 (also called ABS and MPLPF) interacts with several spliceosomal proteins and may recognize the bacterial second messengers cyclic di-GMP and cyclic di-AMP, resulting in the induction of genes involved in the innate immune response.; ncbi-cd \| cd18066 \| 98.5 \| 1.3e-11 \| 2.6e-15 \| 107.7 \| 145 \| (5, 152) \| 471 \| (1, 171) \| 235 \| DEXHc\_RAD54B \| cd18066 DEXHc\_RAD54B; DEXH-box helicase domain of RAD54B. DNA repair and recombination protein RAD54B, also known as RDH54, binds to double-stranded DNA, displays ATPase activity in the presence of DNA, and may have a role in meiotic and mitotic recombination.; ncbi-cd \| cd18034 \| 98.5 \| 1.7e-11 \| 3.3e-15 \| 103.2 \| 144 \| (4, 152) \| 471 \| (2, 174) \| 200 \| DEXHc\_dicer \| cd18034 DEXHc\_dicer; DEXH-box helicase domain of endoribonuclease Dicer. Dicer ribonucleases cleave double-stranded RNA (dsRNA) precursors to generate microRNAs (miRNAs) and small interfering RNAs (siRNAs).; ncbi-cd \| cd00268 \| 98.5 \| 1.7e-11 \| 3.4e-15 \| 103.5 \| 149 \| (4, 156) \| 471 \| (12, 185) \| 196 \| DEADc \| cd00268 DEADc; DEAD-box helicase domain of DEAD box helicases. DEAD-box helicases comprise a diverse family of proteins involved in ATP-dependent RNA unwinding, needed in a variety of cellular processes including splicing, ribosome biogenesis and RNA degradation.; ncbi-cd \| cd18017 \| 98.4 \| 2e-11 \| 4e-15 \| 101.6 \| 144 \| (4, 154) \| 471 \| (12, 174) \| 193 \| DEXHc\_RecQ3 \| cd18017 DEXHc\_RecQ3; DEAH-box helicase domain of RecQ3. DEAD-like helicase RecQ3 (also called Werner syndrome ATP-dependent helicase or WRN) is part of the RecQ family of highly conserved DNA repair helicases that is part of the type II DEAD box helicase superfamily, a diverse family of proteins involved in ATP-dependent RNA or DNA unwinding.; ncbi-cd \| cd18014 \| 98.4 \| 2.1e-11 \| 4.2e-15 \| 102.6 \| 146 \| (5, 156) \| 471 \| (14, 187) \| 205 \| DEXHc\_RecQ5 \| cd18014 DEXHc\_RecQ5; DEAH-box helicase domain of RecQ5. ATP-dependent DNA helicase Q5 (RecQ5) is part of the RecQ family of highly conserved DNA repair helicases that is part of the type II DEAD box helicase superfamily, a diverse family of proteins involved in ATP-dependent RNA or DNA unwinding.; ncbi-cd \| cd18007 \| 98.4 \| 2.1e-11 \| 4.2e-15 \| 106.4 \| 147 \| (5, 152) \| 471 \| (1, 190) \| 239 \| DEXHc\_ATRX-like \| cd18007 DEXHc\_ATRX-like; DEXH-box helicase domain of ATRX-like proteins. This family includes ATRX-like members such as transcriptional regulator ATRX (also called alpha thalassemia/mental retardation syndrome X-linked and X-linked nuclear protein or XNP) which is involved in transcriptional regulation and chromatin remodeling, and ARIP4 (also called androgen receptor-interacting protein 4, RAD54 like 2 or RAD54L2) which modulates androgen receptor (AR)-dependent transactivation in a promoter-dependent manner.; ncbi-cd \| cd18028 \| 98.4 \| 2.5e-11 \| 4.9e-15 \| 98.7 \| 144 \| (4, 150) \| 471 \| (1, 161) \| 177 \| DEXHc\_archSki2 \| cd18028 DEXHc\_archSki2; DEXH-box helicase domain of archaeal Ski2-type helicase. Archaeal Ski2-type RNA helicases play an important role in RNA degradation, processing and splicing pathways.; ncbi-cd \| cd18050 \| 98.4 \| 3.6e-11 \| 7e-15 \| 107.2 \| 149 \| (4, 156) \| 471 \| (84, 258) \| 271 \| DEADc\_DDX17 \| cd18050 DEADc\_DDX17; DEAD-box helicase domain of DEAD box protein 17. DDX17 (also called DEAD Box Protein P72 or DEAD Box Protein P82) has a wide variety of functions including regulating the alternative splicing of exons exhibiting specific features such as the inclusion of AC-rich alternative exons in CD44 transcripts, playing a role in innate immunity, and promoting mRNA degradation mediated by the antiviral zinc-finger protein ZC3HAV1 in an ATPase-dependent manner.; ncbi-cd \| cd18033 \| 98.4 \| 4.3e-11 \| 8.6e-15 \| 97.0 \| 143 \| (4, 151) \| 471 \| (2, 163) \| 182 \| DEXDc\_FANCM \| cd18033 DEXDc\_FANCM; DEAH-box helicase domain of FANCM. Fanconi anemia group M (FANCM) protein is a DNA-dependent ATPase component of the Fanconi anemia (FA) core complex.; ncbi-cd \| cd18016 \| 98.4 \| 4.4e-11 \| 8.8e-15 \| 100.6 \| 145 \| (4, 156) \| 471 \| (17, 191) \| 208 \| DEXHc\_RecQ2\_BLM \| cd18016 DEXHc\_RecQ2\_BLM; DEAH-box helicase domain of RecQ2. ATP-dependent DNA helicase Q2 (RecQ2, also called Bloom syndrome protein homolog or BLM) is part of the RecQ family of highly conserved DNA repair helicases that is part of the type II DEAD box helicase superfamily, a diverse family of proteins involved in ATP-dependent RNA or DNA unwinding.; ncbi-cd \| cd17927 \| 98.4 \| 4.5e-11 \| 9e-15 \| 99.9 \| 145 \| (4, 152) \| 471 \| (2, 175) \| 201 \| DEXHc\_RIG-I \| cd17927 DEXHc\_RIG-I; DEXH-box helicase domain of DEAD-like helicase RIG-I family proteins. Members of the RIG-I family include FANCM, dicer, Hef, and the RIG-I-like receptors.; ncbi-cd \| cd17995 \| 98.4 \| 4.5e-11 \| 9e-15 \| 102.3 \| 149 \| (5, 154) \| 471 \| (1, 176) \| 223 \| DEXHc\_CHD6\_7\_8\_9 \| cd17995 DEXHc\_CHD6\_7\_8\_9; DEXH-box helicase domain of the chromodomain helicase DNA binding protein 6, 7, 8 and 9.; ncbi-cd \| cd18075 \| 98.4 \| 4.8e-11 \| 9.5e-15 \| 100.2 \| 146 \| (4, 153) \| 471 \| (2, 175) \| 200 \| DEXHc\_RLR-3 \| cd18075 DEXHc\_RLR-3; DEXH-box helicase domain of RLR-3. RIG-I-like receptor 3 (RLR-3, also known as laboratory of genetics and physiology 2 or LGP2 and DHX58) appears to positively and negatively regulate MDA5 and RIG-I signaling, respectively.; ncbi-cd \| cd17998 \| 98.4 \| 5.3e-11 \| 1.1e-14 \| 98.4 \| 148 \| (5, 153) \| 471 \| (1, 168) \| 187 \| DEXHc\_SMARCAD1 \| cd17998 DEXHc\_SMARCAD1; DEXH-box helicase domain of SMARCAD1. SWI/SNF-related matrix-associated actin-dependent regulator of chromatin subfamily A containing DEAD/H box 1 (SMARCAD1, also known as ATP-dependent helicase 1 or Hel1) possesses intrinsic ATP-dependent nucleosome-remodeling activity and is required for both DNA repair and heterochromatin organization.; ncbi-cd \| cd18036 \| 98.3 \| 5.4e-11 \| 1.1e-14 \| 99.1 \| 147 \| (4, 155) \| 471 \| (2, 181) \| 204 \| DEXHc\_RLR \| cd18036 DEXHc\_RLR; DEXH-box helicase domain of RIG-I-like receptors. RIG-I-like receptors (RLRs) sense cytoplasmic viral RNA and comprise RIG-I, RLR-2/MDA5 (melanoma differentiation-associated protein 5) and RLR-3/LGP2 (laboratory of genetics and physiology 2).; ncbi-cd \| cd18045 \| 98.3 \| 5.8e-11 \| 1.2e-14 \| 98.2 \| 149 \| (4, 156) \| 471 \| (21, 190) \| 201 \| DEADc\_EIF4AIII\_DDX48 \| cd18045 DEADc\_EIF4AIII\_DDX48; DEAD-box helicase domain of eukaryotic initiation factor 4A-III. Eukaryotic initiation factor 4A-III (EIF4AIII, also known as DDX48) is part of the exon junction complex (EJC) that plays a major role in posttranscriptional regulation of mRNA.; ncbi-cd \| cd18000 \| 98.3 \| 6.5e-11 \| 1.3e-14 \| 97.7 \| 148 \| (5, 153) \| 471 \| (1, 175) \| 193 \| DEXHc\_ERCC6 \| cd18000 DEXHc\_ERCC6; DEXH-box helicase domain of ERCC6. ERCC excision repair 6, chromatin remodeling factor (ERCC6, also known Cockayne syndrome group B (CSB), Rad26 in Saccharomyces cerevisiae, and Rhp26 in Schizosaccharomyces pombe) is a DNA-binding protein that is important in transcription-coupled excision repair.; ncbi-cd \| cd18006 \| 98.3 \| 6.8e-11 \| 1.4e-14 \| 100.0 \| 148 \| (5, 153) \| 471 \| (1, 165) \| 216 \| DEXHc\_CHD1L \| cd18006 DEXHc\_CHD1L; DEAH/Q-box helicase domain of CHD1L. Chromodomain helicase DNA binding protein 1 like (CHD1L, also known as ALC1) is involved in DNA repair by regulating chromatin relaxation following DNA damage.; ncbi-cd \| cd17981 \| 98.3 \| 7.1e-11 \| 1.4e-14 \| 95.9 \| 144 \| (4, 151) \| 471 \| (2, 166) \| 180 \| DEXHc\_DHX36 \| cd17981 DEXHc\_DHX36; DEXH-box helicase domain of DEAH-box helicase 36. DEAH-box helicase 36 (DHX36, also known as G4-resolvase 1 or G4R1, MLE-like protein 1 and RNA helicase associated with AU-rich element or RHAU) unwinds a G4-quadruplex in human telomerase RNA.; ncbi-cd \| cd18015 \| 98.3 \| 7.4e-11 \| 1.5e-14 \| 98.8 \| 144 \| (4, 155) \| 471 \| (18, 191) \| 209 \| DEXHc\_RecQ1 \| cd18015 DEXHc\_RecQ1; DEXH-box helicase domain of RecQ1. ATP-dependent DNA helicase Q1 (RecQ1) is part of the RecQ family of highly conserved DNA repair helicases that is part of the type II DEAD box helicase superfamily, a diverse family of proteins involved in ATP-dependent RNA or DNA unwinding.; ncbi-cd \| cd17975 \| 98.3 \| 9.1e-11 \| 1.8e-14 \| 95.0 \| 144 \| (4, 151) \| 471 \| (2, 169) \| 183 \| DEXHc\_DHX29 \| cd17975 DEXHc\_DHX29; DEXH-box helicase domain of DEAH-box helicase 29. DEAH-box helicase 29 (DHX29) is a part of the 43S pre-initiation complex involved in translation initiation of mRNAs with structured 5'-UTRs.; ncbi-cd \| cd18035 \| 98.3 \| 9.3e-11 \| 1.9e-14 \| 95.9 \| 141 \| (6, 151) \| 471 \| (3, 161) \| 181 \| DEXHc\_Hef \| cd18035 DEXHc\_Hef; DEXH-box helicase domain of Hef. Hef (helicase-associated endonuclease fork-structure) belongs to the XPF/MUS81/FANCM family of endonucleases and is involved in stalled replication fork repair.; ncbi-cd \| cd17966 \| 98.3 \| 9.6e-11 \| 1.9e-14 \| 96.1 \| 149 \| (4, 156) \| 471 \| (12, 186) \| 197 \| DEADc\_DDX5\_DDX17 \| cd17966 DEADc\_DDX5\_DDX17; DEAD-box helicase domain of ATP-dependent RNA helicases DDX5 and DDX17. DDX5 and DDX17 are members of the DEAD-box helicases, a diverse family of proteins involved in ATP-dependent RNA unwinding, needed in a variety of cellular processes including splicing, ribosome biogenesis and RNA degradation.; ncbi-cd \| cd18025 \| 98.3 \| 9.6e-11 \| 1.9e-14 \| 95.5 \| 142 \| (4, 149) \| 471 \| (1, 162) \| 192 \| DEXHc\_DDX60 \| cd18025 DEXHc\_DDX60; DEXH-box helicase domain of DEAD box protein 60. DEAD box protein 60 (DDX60) is an IFN-inducible cytoplasmic helicase that plays a role in RIG-I-mediated type I interferon (IFN) nuclease-mediated viral RNA degradation.; ncbi-cd \| cd17944 \| 98.3 \| 1e-10 \| 2e-14 \| 96.6 \| 149 \| (4, 156) \| 471 \| (12, 190) \| 202 \| DEADc\_DDX21\_DDX50 \| cd17944 DEADc\_DDX21\_DDX50; DEAD-box helicase domain of DEAD box proteins 21 and 50. DDX21 (also called Gu-Alpha and nucleolar RNA helicase 2) is an RNA helicase that acts as a sensor of the transcriptional status of both RNA polymerase (Pol) I and II.; ncbi-cd \| cd18012 \| 98.3 \| 1e-10 \| 2.1e-14 \| 99.1 \| 150 \| (2, 152) \| 471 \| (2, 163) \| 218 \| DEXQc\_arch\_SWI2\_SNF2 \| cd18012 DEXQc\_arch\_SWI2\_SNF2; DEAQ-box helicase domain of archaeal and bacterial SNF2-related proteins. Proteins belonging to SNF2 family of DNA dependent ATPases are important members of the chromatin remodeling complexes that are implicated in epigenetic control of gene expression.; ncbi-cd \| cd17941 \| 98.3 \| 1.1e-10 \| 2.3e-14 \| 95.6 \| 149 \| (4, 156) \| 471 \| (12, 185) \| 198 \| DEADc\_DDX10 \| cd17941 DEADc\_DDX10; DEAD-box helicase domain of DEAD box protein 10. Fusion of the DDX10 gene and the nucleoporin gene, NUP98, by inversion 11 (p15q22) chromosome translocation is found in the patients with de novo or therapy-related myeloid malignancies.; ncbi-cd \| cd17924 \| 98.3 \| 1.1e-10 \| 2.3e-14 \| 95.9 \| 146 \| (2, 151) \| 471 \| (15, 179) \| 189 \| DDXDc\_reverse\_gyrase \| cd17924 DDXDc\_reverse\_gyrase; DDXD-box helicase domain of reverse gyrase. Reverse gyrase modifies the topological state of DNA by introducing positive supercoils in an ATP-dependent process.; ncbi-cd \| cd17942 \| 98.3 \| 1.2e-10 \| 2.4e-14 \| 95.7 \| 149 \| (4, 156) \| 471 \| (12, 186) \| 198 \| DEADc\_DDX18 \| cd17942 DEADc\_DDX18; DEAD-box helicase domain of DEAD box protein 18. This DDX18 gene encodes a DEAD box protein and is activated by Myc protein.; ncbi-cd \| cd17965 \| 98.3 \| 1.2e-10 \| 2.4e-14 \| 101.5 \| 151 \| (4, 156) \| 471 \| (30, 234) \| 251 \| DEADc\_MRH4 \| cd17965 DEADc\_MRH4; DEAD-box helicase domain of ATP-dependent RNA helicase MRH4. Mitochondrial RNA helicase 4 (MRH4) plays an essential role during the late stages of mitochondrial ribosome or mitoribosome assembly by promoting remodeling of the 21S rRNA-protein interactions.; ncbi-cd \| cd18001 \| 98.3 \| 1.3e-10 \| 2.6e-14 \| 99.3 \| 148 \| (5, 153) \| 471 \| (1, 168) \| 232 \| DEXHc\_ERCC6L \| cd18001 DEXHc\_ERCC6L; DEXH-box helicase domain of ERCC6L. ERCC excision repair 6 like, spindle assembly checkpoint helicase (ERCC6L, also known as RAD26L) is an essential component of the mitotic spindle assembly checkpoint, by acting as a tension sensor that associates with catenated DNA which is stretched under tension until it is resolved during anaphase.; ncbi-cd \| cd17963 \| 98.3 \| 1.3e-10 \| 2.6e-14 \| 95.5 \| 151 \| (4, 156) \| 471 \| (16, 185) \| 196 \| DEADc\_DDX19\_DDX25 \| cd17963 DEADc\_DDX19\_DDX25; DEAD-box helicase domain of ATP-dependent RNA helicases DDX19 and DDX25. DDX19 (also called DEAD box RNA helicase DEAD5) and DDX25 (also called gonadotropin-regulated testicular RNA helicase (GRTH)) are members of the DEAD-box helicases, a diverse family of proteins involved in ATP-dependent RNA unwinding, needed in a variety of cellular processes including splicing, ribosome biogenesis and RNA degradation.; ncbi-cd \| cd17997 \| 98.2 \| 1.4e-10 \| 2.7e-14 \| 98.8 \| 151 \| (2, 153) \| 471 \| (1, 168) \| 222 \| DEXHc\_SMARCA1\_SMARCA5 \| cd17997 DEXHc\_SMARCA1\_SMARCA5; DEAH-box helicase domain of SMARCA1 and SMARCA5. SWI/SNF related, matrix associated, actin dependent regulator of chromatin, subfamily a, member 1 and 5 (SMARCA1 and SMARCA5) are members of the DEAD-like helicase superfamily, a diverse family of proteins involved in ATP-dependent RNA or DNA unwinding.; ncbi-cd \| cd17976 \| 98.2 \| 1.4e-10 \| 2.8e-14 \| 94.1 \| 144 \| (5, 151) \| 471 \| (2, 164) \| 178 \| DEXHc\_DHX30 \| cd17976 DEXHc\_DHX30; DEXH-box helicase domain of DEAH-box helicase 30. DEAH-box helicase 30 (DHX30) plays an important role in the assembly of the mitochondrial large ribosomal subunit.; ncbi-cd \| cd17967 \| 98.2 \| 1.4e-10 \| 2.8e-14 \| 97.9 \| 151 \| (4, 158) \| 471 \| (22, 207) \| 221 \| DEADc\_DDX3\_DDX4 \| cd17967 DEADc\_DDX3\_DDX4; DEAD-box helicase domain of ATP-dependent RNA helicases DDX3 and DDX4. This subfamily includes Drosophila melanogaster Vasa, which is essential for development.; ncbi-cd \| cd18027 \| 98.2 \| 1.4e-10 \| 2.9e-14 \| 93.9 \| 144 \| (3, 151) \| 471 \| (7, 158) \| 179 \| DEXHc\_SKIV2L \| cd18027 DEXHc\_SKIV2L; DEXH-box helicase domain of SKIV2L. Superkiller viralicidic activity 2-like (SKIV2L, also called SKI2 or DHX13) plays a role in a number of cellular processes involving alteration of RNA secondary structure such as translation initiation, nuclear and mitochondrial splicing, and ribosome and spliceosome assembly.; ncbi-cd \| cd17923 \| 98.2 \| 1.5e-10 \| 2.9e-14 \| 94.3 \| 141 \| (5, 149) \| 471 \| (1, 173) \| 182 \| DEXHc\_Hrq1-like \| cd17923 DEXHc\_Hrq1-like; DEAH-box helicase domain of Hrq1 and similar proteins. Yeast Hrq1, similar to RecQ4, plays a role in DNA inter-strand crosslink (ICL) repair and in telomere maintenance.; ncbi-cd \| cd17952 \| 98.2 \| 1.5e-10 \| 3e-14 \| 94.7 \| 149 \| (4, 156) \| 471 \| (12, 186) \| 197 \| DEADc\_DDX42 \| cd17952 DEADc\_DDX42; DEAD-box helicase domain of DEAD box protein 42. DDX42 (also called Splicing Factor 3B-Associated 125 kDa Protein, RHELP, or RNAHP) is an NTPase with a preference for ATP, the hydrolysis of which is enhanced by various RNA substrates.; ncbi-cd \| cd18038 \| 98.2 \| 1.6e-10 \| 3.1e-14 \| 98.7 \| 148 \| (4, 152) \| 471 \| (1, 183) \| 229 \| DEXXQc\_Helz-like \| cd18038 DEXXQc\_Helz-like; DEXXQ/H-box helicase domain of Helz-like helicase. This subfamily contains HELZ, Mov10L1, and similar proteins.; ncbi-cd \| cd17991 \| 98.2 \| 1.7e-10 \| 3.4e-14 \| 95.0 \| 147 \| (3, 152) \| 471 \| (14, 177) \| 193 \| DEXHc\_TRCF \| cd17991 DEXHc\_TRCF; DEXH/Q-box helicase domain of the transcription-repair coupling factor. Transcription-repair coupling factor (TrcF) dissociates transcription elongation complexes blocked at nonpairing lesions and mediates recruitment of DNA repair proteins.; ncbi-cd \| cd18032 \| 98.2 \| 1.7e-10 \| 3.4e-14 \| 92.0 \| 146 \| (5, 152) \| 471 \| (1, 155) \| 163 \| DEXHc\_RE\_I\_III\_res \| cd18032 DEXHc\_RE\_I\_III\_res; DEXH-box helicase domain of type III restriction enzyme res subunit. Members of this cd includes both type I and type III restriction enzymes.; ncbi-cd \| cd17945 \| 98.2 \| 1.7e-10 \| 3.5e-14 \| 97.6 \| 149 \| (4, 156) \| 471 \| (12, 209) \| 220 \| DEADc\_DDX23 \| cd17945 DEADc\_DDX23; DEAD-box helicase domain of DEAD box protein 23. DDX23 (also called U5 snRNP 100kD protein and PRP28 homolog) is involved in pre-mRNA splicing and its phosphorylated form (by SRPK2) is required for spliceosomal B complex formation.; ncbi-cd \| cd18047 \| 98.2 \| 1.8e-10 \| 3.6e-14 \| 96.1 \| 151 \| (4, 156) \| 471 \| (23, 194) \| 205 \| DEADc\_DDX19 \| cd18047 DEADc\_DDX19; DEAD-box helicase domain of DEAD box protein 19. DDX19 is an RNA helicase involved in both mRNA (mRNA) export from the nucleus into the cytoplasm and in mRNA translation.; ncbi-cd \| cd17950 \| 98.2 \| 1.9e-10 \| 3.7e-14 \| 95.5 \| 149 \| (4, 156) \| 471 \| (24, 196) \| 208 \| DEADc\_DDX39 \| cd17950 DEADc\_DDX39; DEAD-box helicase domain of DEAD box protein 39. DDX39A is involved in pre-mRNA splicing and is required for the export of mRNA out of the nucleus.; ncbi-cd \| cd17953 \| 98.2 \| 1.9e-10 \| 3.8e-14 \| 96.7 \| 149 \| (4, 156) \| 471 \| (34, 211) \| 222 \| DEADc\_DDX46 \| cd17953 DEADc\_DDX46; DEAD-box helicase domain of DEAD box protein 46. DDX46 (also called Prp5-like DEAD-box protein) is a component of the 17S U2 snRNP complex.; ncbi-cd \| cd17958 \| 98.2 \| 1.9e-10 \| 3.8e-14 \| 94.2 \| 149 \| (4, 156) \| 471 \| (12, 186) \| 197 \| DEADc\_DDX43\_DDX53 \| cd17958 DEADc\_DDX43\_DDX53; DEAD-box helicase domain of DEAD box proteins 43 and 53. DDX43 (also called cancer/testis antigen 13 or helical antigen) displays tumor-specific expression.; ncbi-cd \| cd18019 \| 98.2 \| 2e-10 \| 4e-14 \| 95.9 \| 144 \| (4, 150) \| 471 \| (17, 194) \| 214 \| DEXHc\_Brr2\_1 \| cd18019 DEXHc\_Brr2\_1; N-terminal DEXH-box helicase domain of spliceosomal Brr2 RNA helicase. Brr2 is a type II DEAD box helicase that mediates spliceosome catalytic activation.; ncbi-cd \| cd17947 \| 98.2 \| 2e-10 \| 4e-14 \| 93.6 \| 149 \| (4, 156) \| 471 \| (12, 185) \| 196 \| DEADc\_DDX27 \| cd17947 DEADc\_DDX27; DEAD-box helicase domain of DEAD box protein 27. DDX27 (also called RHLP, deficiency of ribosomal subunits protein 1 homolog, and probable ATP-dependent RNA helicase DDX27) is involved in the processing of 5.; ncbi-cd \| cd18049 \| 98.2 \| 2e-10 \| 4.1e-14 \| 98.4 \| 149 \| (4, 156) \| 471 \| (46, 220) \| 234 \| DEADc\_DDX5 \| cd18049 DEADc\_DDX5; DEAD-box helicase domain of DEAD box protein 5. DDX5 (also called RNA helicase P68, HLR1, G17P1, or HUMP68) is involved in pathways that include the alteration of RNA structures, plays a role as a coregulator of transcription, a regulator of splicing, and in the processing of small noncoding RNAs.; ncbi-cd \| cd17988 \| 98.2 \| 2.1e-10 \| 4.2e-14 \| 92.8 \| 143 \| (5, 151) \| 471 \| (3, 162) \| 180 \| DEXHc\_TDRD9 \| cd17988 DEXHc\_TDRD9; DEXH-box helicase domain of tudor domain containing 9. Tudor domain containing 9 (TDRD9, also known as HIG-1or NET54 or C14orf75) is a part of the nuclear PIWI-interacting RNA (piRNA) pathway essential for transposon silencing and male fertility TDRD9 belongs to the DEAD-like helicase superfamily, a diverse family of proteins involved in ATP-dependent RNA or DNA unwinding.; ncbi-cd \| cd18009 \| 98.2 \| 2.2e-10 \| 4.5e-14 \| 98.0 \| 150 \| (2, 152) \| 471 \| (1, 170) \| 236 \| DEXHc\_HELLS\_SMARCA6 \| cd18009 DEXHc\_HELLS\_SMARCA6; DEXH-box helicase domain of HELLS. HELLS (helicase, lymphoid specific, also known as Lsh or SMARCA6) is a major epigenetic regulator crucial for normal heterochromatin structure and function.; ncbi-cd \| cd17962 \| 98.2 \| 2.3e-10 \| 4.6e-14 \| 93.0 \| 149 \| (4, 156) \| 471 \| (12, 182) \| 193 \| DEADc\_DDX59 \| cd17962 DEADc\_DDX59; DEAD-box helicase domain of DEAD box protein 59. DDX59 plays an important role in lung cancer development by promoting DNA replication.; ncbi-cd \| cd18029 \| 98.2 \| 2.5e-10 \| 5e-14 \| 91.4 \| 143 \| (3, 151) \| 471 \| (7, 166) \| 169 \| DEXHc\_XPB \| cd18029 DEXHc\_XPB; DEXH-box helicase domain of TFIIH XPB subunit and similar proteins. TFIIH basal transcription factor complex helicase XPB subunit (also known as DNA excision repair protein ERCC-3 or TFIIH 89 kDa subunit) is the ATP-dependent 3'-5' DNA helicase component of the core-TFIIH basal transcription factor, involved in nucleotide excision repair (NER) of DNA and, when complexed to CAK, in RNA transcription by RNA polymerase II.; ncbi-cd \| cd18005 \| 98.2 \| 2.6e-10 \| 5.2e-14 \| 100.0 \| 150 \| (5, 156) \| 471 \| (1, 185) \| 245 \| DEXHc\_ERCC6L2 \| cd18005 DEXHc\_ERCC6L2; DEXH-box helicase domain of ERCC6L2. ERCC excision repair 6 like 2 (ERCC6L2, also known as RAD26L) may play a role in DNA repair and mitochondrial function.; ncbi-cd \| cd18026 \| 98.2 \| 2.6e-10 \| 5.2e-14 \| 94.4 \| 145 \| (4, 150) \| 471 \| (16, 183) \| 202 \| DEXHc\_POLQ-like \| cd18026 DEXHc\_POLQ-like; DEXH-box helicase domain of DNA polymerase theta. DNA polymerase theta (POLQ) is important in the repair of genomic double-strand breaks (DSBs).; ncbi-cd \| cd17961 \| 98.2 \| 2.6e-10 \| 5.3e-14 \| 93.8 \| 149 \| (4, 156) \| 471 \| (16, 194) \| 206 \| DEADc\_DDX56 \| cd17961 DEADc\_DDX56; DEAD-box helicase domain of DEAD box protein 56. DDX56 is a helicase required for assembly of infectious West Nile virus particles.; ncbi-cd \| cd17999 \| 98.2 \| 2.7e-10 \| 5.4e-14 \| 97.3 \| 148 \| (5, 153) \| 471 \| (1, 169) \| 232 \| DEXHc\_Mot1 \| cd17999 DEXHc\_Mot1; DEXH-box helicase domain of Mot1. Modifier of transcription 1 (Mot1, also known as TAF172 in eukaryotes) regulates transcription in association with TATA binding protein (TBP).; ncbi-cd \| cd18051 \| 98.2 \| 2.7e-10 \| 5.4e-14 \| 98.8 \| 149 \| (4, 156) \| 471 \| (43, 232) \| 249 \| DEADc\_DDX3 \| cd18051 DEADc\_DDX3; DEAD-box helicase domain of DEAD box protein 3. DDX3 (also called helicase-like protein, DEAD box, X isoform, or DDX14) has been reported to display a high level of RNA-independent ATPase activity stimulated by both RNA and DNA.; ncbi-cd \| cd18046 \| 98.2 \| 2.8e-10 \| 5.6e-14 \| 93.4 \| 149 \| (4, 156) \| 471 \| (21, 190) \| 201 \| DEADc\_EIF4AII\_EIF4AI\_DDX2 \| cd18046 DEADc\_EIF4AII\_EIF4AI\_DDX2; DEAD-box helicase domain of eukaryotic initiation factor 4A-I and 4-II. Eukaryotic initiation factor 4A-I (DDX2A) and eukaryotic initiation factor 4A-II (DDX2B) are involved in cap recognition and are required for mRNA binding to ribosome.; ncbi-cd \| cd17955 \| 98.2 \| 2.8e-10 \| 5.7e-14 \| 93.4 \| 149 \| (4, 156) \| 471 \| (21, 193) \| 204 \| DEADc\_DDX49 \| cd17955 DEADc\_DDX49; DEAD-box helicase domain of DEAD box protein 49. DDX49 (also called Dbp8) is a member of the DEAD-box helicases, a diverse family of proteins involved in ATP-dependent RNA unwinding, needed in a variety of cellular processes including splicing, ribosome biogenesis and RNA degradation.; ncbi-cd \| cd17987 \| 98.2 \| 3e-10 \| 5.9e-14 \| 91.8 \| 144 \| (4, 151) \| 471 \| (2, 162) \| 176 \| DEXHc\_YTHDC2 \| cd17987 DEXHc\_YTHDC2; DEXH-box helicase domain of YTH domain containing 2. YTH domain containing 2 (YTHDC2) regulates mRNA translation and stability via binding to N6-methyladenosine, a modified RNA nucleotide enriched in the stop codons and 3' UTRs of eukaryotic messenger RNAs.; ncbi-cd \| cd17943 \| 98.2 \| 3e-10 \| 6e-14 \| 92.1 \| 148 \| (4, 155) \| 471 \| (12, 180) \| 192 \| DEADc\_DDX20 \| cd17943 DEADc\_DDX20; DEAD-box helicase domain of DEAD box protein 20. DDX20 (also called DEAD Box Protein DP 103, Component Of Gems 3, Gemin-3, and SMN-Interacting Protein) interacts directly with SMN (survival of motor neurons), the spinal muscular atrophy gene product, and may play a catalytic role in the function of the SMN complex on ribonucleoproteins.; ncbi-cd \| cd18024 \| 98.1 \| 3.2e-10 \| 6.5e-14 \| 94.2 \| 143 \| (4, 151) \| 471 \| (32, 182) \| 205 \| DEXHc\_Mtr4-like \| cd18024 DEXHc\_Mtr4-like; DEXH-box helicase domain of ATP-dependent RNA helicase Mtr4. Mtr4 (also known as DOB1 or SKIV2L2) is a type II DEAD box helicase that plays a role in the processing of structured RNAs, including the maturation of 5.; ncbi-cd \| cd17985 \| 98.1 \| 3.6e-10 \| 7.2e-14 \| 91.0 \| 144 \| (4, 151) \| 471 \| (2, 163) \| 177 \| DEXHc\_DHX57 \| cd17985 DEXHc\_DHX57; DEXH-box helicase domain of DEAH-box helicase 57. DEAH-box helicase 57 (DHX57) belongs to the DEAD-like helicase superfamily, a diverse family of proteins involved in ATP-dependent RNA or DNA unwinding.; ncbi-cd \| cd18031 \| 98.1 \| 3.8e-10 \| 7.7e-14 \| 89.3 \| 143 \| (5, 152) \| 471 \| (1, 153) \| 161 \| DEXHc\_UvsW \| cd18031 DEXHc\_UvsW; DEXH-box helicase domain of bacteriophage UvsW. Bacteriophage UvsW is part of the WXY system that repairs DNA damage by a process that involves homologous recombination.; ncbi-cd \| cd17919 \| 98.1 \| 4.1e-10 \| 8.1e-14 \| 91.6 \| 147 \| (5, 152) \| 471 \| (1, 163) \| 182 \| DEXHc\_Snf \| cd17919 DEXHc\_Snf; DEXH/Q-box helicase domain of DEAD-like helicase Snf family proteins. Sucrose Non-Fermenting (SNF) proteins DEAD-like helicases superfamily.; ncbi-cd \| cd18022 \| 98.1 \| 4.2e-10 \| 8.4e-14 \| 91.5 \| 143 \| (5, 150) \| 471 \| (2, 170) \| 189 \| DEXHc\_ASCC3\_2 \| cd18022 DEXHc\_ASCC3\_2; C-terminal DEXH-box helicase domain of Activating signal cointegrator 1 complex subunit 3.; ncbi-cd \| cd17940 \| 98.1 \| 4.4e-10 \| 8.9e-14 \| 91.8 \| 149 \| (4, 156) \| 471 \| (21, 190) \| 201 \| DEADc\_DDX6 \| cd17940 DEADc\_DDX6; DEAD-box helicase domain of DEAD box protein 6. DEAD box protein 6 (DDX6, also known as Rck or p54) participates in mRNA regulation mediated by miRNA-mediated silencing.; ncbi-cd \| cd17964 \| 98.1 \| 4.4e-10 \| 8.9e-14 \| 92.9 \| 150 \| (4, 156) \| 471 \| (16, 199) \| 211 \| DEADc\_MSS116 \| cd17964 DEADc\_MSS116; DEAD-box helicase domain of DEAD-box helicase Mss116. Mss116 is an RNA chaperone important for mitochondrial group I and II intron splicing, translational activation, and RNA end processing.; ncbi-cd \| cd17925 \| 98.1 \| 4.7e-10 \| 9.5e-14 \| 86.9 \| 137 \| (8, 147) \| 471 \| (1, 143) \| 143 \| DEXDc\_ComFA \| cd17925 DEXDc\_ComFA; DEXD-box helicase domain of ComFA. ATP-dependent helicase ComFA (also called ComF operon protein 1) is part of the complex mediating the binding and uptake of single-stranded DNA.; ncbi-cd \| cd18011 \| 98.1 \| 5.3e-10 \| 1.1e-13 \| 93.6 \| 146 \| (5, 153) \| 471 \| (1, 169) \| 207 \| DEXDc\_RapA \| cd18011 DEXDc\_RapA; DEXH-box helicase domain of RapA. In bacteria, RapA is an RNA polymerase (RNAP)-associated SWI2/SNF2 (switch/sucrose non-fermentable) protein that mediates RNAP recycling during transcription.; ncbi-cd \| cd17959 \| 98.1 \| 5.3e-10 \| 1.1e-13 \| 91.6 \| 149 \| (4, 156) \| 471 \| (23, 194) \| 205 \| DEADc\_DDX54 \| cd17959 DEADc\_DDX54; DEAD-box helicase domain of DEAD box protein 54. DDX54 interacts in a hormone-dependent manner with nuclear receptors, and represses their transcriptional activity.; ncbi-cd \| cd17926 \| 98.1 \| 5.5e-10 \| 1.1e-13 \| 86.7 \| 137 \| (5, 148) \| 471 \| (1, 146) \| 146 \| DEXHc\_RE \| cd17926 DEXHc\_RE; DEXH-box helicase domain of DEAD-like helicase restriction enzyme family proteins.; ncbi-cd \| cd18021 \| 98.1 \| 5.7e-10 \| 1.1e-13 \| 91.9 \| 145 \| (4, 151) \| 471 \| (3, 173) \| 191 \| DEXHc\_Brr2\_2 \| cd18021 DEXHc\_Brr2\_2; C-terminal D[D/E]X[H/Q]-box helicase domain of spliceosomal Brr2 RNA helicase. Brr2 is a type II DEAD box helicase that mediates spliceosome catalytic activation.; ncbi-cd \| cd18074 \| 98.1 \| 5.7e-10 \| 1.1e-13 \| 94.2 \| 144 \| (4, 151) \| 471 \| (2, 189) \| 216 \| DEXHc\_RLR-2 \| cd18074 DEXHc\_RLR-2; DEXH-box helicase domain of RLR-2. RIG-I-like receptor 2 (RLR-2, also known as melanoma differentiation-associated protein 5 or Mda5 and IFIH1) is a viral double-stranded RNA (dsRNA) receptor that shares sequence similarity and signaling pathways with RIG-I, yet plays essential functions in antiviral immunity through distinct specificity for viral RNA.; ncbi-cd \| cd18052 \| 98.1 \| 5.8e-10 \| 1.1e-13 \| 97.6 \| 149 \| (4, 156) \| 471 \| (65, 247) \| 264 \| DEADc\_DDX4 \| cd18052 DEADc\_DDX4; DEAD-box helicase domain of DEAD box protein 4. DEAD box protein 4 (DDX4, also known as VASA homolog) is an ATP-dependent RNA helicase required during spermatogenesis and is essential for the germline integrity.; ncbi-cd \| cd18018 \| 98.1 \| 6e-10 \| 1.2e-13 \| 92.9 \| 147 \| (4, 154) \| 471 \| (12, 182) \| 201 \| DEXHc\_RecQ4-like \| cd18018 DEXHc\_RecQ4-like; DEAH-box helicase domain of RecQ4 and similar proteins. ATP-dependent DNA helicase Q4 (RecQ4) is part of the RecQ family of highly conserved DNA repair helicases that is part of the type II DEAD box helicase superfamily, a diverse family of proteins involved in ATP-dependent RNA or DNA unwinding.; ncbi-cd \| cd17921 \| 98.1 \| 6.2e-10 \| 1.2e-13 \| 89.7 \| 142 \| (4, 149) \| 471 \| (1, 163) \| 181 \| DEXHc\_Ski2 \| cd17921 DEXHc\_Ski2; DEXH-box helicase domain of DEAD-like helicase Ski2 family proteins. Ski2-like RNA helicases play an important role in RNA degradation, processing, and splicing pathways.; ncbi-cd \| cd17939 \| 98.1 \| 6.1e-10 \| 1.2e-13 \| 90.3 \| 149 \| (4, 156) \| 471 \| (19, 188) \| 199 \| DEADc\_EIF4A \| cd17939 DEADc\_EIF4A; DEAD-box helicase domain of eukaryotic initiation factor 4A. The eukaryotic initiation factor-4A (eIF4A) family consists of 3 proteins EIF4A1, EIF4A2, and EIF4A3.; ncbi-cd \| cd17960 \| 98.1 \| 6.2e-10 \| 1.2e-13 \| 91.1 \| 147 \| (4, 154) \| 471 \| (12, 189) \| 202 \| DEADc\_DDX55 \| cd17960 DEADc\_DDX55; DEAD-box helicase domain of DEAD box protein 55. DDX55 is a member of the DEAD-box helicases, a diverse family of proteins involved in ATP-dependent RNA unwinding, needed in a variety of cellular processes including splicing, ribosome biogenesis and RNA degradation.; ncbi-cd \| cd17990 \| 98.1 \| 6.5e-10 \| 1.3e-13 \| 88.9 \| 145 \| (5, 152) \| 471 \| (2, 161) \| 174 \| DEXHc\_HrpB \| cd17990 DEXHc\_HrpB; DEXH-box helicase domain of ATP-dependent helicase HrpB. HrpB is part of the HrpB-HrpA two-partner secretion (TPS) system, a secretion pathway important to the secretion of large virulence-associated proteins.; ncbi-cd \| cd17957 \| 98.0 \| 7.2e-10 \| 1.5e-13 \| 90.2 \| 149 \| (4, 156) \| 471 \| (12, 185) \| 198 \| DEADc\_DDX52 \| cd17957 DEADc\_DDX52; DEAD-box helicase domain of DEAD box protein 52. DDX52 (also called ROK1 and HUSSY19) is ubiquitously expressed in testis, endometrium, and other tissues in humans.; ncbi-cd \| cd17948 \| 98.0 \| 7.6e-10 \| 1.5e-13 \| 93.7 \| 150 \| (4, 157) \| 471 \| (12, 202) \| 231 \| DEADc\_DDX28 \| cd17948 DEADc\_DDX28; DEAD-box helicase domain of DEAD box protein 28. DDX28 (also called mitochondrial DEAD-box polypeptide 28) plays an essential role in facilitating the proper assembly of the mitochondrial large ribosomal subunit and its helicase activity is essential for this function.; ncbi-cd \| cd17949 \| 98.0 \| 7.9e-10 \| 1.6e-13 \| 92.3 \| 148 \| (4, 155) \| 471 \| (13, 202) \| 214 \| DEADc\_DDX31 \| cd17949 DEADc\_DDX31; DEAD-box helicase domain of DEAD box protein 31. DDX31 (also called helicain or G2 helicase) plays a role in ribosome biogenesis and TP53/p53 regulation through its interaction with NPM1.; ncbi-cd \| cd17938 \| 98.0 \| 8.3e-10 \| 1.7e-13 \| 91.0 \| 141 \| (4, 152) \| 471 \| (21, 188) \| 204 \| DEADc\_DDX1 \| cd17938 DEADc\_DDX1; DEAD-box helicase domain of DEAD box protein 1. DEAD box protein 1 (DDX1) acts as an ATP-dependent RNA helicase, able to unwind both RNA-RNA and RNA-DNA duplexes.; ncbi-cd \| cd17977 \| 98.0 \| 8.7e-10 \| 1.7e-13 \| 88.0 \| 144 \| (4, 151) \| 471 \| (2, 162) \| 176 \| DEXHc\_DHX32 \| cd17977 DEXHc\_DHX32; DEXH-box helicase domain of DEAH-box helicase 32. DEAH-box helicase 32 (DHX32) belongs to the DEAD-like helicase superfamily, a diverse family of proteins involved in ATP-dependent RNA or DNA unwinding.; ncbi-cd \| cd18008 \| 98.0 \| 9.7e-10 \| 1.9e-13 \| 95.1 \| 142 \| (5, 152) \| 471 \| (1, 189) \| 241 \| DEXDc\_SHPRH-like \| cd18008 DEXDc\_SHPRH-like; DEXH-box helicase domain of SHPRH-like proteins. The SHPRH-like subgroup belongs to the DEAD-like helicase superfamily, a diverse family of proteins involved in ATP-dependent RNA or DNA unwinding.; ncbi-cd \| cd17992 \| 98.0 \| 1e-09 \| 2e-13 \| 93.6 \| 146 \| (4, 152) \| 471 \| (45, 207) \| 225 \| DEXHc\_RecG \| cd17992 DEXHc\_RecG; DEXH/Q-box helicase domain of RecG. ATP-dependent DNA helicase RecG plays a critical role in recombination and DNA repair.; ncbi-cd \| cd18004 \| 98.0 \| 1.1e-09 \| 2.2e-13 \| 93.8 \| 147 \| (5, 152) \| 471 \| (1, 176) \| 240 \| DEXHc\_RAD54 \| cd18004 DEXHc\_RAD54; DEXH-box helicase domain of RAD54. RAD54 proteins play a role in recombination. They are members of the DEAD-like helicase superfamily, a diverse family of proteins involved in ATP-dependent RNA or DNA unwinding.; ncbi-cd \| cd18010 \| 98.0 \| 1.1e-09 \| 2.3e-13 \| 92.0 \| 143 \| (5, 152) \| 471 \| (1, 154) \| 213 \| DEXHc\_HARP\_SMARCAL1 \| cd18010 DEXHc\_HARP\_SMARCAL1; DEXH-box helicase domain of SMARCAL1. SMARCAL1 (SWI/SNF related, matrix associated, actin dependent regulator of chromatin, subfamily a like 1, also known as HARP) is recruited to stalled replication forks to promote repair and helps restart replication.; ncbi-cd \| cd17946 \| 98.0 \| 1.3e-09 \| 2.6e-13 \| 93.2 \| 146 \| (4, 152) \| 471 \| (12, 197) \| 235 \| DEADc\_DDX24 \| cd17946 DEADc\_DDX24; DEAD-box helicase domain of DEAD box protein 24. The human DDX24 gene encodes a DEAD box protein, which shows little similarity to any of the other known human DEAD box proteins, but shows a high similarity to mouse Ddx24 at the amino acid level.; ncbi-cd \| cd17954 \| 98.0 \| 1.5e-09 \| 3e-13 \| 88.4 \| 148 \| (4, 155) \| 471 \| (22, 191) \| 203 \| DEADc\_DDX47 \| cd17954 DEADc\_DDX47; DEAD-box helicase domain of DEAD box protein 47. DDX47 (also called E4-DEAD box protein) can shuttle between the nucleus and the cytoplasm, and has an RNA-independent ATPase activity.; ncbi-cd \| cd18020 \| 98.0 \| 1.5e-09 \| 3e-13 \| 88.9 \| 144 \| (4, 150) \| 471 \| (1, 179) \| 199 \| DEXHc\_ASCC3\_1 \| cd18020 DEXHc\_ASCC3\_1; N-terminal DEXH-box helicase domain of Activating signal cointegrator 1 complex subunit 3.; ncbi-cd \| cd17930 \| 98.0 \| 1.5e-09 \| 3e-13 \| 87.7 \| 131 \| (25, 155) \| 471 \| (3, 179) \| 186 \| DEXHc\_cas3 \| cd17930 DEXHc\_cas3; DEXH/Q-box helicase domain of Cas3. CRISPR-associated (Cas) 3 is a nuclease-helicase responsible for degradation of dsDNA.; ncbi-cd \| cd18068 \| 98.0 \| 1.5e-09 \| 3.1e-13 \| 92.9 \| 147 \| (5, 152) \| 471 \| (1, 197) \| 246 \| DEXHc\_ATRX \| cd18068 DEXHc\_ATRX; DEXH-box helicase domain of ATRX. Transcriptional regulator ATRX (also called alpha thalassemia/mental retardation syndrome X-linked and X-linked nuclear protein or XNP) is involved in transcriptional regulation and chromatin remodeling.; ncbi-cd \| cd18048 \| 97.9 \| 1.7e-09 \| 3.4e-13 \| 92.2 \| 151 \| (4, 156) \| 471 \| (40, 211) \| 229 \| DEADc\_DDX25 \| cd18048 DEADc\_DDX25; DEAD-box helicase domain of DEAD box protein 25. DDX25 (also called gonadotropin-regulated testicular RNA helicase (GRTH) is a testis-specific protein essential for completion of spermatogenesis.; ncbi-cd \| cd17918 \| 97.9 \| 1.7e-09 \| 3.5e-13 \| 87.1 \| 144 \| (4, 151) \| 471 \| (15, 164) \| 180 \| DEXHc\_RecG \| cd17918 DEXHc\_RecG; DEXH/Q-box helicase domain of DEAD-like helicase RecG family proteins. The DEAD-like helicase RecG family is part of the DEAD-like helicases superfamily, a diverse family of proteins involved in ATP-dependent RNA or DNA unwinding.; ncbi-cd \| cd17928 \| 97.9 \| 2e-09 \| 4e-13 \| 92.4 \| 141 \| (5, 151) \| 471 \| (42, 213) \| 230 \| DEXDc\_SecA \| cd17928 DEXDc\_SecA; DEXD-box helicase domain of SecA. SecA is a part of the Sec translocase that transports the vast majority of bacterial and ER-exported proteins.; ncbi-cd \| cd18030 \| 97.9 \| 2.1e-09 \| 4.1e-13 \| 89.3 \| 148 \| (3, 151) \| 471 \| (20, 189) \| 208 \| DEXHc\_RE\_I\_HsdR \| cd18030 DEXHc\_RE\_I\_HsdR; DEXH-box helicase domain of type I restriction enzyme HdsR subunit. The HdsR motor subunit of type I restriction-modification enzymes contains the DNA cleavage and ATP-dependent DNA translocation activities of the heteromeric complex.; ncbi-cd \| cd17974 \| 97.9 \| 2.3e-09 \| 4.7e-13 \| 84.8 \| 144 \| (4, 151) \| 471 \| (2, 160) \| 174 \| DEXHc\_DHX16 \| cd17974 DEXHc\_DHX16; DEXH-box helicase domain of DEAH-box helicase 16. DEAH-box helicase 16 (DHX16) is probably involved in pre-mRNA splicing.; ncbi-cd \| cd00046 \| 97.9 \| 2.4e-09 \| 4.8e-13 \| 82.6 \| 123 \| (25, 147) \| 471 \| (3, 146) \| 146 \| SF2-N \| cd00046 SF2-N; N-terminal DEAD/H-box helicase domain of superfamily 2 helicases. The DEAD/H-like superfamily 2 helicases comprise a diverse family of proteins involved in ATP-dependent RNA or DNA unwinding.; ncbi-cd \| cd18023 \| 97.9 \| 2.6e-09 \| 5.2e-13 \| 87.4 \| 142 \| (5, 149) \| 471 \| (2, 181) \| 206 \| DEXHc\_HFM1 \| cd18023 DEXHc\_HFM1; DEXH-box helicase domain of ATP-dependent DNA helicase HFM1. HFM1 is a type II DEAD box helicase, required for crossover formation and complete synapsis of homologous chromosomes during meiosis.; ncbi-cd \| cd17929 \| 97.9 \| 2.9e-09 \| 5.9e-13 \| 85.0 \| 143 \| (9, 154) \| 471 \| (1, 163) \| 178 \| DEXHc\_priA \| cd17929 DEXHc\_priA; DEXH-box helicase domain of PriA. PriA, also known as replication factor Y or primosomal protein N', is a 3'-5' superfamily 2 DNA helicase that acts to remodel stalled replication forks and as a specificity factor for origin-independent assembly of a new replisome at the stalled fork.; ncbi-cd \| cd18002 \| 97.9 \| 3.1e-09 \| 6.2e-13 \| 88.9 \| 147 \| (5, 152) \| 471 \| (1, 170) \| 229 \| DEXQc\_INO80 \| cd18002 DEXQc\_INO80; DEAQ-box helicase domain of INO80. INO80 is the catalytic ATPase subunit of the INO80 chromatin remodeling complex.; ncbi-cd \| cd18054 \| 97.8 \| 3.3e-09 \| 6.5e-13 \| 89.7 \| 149 \| (3, 152) \| 471 \| (19, 189) \| 237 \| DEXHc\_CHD2 \| cd18054 DEXHc\_CHD2; DEAH-box helicase domain of the chromodomain helicase DNA binding protein 2. Chromodomain-helicase-DNA-binding protein 2 (CHD2) is a DNA-binding helicase that specifically binds to the promoter of target genes, leading to chromatin remodeling, possibly by promoting deposition of histone H3.; ncbi-cd \| cd17956 \| 97.8 \| 3.5e-09 \| 7e-13 \| 89.7 \| 121 \| (4, 124) \| 471 \| (12, 165) \| 231 \| DEADc\_DDX51 \| cd17956 DEADc\_DDX51; DEAD-box helicase domain of DEAD box protein 51. DDX51 aids cell cancer proliferation by regulating multiple signalling pathways.; ncbi-cd \| cd18053 \| 97.8 \| 3.5e-09 \| 7e-13 \| 89.5 \| 149 \| (3, 152) \| 471 \| (19, 189) \| 237 \| DEXHc\_CHD1 \| cd18053 DEXHc\_CHD1; DEAH-box helicase domain of the chromodomain helicase DNA binding protein 1. Chromodomain-helicase-DNA-binding protein 1 (CHD1) is an ATP-dependent chromatin-remodeling factor which functions as substrate recognition component of the transcription regulatory histone acetylation (HAT) complex SAGA.; ncbi-cd \| cd18055 \| 97.8 \| 3.8e-09 \| 7.6e-13 \| 89.0 \| 147 \| (5, 152) \| 471 \| (1, 183) \| 232 \| DEXHc\_CHD3 \| cd18055 DEXHc\_CHD3; DEAH-box helicase domain of the chromodomain helicase DNA binding protein 3. Chromodomain-helicase-DNA-binding protein 3 (CHD3) is a component of the histone deacetylase NuRD complex which participates in the remodeling of chromatin by deacetylating histones.; ncbi-cd \| cd18059 \| 97.8 \| 3.9e-09 \| 7.8e-13 \| 87.7 \| 147 \| (5, 152) \| 471 \| (1, 173) \| 222 \| DEXHc\_CHD7 \| cd18059 DEXHc\_CHD7; DEAH-box helicase domain of the chromodomain helicase DNA binding protein 7. Chromodomain-helicase-DNA-binding protein 7 (CHD7) is a probable transcription regulator.; ncbi-cd \| cd18058 \| 97.8 \| 3.9e-09 \| 7.9e-13 \| 87.9 \| 147 \| (5, 152) \| 471 \| (1, 173) \| 222 \| DEXHc\_CHD6 \| cd18058 DEXHc\_CHD6; DEAH-box helicase domain of the chromodomain helicase DNA binding protein 6. Chromodomain-helicase-DNA-binding protein 6 (CHD6) is a DNA-dependent ATPase that plays a role in chromatin remodeling.; ncbi-cd \| cd17984 \| 97.8 \| 4e-09 \| 8.1e-13 \| 84.4 \| 143 \| (5, 151) \| 471 \| (3, 164) \| 178 \| DEXHc\_DHX40 \| cd17984 DEXHc\_DHX40; DEXH-box helicase domain of DEAH-box helicase 40. DEAH-box helicase 40 (DHX40) belongs to the DEAD-like helicase superfamily, a diverse family of proteins involved in ATP-dependent RNA or DNA unwinding.; ncbi-cd \| cd18057 \| 97.8 \| 4.2e-09 \| 8.5e-13 \| 88.5 \| 147 \| (5, 152) \| 471 \| (1, 183) \| 232 \| DEXHc\_CHD5 \| cd18057 DEXHc\_CHD5; DEAH-box helicase domain of the chromodomain helicase DNA binding protein 5. Chromodomain-helicase-DNA-binding protein 5 (CHD5) is a chromatin-remodeling protein that binds DNA through histones and regulates gene transcription.; ncbi-cd \| cd18060 \| 97.8 \| 4.4e-09 \| 8.8e-13 \| 87.2 \| 147 \| (5, 152) \| 471 \| (1, 173) \| 222 \| DEXHc\_CHD8 \| cd18060 DEXHc\_CHD8; DEAH-box helicase domain of the chromodomain helicase DNA binding protein 8. Chromodomain-helicase-DNA-binding protein 8 (CHD8) is a DNA helicase that acts as a chromatin remodeling factor and regulates transcription.; ncbi-cd \| cd17996 \| 97.8 \| 5e-09 \| 9.9e-13 \| 88.4 \| 149 \| (3, 152) \| 471 \| (2, 167) \| 233 \| DEXHc\_SMARCA2\_SMARCA4 \| cd17996 DEXHc\_SMARCA2\_SMARCA4; DEXH-box helicase domain of SMARCA2 and SMARCA4. SWI/SNF related, matrix associated, actin dependent regulator of chromatin, subfamily a, members 2 and 4 (SMARCA2 and SMARCA4) are members of the DEAD-like helicase superfamily, a diverse family of proteins involved in ATP-dependent RNA or DNA unwinding.; ncbi-cd \| cd17989 \| 97.8 \| 5e-09 \| 1e-12 \| 83.1 \| 143 \| (5, 151) \| 471 \| (3, 159) \| 173 \| DEXHc\_HrpA \| cd17989 DEXHc\_HrpA; DEXH-box helicase domain of ATP-dependent RNA helicase HrpA. HrpA is part of the HrpB-HrpA two-partner secretion (TPS) system, a secretion pathway important to the secretion of large virulence-associated proteins.; ncbi-cd \| cd18072 \| 97.8 \| 5.5e-09 \| 1.1e-12 \| 89.2 \| 147 \| (5, 152) \| 471 \| (1, 193) \| 241 \| DEXHc\_TTF2 \| cd18072 DEXHc\_TTF2; DEAH-box helicase domain of TTF2. Transcription termination factor 2 (TTF2 also called Forkhead-box E1/FOXE1 ) is a transcription termination factor that couples ATP hydrolysis with the removal of RNA polymerase II from the DNA template.; ncbi-cd \| cd17968 \| 97.7 \| 7.4e-09 \| 1.5e-12 \| 78.6 \| 123 \| (25, 150) \| 471 \| (3, 133) \| 134 \| DEAHc\_DDX11\_starthere \| cd17968 DEAHc\_DDX11\_starthere; DEAH-box helicase domain of ATP-dependent DNA helicase DDX11. DDX11 (also called ChlR1) encodes a protein of the conserved family of Iron-Sulfur (Fe-S) cluster DNA helicases and is thought to function in maintaining chromosome transmission fidelity and genome stability.; ncbi-cd \| cd17920 \| 97.7 \| 7.6e-09 \| 1.5e-12 \| 85.5 \| 144 \| (4, 155) \| 471 \| (12, 182) \| 200 \| DEXHc\_RecQ \| cd17920 DEXHc\_RecQ; DEXH-box helicase domain of RecQ family proteins. The RecQ family of the type II DEAD box helicase superfamily is a family of highly conserved DNA repair helicases.; ncbi-cd \| cd18056 \| 97.7 \| 8.1e-09 \| 1.6e-12 \| 86.1 \| 147 \| (5, 152) \| 471 \| (1, 183) \| 232 \| DEXHc\_CHD4 \| cd18056 DEXHc\_CHD4; DEAH-box helicase domain of the chromodomain helicase DNA binding protein 4. Chromodomain-helicase-DNA-binding protein 4 (CHD4) is a component of the histone deacetylase NuRD complex which participates in the remodeling of chromatin by deacetylating histones.; ncbi-cd \| cd18065 \| 97.7 \| 8.8e-09 \| 1.8e-12 \| 87.2 \| 149 \| (3, 152) \| 471 \| (14, 179) \| 233 \| DEXHc\_SMARCA1 \| cd18065 DEXHc\_SMARCA1; DEAH-box helicase domain of SMARCA1. SWI/SNF related, matrix associated, actin dependent regulator of chromatin, subfamily a, member 1 (SMARCA1, also called SNF2L) is a component of NURF (nucleosome-remodeling factor) and CERF (CECR2-containing-remodeling factor) complexes which promote the perturbation of chromatin structure in an ATP-dependent manner.; ncbi-cd \| cd18061 \| 97.7 \| 9.1e-09 \| 1.8e-12 \| 85.5 \| 147 \| (5, 152) \| 471 \| (1, 173) \| 222 \| DEXHc\_CHD9 \| cd18061 DEXHc\_CHD9; DEAH-box helicase domain of the chromodomain helicase DNA binding protein 9. Chromodomain-helicase-DNA-binding protein 9 (CHD9) acts as a transcriptional coactivator for PPARA and possibly other nuclear receptors.; ncbi-cd \| cd18003 \| 97.7 \| 9.6e-09 \| 1.9e-12 \| 85.9 \| 147 \| (5, 152) \| 471 \| (1, 164) \| 223 \| DEXQc\_SRCAP \| cd18003 DEXQc\_SRCAP; DEXH/Q-box helicase domain of SRCAP. Snf2-related CBP activator (SRCAP, also known as SWR1 or DOMO1) is the core catalytic component of the multiprotein chromatin-remodeling SRCAP complex, that is necessary for the incorporation of the histone variant H2A.; ncbi-cd \| cd18063 \| 97.7 \| 1e-08 \| 2e-12 \| 87.3 \| 149 \| (3, 152) \| 471 \| (22, 187) \| 251 \| DEXHc\_SMARCA2 \| cd18063 DEXHc\_SMARCA2; DEXH-box helicase domain of SMARCA2. SWI/SNF related, matrix associated, actin dependent regulator of chromatin, subfamily a, member 2 (SMARCA2, also known as brahma homolog) is a component of the BAF complex.; ncbi-cd \| cd18070 \| 97.7 \| 1.2e-08 \| 2.3e-12 \| 87.8 \| 142 \| (5, 152) \| 471 \| (1, 207) \| 257 \| DEXQc\_SHPRH \| cd18070 DEXQc\_SHPRH; DEXQ-box helicase domain of SHPRH. E3 ubiquitin-protein ligase SHPRH is a ubiquitously expressed protein that contains motifs characteristic of several DNA repair proteins, transcription factors, and helicases.; ncbi-cd \| cd18062 \| 97.7 \| 1.2e-08 \| 2.4e-12 \| 86.8 \| 149 \| (3, 152) \| 471 \| (22, 187) \| 251 \| DEXHc\_SMARCA4 \| cd18062 DEXHc\_SMARCA4; DEXH-box helicase domain of SMARCA4. SWI/SNF related, matrix associated, actin dependent regulator of chromatin, subfamily a, member 4 (SMARCA4, also known as transcription activator BRG1) is a component of the CREST-BRG1 complex that regulates promoter activation by orchestrating a calcium-dependent release of a repressor complex and a recruitment of an activator complex.; ncbi-cd \| cd18064 \| 97.7 \| 1.3e-08 \| 2.5e-12 \| 87.1 \| 149 \| (3, 152) \| 471 \| (14, 179) \| 244 \| DEXHc\_SMARCA5 \| cd18064 DEXHc\_SMARCA5; DEAH-box helicase domain of SMARCA5. SWI/SNF related, matrix associated, actin dependent regulator of chromatin, subfamily a, member 5 (SMARCA5, also called SNF2H) is the catalytic subunit of the four known chromatin-remodeling complexes: CHRAC, RSF, ACF/WCRF, and WICH.; ncbi-cd \| cd17993 \| 97.6 \| 2.3e-08 \| 4.6e-12 \| 81.8 \| 147 \| (4, 151) \| 471 \| (1, 169) \| 218 \| DEXHc\_CHD1\_2 \| cd17993 DEXHc\_CHD1\_2; DEXH-box helicase domain of the chromodomain helicase DNA binding proteins 1 and 2, and similar proteins.; ncbi-cd \| cd18069 \| 97.5 \| 2.9e-08 \| 5.8e-12 \| 83.4 \| 143 \| (5, 152) \| 471 \| (1, 178) \| 227 \| DEXHc\_ARIP4 \| cd18069 DEXHc\_ARIP4; DEXH-box helicase domain of ARIP4. Androgen receptor-interacting protein 4 (ARIP4, also called RAD54 like 2 or RAD54L2 ) modulates androgen receptor (AR)-dependent transactivation in a promoter-dependent manner.; ncbi-cd \| cd17983 \| 97.5 \| 2.9e-08 \| 5.8e-12 \| 78.0 \| 143 \| (5, 151) \| 471 \| (3, 159) \| 173 \| DEXHc\_DHX38 \| cd17983 DEXHc\_DHX38; DEXH-box helicase domain of DEAH-box helicase 38. DEAH-box helicase 38 (DHX38, also known as PRP16) is involved in pre-mRNA splicing.; ncbi-cd \| cd17931 \| 97.5 \| 3.2e-08 \| 6.4e-12 \| 76.1 \| 130 \| (25, 154) \| 471 \| (3, 139) \| 151 \| DEXHc\_viral\_Ns3 \| cd17931 DEXHc\_viral\_Ns3; DEXH-box helicase domain of NS3 protease-helicase. NS3 is a nonstructural multifunctional protein found in pestiviruses that contains an N-terminal protease and a C-terminal helicase.; ncbi-cd \| cd17973 \| 97.5 \| 3.2e-08 \| 6.5e-12 \| 80.2 \| 144 \| (4, 151) \| 471 \| (14, 172) \| 186 \| DEXHc\_DHX15 \| cd17973 DEXHc\_DHX15; DEXH-box helicase domain of DEAH-box helicase 15. DEAH-box helicase 15 (DHX15) is a pre-mRNA processing factor involved in disassembly of spliceosomes after the release of mature mRNA.; ncbi-cd \| cd18044 \| 97.5 \| 3.4e-08 \| 6.7e-12 \| 80.6 \| 139 \| (4, 150) \| 471 \| (1, 151) \| 191 \| DEXXQc\_SMUBP2 \| cd18044 DEXXQc\_SMUBP2; DEXXQ-box helicase domain of SMUBP2. SMUBP2 (also called immunoglobulin mu-binding protein 2, or IGHMBP2) is a 5' to 3' helicase that unwinds RNA and DNA duplexes in an ATP-dependent reaction.; ncbi-cd \| cd17994 \| 97.5 \| 3.9e-08 \| 7.9e-12 \| 79.7 \| 141 \| (5, 151) \| 471 \| (1, 146) \| 196 \| DEXHc\_CHD3\_4\_5 \| cd17994 DEXHc\_CHD3\_4\_5; DEAH-box helicase domain of the chromodomain helicase DNA binding proteins 3, 4 and 5.; ncbi-cd \| cd17972 \| 97.5 \| 4e-08 \| 7.9e-12 \| 82.3 \| 143 \| (4, 150) \| 471 \| (60, 219) \| 234 \| DEXHc\_DHX9 \| cd17972 DEXHc\_DHX9; DEXH-box helicase domain of DEAH-box helicase 9. DEAH-box helicase 9 (DHX9, also known as ATP-dependent RNA helicase A or RHA and leukophysin or LKP) plays an important role in many cellular processes, including regulation of DNA replication, transcription, translation, microRNA biogenesis, RNA processing and transport, and maintenance of genomic stability.; ncbi-cd \| cd18067 \| 97.4 \| 4.6e-08 \| 9.3e-12 \| 82.3 \| 147 \| (5, 152) \| 471 \| (1, 179) \| 243 \| DEXHc\_RAD54A \| cd18067 DEXHc\_RAD54A; DEXH-box helicase domain of RAD54A. DNA repair and recombination protein RAD54A, also known as RAD54L or RAD54, plays a role in homologous recombination related repair of DNA double-strand breaks.; ncbi-cd \| cd17917 \| 97.4 \| 5.5e-08 \| 1.1e-11 \| 74.9 \| 126 \| (25, 150) \| 471 \| (3, 144) \| 159 \| DEXHc\_RHA-like \| cd17917 DEXHc\_RHA-like; DEXH-box helicase domain of DEAD-like helicase RHA family proteins. The RNA helicase A (RHA) family includes RHA, also called DEAH-box helicase 9 (DHX9), DHX8, DHX15-16, DHX32-38, and many others.; ncbi-cd \| cd17979 \| 97.4 \| 5.7e-08 \| 1.1e-11 \| 77.0 \| 142 \| (4, 151) \| 471 \| (2, 156) \| 170 \| DEXHc\_DHX34 \| cd17979 DEXHc\_DHX34; DEXH-box helicase domain of DEAH-box helicase 34. DEAH-box helicase 34 (DHX34) plays a role in the nonsense-mediated decay (NMD), a surveillance mechanism that degrades aberrant mRNAs.; ncbi-cd \| cd18077 \| 97.4 \| 6.2e-08 \| 1.2e-11 \| 80.7 \| 71 \| (5, 75) \| 471 \| (2, 74) \| 226 \| DEXXQc\_HELZ \| cd18077 DEXXQc\_HELZ; DEXXQ-box helicase domain of HELZ. Helicase with zinc finger (HELZ) acts as a helicase that plays a role in RNA metabolism during development.; ncbi-cd \| cd17969 \| 97.4 \| 6.6e-08 \| 1.3e-11 \| 75.1 \| 112 \| (24, 149) \| 471 \| (11, 139) \| 157 \| DEAHc\_XPD \| cd17969 DEAHc\_XPD; DEAH-box helicase domain of TFIIH basal transcription factor complex helicase XPD subunit.; ncbi-cd \| cd17922 \| 97.4 \| 7.2e-08 \| 1.5e-11 \| 74.4 \| 127 \| (25, 151) \| 471 \| (3, 159) \| 166 \| DEXHc\_LHR-like \| cd17922 DEXHc\_LHR-like; DEXH-box helicase domain of LHR. Large helicase-related protein (LHR) is a DNA damage-inducible helicase that uses ATP hydrolysis to drive unidirectional 3'-to-5' translocation along single-stranded DNA (ssDNA) and to unwind RNA:DNA duplexes.; ncbi-cd \| cd17970 \| 97.3 \| 8.4e-08 \| 1.7e-11 \| 76.2 \| 112 \| (25, 150) \| 471 \| (3, 163) \| 181 \| DEAHc\_FancJ \| cd17970 DEAHc\_FancJ; DEAH-box helicase domain of Fanconi anemia group J protein and similar proteins.; ncbi-cd \| cd18071 \| 97.3 \| 8.9e-08 \| 1.8e-11 \| 81.0 \| 140 \| (5, 152) \| 471 \| (1, 187) \| 239 \| DEXHc\_HLTF1\_SMARC3 \| cd18071 DEXHc\_HLTF1\_SMARC3; DEXH-box helicase domain of HLTF1. Helicase like transcription factor (HLTF1, also known as HIP116 or SMARCA3) has both helicase and E3 ubiquitin ligase activities and ATP-dependent nucleosome-remodeling activity.; ncbi-cd \| cd18041 \| 97.3 \| 1.2e-07 \| 2.3e-11 \| 77.7 \| 118 \| (5, 125) \| 471 \| (2, 144) \| 203 \| DEXXQc\_DNA2 \| cd18041 DEXXQc\_DNA2; DEXXQ-box helicase domain of DNA2. DNA2 (DNA Replication Helicase/Nuclease 2) possesses different enzymatic activities, such as single-stranded DNA (ssDNA)-dependent ATPase, 5-3 helicase, and endonuclease activities, and is involved in DNA replication and DNA repair in the nucleus and mitochondrion.; ncbi-cd \| cd18076 \| 97.3 \| 1.3e-07 \| 2.5e-11 \| 78.4 \| 70 \| (5, 74) \| 471 \| (2, 75) \| 230 \| DEXXQc\_HELZ2-N \| cd18076 DEXXQc\_HELZ2-N; N-terminal DEXXQ-box helicase domain of HELZ2. Helicase with zinc finger 2 (HELZ2, also known as PPAR-alpha-interacting complex protein 285 or PRIC285 and PPAR-gamma DBD-interacting protein 1 or PDIP1) acts as a transcriptional coactivator for a number of nuclear receptors including PPARA, PPARG, THRA, THRB, and RXRA.; ncbi-cd \| cd18796 \| 97.2 \| 1.4e-07 \| 2.9e-11 \| 72.1 \| 101 \| (228, 335) \| 471 \| (37, 143) \| 150 \| SF2\_C\_LHR \| cd18796 SF2\_C\_LHR; C-terminal helicase domain of LHR family helicases. Large helicase-related protein (LHR) is a DNA damage-inducible helicase that uses ATP hydrolysis to drive unidirectional 3'-to-5' translocation along single-stranded DNA (ssDNA) and to unwind RNA:DNA duplexes.; ncbi-cd \| cd18793 \| 97.1 \| 2.6e-07 \| 5.3e-11 \| 69.2 \| 100 \| (229, 335) \| 471 \| (27, 128) \| 135 \| SF2\_C\_SNF \| cd18793 SF2\_C\_SNF; C-terminal helicase domain of the SNF family helicases. The Sucrose Non-Fermenting (SNF) family includes chromatin-remodeling factors, such as CHD proteins and SMARCA proteins, recombination proteins Rad54, and many others.; ncbi-cd \| cd17978 \| 97.1 \| 2.7e-07 \| 5.5e-11 \| 71.7 \| 144 \| (4, 151) \| 471 \| (2, 165) \| 179 \| DEXHc\_DHX33 \| cd17978 DEXHc\_DHX33; DEXH-box helicase domain of DEAH-box helicase 33. DEAH-box helicase 33 (DHX33) stimulates RNA polymerase I transcription of the 47S precursor rRNA.; ncbi-cd \| cd17935 \| 97.1 \| 3.3e-07 \| 6.5e-11 \| 76.1 \| 116 \| (3, 125) \| 471 \| (4, 125) \| 207 \| EEXXQc\_AQR \| cd17935 EEXXQc\_AQR; EEXXQ-box helicase domain of AQR. Aquarius (AQR) is a multifunctional RNA helicase that binds precursor-mRNA introns at a defined position and is part of a pentameric intron-binding complex (IBC).; ncbi-cd \| cd18078 \| 97.1 \| 3.9e-07 \| 7.7e-11 \| 76.9 \| 120 \| (5, 124) \| 471 \| (2, 150) \| 230 \| DEXXQc\_Mov10L1 \| cd18078 DEXXQc\_Mov10L1; DEXXQ-box helicase domain of Mov10L1. Moloney leukemia virus 10-like protein 1 (Mov10L1) binds Piwi-interacting RNA (piRNA) precursors to initiate piRNA processing.; ncbi-cd \| cd18799 \| 97.0 \| 4e-07 \| 8.1e-11 \| 64.9 \| 101 \| (228, 335) \| 471 \| (5, 108) \| 116 \| SF2\_C\_EcoAI-like \| cd18799 SF2\_C\_EcoAI-like; C-terminal helicase domain of EcoAI HsdR-like restriction enzyme family helicases.; ncbi-cd \| cd18042 \| 97.0 \| 4.2e-07 \| 8.3e-11 \| 75.9 \| 117 \| (6, 124) \| 471 \| (2, 157) \| 217 \| DEXXQc\_SETX \| cd18042 DEXXQc\_SETX; DEXXQ-box helicase domain of SETX. The RNA/DNA helicase senataxin (SETX) plays a role in transcription, neurogenesis, and antiviral response.; ncbi-cd \| cd17932 \| 97.0 \| 4.4e-07 \| 8.9e-11 \| 72.6 \| 63 \| (7, 75) \| 471 \| (2, 68) \| 189 \| DEXQc\_UvrD \| cd17932 DEXQc\_UvrD; DEXQD-box helicase domain of UvrD. UvrD is a highly conserved helicase involved in mismatch repair, nucleotide excision repair, and recombinational repair.; ncbi-cd \| cd18787 \| 97.0 \| 4.8e-07 \| 9.8e-11 \| 66.4 \| 101 \| (228, 335) \| 471 \| (26, 126) \| 131 \| SF2\_C\_DEAD \| cd18787 SF2\_C\_DEAD; C-terminal helicase domain of the DEAD box helicases. DEAD-box helicases comprise a diverse family of proteins involved in ATP-dependent RNA unwinding, needed in a variety of cellular processes including splicing, ribosome biogenesis, and RNA degradation.; ncbi-cd \| cd18797 \| 97.0 \| 5.1e-07 \| 1e-10 \| 68.1 \| 101 \| (228, 335) \| 471 \| (34, 141) \| 146 \| SF2\_C\_Hrq \| cd18797 SF2\_C\_Hrq; C-terminal helicase domain of HrQ family helicases. Yeast Hrq1, similar to RecQ4, plays a role in DNA inter-strand crosslink (ICL) repair and in telomere maintenance.; ncbi-cd \| cd18802 \| 97.0 \| 6e-07 \| 1.2e-10 \| 67.7 \| 98 \| (229, 333) \| 471 \| (25, 137) \| 142 \| SF2\_C\_dicer \| cd18802 SF2\_C\_dicer; C-terminal helicase domain of the endoribonuclease Dicer. Dicer ribonucleases cleave double-stranded RNA (dsRNA) precursors to generate microRNAs (miRNAs) and small interfering RNAs (siRNAs).; ncbi-cd \| cd18794 \| 96.9 \| 9e-07 \| 1.8e-10 \| 64.7 \| 101 \| (228, 335) \| 471 \| (29, 129) \| 134 \| SF2\_C\_RecQ \| cd18794 SF2\_C\_RecQ; C-terminal helicase domain of the RecQ family helicases. The RecQ helicase family is an evolutionarily conserved class of enzymes, dedicated to preserving genomic integrity by operating in telomere maintenance, DNA repair, and replication.; ncbi-cd \| cd17915 \| 96.9 \| 9.4e-07 \| 1.9e-10 \| 65.7 \| 111 \| (25, 150) \| 471 \| (3, 120) \| 138 \| DEAHc\_XPD-like \| cd17915 DEAHc\_XPD-like; DEAH-box helicase domain of XPD family DEAD-like helicases. The xeroderma pigmentosum group D (XPD)-like family members are DEAD-box helicases, a diverse family of proteins involved in ATP-dependent RNA unwinding, needed in a variety of cellular processes including splicing, ribosome biogenesis and RNA degradation.; ncbi-cd \| cd18810 \| 96.9 \| 9.9e-07 \| 2e-10 \| 67.0 \| 101 \| (228, 335) \| 471 \| (24, 127) \| 151 \| SF2\_C\_TRCF \| cd18810 SF2\_C\_TRCF; C-terminal helicase domain of the transcription-repair coupling factor. Transcription-repair coupling factor (TrcF) dissociates transcription elongation complexes blocked at nonpairing lesions and mediates recruitment of DNA repair proteins.; ncbi-cd \| cd18040 \| 96.8 \| 1.1e-06 \| 2.1e-10 \| 74.7 \| 66 \| (5, 74) \| 471 \| (2, 80) \| 271 \| DEXXc\_HELZ2-C \| cd18040 DEXXc\_HELZ2-C; C-terminal DEXX-box helicase domain of HELZ2. Helicase with zinc finger 2 (HELZ2, also known as PPAR-alpha-interacting complex protein 285 or PRIC285 and PPAR-gamma DBD-interacting protein 1 or PDIP1) acts as a transcriptional coactivator for a number of nuclear receptors including PPARA, PPARG, THRA, THRB and RXRA.; ncbi-cd \| cd18790 \| 96.8 \| 1.2e-06 \| 2.4e-10 \| 69.1 \| 97 \| (228, 331) \| 471 \| (26, 127) \| 171 \| SF2\_C\_UvrB \| cd18790 SF2\_C\_UvrB; C-terminal helicase domain of the UvrB family helicases. Excinuclease ABC subunit B (or UvrB) plays a central role in nucleotide excision repair (NER).; ncbi-cd \| cd18039 \| 96.8 \| 1.3e-06 \| 2.6e-10 \| 71.6 \| 65 \| (5, 73) \| 471 \| (2, 67) \| 234 \| DEXXQc\_UPF1 \| cd18039 DEXXQc\_UPF1; DEXXQ-box helicase domain of UPF1. UPF1 (also called RNA Helicase And ATPase, Regulator Of Nonsense Transcripts, or ATP-Dependent Helicase RENT1) is an RNA-dependent helicase and ATPase required for nonsense-mediated decay (NMD) of mRNAs containing premature stop codons.; ncbi-cd \| cd18803 \| 96.7 \| 1.6e-06 \| 3.3e-10 \| 65.4 \| 99 \| (228, 335) \| 471 \| (29, 135) \| 141 \| SF2\_C\_secA \| cd18803 SF2\_C\_secA; C-terminal helicase domain of the protein translocase subunit secA. SecA is a component of the Sec translocase that transports the vast majority of bacterial and ER-exported proteins.; ncbi-cd \| cd17936 \| 96.7 \| 2.1e-06 \| 4.1e-10 \| 68.2 \| 109 \| (5, 124) \| 471 \| (2, 117) \| 178 \| EEXXEc\_NFX1 \| cd17936 EEXXEc\_NFX1; EEXXE-box helicase domain of NFX1. Human NFX1 protein was identified as a protein that represses class II MHC (major histocompatibility complex) gene expression.; ncbi-cd \| cd18801 \| 96.7 \| 2.3e-06 \| 4.6e-10 \| 63.4 \| 98 \| (229, 333) \| 471 \| (30, 137) \| 143 \| SF2\_C\_FANCM\_Hef \| cd18801 SF2\_C\_FANCM\_Hef; C-terminal helicase domain of Fanconi anemia group M family helicases. Fanconi anemia group M (FANCM) protein is a DNA-dependent ATPase component of the Fanconi anemia (FA) core complex.; ncbi-cd \| cd18792 \| 96.5 \| 4.2e-06 \| 8.4e-10 \| 64.5 \| 101 \| (228, 335) \| 471 \| (25, 136) \| 160 \| SF2\_C\_RecG\_TRCF \| cd18792 SF2\_C\_RecG\_TRCF; C-terminal helicase domain of the RecG family helicases. The DEAD-like helicase RecG family contains recombination factor RecG and transcription-repair coupling factor TrcF.; ncbi-cd \| cd18811 \| 96.5 \| 4.5e-06 \| 9.1e-10 \| 64.1 \| 101 \| (228, 335) \| 471 \| (25, 137) \| 159 \| SF2\_C\_RecG \| cd18811 SF2\_C\_RecG; C-terminal helicase domain of DNA helicase RecG. ATP-dependent DNA helicase RecG plays a critical role in recombination and DNA repair.; ncbi-cd \| cd17913 \| 96.4 \| 5.5e-06 \| 1.1e-09 \| 62.5 \| 92 \| (25, 125) \| 471 \| (3, 96) \| 142 \| DEXQc\_Suv3 \| cd17913 DEXQc\_Suv3; DEXQ-box helicase domain of Suv3. Suppressor of var1 3-like protein (Suv3) is a DNA/RNA unwinding enzyme belonging to the class of DexH-box helicases.; ncbi-cd \| cd18806 \| 96.3 \| 8.1e-06 \| 1.6e-09 \| 62.5 \| 94 \| (228, 333) \| 471 \| (23, 136) \| 145 \| SF2\_C\_viral \| cd18806 SF2\_C\_viral; C-terminal helicase domain of viral helicase. Viral helicases in this family here are DEAD-like helicases belonging to superfamily (SF)2, a diverse family of proteins involved in ATP-dependent RNA or DNA unwinding.; ncbi-cd \| cd21722 \| 96.2 \| 1e-05 \| 2e-09 \| 72.0 \| 109 \| (5, 124) \| 471 \| (11, 131) \| 340 \| betaCoV\_Nsp13-helicase \| cd21722 betaCoV\_Nsp13-helicase; helicase domain of betacoronavirus non-structural protein 13. This model represents the helicase domain of non-structural protein 13 (Nsp13) from betacoronavirus, including pathogenic human viruses such as Severe acute respiratory syndrome coronavirus (SARS-CoV), SARS-CoV2 (also called 2019 novel CoV or 2019-nCoV), and Middle East respiratory syndrome-related (MERS) CoV.; ncbi-cd \| cd17980 \| 96.2 \| 1e-05 \| 2.1e-09 \| 62.8 \| 142 \| (5, 150) \| 471 \| (3, 160) \| 185 \| DEXHc\_DHX35 \| cd17980 DEXHc\_DHX35; DEXH-box helicase domain of DEAH-box helicase 35. DHX35 plays a role in colorectal cancers and seems to be associated with risk to thyroid cancers.; ncbi-cd \| cd18789 \| 96.2 \| 1.1e-05 \| 2.2e-09 \| 61.6 \| 93 \| (229, 333) \| 471 \| (49, 142) \| 153 \| SF2\_C\_XPB \| cd18789 SF2\_C\_XPB; C-terminal helicase domain of XPB-like helicases. TFIIH basal transcription factor complex helicase XPB (xeroderma pigmentosum type B) subunit (also known as DNA excision repair protein ERCC-3 or TFIIH 89 kDa subunit) is the ATP-dependent 3'-5' DNA helicase component of the core-TFIIH basal transcription factor, involved in nucleotide excision repair (NER) of DNA and, when complexed to CAK, in RNA transcription by RNA polymerase II.; ncbi-cd \| cd17982 \| 96.2 \| 1.1e-05 \| 2.3e-09 \| 63.1 \| 144 \| (5, 152) \| 471 \| (3, 174) \| 191 \| DEXHc\_DHX37 \| cd17982 DEXHc\_DHX37; DEXH-box helicase domain of DEAH-box helicase 37. DHX37 plays a role in the development of the human nervous system and has been linked to schizophrenia.; ncbi-cd \| cd17971 \| 96.2 \| 1.1e-05 \| 2.3e-09 \| 61.6 \| 143 \| (4, 150) \| 471 \| (7, 163) \| 179 \| DEXHc\_DHX8 \| cd17971 DEXHc\_DHX8; DEXH-box helicase domain of DEAH-box helicase 8. DEAH-box helicase 8 (DHX8 ,also known as pre-mRNA-splicing factor ATP-dependent RNA helicase PRP22) acts late in the splicing of pre-mRNA and mediates the release of the spliced mRNA from spliceosomes.; ncbi-cd \| cd17986 \| 96.2 \| 1.2e-05 \| 2.4e-09 \| 62.0 \| 40 \| (5, 47) \| 471 \| (3, 42) \| 177 \| DEXQc\_DQX1 \| cd17986 DEXQc\_DQX1; DEXQ-box helicase domain of DEAQ-box RNA dependent ATPase 1. DEAQ-box RNA dependent ATPase 1 (DQX1) belongs to the DEAD-like helicase superfamily, a diverse family of proteins involved in ATP-dependent RNA or DNA unwinding.; ncbi-cd \| cd21718 \| 96.2 \| 1.3e-05 \| 2.6e-09 \| 71.1 \| 108 \| (7, 125) \| 471 \| (13, 132) \| 341 \| CoV\_Nsp13-helicase \| cd21718 CoV\_Nsp13-helicase; helicase domain of coronavirus non-structural protein 13. This model represents the helicase domain of non-structural protein 13 (Nsp13) from alpha-, beta-, gamma-, and deltacoronavirus, including pathogenic human viruses such as Severe acute respiratory syndrome coronavirus (SARS-CoV), SARS-CoV2 (also called 2019 novel CoV or 2019-nCoV), and Middle East respiratory syndrome-related (MERS) CoV.; ncbi-cd \| cd21720 \| 96.0 \| 2e-05 \| 3.9e-09 \| 70.4 \| 109 \| (6, 125) \| 471 \| (12, 132) \| 343 \| gammaCoV\_Nsp13-helicase \| cd21720 gammaCoV\_Nsp13-helicase; helicase domain of gammacoronavirus non-structural protein 13. This model represents the helicase domain of non-structural protein 13 (Nsp13) from gammacoronavirus, including Avian infectious bronchitis virus.; ncbi-cd \| cd18791 \| 95.9 \| 3.1e-05 \| 6.3e-09 \| 59.1 \| 98 \| (228, 332) \| 471 \| (42, 164) \| 171 \| SF2\_C\_RHA \| cd18791 SF2\_C\_RHA; C-terminal helicase domain of the RNA helicase A (RHA) family helicases. The RNA helicase A (RHA) family includes RHA, also called DEAH-box helicase 9 (DHX9), DHX8, DHX15-16, DHX32-38, and many others.; ncbi-cd \| cd21721 \| 95.8 \| 3.6e-05 \| 7.2e-09 \| 68.6 \| 111 \| (4, 125) \| 471 \| (10, 132) \| 342 \| deltaCoV\_Nsp13-helicase \| cd21721 deltaCoV\_Nsp13-helicase; helicase domain of deltacoronavirus non-structural protein 13. This model represents the helicase domain of non-structural protein 13 (Nsp13) from deltacoronavirus, including Bulbul coronavirus (CoV) HKU11 and Common moorhen CoV HKU21.; ncbi-cd \| cd18795 \| 95.8 \| 4.1e-05 \| 8.2e-09 \| 57.9 \| 95 \| (228, 333) \| 471 \| (42, 144) \| 154 \| SF2\_C\_Ski2 \| cd18795 SF2\_C\_Ski2; C-terminal helicase domain of the Ski2 family helicases. Ski2-like RNA helicases play an important role in RNA degradation, processing, and splicing pathways.; ncbi-cd \| cd21723 \| 95.7 \| 5.3e-05 \| 1e-08 \| 67.2 \| 94 \| (24, 124) \| 471 \| (26, 131) \| 340 \| alphaCoV\_Nsp13-helicase \| cd21723 alphaCoV\_Nsp13-helicase; helicase domain of alphacoronavirus non-structural protein 13. This model represents the helicase domain of non-structural protein 13 (Nsp13) from alphacoronavirus, including Porcine epidemic diarrhea virus and Human coronavirus (CoV) NL63.; ncbi-cd \| cd18805 \| 95.6 \| 6.1e-05 \| 1.2e-08 \| 54.7 \| 98 \| (229, 333) \| 471 \| (17, 125) \| 135 \| SF2\_C\_suv3 \| cd18805 SF2\_C\_suv3; C-terminal helicase domain of ATP-dependent RNA helicase. The SUV3 (suppressor of Var 3) gene encodes a DNA and RNA helicase, which is localized in mitochondria and is a subunit of the degradosome complex involved in regulation of RNA surveillance and turnover.; ncbi-cd \| cd18798 \| 95.2 \| 0.00013 \| 2.6e-08 \| 57.1 \| 89 \| (230, 332) \| 471 \| (25, 121) \| 174 \| SF2\_C\_reverse\_gyrase \| cd18798 SF2\_C\_reverse\_gyrase; C-terminal helicase domain of the reverse gyrase. Reverse gyrase modifies the topological state of DNA by introducing positive supercoils in an ATP-dependent process.; ncbi-cd \| cd17933 \| 95.1 \| 0.00016 \| 3.1e-08 \| 53.8 \| 50 \| (9, 62) \| 471 \| (2, 51) \| 155 \| DEXSc\_RecD-like \| cd17933 DEXSc\_RecD-like; DEXS-box helicase domain of RecD and similar proteins. RecD is a member of the RecBCD (EC 3.; ncbi-cd \| cd18804 \| 95.0 \| 0.00021 \| 4.1e-08 \| 59.4 \| 88 \| (241, 335) \| 471 \| (102, 205) \| 238 \| SF2\_C\_priA \| cd18804 SF2\_C\_priA; C-terminal helicase domain of ATP-dependent helicase PriA. PriA, also known as replication factor Y or primosomal protein N', is a 3'-5' DNA helicase that acts to remodel stalled replication forks and as a specificity factor for origin-independent assembly of a new replisome at the stalled fork.; ncbi-cd \| cd01127 \| 94.2 \| 0.00053 \| 1.1e-07 \| 49.5 \| 35 \| (26, 60) \| 471 \| (2, 36) \| 144 \| TrwB\_TraG\_TraD\_VirD4 \| cd01127 TrwB\_TraG\_TraD\_VirD4; TrwB/TraG/TraD/VirD4 family of bacterial conjugation proteins. The TraG/TraD/VirD4 family are bacterial conjugation proteins involved in type IV secretion (T4S) systems, versatile bacterial secretion systems mediating transport of protein and/or DNA.; cath \| 1q0uA00 \| 97.0 \| 5.7e-07 \| 8.1e-11 \| 74.9 \| 165 \| (4, 172) \| 471 \| (26, 215) \| 219 \| Bstdead \| CATHCODE: 3.40.50.300 NAME: Bstdead. Chain: a, b. Fragment: n-terminal domain. Engineered: yes SOURCE: Geobacillus stearothermophilus. Organism\_taxid: 1422. Expressed in: escherichia coli. Expression\_system\_taxid: 562. CLASS: Alpha Beta, ARCH: 3-Layer(aba) Sandwich, TOPOL: Rossmann fold, HOMOL: P-loop containing nucleotide triphosphate hydrolases; cath \| 3h1tA02 \| 96.9 \| 1.1e-06 \| 1.5e-10 \| 71.8 \| 161 \| (3, 165) \| 471 \| (20, 201) \| 202 \| Type i site-specific restriction-modification system, r (restriction) subunit \| CATHCODE: 3.40.50.300 NAME: Type i site-specific restriction-modification system, r (restriction) subunit. Chain: a. Fragment: unp residues 1-590. Engineered: yes SOURCE: Vibrio vulnificus. Organism\_taxid: 196600. Strain: yj016. Gene: vv0265. Expressed in: escherichia coli. Expression\_system\_taxid: 562. CLASS: Alpha Beta, ARCH: 3-Layer(aba) Sandwich, TOPOL: Rossmann fold, HOMOL: P-loop containing nucleotide triphosphate hydrolases; cath \| 2p6rA01 \| 96.9 \| 1.3e-06 \| 1.8e-10 \| 71.4 \| 161 \| (4, 170) \| 471 \| (19, 196) \| 197 \| 25-mer \| CATHCODE: 3.40.50.300 NAME: 25-mer. Chain: x. Engineered: yes. 5'- d(\*cp\*tp\*ap\*gp\*ap\*gp\*ap\*cp\*tp\*ap\*tp\*cp\*gp\*ap\*t)-3'. Chain: y. Engineered: yes. Afuhel308 helicase. Chain: a. Engineered: yes SOURCE: Yes. Yes. CLASS: Alpha Beta, ARCH: 3-Layer(aba) Sandwich, TOPOL: Rossmann fold, HOMOL: P-loop containing nucleotide triphosphate hydrolases; cath \| 1rifA02 \| 96.8 \| 1.8e-06 \| 2.6e-10 \| 69.5 \| 157 \| (3, 164) \| 471 \| (22, 190) \| 192 \| Dna helicase uvsw \| CATHCODE: 3.40.50.300 NAME: Dna helicase uvsw. Chain: a, b. Synonym: dar protein. Engineered: yes.Mutation: yes SOURCE: Enterobacteria phage t4. Organism\_taxid: 10665. Gene: uvsw, dar. Expressed in: escherichia coli bl21(de3). Expression\_system\_taxid: 469008. CLASS: Alpha Beta, ARCH: 3-Layer(aba) Sandwich, TOPOL: Rossmann fold, HOMOL: P-loop containing nucleotide triphosphate hydrolases; cath \| 2fwrA02 \| 96.7 \| 3.2e-06 \| 4.5e-10 \| 67.8 \| 162 \| (3, 174) \| 471 \| (13, 177) \| 178 \| Dna repair protein rad25 \| CATHCODE: 3.40.50.300 NAME: Dna repair protein rad25. Chain: a, b, c, d. Engineered: yes SOURCE: Archaeoglobus fulgidus. Organism\_taxid: 2234. Expressed in: escherichia coli bl21(de3). Expression\_system\_taxid: 469008. CLASS: Alpha Beta, ARCH: 3-Layer(aba) Sandwich, TOPOL: Rossmann fold, HOMOL: P-loop containing nucleotide triphosphate hydrolases; cath \| 1vecA00 \| 96.6 \| 4.2e-06 \| 5.9e-10 \| 67.7 \| 157 \| (4, 164) \| 471 \| (25, 203) \| 206 \| Atp-dependent RNA helicase p54 \| CATHCODE: 3.40.50.300 NAME: Atp-dependent RNA helicase p54. Chain: a, b. Fragment: n-terminal domain. Synonym: rck, dead-box protein 6. Engineered: yes SOURCE: Homo sapiens. Human. Organism\_taxid: 9606. Gene: humrck. Expressed in: escherichia coli. Expression\_system\_taxid: 562. CLASS: Alpha Beta, ARCH: 3-Layer(aba) Sandwich, TOPOL: Rossmann fold, HOMOL: P-loop containing nucleotide triphosphate hydrolases; cath \| 5lstA01 \| 96.6 \| 4.6e-06 \| 6.5e-10 \| 68.8 \| 153 \| (4, 160) \| 471 \| (32, 208) \| 215 \| Atp-dependent DNA helicase q4 \| CATHCODE: 3.40.50.300 NAME: Atp-dependent DNA helicase q4. Chain: a. Synonym: DNA helicase,recq-like type 4,recq4,rts,recq protein-like 4. Engineered: yes SOURCE: Homo sapiens. Human. Organism\_taxid: 9606. Gene: recql4, recq4. Expressed in: escherichia coli. Expression\_system\_taxid: 469008. CLASS: Alpha Beta, ARCH: 3-Layer(aba) Sandwich, TOPOL: Rossmann fold, HOMOL: P-loop containing nucleotide triphosphate hydrolases; cath \| 4a15A01 \| 96.6 \| 4.8e-06 \| 6.8e-10 \| 66.7 \| 155 \| (3, 158) \| 471 \| (2, 174) \| 188 \| Atp-dependent DNA helicase ta0057 \| CATHCODE: 3.40.50.300 NAME: Atp-dependent DNA helicase ta0057. Chain: a. Synonym: xpd helicase. Engineered: yes. 5'-d(\*dtp\*ap\*cp\*gp)-3'. Chain: e SOURCE: Thermoplasma acidophilum. Organism\_taxid: 2303. Expressed in: escherichia coli. Expression\_system\_taxid: 469008. CLASS: Alpha Beta, ARCH: 3-Layer(aba) Sandwich, TOPOL: Rossmann fold, HOMOL: P-loop containing nucleotide triphosphate hydrolases; cath \| 2w00A03 \| 96.5 \| 5.4e-06 \| 7.6e-10 \| 66.3 \| 159 \| (5, 165) \| 471 \| (2, 189) \| 191 \| Hsdr \| CATHCODE: 3.40.50.300 NAME: Hsdr. Chain: a, b. Synonym: r.ecor124i. Engineered: yes SOURCE: Escherichia coli. Organism\_taxid: 562. Strain: b834(de3). Expressed in: escherichia coli. Expression\_system\_taxid: 562. CLASS: Alpha Beta, ARCH: 3-Layer(aba) Sandwich, TOPOL: Rossmann fold, HOMOL: P-loop containing nucleotide triphosphate hydrolases; cath \| 2oxcA00 \| 96.5 \| 7e-06 \| 9.9e-10 \| 66.4 \| 158 \| (4, 165) \| 471 \| (23, 202) \| 207 \| Probable atp-dependent RNA helicase ddx20 \| CATHCODE: 3.40.50.300 NAME: Probable atp-dependent RNA helicase ddx20. Chain: a, b. Fragment: deaddomain. Synonym: dead box protein 20, dead box protein dp 103, component of gems 3, gemin-3. Engineered: yes SOURCE: Homo sapiens. Human. Organism\_taxid: 9606. Gene: ddx20, dp103, gemin3.Expressed in: escherichia coli. Expression\_system\_taxid: 562. CLASS: Alpha Beta, ARCH: 3-Layer(aba) Sandwich, TOPOL: Rossmann fold, HOMOL: P-loop containing nucleotide triphosphate hydrolases; cath \| 3berA00 \| 96.4 \| 7.3e-06 \| 1e-09 \| 67.6 \| 157 \| (4, 165) \| 471 \| (36, 215) \| 220 \| Probable atp-dependent RNA helicase ddx47 \| CATHCODE: 3.40.50.300 NAME: Probable atp-dependent RNA helicase ddx47. Chain: a. Fragment: conserved domain i (dead): residues 5-230. Synonym: dead box protein 47. Engineered: yes SOURCE: Homo sapiens. Human. Organism\_taxid: 9606. Gene: ddx47. Expressed in:escherichia coli. Expression\_system\_taxid: 562. CLASS: Alpha Beta, ARCH: 3-Layer(aba) Sandwich, TOPOL: Rossmann fold, HOMOL: P-loop containing nucleotide triphosphate hydrolases; cath \| 1hv8A01 \| 96.4 \| 7.9e-06 \| 1.1e-09 \| 66.4 \| 157 \| (4, 164) \| 471 \| (28, 204) \| 210 \| Putative atp-dependent RNA helicase mj0669 \| CATHCODE: 3.40.50.300 NAME: Putative atp-dependent RNA helicase mj0669. Chain: a, b. Synonym: deadbox helicase. Engineered: yes SOURCE: Methanocaldococcus jannaschii. Organism\_taxid: 2190. Expressed in: escherichia coli. Expression\_system\_taxid: 562 CLASS: Alpha Beta, ARCH: 3-Layer(aba) Sandwich, TOPOL: Rossmann fold, HOMOL: P-loop containing nucleotide triphosphate hydrolases; cath \| 1xtiA02 \| 96.4 \| 9.1e-06 \| 1.3e-09 \| 64.9 \| 106 \| (222, 334) \| 471 \| (18, 123) \| 166 \| Probable atp-dependent RNA helicase p47 \| CATHCODE: 3.40.50.300 NAME: Probable atp-dependent RNA helicase p47. Chain: a. Fragment: sequence database residues 46-428. Synonym: hla-b associated transcript-1, huap56. Engineered: yes SOURCE: Homo sapiens. Human. Organism\_taxid: 9606. Gene: bat1. Expressed in: escherichia coli bl21(de3). Expression\_system\_taxid: 469008. CLASS: Alpha Beta, ARCH: 3-Layer(aba) Sandwich, TOPOL: Rossmann fold, HOMOL: P-loop containing nucleotide triphosphate hydrolases; cath \| 4q48A01 \| 96.4 \| 9.6e-06 \| 1.4e-09 \| 66.2 \| 147 \| (4, 159) \| 471 \| (24, 194) \| 207 \| Dna helicase recq \| CATHCODE: 3.40.50.300 NAME: Dna helicase recq. Chain: a, b. Fragment: unp residues 1-517. Engineered: yes SOURCE: Deinococcus radiodurans. Organism\_taxid: 243230. Strain: atcc 13939 / dsm 20539 / jcm 16871 / lmg 4051 / nbrc 15346 / ncimb 9279 / r1 / vkm b-1422. Gene: dr\_1289. Expressed in: escherichia coli. Expression\_system\_taxid: 562 CLASS: Alpha Beta, ARCH: 3-Layer(aba) Sandwich, TOPOL: Rossmann fold, HOMOL: P-loop containing nucleotide triphosphate hydrolases; cath \| 1fuuA00 \| 96.3 \| 1e-05 \| 1.5e-09 \| 66.5 \| 156 \| (4, 164) \| 471 \| (43, 219) \| 225 \| Yeast initiation factor 4a \| CATHCODE: 3.40.50.300 NAME: Yeast initiation factor 4a. Chain: a, b. Fragment: mRNA helicase. Synonym: eukaryotic initiation factor 4a. Engineered: yes SOURCE: Saccharomyces cerevisiae. Baker's yeast. Organism\_taxid: 4932. Expressed in: escherichia coli. Expression\_system\_taxid: 562 CLASS: Alpha Beta, ARCH: 3-Layer(aba) Sandwich, TOPOL: Rossmann fold, HOMOL: P-loop containing nucleotide triphosphate hydrolases; cath \| 5gvrA00 \| 96.3 \| 1.4e-05 \| 2e-09 \| 66.5 \| 156 \| (4, 164) \| 471 \| (35, 226) \| 234 \| Probable atp-dependent RNA helicase ddx41 \| CATHCODE: 3.40.50.300 NAME: Probable atp-dependent RNA helicase ddx41. Chain: a. Fragment: unp residues 169-402. Synonym: dead box protein 41,dead box protein abstrakt homolog. Engineered: yes SOURCE: Homo sapiens. Human. Organism\_taxid: 9606. Gene: ddx41, abs. Expressed in: escherichia coli. Expression\_system\_taxid: 562. Expression\_system\_vector\_type: plasmid CLASS: Alpha Beta, ARCH: 3-Layer(aba) Sandwich, TOPOL: Rossmann fold, HOMOL: P-loop containing nucleotide triphosphate hydrolases; cath \| 3dmqA04 \| 96.2 \| 1.6e-05 \| 2.2e-09 \| 70.4 \| 157 \| (1, 160) \| 471 \| (1, 179) \| 302 \| Rna polymerase-associated protein rapa \| CATHCODE: 3.40.50.10810 NAME: Rna polymerase-associated protein rapa. Chain: a, b. Synonym: atp-dependent helicase hepa. Engineered: yes. Mutation: yes SOURCE: Escherichia coli k12. Organism\_taxid: 83333. Strain: k12 / mg1655. Atcc: 47076. Gene: rapa, hepa, yaba, b0059, jw0058. Expressed in: escherichia coli. Expression\_system\_taxid: 562 CLASS: Alpha Beta, ARCH: 3-Layer(aba) Sandwich, TOPOL: Rossmann fold, HOMOL: Tandem AAA-ATPase domain; cath \| 4ljyA01 \| 96.2 \| 1.6e-05 \| 2.2e-09 \| 68.1 \| 157 \| (4, 165) \| 471 \| (73, 261) \| 265 \| Pre-mRNA-processing atp-dependent RNA helicase prp5 \| CATHCODE: 3.40.50.300 NAME: Pre-mRNA-processing atp-dependent RNA helicase prp5. Chain: a. Fragment: unp residues 206-698. Engineered: yes SOURCE: Saccharomyces cerevisiae. Yeast. Organism\_taxid: 559292. Strain: atcc 204508 / s288c. Gene: prp5, RNA5, ybr237w, ybr1603. Expressed in: escherichia coli. Expression\_system\_taxid: 562. Expression\_system\_vector\_type: plasmid CLASS: Alpha Beta, ARCH: 3-Layer(aba) Sandwich, TOPOL: Rossmann fold, HOMOL: P-loop containing nucleotide triphosphate hydrolases; cath \| 4w7sA01 \| 96.2 \| 1.7e-05 \| 2.3e-09 \| 68.8 \| 161 \| (4, 165) \| 471 \| (71, 273) \| 277 \| Pre-mRNA-splicing atp-dependent RNA helicase prp28 \| CATHCODE: 3.40.50.300 NAME: Pre-mRNA-splicing atp-dependent RNA helicase prp28. Chain: a, b. Synonym: helicase ca8. Engineered: yes SOURCE: Saccharomyces cerevisiae. Baker's yeast. Organism\_taxid: 559292. Strain: atcc 204508 / s288c. Gene: prp28, ydr243c, yd8419.10c. Expressed in: escherichia coli. Expression\_system\_taxid: 562. CLASS: Alpha Beta, ARCH: 3-Layer(aba) Sandwich, TOPOL: Rossmann fold, HOMOL: P-loop containing nucleotide triphosphate hydrolases; cath \| 2xgjB01 \| 96.2 \| 1.8e-05 \| 2.6e-09 \| 65.0 \| 143 \| (4, 151) \| 471 \| (33, 183) \| 206 \| Atp-dependent RNA helicase dob1 \| CATHCODE: 3.40.50.300 NAME: Atp-dependent RNA helicase dob1. Chain: a, b. Fragment: residues 81-1073. Synonym: mtr4p, mRNA transport regulator mtr4., 3.6.1.-. Rna (5'-(\*ap\*ap\*ap\*ap\*a)-3'). Chain: c, d SOURCE: Saccharomyces cerevisiae. Baker's yeast. Organism\_taxid: 4932. CLASS: Alpha Beta, ARCH: 3-Layer(aba) Sandwich, TOPOL: Rossmann fold, HOMOL: P-loop containing nucleotide triphosphate hydrolases; cath \| 2db3A01 \| 96.1 \| 2.1e-05 \| 3e-09 \| 67.2 \| 158 \| (4, 165) \| 471 \| (78, 263) \| 267 \| 5'-r(\*up\*up\*up\*up\*up\*up\*up\*up\*up\*u)-3' \| CATHCODE: 3.40.50.300 NAME: 5'-r(\*up\*up\*up\*up\*up\*up\*up\*up\*up\*u)-3'. Chain: e, f, g, h. Engineered:yes. Atp-dependent RNA helicase vasa. Chain: a, b, c, d. Fragment: residues 200-623. Synonym: vasa protein, antigen mab46f11. Engineered: yes SOURCE: Yes. CLASS: Alpha Beta, ARCH: 3-Layer(aba) Sandwich, TOPOL: Rossmann fold, HOMOL: P-loop containing nucleotide triphosphate hydrolases; cath \| 3dkpA00 \| 96.1 \| 2.3e-05 \| 3.2e-09 \| 65.7 \| 158 \| (4, 165) \| 471 \| (51, 237) \| 245 \| Probable atp-dependent RNA helicase ddx52 \| CATHCODE: 3.40.50.300 NAME: Probable atp-dependent RNA helicase ddx52. Chain: a. Fragment: conserved domain i: residues 139-381. Synonym: dead box protein 52, atp-dependent RNA helicase rok1-like. Engineered: yes SOURCE: Homo sapiens. Organism\_taxid: 9606. Gene: ddx52, rok1. Expressed in: escherichia coli. CLASS: Alpha Beta, ARCH: 3-Layer(aba) Sandwich, TOPOL: Rossmann fold, HOMOL: P-loop containing nucleotide triphosphate hydrolases; cath \| 5supC01 \| 96.1 \| 2.4e-05 \| 3.3e-09 \| 64.1 \| 155 \| (4, 163) \| 471 \| (28, 209) \| 220 \| Atp-dependent RNA helicase sub2 \| CATHCODE: 3.40.50.300 NAME: Atp-dependent RNA helicase sub2. Chain: a, b, c. Fragment: residues 61-446. Synonym: suppressor of brr1 protein 2. Engineered: yes. Rna annealing protein yra1. Chain: g, h, i. Fragment: residues 200-226. Engineered: yes. Rna (5'-r(p\*up\*up\*up\*up\*up\*u)-3'). Chain: d, e, f. Engineered: yes SOURCE: Saccharomyces cerevisiae. Baker's yeast. Organism\_taxid: 559292. Strain: atcc 204508 / s288c. Gene: sub2, ydl084w. Expressed in: escherichia coli. Expression\_system\_taxid: 562. Saccharomyces cerevisiae. Baker's yeast. Organism\_taxid: 559292. Strain: atcc 204508 / s288c. Gene: yra1, ydr381w, d9481.2, d9509.1. Expressed in: escherichia coli. Expression\_system\_taxid: 562. CLASS: Alpha Beta, ARCH: 3-Layer(aba) Sandwich, TOPOL: Rossmann fold, HOMOL: P-loop containing nucleotide triphosphate hydrolases; cath \| 1wp9A01 \| 95.9 \| 3.7e-05 \| 5.2e-09 \| 62.0 \| 128 \| (23, 150) \| 471 \| (23, 168) \| 199 \| Atp-dependent RNA helicase, putative \| CATHCODE: 3.40.50.300 NAME: Atp-dependent RNA helicase, putative. Chain: a, b, c, d, e, f. Fragment: residues 2-495. Synonym: hef helicase, nuclease. Engineered: yes SOURCE: Pyrococcus furiosus. Organism\_taxid: 186497. Strain: dsm 3638. Expressed in: escherichia coli bl21(de3). Expression\_system\_taxid: 469008. CLASS: Alpha Beta, ARCH: 3-Layer(aba) Sandwich, TOPOL: Rossmann fold, HOMOL: P-loop containing nucleotide triphosphate hydrolases; cath \| 2hyiC02 \| 95.9 \| 3.8e-05 \| 5.4e-09 \| 60.2 \| 101 \| (228, 335) \| 471 \| (30, 130) \| 167 \| 5'-r(\*up\*up\*up\*up\*up\*u)-3' \| CATHCODE: 3.40.50.300 NAME: 5'-r(\*up\*up\*up\*up\*up\*u)-3'. Chain: f, l. Fragment: mRNA mimick. Engineered: yes. Protein mago nashi homolog. Chain: a, g. Engineered: yes. Rna-binding protein 8a. Chain: b, h. Fragment: n-terminal deletion mutant. Synonym: RNA-binding motif protein 8a, ribonucleoprotein rbm8a, RNA-binding protein y14, binder of ovca1- 1, bov-1. Engineered: yes. Probable atp-dependent RNA helicase ddx48. Chain: c, i. Synonym: dead boxprotein 48, eukaryotic initiation factor 4a-like nuk-34, nuclear matrix protein 265, hnmp 265, eukaryotic translation initiation factor 4a isoform 3. Engineered: yes. Protein casc3. Chain: d, j. Fragment: selor fragment. Synonym: cancer susceptibility candidate gene 3 protein, metastatic lymph node protein 51, mln 51 protein, barentsz protein, btz. Engineered: yes SOURCE: Yes. CLASS: Alpha Beta, ARCH: 3-Layer(aba) Sandwich, TOPOL: Rossmann fold, HOMOL: P-loop containing nucleotide triphosphate hydrolases; cath \| 1gm5A04 \| 95.9 \| 4.3e-05 \| 6.1e-09 \| 62.2 \| 147 \| (3, 152) \| 471 \| (19, 182) \| 204 \| Recg \| CATHCODE: 3.40.50.300 NAME: Recg. Chain: a. Engineered: yes. Dna (5'-(\*cp\*ap\*gp\*cp\*tp\*cp\*cp\*ap\*tp\*gp\*ap\*tp\* cp\*ap\*tp\*tp\*gp\*gp\*cp\*a)-3'). Chain: x. Dna (5'-(\*gp\*cp\*ap\*gp\*tp\*gp\*cp\*tp\*cp\*gp\*cp\*ap\* tp\*gp\*gp\*ap\*gp\*cp\*tp\*g)-3'). Chain: y. Dna(5'-(\*gp\*ap\*gp\*cp\*ap\*cp\*tp\*gp\*c)-3'). Chain: z SOURCE: Thermotoga maritima. Organism\_taxid: 2336. Expressed in: escherichia coli. Expression\_system\_taxid: 562. CLASS: Alpha Beta, ARCH: 3-Layer(aba) Sandwich, TOPOL: Rossmann fold, HOMOL: P-loop containing nucleotide triphosphate hydrolases; cath \| 2eyqA04 \| 95.7 \| 6.2e-05 \| 8.5e-09 \| 64.8 \| 100 \| (229, 335) \| 471 \| (58, 160) \| 249 \| Transcription-repair coupling factor \| CATHCODE: 3.40.50.300 NAME: Transcription-repair coupling factor. Chain: a, b. Synonym: trcf. Engineered: yes SOURCE: Escherichia coli. Organism\_taxid: 562. Gene: mfd. Expressed in: escherichia coli. Expression\_system\_taxid: 562 CLASS: Alpha Beta, ARCH: 3-Layer(aba) Sandwich, TOPOL: Rossmann fold, HOMOL: P-loop containing nucleotide triphosphate hydrolases; cath \| 3i5xA01 \| 95.6 \| 8.5e-05 \| 1.2e-08 \| 62.1 \| 155 \| (4, 160) \| 471 \| (40, 232) \| 247 \| Atp-dependent RNA helicase mss116 \| CATHCODE: 3.40.50.300 NAME: Atp-dependent RNA helicase mss116. Chain: a. Fragment: unp residues 37to 597. Engineered: yes. 5'-r(\*up\*up\*up\*up\*up\*up\*up\*up\*up\*u)-3'. Chain: b. Engineered: yes SOURCE: Saccharomyces cerevisiae. Yeast. Organism\_taxid: 4932. Gene: mss116, yd9346.05c, ydr194c. Expressed in: escherichia coli. Expression\_system\_taxid: 562. CLASS: Alpha Beta, ARCH: 3-Layer(aba) Sandwich, TOPOL: Rossmann fold, HOMOL: P-loop containing nucleotide triphosphate hydrolases; cath \| 2o0jA02 \| 95.6 \| 9.1e-05 \| 1.3e-08 \| 63.1 \| 154 \| (3, 162) \| 471 \| (78, 241) \| 254 \| Dna packaging protein gp17 \| CATHCODE: 3.40.50.300 NAME: Dna packaging protein gp17. Chain: a. Fragment: n-terminal atpase domain. Synonym: terminase. Engineered: yes. Mutation: yes SOURCE: Enterobacteria phage t4. Organism\_taxid: 10665. Gene: 17. Expressed in: escherichia coli. Expression\_system\_taxid: 562. CLASS: Alpha Beta, ARCH: 3-Layer(aba) Sandwich, TOPOL: Rossmann fold, HOMOL: P-loop containing nucleotide triphosphate hydrolases; cath \| 3h1tA03 \| 95.6 \| 9.2e-05 \| 1.3e-08 \| 61.8 \| 107 \| (229, 343) \| 471 \| (80, 200) \| 230 \| Type i site-specific restriction-modification system, r (restriction) subunit \| CATHCODE: 3.40.50.300 NAME: Type i site-specific restriction-modification system, r (restriction) subunit. Chain: a. Fragment: unp residues 1-590. Engineered: yes SOURCE: Vibrio vulnificus. Organism\_taxid: 196600. Strain: yj016. Gene: vv0265. Expressed in: escherichia coli. Expression\_system\_taxid: 562. CLASS: Alpha Beta, ARCH: 3-Layer(aba) Sandwich, TOPOL: Rossmann fold, HOMOL: P-loop containing nucleotide triphosphate hydrolases; cath \| 2pl3A00 \| 95.5 \| 0.00011 \| 1.6e-08 \| 60.1 \| 152 \| (5, 161) \| 471 \| (48, 225) \| 236 \| Probable atp-dependent RNA helicase ddx10 \| CATHCODE: 3.40.50.300 NAME: Probable atp-dependent RNA helicase ddx10. Chain: a. Fragment: dead domain. Synonym: dead box protein 10. Engineered: yes SOURCE: Homo sapiens. Human. Organism\_taxid: 9606. Gene: ddx10. Expressed in:escherichia coli. Expression\_system\_taxid: 562. CLASS: Alpha Beta, ARCH: 3-Layer(aba) Sandwich, TOPOL: Rossmann fold, HOMOL: P-loop containing nucleotide triphosphate hydrolases; cath \| 3dmqA05 \| 95.4 \| 0.00012 \| 1.7e-08 \| 59.7 \| 101 \| (227, 334) \| 471 \| (77, 180) \| 209 \| Rna polymerase-associated protein rapa \| CATHCODE: 3.40.50.300 NAME: Rna polymerase-associated protein rapa. Chain: a, b. Synonym: atp-dependent helicase hepa. Engineered: yes. Mutation: yes SOURCE: Escherichia coli k12. Organism\_taxid: 83333. Strain: k12 / mg1655. Atcc: 47076. Gene: rapa, hepa, yaba, b0059, jw0058. Expressed in: escherichia coli. Expression\_system\_taxid: 562 CLASS: Alpha Beta, ARCH: 3-Layer(aba) Sandwich, TOPOL: Rossmann fold, HOMOL: P-loop containing nucleotide triphosphate hydrolases; cath \| 2ykgA01 \| 95.4 \| 0.00012 \| 1.7e-08 \| 59.4 \| 146 \| (3, 152) \| 471 \| (12, 187) \| 224 \| Probable atp-dependent RNA helicase ddx58 \| CATHCODE: 3.40.50.300 NAME: Probable atp-dependent RNA helicase ddx58. Chain: a. Fragment: residues 230-925. Synonym: dead box protein 58, retinoic acid-inducible gene 1 protein, rig-1, retinoic acid-inducible gene i protein, rig-i. 5'-r(\*gp\*cp\*gp\*cp\*gp\*cp\*gp\*cp\*gp\*cp)-3'. Chain: c, d SOURCE: Homo sapiens. Organism\_taxid: 9606. CLASS: Alpha Beta, ARCH: 3-Layer(aba) Sandwich, TOPOL: Rossmann fold, HOMOL: P-loop containing nucleotide triphosphate hydrolases; cath \| 3llmA01 \| 95.4 \| 0.00013 \| 1.8e-08 \| 60.8 \| 142 \| (4, 151) \| 471 \| (51, 211) \| 225 \| Atp-dependent RNA helicase a \| CATHCODE: 3.40.50.300 NAME: Atp-dependent RNA helicase a. Chain: a, b. Fragment: nucleotide binding domain (unp residues 329-563). Synonym: nuclear DNA helicase ii, ndhii, deah box protein 9. Engineered: yes SOURCE: Homo sapiens. Human. Organism\_taxid: 9606. Gene: ddx9, dhx9, lkp, ndh2. Expressed in: escherichia coli. Expression\_system\_taxid: 562. CLASS: Alpha Beta, ARCH: 3-Layer(aba) Sandwich, TOPOL: Rossmann fold, HOMOL: P-loop containing nucleotide triphosphate hydrolases; cath \| 2p6nA00 \| 95.4 \| 0.00013 \| 1.8e-08 \| 56.7 \| 99 \| (229, 334) \| 471 \| (27, 125) \| 164 \| Atp-dependent RNA helicase ddx41 \| CATHCODE: 3.40.50.300 NAME: Atp-dependent RNA helicase ddx41. Chain: a, b. Fragment: helicase domain. Synonym: dead box protein 41, dead box protein abstrakt homolog. Engineered: yes. Mutation: yes SOURCE: Homo sapiens. Human. Organism\_taxid: 9606. Gene: ddx41, abs. Expressed in: escherichia coli. Expression\_system\_taxid: 562. CLASS: Alpha Beta, ARCH: 3-Layer(aba) Sandwich, TOPOL: Rossmann fold, HOMOL: P-loop containing nucleotide triphosphate hydrolases; cath \| 5lbaD02 \| 95.4 \| 0.00013 \| 1.8e-08 \| 61.0 \| 99 \| (229, 334) \| 471 \| (39, 137) \| 233 \| Atp-dependent DNA helicase q5 \| CATHCODE: 3.40.50.300 NAME: Atp-dependent DNA helicase q5. Chain: b, a, c, d. Synonym: DNA helicase,recq-like type 5,recq5,recq protein-like 5. Engineered: yes SOURCE: Homo sapiens. Human. Organism\_taxid: 9606. Gene: recql5, recq5. Expressed in: escherichia coli bl21. Expression\_system\_taxid: 511693 CLASS: Alpha Beta, ARCH: 3-Layer(aba) Sandwich, TOPOL: Rossmann fold, HOMOL: P-loop containing nucleotide triphosphate hydrolases; cath \| 4nl4H03 \| 95.4 \| 0.00014 \| 1.9e-08 \| 57.8 \| 147 \| (3, 152) \| 471 \| (14, 180) \| 196 \| Primosome assembly protein pria \| CATHCODE: 3.40.50.300 NAME: Primosome assembly protein pria. Chain: h. Engineered: yes SOURCE: Klebsiella pneumoniae subsp. Pneumoniae. Organism\_taxid: 272620. Strain: mgh 78578. Gene: kpn78578\_41850, kpn\_04230, pria. Expressed in: escherichia coli. Expression\_system\_taxid: 562. Expression\_system\_vector\_type: plasmid. CLASS: Alpha Beta, ARCH: 3-Layer(aba) Sandwich, TOPOL: Rossmann fold, HOMOL: P-loop containing nucleotide triphosphate hydrolases; cath \| 3crvA01 \| 95.4 \| 0.00014 \| 2e-08 \| 60.8 \| 67 \| (3, 72) \| 471 \| (2, 68) \| 246 \| Xpd/rad3 related DNA helicase \| CATHCODE: 3.40.50.300 NAME: Xpd/rad3 related DNA helicase. Chain: a. Ec: 3.-.-.-. Engineered: yes SOURCE: Sulfolobus acidocaldarius. Organism\_taxid: 2285. Gene: saci\_0192. Expressed in: escherichia coli. Expression\_system\_taxid: 562. Expression\_system\_vector\_type: plasmid. CLASS: Alpha Beta, ARCH: 3-Layer(aba) Sandwich, TOPOL: Rossmann fold, HOMOL: P-loop containing nucleotide triphosphate hydrolases; cath \| 2yjtD00 \| 95.3 \| 0.00015 \| 2.1e-08 \| 56.5 \| 104 \| (224, 334) \| 471 \| (25, 128) \| 170 \| Regulator of ribonuclease activity a \| CATHCODE: 3.40.50.300 NAME: Regulator of ribonuclease activity a. Chain: a, b, c. Synonym: rraa. Engineered: yes. Atp-dependent RNA helicase srmb. Chain: d. Fragment: residues 219-388. Engineered: yes SOURCE: Escherichia coli. Organism\_taxid: 83333. Strain: k-12. Expressed in: escherichia coli. Expression\_system\_taxid: 511693. CLASS: Alpha Beta, ARCH: 3-Layer(aba) Sandwich, TOPOL: Rossmann fold, HOMOL: P-loop containing nucleotide triphosphate hydrolases; cath \| 4f92B02 \| 95.3 \| 0.00015 \| 2.2e-08 \| 59.3 \| 144 \| (4, 150) \| 471 \| (27, 204) \| 228 \| U5 small nuclear ribonucleoprotein 200 kda helicase \| CATHCODE: 3.40.50.300 NAME: U5 small nuclear ribonucleoprotein 200 kda helicase. Chain: b. Fragment: brr2 helicase region. Synonym: activating signal cointegrator 1 complex subunit 3-like 1, brr2 homolog, u5 snrnp-specific 200 kda protein, u5-200kd. Engineered: yes. Mutation: yes SOURCE: Homo sapiens. Human. Organism\_taxid: 9606. Gene: snrnp200, ascc3l1, helic2, kiaa0788. Expressed in: spodoptera frugiperda. Expression\_system\_taxid: 7108 CLASS: Alpha Beta, ARCH: 3-Layer(aba) Sandwich, TOPOL: Rossmann fold, HOMOL: P-loop containing nucleotide triphosphate hydrolases; cath \| 2v1xA01 \| 95.3 \| 0.00016 \| 2.3e-08 \| 59.5 \| 150 \| (4, 160) \| 471 \| (44, 222) \| 235 \| Atp-dependent DNA helicase q1 \| CATHCODE: 3.40.50.300 NAME: Atp-dependent DNA helicase q1. Chain: a, b. Fragment: residues 49-616.Synonym: DNA-dependent atpase q1, recq DNA helicase. Engineered: yes SOURCE: Homo sapiens. Human. Organism\_taxid: 9606. Expressed in: escherichia coli. Expression\_system\_taxid: 469008. CLASS: Alpha Beta, ARCH: 3-Layer(aba) Sandwich, TOPOL: Rossmann fold, HOMOL: P-loop containing nucleotide triphosphate hydrolases; cath \| 5jajA01 \| 95.2 \| 0.00018 \| 2.5e-08 \| 60.6 \| 154 \| (3, 160) \| 471 \| (8, 189) \| 269 \| Lgp2 \| CATHCODE: 3.40.50.300 NAME: Lgp2. Chain: a. Engineered: yes. Mutation: yes. Rna (5'-r(p\*gp\*gp\*up\*ap\*cp\*gp\*up\*ap\*cp\*cp\*c)-3'). Chain: x. Engineered: yes. Rna (5'-r(p\*gp\*gp\*up\*ap\*cp\*gp\*up\*ap\*cp\*c)-3'). Chain: y. Engineered: yes SOURCE: Gallus gallus. Chicken. Organism\_taxid: 9031. Expressed in: escherichia coli bl21(de3). Expression\_system\_taxid: 469008. Expression\_system\_variant: rosetta 2. CLASS: Alpha Beta, ARCH: 3-Layer(aba) Sandwich, TOPOL: Rossmann fold, HOMOL: P-loop containing nucleotide triphosphate hydrolases; cath \| 1wp9A02 \| 95.2 \| 0.0002 \| 2.8e-08 \| 54.9 \| 108 \| (217, 331) \| 471 \| (19, 134) \| 164 \| Atp-dependent RNA helicase, putative \| CATHCODE: 3.40.50.300 NAME: Atp-dependent RNA helicase, putative. Chain: a, b, c, d, e, f. Fragment: residues 2-495. Synonym: hef helicase, nuclease. Engineered: yes SOURCE: Pyrococcus furiosus. Organism\_taxid: 186497. Strain: dsm 3638. Expressed in: escherichia coli bl21(de3). Expression\_system\_taxid: 469008. CLASS: Alpha Beta, ARCH: 3-Layer(aba) Sandwich, TOPOL: Rossmann fold, HOMOL: P-loop containing nucleotide triphosphate hydrolases; cath \| 2vl7A01 \| 95.2 \| 0.00021 \| 2.9e-08 \| 56.3 \| 143 \| (3, 151) \| 471 \| (6, 165) \| 179 \| Xpd \| CATHCODE: 3.40.50.300 NAME: Xpd. Chain: a. Synonym: uncharacterized protein st1307. Engineered: yes SOURCE: Sulfolobus tokodaii. Organism\_taxid: 111955. Strain: 7. Expressed in:escherichia coli. Expression\_system\_taxid: 562. Expression\_system\_vector\_type: plasmid. CLASS: Alpha Beta, ARCH: 3-Layer(aba) Sandwich, TOPOL: Rossmann fold, HOMOL: P-loop containing nucleotide triphosphate hydrolases; cath \| 2ykgA02 \| 95.1 \| 0.00023 \| 3.2e-08 \| 52.8 \| 98 \| (228, 332) \| 471 \| (34, 144) \| 152 \| Probable atp-dependent RNA helicase ddx58 \| CATHCODE: 3.40.50.300 NAME: Probable atp-dependent RNA helicase ddx58. Chain: a. Fragment: residues 230-925. Synonym: dead box protein 58, retinoic acid-inducible gene 1 protein, rig-1, retinoic acid-inducible gene i protein, rig-i. 5'-r(\*gp\*cp\*gp\*cp\*gp\*cp\*gp\*cp\*gp\*cp)-3'. Chain: c, d SOURCE: Homo sapiens. Organism\_taxid: 9606. CLASS: Alpha Beta, ARCH: 3-Layer(aba) Sandwich, TOPOL: Rossmann fold, HOMOL: P-loop containing nucleotide triphosphate hydrolases; cath \| 3mwyW03 \| 95.1 \| 0.00024 \| 3.4e-08 \| 60.4 \| 152 \| (3, 157) \| 471 \| (33, 212) \| 282 \| Chromo domain-containing protein 1 \| CATHCODE: 3.40.50.10810 NAME: Chromo domain-containing protein 1. Chain: w. Fragment: double chromodomains and atpase motor (unp residues 142- 939). Synonym: atp-dependent helicase chd1. Engineered: yes SOURCE: Saccharomyces cerevisiae. Brewer's yeast,lager beer yeast,yeast. Organism\_taxid: 4932. Gene: chd1, sygp-orf4, yer164w. Expressed in: escherichia coli. Expression\_system\_taxid: 469008. CLASS: Alpha Beta, ARCH: 3-Layer(aba) Sandwich, TOPOL: Rossmann fold, HOMOL: Tandem AAA-ATPase domain; cath \| 3v4rA01 \| 95.0 \| 0.00025 \| 3.5e-08 \| 59.8 \| 69 \| (2, 73) \| 471 \| (16, 85) \| 254 \| Uvrabc system protein b \| CATHCODE: 3.40.50.300 NAME: Uvrabc system protein b. Chain: a, b. Synonym: protein uvrb, excinuclease abc subunit b, protein dina. Engineered: yes. Dna: 5 -tactgttt-3. Chain: c, d. Engineered: yes SOURCE: Bacillus subtilis. Organism\_taxid: 1423. Gene: bsu35170, dina, uvr, uvrb. Expressed in: escherichia coli. Expression\_system\_taxid: 469008. CLASS: Alpha Beta, ARCH: 3-Layer(aba) Sandwich, TOPOL: Rossmann fold, HOMOL: P-loop containing nucleotide triphosphate hydrolases; cath \| 1a1vA01 \| 95.0 \| 0.00025 \| 3.5e-08 \| 53.6 \| 99 \| (23, 124) \| 471 \| (8, 108) \| 135 \| Protein (ns3 protein) \| CATHCODE: 3.40.50.300 NAME: Protein (ns3 protein). Chain: a. Fragment: helicase domain. Engineered: yes. Mutation: yes. Dna (5'-d(\*up\*up\*up\*up\*up\*up\*up\*u)-3'). Chain: b. Engineered: yes. Other\_details: single stranded DNA SOURCE: Hepatitis c virus (isolate h). Organism\_taxid: 11108. Strain: h. Gene:ns3. Expressed in: escherichia coli. Expression\_system\_taxid: 562. CLASS: Alpha Beta, ARCH: 3-Layer(aba) Sandwich, TOPOL: Rossmann fold, HOMOL: P-loop containing nucleotide triphosphate hydrolases; cath \| 1gkuB02 \| 95.0 \| 0.00027 \| 3.8e-08 \| 57.5 \| 117 \| (4, 124) \| 471 \| (24, 157) \| 213 \| Reverse gyrase \| CATHCODE: 3.40.50.300 NAME: Reverse gyrase. Chain: b. Synonym: top-rg. Engineered: yes. Mutation: yes SOURCE: Archaeoglobus fulgidus. Organism\_taxid: 224325. Strain: vc-16. Atcc: 49558. Expressed in: escherichia coli bl21(de3). Expression\_system\_taxid: 469008. Expression\_system\_variant: c41. CLASS: Alpha Beta, ARCH: 3-Layer(aba) Sandwich, TOPOL: Rossmann fold, HOMOL: P-loop containing nucleotide triphosphate hydrolases; cath \| 4w7sA02 \| 95.0 \| 0.00028 \| 4e-08 \| 56.1 \| 98 \| (229, 333) \| 471 \| (34, 132) \| 186 \| Pre-mRNA-splicing atp-dependent RNA helicase prp28 \| CATHCODE: 3.40.50.300 NAME: Pre-mRNA-splicing atp-dependent RNA helicase prp28. Chain: a, b. Synonym: helicase ca8. Engineered: yes SOURCE: Saccharomyces cerevisiae. Baker's yeast. Organism\_taxid: 559292. Strain: atcc 204508 / s288c. Gene: prp28, ydr243c, yd8419.10c. Expressed in: escherichia coli. Expression\_system\_taxid: 562. CLASS: Alpha Beta, ARCH: 3-Layer(aba) Sandwich, TOPOL: Rossmann fold, HOMOL: P-loop containing nucleotide triphosphate hydrolases; cath \| 3i5xA02 \| 94.9 \| 0.00032 \| 4.5e-08 \| 59.5 \| 100 \| (228, 334) \| 471 \| (37, 139) \| 260 \| Atp-dependent RNA helicase mss116 \| CATHCODE: 3.40.50.300 NAME: Atp-dependent RNA helicase mss116. Chain: a. Fragment: unp residues 37to 597. Engineered: yes. 5'-r(\*up\*up\*up\*up\*up\*up\*up\*up\*up\*u)-3'. Chain: b. Engineered: yes SOURCE: Saccharomyces cerevisiae. Yeast. Organism\_taxid: 4932. Gene: mss116, yd9346.05c, ydr194c. Expressed in: escherichia coli. Expression\_system\_taxid: 562. CLASS: Alpha Beta, ARCH: 3-Layer(aba) Sandwich, TOPOL: Rossmann fold, HOMOL: P-loop containing nucleotide triphosphate hydrolases; cath \| 4idhA01 \| 94.9 \| 0.00032 \| 4.5e-08 \| 56.9 \| 137 \| (5, 151) \| 471 \| (2, 149) \| 208 \| Gene 2 protein \| CATHCODE: 3.40.50.300 NAME: Gene 2 protein. Chain: a. Engineered: yes SOURCE: Shigella phage sf6. Shigella flexneri bacteriophage vi. Organism\_taxid: 10761. Expressed in: escherichia coli. Expression\_system\_taxid: 469008. CLASS: Alpha Beta, ARCH: 3-Layer(aba) Sandwich, TOPOL: Rossmann fold, HOMOL: P-loop containing nucleotide triphosphate hydrolases; cath \| 1m6nA01 \| 94.9 \| 0.00033 \| 4.6e-08 \| 60.9 \| 115 \| (4, 124) \| 471 \| (80, 214) \| 286 \| Preprotein translocase seca \| CATHCODE: 3.40.50.300 NAME: Preprotein translocase seca. Chain: a. Synonym: seca. Engineered: yes SOURCE: Bacillus subtilis. Organism\_taxid: 1423. Gene: div. Expressed in: escherichia coli bl21(de3). Expression\_system\_taxid: 469008. CLASS: Alpha Beta, ARCH: 3-Layer(aba) Sandwich, TOPOL: Rossmann fold, HOMOL: P-loop containing nucleotide triphosphate hydrolases; cath \| 4a15A04 \| 94.9 \| 0.00034 \| 4.7e-08 \| 57.8 \| 132 \| (217, 353) \| 471 \| (28, 186) \| 211 \| Atp-dependent DNA helicase ta0057 \| CATHCODE: 3.40.50.300 NAME: Atp-dependent DNA helicase ta0057. Chain: a. Synonym: xpd helicase. Engineered: yes. 5'-d(\*dtp\*ap\*cp\*gp)-3'. Chain: e SOURCE: Thermoplasma acidophilum. Organism\_taxid: 2303. Expressed in: escherichia coli. Expression\_system\_taxid: 469008. CLASS: Alpha Beta, ARCH: 3-Layer(aba) Sandwich, TOPOL: Rossmann fold, HOMOL: P-loop containing nucleotide triphosphate hydrolases; cath \| 5lstA02 \| 94.8 \| 0.00034 \| 4.9e-08 \| 52.1 \| 99 \| (229, 334) \| 471 \| (33, 145) \| 151 \| Atp-dependent DNA helicase q4 \| CATHCODE: 3.40.50.300 NAME: Atp-dependent DNA helicase q4. Chain: a. Synonym: DNA helicase,recq-like type 4,recq4,rts,recq protein-like 4. Engineered: yes SOURCE: Homo sapiens. Human. Organism\_taxid: 9606. Gene: recql4, recq4. Expressed in: escherichia coli. Expression\_system\_taxid: 469008. CLASS: Alpha Beta, ARCH: 3-Layer(aba) Sandwich, TOPOL: Rossmann fold, HOMOL: P-loop containing nucleotide triphosphate hydrolases; cath \| 3b85A00 \| 94.8 \| 0.00035 \| 5e-08 \| 56.2 \| 57 \| (4, 64) \| 471 \| (7, 65) \| 208 \| Phosphate starvation-inducible protein \| CATHCODE: 3.40.50.300 NAME: Phosphate starvation-inducible protein. Chain: a, b. Fragment: residues 116-320. Engineered: yes SOURCE: Corynebacterium glutamicum atcc 13032. Organism\_taxid: 196627. Strain:dsm 20300 / jcm 1318 / lmg 3730 / ncimb 10025. Atcc: 13032. Gene: phoh2, cg2513. Expressed in: escherichia coli bl21(de3). Expression\_system\_taxid: 469008. CLASS: Alpha Beta, ARCH: 3-Layer(aba) Sandwich, TOPOL: Rossmann fold, HOMOL: P-loop containing nucleotide triphosphate hydrolases; cath \| 1c4oA03 \| 94.8 \| 0.00038 \| 5.3e-08 \| 54.4 \| 97 \| (229, 332) \| 471 \| (32, 133) \| 175 \| Dna nucleotide excision repair enzyme uvrb \| CATHCODE: 3.40.50.300 NAME: Dna nucleotide excision repair enzyme uvrb. Chain: a. Engineered: yes SOURCE: Thermus thermophilus. Organism\_taxid: 300852. Strain: hb8. Atcc: 27634. Expressed in: escherichia coli bl21. Expression\_system\_taxid: 511693. CLASS: Alpha Beta, ARCH: 3-Layer(aba) Sandwich, TOPOL: Rossmann fold, HOMOL: P-loop containing nucleotide triphosphate hydrolases; cath \| 1hv8A02 \| 94.8 \| 0.00038 \| 5.4e-08 \| 52.3 \| 99 \| (229, 334) \| 471 \| (25, 123) \| 151 \| Putative atp-dependent RNA helicase mj0669 \| CATHCODE: 3.40.50.300 NAME: Putative atp-dependent RNA helicase mj0669. Chain: a, b. Synonym: deadbox helicase. Engineered: yes SOURCE: Methanocaldococcus jannaschii. Organism\_taxid: 2190. Expressed in: escherichia coli. Expression\_system\_taxid: 562 CLASS: Alpha Beta, ARCH: 3-Layer(aba) Sandwich, TOPOL: Rossmann fold, HOMOL: P-loop containing nucleotide triphosphate hydrolases; cath \| 6b4kB02 \| 94.7 \| 0.00043 \| 6e-08 \| 54.2 \| 100 \| (228, 334) \| 471 \| (29, 134) \| 176 \| Atp-dependent RNA helicase ddx19b \| CATHCODE: 3.40.50.300 NAME: Atp-dependent RNA helicase ddx19b. Chain: a, b. Synonym: dead box RNA helicase dead5,dead box protein 19b. Engineered: yes SOURCE: Homo sapiens. Human. Organism\_taxid: 9606. Gene: ddx19b, dbp5, ddx19, tdbp. Expressed in: escherichia coli. Expression\_system\_taxid: 562 CLASS: Alpha Beta, ARCH: 3-Layer(aba) Sandwich, TOPOL: Rossmann fold, HOMOL: P-loop containing nucleotide triphosphate hydrolases; cath \| 1z63A01 \| 94.7 \| 0.00043 \| 6.1e-08 \| 54.3 \| 144 \| (7, 153) \| 471 \| (2, 154) \| 206 \| 5'-d(\*ap\*ap\*ap\*ap\*ap\*a\*ap\*tp\*tp\*gp\*cp\*cp\*gp\*ap\*ap\*gp\*ap\*cp\* gp\*ap\*ap\*ap\*ap\*ap\*a)-3' \| CATHCODE: 3.40.50.10810 NAME: 5'-d(\*ap\*ap\*ap\*ap\*ap\*a\*ap\*tp\*tp\*gp\*cp\*cp\*gp\*ap\*ap\*gp\*ap\*cp\* gp\*ap\*ap\*ap\*ap\*ap\*a)-3'. Chain: c, e. Engineered: yes. 5'-d(\*tp\*tp\*tp\*tp\*tp\*tp\*tp\*cp\*gp\*tp\*cp\*tp\*tp\*cp\*gp\*gp\*cp\*ap \*ap\*tp\*tp\*tp\*tp\*tp\*t)-3'. Chain: d,f. Engineered: yes. Helicase of the snf2/rad54 family. Chain: a, b. Fragment: residues 407-902. Engineered: yes SOURCE: Yes. Yes. CLASS: Alpha Beta, ARCH: 3-Layer(aba) Sandwich, TOPOL: Rossmann fold, HOMOL: Tandem AAA-ATPase domain; cath \| 1yksA01 \| 94.6 \| 0.00048 \| 6.8e-08 \| 53.5 \| 137 \| (23, 159) \| 471 \| (8, 151) \| 181 \| Genome polyprotein [contains: flavivirin protease ns3 catalytic subunit] \| CATHCODE: 3.40.50.300 NAME: Genome polyprotein [contains: flavivirin protease ns3 catalytic subunit]. Chain: a. Fragment: sequence database residues 1671-2107 (portion of flavivirin protease ns3 catalytic subunit). Engineered: yes SOURCE: Yellow fever virus. Organism\_taxid: 11089. Strain: strain pasteur 17d-204. Expressed in: escherichia coli. Expression\_system\_taxid: 562 CLASS: Alpha Beta, ARCH: 3-Layer(aba) Sandwich, TOPOL: Rossmann fold, HOMOL: P-loop containing nucleotide triphosphate hydrolases; cath \| 4f92B07 \| 94.6 \| 0.0005 \| 7.1e-08 \| 55.9 \| 61 \| (4, 67) \| 471 \| (38, 100) \| 230 \| U5 small nuclear ribonucleoprotein 200 kda helicase \| CATHCODE: 3.40.50.300 NAME: U5 small nuclear ribonucleoprotein 200 kda helicase. Chain: b. Fragment: brr2 helicase region. Synonym: activating signal cointegrator 1 complex subunit 3-like 1, brr2 homolog, u5 snrnp-specific 200 kda protein, u5-200kd. Engineered: yes. Mutation: yes SOURCE: Homo sapiens. Human. Organism\_taxid: 9606. Gene: snrnp200, ascc3l1, helic2, kiaa0788. Expressed in: spodoptera frugiperda. Expression\_system\_taxid: 7108 CLASS: Alpha Beta, ARCH: 3-Layer(aba) Sandwich, TOPOL: Rossmann fold, HOMOL: P-loop containing nucleotide triphosphate hydrolases; cath \| 2v1xA02 \| 94.4 \| 0.0006 \| 8.5e-08 \| 54.1 \| 100 \| (228, 334) \| 471 \| (31, 130) \| 200 \| Atp-dependent DNA helicase q1 \| CATHCODE: 3.40.50.300 NAME: Atp-dependent DNA helicase q1. Chain: a, b. Fragment: residues 49-616.Synonym: DNA-dependent atpase q1, recq DNA helicase. Engineered: yes SOURCE: Homo sapiens. Human. Organism\_taxid: 9606. Expressed in: escherichia coli. Expression\_system\_taxid: 469008. CLASS: Alpha Beta, ARCH: 3-Layer(aba) Sandwich, TOPOL: Rossmann fold, HOMOL: P-loop containing nucleotide triphosphate hydrolases; cath \| 2waxC00 \| 94.3 \| 0.00068 \| 9.6e-08 \| 52.1 \| 99 \| (229, 334) \| 471 \| (28, 126) \| 177 \| Atp-dependent RNA helicase ddx6 \| CATHCODE: 3.40.50.300 NAME: Atp-dependent RNA helicase ddx6. Chain: a, c. Fragment: c-terminal domain, residues 296-483. Synonym: human ddx6, dead box protein 6, atp-dependent RNA helicase p54, oncogene rck. Engineered: yes. Enhancer of mRNA-decapping protein 3. Chain: b, d. Fragment: fdf peptide, residues192-228. Synonym: human edc3, lsm16 homolog, yjef domain-containing protein 1, yjef n-terminal domain-containing protein 2, yjef\_n2, hyjef\_n2. Engineered: yes SOURCE: Homo sapiens. Human. Organism\_taxid: 9606. Expressed in: escherichia coli. Expression\_system\_taxid: 562. CLASS: Alpha Beta, ARCH: 3-Layer(aba) Sandwich, TOPOL: Rossmann fold, HOMOL: P-loop containing nucleotide triphosphate hydrolases; cath \| 3fhtB01 \| 94.3 \| 0.00069 \| 9.7e-08 \| 55.6 \| 147 \| (4, 153) \| 471 \| (47, 215) \| 235 \| Atp-dependent RNA helicase ddx19b \| CATHCODE: 3.40.50.300 NAME: Atp-dependent RNA helicase ddx19b. Chain: a, b. Fragment: helicase atp-binding domain, c-terminal domain, residues 68-479. Synonym: dead-boxhelicase 5, dbp5, dead box protein 19b, dead box RNA helicase dead5. Engineered: yes. Rna (5'-r(\*up\*up\*up\*up\*up\*up\*up\*up\*up\*u)-3'). Chain: c, d. Engineered: yes SOURCE: Homo sapiens. Human. Organism\_taxid: 9606. Gene: ddx19b (dbp5). Expressed in: escherichia coli. Expression\_system\_taxid: 511693. CLASS: Alpha Beta, ARCH: 3-Layer(aba) Sandwich, TOPOL: Rossmann fold, HOMOL: P-loop containing nucleotide triphosphate hydrolases; cath \| 1z3iX01 \| 94.2 \| 0.00084 \| 1.2e-07 \| 56.3 \| 154 \| (2, 158) \| 471 \| (33, 220) \| 277 \| Similar to rad54-like \| CATHCODE: 3.40.50.10810 NAME: Similar to rad54-like. Chain: x. Fragment: proteolytic fragment. Engineered: yes SOURCE: Danio rerio. Zebrafish. Organism\_taxid: 7955. Expressed in: spodoptera frugiperda. CLASS: Alpha Beta, ARCH: 3-Layer(aba) Sandwich, TOPOL: Rossmann fold, HOMOL: Tandem AAA-ATPase domain; cath \| 1z3iX02 \| 94.1 \| 0.00086 \| 1.2e-07 \| 55.8 \| 104 \| (229, 339) \| 471 \| (47, 153) \| 243 \| Similar to rad54-like \| CATHCODE: 3.40.50.300 NAME: Similar to rad54-like. Chain: x. Fragment: proteolytic fragment. Engineered: yes SOURCE: Danio rerio. Zebrafish. Organism\_taxid: 7955. Expressed in: spodoptera frugiperda. CLASS: Alpha Beta, ARCH: 3-Layer(aba) Sandwich, TOPOL: Rossmann fold, HOMOL: P-loop containing nucleotide triphosphate hydrolases; cath \| 2ocaA03 \| 94.1 \| 0.00093 \| 1.3e-07 \| 54.7 \| 107 \| (229, 342) \| 471 \| (52, 160) \| 215 \| Atp-dependent DNA helicase uvsw \| CATHCODE: 3.40.50.300 NAME: Atp-dependent DNA helicase uvsw. Chain: a. Synonym: dar protein. Engineered: yes. Mutation: yes SOURCE: Enterobacteria phage t4. Organism\_taxid: 10665. Gene: uvsw, dar. Expressed in: escherichia coli. Expression\_system\_taxid: 562. CLASS: Alpha Beta, ARCH: 3-Layer(aba) Sandwich, TOPOL: Rossmann fold, HOMOL: P-loop containing nucleotide triphosphate hydrolases; cath \| 2fwrA03 \| 94.0 \| 0.00099 \| 1.4e-07 \| 52.9 \| 94 \| (229, 334) \| 471 \| (92, 185) \| 199 \| Dna repair protein rad25 \| CATHCODE: 3.40.50.300 NAME: Dna repair protein rad25. Chain: a, b, c, d. Engineered: yes SOURCE: Archaeoglobus fulgidus. Organism\_taxid: 2234. Expressed in: escherichia coli bl21(de3). Expression\_system\_taxid: 469008. CLASS: Alpha Beta, ARCH: 3-Layer(aba) Sandwich, TOPOL: Rossmann fold, HOMOL: P-loop containing nucleotide triphosphate hydrolases; cath \| 4ernA00 \| 94.0 \| 0.001 \| 1.4e-07 \| 55.1 \| 92 \| (229, 332) \| 471 \| (62, 155) \| 237 \| Tfiih basal transcription factor complex helicase xpb subunit \| CATHCODE: 3.40.50.300 NAME: Tfiih basal transcription factor complex helicase xpb subunit. Chain: a. Fragment: c-terminal domain (unp residues 494-782). Synonym: basic transcription factor 2 89 kda subunit, btf2 p89, DNA excision repair protein ercc-3, DNA repair protein complementing xp-b cells, tfiih basal transcription factor complex 89 kda subunit, tfiih 89 kda subunit, tfiih p89, xeroderma pigmentosum group b- complementing protein. Engineered: yes SOURCE: Homo sapiens. Human. Organism\_taxid: 9606. Gene: ercc3, xpb, xpbc. Expressed in: escherichia coli. Expression\_system\_taxid: 562. CLASS: Alpha Beta, ARCH: 3-Layer(aba) Sandwich, TOPOL: Rossmann fold, HOMOL: P-loop containing nucleotide triphosphate hydrolases; phrogs \| 62 \| 99.9 \| 6.2e-30 \| 8.3e-34 \| 262.7 \| 320 \| (2, 359) \| 471 \| (9, 417) \| 434 \| DNA helicase \| DNA helicase; Category: DNA, RNA and nucleotide metabolism; NC\_028990\_p38; phrogs \| 16 \| 99.8 \| 4.8e-24 \| 6.2e-28 \| 224.2 \| 307 \| (3, 346) \| 471 \| (119, 501) \| 568 \| DNA helicase \| DNA helicase; Category: DNA, RNA and nucleotide metabolism; NC\_027402\_p36; phrogs \| 9690 \| 99.7 \| 1.4e-22 \| 1.6e-26 \| 186.5 \| 213 \| (113, 344) \| 471 \| (30, 261) \| 306 \| DNA helicase \| DNA helicase; Category: DNA, RNA and nucleotide metabolism; MF001355\_p164; phrogs \| 23807 \| 99.6 \| 4.4e-21 \| 5e-25 \| 193.0 \| 333 \| (3, 349) \| 471 \| (121, 643) \| 1044 \| NA \| NA; Category: unknown function; p54525 VI\_01114; phrogs \| 6819 \| 99.6 \| 5.1e-21 \| 5.9e-25 \| 187.7 \| 286 \| (3, 334) \| 471 \| (2, 324) \| 593 \| Holliday junction branch migration helicase \| Holliday junction branch migration helicase; Category: DNA, RNA and nucleotide metabolism; NC\_010155\_p9; phrogs \| 16694 \| 99.5 \| 2.1e-19 \| 2.3e-23 \| 178.7 \| 316 \| (2, 342) \| 471 \| (271, 675) \| 977 \| NA \| NA; Category: unknown function; p110125 VI\_04313; phrogs \| 29996 \| 99.4 \| 5.7e-18 \| 6.4e-22 \| 166.4 \| 324 \| (3, 345) \| 471 \| (711, 1132) \| 1178 \| NA \| NA; Category: unknown function; p435648 VI\_04262; phrogs \| 3830 \| 99.3 \| 1.7e-16 \| 2e-20 \| 168.7 \| 326 \| (1, 341) \| 471 \| (265, 688) \| 1038 \| NA \| NA; Category: unknown function; p169260 VI\_00445; phrogs \| 26785 \| 99.2 \| 2.7e-15 \| 3.1e-19 \| 149.3 \| 108 \| (227, 341) \| 471 \| (705, 841) \| 1087 \| NA \| NA; Category: unknown function; MF172979\_p4; phrogs \| 170 \| 99.1 \| 6.8e-15 \| 8.5e-19 \| 136.1 \| 141 \| (3, 154) \| 471 \| (7, 171) \| 239 \| PhoH-like phosphate starvation-inducible \| PhoH-like phosphate starvation-inducible; Category: other; KT995480\_p45; phrogs \| 24008 \| 99.1 \| 1.8e-14 \| 2e-18 \| 142.2 \| 320 \| (2, 341) \| 471 \| (350, 811) \| 879 \| helicase \| helicase; Category: DNA, RNA and nucleotide metabolism; p22513 VI\_12484; phrogs \| 1685 \| 98.8 \| 3.5e-13 \| 4.1e-17 \| 144.6 \| 115 \| (227, 347) \| 471 \| (1366, 1534) \| 2255 \| NA \| NA; Category: unknown function; FO818745\_p8; phrogs \| 4913 \| 98.8 \| 5.1e-13 \| 6.1e-17 \| 140.8 \| 272 \| (25, 332) \| 471 \| (351, 639) \| 1099 \| replication \| replication; Category: DNA, RNA and nucleotide metabolism; NC\_019722\_p45; phrogs \| 325 \| 98.7 \| 1.1e-12 \| 1.5e-16 \| 130.7 \| 136 \| (1, 153) \| 471 \| (2, 159) \| 405 \| Dda-like helicase \| Dda-like helicase; Category: DNA, RNA and nucleotide metabolism; MF360957\_p200; phrogs \| 34613 \| 98.7 \| 4e-12 \| 4.5e-16 \| 118.7 \| 287 \| (2, 334) \| 471 \| (250, 570) \| 666 \| NA \| NA; Category: unknown function; p298842 VI\_05329; phrogs \| 11801 \| 98.4 \| 4.6e-11 \| 5.3e-15 \| 126.6 \| 94 \| (228, 334) \| 471 \| (1311, 1408) \| 1493 \| DNA methyltransferase \| DNA methyltransferase; Category: other; NC\_031129\_p54; phrogs \| 16259 \| 98.4 \| 8.7e-11 \| 1e-14 \| 106.6 \| 145 \| (2, 155) \| 471 \| (111, 264) \| 326 \| DNA helicase \| DNA helicase; Category: DNA, RNA and nucleotide metabolism; NC\_026607\_p174; phrogs \| 296 \| 98.1 \| 8.7e-10 \| 1.1e-13 \| 105.4 \| 110 \| (25, 152) \| 471 \| (95, 207) \| 312 \| DNA transposition protein \| DNA transposition protein; Category: integration and excision; p199288 VI\_08191; phrogs \| 4412 \| 98.0 \| 1.6e-09 \| 1.9e-13 \| 110.9 \| 139 \| (2, 158) \| 471 \| (338, 490) \| 869 \| exonuclease V \| exonuclease V; Category: DNA, RNA and nucleotide metabolism; p152729 VI\_00586; phrogs \| 2765 \| 98.0 \| 2.5e-09 \| 3e-13 \| 102.7 \| 140 \| (4, 155) \| 471 \| (204, 364) \| 405 \| PhoH-like phosphate starvation-inducible \| PhoH-like phosphate starvation-inducible; Category: other; KT070867\_p233; phrogs \| 3712 \| 97.4 \| 8.2e-08 \| 9.5e-12 \| 96.3 \| 267 \| (24, 331) \| 471 \| (393, 683) \| 928 \| replication origin binding \| replication origin binding; Category: DNA, RNA and nucleotide metabolism; NC\_005859\_p108; phrogs \| 5173 \| 97.3 \| 2.3e-07 \| 2.6e-11 \| 84.1 \| 255 \| (25, 332) \| 471 \| (59, 322) \| 449 \| NA \| NA; Category: unknown function; NC\_029001\_p35; phrogs \| 25471 \| 97.2 \| 2.6e-07 \| 2.9e-11 \| 73.2 \| 122 \| (3, 129) \| 471 \| (1, 139) \| 162 \| NA \| NA; Category: unknown function; KT876724\_p67; phrogs \| 25411 \| 97.2 \| 3.7e-07 \| 4.3e-11 \| 87.2 \| 139 \| (6, 153) \| 471 \| (206, 372) \| 600 \| NA \| NA; Category: unknown function; p367370 VI\_12217; phrogs \| 35989 \| 97.0 \| 9.3e-07 \| 1e-10 \| 83.5 \| 78 \| (255, 341) \| 471 \| (445, 529) \| 860 \| NA \| NA; Category: unknown function; p71570 VI\_01674; phrogs \| 11354 \| 96.9 \| 1.2e-06 \| 1.4e-10 \| 66.1 \| 63 \| (273, 341) \| 471 \| (16, 79) \| 115 \| NA \| NA; Category: unknown function; MG029509\_p47; phrogs \| 10089 \| 96.9 \| 1.4e-06 \| 1.6e-10 \| 95.4 \| 80 \| (255, 341) \| 471 \| (3683, 3765) \| 5483 \| DarB-like antirestriction \| DarB-like antirestriction; Category: moron, auxiliary metabolic gene and host takeover; p120769 VI\_04205; phrogs \| 21391 \| 96.8 \| 1.9e-06 \| 2.2e-10 \| 90.0 \| 99 \| (227, 332) \| 471 \| (1290, 1393) \| 1951 \| DNA helicase \| DNA helicase; Category: DNA, RNA and nucleotide metabolism; KY984068\_p176; phrogs \| 1967 \| 96.2 \| 1.7e-05 \| 2.1e-09 \| 79.0 \| 75 \| (3, 83) \| 471 \| (1, 79) \| 499 \| DNA helicase \| DNA helicase; Category: DNA, RNA and nucleotide metabolism; KT876724\_p41; phrogs \| 33220 \| 96.2 \| 2e-05 \| 2.2e-09 \| 77.2 \| 99 \| (227, 332) \| 471 \| (1041, 1153) \| 1558 \| NA \| NA; Category: unknown function; p329573 VI\_10960; phrogs \| 5846 \| 96.2 \| 1.9e-05 \| 2.4e-09 \| 72.9 \| 98 \| (24, 151) \| 471 \| (37, 136) \| 306 \| clamp loader of DNA polymerase \| clamp loader of DNA polymerase; Category: DNA, RNA and nucleotide metabolism; KY606587\_p99; phrogs \| 27849 \| 96.2 \| 2.2e-05 \| 2.4e-09 \| 73.3 \| 100 \| (227, 334) \| 471 \| (444, 549) \| 664 \| NA \| NA; Category: unknown function; p108771 VI\_06683; phrogs \| 168 \| 95.9 \| 4.5e-05 \| 5.6e-09 \| 71.4 \| 23 \| (25, 47) \| 471 \| (44, 66) \| 318 \| clamp loader of DNA polymerase \| clamp loader of DNA polymerase; Category: DNA, RNA and nucleotide metabolism; p26368 VI\_12322; phrogs \| 17901 \| 95.9 \| 5.3e-05 \| 5.9e-09 \| 55.2 \| 59 \| (278, 344) \| 471 \| (10, 70) \| 109 \| DNA helicase \| DNA helicase; Category: DNA, RNA and nucleotide metabolism; KY942057\_p118; phrogs \| 22930 \| 95.8 \| 6.1e-05 \| 6.8e-09 \| 71.1 \| 306 \| (3, 341) \| 471 \| (115, 466) \| 704 \| NA \| NA; Category: unknown function; NC\_031927\_p55; phrogs \| 10417 \| 95.8 \| 6.5e-05 \| 7.3e-09 \| 71.1 \| 99 \| (227, 332) \| 471 \| (496, 600) \| 728 \| helicase \| helicase; Category: DNA, RNA and nucleotide metabolism; NC\_027997\_p153; phrogs \| 6891 \| 95.6 \| 9.4e-05 \| 1.1e-08 \| 64.1 \| 92 \| (25, 125) \| 471 \| (36, 127) \| 205 \| ATPase \| ATPase; Category: other; p126525 VI\_01034; phrogs \| 10691 \| 95.4 \| 0.00018 \| 2e-08 \| 72.6 \| 115 \| (5, 125) \| 471 \| (328, 453) \| 1276 \| NA \| NA; Category: unknown function; p118307 VI\_06669; phrogs \| 2488 \| 94.2 \| 0.00089 \| 1.1e-07 \| 64.9 \| 59 \| (3, 69) \| 471 \| (2, 63) \| 494 \| DNA helicase \| DNA helicase; Category: DNA, RNA and nucleotide metabolism; NC\_005361\_p10 |
| Top keywords  (threshold 1.00e-03 (evalue)) | **helicase, a, and, in, the, DNA, RNA, i, yes, ATP\_dependent** |
| Output files | ../../domain\_architecture/44\_FANPEZAQ\_CDS\_0044\_cath.hhr ../../domain\_architecture/44\_FANPEZAQ\_CDS\_0044\_merged.svg ../../domain\_architecture/44\_FANPEZAQ\_CDS\_0044\_ncbi-cd.hhr ../../domain\_architecture/44\_FANPEZAQ\_CDS\_0044\_pfam.hhr ../../domain\_architecture/44\_FANPEZAQ\_CDS\_0044\_phrogs.hhr |

### Identical protein sequences/structures

#### Search results

|  |  |
| --- | --- |
| Protein sequence databases searched | Pdb, Swissprot, Refseq |
| Identical proteins found | -- |
| Top keywords | -- |
| Output files | -- |

### Similar protein sequences/structures

#### Sequence similarity search results (HHblits)^1^

|  |  |
| --- | --- |
| Sequence databases searched | Uniclust, Pdb70 |
| Results, scheme(s)  (Top layers only, threshold 1.00e-03 (evalue)) | xml version="1.0" encoding="utf-8" standalone="no"?       2024-09-02T21:08:49.262162 image/svg+xml   Matplotlib v3.7.2, https://matplotlib.org/ |
| Results, table(s)  (threshold 1.00e-03 (evalue)) | db \| id \| prob \| evalue \| pvalue \| score \| cols \| query \| query\_len \| template \| template\_len \| name \| description; uniclust \| UniRef100\_A0A017TJA4 \| 100.0 \| 2e-108 \| 5e-114 \| 792.8 \| 451 \| (1, 469) \| 471 \| (41, 493) \| 567 \| Uncharacterized protein \| Uncharacterized protein; uniclust \| UniRef100\_A0A062XM93 \| 100.0 \| 4e-105 \| 8e-111 \| 745.1 \| 444 \| (1, 470) \| 471 \| (19, 472) \| 501 \| Putative helicase \| Putative helicase; uniclust \| UniRef100\_A0A0F9LT42 \| 100.0 \| 4e-104 \| 8e-110 \| 749.9 \| 452 \| (1, 469) \| 471 \| (46, 501) \| 550 \| Helicase ATP-binding domain-containing protein \| Helicase ATP-binding domain-containing protein; uniclust \| UniRef100\_A0A011PS19 \| 100.0 \| 2.4e-99 \| 5e-105 \| 752.5 \| 447 \| (1, 467) \| 471 \| (113, 561) \| 656 \| Hef nuclease \| Hef nuclease; uniclust \| UniRef100\_A0A158DXS0 \| 100.0 \| 6.9e-99 \| 1e-104 \| 701.4 \| 455 \| (1, 470) \| 471 \| (12, 472) \| 581 \| Type III restriction enzyme, res subunit \| Type III restriction enzyme, res subunit; uniclust \| UniRef100\_A0A0C9NWV0 \| 100.0 \| 5.9e-96 \| 1e-101 \| 667.6 \| 443 \| (1, 469) \| 471 \| (8, 455) \| 489 \| Putative helicase \| Putative helicase; uniclust \| UniRef100\_A0A0F8ZEC1 \| 100.0 \| 8.9e-95 \| 2e-100 \| 675.4 \| 451 \| (2, 467) \| 471 \| (1, 462) \| 547 \| Helicase C-terminal domain-containing protein (Fragment) \| Helicase C-terminal domain-containing protein (Fragment); uniclust \| UniRef100\_A0A011TAH4 \| 100.0 \| 2.7e-92 \| 5.3e-98 \| 682.0 \| 444 \| (1, 467) \| 471 \| (60, 516) \| 603 \| Helicase \| Helicase; uniclust \| UniRef100\_A0A060DLC1 \| 100.0 \| 2.3e-91 \| 4.5e-97 \| 664.3 \| 434 \| (1, 466) \| 471 \| (33, 474) \| 595 \| DEAD/DEAH box helicase \| DEAD/DEAH box helicase; uniclust \| UniRef100\_A0A178MUY0 \| 100.0 \| 8.7e-89 \| 1.7e-94 \| 623.7 \| 449 \| (1, 469) \| 471 \| (2, 453) \| 468 \| Helicase C-terminal domain-containing protein \| Helicase C-terminal domain-containing protein; uniclust \| UniRef100\_A0A1M3E4Z4 \| 100.0 \| 1.1e-87 \| 2.2e-93 \| 635.2 \| 402 \| (1, 419) \| 471 \| (31, 440) \| 533 \| Helicase ATP-binding domain-containing protein \| Helicase ATP-binding domain-containing protein; uniclust \| UniRef100\_A0A1D8A331 \| 100.0 \| 2.4e-84 \| 4.6e-90 \| 600.5 \| 450 \| (4, 466) \| 471 \| (39, 499) \| 547 \| Helicase \| Helicase; uniclust \| UniRef100\_A0A060D1N3 \| 100.0 \| 6.8e-83 \| 1.3e-88 \| 619.2 \| 407 \| (1, 422) \| 471 \| (68, 504) \| 620 \| Putative helicase DEXDc superfamily protein \| Putative helicase DEXDc superfamily protein; uniclust \| UniRef100\_A0A2E2RSB8 \| 100.0 \| 1.1e-82 \| 2.1e-88 \| 592.9 \| 403 \| (1, 420) \| 471 \| (3, 406) \| 664 \| Helicase \| Helicase; uniclust \| UniRef100\_A0A965K7S0 \| 100.0 \| 1.3e-80 \| 2.5e-86 \| 563.1 \| 450 \| (2, 470) \| 471 \| (3, 475) \| 562 \| DEAD/DEAH box helicase \| DEAD/DEAH box helicase; uniclust \| UniRef100\_A0A021X9K7 \| 100.0 \| 1.8e-78 \| 3.4e-84 \| 601.0 \| 387 \| (1, 404) \| 471 \| (110, 514) \| 763 \| DNA or RNA helicase of superfamily II \| DNA or RNA helicase of superfamily II; uniclust \| UniRef100\_A0A2I7QJN8 \| 100.0 \| 2.8e-78 \| 5.2e-84 \| 587.8 \| 426 \| (24, 468) \| 471 \| (386, 817) \| 878 \| Homing endonuclease \| Homing endonuclease; uniclust \| UniRef100\_A0A0R1ZNP6 \| 100.0 \| 2.9e-77 \| 5.5e-83 \| 546.3 \| 376 \| (1, 405) \| 471 \| (1, 385) \| 538 \| Helicase \| Helicase; uniclust \| UniRef100\_A0A1F9YLJ7 \| 100.0 \| 4.6e-77 \| 8.8e-83 \| 565.0 \| 392 \| (1, 405) \| 471 \| (2, 415) \| 560 \| Helicase \| Helicase; uniclust \| UniRef100\_A0A2D8IZ10 \| 100.0 \| 5.9e-77 \| 1.1e-82 \| 569.8 \| 326 \| (1, 337) \| 471 \| (9, 340) \| 856 \| Helicase \| Helicase; uniclust \| UniRef100\_A0A011PZD2 \| 100.0 \| 1.3e-76 \| 2.5e-82 \| 567.5 \| 401 \| (2, 420) \| 471 \| (102, 522) \| 677 \| UvsW helicase \| UvsW helicase; uniclust \| UniRef100\_A0A0C1NLT0 \| 100.0 \| 1.6e-76 \| 3e-82 \| 564.8 \| 389 \| (3, 407) \| 471 \| (53, 452) \| 734 \| Helicase \| Helicase; uniclust \| UniRef100\_A0A258YNK0 \| 100.0 \| 4.3e-76 \| 8.2e-82 \| 538.1 \| 355 \| (1, 366) \| 471 \| (18, 377) \| 514 \| DEAD/DEAH box helicase \| DEAD/DEAH box helicase; uniclust \| UniRef100\_A0A0A0M1Y7 \| 100.0 \| 4.7e-76 \| 8.9e-82 \| 521.5 \| 345 \| (1, 366) \| 471 \| (6, 351) \| 366 \| Helicase ATP-binding domain-containing protein (Fragment) \| Helicase ATP-binding domain-containing protein (Fragment); uniclust \| UniRef100\_A0A257JGZ3 \| 100.0 \| 7.8e-76 \| 1.5e-81 \| 530.0 \| 404 \| (1, 420) \| 471 \| (9, 430) \| 455 \| DNA helicase (Fragment) \| DNA helicase (Fragment); uniclust \| UniRef100\_A0A2D0IPH5 \| 100.0 \| 1.1e-74 \| 2.1e-80 \| 516.0 \| 430 \| (1, 452) \| 471 \| (10, 456) \| 463 \| Helicase \| Helicase; uniclust \| UniRef100\_A0A3M1TKW1 \| 100.0 \| 7.9e-73 \| 1.5e-78 \| 503.1 \| 389 \| (3, 404) \| 471 \| (42, 439) \| 440 \| DEAD/DEAH box helicase (Fragment) \| DEAD/DEAH box helicase (Fragment); uniclust \| UniRef100\_A0A179S5I4 \| 100.0 \| 2.4e-72 \| 4.4e-78 \| 522.9 \| 443 \| (1, 467) \| 471 \| (12, 714) \| 759 \| Helicase \| Helicase; uniclust \| UniRef100\_A0A078LFY8 \| 100.0 \| 2.5e-72 \| 4.6e-78 \| 511.9 \| 442 \| (5, 467) \| 471 \| (13, 473) \| 644 \| Helicase \| Helicase; uniclust \| UniRef100\_A0A1Y2K431 \| 100.0 \| 4e-72 \| 7.7e-78 \| 521.8 \| 384 \| (3, 405) \| 471 \| (1, 411) \| 556 \| Putative superfamily II DNA/RNA helicase \| Putative superfamily II DNA/RNA helicase; uniclust \| UniRef100\_A0A1E2WND8 \| 100.0 \| 1.1e-71 \| 2e-77 \| 515.6 \| 387 \| (2, 402) \| 471 \| (9, 421) \| 682 \| DEAD/DEAH box helicase \| DEAD/DEAH box helicase; uniclust \| UniRef100\_A0A1H4CGZ5 \| 100.0 \| 1.9e-71 \| 3.7e-77 \| 518.1 \| 406 \| (1, 420) \| 471 \| (6, 417) \| 556 \| Superfamily II DNA or RNA helicase \| Superfamily II DNA or RNA helicase; uniclust \| UniRef100\_A0A0C6F7K9 \| 100.0 \| 2.8e-71 \| 5.3e-77 \| 510.4 \| 392 \| (1, 405) \| 471 \| (1, 412) \| 573 \| Helicase \| Helicase; uniclust \| UniRef100\_A0A357XW32 \| 100.0 \| 3.5e-71 \| 6.7e-77 \| 494.4 \| 389 \| (2, 407) \| 471 \| (4, 409) \| 443 \| DNA helicase \| DNA helicase; uniclust \| UniRef100\_A0A0C1N4V4 \| 100.0 \| 3.9e-71 \| 7.4e-77 \| 509.1 \| 382 \| (2, 403) \| 471 \| (33, 438) \| 666 \| DEAD/DEAH box helicase \| DEAD/DEAH box helicase; uniclust \| UniRef100\_A0A968H947 \| 100.0 \| 6e-70 \| 1.1e-75 \| 494.2 \| 449 \| (3, 468) \| 471 \| (304, 761) \| 804 \| Toprim domain-containing protein \| Toprim domain-containing protein; uniclust \| UniRef100\_A0A1Q7G082 \| 100.0 \| 1e-69 \| 2e-75 \| 508.7 \| 350 \| (1, 357) \| 471 \| (20, 380) \| 445 \| Helicase (Fragment) \| Helicase (Fragment); uniclust \| UniRef100\_A0A1F9KU86 \| 100.0 \| 3.2e-69 \| 6.2e-75 \| 515.2 \| 346 \| (1, 355) \| 471 \| (24, 382) \| 600 \| DEAD/DEAH box helicase \| DEAD/DEAH box helicase; uniclust \| UniRef100\_A0A1X1FC89 \| 100.0 \| 3.4e-69 \| 6.4e-75 \| 478.0 \| 332 \| (125, 469) \| 471 \| (3, 347) \| 371 \| ATP-dependent RNA helicase RhlE \| ATP-dependent RNA helicase RhlE; uniclust \| UniRef100\_A0A2H2YQX1 \| 100.0 \| 3.4e-69 \| 6.5e-75 \| 486.3 \| 389 \| (3, 407) \| 471 \| (30, 424) \| 504 \| Putative helicase \| Putative helicase; uniclust \| UniRef100\_A0A5C7M4H4 \| 100.0 \| 5.9e-69 \| 1.1e-74 \| 486.4 \| 346 \| (1, 356) \| 471 \| (5, 362) \| 416 \| DEAD/DEAH box helicase \| DEAD/DEAH box helicase; uniclust \| UniRef100\_A0A059VBC1 \| 100.0 \| 2.3e-68 \| 4.5e-74 \| 527.0 \| 343 \| (1, 351) \| 471 \| (55, 408) \| 640 \| DNA helicase \| DNA helicase; uniclust \| UniRef100\_A0A3M1T4P2 \| 100.0 \| 5.4e-68 \| 1e-73 \| 485.6 \| 384 \| (3, 404) \| 471 \| (15, 424) \| 674 \| DEAD/DEAH box helicase \| DEAD/DEAH box helicase; uniclust \| UniRef100\_A0A218QTQ5 \| 100.0 \| 5.6e-67 \| 1.1e-72 \| 464.5 \| 352 \| (2, 362) \| 471 \| (27, 382) \| 407 \| DEAD/DEAH box helicase-like protein \| DEAD/DEAH box helicase-like protein; uniclust \| UniRef100\_A0A2E5HBX7 \| 100.0 \| 1.2e-66 \| 2.2e-72 \| 468.0 \| 406 \| (3, 422) \| 471 \| (1, 409) \| 712 \| Helicase \| Helicase; uniclust \| UniRef100\_A0A2J9QM80 \| 100.0 \| 1.4e-66 \| 2.6e-72 \| 488.4 \| 395 \| (2, 420) \| 471 \| (1, 438) \| 559 \| Helicase \| Helicase; uniclust \| UniRef100\_A0A0F9GHN0 \| 100.0 \| 2e-66 \| 3.7e-72 \| 466.2 \| 384 \| (3, 404) \| 471 \| (1, 400) \| 475 \| Helicase ATP-binding domain-containing protein (Fragment) \| Helicase ATP-binding domain-containing protein (Fragment); uniclust \| UniRef100\_A0A2I7QZ94 \| 100.0 \| 2.1e-66 \| 3.9e-72 \| 504.1 \| 340 \| (114, 467) \| 471 \| (493, 877) \| 918 \| P-loop containing nucleoside triphosphate hydrolase \| P-loop containing nucleoside triphosphate hydrolase; uniclust \| UniRef100\_A0A6B0YY87 \| 100.0 \| 2.2e-66 \| 4.1e-72 \| 490.5 \| 386 \| (2, 403) \| 471 \| (5, 407) \| 780 \| DEAD/DEAH box helicase \| DEAD/DEAH box helicase; uniclust \| UniRef100\_A0A934JM14 \| 100.0 \| 4.5e-66 \| 8.3e-72 \| 458.0 \| 403 \| (2, 420) \| 471 \| (1, 414) \| 617 \| DEAD/DEAH box helicase family protein \| DEAD/DEAH box helicase family protein; uniclust \| UniRef100\_A0A0N8GP87 \| 100.0 \| 8.5e-66 \| 1.6e-71 \| 488.6 \| 338 \| (2, 350) \| 471 \| (14, 369) \| 622 \| DEAD/DEAH box helicase \| DEAD/DEAH box helicase; uniclust \| UniRef100\_A0A0F9S4V7 \| 100.0 \| 4.2e-65 \| 8.2e-71 \| 497.6 \| 343 \| (2, 352) \| 471 \| (54, 432) \| 648 \| Helicase ATP-binding domain-containing protein \| Helicase ATP-binding domain-containing protein; uniclust \| UniRef100\_A0A2W4XRR0 \| 100.0 \| 5.1e-65 \| 9.5e-71 \| 458.3 \| 352 \| (2, 362) \| 471 \| (1, 360) \| 540 \| Type III restriction endonuclease subunit R \| Type III restriction endonuclease subunit R; uniclust \| UniRef100\_A0A374CI74 \| 100.0 \| 5.3e-65 \| 9.9e-71 \| 468.2 \| 349 \| (3, 360) \| 471 \| (38, 405) \| 575 \| ATP-dependent helicase \| ATP-dependent helicase; uniclust \| UniRef100\_A0A1Q4RMU9 \| 100.0 \| 5.5e-65 \| 1e-70 \| 485.9 \| 401 \| (3, 419) \| 471 \| (20, 434) \| 837 \| Helicase \| Helicase; uniclust \| UniRef100\_A0A023WZC6 \| 100.0 \| 9.3e-65 \| 1.8e-70 \| 459.5 \| 318 \| (1, 330) \| 471 \| (21, 346) \| 376 \| Helicase \| Helicase; uniclust \| UniRef100\_A0A069SFB8 \| 100.0 \| 2.4e-64 \| 4.5e-70 \| 453.8 \| 340 \| (3, 361) \| 471 \| (22, 371) \| 526 \| Type III restriction enzyme, res subunit \| Type III restriction enzyme, res subunit; uniclust \| UniRef100\_A0A2T1EBN6 \| 100.0 \| 2.4e-64 \| 4.5e-70 \| 465.5 \| 391 \| (1, 407) \| 471 \| (17, 420) \| 601 \| Helicase \| Helicase; uniclust \| UniRef100\_A0A1V5YWC8 \| 100.0 \| 2.8e-64 \| 5.3e-70 \| 471.2 \| 351 \| (2, 361) \| 471 \| (13, 377) \| 710 \| ATP-dependent RNA helicase SrmB \| ATP-dependent RNA helicase SrmB; uniclust \| UniRef100\_A0A7W1N8N9 \| 100.0 \| 5.7e-64 \| 1e-69 \| 439.7 \| 399 \| (2, 419) \| 471 \| (1, 422) \| 459 \| DEAD/DEAH box helicase \| DEAD/DEAH box helicase; uniclust \| UniRef100\_A0A2G0ECK2 \| 100.0 \| 9.9e-64 \| 1.8e-69 \| 439.3 \| 398 \| (2, 422) \| 471 \| (20, 428) \| 526 \| ATP-dependent helicase \| ATP-dependent helicase; uniclust \| UniRef100\_A0A2H6CIW6 \| 100.0 \| 1.3e-63 \| 2.4e-69 \| 439.6 \| 380 \| (2, 406) \| 471 \| (3, 387) \| 504 \| Helicase \| Helicase; uniclust \| UniRef100\_A0A060AZZ9 \| 100.0 \| 1.8e-63 \| 3.4e-69 \| 472.9 \| 390 \| (1, 403) \| 471 \| (63, 518) \| 676 \| Carboxylate--amine ligase \| Carboxylate--amine ligase; uniclust \| UniRef100\_A0A081GLT7 \| 100.0 \| 2.2e-63 \| 4.2e-69 \| 437.2 \| 375 \| (1, 389) \| 471 \| (1, 381) \| 405 \| Helicase \| Helicase; uniclust \| UniRef100\_A0A096DF57 \| 100.0 \| 2.3e-63 \| 4.4e-69 \| 471.6 \| 338 \| (1, 351) \| 471 \| (20, 366) \| 598 \| DEAD/DEAH box helicase \| DEAD/DEAH box helicase; uniclust \| UniRef100\_A0A1W1ZVY3 \| 100.0 \| 4.9e-63 \| 9.3e-69 \| 450.7 \| 443 \| (1, 467) \| 471 \| (15, 488) \| 501 \| Superfamily II DNA or RNA helicase \| Superfamily II DNA or RNA helicase; uniclust \| UniRef100\_UPI0019D07770 \| 100.0 \| 9.5e-63 \| 1.7e-68 \| 463.8 \| 388 \| (2, 401) \| 471 \| (1, 427) \| 1277 \| VapE family protein \| VapE family protein; uniclust \| UniRef100\_A0A1F8QJA7 \| 100.0 \| 9.4e-63 \| 1.8e-68 \| 436.4 \| 404 \| (3, 466) \| 471 \| (1, 440) \| 451 \| Helicase C-terminal domain-containing protein \| Helicase C-terminal domain-containing protein; uniclust \| UniRef100\_A0A255SWQ4 \| 100.0 \| 2.3e-62 \| 4.4e-68 \| 478.0 \| 344 \| (3, 361) \| 471 \| (96, 458) \| 905 \| Helicase \| Helicase; uniclust \| UniRef100\_A0A069S9L0 \| 100.0 \| 8e-62 \| 1.5e-67 \| 468.7 \| 339 \| (3, 360) \| 471 \| (56, 404) \| 818 \| Helicase conserved C-terminal domain protein (Fragment) \| Helicase conserved C-terminal domain protein (Fragment); uniclust \| UniRef100\_A0A014PW86 \| 100.0 \| 1.4e-61 \| 2.7e-67 \| 452.0 \| 339 \| (1, 351) \| 471 \| (52, 417) \| 658 \| Carboxylate--amine ligase \| Carboxylate--amine ligase; uniclust \| UniRef100\_A0A099P4G6 \| 100.0 \| 1.7e-61 \| 3.3e-67 \| 486.3 \| 337 \| (2, 345) \| 471 \| (126, 483) \| 847 \| ATP-dependent helicase IRC3 \| ATP-dependent helicase IRC3; uniclust \| UniRef100\_A0A345BQ98 \| 100.0 \| 3e-61 \| 5.6e-67 \| 447.8 \| 389 \| (1, 406) \| 471 \| (14, 424) \| 544 \| DNA or RNA helicase of superfamily II \| DNA or RNA helicase of superfamily II; uniclust \| UniRef100\_A0A1I3PAS7 \| 100.0 \| 4.4e-61 \| 8.4e-67 \| 433.2 \| 346 \| (1, 354) \| 471 \| (29, 392) \| 400 \| Superfamily II DNA or RNA helicase \| Superfamily II DNA or RNA helicase; uniclust \| UniRef100\_A0A0A2U5K4 \| 100.0 \| 6.3e-61 \| 1.2e-66 \| 442.0 \| 341 \| (1, 350) \| 471 \| (9, 360) \| 596 \| DEAD/DEAH box helicase \| DEAD/DEAH box helicase; uniclust \| UniRef100\_A0A015LNM1 \| 100.0 \| 6.1e-61 \| 1.2e-66 \| 481.9 \| 339 \| (2, 347) \| 471 \| (143, 511) \| 862 \| Irc3p \| Irc3p; uniclust \| UniRef100\_A0A258JF07 \| 100.0 \| 9.1e-61 \| 1.7e-66 \| 430.8 \| 366 \| (3, 387) \| 471 \| (19, 397) \| 413 \| DEAD/DEAH box helicase (Fragment) \| DEAD/DEAH box helicase (Fragment); uniclust \| UniRef100\_A0A068NS99 \| 100.0 \| 1.3e-60 \| 2.5e-66 \| 458.4 \| 344 \| (2, 352) \| 471 \| (68, 441) \| 665 \| DEAD box family helicase \| DEAD box family helicase; uniclust \| UniRef100\_A0A124IPL3 \| 100.0 \| 1.6e-60 \| 3.1e-66 \| 446.7 \| 342 \| (1, 359) \| 471 \| (12, 365) \| 558 \| DEAD/DEAH box helicase \| DEAD/DEAH box helicase; uniclust \| UniRef100\_A0A088C524 \| 100.0 \| 1.7e-60 \| 3.2e-66 \| 459.6 \| 369 \| (1, 384) \| 471 \| (29, 425) \| 660 \| Helicase \| Helicase; uniclust \| UniRef100\_A0A0F9DTA3 \| 100.0 \| 4.2e-60 \| 7.9e-66 \| 419.2 \| 388 \| (3, 405) \| 471 \| (1, 405) \| 435 \| Uncharacterized protein (Fragment) \| Uncharacterized protein (Fragment); uniclust \| UniRef100\_A0A068S0Z1 \| 100.0 \| 4.9e-60 \| 9.5e-66 \| 467.4 \| 341 \| (3, 350) \| 471 \| (126, 488) \| 850 \| p-loop containing nucleoside triphosphatehydrolase protein \| p-loop containing nucleoside triphosphatehydrolase protein; uniclust \| UniRef100\_A0A7C9U239 \| 100.0 \| 5.7e-60 \| 1e-65 \| 425.2 \| 384 \| (2, 402) \| 471 \| (13, 407) \| 707 \| DEAD/DEAH box helicase \| DEAD/DEAH box helicase; uniclust \| UniRef100\_A0A3B1AZQ2 \| 100.0 \| 6.6e-60 \| 1.2e-65 \| 438.3 \| 335 \| (1, 344) \| 471 \| (15, 362) \| 598 \| DNA helicase, phage-associated Type III restriction enzme \| DNA helicase, phage-associated Type III restriction enzme; uniclust \| UniRef100\_A0A1Y5RRB6 \| 100.0 \| 8.2e-60 \| 1.5e-65 \| 456.4 \| 371 \| (32, 420) \| 471 \| (374, 763) \| 902 \| ATP-dependent RNA helicase DbpA \| ATP-dependent RNA helicase DbpA; uniclust \| UniRef100\_A0A016FQE1 \| 100.0 \| 1.5e-59 \| 2.8e-65 \| 432.5 \| 338 \| (1, 355) \| 471 \| (22, 384) \| 499 \| Helicase conserved C-terminal domain protein \| Helicase conserved C-terminal domain protein; uniclust \| UniRef100\_A0A077ND82 \| 100.0 \| 3.2e-59 \| 6e-65 \| 412.1 \| 372 \| (75, 467) \| 471 \| (7, 394) \| 472 \| Helicase C-terminal domain-containing protein \| Helicase C-terminal domain-containing protein; uniclust \| UniRef100\_A0A6M3II10 \| 100.0 \| 5.7e-59 \| 1.1e-64 \| 416.8 \| 401 \| (3, 422) \| 471 \| (7, 431) \| 531 \| Putative type III restriction enzyme \| Putative type III restriction enzyme; uniclust \| UniRef100\_A0A965PM57 \| 100.0 \| 5.8e-59 \| 1.1e-64 \| 408.4 \| 398 \| (42, 468) \| 471 \| (4, 411) \| 439 \| Helicase ATP-binding domain-containing protein \| Helicase ATP-binding domain-containing protein; uniclust \| UniRef100\_A0A4P7F6U5 \| 100.0 \| 8.7e-59 \| 1.6e-64 \| 417.4 \| 446 \| (3, 468) \| 471 \| (213, 664) \| 703 \| Helicase ATP-binding domain-containing protein \| Helicase ATP-binding domain-containing protein; uniclust \| UniRef100\_A0A327L2Q1 \| 100.0 \| 1.1e-58 \| 2e-64 \| 419.4 \| 422 \| (3, 448) \| 471 \| (264, 692) \| 754 \| Helicase ATP-binding domain-containing protein \| Helicase ATP-binding domain-containing protein; uniclust \| UniRef100\_A0A7V2IN70 \| 100.0 \| 3e-58 \| 5.5e-64 \| 410.8 \| 387 \| (2, 405) \| 471 \| (15, 419) \| 577 \| DEAD/DEAH box helicase \| DEAD/DEAH box helicase; uniclust \| UniRef100\_UPI000BAF482B \| 100.0 \| 4e-58 \| 7.5e-64 \| 384.1 \| 284 \| (94, 388) \| 471 \| (1, 285) \| 289 \| helicase-related protein \| helicase-related protein; uniclust \| UniRef100\_A0A432VA30 \| 100.0 \| 5e-58 \| 9.5e-64 \| 448.9 \| 358 \| (32, 406) \| 471 \| (388, 762) \| 908 \| DEAD/DEAH box helicase \| DEAD/DEAH box helicase; uniclust \| UniRef100\_A0A0P0M3T6 \| 100.0 \| 6e-58 \| 1.1e-63 \| 424.3 \| 340 \| (2, 359) \| 471 \| (31, 380) \| 684 \| Putative helicase \| Putative helicase; uniclust \| UniRef100\_A0A0C2X884 \| 100.0 \| 6.8e-58 \| 1.3e-63 \| 437.7 \| 339 \| (2, 347) \| 471 \| (62, 424) \| 681 \| P-loop containing nucleoside triphosphate hydrolase protein \| P-loop containing nucleoside triphosphate hydrolase protein; uniclust \| UniRef100\_A0A2T1LW14 \| 100.0 \| 7.6e-58 \| 1.4e-63 \| 420.2 \| 353 \| (2, 363) \| 471 \| (107, 463) \| 745 \| DEAD/DEAH box helicase \| DEAD/DEAH box helicase; uniclust \| UniRef100\_A0A090S396 \| 100.0 \| 7.7e-58 \| 1.5e-63 \| 433.6 \| 342 \| (1, 354) \| 471 \| (36, 404) \| 634 \| ATP-dependent RNA helicase YejH \| ATP-dependent RNA helicase YejH; uniclust \| UniRef100\_A0A6M2AY28 \| 100.0 \| 1.1e-57 \| 2e-63 \| 404.5 \| 353 \| (1, 362) \| 471 \| (1, 358) \| 515 \| DEAD/DEAH box helicase family protein \| DEAD/DEAH box helicase family protein; uniclust \| UniRef100\_A0A1U7P314 \| 100.0 \| 1.2e-57 \| 2.2e-63 \| 421.0 \| 451 \| (2, 466) \| 471 \| (134, 652) \| 749 \| DNA helicase, phage-associated \| DNA helicase, phage-associated; uniclust \| UniRef100\_A0A162TW98 \| 100.0 \| 1.2e-57 \| 2.3e-63 \| 417.6 \| 336 \| (3, 345) \| 471 \| (80, 436) \| 501 \| P-loop containing nucleoside triphosphate hydrolase protein (Fragment) \| P-loop containing nucleoside triphosphate hydrolase protein (Fragment); uniclust \| UniRef100\_A0A1H7KEX0 \| 100.0 \| 2.4e-57 \| 4.4e-63 \| 412.3 \| 349 \| (3, 360) \| 471 \| (12, 377) \| 565 \| Helicase conserved C-terminal domain-containing protein \| Helicase conserved C-terminal domain-containing protein; uniclust \| UniRef100\_A0A1G3ADG1 \| 100.0 \| 3.1e-57 \| 5.9e-63 \| 418.5 \| 433 \| (3, 451) \| 471 \| (12, 477) \| 529 \| Helicase ATP-binding domain-containing protein \| Helicase ATP-binding domain-containing protein; uniclust \| UniRef100\_A0A517T222 \| 100.0 \| 6.3e-57 \| 1.2e-62 \| 432.5 \| 279 \| (112, 406) \| 471 \| (441, 741) \| 884 \| Type I restriction enzyme EcoKI subunit R \| Type I restriction enzyme EcoKI subunit R; uniclust \| UniRef100\_A0A5C7PI05 \| 100.0 \| 6.9e-57 \| 1.3e-62 \| 395.1 \| 413 \| (3, 455) \| 471 \| (1, 414) \| 541 \| DEAD/DEAH box helicase \| DEAD/DEAH box helicase; uniclust \| UniRef100\_A0A858NNU1 \| 100.0 \| 9.1e-57 \| 1.7e-62 \| 408.7 \| 349 \| (114, 466) \| 471 \| (417, 769) \| 788 \| Putative DNA helicase \| Putative DNA helicase; uniclust \| UniRef100\_A0A010S8Y7 \| 100.0 \| 9.5e-57 \| 1.8e-62 \| 434.1 \| 338 \| (2, 346) \| 471 \| (86, 442) \| 789 \| DEAD/DEAH box helicase \| DEAD/DEAH box helicase; uniclust \| UniRef100\_UPI00201384D1 \| 100.0 \| 1.3e-56 \| 2.4e-62 \| 407.9 \| 367 \| (3, 387) \| 471 \| (12, 383) \| 793 \| DEAD/DEAH box helicase \| DEAD/DEAH box helicase; uniclust \| UniRef100\_A0A951QK77 \| 100.0 \| 1.6e-56 \| 3e-62 \| 423.0 \| 312 \| (2, 325) \| 471 \| (73, 387) \| 998 \| DEAD/DEAH box helicase family protein \| DEAD/DEAH box helicase family protein; uniclust \| UniRef100\_A0A011V5Z9 \| 100.0 \| 2e-56 \| 3.7e-62 \| 428.5 \| 325 \| (1, 352) \| 471 \| (118, 446) \| 748 \| DEAD/DEAH box helicase \| DEAD/DEAH box helicase; uniclust \| UniRef100\_A0A1V1UMG2 \| 100.0 \| 2.5e-56 \| 4.7e-62 \| 404.4 \| 373 \| (1, 405) \| 471 \| (17, 394) \| 501 \| Type III restriction enzyme, res subunit \| Type III restriction enzyme, res subunit; uniclust \| UniRef100\_A0A060T456 \| 100.0 \| 3.6e-56 \| 6.8e-62 \| 419.7 \| 338 \| (3, 347) \| 471 \| (35, 391) \| 699 \| ARAD1C38984p \| ARAD1C38984p; uniclust \| UniRef100\_T2JKH3 \| 100.0 \| 8.1e-56 \| 1.5e-61 \| 384.4 \| 353 \| (2, 363) \| 471 \| (107, 463) \| 491 \| D-alanine-D-alanine ligase \| D-alanine-D-alanine ligase; uniclust \| UniRef100\_A0A1H3CT34 \| 100.0 \| 9.5e-56 \| 1.8e-61 \| 409.5 \| 315 \| (76, 406) \| 471 \| (15, 342) \| 519 \| DNA repair protein RadD \| DNA repair protein RadD; uniclust \| UniRef100\_A0A2W4YBE5 \| 100.0 \| 9.8e-56 \| 1.8e-61 \| 395.1 \| 354 \| (2, 364) \| 471 \| (1, 360) \| 651 \| Restriction endonuclease subunit R \| Restriction endonuclease subunit R; uniclust \| UniRef100\_A0A023BZQ2 \| 100.0 \| 9.9e-56 \| 1.9e-61 \| 415.9 \| 345 \| (3, 360) \| 471 \| (65, 423) \| 583 \| DEAD/DEAH box helicase \| DEAD/DEAH box helicase; uniclust \| UniRef100\_A0A3M1Q2S6 \| 100.0 \| 1.4e-55 \| 2.5e-61 \| 402.8 \| 388 \| (2, 406) \| 471 \| (265, 671) \| 827 \| DEAD/DEAH box helicase \| DEAD/DEAH box helicase; uniclust \| UniRef100\_Q6VSX2 \| 100.0 \| 1.9e-55 \| 3.5e-61 \| 396.9 \| 404 \| (2, 420) \| 471 \| (1, 437) \| 718 \| Putative helicase \| Putative helicase; uniclust \| UniRef100\_A0A1B9GD36 \| 100.0 \| 2.3e-55 \| 4.3e-61 \| 416.2 \| 338 \| (3, 348) \| 471 \| (51, 404) \| 673 \| Helicase ATP-binding domain-containing protein \| Helicase ATP-binding domain-containing protein; uniclust \| UniRef100\_A0A191SAQ9 \| 100.0 \| 3.1e-55 \| 5.9e-61 \| 395.4 \| 378 \| (1, 402) \| 471 \| (1, 385) \| 473 \| Helicase \| Helicase; uniclust \| UniRef100\_UPI0015E48027 \| 100.0 \| 4.1e-55 \| 7.7e-61 \| 392.7 \| 369 \| (1, 405) \| 471 \| (8, 386) \| 469 \| DEAD/DEAH box helicase \| DEAD/DEAH box helicase; uniclust \| UniRef100\_A0A218QTQ8 \| 100.0 \| 4.2e-55 \| 7.8e-61 \| 394.4 \| 405 \| (2, 420) \| 471 \| (1, 421) \| 712 \| Putative helicase \| Putative helicase; uniclust \| UniRef100\_L7U5Z6 \| 100.0 \| 4.7e-55 \| 8.6e-61 \| 384.0 \| 379 \| (3, 403) \| 471 \| (4, 394) \| 545 \| DEAD/DEAH box helicase \| DEAD/DEAH box helicase; uniclust \| UniRef100\_A0A356U8J2 \| 100.0 \| 4.9e-55 \| 9.3e-61 \| 419.2 \| 335 \| (3, 350) \| 471 \| (157, 498) \| 775 \| DEAD/DEAH box helicase \| DEAD/DEAH box helicase; uniclust \| UniRef100\_F3MW75 \| 100.0 \| 6.5e-55 \| 1.2e-60 \| 380.3 \| 343 \| (2, 366) \| 471 \| (126, 470) \| 509 \| Phage helicase \| Phage helicase; uniclust \| UniRef100\_A0A6L7TM14 \| 100.0 \| 7.8e-55 \| 1.4e-60 \| 381.9 \| 385 \| (2, 402) \| 471 \| (1, 402) \| 536 \| DEAD/DEAH box helicase \| DEAD/DEAH box helicase; uniclust \| UniRef100\_A0A9C9DIN2 \| 100.0 \| 8.4e-55 \| 1.5e-60 \| 396.3 \| 316 \| (139, 467) \| 471 \| (433, 754) \| 793 \| Uncharacterized protein \| Uncharacterized protein; uniclust \| UniRef100\_A0A3D4X7E0 \| 100.0 \| 8.5e-55 \| 1.6e-60 \| 379.8 \| 333 \| (2, 347) \| 471 \| (4, 345) \| 392 \| DEAD/DEAH box helicase (Fragment) \| DEAD/DEAH box helicase (Fragment); uniclust \| UniRef100\_A0A2C0F336 \| 100.0 \| 1.1e-54 \| 2e-60 \| 373.0 \| 240 \| (215, 469) \| 471 \| (8, 248) \| 267 \| Helicase (Fragment) \| Helicase (Fragment); uniclust \| UniRef100\_A0A646HI40 \| 100.0 \| 1.4e-54 \| 2.6e-60 \| 381.5 \| 330 \| (2, 344) \| 471 \| (21, 361) \| 364 \| DEAD/DEAH box helicase (Fragment) \| DEAD/DEAH box helicase (Fragment); uniclust \| UniRef100\_A0A353A170 \| 100.0 \| 2.5e-54 \| 4.7e-60 \| 387.6 \| 347 \| (2, 359) \| 471 \| (3, 381) \| 527 \| DEAD/DEAH box helicase \| DEAD/DEAH box helicase; uniclust \| UniRef100\_UPI00146F4154 \| 100.0 \| 3.7e-54 \| 6.8e-60 \| 390.8 \| 352 \| (2, 362) \| 471 \| (10, 369) \| 762 \| DEAD/DEAH box helicase family protein \| DEAD/DEAH box helicase family protein; uniclust \| UniRef100\_A0A0U2VII8 \| 100.0 \| 6.4e-54 \| 1.2e-59 \| 377.9 \| 388 \| (2, 404) \| 471 \| (1, 409) \| 445 \| ATP-dependent helicase IRC3 \| ATP-dependent helicase IRC3; uniclust \| UniRef100\_A0A3N4KIA0 \| 100.0 \| 8.5e-54 \| 1.6e-59 \| 410.2 \| 339 \| (2, 347) \| 471 \| (145, 507) \| 869 \| P-loop containing nucleoside triphosphate hydrolase protein \| P-loop containing nucleoside triphosphate hydrolase protein; uniclust \| UniRef100\_A0A021W1L9 \| 100.0 \| 1e-53 \| 2e-59 \| 415.7 \| 337 \| (3, 349) \| 471 \| (171, 522) \| 630 \| Restriction endonuclease subunit R \| Restriction endonuclease subunit R; uniclust \| UniRef100\_A0A1B8P5S8 \| 100.0 \| 1.1e-53 \| 2e-59 \| 390.0 \| 363 \| (2, 387) \| 471 \| (1, 390) \| 584 \| Type I restriction enzyme EcoKI subunit R \| Type I restriction enzyme EcoKI subunit R; uniclust \| UniRef100\_A0A1W2H7H3 \| 100.0 \| 1.2e-53 \| 2.2e-59 \| 391.8 \| 347 \| (3, 359) \| 471 \| (149, 505) \| 742 \| Helicase conserved C-terminal domain-containing protein \| Helicase conserved C-terminal domain-containing protein; uniclust \| UniRef100\_A0A497BHL8 \| 100.0 \| 1.3e-53 \| 2.5e-59 \| 383.8 \| 386 \| (1, 406) \| 471 \| (34, 437) \| 557 \| DNA helicase \| DNA helicase; uniclust \| UniRef100\_UPI0011841A0E \| 100.0 \| 1.4e-53 \| 2.5e-59 \| 392.7 \| 348 \| (3, 359) \| 471 \| (264, 624) \| 829 \| DEAD/DEAH box helicase \| DEAD/DEAH box helicase; uniclust \| UniRef100\_A0A4R2VIA0 \| 100.0 \| 1.6e-53 \| 2.9e-59 \| 384.6 \| 384 \| (1, 404) \| 471 \| (3, 400) \| 538 \| DNA repair protein RadD \| DNA repair protein RadD; uniclust \| UniRef100\_A0A061BDT3 \| 100.0 \| 1.7e-53 \| 3.3e-59 \| 426.2 \| 350 \| (3, 360) \| 471 \| (102, 551) \| 938 \| RHTO0S14e05072g1\_1 \| RHTO0S14e05072g1\_1; uniclust \| UniRef100\_A0A1C5NVQ0 \| 100.0 \| 2e-53 \| 3.7e-59 \| 401.3 \| 359 \| (31, 406) \| 471 \| (383, 756) \| 820 \| Predicted HKD family nuclease \| Predicted HKD family nuclease; uniclust \| UniRef100\_UPI000C287AAC \| 100.0 \| 2.1e-53 \| 3.8e-59 \| 383.9 \| 354 \| (2, 363) \| 471 \| (1, 360) \| 717 \| DEAD/DEAH box helicase family protein \| DEAD/DEAH box helicase family protein; uniclust \| UniRef100\_A0A2N1TNP0 \| 100.0 \| 2.7e-53 \| 5.1e-59 \| 375.6 \| 407 \| (2, 422) \| 471 \| (13, 434) \| 502 \| Helicase \| Helicase; uniclust \| UniRef100\_A0A936VSQ7 \| 100.0 \| 3.7e-53 \| 6.8e-59 \| 379.8 \| 388 \| (2, 405) \| 471 \| (1, 391) \| 667 \| DEAD/DEAH box helicase \| DEAD/DEAH box helicase; uniclust \| UniRef100\_UPI0005708FED \| 100.0 \| 4.7e-53 \| 8.8e-59 \| 375.3 \| 339 \| (109, 467) \| 471 \| (22, 376) \| 417 \| helicase-related protein \| helicase-related protein; uniclust \| UniRef100\_UPI000C1E5455 \| 100.0 \| 5.7e-53 \| 1e-58 \| 379.3 \| 372 \| (3, 401) \| 471 \| (38, 427) \| 679 \| DEAD/DEAH box helicase \| DEAD/DEAH box helicase; uniclust \| UniRef100\_A0A0Q9ZBC2 \| 100.0 \| 6.7e-53 \| 1.3e-58 \| 387.1 \| 346 \| (3, 361) \| 471 \| (37, 396) \| 542 \| DEAD/DEAH box helicase \| DEAD/DEAH box helicase; uniclust \| UniRef100\_A0A1C5X7B1 \| 100.0 \| 7.5e-53 \| 1.4e-58 \| 402.7 \| 338 \| (3, 352) \| 471 \| (47, 395) \| 675 \| Probable DEAD-box ATP-dependent RNA helicase SA1885 \| Probable DEAD-box ATP-dependent RNA helicase SA1885; uniclust \| UniRef100\_A0A023DJ00 \| 100.0 \| 9.3e-53 \| 1.8e-58 \| 424.1 \| 330 \| (3, 347) \| 471 \| (283, 622) \| 1048 \| DNA helicase \| DNA helicase; uniclust \| UniRef100\_A0A0C9LV95 \| 100.0 \| 9.7e-53 \| 1.8e-58 \| 380.8 \| 330 \| (2, 347) \| 471 \| (52, 402) \| 503 \| P-loop containing nucleoside triphosphate hydrolase protein \| P-loop containing nucleoside triphosphate hydrolase protein; uniclust \| UniRef100\_A0A084AJT0 \| 100.0 \| 1.4e-52 \| 2.7e-58 \| 392.3 \| 338 \| (2, 346) \| 471 \| (106, 462) \| 745 \| Helicase ATP-binding domain-containing protein \| Helicase ATP-binding domain-containing protein; uniclust \| UniRef100\_A0A0B8Q733 \| 100.0 \| 1.5e-52 \| 2.9e-58 \| 423.0 \| 340 \| (2, 361) \| 471 \| (269, 622) \| 1060 \| Helicase \| Helicase; uniclust \| UniRef100\_UPI0021062436 \| 100.0 \| 1.9e-52 \| 3.5e-58 \| 369.7 \| 351 \| (2, 366) \| 471 \| (69, 426) \| 573 \| DEAD/DEAH box helicase \| DEAD/DEAH box helicase; uniclust \| UniRef100\_A0A0J6WV84 \| 100.0 \| 2.9e-52 \| 5.6e-58 \| 410.7 \| 307 \| (33, 352) \| 471 \| (400, 714) \| 910 \| Helicase \| Helicase; uniclust \| UniRef100\_A0A1K1P3W3 \| 100.0 \| 7.5e-52 \| 1.4e-57 \| 396.8 \| 349 \| (3, 360) \| 471 \| (14, 379) \| 1249 \| Superfamily II DNA or RNA helicase \| Superfamily II DNA or RNA helicase; uniclust \| UniRef100\_A0A482MSI2 \| 100.0 \| 1e-51 \| 1.9e-57 \| 368.2 \| 410 \| (4, 422) \| 471 \| (5, 490) \| 622 \| Putative helicase \| Putative helicase; uniclust \| UniRef100\_A0A3G6VAB1 \| 100.0 \| 1.8e-51 \| 3.3e-57 \| 341.4 \| 287 \| (164, 471) \| 471 \| (2, 289) \| 299 \| Helicase C-terminal domain-containing protein \| Helicase C-terminal domain-containing protein; uniclust \| UniRef100\_A0A059ZRC3 \| 100.0 \| 1.9e-51 \| 3.6e-57 \| 396.7 \| 344 \| (1, 356) \| 471 \| (19, 381) \| 782 \| ATP-dependent RNA helicase YejH \| ATP-dependent RNA helicase YejH; uniclust \| UniRef100\_A0A2W5D8E7 \| 100.0 \| 2.1e-51 \| 3.9e-57 \| 403.4 \| 309 \| (2, 321) \| 471 \| (88, 404) \| 980 \| Helicase \| Helicase; uniclust \| UniRef100\_UPI000BE17027 \| 100.0 \| 2.3e-51 \| 4.2e-57 \| 356.6 \| 346 \| (2, 361) \| 471 \| (1, 357) \| 485 \| DEAD/DEAH box helicase \| DEAD/DEAH box helicase; uniclust \| UniRef100\_A0A4Q6F0N9 \| 100.0 \| 4.7e-51 \| 8.9e-57 \| 352.3 \| 308 \| (1, 320) \| 471 \| (20, 332) \| 333 \| DEAD/DEAH box helicase (Fragment) \| DEAD/DEAH box helicase (Fragment); uniclust \| UniRef100\_A0A1Y4D436 \| 100.0 \| 6.6e-51 \| 1.2e-56 \| 371.9 \| 393 \| (2, 403) \| 471 \| (86, 510) \| 717 \| Helicase \| Helicase; uniclust \| UniRef100\_A0A151CDR3 \| 100.0 \| 6.9e-51 \| 1.3e-56 \| 376.5 \| 337 \| (2, 355) \| 471 \| (104, 443) \| 732 \| Helicase \| Helicase; uniclust \| UniRef100\_UPI001F42B10E \| 100.0 \| 7.3e-51 \| 1.3e-56 \| 368.8 \| 436 \| (3, 455) \| 471 \| (223, 671) \| 740 \| DEAD/DEAH box helicase family protein \| DEAD/DEAH box helicase family protein; uniclust \| UniRef100\_A0A3S0C816 \| 100.0 \| 1.3e-50 \| 2.4e-56 \| 356.5 \| 405 \| (2, 420) \| 471 \| (20, 434) \| 547 \| Helicase ATP-binding domain-containing protein \| Helicase ATP-binding domain-containing protein; uniclust \| UniRef100\_A0A662AKH4 \| 100.0 \| 2.1e-50 \| 3.9e-56 \| 335.7 \| 284 \| (50, 342) \| 471 \| (6, 293) \| 293 \| ATP-dependent helicase (Fragment) \| ATP-dependent helicase (Fragment); uniclust \| UniRef100\_A0A3B8W691 \| 100.0 \| 2.2e-50 \| 4.1e-56 \| 362.8 \| 348 \| (3, 359) \| 471 \| (146, 510) \| 678 \| DEAD/DEAH box helicase \| DEAD/DEAH box helicase; uniclust \| UniRef100\_A0A0C3BHF4 \| 100.0 \| 2.4e-50 \| 4.5e-56 \| 353.3 \| 335 \| (3, 344) \| 471 \| (38, 399) \| 435 \| P-loop containing nucleoside triphosphate hydrolase protein \| P-loop containing nucleoside triphosphate hydrolase protein; uniclust \| UniRef100\_A0A086WDS7 \| 100.0 \| 3.2e-50 \| 6.2e-56 \| 409.0 \| 336 \| (3, 347) \| 471 \| (428, 778) \| 1146 \| ATP-dependent helicase (Fragment) \| ATP-dependent helicase (Fragment); uniclust \| UniRef100\_A0A069RED1 \| 100.0 \| 3.7e-50 \| 7.1e-56 \| 406.8 \| 334 \| (3, 351) \| 471 \| (406, 754) \| 1213 \| 51.5 kDa protein \| 51.5 kDa protein; uniclust \| UniRef100\_A0A853IXQ9 \| 100.0 \| 4.2e-50 \| 7.8e-56 \| 362.0 \| 420 \| (3, 450) \| 471 \| (16, 451) \| 698 \| DEAD/DEAH box helicase \| DEAD/DEAH box helicase; uniclust \| UniRef100\_A0A0B2Y537 \| 100.0 \| 4.3e-50 \| 8.5e-56 \| 396.9 \| 342 \| (3, 351) \| 471 \| (214, 600) \| 621 \| Restriction endonuclease subunit R \| Restriction endonuclease subunit R; uniclust \| UniRef100\_A0A086WDP1 \| 100.0 \| 5.5e-50 \| 1e-55 \| 404.5 \| 336 \| (3, 347) \| 471 \| (274, 625) \| 1048 \| ATP-dependent helicase (Fragment) \| ATP-dependent helicase (Fragment); uniclust \| UniRef100\_A0A256SP93 \| 100.0 \| 5.5e-50 \| 1e-55 \| 393.3 \| 333 \| (3, 350) \| 471 \| (168, 512) \| 920 \| DEAD/DEAH box helicase \| DEAD/DEAH box helicase; uniclust \| UniRef100\_A0A017RRU4 \| 100.0 \| 6.9e-50 \| 1.3e-55 \| 365.4 \| 337 \| (3, 352) \| 471 \| (153, 496) \| 602 \| DEAD/DEAH box helicase \| DEAD/DEAH box helicase; uniclust \| UniRef100\_A0A6J5PQM8 \| 100.0 \| 7.7e-50 \| 1.4e-55 \| 346.1 \| 397 \| (3, 420) \| 471 \| (1, 401) \| 470 \| SSL2 DNA or RNA helicases of superfamily II \| SSL2 DNA or RNA helicases of superfamily II; uniclust \| UniRef100\_A0A017TGD1 \| 100.0 \| 8.6e-50 \| 1.7e-55 \| 416.7 \| 336 \| (3, 347) \| 471 \| (430, 782) \| 1219 \| Helicase/Type III restriction enzyme \| Helicase/Type III restriction enzyme; uniclust \| UniRef100\_A0A0F9KYB5 \| 100.0 \| 1.3e-49 \| 2.4e-55 \| 339.9 \| 323 \| (3, 333) \| 471 \| (1, 339) \| 342 \| Helicase ATP-binding domain-containing protein \| Helicase ATP-binding domain-containing protein; uniclust \| UniRef100\_A0A661MM81 \| 100.0 \| 1.6e-49 \| 2.9e-55 \| 347.8 \| 343 \| (2, 352) \| 471 \| (75, 432) \| 517 \| DEAD/DEAH box helicase (Fragment) \| DEAD/DEAH box helicase (Fragment); uniclust \| UniRef100\_A0A8F2W1H3 \| 100.0 \| 1.7e-49 \| 3.2e-55 \| 397.9 \| 342 \| (2, 351) \| 471 \| (673, 1034) \| 1643 \| DNA polymerase alpha subunit B \| DNA polymerase alpha subunit B; uniclust \| UniRef100\_A0A518BMG2 \| 100.0 \| 4.1e-49 \| 7.6e-55 \| 358.9 \| 339 \| (2, 353) \| 471 \| (144, 496) \| 657 \| Type I restriction enzyme EcoKI subunit R \| Type I restriction enzyme EcoKI subunit R; uniclust \| UniRef100\_A0A2D6MRW8 \| 100.0 \| 5.9e-49 \| 1.1e-54 \| 385.2 \| 336 \| (3, 351) \| 471 \| (338, 683) \| 767 \| HIT domain-containing protein \| HIT domain-containing protein; uniclust \| UniRef100\_A0A450Z507 \| 100.0 \| 6.8e-49 \| 1.3e-54 \| 359.2 \| 339 \| (3, 352) \| 471 \| (1, 429) \| 614 \| Helicase conserved C-terminal domain-containing protein \| Helicase conserved C-terminal domain-containing protein; uniclust \| UniRef100\_A0A2N3F2A2 \| 100.0 \| 8.3e-49 \| 1.6e-54 \| 383.2 \| 329 \| (4, 347) \| 471 \| (327, 665) \| 712 \| DUF3427 domain-containing protein (Fragment) \| DUF3427 domain-containing protein (Fragment); uniclust \| UniRef100\_A0A1G0KS28 \| 100.0 \| 1.1e-48 \| 2e-54 \| 375.0 \| 320 \| (3, 344) \| 471 \| (156, 487) \| 527 \| Helicase ATP-binding domain-containing protein \| Helicase ATP-binding domain-containing protein; uniclust \| UniRef100\_A0A1V6HEI4 \| 100.0 \| 1.5e-48 \| 2.9e-54 \| 362.0 \| 320 \| (2, 343) \| 471 \| (106, 435) \| 487 \| ATP-dependent RNA helicase DbpA \| ATP-dependent RNA helicase DbpA; uniclust \| UniRef100\_A0A2N2IT31 \| 100.0 \| 1.6e-48 \| 3e-54 \| 390.8 \| 334 \| (3, 347) \| 471 \| (342, 688) \| 989 \| DEAD/DEAH box helicase (Fragment) \| DEAD/DEAH box helicase (Fragment); uniclust \| UniRef100\_A0A068EU29 \| 100.0 \| 2.3e-48 \| 4.4e-54 \| 383.8 \| 320 \| (2, 345) \| 471 \| (217, 564) \| 617 \| Putative helicase \| Putative helicase; uniclust \| UniRef100\_A0A075LV11 \| 100.0 \| 2.3e-48 \| 4.5e-54 \| 368.0 \| 319 \| (3, 344) \| 471 \| (115, 442) \| 492 \| DNA repair protein Rad25 \| DNA repair protein Rad25; uniclust \| UniRef100\_A0A014LC28 \| 100.0 \| 2.7e-48 \| 5.2e-54 \| 402.7 \| 336 \| (3, 347) \| 471 \| (350, 702) \| 1318 \| Helicase \| Helicase; uniclust \| UniRef100\_A0A1I0IRV7 \| 100.0 \| 3.6e-48 \| 6.8e-54 \| 349.5 \| 339 \| (2, 350) \| 471 \| (8, 361) \| 427 \| Type III restriction enzyme, res subunit \| Type III restriction enzyme, res subunit; uniclust \| UniRef100\_A0A6N3G5E7 \| 100.0 \| 4.2e-48 \| 7.7e-54 \| 345.8 \| 346 \| (3, 359) \| 471 \| (9, 364) \| 628 \| ATP-dependent RNA helicase SrmB \| ATP-dependent RNA helicase SrmB; uniclust \| UniRef100\_A0A1G0H321 \| 100.0 \| 4.3e-48 \| 7.9e-54 \| 340.6 \| 334 \| (2, 347) \| 471 \| (4, 343) \| 460 \| DEAD/DEAH box helicase \| DEAD/DEAH box helicase; uniclust \| UniRef100\_A0A011NKT6 \| 100.0 \| 6.3e-48 \| 1.2e-53 \| 399.9 \| 332 \| (3, 348) \| 471 \| (469, 811) \| 1419 \| Type I restriction enzyme EcoKI subunit R \| Type I restriction enzyme EcoKI subunit R; uniclust \| UniRef100\_A0A0F9JQ39 \| 100.0 \| 7.8e-48 \| 1.4e-53 \| 335.2 \| 345 \| (112, 468) \| 471 \| (135, 486) \| 488 \| Intein C-terminal splicing domain-containing protein (Fragment) \| Intein C-terminal splicing domain-containing protein (Fragment); uniclust \| UniRef100\_C5B0S1 \| 100.0 \| 7.9e-48 \| 1.5e-53 \| 341.0 \| 334 \| (112, 467) \| 471 \| (189, 530) \| 575 \| Helicase domain protein \| Helicase domain protein; uniclust \| UniRef100\_A0A0H3XKY0 \| 100.0 \| 7.9e-48 \| 1.5e-53 \| 392.0 \| 333 \| (2, 346) \| 471 \| (293, 640) \| 1003 \| DNA/RNA helicase \| DNA/RNA helicase; uniclust \| UniRef100\_A0A455V0T9 \| 100.0 \| 8.1e-48 \| 1.5e-53 \| 344.8 \| 317 \| (1, 329) \| 471 \| (18, 360) \| 377 \| Helicase ATP-binding domain-containing protein \| Helicase ATP-binding domain-containing protein; uniclust \| UniRef100\_A0A1D2YTV2 \| 100.0 \| 8.9e-48 \| 1.7e-53 \| 364.6 \| 336 \| (3, 347) \| 471 \| (128, 480) \| 640 \| DNA helicase \| DNA helicase; uniclust \| UniRef100\_A0A031LPW7 \| 100.0 \| 9.6e-48 \| 1.8e-53 \| 364.9 \| 317 \| (3, 341) \| 471 \| (167, 524) \| 567 \| DNA helicase \| DNA helicase; uniclust \| UniRef100\_A0A0C2D3F5 \| 100.0 \| 1e-47 \| 2e-53 \| 389.7 \| 344 \| (3, 354) \| 471 \| (196, 583) \| 987 \| DNA helicase \| DNA helicase; uniclust \| UniRef100\_A0A8X8SQH6 \| 100.0 \| 1.4e-47 \| 2.6e-53 \| 349.7 \| 385 \| (2, 404) \| 471 \| (8, 423) \| 583 \| DEAD/DEAH box helicase family protein \| DEAD/DEAH box helicase family protein; uniclust \| UniRef100\_A0A0C1ZME6 \| 100.0 \| 1.5e-47 \| 2.8e-53 \| 388.5 \| 334 \| (3, 347) \| 471 \| (376, 722) \| 1455 \| HIT family hydrolase \| HIT family hydrolase; uniclust \| UniRef100\_A0A6L4ZR19 \| 100.0 \| 1.9e-47 \| 3.5e-53 \| 340.7 \| 348 \| (3, 359) \| 471 \| (149, 507) \| 611 \| DEAD/DEAH box helicase (Fragment) \| DEAD/DEAH box helicase (Fragment); uniclust \| UniRef100\_A0A936H9W6 \| 100.0 \| 1.9e-47 \| 3.5e-53 \| 346.9 \| 395 \| (3, 420) \| 471 \| (1, 441) \| 739 \| site-specific DNA-methyltransferase (adenine-specific) \| site-specific DNA-methyltransferase (adenine-specific); uniclust \| UniRef100\_A0A094WFY6 \| 100.0 \| 2.8e-47 \| 5.4e-53 \| 368.3 \| 346 \| (3, 355) \| 471 \| (193, 589) \| 623 \| DEAD/DEAH box helicase \| DEAD/DEAH box helicase; uniclust \| UniRef100\_A0A2R5GN63 \| 100.0 \| 3.6e-47 \| 6.6e-53 \| 336.4 \| 335 \| (4, 358) \| 471 \| (113, 466) \| 566 \| ATP-dependent RNA helicase ded1 \| ATP-dependent RNA helicase ded1; uniclust \| UniRef100\_A0A1B7X2J6 \| 100.0 \| 4.2e-47 \| 7.7e-53 \| 339.4 \| 390 \| (4, 403) \| 471 \| (20, 443) \| 625 \| DEAD/DEAH box helicase \| DEAD/DEAH box helicase; uniclust \| UniRef100\_A0A2H0MXI2 \| 100.0 \| 4.3e-47 \| 7.9e-53 \| 321.0 \| 296 \| (159, 467) \| 471 \| (2, 300) \| 342 \| Helicase C-terminal domain-containing protein \| Helicase C-terminal domain-containing protein; uniclust \| UniRef100\_A0A1A0HBL8 \| 100.0 \| 4.5e-47 \| 8.4e-53 \| 347.2 \| 340 \| (3, 350) \| 471 \| (1, 360) \| 613 \| P-loop containing nucleoside triphosphate hydrolase protein (Fragment) \| P-loop containing nucleoside triphosphate hydrolase protein (Fragment); uniclust \| UniRef100\_A0A7C6KF04 \| 100.0 \| 5.5e-47 \| 1e-52 \| 340.5 \| 387 \| (3, 403) \| 471 \| (75, 482) \| 662 \| DEAD/DEAH box helicase \| DEAD/DEAH box helicase; uniclust \| UniRef100\_A0A1Y5SXW7 \| 100.0 \| 6e-47 \| 1.1e-52 \| 341.7 \| 296 \| (109, 420) \| 471 \| (2, 311) \| 465 \| UvrABC system protein B \| UvrABC system protein B; uniclust \| UniRef100\_A0A1W9U5Q2 \| 100.0 \| 6.2e-47 \| 1.1e-52 \| 334.8 \| 384 \| (2, 405) \| 471 \| (1, 419) \| 564 \| Helicase \| Helicase; uniclust \| UniRef100\_A0A1C3IJV1 \| 100.0 \| 6.7e-47 \| 1.3e-52 \| 364.0 \| 341 \| (3, 351) \| 471 \| (142, 534) \| 554 \| UvrABC system protein B \| UvrABC system protein B; uniclust \| UniRef100\_A0A379T0C9 \| 100.0 \| 7.6e-47 \| 1.4e-52 \| 345.1 \| 373 \| (75, 468) \| 471 \| (5, 394) \| 790 \| ATP-dependent helicase \| ATP-dependent helicase; uniclust \| UniRef100\_A0A1F5P0I6 \| 100.0 \| 8.5e-47 \| 1.6e-52 \| 357.8 \| 337 \| (3, 352) \| 471 \| (7, 357) \| 672 \| DEAD/DEAH box helicase \| DEAD/DEAH box helicase; uniclust \| UniRef100\_A0A1A8TF80 \| 100.0 \| 1e-46 \| 1.9e-52 \| 366.2 \| 336 \| (3, 348) \| 471 \| (364, 716) \| 854 \| Type I restriction enzyme EcoKI subunit R \| Type I restriction enzyme EcoKI subunit R; uniclust \| UniRef100\_A0A2S9TD02 \| 100.0 \| 1.2e-46 \| 2.2e-52 \| 345.2 \| 340 \| (2, 355) \| 471 \| (3, 350) \| 792 \| DEAD/DEAH box helicase \| DEAD/DEAH box helicase; uniclust \| UniRef100\_A0A1M5ATC4 \| 100.0 \| 1.2e-46 \| 2.3e-52 \| 345.8 \| 348 \| (4, 360) \| 471 \| (60, 421) \| 845 \| Superfamily II DNA or RNA helicase \| Superfamily II DNA or RNA helicase; uniclust \| UniRef100\_A0A0G1U7D7 \| 100.0 \| 1.3e-46 \| 2.4e-52 \| 366.3 \| 333 \| (1, 354) \| 471 \| (4, 356) \| 798 \| Type III restriction protein res subunit \| Type III restriction protein res subunit; uniclust \| UniRef100\_A0A090R115 \| 100.0 \| 1.6e-46 \| 3e-52 \| 360.9 \| 332 \| (3, 348) \| 471 \| (219, 561) \| 676 \| Helicase-related protein \| Helicase-related protein; uniclust \| UniRef100\_A0A954J9P4 \| 100.0 \| 2e-46 \| 3.6e-52 \| 346.8 \| 312 \| (3, 321) \| 471 \| (21, 356) \| 919 \| DEAD/DEAH box helicase family protein \| DEAD/DEAH box helicase family protein; uniclust \| UniRef100\_A0A0F9KM15 \| 100.0 \| 1.9e-46 \| 3.7e-52 \| 339.1 \| 252 \| (139, 406) \| 471 \| (19, 277) \| 455 \| Helicase C-terminal domain-containing protein (Fragment) \| Helicase C-terminal domain-containing protein (Fragment); uniclust \| UniRef100\_A0A069IGA8 \| 100.0 \| 2.7e-46 \| 5.3e-52 \| 373.5 \| 320 \| (4, 344) \| 471 \| (426, 758) \| 822 \| DEAD/DEAH box helicase \| DEAD/DEAH box helicase; uniclust \| UniRef100\_A0A1Q4RSB3 \| 100.0 \| 3.2e-46 \| 6e-52 \| 346.7 \| 387 \| (2, 403) \| 471 \| (7, 404) \| 824 \| Helicase \| Helicase; uniclust \| UniRef100\_A0A0B7NDC9 \| 100.0 \| 3.5e-46 \| 6.4e-52 \| 333.1 \| 321 \| (3, 348) \| 471 \| (6, 352) \| 463 \| P-loop containing nucleoside triphosphate hydrolase protein \| P-loop containing nucleoside triphosphate hydrolase protein; uniclust \| UniRef100\_A0A2P2DIA8 \| 100.0 \| 3.6e-46 \| 6.8e-52 \| 334.9 \| 341 \| (3, 352) \| 471 \| (15, 363) \| 531 \| Helicase C-terminal domain protein \| Helicase C-terminal domain protein; uniclust \| UniRef100\_A0A0K1QBS4 \| 100.0 \| 3.6e-46 \| 6.9e-52 \| 346.3 \| 318 \| (2, 342) \| 471 \| (54, 415) \| 451 \| DNA helicase \| DNA helicase; uniclust \| UniRef100\_A0A7X7MWV3 \| 100.0 \| 3.7e-46 \| 7e-52 \| 335.6 \| 342 \| (3, 361) \| 471 \| (24, 379) \| 529 \| DEAD/DEAH box helicase \| DEAD/DEAH box helicase; uniclust \| UniRef100\_A0A174VXQ6 \| 100.0 \| 5.3e-46 \| 9.9e-52 \| 358.9 \| 322 \| (1, 342) \| 471 \| (87, 421) \| 846 \| Type I restriction enzyme EcoKI subunit R \| Type I restriction enzyme EcoKI subunit R; uniclust \| UniRef100\_A0A1C3RE97 \| 100.0 \| 6.2e-46 \| 1.2e-51 \| 346.1 \| 351 \| (2, 361) \| 471 \| (49, 416) \| 666 \| DNA/RNA helicase, superfamily II \| DNA/RNA helicase, superfamily II; uniclust \| UniRef100\_A0A090QT69 \| 100.0 \| 6.3e-46 \| 1.2e-51 \| 368.9 \| 331 \| (3, 347) \| 471 \| (221, 562) \| 1121 \| ATP-dependent RNA helicase YejH \| ATP-dependent RNA helicase YejH; uniclust \| UniRef100\_A0A023X515 \| 100.0 \| 6.9e-46 \| 1.3e-51 \| 361.8 \| 318 \| (2, 342) \| 471 \| (154, 515) \| 656 \| DNA helicase \| DNA helicase; uniclust \| UniRef100\_A0A0A8WX06 \| 100.0 \| 9e-46 \| 1.7e-51 \| 361.1 \| 332 \| (3, 348) \| 471 \| (185, 526) \| 880 \| DNA/RNA helicase \| DNA/RNA helicase; uniclust \| UniRef100\_A0A256ZIB2 \| 100.0 \| 9.7e-46 \| 1.8e-51 \| 351.9 \| 316 \| (3, 340) \| 471 \| (262, 618) \| 657 \| DNA methylase (Fragment) \| DNA methylase (Fragment); uniclust \| UniRef100\_A0A256Y0L1 \| 100.0 \| 1.1e-45 \| 2e-51 \| 342.4 \| 325 \| (2, 344) \| 471 \| (64, 426) \| 447 \| Helicase \| Helicase; uniclust \| UniRef100\_A0A061AZP5 \| 100.0 \| 1.1e-45 \| 2.1e-51 \| 343.5 \| 338 \| (1, 345) \| 471 \| (52, 408) \| 770 \| CYFA0S11e00452g1\_1 \| CYFA0S11e00452g1\_1; uniclust \| UniRef100\_A0A011QTS1 \| 100.0 \| 1.2e-45 \| 2.3e-51 \| 392.1 \| 328 \| (3, 347) \| 471 \| (379, 722) \| 1304 \| DNA/RNA helicase of DEAD/DEAH box family \| DNA/RNA helicase of DEAD/DEAH box family; uniclust \| UniRef100\_A0A920AJR7 \| 100.0 \| 1.4e-45 \| 2.5e-51 \| 330.5 \| 378 \| (3, 404) \| 471 \| (4, 394) \| 639 \| Uncharacterized protein \| Uncharacterized protein; uniclust \| UniRef100\_A0A0M6WTG7 \| 100.0 \| 1.4e-45 \| 2.6e-51 \| 349.9 \| 336 \| (5, 351) \| 471 \| (25, 377) \| 653 \| Type I restriction enzyme EcoKI subunit R \| Type I restriction enzyme EcoKI subunit R; uniclust \| UniRef100\_A0A0F2L4X5 \| 100.0 \| 1.4e-45 \| 2.6e-51 \| 358.9 \| 321 \| (2, 341) \| 471 \| (389, 721) \| 808 \| DEAD/DEAH box helicase \| DEAD/DEAH box helicase; uniclust \| UniRef100\_A0A968RBZ8 \| 100.0 \| 1.5e-45 \| 2.7e-51 \| 323.6 \| 408 \| (2, 420) \| 471 \| (1, 437) \| 521 \| DEAD/DEAH box helicase family protein \| DEAD/DEAH box helicase family protein; uniclust \| UniRef100\_UPI0013D759F8 \| 100.0 \| 1.5e-45 \| 2.8e-51 \| 334.9 \| 384 \| (3, 404) \| 471 \| (1, 401) \| 741 \| DEAD/DEAH box helicase \| DEAD/DEAH box helicase; uniclust \| UniRef100\_A0A0W0F5M0 \| 100.0 \| 2.1e-45 \| 3.9e-51 \| 354.0 \| 328 \| (14, 348) \| 471 \| (313, 675) \| 950 \| P-loop containing nucleoside triphosphate hydrolase protein \| P-loop containing nucleoside triphosphate hydrolase protein; uniclust \| UniRef100\_A0A4R8ZEZ6 \| 100.0 \| 3.5e-45 \| 6.6e-51 \| 340.6 \| 343 \| (3, 352) \| 471 \| (84, 470) \| 487 \| DEAD/DEAH box helicase \| DEAD/DEAH box helicase; uniclust \| UniRef100\_A0A0D2VP34 \| 100.0 \| 3.9e-45 \| 7.3e-51 \| 345.3 \| 342 \| (2, 351) \| 471 \| (243, 610) \| 884 \| ATP-dependent DNA helicase \| ATP-dependent DNA helicase; uniclust \| UniRef100\_A0A023ZUD9 \| 100.0 \| 3.9e-45 \| 7.5e-51 \| 361.0 \| 381 \| (3, 397) \| 471 \| (60, 504) \| 743 \| Helicase ATP-binding domain-containing protein \| Helicase ATP-binding domain-containing protein; uniclust \| UniRef100\_A0A521UH50 \| 100.0 \| 4.4e-45 \| 8.3e-51 \| 359.5 \| 332 \| (3, 347) \| 471 \| (299, 642) \| 1187 \| DEAD/DEAH box helicase \| DEAD/DEAH box helicase; uniclust \| UniRef100\_UPI0022835FE8 \| 100.0 \| 4.9e-45 \| 8.9e-51 \| 313.7 \| 336 \| (1, 364) \| 471 \| (1, 343) \| 432 \| DEAD/DEAH box helicase \| DEAD/DEAH box helicase; uniclust \| UniRef100\_A0A2E8DZC4 \| 100.0 \| 6e-45 \| 1.1e-50 \| 323.3 \| 391 \| (2, 404) \| 471 \| (7, 445) \| 578 \| ATP-dependent helicase \| ATP-dependent helicase; uniclust \| UniRef100\_A0A1I4DLJ8 \| 100.0 \| 6.5e-45 \| 1.2e-50 \| 356.5 \| 331 \| (3, 348) \| 471 \| (143, 483) \| 905 \| PLD-like domain-containing protein \| PLD-like domain-containing protein; uniclust \| UniRef100\_A0A059G1H1 \| 100.0 \| 6.5e-45 \| 1.3e-50 \| 353.6 \| 334 \| (3, 343) \| 471 \| (191, 571) \| 600 \| Type III restriction protein res subunit \| Type III restriction protein res subunit; uniclust \| UniRef100\_A0A935XXU2 \| 100.0 \| 7e-45 \| 1.3e-50 \| 320.9 \| 300 \| (3, 405) \| 471 \| (1, 311) \| 470 \| DEAD/DEAH box helicase family protein \| DEAD/DEAH box helicase family protein; uniclust \| UniRef100\_A0A1W1HFH2 \| 100.0 \| 7.2e-45 \| 1.4e-50 \| 372.2 \| 334 \| (3, 347) \| 471 \| (343, 693) \| 1161 \| Type III restriction protein res subunit \| Type III restriction protein res subunit; uniclust \| UniRef100\_A0A660T5X1 \| 100.0 \| 7.8e-45 \| 1.4e-50 \| 327.8 \| 345 \| (2, 358) \| 471 \| (1, 347) \| 681 \| DEAD/DEAH box helicase \| DEAD/DEAH box helicase; uniclust \| UniRef100\_A0A2N5YN75 \| 100.0 \| 9.1e-45 \| 1.7e-50 \| 360.4 \| 333 \| (5, 347) \| 471 \| (103, 453) \| 833 \| Helicase C-terminal domain-containing protein \| Helicase C-terminal domain-containing protein; uniclust \| UniRef100\_A0A3N5GNH0 \| 100.0 \| 9.6e-45 \| 1.8e-50 \| 333.6 \| 274 \| (2, 297) \| 471 \| (64, 340) \| 847 \| DEAD/DEAH box helicase \| DEAD/DEAH box helicase; uniclust \| UniRef100\_A0A150TWR4 \| 100.0 \| 9.9e-45 \| 1.9e-50 \| 368.3 \| 334 \| (3, 347) \| 471 \| (348, 698) \| 1590 \| DEAD/DEAH box helicase \| DEAD/DEAH box helicase; uniclust \| UniRef100\_A0A965URS7 \| 100.0 \| 1.2e-44 \| 2.2e-50 \| 333.5 \| 358 \| (1, 468) \| 471 \| (31, 664) \| 708 \| Uncharacterized protein \| Uncharacterized protein; uniclust \| UniRef100\_A0A2P2BZ99 \| 100.0 \| 1.2e-44 \| 2.3e-50 \| 343.7 \| 335 \| (3, 347) \| 471 \| (163, 515) \| 720 \| Type III restriction protein res subunit \| Type III restriction protein res subunit; uniclust \| UniRef100\_A0A661TPG1 \| 100.0 \| 1.3e-44 \| 2.5e-50 \| 311.6 \| 303 \| (108, 421) \| 471 \| (14, 325) \| 438 \| ATP-dependent helicase (Fragment) \| ATP-dependent helicase (Fragment); uniclust \| UniRef100\_A0A3B0VE26 \| 100.0 \| 1.6e-44 \| 2.9e-50 \| 319.7 \| 447 \| (2, 468) \| 471 \| (23, 515) \| 562 \| Helicase \| Helicase; uniclust \| UniRef100\_A0A2P6V7A7 \| 100.0 \| 1.7e-44 \| 3.2e-50 \| 341.9 \| 332 \| (2, 344) \| 471 \| (561, 926) \| 1254 \| Metal transporter Nramp2-like \| Metal transporter Nramp2-like; uniclust \| UniRef100\_A0A0F9PA38 \| 100.0 \| 1.8e-44 \| 3.5e-50 \| 342.1 \| 325 \| (2, 344) \| 471 \| (105, 440) \| 492 \| Helicase ATP-binding domain-containing protein \| Helicase ATP-binding domain-containing protein; uniclust \| UniRef100\_A0A421JNJ2 \| 100.0 \| 2.1e-44 \| 3.9e-50 \| 341.6 \| 341 \| (2, 350) \| 471 \| (21, 381) \| 996 \| GPI-anchor transamidase \| GPI-anchor transamidase; uniclust \| UniRef100\_A0A816ZT36 \| 100.0 \| 2.3e-44 \| 4.3e-50 \| 330.8 \| 379 \| (3, 403) \| 471 \| (1, 400) \| 837 \| Uncharacterized protein \| Uncharacterized protein; uniclust \| UniRef100\_A0A2W6ZLK6 \| 100.0 \| 2.6e-44 \| 4.9e-50 \| 304.8 \| 255 \| (199, 469) \| 471 \| (2, 269) \| 293 \| Helicase C-terminal domain-containing protein \| Helicase C-terminal domain-containing protein; uniclust \| UniRef100\_A0A0H3JNJ1 \| 100.0 \| 2.6e-44 \| 4.9e-50 \| 357.3 \| 328 \| (3, 347) \| 471 \| (39, 380) \| 787 \| Helicase C-terminal domain-containing protein \| Helicase C-terminal domain-containing protein; uniclust \| UniRef100\_A0A812NDJ4 \| 100.0 \| 3.2e-44 \| 5.8e-50 \| 326.9 \| 338 \| (2, 352) \| 471 \| (103, 454) \| 753 \| acylphosphatase \| acylphosphatase; uniclust \| UniRef100\_A0A0D6AWB2 \| 100.0 \| 3.2e-44 \| 6.1e-50 \| 342.8 \| 343 \| (3, 352) \| 471 \| (166, 557) \| 575 \| Type I restriction-modification system \| Type I restriction-modification system; uniclust \| UniRef100\_UPI0003496CD4 \| 100.0 \| 3.6e-44 \| 6.5e-50 \| 318.2 \| 346 \| (3, 357) \| 471 \| (1, 353) \| 574 \| DEAD/DEAH box helicase \| DEAD/DEAH box helicase; uniclust \| UniRef100\_A0A965L6M0 \| 100.0 \| 4.4e-44 \| 8.2e-50 \| 310.8 \| 399 \| (5, 466) \| 471 \| (54, 458) \| 469 \| Helicase ATP-binding domain-containing protein \| Helicase ATP-binding domain-containing protein; uniclust \| UniRef100\_A0A0F6W5J8 \| 100.0 \| 4.3e-44 \| 8.3e-50 \| 343.7 \| 318 \| (3, 343) \| 471 \| (156, 515) \| 573 \| DNA helicase \| DNA helicase; uniclust \| UniRef100\_A0A016QL41 \| 100.0 \| 4.7e-44 \| 9e-50 \| 336.3 \| 318 \| (3, 343) \| 471 \| (102, 462) \| 549 \| DNA helicase \| DNA helicase; uniclust \| UniRef100\_A0A950F2N3 \| 100.0 \| 5.3e-44 \| 9.9e-50 \| 325.5 \| 350 \| (3, 359) \| 471 \| (31, 416) \| 642 \| DEAD/DEAH box helicase \| DEAD/DEAH box helicase; uniclust \| UniRef100\_A0A367XRL6 \| 100.0 \| 6.2e-44 \| 1.1e-49 \| 328.0 \| 335 \| (2, 343) \| 471 \| (24, 376) \| 836 \| Putative ATP-dependent helicase IRC3 \| Putative ATP-dependent helicase IRC3; uniclust \| UniRef100\_A0A962CYC4 \| 100.0 \| 6.7e-44 \| 1.2e-49 \| 286.9 \| 198 \| (259, 469) \| 471 \| (2, 200) \| 208 \| Uncharacterized protein \| Uncharacterized protein; uniclust \| UniRef100\_A0A093YJB7 \| 100.0 \| 8e-44 \| 1.5e-49 \| 345.6 \| 338 \| (2, 346) \| 471 \| (86, 442) \| 1272 \| Helicase ATP-binding domain-containing protein \| Helicase ATP-binding domain-containing protein; uniclust \| UniRef100\_A0A6L4ZSK1 \| 100.0 \| 1e-43 \| 1.9e-49 \| 328.6 \| 348 \| (3, 361) \| 471 \| (32, 391) \| 898 \| Type III restriction protein res subunit \| Type III restriction protein res subunit; uniclust \| UniRef100\_A0A5N5XQV3 \| 100.0 \| 1.3e-43 \| 2.4e-49 \| 322.9 \| 336 \| (1, 344) \| 471 \| (5, 356) \| 578 \| DEAD/DEAH box helicase \| DEAD/DEAH box helicase; uniclust \| UniRef100\_A0A956GY33 \| 100.0 \| 1.3e-43 \| 2.5e-49 \| 315.1 \| 338 \| (3, 351) \| 471 \| (1, 345) \| 582 \| DEAD/DEAH box helicase \| DEAD/DEAH box helicase; uniclust \| UniRef100\_A0A061HBV4 \| 100.0 \| 1.4e-43 \| 2.5e-49 \| 329.7 \| 339 \| (2, 347) \| 471 \| (99, 508) \| 835 \| Uncharacterized protein \| Uncharacterized protein; uniclust \| UniRef100\_A9UTX7 \| 100.0 \| 1.9e-43 \| 3.5e-49 \| 306.1 \| 336 \| (3, 346) \| 471 \| (65, 423) \| 457 \| Uncharacterized protein \| Uncharacterized protein; uniclust \| UniRef100\_A0A0A0J1H6 \| 100.0 \| 1.8e-43 \| 3.6e-49 \| 385.6 \| 343 \| (3, 352) \| 471 \| (1223, 1612) \| 1644 \| Helicase \| Helicase; uniclust \| UniRef100\_A0A0R3EQW8 \| 100.0 \| 2e-43 \| 3.6e-49 \| 324.9 \| 341 \| (3, 351) \| 471 \| (28, 387) \| 657 \| Helicase \| Helicase; uniclust \| UniRef100\_A0A4P7WCN2 \| 100.0 \| 2.2e-43 \| 4e-49 \| 320.8 \| 385 \| (2, 405) \| 471 \| (7, 412) \| 734 \| DEAD/DEAH box helicase \| DEAD/DEAH box helicase; uniclust \| UniRef100\_A0A086ZGU5 \| 100.0 \| 2.4e-43 \| 4.5e-49 \| 355.1 \| 330 \| (5, 347) \| 471 \| (151, 492) \| 907 \| DNA/RNA helicase \| DNA/RNA helicase; uniclust \| UniRef100\_A0A538Q2V0 \| 100.0 \| 2.5e-43 \| 4.6e-49 \| 313.4 \| 366 \| (3, 407) \| 471 \| (143, 514) \| 582 \| Helicase ATP-binding domain-containing protein \| Helicase ATP-binding domain-containing protein; uniclust \| UniRef100\_A0A2B4QYV1 \| 100.0 \| 2.7e-43 \| 5e-49 \| 325.0 \| 364 \| (20, 404) \| 471 \| (194, 578) \| 871 \| Putative ATP-dependent helicase IRC3 (Fragment) \| Putative ATP-dependent helicase IRC3 (Fragment); uniclust \| UniRef100\_A0A1L8MY55 \| 100.0 \| 3.6e-43 \| 6.6e-49 \| 315.3 \| 344 \| (2, 362) \| 471 \| (4, 371) \| 548 \| Helicase \| Helicase; uniclust \| UniRef100\_A0A011QEE7 \| 100.0 \| 4.2e-43 \| 8e-49 \| 349.0 \| 339 \| (3, 353) \| 471 \| (216, 601) \| 924 \| Type-1 restriction enzyme R protein \| Type-1 restriction enzyme R protein; uniclust \| UniRef100\_D3SGN6 \| 100.0 \| 4.5e-43 \| 8.3e-49 \| 327.3 \| 373 \| (4, 387) \| 471 \| (28, 442) \| 696 \| Type III restriction protein res subunit \| Type III restriction protein res subunit; uniclust \| UniRef100\_UPI0021B4E6CE \| 100.0 \| 4.8e-43 \| 8.8e-49 \| 309.8 \| 338 \| (1, 355) \| 471 \| (1, 344) \| 550 \| DEAD/DEAH box helicase \| DEAD/DEAH box helicase; uniclust \| UniRef100\_A0A2W0H9H5 \| 100.0 \| 4.8e-43 \| 9.1e-49 \| 331.8 \| 328 \| (3, 348) \| 471 \| (84, 426) \| 628 \| Restriction endonuclease subunit R \| Restriction endonuclease subunit R; uniclust \| UniRef100\_A0A261EUJ9 \| 100.0 \| 4.9e-43 \| 9.4e-49 \| 343.4 \| 328 \| (3, 347) \| 471 \| (262, 606) \| 735 \| RNA helicase \| RNA helicase; uniclust \| UniRef100\_A0A088QJ79 \| 100.0 \| 5.5e-43 \| 1e-48 \| 353.4 \| 329 \| (3, 347) \| 471 \| (248, 595) \| 1030 \| DEAD/DEAH box helicase family protein \| DEAD/DEAH box helicase family protein; uniclust \| UniRef100\_A0A3A4WK42 \| 100.0 \| 5.6e-43 \| 1.1e-48 \| 322.3 \| 321 \| (4, 344) \| 471 \| (141, 469) \| 491 \| DEAD/DEAH box helicase (Fragment) \| DEAD/DEAH box helicase (Fragment); uniclust \| UniRef100\_A0A1M3B859 \| 100.0 \| 6.2e-43 \| 1.2e-48 \| 322.2 \| 240 \| (109, 355) \| 471 \| (7, 252) \| 473 \| Helicase C-terminal domain-containing protein \| Helicase C-terminal domain-containing protein; uniclust \| UniRef100\_A0A2E8RD89 \| 100.0 \| 6.8e-43 \| 1.3e-48 \| 321.2 \| 330 \| (2, 344) \| 471 \| (31, 381) \| 553 \| DEAD/DEAH box helicase \| DEAD/DEAH box helicase; uniclust \| UniRef100\_A0A150K7H9 \| 100.0 \| 7.6e-43 \| 1.4e-48 \| 338.8 \| 328 \| (3, 348) \| 471 \| (256, 598) \| 825 \| Uncharacterized protein \| Uncharacterized protein; uniclust \| UniRef100\_A0A0X1RVT1 \| 100.0 \| 7.7e-43 \| 1.5e-48 \| 347.2 \| 341 \| (2, 349) \| 471 \| (111, 493) \| 812 \| DEAD/DEAH box helicase family protein \| DEAD/DEAH box helicase family protein; uniclust \| UniRef100\_A0A060LX62 \| 100.0 \| 9.5e-43 \| 1.8e-48 \| 361.2 \| 340 \| (2, 348) \| 471 \| (232, 629) \| 1262 \| Helicase \| Helicase; uniclust \| UniRef100\_A0A010SLW4 \| 100.0 \| 1.2e-42 \| 2.4e-48 \| 341.8 \| 310 \| (4, 348) \| 471 \| (124, 459) \| 620 \| RNA helicase \| RNA helicase; uniclust \| UniRef100\_A0A496A9P6 \| 100.0 \| 1.3e-42 \| 2.5e-48 \| 307.2 \| 331 \| (2, 343) \| 471 \| (8, 367) \| 553 \| DEAD/DEAH box helicase \| DEAD/DEAH box helicase; uniclust \| UniRef100\_A0A2N2IYY4 \| 100.0 \| 1.4e-42 \| 2.6e-48 \| 305.8 \| 316 \| (4, 343) \| 471 \| (8, 332) \| 362 \| Helicase (Fragment) \| Helicase (Fragment); uniclust \| UniRef100\_A0A2T2WR37 \| 100.0 \| 1.5e-42 \| 2.7e-48 \| 318.5 \| 339 \| (2, 353) \| 471 \| (3, 478) \| 696 \| Restriction endonuclease subunit R (Fragment) \| Restriction endonuclease subunit R (Fragment); uniclust \| UniRef100\_UPI00232C099B \| 100.0 \| 1.6e-42 \| 2.9e-48 \| 299.8 \| 343 \| (3, 361) \| 471 \| (1, 357) \| 448 \| DEAD/DEAH box helicase \| DEAD/DEAH box helicase; uniclust \| UniRef100\_A0A058ZJS0 \| 100.0 \| 1.7e-42 \| 3.1e-48 \| 345.0 \| 406 \| (3, 419) \| 471 \| (308, 773) \| 1071 \| Type I site-specific deoxyribonuclease \| Type I site-specific deoxyribonuclease; uniclust \| UniRef100\_A0A8H6BUF5 \| 100.0 \| 1.7e-42 \| 3.1e-48 \| 316.6 \| 337 \| (2, 345) \| 471 \| (33, 384) \| 738 \| Type III restriction enzyme, res subunit family protein \| Type III restriction enzyme, res subunit family protein; uniclust \| UniRef100\_UPI00055AE175 \| 100.0 \| 2e-42 \| 3.7e-48 \| 289.9 \| 295 \| (4, 310) \| 471 \| (5, 308) \| 309 \| DEAD/DEAH box helicase \| DEAD/DEAH box helicase; uniclust \| UniRef100\_A0A0D0RIZ1 \| 100.0 \| 2.2e-42 \| 4.2e-48 \| 338.0 \| 326 \| (4, 347) \| 471 \| (65, 412) \| 834 \| DNA/RNA helicase of DEAD/DEAH box family \| DNA/RNA helicase of DEAD/DEAH box family; uniclust \| UniRef100\_A0A0F9HEE8 \| 100.0 \| 2.4e-42 \| 4.6e-48 \| 321.7 \| 320 \| (3, 345) \| 471 \| (108, 445) \| 479 \| Helicase ATP-binding domain-containing protein (Fragment) \| Helicase ATP-binding domain-containing protein (Fragment); uniclust \| UniRef100\_A0A1N6QDG8 \| 100.0 \| 3.3e-42 \| 6.1e-48 \| 335.5 \| 331 \| (3, 347) \| 471 \| (398, 739) \| 985 \| Superfamily II DNA or RNA helicase \| Superfamily II DNA or RNA helicase; uniclust \| UniRef100\_A0A011PF13 \| 100.0 \| 3.7e-42 \| 7.2e-48 \| 339.6 \| 304 \| (4, 343) \| 471 \| (102, 438) \| 672 \| ATP-dependent RNA helicase RhlE \| ATP-dependent RNA helicase RhlE; uniclust \| UniRef100\_A0A2A4UZJ5 \| 100.0 \| 3.9e-42 \| 7.5e-48 \| 323.8 \| 308 \| (4, 346) \| 471 \| (30, 361) \| 452 \| RNA helicase \| RNA helicase; uniclust \| UniRef100\_A0A197K8K7 \| 100.0 \| 5.4e-42 \| 1e-47 \| 322.7 \| 345 \| (2, 353) \| 471 \| (233, 596) \| 930 \| p-loop containing nucleoside triphosphate hydrolase protein \| p-loop containing nucleoside triphosphate hydrolase protein; uniclust \| UniRef100\_A0A929LPA0 \| 100.0 \| 5.5e-42 \| 1e-47 \| 316.5 \| 336 \| (3, 351) \| 471 \| (146, 518) \| 713 \| DEAD/DEAH box helicase \| DEAD/DEAH box helicase; uniclust \| UniRef100\_A0A0C2WKU0 \| 100.0 \| 5.3e-42 \| 1e-47 \| 332.0 \| 304 \| (4, 345) \| 471 \| (85, 411) \| 647 \| Helicase \| Helicase; uniclust \| UniRef100\_A0A2T3GEM9 \| 100.0 \| 5.6e-42 \| 1.1e-47 \| 334.3 \| 329 \| (3, 348) \| 471 \| (237, 579) \| 783 \| DEAD/DEAH box helicase \| DEAD/DEAH box helicase; uniclust \| UniRef100\_A0A3G1GLB6 \| 100.0 \| 6.1e-42 \| 1.1e-47 \| 316.6 \| 343 \| (3, 354) \| 471 \| (25, 394) \| 635 \| DEAD/DEAH box helicase family protein \| DEAD/DEAH box helicase family protein; uniclust \| UniRef100\_UPI000834A4D0 \| 100.0 \| 6.7e-42 \| 1.2e-47 \| 313.2 \| 337 \| (2, 350) \| 471 \| (1, 369) \| 636 \| DEAD/DEAH box helicase \| DEAD/DEAH box helicase; uniclust \| UniRef100\_A0A010SCV3 \| 100.0 \| 7e-42 \| 1.3e-47 \| 326.6 \| 336 \| (3, 349) \| 471 \| (153, 507) \| 755 \| Restriction endonuclease \| Restriction endonuclease; uniclust \| UniRef100\_A0A955J4R2 \| 100.0 \| 8e-42 \| 1.5e-47 \| 289.2 \| 280 \| (3, 282) \| 471 \| (1, 296) \| 297 \| DEAD/DEAH box helicase family protein (Fragment) \| DEAD/DEAH box helicase family protein (Fragment); uniclust \| UniRef100\_A0A956CHY4 \| 100.0 \| 7.9e-42 \| 1.5e-47 \| 337.1 \| 333 \| (3, 348) \| 471 \| (330, 670) \| 1325 \| DUF3427 domain-containing protein \| DUF3427 domain-containing protein; uniclust \| UniRef100\_A0A109RM47 \| 100.0 \| 8.5e-42 \| 1.6e-47 \| 326.5 \| 338 \| (3, 347) \| 471 \| (110, 497) \| 741 \| Helicase \| Helicase; uniclust \| UniRef100\_A0A255TER8 \| 100.0 \| 9.4e-42 \| 1.7e-47 \| 318.1 \| 350 \| (2, 360) \| 471 \| (14, 592) \| 984 \| Helicase ATP-binding domain-containing protein \| Helicase ATP-binding domain-containing protein; uniclust \| UniRef100\_A0A538HN94 \| 100.0 \| 1e-41 \| 1.9e-47 \| 300.3 \| 335 \| (4, 347) \| 471 \| (95, 449) \| 459 \| DEAD/DEAH box helicase (Fragment) \| DEAD/DEAH box helicase (Fragment); uniclust \| UniRef100\_A0A1E3QMQ9 \| 100.0 \| 1.2e-41 \| 2.3e-47 \| 316.5 \| 335 \| (4, 345) \| 471 \| (23, 372) \| 951 \| ATP-dependent helicase IRC3 \| ATP-dependent helicase IRC3; uniclust \| UniRef100\_A0A1Y2HFK6 \| 100.0 \| 1.3e-41 \| 2.5e-47 \| 302.1 \| 336 \| (3, 345) \| 471 \| (30, 398) \| 572 \| p-loop containing nucleoside triphosphate hydrolase protein \| p-loop containing nucleoside triphosphate hydrolase protein; uniclust \| UniRef100\_A0A7C5N2Q9 \| 100.0 \| 1.4e-41 \| 2.6e-47 \| 284.1 \| 252 \| (139, 406) \| 471 \| (16, 274) \| 300 \| Helicase C-terminal domain-containing protein \| Helicase C-terminal domain-containing protein; uniclust \| UniRef100\_A0A0G4KNW0 \| 100.0 \| 1.5e-41 \| 2.8e-47 \| 315.5 \| 337 \| (3, 346) \| 471 \| (44, 399) \| 892 \| Helicase C-terminal domain-containing protein \| Helicase C-terminal domain-containing protein; uniclust \| UniRef100\_A0A011RQJ8 \| 100.0 \| 1.5e-41 \| 2.9e-47 \| 343.1 \| 305 \| (4, 345) \| 471 \| (91, 416) \| 728 \| DEAD-box ATP-dependent RNA helicase CshA \| DEAD-box ATP-dependent RNA helicase CshA; uniclust \| UniRef100\_A0A0D8FYE7 \| 100.0 \| 1.6e-41 \| 3e-47 \| 332.6 \| 320 \| (2, 334) \| 471 \| (184, 562) \| 747 \| UvrABC system protein B \| UvrABC system protein B; uniclust \| UniRef100\_A0A081CC42 \| 100.0 \| 1.8e-41 \| 3.3e-47 \| 320.6 \| 339 \| (2, 347) \| 471 \| (90, 462) \| 1183 \| p-loop containing nucleoside triphosphate hydrolase protein \| p-loop containing nucleoside triphosphate hydrolase protein; uniclust \| UniRef100\_A0A0F8YP13 \| 100.0 \| 2.3e-41 \| 4.3e-47 \| 279.7 \| 187 \| (1, 189) \| 471 \| (16, 203) \| 219 \| Helicase ATP-binding domain-containing protein (Fragment) \| Helicase ATP-binding domain-containing protein (Fragment); uniclust \| UniRef100\_A0A069EZS9 \| 100.0 \| 2.3e-41 \| 4.5e-47 \| 331.2 \| 329 \| (2, 343) \| 471 \| (84, 474) \| 552 \| Type III restriction protein res subunit \| Type III restriction protein res subunit; uniclust \| UniRef100\_A0A0G2ZBN3 \| 100.0 \| 2.6e-41 \| 4.9e-47 \| 334.9 \| 330 \| (4, 345) \| 471 \| (213, 555) \| 944 \| Superfamily II DNA or RNA helicase \| Superfamily II DNA or RNA helicase; uniclust \| UniRef100\_A0A0F9FJE3 \| 100.0 \| 2.8e-41 \| 5.2e-47 \| 305.4 \| 346 \| (4, 362) \| 471 \| (24, 373) \| 685 \| Helicase ATP-binding domain-containing protein \| Helicase ATP-binding domain-containing protein; uniclust \| UniRef100\_A0A510LBK3 \| 100.0 \| 3e-41 \| 5.6e-47 \| 319.6 \| 329 \| (4, 348) \| 471 \| (264, 609) \| 749 \| Type III restriction protein Res subunit \| Type III restriction protein Res subunit; uniclust \| UniRef100\_A0A421J579 \| 100.0 \| 3.1e-41 \| 5.7e-47 \| 327.7 \| 343 \| (2, 352) \| 471 \| (672, 1033) \| 1339 \| DNA polymerase alpha subunit B \| DNA polymerase alpha subunit B; uniclust \| UniRef100\_A0A0R3DQ16 \| 100.0 \| 3.1e-41 \| 5.8e-47 \| 302.8 \| 391 \| (2, 404) \| 471 \| (1, 425) \| 548 \| Helicase C-terminal domain-containing protein \| Helicase C-terminal domain-containing protein; uniclust \| UniRef100\_A0A084SIF9 \| 100.0 \| 3.1e-41 \| 6e-47 \| 321.0 \| 311 \| (4, 353) \| 471 \| (69, 402) \| 533 \| DEAD/DEAH box helicase \| DEAD/DEAH box helicase; uniclust \| UniRef100\_A0A059ELD1 \| 100.0 \| 3.2e-41 \| 6.2e-47 \| 318.8 \| 299 \| (4, 341) \| 471 \| (68, 392) \| 515 \| Uncharacterized protein \| Uncharacterized protein; uniclust \| UniRef100\_A0A0N0XGX0 \| 100.0 \| 3.3e-41 \| 6.2e-47 \| 330.2 \| 304 \| (4, 344) \| 471 \| (82, 412) \| 720 \| ATP-dependent RNA helicase RhlE \| ATP-dependent RNA helicase RhlE; uniclust \| UniRef100\_A0A014M0H5 \| 100.0 \| 3.7e-41 \| 7.3e-47 \| 339.7 \| 305 \| (4, 347) \| 471 \| (149, 477) \| 733 \| ATP-dependent RNA helicase \| ATP-dependent RNA helicase; uniclust \| UniRef100\_A0A021VTQ4 \| 100.0 \| 4.1e-41 \| 7.9e-47 \| 311.7 \| 328 \| (2, 343) \| 471 \| (25, 387) \| 423 \| Helicase \| Helicase; uniclust \| UniRef100\_A0A6S6SZY5 \| 100.0 \| 4.3e-41 \| 7.9e-47 \| 287.6 \| 336 \| (3, 355) \| 471 \| (6, 345) \| 404 \| Helicase \| Helicase; uniclust \| UniRef100\_A0A014MMG9 \| 100.0 \| 4.5e-41 \| 8.7e-47 \| 323.6 \| 313 \| (4, 355) \| 471 \| (114, 449) \| 623 \| DEAD/DEAH box helicase \| DEAD/DEAH box helicase; uniclust \| UniRef100\_A0A021XEI8 \| 100.0 \| 4.7e-41 \| 9.1e-47 \| 331.7 \| 307 \| (4, 348) \| 471 \| (94, 427) \| 682 \| ATP-dependent RNA helicase RhlE \| ATP-dependent RNA helicase RhlE; uniclust \| UniRef100\_A0A1I0HAW8 \| 100.0 \| 5.9e-41 \| 1.1e-46 \| 305.8 \| 291 \| (4, 331) \| 471 \| (73, 388) \| 389 \| ATP-dependent RNA helicase RhlE (Fragment) \| ATP-dependent RNA helicase RhlE (Fragment); uniclust \| UniRef100\_A0A011N3G2 \| 100.0 \| 6.2e-41 \| 1.2e-46 \| 334.2 \| 301 \| (4, 344) \| 471 \| (142, 472) \| 718 \| ATP-dependent RNA helicase RhlE \| ATP-dependent RNA helicase RhlE; uniclust \| UniRef100\_A0A2S6CGS2 \| 100.0 \| 7.1e-41 \| 1.3e-46 \| 312.4 \| 337 \| (3, 346) \| 471 \| (311, 667) \| 928 \| Helicase ATP-binding domain-containing protein \| Helicase ATP-binding domain-containing protein; uniclust \| UniRef100\_A0A2J6XMU8 \| 100.0 \| 8.4e-41 \| 1.6e-46 \| 318.1 \| 334 \| (4, 348) \| 471 \| (167, 557) \| 605 \| type I site-specific deoxyribonuclease (Fragment) \| type I site-specific deoxyribonuclease (Fragment); uniclust \| UniRef100\_A0A011SEZ8 \| 100.0 \| 8.9e-41 \| 1.7e-46 \| 336.7 \| 305 \| (4, 345) \| 471 \| (265, 591) \| 873 \| DEAD-box ATP-dependent RNA helicase CshB \| DEAD-box ATP-dependent RNA helicase CshB; uniclust \| UniRef100\_A0A060T2W9 \| 100.0 \| 9.1e-41 \| 1.7e-46 \| 323.4 \| 304 \| (4, 345) \| 471 \| (212, 546) \| 659 \| ARAD1C28336p \| ARAD1C28336p; uniclust \| UniRef100\_A0A031GUF1 \| 100.0 \| 9.1e-41 \| 1.8e-46 \| 333.4 \| 308 \| (4, 356) \| 471 \| (99, 432) \| 786 \| RecQ family ATP-dependent DNA helicase \| RecQ family ATP-dependent DNA helicase; uniclust \| UniRef100\_A0A070A925 \| 100.0 \| 9.9e-41 \| 1.9e-46 \| 325.0 \| 306 \| (4, 346) \| 471 \| (101, 427) \| 629 \| RNA helicase \| RNA helicase; uniclust \| UniRef100\_A0A0D6MW56 \| 100.0 \| 1e-40 \| 1.9e-46 \| 308.4 \| 298 \| (4, 340) \| 471 \| (60, 381) \| 443 \| RNA helicase \| RNA helicase; uniclust \| UniRef100\_A0A3M0ZKB0 \| 100.0 \| 1e-40 \| 1.9e-46 \| 308.5 \| 332 \| (3, 356) \| 471 \| (77, 434) \| 443 \| DEAD/DEAH box helicase \| DEAD/DEAH box helicase; uniclust \| UniRef100\_A0A0F9CQ82 \| 100.0 \| 1.1e-40 \| 2e-46 \| 276.7 \| 311 \| (2, 322) \| 471 \| (1, 318) \| 318 \| Helicase ATP-binding domain-containing protein (Fragment) \| Helicase ATP-binding domain-containing protein (Fragment); uniclust \| UniRef100\_UPI0023500254 \| 100.0 \| 1.1e-40 \| 2e-46 \| 302.6 \| 334 \| (2, 353) \| 471 \| (22, 376) \| 709 \| DEAD/DEAH box helicase family protein \| DEAD/DEAH box helicase family protein; uniclust \| UniRef100\_A0A1C6IYF2 \| 100.0 \| 1.1e-40 \| 2.2e-46 \| 315.6 \| 310 \| (4, 347) \| 471 \| (74, 410) \| 486 \| ATP-dependent DNA helicase recQ \| ATP-dependent DNA helicase recQ; uniclust \| UniRef100\_A0A174C369 \| 100.0 \| 1.5e-40 \| 2.9e-46 \| 330.2 \| 337 \| (3, 346) \| 471 \| (26, 415) \| 1025 \| Restriction/helicase domain-containing protein \| Restriction/helicase domain-containing protein; uniclust \| UniRef100\_UPI001E5A363B \| 100.0 \| 1.6e-40 \| 3e-46 \| 268.1 \| 243 \| (111, 362) \| 471 \| (3, 247) \| 259 \| DEAD/DEAH box helicase \| DEAD/DEAH box helicase; uniclust \| UniRef100\_A0A355M695 \| 100.0 \| 1.6e-40 \| 3e-46 \| 290.2 \| 327 \| (2, 353) \| 471 \| (8, 340) \| 486 \| DEAD/DEAH box helicase \| DEAD/DEAH box helicase; uniclust \| UniRef100\_A0A061N506 \| 100.0 \| 1.6e-40 \| 3.1e-46 \| 328.3 \| 300 \| (4, 340) \| 471 \| (143, 463) \| 707 \| ATP-dependent RNA helicase YqfR \| ATP-dependent RNA helicase YqfR; uniclust \| UniRef100\_A0A099I5T5 \| 100.0 \| 1.7e-40 \| 3.3e-46 \| 334.8 \| 312 \| (4, 353) \| 471 \| (178, 511) \| 851 \| Helicase \| Helicase; uniclust \| UniRef100\_A0A024C7J1 \| 100.0 \| 1.9e-40 \| 3.6e-46 \| 315.7 \| 304 \| (4, 345) \| 471 \| (91, 415) \| 602 \| DEAD/DEAH box helicase \| DEAD/DEAH box helicase; uniclust \| UniRef100\_A0A0F8ZGZ7 \| 100.0 \| 1.9e-40 \| 3.6e-46 \| 300.7 \| 321 \| (2, 346) \| 471 \| (94, 430) \| 465 \| Helicase C-terminal domain-containing protein (Fragment) \| Helicase C-terminal domain-containing protein (Fragment); uniclust \| UniRef100\_A0A5C7Q8J0 \| 100.0 \| 2.4e-40 \| 4.3e-46 \| 287.4 \| 393 \| (2, 401) \| 471 \| (3, 450) \| 459 \| Helicase ATP-binding domain-containing protein (Fragment) \| Helicase ATP-binding domain-containing protein (Fragment); uniclust \| UniRef100\_A0A369S3R1 \| 100.0 \| 2.6e-40 \| 4.7e-46 \| 299.7 \| 337 \| (4, 347) \| 471 \| (48, 415) \| 695 \| Putative mitochondrial ATP-dependent helicase irc3 \| Putative mitochondrial ATP-dependent helicase irc3; uniclust \| UniRef100\_A0A2H6AK24 \| 100.0 \| 2.5e-40 \| 4.8e-46 \| 317.8 \| 333 \| (4, 345) \| 471 \| (147, 517) \| 765 \| DNA repair helicase RadD \| DNA repair helicase RadD; uniclust \| UniRef100\_A0A093Z2C4 \| 100.0 \| 2.7e-40 \| 5e-46 \| 323.6 \| 338 \| (2, 346) \| 471 \| (597, 953) \| 1718 \| Helicase ATP-binding domain-containing protein \| Helicase ATP-binding domain-containing protein; uniclust \| UniRef100\_A0A0G0CC59 \| 100.0 \| 3.2e-40 \| 5.9e-46 \| 301.4 \| 351 \| (3, 361) \| 471 \| (108, 473) \| 757 \| Or RNA helicase of superfamily II protein \| Or RNA helicase of superfamily II protein; uniclust \| UniRef100\_A0A085LEP0 \| 100.0 \| 3.2e-40 \| 6.1e-46 \| 351.0 \| 345 \| (4, 355) \| 471 \| (1241, 1632) \| 1706 \| Helicase \| Helicase; uniclust \| UniRef100\_A0A059IV36 \| 100.0 \| 3.2e-40 \| 6.3e-46 \| 325.8 \| 297 \| (4, 339) \| 471 \| (128, 444) \| 681 \| DEAD-box ATP dependent DNA helicase \| DEAD-box ATP dependent DNA helicase; uniclust \| UniRef100\_A0A062VDK9 \| 100.0 \| 3.3e-40 \| 6.4e-46 \| 312.4 \| 297 \| (4, 339) \| 471 \| (67, 383) \| 574 \| DNA/RNA helicase, superfamily II \| DNA/RNA helicase, superfamily II; uniclust \| UniRef100\_A0A090VJF3 \| 100.0 \| 3.4e-40 \| 6.6e-46 \| 315.9 \| 301 \| (4, 344) \| 471 \| (103, 430) \| 536 \| ATP-dependent RNA helicase RhlE \| ATP-dependent RNA helicase RhlE; uniclust \| UniRef100\_A0A1G4A0W1 \| 100.0 \| 3.8e-40 \| 7e-46 \| 301.8 \| 332 \| (5, 347) \| 471 \| (197, 536) \| 658 \| DEAD/DEAH box helicase \| DEAD/DEAH box helicase; uniclust \| UniRef100\_A0A0L0RYB0 \| 100.0 \| 3.8e-40 \| 7e-46 \| 304.0 \| 340 \| (3, 349) \| 471 \| (186, 558) \| 849 \| P-loop containing nucleoside triphosphate hydrolase protein \| P-loop containing nucleoside triphosphate hydrolase protein; uniclust \| UniRef100\_A0A068N2T0 \| 100.0 \| 4e-40 \| 7.7e-46 \| 314.7 \| 304 \| (4, 345) \| 471 \| (122, 447) \| 578 \| DEAD/DEAH box helicase \| DEAD/DEAH box helicase; uniclust \| UniRef100\_F9FE71 \| 100.0 \| 4.3e-40 \| 7.8e-46 \| 311.5 \| 336 \| (3, 345) \| 471 \| (17, 377) \| 1182 \| ATP-dependent helicase IRC3 \| ATP-dependent helicase IRC3; uniclust \| UniRef100\_A0A014KUF5 \| 100.0 \| 4.4e-40 \| 8.4e-46 \| 335.1 \| 338 \| (3, 352) \| 471 \| (340, 723) \| 1077 \| Type I restriction modification system endonuclease (R) subunit, HsdR \| Type I restriction modification system endonuclease (R) subunit, HsdR; uniclust \| UniRef100\_A0A011QU62 \| 100.0 \| 4.5e-40 \| 8.6e-46 \| 318.3 \| 302 \| (4, 343) \| 471 \| (82, 405) \| 561 \| ATP-dependent RNA helicase YxiN \| ATP-dependent RNA helicase YxiN; uniclust \| UniRef100\_A0A357XX89 \| 100.0 \| 4.7e-40 \| 8.9e-46 \| 297.8 \| 229 \| (163, 406) \| 471 \| (5, 236) \| 417 \| Helicase C-terminal domain-containing protein (Fragment) \| Helicase C-terminal domain-containing protein (Fragment); uniclust \| UniRef100\_A0A059DMQ1 \| 100.0 \| 5.3e-40 \| 1e-45 \| 325.3 \| 300 \| (4, 341) \| 471 \| (95, 422) \| 701 \| DEAD/DEAH box helicase \| DEAD/DEAH box helicase; uniclust \| UniRef100\_A0A373MJZ5 \| 100.0 \| 6.8e-40 \| 1.3e-45 \| 307.1 \| 325 \| (2, 348) \| 471 \| (252, 588) \| 662 \| ATP-dependent helicase \| ATP-dependent helicase; uniclust \| UniRef100\_A0A059XBL3 \| 100.0 \| 7.2e-40 \| 1.4e-45 \| 314.7 \| 297 \| (4, 339) \| 471 \| (191, 509) \| 592 \| DEAD/DEAH box helicase (Fragment) \| DEAD/DEAH box helicase (Fragment); uniclust \| UniRef100\_A0A059X5H7 \| 100.0 \| 8.6e-40 \| 1.6e-45 \| 311.6 \| 295 \| (4, 337) \| 471 \| (113, 428) \| 536 \| DEAD/DEAH box helicase (Fragment) \| DEAD/DEAH box helicase (Fragment); uniclust \| UniRef100\_A0A873WKQ9 \| 100.0 \| 9.2e-40 \| 1.7e-45 \| 266.3 \| 265 \| (199, 463) \| 471 \| (2, 266) \| 277 \| DNA helicase \| DNA helicase; uniclust \| UniRef100\_A0A177QN03 \| 100.0 \| 8.8e-40 \| 1.7e-45 \| 312.4 \| 300 \| (4, 339) \| 471 \| (24, 346) \| 562 \| DEAD/DEAH box helicase \| DEAD/DEAH box helicase; uniclust \| UniRef100\_A0A133YXG2 \| 100.0 \| 9.8e-40 \| 1.9e-45 \| 314.0 \| 306 \| (4, 347) \| 471 \| (91, 422) \| 626 \| DEAD/DEAH box helicase \| DEAD/DEAH box helicase; uniclust \| UniRef100\_A0A2E8T8A0 \| 100.0 \| 1.1e-39 \| 2e-45 \| 282.5 \| 350 \| (3, 361) \| 471 \| (1, 367) \| 402 \| DEAD/DEAH box helicase (Fragment) \| DEAD/DEAH box helicase (Fragment); uniclust \| UniRef100\_A0A096KUJ1 \| 100.0 \| 1.1e-39 \| 2e-45 \| 320.4 \| 296 \| (4, 337) \| 471 \| (143, 469) \| 802 \| DEAD/DEAH box helicase \| DEAD/DEAH box helicase; uniclust \| UniRef100\_A0A3S0BS24 \| 100.0 \| 1.1e-39 \| 2.1e-45 \| 281.2 \| 389 \| (3, 466) \| 471 \| (4, 401) \| 431 \| Helicase C-terminal domain-containing protein \| Helicase C-terminal domain-containing protein; uniclust \| UniRef100\_A0A090PWV5 \| 100.0 \| 1.1e-39 \| 2.1e-45 \| 307.1 \| 304 \| (4, 345) \| 471 \| (50, 376) \| 503 \| ATP-dependent RNA helicase \| ATP-dependent RNA helicase; uniclust \| UniRef100\_A0A6J5LKN8 \| 100.0 \| 1.2e-39 \| 2.2e-45 \| 282.1 \| 349 \| (50, 420) \| 471 \| (7, 368) \| 446 \| SSL2 DNA or RNA helicases of superfamily II \| SSL2 DNA or RNA helicases of superfamily II; uniclust \| UniRef100\_A0A062XLX7 \| 100.0 \| 1.2e-39 \| 2.3e-45 \| 338.2 \| 328 \| (3, 343) \| 471 \| (305, 692) \| 776 \| Putative DNA-repair helicase \| Putative DNA-repair helicase; uniclust \| UniRef100\_A0A2G0Q3E4 \| 100.0 \| 1.4e-39 \| 2.6e-45 \| 287.9 \| 311 \| (4, 326) \| 471 \| (5, 324) \| 543 \| Helicase \| Helicase; uniclust \| UniRef100\_A0A3N4IK77 \| 100.0 \| 1.4e-39 \| 2.6e-45 \| 299.4 \| 336 \| (3, 345) \| 471 \| (75, 432) \| 819 \| P-loop containing nucleoside triphosphate hydrolase protein \| P-loop containing nucleoside triphosphate hydrolase protein; uniclust \| UniRef100\_A0A0B7NDA0 \| 100.0 \| 1.5e-39 \| 2.7e-45 \| 311.4 \| 316 \| (24, 349) \| 471 \| (375, 711) \| 970 \| Uncharacterized protein \| Uncharacterized protein; uniclust \| UniRef100\_A0A235B1F3 \| 100.0 \| 1.5e-39 \| 2.8e-45 \| 290.9 \| 247 \| (108, 361) \| 471 \| (14, 265) \| 429 \| Helicase C-terminal domain-containing protein (Fragment) \| Helicase C-terminal domain-containing protein (Fragment); uniclust \| UniRef100\_UPI001E5A22BD \| 100.0 \| 1.5e-39 \| 2.8e-45 \| 293.0 \| 342 \| (2, 351) \| 471 \| (4, 373) \| 652 \| DEAD/DEAH box helicase \| DEAD/DEAH box helicase; uniclust \| UniRef100\_A0A0R2HE16 \| 100.0 \| 1.5e-39 \| 2.9e-45 \| 314.9 \| 296 \| (4, 339) \| 471 \| (93, 415) \| 691 \| ATP-dependent RNA helicase \| ATP-dependent RNA helicase; uniclust \| UniRef100\_A0A015S7S3 \| 100.0 \| 1.5e-39 \| 3e-45 \| 328.9 \| 338 \| (3, 352) \| 471 \| (253, 636) \| 928 \| type I site-specific deoxyribonuclease \| type I site-specific deoxyribonuclease; uniclust \| UniRef100\_A0A1Q6KXU7 \| 100.0 \| 1.8e-39 \| 3.4e-45 \| 305.2 \| 331 \| (3, 349) \| 471 \| (205, 547) \| 686 \| Uncharacterized protein \| Uncharacterized protein; uniclust \| UniRef100\_A0A947JEL1 \| 100.0 \| 2e-39 \| 3.7e-45 \| 267.0 \| 254 \| (2, 258) \| 471 \| (3, 257) \| 258 \| DEAD/DEAH box helicase family protein (Fragment) \| DEAD/DEAH box helicase family protein (Fragment); uniclust \| UniRef100\_A0A017TC97 \| 100.0 \| 2.1e-39 \| 4e-45 \| 317.5 \| 312 \| (4, 352) \| 471 \| (108, 441) \| 713 \| Cold-shock DEAD-box protein A \| Cold-shock DEAD-box protein A; uniclust \| UniRef100\_A0A0C9NPH8 \| 100.0 \| 2.3e-39 \| 4.3e-45 \| 317.6 \| 323 \| (3, 345) \| 471 \| (477, 815) \| 867 \| DNA or RNA helicases of superfamily II \| DNA or RNA helicases of superfamily II; uniclust \| UniRef100\_A0A090AD17 \| 100.0 \| 2.4e-39 \| 4.5e-45 \| 301.7 \| 303 \| (4, 344) \| 471 \| (34, 362) \| 481 \| ATP-dependent RNA helicase RhlE \| ATP-dependent RNA helicase RhlE; uniclust \| UniRef100\_A0A031GEQ0 \| 100.0 \| 2.6e-39 \| 5e-45 \| 338.6 \| 333 \| (5, 350) \| 471 \| (363, 752) \| 1283 \| Uncharacterized protein \| Uncharacterized protein; uniclust \| UniRef100\_A0A1G5X3X8 \| 100.0 \| 2.8e-39 \| 5.4e-45 \| 320.7 \| 327 \| (3, 343) \| 471 \| (302, 692) \| 770 \| Superfamily II DNA or RNA helicase \| Superfamily II DNA or RNA helicase; uniclust \| UniRef100\_A0A0B8PE49 \| 100.0 \| 3e-39 \| 5.8e-45 \| 314.6 \| 339 \| (3, 352) \| 471 \| (192, 581) \| 733 \| Type I restriction-modification system \| Type I restriction-modification system; uniclust \| UniRef100\_A0A844Y7C8 \| 100.0 \| 3.2e-39 \| 5.9e-45 \| 270.0 \| 225 \| (2, 234) \| 471 \| (11, 238) \| 251 \| DEAD/DEAH box helicase \| DEAD/DEAH box helicase; uniclust \| UniRef100\_A0A1C4TDC2 \| 100.0 \| 3.1e-39 \| 6e-45 \| 324.1 \| 344 \| (3, 353) \| 471 \| (677, 1066) \| 1087 \| Superfamily II DNA or RNA helicase \| Superfamily II DNA or RNA helicase; uniclust \| UniRef100\_A0A0R2SWX3 \| 100.0 \| 3.1e-39 \| 6e-45 \| 307.0 \| 312 \| (4, 353) \| 471 \| (65, 399) \| 556 \| ATP-dependent RNA helicase \| ATP-dependent RNA helicase; uniclust \| UniRef100\_A0A010RIX9 \| 100.0 \| 3.2e-39 \| 6.3e-45 \| 334.1 \| 312 \| (4, 352) \| 471 \| (311, 644) \| 1043 \| ATP-dependent RNA helicase DeaD \| ATP-dependent RNA helicase DeaD; uniclust \| UniRef100\_A0A0P6YGE4 \| 100.0 \| 3.4e-39 \| 6.5e-45 \| 322.2 \| 304 \| (4, 344) \| 471 \| (152, 476) \| 780 \| RNA helicase \| RNA helicase; uniclust \| UniRef100\_A0A060HM29 \| 100.0 \| 3.4e-39 \| 6.5e-45 \| 307.5 \| 303 \| (4, 344) \| 471 \| (133, 456) \| 624 \| DEAD-box ATP-dependent RNA helicase \| DEAD-box ATP-dependent RNA helicase; uniclust \| UniRef100\_A0A094NXF7 \| 100.0 \| 3.6e-39 \| 6.8e-45 \| 301.3 \| 314 \| (3, 339) \| 471 \| (135, 497) \| 561 \| DNA helicase (Fragment) \| DNA helicase (Fragment); uniclust \| UniRef100\_A0A1B6Y742 \| 100.0 \| 3.8e-39 \| 7.2e-45 \| 304.5 \| 295 \| (4, 337) \| 471 \| (23, 340) \| 544 \| DEAD/DEAH box helicase \| DEAD/DEAH box helicase; uniclust \| UniRef100\_A0A1E4ZY87 \| 100.0 \| 4e-39 \| 7.4e-45 \| 283.5 \| 340 \| (4, 355) \| 471 \| (2, 349) \| 516 \| DEAD/DEAH box helicase \| DEAD/DEAH box helicase; uniclust \| UniRef100\_A0A0R1EYB2 \| 100.0 \| 4.1e-39 \| 7.9e-45 \| 298.7 \| 305 \| (4, 352) \| 471 \| (84, 412) \| 485 \| ATP-dependent DNA helicase (RecQ) \| ATP-dependent DNA helicase (RecQ); uniclust \| UniRef100\_A0A059X7D8 \| 100.0 \| 4.2e-39 \| 8.1e-45 \| 312.4 \| 304 \| (4, 349) \| 471 \| (93, 420) \| 665 \| ATP-dependent DNA helicase, RecQ family (Fragment) \| ATP-dependent DNA helicase, RecQ family (Fragment); uniclust \| UniRef100\_A0A094FXL0 \| 100.0 \| 4.4e-39 \| 8.1e-45 \| 305.6 \| 338 \| (2, 346) \| 471 \| (615, 971) \| 1228 \| Helicase ATP-binding domain-containing protein \| Helicase ATP-binding domain-containing protein; uniclust \| UniRef100\_A0A5B9MCL1 \| 100.0 \| 4.4e-39 \| 8.2e-45 \| 308.0 \| 333 \| (3, 349) \| 471 \| (37, 393) \| 769 \| ATP-dependent RNA helicase RhlE \| ATP-dependent RNA helicase RhlE; uniclust \| UniRef100\_A0A022N5P9 \| 100.0 \| 4.4e-39 \| 8.6e-45 \| 316.2 \| 301 \| (4, 343) \| 471 \| (78, 399) \| 588 \| RNA helicase \| RNA helicase; uniclust \| UniRef100\_A0A5C7P768 \| 100.0 \| 4.7e-39 \| 8.6e-45 \| 291.7 \| 452 \| (1, 467) \| 471 \| (1, 577) \| 598 \| DEAD/DEAH box helicase \| DEAD/DEAH box helicase; uniclust \| UniRef100\_A0A0B4N044 \| 100.0 \| 5.1e-39 \| 9.6e-45 \| 315.1 \| 301 \| (4, 344) \| 471 \| (157, 482) \| 893 \| Putative DEAD/DEAH box helicase \| Putative DEAD/DEAH box helicase; uniclust \| UniRef100\_A0A1C6H4Y0 \| 100.0 \| 5.1e-39 \| 9.8e-45 \| 307.9 \| 339 \| (3, 348) \| 471 \| (146, 560) \| 587 \| Type I restriction enzyme EcoKI subunit R \| Type I restriction enzyme EcoKI subunit R; uniclust \| UniRef100\_A0A015TVL0 \| 100.0 \| 5.1e-39 \| 9.8e-45 \| 311.1 \| 300 \| (4, 342) \| 471 \| (101, 419) \| 572 \| Helicase conserved C-terminal domain protein \| Helicase conserved C-terminal domain protein; uniclust \| UniRef100\_A0A139KGF0 \| 100.0 \| 5.6e-39 \| 1.1e-44 \| 309.3 \| 303 \| (4, 342) \| 471 \| (85, 412) \| 654 \| Putative cold-shock DEAD-box protein A \| Putative cold-shock DEAD-box protein A; uniclust \| UniRef100\_A0A521UGQ1 \| 100.0 \| 6.1e-39 \| 1.1e-44 \| 278.5 \| 375 \| (3, 416) \| 471 \| (6, 388) \| 455 \| Helicase ATP-binding domain-containing protein \| Helicase ATP-binding domain-containing protein; uniclust \| UniRef100\_A0A031LM96 \| 100.0 \| 6.1e-39 \| 1.2e-44 \| 294.5 \| 291 \| (4, 340) \| 471 \| (66, 374) \| 466 \| RNA helicase \| RNA helicase; uniclust \| UniRef100\_A0A0G0WNU3 \| 100.0 \| 6.2e-39 \| 1.2e-44 \| 319.7 \| 333 \| (4, 348) \| 471 \| (181, 574) \| 884 \| Type I site-specific deoxyribonuclease \| Type I site-specific deoxyribonuclease; uniclust \| UniRef100\_A0A061QIW8 \| 100.0 \| 6.2e-39 \| 1.2e-44 \| 316.1 \| 302 \| (4, 343) \| 471 \| (97, 425) \| 735 \| ATP-dependent RNA helicase RhlE \| ATP-dependent RNA helicase RhlE; uniclust \| UniRef100\_A0A0G0GL02 \| 100.0 \| 6.3e-39 \| 1.2e-44 \| 308.5 \| 296 \| (4, 339) \| 471 \| (105, 421) \| 682 \| Dead-box ATP-dependent RNA helicase \| Dead-box ATP-dependent RNA helicase; uniclust \| UniRef100\_A0A074LXA7 \| 100.0 \| 6.1e-39 \| 1.2e-44 \| 330.5 \| 329 \| (3, 344) \| 471 \| (288, 678) \| 766 \| DNA repair helicase \| DNA repair helicase; uniclust \| UniRef100\_A0A497HKE1 \| 100.0 \| 6.8e-39 \| 1.3e-44 \| 292.8 \| 327 \| (2, 343) \| 471 \| (14, 416) \| 479 \| DEAD/DEAH box helicase \| DEAD/DEAH box helicase; uniclust \| UniRef100\_A0A090Q5R9 \| 100.0 \| 6.9e-39 \| 1.3e-44 \| 319.1 \| 340 \| (5, 356) \| 471 \| (368, 762) \| 902 \| Type I restriction-modification system restriction subunit R \| Type I restriction-modification system restriction subunit R; uniclust \| UniRef100\_A0A011M776 \| 100.0 \| 7.4e-39 \| 1.4e-44 \| 314.6 \| 303 \| (4, 344) \| 471 \| (150, 481) \| 732 \| ATP-dependent RNA helicase RhlE \| ATP-dependent RNA helicase RhlE; uniclust \| UniRef100\_A0A938MTN5 \| 100.0 \| 8.1e-39 \| 1.5e-44 \| 283.1 \| 343 \| (2, 358) \| 471 \| (1, 348) \| 541 \| DEAD/DEAH box helicase \| DEAD/DEAH box helicase; uniclust \| UniRef100\_A0A1H9P6L4 \| 100.0 \| 8.2e-39 \| 1.6e-44 \| 303.1 \| 307 \| (4, 351) \| 471 \| (64, 393) \| 543 \| ATP-dependent DNA helicase RecQ \| ATP-dependent DNA helicase RecQ; uniclust \| UniRef100\_A0A1Q9NU84 \| 100.0 \| 8.2e-39 \| 1.6e-44 \| 304.3 \| 317 \| (2, 341) \| 471 \| (165, 528) \| 555 \| DNA helicase \| DNA helicase; uniclust \| UniRef100\_A0A090VRU3 \| 100.0 \| 8.4e-39 \| 1.6e-44 \| 300.4 \| 305 \| (4, 346) \| 471 \| (80, 408) \| 527 \| DEAD-box ATP-dependent RNA helicase CshA \| DEAD-box ATP-dependent RNA helicase CshA; uniclust \| UniRef100\_A0A059LDH5 \| 100.0 \| 8.4e-39 \| 1.6e-44 \| 305.3 \| 294 \| (4, 334) \| 471 \| (96, 425) \| 583 \| RNA helicase (Fragment) \| RNA helicase (Fragment); uniclust \| UniRef100\_UPI0021B548A4 \| 100.0 \| 9.3e-39 \| 1.7e-44 \| 282.3 \| 333 \| (4, 355) \| 471 \| (8, 344) \| 534 \| DEAD/DEAH box helicase \| DEAD/DEAH box helicase; uniclust \| UniRef100\_A0A6A5YFT1 \| 100.0 \| 9.7e-39 \| 1.8e-44 \| 314.0 \| 335 \| (3, 345) \| 471 \| (50, 403) \| 1677 \| GPI inositol-deacylase \| GPI inositol-deacylase; uniclust \| UniRef100\_A0A0T6A863 \| 100.0 \| 9.4e-39 \| 1.8e-44 \| 309.4 \| 299 \| (4, 340) \| 471 \| (108, 429) \| 710 \| ATP-dependent RNA helicase \| ATP-dependent RNA helicase; uniclust \| UniRef100\_A0A011TJA1 \| 100.0 \| 9.7e-39 \| 1.9e-44 \| 315.8 \| 302 \| (4, 343) \| 471 \| (130, 458) \| 720 \| RNA helicase \| RNA helicase; uniclust \| UniRef100\_A0A0P7WKD9 \| 100.0 \| 1.1e-38 \| 2.1e-44 \| 311.1 \| 299 \| (4, 341) \| 471 \| (55, 381) \| 683 \| ATP-dependent RNA helicase RhlE \| ATP-dependent RNA helicase RhlE; uniclust \| UniRef100\_A0A011QWJ0 \| 100.0 \| 1.1e-38 \| 2.1e-44 \| 316.0 \| 301 \| (4, 345) \| 471 \| (56, 378) \| 569 \| ATP-dependent DNA helicase RecQ \| ATP-dependent DNA helicase RecQ; uniclust \| UniRef100\_A0A022N706 \| 100.0 \| 1.1e-38 \| 2.3e-44 \| 329.7 \| 306 \| (4, 352) \| 471 \| (149, 478) \| 894 \| DNA helicase \| DNA helicase; uniclust \| UniRef100\_A0A0P9UNN4 \| 100.0 \| 1.2e-38 \| 2.3e-44 \| 315.3 \| 304 \| (4, 348) \| 471 \| (178, 503) \| 767 \| ATP-dependent DNA helicase RecQ \| ATP-dependent DNA helicase RecQ; uniclust \| UniRef100\_A0A7C8NB46 \| 100.0 \| 1.3e-38 \| 2.4e-44 \| 316.1 \| 336 \| (3, 345) \| 471 \| (23, 377) \| 2893 \| Lon protease homolog, mitochondrial \| Lon protease homolog, mitochondrial; uniclust \| UniRef100\_A0A0F8YQH4 \| 100.0 \| 1.2e-38 \| 2.4e-44 \| 298.8 \| 301 \| (4, 346) \| 471 \| (84, 408) \| 548 \| Helicase ATP-binding domain-containing protein (Fragment) \| Helicase ATP-binding domain-containing protein (Fragment); uniclust \| UniRef100\_A0A009N196 \| 100.0 \| 1.3e-38 \| 2.5e-44 \| 337.8 \| 333 \| (5, 349) \| 471 \| (586, 979) \| 1481 \| type I site-specific deoxyribonuclease \| type I site-specific deoxyribonuclease; uniclust \| UniRef100\_A0A177R869 \| 100.0 \| 1.4e-38 \| 2.7e-44 \| 301.1 \| 293 \| (4, 335) \| 471 \| (74, 387) \| 537 \| RNA helicase \| RNA helicase; uniclust \| UniRef100\_A0A8H6ACR5 \| 100.0 \| 1.5e-38 \| 2.8e-44 \| 304.5 \| 337 \| (2, 345) \| 471 \| (50, 405) \| 1390 \| P-loop containing nucleoside triphosphate hydrolase protein \| P-loop containing nucleoside triphosphate hydrolase protein; uniclust \| UniRef100\_UPI00141F429B \| 100.0 \| 1.7e-38 \| 3e-44 \| 290.0 \| 372 \| (70, 467) \| 471 \| (318, 702) \| 744 \| helicase-related protein \| helicase-related protein; uniclust \| UniRef100\_UPI00221F344C \| 100.0 \| 1.7e-38 \| 3.1e-44 \| 273.4 \| 321 \| (4, 343) \| 471 \| (56, 400) \| 422 \| P-loop containing nucleoside triphosphate hydrolase protein \| P-loop containing nucleoside triphosphate hydrolase protein; uniclust \| UniRef100\_A0A150QSB4 \| 100.0 \| 1.7e-38 \| 3.2e-44 \| 306.0 \| 338 \| (3, 347) \| 471 \| (267, 673) \| 703 \| DEAD/DEAH box helicase \| DEAD/DEAH box helicase; uniclust \| UniRef100\_A0A0B5B838 \| 100.0 \| 1.8e-38 \| 3.4e-44 \| 309.9 \| 300 \| (4, 342) \| 471 \| (119, 441) \| 704 \| RNA helicase \| RNA helicase; uniclust \| UniRef100\_A0A139DPM2 \| 100.0 \| 1.9e-38 \| 3.6e-44 \| 299.6 \| 334 \| (4, 347) \| 471 \| (51, 425) \| 871 \| Type III restriction protein res subunit \| Type III restriction protein res subunit; uniclust \| UniRef100\_A0A0A8X3C0 \| 100.0 \| 1.9e-38 \| 3.6e-44 \| 305.1 \| 308 \| (4, 351) \| 471 \| (87, 419) \| 677 \| ATP-dependent RNA helicase BA2475 \| ATP-dependent RNA helicase BA2475; uniclust \| UniRef100\_A0A075HY00 \| 100.0 \| 2.1e-38 \| 3.9e-44 \| 290.0 \| 311 \| (4, 353) \| 471 \| (37, 367) \| 404 \| Dead/deah box helicase domain-containing protein (DeaD) \| Dead/deah box helicase domain-containing protein (DeaD); uniclust \| UniRef100\_A0A059DQS2 \| 100.0 \| 2.2e-38 \| 4.1e-44 \| 308.2 \| 304 \| (4, 345) \| 471 \| (150, 478) \| 843 \| RNA helicase \| RNA helicase; uniclust \| UniRef100\_A0A1Q7B0N5 \| 100.0 \| 2.4e-38 \| 4.5e-44 \| 290.6 \| 303 \| (4, 346) \| 471 \| (96, 423) \| 531 \| DEAD/DEAH box helicase \| DEAD/DEAH box helicase; uniclust \| UniRef100\_A0A0C2IER9 \| 100.0 \| 2.5e-38 \| 4.8e-44 \| 299.5 \| 307 \| (4, 347) \| 471 \| (87, 417) \| 512 \| RNA helicase \| RNA helicase; uniclust \| UniRef100\_A0A1J4UEU0 \| 100.0 \| 2.6e-38 \| 5.1e-44 \| 304.9 \| 340 \| (2, 363) \| 471 \| (48, 507) \| 576 \| DEAD/DEAH box helicase \| DEAD/DEAH box helicase; uniclust \| UniRef100\_A0A015V834 \| 100.0 \| 2.8e-38 \| 5.4e-44 \| 326.8 \| 304 \| (4, 352) \| 471 \| (190, 517) \| 992 \| DNA helicase \| DNA helicase; uniclust \| UniRef100\_A0A0F9ZSQ4 \| 100.0 \| 3e-38 \| 5.6e-44 \| 298.2 \| 330 \| (3, 345) \| 471 \| (104, 446) \| 859 \| Or RNA helicase of superfamily II-like protein \| Or RNA helicase of superfamily II-like protein; uniclust \| UniRef100\_A0A010YCY8 \| 100.0 \| 3.1e-38 \| 5.9e-44 \| 317.8 \| 306 \| (4, 353) \| 471 \| (109, 438) \| 841 \| DNA helicase \| DNA helicase; uniclust \| UniRef100\_A0A2E6BRS8 \| 100.0 \| 3.2e-38 \| 5.9e-44 \| 304.2 \| 336 \| (3, 353) \| 471 \| (48, 402) \| 934 \| Restriction endonuclease subunit R \| Restriction endonuclease subunit R; uniclust \| UniRef100\_A0A0B3B2U7 \| 100.0 \| 3.2e-38 \| 6.1e-44 \| 296.0 \| 297 \| (4, 339) \| 471 \| (86, 402) \| 561 \| DEAD/DEAH box helicase-like protein \| DEAD/DEAH box helicase-like protein; uniclust \| UniRef100\_A0A061QCN4 \| 100.0 \| 3.3e-38 \| 6.4e-44 \| 308.3 \| 300 \| (4, 340) \| 471 \| (110, 439) \| 671 \| ATP-dependent RNA helicase RhlB \| ATP-dependent RNA helicase RhlB; uniclust \| UniRef100\_A0A5N7MB36 \| 100.0 \| 3.4e-38 \| 6.4e-44 \| 295.5 \| 345 \| (4, 355) \| 471 \| (16, 388) \| 580 \| DEAD/DEAH box helicase \| DEAD/DEAH box helicase; uniclust \| UniRef100\_A0A017RY23 \| 100.0 \| 3.3e-38 \| 6.4e-44 \| 327.2 \| 333 \| (6, 350) \| 471 \| (468, 856) \| 1252 \| Type III restriction endonuclease subunit R \| Type III restriction endonuclease subunit R; uniclust \| UniRef100\_A0A0Q9YTK0 \| 100.0 \| 3.5e-38 \| 6.8e-44 \| 309.2 \| 302 \| (4, 344) \| 471 \| (97, 420) \| 714 \| DEAD-box ATP-dependent RNA helicase CshA \| DEAD-box ATP-dependent RNA helicase CshA; uniclust \| UniRef100\_A0A4T0FKT3 \| 100.0 \| 3.9e-38 \| 7.1e-44 \| 308.0 \| 340 \| (3, 349) \| 471 \| (1177, 1543) \| 1791 \| Mitochondrial presequence protease \| Mitochondrial presequence protease; uniclust \| UniRef100\_A0A1G3A7P0 \| 100.0 \| 3.8e-38 \| 7.2e-44 \| 298.3 \| 299 \| (4, 340) \| 471 \| (226, 547) \| 632 \| DEAD/DEAH box helicase \| DEAD/DEAH box helicase; uniclust \| UniRef100\_A0A097IEA7 \| 100.0 \| 4.2e-38 \| 8e-44 \| 305.0 \| 298 \| (4, 337) \| 471 \| (154, 479) \| 633 \| DEAD/DEAH box helicase \| DEAD/DEAH box helicase; uniclust \| UniRef100\_A0A084SY92 \| 100.0 \| 4.3e-38 \| 8.2e-44 \| 311.4 \| 299 \| (4, 342) \| 471 \| (56, 378) \| 786 \| DEAD/DEAH box helicase \| DEAD/DEAH box helicase; uniclust \| UniRef100\_A5A3T1 \| 100.0 \| 4.5e-38 \| 8.2e-44 \| 281.5 \| 389 \| (2, 405) \| 471 \| (1, 467) \| 600 \| BcepGomrgp42 \| BcepGomrgp42; uniclust \| UniRef100\_A0A2C8EFP9 \| 100.0 \| 4.4e-38 \| 8.3e-44 \| 283.4 \| 328 \| (4, 348) \| 471 \| (43, 387) \| 422 \| Helicase ATP-binding domain-containing protein \| Helicase ATP-binding domain-containing protein; uniclust \| UniRef100\_A0A016VEN6 \| 100.0 \| 4.5e-38 \| 8.7e-44 \| 306.9 \| 309 \| (4, 345) \| 471 \| (192, 526) \| 697 \| RNA helicase \| RNA helicase; uniclust \| UniRef100\_A0A089JXU6 \| 100.0 \| 4.6e-38 \| 8.8e-44 \| 311.7 \| 338 \| (3, 352) \| 471 \| (198, 581) \| 908 \| Restriction endonuclease subunit R \| Restriction endonuclease subunit R; uniclust \| UniRef100\_A0A063BKQ2 \| 100.0 \| 4.7e-38 \| 9e-44 \| 305.1 \| 304 \| (4, 339) \| 471 \| (368, 701) \| 765 \| Uncharacterized protein \| Uncharacterized protein; uniclust \| UniRef100\_A0A0F9YWH9 \| 100.0 \| 4.8e-38 \| 9.3e-44 \| 306.8 \| 308 \| (4, 351) \| 471 \| (56, 388) \| 638 \| DEAD/DEAH box helicase \| DEAD/DEAH box helicase; uniclust \| UniRef100\_A0A098QYC1 \| 100.0 \| 4.9e-38 \| 9.4e-44 \| 308.1 \| 302 \| (4, 342) \| 471 \| (69, 395) \| 673 \| DEAD/DEAH box helicase \| DEAD/DEAH box helicase; uniclust \| UniRef100\_A0A143ZV74 \| 100.0 \| 5e-38 \| 9.5e-44 \| 300.3 \| 305 \| (4, 352) \| 471 \| (54, 382) \| 635 \| ATP-dependent DNA helicase RecQ \| ATP-dependent DNA helicase RecQ; uniclust \| UniRef100\_A0A3M7N0V4 \| 100.0 \| 5.3e-38 \| 9.8e-44 \| 298.2 \| 338 \| (2, 346) \| 471 \| (542, 923) \| 1220 \| Uncharacterized protein (Fragment) \| Uncharacterized protein (Fragment); uniclust \| UniRef100\_A0A0F9S804 \| 100.0 \| 5.1e-38 \| 9.9e-44 \| 307.9 \| 314 \| (3, 344) \| 471 \| (95, 435) \| 536 \| Helicase ATP-binding domain-containing protein \| Helicase ATP-binding domain-containing protein; uniclust \| UniRef100\_A0A069SX40 \| 100.0 \| 5.2e-38 \| 9.9e-44 \| 307.7 \| 328 \| (3, 345) \| 471 \| (97, 477) \| 707 \| DEAD/DEAH box helicase \| DEAD/DEAH box helicase; uniclust \| UniRef100\_A0A010QKI1 \| 100.0 \| 5.3e-38 \| 1e-43 \| 305.1 \| 304 \| (4, 344) \| 471 \| (271, 604) \| 750 \| DEAD/DEAH box helicase \| DEAD/DEAH box helicase; uniclust \| UniRef100\_A0A2Z6IA22 \| 100.0 \| 5.8e-38 \| 1.1e-43 \| 298.4 \| 330 \| (5, 348) \| 471 \| (246, 603) \| 790 \| Helicase \| Helicase; uniclust \| UniRef100\_A0A015KPS7 \| 100.0 \| 6.6e-38 \| 1.3e-43 \| 308.0 \| 304 \| (3, 343) \| 471 \| (116, 441) \| 599 \| DEAD/DEAH box helicase \| DEAD/DEAH box helicase; uniclust \| UniRef100\_UPI0014432E6F \| 100.0 \| 7e-38 \| 1.3e-43 \| 281.0 \| 329 \| (6, 349) \| 471 \| (145, 488) \| 616 \| DEAD/DEAH box helicase \| DEAD/DEAH box helicase; uniclust \| UniRef100\_A0A0E3UHL8 \| 100.0 \| 7e-38 \| 1.3e-43 \| 295.5 \| 304 \| (4, 345) \| 471 \| (185, 509) \| 576 \| DEAD/DEAH box helicase \| DEAD/DEAH box helicase; pdb70 \| 2YKG\_A \| 99.9 \| 2.6e-29 \| 2e-33 \| 258.0 \| 190 \| (213, 410) \| 471 \| (379, 612) \| 696 \| PROBABLE ATP-DEPENDENT RNA HELICASE DDX58 \| 2YKG\_A PROBABLE ATP-DEPENDENT RNA HELICASE DDX58 HYDROLASE, INNATE IMMUNITY; pdb70 \| 4BPB\_A \| 99.9 \| 3.1e-29 \| 2.3e-33 \| 257.5 \| 189 \| (213, 409) \| 471 \| (379, 613) \| 698 \| PROBABLE ATP-DEPENDENT RNA HELICASE DDX58 \| 4BPB\_A PROBABLE ATP-DEPENDENT RNA HELICASE DDX58 HYDROLASE-RNA COMPLEX, ADENOSINE TRIPHOSPHATE, DEAD-BOX; pdb70 \| 5F9F\_I \| 99.9 \| 5.5e-28 \| 4.1e-32 \| 248.0 \| 191 \| (213, 411) \| 471 \| (378, 612) \| 695 \| Probable ATP-dependent RNA helicase DDX58/RNA \| 5F9F\_I Probable ATP-dependent RNA helicase DDX58/RNA Complex, RIG-I, capped RNA, self HET: BU3, ETF; pdb70 \| 5JB2\_A \| 99.9 \| 2.6e-27 \| 1.9e-31 \| 242.5 \| 189 \| (213, 409) \| 471 \| (352, 590) \| 680 \| LGP2/RNA Complex \| 5JB2\_A LGP2/RNA Complex Innate immune pattern recognition receptor HET: GTP, ADP; pdb70 \| 5JAJ\_A \| 99.9 \| 3e-27 \| 2.2e-31 \| 242.0 \| 189 \| (213, 409) \| 471 \| (353, 591) \| 681 \| LGP2/RNA Complex \| 5JAJ\_A LGP2/RNA Complex Innate immune pattern recognition receptor HET: EDO, ADP; pdb70 \| 6JDE\_B \| 99.9 \| 7.2e-27 \| 5.3e-31 \| 234.0 \| 398 \| (1, 412) \| 471 \| (1, 465) \| 586 \| Putative DNA repair helicase RadD \| 6JDE\_B Putative DNA repair helicase RadD DNA repair protein, HYDROLASE; pdb70 \| 6GKM\_A \| 99.8 \| 3.3e-26 \| 2.4e-30 \| 242.4 \| 192 \| (214, 413) \| 471 \| (681, 923) \| 1007 \| Interferon-induced helicase C domain-containing protein \| 6GKM\_A Interferon-induced helicase C domain-containing protein Protein-RNA complex, helical filament, ATPase HET: ATP; pdb70 \| 5JC3\_A \| 99.8 \| 8.4e-25 \| 6.3e-29 \| 224.4 \| 192 \| (213, 412) \| 471 \| (380, 621) \| 701 \| LGP2/RNA Complex \| 5JC3\_A LGP2/RNA Complex Innate immune pattern recognition receptor HET: ADP; pdb70 \| 5JCH\_A \| 99.8 \| 8.4e-25 \| 6.3e-29 \| 224.4 \| 192 \| (213, 412) \| 471 \| (380, 621) \| 701 \| LGP2/RNA Complex \| 5JCH\_A LGP2/RNA Complex Innate immune pattern recognition receptor HET: ADP; pdb70 \| 4A2W\_B \| 99.8 \| 1.3e-24 \| 9.7e-29 \| 228.0 \| 189 \| (213, 409) \| 471 \| (612, 845) \| 936 \| RETINOIC ACID INDUCIBLE PROTEIN I \| 4A2W\_B RETINOIC ACID INDUCIBLE PROTEIN I HYDROLASE, SUPERFAMILY 2 RNA HELICASE; pdb70 \| 4A2W\_A \| 99.8 \| 2.3e-24 \| 1.7e-28 \| 226.1 \| 189 \| (213, 409) \| 471 \| (612, 845) \| 936 \| RETINOIC ACID INDUCIBLE PROTEIN I \| 4A2W\_A RETINOIC ACID INDUCIBLE PROTEIN I HYDROLASE, SUPERFAMILY 2 RNA HELICASE; pdb70 \| 6G19\_A \| 99.8 \| 4.5e-24 \| 3.4e-28 \| 218.8 \| 194 \| (214, 415) \| 471 \| (375, 619) \| 696 \| Interferon-induced helicase C domain-containing protein \| 6G19\_A Interferon-induced helicase C domain-containing protein Protein-RNA complex, helical filament, ATPase HET: ANP; pdb70 \| 2OCA\_A \| 99.8 \| 1.2e-23 \| 8.5e-28 \| 207.6 \| 326 \| (3, 343) \| 471 \| (112, 456) \| 510 \| ATP-dependent DNA helicase uvsW (E.C.3.6.1.8) \| 2OCA\_A ATP-dependent DNA helicase uvsW (E.C.3.6.1.8) ATP-dependant helicase, T4-bacteriophage, Recombination, HYDROLASE; pdb70 \| 5SUQ\_C \| 99.6 \| 3.4e-20 \| 2.5e-24 \| 179.1 \| 299 \| (3, 338) \| 471 \| (83, 408) \| 446 \| ATP-dependent RNA helicase SUB2 (E.C.3.6.4.13) \| 5SUQ\_C ATP-dependent RNA helicase SUB2 (E.C.3.6.4.13) mRNA export, HYDROLASE HET: KEG; pdb70 \| 4A2Q\_B \| 99.6 \| 1.3e-19 \| 9.9e-24 \| 187.3 \| 113 \| (213, 332) \| 471 \| (612, 740) \| 797 \| RETINOIC ACID INDUCIBLE PROTEIN I \| 4A2Q\_B RETINOIC ACID INDUCIBLE PROTEIN I HYDROLASE, SUPERFAMILY 2 RNA HELICASE; pdb70 \| 2V1X\_B \| 99.6 \| 1.6e-19 \| 1.2e-23 \| 180.4 \| 295 \| (4, 337) \| 471 \| (44, 368) \| 591 \| ATP-DEPENDENT DNA HELICASE Q1 (E.C.3.6.1.-) \| 2V1X\_B ATP-DEPENDENT DNA HELICASE Q1 (E.C.3.6.1.-) DNA STRAND ANNEALING, MISMATCH REPAIR HET: ADP; pdb70 \| 2WWY\_B \| 99.6 \| 1.6e-19 \| 1.2e-23 \| 180.4 \| 295 \| (4, 337) \| 471 \| (44, 368) \| 591 \| ATP-DEPENDENT DNA HELICASE Q1 (E.C.3.6.1.-) \| 2WWY\_B ATP-DEPENDENT DNA HELICASE Q1 (E.C.3.6.1.-) HYDROLASE-DNA COMPLEX, NUCLEAR PROTEIN, HYDROLASE HET: SO4; pdb70 \| 4U7D\_C \| 99.6 \| 1.8e-19 \| 1.3e-23 \| 180.1 \| 295 \| (4, 337) \| 471 \| (44, 368) \| 591 \| ATP-DEPENDENT DNA HELICASE Q1/DNA Complex \| 4U7D\_C ATP-DEPENDENT DNA HELICASE Q1/DNA Complex HYDROLASE-DNA COMPLEX, NUCLEAR PROTEIN, HYDROLASE; pdb70 \| 4Q48\_A \| 99.6 \| 2e-19 \| 1.5e-23 \| 177.5 \| 288 \| (4, 334) \| 471 \| (24, 334) \| 525 \| DNA helicase RecQ \| 4Q48\_A DNA helicase RecQ DNA unwinding, helicase, DNA BINDING; pdb70 \| 4Q47\_A \| 99.6 \| 2.1e-19 \| 1.5e-23 \| 177.4 \| 289 \| (4, 334) \| 471 \| (24, 334) \| 525 \| DNA helicase RecQ \| 4Q47\_A DNA helicase RecQ DNA unwinding, Topoisomerase, helicase, DNA HET: ADP; pdb70 \| 5Z3G\_Y \| 99.6 \| 2.6e-19 \| 1.9e-23 \| 175.9 \| 294 \| (4, 332) \| 471 \| (64, 385) \| 505 \| Ribosome biogenesis protein RLP7, Ribosome \| 5Z3G\_Y Ribosome biogenesis protein RLP7, Ribosome ribosome, pre-60S, pre-ribosome, protein-RNA complex; pdb70 \| 6EM3\_D \| 99.6 \| 2.6e-19 \| 1.9e-23 \| 175.9 \| 294 \| (4, 332) \| 471 \| (64, 385) \| 505 \| Ribosome production factor 1, 60S \| 6EM3\_D Ribosome production factor 1, 60S Large Subunit Biogenesis Nucleolus, RIBOSOME; pdb70 \| 6QW6\_5X \| 99.6 \| 2.7e-19 \| 2e-23 \| 185.9 \| 99 \| (229, 334) \| 471 \| (664, 762) \| 820 \| Small nuclear ribonucleoprotein Sm D1 \| 6QW6\_5X Small nuclear ribonucleoprotein Sm D1 RNP complex, splicing, RNA, protein HET: IHP, M7M, GTP; pdb70 \| 5ZWN\_y \| 99.6 \| 2.8e-19 \| 2e-23 \| 178.5 \| 296 \| (4, 334) \| 471 \| (196, 535) \| 588 \| U1 small nuclear ribonucleoprotein 70 \| 5ZWN\_y U1 small nuclear ribonucleoprotein 70 spliceosme, assemply, pre-B complex, U1; pdb70 \| 5VVR\_M \| 99.5 \| 3.1e-19 \| 2.3e-23 \| 187.9 \| 122 \| (213, 341) \| 471 \| (650, 778) \| 1085 \| DNA-directed RNA polymerase II subunit \| 5VVR\_M DNA-directed RNA polymerase II subunit complex, RNA polymerase, CSB, transcription; pdb70 \| 4TMU\_A \| 99.5 \| 3.4e-19 \| 2.5e-23 \| 176.5 \| 290 \| (4, 337) \| 471 \| (45, 357) \| 541 \| Protein/DNA Complex \| 4TMU\_A Protein/DNA Complex RecQ, helicase, Winged helix, ATP; pdb70 \| 5JXR\_A \| 99.5 \| 3.6e-19 \| 2.7e-23 \| 181.1 \| 329 \| (3, 341) \| 471 \| (104, 534) \| 723 \| Chromatin-remodeling complex ATPase-like protein \| 5JXR\_A Chromatin-remodeling complex ATPase-like protein chromatin remodeler, ISWI, TRANSCRIPTION; pdb70 \| 1OYW\_A \| 99.5 \| 3.9e-19 \| 2.9e-23 \| 175.4 \| 291 \| (4, 338) \| 471 \| (25, 338) \| 523 \| ATP-dependent DNA helicase (E.C.3.6.1.-) \| 1OYW\_A ATP-dependent DNA helicase (E.C.3.6.1.-) RecQ, helicase, Winged helix, Helix-turn-helix; pdb70 \| 6UV1\_A \| 99.5 \| 4e-19 \| 2.9e-23 \| 171.4 \| 295 \| (4, 334) \| 471 \| (85, 408) \| 448 \| Probable ATP-dependent RNA helicase DDX17/RNA \| 6UV1\_A Probable ATP-dependent RNA helicase DDX17/RNA DEAD-box ATPase, RNA helicase, RNA HET: MG, ADP; pdb70 \| 6UV2\_A \| 99.5 \| 4e-19 \| 2.9e-23 \| 171.4 \| 295 \| (4, 334) \| 471 \| (85, 408) \| 448 \| Probable ATP-dependent RNA helicase DDX17/RNA \| 6UV2\_A Probable ATP-dependent RNA helicase DDX17/RNA DEAD-box ATPase, RNA helicase, RNA HET: ADP; pdb70 \| 6UV3\_A \| 99.5 \| 4e-19 \| 3e-23 \| 171.3 \| 294 \| (4, 334) \| 471 \| (85, 408) \| 448 \| Probable ATP-dependent RNA helicase DDX17/RNA \| 6UV3\_A Probable ATP-dependent RNA helicase DDX17/RNA DEAD-box ATPase, RNA helicase, RNA HET: ADP; pdb70 \| 4LJY\_A \| 99.5 \| 4.3e-19 \| 3.2e-23 \| 173.6 \| 308 \| (4, 334) \| 471 \| (73, 418) \| 493 \| Pre-mRNA-processing ATP-dependent RNA helicase PRP5 \| 4LJY\_A Pre-mRNA-processing ATP-dependent RNA helicase PRP5 Prp5, DEAD box, RNA splicing HET: MRD, ADP; pdb70 \| 6IY2\_O \| 99.5 \| 4.5e-19 \| 3.3e-23 \| 179.8 \| 121 \| (214, 341) \| 471 \| (418, 542) \| 679 \| Transcription regulatory protein SNF2, Histone \| 6IY2\_O Transcription regulatory protein SNF2, Histone complex, nucleosome, chromatin remodeling, gene HET: ADP; pdb70 \| 5MC6\_h \| 99.5 \| 5.4e-19 \| 3.9e-23 \| 189.9 \| 318 \| (2, 334) \| 471 \| (327, 771) \| 1287 \| 40S ribosomal protein S3, 40S \| 5MC6\_h 40S ribosomal protein S3, 40S cryo-EM, ribosome, RNA, helicase HET: 5CT; pdb70 \| 5OF4\_A \| 99.5 \| 5.4e-19 \| 4e-23 \| 175.1 \| 309 \| (3, 332) \| 471 \| (85, 419) \| 553 \| TFIIH basal transcription factor complex \| 5OF4\_A TFIIH basal transcription factor complex transcription initiation, DNA repair, multiprotein; pdb70 \| 5E7I\_B \| 99.5 \| 5.6e-19 \| 4.1e-23 \| 170.6 \| 296 \| (3, 335) \| 471 \| (69, 408) \| 452 \| DEAD (Asp-Glu-Ala-Asp) box helicase 3 \| 5E7I\_B DEAD (Asp-Glu-Ala-Asp) box helicase 3 DEAD-box protein, RNA helicase, RecA; pdb70 \| 5HZR\_A \| 99.5 \| 5.7e-19 \| 4.2e-23 \| 181.2 \| 122 \| (213, 341) \| 471 \| (428, 553) \| 732 \| SNF2-family ATP dependent chromatin remodeling \| 5HZR\_A SNF2-family ATP dependent chromatin remodeling Swi2/Snf2, chromatin remodeling, TRANSCRIPTION HET: KH2, SO4; pdb70 \| 5E7J\_A \| 99.5 \| 6.1e-19 \| 4.6e-23 \| 170.3 \| 297 \| (3, 336) \| 471 \| (69, 409) \| 452 \| DEAD (Asp-Glu-Ala-Asp) box helicase 3 \| 5E7J\_A DEAD (Asp-Glu-Ala-Asp) box helicase 3 DEAD-box protein, RNA helicase, RecA HET: AMP; pdb70 \| 6O9M\_7 \| 99.5 \| 7e-19 \| 5.2e-23 \| 179.9 \| 309 \| (3, 332) \| 471 \| (314, 648) \| 782 \| TFIIH basal transcription factor complex \| 6O9M\_7 TFIIH basal transcription factor complex Transcription initiation, Molecular dynamics, Gene; pdb70 \| 6RO4\_A \| 99.5 \| 7e-19 \| 5.2e-23 \| 179.9 \| 309 \| (3, 332) \| 471 \| (314, 648) \| 782 \| General transcription and DNA repair \| 6RO4\_A General transcription and DNA repair Complex, Helicase, Translocase, DNA repair; pdb70 \| 5OQJ\_7 \| 99.5 \| 7.3e-19 \| 5.3e-23 \| 180.3 \| 309 \| (3, 332) \| 471 \| (360, 695) \| 843 \| DNA-directed RNA polymerase II subunit \| 5OQJ\_7 DNA-directed RNA polymerase II subunit transcription initiation, TRANSCRIPTION, macromolecular complex HET: ZN; pdb70 \| 4KBF\_A \| 99.5 \| 7.3e-19 \| 5.5e-23 \| 163.8 \| 292 \| (3, 334) \| 471 \| (22, 336) \| 365 \| Heat resistant RNA dependent ATPase \| 4KBF\_A Heat resistant RNA dependent ATPase DEAD BOX RNA HELICASE, DIMER HET: AMP; pdb70 \| 5SUP\_B \| 99.5 \| 8.6e-19 \| 6.4e-23 \| 165.5 \| 295 \| (3, 334) \| 471 \| (27, 348) \| 390 \| ATP-dependent RNA helicase SUB2, YRA1/RNA \| 5SUP\_B ATP-dependent RNA helicase SUB2, YRA1/RNA mRNA export, HYDROLASE-RNA complex HET: ADP, BEF; pdb70 \| 6PWF\_K \| 99.5 \| 8.8e-19 \| 6.5e-23 \| 176.7 \| 121 \| (214, 341) \| 471 \| (407, 531) \| 640 \| Histone H3, Histone H4, Histone \| 6PWF\_K Histone H3, Histone H4, Histone nucleosome, DNA-binding protein, ATP-dependent chromatin; pdb70 \| 6O5F\_A \| 99.5 \| 8.7e-19 \| 6.5e-23 \| 170.5 \| 297 \| (4, 337) \| 471 \| (71, 411) \| 476 \| Heterogeneous nuclear ribonucleoprotein A1/RNA Complex \| 6O5F\_A Heterogeneous nuclear ribonucleoprotein A1/RNA Complex DDX3X, DEAD-box, RNA helicase, HYDROLASE; pdb70 \| 1XTK\_A \| 99.5 \| 9.4e-19 \| 7e-23 \| 165.1 \| 297 \| (4, 338) \| 471 \| (29, 351) \| 390 \| Probable ATP-dependent RNA helicase p47 \| 1XTK\_A Probable ATP-dependent RNA helicase p47 alpha-beta fold, GENE REGULATION; pdb70 \| 6FTX\_W \| 99.5 \| 1.1e-18 \| 7.9e-23 \| 182.2 \| 122 \| (213, 341) \| 471 \| (499, 624) \| 878 \| Histone H3.3C, Histone H4, Histone \| 6FTX\_W Histone H3.3C, Histone H4, Histone Chromatin remodellers, MOTOR PROTEIN HET: ADP; pdb70 \| 5IVL\_B \| 99.5 \| 1.1e-18 \| 8.5e-23 \| 167.1 \| 297 \| (4, 338) \| 471 \| (24, 342) \| 429 \| DEAD-box ATP-dependent RNA helicase CshA \| 5IVL\_B DEAD-box ATP-dependent RNA helicase CshA Rec-A like domain DEAD-box helicase HET: SO4; pdb70 \| 4D26\_A \| 99.5 \| 1.1e-18 \| 8.5e-23 \| 167.2 \| 294 \| (3, 334) \| 471 \| (71, 399) \| 434 \| BMVLG PROTEIN \| 4D26\_A BMVLG PROTEIN HYDROLASE, PIRNA, AMPLIFIER COMPLEX, TRANSPOSON HET: ADP; pdb70 \| 4D25\_A \| 99.5 \| 1.2e-18 \| 8.6e-23 \| 167.2 \| 294 \| (3, 334) \| 471 \| (71, 399) \| 434 \| BMVLG PROTEIN \| 4D25\_A BMVLG PROTEIN HYDROLASE, PIRNA, AMPLIFIER COMPLEX, TRANSPOSON HET: ANP; pdb70 \| 1XTI\_A \| 99.5 \| 1.2e-18 \| 8.7e-23 \| 164.5 \| 298 \| (3, 338) \| 471 \| (29, 352) \| 391 \| Probable ATP-dependent RNA helicase p47 \| 1XTI\_A Probable ATP-dependent RNA helicase p47 alpha-beta fold, GENE REGULATION HET: IPA; pdb70 \| 2I4I\_A \| 99.5 \| 1.2e-18 \| 8.7e-23 \| 165.9 \| 298 \| (3, 337) \| 471 \| (36, 377) \| 417 \| ATP-dependent RNA helicase DDX3X (E.C.3.6.1.-) \| 2I4I\_A ATP-dependent RNA helicase DDX3X (E.C.3.6.1.-) RNA, HELICASE, DEAD, STRUCTURAL GENOMICS HET: AMP; pdb70 \| 6JYL\_K \| 99.5 \| 1.2e-18 \| 8.9e-23 \| 181.6 \| 122 \| (213, 341) \| 471 \| (433, 558) \| 1061 \| Histone H3, Histone H4, Histone \| 6JYL\_K Histone H3, Histone H4, Histone chromatin remodelling, single particle Cryo-EM HET: ADP; pdb70 \| 2VSO\_B \| 99.5 \| 1.2e-18 \| 9.2e-23 \| 164.7 \| 296 \| (3, 336) \| 471 \| (43, 360) \| 395 \| ATP-DEPENDENT RNA HELICASE EIF4A (E.C.3.6.1.-) \| 2VSO\_B ATP-DEPENDENT RNA HELICASE EIF4A (E.C.3.6.1.-) ACETYLATION, ATP-BINDING, PHOSPHOPROTEIN, PROTEIN BIOSYNTHESIS HET: AMP; pdb70 \| 4BRW\_A \| 99.5 \| 1.3e-18 \| 9.5e-23 \| 163.1 \| 294 \| (3, 334) \| 471 \| (22, 336) \| 377 \| ATP-DEPENDENT RNA HELICASE DHH1 (E.C.3.6.4.13) \| 4BRW\_A ATP-DEPENDENT RNA HELICASE DHH1 (E.C.3.6.4.13) HYDROLASE, TRANSLATIONAL REPRESSION, MRNP REMODELING HET: 1PE; pdb70 \| 4NHO\_A \| 99.5 \| 1.5e-18 \| 1.1e-22 \| 169.4 \| 99 \| (229, 334) \| 471 \| (332, 430) \| 488 \| Probable ATP-dependent RNA helicase DDX23 \| 4NHO\_A Probable ATP-dependent RNA helicase DDX23 DEAD-BOX, HELICASE, HYDROLASE HET: GOL, CXS, SO4; pdb70 \| 1S2M\_A \| 99.5 \| 1.5e-18 \| 1.1e-22 \| 164.4 \| 292 \| (4, 334) \| 471 \| (43, 356) \| 400 \| Putative ATP-dependent RNA helicase DHH1 \| 1S2M\_A Putative ATP-dependent RNA helicase DHH1 ATP-binding, RNA-binding, Helicase, RNA BINDING; pdb70 \| 6GEJ\_M \| 99.5 \| 1.5e-18 \| 1.1e-22 \| 186.9 \| 122 \| (214, 342) \| 471 \| (1246, 1370) \| 1514 \| Vacuolar protein sorting-associated protein 72 \| 6GEJ\_M Vacuolar protein sorting-associated protein 72 Chromatin, Remodeller, ATPase, Histone, NUCLEAR HET: ADP; pdb70 \| 4CT4\_B \| 99.5 \| 1.5e-18 \| 1.1e-22 \| 162.8 \| 292 \| (4, 334) \| 471 \| (27, 341) \| 378 \| CCR4-NOT TRANSCRIPTION COMPLEX SUBUNIT 1 \| 4CT4\_B CCR4-NOT TRANSCRIPTION COMPLEX SUBUNIT 1 RNA BINDING PROTEIN, DEADENYLATION, TRANSCRIPTION; pdb70 \| 6NE3\_W \| 99.5 \| 1.5e-18 \| 1.1e-22 \| 167.4 \| 119 \| (214, 339) \| 471 \| (316, 438) \| 467 \| Histone H3.2, Histone H4, Histone \| 6NE3\_W Histone H3.2, Histone H4, Histone ISWI, Chromatin, Nucleosome, DNA, SNF2h HET: ADP; pdb70 \| 1Z3I\_X \| 99.5 \| 1.5e-18 \| 1.1e-22 \| 175.6 \| 106 \| (229, 341) \| 471 \| (416, 524) \| 644 \| similar to RAD54-like \| 1Z3I\_X similar to RAD54-like Recombination ATPase helicase, Recombination-DNA binding HET: SO4; pdb70 \| 4BUJ\_A \| 99.5 \| 1.6e-18 \| 1.2e-22 \| 183.9 \| 317 \| (2, 333) \| 471 \| (331, 774) \| 1044 \| ANTIVIRAL HELICASE SKI2 (E.C.3.6.4.13), SUPERKILLER \| 4BUJ\_A ANTIVIRAL HELICASE SKI2 (E.C.3.6.4.13), SUPERKILLER HYDROLASE, DEXH BOX HELICASE, RNA HET: SO4; pdb70 \| 5E02\_A \| 99.5 \| 1.6e-18 \| 1.2e-22 \| 185.6 \| 314 \| (3, 332) \| 471 \| (179, 583) \| 1106 \| FRQ-interacting RNA helicase/RNA Complex \| 5E02\_A FRQ-interacting RNA helicase/RNA Complex circadian clock, RNA BINDING PROTEIN-RNA HET: ADP; pdb70 \| 6BB8\_A \| 99.5 \| 1.6e-18 \| 1.2e-22 \| 185.6 \| 314 \| (3, 332) \| 471 \| (179, 583) \| 1106 \| FRQ-interacting RNA helicase \| 6BB8\_A FRQ-interacting RNA helicase RNA helicase, RNA BINDING PROTEIN; pdb70 \| 6UXW\_A \| 99.5 \| 1.7e-18 \| 1.2e-22 \| 184.0 \| 121 \| (214, 341) \| 471 \| (1087, 1211) \| 1703 \| Histone H3.2, Histone H4, Histone \| 6UXW\_A Histone H3.2, Histone H4, Histone SWI/SNF, chromatin remodeler, TRANSCRIPTION, TRANSCRIPTION-DNA HET: ADP, PO4; pdb70 \| 5ANR\_B \| 99.5 \| 1.7e-18 \| 1.3e-22 \| 162.4 \| 293 \| (4, 335) \| 471 \| (27, 342) \| 378 \| CCR4-NOT TRANSCRIPTION COMPLEX SUBUNIT 1 \| 5ANR\_B CCR4-NOT TRANSCRIPTION COMPLEX SUBUNIT 1 RNA BINDING PROTEIN; pdb70 \| 1FUU\_A \| 99.5 \| 1.8e-18 \| 1.3e-22 \| 163.5 \| 294 \| (3, 334) \| 471 \| (42, 357) \| 394 \| INITIATION FACTOR 4A \| 1FUU\_A INITIATION FACTOR 4A IF4A, HELICASE, DEAD-BOX PROTEIN, TRANSLATION HET: MSE; pdb70 \| 4BUJ\_E \| 99.5 \| 1.8e-18 \| 1.3e-22 \| 183.4 \| 317 \| (2, 333) \| 471 \| (331, 774) \| 1044 \| ANTIVIRAL HELICASE SKI2 (E.C.3.6.4.13), SUPERKILLER \| 4BUJ\_E ANTIVIRAL HELICASE SKI2 (E.C.3.6.4.13), SUPERKILLER HYDROLASE, DEXH BOX HELICASE, RNA HET: SO4; pdb70 \| 6G0L\_W \| 99.5 \| 1.8e-18 \| 1.4e-22 \| 187.8 \| 123 \| (213, 342) \| 471 \| (694, 820) \| 1468 \| Histone H3.3C, Histone H4, Histone \| 6G0L\_W Histone H3.3C, Histone H4, Histone Chromatin remodellers, MOTOR PROTEIN HET: ADP; pdb70 \| 5SVA\_Z \| 99.5 \| 1.9e-18 \| 1.4e-22 \| 177.0 \| 310 \| (3, 333) \| 471 \| (360, 696) \| 843 \| DNA-directed RNA polymerase II subunit \| 5SVA\_Z DNA-directed RNA polymerase II subunit Transcriptional Initiation, Mediator, Pre-Initiation Complex; pdb70 \| 5O9G\_W \| 99.5 \| 1.9e-18 \| 1.4e-22 \| 187.7 \| 123 \| (213, 342) \| 471 \| (694, 820) \| 1468 \| Histone H3.2, Histone H4, Histone \| 5O9G\_W Histone H3.2, Histone H4, Histone ATPase, Complex, Nucleosome, DNA, DNA HET: ADP; pdb70 \| 6TDA\_S \| 99.5 \| 2e-18 \| 1.5e-22 \| 182.6 \| 120 \| (214, 340) \| 471 \| (791, 914) \| 1359 \| Histone H3.2, Histone H4, Histone \| 6TDA\_S Histone H3.2, Histone H4, Histone Chromatin remodeler DNA binding Nucleosome HET: MSE; pdb70 \| 6K15\_J \| 99.5 \| 2.1e-18 \| 1.5e-22 \| 182.5 \| 120 \| (214, 340) \| 471 \| (791, 914) \| 1359 \| RSC7, Chromatin structure-remodeling complex protein \| 6K15\_J RSC7, Chromatin structure-remodeling complex protein chromatin remodeler, SWI/SNF family, DNA; pdb70 \| 3FMP\_B \| 99.5 \| 2.2e-18 \| 1.6e-22 \| 167.9 \| 297 \| (4, 336) \| 471 \| (114, 439) \| 479 \| Nuclear pore complex protein Nup214 \| 3FMP\_B Nuclear pore complex protein Nup214 nuclear porin, nuclear pore complex HET: ADP; pdb70 \| 3MWY\_W \| 99.5 \| 2.3e-18 \| 1.7e-22 \| 178.7 \| 122 \| (213, 341) \| 471 \| (555, 680) \| 800 \| --REMARK 3 \| 3MWY\_W --REMARK 3 SWI2/SNF2 ATPase, double chromodomains, HYDROLASE HET: AGS; pdb70 \| 5ZC9\_A \| 99.5 \| 2.3e-18 \| 1.7e-22 \| 162.6 \| 295 \| (3, 335) \| 471 \| (41, 359) \| 394 \| Eukaryotic initiation factor 4A-I/RNA Complex \| 5ZC9\_A Eukaryotic initiation factor 4A-I/RNA Complex INITIATION FACTOR, DEAD-BOX, HELICASE, PROTEIN-RNA HET: ANP, RCG; pdb70 \| 4W7S\_A \| 99.5 \| 2.6e-18 \| 1.9e-22 \| 166.4 \| 296 \| (4, 334) \| 471 \| (71, 410) \| 463 \| S.cerevisiae Prp28 (127-588 aa) \| 4W7S\_A S.cerevisiae Prp28 (127-588 aa) splicing factor, DEAD-box protein, ATPase HET: P6G, ANP, MSE; pdb70 \| 3I62\_A \| 99.5 \| 2.7e-18 \| 2e-22 \| 170.9 \| 306 \| (4, 334) \| 471 \| (94, 440) \| 563 \| ATP-dependent RNA helicase MSS116 (E.C.3.6.1.-)/RNA \| 3I62\_A ATP-dependent RNA helicase MSS116 (E.C.3.6.1.-)/RNA Protein-RNA complex, RNA helicase, DEAD-box HET: ADP; pdb70 \| 4W7S\_B \| 99.5 \| 2.7e-18 \| 2e-22 \| 166.2 \| 296 \| (4, 334) \| 471 \| (71, 410) \| 463 \| S.cerevisiae Prp28 (127-588 aa) \| 4W7S\_B S.cerevisiae Prp28 (127-588 aa) splicing factor, DEAD-box protein, ATPase HET: ANP, MSE, P6G; pdb70 \| 3I5X\_A \| 99.5 \| 2.8e-18 \| 2.1e-22 \| 170.7 \| 306 \| (4, 334) \| 471 \| (94, 440) \| 563 \| ATP-dependent RNA helicase MSS116 (E.C.3.6.1.-)/RNA \| 3I5X\_A ATP-dependent RNA helicase MSS116 (E.C.3.6.1.-)/RNA Protein-RNA complex, RNA helicase, DEAD-box HET: ANP; pdb70 \| 1FUU\_B \| 99.5 \| 2.9e-18 \| 2.1e-22 \| 162.0 \| 294 \| (3, 334) \| 471 \| (42, 357) \| 394 \| INITIATION FACTOR 4A \| 1FUU\_B INITIATION FACTOR 4A IF4A, HELICASE, DEAD-BOX PROTEIN, TRANSLATION; pdb70 \| 5FMF\_1 \| 99.5 \| 2.9e-18 \| 2.2e-22 \| 167.7 \| 310 \| (3, 333) \| 471 \| (67, 403) \| 492 \| DNA REPAIR HELICASE RAD25, SSL2 \| 5FMF\_1 DNA REPAIR HELICASE RAD25, SSL2 TRANSCRIPTION, PRE-INITIATION COMPLEX, RNA POLYMERASE; pdb70 \| 2DB3\_D \| 99.5 \| 3e-18 \| 2.2e-22 \| 164.4 \| 293 \| (4, 334) \| 471 \| (78, 398) \| 434 \| ATP-dependent RNA helicase vasa(E.C.3.6.1.3)/RNA complex \| 2DB3\_D ATP-dependent RNA helicase vasa(E.C.3.6.1.3)/RNA complex DEAD-BOX, HELICASE, PROTEIN-RNA COMPLEX, ATPase HET: ANP; pdb70 \| 3H1T\_A \| 99.5 \| 3.1e-18 \| 2.3e-22 \| 171.4 \| 331 \| (3, 345) \| 471 \| (177, 561) \| 590 \| Type I site-specific restriction-modification system \| 3H1T\_A Type I site-specific restriction-modification system hydrolase, restriction enzyme hsdR, ATP-binding; pdb70 \| 3FHO\_B \| 99.5 \| 3.3e-18 \| 2.4e-22 \| 167.6 \| 298 \| (4, 337) \| 471 \| (141, 464) \| 508 \| ATP-dependent RNA helicase dbp5 (E.C.3.6.1.-) \| 3FHO\_B ATP-dependent RNA helicase dbp5 (E.C.3.6.1.-) RNA helicase, mRNA export, ATPase; pdb70 \| 6QDV\_7 \| 99.5 \| 3.8e-18 \| 2.8e-22 \| 160.6 \| 295 \| (4, 336) \| 471 \| (39, 356) \| 390 \| Eukaryotic initiation factor 4A-III, RNA-binding \| 6QDV\_7 Eukaryotic initiation factor 4A-III, RNA-binding spliceosome, RNA, complex, SPLICING HET: GTP, I6P, ATP, SEP; pdb70 \| 4A36\_A \| 99.5 \| 3.9e-18 \| 2.9e-22 \| 169.2 \| 113 \| (213, 332) \| 471 \| (371, 499) \| 556 \| RETINOIC ACID INDUCIBLE PROTEIN I \| 4A36\_A RETINOIC ACID INDUCIBLE PROTEIN I RNA BINDING PROTEIN-RNA COMPLEX, SUPERFAMILY HET: ADP; pdb70 \| 3SQW\_A \| 99.5 \| 4e-18 \| 3e-22 \| 170.0 \| 306 \| (4, 334) \| 471 \| (43, 389) \| 579 \| ATP-dependent RNA helicase MSS116, mitochondrial \| 3SQW\_A ATP-dependent RNA helicase MSS116, mitochondrial RecA fold, RNA dependent ATPase HET: ANP; pdb70 \| 2J0S\_A \| 99.5 \| 4.2e-18 \| 3.2e-22 \| 161.9 \| 298 \| (4, 339) \| 471 \| (59, 379) \| 410 \| ATP-DEPENDENT RNA HELICASE DDX48, PROTEIN \| 2J0S\_A ATP-DEPENDENT RNA HELICASE DDX48, PROTEIN MRNA PROCESSING, PHOSPHORYLATION, RRNA PROCESSING HET: ANP; pdb70 \| 5ELX\_A \| 99.5 \| 4.3e-18 \| 3.2e-22 \| 160.3 \| 296 \| (3, 334) \| 471 \| (23, 344) \| 391 \| ATP-dependent RNA helicase DBP5/RNA Complex \| 5ELX\_A ATP-dependent RNA helicase DBP5/RNA Complex Fluorescent, Nucleotide, Mant, ADP, RNA HET: M2A; pdb70 \| 5LST\_A \| 99.5 \| 5.2e-18 \| 3.8e-22 \| 172.7 \| 294 \| (4, 336) \| 471 \| (57, 391) \| 693 \| ATP-dependent DNA helicase Q4 (E.C.3.6.4.12) \| 5LST\_A ATP-dependent DNA helicase Q4 (E.C.3.6.4.12) RecQ4, helicase, Rothmund-Thomson-Syndrome, RAPADILINO-Syndrome, hydrolase; pdb70 \| 5LB8\_A \| 99.5 \| 5.8e-18 \| 4.3e-22 \| 166.7 \| 103 \| (229, 338) \| 471 \| (251, 353) \| 518 \| ATP-dependent DNA helicase Q5 (E.C.3.6.4.12) \| 5LB8\_A ATP-dependent DNA helicase Q5 (E.C.3.6.4.12) Helicase, RecQ, Transcription, DNA repair; pdb70 \| 4A2P\_A \| 99.4 \| 6.2e-18 \| 4.6e-22 \| 167.8 \| 113 \| (213, 332) \| 471 \| (371, 499) \| 556 \| RETINOIC ACID INDUCIBLE PROTEIN I \| 4A2P\_A RETINOIC ACID INDUCIBLE PROTEIN I HYDROLASE, SUPERFAMILY 2 RNA HELICASE; pdb70 \| 6LTJ\_I \| 99.4 \| 6.7e-18 \| 4.9e-22 \| 184.5 \| 120 \| (215, 341) \| 471 \| (1081, 1204) \| 1647 \| Histone H3.2, Histone H4, Histone \| 6LTJ\_I Histone H3.2, Histone H4, Histone Chromatin remodeler, Complex, GENE REGULATION; pdb70 \| 5DTU\_A \| 99.4 \| 6.8e-18 \| 5e-22 \| 163.7 \| 102 \| (229, 337) \| 471 \| (315, 416) \| 475 \| Putative uncharacterized protein \| 5DTU\_A Putative uncharacterized protein DEAD-box Protein, ATPase, RNA-helicase, DDX23 HET: ADP; pdb70 \| 1HV8\_A \| 99.4 \| 6.9e-18 \| 5.1e-22 \| 157.2 \| 288 \| (4, 334) \| 471 \| (28, 336) \| 367 \| DEAD BOX HELICASE \| 1HV8\_A DEAD BOX HELICASE Helicase, RNA-binding Protein, ATPase, RNA HET: MSE, SO4; pdb70 \| 1Z6A\_A \| 99.4 \| 7.1e-18 \| 5.3e-22 \| 165.1 \| 119 \| (214, 339) \| 471 \| (325, 447) \| 500 \| Helicase of the snf2/rad54 family \| 1Z6A\_A Helicase of the snf2/rad54 family HYDROLASE, RECOMBINATION, HYDROLASE-RECOMBINATION COMPLEX HET: PO4; pdb70 \| 4TYN\_A \| 99.4 \| 7.2e-18 \| 5.3e-22 \| 165.6 \| 307 \| (3, 334) \| 471 \| (40, 387) \| 509 \| Mss116 catalytic core/DNA Complex \| 4TYN\_A Mss116 catalytic core/DNA Complex DEAD-box, RNA helicase, hydrolase, RNA HET: ADP; pdb70 \| 5FFJ\_B \| 99.4 \| 7.4e-18 \| 5.4e-22 \| 183.9 \| 323 \| (2, 334) \| 471 \| (11, 440) \| 1406 \| LlaGI/DNA Complex \| 5FFJ\_B LlaGI/DNA Complex Helicase-like ATPase, methyltransferase, DNA-binding protein; pdb70 \| 3TBK\_A \| 99.4 \| 7.7e-18 \| 5.7e-22 \| 167.0 \| 112 \| (213, 331) \| 471 \| (370, 497) \| 555 \| RIG-I Helicase Domain (E.C.3.6.4.13) \| 3TBK\_A RIG-I Helicase Domain (E.C.3.6.4.13) DECH Helicase, Helicase, ATP binding HET: ANP; pdb70 \| 4CGZ\_A \| 99.4 \| 8e-18 \| 5.9e-22 \| 170.8 \| 291 \| (4, 334) \| 471 \| (34, 354) \| 665 \| BLOOM'S SYNDROME HELICASE (E.C.3.6.4.12) \| 4CGZ\_A BLOOM'S SYNDROME HELICASE (E.C.3.6.4.12) HYDROLASE-DNA COMPLEX HET: ADP; pdb70 \| 6NMI\_A \| 99.4 \| 8.1e-18 \| 5.9e-22 \| 170.1 \| 309 \| (3, 332) \| 471 \| (237, 571) \| 653 \| General transcription and DNA repair \| 6NMI\_A General transcription and DNA repair transcription, DNA repair, helicase, multiprotein; pdb70 \| 3PEY\_A \| 99.4 \| 8.2e-18 \| 6.1e-22 \| 158.7 \| 298 \| (3, 336) \| 471 \| (26, 349) \| 395 \| ATP-dependent RNA helicase DBP5 (E.C.3.6.4.13) \| 3PEY\_A ATP-dependent RNA helicase DBP5 (E.C.3.6.4.13) RecA, DEAD-box, ATPase, Helicase, mRNA-export HET: ADP, NO3; pdb70 \| 6FT6\_MM \| 99.4 \| 8.4e-18 \| 6.2e-22 \| 178.2 \| 317 \| (3, 334) \| 471 \| (86, 489) \| 1011 \| Ribosomal 60S subunit protein L2B \| 6FT6\_MM Ribosomal 60S subunit protein L2B RNA exosome, Ribosome, pre-ribosome, Mtr4 HET: GTP; pdb70 \| 4XQK\_A \| 99.4 \| 8.6e-18 \| 6.3e-22 \| 184.2 \| 322 \| (3, 334) \| 471 \| (176, 604) \| 1578 \| LlaBIII/DNA Complex \| 4XQK\_A LlaBIII/DNA Complex ATP-dependent restriction-modification enzyme, Type ISP; pdb70 \| 5V9X\_A \| 99.4 \| 1.1e-17 \| 8e-22 \| 174.4 \| 296 \| (4, 331) \| 471 \| (47, 420) \| 877 \| ATP-dependent DNA helicase/DNA Complex \| 5V9X\_A ATP-dependent DNA helicase/DNA Complex Helicase, HYDROLASE-DNA complex HET: GOL, ANP; pdb70 \| 2FWR\_A \| 99.4 \| 1.1e-17 \| 8.3e-22 \| 162.6 \| 314 \| (3, 341) \| 471 \| (92, 450) \| 472 \| DNA repair protein RAD25 \| 2FWR\_A DNA repair protein RAD25 DNA Unwinding, DNA Repair, XPB HET: IPA, PO4; pdb70 \| 4O3M\_A \| 99.4 \| 1.1e-17 \| 8.4e-22 \| 169.5 \| 291 \| (4, 334) \| 471 \| (28, 348) \| 659 \| Bloom syndrome protein (E.C.3.6.4.12) \| 4O3M\_A Bloom syndrome protein (E.C.3.6.4.12) Winged Helix, Helicase, Hydrolase-DNA complex HET: EDO, ADP; pdb70 \| 6HTS\_G \| 99.4 \| 1.2e-17 \| 9e-22 \| 177.1 \| 122 \| (213, 341) \| 471 \| (834, 958) \| 1290 \| RuvB-like 1, RuvB-like 2, DNA \| 6HTS\_G RuvB-like 1, RuvB-like 2, DNA Chromatin, Remodeller, Nucleosome, DNA Binding HET: ADP; pdb70 \| 4CBG\_B \| 99.4 \| 1.3e-17 \| 9.3e-22 \| 164.1 \| 282 \| (5, 339) \| 471 \| (30, 344) \| 516 \| SERINE PROTEASE NS3 (E.C.3.4.21.113, 3.6.1.15 \| 4CBG\_B SERINE PROTEASE NS3 (E.C.3.4.21.113, 3.6.1.15 HYDROLASE, FLAVIVIRIDAE NS3, SAXS HET: MSE; pdb70 \| 4CBH\_A \| 99.4 \| 1.3e-17 \| 9.3e-22 \| 164.1 \| 282 \| (5, 339) \| 471 \| (30, 344) \| 516 \| SERINE PROTEASE NS3 (E.C.3.4.21.113, 3.6.1.15 \| 4CBH\_A SERINE PROTEASE NS3 (E.C.3.4.21.113, 3.6.1.15 HYDROLASE, SF2 HELICASES, FLAVIVIRIDAE NS3; pdb70 \| 6FHS\_G \| 99.4 \| 1.3e-17 \| 9.7e-22 \| 176.3 \| 122 \| (213, 341) \| 471 \| (803, 927) \| 1107 \| RuvB-like helicase (E.C.3.6.4.12), Ino80, Uncharacterized \| 6FHS\_G RuvB-like helicase (E.C.3.6.4.12), Ino80, Uncharacterized DNA BINDING PROTEIN HET: ADP, ATP; pdb70 \| 1C4O\_A \| 99.4 \| 1.4e-17 \| 1e-21 \| 168.9 \| 117 \| (217, 340) \| 471 \| (426, 548) \| 664 \| DNA NUCLEOTIDE EXCISION REPAIR ENZYME \| 1C4O\_A DNA NUCLEOTIDE EXCISION REPAIR ENZYME DNA NUCLEOTIDE EXCISION REPAIR, UVRABC HET: BOG, SO4; pdb70 \| 4U4C\_A \| 99.4 \| 1.4e-17 \| 1e-21 \| 176.5 \| 317 \| (3, 334) \| 471 \| (73, 476) \| 998 \| Mtr4, Dob1 (E.C.3.6.4.13), Air2-Trf4 Fusion \| 4U4C\_A Mtr4, Dob1 (E.C.3.6.4.13), Air2-Trf4 Fusion helicase, ATPase, poly(A)polymerase, RNA degradation HET: PG4, SO4; pdb70 \| 5LB3\_B \| 99.4 \| 1.5e-17 \| 1.1e-21 \| 160.2 \| 102 \| (229, 337) \| 471 \| (251, 352) \| 445 \| ATP-dependent DNA helicase Q5 (E.C.3.6.4.12) \| 5LB3\_B ATP-dependent DNA helicase Q5 (E.C.3.6.4.12) Helicase, RecQ, Transcription, DNA repair HET: ADP; pdb70 \| 1Z63\_A \| 99.4 \| 1.7e-17 \| 1.2e-21 \| 162.5 \| 119 \| (214, 339) \| 471 \| (325, 447) \| 500 \| Helicase of the snf2/rad54 family/DNA \| 1Z63\_A Helicase of the snf2/rad54 family/DNA protein-DNA complex, HYDROLASE-DNA complex COMPLEX HET: MSE; pdb70 \| 6FF7\_q \| 99.4 \| 2e-17 \| 1.5e-21 \| 174.3 \| 298 \| (3, 339) \| 471 \| (399, 738) \| 1041 \| RNA-binding motif protein, X-linked 2 \| 6FF7\_q RNA-binding motif protein, X-linked 2 spliceosome, human, HELA, BACT, dynamics HET: GTP, IHP, ADP, ZN; pdb70 \| 3FHT\_A \| 99.4 \| 2e-17 \| 1.5e-21 \| 157.3 \| 295 \| (4, 334) \| 471 \| (47, 370) \| 412 \| ATP-dependent RNA helicase DDX19B (E.C.3.6.1.-)/RNA \| 3FHT\_A ATP-dependent RNA helicase DDX19B (E.C.3.6.1.-)/RNA Dbp5, DEAD-box helicase, RNA dependent HET: GOL, ANP; pdb70 \| 2Z0M\_A \| 99.4 \| 2.1e-17 \| 1.6e-21 \| 151.7 \| 282 \| (3, 334) \| 471 \| (15, 314) \| 337 \| 337aa long hypothetical ATP-dependent RNA \| 2Z0M\_A 337aa long hypothetical ATP-dependent RNA helicase, ATP-binding, Hydrolase, Nucleotide-binding, RNA; pdb70 \| 1D2M\_A \| 99.4 \| 2.3e-17 \| 1.7e-21 \| 167.2 \| 117 \| (217, 340) \| 471 \| (427, 549) \| 665 \| EXCINUCLEASE ABC SUBUNIT B \| 1D2M\_A EXCINUCLEASE ABC SUBUNIT B MULTIDOMAIN PROTEIN, RIKEN Structural Genomics/Proteomics HET: BOG; pdb70 \| 5TNU\_A \| 99.4 \| 2.3e-17 \| 1.7e-21 \| 159.6 \| 304 \| (4, 332) \| 471 \| (88, 435) \| 459 \| DNA-dependent ATPase XPBII \| 5TNU\_A DNA-dependent ATPase XPBII Helicase, NER, transcription HET: GOL, SO4; pdb70 \| 6RFL\_Y \| 99.4 \| 2.9e-17 \| 2.2e-21 \| 165.5 \| 109 \| (216, 332) \| 471 \| (365, 482) \| 631 \| 15 kDa core protein \| 6RFL\_Y 15 kDa core protein Vaccinia, RNA polymerase, RNA Polymerase HET: SEP; pdb70 \| 2EYQ\_A \| 99.4 \| 3e-17 \| 2.2e-21 \| 174.9 \| 297 \| (3, 339) \| 471 \| (602, 918) \| 1151 \| Transcription-repair coupling factor \| 2EYQ\_A Transcription-repair coupling factor Mfd, SF2 ATPase, HYDROLASE HET: EPE, SO4; pdb70 \| 2Z83\_A \| 99.4 \| 3e-17 \| 2.2e-21 \| 158.9 \| 277 \| (3, 339) \| 471 \| (5, 309) \| 459 \| Helicase/Nucleoside Triphosphatase \| 2Z83\_A Helicase/Nucleoside Triphosphatase HELICASE, Hydrolase, Membrane, Nucleotide-binding, RNA; pdb70 \| 3P4X\_B \| 99.4 \| 3e-17 \| 2.2e-21 \| 156.6 \| 283 \| (2, 332) \| 471 \| (18, 349) \| 413 \| reverse gyrase helicase-like domain \| 3P4X\_B reverse gyrase helicase-like domain TOPOISOMERASE, DNA SUPERCOILING, ARCHAEA, HELICASE HET: ADP; pdb70 \| 6C90\_A \| 99.4 \| 3.1e-17 \| 2.3e-21 \| 168.3 \| 316 \| (3, 333) \| 471 \| (71, 463) \| 734 \| Superkiller viralicidic activity 2-like 2 \| 6C90\_A Superkiller viralicidic activity 2-like 2 HYDROLASE, NEXT, EXOSOME, NUCLEOTIDE-BINDING, ATPASE HET: ADP, TLA, MLI; pdb70 \| 4XGT\_A \| 99.4 \| 3.2e-17 \| 2.4e-21 \| 173.5 \| 316 \| (3, 334) \| 471 \| (66, 472) \| 993 \| FRQ-interacting RNA helicase \| 4XGT\_A FRQ-interacting RNA helicase ATPase, circadian clock, HYDROLASE; pdb70 \| 6FML\_G \| 99.4 \| 3.6e-17 \| 2.6e-21 \| 173.4 \| 121 \| (214, 341) \| 471 \| (1553, 1676) \| 1856 \| RuvB-like helicase (E.C.3.6.4.12), Ino80, Uncharacterized \| 6FML\_G RuvB-like helicase (E.C.3.6.4.12), Ino80, Uncharacterized INO80, Nucleosome, ATP dependent Chromatin HET: ATP, ADP; pdb70 \| 3OIY\_A \| 99.4 \| 3.7e-17 \| 2.8e-21 \| 156.0 \| 283 \| (2, 332) \| 471 \| (19, 350) \| 414 \| reverse gyrase helicase domain \| 3OIY\_A reverse gyrase helicase domain TOPOISOMERASE, DNA SUPERCOILING, ARCHAEA, HELICASE; pdb70 \| 6B4J\_F \| 99.4 \| 4.4e-17 \| 3.2e-21 \| 156.0 \| 298 \| (4, 337) \| 471 \| (65, 391) \| 430 \| Nucleoporin GLE1, Nucleoporin like 2 \| 6B4J\_F Nucleoporin GLE1, Nucleoporin like 2 Complex, Nuclear Pore Complex, mRNA HET: PO4, ANP; pdb70 \| 6BOG\_A \| 99.4 \| 4.4e-17 \| 3.3e-21 \| 170.8 \| 114 \| (215, 336) \| 471 \| (490, 606) \| 968 \| RNA polymerase-associated protein RapA (E.C.3.6.4.-) \| 6BOG\_A RNA polymerase-associated protein RapA (E.C.3.6.4.-) hydrolase, TRANSCRIPTION HET: SO4, MSE; pdb70 \| 2EYQ\_B \| 99.4 \| 4.7e-17 \| 3.4e-21 \| 173.4 \| 297 \| (3, 339) \| 471 \| (602, 918) \| 1151 \| Transcription-repair coupling factor \| 2EYQ\_B Transcription-repair coupling factor Mfd, SF2 ATPase, HYDROLASE HET: EPE, SO4; pdb70 \| 5AGA\_A \| 99.4 \| 4.9e-17 \| 3.6e-21 \| 169.1 \| 306 \| (3, 332) \| 471 \| (26, 422) \| 830 \| DNA POLYMERASE THETA (E.C.2.7.7.7) \| 5AGA\_A DNA POLYMERASE THETA (E.C.2.7.7.7) TRANSFERASE, POLQ, DNA REPAIR HET: ANP, FLC; pdb70 \| 5YZG\_Z \| 99.4 \| 5.9e-17 \| 4.3e-21 \| 172.4 \| 299 \| (3, 340) \| 471 \| (532, 870) \| 1227 \| Pre-mRNA-processing-splicing factor 8, 116 kDa \| 5YZG\_Z Pre-mRNA-processing-splicing factor 8, 116 kDa Structure of a Human Catalytic HET: GTP, I6P, ATP, ADP, SEP; pdb70 \| 6AC8\_A \| 99.4 \| 6e-17 \| 4.4e-21 \| 172.5 \| 293 \| (3, 336) \| 471 \| (661, 974) \| 1235 \| Mycobacterium smegmatis Mfd (E.C.3.6.4.-) \| 6AC8\_A Mycobacterium smegmatis Mfd (E.C.3.6.4.-) Transcription repair coupling factor, Mfd HET: SO4; pdb70 \| 6ACX\_A \| 99.4 \| 6e-17 \| 4.4e-21 \| 172.5 \| 293 \| (3, 336) \| 471 \| (661, 974) \| 1235 \| Mycobacterium smegmatis Mfd (E.C.3.6.4.-) \| 6ACX\_A Mycobacterium smegmatis Mfd (E.C.3.6.4.-) Transcription repair coupling factor, Mfd HET: ADP, SO4; pdb70 \| 6ID1\_V \| 99.4 \| 6.1e-17 \| 4.5e-21 \| 167.5 \| 291 \| (3, 331) \| 471 \| (137, 477) \| 795 \| Pre-mRNA-processing-splicing factor 8, 116 kDa \| 6ID1\_V Pre-mRNA-processing-splicing factor 8, 116 kDa Human Intron Lariat Spliceosome, SPLICING HET: GTP, SEP, I6P; pdb70 \| 5XC7\_A \| 99.4 \| 6.6e-17 \| 4.9e-21 \| 156.1 \| 272 \| (3, 333) \| 471 \| (3, 300) \| 451 \| NS3 Helicase \| 5XC7\_A NS3 Helicase Helicase, Dengue NS3, Hydrolase HET: GOL; pdb70 \| 3BXZ\_B \| 99.4 \| 7.5e-17 \| 5.6e-21 \| 156.6 \| 119 \| (216, 343) \| 471 \| (297, 454) \| 471 \| Preprotein translocase subunit secA \| 3BXZ\_B Preprotein translocase subunit secA Protein transport, Translocation, DEAD motor HET: SPD, ADP; pdb70 \| 6ICZ\_Y \| 99.3 \| 8.2e-17 \| 6.1e-21 \| 173.0 \| 298 \| (3, 339) \| 471 \| (565, 903) \| 1220 \| Protein mago nashi homolog 2 \| 6ICZ\_Y Protein mago nashi homolog 2 Human Post-catalytic Spliceosome, SPLICING HET: GTP, ATP, SEP, I6P; pdb70 \| 5DZR\_A \| 99.3 \| 8.5e-17 \| 6.3e-21 \| 170.2 \| 316 \| (3, 334) \| 471 \| (66, 472) \| 993 \| FRQ-interacting RNA helicase \| 5DZR\_A FRQ-interacting RNA helicase helicase, circadian clock, RNA BINDING; pdb70 \| 5GM6\_Y \| 99.3 \| 1e-16 \| 7.4e-21 \| 166.8 \| 298 \| (5, 340) \| 471 \| (225, 566) \| 876 \| Prp8, Brr2, Snu114, Rse1, Cus1 \| 5GM6\_Y Prp8, Brr2, Snu114, Rse1, Cus1 spliceosome, RNA splicing, Bact, Catalytically HET: GTP, ADP, ZN; pdb70 \| 5LQW\_O \| 99.3 \| 1e-16 \| 7.4e-21 \| 166.8 \| 298 \| (5, 340) \| 471 \| (225, 566) \| 876 \| Pre-mRNA-splicing factor 8, Pre-mRNA-splicing factor \| 5LQW\_O Pre-mRNA-splicing factor 8, Pre-mRNA-splicing factor activated spliceosome, spliceosome, pre-mRNA splicing; pdb70 \| 1GM5\_A \| 99.3 \| 1.1e-16 \| 7.7e-21 \| 164.0 \| 298 \| (2, 339) \| 471 \| (366, 693) \| 780 \| RECG \| 1GM5\_A RECG HELICASE, REPLICATION RESTART HET: ADP; pdb70 \| 2P6R\_A \| 99.3 \| 1.1e-16 \| 7.8e-21 \| 163.5 \| 296 \| (4, 332) \| 471 \| (25, 375) \| 702 \| afUHEL308 HELICASE/DNA Complex \| 2P6R\_A afUHEL308 HELICASE/DNA Complex PROTEIN-DNA COMPLEX, SF2 HELICASE, ARCHAEAL; pdb70 \| 1WP9\_C \| 99.3 \| 1.1e-16 \| 8.2e-21 \| 154.6 \| 96 \| (229, 331) \| 471 \| (361, 464) \| 494 \| Hef helicase/nuclease \| 1WP9\_C Hef helicase/nuclease Helicase, ATPase, DNA replication, DNA HET: PO4; pdb70 \| 5WX1\_A \| 99.3 \| 1.1e-16 \| 8.3e-21 \| 163.3 \| 286 \| (5, 339) \| 471 \| (248, 569) \| 733 \| Serine protease NS3 \| 5WX1\_A Serine protease NS3 protease, RNA helicase, HYDROLASE; pdb70 \| 5ZAM\_A \| 99.3 \| 1.3e-16 \| 9.7e-21 \| 176.7 \| 105 \| (229, 341) \| 471 \| (441, 566) \| 1922 \| Endoribonuclease Dicer, RISC-loading complex subunit \| 5ZAM\_A Endoribonuclease Dicer, RISC-loading complex subunit Dicer, TRBP, Cryo-EM, RNA interference; pdb70 \| 2V8O\_A \| 99.3 \| 1.4e-16 \| 1e-20 \| 152.7 \| 254 \| (23, 332) \| 471 \| (12, 292) \| 444 \| FLAVIVIRIN PROTEASE NS3 (E.C.3.4.21.91) \| 2V8O\_A FLAVIVIRIN PROTEASE NS3 (E.C.3.4.21.91) MURRAY VALLEY ENCEPHALITIS VIRUS, GLYCOPROTEIN; pdb70 \| 1WP9\_A \| 99.3 \| 1.4e-16 \| 1.1e-20 \| 153.7 \| 96 \| (229, 331) \| 471 \| (361, 464) \| 494 \| Hef helicase/nuclease \| 1WP9\_A Hef helicase/nuclease Helicase, ATPase, DNA replication, DNA HET: PO4; pdb70 \| 5LJ5\_Q \| 99.3 \| 1.9e-16 \| 1.4e-20 \| 167.0 \| 300 \| (3, 340) \| 471 \| (350, 703) \| 1071 \| Pre-mRNA-splicing factor 8, Pre-mRNA-splicing helicase \| 5LJ5\_Q Pre-mRNA-splicing factor 8, Pre-mRNA-splicing helicase spliceosome, snRNP, pre-mRNA splicing, trans-esterification HET: GTP; pdb70 \| 5WSG\_e \| 99.3 \| 1.9e-16 \| 1.4e-20 \| 167.0 \| 300 \| (3, 340) \| 471 \| (350, 703) \| 1071 \| Pre-mRNA-splicing factor 8, Pre-mRNA-splicing factor \| 5WSG\_e Pre-mRNA-splicing factor 8, Pre-mRNA-splicing factor Catalytic Step II spliceosome, C\* HET: GTP; pdb70 \| 5MZ4\_B \| 99.3 \| 2e-16 \| 1.5e-20 \| 161.8 \| 268 \| (23, 339) \| 471 \| (280, 580) \| 752 \| NS3/NS4A (E.C.3.4.22.-,3.4.21.113,3.6.1.15,3.6.4.13,2.7.7.48) \| 5MZ4\_B NS3/NS4A (E.C.3.4.22.-,3.4.21.113,3.6.1.15,3.6.4.13,2.7.7.48) Flaviviridae, NS3/NS4A, Protease/Helicase, viral protein; pdb70 \| 2JLQ\_A \| 99.3 \| 2.1e-16 \| 1.6e-20 \| 152.5 \| 278 \| (4, 333) \| 471 \| (4, 300) \| 451 \| SERINE PROTEASE SUBUNIT NS3 (E.C.3.4.21.91) \| 2JLQ\_A SERINE PROTEASE SUBUNIT NS3 (E.C.3.4.21.91) RIBONUCLEOPROTEIN, NUCLEOTIDE-BINDING, VIRAL NUCLEOPROTEIN, ENDOPLASMIC; pdb70 \| 6HYS\_A \| 99.3 \| 2.3e-16 \| 1.7e-20 \| 160.1 \| 298 \| (3, 339) \| 471 \| (18, 356) \| 673 \| ATP-dependent RNA helicase DHX8 (E.C.3.6.4.13) \| 6HYS\_A ATP-dependent RNA helicase DHX8 (E.C.3.6.4.13) Helicase, Splicing, RNA, RNA BINDING HET: ADP, DMS, EDO, ACT; pdb70 \| 6HYT\_A \| 99.3 \| 2.3e-16 \| 1.7e-20 \| 160.1 \| 298 \| (3, 339) \| 471 \| (18, 356) \| 673 \| ATP-dependent RNA helicase DHX8 (E.C.3.6.4.13) \| 6HYT\_A ATP-dependent RNA helicase DHX8 (E.C.3.6.4.13) helicase, splicing, RNA, RNA BINDING HET: ADP, DMS, EDO; pdb70 \| 2V6I\_A \| 99.3 \| 2.4e-16 \| 1.8e-20 \| 151.1 \| 254 \| (23, 332) \| 471 \| (2, 279) \| 431 \| RNA HELICASE \| 2V6I\_A RNA HELICASE MEMBRANE, HELICASE, HYDROLASE, RNA HELICASE; pdb70 \| 6HYT\_B \| 99.3 \| 2.4e-16 \| 1.8e-20 \| 160.0 \| 298 \| (3, 339) \| 471 \| (18, 356) \| 673 \| ATP-dependent RNA helicase DHX8 (E.C.3.6.4.13) \| 6HYT\_B ATP-dependent RNA helicase DHX8 (E.C.3.6.4.13) helicase, splicing, RNA, RNA BINDING HET: ADP, EDO, DMS; pdb70 \| 6IEH\_B \| 99.3 \| 2.5e-16 \| 1.9e-20 \| 166.2 \| 316 \| (3, 333) \| 471 \| (75, 467) \| 979 \| Exosome RNA helicase MTR4 (E.C.3.6.4.13) \| 6IEH\_B Exosome RNA helicase MTR4 (E.C.3.6.4.13) RNA helicase, MTR4, NRDE2, Complex HET: ATP; pdb70 \| 2ZJ8\_A \| 99.3 \| 2.7e-16 \| 2e-20 \| 160.9 \| 294 \| (3, 332) \| 471 \| (22, 374) \| 720 \| Putative ski2-type helicase (E.C.3.6.1.-) \| 2ZJ8\_A Putative ski2-type helicase (E.C.3.6.1.-) RecA fold, ATP-binding, Helicase, Hydrolase; pdb70 \| 2ZJA\_A \| 99.3 \| 2.7e-16 \| 2e-20 \| 160.9 \| 294 \| (3, 332) \| 471 \| (22, 374) \| 720 \| Putative ski2-type helicase (E.C.3.6.1.-) \| 2ZJA\_A Putative ski2-type helicase (E.C.3.6.1.-) RecA fold, ATP-binding, Helicase, Hydrolase HET: ACP; pdb70 \| 2WV9\_A \| 99.3 \| 2.9e-16 \| 2.2e-20 \| 158.7 \| 277 \| (4, 339) \| 471 \| (225, 529) \| 673 \| FLAVIVIRIN PROTEASE NS2B REGULATORY SUBUNIT \| 2WV9\_A FLAVIVIRIN PROTEASE NS2B REGULATORY SUBUNIT NUCLEOTIDE-BINDING, CAPSID PROTEIN, RNA REPLICATION; pdb70 \| 6FAC\_A \| 99.3 \| 3.4e-16 \| 2.6e-20 \| 158.4 \| 297 \| (4, 339) \| 471 \| (32, 372) \| 655 \| Putative mRNA splicing factor \| 6FAC\_A Putative mRNA splicing factor SPLICING, ATPASE, HELICASE, G-PATCH, HYDROLASE HET: ADP; pdb70 \| 2V6J\_A \| 99.3 \| 3.6e-16 \| 2.7e-20 \| 149.8 \| 254 \| (23, 332) \| 471 \| (2, 279) \| 431 \| RNA HELICASE \| 2V6J\_A RNA HELICASE MEMBRANE, HELICASE, HYDROLASE, RNA HELICASE; pdb70 \| 6EUD\_A \| 99.3 \| 4.3e-16 \| 3.1e-20 \| 161.1 \| 298 \| (4, 340) \| 471 \| (8, 340) \| 812 \| ATP-dependent RNA helicase HrpB (E.C.3.6.4.13) \| 6EUD\_A ATP-dependent RNA helicase HrpB (E.C.3.6.4.13) HrpB, DEAH/RHA helicase, bacterial helicase; pdb70 \| 5XDR\_A \| 99.3 \| 4.3e-16 \| 3.2e-20 \| 158.6 \| 299 \| (3, 339) \| 471 \| (32, 380) \| 690 \| Pre-mRNA-splicing factor ATP-dependent RNA helicase \| 5XDR\_A Pre-mRNA-splicing factor ATP-dependent RNA helicase RNA helicase, DEAH-box, DHX15, Prp43 HET: ADP, SO4; pdb70 \| 5WSO\_A \| 99.3 \| 4.9e-16 \| 3.6e-20 \| 149.7 \| 278 \| (9, 339) \| 471 \| (2, 312) \| 476 \| NS3 helicase \| 5WSO\_A NS3 helicase BVDV, Bovine viral diarrhea virus; pdb70 \| 6G7E\_B \| 99.3 \| 5.7e-16 \| 4.3e-20 \| 174.1 \| 106 \| (229, 341) \| 471 \| (1664, 1774) \| 1852 \| Helicase-like protein \| 6G7E\_B Helicase-like protein ATPase, Hydrolase, TRANSCRIPTION HET: MSE; pdb70 \| 1YKS\_A \| 99.3 \| 6.7e-16 \| 5e-20 \| 148.4 \| 254 \| (23, 331) \| 471 \| (8, 286) \| 440 \| Genome polyprotein [contains: Flavivirin protease \| 1YKS\_A Genome polyprotein [contains: Flavivirin protease helicase, flavivirus, DEAD-box, ATPase, RTPase; pdb70 \| 6BU9\_A \| 99.2 \| 7.9e-16 \| 5.8e-20 \| 170.3 \| 119 \| (214, 340) \| 471 \| (367, 509) \| 1724 \| Dicer-2, isoform A/RNA Complex \| 6BU9\_A Dicer-2, isoform A/RNA Complex Dicer, Dcr2, Dcr-2, dmDcr-2, Dicer-2; pdb70 \| 6BUA\_A \| 99.2 \| 7.9e-16 \| 5.8e-20 \| 170.3 \| 119 \| (214, 340) \| 471 \| (367, 509) \| 1724 \| Drosophila Dicer-2 \| 6BUA\_A Drosophila Dicer-2 Dicer, dmDcr2, Dicer-2, helicase, platform; pdb70 \| 5N8R\_B \| 99.2 \| 7.9e-16 \| 5.8e-20 \| 162.0 \| 104 \| (229, 339) \| 471 \| (422, 553) \| 944 \| CG9323, isoform A/DNA Complex \| 5N8R\_B CG9323, isoform A/DNA Complex Helicase, DExH, ssDNA, hydrolase; pdb70 \| 5VHE\_A \| 99.2 \| 8.4e-16 \| 6.2e-20 \| 161.4 \| 104 \| (229, 339) \| 471 \| (404, 531) \| 933 \| DEAH (Asp-Glu-Ala-His) box polypeptide 36/DNA \| 5VHE\_A DEAH (Asp-Glu-Ala-His) box polypeptide 36/DNA HYDROLASE; pdb70 \| 5N9D\_B \| 99.2 \| 8.7e-16 \| 6.5e-20 \| 161.6 \| 104 \| (229, 339) \| 471 \| (422, 553) \| 944 \| CG9323, isoform A/DNA Complex \| 5N9D\_B CG9323, isoform A/DNA Complex Helicase DExH ssDNA, hydrolase; pdb70 \| 6RM9\_A \| 99.2 \| 8.8e-16 \| 6.6e-20 \| 155.5 \| 297 \| (4, 339) \| 471 \| (37, 377) \| 660 \| Putative mRNA splicing factor, Putative \| 6RM9\_A Putative mRNA splicing factor, Putative Prp2, DEAH-box ATPase, G-patch, spliceosome HET: GOL, ADP; pdb70 \| 4A4Z\_A \| 99.2 \| 9.3e-16 \| 6.9e-20 \| 161.6 \| 143 \| (2, 149) \| 471 \| (37, 189) \| 997 \| ANTIVIRAL HELICASE SKI2 (E.C.3.6.4.13) \| 4A4Z\_A ANTIVIRAL HELICASE SKI2 (E.C.3.6.4.13) HYDROLASE, ATPASE, MRNA DEGRADATION, EXOSOME HET: ANP, EDO; pdb70 \| 5VHA\_A \| 99.2 \| 1e-15 \| 7.5e-20 \| 159.9 \| 104 \| (229, 339) \| 471 \| (341, 468) \| 870 \| DEAH (Asp-Glu-Ala-His) box polypeptide 36 \| 5VHA\_A DEAH (Asp-Glu-Ala-His) box polypeptide 36 HYDROLASE; pdb70 \| 2IBM\_A \| 99.2 \| 1.2e-15 \| 8.7e-20 \| 157.5 \| 113 \| (213, 334) \| 471 \| (412, 533) \| 780 \| Preprotein translocase secA subunit \| 2IBM\_A Preprotein translocase secA subunit protein translocation, SecA, signal peptide HET: ADP; pdb70 \| 1TF5\_A \| 99.2 \| 1.2e-15 \| 9.2e-20 \| 158.8 \| 112 \| (214, 334) \| 471 \| (416, 536) \| 844 \| Preprotein translocase secA subunit \| 1TF5\_A Preprotein translocase secA subunit ATPase, helicase, translocation, secretion, PROTEIN; pdb70 \| 5VHD\_D \| 99.2 \| 1.3e-15 \| 9.7e-20 \| 159.1 \| 104 \| (229, 339) \| 471 \| (341, 468) \| 870 \| DEAH (Asp-Glu-Ala-His) box polypeptide 36 \| 5VHD\_D DEAH (Asp-Glu-Ala-His) box polypeptide 36 HYDROLASE HET: ADP; pdb70 \| 6H57\_A \| 99.2 \| 1.4e-15 \| 1e-19 \| 161.6 \| 79 \| (255, 340) \| 471 \| (730, 826) \| 1267 \| Probable ATP-dependent RNA helicase DHR1 \| 6H57\_A Probable ATP-dependent RNA helicase DHR1 protein, HYDROLASE HET: EDO; pdb70 \| 5Y4Z\_A \| 99.2 \| 1.4e-15 \| 1.1e-19 \| 145.9 \| 92 \| (229, 331) \| 471 \| (179, 287) \| 440 \| NS3 helicase (E.C.3.4.21.91,3.6.1.15,3.6.4.13) \| 5Y4Z\_A NS3 helicase (E.C.3.4.21.91,3.6.1.15,3.6.4.13) ZIKA virus, NS3 helicase, AMPPNP HET: ANP; pdb70 \| 5YVW\_B \| 99.2 \| 1.5e-15 \| 1.1e-19 \| 151.7 \| 255 \| (23, 333) \| 471 \| (167, 448) \| 599 \| Genome polyprotein \| 5YVW\_B Genome polyprotein SERINE PROTEASE, NON-STRUCTURAL PROTEIN 3 HET: EPE; pdb70 \| 3JV2\_A \| 99.2 \| 1.7e-15 \| 1.2e-19 \| 156.4 \| 113 \| (213, 334) \| 471 \| (415, 536) \| 783 \| Protein translocase subunit secA \| 3JV2\_A Protein translocase subunit secA protein translocation, ATPase, conformational change HET: ADP; pdb70 \| 5MFX\_A \| 99.2 \| 2.2e-15 \| 1.6e-19 \| 145.2 \| 101 \| (229, 340) \| 471 \| (184, 302) \| 451 \| NS3 helicase/RNA Complex \| 5MFX\_A NS3 helicase/RNA Complex Helicase, RNA, hydrolase HET: FLC; pdb70 \| 1GL9\_B \| 99.2 \| 2.3e-15 \| 1.7e-19 \| 159.5 \| 262 \| (3, 315) \| 471 \| (55, 353) \| 1054 \| REVERSE GYRASE \| 1GL9\_B REVERSE GYRASE TOPOISOMERASE, DNA SUPERCOILING, ARCHAEA, HELICASE HET: ANP; pdb70 \| 3RC3\_A \| 99.2 \| 2.4e-15 \| 1.8e-19 \| 151.1 \| 96 \| (229, 332) \| 471 \| (320, 431) \| 677 \| ATP-dependent RNA helicase SUPV3L1, mitochondrial \| 3RC3\_A ATP-dependent RNA helicase SUPV3L1, mitochondrial helicase, mitochondria, Suv3, Nucleus, HYDROLASE HET: ANP; pdb70 \| 2VA8\_B \| 99.2 \| 2.5e-15 \| 1.9e-19 \| 153.5 \| 304 \| (3, 332) \| 471 \| (29, 395) \| 715 \| SKI2-TYPE HELICASE \| 2VA8\_B SKI2-TYPE HELICASE HEL308, SSO2462, HELICASE, HYDROLASE, DNA HET: SO4; pdb70 \| 3RC8\_A \| 99.2 \| 2.9e-15 \| 2.1e-19 \| 150.5 \| 96 \| (229, 332) \| 471 \| (320, 431) \| 677 \| ATP-dependent RNA helicase SUPV3L1, mitochondrial \| 3RC8\_A ATP-dependent RNA helicase SUPV3L1, mitochondrial Helicaase, Suv3, Mitochondria, RNA, Helicase; pdb70 \| 5YW1\_B \| 99.2 \| 3.1e-15 \| 2.3e-19 \| 149.8 \| 256 \| (23, 333) \| 471 \| (186, 467) \| 618 \| Genome polyprotein, PTI protein \| 5YW1\_B Genome polyprotein, PTI protein SERINE PROTEASE, NON-STRUCTURAL PROTEIN 3; pdb70 \| 1GKU\_B \| 99.2 \| 3.8e-15 \| 2.8e-19 \| 157.6 \| 262 \| (4, 315) \| 471 \| (56, 353) \| 1054 \| REVERSE GYRASE \| 1GKU\_B REVERSE GYRASE TOPOISOMERASE, DNA SUPERCOILING, ARCHAEA, HELICASE; pdb70 \| 6QV4\_A \| 99.2 \| 4e-15 \| 3e-19 \| 166.3 \| 300 \| (4, 332) \| 471 \| (910, 1280) \| 1725 \| Pre-mRNA splicing helicase-like protein \| 6QV4\_A Pre-mRNA splicing helicase-like protein Helicase ATP-gamma-S Brr2 Ski2 Chaetomium HET: AGS; pdb70 \| 6H2J\_B \| 99.2 \| 4.8e-15 \| 3.5e-19 \| 156.6 \| 330 \| (3, 342) \| 471 \| (270, 706) \| 1038 \| Type I restriction enzyme R \| 6H2J\_B Type I restriction enzyme R EcoR124, HsdR, C-terminal domain, restriction-modification HET: MSE, ATP; pdb70 \| 5AOR\_B \| 99.1 \| 5.2e-15 \| 3.9e-19 \| 160.3 \| 104 \| (229, 339) \| 471 \| (653, 781) \| 1293 \| DOSAGE COMPENSATION REGULATOR (E.C.3.6.4.13) \| 5AOR\_B DOSAGE COMPENSATION REGULATOR (E.C.3.6.4.13) HYDROLASE-RNA COMPLEX, HELICASE, DOSAGE COMPENSATION HET: ADP, ALF, GOL; pdb70 \| 5M59\_E \| 99.1 \| 5.3e-15 \| 4e-19 \| 165.6 \| 300 \| (4, 332) \| 471 \| (957, 1327) \| 1772 \| Putative pre-mRNA splicing factor, Pre-mRNA \| 5M59\_E Putative pre-mRNA splicing factor, Pre-mRNA Brr2, pre-mRNA splicing, RNA-helicase, Prp8 HET: ACT; pdb70 \| 5AOR\_A \| 99.1 \| 5.6e-15 \| 4.1e-19 \| 160.1 \| 104 \| (229, 339) \| 471 \| (653, 781) \| 1293 \| DOSAGE COMPENSATION REGULATOR (E.C.3.6.4.13) \| 5AOR\_A DOSAGE COMPENSATION REGULATOR (E.C.3.6.4.13) HYDROLASE-RNA COMPLEX, HELICASE, DOSAGE COMPENSATION HET: ADP, ALF, GOL; pdb70 \| 4DDU\_A \| 99.1 \| 6.1e-15 \| 4.5e-19 \| 157.4 \| 292 \| (3, 341) \| 471 \| (77, 427) \| 1104 \| Reverse gyrase (E.C.3.6.4.12, 5.99.1.3) \| 4DDU\_A Reverse gyrase (E.C.3.6.4.12, 5.99.1.3) TOPOISOMERASE, DNA SUPERCOILING, ARCHAEA, HELICASE; pdb70 \| 2WHX\_A \| 99.1 \| 6.3e-15 \| 4.7e-19 \| 147.6 \| 256 \| (23, 334) \| 471 \| (186, 468) \| 618 \| SERINE PROTEASE/NTPASE/HELICASE NS3 (E.C.3.4.21.91, 3.6.1.15 \| 2WHX\_A SERINE PROTEASE/NTPASE/HELICASE NS3 (E.C.3.4.21.91, 3.6.1.15 TRANSCRIPTION, HYDROLASE, ATP-BINDING, RETICULUM, NUCLEOTIDYLTRANSFERASE HET: ADP; pdb70 \| 4XJX\_A \| 99.1 \| 6.8e-15 \| 5e-19 \| 155.4 \| 330 \| (3, 342) \| 471 \| (270, 706) \| 1038 \| HSDR subunit of type I \| 4XJX\_A HSDR subunit of type I Restriction enzyme, ATP, hydrolase HET: ATP; pdb70 \| 6QX9\_5B \| 99.1 \| 7e-15 \| 5.1e-19 \| 166.2 \| 300 \| (4, 331) \| 471 \| (480, 860) \| 2136 \| Splicing factor 3B subunit 4 \| 6QX9\_5B Splicing factor 3B subunit 4 RNP complex, splicing, RNA, protein HET: M7M, IHP, GTP; pdb70 \| 4DDT\_A \| 99.1 \| 7.1e-15 \| 5.3e-19 \| 156.9 \| 290 \| (3, 340) \| 471 \| (77, 426) \| 1104 \| Reverse gyrase (E.C.3.6.4.12, 5.99.1.3) \| 4DDT\_A Reverse gyrase (E.C.3.6.4.12, 5.99.1.3) TOPOISOMERASE, DNA SUPERCOILING, ARCHAEA, HELICASE; pdb70 \| 4DDW\_A \| 99.1 \| 7.1e-15 \| 5.3e-19 \| 156.9 \| 290 \| (3, 340) \| 471 \| (77, 426) \| 1104 \| Reverse gyrase (E.C.3.6.4.12, 5.99.1.3) \| 4DDW\_A Reverse gyrase (E.C.3.6.4.12, 5.99.1.3) TOPOISOMERASE, DNA SUPERCOILING, ARCHAEA, HELICASE; pdb70 \| 6QW6\_5B \| 99.1 \| 7.2e-15 \| 5.3e-19 \| 166.1 \| 300 \| (4, 331) \| 471 \| (480, 860) \| 2136 \| Small nuclear ribonucleoprotein Sm D1 \| 6QW6\_5B Small nuclear ribonucleoprotein Sm D1 RNP complex, splicing, RNA, protein HET: IHP, M7M, GTP; pdb70 \| 3KQN\_A \| 99.1 \| 7.7e-15 \| 5.7e-19 \| 140.7 \| 255 \| (23, 341) \| 471 \| (9, 290) \| 437 \| Serine protease/NTPase/helicase NS3 (E.C.3.4.21.98, 3.6.1.15 \| 3KQN\_A Serine protease/NTPase/helicase NS3 (E.C.3.4.21.98, 3.6.1.15 helicase-substrate transition-state complex, HCV, NS3 HET: ADP; pdb70 \| 3KQL\_A \| 99.1 \| 8e-15 \| 5.9e-19 \| 140.6 \| 254 \| (23, 340) \| 471 \| (9, 289) \| 437 \| Serine protease/NTPase/helicase NS3 (E.C.3.4.21.98, 3.6.1.15 \| 3KQL\_A Serine protease/NTPase/helicase NS3 (E.C.3.4.21.98, 3.6.1.15 helicase-substrate transition-state complex, HCV, NS3 HET: ADP, ALF; pdb70 \| 3KQU\_D \| 99.1 \| 8e-15 \| 5.9e-19 \| 140.6 \| 254 \| (23, 340) \| 471 \| (9, 289) \| 437 \| Serine protease/NTPase/helicase NS3 (E.C.3.4.21.98, 3.6.1.15 \| 3KQU\_D Serine protease/NTPase/helicase NS3 (E.C.3.4.21.98, 3.6.1.15 helicase-substrate ground-state complex, HCV, NS3 HET: ADP, BEF; pdb70 \| 4KIT\_B \| 99.1 \| 8.4e-15 \| 6.3e-19 \| 163.7 \| 300 \| (4, 331) \| 471 \| (90, 470) \| 1739 \| U5 small nuclear ribonucleoprotein 200 \| 4KIT\_B U5 small nuclear ribonucleoprotein 200 RecA domain, winged helix domain HET: ADP; pdb70 \| 1CU1\_B \| 99.1 \| 8.8e-15 \| 6.5e-19 \| 146.8 \| 254 \| (23, 340) \| 471 \| (211, 491) \| 645 \| PROTEIN (HEPATITIS C VIRUS) \| 1CU1\_B PROTEIN (HEPATITIS C VIRUS) hepatitis C Virus, bifunctional, protease-helicase HET: PO4; pdb70 \| 4UAQ\_A \| 99.1 \| 8.9e-15 \| 6.6e-19 \| 150.5 \| 119 \| (213, 340) \| 471 \| (425, 560) \| 778 \| SecA2 \| 4UAQ\_A SecA2 Protein Transport, DEAD/DEAH box helicase HET: MSE; pdb70 \| 4Q2C\_A \| 99.1 \| 1e-14 \| 7.5e-19 \| 152.9 \| 94 \| (229, 332) \| 471 \| (595, 704) \| 949 \| CRISPR-associated helicase Cas3 \| 4Q2C\_A CRISPR-associated helicase Cas3 RecA, HD nuclease, Hydrolase; pdb70 \| 4Q2D\_A \| 99.1 \| 1e-14 \| 7.5e-19 \| 152.9 \| 94 \| (229, 332) \| 471 \| (595, 704) \| 949 \| CRISPR-associated helicase Cas3 \| 4Q2D\_A CRISPR-associated helicase Cas3 RecA, HD nuclease, Hydrolase HET: DTP; pdb70 \| 2FSF\_A \| 99.1 \| 1e-14 \| 7.6e-19 \| 151.5 \| 113 \| (213, 334) \| 471 \| (424, 574) \| 853 \| Preprotein translocase secA subunit \| 2FSF\_A Preprotein translocase secA subunit ATPase, DNA-RNA helicase, Protein translocation; pdb70 \| 2FSH\_A \| 99.1 \| 1e-14 \| 7.6e-19 \| 151.5 \| 113 \| (213, 334) \| 471 \| (424, 574) \| 853 \| Preprotein translocase secA subunit \| 2FSH\_A Preprotein translocase secA subunit ATPase, DNA-RNA helicase, Protein translocation HET: ANP; pdb70 \| 6S8Q\_B \| 99.1 \| 1.1e-14 \| 7.9e-19 \| 163.0 \| 300 \| (4, 331) \| 471 \| (91, 471) \| 1747 \| U5 small nuclear ribonucleoprotein 200 \| 6S8Q\_B U5 small nuclear ribonucleoprotein 200 RNP REMODELING, PRE-MRNA SPLICING, SPLICEOSOME HET: SO4; pdb70 \| 3O8B\_B \| 99.1 \| 1.3e-14 \| 9.9e-19 \| 145.7 \| 254 \| (23, 340) \| 471 \| (232, 512) \| 666 \| HCV NS3 protease/helicase (E.C.3.4.21.98, 3.6.1.15 \| 3O8B\_B HCV NS3 protease/helicase (E.C.3.4.21.98, 3.6.1.15 helicase, NTPase, HCV, RNA, translocation HET: SO4; pdb70 \| 6QDV\_B \| 99.1 \| 1.3e-14 \| 9.9e-19 \| 161.9 \| 300 \| (4, 331) \| 471 \| (77, 457) \| 1722 \| Eukaryotic initiation factor 4A-III, RNA-binding \| 6QDV\_B Eukaryotic initiation factor 4A-III, RNA-binding spliceosome, RNA, complex, SPLICING HET: GTP, I6P, ATP, SEP; pdb70 \| 6O16\_A \| 99.1 \| 1.4e-14 \| 1e-18 \| 152.5 \| 78 \| (255, 339) \| 471 \| (407, 502) \| 975 \| DEAH (Asp-Glu-Ala-His) box polypeptide 37/RNA \| 6O16\_A DEAH (Asp-Glu-Ala-His) box polypeptide 37/RNA RNA helicase, ribosome biogenesis, RNA-dependent; pdb70 \| 5DCA\_A \| 99.1 \| 1.5e-14 \| 1.1e-18 \| 161.8 \| 296 \| (4, 331) \| 471 \| (1144, 1511) \| 1948 \| Pre-mRNA-splicing helicase BRR2 (E.C.3.6.4.13), Pre-mRNA-splicing \| 5DCA\_A Pre-mRNA-splicing helicase BRR2 (E.C.3.6.4.13), Pre-mRNA-splicing protein complex, helicase, RNP remodeling; pdb70 \| 4QQW\_A \| 99.1 \| 1.6e-14 \| 1.2e-18 \| 151.7 \| 94 \| (229, 332) \| 471 \| (602, 715) \| 964 \| CRISPR-associated helicase, Cas3 family/dna \| 4QQW\_A CRISPR-associated helicase, Cas3 family/dna CRISPR, Cas3, helicase, Hydrolase-DNA complex; pdb70 \| 5DCA\_A \| 99.1 \| 2e-14 \| 1.5e-18 \| 160.8 \| 300 \| (3, 331) \| 471 \| (294, 676) \| 1948 \| Pre-mRNA-splicing helicase BRR2 (E.C.3.6.4.13), Pre-mRNA-splicing \| 5DCA\_A Pre-mRNA-splicing helicase BRR2 (E.C.3.6.4.13), Pre-mRNA-splicing protein complex, helicase, RNP remodeling; pdb70 \| 2VDA\_A \| 99.1 \| 2.3e-14 \| 1.7e-18 \| 148.3 \| 113 \| (213, 334) \| 471 \| (424, 574) \| 828 \| TRANSLOCASE SUBUNIT SECA, MALTOPORIN \| 2VDA\_A TRANSLOCASE SUBUNIT SECA, MALTOPORIN SUGAR TRANSPORT, PROTEIN TRANSPORT, PROTEIN; pdb70 \| 3JUX\_A \| 99.1 \| 2.3e-14 \| 1.7e-18 \| 147.6 \| 112 \| (214, 334) \| 471 \| (458, 578) \| 822 \| Protein translocase subunit secA \| 3JUX\_A Protein translocase subunit secA protein translocation, ATPase, conformational change HET: ADP; pdb70 \| 4BGD\_A \| 99.1 \| 2.3e-14 \| 1.7e-18 \| 159.6 \| 300 \| (3, 331) \| 471 \| (56, 438) \| 1722 \| PRE-MRNA-SPLICING HELICASE BRR2 (E.C.3.6.4.13), PRE-MRNA-SPLICING \| 4BGD\_A PRE-MRNA-SPLICING HELICASE BRR2 (E.C.3.6.4.13), PRE-MRNA-SPLICING TRANSCRIPTION, SPLICEOSOME, RNA HELICASE, U5 HET: ADP, PE5; pdb70 \| 5K9T\_A \| 99.1 \| 2.4e-14 \| 1.8e-18 \| 142.4 \| 112 \| (214, 334) \| 471 \| (421, 570) \| 578 \| Protein translocase subunit SecA \| 5K9T\_A Protein translocase subunit SecA preprotein translocase, SecA-N68, ATPase, C-terminal HET: ADP; pdb70 \| 6F4A\_B \| 99.1 \| 2.6e-14 \| 1.9e-18 \| 143.4 \| 252 \| (24, 332) \| 471 \| (160, 431) \| 644 \| Exoribonuclease II, mitochondrial, Suv3 helicase/RNA \| 6F4A\_B Exoribonuclease II, mitochondrial, Suv3 helicase/RNA RNA degradation, mitochondria, nuclease, helicase; pdb70 \| 4YS0\_A \| 99.1 \| 2.7e-14 \| 2e-18 \| 147.3 \| 113 \| (213, 334) \| 471 \| (457, 578) \| 824 \| Protein translocase subunit SecA \| 4YS0\_A Protein translocase subunit SecA Protein translocation, SecA, ATPase, PROTEIN HET: ADP; pdb70 \| 5B7I\_A \| 99.0 \| 3.4e-14 \| 2.5e-18 \| 148.7 \| 94 \| (229, 332) \| 471 \| (733, 863) \| 1082 \| CRISPR-associated nuclease/helicase Cas3 subtype I-F/YPEST \| 5B7I\_A CRISPR-associated nuclease/helicase Cas3 subtype I-F/YPEST DNA nuclease, phagy protein, Anti-CRISPR HET: ADP, MSE; pdb70 \| 4BGD\_A \| 99.0 \| 3.4e-14 \| 2.5e-18 \| 158.3 \| 298 \| (4, 332) \| 471 \| (906, 1274) \| 1722 \| PRE-MRNA-SPLICING HELICASE BRR2 (E.C.3.6.4.13), PRE-MRNA-SPLICING \| 4BGD\_A PRE-MRNA-SPLICING HELICASE BRR2 (E.C.3.6.4.13), PRE-MRNA-SPLICING TRANSCRIPTION, SPLICEOSOME, RNA HELICASE, U5 HET: ADP, PE5; pdb70 \| 6QDV\_B \| 99.0 \| 4.4e-14 \| 3.2e-18 \| 157.7 \| 298 \| (4, 331) \| 471 \| (924, 1292) \| 1722 \| Eukaryotic initiation factor 4A-III, RNA-binding \| 6QDV\_B Eukaryotic initiation factor 4A-III, RNA-binding spliceosome, RNA, complex, SPLICING HET: GTP, I6P, ATP, SEP; pdb70 \| 6QW6\_5B \| 99.0 \| 4.8e-14 \| 3.5e-18 \| 159.3 \| 297 \| (4, 331) \| 471 \| (1327, 1695) \| 2136 \| Small nuclear ribonucleoprotein Sm D1 \| 6QW6\_5B Small nuclear ribonucleoprotein Sm D1 RNP complex, splicing, RNA, protein HET: IHP, M7M, GTP; pdb70 \| 4KIT\_B \| 99.0 \| 5e-14 \| 3.7e-18 \| 157.4 \| 298 \| (4, 331) \| 471 \| (937, 1305) \| 1739 \| U5 small nuclear ribonucleoprotein 200 \| 4KIT\_B U5 small nuclear ribonucleoprotein 200 RecA domain, winged helix domain HET: ADP; pdb70 \| 6QX9\_5B \| 99.0 \| 5e-14 \| 3.7e-18 \| 159.2 \| 297 \| (4, 331) \| 471 \| (1327, 1695) \| 2136 \| Splicing factor 3B subunit 4 \| 6QX9\_5B Splicing factor 3B subunit 4 RNP complex, splicing, RNA, protein HET: M7M, IHP, GTP; pdb70 \| 6DCR\_B \| 99.0 \| 5.8e-14 \| 4.3e-18 \| 142.7 \| 78 \| (255, 339) \| 471 \| (468, 559) \| 694 \| Primosomal protein N' (E.C.3.6.4.-) \| 6DCR\_B Primosomal protein N' (E.C.3.6.4.-) PriA, Helicase, DNA replication restart HET: SO4; pdb70 \| 5M59\_E \| 99.0 \| 5.8e-14 \| 4.3e-18 \| 157.1 \| 304 \| (3, 332) \| 471 \| (106, 490) \| 1772 \| Putative pre-mRNA splicing factor, Pre-mRNA \| 5M59\_E Putative pre-mRNA splicing factor, Pre-mRNA Brr2, pre-mRNA splicing, RNA-helicase, Prp8 HET: ACT; pdb70 \| 6DGD\_A \| 99.0 \| 6.3e-14 \| 4.7e-18 \| 143.7 \| 144 \| (3, 149) \| 471 \| (216, 379) \| 751 \| Primosomal protein N'/DNA Complex \| 6DGD\_A Primosomal protein N'/DNA Complex DNA replication restart, PriA helicase HET: SO4; pdb70 \| 6S8Q\_B \| 99.0 \| 7e-14 \| 5.2e-18 \| 156.3 \| 298 \| (4, 331) \| 471 \| (938, 1306) \| 1747 \| U5 small nuclear ribonucleoprotein 200 \| 6S8Q\_B U5 small nuclear ribonucleoprotein 200 RNP REMODELING, PRE-MRNA SPLICING, SPLICEOSOME HET: SO4; pdb70 \| 5YLZ\_W \| 99.0 \| 9.4e-14 \| 6.9e-18 \| 146.9 \| 105 \| (229, 340) \| 471 \| (691, 822) \| 1145 \| Pre-mRNA-splicing factor 8, Pre-mRNA-splicing factor \| 5YLZ\_W Pre-mRNA-splicing factor 8, Pre-mRNA-splicing factor Post-catalytic Spliceosome (P complex), RNA HET: I6P, GTP; pdb70 \| 6BK8\_P \| 99.0 \| 9.4e-14 \| 6.9e-18 \| 146.9 \| 105 \| (229, 340) \| 471 \| (691, 822) \| 1145 \| Pre-mRNA-splicing factor 8, Pre-mRNA-splicing factor \| 6BK8\_P Pre-mRNA-splicing factor 8, Pre-mRNA-splicing factor pre-mRNA splicing, spliceosome, post-catalytic, P HET: IHP, GTP; pdb70 \| 5LTJ\_A \| 99.0 \| 1.1e-13 \| 8.3e-18 \| 140.9 \| 104 \| (229, 339) \| 471 \| (248, 387) \| 714 \| Pre-mRNA-splicing factor ATP-dependent RNA helicase \| 5LTJ\_A Pre-mRNA-splicing factor ATP-dependent RNA helicase Spliceosome, RNA HELICASE, DEAH-BOX PROTEIN HET: MPD, ADP; pdb70 \| 6QV4\_A \| 99.0 \| 1.2e-13 \| 8.7e-18 \| 154.3 \| 302 \| (3, 331) \| 471 \| (59, 442) \| 1725 \| Pre-mRNA splicing helicase-like protein \| 6QV4\_A Pre-mRNA splicing helicase-like protein Helicase ATP-gamma-S Brr2 Ski2 Chaetomium HET: AGS; pdb70 \| 6QID\_A \| 98.9 \| 2e-13 \| 1.5e-17 \| 138.8 \| 104 \| (229, 339) \| 471 \| (247, 386) \| 705 \| Putative pre-mRNA splicing factor/RNA Complex \| 6QID\_A Putative pre-mRNA splicing factor/RNA Complex Splicing, DEAH, ATPase, helicase, HYDROLASE HET: MPD, PEG, ADP, SO4; pdb70 \| 1NKT\_A \| 98.9 \| 2.1e-13 \| 1.5e-17 \| 141.4 \| 123 \| (213, 341) \| 471 \| (443, 616) \| 922 \| PREPROTEIN TRANSLOCASE SECA 1 SUBUNIT \| 1NKT\_A PREPROTEIN TRANSLOCASE SECA 1 SUBUNIT PREPROTEIN TRANSLOCATION, ATPASE, TRANSMEMBRANE TRANSPORT HET: ADP; pdb70 \| 1NL3\_A \| 98.9 \| 2.1e-13 \| 1.5e-17 \| 141.4 \| 123 \| (213, 341) \| 471 \| (443, 616) \| 922 \| PREPROTEIN TRANSLOCASE SECA 1 SUBUNIT \| 1NL3\_A PREPROTEIN TRANSLOCASE SECA 1 SUBUNIT PREPROTEIN TRANSLOCATION, ATPASE, TRANSMEMBRANE TRANSPORT; pdb70 \| 1NL3\_B \| 98.9 \| 2.1e-13 \| 1.5e-17 \| 141.4 \| 123 \| (213, 341) \| 471 \| (443, 616) \| 922 \| PREPROTEIN TRANSLOCASE SECA 1 SUBUNIT \| 1NL3\_B PREPROTEIN TRANSLOCASE SECA 1 SUBUNIT PREPROTEIN TRANSLOCATION, ATPASE, TRANSMEMBRANE TRANSPORT; pdb70 \| 2IPC\_A \| 98.9 \| 4.5e-13 \| 3.3e-17 \| 138.3 \| 64 \| (3, 72) \| 471 \| (78, 141) \| 997 \| Preprotein translocase SecA subunit \| 2IPC\_A Preprotein translocase SecA subunit nucleotide binding fold, ATPase, parallel; pdb70 \| 4ZCF\_C \| 98.8 \| 1.6e-12 \| 1.2e-16 \| 135.8 \| 47 \| (278, 331) \| 471 \| (493, 539) \| 970 \| EcoP15I, ModA, EcoP15I, Res/RNA Complex \| 4ZCF\_C EcoP15I, ModA, EcoP15I, Res/RNA Complex Hydrolase/DNA, ATP motor, DNA methyltransferase HET: AMP; pdb70 \| 5GI4\_B \| 98.7 \| 3.9e-12 \| 2.9e-16 \| 110.0 \| 112 \| (215, 334) \| 471 \| (15, 126) \| 228 \| ATP-dependent RNA helicase DeaD (E.C.3.6.4.13) \| 5GI4\_B ATP-dependent RNA helicase DeaD (E.C.3.6.4.13) Dimer, RecA-like, Wild type, RNA; pdb70 \| 3EAS\_B \| 98.6 \| 1.4e-11 \| 1e-15 \| 105.2 \| 113 \| (214, 334) \| 471 \| (17, 129) \| 212 \| Hera \| 3EAS\_B Hera DEAD box RNA helicase, dimer; pdb70 \| 3I32\_A \| 98.6 \| 1.5e-11 \| 1.1e-15 \| 110.6 \| 113 \| (214, 334) \| 471 \| (14, 126) \| 300 \| Heat resistant RNA dependent ATPase \| 3I32\_A Heat resistant RNA dependent ATPase RNA helicase, ATPase, Dimer, RNA; pdb70 \| 3EAQ\_A \| 98.5 \| 5.3e-11 \| 3.9e-15 \| 101.4 \| 113 \| (214, 334) \| 471 \| (17, 129) \| 212 \| Heat resistant RNA dependent ATPase \| 3EAQ\_A Heat resistant RNA dependent ATPase DEAD box RNA helicase, dimer; pdb70 \| 3EAR\_A \| 98.5 \| 5.3e-11 \| 3.9e-15 \| 101.4 \| 113 \| (214, 334) \| 471 \| (17, 129) \| 212 \| Hera \| 3EAR\_A Hera DEAD box RNA helicase, dimer; pdb70 \| 2JGN\_B \| 98.4 \| 9e-11 \| 6.6e-15 \| 97.6 \| 114 \| (214, 334) \| 471 \| (31, 144) \| 185 \| ATP-DEPENDENT RNA HELICASE DDX3X (E.C.3.6.1.-) \| 2JGN\_B ATP-DEPENDENT RNA HELICASE DDX3X (E.C.3.6.1.-) PHOSPHORYLATION, NUCLEOTIDE-BINDING, HELICASE, HYDROLASE, RNA-BINDING; pdb70 \| 5JXT\_B \| 98.4 \| 1.2e-10 \| 8.7e-15 \| 107.3 \| 121 \| (213, 340) \| 471 \| (84, 208) \| 349 \| Chromatin-remodeling complex ATPase-like protein, Histone \| 5JXT\_B Chromatin-remodeling complex ATPase-like protein, Histone chromatin remodeler, ISWI, TRANSCRIPTION; pdb70 \| 5JXT\_D \| 98.4 \| 1.2e-10 \| 8.7e-15 \| 107.3 \| 121 \| (213, 340) \| 471 \| (84, 208) \| 349 \| Chromatin-remodeling complex ATPase-like protein, Histone \| 5JXT\_D Chromatin-remodeling complex ATPase-like protein, Histone chromatin remodeler, ISWI, TRANSCRIPTION; pdb70 \| 4CRW\_B \| 98.4 \| 1.2e-10 \| 8.8e-15 \| 96.5 \| 112 \| (215, 334) \| 471 \| (20, 131) \| 182 \| CCR4-NOT TRANSCRIPTION COMPLEX SUBUNIT 1 \| 4CRW\_B CCR4-NOT TRANSCRIPTION COMPLEX SUBUNIT 1 GENE REGULATION, CCR4-NOT, TRANSLATIONAL REPRESSION HET: GOL; pdb70 \| 5NT7\_D \| 98.3 \| 1.5e-10 \| 1.1e-14 \| 93.6 \| 112 \| (214, 334) \| 471 \| (16, 127) \| 163 \| ATP-dependent RNA helicase vasa, isoform \| 5NT7\_D ATP-dependent RNA helicase vasa, isoform DEAD-box RNA helicase, ATPase, RNA; pdb70 \| 6S8S\_C \| 98.3 \| 1.6e-10 \| 1.2e-14 \| 96.7 \| 112 \| (215, 334) \| 471 \| (31, 142) \| 193 \| Probable ATP-dependent RNA helicase DDX6 \| 6S8S\_C Probable ATP-dependent RNA helicase DDX6 Translational control, mRNA decay, miRNA HET: PO4, CXS; pdb70 \| 6IGM\_H \| 98.3 \| 2.4e-10 \| 1.7e-14 \| 123.9 \| 146 \| (3, 151) \| 471 \| (616, 780) \| 3230 \| RuvB-like 1 (E.C.3.6.4.12), RuvB-like 2 \| 6IGM\_H RuvB-like 1 (E.C.3.6.4.12), RuvB-like 2 SRCAP complex, TRANSCRIPTION; pdb70 \| 1FUK\_A \| 98.3 \| 2.7e-10 \| 2e-14 \| 92.3 \| 112 \| (215, 334) \| 471 \| (17, 128) \| 165 \| EUKARYOTIC INITIATION FACTOR 4A \| 1FUK\_A EUKARYOTIC INITIATION FACTOR 4A Helicase, Initiation Factor 4A, DEAD-Box; pdb70 \| 2JGN\_C \| 98.3 \| 2.9e-10 \| 2.2e-14 \| 94.4 \| 114 \| (214, 334) \| 471 \| (31, 144) \| 185 \| ATP-DEPENDENT RNA HELICASE DDX3X (E.C.3.6.1.-) \| 2JGN\_C ATP-DEPENDENT RNA HELICASE DDX3X (E.C.3.6.1.-) PHOSPHORYLATION, NUCLEOTIDE-BINDING, HELICASE, HYDROLASE, RNA-BINDING; pdb70 \| 4DB2\_D \| 98.3 \| 3.2e-10 \| 2.3e-14 \| 99.6 \| 99 \| (229, 334) \| 471 \| (33, 134) \| 257 \| ATP-dependent RNA helicase MSS116, mitochondrial \| 4DB2\_D ATP-dependent RNA helicase MSS116, mitochondrial DEAD-box, RNA helicase, hydrolase, RNA; pdb70 \| 1T5I\_A \| 98.2 \| 3.6e-10 \| 2.7e-14 \| 92.5 \| 99 \| (229, 334) \| 471 \| (31, 129) \| 172 \| Probable ATP-dependent RNA helicase p47 \| 1T5I\_A Probable ATP-dependent RNA helicase p47 RecA-like fold, PRE-MRNA PROCESSING PROTEIN; pdb70 \| 4ERN\_A \| 98.2 \| 4.8e-10 \| 3.5e-14 \| 100.0 \| 108 \| (212, 331) \| 471 \| (44, 154) \| 289 \| TFIIH basal transcription factor complex \| 4ERN\_A TFIIH basal transcription factor complex Helicase domain 2, general transcription; pdb70 \| 2YJT\_D \| 98.2 \| 7.1e-10 \| 5.3e-14 \| 90.3 \| 112 \| (215, 334) \| 471 \| (17, 128) \| 170 \| REGULATOR OF RIBONUCLEASE ACTIVITY A \| 2YJT\_D REGULATOR OF RIBONUCLEASE ACTIVITY A HYDROLASE INHIBITOR-HYDROLASE COMPLEX, DEAD BOX; pdb70 \| 4DB4\_A \| 98.2 \| 7.7e-10 \| 5.7e-14 \| 97.0 \| 99 \| (229, 334) \| 471 \| (33, 134) \| 256 \| ATP-dependent RNA helicase MSS116, mitochondrial \| 4DB4\_A ATP-dependent RNA helicase MSS116, mitochondrial DEAD-box, RNA helicase, hydrolase, RNA-BINDING; pdb70 \| 2HJV\_B \| 98.2 \| 7.9e-10 \| 5.9e-14 \| 89.2 \| 99 \| (229, 334) \| 471 \| (35, 133) \| 163 \| ATP-dependent RNA helicase dbpA (E.C.3.6.1.-) \| 2HJV\_B ATP-dependent RNA helicase dbpA (E.C.3.6.1.-) parallel alpha-beta, HYDROLASE; pdb70 \| 2HJV\_A \| 98.2 \| 8.4e-10 \| 6.3e-14 \| 89.0 \| 112 \| (215, 334) \| 471 \| (22, 133) \| 163 \| ATP-dependent RNA helicase dbpA (E.C.3.6.1.-) \| 2HJV\_A ATP-dependent RNA helicase dbpA (E.C.3.6.1.-) parallel alpha-beta, HYDROLASE; pdb70 \| 2P6N\_B \| 98.1 \| 8.7e-10 \| 6.5e-14 \| 91.9 \| 112 \| (214, 334) \| 471 \| (41, 152) \| 191 \| ATP-dependent RNA helicase DDX41 (E.C.3.6.1.-) \| 2P6N\_B ATP-dependent RNA helicase DDX41 (E.C.3.6.1.-) RNA, HELICASE, DEAD, STRUCTURAL GENOMICS; pdb70 \| 6SXA\_F \| 98.1 \| 8.7e-10 \| 6.5e-14 \| 114.9 \| 143 \| (2, 150) \| 471 \| (12, 170) \| 916 \| DNA repair endonuclease XPF (E.C.3.1.-.-) \| 6SXA\_F DNA repair endonuclease XPF (E.C.3.1.-.-) DNA Repair enzyme. Nucleotide excision; pdb70 \| 6F9S\_A \| 98.1 \| 9.2e-10 \| 6.8e-14 \| 89.9 \| 112 \| (215, 334) \| 471 \| (24, 135) \| 172 \| Probable ATP-dependent RNA helicase DDX6 \| 6F9S\_A Probable ATP-dependent RNA helicase DDX6 mRNA turnover, translational repression, decapping HET: SO4; pdb70 \| 6S8R\_A \| 98.1 \| 1.4e-09 \| 1e-13 \| 88.8 \| 114 \| (214, 335) \| 471 \| (22, 135) \| 172 \| ATP-dependent RNA helicase me31b (E.C.3.6.4.13) \| 6S8R\_A ATP-dependent RNA helicase me31b (E.C.3.6.4.13) Translation, Translational control, mRNA decay HET: ACT; pdb70 \| 2RB4\_A \| 98.1 \| 1.5e-09 \| 1.1e-13 \| 88.8 \| 112 \| (215, 334) \| 471 \| (21, 138) \| 175 \| ATP-dependent RNA helicase DDX25 (E.C.3.6.1.-) \| 2RB4\_A ATP-dependent RNA helicase DDX25 (E.C.3.6.1.-) RNA helicase, Rossmann fold, Structural HET: SO4; pdb70 \| 2VL7\_A \| 98.1 \| 1.6e-09 \| 1.2e-13 \| 105.9 \| 64 \| (3, 69) \| 471 \| (6, 69) \| 540 \| XPD \| 2VL7\_A XPD HELICASE, UNKNOWN FUNCTION; pdb70 \| 2P6N\_A \| 98.1 \| 1.7e-09 \| 1.2e-13 \| 90.2 \| 112 \| (214, 334) \| 471 \| (41, 152) \| 191 \| ATP-dependent RNA helicase DDX41 (E.C.3.6.1.-) \| 2P6N\_A ATP-dependent RNA helicase DDX41 (E.C.3.6.1.-) RNA, HELICASE, DEAD, STRUCTURAL GENOMICS; pdb70 \| 3GFP\_A \| 98.1 \| 1.7e-09 \| 1.3e-13 \| 89.8 \| 112 \| (215, 334) \| 471 \| (24, 141) \| 189 \| DEAD box protein 5 (E.C.3.6.1.-) \| 3GFP\_A DEAD box protein 5 (E.C.3.6.1.-) mRNA export, ATPase, RecA-fold, ATP-binding; pdb70 \| 2KBF\_A \| 97.9 \| 4.3e-09 \| 3.2e-13 \| 87.2 \| 113 \| (214, 334) \| 471 \| (21, 139) \| 187 \| ATP-dependent RNA helicase DBP5 (E.C.3.6.1.-) \| 2KBF\_A ATP-dependent RNA helicase DBP5 (E.C.3.6.1.-) Dbp5p, ATP-binding, Helicase, Hydrolase, Membrane; pdb70 \| 3PEU\_A \| 97.9 \| 5.1e-09 \| 3.8e-13 \| 86.7 \| 113 \| (214, 334) \| 471 \| (22, 140) \| 188 \| ATP-dependent RNA helicase DBP5 (E.C.3.6.4.13) \| 3PEU\_A ATP-dependent RNA helicase DBP5 (E.C.3.6.4.13) RecA, HEAT, DEAD-box, ATPase, Helicase HET: GOL, IHP; pdb70 \| 2FZL\_A \| 97.9 \| 6.5e-09 \| 4.8e-13 \| 88.4 \| 114 \| (214, 340) \| 471 \| (100, 214) \| 219 \| DNA repair protein RAD25 \| 2FZL\_A DNA repair protein RAD25 XPB, nucleotide excision repair, dna; pdb70 \| 1Z5Z\_A \| 97.9 \| 8e-09 \| 5.9e-13 \| 91.1 \| 118 \| (214, 338) \| 471 \| (96, 217) \| 271 \| Helicase of the snf2/rad54 family \| 1Z5Z\_A Helicase of the snf2/rad54 family HYDROLASE, RECOMBINATION, HYDROLASE-RECOMBINATION COMPLEX; pdb70 \| 5GN1\_C \| 97.8 \| 1.4e-08 \| 1e-12 \| 93.8 \| 117 \| (213, 336) \| 471 \| (184, 304) \| 366 \| ATP-dependent helicase FUN30 (E.C.3.6.4.12) \| 5GN1\_C ATP-dependent helicase FUN30 (E.C.3.6.4.12) helicase, recA, remodeler, HYDROLASE; pdb70 \| 5GN1\_D \| 97.7 \| 1.9e-08 \| 1.4e-12 \| 92.9 \| 118 \| (212, 336) \| 471 \| (183, 304) \| 366 \| ATP-dependent helicase FUN30 (E.C.3.6.4.12) \| 5GN1\_D ATP-dependent helicase FUN30 (E.C.3.6.4.12) helicase, recA, remodeler, HYDROLASE; pdb70 \| 1RIF\_A \| 97.7 \| 2.4e-08 \| 1.8e-12 \| 88.5 \| 142 \| (3, 149) \| 471 \| (112, 263) \| 282 \| DNA helicase uvsW \| 1RIF\_A DNA helicase uvsW Bacteriophage, T4, Helicase, UvsW, RecG; pdb70 \| 1ONB\_A \| 97.7 \| 2.6e-08 \| 1.9e-12 \| 77.8 \| 91 \| (229, 335) \| 471 \| (39, 133) \| 142 \| helicase NS3 (E.C.3.4.21.98) \| 1ONB\_A helicase NS3 (E.C.3.4.21.98) alpha-beta-alpha, HYDROLASE; pdb70 \| 3V4R\_A \| 97.4 \| 1.3e-07 \| 9.9e-12 \| 94.4 \| 100 \| (229, 335) \| 471 \| (451, 555) \| 667 \| UvrABC system protein B (E.C.3.1.21.5)/DNA \| 3V4R\_A UvrABC system protein B (E.C.3.1.21.5)/DNA Helicase motifs and a beta-hairpin HET: ADP; pdb70 \| 6O8H\_A \| 97.4 \| 1.5e-07 \| 1.1e-11 \| 92.8 \| 96 \| (229, 331) \| 471 \| (445, 545) \| 593 \| UvrABC system protein B/DNA Complex \| 6O8H\_A UvrABC system protein B/DNA Complex DNA repair, Nucleotide excision repair HET: SO4; pdb70 \| 6O8E\_A \| 97.3 \| 2.2e-07 \| 1.6e-11 \| 91.5 \| 96 \| (229, 331) \| 471 \| (445, 545) \| 593 \| UvrABC system protein B/DNA Complex \| 6O8E\_A UvrABC system protein B/DNA Complex Nucleotide excision repair, DNA repair HET: ADP, PO4; pdb70 \| 6O8F\_A \| 97.3 \| 2.2e-07 \| 1.6e-11 \| 91.5 \| 96 \| (229, 331) \| 471 \| (445, 545) \| 593 \| UvrABC system protein B/DNA Complex \| 6O8F\_A UvrABC system protein B/DNA Complex Nucleotide excision repair, DNA repair; pdb70 \| 3HGQ\_A \| 97.3 \| 3.1e-07 \| 2.3e-11 \| 83.1 \| 116 \| (212, 339) \| 471 \| (107, 231) \| 328 \| HDA1 complex subunit 3 \| 3HGQ\_A HDA1 complex subunit 3 RecA-like domain, SWI2/SNF2 helical domain; pdb70 \| 3HGT\_B \| 97.3 \| 3.2e-07 \| 2.4e-11 \| 83.0 \| 116 \| (212, 339) \| 471 \| (107, 231) \| 328 \| HDA1 complex subunit 3 \| 3HGT\_B HDA1 complex subunit 3 RecA-like domain, SWI2/SNF2 helical domain; pdb70 \| 2NMV\_A \| 97.2 \| 4e-07 \| 2.9e-11 \| 90.9 \| 96 \| (229, 331) \| 471 \| (445, 545) \| 661 \| UvrABC system protein B (E.C.3.1.-.-)/DNA \| 2NMV\_A UvrABC system protein B (E.C.3.1.-.-)/DNA Protein-DNA complex, T-fluorescein, hairpin, HYDROLASE-DNA HET: ADP, FLU; pdb70 \| 2D7D\_A \| 97.2 \| 4e-07 \| 3e-11 \| 90.8 \| 96 \| (229, 331) \| 471 \| (445, 545) \| 661 \| UvrABC system protein B/DNA Complex \| 2D7D\_A UvrABC system protein B/DNA Complex Helicase, Protein-DNA-ADP ternary complex, HYDROLASE-DNA HET: ADP; pdb70 \| 3GA3\_A \| 97.1 \| 6.9e-07 \| 5e-11 \| 68.4 \| 46 \| (368, 413) \| 471 \| (3, 49) \| 133 \| Interferon-induced helicase C domain-containing protein \| 3GA3\_A Interferon-induced helicase C domain-containing protein Innate immune receptor, RNA biniding; pdb70 \| 2RQA\_A \| 97.1 \| 1.1e-06 \| 7.8e-11 \| 67.5 \| 45 \| (369, 413) \| 471 \| (4, 49) \| 137 \| ATP-dependent RNA helicase DHX58 \| 2RQA\_A ATP-dependent RNA helicase DHX58 RNA binding protein, ATP-binding, Helicase HET: ZN; pdb70 \| 3EQT\_B \| 97.0 \| 1.3e-06 \| 9.8e-11 \| 67.5 \| 47 \| (368, 414) \| 471 \| (4, 51) \| 145 \| ATP-dependent RNA helicase DHX58 /RNA \| 3EQT\_B ATP-dependent RNA helicase DHX58 /RNA innate immunity, RIG-I-like helicases, viral; pdb70 \| 4I1S\_A \| 97.0 \| 1.5e-06 \| 1.1e-10 \| 74.5 \| 79 \| (229, 314) \| 471 \| (150, 242) \| 243 \| Melanoma differentiation associated protein-5, Non-structural \| 4I1S\_A Melanoma differentiation associated protein-5, Non-structural SF2-ATPase, Helicase, Hydrolase-Hydrolase Inhibitor complex; pdb70 \| 3NCU\_B \| 97.0 \| 1.7e-06 \| 1.2e-10 \| 66.2 \| 44 \| (369, 412) \| 471 \| (8, 52) \| 134 \| RIG-I (E.C.3.6.1.-)/RNA complex \| 3NCU\_B RIG-I (E.C.3.6.1.-)/RNA complex innate immune receptor, RIG-I c-terminal HET: GDP; pdb70 \| 2RMJ\_A \| 96.9 \| 1.8e-06 \| 1.3e-10 \| 66.1 \| 44 \| (369, 412) \| 471 \| (8, 52) \| 134 \| Probable ATP-dependent RNA helicase DDX58 \| 2RMJ\_A Probable ATP-dependent RNA helicase DDX58 RNA binding protein, Antiviral defense; pdb70 \| 2FZ4\_A \| 96.9 \| 2e-06 \| 1.4e-10 \| 73.6 \| 136 \| (4, 150) \| 471 \| (93, 230) \| 237 \| DNA repair protein RAD25 \| 2FZ4\_A DNA repair protein RAD25 RecA-like domain, DNA damage recognition; pdb70 \| 3LRR\_A \| 96.9 \| 2.7e-06 \| 2e-10 \| 63.9 \| 38 \| (374, 411) \| 471 \| (2, 40) \| 121 \| Probable ATP-dependent RNA helicase DDX58 \| 3LRR\_A Probable ATP-dependent RNA helicase DDX58 Innate immunity, viral RNA, RIG-I HET: ATP; pdb70 \| 4A2V\_A \| 96.8 \| 2.8e-06 \| 2.1e-10 \| 64.8 \| 38 \| (373, 410) \| 471 \| (3, 41) \| 131 \| RETINOIC ACID INDUCIBLE PROTEIN I \| 4A2V\_A RETINOIC ACID INDUCIBLE PROTEIN I HYDROLASE, SUPERFAMILY 2 RNA HELICASE HET: GOL; pdb70 \| 3OG8\_B \| 96.8 \| 2.9e-06 \| 2.1e-10 \| 64.4 \| 39 \| (373, 411) \| 471 \| (3, 42) \| 128 \| ATP-dependent RNA helicase DDX58 (E.C.3.6.4.13) \| 3OG8\_B ATP-dependent RNA helicase DDX58 (E.C.3.6.4.13) Innate immunity, viral RNA sensing; pdb70 \| 3BER\_A \| 96.8 \| 3.2e-06 \| 2.3e-10 \| 72.8 \| 142 \| (4, 150) \| 471 \| (65, 229) \| 249 \| Probable ATP-dependent RNA helicase DDX47 \| 3BER\_A Probable ATP-dependent RNA helicase DDX47 RNA HELICASE, DEAD, AMP, Structural HET: PGE, AMP; pdb70 \| 3MWL\_A \| 96.7 \| 4.8e-06 \| 3.6e-10 \| 68.7 \| 61 \| (4, 68) \| 471 \| (23, 89) \| 207 \| Heat resistant RNA dependent ATPase \| 3MWL\_A Heat resistant RNA dependent ATPase RNA HELICASE, RIBOSOME BIOGENESIS, THERMOPHILIC HET: SO4, 8OX; pdb70 \| 2GXQ\_A \| 96.6 \| 8.2e-06 \| 6.1e-10 \| 67.1 \| 143 \| (4, 150) \| 471 \| (23, 187) \| 207 \| heat resistant RNA dependent ATPase \| 2GXQ\_A heat resistant RNA dependent ATPase RNA helicase, atomic resolution, AMP HET: AMP; pdb70 \| 3FMO\_B \| 96.6 \| 8.5e-06 \| 6.3e-10 \| 72.3 \| 144 \| (4, 150) \| 471 \| (114, 279) \| 300 \| Nuclear pore complex protein Nup214 \| 3FMO\_B Nuclear pore complex protein Nup214 nuclear porin, nuclear pore complex HET: ADP; pdb70 \| 6AIB\_A \| 96.6 \| 8.6e-06 \| 6.4e-10 \| 67.6 \| 61 \| (4, 68) \| 471 \| (24, 86) \| 214 \| DEAD-box ATP-dependent RNA helicase CshA \| 6AIB\_A DEAD-box ATP-dependent RNA helicase CshA STRUCTURAL PROTEIN; pdb70 \| 6AIC\_A \| 96.5 \| 9.5e-06 \| 7.1e-10 \| 67.4 \| 62 \| (4, 69) \| 471 \| (24, 87) \| 214 \| DEAD-box ATP-dependent RNA helicase CshA \| 6AIC\_A DEAD-box ATP-dependent RNA helicase CshA STRUCTURAL PROTEIN HET: AMP; pdb70 \| 1Q0U\_B \| 96.5 \| 1e-05 \| 7.6e-10 \| 67.1 \| 63 \| (3, 69) \| 471 \| (25, 90) \| 219 \| BstDEAD \| 1Q0U\_B BstDEAD DEAD PROTEIN, RNA BINDING PROTEIN; pdb70 \| 1Q0U\_A \| 96.5 \| 1e-05 \| 7.8e-10 \| 67.1 \| 63 \| (3, 69) \| 471 \| (25, 90) \| 219 \| BstDEAD \| 1Q0U\_A BstDEAD DEAD PROTEIN, RNA BINDING PROTEIN HET: MSE; pdb70 \| 1VEC\_B \| 96.5 \| 1.2e-05 \| 9.1e-10 \| 66.5 \| 143 \| (4, 150) \| 471 \| (25, 189) \| 206 \| ATP-dependent RNA helicase p54 \| 1VEC\_B ATP-dependent RNA helicase p54 RNA HELICASE, DEAD-BOX PROTEIN, RNA HET: TLA; pdb70 \| 3LY5\_A \| 96.5 \| 1.3e-05 \| 9.9e-10 \| 69.1 \| 142 \| (4, 150) \| 471 \| (76, 244) \| 262 \| ATP-dependent RNA helicase DDX18 (E.C.3.6.1.-) \| 3LY5\_A ATP-dependent RNA helicase DDX18 (E.C.3.6.1.-) alpha-beta, Structural Genomics, Structural Genomics; pdb70 \| 3LY5\_B \| 96.4 \| 1.5e-05 \| 1.1e-09 \| 68.8 \| 142 \| (4, 150) \| 471 \| (76, 244) \| 262 \| ATP-dependent RNA helicase DDX18 (E.C.3.6.1.-) \| 3LY5\_B ATP-dependent RNA helicase DDX18 (E.C.3.6.1.-) alpha-beta, Structural Genomics, Structural Genomics; pdb70 \| 3FE2\_B \| 96.4 \| 1.9e-05 \| 1.4e-09 \| 66.8 \| 61 \| (4, 68) \| 471 \| (51, 119) \| 242 \| Probable ATP-dependent RNA helicase DDX5 \| 3FE2\_B Probable ATP-dependent RNA helicase DDX5 RNA HELICASE, DEAD, ADP, ATP-binding HET: SO4, ADP; pdb70 \| 5ZBZ\_A \| 96.3 \| 2e-05 \| 1.5e-09 \| 65.7 \| 61 \| (4, 68) \| 471 \| (36, 99) \| 220 \| Eukaryotic initiation factor 4A-I (E.C.3.6.4.13) \| 5ZBZ\_A Eukaryotic initiation factor 4A-I (E.C.3.6.4.13) eukaryotic translation initiation factor 4A HET: MLY, SAU; pdb70 \| 3B6E\_A \| 96.3 \| 2e-05 \| 1.5e-09 \| 65.3 \| 59 \| (3, 65) \| 471 \| (32, 96) \| 216 \| Interferon-induced helicase C domain-containing protein \| 3B6E\_A Interferon-induced helicase C domain-containing protein DECH, DExD/H RNA-binding helicase, innate; pdb70 \| 2PL3\_A \| 96.3 \| 2.1e-05 \| 1.6e-09 \| 66.5 \| 61 \| (4, 68) \| 471 \| (47, 114) \| 236 \| Probable ATP-dependent RNA helicase DDX10 \| 2PL3\_A Probable ATP-dependent RNA helicase DDX10 RNA, HELICASE, DEAD, STRUCTURAL GENOMICS HET: EOH, ADP; pdb70 \| 5GVR\_A \| 96.3 \| 2.2e-05 \| 1.6e-09 \| 66.0 \| 61 \| (4, 68) \| 471 \| (35, 106) \| 234 \| Probable ATP-dependent RNA helicase DDX41 \| 5GVR\_A Probable ATP-dependent RNA helicase DDX41 ATPase, DEAD box protein, HYDROLASE; pdb70 \| 1QVA\_A \| 96.3 \| 2.3e-05 \| 1.7e-09 \| 65.6 \| 142 \| (4, 150) \| 471 \| (43, 205) \| 223 \| INITIATION FACTOR 4A \| 1QVA\_A INITIATION FACTOR 4A RNA HELICASE, DEAD BOX, EIF4A; pdb70 \| 6FWR\_A \| 96.3 \| 2.6e-05 \| 1.9e-09 \| 78.1 \| 104 \| (229, 341) \| 471 \| (534, 672) \| 716 \| ATP-dependent DNA helicase DinG/DNA Complex \| 6FWR\_A ATP-dependent DNA helicase DinG/DNA Complex ATP, helicase, translocase, DNA binding; pdb70 \| 6FWS\_B \| 96.3 \| 2.6e-05 \| 1.9e-09 \| 78.1 \| 104 \| (229, 341) \| 471 \| (534, 672) \| 716 \| ATP-dependent DNA helicase DinG/DNA Complex \| 6FWS\_B ATP-dependent DNA helicase DinG/DNA Complex ATP, helicase, translocase, DNA binding HET: ADP, SF4; pdb70 \| 3DKP\_A \| 96.2 \| 3.4e-05 \| 2.5e-09 \| 65.7 \| 64 \| (4, 71) \| 471 \| (51, 118) \| 245 \| Probable ATP-dependent RNA helicase DDX52 \| 3DKP\_A Probable ATP-dependent RNA helicase DDX52 RNA HELICASE, DEAD, ADP, Structural HET: ADP; pdb70 \| 5H1Y\_B \| 96.2 \| 3.5e-05 \| 2.6e-09 \| 67.3 \| 141 \| (4, 149) \| 471 \| (74, 250) \| 281 \| Probable ATP-dependent RNA helicase DDX41 \| 5H1Y\_B Probable ATP-dependent RNA helicase DDX41 DNA sensor CDNs receptor ATPase HET: SO4; pdb70 \| 5GJU\_A \| 96.2 \| 3.5e-05 \| 2.6e-09 \| 62.8 \| 60 \| (4, 67) \| 471 \| (23, 85) \| 205 \| ATP-dependent RNA helicase DeaD (E.C.3.6.4.13) \| 5GJU\_A ATP-dependent RNA helicase DeaD (E.C.3.6.4.13) RecA-like, RNA helicase, HYDROLASE HET: AMP; pdb70 \| 3FHC\_B \| 96.2 \| 3.7e-05 \| 2.7e-09 \| 64.6 \| 144 \| (4, 150) \| 471 \| (47, 212) \| 235 \| Nuclear pore complex protein Nup214 \| 3FHC\_B Nuclear pore complex protein Nup214 DEAD-box helicase, mRNA export, Nucleoporin; pdb70 \| 4A4D\_A \| 96.1 \| 3.8e-05 \| 2.8e-09 \| 65.5 \| 61 \| (4, 68) \| 471 \| (65, 133) \| 253 \| PROBABLE ATP-DEPENDENT RNA HELICASE DDX5 \| 4A4D\_A PROBABLE ATP-DEPENDENT RNA HELICASE DDX5 ATP-BINDING, HYDROLASE, RNA-BINDING; pdb70 \| 1QDE\_A \| 96.1 \| 3.8e-05 \| 2.9e-09 \| 64.2 \| 62 \| (4, 69) \| 471 \| (36, 100) \| 224 \| TRANSLATION INITIATION FACTOR 4A \| 1QDE\_A TRANSLATION INITIATION FACTOR 4A TRANSLATION INITIATION, SACCHAROMYCES CEREVISIAE, DEAD; pdb70 \| 2KBE\_A \| 96.1 \| 4.5e-05 \| 3.4e-09 \| 63.5 \| 145 \| (3, 150) \| 471 \| (43, 206) \| 226 \| ATP-dependent RNA helicase DBP5 (E.C.3.6.1.-) \| 2KBE\_A ATP-dependent RNA helicase DBP5 (E.C.3.6.1.-) dbp5p, ATP-binding, Helicase, Hydrolase, Membrane; pdb70 \| 5H8W\_A \| 96.1 \| 4.6e-05 \| 3.4e-09 \| 74.5 \| 105 \| (229, 341) \| 471 \| (430, 565) \| 597 \| ATP-dependent DNA helicase Ta0057/DNA Complex \| 5H8W\_A ATP-dependent DNA helicase Ta0057/DNA Complex helicase, hydrolase; pdb70 \| 2OXC\_A \| 96.1 \| 4.7e-05 \| 3.5e-09 \| 63.5 \| 60 \| (4, 67) \| 471 \| (46, 108) \| 230 \| Probable ATP-dependent RNA helicase DDX20 \| 2OXC\_A Probable ATP-dependent RNA helicase DDX20 RNA, HELICASE, DEAD, STRUCTURAL GENOMICS HET: ADP; pdb70 \| 3B7G\_A \| 96.1 \| 4.7e-05 \| 3.5e-09 \| 63.5 \| 60 \| (4, 67) \| 471 \| (46, 108) \| 230 \| Probable ATP-dependent RNA helicase DDX20 \| 3B7G\_A Probable ATP-dependent RNA helicase DDX20 RNA, HELICASE, DEAD, CONSERVED DOMAIN HET: ANP; pdb70 \| 4PX9\_A \| 96.0 \| 5.7e-05 \| 4.2e-09 \| 66.4 \| 117 \| (4, 124) \| 471 \| (87, 239) \| 292 \| ATP-dependent RNA helicase DDX3X (E.C.3.6.4.13) \| 4PX9\_A ATP-dependent RNA helicase DDX3X (E.C.3.6.4.13) DEAD-box helicase, HYDROLASE, TRANSLATION, RNA HET: ADP; pdb70 \| 1T6N\_A \| 95.9 \| 7.5e-05 \| 5.5e-09 \| 62.2 \| 60 \| (4, 67) \| 471 \| (36, 98) \| 220 \| Probable ATP-dependent RNA helicase p47 \| 1T6N\_A Probable ATP-dependent RNA helicase p47 RecA-like fold, pre-mRNA processing protein HET: FLC; pdb70 \| 4PX9\_C \| 95.9 \| 7.6e-05 \| 5.6e-09 \| 65.5 \| 117 \| (4, 124) \| 471 \| (87, 239) \| 292 \| ATP-dependent RNA helicase DDX3X (E.C.3.6.4.13) \| 4PX9\_C ATP-dependent RNA helicase DDX3X (E.C.3.6.4.13) DEAD-box helicase, HYDROLASE, TRANSLATION, RNA HET: ADP; pdb70 \| 3IUY\_B \| 95.8 \| 9.4e-05 \| 7e-09 \| 61.8 \| 61 \| (4, 68) \| 471 \| (42, 111) \| 228 \| Probable ATP-dependent RNA helicase DDX53 \| 3IUY\_B Probable ATP-dependent RNA helicase DDX53 REC-A-like, DEAD-box, Structural Genomics, Structural HET: AMP; pdb70 \| 3IUY\_A \| 95.8 \| 0.00011 \| 7.9e-09 \| 61.5 \| 62 \| (3, 68) \| 471 \| (41, 111) \| 228 \| Probable ATP-dependent RNA helicase DDX53 \| 3IUY\_A Probable ATP-dependent RNA helicase DDX53 REC-A-like, DEAD-box, Structural Genomics, Structural HET: AMP; pdb70 \| 3B85\_A \| 95.7 \| 0.00013 \| 9.4e-09 \| 59.4 \| 55 \| (3, 61) \| 471 \| (6, 62) \| 208 \| Phosphate starvation-inducible protein \| 3B85\_A Phosphate starvation-inducible protein Corynebacterium glutamicum, PhoH2, ATPase, PFAM: HET: MSE, SO4; pdb70 \| 3LLM\_B \| 95.3 \| 0.00027 \| 2e-08 \| 59.0 \| 60 \| (4, 67) \| 471 \| (61, 125) \| 235 \| ATP-dependent RNA helicase A (E.C.3.6.1.-) \| 3LLM\_B ATP-dependent RNA helicase A (E.C.3.6.1.-) alpha-beta-alpha, Structural Genomics, Structural Genomics HET: ADP, MSE; pdb70 \| 6EG2\_A \| 95.3 \| 0.00028 \| 2.1e-08 \| 68.6 \| 147 \| (3, 151) \| 471 \| (388, 552) \| 621 \| Maltose/maltodextrin-binding periplasmic protein,Probable global transcription \| 6EG2\_A Maltose/maltodextrin-binding periplasmic protein,Probable global transcription Helicase, ATPase, Chromatin remodeling, inhibitor HET: J7J; pdb70 \| 6EG3\_A \| 95.3 \| 0.00028 \| 2.1e-08 \| 68.6 \| 147 \| (3, 151) \| 471 \| (388, 552) \| 621 \| Maltose/maltodextrin-binding periplasmic protein,Probable global transcription \| 6EG3\_A Maltose/maltodextrin-binding periplasmic protein,Probable global transcription Helicase, ATPase, Chromatin remodeling, inhibitor HET: EOH, J7G; pdb70 \| 6RO4\_B \| 95.3 \| 0.00033 \| 2.4e-08 \| 70.3 \| 83 \| (229, 316) \| 471 \| (532, 626) \| 760 \| General transcription and DNA repair \| 6RO4\_B General transcription and DNA repair Complex, Helicase, Translocase, DNA repair; pdb70 \| 1WRB\_B \| 95.1 \| 0.00046 \| 3.4e-08 \| 58.7 \| 61 \| (4, 68) \| 471 \| (45, 117) \| 253 \| DjVLGB \| 1WRB\_B DjVLGB RNA HELICASE, DEAD BOX, VASA HET: SO4; pdb70 \| 6NMI\_B \| 95.0 \| 0.00049 \| 3.6e-08 \| 69.0 \| 83 \| (229, 316) \| 471 \| (532, 626) \| 760 \| General transcription and DNA repair \| 6NMI\_B General transcription and DNA repair transcription, DNA repair, helicase, multiprotein; pdb70 \| 2L8B\_A \| 94.9 \| 0.00062 \| 4.6e-08 \| 55.0 \| 54 \| (4, 59) \| 471 \| (34, 88) \| 189 \| Protein traI (E.C.3.6.4.12) \| 2L8B\_A Protein traI (E.C.3.6.4.12) RecD, HYDROLASE; pdb70 \| 5WWP\_A \| 94.7 \| 0.00081 \| 5.9e-08 \| 65.1 \| 60 \| (4, 66) \| 471 \| (259, 318) \| 600 \| ORF1ab \| 5WWP\_A ORF1ab Middle East respiratory syndrome coronavirus HET: SO4; pdb70 \| 3E1S\_A \| 94.6 \| 0.00092 \| 6.8e-08 \| 64.5 \| 55 \| (4, 62) \| 471 \| (189, 243) \| 574 \| Exodeoxyribonuclease V, subunit RecD \| 3E1S\_A Exodeoxyribonuclease V, subunit RecD alpha and beta protein, ATP-binding; pdb70 \| 3GP8\_A \| 94.6 \| 0.00092 \| 6.8e-08 \| 64.5 \| 55 \| (4, 62) \| 471 \| (189, 243) \| 574 \| Exodeoxyribonuclease V, subunit RecD, putative \| 3GP8\_A Exodeoxyribonuclease V, subunit RecD, putative ALPHA AND BETA PROTEIN, ATP-BINDING |
| Top keywords  (threshold 1.00e-03 (evalue)) | **helicase, RNA, DEAD, DNA, box, ATP\_dependent, DEAH, complex, C, HYDROLASE** |
| Output files | ../../similar\_sequences/44\_FANPEZAQ\_CDS\_0044\_merged.svg ../../similar\_sequences/44\_FANPEZAQ\_CDS\_0044\_pdb70.a3m ../../similar\_sequences/44\_FANPEZAQ\_CDS\_0044\_pdb70.hhr ../../similar\_sequences/44\_FANPEZAQ\_CDS\_0044\_uniclust.a3m ../../similar\_sequences/44\_FANPEZAQ\_CDS\_0044\_uniclust.hhr |

#### Structure prediction (AlphaFold)^2^

|  |  |
| --- | --- |
| Stats | **Files missing!** |
| Predicted structure | **NGL Viewer Controls:**  - Center: *Left-Click* - Rotate: *Left-Click + Drag* - Translate: *Right-Click + Drag* - Zoom: *Shift + Left-Click + Drag* |
| Output files | ../../predicted\_structures/44\_FANPEZAQ\_CDS\_0044/features.pkl ../../predicted\_structures/44\_FANPEZAQ\_CDS\_0044/ranked\_0.pdb ../../predicted\_structures/44\_FANPEZAQ\_CDS\_0044/result\_model\_1\_ptm\_pred\_0.pkl |

#### Structure similarity search results (Foldseek)^3^

|  |  |
| --- | --- |
| Structure databases searched | Pdb, Afdb-proteome, Afdb-uniprot50 |
| Results, scheme(s)  (Top layers only, threshold 1.00e-02 (evalue)) | xml version="1.0" encoding="utf-8" standalone="no"?       2024-09-02T21:11:22.137045 image/svg+xml   Matplotlib v3.7.2, https://matplotlib.org/ |
| Results, table  (threshold 1.00e-02 (evalue)) | -- |
| Top keywords  (threshold 1.00e-02 (evalue)) | -- |
| Output files | ../../similar\_structures/43\_FANPEZAQ\_CDS\_0043\_merged.svg |

  
  
  

Return to summary | Go to previous | Go to next

  


---

**Sequence/structure alignments coloring**  
Each object in the alignment figures is colored according to its E-value following this color coding:

1e-100
10
