## Supplementary material for "Completing the BASEL phage collection to unlock hidden diversity for systematic exploration of phage-host interactions": Data S2: 45.html

FANPEZAQ\_CDS\_0045


Return to summary | Go to previous | Go to next

|  |  |
| --- | --- |
| FANPEZAQ\_CDS\_0045  Page creation date: 02 Sep 2024, 12:00  Project folder: n/a  Input sequences file: Escherichia\_virus\_HeidiAbel.gb | duf1391 domain\_containing prophage putative phage fragment p287170 vi\_07176 p274931 vi\_10756 pf07151 ydaf ydfa |

### Sequence information

|  |  |
| --- | --- |
| Name | FANPEZAQ\_CDS\_0045  45\_FANPEZAQ\_CDS\_0045 (pipeline id) |
| Imported annotations | Escherichia\_virus\_HeidiAbel Bas97 |
| Protein sequence | MNTTHDMGNNETIKTGVFPNTDGTFTALTFTKSKTFKTEAGALKWFNRQMAD |
| Number of residues | 52 |
| Molecular weight (Da) | 5817.46 |
| Output files | ../../query\_sequences/45\_FANPEZAQ\_CDS\_0045.fasta |

### Putative domain architecture and protein family

#### Search results (HHblits)^1^

|  |  |
| --- | --- |
| Domain family databases searched | Pfam, Ncbi-cd, Cath, Phrogs |
| Results, scheme(s)  (Top layers only; threshold 1.00e-03 (evalue)) | xml version="1.0" encoding="utf-8" standalone="no"?       2024-09-02T21:08:22.690407 image/svg+xml   Matplotlib v3.7.2, https://matplotlib.org/ |
| Results, table  (E-value ≤ 1.00e-03 (evalue)) | db \| id \| prob \| evalue \| pvalue \| score \| cols \| query \| query\_len \| template \| template\_len \| name \| description; pfam \| PF07151 \| 100.0 \| 5.3e-39 \| 9.8e-43 \| 179.2 \| 47 \| (4, 50) \| 52 \| (2, 48) \| 48 \| DUF1391 \| Protein of unknown function (DUF1391); phrogs \| 751 \| 100.0 \| 1e-37 \| 1.2e-41 \| 190.6 \| 49 \| (2, 50) \| 52 \| (28, 76) \| 77 \| NA \| NA; Category: unknown function; p287170 VI\_07176; phrogs \| 37620 \| 97.9 \| 5.6e-09 \| 6.3e-13 \| 57.4 \| 18 \| (3, 20) \| 52 \| (29, 46) \| 46 \| NA \| NA; Category: unknown function; p274931 VI\_10756 |
| Top keywords  (threshold 1.00e-03 (evalue)) | **DUF1391, p287170, VI\_07176, p274931, VI\_10756** |
| Output files | ../../domain\_architecture/45\_FANPEZAQ\_CDS\_0045\_cath.hhr ../../domain\_architecture/45\_FANPEZAQ\_CDS\_0045\_merged.svg ../../domain\_architecture/45\_FANPEZAQ\_CDS\_0045\_ncbi-cd.hhr ../../domain\_architecture/45\_FANPEZAQ\_CDS\_0045\_pfam.hhr ../../domain\_architecture/45\_FANPEZAQ\_CDS\_0045\_phrogs.hhr |

### Identical protein sequences/structures

#### Search results

|  |  |
| --- | --- |
| Protein sequence databases searched | Pdb, Swissprot, Refseq |
| Identical proteins found | -- |
| Top keywords | -- |
| Output files | -- |

### Similar protein sequences/structures

#### Sequence similarity search results (HHblits)^1^

|  |  |
| --- | --- |
| Sequence databases searched | Uniclust, Pdb70 |
| Results, scheme(s)  (Top layers only, threshold 1.00e-03 (evalue)) | xml version="1.0" encoding="utf-8" standalone="no"?       2024-09-02T21:08:49.514118 image/svg+xml   Matplotlib v3.7.2, https://matplotlib.org/ |
| Results, table(s)  (threshold 1.00e-03 (evalue)) | db \| id \| prob \| evalue \| pvalue \| score \| cols \| query \| query\_len \| template \| template\_len \| name \| description; uniclust \| UniRef100\_A0A0A0G5S2 \| 100.0 \| 1.2e-40 \| 2.7e-46 \| 208.6 \| 51 \| (2, 52) \| 52 \| (27, 77) \| 77 \| Uncharacterized protein \| Uncharacterized protein; uniclust \| UniRef100\_A0A0A2RM88 \| 100.0 \| 3.7e-40 \| 8.7e-46 \| 203.0 \| 52 \| (1, 52) \| 52 \| (8, 59) \| 69 \| DUF1391 domain-containing protein \| DUF1391 domain-containing protein; uniclust \| UniRef100\_A0A0A1A890 \| 100.0 \| 5.5e-38 \| 1.1e-43 \| 190.2 \| 51 \| (2, 52) \| 52 \| (5, 55) \| 61 \| DUF1391 domain-containing protein \| DUF1391 domain-containing protein; uniclust \| UniRef100\_A0A0B0VS76 \| 100.0 \| 2.7e-37 \| 5.2e-43 \| 203.0 \| 51 \| (2, 52) \| 52 \| (58, 108) \| 108 \| Phage protein \| Phage protein; uniclust \| UniRef100\_A0A376Y8S3 \| 99.9 \| 2e-30 \| 3.8e-36 \| 173.1 \| 49 \| (2, 50) \| 52 \| (1, 49) \| 119 \| Putative prophage protein \| Putative prophage protein; uniclust \| UniRef100\_A8GLQ0 \| 99.8 \| 1.8e-23 \| 3.4e-29 \| 131.9 \| 51 \| (2, 52) \| 52 \| (24, 74) \| 74 \| DUF1391 domain-containing protein \| DUF1391 domain-containing protein; uniclust \| UniRef100\_A0A376ZVH1 \| 99.8 \| 8e-23 \| 1.5e-28 \| 138.0 \| 38 \| (2, 39) \| 52 \| (28, 65) \| 121 \| Putative prophage protein \| Putative prophage protein; uniclust \| UniRef100\_UPI002035C54D \| 99.8 \| 4.3e-22 \| 8.1e-28 \| 121.8 \| 33 \| (2, 34) \| 52 \| (27, 59) \| 59 \| DUF1391 family protein \| DUF1391 family protein; uniclust \| UniRef100\_F4V2C7 \| 99.7 \| 4.2e-21 \| 7.7e-27 \| 143.2 \| 38 \| (2, 39) \| 52 \| (53, 90) \| 295 \| Prophage protein \| Prophage protein; uniclust \| UniRef100\_A0A3S4MIN5 \| 99.7 \| 6.1e-20 \| 1.1e-25 \| 103.4 \| 32 \| (2, 33) \| 52 \| (1, 32) \| 35 \| Putative prophage protein \| Putative prophage protein; uniclust \| UniRef100\_UPI0020335C76 \| 99.7 \| 7e-20 \| 1.3e-25 \| 114.1 \| 30 \| (2, 31) \| 52 \| (36, 65) \| 65 \| DUF1391 family protein \| DUF1391 family protein; uniclust \| UniRef100\_A0A0Q5E3H7 \| 99.6 \| 5.8e-18 \| 1.1e-23 \| 108.6 \| 49 \| (1, 50) \| 52 \| (22, 70) \| 78 \| DUF1391 domain-containing protein \| DUF1391 domain-containing protein; uniclust \| UniRef100\_A0A827QYV4 \| 99.5 \| 1.2e-17 \| 2.4e-23 \| 103.8 \| 34 \| (2, 35) \| 52 \| (28, 61) \| 62 \| DUF1391 domain-containing protein (Fragment) \| DUF1391 domain-containing protein (Fragment); uniclust \| UniRef100\_A0A8E0FTF2 \| 99.5 \| 1.7e-17 \| 3.1e-23 \| 97.6 \| 38 \| (2, 39) \| 52 \| (1, 38) \| 44 \| PF07151 family protein \| PF07151 family protein; uniclust \| UniRef100\_A0A1B1UD71 \| 99.4 \| 1e-15 \| 1.9e-21 \| 96.3 \| 48 \| (2, 50) \| 52 \| (1, 48) \| 66 \| Uncharacterized protein \| Uncharacterized protein; uniclust \| UniRef100\_A0A6C9QHP0 \| 98.2 \| 1.7e-08 \| 3.1e-14 \| 58.8 \| 17 \| (2, 18) \| 52 \| (19, 35) \| 35 \| DUF1391 domain-containing protein (Fragment) \| DUF1391 domain-containing protein (Fragment); uniclust \| UniRef100\_A0A6J5LVG2 \| 97.6 \| 1.8e-06 \| 3.2e-12 \| 56.0 \| 33 \| (20, 52) \| 52 \| (21, 53) \| 62 \| Uncharacterized protein \| Uncharacterized protein |
| Top keywords  (threshold 1.00e-03 (evalue)) | **DUF1391, domain\_containing, prophage, Putative, Fragment, Phage, PF07151** |
| Output files | ../../similar\_sequences/45\_FANPEZAQ\_CDS\_0045\_merged.svg ../../similar\_sequences/45\_FANPEZAQ\_CDS\_0045\_pdb70.a3m ../../similar\_sequences/45\_FANPEZAQ\_CDS\_0045\_pdb70.hhr ../../similar\_sequences/45\_FANPEZAQ\_CDS\_0045\_uniclust.a3m ../../similar\_sequences/45\_FANPEZAQ\_CDS\_0045\_uniclust.hhr |

#### Structure prediction (AlphaFold)^2^

|  |  |
| --- | --- |
| Stats | xml version="1.0" encoding="utf-8" standalone="no"?       2024-09-02T21:09:42.611819 image/svg+xml   Matplotlib v3.7.2, https://matplotlib.org/ |
| Predicted structure | **NGL Viewer Controls:**  - Center: *Left-Click* - Rotate: *Left-Click + Drag* - Translate: *Right-Click + Drag* - Zoom: *Shift + Left-Click + Drag* |
| Output files | ../../predicted\_structures/45\_FANPEZAQ\_CDS\_0045/features.pkl ../../predicted\_structures/45\_FANPEZAQ\_CDS\_0045/ranked\_0.pdb ../../predicted\_structures/45\_FANPEZAQ\_CDS\_0045/ranked\_0\_plots.svg ../../predicted\_structures/45\_FANPEZAQ\_CDS\_0045/result\_model\_1\_ptm\_pred\_0.pkl |

#### Structure similarity search results (Foldseek)^3^

|  |  |
| --- | --- |
| Structure databases searched | Pdb, Afdb-proteome, Afdb-uniprot50 |
| Results, scheme(s)  (Top layers only, threshold 1.00e-02 (evalue)) | xml version="1.0" encoding="utf-8" standalone="no"?       2024-09-02T21:11:22.327438 image/svg+xml   Matplotlib v3.7.2, https://matplotlib.org/ |
| Results, table  (threshold 1.00e-02 (evalue)) | db \| id \| prob \| evalue \| bits \| fident \| alnlen \| mismatch \| gapopen \| qstart \| qend \| tstart \| tend \| name \| description; afdb-proteome \| AF-P0ACW0-F1-MODEL\_V4 \| 1.0 \| 7.805e-05 \| 228 \| 0.63 \| 46 \| 17 \| 0 \| 3 \| 48 \| 2 \| 47 \| Uncharacterized protein YdaF \| Uncharacterized protein YdaF; afdb-proteome \| AF-P0ACW8-F1-MODEL\_V4 \| 1.0 \| 9.4e-05 \| 211 \| 0.62 \| 50 \| 19 \| 0 \| 3 \| 52 \| 2 \| 51 \| Uncharacterized protein YdfA \| Uncharacterized protein YdfA; afdb-uniprot50 \| AF-A0A1B3FBA6-F1-MODEL\_V4 \| 1.0 \| 2.142e-06 \| 320 \| 0.826 \| 52 \| 9 \| 0 \| 1 \| 52 \| 1 \| 52 \| DUF1391 domain-containing protein \| DUF1391 domain-containing protein; afdb-uniprot50 \| AF-A0A7W4F9P6-F1-MODEL\_V4 \| 1.0 \| 1.463e-05 \| 282 \| 0.711 \| 52 \| 15 \| 0 \| 1 \| 52 \| 1 \| 52 \| DUF1391 family protein \| DUF1391 family protein; afdb-uniprot50 \| AF-D3QX46-F1-MODEL\_V4 \| 1.0 \| 5.724e-05 \| 265 \| 0.62 \| 50 \| 19 \| 0 \| 1 \| 50 \| 52 \| 101 \| Uncharacterized protein \| Uncharacterized protein; afdb-uniprot50 \| AF-A8GLQ0-F1-MODEL\_V4 \| 1.0 \| 6.479e-05 \| 259 \| 0.686 \| 51 \| 16 \| 0 \| 2 \| 52 \| 24 \| 74 \| Uncharacterized protein \| Uncharacterized protein; afdb-uniprot50 \| AF-A0A6H2VLD3-F1-MODEL\_V4 \| 1.0 \| 8.303e-05 \| 256 \| 0.557 \| 52 \| 23 \| 0 \| 1 \| 52 \| 1 \| 52 \| DUF1391 domain-containing protein \| DUF1391 domain-containing protein; afdb-uniprot50 \| AF-A0A749PJ15-F1-MODEL\_V4 \| 1.0 \| 7.335e-05 \| 253 \| 0.588 \| 51 \| 21 \| 0 \| 2 \| 52 \| 9 \| 59 \| DUF1391 domain-containing protein \| DUF1391 domain-containing protein; afdb-uniprot50 \| AF-A0A6C7C4R7-F1-MODEL\_V4 \| 1.0 \| 0.0001132 \| 251 \| 0.66 \| 50 \| 17 \| 0 \| 3 \| 52 \| 2 \| 51 \| DUF1391 domain-containing protein \| DUF1391 domain-containing protein; afdb-uniprot50 \| AF-A0A5U8J6R4-F1-MODEL\_V4 \| 1.0 \| 0.0001543 \| 244 \| 0.632 \| 49 \| 18 \| 0 \| 3 \| 51 \| 2 \| 50 \| DUF1391 domain-containing protein \| DUF1391 domain-containing protein; afdb-uniprot50 \| AF-A0A3V2NXN8-F1-MODEL\_V4 \| 1.0 \| 0.0001451 \| 240 \| 0.6 \| 50 \| 20 \| 0 \| 3 \| 52 \| 2 \| 51 \| DUF1391 domain-containing protein \| DUF1391 domain-containing protein; afdb-uniprot50 \| AF-D6I777-F1-MODEL\_V4 \| 1.0 \| 0.0001132 \| 240 \| 0.596 \| 52 \| 21 \| 0 \| 1 \| 52 \| 57 \| 108 \| Uncharacterized protein \| Uncharacterized protein; afdb-uniprot50 \| AF-K8A7Y3-F1-MODEL\_V4 \| 1.0 \| 0.0001859 \| 239 \| 0.68 \| 47 \| 15 \| 0 \| 3 \| 49 \| 2 \| 48 \| Uncharacterized protein \| Uncharacterized protein; afdb-uniprot50 \| AF-Q3Z1T6-F1-MODEL\_V4 \| 1.0 \| 0.0001543 \| 237 \| 0.632 \| 49 \| 18 \| 0 \| 2 \| 50 \| 19 \| 67 \| Uncharacterized protein \| Uncharacterized protein; afdb-uniprot50 \| AF-A0A376P4T6-F1-MODEL\_V4 \| 1.0 \| 0.0001642 \| 237 \| 0.568 \| 51 \| 22 \| 0 \| 2 \| 52 \| 49 \| 99 \| Uncharacterized protein \| Uncharacterized protein; afdb-uniprot50 \| AF-A0A376Y8S3-F1-MODEL\_V4 \| 1.0 \| 0.0001363 \| 237 \| 0.612 \| 49 \| 19 \| 0 \| 3 \| 51 \| 2 \| 50 \| Putative prophage protein \| Putative prophage protein; afdb-uniprot50 \| AF-W1Y735-F1-MODEL\_V4 \| 1.0 \| 0.0001543 \| 236 \| 0.591 \| 49 \| 20 \| 0 \| 2 \| 50 \| 18 \| 66 \| Uncharacterized protein \| Uncharacterized protein; afdb-uniprot50 \| AF-A0A826I354-F1-MODEL\_V4 \| 1.0 \| 0.0001978 \| 234 \| 0.63 \| 46 \| 17 \| 0 \| 3 \| 48 \| 2 \| 47 \| DUF1391 domain-containing protein \| DUF1391 domain-containing protein; afdb-uniprot50 \| AF-A0A4C3QBM3-F1-MODEL\_V4 \| 1.0 \| 0.0002104 \| 231 \| 0.617 \| 47 \| 18 \| 0 \| 2 \| 48 \| 15 \| 61 \| Uncharacterized protein \| Uncharacterized protein; afdb-uniprot50 \| AF-W0AQM8-F1-MODEL\_V4 \| 1.0 \| 0.0001642 \| 230 \| 0.632 \| 49 \| 18 \| 0 \| 4 \| 52 \| 29 \| 77 \| Phage protein \| Phage protein; afdb-uniprot50 \| AF-E9XM46-F1-MODEL\_V4 \| 1.0 \| 0.0002239 \| 225 \| 0.596 \| 52 \| 21 \| 0 \| 1 \| 52 \| 35 \| 86 \| Uncharacterized protein \| Uncharacterized protein; afdb-uniprot50 \| AF-B7MV19-F1-MODEL\_V4 \| 1.0 \| 0.0003052 \| 222 \| 0.617 \| 47 \| 18 \| 0 \| 4 \| 50 \| 67 \| 113 \| Uncharacterized protein \| Uncharacterized protein; afdb-uniprot50 \| AF-A0A0B1MRQ9-F1-MODEL\_V4 \| 1.0 \| 0.0002869 \| 221 \| 0.645 \| 48 \| 17 \| 0 \| 2 \| 49 \| 19 \| 66 \| Phage protein \| Phage protein; afdb-uniprot50 \| AF-A0A729FUV0-F1-MODEL\_V4 \| 1.0 \| 0.0003247 \| 219 \| 0.652 \| 46 \| 16 \| 0 \| 3 \| 48 \| 2 \| 47 \| DUF1391 domain-containing protein \| DUF1391 domain-containing protein; afdb-uniprot50 \| AF-A0A447XGR3-F1-MODEL\_V4 \| 1.0 \| 0.0002696 \| 218 \| 0.604 \| 48 \| 19 \| 0 \| 4 \| 51 \| 30 \| 77 \| Putative prophage protein \| Putative prophage protein; afdb-uniprot50 \| AF-A0A828FR55-F1-MODEL\_V4 \| 1.0 \| 0.0005673 \| 217 \| 0.608 \| 46 \| 18 \| 0 \| 3 \| 48 \| 2 \| 47 \| DUF1391 domain-containing protein \| DUF1391 domain-containing protein; afdb-uniprot50 \| AF-A0A376K109-F1-MODEL\_V4 \| 1.0 \| 0.0004161 \| 214 \| 0.62 \| 50 \| 19 \| 0 \| 3 \| 52 \| 2 \| 51 \| DUF1391 family protein \| DUF1391 family protein; afdb-uniprot50 \| AF-A0A3P6LQE1-F1-MODEL\_V4 \| 1.0 \| 0.0005012 \| 202 \| 0.659 \| 47 \| 16 \| 0 \| 4 \| 50 \| 44 \| 90 \| Prophage protein \| Prophage protein; afdb-uniprot50 \| AF-A0A243UXC8-F1-MODEL\_V4 \| 1.0 \| 0.001194 \| 197 \| 0.75 \| 40 \| 10 \| 0 \| 3 \| 42 \| 2 \| 41 \| Phage protein \| Phage protein; afdb-uniprot50 \| AF-A0A6C6Z2M3-F1-MODEL\_V4 \| 1.0 \| 0.003644 \| 192 \| 0.684 \| 38 \| 12 \| 0 \| 15 \| 52 \| 3 \| 40 \| Uncharacterized protein \| Uncharacterized protein; afdb-uniprot50 \| AF-A0A1B1UD71-F1-MODEL\_V4 \| 1.0 \| 0.005983 \| 169 \| 0.531 \| 47 \| 21 \| 1 \| 3 \| 49 \| 2 \| 47 \| Uncharacterized protein \| Uncharacterized protein |
| Top keywords  (threshold 1.00e-02 (evalue)) | **DUF1391, domain\_containing, prophage, Phage, Putative, YdaF, YdfA** |
| Output files | ../../similar\_structures/45\_FANPEZAQ\_CDS\_0045\_afdb-proteome\_foldseek.tsv ../../similar\_structures/45\_FANPEZAQ\_CDS\_0045\_afdb-uniprot50\_foldseek.tsv ../../similar\_structures/45\_FANPEZAQ\_CDS\_0045\_merged.svg ../../similar\_structures/45\_FANPEZAQ\_CDS\_0045\_pdb\_foldseek.tsv |

  
  
  

Return to summary | Go to previous | Go to next

  


---

**Sequence/structure alignments coloring**  
Each object in the alignment figures is colored according to its E-value following this color coding:

1e-100
10
