## Supplementary material for "Completing the BASEL phage collection to unlock hidden diversity for systematic exploration of phage-host interactions": Data S2: 46.html

FANPEZAQ\_CDS\_0046


Return to summary | Go to previous | Go to next

|  |  |
| --- | --- |
| FANPEZAQ\_CDS\_0046  Page creation date: 02 Sep 2024, 12:00  Project folder: n/a  Input sequences file: Escherichia\_virus\_HeidiAbel.gb | domain\_containing nucleotide modification nmad5 associated putative fragment hypothetical nc\_028980\_p51 gp23 dna\_directed dna polymerase a palm contig\_80 whole genome shotgun sequence |

### Sequence information

|  |  |
| --- | --- |
| Name | FANPEZAQ\_CDS\_0046  46\_FANPEZAQ\_CDS\_0046 (pipeline id) |
| Imported annotations | Escherichia\_virus\_HeidiAbel Bas97 |
| Protein sequence | MARLTAAIRDGIIANAIKTKDFAGRDLAIVQKRADFAERLRKYALAQYGLTDARLTEIKK QIEELQNEVKHEGARFVRITFDSTNRAWDVNLAGQTRRLYLDGTCSGESVKPLYGDSRVE YSDDKITPHFHESDFTVRDPKWREELDAIDLEAAQLREEYNTLKSTLRATISAFTTVEKL LEAWPDVKELIPETTPIAKQPGTGIALSVADLNAL |
| Number of residues | 215 |
| Molecular weight (Da) | 24239.10 |
| Output files | ../../query\_sequences/46\_FANPEZAQ\_CDS\_0046.fasta |

### Putative domain architecture and protein family

#### Search results (HHblits)^1^

|  |  |
| --- | --- |
| Domain family databases searched | Pfam, Ncbi-cd, Cath, Phrogs |
| Results, scheme(s)  (Top layers only; threshold 1.00e-03 (evalue)) | xml version="1.0" encoding="utf-8" standalone="no"?       2024-09-02T21:08:22.899328 image/svg+xml   Matplotlib v3.7.2, https://matplotlib.org/ |
| Results, table  (E-value ≤ 1.00e-03 (evalue)) | db \| id \| prob \| evalue \| pvalue \| score \| cols \| query \| query\_len \| template \| template\_len \| name \| description; pfam \| PF18757 \| 100.0 \| 1.7e-46 \| 3.4e-50 \| 309.2 \| 200 \| (2, 215) \| 215 \| (1, 200) \| 204 \| Nmad5 \| Nucleotide modification associated domain 5; phrogs \| 1965 \| 100.0 \| 5.6e-54 \| 7.4e-58 \| 359.1 \| 175 \| (1, 215) \| 215 \| (2, 176) \| 186 \| NA \| NA; Category: unknown function; NC\_028980\_p51 |
| Top keywords  (threshold 1.00e-03 (evalue)) | **Nucleotide, modification, associated, NC\_028980\_p51** |
| Output files | ../../domain\_architecture/46\_FANPEZAQ\_CDS\_0046\_cath.hhr ../../domain\_architecture/46\_FANPEZAQ\_CDS\_0046\_merged.svg ../../domain\_architecture/46\_FANPEZAQ\_CDS\_0046\_ncbi-cd.hhr ../../domain\_architecture/46\_FANPEZAQ\_CDS\_0046\_pfam.hhr ../../domain\_architecture/46\_FANPEZAQ\_CDS\_0046\_phrogs.hhr |

### Identical protein sequences/structures

#### Search results

|  |  |
| --- | --- |
| Protein sequence databases searched | Pdb, Swissprot, Refseq |
| Identical proteins found | -- |
| Top keywords | -- |
| Output files | -- |

### Similar protein sequences/structures

#### Sequence similarity search results (HHblits)^1^

|  |  |
| --- | --- |
| Sequence databases searched | Uniclust, Pdb70 |
| Results, scheme(s)  (Top layers only, threshold 1.00e-03 (evalue)) | xml version="1.0" encoding="utf-8" standalone="no"?       2024-09-02T21:08:49.713837 image/svg+xml   Matplotlib v3.7.2, https://matplotlib.org/ |
| Results, table(s)  (threshold 1.00e-03 (evalue)) | db \| id \| prob \| evalue \| pvalue \| score \| cols \| query \| query\_len \| template \| template\_len \| name \| description; uniclust \| UniRef100\_A0A0A0YU75 \| 100.0 \| 1.7e-51 \| 3.9e-57 \| 347.0 \| 196 \| (1, 215) \| 215 \| (31, 226) \| 236 \| Nucleotide modification associated domain-containing protein \| Nucleotide modification associated domain-containing protein; uniclust \| UniRef100\_A0A068R047 \| 100.0 \| 2.1e-48 \| 4.9e-54 \| 333.3 \| 196 \| (1, 215) \| 215 \| (48, 246) \| 262 \| Nucleotide modification associated domain-containing protein \| Nucleotide modification associated domain-containing protein; uniclust \| UniRef100\_A0A371YIE3 \| 100.0 \| 2.7e-38 \| 5.7e-44 \| 256.5 \| 175 \| (1, 214) \| 215 \| (1, 175) \| 189 \| Nucleotide modification associated domain-containing protein (Fragment) \| Nucleotide modification associated domain-containing protein (Fragment); uniclust \| UniRef100\_A0A2Z3I5W0 \| 100.0 \| 1.9e-35 \| 4.4e-41 \| 248.3 \| 177 \| (1, 215) \| 215 \| (4, 183) \| 209 \| Nucleotide modification associated domain-containing protein \| Nucleotide modification associated domain-containing protein; uniclust \| UniRef100\_A0A1I3TCD5 \| 100.0 \| 2.2e-35 \| 4.8e-41 \| 244.7 \| 177 \| (1, 215) \| 215 \| (9, 191) \| 203 \| Nucleotide modification associated domain-containing protein \| Nucleotide modification associated domain-containing protein; uniclust \| UniRef100\_A0A0H3EFP6 \| 100.0 \| 3.4e-34 \| 6.3e-40 \| 232.7 \| 196 \| (1, 215) \| 215 \| (26, 222) \| 232 \| Nucleotide modification associated domain-containing protein \| Nucleotide modification associated domain-containing protein; uniclust \| UniRef100\_A0A0N7EMH6 \| 100.0 \| 2.6e-33 \| 5.5e-39 \| 229.9 \| 174 \| (1, 215) \| 215 \| (6, 180) \| 200 \| Nucleotide modification associated domain-containing protein \| Nucleotide modification associated domain-containing protein; uniclust \| UniRef100\_A0A0M0T5J9 \| 99.9 \| 4.1e-28 \| 7.9e-34 \| 193.3 \| 162 \| (35, 215) \| 215 \| (6, 170) \| 181 \| Nucleotide modification associated domain-containing protein (Fragment) \| Nucleotide modification associated domain-containing protein (Fragment); uniclust \| UniRef100\_Q5DN82 \| 99.9 \| 1.9e-27 \| 3.9e-33 \| 195.5 \| 186 \| (1, 214) \| 215 \| (4, 198) \| 209 \| Gp23 \| Gp23; uniclust \| UniRef100\_A0A329VKA5 \| 99.9 \| 2.9e-27 \| 5.6e-33 \| 182.3 \| 116 \| (84, 215) \| 215 \| (14, 130) \| 140 \| Nucleotide modification associated domain-containing protein \| Nucleotide modification associated domain-containing protein; uniclust \| UniRef100\_A0A0P7PEB3 \| 99.9 \| 4.8e-27 \| 9.6e-33 \| 197.8 \| 203 \| (1, 215) \| 215 \| (39, 253) \| 263 \| Nucleotide modification associated domain-containing protein \| Nucleotide modification associated domain-containing protein; uniclust \| UniRef100\_UPI0012FF825C \| 99.9 \| 9.2e-27 \| 2e-32 \| 194.0 \| 163 \| (1, 180) \| 215 \| (26, 189) \| 196 \| Nmad5 family putative nucleotide modification protein \| Nmad5 family putative nucleotide modification protein; uniclust \| UniRef100\_A0A345MLV4 \| 99.9 \| 1e-25 \| 1.9e-31 \| 182.2 \| 184 \| (1, 215) \| 215 \| (2, 188) \| 199 \| Nucleotide modification associated domain-containing protein \| Nucleotide modification associated domain-containing protein; uniclust \| UniRef100\_A0A6M9Z847 \| 99.8 \| 1.3e-23 \| 2.4e-29 \| 172.1 \| 172 \| (1, 214) \| 215 \| (17, 188) \| 243 \| Nucleotide modification associated domain-containing protein \| Nucleotide modification associated domain-containing protein; uniclust \| UniRef100\_A0A5Z4LDL4 \| 99.8 \| 1.4e-23 \| 2.6e-29 \| 163.0 \| 148 \| (2, 165) \| 215 \| (3, 153) \| 154 \| Nucleotide modification associated domain-containing protein (Fragment) \| Nucleotide modification associated domain-containing protein (Fragment); uniclust \| UniRef100\_A0A076YQL7 \| 99.8 \| 4.5e-23 \| 8.6e-29 \| 173.0 \| 202 \| (2, 215) \| 215 \| (16, 264) \| 269 \| Nucleotide modification associated domain-containing protein \| Nucleotide modification associated domain-containing protein; uniclust \| UniRef100\_A0A6G5XY22 \| 99.8 \| 1.2e-21 \| 2.2e-27 \| 157.9 \| 190 \| (1, 215) \| 215 \| (1, 200) \| 209 \| Nucleotide modification associated domain-containing protein \| Nucleotide modification associated domain-containing protein; uniclust \| UniRef100\_A0A455UAL4 \| 99.7 \| 4e-20 \| 7.7e-26 \| 153.5 \| 206 \| (2, 214) \| 215 \| (10, 221) \| 230 \| Nucleotide modification associated domain-containing protein \| Nucleotide modification associated domain-containing protein; uniclust \| UniRef100\_A0A2E3MWD4 \| 99.7 \| 4.3e-20 \| 7.9e-26 \| 147.6 \| 179 \| (1, 214) \| 215 \| (3, 183) \| 192 \| Nucleotide modification associated domain-containing protein \| Nucleotide modification associated domain-containing protein; uniclust \| UniRef100\_A0A1G3M2G9 \| 99.6 \| 4.6e-19 \| 8.4e-25 \| 140.5 \| 169 \| (1, 215) \| 215 \| (1, 169) \| 178 \| Nucleotide modification associated domain-containing protein \| Nucleotide modification associated domain-containing protein; uniclust \| UniRef100\_A0A1B9LQJ1 \| 99.5 \| 2.3e-17 \| 4.2e-23 \| 119.7 \| 82 \| (132, 215) \| 215 \| (7, 89) \| 95 \| Nucleotide modification associated domain-containing protein \| Nucleotide modification associated domain-containing protein; uniclust \| UniRef100\_A0A174S3N4 \| 99.5 \| 3.3e-17 \| 7e-23 \| 134.8 \| 165 \| (2, 214) \| 215 \| (1, 165) \| 175 \| Uncharacterized protein \| Uncharacterized protein; uniclust \| UniRef100\_A0A658IDF4 \| 99.4 \| 2.3e-15 \| 4.4e-21 \| 109.1 \| 76 \| (137, 215) \| 215 \| (2, 77) \| 88 \| Nucleotide modification associated domain-containing protein \| Nucleotide modification associated domain-containing protein; uniclust \| UniRef100\_A0A384ZRN4 \| 99.4 \| 2.7e-15 \| 4.9e-21 \| 124.2 \| 127 \| (84, 215) \| 215 \| (95, 227) \| 232 \| Nucleotide modification associated domain-containing protein \| Nucleotide modification associated domain-containing protein; uniclust \| UniRef100\_UPI001F614FAB \| 99.3 \| 5.8e-15 \| 1.1e-20 \| 102.3 \| 60 \| (154, 215) \| 215 \| (3, 62) \| 70 \| Nmad5 family putative nucleotide modification protein \| Nmad5 family putative nucleotide modification protein; uniclust \| UniRef100\_A0A497QQK1 \| 99.3 \| 1.9e-14 \| 3.5e-20 \| 116.3 \| 172 \| (1, 215) \| 215 \| (3, 177) \| 190 \| Nucleotide modification associated domain-containing protein \| Nucleotide modification associated domain-containing protein; uniclust \| UniRef100\_G1V8X6 \| 99.3 \| 2.1e-14 \| 3.8e-20 \| 116.9 \| 170 \| (1, 208) \| 215 \| (3, 176) \| 199 \| Nucleotide modification associated domain-containing protein \| Nucleotide modification associated domain-containing protein; uniclust \| UniRef100\_A0A118DTA8 \| 99.3 \| 2.2e-14 \| 5e-20 \| 120.9 \| 81 \| (132, 214) \| 215 \| (80, 162) \| 179 \| Nucleotide modification associated domain-containing protein \| Nucleotide modification associated domain-containing protein; uniclust \| UniRef100\_UPI001B8ABC59 \| 99.2 \| 3e-14 \| 5.4e-20 \| 114.7 \| 165 \| (2, 214) \| 215 \| (1, 166) \| 183 \| Nmad5 family putative nucleotide modification protein \| Nmad5 family putative nucleotide modification protein; uniclust \| UniRef100\_UPI00056EFD59 \| 99.2 \| 1.1e-13 \| 2.1e-19 \| 116.2 \| 175 \| (14, 214) \| 215 \| (18, 208) \| 218 \| Nmad5 family putative nucleotide modification protein \| Nmad5 family putative nucleotide modification protein; uniclust \| UniRef100\_A0A076G521 \| 99.2 \| 1.6e-13 \| 3.4e-19 \| 117.2 \| 69 \| (144, 214) \| 215 \| (135, 204) \| 211 \| Nucleotide modification associated domain-containing protein \| Nucleotide modification associated domain-containing protein; uniclust \| UniRef100\_A0A0U2AVF2 \| 99.1 \| 2.4e-13 \| 4.3e-19 \| 110.5 \| 171 \| (1, 208) \| 215 \| (1, 174) \| 192 \| Nucleotide modification associated domain-containing protein \| Nucleotide modification associated domain-containing protein; uniclust \| UniRef100\_UPI00037C3BEE \| 99.1 \| 2.4e-13 \| 4.3e-19 \| 111.2 \| 187 \| (4, 214) \| 215 \| (1, 191) \| 201 \| Nmad5 family putative nucleotide modification protein \| Nmad5 family putative nucleotide modification protein; uniclust \| UniRef100\_UPI001FC9B87A \| 99.1 \| 2.8e-13 \| 5.4e-19 \| 93.1 \| 50 \| (163, 215) \| 215 \| (1, 50) \| 62 \| Nmad5 family putative nucleotide modification protein \| Nmad5 family putative nucleotide modification protein; uniclust \| UniRef100\_A0A142BA81 \| 99.1 \| 2.8e-13 \| 5.5e-19 \| 114.4 \| 185 \| (2, 215) \| 215 \| (1, 205) \| 215 \| Nucleotide modification associated domain-containing protein \| Nucleotide modification associated domain-containing protein; uniclust \| UniRef100\_UPI00234176EE \| 99.1 \| 4.1e-13 \| 7.5e-19 \| 111.3 \| 189 \| (2, 215) \| 215 \| (7, 208) \| 221 \| Nmad5 family putative nucleotide modification protein \| Nmad5 family putative nucleotide modification protein; uniclust \| UniRef100\_UPI001EF750B4 \| 99.1 \| 5.3e-13 \| 9.8e-19 \| 110.8 \| 199 \| (1, 215) \| 215 \| (1, 214) \| 224 \| Nmad5 family putative nucleotide modification protein \| Nmad5 family putative nucleotide modification protein; uniclust \| UniRef100\_A0A1I3RUC2 \| 99.1 \| 6.4e-13 \| 1.2e-18 \| 109.2 \| 189 \| (1, 214) \| 215 \| (1, 196) \| 207 \| Nucleotide modification associated domain-containing protein \| Nucleotide modification associated domain-containing protein; uniclust \| UniRef100\_A0A1B9MFZ6 \| 99.0 \| 2.4e-12 \| 4.5e-18 \| 99.7 \| 128 \| (1, 159) \| 215 \| (1, 128) \| 134 \| Nucleotide modification associated domain-containing protein \| Nucleotide modification associated domain-containing protein; uniclust \| UniRef100\_UPI001EE816BC \| 99.0 \| 2.7e-12 \| 4.9e-18 \| 104.2 \| 167 \| (2, 213) \| 215 \| (9, 175) \| 185 \| Nmad5 family putative nucleotide modification protein \| Nmad5 family putative nucleotide modification protein; uniclust \| UniRef100\_A0A9E7DYX3 \| 99.0 \| 4.5e-12 \| 8.2e-18 \| 106.0 \| 194 \| (2, 214) \| 215 \| (5, 223) \| 231 \| Nucleotide modification associated protein \| Nucleotide modification associated protein; uniclust \| UniRef100\_UPI002107D559 \| 98.8 \| 5.5e-11 \| 1e-16 \| 90.9 \| 102 \| (2, 109) \| 215 \| (4, 108) \| 119 \| Nmad5 family putative nucleotide modification protein \| Nmad5 family putative nucleotide modification protein; uniclust \| UniRef100\_UPI001FCF116F \| 98.7 \| 8.6e-11 \| 1.6e-16 \| 86.3 \| 80 \| (2, 82) \| 215 \| (4, 86) \| 88 \| Nmad5 family putative nucleotide modification protein \| Nmad5 family putative nucleotide modification protein; uniclust \| UniRef100\_A0A369WST6 \| 98.7 \| 1.8e-10 \| 3.4e-16 \| 95.6 \| 183 \| (2, 214) \| 215 \| (4, 195) \| 207 \| Nucleotide modification associated domain-containing protein \| Nucleotide modification associated domain-containing protein; uniclust \| UniRef100\_UPI0018EAAC65 \| 98.6 \| 3e-10 \| 5.5e-16 \| 96.2 \| 198 \| (2, 214) \| 215 \| (1, 228) \| 240 \| Nmad5 family putative nucleotide modification protein \| Nmad5 family putative nucleotide modification protein; uniclust \| UniRef100\_UPI002001BDC6 \| 98.6 \| 3.9e-10 \| 7.1e-16 \| 87.5 \| 87 \| (126, 215) \| 215 \| (35, 121) \| 128 \| Nmad5 family putative nucleotide modification protein \| Nmad5 family putative nucleotide modification protein; uniclust \| UniRef100\_UPI00201A8D73 \| 98.5 \| 9.8e-10 \| 1.8e-15 \| 87.1 \| 104 \| (1, 107) \| 215 \| (26, 130) \| 145 \| Nmad5 family putative nucleotide modification protein \| Nmad5 family putative nucleotide modification protein; uniclust \| UniRef100\_UPI0012EC2E42 \| 98.4 \| 3.1e-09 \| 5.6e-15 \| 89.5 \| 172 \| (2, 212) \| 215 \| (28, 205) \| 220 \| hypothetical protein \| hypothetical protein; uniclust \| UniRef100\_A0A258K0Y5 \| 98.4 \| 2.9e-09 \| 5.8e-15 \| 89.0 \| 151 \| (1, 196) \| 215 \| (1, 151) \| 169 \| Uncharacterized protein \| Uncharacterized protein; uniclust \| UniRef100\_A0A853N6U5 \| 98.4 \| 3.7e-09 \| 6.7e-15 \| 75.6 \| 70 \| (1, 72) \| 215 \| (1, 70) \| 76 \| Nucleotide modification associated domain-containing protein \| Nucleotide modification associated domain-containing protein; uniclust \| UniRef100\_A0A0S3UFR8 \| 98.4 \| 3.8e-09 \| 7.9e-15 \| 93.8 \| 67 \| (132, 199) \| 215 \| (153, 219) \| 246 \| Uncharacterized protein \| Uncharacterized protein; uniclust \| UniRef100\_UPI0004740F64 \| 98.2 \| 3.2e-08 \| 5.8e-14 \| 82.4 \| 162 \| (2, 194) \| 215 \| (1, 164) \| 194 \| Nmad5 family putative nucleotide modification protein \| Nmad5 family putative nucleotide modification protein; uniclust \| UniRef100\_A0A0L0CE49 \| 98.2 \| 3.3e-08 \| 6.1e-14 \| 103.5 \| 204 \| (1, 212) \| 215 \| (3403, 3634) \| 3640 \| DNA-directed DNA polymerase family A palm domain-containing protein \| DNA-directed DNA polymerase family A palm domain-containing protein; uniclust \| UniRef100\_Q31HW8 \| 98.0 \| 1.5e-07 \| 2.7e-13 \| 80.3 \| 193 \| (2, 212) \| 215 \| (6, 211) \| 225 \| Nucleotide modification associated domain-containing protein \| Nucleotide modification associated domain-containing protein; uniclust \| UniRef100\_UPI000442AFC4 \| 97.9 \| 2.2e-07 \| 4.1e-13 \| 76.5 \| 77 \| (133, 213) \| 215 \| (89, 165) \| 172 \| Nmad5 family putative nucleotide modification protein \| Nmad5 family putative nucleotide modification protein; uniclust \| UniRef100\_UPI00167604E6 \| 97.8 \| 3.6e-07 \| 6.6e-13 \| 62.5 \| 47 \| (2, 48) \| 215 \| (1, 47) \| 56 \| Nmad5 family putative nucleotide modification protein \| Nmad5 family putative nucleotide modification protein; uniclust \| UniRef100\_W1J8V1 \| 97.8 \| 3.7e-07 \| 6.8e-13 \| 68.0 \| 55 \| (3, 57) \| 215 \| (5, 59) \| 90 \| Nucleotide modification associated domain-containing protein \| Nucleotide modification associated domain-containing protein; uniclust \| UniRef100\_UPI0003EE2D2C \| 97.8 \| 6e-07 \| 1.2e-12 \| 68.8 \| 57 \| (1, 57) \| 215 \| (3, 59) \| 98 \| Nmad5 family putative nucleotide modification protein \| Nmad5 family putative nucleotide modification protein; uniclust \| UniRef100\_A0A516M5Q1 \| 97.8 \| 6.8e-07 \| 1.2e-12 \| 74.4 \| 157 \| (2, 198) \| 215 \| (1, 164) \| 181 \| Uncharacterized protein \| Uncharacterized protein; uniclust \| UniRef100\_UPI001F03F4A1 \| 97.7 \| 7.1e-07 \| 1.3e-12 \| 70.0 \| 95 \| (1, 104) \| 215 \| (1, 95) \| 118 \| Nmad5 family putative nucleotide modification protein \| Nmad5 family putative nucleotide modification protein; uniclust \| UniRef100\_A0A2I7R0Q1 \| 97.3 \| 9.5e-06 \| 1.8e-11 \| 69.0 \| 81 \| (133, 214) \| 215 \| (105, 195) \| 198 \| Nucleotide modification associated domain-containing protein \| Nucleotide modification associated domain-containing protein; uniclust \| UniRef100\_UPI001FCEE419 \| 97.3 \| 1.4e-05 \| 2.6e-11 \| 60.2 \| 60 \| (84, 157) \| 215 \| (28, 88) \| 89 \| Nmad5 family putative nucleotide modification protein \| Nmad5 family putative nucleotide modification protein; uniclust \| UniRef100\_A0A2E3DFL3 \| 97.3 \| 1.5e-05 \| 2.7e-11 \| 71.6 \| 65 \| (132, 197) \| 215 \| (179, 250) \| 286 \| Nucleotide modification associated domain-containing protein \| Nucleotide modification associated domain-containing protein; uniclust \| UniRef100\_A0A8S5MMN6 \| 97.2 \| 2.2e-05 \| 4.3e-11 \| 68.1 \| 155 \| (1, 191) \| 215 \| (3, 158) \| 200 \| Nucleotide modification associated domain 5 \| Nucleotide modification associated domain 5; uniclust \| UniRef100\_A0A377D698 \| 97.1 \| 4e-05 \| 7.3e-11 \| 54.5 \| 49 \| (1, 49) \| 215 \| (3, 51) \| 63 \| Nucleotide modification associated domain-containing protein \| Nucleotide modification associated domain-containing protein; uniclust \| UniRef100\_UPI0020CC0AC4 \| 96.7 \| 0.00026 \| 4.7e-10 \| 59.7 \| 150 \| (2, 198) \| 215 \| (1, 151) \| 168 \| hypothetical protein \| hypothetical protein; uniclust \| UniRef100\_A0A5P9EY15 \| 96.6 \| 0.00028 \| 5.2e-10 \| 62.3 \| 61 \| (139, 199) \| 215 \| (148, 208) \| 236 \| Nucleotide modification associated domain-containing protein \| Nucleotide modification associated domain-containing protein; uniclust \| UniRef100\_A0A384WWP2 \| 96.3 \| 0.00086 \| 1.6e-09 \| 58.7 \| 74 \| (123, 198) \| 215 \| (100, 180) \| 191 \| Uncharacterized protein \| Uncharacterized protein; uniclust \| UniRef100\_A0A381XKK1 \| 96.3 \| 0.00092 \| 1.8e-09 \| 60.1 \| 164 \| (1, 198) \| 215 \| (1, 174) \| 206 \| Uncharacterized protein \| Uncharacterized protein |
| Top keywords  (threshold 1.00e-03 (evalue)) | **Nucleotide, modification, associated, domain\_containing, Nmad5, putative, Fragment, hypothetical, Gp23, DNA\_directed** |
| Output files | ../../similar\_sequences/46\_FANPEZAQ\_CDS\_0046\_merged.svg ../../similar\_sequences/46\_FANPEZAQ\_CDS\_0046\_pdb70.a3m ../../similar\_sequences/46\_FANPEZAQ\_CDS\_0046\_pdb70.hhr ../../similar\_sequences/46\_FANPEZAQ\_CDS\_0046\_uniclust.a3m ../../similar\_sequences/46\_FANPEZAQ\_CDS\_0046\_uniclust.hhr |

#### Structure prediction (AlphaFold)^2^

|  |  |
| --- | --- |
| Stats | xml version="1.0" encoding="utf-8" standalone="no"?       2024-09-02T21:09:43.386372 image/svg+xml   Matplotlib v3.7.2, https://matplotlib.org/ |
| Predicted structure | **NGL Viewer Controls:**  - Center: *Left-Click* - Rotate: *Left-Click + Drag* - Translate: *Right-Click + Drag* - Zoom: *Shift + Left-Click + Drag* |
| Output files | ../../predicted\_structures/46\_FANPEZAQ\_CDS\_0046/features.pkl ../../predicted\_structures/46\_FANPEZAQ\_CDS\_0046/ranked\_0.pdb ../../predicted\_structures/46\_FANPEZAQ\_CDS\_0046/ranked\_0\_plots.svg ../../predicted\_structures/46\_FANPEZAQ\_CDS\_0046/result\_model\_1\_ptm\_pred\_0.pkl |

#### Structure similarity search results (Foldseek)^3^

|  |  |
| --- | --- |
| Structure databases searched | Pdb, Afdb-proteome, Afdb-uniprot50 |
| Results, scheme(s)  (Top layers only, threshold 1.00e-02 (evalue)) | xml version="1.0" encoding="utf-8" standalone="no"?       2024-09-02T21:11:22.487307 image/svg+xml   Matplotlib v3.7.2, https://matplotlib.org/ |
| Results, table  (threshold 1.00e-02 (evalue)) | db \| id \| prob \| evalue \| bits \| fident \| alnlen \| mismatch \| gapopen \| qstart \| qend \| tstart \| tend \| name \| description; afdb-uniprot50 \| AF-A0A0A2RQ45-F1-MODEL\_V4 \| 1.0 \| 1.222e-09 \| 330 \| 0.226 \| 221 \| 139 \| 10 \| 1 \| 214 \| 1 \| 196 \| Nmad5 domain-containing protein \| Nmad5 domain-containing protein; afdb-uniprot50 \| AF-A0A6N8JYJ7-F1-MODEL\_V4 \| 1.0 \| 6.971e-09 \| 301 \| 0.193 \| 227 \| 157 \| 6 \| 1 \| 214 \| 1 \| 214 \| Nmad5 domain-containing protein \| Nmad5 domain-containing protein; afdb-uniprot50 \| AF-A0A358RVT3-F1-MODEL\_V4 \| 1.0 \| 1.665e-08 \| 297 \| 0.243 \| 218 \| 131 \| 9 \| 1 \| 214 \| 1 \| 188 \| Nmad5 domain-containing protein \| Nmad5 domain-containing protein; afdb-uniprot50 \| AF-A0A756K3A0-F1-MODEL\_V4 \| 1.0 \| 1.982e-08 \| 292 \| 0.288 \| 222 \| 137 \| 8 \| 2 \| 214 \| 7 \| 216 \| Nmad5 domain-containing protein \| Nmad5 domain-containing protein; afdb-uniprot50 \| AF-A0A379DND9-F1-MODEL\_V4 \| 1.0 \| 2.5e-08 \| 275 \| 0.244 \| 217 \| 136 \| 10 \| 3 \| 214 \| 5 \| 198 \| Nmad5 domain-containing protein \| Nmad5 domain-containing protein; afdb-uniprot50 \| AF-A0A8B2F7G3-F1-MODEL\_V4 \| 1.0 \| 7.531e-08 \| 274 \| 0.19 \| 221 \| 147 \| 10 \| 1 \| 214 \| 1 \| 196 \| Uncharacterized protein \| Uncharacterized protein; afdb-uniprot50 \| AF-A0A068R047-F1-MODEL\_V4 \| 1.0 \| 1.067e-07 \| 264 \| 0.232 \| 219 \| 138 \| 9 \| 2 \| 214 \| 4 \| 198 \| Nmad5 domain-containing protein \| Nmad5 domain-containing protein; afdb-uniprot50 \| AF-A0A759RC34-F1-MODEL\_V4 \| 1.0 \| 1.198e-07 \| 262 \| 0.246 \| 211 \| 129 \| 11 \| 1 \| 202 \| 1 \| 190 \| Nmad5 domain-containing protein \| Nmad5 domain-containing protein; afdb-uniprot50 \| AF-A0A653KZM3-F1-MODEL\_V4 \| 1.0 \| 1.27e-07 \| 251 \| 0.235 \| 217 \| 136 \| 10 \| 3 \| 214 \| 2 \| 193 \| Contig\_80, whole genome shotgun sequence \| Contig\_80, whole genome shotgun sequence; afdb-uniprot50 \| AF-A0A377PCD9-F1-MODEL\_V4 \| 1.0 \| 9.499e-08 \| 245 \| 0.253 \| 225 \| 126 \| 10 \| 2 \| 214 \| 4 \| 198 \| Nmad5 domain-containing protein \| Nmad5 domain-containing protein; afdb-uniprot50 \| AF-A0A3Y4Y6V3-F1-MODEL\_V4 \| 1.0 \| 1.764e-05 \| 208 \| 0.175 \| 200 \| 133 \| 8 \| 2 \| 193 \| 4 \| 179 \| Nmad5 domain-containing protein \| Nmad5 domain-containing protein; afdb-uniprot50 \| AF-A0A242NEU4-F1-MODEL\_V4 \| 1.0 \| 1.398e-05 \| 202 \| 0.19 \| 215 \| 144 \| 6 \| 1 \| 214 \| 1 \| 186 \| Nmad5 domain-containing protein \| Nmad5 domain-containing protein; afdb-uniprot50 \| AF-A0A3T7E4H6-F1-MODEL\_V4 \| 1.0 \| 1.175e-05 \| 201 \| 0.18 \| 222 \| 146 \| 9 \| 2 \| 214 \| 4 \| 198 \| Nmad5 domain-containing protein \| Nmad5 domain-containing protein; afdb-uniprot50 \| AF-U9Y7X5-F1-MODEL\_V4 \| 1.0 \| 3.975e-05 \| 187 \| 0.175 \| 211 \| 146 \| 10 \| 9 \| 214 \| 2 \| 189 \| Nmad5 domain-containing protein \| Nmad5 domain-containing protein; afdb-uniprot50 \| AF-A0A1B9NL72-F1-MODEL\_V4 \| 1.0 \| 6.325e-05 \| 185 \| 0.172 \| 215 \| 144 \| 7 \| 1 \| 214 \| 1 \| 182 \| Nmad5 domain-containing protein \| Nmad5 domain-containing protein; afdb-uniprot50 \| AF-H8FYC4-F1-MODEL\_V4 \| 1.0 \| 3.975e-05 \| 179 \| 0.195 \| 220 \| 122 \| 11 \| 1 \| 214 \| 1 \| 171 \| Nmad5 domain-containing protein \| Nmad5 domain-containing protein; afdb-uniprot50 \| AF-A0A556RGF3-F1-MODEL\_V4 \| 1.0 \| 0.0001601 \| 178 \| 0.185 \| 216 \| 148 \| 8 \| 1 \| 214 \| 1 \| 190 \| Nmad5 domain-containing protein \| Nmad5 domain-containing protein; afdb-uniprot50 \| AF-A0A1M3PIE9-F1-MODEL\_V4 \| 1.0 \| 9.315e-06 \| 177 \| 0.202 \| 222 \| 137 \| 13 \| 1 \| 214 \| 1 \| 190 \| Nmad5 domain-containing protein \| Nmad5 domain-containing protein; afdb-uniprot50 \| AF-G1V8X6-F1-MODEL\_V4 \| 1.0 \| 0.0001697 \| 161 \| 0.162 \| 216 \| 140 \| 10 \| 2 \| 214 \| 4 \| 181 \| Nmad5 domain-containing protein \| Nmad5 domain-containing protein; afdb-uniprot50 \| AF-A0A8B3IJ52-F1-MODEL\_V4 \| 1.0 \| 0.0001006 \| 160 \| 0.168 \| 225 \| 142 \| 11 \| 1 \| 214 \| 1 \| 191 \| Uncharacterized protein \| Uncharacterized protein; afdb-uniprot50 \| AF-A0A1I3RUC2-F1-MODEL\_V4 \| 1.0 \| 0.0001601 \| 147 \| 0.184 \| 201 \| 125 \| 10 \| 1 \| 190 \| 1 \| 173 \| Nmad5 domain-containing protein \| Nmad5 domain-containing protein; afdb-uniprot50 \| AF-A0A3N8QQM7-F1-MODEL\_V4 \| 1.0 \| 0.0002268 \| 143 \| 0.192 \| 223 \| 123 \| 15 \| 2 \| 214 \| 5 \| 180 \| Uncharacterized protein \| Uncharacterized protein; afdb-uniprot50 \| AF-A0A2E3MWD4-F1-MODEL\_V4 \| 1.0 \| 0.0004295 \| 141 \| 0.165 \| 218 \| 139 \| 8 \| 2 \| 214 \| 4 \| 183 \| Nmad5 domain-containing protein \| Nmad5 domain-containing protein; afdb-uniprot50 \| AF-A0A5E4SM04-F1-MODEL\_V4 \| 1.0 \| 0.002182 \| 126 \| 0.187 \| 224 \| 122 \| 12 \| 3 \| 214 \| 2 \| 177 \| Nmad5 domain-containing protein \| Nmad5 domain-containing protein; afdb-uniprot50 \| AF-A0A2Z3I5W0-F1-MODEL\_V4 \| 1.0 \| 0.002312 \| 121 \| 0.2 \| 224 \| 120 \| 12 \| 3 \| 214 \| 2 \| 178 \| Nmad5 domain-containing protein \| Nmad5 domain-containing protein; afdb-uniprot50 \| AF-A0A378XRC0-F1-MODEL\_V4 \| 1.0 \| 0.003275 \| 117 \| 0.171 \| 216 \| 117 \| 11 \| 3 \| 201 \| 2 \| 172 \| Nmad5 domain-containing protein \| Nmad5 domain-containing protein; afdb-uniprot50 \| AF-A0A369WST6-F1-MODEL\_V4 \| 1.0 \| 0.00464 \| 109 \| 0.188 \| 218 \| 144 \| 10 \| 3 \| 214 \| 5 \| 195 \| Nmad5 domain-containing protein \| Nmad5 domain-containing protein; afdb-uniprot50 \| AF-A0A6H3FZV3-F1-MODEL\_V4 \| 0.998 \| 0.00464 \| 95 \| 0.15 \| 252 \| 150 \| 16 \| 1 \| 214 \| 1 \| 226 \| Uncharacterized protein \| Uncharacterized protein; afdb-uniprot50 \| AF-A0A545THF2-F1-MODEL\_V4 \| 0.993 \| 0.008786 \| 86 \| 0.172 \| 220 \| 145 \| 10 \| 1 \| 214 \| 1 \| 189 \| Nmad5 domain-containing protein \| Nmad5 domain-containing protein |
| Top keywords  (threshold 1.00e-02 (evalue)) | **Nmad5, domain\_containing, Contig\_80, whole, genome, shotgun, sequence** |
| Output files | ../../similar\_structures/46\_FANPEZAQ\_CDS\_0046\_afdb-proteome\_foldseek.tsv ../../similar\_structures/46\_FANPEZAQ\_CDS\_0046\_afdb-uniprot50\_foldseek.tsv ../../similar\_structures/46\_FANPEZAQ\_CDS\_0046\_merged.svg ../../similar\_structures/46\_FANPEZAQ\_CDS\_0046\_pdb\_foldseek.tsv |

  
  
  

Return to summary | Go to previous | Go to next

  


---

**Sequence/structure alignments coloring**  
Each object in the alignment figures is colored according to its E-value following this color coding:

1e-100
10
