## Supplementary material for "Completing the BASEL phage collection to unlock hidden diversity for systematic exploration of phage-host interactions": Data S2: 48.html

FANPEZAQ\_CDS\_0048


Return to summary | Go to previous | Go to next

|  |  |
| --- | --- |
| FANPEZAQ\_CDS\_0048  Page creation date: 02 Sep 2024, 12:00  Project folder: n/a  Input sequences file: Escherichia\_virus\_HeidiAbel.gb | vrr\_nuc domain\_containing nuclease fragment endonuclease a containing hypothetical putative trna dna hydrolase engineered yes organism\_taxid expressed in escherichia coli expression\_system\_taxid alpha beta arch 3\_layer aba sandwich holliday junction resolvase phage e c fanconi\_associated and b factor recombinase recb complex \_ repair this model characterizes set that resemble holliday\_junction resolving enzymes |

### Sequence information

|  |  |
| --- | --- |
| Name | FANPEZAQ\_CDS\_0048  48\_FANPEZAQ\_CDS\_0048 (pipeline id) |
| Imported annotations | Escherichia\_virus\_HeidiAbel Bas97 |
| Protein sequence | MNYSGCWWFWLLALSWWGLSMANKEHALQNAIRNALAGRCLLFRANVGKAYTSNDVVKVP RQMPVVMGPRDILLKNARPFDTGLPPGFSDLFGMVSVEITPDMVGQKVAIFTGLEVKDGA RVSPLQRNFINAVNDNGGRAGVVRSVDDAEKLVFGK |
| Number of residues | 156 |
| Molecular weight (Da) | 17167.72 |
| Output files | ../../query\_sequences/48\_FANPEZAQ\_CDS\_0048.fasta |

### Putative domain architecture and protein family

#### Search results (HHblits)^1^

|  |  |
| --- | --- |
| Domain family databases searched | Pfam, Ncbi-cd, Cath, Phrogs |
| Results, scheme(s)  (Top layers only; threshold 1.00e-03 (evalue)) | xml version="1.0" encoding="utf-8" standalone="no"?       2024-09-02T21:08:23.317903 image/svg+xml   Matplotlib v3.7.2, https://matplotlib.org/ |
| Results, table  (E-value ≤ 1.00e-03 (evalue)) | db \| id \| prob \| evalue \| pvalue \| score \| cols \| query \| query\_len \| template \| template\_len \| name \| description; pfam \| PF08774 \| 98.3 \| 5.5e-11 \| 1.3e-14 \| 80.1 \| 65 \| (81, 146) \| 156 \| (57, 126) \| 127 \| VRR\_NUC \| VRR-NUC domain; ncbi-cd \| cd22326 \| 97.8 \| 2.9e-09 \| 7.1e-13 \| 87.9 \| 56 \| (82, 147) \| 156 \| (588, 644) \| 652 \| FAN1-like \| cd22326 FAN1-like; repair nuclease FAN1. This model characterizes a set of nucleases that resemble Holliday-junction resolving enzymes.; ncbi-cd \| cd22354 \| 97.6 \| 1.5e-08 \| 3.4e-12 \| 72.0 \| 59 \| (82, 152) \| 156 \| (42, 107) \| 157 \| RecU-like \| cd22354 RecU-like; Holliday junction resolvase RecU (recombination protein U) and similar nucleases.; ncbi-cd \| cd22365 \| 97.5 \| 2.4e-08 \| 5.9e-12 \| 61.6 \| 59 \| (85, 154) \| 156 \| (29, 89) \| 92 \| VRR-NUC-like \| cd22365 VRR-NUC-like; Virus-type replication repair nuclease. This model characterizes a set of nucleases that resemble Holliday-junction resolving enzymes.; cath \| 4qbnA00 \| 99.0 \| 2.3e-14 \| 3.8e-18 \| 91.6 \| 88 \| (22, 155) \| 156 \| (1, 91) \| 93 \| Nuclease \| CATHCODE: 3.40.1350.10 NAME: Nuclease. Chain: a, b. Engineered: yes SOURCE: Salmonella phage setp3. Organism\_taxid: 424944. Expressed in: escherichia coli. Expression\_system\_taxid: 562 CLASS: Alpha Beta, ARCH: 3-Layer(aba) Sandwich, TOPOL: Trna Endonuclease; Chain: A, domain 1, HOMOL: Trna Endonuclease; Chain: A, domain 1; cath \| 4qblA00 \| 98.9 \| 1.3e-13 \| 2e-17 \| 97.4 \| 116 \| (21, 155) \| 156 \| (3, 128) \| 145 \| Vrr-nuc \| CATHCODE: 3.40.1350.10 NAME: Vrr-nuc. Chain: a, b, d, e, f, c. Engineered: yes SOURCE: Psychrobacter sp.. Organism\_taxid: 349106. Strain: prwf-1. Gene: psycprwf\_1331. Expressed in: escherichia coli. Expression\_system\_taxid: 562 CLASS: Alpha Beta, ARCH: 3-Layer(aba) Sandwich, TOPOL: Trna Endonuclease; Chain: A, domain 1, HOMOL: Trna Endonuclease; Chain: A, domain 1; cath \| 4qboA00 \| 98.8 \| 6.7e-13 \| 1.1e-16 \| 84.7 \| 87 \| (20, 155) \| 156 \| (1, 88) \| 92 \| Nuclease \| CATHCODE: 3.40.1350.10 NAME: Nuclease. Chain: a. Engineered: yes SOURCE: Streptococcus phage p9. Organism\_taxid: 403905. Expressed in: escherichia coli. Expression\_system\_taxid: 562 CLASS: Alpha Beta, ARCH: 3-Layer(aba) Sandwich, TOPOL: Trna Endonuclease; Chain: A, domain 1, HOMOL: Trna Endonuclease; Chain: A, domain 1; cath \| 1rznA00 \| 98.1 \| 8.7e-10 \| 1.3e-13 \| 79.6 \| 98 \| (24, 147) \| 156 \| (2, 108) \| 173 \| Recombination protein u \| CATHCODE: 3.40.1350.10 NAME: Recombination protein u. Chain: a, b. Synonym: penicillin-binding protein-related factor a, pbp related factor a. Engineered: yes SOURCE: Bacillus subtilis. Organism\_taxid: 1423. Gene: recu, prfa, bsu22310. Expressed in: escherichia coli. Expression\_system\_taxid: 562 CLASS: Alpha Beta, ARCH: 3-Layer(aba) Sandwich, TOPOL: Trna Endonuclease; Chain: A, domain 1, HOMOL: Trna Endonuclease; Chain: A, domain 1; phrogs \| 824 \| 99.9 \| 1.9e-31 \| 2.7e-35 \| 201.5 \| 110 \| (19, 155) \| 156 \| (16, 129) \| 136 \| NA \| NA; Category: unknown function; NC\_025471\_p57; phrogs \| 5702 \| 99.4 \| 1.1e-17 \| 1.3e-21 \| 116.7 \| 70 \| (76, 156) \| 156 \| (14, 84) \| 86 \| Holliday junction resolvase \| Holliday junction resolvase; Category: DNA, RNA and nucleotide metabolism; NC\_004303\_p15; phrogs \| 2363 \| 99.3 \| 5.4e-17 \| 6.1e-21 \| 124.6 \| 111 \| (18, 154) \| 156 \| (54, 168) \| 192 \| NA \| NA; Category: unknown function; p68790 VI\_09761; phrogs \| 93 \| 98.5 \| 1.4e-11 \| 2e-15 \| 88.2 \| 58 \| (84, 155) \| 156 \| (30, 91) \| 116 \| endonuclease \| endonuclease; Category: DNA, RNA and nucleotide metabolism; NC\_028826\_p5 |
| Top keywords  (threshold 1.00e-03 (evalue)) | **a, Endonuclease, Trna, nuclease, Engineered, yes, Organism\_taxid, Expressed, in, escherichia** |
| Output files | ../../domain\_architecture/48\_FANPEZAQ\_CDS\_0048\_cath.hhr ../../domain\_architecture/48\_FANPEZAQ\_CDS\_0048\_merged.svg ../../domain\_architecture/48\_FANPEZAQ\_CDS\_0048\_ncbi-cd.hhr ../../domain\_architecture/48\_FANPEZAQ\_CDS\_0048\_pfam.hhr ../../domain\_architecture/48\_FANPEZAQ\_CDS\_0048\_phrogs.hhr |

### Identical protein sequences/structures

#### Search results

|  |  |
| --- | --- |
| Protein sequence databases searched | Pdb, Swissprot, Refseq |
| Identical proteins found | -- |
| Top keywords | -- |
| Output files | -- |

### Similar protein sequences/structures

#### Sequence similarity search results (HHblits)^1^

|  |  |
| --- | --- |
| Sequence databases searched | Uniclust, Pdb70 |
| Results, scheme(s)  (Top layers only, threshold 1.00e-03 (evalue)) | xml version="1.0" encoding="utf-8" standalone="no"?       2024-09-02T21:08:50.496807 image/svg+xml   Matplotlib v3.7.2, https://matplotlib.org/ |
| Results, table(s)  (threshold 1.00e-03 (evalue)) | db \| id \| prob \| evalue \| pvalue \| score \| cols \| query \| query\_len \| template \| template\_len \| name \| description; uniclust \| UniRef100\_A0A088FQI8 \| 100.0 \| 2.5e-41 \| 5.4e-47 \| 251.0 \| 147 \| (1, 155) \| 156 \| (2, 149) \| 164 \| VRR-NUC domain-containing protein \| VRR-NUC domain-containing protein; uniclust \| UniRef100\_A0A0F9MG75 \| 99.9 \| 2.9e-31 \| 6.3e-37 \| 198.9 \| 136 \| (19, 155) \| 156 \| (1, 138) \| 162 \| VRR-NUC domain-containing protein \| VRR-NUC domain-containing protein; uniclust \| UniRef100\_A0A0F9LJW8 \| 99.9 \| 9.8e-31 \| 2.1e-36 \| 194.5 \| 117 \| (16, 154) \| 156 \| (16, 134) \| 153 \| Uncharacterized protein \| Uncharacterized protein; uniclust \| UniRef100\_A0A0U5E8J8 \| 99.9 \| 9.2e-28 \| 1.8e-33 \| 174.6 \| 126 \| (20, 154) \| 156 \| (18, 145) \| 149 \| VRR-NUC domain protein \| VRR-NUC domain protein; uniclust \| UniRef100\_A0A091AJ71 \| 99.9 \| 1.3e-27 \| 3e-33 \| 176.8 \| 109 \| (16, 155) \| 156 \| (8, 118) \| 138 \| VRR-NUC domain-containing protein \| VRR-NUC domain-containing protein; uniclust \| UniRef100\_A0A0A0X031 \| 99.9 \| 3.1e-26 \| 7.2e-32 \| 169.5 \| 107 \| (18, 155) \| 156 \| (13, 121) \| 137 \| VRR-NUC domain-containing protein \| VRR-NUC domain-containing protein; uniclust \| UniRef100\_A0A0F9IJ46 \| 99.9 \| 4e-26 \| 7.9e-32 \| 161.0 \| 111 \| (22, 154) \| 156 \| (1, 112) \| 122 \| VRR-NUC domain-containing protein \| VRR-NUC domain-containing protein; uniclust \| UniRef100\_A0A084AC97 \| 99.9 \| 9.9e-26 \| 2.1e-31 \| 166.0 \| 105 \| (18, 155) \| 156 \| (18, 124) \| 148 \| VRR-NUC domain-containing protein \| VRR-NUC domain-containing protein; uniclust \| UniRef100\_A0A069QR83 \| 99.9 \| 1.9e-25 \| 4.3e-31 \| 172.0 \| 115 \| (12, 155) \| 156 \| (35, 154) \| 178 \| VRR-NUC domain protein \| VRR-NUC domain protein; uniclust \| UniRef100\_A0A090JN26 \| 99.9 \| 2.3e-25 \| 4.7e-31 \| 163.6 \| 107 \| (17, 155) \| 156 \| (21, 129) \| 146 \| VRR-NUC domain-containing protein \| VRR-NUC domain-containing protein; uniclust \| UniRef100\_A0A0F9HTW1 \| 99.8 \| 4.8e-25 \| 1.1e-30 \| 162.7 \| 102 \| (19, 155) \| 156 \| (13, 117) \| 136 \| VRR-NUC domain-containing protein \| VRR-NUC domain-containing protein; uniclust \| UniRef100\_A0A0F3IMV8 \| 99.8 \| 6.6e-25 \| 1.5e-30 \| 168.5 \| 109 \| (18, 155) \| 156 \| (31, 145) \| 175 \| VRR-NUC domain-containing protein \| VRR-NUC domain-containing protein; uniclust \| UniRef100\_A0A059T805 \| 99.8 \| 8e-25 \| 1.7e-30 \| 165.8 \| 107 \| (17, 155) \| 156 \| (32, 140) \| 170 \| VRR-NUC domain-containing protein \| VRR-NUC domain-containing protein; uniclust \| UniRef100\_A0A0B5A593 \| 99.8 \| 1e-24 \| 2.4e-30 \| 170.6 \| 107 \| (18, 155) \| 156 \| (50, 166) \| 197 \| Vsr endonuclease \| Vsr endonuclease; uniclust \| UniRef100\_A0A0F8ZZ41 \| 99.8 \| 1.2e-24 \| 2.6e-30 \| 159.5 \| 106 \| (18, 155) \| 156 \| (9, 119) \| 139 \| VRR-NUC domain-containing protein \| VRR-NUC domain-containing protein; uniclust \| UniRef100\_A0A009Z9C9 \| 99.8 \| 1.7e-24 \| 3.6e-30 \| 162.5 \| 110 \| (17, 155) \| 156 \| (22, 140) \| 157 \| VRR-NUC domain protein \| VRR-NUC domain protein; uniclust \| UniRef100\_A0A134B5D7 \| 99.8 \| 1.8e-24 \| 3.9e-30 \| 163.6 \| 109 \| (17, 155) \| 156 \| (30, 145) \| 164 \| VRR-NUC domain protein \| VRR-NUC domain protein; uniclust \| UniRef100\_A0A076G6E5 \| 99.8 \| 1.8e-24 \| 4.2e-30 \| 161.4 \| 116 \| (18, 155) \| 156 \| (1, 133) \| 140 \| VRR-NUC domain-containing protein \| VRR-NUC domain-containing protein; uniclust \| UniRef100\_A0A010YDT0 \| 99.8 \| 6.4e-24 \| 1.5e-29 \| 163.5 \| 109 \| (18, 155) \| 156 \| (26, 143) \| 176 \| VRR-NUC domain-containing protein \| VRR-NUC domain-containing protein; uniclust \| UniRef100\_A0A0A1Q2B7 \| 99.8 \| 1e-23 \| 2.2e-29 \| 163.8 \| 108 \| (17, 155) \| 156 \| (29, 140) \| 197 \| VRR-NUC domain protein \| VRR-NUC domain protein; uniclust \| UniRef100\_A0A0F9A7G8 \| 99.8 \| 3.4e-23 \| 7.5e-29 \| 152.3 \| 110 \| (1, 155) \| 156 \| (4, 115) \| 132 \| VRR-NUC domain-containing protein \| VRR-NUC domain-containing protein; uniclust \| UniRef100\_A0A0R3MLZ8 \| 99.8 \| 3.5e-23 \| 7.7e-29 \| 157.0 \| 102 \| (23, 155) \| 156 \| (33, 139) \| 161 \| VRR-NUC domain-containing protein \| VRR-NUC domain-containing protein; uniclust \| UniRef100\_A0A0S2ZD03 \| 99.8 \| 7.5e-23 \| 1.7e-28 \| 150.2 \| 108 \| (18, 155) \| 156 \| (9, 122) \| 131 \| VRR-NUC domain-containing protein \| VRR-NUC domain-containing protein; uniclust \| UniRef100\_A0A011VZ13 \| 99.8 \| 1.1e-22 \| 2.3e-28 \| 156.5 \| 108 \| (19, 155) \| 156 \| (41, 155) \| 182 \| Nuclease \| Nuclease; uniclust \| UniRef100\_A0A0C5XU96 \| 99.8 \| 1.5e-22 \| 3e-28 \| 149.1 \| 113 \| (19, 155) \| 156 \| (25, 140) \| 145 \| VRR-NUC domain-containing protein \| VRR-NUC domain-containing protein; uniclust \| UniRef100\_A0A0C7SLG5 \| 99.8 \| 2.5e-22 \| 5.1e-28 \| 146.1 \| 110 \| (17, 155) \| 156 \| (3, 120) \| 134 \| VRR-NUC domain-containing protein \| VRR-NUC domain-containing protein; uniclust \| UniRef100\_A0A015SVT4 \| 99.8 \| 2.3e-22 \| 5.1e-28 \| 155.9 \| 109 \| (18, 155) \| 156 \| (50, 165) \| 190 \| VRR-NUC domain protein \| VRR-NUC domain protein; uniclust \| UniRef100\_A0A0F9GCV1 \| 99.8 \| 4.7e-22 \| 1e-27 \| 147.4 \| 106 \| (17, 155) \| 156 \| (18, 128) \| 141 \| VRR-NUC domain-containing protein \| VRR-NUC domain-containing protein; uniclust \| UniRef100\_A0A149SVC6 \| 99.8 \| 5.4e-22 \| 1.1e-27 \| 142.7 \| 129 \| (23, 154) \| 156 \| (1, 130) \| 133 \| VRR-NUC domain-containing protein \| VRR-NUC domain-containing protein; uniclust \| UniRef100\_A0A081N321 \| 99.8 \| 6.3e-22 \| 1.4e-27 \| 150.0 \| 109 \| (18, 155) \| 156 \| (18, 137) \| 161 \| VRR-NUC domain-containing protein \| VRR-NUC domain-containing protein; uniclust \| UniRef100\_A0A017HBU9 \| 99.8 \| 7e-22 \| 1.5e-27 \| 151.4 \| 110 \| (18, 155) \| 156 \| (26, 140) \| 175 \| Uncharacterized protein \| Uncharacterized protein; uniclust \| UniRef100\_A0A0F7L4G6 \| 99.7 \| 1.4e-21 \| 3e-27 \| 147.1 \| 108 \| (19, 155) \| 156 \| (22, 133) \| 153 \| VRR-NUC domain-containing protein \| VRR-NUC domain-containing protein; uniclust \| UniRef100\_A0A0F9HWT1 \| 99.7 \| 1.7e-21 \| 3.7e-27 \| 143.4 \| 108 \| (13, 154) \| 156 \| (3, 114) \| 133 \| VRR-NUC domain-containing protein \| VRR-NUC domain-containing protein; uniclust \| UniRef100\_A0A0F9KD86 \| 99.7 \| 2.5e-21 \| 5.1e-27 \| 138.3 \| 96 \| (15, 154) \| 156 \| (8, 105) \| 119 \| VRR-NUC domain-containing protein \| VRR-NUC domain-containing protein; uniclust \| UniRef100\_A0A0F5ALC1 \| 99.7 \| 2.9e-21 \| 6.2e-27 \| 151.5 \| 110 \| (17, 155) \| 156 \| (40, 157) \| 208 \| Uncharacterized protein \| Uncharacterized protein; uniclust \| UniRef100\_A0A1V5V041 \| 99.7 \| 3.4e-21 \| 7.7e-27 \| 142.3 \| 108 \| (14, 154) \| 156 \| (8, 121) \| 130 \| VRR-NUC domain protein \| VRR-NUC domain protein; uniclust \| UniRef100\_A0A0A2TKQ8 \| 99.7 \| 4.9e-21 \| 1.1e-26 \| 146.1 \| 93 \| (17, 154) \| 156 \| (33, 127) \| 159 \| Nuclease \| Nuclease; uniclust \| UniRef100\_A0A1Q3W833 \| 99.7 \| 8.2e-21 \| 1.7e-26 \| 141.1 \| 107 \| (20, 155) \| 156 \| (32, 146) \| 151 \| VRR-NUC domain-containing protein \| VRR-NUC domain-containing protein; uniclust \| UniRef100\_A0A0F8VVC6 \| 99.7 \| 1.3e-20 \| 2.6e-26 \| 135.8 \| 106 \| (19, 155) \| 156 \| (4, 114) \| 125 \| VRR-NUC domain-containing protein \| VRR-NUC domain-containing protein; uniclust \| UniRef100\_A0A1C6FI89 \| 99.7 \| 1.9e-20 \| 3.9e-26 \| 135.9 \| 108 \| (19, 155) \| 156 \| (1, 118) \| 129 \| VRR-NUC domain \| VRR-NUC domain; uniclust \| UniRef100\_A0A3M2B9U5 \| 99.7 \| 1.9e-20 \| 4e-26 \| 136.8 \| 109 \| (18, 155) \| 156 \| (3, 124) \| 132 \| VRR-NUC domain-containing protein (Fragment) \| VRR-NUC domain-containing protein (Fragment); uniclust \| UniRef100\_A0A1V5XRF5 \| 99.7 \| 2.2e-20 \| 4.8e-26 \| 137.3 \| 96 \| (15, 155) \| 156 \| (5, 102) \| 130 \| VRR-NUC domain protein \| VRR-NUC domain protein; uniclust \| UniRef100\_A0A2S2F862 \| 99.7 \| 3.7e-20 \| 7.6e-26 \| 141.6 \| 104 \| (23, 155) \| 156 \| (41, 153) \| 182 \| VRR-NUC domain-containing protein \| VRR-NUC domain-containing protein; uniclust \| UniRef100\_A0A328AAM2 \| 99.7 \| 4.3e-20 \| 8.8e-26 \| 137.4 \| 104 \| (21, 155) \| 156 \| (26, 134) \| 153 \| VRR-NUC domain-containing protein \| VRR-NUC domain-containing protein; uniclust \| UniRef100\_A0A0F9BCA0 \| 99.7 \| 5.3e-20 \| 1.1e-25 \| 138.1 \| 106 \| (17, 154) \| 156 \| (22, 136) \| 152 \| VRR-NUC domain-containing protein \| VRR-NUC domain-containing protein; uniclust \| UniRef100\_A0A8K2ATK7 \| 99.7 \| 7e-20 \| 1.3e-25 \| 136.9 \| 135 \| (21, 155) \| 156 \| (5, 141) \| 185 \| Endonuclease \| Endonuclease; uniclust \| UniRef100\_A0A5C8V1Q9 \| 99.7 \| 6.7e-20 \| 1.4e-25 \| 135.0 \| 106 \| (18, 154) \| 156 \| (1, 109) \| 140 \| VRR-NUC domain-containing protein \| VRR-NUC domain-containing protein; uniclust \| UniRef100\_A0A2N2E106 \| 99.7 \| 7.5e-20 \| 1.6e-25 \| 135.9 \| 117 \| (16, 155) \| 156 \| (12, 132) \| 142 \| VRR-NUC domain-containing protein \| VRR-NUC domain-containing protein; uniclust \| UniRef100\_A0A1V6B667 \| 99.7 \| 1e-19 \| 2.1e-25 \| 129.5 \| 101 \| (22, 155) \| 156 \| (1, 103) \| 113 \| VRR-NUC domain protein \| VRR-NUC domain protein; uniclust \| UniRef100\_A0A1Q8Y988 \| 99.6 \| 3.3e-19 \| 6.6e-25 \| 134.9 \| 108 \| (19, 155) \| 156 \| (41, 156) \| 175 \| VRR-NUC domain-containing protein \| VRR-NUC domain-containing protein; uniclust \| UniRef100\_A0A1D9FH02 \| 99.6 \| 4.3e-19 \| 9.1e-25 \| 128.3 \| 94 \| (17, 155) \| 156 \| (3, 99) \| 120 \| Nuclease \| Nuclease; uniclust \| UniRef100\_A0A2E5G474 \| 99.6 \| 4.7e-19 \| 9.4e-25 \| 132.0 \| 113 \| (17, 155) \| 156 \| (21, 138) \| 159 \| VRR-NUC domain-containing protein \| VRR-NUC domain-containing protein; uniclust \| UniRef100\_A0A966V8T8 \| 99.6 \| 5e-19 \| 9.7e-25 \| 128.3 \| 105 \| (18, 153) \| 156 \| (15, 121) \| 135 \| VRR-NUC domain-containing protein \| VRR-NUC domain-containing protein; uniclust \| UniRef100\_A0A2D6ZZN4 \| 99.6 \| 5.1e-19 \| 1e-24 \| 129.0 \| 102 \| (18, 154) \| 156 \| (12, 116) \| 131 \| VRR-NUC domain-containing protein \| VRR-NUC domain-containing protein; uniclust \| UniRef100\_A0A096P758 \| 99.6 \| 5e-19 \| 1.1e-24 \| 138.3 \| 112 \| (22, 155) \| 156 \| (54, 181) \| 200 \| Uncharacterized protein \| Uncharacterized protein; uniclust \| UniRef100\_A0A383X5C3 \| 99.6 \| 7.6e-19 \| 1.6e-24 \| 134.2 \| 107 \| (20, 155) \| 156 \| (39, 153) \| 179 \| VRR-NUC domain-containing protein \| VRR-NUC domain-containing protein; uniclust \| UniRef100\_A0A0R1QJF2 \| 99.6 \| 1.2e-18 \| 2.5e-24 \| 122.3 \| 92 \| (20, 155) \| 156 \| (1, 93) \| 100 \| VRR-NUC domain-containing protein \| VRR-NUC domain-containing protein; uniclust \| UniRef100\_A0A0F9VL45 \| 99.6 \| 1.3e-18 \| 3e-24 \| 125.3 \| 91 \| (19, 155) \| 156 \| (4, 96) \| 109 \| VRR-NUC domain-containing protein \| VRR-NUC domain-containing protein; uniclust \| UniRef100\_A0A3M1NGK4 \| 99.6 \| 1.5e-18 \| 3e-24 \| 125.3 \| 104 \| (20, 155) \| 156 \| (8, 113) \| 124 \| VRR-NUC domain-containing protein \| VRR-NUC domain-containing protein; uniclust \| UniRef100\_A0A1Z7ZMK4 \| 99.6 \| 1.6e-18 \| 3.3e-24 \| 133.0 \| 106 \| (21, 155) \| 156 \| (56, 169) \| 185 \| VRR-NUC domain-containing protein \| VRR-NUC domain-containing protein; uniclust \| UniRef100\_A0A142XR66 \| 99.6 \| 2.1e-18 \| 4.4e-24 \| 126.4 \| 105 \| (16, 154) \| 156 \| (2, 113) \| 135 \| VRR-NUC domain protein \| VRR-NUC domain protein; uniclust \| UniRef100\_UPI000EF1452C \| 99.6 \| 3.1e-18 \| 6e-24 \| 130.8 \| 106 \| (21, 155) \| 156 \| (73, 187) \| 195 \| VRR-NUC domain-containing protein \| VRR-NUC domain-containing protein; uniclust \| UniRef100\_A0A0A6PYH0 \| 99.6 \| 3.1e-18 \| 6.6e-24 \| 124.6 \| 94 \| (16, 154) \| 156 \| (1, 99) \| 123 \| VRR-NUC domain-containing protein \| VRR-NUC domain-containing protein; uniclust \| UniRef100\_A0A086MIG4 \| 99.6 \| 3.7e-18 \| 7.8e-24 \| 130.0 \| 105 \| (20, 155) \| 156 \| (32, 141) \| 165 \| VRR-NUC domain-containing protein \| VRR-NUC domain-containing protein; uniclust \| UniRef100\_A0A2X1LNR6 \| 99.6 \| 3.7e-18 \| 8e-24 \| 116.9 \| 68 \| (79, 155) \| 156 \| (2, 70) \| 83 \| Putative norphogenetic protein \| Putative norphogenetic protein; uniclust \| UniRef100\_A0A4Q2XQZ3 \| 99.6 \| 4.3e-18 \| 8.5e-24 \| 123.0 \| 105 \| (20, 155) \| 156 \| (7, 114) \| 131 \| VRR-NUC domain-containing protein \| VRR-NUC domain-containing protein; uniclust \| UniRef100\_A0A323U8N5 \| 99.6 \| 4.1e-18 \| 8.6e-24 \| 125.3 \| 99 \| (18, 155) \| 156 \| (13, 117) \| 133 \| VRR-NUC domain-containing protein \| VRR-NUC domain-containing protein; uniclust \| UniRef100\_A0A6H1Z980 \| 99.6 \| 5e-18 \| 9.9e-24 \| 124.5 \| 109 \| (19, 155) \| 156 \| (23, 134) \| 141 \| VRR-NUC domain-containing protein \| VRR-NUC domain-containing protein; uniclust \| UniRef100\_A0A661RNB2 \| 99.6 \| 5.9e-18 \| 1.1e-23 \| 117.9 \| 99 \| (21, 154) \| 156 \| (3, 107) \| 109 \| VRR-NUC domain-containing protein \| VRR-NUC domain-containing protein; uniclust \| UniRef100\_A0A1C6I8N5 \| 99.6 \| 7.9e-18 \| 1.6e-23 \| 127.8 \| 108 \| (19, 155) \| 156 \| (10, 125) \| 180 \| VRR-NUC domain \| VRR-NUC domain; uniclust \| UniRef100\_A0A0F9B8D5 \| 99.5 \| 1.1e-17 \| 2.4e-23 \| 120.6 \| 92 \| (19, 155) \| 156 \| (13, 106) \| 119 \| VRR-NUC domain-containing protein \| VRR-NUC domain-containing protein; uniclust \| UniRef100\_A0A059N063 \| 99.5 \| 1.2e-17 \| 2.7e-23 \| 127.0 \| 94 \| (16, 155) \| 156 \| (16, 111) \| 151 \| VRR-NUC domain protein \| VRR-NUC domain protein; uniclust \| UniRef100\_A0A150J612 \| 99.5 \| 1.4e-17 \| 2.7e-23 \| 124.9 \| 107 \| (19, 155) \| 156 \| (3, 115) \| 165 \| VRR-NUC domain protein \| VRR-NUC domain protein; uniclust \| UniRef100\_A0A095WF53 \| 99.5 \| 1.3e-17 \| 2.8e-23 \| 123.5 \| 98 \| (12, 155) \| 156 \| (9, 109) \| 134 \| VRR-NUC domain-containing protein \| VRR-NUC domain-containing protein; uniclust \| UniRef100\_A0A2W6ZLY5 \| 99.5 \| 1.6e-17 \| 3e-23 \| 122.6 \| 111 \| (21, 153) \| 156 \| (3, 115) \| 174 \| VRR-NUC domain-containing protein \| VRR-NUC domain-containing protein; uniclust \| UniRef100\_A0A0N1CD04 \| 99.5 \| 1.6e-17 \| 3.3e-23 \| 128.3 \| 103 \| (21, 155) \| 156 \| (51, 159) \| 188 \| Nuclease \| Nuclease; uniclust \| UniRef100\_A0A2M8G080 \| 99.5 \| 2e-17 \| 3.9e-23 \| 119.3 \| 107 \| (19, 155) \| 156 \| (3, 111) \| 126 \| VRR-NUC domain-containing protein \| VRR-NUC domain-containing protein; uniclust \| UniRef100\_G4Q3Y9 \| 99.5 \| 2.5e-17 \| 4.8e-23 \| 128.5 \| 103 \| (21, 155) \| 156 \| (19, 123) \| 223 \| VRR-NUC domain protein \| VRR-NUC domain protein; uniclust \| UniRef100\_A0A1V5MV66 \| 99.5 \| 2.6e-17 \| 5.1e-23 \| 117.0 \| 107 \| (20, 155) \| 156 \| (1, 109) \| 115 \| VRR-NUC domain protein \| VRR-NUC domain protein; uniclust \| UniRef100\_A0A1I3QHP0 \| 99.5 \| 2.6e-17 \| 5.4e-23 \| 122.0 \| 108 \| (19, 155) \| 156 \| (5, 120) \| 139 \| VRR-NUC domain-containing protein \| VRR-NUC domain-containing protein; uniclust \| UniRef100\_A0A1E3LZS1 \| 99.5 \| 2.5e-17 \| 5.5e-23 \| 124.8 \| 102 \| (22, 155) \| 156 \| (30, 135) \| 152 \| VRR-NUC domain-containing protein \| VRR-NUC domain-containing protein; uniclust \| UniRef100\_A0A0S7XPN7 \| 99.5 \| 4.6e-17 \| 9.5e-23 \| 118.5 \| 90 \| (18, 154) \| 156 \| (19, 110) \| 126 \| VRR-NUC domain-containing protein \| VRR-NUC domain-containing protein; uniclust \| UniRef100\_A0A127SDT5 \| 99.5 \| 5.3e-17 \| 1.1e-22 \| 122.5 \| 100 \| (11, 155) \| 156 \| (15, 118) \| 149 \| VRR-NUC domain protein \| VRR-NUC domain protein; uniclust \| UniRef100\_UPI00223A9835 \| 99.5 \| 6.6e-17 \| 1.2e-22 \| 119.6 \| 112 \| (21, 154) \| 156 \| (55, 168) \| 175 \| VRR-NUC domain-containing protein \| VRR-NUC domain-containing protein; uniclust \| UniRef100\_A0A015ZPF0 \| 99.5 \| 5.8e-17 \| 1.3e-22 \| 122.1 \| 94 \| (16, 155) \| 156 \| (17, 112) \| 141 \| VRR-NUC domain protein \| VRR-NUC domain protein; uniclust \| UniRef100\_A0A094Z4S5 \| 99.5 \| 7.6e-17 \| 1.6e-22 \| 115.6 \| 91 \| (19, 155) \| 156 \| (5, 97) \| 111 \| Phage protein \| Phage protein; uniclust \| UniRef100\_UPI0009B54E9D \| 99.5 \| 8.5e-17 \| 1.6e-22 \| 117.2 \| 101 \| (22, 154) \| 156 \| (29, 131) \| 143 \| VRR-NUC domain-containing protein \| VRR-NUC domain-containing protein; uniclust \| UniRef100\_A0A060DAM7 \| 99.5 \| 8.1e-17 \| 1.8e-22 \| 124.4 \| 113 \| (21, 155) \| 156 \| (40, 169) \| 173 \| Nuclease \| Nuclease; uniclust \| UniRef100\_A0A0F8X2W8 \| 99.5 \| 9.4e-17 \| 2.1e-22 \| 120.8 \| 85 \| (20, 147) \| 156 \| (29, 115) \| 144 \| VRR-NUC domain-containing protein \| VRR-NUC domain-containing protein; uniclust \| UniRef100\_A0A5P1RAG0 \| 99.5 \| 1e-16 \| 2.1e-22 \| 119.8 \| 108 \| (19, 155) \| 156 \| (18, 134) \| 151 \| VRR-NUC domain-containing protein \| VRR-NUC domain-containing protein; uniclust \| UniRef100\_A0A165PXT7 \| 99.5 \| 1e-16 \| 2.1e-22 \| 119.5 \| 106 \| (19, 155) \| 156 \| (17, 127) \| 155 \| Uncharacterized protein \| Uncharacterized protein; uniclust \| UniRef100\_A0A089WVF8 \| 99.5 \| 1e-16 \| 2.1e-22 \| 125.6 \| 106 \| (21, 155) \| 156 \| (82, 196) \| 213 \| Nuclease \| Nuclease; uniclust \| UniRef100\_A0A072NS31 \| 99.5 \| 1.1e-16 \| 2.5e-22 \| 121.0 \| 106 \| (4, 155) \| 156 \| (10, 117) \| 147 \| VRR-NUC domain-containing protein \| VRR-NUC domain-containing protein; uniclust \| UniRef100\_A0A0F9EIV8 \| 99.5 \| 1.4e-16 \| 2.7e-22 \| 109.2 \| 80 \| (76, 155) \| 156 \| (5, 86) \| 99 \| VRR-NUC domain-containing protein (Fragment) \| VRR-NUC domain-containing protein (Fragment); uniclust \| UniRef100\_A0A1R4EF88 \| 99.5 \| 1.4e-16 \| 2.7e-22 \| 118.9 \| 111 \| (16, 155) \| 156 \| (15, 135) \| 154 \| VRR-NUC domain protein \| VRR-NUC domain protein; uniclust \| UniRef100\_A0A1H4CGG7 \| 99.5 \| 1.6e-16 \| 2.9e-22 \| 115.1 \| 101 \| (22, 154) \| 156 \| (39, 145) \| 150 \| VRR-NUC domain-containing protein \| VRR-NUC domain-containing protein; uniclust \| UniRef100\_A0A352Z8N4 \| 99.5 \| 1.5e-16 \| 3.1e-22 \| 114.4 \| 100 \| (19, 154) \| 156 \| (8, 112) \| 122 \| Recombinase RecB \| Recombinase RecB; uniclust \| UniRef100\_A0A011PYA4 \| 99.5 \| 1.4e-16 \| 3.1e-22 \| 116.1 \| 94 \| (16, 155) \| 156 \| (4, 99) \| 114 \| VRR-NUC domain protein \| VRR-NUC domain protein; uniclust \| UniRef100\_A0A222YXP1 \| 99.5 \| 1.6e-16 \| 3.2e-22 \| 121.7 \| 105 \| (22, 155) \| 156 \| (56, 166) \| 179 \| Putative morphogenetic function protein \| Putative morphogenetic function protein; uniclust \| UniRef100\_A0A3M1AH50 \| 99.5 \| 2.1e-16 \| 4.1e-22 \| 106.4 \| 71 \| (73, 154) \| 156 \| (9, 80) \| 87 \| VRR-NUC domain-containing protein \| VRR-NUC domain-containing protein; uniclust \| UniRef100\_UPI001FA97645 \| 99.5 \| 2.4e-16 \| 4.5e-22 \| 115.8 \| 118 \| (9, 155) \| 156 \| (36, 161) \| 165 \| VRR-NUC domain-containing protein \| VRR-NUC domain-containing protein; uniclust \| UniRef100\_A0A1F2RUT7 \| 99.4 \| 2.2e-16 \| 4.6e-22 \| 115.0 \| 97 \| (15, 155) \| 156 \| (5, 103) \| 121 \| VRR-NUC domain-containing protein \| VRR-NUC domain-containing protein; uniclust \| UniRef100\_A0A1V6K0J2 \| 99.4 \| 2.6e-16 \| 5e-22 \| 118.7 \| 118 \| (18, 155) \| 156 \| (23, 146) \| 174 \| VRR-NUC domain protein \| VRR-NUC domain protein; uniclust \| UniRef100\_A0A0I9SDG8 \| 99.4 \| 2.6e-16 \| 5.5e-22 \| 113.6 \| 92 \| (16, 154) \| 156 \| (16, 109) \| 116 \| VRR-NUC domain-containing protein \| VRR-NUC domain-containing protein; uniclust \| UniRef100\_A0A1G2ZDQ9 \| 99.4 \| 2.8e-16 \| 5.8e-22 \| 115.8 \| 111 \| (15, 154) \| 156 \| (4, 117) \| 135 \| VRR-NUC domain-containing protein \| VRR-NUC domain-containing protein; uniclust \| UniRef100\_A0A1E4ZHT9 \| 99.4 \| 3.2e-16 \| 6.8e-22 \| 116.2 \| 91 \| (18, 155) \| 156 \| (24, 116) \| 131 \| VRR-NUC domain-containing protein \| VRR-NUC domain-containing protein; uniclust \| UniRef100\_A0A927XP81 \| 99.4 \| 4e-16 \| 7.7e-22 \| 116.2 \| 105 \| (22, 155) \| 156 \| (42, 154) \| 164 \| VRR-NUC domain-containing protein \| VRR-NUC domain-containing protein; uniclust \| UniRef100\_A0A093UG92 \| 99.4 \| 4.3e-16 \| 8.5e-22 \| 116.3 \| 107 \| (20, 155) \| 156 \| (29, 144) \| 156 \| VRR-NUC domain-containing protein \| VRR-NUC domain-containing protein; uniclust \| UniRef100\_A0A1F8N6W9 \| 99.4 \| 4.4e-16 \| 9.6e-22 \| 114.2 \| 95 \| (17, 154) \| 156 \| (15, 113) \| 122 \| VRR-NUC domain-containing protein (Fragment) \| VRR-NUC domain-containing protein (Fragment); uniclust \| UniRef100\_A0A6J5PBK3 \| 99.4 \| 4.9e-16 \| 9.8e-22 \| 116.8 \| 103 \| (21, 155) \| 156 \| (1, 108) \| 161 \| VRR-NUC domain containing protein \| VRR-NUC domain containing protein; uniclust \| UniRef100\_A0A823KYF0 \| 99.4 \| 6.6e-16 \| 1.3e-21 \| 103.9 \| 79 \| (22, 132) \| 156 \| (1, 81) \| 82 \| VRR-NUC domain-containing protein (Fragment) \| VRR-NUC domain-containing protein (Fragment); uniclust \| UniRef100\_A0A0F9AK81 \| 99.4 \| 7.1e-16 \| 1.5e-21 \| 114.3 \| 90 \| (20, 155) \| 156 \| (16, 107) \| 134 \| VRR-NUC domain-containing protein \| VRR-NUC domain-containing protein; uniclust \| UniRef100\_A0A179DMZ1 \| 99.4 \| 8.1e-16 \| 1.7e-21 \| 119.3 \| 99 \| (24, 155) \| 156 \| (48, 148) \| 181 \| VRR-NUC domain-containing protein \| VRR-NUC domain-containing protein; uniclust \| UniRef100\_A0A0T9UVH8 \| 99.4 \| 8.9e-16 \| 1.7e-21 \| 118.3 \| 106 \| (21, 155) \| 156 \| (47, 161) \| 197 \| VRR-NUC domain \| VRR-NUC domain; uniclust \| UniRef100\_A0A1X0Y852 \| 99.4 \| 9.6e-16 \| 1.8e-21 \| 111.0 \| 98 \| (22, 154) \| 156 \| (28, 127) \| 139 \| VRR-NUC domain-containing protein \| VRR-NUC domain-containing protein; uniclust \| UniRef100\_A0A133ZYW0 \| 99.4 \| 1e-15 \| 2.1e-21 \| 117.6 \| 92 \| (19, 155) \| 156 \| (60, 156) \| 176 \| VRR-NUC domain protein \| VRR-NUC domain protein; uniclust \| UniRef100\_A0A1E5GIK2 \| 99.4 \| 1.1e-15 \| 2.2e-21 \| 110.5 \| 91 \| (19, 155) \| 156 \| (13, 105) \| 117 \| VRR-NUC domain-containing protein \| VRR-NUC domain-containing protein; uniclust \| UniRef100\_A0A2A4V727 \| 99.4 \| 1.2e-15 \| 2.4e-21 \| 112.2 \| 104 \| (20, 155) \| 156 \| (17, 126) \| 140 \| VRR-NUC domain-containing protein \| VRR-NUC domain-containing protein; uniclust \| UniRef100\_A0A0F9LX93 \| 99.4 \| 1.2e-15 \| 2.7e-21 \| 111.2 \| 90 \| (21, 155) \| 156 \| (4, 97) \| 115 \| VRR-NUC domain-containing protein \| VRR-NUC domain-containing protein; uniclust \| UniRef100\_A0A015XBN2 \| 99.4 \| 1.5e-15 \| 3e-21 \| 108.4 \| 86 \| (40, 155) \| 156 \| (19, 106) \| 115 \| VRR-NUC domain protein \| VRR-NUC domain protein; uniclust \| UniRef100\_A0A858NP77 \| 99.4 \| 1.7e-15 \| 3.2e-21 \| 105.8 \| 87 \| (69, 155) \| 156 \| (3, 90) \| 116 \| VRR-NUC domain-containing protein \| VRR-NUC domain-containing protein; uniclust \| UniRef100\_A0A1L7AL59 \| 99.4 \| 2.2e-15 \| 4.2e-21 \| 113.7 \| 111 \| (20, 154) \| 156 \| (21, 135) \| 171 \| VRR-NUC domain-containing protein \| VRR-NUC domain-containing protein; uniclust \| UniRef100\_A0A0U4JU24 \| 99.4 \| 2.1e-15 \| 4.5e-21 \| 112.1 \| 85 \| (20, 147) \| 156 \| (20, 106) \| 135 \| VRR-NUC domain-containing protein \| VRR-NUC domain-containing protein; uniclust \| UniRef100\_A0A1V5TXC5 \| 99.4 \| 2.4e-15 \| 4.8e-21 \| 110.3 \| 104 \| (20, 155) \| 156 \| (16, 125) \| 132 \| VRR-NUC domain protein \| VRR-NUC domain protein; uniclust \| UniRef100\_A0A5B8I858 \| 99.4 \| 2.6e-15 \| 4.9e-21 \| 109.0 \| 104 \| (20, 155) \| 156 \| (7, 115) \| 134 \| VRR-NUC domain-containing protein \| VRR-NUC domain-containing protein; uniclust \| UniRef100\_UPI002103C52B \| 99.4 \| 2.9e-15 \| 5.4e-21 \| 98.7 \| 70 \| (85, 154) \| 156 \| (2, 72) \| 81 \| VRR-NUC domain-containing protein \| VRR-NUC domain-containing protein; uniclust \| UniRef100\_A0A068YHT1 \| 99.3 \| 3.6e-15 \| 7e-21 \| 114.7 \| 99 \| (21, 154) \| 156 \| (65, 167) \| 195 \| Uncharacterized protein \| Uncharacterized protein; uniclust \| UniRef100\_A0A2E5G178 \| 99.3 \| 3.5e-15 \| 7.4e-21 \| 122.9 \| 103 \| (24, 155) \| 156 \| (142, 250) \| 277 \| VRR-NUC domain-containing protein \| VRR-NUC domain-containing protein; uniclust \| UniRef100\_A0A024YQ59 \| 99.3 \| 4.4e-15 \| 9.8e-21 \| 112.3 \| 88 \| (22, 154) \| 156 \| (32, 124) \| 143 \| VRR-NUC domain-containing protein \| VRR-NUC domain-containing protein; uniclust \| UniRef100\_A0A084EGU8 \| 99.3 \| 5.6e-15 \| 1.2e-20 \| 106.4 \| 100 \| (21, 154) \| 156 \| (1, 102) \| 112 \| VRR-NUC domain-containing protein \| VRR-NUC domain-containing protein; uniclust \| UniRef100\_A0A6B9L9E1 \| 99.3 \| 6e-15 \| 1.2e-20 \| 118.2 \| 103 \| (20, 154) \| 156 \| (56, 164) \| 271 \| VRR-NUC domain-containing protein \| VRR-NUC domain-containing protein; uniclust \| UniRef100\_A0A5C7QIV6 \| 99.3 \| 6.4e-15 \| 1.2e-20 \| 108.9 \| 122 \| (20, 151) \| 156 \| (36, 160) \| 168 \| VRR-NUC domain-containing protein \| VRR-NUC domain-containing protein; uniclust \| UniRef100\_A0A0U3SL53 \| 99.3 \| 6.2e-15 \| 1.2e-20 \| 112.0 \| 102 \| (23, 154) \| 156 \| (35, 138) \| 172 \| VRR-NUC domain-containing protein \| VRR-NUC domain-containing protein; uniclust \| UniRef100\_A0A0B6XGD4 \| 99.3 \| 5.6e-15 \| 1.2e-20 \| 120.1 \| 105 \| (22, 155) \| 156 \| (95, 205) \| 244 \| Putative norphogenetic protein \| Putative norphogenetic protein; uniclust \| UniRef100\_A0A1Z3CMV2 \| 99.3 \| 7e-15 \| 1.5e-20 \| 101.0 \| 61 \| (83, 155) \| 156 \| (12, 73) \| 82 \| Nuclease \| Nuclease; uniclust \| UniRef100\_A0A662KWK5 \| 99.3 \| 8.7e-15 \| 1.6e-20 \| 107.4 \| 100 \| (22, 155) \| 156 \| (49, 150) \| 159 \| VRR-NUC domain-containing protein \| VRR-NUC domain-containing protein; uniclust \| UniRef100\_A0A151A655 \| 99.3 \| 8.2e-15 \| 1.6e-20 \| 108.2 \| 90 \| (20, 153) \| 156 \| (17, 109) \| 140 \| VRR-NUC domain-containing protein \| VRR-NUC domain-containing protein; uniclust \| UniRef100\_A0A6G5Y5C2 \| 99.3 \| 9.2e-15 \| 1.7e-20 \| 104.3 \| 112 \| (21, 155) \| 156 \| (13, 126) \| 131 \| VRR-NUC domain-containing protein \| VRR-NUC domain-containing protein; uniclust \| UniRef100\_A0A2D9X072 \| 99.3 \| 1e-14 \| 1.9e-20 \| 103.0 \| 104 \| (19, 155) \| 156 \| (3, 111) \| 116 \| VRR-NUC domain-containing protein \| VRR-NUC domain-containing protein; uniclust \| UniRef100\_A0A0R3DPT5 \| 99.3 \| 9.7e-15 \| 1.9e-20 \| 106.0 \| 87 \| (20, 138) \| 156 \| (6, 99) \| 126 \| VRR-NUC domain-containing protein \| VRR-NUC domain-containing protein; uniclust \| UniRef100\_UPI001903D17B \| 99.3 \| 1e-14 \| 2e-20 \| 109.7 \| 111 \| (19, 155) \| 156 \| (9, 123) \| 165 \| hypothetical protein \| hypothetical protein; uniclust \| UniRef100\_A0A3D8IYP5 \| 99.3 \| 1e-14 \| 2e-20 \| 110.5 \| 108 \| (19, 155) \| 156 \| (25, 140) \| 160 \| VRR-NUC domain-containing protein \| VRR-NUC domain-containing protein; uniclust \| UniRef100\_A0A7U9NG49 \| 99.3 \| 1.1e-14 \| 2.1e-20 \| 114.0 \| 111 \| (16, 155) \| 156 \| (74, 192) \| 239 \| VRR-NUC domain-containing protein \| VRR-NUC domain-containing protein; uniclust \| UniRef100\_A0A3A9F823 \| 99.3 \| 1.2e-14 \| 2.3e-20 \| 112.6 \| 105 \| (22, 155) \| 156 \| (6, 118) \| 216 \| VRR-NUC domain-containing protein \| VRR-NUC domain-containing protein; uniclust \| UniRef100\_A0A839GEX4 \| 99.3 \| 1.4e-14 \| 2.7e-20 \| 105.5 \| 105 \| (22, 155) \| 156 \| (7, 117) \| 132 \| VRR-NUC domain-containing protein \| VRR-NUC domain-containing protein; uniclust \| UniRef100\_L1PCC8 \| 99.3 \| 1.5e-14 \| 2.7e-20 \| 109.9 \| 103 \| (22, 155) \| 156 \| (2, 108) \| 202 \| VRR-NUC domain protein \| VRR-NUC domain protein; uniclust \| UniRef100\_A0A5C7JC15 \| 99.3 \| 1.5e-14 \| 2.7e-20 \| 106.6 \| 107 \| (20, 155) \| 156 \| (32, 153) \| 163 \| VRR-NUC domain-containing protein \| VRR-NUC domain-containing protein; uniclust \| UniRef100\_A0A1B7X3P3 \| 99.3 \| 1.3e-14 \| 2.8e-20 \| 106.2 \| 94 \| (19, 149) \| 156 \| (1, 96) \| 121 \| VRR-NUC domain-containing protein \| VRR-NUC domain-containing protein; uniclust \| UniRef100\_A0A212T8L3 \| 99.3 \| 1.7e-14 \| 3.3e-20 \| 110.5 \| 102 \| (22, 154) \| 156 \| (61, 164) \| 177 \| VRR-NUC domain-containing protein \| VRR-NUC domain-containing protein; uniclust \| UniRef100\_A0A2T0YJ69 \| 99.3 \| 1.6e-14 \| 3.4e-20 \| 107.3 \| 85 \| (19, 147) \| 156 \| (19, 105) \| 134 \| VRR-NUC domain-containing protein \| VRR-NUC domain-containing protein; uniclust \| UniRef100\_A0A354H0Y4 \| 99.3 \| 1.9e-14 \| 3.7e-20 \| 104.4 \| 113 \| (13, 154) \| 156 \| (6, 120) \| 126 \| VRR-NUC domain-containing protein \| VRR-NUC domain-containing protein; uniclust \| UniRef100\_A0A3N5V863 \| 99.3 \| 2.1e-14 \| 3.8e-20 \| 105.4 \| 103 \| (22, 153) \| 156 \| (51, 157) \| 158 \| VRR-NUC domain-containing protein \| VRR-NUC domain-containing protein; uniclust \| UniRef100\_A0A2D6XBS1 \| 99.3 \| 2.6e-14 \| 4.9e-20 \| 108.3 \| 105 \| (22, 154) \| 156 \| (62, 169) \| 180 \| VRR-NUC domain-containing protein \| VRR-NUC domain-containing protein; uniclust \| UniRef100\_A0A349HTR8 \| 99.3 \| 2.9e-14 \| 5.3e-20 \| 111.0 \| 102 \| (22, 154) \| 156 \| (2, 109) \| 245 \| VRR-NUC domain-containing protein \| VRR-NUC domain-containing protein; uniclust \| UniRef100\_A0A501W5S5 \| 99.2 \| 2.9e-14 \| 5.6e-20 \| 106.4 \| 110 \| (19, 155) \| 156 \| (26, 141) \| 161 \| VRR-NUC domain-containing protein \| VRR-NUC domain-containing protein; uniclust \| UniRef100\_A0A174X9Y2 \| 99.2 \| 3e-14 \| 5.8e-20 \| 106.9 \| 106 \| (19, 155) \| 156 \| (6, 119) \| 167 \| VRR-NUC domain-containing protein \| VRR-NUC domain-containing protein; uniclust \| UniRef100\_A0A8S5PT49 \| 99.2 \| 4.4e-14 \| 8.4e-20 \| 104.1 \| 89 \| (22, 155) \| 156 \| (47, 138) \| 145 \| Nuclease \| Nuclease; uniclust \| UniRef100\_A0A2D8XMR4 \| 99.2 \| 5.2e-14 \| 9.6e-20 \| 102.9 \| 109 \| (21, 155) \| 156 \| (35, 148) \| 152 \| VRR-NUC domain-containing protein \| VRR-NUC domain-containing protein; uniclust \| UniRef100\_A0A088GMG7 \| 99.2 \| 5.2e-14 \| 1e-19 \| 98.0 \| 90 \| (22, 154) \| 156 \| (1, 94) \| 100 \| VRR-NUC domain-containing protein \| VRR-NUC domain-containing protein; uniclust \| UniRef100\_UPI0009BC488B \| 99.2 \| 5.4e-14 \| 1e-19 \| 104.9 \| 105 \| (19, 155) \| 156 \| (43, 153) \| 161 \| VRR-NUC domain-containing protein \| VRR-NUC domain-containing protein; uniclust \| UniRef100\_UPI00223C1613 \| 99.2 \| 5.7e-14 \| 1e-19 \| 103.3 \| 107 \| (20, 155) \| 156 \| (4, 113) \| 159 \| VRR-NUC domain-containing protein \| VRR-NUC domain-containing protein; uniclust \| UniRef100\_A0A3M1MRP5 \| 99.2 \| 6.3e-14 \| 1.2e-19 \| 100.1 \| 102 \| (17, 155) \| 156 \| (22, 125) \| 130 \| VRR-NUC domain-containing protein (Fragment) \| VRR-NUC domain-containing protein (Fragment); uniclust \| UniRef100\_A0A8S5TAK1 \| 99.2 \| 6.4e-14 \| 1.2e-19 \| 106.2 \| 106 \| (21, 155) \| 156 \| (68, 182) \| 198 \| Nuclease \| Nuclease; uniclust \| UniRef100\_A0A6J5SJ08 \| 99.2 \| 6.3e-14 \| 1.2e-19 \| 103.7 \| 105 \| (21, 154) \| 156 \| (28, 137) \| 149 \| VRR-NUC domain containing protein \| VRR-NUC domain containing protein; uniclust \| UniRef100\_A0A023ZVH9 \| 99.2 \| 5.7e-14 \| 1.2e-19 \| 109.3 \| 106 \| (22, 155) \| 156 \| (47, 160) \| 181 \| VRR-NUC domain-containing protein \| VRR-NUC domain-containing protein; uniclust \| UniRef100\_A0A2T0RG28 \| 99.2 \| 6.7e-14 \| 1.3e-19 \| 101.3 \| 107 \| (21, 155) \| 156 \| (4, 115) \| 136 \| VRR-NUC domain-containing protein \| VRR-NUC domain-containing protein; uniclust \| UniRef100\_A0A104Z6Z7 \| 99.2 \| 7.1e-14 \| 1.4e-19 \| 106.2 \| 105 \| (22, 155) \| 156 \| (31, 145) \| 166 \| VRR-NUC domain-containing protein \| VRR-NUC domain-containing protein; uniclust \| UniRef100\_A0A965PCU2 \| 99.2 \| 9e-14 \| 1.7e-19 \| 105.6 \| 106 \| (19, 154) \| 156 \| (71, 185) \| 199 \| VRR-NUC domain-containing protein \| VRR-NUC domain-containing protein; uniclust \| UniRef100\_A0A016QSZ2 \| 99.2 \| 9e-14 \| 1.7e-19 \| 99.4 \| 93 \| (22, 153) \| 156 \| (14, 109) \| 121 \| VRR-NUC domain-containing protein \| VRR-NUC domain-containing protein; uniclust \| UniRef100\_A0A6C2YRG2 \| 99.2 \| 9.1e-14 \| 1.7e-19 \| 97.6 \| 99 \| (23, 154) \| 156 \| (1, 101) \| 110 \| VRR-NUC domain-containing protein \| VRR-NUC domain-containing protein; uniclust \| UniRef100\_A0A1M5CEU7 \| 99.2 \| 8.8e-14 \| 1.7e-19 \| 100.6 \| 88 \| (23, 155) \| 156 \| (1, 93) \| 122 \| VRR-NUC domain-containing protein \| VRR-NUC domain-containing protein; uniclust \| UniRef100\_A0A516M9P1 \| 99.2 \| 1.1e-13 \| 1.9e-19 \| 97.3 \| 101 \| (22, 153) \| 156 \| (1, 105) \| 116 \| VRR-NUC domain-containing protein \| VRR-NUC domain-containing protein; uniclust \| UniRef100\_A0A3E0Q3U7 \| 99.2 \| 8.7e-14 \| 1.9e-19 \| 104.0 \| 99 \| (21, 154) \| 156 \| (15, 118) \| 129 \| Uncharacterized protein \| Uncharacterized protein; uniclust \| UniRef100\_A0A1C7G917 \| 99.2 \| 1.1e-13 \| 2e-19 \| 104.3 \| 106 \| (21, 155) \| 156 \| (10, 123) \| 187 \| Nuclease \| Nuclease; uniclust \| UniRef100\_A0A1W1ZWA3 \| 99.2 \| 1.3e-13 \| 2.5e-19 \| 105.1 \| 106 \| (17, 154) \| 156 \| (2, 113) \| 171 \| VRR-NUC domain-containing protein \| VRR-NUC domain-containing protein; uniclust \| UniRef100\_A0A0F8XH15 \| 99.2 \| 1.2e-13 \| 2.6e-19 \| 107.7 \| 100 \| (21, 154) \| 156 \| (61, 163) \| 180 \| VRR-NUC domain-containing protein (Fragment) \| VRR-NUC domain-containing protein (Fragment); uniclust \| UniRef100\_A0A2E6C829 \| 99.2 \| 1.5e-13 \| 2.7e-19 \| 99.6 \| 101 \| (21, 154) \| 156 \| (16, 121) \| 142 \| VRR-NUC domain-containing protein \| VRR-NUC domain-containing protein; uniclust \| UniRef100\_A0A3A9C013 \| 99.2 \| 1.4e-13 \| 2.8e-19 \| 105.6 \| 109 \| (18, 155) \| 156 \| (8, 124) \| 180 \| VRR-NUC domain-containing protein \| VRR-NUC domain-containing protein; uniclust \| UniRef100\_A0A965AG40 \| 99.2 \| 1.6e-13 \| 2.9e-19 \| 103.2 \| 98 \| (22, 154) \| 156 \| (76, 176) \| 184 \| VRR-NUC domain-containing protein \| VRR-NUC domain-containing protein; uniclust \| UniRef100\_A0A3B8HLD6 \| 99.2 \| 1.6e-13 \| 2.9e-19 \| 98.8 \| 109 \| (20, 155) \| 156 \| (23, 134) \| 136 \| VRR-NUC domain-containing protein \| VRR-NUC domain-containing protein; uniclust \| UniRef100\_A0A661JDB3 \| 99.2 \| 1.7e-13 \| 3.2e-19 \| 101.8 \| 102 \| (24, 155) \| 156 \| (34, 137) \| 154 \| VRR-NUC domain-containing protein \| VRR-NUC domain-containing protein; uniclust \| UniRef100\_A0A969LIU0 \| 99.2 \| 1.8e-13 \| 3.3e-19 \| 98.0 \| 103 \| (21, 153) \| 156 \| (23, 127) \| 131 \| VRR-NUC domain-containing protein \| VRR-NUC domain-containing protein; uniclust \| UniRef100\_A0A076YP05 \| 99.2 \| 1.6e-13 \| 3.3e-19 \| 101.1 \| 97 \| (13, 155) \| 156 \| (12, 110) \| 129 \| VRR-NUC domain-containing protein \| VRR-NUC domain-containing protein; uniclust \| UniRef100\_A0A3B8HH35 \| 99.2 \| 1.7e-13 \| 3.4e-19 \| 101.4 \| 93 \| (23, 155) \| 156 \| (30, 124) \| 134 \| VRR-NUC domain-containing protein \| VRR-NUC domain-containing protein; uniclust \| UniRef100\_A0A7C2EKI6 \| 99.2 \| 1.8e-13 \| 3.6e-19 \| 103.0 \| 111 \| (20, 156) \| 156 \| (29, 147) \| 158 \| Uncharacterized protein \| Uncharacterized protein; uniclust \| UniRef100\_A0A2S1GLC8 \| 99.2 \| 1.9e-13 \| 3.6e-19 \| 105.2 \| 105 \| (21, 154) \| 156 \| (27, 140) \| 187 \| VRR-NUC domain-containing protein \| VRR-NUC domain-containing protein; uniclust \| UniRef100\_A0A060BFY8 \| 99.1 \| 1.9e-13 \| 3.8e-19 \| 107.8 \| 105 \| (23, 156) \| 156 \| (46, 165) \| 210 \| VRR-NUC domain-containing protein \| VRR-NUC domain-containing protein; uniclust \| UniRef100\_A0A1C5RXP7 \| 99.1 \| 2e-13 \| 3.9e-19 \| 109.0 \| 106 \| (18, 154) \| 156 \| (3, 116) \| 249 \| VRR-NUC domain \| VRR-NUC domain; uniclust \| UniRef100\_A0A511FFQ3 \| 99.1 \| 2e-13 \| 4e-19 \| 102.1 \| 87 \| (20, 149) \| 156 \| (40, 128) \| 145 \| VRR-NUC domain-containing protein \| VRR-NUC domain-containing protein; uniclust \| UniRef100\_A0A1V5WI66 \| 99.1 \| 2.4e-13 \| 4.3e-19 \| 98.7 \| 98 \| (23, 154) \| 156 \| (1, 100) \| 143 \| VRR-NUC domain protein \| VRR-NUC domain protein; uniclust \| UniRef100\_UPI00130D683B \| 99.1 \| 2.4e-13 \| 4.4e-19 \| 101.0 \| 108 \| (22, 155) \| 156 \| (29, 139) \| 168 \| VRR-NUC domain-containing protein \| VRR-NUC domain-containing protein; uniclust \| UniRef100\_A0A1V5T3V5 \| 99.1 \| 2.4e-13 \| 4.4e-19 \| 85.8 \| 56 \| (85, 151) \| 156 \| (3, 59) \| 62 \| VRR-NUC domain protein \| VRR-NUC domain protein; uniclust \| UniRef100\_A0A3S0DF79 \| 99.1 \| 2.4e-13 \| 4.4e-19 \| 101.3 \| 104 \| (21, 153) \| 156 \| (47, 155) \| 172 \| VRR-NUC domain-containing protein \| VRR-NUC domain-containing protein; uniclust \| UniRef100\_A0A1F8V0P7 \| 99.1 \| 2.5e-13 \| 4.5e-19 \| 102.5 \| 105 \| (21, 155) \| 156 \| (7, 120) \| 188 \| VRR-NUC domain-containing protein \| VRR-NUC domain-containing protein; uniclust \| UniRef100\_A0A285X397 \| 99.1 \| 2.1e-13 \| 4.6e-19 \| 103.1 \| 105 \| (20, 155) \| 156 \| (3, 125) \| 143 \| VRR-NUC domain-containing protein \| VRR-NUC domain-containing protein; uniclust \| UniRef100\_A0A9E6MYT2 \| 99.1 \| 2.5e-13 \| 4.6e-19 \| 106.3 \| 107 \| (20, 155) \| 156 \| (14, 128) \| 250 \| VRR-NUC domain-containing protein \| VRR-NUC domain-containing protein; uniclust \| UniRef100\_A0A0R1VXF8 \| 99.1 \| 2.6e-13 \| 4.7e-19 \| 97.7 \| 103 \| (20, 155) \| 156 \| (30, 134) \| 135 \| VRR-NUC domain protein \| VRR-NUC domain protein; uniclust \| UniRef100\_A0A1M7G6C6 \| 99.1 \| 2.4e-13 \| 4.8e-19 \| 104.2 \| 101 \| (24, 154) \| 156 \| (22, 124) \| 177 \| VRR-NUC domain-containing protein \| VRR-NUC domain-containing protein; uniclust \| UniRef100\_A0A645G019 \| 99.1 \| 2.5e-13 \| 4.8e-19 \| 96.9 \| 91 \| (21, 155) \| 156 \| (9, 103) \| 116 \| VRR-NUC domain-containing protein \| VRR-NUC domain-containing protein; uniclust \| UniRef100\_E5Y6D9 \| 99.1 \| 2.7e-13 \| 5e-19 \| 101.9 \| 107 \| (20, 155) \| 156 \| (30, 143) \| 178 \| VRR-NUC domain-containing protein \| VRR-NUC domain-containing protein; uniclust \| UniRef100\_A0A512B9U6 \| 99.1 \| 2.7e-13 \| 5.2e-19 \| 98.5 \| 103 \| (19, 154) \| 156 \| (21, 127) \| 133 \| VRR-NUC domain-containing protein \| VRR-NUC domain-containing protein; uniclust \| UniRef100\_A0A351TKZ0 \| 99.1 \| 2.6e-13 \| 5.2e-19 \| 90.5 \| 59 \| (85, 155) \| 156 \| (7, 66) \| 73 \| VRR-NUC domain-containing protein (Fragment) \| VRR-NUC domain-containing protein (Fragment); uniclust \| UniRef100\_A0A1Y1QLN9 \| 99.1 \| 2.4e-13 \| 5.4e-19 \| 98.7 \| 92 \| (18, 155) \| 156 \| (3, 96) \| 108 \| VRR-NUC domain-containing protein \| VRR-NUC domain-containing protein; uniclust \| UniRef100\_A0A431K292 \| 99.1 \| 2.9e-13 \| 5.4e-19 \| 100.2 \| 128 \| (1, 151) \| 156 \| (1, 154) \| 164 \| VRR-NUC domain-containing protein \| VRR-NUC domain-containing protein; uniclust \| UniRef100\_UPI001FFBE019 \| 99.1 \| 3.4e-13 \| 6.2e-19 \| 99.5 \| 103 \| (21, 154) \| 156 \| (16, 123) \| 159 \| VRR-NUC domain-containing protein \| VRR-NUC domain-containing protein; uniclust \| UniRef100\_A0A7C6QZC2 \| 99.1 \| 3.4e-13 \| 6.2e-19 \| 95.0 \| 86 \| (23, 153) \| 156 \| (1, 88) \| 117 \| VRR-NUC domain-containing protein \| VRR-NUC domain-containing protein; uniclust \| UniRef100\_A0A2E6PX40 \| 99.1 \| 3.4e-13 \| 6.4e-19 \| 101.0 \| 104 \| (21, 155) \| 156 \| (4, 112) \| 166 \| VRR-NUC domain-containing protein \| VRR-NUC domain-containing protein; uniclust \| UniRef100\_A0A0A0P383 \| 99.1 \| 3.1e-13 \| 6.6e-19 \| 105.0 \| 108 \| (21, 155) \| 156 \| (39, 153) \| 173 \| Type III restriction endonuclease \| Type III restriction endonuclease; uniclust \| UniRef100\_A0A6N2ZXV1 \| 99.1 \| 3.6e-13 \| 7e-19 \| 96.5 \| 90 \| (20, 155) \| 156 \| (3, 94) \| 114 \| VRR-NUC domain protein \| VRR-NUC domain protein; uniclust \| UniRef100\_A0A3N5H338 \| 99.1 \| 3.8e-13 \| 7e-19 \| 89.3 \| 67 \| (77, 155) \| 156 \| (7, 75) \| 82 \| VRR-NUC domain-containing protein \| VRR-NUC domain-containing protein; uniclust \| UniRef100\_A0A060AHI1 \| 99.1 \| 3.6e-13 \| 7.3e-19 \| 99.4 \| 90 \| (19, 153) \| 156 \| (2, 97) \| 128 \| Endonuclease \| Endonuclease; uniclust \| UniRef100\_A0A1B1IQE5 \| 99.1 \| 3.4e-13 \| 7.3e-19 \| 99.9 \| 85 \| (18, 147) \| 156 \| (11, 97) \| 124 \| VRR-NUC domain-containing protein \| VRR-NUC domain-containing protein; uniclust \| UniRef100\_A0A6J5NKG2 \| 99.1 \| 4.5e-13 \| 8.2e-19 \| 98.9 \| 102 \| (22, 154) \| 156 \| (2, 109) \| 159 \| VRR-NUC domain containing protein \| VRR-NUC domain containing protein; uniclust \| UniRef100\_A0A4Q2Y3F1 \| 99.1 \| 4.9e-13 \| 9.1e-19 \| 103.2 \| 111 \| (19, 154) \| 156 \| (98, 216) \| 222 \| VRR-NUC domain-containing protein \| VRR-NUC domain-containing protein; uniclust \| UniRef100\_A0A2T3K2K6 \| 99.1 \| 5.3e-13 \| 1e-18 \| 98.9 \| 108 \| (19, 155) \| 156 \| (27, 143) \| 145 \| VRR-NUC domain-containing protein \| VRR-NUC domain-containing protein; uniclust \| UniRef100\_A0A3G2KI64 \| 99.1 \| 6.7e-13 \| 1.3e-18 \| 97.4 \| 85 \| (20, 147) \| 156 \| (27, 114) \| 128 \| Nuclease \| Nuclease; uniclust \| UniRef100\_A0A350UDK1 \| 99.1 \| 7.7e-13 \| 1.4e-18 \| 100.4 \| 100 \| (22, 154) \| 156 \| (81, 185) \| 194 \| VRR-NUC domain-containing protein \| VRR-NUC domain-containing protein; uniclust \| UniRef100\_A0A6J5PR03 \| 99.1 \| 7e-13 \| 1.4e-18 \| 99.9 \| 94 \| (17, 148) \| 156 \| (25, 123) \| 145 \| VRR-NUC domain containing protein \| VRR-NUC domain containing protein; uniclust \| UniRef100\_A0A250JC22 \| 99.1 \| 8e-13 \| 1.6e-18 \| 97.5 \| 58 \| (86, 155) \| 156 \| (35, 93) \| 132 \| Nuclease \| Nuclease; uniclust \| UniRef100\_A0A6J5LZW8 \| 99.1 \| 9.1e-13 \| 1.7e-18 \| 99.7 \| 104 \| (22, 154) \| 156 \| (9, 120) \| 171 \| VRR-NUC domain containing protein \| VRR-NUC domain containing protein; uniclust \| UniRef100\_A0A068CGD0 \| 99.1 \| 8.1e-13 \| 1.8e-18 \| 100.4 \| 92 \| (12, 154) \| 156 \| (21, 119) \| 143 \| VRR-NUC domain-containing protein \| VRR-NUC domain-containing protein; uniclust \| UniRef100\_A0A0Q5KUS2 \| 99.1 \| 8.6e-13 \| 1.8e-18 \| 102.3 \| 93 \| (23, 153) \| 156 \| (64, 158) \| 173 \| VRR-NUC domain-containing protein \| VRR-NUC domain-containing protein; uniclust \| UniRef100\_A0A0F8W6U3 \| 99.1 \| 9.5e-13 \| 1.8e-18 \| 97.0 \| 109 \| (21, 154) \| 156 \| (15, 125) \| 139 \| Uncharacterized protein (Fragment) \| Uncharacterized protein (Fragment); uniclust \| UniRef100\_A0A317H910 \| 99.1 \| 1e-12 \| 1.9e-18 \| 96.3 \| 101 \| (22, 153) \| 156 \| (5, 110) \| 150 \| VRR-NUC domain-containing protein \| VRR-NUC domain-containing protein; uniclust \| UniRef100\_UPI001FFBBF5C \| 99.1 \| 1e-12 \| 1.9e-18 \| 95.5 \| 106 \| (21, 155) \| 156 \| (13, 123) \| 134 \| hypothetical protein \| hypothetical protein; uniclust \| UniRef100\_A0A6J5LIH6 \| 99.0 \| 1.3e-12 \| 2.4e-18 \| 94.4 \| 110 \| (23, 155) \| 156 \| (3, 118) \| 137 \| VRR-NUC domain containing protein \| VRR-NUC domain containing protein; uniclust \| UniRef100\_A0A139CFL3 \| 99.0 \| 1.4e-12 \| 2.6e-18 \| 96.5 \| 105 \| (22, 155) \| 156 \| (26, 135) \| 140 \| VRR-NUC domain-containing protein (Fragment) \| VRR-NUC domain-containing protein (Fragment); uniclust \| UniRef100\_A0A1A0IRN8 \| 99.0 \| 1.4e-12 \| 2.9e-18 \| 99.4 \| 96 \| (16, 153) \| 156 \| (1, 98) \| 151 \| Nuclease \| Nuclease; uniclust \| UniRef100\_A0A838SYU0 \| 99.0 \| 1.7e-12 \| 3.1e-18 \| 97.0 \| 104 \| (22, 154) \| 156 \| (10, 124) \| 171 \| VRR-NUC domain-containing protein \| VRR-NUC domain-containing protein; uniclust \| UniRef100\_A0A7K0AAW2 \| 99.0 \| 1.7e-12 \| 3.3e-18 \| 99.6 \| 115 \| (20, 155) \| 156 \| (25, 153) \| 191 \| VRR-NUC domain-containing protein \| VRR-NUC domain-containing protein; uniclust \| UniRef100\_UPI0022E3BF2E \| 99.0 \| 2.1e-12 \| 3.9e-18 \| 99.9 \| 105 \| (21, 155) \| 156 \| (94, 207) \| 222 \| VRR-NUC domain-containing protein \| VRR-NUC domain-containing protein; uniclust \| UniRef100\_A0A0F9IFS0 \| 99.0 \| 1.9e-12 \| 3.9e-18 \| 91.0 \| 81 \| (23, 154) \| 156 \| (7, 89) \| 94 \| VRR-NUC domain-containing protein \| VRR-NUC domain-containing protein; uniclust \| UniRef100\_A0A060RIT6 \| 99.0 \| 2.1e-12 \| 4.1e-18 \| 100.6 \| 111 \| (22, 155) \| 156 \| (17, 167) \| 187 \| VRR-NUC domain-containing protein \| VRR-NUC domain-containing protein; uniclust \| UniRef100\_A0A3S4F827 \| 99.0 \| 2.3e-12 \| 4.2e-18 \| 93.8 \| 94 \| (29, 154) \| 156 \| (2, 100) \| 142 \| VRR-NUC domain-containing protein \| VRR-NUC domain-containing protein; uniclust \| UniRef100\_A0A1T2BEL0 \| 99.0 \| 2.6e-12 \| 4.7e-18 \| 85.2 \| 62 \| (83, 155) \| 156 \| (12, 75) \| 80 \| VRR-NUC domain-containing protein \| VRR-NUC domain-containing protein; uniclust \| UniRef100\_A0A086D0Z9 \| 99.0 \| 2.7e-12 \| 5.3e-18 \| 97.9 \| 106 \| (22, 156) \| 156 \| (22, 142) \| 167 \| VRR-NUC domain-containing protein \| VRR-NUC domain-containing protein; uniclust \| UniRef100\_A0A0B4Y550 \| 99.0 \| 2.5e-12 \| 5.3e-18 \| 99.1 \| 106 \| (22, 155) \| 156 \| (36, 149) \| 163 \| VRR-NUC domain-containing protein \| VRR-NUC domain-containing protein; uniclust \| UniRef100\_J5UZX6 \| 99.0 \| 3.4e-12 \| 6.2e-18 \| 90.2 \| 88 \| (23, 155) \| 156 \| (1, 91) \| 117 \| VRR-NUC domain protein \| VRR-NUC domain protein; uniclust \| UniRef100\_A0A0S7CQU9 \| 99.0 \| 3e-12 \| 6.4e-18 \| 98.8 \| 112 \| (16, 155) \| 156 \| (12, 139) \| 161 \| Uncharacterized protein \| Uncharacterized protein; uniclust \| UniRef100\_A0A0Q5IHK8 \| 99.0 \| 3.7e-12 \| 7.4e-18 \| 99.0 \| 113 \| (23, 155) \| 156 \| (33, 154) \| 189 \| VRR-NUC domain-containing protein \| VRR-NUC domain-containing protein; uniclust \| UniRef100\_A0A0F8Y460 \| 99.0 \| 3.8e-12 \| 7.6e-18 \| 95.5 \| 102 \| (21, 155) \| 156 \| (39, 143) \| 147 \| VRR-NUC domain-containing protein (Fragment) \| VRR-NUC domain-containing protein (Fragment); uniclust \| UniRef100\_A0A6B1DRS4 \| 99.0 \| 4.3e-12 \| 7.8e-18 \| 97.4 \| 101 \| (23, 155) \| 156 \| (100, 204) \| 207 \| VRR-NUC domain-containing protein \| VRR-NUC domain-containing protein; uniclust \| UniRef100\_A0A965X540 \| 99.0 \| 4.3e-12 \| 8e-18 \| 93.8 \| 102 \| (22, 154) \| 156 \| (35, 146) \| 157 \| VRR-NUC domain-containing protein \| VRR-NUC domain-containing protein; uniclust \| UniRef100\_A0A3M1SPX9 \| 99.0 \| 4.4e-12 \| 8e-18 \| 90.0 \| 96 \| (23, 155) \| 156 \| (1, 100) \| 120 \| VRR-NUC domain-containing protein \| VRR-NUC domain-containing protein; uniclust \| UniRef100\_UPI001D0E449F \| 99.0 \| 4.6e-12 \| 8.5e-18 \| 86.3 \| 69 \| (75, 155) \| 156 \| (14, 83) \| 93 \| VRR-NUC domain-containing protein \| VRR-NUC domain-containing protein; uniclust \| UniRef100\_A0A0G3EVL4 \| 99.0 \| 4.5e-12 \| 8.8e-18 \| 100.2 \| 112 \| (21, 154) \| 156 \| (61, 193) \| 217 \| Endonuclease \| Endonuclease; uniclust \| UniRef100\_A0A3N9NZB7 \| 99.0 \| 4.7e-12 \| 9.1e-18 \| 89.0 \| 87 \| (18, 133) \| 156 \| (7, 102) \| 102 \| VRR-NUC domain-containing protein (Fragment) \| VRR-NUC domain-containing protein (Fragment); uniclust \| UniRef100\_A0A371YWW9 \| 98.9 \| 5.5e-12 \| 1e-17 \| 93.6 \| 69 \| (75, 155) \| 156 \| (81, 150) \| 161 \| VRR-NUC domain-containing protein \| VRR-NUC domain-containing protein; uniclust \| UniRef100\_A0A2K4QBL7 \| 98.9 \| 7.3e-12 \| 1.3e-17 \| 93.5 \| 102 \| (21, 155) \| 156 \| (64, 167) \| 168 \| VRR-NUC domain-containing protein (Modular protein) \| VRR-NUC domain-containing protein (Modular protein); uniclust \| UniRef100\_UPI0022E05DE5 \| 98.9 \| 8.1e-12 \| 1.5e-17 \| 101.1 \| 106 \| (21, 155) \| 156 \| (169, 281) \| 316 \| VRR-NUC domain-containing protein \| VRR-NUC domain-containing protein; uniclust \| UniRef100\_A0A2K9NFU2 \| 98.9 \| 8.2e-12 \| 1.5e-17 \| 92.9 \| 102 \| (21, 155) \| 156 \| (37, 152) \| 163 \| VRR-NUC domain-containing protein \| VRR-NUC domain-containing protein; uniclust \| UniRef100\_A0A451GGP2 \| 98.9 \| 8.2e-12 \| 1.6e-17 \| 86.3 \| 69 \| (75, 155) \| 156 \| (9, 78) \| 94 \| VRR-NUC domain-containing protein \| VRR-NUC domain-containing protein; uniclust \| UniRef100\_A0A7X5Y9R6 \| 98.9 \| 8.6e-12 \| 1.6e-17 \| 85.3 \| 70 \| (74, 155) \| 156 \| (15, 85) \| 95 \| VRR-NUC domain-containing protein \| VRR-NUC domain-containing protein; uniclust \| UniRef100\_UPI001305397F \| 98.9 \| 9.3e-12 \| 1.7e-17 \| 83.3 \| 68 \| (75, 154) \| 156 \| (14, 82) \| 83 \| VRR-NUC domain-containing protein \| VRR-NUC domain-containing protein; uniclust \| UniRef100\_UPI000FCC612F \| 98.9 \| 9.5e-12 \| 1.7e-17 \| 84.1 \| 76 \| (79, 154) \| 156 \| (6, 82) \| 88 \| hypothetical protein \| hypothetical protein; uniclust \| UniRef100\_A0A0R1IE25 \| 98.9 \| 8.9e-12 \| 1.7e-17 \| 91.8 \| 86 \| (17, 146) \| 156 \| (1, 88) \| 130 \| VRR-NUC domain-containing protein \| VRR-NUC domain-containing protein; uniclust \| UniRef100\_A0A2L0HNZ2 \| 98.9 \| 1.1e-11 \| 2.1e-17 \| 86.5 \| 88 \| (22, 154) \| 156 \| (1, 94) \| 99 \| Nuclease \| Nuclease; uniclust \| UniRef100\_A0A143XCJ5 \| 98.9 \| 1.1e-11 \| 2.2e-17 \| 92.6 \| 88 \| (22, 155) \| 156 \| (46, 135) \| 139 \| VRR-NUC domain protein \| VRR-NUC domain protein; uniclust \| UniRef100\_A0A7C3YQK5 \| 98.9 \| 1.2e-11 \| 2.2e-17 \| 91.9 \| 111 \| (19, 155) \| 156 \| (18, 135) \| 160 \| VRR-NUC domain-containing protein \| VRR-NUC domain-containing protein; uniclust \| UniRef100\_A0A6J5QEF0 \| 98.9 \| 1.1e-11 \| 2.2e-17 \| 91.7 \| 89 \| (21, 154) \| 156 \| (2, 93) \| 143 \| VRR-NUC domain containing protein \| VRR-NUC domain containing protein; uniclust \| UniRef100\_A0A2S0JKP9 \| 98.9 \| 1.2e-11 \| 2.3e-17 \| 81.7 \| 60 \| (84, 155) \| 156 \| (9, 69) \| 73 \| Recombinase RecB \| Recombinase RecB; uniclust \| UniRef100\_A0A0F9JDU4 \| 98.9 \| 1.3e-11 \| 2.3e-17 \| 92.9 \| 85 \| (26, 155) \| 156 \| (86, 172) \| 175 \| Uncharacterized protein \| Uncharacterized protein; uniclust \| UniRef100\_A0A1V5WR11 \| 98.9 \| 1.2e-11 \| 2.4e-17 \| 95.6 \| 104 \| (19, 154) \| 156 \| (28, 135) \| 171 \| VRR-NUC domain protein \| VRR-NUC domain protein; uniclust \| UniRef100\_R9KAU3 \| 98.9 \| 1.4e-11 \| 2.6e-17 \| 97.6 \| 106 \| (21, 155) \| 156 \| (10, 123) \| 261 \| VRR-NUC domain-containing protein \| VRR-NUC domain-containing protein; uniclust \| UniRef100\_A0A832HUD1 \| 98.9 \| 1.5e-11 \| 2.8e-17 \| 87.7 \| 91 \| (21, 153) \| 156 \| (28, 120) \| 122 \| VRR-NUC domain-containing protein \| VRR-NUC domain-containing protein; uniclust \| UniRef100\_A0A383C7W8 \| 98.9 \| 1.6e-11 \| 2.9e-17 \| 84.4 \| 68 \| (76, 155) \| 156 \| (4, 77) \| 97 \| VRR-NUC domain-containing protein \| VRR-NUC domain-containing protein; uniclust \| UniRef100\_A0A7H5F1N7 \| 98.9 \| 1.6e-11 \| 3e-17 \| 94.1 \| 112 \| (21, 155) \| 156 \| (58, 176) \| 201 \| VRR-NUC domain-containing protein \| VRR-NUC domain-containing protein; uniclust \| UniRef100\_A0A6M5C9Y6 \| 98.9 \| 1.7e-11 \| 3.1e-17 \| 90.5 \| 108 \| (21, 155) \| 156 \| (12, 126) \| 149 \| Endonuclease \| Endonuclease; uniclust \| UniRef100\_A0A3C0STG0 \| 98.9 \| 1.6e-11 \| 3.2e-17 \| 85.6 \| 58 \| (85, 154) \| 156 \| (18, 78) \| 87 \| VRR-NUC domain-containing protein \| VRR-NUC domain-containing protein; uniclust \| UniRef100\_A0A2S7YZ17 \| 98.9 \| 1.7e-11 \| 3.5e-17 \| 88.9 \| 89 \| (20, 154) \| 156 \| (13, 103) \| 115 \| VRR-NUC domain-containing protein \| VRR-NUC domain-containing protein; uniclust \| UniRef100\_A0A017N0S4 \| 98.9 \| 1.7e-11 \| 3.5e-17 \| 86.4 \| 60 \| (85, 155) \| 156 \| (25, 85) \| 93 \| VRR-NUC domain protein \| VRR-NUC domain protein; uniclust \| UniRef100\_A0A317HKU5 \| 98.9 \| 2.1e-11 \| 3.8e-17 \| 90.1 \| 103 \| (21, 155) \| 156 \| (42, 149) \| 154 \| VRR-NUC domain-containing protein \| VRR-NUC domain-containing protein; uniclust \| UniRef100\_UPI000A0099DC \| 98.8 \| 2.4e-11 \| 4.3e-17 \| 94.0 \| 104 \| (20, 153) \| 156 \| (70, 182) \| 214 \| VRR-NUC domain-containing protein \| VRR-NUC domain-containing protein; uniclust \| UniRef100\_A0A6H2A0C5 \| 98.8 \| 2.2e-11 \| 4.5e-17 \| 93.8 \| 90 \| (22, 154) \| 156 \| (52, 143) \| 173 \| Putative VRR-NUC domain-containing protein \| Putative VRR-NUC domain-containing protein; uniclust \| UniRef100\_A0A7Y5S375 \| 98.8 \| 2.7e-11 \| 4.9e-17 \| 81.8 \| 68 \| (75, 154) \| 156 \| (12, 80) \| 87 \| VRR-NUC domain-containing protein \| VRR-NUC domain-containing protein; uniclust \| UniRef100\_A0A2K1P5E6 \| 98.8 \| 2.7e-11 \| 5e-17 \| 89.4 \| 87 \| (23, 154) \| 156 \| (1, 90) \| 149 \| VRR-NUC domain-containing protein \| VRR-NUC domain-containing protein; uniclust \| UniRef100\_A0A2V2CT51 \| 98.8 \| 3.2e-11 \| 6.2e-17 \| 85.3 \| 100 \| (22, 154) \| 156 \| (1, 102) \| 106 \| VRR-NUC domain-containing protein \| VRR-NUC domain-containing protein; uniclust \| UniRef100\_UPI001ED9FD3A \| 98.8 \| 3.5e-11 \| 6.4e-17 \| 90.2 \| 115 \| (21, 155) \| 156 \| (1, 126) \| 169 \| hypothetical protein \| hypothetical protein; uniclust \| UniRef100\_A0A965URT5 \| 98.8 \| 3.6e-11 \| 6.6e-17 \| 82.3 \| 66 \| (80, 154) \| 156 \| (2, 68) \| 94 \| VRR-NUC domain-containing protein \| VRR-NUC domain-containing protein; uniclust \| UniRef100\_A0A7C4LPF3 \| 98.8 \| 3.6e-11 \| 6.8e-17 \| 86.9 \| 112 \| (1, 154) \| 156 \| (1, 118) \| 120 \| VRR-NUC domain-containing protein \| VRR-NUC domain-containing protein; uniclust \| UniRef100\_A0A1D7TSS5 \| 98.8 \| 3.4e-11 \| 6.8e-17 \| 90.4 \| 58 \| (85, 154) \| 156 \| (49, 109) \| 141 \| VRR-NUC domain-containing protein \| VRR-NUC domain-containing protein; uniclust \| UniRef100\_A0A7U9N3H7 \| 98.8 \| 3.9e-11 \| 7.2e-17 \| 92.9 \| 117 \| (10, 155) \| 156 \| (72, 196) \| 214 \| VRR-NUC domain-containing protein \| VRR-NUC domain-containing protein; uniclust \| UniRef100\_A0A660V061 \| 98.8 \| 4.2e-11 \| 7.8e-17 \| 85.3 \| 87 \| (22, 153) \| 156 \| (1, 89) \| 120 \| VRR-NUC domain-containing protein \| VRR-NUC domain-containing protein; uniclust \| UniRef100\_UPI000558DD9B \| 98.8 \| 4.3e-11 \| 8e-17 \| 89.3 \| 93 \| (23, 154) \| 156 \| (65, 159) \| 164 \| VRR-NUC domain-containing protein \| VRR-NUC domain-containing protein; uniclust \| UniRef100\_UPI0021F2D242 \| 98.8 \| 4.5e-11 \| 8.3e-17 \| 78.4 \| 63 \| (82, 155) \| 156 \| (4, 67) \| 73 \| hypothetical protein \| hypothetical protein; uniclust \| UniRef100\_A0A2E0FGH7 \| 98.8 \| 4.6e-11 \| 8.4e-17 \| 85.1 \| 102 \| (22, 154) \| 156 \| (10, 117) \| 120 \| VRR-NUC domain-containing protein \| VRR-NUC domain-containing protein; uniclust \| UniRef100\_A0A7T5QUK7 \| 98.8 \| 4.8e-11 \| 8.8e-17 \| 89.5 \| 106 \| (22, 154) \| 156 \| (2, 112) \| 169 \| VRR-NUC domain-containing protein \| VRR-NUC domain-containing protein; uniclust \| UniRef100\_R5J1V9 \| 98.8 \| 4.8e-11 \| 9.2e-17 \| 87.0 \| 59 \| (86, 155) \| 156 \| (35, 94) \| 126 \| VRR-NUC domain protein \| VRR-NUC domain protein; uniclust \| UniRef100\_A0A949GBL7 \| 98.8 \| 5.3e-11 \| 9.7e-17 \| 84.3 \| 70 \| (73, 154) \| 156 \| (21, 91) \| 115 \| VRR-NUC domain-containing protein \| VRR-NUC domain-containing protein; uniclust \| UniRef100\_A0A6J5SIW3 \| 98.8 \| 5.5e-11 \| 1e-16 \| 91.7 \| 105 \| (22, 155) \| 156 \| (58, 169) \| 183 \| VRR-NUC domain containing protein \| VRR-NUC domain containing protein; uniclust \| UniRef100\_B0N9W1 \| 98.8 \| 5.5e-11 \| 1e-16 \| 86.8 \| 92 \| (17, 154) \| 156 \| (28, 121) \| 129 \| VRR-NUC domain protein \| VRR-NUC domain protein; uniclust \| UniRef100\_A0A1E7I3F0 \| 98.8 \| 5.5e-11 \| 1e-16 \| 86.1 \| 89 \| (23, 155) \| 156 \| (25, 115) \| 124 \| VRR-NUC domain-containing protein \| VRR-NUC domain-containing protein; uniclust \| UniRef100\_A0A0F9BWP4 \| 98.8 \| 5.6e-11 \| 1.1e-16 \| 82.3 \| 84 \| (20, 138) \| 156 \| (2, 89) \| 90 \| VRR-NUC domain-containing protein (Fragment) \| VRR-NUC domain-containing protein (Fragment); uniclust \| UniRef100\_A0A0F9FVZ7 \| 98.8 \| 6.2e-11 \| 1.1e-16 \| 78.0 \| 69 \| (22, 122) \| 156 \| (2, 72) \| 74 \| VRR-NUC domain-containing protein (Fragment) \| VRR-NUC domain-containing protein (Fragment); uniclust \| UniRef100\_A0A6P2FZW6 \| 98.8 \| 6.2e-11 \| 1.1e-16 \| 91.7 \| 107 \| (22, 155) \| 156 \| (83, 197) \| 203 \| VRR-NUC domain protein \| VRR-NUC domain protein; uniclust \| UniRef100\_A0A016QS04 \| 98.8 \| 5.7e-11 \| 1.2e-16 \| 91.9 \| 108 \| (20, 155) \| 156 \| (17, 127) \| 160 \| VRR-NUC domain-containing protein \| VRR-NUC domain-containing protein; uniclust \| UniRef100\_A0A7V0Q709 \| 98.8 \| 6.8e-11 \| 1.2e-16 \| 89.0 \| 110 \| (22, 155) \| 156 \| (53, 165) \| 172 \| VRR-NUC domain-containing protein \| VRR-NUC domain-containing protein; uniclust \| UniRef100\_A0A965UUD2 \| 98.8 \| 7e-11 \| 1.3e-16 \| 92.0 \| 108 \| (21, 155) \| 156 \| (89, 203) \| 223 \| VRR-NUC domain-containing protein \| VRR-NUC domain-containing protein; uniclust \| UniRef100\_A0A2P6BVS4 \| 98.7 \| 7.9e-11 \| 1.5e-16 \| 88.0 \| 113 \| (22, 154) \| 156 \| (32, 152) \| 164 \| VRR-NUC domain-containing protein \| VRR-NUC domain-containing protein; uniclust \| UniRef100\_A0A0S7BVS5 \| 98.7 \| 7.8e-11 \| 1.7e-16 \| 95.3 \| 114 \| (23, 155) \| 156 \| (69, 185) \| 205 \| VRR-NUC domain-containing protein \| VRR-NUC domain-containing protein; uniclust \| UniRef100\_A0A1Q6RAF5 \| 98.7 \| 9.5e-11 \| 1.7e-16 \| 85.9 \| 111 \| (22, 154) \| 156 \| (2, 118) \| 143 \| VRR-NUC domain-containing protein \| VRR-NUC domain-containing protein; uniclust \| UniRef100\_A0A8J3EF72 \| 98.7 \| 9.8e-11 \| 1.8e-16 \| 87.0 \| 108 \| (20, 154) \| 156 \| (27, 142) \| 157 \| VRR-NUC domain-containing protein \| VRR-NUC domain-containing protein; uniclust \| UniRef100\_A0A2L0XD71 \| 98.7 \| 9.7e-11 \| 1.8e-16 \| 89.0 \| 104 \| (20, 154) \| 156 \| (42, 149) \| 171 \| VRR-NUC domain-containing protein \| VRR-NUC domain-containing protein; uniclust \| UniRef100\_A0A6M3Y0W3 \| 98.7 \| 1e-10 \| 1.9e-16 \| 85.0 \| 101 \| (16, 155) \| 156 \| (31, 133) \| 135 \| Putative VRR-NUC domain-containing protein \| Putative VRR-NUC domain-containing protein; uniclust \| UniRef100\_A0A7C5WRI5 \| 98.7 \| 1.1e-10 \| 2e-16 \| 88.0 \| 106 \| (23, 155) \| 156 \| (56, 163) \| 173 \| VRR-NUC domain-containing protein \| VRR-NUC domain-containing protein; uniclust \| UniRef100\_A0A2W5ZUK3 \| 98.7 \| 1.1e-10 \| 2e-16 \| 80.4 \| 71 \| (74, 155) \| 156 \| (10, 81) \| 96 \| VRR-NUC domain-containing protein \| VRR-NUC domain-containing protein; uniclust \| UniRef100\_A0A0F9H755 \| 98.7 \| 1.1e-10 \| 2e-16 \| 82.9 \| 99 \| (23, 154) \| 156 \| (1, 104) \| 116 \| VRR-NUC domain-containing protein \| VRR-NUC domain-containing protein; uniclust \| UniRef100\_A0A522SU77 \| 98.7 \| 1.1e-10 \| 2.1e-16 \| 88.3 \| 103 \| (21, 155) \| 156 \| (43, 167) \| 178 \| VRR-NUC domain-containing protein \| VRR-NUC domain-containing protein; uniclust \| UniRef100\_A0A142KBX3 \| 98.7 \| 1.2e-10 \| 2.2e-16 \| 76.5 \| 59 \| (85, 155) \| 156 \| (11, 70) \| 73 \| Holliday junction resolvase \| Holliday junction resolvase; uniclust \| UniRef100\_A0A6J5PJ07 \| 98.7 \| 1.2e-10 \| 2.2e-16 \| 85.9 \| 87 \| (14, 145) \| 156 \| (18, 106) \| 134 \| VRR-NUC domain containing protein \| VRR-NUC domain containing protein; uniclust \| UniRef100\_UPI001F5E1DBC \| 98.7 \| 1.2e-10 \| 2.3e-16 \| 76.1 \| 60 \| (83, 155) \| 156 \| (7, 68) \| 71 \| VRR-NUC domain-containing protein \| VRR-NUC domain-containing protein; uniclust \| UniRef100\_A0A0G1CCZ0 \| 98.7 \| 1.2e-10 \| 2.3e-16 \| 82.1 \| 87 \| (21, 152) \| 156 \| (4, 92) \| 111 \| VRR-NUC domain-containing protein \| VRR-NUC domain-containing protein; uniclust \| UniRef100\_A0A0F9TBD8 \| 98.7 \| 1.2e-10 \| 2.3e-16 \| 82.1 \| 57 \| (85, 151) \| 156 \| (31, 88) \| 102 \| VRR-NUC domain-containing protein \| VRR-NUC domain-containing protein; uniclust \| UniRef100\_A0A0F9NEB0 \| 98.7 \| 1.3e-10 \| 2.4e-16 \| 83.5 \| 95 \| (22, 147) \| 156 \| (5, 106) \| 125 \| VRR-NUC domain-containing protein \| VRR-NUC domain-containing protein; uniclust \| UniRef100\_A0A2V8RVF6 \| 98.7 \| 1.2e-10 \| 2.4e-16 \| 86.6 \| 92 \| (22, 155) \| 156 \| (27, 120) \| 142 \| VRR-NUC domain-containing protein \| VRR-NUC domain-containing protein; uniclust \| UniRef100\_A0A497BB15 \| 98.7 \| 1.3e-10 \| 2.4e-16 \| 85.0 \| 91 \| (20, 155) \| 156 \| (26, 118) \| 140 \| VRR-NUC domain-containing protein \| VRR-NUC domain-containing protein; uniclust \| UniRef100\_A0A6M3K5H3 \| 98.7 \| 1.3e-10 \| 2.4e-16 \| 84.3 \| 93 \| (19, 155) \| 156 \| (3, 101) \| 119 \| VRR-NUC domain-containing protein \| VRR-NUC domain-containing protein; uniclust \| UniRef100\_A0A6J5L3D2 \| 98.7 \| 1.3e-10 \| 2.5e-16 \| 86.6 \| 106 \| (21, 155) \| 156 \| (48, 157) \| 160 \| VRR-NUC domain containing protein \| VRR-NUC domain containing protein; uniclust \| UniRef100\_A0A1V3QA50 \| 98.7 \| 1.3e-10 \| 2.5e-16 \| 88.3 \| 104 \| (21, 153) \| 156 \| (42, 157) \| 170 \| VRR-NUC domain-containing protein \| VRR-NUC domain-containing protein; uniclust \| UniRef100\_A3JK17 \| 98.7 \| 1.5e-10 \| 2.7e-16 \| 89.6 \| 104 \| (23, 154) \| 156 \| (80, 192) \| 209 \| VRR-NUC domain-containing protein \| VRR-NUC domain-containing protein; uniclust \| UniRef100\_A0A2N5RCI0 \| 98.7 \| 1.4e-10 \| 2.7e-16 \| 90.8 \| 103 \| (22, 153) \| 156 \| (43, 148) \| 187 \| Nuclease \| Nuclease; uniclust \| UniRef100\_A0A6J5NIF6 \| 98.7 \| 1.5e-10 \| 2.9e-16 \| 87.2 \| 112 \| (22, 155) \| 156 \| (29, 149) \| 154 \| VRR-NUC domain containing protein \| VRR-NUC domain containing protein; uniclust \| UniRef100\_A0A6J5LH34 \| 98.7 \| 1.6e-10 \| 2.9e-16 \| 83.6 \| 105 \| (21, 155) \| 156 \| (5, 115) \| 130 \| VRR-NUC domain containing protein \| VRR-NUC domain containing protein; uniclust \| UniRef100\_A0A6H1ZUM5 \| 98.7 \| 1.6e-10 \| 2.9e-16 \| 86.7 \| 111 \| (21, 153) \| 156 \| (49, 165) \| 166 \| Putative VRR-NUC domain-containing protein \| Putative VRR-NUC domain-containing protein; uniclust \| UniRef100\_A0A843H219 \| 98.7 \| 1.6e-10 \| 2.9e-16 \| 80.7 \| 101 \| (22, 153) \| 156 \| (2, 104) \| 104 \| VRR-NUC domain-containing protein \| VRR-NUC domain-containing protein; uniclust \| UniRef100\_A0A948NR45 \| 98.7 \| 1.5e-10 \| 3e-16 \| 86.4 \| 82 \| (21, 147) \| 156 \| (24, 108) \| 129 \| VRR-NUC domain-containing protein \| VRR-NUC domain-containing protein; uniclust \| UniRef100\_UPI0002D4C647 \| 98.7 \| 1.7e-10 \| 3.1e-16 \| 85.3 \| 103 \| (22, 153) \| 156 \| (2, 112) \| 150 \| VRR-NUC domain-containing protein \| VRR-NUC domain-containing protein; uniclust \| UniRef100\_A0A7C1JCQ0 \| 98.7 \| 1.7e-10 \| 3.1e-16 \| 80.7 \| 65 \| (75, 151) \| 156 \| (37, 102) \| 105 \| VRR-NUC domain-containing protein \| VRR-NUC domain-containing protein; uniclust \| UniRef100\_A0A1J5SCT5 \| 98.7 \| 1.7e-10 \| 3.1e-16 \| 86.3 \| 105 \| (26, 155) \| 156 \| (46, 158) \| 164 \| VRR-NUC domain protein \| VRR-NUC domain protein; uniclust \| UniRef100\_UPI001D07F940 \| 98.7 \| 1.7e-10 \| 3.2e-16 \| 90.3 \| 90 \| (20, 154) \| 156 \| (60, 152) \| 230 \| VRR-NUC domain-containing protein \| VRR-NUC domain-containing protein; uniclust \| UniRef100\_A0A373ZU69 \| 98.7 \| 1.7e-10 \| 3.2e-16 \| 86.4 \| 103 \| (23, 154) \| 156 \| (4, 114) \| 165 \| VRR-NUC domain-containing protein \| VRR-NUC domain-containing protein; uniclust \| UniRef100\_A0A958CYF5 \| 98.7 \| 1.8e-10 \| 3.3e-16 \| 84.4 \| 101 \| (21, 153) \| 156 \| (10, 125) \| 141 \| Uncharacterized protein \| Uncharacterized protein; uniclust \| UniRef100\_A0A7X7LEV0 \| 98.7 \| 1.8e-10 \| 3.3e-16 \| 86.3 \| 88 \| (23, 152) \| 156 \| (22, 111) \| 165 \| VRR-NUC domain-containing protein \| VRR-NUC domain-containing protein; uniclust \| UniRef100\_A0A6J5L990 \| 98.7 \| 2e-10 \| 3.6e-16 \| 89.6 \| 100 \| (22, 153) \| 156 \| (1, 107) \| 220 \| VRR-NUC domain-containing protein \| VRR-NUC domain-containing protein; uniclust \| UniRef100\_A0A965UQW0 \| 98.7 \| 2e-10 \| 3.6e-16 \| 84.4 \| 68 \| (76, 153) \| 156 \| (63, 131) \| 144 \| VRR-NUC domain-containing protein \| VRR-NUC domain-containing protein; uniclust \| UniRef100\_A0A3E2TD32 \| 98.7 \| 2e-10 \| 3.7e-16 \| 89.3 \| 108 \| (18, 154) \| 156 \| (36, 151) \| 217 \| VRR-NUC domain-containing protein \| VRR-NUC domain-containing protein; uniclust \| UniRef100\_A0A6G6YUN2 \| 98.7 \| 2e-10 \| 3.8e-16 \| 80.2 \| 70 \| (75, 155) \| 156 \| (22, 93) \| 104 \| VRR-NUC domain-containing protein \| VRR-NUC domain-containing protein; uniclust \| UniRef100\_UPI001391A1F6 \| 98.7 \| 2.1e-10 \| 3.9e-16 \| 85.2 \| 82 \| (40, 152) \| 156 \| (26, 111) \| 156 \| VRR-NUC domain-containing protein \| VRR-NUC domain-containing protein; uniclust \| UniRef100\_A0A925VSB9 \| 98.7 \| 2.2e-10 \| 4e-16 \| 88.8 \| 103 \| (22, 154) \| 156 \| (92, 196) \| 210 \| VRR-NUC domain-containing protein \| VRR-NUC domain-containing protein; uniclust \| UniRef100\_A0A0F9NEN3 \| 98.7 \| 2.2e-10 \| 4.1e-16 \| 82.3 \| 102 \| (21, 153) \| 156 \| (6, 113) \| 124 \| VRR-NUC domain-containing protein \| VRR-NUC domain-containing protein; uniclust \| UniRef100\_A0A0F9BLX2 \| 98.7 \| 2.3e-10 \| 4.2e-16 \| 83.0 \| 104 \| (21, 155) \| 156 \| (23, 129) \| 132 \| VRR-NUC domain-containing protein \| VRR-NUC domain-containing protein; uniclust \| UniRef100\_A0A7V8NUZ0 \| 98.7 \| 2.4e-10 \| 4.3e-16 \| 86.6 \| 105 \| (20, 155) \| 156 \| (36, 145) \| 178 \| VRR-NUC domain-containing protein \| VRR-NUC domain-containing protein; uniclust \| UniRef100\_A0A7U3ZN37 \| 98.7 \| 2.3e-10 \| 4.4e-16 \| 86.0 \| 97 \| (25, 154) \| 156 \| (32, 130) \| 143 \| VRR-NUC domain-containing protein \| VRR-NUC domain-containing protein; uniclust \| UniRef100\_UPI001C60F457 \| 98.6 \| 2.7e-10 \| 4.9e-16 \| 85.0 \| 112 \| (21, 155) \| 156 \| (23, 143) \| 159 \| VRR-NUC domain-containing protein \| VRR-NUC domain-containing protein; uniclust \| UniRef100\_A0A7C6EBY8 \| 98.6 \| 2.7e-10 \| 4.9e-16 \| 83.0 \| 89 \| (23, 155) \| 156 \| (2, 92) \| 135 \| VRR-NUC domain-containing protein \| VRR-NUC domain-containing protein; uniclust \| UniRef100\_A0A101GVW2 \| 98.6 \| 2.8e-10 \| 5.1e-16 \| 86.7 \| 91 \| (20, 155) \| 156 \| (88, 181) \| 184 \| VRR-NUC domain-containing protein \| VRR-NUC domain-containing protein; uniclust \| UniRef100\_A0A225DD89 \| 98.6 \| 2.7e-10 \| 5.1e-16 \| 84.5 \| 110 \| (20, 151) \| 156 \| (28, 139) \| 149 \| VRR-NUC domain-containing protein \| VRR-NUC domain-containing protein; uniclust \| UniRef100\_A0A497C3L4 \| 98.6 \| 2.8e-10 \| 5.1e-16 \| 86.3 \| 102 \| (23, 153) \| 156 \| (55, 165) \| 179 \| VRR-NUC domain-containing protein \| VRR-NUC domain-containing protein; uniclust \| UniRef100\_A0A1B4G129 \| 98.6 \| 2.6e-10 \| 5.2e-16 \| 90.0 \| 104 \| (21, 155) \| 156 \| (54, 169) \| 188 \| VRR-NUC domain-containing protein \| VRR-NUC domain-containing protein; uniclust \| UniRef100\_A0A966WJ63 \| 98.6 \| 2.9e-10 \| 5.4e-16 \| 79.8 \| 85 \| (21, 134) \| 156 \| (14, 106) \| 107 \| VRR-NUC domain-containing protein (Fragment) \| VRR-NUC domain-containing protein (Fragment); uniclust \| UniRef100\_A0A2D8XCT8 \| 98.6 \| 2.8e-10 \| 5.5e-16 \| 75.3 \| 57 \| (85, 153) \| 156 \| (9, 66) \| 68 \| VRR-NUC domain-containing protein \| VRR-NUC domain-containing protein; uniclust \| UniRef100\_A0A511MZ74 \| 98.6 \| 3.3e-10 \| 6e-16 \| 86.8 \| 94 \| (21, 153) \| 156 \| (82, 179) \| 192 \| VRR-NUC domain-containing protein \| VRR-NUC domain-containing protein; uniclust \| UniRef100\_A0A1Q3M5C5 \| 98.6 \| 3.3e-10 \| 6.1e-16 \| 80.8 \| 82 \| (21, 131) \| 156 \| (23, 113) \| 114 \| Nuclease (Fragment) \| Nuclease (Fragment); uniclust \| UniRef100\_A0A1W9LSN6 \| 98.6 \| 3.1e-10 \| 6.2e-16 \| 81.7 \| 90 \| (19, 154) \| 156 \| (4, 95) \| 108 \| VRR-NUC domain-containing protein \| VRR-NUC domain-containing protein; uniclust \| UniRef100\_UPI000C5CBA07 \| 98.6 \| 3.4e-10 \| 6.2e-16 \| 85.6 \| 102 \| (22, 151) \| 156 \| (34, 143) \| 174 \| VRR-NUC domain-containing protein \| VRR-NUC domain-containing protein; uniclust \| UniRef100\_A0A6S4IR19 \| 98.6 \| 3.6e-10 \| 6.7e-16 \| 79.5 \| 97 \| (21, 154) \| 156 \| (2, 100) \| 108 \| VRR-NUC domain-containing protein \| VRR-NUC domain-containing protein; uniclust \| UniRef100\_A0A561PL50 \| 98.6 \| 3.6e-10 \| 6.9e-16 \| 75.0 \| 64 \| (79, 154) \| 156 \| (3, 67) \| 71 \| VRR-NUC domain-containing protein \| VRR-NUC domain-containing protein; uniclust \| UniRef100\_A0A6J5M4C0 \| 98.6 \| 4e-10 \| 7.3e-16 \| 85.1 \| 106 \| (21, 154) \| 156 \| (64, 171) \| 173 \| VRR-NUC domain containing protein \| VRR-NUC domain containing protein; uniclust \| UniRef100\_A0A0N8HQ47 \| 98.6 \| 4.1e-10 \| 7.5e-16 \| 85.8 \| 104 \| (22, 154) \| 156 \| (53, 162) \| 178 \| VRR-NUC domain-containing protein \| VRR-NUC domain-containing protein; uniclust \| UniRef100\_A0A524RQI1 \| 98.6 \| 4.2e-10 \| 7.8e-16 \| 71.9 \| 51 \| (103, 153) \| 156 \| (1, 52) \| 62 \| VRR-NUC domain-containing protein \| VRR-NUC domain-containing protein; uniclust \| UniRef100\_A0A6J5LGZ3 \| 98.6 \| 4.3e-10 \| 7.9e-16 \| 82.9 \| 105 \| (22, 155) \| 156 \| (28, 134) \| 146 \| VRR-NUC domain containing protein \| VRR-NUC domain containing protein; uniclust \| UniRef100\_A0A7C3EGK7 \| 98.6 \| 4.6e-10 \| 8.4e-16 \| 80.9 \| 85 \| (23, 154) \| 156 \| (2, 88) \| 125 \| VRR-NUC domain-containing protein \| VRR-NUC domain-containing protein; uniclust \| UniRef100\_A0A7C6KEH1 \| 98.6 \| 4.6e-10 \| 8.8e-16 \| 78.0 \| 57 \| (85, 153) \| 156 \| (33, 90) \| 91 \| VRR-NUC domain-containing protein \| VRR-NUC domain-containing protein; uniclust \| UniRef100\_UPI00164113D9 \| 98.6 \| 4.5e-10 \| 9.6e-16 \| 101.1 \| 87 \| (23, 155) \| 156 \| (1, 89) \| 512 \| VRR-NUC domain-containing protein \| VRR-NUC domain-containing protein; uniclust \| UniRef100\_UPI001B800FDB \| 98.6 \| 5.5e-10 \| 1e-15 \| 84.6 \| 104 \| (20, 155) \| 156 \| (25, 140) \| 175 \| hypothetical protein \| hypothetical protein; uniclust \| UniRef100\_A0A519WTD9 \| 98.6 \| 6e-10 \| 1.1e-15 \| 74.3 \| 69 \| (75, 155) \| 156 \| (4, 73) \| 78 \| VRR-NUC domain-containing protein (Fragment) \| VRR-NUC domain-containing protein (Fragment); uniclust \| UniRef100\_A0A0G1U713 \| 98.6 \| 6.2e-10 \| 1.1e-15 \| 79.0 \| 71 \| (76, 154) \| 156 \| (36, 107) \| 113 \| VRR-NUC domain-containing protein \| VRR-NUC domain-containing protein; uniclust \| UniRef100\_A0A0F9QS03 \| 98.6 \| 7.1e-10 \| 1.3e-15 \| 82.7 \| 105 \| (21, 154) \| 156 \| (19, 150) \| 157 \| VRR-NUC domain-containing protein \| VRR-NUC domain-containing protein; uniclust \| UniRef100\_UPI000B7945AE \| 98.6 \| 7.1e-10 \| 1.4e-15 \| 84.0 \| 101 \| (21, 152) \| 156 \| (14, 123) \| 149 \| hypothetical protein \| hypothetical protein; uniclust \| UniRef100\_A0A2H5X7U0 \| 98.5 \| 8.1e-10 \| 1.6e-15 \| 78.9 \| 75 \| (25, 144) \| 156 \| (7, 83) \| 105 \| VRR-NUC domain-containing protein \| VRR-NUC domain-containing protein; uniclust \| UniRef100\_A0A1F2WNL4 \| 98.5 \| 9.5e-10 \| 1.7e-15 \| 83.6 \| 115 \| (21, 155) \| 156 \| (31, 149) \| 179 \| VRR-NUC domain-containing protein \| VRR-NUC domain-containing protein; uniclust \| UniRef100\_A0A4Q0T6H7 \| 98.5 \| 1e-09 \| 1.9e-15 \| 78.3 \| 64 \| (79, 154) \| 156 \| (40, 104) \| 116 \| VRR-NUC domain-containing protein \| VRR-NUC domain-containing protein; uniclust \| UniRef100\_A0A2H0NCQ0 \| 98.5 \| 1e-09 \| 1.9e-15 \| 78.8 \| 86 \| (23, 155) \| 156 \| (8, 96) \| 111 \| VRR-NUC domain-containing protein \| VRR-NUC domain-containing protein; uniclust \| UniRef100\_A0A1L8TFK0 \| 98.5 \| 1.1e-09 \| 1.9e-15 \| 74.6 \| 64 \| (82, 155) \| 156 \| (14, 78) \| 87 \| VRR-NUC domain-containing protein \| VRR-NUC domain-containing protein; uniclust \| UniRef100\_A0A0A0BLP2 \| 98.5 \| 9.8e-10 \| 2e-15 \| 83.7 \| 97 \| (22, 153) \| 156 \| (1, 108) \| 150 \| Recombinase RecB \| Recombinase RecB; uniclust \| UniRef100\_E4LIQ7 \| 98.5 \| 1.1e-09 \| 2e-15 \| 78.7 \| 88 \| (21, 154) \| 156 \| (31, 120) \| 122 \| VRR-NUC domain protein \| VRR-NUC domain protein; uniclust \| UniRef100\_A0A316S5X2 \| 98.5 \| 1.1e-09 \| 2.2e-15 \| 84.8 \| 93 \| (18, 155) \| 156 \| (34, 131) \| 170 \| Nuclease \| Nuclease; uniclust \| UniRef100\_UPI001F17BD1E \| 98.5 \| 1.2e-09 \| 2.2e-15 \| 81.7 \| 94 \| (21, 144) \| 156 \| (24, 124) \| 144 \| VRR-NUC domain-containing protein \| VRR-NUC domain-containing protein; uniclust \| UniRef100\_A0A7X9Q8X8 \| 98.5 \| 1.2e-09 \| 2.3e-15 \| 85.6 \| 103 \| (20, 151) \| 156 \| (105, 216) \| 224 \| VRR-NUC domain-containing protein \| VRR-NUC domain-containing protein; uniclust \| UniRef100\_UPI0018D3F871 \| 98.5 \| 1.2e-09 \| 2.3e-15 \| 82.8 \| 118 \| (23, 153) \| 156 \| (1, 128) \| 158 \| hypothetical protein \| hypothetical protein; uniclust \| UniRef100\_L7U9Y9 \| 98.5 \| 1.3e-09 \| 2.4e-15 \| 80.3 \| 111 \| (21, 154) \| 156 \| (25, 138) \| 144 \| VRR-NUC domain-containing protein \| VRR-NUC domain-containing protein; uniclust \| UniRef100\_A0A1V5LEE2 \| 98.5 \| 1.3e-09 \| 2.4e-15 \| 77.7 \| 108 \| (19, 151) \| 156 \| (4, 115) \| 116 \| VRR-NUC domain protein \| VRR-NUC domain protein; uniclust \| UniRef100\_A0A0F9VBY1 \| 98.5 \| 1.3e-09 \| 2.5e-15 \| 79.8 \| 112 \| (22, 155) \| 156 \| (3, 117) \| 127 \| VRR-NUC domain-containing protein \| VRR-NUC domain-containing protein; uniclust \| UniRef100\_A0A6G7X067 \| 98.5 \| 1.3e-09 \| 2.5e-15 \| 79.1 \| 89 \| (23, 155) \| 156 \| (1, 94) \| 119 \| VRR-NUC domain-containing protein \| VRR-NUC domain-containing protein; uniclust \| UniRef100\_A0A3N5E8N3 \| 98.5 \| 1.4e-09 \| 2.6e-15 \| 76.3 \| 90 \| (14, 144) \| 156 \| (13, 104) \| 104 \| VRR-NUC domain-containing protein (Fragment) \| VRR-NUC domain-containing protein (Fragment); uniclust \| UniRef100\_A0A177R4V3 \| 98.5 \| 1.4e-09 \| 2.6e-15 \| 83.6 \| 110 \| (22, 154) \| 156 \| (77, 191) \| 194 \| Uncharacterized protein \| Uncharacterized protein; uniclust \| UniRef100\_A0A0L6JT85 \| 98.5 \| 1.5e-09 \| 2.8e-15 \| 81.6 \| 60 \| (83, 154) \| 156 \| (83, 143) \| 148 \| VRR-NUC domain-containing protein \| VRR-NUC domain-containing protein; uniclust \| UniRef100\_A0A1G4TXQ0 \| 98.5 \| 1.5e-09 \| 2.9e-15 \| 83.1 \| 89 \| (21, 155) \| 156 \| (70, 160) \| 166 \| YozE SAM-like fold \| YozE SAM-like fold; uniclust \| UniRef100\_A0A537MDI3 \| 98.5 \| 1.6e-09 \| 2.9e-15 \| 74.3 \| 67 \| (77, 155) \| 156 \| (13, 80) \| 91 \| VRR-NUC domain-containing protein \| VRR-NUC domain-containing protein; uniclust \| UniRef100\_A0A2V5PQQ7 \| 98.5 \| 1.7e-09 \| 3e-15 \| 82.1 \| 59 \| (86, 155) \| 156 \| (114, 173) \| 174 \| VRR-NUC domain-containing protein \| VRR-NUC domain-containing protein; uniclust \| UniRef100\_A0A2T3ITI0 \| 98.5 \| 1.6e-09 \| 3.2e-15 \| 88.0 \| 105 \| (22, 155) \| 156 \| (78, 198) \| 231 \| VRR-NUC domain-containing protein \| VRR-NUC domain-containing protein; uniclust \| UniRef100\_A0A1H0PBQ8 \| 98.5 \| 1.8e-09 \| 3.2e-15 \| 79.9 \| 57 \| (86, 154) \| 156 \| (34, 91) \| 146 \| VRR-NUC domain-containing protein \| VRR-NUC domain-containing protein; uniclust \| UniRef100\_A0A239DNY6 \| 98.5 \| 1.9e-09 \| 3.4e-15 \| 81.3 \| 67 \| (75, 153) \| 156 \| (90, 157) \| 166 \| VRR-NUC domain-containing protein \| VRR-NUC domain-containing protein; uniclust \| UniRef100\_A0A0H3ZPV8 \| 98.5 \| 1.8e-09 \| 3.5e-15 \| 83.2 \| 102 \| (22, 152) \| 156 \| (52, 159) \| 175 \| Phage protein \| Phage protein; uniclust \| UniRef100\_A0A968LFM5 \| 98.5 \| 1.9e-09 \| 3.5e-15 \| 78.1 \| 63 \| (76, 150) \| 156 \| (55, 118) \| 128 \| VRR-NUC domain-containing protein \| VRR-NUC domain-containing protein; uniclust \| UniRef100\_A0A1C6CWA0 \| 98.5 \| 2.1e-09 \| 3.8e-15 \| 84.1 \| 107 \| (21, 154) \| 156 \| (12, 130) \| 219 \| VRR-NUC domain \| VRR-NUC domain; uniclust \| UniRef100\_A0A7C5QET6 \| 98.4 \| 2.2e-09 \| 4.1e-15 \| 78.9 \| 100 \| (21, 155) \| 156 \| (37, 138) \| 140 \| VRR-NUC domain-containing protein \| VRR-NUC domain-containing protein; uniclust \| UniRef100\_A0A370CU33 \| 98.4 \| 2.3e-09 \| 4.2e-15 \| 80.0 \| 112 \| (20, 154) \| 156 \| (25, 142) \| 154 \| VRR-NUC domain-containing protein \| VRR-NUC domain-containing protein; uniclust \| UniRef100\_A0A418MBC5 \| 98.4 \| 2.4e-09 \| 4.6e-15 \| 80.4 \| 101 \| (21, 154) \| 156 \| (17, 123) \| 149 \| VRR-NUC domain-containing protein \| VRR-NUC domain-containing protein; uniclust \| UniRef100\_A0A257TCZ7 \| 98.4 \| 2.5e-09 \| 4.6e-15 \| 69.2 \| 56 \| (87, 154) \| 156 \| (1, 57) \| 65 \| Uncharacterized protein (Fragment) \| Uncharacterized protein (Fragment); uniclust \| UniRef100\_A0A7X3R873 \| 98.4 \| 2.6e-09 \| 4.8e-15 \| 72.0 \| 57 \| (85, 153) \| 156 \| (23, 80) \| 82 \| VRR-NUC domain-containing protein \| VRR-NUC domain-containing protein; uniclust \| UniRef100\_A0A2A1K7C6 \| 98.4 \| 2.9e-09 \| 5.4e-15 \| 72.0 \| 80 \| (22, 134) \| 156 \| (1, 82) \| 83 \| VRR-NUC domain-containing protein (Fragment) \| VRR-NUC domain-containing protein (Fragment); uniclust \| UniRef100\_UPI0007DC280A \| 98.4 \| 2.7e-09 \| 5.4e-15 \| 79.7 \| 86 \| (21, 154) \| 156 \| (9, 96) \| 132 \| VRR-NUC domain-containing protein \| VRR-NUC domain-containing protein; uniclust \| UniRef100\_A0A4Q5NWE0 \| 98.4 \| 3e-09 \| 5.6e-15 \| 84.1 \| 100 \| (21, 152) \| 156 \| (77, 184) \| 236 \| VRR-NUC domain-containing protein \| VRR-NUC domain-containing protein; uniclust \| UniRef100\_A0A3C1HPS2 \| 98.4 \| 3.2e-09 \| 5.9e-15 \| 73.1 \| 84 \| (22, 128) \| 156 \| (2, 92) \| 92 \| Uncharacterized protein \| Uncharacterized protein; uniclust \| UniRef100\_UPI001FFCBC52 \| 98.4 \| 3.3e-09 \| 6.2e-15 \| 74.9 \| 51 \| (85, 147) \| 156 \| (12, 63) \| 99 \| VRR-NUC domain-containing protein \| VRR-NUC domain-containing protein; uniclust \| UniRef100\_A0A7C1C9X5 \| 98.4 \| 3.5e-09 \| 6.5e-15 \| 78.6 \| 89 \| (23, 155) \| 156 \| (53, 143) \| 149 \| VRR-NUC domain-containing protein \| VRR-NUC domain-containing protein; uniclust \| UniRef100\_A0A2S7ZHM1 \| 98.4 \| 3.7e-09 \| 6.9e-15 \| 77.9 \| 58 \| (85, 153) \| 156 \| (50, 108) \| 132 \| VRR-NUC domain-containing protein \| VRR-NUC domain-containing protein; uniclust \| UniRef100\_A0A661DS26 \| 98.4 \| 3.8e-09 \| 7e-15 \| 75.7 \| 85 \| (23, 154) \| 156 \| (1, 88) \| 118 \| VRR-NUC domain-containing protein \| VRR-NUC domain-containing protein; uniclust \| UniRef100\_UPI0018E07DFE \| 98.4 \| 3.6e-09 \| 7.1e-15 \| 85.3 \| 58 \| (86, 154) \| 156 \| (30, 88) \| 219 \| VRR-NUC domain-containing protein \| VRR-NUC domain-containing protein; uniclust \| UniRef100\_A0A2I7QVY9 \| 98.4 \| 4.1e-09 \| 7.6e-15 \| 79.0 \| 72 \| (75, 155) \| 156 \| (68, 143) \| 150 \| VRR-NUC domain protein \| VRR-NUC domain protein; uniclust \| UniRef100\_A0A0P0J010 \| 98.4 \| 3.9e-09 \| 7.6e-15 \| 80.4 \| 105 \| (22, 153) \| 156 \| (25, 136) \| 148 \| VRR-NUC domain-containing protein \| VRR-NUC domain-containing protein; uniclust \| UniRef100\_A0A6M3TDT4 \| 98.4 \| 4.2e-09 \| 8e-15 \| 78.2 \| 107 \| (18, 145) \| 156 \| (9, 132) \| 136 \| Uncharacterized protein \| Uncharacterized protein; uniclust \| UniRef100\_A0A0F9HQT7 \| 98.4 \| 4.5e-09 \| 8.2e-15 \| 74.7 \| 95 \| (20, 153) \| 156 \| (6, 102) \| 111 \| VRR-NUC domain-containing protein \| VRR-NUC domain-containing protein; uniclust \| UniRef100\_A0A951MTT0 \| 98.4 \| 4.5e-09 \| 8.2e-15 \| 81.4 \| 106 \| (21, 155) \| 156 \| (63, 190) \| 200 \| VRR-NUC domain-containing protein \| VRR-NUC domain-containing protein; uniclust \| UniRef100\_A0A6J7W6B7 \| 98.4 \| 4.5e-09 \| 8.3e-15 \| 73.3 \| 62 \| (82, 155) \| 156 \| (26, 88) \| 99 \| VRR-NUC domain containing protein \| VRR-NUC domain containing protein; uniclust \| UniRef100\_A0A117IRP9 \| 98.4 \| 4.4e-09 \| 8.4e-15 \| 77.3 \| 94 \| (15, 155) \| 156 \| (10, 105) \| 123 \| VRR-NUC domain-containing protein \| VRR-NUC domain-containing protein; uniclust \| UniRef100\_A0A249XQD2 \| 98.4 \| 4.8e-09 \| 8.9e-15 \| 75.2 \| 60 \| (85, 155) \| 156 \| (52, 112) \| 117 \| Nuclease \| Nuclease; uniclust \| UniRef100\_A0A2D6X4D2 \| 98.4 \| 4.9e-09 \| 9e-15 \| 79.0 \| 106 \| (23, 155) \| 156 \| (46, 159) \| 164 \| Uncharacterized protein \| Uncharacterized protein; uniclust \| UniRef100\_A0A6L9GF55 \| 98.4 \| 5.3e-09 \| 9.8e-15 \| 69.3 \| 48 \| (108, 155) \| 156 \| (7, 55) \| 71 \| VRR-NUC domain-containing protein (Fragment) \| VRR-NUC domain-containing protein (Fragment); uniclust \| UniRef100\_UPI0003B37613 \| 98.4 \| 5.4e-09 \| 1e-14 \| 77.3 \| 106 \| (23, 154) \| 156 \| (15, 124) \| 143 \| hypothetical protein \| hypothetical protein; uniclust \| UniRef100\_UPI002240A71F \| 98.4 \| 5.5e-09 \| 1e-14 \| 79.5 \| 104 \| (22, 153) \| 156 \| (41, 148) \| 174 \| VRR-NUC domain-containing protein \| VRR-NUC domain-containing protein; uniclust \| UniRef100\_A0A1E7Y0A1 \| 98.4 \| 5.6e-09 \| 1e-14 \| 77.5 \| 57 \| (86, 153) \| 156 \| (30, 87) \| 147 \| VRR-NUC domain-containing protein \| VRR-NUC domain-containing protein; uniclust \| UniRef100\_Q45PU7 \| 98.4 \| 5.8e-09 \| 1.1e-14 \| 71.8 \| 67 \| (76, 154) \| 156 \| (12, 82) \| 91 \| JK\_69P \| JK\_69P; uniclust \| UniRef100\_A0A9E9GX09 \| 98.3 \| 5.7e-09 \| 1.1e-14 \| 81.2 \| 55 \| (86, 152) \| 156 \| (31, 87) \| 184 \| Uncharacterized protein \| Uncharacterized protein; uniclust \| UniRef100\_A0A1V6G008 \| 98.3 \| 6e-09 \| 1.1e-14 \| 75.2 \| 92 \| (22, 154) \| 156 \| (1, 94) \| 122 \| VRR-NUC domain protein \| VRR-NUC domain protein; uniclust \| UniRef100\_UPI001C6961CB \| 98.3 \| 6.1e-09 \| 1.1e-14 \| 68.4 \| 62 \| (82, 155) \| 156 \| (1, 65) \| 70 \| hypothetical protein \| hypothetical protein; uniclust \| UniRef100\_A0A7C6T0D3 \| 98.3 \| 6.1e-09 \| 1.1e-14 \| 76.3 \| 90 \| (19, 153) \| 156 \| (38, 129) \| 134 \| VRR-NUC domain-containing protein \| VRR-NUC domain-containing protein; uniclust \| UniRef100\_A0A3C1SW60 \| 98.3 \| 6e-09 \| 1.1e-14 \| 77.1 \| 112 \| (20, 155) \| 156 \| (12, 131) \| 134 \| VRR-NUC domain-containing protein \| VRR-NUC domain-containing protein; uniclust \| UniRef100\_UPI00164128D5 \| 98.3 \| 5.8e-09 \| 1.1e-14 \| 90.5 \| 87 \| (23, 155) \| 156 \| (1, 89) \| 403 \| hypothetical protein \| hypothetical protein; uniclust \| UniRef100\_A0A7T4PUE8 \| 98.3 \| 6.3e-09 \| 1.1e-14 \| 76.1 \| 90 \| (22, 155) \| 156 \| (2, 103) \| 133 \| VRR-NUC domain-containing protein \| VRR-NUC domain-containing protein; uniclust \| UniRef100\_E3GF55 \| 98.3 \| 6.3e-09 \| 1.2e-14 \| 79.7 \| 57 \| (86, 154) \| 156 \| (34, 91) \| 183 \| VRR-NUC domain protein \| VRR-NUC domain protein; uniclust \| UniRef100\_A0A5C7JER8 \| 98.3 \| 6.2e-09 \| 1.2e-14 \| 77.6 \| 97 \| (23, 148) \| 156 \| (31, 131) \| 142 \| VRR-NUC domain-containing protein \| VRR-NUC domain-containing protein; uniclust \| UniRef100\_A0A966K7I6 \| 98.3 \| 6.4e-09 \| 1.2e-14 \| 78.8 \| 104 \| (22, 155) \| 156 \| (51, 159) \| 169 \| VRR-NUC domain-containing protein \| VRR-NUC domain-containing protein; uniclust \| UniRef100\_A0A952PDW2 \| 98.3 \| 6.8e-09 \| 1.3e-14 \| 77.0 \| 115 \| (21, 155) \| 156 \| (14, 130) \| 132 \| Uncharacterized protein \| Uncharacterized protein; uniclust \| UniRef100\_UPI001F37818D \| 98.3 \| 7.1e-09 \| 1.3e-14 \| 76.4 \| 89 \| (21, 155) \| 156 \| (42, 132) \| 139 \| VRR-NUC domain-containing protein \| VRR-NUC domain-containing protein; uniclust \| UniRef100\_A0A6M3JW76 \| 98.3 \| 6.9e-09 \| 1.3e-14 \| 80.1 \| 88 \| (22, 154) \| 156 \| (15, 106) \| 176 \| VRR-NUC domain-containing protein \| VRR-NUC domain-containing protein; uniclust \| UniRef100\_A0A0E4BRS8 \| 98.3 \| 6.7e-09 \| 1.3e-14 \| 80.0 \| 121 \| (4, 152) \| 156 \| (11, 137) \| 161 \| VRR-NUC domain-containing protein \| VRR-NUC domain-containing protein; uniclust \| UniRef100\_A0A965H4R9 \| 98.3 \| 7.2e-09 \| 1.4e-14 \| 67.1 \| 52 \| (103, 154) \| 156 \| (1, 53) \| 61 \| VRR-NUC domain-containing protein \| VRR-NUC domain-containing protein; uniclust \| UniRef100\_A0A849N9L9 \| 98.3 \| 7.5e-09 \| 1.4e-14 \| 76.1 \| 90 \| (23, 144) \| 156 \| (16, 107) \| 137 \| VRR-NUC domain-containing protein \| VRR-NUC domain-containing protein; uniclust \| UniRef100\_A0A6J5SRX6 \| 98.3 \| 7.7e-09 \| 1.4e-14 \| 69.3 \| 71 \| (76, 155) \| 156 \| (4, 75) \| 78 \| Uncharacterized protein \| Uncharacterized protein; uniclust \| UniRef100\_A0A2E0R731 \| 98.3 \| 8.1e-09 \| 1.5e-14 \| 69.5 \| 68 \| (85, 155) \| 156 \| (2, 70) \| 80 \| VRR-NUC domain-containing protein \| VRR-NUC domain-containing protein; uniclust \| UniRef100\_A0A2A4V7W8 \| 98.3 \| 8e-09 \| 1.5e-14 \| 76.5 \| 99 \| (21, 154) \| 156 \| (2, 110) \| 131 \| VRR-NUC domain-containing protein \| VRR-NUC domain-containing protein; uniclust \| UniRef100\_A0A661MNW6 \| 98.3 \| 9e-09 \| 1.6e-14 \| 76.0 \| 115 \| (23, 154) \| 156 \| (1, 124) \| 141 \| VRR-NUC domain-containing protein \| VRR-NUC domain-containing protein; uniclust \| UniRef100\_A0A1A8XYT3 \| 98.3 \| 9.1e-09 \| 1.7e-14 \| 79.1 \| 98 \| (21, 153) \| 156 \| (27, 128) \| 187 \| VRR-NUC domain-containing protein \| VRR-NUC domain-containing protein; uniclust \| UniRef100\_A0A224AIU8 \| 98.3 \| 8.8e-09 \| 1.7e-14 \| 77.8 \| 103 \| (23, 155) \| 156 \| (30, 137) \| 147 \| VRR-NUC domain-containing protein \| VRR-NUC domain-containing protein; uniclust \| UniRef100\_UPI00204095D6 \| 98.3 \| 9.3e-09 \| 1.7e-14 \| 74.9 \| 92 \| (21, 146) \| 156 \| (2, 98) \| 129 \| hypothetical protein \| hypothetical protein; uniclust \| UniRef100\_A0A516LPK8 \| 98.3 \| 9.5e-09 \| 1.7e-14 \| 73.3 \| 59 \| (86, 155) \| 156 \| (50, 109) \| 113 \| Putative nuclease \| Putative nuclease; uniclust \| UniRef100\_A0A6M3T9R2 \| 98.3 \| 1e-08 \| 1.9e-14 \| 81.6 \| 107 \| (21, 149) \| 156 \| (54, 177) \| 212 \| VRR-NUC domain endonuclease \| VRR-NUC domain endonuclease; uniclust \| UniRef100\_A0A7C3D595 \| 98.3 \| 1.1e-08 \| 2e-14 \| 76.8 \| 98 \| (25, 153) \| 156 \| (22, 129) \| 156 \| VRR-NUC domain-containing protein \| VRR-NUC domain-containing protein; uniclust \| UniRef100\_A0A516L0Z0 \| 98.3 \| 1.1e-08 \| 2e-14 \| 82.5 \| 104 \| (23, 155) \| 156 \| (108, 215) \| 226 \| Chromosomal replication initiator DnaA C-terminal domain-containing protein \| Chromosomal replication initiator DnaA C-terminal domain-containing protein; uniclust \| UniRef100\_A0A088C3V3 \| 98.3 \| 1e-08 \| 2.3e-14 \| 80.8 \| 101 \| (22, 155) \| 156 \| (32, 142) \| 163 \| Holliday junction resolvase \| Holliday junction resolvase; uniclust \| UniRef100\_A0A3D6BPK0 \| 98.3 \| 1.3e-08 \| 2.3e-14 \| 74.0 \| 102 \| (22, 154) \| 156 \| (18, 121) \| 126 \| VRR-NUC domain-containing protein \| VRR-NUC domain-containing protein; uniclust \| UniRef100\_UPI00068F67A1 \| 98.3 \| 1.4e-08 \| 2.6e-14 \| 75.5 \| 112 \| (23, 154) \| 156 \| (1, 121) \| 147 \| VRR-NUC domain-containing protein \| VRR-NUC domain-containing protein; uniclust \| UniRef100\_UPI001E6F27A4 \| 98.2 \| 1.5e-08 \| 2.8e-14 \| 78.8 \| 101 \| (21, 143) \| 156 \| (81, 197) \| 203 \| putative nuclease \| putative nuclease; uniclust \| UniRef100\_A0A5C7LMD1 \| 98.2 \| 1.5e-08 \| 2.9e-14 \| 75.5 \| 91 \| (23, 155) \| 156 \| (44, 136) \| 145 \| VRR-NUC domain-containing protein \| VRR-NUC domain-containing protein; uniclust \| UniRef100\_A0A011PI35 \| 98.2 \| 1.6e-08 \| 2.9e-14 \| 76.7 \| 110 \| (19, 155) \| 156 \| (38, 150) \| 152 \| VRR-NUC domain protein \| VRR-NUC domain protein; uniclust \| UniRef100\_A0A1H0RUY3 \| 98.2 \| 1.6e-08 \| 3.3e-14 \| 77.3 \| 116 \| (17, 155) \| 156 \| (1, 131) \| 144 \| VRR-NUC domain-containing protein \| VRR-NUC domain-containing protein; uniclust \| UniRef100\_A0A0R1XFK4 \| 98.2 \| 1.7e-08 \| 3.3e-14 \| 67.2 \| 55 \| (89, 155) \| 156 \| (2, 57) \| 64 \| VRR-NUC domain-containing protein \| VRR-NUC domain-containing protein; uniclust \| UniRef100\_A0A1I2LF52 \| 98.2 \| 1.9e-08 \| 3.4e-14 \| 74.9 \| 110 \| (20, 154) \| 156 \| (20, 134) \| 146 \| VRR-NUC domain-containing protein \| VRR-NUC domain-containing protein; uniclust \| UniRef100\_X0S3S5 \| 98.2 \| 2e-08 \| 3.6e-14 \| 65.7 \| 59 \| (85, 154) \| 156 \| (2, 61) \| 67 \| VRR-NUC domain-containing protein (Fragment) \| VRR-NUC domain-containing protein (Fragment); uniclust \| UniRef100\_A0A3C1F314 \| 98.2 \| 2e-08 \| 3.7e-14 \| 69.5 \| 59 \| (85, 151) \| 156 \| (30, 89) \| 93 \| VRR-NUC domain-containing protein \| VRR-NUC domain-containing protein; uniclust \| UniRef100\_A0A0U4C7T5 \| 98.2 \| 2.1e-08 \| 3.8e-14 \| 80.2 \| 102 \| (19, 153) \| 156 \| (138, 243) \| 248 \| VRR-NUC domain-containing protein \| VRR-NUC domain-containing protein; uniclust \| UniRef100\_UPI00200E4F20 \| 98.2 \| 2.1e-08 \| 3.8e-14 \| 66.9 \| 61 \| (83, 155) \| 156 \| (2, 64) \| 75 \| hypothetical protein \| hypothetical protein; uniclust \| UniRef100\_A0A3A8JIG7 \| 98.2 \| 2.1e-08 \| 3.8e-14 \| 76.8 \| 58 \| (85, 154) \| 156 \| (80, 138) \| 178 \| VRR-NUC domain-containing protein \| VRR-NUC domain-containing protein; uniclust \| UniRef100\_A0A2A7AQM9 \| 98.2 \| 2.1e-08 \| 4e-14 \| 75.7 \| 58 \| (85, 154) \| 156 \| (80, 139) \| 147 \| Nuclease \| Nuclease; uniclust \| UniRef100\_A0A4U0RCC9 \| 98.2 \| 2.3e-08 \| 4.1e-14 \| 79.0 \| 105 \| (22, 153) \| 156 \| (91, 200) \| 225 \| VRR-NUC domain-containing protein \| VRR-NUC domain-containing protein; uniclust \| UniRef100\_UPI000A67298D \| 98.2 \| 2.3e-08 \| 4.2e-14 \| 78.3 \| 79 \| (23, 146) \| 156 \| (2, 83) \| 210 \| VRR-NUC domain-containing protein \| VRR-NUC domain-containing protein; uniclust \| UniRef100\_A0A5C0CFL2 \| 98.2 \| 2.3e-08 \| 4.2e-14 \| 65.9 \| 60 \| (85, 155) \| 156 \| (6, 67) \| 70 \| Restriction endonuclease \| Restriction endonuclease; uniclust \| UniRef100\_A0A2G2PNU2 \| 98.2 \| 2.3e-08 \| 4.2e-14 \| 70.9 \| 59 \| (85, 154) \| 156 \| (24, 83) \| 107 \| VRR-NUC domain-containing protein \| VRR-NUC domain-containing protein; uniclust \| UniRef100\_A0A1G0QSC3 \| 98.2 \| 2.5e-08 \| 4.5e-14 \| 70.1 \| 84 \| (22, 149) \| 156 \| (3, 88) \| 101 \| VRR-NUC domain-containing protein \| VRR-NUC domain-containing protein; uniclust \| UniRef100\_UPI001ABAAAB1 \| 98.2 \| 2.4e-08 \| 4.7e-14 \| 82.8 \| 60 \| (85, 155) \| 156 \| (29, 89) \| 288 \| hypothetical protein \| hypothetical protein; uniclust \| UniRef100\_UPI0019D2A3A2 \| 98.2 \| 2.4e-08 \| 4.7e-14 \| 86.7 \| 87 \| (23, 155) \| 156 \| (1, 89) \| 393 \| VRR-NUC domain-containing protein \| VRR-NUC domain-containing protein; uniclust \| UniRef100\_A0A966D9Q4 \| 98.2 \| 2.8e-08 \| 5.1e-14 \| 74.6 \| 116 \| (21, 155) \| 156 \| (15, 141) \| 154 \| Uncharacterized protein \| Uncharacterized protein; uniclust \| UniRef100\_A0A257PP85 \| 98.2 \| 2.6e-08 \| 5.1e-14 \| 78.5 \| 108 \| (20, 154) \| 156 \| (44, 155) \| 188 \| Uncharacterized protein \| Uncharacterized protein; uniclust \| UniRef100\_UPI000B1D41EA \| 98.2 \| 2.9e-08 \| 5.4e-14 \| 73.3 \| 58 \| (85, 153) \| 156 \| (35, 93) \| 139 \| VRR-NUC domain-containing protein \| VRR-NUC domain-containing protein; uniclust \| UniRef100\_A0A7S2FV51 \| 98.2 \| 2.9e-08 \| 5.4e-14 \| 81.0 \| 108 \| (22, 155) \| 156 \| (17, 128) \| 293 \| VRR-NUC domain-containing protein \| VRR-NUC domain-containing protein; uniclust \| UniRef100\_A0A2N4UWG8 \| 98.2 \| 3e-08 \| 6e-14 \| 81.0 \| 107 \| (19, 154) \| 156 \| (87, 209) \| 238 \| VRR-NUC domain-containing protein \| VRR-NUC domain-containing protein; uniclust \| UniRef100\_A0A021X9L9 \| 98.1 \| 3.2e-08 \| 6.6e-14 \| 79.9 \| 94 \| (21, 145) \| 156 \| (57, 162) \| 206 \| VRR-NUC domain-containing protein \| VRR-NUC domain-containing protein; uniclust \| UniRef100\_A0A7Z9G7R6 \| 98.1 \| 3.6e-08 \| 6.7e-14 \| 71.3 \| 102 \| (23, 154) \| 156 \| (1, 108) \| 121 \| VRR-NUC domain-containing protein \| VRR-NUC domain-containing protein; uniclust \| UniRef100\_R1BRB5 \| 98.1 \| 3.7e-08 \| 6.7e-14 \| 83.1 \| 111 \| (21, 155) \| 156 \| (5, 120) \| 392 \| VRR-NUC domain-containing protein \| VRR-NUC domain-containing protein; uniclust \| UniRef100\_A0A6G5Y0E8 \| 98.1 \| 3.7e-08 \| 6.8e-14 \| 70.8 \| 82 \| (23, 147) \| 156 \| (5, 88) \| 116 \| VRR-NUC domain-containing protein \| VRR-NUC domain-containing protein; uniclust \| UniRef100\_A0A1F8QJ77 \| 98.1 \| 3.9e-08 \| 7.2e-14 \| 71.9 \| 101 \| (22, 154) \| 156 \| (1, 128) \| 129 \| VRR-NUC domain-containing protein \| VRR-NUC domain-containing protein; uniclust \| UniRef100\_A0A4Q5VAH5 \| 98.1 \| 3.8e-08 \| 7.4e-14 \| 79.6 \| 104 \| (22, 154) \| 156 \| (75, 186) \| 221 \| Uncharacterized protein \| Uncharacterized protein; uniclust \| UniRef100\_UPI001D0CB0F5 \| 98.1 \| 4.1e-08 \| 7.6e-14 \| 65.1 \| 48 \| (108, 155) \| 156 \| (4, 52) \| 72 \| hypothetical protein \| hypothetical protein; uniclust \| UniRef100\_UPI00227ADC10 \| 98.1 \| 4.2e-08 \| 7.7e-14 \| 72.0 \| 95 \| (40, 155) \| 156 \| (5, 102) \| 132 \| VRR-NUC domain-containing protein \| VRR-NUC domain-containing protein; uniclust \| UniRef100\_A0A4Z0F6A1 \| 98.1 \| 4.3e-08 \| 7.9e-14 \| 68.5 \| 61 \| (84, 155) \| 156 \| (31, 92) \| 97 \| VRR-NUC domain-containing protein \| VRR-NUC domain-containing protein; uniclust \| UniRef100\_A0A1Q4FJ35 \| 98.1 \| 4.4e-08 \| 8.1e-14 \| 76.4 \| 104 \| (22, 154) \| 156 \| (40, 156) \| 202 \| VRR-NUC domain-containing protein \| VRR-NUC domain-containing protein; uniclust \| UniRef100\_A0A0F9G0T4 \| 98.1 \| 4.5e-08 \| 8.3e-14 \| 69.5 \| 58 \| (85, 154) \| 156 \| (45, 103) \| 107 \| VRR-NUC domain-containing protein \| VRR-NUC domain-containing protein; uniclust \| UniRef100\_A0A965PF78 \| 98.1 \| 4.6e-08 \| 8.5e-14 \| 70.8 \| 80 \| (23, 149) \| 156 \| (1, 82) \| 121 \| VRR-NUC domain-containing protein \| VRR-NUC domain-containing protein; uniclust \| UniRef100\_A0A522RZJ2 \| 98.1 \| 4.7e-08 \| 8.7e-14 \| 76.6 \| 104 \| (22, 154) \| 156 \| (33, 148) \| 210 \| VRR-NUC domain-containing protein \| VRR-NUC domain-containing protein; uniclust \| UniRef100\_A0A4Z0MR71 \| 98.1 \| 4.8e-08 \| 8.9e-14 \| 63.3 \| 48 \| (107, 154) \| 156 \| (7, 55) \| 63 \| VRR-NUC domain-containing protein \| VRR-NUC domain-containing protein; uniclust \| UniRef100\_A0A0X7A620 \| 98.1 \| 4.3e-08 \| 9e-14 \| 73.9 \| 89 \| (21, 155) \| 156 \| (11, 106) \| 124 \| VRR-NUC domain-containing protein \| VRR-NUC domain-containing protein; uniclust \| UniRef100\_UPI001FA731F7 \| 98.1 \| 4.9e-08 \| 9.1e-14 \| 75.1 \| 87 \| (22, 153) \| 156 \| (28, 119) \| 182 \| VRR-NUC domain-containing protein \| VRR-NUC domain-containing protein; uniclust \| UniRef100\_A0A7C3U5D4 \| 98.1 \| 5.1e-08 \| 9.4e-14 \| 72.0 \| 80 \| (21, 145) \| 156 \| (41, 122) \| 137 \| VRR-NUC domain-containing protein \| VRR-NUC domain-containing protein; uniclust \| UniRef100\_A0A8S5M5J1 \| 98.1 \| 4.9e-08 \| 9.5e-14 \| 72.3 \| 100 \| (22, 154) \| 156 \| (3, 110) \| 122 \| Hydrolase \| Hydrolase; uniclust \| UniRef100\_UPI001C8DF2A0 \| 98.1 \| 5.3e-08 \| 9.8e-14 \| 75.3 \| 96 \| (11, 153) \| 156 \| (83, 180) \| 189 \| VRR-NUC domain-containing protein \| VRR-NUC domain-containing protein; uniclust \| UniRef100\_A0A7K1CW68 \| 98.1 \| 5.7e-08 \| 1.1e-13 \| 70.2 \| 82 \| (28, 144) \| 156 \| (10, 93) \| 119 \| VRR-NUC domain-containing protein \| VRR-NUC domain-containing protein; uniclust \| UniRef100\_A0A6J5LHN5 \| 98.1 \| 6.1e-08 \| 1.1e-13 \| 63.4 \| 60 \| (86, 155) \| 156 \| (2, 63) \| 66 \| VRR-NUC domain containing protein \| VRR-NUC domain containing protein; uniclust \| UniRef100\_A0A965GSE7 \| 98.1 \| 6.4e-08 \| 1.2e-13 \| 70.7 \| 78 \| (23, 145) \| 156 \| (2, 81) \| 127 \| Uncharacterized protein \| Uncharacterized protein; pdb70 \| 4QBL\_B \| 99.7 \| 1.5e-21 \| 1.5e-25 \| 145.1 \| 116 \| (20, 155) \| 156 \| (2, 128) \| 145 \| VRR-NUC \| 4QBL\_B VRR-NUC Nuclease, HYDROLASE HET: MSE; pdb70 \| 4QBL\_E \| 99.7 \| 1.5e-21 \| 1.5e-25 \| 145.1 \| 116 \| (20, 155) \| 156 \| (2, 128) \| 145 \| VRR-NUC \| 4QBL\_E VRR-NUC Nuclease, HYDROLASE; pdb70 \| 4QBN\_A \| 99.4 \| 7.9e-18 \| 7.8e-22 \| 115.0 \| 88 \| (22, 155) \| 156 \| (1, 91) \| 93 \| Uncharacterized protein \| 4QBN\_A Uncharacterized protein Nuclease, HYDROLASE HET: SO4; pdb70 \| 4QBO\_A \| 99.4 \| 5.1e-17 \| 4.9e-21 \| 111.2 \| 85 \| (22, 155) \| 156 \| (3, 88) \| 92 \| Putative uncharacterized protein \| 4QBO\_A Putative uncharacterized protein nuclease, HYDROLASE; pdb70 \| 4R8A\_A \| 98.8 \| 6.4e-13 \| 6.2e-17 \| 115.8 \| 65 \| (76, 150) \| 156 \| (493, 558) \| 559 \| Uncharacterized protein, DNA \| 4R8A\_A Uncharacterized protein, DNA DNA binding, metal binding nuclease; pdb70 \| 5Y7Q\_A \| 98.8 \| 7.7e-13 \| 7.3e-17 \| 115.9 \| 66 \| (75, 150) \| 156 \| (513, 579) \| 580 \| Fanconi-associated nuclease 1 homolog/DNA Complex \| 5Y7Q\_A Fanconi-associated nuclease 1 homolog/DNA Complex Nuclease, HYDROLASE-DNA complex; pdb70 \| 4REC\_A \| 98.7 \| 1.7e-12 \| 1.7e-16 \| 114.4 \| 66 \| (76, 151) \| 156 \| (576, 642) \| 647 \| Fanconi-associated nuclease 1 (E.C.3.1.21.-, 3.1.4.1) \| 4REC\_A Fanconi-associated nuclease 1 (E.C.3.1.21.-, 3.1.4.1) HJC, TPR, SAP, structure specific; pdb70 \| 4RY3\_A \| 98.7 \| 1.9e-12 \| 1.8e-16 \| 114.2 \| 66 \| (76, 151) \| 156 \| (576, 642) \| 648 \| Fanconi-associated nuclease 1 (E.C.3.1.21.-, 3.1.4.1) \| 4RY3\_A Fanconi-associated nuclease 1 (E.C.3.1.21.-, 3.1.4.1) Endonuclease 5'-3'exonulease, FANCD2, FAN1, HYDROLASE; pdb70 \| 4RIC\_B \| 98.6 \| 6.9e-12 \| 6.7e-16 \| 110.1 \| 61 \| (77, 147) \| 156 \| (569, 630) \| 631 \| Fanconi-associated nuclease 1 (E.C.3.1.21.-, 3.1.4.1)/dna \| 4RIC\_B Fanconi-associated nuclease 1 (E.C.3.1.21.-, 3.1.4.1)/dna nuclease, hydrolase-dna complex |
| Top keywords  (threshold 1.00e-03 (evalue)) | **VRR\_NUC, domain\_containing, Nuclease, Fragment, containing, hypothetical, Endonuclease, Putative, HYDROLASE, DNA** |
| Output files | ../../similar\_sequences/48\_FANPEZAQ\_CDS\_0048\_merged.svg ../../similar\_sequences/48\_FANPEZAQ\_CDS\_0048\_pdb70.a3m ../../similar\_sequences/48\_FANPEZAQ\_CDS\_0048\_pdb70.hhr ../../similar\_sequences/48\_FANPEZAQ\_CDS\_0048\_uniclust.a3m ../../similar\_sequences/48\_FANPEZAQ\_CDS\_0048\_uniclust.hhr |

#### Structure prediction (AlphaFold)^2^

|  |  |
| --- | --- |
| Stats | xml version="1.0" encoding="utf-8" standalone="no"?       2024-09-02T21:09:45.365222 image/svg+xml   Matplotlib v3.7.2, https://matplotlib.org/ |
| Predicted structure | **NGL Viewer Controls:**  - Center: *Left-Click* - Rotate: *Left-Click + Drag* - Translate: *Right-Click + Drag* - Zoom: *Shift + Left-Click + Drag* |
| Output files | ../../predicted\_structures/48\_FANPEZAQ\_CDS\_0048/features.pkl ../../predicted\_structures/48\_FANPEZAQ\_CDS\_0048/ranked\_0.pdb ../../predicted\_structures/48\_FANPEZAQ\_CDS\_0048/ranked\_0\_plots.svg ../../predicted\_structures/48\_FANPEZAQ\_CDS\_0048/result\_model\_1\_ptm\_pred\_0.pkl |

#### Structure similarity search results (Foldseek)^3^

|  |  |
| --- | --- |
| Structure databases searched | Pdb, Afdb-proteome, Afdb-uniprot50 |
| Results, scheme(s)  (Top layers only, threshold 1.00e-02 (evalue)) | xml version="1.0" encoding="utf-8" standalone="no"?       2024-09-02T21:11:23.045883 image/svg+xml   Matplotlib v3.7.2, https://matplotlib.org/ |
| Results, table  (threshold 1.00e-02 (evalue)) | db \| id \| prob \| evalue \| bits \| fident \| alnlen \| mismatch \| gapopen \| qstart \| qend \| tstart \| tend \| name \| description; pdb \| 1Y1O\_D \| 0.975 \| 0.002521 \| 77 \| 0.105 \| 151 \| 92 \| 8 \| 20 \| 154 \| 1 \| 124 \| Penicillin-binding protein-related factor A \| Penicillin-binding protein-related factor A; afdb-uniprot50 \| AF-A0A2W5MTV2-F1-MODEL\_V4 \| 1.0 \| 7.381e-17 \| 592 \| 0.526 \| 131 \| 53 \| 2 \| 25 \| 154 \| 3 \| 125 \| VRR-NUC domain-containing protein \| VRR-NUC domain-containing protein; afdb-uniprot50 \| AF-A0A7W3KP85-F1-MODEL\_V4 \| 1.0 \| 3.895e-14 \| 490 \| 0.402 \| 134 \| 76 \| 4 \| 25 \| 156 \| 3 \| 134 \| VRR-NUC domain-containing protein \| VRR-NUC domain-containing protein; afdb-uniprot50 \| AF-A0A560BMV8-F1-MODEL\_V4 \| 1.0 \| 9.269e-14 \| 466 \| 0.39 \| 133 \| 77 \| 3 \| 25 \| 154 \| 5 \| 136 \| VRR-NUC domain-containing protein \| VRR-NUC domain-containing protein; afdb-uniprot50 \| AF-I8TQK1-F1-MODEL\_V4 \| 1.0 \| 4.593e-13 \| 446 \| 0.428 \| 133 \| 65 \| 3 \| 25 \| 154 \| 22 \| 146 \| VRR-NUC domain-containing protein \| VRR-NUC domain-containing protein; afdb-uniprot50 \| AF-A0A1B1I055-F1-MODEL\_V4 \| 1.0 \| 1.863e-12 \| 434 \| 0.393 \| 132 \| 68 \| 3 \| 25 \| 153 \| 5 \| 127 \| Nuclease \| Nuclease; afdb-uniprot50 \| AF-A0A853YK68-F1-MODEL\_V4 \| 1.0 \| 5.997e-13 \| 414 \| 0.388 \| 126 \| 73 \| 3 \| 29 \| 154 \| 4 \| 125 \| Uncharacterized protein \| Uncharacterized protein; afdb-uniprot50 \| AF-A0A4P5NU28-F1-MODEL\_V4 \| 1.0 \| 8.081e-12 \| 409 \| 0.36 \| 133 \| 73 \| 4 \| 25 \| 154 \| 3 \| 126 \| VRR-NUC domain-containing protein \| VRR-NUC domain-containing protein; afdb-uniprot50 \| AF-A0A0D6PDS2-F1-MODEL\_V4 \| 1.0 \| 9.871e-12 \| 399 \| 0.333 \| 135 \| 78 \| 4 \| 25 \| 156 \| 4 \| 129 \| Uncharacterized protein \| Uncharacterized protein; afdb-uniprot50 \| AF-A0A1I4Q9V7-F1-MODEL\_V4 \| 1.0 \| 3.396e-12 \| 398 \| 0.416 \| 132 \| 66 \| 3 \| 25 \| 153 \| 6 \| 129 \| VRR-NUC domain-containing protein \| VRR-NUC domain-containing protein; afdb-uniprot50 \| AF-A0A7X6FPB3-F1-MODEL\_V4 \| 1.0 \| 2.869e-11 \| 398 \| 0.31 \| 129 \| 86 \| 3 \| 23 \| 149 \| 2 \| 129 \| VRR-NUC domain-containing protein \| VRR-NUC domain-containing protein; afdb-uniprot50 \| AF-A0A662MG58-F1-MODEL\_V4 \| 1.0 \| 1.473e-11 \| 381 \| 0.33 \| 133 \| 85 \| 4 \| 25 \| 155 \| 3 \| 133 \| Uncharacterized protein \| Uncharacterized protein; afdb-uniprot50 \| AF-A0A2G2PN72-F1-MODEL\_V4 \| 1.0 \| 1.799e-11 \| 368 \| 0.326 \| 141 \| 82 \| 4 \| 25 \| 153 \| 3 \| 142 \| Uncharacterized protein \| Uncharacterized protein; afdb-uniprot50 \| AF-A0A424NXD6-F1-MODEL\_V4 \| 1.0 \| 2.684e-11 \| 365 \| 0.292 \| 157 \| 80 \| 5 \| 1 \| 154 \| 1 \| 129 \| VRR-NUC domain-containing protein \| VRR-NUC domain-containing protein; afdb-uniprot50 \| AF-A0A839H9H4-F1-MODEL\_V4 \| 1.0 \| 2.424e-10 \| 355 \| 0.385 \| 127 \| 68 \| 3 \| 25 \| 149 \| 5 \| 123 \| VRR-NUC domain-containing protein \| VRR-NUC domain-containing protein; afdb-uniprot50 \| AF-A0A1S1SLT7-F1-MODEL\_V4 \| 1.0 \| 4.891e-11 \| 355 \| 0.403 \| 129 \| 68 \| 3 \| 23 \| 149 \| 2 \| 123 \| Uncharacterized protein \| Uncharacterized protein; afdb-uniprot50 \| AF-A0A838BWW6-F1-MODEL\_V4 \| 1.0 \| 1.574e-11 \| 348 \| 0.398 \| 128 \| 72 \| 5 \| 25 \| 148 \| 2 \| 128 \| Uncharacterized protein \| Uncharacterized protein; afdb-uniprot50 \| AF-A0A1H3DQD1-F1-MODEL\_V4 \| 1.0 \| 3.866e-10 \| 344 \| 0.338 \| 130 \| 75 \| 4 \| 25 \| 152 \| 3 \| 123 \| VRR-NUC domain-containing protein \| VRR-NUC domain-containing protein; afdb-uniprot50 \| AF-A0A149SVC6-F1-MODEL\_V4 \| 1.0 \| 6.801e-09 \| 305 \| 0.478 \| 92 \| 47 \| 1 \| 63 \| 153 \| 1 \| 92 \| VRR-NUC domain-containing protein \| VRR-NUC domain-containing protein; afdb-uniprot50 \| AF-A0A1L7AG69-F1-MODEL\_V4 \| 1.0 \| 1.124e-09 \| 304 \| 0.346 \| 130 \| 69 \| 4 \| 28 \| 153 \| 4 \| 121 \| Uncharacterized protein \| Uncharacterized protein; afdb-uniprot50 \| AF-A0A836VDS1-F1-MODEL\_V4 \| 1.0 \| 4.873e-09 \| 302 \| 0.284 \| 137 \| 83 \| 4 \| 25 \| 155 \| 3 \| 130 \| Uncharacterized protein \| Uncharacterized protein; afdb-uniprot50 \| AF-A0A4P5VJF4-F1-MODEL\_V4 \| 1.0 \| 7.27e-09 \| 285 \| 0.286 \| 136 \| 71 \| 4 \| 25 \| 154 \| 8 \| 123 \| Uncharacterized protein \| Uncharacterized protein; afdb-uniprot50 \| AF-A0A5C7QIV6-F1-MODEL\_V4 \| 1.0 \| 6.362e-09 \| 275 \| 0.22 \| 159 \| 107 \| 4 \| 1 \| 156 \| 21 \| 165 \| VRR-NUC domain-containing protein \| VRR-NUC domain-containing protein; afdb-uniprot50 \| AF-A0A679JRI6-F1-MODEL\_V4 \| 1.0 \| 3.974e-07 \| 223 \| 0.421 \| 102 \| 54 \| 3 \| 56 \| 153 \| 4 \| 104 \| VRR-NUC domain-containing protein \| VRR-NUC domain-containing protein |
| Top keywords  (threshold 1.00e-02 (evalue)) | **VRR\_NUC, domain\_containing, Penicillin\_binding, protein\_related, factor, A, Nuclease** |
| Output files | ../../similar\_structures/48\_FANPEZAQ\_CDS\_0048\_afdb-proteome\_foldseek.tsv ../../similar\_structures/48\_FANPEZAQ\_CDS\_0048\_afdb-uniprot50\_foldseek.tsv ../../similar\_structures/48\_FANPEZAQ\_CDS\_0048\_merged.svg ../../similar\_structures/48\_FANPEZAQ\_CDS\_0048\_pdb\_foldseek.tsv |

  
  
  

Return to summary | Go to previous | Go to next

  


---

**Sequence/structure alignments coloring**  
Each object in the alignment figures is colored according to its E-value following this color coding:

1e-100
10
