## Supplementary material for "Completing the BASEL phage collection to unlock hidden diversity for systematic exploration of phage-host interactions": Data S2: 51.html

FANPEZAQ\_CDS\_0051


Return to summary | Go to previous | Go to next

|  |  |
| --- | --- |
| FANPEZAQ\_CDS\_0051  Page creation date: 02 Sep 2024, 12:00  Project folder: n/a  Input sequences file: Escherichia\_virus\_HeidiAbel.gb | hypothetical phage transcriptional regulator p119269 vi\_03293 cell envelope biogenesis ompa membrane lipoprotein duf4222 domain\_containing cro ci coil containing lysis system i\_spanin rz |

### Sequence information

|  |  |
| --- | --- |
| Name | FANPEZAQ\_CDS\_0051  51\_FANPEZAQ\_CDS\_0051 (pipeline id) |
| Imported annotations | Escherichia\_virus\_HeidiAbel Bas97 |
| Protein sequence | MNPVDFIKKQVMAELVKQGASPGTAQHAADYAAQQYRTTSAIGKDPFAELIRLAGAMAMK HQAGFRFVMPQSSR |
| Number of residues | 74 |
| Molecular weight (Da) | 8068.24 |
| Output files | ../../query\_sequences/51\_FANPEZAQ\_CDS\_0051.fasta |

### Putative domain architecture and protein family

#### Search results (HHblits)^1^

|  |  |
| --- | --- |
| Domain family databases searched | Pfam, Ncbi-cd, Cath, Phrogs |
| Results, scheme(s)  (Top layers only; threshold 1.00e-03 (evalue)) | xml version="1.0" encoding="utf-8" standalone="no"?       2024-09-02T21:08:23.865001 image/svg+xml   Matplotlib v3.7.2, https://matplotlib.org/ |
| Results, table  (E-value ≤ 1.00e-03 (evalue)) | db \| id \| prob \| evalue \| pvalue \| score \| cols \| query \| query\_len \| template \| template\_len \| name \| description; phrogs \| 4439 \| 99.9 \| 6.7e-34 \| 8.8e-38 \| 173.2 \| 58 \| (1, 58) \| 74 \| (1, 60) \| 60 \| NA \| NA; Category: unknown function; p119269 VI\_03293 |
| Top keywords  (threshold 1.00e-03 (evalue)) | **p119269, VI\_03293** |
| Output files | ../../domain\_architecture/51\_FANPEZAQ\_CDS\_0051\_cath.hhr ../../domain\_architecture/51\_FANPEZAQ\_CDS\_0051\_merged.svg ../../domain\_architecture/51\_FANPEZAQ\_CDS\_0051\_ncbi-cd.hhr ../../domain\_architecture/51\_FANPEZAQ\_CDS\_0051\_pfam.hhr ../../domain\_architecture/51\_FANPEZAQ\_CDS\_0051\_phrogs.hhr |

### Identical protein sequences/structures

#### Search results

|  |  |
| --- | --- |
| Protein sequence databases searched | Pdb, Swissprot, Refseq |
| Identical proteins found | -- |
| Top keywords | -- |
| Output files | -- |

### Similar protein sequences/structures

#### Sequence similarity search results (HHblits)^1^

|  |  |
| --- | --- |
| Sequence databases searched | Uniclust, Pdb70 |
| Results, scheme(s)  (Top layers only, threshold 1.00e-03 (evalue)) | xml version="1.0" encoding="utf-8" standalone="no"?       2024-09-02T21:08:51.007845 image/svg+xml   Matplotlib v3.7.2, https://matplotlib.org/ |
| Results, table(s)  (threshold 1.00e-03 (evalue)) | db \| id \| prob \| evalue \| pvalue \| score \| cols \| query \| query\_len \| template \| template\_len \| name \| description; uniclust \| UniRef100\_A0A094U7C3 \| 100.0 \| 1.2e-33 \| 2.6e-39 \| 175.5 \| 69 \| (1, 69) \| 74 \| (3, 73) \| 90 \| Uncharacterized protein \| Uncharacterized protein; uniclust \| UniRef100\_A0A0T9RNR7 \| 99.9 \| 6.7e-32 \| 1.4e-37 \| 168.6 \| 73 \| (1, 73) \| 74 \| (1, 75) \| 93 \| Uncharacterized protein \| Uncharacterized protein; uniclust \| UniRef100\_A0A085ADJ9 \| 99.9 \| 3.7e-31 \| 8e-37 \| 171.7 \| 72 \| (1, 72) \| 74 \| (18, 91) \| 119 \| Phage protein \| Phage protein; uniclust \| UniRef100\_A0A077NE64 \| 99.9 \| 6.1e-31 \| 1.4e-36 \| 167.0 \| 72 \| (1, 72) \| 74 \| (15, 88) \| 99 \| Phage protein \| Phage protein; uniclust \| UniRef100\_A0A2S1BH24 \| 99.8 \| 6.9e-22 \| 1.3e-27 \| 121.7 \| 72 \| (1, 72) \| 74 \| (1, 74) \| 85 \| Uncharacterized protein \| Uncharacterized protein; uniclust \| UniRef100\_UPI0020173D3F \| 99.8 \| 1.2e-21 \| 2.2e-27 \| 125.1 \| 73 \| (1, 73) \| 74 \| (30, 104) \| 114 \| hypothetical protein \| hypothetical protein; uniclust \| UniRef100\_A0A2U1TMP1 \| 99.7 \| 1.2e-20 \| 2.4e-26 \| 115.9 \| 70 \| (1, 70) \| 74 \| (3, 74) \| 82 \| Uncharacterized protein \| Uncharacterized protein; uniclust \| UniRef100\_UPI0019099C25 \| 99.7 \| 2.3e-19 \| 4.2e-25 \| 118.9 \| 72 \| (1, 72) \| 74 \| (62, 135) \| 145 \| hypothetical protein \| hypothetical protein; uniclust \| UniRef100\_A0A0P6ZIQ1 \| 99.6 \| 7.3e-19 \| 1.5e-24 \| 113.0 \| 71 \| (2, 72) \| 74 \| (20, 93) \| 102 \| Uncharacterized protein \| Uncharacterized protein; uniclust \| UniRef100\_A0A2V4G0H8 \| 99.6 \| 1.6e-18 \| 3.3e-24 \| 108.5 \| 71 \| (1, 71) \| 74 \| (4, 76) \| 89 \| Uncharacterized protein \| Uncharacterized protein; uniclust \| UniRef100\_UPI00090789F6 \| 99.6 \| 2e-18 \| 3.8e-24 \| 111.8 \| 69 \| (2, 70) \| 74 \| (4, 74) \| 113 \| hypothetical protein \| hypothetical protein; uniclust \| UniRef100\_A0A2L0TW67 \| 99.6 \| 5.6e-18 \| 1.1e-23 \| 105.9 \| 67 \| (1, 67) \| 74 \| (1, 68) \| 88 \| Uncharacterized protein \| Uncharacterized protein; uniclust \| UniRef100\_A0A078L883 \| 99.6 \| 8.1e-18 \| 1.6e-23 \| 99.1 \| 54 \| (1, 54) \| 74 \| (1, 56) \| 60 \| Uncharacterized protein \| Uncharacterized protein; uniclust \| UniRef100\_A0A379D1S2 \| 99.5 \| 1.5e-17 \| 3.2e-23 \| 99.7 \| 51 \| (23, 73) \| 74 \| (2, 54) \| 65 \| Uncharacterized protein \| Uncharacterized protein; uniclust \| UniRef100\_A0A090IT71 \| 99.5 \| 2.2e-17 \| 4.4e-23 \| 103.0 \| 71 \| (2, 72) \| 74 \| (9, 80) \| 84 \| Uncharacterized phage protein \| Uncharacterized phage protein; uniclust \| UniRef100\_A0A173GC14 \| 99.4 \| 4.6e-16 \| 8.9e-22 \| 91.1 \| 56 \| (1, 62) \| 74 \| (1, 57) \| 58 \| Uncharacterized protein \| Uncharacterized protein; uniclust \| UniRef100\_A0A087KTT2 \| 99.4 \| 4.5e-16 \| 9.8e-22 \| 97.6 \| 66 \| (4, 72) \| 74 \| (6, 71) \| 83 \| Cell envelope biogenesis protein OmpA \| Cell envelope biogenesis protein OmpA; uniclust \| UniRef100\_A0A241XEA3 \| 99.4 \| 1.3e-15 \| 2.5e-21 \| 92.3 \| 67 \| (1, 67) \| 74 \| (1, 67) \| 71 \| Uncharacterized protein \| Uncharacterized protein; uniclust \| UniRef100\_UPI000A4DD1B9 \| 99.4 \| 1.5e-15 \| 2.8e-21 \| 103.8 \| 71 \| (1, 71) \| 74 \| (88, 161) \| 173 \| hypothetical protein \| hypothetical protein; uniclust \| UniRef100\_UPI00226A8289 \| 99.4 \| 2.2e-15 \| 4e-21 \| 99.2 \| 63 \| (1, 63) \| 74 \| (63, 127) \| 130 \| hypothetical protein \| hypothetical protein; uniclust \| UniRef100\_A0A209B2B0 \| 99.3 \| 1.7e-14 \| 3.2e-20 \| 85.8 \| 63 \| (1, 64) \| 74 \| (1, 65) \| 66 \| Uncharacterized protein \| Uncharacterized protein; uniclust \| UniRef100\_A0A0E1SXQ1 \| 99.2 \| 3.2e-14 \| 5.9e-20 \| 79.2 \| 38 \| (27, 64) \| 74 \| (2, 41) \| 45 \| Uncharacterized protein \| Uncharacterized protein; uniclust \| UniRef100\_UPI0002F40243 \| 99.2 \| 5.2e-14 \| 9.5e-20 \| 86.1 \| 52 \| (12, 63) \| 74 \| (2, 55) \| 77 \| hypothetical protein \| hypothetical protein; uniclust \| UniRef100\_A0A5M7L259 \| 99.2 \| 9e-14 \| 1.7e-19 \| 87.0 \| 70 \| (1, 74) \| 74 \| (8, 80) \| 85 \| Uncharacterized protein \| Uncharacterized protein; uniclust \| UniRef100\_A0A0A7KY33 \| 99.2 \| 1.2e-13 \| 2.2e-19 \| 90.7 \| 67 \| (1, 67) \| 74 \| (42, 109) \| 123 \| Uncharacterized protein \| Uncharacterized protein; uniclust \| UniRef100\_A0A8I1W9T1 \| 99.1 \| 2.1e-13 \| 3.8e-19 \| 84.5 \| 62 \| (1, 62) \| 74 \| (1, 63) \| 83 \| Uncharacterized protein \| Uncharacterized protein; uniclust \| UniRef100\_A0A1Q8CZL4 \| 99.1 \| 3.4e-13 \| 6.3e-19 \| 84.1 \| 66 \| (1, 66) \| 74 \| (12, 78) \| 84 \| Uncharacterized protein \| Uncharacterized protein; uniclust \| UniRef100\_A0A1V3K6Q6 \| 99.1 \| 9.1e-13 \| 1.8e-18 \| 82.3 \| 66 \| (1, 66) \| 74 \| (1, 66) \| 80 \| Uncharacterized protein \| Uncharacterized protein; uniclust \| UniRef100\_A0A655UTX9 \| 99.1 \| 1e-12 \| 2e-18 \| 83.3 \| 71 \| (2, 72) \| 74 \| (9, 81) \| 89 \| Uncharacterized protein \| Uncharacterized protein; uniclust \| UniRef100\_UPI001FB599ED \| 99.1 \| 1.1e-12 \| 2e-18 \| 77.5 \| 43 \| (23, 65) \| 74 \| (2, 46) \| 61 \| hypothetical protein \| hypothetical protein; uniclust \| UniRef100\_A0A0F4NNX2 \| 99.0 \| 2e-12 \| 4e-18 \| 83.5 \| 71 \| (1, 72) \| 74 \| (1, 71) \| 97 \| Uncharacterized protein \| Uncharacterized protein; uniclust \| UniRef100\_UPI00198254EA \| 99.0 \| 3e-12 \| 5.5e-18 \| 78.0 \| 47 \| (18, 64) \| 74 \| (23, 71) \| 73 \| hypothetical protein \| hypothetical protein; uniclust \| UniRef100\_A0A089Q372 \| 99.0 \| 3.1e-12 \| 7e-18 \| 81.7 \| 63 \| (1, 66) \| 74 \| (2, 64) \| 86 \| Uncharacterized protein \| Uncharacterized protein; uniclust \| UniRef100\_H2BDF4 \| 98.9 \| 1e-11 \| 1.9e-17 \| 83.4 \| 66 \| (1, 66) \| 74 \| (71, 136) \| 138 \| Uncharacterized protein \| Uncharacterized protein; uniclust \| UniRef100\_UPI0019D9CF8F \| 98.9 \| 1.3e-11 \| 2.3e-17 \| 75.5 \| 57 \| (1, 57) \| 74 \| (1, 60) \| 73 \| hypothetical protein \| hypothetical protein; uniclust \| UniRef100\_UPI001F319DAE \| 98.9 \| 1.7e-11 \| 3.3e-17 \| 79.2 \| 55 \| (1, 55) \| 74 \| (29, 83) \| 98 \| hypothetical protein \| hypothetical protein; uniclust \| UniRef100\_A0A0N7JPH6 \| 98.9 \| 1.7e-11 \| 3.3e-17 \| 77.4 \| 68 \| (1, 71) \| 74 \| (3, 70) \| 82 \| Uncharacterized protein \| Uncharacterized protein; uniclust \| UniRef100\_UPI001E649674 \| 98.8 \| 4.5e-11 \| 8.2e-17 \| 69.0 \| 40 \| (1, 40) \| 74 \| (1, 41) \| 52 \| hypothetical protein \| hypothetical protein; uniclust \| UniRef100\_UPI000573BB50 \| 98.8 \| 5.1e-11 \| 9.4e-17 \| 72.4 \| 61 \| (1, 66) \| 74 \| (1, 61) \| 70 \| hypothetical protein \| hypothetical protein; uniclust \| UniRef100\_A0A4V1DBZ5 \| 98.8 \| 6.6e-11 \| 1.2e-16 \| 73.5 \| 63 \| (1, 63) \| 74 \| (1, 63) \| 80 \| Uncharacterized protein \| Uncharacterized protein; uniclust \| UniRef100\_A0A0Q1CBT0 \| 98.7 \| 1e-10 \| 2.2e-16 \| 77.9 \| 35 \| (1, 35) \| 74 \| (1, 35) \| 110 \| Uncharacterized protein \| Uncharacterized protein; uniclust \| UniRef100\_A0A513SPV1 \| 98.7 \| 1.4e-10 \| 2.6e-16 \| 77.2 \| 67 \| (2, 68) \| 74 \| (45, 115) \| 125 \| Uncharacterized protein \| Uncharacterized protein; uniclust \| UniRef100\_A0A516MMM3 \| 98.7 \| 2.4e-10 \| 4.4e-16 \| 68.3 \| 60 \| (1, 60) \| 74 \| (1, 60) \| 62 \| Uncharacterized protein \| Uncharacterized protein; uniclust \| UniRef100\_A0A7I6Q891 \| 98.6 \| 2.7e-10 \| 5e-16 \| 62.3 \| 29 \| (1, 29) \| 74 \| (1, 29) \| 38 \| Uncharacterized protein \| Uncharacterized protein; uniclust \| UniRef100\_A0A4R3Z3M5 \| 98.6 \| 2.7e-10 \| 5.2e-16 \| 60.9 \| 29 \| (1, 29) \| 74 \| (1, 29) \| 33 \| Uncharacterized protein \| Uncharacterized protein; uniclust \| UniRef100\_UPI00211A06C2 \| 98.6 \| 4.4e-10 \| 8.1e-16 \| 72.4 \| 66 \| (1, 66) \| 74 \| (1, 66) \| 99 \| hypothetical protein \| hypothetical protein; uniclust \| UniRef100\_A0A9E7LPW0 \| 98.6 \| 7.3e-10 \| 1.3e-15 \| 77.3 \| 54 \| (1, 60) \| 74 \| (118, 172) \| 174 \| Uncharacterized protein \| Uncharacterized protein; uniclust \| UniRef100\_A0A066RYV4 \| 98.5 \| 1.2e-09 \| 2.3e-15 \| 67.7 \| 59 \| (2, 60) \| 74 \| (7, 65) \| 70 \| Transcriptional regulator \| Transcriptional regulator; uniclust \| UniRef100\_A0A2I7S8J3 \| 98.5 \| 1.6e-09 \| 3.4e-15 \| 70.5 \| 70 \| (1, 70) \| 74 \| (1, 70) \| 92 \| Membrane lipoprotein \| Membrane lipoprotein; uniclust \| UniRef100\_UPI0021CBB242 \| 98.5 \| 2.1e-09 \| 3.9e-15 \| 70.8 \| 68 \| (2, 69) \| 74 \| (9, 78) \| 112 \| hypothetical protein \| hypothetical protein; uniclust \| UniRef100\_UPI0003BEDC01 \| 98.4 \| 3.2e-09 \| 6.1e-15 \| 62.1 \| 33 \| (1, 33) \| 74 \| (1, 33) \| 52 \| hypothetical protein \| hypothetical protein; uniclust \| UniRef100\_A0A1I3XT93 \| 98.4 \| 4e-09 \| 7.3e-15 \| 55.0 \| 27 \| (1, 27) \| 74 \| (1, 27) \| 29 \| Uncharacterized protein \| Uncharacterized protein; uniclust \| UniRef100\_UPI0018F0D1C9 \| 98.4 \| 4.3e-09 \| 7.8e-15 \| 63.2 \| 59 \| (1, 60) \| 74 \| (1, 59) \| 62 \| hypothetical protein \| hypothetical protein; uniclust \| UniRef100\_UPI000AF57BD8 \| 98.3 \| 9.6e-09 \| 1.9e-14 \| 69.7 \| 30 \| (1, 30) \| 74 \| (1, 30) \| 122 \| hypothetical protein \| hypothetical protein; uniclust \| UniRef100\_UPI000988BEDA \| 98.3 \| 1.1e-08 \| 2e-14 \| 64.8 \| 70 \| (2, 71) \| 74 \| (8, 77) \| 84 \| hypothetical protein \| hypothetical protein; uniclust \| UniRef100\_UPI0020A36E27 \| 98.3 \| 1.3e-08 \| 2.3e-14 \| 62.3 \| 35 \| (1, 35) \| 74 \| (1, 35) \| 68 \| hypothetical protein \| hypothetical protein; uniclust \| UniRef100\_A0A2N7JZE4 \| 98.2 \| 2.9e-08 \| 5.4e-14 \| 61.8 \| 57 \| (14, 70) \| 74 \| (12, 71) \| 75 \| Uncharacterized protein \| Uncharacterized protein; uniclust \| UniRef100\_A0A127MXN3 \| 98.1 \| 3.9e-08 \| 7.2e-14 \| 57.1 \| 44 \| (1, 44) \| 74 \| (1, 44) \| 50 \| Uncharacterized protein \| Uncharacterized protein; uniclust \| UniRef100\_A0A087L1N4 \| 98.1 \| 3.8e-08 \| 7.9e-14 \| 64.5 \| 33 \| (1, 33) \| 74 \| (1, 33) \| 92 \| Uncharacterized protein \| Uncharacterized protein; uniclust \| UniRef100\_A0A1Q8Z4M5 \| 98.0 \| 8.2e-08 \| 1.6e-13 \| 63.8 \| 33 \| (1, 33) \| 74 \| (1, 33) \| 107 \| Uncharacterized protein \| Uncharacterized protein; uniclust \| UniRef100\_A0A927DSF2 \| 98.0 \| 9.3e-08 \| 1.8e-13 \| 66.8 \| 33 \| (1, 33) \| 74 \| (17, 49) \| 150 \| Uncharacterized protein \| Uncharacterized protein; uniclust \| UniRef100\_A0A380QAC9 \| 97.9 \| 2.1e-07 \| 3.9e-13 \| 55.0 \| 33 \| (1, 33) \| 74 \| (1, 33) \| 53 \| Uncharacterized protein \| Uncharacterized protein; uniclust \| UniRef100\_UPI001FCE5DAA \| 97.9 \| 2.2e-07 \| 4e-13 \| 63.4 \| 34 \| (1, 34) \| 74 \| (1, 34) \| 132 \| DUF4222 domain-containing protein \| DUF4222 domain-containing protein; uniclust \| UniRef100\_A0A1X3IVQ3 \| 97.9 \| 2.3e-07 \| 4.2e-13 \| 57.5 \| 32 \| (1, 32) \| 74 \| (1, 32) \| 70 \| Uncharacterized protein \| Uncharacterized protein; uniclust \| UniRef100\_UPI000791D832 \| 97.9 \| 2.6e-07 \| 5e-13 \| 62.0 \| 28 \| (1, 28) \| 74 \| (1, 28) \| 109 \| hypothetical protein \| hypothetical protein; uniclust \| UniRef100\_UPI0004ADB128 \| 97.8 \| 4e-07 \| 7.8e-13 \| 55.3 \| 32 \| (1, 32) \| 74 \| (1, 32) \| 58 \| hypothetical protein \| hypothetical protein; uniclust \| UniRef100\_UPI000B237913 \| 97.7 \| 8.5e-07 \| 1.6e-12 \| 56.2 \| 53 \| (13, 65) \| 74 \| (3, 56) \| 78 \| hypothetical protein \| hypothetical protein; uniclust \| UniRef100\_A0A7Y3Z2V1 \| 97.7 \| 1.2e-06 \| 2.2e-12 \| 53.5 \| 43 \| (2, 44) \| 74 \| (9, 51) \| 63 \| Uncharacterized protein \| Uncharacterized protein; uniclust \| UniRef100\_A0A927HPP5 \| 97.7 \| 1.3e-06 \| 2.4e-12 \| 55.9 \| 26 \| (1, 26) \| 74 \| (58, 83) \| 84 \| Uncharacterized protein \| Uncharacterized protein; uniclust \| UniRef100\_UPI0021BD29FD \| 97.6 \| 1.6e-06 \| 3e-12 \| 59.5 \| 35 \| (1, 35) \| 74 \| (84, 118) \| 133 \| Cro/CI family transcriptional regulator \| Cro/CI family transcriptional regulator; uniclust \| UniRef100\_UPI001CBDE202 \| 97.5 \| 3.8e-06 \| 7e-12 \| 49.4 \| 37 \| (1, 37) \| 74 \| (1, 37) \| 50 \| hypothetical protein \| hypothetical protein; uniclust \| UniRef100\_UPI001F40ADE3 \| 97.5 \| 4e-06 \| 7.3e-12 \| 49.7 \| 32 \| (1, 32) \| 74 \| (1, 32) \| 52 \| hypothetical protein \| hypothetical protein; uniclust \| UniRef100\_UPI000A1EE641 \| 97.5 \| 4.2e-06 \| 7.9e-12 \| 52.0 \| 56 \| (3, 58) \| 74 \| (4, 59) \| 66 \| hypothetical protein \| hypothetical protein; uniclust \| UniRef100\_UPI0022CE1587 \| 97.4 \| 5e-06 \| 9.1e-12 \| 53.0 \| 58 \| (2, 59) \| 74 \| (9, 66) \| 79 \| hypothetical protein \| hypothetical protein; uniclust \| UniRef100\_UPI000B35E1F4 \| 97.4 \| 4.8e-06 \| 9.2e-12 \| 55.0 \| 67 \| (1, 67) \| 74 \| (1, 69) \| 94 \| hypothetical protein \| hypothetical protein; uniclust \| UniRef100\_A0A927HJ22 \| 97.4 \| 6e-06 \| 1.1e-11 \| 58.3 \| 32 \| (1, 32) \| 74 \| (58, 89) \| 158 \| Uncharacterized protein \| Uncharacterized protein; uniclust \| UniRef100\_A0A2I7R4F3 \| 97.4 \| 8.2e-06 \| 1.5e-11 \| 49.6 \| 56 \| (4, 59) \| 74 \| (2, 57) \| 59 \| Coil containing protein \| Coil containing protein; uniclust \| UniRef100\_A0A7Y6JDD3 \| 97.3 \| 1.3e-05 \| 2.5e-11 \| 49.8 \| 55 \| (3, 57) \| 74 \| (6, 61) \| 67 \| Uncharacterized protein \| Uncharacterized protein; uniclust \| UniRef100\_A0A1M7YP10 \| 97.3 \| 1.5e-05 \| 2.7e-11 \| 48.3 \| 38 \| (3, 40) \| 74 \| (9, 46) \| 57 \| Uncharacterized protein \| Uncharacterized protein; uniclust \| UniRef100\_UPI000A2E50F0 \| 97.2 \| 2e-05 \| 3.7e-11 \| 50.2 \| 38 \| (1, 40) \| 74 \| (1, 38) \| 75 \| hypothetical protein \| hypothetical protein; uniclust \| UniRef100\_UPI00053E6714 \| 97.2 \| 2.2e-05 \| 4.1e-11 \| 55.0 \| 40 \| (1, 40) \| 74 \| (1, 40) \| 144 \| hypothetical protein \| hypothetical protein; uniclust \| UniRef100\_A0A7D5XSG2 \| 97.1 \| 3e-05 \| 5.5e-11 \| 45.3 \| 33 \| (1, 33) \| 74 \| (1, 33) \| 46 \| Uncharacterized protein \| Uncharacterized protein; uniclust \| UniRef100\_A0A6S4Y2E2 \| 97.0 \| 5.9e-05 \| 1.1e-10 \| 40.6 \| 28 \| (1, 28) \| 74 \| (1, 28) \| 30 \| Uncharacterized protein \| Uncharacterized protein; uniclust \| UniRef100\_A0A2G2PXP6 \| 96.7 \| 0.00023 \| 4.4e-10 \| 50.9 \| 58 \| (1, 58) \| 74 \| (1, 58) \| 150 \| Uncharacterized protein \| Uncharacterized protein; uniclust \| UniRef100\_UPI0012B549EA \| 96.6 \| 0.00029 \| 5.4e-10 \| 43.0 \| 21 \| (1, 21) \| 74 \| (1, 21) \| 55 \| hypothetical protein \| hypothetical protein; uniclust \| UniRef100\_UPI0020C6DD36 \| 96.5 \| 0.0004 \| 7.4e-10 \| 44.1 \| 26 \| (41, 66) \| 74 \| (43, 68) \| 70 \| lysis system i-spanin subunit Rz \| lysis system i-spanin subunit Rz; uniclust \| UniRef100\_Q9MC75 \| 96.4 \| 0.00055 \| 1e-09 \| 39.9 \| 38 \| (29, 66) \| 74 \| (4, 41) \| 43 \| Uncharacterized protein \| Uncharacterized protein |
| Top keywords  (threshold 1.00e-03 (evalue)) | **hypothetical, Phage, Transcriptional, regulator, Cell, envelope, biogenesis, OmpA, Membrane, lipoprotein** |
| Output files | ../../similar\_sequences/51\_FANPEZAQ\_CDS\_0051\_merged.svg ../../similar\_sequences/51\_FANPEZAQ\_CDS\_0051\_pdb70.a3m ../../similar\_sequences/51\_FANPEZAQ\_CDS\_0051\_pdb70.hhr ../../similar\_sequences/51\_FANPEZAQ\_CDS\_0051\_uniclust.a3m ../../similar\_sequences/51\_FANPEZAQ\_CDS\_0051\_uniclust.hhr |

#### Structure prediction (AlphaFold)^2^

|  |  |
| --- | --- |
| Stats | xml version="1.0" encoding="utf-8" standalone="no"?       2024-09-02T21:09:47.360087 image/svg+xml   Matplotlib v3.7.2, https://matplotlib.org/ |
| Predicted structure | **NGL Viewer Controls:**  - Center: *Left-Click* - Rotate: *Left-Click + Drag* - Translate: *Right-Click + Drag* - Zoom: *Shift + Left-Click + Drag* |
| Output files | ../../predicted\_structures/51\_FANPEZAQ\_CDS\_0051/features.pkl ../../predicted\_structures/51\_FANPEZAQ\_CDS\_0051/ranked\_0.pdb ../../predicted\_structures/51\_FANPEZAQ\_CDS\_0051/ranked\_0\_plots.svg ../../predicted\_structures/51\_FANPEZAQ\_CDS\_0051/result\_model\_1\_ptm\_pred\_0.pkl |

#### Structure similarity search results (Foldseek)^3^

|  |  |
| --- | --- |
| Structure databases searched | Pdb, Afdb-proteome, Afdb-uniprot50 |
| Results, scheme(s)  (Top layers only, threshold 1.00e-02 (evalue)) | xml version="1.0" encoding="utf-8" standalone="no"?       2024-09-02T21:11:23.410408 image/svg+xml   Matplotlib v3.7.2, https://matplotlib.org/ |
| Results, table  (threshold 1.00e-02 (evalue)) | -- |
| Top keywords  (threshold 1.00e-02 (evalue)) | -- |
| Output files | ../../similar\_structures/51\_FANPEZAQ\_CDS\_0051\_afdb-proteome\_foldseek.tsv ../../similar\_structures/51\_FANPEZAQ\_CDS\_0051\_afdb-uniprot50\_foldseek.tsv ../../similar\_structures/51\_FANPEZAQ\_CDS\_0051\_merged.svg ../../similar\_structures/51\_FANPEZAQ\_CDS\_0051\_pdb\_foldseek.tsv |

  
  
  

Return to summary | Go to previous | Go to next

  


---

**Sequence/structure alignments coloring**  
Each object in the alignment figures is colored according to its E-value following this color coding:

1e-100
10
