## Supplementary material for "Completing the BASEL phage collection to unlock hidden diversity for systematic exploration of phage-host interactions": Data S2: 52.html

FANPEZAQ\_CDS\_0052


Return to summary | Go to previous | Go to next

|  |  |
| --- | --- |
| FANPEZAQ\_CDS\_0052  Page creation date: 02 Sep 2024, 12:00  Project folder: n/a  Input sequences file: Escherichia\_virus\_HeidiAbel.gb | transcriptional regulator hth domain\_containing cro c1\_type helix\_turn\_helix xre dna\_binding repressor hth\_type transcription dna regulation and xre\_family immr dica binding rna nucleotide metabolism putative contains antitoxin phage ci lexa integration excision terminase small head packaging |

### Sequence information

|  |  |
| --- | --- |
| Name | FANPEZAQ\_CDS\_0052  52\_FANPEZAQ\_CDS\_0052 (pipeline id) |
| Imported annotations | Escherichia\_virus\_HeidiAbel Bas97 |
| Protein sequence | MRPVLWAHCYYEAITMDERTERLHNIMAEHGLTCRDVGRLLGRTEQTVLIWRCKGGKVIP THQLELLELKLNAQAAQ |
| Number of residues | 77 |
| Molecular weight (Da) | 8927.33 |
| Output files | ../../query\_sequences/52\_FANPEZAQ\_CDS\_0052.fasta |

### Putative domain architecture and protein family

#### Search results (HHblits)^1^

|  |  |
| --- | --- |
| Domain family databases searched | Pfam, Ncbi-cd, Cath, Phrogs |
| Results, scheme(s)  (Top layers only; threshold 1.00e-03 (evalue)) | xml version="1.0" encoding="utf-8" standalone="no"?       2024-09-02T21:08:24.052090 image/svg+xml   Matplotlib v3.7.2, https://matplotlib.org/ |
| Results, table  (E-value ≤ 1.00e-03 (evalue)) | db \| id \| prob \| evalue \| pvalue \| score \| cols \| query \| query\_len \| template \| template\_len \| name \| description; phrogs \| 14341 \| 99.6 \| 7.7e-20 \| 9.3e-24 \| 108.3 \| 62 \| (16, 77) \| 77 \| (1, 63) \| 63 \| transcriptional repressor \| transcriptional repressor; Category: transcription regulation; NC\_023006\_p49; phrogs \| 4779 \| 99.5 \| 1.1e-18 \| 1.4e-22 \| 106.6 \| 56 \| (18, 73) \| 77 \| (1, 61) \| 78 \| transcriptional regulator \| transcriptional regulator; Category: transcription regulation; p225361 VI\_06249; phrogs \| 8898 \| 99.1 \| 1.5e-14 \| 1.9e-18 \| 86.4 \| 50 \| (18, 69) \| 77 \| (1, 50) \| 69 \| NA \| NA; Category: unknown function; p78052 VI\_06244; phrogs \| 7950 \| 98.8 \| 7.6e-13 \| 9.4e-17 \| 75.4 \| 37 \| (19, 55) \| 77 \| (1, 37) \| 53 \| transcriptional repressor \| transcriptional repressor; Category: transcription regulation; p141193 VI\_04895; phrogs \| 2104 \| 98.0 \| 1.3e-09 \| 1.6e-13 \| 67.2 \| 38 \| (18, 55) \| 77 \| (30, 67) \| 92 \| NA \| NA; Category: unknown function; p72070 VI\_03178; phrogs \| 14076 \| 97.7 \| 1.6e-08 \| 1.9e-12 \| 61.1 \| 51 \| (17, 68) \| 77 \| (13, 68) \| 81 \| transcriptional regulator \| transcriptional regulator; Category: transcription regulation; p109717 VI\_00077; phrogs \| 5119 \| 97.4 \| 1.1e-07 \| 1.3e-11 \| 62.3 \| 41 \| (16, 56) \| 77 \| (51, 91) \| 152 \| NA \| NA; Category: unknown function; p324576 VI\_09304; phrogs \| 2328 \| 97.3 \| 2e-07 \| 2.6e-11 \| 59.3 \| 36 \| (19, 54) \| 77 \| (19, 54) \| 123 \| transcriptional repressor \| transcriptional repressor; Category: transcription regulation; MG099936\_p34; phrogs \| 1373 \| 97.2 \| 2.1e-07 \| 2.8e-11 \| 56.4 \| 48 \| (19, 66) \| 77 \| (34, 81) \| 95 \| transcriptional regulator \| transcriptional regulator; Category: transcription regulation; NC\_005857\_p48; phrogs \| 25249 \| 96.9 \| 1.8e-06 \| 2e-10 \| 59.7 \| 51 \| (18, 71) \| 77 \| (1, 51) \| 263 \| NA \| NA; Category: unknown function; p347442 VI\_06291; phrogs \| 35 \| 96.7 \| 2.4e-06 \| 3.6e-10 \| 47.4 \| 38 \| (16, 54) \| 77 \| (2, 39) \| 75 \| transcriptional repressor \| transcriptional repressor; Category: transcription regulation; p323300 VI\_10199; phrogs \| 2339 \| 96.7 \| 3.1e-06 \| 4e-10 \| 51.4 \| 34 \| (22, 55) \| 77 \| (32, 65) \| 93 \| HTH DNA binding protein \| HTH DNA binding protein; Category: DNA, RNA and nucleotide metabolism; p425338 VI\_07226; phrogs \| 3 \| 96.7 \| 3e-06 \| 4.6e-10 \| 47.4 \| 37 \| (16, 54) \| 77 \| (3, 41) \| 80 \| transcriptional repressor \| transcriptional repressor; Category: transcription regulation; p29899 VI\_00998; phrogs \| 2967 \| 96.6 \| 3.6e-06 \| 4.8e-10 \| 55.4 \| 52 \| (18, 69) \| 77 \| (6, 60) \| 167 \| transcriptional regulator \| transcriptional regulator; Category: transcription regulation; p123578 VI\_00485; phrogs \| 147 \| 96.6 \| 5.2e-06 \| 6.7e-10 \| 51.9 \| 46 \| (10, 55) \| 77 \| (4, 49) \| 112 \| transcriptional regulator \| transcriptional regulator; Category: transcription regulation; p315289 VI\_07968; phrogs \| 7241 \| 96.4 \| 8.8e-06 \| 1.1e-09 \| 52.1 \| 41 \| (14, 55) \| 77 \| (58, 98) \| 128 \| transcriptional regulator \| transcriptional regulator; Category: transcription regulation; p63620 VI\_05511; phrogs \| 1468 \| 96.4 \| 8.8e-06 \| 1.1e-09 \| 47.0 \| 52 \| (19, 70) \| 77 \| (11, 64) \| 69 \| transcriptional repressor \| transcriptional repressor; Category: transcription regulation; p307668 VI\_06058; phrogs \| 6831 \| 96.4 \| 1.1e-05 \| 1.4e-09 \| 46.5 \| 34 \| (21, 54) \| 77 \| (5, 38) \| 66 \| transcriptional regulator \| transcriptional regulator; Category: transcription regulation; p431926 VI\_12195; phrogs \| 1337 \| 96.3 \| 1.2e-05 \| 1.5e-09 \| 45.2 \| 37 \| (19, 55) \| 77 \| (3, 39) \| 61 \| DNA binding protein \| DNA binding protein; Category: DNA, RNA and nucleotide metabolism; p177899 VI\_00920; phrogs \| 409 \| 96.2 \| 1.5e-05 \| 2.1e-09 \| 45.2 \| 28 \| (25, 54) \| 77 \| (5, 32) \| 72 \| transcriptional regulator \| transcriptional regulator; Category: transcription regulation; p88852 VI\_07592; phrogs \| 37751 \| 96.2 \| 2.2e-05 \| 2.5e-09 \| 52.3 \| 38 \| (18, 55) \| 77 \| (152, 191) \| 205 \| NA \| NA; Category: unknown function; KR093631\_p2; phrogs \| 6962 \| 96.2 \| 2.2e-05 \| 2.6e-09 \| 44.3 \| 52 \| (19, 70) \| 77 \| (2, 53) \| 59 \| NA \| NA; Category: unknown function; p75562 VI\_02384; phrogs \| 4151 \| 96.1 \| 2.6e-05 \| 3.3e-09 \| 51.5 \| 30 \| (25, 54) \| 77 \| (14, 43) \| 168 \| DNA binding protein \| DNA binding protein; Category: DNA, RNA and nucleotide metabolism; p284557 VI\_05952; phrogs \| 296 \| 96.1 \| 2.4e-05 \| 3.5e-09 \| 54.0 \| 41 \| (16, 56) \| 77 \| (3, 43) \| 312 \| DNA transposition protein \| DNA transposition protein; Category: integration and excision; p199288 VI\_08191; phrogs \| 4836 \| 96.1 \| 3.1e-05 \| 3.7e-09 \| 53.3 \| 37 \| (20, 56) \| 77 \| (84, 120) \| 233 \| CII-like regulator \| CII-like regulator; Category: transcription regulation; p13677 VI\_07483; phrogs \| 6223 \| 95.9 \| 4.6e-05 \| 5.6e-09 \| 44.9 \| 34 \| (22, 55) \| 77 \| (6, 41) \| 77 \| transcriptional repressor \| transcriptional repressor; Category: transcription regulation; p192116 VI\_09192; phrogs \| 8008 \| 95.8 \| 5.3e-05 \| 6.5e-09 \| 43.5 \| 47 \| (23, 73) \| 77 \| (3, 49) \| 66 \| DNA binding protein \| DNA binding protein; Category: DNA, RNA and nucleotide metabolism; p180387 VI\_11803; phrogs \| 2439 \| 95.8 \| 5.1e-05 \| 7e-09 \| 54.9 \| 39 \| (16, 54) \| 77 \| (5, 43) \| 385 \| transcriptional regulator \| transcriptional regulator; Category: transcription regulation; p298790 VI\_12393; phrogs \| 7676 \| 95.8 \| 6.4e-05 \| 7.5e-09 \| 44.0 \| 29 \| (26, 54) \| 77 \| (11, 39) \| 75 \| NA \| NA; Category: unknown function; NC\_023688\_p227; phrogs \| 3642 \| 95.7 \| 6.9e-05 \| 8.7e-09 \| 45.1 \| 38 \| (19, 56) \| 77 \| (6, 43) \| 90 \| HTH DNA binding protein \| HTH DNA binding protein; Category: DNA, RNA and nucleotide metabolism; p184909 VI\_05451; phrogs \| 2538 \| 95.7 \| 7.9e-05 \| 1.1e-08 \| 46.0 \| 33 \| (22, 54) \| 77 \| (7, 39) \| 113 \| terminase small subunit \| terminase small subunit; Category: head and packaging; p133796 VI\_05051; phrogs \| 3626 \| 95.6 \| 0.00011 \| 1.2e-08 \| 43.6 \| 49 \| (16, 65) \| 77 \| (8, 58) \| 82 \| NA \| NA; Category: unknown function; p164942 VI\_09747; phrogs \| 664 \| 95.6 \| 8.9e-05 \| 1.3e-08 \| 45.1 \| 40 \| (16, 55) \| 77 \| (57, 96) \| 117 \| DNA binding protein \| DNA binding protein; Category: DNA, RNA and nucleotide metabolism; MF072690\_p140; phrogs \| 1124 \| 95.5 \| 0.00011 \| 1.5e-08 \| 48.3 \| 44 \| (22, 67) \| 77 \| (8, 51) \| 182 \| transcriptional regulator \| transcriptional regulator; Category: transcription regulation; p275302 VI\_05197; phrogs \| 4583 \| 95.5 \| 0.00013 \| 1.6e-08 \| 42.7 \| 37 \| (19, 55) \| 77 \| (11, 47) \| 77 \| transcriptional repressor \| transcriptional repressor; Category: transcription regulation; NC\_031243\_p35; phrogs \| 34341 \| 95.4 \| 0.00015 \| 1.7e-08 \| 44.9 \| 35 \| (20, 54) \| 77 \| (73, 107) \| 122 \| transcriptional regulator \| transcriptional regulator; Category: transcription regulation; MF360957\_p141; phrogs \| 925 \| 95.4 \| 0.00014 \| 1.9e-08 \| 42.1 \| 43 \| (12, 54) \| 77 \| (7, 49) \| 76 \| HTH DNA binding protein \| HTH DNA binding protein; Category: DNA, RNA and nucleotide metabolism; p18585 VI\_03080; phrogs \| 66 \| 95.4 \| 0.00012 \| 1.9e-08 \| 39.7 \| 29 \| (27, 55) \| 77 \| (13, 41) \| 81 \| excisionase and transcriptional regulator \| excisionase and transcriptional regulator; Category: integration and excision; NC\_023692\_p35; phrogs \| 725 \| 95.4 \| 0.00016 \| 2e-08 \| 44.5 \| 25 \| (30, 54) \| 77 \| (28, 52) \| 106 \| CII-like regulator \| CII-like regulator; Category: transcription regulation; p72240 VI\_06449; phrogs \| 8 \| 95.2 \| 0.00016 \| 2.4e-08 \| 41.3 \| 23 \| (32, 54) \| 77 \| (1, 23) \| 95 \| transcriptional regulator \| transcriptional regulator; Category: transcription regulation; p309030 VI\_07697; phrogs \| 1668 \| 95.2 \| 0.00021 \| 2.5e-08 \| 45.3 \| 42 \| (28, 71) \| 77 \| (18, 59) \| 126 \| tail assembly protein \| tail assembly protein; Category: tail; p440314 VI\_06404; phrogs \| 38149 \| 95.2 \| 0.00024 \| 2.7e-08 \| 46.9 \| 57 \| (18, 74) \| 77 \| (8, 64) \| 184 \| NA \| NA; Category: unknown function; p54375 VI\_01307; phrogs \| 597 \| 95.0 \| 0.00028 \| 3.8e-08 \| 39.1 \| 21 \| (31, 51) \| 77 \| (4, 24) \| 63 \| DNA binding protein \| DNA binding protein; Category: DNA, RNA and nucleotide metabolism; KU160644\_p26; phrogs \| 151 \| 94.9 \| 0.0003 \| 4.4e-08 \| 40.4 \| 26 \| (29, 54) \| 77 \| (20, 45) \| 91 \| transposase \| transposase; Category: integration and excision; p268658 VI\_04551; phrogs \| 769 \| 94.8 \| 0.00039 \| 5.1e-08 \| 41.4 \| 37 \| (18, 54) \| 77 \| (1, 37) \| 90 \| transcriptional repressor \| transcriptional repressor; Category: transcription regulation; p166433 VI\_07600; phrogs \| 1381 \| 94.8 \| 0.00039 \| 5.1e-08 \| 41.8 \| 25 \| (31, 55) \| 77 \| (19, 43) \| 99 \| plasmid antitoxin with HTH domain \| plasmid antitoxin with HTH domain; Category: moron, auxiliary metabolic gene and host takeover; p79485 VI\_11488; phrogs \| 7061 \| 94.8 \| 0.00045 \| 5.5e-08 \| 40.1 \| 27 \| (28, 54) \| 77 \| (44, 70) \| 73 \| HTH DNA binding protein \| HTH DNA binding protein; Category: DNA, RNA and nucleotide metabolism; KU160668\_p38; phrogs \| 1889 \| 94.7 \| 0.00046 \| 6.1e-08 \| 46.9 \| 36 \| (19, 54) \| 77 \| (8, 45) \| 233 \| terminase small subunit \| terminase small subunit; Category: head and packaging; p363276 VI\_05842; phrogs \| 517 \| 94.7 \| 0.0005 \| 6.2e-08 \| 42.3 \| 28 \| (27, 54) \| 77 \| (76, 103) \| 107 \| HTH DNA binding protein \| HTH DNA binding protein; Category: DNA, RNA and nucleotide metabolism; MF668278\_p45; phrogs \| 955 \| 94.5 \| 0.00062 \| 7.5e-08 \| 44.9 \| 38 \| (18, 55) \| 77 \| (55, 92) \| 174 \| HTH DNA binding protein \| HTH DNA binding protein; Category: DNA, RNA and nucleotide metabolism; JN699010\_p46; phrogs \| 10045 \| 94.4 \| 0.00073 \| 8.7e-08 \| 38.1 \| 29 \| (32, 62) \| 77 \| (10, 38) \| 62 \| HTH DNA binding protein \| HTH DNA binding protein; Category: DNA, RNA and nucleotide metabolism; KU963258\_p53; phrogs \| 57 \| 94.4 \| 0.00059 \| 9e-08 \| 42.4 \| 25 \| (30, 54) \| 77 \| (3, 27) \| 173 \| terminase small subunit \| terminase small subunit; Category: head and packaging; p277333 VI\_02593; phrogs \| 18342 \| 94.4 \| 0.00078 \| 9.2e-08 \| 42.3 \| 34 \| (21, 54) \| 77 \| (21, 54) \| 123 \| transcriptional repressor \| transcriptional repressor; Category: transcription regulation; p158151 VI\_11365; phrogs \| 7344 \| 94.4 \| 0.00074 \| 9.3e-08 \| 43.9 \| 25 \| (29, 53) \| 77 \| (26, 50) \| 158 \| terminase small subunit \| terminase small subunit; Category: head and packaging; p355407 VI\_04507; phrogs \| 11187 \| 94.3 \| 0.00081 \| 9.7e-08 \| 38.0 \| 35 \| (21, 55) \| 77 \| (10, 44) \| 63 \| transcriptional regulator \| transcriptional regulator; Category: transcription regulation; p82712 VI\_00440; phrogs \| 309 \| 94.3 \| 0.00074 \| 1.1e-07 \| 42.3 \| 32 \| (22, 54) \| 77 \| (15, 46) \| 153 \| terminase small subunit \| terminase small subunit; Category: head and packaging; p301521 VI\_00938; phrogs \| 2907 \| 94.2 \| 0.00086 \| 1.1e-07 \| 40.9 \| 28 \| (26, 53) \| 77 \| (4, 31) \| 103 \| excisionase \| excisionase; Category: integration and excision; p256170 VI\_09967; phrogs \| 6620 \| 94.2 \| 0.00093 \| 1.1e-07 \| 43.9 \| 25 \| (31, 55) \| 77 \| (5, 29) \| 170 \| transposase \| transposase; Category: integration and excision; p336225 VI\_06314 |
| Top keywords  (threshold 1.00e-03 (evalue)) | **transcription, regulation, DNA, transcriptional, and, regulator, binding, RNA, nucleotide, metabolism** |
| Output files | ../../domain\_architecture/52\_FANPEZAQ\_CDS\_0052\_cath.hhr ../../domain\_architecture/52\_FANPEZAQ\_CDS\_0052\_merged.svg ../../domain\_architecture/52\_FANPEZAQ\_CDS\_0052\_ncbi-cd.hhr ../../domain\_architecture/52\_FANPEZAQ\_CDS\_0052\_pfam.hhr ../../domain\_architecture/52\_FANPEZAQ\_CDS\_0052\_phrogs.hhr |

### Identical protein sequences/structures

#### Search results

|  |  |
| --- | --- |
| Protein sequence databases searched | Pdb, Swissprot, Refseq |
| Identical proteins found | -- |
| Top keywords | -- |
| Output files | -- |

### Similar protein sequences/structures

#### Sequence similarity search results (HHblits)^1^

|  |  |
| --- | --- |
| Sequence databases searched | Uniclust, Pdb70 |
| Results, scheme(s)  (Top layers only, threshold 1.00e-03 (evalue)) | xml version="1.0" encoding="utf-8" standalone="no"?       2024-09-02T21:08:51.590618 image/svg+xml   Matplotlib v3.7.2, https://matplotlib.org/ |
| Results, table(s)  (threshold 1.00e-03 (evalue)) | db \| id \| prob \| evalue \| pvalue \| score \| cols \| query \| query\_len \| template \| template\_len \| name \| description; uniclust \| UniRef100\_A0A060M6C1 \| 99.6 \| 1e-18 \| 2.4e-24 \| 106.8 \| 71 \| (1, 72) \| 77 \| (1, 71) \| 113 \| Cryptic phage CTXphi transcriptional repressor rstR \| Cryptic phage CTXphi transcriptional repressor rstR; uniclust \| UniRef100\_A0A0C1PUG7 \| 99.6 \| 1.2e-18 \| 2.7e-24 \| 110.2 \| 61 \| (11, 72) \| 77 \| (3, 63) \| 143 \| Transcriptional regulator, XRE family \| Transcriptional regulator, XRE family; uniclust \| UniRef100\_A0A329KDZ8 \| 99.5 \| 1.5e-17 \| 3.1e-23 \| 95.1 \| 56 \| (16, 72) \| 77 \| (1, 56) \| 78 \| XRE family transcriptional regulator \| XRE family transcriptional regulator; uniclust \| UniRef100\_A0A125W4T6 \| 99.5 \| 2.3e-17 \| 5e-23 \| 97.0 \| 60 \| (12, 72) \| 77 \| (10, 69) \| 91 \| DNA-binding helix-turn-helix protein \| DNA-binding helix-turn-helix protein; uniclust \| UniRef100\_A0A074JBZ5 \| 99.5 \| 2.6e-17 \| 5.7e-23 \| 108.1 \| 61 \| (11, 72) \| 77 \| (2, 62) \| 188 \| Cro/Cl family transcriptional regulator \| Cro/Cl family transcriptional regulator; uniclust \| UniRef100\_A0A0U5NTU9 \| 99.5 \| 3e-17 \| 6.3e-23 \| 106.6 \| 60 \| (14, 74) \| 77 \| (22, 81) \| 177 \| HTH-type transcriptional regulator ImmR \| HTH-type transcriptional regulator ImmR; uniclust \| UniRef100\_A0A023CY80 \| 99.5 \| 3.1e-17 \| 7.5e-23 \| 106.6 \| 70 \| (2, 72) \| 77 \| (13, 82) \| 169 \| HTH cro/C1-type domain-containing protein \| HTH cro/C1-type domain-containing protein; uniclust \| UniRef100\_A0A0H5SX81 \| 99.5 \| 4.5e-17 \| 9.5e-23 \| 98.3 \| 59 \| (13, 72) \| 77 \| (21, 79) \| 107 \| HTH cro/C1-type domain-containing protein \| HTH cro/C1-type domain-containing protein; uniclust \| UniRef100\_A0A1C6GRS4 \| 99.5 \| 4.5e-17 \| 9.7e-23 \| 103.3 \| 64 \| (8, 72) \| 77 \| (21, 84) \| 149 \| Helix-turn-helix \| Helix-turn-helix; uniclust \| UniRef100\_A0A0B3W1R4 \| 99.5 \| 4.6e-17 \| 1.1e-22 \| 102.7 \| 64 \| (8, 72) \| 77 \| (4, 67) \| 138 \| Repressor \| Repressor; uniclust \| UniRef100\_A0A0F0C6Y2 \| 99.5 \| 4.7e-17 \| 1.1e-22 \| 103.4 \| 61 \| (11, 72) \| 77 \| (18, 78) \| 146 \| HTH-type transcriptional regulator Xre \| HTH-type transcriptional regulator Xre; uniclust \| UniRef100\_A0A0D6XQB1 \| 99.5 \| 5.9e-17 \| 1.2e-22 \| 99.5 \| 58 \| (14, 72) \| 77 \| (1, 58) \| 120 \| DNA-binding protein \| DNA-binding protein; uniclust \| UniRef100\_A0A0A2SKT0 \| 99.5 \| 6.6e-17 \| 1.5e-22 \| 103.7 \| 67 \| (5, 72) \| 77 \| (5, 71) \| 154 \| Transcriptional regulator \| Transcriptional regulator; uniclust \| UniRef100\_A0A173SJ57 \| 99.5 \| 7.3e-17 \| 1.7e-22 \| 104.2 \| 62 \| (11, 73) \| 77 \| (4, 65) \| 163 \| Anaerobic benzoate catabolism transcriptional regulator \| Anaerobic benzoate catabolism transcriptional regulator; uniclust \| UniRef100\_A0A173URG9 \| 99.5 \| 7.4e-17 \| 1.7e-22 \| 97.3 \| 61 \| (11, 72) \| 77 \| (12, 72) \| 102 \| Helix-turn-helix \| Helix-turn-helix; uniclust \| UniRef100\_A0A031WE04 \| 99.5 \| 6.9e-17 \| 1.7e-22 \| 105.7 \| 59 \| (13, 72) \| 77 \| (11, 69) \| 177 \| Transcriptional regulator, HTH-type \| Transcriptional regulator, HTH-type; uniclust \| UniRef100\_A0A0C3LWL5 \| 99.5 \| 7.3e-17 \| 1.7e-22 \| 105.5 \| 70 \| (2, 72) \| 77 \| (4, 81) \| 176 \| HTH cro/C1-type domain-containing protein \| HTH cro/C1-type domain-containing protein; uniclust \| UniRef100\_A0A087N614 \| 99.5 \| 7.2e-17 \| 1.7e-22 \| 104.0 \| 62 \| (10, 72) \| 77 \| (22, 83) \| 158 \| XRE family transcriptional regulator \| XRE family transcriptional regulator; uniclust \| UniRef100\_A0A075KGJ7 \| 99.5 \| 8.4e-17 \| 1.8e-22 \| 96.3 \| 57 \| (15, 72) \| 77 \| (6, 62) \| 100 \| Transcriptional regulator, XRE family \| Transcriptional regulator, XRE family; uniclust \| UniRef100\_A0A173TQ57 \| 99.5 \| 1.1e-16 \| 2.3e-22 \| 93.1 \| 58 \| (15, 73) \| 77 \| (1, 58) \| 86 \| Helix-turn-helix \| Helix-turn-helix; uniclust \| UniRef100\_A0A062XL10 \| 99.5 \| 1e-16 \| 2.4e-22 \| 102.8 \| 64 \| (8, 72) \| 77 \| (13, 76) \| 154 \| Transcriptional regulator, xre family \| Transcriptional regulator, xre family; uniclust \| UniRef100\_A0A1D2YT19 \| 99.5 \| 1.2e-16 \| 2.6e-22 \| 104.1 \| 61 \| (11, 72) \| 77 \| (29, 89) \| 177 \| HTH cro/C1-type domain-containing protein \| HTH cro/C1-type domain-containing protein; uniclust \| UniRef100\_A0A061PBD1 \| 99.5 \| 1.1e-16 \| 2.7e-22 \| 104.8 \| 58 \| (14, 72) \| 77 \| (21, 78) \| 178 \| Transcriptional regulator \| Transcriptional regulator; uniclust \| UniRef100\_A0A0A2HHH8 \| 99.5 \| 1.2e-16 \| 2.7e-22 \| 101.6 \| 64 \| (8, 72) \| 77 \| (2, 65) \| 145 \| DNA-binding protein \| DNA-binding protein; uniclust \| UniRef100\_A0A075JR98 \| 99.5 \| 1.2e-16 \| 2.9e-22 \| 100.7 \| 60 \| (12, 72) \| 77 \| (4, 63) \| 135 \| XRE family transcriptional regulator \| XRE family transcriptional regulator; uniclust \| UniRef100\_A0A099I4W4 \| 99.5 \| 1.3e-16 \| 2.9e-22 \| 108.3 \| 61 \| (11, 72) \| 77 \| (18, 78) \| 242 \| DNA-binding protein \| DNA-binding protein; uniclust \| UniRef100\_A0A0A8WJW3 \| 99.5 \| 1.4e-16 \| 3.3e-22 \| 101.6 \| 57 \| (15, 72) \| 77 \| (13, 69) \| 148 \| Putative repressor \| Putative repressor; uniclust \| UniRef100\_A0A090HZD4 \| 99.5 \| 1.8e-16 \| 3.7e-22 \| 84.8 \| 43 \| (12, 54) \| 77 \| (5, 47) \| 54 \| Helix-turn-helix XRE-family like proteins \| Helix-turn-helix XRE-family like proteins; uniclust \| UniRef100\_A0A173T283 \| 99.5 \| 1.8e-16 \| 3.9e-22 \| 104.2 \| 62 \| (10, 72) \| 77 \| (23, 84) \| 190 \| Transcriptional repressor DicA \| Transcriptional repressor DicA; uniclust \| UniRef100\_A0A089I2J7 \| 99.5 \| 2.1e-16 \| 4.5e-22 \| 108.7 \| 60 \| (12, 72) \| 77 \| (13, 72) \| 282 \| HTH cro/C1-type domain-containing protein \| HTH cro/C1-type domain-containing protein; uniclust \| UniRef100\_A0A086YZY1 \| 99.5 \| 2e-16 \| 4.5e-22 \| 92.5 \| 62 \| (10, 72) \| 77 \| (3, 64) \| 85 \| HTH cro/C1-type domain-containing protein \| HTH cro/C1-type domain-containing protein; uniclust \| UniRef100\_A0A090IVJ4 \| 99.4 \| 2e-16 \| 4.7e-22 \| 103.3 \| 63 \| (9, 72) \| 77 \| (24, 86) \| 173 \| HTH cro/C1-type domain-containing protein \| HTH cro/C1-type domain-containing protein; uniclust \| UniRef100\_A0A150BTM0 \| 99.4 \| 2.1e-16 \| 4.9e-22 \| 100.8 \| 61 \| (11, 72) \| 77 \| (5, 65) \| 147 \| Helix-turn-helix domain-containing protein \| Helix-turn-helix domain-containing protein; uniclust \| UniRef100\_A0A059N6C6 \| 99.4 \| 2.4e-16 \| 5.3e-22 \| 94.8 \| 64 \| (8, 72) \| 77 \| (5, 68) \| 100 \| DNA-binding helix-turn-helix protein \| DNA-binding helix-turn-helix protein; uniclust \| UniRef100\_A0A061BX60 \| 99.4 \| 2.6e-16 \| 5.5e-22 \| 97.1 \| 57 \| (15, 72) \| 77 \| (2, 58) \| 124 \| Helix-turn-helix transcriptional regulator \| Helix-turn-helix transcriptional regulator; uniclust \| UniRef100\_A0A0R1F3Q5 \| 99.4 \| 2.6e-16 \| 5.5e-22 \| 100.2 \| 60 \| (12, 72) \| 77 \| (3, 62) \| 150 \| Transcriptional regulator \| Transcriptional regulator; uniclust \| UniRef100\_A0A074LR75 \| 99.4 \| 2.5e-16 \| 5.6e-22 \| 101.6 \| 66 \| (6, 72) \| 77 \| (6, 71) \| 161 \| HTH cro/C1-type domain-containing protein \| HTH cro/C1-type domain-containing protein; uniclust \| UniRef100\_A0A0R2G103 \| 99.4 \| 2.7e-16 \| 5.6e-22 \| 91.1 \| 59 \| (15, 74) \| 77 \| (10, 68) \| 84 \| HTH cro/C1-type domain-containing protein \| HTH cro/C1-type domain-containing protein; uniclust \| UniRef100\_A0A0R2GF18 \| 99.4 \| 2.6e-16 \| 6e-22 \| 101.7 \| 60 \| (12, 72) \| 77 \| (24, 83) \| 161 \| HTH cro/C1-type domain-containing protein \| HTH cro/C1-type domain-containing protein; uniclust \| UniRef100\_A0A0M2NH40 \| 99.4 \| 2.9e-16 \| 6.2e-22 \| 104.0 \| 62 \| (10, 72) \| 77 \| (4, 65) \| 203 \| Transcriptional regulator, XRE family \| Transcriptional regulator, XRE family; uniclust \| UniRef100\_A0A0A0M2F1 \| 99.4 \| 3e-16 \| 6.2e-22 \| 100.0 \| 65 \| (7, 72) \| 77 \| (15, 79) \| 153 \| HTH cro/C1-type domain-containing protein \| HTH cro/C1-type domain-containing protein; uniclust \| UniRef100\_A0A096AZ92 \| 99.4 \| 3.7e-16 \| 7.8e-22 \| 93.0 \| 58 \| (14, 72) \| 77 \| (10, 67) \| 96 \| HTH cro/C1-type domain-containing protein \| HTH cro/C1-type domain-containing protein; uniclust \| UniRef100\_A0A073KEL8 \| 99.4 \| 3.6e-16 \| 8.2e-22 \| 102.7 \| 61 \| (11, 72) \| 77 \| (25, 85) \| 183 \| XRE family transcriptional regulator \| XRE family transcriptional regulator; uniclust \| UniRef100\_A0A0C2YFC1 \| 99.4 \| 4.1e-16 \| 9.3e-22 \| 97.4 \| 57 \| (15, 72) \| 77 \| (2, 58) \| 127 \| Transcriptional regulator, XRE family \| Transcriptional regulator, XRE family; uniclust \| UniRef100\_A0A011VZ04 \| 99.4 \| 4e-16 \| 9.5e-22 \| 106.0 \| 66 \| (7, 73) \| 77 \| (2, 67) \| 234 \| DNA-binding protein \| DNA-binding protein; uniclust \| UniRef100\_A0A0H1RPT5 \| 99.4 \| 5e-16 \| 1.1e-21 \| 91.6 \| 65 \| (8, 73) \| 77 \| (7, 71) \| 91 \| Transcriptional regulator, y4mF family \| Transcriptional regulator, y4mF family; uniclust \| UniRef100\_A0A1C6J0G0 \| 99.4 \| 5e-16 \| 1.1e-21 \| 106.8 \| 61 \| (14, 75) \| 77 \| (15, 75) \| 270 \| Transcriptional repressor DicA \| Transcriptional repressor DicA; uniclust \| UniRef100\_A0A074IXR9 \| 99.4 \| 4.8e-16 \| 1.1e-21 \| 104.8 \| 60 \| (12, 72) \| 77 \| (4, 63) \| 220 \| HTH cro/C1-type domain-containing protein \| HTH cro/C1-type domain-containing protein; uniclust \| UniRef100\_A0A0C1W593 \| 99.4 \| 5.2e-16 \| 1.1e-21 \| 105.0 \| 65 \| (8, 73) \| 77 \| (5, 69) \| 233 \| Phage repressor protein \| Phage repressor protein; uniclust \| UniRef100\_A0A0E1KYC3 \| 99.4 \| 5.4e-16 \| 1.2e-21 \| 101.6 \| 66 \| (6, 72) \| 77 \| (2, 67) \| 180 \| Helix-turn-helix family protein \| Helix-turn-helix family protein; uniclust \| UniRef100\_A0A0D0QRS3 \| 99.4 \| 5.6e-16 \| 1.2e-21 \| 97.4 \| 61 \| (11, 72) \| 77 \| (7, 67) \| 137 \| MerR family transcriptional regulator \| MerR family transcriptional regulator; uniclust \| UniRef100\_A0A023BMV4 \| 99.4 \| 5.4e-16 \| 1.2e-21 \| 100.8 \| 61 \| (11, 72) \| 77 \| (26, 86) \| 169 \| HTH cro/C1-type domain-containing protein \| HTH cro/C1-type domain-containing protein; uniclust \| UniRef100\_A0A0A8WJE4 \| 99.4 \| 5.3e-16 \| 1.2e-21 \| 100.1 \| 59 \| (13, 72) \| 77 \| (8, 66) \| 158 \| DNA-binding protein Putative repressor RepR \| DNA-binding protein Putative repressor RepR; uniclust \| UniRef100\_A0A143ZRN3 \| 99.4 \| 5.9e-16 \| 1.2e-21 \| 93.4 \| 63 \| (9, 72) \| 77 \| (8, 72) \| 105 \| HTH cro/C1-type domain-containing protein \| HTH cro/C1-type domain-containing protein; uniclust \| UniRef100\_A0A0C7NN68 \| 99.4 \| 6.3e-16 \| 1.3e-21 \| 101.1 \| 65 \| (7, 72) \| 77 \| (26, 90) \| 187 \| Uncharacterized \| Uncharacterized; uniclust \| UniRef100\_A0A0V8QIR2 \| 99.4 \| 6.1e-16 \| 1.4e-21 \| 102.0 \| 60 \| (12, 72) \| 77 \| (16, 75) \| 188 \| HTH cro/C1-type domain-containing protein \| HTH cro/C1-type domain-containing protein; uniclust \| UniRef100\_A0A0C7G2Y4 \| 99.4 \| 8.2e-16 \| 1.7e-21 \| 99.5 \| 57 \| (15, 72) \| 77 \| (5, 61) \| 172 \| Transcriptional regulator \| Transcriptional regulator; uniclust \| UniRef100\_A0A059NZ33 \| 99.4 \| 7.8e-16 \| 1.8e-21 \| 100.6 \| 63 \| (9, 72) \| 77 \| (15, 77) \| 174 \| HTH-type transcriptional regulator ImmR \| HTH-type transcriptional regulator ImmR; uniclust \| UniRef100\_A0A088T2X6 \| 99.4 \| 8.5e-16 \| 1.9e-21 \| 102.9 \| 58 \| (14, 72) \| 77 \| (4, 61) \| 217 \| HTH-type transcriptional regulator ImmR \| HTH-type transcriptional regulator ImmR; uniclust \| UniRef100\_A0A084J9J4 \| 99.4 \| 9.5e-16 \| 1.9e-21 \| 98.5 \| 58 \| (14, 72) \| 77 \| (1, 58) \| 170 \| Transcriptional regulator \| Transcriptional regulator; uniclust \| UniRef100\_A0A062X6M5 \| 99.4 \| 8.9e-16 \| 2e-21 \| 100.1 \| 60 \| (12, 72) \| 77 \| (27, 86) \| 172 \| Transcriptional regulator, y4mF family \| Transcriptional regulator, y4mF family; uniclust \| UniRef100\_A0A354UQ85 \| 99.4 \| 1e-15 \| 2e-21 \| 86.5 \| 59 \| (14, 73) \| 77 \| (11, 69) \| 76 \| Transcriptional regulator \| Transcriptional regulator; uniclust \| UniRef100\_A0A0R1HYU3 \| 99.4 \| 9.3e-16 \| 2.1e-21 \| 102.8 \| 59 \| (13, 72) \| 77 \| (31, 89) \| 215 \| HTH cro/C1-type domain-containing protein \| HTH cro/C1-type domain-containing protein; uniclust \| UniRef100\_A0A0V8QD66 \| 99.4 \| 1.1e-15 \| 2.2e-21 \| 83.3 \| 55 \| (13, 68) \| 77 \| (6, 60) \| 63 \| Transcriptional regulator \| Transcriptional regulator; uniclust \| UniRef100\_A0A0K9GGF8 \| 99.4 \| 1e-15 \| 2.2e-21 \| 96.4 \| 59 \| (14, 73) \| 77 \| (13, 71) \| 139 \| HTH cro/C1-type domain-containing protein \| HTH cro/C1-type domain-containing protein; uniclust \| UniRef100\_A0A0R1I3I0 \| 99.4 \| 1.1e-15 \| 2.2e-21 \| 87.5 \| 55 \| (16, 71) \| 77 \| (1, 55) \| 76 \| HTH cro/C1-type domain-containing protein \| HTH cro/C1-type domain-containing protein; uniclust \| UniRef100\_A0A0E3ZTG7 \| 99.4 \| 1.1e-15 \| 2.3e-21 \| 90.1 \| 63 \| (10, 73) \| 77 \| (7, 69) \| 89 \| XRE family transcriptional regulator \| XRE family transcriptional regulator; uniclust \| UniRef100\_A0A0E2BX49 \| 99.4 \| 1.1e-15 \| 2.3e-21 \| 90.2 \| 58 \| (14, 72) \| 77 \| (13, 70) \| 89 \| HTH cro/C1-type domain-containing protein \| HTH cro/C1-type domain-containing protein; uniclust \| UniRef100\_A0A090ZIW3 \| 99.4 \| 1.1e-15 \| 2.5e-21 \| 97.1 \| 65 \| (8, 73) \| 77 \| (5, 69) \| 142 \| Helix-turn-helix domain-containing protein \| Helix-turn-helix domain-containing protein; uniclust \| UniRef100\_A0A095X2P6 \| 99.4 \| 1.2e-15 \| 2.5e-21 \| 102.4 \| 63 \| (8, 71) \| 77 \| (36, 98) \| 223 \| XRE family transcriptional regulator \| XRE family transcriptional regulator; uniclust \| UniRef100\_A0A0R1KZW4 \| 99.4 \| 1.2e-15 \| 2.6e-21 \| 100.0 \| 67 \| (5, 72) \| 77 \| (16, 82) \| 190 \| Helix-turn-helix domain-containing protein \| Helix-turn-helix domain-containing protein; uniclust \| UniRef100\_A0A024QFL1 \| 99.4 \| 1.2e-15 \| 2.6e-21 \| 98.7 \| 65 \| (8, 73) \| 77 \| (9, 73) \| 163 \| Helix-turn-helix protein \| Helix-turn-helix protein; uniclust \| UniRef100\_A0A075SB65 \| 99.4 \| 1.5e-15 \| 3e-21 \| 97.4 \| 60 \| (11, 71) \| 77 \| (28, 87) \| 169 \| HTH cro/C1-type domain-containing protein \| HTH cro/C1-type domain-containing protein; uniclust \| UniRef100\_A0A2K9P5D5 \| 99.4 \| 1.6e-15 \| 3.2e-21 \| 88.2 \| 58 \| (14, 72) \| 77 \| (4, 61) \| 84 \| Helix-turn-helix domain protein \| Helix-turn-helix domain protein; uniclust \| UniRef100\_A0A0F6CKR7 \| 99.4 \| 1.6e-15 \| 3.3e-21 \| 86.5 \| 58 \| (15, 73) \| 77 \| (1, 58) \| 75 \| XRE family transcriptional regulator \| XRE family transcriptional regulator; uniclust \| UniRef100\_A0A0F2Q257 \| 99.4 \| 1.6e-15 \| 3.4e-21 \| 89.0 \| 59 \| (13, 72) \| 77 \| (5, 63) \| 88 \| HTH cro/C1-type domain-containing protein \| HTH cro/C1-type domain-containing protein; uniclust \| UniRef100\_A0A0R2JC93 \| 99.4 \| 1.7e-15 \| 3.4e-21 \| 96.6 \| 58 \| (14, 72) \| 77 \| (8, 65) \| 156 \| Helix-turn-helix domain-containing protein \| Helix-turn-helix domain-containing protein; uniclust \| UniRef100\_A0A077EIM8 \| 99.4 \| 1.6e-15 \| 3.5e-21 \| 98.1 \| 59 \| (13, 72) \| 77 \| (21, 79) \| 165 \| BcrR \| BcrR; uniclust \| UniRef100\_A0A0R2F9F5 \| 99.4 \| 1.8e-15 \| 3.7e-21 \| 88.4 \| 59 \| (13, 72) \| 77 \| (3, 61) \| 85 \| HTH cro/C1-type domain-containing protein \| HTH cro/C1-type domain-containing protein; uniclust \| UniRef100\_A0A173SRG9 \| 99.4 \| 1.8e-15 \| 3.8e-21 \| 95.2 \| 66 \| (6, 72) \| 77 \| (2, 67) \| 137 \| HTH-type transcriptional regulator immR \| HTH-type transcriptional regulator immR; uniclust \| UniRef100\_A0A162KR50 \| 99.4 \| 1.9e-15 \| 3.8e-21 \| 96.1 \| 58 \| (14, 72) \| 77 \| (11, 68) \| 158 \| HTH-type transcriptional regulator ImmR \| HTH-type transcriptional regulator ImmR; uniclust \| UniRef100\_A0A062XC28 \| 99.4 \| 1.7e-15 \| 3.9e-21 \| 97.4 \| 64 \| (8, 72) \| 77 \| (10, 73) \| 152 \| Transcriptional regulator, y4mF family \| Transcriptional regulator, y4mF family; uniclust \| UniRef100\_A0A1G9NPQ0 \| 99.4 \| 1.9e-15 \| 3.9e-21 \| 89.9 \| 59 \| (13, 72) \| 77 \| (8, 66) \| 97 \| DNA-binding transcriptional regulator, XRE-family HTH domain \| DNA-binding transcriptional regulator, XRE-family HTH domain; uniclust \| UniRef100\_A0A352BGN2 \| 99.4 \| 2e-15 \| 3.9e-21 \| 86.6 \| 57 \| (15, 72) \| 77 \| (2, 58) \| 80 \| HTH cro/C1-type domain-containing protein \| HTH cro/C1-type domain-containing protein; uniclust \| UniRef100\_A0A068EMF1 \| 99.4 \| 1.9e-15 \| 4.2e-21 \| 96.3 \| 63 \| (9, 72) \| 77 \| (8, 70) \| 147 \| Transcriptional regulator \| Transcriptional regulator; uniclust \| UniRef100\_A0A0R2D4N1 \| 99.4 \| 1.9e-15 \| 4.2e-21 \| 94.3 \| 62 \| (11, 73) \| 77 \| (3, 64) \| 124 \| HTH cro/C1-type domain-containing protein \| HTH cro/C1-type domain-containing protein; uniclust \| UniRef100\_A0A0A1SME0 \| 99.4 \| 2e-15 \| 4.2e-21 \| 97.0 \| 55 \| (17, 72) \| 77 \| (1, 55) \| 159 \| DNA-binding protein \| DNA-binding protein; uniclust \| UniRef100\_A0A088GHG5 \| 99.4 \| 2e-15 \| 4.3e-21 \| 88.9 \| 57 \| (16, 73) \| 77 \| (3, 59) \| 90 \| Transcriptional regulator, Cro/CI family protein \| Transcriptional regulator, Cro/CI family protein; uniclust \| UniRef100\_A0A098ELK1 \| 99.4 \| 1.8e-15 \| 4.3e-21 \| 104.1 \| 64 \| (8, 72) \| 77 \| (31, 94) \| 255 \| LexA repressor \| LexA repressor; uniclust \| UniRef100\_A0A063Y955 \| 99.4 \| 2.1e-15 \| 4.5e-21 \| 89.2 \| 62 \| (11, 73) \| 77 \| (6, 67) \| 90 \| HTH cro/C1-type domain-containing protein \| HTH cro/C1-type domain-containing protein; uniclust \| UniRef100\_A0A242B388 \| 99.4 \| 2.2e-15 \| 4.5e-21 \| 91.3 \| 56 \| (16, 72) \| 77 \| (3, 58) \| 111 \| HTH cro/C1-type domain-containing protein \| HTH cro/C1-type domain-containing protein; uniclust \| UniRef100\_A0A073IPP6 \| 99.4 \| 2e-15 \| 4.5e-21 \| 103.1 \| 58 \| (14, 72) \| 77 \| (17, 74) \| 252 \| HTH cro/C1-type domain-containing protein \| HTH cro/C1-type domain-containing protein; uniclust \| UniRef100\_A0A1C6GXN4 \| 99.4 \| 2.2e-15 \| 4.5e-21 \| 90.8 \| 58 \| (14, 72) \| 77 \| (6, 63) \| 106 \| HTH-type transcriptional regulator immR \| HTH-type transcriptional regulator immR; uniclust \| UniRef100\_A0A0C7N6M6 \| 99.4 \| 2.1e-15 \| 4.6e-21 \| 89.8 \| 62 \| (10, 72) \| 77 \| (3, 64) \| 93 \| Cro/C1-type helix-turn-helix domain \| Cro/C1-type helix-turn-helix domain; uniclust \| UniRef100\_A0A0R1K8U9 \| 99.4 \| 2.1e-15 \| 4.6e-21 \| 97.6 \| 59 \| (13, 72) \| 77 \| (29, 87) \| 166 \| HTH cro/C1-type domain-containing protein \| HTH cro/C1-type domain-containing protein; uniclust \| UniRef100\_A0A0R1Q4I5 \| 99.4 \| 2.2e-15 \| 5.1e-21 \| 94.8 \| 57 \| (15, 72) \| 77 \| (1, 57) \| 132 \| HTH cro/C1-type domain-containing protein \| HTH cro/C1-type domain-containing protein; uniclust \| UniRef100\_A0A0R1XFR0 \| 99.4 \| 2.5e-15 \| 5.1e-21 \| 96.8 \| 57 \| (15, 72) \| 77 \| (15, 71) \| 177 \| HTH cro/C1-type domain-containing protein \| HTH cro/C1-type domain-containing protein; uniclust \| UniRef100\_A0A0D8IC64 \| 99.4 \| 2.3e-15 \| 5.2e-21 \| 99.6 \| 63 \| (9, 72) \| 77 \| (26, 88) \| 188 \| Putative phage repressor \| Putative phage repressor; uniclust \| UniRef100\_A0A098F0D7 \| 99.4 \| 2.3e-15 \| 5.4e-21 \| 99.7 \| 65 \| (7, 72) \| 77 \| (18, 82) \| 189 \| SPBc2 prophage-derived uncharacterized HTH-type transcriptional regulator yonR \| SPBc2 prophage-derived uncharacterized HTH-type transcriptional regulator yonR; uniclust \| UniRef100\_A0A098APU9 \| 99.4 \| 2.4e-15 \| 5.5e-21 \| 100.6 \| 57 \| (15, 72) \| 77 \| (4, 60) \| 205 \| Helix-turn-helix domain-containing protein \| Helix-turn-helix domain-containing protein; uniclust \| UniRef100\_A0A3D4S5S0 \| 99.4 \| 2.7e-15 \| 5.5e-21 \| 86.5 \| 55 \| (16, 71) \| 77 \| (2, 56) \| 80 \| XRE family transcriptional regulator \| XRE family transcriptional regulator; uniclust \| UniRef100\_A0A1M5U3U8 \| 99.4 \| 2.7e-15 \| 5.6e-21 \| 86.6 \| 56 \| (16, 72) \| 77 \| (3, 58) \| 80 \| Helix-turn-helix \| Helix-turn-helix; uniclust \| UniRef100\_A0A068A5Q8 \| 99.4 \| 2.6e-15 \| 5.6e-21 \| 90.1 \| 66 \| (7, 73) \| 77 \| (6, 71) \| 98 \| Helix-turn-helix domain-containing protein \| Helix-turn-helix domain-containing protein; uniclust \| UniRef100\_A0A081PXA1 \| 99.4 \| 2.6e-15 \| 5.6e-21 \| 98.5 \| 56 \| (16, 72) \| 77 \| (2, 57) \| 185 \| Helix-turn-helix family protein \| Helix-turn-helix family protein; uniclust \| UniRef100\_A0A1B3WME1 \| 99.4 \| 2.8e-15 \| 5.8e-21 \| 87.3 \| 55 \| (17, 72) \| 77 \| (6, 60) \| 83 \| HTH cro/C1-type domain-containing protein \| HTH cro/C1-type domain-containing protein; uniclust \| UniRef100\_A0A173YT44 \| 99.3 \| 3e-15 \| 6.1e-21 \| 91.1 \| 56 \| (16, 72) \| 77 \| (5, 60) \| 115 \| HTH-type transcriptional regulator immR \| HTH-type transcriptional regulator immR; uniclust \| UniRef100\_A0A099Y982 \| 99.3 \| 3.1e-15 \| 6.6e-21 \| 88.5 \| 53 \| (19, 72) \| 77 \| (8, 60) \| 91 \| HTH cro/C1-type domain-containing protein \| HTH cro/C1-type domain-containing protein; uniclust \| UniRef100\_A0A1G6DIB8 \| 99.3 \| 3.1e-15 \| 6.6e-21 \| 92.9 \| 60 \| (15, 75) \| 77 \| (5, 64) \| 125 \| DNA-binding transcriptional regulator, XRE-family HTH domain \| DNA-binding transcriptional regulator, XRE-family HTH domain; uniclust \| UniRef100\_A0A1Y4S908 \| 99.3 \| 3.3e-15 \| 6.8e-21 \| 95.9 \| 58 \| (13, 71) \| 77 \| (5, 62) \| 161 \| HTH cro/C1-type domain-containing protein \| HTH cro/C1-type domain-containing protein; uniclust \| UniRef100\_A0A1G8HQ68 \| 99.3 \| 3.2e-15 \| 6.8e-21 \| 100.0 \| 61 \| (11, 72) \| 77 \| (20, 80) \| 220 \| Helix-turn-helix \| Helix-turn-helix; uniclust \| UniRef100\_A0A198AE04 \| 99.3 \| 3.5e-15 \| 7.5e-21 \| 93.2 \| 58 \| (14, 72) \| 77 \| (1, 58) \| 129 \| HTH cro/C1-type domain-containing protein \| HTH cro/C1-type domain-containing protein; uniclust \| UniRef100\_A0A416ZRR7 \| 99.3 \| 3.8e-15 \| 7.6e-21 \| 86.8 \| 56 \| (16, 72) \| 77 \| (3, 58) \| 88 \| XRE family transcriptional regulator \| XRE family transcriptional regulator; uniclust \| UniRef100\_A0A0R1RXY9 \| 99.3 \| 3.7e-15 \| 7.7e-21 \| 94.5 \| 58 \| (14, 72) \| 77 \| (1, 58) \| 147 \| Transcriptional regulator, XRE family \| Transcriptional regulator, XRE family; uniclust \| UniRef100\_A0A0H2UUS6 \| 99.3 \| 3.8e-15 \| 7.7e-21 \| 91.3 \| 53 \| (19, 72) \| 77 \| (3, 55) \| 119 \| HTH cro/C1-type domain-containing protein \| HTH cro/C1-type domain-containing protein; uniclust \| UniRef100\_A0A1G7QJ77 \| 99.3 \| 3.7e-15 \| 7.7e-21 \| 92.2 \| 55 \| (17, 72) \| 77 \| (1, 55) \| 125 \| DNA-binding transcriptional regulator, XRE-family HTH domain \| DNA-binding transcriptional regulator, XRE-family HTH domain; uniclust \| UniRef100\_A0A8S5MSF4 \| 99.3 \| 3.9e-15 \| 7.7e-21 \| 81.3 \| 56 \| (17, 73) \| 77 \| (1, 56) \| 62 \| Helix-turn-helix domain protein \| Helix-turn-helix domain protein; uniclust \| UniRef100\_A0A1A5Y9X4 \| 99.3 \| 3.7e-15 \| 8.2e-21 \| 93.8 \| 57 \| (16, 73) \| 77 \| (1, 57) \| 133 \| HTH cro/C1-type domain-containing protein \| HTH cro/C1-type domain-containing protein; uniclust \| UniRef100\_A0A0A2HEB2 \| 99.3 \| 3.9e-15 \| 8.3e-21 \| 96.2 \| 59 \| (13, 72) \| 77 \| (1, 59) \| 166 \| XRE family transcriptional regulator \| XRE family transcriptional regulator; uniclust \| UniRef100\_A0A069AZ01 \| 99.3 \| 4.1e-15 \| 8.4e-21 \| 80.0 \| 41 \| (14, 54) \| 77 \| (2, 42) \| 55 \| Antitoxin PezA \| Antitoxin PezA; uniclust \| UniRef100\_A0A0B2JZ21 \| 99.3 \| 4e-15 \| 8.6e-21 \| 89.6 \| 60 \| (12, 72) \| 77 \| (14, 73) \| 102 \| HTH cro/C1-type domain-containing protein \| HTH cro/C1-type domain-containing protein; uniclust \| UniRef100\_A0A0R1H4Y4 \| 99.3 \| 4.3e-15 \| 8.8e-21 \| 92.8 \| 59 \| (13, 72) \| 77 \| (4, 62) \| 136 \| Helix-turn-helix XRE-family transcriptional regulator \| Helix-turn-helix XRE-family transcriptional regulator; uniclust \| UniRef100\_A0A1G6DJ03 \| 99.3 \| 4.2e-15 \| 8.9e-21 \| 93.2 \| 63 \| (10, 73) \| 77 \| (3, 66) \| 134 \| Helix-turn-helix domain-containing protein \| Helix-turn-helix domain-containing protein; uniclust \| UniRef100\_A0A0D0SD34 \| 99.3 \| 4.2e-15 \| 9.1e-21 \| 90.7 \| 64 \| (8, 72) \| 77 \| (10, 73) \| 110 \| Transcriptional regulator, XRE family \| Transcriptional regulator, XRE family; uniclust \| UniRef100\_A0A099I752 \| 99.3 \| 4.5e-15 \| 9.1e-21 \| 80.4 \| 44 \| (11, 54) \| 77 \| (7, 50) \| 57 \| XRE family transcriptional regulator \| XRE family transcriptional regulator; uniclust \| UniRef100\_A0A347WIN2 \| 99.3 \| 4.4e-15 \| 9.2e-21 \| 86.7 \| 58 \| (15, 73) \| 77 \| (11, 68) \| 86 \| HTH cro/C1-type domain-containing protein \| HTH cro/C1-type domain-containing protein; uniclust \| UniRef100\_A0A0D6DZ12 \| 99.3 \| 4.1e-15 \| 9.2e-21 \| 99.8 \| 57 \| (15, 72) \| 77 \| (7, 63) \| 211 \| HTH-type DNA-binding domain \| HTH-type DNA-binding domain; uniclust \| UniRef100\_A0A0R1HZP7 \| 99.3 \| 4.4e-15 \| 9.6e-21 \| 92.9 \| 60 \| (12, 72) \| 77 \| (5, 64) \| 128 \| HTH cro/C1-type domain-containing protein \| HTH cro/C1-type domain-containing protein; uniclust \| UniRef100\_A0A0C9PMQ7 \| 99.3 \| 4.3e-15 \| 9.7e-21 \| 100.9 \| 64 \| (8, 72) \| 77 \| (28, 91) \| 235 \| XRE family transcriptional regulator \| XRE family transcriptional regulator; uniclust \| UniRef100\_A0A084JB70 \| 99.3 \| 4.8e-15 \| 1e-20 \| 96.5 \| 65 \| (6, 71) \| 77 \| (10, 74) \| 178 \| DNA-binding protein \| DNA-binding protein; uniclust \| UniRef100\_A0A096DH56 \| 99.3 \| 5e-15 \| 1.1e-20 \| 104.0 \| 58 \| (14, 72) \| 77 \| (12, 69) \| 309 \| HTH cro/C1-type domain-containing protein \| HTH cro/C1-type domain-containing protein; uniclust \| UniRef100\_A0A0Q5D1Z3 \| 99.3 \| 5.5e-15 \| 1.2e-20 \| 85.1 \| 62 \| (8, 71) \| 77 \| (4, 65) \| 78 \| HTH cro/C1-type domain-containing protein \| HTH cro/C1-type domain-containing protein; uniclust \| UniRef100\_A0A069D8G9 \| 99.3 \| 5.4e-15 \| 1.2e-20 \| 91.9 \| 60 \| (13, 73) \| 77 \| (3, 62) \| 121 \| Putative transcriptional regulator \| Putative transcriptional regulator; uniclust \| UniRef100\_A0A0A3J3C4 \| 99.3 \| 5.5e-15 \| 1.2e-20 \| 94.9 \| 57 \| (15, 72) \| 77 \| (20, 76) \| 153 \| HTH cro/C1-type domain-containing protein \| HTH cro/C1-type domain-containing protein; uniclust \| UniRef100\_A0A356ZA25 \| 99.3 \| 6.2e-15 \| 1.2e-20 \| 92.0 \| 67 \| (5, 72) \| 77 \| (5, 71) \| 138 \| XRE family transcriptional regulator \| XRE family transcriptional regulator; uniclust \| UniRef100\_A0A084A723 \| 99.3 \| 5.4e-15 \| 1.2e-20 \| 96.2 \| 68 \| (3, 72) \| 77 \| (8, 75) \| 166 \| HTH-type transcriptional regulator \| HTH-type transcriptional regulator; uniclust \| UniRef100\_A0A023CJ53 \| 99.3 \| 6.3e-15 \| 1.3e-20 \| 94.6 \| 60 \| (12, 72) \| 77 \| (23, 82) \| 162 \| XRE family transcriptional regulator \| XRE family transcriptional regulator; uniclust \| UniRef100\_A0A1C5QRJ2 \| 99.3 \| 6.5e-15 \| 1.3e-20 \| 95.4 \| 63 \| (13, 76) \| 77 \| (21, 83) \| 178 \| Transcriptional repressor DicA \| Transcriptional repressor DicA; uniclust \| UniRef100\_A0A0V8DXP7 \| 99.3 \| 6.6e-15 \| 1.3e-20 \| 86.0 \| 59 \| (13, 72) \| 77 \| (11, 69) \| 87 \| HTH cro/C1-type domain-containing protein \| HTH cro/C1-type domain-containing protein; uniclust \| UniRef100\_A0A031WFF9 \| 99.3 \| 6e-15 \| 1.4e-20 \| 96.7 \| 63 \| (9, 72) \| 77 \| (25, 87) \| 175 \| DNA-binding helix-turn-helix protein \| DNA-binding helix-turn-helix protein; uniclust \| UniRef100\_A0A011UYL9 \| 99.3 \| 6.7e-15 \| 1.4e-20 \| 101.2 \| 59 \| (13, 72) \| 77 \| (43, 101) \| 280 \| DNA-binding protein \| DNA-binding protein; uniclust \| UniRef100\_A0A1C7GW64 \| 99.3 \| 7.1e-15 \| 1.4e-20 \| 89.7 \| 59 \| (15, 74) \| 77 \| (1, 59) \| 116 \| Transcriptional regulator \| Transcriptional regulator; uniclust \| UniRef100\_A0A1F8UDN2 \| 99.3 \| 6.7e-15 \| 1.5e-20 \| 95.0 \| 58 \| (14, 72) \| 77 \| (1, 58) \| 159 \| Transcriptional regulator \| Transcriptional regulator; uniclust \| UniRef100\_A0A0G3WDU7 \| 99.3 \| 7.4e-15 \| 1.5e-20 \| 82.8 \| 59 \| (13, 72) \| 77 \| (2, 60) \| 72 \| Putative transcriptional regulator \| Putative transcriptional regulator; uniclust \| UniRef100\_A0A1E9IL17 \| 99.3 \| 7.5e-15 \| 1.6e-20 \| 96.5 \| 57 \| (15, 72) \| 77 \| (13, 69) \| 190 \| Transcriptional regulator \| Transcriptional regulator; uniclust \| UniRef100\_A0A011UWQ9 \| 99.3 \| 7e-15 \| 1.6e-20 \| 88.9 \| 63 \| (10, 73) \| 77 \| (9, 71) \| 101 \| ABC transporter substrate-binding protein \| ABC transporter substrate-binding protein; uniclust \| UniRef100\_A0A024QBX0 \| 99.3 \| 7.3e-15 \| 1.6e-20 \| 91.5 \| 62 \| (11, 73) \| 77 \| (31, 92) \| 125 \| Helix-turn-helix \| Helix-turn-helix; uniclust \| UniRef100\_A0A0Z8ISK9 \| 99.3 \| 8e-15 \| 1.6e-20 \| 97.1 \| 59 \| (13, 72) \| 77 \| (5, 63) \| 207 \| XRE family transcriptional regulator \| XRE family transcriptional regulator; uniclust \| UniRef100\_A0A1C5QEW4 \| 99.3 \| 8e-15 \| 1.7e-20 \| 99.7 \| 57 \| (15, 72) \| 77 \| (28, 84) \| 248 \| HTH-type transcriptional regulator immR \| HTH-type transcriptional regulator immR; uniclust \| UniRef100\_A0A0F5R9Z3 \| 99.3 \| 7.8e-15 \| 1.7e-20 \| 91.7 \| 61 \| (11, 72) \| 77 \| (7, 67) \| 129 \| HTH cro/C1-type domain-containing protein \| HTH cro/C1-type domain-containing protein; uniclust \| UniRef100\_A0A1C6D7B9 \| 99.3 \| 8.7e-15 \| 1.7e-20 \| 75.3 \| 38 \| (17, 54) \| 77 \| (1, 38) \| 45 \| Antitoxin PezA \| Antitoxin PezA; uniclust \| UniRef100\_A0A0V8C4S1 \| 99.3 \| 7.5e-15 \| 1.7e-20 \| 96.8 \| 64 \| (8, 72) \| 77 \| (16, 79) \| 182 \| Transcriptional regulator Cro/CI family \| Transcriptional regulator Cro/CI family; uniclust \| UniRef100\_A0A927NWC2 \| 99.3 \| 8.7e-15 \| 1.7e-20 \| 91.3 \| 58 \| (14, 72) \| 77 \| (6, 63) \| 139 \| Helix-turn-helix transcriptional regulator \| Helix-turn-helix transcriptional regulator; uniclust \| UniRef100\_A0A085ZCW4 \| 99.3 \| 8.1e-15 \| 1.7e-20 \| 94.5 \| 62 \| (10, 72) \| 77 \| (25, 86) \| 161 \| HTH cro/C1-type domain-containing protein \| HTH cro/C1-type domain-containing protein; uniclust \| UniRef100\_A0A062X1K8 \| 99.3 \| 8.4e-15 \| 1.7e-20 \| 86.7 \| 54 \| (19, 73) \| 77 \| (3, 56) \| 93 \| Zinc finger/helix-turn-helix protein, YgiT family \| Zinc finger/helix-turn-helix protein, YgiT family; uniclust \| UniRef100\_A0A061K0S8 \| 99.3 \| 8.7e-15 \| 1.9e-20 \| 94.9 \| 64 \| (8, 72) \| 77 \| (12, 75) \| 165 \| Helix-turn-helix domain-containing protein \| Helix-turn-helix domain-containing protein; uniclust \| UniRef100\_A0A1M4ZB90 \| 99.3 \| 9.2e-15 \| 1.9e-20 \| 92.1 \| 58 \| (15, 73) \| 77 \| (6, 63) \| 139 \| Helix-turn-helix domain-containing protein \| Helix-turn-helix domain-containing protein; uniclust \| UniRef100\_A0A0R1MYT8 \| 99.3 \| 9.2e-15 \| 2e-20 \| 97.4 \| 57 \| (15, 72) \| 77 \| (1, 57) \| 207 \| HTH cro/C1-type domain-containing protein \| HTH cro/C1-type domain-containing protein; uniclust \| UniRef100\_A0A174CCW1 \| 99.3 \| 9.1e-15 \| 2e-20 \| 98.4 \| 58 \| (14, 72) \| 77 \| (4, 61) \| 218 \| HTH-type transcriptional regulator sinR \| HTH-type transcriptional regulator sinR; uniclust \| UniRef100\_A0A1C5UWY8 \| 99.3 \| 1e-14 \| 2e-20 \| 78.0 \| 41 \| (14, 54) \| 77 \| (4, 44) \| 53 \| Putative zinc finger/helix-turn-helix protein, YgiT family \| Putative zinc finger/helix-turn-helix protein, YgiT family; uniclust \| UniRef100\_A0A1C5P1N9 \| 99.3 \| 9.4e-15 \| 2e-20 \| 97.9 \| 60 \| (13, 73) \| 77 \| (3, 62) \| 213 \| HTH-type transcriptional regulator immR \| HTH-type transcriptional regulator immR; uniclust \| UniRef100\_A0A0C7N4G8 \| 99.3 \| 9.8e-15 \| 2e-20 \| 91.8 \| 60 \| (12, 72) \| 77 \| (2, 61) \| 140 \| Cro/C1-type helix-turn-helix domain \| Cro/C1-type helix-turn-helix domain; uniclust \| UniRef100\_A0A0F0CL80 \| 99.3 \| 9.6e-15 \| 2.1e-20 \| 105.2 \| 59 \| (13, 72) \| 77 \| (33, 91) \| 396 \| Transcriptional repressor DicA \| Transcriptional repressor DicA; uniclust \| UniRef100\_A0A1Y3WG13 \| 99.3 \| 1.1e-14 \| 2.2e-20 \| 77.7 \| 44 \| (12, 55) \| 77 \| (3, 46) \| 52 \| Transcriptional regulator \| Transcriptional regulator; uniclust \| UniRef100\_A0A7C6WH00 \| 99.3 \| 1e-14 \| 2.2e-20 \| 95.1 \| 61 \| (11, 72) \| 77 \| (14, 74) \| 181 \| Helix-turn-helix transcriptional regulator \| Helix-turn-helix transcriptional regulator; uniclust \| UniRef100\_A0A2K9ADE1 \| 99.3 \| 1.1e-14 \| 2.2e-20 \| 85.6 \| 59 \| (15, 74) \| 77 \| (2, 60) \| 88 \| HTH cro/C1-type domain-containing protein \| HTH cro/C1-type domain-containing protein; uniclust \| UniRef100\_A0A1D2LGT8 \| 99.3 \| 1e-14 \| 2.2e-20 \| 90.1 \| 58 \| (14, 72) \| 77 \| (3, 60) \| 120 \| XRE family transcriptional regulator \| XRE family transcriptional regulator; uniclust \| UniRef100\_A0A086BKQ7 \| 99.3 \| 9.6e-15 \| 2.3e-20 \| 95.4 \| 60 \| (12, 72) \| 77 \| (30, 89) \| 169 \| DNA-binding protein \| DNA-binding protein; uniclust \| UniRef100\_A0A316QXQ5 \| 99.3 \| 1.1e-14 \| 2.3e-20 \| 95.2 \| 57 \| (15, 72) \| 77 \| (2, 58) \| 178 \| HTH cro/C1-type domain-containing protein \| HTH cro/C1-type domain-containing protein; uniclust \| UniRef100\_A0A845JTS9 \| 99.3 \| 1.1e-14 \| 2.3e-20 \| 90.1 \| 57 \| (15, 72) \| 77 \| (4, 60) \| 121 \| Helix-turn-helix transcriptional regulator \| Helix-turn-helix transcriptional regulator; uniclust \| UniRef100\_A0A011RNT1 \| 99.3 \| 1.1e-14 \| 2.3e-20 \| 96.9 \| 60 \| (12, 72) \| 77 \| (20, 79) \| 215 \| Transcriptional regulator, XRE family \| Transcriptional regulator, XRE family; uniclust \| UniRef100\_A0A060M4E5 \| 99.3 \| 1e-14 \| 2.3e-20 \| 91.8 \| 61 \| (11, 72) \| 77 \| (6, 66) \| 134 \| HTH-type transcriptional regulator \| HTH-type transcriptional regulator; uniclust \| UniRef100\_A0A1R0XIP9 \| 99.3 \| 1.1e-14 \| 2.3e-20 \| 94.5 \| 58 \| (16, 74) \| 77 \| (1, 58) \| 176 \| HTH cro/C1-type domain-containing protein \| HTH cro/C1-type domain-containing protein; uniclust \| UniRef100\_A0A0K1F1X2 \| 99.3 \| 1.1e-14 \| 2.3e-20 \| 99.1 \| 63 \| (10, 73) \| 77 \| (14, 76) \| 240 \| HTH cro/C1-type domain-containing protein \| HTH cro/C1-type domain-containing protein; uniclust \| UniRef100\_A0A069D9R6 \| 99.3 \| 1e-14 \| 2.4e-20 \| 95.8 \| 63 \| (10, 72) \| 77 \| (20, 82) \| 177 \| HTH cro/C1-type domain-containing protein \| HTH cro/C1-type domain-containing protein; uniclust \| UniRef100\_A0A0M2NJ60 \| 99.3 \| 1.1e-14 \| 2.4e-20 \| 89.5 \| 56 \| (16, 72) \| 77 \| (2, 57) \| 117 \| HTH cro/C1-type domain-containing protein \| HTH cro/C1-type domain-containing protein; uniclust \| UniRef100\_A0A351SRF1 \| 99.3 \| 1.2e-14 \| 2.4e-20 \| 98.4 \| 63 \| (9, 72) \| 77 \| (23, 85) \| 243 \| HTH cro/C1-type domain-containing protein \| HTH cro/C1-type domain-containing protein; uniclust \| UniRef100\_A0A149UEU3 \| 99.3 \| 1.1e-14 \| 2.4e-20 \| 91.5 \| 62 \| (10, 72) \| 77 \| (15, 76) \| 132 \| HTH cro/C1-type domain-containing protein \| HTH cro/C1-type domain-containing protein; uniclust \| UniRef100\_A0A0D8J1P7 \| 99.3 \| 1.2e-14 \| 2.4e-20 \| 96.9 \| 60 \| (12, 72) \| 77 \| (5, 64) \| 219 \| Helix-turn-helix domain-containing protein \| Helix-turn-helix domain-containing protein; uniclust \| UniRef100\_A0A0E1Y5F5 \| 99.3 \| 1.2e-14 \| 2.5e-20 \| 91.0 \| 55 \| (18, 73) \| 77 \| (3, 57) \| 130 \| DNA-binding protein (Fragment) \| DNA-binding protein (Fragment); uniclust \| UniRef100\_A0A0H4P7Y6 \| 99.3 \| 1.2e-14 \| 2.5e-20 \| 87.7 \| 60 \| (12, 72) \| 77 \| (5, 64) \| 101 \| Prophage LambdaBa02repressor protein \| Prophage LambdaBa02repressor protein; uniclust \| UniRef100\_A0A099WFI4 \| 99.3 \| 1.3e-14 \| 2.6e-20 \| 91.4 \| 62 \| (10, 72) \| 77 \| (12, 73) \| 143 \| DNA-binding protein \| DNA-binding protein; uniclust \| UniRef100\_A0A0H5Q6F9 \| 99.3 \| 1.3e-14 \| 2.7e-20 \| 92.4 \| 57 \| (15, 72) \| 77 \| (8, 64) \| 150 \| HTH cro/C1-type domain-containing protein \| HTH cro/C1-type domain-containing protein; uniclust \| UniRef100\_A0A015LSZ1 \| 99.3 \| 1.2e-14 \| 2.7e-20 \| 93.3 \| 56 \| (16, 72) \| 77 \| (16, 71) \| 151 \| XRE family transcriptional regulator \| XRE family transcriptional regulator; uniclust \| UniRef100\_A0A1B1YAB2 \| 99.3 \| 1.3e-14 \| 2.8e-20 \| 91.8 \| 60 \| (13, 73) \| 77 \| (3, 62) \| 145 \| HTH cro/C1-type domain-containing protein \| HTH cro/C1-type domain-containing protein; uniclust \| UniRef100\_A0A023CPN5 \| 99.3 \| 1.3e-14 \| 2.9e-20 \| 90.4 \| 60 \| (13, 73) \| 77 \| (2, 61) \| 121 \| HTH-type transcriptional regulator SlrR \| HTH-type transcriptional regulator SlrR; uniclust \| UniRef100\_A0A1M6XC60 \| 99.3 \| 1.4e-14 \| 2.9e-20 \| 93.5 \| 57 \| (15, 72) \| 77 \| (10, 66) \| 165 \| Helix-turn-helix \| Helix-turn-helix; uniclust \| UniRef100\_A0A084JI99 \| 99.3 \| 1.4e-14 \| 2.9e-20 \| 93.1 \| 63 \| (9, 72) \| 77 \| (7, 69) \| 158 \| HTH cro/C1-type domain-containing protein \| HTH cro/C1-type domain-containing protein; uniclust \| UniRef100\_A0A095XNQ4 \| 99.3 \| 1.4e-14 \| 3e-20 \| 92.9 \| 57 \| (15, 72) \| 77 \| (2, 58) \| 153 \| HTH cro/C1-type domain-containing protein \| HTH cro/C1-type domain-containing protein; uniclust \| UniRef100\_A0A0F8X4V8 \| 99.3 \| 1.4e-14 \| 3e-20 \| 80.7 \| 56 \| (18, 74) \| 77 \| (4, 59) \| 64 \| HTH cro/C1-type domain-containing protein \| HTH cro/C1-type domain-containing protein; uniclust \| UniRef100\_A0A081P4C3 \| 99.3 \| 1.4e-14 \| 3e-20 \| 92.0 \| 58 \| (14, 72) \| 77 \| (2, 59) \| 139 \| HTH cro/C1-type domain-containing protein \| HTH cro/C1-type domain-containing protein; uniclust \| UniRef100\_A0A024P3C3 \| 99.3 \| 1.5e-14 \| 3.1e-20 \| 91.5 \| 57 \| (15, 72) \| 77 \| (9, 65) \| 142 \| HTH-type transcriptional regulator ImmR \| HTH-type transcriptional regulator ImmR; uniclust \| UniRef100\_A0A0J1F8E5 \| 99.3 \| 1.5e-14 \| 3.2e-20 \| 100.5 \| 62 \| (10, 72) \| 77 \| (12, 73) \| 291 \| HTH-type transcriptional regulator Xre \| HTH-type transcriptional regulator Xre; uniclust \| UniRef100\_A0A084ACG0 \| 99.3 \| 1.4e-14 \| 3.3e-20 \| 98.2 \| 62 \| (10, 72) \| 77 \| (11, 72) \| 224 \| HTH cro/C1-type domain-containing protein \| HTH cro/C1-type domain-containing protein; uniclust \| UniRef100\_A0A1W2ASU0 \| 99.3 \| 1.5e-14 \| 3.3e-20 \| 95.6 \| 55 \| (17, 72) \| 77 \| (1, 55) \| 193 \| Transcriptional regulator, contains XRE-family HTH domain \| Transcriptional regulator, contains XRE-family HTH domain; uniclust \| UniRef100\_A0A069DDP2 \| 99.3 \| 1.6e-14 \| 3.3e-20 \| 96.2 \| 58 \| (14, 72) \| 77 \| (55, 112) \| 214 \| Transcriptional regulator \| Transcriptional regulator; uniclust \| UniRef100\_A0A174HLJ4 \| 99.3 \| 1.5e-14 \| 3.4e-20 \| 94.7 \| 55 \| (17, 72) \| 77 \| (1, 55) \| 172 \| MerR family transcriptional regulator \| MerR family transcriptional regulator; uniclust \| UniRef100\_A0A150LHC3 \| 99.3 \| 1.6e-14 \| 3.4e-20 \| 90.4 \| 61 \| (11, 72) \| 77 \| (5, 65) \| 128 \| HTH cro/C1-type domain-containing protein \| HTH cro/C1-type domain-containing protein; uniclust \| UniRef100\_A0A084JPR8 \| 99.3 \| 1.6e-14 \| 3.4e-20 \| 97.9 \| 60 \| (11, 71) \| 77 \| (14, 73) \| 233 \| XRE family transcriptional regulator \| XRE family transcriptional regulator; uniclust \| UniRef100\_A0A0V8DXH0 \| 99.3 \| 1.9e-14 \| 3.8e-20 \| 81.7 \| 57 \| (15, 72) \| 77 \| (4, 60) \| 75 \| Transcriptional regulator XRE family \| Transcriptional regulator XRE family; uniclust \| UniRef100\_A0A143HDQ9 \| 99.3 \| 1.8e-14 \| 4e-20 \| 96.4 \| 58 \| (14, 72) \| 77 \| (20, 77) \| 209 \| HTH cro/C1-type domain-containing protein \| HTH cro/C1-type domain-containing protein; uniclust \| UniRef100\_A0A1Q9YGQ7 \| 99.3 \| 1.9e-14 \| 4.2e-20 \| 90.6 \| 61 \| (13, 73) \| 77 \| (6, 66) \| 134 \| HTH cro/C1-type domain-containing protein \| HTH cro/C1-type domain-containing protein; uniclust \| UniRef100\_A0A1C5MER2 \| 99.3 \| 2e-14 \| 4.2e-20 \| 89.2 \| 60 \| (11, 71) \| 77 \| (4, 64) \| 123 \| Helix-turn-helix \| Helix-turn-helix; uniclust \| UniRef100\_A0A0U2MC24 \| 99.3 \| 1.9e-14 \| 4.4e-20 \| 93.5 \| 63 \| (11, 74) \| 77 \| (5, 67) \| 161 \| HTH domain-containing protein \| HTH domain-containing protein; uniclust \| UniRef100\_A0A3D0XZX5 \| 99.3 \| 2.3e-14 \| 4.5e-20 \| 80.0 \| 54 \| (17, 71) \| 77 \| (1, 54) \| 70 \| Transcriptional regulator (Fragment) \| Transcriptional regulator (Fragment); uniclust \| UniRef100\_A0A0J5PB61 \| 99.3 \| 2.2e-14 \| 4.5e-20 \| 95.7 \| 57 \| (14, 71) \| 77 \| (2, 58) \| 211 \| HTH-type transcriptional regulator Xre \| HTH-type transcriptional regulator Xre; uniclust \| UniRef100\_A0A1D2M0A2 \| 99.3 \| 2.1e-14 \| 4.7e-20 \| 97.8 \| 59 \| (13, 72) \| 77 \| (4, 62) \| 236 \| XRE family transcriptional regulator \| XRE family transcriptional regulator; uniclust \| UniRef100\_A0A024QIL8 \| 99.3 \| 2.2e-14 \| 4.8e-20 \| 92.0 \| 61 \| (11, 72) \| 77 \| (6, 66) \| 152 \| Antitoxin HipB \| Antitoxin HipB; uniclust \| UniRef100\_A0A031WFK4 \| 99.3 \| 2.2e-14 \| 4.8e-20 \| 92.0 \| 63 \| (9, 72) \| 77 \| (5, 67) \| 151 \| Helix-turn-helix domain protein \| Helix-turn-helix domain protein; uniclust \| UniRef100\_A0A081NRH7 \| 99.3 \| 2.1e-14 \| 4.9e-20 \| 99.3 \| 61 \| (10, 71) \| 77 \| (23, 83) \| 262 \| XRE family transcriptional regulator \| XRE family transcriptional regulator; uniclust \| UniRef100\_A0A095X9F7 \| 99.3 \| 2.4e-14 \| 5e-20 \| 98.9 \| 64 \| (8, 72) \| 77 \| (33, 96) \| 285 \| DNA-binding protein \| DNA-binding protein; uniclust \| UniRef100\_A0A179ESE4 \| 99.3 \| 2.4e-14 \| 5e-20 \| 92.8 \| 58 \| (14, 72) \| 77 \| (5, 62) \| 170 \| XRE family transcriptional regulator \| XRE family transcriptional regulator; uniclust \| UniRef100\_A0A075RC30 \| 99.3 \| 2.5e-14 \| 5.1e-20 \| 89.7 \| 62 \| (12, 74) \| 77 \| (2, 63) \| 134 \| Transcriptional repressor DicA \| Transcriptional repressor DicA; uniclust \| UniRef100\_A0A075WVE9 \| 99.3 \| 2.3e-14 \| 5.1e-20 \| 98.0 \| 57 \| (15, 72) \| 77 \| (33, 89) \| 244 \| HTH cro/C1-type domain-containing protein \| HTH cro/C1-type domain-containing protein; uniclust \| UniRef100\_A0A139QKZ0 \| 99.3 \| 2.7e-14 \| 5.4e-20 \| 81.5 \| 56 \| (16, 72) \| 77 \| (5, 60) \| 76 \| HTH cro/C1-type domain-containing protein \| HTH cro/C1-type domain-containing protein; uniclust \| UniRef100\_A0A1L7GTF0 \| 99.2 \| 2.6e-14 \| 5.4e-20 \| 88.8 \| 57 \| (15, 72) \| 77 \| (2, 58) \| 125 \| Transcriptional regulator \| Transcriptional regulator; uniclust \| UniRef100\_A0A1Y4LJ54 \| 99.2 \| 2.6e-14 \| 5.4e-20 \| 93.0 \| 55 \| (17, 72) \| 77 \| (1, 55) \| 172 \| HTH cro/C1-type domain-containing protein \| HTH cro/C1-type domain-containing protein; uniclust \| UniRef100\_A0A084JDN7 \| 99.2 \| 2.7e-14 \| 5.4e-20 \| 91.8 \| 57 \| (14, 71) \| 77 \| (11, 67) \| 170 \| DNA-binding protein \| DNA-binding protein; uniclust \| UniRef100\_A0A0Q4FSA8 \| 99.2 \| 2.5e-14 \| 5.4e-20 \| 82.7 \| 56 \| (16, 72) \| 77 \| (2, 57) \| 78 \| HTH cro/C1-type domain-containing protein \| HTH cro/C1-type domain-containing protein; uniclust \| UniRef100\_A0A0F9I898 \| 99.2 \| 2.7e-14 \| 5.6e-20 \| 83.1 \| 63 \| (8, 71) \| 77 \| (5, 67) \| 83 \| HTH cro/C1-type domain-containing protein \| HTH cro/C1-type domain-containing protein; uniclust \| UniRef100\_A0A140L514 \| 99.2 \| 2.7e-14 \| 5.6e-20 \| 87.6 \| 59 \| (13, 72) \| 77 \| (1, 59) \| 117 \| HTH-type transcriptional regulator ImmR \| HTH-type transcriptional regulator ImmR; uniclust \| UniRef100\_A0A1E3A252 \| 99.2 \| 2.5e-14 \| 5.6e-20 \| 90.1 \| 60 \| (12, 72) \| 77 \| (12, 71) \| 133 \| Transcriptional repressor DicA \| Transcriptional repressor DicA; uniclust \| UniRef100\_A0A174NGD7 \| 99.2 \| 2.8e-14 \| 5.7e-20 \| 75.7 \| 39 \| (17, 55) \| 77 \| (1, 39) \| 51 \| Transcriptional repressor DicA \| Transcriptional repressor DicA; uniclust \| UniRef100\_A0A134AB19 \| 99.2 \| 2.8e-14 \| 5.7e-20 \| 89.6 \| 68 \| (4, 72) \| 77 \| (4, 71) \| 137 \| DNA-binding helix-turn-helix protein \| DNA-binding helix-turn-helix protein; uniclust \| UniRef100\_A0A024QB89 \| 99.2 \| 2.7e-14 \| 5.9e-20 \| 83.8 \| 57 \| (16, 73) \| 77 \| (2, 58) \| 85 \| Anaerobic benzoate catabolism transcriptional regulator \| Anaerobic benzoate catabolism transcriptional regulator; uniclust \| UniRef100\_A0A098N141 \| 99.2 \| 2.7e-14 \| 5.9e-20 \| 91.3 \| 59 \| (13, 72) \| 77 \| (18, 76) \| 148 \| DNA-binding protein \| DNA-binding protein; uniclust \| UniRef100\_A0A0A7FXJ6 \| 99.2 \| 3e-14 \| 6.1e-20 \| 90.7 \| 61 \| (11, 72) \| 77 \| (16, 76) \| 154 \| Helix-turn-helix family protein \| Helix-turn-helix family protein; uniclust \| UniRef100\_A0A061PBI6 \| 99.2 \| 2.9e-14 \| 6.2e-20 \| 84.7 \| 56 \| (17, 73) \| 77 \| (10, 65) \| 91 \| HTH cro/C1-type domain-containing protein \| HTH cro/C1-type domain-containing protein; uniclust \| UniRef100\_A0A090Y2E7 \| 99.2 \| 2.9e-14 \| 6.4e-20 \| 90.8 \| 59 \| (13, 72) \| 77 \| (3, 61) \| 145 \| Helix-turn-helix family protein \| Helix-turn-helix family protein; uniclust \| UniRef100\_A0A0R1ZQ19 \| 99.2 \| 2.7e-14 \| 6.4e-20 \| 97.4 \| 62 \| (10, 72) \| 77 \| (3, 64) \| 235 \| Prophage lsa1 XRE family DNA-binding protein \| Prophage lsa1 XRE family DNA-binding protein; uniclust \| UniRef100\_A0A364K8X3 \| 99.2 \| 3.2e-14 \| 6.6e-20 \| 93.0 \| 56 \| (16, 72) \| 77 \| (1, 56) \| 184 \| Transcriptional regulator \| Transcriptional regulator; uniclust \| UniRef100\_A0A0P9FPS6 \| 99.2 \| 3.1e-14 \| 6.7e-20 \| 90.7 \| 59 \| (13, 72) \| 77 \| (7, 65) \| 147 \| HTH cro/C1-type domain-containing protein \| HTH cro/C1-type domain-containing protein; uniclust \| UniRef100\_A0A0H5SHR8 \| 99.2 \| 3.3e-14 \| 6.9e-20 \| 85.8 \| 60 \| (12, 72) \| 77 \| (16, 75) \| 102 \| HTH cro/C1-type domain-containing protein \| HTH cro/C1-type domain-containing protein; uniclust \| UniRef100\_A0A024QF78 \| 99.2 \| 3.5e-14 \| 7e-20 \| 92.1 \| 59 \| (13, 72) \| 77 \| (24, 82) \| 183 \| Helix-turn-helix protein \| Helix-turn-helix protein; uniclust \| UniRef100\_A0A072MV23 \| 99.2 \| 3.2e-14 \| 7e-20 \| 90.7 \| 63 \| (10, 73) \| 77 \| (5, 67) \| 146 \| XRE family transcriptional regulator \| XRE family transcriptional regulator; uniclust \| UniRef100\_A0A1W1Y945 \| 99.2 \| 3.1e-14 \| 7e-20 \| 89.1 \| 60 \| (12, 72) \| 77 \| (4, 63) \| 125 \| Transcriptional regulator, contains XRE-family HTH domain \| Transcriptional regulator, contains XRE-family HTH domain; uniclust \| UniRef100\_A0A096ALY3 \| 99.2 \| 3e-14 \| 7e-20 \| 97.8 \| 60 \| (12, 72) \| 77 \| (27, 86) \| 248 \| HTH cro/C1-type domain-containing protein \| HTH cro/C1-type domain-containing protein; uniclust \| UniRef100\_A0A0H5SUC8 \| 99.2 \| 3.3e-14 \| 7e-20 \| 88.2 \| 56 \| (17, 73) \| 77 \| (3, 58) \| 122 \| HTH cro/C1-type domain-containing protein \| HTH cro/C1-type domain-containing protein; uniclust \| UniRef100\_A0A2S7FD02 \| 99.2 \| 3.3e-14 \| 7.3e-20 \| 85.5 \| 58 \| (14, 72) \| 77 \| (13, 70) \| 98 \| HTH cro/C1-type domain-containing protein \| HTH cro/C1-type domain-containing protein; uniclust \| UniRef100\_A0A1C5M5K7 \| 99.2 \| 3.5e-14 \| 7.3e-20 \| 95.2 \| 55 \| (17, 72) \| 77 \| (1, 55) \| 218 \| Transcriptional repressor DicA \| Transcriptional repressor DicA; uniclust \| UniRef100\_A0A010NY17 \| 99.2 \| 3.5e-14 \| 7.5e-20 \| 85.2 \| 63 \| (11, 74) \| 77 \| (7, 69) \| 98 \| DNA-binding protein \| DNA-binding protein; uniclust \| UniRef100\_A0A0H5Q551 \| 99.2 \| 3.6e-14 \| 7.7e-20 \| 97.1 \| 58 \| (14, 72) \| 77 \| (31, 88) \| 254 \| HTH cro/C1-type domain-containing protein \| HTH cro/C1-type domain-containing protein; uniclust \| UniRef100\_A0A0D8I6J6 \| 99.2 \| 3.5e-14 \| 8e-20 \| 89.8 \| 59 \| (15, 73) \| 77 \| (1, 59) \| 134 \| Putative phage repressor \| Putative phage repressor; uniclust \| UniRef100\_A0A1V4GWD0 \| 99.2 \| 3.5e-14 \| 8.2e-20 \| 97.2 \| 65 \| (9, 74) \| 77 \| (13, 77) \| 241 \| HTH cro/C1-type domain-containing protein \| HTH cro/C1-type domain-containing protein; uniclust \| UniRef100\_A0A0S2W8V1 \| 99.2 \| 4e-14 \| 8.2e-20 \| 89.4 \| 58 \| (14, 72) \| 77 \| (14, 71) \| 144 \| HTH-type transcriptional regulator xre \| HTH-type transcriptional regulator xre; uniclust \| UniRef100\_A0A0S2W0A6 \| 99.2 \| 3.9e-14 \| 8.3e-20 \| 95.3 \| 56 \| (16, 72) \| 77 \| (1, 56) \| 222 \| Transcriptional regulator, XRE family \| Transcriptional regulator, XRE family; uniclust \| UniRef100\_A0A0R1EZ08 \| 99.2 \| 4e-14 \| 8.4e-20 \| 94.2 \| 55 \| (16, 71) \| 77 \| (1, 55) \| 208 \| XRE family transcriptional regulator \| XRE family transcriptional regulator; uniclust \| UniRef100\_A0A323UEM4 \| 99.2 \| 4.1e-14 \| 8.5e-20 \| 86.4 \| 59 \| (13, 72) \| 77 \| (24, 82) \| 113 \| XRE family transcriptional regulator \| XRE family transcriptional regulator; uniclust \| UniRef100\_A0A1W1IH47 \| 99.2 \| 4.2e-14 \| 8.5e-20 \| 84.1 \| 55 \| (16, 71) \| 77 \| (1, 55) \| 96 \| HTH cro/C1-type domain-containing protein \| HTH cro/C1-type domain-containing protein; uniclust \| UniRef100\_A0A0A0IG66 \| 99.2 \| 3.9e-14 \| 8.5e-20 \| 89.2 \| 64 \| (8, 72) \| 77 \| (8, 71) \| 132 \| HTH cro/C1-type domain-containing protein \| HTH cro/C1-type domain-containing protein; uniclust \| UniRef100\_A0A1S9BXS8 \| 99.2 \| 4.4e-14 \| 8.6e-20 \| 81.5 \| 56 \| (15, 71) \| 77 \| (2, 57) \| 85 \| HTH cro/C1-type domain-containing protein \| HTH cro/C1-type domain-containing protein; uniclust \| UniRef100\_A0A124IPR8 \| 99.2 \| 4e-14 \| 8.8e-20 \| 88.6 \| 55 \| (17, 72) \| 77 \| (1, 55) \| 127 \| HTH cro/C1-type domain-containing protein \| HTH cro/C1-type domain-containing protein; uniclust \| UniRef100\_A0A174NJ84 \| 99.2 \| 4.5e-14 \| 9.3e-20 \| 91.4 \| 58 \| (14, 72) \| 77 \| (10, 67) \| 169 \| HTH-type transcriptional regulator immR \| HTH-type transcriptional regulator immR; uniclust \| UniRef100\_A0A0C2PWR1 \| 99.2 \| 4.4e-14 \| 9.3e-20 \| 85.3 \| 69 \| (5, 74) \| 77 \| (5, 73) \| 103 \| XRE family transcriptional regulator \| XRE family transcriptional regulator; uniclust \| UniRef100\_A0A099I8V5 \| 99.2 \| 4.2e-14 \| 9.4e-20 \| 99.1 \| 58 \| (13, 72) \| 77 \| (1, 58) \| 298 \| HTH cro/C1-type domain-containing protein \| HTH cro/C1-type domain-containing protein; uniclust \| UniRef100\_A0A0K9N8I6 \| 99.2 \| 4.8e-14 \| 9.6e-20 \| 81.0 \| 58 \| (14, 72) \| 77 \| (2, 59) \| 80 \| Toxin-antitoxin system, antitoxin component, Xre family \| Toxin-antitoxin system, antitoxin component, Xre family; uniclust \| UniRef100\_A0A0S2W086 \| 99.2 \| 4.6e-14 \| 9.6e-20 \| 87.0 \| 58 \| (14, 72) \| 77 \| (1, 58) \| 118 \| Transcriptional regulator, XRE family \| Transcriptional regulator, XRE family; uniclust \| UniRef100\_A0A096B2S2 \| 99.2 \| 4.6e-14 \| 9.8e-20 \| 89.3 \| 65 \| (6, 72) \| 77 \| (2, 66) \| 140 \| HTH cro/C1-type domain-containing protein \| HTH cro/C1-type domain-containing protein; uniclust \| UniRef100\_A0A174DE52 \| 99.2 \| 4.5e-14 \| 9.8e-20 \| 95.5 \| 62 \| (10, 72) \| 77 \| (6, 67) \| 224 \| Antitoxin HipB \| Antitoxin HipB; uniclust \| UniRef100\_A0A1R0Z6W7 \| 99.2 \| 5e-14 \| 1e-19 \| 87.3 \| 59 \| (14, 73) \| 77 \| (1, 59) \| 124 \| HTH cro/C1-type domain-containing protein \| HTH cro/C1-type domain-containing protein; uniclust \| UniRef100\_A0A0K8JHW7 \| 99.2 \| 4.7e-14 \| 1e-19 \| 89.3 \| 60 \| (12, 72) \| 77 \| (4, 63) \| 138 \| HTH cro/C1-type domain-containing protein \| HTH cro/C1-type domain-containing protein; uniclust \| UniRef100\_A0A101ETY0 \| 99.2 \| 5e-14 \| 1e-19 \| 86.5 \| 57 \| (16, 73) \| 77 \| (5, 61) \| 117 \| HTH cro/C1-type domain-containing protein \| HTH cro/C1-type domain-containing protein; uniclust \| UniRef100\_A0A354UQS3 \| 99.2 \| 5.4e-14 \| 1.1e-19 \| 92.5 \| 58 \| (15, 73) \| 77 \| (28, 85) \| 204 \| HTH cro/C1-type domain-containing protein \| HTH cro/C1-type domain-containing protein; uniclust \| UniRef100\_A0A1X7I7X0 \| 99.2 \| 4.8e-14 \| 1.1e-19 \| 89.0 \| 58 \| (14, 72) \| 77 \| (3, 60) \| 133 \| DNA-binding transcriptional regulator, XRE-family HTH domain \| DNA-binding transcriptional regulator, XRE-family HTH domain; uniclust \| UniRef100\_A0A022N9X9 \| 99.2 \| 4.7e-14 \| 1.1e-19 \| 98.2 \| 66 \| (7, 73) \| 77 \| (28, 93) \| 275 \| Peptidase S24 \| Peptidase S24; uniclust \| UniRef100\_A0A4U9R0I1 \| 99.2 \| 5.8e-14 \| 1.1e-19 \| 74.0 \| 46 \| (17, 63) \| 77 \| (1, 46) \| 52 \| XRE family transcriptional regulator \| XRE family transcriptional regulator; uniclust \| UniRef100\_A0A0R1KF97 \| 99.2 \| 5.3e-14 \| 1.1e-19 \| 88.6 \| 59 \| (13, 72) \| 77 \| (12, 70) \| 134 \| Phage-related Cro CI family transcription regulator \| Phage-related Cro CI family transcription regulator; uniclust \| UniRef100\_A0A1J1CTL4 \| 99.2 \| 5.8e-14 \| 1.1e-19 \| 79.7 \| 54 \| (17, 71) \| 77 \| (1, 54) \| 78 \| Helix-turn-helix family protein \| Helix-turn-helix family protein; uniclust \| UniRef100\_A0A0B5QNQ3 \| 99.2 \| 5.5e-14 \| 1.2e-19 \| 88.0 \| 65 \| (12, 76) \| 77 \| (3, 67) \| 130 \| Helix-turn-helix domain protein \| Helix-turn-helix domain protein; uniclust \| UniRef100\_A0A1C6HMP3 \| 99.2 \| 5.6e-14 \| 1.2e-19 \| 87.4 \| 59 \| (13, 72) \| 77 \| (20, 78) \| 123 \| Transcriptional regulator, y4mF family \| Transcriptional regulator, y4mF family; uniclust \| UniRef100\_A0A011WSF5 \| 99.2 \| 5.9e-14 \| 1.2e-19 \| 93.8 \| 59 \| (13, 72) \| 77 \| (46, 104) \| 214 \| AbrB family transcriptional regulator \| AbrB family transcriptional regulator; uniclust \| UniRef100\_A0A084A7B7 \| 99.2 \| 5.8e-14 \| 1.2e-19 \| 94.7 \| 58 \| (14, 72) \| 77 \| (20, 77) \| 225 \| HTH cro/C1-type domain-containing protein \| HTH cro/C1-type domain-containing protein; uniclust \| UniRef100\_A0A161S4G1 \| 99.2 \| 5.9e-14 \| 1.2e-19 \| 80.5 \| 56 \| (16, 72) \| 77 \| (7, 62) \| 74 \| DNA-binding protein \| DNA-binding protein; uniclust \| UniRef100\_A0A023UJ95 \| 99.2 \| 5.7e-14 \| 1.3e-19 \| 91.8 \| 60 \| (12, 72) \| 77 \| (18, 77) \| 171 \| Repressor-like protein \| Repressor-like protein; uniclust \| UniRef100\_A0A168PAQ4 \| 99.2 \| 5.6e-14 \| 1.3e-19 \| 89.5 \| 67 \| (5, 72) \| 77 \| (9, 77) \| 141 \| HTH cro/C1-type domain-containing protein \| HTH cro/C1-type domain-containing protein; uniclust \| UniRef100\_A0A1Y3SKS7 \| 99.2 \| 5.7e-14 \| 1.3e-19 \| 90.6 \| 56 \| (16, 72) \| 77 \| (4, 59) \| 155 \| HTH cro/C1-type domain-containing protein \| HTH cro/C1-type domain-containing protein; uniclust \| UniRef100\_A0A094XY50 \| 99.2 \| 5.7e-14 \| 1.3e-19 \| 97.5 \| 57 \| (14, 71) \| 77 \| (46, 102) \| 270 \| XRE family transcriptional regulator \| XRE family transcriptional regulator; uniclust \| UniRef100\_A0A1C7G621 \| 99.2 \| 6.4e-14 \| 1.3e-19 \| 89.1 \| 55 \| (17, 72) \| 77 \| (1, 55) \| 149 \| Transcriptional regulator \| Transcriptional regulator; uniclust \| UniRef100\_A0A7W2AJ58 \| 99.2 \| 6.3e-14 \| 1.3e-19 \| 90.9 \| 58 \| (14, 72) \| 77 \| (1, 58) \| 168 \| Helix-turn-helix domain-containing protein \| Helix-turn-helix domain-containing protein; uniclust \| UniRef100\_A0A0E2UAE2 \| 99.2 \| 6.5e-14 \| 1.4e-19 \| 90.4 \| 60 \| (12, 72) \| 77 \| (5, 64) \| 164 \| DNA-binding helix-turn-helix protein \| DNA-binding helix-turn-helix protein; uniclust \| UniRef100\_A0A061PBZ7 \| 99.2 \| 6.6e-14 \| 1.4e-19 \| 84.2 \| 62 \| (11, 73) \| 77 \| (16, 77) \| 101 \| Transcriptional regulator, Cro/CI family \| Transcriptional regulator, Cro/CI family; uniclust \| UniRef100\_A0A125W4S8 \| 99.2 \| 6.8e-14 \| 1.4e-19 \| 87.4 \| 59 \| (13, 72) \| 77 \| (2, 60) \| 133 \| DNA-binding helix-turn-helix protein \| DNA-binding helix-turn-helix protein; uniclust \| UniRef100\_A0A5F0MF59 \| 99.2 \| 6.7e-14 \| 1.4e-19 \| 86.7 \| 61 \| (11, 72) \| 77 \| (12, 72) \| 122 \| HTH cro/C1-type domain-containing protein \| HTH cro/C1-type domain-containing protein; uniclust \| UniRef100\_A0A099I7D2 \| 99.2 \| 6.9e-14 \| 1.4e-19 \| 88.8 \| 61 \| (11, 72) \| 77 \| (18, 78) \| 146 \| HTH cro/C1-type domain-containing protein \| HTH cro/C1-type domain-containing protein; uniclust \| UniRef100\_A0A0R2CQI8 \| 99.2 \| 6.9e-14 \| 1.4e-19 \| 88.3 \| 61 \| (11, 72) \| 77 \| (9, 69) \| 139 \| HTH cro/C1-type domain-containing protein \| HTH cro/C1-type domain-containing protein; uniclust \| UniRef100\_A0A084J9P7 \| 99.2 \| 6.9e-14 \| 1.4e-19 \| 94.9 \| 58 \| (14, 72) \| 77 \| (8, 65) \| 239 \| XRE family transcriptional regulator \| XRE family transcriptional regulator; uniclust \| UniRef100\_A0A1Z4JR02 \| 99.2 \| 6.6e-14 \| 1.5e-19 \| 84.2 \| 62 \| (10, 72) \| 77 \| (6, 69) \| 97 \| Helix-turn-helix domain protein \| Helix-turn-helix domain protein; uniclust \| UniRef100\_A0A178GS71 \| 99.2 \| 6.8e-14 \| 1.5e-19 \| 89.4 \| 64 \| (8, 72) \| 77 \| (16, 79) \| 147 \| Transcriptional regulator \| Transcriptional regulator; uniclust \| UniRef100\_A0A172S040 \| 99.2 \| 7.3e-14 \| 1.5e-19 \| 88.3 \| 56 \| (16, 72) \| 77 \| (3, 58) \| 147 \| Looped-hinge helix DNA binding domain-containing protein, AbrB family \| Looped-hinge helix DNA binding domain-containing protein, AbrB family; uniclust \| UniRef100\_A0A0F2RS12 \| 99.2 \| 7.1e-14 \| 1.5e-19 \| 83.2 \| 70 \| (4, 74) \| 77 \| (4, 73) \| 94 \| XRE family transcriptional regulator \| XRE family transcriptional regulator; uniclust \| UniRef100\_A0A0R3JXL2 \| 99.2 \| 7.1e-14 \| 1.5e-19 \| 97.9 \| 58 \| (14, 72) \| 77 \| (15, 72) \| 311 \| HTH-type transcriptional regulator Xre \| HTH-type transcriptional regulator Xre; uniclust \| UniRef100\_A0A0K9NFW5 \| 99.2 \| 6.5e-14 \| 1.5e-19 \| 92.3 \| 61 \| (11, 72) \| 77 \| (22, 82) \| 178 \| Toxin-antitoxin system, antitoxin component, Xre family \| Toxin-antitoxin system, antitoxin component, Xre family; uniclust \| UniRef100\_A0A0C3HC21 \| 99.2 \| 7.2e-14 \| 1.5e-19 \| 90.3 \| 61 \| (11, 72) \| 77 \| (10, 70) \| 163 \| Immunity protein \| Immunity protein; uniclust \| UniRef100\_A0A1C6BIG0 \| 99.2 \| 7.5e-14 \| 1.6e-19 \| 89.1 \| 55 \| (17, 72) \| 77 \| (1, 55) \| 151 \| Transcriptional repressor DicA \| Transcriptional repressor DicA; uniclust \| UniRef100\_A0A116KFN5 \| 99.2 \| 7.6e-14 \| 1.6e-19 \| 89.1 \| 55 \| (18, 73) \| 77 \| (2, 56) \| 152 \| ICESt1 APR2 Cro/CI family transcriptional regulator \| ICESt1 APR2 Cro/CI family transcriptional regulator; uniclust \| UniRef100\_A0A1I2JTK0 \| 99.2 \| 7.5e-14 \| 1.6e-19 \| 85.5 \| 57 \| (15, 72) \| 77 \| (15, 71) \| 113 \| Helix-turn-helix \| Helix-turn-helix; uniclust \| UniRef100\_A0A1Y4IDQ4 \| 99.2 \| 7.5e-14 \| 1.6e-19 \| 95.1 \| 59 \| (14, 73) \| 77 \| (5, 63) \| 245 \| HTH cro/C1-type domain-containing protein \| HTH cro/C1-type domain-containing protein; uniclust \| UniRef100\_A0A091BSN2 \| 99.2 \| 7.6e-14 \| 1.6e-19 \| 91.8 \| 54 \| (17, 71) \| 77 \| (1, 54) \| 191 \| Transcriptional regulator \| Transcriptional regulator; uniclust \| UniRef100\_A0A0R1JS02 \| 99.2 \| 7.4e-14 \| 1.6e-19 \| 98.1 \| 60 \| (11, 71) \| 77 \| (38, 97) \| 313 \| HTH cro/C1-type domain-containing protein \| HTH cro/C1-type domain-containing protein; uniclust \| UniRef100\_A0A089LK22 \| 99.2 \| 7.6e-14 \| 1.6e-19 \| 87.3 \| 57 \| (15, 72) \| 77 \| (1, 57) \| 130 \| HTH cro/C1-type domain-containing protein \| HTH cro/C1-type domain-containing protein; uniclust \| UniRef100\_A0A011VZ09 \| 99.2 \| 6.9e-14 \| 1.6e-19 \| 97.3 \| 64 \| (9, 73) \| 77 \| (46, 109) \| 272 \| LexA family transcriptional regulator \| LexA family transcriptional regulator; uniclust \| UniRef100\_A0A1Y3RF54 \| 99.2 \| 8e-14 \| 1.6e-19 \| 90.8 \| 59 \| (13, 72) \| 77 \| (14, 72) \| 179 \| Transcriptional regulator \| Transcriptional regulator; uniclust \| UniRef100\_A0A154BQ77 \| 99.2 \| 8e-14 \| 1.7e-19 \| 80.4 \| 58 \| (14, 72) \| 77 \| (1, 58) \| 78 \| HTH cro/C1-type domain-containing protein \| HTH cro/C1-type domain-containing protein; uniclust \| UniRef100\_A0A024QAL7 \| 99.2 \| 7.8e-14 \| 1.7e-19 \| 82.1 \| 60 \| (12, 72) \| 77 \| (5, 64) \| 86 \| Helix-turn-helix \| Helix-turn-helix; uniclust \| UniRef100\_A0A0H2T341 \| 99.2 \| 8e-14 \| 1.7e-19 \| 84.9 \| 68 \| (5, 73) \| 77 \| (14, 81) \| 108 \| Cro/Cl family transcriptional regulator \| Cro/Cl family transcriptional regulator; uniclust \| UniRef100\_A0A1E3A954 \| 99.2 \| 8.4e-14 \| 1.7e-19 \| 95.0 \| 66 \| (6, 72) \| 77 \| (6, 71) \| 265 \| Transcriptional repressor DicA \| Transcriptional repressor DicA; uniclust \| UniRef100\_A0A0J1J383 \| 99.2 \| 8.1e-14 \| 1.7e-19 \| 90.6 \| 57 \| (16, 73) \| 77 \| (21, 78) \| 167 \| HTH cro/C1-type domain-containing protein \| HTH cro/C1-type domain-containing protein; uniclust \| UniRef100\_A0A015KU58 \| 99.2 \| 8.9e-14 \| 1.8e-19 \| 88.9 \| 58 \| (14, 72) \| 77 \| (14, 71) \| 155 \| XRE family transcriptional regulator \| XRE family transcriptional regulator; uniclust \| UniRef100\_A0A095YLK0 \| 99.2 \| 8.7e-14 \| 1.8e-19 \| 94.0 \| 67 \| (5, 72) \| 77 \| (43, 110) \| 227 \| HTH cro/C1-type domain-containing protein \| HTH cro/C1-type domain-containing protein; uniclust \| UniRef100\_A0A174AVY8 \| 99.2 \| 8.7e-14 \| 1.9e-19 \| 99.0 \| 58 \| (14, 72) \| 77 \| (33, 90) \| 355 \| Transcription factor \| Transcription factor; uniclust \| UniRef100\_A0A1S8SA91 \| 99.2 \| 8.7e-14 \| 1.9e-19 \| 92.5 \| 58 \| (14, 72) \| 77 \| (11, 68) \| 196 \| Transcriptional repressor DicA \| Transcriptional repressor DicA; uniclust \| UniRef100\_A0A2V2EQU8 \| 99.2 \| 9.3e-14 \| 1.9e-19 \| 83.1 \| 62 \| (12, 74) \| 77 \| (15, 76) \| 97 \| HTH cro/C1-type domain-containing protein \| HTH cro/C1-type domain-containing protein; uniclust \| UniRef100\_A0A431UEA1 \| 99.2 \| 1e-13 \| 2e-19 \| 77.4 \| 56 \| (16, 72) \| 77 \| (1, 56) \| 69 \| XRE family transcriptional regulator \| XRE family transcriptional regulator; uniclust \| UniRef100\_A0A4Y9K6V4 \| 99.2 \| 9.7e-14 \| 2e-19 \| 84.3 \| 56 \| (16, 72) \| 77 \| (1, 56) \| 106 \| XRE family transcriptional regulator \| XRE family transcriptional regulator; uniclust \| UniRef100\_A0A1Y3S0A0 \| 99.2 \| 1e-13 \| 2.1e-19 \| 99.0 \| 59 \| (13, 72) \| 77 \| (17, 75) \| 393 \| HTH cro/C1-type domain-containing protein \| HTH cro/C1-type domain-containing protein; uniclust \| UniRef100\_A0A926NBJ0 \| 99.2 \| 1.1e-13 \| 2.1e-19 \| 86.0 \| 55 \| (17, 72) \| 77 \| (1, 55) \| 135 \| Helix-turn-helix transcriptional regulator \| Helix-turn-helix transcriptional regulator; uniclust \| UniRef100\_A0A1G4SNC0 \| 99.2 \| 1e-13 \| 2.1e-19 \| 94.4 \| 60 \| (13, 73) \| 77 \| (20, 79) \| 252 \| Stage 0 sporulation protein A homolog \| Stage 0 sporulation protein A homolog; uniclust \| UniRef100\_A0A091BL71 \| 99.2 \| 1e-13 \| 2.1e-19 \| 92.0 \| 56 \| (16, 72) \| 77 \| (1, 56) \| 215 \| Transcriptional regulator \| Transcriptional regulator; uniclust \| UniRef100\_A0A3A4JVA9 \| 99.2 \| 9.8e-14 \| 2.1e-19 \| 81.1 \| 59 \| (15, 74) \| 77 \| (8, 66) \| 83 \| XRE family transcriptional regulator \| XRE family transcriptional regulator; uniclust \| UniRef100\_A0A160HBR0 \| 99.2 \| 9.8e-14 \| 2.1e-19 \| 88.9 \| 60 \| (12, 72) \| 77 \| (24, 83) \| 150 \| Uncharacterized protein \| Uncharacterized protein; uniclust \| UniRef100\_A0A0A8B2Q9 \| 99.2 \| 1.1e-13 \| 2.2e-19 \| 81.8 \| 58 \| (15, 72) \| 77 \| (5, 62) \| 90 \| HTH cro/C1-type domain-containing protein \| HTH cro/C1-type domain-containing protein; uniclust \| UniRef100\_A0A174WG95 \| 99.2 \| 1.1e-13 \| 2.2e-19 \| 82.1 \| 66 \| (7, 73) \| 77 \| (10, 75) \| 93 \| Helix-turn-helix domain-containing protein \| Helix-turn-helix domain-containing protein; uniclust \| UniRef100\_A0A023PA98 \| 99.2 \| 1e-13 \| 2.2e-19 \| 98.5 \| 57 \| (14, 71) \| 77 \| (30, 86) \| 335 \| XRE family transcriptional regulator \| XRE family transcriptional regulator; uniclust \| UniRef100\_A0A098TGA3 \| 99.2 \| 1.1e-13 \| 2.3e-19 \| 84.7 \| 54 \| (18, 72) \| 77 \| (21, 76) \| 109 \| HTH cro/C1-type domain-containing protein \| HTH cro/C1-type domain-containing protein; uniclust \| UniRef100\_A0A0R2D5X0 \| 99.2 \| 1.1e-13 \| 2.3e-19 \| 81.7 \| 52 \| (20, 72) \| 77 \| (11, 62) \| 90 \| HTH cro/C1-type domain-containing protein \| HTH cro/C1-type domain-containing protein; uniclust \| UniRef100\_A0A0B3BRB6 \| 99.2 \| 1.1e-13 \| 2.3e-19 \| 88.6 \| 62 \| (10, 72) \| 77 \| (17, 78) \| 147 \| XRE family transcriptional regulator \| XRE family transcriptional regulator; uniclust \| UniRef100\_A0A1C6G8A2 \| 99.2 \| 1.1e-13 \| 2.3e-19 \| 82.9 \| 57 \| (15, 72) \| 77 \| (2, 58) \| 99 \| Plasmid maintenance system antidote protein \| Plasmid maintenance system antidote protein; uniclust \| UniRef100\_A0A2T4LBR6 \| 99.2 \| 1.3e-13 \| 2.5e-19 \| 80.2 \| 60 \| (13, 73) \| 77 \| (3, 62) \| 84 \| XRE family transcriptional regulator \| XRE family transcriptional regulator; uniclust \| UniRef100\_A0A059NYG7 \| 99.2 \| 1.2e-13 \| 2.5e-19 \| 82.8 \| 61 \| (12, 72) \| 77 \| (3, 63) \| 94 \| HTH-type transcriptional regulator SinR \| HTH-type transcriptional regulator SinR; uniclust \| UniRef100\_A0A0F2DXV9 \| 99.2 \| 1.2e-13 \| 2.6e-19 \| 92.9 \| 56 \| (16, 72) \| 77 \| (28, 83) \| 224 \| DNA-binding phage protein \| DNA-binding phage protein; uniclust \| UniRef100\_A0A095YT12 \| 99.2 \| 1.1e-13 \| 2.6e-19 \| 94.7 \| 62 \| (10, 72) \| 77 \| (14, 75) \| 236 \| HTH cro/C1-type domain-containing protein \| HTH cro/C1-type domain-containing protein; uniclust \| UniRef100\_A0A0R2FDG7 \| 99.2 \| 1.2e-13 \| 2.6e-19 \| 93.8 \| 60 \| (12, 72) \| 77 \| (18, 77) \| 244 \| Transcriptional regulator \| Transcriptional regulator; uniclust \| UniRef100\_A0A1C5XDG0 \| 99.2 \| 1.2e-13 \| 2.7e-19 \| 96.6 \| 57 \| (15, 72) \| 77 \| (4, 60) \| 296 \| HTH-type transcriptional regulator immR \| HTH-type transcriptional regulator immR; uniclust \| UniRef100\_A0A059NUZ0 \| 99.2 \| 1.3e-13 \| 2.7e-19 \| 87.7 \| 56 \| (17, 73) \| 77 \| (13, 68) \| 142 \| Transcriptional repressor DicA \| Transcriptional repressor DicA; uniclust \| UniRef100\_A0A174BT49 \| 99.2 \| 1.4e-13 \| 2.8e-19 \| 89.2 \| 57 \| (15, 72) \| 77 \| (30, 86) \| 184 \| Predicted transcriptional regulator \| Predicted transcriptional regulator; uniclust \| UniRef100\_A0A926F381 \| 99.2 \| 1.3e-13 \| 2.8e-19 \| 91.0 \| 54 \| (18, 72) \| 77 \| (1, 54) \| 187 \| Helix-turn-helix transcriptional regulator \| Helix-turn-helix transcriptional regulator; uniclust \| UniRef100\_A0A1C5P4I0 \| 99.2 \| 1.3e-13 \| 2.9e-19 \| 84.7 \| 58 \| (14, 72) \| 77 \| (1, 58) \| 111 \| HTH-type transcriptional regulator immR \| HTH-type transcriptional regulator immR; uniclust \| UniRef100\_A0A2V2GP97 \| 99.2 \| 1.3e-13 \| 2.9e-19 \| 100.3 \| 68 \| (3, 72) \| 77 \| (53, 120) \| 431 \| Uncharacterized protein \| Uncharacterized protein; uniclust \| UniRef100\_A0A098ESC9 \| 99.2 \| 1.4e-13 \| 2.9e-19 \| 82.3 \| 58 \| (14, 72) \| 77 \| (8, 65) \| 97 \| Helix-turn-helix domain protein \| Helix-turn-helix domain protein; uniclust \| UniRef100\_A0A0B7GNK0 \| 99.2 \| 1.5e-13 \| 2.9e-19 \| 89.3 \| 56 \| (15, 71) \| 77 \| (2, 57) \| 178 \| HTH-type transcriptional regulator \| HTH-type transcriptional regulator; uniclust \| UniRef100\_A0A075SKL1 \| 99.2 \| 1.5e-13 \| 3e-19 \| 91.7 \| 59 \| (12, 71) \| 77 \| (45, 103) \| 216 \| XRE family transcriptional regulator \| XRE family transcriptional regulator; uniclust \| UniRef100\_A0A1F8V9K7 \| 99.2 \| 1.4e-13 \| 3e-19 \| 98.7 \| 60 \| (12, 72) \| 77 \| (8, 67) \| 365 \| HTH cro/C1-type domain-containing protein \| HTH cro/C1-type domain-containing protein; uniclust \| UniRef100\_A0A135YRB4 \| 99.2 \| 1.5e-13 \| 3e-19 \| 85.7 \| 52 \| (20, 72) \| 77 \| (2, 53) \| 128 \| DNA-binding helix-turn-helix protein \| DNA-binding helix-turn-helix protein; uniclust \| UniRef100\_A0A098AP68 \| 99.2 \| 1.4e-13 \| 3.1e-19 \| 93.2 \| 59 \| (13, 72) \| 77 \| (20, 78) \| 230 \| Helix-turn-helix domain-containing protein \| Helix-turn-helix domain-containing protein; uniclust \| UniRef100\_A0A094J0N9 \| 99.2 \| 1.5e-13 \| 3.2e-19 \| 78.4 \| 54 \| (18, 72) \| 77 \| (5, 58) \| 73 \| Transcriptional repressor DicA \| Transcriptional repressor DicA; uniclust \| UniRef100\_A0A173SZC8 \| 99.2 \| 1.4e-13 \| 3.2e-19 \| 90.8 \| 66 \| (9, 74) \| 77 \| (16, 81) \| 183 \| Helix-turn-helix \| Helix-turn-helix; uniclust \| UniRef100\_A0A095XIG0 \| 99.2 \| 1.5e-13 \| 3.2e-19 \| 80.9 \| 62 \| (11, 73) \| 77 \| (4, 65) \| 87 \| HTH cro/C1-type domain-containing protein \| HTH cro/C1-type domain-containing protein; uniclust \| UniRef100\_A0A134A0B4 \| 99.2 \| 1.4e-13 \| 3.3e-19 \| 95.5 \| 62 \| (11, 73) \| 77 \| (9, 70) \| 267 \| Peptidase S24-like protein \| Peptidase S24-like protein; uniclust \| UniRef100\_A0A1I0R8C4 \| 99.2 \| 1.6e-13 \| 3.3e-19 \| 86.2 \| 57 \| (15, 72) \| 77 \| (1, 57) \| 132 \| Helix-turn-helix \| Helix-turn-helix; uniclust \| UniRef100\_A0A022N9C9 \| 99.2 \| 1.6e-13 \| 3.3e-19 \| 91.3 \| 56 \| (16, 72) \| 77 \| (1, 56) \| 208 \| XRE family transcriptional regulator \| XRE family transcriptional regulator; uniclust \| UniRef100\_A0A167V013 \| 99.2 \| 1.5e-13 \| 3.3e-19 \| 88.8 \| 58 \| (15, 73) \| 77 \| (16, 73) \| 159 \| Helix-turn-helix family protein \| Helix-turn-helix family protein; uniclust \| UniRef100\_A0A1C5ZA08 \| 99.2 \| 1.6e-13 \| 3.4e-19 \| 92.5 \| 63 \| (11, 73) \| 77 \| (4, 66) \| 222 \| LexA repressor \| LexA repressor; uniclust \| UniRef100\_A0A264W7S8 \| 99.2 \| 1.8e-13 \| 3.5e-19 \| 80.3 \| 57 \| (16, 73) \| 77 \| (21, 77) \| 89 \| HTH cro/C1-type domain-containing protein \| HTH cro/C1-type domain-containing protein; uniclust \| UniRef100\_A0A0W7TL75 \| 99.2 \| 1.8e-13 \| 3.6e-19 \| 75.4 \| 44 \| (11, 54) \| 77 \| (7, 50) \| 62 \| XRE family transcriptional regulator \| XRE family transcriptional regulator; uniclust \| UniRef100\_A0A0Q6VM94 \| 99.2 \| 1.7e-13 \| 3.7e-19 \| 87.3 \| 60 \| (13, 73) \| 77 \| (16, 75) \| 145 \| HTH cro/C1-type domain-containing protein \| HTH cro/C1-type domain-containing protein; uniclust \| UniRef100\_A0A173STR4 \| 99.1 \| 1.8e-13 \| 3.8e-19 \| 74.4 \| 45 \| (11, 55) \| 77 \| (4, 48) \| 56 \| Transcriptional repressor DicA \| Transcriptional repressor DicA; uniclust \| UniRef100\_A0A095XQU1 \| 99.1 \| 1.8e-13 \| 3.8e-19 \| 80.9 \| 60 \| (12, 72) \| 77 \| (3, 62) \| 89 \| HTH cro/C1-type domain-containing protein \| HTH cro/C1-type domain-containing protein; uniclust \| UniRef100\_A0A3A6HEE8 \| 99.1 \| 1.9e-13 \| 3.9e-19 \| 85.9 \| 61 \| (12, 73) \| 77 \| (12, 72) \| 136 \| XRE family transcriptional regulator \| XRE family transcriptional regulator; uniclust \| UniRef100\_A0A0A0HWB2 \| 99.1 \| 1.8e-13 \| 3.9e-19 \| 93.2 \| 65 \| (7, 72) \| 77 \| (24, 88) \| 237 \| Transcriptional regulator \| Transcriptional regulator; uniclust \| UniRef100\_A0A0E2B8K5 \| 99.1 \| 1.9e-13 \| 4e-19 \| 88.6 \| 61 \| (11, 72) \| 77 \| (29, 89) \| 161 \| DNA-binding helix-turn-helix protein \| DNA-binding helix-turn-helix protein; uniclust \| UniRef100\_A0A090HXE6 \| 99.1 \| 1.8e-13 \| 4e-19 \| 93.7 \| 62 \| (11, 73) \| 77 \| (15, 76) \| 242 \| LexA protein \| LexA protein; uniclust \| UniRef100\_A0A061CEQ4 \| 99.1 \| 2e-13 \| 4e-19 \| 86.9 \| 58 \| (14, 72) \| 77 \| (2, 59) \| 157 \| XRE family transcriptional regulator \| XRE family transcriptional regulator; uniclust \| UniRef100\_A0A084JP97 \| 99.1 \| 1.8e-13 \| 4.1e-19 \| 99.8 \| 61 \| (11, 72) \| 77 \| (55, 115) \| 427 \| HTH cro/C1-type domain-containing protein \| HTH cro/C1-type domain-containing protein; uniclust \| UniRef100\_A0A0L6Z9N2 \| 99.1 \| 2e-13 \| 4.2e-19 \| 91.4 \| 58 \| (14, 72) \| 77 \| (1, 58) \| 220 \| HTH cro/C1-type domain-containing protein \| HTH cro/C1-type domain-containing protein; uniclust \| UniRef100\_A0A064C1R1 \| 99.1 \| 2.1e-13 \| 4.2e-19 \| 83.4 \| 57 \| (15, 72) \| 77 \| (15, 71) \| 113 \| Helix-turn-helix transcriptional regulator \| Helix-turn-helix transcriptional regulator; uniclust \| UniRef100\_A0A0F9HIX3 \| 99.1 \| 2e-13 \| 4.3e-19 \| 80.6 \| 58 \| (15, 73) \| 77 \| (2, 59) \| 86 \| HTH cro/C1-type domain-containing protein \| HTH cro/C1-type domain-containing protein; uniclust \| UniRef100\_A0A1W1XSD5 \| 99.1 \| 2.1e-13 \| 4.4e-19 \| 85.9 \| 57 \| (15, 72) \| 77 \| (1, 57) \| 133 \| DNA-binding transcriptional regulator, XRE-family HTH domain \| DNA-binding transcriptional regulator, XRE-family HTH domain; uniclust \| UniRef100\_A0A011VY25 \| 99.1 \| 2.2e-13 \| 4.4e-19 \| 86.1 \| 54 \| (17, 71) \| 77 \| (1, 54) \| 144 \| DNA-binding protein \| DNA-binding protein; uniclust \| UniRef100\_A0A1M6I385 \| 99.1 \| 2.1e-13 \| 4.5e-19 \| 90.5 \| 55 \| (17, 72) \| 77 \| (1, 55) \| 192 \| Helix-turn-helix domain-containing protein \| Helix-turn-helix domain-containing protein; uniclust \| UniRef100\_A0A1Y4L638 \| 99.1 \| 2.1e-13 \| 4.5e-19 \| 82.3 \| 57 \| (15, 72) \| 77 \| (4, 60) \| 101 \| DNA-binding protein \| DNA-binding protein; uniclust \| UniRef100\_A0A081BKJ0 \| 99.1 \| 2.1e-13 \| 4.6e-19 \| 91.3 \| 55 \| (16, 71) \| 77 \| (3, 57) \| 203 \| HTH cro/C1-type domain-containing protein \| HTH cro/C1-type domain-containing protein; uniclust \| UniRef100\_A0A3A8YLP0 \| 99.1 \| 2.4e-13 \| 4.7e-19 \| 85.7 \| 54 \| (18, 72) \| 77 \| (2, 55) \| 147 \| XRE family transcriptional regulator \| XRE family transcriptional regulator; uniclust \| UniRef100\_A0A0V8CR97 \| 99.1 \| 2.3e-13 \| 4.7e-19 \| 88.8 \| 60 \| (12, 72) \| 77 \| (2, 61) \| 184 \| Phage repressor \| Phage repressor; uniclust \| UniRef100\_A0A0M2ZP61 \| 99.1 \| 2.2e-13 \| 4.7e-19 \| 93.9 \| 56 \| (16, 72) \| 77 \| (9, 64) \| 255 \| Zinc finger/helix-turn-helix protein, YgiT family \| Zinc finger/helix-turn-helix protein, YgiT family; uniclust \| UniRef100\_A0A1R1MJE6 \| 99.1 \| 2.2e-13 \| 4.7e-19 \| 91.5 \| 57 \| (15, 72) \| 77 \| (17, 73) \| 216 \| HTH cro/C1-type domain-containing protein \| HTH cro/C1-type domain-containing protein; uniclust \| UniRef100\_A0A1I5QA44 \| 99.1 \| 2.4e-13 \| 4.8e-19 \| 71.1 \| 41 \| (14, 54) \| 77 \| (3, 43) \| 47 \| Helix-turn-helix \| Helix-turn-helix; uniclust \| UniRef100\_A0A379DAL3 \| 99.1 \| 2.4e-13 \| 4.8e-19 \| 85.3 \| 56 \| (16, 72) \| 77 \| (1, 56) \| 134 \| HTH-type transcriptional regulator immR \| HTH-type transcriptional regulator immR; uniclust \| UniRef100\_A0A1H8CHK5 \| 99.1 \| 2.4e-13 \| 4.9e-19 \| 80.3 \| 58 \| (14, 72) \| 77 \| (7, 64) \| 91 \| DNA-binding transcriptional regulator, XRE-family HTH domain \| DNA-binding transcriptional regulator, XRE-family HTH domain; uniclust \| UniRef100\_A0A328UKW6 \| 99.1 \| 2.5e-13 \| 4.9e-19 \| 72.0 \| 39 \| (16, 54) \| 77 \| (6, 44) \| 51 \| XRE family transcriptional regulator \| XRE family transcriptional regulator; uniclust \| UniRef100\_A0A028ZGG7 \| 99.1 \| 2.3e-13 \| 4.9e-19 \| 91.8 \| 57 \| (14, 71) \| 77 \| (18, 74) \| 217 \| Transcriptional regulator \| Transcriptional regulator; uniclust \| UniRef100\_A0A0M6WJ36 \| 99.1 \| 2.3e-13 \| 5e-19 \| 85.3 \| 61 \| (15, 75) \| 77 \| (5, 65) \| 129 \| HTH cro/C1-type domain-containing protein \| HTH cro/C1-type domain-containing protein; uniclust \| UniRef100\_A0A062XFL7 \| 99.1 \| 2.2e-13 \| 5e-19 \| 94.9 \| 58 \| (15, 73) \| 77 \| (33, 90) \| 275 \| HTH cro/C1-type domain-containing protein \| HTH cro/C1-type domain-containing protein; uniclust \| UniRef100\_A0A0R1L3N4 \| 99.1 \| 2.3e-13 \| 5.1e-19 \| 85.0 \| 55 \| (19, 74) \| 77 \| (49, 103) \| 123 \| HTH cro/C1-type domain-containing protein \| HTH cro/C1-type domain-containing protein; uniclust \| UniRef100\_A0A0H0YL11 \| 99.1 \| 2.4e-13 \| 5.1e-19 \| 86.7 \| 58 \| (14, 72) \| 77 \| (8, 65) \| 146 \| HTH cro/C1-type domain-containing protein \| HTH cro/C1-type domain-containing protein; uniclust \| UniRef100\_A0A0V8QET0 \| 99.1 \| 2.4e-13 \| 5.2e-19 \| 81.4 \| 57 \| (15, 72) \| 77 \| (5, 61) \| 96 \| XRE family transcriptional regulator \| XRE family transcriptional regulator; uniclust \| UniRef100\_A0A1C0TYE9 \| 99.1 \| 2.5e-13 \| 5.2e-19 \| 85.4 \| 63 \| (10, 73) \| 77 \| (10, 72) \| 135 \| Transcriptional repressor DicA \| Transcriptional repressor DicA; uniclust \| UniRef100\_A0A0A1GU27 \| 99.1 \| 2.5e-13 \| 5.2e-19 \| 81.8 \| 56 \| (16, 72) \| 77 \| (15, 70) \| 102 \| Transcriptional regulator \| Transcriptional regulator; uniclust \| UniRef100\_A0A239QG85 \| 99.1 \| 2.5e-13 \| 5.3e-19 \| 93.0 \| 57 \| (15, 72) \| 77 \| (32, 88) \| 256 \| Transcriptional regulator, contains XRE-family HTH domain \| Transcriptional regulator, contains XRE-family HTH domain; uniclust \| UniRef100\_A0A1C5NC58 \| 99.1 \| 2.6e-13 \| 5.3e-19 \| 94.7 \| 59 \| (13, 72) \| 77 \| (35, 93) \| 308 \| HTH-type transcriptional regulator immR \| HTH-type transcriptional regulator immR; uniclust \| UniRef100\_A0A089I0P5 \| 99.1 \| 2.6e-13 \| 5.4e-19 \| 95.2 \| 60 \| (12, 72) \| 77 \| (34, 93) \| 323 \| XRE family transcriptional regulator \| XRE family transcriptional regulator; uniclust \| UniRef100\_A0A1C3T4X4 \| 99.1 \| 2.7e-13 \| 5.4e-19 \| 81.6 \| 57 \| (15, 72) \| 77 \| (9, 65) \| 104 \| HTH cro/C1-type domain-containing protein \| HTH cro/C1-type domain-containing protein; uniclust \| UniRef100\_A0A0U9HDI7 \| 99.1 \| 2.6e-13 \| 5.4e-19 \| 84.7 \| 55 \| (17, 72) \| 77 \| (1, 55) \| 129 \| Helix-turn-helix domain-containing protein \| Helix-turn-helix domain-containing protein; uniclust \| UniRef100\_A0A430AU27 \| 99.1 \| 2.7e-13 \| 5.5e-19 \| 85.3 \| 54 \| (19, 73) \| 77 \| (4, 57) \| 140 \| HTH cro/C1-type domain-containing protein \| HTH cro/C1-type domain-containing protein; uniclust \| UniRef100\_A0A1G9IKI2 \| 99.1 \| 2.7e-13 \| 5.6e-19 \| 85.2 \| 57 \| (15, 72) \| 77 \| (5, 61) \| 131 \| DNA-binding transcriptional regulator, XRE-family HTH domain \| DNA-binding transcriptional regulator, XRE-family HTH domain; uniclust \| UniRef100\_A0A1M2Z9A3 \| 99.1 \| 2.5e-13 \| 5.7e-19 \| 98.3 \| 59 \| (13, 72) \| 77 \| (17, 75) \| 388 \| HTH cro/C1-type domain-containing protein \| HTH cro/C1-type domain-containing protein; uniclust \| UniRef100\_A0A0G0K4J7 \| 99.1 \| 2.6e-13 \| 5.9e-19 \| 82.3 \| 59 \| (14, 73) \| 77 \| (16, 74) \| 101 \| Transcriptional regulator, XRE family \| Transcriptional regulator, XRE family; uniclust \| UniRef100\_A0A358AQ13 \| 99.1 \| 2.8e-13 \| 5.9e-19 \| 83.1 \| 56 \| (16, 72) \| 77 \| (1, 56) \| 113 \| XRE family transcriptional regulator \| XRE family transcriptional regulator; uniclust \| UniRef100\_A0A173TGI2 \| 99.1 \| 3e-13 \| 5.9e-19 \| 88.1 \| 56 \| (16, 72) \| 77 \| (5, 60) \| 180 \| HTH-type transcriptional regulator immR \| HTH-type transcriptional regulator immR; uniclust \| UniRef100\_A0A061NRM5 \| 99.1 \| 2.8e-13 \| 6e-19 \| 80.9 \| 58 \| (14, 72) \| 77 \| (2, 59) \| 94 \| HTH cro/C1-type domain-containing protein \| HTH cro/C1-type domain-containing protein; uniclust \| UniRef100\_A0A1I3H5B9 \| 99.1 \| 2.8e-13 \| 6e-19 \| 78.7 \| 57 \| (16, 73) \| 77 \| (2, 58) \| 79 \| DNA-binding transcriptional regulator, XRE-family HTH domain \| DNA-binding transcriptional regulator, XRE-family HTH domain; uniclust \| UniRef100\_A0A2S0M7U0 \| 99.1 \| 3e-13 \| 6.1e-19 \| 77.4 \| 53 \| (19, 72) \| 77 \| (4, 56) \| 76 \| XRE family transcriptional regulator \| XRE family transcriptional regulator; uniclust \| UniRef100\_A0A135YNR8 \| 99.1 \| 3.3e-13 \| 6.4e-19 \| 87.2 \| 52 \| (20, 72) \| 77 \| (2, 53) \| 175 \| DNA-binding helix-turn-helix protein \| DNA-binding helix-turn-helix protein; uniclust \| UniRef100\_A0A075R1N1 \| 99.1 \| 3e-13 \| 6.5e-19 \| 88.9 \| 60 \| (13, 73) \| 77 \| (31, 90) \| 180 \| HTH-type transcriptional regulator ImmR \| HTH-type transcriptional regulator ImmR; uniclust \| UniRef100\_A0A0C1R042 \| 99.1 \| 2.9e-13 \| 6.5e-19 \| 92.0 \| 64 \| (8, 72) \| 77 \| (16, 79) \| 228 \| Cro/Cl family transcriptional regulator \| Cro/Cl family transcriptional regulator; uniclust \| UniRef100\_A0A078KP50 \| 99.1 \| 3.1e-13 \| 6.5e-19 \| 96.9 \| 59 \| (13, 72) \| 77 \| (27, 85) \| 387 \| Helix-turn-helix domain-containing protein \| Helix-turn-helix domain-containing protein; uniclust \| UniRef100\_A0A0K0MX67 \| 99.1 \| 3.2e-13 \| 6.6e-19 \| 83.3 \| 54 \| (17, 71) \| 77 \| (15, 68) \| 119 \| Putative DNA binding protein \| Putative DNA binding protein; uniclust \| UniRef100\_A0A1G6FFN7 \| 99.1 \| 3.2e-13 \| 6.6e-19 \| 90.9 \| 55 \| (16, 71) \| 77 \| (3, 57) \| 229 \| Transcriptional regulator, contains XRE-family HTH domain \| Transcriptional regulator, contains XRE-family HTH domain; uniclust \| UniRef100\_A0A4U4D9H3 \| 99.1 \| 3.2e-13 \| 6.6e-19 \| 88.8 \| 55 \| (16, 71) \| 77 \| (17, 71) \| 190 \| XRE family transcriptional regulator \| XRE family transcriptional regulator; uniclust \| UniRef100\_A0A3D2G1G4 \| 99.1 \| 3.3e-13 \| 6.6e-19 \| 84.2 \| 56 \| (16, 72) \| 77 \| (1, 56) \| 133 \| HTH cro/C1-type domain-containing protein \| HTH cro/C1-type domain-containing protein; uniclust \| UniRef100\_A0A0M6WHG0 \| 99.1 \| 2.9e-13 \| 6.6e-19 \| 94.5 \| 59 \| (13, 72) \| 77 \| (20, 78) \| 281 \| Putative transcriptional regulator \| Putative transcriptional regulator; uniclust \| UniRef100\_A0A239PSQ2 \| 99.1 \| 3.3e-13 \| 7e-19 \| 92.8 \| 59 \| (15, 74) \| 77 \| (55, 113) \| 265 \| Transcriptional regulator, contains XRE-family HTH domain \| Transcriptional regulator, contains XRE-family HTH domain; uniclust \| UniRef100\_A0A1C0BS68 \| 99.1 \| 3.4e-13 \| 7e-19 \| 86.5 \| 54 \| (15, 69) \| 77 \| (7, 60) \| 155 \| HTH cro/C1-type domain-containing protein \| HTH cro/C1-type domain-containing protein; uniclust \| UniRef100\_A0A1M6UBA3 \| 99.1 \| 3.3e-13 \| 7e-19 \| 95.1 \| 67 \| (5, 73) \| 77 \| (33, 99) \| 324 \| FHIPEP family protein \| FHIPEP family protein; uniclust \| UniRef100\_A0A023BQT6 \| 99.1 \| 3.2e-13 \| 7e-19 \| 83.5 \| 61 \| (12, 73) \| 77 \| (31, 91) \| 116 \| HTH cro/C1-type domain-containing protein \| HTH cro/C1-type domain-containing protein; uniclust \| UniRef100\_A0A0X1TLQ8 \| 99.1 \| 3.2e-13 \| 7e-19 \| 90.7 \| 56 \| (16, 72) \| 77 \| (3, 58) \| 210 \| HTH cro/C1-type domain-containing protein \| HTH cro/C1-type domain-containing protein; uniclust \| UniRef100\_A0A1C5Y1I1 \| 99.1 \| 3.3e-13 \| 7.1e-19 \| 91.6 \| 55 \| (17, 72) \| 77 \| (1, 55) \| 232 \| HTH-type transcriptional regulator immR \| HTH-type transcriptional regulator immR; uniclust \| UniRef100\_A0A0C2URG5 \| 99.1 \| 3.3e-13 \| 7.1e-19 \| 82.1 \| 60 \| (12, 72) \| 77 \| (7, 66) \| 104 \| HTH cro/C1-type domain-containing protein \| HTH cro/C1-type domain-containing protein; uniclust \| UniRef100\_A0A0R2I9Z4 \| 99.1 \| 3.5e-13 \| 7.2e-19 \| 90.4 \| 57 \| (15, 72) \| 77 \| (8, 64) \| 215 \| HTH cro/C1-type domain-containing protein \| HTH cro/C1-type domain-containing protein; uniclust \| UniRef100\_A0A1G9K086 \| 99.1 \| 3.6e-13 \| 7.4e-19 \| 91.6 \| 55 \| (15, 71) \| 77 \| (4, 58) \| 245 \| Transcriptional regulator, contains XRE-family HTH domain \| Transcriptional regulator, contains XRE-family HTH domain; uniclust \| UniRef100\_A0A1V4SGG0 \| 99.1 \| 3.6e-13 \| 7.4e-19 \| 85.8 \| 59 \| (13, 72) \| 77 \| (8, 66) \| 150 \| HTH-type transcriptional regulator ImmR \| HTH-type transcriptional regulator ImmR; uniclust \| UniRef100\_A0A0Q6ASP6 \| 99.1 \| 3.6e-13 \| 7.4e-19 \| 83.6 \| 58 \| (14, 72) \| 77 \| (9, 66) \| 123 \| HTH cro/C1-type domain-containing protein \| HTH cro/C1-type domain-containing protein; uniclust \| UniRef100\_A0A1C0BS96 \| 99.1 \| 3.5e-13 \| 7.5e-19 \| 88.1 \| 66 \| (7, 73) \| 77 \| (9, 74) \| 173 \| HTH cro/C1-type domain-containing protein \| HTH cro/C1-type domain-containing protein; uniclust \| UniRef100\_A0A1C6HMS0 \| 99.1 \| 3.7e-13 \| 7.5e-19 \| 79.2 \| 55 \| (17, 72) \| 77 \| (4, 58) \| 88 \| Predicted transcriptional regulator \| Predicted transcriptional regulator; uniclust \| UniRef100\_A0A0D5LVP1 \| 99.1 \| 3.5e-13 \| 7.5e-19 \| 82.5 \| 60 \| (13, 73) \| 77 \| (23, 82) \| 108 \| XRE family transcriptional regulator \| XRE family transcriptional regulator; uniclust \| UniRef100\_A0A066Q0N8 \| 99.1 \| 3.6e-13 \| 7.7e-19 \| 87.2 \| 58 \| (15, 73) \| 77 \| (23, 80) \| 159 \| DNA-binding protein \| DNA-binding protein; uniclust \| UniRef100\_A0A1A7QK35 \| 99.1 \| 3.5e-13 \| 7.7e-19 \| 85.9 \| 63 \| (11, 74) \| 77 \| (16, 78) \| 141 \| HTH cro/C1-type domain-containing protein \| HTH cro/C1-type domain-containing protein; uniclust \| UniRef100\_A0A2S6HCI4 \| 99.1 \| 4e-13 \| 7.7e-19 \| 76.4 \| 56 \| (16, 72) \| 77 \| (1, 56) \| 77 \| Helix-turn-helix protein \| Helix-turn-helix protein; uniclust \| UniRef100\_A0A1G7NVX8 \| 99.1 \| 3.8e-13 \| 7.7e-19 \| 84.1 \| 55 \| (17, 72) \| 77 \| (1, 55) \| 131 \| Transcriptional regulator, contains XRE-family HTH domain \| Transcriptional regulator, contains XRE-family HTH domain; uniclust \| UniRef100\_A0A073CX64 \| 99.1 \| 3.4e-13 \| 7.7e-19 \| 86.7 \| 72 \| (3, 74) \| 77 \| (8, 80) \| 148 \| HTH cro/C1-type domain-containing protein \| HTH cro/C1-type domain-containing protein; uniclust \| UniRef100\_A0A0R2IDY0 \| 99.1 \| 3.8e-13 \| 7.8e-19 \| 85.4 \| 55 \| (17, 72) \| 77 \| (14, 68) \| 145 \| HTH cro/C1-type domain-containing protein \| HTH cro/C1-type domain-containing protein; uniclust \| UniRef100\_A0A357QYW2 \| 99.1 \| 4e-13 \| 8e-19 \| 86.5 \| 62 \| (10, 72) \| 77 \| (7, 68) \| 164 \| HTH cro/C1-type domain-containing protein \| HTH cro/C1-type domain-containing protein; uniclust \| UniRef100\_A0A173UEQ1 \| 99.1 \| 3.8e-13 \| 8.1e-19 \| 87.7 \| 59 \| (14, 73) \| 77 \| (1, 59) \| 168 \| Helix-turn-helix domain \| Helix-turn-helix domain; uniclust \| UniRef100\_A0A0H4VVM8 \| 99.1 \| 3.8e-13 \| 8.1e-19 \| 86.0 \| 59 \| (12, 72) \| 77 \| (7, 65) \| 148 \| Transcriptional repressor DicA \| Transcriptional repressor DicA; uniclust \| UniRef100\_A0A1G5DD51 \| 99.1 \| 4e-13 \| 8.2e-19 \| 78.1 \| 57 \| (17, 74) \| 77 \| (4, 60) \| 81 \| Helix-turn-helix \| Helix-turn-helix; uniclust \| UniRef100\_A0A0N0DL77 \| 99.1 \| 4.1e-13 \| 8.4e-19 \| 79.3 \| 58 \| (14, 72) \| 77 \| (12, 69) \| 90 \| Transcriptional regulator \| Transcriptional regulator; uniclust \| UniRef100\_A0A1Q6U0U5 \| 99.1 \| 4e-13 \| 8.4e-19 \| 83.9 \| 59 \| (14, 73) \| 77 \| (13, 71) \| 126 \| HTH cro/C1-type domain-containing protein \| HTH cro/C1-type domain-containing protein; uniclust \| UniRef100\_A0A173WE40 \| 99.1 \| 4e-13 \| 8.5e-19 \| 96.3 \| 56 \| (16, 72) \| 77 \| (33, 88) \| 384 \| HTH-type transcriptional regulator immR \| HTH-type transcriptional regulator immR; uniclust \| UniRef100\_A0A023PEX3 \| 99.1 \| 3.9e-13 \| 8.5e-19 \| 83.5 \| 60 \| (12, 72) \| 77 \| (19, 78) \| 118 \| Transcriptional regulator \| Transcriptional regulator; uniclust \| UniRef100\_A0A2G3DYU1 \| 99.1 \| 4e-13 \| 8.7e-19 \| 85.3 \| 59 \| (14, 73) \| 77 \| (17, 75) \| 138 \| HTH cro/C1-type domain-containing protein \| HTH cro/C1-type domain-containing protein; uniclust \| UniRef100\_A0A1D3TPS8 \| 99.1 \| 4.3e-13 \| 8.7e-19 \| 86.0 \| 56 \| (16, 72) \| 77 \| (1, 56) \| 158 \| Transcriptional regulator, contains XRE-family HTH domain \| Transcriptional regulator, contains XRE-family HTH domain; uniclust \| UniRef100\_A0A350BBX8 \| 99.1 \| 4.3e-13 \| 8.9e-19 \| 86.6 \| 56 \| (16, 72) \| 77 \| (4, 59) \| 160 \| HTH cro/C1-type domain-containing protein \| HTH cro/C1-type domain-containing protein; uniclust \| UniRef100\_A0A059NZG4 \| 99.1 \| 4.6e-13 \| 9.2e-19 \| 79.3 \| 58 \| (15, 73) \| 77 \| (6, 64) \| 94 \| Helix-turn-helix \| Helix-turn-helix; uniclust \| UniRef100\_A0A096B436 \| 99.1 \| 4.3e-13 \| 9.2e-19 \| 86.3 \| 64 \| (8, 72) \| 77 \| (12, 75) \| 154 \| HTH cro/C1-type domain-containing protein \| HTH cro/C1-type domain-containing protein; uniclust \| UniRef100\_A0A1Y3PDK6 \| 99.1 \| 4.8e-13 \| 9.3e-19 \| 80.7 \| 57 \| (15, 72) \| 77 \| (2, 58) \| 111 \| HTH cro/C1-type domain-containing protein \| HTH cro/C1-type domain-containing protein; uniclust \| UniRef100\_A0A4R1RIV0 \| 99.1 \| 4.6e-13 \| 9.6e-19 \| 82.8 \| 57 \| (15, 72) \| 77 \| (1, 57) \| 119 \| DNA-binding XRE family transcriptional regulator \| DNA-binding XRE family transcriptional regulator; uniclust \| UniRef100\_A0A1Y4G4T6 \| 99.1 \| 4.5e-13 \| 9.6e-19 \| 78.0 \| 61 \| (12, 73) \| 77 \| (4, 64) \| 80 \| HTH cro/C1-type domain-containing protein \| HTH cro/C1-type domain-containing protein; uniclust \| UniRef100\_A0A073IMZ9 \| 99.1 \| 4.5e-13 \| 9.7e-19 \| 80.3 \| 51 \| (21, 72) \| 77 \| (8, 58) \| 95 \| HTH cro/C1-type domain-containing protein \| HTH cro/C1-type domain-containing protein; uniclust \| UniRef100\_A0A173Z0X1 \| 99.1 \| 4.9e-13 \| 9.8e-19 \| 77.2 \| 63 \| (11, 74) \| 77 \| (5, 67) \| 80 \| DNA-binding transcriptional regulator, XRE-family HTH domain \| DNA-binding transcriptional regulator, XRE-family HTH domain; uniclust \| UniRef100\_A0A0B2JZ35 \| 99.1 \| 4.5e-13 \| 9.9e-19 \| 80.8 \| 57 \| (15, 72) \| 77 \| (14, 70) \| 98 \| XRE family transcriptional regulator \| XRE family transcriptional regulator; uniclust \| UniRef100\_A0A064ABB6 \| 99.1 \| 4.5e-13 \| 9.9e-19 \| 85.3 \| 58 \| (14, 72) \| 77 \| (3, 66) \| 139 \| HTH cro/C1-type domain-containing protein \| HTH cro/C1-type domain-containing protein; uniclust \| UniRef100\_A0A089HUS8 \| 99.1 \| 4.5e-13 \| 9.9e-19 \| 93.8 \| 59 \| (13, 72) \| 77 \| (55, 113) \| 289 \| HTH cro/C1-type domain-containing protein \| HTH cro/C1-type domain-containing protein; uniclust \| UniRef100\_A0A1C5L5I0 \| 99.1 \| 4.8e-13 \| 1e-18 \| 90.0 \| 62 \| (11, 73) \| 77 \| (4, 65) \| 219 \| LexA repressor \| LexA repressor; uniclust \| UniRef100\_A0A140C2Z8 \| 99.1 \| 5e-13 \| 1e-18 \| 83.9 \| 59 \| (13, 72) \| 77 \| (3, 61) \| 134 \| Helix-turn-helix domain protein \| Helix-turn-helix domain protein; uniclust \| UniRef100\_A0A0R2CGT4 \| 99.1 \| 5.1e-13 \| 1e-18 \| 80.2 \| 54 \| (17, 71) \| 77 \| (1, 54) \| 101 \| XRE family transcriptional regulator \| XRE family transcriptional regulator; uniclust \| UniRef100\_A0A0C1K7L7 \| 99.1 \| 5.3e-13 \| 1e-18 \| 78.0 \| 54 \| (17, 71) \| 77 \| (3, 56) \| 91 \| XRE family transcriptional regulator \| XRE family transcriptional regulator; uniclust \| UniRef100\_A0A927PAF1 \| 99.1 \| 5.3e-13 \| 1e-18 \| 81.5 \| 54 \| (18, 72) \| 77 \| (5, 58) \| 120 \| Helix-turn-helix transcriptional regulator \| Helix-turn-helix transcriptional regulator; uniclust \| UniRef100\_A0A0F2S7G2 \| 99.1 \| 4.7e-13 \| 1e-18 \| 83.9 \| 57 \| (15, 72) \| 77 \| (7, 63) \| 125 \| HTH cro/C1-type domain-containing protein \| HTH cro/C1-type domain-containing protein; uniclust \| UniRef100\_A0A075U1V1 \| 99.1 \| 5.1e-13 \| 1.1e-18 \| 87.4 \| 57 \| (15, 72) \| 77 \| (1, 57) \| 179 \| HTH cro/C1-type domain-containing protein \| HTH cro/C1-type domain-containing protein; uniclust \| UniRef100\_A0A0R2DDB6 \| 99.1 \| 5e-13 \| 1.1e-18 \| 86.8 \| 62 \| (12, 74) \| 77 \| (12, 73) \| 163 \| HTH cro/C1-type domain-containing protein \| HTH cro/C1-type domain-containing protein; uniclust \| UniRef100\_A0A0Q2USQ7 \| 99.1 \| 5e-13 \| 1.1e-18 \| 85.2 \| 58 \| (15, 73) \| 77 \| (12, 69) \| 141 \| HTH cro/C1-type domain-containing protein \| HTH cro/C1-type domain-containing protein; uniclust \| UniRef100\_A0A0D6DWE9 \| 99.1 \| 5.5e-13 \| 1.1e-18 \| 82.2 \| 56 \| (16, 72) \| 77 \| (1, 56) \| 123 \| XRE-family transcriptional regulator \| XRE-family transcriptional regulator; uniclust \| UniRef100\_A0A0M6WVC9 \| 99.1 \| 5e-13 \| 1.1e-18 \| 87.3 \| 65 \| (8, 73) \| 77 \| (13, 77) \| 168 \| Helix-turn-helix domain-containing protein \| Helix-turn-helix domain-containing protein; uniclust \| UniRef100\_A0A0A8VZS6 \| 99.1 \| 5.2e-13 \| 1.1e-18 \| 84.9 \| 58 \| (14, 72) \| 77 \| (1, 58) \| 142 \| Helix-turn-helix transcriptional regulator \| Helix-turn-helix transcriptional regulator; uniclust \| UniRef100\_A0A174ZKR9 \| 99.1 \| 5e-13 \| 1.1e-18 \| 96.7 \| 57 \| (15, 72) \| 77 \| (12, 68) \| 385 \| Antitoxin PezA \| Antitoxin PezA; uniclust \| UniRef100\_A0A0J6HEI8 \| 99.1 \| 5.2e-13 \| 1.1e-18 \| 84.2 \| 57 \| (15, 72) \| 77 \| (5, 61) \| 132 \| XRE family transcriptional regulator \| XRE family transcriptional regulator; uniclust \| UniRef100\_A0A074LWD2 \| 99.1 \| 5.3e-13 \| 1.1e-18 \| 81.4 \| 57 \| (15, 72) \| 77 \| (21, 77) \| 107 \| HTH cro/C1-type domain-containing protein \| HTH cro/C1-type domain-containing protein; uniclust \| UniRef100\_A0A0U4X1Q9 \| 99.1 \| 5.8e-13 \| 1.1e-18 \| 85.5 \| 56 \| (17, 73) \| 77 \| (3, 58) \| 168 \| HTH cro/C1-type domain-containing protein \| HTH cro/C1-type domain-containing protein; uniclust \| UniRef100\_A0A087BBT1 \| 99.1 \| 5.6e-13 \| 1.1e-18 \| 88.2 \| 59 \| (13, 72) \| 77 \| (18, 76) \| 198 \| Putative transcriptional regulator \| Putative transcriptional regulator; uniclust \| UniRef100\_A0A1B1LIU2 \| 99.1 \| 5.8e-13 \| 1.1e-18 \| 81.3 \| 58 \| (14, 72) \| 77 \| (1, 58) \| 117 \| HTH cro/C1-type domain-containing protein \| HTH cro/C1-type domain-containing protein; uniclust \| UniRef100\_A0A1I7IJU0 \| 99.1 \| 5.7e-13 \| 1.2e-18 \| 86.7 \| 55 \| (17, 72) \| 77 \| (1, 55) \| 179 \| Transcriptional regulator, contains XRE-family HTH domain \| Transcriptional regulator, contains XRE-family HTH domain; uniclust \| UniRef100\_A0A1H0KR93 \| 99.1 \| 5.5e-13 \| 1.2e-18 \| 80.4 \| 57 \| (15, 72) \| 77 \| (14, 70) \| 99 \| DNA-binding transcriptional regulator, XRE-family HTH domain \| DNA-binding transcriptional regulator, XRE-family HTH domain; uniclust \| UniRef100\_A0A0B7IC89 \| 99.1 \| 5.7e-13 \| 1.2e-18 \| 77.6 \| 58 \| (14, 72) \| 77 \| (7, 64) \| 84 \| Anaerobic benzoate catabolism transcriptional regulator \| Anaerobic benzoate catabolism transcriptional regulator; uniclust \| UniRef100\_A0A3Q9V8T4 \| 99.1 \| 5.8e-13 \| 1.2e-18 \| 81.3 \| 53 \| (19, 72) \| 77 \| (2, 54) \| 112 \| XRE family transcriptional regulator \| XRE family transcriptional regulator; uniclust \| UniRef100\_A0A235IZF3 \| 99.1 \| 5.9e-13 \| 1.2e-18 \| 81.4 \| 56 \| (15, 71) \| 77 \| (21, 78) \| 115 \| Transcriptional regulator \| Transcriptional regulator; uniclust \| UniRef100\_A0A1Y4TPK4 \| 99.1 \| 5.9e-13 \| 1.2e-18 \| 88.9 \| 56 \| (16, 72) \| 77 \| (3, 58) \| 214 \| HTH cro/C1-type domain-containing protein \| HTH cro/C1-type domain-containing protein; uniclust \| UniRef100\_A0A073JPW0 \| 99.1 \| 5.5e-13 \| 1.2e-18 \| 85.3 \| 56 \| (16, 72) \| 77 \| (17, 72) \| 144 \| Helix-turn-helix domain-containing protein \| Helix-turn-helix domain-containing protein; uniclust \| UniRef100\_A0A173WDM4 \| 99.1 \| 6e-13 \| 1.2e-18 \| 81.2 \| 58 \| (14, 72) \| 77 \| (3, 60) \| 116 \| DNA-binding transcriptional repressor PuuR \| DNA-binding transcriptional repressor PuuR; uniclust \| UniRef100\_A0A084J761 \| 99.1 \| 5.3e-13 \| 1.2e-18 \| 95.2 \| 66 \| (7, 73) \| 77 \| (40, 105) \| 335 \| HTH cro/C1-type domain-containing protein \| HTH cro/C1-type domain-containing protein; uniclust \| UniRef100\_A0A143X185 \| 99.1 \| 5.9e-13 \| 1.2e-18 \| 87.1 \| 55 \| (16, 71) \| 77 \| (10, 64) \| 182 \| DNA-binding transcriptional repressor PuuR \| DNA-binding transcriptional repressor PuuR; uniclust \| UniRef100\_A0A2G5W151 \| 99.1 \| 6.2e-13 \| 1.2e-18 \| 77.8 \| 55 \| (17, 72) \| 77 \| (1, 55) \| 88 \| HTH cro/C1-type domain-containing protein \| HTH cro/C1-type domain-containing protein; uniclust \| UniRef100\_A0A069RGK5 \| 99.1 \| 5.7e-13 \| 1.2e-18 \| 85.9 \| 59 \| (13, 72) \| 77 \| (13, 71) \| 152 \| XRE family transcriptional regulator \| XRE family transcriptional regulator; uniclust \| UniRef100\_A0A351SPC4 \| 99.1 \| 6.3e-13 \| 1.2e-18 \| 78.1 \| 58 \| (14, 72) \| 77 \| (6, 63) \| 91 \| Transcriptional regulator (Fragment) \| Transcriptional regulator (Fragment); uniclust \| UniRef100\_A0A1V5P064 \| 99.1 \| 5.8e-13 \| 1.2e-18 \| 92.0 \| 61 \| (12, 73) \| 77 \| (32, 92) \| 265 \| LexA repressor \| LexA repressor; uniclust \| UniRef100\_A0A090Y881 \| 99.1 \| 5.8e-13 \| 1.3e-18 \| 96.5 \| 60 \| (12, 72) \| 77 \| (54, 113) \| 412 \| Helix-turn-helix family protein \| Helix-turn-helix family protein; uniclust \| UniRef100\_A0A0B0CZI8 \| 99.1 \| 6e-13 \| 1.3e-18 \| 78.3 \| 63 \| (10, 73) \| 77 \| (2, 64) \| 85 \| HTH cro/C1-type domain-containing protein \| HTH cro/C1-type domain-containing protein; uniclust \| UniRef100\_A0A133YTS9 \| 99.1 \| 6e-13 \| 1.3e-18 \| 88.4 \| 65 \| (8, 73) \| 77 \| (32, 96) \| 196 \| HTH cro/C1-type domain-containing protein \| HTH cro/C1-type domain-containing protein; uniclust \| UniRef100\_A0A0R1N1Y1 \| 99.1 \| 5.7e-13 \| 1.3e-18 \| 96.9 \| 63 \| (9, 72) \| 77 \| (45, 107) \| 414 \| HTH cro/C1-type domain-containing protein \| HTH cro/C1-type domain-containing protein; uniclust \| UniRef100\_A0A090JLT7 \| 99.1 \| 6.1e-13 \| 1.3e-18 \| 85.1 \| 61 \| (11, 72) \| 77 \| (3, 63) \| 150 \| HTH cro/C1-type domain-containing protein \| HTH cro/C1-type domain-containing protein; uniclust \| UniRef100\_A0A173SEW4 \| 99.1 \| 6.4e-13 \| 1.3e-18 \| 88.6 \| 59 \| (13, 72) \| 77 \| (19, 77) \| 220 \| Predicted transcriptional regulator \| Predicted transcriptional regulator; uniclust \| UniRef100\_A0A0R1TWH7 \| 99.1 \| 6.6e-13 \| 1.3e-18 \| 83.7 \| 56 \| (16, 72) \| 77 \| (1, 56) \| 147 \| Cro CI family phage transcriptional regulator \| Cro CI family phage transcriptional regulator; uniclust \| UniRef100\_A0A0R2G5H7 \| 99.1 \| 6.1e-13 \| 1.3e-18 \| 85.9 \| 59 \| (15, 73) \| 77 \| (6, 64) \| 158 \| HTH cro/C1-type domain-containing protein \| HTH cro/C1-type domain-containing protein; uniclust \| UniRef100\_A0A5M9NFB2 \| 99.1 \| 6.2e-13 \| 1.3e-18 \| 85.8 \| 57 \| (15, 72) \| 77 \| (4, 60) \| 158 \| Helix-turn-helix transcriptional regulator \| Helix-turn-helix transcriptional regulator; uniclust \| UniRef100\_A0A162QEW9 \| 99.1 \| 6.3e-13 \| 1.3e-18 \| 90.8 \| 58 \| (14, 72) \| 77 \| (1, 58) \| 248 \| HTH-type transcriptional regulator Xre \| HTH-type transcriptional regulator Xre; uniclust \| UniRef100\_A0A1F9VE86 \| 99.1 \| 6.3e-13 \| 1.3e-18 \| 75.5 \| 49 \| (16, 65) \| 77 \| (5, 53) \| 69 \| HTH cro/C1-type domain-containing protein \| HTH cro/C1-type domain-containing protein; uniclust \| UniRef100\_A0A7X6XEI5 \| 99.1 \| 6.8e-13 \| 1.3e-18 \| 76.4 \| 53 \| (18, 71) \| 77 \| (2, 54) \| 80 \| Helix-turn-helix transcriptional regulator \| Helix-turn-helix transcriptional regulator; uniclust \| UniRef100\_A0A072YE13 \| 99.1 \| 6.4e-13 \| 1.3e-18 \| 79.0 \| 59 \| (13, 72) \| 77 \| (8, 68) \| 92 \| Putative DNA-binding protein \| Putative DNA-binding protein; uniclust \| UniRef100\_A0A1C5PFS1 \| 99.1 \| 6.2e-13 \| 1.3e-18 \| 83.9 \| 63 \| (11, 73) \| 77 \| (8, 70) \| 131 \| Transcriptional repressor DicA \| Transcriptional repressor DicA; uniclust \| UniRef100\_A0A0V8QH06 \| 99.1 \| 6.9e-13 \| 1.4e-18 \| 79.2 \| 62 \| (9, 71) \| 77 \| (9, 70) \| 101 \| HTH cro/C1-type domain-containing protein \| HTH cro/C1-type domain-containing protein; uniclust \| UniRef100\_A0A413LW35 \| 99.1 \| 7.1e-13 \| 1.4e-18 \| 75.7 \| 59 \| (16, 74) \| 77 \| (1, 59) \| 79 \| XRE family transcriptional regulator \| XRE family transcriptional regulator; uniclust \| UniRef100\_A0A069DTQ8 \| 99.1 \| 6.5e-13 \| 1.4e-18 \| 81.0 \| 62 \| (12, 73) \| 77 \| (15, 76) \| 107 \| HTH cro/C1-type domain-containing protein \| HTH cro/C1-type domain-containing protein; uniclust \| UniRef100\_A0A076Z3U0 \| 99.1 \| 6.6e-13 \| 1.4e-18 \| 88.4 \| 54 \| (18, 72) \| 77 \| (3, 56) \| 201 \| Helix-turn-helix domain-containing protein \| Helix-turn-helix domain-containing protein |
| Top keywords  (threshold 1.00e-03 (evalue)) | **transcriptional, regulator, domain\_containing, HTH, cro, C1\_type, Helix\_turn\_helix, XRE, DNA\_binding, HTH\_type** |
| Output files | ../../similar\_sequences/52\_FANPEZAQ\_CDS\_0052\_merged.svg ../../similar\_sequences/52\_FANPEZAQ\_CDS\_0052\_pdb70.a3m ../../similar\_sequences/52\_FANPEZAQ\_CDS\_0052\_pdb70.hhr ../../similar\_sequences/52\_FANPEZAQ\_CDS\_0052\_uniclust.a3m ../../similar\_sequences/52\_FANPEZAQ\_CDS\_0052\_uniclust.hhr |

#### Structure prediction (AlphaFold)^2^

|  |  |
| --- | --- |
| Stats | xml version="1.0" encoding="utf-8" standalone="no"?       2024-09-02T21:09:47.994922 image/svg+xml   Matplotlib v3.7.2, https://matplotlib.org/ |
| Predicted structure | **NGL Viewer Controls:**  - Center: *Left-Click* - Rotate: *Left-Click + Drag* - Translate: *Right-Click + Drag* - Zoom: *Shift + Left-Click + Drag* |
| Output files | ../../predicted\_structures/52\_FANPEZAQ\_CDS\_0052/features.pkl ../../predicted\_structures/52\_FANPEZAQ\_CDS\_0052/ranked\_0.pdb ../../predicted\_structures/52\_FANPEZAQ\_CDS\_0052/ranked\_0\_plots.svg ../../predicted\_structures/52\_FANPEZAQ\_CDS\_0052/result\_model\_1\_ptm\_pred\_0.pkl |

#### Structure similarity search results (Foldseek)^3^

|  |  |
| --- | --- |
| Structure databases searched | Pdb, Afdb-proteome, Afdb-uniprot50 |
| Results, scheme(s)  (Top layers only, threshold 1.00e-02 (evalue)) | xml version="1.0" encoding="utf-8" standalone="no"?       2024-09-02T21:11:23.614800 image/svg+xml   Matplotlib v3.7.2, https://matplotlib.org/ |
| Results, table  (threshold 1.00e-02 (evalue)) | db \| id \| prob \| evalue \| bits \| fident \| alnlen \| mismatch \| gapopen \| qstart \| qend \| tstart \| tend \| name \| description; afdb-uniprot50 \| AF-A0A2Z3I919-F1-MODEL\_V4 \| 1.0 \| 0.0006904 \| 217 \| 0.5 \| 58 \| 29 \| 0 \| 16 \| 73 \| 1 \| 58 \| Uncharacterized protein \| Uncharacterized protein |
| Top keywords  (threshold 1.00e-02 (evalue)) | -- |
| Output files | ../../similar\_structures/52\_FANPEZAQ\_CDS\_0052\_afdb-proteome\_foldseek.tsv ../../similar\_structures/52\_FANPEZAQ\_CDS\_0052\_afdb-uniprot50\_foldseek.tsv ../../similar\_structures/52\_FANPEZAQ\_CDS\_0052\_merged.svg ../../similar\_structures/52\_FANPEZAQ\_CDS\_0052\_pdb\_foldseek.tsv |

  
  
  

Return to summary | Go to previous | Go to next

  


---

**Sequence/structure alignments coloring**  
Each object in the alignment figures is colored according to its E-value following this color coding:

1e-100
10
