## Supplementary material for "Completing the BASEL phage collection to unlock hidden diversity for systematic exploration of phage-host interactions": Data S2: 53.html

FANPEZAQ\_CDS\_0053


Return to summary | Go to previous | Go to next

|  |  |
| --- | --- |
| FANPEZAQ\_CDS\_0053  Page creation date: 02 Sep 2024, 12:00  Project folder: n/a  Input sequences file: Escherichia\_virus\_HeidiAbel.gb | hypothetical ep3\_0015 domain\_containing ky705409\_p11 p222118 vi\_08569 duf1289 kfra n\_terminal dna\_binding |

### Sequence information

|  |  |
| --- | --- |
| Name | FANPEZAQ\_CDS\_0053  53\_FANPEZAQ\_CDS\_0053 (pipeline id) |
| Imported annotations | Escherichia\_virus\_HeidiAbel Bas97 |
| Protein sequence | MNACTWQPTTVTLQDGREVLSDSREWLLECEAKYVLNQPTIEARRTLLAAIEKRRGATAR EELEQRALAIWRANRP |
| Number of residues | 76 |
| Molecular weight (Da) | 8795.92 |
| Output files | ../../query\_sequences/53\_FANPEZAQ\_CDS\_0053.fasta |

### Putative domain architecture and protein family

#### Search results (HHblits)^1^

|  |  |
| --- | --- |
| Domain family databases searched | Pfam, Ncbi-cd, Cath, Phrogs |
| Results, scheme(s)  (Top layers only; threshold 1.00e-03 (evalue)) | xml version="1.0" encoding="utf-8" standalone="no"?       2024-09-02T21:08:24.222966 image/svg+xml   Matplotlib v3.7.2, https://matplotlib.org/ |
| Results, table  (E-value ≤ 1.00e-03 (evalue)) | db \| id \| prob \| evalue \| pvalue \| score \| cols \| query \| query\_len \| template \| template\_len \| name \| description; phrogs \| 11642 \| 100.0 \| 9.5e-42 \| 1.1e-45 \| 219.3 \| 76 \| (1, 76) \| 76 \| (1, 76) \| 76 \| NA \| NA; Category: unknown function; KY705409\_p11; phrogs \| 14931 \| 99.6 \| 3.4e-20 \| 3.9e-24 \| 116.3 \| 52 \| (24, 75) \| 76 \| (8, 61) \| 69 \| NA \| NA; Category: unknown function; p222118 VI\_08569 |
| Top keywords  (threshold 1.00e-03 (evalue)) | **KY705409\_p11, p222118, VI\_08569** |
| Output files | ../../domain\_architecture/53\_FANPEZAQ\_CDS\_0053\_cath.hhr ../../domain\_architecture/53\_FANPEZAQ\_CDS\_0053\_merged.svg ../../domain\_architecture/53\_FANPEZAQ\_CDS\_0053\_ncbi-cd.hhr ../../domain\_architecture/53\_FANPEZAQ\_CDS\_0053\_pfam.hhr ../../domain\_architecture/53\_FANPEZAQ\_CDS\_0053\_phrogs.hhr |

### Identical protein sequences/structures

#### Search results

|  |  |
| --- | --- |
| Protein sequence databases searched | Pdb, Swissprot, Refseq |
| Identical proteins found | Refseq  - YP\_009100010.1: Hypothetical protein ep3\_0015 (Escherichia phage vB\_EcoM-ep3) - AIM50545.1: Hypothetical protein ep3\_0015 (Escherichia phage vB\_EcoM-ep3) |
| Top keywords | **Hypothetical, ep3\_0015** |
| Output files | ../../identical\_sequences/53\_FANPEZAQ\_CDS\_0053\_refseq.fasta |

### Similar protein sequences/structures

#### Sequence similarity search results (HHblits)^1^

|  |  |
| --- | --- |
| Sequence databases searched | Uniclust, Pdb70 |
| Results, scheme(s)  (Top layers only, threshold 1.00e-03 (evalue)) | xml version="1.0" encoding="utf-8" standalone="no"?       2024-09-02T21:08:51.793827 image/svg+xml   Matplotlib v3.7.2, https://matplotlib.org/ |
| Results, table(s)  (threshold 1.00e-03 (evalue)) | db \| id \| prob \| evalue \| pvalue \| score \| cols \| query \| query\_len \| template \| template\_len \| name \| description; uniclust \| UniRef100\_A0A088FQJ3 \| 99.9 \| 4e-30 \| 8.7e-36 \| 167.6 \| 75 \| (1, 75) \| 76 \| (11, 85) \| 97 \| Uncharacterized protein \| Uncharacterized protein; uniclust \| UniRef100\_A0A1B2ECS2 \| 99.9 \| 4.6e-29 \| 1.1e-34 \| 163.7 \| 75 \| (1, 75) \| 76 \| (5, 90) \| 98 \| Uncharacterized protein \| Uncharacterized protein; uniclust \| UniRef100\_A0A7G8LJG8 \| 99.8 \| 7.4e-23 \| 1.6e-28 \| 139.2 \| 68 \| (1, 69) \| 76 \| (2, 69) \| 126 \| Uncharacterized protein \| Uncharacterized protein; uniclust \| UniRef100\_A0A086MEE7 \| 99.6 \| 6.2e-19 \| 1.3e-24 \| 118.8 \| 67 \| (8, 75) \| 76 \| (23, 101) \| 112 \| Uncharacterized protein \| Uncharacterized protein; uniclust \| UniRef100\_A0A2W5MTW2 \| 99.6 \| 1.2e-18 \| 2.7e-24 \| 116.2 \| 69 \| (7, 75) \| 76 \| (19, 96) \| 103 \| Uncharacterized protein \| Uncharacterized protein; uniclust \| UniRef100\_A0A061JNG3 \| 99.6 \| 1.5e-18 \| 3.3e-24 \| 111.8 \| 67 \| (9, 75) \| 76 \| (3, 71) \| 81 \| Uncharacterized protein \| Uncharacterized protein; uniclust \| UniRef100\_A0A085FCV0 \| 99.5 \| 3.7e-17 \| 7.8e-23 \| 107.0 \| 62 \| (10, 72) \| 76 \| (12, 84) \| 90 \| Uncharacterized protein \| Uncharacterized protein; uniclust \| UniRef100\_A0A2I8DIP3 \| 99.5 \| 1.5e-16 \| 3.2e-22 \| 98.5 \| 58 \| (17, 75) \| 76 \| (3, 60) \| 63 \| Uncharacterized protein \| Uncharacterized protein; uniclust \| UniRef100\_A0A0Q9HPA0 \| 99.4 \| 2.5e-16 \| 5e-22 \| 102.1 \| 62 \| (8, 70) \| 76 \| (9, 77) \| 86 \| Uncharacterized protein \| Uncharacterized protein; uniclust \| UniRef100\_A0A375IY35 \| 99.4 \| 1.7e-15 \| 3.4e-21 \| 97.6 \| 53 \| (23, 75) \| 76 \| (16, 68) \| 82 \| Uncharacterized protein \| Uncharacterized protein; uniclust \| UniRef100\_A0A5C7JBT4 \| 99.4 \| 2.6e-15 \| 5.1e-21 \| 93.5 \| 59 \| (17, 75) \| 76 \| (2, 60) \| 66 \| Uncharacterized protein \| Uncharacterized protein; uniclust \| UniRef100\_A0A1D7U039 \| 99.3 \| 3.4e-15 \| 6.8e-21 \| 98.6 \| 63 \| (12, 75) \| 76 \| (14, 87) \| 95 \| Uncharacterized protein \| Uncharacterized protein; uniclust \| UniRef100\_A0A6J5QN02 \| 99.3 \| 4.9e-15 \| 8.9e-21 \| 92.8 \| 65 \| (7, 72) \| 76 \| (3, 67) \| 72 \| Uncharacterized protein \| Uncharacterized protein; uniclust \| UniRef100\_A0A2Z3I5Z9 \| 99.3 \| 1.9e-14 \| 3.4e-20 \| 92.7 \| 70 \| (7, 76) \| 76 \| (8, 78) \| 85 \| DUF1289 domain-containing protein \| DUF1289 domain-containing protein; uniclust \| UniRef100\_UPI0021E4EEF5 \| 99.2 \| 4.3e-14 \| 7.9e-20 \| 95.1 \| 59 \| (17, 75) \| 76 \| (4, 62) \| 114 \| hypothetical protein \| hypothetical protein; uniclust \| UniRef100\_A0A6J5NE18 \| 99.1 \| 1.9e-13 \| 4.1e-19 \| 87.7 \| 57 \| (19, 75) \| 76 \| (7, 65) \| 75 \| Uncharacterized protein \| Uncharacterized protein; uniclust \| UniRef100\_A0A2Z3HWP4 \| 99.1 \| 9.4e-13 \| 1.9e-18 \| 82.1 \| 55 \| (20, 74) \| 76 \| (4, 60) \| 64 \| Uncharacterized protein \| Uncharacterized protein; uniclust \| UniRef100\_UPI0007826AF5 \| 98.9 \| 5.1e-12 \| 9.9e-18 \| 83.7 \| 72 \| (4, 75) \| 76 \| (6, 82) \| 93 \| hypothetical protein \| hypothetical protein; uniclust \| UniRef100\_A0A2G7RZZ6 \| 98.9 \| 7.6e-12 \| 1.5e-17 \| 83.2 \| 65 \| (10, 74) \| 76 \| (1, 80) \| 93 \| Uncharacterized protein \| Uncharacterized protein; uniclust \| UniRef100\_A0A1G8AZ57 \| 98.9 \| 1.3e-11 \| 2.5e-17 \| 77.5 \| 57 \| (19, 75) \| 76 \| (1, 58) \| 66 \| Uncharacterized protein \| Uncharacterized protein; uniclust \| UniRef100\_A0A5B2VSF5 \| 98.9 \| 1.9e-11 \| 3.4e-17 \| 86.8 \| 66 \| (9, 75) \| 76 \| (78, 154) \| 158 \| Uncharacterized protein \| Uncharacterized protein; uniclust \| UniRef100\_A0A0R0CJT0 \| 98.8 \| 2.5e-11 \| 5.5e-17 \| 77.4 \| 58 \| (18, 75) \| 76 \| (2, 61) \| 69 \| KfrA N-terminal DNA-binding domain-containing protein \| KfrA N-terminal DNA-binding domain-containing protein; uniclust \| UniRef100\_A0A1H7WFY8 \| 98.8 \| 3.8e-11 \| 7.6e-17 \| 80.5 \| 64 \| (10, 74) \| 76 \| (12, 86) \| 97 \| Uncharacterized protein \| Uncharacterized protein; uniclust \| UniRef100\_A0A6J7X307 \| 98.7 \| 8.1e-11 \| 1.5e-16 \| 75.0 \| 62 \| (11, 72) \| 76 \| (5, 71) \| 74 \| Uncharacterized protein \| Uncharacterized protein; uniclust \| UniRef100\_A0A1E4HKI7 \| 98.7 \| 8.3e-11 \| 1.6e-16 \| 75.4 \| 54 \| (22, 75) \| 76 \| (12, 67) \| 74 \| Uncharacterized protein \| Uncharacterized protein; uniclust \| UniRef100\_A0A6J5NVJ8 \| 98.6 \| 3.4e-10 \| 6.3e-16 \| 73.8 \| 65 \| (8, 74) \| 76 \| (11, 80) \| 84 \| Uncharacterized protein \| Uncharacterized protein; uniclust \| UniRef100\_A0A1E4FVV8 \| 98.6 \| 4.1e-10 \| 8e-16 \| 70.9 \| 52 \| (20, 72) \| 76 \| (2, 53) \| 64 \| Uncharacterized protein \| Uncharacterized protein; uniclust \| UniRef100\_A0A5B2VQR8 \| 98.6 \| 4.5e-10 \| 8.3e-16 \| 69.4 \| 44 \| (9, 53) \| 76 \| (5, 48) \| 60 \| Uncharacterized protein \| Uncharacterized protein; uniclust \| UniRef100\_A0A1V5QQ95 \| 98.6 \| 4.8e-10 \| 9.3e-16 \| 71.7 \| 59 \| (16, 75) \| 76 \| (8, 66) \| 71 \| Uncharacterized protein \| Uncharacterized protein; uniclust \| UniRef100\_A0A9E5K5N7 \| 98.5 \| 8.5e-10 \| 1.6e-15 \| 73.1 \| 68 \| (2, 70) \| 76 \| (8, 75) \| 92 \| Uncharacterized protein \| Uncharacterized protein; uniclust \| UniRef100\_UPI00135B86F6 \| 98.5 \| 1.1e-09 \| 2e-15 \| 73.1 \| 63 \| (13, 75) \| 76 \| (9, 88) \| 95 \| hypothetical protein \| hypothetical protein; uniclust \| UniRef100\_A0A069P165 \| 98.5 \| 1e-09 \| 2.1e-15 \| 77.7 \| 53 \| (17, 70) \| 76 \| (54, 106) \| 133 \| Uncharacterized protein \| Uncharacterized protein; uniclust \| UniRef100\_A0A1B4G289 \| 98.4 \| 2.4e-09 \| 4.8e-15 \| 74.5 \| 52 \| (18, 70) \| 76 \| (56, 107) \| 120 \| Uncharacterized protein \| Uncharacterized protein; uniclust \| UniRef100\_A0A4R3UNM8 \| 98.4 \| 3e-09 \| 5.6e-15 \| 65.5 \| 51 \| (23, 73) \| 76 \| (3, 53) \| 58 \| Uncharacterized protein \| Uncharacterized protein; uniclust \| UniRef100\_A0A9E3YW76 \| 98.4 \| 3.2e-09 \| 5.8e-15 \| 66.6 \| 55 \| (20, 74) \| 76 \| (5, 61) \| 64 \| Uncharacterized protein \| Uncharacterized protein; uniclust \| UniRef100\_A0A4Q8M7C5 \| 98.4 \| 3.4e-09 \| 6.7e-15 \| 73.8 \| 58 \| (18, 75) \| 76 \| (51, 110) \| 118 \| Uncharacterized protein \| Uncharacterized protein; uniclust \| UniRef100\_UPI001F0C0D3A \| 98.2 \| 1.7e-08 \| 3e-14 \| 66.8 \| 50 \| (26, 75) \| 76 \| (28, 79) \| 87 \| hypothetical protein \| hypothetical protein; uniclust \| UniRef100\_UPI001FC80872 \| 98.2 \| 2.7e-08 \| 4.9e-14 \| 71.5 \| 53 \| (17, 70) \| 76 \| (91, 143) \| 156 \| hypothetical protein \| hypothetical protein; uniclust \| UniRef100\_A0A0Q9J9L1 \| 98.1 \| 4.1e-08 \| 7.5e-14 \| 68.8 \| 52 \| (14, 66) \| 76 \| (38, 100) \| 128 \| Uncharacterized protein \| Uncharacterized protein; uniclust \| UniRef100\_UPI00054B42C1 \| 98.0 \| 8.9e-08 \| 1.6e-13 \| 63.0 \| 52 \| (24, 75) \| 76 \| (11, 64) \| 82 \| hypothetical protein \| hypothetical protein; uniclust \| UniRef100\_A0A0H1RC28 \| 98.0 \| 1.6e-07 \| 3e-13 \| 64.7 \| 59 \| (9, 68) \| 76 \| (8, 78) \| 111 \| Uncharacterized protein \| Uncharacterized protein; uniclust \| UniRef100\_A0A2S5MC58 \| 98.0 \| 1.6e-07 \| 3e-13 \| 66.4 \| 50 \| (18, 68) \| 76 \| (29, 78) \| 133 \| Uncharacterized protein \| Uncharacterized protein; uniclust \| UniRef100\_I4YKC4 \| 97.9 \| 2.3e-07 \| 4.4e-13 \| 61.4 \| 47 \| (9, 56) \| 76 \| (17, 63) \| 82 \| Uncharacterized protein \| Uncharacterized protein; uniclust \| UniRef100\_UPI0011C05C5A \| 97.8 \| 4.3e-07 \| 7.9e-13 \| 63.9 \| 70 \| (5, 75) \| 76 \| (39, 108) \| 126 \| hypothetical protein \| hypothetical protein; uniclust \| UniRef100\_A0A9E8CZF4 \| 97.8 \| 4.4e-07 \| 8.1e-13 \| 63.2 \| 49 \| (15, 64) \| 76 \| (42, 97) \| 117 \| Uncharacterized protein \| Uncharacterized protein; uniclust \| UniRef100\_A0A4Q8LD98 \| 97.8 \| 4.5e-07 \| 8.2e-13 \| 59.1 \| 57 \| (19, 75) \| 76 \| (9, 67) \| 75 \| Uncharacterized protein \| Uncharacterized protein; uniclust \| UniRef100\_A0A202B296 \| 97.6 \| 1.7e-06 \| 3.6e-12 \| 57.9 \| 56 \| (20, 75) \| 76 \| (20, 77) \| 80 \| Uncharacterized protein \| Uncharacterized protein; uniclust \| UniRef100\_A0A6J5PKL6 \| 97.4 \| 6.7e-06 \| 1.2e-11 \| 56.7 \| 64 \| (5, 70) \| 76 \| (35, 100) \| 103 \| Uncharacterized protein \| Uncharacterized protein; uniclust \| UniRef100\_A0A0Q7F577 \| 97.3 \| 1.3e-05 \| 2.3e-11 \| 56.9 \| 58 \| (16, 73) \| 76 \| (5, 62) \| 122 \| Uncharacterized protein \| Uncharacterized protein; uniclust \| UniRef100\_A0A0S4U452 \| 97.2 \| 1.4e-05 \| 2.9e-11 \| 57.1 \| 47 \| (23, 70) \| 76 \| (43, 89) \| 120 \| Uncharacterized protein \| Uncharacterized protein; uniclust \| UniRef100\_UPI0020796AD7 \| 97.2 \| 2.5e-05 \| 4.7e-11 \| 50.2 \| 51 \| (22, 73) \| 76 \| (3, 53) \| 64 \| hypothetical protein \| hypothetical protein; uniclust \| UniRef100\_UPI00036E2260 \| 97.0 \| 5.9e-05 \| 1.1e-10 \| 50.4 \| 40 \| (19, 58) \| 76 \| (3, 42) \| 77 \| hypothetical protein \| hypothetical protein; uniclust \| UniRef100\_H6BJ70 \| 96.9 \| 7.7e-05 \| 1.4e-10 \| 48.5 \| 64 \| (6, 73) \| 76 \| (1, 64) \| 67 \| Uncharacterized protein \| Uncharacterized protein; uniclust \| UniRef100\_UPI001E405AA9 \| 96.6 \| 0.00033 \| 6e-10 \| 51.0 \| 46 \| (23, 69) \| 76 \| (59, 104) \| 135 \| hypothetical protein \| hypothetical protein; uniclust \| UniRef100\_A0A6J5KQ13 \| 96.5 \| 0.00038 \| 7.7e-10 \| 46.8 \| 56 \| (15, 73) \| 76 \| (4, 62) \| 74 \| Uncharacterized protein \| Uncharacterized protein; uniclust \| UniRef100\_C1D867 \| 96.5 \| 0.00052 \| 9.7e-10 \| 51.8 \| 55 \| (21, 75) \| 76 \| (101, 155) \| 169 \| Uncharacterized protein \| Uncharacterized protein; uniclust \| UniRef100\_A0A6S7BPC1 \| 96.5 \| 0.00055 \| 1e-09 \| 51.2 \| 53 \| (17, 70) \| 76 \| (81, 133) \| 162 \| Uncharacterized protein \| Uncharacterized protein; uniclust \| UniRef100\_UPI00131C9C71 \| 96.4 \| 0.00069 \| 1.3e-09 \| 43.9 \| 43 \| (31, 74) \| 76 \| (13, 55) \| 62 \| hypothetical protein \| hypothetical protein |
| Top keywords  (threshold 1.00e-03 (evalue)) | **hypothetical, domain\_containing, DUF1289, KfrA, N\_terminal, DNA\_binding** |
| Output files | ../../similar\_sequences/53\_FANPEZAQ\_CDS\_0053\_merged.svg ../../similar\_sequences/53\_FANPEZAQ\_CDS\_0053\_pdb70.a3m ../../similar\_sequences/53\_FANPEZAQ\_CDS\_0053\_pdb70.hhr ../../similar\_sequences/53\_FANPEZAQ\_CDS\_0053\_uniclust.a3m ../../similar\_sequences/53\_FANPEZAQ\_CDS\_0053\_uniclust.hhr |

#### Structure prediction (AlphaFold)^2^

|  |  |
| --- | --- |
| Stats | xml version="1.0" encoding="utf-8" standalone="no"?       2024-09-02T21:09:48.666323 image/svg+xml   Matplotlib v3.7.2, https://matplotlib.org/ |
| Predicted structure | **NGL Viewer Controls:**  - Center: *Left-Click* - Rotate: *Left-Click + Drag* - Translate: *Right-Click + Drag* - Zoom: *Shift + Left-Click + Drag* |
| Output files | ../../predicted\_structures/53\_FANPEZAQ\_CDS\_0053/features.pkl ../../predicted\_structures/53\_FANPEZAQ\_CDS\_0053/ranked\_0.pdb ../../predicted\_structures/53\_FANPEZAQ\_CDS\_0053/ranked\_0\_plots.svg ../../predicted\_structures/53\_FANPEZAQ\_CDS\_0053/result\_model\_1\_ptm\_pred\_0.pkl |

#### Structure similarity search results (Foldseek)^3^

|  |  |
| --- | --- |
| Structure databases searched | Pdb, Afdb-proteome, Afdb-uniprot50 |
| Results, scheme(s)  (Top layers only, threshold 1.00e-02 (evalue)) | xml version="1.0" encoding="utf-8" standalone="no"?       2024-09-02T21:11:23.792508 image/svg+xml   Matplotlib v3.7.2, https://matplotlib.org/ |
| Results, table  (threshold 1.00e-02 (evalue)) | db \| id \| prob \| evalue \| bits \| fident \| alnlen \| mismatch \| gapopen \| qstart \| qend \| tstart \| tend \| name \| description; afdb-uniprot50 \| AF-A0A2W5FN44-F1-MODEL\_V4 \| 1.0 \| 1.574e-06 \| 311 \| 0.479 \| 73 \| 38 \| 0 \| 1 \| 73 \| 11 \| 83 \| Uncharacterized protein \| Uncharacterized protein; afdb-uniprot50 \| AF-A0A2W5MTW2-F1-MODEL\_V4 \| 1.0 \| 3.056e-05 \| 247 \| 0.364 \| 74 \| 42 \| 1 \| 8 \| 76 \| 20 \| 93 \| Uncharacterized protein \| Uncharacterized protein; afdb-uniprot50 \| AF-A0A7R7YH54-F1-MODEL\_V4 \| 1.0 \| 6.941e-05 \| 238 \| 0.422 \| 71 \| 40 \| 1 \| 1 \| 71 \| 1 \| 70 \| Uncharacterized protein \| Uncharacterized protein; afdb-uniprot50 \| AF-A0A0Q7SHZ6-F1-MODEL\_V4 \| 1.0 \| 0.0008133 \| 204 \| 0.476 \| 65 \| 34 \| 0 \| 9 \| 73 \| 2 \| 66 \| Uncharacterized protein \| Uncharacterized protein; afdb-uniprot50 \| AF-A0A2Z3I5Z9-F1-MODEL\_V4 \| 1.0 \| 0.001435 \| 192 \| 0.402 \| 72 \| 41 \| 2 \| 2 \| 72 \| 4 \| 74 \| Uncharacterized protein \| Uncharacterized protein; afdb-uniprot50 \| AF-A0A246JHM4-F1-MODEL\_V4 \| 1.0 \| 0.0002302 \| 169 \| 0.375 \| 72 \| 40 \| 1 \| 9 \| 75 \| 2 \| 73 \| Uncharacterized protein \| Uncharacterized protein |
| Top keywords  (threshold 1.00e-02 (evalue)) | -- |
| Output files | ../../similar\_structures/53\_FANPEZAQ\_CDS\_0053\_afdb-proteome\_foldseek.tsv ../../similar\_structures/53\_FANPEZAQ\_CDS\_0053\_afdb-uniprot50\_foldseek.tsv ../../similar\_structures/53\_FANPEZAQ\_CDS\_0053\_merged.svg ../../similar\_structures/53\_FANPEZAQ\_CDS\_0053\_pdb\_foldseek.tsv |

  
  
  

Return to summary | Go to previous | Go to next

  


---

**Sequence/structure alignments coloring**  
Each object in the alignment figures is colored according to its E-value following this color coding:

1e-100
10
