## Supplementary material for "Completing the BASEL phage collection to unlock hidden diversity for systematic exploration of phage-host interactions": Data S2: 54.html

FANPEZAQ\_CDS\_0054


Return to summary | Go to previous | Go to next

|  |  |
| --- | --- |
| FANPEZAQ\_CDS\_0054  Page creation date: 02 Sep 2024, 12:00  Project folder: n/a  Input sequences file: Escherichia\_virus\_HeidiAbel.gb | dna domain\_containing primase virulence\_associated e replication factor licensing toprim helicase putative e\_like atpase p\_loop and virulence mcm2 trac mcm3 prict\_2 fragment rna phage nucleotide metabolism inactivated duf3874 predicted derivatives mcm6 polymerase n\_terminal prim\_pol c plasmid mcm7 vire\_n vape vire mcm4 sf3 duf3854 bifunctional associated maintenance \_ duf927 minichromosome p4 terminal |

### Sequence information

|  |  |
| --- | --- |
| Name | FANPEZAQ\_CDS\_0054  54\_FANPEZAQ\_CDS\_0054 (pipeline id) |
| Imported annotations | Escherichia\_virus\_HeidiAbel Bas97 |
| Protein sequence | MSAQQNYKDLTEQEIAEALSYIDAGCDREAWVRMAMAVKSELGDGGFTVWNDWSRQSDKY NSKDARDTWKSVKRHGGIGIGTLIGEAQQFGFSLNDEERTPLSAEEIEARKRKREAEEKA EQQRRAERMAQAAEQARNIWESATELEGDEHPYLQRKKVQAFGLRVGRFWTENRSIDGAL LVPVRNIDGQITSLQAIFPNENPQLGRDRDYLPGGQKRGCFHVIGGKPTGANPVIVICEG YSTGASIHQATGYCVVVAFDAGNVPTVAKLMRQQFGRATIVVAADNDQWHEDGKQNDGVH YARQAATTCGGLLVVPKFAALSDKPTDFNDLHNLQGLAEVKRQINAAIPEPANDNFLPLD ATVNPFMYPHMSHQQKPLSTWENLEWLLDQYGITARYNEISKDVVVTIPGRDYGVDASAN CSLAEVSSLCARNGMPKGDVGDYIKLIGVANRYNPAAEFITSRPWDGVSRIYDLVDTLAT PAGFDRGLVLMMVRRWLISAVAAVMKPTGFWSKGVLVLQGDQSLGKTSWFKALLPPTMRD LVKDGATIDPSNKDSVTTAIGHWMVELGELDATFRKSDIARLKSFISSDVDMLRRPYDRL ESKYQRRTVFFASVNPKHFLADDTGNVRWWTIPVTGVNYAHDIDTQQLWAEVAQLYRDGE RWWLDRDEEAMLEGVNKEHESIDPVEEMILARFEWGSERISAYREHTASQVLQEIGFDKP NKSQATHCSNVLRKLTGQEPRKTKNGRFFNLPPKVLQPDYRRPDDDKPF |
| Number of residues | 769 |
| Molecular weight (Da) | 86447.26 |
| Output files | ../../query\_sequences/54\_FANPEZAQ\_CDS\_0054.fasta |

### Putative domain architecture and protein family

#### Search results (HHblits)^1^

|  |  |
| --- | --- |
| Domain family databases searched | Pfam, Ncbi-cd, Cath, Phrogs |
| Results, scheme(s)  (Top layers only; threshold 1.00e-03 (evalue)) | xml version="1.0" encoding="utf-8" standalone="no"?       2024-09-02T21:08:24.476812 image/svg+xml   Matplotlib v3.7.2, https://matplotlib.org/ |
| Results, table  (E-value ≤ 1.00e-03 (evalue)) | db \| id \| prob \| evalue \| pvalue \| score \| cols \| query \| query\_len \| template \| template\_len \| name \| description; pfam \| PF05272 \| 99.4 \| 1.8e-17 \| 4.5e-21 \| 150.8 \| 214 \| (460, 680) \| 769 \| (1, 220) \| 221 \| VirE \| Virulence-associated protein E; pfam \| PF04735 \| 97.9 \| 2.5e-09 \| 5e-13 \| 124.9 \| 138 \| (490, 634) \| 769 \| (884, 1028) \| 1217 \| Baculo\_helicase \| Baculovirus DNA helicase; pfam \| PF00519 \| 97.8 \| 3.6e-09 \| 8.3e-13 \| 102.0 \| 120 \| (509, 634) \| 769 \| (116, 237) \| 289 \| PPV\_E1\_C \| Papillomavirus helicase; pfam \| PF01057 \| 97.8 \| 4.2e-09 \| 1e-12 \| 99.7 \| 129 \| (497, 634) \| 769 \| (103, 241) \| 284 \| Parvo\_NS1 \| Parvovirus non-structural protein NS1; pfam \| PF06431 \| 97.3 \| 8.9e-08 \| 2e-11 \| 98.9 \| 119 \| (510, 634) \| 769 \| (153, 283) \| 417 \| Polyoma\_lg\_T\_C \| Polyomavirus large T antigen C-terminus; pfam \| PF11868 \| 96.4 \| 6.6e-06 \| 1.3e-09 \| 76.9 \| 76 \| (495, 574) \| 769 \| (39, 118) \| 190 \| DUF3388 \| Protein of unknown function (DUF3388); pfam \| PF00910 \| 95.5 \| 5.9e-05 \| 1.5e-08 \| 54.6 \| 100 \| (516, 616) \| 769 \| (1, 100) \| 100 \| RNA\_helicase \| RNA helicase; pfam \| PF20030 \| 94.7 \| 0.00023 \| 5.6e-08 \| 60.0 \| 132 \| (514, 654) \| 769 \| (32, 183) \| 204 \| bpMoxR \| MoxR domain in the MoxR-vWA-beta-propeller ternary systems; pfam \| PF01078 \| 94.6 \| 0.00028 \| 6.7e-08 \| 60.5 \| 24 \| (514, 537) \| 769 \| (23, 46) \| 208 \| Mg\_chelatase \| Magnesium chelatase, subunit ChlI; pfam \| PF07726 \| 94.6 \| 0.00029 \| 7.3e-08 \| 53.8 \| 23 \| (516, 538) \| 769 \| (2, 24) \| 134 \| AAA\_3 \| ATPase family associated with various cellular activities (AAA); pfam \| PF05496 \| 94.5 \| 0.00032 \| 8e-08 \| 55.0 \| 116 \| (514, 632) \| 769 \| (34, 156) \| 159 \| RuvB\_N \| Holliday junction DNA helicase RuvB P-loop domain; pfam \| PF19263 \| 94.2 \| 0.00042 \| 1.1e-07 \| 49.9 \| 95 \| (518, 618) \| 769 \| (1, 98) \| 110 \| DUF5906 \| Family of unknown function (DUF5906); pfam \| PF00493 \| 94.2 \| 0.00044 \| 1.1e-07 \| 56.9 \| 25 \| (514, 538) \| 769 \| (53, 77) \| 216 \| MCM \| MCM P-loop domain; cath \| 1q57G02 \| 94.6 \| 0.00026 \| 6.8e-08 \| 46.8 \| 59 \| (231, 289) \| 769 \| (13, 75) \| 126 \| Dna primase/helicase \| CATHCODE: 3.40.1360.10 NAME: Dna primase/helicase. Chain: a, b, c, d, e, f, g. Engineered: yes. Mutation: yes SOURCE: Enterobacteria phage t7. Organism\_taxid: 10760. Gene: 4. Expressed in: escherichia coli. Expression\_system\_taxid: 562 CLASS: Alpha Beta, ARCH: 3-Layer(aba) Sandwich, TOPOL: Dna Topoisomerase Vi A Subunit; Chain: A, domain 2, HOMOL: Dna Topoisomerase Vi A Subunit; Chain: A, domain 2; phrogs \| 2249 \| 100.0 \| 2.4e-76 \| 2.8e-80 \| 651.9 \| 633 \| (6, 749) \| 769 \| (46, 797) \| 814 \| DNA primase \| DNA primase; Category: DNA, RNA and nucleotide metabolism; p352003 VI\_04181; phrogs \| 256 \| 100.0 \| 9.7e-49 \| 1.2e-52 \| 437.6 \| 294 \| (439, 741) \| 769 \| (460, 789) \| 808 \| DNA helicase \| DNA helicase; Category: DNA, RNA and nucleotide metabolism; NC\_028991\_p19; phrogs \| 1313 \| 100.0 \| 2.6e-45 \| 3.2e-49 \| 394.4 \| 295 \| (443, 746) \| 769 \| (104, 436) \| 460 \| DNA helicase \| DNA helicase; Category: DNA, RNA and nucleotide metabolism; p88668 VI\_00162; phrogs \| 6088 \| 100.0 \| 1.7e-39 \| 2e-43 \| 353.0 \| 308 \| (424, 758) \| 769 \| (413, 753) \| 754 \| DNA primase \| DNA primase; Category: DNA, RNA and nucleotide metabolism; p124025 VI\_05756; phrogs \| 4913 \| 99.9 \| 2.8e-32 \| 3.3e-36 \| 305.1 \| 323 \| (1, 350) \| 769 \| (1, 332) \| 1099 \| replication \| replication; Category: DNA, RNA and nucleotide metabolism; NC\_019722\_p45; phrogs \| 7601 \| 99.9 \| 8.4e-32 \| 9.8e-36 \| 259.8 \| 264 \| (494, 765) \| 769 \| (8, 286) \| 304 \| NA \| NA; Category: unknown function; p167392 VI\_00748; phrogs \| 1156 \| 99.9 \| 6e-31 \| 7.6e-35 \| 307.2 \| 304 \| (426, 758) \| 769 \| (426, 777) \| 790 \| DNA primase \| DNA primase; Category: DNA, RNA and nucleotide metabolism; NC\_019550\_p28; phrogs \| 15962 \| 99.9 \| 1.3e-30 \| 1.5e-34 \| 273.3 \| 303 \| (440, 752) \| 769 \| (397, 722) \| 737 \| DNA primase \| DNA primase; Category: DNA, RNA and nucleotide metabolism; NC\_021800\_p32; phrogs \| 16593 \| 99.9 \| 1.6e-30 \| 1.8e-34 \| 269.5 \| 358 \| (380, 749) \| 769 \| (423, 795) \| 824 \| DNA primase \| DNA primase; Category: DNA, RNA and nucleotide metabolism; NC\_009604\_p54; phrogs \| 6444 \| 99.9 \| 6.2e-30 \| 7.2e-34 \| 271.5 \| 283 \| (6, 346) \| 769 \| (41, 368) \| 600 \| DNA primase \| DNA primase; Category: DNA, RNA and nucleotide metabolism; p94528 VI\_00468; phrogs \| 864 \| 99.9 \| 1.3e-29 \| 1.6e-33 \| 274.5 \| 274 \| (427, 735) \| 769 \| (169, 489) \| 526 \| DNA primase \| DNA primase; Category: DNA, RNA and nucleotide metabolism; NC\_025465\_p48; phrogs \| 510 \| 99.9 \| 1.5e-29 \| 1.8e-33 \| 266.0 \| 296 \| (6, 345) \| 769 \| (35, 367) \| 389 \| DNA primase \| DNA primase; Category: DNA, RNA and nucleotide metabolism; p276824 VI\_04255; phrogs \| 1139 \| 99.9 \| 4.2e-29 \| 5.1e-33 \| 281.8 \| 279 \| (6, 350) \| 769 \| (33, 318) \| 890 \| DNA primase \| DNA primase; Category: DNA, RNA and nucleotide metabolism; p412832 VI\_12129; phrogs \| 1883 \| 99.9 \| 4e-28 \| 4.6e-32 \| 255.9 \| 310 \| (425, 761) \| 769 \| (418, 760) \| 761 \| DNA helicase \| DNA helicase; Category: DNA, RNA and nucleotide metabolism; p89953 VI\_04942; phrogs \| 2478 \| 99.9 \| 2.9e-27 \| 3.3e-31 \| 253.7 \| 310 \| (424, 759) \| 769 \| (511, 853) \| 873 \| DNA polymerase/primase \| DNA polymerase/primase; Category: DNA, RNA and nucleotide metabolism; KY087992\_p73; phrogs \| 1829 \| 99.8 \| 1.1e-26 \| 1.3e-30 \| 261.9 \| 270 \| (423, 734) \| 769 \| (413, 698) \| 785 \| DNA primase \| DNA primase; Category: DNA, RNA and nucleotide metabolism; MG432137\_p16; phrogs \| 35358 \| 99.8 \| 7.4e-26 \| 8.3e-30 \| 233.4 \| 220 \| (135, 358) \| 769 \| (116, 354) \| 947 \| DNA primase \| DNA primase; Category: DNA, RNA and nucleotide metabolism; MF663786\_p98; phrogs \| 25777 \| 99.8 \| 2.2e-23 \| 2.5e-27 \| 214.1 \| 281 \| (6, 350) \| 769 \| (68, 354) \| 905 \| DNA primase \| DNA primase; Category: DNA, RNA and nucleotide metabolism; NC\_025458\_p2; phrogs \| 36873 \| 99.7 \| 3.1e-22 \| 3.4e-26 \| 203.5 \| 309 \| (429, 751) \| 769 \| (442, 757) \| 1062 \| DNA helicase \| DNA helicase; Category: DNA, RNA and nucleotide metabolism; NC\_029032\_p60; phrogs \| 16950 \| 99.7 \| 3.6e-21 \| 4.1e-25 \| 194.9 \| 280 \| (427, 734) \| 769 \| (480, 808) \| 844 \| DNA primase \| DNA primase; Category: DNA, RNA and nucleotide metabolism; p371263 VI\_11800; phrogs \| 47 \| 99.6 \| 1.4e-20 \| 1.8e-24 \| 198.5 \| 256 \| (8, 339) \| 769 \| (44, 313) \| 327 \| DNA primase \| DNA primase; Category: DNA, RNA and nucleotide metabolism; KX925554\_p102; phrogs \| 4630 \| 99.6 \| 2.8e-20 \| 3.2e-24 \| 188.3 \| 299 \| (444, 751) \| 769 \| (173, 522) \| 579 \| DNA primase \| DNA primase; Category: DNA, RNA and nucleotide metabolism; KP340288\_p74; phrogs \| 16213 \| 99.3 \| 4.6e-16 \| 5.1e-20 \| 155.3 \| 325 \| (428, 768) \| 769 \| (315, 668) \| 732 \| DNA primase \| DNA primase; Category: DNA, RNA and nucleotide metabolism; KU530220\_p53; phrogs \| 849 \| 99.1 \| 3.6e-15 \| 4.2e-19 \| 140.0 \| 177 \| (149, 346) \| 769 \| (29, 220) \| 223 \| DNA primase \| DNA primase; Category: DNA, RNA and nucleotide metabolism; NC\_027625\_p30; phrogs \| 168 \| 99.1 \| 7.4e-15 \| 9.1e-19 \| 150.1 \| 173 \| (497, 680) \| 769 \| (28, 216) \| 318 \| clamp loader of DNA polymerase \| clamp loader of DNA polymerase; Category: DNA, RNA and nucleotide metabolism; p26368 VI\_12322; phrogs \| 24712 \| 98.7 \| 2.1e-12 \| 2.4e-16 \| 131.2 \| 220 \| (376, 638) \| 769 \| (369, 599) \| 801 \| NA \| NA; Category: unknown function; LC102729\_p32; phrogs \| 4210 \| 98.5 \| 1.4e-11 \| 1.6e-15 \| 133.4 \| 245 \| (466, 734) \| 769 \| (137, 409) \| 1350 \| DNA polymerase \| DNA polymerase; Category: DNA, RNA and nucleotide metabolism; MG757154\_p64; phrogs \| 5629 \| 98.5 \| 3.3e-11 \| 3.7e-15 \| 122.7 \| 274 \| (10, 347) \| 769 \| (35, 351) \| 930 \| DNA primase \| DNA primase; Category: DNA, RNA and nucleotide metabolism; DQ087285\_p11; phrogs \| 4762 \| 97.7 \| 1.5e-08 \| 1.7e-12 \| 99.8 \| 212 \| (382, 632) \| 769 \| (404, 624) \| 822 \| NA \| NA; Category: unknown function; p277771 VI\_05561; phrogs \| 239 \| 97.4 \| 8.5e-08 \| 1.1e-11 \| 104.5 \| 164 \| (148, 342) \| 769 \| (47, 218) \| 543 \| DNA primase/helicase \| DNA primase/helicase; Category: DNA, RNA and nucleotide metabolism; NC\_031029\_p115; phrogs \| 5846 \| 97.4 \| 9.1e-08 \| 1.1e-11 \| 94.3 \| 119 \| (495, 638) \| 769 \| (25, 154) \| 306 \| clamp loader of DNA polymerase \| clamp loader of DNA polymerase; Category: DNA, RNA and nucleotide metabolism; KY606587\_p99; phrogs \| 36509 \| 97.2 \| 2.6e-07 \| 3e-11 \| 94.6 \| 259 \| (426, 706) \| 769 \| (516, 829) \| 1021 \| DNA primase \| DNA primase; Category: DNA, RNA and nucleotide metabolism; KU234533\_p69; phrogs \| 3757 \| 97.1 \| 6.3e-07 \| 7.5e-11 \| 89.6 \| 129 \| (509, 652) \| 769 \| (128, 274) \| 379 \| NA \| NA; Category: unknown function; p40711 VI\_00882; phrogs \| 11992 \| 97.1 \| 6.5e-07 \| 7.6e-11 \| 78.3 \| 60 \| (6, 81) \| 769 \| (33, 93) \| 166 \| NA \| NA; Category: unknown function; p110551 VI\_01281; phrogs \| 19785 \| 97.0 \| 9.3e-07 \| 1e-10 \| 88.2 \| 156 \| (468, 638) \| 769 \| (428, 588) \| 753 \| DNA polymerase/primase \| DNA polymerase/primase; Category: DNA, RNA and nucleotide metabolism; NC\_019420\_p20; phrogs \| 7485 \| 96.7 \| 3.4e-06 \| 3.9e-10 \| 84.3 \| 62 \| (509, 572) \| 769 \| (215, 280) \| 408 \| NA \| NA; Category: unknown function; NC\_019406\_p235; phrogs \| 11838 \| 96.3 \| 1.3e-05 \| 1.5e-09 \| 77.7 \| 62 \| (510, 572) \| 769 \| (42, 114) \| 413 \| clamp loader of DNA polymerase \| clamp loader of DNA polymerase; Category: DNA, RNA and nucleotide metabolism; p389573 VI\_12493; phrogs \| 9172 \| 95.5 \| 0.00013 \| 1.5e-08 \| 73.4 \| 134 \| (495, 636) \| 769 \| (82, 239) \| 518 \| NA \| NA; Category: unknown function; KX607102\_p48; phrogs \| 30773 \| 94.9 \| 0.00038 \| 4.3e-08 \| 67.7 \| 102 \| (509, 622) \| 769 \| (176, 300) \| 647 \| NA \| NA; Category: unknown function; p438992 VI\_07698; phrogs \| 25243 \| 94.9 \| 0.0004 \| 4.6e-08 \| 68.9 \| 118 \| (513, 632) \| 769 \| (230, 354) \| 619 \| DNA primase \| DNA primase; Category: DNA, RNA and nucleotide metabolism; NC\_019516\_p159; phrogs \| 16448 \| 94.8 \| 0.00045 \| 5.2e-08 \| 62.5 \| 121 \| (512, 636) \| 769 \| (26, 169) \| 246 \| ATPase \| ATPase; Category: other; KU886274\_p16; phrogs \| 10401 \| 94.8 \| 0.00046 \| 5.4e-08 \| 67.8 \| 115 \| (512, 638) \| 769 \| (198, 330) \| 385 \| ATPase \| ATPase; Category: other; MG189906\_p41; phrogs \| 14322 \| 94.6 \| 0.00062 \| 7.1e-08 \| 64.6 \| 114 \| (512, 638) \| 769 \| (194, 321) \| 366 \| NA \| NA; Category: unknown function; KU521356\_p250 |
| Top keywords  (threshold 1.00e-03 (evalue)) | **DNA, RNA, and, nucleotide, metabolism, primase, helicase, polymerase, A, ATPase** |
| Output files | ../../domain\_architecture/54\_FANPEZAQ\_CDS\_0054\_cath.hhr ../../domain\_architecture/54\_FANPEZAQ\_CDS\_0054\_merged.svg ../../domain\_architecture/54\_FANPEZAQ\_CDS\_0054\_ncbi-cd.hhr ../../domain\_architecture/54\_FANPEZAQ\_CDS\_0054\_pfam.hhr ../../domain\_architecture/54\_FANPEZAQ\_CDS\_0054\_phrogs.hhr |

### Identical protein sequences/structures

#### Search results

|  |  |
| --- | --- |
| Protein sequence databases searched | Pdb, Swissprot, Refseq |
| Identical proteins found | -- |
| Top keywords | -- |
| Output files | -- |

### Similar protein sequences/structures

#### Sequence similarity search results (HHblits)^1^

|  |  |
| --- | --- |
| Sequence databases searched | Uniclust, Pdb70 |
| Results, scheme(s)  (Top layers only, threshold 1.00e-03 (evalue)) | xml version="1.0" encoding="utf-8" standalone="no"?       2024-09-02T21:08:52.456549 image/svg+xml   Matplotlib v3.7.2, https://matplotlib.org/ |
| Results, table(s)  (threshold 1.00e-03 (evalue)) | db \| id \| prob \| evalue \| pvalue \| score \| cols \| query \| query\_len \| template \| template\_len \| name \| description; uniclust \| UniRef100\_A0A073ISL6 \| 100.0 \| 2e-108 \| 5e-114 \| 879.9 \| 728 \| (14, 762) \| 769 \| (49, 798) \| 813 \| Uncharacterized protein \| Uncharacterized protein; uniclust \| UniRef100\_A0A158S187 \| 100.0 \| 2.6e-96 \| 5e-102 \| 756.6 \| 728 \| (14, 754) \| 769 \| (26, 767) \| 779 \| DNA primase domain protein \| DNA primase domain protein; uniclust \| UniRef100\_A0A024HDH8 \| 100.0 \| 1.9e-92 \| 3.6e-98 \| 754.5 \| 687 \| (5, 748) \| 769 \| (10, 733) \| 767 \| Virulence-associated E family protein \| Virulence-associated E family protein; uniclust \| UniRef100\_A0A1G3JY92 \| 100.0 \| 5.4e-92 \| 1e-97 \| 731.9 \| 679 \| (16, 739) \| 769 \| (19, 737) \| 778 \| Uncharacterized protein \| Uncharacterized protein; uniclust \| UniRef100\_A0A2G2PN86 \| 100.0 \| 7.3e-90 \| 1.4e-95 \| 705.2 \| 702 \| (5, 756) \| 769 \| (4, 753) \| 769 \| Toprim domain-containing protein \| Toprim domain-containing protein; uniclust \| UniRef100\_A0A4V2HU15 \| 100.0 \| 7.9e-82 \| 1.5e-87 \| 650.5 \| 711 \| (14, 750) \| 769 \| (29, 994) \| 1004 \| Uncharacterized protein \| Uncharacterized protein; uniclust \| UniRef100\_A0A349EG17 \| 100.0 \| 4.2e-77 \| 7.8e-83 \| 593.8 \| 669 \| (9, 754) \| 769 \| (3, 689) \| 694 \| Toprim domain-containing protein \| Toprim domain-containing protein; uniclust \| UniRef100\_A0A080LT75 \| 100.0 \| 4.7e-73 \| 9.1e-79 \| 615.4 \| 683 \| (14, 755) \| 769 \| (32, 804) \| 831 \| DNA primase TraC \| DNA primase TraC; uniclust \| UniRef100\_A0A345ANS2 \| 100.0 \| 2e-66 \| 3.7e-72 \| 543.8 \| 693 \| (37, 769) \| 769 \| (60, 936) \| 936 \| Putative primase \| Putative primase; uniclust \| UniRef100\_A0A159Z405 \| 100.0 \| 2.3e-66 \| 4.4e-72 \| 534.6 \| 582 \| (135, 742) \| 769 \| (173, 868) \| 902 \| Virulence-associated protein E \| Virulence-associated protein E; uniclust \| UniRef100\_A0A016XJH4 \| 100.0 \| 1.2e-64 \| 2.3e-70 \| 546.5 \| 652 \| (55, 744) \| 769 \| (77, 804) \| 837 \| Conjugal transfer protein TraC \| Conjugal transfer protein TraC; uniclust \| UniRef100\_A0A151FI48 \| 100.0 \| 1.1e-63 \| 2.2e-69 \| 524.7 \| 684 \| (10, 752) \| 769 \| (17, 804) \| 828 \| Zinc finger CHC2-type domain-containing protein \| Zinc finger CHC2-type domain-containing protein; uniclust \| UniRef100\_A0A158EA01 \| 100.0 \| 6.2e-61 \| 1.2e-66 \| 487.7 \| 517 \| (233, 768) \| 769 \| (191, 726) \| 727 \| Virulence-associated protein E \| Virulence-associated protein E; uniclust \| UniRef100\_A0A094ZFQ5 \| 100.0 \| 9.8e-61 \| 1.9e-66 \| 494.3 \| 496 \| (231, 756) \| 769 \| (164, 681) \| 699 \| Virulence-associated E-like domain-containing protein \| Virulence-associated E-like domain-containing protein; uniclust \| UniRef100\_A0A1U9VG46 \| 100.0 \| 1.4e-60 \| 2.6e-66 \| 497.4 \| 559 \| (180, 751) \| 769 \| (150, 772) \| 810 \| Virulence-associated E-like domain-containing protein \| Virulence-associated E-like domain-containing protein; uniclust \| UniRef100\_A0A2G4RGB3 \| 100.0 \| 1.5e-58 \| 2.8e-64 \| 469.5 \| 436 \| (277, 741) \| 769 \| (28, 482) \| 516 \| Virulence-associated E family protein \| Virulence-associated E family protein; uniclust \| UniRef100\_A0A962X8E3 \| 100.0 \| 1.9e-58 \| 3.5e-64 \| 456.6 \| 550 \| (177, 752) \| 769 \| (13, 612) \| 633 \| Toprim domain-containing protein (Fragment) \| Toprim domain-containing protein (Fragment); uniclust \| UniRef100\_A0A1D2U957 \| 100.0 \| 3e-58 \| 5.8e-64 \| 463.7 \| 384 \| (368, 754) \| 769 \| (78, 461) \| 505 \| Virulence-associated E-like domain-containing protein \| Virulence-associated E-like domain-containing protein; uniclust \| UniRef100\_A0A059N520 \| 100.0 \| 2.2e-57 \| 4.3e-63 \| 482.4 \| 359 \| (372, 741) \| 769 \| (217, 590) \| 642 \| Virulence-associated protein E \| Virulence-associated protein E; uniclust \| UniRef100\_A0A096HFV3 \| 100.0 \| 2.5e-57 \| 4.8e-63 \| 484.6 \| 665 \| (2, 735) \| 769 \| (2, 748) \| 863 \| DUF5906 domain-containing protein \| DUF5906 domain-containing protein; uniclust \| UniRef100\_A0A0J6VXK3 \| 100.0 \| 2e-56 \| 3.9e-62 \| 461.8 \| 367 \| (371, 752) \| 769 \| (139, 522) \| 563 \| Virulence-associated E family protein \| Virulence-associated E family protein; uniclust \| UniRef100\_UPI00196A8CA5 \| 100.0 \| 1.1e-55 \| 2e-61 \| 445.8 \| 676 \| (48, 755) \| 769 \| (36, 766) \| 771 \| VapE family protein \| VapE family protein; uniclust \| UniRef100\_A0A0E3K4D7 \| 100.0 \| 1e-54 \| 1.9e-60 \| 426.2 \| 368 \| (378, 753) \| 769 \| (48, 448) \| 464 \| Virulence-associated E family protein \| Virulence-associated E family protein; uniclust \| UniRef100\_A0A094YPP6 \| 100.0 \| 3.5e-54 \| 6.8e-60 \| 470.0 \| 359 \| (373, 746) \| 769 \| (380, 754) \| 775 \| Virulence-associated E \| Virulence-associated E; uniclust \| UniRef100\_A0A2M8YCJ3 \| 100.0 \| 5.5e-54 \| 1e-59 \| 425.7 \| 390 \| (365, 754) \| 769 \| (156, 545) \| 575 \| Virulence-associated protein E \| Virulence-associated protein E; uniclust \| UniRef100\_A0A551Z1W1 \| 100.0 \| 7e-54 \| 1.3e-59 \| 444.9 \| 481 \| (233, 753) \| 769 \| (171, 673) \| 804 \| Virulence-associated E-like domain-containing protein \| Virulence-associated E-like domain-containing protein; uniclust \| UniRef100\_A0A0A2WCR2 \| 100.0 \| 8.3e-54 \| 1.6e-59 \| 433.2 \| 353 \| (372, 741) \| 769 \| (82, 443) \| 498 \| Phage associated DNA primase \| Phage associated DNA primase; uniclust \| UniRef100\_UPI000DE53851 \| 100.0 \| 1.3e-53 \| 2.4e-59 \| 398.0 \| 482 \| (13, 503) \| 769 \| (13, 494) \| 496 \| PriCT-2 domain-containing protein \| PriCT-2 domain-containing protein; uniclust \| UniRef100\_A0A1H1ILW6 \| 100.0 \| 3.2e-53 \| 5.9e-59 \| 402.3 \| 542 \| (31, 619) \| 769 \| (19, 579) \| 581 \| Putative DNA primase/helicase \| Putative DNA primase/helicase; uniclust \| UniRef100\_A0A258SSH2 \| 100.0 \| 3.2e-53 \| 6e-59 \| 411.6 \| 386 \| (371, 757) \| 769 \| (15, 400) \| 420 \| Virulence-associated E-like domain-containing protein (Fragment) \| Virulence-associated E-like domain-containing protein (Fragment); uniclust \| UniRef100\_A0A1E5XQN8 \| 100.0 \| 3.3e-53 \| 6.2e-59 \| 430.5 \| 466 \| (233, 741) \| 769 \| (147, 623) \| 679 \| Virulence-associated E-like domain-containing protein \| Virulence-associated E-like domain-containing protein; uniclust \| UniRef100\_A0A0J7XW03 \| 100.0 \| 3.3e-53 \| 6.3e-59 \| 431.5 \| 381 \| (373, 755) \| 769 \| (147, 529) \| 558 \| Virulence-associated E-like domain-containing protein \| Virulence-associated E-like domain-containing protein; uniclust \| UniRef100\_A0A023WUM9 \| 100.0 \| 1.1e-52 \| 2e-58 \| 440.4 \| 677 \| (10, 735) \| 769 \| (52, 877) \| 956 \| DNA primase \| DNA primase; uniclust \| UniRef100\_A0A090TZ23 \| 100.0 \| 1.2e-52 \| 2.2e-58 \| 436.5 \| 687 \| (14, 749) \| 769 \| (14, 816) \| 834 \| DNA primase phage-associated \| DNA primase phage-associated; uniclust \| UniRef100\_A0A2Z5ME49 \| 100.0 \| 1.7e-52 \| 3.3e-58 \| 416.8 \| 373 \| (378, 758) \| 769 \| (160, 533) \| 559 \| Virulence-associated E family protein \| Virulence-associated E family protein; uniclust \| UniRef100\_A0A1Z4GW22 \| 100.0 \| 2.6e-52 \| 4.9e-58 \| 442.7 \| 477 \| (232, 742) \| 769 \| (159, 650) \| 933 \| Virulence-associated E family protein \| Virulence-associated E family protein; uniclust \| UniRef100\_A0A013UG67 \| 100.0 \| 3.4e-52 \| 6.7e-58 \| 461.8 \| 600 \| (100, 736) \| 769 \| (101, 815) \| 888 \| Toprim domain protein \| Toprim domain protein; uniclust \| UniRef100\_A0A0T7BQB0 \| 100.0 \| 4.4e-52 \| 8.4e-58 \| 442.7 \| 477 \| (231, 743) \| 769 \| (173, 666) \| 908 \| Uncharacterized protein \| Uncharacterized protein; uniclust \| UniRef100\_A0A0N8K9E5 \| 100.0 \| 4.6e-52 \| 8.7e-58 \| 415.6 \| 371 \| (372, 751) \| 769 \| (62, 458) \| 497 \| Putative P-loop ATPase \| Putative P-loop ATPase; uniclust \| UniRef100\_A0A0Q6QE44 \| 100.0 \| 5.2e-52 \| 9.8e-58 \| 425.5 \| 374 \| (374, 756) \| 769 \| (195, 590) \| 667 \| Virulence protein E \| Virulence protein E; uniclust \| UniRef100\_A0A099GLM4 \| 100.0 \| 6.4e-52 \| 1.2e-57 \| 416.5 \| 582 \| (136, 743) \| 769 \| (192, 903) \| 944 \| p-loop ATPase \| p-loop ATPase; uniclust \| UniRef100\_A0A2N3DYE7 \| 100.0 \| 9.3e-52 \| 1.8e-57 \| 444.1 \| 343 \| (393, 749) \| 769 \| (345, 699) \| 726 \| DNA primase/polymerase bifunctional N-terminal domain-containing protein \| DNA primase/polymerase bifunctional N-terminal domain-containing protein; uniclust \| UniRef100\_A0A2T1EHM2 \| 100.0 \| 2.2e-51 \| 4.2e-57 \| 417.9 \| 488 \| (231, 758) \| 769 \| (150, 661) \| 676 \| Virulence-associated E-like domain-containing protein \| Virulence-associated E-like domain-containing protein; uniclust \| UniRef100\_A0A1M5W1E5 \| 100.0 \| 5.7e-51 \| 1.1e-56 \| 430.7 \| 637 \| (14, 697) \| 769 \| (24, 767) \| 933 \| Phage/plasmid primase, P4 family, C-terminal domain-containing protein \| Phage/plasmid primase, P4 family, C-terminal domain-containing protein; uniclust \| UniRef100\_A0A0S7XZ70 \| 100.0 \| 2.1e-50 \| 4e-56 \| 431.2 \| 564 \| (14, 637) \| 769 \| (61, 727) \| 965 \| Toprim domain-containing protein \| Toprim domain-containing protein; uniclust \| UniRef100\_A0A068MV01 \| 100.0 \| 2.3e-50 \| 4.4e-56 \| 443.9 \| 483 \| (233, 753) \| 769 \| (160, 666) \| 903 \| Uncharacterized protein \| Uncharacterized protein; uniclust \| UniRef100\_A0A060H485 \| 100.0 \| 2.8e-50 \| 5.3e-56 \| 402.1 \| 360 \| (372, 742) \| 769 \| (105, 495) \| 535 \| Virulence factor \| Virulence factor; uniclust \| UniRef100\_A0A317J441 \| 100.0 \| 3.6e-50 \| 6.8e-56 \| 402.6 \| 501 \| (233, 760) \| 769 \| (158, 716) \| 731 \| Uncharacterized protein \| Uncharacterized protein; uniclust \| UniRef100\_A0A0F9U4J5 \| 100.0 \| 3.9e-50 \| 7.4e-56 \| 419.5 \| 379 \| (371, 755) \| 769 \| (355, 756) \| 777 \| DNA primase/polymerase bifunctional N-terminal domain-containing protein \| DNA primase/polymerase bifunctional N-terminal domain-containing protein; uniclust \| UniRef100\_A0A0W7WDM8 \| 100.0 \| 5.7e-50 \| 1.1e-55 \| 419.7 \| 370 \| (372, 753) \| 769 \| (383, 764) \| 771 \| Virulence-associated E-like domain-containing protein \| Virulence-associated E-like domain-containing protein; uniclust \| UniRef100\_A0A014ML09 \| 100.0 \| 8.4e-50 \| 1.6e-55 \| 419.9 \| 516 \| (219, 742) \| 769 \| (223, 837) \| 903 \| Virulence protein E \| Virulence protein E; uniclust \| UniRef100\_A0A031IMY7 \| 100.0 \| 9.2e-50 \| 1.8e-55 \| 423.1 \| 372 \| (373, 753) \| 769 \| (380, 773) \| 824 \| Phage/plasmid primase, P4 family \| Phage/plasmid primase, P4 family; uniclust \| UniRef100\_A0A0D0QS22 \| 100.0 \| 1.3e-49 \| 2.4e-55 \| 406.5 \| 357 \| (372, 739) \| 769 \| (193, 595) \| 639 \| DNA primase, phage associated \| DNA primase, phage associated; uniclust \| UniRef100\_A0A0H4XB71 \| 100.0 \| 1.6e-49 \| 3.2e-55 \| 394.5 \| 308 \| (437, 753) \| 769 \| (23, 338) \| 380 \| DNA primase domain protein \| DNA primase domain protein; uniclust \| UniRef100\_A0A017N0X3 \| 100.0 \| 1.7e-49 \| 3.4e-55 \| 441.4 \| 356 \| (371, 738) \| 769 \| (408, 808) \| 878 \| Virulence-associated E family protein \| Virulence-associated E family protein; uniclust \| UniRef100\_UPI001FCAD0C1 \| 100.0 \| 2.2e-49 \| 4e-55 \| 389.3 \| 683 \| (14, 750) \| 769 \| (15, 805) \| 826 \| phage/plasmid primase, P4 family \| phage/plasmid primase, P4 family; uniclust \| UniRef100\_A0A4P7L7L6 \| 100.0 \| 2.4e-49 \| 4.3e-55 \| 387.3 \| 575 \| (130, 743) \| 769 \| (107, 701) \| 713 \| DNA primase TraC \| DNA primase TraC; uniclust \| UniRef100\_A0A078S3U0 \| 100.0 \| 2.5e-49 \| 4.7e-55 \| 394.7 \| 368 \| (377, 755) \| 769 \| (92, 472) \| 511 \| Virulence-associated E family protein (Fragment) \| Virulence-associated E family protein (Fragment); uniclust \| UniRef100\_A0A158DR21 \| 100.0 \| 3.6e-49 \| 6.9e-55 \| 426.9 \| 478 \| (14, 589) \| 769 \| (46, 535) \| 858 \| TOPRIM domain-containing protein \| TOPRIM domain-containing protein; uniclust \| UniRef100\_A0A109LYF1 \| 100.0 \| 3.7e-49 \| 7.1e-55 \| 424.8 \| 365 \| (367, 742) \| 769 \| (390, 759) \| 817 \| Virulence-associated E-like domain-containing protein \| Virulence-associated E-like domain-containing protein; uniclust \| UniRef100\_A0A061P292 \| 100.0 \| 3.7e-49 \| 7.2e-55 \| 404.7 \| 353 \| (375, 737) \| 769 \| (43, 451) \| 502 \| DNA primase \| DNA primase; uniclust \| UniRef100\_A0A0D6MP65 \| 100.0 \| 3.8e-49 \| 7.3e-55 \| 409.6 \| 366 \| (370, 750) \| 769 \| (333, 714) \| 721 \| Virulence-associated E family protein \| Virulence-associated E family protein; uniclust \| UniRef100\_A0A945RBT7 \| 100.0 \| 4e-49 \| 7.4e-55 \| 386.6 \| 653 \| (14, 747) \| 769 \| (44, 785) \| 805 \| Toprim domain-containing protein \| Toprim domain-containing protein; uniclust \| UniRef100\_A0A096BB79 \| 100.0 \| 4.9e-49 \| 9.4e-55 \| 409.8 \| 355 \| (370, 738) \| 769 \| (129, 508) \| 618 \| Virulence-associated E-like domain-containing protein \| Virulence-associated E-like domain-containing protein; uniclust \| UniRef100\_A0A0F9W6J4 \| 100.0 \| 8.3e-49 \| 1.6e-54 \| 382.9 \| 355 \| (372, 741) \| 769 \| (85, 455) \| 475 \| Virulence-associated E-like domain-containing protein \| Virulence-associated E-like domain-containing protein; uniclust \| UniRef100\_A0A0D6Q293 \| 100.0 \| 8.6e-49 \| 1.6e-54 \| 393.5 \| 359 \| (367, 737) \| 769 \| (58, 440) \| 486 \| DNA primase \| DNA primase; uniclust \| UniRef100\_A0A651E934 \| 100.0 \| 9.3e-49 \| 1.8e-54 \| 403.6 \| 470 \| (233, 741) \| 769 \| (134, 617) \| 803 \| DUF3854 domain-containing protein \| DUF3854 domain-containing protein; uniclust \| UniRef100\_A0A084J7F2 \| 100.0 \| 1.3e-48 \| 2.5e-54 \| 401.9 \| 488 \| (231, 757) \| 769 \| (187, 693) \| 730 \| Zinc finger CHC2-type domain-containing protein \| Zinc finger CHC2-type domain-containing protein; uniclust \| UniRef100\_A0A0T2QCS5 \| 100.0 \| 1.8e-48 \| 3.4e-54 \| 381.3 \| 369 \| (374, 755) \| 769 \| (39, 408) \| 431 \| Virulence-associated E-like domain-containing protein \| Virulence-associated E-like domain-containing protein; uniclust \| UniRef100\_A0A1U7IAQ8 \| 100.0 \| 2e-48 \| 3.8e-54 \| 409.5 \| 483 \| (231, 756) \| 769 \| (159, 665) \| 933 \| Virulence-associated E-like domain-containing protein \| Virulence-associated E-like domain-containing protein; uniclust \| UniRef100\_A0A0K0WP96 \| 100.0 \| 3.3e-48 \| 6.1e-54 \| 386.0 \| 525 \| (177, 740) \| 769 \| (111, 661) \| 773 \| Uncharacterized protein \| Uncharacterized protein; uniclust \| UniRef100\_A0A076Z3T2 \| 100.0 \| 3.2e-48 \| 6.3e-54 \| 400.0 \| 317 \| (372, 696) \| 769 \| (52, 383) \| 500 \| Predicted P-loop ATPase and inactivated derivatives \| Predicted P-loop ATPase and inactivated derivatives; uniclust \| UniRef100\_A0A0P1IL64 \| 100.0 \| 3.4e-48 \| 6.6e-54 \| 415.6 \| 381 \| (366, 756) \| 769 \| (408, 799) \| 829 \| Putative P-loop ATPase \| Putative P-loop ATPase; uniclust \| UniRef100\_A0A1I1CGG0 \| 100.0 \| 1.1e-47 \| 2.1e-53 \| 374.2 \| 366 \| (377, 754) \| 769 \| (8, 389) \| 439 \| Virulence-associated protein E (Fragment) \| Virulence-associated protein E (Fragment); uniclust \| UniRef100\_A0A2V5PJC8 \| 100.0 \| 1.7e-47 \| 3.2e-53 \| 401.6 \| 656 \| (12, 735) \| 769 \| (8, 763) \| 837 \| DUF5906 domain-containing protein \| DUF5906 domain-containing protein; uniclust \| UniRef100\_A0A017RTM7 \| 100.0 \| 3.1e-47 \| 6.1e-53 \| 419.7 \| 356 \| (371, 737) \| 769 \| (421, 824) \| 891 \| Virulence-associated protein E \| Virulence-associated protein E; uniclust \| UniRef100\_A0A2H9UPD3 \| 100.0 \| 4.7e-47 \| 8.9e-53 \| 386.4 \| 390 \| (367, 758) \| 769 \| (357, 753) \| 760 \| Primase C-terminal 2 domain-containing protein \| Primase C-terminal 2 domain-containing protein; uniclust \| UniRef100\_J9ZDI3 \| 100.0 \| 5.6e-47 \| 1e-52 \| 375.7 \| 528 \| (177, 741) \| 769 \| (95, 646) \| 747 \| Putative P-loop ATPase \| Putative P-loop ATPase; uniclust \| UniRef100\_A0A094SVM7 \| 100.0 \| 6.8e-47 \| 1.3e-52 \| 392.9 \| 556 \| (150, 757) \| 769 \| (159, 774) \| 802 \| Helicase \| Helicase; uniclust \| UniRef100\_A0A015SLF0 \| 100.0 \| 7e-47 \| 1.4e-52 \| 416.2 \| 366 \| (378, 755) \| 769 \| (408, 783) \| 822 \| Virulence-associated E family protein \| Virulence-associated E family protein; uniclust \| UniRef100\_A0A0C1QT32 \| 100.0 \| 8.8e-47 \| 1.7e-52 \| 365.0 \| 288 \| (14, 345) \| 769 \| (18, 305) \| 314 \| Toprim domain-containing protein \| Toprim domain-containing protein; uniclust \| UniRef100\_UPI0003678CFF \| 100.0 \| 1.2e-46 \| 2.1e-52 \| 378.5 \| 685 \| (14, 760) \| 769 \| (14, 845) \| 867 \| DUF5906 domain-containing protein \| DUF5906 domain-containing protein; uniclust \| UniRef100\_A0A1H8FZ14 \| 100.0 \| 1.2e-46 \| 2.3e-52 \| 388.3 \| 369 \| (376, 753) \| 769 \| (456, 842) \| 903 \| Virulence-associated protein E \| Virulence-associated protein E; uniclust \| UniRef100\_A0A0R3NCG4 \| 100.0 \| 1.3e-46 \| 2.4e-52 \| 383.1 \| 346 \| (384, 741) \| 769 \| (196, 549) \| 594 \| Virulence-associated E-like domain-containing protein \| Virulence-associated E-like domain-containing protein; uniclust \| UniRef100\_A0A6J5LA04 \| 100.0 \| 1.3e-46 \| 2.5e-52 \| 383.6 \| 396 \| (366, 767) \| 769 \| (388, 784) \| 786 \| Virulence-associated E \| Virulence-associated E; uniclust \| UniRef100\_A0A076P157 \| 100.0 \| 1.9e-46 \| 3.7e-52 \| 395.1 \| 363 \| (379, 755) \| 769 \| (108, 475) \| 518 \| Virulence-associated protein E \| Virulence-associated protein E; uniclust \| UniRef100\_A0A3G9FYK3 \| 100.0 \| 2e-46 \| 3.8e-52 \| 370.3 \| 387 \| (366, 756) \| 769 \| (193, 585) \| 591 \| Putative virulence-associated protein E \| Putative virulence-associated protein E; uniclust \| UniRef100\_A0A132Z4M9 \| 100.0 \| 2.1e-46 \| 4.1e-52 \| 377.4 \| 313 \| (378, 697) \| 769 \| (40, 366) \| 508 \| Helicase \| Helicase; uniclust \| UniRef100\_A0A1F9M777 \| 100.0 \| 2.2e-46 \| 4.3e-52 \| 375.1 \| 296 \| (14, 349) \| 769 \| (86, 382) \| 429 \| Toprim domain-containing protein \| Toprim domain-containing protein; uniclust \| UniRef100\_A0A251W7E6 \| 100.0 \| 2.4e-46 \| 4.6e-52 \| 380.2 \| 336 \| (386, 741) \| 769 \| (228, 571) \| 612 \| Virulence-associated E-like domain-containing protein \| Virulence-associated E-like domain-containing protein; uniclust \| UniRef100\_A0A7C2DBA8 \| 100.0 \| 2.7e-46 \| 5e-52 \| 374.7 \| 345 \| (391, 752) \| 769 \| (557, 917) \| 920 \| Zinc finger CHC2-type domain-containing protein \| Zinc finger CHC2-type domain-containing protein; uniclust \| UniRef100\_A0A2H2X2H4 \| 100.0 \| 3.1e-46 \| 5.9e-52 \| 386.0 \| 470 \| (233, 741) \| 769 \| (187, 675) \| 869 \| Virulence-associated E family protein \| Virulence-associated E family protein; uniclust \| UniRef100\_A0A1W7AEU1 \| 100.0 \| 5e-46 \| 9.4e-52 \| 388.7 \| 356 \| (374, 741) \| 769 \| (324, 721) \| 742 \| Virulence-associated protein E \| Virulence-associated protein E; uniclust \| UniRef100\_A0A085TW06 \| 100.0 \| 5.3e-46 \| 9.9e-52 \| 382.0 \| 365 \| (373, 752) \| 769 \| (414, 795) \| 805 \| Uncharacterized protein \| Uncharacterized protein; uniclust \| UniRef100\_A0A1N7PIB5 \| 100.0 \| 5.4e-46 \| 1e-51 \| 381.9 \| 365 \| (374, 753) \| 769 \| (339, 720) \| 728 \| Primase C terminal 2 (PriCT-2) \| Primase C terminal 2 (PriCT-2); uniclust \| UniRef100\_A0A031JL01 \| 100.0 \| 7.1e-46 \| 1.4e-51 \| 377.8 \| 285 \| (14, 340) \| 769 \| (69, 354) \| 402 \| Toprim domain-containing protein \| Toprim domain-containing protein; uniclust \| UniRef100\_A0A6L3T504 \| 100.0 \| 7.8e-46 \| 1.4e-51 \| 366.4 \| 698 \| (14, 750) \| 769 \| (16, 863) \| 871 \| SF3 helicase domain-containing protein \| SF3 helicase domain-containing protein; uniclust \| UniRef100\_A0A0P1I0V3 \| 100.0 \| 1.2e-45 \| 2.3e-51 \| 383.1 \| 340 \| (380, 738) \| 769 \| (406, 767) \| 815 \| Putative P-loop ATPase \| Putative P-loop ATPase; uniclust \| UniRef100\_A0A838CHG7 \| 100.0 \| 1.3e-45 \| 2.5e-51 \| 353.7 \| 410 \| (324, 751) \| 769 \| (12, 469) \| 498 \| Virulence protein E \| Virulence protein E; uniclust \| UniRef100\_A0A015VAS0 \| 100.0 \| 1.5e-45 \| 2.9e-51 \| 380.4 \| 356 \| (379, 741) \| 769 \| (80, 443) \| 497 \| Virulence-associated E family protein \| Virulence-associated E family protein; uniclust \| UniRef100\_A0A014P2B8 \| 100.0 \| 1.5e-45 \| 2.9e-51 \| 396.9 \| 332 \| (3, 349) \| 769 \| (12, 347) \| 991 \| Membrane protein \| Membrane protein; uniclust \| UniRef100\_A0A374DQX8 \| 100.0 \| 1.7e-45 \| 3.2e-51 \| 387.0 \| 348 \| (379, 739) \| 769 \| (315, 683) \| 731 \| Virulence-associated E-like domain-containing protein \| Virulence-associated E-like domain-containing protein; uniclust \| UniRef100\_A0A0N7LN66 \| 100.0 \| 2e-45 \| 3.9e-51 \| 387.0 \| 357 \| (371, 739) \| 769 \| (400, 775) \| 812 \| Putative P-loop ATPase \| Putative P-loop ATPase; uniclust \| UniRef100\_A0A317N8C0 \| 100.0 \| 2.1e-45 \| 4.1e-51 \| 388.9 \| 599 \| (14, 637) \| 769 \| (27, 693) \| 870 \| Phage/plasmid primase-like uncharacterized protein \| Phage/plasmid primase-like uncharacterized protein; uniclust \| UniRef100\_A0A158DVE4 \| 100.0 \| 2.2e-45 \| 4.3e-51 \| 409.4 \| 316 \| (14, 351) \| 769 \| (57, 382) \| 994 \| Inner membrane protein \| Inner membrane protein; uniclust \| UniRef100\_A0A4R5Q8H9 \| 100.0 \| 2.6e-45 \| 4.8e-51 \| 373.1 \| 361 \| (372, 750) \| 769 \| (350, 729) \| 753 \| Virulence-associated E-like domain-containing protein \| Virulence-associated E-like domain-containing protein; uniclust \| UniRef100\_A0A0Q9YMA6 \| 100.0 \| 2.5e-45 \| 4.9e-51 \| 404.9 \| 290 \| (14, 350) \| 769 \| (47, 336) \| 946 \| DNA primase TraC \| DNA primase TraC; uniclust \| UniRef100\_A0A2M7EZU9 \| 100.0 \| 2.6e-45 \| 5e-51 \| 361.9 \| 350 \| (378, 740) \| 769 \| (68, 444) \| 469 \| Virulence-associated E-like domain-containing protein \| Virulence-associated E-like domain-containing protein; uniclust \| UniRef100\_A0A078R7F8 \| 100.0 \| 4.2e-45 \| 8.2e-51 \| 397.6 \| 372 \| (377, 756) \| 769 \| (337, 717) \| 748 \| Virulence-associated E family protein \| Virulence-associated E family protein; uniclust \| UniRef100\_A0A034T3J5 \| 100.0 \| 4.5e-45 \| 8.6e-51 \| 393.3 \| 479 \| (132, 656) \| 769 \| (139, 662) \| 829 \| Toprim domain protein \| Toprim domain protein; uniclust \| UniRef100\_A0A0C9PJH3 \| 100.0 \| 5.6e-45 \| 1.1e-50 \| 406.9 \| 483 \| (233, 754) \| 769 \| (195, 759) \| 898 \| ATPase \| ATPase; uniclust \| UniRef100\_A0A442XE73 \| 100.0 \| 6e-45 \| 1.1e-50 \| 348.3 \| 295 \| (434, 737) \| 769 \| (59, 388) \| 420 \| Virulence-associated E-like domain-containing protein \| Virulence-associated E-like domain-containing protein; uniclust \| UniRef100\_A0A1G3DUL0 \| 100.0 \| 9.6e-45 \| 1.9e-50 \| 361.6 \| 283 \| (14, 343) \| 769 \| (44, 326) \| 361 \| Toprim domain-containing protein (Fragment) \| Toprim domain-containing protein (Fragment); uniclust \| UniRef100\_A0A139JUI2 \| 100.0 \| 1.1e-44 \| 2e-50 \| 363.4 \| 370 \| (377, 755) \| 769 \| (111, 492) \| 525 \| Virulence-associated E-like domain-containing protein \| Virulence-associated E-like domain-containing protein; uniclust \| UniRef100\_A0A1B0Z1F4 \| 100.0 \| 1.1e-44 \| 2e-50 \| 384.0 \| 357 \| (376, 735) \| 769 \| (528, 897) \| 962 \| Virulence-associated E-like domain-containing protein \| Virulence-associated E-like domain-containing protein; uniclust \| UniRef100\_A0A078BQ56 \| 100.0 \| 1.5e-44 \| 2.9e-50 \| 357.8 \| 290 \| (14, 349) \| 769 \| (46, 335) \| 393 \| Toprim domain-containing protein \| Toprim domain-containing protein; uniclust \| UniRef100\_A0A010SRZ9 \| 100.0 \| 3.2e-44 \| 6.1e-50 \| 349.2 \| 264 \| (31, 335) \| 769 \| (60, 324) \| 354 \| Topoisomerase \| Topoisomerase; uniclust \| UniRef100\_UPI000CFF6687 \| 100.0 \| 5.2e-44 \| 9.6e-50 \| 360.8 \| 590 \| (14, 638) \| 769 \| (16, 650) \| 817 \| DUF5906 domain-containing protein \| DUF5906 domain-containing protein; uniclust \| UniRef100\_A0A5C7M7G3 \| 100.0 \| 5.6e-44 \| 1.1e-49 \| 370.6 \| 355 \| (375, 741) \| 769 \| (293, 654) \| 720 \| Uncharacterized protein \| Uncharacterized protein; uniclust \| UniRef100\_A0A6M4GU76 \| 100.0 \| 6.1e-44 \| 1.1e-49 \| 348.5 \| 668 \| (16, 737) \| 769 \| (22, 717) \| 745 \| Uncharacterized protein \| Uncharacterized protein; uniclust \| UniRef100\_A0A1U7MHB1 \| 100.0 \| 8.2e-44 \| 1.6e-49 \| 371.1 \| 382 \| (371, 759) \| 769 \| (319, 716) \| 738 \| Virulence-associated protein E \| Virulence-associated protein E; uniclust \| UniRef100\_A0A5K7Y2P6 \| 100.0 \| 9.8e-44 \| 1.8e-49 \| 333.8 \| 372 \| (367, 741) \| 769 \| (121, 493) \| 505 \| Virulence-associated E-like domain-containing protein \| Virulence-associated E-like domain-containing protein; uniclust \| UniRef100\_A0A015V8N0 \| 100.0 \| 9.5e-44 \| 1.9e-49 \| 390.0 \| 367 \| (382, 759) \| 769 \| (425, 801) \| 826 \| VirE N-terminal domain protein \| VirE N-terminal domain protein; uniclust \| UniRef100\_A0A176F820 \| 100.0 \| 1e-43 \| 2e-49 \| 389.4 \| 362 \| (370, 740) \| 769 \| (448, 875) \| 925 \| Virulence-associated E-like domain-containing protein \| Virulence-associated E-like domain-containing protein; uniclust \| UniRef100\_A0A3A8HXS5 \| 100.0 \| 1.5e-43 \| 2.9e-49 \| 358.2 \| 370 \| (376, 754) \| 769 \| (351, 739) \| 775 \| DNA primase \| DNA primase; uniclust \| UniRef100\_A0A3G8X3W9 \| 100.0 \| 1.7e-43 \| 3.3e-49 \| 355.3 \| 366 \| (378, 755) \| 769 \| (156, 529) \| 561 \| Virulence-associated e family protein \| Virulence-associated e family protein; uniclust \| UniRef100\_A0A0A1VBE9 \| 100.0 \| 2e-43 \| 4e-49 \| 359.6 \| 273 \| (14, 333) \| 769 \| (76, 348) \| 413 \| Toprim domain-containing protein \| Toprim domain-containing protein; uniclust \| UniRef100\_A0A6N6T5K1 \| 100.0 \| 2.2e-43 \| 4.3e-49 \| 375.2 \| 311 \| (14, 351) \| 769 \| (27, 353) \| 903 \| Toprim domain-containing protein \| Toprim domain-containing protein; uniclust \| UniRef100\_UPI001C268BC4 \| 100.0 \| 2.8e-43 \| 5.2e-49 \| 335.7 \| 355 \| (391, 754) \| 769 \| (32, 402) \| 452 \| virulence-associated E family protein \| virulence-associated E family protein; uniclust \| UniRef100\_A0A011VM22 \| 100.0 \| 4e-43 \| 7.6e-49 \| 361.7 \| 359 \| (366, 739) \| 769 \| (344, 709) \| 765 \| Virulence protein E \| Virulence protein E; uniclust \| UniRef100\_A0A1A5HIC1 \| 100.0 \| 4.3e-43 \| 8.3e-49 \| 367.3 \| 476 \| (233, 755) \| 769 \| (240, 724) \| 763 \| Toprim domain-containing protein \| Toprim domain-containing protein; uniclust \| UniRef100\_A0A374J606 \| 100.0 \| 6.1e-43 \| 1.1e-48 \| 350.7 \| 464 \| (218, 714) \| 769 \| (147, 646) \| 725 \| Virulence-associated E-like domain-containing protein \| Virulence-associated E-like domain-containing protein; uniclust \| UniRef100\_A0A241VC66 \| 100.0 \| 6.7e-43 \| 1.2e-48 \| 336.3 \| 432 \| (324, 756) \| 769 \| (194, 629) \| 636 \| Virulence-associated E-like domain-containing protein \| Virulence-associated E-like domain-containing protein; uniclust \| UniRef100\_A0A011PNF2 \| 100.0 \| 6.6e-43 \| 1.3e-48 \| 382.3 \| 292 \| (14, 350) \| 769 \| (70, 362) \| 877 \| DNA primase TraC \| DNA primase TraC; uniclust \| UniRef100\_A0A059XSZ9 \| 100.0 \| 7.4e-43 \| 1.4e-48 \| 383.9 \| 485 \| (234, 756) \| 769 \| (187, 753) \| 869 \| DNA primase \| DNA primase; uniclust \| UniRef100\_A0A1G7N5S7 \| 100.0 \| 1e-42 \| 1.9e-48 \| 346.7 \| 300 \| (3, 319) \| 769 \| (29, 370) \| 411 \| Putative DNA primase/helicase \| Putative DNA primase/helicase; uniclust \| UniRef100\_A0A0Q4IXU5 \| 100.0 \| 1.1e-42 \| 2.1e-48 \| 368.4 \| 362 \| (369, 739) \| 769 \| (432, 859) \| 897 \| Uncharacterized protein \| Uncharacterized protein; uniclust \| UniRef100\_A0A5Q0BKD1 \| 100.0 \| 1.1e-42 \| 2.1e-48 \| 335.6 \| 316 \| (8, 348) \| 769 \| (15, 337) \| 367 \| Toprim domain-containing protein \| Toprim domain-containing protein; uniclust \| UniRef100\_A0A071MGH5 \| 100.0 \| 1.7e-42 \| 3.4e-48 \| 387.0 \| 322 \| (12, 350) \| 769 \| (64, 390) \| 1076 \| DUF927 domain-containing protein \| DUF927 domain-containing protein; uniclust \| UniRef100\_A0A0C3IBC7 \| 100.0 \| 2.1e-42 \| 4e-48 \| 355.8 \| 370 \| (375, 757) \| 769 \| (126, 519) \| 537 \| Virulence-associated E-like domain-containing protein \| Virulence-associated E-like domain-containing protein; uniclust \| UniRef100\_A0A2S7FDM4 \| 100.0 \| 2.4e-42 \| 4.7e-48 \| 349.3 \| 373 \| (375, 756) \| 769 \| (85, 470) \| 509 \| Virulence-associated E-like domain-containing protein \| Virulence-associated E-like domain-containing protein; uniclust \| UniRef100\_A0A0P6WTJ1 \| 100.0 \| 3.3e-42 \| 6.2e-48 \| 341.4 \| 339 \| (386, 742) \| 769 \| (132, 480) \| 501 \| Virulence-associated E-like domain-containing protein \| Virulence-associated E-like domain-containing protein; uniclust \| UniRef100\_A0A069D3U0 \| 100.0 \| 3.4e-42 \| 6.5e-48 \| 338.1 \| 365 \| (382, 758) \| 769 \| (50, 424) \| 434 \| Putative helicase \| Putative helicase; uniclust \| UniRef100\_A0A1C4C0F2 \| 100.0 \| 3.7e-42 \| 6.9e-48 \| 353.8 \| 465 \| (127, 637) \| 769 \| (110, 619) \| 698 \| Putative DNA primase/helicase \| Putative DNA primase/helicase; uniclust \| UniRef100\_A0A6H1Z9Z7 \| 100.0 \| 3.9e-42 \| 7.4e-48 \| 351.8 \| 356 \| (377, 742) \| 769 \| (378, 746) \| 778 \| Putative VirE domain containing protein \| Putative VirE domain containing protein; uniclust \| UniRef100\_A0A1I9YDF5 \| 100.0 \| 4.7e-42 \| 8.6e-48 \| 334.7 \| 674 \| (12, 749) \| 769 \| (6, 716) \| 725 \| PriCT-2 domain-containing protein \| PriCT-2 domain-containing protein; uniclust \| UniRef100\_A0A7C5A2E1 \| 100.0 \| 4.7e-42 \| 8.8e-48 \| 339.3 \| 462 \| (231, 741) \| 769 \| (172, 643) \| 665 \| Virulence-associated E-like domain-containing protein \| Virulence-associated E-like domain-containing protein; uniclust \| UniRef100\_A0A2G0Q1H7 \| 100.0 \| 5.7e-42 \| 1.1e-47 \| 339.8 \| 371 \| (371, 752) \| 769 \| (179, 567) \| 583 \| ATPase \| ATPase; uniclust \| UniRef100\_A0A127SEK4 \| 100.0 \| 7.6e-42 \| 1.4e-47 \| 307.8 \| 315 \| (437, 755) \| 769 \| (3, 317) \| 321 \| Virulence-associated protein E (Fragment) \| Virulence-associated protein E (Fragment); uniclust \| UniRef100\_A0A068R0U5 \| 100.0 \| 7.3e-42 \| 1.4e-47 \| 369.0 \| 363 \| (370, 742) \| 769 \| (360, 736) \| 776 \| Predicted P-loop ATPase and inactivated derivatives \| Predicted P-loop ATPase and inactivated derivatives; uniclust \| UniRef100\_A0A1H4GIG7 \| 100.0 \| 9.1e-42 \| 1.7e-47 \| 342.7 \| 365 \| (383, 758) \| 769 \| (179, 553) \| 574 \| Virulence-associated protein E \| Virulence-associated protein E; uniclust \| UniRef100\_A0A0B2JZB9 \| 100.0 \| 1.4e-41 \| 2.7e-47 \| 348.7 \| 371 \| (379, 758) \| 769 \| (332, 713) \| 728 \| Virulence-associated E-like domain-containing protein \| Virulence-associated E-like domain-containing protein; uniclust \| UniRef100\_A0A0R1VXQ4 \| 100.0 \| 1.8e-41 \| 3.4e-47 \| 319.1 \| 310 \| (379, 698) \| 769 \| (40, 370) \| 375 \| Prophage Lp3 protein 8, helicase \| Prophage Lp3 protein 8, helicase; uniclust \| UniRef100\_A0A255MSC6 \| 100.0 \| 1.9e-41 \| 3.6e-47 \| 333.8 \| 371 \| (372, 751) \| 769 \| (73, 453) \| 460 \| Virulence-associated E-like domain-containing protein \| Virulence-associated E-like domain-containing protein; uniclust \| UniRef100\_A0A0P1FRF1 \| 100.0 \| 2e-41 \| 3.8e-47 \| 353.0 \| 356 \| (375, 738) \| 769 \| (351, 748) \| 776 \| Putative P-loop ATPase \| Putative P-loop ATPase; uniclust \| UniRef100\_A0A433VJU3 \| 100.0 \| 2e-41 \| 3.8e-47 \| 347.1 \| 357 \| (385, 761) \| 769 \| (327, 694) \| 812 \| Virulence-associated E-like domain-containing protein \| Virulence-associated E-like domain-containing protein; uniclust \| UniRef100\_A0A3M0XHA9 \| 100.0 \| 2.5e-41 \| 4.7e-47 \| 318.3 \| 266 \| (466, 741) \| 769 \| (1, 273) \| 344 \| Virulence-associated E-like domain-containing protein (Fragment) \| Virulence-associated E-like domain-containing protein (Fragment); uniclust \| UniRef100\_A0A6J5P9W1 \| 100.0 \| 2.7e-41 \| 5e-47 \| 324.8 \| 488 \| (233, 742) \| 769 \| (68, 590) \| 625 \| Virulence-associated E \| Virulence-associated E; uniclust \| UniRef100\_UPI0007164CDC \| 100.0 \| 3.1e-41 \| 5.6e-47 \| 330.7 \| 663 \| (16, 740) \| 769 \| (56, 746) \| 763 \| VapE family protein \| VapE family protein; uniclust \| UniRef100\_A0A0X3TJ34 \| 100.0 \| 2.9e-41 \| 5.6e-47 \| 337.1 \| 306 \| (14, 334) \| 769 \| (28, 343) \| 363 \| Toprim domain-containing protein \| Toprim domain-containing protein; uniclust \| UniRef100\_A0A5C7W4R6 \| 100.0 \| 3.2e-41 \| 6e-47 \| 319.1 \| 387 \| (366, 755) \| 769 \| (140, 529) \| 535 \| Virulence-associated E family protein \| Virulence-associated E family protein; uniclust \| UniRef100\_A0A1F4HLP3 \| 100.0 \| 4.8e-41 \| 9e-47 \| 349.2 \| 321 \| (14, 350) \| 769 \| (13, 336) \| 747 \| Toprim domain-containing protein (Fragment) \| Toprim domain-containing protein (Fragment); uniclust \| UniRef100\_A0A095Y5Z3 \| 100.0 \| 5.2e-41 \| 9.9e-47 \| 339.4 \| 366 \| (383, 760) \| 769 \| (101, 486) \| 522 \| Virulence-associated E-like domain-containing protein \| Virulence-associated E-like domain-containing protein; uniclust \| UniRef100\_A0A378IRY1 \| 100.0 \| 6.6e-41 \| 1.2e-46 \| 346.1 \| 682 \| (14, 742) \| 769 \| (16, 792) \| 812 \| 5' DNA primase TraC \| 5' DNA primase TraC; uniclust \| UniRef100\_A0A059PAJ6 \| 100.0 \| 6.8e-41 \| 1.3e-46 \| 346.5 \| 353 \| (372, 737) \| 769 \| (104, 479) \| 519 \| Putative DNA helicase \| Putative DNA helicase; uniclust \| UniRef100\_A0A097EN21 \| 100.0 \| 7.2e-41 \| 1.4e-46 \| 327.8 \| 268 \| (14, 337) \| 769 \| (49, 316) \| 330 \| DNA primase \| DNA primase; uniclust \| UniRef100\_A0A679K914 \| 100.0 \| 8.3e-41 \| 1.6e-46 \| 332.0 \| 352 \| (373, 738) \| 769 \| (108, 479) \| 525 \| Virulence-associated E-like domain-containing protein \| Virulence-associated E-like domain-containing protein; uniclust \| UniRef100\_A0A0D0NW01 \| 100.0 \| 8.8e-41 \| 1.7e-46 \| 358.5 \| 530 \| (177, 742) \| 769 \| (233, 862) \| 914 \| DNA primase \| DNA primase; uniclust \| UniRef100\_A0A1M6YKQ7 \| 100.0 \| 1.1e-40 \| 2.1e-46 \| 347.7 \| 290 \| (14, 348) \| 769 \| (111, 401) \| 804 \| Uncharacterized domain associated with phage/plasmid primase \| Uncharacterized domain associated with phage/plasmid primase; uniclust \| UniRef100\_A0A1I2NF75 \| 100.0 \| 1.2e-40 \| 2.2e-46 \| 339.1 \| 336 \| (391, 741) \| 769 \| (364, 713) \| 733 \| Primase C terminal 2 (PriCT-2) \| Primase C terminal 2 (PriCT-2); uniclust \| UniRef100\_A0A0B6RPX2 \| 100.0 \| 1.1e-40 \| 2.3e-46 \| 380.7 \| 324 \| (11, 350) \| 769 \| (130, 454) \| 1165 \| Uncharacterized protein \| Uncharacterized protein; uniclust \| UniRef100\_A0A0B3W190 \| 100.0 \| 1.3e-40 \| 2.4e-46 \| 338.8 \| 375 \| (366, 757) \| 769 \| (154, 536) \| 575 \| Zinc finger CHC2-type domain-containing protein \| Zinc finger CHC2-type domain-containing protein; uniclust \| UniRef100\_A0A0R1TQE1 \| 100.0 \| 1.7e-40 \| 3.2e-46 \| 326.9 \| 309 \| (380, 697) \| 769 \| (73, 394) \| 489 \| Virulence-associated E-like domain-containing protein \| Virulence-associated E-like domain-containing protein; uniclust \| UniRef100\_A0A086PE93 \| 100.0 \| 1.8e-40 \| 3.5e-46 \| 351.8 \| 691 \| (4, 740) \| 769 \| (2, 1020) \| 1075 \| DNA primase \| DNA primase; uniclust \| UniRef100\_A0A846FJ73 \| 100.0 \| 3.2e-40 \| 5.9e-46 \| 333.5 \| 478 \| (233, 754) \| 769 \| (154, 652) \| 826 \| DUF3854 domain-containing protein \| DUF3854 domain-containing protein; uniclust \| UniRef100\_A0A174SAF4 \| 100.0 \| 3.3e-40 \| 6.3e-46 \| 316.5 \| 304 \| (370, 682) \| 769 \| (47, 370) \| 403 \| Predicted P-loop ATPase and inactivated derivatives \| Predicted P-loop ATPase and inactivated derivatives; uniclust \| UniRef100\_A0A069SD21 \| 100.0 \| 3.9e-40 \| 7.5e-46 \| 339.2 \| 367 \| (379, 757) \| 769 \| (121, 497) \| 534 \| Uncharacterized protein \| Uncharacterized protein; uniclust \| UniRef100\_A0A7C1H7I6 \| 100.0 \| 4.3e-40 \| 7.9e-46 \| 316.9 \| 292 \| (14, 350) \| 769 \| (53, 344) \| 523 \| Toprim domain-containing protein \| Toprim domain-containing protein; uniclust \| UniRef100\_A0A0C1Y5L2 \| 100.0 \| 4.1e-40 \| 8e-46 \| 343.2 \| 278 \| (469, 755) \| 769 \| (223, 511) \| 546 \| Virulence-associated E-like domain-containing protein \| Virulence-associated E-like domain-containing protein; uniclust \| UniRef100\_A0A1X7JKC4 \| 100.0 \| 4.5e-40 \| 8.5e-46 \| 321.5 \| 352 \| (371, 741) \| 769 \| (161, 521) \| 546 \| Virulence-associated protein E \| Virulence-associated protein E; uniclust \| UniRef100\_A0A1Q2T5M1 \| 100.0 \| 4.8e-40 \| 9.1e-46 \| 315.8 \| 255 \| (435, 697) \| 769 \| (13, 273) \| 379 \| Virulence-associated protein E \| Virulence-associated protein E; uniclust \| UniRef100\_A0A0B6RVH3 \| 100.0 \| 7.5e-40 \| 1.4e-45 \| 346.7 \| 314 \| (11, 350) \| 769 \| (12, 328) \| 814 \| Uncharacterized protein \| Uncharacterized protein; uniclust \| UniRef100\_A0A1H0TMV8 \| 100.0 \| 8.1e-40 \| 1.5e-45 \| 353.1 \| 567 \| (14, 636) \| 769 \| (26, 676) \| 917 \| Putative DNA primase/helicase \| Putative DNA primase/helicase; uniclust \| UniRef100\_A0A142BA72 \| 100.0 \| 1.2e-39 \| 2.4e-45 \| 339.7 \| 287 \| (14, 346) \| 769 \| (32, 318) \| 693 \| Putative primase-helicase \| Putative primase-helicase; uniclust \| UniRef100\_A0A0F9PAW5 \| 100.0 \| 1.4e-39 \| 2.6e-45 \| 324.3 \| 365 \| (375, 752) \| 769 \| (306, 687) \| 696 \| DNA primase/polymerase bifunctional N-terminal domain-containing protein \| DNA primase/polymerase bifunctional N-terminal domain-containing protein; uniclust \| UniRef100\_A0A0M7MF94 \| 100.0 \| 2.1e-39 \| 3.9e-45 \| 340.3 \| 375 \| (370, 754) \| 769 \| (377, 771) \| 792 \| Predicted P-loop ATPase and inactivated derivatives \| Predicted P-loop ATPase and inactivated derivatives; uniclust \| UniRef100\_A0A148KLT5 \| 100.0 \| 2.2e-39 \| 4.1e-45 \| 328.5 \| 378 \| (376, 756) \| 769 \| (464, 843) \| 849 \| Virulence-associated E-like domain-containing protein \| Virulence-associated E-like domain-containing protein; uniclust \| UniRef100\_A0A174V3L9 \| 100.0 \| 2.4e-39 \| 4.5e-45 \| 308.6 \| 352 \| (372, 734) \| 769 \| (23, 394) \| 457 \| Predicted P-loop ATPase and inactivated derivatives \| Predicted P-loop ATPase and inactivated derivatives; uniclust \| UniRef100\_A0A1E2USX5 \| 100.0 \| 2.4e-39 \| 4.7e-45 \| 348.4 \| 285 \| (14, 344) \| 769 \| (31, 316) \| 721 \| Toprim domain-containing protein \| Toprim domain-containing protein; uniclust \| UniRef100\_A0A0D5Y4M5 \| 100.0 \| 2.6e-39 \| 5e-45 \| 306.0 \| 264 \| (32, 333) \| 769 \| (22, 286) \| 301 \| Topoisomerase \| Topoisomerase; uniclust \| UniRef100\_A0A081B651 \| 100.0 \| 3.3e-39 \| 6.3e-45 \| 350.4 \| 299 \| (14, 349) \| 769 \| (420, 723) \| 811 \| DNA primase TraC \| DNA primase TraC; uniclust \| UniRef100\_A0A095ZAT3 \| 100.0 \| 3.8e-39 \| 7.1e-45 \| 319.7 \| 320 \| (371, 697) \| 769 \| (141, 474) \| 612 \| Virulence-associated E family protein \| Virulence-associated E family protein; uniclust \| UniRef100\_A0A2T7SVQ2 \| 100.0 \| 4.2e-39 \| 7.8e-45 \| 314.5 \| 366 \| (366, 737) \| 769 \| (342, 707) \| 725 \| Virulence-associated E-like domain-containing protein \| Virulence-associated E-like domain-containing protein; uniclust \| UniRef100\_A0A239KTG6 \| 100.0 \| 4.4e-39 \| 8.2e-45 \| 317.3 \| 350 \| (372, 740) \| 769 \| (288, 653) \| 672 \| Predicted P-loop ATPase and inactivated derivatives \| Predicted P-loop ATPase and inactivated derivatives; uniclust \| UniRef100\_A0A2E5HC47 \| 100.0 \| 6e-39 \| 1.1e-44 \| 338.0 \| 344 \| (383, 742) \| 769 \| (405, 759) \| 783 \| Virulence-associated E-like domain-containing protein \| Virulence-associated E-like domain-containing protein; uniclust \| UniRef100\_A0A1Z8N2Q3 \| 100.0 \| 6.6e-39 \| 1.2e-44 \| 317.9 \| 347 \| (377, 740) \| 769 \| (360, 714) \| 731 \| Uncharacterized protein \| Uncharacterized protein; uniclust \| UniRef100\_A0A1G9GZG2 \| 100.0 \| 7.5e-39 \| 1.4e-44 \| 309.6 \| 348 \| (379, 741) \| 769 \| (124, 487) \| 514 \| Virulence-associated protein E \| Virulence-associated protein E; uniclust \| UniRef100\_A0A1C5MBB1 \| 100.0 \| 8.2e-39 \| 1.6e-44 \| 328.5 \| 373 \| (376, 758) \| 769 \| (302, 689) \| 717 \| Predicted P-loop ATPase and inactivated derivatives \| Predicted P-loop ATPase and inactivated derivatives; uniclust \| UniRef100\_A0A095YT29 \| 100.0 \| 8.8e-39 \| 1.7e-44 \| 340.3 \| 321 \| (4, 347) \| 769 \| (3, 346) \| 921 \| Toprim domain-containing protein \| Toprim domain-containing protein; uniclust \| UniRef100\_A0A934ECD9 \| 100.0 \| 9e-39 \| 1.7e-44 \| 314.7 \| 293 \| (14, 350) \| 769 \| (12, 305) \| 551 \| AAA family ATPase \| AAA family ATPase; uniclust \| UniRef100\_A0A968S1F8 \| 100.0 \| 9.2e-39 \| 1.7e-44 \| 327.4 \| 473 \| (233, 739) \| 769 \| (212, 698) \| 961 \| Virulence-associated E-like domain-containing protein \| Virulence-associated E-like domain-containing protein; uniclust \| UniRef100\_R6WKB3 \| 100.0 \| 9.4e-39 \| 1.8e-44 \| 311.0 \| 364 \| (381, 755) \| 769 \| (44, 417) \| 448 \| VirE N-terminal domain protein \| VirE N-terminal domain protein; uniclust \| UniRef100\_A0A2N2S409 \| 100.0 \| 9.8e-39 \| 1.8e-44 \| 301.8 \| 350 \| (6, 402) \| 769 \| (3, 354) \| 395 \| Toprim domain-containing protein (Fragment) \| Toprim domain-containing protein (Fragment); uniclust \| UniRef100\_A0A2Z5WWH0 \| 100.0 \| 9.9e-39 \| 1.9e-44 \| 314.1 \| 333 \| (390, 742) \| 769 \| (60, 403) \| 431 \| Virulence-associated E \| Virulence-associated E; uniclust \| UniRef100\_A0A0A2B4H4 \| 100.0 \| 1e-38 \| 1.9e-44 \| 312.6 \| 357 \| (381, 755) \| 769 \| (66, 437) \| 455 \| Virulence-associated E-like domain-containing protein \| Virulence-associated E-like domain-containing protein; uniclust \| UniRef100\_A0A089PBP6 \| 100.0 \| 1.1e-38 \| 2e-44 \| 320.3 \| 361 \| (380, 756) \| 769 \| (88, 464) \| 561 \| DNA primase \| DNA primase; uniclust \| UniRef100\_A0A081RVY4 \| 100.0 \| 1.2e-38 \| 2.2e-44 \| 329.8 \| 362 \| (370, 742) \| 769 \| (377, 753) \| 780 \| Putative P-loop ATPase \| Putative P-loop ATPase; uniclust \| UniRef100\_A0A0G0A8B1 \| 100.0 \| 1.1e-38 \| 2.2e-44 \| 342.6 \| 363 \| (382, 757) \| 769 \| (339, 711) \| 731 \| Virulence-associated E family protein \| Virulence-associated E family protein; uniclust \| UniRef100\_A0A0B7MHW3 \| 100.0 \| 1.1e-38 \| 2.3e-44 \| 346.4 \| 322 \| (2, 349) \| 769 \| (20, 383) \| 638 \| DNA primase \| DNA primase; uniclust \| UniRef100\_A0A069S6V9 \| 100.0 \| 1.5e-38 \| 2.8e-44 \| 301.1 \| 298 \| (450, 757) \| 769 \| (11, 315) \| 322 \| Uncharacterized protein \| Uncharacterized protein; uniclust \| UniRef100\_A0A059ZR08 \| 100.0 \| 1.8e-38 \| 3.5e-44 \| 339.3 \| 484 \| (231, 755) \| 769 \| (206, 762) \| 805 \| DNA primase/helicase, phage-associated \| DNA primase/helicase, phage-associated; uniclust \| UniRef100\_A0A088F8A2 \| 100.0 \| 1.8e-38 \| 3.5e-44 \| 331.9 \| 273 \| (14, 336) \| 769 \| (31, 304) \| 706 \| Putative primase-helicase \| Putative primase-helicase; uniclust \| UniRef100\_A0A6M8VRM0 \| 100.0 \| 1.9e-38 \| 3.6e-44 \| 285.5 \| 276 \| (448, 739) \| 769 \| (3, 282) \| 300 \| Virulence-associated E family protein \| Virulence-associated E family protein; uniclust \| UniRef100\_A0A1F9Y098 \| 100.0 \| 2e-38 \| 3.8e-44 \| 330.4 \| 287 \| (14, 348) \| 769 \| (12, 300) \| 878 \| Toprim domain-containing protein \| Toprim domain-containing protein; uniclust \| UniRef100\_A0A2I8A4T1 \| 100.0 \| 2.4e-38 \| 4.5e-44 \| 332.9 \| 344 \| (383, 743) \| 769 \| (353, 714) \| 754 \| Virulence-associated E \| Virulence-associated E; uniclust \| UniRef100\_A0A1B9LY80 \| 100.0 \| 2.5e-38 \| 4.6e-44 \| 318.5 \| 456 \| (129, 625) \| 769 \| (115, 607) \| 610 \| Bacteriophage T7 Gp4 DNA primase/helicase N-terminal domain-containing protein \| Bacteriophage T7 Gp4 DNA primase/helicase N-terminal domain-containing protein; uniclust \| UniRef100\_A0A0H5Q210 \| 100.0 \| 2.8e-38 \| 5.4e-44 \| 333.5 \| 289 \| (14, 348) \| 769 \| (37, 326) \| 661 \| Toprim domain-containing protein \| Toprim domain-containing protein; uniclust \| UniRef100\_UPI000469875C \| 100.0 \| 3.3e-38 \| 6.2e-44 \| 283.9 \| 227 \| (451, 686) \| 769 \| (4, 236) \| 247 \| virulence-associated E family protein \| virulence-associated E family protein; uniclust \| UniRef100\_A0A1I5CW14 \| 100.0 \| 3.8e-38 \| 7.2e-44 \| 317.1 \| 366 \| (371, 746) \| 769 \| (218, 598) \| 614 \| Predicted P-loop ATPase and inactivated derivatives \| Predicted P-loop ATPase and inactivated derivatives; uniclust \| UniRef100\_A0A1K1LFZ9 \| 100.0 \| 3.9e-38 \| 7.4e-44 \| 337.6 \| 297 \| (14, 350) \| 769 \| (53, 353) \| 971 \| DNA primase, phage-associated \| DNA primase, phage-associated; uniclust \| UniRef100\_A0A2C8EXD6 \| 100.0 \| 4.4e-38 \| 8.2e-44 \| 315.5 \| 354 \| (377, 741) \| 769 \| (380, 755) \| 779 \| DNA primase/polymerase bifunctional N-terminal domain-containing protein \| DNA primase/polymerase bifunctional N-terminal domain-containing protein; uniclust \| UniRef100\_A0A6H9GBR6 \| 100.0 \| 5.9e-38 \| 1.1e-43 \| 311.3 \| 327 \| (393, 742) \| 769 \| (177, 517) \| 550 \| Virulence-associated E-like domain-containing protein \| Virulence-associated E-like domain-containing protein; uniclust \| UniRef100\_A0A1F3BA57 \| 100.0 \| 6.3e-38 \| 1.2e-43 \| 343.8 \| 322 \| (1, 350) \| 769 \| (54, 419) \| 676 \| DNA primase \| DNA primase; uniclust \| UniRef100\_A0A075KDN3 \| 100.0 \| 6.4e-38 \| 1.3e-43 \| 342.0 \| 324 \| (2, 350) \| 769 \| (51, 417) \| 676 \| DNA primase \| DNA primase; uniclust \| UniRef100\_A0A024GWJ1 \| 100.0 \| 7.9e-38 \| 1.5e-43 \| 339.7 \| 558 \| (150, 742) \| 769 \| (166, 826) \| 886 \| Phage/plasmid primase, P4 family, C-terminal domain protein \| Phage/plasmid primase, P4 family, C-terminal domain protein; uniclust \| UniRef100\_A0A1Z4I5E7 \| 100.0 \| 8.3e-38 \| 1.6e-43 \| 321.6 \| 343 \| (391, 756) \| 769 \| (337, 690) \| 814 \| Virulence-associated E \| Virulence-associated E; uniclust \| UniRef100\_A0A011NP97 \| 100.0 \| 1e-37 \| 1.9e-43 \| 335.9 \| 291 \| (14, 348) \| 769 \| (42, 338) \| 860 \| DNA primase TraC \| DNA primase TraC; uniclust \| UniRef100\_UPI001C2DDBBB \| 100.0 \| 1.1e-37 \| 2e-43 \| 282.9 \| 255 \| (433, 696) \| 769 \| (32, 294) \| 312 \| virulence-associated E family protein \| virulence-associated E family protein; uniclust \| UniRef100\_A0A0B8WUT7 \| 100.0 \| 1.3e-37 \| 2.6e-43 \| 340.4 \| 323 \| (2, 349) \| 769 \| (25, 391) \| 644 \| DNA primase \| DNA primase; uniclust \| UniRef100\_A0A1V5XFC1 \| 100.0 \| 1.4e-37 \| 2.7e-43 \| 305.6 \| 351 \| (371, 738) \| 769 \| (115, 483) \| 520 \| Virulence-associated protein E \| Virulence-associated protein E; uniclust \| UniRef100\_UPI00163E2698 \| 100.0 \| 1.5e-37 \| 2.7e-43 \| 304.5 \| 521 \| (16, 585) \| 769 \| (11, 648) \| 648 \| VapE family protein \| VapE family protein; uniclust \| UniRef100\_A0A135YTI2 \| 100.0 \| 1.8e-37 \| 3.4e-43 \| 285.3 \| 281 \| (373, 659) \| 769 \| (40, 336) \| 340 \| Virulence-associated protein E \| Virulence-associated protein E; uniclust \| UniRef100\_A0A916EP12 \| 100.0 \| 1.8e-37 \| 3.4e-43 \| 318.4 \| 383 \| (372, 759) \| 769 \| (539, 934) \| 962 \| Virulence-associated E-like domain-containing protein \| Virulence-associated E-like domain-containing protein; uniclust \| UniRef100\_A0A1F5A0V8 \| 100.0 \| 1.7e-37 \| 3.4e-43 \| 338.0 \| 322 \| (2, 349) \| 769 \| (54, 418) \| 681 \| DNA primase \| DNA primase; uniclust \| UniRef100\_UPI0022AFBBF4 \| 100.0 \| 2.1e-37 \| 3.8e-43 \| 302.3 \| 384 \| (369, 757) \| 769 \| (316, 700) \| 707 \| VapE family protein \| VapE family protein; uniclust \| UniRef100\_A0A1H8DHD7 \| 100.0 \| 2.2e-37 \| 4.1e-43 \| 324.0 \| 369 \| (379, 756) \| 769 \| (277, 657) \| 662 \| Virulence-associated protein E \| Virulence-associated protein E; uniclust \| UniRef100\_A0A6M0JXK6 \| 100.0 \| 2.4e-37 \| 4.4e-43 \| 317.1 \| 658 \| (14, 695) \| 769 \| (92, 869) \| 995 \| SF3 helicase domain-containing protein \| SF3 helicase domain-containing protein; uniclust \| UniRef100\_A0A4Y4D005 \| 100.0 \| 2.5e-37 \| 4.6e-43 \| 312.3 \| 382 \| (377, 759) \| 769 \| (440, 822) \| 836 \| DNA primase/polymerase bifunctional N-terminal domain-containing protein \| DNA primase/polymerase bifunctional N-terminal domain-containing protein; uniclust \| UniRef100\_A0A1H0PAA2 \| 100.0 \| 2.5e-37 \| 4.8e-43 \| 325.2 \| 297 \| (14, 348) \| 769 \| (437, 734) \| 790 \| Antirestriction protein ArdC \| Antirestriction protein ArdC; uniclust \| UniRef100\_A0A1H4CGI7 \| 100.0 \| 2.5e-37 \| 4.9e-43 \| 341.1 \| 359 \| (381, 753) \| 769 \| (378, 742) \| 790 \| Primase C terminal 2 (PriCT-2) \| Primase C terminal 2 (PriCT-2); uniclust \| UniRef100\_A0A136K0K7 \| 100.0 \| 2.7e-37 \| 5.1e-43 \| 307.0 \| 322 \| (2, 350) \| 769 \| (1, 366) \| 441 \| DNA primase catalytic subunit \| DNA primase catalytic subunit; uniclust \| UniRef100\_A0A081C5E7 \| 100.0 \| 3e-37 \| 5.9e-43 \| 342.1 \| 322 \| (2, 350) \| 769 \| (64, 427) \| 711 \| DNA primase \| DNA primase; uniclust \| UniRef100\_A0A1E1GE69 \| 100.0 \| 3.3e-37 \| 6.1e-43 \| 310.7 \| 387 \| (368, 764) \| 769 \| (399, 787) \| 794 \| Putative primase \| Putative primase; uniclust \| UniRef100\_A0A1H7JV05 \| 100.0 \| 4.3e-37 \| 8.2e-43 \| 294.7 \| 243 \| (487, 736) \| 769 \| (2, 278) \| 332 \| Virulence-associated protein E \| Virulence-associated protein E; uniclust \| UniRef100\_A0A0K2WEP3 \| 100.0 \| 4.5e-37 \| 8.8e-43 \| 332.2 \| 310 \| (14, 350) \| 769 \| (29, 383) \| 647 \| DNA primase \| DNA primase; uniclust \| UniRef100\_A0A1L5KF83 \| 100.0 \| 4.8e-37 \| 9.3e-43 \| 289.3 \| 240 \| (450, 695) \| 769 \| (4, 254) \| 280 \| NHLM\_micro\_ABC2: NHLM bacteriocin system ABC transporter, ATP-binding (Fragment) \| NHLM\_micro\_ABC2: NHLM bacteriocin system ABC transporter, ATP-binding (Fragment); uniclust \| UniRef100\_A0A0F9HLD4 \| 100.0 \| 5.2e-37 \| 1e-42 \| 329.6 \| 322 \| (3, 350) \| 769 \| (52, 417) \| 657 \| Toprim domain-containing protein (Fragment) \| Toprim domain-containing protein (Fragment); uniclust \| UniRef100\_A0A315D5W5 \| 100.0 \| 5.7e-37 \| 1.1e-42 \| 304.1 \| 374 \| (377, 754) \| 769 \| (350, 726) \| 728 \| DNA primase/polymerase bifunctional N-terminal domain-containing protein \| DNA primase/polymerase bifunctional N-terminal domain-containing protein; uniclust \| UniRef100\_A0A1G7B4P0 \| 100.0 \| 5.8e-37 \| 1.1e-42 \| 314.6 \| 287 \| (14, 348) \| 769 \| (31, 321) \| 686 \| Uncharacterized domain associated with phage/plasmid primase \| Uncharacterized domain associated with phage/plasmid primase; uniclust \| UniRef100\_A0A062Y2K0 \| 100.0 \| 5.6e-37 \| 1.1e-42 \| 335.8 \| 311 \| (13, 350) \| 769 \| (101, 455) \| 732 \| DNA primase \| DNA primase; uniclust \| UniRef100\_A0A0F5Y7I5 \| 100.0 \| 6.7e-37 \| 1.2e-42 \| 312.8 \| 331 \| (391, 741) \| 769 \| (326, 673) \| 866 \| Virulence-associated E-like domain-containing protein \| Virulence-associated E-like domain-containing protein; uniclust \| UniRef100\_A0A8S0YIP5 \| 100.0 \| 7e-37 \| 1.3e-42 \| 299.6 \| 508 \| (230, 757) \| 769 \| (169, 699) \| 732 \| Virulence-associated E-like domain-containing protein \| Virulence-associated E-like domain-containing protein; uniclust \| UniRef100\_A0A1E7I3V3 \| 100.0 \| 6.6e-37 \| 1.3e-42 \| 333.1 \| 318 \| (7, 349) \| 769 \| (60, 420) \| 670 \| DNA primase \| DNA primase; uniclust \| UniRef100\_A0A352X4W4 \| 100.0 \| 8.1e-37 \| 1.5e-42 \| 307.3 \| 486 \| (233, 750) \| 769 \| (283, 823) \| 1004 \| Toprim domain-containing protein \| Toprim domain-containing protein; uniclust \| UniRef100\_A0A1G0RDT0 \| 100.0 \| 7.9e-37 \| 1.5e-42 \| 296.6 \| 373 \| (379, 755) \| 769 \| (55, 450) \| 471 \| Virulence-associated E-like domain-containing protein \| Virulence-associated E-like domain-containing protein; uniclust \| UniRef100\_A0A2N1CUR9 \| 100.0 \| 8.5e-37 \| 1.6e-42 \| 289.4 \| 384 \| (368, 754) \| 769 \| (107, 496) \| 528 \| Virulence-associated E family protein \| Virulence-associated E family protein; uniclust \| UniRef100\_A0A0P9CJD9 \| 100.0 \| 8.8e-37 \| 1.8e-42 \| 338.7 \| 322 \| (3, 349) \| 769 \| (47, 411) \| 668 \| DNA primase \| DNA primase; uniclust \| UniRef100\_A0A1F4ANV0 \| 100.0 \| 1e-36 \| 1.9e-42 \| 316.8 \| 380 \| (372, 756) \| 769 \| (364, 789) \| 814 \| Virulence-associated E-like domain-containing protein \| Virulence-associated E-like domain-containing protein; uniclust \| UniRef100\_A0A5B2V8U9 \| 100.0 \| 1.1e-36 \| 2e-42 \| 290.1 \| 294 \| (434, 742) \| 769 \| (243, 541) \| 553 \| Virulence-associated E family protein \| Virulence-associated E family protein; uniclust \| UniRef100\_A0A2H2XUZ8 \| 100.0 \| 1.1e-36 \| 2.1e-42 \| 308.9 \| 337 \| (383, 741) \| 769 \| (233, 579) \| 755 \| Virulence-associated E-like domain-containing protein \| Virulence-associated E-like domain-containing protein; uniclust \| UniRef100\_A0A061N9C7 \| 100.0 \| 1.1e-36 \| 2.2e-42 \| 338.3 \| 323 \| (2, 350) \| 769 \| (73, 441) \| 723 \| DNA primase \| DNA primase; uniclust \| UniRef100\_A0A1Y4HHS4 \| 100.0 \| 1.2e-36 \| 2.3e-42 \| 302.0 \| 364 \| (380, 755) \| 769 \| (147, 529) \| 547 \| Uncharacterized protein \| Uncharacterized protein; uniclust \| UniRef100\_A0A269PIH2 \| 100.0 \| 1.3e-36 \| 2.4e-42 \| 276.2 \| 274 \| (376, 650) \| 769 \| (43, 316) \| 316 \| Virulence-associated E-like domain-containing protein \| Virulence-associated E-like domain-containing protein; uniclust \| UniRef100\_A0A2X3IEU8 \| 100.0 \| 1.4e-36 \| 2.6e-42 \| 318.6 \| 481 \| (231, 753) \| 769 \| (193, 765) \| 792 \| Phage/plasmid primase, P4 family \| Phage/plasmid primase, P4 family; uniclust \| UniRef100\_A0A357A780 \| 100.0 \| 1.9e-36 \| 3.5e-42 \| 302.6 \| 360 \| (377, 757) \| 769 \| (327, 700) \| 918 \| Uncharacterized protein \| Uncharacterized protein; uniclust \| UniRef100\_UPI0009B50754 \| 100.0 \| 2e-36 \| 3.7e-42 \| 299.1 \| 486 \| (232, 738) \| 769 \| (229, 785) \| 808 \| virulence-associated E family protein \| virulence-associated E family protein; uniclust \| UniRef100\_A0A3B0ME56 \| 100.0 \| 2e-36 \| 3.8e-42 \| 295.4 \| 352 \| (373, 740) \| 769 \| (321, 685) \| 705 \| Virulence-associated E-like domain-containing protein \| Virulence-associated E-like domain-containing protein; uniclust \| UniRef100\_A0A133YEF9 \| 100.0 \| 2e-36 \| 3.9e-42 \| 324.6 \| 315 \| (8, 350) \| 769 \| (60, 417) \| 681 \| DNA primase \| DNA primase; uniclust \| UniRef100\_UPI00203AF1B9 \| 100.0 \| 2.4e-36 \| 4.3e-42 \| 295.4 \| 468 \| (231, 734) \| 769 \| (156, 654) \| 716 \| VapE family protein \| VapE family protein; uniclust \| UniRef100\_A0A0E3MC62 \| 100.0 \| 2.3e-36 \| 4.4e-42 \| 315.1 \| 274 \| (370, 655) \| 769 \| (382, 672) \| 673 \| Virulence-associated E family protein \| Virulence-associated E family protein; uniclust \| UniRef100\_A0A0R1U9K6 \| 100.0 \| 2.6e-36 \| 4.9e-42 \| 300.2 \| 309 \| (378, 694) \| 769 \| (139, 462) \| 584 \| Prophage Lp3 protein 8, helicase \| Prophage Lp3 protein 8, helicase; uniclust \| UniRef100\_A0A073IRN6 \| 100.0 \| 2.7e-36 \| 5.2e-42 \| 320.3 \| 316 \| (5, 351) \| 769 \| (22, 379) \| 617 \| DNA primase \| DNA primase; uniclust \| UniRef100\_A0A095XT20 \| 100.0 \| 2.7e-36 \| 5.2e-42 \| 288.4 \| 282 \| (470, 755) \| 769 \| (58, 345) \| 376 \| Virulence-associated protein E (Fragment) \| Virulence-associated protein E (Fragment); uniclust \| UniRef100\_A0A010NL17 \| 100.0 \| 2.6e-36 \| 5.2e-42 \| 328.8 \| 320 \| (3, 350) \| 769 \| (39, 398) \| 644 \| DNA primase \| DNA primase; uniclust \| UniRef100\_A0A2E0AGP3 \| 100.0 \| 2.8e-36 \| 5.3e-42 \| 289.9 \| 351 \| (389, 753) \| 769 \| (51, 413) \| 424 \| Virulence-associated E-like domain-containing protein \| Virulence-associated E-like domain-containing protein; uniclust \| UniRef100\_A0A418RT30 \| 100.0 \| 3e-36 \| 5.6e-42 \| 279.8 \| 269 \| (14, 336) \| 769 \| (38, 306) \| 312 \| Toprim domain-containing protein \| Toprim domain-containing protein; uniclust \| UniRef100\_A0A072Y8M3 \| 100.0 \| 2.9e-36 \| 5.7e-42 \| 334.8 \| 317 \| (3, 349) \| 769 \| (71, 429) \| 684 \| DNA primase \| DNA primase; uniclust \| UniRef100\_A0A2P8WAJ2 \| 100.0 \| 3.2e-36 \| 6e-42 \| 277.8 \| 286 \| (445, 741) \| 769 \| (4, 297) \| 356 \| Virulence-associated E-like domain-containing protein \| Virulence-associated E-like domain-containing protein; uniclust \| UniRef100\_UPI0019D07770 \| 100.0 \| 3.8e-36 \| 6.9e-42 \| 308.0 \| 349 \| (392, 755) \| 769 \| (905, 1269) \| 1277 \| VapE family protein \| VapE family protein; uniclust \| UniRef100\_A0A374UKX2 \| 100.0 \| 4.2e-36 \| 7.8e-42 \| 289.6 \| 340 \| (380, 735) \| 769 \| (25, 385) \| 415 \| Virulence-associated E-like domain-containing protein \| Virulence-associated E-like domain-containing protein; uniclust \| UniRef100\_UPI001CDC2532 \| 100.0 \| 4.6e-36 \| 8.4e-42 \| 282.6 \| 312 \| (376, 696) \| 769 \| (25, 351) \| 495 \| virulence-associated E family protein \| virulence-associated E family protein; uniclust \| UniRef100\_A0A016QSH3 \| 100.0 \| 4.4e-36 \| 8.9e-42 \| 340.1 \| 323 \| (2, 349) \| 769 \| (77, 441) \| 758 \| DNA primase \| DNA primase; uniclust \| UniRef100\_A0A011UA69 \| 100.0 \| 4.5e-36 \| 9e-42 \| 334.6 \| 320 \| (2, 348) \| 769 \| (30, 391) \| 651 \| DNA primase \| DNA primase; uniclust \| UniRef100\_A0A0G0HEE0 \| 100.0 \| 4.9e-36 \| 9.5e-42 \| 326.3 \| 322 \| (1, 350) \| 769 \| (42, 408) \| 684 \| Primase protein (Fragment) \| Primase protein (Fragment); uniclust \| UniRef100\_A0A2S5DRL8 \| 100.0 \| 5.2e-36 \| 9.6e-42 \| 304.9 \| 362 \| (370, 741) \| 769 \| (357, 734) \| 763 \| Replication protein \| Replication protein; uniclust \| UniRef100\_A0A1F4YB42 \| 100.0 \| 5.4e-36 \| 1.1e-41 \| 325.5 \| 316 \| (8, 350) \| 769 \| (25, 387) \| 642 \| DNA primase \| DNA primase; uniclust \| UniRef100\_A0A1I1YQB9 \| 100.0 \| 6.4e-36 \| 1.2e-41 \| 307.2 \| 386 \| (370, 764) \| 769 \| (328, 738) \| 758 \| Virulence-associated protein E \| Virulence-associated protein E; uniclust \| UniRef100\_A0A7W0X7N0 \| 100.0 \| 6.9e-36 \| 1.3e-41 \| 296.0 \| 292 \| (14, 350) \| 769 \| (11, 303) \| 680 \| DUF3987 domain-containing protein (Fragment) \| DUF3987 domain-containing protein (Fragment); uniclust \| UniRef100\_A0A2D9N1G6 \| 100.0 \| 6.9e-36 \| 1.3e-41 \| 277.7 \| 308 \| (376, 692) \| 769 \| (38, 363) \| 369 \| Virulence-associated E-like domain-containing protein (Fragment) \| Virulence-associated E-like domain-containing protein (Fragment); uniclust \| UniRef100\_A0A928SS04 \| 100.0 \| 7.2e-36 \| 1.3e-41 \| 282.5 \| 357 \| (380, 750) \| 769 \| (126, 492) \| 515 \| Virulence-associated E-like domain-containing protein \| Virulence-associated E-like domain-containing protein; uniclust \| UniRef100\_A0A0H3A991 \| 100.0 \| 7.4e-36 \| 1.4e-41 \| 318.1 \| 640 \| (43, 735) \| 769 \| (28, 758) \| 812 \| Plasmid/phage primase, P4 family \| Plasmid/phage primase, P4 family; uniclust \| UniRef100\_A0A068F471 \| 100.0 \| 7.4e-36 \| 1.5e-41 \| 335.0 \| 321 \| (3, 350) \| 769 \| (96, 459) \| 736 \| DNA primase \| DNA primase; uniclust \| UniRef100\_A0A101PRR1 \| 100.0 \| 8.1e-36 \| 1.6e-41 \| 318.9 \| 477 \| (233, 752) \| 769 \| (17, 573) \| 625 \| DNA primase (Fragment) \| DNA primase (Fragment); uniclust \| UniRef100\_A0A0G1XGQ8 \| 100.0 \| 8.9e-36 \| 1.7e-41 \| 310.3 \| 320 \| (3, 350) \| 769 \| (27, 389) \| 624 \| DNA primase \| DNA primase; uniclust \| UniRef100\_A0A0A8WPJ3 \| 100.0 \| 8.7e-36 \| 1.7e-41 \| 332.3 \| 320 \| (2, 349) \| 769 \| (84, 447) \| 710 \| DNA primase \| DNA primase; uniclust \| UniRef100\_A0A8J6XJ06 \| 100.0 \| 9.4e-36 \| 1.8e-41 \| 302.0 \| 490 \| (217, 740) \| 769 \| (146, 655) \| 874 \| DUF3854 domain-containing protein \| DUF3854 domain-containing protein; uniclust \| UniRef100\_A0A011PBN5 \| 100.0 \| 9.9e-36 \| 1.9e-41 \| 277.9 \| 289 \| (447, 743) \| 769 \| (4, 299) \| 316 \| Putative P-loop ATPase \| Putative P-loop ATPase; uniclust \| UniRef100\_A0A0H4B4X1 \| 100.0 \| 1e-35 \| 1.9e-41 \| 287.8 \| 359 \| (382, 755) \| 769 \| (28, 398) \| 438 \| Virulence-associated E-like domain-containing protein \| Virulence-associated E-like domain-containing protein; uniclust \| UniRef100\_A0A0Q7CC84 \| 100.0 \| 1.1e-35 \| 2e-41 \| 276.7 \| 217 \| (99, 339) \| 769 \| (101, 319) \| 322 \| Toprim domain-containing protein \| Toprim domain-containing protein; uniclust \| UniRef100\_A0A0B4XUC8 \| 100.0 \| 1.1e-35 \| 2e-41 \| 314.9 \| 293 \| (14, 348) \| 769 \| (658, 953) \| 990 \| Toprim domain-containing protein \| Toprim domain-containing protein; uniclust \| UniRef100\_A0A069RDL3 \| 100.0 \| 1.1e-35 \| 2.2e-41 \| 329.5 \| 321 \| (3, 350) \| 769 \| (79, 441) \| 687 \| DNA primase \| DNA primase; uniclust \| UniRef100\_A0A0D6QLL8 \| 100.0 \| 1.2e-35 \| 2.4e-41 \| 316.2 \| 322 \| (3, 349) \| 769 \| (13, 381) \| 633 \| DNA primase \| DNA primase; uniclust \| UniRef100\_A0A2E8UJR5 \| 100.0 \| 1.3e-35 \| 2.5e-41 \| 295.4 \| 344 \| (381, 741) \| 769 \| (28, 394) \| 571 \| Virulence-associated E-like domain-containing protein \| Virulence-associated E-like domain-containing protein; uniclust \| UniRef100\_A0A923AMT9 \| 100.0 \| 1.4e-35 \| 2.5e-41 \| 296.0 \| 357 \| (373, 737) \| 769 \| (500, 873) \| 895 \| Uncharacterized protein \| Uncharacterized protein; uniclust \| UniRef100\_A0A066ZPD8 \| 100.0 \| 1.3e-35 \| 2.6e-41 \| 319.3 \| 319 \| (9, 350) \| 769 \| (41, 402) \| 644 \| DNA primase \| DNA primase; uniclust \| UniRef100\_A0A178MKK8 \| 100.0 \| 1.4e-35 \| 2.6e-41 \| 289.0 \| 368 \| (373, 751) \| 769 \| (105, 511) \| 535 \| Virulence-associated E-like domain-containing protein \| Virulence-associated E-like domain-containing protein; uniclust \| UniRef100\_A0A0G1KEK0 \| 100.0 \| 1.4e-35 \| 2.7e-41 \| 309.4 \| 321 \| (2, 349) \| 769 \| (38, 404) \| 515 \| Primase protein \| Primase protein; uniclust \| UniRef100\_A0A2P8WI05 \| 100.0 \| 1.7e-35 \| 3.1e-41 \| 287.0 \| 341 \| (390, 750) \| 769 \| (127, 475) \| 652 \| Virulence-associated E-like domain-containing protein (Fragment) \| Virulence-associated E-like domain-containing protein (Fragment); uniclust \| UniRef100\_A0A2E8UEW4 \| 100.0 \| 1.7e-35 \| 3.2e-41 \| 305.6 \| 318 \| (2, 349) \| 769 \| (19, 379) \| 497 \| DNA primase (Fragment) \| DNA primase (Fragment); uniclust \| UniRef100\_A0A011MS96 \| 100.0 \| 1.7e-35 \| 3.4e-41 \| 317.7 \| 321 \| (3, 349) \| 769 \| (51, 412) \| 648 \| DNA primase \| DNA primase; uniclust \| UniRef100\_A0A0F5MPZ0 \| 100.0 \| 2.1e-35 \| 4e-41 \| 302.8 \| 317 \| (6, 349) \| 769 \| (38, 399) \| 513 \| DNA primase (Fragment) \| DNA primase (Fragment); uniclust \| UniRef100\_A0A1V5VAP1 \| 100.0 \| 2.1e-35 \| 4e-41 \| 317.6 \| 322 \| (4, 350) \| 769 \| (69, 434) \| 735 \| DNA primase \| DNA primase; uniclust \| UniRef100\_A0A6J5MWU5 \| 100.0 \| 2.3e-35 \| 4.2e-41 \| 294.6 \| 349 \| (381, 741) \| 769 \| (313, 678) \| 697 \| COG5545 Predicted P-loop ATPase and inactivated derivatives \| COG5545 Predicted P-loop ATPase and inactivated derivatives; uniclust \| UniRef100\_A0A011VIC8 \| 100.0 \| 2.1e-35 \| 4.2e-41 \| 335.4 \| 322 \| (1, 349) \| 769 \| (85, 453) \| 830 \| DNA primase \| DNA primase; uniclust \| UniRef100\_A0A0P7YWN9 \| 100.0 \| 2.2e-35 \| 4.2e-41 \| 315.0 \| 321 \| (2, 348) \| 769 \| (3, 369) \| 682 \| DNA primase \| DNA primase; uniclust \| UniRef100\_A0A7Z0LRG9 \| 100.0 \| 2.4e-35 \| 4.3e-41 \| 292.4 \| 377 \| (378, 756) \| 769 \| (442, 820) \| 827 \| Virulence-associated E-like domain-containing protein \| Virulence-associated E-like domain-containing protein; uniclust \| UniRef100\_A0A3P1ZAJ6 \| 100.0 \| 2.4e-35 \| 4.6e-41 \| 296.4 \| 363 \| (382, 757) \| 769 \| (94, 467) \| 473 \| Helicase \| Helicase; uniclust \| UniRef100\_A0A088GFL0 \| 100.0 \| 2.4e-35 \| 4.8e-41 \| 319.6 \| 320 \| (3, 349) \| 769 \| (48, 411) \| 657 \| DNA primase \| DNA primase; uniclust \| UniRef100\_A0A2T4F5R5 \| 100.0 \| 2.6e-35 \| 4.9e-41 \| 272.0 \| 274 \| (453, 740) \| 769 \| (9, 286) \| 318 \| Virulence-associated E-like domain-containing protein \| Virulence-associated E-like domain-containing protein; uniclust \| UniRef100\_A0A011PSP2 \| 100.0 \| 2.6e-35 \| 4.9e-41 \| 319.0 \| 302 \| (14, 349) \| 769 \| (648, 958) \| 1044 \| DNA primase TraC \| DNA primase TraC; uniclust \| UniRef100\_A0A0G1C3E6 \| 100.0 \| 2.6e-35 \| 5e-41 \| 297.8 \| 320 \| (2, 349) \| 769 \| (4, 366) \| 424 \| Primase protein \| Primase protein; uniclust \| UniRef100\_A0A1M3GZV5 \| 100.0 \| 2.8e-35 \| 5.3e-41 \| 316.0 \| 286 \| (14, 348) \| 769 \| (57, 342) \| 849 \| Toprim domain-containing protein \| Toprim domain-containing protein; uniclust \| UniRef100\_A0A7H0HGN8 \| 100.0 \| 3.1e-35 \| 5.6e-41 \| 287.0 \| 290 \| (14, 349) \| 769 \| (12, 302) \| 612 \| DUF3987 domain-containing protein \| DUF3987 domain-containing protein; uniclust \| UniRef100\_A0A6N6T496 \| 100.0 \| 3.4e-35 \| 6.2e-41 \| 275.9 \| 329 \| (380, 718) \| 769 \| (73, 411) \| 480 \| Virulence-associated E-like domain-containing protein \| Virulence-associated E-like domain-containing protein; uniclust \| UniRef100\_A0A1M6YK98 \| 100.0 \| 3.3e-35 \| 6.3e-41 \| 278.2 \| 261 \| (492, 756) \| 769 \| (6, 268) \| 304 \| Virulence-associated protein E \| Virulence-associated protein E; uniclust \| UniRef100\_A0A9D5UT42 \| 100.0 \| 3.4e-35 \| 6.3e-41 \| 295.7 \| 563 \| (127, 738) \| 769 \| (126, 787) \| 884 \| Uncharacterized protein \| Uncharacterized protein; uniclust \| UniRef100\_A0A5C7LJI8 \| 100.0 \| 3.6e-35 \| 6.6e-41 \| 287.3 \| 367 \| (372, 749) \| 769 \| (317, 705) \| 714 \| Virulence-associated E-like domain-containing protein \| Virulence-associated E-like domain-containing protein; uniclust \| UniRef100\_A0A017T7Q0 \| 100.0 \| 3.5e-35 \| 6.8e-41 \| 312.6 \| 324 \| (3, 351) \| 769 \| (13, 381) \| 628 \| DNA primase \| DNA primase; uniclust \| UniRef100\_A0A099UD78 \| 100.0 \| 3.9e-35 \| 7.4e-41 \| 297.3 \| 299 \| (14, 349) \| 769 \| (100, 419) \| 500 \| DNA primase \| DNA primase; uniclust \| UniRef100\_A0A1F2X0U7 \| 100.0 \| 3.9e-35 \| 7.5e-41 \| 293.8 \| 317 \| (2, 348) \| 769 \| (29, 388) \| 421 \| DNA primase (Fragment) \| DNA primase (Fragment); uniclust \| UniRef100\_A0A094K1B3 \| 100.0 \| 3.9e-35 \| 7.6e-41 \| 285.2 \| 276 \| (14, 335) \| 769 \| (22, 304) \| 316 \| Toprim domain-containing protein \| Toprim domain-containing protein; uniclust \| UniRef100\_A0A969EVW1 \| 100.0 \| 4.5e-35 \| 8.4e-41 \| 280.1 \| 294 \| (14, 346) \| 769 \| (18, 313) \| 487 \| DUF1738 domain-containing protein (Fragment) \| DUF1738 domain-containing protein (Fragment); uniclust \| UniRef100\_A0A077XNU5 \| 100.0 \| 4.4e-35 \| 8.5e-41 \| 304.6 \| 321 \| (3, 348) \| 769 \| (21, 387) \| 535 \| DNA primase \| DNA primase; uniclust \| UniRef100\_A0A0B0HLY2 \| 100.0 \| 4.6e-35 \| 8.9e-41 \| 308.6 \| 313 \| (10, 348) \| 769 \| (50, 407) \| 638 \| DNA primase \| DNA primase; uniclust \| UniRef100\_A0A0A2HRP0 \| 100.0 \| 4.7e-35 \| 9.2e-41 \| 315.4 \| 312 \| (14, 350) \| 769 \| (21, 373) \| 618 \| DNA primase \| DNA primase; uniclust \| UniRef100\_A0A4P7IPZ1 \| 100.0 \| 5.2e-35 \| 9.7e-41 \| 300.7 \| 350 \| (373, 735) \| 769 \| (324, 692) \| 712 \| ATPase \| ATPase; uniclust \| UniRef100\_A0A8J2YXK5 \| 100.0 \| 5.4e-35 \| 1e-40 \| 281.0 \| 291 \| (436, 742) \| 769 \| (276, 578) \| 595 \| Virulence-associated E-like domain-containing protein \| Virulence-associated E-like domain-containing protein; uniclust \| UniRef100\_A0A2E5G1Q1 \| 100.0 \| 5.6e-35 \| 1e-40 \| 286.8 \| 364 \| (372, 751) \| 769 \| (349, 731) \| 738 \| Virulence-associated E-like domain-containing protein \| Virulence-associated E-like domain-containing protein; uniclust \| UniRef100\_A0A229I531 \| 100.0 \| 5.6e-35 \| 1.1e-40 \| 309.0 \| 372 \| (378, 755) \| 769 \| (355, 755) \| 794 \| Uncharacterized protein \| Uncharacterized protein; uniclust \| UniRef100\_A0A0C3IN80 \| 100.0 \| 6.1e-35 \| 1.2e-40 \| 288.7 \| 280 \| (30, 335) \| 769 \| (74, 373) \| 418 \| DNA primase \| DNA primase; uniclust \| UniRef100\_A0A031WEN8 \| 100.0 \| 6e-35 \| 1.2e-40 \| 323.4 \| 320 \| (2, 350) \| 769 \| (118, 481) \| 727 \| DNA primase \| DNA primase; uniclust \| UniRef100\_A0A010SLU5 \| 100.0 \| 6.2e-35 \| 1.2e-40 \| 319.2 \| 319 \| (6, 349) \| 769 \| (36, 402) \| 693 \| DNA primase \| DNA primase; uniclust \| UniRef100\_A0A1Z4QLR4 \| 100.0 \| 6.7e-35 \| 1.2e-40 \| 295.8 \| 473 \| (233, 741) \| 769 \| (134, 627) \| 1094 \| Virulence-associated E family protein \| Virulence-associated E family protein; uniclust \| UniRef100\_A0A0S3U510 \| 100.0 \| 6.6e-35 \| 1.2e-40 \| 298.0 \| 338 \| (386, 742) \| 769 \| (486, 831) \| 929 \| Virulence-associated E-like domain-containing protein \| Virulence-associated E-like domain-containing protein; uniclust \| UniRef100\_A0A069D3P6 \| 100.0 \| 6.8e-35 \| 1.3e-40 \| 301.8 \| 359 \| (384, 753) \| 769 \| (335, 703) \| 705 \| Putative helicase \| Putative helicase; uniclust \| UniRef100\_A0A1Y6D883 \| 100.0 \| 7.2e-35 \| 1.4e-40 \| 298.0 \| 327 \| (6, 350) \| 769 \| (3, 343) \| 660 \| Primase C terminal 2 (PriCT-2) \| Primase C terminal 2 (PriCT-2); uniclust \| UniRef100\_A0A060AZD4 \| 100.0 \| 7.4e-35 \| 1.4e-40 \| 310.2 \| 323 \| (2, 350) \| 769 \| (7, 376) \| 645 \| DNA primase \| DNA primase; uniclust \| UniRef100\_UPI000BFFB864 \| 100.0 \| 8e-35 \| 1.5e-40 \| 297.3 \| 468 \| (235, 739) \| 769 \| (150, 633) \| 1203 \| VapE family protein \| VapE family protein; uniclust \| UniRef100\_A0A419H4F3 \| 100.0 \| 8.2e-35 \| 1.5e-40 \| 285.7 \| 378 \| (385, 769) \| 769 \| (178, 562) \| 562 \| Virulence-associated E-like domain-containing protein \| Virulence-associated E-like domain-containing protein; uniclust \| UniRef100\_A0A261S325 \| 100.0 \| 8.2e-35 \| 1.6e-40 \| 290.6 \| 307 \| (8, 348) \| 769 \| (40, 352) \| 401 \| Primase C-terminal 2 domain-containing protein \| Primase C-terminal 2 domain-containing protein; uniclust \| UniRef100\_A0A424HP40 \| 100.0 \| 8.8e-35 \| 1.7e-40 \| 298.9 \| 341 \| (384, 742) \| 769 \| (300, 647) \| 670 \| Virulence-associated E family protein \| Virulence-associated E family protein; uniclust \| UniRef100\_A0A022GRU9 \| 100.0 \| 8.9e-35 \| 1.7e-40 \| 310.3 \| 323 \| (3, 350) \| 769 \| (173, 544) \| 790 \| DNA primase \| DNA primase; uniclust \| UniRef100\_A0A078LXT8 \| 100.0 \| 8.7e-35 \| 1.7e-40 \| 323.9 \| 316 \| (3, 349) \| 769 \| (50, 411) \| 666 \| DNA primase \| DNA primase; uniclust \| UniRef100\_UPI000BE33B30 \| 100.0 \| 9.8e-35 \| 1.8e-40 \| 283.7 \| 462 \| (232, 735) \| 769 \| (181, 672) \| 698 \| VapE family protein \| VapE family protein; uniclust \| UniRef100\_A0A2D4SRW8 \| 100.0 \| 9.5e-35 \| 1.8e-40 \| 307.6 \| 320 \| (3, 350) \| 769 \| (30, 393) \| 683 \| DNA primase \| DNA primase; uniclust \| UniRef100\_A0A1V8RP15 \| 100.0 \| 1e-34 \| 1.9e-40 \| 301.2 \| 364 \| (370, 741) \| 769 \| (415, 837) \| 879 \| Uncharacterized protein \| Uncharacterized protein; uniclust \| UniRef100\_A0A1H6V3P5 \| 100.0 \| 1.1e-34 \| 2e-40 \| 294.7 \| 354 \| (375, 742) \| 769 \| (486, 866) \| 885 \| Primase C terminal 2 (PriCT-2) \| Primase C terminal 2 (PriCT-2); uniclust \| UniRef100\_A0A238HHW5 \| 100.0 \| 1.2e-34 \| 2.3e-40 \| 308.2 \| 317 \| (9, 345) \| 769 \| (45, 376) \| 1258 \| DNA primase TraC \| DNA primase TraC; uniclust \| UniRef100\_A0A031JRC8 \| 100.0 \| 1.3e-34 \| 2.5e-40 \| 290.6 \| 278 \| (33, 335) \| 769 \| (42, 331) \| 407 \| DNA primase \| DNA primase; uniclust \| UniRef100\_A0A1G3FM87 \| 100.0 \| 1.3e-34 \| 2.5e-40 \| 305.4 \| 658 \| (38, 752) \| 769 \| (63, 838) \| 860 \| SF3 helicase domain-containing protein \| SF3 helicase domain-containing protein; uniclust \| UniRef100\_A0A1H6L3X8 \| 100.0 \| 1.5e-34 \| 2.8e-40 \| 270.2 \| 264 \| (487, 755) \| 769 \| (27, 293) \| 325 \| Virulence-associated E-like domain-containing protein \| Virulence-associated E-like domain-containing protein; uniclust \| UniRef100\_A0A014N801 \| 100.0 \| 1.4e-34 \| 2.8e-40 \| 316.4 \| 322 \| (3, 350) \| 769 \| (36, 402) \| 644 \| DNA primase \| DNA primase; uniclust \| UniRef100\_A0A0F9NPG2 \| 100.0 \| 1.5e-34 \| 2.9e-40 \| 310.3 \| 272 \| (467, 755) \| 769 \| (274, 565) \| 624 \| SF3 helicase domain-containing protein (Fragment) \| SF3 helicase domain-containing protein (Fragment); uniclust \| UniRef100\_A0A7T2PJJ0 \| 100.0 \| 1.6e-34 \| 3.1e-40 \| 280.4 \| 377 \| (366, 755) \| 769 \| (23, 424) \| 434 \| ATPase \| ATPase; uniclust \| UniRef100\_A0A096AAC7 \| 100.0 \| 1.6e-34 \| 3.1e-40 \| 280.5 \| 365 \| (382, 755) \| 769 \| (58, 432) \| 445 \| Helicase \| Helicase; uniclust \| UniRef100\_UPI002277702C \| 100.0 \| 1.7e-34 \| 3.1e-40 \| 274.7 \| 410 \| (131, 565) \| 769 \| (20, 538) \| 538 \| VapE family protein \| VapE family protein; uniclust \| UniRef100\_A0A0H5DU62 \| 100.0 \| 1.7e-34 \| 3.3e-40 \| 311.6 \| 316 \| (4, 350) \| 769 \| (1, 360) \| 591 \| DNA primase \| DNA primase; uniclust \| UniRef100\_A0A966PIM5 \| 100.0 \| 1.9e-34 \| 3.5e-40 \| 262.6 \| 317 \| (376, 693) \| 769 \| (48, 366) \| 369 \| Virulence-associated E family protein (Fragment) \| Virulence-associated E family protein (Fragment); uniclust \| UniRef100\_A0A1F5RLM4 \| 100.0 \| 1.8e-34 \| 3.6e-40 \| 312.4 \| 312 \| (14, 350) \| 769 \| (111, 465) \| 738 \| DNA primase \| DNA primase; uniclust \| UniRef100\_A0A9D7D5W3 \| 100.0 \| 2e-34 \| 3.8e-40 \| 252.0 \| 234 \| (450, 689) \| 769 \| (5, 239) \| 240 \| Uncharacterized protein \| Uncharacterized protein; uniclust \| UniRef100\_A0A099I9C9 \| 100.0 \| 1.9e-34 \| 3.8e-40 \| 324.1 \| 320 \| (4, 349) \| 769 \| (52, 414) \| 665 \| DNA primase \| DNA primase; uniclust \| UniRef100\_A0A1F5V2M0 \| 100.0 \| 2e-34 \| 3.8e-40 \| 297.8 \| 312 \| (6, 350) \| 769 \| (19, 368) \| 608 \| DNA primase \| DNA primase; uniclust \| UniRef100\_A0A7Z9GAF4 \| 100.0 \| 2.2e-34 \| 4e-40 \| 281.5 \| 350 \| (375, 742) \| 769 \| (317, 673) \| 704 \| Virulence-associated E-like domain-containing protein \| Virulence-associated E-like domain-containing protein; uniclust \| UniRef100\_A0A937KUZ1 \| 100.0 \| 2.7e-34 \| 5.1e-40 \| 259.7 \| 244 \| (484, 740) \| 769 \| (23, 271) \| 304 \| Virulence-associated E family protein \| Virulence-associated E family protein; uniclust \| UniRef100\_A0A139C8D1 \| 100.0 \| 2.9e-34 \| 5.4e-40 \| 264.8 \| 258 \| (31, 336) \| 769 \| (46, 304) \| 308 \| Toprim domain-containing protein \| Toprim domain-containing protein; uniclust \| UniRef100\_UPI0010F7918A \| 100.0 \| 3.3e-34 \| 6.1e-40 \| 271.1 \| 356 \| (367, 737) \| 769 \| (10, 372) \| 509 \| VapE family protein \| VapE family protein; uniclust \| UniRef100\_A0A955RF62 \| 100.0 \| 3.8e-34 \| 6.9e-40 \| 275.4 \| 351 \| (376, 740) \| 769 \| (215, 581) \| 596 \| Virulence-associated E-like domain-containing protein \| Virulence-associated E-like domain-containing protein; uniclust \| UniRef100\_UPI001F548CF2 \| 100.0 \| 3.9e-34 \| 7.2e-40 \| 277.3 \| 346 \| (386, 755) \| 769 \| (205, 565) \| 641 \| virulence-associated E family protein \| virulence-associated E family protein; uniclust \| UniRef100\_A0A077MUW8 \| 100.0 \| 4e-34 \| 7.8e-40 \| 331.3 \| 300 \| (14, 348) \| 769 \| (976, 1311) \| 1402 \| Toprim domain-containing protein \| Toprim domain-containing protein; uniclust \| UniRef100\_A0A2N7R2N6 \| 100.0 \| 4.3e-34 \| 8e-40 \| 270.3 \| 423 \| (17, 482) \| 769 \| (10, 451) \| 480 \| DNA primase TraC \| DNA primase TraC; uniclust \| UniRef100\_A0A011RLF2 \| 100.0 \| 4.1e-34 \| 8.2e-40 \| 318.1 \| 311 \| (12, 347) \| 769 \| (82, 437) \| 717 \| DNA primase \| DNA primase; uniclust \| UniRef100\_A0A1E4C277 \| 100.0 \| 4.6e-34 \| 8.4e-40 \| 279.1 \| 359 \| (379, 751) \| 769 \| (166, 534) \| 595 \| Virulence-associated E-like domain-containing protein \| Virulence-associated E-like domain-containing protein; uniclust \| UniRef100\_A0A013VTI8 \| 100.0 \| 4.3e-34 \| 8.5e-40 \| 312.9 \| 320 \| (3, 349) \| 769 \| (92, 456) \| 733 \| DNA primase \| DNA primase; uniclust \| UniRef100\_A0A0D6JI60 \| 100.0 \| 4.5e-34 \| 8.7e-40 \| 305.0 \| 322 \| (2, 350) \| 769 \| (78, 446) \| 739 \| DNA primase \| DNA primase; uniclust \| UniRef100\_A0A350YX52 \| 100.0 \| 4.8e-34 \| 9e-40 \| 271.8 \| 345 \| (384, 741) \| 769 \| (13, 374) \| 400 \| Virulence-associated E-like domain-containing protein (Fragment) \| Virulence-associated E-like domain-containing protein (Fragment); uniclust \| UniRef100\_A0A068NYD4 \| 100.0 \| 4.9e-34 \| 9.4e-40 \| 302.9 \| 319 \| (4, 350) \| 769 \| (3, 361) \| 621 \| DNA primase \| DNA primase; uniclust \| UniRef100\_A0A2H0MZN7 \| 100.0 \| 5.5e-34 \| 1e-39 \| 296.9 \| 287 \| (14, 350) \| 769 \| (31, 319) \| 807 \| Toprim domain-containing protein \| Toprim domain-containing protein; uniclust \| UniRef100\_A0A948T5F1 \| 100.0 \| 5.7e-34 \| 1e-39 \| 271.1 \| 379 \| (370, 755) \| 769 \| (75, 468) \| 474 \| Virulence-associated E family protein \| Virulence-associated E family protein; uniclust \| UniRef100\_A0A163VSB7 \| 100.0 \| 5.9e-34 \| 1.1e-39 \| 297.2 \| 344 \| (382, 741) \| 769 \| (400, 755) \| 928 \| Virulence-associated protein E \| Virulence-associated protein E; uniclust \| UniRef100\_A0A6G7VDW5 \| 100.0 \| 6e-34 \| 1.1e-39 \| 272.3 \| 314 \| (12, 348) \| 769 \| (7, 323) \| 462 \| AAA family ATPase \| AAA family ATPase; uniclust \| UniRef100\_A0A965PG93 \| 100.0 \| 6.3e-34 \| 1.2e-39 \| 277.3 \| 486 \| (232, 751) \| 769 \| (163, 673) \| 676 \| Bacteriophage T7 Gp4 DNA primase/helicase N-terminal domain-containing protein \| Bacteriophage T7 Gp4 DNA primase/helicase N-terminal domain-containing protein; uniclust \| UniRef100\_A0A0R2W862 \| 100.0 \| 6.4e-34 \| 1.2e-39 \| 272.9 \| 346 \| (378, 740) \| 769 \| (176, 554) \| 576 \| Virulence-associated E-like domain-containing protein (Fragment) \| Virulence-associated E-like domain-containing protein (Fragment); uniclust \| UniRef100\_A0A081SGQ8 \| 100.0 \| 6.1e-34 \| 1.2e-39 \| 317.0 \| 320 \| (4, 349) \| 769 \| (80, 447) \| 777 \| DNA primase \| DNA primase; uniclust \| UniRef100\_A0A2N2U4N7 \| 100.0 \| 6.5e-34 \| 1.2e-39 \| 266.0 \| 332 \| (6, 349) \| 769 \| (3, 347) \| 404 \| Toprim domain-containing protein \| Toprim domain-containing protein; uniclust \| UniRef100\_A0A6L3SS05 \| 100.0 \| 7.1e-34 \| 1.3e-39 \| 285.1 \| 313 \| (14, 345) \| 769 \| (53, 377) \| 936 \| Toprim domain-containing protein \| Toprim domain-containing protein; uniclust \| UniRef100\_A0A0G1WVT7 \| 100.0 \| 6.9e-34 \| 1.4e-39 \| 317.4 \| 320 \| (3, 350) \| 769 \| (96, 481) \| 777 \| DNA primase \| DNA primase; uniclust \| UniRef100\_A0A1M6R249 \| 100.0 \| 7.3e-34 \| 1.4e-39 \| 315.1 \| 298 \| (14, 349) \| 769 \| (1089, 1389) \| 1483 \| Conjugative relaxase domain-containing protein, TrwC/TraI family \| Conjugative relaxase domain-containing protein, TrwC/TraI family; uniclust \| UniRef100\_A0A371XDT2 \| 100.0 \| 8e-34 \| 1.5e-39 \| 280.2 \| 336 \| (391, 742) \| 769 \| (406, 754) \| 777 \| Virulence-associated E-like domain-containing protein \| Virulence-associated E-like domain-containing protein; uniclust \| UniRef100\_A0A0B6WZ92 \| 100.0 \| 7.8e-34 \| 1.5e-39 \| 302.0 \| 323 \| (3, 350) \| 769 \| (7, 372) \| 617 \| DNA primase \| DNA primase; uniclust \| UniRef100\_A0A6N9AAB1 \| 100.0 \| 8.5e-34 \| 1.6e-39 \| 282.5 \| 573 \| (14, 639) \| 769 \| (11, 674) \| 810 \| Toprim domain-containing protein \| Toprim domain-containing protein; uniclust \| UniRef100\_UPI00226D0A03 \| 100.0 \| 8.6e-34 \| 1.6e-39 \| 270.8 \| 350 \| (372, 739) \| 769 \| (178, 532) \| 553 \| virulence-associated E family protein \| virulence-associated E family protein; uniclust \| UniRef100\_Q1YDH5 \| 100.0 \| 8.9e-34 \| 1.6e-39 \| 262.8 \| 280 \| (383, 673) \| 769 \| (123, 414) \| 424 \| Putative virulence-associated protein E \| Putative virulence-associated protein E; uniclust \| UniRef100\_A0A0G1YTK1 \| 100.0 \| 1e-33 \| 1.9e-39 \| 284.6 \| 371 \| (371, 749) \| 769 \| (543, 923) \| 962 \| Virulence-associated protein E domain protein \| Virulence-associated protein E domain protein; uniclust \| UniRef100\_A0A0F8W6I5 \| 100.0 \| 1e-33 \| 2e-39 \| 289.6 \| 314 \| (6, 350) \| 769 \| (48, 408) \| 510 \| Toprim domain-containing protein (Fragment) \| Toprim domain-containing protein (Fragment); uniclust \| UniRef100\_A0A969Q8P7 \| 100.0 \| 1.1e-33 \| 2e-39 \| 261.7 \| 280 \| (446, 741) \| 769 \| (3, 287) \| 418 \| Virulence-associated E-like domain-containing protein \| Virulence-associated E-like domain-containing protein; uniclust \| UniRef100\_A0A6J5KG32 \| 100.0 \| 1.1e-33 \| 2.1e-39 \| 292.7 \| 302 \| (437, 742) \| 769 \| (482, 811) \| 859 \| Virulence-associated E \| Virulence-associated E; uniclust \| UniRef100\_A0A063Y669 \| 100.0 \| 1.1e-33 \| 2.1e-39 \| 299.4 \| 318 \| (7, 350) \| 769 \| (10, 374) \| 674 \| DNA primase \| DNA primase; uniclust \| UniRef100\_A0A081N1X8 \| 100.0 \| 1.1e-33 \| 2.1e-39 \| 277.9 \| 245 \| (67, 336) \| 769 \| (76, 331) \| 338 \| Bacteriophage T7 Gp4 DNA primase/helicase N-terminal domain-containing protein \| Bacteriophage T7 Gp4 DNA primase/helicase N-terminal domain-containing protein; uniclust \| UniRef100\_UPI0005DFE532 \| 100.0 \| 1.1e-33 \| 2.1e-39 \| 271.3 \| 288 \| (435, 736) \| 769 \| (58, 357) \| 384 \| virulence-associated E family protein \| virulence-associated E family protein; uniclust \| UniRef100\_L8N3H0 \| 100.0 \| 1.2e-33 \| 2.1e-39 \| 272.1 \| 333 \| (391, 749) \| 769 \| (122, 463) \| 597 \| Virulence-associated E family protein \| Virulence-associated E family protein; uniclust \| UniRef100\_A0A7C9NE92 \| 100.0 \| 1.2e-33 \| 2.2e-39 \| 279.1 \| 356 \| (375, 742) \| 769 \| (402, 769) \| 780 \| Virulence-associated E-like domain-containing protein \| Virulence-associated E-like domain-containing protein; uniclust \| UniRef100\_A0A0B1U8J8 \| 100.0 \| 1.1e-33 \| 2.2e-39 \| 299.5 \| 321 \| (4, 349) \| 769 \| (18, 380) \| 607 \| DNA primase \| DNA primase; uniclust \| UniRef100\_A0A1F6PUQ3 \| 100.0 \| 1.2e-33 \| 2.2e-39 \| 288.6 \| 318 \| (3, 349) \| 769 \| (17, 378) \| 487 \| DNA primase (Fragment) \| DNA primase (Fragment); uniclust \| UniRef100\_A0A011NMP9 \| 100.0 \| 1.3e-33 \| 2.5e-39 \| 282.3 \| 301 \| (14, 348) \| 769 \| (159, 468) \| 505 \| DNA primase TraC \| DNA primase TraC; uniclust \| UniRef100\_A0A022MEH4 \| 100.0 \| 1.3e-33 \| 2.5e-39 \| 305.5 \| 323 \| (4, 349) \| 769 \| (113, 480) \| 768 \| DNA primase (Fragment) \| DNA primase (Fragment); uniclust \| UniRef100\_A0A4P2QP84 \| 100.0 \| 1.4e-33 \| 2.5e-39 \| 280.6 \| 295 \| (435, 740) \| 769 \| (460, 758) \| 841 \| Uncharacterized protein \| Uncharacterized protein; uniclust \| UniRef100\_A0A0A2G2K2 \| 100.0 \| 1.4e-33 \| 2.5e-39 \| 267.7 \| 353 \| (381, 741) \| 769 \| (51, 413) \| 443 \| Virulence-associated protein E (Fragment) \| Virulence-associated protein E (Fragment); uniclust \| UniRef100\_A0A0G2ZH66 \| 100.0 \| 1.3e-33 \| 2.6e-39 \| 302.9 \| 321 \| (3, 350) \| 769 \| (19, 382) \| 634 \| DNA primase \| DNA primase; uniclust \| UniRef100\_A0A2E4HX51 \| 100.0 \| 1.3e-33 \| 2.6e-39 \| 284.8 \| 320 \| (3, 348) \| 769 \| (35, 398) \| 412 \| DNA primase (Fragment) \| DNA primase (Fragment); uniclust \| UniRef100\_A0A0F6W4Z0 \| 100.0 \| 1.5e-33 \| 2.8e-39 \| 298.7 \| 325 \| (3, 350) \| 769 \| (65, 441) \| 702 \| DNA primase \| DNA primase; uniclust \| UniRef100\_A0A060H525 \| 100.0 \| 1.5e-33 \| 2.9e-39 \| 273.7 \| 301 \| (14, 348) \| 769 \| (59, 368) \| 406 \| DNA primase \| DNA primase; uniclust \| UniRef100\_A0A3N5GQR2 \| 100.0 \| 1.6e-33 \| 2.9e-39 \| 275.9 \| 354 \| (380, 751) \| 769 \| (336, 697) \| 710 \| Virulence-associated E-like domain-containing protein \| Virulence-associated E-like domain-containing protein; uniclust \| UniRef100\_A0A2P5L752 \| 100.0 \| 1.6e-33 \| 3.1e-39 \| 266.8 \| 201 \| (127, 349) \| 769 \| (148, 348) \| 375 \| Toprim domain-containing protein \| Toprim domain-containing protein; uniclust \| UniRef100\_A0A838WBL7 \| 100.0 \| 1.7e-33 \| 3.1e-39 \| 270.1 \| 290 \| (14, 348) \| 769 \| (14, 324) \| 576 \| Toprim domain-containing protein (Fragment) \| Toprim domain-containing protein (Fragment); uniclust \| UniRef100\_A0A1F8DMI8 \| 100.0 \| 1.9e-33 \| 3.6e-39 \| 277.6 \| 319 \| (3, 349) \| 769 \| (1, 366) \| 413 \| DNA primase (Fragment) \| DNA primase (Fragment); uniclust \| UniRef100\_A0A0R1FLZ9 \| 100.0 \| 2e-33 \| 3.7e-39 \| 282.6 \| 298 \| (389, 696) \| 769 \| (322, 636) \| 741 \| Prophage lp3 protein 8, helicase \| Prophage lp3 protein 8, helicase; uniclust \| UniRef100\_UPI00109CB367 \| 100.0 \| 2e-33 \| 3.7e-39 \| 286.7 \| 357 \| (391, 755) \| 769 \| (380, 751) \| 759 \| VapE family protein \| VapE family protein; uniclust \| UniRef100\_A0A9E1U8H1 \| 100.0 \| 2.1e-33 \| 3.9e-39 \| 281.5 \| 351 \| (376, 741) \| 769 \| (415, 775) \| 926 \| Uncharacterized protein \| Uncharacterized protein; uniclust \| UniRef100\_A0A078KUP1 \| 100.0 \| 2e-33 \| 4e-39 \| 305.9 \| 315 \| (2, 349) \| 769 \| (54, 407) \| 626 \| DNA primase \| DNA primase; uniclust \| UniRef100\_A0A956I739 \| 100.0 \| 2.2e-33 \| 4e-39 \| 278.4 \| 316 \| (373, 696) \| 769 \| (402, 728) \| 812 \| Virulence-associated E-like domain-containing protein \| Virulence-associated E-like domain-containing protein; uniclust \| UniRef100\_UPI00211665FF \| 100.0 \| 2.2e-33 \| 4e-39 \| 266.3 \| 372 \| (31, 416) \| 769 \| (36, 481) \| 518 \| toprim domain-containing protein \| toprim domain-containing protein; uniclust \| UniRef100\_A0A0A7UW42 \| 100.0 \| 2e-33 \| 4e-39 \| 303.5 \| 318 \| (3, 349) \| 769 \| (23, 380) \| 658 \| DNA primase \| DNA primase; uniclust \| UniRef100\_A0A1C5YKD3 \| 100.0 \| 2.2e-33 \| 4.1e-39 \| 280.7 \| 472 \| (233, 737) \| 769 \| (197, 683) \| 780 \| DNA primase \| DNA primase; uniclust \| UniRef100\_A0A930XU13 \| 100.0 \| 2.3e-33 \| 4.2e-39 \| 265.2 \| 295 \| (436, 744) \| 769 \| (47, 349) \| 501 \| Virulence-associated E-like domain-containing protein \| Virulence-associated E-like domain-containing protein; uniclust \| UniRef100\_UPI000781351A \| 100.0 \| 2.3e-33 \| 4.2e-39 \| 270.3 \| 406 \| (324, 756) \| 769 \| (38, 455) \| 599 \| VapE family protein \| VapE family protein; uniclust \| UniRef100\_A0A1F8JGT0 \| 100.0 \| 2.3e-33 \| 4.4e-39 \| 290.1 \| 321 \| (2, 349) \| 769 \| (9, 370) \| 551 \| DNA primase (Fragment) \| DNA primase (Fragment); uniclust \| UniRef100\_UPI001A9764D7 \| 100.0 \| 2.6e-33 \| 4.9e-39 \| 283.1 \| 382 \| (374, 759) \| 769 \| (352, 742) \| 759 \| VapE family protein \| VapE family protein; uniclust \| UniRef100\_A0A946HIV3 \| 100.0 \| 3e-33 \| 5.6e-39 \| 273.7 \| 350 \| (377, 740) \| 769 \| (305, 684) \| 703 \| Virulence-associated E-like domain-containing protein \| Virulence-associated E-like domain-containing protein; uniclust \| UniRef100\_A0A0P0FR19 \| 100.0 \| 3.3e-33 \| 6.2e-39 \| 293.2 \| 321 \| (4, 349) \| 769 \| (82, 453) \| 720 \| DNA primase \| DNA primase; uniclust \| UniRef100\_A0A926WX44 \| 100.0 \| 3.5e-33 \| 6.4e-39 \| 282.2 \| 481 \| (234, 740) \| 769 \| (128, 623) \| 1023 \| DUF3854 domain-containing protein \| DUF3854 domain-containing protein; uniclust \| UniRef100\_A0A516M9R7 \| 100.0 \| 3.4e-33 \| 6.5e-39 \| 297.8 \| 285 \| (14, 345) \| 769 \| (46, 332) \| 777 \| Toprim domain-containing protein \| Toprim domain-containing protein; uniclust \| UniRef100\_A0A1Y6D1M0 \| 100.0 \| 3.4e-33 \| 6.6e-39 \| 294.8 \| 311 \| (12, 350) \| 769 \| (32, 386) \| 617 \| DNA primase \| DNA primase; uniclust \| UniRef100\_UPI001E4C4C0F \| 100.0 \| 3.9e-33 \| 7.2e-39 \| 250.3 \| 290 \| (375, 666) \| 769 \| (12, 303) \| 327 \| virulence-associated E family protein \| virulence-associated E family protein; uniclust \| UniRef100\_A0A0D2JMB5 \| 100.0 \| 3.8e-33 \| 7.5e-39 \| 309.9 \| 319 \| (1, 349) \| 769 \| (26, 389) \| 642 \| DNA primase \| DNA primase; uniclust \| UniRef100\_A0A1H2VE35 \| 100.0 \| 4.1e-33 \| 7.7e-39 \| 279.1 \| 368 \| (381, 757) \| 769 \| (248, 632) \| 643 \| Virulence-associated protein E \| Virulence-associated protein E; uniclust \| UniRef100\_A0A969AB06 \| 100.0 \| 4.4e-33 \| 8.1e-39 \| 270.2 \| 434 \| (231, 695) \| 769 \| (144, 607) \| 641 \| Virulence-associated E-like domain-containing protein \| Virulence-associated E-like domain-containing protein; uniclust \| UniRef100\_A0A1U7H6I1 \| 100.0 \| 4.7e-33 \| 8.7e-39 \| 283.2 \| 349 \| (386, 754) \| 769 \| (335, 697) \| 859 \| Virulence-associated E-like domain-containing protein \| Virulence-associated E-like domain-containing protein; uniclust \| UniRef100\_A0A537Y110 \| 100.0 \| 4.7e-33 \| 8.9e-39 \| 292.9 \| 318 \| (5, 350) \| 769 \| (3, 363) \| 697 \| DNA primase \| DNA primase; uniclust \| UniRef100\_A0A0F9YUN7 \| 100.0 \| 5e-33 \| 9.8e-39 \| 305.5 \| 315 \| (4, 349) \| 769 \| (36, 392) \| 661 \| DNA primase \| DNA primase; uniclust \| UniRef100\_A0A164AHS2 \| 100.0 \| 5.3e-33 \| 1e-38 \| 269.5 \| 335 \| (390, 739) \| 769 \| (62, 406) \| 443 \| Virulence-associated protein E \| Virulence-associated protein E; uniclust \| UniRef100\_A0A1D2SKX3 \| 100.0 \| 5.2e-33 \| 1e-38 \| 298.2 \| 302 \| (14, 351) \| 769 \| (28, 333) \| 724 \| Uncharacterized protein \| Uncharacterized protein; uniclust \| UniRef100\_A0A023Y828 \| 100.0 \| 5.7e-33 \| 1.1e-38 \| 290.3 \| 323 \| (2, 350) \| 769 \| (43, 409) \| 632 \| DNA primase \| DNA primase; uniclust \| UniRef100\_A0A2E1R907 \| 100.0 \| 6.1e-33 \| 1.1e-38 \| 281.5 \| 343 \| (392, 751) \| 769 \| (379, 737) \| 741 \| Primase C-terminal 2 domain-containing protein \| Primase C-terminal 2 domain-containing protein; uniclust \| UniRef100\_A0A1F9TV27 \| 100.0 \| 6e-33 \| 1.1e-38 \| 290.7 \| 320 \| (2, 349) \| 769 \| (2, 365) \| 596 \| DNA primase \| DNA primase; uniclust \| UniRef100\_A0A059KN83 \| 100.0 \| 6.1e-33 \| 1.2e-38 \| 271.7 \| 287 \| (14, 339) \| 769 \| (41, 342) \| 364 \| Toprim domain-containing protein \| Toprim domain-containing protein; uniclust \| UniRef100\_A0A8S5MVS7 \| 100.0 \| 6.7e-33 \| 1.3e-38 \| 286.7 \| 311 \| (372, 696) \| 769 \| (403, 728) \| 834 \| Virulence associated protein E \| Virulence associated protein E; uniclust \| UniRef100\_A0A926PMV0 \| 100.0 \| 6.8e-33 \| 1.3e-38 \| 275.0 \| 357 \| (389, 769) \| 769 \| (448, 815) \| 815 \| Virulence-associated E-like domain-containing protein \| Virulence-associated E-like domain-containing protein; uniclust \| UniRef100\_A0A0A1H6W6 \| 100.0 \| 7.1e-33 \| 1.4e-38 \| 288.9 \| 320 \| (5, 352) \| 769 \| (1, 361) \| 624 \| DNA primase \| DNA primase; uniclust \| UniRef100\_A0A017H3G2 \| 100.0 \| 6.9e-33 \| 1.4e-38 \| 313.7 \| 319 \| (1, 346) \| 769 \| (30, 392) \| 717 \| DNA primase \| DNA primase; uniclust \| UniRef100\_A0A382QEF6 \| 100.0 \| 8.3e-33 \| 1.6e-38 \| 265.4 \| 286 \| (13, 343) \| 769 \| (37, 328) \| 366 \| Toprim domain-containing protein (Fragment) \| Toprim domain-containing protein (Fragment); uniclust \| UniRef100\_UPI001C0AE9DC \| 100.0 \| 8.8e-33 \| 1.6e-38 \| 267.1 \| 377 \| (366, 757) \| 769 \| (228, 609) \| 616 \| VapE family protein \| VapE family protein; uniclust \| UniRef100\_A0A1F5F2Q7 \| 100.0 \| 9.1e-33 \| 1.8e-38 \| 300.2 \| 320 \| (4, 348) \| 769 \| (107, 473) \| 751 \| DNA primase \| DNA primase; uniclust \| UniRef100\_A0A014NIL8 \| 100.0 \| 9.4e-33 \| 1.8e-38 \| 289.0 \| 322 \| (4, 350) \| 769 \| (60, 430) \| 722 \| DNA primase \| DNA primase; uniclust \| UniRef100\_A0A078BNX3 \| 100.0 \| 9.5e-33 \| 1.8e-38 \| 284.1 \| 322 \| (3, 349) \| 769 \| (1, 367) \| 652 \| DNA primase \| DNA primase; uniclust \| UniRef100\_A0A2D9A9Z1 \| 100.0 \| 9.5e-33 \| 1.8e-38 \| 287.8 \| 320 \| (3, 350) \| 769 \| (41, 407) \| 642 \| DNA primase (Fragment) \| DNA primase (Fragment); uniclust \| UniRef100\_A0A134A587 \| 100.0 \| 9.9e-33 \| 1.8e-38 \| 280.4 \| 309 \| (376, 694) \| 769 \| (318, 632) \| 746 \| Virulence-associated protein E \| Virulence-associated protein E; uniclust \| UniRef100\_A0A101I1T0 \| 100.0 \| 1.1e-32 \| 2.1e-38 \| 295.9 \| 313 \| (14, 350) \| 769 \| (45, 401) \| 629 \| DNA primase \| DNA primase; uniclust \| UniRef100\_A0A0F9MKQ4 \| 100.0 \| 1.1e-32 \| 2.1e-38 \| 298.0 \| 282 \| (457, 755) \| 769 \| (225, 532) \| 595 \| SF3 helicase domain-containing protein \| SF3 helicase domain-containing protein; uniclust \| UniRef100\_A0A1F8RM87 \| 100.0 \| 1.1e-32 \| 2.1e-38 \| 293.4 \| 313 \| (10, 351) \| 769 \| (2, 373) \| 649 \| DNA primase \| DNA primase; uniclust \| UniRef100\_A0A936HA23 \| 100.0 \| 1.2e-32 \| 2.2e-38 \| 272.7 \| 493 \| (230, 741) \| 769 \| (143, 667) \| 797 \| Virulence-associated E-like domain-containing protein \| Virulence-associated E-like domain-containing protein; uniclust \| UniRef100\_A0A0R1XGL3 \| 100.0 \| 1.2e-32 \| 2.2e-38 \| 285.8 \| 270 \| (467, 754) \| 769 \| (287, 577) \| 622 \| Primase \| Primase; uniclust \| UniRef100\_UPI0019D1F5EF \| 100.0 \| 1.4e-32 \| 2.6e-38 \| 279.3 \| 286 \| (14, 350) \| 769 \| (64, 350) \| 747 \| AAA family ATPase \| AAA family ATPase; uniclust \| UniRef100\_A0A1F2W6T8 \| 100.0 \| 1.4e-32 \| 2.6e-38 \| 291.8 \| 321 \| (2, 350) \| 769 \| (31, 394) \| 630 \| DNA primase \| DNA primase; uniclust \| UniRef100\_A0A0X1KQ28 \| 100.0 \| 1.5e-32 \| 2.8e-38 \| 283.5 \| 312 \| (8, 350) \| 769 \| (3, 354) \| 574 \| DNA primase \| DNA primase; uniclust \| UniRef100\_A0A0E3MU70 \| 100.0 \| 1.5e-32 \| 2.8e-38 \| 277.4 \| 296 \| (14, 349) \| 769 \| (87, 385) \| 599 \| Pri \| Pri; uniclust \| UniRef100\_UPI0020CC0711 \| 100.0 \| 1.5e-32 \| 2.9e-38 \| 286.8 \| 337 \| (393, 740) \| 769 \| (470, 829) \| 995 \| virulence-associated E family protein \| virulence-associated E family protein; uniclust \| UniRef100\_A0A1G5H383 \| 100.0 \| 1.7e-32 \| 3.1e-38 \| 270.8 \| 519 \| (131, 697) \| 769 \| (113, 713) \| 767 \| Phage/plasmid primase, P4 family, C-terminal domain-containing protein \| Phage/plasmid primase, P4 family, C-terminal domain-containing protein; uniclust \| UniRef100\_A0A068Z0K6 \| 100.0 \| 1.7e-32 \| 3.4e-38 \| 314.0 \| 298 \| (14, 348) \| 769 \| (283, 583) \| 1434 \| DNA primase \| DNA primase; uniclust \| UniRef100\_A0A3B8XX17 \| 100.0 \| 1.9e-32 \| 3.5e-38 \| 267.6 \| 314 \| (372, 695) \| 769 \| (249, 588) \| 684 \| Virulence-associated E-like domain-containing protein \| Virulence-associated E-like domain-containing protein; uniclust \| UniRef100\_A0A158KPG1 \| 100.0 \| 1.9e-32 \| 3.5e-38 \| 274.0 \| 347 \| (393, 753) \| 769 \| (517, 887) \| 893 \| Virulence-associated E family protein \| Virulence-associated E family protein; uniclust \| UniRef100\_A0A177R8F8 \| 100.0 \| 2e-32 \| 3.9e-38 \| 293.6 \| 321 \| (3, 350) \| 769 \| (25, 391) \| 641 \| DNA primase \| DNA primase; uniclust \| UniRef100\_UPI001CD65D62 \| 100.0 \| 2.2e-32 \| 4.1e-38 \| 261.6 \| 307 \| (380, 695) \| 769 \| (155, 468) \| 557 \| virulence-associated E family protein \| virulence-associated E family protein; uniclust \| UniRef100\_A0A645CY77 \| 100.0 \| 2.4e-32 \| 4.4e-38 \| 244.4 \| 229 \| (436, 672) \| 769 \| (65, 299) \| 318 \| Virulence-associated E-like domain-containing protein \| Virulence-associated E-like domain-containing protein; uniclust \| UniRef100\_A0A1G3LYT4 \| 100.0 \| 2.4e-32 \| 4.5e-38 \| 285.3 \| 320 \| (4, 349) \| 769 \| (17, 378) \| 611 \| DNA primase \| DNA primase; uniclust \| UniRef100\_A0A143Y5P7 \| 100.0 \| 2.6e-32 \| 4.8e-38 \| 269.3 \| 309 \| (378, 695) \| 769 \| (340, 662) \| 760 \| Virulence-associated E-like domain-containing protein \| Virulence-associated E-like domain-containing protein; uniclust \| UniRef100\_A0A2V5PV48 \| 99.9 \| 2.7e-32 \| 5e-38 \| 276.4 \| 340 \| (390, 742) \| 769 \| (405, 759) \| 780 \| DNA primase/polymerase bifunctional N-terminal domain-containing protein \| DNA primase/polymerase bifunctional N-terminal domain-containing protein; uniclust \| UniRef100\_A0A521XCL3 \| 99.9 \| 2.7e-32 \| 5e-38 \| 276.0 \| 352 \| (381, 740) \| 769 \| (385, 754) \| 780 \| DNA primase/polymerase bifunctional N-terminal domain-containing protein \| DNA primase/polymerase bifunctional N-terminal domain-containing protein; uniclust \| UniRef100\_A0A2N5YUX1 \| 99.9 \| 2.6e-32 \| 5e-38 \| 255.5 \| 269 \| (476, 755) \| 769 \| (5, 279) \| 286 \| Virulence-associated E-like domain-containing protein \| Virulence-associated E-like domain-containing protein; uniclust \| UniRef100\_A0A031K4Q8 \| 99.9 \| 2.9e-32 \| 5.5e-38 \| 292.0 \| 319 \| (4, 349) \| 769 \| (19, 383) \| 666 \| DNA primase \| DNA primase; uniclust \| UniRef100\_A0A1Y6ISW7 \| 99.9 \| 3.2e-32 \| 6e-38 \| 258.6 \| 312 \| (435, 755) \| 769 \| (6, 341) \| 348 \| Virulence-associated protein E \| Virulence-associated protein E; uniclust \| UniRef100\_A0A6J5S9L8 \| 99.9 \| 3.5e-32 \| 6.4e-38 \| 267.9 \| 372 \| (376, 755) \| 769 \| (355, 740) \| 745 \| Virulence-associated E \| Virulence-associated E; uniclust \| UniRef100\_A0A072NKF5 \| 99.9 \| 3.3e-32 \| 6.5e-38 \| 272.5 \| 281 \| (3, 307) \| 769 \| (26, 347) \| 362 \| DNA primase, catalytic core \| DNA primase, catalytic core; uniclust \| UniRef100\_UPI0016899E5E \| 99.9 \| 3.5e-32 \| 6.5e-38 \| 275.7 \| 304 \| (438, 755) \| 769 \| (662, 976) \| 1051 \| VapE family protein \| VapE family protein; uniclust \| UniRef100\_UPI001E53A1EB \| 99.9 \| 3.8e-32 \| 7.1e-38 \| 273.7 \| 563 \| (150, 739) \| 769 \| (51, 704) \| 796 \| VapE family protein \| VapE family protein; uniclust \| UniRef100\_UPI00096A411E \| 99.9 \| 4e-32 \| 7.4e-38 \| 258.6 \| 320 \| (14, 350) \| 769 \| (23, 360) \| 436 \| toprim domain-containing protein \| toprim domain-containing protein; uniclust \| UniRef100\_A0A528WAA4 \| 99.9 \| 4.1e-32 \| 7.6e-38 \| 270.2 \| 358 \| (371, 737) \| 769 \| (208, 615) \| 657 \| Virulence-associated E-like domain-containing protein \| Virulence-associated E-like domain-containing protein; uniclust \| UniRef100\_A0A101H196 \| 99.9 \| 4e-32 \| 7.9e-38 \| 290.9 \| 319 \| (1, 351) \| 769 \| (1, 360) \| 576 \| DNA primase \| DNA primase; uniclust \| UniRef100\_A0A068YJZ3 \| 99.9 \| 4.5e-32 \| 8.6e-38 \| 258.2 \| 277 \| (10, 338) \| 769 \| (5, 282) \| 305 \| DNA primase TraC \| DNA primase TraC; uniclust \| UniRef100\_A0A0G0BXR0 \| 99.9 \| 4.6e-32 \| 8.8e-38 \| 285.3 \| 315 \| (4, 349) \| 769 \| (20, 382) \| 628 \| DNA primase \| DNA primase; uniclust \| UniRef100\_A0A2A4ZUQ6 \| 99.9 \| 4.7e-32 \| 8.9e-38 \| 252.4 \| 240 \| (12, 288) \| 769 \| (27, 266) \| 305 \| Toprim domain-containing protein (Fragment) \| Toprim domain-containing protein (Fragment); pdb70 \| 3B39\_A \| 99.6 \| 6e-20 \| 7.4e-24 \| 180.8 \| 161 \| (174, 350) \| 769 \| (91, 261) \| 322 \| DNA primase/DNA Complex \| 3B39\_A DNA primase/DNA Complex Protein-DNA complex, TOPRIM fold, DNA; pdb70 \| 3B39\_B \| 99.6 \| 6e-20 \| 7.4e-24 \| 180.8 \| 161 \| (174, 350) \| 769 \| (91, 261) \| 322 \| DNA primase/DNA Complex \| 3B39\_B DNA primase/DNA Complex Protein-DNA complex, TOPRIM fold, DNA; pdb70 \| 1DD9\_A \| 99.6 \| 1.6e-19 \| 1.9e-23 \| 180.3 \| 161 \| (174, 350) \| 769 \| (103, 273) \| 338 \| DNA PRIMASE (2.7.7.-) \| 1DD9\_A DNA PRIMASE (2.7.7.-) TOPRIM, 3-HELIX BUNDLE, DNA-BINDING PROTEIN; pdb70 \| 2AU3\_A \| 99.6 \| 1.5e-19 \| 2e-23 \| 179.3 \| 162 \| (175, 351) \| 769 \| (184, 352) \| 407 \| DNA primase (E.C.2.7.7.-) \| 2AU3\_A DNA primase (E.C.2.7.7.-) Zinc Ribbon, TOPRIM, RNA POLYMERASE; pdb70 \| 5VAZ\_A \| 99.5 \| 2.3e-19 \| 2.7e-23 \| 183.4 \| 161 \| (173, 349) \| 769 \| (103, 274) \| 389 \| DNA primase (E.C.2.7.7.-) \| 5VAZ\_A DNA primase (E.C.2.7.7.-) structural genomics, DNA-binding, primase, truncation HET: SO4; pdb70 \| 5VAZ\_B \| 99.5 \| 2.8e-19 \| 3.3e-23 \| 182.9 \| 161 \| (173, 349) \| 769 \| (103, 274) \| 389 \| DNA primase (E.C.2.7.7.-) \| 5VAZ\_B DNA primase (E.C.2.7.7.-) structural genomics, DNA-binding, primase, truncation HET: SO4; pdb70 \| 5W35\_B \| 99.5 \| 3.5e-19 \| 4.4e-23 \| 173.8 \| 167 \| (173, 350) \| 769 \| (90, 268) \| 325 \| DNA primase/DNA Complex \| 5W35\_B DNA primase/DNA Complex DNA replication, replisome, TOPRIM fold; pdb70 \| 5W36\_B \| 99.5 \| 3.5e-19 \| 4.4e-23 \| 173.8 \| 167 \| (173, 350) \| 769 \| (90, 268) \| 325 \| DNA primase/DNA Complex \| 5W36\_B DNA primase/DNA Complex DNA replication, replisome, TOPRIM fold; pdb70 \| 4EDG\_A \| 99.5 \| 4.5e-19 \| 5.6e-23 \| 175.2 \| 160 \| (173, 348) \| 769 \| (91, 257) \| 329 \| DNA primase (E.C.2.7.7.-) \| 4EDG\_A DNA primase (E.C.2.7.7.-) Catalytic Domain, nucleoside triphosphate, nucleoside HET: BEN, ATP; pdb70 \| 4EDK\_A \| 99.5 \| 4.5e-19 \| 5.6e-23 \| 175.2 \| 160 \| (173, 348) \| 769 \| (91, 257) \| 329 \| DNA primase (E.C.2.7.7.-) \| 4EDK\_A DNA primase (E.C.2.7.7.-) Catalytic Domain, nucleoside triphosphate, nucleoside HET: BEN, GTP; pdb70 \| 4EDR\_A \| 99.5 \| 4.5e-19 \| 5.6e-23 \| 175.2 \| 160 \| (173, 348) \| 769 \| (91, 257) \| 329 \| DNA primase (E.C.2.7.7.-) \| 4EDR\_A DNA primase (E.C.2.7.7.-) Catalytic Domain, nucleoside triphosphate, nucleoside HET: BEN, UTP; pdb70 \| 5GUJ\_A \| 99.5 \| 4.5e-19 \| 5.7e-23 \| 174.5 \| 157 \| (173, 345) \| 769 \| (89, 252) \| 324 \| DNA primase (E.C.2.7.7.-) \| 5GUJ\_A DNA primase (E.C.2.7.7.-) TRANSFERASE; pdb70 \| 1NUI\_A \| 98.9 \| 7.2e-14 \| 9.9e-18 \| 128.0 \| 139 \| (178, 343) \| 769 \| (105, 253) \| 255 \| DNA primase/helicase (E.C.2.7.7.-) \| 1NUI\_A DNA primase/helicase (E.C.2.7.7.-) zinc-biding domain, TOPRIM fold, DNA; pdb70 \| 6N9X\_B \| 98.9 \| 8.3e-14 \| 1.2e-17 \| 139.3 \| 170 \| (148, 349) \| 769 \| (75, 259) \| 566 \| DNA-directed DNA polymerase, TrxA/DNA Complex \| 6N9X\_B DNA-directed DNA polymerase, TrxA/DNA Complex DNA polymerase, primase, helicase, DNA HET: DOC, TTP; pdb70 \| 6N7I\_D \| 98.9 \| 9.7e-14 \| 1.4e-17 \| 138.8 \| 170 \| (148, 349) \| 769 \| (75, 259) \| 566 \| DNA primase/helicase/DNA Complex \| 6N7I\_D DNA primase/helicase/DNA Complex helicase, ATPase, hexamer, DNA replication HET: TTP; pdb70 \| 1Q57\_G \| 98.8 \| 2.9e-13 \| 4e-17 \| 134.8 \| 148 \| (178, 349) \| 769 \| (42, 196) \| 503 \| DNA primase/helicase (E.C.2.7.7.-) \| 1Q57\_G DNA primase/helicase (E.C.2.7.7.-) primase, helicase, dNTPase, DNA replication; pdb70 \| 5IKN\_E \| 98.8 \| 6.5e-13 \| 9.2e-17 \| 129.8 \| 141 \| (178, 345) \| 769 \| (42, 192) \| 486 \| DNA-directed DNA polymerase (E.C.2.7.7.7,3.1.11.-), DNA \| 5IKN\_E DNA-directed DNA polymerase (E.C.2.7.7.7,3.1.11.-), DNA Replisome, TRANSFERASE; pdb70 \| 5IKN\_H \| 98.8 \| 6.5e-13 \| 9.2e-17 \| 129.8 \| 141 \| (178, 345) \| 769 \| (42, 192) \| 486 \| DNA-directed DNA polymerase (E.C.2.7.7.7,3.1.11.-), DNA \| 5IKN\_H DNA-directed DNA polymerase (E.C.2.7.7.7,3.1.11.-), DNA Replisome, TRANSFERASE; pdb70 \| 2I5R\_C \| 98.3 \| 1.3e-10 \| 1.6e-14 \| 99.1 \| 93 \| (231, 336) \| 769 \| (6, 100) \| 122 \| Hypothetical conserved protein \| 2I5R\_C Hypothetical conserved protein Toprim domain, Structural Genomics, PSI-2 HET: GOL, SO4, MES; pdb70 \| 2FCJ\_B \| 98.3 \| 1.9e-10 \| 2.4e-14 \| 97.2 \| 92 \| (232, 336) \| 769 \| (4, 97) \| 119 \| small TOPRIM domain protein \| 2FCJ\_B small TOPRIM domain protein TOPRIM domain, Structural Genomics, PSI HET: MES, SO4, GOL; pdb70 \| 2FCJ\_C \| 98.3 \| 1.9e-10 \| 2.4e-14 \| 97.2 \| 92 \| (232, 336) \| 769 \| (4, 97) \| 119 \| small TOPRIM domain protein \| 2FCJ\_C small TOPRIM domain protein TOPRIM domain, Structural Genomics, PSI HET: GOL, SO4, MES; pdb70 \| 1T6T\_1 \| 97.4 \| 9.1e-08 \| 1e-11 \| 82.4 \| 70 \| (232, 312) \| 769 \| (20, 89) \| 118 \| putative protein \| 1T6T\_1 putative protein structural genomics, Aquifex aeolicus, PSI HET: MSE; pdb70 \| 1T6T\_2 \| 97.4 \| 9.1e-08 \| 1e-11 \| 82.4 \| 70 \| (232, 312) \| 769 \| (20, 89) \| 118 \| putative protein \| 1T6T\_2 putative protein structural genomics, Aquifex aeolicus, PSI |
| Top keywords  (threshold 1.00e-03 (evalue)) | **primase, domain\_containing, DNA, Virulence\_associated, E, E\_like, Toprim, Fragment, helicase, ATPase** |
| Output files | ../../similar\_sequences/54\_FANPEZAQ\_CDS\_0054\_merged.svg ../../similar\_sequences/54\_FANPEZAQ\_CDS\_0054\_pdb70.a3m ../../similar\_sequences/54\_FANPEZAQ\_CDS\_0054\_pdb70.hhr ../../similar\_sequences/54\_FANPEZAQ\_CDS\_0054\_uniclust.a3m ../../similar\_sequences/54\_FANPEZAQ\_CDS\_0054\_uniclust.hhr |

#### Structure prediction (AlphaFold)^2^

|  |  |
| --- | --- |
| Stats | xml version="1.0" encoding="utf-8" standalone="no"?       2024-09-02T21:09:50.850261 image/svg+xml   Matplotlib v3.7.2, https://matplotlib.org/ |
| Predicted structure | **NGL Viewer Controls:**  - Center: *Left-Click* - Rotate: *Left-Click + Drag* - Translate: *Right-Click + Drag* - Zoom: *Shift + Left-Click + Drag* |
| Output files | ../../predicted\_structures/54\_FANPEZAQ\_CDS\_0054/features.pkl ../../predicted\_structures/54\_FANPEZAQ\_CDS\_0054/ranked\_0.pdb ../../predicted\_structures/54\_FANPEZAQ\_CDS\_0054/ranked\_0\_plots.svg ../../predicted\_structures/54\_FANPEZAQ\_CDS\_0054/result\_model\_1\_ptm\_pred\_0.pkl |

#### Structure similarity search results (Foldseek)^3^

|  |  |
| --- | --- |
| Structure databases searched | Pdb, Afdb-proteome, Afdb-uniprot50 |
| Results, scheme(s)  (Top layers only, threshold 1.00e-02 (evalue)) | xml version="1.0" encoding="utf-8" standalone="no"?       2024-09-02T21:11:28.495615 image/svg+xml   Matplotlib v3.7.2, https://matplotlib.org/ |
| Results, table  (threshold 1.00e-02 (evalue)) | db \| id \| prob \| evalue \| bits \| fident \| alnlen \| mismatch \| gapopen \| qstart \| qend \| tstart \| tend \| name \| description; pdb \| 7PDS\_C \| 1.0 \| 7.1e-19 \| 583 \| 0.207 \| 449 \| 278 \| 22 \| 370 \| 751 \| 8 \| 445 \| Similar to D. nodosus vapE \| Similar to D. nodosus vapE; pdb \| 7PDS\_F \| 1.0 \| 5.486e-19 \| 577 \| 0.205 \| 452 \| 275 \| 26 \| 370 \| 751 \| 8 \| 445 \| Similar to D. nodosus vapE \| Similar to D. nodosus vapE; pdb \| 7PDS\_B \| 1.0 \| 2.326e-18 \| 575 \| 0.204 \| 445 \| 282 \| 22 \| 370 \| 752 \| 8 \| 442 \| Similar to D. nodosus vapE \| Similar to D. nodosus vapE; pdb \| 7PDS\_A \| 1.0 \| 5.593e-18 \| 567 \| 0.197 \| 456 \| 283 \| 24 \| 368 \| 752 \| 7 \| 450 \| Similar to D. nodosus vapE \| Similar to D. nodosus vapE; pdb \| 7PDS\_E \| 1.0 \| 2.326e-18 \| 555 \| 0.202 \| 459 \| 286 \| 24 \| 370 \| 761 \| 8 \| 453 \| Similar to D. nodosus vapE \| Similar to D. nodosus vapE; pdb \| 7PDS\_D \| 1.0 \| 1.416e-17 \| 544 \| 0.202 \| 450 \| 280 \| 23 \| 370 \| 751 \| 8 \| 446 \| Similar to D. nodosus vapE \| Similar to D. nodosus vapE; pdb \| 6K9E\_A \| 1.0 \| 1.631e-07 \| 175 \| 0.13 \| 443 \| 267 \| 32 \| 384 \| 755 \| 7 \| 402 \| Primase \| Primase; pdb \| 6K9E\_C \| 1.0 \| 2.341e-07 \| 174 \| 0.133 \| 442 \| 262 \| 30 \| 384 \| 751 \| 6 \| 400 \| Primase \| Primase; pdb \| 6LRB\_C \| 1.0 \| 7.285e-07 \| 159 \| 0.128 \| 450 \| 269 \| 31 \| 384 \| 755 \| 6 \| 410 \| Primase \| Primase; pdb \| 6LRB\_D \| 1.0 \| 2.786e-06 \| 158 \| 0.125 \| 447 \| 271 \| 31 \| 384 \| 755 \| 8 \| 409 \| Primase \| Primase; pdb \| 6K9C\_A \| 1.0 \| 4.348e-07 \| 157 \| 0.138 \| 447 \| 272 \| 26 \| 384 \| 755 \| 8 \| 416 \| Primase \| Primase; pdb \| 6K9E\_F \| 1.0 \| 7.67e-07 \| 153 \| 0.128 \| 444 \| 262 \| 30 \| 384 \| 751 \| 7 \| 401 \| Primase \| Primase; pdb \| 6LRB\_F \| 1.0 \| 2.153e-06 \| 152 \| 0.138 \| 442 \| 272 \| 29 \| 384 \| 755 \| 6 \| 408 \| Primase \| Primase; pdb \| 5W33\_A \| 1.0 \| 2.698e-05 \| 145 \| 0.154 \| 279 \| 157 \| 15 \| 123 \| 345 \| 5 \| 260 \| DNA primase \| DNA primase; pdb \| 6LRB\_A \| 1.0 \| 2.513e-06 \| 145 \| 0.137 \| 415 \| 263 \| 26 \| 384 \| 745 \| 7 \| 379 \| Primase \| Primase; pdb \| 7OLA\_B \| 1.0 \| 2.153e-06 \| 141 \| 0.115 \| 432 \| 271 \| 29 \| 372 \| 731 \| 26 \| 418 \| DNA primase \| DNA primase; pdb \| 7OLA\_D \| 1.0 \| 1.122e-05 \| 138 \| 0.12 \| 375 \| 238 \| 25 \| 374 \| 685 \| 28 \| 373 \| DNA primase \| DNA primase; pdb \| 2AU3\_A \| 1.0 \| 2.991e-05 \| 138 \| 0.131 \| 326 \| 189 \| 20 \| 70 \| 347 \| 67 \| 346 \| DNA primase \| DNA primase; pdb \| 5W36\_B \| 1.0 \| 6.83e-05 \| 134 \| 0.16 \| 287 \| 154 \| 14 \| 123 \| 349 \| 6 \| 265 \| DNA primase \| DNA primase; pdb \| 7OLA\_A \| 1.0 \| 1.98e-05 \| 131 \| 0.105 \| 434 \| 281 \| 31 \| 372 \| 735 \| 26 \| 422 \| DNA primase \| DNA primase; pdb \| 8IQI\_E \| 1.0 \| 4.348e-07 \| 128 \| 0.105 \| 856 \| 433 \| 58 \| 13 \| 766 \| 283 \| 907 \| Putative primase C962R \| Putative primase C962R; pdb \| 3B39\_A \| 1.0 \| 5.85e-05 \| 127 \| 0.157 \| 247 \| 133 \| 18 \| 143 \| 352 \| 52 \| 260 \| DNA primase \| DNA primase; pdb \| 7OLA\_C \| 1.0 \| 1.786e-05 \| 127 \| 0.109 \| 431 \| 275 \| 29 \| 370 \| 730 \| 24 \| 415 \| DNA primase \| DNA primase; pdb \| 5GUJ\_A \| 1.0 \| 2.311e-05 \| 126 \| 0.084 \| 333 \| 184 \| 20 \| 154 \| 447 \| 73 \| 323 \| DNA primase \| DNA primase; pdb \| 8IQI\_B \| 1.0 \| 9.429e-07 \| 125 \| 0.104 \| 845 \| 430 \| 64 \| 13 \| 758 \| 283 \| 899 \| Putative primase C962R \| Putative primase C962R; pdb \| 3B39\_B \| 1.0 \| 9.801e-05 \| 124 \| 0.124 \| 387 \| 191 \| 24 \| 124 \| 446 \| 3 \| 305 \| DNA primase \| DNA primase; pdb \| 1DD9\_A \| 1.0 \| 4.52e-05 \| 124 \| 0.121 \| 347 \| 177 \| 19 \| 153 \| 444 \| 28 \| 301 \| DNA PRIMASE \| DNA PRIMASE; pdb \| 7OM0\_F \| 1.0 \| 3.15e-05 \| 124 \| 0.125 \| 423 \| 261 \| 30 \| 384 \| 735 \| 38 \| 422 \| DNA primase \| DNA primase; pdb \| 1DDE\_A \| 1.0 \| 5.011e-05 \| 122 \| 0.133 \| 359 \| 166 \| 22 \| 153 \| 447 \| 28 \| 305 \| DNA PRIMASE \| DNA PRIMASE; pdb \| 5W34\_A \| 1.0 \| 0.0001336 \| 122 \| 0.125 \| 384 \| 197 \| 20 \| 126 \| 444 \| 5 \| 314 \| DNA primase \| DNA primase; pdb \| 7OLA\_E \| 1.0 \| 0.0004608 \| 116 \| 0.151 \| 185 \| 127 \| 14 \| 492 \| 665 \| 174 \| 339 \| DNA primase \| DNA primase; pdb \| 1EQN\_E \| 1.0 \| 0.0001144 \| 114 \| 0.136 \| 264 \| 136 \| 20 \| 153 \| 350 \| 28 \| 265 \| DNA PRIMASE \| DNA PRIMASE; pdb \| 7OLA\_F \| 1.0 \| 0.0001144 \| 112 \| 0.095 \| 344 \| 214 \| 23 \| 372 \| 653 \| 26 \| 334 \| DNA primase \| DNA primase; pdb \| 8IQI\_A \| 1.0 \| 9.13e-06 \| 104 \| 0.103 \| 825 \| 432 \| 56 \| 13 \| 750 \| 284 \| 887 \| Putative primase C962R \| Putative primase C962R; pdb \| 4EDV\_A \| 1.0 \| 9.801e-05 \| 103 \| 0.121 \| 347 \| 182 \| 20 \| 139 \| 444 \| 46 \| 310 \| DNA primase \| DNA primase; pdb \| 8APL\_A \| 1.0 \| 0.000305 \| 101 \| 0.107 \| 473 \| 286 \| 28 \| 381 \| 752 \| 7 \| 444 \| Primase D5 \| Primase D5; pdb \| 8IQH\_D \| 0.999 \| 6.362e-06 \| 98 \| 0.105 \| 837 \| 417 \| 61 \| 12 \| 762 \| 278 \| 868 \| Putative primase C962R \| Putative primase C962R; pdb \| 4E2K\_A \| 0.999 \| 0.0001729 \| 97 \| 0.122 \| 342 \| 185 \| 21 \| 139 \| 444 \| 46 \| 308 \| DNA primase \| DNA primase; pdb \| 8APM\_F \| 0.998 \| 0.0005664 \| 95 \| 0.108 \| 369 \| 224 \| 24 \| 370 \| 657 \| 1 \| 345 \| Primase D5 \| Primase D5; pdb \| 8B9A\_6 \| 0.996 \| 6.83e-05 \| 90 \| 0.117 \| 333 \| 212 \| 20 \| 456 \| 735 \| 319 \| 622 \| DNA replication licensing factor MCM6 \| DNA replication licensing factor MCM6; pdb \| 8IQH\_G \| 0.996 \| 4.293e-05 \| 90 \| 0.094 \| 806 \| 434 \| 54 \| 12 \| 762 \| 276 \| 840 \| Putative primase C962R \| Putative primase C962R; pdb \| 8IQH\_B \| 0.996 \| 2.698e-05 \| 90 \| 0.105 \| 824 \| 430 \| 57 \| 12 \| 758 \| 280 \| 873 \| Putative primase C962R \| Putative primase C962R; pdb \| 5VAZ\_A \| 0.995 \| 0.0008559 \| 88 \| 0.136 \| 315 \| 156 \| 19 \| 169 \| 442 \| 80 \| 319 \| DNA primase \| DNA primase; pdb \| 7PT7\_F \| 0.994 \| 0.0002125 \| 87 \| 0.116 \| 344 \| 212 \| 19 \| 456 \| 735 \| 298 \| 613 \| DNA replication licensing factor MCM6 \| DNA replication licensing factor MCM6; pdb \| 8IQH\_M \| 0.994 \| 3.872e-05 \| 87 \| 0.097 \| 811 \| 420 \| 54 \| 13 \| 755 \| 275 \| 841 \| Putative primase C962R \| Putative primase C962R; pdb \| 7PT6\_F \| 0.993 \| 0.0001336 \| 86 \| 0.123 \| 339 \| 218 \| 20 \| 456 \| 735 \| 315 \| 633 \| DNA replication licensing factor MCM6 \| DNA replication licensing factor MCM6; pdb \| 6SKL\_3 \| 0.991 \| 0.0001917 \| 84 \| 0.112 \| 339 \| 209 \| 23 \| 468 \| 738 \| 302 \| 616 \| DNA replication licensing factor MCM3 \| DNA replication licensing factor MCM3; pdb \| 2V9P\_E \| 0.99 \| 0.003109 \| 83 \| 0.109 \| 265 \| 193 \| 17 \| 416 \| 657 \| 21 \| 265 \| REPLICATION PROTEIN E1 \| REPLICATION PROTEIN E1; pdb \| 7QHS\_3 \| 0.99 \| 0.0002612 \| 83 \| 0.097 \| 360 \| 220 \| 21 \| 456 \| 735 \| 294 \| 628 \| DNA replication licensing factor MCM3 \| DNA replication licensing factor MCM3; pdb \| 3JC7\_2 \| 0.988 \| 0.0003749 \| 82 \| 0.098 \| 326 \| 206 \| 20 \| 470 \| 736 \| 261 \| 557 \| DNA replication licensing factor MCM2 \| DNA replication licensing factor MCM2; pdb \| 7W1Y\_2 \| 0.988 \| 0.0002238 \| 82 \| 0.092 \| 346 \| 225 \| 24 \| 456 \| 735 \| 281 \| 603 \| DNA replication licensing factor MCM2 \| DNA replication licensing factor MCM2; pdb \| 8IQH\_C \| 0.988 \| 9.801e-05 \| 82 \| 0.108 \| 826 \| 425 \| 63 \| 12 \| 758 \| 279 \| 871 \| Putative primase C962R \| Putative primase C962R; pdb \| 8B9A\_4 \| 0.986 \| 0.0002751 \| 81 \| 0.118 \| 337 \| 210 \| 25 \| 457 \| 735 \| 303 \| 610 \| DNA replication licensing factor MCM4 \| DNA replication licensing factor MCM4; pdb \| 7W1Y\_E \| 0.986 \| 5.011e-05 \| 81 \| 0.095 \| 409 \| 247 \| 24 \| 402 \| 735 \| 277 \| 637 \| DNA replication licensing factor MCM6 \| DNA replication licensing factor MCM6; pdb \| 7W1Y\_A \| 0.984 \| 0.0002896 \| 80 \| 0.102 \| 350 \| 220 \| 26 \| 453 \| 735 \| 277 \| 599 \| DNA replication licensing factor MCM2 \| DNA replication licensing factor MCM2; pdb \| 6MII\_A \| 0.978 \| 0.0002896 \| 78 \| 0.111 \| 342 \| 217 \| 23 \| 456 \| 738 \| 267 \| 580 \| Minichromosome maintenance protein MCM \| Minichromosome maintenance protein MCM; pdb \| 6WGI\_3 \| 0.975 \| 0.001108 \| 77 \| 0.107 \| 344 \| 206 \| 19 \| 456 \| 735 \| 260 \| 566 \| DNA replication licensing factor MCM3 \| DNA replication licensing factor MCM3; pdb \| 7PMN\_5 \| 0.975 \| 0.0009992 \| 77 \| 0.128 \| 272 \| 160 \| 19 \| 457 \| 682 \| 259 \| 499 \| Minichromosome maintenance protein 5 \| Minichromosome maintenance protein 5; pdb \| 7PLO\_2 \| 0.975 \| 0.001293 \| 77 \| 0.095 \| 335 \| 215 \| 21 \| 456 \| 737 \| 292 \| 591 \| DNA replication licensing factor MCM2 \| DNA replication licensing factor MCM2; pdb \| 1SVM\_C \| 0.971 \| 0.003109 \| 76 \| 0.134 \| 313 \| 180 \| 21 \| 415 \| 677 \| 66 \| 337 \| large T antigen \| large T antigen; pdb \| 6PTN\_M \| 0.971 \| 0.000356 \| 76 \| 0.114 \| 358 \| 198 \| 23 \| 456 \| 734 \| 290 \| 607 \| DNA replication licensing factor MCM6 \| DNA replication licensing factor MCM6; pdb \| 3JC5\_2 \| 0.967 \| 0.0006963 \| 75 \| 0.102 \| 333 \| 205 \| 20 \| 470 \| 736 \| 261 \| 565 \| DNA replication licensing factor MCM2 \| DNA replication licensing factor MCM2; pdb \| 5U8T\_3 \| 0.967 \| 0.001674 \| 75 \| 0.121 \| 395 \| 214 \| 27 \| 405 \| 736 \| 264 \| 588 \| DNA replication licensing factor MCM3 \| DNA replication licensing factor MCM3; pdb \| 5U8S\_3 \| 0.967 \| 0.001856 \| 75 \| 0.112 \| 354 \| 197 \| 25 \| 456 \| 735 \| 279 \| 589 \| DNA replication licensing factor MCM3 \| DNA replication licensing factor MCM3; pdb \| 6XTY\_3 \| 0.967 \| 0.0003947 \| 75 \| 0.092 \| 378 \| 210 \| 28 \| 456 \| 757 \| 288 \| 608 \| DNA replication licensing factor MCM3 \| DNA replication licensing factor MCM3; pdb \| 8OUW\_6 \| 0.967 \| 0.001052 \| 75 \| 0.093 \| 353 \| 222 \| 24 \| 455 \| 735 \| 309 \| 635 \| DNA replication licensing factor mcm-6 \| DNA replication licensing factor mcm-6; pdb \| 6SKO\_3 \| 0.961 \| 0.0008129 \| 74 \| 0.11 \| 352 \| 208 \| 23 \| 456 \| 735 \| 13 \| 331 \| DNA replication licensing factor MCM3 \| DNA replication licensing factor MCM3; pdb \| 7PMN\_6 \| 0.961 \| 0.0002481 \| 74 \| 0.102 \| 399 \| 238 \| 25 \| 403 \| 735 \| 293 \| 637 \| DNA replication licensing factor MCM6 \| DNA replication licensing factor MCM6; pdb \| 8IQH\_E \| 0.961 \| 0.0001642 \| 74 \| 0.116 \| 671 \| 334 \| 51 \| 243 \| 762 \| 298 \| 860 \| Putative primase C962R \| Putative primase C962R; pdb \| 8IQH\_A \| 0.956 \| 0.0004608 \| 73 \| 0.102 \| 830 \| 423 \| 56 \| 12 \| 762 \| 277 \| 863 \| Putative primase C962R \| Putative primase C962R; pdb \| 7PMK\_3 \| 0.949 \| 0.0009992 \| 72 \| 0.103 \| 358 \| 209 \| 25 \| 456 \| 738 \| 287 \| 607 \| DNA replication licensing factor MCM3 \| DNA replication licensing factor MCM3; pdb \| 8OUW\_3 \| 0.949 \| 0.0009992 \| 72 \| 0.103 \| 376 \| 222 \| 22 \| 456 \| 752 \| 276 \| 615 \| DNA replication licensing factor MCM3 \| DNA replication licensing factor MCM3; pdb \| 7V3V\_C \| 0.949 \| 0.001293 \| 72 \| 0.105 \| 369 \| 213 \| 22 \| 456 \| 735 \| 295 \| 635 \| DNA replication licensing factor MCM3 \| DNA replication licensing factor MCM3; pdb \| 6WGG\_3 \| 0.949 \| 0.002663 \| 72 \| 0.103 \| 347 \| 204 \| 20 \| 456 \| 735 \| 265 \| 571 \| DNA replication licensing factor MCM3 \| DNA replication licensing factor MCM3; pdb \| 7PMK\_2 \| 0.949 \| 0.0006613 \| 72 \| 0.097 \| 369 \| 215 \| 22 \| 457 \| 737 \| 296 \| 634 \| DNA replication licensing factor MCM2 \| DNA replication licensing factor MCM2; pdb \| 6SKL\_4 \| 0.949 \| 0.002663 \| 72 \| 0.093 \| 407 \| 223 \| 28 \| 456 \| 747 \| 301 \| 676 \| DNA replication licensing factor MCM4 \| DNA replication licensing factor MCM4; pdb \| 8IQH\_J \| 0.949 \| 0.0005964 \| 72 \| 0.088 \| 827 \| 437 \| 53 \| 12 \| 762 \| 273 \| 858 \| Putative primase C962R \| Putative primase C962R; pdb \| 7PT6\_2 \| 0.941 \| 0.000538 \| 71 \| 0.107 \| 344 \| 196 \| 26 \| 457 \| 735 \| 291 \| 588 \| DNA replication licensing factor MCM2 \| DNA replication licensing factor MCM2; pdb \| 6XTX\_6 \| 0.941 \| 0.001434 \| 71 \| 0.107 \| 373 \| 224 \| 26 \| 456 \| 760 \| 280 \| 611 \| DNA replication licensing factor MCM6 \| DNA replication licensing factor MCM6; pdb \| 7V3U\_G \| 0.941 \| 0.001108 \| 71 \| 0.129 \| 356 \| 201 \| 27 \| 456 \| 735 \| 370 \| 692 \| DNA replication licensing factor MCM7 \| DNA replication licensing factor MCM7; pdb \| 4GDF\_A \| 0.933 \| 0.005485 \| 70 \| 0.118 \| 322 \| 195 \| 22 \| 415 \| 677 \| 180 \| 471 \| Large T antigen \| Large T antigen; pdb \| 6WGG\_2 \| 0.933 \| 0.003274 \| 70 \| 0.112 \| 347 \| 208 \| 19 \| 457 \| 737 \| 251 \| 563 \| DNA replication licensing factor MCM2 \| DNA replication licensing factor MCM2; pdb \| 5XF8\_3 \| 0.933 \| 0.0009992 \| 70 \| 0.117 \| 348 \| 200 \| 23 \| 456 \| 736 \| 293 \| 600 \| DNA replication licensing factor MCM3 \| DNA replication licensing factor MCM3; pdb \| 7PFO\_5 \| 0.933 \| 0.002663 \| 70 \| 0.116 \| 267 \| 164 \| 19 \| 457 \| 675 \| 273 \| 515 \| DNA replication licensing factor MCM5 \| DNA replication licensing factor MCM5; pdb \| 7PLO\_4 \| 0.933 \| 0.001362 \| 70 \| 0.145 \| 358 \| 210 \| 26 \| 456 \| 746 \| 273 \| 601 \| DNA replication licensing factor MCM4 \| DNA replication licensing factor MCM4; pdb \| 7QHS\_6 \| 0.933 \| 0.000356 \| 70 \| 0.131 \| 351 \| 196 \| 24 \| 456 \| 735 \| 316 \| 628 \| DNA replication licensing factor MCM6 \| DNA replication licensing factor MCM6; pdb \| 7W1Y\_4 \| 0.933 \| 0.0005109 \| 70 \| 0.11 \| 362 \| 216 \| 22 \| 445 \| 737 \| 316 \| 640 \| DNA replication licensing factor MCM4 \| DNA replication licensing factor MCM4; pdb \| 7PMN\_2 \| 0.933 \| 0.0009489 \| 70 \| 0.118 \| 362 \| 210 \| 25 \| 457 \| 735 \| 297 \| 632 \| DNA replication licensing factor MCM2 \| DNA replication licensing factor MCM2; pdb \| 7V3U\_E \| 0.933 \| 0.001108 \| 70 \| 0.119 \| 367 \| 205 \| 26 \| 457 \| 737 \| 313 \| 647 \| Minichromosome maintenance protein 5 \| Minichromosome maintenance protein 5; pdb \| 6SKL\_2 \| 0.933 \| 0.001674 \| 70 \| 0.101 \| 366 \| 212 \| 24 \| 457 \| 735 \| 310 \| 645 \| DNA replication licensing factor MCM2 \| DNA replication licensing factor MCM2; pdb \| 1QVR\_C \| 0.933 \| 0.006081 \| 70 \| 0.107 \| 288 \| 177 \| 20 \| 420 \| 684 \| 465 \| 695 \| ClpB protein \| ClpB protein; pdb \| 6WGF\_3 \| 0.923 \| 0.005485 \| 69 \| 0.1 \| 347 \| 205 \| 19 \| 456 \| 735 \| 258 \| 564 \| DNA replication licensing factor MCM3 \| DNA replication licensing factor MCM3; pdb \| 8EAF\_B \| 0.923 \| 0.001166 \| 69 \| 0.101 \| 353 \| 210 \| 27 \| 456 \| 737 \| 267 \| 583 \| Minichromosome maintenance protein MCM \| Minichromosome maintenance protein MCM; pdb \| 6XTY\_6 \| 0.923 \| 0.00159 \| 69 \| 0.1 \| 369 \| 217 \| 27 \| 456 \| 751 \| 280 \| 606 \| DNA replication licensing factor MCM6 \| DNA replication licensing factor MCM6; pdb \| 3JA8\_5 \| 0.923 \| 0.0008559 \| 69 \| 0.116 \| 413 \| 238 \| 25 \| 402 \| 737 \| 267 \| 629 \| Minichromosome Maintenance 5 \| Minichromosome Maintenance 5; pdb \| 7T8C\_B \| 0.912 \| 0.0003381 \| 68 \| 0.101 \| 564 \| 272 \| 37 \| 137 \| 625 \| 73 \| 476 \| Twinkle mtDNA helicase \| Twinkle mtDNA helicase; pdb \| 5J47\_A \| 0.9 \| 0.004237 \| 67 \| 0.114 \| 332 \| 194 \| 21 \| 415 \| 683 \| 66 \| 360 \| Large T antigen \| Large T antigen; pdb \| 8E2L\_A \| 0.9 \| 0.002804 \| 67 \| 0.098 \| 526 \| 277 \| 25 \| 143 \| 616 \| 7 \| 387 \| Twinkle mtDNA helicase \| Twinkle mtDNA helicase; pdb \| 3JC5\_3 \| 0.9 \| 0.00151 \| 67 \| 0.106 \| 396 \| 218 \| 21 \| 405 \| 736 \| 264 \| 587 \| DNA replication licensing factor MCM3 \| DNA replication licensing factor MCM3; pdb \| 7PT7\_C \| 0.9 \| 0.002402 \| 67 \| 0.102 \| 382 \| 210 \| 22 \| 456 \| 738 \| 284 \| 631 \| DNA replication licensing factor MCM3 \| DNA replication licensing factor MCM3; pdb \| 7W1Y\_B \| 0.9 \| 0.001052 \| 67 \| 0.102 \| 401 \| 217 \| 24 \| 456 \| 749 \| 283 \| 647 \| Isoform 2 of DNA replication licensing factor MCM3 \| Isoform 2 of DNA replication licensing factor MCM3; pdb \| 3JC7\_5 \| 0.9 \| 0.001166 \| 67 \| 0.106 \| 412 \| 238 \| 26 \| 402 \| 737 \| 238 \| 595 \| Minichromosome maintenance protein 5 \| Minichromosome maintenance protein 5; pdb \| 7QHS\_2 \| 0.9 \| 0.00151 \| 67 \| 0.095 \| 368 \| 211 \| 25 \| 457 \| 735 \| 303 \| 637 \| DNA replication licensing factor MCM2 \| DNA replication licensing factor MCM2; pdb \| 5V8F\_7 \| 0.9 \| 0.001856 \| 67 \| 0.139 \| 336 \| 201 \| 28 \| 456 \| 736 \| 335 \| 637 \| DNA replication licensing factor MCM7 \| DNA replication licensing factor MCM7; pdb \| 5V8F\_4 \| 0.9 \| 0.0005109 \| 67 \| 0.089 \| 505 \| 260 \| 32 \| 403 \| 750 \| 278 \| 739 \| DNA replication licensing factor MCM4 \| DNA replication licensing factor MCM4; pdb \| 6WGI\_2 \| 0.887 \| 0.002804 \| 66 \| 0.108 \| 351 \| 205 \| 21 \| 457 \| 737 \| 243 \| 555 \| DNA replication licensing factor MCM2 \| DNA replication licensing factor MCM2; pdb \| 7PFO\_4 \| 0.887 \| 0.001108 \| 66 \| 0.119 \| 403 \| 211 \| 28 \| 402 \| 735 \| 263 \| 590 \| DNA replication licensing factor MCM4 \| DNA replication licensing factor MCM4; pdb \| 7W1Y\_F \| 0.887 \| 0.002167 \| 66 \| 0.115 \| 339 \| 199 \| 25 \| 471 \| 737 \| 331 \| 640 \| DNA replication licensing factor MCM7 \| DNA replication licensing factor MCM7; pdb \| 5V8F\_3 \| 0.887 \| 0.00159 \| 66 \| 0.105 \| 407 \| 219 \| 26 \| 402 \| 736 \| 251 \| 584 \| DNA replication licensing factor MCM3 \| DNA replication licensing factor MCM3; pdb \| 5V8F\_6 \| 0.887 \| 0.00159 \| 66 \| 0.108 \| 359 \| 217 \| 22 \| 456 \| 745 \| 297 \| 621 \| DNA replication licensing factor MCM6 \| DNA replication licensing factor MCM6; pdb \| 8OUW\_4 \| 0.872 \| 0.003109 \| 65 \| 0.113 \| 351 \| 211 \| 22 \| 456 \| 738 \| 301 \| 619 \| DNA replication licensing factor mcm-4 \| DNA replication licensing factor mcm-4; pdb \| 7PT6\_C \| 0.872 \| 0.006742 \| 65 \| 0.108 \| 359 \| 223 \| 20 \| 456 \| 735 \| 291 \| 631 \| DNA replication licensing factor MCM3 \| DNA replication licensing factor MCM3; pdb \| 6EYC\_7 \| 0.872 \| 0.001954 \| 65 \| 0.129 \| 341 \| 209 \| 26 \| 453 \| 736 \| 354 \| 663 \| DNA replication licensing factor MCM7 \| DNA replication licensing factor MCM7; pdb \| 6PTN\_J \| 0.855 \| 0.002953 \| 64 \| 0.122 \| 374 \| 206 \| 26 \| 430 \| 735 \| 273 \| 592 \| DNA replication licensing factor MCM3 \| DNA replication licensing factor MCM3; pdb \| 5BK4\_B \| 0.855 \| 0.001293 \| 64 \| 0.102 \| 341 \| 195 \| 23 \| 467 \| 735 \| 301 \| 602 \| DNA replication licensing factor MCM3 \| DNA replication licensing factor MCM3; pdb \| 7PLO\_7 \| 0.855 \| 0.002953 \| 64 \| 0.112 \| 339 \| 201 \| 23 \| 465 \| 737 \| 300 \| 604 \| DNA replication licensing factor MCM7 \| DNA replication licensing factor MCM7; pdb \| 7V3V\_B \| 0.855 \| 0.002529 \| 64 \| 0.115 \| 345 \| 214 \| 25 \| 457 \| 736 \| 289 \| 607 \| DNA replication licensing factor MCM2 \| DNA replication licensing factor MCM2; pdb \| 7PMK\_7 \| 0.855 \| 0.003447 \| 64 \| 0.122 \| 344 \| 205 \| 25 \| 456 \| 737 \| 317 \| 625 \| DNA replication licensing factor MCM7 \| DNA replication licensing factor MCM7; pdb \| 8OUW\_7 \| 0.855 \| 0.001954 \| 64 \| 0.125 \| 358 \| 201 \| 30 \| 457 \| 737 \| 304 \| 626 \| DNA replication licensing factor MCM7 \| DNA replication licensing factor MCM7; pdb \| 7W1Y\_6 \| 0.855 \| 0.001362 \| 64 \| 0.099 \| 413 \| 241 \| 29 \| 402 \| 735 \| 278 \| 638 \| DNA replication licensing factor MCM6 \| DNA replication licensing factor MCM6; pdb \| 6WGF\_7 \| 0.837 \| 0.00363 \| 63 \| 0.123 \| 364 \| 218 \| 28 \| 449 \| 737 \| 222 \| 559 \| DNA replication licensing factor MCM7 \| DNA replication licensing factor MCM7; pdb \| 8B9D\_6 \| 0.837 \| 0.002953 \| 63 \| 0.116 \| 343 \| 205 \| 24 \| 456 \| 736 \| 296 \| 602 \| DNA replication licensing factor MCM6 \| DNA replication licensing factor MCM6; pdb \| 7QHS\_4 \| 0.837 \| 0.002663 \| 63 \| 0.108 \| 350 \| 202 \| 24 \| 457 \| 738 \| 302 \| 609 \| DNA replication licensing factor MCM4 \| DNA replication licensing factor MCM4; pdb \| 7P30\_2 \| 0.837 \| 0.002663 \| 63 \| 0.115 \| 347 \| 211 \| 21 \| 457 \| 736 \| 287 \| 604 \| DNA replication licensing factor MCM2 \| DNA replication licensing factor MCM2; pdb \| 7PMK\_5 \| 0.837 \| 0.003822 \| 63 \| 0.108 \| 361 \| 216 \| 24 \| 457 \| 737 \| 272 \| 606 \| DNA helicase \| DNA helicase; pdb \| 7PT6\_7 \| 0.837 \| 0.0009992 \| 63 \| 0.146 \| 362 \| 182 \| 30 \| 457 \| 733 \| 357 \| 676 \| DNA replication licensing factor MCM7 \| DNA replication licensing factor MCM7; pdb \| 1G8P\_A \| 0.817 \| 0.006081 \| 62 \| 0.138 \| 317 \| 167 \| 24 \| 495 \| 735 \| 15 \| 301 \| MAGNESIUM-CHELATASE 38 KDA SUBUNIT \| MAGNESIUM-CHELATASE 38 KDA SUBUNIT; pdb \| 7WI7\_A \| 0.817 \| 0.008287 \| 62 \| 0.127 \| 321 \| 192 \| 21 \| 456 \| 736 \| 288 \| 560 \| DNA helicase MCM8 \| DNA helicase MCM8; pdb \| 8B9D\_3 \| 0.817 \| 0.005209 \| 62 \| 0.106 \| 356 \| 215 \| 20 \| 456 \| 735 \| 237 \| 565 \| DNA replication licensing factor MCM3 \| DNA replication licensing factor MCM3; pdb \| 6WGG\_7 \| 0.817 \| 0.004237 \| 62 \| 0.123 \| 364 \| 218 \| 28 \| 449 \| 737 \| 248 \| 585 \| DNA replication licensing factor MCM7 \| DNA replication licensing factor MCM7; pdb \| 3JC5\_5 \| 0.817 \| 0.003274 \| 62 \| 0.109 \| 357 \| 211 \| 24 \| 457 \| 737 \| 269 \| 594 \| Minichromosome maintenance protein 5 \| Minichromosome maintenance protein 5; pdb \| 7P30\_F \| 0.817 \| 0.002058 \| 62 \| 0.143 \| 342 \| 197 \| 24 \| 456 \| 738 \| 361 \| 665 \| DNA replication licensing factor MCM7 \| DNA replication licensing factor MCM7; pdb \| 7PMK\_4 \| 0.795 \| 0.004462 \| 61 \| 0.127 \| 337 \| 201 \| 23 \| 457 \| 738 \| 299 \| 597 \| DNA replication licensing factor MCM4 \| DNA replication licensing factor MCM4; pdb \| 7P30\_E \| 0.795 \| 0.009188 \| 61 \| 0.127 \| 345 \| 203 \| 21 \| 457 \| 735 \| 289 \| 601 \| DNA replication licensing factor MCM6 \| DNA replication licensing factor MCM6; pdb \| 7W1Y\_3 \| 0.795 \| 0.001166 \| 61 \| 0.093 \| 479 \| 238 \| 26 \| 404 \| 751 \| 235 \| 648 \| Isoform 2 of DNA replication licensing factor MCM3 \| Isoform 2 of DNA replication licensing factor MCM3; pdb \| 8A8W\_D \| 0.772 \| 0.001434 \| 60 \| 0.125 \| 479 \| 249 \| 32 \| 232 \| 618 \| 44 \| 444 \| ATP-dependent Clp protease ATP-binding subunit ClpC1 \| ATP-dependent Clp protease ATP-binding subunit ClpC1; pdb \| 7PT7\_3 \| 0.747 \| 0.004698 \| 59 \| 0.099 \| 392 \| 204 \| 30 \| 456 \| 739 \| 284 \| 634 \| DNA replication licensing factor MCM3 \| DNA replication licensing factor MCM3; pdb \| 6F0L\_4 \| 0.747 \| 0.002167 \| 59 \| 0.096 \| 416 \| 244 \| 28 \| 403 \| 737 \| 274 \| 638 \| DNA replication licensing factor MCM4 \| DNA replication licensing factor MCM4; pdb \| 7W1Y\_5 \| 0.747 \| 0.004947 \| 59 \| 0.104 \| 343 \| 208 \| 29 \| 457 \| 738 \| 306 \| 610 \| DNA replication licensing factor MCM5 \| DNA replication licensing factor MCM5; pdb \| 3JC7\_3 \| 0.72 \| 0.004462 \| 58 \| 0.123 \| 371 \| 198 \| 25 \| 446 \| 736 \| 253 \| 576 \| DNA replication licensing factor MCM3 \| DNA replication licensing factor MCM3; pdb \| 7PFO\_7 \| 0.692 \| 0.008726 \| 57 \| 0.116 \| 342 \| 207 \| 25 \| 458 \| 737 \| 284 \| 592 \| DNA replication licensing factor MCM7 \| DNA replication licensing factor MCM7; pdb \| 7PT7\_E \| 0.692 \| 0.003274 \| 57 \| 0.109 \| 348 \| 198 \| 28 \| 457 \| 735 \| 294 \| 598 \| Minichromosome maintenance protein 5 \| Minichromosome maintenance protein 5; pdb \| 7QHS\_5 \| 0.663 \| 0.007871 \| 56 \| 0.086 \| 449 \| 229 \| 27 \| 457 \| 753 \| 270 \| 689 \| DNA replication licensing factor MCM5 \| DNA replication licensing factor MCM5; pdb \| 7PLO\_5 \| 0.632 \| 0.006742 \| 55 \| 0.109 \| 357 \| 213 \| 30 \| 457 \| 736 \| 282 \| 610 \| DNA replication licensing factor MCM5 \| DNA replication licensing factor MCM5; pdb \| 7PT6\_E \| 0.632 \| 0.003274 \| 55 \| 0.109 \| 365 \| 211 \| 26 \| 457 \| 737 \| 299 \| 633 \| Minichromosome maintenance protein 5 \| Minichromosome maintenance protein 5; pdb \| 7W1Y\_C \| 0.632 \| 0.004462 \| 55 \| 0.107 \| 411 \| 224 \| 25 \| 402 \| 737 \| 296 \| 638 \| DNA replication licensing factor MCM4 \| DNA replication licensing factor MCM4; pdb \| 7P5Z\_D \| 0.632 \| 0.005485 \| 55 \| 0.107 \| 372 \| 204 \| 28 \| 457 \| 737 \| 308 \| 642 \| Minichromosome maintenance protein 5 \| Minichromosome maintenance protein 5; pdb \| 7PT6\_4 \| 0.632 \| 0.009188 \| 55 \| 0.113 \| 413 \| 243 \| 28 \| 403 \| 737 \| 282 \| 649 \| DNA replication licensing factor MCM4 \| DNA replication licensing factor MCM4; pdb \| 7W8G\_D \| 0.632 \| 0.004698 \| 55 \| 0.114 \| 374 \| 222 \| 26 \| 439 \| 737 \| 312 \| 651 \| DNA replication licensing factor MCM4 \| DNA replication licensing factor MCM4; pdb \| 6XTY\_4 \| 0.601 \| 0.006081 \| 54 \| 0.128 \| 405 \| 219 \| 29 \| 403 \| 735 \| 254 \| 596 \| DNA replication licensing factor MCM4 \| DNA replication licensing factor MCM4; pdb \| 7UIV\_F \| 0.505 \| 0.008287 \| 51 \| 0.101 \| 474 \| 243 \| 30 \| 235 \| 615 \| 41 \| 424 \| ATP-dependent Clp protease ATP-binding subunit ClpA \| ATP-dependent Clp protease ATP-binding subunit ClpA; pdb \| 7MGM\_A \| 0.505 \| 0.005209 \| 51 \| 0.09 \| 496 \| 239 \| 31 \| 381 \| 739 \| 164 \| 584 \| dynein AAA3-WalkerB mutant (E2488Q) \| dynein AAA3-WalkerB mutant (E2488Q); afdb-proteome \| AF-A0A077ZFG4-F1-MODEL\_V4 \| 1.0 \| 5.739e-14 \| 550 \| 0.289 \| 235 \| 145 \| 9 \| 115 \| 346 \| 157 \| 372 \| Toprim 4 domain containing protein \| Toprim 4 domain containing protein; afdb-proteome \| AF-A0A0H3GIU0-F1-MODEL\_V4 \| 1.0 \| 1.237e-24 \| 472 \| 0.174 \| 812 \| 456 \| 47 \| 64 \| 761 \| 67 \| 777 \| Bacteriophage P4 DNA primase \| Bacteriophage P4 DNA primase; afdb-proteome \| AF-Q2FYB7-F1-MODEL\_V4 \| 1.0 \| 3.03e-15 \| 465 \| 0.222 \| 360 \| 237 \| 19 \| 317 \| 651 \| 323 \| 664 \| Conserved hypothetical phage protein \| Conserved hypothetical phage protein; afdb-proteome \| AF-A0A0H3GLV7-F1-MODEL\_V4 \| 1.0 \| 1.718e-23 \| 436 \| 0.176 \| 812 \| 454 \| 50 \| 64 \| 761 \| 67 \| 777 \| Nucleoside triphosphatase, D5 family \| Nucleoside triphosphatase, D5 family; afdb-proteome \| AF-A0A0H3H6B1-F1-MODEL\_V4 \| 1.0 \| 2.296e-08 \| 255 \| 0.176 \| 301 \| 180 \| 21 \| 91 \| 346 \| 289 \| 566 \| Prim\_Zn\_Ribbon domain-containing protein \| Prim\_Zn\_Ribbon domain-containing protein; afdb-proteome \| AF-A0A0H3GM72-F1-MODEL\_V4 \| 1.0 \| 1.045e-06 \| 173 \| 0.143 \| 292 \| 163 \| 19 \| 71 \| 331 \| 72 \| 307 \| Putative phage DNA primase \| Putative phage DNA primase; afdb-proteome \| AF-O06608-F1-MODEL\_V4 \| 1.0 \| 2.465e-07 \| 165 \| 0.135 \| 510 \| 292 \| 36 \| 348 \| 762 \| 16 \| 471 \| Probable PhiRv1 phage protein \| Probable PhiRv1 phage protein; afdb-proteome \| AF-Q8ZQI0-F1-MODEL\_V4 \| 1.0 \| 1.182e-05 \| 151 \| 0.105 \| 304 \| 180 \| 19 \| 60 \| 331 \| 60 \| 303 \| Fels-1 putative prophage DNA primase \| Fels-1 putative prophage DNA primase; afdb-proteome \| AF-Q8DPW1-F1-MODEL\_V4 \| 1.0 \| 2.646e-06 \| 128 \| 0.103 \| 485 \| 295 \| 30 \| 70 \| 503 \| 74 \| 469 \| DNA primase \| DNA primase; afdb-proteome \| AF-A0A132Z8R3-F1-MODEL\_V4 \| 1.0 \| 5.449e-06 \| 122 \| 0.109 \| 504 \| 282 \| 33 \| 77 \| 505 \| 79 \| 490 \| DNA primase \| DNA primase; afdb-proteome \| AF-A0A0H3GVT7-F1-MODEL\_V4 \| 1.0 \| 1.045e-06 \| 115 \| 0.105 \| 765 \| 346 \| 40 \| 61 \| 738 \| 69 \| 581 \| DNA primase \| DNA primase; afdb-proteome \| AF-P07362-F1-MODEL\_V4 \| 1.0 \| 1.353e-06 \| 114 \| 0.106 \| 750 \| 345 \| 46 \| 69 \| 738 \| 77 \| 581 \| DNA primase \| DNA primase; afdb-proteome \| AF-P0ABS5-F1-MODEL\_V4 \| 1.0 \| 1.663e-06 \| 113 \| 0.099 \| 753 \| 363 \| 43 \| 61 \| 738 \| 69 \| 581 \| DNA primase \| DNA primase; afdb-proteome \| AF-Q54Q10-F1-MODEL\_V4 \| 1.0 \| 0.000182 \| 113 \| 0.107 \| 354 \| 232 \| 22 \| 470 \| 760 \| 284 \| 616 \| Uncharacterized protein \| Uncharacterized protein; afdb-proteome \| AF-Q5F806-F1-MODEL\_V4 \| 1.0 \| 9.129e-06 \| 109 \| 0.146 \| 519 \| 242 \| 34 \| 77 \| 492 \| 76 \| 496 \| DNA primase \| DNA primase; afdb-proteome \| AF-Q32BQ1-F1-MODEL\_V4 \| 1.0 \| 2.646e-06 \| 108 \| 0.099 \| 761 \| 355 \| 46 \| 61 \| 738 \| 69 \| 581 \| DNA primase \| DNA primase; afdb-proteome \| AF-Q08346-F1-MODEL\_V4 \| 1.0 \| 6.699e-06 \| 104 \| 0.111 \| 521 \| 275 \| 34 \| 68 \| 510 \| 76 \| 486 \| DNA primase \| DNA primase; afdb-proteome \| AF-A0A0R4ICC1-F1-MODEL\_V4 \| 1.0 \| 7.82e-06 \| 104 \| 0.112 \| 649 \| 286 \| 34 \| 60 \| 616 \| 150 \| 600 \| Twinkle mtDNA helicase \| Twinkle mtDNA helicase; afdb-proteome \| AF-Q2FY11-F1-MODEL\_V4 \| 0.999 \| 1.529e-05 \| 99 \| 0.1 \| 747 \| 373 \| 42 \| 67 \| 737 \| 73 \| 596 \| DNA primase \| DNA primase; afdb-proteome \| AF-I6Y1F0-F1-MODEL\_V4 \| 0.999 \| 0.0005664 \| 98 \| 0.109 \| 373 \| 218 \| 29 \| 456 \| 751 \| 27 \| 362 \| Possible PhiRv2 prophage protein \| Possible PhiRv2 prophage protein; afdb-proteome \| AF-K0EWG0-F1-MODEL\_V4 \| 0.999 \| 1.012e-05 \| 97 \| 0.125 \| 533 \| 263 \| 30 \| 143 \| 595 \| 161 \| 570 \| DNA primase \| DNA primase; afdb-proteome \| AF-Q9PM37-F1-MODEL\_V4 \| 0.999 \| 5.556e-05 \| 96 \| 0.1 \| 539 \| 266 \| 30 \| 46 \| 493 \| 67 \| 477 \| DNA primase \| DNA primase; afdb-proteome \| AF-Q9CCG2-F1-MODEL\_V4 \| 0.999 \| 0.0001205 \| 96 \| 0.135 \| 525 \| 264 \| 35 \| 67 \| 503 \| 76 \| 498 \| DNA primase \| DNA primase; afdb-proteome \| AF-K7MB62-F1-MODEL\_V4 \| 0.998 \| 4.52e-05 \| 95 \| 0.112 \| 543 \| 254 \| 30 \| 154 \| 616 \| 216 \| 610 \| Uncharacterized protein \| Uncharacterized protein; afdb-proteome \| AF-F1R5P3-F1-MODEL\_V4 \| 0.998 \| 3.492e-05 \| 95 \| 0.117 \| 393 \| 248 \| 23 \| 403 \| 735 \| 300 \| 653 \| DNA helicase \| DNA helicase; afdb-proteome \| AF-U7PYU5-F1-MODEL\_V4 \| 0.998 \| 3.872e-05 \| 95 \| 0.127 \| 423 \| 217 \| 25 \| 407 \| 742 \| 459 \| 816 \| DNA replication licensing factor MCM6 \| DNA replication licensing factor MCM6; afdb-proteome \| AF-K7LBI0-F1-MODEL\_V4 \| 0.998 \| 2.433e-05 \| 94 \| 0.106 \| 547 \| 254 \| 32 \| 154 \| 616 \| 217 \| 612 \| Uncharacterized protein \| Uncharacterized protein; afdb-proteome \| AF-B5X582-F1-MODEL\_V4 \| 0.998 \| 8.395e-05 \| 93 \| 0.105 \| 576 \| 270 \| 33 \| 142 \| 623 \| 200 \| 623 \| Twinkle homolog protein, chloroplastic/mitochondrial \| Twinkle homolog protein, chloroplastic/mitochondrial; afdb-proteome \| AF-A0A3Q0KJA1-F1-MODEL\_V4 \| 0.998 \| 8.395e-05 \| 93 \| 0.103 \| 347 \| 222 \| 23 \| 456 \| 735 \| 331 \| 655 \| DNA replication licensing factor MCM6 \| DNA replication licensing factor MCM6; afdb-proteome \| AF-P53091-F1-MODEL\_V4 \| 0.998 \| 1.122e-05 \| 93 \| 0.118 \| 423 \| 263 \| 29 \| 403 \| 742 \| 449 \| 844 \| DNA replication licensing factor MCM6 \| DNA replication licensing factor MCM6; afdb-proteome \| AF-Q9I5W0-F1-MODEL\_V4 \| 0.996 \| 0.0004376 \| 90 \| 0.12 \| 458 \| 229 \| 31 \| 68 \| 447 \| 76 \| 437 \| DNA primase \| DNA primase; afdb-proteome \| AF-A0A1C1CIE0-F1-MODEL\_V4 \| 0.996 \| 5.85e-05 \| 90 \| 0.109 \| 528 \| 275 \| 34 \| 393 \| 757 \| 510 \| 1005 \| DNA helicase \| DNA helicase; afdb-proteome \| AF-A0A1D6M0G4-F1-MODEL\_V4 \| 0.996 \| 0.0001559 \| 89 \| 0.078 \| 332 \| 222 \| 22 \| 456 \| 736 \| 289 \| 587 \| DNA helicase \| DNA helicase; afdb-proteome \| AF-P30665-F1-MODEL\_V4 \| 0.996 \| 6.829e-05 \| 89 \| 0.092 \| 528 \| 271 \| 32 \| 393 \| 751 \| 441 \| 929 \| DNA replication licensing factor MCM4 \| DNA replication licensing factor MCM4; afdb-proteome \| AF-A0A0E4AYA7-F1-MODEL\_V4 \| 0.995 \| 6.486e-05 \| 88 \| 0.107 \| 401 \| 245 \| 24 \| 402 \| 735 \| 295 \| 649 \| DNA helicase \| DNA helicase; afdb-proteome \| AF-P97311-F1-MODEL\_V4 \| 0.994 \| 7.191e-05 \| 87 \| 0.11 \| 398 \| 245 \| 24 \| 403 \| 735 \| 302 \| 655 \| DNA replication licensing factor MCM6 \| DNA replication licensing factor MCM6; afdb-proteome \| AF-A0A1C1D2E4-F1-MODEL\_V4 \| 0.994 \| 3.677e-05 \| 87 \| 0.096 \| 435 \| 258 \| 27 \| 402 \| 742 \| 402 \| 795 \| DNA replication licensing factor MCM6 \| DNA replication licensing factor MCM6; afdb-proteome \| AF-P29496-F1-MODEL\_V4 \| 0.993 \| 0.0001087 \| 86 \| 0.116 \| 342 \| 209 \| 27 \| 457 \| 735 \| 355 \| 666 \| Minichromosome maintenance protein 5 \| Minichromosome maintenance protein 5; afdb-proteome \| AF-A0A077ZEI1-F1-MODEL\_V4 \| 0.993 \| 4.077e-05 \| 86 \| 0.076 \| 749 \| 398 \| 43 \| 67 \| 738 \| 273 \| 804 \| DNA primase \| DNA primase; afdb-proteome \| AF-A0A0K0ECF1-F1-MODEL\_V4 \| 0.993 \| 0.0001481 \| 86 \| 0.112 \| 356 \| 221 \| 21 \| 456 \| 735 \| 323 \| 659 \| DNA replication licensing factor MCM6 \| DNA replication licensing factor MCM6; afdb-proteome \| AF-A0A0R0GCD2-F1-MODEL\_V4 \| 0.993 \| 0.0002125 \| 86 \| 0.107 \| 345 \| 221 \| 28 \| 456 \| 735 \| 340 \| 662 \| DNA helicase \| DNA helicase; afdb-proteome \| AF-Q95XQ8-F1-MODEL\_V4 \| 0.992 \| 0.0003211 \| 85 \| 0.111 \| 438 \| 221 \| 24 \| 457 \| 750 \| 406 \| 819 \| DNA replication licensing factor mcm-4 \| DNA replication licensing factor mcm-4; afdb-proteome \| AF-Q4CZ61-F1-MODEL\_V4 \| 0.992 \| 0.0002125 \| 85 \| 0.121 \| 380 \| 227 \| 25 \| 438 \| 737 \| 334 \| 686 \| DNA helicase \| DNA helicase; afdb-proteome \| AF-Q86B14-F1-MODEL\_V4 \| 0.992 \| 6.829e-05 \| 85 \| 0.119 \| 393 \| 236 \| 26 \| 407 \| 735 \| 383 \| 729 \| DNA replication licensing factor mcm6 \| DNA replication licensing factor mcm6; afdb-proteome \| AF-A4I3W9-F1-MODEL\_V4 \| 0.992 \| 0.0001032 \| 85 \| 0.131 \| 366 \| 213 \| 29 \| 456 \| 744 \| 378 \| 715 \| DNA helicase \| DNA helicase; afdb-proteome \| AF-Q9HZQ5-F1-MODEL\_V4 \| 0.991 \| 0.0006963 \| 84 \| 0.123 \| 291 \| 189 \| 18 \| 474 \| 737 \| 9 \| 260 \| Probable magnesium chelatase \| Probable magnesium chelatase; afdb-proteome \| AF-P56064-F1-MODEL\_V4 \| 0.991 \| 0.0005664 \| 84 \| 0.108 \| 388 \| 207 \| 24 \| 154 \| 503 \| 168 \| 454 \| DNA primase \| DNA primase; afdb-proteome \| AF-I1JQR6-F1-MODEL\_V4 \| 0.991 \| 0.0001406 \| 84 \| 0.117 \| 433 \| 215 \| 29 \| 456 \| 752 \| 315 \| 716 \| DNA replication licensing factor MCM7 \| DNA replication licensing factor MCM7; afdb-proteome \| AF-Q9XYU0-F1-MODEL\_V4 \| 0.991 \| 0.0002612 \| 84 \| 0.106 \| 412 \| 212 \| 26 \| 471 \| 751 \| 333 \| 719 \| DNA replication licensing factor Mcm7 \| DNA replication licensing factor Mcm7; afdb-proteome \| AF-A0A0K0EDS2-F1-MODEL\_V4 \| 0.991 \| 0.0001144 \| 84 \| 0.138 \| 355 \| 204 \| 28 \| 457 \| 736 \| 324 \| 651 \| DNA replication licensing factor MCM5 \| DNA replication licensing factor MCM5; afdb-proteome \| AF-Q6F353-F1-MODEL\_V4 \| 0.991 \| 5.85e-05 \| 84 \| 0.104 \| 410 \| 247 \| 32 \| 402 \| 735 \| 293 \| 658 \| DNA replication licensing factor MCM6 \| DNA replication licensing factor MCM6; afdb-proteome \| AF-A0A3Q0KG44-F1-MODEL\_V4 \| 0.991 \| 0.0003749 \| 84 \| 0.113 \| 344 \| 222 \| 21 \| 456 \| 737 \| 445 \| 767 \| DNA replication licensing factor MCM2 \| DNA replication licensing factor MCM2; afdb-proteome \| AF-Q57809-F1-MODEL\_V4 \| 0.99 \| 0.000305 \| 83 \| 0.105 \| 369 \| 218 \| 26 \| 455 \| 735 \| 327 \| 671 \| Uncharacterized MCM-type protein MJ0363 \| Uncharacterized MCM-type protein MJ0363; afdb-proteome \| AF-A0A0K0JSP7-F1-MODEL\_V4 \| 0.99 \| 9.8e-05 \| 83 \| 0.101 \| 403 \| 246 \| 24 \| 403 \| 735 \| 301 \| 657 \| DNA replication licensing factor MCM6 \| DNA replication licensing factor MCM6; afdb-proteome \| AF-P30666-F1-MODEL\_V4 \| 0.99 \| 0.0004156 \| 83 \| 0.114 \| 314 \| 169 \| 20 \| 402 \| 666 \| 268 \| 521 \| DNA replication licensing factor mcm3 \| DNA replication licensing factor mcm3; afdb-proteome \| AF-P49731-F1-MODEL\_V4 \| 0.99 \| 0.0001406 \| 83 \| 0.103 \| 406 \| 251 \| 29 \| 403 \| 735 \| 370 \| 735 \| DNA replication licensing factor mcm6 \| DNA replication licensing factor mcm6; afdb-proteome \| AF-P41389-F1-MODEL\_V4 \| 0.988 \| 0.0003211 \| 82 \| 0.127 \| 353 \| 213 \| 26 \| 457 \| 736 \| 311 \| 641 \| DNA replication licensing factor mcm5 \| DNA replication licensing factor mcm5; afdb-proteome \| AF-A0A077Z6A7-F1-MODEL\_V4 \| 0.988 \| 7.572e-05 \| 82 \| 0.118 \| 405 \| 242 \| 26 \| 403 \| 735 \| 261 \| 622 \| DNA replication licensing factor MCM6 \| DNA replication licensing factor MCM6; afdb-proteome \| AF-A0A0N4UBH7-F1-MODEL\_V4 \| 0.988 \| 0.002663 \| 82 \| 0.128 \| 234 \| 143 \| 16 \| 467 \| 657 \| 282 \| 497 \| DNA replication licensing factor MCM3 \| DNA replication licensing factor MCM3; afdb-proteome \| AF-A0A1D6FQT3-F1-MODEL\_V4 \| 0.988 \| 0.0001559 \| 82 \| 0.087 \| 389 \| 239 \| 27 \| 406 \| 737 \| 306 \| 635 \| DNA helicase \| DNA helicase; afdb-proteome \| AF-Q4DCB3-F1-MODEL\_V4 \| 0.988 \| 0.0003749 \| 82 \| 0.117 \| 332 \| 197 \| 24 \| 455 \| 735 \| 452 \| 738 \| DNA replication licensing factor MCM2 \| DNA replication licensing factor MCM2; afdb-proteome \| AF-Q53RM0-F1-MODEL\_V4 \| 0.986 \| 0.0004852 \| 81 \| 0.157 \| 311 \| 170 \| 19 \| 492 \| 738 \| 90 \| 372 \| Magnesium-chelatase subunit ChlI, chloroplastic \| Magnesium-chelatase subunit ChlI, chloroplastic; afdb-proteome \| AF-A0A0K0E3B8-F1-MODEL\_V4 \| 0.986 \| 0.0001336 \| 81 \| 0.135 \| 355 \| 200 \| 27 \| 456 \| 737 \| 332 \| 652 \| DNA replication licensing factor MCM7 \| DNA replication licensing factor MCM7; afdb-proteome \| AF-P34647-F1-MODEL\_V4 \| 0.986 \| 0.0001205 \| 81 \| 0.111 \| 404 \| 240 \| 25 \| 405 \| 735 \| 301 \| 658 \| DNA replication licensing factor mcm-6 \| DNA replication licensing factor mcm-6; afdb-proteome \| AF-A0A077ZGF4-F1-MODEL\_V4 \| 0.986 \| 0.0001481 \| 81 \| 0.114 \| 411 \| 229 \| 29 \| 402 \| 738 \| 359 \| 708 \| DNA replication licensing factor MCM4 \| DNA replication licensing factor MCM4; afdb-proteome \| AF-I1L122-F1-MODEL\_V4 \| 0.986 \| 0.0002125 \| 81 \| 0.108 \| 404 \| 251 \| 30 \| 402 \| 735 \| 289 \| 653 \| DNA helicase \| DNA helicase; afdb-proteome \| AF-C1GVN3-F1-MODEL\_V4 \| 0.986 \| 6.829e-05 \| 81 \| 0.122 \| 425 \| 232 \| 36 \| 405 \| 735 \| 415 \| 792 \| DNA replication licensing factor MCM6 \| DNA replication licensing factor MCM6; afdb-proteome \| AF-I1KLS0-F1-MODEL\_V4 \| 0.984 \| 0.0004376 \| 80 \| 0.142 \| 316 \| 169 \| 23 \| 492 \| 738 \| 89 \| 371 \| Mg-protoporphyrin IX chelatase \| Mg-protoporphyrin IX chelatase; afdb-proteome \| AF-P9WNW1-F1-MODEL\_V4 \| 0.984 \| 0.0001481 \| 80 \| 0.11 \| 725 \| 334 \| 46 \| 123 \| 739 \| 115 \| 636 \| DNA primase \| DNA primase; afdb-proteome \| AF-Q54RU0-F1-MODEL\_V4 \| 0.984 \| 0.0002896 \| 80 \| 0.123 \| 356 \| 208 \| 24 \| 456 \| 737 \| 390 \| 715 \| DNA replication licensing factor MCM7 \| DNA replication licensing factor MCM7; afdb-proteome \| AF-Q14566-F1-MODEL\_V4 \| 0.984 \| 0.0001917 \| 80 \| 0.116 \| 402 \| 238 \| 27 \| 403 \| 735 \| 302 \| 655 \| DNA replication licensing factor MCM6 \| DNA replication licensing factor MCM6; afdb-proteome \| AF-A0A175W509-F1-MODEL\_V4 \| 0.984 \| 0.0001729 \| 80 \| 0.106 \| 534 \| 271 \| 35 \| 393 \| 757 \| 503 \| 999 \| DNA helicase \| DNA helicase; afdb-proteome \| AF-Q2QNM1-F1-MODEL\_V4 \| 0.981 \| 0.0001729 \| 79 \| 0.095 \| 482 \| 239 \| 31 \| 405 \| 751 \| 303 \| 722 \| DNA replication licensing factor MCM7 \| DNA replication licensing factor MCM7; afdb-proteome \| AF-Q8I5T4-F1-MODEL\_V4 \| 0.981 \| 0.0003749 \| 79 \| 0.089 \| 434 \| 225 \| 28 \| 457 \| 750 \| 354 \| 757 \| DNA helicase \| DNA helicase; afdb-proteome \| AF-J9F6T9-F1-MODEL\_V4 \| 0.981 \| 0.0008559 \| 79 \| 0.107 \| 352 \| 229 \| 25 \| 456 \| 743 \| 466 \| 796 \| DNA replication licensing factor MCM4 \| DNA replication licensing factor MCM4; afdb-proteome \| AF-P9WPR3-F1-MODEL\_V4 \| 0.978 \| 0.0005379 \| 78 \| 0.123 \| 323 \| 192 \| 23 \| 475 \| 743 \| 6 \| 291 \| Uncharacterized protein Rv2850c \| Uncharacterized protein Rv2850c; afdb-proteome \| AF-A0A1D6MX99-F1-MODEL\_V4 \| 0.978 \| 7.973e-05 \| 78 \| 0.13 \| 397 \| 214 \| 30 \| 403 \| 735 \| 291 \| 620 \| DNA replication licensing factor MCM7 \| DNA replication licensing factor MCM7; afdb-proteome \| AF-Q9VGW6-F1-MODEL\_V4 \| 0.978 \| 0.0006963 \| 78 \| 0.085 \| 443 \| 228 \| 29 \| 457 \| 750 \| 317 \| 731 \| DNA replication licensing factor Mcm5 \| DNA replication licensing factor Mcm5; afdb-proteome \| AF-A0A044QUL7-F1-MODEL\_V4 \| 0.978 \| 0.0005664 \| 78 \| 0.117 \| 358 \| 223 \| 24 \| 457 \| 746 \| 468 \| 800 \| DNA replication licensing factor MCM4 \| DNA replication licensing factor MCM4; afdb-proteome \| AF-Q54LI2-F1-MODEL\_V4 \| 0.978 \| 0.0003381 \| 78 \| 0.101 \| 452 \| 230 \| 30 \| 456 \| 751 \| 574 \| 1005 \| DNA replication licensing factor MCM2 \| DNA replication licensing factor MCM2; afdb-proteome \| AF-Q58321-F1-MODEL\_V4 \| 0.975 \| 0.0009992 \| 77 \| 0.112 \| 330 \| 186 \| 21 \| 475 \| 738 \| 10 \| 298 \| Magnesium-chelatase subunit ChlI homolog \| Magnesium-chelatase subunit ChlI homolog; afdb-proteome \| AF-I1M038-F1-MODEL\_V4 \| 0.975 \| 0.0001917 \| 77 \| 0.126 \| 403 \| 208 \| 28 \| 407 \| 738 \| 49 \| 378 \| Mg-protoporphyrin IX chelatase \| Mg-protoporphyrin IX chelatase; afdb-proteome \| AF-I1NBC2-F1-MODEL\_V4 \| 0.975 \| 0.000182 \| 77 \| 0.098 \| 486 \| 233 \| 31 \| 405 \| 752 \| 298 \| 716 \| DNA replication licensing factor MCM7 \| DNA replication licensing factor MCM7; afdb-proteome \| AF-A0A0J9XQV4-F1-MODEL\_V4 \| 0.975 \| 0.0004852 \| 77 \| 0.118 \| 355 \| 206 \| 25 \| 456 \| 737 \| 331 \| 651 \| DNA replication licensing factor MCM7 \| DNA replication licensing factor MCM7; afdb-proteome \| AF-A0A1D6LPW0-F1-MODEL\_V4 \| 0.975 \| 0.0003381 \| 77 \| 0.11 \| 545 \| 252 \| 36 \| 154 \| 616 \| 229 \| 622 \| Twinkle homolog protein chloroplastic/mitochondrial \| Twinkle homolog protein chloroplastic/mitochondrial; afdb-proteome \| AF-C1H7I1-F1-MODEL\_V4 \| 0.975 \| 0.0005664 \| 77 \| 0.121 \| 444 \| 208 \| 25 \| 471 \| 767 \| 401 \| 809 \| DNA replication licensing factor MCM7 \| DNA replication licensing factor MCM7; afdb-proteome \| AF-A0A5S6PT53-F1-MODEL\_V4 \| 0.975 \| 0.001228 \| 77 \| 0.109 \| 357 \| 229 \| 22 \| 457 \| 747 \| 396 \| 729 \| DNA replication licensing factor MCM4 \| DNA replication licensing factor MCM4; afdb-proteome \| AF-A0A077YZA0-F1-MODEL\_V4 \| 0.975 \| 0.000356 \| 77 \| 0.11 \| 644 \| 328 \| 41 \| 129 \| 714 \| 20 \| 476 \| Alpha:beta hydrolase fold protein \| Alpha:beta hydrolase fold protein; afdb-proteome \| AF-Q8ILR7-F1-MODEL\_V4 \| 0.975 \| 0.0007331 \| 77 \| 0.099 \| 371 \| 222 \| 24 \| 456 \| 737 \| 473 \| 820 \| DNA replication licensing factor MCM2 \| DNA replication licensing factor MCM2; afdb-proteome \| AF-A0A0D2EQJ7-F1-MODEL\_V4 \| 0.975 \| 0.0002125 \| 77 \| 0.123 \| 430 \| 256 \| 25 \| 393 \| 737 \| 509 \| 902 \| DNA helicase \| DNA helicase; afdb-proteome \| AF-Q6KAJ4-F1-MODEL\_V4 \| 0.971 \| 0.000628 \| 76 \| 0.104 \| 442 \| 218 \| 28 \| 456 \| 750 \| 317 \| 727 \| DNA replication licensing factor MCM5 \| DNA replication licensing factor MCM5; afdb-proteome \| AF-Q5RIC5-F1-MODEL\_V4 \| 0.971 \| 0.0006613 \| 76 \| 0.123 \| 340 \| 210 \| 24 \| 456 \| 745 \| 283 \| 584 \| DNA helicase \| DNA helicase; afdb-proteome \| AF-D3ZFP4-F1-MODEL\_V4 \| 0.971 \| 0.0003947 \| 76 \| 0.096 \| 424 \| 239 \| 28 \| 456 \| 769 \| 284 \| 673 \| DNA helicase \| DNA helicase; afdb-proteome \| AF-A0A0K0EHC7-F1-MODEL\_V4 \| 0.971 \| 0.0005664 \| 76 \| 0.116 \| 370 \| 203 \| 24 \| 456 \| 735 \| 450 \| 785 \| DNA replication licensing factor MCM2 \| DNA replication licensing factor MCM2; afdb-proteome \| AF-A0A183XGH6-F1-MODEL\_V4 \| 0.971 \| 0.0005109 \| 76 \| 0.096 \| 333 \| 208 \| 20 \| 471 \| 735 \| 350 \| 657 \| DNA replication licensing factor MCM6 \| DNA replication licensing factor MCM6; afdb-proteome \| AF-K7TMN0-F1-MODEL\_V4 \| 0.967 \| 0.0007331 \| 75 \| 0.15 \| 320 \| 166 \| 23 \| 492 \| 738 \| 130 \| 416 \| Mg-protoporphyrin IX chelatase \| Mg-protoporphyrin IX chelatase; afdb-proteome \| AF-C0HFI8-F1-MODEL\_V4 \| 0.967 \| 0.0001729 \| 75 \| 0.098 \| 485 \| 234 \| 33 \| 405 \| 751 \| 298 \| 717 \| DNA replication licensing factor MCM7 \| DNA replication licensing factor MCM7; afdb-proteome \| AF-Q9CWV1-F1-MODEL\_V4 \| 0.967 \| 0.002953 \| 75 \| 0.16 \| 249 \| 138 \| 21 \| 456 \| 659 \| 386 \| 608 \| DNA helicase MCM8 \| DNA helicase MCM8; afdb-proteome \| AF-P38132-F1-MODEL\_V4 \| 0.967 \| 0.0003381 \| 75 \| 0.132 \| 347 \| 191 \| 27 \| 456 \| 736 \| 401 \| 703 \| DNA replication licensing factor MCM7 \| DNA replication licensing factor MCM7; afdb-proteome \| AF-P29469-F1-MODEL\_V4 \| 0.967 \| 0.001362 \| 75 \| 0.119 \| 342 \| 203 \| 22 \| 456 \| 744 \| 484 \| 780 \| DNA replication licensing factor MCM2 \| DNA replication licensing factor MCM2; afdb-proteome \| AF-A0A0R4IF65-F1-MODEL\_V4 \| 0.967 \| 0.0007331 \| 75 \| 0.095 \| 463 \| 224 \| 29 \| 456 \| 748 \| 448 \| 885 \| DNA replication licensing factor MCM2 \| DNA replication licensing factor MCM2; afdb-proteome \| AF-D3ZP96-F1-MODEL\_V4 \| 0.967 \| 0.0006613 \| 75 \| 0.093 \| 352 \| 216 \| 21 \| 456 \| 737 \| 461 \| 779 \| DNA replication licensing factor MCM2 \| DNA replication licensing factor MCM2; afdb-proteome \| AF-C0NCM7-F1-MODEL\_V4 \| 0.967 \| 0.0002751 \| 75 \| 0.127 \| 430 \| 256 \| 26 \| 393 \| 738 \| 521 \| 915 \| DNA helicase \| DNA helicase; afdb-proteome \| AF-P49735-F1-MODEL\_V4 \| 0.961 \| 0.001108 \| 74 \| 0.109 \| 373 \| 212 \| 28 \| 456 \| 735 \| 434 \| 779 \| DNA replication licensing factor Mcm2 \| DNA replication licensing factor Mcm2; afdb-proteome \| AF-I1MRS5-F1-MODEL\_V4 \| 0.961 \| 0.00151 \| 74 \| 0.145 \| 310 \| 191 \| 18 \| 428 \| 684 \| 451 \| 739 \| DNA replication licensing factor MCM2 \| DNA replication licensing factor MCM2; afdb-proteome \| AF-Q384N2-F1-MODEL\_V4 \| 0.961 \| 0.001052 \| 74 \| 0.129 \| 340 \| 188 \| 27 \| 455 \| 737 \| 447 \| 735 \| DNA replication licensing factor MCM2 \| DNA replication licensing factor MCM2; afdb-proteome \| AF-A0A3P7GBU4-F1-MODEL\_V4 \| 0.956 \| 0.00159 \| 73 \| 0.068 \| 466 \| 237 \| 27 \| 456 \| 748 \| 446 \| 887 \| DNA replication licensing factor MCM2 \| DNA replication licensing factor MCM2; afdb-proteome \| AF-I1KMP8-F1-MODEL\_V4 \| 0.956 \| 0.001166 \| 73 \| 0.14 \| 328 \| 198 \| 22 \| 456 \| 735 \| 480 \| 771 \| DNA replication licensing factor MCM2 \| DNA replication licensing factor MCM2; afdb-proteome \| AF-Q2R482-F1-MODEL\_V4 \| 0.956 \| 0.0005964 \| 73 \| 0.108 \| 514 \| 245 \| 32 \| 428 \| 756 \| 476 \| 961 \| DNA replication licensing factor MCM2 \| DNA replication licensing factor MCM2; afdb-proteome \| AF-A0A1D6PT01-F1-MODEL\_V4 \| 0.956 \| 0.0008559 \| 73 \| 0.104 \| 471 \| 222 \| 29 \| 456 \| 753 \| 548 \| 991 \| DNA replication licensing factor MCM2 \| DNA replication licensing factor MCM2; afdb-proteome \| AF-U7Q2I4-F1-MODEL\_V4 \| 0.956 \| 0.0001917 \| 73 \| 0.111 \| 457 \| 267 \| 31 \| 393 \| 747 \| 558 \| 977 \| DNA helicase \| DNA helicase; afdb-proteome \| AF-C1GR90-F1-MODEL\_V4 \| 0.956 \| 0.0004156 \| 73 \| 0.138 \| 411 \| 244 \| 27 \| 402 \| 738 \| 556 \| 930 \| DNA helicase \| DNA helicase; afdb-proteome \| AF-P16127-F1-MODEL\_V4 \| 0.949 \| 0.001362 \| 72 \| 0.141 \| 326 \| 165 \| 22 \| 487 \| 738 \| 97 \| 381 \| Magnesium-chelatase subunit ChlI-1, chloroplastic \| Magnesium-chelatase subunit ChlI-1, chloroplastic; afdb-proteome \| AF-Q58371-F1-MODEL\_V4 \| 0.949 \| 0.0003381 \| 72 \| 0.096 \| 539 \| 228 \| 31 \| 403 \| 751 \| 293 \| 762 \| Uncharacterized MCM-type protein MJ0961 \| Uncharacterized MCM-type protein MJ0961; afdb-proteome \| AF-P49717-F1-MODEL\_V4 \| 0.949 \| 0.0009992 \| 72 \| 0.125 \| 407 \| 232 \| 26 \| 402 \| 738 \| 418 \| 770 \| DNA replication licensing factor MCM4 \| DNA replication licensing factor MCM4; afdb-proteome \| AF-A0A158Q488-F1-MODEL\_V4 \| 0.949 \| 0.004698 \| 72 \| 0.098 \| 285 \| 184 \| 20 \| 456 \| 697 \| 475 \| 729 \| DNA replication licensing factor MCM2 \| DNA replication licensing factor MCM2; afdb-proteome \| AF-Q4E4H4-F1-MODEL\_V4 \| 0.941 \| 0.001856 \| 71 \| 0.114 \| 350 \| 211 \| 27 \| 456 \| 735 \| 287 \| 607 \| DNA replication factor, putative \| DNA replication factor, putative; afdb-proteome \| AF-A0A1C1CC10-F1-MODEL\_V4 \| 0.941 \| 0.0009012 \| 71 \| 0.1 \| 447 \| 222 \| 27 \| 471 \| 769 \| 394 \| 808 \| DNA replication licensing factor MCM7 \| DNA replication licensing factor MCM7; afdb-proteome \| AF-A0A044UXW5-F1-MODEL\_V4 \| 0.941 \| 0.0004376 \| 71 \| 0.107 \| 454 \| 276 \| 26 \| 405 \| 769 \| 301 \| 714 \| DNA replication licensing factor MCM6 \| DNA replication licensing factor MCM6; afdb-proteome \| AF-G3V681-F1-MODEL\_V4 \| 0.941 \| 0.0006613 \| 71 \| 0.121 \| 429 \| 241 \| 27 \| 393 \| 738 \| 395 \| 770 \| DNA helicase \| DNA helicase; afdb-proteome \| AF-Q383B6-F1-MODEL\_V4 \| 0.941 \| 0.000356 \| 71 \| 0.104 \| 420 \| 257 \| 29 \| 403 \| 743 \| 322 \| 701 \| DNA replication licensing factor MCM6 \| DNA replication licensing factor MCM6; afdb-proteome \| AF-A0A044VIL0-F1-MODEL\_V4 \| 0.941 \| 0.003822 \| 71 \| 0.106 \| 272 \| 173 \| 20 \| 456 \| 683 \| 449 \| 694 \| DNA replication licensing factor MCM2 \| DNA replication licensing factor MCM2; afdb-proteome \| AF-Q8IDF0-F1-MODEL\_V4 \| 0.941 \| 0.0004156 \| 71 \| 0.107 \| 418 \| 248 \| 30 \| 402 \| 743 \| 391 \| 759 \| DNA helicase \| DNA helicase; afdb-proteome \| AF-Q9LPD9-F1-MODEL\_V4 \| 0.941 \| 0.000772 \| 71 \| 0.096 \| 507 \| 242 \| 30 \| 428 \| 748 \| 451 \| 927 \| DNA replication licensing factor MCM2 \| DNA replication licensing factor MCM2; afdb-proteome \| AF-I1MEQ1-F1-MODEL\_V4 \| 0.933 \| 0.001052 \| 70 \| 0.12 \| 356 \| 190 \| 25 \| 457 \| 738 \| 72 \| 378 \| Mg-protoporphyrin IX chelatase \| Mg-protoporphyrin IX chelatase; afdb-proteome \| AF-P93162-F1-MODEL\_V4 \| 0.933 \| 0.001674 \| 70 \| 0.14 \| 321 \| 177 \| 23 \| 487 \| 738 \| 88 \| 378 \| Magnesium-chelatase subunit ChlI, chloroplastic \| Magnesium-chelatase subunit ChlI, chloroplastic; afdb-proteome \| AF-P40377-F1-MODEL\_V4 \| 0.933 \| 0.0008559 \| 70 \| 0.096 \| 416 \| 238 \| 29 \| 405 \| 736 \| 448 \| 809 \| DNA replication licensing factor mcm2 \| DNA replication licensing factor mcm2; afdb-proteome \| AF-Q6NZV2-F1-MODEL\_V4 \| 0.933 \| 0.001954 \| 70 \| 0.097 \| 449 \| 218 \| 29 \| 456 \| 751 \| 431 \| 845 \| DNA helicase \| DNA helicase; afdb-proteome \| AF-K0F2Y6-F1-MODEL\_V4 \| 0.923 \| 0.004024 \| 69 \| 0.144 \| 298 \| 180 \| 20 \| 482 \| 737 \| 24 \| 288 \| Putative magnesium chelatase subunit \| Putative magnesium chelatase subunit; afdb-proteome \| AF-A0A3Q0KEL0-F1-MODEL\_V4 \| 0.923 \| 0.0009992 \| 69 \| 0.122 \| 400 \| 224 \| 27 \| 405 \| 744 \| 254 \| 586 \| DNA replication licensing factor MCM3 \| DNA replication licensing factor MCM3; afdb-proteome \| AF-A0A0A2V2X0-F1-MODEL\_V4 \| 0.923 \| 0.002167 \| 69 \| 0.123 \| 349 \| 191 \| 24 \| 456 \| 744 \| 285 \| 578 \| DNA replication licensing factor MCM3 \| DNA replication licensing factor MCM3; afdb-proteome \| AF-A0A0D2GY10-F1-MODEL\_V4 \| 0.923 \| 0.002281 \| 69 \| 0.101 \| 353 \| 206 \| 26 \| 456 \| 744 \| 295 \| 600 \| DNA replication licensing factor MCM3 \| DNA replication licensing factor MCM3; afdb-proteome \| AF-P97310-F1-MODEL\_V4 \| 0.923 \| 0.001954 \| 69 \| 0.092 \| 467 \| 224 \| 28 \| 456 \| 748 \| 460 \| 900 \| DNA replication licensing factor MCM2 \| DNA replication licensing factor MCM2; afdb-proteome \| AF-P49736-F1-MODEL\_V4 \| 0.923 \| 0.00159 \| 69 \| 0.093 \| 472 \| 226 \| 28 \| 456 \| 753 \| 460 \| 903 \| DNA replication licensing factor MCM2 \| DNA replication licensing factor MCM2; afdb-proteome \| AF-U7Q723-F1-MODEL\_V4 \| 0.923 \| 0.001674 \| 69 \| 0.115 \| 354 \| 207 \| 25 \| 456 \| 744 \| 292 \| 604 \| DNA replication licensing factor MCM3 \| DNA replication licensing factor MCM3; afdb-proteome \| AF-Q5XF33-F1-MODEL\_V4 \| 0.912 \| 0.001763 \| 68 \| 0.122 \| 334 \| 182 \| 24 \| 478 \| 738 \| 80 \| 375 \| Magnesium-chelatase subunit ChlI-2, chloroplastic \| Magnesium-chelatase subunit ChlI-2, chloroplastic; afdb-proteome \| AF-A0A077Z5J5-F1-MODEL\_V4 \| 0.912 \| 0.001674 \| 68 \| 0.102 \| 418 \| 202 \| 27 \| 471 \| 751 \| 344 \| 725 \| DNA replication licensing factor MCM7 \| DNA replication licensing factor MCM7; afdb-proteome \| AF-Q54VI9-F1-MODEL\_V4 \| 0.912 \| 0.001856 \| 68 \| 0.138 \| 347 \| 195 \| 26 \| 456 \| 744 \| 300 \| 600 \| DNA helicase \| DNA helicase; afdb-proteome \| AF-A0A175WE06-F1-MODEL\_V4 \| 0.912 \| 0.006742 \| 68 \| 0.1 \| 339 \| 223 \| 22 \| 455 \| 744 \| 465 \| 770 \| DNA replication licensing factor MCM2 \| DNA replication licensing factor MCM2; afdb-proteome \| AF-C0NSR0-F1-MODEL\_V4 \| 0.912 \| 0.001166 \| 68 \| 0.094 \| 434 \| 226 \| 28 \| 456 \| 767 \| 295 \| 683 \| DNA replication licensing factor MCM3 \| DNA replication licensing factor MCM3; afdb-proteome \| AF-P33992-F1-MODEL\_V4 \| 0.9 \| 0.002058 \| 67 \| 0.11 \| 444 \| 216 \| 27 \| 457 \| 751 \| 320 \| 733 \| DNA replication licensing factor MCM5 \| DNA replication licensing factor MCM5; afdb-proteome \| AF-I1KRF9-F1-MODEL\_V4 \| 0.9 \| 0.002663 \| 67 \| 0.11 \| 317 \| 188 \| 21 \| 456 \| 725 \| 280 \| 549 \| DNA helicase \| DNA helicase; afdb-proteome \| AF-A0A1D8PHW4-F1-MODEL\_V4 \| 0.9 \| 0.001763 \| 67 \| 0.134 \| 393 \| 224 \| 30 \| 402 \| 744 \| 285 \| 611 \| DNA helicase \| DNA helicase; afdb-proteome \| AF-Q9XXI9-F1-MODEL\_V4 \| 0.9 \| 0.0008559 \| 67 \| 0.092 \| 456 \| 205 \| 33 \| 456 \| 730 \| 439 \| 866 \| DNA replication licensing factor MCM2 \| DNA replication licensing factor MCM2; afdb-proteome \| AF-C0NPQ6-F1-MODEL\_V4 \| 0.9 \| 0.002058 \| 67 \| 0.112 \| 357 \| 216 \| 28 \| 456 \| 736 \| 474 \| 805 \| DNA replication licensing factor MCM2 \| DNA replication licensing factor MCM2; afdb-proteome \| AF-U7PSF8-F1-MODEL\_V4 \| 0.9 \| 0.002058 \| 67 \| 0.098 \| 384 \| 212 \| 24 \| 456 \| 737 \| 488 \| 839 \| DNA replication licensing factor MCM2 \| DNA replication licensing factor MCM2; afdb-proteome \| AF-A0A3P7EEJ1-F1-MODEL\_V4 \| 0.887 \| 0.008287 \| 66 \| 0.137 \| 320 \| 186 \| 27 \| 471 \| 738 \| 258 \| 539 \| Mini-chromosome maintenance complex-binding protein \| Mini-chromosome maintenance complex-binding protein; afdb-proteome \| AF-Q0DHC4-F1-MODEL\_V4 \| 0.887 \| 0.004024 \| 66 \| 0.12 \| 341 \| 202 \| 21 \| 456 \| 744 \| 281 \| 575 \| DNA replication licensing factor MCM3 \| DNA replication licensing factor MCM3; afdb-proteome \| AF-C1GZP9-F1-MODEL\_V4 \| 0.887 \| 0.002663 \| 66 \| 0.087 \| 341 \| 208 \| 23 \| 456 \| 744 \| 483 \| 772 \| DNA replication licensing factor MCM2 \| DNA replication licensing factor MCM2; afdb-proteome \| AF-Q5A034-F1-MODEL\_V4 \| 0.887 \| 0.004024 \| 66 \| 0.124 \| 330 \| 196 \| 22 \| 457 \| 735 \| 512 \| 799 \| DNA replication licensing factor MCM2 \| DNA replication licensing factor MCM2; afdb-proteome \| AF-A0A150ASS4-F1-MODEL\_V4 \| 0.887 \| 0.001108 \| 66 \| 0.115 \| 459 \| 220 \| 31 \| 402 \| 735 \| 266 \| 663 \| DNA replication licensing factor MCM3 \| DNA replication licensing factor MCM3; afdb-proteome \| AF-A4IC27-F1-MODEL\_V4 \| 0.887 \| 0.00363 \| 66 \| 0.12 \| 357 \| 203 \| 26 \| 456 \| 735 \| 298 \| 620 \| DNA\_replication\_licensing\_factor\_MCM9\_-\_putative \| DNA\_replication\_licensing\_factor\_MCM9\_-\_putative; afdb-proteome \| AF-Q38E36-F1-MODEL\_V4 \| 0.872 \| 0.002402 \| 65 \| 0.101 \| 425 \| 250 \| 26 \| 393 \| 735 \| 220 \| 594 \| Minichromosome maintenance (MCM) complex subunit, putative \| Minichromosome maintenance (MCM) complex subunit, putative; afdb-proteome \| AF-I1K328-F1-MODEL\_V4 \| 0.872 \| 0.003109 \| 65 \| 0.095 \| 344 \| 207 \| 22 \| 456 \| 744 \| 280 \| 574 \| DNA helicase \| DNA helicase; afdb-proteome \| AF-P25205-F1-MODEL\_V4 \| 0.872 \| 0.002281 \| 65 \| 0.099 \| 431 \| 237 \| 26 \| 428 \| 744 \| 269 \| 662 \| DNA replication licensing factor MCM3 \| DNA replication licensing factor MCM3; afdb-proteome \| AF-A0A0D2DM29-F1-MODEL\_V4 \| 0.872 \| 0.001763 \| 65 \| 0.123 \| 381 \| 204 \| 30 \| 456 \| 737 \| 471 \| 820 \| DNA replication licensing factor MCM2 \| DNA replication licensing factor MCM2; afdb-proteome \| AF-A0A1C1CXS2-F1-MODEL\_V4 \| 0.855 \| 0.002953 \| 64 \| 0.123 \| 340 \| 196 \| 25 \| 456 \| 744 \| 472 \| 760 \| DNA replication licensing factor MCM2 \| DNA replication licensing factor MCM2; afdb-proteome \| AF-Q9SX03-F1-MODEL\_V4 \| 0.817 \| 0.002402 \| 62 \| 0.124 \| 411 \| 229 \| 26 \| 393 \| 744 \| 237 \| 575 \| DNA replication licensing factor MCM3 homolog 3 \| DNA replication licensing factor MCM3 homolog 3; afdb-proteome \| AF-Q9SX04-F1-MODEL\_V4 \| 0.817 \| 0.004237 \| 62 \| 0.093 \| 504 \| 231 \| 23 \| 456 \| 750 \| 281 \| 767 \| DNA replication licensing factor MCM3 homolog 2 \| DNA replication licensing factor MCM3 homolog 2; afdb-proteome \| AF-Q9SF37-F1-MODEL\_V4 \| 0.817 \| 0.003822 \| 62 \| 0.099 \| 391 \| 215 \| 27 \| 456 \| 737 \| 340 \| 702 \| Probable DNA helicase MCM8 \| Probable DNA helicase MCM8; afdb-proteome \| AF-D3ZVK1-F1-MODEL\_V4 \| 0.817 \| 0.001166 \| 62 \| 0.112 \| 446 \| 228 \| 32 \| 402 \| 736 \| 350 \| 738 \| DNA helicase MCM8 \| DNA helicase MCM8; afdb-proteome \| AF-Q38AM6-F1-MODEL\_V4 \| 0.795 \| 0.005484 \| 61 \| 0.096 \| 373 \| 203 \| 26 \| 471 \| 737 \| 246 \| 590 \| Uncharacterized protein \| Uncharacterized protein; afdb-proteome \| AF-Q9FL33-F1-MODEL\_V4 \| 0.747 \| 0.002529 \| 59 \| 0.095 \| 461 \| 240 \| 29 \| 393 \| 735 \| 232 \| 633 \| DNA replication licensing factor MCM3 \| DNA replication licensing factor MCM3; afdb-proteome \| AF-A0A1C1D0R7-F1-MODEL\_V4 \| 0.747 \| 0.002281 \| 59 \| 0.104 \| 401 \| 235 \| 26 \| 402 \| 744 \| 269 \| 603 \| DNA replication licensing factor MCM3 \| DNA replication licensing factor MCM3; afdb-proteome \| AF-E7F643-F1-MODEL\_V4 \| 0.72 \| 0.004024 \| 58 \| 0.108 \| 444 \| 232 \| 33 \| 402 \| 736 \| 372 \| 760 \| DNA helicase \| DNA helicase; afdb-proteome \| AF-A0A0N4U787-F1-MODEL\_V4 \| 0.692 \| 0.006403 \| 57 \| 0.123 \| 445 \| 235 \| 30 \| 393 \| 747 \| 295 \| 674 \| DNA replication licensing factor MCM4 \| DNA replication licensing factor MCM4; afdb-proteome \| AF-Q54RN8-F1-MODEL\_V4 \| 0.632 \| 0.004947 \| 55 \| 0.103 \| 407 \| 225 \| 26 \| 402 \| 744 \| 329 \| 659 \| MCM domain-containing protein \| MCM domain-containing protein; afdb-proteome \| AF-A0A1D6NMY4-F1-MODEL\_V4 \| 0.632 \| 0.004947 \| 55 \| 0.113 \| 466 \| 233 \| 33 \| 393 \| 738 \| 370 \| 775 \| DNA helicase \| DNA helicase; afdb-proteome \| AF-P24279-F1-MODEL\_V4 \| 0.632 \| 0.004462 \| 55 \| 0.098 \| 489 \| 246 \| 30 \| 404 \| 738 \| 293 \| 740 \| DNA replication licensing factor MCM3 \| DNA replication licensing factor MCM3; afdb-proteome \| AF-C0PDH6-F1-MODEL\_V4 \| 0.601 \| 0.008726 \| 54 \| 0.088 \| 541 \| 255 \| 38 \| 393 \| 755 \| 371 \| 851 \| DNA helicase \| DNA helicase; afdb-proteome \| AF-A0A077ZCX8-F1-MODEL\_V4 \| 0.505 \| 0.008287 \| 51 \| 0.104 \| 468 \| 239 \| 30 \| 393 \| 738 \| 346 \| 755 \| DNA replication licensing factor MCM8 \| DNA replication licensing factor MCM8; afdb-proteome \| AF-Q8ILY1-F1-MODEL\_V4 \| 0.382 \| 0.006742 \| 47 \| 0.096 \| 693 \| 315 \| 41 \| 153 \| 616 \| 249 \| 859 \| Plastid replication-repair enzyme \| Plastid replication-repair enzyme; afdb-uniprot50 \| AF-A0A2Z3IKG0-F1-MODEL\_V4 \| 1.0 \| 3.972e-71 \| 2233 \| 0.449 \| 685 \| 347 \| 13 \| 102 \| 769 \| 88 \| 759 \| Integrase \| Integrase; afdb-uniprot50 \| AF-A0A7L6EDN8-F1-MODEL\_V4 \| 1.0 \| 2.18e-54 \| 2226 \| 0.6 \| 443 \| 168 \| 5 \| 330 \| 769 \| 8 \| 444 \| Virulence-associated E family protein \| Virulence-associated E family protein; afdb-uniprot50 \| AF-A0A6G8CKG5-F1-MODEL\_V4 \| 1.0 \| 9.011e-74 \| 2189 \| 0.378 \| 929 \| 373 \| 19 \| 22 \| 757 \| 1 \| 917 \| PriCT\_2 domain-containing protein \| PriCT\_2 domain-containing protein; afdb-uniprot50 \| AF-B4EME0-F1-MODEL\_V4 \| 1.0 \| 2.545e-54 \| 2167 \| 0.579 \| 461 \| 169 \| 8 \| 314 \| 757 \| 7 \| 459 \| Hypothetical phage protein \| Hypothetical phage protein; afdb-uniprot50 \| AF-A0A8A6KD37-F1-MODEL\_V4 \| 1.0 \| 1.892e-72 \| 2132 \| 0.396 \| 789 \| 428 \| 21 \| 3 \| 769 \| 1 \| 762 \| PriCT-2 domain-containing protein \| PriCT-2 domain-containing protein; afdb-uniprot50 \| AF-A0A370WXM5-F1-MODEL\_V4 \| 1.0 \| 2.714e-64 \| 2083 \| 0.45 \| 671 \| 321 \| 14 \| 100 \| 757 \| 1 \| 636 \| Toprim domain-containing protein \| Toprim domain-containing protein; afdb-uniprot50 \| AF-A0A0D7EBT5-F1-MODEL\_V4 \| 1.0 \| 6.079e-65 \| 1947 \| 0.432 \| 721 \| 331 \| 23 \| 63 \| 757 \| 97 \| 764 \| Toprim domain-containing protein \| Toprim domain-containing protein; afdb-uniprot50 \| AF-A0A3R8TDC4-F1-MODEL\_V4 \| 1.0 \| 4.023e-65 \| 1940 \| 0.407 \| 702 \| 372 \| 15 \| 102 \| 769 \| 87 \| 778 \| Toprim domain-containing protein \| Toprim domain-containing protein; afdb-uniprot50 \| AF-A0A4R6JMW9-F1-MODEL\_V4 \| 1.0 \| 1.509e-65 \| 1919 \| 0.408 \| 724 \| 357 \| 19 \| 98 \| 761 \| 84 \| 795 \| Putative DNA primase/helicase \| Putative DNA primase/helicase; afdb-uniprot50 \| AF-A0A8B2Q819-F1-MODEL\_V4 \| 1.0 \| 5.342e-45 \| 1878 \| 0.449 \| 414 \| 222 \| 4 \| 356 \| 769 \| 51 \| 458 \| Virulence-associated E family protein \| Virulence-associated E family protein; afdb-uniprot50 \| AF-A0A810TDZ1-F1-MODEL\_V4 \| 1.0 \| 2.138e-55 \| 1874 \| 0.491 \| 576 \| 232 \| 15 \| 228 \| 769 \| 11 \| 559 \| Uncharacterized protein \| Uncharacterized protein; afdb-uniprot50 \| AF-A0A5K7Y2P6-F1-MODEL\_V4 \| 1.0 \| 2.355e-42 \| 1805 \| 0.453 \| 390 \| 204 \| 6 \| 369 \| 756 \| 123 \| 505 \| Uncharacterized protein \| Uncharacterized protein; afdb-uniprot50 \| AF-A0A5C7W4R6-F1-MODEL\_V4 \| 1.0 \| 8.336e-46 \| 1706 \| 0.448 \| 484 \| 236 \| 10 \| 287 \| 762 \| 74 \| 534 \| Virulence-associated E family protein \| Virulence-associated E family protein; afdb-uniprot50 \| AF-A0A550ECX4-F1-MODEL\_V4 \| 1.0 \| 3.048e-58 \| 1682 \| 0.35 \| 728 \| 374 \| 25 \| 48 \| 757 \| 99 \| 745 \| Toprim domain-containing protein \| Toprim domain-containing protein; afdb-uniprot50 \| AF-A0A2N1CUR9-F1-MODEL\_V4 \| 1.0 \| 2.03e-39 \| 1607 \| 0.412 \| 417 \| 236 \| 7 \| 346 \| 756 \| 85 \| 498 \| Virulence-associated E family protein \| Virulence-associated E family protein; afdb-uniprot50 \| AF-A0A7Y9SYN6-F1-MODEL\_V4 \| 1.0 \| 7.233e-40 \| 1602 \| 0.407 \| 400 \| 232 \| 5 \| 369 \| 766 \| 10 \| 406 \| Putative DNA primase/helicase \| Putative DNA primase/helicase; afdb-uniprot50 \| AF-Q1NPY4-F1-MODEL\_V4 \| 1.0 \| 3.401e-39 \| 1602 \| 0.416 \| 406 \| 228 \| 6 \| 354 \| 757 \| 49 \| 447 \| Virulence-associated E \| Virulence-associated E; afdb-uniprot50 \| AF-A0A2T1K8Q3-F1-MODEL\_V4 \| 1.0 \| 3.148e-43 \| 1599 \| 0.392 \| 482 \| 260 \| 11 \| 298 \| 769 \| 331 \| 789 \| RepB\_primase domain-containing protein \| RepB\_primase domain-containing protein; afdb-uniprot50 \| AF-A0A370DD10-F1-MODEL\_V4 \| 1.0 \| 6.869e-40 \| 1594 \| 0.428 \| 411 \| 221 \| 7 \| 356 \| 762 \| 446 \| 846 \| Uncharacterized protein \| Uncharacterized protein; afdb-uniprot50 \| AF-A0A241VC66-F1-MODEL\_V4 \| 1.0 \| 4.154e-50 \| 1592 \| 0.334 \| 660 \| 381 \| 18 \| 136 \| 769 \| 10 \| 636 \| Uncharacterized protein \| Uncharacterized protein; afdb-uniprot50 \| AF-A0A2D7BYY9-F1-MODEL\_V4 \| 1.0 \| 3.029e-53 \| 1587 \| 0.331 \| 679 \| 371 \| 24 \| 92 \| 757 \| 89 \| 697 \| Toprim domain-containing protein \| Toprim domain-containing protein; afdb-uniprot50 \| AF-A0A329BI33-F1-MODEL\_V4 \| 1.0 \| 6.196e-40 \| 1586 \| 0.436 \| 417 \| 219 \| 7 \| 354 \| 769 \| 2 \| 403 \| Virulence-associated protein E \| Virulence-associated protein E; afdb-uniprot50 \| AF-A0A126RQW2-F1-MODEL\_V4 \| 1.0 \| 2.03e-39 \| 1586 \| 0.41 \| 414 \| 230 \| 8 \| 368 \| 769 \| 35 \| 446 \| Uncharacterized protein \| Uncharacterized protein; afdb-uniprot50 \| AF-A0A2N2RJ59-F1-MODEL\_V4 \| 1.0 \| 6.196e-40 \| 1570 \| 0.413 \| 433 \| 236 \| 9 \| 346 \| 769 \| 92 \| 515 \| Virulence-associated E family protein \| Virulence-associated E family protein; afdb-uniprot50 \| AF-W0DSD8-F1-MODEL\_V4 \| 1.0 \| 2.48e-50 \| 1563 \| 0.382 \| 638 \| 325 \| 20 \| 169 \| 769 \| 88 \| 693 \| Virulence-associated E family protein \| Virulence-associated E family protein; afdb-uniprot50 \| AF-A0A8A7VX14-F1-MODEL\_V4 \| 1.0 \| 1.63e-37 \| 1553 \| 0.39 \| 394 \| 233 \| 6 \| 368 \| 757 \| 143 \| 533 \| Uncharacterized protein \| Uncharacterized protein; afdb-uniprot50 \| AF-A0A1B2I2L3-F1-MODEL\_V4 \| 1.0 \| 4.403e-55 \| 1551 \| 0.311 \| 755 \| 401 \| 24 \| 68 \| 762 \| 69 \| 764 \| Toprim domain-containing protein \| Toprim domain-containing protein; afdb-uniprot50 \| AF-A0A148KLT5-F1-MODEL\_V4 \| 1.0 \| 6.651e-39 \| 1550 \| 0.382 \| 410 \| 240 \| 8 \| 351 \| 756 \| 403 \| 803 \| Uncharacterized protein \| Uncharacterized protein; afdb-uniprot50 \| AF-A0A2D9C8Q1-F1-MODEL\_V4 \| 1.0 \| 3.87e-51 \| 1521 \| 0.306 \| 685 \| 407 \| 21 \| 92 \| 767 \| 92 \| 717 \| Toprim domain-containing protein \| Toprim domain-containing protein; afdb-uniprot50 \| AF-A0A4P5WEG0-F1-MODEL\_V4 \| 1.0 \| 5.624e-37 \| 1517 \| 0.373 \| 415 \| 247 \| 8 \| 362 \| 769 \| 74 \| 482 \| Uncharacterized protein \| Uncharacterized protein; afdb-uniprot50 \| AF-A0A7X1XSH3-F1-MODEL\_V4 \| 1.0 \| 1.025e-37 \| 1492 \| 0.39 \| 433 \| 242 \| 10 \| 354 \| 769 \| 84 \| 511 \| Virulence-associated E family protein \| Virulence-associated E family protein; afdb-uniprot50 \| AF-A0A4P6X2C8-F1-MODEL\_V4 \| 1.0 \| 4.725e-38 \| 1487 \| 0.378 \| 410 \| 249 \| 6 \| 363 \| 769 \| 70 \| 476 \| Virulence-associated E family protein \| Virulence-associated E family protein; afdb-uniprot50 \| AF-A0A838L9S2-F1-MODEL\_V4 \| 1.0 \| 1.158e-36 \| 1486 \| 0.363 \| 410 \| 254 \| 7 \| 363 \| 769 \| 42 \| 447 \| Uncharacterized protein \| Uncharacterized protein; afdb-uniprot50 \| AF-A0A6N9HMH7-F1-MODEL\_V4 \| 1.0 \| 7.665e-37 \| 1486 \| 0.379 \| 427 \| 240 \| 10 \| 349 \| 756 \| 4 \| 424 \| Virulence-associated E family protein \| Virulence-associated E family protein; afdb-uniprot50 \| AF-A0A2X1BCW5-F1-MODEL\_V4 \| 1.0 \| 3.604e-36 \| 1455 \| 0.426 \| 387 \| 208 \| 10 \| 389 \| 769 \| 2 \| 380 \| Predicted P-loop ATPase and inactivated derivatives \| Predicted P-loop ATPase and inactivated derivatives; afdb-uniprot50 \| AF-A0A0Q4G1V4-F1-MODEL\_V4 \| 1.0 \| 7.048e-36 \| 1454 \| 0.379 \| 387 \| 232 \| 5 \| 388 \| 769 \| 2 \| 385 \| Uncharacterized protein \| Uncharacterized protein; afdb-uniprot50 \| AF-A0A2D4Y2B1-F1-MODEL\_V4 \| 1.0 \| 9.422e-37 \| 1447 \| 0.361 \| 409 \| 251 \| 6 \| 366 \| 769 \| 9 \| 412 \| Uncharacterized protein \| Uncharacterized protein; afdb-uniprot50 \| AF-A0A4P7QW42-F1-MODEL\_V4 \| 1.0 \| 7.094e-41 \| 1444 \| 0.366 \| 461 \| 271 \| 9 \| 315 \| 769 \| 30 \| 475 \| Virulence-associated E family protein \| Virulence-associated E family protein; afdb-uniprot50 \| AF-A0A848H5Z6-F1-MODEL\_V4 \| 1.0 \| 5.922e-37 \| 1444 \| 0.371 \| 406 \| 232 \| 9 \| 368 \| 753 \| 82 \| 484 \| Uncharacterized protein \| Uncharacterized protein; afdb-uniprot50 \| AF-A0A1H9PLQ6-F1-MODEL\_V4 \| 1.0 \| 4.757e-51 \| 1429 \| 0.391 \| 616 \| 297 \| 21 \| 81 \| 671 \| 105 \| 667 \| Putative DNA primase/helicase \| Putative DNA primase/helicase; afdb-uniprot50 \| AF-A0A3N5V9B1-F1-MODEL\_V4 \| 1.0 \| 2.138e-47 \| 1428 \| 0.333 \| 638 \| 319 \| 25 \| 181 \| 763 \| 138 \| 723 \| Uncharacterized protein \| Uncharacterized protein; afdb-uniprot50 \| AF-A0A162LR67-F1-MODEL\_V4 \| 1.0 \| 6.357e-36 \| 1422 \| 0.362 \| 408 \| 250 \| 7 \| 366 \| 767 \| 31 \| 434 \| Uncharacterized protein \| Uncharacterized protein; afdb-uniprot50 \| AF-A0A0W1I738-F1-MODEL\_V4 \| 1.0 \| 1.309e-35 \| 1418 \| 0.362 \| 405 \| 245 \| 6 \| 362 \| 764 \| 4 \| 397 \| Uncharacterized protein \| Uncharacterized protein; afdb-uniprot50 \| AF-A0A257G4S2-F1-MODEL\_V4 \| 1.0 \| 8.389e-35 \| 1408 \| 0.369 \| 398 \| 238 \| 9 \| 377 \| 769 \| 71 \| 460 \| Uncharacterized protein \| Uncharacterized protein; afdb-uniprot50 \| AF-A0A6I5YGU0-F1-MODEL\_V4 \| 1.0 \| 1.235e-38 \| 1405 \| 0.378 \| 433 \| 254 \| 8 \| 340 \| 769 \| 23 \| 443 \| Virulence-associated E family protein \| Virulence-associated E family protein; afdb-uniprot50 \| AF-A0A1B6Z6D8-F1-MODEL\_V4 \| 1.0 \| 1.268e-34 \| 1397 \| 0.376 \| 393 \| 241 \| 4 \| 378 \| 769 \| 5 \| 394 \| Uncharacterized protein \| Uncharacterized protein; afdb-uniprot50 \| AF-A0A315BDU7-F1-MODEL\_V4 \| 1.0 \| 4.575e-37 \| 1387 \| 0.368 \| 450 \| 252 \| 13 \| 328 \| 758 \| 2 \| 438 \| Uncharacterized protein \| Uncharacterized protein; afdb-uniprot50 \| AF-A0A6P1J3W7-F1-MODEL\_V4 \| 1.0 \| 7.048e-36 \| 1379 \| 0.378 \| 391 \| 232 \| 8 \| 369 \| 757 \| 44 \| 425 \| Uncharacterized protein \| Uncharacterized protein; afdb-uniprot50 \| AF-A0A1B6Z6F3-F1-MODEL\_V4 \| 1.0 \| 5.007e-35 \| 1364 \| 0.36 \| 419 \| 249 \| 10 \| 348 \| 758 \| 7 \| 414 \| Uncharacterized protein \| Uncharacterized protein; afdb-uniprot50 \| AF-A0A158EA01-F1-MODEL\_V4 \| 1.0 \| 1.284e-44 \| 1360 \| 0.293 \| 654 \| 370 \| 22 \| 181 \| 766 \| 147 \| 776 \| Virulence-associated protein E \| Virulence-associated protein E; afdb-uniprot50 \| AF-A0A6G6X8A9-F1-MODEL\_V4 \| 1.0 \| 1.855e-33 \| 1336 \| 0.379 \| 398 \| 230 \| 7 \| 368 \| 762 \| 6 \| 389 \| Uncharacterized protein \| Uncharacterized protein; afdb-uniprot50 \| AF-A0A7Z0LRG9-F1-MODEL\_V4 \| 1.0 \| 7.715e-42 \| 1321 \| 0.366 \| 521 \| 274 \| 12 \| 246 \| 756 \| 346 \| 820 \| Uncharacterized protein \| Uncharacterized protein; afdb-uniprot50 \| AF-A0A7Y8YLU9-F1-MODEL\_V4 \| 1.0 \| 9.793e-35 \| 1316 \| 0.355 \| 430 \| 251 \| 9 \| 356 \| 769 \| 8 \| 427 \| P-loop ATPase \| P-loop ATPase; afdb-uniprot50 \| AF-A0A2U2MYE5-F1-MODEL\_V4 \| 1.0 \| 1.065e-51 \| 1316 \| 0.257 \| 855 \| 498 \| 34 \| 1 \| 757 \| 1 \| 816 \| PriCT\_2 domain-containing protein \| PriCT\_2 domain-containing protein; afdb-uniprot50 \| AF-A0A3B9WZT9-F1-MODEL\_V4 \| 1.0 \| 9.423e-45 \| 1307 \| 0.336 \| 561 \| 301 \| 18 \| 102 \| 651 \| 102 \| 601 \| Toprim domain-containing protein \| Toprim domain-containing protein; afdb-uniprot50 \| AF-S6SGC4-F1-MODEL\_V4 \| 1.0 \| 9.124e-44 \| 1278 \| 0.287 \| 675 \| 407 \| 32 \| 128 \| 769 \| 2 \| 635 \| DNA primase domain protein \| DNA primase domain protein; afdb-uniprot50 \| AF-A0A4Y4D005-F1-MODEL\_V4 \| 1.0 \| 5.922e-45 \| 1256 \| 0.284 \| 786 \| 352 \| 19 \| 1 \| 756 \| 215 \| 819 \| Uncharacterized protein \| Uncharacterized protein; afdb-uniprot50 \| AF-A0A450WC63-F1-MODEL\_V4 \| 1.0 \| 3.445e-49 \| 1255 \| 0.29 \| 825 \| 471 \| 38 \| 10 \| 750 \| 4 \| 797 \| Putative DNA primase/helicase \| Putative DNA primase/helicase; afdb-uniprot50 \| AF-A0A4Q9VPA0-F1-MODEL\_V4 \| 1.0 \| 7.141e-46 \| 1241 \| 0.291 \| 680 \| 416 \| 30 \| 100 \| 752 \| 75 \| 715 \| Toprim domain-containing protein \| Toprim domain-containing protein; afdb-uniprot50 \| AF-L0GU11-F1-MODEL\_V4 \| 1.0 \| 4.373e-34 \| 1232 \| 0.325 \| 452 \| 267 \| 11 \| 328 \| 762 \| 137 \| 567 \| Putative P-loop ATPase \| Putative P-loop ATPase; afdb-uniprot50 \| AF-A0A3G9FYK3-F1-MODEL\_V4 \| 1.0 \| 1.343e-39 \| 1224 \| 0.275 \| 632 \| 361 \| 26 \| 161 \| 762 \| 25 \| 589 \| Putative virulence-associated protein E \| Putative virulence-associated protein E; afdb-uniprot50 \| AF-A0A1D2U957-F1-MODEL\_V4 \| 1.0 \| 2.137e-31 \| 1211 \| 0.326 \| 404 \| 238 \| 7 \| 376 \| 750 \| 22 \| 420 \| Uncharacterized protein \| Uncharacterized protein; afdb-uniprot50 \| AF-A0A2T7SVQ2-F1-MODEL\_V4 \| 1.0 \| 3.722e-37 \| 1211 \| 0.293 \| 538 \| 313 \| 16 \| 239 \| 752 \| 229 \| 723 \| Uncharacterized protein \| Uncharacterized protein; afdb-uniprot50 \| AF-A0A7W7NUF6-F1-MODEL\_V4 \| 1.0 \| 4.047e-30 \| 1208 \| 0.424 \| 325 \| 181 \| 6 \| 435 \| 757 \| 1 \| 321 \| Putative DNA primase/helicase \| Putative DNA primase/helicase; afdb-uniprot50 \| AF-A0A258QD56-F1-MODEL\_V4 \| 1.0 \| 4.43e-36 \| 1189 \| 0.263 \| 547 \| 324 \| 14 \| 242 \| 755 \| 293 \| 793 \| Uncharacterized protein \| Uncharacterized protein; afdb-uniprot50 \| AF-A0A662BLW6-F1-MODEL\_V4 \| 1.0 \| 4.317e-32 \| 1164 \| 0.307 \| 459 \| 281 \| 13 \| 329 \| 766 \| 363 \| 805 \| Prim-Pol domain-containing protein \| Prim-Pol domain-containing protein; afdb-uniprot50 \| AF-A0A127SEK4-F1-MODEL\_V4 \| 1.0 \| 5.171e-28 \| 1161 \| 0.474 \| 270 \| 140 \| 2 \| 488 \| 757 \| 4 \| 271 \| Virulence-associated protein E \| Virulence-associated protein E; afdb-uniprot50 \| AF-A0A269PIH2-F1-MODEL\_V4 \| 1.0 \| 1.978e-27 \| 1146 \| 0.439 \| 287 \| 157 \| 2 \| 355 \| 638 \| 19 \| 304 \| Uncharacterized protein \| Uncharacterized protein; afdb-uniprot50 \| AF-A0A2K4XP73-F1-MODEL\_V4 \| 1.0 \| 9.544e-39 \| 1142 \| 0.285 \| 652 \| 371 \| 24 \| 153 \| 756 \| 79 \| 683 \| Uncharacterized protein \| Uncharacterized protein; afdb-uniprot50 \| AF-A0A363ULB6-F1-MODEL\_V4 \| 1.0 \| 1.916e-42 \| 1130 \| 0.241 \| 725 \| 455 \| 30 \| 92 \| 769 \| 82 \| 758 \| Uncharacterized protein \| Uncharacterized protein; afdb-uniprot50 \| AF-A0A059KRR5-F1-MODEL\_V4 \| 1.0 \| 4.262e-46 \| 1125 \| 0.246 \| 872 \| 518 \| 37 \| 1 \| 769 \| 1 \| 836 \| Uncharacterized protein \| Uncharacterized protein; afdb-uniprot50 \| AF-A0A5C7PWX5-F1-MODEL\_V4 \| 1.0 \| 2.696e-43 \| 1121 \| 0.265 \| 742 \| 421 \| 30 \| 92 \| 757 \| 81 \| 774 \| Toprim domain-containing protein \| Toprim domain-containing protein; afdb-uniprot50 \| AF-A0A5C7Q9V4-F1-MODEL\_V4 \| 1.0 \| 1.335e-26 \| 1117 \| 0.423 \| 274 \| 151 \| 4 \| 380 \| 649 \| 9 \| 279 \| Uncharacterized protein \| Uncharacterized protein; afdb-uniprot50 \| AF-A0A844Y3H4-F1-MODEL\_V4 \| 1.0 \| 3.422e-28 \| 1097 \| 0.38 \| 352 \| 206 \| 7 \| 423 \| 767 \| 2 \| 348 \| Uncharacterized protein \| Uncharacterized protein; afdb-uniprot50 \| AF-A0A5B9W769-F1-MODEL\_V4 \| 1.0 \| 2.124e-42 \| 1088 \| 0.294 \| 710 \| 409 \| 28 \| 91 \| 750 \| 75 \| 742 \| DNA primase TraC \| DNA primase TraC; afdb-uniprot50 \| AF-A0A315D5W5-F1-MODEL\_V4 \| 1.0 \| 1.47e-37 \| 1074 \| 0.226 \| 764 \| 337 \| 17 \| 2 \| 752 \| 196 \| 718 \| Prim-Pol domain-containing protein \| Prim-Pol domain-containing protein; afdb-uniprot50 \| AF-A0A848EXA0-F1-MODEL\_V4 \| 1.0 \| 2.875e-29 \| 1064 \| 0.28 \| 421 \| 272 \| 14 \| 370 \| 768 \| 19 \| 430 \| Uncharacterized protein \| Uncharacterized protein; afdb-uniprot50 \| AF-A0A1C6F2F8-F1-MODEL\_V4 \| 1.0 \| 1.151e-31 \| 1062 \| 0.277 \| 486 \| 300 \| 18 \| 298 \| 761 \| 78 \| 534 \| DNA primase \| DNA primase; afdb-uniprot50 \| AF-A0A4V2HU15-F1-MODEL\_V4 \| 1.0 \| 5.961e-42 \| 1053 \| 0.231 \| 937 \| 422 \| 38 \| 98 \| 752 \| 95 \| 1014 \| Toprim domain-containing protein \| Toprim domain-containing protein; afdb-uniprot50 \| AF-A0A2S5TKJ6-F1-MODEL\_V4 \| 1.0 \| 1.093e-39 \| 1037 \| 0.262 \| 698 \| 432 \| 33 \| 102 \| 762 \| 90 \| 741 \| Uncharacterized protein \| Uncharacterized protein; afdb-uniprot50 \| AF-A0A1U7MHB1-F1-MODEL\_V4 \| 1.0 \| 4.545e-32 \| 1034 \| 0.247 \| 545 \| 333 \| 20 \| 260 \| 769 \| 202 \| 704 \| Virulence-associated protein E \| Virulence-associated protein E; afdb-uniprot50 \| AF-A0A1M6WA62-F1-MODEL\_V4 \| 1.0 \| 6.737e-33 \| 1016 \| 0.263 \| 577 \| 320 \| 20 \| 242 \| 762 \| 33 \| 560 \| Virulence-associated protein E \| Virulence-associated protein E; afdb-uniprot50 \| AF-A0A661CNY3-F1-MODEL\_V4 \| 1.0 \| 2.593e-29 \| 1001 \| 0.234 \| 457 \| 304 \| 15 \| 330 \| 758 \| 299 \| 737 \| Uncharacterized protein \| Uncharacterized protein; afdb-uniprot50 \| AF-A0A7Y8ZYJ0-F1-MODEL\_V4 \| 1.0 \| 7.967e-35 \| 984 \| 0.213 \| 646 \| 407 \| 31 \| 168 \| 768 \| 103 \| 692 \| Putative P-loop ATPase \| Putative P-loop ATPase; afdb-uniprot50 \| AF-A0A349EG17-F1-MODEL\_V4 \| 1.0 \| 1.196e-37 \| 982 \| 0.263 \| 686 \| 414 \| 34 \| 99 \| 757 \| 74 \| 694 \| Toprim domain-containing protein \| Toprim domain-containing protein; afdb-uniprot50 \| AF-A0A7Z8QIE3-F1-MODEL\_V4 \| 1.0 \| 9.605e-28 \| 979 \| 0.268 \| 410 \| 270 \| 12 \| 370 \| 762 \| 9 \| 405 \| Uncharacterized protein \| Uncharacterized protein; afdb-uniprot50 \| AF-A0A431IWX9-F1-MODEL\_V4 \| 1.0 \| 8.947e-29 \| 975 \| 0.244 \| 482 \| 324 \| 15 \| 291 \| 762 \| 280 \| 731 \| PriCT\_2 domain-containing protein \| PriCT\_2 domain-containing protein; afdb-uniprot50 \| AF-A0A7T4N249-F1-MODEL\_V4 \| 1.0 \| 2.447e-24 \| 964 \| 0.335 \| 349 \| 200 \| 10 \| 13 \| 347 \| 9 \| 339 \| DUF927 domain-containing protein \| DUF927 domain-containing protein; afdb-uniprot50 \| AF-A0A2N2U4N7-F1-MODEL\_V4 \| 1.0 \| 2.265e-28 \| 957 \| 0.366 \| 360 \| 200 \| 11 \| 1 \| 347 \| 1 \| 345 \| Toprim domain-containing protein \| Toprim domain-containing protein; afdb-uniprot50 \| AF-A0A412CFG8-F1-MODEL\_V4 \| 1.0 \| 1.301e-30 \| 952 \| 0.235 \| 557 \| 341 \| 21 \| 243 \| 769 \| 241 \| 742 \| Uncharacterized protein \| Uncharacterized protein; afdb-uniprot50 \| AF-A0A2G2PN86-F1-MODEL\_V4 \| 1.0 \| 3.401e-39 \| 948 \| 0.232 \| 788 \| 454 \| 33 \| 93 \| 769 \| 54 \| 801 \| Uncharacterized protein \| Uncharacterized protein; afdb-uniprot50 \| AF-A0A6A4U5X7-F1-MODEL\_V4 \| 1.0 \| 5.105e-26 \| 947 \| 0.246 \| 386 \| 276 \| 13 \| 379 \| 756 \| 310 \| 688 \| Virulence-associated E family protein \| Virulence-associated E family protein; afdb-uniprot50 \| AF-A0A2S7FDM4-F1-MODEL\_V4 \| 1.0 \| 8.28e-25 \| 929 \| 0.246 \| 414 \| 270 \| 19 \| 366 \| 757 \| 18 \| 411 \| Uncharacterized protein \| Uncharacterized protein; afdb-uniprot50 \| AF-A0A849UAZ0-F1-MODEL\_V4 \| 1.0 \| 5.66e-34 \| 925 \| 0.228 \| 652 \| 416 \| 32 \| 127 \| 754 \| 49 \| 637 \| Uncharacterized protein \| Uncharacterized protein; afdb-uniprot50 \| AF-A0A193CM65-F1-MODEL\_V4 \| 1.0 \| 1.694e-27 \| 920 \| 0.277 \| 480 \| 300 \| 16 \| 286 \| 757 \| 292 \| 732 \| Uncharacterized protein \| Uncharacterized protein; afdb-uniprot50 \| AF-A0A661DIC7-F1-MODEL\_V4 \| 1.0 \| 7.278e-29 \| 920 \| 0.209 \| 496 \| 326 \| 16 \| 291 \| 758 \| 376 \| 833 \| Uncharacterized protein \| Uncharacterized protein; afdb-uniprot50 \| AF-C9LQR2-F1-MODEL\_V4 \| 1.0 \| 4.126e-29 \| 917 \| 0.238 \| 537 \| 341 \| 23 \| 240 \| 757 \| 223 \| 710 \| Virulence-associated protein E \| Virulence-associated protein E; afdb-uniprot50 \| AF-A0A4P8XWJ9-F1-MODEL\_V4 \| 1.0 \| 8.719e-25 \| 910 \| 0.253 \| 414 \| 273 \| 16 \| 365 \| 757 \| 133 \| 531 \| Uncharacterized protein \| Uncharacterized protein; afdb-uniprot50 \| AF-A0A354KW19-F1-MODEL\_V4 \| 1.0 \| 3.995e-28 \| 902 \| 0.275 \| 497 \| 295 \| 17 \| 299 \| 757 \| 28 \| 497 \| ZnF\_CHCC domain-containing protein \| ZnF\_CHCC domain-containing protein; afdb-uniprot50 \| AF-A0A419U224-F1-MODEL\_V4 \| 1.0 \| 5.734e-36 \| 902 \| 0.218 \| 829 \| 456 \| 36 \| 92 \| 769 \| 62 \| 849 \| Putative P-loop ATPase \| Putative P-loop ATPase; afdb-uniprot50 \| AF-A0A3M9MMY6-F1-MODEL\_V4 \| 1.0 \| 1.854e-25 \| 901 \| 0.252 \| 404 \| 271 \| 17 \| 384 \| 769 \| 31 \| 421 \| Virulence-associated e family protein \| Virulence-associated e family protein; afdb-uniprot50 \| AF-A0A0B2JZB9-F1-MODEL\_V4 \| 1.0 \| 8.227e-28 \| 897 \| 0.232 \| 521 \| 320 \| 21 \| 261 \| 756 \| 230 \| 695 \| Uncharacterized protein \| Uncharacterized protein; afdb-uniprot50 \| AF-A0A450WD24-F1-MODEL\_V4 \| 1.0 \| 2.03e-39 \| 892 \| 0.237 \| 870 \| 460 \| 50 \| 10 \| 753 \| 4 \| 795 \| Putative DNA primase/helicase \| Putative DNA primase/helicase; afdb-uniprot50 \| AF-A0A7Z0SE33-F1-MODEL\_V4 \| 1.0 \| 4.546e-40 \| 890 \| 0.247 \| 876 \| 483 \| 48 \| 12 \| 757 \| 9 \| 837 \| PriCT-2 domain-containing protein \| PriCT-2 domain-containing protein; afdb-uniprot50 \| AF-A0A1S8TCW1-F1-MODEL\_V4 \| 1.0 \| 1.335e-34 \| 887 \| 0.224 \| 744 \| 428 \| 35 \| 64 \| 769 \| 67 \| 699 \| DNA primase \| DNA primase; afdb-uniprot50 \| AF-A0A6M4GU76-F1-MODEL\_V4 \| 1.0 \| 1.326e-37 \| 877 \| 0.204 \| 816 \| 508 \| 36 \| 1 \| 757 \| 6 \| 738 \| PriCT\_2 domain-containing protein \| PriCT\_2 domain-containing protein; afdb-uniprot50 \| AF-A0A5B3G911-F1-MODEL\_V4 \| 1.0 \| 2.577e-24 \| 873 \| 0.248 \| 383 \| 264 \| 13 \| 381 \| 755 \| 54 \| 420 \| Uncharacterized protein \| Uncharacterized protein; afdb-uniprot50 \| AF-A0A258SSH2-F1-MODEL\_V4 \| 1.0 \| 3.335e-24 \| 867 \| 0.461 \| 286 \| 140 \| 7 \| 493 \| 769 \| 2 \| 282 \| Uncharacterized protein \| Uncharacterized protein; afdb-uniprot50 \| AF-A0A1Q2M5G7-F1-MODEL\_V4 \| 1.0 \| 9.855e-24 \| 866 \| 0.344 \| 354 \| 177 \| 11 \| 10 \| 324 \| 5 \| 342 \| Uncharacterized protein \| Uncharacterized protein; afdb-uniprot50 \| AF-A0A158KF25-F1-MODEL\_V4 \| 1.0 \| 1.83e-23 \| 866 \| 0.343 \| 361 \| 200 \| 14 \| 3 \| 347 \| 2 \| 341 \| DNA replication protein \| DNA replication protein; afdb-uniprot50 \| AF-A0A1G0RDT0-F1-MODEL\_V4 \| 1.0 \| 7.231e-24 \| 864 \| 0.215 \| 403 \| 282 \| 15 \| 381 \| 756 \| 31 \| 426 \| Uncharacterized protein \| Uncharacterized protein; afdb-uniprot50 \| AF-D7K6T8-F1-MODEL\_V4 \| 1.0 \| 1.461e-24 \| 863 \| 0.241 \| 389 \| 261 \| 16 \| 384 \| 757 \| 19 \| 388 \| Putative helicase \| Putative helicase; afdb-uniprot50 \| AF-A0A1I9YDF5-F1-MODEL\_V4 \| 1.0 \| 3.558e-34 \| 859 \| 0.193 \| 818 \| 493 \| 39 \| 13 \| 762 \| 7 \| 725 \| PriCT\_2 domain-containing protein \| PriCT\_2 domain-containing protein; afdb-uniprot50 \| AF-R5CCA7-F1-MODEL\_V4 \| 1.0 \| 3.511e-24 \| 855 \| 0.239 \| 389 \| 272 \| 16 \| 378 \| 756 \| 35 \| 409 \| Uncharacterized protein \| Uncharacterized protein; afdb-uniprot50 \| AF-A0A7W8FU46-F1-MODEL\_V4 \| 1.0 \| 2.193e-27 \| 852 \| 0.238 \| 528 \| 335 \| 19 \| 261 \| 769 \| 216 \| 695 \| PriCT\_1 domain-containing protein \| PriCT\_1 domain-containing protein; afdb-uniprot50 \| AF-A0A4P7L7L6-F1-MODEL\_V4 \| 1.0 \| 2.124e-34 \| 850 \| 0.21 \| 723 \| 467 \| 37 \| 60 \| 755 \| 61 \| 706 \| DNA primase TraC \| DNA primase TraC; afdb-uniprot50 \| AF-A0A3G9IPK2-F1-MODEL\_V4 \| 1.0 \| 5.515e-22 \| 848 \| 0.28 \| 339 \| 212 \| 9 \| 366 \| 682 \| 41 \| 369 \| Uncharacterized protein \| Uncharacterized protein; afdb-uniprot50 \| AF-A0A4Q3Q5F2-F1-MODEL\_V4 \| 1.0 \| 8.069e-21 \| 843 \| 0.388 \| 270 \| 148 \| 8 \| 6 \| 272 \| 1 \| 256 \| Toprim domain-containing protein \| Toprim domain-containing protein; afdb-uniprot50 \| AF-A0A7X6JZJ4-F1-MODEL\_V4 \| 1.0 \| 5.998e-31 \| 843 \| 0.235 \| 658 \| 348 \| 22 \| 183 \| 755 \| 351 \| 938 \| Uncharacterized protein \| Uncharacterized protein; afdb-uniprot50 \| AF-A0A6M3KDE2-F1-MODEL\_V4 \| 1.0 \| 2.029e-23 \| 842 \| 0.221 \| 389 \| 272 \| 16 \| 384 \| 751 \| 51 \| 429 \| Putative VirE domain containing protein \| Putative VirE domain containing protein; afdb-uniprot50 \| AF-A0A6N4AQ20-F1-MODEL\_V4 \| 1.0 \| 8.017e-24 \| 841 \| 0.301 \| 398 \| 247 \| 15 \| 377 \| 757 \| 28 \| 411 \| Uncharacterized protein \| Uncharacterized protein; afdb-uniprot50 \| AF-A0A6I7WDH6-F1-MODEL\_V4 \| 1.0 \| 2.543e-22 \| 840 \| 0.298 \| 335 \| 217 \| 7 \| 376 \| 699 \| 19 \| 346 \| Uncharacterized protein \| Uncharacterized protein; afdb-uniprot50 \| AF-A0A6L4NDA3-F1-MODEL\_V4 \| 1.0 \| 9.855e-24 \| 840 \| 0.232 \| 387 \| 271 \| 14 \| 384 \| 755 \| 18 \| 393 \| Virulence protein E \| Virulence protein E; afdb-uniprot50 \| AF-A0A5B5U8H6-F1-MODEL\_V4 \| 1.0 \| 2.627e-23 \| 838 \| 0.24 \| 387 \| 267 \| 16 \| 382 \| 757 \| 67 \| 437 \| Virulence-associated e family protein \| Virulence-associated e family protein; afdb-uniprot50 \| AF-A0A6M3J7V5-F1-MODEL\_V4 \| 1.0 \| 4.179e-23 \| 836 \| 0.266 \| 401 \| 262 \| 16 \| 370 \| 753 \| 43 \| 428 \| Putative VirE domain containing protein \| Putative VirE domain containing protein; afdb-uniprot50 \| AF-A0A078S3U0-F1-MODEL\_V4 \| 1.0 \| 1.651e-23 \| 835 \| 0.23 \| 391 \| 274 \| 12 \| 381 \| 757 \| 15 \| 392 \| Virulence-associated E family protein \| Virulence-associated E family protein; afdb-uniprot50 \| AF-A0A7Z8VJ41-F1-MODEL\_V4 \| 1.0 \| 2.003e-21 \| 833 \| 0.305 \| 311 \| 196 \| 7 \| 370 \| 665 \| 4 \| 309 \| Uncharacterized protein \| Uncharacterized protein; afdb-uniprot50 \| AF-A0A7J5V6L9-F1-MODEL\_V4 \| 1.0 \| 2.017e-26 \| 831 \| 0.221 \| 479 \| 317 \| 20 \| 297 \| 756 \| 151 \| 592 \| Virulence-associated E family protein \| Virulence-associated E family protein; afdb-uniprot50 \| AF-A0A4R5B4A1-F1-MODEL\_V4 \| 1.0 \| 7.763e-23 \| 826 \| 0.229 \| 387 \| 268 \| 17 \| 384 \| 757 \| 2 \| 371 \| Virulence-associated e family protein \| Virulence-associated e family protein; afdb-uniprot50 \| AF-A0A1H1CRS4-F1-MODEL\_V4 \| 1.0 \| 1.414e-23 \| 825 \| 0.202 \| 404 \| 282 \| 15 \| 379 \| 751 \| 21 \| 415 \| Virulence-associated protein E \| Virulence-associated protein E; afdb-uniprot50 \| AF-A0A6B2H5H0-F1-MODEL\_V4 \| 1.0 \| 1.114e-22 \| 821 \| 0.244 \| 385 \| 266 \| 15 \| 384 \| 756 \| 20 \| 391 \| Uncharacterized protein \| Uncharacterized protein; afdb-uniprot50 \| AF-A0A1I0LZP8-F1-MODEL\_V4 \| 1.0 \| 1.005e-22 \| 817 \| 0.196 \| 392 \| 292 \| 15 \| 378 \| 756 \| 181 \| 562 \| VirE N-terminal domain-containing protein \| VirE N-terminal domain-containing protein; afdb-uniprot50 \| AF-A0A3A5S119-F1-MODEL\_V4 \| 1.0 \| 8.607e-23 \| 816 \| 0.231 \| 389 \| 276 \| 12 \| 381 \| 757 \| 330 \| 707 \| VirE protein \| VirE protein; afdb-uniprot50 \| AF-A0A431MH39-F1-MODEL\_V4 \| 1.0 \| 3.969e-23 \| 815 \| 0.215 \| 395 \| 289 \| 14 \| 372 \| 757 \| 23 \| 405 \| Uncharacterized protein \| Uncharacterized protein; afdb-uniprot50 \| AF-A0A6I0LF08-F1-MODEL\_V4 \| 1.0 \| 9.062e-23 \| 815 \| 0.231 \| 388 \| 275 \| 11 \| 381 \| 756 \| 330 \| 706 \| VirE protein \| VirE protein; afdb-uniprot50 \| AF-A0A1Y0EKF6-F1-MODEL\_V4 \| 1.0 \| 1.309e-35 \| 814 \| 0.221 \| 940 \| 452 \| 49 \| 12 \| 731 \| 9 \| 888 \| Uncharacterized protein \| Uncharacterized protein; afdb-uniprot50 \| AF-A0A7K3MY39-F1-MODEL\_V4 \| 1.0 \| 9.481e-26 \| 813 \| 0.217 \| 515 \| 320 \| 22 \| 264 \| 756 \| 4 \| 457 \| Virulence-associated E family protein \| Virulence-associated E family protein; afdb-uniprot50 \| AF-A0A7D6XXE7-F1-MODEL\_V4 \| 1.0 \| 3.77e-23 \| 813 \| 0.23 \| 438 \| 284 \| 17 \| 363 \| 757 \| 76 \| 503 \| Uncharacterized protein \| Uncharacterized protein; afdb-uniprot50 \| AF-A0A7T1WPK4-F1-MODEL\_V4 \| 1.0 \| 8.442e-24 \| 813 \| 0.234 \| 426 \| 270 \| 16 \| 370 \| 751 \| 532 \| 945 \| Uncharacterized protein \| Uncharacterized protein; afdb-uniprot50 \| AF-A0A414KVM2-F1-MODEL\_V4 \| 1.0 \| 2.627e-23 \| 812 \| 0.215 \| 389 \| 276 \| 13 \| 384 \| 756 \| 100 \| 475 \| Virulence protein E \| Virulence protein E; afdb-uniprot50 \| AF-D0X1Z3-F1-MODEL\_V4 \| 1.0 \| 1.772e-22 \| 808 \| 0.271 \| 350 \| 218 \| 10 \| 435 \| 756 \| 6 \| 346 \| Uncharacterized protein \| Uncharacterized protein; afdb-uniprot50 \| AF-A0A2T4J627-F1-MODEL\_V4 \| 1.0 \| 7.325e-26 \| 808 \| 0.245 \| 518 \| 333 \| 23 \| 259 \| 752 \| 3 \| 486 \| Uncharacterized protein \| Uncharacterized protein; afdb-uniprot50 \| AF-A0A1Y6D883-F1-MODEL\_V4 \| 1.0 \| 1.251e-24 \| 808 \| 0.326 \| 361 \| 201 \| 14 \| 11 \| 347 \| 8 \| 350 \| Uncharacterized protein \| Uncharacterized protein; afdb-uniprot50 \| AF-R1ISA0-F1-MODEL\_V4 \| 1.0 \| 2.385e-28 \| 808 \| 0.217 \| 562 \| 334 \| 25 \| 260 \| 757 \| 12 \| 531 \| Virulence-associated E family protein \| Virulence-associated E family protein; afdb-uniprot50 \| AF-A0A1H0G3S0-F1-MODEL\_V4 \| 1.0 \| 6.48e-27 \| 808 \| 0.203 \| 502 \| 351 \| 20 \| 264 \| 751 \| 243 \| 709 \| Virulence-associated protein E \| Virulence-associated protein E; afdb-uniprot50 \| AF-R5JFQ8-F1-MODEL\_V4 \| 1.0 \| 7.713e-26 \| 807 \| 0.233 \| 527 \| 308 \| 19 \| 258 \| 756 \| 275 \| 733 \| Virulence-associated protein E \| Virulence-associated protein E; afdb-uniprot50 \| AF-A0A6V6ZI02-F1-MODEL\_V4 \| 1.0 \| 7.372e-23 \| 804 \| 0.259 \| 405 \| 262 \| 17 \| 370 \| 751 \| 36 \| 425 \| Putative P-loop ATPase \| Putative P-loop ATPase; afdb-uniprot50 \| AF-A0A252F7E8-F1-MODEL\_V4 \| 1.0 \| 3.008e-24 \| 804 \| 0.245 \| 488 \| 294 \| 23 \| 321 \| 769 \| 97 \| 549 \| Uncharacterized protein \| Uncharacterized protein; afdb-uniprot50 \| AF-A0A240U2T6-F1-MODEL\_V4 \| 1.0 \| 5.445e-36 \| 802 \| 0.218 \| 930 \| 480 \| 52 \| 1 \| 750 \| 10 \| 872 \| Uncharacterized protein \| Uncharacterized protein; afdb-uniprot50 \| AF-A0A3A9UXP7-F1-MODEL\_V4 \| 1.0 \| 4.487e-22 \| 800 \| 0.238 \| 385 \| 272 \| 14 \| 384 \| 759 \| 47 \| 419 \| Virulence-associated e family protein \| Virulence-associated e family protein; afdb-uniprot50 \| AF-A0A173MFG2-F1-MODEL\_V4 \| 1.0 \| 4.974e-22 \| 797 \| 0.184 \| 384 \| 289 \| 15 \| 381 \| 752 \| 79 \| 450 \| Virulence-associated protein E \| Virulence-associated protein E; afdb-uniprot50 \| AF-A0A315Z5E8-F1-MODEL\_V4 \| 1.0 \| 3.77e-23 \| 797 \| 0.221 \| 401 \| 272 \| 16 \| 382 \| 751 \| 59 \| 450 \| Virulence-associated protein E \| Virulence-associated protein E; afdb-uniprot50 \| AF-A0A3D3MX14-F1-MODEL\_V4 \| 1.0 \| 2.179e-22 \| 797 \| 0.227 \| 395 \| 272 \| 19 \| 381 \| 757 \| 309 \| 688 \| VirE\_N domain-containing protein \| VirE\_N domain-containing protein; afdb-uniprot50 \| AF-A0A1H0Y2A4-F1-MODEL\_V4 \| 1.0 \| 2.447e-24 \| 796 \| 0.241 \| 427 \| 292 \| 17 \| 339 \| 755 \| 164 \| 568 \| VirE N-terminal domain-containing protein \| VirE N-terminal domain-containing protein; afdb-uniprot50 \| AF-D4VNM0-F1-MODEL\_V4 \| 1.0 \| 4.487e-22 \| 795 \| 0.206 \| 383 \| 281 \| 16 \| 385 \| 757 \| 4 \| 373 \| Putative helicase \| Putative helicase; afdb-uniprot50 \| AF-A0A521XCL3-F1-MODEL\_V4 \| 1.0 \| 2.627e-23 \| 793 \| 0.2 \| 433 \| 301 \| 18 \| 333 \| 755 \| 327 \| 724 \| Prim-Pol domain-containing protein \| Prim-Pol domain-containing protein; afdb-uniprot50 \| AF-W1YBE3-F1-MODEL\_V4 \| 1.0 \| 9.73e-30 \| 792 \| 0.204 \| 651 \| 410 \| 41 \| 153 \| 754 \| 113 \| 704 \| Sigma 54 interacting protein \| Sigma 54 interacting protein; afdb-uniprot50 \| AF-A0A316MW33-F1-MODEL\_V4 \| 1.0 \| 3.422e-20 \| 791 \| 0.379 \| 258 \| 142 \| 8 \| 93 \| 347 \| 70 \| 312 \| Uncharacterized protein \| Uncharacterized protein; afdb-uniprot50 \| AF-A0A382QEF6-F1-MODEL\_V4 \| 1.0 \| 6.48e-19 \| 789 \| 0.372 \| 247 \| 130 \| 6 \| 102 \| 347 \| 70 \| 292 \| Toprim domain-containing protein \| Toprim domain-containing protein; afdb-uniprot50 \| AF-A0A316F066-F1-MODEL\_V4 \| 1.0 \| 2.416e-22 \| 786 \| 0.236 \| 411 \| 284 \| 17 \| 369 \| 762 \| 2 \| 399 \| Virulence-associated protein E \| Virulence-associated protein E; afdb-uniprot50 \| AF-A0A2E3H3V5-F1-MODEL\_V4 \| 1.0 \| 8.607e-23 \| 786 \| 0.267 \| 411 \| 255 \| 17 \| 370 \| 757 \| 32 \| 419 \| Virulence-associated E family protein \| Virulence-associated E family protein; afdb-uniprot50 \| AF-A0A5C7J8I3-F1-MODEL\_V4 \| 1.0 \| 2.678e-22 \| 786 \| 0.252 \| 392 \| 261 \| 15 \| 385 \| 760 \| 125 \| 500 \| Uncharacterized protein \| Uncharacterized protein; afdb-uniprot50 \| AF-A0A0F9B953-F1-MODEL\_V4 \| 1.0 \| 8.174e-23 \| 785 \| 0.266 \| 405 \| 255 \| 21 \| 370 \| 753 \| 53 \| 436 \| Uncharacterized protein \| Uncharacterized protein; afdb-uniprot50 \| AF-A0A095YT29-F1-MODEL\_V4 \| 1.0 \| 2.221e-21 \| 785 \| 0.269 \| 371 \| 215 \| 14 \| 1 \| 350 \| 1 \| 336 \| Uncharacterized protein \| Uncharacterized protein; afdb-uniprot50 \| AF-A0A7W7EQK1-F1-MODEL\_V4 \| 1.0 \| 4.288e-19 \| 784 \| 0.422 \| 251 \| 139 \| 5 \| 363 \| 609 \| 9 \| 257 \| Putative P-loop ATPase \| Putative P-loop ATPase; afdb-uniprot50 \| AF-A0A5Q0BL22-F1-MODEL\_V4 \| 1.0 \| 1.211e-23 \| 783 \| 0.335 \| 340 \| 169 \| 12 \| 1 \| 333 \| 1 \| 290 \| Toprim domain-containing protein \| Toprim domain-containing protein; afdb-uniprot50 \| AF-A0A7Z1L1I9-F1-MODEL\_V4 \| 1.0 \| 5.807e-22 \| 783 \| 0.233 \| 385 \| 275 \| 15 \| 381 \| 757 \| 15 \| 387 \| Uncharacterized protein \| Uncharacterized protein; afdb-uniprot50 \| AF-A0A1Y6CVM3-F1-MODEL\_V4 \| 1.0 \| 6.736e-25 \| 783 \| 0.263 \| 474 \| 278 \| 20 \| 2 \| 447 \| 4 \| 434 \| Uncharacterized domain associated with phage/plasmid primase \| Uncharacterized domain associated with phage/plasmid primase; afdb-uniprot50 \| AF-A0A7L5DSN9-F1-MODEL\_V4 \| 1.0 \| 4.879e-23 \| 780 \| 0.248 \| 399 \| 266 \| 16 \| 369 \| 751 \| 28 \| 408 \| Virulence protein \| Virulence protein; afdb-uniprot50 \| AF-A0A246G7T7-F1-MODEL\_V4 \| 1.0 \| 1.991e-24 \| 780 \| 0.195 \| 506 \| 308 \| 22 \| 261 \| 750 \| 28 \| 450 \| Virulence-associated e family protein \| Virulence-associated e family protein; afdb-uniprot50 \| AF-A0A0X8JSM9-F1-MODEL\_V4 \| 1.0 \| 1.878e-19 \| 780 \| 0.37 \| 251 \| 138 \| 6 \| 101 \| 349 \| 455 \| 687 \| Toprim domain-containing protein \| Toprim domain-containing protein; afdb-uniprot50 \| AF-A0A149STJ7-F1-MODEL\_V4 \| 1.0 \| 6.115e-22 \| 778 \| 0.227 \| 408 \| 265 \| 17 \| 370 \| 752 \| 3 \| 385 \| Virulence-associated E family protein \| Virulence-associated E family protein; afdb-uniprot50 \| AF-A0A2C8Z209-F1-MODEL\_V4 \| 1.0 \| 1.784e-35 \| 778 \| 0.222 \| 850 \| 461 \| 43 \| 13 \| 737 \| 5 \| 779 \| Putative DNA primase/helicase \| Putative DNA primase/helicase; afdb-uniprot50 \| AF-A0A7D7QZK8-F1-MODEL\_V4 \| 1.0 \| 4.816e-21 \| 777 \| 0.23 \| 351 \| 248 \| 12 \| 414 \| 756 \| 1 \| 337 \| Virulence-associated e family protein \| Virulence-associated e family protein; afdb-uniprot50 \| AF-A0A1F3A5U3-F1-MODEL\_V4 \| 1.0 \| 1.058e-22 \| 776 \| 0.275 \| 410 \| 248 \| 19 \| 370 \| 751 \| 18 \| 406 \| Uncharacterized protein \| Uncharacterized protein; afdb-uniprot50 \| AF-A0A6G1V777-F1-MODEL\_V4 \| 1.0 \| 1.3e-22 \| 773 \| 0.215 \| 417 \| 284 \| 16 \| 377 \| 769 \| 4 \| 401 \| Uncharacterized protein \| Uncharacterized protein; afdb-uniprot50 \| AF-A0A1I1YQB9-F1-MODEL\_V4 \| 1.0 \| 3.557e-26 \| 773 \| 0.213 \| 524 \| 319 \| 20 \| 261 \| 754 \| 263 \| 723 \| Virulence-associated protein E \| Virulence-associated protein E; afdb-uniprot50 \| AF-A0A7Z8R0X3-F1-MODEL\_V4 \| 1.0 \| 3.969e-23 \| 772 \| 0.263 \| 410 \| 260 \| 16 \| 376 \| 762 \| 5 \| 395 \| Uncharacterized protein \| Uncharacterized protein; afdb-uniprot50 \| AF-A0A357CEQ6-F1-MODEL\_V4 \| 1.0 \| 2.73e-21 \| 772 \| 0.255 \| 392 \| 258 \| 19 \| 377 \| 753 \| 63 \| 435 \| Virulence-associated E family protein \| Virulence-associated E family protein; afdb-uniprot50 \| AF-A0A2R4KET0-F1-MODEL\_V4 \| 1.0 \| 1.235e-22 \| 772 \| 0.217 \| 405 \| 287 \| 17 \| 370 \| 756 \| 39 \| 431 \| Virulence protein \| Virulence protein; afdb-uniprot50 \| AF-A0A8B3TM54-F1-MODEL\_V4 \| 1.0 \| 4.207e-28 \| 771 \| 0.23 \| 624 \| 345 \| 28 \| 183 \| 752 \| 76 \| 618 \| Uncharacterized protein \| Uncharacterized protein; afdb-uniprot50 \| AF-A0A4Q6JDT2-F1-MODEL\_V4 \| 1.0 \| 1.276e-23 \| 770 \| 0.315 \| 364 \| 188 \| 13 \| 3 \| 324 \| 1 \| 345 \| Toprim domain-containing protein \| Toprim domain-containing protein; afdb-uniprot50 \| AF-A0A4R5Q8H9-F1-MODEL\_V4 \| 1.0 \| 2.593e-21 \| 770 \| 0.24 \| 400 \| 271 \| 18 \| 370 \| 752 \| 319 \| 702 \| Uncharacterized protein \| Uncharacterized protein; afdb-uniprot50 \| AF-A0A7X6HWL2-F1-MODEL\_V4 \| 1.0 \| 1.952e-25 \| 770 \| 0.23 \| 537 \| 328 \| 24 \| 261 \| 762 \| 261 \| 747 \| PriCT\_1 domain-containing protein \| PriCT\_1 domain-containing protein; afdb-uniprot50 \| AF-A0A7X7UY95-F1-MODEL\_V4 \| 1.0 \| 1.251e-24 \| 767 \| 0.203 \| 482 \| 335 \| 21 \| 297 \| 762 \| 257 \| 705 \| Prim-Pol domain-containing protein \| Prim-Pol domain-containing protein; afdb-uniprot50 \| AF-A0A1Z5HBM1-F1-MODEL\_V4 \| 1.0 \| 1.219e-28 \| 766 \| 0.178 \| 777 \| 339 \| 15 \| 1 \| 762 \| 252 \| 743 \| Uncharacterized protein \| Uncharacterized protein; afdb-uniprot50 \| AF-A0A847QTI9-F1-MODEL\_V4 \| 1.0 \| 2.495e-23 \| 765 \| 0.197 \| 496 \| 337 \| 22 \| 292 \| 757 \| 279 \| 743 \| Uncharacterized protein \| Uncharacterized protein; afdb-uniprot50 \| AF-A0A2E4PDU5-F1-MODEL\_V4 \| 1.0 \| 8.333e-22 \| 764 \| 0.223 \| 426 \| 269 \| 16 \| 381 \| 759 \| 4 \| 414 \| Virulence-associated e family protein \| Virulence-associated e family protein; afdb-uniprot50 \| AF-A0A3M3TAF3-F1-MODEL\_V4 \| 1.0 \| 1.024e-29 \| 764 \| 0.308 \| 509 \| 304 \| 25 \| 102 \| 594 \| 78 \| 554 \| DNA primase domain protein \| DNA primase domain protein; afdb-uniprot50 \| AF-A0A1M4N4I5-F1-MODEL\_V4 \| 1.0 \| 2.179e-22 \| 763 \| 0.262 \| 412 \| 249 \| 20 \| 370 \| 753 \| 49 \| 433 \| Death domain-containing protein \| Death domain-containing protein; afdb-uniprot50 \| AF-A0A7Z9GAF4-F1-MODEL\_V4 \| 1.0 \| 7.516e-22 \| 760 \| 0.235 \| 404 \| 263 \| 19 \| 370 \| 753 \| 308 \| 685 \| Uncharacterized protein \| Uncharacterized protein; afdb-uniprot50 \| AF-A0A641RMU6-F1-MODEL\_V4 \| 1.0 \| 2.109e-21 \| 758 \| 0.241 \| 360 \| 244 \| 12 \| 408 \| 755 \| 5 \| 347 \| Virulence protein E \| Virulence protein E; afdb-uniprot50 \| AF-A0A2E7IPX4-F1-MODEL\_V4 \| 1.0 \| 7.614e-24 \| 758 \| 0.239 \| 493 \| 304 \| 26 \| 317 \| 762 \| 1 \| 469 \| Uncharacterized protein \| Uncharacterized protein; afdb-uniprot50 \| AF-A0A5C7NHB2-F1-MODEL\_V4 \| 1.0 \| 2.875e-29 \| 758 \| 0.253 \| 671 \| 334 \| 34 \| 8 \| 563 \| 2 \| 620 \| DUF927 domain-containing protein \| DUF927 domain-containing protein; afdb-uniprot50 \| AF-A0A645B275-F1-MODEL\_V4 \| 1.0 \| 2.73e-21 \| 757 \| 0.21 \| 399 \| 282 \| 17 \| 375 \| 755 \| 53 \| 436 \| DUF3874 domain-containing protein \| DUF3874 domain-containing protein; afdb-uniprot50 \| AF-A0A7V9SQL9-F1-MODEL\_V4 \| 1.0 \| 1.903e-21 \| 757 \| 0.221 \| 401 \| 280 \| 16 \| 369 \| 754 \| 37 \| 420 \| Virulence protein \| Virulence protein; afdb-uniprot50 \| AF-A0A2G5L038-F1-MODEL\_V4 \| 1.0 \| 2.73e-21 \| 755 \| 0.203 \| 394 \| 286 \| 15 \| 381 \| 757 \| 19 \| 401 \| Uncharacterized protein \| Uncharacterized protein; afdb-uniprot50 \| AF-Q1QLE7-F1-MODEL\_V4 \| 1.0 \| 5.34e-21 \| 755 \| 0.233 \| 394 \| 268 \| 14 \| 370 \| 752 \| 317 \| 687 \| Virulence-associated E \| Virulence-associated E; afdb-uniprot50 \| AF-A0A414PMJ6-F1-MODEL\_V4 \| 1.0 \| 2.003e-21 \| 753 \| 0.19 \| 393 \| 292 \| 15 \| 378 \| 757 \| 7 \| 386 \| Virulence protein E \| Virulence protein E; afdb-uniprot50 \| AF-A0A2E2PC01-F1-MODEL\_V4 \| 1.0 \| 2.339e-21 \| 751 \| 0.215 \| 380 \| 280 \| 13 \| 379 \| 750 \| 2 \| 371 \| Uncharacterized protein \| Uncharacterized protein; afdb-uniprot50 \| AF-A0A1I4P1M4-F1-MODEL\_V4 \| 1.0 \| 2.25e-23 \| 750 \| 0.237 \| 468 \| 304 \| 18 \| 329 \| 755 \| 307 \| 762 \| Predicted P-loop ATPase and inactivated derivatives \| Predicted P-loop ATPase and inactivated derivatives; afdb-uniprot50 \| AF-A0A2Z6UA34-F1-MODEL\_V4 \| 1.0 \| 2.221e-21 \| 749 \| 0.288 \| 371 \| 199 \| 14 \| 1 \| 323 \| 1 \| 354 \| PriCT\_2 domain-containing protein \| PriCT\_2 domain-containing protein; afdb-uniprot50 \| AF-A0A1F3KBL0-F1-MODEL\_V4 \| 1.0 \| 1.716e-21 \| 749 \| 0.237 \| 388 \| 271 \| 16 \| 381 \| 755 \| 104 \| 479 \| Uncharacterized protein \| Uncharacterized protein; afdb-uniprot50 \| AF-A0A3B9IYP5-F1-MODEL\_V4 \| 1.0 \| 7.093e-25 \| 749 \| 0.226 \| 534 \| 319 \| 26 \| 263 \| 757 \| 243 \| 721 \| Uncharacterized protein \| Uncharacterized protein; afdb-uniprot50 \| AF-A0A1Q3GLM4-F1-MODEL\_V4 \| 1.0 \| 7.138e-22 \| 748 \| 0.228 \| 390 \| 266 \| 14 \| 381 \| 751 \| 43 \| 416 \| Uncharacterized protein \| Uncharacterized protein; afdb-uniprot50 \| AF-A0A6F8PR07-F1-MODEL\_V4 \| 1.0 \| 1.442e-22 \| 743 \| 0.303 \| 362 \| 216 \| 11 \| 1 \| 346 \| 3 \| 344 \| Uncharacterized protein \| Uncharacterized protein; afdb-uniprot50 \| AF-A0A350YX52-F1-MODEL\_V4 \| 1.0 \| 4.663e-20 \| 742 \| 0.236 \| 377 \| 260 \| 12 \| 380 \| 748 \| 7 \| 363 \| Uncharacterized protein \| Uncharacterized protein; afdb-uniprot50 \| AF-A0A7C5FUS1-F1-MODEL\_V4 \| 1.0 \| 6.564e-21 \| 741 \| 0.251 \| 418 \| 270 \| 19 \| 370 \| 769 \| 70 \| 462 \| Virulence-associated E family protein \| Virulence-associated E family protein; afdb-uniprot50 \| AF-A0A7W6EUR7-F1-MODEL\_V4 \| 1.0 \| 7.184e-19 \| 740 \| 0.38 \| 260 \| 149 \| 4 \| 358 \| 614 \| 24 \| 274 \| Putative P-loop ATPase \| Putative P-loop ATPase; afdb-uniprot50 \| AF-A0A7T9JLM2-F1-MODEL\_V4 \| 1.0 \| 2.73e-21 \| 740 \| 0.221 \| 398 \| 270 \| 18 \| 381 \| 756 \| 27 \| 406 \| Virulence-associated E family protein \| Virulence-associated E family protein; afdb-uniprot50 \| AF-A0A512HAX8-F1-MODEL\_V4 \| 1.0 \| 6.912e-21 \| 740 \| 0.223 \| 408 \| 274 \| 15 \| 370 \| 752 \| 40 \| 429 \| Uncharacterized protein \| Uncharacterized protein; afdb-uniprot50 \| AF-A0A4S0QEA6-F1-MODEL\_V4 \| 1.0 \| 1.079e-21 \| 739 \| 0.261 \| 409 \| 259 \| 17 \| 378 \| 753 \| 25 \| 423 \| Uncharacterized protein \| Uncharacterized protein; afdb-uniprot50 \| AF-A0A0Q5HQG4-F1-MODEL\_V4 \| 1.0 \| 3.25e-20 \| 738 \| 0.203 \| 407 \| 282 \| 20 \| 370 \| 755 \| 1 \| 386 \| Uncharacterized protein \| Uncharacterized protein; afdb-uniprot50 \| AF-A0A078BCE6-F1-MODEL\_V4 \| 1.0 \| 4.234e-33 \| 738 \| 0.214 \| 855 \| 471 \| 47 \| 1 \| 737 \| 1 \| 772 \| Primase C 2 (PriCT-2) family \| Primase C 2 (PriCT-2) family; afdb-uniprot50 \| AF-A0A2W6XRY1-F1-MODEL\_V4 \| 1.0 \| 1.158e-36 \| 738 \| 0.214 \| 1029 \| 451 \| 58 \| 12 \| 751 \| 13 \| 972 \| Uncharacterized protein \| Uncharacterized protein; afdb-uniprot50 \| AF-E4MB64-F1-MODEL\_V4 \| 1.0 \| 2.109e-21 \| 737 \| 0.227 \| 400 \| 278 \| 19 \| 369 \| 756 \| 13 \| 393 \| Uncharacterized protein \| Uncharacterized protein; afdb-uniprot50 \| AF-A0A1Y2RF59-F1-MODEL\_V4 \| 1.0 \| 8.775e-22 \| 737 \| 0.231 \| 402 \| 282 \| 16 \| 370 \| 757 \| 26 \| 414 \| Uncharacterized protein \| Uncharacterized protein; afdb-uniprot50 \| AF-A0A2T1HTE2-F1-MODEL\_V4 \| 1.0 \| 9.239e-22 \| 737 \| 0.257 \| 416 \| 263 \| 16 \| 366 \| 762 \| 36 \| 424 \| Virulence-associated E family protein \| Virulence-associated E family protein; afdb-uniprot50 \| AF-A0A528UG59-F1-MODEL\_V4 \| 1.0 \| 8.496e-21 \| 736 \| 0.237 \| 387 \| 268 \| 14 \| 379 \| 751 \| 41 \| 414 \| Virulence protein E \| Virulence protein E; afdb-uniprot50 \| AF-A0A318SE39-F1-MODEL\_V4 \| 1.0 \| 4.344e-21 \| 735 \| 0.353 \| 297 \| 165 \| 12 \| 26 \| 313 \| 1 \| 279 \| Toprim domain-containing protein \| Toprim domain-containing protein; afdb-uniprot50 \| AF-A0A7D4NP15-F1-MODEL\_V4 \| 1.0 \| 6.912e-21 \| 734 \| 0.299 \| 364 \| 198 \| 15 \| 3 \| 334 \| 2 \| 340 \| PriCT-2 domain-containing protein \| PriCT-2 domain-containing protein; afdb-uniprot50 \| AF-A0A1M3LBF8-F1-MODEL\_V4 \| 1.0 \| 2.593e-21 \| 734 \| 0.219 \| 406 \| 282 \| 19 \| 370 \| 756 \| 4 \| 393 \| Virulence protein \| Virulence protein; afdb-uniprot50 \| AF-A0A1M6SXZ2-F1-MODEL\_V4 \| 1.0 \| 1.024e-21 \| 734 \| 0.209 \| 387 \| 276 \| 16 \| 381 \| 751 \| 69 \| 441 \| Virulence-associated protein E \| Virulence-associated protein E; afdb-uniprot50 \| AF-A0A836NRY6-F1-MODEL\_V4 \| 1.0 \| 7.093e-25 \| 734 \| 0.292 \| 404 \| 249 \| 18 \| 102 \| 489 \| 78 \| 460 \| Conjugal transfer protein TraC \| Conjugal transfer protein TraC; afdb-uniprot50 \| AF-A0A419H4F3-F1-MODEL\_V4 \| 1.0 \| 2.207e-24 \| 734 \| 0.193 \| 528 \| 349 \| 19 \| 239 \| 751 \| 42 \| 507 \| Uncharacterized protein \| Uncharacterized protein; afdb-uniprot50 \| AF-A0A2A5B9Y8-F1-MODEL\_V4 \| 1.0 \| 3.167e-24 \| 734 \| 0.201 \| 506 \| 319 \| 23 \| 298 \| 768 \| 256 \| 711 \| Uncharacterized protein \| Uncharacterized protein; afdb-uniprot50 \| AF-A0A7W6CYB3-F1-MODEL\_V4 \| 1.0 \| 1.219e-20 \| 733 \| 0.216 \| 411 \| 274 \| 17 \| 370 \| 752 \| 336 \| 726 \| Putative P-loop ATPase \| Putative P-loop ATPase; afdb-uniprot50 \| AF-A0A558BMX0-F1-MODEL\_V4 \| 1.0 \| 5.92e-21 \| 732 \| 0.224 \| 387 \| 263 \| 15 \| 384 \| 751 \| 32 \| 400 \| Uncharacterized protein \| Uncharacterized protein; afdb-uniprot50 \| AF-A0A845B9K8-F1-MODEL\_V4 \| 1.0 \| 6.115e-22 \| 730 \| 0.217 \| 459 \| 313 \| 20 \| 316 \| 753 \| 18 \| 451 \| Virulence-associated E family protein \| Virulence-associated E family protein; afdb-uniprot50 \| AF-A0A0T7AN15-F1-MODEL\_V4 \| 1.0 \| 1.079e-21 \| 729 \| 0.222 \| 423 \| 295 \| 14 \| 370 \| 767 \| 62 \| 475 \| Virulence-associated E family protein \| Virulence-associated E family protein; afdb-uniprot50 \| AF-A0A7D4FWX9-F1-MODEL\_V4 \| 1.0 \| 7.864e-25 \| 729 \| 0.205 \| 530 \| 324 \| 18 \| 239 \| 757 \| 255 \| 698 \| Virulence protein E \| Virulence protein E; afdb-uniprot50 \| AF-A0A150TWT5-F1-MODEL\_V4 \| 1.0 \| 1.47e-21 \| 729 \| 0.23 \| 439 \| 277 \| 19 \| 370 \| 762 \| 423 \| 846 \| Uncharacterized protein \| Uncharacterized protein; afdb-uniprot50 \| AF-A0A8B2CHX5-F1-MODEL\_V4 \| 1.0 \| 5.071e-21 \| 728 \| 0.18 \| 398 \| 297 \| 15 \| 384 \| 762 \| 27 \| 414 \| DUF3874 domain-containing protein \| DUF3874 domain-containing protein; afdb-uniprot50 \| AF-A0A126ZEG5-F1-MODEL\_V4 \| 1.0 \| 1.432e-17 \| 726 \| 0.372 \| 228 \| 118 \| 8 \| 113 \| 338 \| 2 \| 206 \| Toprim domain-containing protein \| Toprim domain-containing protein; afdb-uniprot50 \| AF-A0A2R3MUY2-F1-MODEL\_V4 \| 1.0 \| 7.278e-21 \| 726 \| 0.211 \| 407 \| 286 \| 21 \| 371 \| 757 \| 1 \| 392 \| Virulence-associated protein E \| Virulence-associated protein E; afdb-uniprot50 \| AF-R6WKB3-F1-MODEL\_V4 \| 1.0 \| 1.065e-19 \| 725 \| 0.25 \| 311 \| 216 \| 12 \| 453 \| 756 \| 25 \| 325 \| VirE N-terminal domain protein \| VirE N-terminal domain protein; afdb-uniprot50 \| AF-A0A1I5LS54-F1-MODEL\_V4 \| 1.0 \| 1.423e-20 \| 725 \| 0.21 \| 394 \| 278 \| 17 \| 381 \| 757 \| 237 \| 614 \| VirE N-terminal domain-containing protein \| VirE N-terminal domain-containing protein; afdb-uniprot50 \| AF-A0A1Y1QLP3-F1-MODEL\_V4 \| 1.0 \| 2.748e-18 \| 724 \| 0.405 \| 237 \| 121 \| 8 \| 100 \| 333 \| 75 \| 294 \| Toprim domain-containing protein \| Toprim domain-containing protein; afdb-uniprot50 \| AF-A0A1M7DJB2-F1-MODEL\_V4 \| 1.0 \| 1.211e-23 \| 724 \| 0.191 \| 497 \| 324 \| 22 \| 327 \| 766 \| 271 \| 746 \| Virulence-associated protein E \| Virulence-associated protein E; afdb-uniprot50 \| AF-A0A3C0IVH4-F1-MODEL\_V4 \| 1.0 \| 4.847e-18 \| 723 \| 0.294 \| 251 \| 169 \| 6 \| 453 \| 701 \| 15 \| 259 \| Virulence-associated E family protein \| Virulence-associated E family protein; afdb-uniprot50 \| AF-G8UM15-F1-MODEL\_V4 \| 1.0 \| 2.109e-21 \| 723 \| 0.231 \| 414 \| 268 \| 18 \| 379 \| 756 \| 22 \| 421 \| Virulence-associated protein E \| Virulence-associated protein E; afdb-uniprot50 \| AF-A0A2D0MXJ1-F1-MODEL\_V4 \| 1.0 \| 2.511e-20 \| 723 \| 0.206 \| 388 \| 289 \| 9 \| 381 \| 755 \| 27 \| 408 \| Virulence protein \| Virulence protein; afdb-uniprot50 \| AF-A0A1Y6ELX1-F1-MODEL\_V4 \| 1.0 \| 3.047e-26 \| 723 \| 0.194 \| 608 \| 416 \| 31 \| 167 \| 752 \| 26 \| 581 \| Virulence-associated protein E \| Virulence-associated protein E; afdb-uniprot50 \| AF-A0A4Z1A268-F1-MODEL\_V4 \| 1.0 \| 1.065e-27 \| 723 \| 0.188 \| 663 \| 433 \| 36 \| 153 \| 759 \| 83 \| 696 \| Uncharacterized protein \| Uncharacterized protein; afdb-uniprot50 \| AF-A0A5C6JXI2-F1-MODEL\_V4 \| 1.0 \| 1.499e-20 \| 722 \| 0.173 \| 387 \| 292 \| 15 \| 381 \| 755 \| 13 \| 383 \| Virulence-associated protein E \| Virulence-associated protein E; afdb-uniprot50 \| AF-A0A2N2YSH7-F1-MODEL\_V4 \| 1.0 \| 2.339e-21 \| 722 \| 0.189 \| 416 \| 298 \| 13 \| 370 \| 755 \| 13 \| 419 \| Virulence protein \| Virulence protein; afdb-uniprot50 \| AF-A0A373NTM4-F1-MODEL\_V4 \| 1.0 \| 4.344e-21 \| 722 \| 0.231 \| 428 \| 286 \| 19 \| 370 \| 769 \| 15 \| 427 \| Virulence-associated protein E \| Virulence-associated protein E; afdb-uniprot50 \| AF-A0A094ZFQ5-F1-MODEL\_V4 \| 1.0 \| 2.193e-27 \| 722 \| 0.199 \| 628 \| 414 \| 32 \| 170 \| 752 \| 89 \| 672 \| Uncharacterized protein \| Uncharacterized protein; afdb-uniprot50 \| AF-A0A1T5CXN1-F1-MODEL\_V4 \| 1.0 \| 1.018e-24 \| 722 \| 0.197 \| 538 \| 341 \| 23 \| 261 \| 768 \| 366 \| 842 \| Uncharacterized protein \| Uncharacterized protein; afdb-uniprot50 \| AF-A0A177N064-F1-MODEL\_V4 \| 1.0 \| 1.479e-18 \| 721 \| 0.356 \| 250 \| 138 \| 6 \| 92 \| 332 \| 68 \| 303 \| Toprim domain-containing protein \| Toprim domain-containing protein; afdb-uniprot50 \| AF-A0A0E4FNC5-F1-MODEL\_V4 \| 1.0 \| 4.974e-22 \| 721 \| 0.23 \| 404 \| 280 \| 19 \| 370 \| 756 \| 4 \| 393 \| Virulence-associated protein E \| Virulence-associated protein E; afdb-uniprot50 \| AF-A0A413DD07-F1-MODEL\_V4 \| 1.0 \| 2.784e-20 \| 721 \| 0.181 \| 386 \| 290 \| 14 \| 384 \| 755 \| 237 \| 610 \| Helicase \| Helicase; afdb-uniprot50 \| AF-A0A1I5I2J5-F1-MODEL\_V4 \| 1.0 \| 3.511e-24 \| 721 \| 0.189 \| 516 \| 341 \| 21 \| 261 \| 756 \| 244 \| 702 \| Virulence-associated protein E \| Virulence-associated protein E; afdb-uniprot50 \| AF-A0A3E5DTG1-F1-MODEL\_V4 \| 1.0 \| 2.384e-20 \| 720 \| 0.191 \| 381 \| 283 \| 13 \| 385 \| 751 \| 2 \| 371 \| Helicase \| Helicase; afdb-uniprot50 \| AF-A0A2N7R2N6-F1-MODEL\_V4 \| 1.0 \| 3.4e-23 \| 720 \| 0.283 \| 370 \| 226 \| 16 \| 118 \| 471 \| 6 \| 352 \| DNA primase TraC \| DNA primase TraC; afdb-uniprot50 \| AF-T0GQL6-F1-MODEL\_V4 \| 1.0 \| 1.326e-21 \| 720 \| 0.26 \| 407 \| 268 \| 22 \| 370 \| 762 \| 15 \| 402 \| Uncharacterized protein \| Uncharacterized protein; afdb-uniprot50 \| AF-A0A1G0X7P3-F1-MODEL\_V4 \| 1.0 \| 2.003e-21 \| 720 \| 0.188 \| 484 \| 330 \| 24 \| 316 \| 766 \| 6 \| 459 \| Uncharacterized protein \| Uncharacterized protein; afdb-uniprot50 \| AF-A0A5F0UFE8-F1-MODEL\_V4 \| 1.0 \| 5.17e-20 \| 719 \| 0.201 \| 392 \| 272 \| 22 \| 384 \| 756 \| 4 \| 373 \| Helicase \| Helicase; afdb-uniprot50 \| AF-A0A7C4P9F5-F1-MODEL\_V4 \| 1.0 \| 3.229e-23 \| 719 \| 0.213 \| 486 \| 305 \| 24 \| 287 \| 757 \| 280 \| 703 \| Uncharacterized protein \| Uncharacterized protein; afdb-uniprot50 \| AF-A0A847L830-F1-MODEL\_V4 \| 1.0 \| 5.071e-21 \| 718 \| 0.212 \| 405 \| 293 \| 13 \| 367 \| 757 \| 69 \| 461 \| Uncharacterized protein \| Uncharacterized protein; afdb-uniprot50 \| AF-A0A644UL52-F1-MODEL\_V4 \| 1.0 \| 5.997e-23 \| 718 \| 0.2 \| 480 \| 339 \| 17 \| 291 \| 755 \| 217 \| 666 \| VirE\_N domain-containing protein \| VirE\_N domain-containing protein; afdb-uniprot50 \| AF-Q01RE1-F1-MODEL\_V4 \| 1.0 \| 1.181e-19 \| 716 \| 0.225 \| 408 \| 269 \| 17 \| 370 \| 755 \| 1 \| 383 \| Virulence-associated E family protein \| Virulence-associated E family protein; afdb-uniprot50 \| AF-A0A0F5JLY6-F1-MODEL\_V4 \| 1.0 \| 4.099e-24 \| 716 \| 0.194 \| 530 \| 335 \| 25 \| 243 \| 755 \| 212 \| 666 \| Uncharacterized protein \| Uncharacterized protein; afdb-uniprot50 \| AF-A0A2S5R1A1-F1-MODEL\_V4 \| 1.0 \| 1.672e-17 \| 716 \| 0.363 \| 231 \| 129 \| 6 \| 118 \| 347 \| 503 \| 716 \| Toprim domain-containing protein \| Toprim domain-containing protein; afdb-uniprot50 \| AF-A0A1G8MPZ8-F1-MODEL\_V4 \| 1.0 \| 5.844e-19 \| 715 \| 0.284 \| 330 \| 211 \| 12 \| 370 \| 684 \| 58 \| 377 \| Virulence-associated protein E \| Virulence-associated protein E; afdb-uniprot50 \| AF-B6VV90-F1-MODEL\_V4 \| 1.0 \| 1.75e-20 \| 715 \| 0.175 \| 387 \| 291 \| 16 \| 381 \| 755 \| 70 \| 440 \| DUF3874 domain-containing protein \| DUF3874 domain-containing protein; afdb-uniprot50 \| AF-R6W8X3-F1-MODEL\_V4 \| 1.0 \| 2.384e-20 \| 713 \| 0.206 \| 392 \| 283 \| 15 \| 381 \| 757 \| 57 \| 435 \| DUF3874 domain-containing protein \| DUF3874 domain-containing protein; afdb-uniprot50 \| AF-B7BC91-F1-MODEL\_V4 \| 1.0 \| 2.678e-22 \| 712 \| 0.22 \| 454 \| 302 \| 23 \| 318 \| 757 \| 161 \| 576 \| VirE N-terminal domain protein \| VirE N-terminal domain protein; afdb-uniprot50 \| AF-A0A257L5C6-F1-MODEL\_V4 \| 1.0 \| 2.931e-20 \| 711 \| 0.217 \| 400 \| 273 \| 18 \| 371 \| 756 \| 6 \| 379 \| Virulence protein E \| Virulence protein E; afdb-uniprot50 \| AF-A0A2G4RGB3-F1-MODEL\_V4 \| 1.0 \| 3.58e-23 \| 711 \| 0.197 \| 511 \| 352 \| 22 \| 268 \| 752 \| 20 \| 498 \| Virulence-associated E family protein \| Virulence-associated E family protein; afdb-uniprot50 \| AF-A0A442KKS8-F1-MODEL\_V4 \| 1.0 \| 8.388e-27 \| 711 \| 0.173 \| 644 \| 426 \| 33 \| 167 \| 752 \| 19 \| 613 \| Uncharacterized protein \| Uncharacterized protein; afdb-uniprot50 \| AF-A0A0J6CHU3-F1-MODEL\_V4 \| 1.0 \| 6.194e-24 \| 711 \| 0.224 \| 513 \| 304 \| 23 \| 261 \| 755 \| 249 \| 685 \| Uncharacterized protein \| Uncharacterized protein; afdb-uniprot50 \| AF-A0A497G7H7-F1-MODEL\_V4 \| 1.0 \| 8.497e-29 \| 711 \| 0.192 \| 783 \| 379 \| 26 \| 2 \| 752 \| 193 \| 753 \| PriCT\_2 domain-containing protein \| PriCT\_2 domain-containing protein; afdb-uniprot50 \| AF-A0A413HE39-F1-MODEL\_V4 \| 1.0 \| 4.724e-22 \| 710 \| 0.199 \| 466 \| 315 \| 18 \| 328 \| 769 \| 22 \| 453 \| Uncharacterized protein \| Uncharacterized protein; afdb-uniprot50 \| AF-R6SXJ8-F1-MODEL\_V4 \| 1.0 \| 2.416e-22 \| 710 \| 0.187 \| 443 \| 313 \| 22 \| 330 \| 757 \| 54 \| 464 \| DUF3874 domain-containing protein \| DUF3874 domain-containing protein; afdb-uniprot50 \| AF-U4QVN3-F1-MODEL\_V4 \| 1.0 \| 2.431e-27 \| 710 \| 0.223 \| 621 \| 401 \| 36 \| 177 \| 762 \| 111 \| 684 \| Putative virulence-associated protein E \| Putative virulence-associated protein E; afdb-uniprot50 \| AF-A0A7Y9W3V7-F1-MODEL\_V4 \| 1.0 \| 9.605e-28 \| 710 \| 0.208 \| 738 \| 380 \| 28 \| 216 \| 769 \| 2 \| 719 \| Putative P-loop ATPase \| Putative P-loop ATPase; afdb-uniprot50 \| AF-A0A7X7X0D4-F1-MODEL\_V4 \| 1.0 \| 1.75e-20 \| 708 \| 0.22 \| 385 \| 275 \| 16 \| 381 \| 755 \| 23 \| 392 \| DUF3874 domain-containing protein \| DUF3874 domain-containing protein; afdb-uniprot50 \| AF-A0A255T6X0-F1-MODEL\_V4 \| 1.0 \| 1.842e-20 \| 708 \| 0.194 \| 390 \| 279 \| 14 \| 384 \| 753 \| 244 \| 618 \| Uncharacterized protein \| Uncharacterized protein; afdb-uniprot50 \| AF-G1UM92-F1-MODEL\_V4 \| 1.0 \| 5.34e-21 \| 707 \| 0.22 \| 430 \| 278 \| 21 \| 384 \| 769 \| 2 \| 418 \| Uncharacterized protein \| Uncharacterized protein; afdb-uniprot50 \| AF-A0A659V3T7-F1-MODEL\_V4 \| 1.0 \| 1.158e-20 \| 707 \| 0.251 \| 410 \| 265 \| 18 \| 381 \| 766 \| 70 \| 461 \| Uncharacterized protein \| Uncharacterized protein; afdb-uniprot50 \| AF-Q5P8W9-F1-MODEL\_V4 \| 1.0 \| 3.626e-17 \| 707 \| 0.327 \| 247 \| 140 \| 9 \| 102 \| 347 \| 77 \| 298 \| Toprim domain-containing protein \| Toprim domain-containing protein; afdb-uniprot50 \| AF-A0A7Y8NQQ4-F1-MODEL\_V4 \| 1.0 \| 9.18e-25 \| 707 \| 0.189 \| 550 \| 347 \| 24 \| 263 \| 755 \| 231 \| 738 \| Primase C-terminal domain-containing protein \| Primase C-terminal domain-containing protein; afdb-uniprot50 \| AF-U6RKE9-F1-MODEL\_V4 \| 1.0 \| 1.423e-20 \| 706 \| 0.209 \| 386 \| 278 \| 19 \| 381 \| 755 \| 35 \| 404 \| DUF3874 domain-containing protein \| DUF3874 domain-containing protein; afdb-uniprot50 \| AF-A0A174QPU8-F1-MODEL\_V4 \| 1.0 \| 1.738e-23 \| 706 \| 0.165 \| 508 \| 347 \| 23 \| 261 \| 756 \| 237 \| 679 \| Predicted P-loop ATPase and inactivated derivatives \| Predicted P-loop ATPase and inactivated derivatives; afdb-uniprot50 \| AF-A0A255TQL6-F1-MODEL\_V4 \| 1.0 \| 4.943e-25 \| 706 \| 0.212 \| 545 \| 344 \| 28 \| 260 \| 769 \| 214 \| 708 \| VirE\_N domain-containing protein \| VirE\_N domain-containing protein; afdb-uniprot50 \| AF-A0A6M0S7I4-F1-MODEL\_V4 \| 1.0 \| 1.1e-20 \| 705 \| 0.276 \| 401 \| 257 \| 16 \| 369 \| 757 \| 213 \| 592 \| Uncharacterized protein \| Uncharacterized protein; afdb-uniprot50 \| AF-A0A3E5A259-F1-MODEL\_V4 \| 1.0 \| 1.423e-20 \| 704 \| 0.193 \| 424 \| 291 \| 21 \| 370 \| 755 \| 17 \| 427 \| Uncharacterized protein \| Uncharacterized protein; afdb-uniprot50 \| AF-A0A255TBA5-F1-MODEL\_V4 \| 1.0 \| 9.359e-24 \| 704 \| 0.177 \| 530 \| 341 \| 24 \| 263 \| 769 \| 245 \| 702 \| VirE\_N domain-containing protein \| VirE\_N domain-containing protein; afdb-uniprot50 \| AF-A0A5B0WFK2-F1-MODEL\_V4 \| 1.0 \| 1.662e-20 \| 702 \| 0.218 \| 411 \| 288 \| 19 \| 370 \| 762 \| 27 \| 422 \| Uncharacterized protein \| Uncharacterized protein; afdb-uniprot50 \| AF-E6SUT3-F1-MODEL\_V4 \| 1.0 \| 5.137e-23 \| 702 \| 0.18 \| 505 \| 320 \| 18 \| 264 \| 756 \| 273 \| 695 \| VirE\_N domain-containing protein \| VirE\_N domain-containing protein; afdb-uniprot50 \| AF-A0A840L725-F1-MODEL\_V4 \| 1.0 \| 6.275e-34 \| 702 \| 0.214 \| 935 \| 454 \| 53 \| 1 \| 740 \| 1 \| 849 \| Putative DNA primase/helicase \| Putative DNA primase/helicase; afdb-uniprot50 \| AF-A0A0Q7CC84-F1-MODEL\_V4 \| 1.0 \| 5.769e-17 \| 701 \| 0.375 \| 237 \| 121 \| 8 \| 107 \| 340 \| 4 \| 216 \| Toprim domain-containing protein \| Toprim domain-containing protein; afdb-uniprot50 \| AF-A0A7T0JTE9-F1-MODEL\_V4 \| 1.0 \| 2.123e-18 \| 701 \| 0.392 \| 247 \| 119 \| 11 \| 99 \| 333 \| 78 \| 305 \| Toprim domain-containing protein \| Toprim domain-containing protein; afdb-uniprot50 \| AF-R7J768-F1-MODEL\_V4 \| 1.0 \| 1.219e-20 \| 701 \| 0.195 \| 388 \| 283 \| 16 \| 381 \| 755 \| 45 \| 416 \| DUF3874 domain-containing protein \| DUF3874 domain-containing protein; afdb-uniprot50 \| AF-A0A3C1P7C6-F1-MODEL\_V4 \| 1.0 \| 2.384e-20 \| 701 \| 0.239 \| 426 \| 277 \| 19 \| 344 \| 750 \| 36 \| 433 \| Uncharacterized protein \| Uncharacterized protein; afdb-uniprot50 \| AF-A0A0F5JAQ5-F1-MODEL\_V4 \| 1.0 \| 4.206e-20 \| 701 \| 0.199 \| 391 \| 274 \| 22 \| 384 \| 756 \| 83 \| 452 \| DUF3874 domain-containing protein \| DUF3874 domain-containing protein; afdb-uniprot50 \| AF-A0A158A108-F1-MODEL\_V4 \| 1.0 \| 1.538e-32 \| 701 \| 0.192 \| 981 \| 474 \| 61 \| 3 \| 751 \| 1 \| 894 \| Inner membrane protein \| Inner membrane protein; afdb-uniprot50 \| AF-A0A1Y4HDZ0-F1-MODEL\_V4 \| 1.0 \| 3.086e-20 \| 700 \| 0.198 \| 388 \| 280 \| 17 \| 381 \| 752 \| 33 \| 405 \| DUF3874 domain-containing protein \| DUF3874 domain-containing protein; afdb-uniprot50 \| AF-A0A2U3I7B7-F1-MODEL\_V4 \| 1.0 \| 5.444e-20 \| 699 \| 0.221 \| 411 \| 266 \| 22 \| 382 \| 769 \| 37 \| 416 \| Virulence protein E \| Virulence protein E; afdb-uniprot50 \| AF-A0A646HEP4-F1-MODEL\_V4 \| 1.0 \| 5.444e-20 \| 699 \| 0.183 \| 386 \| 288 \| 14 \| 381 \| 751 \| 139 \| 512 \| Helicase \| Helicase; afdb-uniprot50 \| AF-A0A2N7VR70-F1-MODEL\_V4 \| 1.0 \| 3.229e-23 \| 699 \| 0.318 \| 380 \| 202 \| 16 \| 3 \| 354 \| 1 \| 351 \| DNA replication protein \| DNA replication protein; afdb-uniprot50 \| AF-A0A4P7YXX2-F1-MODEL\_V4 \| 1.0 \| 8.279e-17 \| 699 \| 0.334 \| 233 \| 136 \| 7 \| 116 \| 347 \| 489 \| 703 \| DUF1738 domain-containing protein \| DUF1738 domain-containing protein; afdb-uniprot50 \| AF-A0A4P7YZT8-F1-MODEL\_V4 \| 1.0 \| 2.802e-17 \| 698 \| 0.332 \| 250 \| 145 \| 9 \| 102 \| 347 \| 1 \| 232 \| Toprim domain-containing protein \| Toprim domain-containing protein; afdb-uniprot50 \| AF-A0A257ZQW8-F1-MODEL\_V4 \| 1.0 \| 9.792e-27 \| 698 \| 0.188 \| 723 \| 403 \| 38 \| 179 \| 767 \| 3 \| 675 \| P-loop ATPase \| P-loop ATPase; afdb-uniprot50 \| AF-A0A1H5S3J8-F1-MODEL\_V4 \| 1.0 \| 7.565e-19 \| 696 \| 0.279 \| 358 \| 200 \| 14 \| 8 \| 320 \| 3 \| 347 \| Putative DNA primase/helicase \| Putative DNA primase/helicase; afdb-uniprot50 \| AF-D5EXB0-F1-MODEL\_V4 \| 1.0 \| 4.344e-21 \| 696 \| 0.217 \| 390 \| 278 \| 16 \| 384 \| 757 \| 21 \| 399 \| DUF3874 domain-containing protein \| DUF3874 domain-containing protein; afdb-uniprot50 \| AF-A0A3E5B532-F1-MODEL\_V4 \| 1.0 \| 6.522e-24 \| 696 \| 0.192 \| 535 \| 326 \| 22 \| 240 \| 757 \| 260 \| 705 \| Virulence protein E \| Virulence protein E; afdb-uniprot50 \| AF-A0A2L1GL73-F1-MODEL\_V4 \| 1.0 \| 1.165e-17 \| 695 \| 0.352 \| 250 \| 138 \| 11 \| 102 \| 346 \| 506 \| 736 \| Uncharacterized protein \| Uncharacterized protein; afdb-uniprot50 \| AF-A0A6G6YPV6-F1-MODEL\_V4 \| 1.0 \| 6.356e-20 \| 694 \| 0.216 \| 406 \| 279 \| 20 \| 370 \| 755 \| 18 \| 404 \| Virulence-associated E family protein \| Virulence-associated E family protein; afdb-uniprot50 \| AF-A0A239FXE1-F1-MODEL\_V4 \| 1.0 \| 3.697e-24 \| 694 \| 0.216 \| 537 \| 359 \| 26 \| 261 \| 769 \| 252 \| 754 \| Uncharacterized protein \| Uncharacterized protein; afdb-uniprot50 \| AF-A0A4U1AJI8-F1-MODEL\_V4 \| 1.0 \| 2.527e-17 \| 694 \| 0.322 \| 257 \| 142 \| 9 \| 102 \| 346 \| 587 \| 823 \| DUF1738 domain-containing protein \| DUF1738 domain-containing protein; afdb-uniprot50 \| AF-A0A0F9XQ22-F1-MODEL\_V4 \| 1.0 \| 2.56e-19 \| 693 \| 0.211 \| 406 \| 280 \| 18 \| 370 \| 755 \| 31 \| 416 \| Uncharacterized protein \| Uncharacterized protein; afdb-uniprot50 \| AF-A0A0A2WCR2-F1-MODEL\_V4 \| 1.0 \| 1.578e-20 \| 693 \| 0.204 \| 415 \| 293 \| 18 \| 370 \| 769 \| 42 \| 434 \| Phage associated DNA primase \| Phage associated DNA primase; afdb-uniprot50 \| AF-A0A844FYD8-F1-MODEL\_V4 \| 1.0 \| 2.644e-28 \| 693 \| 0.178 \| 848 \| 469 \| 45 \| 1 \| 757 \| 14 \| 724 \| Uncharacterized protein \| Uncharacterized protein; afdb-uniprot50 \| AF-A0A1X7F2Z1-F1-MODEL\_V4 \| 1.0 \| 1.432e-17 \| 692 \| 0.319 \| 257 \| 154 \| 7 \| 98 \| 351 \| 48 \| 286 \| Uncharacterized domain associated with phage/plasmid primase \| Uncharacterized domain associated with phage/plasmid primase; afdb-uniprot50 \| AF-A0A7Y5I7S5-F1-MODEL\_V4 \| 1.0 \| 5.732e-20 \| 692 \| 0.264 \| 378 \| 244 \| 15 \| 384 \| 753 \| 119 \| 470 \| Uncharacterized protein \| Uncharacterized protein; afdb-uniprot50 \| AF-A0A1H1AY48-F1-MODEL\_V4 \| 1.0 \| 1.259e-21 \| 691 \| 0.197 \| 435 \| 308 \| 22 \| 335 \| 755 \| 278 \| 685 \| Uncharacterized protein \| Uncharacterized protein; afdb-uniprot50 \| AF-A0A0A2B4H4-F1-MODEL\_V4 \| 1.0 \| 4.429e-20 \| 689 \| 0.238 \| 398 \| 262 \| 20 \| 376 \| 756 \| 20 \| 393 \| Uncharacterized protein \| Uncharacterized protein; afdb-uniprot50 \| AF-R6SB64-F1-MODEL\_V4 \| 1.0 \| 2.179e-22 \| 689 \| 0.183 \| 517 \| 327 \| 24 \| 260 \| 758 \| 254 \| 693 \| VirE domain protein \| VirE domain protein; afdb-uniprot50 \| AF-A0A412XPF4-F1-MODEL\_V4 \| 1.0 \| 3.995e-20 \| 687 \| 0.233 \| 390 \| 271 \| 14 \| 379 \| 751 \| 5 \| 383 \| Virulence protein \| Virulence protein; afdb-uniprot50 \| AF-A7V3I1-F1-MODEL\_V4 \| 1.0 \| 2.137e-23 \| 687 \| 0.17 \| 510 \| 341 \| 23 \| 259 \| 755 \| 48 \| 488 \| Uncharacterized protein \| Uncharacterized protein; afdb-uniprot50 \| AF-A0A0S3U510-F1-MODEL\_V4 \| 1.0 \| 2.151e-20 \| 687 \| 0.243 \| 395 \| 267 \| 16 \| 384 \| 769 \| 448 \| 819 \| Uncharacterized protein \| Uncharacterized protein; afdb-uniprot50 \| AF-A0A2W4V1A4-F1-MODEL\_V4 \| 1.0 \| 6.65e-31 \| 687 \| 0.194 \| 904 \| 490 \| 48 \| 1 \| 737 \| 1 \| 832 \| Uncharacterized protein \| Uncharacterized protein; afdb-uniprot50 \| AF-E2N8F3-F1-MODEL\_V4 \| 1.0 \| 1.405e-18 \| 686 \| 0.261 \| 295 \| 205 \| 8 \| 467 \| 757 \| 2 \| 287 \| Uncharacterized protein \| Uncharacterized protein; afdb-uniprot50 \| AF-A0A412LCT6-F1-MODEL\_V4 \| 1.0 \| 5.271e-19 \| 686 \| 0.256 \| 374 \| 249 \| 13 \| 398 \| 752 \| 2 \| 365 \| Uncharacterized protein \| Uncharacterized protein; afdb-uniprot50 \| AF-A0A1B3W9W8-F1-MODEL\_V4 \| 1.0 \| 1.499e-20 \| 686 \| 0.241 \| 431 \| 266 \| 23 \| 370 \| 757 \| 29 \| 441 \| Uncharacterized protein \| Uncharacterized protein; afdb-uniprot50 \| AF-A0A7C1RII9-F1-MODEL\_V4 \| 1.0 \| 8.226e-20 \| 686 \| 0.215 \| 408 \| 276 \| 16 \| 370 \| 755 \| 95 \| 480 \| Virulence-associated E family protein \| Virulence-associated E family protein; afdb-uniprot50 \| AF-A0A239KTG6-F1-MODEL\_V4 \| 1.0 \| 6.115e-22 \| 686 \| 0.235 \| 492 \| 303 \| 24 \| 295 \| 755 \| 222 \| 671 \| Predicted P-loop ATPase and inactivated derivatives \| Predicted P-loop ATPase and inactivated derivatives; afdb-uniprot50 \| AF-A0A6A7WDK3-F1-MODEL\_V4 \| 1.0 \| 7.914e-22 \| 686 \| 0.194 \| 489 \| 320 \| 26 \| 330 \| 769 \| 293 \| 756 \| VirE\_N domain-containing protein \| VirE\_N domain-containing protein; afdb-uniprot50 \| AF-A0A1Y4IN65-F1-MODEL\_V4 \| 1.0 \| 8.226e-20 \| 685 \| 0.208 \| 398 \| 281 \| 20 \| 381 \| 762 \| 43 \| 422 \| Helicase \| Helicase; afdb-uniprot50 \| AF-A0A2J4JNZ5-F1-MODEL\_V4 \| 1.0 \| 6.692e-20 \| 685 \| 0.206 \| 422 \| 283 \| 20 \| 370 \| 756 \| 3 \| 407 \| Virulence-associated protein E \| Virulence-associated protein E; afdb-uniprot50 \| AF-A0A843HRX9-F1-MODEL\_V4 \| 1.0 \| 1.3e-22 \| 685 \| 0.185 \| 523 \| 339 \| 20 \| 264 \| 760 \| 6 \| 467 \| Uncharacterized protein \| Uncharacterized protein; afdb-uniprot50 \| AF-A0A7X5RAN1-F1-MODEL\_V4 \| 1.0 \| 4.261e-22 \| 685 \| 0.209 \| 521 \| 335 \| 22 \| 260 \| 761 \| 170 \| 632 \| Uncharacterized protein \| Uncharacterized protein; afdb-uniprot50 \| AF-A0A522TUF4-F1-MODEL\_V4 \| 1.0 \| 1.284e-20 \| 685 \| 0.215 \| 426 \| 285 \| 20 \| 349 \| 751 \| 345 \| 744 \| Uncharacterized protein \| Uncharacterized protein; afdb-uniprot50 \| AF-A0A3E1EZ76-F1-MODEL\_V4 \| 1.0 \| 2.265e-20 \| 684 \| 0.215 \| 409 \| 268 \| 20 \| 376 \| 755 \| 11 \| 395 \| Virulence protein E \| Virulence protein E; afdb-uniprot50 \| AF-A0A7C9G0C5-F1-MODEL\_V4 \| 1.0 \| 8.226e-20 \| 684 \| 0.221 \| 401 \| 281 \| 15 \| 370 \| 759 \| 34 \| 414 \| Uncharacterized protein \| Uncharacterized protein; afdb-uniprot50 \| AF-A0A1T5AQ44-F1-MODEL\_V4 \| 1.0 \| 1.518e-22 \| 684 \| 0.199 \| 506 \| 327 \| 21 \| 261 \| 755 \| 255 \| 693 \| Uncharacterized protein \| Uncharacterized protein; afdb-uniprot50 \| AF-A0A5N7RS25-F1-MODEL\_V4 \| 1.0 \| 1.819e-26 \| 684 \| 0.2 \| 668 \| 419 \| 36 \| 181 \| 769 \| 144 \| 775 \| Uncharacterized protein \| Uncharacterized protein; afdb-uniprot50 \| AF-A0A7H9BIY7-F1-MODEL\_V4 \| 1.0 \| 4.605e-34 \| 684 \| 0.196 \| 932 \| 485 \| 51 \| 1 \| 750 \| 1 \| 850 \| DUF3631 domain-containing protein \| DUF3631 domain-containing protein; afdb-uniprot50 \| AF-A0A522TWI5-F1-MODEL\_V4 \| 1.0 \| 4.126e-21 \| 683 \| 0.371 \| 291 \| 167 \| 8 \| 1 \| 280 \| 12 \| 297 \| Toprim domain-containing protein \| Toprim domain-containing protein; afdb-uniprot50 \| AF-A0A7X8AE49-F1-MODEL\_V4 \| 1.0 \| 6.692e-20 \| 683 \| 0.194 \| 395 \| 283 \| 18 \| 381 \| 755 \| 16 \| 395 \| DUF3874 domain-containing protein \| DUF3874 domain-containing protein; afdb-uniprot50 \| AF-A0A414I4A7-F1-MODEL\_V4 \| 1.0 \| 1.1e-20 \| 683 \| 0.211 \| 444 \| 288 \| 24 \| 364 \| 769 \| 9 \| 428 \| Virulence-associated protein E \| Virulence-associated protein E; afdb-uniprot50 \| AF-A0A809PTW9-F1-MODEL\_V4 \| 1.0 \| 2.369e-23 \| 683 \| 0.267 \| 486 \| 261 \| 24 \| 10 \| 457 \| 4 \| 432 \| Uncharacterized protein \| Uncharacterized protein; afdb-uniprot50 \| AF-A0A4P6RG19-F1-MODEL\_V4 \| 1.0 \| 2.416e-22 \| 683 \| 0.203 \| 505 \| 322 \| 25 \| 264 \| 757 \| 262 \| 697 \| Helicase \| Helicase; afdb-uniprot50 \| AF-S8GYB6-F1-MODEL\_V4 \| 1.0 \| 6.564e-21 \| 682 \| 0.218 \| 440 \| 275 \| 19 \| 369 \| 757 \| 16 \| 437 \| Virulence-associated protein E \| Virulence-associated protein E; afdb-uniprot50 \| AF-A0A3R6BK15-F1-MODEL\_V4 \| 1.0 \| 2.016e-18 \| 681 \| 0.191 \| 344 \| 245 \| 14 \| 453 \| 768 \| 23 \| 361 \| Uncharacterized protein \| Uncharacterized protein; afdb-uniprot50 \| AF-A0A2H6BA53-F1-MODEL\_V4 \| 1.0 \| 8.069e-21 \| 681 \| 0.241 \| 459 \| 285 \| 23 \| 335 \| 769 \| 178 \| 597 \| Uncharacterized protein \| Uncharacterized protein; afdb-uniprot50 \| AF-A0A1F3KP86-F1-MODEL\_V4 \| 1.0 \| 3.819e-25 \| 681 \| 0.158 \| 776 \| 366 \| 26 \| 12 \| 769 \| 189 \| 695 \| Uncharacterized protein \| Uncharacterized protein; afdb-uniprot50 \| AF-A0A3N8QT28-F1-MODEL\_V4 \| 1.0 \| 6.608e-34 \| 681 \| 0.209 \| 971 \| 461 \| 45 \| 6 \| 752 \| 1 \| 888 \| DUF927 domain-containing protein \| DUF927 domain-containing protein; afdb-uniprot50 \| AF-A0A0Q6MHX9-F1-MODEL\_V4 \| 1.0 \| 1.538e-32 \| 681 \| 0.201 \| 898 \| 480 \| 48 \| 13 \| 738 \| 2 \| 833 \| Toprim domain-containing protein \| Toprim domain-containing protein; afdb-uniprot50 \| AF-A0A2T3MTH8-F1-MODEL\_V4 \| 1.0 \| 2.016e-18 \| 680 \| 0.329 \| 267 \| 126 \| 8 \| 80 \| 346 \| 82 \| 295 \| Toprim domain-containing protein \| Toprim domain-containing protein; afdb-uniprot50 \| AF-A0A416XEW5-F1-MODEL\_V4 \| 1.0 \| 1.326e-21 \| 680 \| 0.188 \| 536 \| 323 \| 25 \| 239 \| 755 \| 37 \| 479 \| DUF3874 domain-containing protein \| DUF3874 domain-containing protein; afdb-uniprot50 \| AF-A0A641ULV4-F1-MODEL\_V4 \| 1.0 \| 2.416e-22 \| 680 \| 0.193 \| 528 \| 331 \| 25 \| 242 \| 751 \| 28 \| 478 \| Virulence protein E \| Virulence protein E; afdb-uniprot50 \| AF-A0A8A6EWM4-F1-MODEL\_V4 \| 1.0 \| 9.42e-21 \| 680 \| 0.206 \| 411 \| 279 \| 22 \| 370 \| 753 \| 348 \| 738 \| Bifunctional DNA primase/polymerase \| Bifunctional DNA primase/polymerase; afdb-uniprot50 \| AF-A0A1I5LX50-F1-MODEL\_V4 \| 1.0 \| 1.568e-23 \| 680 \| 0.196 \| 530 \| 322 \| 25 \| 261 \| 751 \| 253 \| 717 \| Virulence-associated protein E \| Virulence-associated protein E; afdb-uniprot50 \| AF-A0A4R8FHG7-F1-MODEL\_V4 \| 1.0 \| 2.784e-20 \| 680 \| 0.232 \| 426 \| 277 \| 16 \| 370 \| 759 \| 428 \| 839 \| Virulence-associated protein E \| Virulence-associated protein E; afdb-uniprot50 \| AF-A0A6M8VRM0-F1-MODEL\_V4 \| 1.0 \| 6.956e-18 \| 679 \| 0.285 \| 308 \| 195 \| 12 \| 453 \| 753 \| 8 \| 297 \| Virulence-associated E family protein \| Virulence-associated E family protein; afdb-uniprot50 \| AF-K9E0U4-F1-MODEL\_V4 \| 1.0 \| 2.309e-19 \| 679 \| 0.184 \| 385 \| 284 \| 17 \| 381 \| 751 \| 50 \| 418 \| DUF3874 domain-containing protein \| DUF3874 domain-containing protein; afdb-uniprot50 \| AF-A0A7G8F7P4-F1-MODEL\_V4 \| 1.0 \| 1.121e-19 \| 679 \| 0.205 \| 409 \| 284 \| 16 \| 370 \| 757 \| 52 \| 440 \| Virulence-associated E family protein \| Virulence-associated E family protein; afdb-uniprot50 \| AF-A0A1K1M310-F1-MODEL\_V4 \| 1.0 \| 1.309e-19 \| 679 \| 0.194 \| 390 \| 287 \| 14 \| 384 \| 756 \| 228 \| 607 \| Uncharacterized protein \| Uncharacterized protein; afdb-uniprot50 \| AF-A0A508X032-F1-MODEL\_V4 \| 1.0 \| 3.356e-21 \| 678 \| 0.214 \| 461 \| 306 \| 20 \| 331 \| 769 \| 112 \| 538 \| Uncharacterized protein \| Uncharacterized protein; afdb-uniprot50 \| AF-A0A843BCM8-F1-MODEL\_V4 \| 1.0 \| 7.467e-17 \| 677 \| 0.362 \| 248 \| 132 \| 9 \| 102 \| 347 \| 120 \| 343 \| Toprim domain-containing protein \| Toprim domain-containing protein; afdb-uniprot50 \| AF-A0A089P3D8-F1-MODEL\_V4 \| 1.0 \| 9.604e-20 \| 677 \| 0.231 \| 410 \| 277 \| 17 \| 378 \| 769 \| 23 \| 412 \| Uncharacterized protein \| Uncharacterized protein; afdb-uniprot50 \| AF-A0A374W536-F1-MODEL\_V4 \| 1.0 \| 3.969e-23 \| 677 \| 0.203 \| 515 \| 334 \| 29 \| 264 \| 756 \| 215 \| 675 \| VirE\_N domain-containing protein \| VirE\_N domain-containing protein; afdb-uniprot50 \| AF-A0A2D6XDQ3-F1-MODEL\_V4 \| 1.0 \| 3.794e-20 \| 676 \| 0.209 \| 415 \| 285 \| 19 \| 370 \| 768 \| 47 \| 434 \| Uncharacterized protein \| Uncharacterized protein; afdb-uniprot50 \| AF-A0A558E116-F1-MODEL\_V4 \| 1.0 \| 3.292e-22 \| 676 \| 0.25 \| 512 \| 303 \| 26 \| 316 \| 768 \| 2 \| 491 \| Uncharacterized protein \| Uncharacterized protein; afdb-uniprot50 \| AF-A0A4Q3PJ79-F1-MODEL\_V4 \| 1.0 \| 1.928e-31 \| 676 \| 0.193 \| 964 \| 453 \| 49 \| 6 \| 753 \| 1 \| 855 \| DUF3987 domain-containing protein \| DUF3987 domain-containing protein; afdb-uniprot50 \| AF-A0A7Z8VSS1-F1-MODEL\_V4 \| 1.0 \| 3.378e-18 \| 675 \| 0.231 \| 307 \| 203 \| 10 \| 370 \| 659 \| 7 \| 297 \| Uncharacterized protein \| Uncharacterized protein; afdb-uniprot50 \| AF-A0A5P2FXH3-F1-MODEL\_V4 \| 1.0 \| 1.011e-19 \| 675 \| 0.21 \| 390 \| 279 \| 18 \| 381 \| 757 \| 17 \| 390 \| Virulence protein E \| Virulence protein E; afdb-uniprot50 \| AF-A0A7Y9FR30-F1-MODEL\_V4 \| 1.0 \| 1.672e-25 \| 675 \| 0.181 \| 635 \| 403 \| 35 \| 181 \| 753 \| 61 \| 640 \| Uncharacterized protein \| Uncharacterized protein; afdb-uniprot50 \| AF-R6FMS2-F1-MODEL\_V4 \| 1.0 \| 3.126e-22 \| 675 \| 0.171 \| 506 \| 348 \| 21 \| 259 \| 755 \| 230 \| 673 \| VirE domain-containing protein \| VirE domain-containing protein; afdb-uniprot50 \| AF-A0A1I3TR65-F1-MODEL\_V4 \| 1.0 \| 2.069e-22 \| 675 \| 0.209 \| 531 \| 319 \| 24 \| 265 \| 750 \| 268 \| 742 \| Predicted P-loop ATPase and inactivated derivatives \| Predicted P-loop ATPase and inactivated derivatives; afdb-uniprot50 \| AF-A0A2S7IR16-F1-MODEL\_V4 \| 1.0 \| 2.043e-20 \| 675 \| 0.232 \| 439 \| 274 \| 23 \| 370 \| 755 \| 369 \| 797 \| Uncharacterized protein \| Uncharacterized protein; afdb-uniprot50 \| AF-G0AAH9-F1-MODEL\_V4 \| 1.0 \| 1.093e-31 \| 675 \| 0.199 \| 893 \| 485 \| 53 \| 6 \| 751 \| 1 \| 810 \| PriCT\_2 domain-containing protein \| PriCT\_2 domain-containing protein; afdb-uniprot50 \| AF-A0A1H6TNE7-F1-MODEL\_V4 \| 1.0 \| 5.273e-35 \| 675 \| 0.211 \| 879 \| 486 \| 45 \| 1 \| 730 \| 10 \| 829 \| Putative DNA primase/helicase \| Putative DNA primase/helicase; afdb-uniprot50 \| AF-Q0AG08-F1-MODEL\_V4 \| 1.0 \| 1.309e-19 \| 673 \| 0.243 \| 419 \| 280 \| 19 \| 370 \| 769 \| 1 \| 401 \| Virulence-associated E family protein \| Virulence-associated E family protein; afdb-uniprot50 \| AF-A0A839HPN6-F1-MODEL\_V4 \| 1.0 \| 5.771e-33 \| 673 \| 0.214 \| 884 \| 477 \| 43 \| 8 \| 737 \| 2 \| 821 \| DUF927 domain-containing protein \| DUF927 domain-containing protein; afdb-uniprot50 \| AF-A0A3A6KL80-F1-MODEL\_V4 \| 1.0 \| 1.158e-20 \| 672 \| 0.209 \| 444 \| 289 \| 24 \| 364 \| 769 \| 9 \| 428 \| Virulence-associated protein E \| Virulence-associated protein E; afdb-uniprot50 \| AF-B6ATL0-F1-MODEL\_V4 \| 1.0 \| 2.593e-21 \| 672 \| 0.232 \| 431 \| 272 \| 21 \| 368 \| 757 \| 28 \| 440 \| Virulence-associated E family protein \| Virulence-associated E family protein; afdb-uniprot50 \| AF-J9GG52-F1-MODEL\_V4 \| 1.0 \| 6.912e-21 \| 672 \| 0.215 \| 422 \| 282 \| 20 \| 369 \| 755 \| 107 \| 514 \| Helicase \| Helicase; afdb-uniprot50 \| AF-A0A1M3H1Y9-F1-MODEL\_V4 \| 1.0 \| 5.959e-18 \| 672 \| 0.329 \| 270 \| 155 \| 8 \| 92 \| 361 \| 87 \| 330 \| AAA domain-containing protein \| AAA domain-containing protein; afdb-uniprot50 \| AF-A0A496KVV9-F1-MODEL\_V4 \| 1.0 \| 9.121e-20 \| 671 \| 0.198 \| 403 \| 289 \| 19 \| 369 \| 757 \| 33 \| 415 \| Helicase \| Helicase; afdb-uniprot50 \| AF-A0A3B0ME56-F1-MODEL\_V4 \| 1.0 \| 3.969e-23 \| 671 \| 0.228 \| 534 \| 308 \| 28 \| 261 \| 756 \| 238 \| 705 \| Uncharacterized protein \| Uncharacterized protein; afdb-uniprot50 \| AF-J9GP52-F1-MODEL\_V4 \| 1.0 \| 1.36e-17 \| 670 \| 0.258 \| 286 \| 199 \| 9 \| 475 \| 756 \| 2 \| 278 \| Virulence-associated E family protein \| Virulence-associated E family protein; afdb-uniprot50 \| AF-A0A0F9EF48-F1-MODEL\_V4 \| 1.0 \| 1.978e-19 \| 670 \| 0.199 \| 406 \| 284 \| 20 \| 370 \| 757 \| 4 \| 386 \| Uncharacterized protein \| Uncharacterized protein; afdb-uniprot50 \| AF-A0A6G9HJR3-F1-MODEL\_V4 \| 1.0 \| 5.479e-17 \| 670 \| 0.293 \| 252 \| 155 \| 9 \| 100 \| 347 \| 143 \| 375 \| DNA primase TraC \| DNA primase TraC; afdb-uniprot50 \| AF-A0A7G7KL89-F1-MODEL\_V4 \| 1.0 \| 1.121e-19 \| 670 \| 0.217 \| 418 \| 276 \| 20 \| 369 \| 757 \| 12 \| 407 \| Uncharacterized protein \| Uncharacterized protein; afdb-uniprot50 \| AF-A0A0B4XZW9-F1-MODEL\_V4 \| 1.0 \| 6.075e-17 \| 670 \| 0.329 \| 249 \| 144 \| 8 \| 100 \| 346 \| 466 \| 693 \| Uncharacterized protein \| Uncharacterized protein; afdb-uniprot50 \| AF-A0A4T2A6U5-F1-MODEL\_V4 \| 1.0 \| 5.479e-17 \| 670 \| 0.326 \| 257 \| 142 \| 10 \| 99 \| 346 \| 513 \| 747 \| DUF1738 domain-containing protein \| DUF1738 domain-containing protein; afdb-uniprot50 \| AF-A0A0R3NCG4-F1-MODEL\_V4 \| 1.0 \| 1.018e-24 \| 669 \| 0.208 \| 567 \| 367 \| 26 \| 234 \| 751 \| 9 \| 542 \| Uncharacterized protein \| Uncharacterized protein; afdb-uniprot50 \| AF-A0A4Q3L9Q7-F1-MODEL\_V4 \| 1.0 \| 1.018e-16 \| 668 \| 0.357 \| 238 \| 149 \| 4 \| 368 \| 602 \| 60 \| 296 \| Uncharacterized protein \| Uncharacterized protein; afdb-uniprot50 \| AF-R5CMD4-F1-MODEL\_V4 \| 1.0 \| 3.65e-22 \| 668 \| 0.179 \| 519 \| 339 \| 22 \| 261 \| 757 \| 241 \| 694 \| VirE N-terminal domain protein \| VirE N-terminal domain protein; afdb-uniprot50 \| AF-A0A412M1I6-F1-MODEL\_V4 \| 1.0 \| 1.599e-22 \| 668 \| 0.195 \| 523 \| 332 \| 26 \| 261 \| 755 \| 233 \| 694 \| Helicase \| Helicase; afdb-uniprot50 \| AF-A0A1X6Y4N3-F1-MODEL\_V4 \| 1.0 \| 6.275e-26 \| 668 \| 0.199 \| 661 \| 398 \| 37 \| 176 \| 753 \| 64 \| 676 \| Uncharacterized protein \| Uncharacterized protein; afdb-uniprot50 \| AF-K9Z0R0-F1-MODEL\_V4 \| 1.0 \| 3.868e-19 \| 668 \| 0.253 \| 379 \| 247 \| 18 \| 384 \| 751 \| 343 \| 696 \| Virulence-associated E family protein \| Virulence-associated E family protein; afdb-uniprot50 \| AF-A0A355VT45-F1-MODEL\_V4 \| 1.0 \| 1.378e-19 \| 667 \| 0.215 \| 417 \| 273 \| 16 \| 379 \| 755 \| 11 \| 413 \| Uncharacterized protein \| Uncharacterized protein; afdb-uniprot50 \| AF-A0A7T1I9T7-F1-MODEL\_V4 \| 1.0 \| 4.91e-20 \| 667 \| 0.226 \| 424 \| 267 \| 21 \| 371 \| 757 \| 177 \| 576 \| Uncharacterized protein \| Uncharacterized protein; afdb-uniprot50 \| AF-A0A1M6MNJ7-F1-MODEL\_V4 \| 1.0 \| 1.243e-19 \| 666 \| 0.216 \| 429 \| 292 \| 20 \| 370 \| 769 \| 9 \| 422 \| Virulence-associated protein E \| Virulence-associated protein E; afdb-uniprot50 \| AF-A0A5B2V8U9-F1-MODEL\_V4 \| 1.0 \| 9.542e-23 \| 666 \| 0.195 \| 532 \| 344 \| 24 \| 261 \| 753 \| 65 \| 551 \| Virulence-associated E family protein \| Virulence-associated E family protein; afdb-uniprot50 \| AF-A0A7W0BSW7-F1-MODEL\_V4 \| 1.0 \| 5.696e-23 \| 666 \| 0.223 \| 536 \| 342 \| 28 \| 245 \| 756 \| 259 \| 744 \| Uncharacterized protein \| Uncharacterized protein; afdb-uniprot50 \| AF-A0A6S6SDG9-F1-MODEL\_V4 \| 1.0 \| 6.735e-17 \| 665 \| 0.315 \| 260 \| 155 \| 8 \| 91 \| 347 \| 48 \| 287 \| Cpp22 \| Cpp22; afdb-uniprot50 \| AF-A0A841EPH5-F1-MODEL\_V4 \| 1.0 \| 2.56e-19 \| 665 \| 0.223 \| 399 \| 270 \| 20 \| 381 \| 756 \| 42 \| 423 \| Putative P-loop ATPase \| Putative P-loop ATPase; afdb-uniprot50 \| AF-A0A7W9AWL7-F1-MODEL\_V4 \| 1.0 \| 3.147e-19 \| 665 \| 0.222 \| 414 \| 291 \| 17 \| 370 \| 767 \| 61 \| 459 \| Putative DNA primase/helicase \| Putative DNA primase/helicase; afdb-uniprot50 \| AF-A0A7C9KA34-F1-MODEL\_V4 \| 1.0 \| 1.978e-19 \| 665 \| 0.213 \| 399 \| 282 \| 19 \| 377 \| 760 \| 185 \| 566 \| Uncharacterized protein \| Uncharacterized protein; afdb-uniprot50 \| AF-A0A1C3X7K5-F1-MODEL\_V4 \| 1.0 \| 2.043e-20 \| 665 \| 0.208 \| 451 \| 306 \| 18 \| 336 \| 767 \| 239 \| 657 \| Virulence-associated protein E \| Virulence-associated protein E; afdb-uniprot50 \| AF-A0A553EPP8-F1-MODEL\_V4 \| 1.0 \| 1.173e-22 \| 665 \| 0.197 \| 537 \| 328 \| 26 \| 241 \| 751 \| 215 \| 674 \| Virulence protein E \| Virulence protein E; afdb-uniprot50 \| AF-A0A6J4F9N4-F1-MODEL\_V4 \| 1.0 \| 5.921e-29 \| 665 \| 0.175 \| 918 \| 501 \| 52 \| 2 \| 752 \| 6 \| 834 \| Putative Virulence-associated protein E \| Putative Virulence-associated protein E; afdb-uniprot50 \| AF-A0A0N7LN66-F1-MODEL\_V4 \| 1.0 \| 4.207e-28 \| 665 \| 0.18 \| 809 \| 410 \| 38 \| 1 \| 764 \| 244 \| 844 \| Putative P-loop ATPase \| Putative P-loop ATPase; afdb-uniprot50 \| AF-A0A1X7JKC4-F1-MODEL\_V4 \| 1.0 \| 8.226e-20 \| 664 \| 0.211 \| 417 \| 279 \| 22 \| 370 \| 769 \| 87 \| 470 \| Virulence-associated protein E \| Virulence-associated protein E; afdb-uniprot50 \| AF-A0A5E4VTX6-F1-MODEL\_V4 \| 1.0 \| 4.694e-33 \| 664 \| 0.186 \| 940 \| 510 \| 53 \| 1 \| 750 \| 18 \| 892 \| DNA primase \| DNA primase; afdb-uniprot50 \| AF-A0A416T296-F1-MODEL\_V4 \| 1.0 \| 1.219e-20 \| 663 \| 0.214 \| 434 \| 295 \| 19 \| 328 \| 744 \| 61 \| 465 \| DUF3874 domain-containing protein \| DUF3874 domain-containing protein; afdb-uniprot50 \| AF-A0A6G1UCW6-F1-MODEL\_V4 \| 1.0 \| 1.326e-21 \| 663 \| 0.174 \| 527 \| 360 \| 28 \| 286 \| 768 \| 81 \| 576 \| Uncharacterized protein \| Uncharacterized protein; afdb-uniprot50 \| AF-A0A2W4XDU3-F1-MODEL\_V4 \| 1.0 \| 4.574e-21 \| 663 \| 0.24 \| 450 \| 281 \| 22 \| 321 \| 757 \| 194 \| 595 \| Uncharacterized protein \| Uncharacterized protein; afdb-uniprot50 \| AF-A0A0F5INE2-F1-MODEL\_V4 \| 1.0 \| 2.543e-22 \| 663 \| 0.193 \| 516 \| 316 \| 26 \| 261 \| 755 \| 253 \| 689 \| Uncharacterized protein \| Uncharacterized protein; afdb-uniprot50 \| AF-A0A6G1U079-F1-MODEL\_V4 \| 1.0 \| 2.495e-23 \| 663 \| 0.187 \| 559 \| 335 \| 20 \| 259 \| 769 \| 229 \| 716 \| Virulence protein E \| Virulence protein E; afdb-uniprot50 \| AF-A0A3M1WWN1-F1-MODEL\_V4 \| 1.0 \| 2.838e-19 \| 662 \| 0.221 \| 406 \| 272 \| 20 \| 380 \| 769 \| 352 \| 729 \| Uncharacterized protein \| Uncharacterized protein; afdb-uniprot50 \| AF-A0A0R1T496-F1-MODEL\_V4 \| 1.0 \| 3.25e-20 \| 661 \| 0.227 \| 404 \| 265 \| 16 \| 367 \| 750 \| 23 \| 399 \| Prophage lp3 protein 8, helicase \| Prophage lp3 protein 8, helicase; afdb-uniprot50 \| AF-A0A352JCJ9-F1-MODEL\_V4 \| 1.0 \| 2.082e-19 \| 661 \| 0.236 \| 394 \| 261 \| 17 \| 382 \| 760 \| 104 \| 472 \| Uncharacterized protein \| Uncharacterized protein; afdb-uniprot50 \| AF-Q027Z8-F1-MODEL\_V4 \| 1.0 \| 1.509e-25 \| 661 \| 0.183 \| 669 \| 430 \| 33 \| 178 \| 757 \| 59 \| 699 \| Virulence-associated E family protein \| Virulence-associated E family protein; afdb-uniprot50 \| AF-A0A1F4EP90-F1-MODEL\_V4 \| 1.0 \| 1.558e-18 \| 660 \| 0.273 \| 347 \| 192 \| 18 \| 11 \| 346 \| 1 \| 298 \| PriCT\_2 domain-containing protein \| PriCT\_2 domain-containing protein; afdb-uniprot50 \| AF-A0A3P1ZAJ6-F1-MODEL\_V4 \| 1.0 \| 3.868e-19 \| 660 \| 0.184 \| 396 \| 281 \| 20 \| 381 \| 757 \| 99 \| 471 \| Helicase \| Helicase; afdb-uniprot50 \| AF-G7GJU6-F1-MODEL\_V4 \| 1.0 \| 3.534e-21 \| 660 \| 0.209 \| 443 \| 286 \| 19 \| 380 \| 769 \| 38 \| 469 \| Uncharacterized protein \| Uncharacterized protein; afdb-uniprot50 \| AF-A0A1G7G4Y5-F1-MODEL\_V4 \| 1.0 \| 5.17e-20 \| 659 \| 0.195 \| 445 \| 297 \| 18 \| 370 \| 762 \| 371 \| 806 \| Virulence-associated protein E \| Virulence-associated protein E; afdb-uniprot50 \| AF-A0A5C7BZC7-F1-MODEL\_V4 \| 1.0 \| 2.856e-16 \| 659 \| 0.312 \| 253 \| 155 \| 7 \| 98 \| 347 \| 139 \| 375 \| Uncharacterized protein \| Uncharacterized protein; afdb-uniprot50 \| AF-A0A0M3Q0Y7-F1-MODEL\_V4 \| 1.0 \| 4.515e-19 \| 658 \| 0.23 \| 394 \| 268 \| 17 \| 393 \| 766 \| 5 \| 383 \| Virulence associated protein \| Virulence associated protein; afdb-uniprot50 \| AF-A0A174RE62-F1-MODEL\_V4 \| 1.0 \| 3.868e-19 \| 658 \| 0.182 \| 390 \| 280 \| 18 \| 381 \| 751 \| 68 \| 437 \| Predicted P-loop ATPase and inactivated derivatives \| Predicted P-loop ATPase and inactivated derivatives; afdb-uniprot50 \| AF-A0A7W1X8F7-F1-MODEL\_V4 \| 1.0 \| 4.755e-19 \| 658 \| 0.221 \| 424 \| 272 \| 20 \| 379 \| 755 \| 376 \| 788 \| Uncharacterized protein \| Uncharacterized protein; afdb-uniprot50 \| AF-A0A068MV01-F1-MODEL\_V4 \| 1.0 \| 1.031e-26 \| 658 \| 0.19 \| 645 \| 389 \| 37 \| 178 \| 752 \| 75 \| 656 \| DUF3854 domain-containing protein \| DUF3854 domain-containing protein; afdb-uniprot50 \| AF-A0A7U5WHF4-F1-MODEL\_V4 \| 1.0 \| 1.243e-19 \| 657 \| 0.233 \| 424 \| 262 \| 23 \| 370 \| 749 \| 35 \| 439 \| Virulence factor \| Virulence factor; afdb-uniprot50 \| AF-A0A0J7L464-F1-MODEL\_V4 \| 1.0 \| 7.965e-19 \| 657 \| 0.165 \| 406 \| 288 \| 18 \| 370 \| 749 \| 333 \| 713 \| Virulence-associated e family protein \| Virulence-associated e family protein; afdb-uniprot50 \| AF-U4QRR7-F1-MODEL\_V4 \| 1.0 \| 1.405e-26 \| 656 \| 0.212 \| 674 \| 434 \| 38 \| 133 \| 756 \| 31 \| 657 \| Putative Virulence-associated E \| Putative Virulence-associated E; afdb-uniprot50 \| AF-A0A1M4WF05-F1-MODEL\_V4 \| 1.0 \| 1.716e-21 \| 656 \| 0.173 \| 519 \| 320 \| 22 \| 261 \| 755 \| 268 \| 701 \| VirE N-terminal domain-containing protein \| VirE N-terminal domain-containing protein; afdb-uniprot50 \| AF-A0A1V0GS15-F1-MODEL\_V4 \| 1.0 \| 1.005e-22 \| 655 \| 0.211 \| 502 \| 332 \| 19 \| 300 \| 754 \| 180 \| 664 \| RepB\_primase domain-containing protein \| RepB\_primase domain-containing protein; afdb-uniprot50 \| AF-A0A1M5BFH6-F1-MODEL\_V4 \| 1.0 \| 3.969e-23 \| 655 \| 0.203 \| 536 \| 332 \| 25 \| 239 \| 755 \| 212 \| 671 \| Uncharacterized protein \| Uncharacterized protein; afdb-uniprot50 \| AF-A0A6G1U056-F1-MODEL\_V4 \| 1.0 \| 1.196e-21 \| 655 \| 0.155 \| 539 \| 365 \| 29 \| 260 \| 756 \| 245 \| 735 \| VirE\_N domain-containing protein \| VirE\_N domain-containing protein; afdb-uniprot50 \| AF-A0A1N7MGC5-F1-MODEL\_V4 \| 1.0 \| 4.099e-16 \| 655 \| 0.336 \| 238 \| 131 \| 7 \| 120 \| 346 \| 547 \| 768 \| Antirestriction protein ArdC \| Antirestriction protein ArdC; afdb-uniprot50 \| AF-A0A1Q4RMJ7-F1-MODEL\_V4 \| 1.0 \| 2.56e-27 \| 655 \| 0.19 \| 682 \| 442 \| 33 \| 138 \| 769 \| 56 \| 677 \| DUF3854 domain-containing protein \| DUF3854 domain-containing protein; afdb-uniprot50 \| AF-A0A448QER1-F1-MODEL\_V4 \| 1.0 \| 1.212e-31 \| 655 \| 0.19 \| 934 \| 507 \| 57 \| 1 \| 759 \| 1 \| 859 \| Bifunctional DNA primase/polymerase \| Bifunctional DNA primase/polymerase; afdb-uniprot50 \| AF-A0A4P6HIC1-F1-MODEL\_V4 \| 1.0 \| 1.018e-16 \| 654 \| 0.337 \| 252 \| 137 \| 11 \| 100 \| 347 \| 74 \| 299 \| Toprim domain-containing protein \| Toprim domain-containing protein; afdb-uniprot50 \| AF-A0A7U4E6T9-F1-MODEL\_V4 \| 1.0 \| 5.844e-19 \| 654 \| 0.217 \| 400 \| 270 \| 17 \| 381 \| 755 \| 12 \| 393 \| Uncharacterized protein \| Uncharacterized protein; afdb-uniprot50 \| AF-A0A373IIN3-F1-MODEL\_V4 \| 1.0 \| 4.073e-19 \| 654 \| 0.17 \| 410 \| 291 \| 17 \| 381 \| 755 \| 92 \| 487 \| DUF3874 domain-containing protein \| DUF3874 domain-containing protein; afdb-uniprot50 \| AF-A0A7W5TKY8-F1-MODEL\_V4 \| 1.0 \| 3.086e-20 \| 654 \| 0.228 \| 442 \| 284 \| 26 \| 360 \| 769 \| 8 \| 424 \| Uncharacterized protein \| Uncharacterized protein; afdb-uniprot50 \| AF-A0A3R5Z023-F1-MODEL\_V4 \| 1.0 \| 3.843e-22 \| 654 \| 0.175 \| 535 \| 345 \| 19 \| 259 \| 769 \| 86 \| 548 \| Virulence protein E \| Virulence protein E; afdb-uniprot50 \| AF-A0A5C7LJI8-F1-MODEL\_V4 \| 1.0 \| 5.807e-22 \| 654 \| 0.188 \| 546 \| 348 \| 23 \| 268 \| 757 \| 206 \| 712 \| Uncharacterized protein \| Uncharacterized protein; afdb-uniprot50 \| AF-A0A0D6QBU6-F1-MODEL\_V4 \| 1.0 \| 3.313e-19 \| 653 \| 0.214 \| 415 \| 276 \| 21 \| 370 \| 753 \| 18 \| 413 \| DNA primase \| DNA primase; afdb-uniprot50 \| AF-A0A0P0FP05-F1-MODEL\_V4 \| 1.0 \| 2.478e-18 \| 653 \| 0.172 \| 389 \| 296 \| 16 \| 381 \| 757 \| 47 \| 421 \| DUF3874 domain-containing protein \| DUF3874 domain-containing protein; afdb-uniprot50 \| AF-A0A5P0X0N2-F1-MODEL\_V4 \| 1.0 \| 4.816e-21 \| 653 \| 0.18 \| 510 \| 346 \| 27 \| 286 \| 755 \| 178 \| 655 \| VirE\_N domain-containing protein \| VirE\_N domain-containing protein; afdb-uniprot50 \| AF-A0A5P0XUM3-F1-MODEL\_V4 \| 1.0 \| 2.069e-22 \| 653 \| 0.178 \| 537 \| 341 \| 19 \| 259 \| 769 \| 219 \| 681 \| Virulence protein E \| Virulence protein E; afdb-uniprot50 \| AF-A0A2S5P5D7-F1-MODEL\_V4 \| 1.0 \| 5.623e-21 \| 653 \| 0.216 \| 489 \| 325 \| 21 \| 287 \| 751 \| 303 \| 757 \| Uncharacterized protein \| Uncharacterized protein; afdb-uniprot50 \| AF-A0A139K8J3-F1-MODEL\_V4 \| 1.0 \| 8.831e-19 \| 652 \| 0.184 \| 396 \| 296 \| 18 \| 384 \| 768 \| 63 \| 442 \| DUF3874 domain-containing protein \| DUF3874 domain-containing protein; afdb-uniprot50 \| AF-A0A4Q7LBG2-F1-MODEL\_V4 \| 1.0 \| 1.831e-31 \| 652 \| 0.199 \| 914 \| 483 \| 47 \| 1 \| 737 \| 3 \| 844 \| Putative DNA primase/helicase \| Putative DNA primase/helicase; afdb-uniprot50 \| AF-A0A7W6QX48-F1-MODEL\_V4 \| 1.0 \| 3.444e-17 \| 651 \| 0.234 \| 303 \| 202 \| 12 \| 457 \| 750 \| 53 \| 334 \| Putative P-loop ATPase \| Putative P-loop ATPase; afdb-uniprot50 \| AF-A0A3R6QWH6-F1-MODEL\_V4 \| 1.0 \| 1.309e-19 \| 651 \| 0.202 \| 429 \| 295 \| 21 \| 370 \| 767 \| 18 \| 430 \| Virulence-associated protein E \| Virulence-associated protein E; afdb-uniprot50 \| AF-A0A4R3GMS2-F1-MODEL\_V4 \| 1.0 \| 1.011e-19 \| 651 \| 0.239 \| 439 \| 279 \| 24 \| 370 \| 769 \| 18 \| 440 \| Virulence-associated protein E \| Virulence-associated protein E; afdb-uniprot50 \| AF-A0A357A780-F1-MODEL\_V4 \| 1.0 \| 8.552e-26 \| 651 \| 0.216 \| 652 \| 392 \| 32 \| 180 \| 757 \| 89 \| 695 \| DUF3854 domain-containing protein \| DUF3854 domain-containing protein; afdb-uniprot50 \| AF-A0A037USG8-F1-MODEL\_V4 \| 1.0 \| 8.718e-17 \| 650 \| 0.338 \| 260 \| 149 \| 10 \| 100 \| 345 \| 68 \| 318 \| Toprim domain-containing protein \| Toprim domain-containing protein; afdb-uniprot50 \| AF-A0A1I6WIK4-F1-MODEL\_V4 \| 1.0 \| 2.431e-19 \| 650 \| 0.213 \| 407 \| 276 \| 18 \| 394 \| 769 \| 13 \| 406 \| Virulence-associated protein E \| Virulence-associated protein E; afdb-uniprot50 \| AF-A0A1Y3RIE9-F1-MODEL\_V4 \| 1.0 \| 4.073e-19 \| 650 \| 0.217 \| 418 \| 278 \| 21 \| 370 \| 755 \| 19 \| 419 \| Virulence-associated protein E \| Virulence-associated protein E; afdb-uniprot50 \| AF-Q3AX57-F1-MODEL\_V4 \| 1.0 \| 6.823e-19 \| 649 \| 0.206 \| 397 \| 279 \| 15 \| 378 \| 756 \| 18 \| 396 \| p-loop ATPase and inactivated derivatives-like \| p-loop ATPase and inactivated derivatives-like; afdb-uniprot50 \| AF-J9GWN3-F1-MODEL\_V4 \| 1.0 \| 1.181e-19 \| 649 \| 0.202 \| 414 \| 281 \| 18 \| 369 \| 755 \| 82 \| 473 \| Helicase \| Helicase; afdb-uniprot50 \| AF-A0A7W5Y0N8-F1-MODEL\_V4 \| 1.0 \| 1.326e-21 \| 648 \| 0.17 \| 516 \| 340 \| 27 \| 260 \| 761 \| 258 \| 699 \| Uncharacterized protein \| Uncharacterized protein; afdb-uniprot50 \| AF-A0A4V2AU08-F1-MODEL\_V4 \| 1.0 \| 2.137e-23 \| 648 \| 0.228 \| 533 \| 330 \| 29 \| 289 \| 769 \| 255 \| 758 \| DNA primase \| DNA primase; afdb-uniprot50 \| AF-A0A318GP25-F1-MODEL\_V4 \| 1.0 \| 7.278e-21 \| 648 \| 0.21 \| 479 \| 297 \| 26 \| 332 \| 752 \| 326 \| 781 \| Virulence-associated protein E \| Virulence-associated protein E; afdb-uniprot50 \| AF-A0A5C8NRV2-F1-MODEL\_V4 \| 1.0 \| 9.792e-27 \| 648 \| 0.172 \| 789 \| 425 \| 35 \| 1 \| 755 \| 243 \| 837 \| PriCT\_2 domain-containing protein \| PriCT\_2 domain-containing protein; afdb-uniprot50 \| AF-A0A6L3T504-F1-MODEL\_V4 \| 1.0 \| 6.076e-33 \| 648 \| 0.213 \| 820 \| 450 \| 41 \| 100 \| 757 \| 84 \| 870 \| SF3 helicase domain-containing protein \| SF3 helicase domain-containing protein; afdb-uniprot50 \| AF-A0A3R8ZF92-F1-MODEL\_V4 \| 1.0 \| 1.165e-17 \| 647 \| 0.297 \| 299 \| 160 \| 11 \| 93 \| 355 \| 75 \| 359 \| Toprim domain-containing protein \| Toprim domain-containing protein; afdb-uniprot50 \| AF-A0A7Y3JLU6-F1-MODEL\_V4 \| 1.0 \| 9.122e-28 \| 647 \| 0.211 \| 767 \| 447 \| 49 \| 76 \| 769 \| 80 \| 761 \| Uncharacterized protein \| Uncharacterized protein; afdb-uniprot50 \| AF-Q0BD11-F1-MODEL\_V4 \| 1.0 \| 4.126e-29 \| 647 \| 0.197 \| 864 \| 411 \| 45 \| 12 \| 769 \| 7 \| 693 \| TOPRIM domain protein \| TOPRIM domain protein; afdb-uniprot50 \| AF-A0A1U9VG46-F1-MODEL\_V4 \| 1.0 \| 1.761e-25 \| 647 \| 0.199 \| 666 \| 418 \| 39 \| 164 \| 753 \| 153 \| 779 \| Uncharacterized protein \| Uncharacterized protein; afdb-uniprot50 \| AF-D4WHI7-F1-MODEL\_V4 \| 1.0 \| 9.729e-22 \| 646 \| 0.177 \| 513 \| 330 \| 23 \| 261 \| 756 \| 261 \| 698 \| VirE N-terminal domain protein \| VirE N-terminal domain protein; afdb-uniprot50 \| AF-A0A2M8UBK7-F1-MODEL\_V4 \| 1.0 \| 7.467e-17 \| 646 \| 0.344 \| 264 \| 139 \| 8 \| 102 \| 347 \| 460 \| 707 \| Uncharacterized protein \| Uncharacterized protein; afdb-uniprot50 \| AF-A0A3C1F3D1-F1-MODEL\_V4 \| 1.0 \| 2.644e-20 \| 645 \| 0.209 \| 449 \| 309 \| 19 \| 338 \| 769 \| 19 \| 438 \| Uncharacterized protein \| Uncharacterized protein; afdb-uniprot50 \| AF-B8IT86-F1-MODEL\_V4 \| 1.0 \| 5.623e-21 \| 645 \| 0.199 \| 547 \| 331 \| 24 \| 261 \| 752 \| 10 \| 504 \| Virulence-associated E family protein \| Virulence-associated E family protein; afdb-uniprot50 \| AF-A0A4R1QM10-F1-MODEL\_V4 \| 1.0 \| 1.878e-19 \| 644 \| 0.208 \| 431 \| 294 \| 21 \| 370 \| 769 \| 18 \| 432 \| Virulence-associated protein E \| Virulence-associated protein E; afdb-uniprot50 \| AF-A0A1H1AAV9-F1-MODEL\_V4 \| 1.0 \| 1.251e-16 \| 644 \| 0.286 \| 265 \| 157 \| 9 \| 102 \| 352 \| 276 \| 522 \| Uncharacterized domain associated with phage/plasmid primase \| Uncharacterized domain associated with phage/plasmid primase; afdb-uniprot50 \| AF-A0A351YBI0-F1-MODEL\_V4 \| 1.0 \| 1.005e-22 \| 644 \| 0.187 \| 539 \| 327 \| 30 \| 261 \| 767 \| 244 \| 703 \| Uncharacterized protein \| Uncharacterized protein; afdb-uniprot50 \| AF-A0A349MYJ9-F1-MODEL\_V4 \| 1.0 \| 1.866e-22 \| 644 \| 0.206 \| 537 \| 322 \| 26 \| 263 \| 751 \| 242 \| 722 \| Uncharacterized protein \| Uncharacterized protein; afdb-uniprot50 \| AF-E5UXW1-F1-MODEL\_V4 \| 1.0 \| 6.48e-19 \| 643 \| 0.195 \| 405 \| 297 \| 19 \| 370 \| 761 \| 56 \| 444 \| DUF3874 domain-containing protein \| DUF3874 domain-containing protein; afdb-uniprot50 \| AF-A0A2T1LQX5-F1-MODEL\_V4 \| 1.0 \| 6.275e-26 \| 643 \| 0.191 \| 653 \| 407 \| 34 \| 177 \| 769 \| 85 \| 676 \| DUF3854 domain-containing protein \| DUF3854 domain-containing protein; afdb-uniprot50 \| AF-A0A089PBP6-F1-MODEL\_V4 \| 1.0 \| 4.755e-19 \| 642 \| 0.223 \| 389 \| 268 \| 16 \| 384 \| 756 \| 51 \| 421 \| DNA primase \| DNA primase; afdb-uniprot50 \| AF-A0A2N5YK98-F1-MODEL\_V4 \| 1.0 \| 6.438e-22 \| 642 \| 0.181 \| 513 \| 353 \| 25 \| 260 \| 757 \| 193 \| 653 \| VirE\_N domain-containing protein \| VirE\_N domain-containing protein; afdb-uniprot50 \| AF-A0A0B1RHV7-F1-MODEL\_V4 \| 1.0 \| 1.352e-20 \| 642 \| 0.223 \| 434 \| 282 \| 23 \| 379 \| 768 \| 49 \| 471 \| Uncharacterized protein \| Uncharacterized protein; afdb-uniprot50 \| AF-A0A3R6QTT6-F1-MODEL\_V4 \| 1.0 \| 3.466e-22 \| 642 \| 0.188 \| 536 \| 353 \| 27 \| 246 \| 757 \| 216 \| 693 \| Uncharacterized protein \| Uncharacterized protein; afdb-uniprot50 \| AF-A0A396NE55-F1-MODEL\_V4 \| 1.0 \| 1.031e-18 \| 641 \| 0.204 \| 421 \| 286 \| 23 \| 370 \| 759 \| 2 \| 404 \| Virulence-associated protein E \| Virulence-associated protein E; afdb-uniprot50 \| AF-A0A2S7YZ26-F1-MODEL\_V4 \| 1.0 \| 5.844e-19 \| 641 \| 0.214 \| 442 \| 276 \| 23 \| 370 \| 756 \| 394 \| 819 \| Virulence-associated protein E \| Virulence-associated protein E; afdb-uniprot50 \| AF-A0A1M5AIN5-F1-MODEL\_V4 \| 1.0 \| 6.154e-19 \| 640 \| 0.169 \| 390 \| 294 \| 17 \| 381 \| 757 \| 58 \| 430 \| Virulence-associated protein E \| Virulence-associated protein E; afdb-uniprot50 \| AF-A0A646FJ01-F1-MODEL\_V4 \| 1.0 \| 1.842e-20 \| 640 \| 0.172 \| 528 \| 358 \| 27 \| 261 \| 751 \| 14 \| 499 \| Uncharacterized protein \| Uncharacterized protein; afdb-uniprot50 \| AF-A0A0P1IHX5-F1-MODEL\_V4 \| 1.0 \| 5.375e-26 \| 640 \| 0.173 \| 782 \| 364 \| 29 \| 2 \| 752 \| 234 \| 763 \| Putative P-loop ATPase \| Putative P-loop ATPase; afdb-uniprot50 \| AF-A0A521ZVA1-F1-MODEL\_V4 \| 1.0 \| 3.378e-26 \| 640 \| 0.198 \| 707 \| 430 \| 44 \| 160 \| 769 \| 180 \| 846 \| Uncharacterized protein \| Uncharacterized protein; afdb-uniprot50 \| AF-A0A6L3SS05-F1-MODEL\_V4 \| 1.0 \| 1.953e-33 \| 640 \| 0.223 \| 827 \| 443 \| 42 \| 100 \| 762 \| 121 \| 911 \| Toprim domain-containing protein \| Toprim domain-containing protein; afdb-uniprot50 \| AF-A0A5B5VUS0-F1-MODEL\_V4 \| 1.0 \| 2.802e-17 \| 638 \| 0.25 \| 300 \| 207 \| 11 \| 463 \| 755 \| 5 \| 293 \| Helicase \| Helicase; afdb-uniprot50 \| AF-A0A3E4JMT1-F1-MODEL\_V4 \| 1.0 \| 3.868e-19 \| 638 \| 0.19 \| 394 \| 281 \| 21 \| 381 \| 756 \| 45 \| 418 \| Helicase \| Helicase; afdb-uniprot50 \| AF-A0A7G8HY62-F1-MODEL\_V4 \| 1.0 \| 5.006e-19 \| 638 \| 0.206 \| 406 \| 280 \| 18 \| 370 \| 757 \| 75 \| 456 \| Virulence-associated E family protein \| Virulence-associated E family protein; afdb-uniprot50 \| AF-J9D1I1-F1-MODEL\_V4 \| 1.0 \| 2.695e-19 \| 638 \| 0.194 \| 412 \| 287 \| 18 \| 369 \| 755 \| 180 \| 571 \| Helicase \| Helicase; afdb-uniprot50 \| AF-A0A2D8XE83-F1-MODEL\_V4 \| 1.0 \| 6.154e-19 \| 638 \| 0.201 \| 397 \| 293 \| 16 \| 368 \| 753 \| 346 \| 729 \| Uncharacterized protein \| Uncharacterized protein; afdb-uniprot50 \| AF-A0A7Y0G5D6-F1-MODEL\_V4 \| 1.0 \| 3.356e-21 \| 638 \| 0.207 \| 481 \| 325 \| 26 \| 302 \| 756 \| 328 \| 778 \| PriCT\_2 domain-containing protein \| PriCT\_2 domain-containing protein; afdb-uniprot50 \| AF-A0A328I9Q7-F1-MODEL\_V4 \| 1.0 \| 1.107e-25 \| 638 \| 0.194 \| 658 \| 411 \| 35 \| 167 \| 769 \| 52 \| 645 \| DUF3854 domain-containing protein \| DUF3854 domain-containing protein; afdb-uniprot50 \| AF-A0A1Y6CYW9-F1-MODEL\_V4 \| 1.0 \| 3.028e-29 \| 638 \| 0.166 \| 992 \| 493 \| 57 \| 1 \| 735 \| 1 \| 915 \| Primase C terminal 2 (PriCT-2) \| Primase C terminal 2 (PriCT-2); afdb-uniprot50 \| AF-A0A1C5XKF3-F1-MODEL\_V4 \| 1.0 \| 9.299e-19 \| 637 \| 0.194 \| 427 \| 294 \| 23 \| 365 \| 757 \| 17 \| 427 \| Predicted P-loop ATPase and inactivated derivatives \| Predicted P-loop ATPase and inactivated derivatives; afdb-uniprot50 \| AF-A0A5A9EX48-F1-MODEL\_V4 \| 1.0 \| 1.75e-20 \| 637 \| 0.209 \| 468 \| 266 \| 21 \| 370 \| 751 \| 140 \| 589 \| Uncharacterized protein \| Uncharacterized protein; afdb-uniprot50 \| AF-A0A857J6L1-F1-MODEL\_V4 \| 1.0 \| 2.695e-19 \| 637 \| 0.243 \| 444 \| 275 \| 26 \| 369 \| 769 \| 365 \| 790 \| Uncharacterized protein \| Uncharacterized protein; afdb-uniprot50 \| AF-A0A6N2WVJ0-F1-MODEL\_V4 \| 1.0 \| 1.819e-18 \| 636 \| 0.184 \| 396 \| 284 \| 18 \| 384 \| 755 \| 157 \| 537 \| DUF based on B. Theta Gene description \| DUF based on B. Theta Gene description; afdb-uniprot50 \| AF-A0A251WJM1-F1-MODEL\_V4 \| 1.0 \| 3.25e-20 \| 636 \| 0.238 \| 444 \| 282 \| 21 \| 328 \| 755 \| 136 \| 539 \| Uncharacterized protein \| Uncharacterized protein; afdb-uniprot50 \| AF-A0A6S6TV87-F1-MODEL\_V4 \| 1.0 \| 3.893e-16 \| 636 \| 0.301 \| 265 \| 152 \| 9 \| 91 \| 347 \| 406 \| 645 \| DNA primase (EC) \| DNA primase (EC); afdb-uniprot50 \| AF-A0A2C6A5S6-F1-MODEL\_V4 \| 1.0 \| 1.978e-19 \| 636 \| 0.224 \| 450 \| 271 \| 25 \| 370 \| 757 \| 384 \| 817 \| Virulence-associated protein E \| Virulence-associated protein E; afdb-uniprot50 \| AF-A0A374WBI8-F1-MODEL\_V4 \| 1.0 \| 1.086e-18 \| 635 \| 0.195 \| 389 \| 282 \| 16 \| 384 \| 756 \| 34 \| 407 \| DUF3874 domain-containing protein \| DUF3874 domain-containing protein; afdb-uniprot50 \| AF-A0A255TI73-F1-MODEL\_V4 \| 1.0 \| 3.292e-22 \| 635 \| 0.147 \| 801 \| 396 \| 30 \| 1 \| 769 \| 34 \| 579 \| PriCT\_2 domain-containing protein \| PriCT\_2 domain-containing protein; afdb-uniprot50 \| AF-A0A258ZJK0-F1-MODEL\_V4 \| 1.0 \| 5.375e-26 \| 635 \| 0.196 \| 672 \| 422 \| 41 \| 181 \| 769 \| 43 \| 679 \| Uncharacterized protein \| Uncharacterized protein; afdb-uniprot50 \| AF-A0A7R8AWV8-F1-MODEL\_V4 \| 1.0 \| 6.356e-20 \| 635 \| 0.203 \| 438 \| 317 \| 20 \| 332 \| 753 \| 312 \| 733 \| Prim-Pol domain-containing protein \| Prim-Pol domain-containing protein; afdb-uniprot50 \| AF-I3YBE5-F1-MODEL\_V4 \| 1.0 \| 8.175e-31 \| 635 \| 0.194 \| 945 \| 523 \| 56 \| 1 \| 763 \| 72 \| 959 \| Putative ATPase \| Putative ATPase; afdb-uniprot50 \| AF-A0A5C7R6K8-F1-MODEL\_V4 \| 1.0 \| 2.165e-17 \| 634 \| 0.363 \| 253 \| 136 \| 11 \| 99 \| 346 \| 78 \| 310 \| Toprim domain-containing protein \| Toprim domain-containing protein; afdb-uniprot50 \| AF-A0A849UD71-F1-MODEL\_V4 \| 1.0 \| 2.988e-19 \| 634 \| 0.277 \| 331 \| 214 \| 10 \| 101 \| 431 \| 69 \| 374 \| Toprim domain-containing protein \| Toprim domain-containing protein; afdb-uniprot50 \| AF-X5UMY1-F1-MODEL\_V4 \| 1.0 \| 4.288e-19 \| 634 \| 0.219 \| 428 \| 291 \| 19 \| 350 \| 757 \| 64 \| 468 \| Virulence protein E \| Virulence protein E; afdb-uniprot50 \| AF-E3PGT5-F1-MODEL\_V4 \| 1.0 \| 1.352e-20 \| 634 \| 0.219 \| 464 \| 298 \| 26 \| 319 \| 753 \| 139 \| 567 \| Putative virulence associated protein virE \| Putative virulence associated protein virE; afdb-uniprot50 \| AF-A0A7Y6YZ19-F1-MODEL\_V4 \| 1.0 \| 5.305e-16 \| 634 \| 0.305 \| 255 \| 148 \| 8 \| 100 \| 344 \| 715 \| 950 \| DUF1738 domain-containing protein \| DUF1738 domain-containing protein; afdb-uniprot50 \| AF-F0F4L8-F1-MODEL\_V4 \| 1.0 \| 1.188e-16 \| 633 \| 0.237 \| 303 \| 215 \| 10 \| 466 \| 758 \| 4 \| 300 \| Virulence-associated protein E \| Virulence-associated protein E; afdb-uniprot50 \| AF-A0A212JQE2-F1-MODEL\_V4 \| 1.0 \| 4.604e-18 \| 633 \| 0.218 \| 366 \| 239 \| 14 \| 370 \| 723 \| 4 \| 334 \| Uncharacterized protein \| Uncharacterized protein; afdb-uniprot50 \| AF-A0A164AHS2-F1-MODEL\_V4 \| 1.0 \| 1.451e-19 \| 633 \| 0.251 \| 398 \| 247 \| 19 \| 381 \| 750 \| 51 \| 425 \| Virulence-associated protein E \| Virulence-associated protein E; afdb-uniprot50 \| AF-A0A416H079-F1-MODEL\_V4 \| 1.0 \| 3.027e-21 \| 633 \| 0.2 \| 485 \| 322 \| 26 \| 333 \| 769 \| 241 \| 707 \| Uncharacterized protein \| Uncharacterized protein; afdb-uniprot50 \| AF-A0A068R0U5-F1-MODEL\_V4 \| 1.0 \| 4.126e-21 \| 633 \| 0.214 \| 457 \| 301 \| 25 \| 327 \| 753 \| 294 \| 722 \| Predicted P-loop ATPase and inactivated derivatives \| Predicted P-loop ATPase and inactivated derivatives; afdb-uniprot50 \| AF-A0A844YF13-F1-MODEL\_V4 \| 1.0 \| 6.912e-21 \| 633 \| 0.207 \| 511 \| 348 \| 21 \| 265 \| 758 \| 247 \| 717 \| Uncharacterized protein \| Uncharacterized protein; afdb-uniprot50 \| AF-A0A255T784-F1-MODEL\_V4 \| 1.0 \| 4.515e-19 \| 632 \| 0.219 \| 405 \| 277 \| 14 \| 371 \| 752 \| 213 \| 601 \| Uncharacterized protein \| Uncharacterized protein; afdb-uniprot50 \| AF-A0A7Z0SBX3-F1-MODEL\_V4 \| 1.0 \| 3.65e-22 \| 632 \| 0.194 \| 525 \| 340 \| 25 \| 262 \| 753 \| 146 \| 620 \| Uncharacterized protein \| Uncharacterized protein; afdb-uniprot50 \| AF-A0A7G8FP95-F1-MODEL\_V4 \| 1.0 \| 8.831e-19 \| 632 \| 0.227 \| 400 \| 273 \| 20 \| 370 \| 753 \| 278 \| 657 \| Virulence-associated E family protein \| Virulence-associated E family protein; afdb-uniprot50 \| AF-A0A2T0NTU5-F1-MODEL\_V4 \| 1.0 \| 6.912e-21 \| 632 \| 0.2 \| 485 \| 326 \| 26 \| 295 \| 757 \| 215 \| 659 \| Primase-like protein \| Primase-like protein; afdb-uniprot50 \| AF-A0A255ZEU1-F1-MODEL\_V4 \| 1.0 \| 2.207e-32 \| 632 \| 0.218 \| 877 \| 455 \| 51 \| 8 \| 737 \| 2 \| 794 \| Uncharacterized protein \| Uncharacterized protein; afdb-uniprot50 \| AF-U2QHM6-F1-MODEL\_V4 \| 1.0 \| 1.819e-18 \| 631 \| 0.199 \| 427 \| 292 \| 23 \| 365 \| 757 \| 17 \| 427 \| Virulence-associated protein E \| Virulence-associated protein E; afdb-uniprot50 \| AF-A0A0S6UW49-F1-MODEL\_V4 \| 1.0 \| 3.147e-19 \| 631 \| 0.205 \| 433 \| 278 \| 22 \| 370 \| 756 \| 11 \| 423 \| Uncharacterized protein \| Uncharacterized protein; afdb-uniprot50 \| AF-A0A1M4VDX6-F1-MODEL\_V4 \| 1.0 \| 4.574e-21 \| 631 \| 0.195 \| 512 \| 314 \| 26 \| 264 \| 755 \| 259 \| 692 \| Uncharacterized protein \| Uncharacterized protein; afdb-uniprot50 \| AF-Q6AIG1-F1-MODEL\_V4 \| 1.0 \| 2.494e-15 \| 631 \| 0.32 \| 250 \| 152 \| 6 \| 100 \| 346 \| 449 \| 683 \| Related to DNA primase TraC \| Related to DNA primase TraC; afdb-uniprot50 \| AF-A0A252DZZ4-F1-MODEL\_V4 \| 1.0 \| 6.607e-26 \| 631 \| 0.197 \| 678 \| 422 \| 40 \| 151 \| 762 \| 25 \| 646 \| DUF3854 domain-containing protein \| DUF3854 domain-containing protein; afdb-uniprot50 \| AF-A0A2S0V1Y7-F1-MODEL\_V4 \| 1.0 \| 2.447e-16 \| 630 \| 0.291 \| 257 \| 153 \| 9 \| 92 \| 347 \| 67 \| 295 \| Toprim domain-containing protein \| Toprim domain-containing protein; afdb-uniprot50 \| AF-A0A2D9N1G6-F1-MODEL\_V4 \| 1.0 \| 1.165e-17 \| 630 \| 0.224 \| 339 \| 237 \| 16 \| 370 \| 690 \| 27 \| 357 \| Uncharacterized protein \| Uncharacterized protein; afdb-uniprot50 \| AF-A0A126UX24-F1-MODEL\_V4 \| 1.0 \| 3.603e-20 \| 630 \| 0.197 \| 450 \| 312 \| 18 \| 329 \| 758 \| 183 \| 603 \| Uncharacterized protein \| Uncharacterized protein; afdb-uniprot50 \| AF-A0A1Q6HTS3-F1-MODEL\_V4 \| 1.0 \| 7.763e-23 \| 630 \| 0.188 \| 572 \| 350 \| 31 \| 213 \| 756 \| 212 \| 697 \| VirE\_N domain-containing protein \| VirE\_N domain-containing protein; afdb-uniprot50 \| AF-A0A1H1JG68-F1-MODEL\_V4 \| 1.0 \| 2.856e-16 \| 630 \| 0.274 \| 270 \| 164 \| 8 \| 100 \| 352 \| 459 \| 713 \| Antirestriction protein ArdC \| Antirestriction protein ArdC; afdb-uniprot50 \| AF-A0A2D8CJQ3-F1-MODEL\_V4 \| 1.0 \| 7.863e-17 \| 629 \| 0.282 \| 262 \| 181 \| 5 \| 498 \| 756 \| 2 \| 259 \| Uncharacterized protein \| Uncharacterized protein; afdb-uniprot50 \| AF-E6K9V5-F1-MODEL\_V4 \| 1.0 \| 5.374e-18 \| 629 \| 0.234 \| 350 \| 229 \| 11 \| 379 \| 699 \| 23 \| 362 \| Virulence-associated protein E \| Virulence-associated protein E; afdb-uniprot50 \| AF-A0A2E0B660-F1-MODEL\_V4 \| 1.0 \| 1.558e-18 \| 629 \| 0.205 \| 395 \| 277 \| 18 \| 394 \| 769 \| 3 \| 379 \| Uncharacterized protein \| Uncharacterized protein; afdb-uniprot50 \| AF-A0A496KNL5-F1-MODEL\_V4 \| 1.0 \| 7.42e-20 \| 629 \| 0.206 \| 455 \| 299 \| 21 \| 329 \| 752 \| 4 \| 427 \| Helicase \| Helicase; afdb-uniprot50 \| AF-A0A345REX8-F1-MODEL\_V4 \| 1.0 \| 3.313e-19 \| 629 \| 0.2 \| 433 \| 286 \| 24 \| 370 \| 757 \| 39 \| 456 \| Uncharacterized protein \| Uncharacterized protein; afdb-uniprot50 \| AF-A0A7Y5C283-F1-MODEL\_V4 \| 1.0 \| 9.666e-25 \| 629 \| 0.172 \| 636 \| 435 \| 34 \| 153 \| 757 \| 106 \| 680 \| Uncharacterized protein \| Uncharacterized protein; afdb-uniprot50 \| AF-B7UEZ2-F1-MODEL\_V4 \| 1.0 \| 2.029e-15 \| 629 \| 0.276 \| 239 \| 145 \| 7 \| 117 \| 346 \| 164 \| 383 \| DNA primase \| DNA primase; afdb-uniprot50 \| AF-A0A6F8PSK9-F1-MODEL\_V4 \| 1.0 \| 7.813e-20 \| 628 \| 0.26 \| 395 \| 220 \| 13 \| 3 \| 357 \| 1 \| 363 \| Uncharacterized protein \| Uncharacterized protein; afdb-uniprot50 \| AF-A0A6L9HKV5-F1-MODEL\_V4 \| 1.0 \| 2.893e-18 \| 628 \| 0.199 \| 436 \| 287 \| 17 \| 370 \| 755 \| 6 \| 429 \| Virulence-associated protein E \| Virulence-associated protein E; afdb-uniprot50 \| AF-A0A1C6HAC9-F1-MODEL\_V4 \| 1.0 \| 3.868e-19 \| 628 \| 0.19 \| 410 \| 291 \| 21 \| 370 \| 757 \| 44 \| 434 \| Predicted P-loop ATPase and inactivated derivatives \| Predicted P-loop ATPase and inactivated derivatives; afdb-uniprot50 \| AF-G5SV99-F1-MODEL\_V4 \| 1.0 \| 4.073e-19 \| 628 \| 0.184 \| 391 \| 273 \| 21 \| 384 \| 751 \| 102 \| 469 \| DUF3874 domain-containing protein \| DUF3874 domain-containing protein; afdb-uniprot50 \| AF-A0A7G8C997-F1-MODEL\_V4 \| 1.0 \| 2.431e-19 \| 628 \| 0.2 \| 428 \| 283 \| 21 \| 363 \| 752 \| 81 \| 487 \| Virulence-associated E family protein \| Virulence-associated E family protein; afdb-uniprot50 \| AF-A0A1X9YSK3-F1-MODEL\_V4 \| 1.0 \| 5.271e-19 \| 628 \| 0.2 \| 413 \| 289 \| 20 \| 372 \| 755 \| 56 \| 456 \| Uncharacterized protein \| Uncharacterized protein; afdb-uniprot50 \| AF-A0A7T1MDW8-F1-MODEL\_V4 \| 1.0 \| 6.823e-19 \| 628 \| 0.202 \| 410 \| 275 \| 18 \| 379 \| 757 \| 245 \| 633 \| Uncharacterized protein \| Uncharacterized protein; afdb-uniprot50 \| AF-A0A846FJ73-F1-MODEL\_V4 \| 1.0 \| 5.77e-25 \| 628 \| 0.208 \| 629 \| 389 \| 35 \| 178 \| 757 \| 70 \| 638 \| DUF3854 domain-containing protein \| DUF3854 domain-containing protein; afdb-uniprot50 \| AF-A0A6I2N1Y2-F1-MODEL\_V4 \| 1.0 \| 1.196e-21 \| 627 \| 0.178 \| 526 \| 357 \| 23 \| 260 \| 756 \| 236 \| 715 \| VirE\_N domain-containing protein \| VirE\_N domain-containing protein; afdb-uniprot50 \| AF-A0A7J4XCA9-F1-MODEL\_V4 \| 1.0 \| 7.138e-22 \| 627 \| 0.18 \| 531 \| 341 \| 25 \| 242 \| 755 \| 252 \| 705 \| VirE\_N domain-containing protein \| VirE\_N domain-containing protein; afdb-uniprot50 \| AF-A0A0X3TJ34-F1-MODEL\_V4 \| 1.0 \| 2.123e-18 \| 626 \| 0.292 \| 390 \| 196 \| 17 \| 3 \| 336 \| 1 \| 366 \| Uncharacterized protein \| Uncharacterized protein; afdb-uniprot50 \| AF-A0A2W6Z2Y3-F1-MODEL\_V4 \| 1.0 \| 1.479e-18 \| 626 \| 0.237 \| 408 \| 262 \| 21 \| 369 \| 755 \| 20 \| 399 \| Uncharacterized protein \| Uncharacterized protein; afdb-uniprot50 \| AF-A0A1M6CUR8-F1-MODEL\_V4 \| 1.0 \| 5.659e-18 \| 626 \| 0.187 \| 385 \| 277 \| 17 \| 384 \| 751 \| 50 \| 415 \| Virulence-associated protein E \| Virulence-associated protein E; afdb-uniprot50 \| AF-A0A5P0WS35-F1-MODEL\_V4 \| 1.0 \| 2.838e-19 \| 626 \| 0.204 \| 386 \| 274 \| 19 \| 384 \| 755 \| 75 \| 441 \| DUF3874 domain-containing protein \| DUF3874 domain-containing protein; afdb-uniprot50 \| AF-A0A239KSC1-F1-MODEL\_V4 \| 1.0 \| 1.94e-20 \| 626 \| 0.201 \| 501 \| 324 \| 26 \| 292 \| 753 \| 35 \| 498 \| Predicted P-loop ATPase and inactivated derivatives \| Predicted P-loop ATPase and inactivated derivatives; afdb-uniprot50 \| AF-B2J1H6-F1-MODEL\_V4 \| 1.0 \| 8.28e-25 \| 626 \| 0.203 \| 644 \| 398 \| 39 \| 171 \| 756 \| 53 \| 639 \| Virulence-associated E family protein \| Virulence-associated E family protein; afdb-uniprot50 \| AF-K1JGA8-F1-MODEL\_V4 \| 1.0 \| 1.387e-32 \| 626 \| 0.2 \| 888 \| 431 \| 50 \| 100 \| 752 \| 92 \| 935 \| Toprim domain-containing protein \| Toprim domain-containing protein; afdb-uniprot50 \| AF-F9D560-F1-MODEL\_V4 \| 1.0 \| 2.207e-16 \| 625 \| 0.267 \| 277 \| 190 \| 8 \| 471 \| 742 \| 12 \| 280 \| Virulence-associated protein E \| Virulence-associated protein E; afdb-uniprot50 \| AF-A0A3E5E5N8-F1-MODEL\_V4 \| 1.0 \| 1.784e-19 \| 625 \| 0.24 \| 440 \| 257 \| 21 \| 370 \| 747 \| 10 \| 434 \| Virulence-associated E family protein \| Virulence-associated E family protein; afdb-uniprot50 \| AF-M4UF70-F1-MODEL\_V4 \| 1.0 \| 1.538e-24 \| 625 \| 0.206 \| 643 \| 396 \| 37 \| 181 \| 769 \| 101 \| 682 \| Uncharacterized protein \| Uncharacterized protein; afdb-uniprot50 \| AF-A0A7R8BVV0-F1-MODEL\_V4 \| 1.0 \| 7.047e-20 \| 625 \| 0.202 \| 478 \| 318 \| 26 \| 328 \| 757 \| 218 \| 680 \| Uncharacterized protein \| Uncharacterized protein; afdb-uniprot50 \| AF-A0A5P3VSJ1-F1-MODEL\_V4 \| 1.0 \| 4.785e-16 \| 624 \| 0.317 \| 252 \| 151 \| 9 \| 99 \| 347 \| 75 \| 308 \| Toprim domain-containing protein \| Toprim domain-containing protein; afdb-uniprot50 \| AF-A0A3G3GKY0-F1-MODEL\_V4 \| 1.0 \| 1.558e-18 \| 624 \| 0.203 \| 402 \| 281 \| 14 \| 382 \| 756 \| 9 \| 398 \| Uncharacterized protein \| Uncharacterized protein; afdb-uniprot50 \| AF-R6LH63-F1-MODEL\_V4 \| 1.0 \| 4.604e-18 \| 624 \| 0.189 \| 386 \| 275 \| 19 \| 384 \| 751 \| 53 \| 418 \| DUF3874 domain-containing protein \| DUF3874 domain-containing protein; afdb-uniprot50 \| AF-A0A3S4WPJ7-F1-MODEL\_V4 \| 1.0 \| 4.206e-20 \| 624 \| 0.213 \| 460 \| 308 \| 27 \| 319 \| 755 \| 293 \| 721 \| Predicted P-loop ATPase and inactivated derivatives \| Predicted P-loop ATPase and inactivated derivatives; afdb-uniprot50 \| AF-A0A7U3BA69-F1-MODEL\_V4 \| 1.0 \| 2.931e-20 \| 624 \| 0.214 \| 461 \| 299 \| 25 \| 319 \| 751 \| 293 \| 718 \| Replication protein \| Replication protein; afdb-uniprot50 \| AF-A0A1Z4R883-F1-MODEL\_V4 \| 1.0 \| 3.627e-25 \| 624 \| 0.204 \| 655 \| 409 \| 33 \| 168 \| 769 \| 82 \| 677 \| Virulence-associated E family protein \| Virulence-associated E family protein; afdb-uniprot50 \| AF-A0A2C6W764-F1-MODEL\_V4 \| 1.0 \| 4.847e-18 \| 623 \| 0.199 \| 396 \| 272 \| 21 \| 379 \| 753 \| 2 \| 373 \| Uncharacterized protein \| Uncharacterized protein; afdb-uniprot50 \| AF-A0A7C9NE92-F1-MODEL\_V4 \| 1.0 \| 5.34e-21 \| 623 \| 0.204 \| 513 \| 330 \| 25 \| 263 \| 751 \| 321 \| 779 \| Uncharacterized protein \| Uncharacterized protein; afdb-uniprot50 \| AF-A0A6S6U6M2-F1-MODEL\_V4 \| 1.0 \| 1.343e-15 \| 623 \| 0.311 \| 260 \| 156 \| 7 \| 91 \| 347 \| 560 \| 799 \| DNA primase (EC) \| DNA primase (EC); afdb-uniprot50 \| AF-A0A2A4P7I2-F1-MODEL\_V4 \| 1.0 \| 8.831e-19 \| 622 \| 0.235 \| 390 \| 259 \| 18 \| 385 \| 752 \| 38 \| 410 \| Uncharacterized protein \| Uncharacterized protein; afdb-uniprot50 \| AF-S9S807-F1-MODEL\_V4 \| 1.0 \| 8.016e-16 \| 622 \| 0.348 \| 241 \| 124 \| 10 \| 120 \| 347 \| 610 \| 830 \| Toprim domain-containing protein \| Toprim domain-containing protein; afdb-uniprot50 \| AF-A0A2S6VC66-F1-MODEL\_V4 \| 1.0 \| 1.387e-16 \| 621 \| 0.263 \| 307 \| 203 \| 12 \| 460 \| 757 \| 2 \| 294 \| Uncharacterized protein \| Uncharacterized protein; afdb-uniprot50 \| AF-A0A858Q8N5-F1-MODEL\_V4 \| 1.0 \| 5.697e-31 \| 621 \| 0.186 \| 928 \| 516 \| 52 \| 12 \| 755 \| 7 \| 879 \| Uncharacterized protein \| Uncharacterized protein; afdb-uniprot50 \| AF-A0A6L7WJY2-F1-MODEL\_V4 \| 1.0 \| 1.188e-16 \| 620 \| 0.277 \| 332 \| 211 \| 14 \| 379 \| 687 \| 21 \| 346 \| Uncharacterized protein \| Uncharacterized protein; afdb-uniprot50 \| AF-A0A3D5PJD2-F1-MODEL\_V4 \| 1.0 \| 9.299e-19 \| 620 \| 0.224 \| 405 \| 247 \| 16 \| 370 \| 737 \| 20 \| 394 \| Uncharacterized protein \| Uncharacterized protein; afdb-uniprot50 \| AF-A0A7W3DK48-F1-MODEL\_V4 \| 1.0 \| 7.965e-19 \| 620 \| 0.203 \| 422 \| 273 \| 24 \| 372 \| 755 \| 35 \| 431 \| Uncharacterized protein \| Uncharacterized protein; afdb-uniprot50 \| AF-A0A660LQS4-F1-MODEL\_V4 \| 1.0 \| 9.791e-19 \| 620 \| 0.203 \| 408 \| 287 \| 18 \| 369 \| 757 \| 85 \| 473 \| Helicase \| Helicase; afdb-uniprot50 \| AF-A0A1H2RHG9-F1-MODEL\_V4 \| 1.0 \| 7.663e-21 \| 620 \| 0.208 \| 504 \| 310 \| 24 \| 327 \| 762 \| 2 \| 484 \| Predicted P-loop ATPase and inactivated derivatives \| Predicted P-loop ATPase and inactivated derivatives; afdb-uniprot50 \| AF-A0A857L932-F1-MODEL\_V4 \| 1.0 \| 9.299e-19 \| 620 \| 0.212 \| 419 \| 271 \| 24 \| 380 \| 757 \| 110 \| 510 \| Uncharacterized protein \| Uncharacterized protein; afdb-uniprot50 \| AF-I5B0X5-F1-MODEL\_V4 \| 1.0 \| 9.854e-16 \| 620 \| 0.301 \| 239 \| 135 \| 11 \| 116 \| 346 \| 551 \| 765 \| Antirestriction protein \| Antirestriction protein; afdb-uniprot50 \| AF-A0A178MCH8-F1-MODEL\_V4 \| 1.0 \| 1.317e-16 \| 620 \| 0.34 \| 244 \| 123 \| 10 \| 120 \| 347 \| 567 \| 788 \| Toprim domain-containing protein \| Toprim domain-containing protein; afdb-uniprot50 \| AF-A0A661E6V8-F1-MODEL\_V4 \| 1.0 \| 1.795e-24 \| 620 \| 0.161 \| 918 \| 350 \| 29 \| 195 \| 757 \| 348 \| 1200 \| Uncharacterized protein \| Uncharacterized protein; afdb-uniprot50 \| AF-C9MTT5-F1-MODEL\_V4 \| 1.0 \| 6.606e-18 \| 619 \| 0.252 \| 329 \| 207 \| 13 \| 458 \| 756 \| 3 \| 322 \| Virulence-associated protein E \| Virulence-associated protein E; afdb-uniprot50 \| AF-A0A661CSP6-F1-MODEL\_V4 \| 1.0 \| 7.565e-19 \| 619 \| 0.212 \| 429 \| 288 \| 21 \| 354 \| 757 \| 6 \| 409 \| Uncharacterized protein \| Uncharacterized protein; afdb-uniprot50 \| AF-R5KUI2-F1-MODEL\_V4 \| 1.0 \| 3.25e-20 \| 619 \| 0.195 \| 444 \| 317 \| 20 \| 329 \| 757 \| 281 \| 699 \| Uncharacterized protein \| Uncharacterized protein; afdb-uniprot50 \| AF-A0A1I2KRS2-F1-MODEL\_V4 \| 1.0 \| 1.044e-20 \| 619 \| 0.192 \| 534 \| 343 \| 26 \| 260 \| 755 \| 281 \| 764 \| Virulence-associated protein E \| Virulence-associated protein E; afdb-uniprot50 \| AF-A0A3M0XHA9-F1-MODEL\_V4 \| 1.0 \| 8.551e-18 \| 618 \| 0.318 \| 308 \| 189 \| 11 \| 466 \| 766 \| 1 \| 294 \| Uncharacterized protein \| Uncharacterized protein; afdb-uniprot50 \| AF-A0A4P2SPK5-F1-MODEL\_V4 \| 1.0 \| 1.46e-16 \| 618 \| 0.308 \| 256 \| 145 \| 12 \| 107 \| 358 \| 155 \| 382 \| Uncharacterized protein \| Uncharacterized protein; afdb-uniprot50 \| AF-A0A7K3KGX5-F1-MODEL\_V4 \| 1.0 \| 1.807e-21 \| 618 \| 0.19 \| 547 \| 338 \| 28 \| 261 \| 768 \| 123 \| 603 \| VirE\_N domain-containing protein \| VirE\_N domain-containing protein; afdb-uniprot50 \| AF-A0A3N5GQR2-F1-MODEL\_V4 \| 1.0 \| 1.079e-21 \| 618 \| 0.208 \| 546 \| 354 \| 27 \| 231 \| 757 \| 218 \| 704 \| Uncharacterized protein \| Uncharacterized protein; afdb-uniprot50 \| AF-A0A127SE37-F1-MODEL\_V4 \| 1.0 \| 2.324e-16 \| 617 \| 0.469 \| 196 \| 103 \| 1 \| 562 \| 757 \| 6 \| 200 \| Virulence-associated protein E \| Virulence-associated protein E; afdb-uniprot50 \| AF-A0A522RNX4-F1-MODEL\_V4 \| 1.0 \| 8.718e-17 \| 617 \| 0.301 \| 252 \| 139 \| 11 \| 92 \| 333 \| 79 \| 303 \| Toprim domain-containing protein \| Toprim domain-containing protein; afdb-uniprot50 \| AF-A0A134BGQ9-F1-MODEL\_V4 \| 1.0 \| 4.693e-17 \| 617 \| 0.268 \| 298 \| 195 \| 10 \| 452 \| 735 \| 35 \| 323 \| Virulence-associated protein E \| Virulence-associated protein E; afdb-uniprot50 \| AF-A0A1G2ZMJ1-F1-MODEL\_V4 \| 1.0 \| 1.508e-17 \| 617 \| 0.184 \| 402 \| 291 \| 16 \| 370 \| 755 \| 37 \| 417 \| Uncharacterized protein \| Uncharacterized protein; afdb-uniprot50 \| AF-A0A1G8LAT9-F1-MODEL\_V4 \| 1.0 \| 5.92e-21 \| 617 \| 0.217 \| 473 \| 304 \| 26 \| 335 \| 769 \| 13 \| 457 \| Virulence-associated protein E \| Virulence-associated protein E; afdb-uniprot50 \| AF-A0A3C0D4T8-F1-MODEL\_V4 \| 1.0 \| 1.158e-20 \| 617 \| 0.167 \| 548 \| 371 \| 30 \| 265 \| 768 \| 259 \| 765 \| VirE\_N domain-containing protein \| VirE\_N domain-containing protein; afdb-uniprot50 \| AF-A0A252DL02-F1-MODEL\_V4 \| 1.0 \| 2.527e-25 \| 617 \| 0.195 \| 689 \| 403 \| 37 \| 153 \| 757 \| 26 \| 646 \| KOW domain-containing protein \| KOW domain-containing protein; afdb-uniprot50 \| AF-A6GUP7-F1-MODEL\_V4 \| 1.0 \| 5.586e-16 \| 617 \| 0.386 \| 233 \| 121 \| 8 \| 116 \| 347 \| 780 \| 991 \| Toprim domain-containing protein \| Toprim domain-containing protein; afdb-uniprot50 \| AF-A0A0F9W6J4-F1-MODEL\_V4 \| 1.0 \| 1.46e-16 \| 616 \| 0.241 \| 306 \| 206 \| 12 \| 370 \| 659 \| 33 \| 328 \| Uncharacterized protein \| Uncharacterized protein; afdb-uniprot50 \| AF-A0A414JB35-F1-MODEL\_V4 \| 1.0 \| 5.17e-20 \| 616 \| 0.215 \| 440 \| 284 \| 23 \| 370 \| 769 \| 25 \| 443 \| Uncharacterized protein \| Uncharacterized protein; afdb-uniprot50 \| AF-A0A1T4ZJ82-F1-MODEL\_V4 \| 1.0 \| 9.666e-25 \| 616 \| 0.203 \| 673 \| 387 \| 34 \| 181 \| 769 \| 18 \| 625 \| DUF3854 domain-containing protein \| DUF3854 domain-containing protein; afdb-uniprot50 \| AF-A0A3C0FNH6-F1-MODEL\_V4 \| 1.0 \| 9.358e-16 \| 615 \| 0.342 \| 228 \| 123 \| 9 \| 128 \| 349 \| 3 \| 209 \| Toprim domain-containing protein \| Toprim domain-containing protein; afdb-uniprot50 \| AF-A0A417P6N5-F1-MODEL\_V4 \| 1.0 \| 2.661e-17 \| 615 \| 0.207 \| 386 \| 259 \| 19 \| 370 \| 727 \| 2 \| 368 \| Virulence-associated protein E \| Virulence-associated protein E; afdb-uniprot50 \| AF-A0A1S1NXN9-F1-MODEL\_V4 \| 1.0 \| 4.574e-21 \| 615 \| 0.194 \| 530 \| 340 \| 27 \| 285 \| 750 \| 337 \| 843 \| Uncharacterized protein \| Uncharacterized protein; afdb-uniprot50 \| AF-A0A353E5I1-F1-MODEL\_V4 \| 1.0 \| 5.071e-21 \| 614 \| 0.18 \| 514 \| 331 \| 23 \| 261 \| 755 \| 249 \| 691 \| Helicase \| Helicase; afdb-uniprot50 \| AF-A0A662MET1-F1-MODEL\_V4 \| 1.0 \| 5.204e-25 \| 614 \| 0.18 \| 663 \| 411 \| 34 \| 181 \| 757 \| 149 \| 764 \| Uncharacterized protein \| Uncharacterized protein; afdb-uniprot50 \| AF-I5D2C8-F1-MODEL\_V4 \| 1.0 \| 3.919e-29 \| 614 \| 0.189 \| 854 \| 484 \| 47 \| 6 \| 736 \| 2 \| 769 \| Uncharacterized protein \| Uncharacterized protein; afdb-uniprot50 \| AF-A0A6B2KQS6-F1-MODEL\_V4 \| 1.0 \| 2.593e-29 \| 614 \| 0.183 \| 936 \| 464 \| 54 \| 10 \| 737 \| 4 \| 846 \| DUF927 domain-containing protein \| DUF927 domain-containing protein; afdb-uniprot50 \| AF-A0A3Q9MY88-F1-MODEL\_V4 \| 1.0 \| 1.489e-15 \| 614 \| 0.295 \| 244 \| 140 \| 8 \| 120 \| 350 \| 170 \| 394 \| Uncharacterized protein \| Uncharacterized protein; afdb-uniprot50 \| AF-A0A0D6MP65-F1-MODEL\_V4 \| 1.0 \| 1.662e-20 \| 613 \| 0.206 \| 522 \| 327 \| 25 \| 262 \| 752 \| 224 \| 689 \| Virulence-associated E family protein \| Virulence-associated E family protein; afdb-uniprot50 \| AF-A0A2I9D7G1-F1-MODEL\_V4 \| 1.0 \| 3.819e-25 \| 613 \| 0.196 \| 635 \| 401 \| 35 \| 178 \| 757 \| 42 \| 621 \| Virulence protein E \| Virulence protein E; afdb-uniprot50 \| AF-A0A4R3GWI1-F1-MODEL\_V4 \| 1.0 \| 8.552e-26 \| 613 \| 0.222 \| 696 \| 395 \| 39 \| 174 \| 769 \| 157 \| 806 \| CHC2-type zinc finger protein \| CHC2-type zinc finger protein; afdb-uniprot50 \| AF-A0A1Y6ISW7-F1-MODEL\_V4 \| 1.0 \| 5.769e-17 \| 612 \| 0.292 \| 291 \| 170 \| 11 \| 494 \| 756 \| 2 \| 284 \| Virulence-associated protein E \| Virulence-associated protein E; afdb-uniprot50 \| AF-A0A285D5W0-F1-MODEL\_V4 \| 1.0 \| 3.745e-18 \| 612 \| 0.236 \| 384 \| 246 \| 18 \| 407 \| 757 \| 5 \| 374 \| Virulence-associated protein E \| Virulence-associated protein E; afdb-uniprot50 \| AF-A0A2Z5WWH0-F1-MODEL\_V4 \| 1.0 \| 2.695e-19 \| 612 \| 0.223 \| 412 \| 257 \| 19 \| 368 \| 751 \| 36 \| 412 \| Virulence-associated E \| Virulence-associated E; afdb-uniprot50 \| AF-A0A522ISJ4-F1-MODEL\_V4 \| 1.0 \| 8.175e-31 \| 612 \| 0.212 \| 830 \| 438 \| 43 \| 98 \| 750 \| 95 \| 885 \| Toprim domain-containing protein \| Toprim domain-containing protein; afdb-uniprot50 \| AF-A0A1Z5HMC9-F1-MODEL\_V4 \| 1.0 \| 4.126e-21 \| 611 \| 0.22 \| 472 \| 319 \| 17 \| 260 \| 715 \| 204 \| 642 \| VirE\_N domain-containing protein \| VirE\_N domain-containing protein; afdb-uniprot50 \| AF-A0A1Z4U8L3-F1-MODEL\_V4 \| 1.0 \| 3.008e-24 \| 611 \| 0.18 \| 660 \| 430 \| 40 \| 153 \| 756 \| 21 \| 625 \| Virulence-associated E family protein \| Virulence-associated E family protein; afdb-uniprot50 \| AF-A0A844A8E2-F1-MODEL\_V4 \| 1.0 \| 4.786e-24 \| 611 \| 0.176 \| 780 \| 367 \| 29 \| 1 \| 761 \| 285 \| 807 \| Uncharacterized protein \| Uncharacterized protein; afdb-uniprot50 \| AF-A0A286GSM2-F1-MODEL\_V4 \| 1.0 \| 1.83e-15 \| 611 \| 0.305 \| 255 \| 153 \| 9 \| 102 \| 347 \| 689 \| 928 \| Antirestriction protein ArdC \| Antirestriction protein ArdC; afdb-uniprot50 \| AF-A0A7C9LWM4-F1-MODEL\_V4 \| 1.0 \| 1.819e-18 \| 610 \| 0.225 \| 386 \| 250 \| 16 \| 369 \| 721 \| 61 \| 430 \| Virulence protein \| Virulence protein; afdb-uniprot50 \| AF-A0A702EU54-F1-MODEL\_V4 \| 1.0 \| 1.058e-14 \| 610 \| 0.288 \| 232 \| 146 \| 7 \| 118 \| 346 \| 162 \| 377 \| DUF5710 domain-containing protein \| DUF5710 domain-containing protein; afdb-uniprot50 \| AF-A0A5C9BA17-F1-MODEL\_V4 \| 1.0 \| 2.857e-24 \| 610 \| 0.164 \| 790 \| 361 \| 32 \| 1 \| 763 \| 194 \| 711 \| Virulence-associated E family protein \| Virulence-associated E family protein; afdb-uniprot50 \| AF-A0A651E934-F1-MODEL\_V4 \| 1.0 \| 5.48e-25 \| 610 \| 0.205 \| 675 \| 416 \| 38 \| 149 \| 769 \| 34 \| 641 \| DUF3854 domain-containing protein \| DUF3854 domain-containing protein; afdb-uniprot50 \| AF-A0A7T8TC66-F1-MODEL\_V4 \| 1.0 \| 4.91e-20 \| 609 \| 0.221 \| 483 \| 299 \| 28 \| 321 \| 769 \| 107 \| 546 \| Uncharacterized protein \| Uncharacterized protein; afdb-uniprot50 \| AF-K9PF86-F1-MODEL\_V4 \| 1.0 \| 5.48e-25 \| 609 \| 0.186 \| 675 \| 423 \| 37 \| 139 \| 757 \| 36 \| 640 \| Virulence-associated E family protein \| Virulence-associated E family protein; afdb-uniprot50 \| AF-A0A2K8T0V0-F1-MODEL\_V4 \| 1.0 \| 4.694e-25 \| 609 \| 0.204 \| 642 \| 411 \| 34 \| 171 \| 762 \| 53 \| 644 \| DNA primase \| DNA primase; afdb-uniprot50 \| AF-W6M4H4-F1-MODEL\_V4 \| 1.0 \| 3.127e-30 \| 609 \| 0.191 \| 823 \| 463 \| 44 \| 102 \| 757 \| 76 \| 862 \| Putative Phage/plasmid primase, P4 family \| Putative Phage/plasmid primase, P4 family; afdb-uniprot50 \| AF-A0A3P1YTN5-F1-MODEL\_V4 \| 1.0 \| 1.317e-16 \| 608 \| 0.21 \| 313 \| 222 \| 14 \| 384 \| 684 \| 58 \| 357 \| Helicase \| Helicase; afdb-uniprot50 \| AF-A0A6H0KWD7-F1-MODEL\_V4 \| 1.0 \| 1.508e-17 \| 608 \| 0.173 \| 393 \| 282 \| 19 \| 381 \| 751 \| 37 \| 408 \| DUF3874 domain-containing protein \| DUF3874 domain-containing protein; afdb-uniprot50 \| AF-A0A354WYX5-F1-MODEL\_V4 \| 1.0 \| 3.603e-20 \| 607 \| 0.145 \| 528 \| 359 \| 26 \| 246 \| 755 \| 233 \| 686 \| Helicase \| Helicase; afdb-uniprot50 \| AF-A0A562ZHG6-F1-MODEL\_V4 \| 1.0 \| 1.878e-19 \| 607 \| 0.206 \| 469 \| 315 \| 29 \| 325 \| 757 \| 280 \| 727 \| Prim-Pol domain-containing protein \| Prim-Pol domain-containing protein; afdb-uniprot50 \| AF-A0A4Q6HNM1-F1-MODEL\_V4 \| 1.0 \| 6.314e-15 \| 606 \| 0.277 \| 234 \| 147 \| 8 \| 116 \| 347 \| 165 \| 378 \| DUF5710 domain-containing protein \| DUF5710 domain-containing protein; afdb-uniprot50 \| AF-A0A562Q1D0-F1-MODEL\_V4 \| 1.0 \| 7.42e-20 \| 606 \| 0.202 \| 459 \| 318 \| 20 \| 331 \| 769 \| 61 \| 491 \| Virulence-associated protein E \| Virulence-associated protein E; afdb-uniprot50 \| AF-A0A6A7VW87-F1-MODEL\_V4 \| 1.0 \| 7.516e-22 \| 606 \| 0.164 \| 578 \| 356 \| 34 \| 226 \| 756 \| 61 \| 558 \| Uncharacterized protein \| Uncharacterized protein; afdb-uniprot50 \| AF-A0A6M0RZD1-F1-MODEL\_V4 \| 1.0 \| 3.511e-24 \| 606 \| 0.196 \| 632 \| 396 \| 35 \| 181 \| 765 \| 104 \| 670 \| Uncharacterized protein \| Uncharacterized protein; afdb-uniprot50 \| AF-A0A515L9P3-F1-MODEL\_V4 \| 1.0 \| 6.606e-18 \| 606 \| 0.203 \| 407 \| 285 \| 21 \| 370 \| 752 \| 345 \| 736 \| Uncharacterized protein \| Uncharacterized protein; afdb-uniprot50 \| AF-A0A5C7PXR7-F1-MODEL\_V4 \| 1.0 \| 3.313e-19 \| 606 \| 0.21 \| 460 \| 313 \| 24 \| 328 \| 757 \| 393 \| 832 \| Uncharacterized protein \| Uncharacterized protein; afdb-uniprot50 \| AF-A0A1M5F3B9-F1-MODEL\_V4 \| 1.0 \| 1.219e-28 \| 606 \| 0.199 \| 861 \| 414 \| 50 \| 10 \| 735 \| 2 \| 721 \| Putative DNA primase/helicase \| Putative DNA primase/helicase; afdb-uniprot50 \| AF-A0A4P7CQ19-F1-MODEL\_V4 \| 1.0 \| 1.683e-30 \| 606 \| 0.193 \| 841 \| 495 \| 48 \| 13 \| 750 \| 19 \| 778 \| DUF3631 domain-containing protein \| DUF3631 domain-containing protein; afdb-uniprot50 \| AF-A0A370V047-F1-MODEL\_V4 \| 1.0 \| 7.001e-15 \| 606 \| 0.322 \| 236 \| 139 \| 8 \| 114 \| 346 \| 160 \| 377 \| DNA primase TraC \| DNA primase TraC; afdb-uniprot50 \| AF-A0A450TW60-F1-MODEL\_V4 \| 1.0 \| 2.016e-18 \| 605 \| 0.325 \| 323 \| 156 \| 12 \| 76 \| 351 \| 8 \| 315 \| Toprim domain-containing protein \| Toprim domain-containing protein; afdb-uniprot50 \| AF-A0A1C5VVZ5-F1-MODEL\_V4 \| 1.0 \| 1.64e-18 \| 605 \| 0.207 \| 433 \| 280 \| 20 \| 370 \| 753 \| 387 \| 805 \| Predicted P-loop ATPase and inactivated derivatives \| Predicted P-loop ATPase and inactivated derivatives; afdb-uniprot50 \| AF-A0A2M8JR34-F1-MODEL\_V4 \| 1.0 \| 1.903e-21 \| 605 \| 0.229 \| 531 \| 335 \| 28 \| 266 \| 769 \| 292 \| 775 \| Uncharacterized protein \| Uncharacterized protein; afdb-uniprot50 \| AF-A0A1Z4QLR4-F1-MODEL\_V4 \| 1.0 \| 8.719e-25 \| 605 \| 0.18 \| 693 \| 443 \| 40 \| 145 \| 762 \| 9 \| 651 \| Virulence-associated E family protein \| Virulence-associated E family protein; afdb-uniprot50 \| AF-S2YJ98-F1-MODEL\_V4 \| 1.0 \| 2.235e-18 \| 604 \| 0.192 \| 420 \| 278 \| 22 \| 384 \| 769 \| 5 \| 397 \| Uncharacterized protein \| Uncharacterized protein; afdb-uniprot50 \| AF-A0A011VM22-F1-MODEL\_V4 \| 1.0 \| 1.121e-19 \| 604 \| 0.203 \| 492 \| 317 \| 20 \| 321 \| 769 \| 102 \| 561 \| Virulence protein E \| Virulence protein E; afdb-uniprot50 \| AF-A0A1Q3JRW3-F1-MODEL\_V4 \| 1.0 \| 5.104e-18 \| 603 \| 0.19 \| 393 \| 277 \| 21 \| 384 \| 758 \| 38 \| 407 \| Uncharacterized protein \| Uncharacterized protein; afdb-uniprot50 \| AF-A0A1C7HSV1-F1-MODEL\_V4 \| 1.0 \| 6.234e-21 \| 603 \| 0.2 \| 574 \| 340 \| 28 \| 252 \| 769 \| 290 \| 800 \| Uncharacterized protein \| Uncharacterized protein; afdb-uniprot50 \| AF-A0A1B1TQM4-F1-MODEL\_V4 \| 1.0 \| 9.3e-27 \| 603 \| 0.205 \| 685 \| 412 \| 36 \| 177 \| 769 \| 66 \| 709 \| DUF3854 domain-containing protein \| DUF3854 domain-containing protein; afdb-uniprot50 \| AF-A0A3M1BAH4-F1-MODEL\_V4 \| 1.0 \| 1.75e-20 \| 601 \| 0.206 \| 490 \| 312 \| 18 \| 261 \| 737 \| 35 \| 460 \| Uncharacterized protein \| Uncharacterized protein; afdb-uniprot50 \| AF-Q7U784-F1-MODEL\_V4 \| 1.0 \| 1.978e-19 \| 601 \| 0.232 \| 418 \| 269 \| 23 \| 371 \| 763 \| 298 \| 688 \| Uncharacterized protein \| Uncharacterized protein; afdb-uniprot50 \| AF-A0A3S0D2I0-F1-MODEL\_V4 \| 1.0 \| 6.115e-22 \| 601 \| 0.199 \| 608 \| 347 \| 28 \| 240 \| 753 \| 215 \| 776 \| Uncharacterized protein \| Uncharacterized protein; afdb-uniprot50 \| AF-R4MXK3-F1-MODEL\_V4 \| 1.0 \| 4.88e-31 \| 601 \| 0.206 \| 880 \| 484 \| 45 \| 1 \| 738 \| 5 \| 811 \| DNA primase \| DNA primase; afdb-uniprot50 \| AF-A0A3M3AN02-F1-MODEL\_V4 \| 1.0 \| 5.586e-16 \| 600 \| 0.306 \| 261 \| 149 \| 10 \| 107 \| 355 \| 12 \| 252 \| DNA primase TraC \| DNA primase TraC; afdb-uniprot50 \| AF-R9IAU2-F1-MODEL\_V4 \| 1.0 \| 1.031e-18 \| 600 \| 0.16 \| 398 \| 291 \| 20 \| 382 \| 757 \| 40 \| 416 \| DUF3874 domain-containing protein \| DUF3874 domain-containing protein; afdb-uniprot50 \| AF-F8L0K0-F1-MODEL\_V4 \| 1.0 \| 3.868e-19 \| 600 \| 0.274 \| 353 \| 190 \| 16 \| 102 \| 442 \| 82 \| 380 \| Toprim domain-containing protein \| Toprim domain-containing protein; afdb-uniprot50 \| AF-A0A7X5TDU8-F1-MODEL\_V4 \| 1.0 \| 1.609e-19 \| 600 \| 0.257 \| 349 \| 213 \| 17 \| 95 \| 431 \| 115 \| 429 \| DNA primase TraC \| DNA primase TraC; afdb-uniprot50 \| AF-A0A353YA43-F1-MODEL\_V4 \| 1.0 \| 2.056e-25 \| 600 \| 0.184 \| 742 \| 418 \| 48 \| 159 \| 768 \| 172 \| 858 \| Virulence protein E \| Virulence protein E; afdb-uniprot50 \| AF-A0A1M7E7Q2-F1-MODEL\_V4 \| 1.0 \| 1.548e-21 \| 599 \| 0.171 \| 512 \| 336 \| 24 \| 261 \| 751 \| 257 \| 701 \| Virulence-associated protein E \| Virulence-associated protein E; afdb-uniprot50 \| AF-A0A269PII8-F1-MODEL\_V4 \| 1.0 \| 1.651e-15 \| 598 \| 0.296 \| 273 \| 160 \| 9 \| 92 \| 346 \| 72 \| 330 \| Toprim domain-containing protein \| Toprim domain-containing protein; afdb-uniprot50 \| AF-A0A3N2MCU4-F1-MODEL\_V4 \| 1.0 \| 7.324e-18 \| 598 \| 0.242 \| 383 \| 252 \| 14 \| 317 \| 680 \| 92 \| 455 \| Uncharacterized protein \| Uncharacterized protein; afdb-uniprot50 \| AF-A0A433UZH8-F1-MODEL\_V4 \| 1.0 \| 2.207e-24 \| 598 \| 0.199 \| 642 \| 396 \| 36 \| 178 \| 762 \| 20 \| 600 \| Uncharacterized protein \| Uncharacterized protein; afdb-uniprot50 \| AF-A0A7W8TA26-F1-MODEL\_V4 \| 1.0 \| 5.204e-25 \| 598 \| 0.223 \| 719 \| 358 \| 40 \| 6 \| 602 \| 2 \| 641 \| Putative DNA primase/helicase \| Putative DNA primase/helicase; afdb-uniprot50 \| AF-A0A3D4KD37-F1-MODEL\_V4 \| 1.0 \| 8.888e-16 \| 597 \| 0.306 \| 261 \| 147 \| 10 \| 92 \| 346 \| 59 \| 291 \| Toprim domain-containing protein \| Toprim domain-containing protein; afdb-uniprot50 \| AF-A0A522W194-F1-MODEL\_V4 \| 1.0 \| 3.167e-16 \| 597 \| 0.304 \| 263 \| 143 \| 12 \| 100 \| 346 \| 454 \| 692 \| DUF1738 domain-containing protein \| DUF1738 domain-containing protein; afdb-uniprot50 \| AF-A0A442FIW1-F1-MODEL\_V4 \| 1.0 \| 5.883e-24 \| 597 \| 0.181 \| 779 \| 359 \| 32 \| 1 \| 752 \| 182 \| 708 \| Prim-Pol domain-containing protein \| Prim-Pol domain-containing protein; afdb-uniprot50 \| AF-A0A412ETP2-F1-MODEL\_V4 \| 1.0 \| 1.072e-24 \| 597 \| 0.177 \| 715 \| 448 \| 44 \| 145 \| 769 \| 97 \| 761 \| ZnF\_CHCC domain-containing protein \| ZnF\_CHCC domain-containing protein; afdb-uniprot50 \| AF-A0A6N3GAZ3-F1-MODEL\_V4 \| 1.0 \| 2.627e-15 \| 596 \| 0.206 \| 315 \| 226 \| 14 \| 453 \| 757 \| 22 \| 322 \| Virulence-associated protein E \| Virulence-associated protein E; afdb-uniprot50 \| AF-A0A173U6S3-F1-MODEL\_V4 \| 1.0 \| 9.179e-17 \| 596 \| 0.206 \| 330 \| 237 \| 14 \| 381 \| 699 \| 43 \| 358 \| Predicted P-loop ATPase and inactivated derivatives \| Predicted P-loop ATPase and inactivated derivatives; afdb-uniprot50 \| AF-A0A1V5XFC1-F1-MODEL\_V4 \| 1.0 \| 5.104e-18 \| 596 \| 0.192 \| 432 \| 299 \| 18 \| 368 \| 769 \| 36 \| 447 \| Virulence-associated protein E \| Virulence-associated protein E; afdb-uniprot50 \| AF-A0A0D7K6F3-F1-MODEL\_V4 \| 1.0 \| 1.978e-19 \| 596 \| 0.263 \| 436 \| 247 \| 22 \| 370 \| 753 \| 217 \| 630 \| Uncharacterized protein \| Uncharacterized protein; afdb-uniprot50 \| AF-A0A5R1X1K9-F1-MODEL\_V4 \| 1.0 \| 2.713e-24 \| 596 \| 0.179 \| 706 \| 432 \| 42 \| 164 \| 769 \| 179 \| 836 \| Uncharacterized protein \| Uncharacterized protein; afdb-uniprot50 \| AF-A0A5C7RLC4-F1-MODEL\_V4 \| 1.0 \| 2.137e-15 \| 595 \| 0.348 \| 238 \| 128 \| 10 \| 99 \| 333 \| 82 \| 295 \| Toprim domain-containing protein \| Toprim domain-containing protein; afdb-uniprot50 \| AF-A0A7J5Q366-F1-MODEL\_V4 \| 1.0 \| 3.444e-17 \| 595 \| 0.178 \| 381 \| 274 \| 18 \| 393 \| 755 \| 3 \| 362 \| DUF3874 domain-containing protein \| DUF3874 domain-containing protein; afdb-uniprot50 \| AF-A0A251WKU0-F1-MODEL\_V4 \| 1.0 \| 1.479e-18 \| 595 \| 0.223 \| 442 \| 281 \| 18 \| 328 \| 750 \| 175 \| 573 \| Uncharacterized protein \| Uncharacterized protein; afdb-uniprot50 \| AF-A0A3R5ZLK0-F1-MODEL\_V4 \| 1.0 \| 8.946e-21 \| 595 \| 0.179 \| 485 \| 308 \| 17 \| 259 \| 725 \| 220 \| 632 \| Virulence protein E \| Virulence protein E; afdb-uniprot50 \| AF-A0A2V5PV48-F1-MODEL\_V4 \| 1.0 \| 1.196e-21 \| 595 \| 0.204 \| 548 \| 341 \| 26 \| 243 \| 762 \| 248 \| 728 \| Prim-Pol domain-containing protein \| Prim-Pol domain-containing protein; afdb-uniprot50 \| AF-A0A6I5P4Y0-F1-MODEL\_V4 \| 1.0 \| 3.627e-25 \| 595 \| 0.202 \| 677 \| 409 \| 39 \| 144 \| 751 \| 2 \| 616 \| DUF3854 domain-containing protein \| DUF3854 domain-containing protein; afdb-uniprot50 \| AF-Q7NEJ1-F1-MODEL\_V4 \| 1.0 \| 5.039e-24 \| 595 \| 0.203 \| 663 \| 395 \| 42 \| 179 \| 769 \| 72 \| 673 \| Glr3888 protein \| Glr3888 protein; afdb-uniprot50 \| AF-A0A3M4XFL8-F1-MODEL\_V4 \| 1.0 \| 2.766e-15 \| 595 \| 0.317 \| 255 \| 144 \| 11 \| 114 \| 358 \| 162 \| 396 \| Putative DNA primase \| Putative DNA primase; afdb-uniprot50 \| AF-A0A095ZZV9-F1-MODEL\_V4 \| 1.0 \| 1.093e-15 \| 594 \| 0.246 \| 264 \| 188 \| 6 \| 467 \| 725 \| 57 \| 314 \| Virulence-associated protein E \| Virulence-associated protein E; afdb-uniprot50 \| AF-A0A433VS18-F1-MODEL\_V4 \| 1.0 \| 6.956e-18 \| 593 \| 0.208 \| 394 \| 272 \| 19 \| 381 \| 758 \| 88 \| 457 \| Uncharacterized protein \| Uncharacterized protein; afdb-uniprot50 \| AF-A0A1H2VE35-F1-MODEL\_V4 \| 1.0 \| 3.77e-23 \| 593 \| 0.181 \| 688 \| 386 \| 38 \| 142 \| 756 \| 46 \| 629 \| Virulence-associated protein E \| Virulence-associated protein E; afdb-uniprot50 \| AF-A0A3R6S2F5-F1-MODEL\_V4 \| 1.0 \| 5.17e-20 \| 593 \| 0.188 \| 521 \| 342 \| 26 \| 261 \| 757 \| 231 \| 694 \| Uncharacterized protein \| Uncharacterized protein; afdb-uniprot50 \| AF-A0A7G8IZC6-F1-MODEL\_V4 \| 1.0 \| 9.003e-18 \| 592 \| 0.212 \| 414 \| 273 \| 20 \| 384 \| 769 \| 4 \| 392 \| Virulence-associated E family protein \| Virulence-associated E family protein; afdb-uniprot50 \| AF-A0A4Q3P7C9-F1-MODEL\_V4 \| 1.0 \| 3.229e-23 \| 592 \| 0.162 \| 792 \| 389 \| 35 \| 1 \| 761 \| 94 \| 641 \| PriCT\_2 domain-containing protein \| PriCT\_2 domain-containing protein; afdb-uniprot50 \| AF-A0A0D0QDV7-F1-MODEL\_V4 \| 1.0 \| 1.396e-21 \| 592 \| 0.214 \| 584 \| 355 \| 30 \| 206 \| 762 \| 188 \| 694 \| Putative P-loop ATPase \| Putative P-loop ATPase; afdb-uniprot50 \| AF-C7GYW9-F1-MODEL\_V4 \| 1.0 \| 8.069e-21 \| 592 \| 0.208 \| 562 \| 324 \| 32 \| 257 \| 755 \| 285 \| 788 \| Virulence-associated protein E \| Virulence-associated protein E; afdb-uniprot50 \| AF-A5Z5J5-F1-MODEL\_V4 \| 1.0 \| 1.292e-17 \| 591 \| 0.217 \| 377 \| 250 \| 18 \| 420 \| 769 \| 4 \| 362 \| Virulence-associated protein E \| Virulence-associated protein E; afdb-uniprot50 \| AF-A0A3L8PZN3-F1-MODEL\_V4 \| 1.0 \| 1.651e-15 \| 591 \| 0.309 \| 255 \| 142 \| 7 \| 118 \| 355 \| 163 \| 400 \| Uncharacterized protein \| Uncharacterized protein; afdb-uniprot50 \| AF-A0A1M6Q060-F1-MODEL\_V4 \| 1.0 \| 1.196e-21 \| 591 \| 0.214 \| 536 \| 321 \| 26 \| 260 \| 756 \| 246 \| 720 \| Predicted P-loop ATPase and inactivated derivatives \| Predicted P-loop ATPase and inactivated derivatives; afdb-uniprot50 \| AF-K9Q2E5-F1-MODEL\_V4 \| 1.0 \| 2.017e-26 \| 591 \| 0.185 \| 726 \| 409 \| 39 \| 178 \| 762 \| 78 \| 761 \| Virulence-associated E family protein \| Virulence-associated E family protein; afdb-uniprot50 \| AF-D7I880-F1-MODEL\_V4 \| 1.0 \| 3.008e-16 \| 590 \| 0.25 \| 272 \| 190 \| 10 \| 491 \| 755 \| 10 \| 274 \| Helicase \| Helicase; afdb-uniprot50 \| AF-A0A365U406-F1-MODEL\_V4 \| 1.0 \| 8.441e-16 \| 590 \| 0.377 \| 236 \| 120 \| 9 \| 128 \| 358 \| 140 \| 353 \| Toprim domain-containing protein \| Toprim domain-containing protein; afdb-uniprot50 \| AF-A0A8B3N3Z6-F1-MODEL\_V4 \| 1.0 \| 1.309e-19 \| 590 \| 0.193 \| 491 \| 327 \| 24 \| 306 \| 751 \| 22 \| 488 \| Uncharacterized protein \| Uncharacterized protein; afdb-uniprot50 \| AF-A0A5C7PH59-F1-MODEL\_V4 \| 1.0 \| 6.522e-24 \| 590 \| 0.188 \| 691 \| 405 \| 41 \| 164 \| 756 \| 33 \| 665 \| Uncharacterized protein \| Uncharacterized protein; afdb-uniprot50 \| AF-A0A2T1G467-F1-MODEL\_V4 \| 1.0 \| 3.721e-21 \| 590 \| 0.208 \| 546 \| 340 \| 28 \| 260 \| 769 \| 334 \| 823 \| Uncharacterized protein \| Uncharacterized protein; afdb-uniprot50 \| AF-A0A826KI04-F1-MODEL\_V4 \| 1.0 \| 2.069e-14 \| 589 \| 0.323 \| 235 \| 138 \| 8 \| 115 \| 346 \| 161 \| 377 \| Uncharacterized protein \| Uncharacterized protein; afdb-uniprot50 \| AF-A0A353UWV3-F1-MODEL\_V4 \| 1.0 \| 4.261e-22 \| 589 \| 0.199 \| 626 \| 365 \| 32 \| 226 \| 752 \| 19 \| 607 \| P-loop ATPase \| P-loop ATPase; afdb-uniprot50 \| AF-A0A2P8WI05-F1-MODEL\_V4 \| 1.0 \| 3.25e-20 \| 589 \| 0.21 \| 512 \| 336 \| 21 \| 275 \| 760 \| 16 \| 485 \| Uncharacterized protein \| Uncharacterized protein; afdb-uniprot50 \| AF-A0A4U6M2C1-F1-MODEL\_V4 \| 1.0 \| 1.773e-30 \| 589 \| 0.198 \| 872 \| 447 \| 51 \| 91 \| 751 \| 77 \| 907 \| Bifunctional DNA primase/helicase \| Bifunctional DNA primase/helicase; afdb-uniprot50 \| AF-A0A1U7IM20-F1-MODEL\_V4 \| 1.0 \| 1.489e-23 \| 589 \| 0.186 \| 648 \| 425 \| 38 \| 159 \| 759 \| 62 \| 654 \| Uncharacterized protein \| Uncharacterized protein; afdb-uniprot50 \| AF-G2KRX2-F1-MODEL\_V4 \| 1.0 \| 1.3e-14 \| 588 \| 0.303 \| 237 \| 143 \| 7 \| 100 \| 335 \| 73 \| 288 \| Toprim domain-containing protein \| Toprim domain-containing protein; afdb-uniprot50 \| AF-A0A0D6Q293-F1-MODEL\_V4 \| 1.0 \| 8.441e-16 \| 588 \| 0.213 \| 342 \| 241 \| 12 \| 370 \| 691 \| 22 \| 355 \| DNA primase \| DNA primase; afdb-uniprot50 \| AF-A0A1E4C277-F1-MODEL\_V4 \| 1.0 \| 3.147e-19 \| 588 \| 0.208 \| 495 \| 329 \| 26 \| 292 \| 756 \| 7 \| 468 \| Uncharacterized protein \| Uncharacterized protein; afdb-uniprot50 \| AF-A0A016XJH4-F1-MODEL\_V4 \| 1.0 \| 6.076e-25 \| 588 \| 0.179 \| 742 \| 428 \| 44 \| 160 \| 769 \| 50 \| 742 \| Conjugal transfer protein TraC \| Conjugal transfer protein TraC; afdb-uniprot50 \| AF-A0A2T5NFF6-F1-MODEL\_V4 \| 1.0 \| 2.294e-30 \| 588 \| 0.197 \| 856 \| 478 \| 46 \| 12 \| 737 \| 5 \| 781 \| Uncharacterized protein \| Uncharacterized protein; afdb-uniprot50 \| AF-A0A6L9H472-F1-MODEL\_V4 \| 1.0 \| 1.99e-16 \| 586 \| 0.254 \| 287 \| 195 \| 11 \| 494 \| 769 \| 2 \| 280 \| Uncharacterized protein \| Uncharacterized protein; afdb-uniprot50 \| AF-A0A2N0AT84-F1-MODEL\_V4 \| 1.0 \| 1.378e-19 \| 586 \| 0.174 \| 481 \| 336 \| 25 \| 311 \| 755 \| 30 \| 485 \| Uncharacterized protein \| Uncharacterized protein; afdb-uniprot50 \| AF-A0A846FT90-F1-MODEL\_V4 \| 1.0 \| 9.855e-24 \| 586 \| 0.202 \| 636 \| 384 \| 32 \| 170 \| 743 \| 72 \| 645 \| DUF3854 domain-containing protein \| DUF3854 domain-containing protein; afdb-uniprot50 \| AF-A0A807K447-F1-MODEL\_V4 \| 1.0 \| 1.903e-21 \| 586 \| 0.191 \| 573 \| 347 \| 30 \| 239 \| 769 \| 201 \| 698 \| Uncharacterized protein \| Uncharacterized protein; afdb-uniprot50 \| AF-A0A1M6RY44-F1-MODEL\_V4 \| 1.0 \| 3.603e-20 \| 586 \| 0.157 \| 519 \| 362 \| 22 \| 260 \| 752 \| 253 \| 721 \| VirE N-terminal domain-containing protein \| VirE N-terminal domain-containing protein; afdb-uniprot50 \| AF-A0A239LDP2-F1-MODEL\_V4 \| 1.0 \| 1.038e-23 \| 586 \| 0.18 \| 799 \| 374 \| 31 \| 2 \| 769 \| 197 \| 745 \| Predicted P-loop ATPase and inactivated derivatives \| Predicted P-loop ATPase and inactivated derivatives; afdb-uniprot50 \| AF-G5SU78-F1-MODEL\_V4 \| 1.0 \| 2.016e-18 \| 585 \| 0.187 \| 447 \| 320 \| 24 \| 327 \| 755 \| 2 \| 423 \| DUF3874 domain-containing protein \| DUF3874 domain-containing protein; afdb-uniprot50 \| AF-A0A7X5C8R8-F1-MODEL\_V4 \| 1.0 \| 9.604e-20 \| 585 \| 0.179 \| 518 \| 359 \| 25 \| 288 \| 769 \| 23 \| 510 \| Uncharacterized protein \| Uncharacterized protein; afdb-uniprot50 \| AF-A0A1Z4SI32-F1-MODEL\_V4 \| 1.0 \| 2.447e-24 \| 585 \| 0.189 \| 692 \| 416 \| 37 \| 139 \| 756 \| 37 \| 657 \| Virulence-associated E family protein \| Virulence-associated E family protein; afdb-uniprot50 \| AF-A0A2D3TGD8-F1-MODEL\_V4 \| 1.0 \| 1.005e-14 \| 585 \| 0.335 \| 200 \| 114 \| 6 \| 149 \| 346 \| 6 \| 188 \| LPD7 domain-containing protein \| LPD7 domain-containing protein; afdb-uniprot50 \| AF-A0A1I7KD97-F1-MODEL\_V4 \| 1.0 \| 6.356e-20 \| 585 \| 0.201 \| 502 \| 332 \| 26 \| 311 \| 761 \| 295 \| 778 \| Bifunctional DNA primase/polymerase, N-terminal \| Bifunctional DNA primase/polymerase, N-terminal; afdb-uniprot50 \| AF-A0A838BUZ2-F1-MODEL\_V4 \| 1.0 \| 2.4e-25 \| 585 \| 0.178 \| 773 \| 393 \| 46 \| 167 \| 759 \| 137 \| 847 \| Uncharacterized protein \| Uncharacterized protein; afdb-uniprot50 \| AF-A0A2T5L4M6-F1-MODEL\_V4 \| 1.0 \| 7.002e-31 \| 585 \| 0.204 \| 927 \| 445 \| 51 \| 100 \| 769 \| 78 \| 968 \| Putative DNA primase/helicase \| Putative DNA primase/helicase; afdb-uniprot50 \| AF-A0A0F2P8W8-F1-MODEL\_V4 \| 1.0 \| 1.005e-14 \| 584 \| 0.297 \| 235 \| 143 \| 6 \| 106 \| 338 \| 61 \| 275 \| Toprim domain-containing protein \| Toprim domain-containing protein; afdb-uniprot50 \| AF-K1V3K5-F1-MODEL\_V4 \| 1.0 \| 1.538e-16 \| 584 \| 0.23 \| 339 \| 228 \| 15 \| 436 \| 753 \| 9 \| 335 \| Virulence-associated E \| Virulence-associated E; afdb-uniprot50 \| AF-A0A556QUB3-F1-MODEL\_V4 \| 1.0 \| 1.011e-19 \| 584 \| 0.181 \| 517 \| 345 \| 30 \| 267 \| 755 \| 25 \| 491 \| Uncharacterized protein \| Uncharacterized protein; afdb-uniprot50 \| AF-X5J840-F1-MODEL\_V4 \| 1.0 \| 1.738e-23 \| 584 \| 0.159 \| 658 \| 433 \| 41 \| 154 \| 755 \| 162 \| 755 \| Virulence-associated E family protein \| Virulence-associated E family protein; afdb-uniprot50 \| AF-A0A6J4IEE4-F1-MODEL\_V4 \| 1.0 \| 1.651e-15 \| 583 \| 0.223 \| 300 \| 211 \| 10 \| 463 \| 757 \| 17 \| 299 \| Virulence-associated E \| Virulence-associated E; afdb-uniprot50 \| AF-A0A2N5C915-F1-MODEL\_V4 \| 1.0 \| 1.275e-15 \| 583 \| 0.337 \| 240 \| 124 \| 9 \| 122 \| 347 \| 78 \| 296 \| Toprim domain-containing protein \| Toprim domain-containing protein; afdb-uniprot50 \| AF-A0A430HF59-F1-MODEL\_V4 \| 1.0 \| 1.204e-18 \| 583 \| 0.292 \| 352 \| 193 \| 15 \| 10 \| 347 \| 1 \| 310 \| Uncharacterized protein \| Uncharacterized protein; afdb-uniprot50 \| AF-A0A4S2GDP1-F1-MODEL\_V4 \| 1.0 \| 4.515e-19 \| 583 \| 0.217 \| 478 \| 278 \| 23 \| 370 \| 769 \| 35 \| 494 \| Virulence-associated E family protein \| Virulence-associated E family protein; afdb-uniprot50 \| AF-A0A0U2U1R7-F1-MODEL\_V4 \| 1.0 \| 1.204e-18 \| 583 \| 0.203 \| 467 \| 282 \| 25 \| 366 \| 761 \| 50 \| 497 \| Uncharacterized protein \| Uncharacterized protein; afdb-uniprot50 \| AF-A0A2S5MV44-F1-MODEL\_V4 \| 1.0 \| 2.56e-19 \| 583 \| 0.178 \| 505 \| 332 \| 20 \| 261 \| 750 \| 15 \| 451 \| Uncharacterized protein \| Uncharacterized protein; afdb-uniprot50 \| AF-A0A317MT75-F1-MODEL\_V4 \| 1.0 \| 8.496e-21 \| 583 \| 0.18 \| 538 \| 348 \| 28 \| 268 \| 763 \| 251 \| 737 \| DNA primase RepB-like protein \| DNA primase RepB-like protein; afdb-uniprot50 \| AF-A0A255TPW4-F1-MODEL\_V4 \| 1.0 \| 3.534e-21 \| 583 \| 0.181 \| 572 \| 342 \| 26 \| 241 \| 755 \| 225 \| 727 \| VirE\_N domain-containing protein \| VirE\_N domain-containing protein; afdb-uniprot50 \| AF-A0A353X8G7-F1-MODEL\_V4 \| 1.0 \| 5.997e-15 \| 582 \| 0.296 \| 243 \| 141 \| 6 \| 91 \| 333 \| 67 \| 279 \| Toprim domain-containing protein \| Toprim domain-containing protein; afdb-uniprot50 \| AF-A0A433HYK3-F1-MODEL\_V4 \| 1.0 \| 5.374e-18 \| 582 \| 0.22 \| 421 \| 259 \| 18 \| 395 \| 755 \| 4 \| 415 \| Uncharacterized protein \| Uncharacterized protein; afdb-uniprot50 \| AF-A0A844MXR4-F1-MODEL\_V4 \| 1.0 \| 2.61e-18 \| 582 \| 0.231 \| 415 \| 260 \| 21 \| 367 \| 753 \| 430 \| 813 \| Uncharacterized protein \| Uncharacterized protein; afdb-uniprot50 \| AF-A0A6P0YU45-F1-MODEL\_V4 \| 1.0 \| 1.317e-16 \| 581 \| 0.249 \| 341 \| 223 \| 17 \| 438 \| 762 \| 43 \| 366 \| Uncharacterized protein \| Uncharacterized protein; afdb-uniprot50 \| AF-A0A551Z1W1-F1-MODEL\_V4 \| 1.0 \| 3.271e-25 \| 581 \| 0.169 \| 698 \| 410 \| 37 \| 163 \| 762 \| 50 \| 675 \| Uncharacterized protein \| Uncharacterized protein; afdb-uniprot50 \| AF-A0A1F4ANV0-F1-MODEL\_V4 \| 1.0 \| 2.043e-20 \| 580 \| 0.19 \| 567 \| 347 \| 30 \| 246 \| 753 \| 221 \| 734 \| Uncharacterized protein \| Uncharacterized protein; afdb-uniprot50 \| AF-A0A2T5K521-F1-MODEL\_V4 \| 1.0 \| 2.369e-23 \| 580 \| 0.195 \| 664 \| 417 \| 38 \| 181 \| 762 \| 143 \| 770 \| Putative P-loop ATPase \| Putative P-loop ATPase; afdb-uniprot50 \| AF-A0A1Y6D7X7-F1-MODEL\_V4 \| 1.0 \| 1.965e-30 \| 580 \| 0.179 \| 1009 \| 512 \| 53 \| 11 \| 756 \| 8 \| 963 \| Putative DNA primase/helicase \| Putative DNA primase/helicase; afdb-uniprot50 \| AF-A0A374J606-F1-MODEL\_V4 \| 1.0 \| 4.316e-24 \| 579 \| 0.189 \| 690 \| 428 \| 43 \| 149 \| 769 \| 81 \| 707 \| Uncharacterized protein \| Uncharacterized protein; afdb-uniprot50 \| AF-A0A2T4IDI9-F1-MODEL\_V4 \| 1.0 \| 6.316e-31 \| 579 \| 0.186 \| 998 \| 495 \| 53 \| 1 \| 751 \| 1 \| 928 \| Uncharacterized protein \| Uncharacterized protein; afdb-uniprot50 \| AF-A0A5U2TXN0-F1-MODEL\_V4 \| 1.0 \| 1.173e-14 \| 579 \| 0.28 \| 250 \| 157 \| 8 \| 116 \| 358 \| 4 \| 237 \| LPD7 domain-containing protein \| LPD7 domain-containing protein; afdb-uniprot50 \| AF-A0A6G3Z197-F1-MODEL\_V4 \| 1.0 \| 1.568e-23 \| 578 \| 0.201 \| 649 \| 405 \| 35 \| 169 \| 766 \| 74 \| 659 \| Uncharacterized protein \| Uncharacterized protein; afdb-uniprot50 \| AF-U7QLA3-F1-MODEL\_V4 \| 1.0 \| 4.316e-24 \| 578 \| 0.17 \| 687 \| 410 \| 40 \| 166 \| 769 \| 99 \| 708 \| Virulence-associated E family protein \| Virulence-associated E family protein; afdb-uniprot50 \| AF-A0A317N8C0-F1-MODEL\_V4 \| 1.0 \| 1.079e-29 \| 578 \| 0.205 \| 798 \| 452 \| 47 \| 92 \| 760 \| 87 \| 831 \| Phage/plasmid primase-like uncharacterized protein \| Phage/plasmid primase-like uncharacterized protein; afdb-uniprot50 \| AF-A0A1M4ZAH0-F1-MODEL\_V4 \| 1.0 \| 2.056e-17 \| 577 \| 0.184 \| 391 \| 284 \| 17 \| 381 \| 755 \| 41 \| 412 \| Virulence-associated protein E \| Virulence-associated protein E; afdb-uniprot50 \| AF-A0A840ASX3-F1-MODEL\_V4 \| 1.0 \| 2.431e-19 \| 577 \| 0.198 \| 525 \| 341 \| 22 \| 261 \| 769 \| 257 \| 717 \| Putative P-loop ATPase \| Putative P-loop ATPase; afdb-uniprot50 \| AF-A0A522CQB3-F1-MODEL\_V4 \| 1.0 \| 1.672e-17 \| 577 \| 0.195 \| 414 \| 276 \| 25 \| 370 \| 750 \| 378 \| 767 \| Prim-Pol domain-containing protein \| Prim-Pol domain-containing protein; afdb-uniprot50 \| AF-A0A840RUQ5-F1-MODEL\_V4 \| 1.0 \| 6.956e-18 \| 576 \| 0.279 \| 354 \| 195 \| 17 \| 10 \| 347 \| 1 \| 310 \| Putative DNA primase/helicase \| Putative DNA primase/helicase; afdb-uniprot50 \| AF-A8SSN2-F1-MODEL\_V4 \| 1.0 \| 2.527e-17 \| 576 \| 0.227 \| 378 \| 245 \| 19 \| 420 \| 769 \| 4 \| 362 \| Virulence-associated protein E \| Virulence-associated protein E; afdb-uniprot50 \| AF-A0A828JM73-F1-MODEL\_V4 \| 1.0 \| 1.3e-14 \| 576 \| 0.306 \| 251 \| 152 \| 9 \| 100 \| 346 \| 83 \| 315 \| Uncharacterized protein \| Uncharacterized protein; afdb-uniprot50 \| AF-A0A1M5CP24-F1-MODEL\_V4 \| 1.0 \| 1.578e-20 \| 576 \| 0.172 \| 539 \| 348 \| 27 \| 240 \| 751 \| 220 \| 687 \| Uncharacterized protein \| Uncharacterized protein; afdb-uniprot50 \| AF-A0A218QNW9-F1-MODEL\_V4 \| 1.0 \| 9.18e-25 \| 575 \| 0.177 \| 756 \| 457 \| 36 \| 89 \| 757 \| 8 \| 685 \| Virulence-associated E family protein \| Virulence-associated E family protein; afdb-uniprot50 \| AF-A0A414TCF4-F1-MODEL\_V4 \| 1.0 \| 1.705e-16 \| 574 \| 0.21 \| 346 \| 231 \| 16 \| 457 \| 769 \| 2 \| 338 \| Uncharacterized protein \| Uncharacterized protein; afdb-uniprot50 \| AF-A0A2N7QRK2-F1-MODEL\_V4 \| 1.0 \| 5.238e-30 \| 574 \| 0.197 \| 779 \| 435 \| 42 \| 100 \| 739 \| 99 \| 825 \| DNA primase TraC \| DNA primase TraC; afdb-uniprot50 \| AF-A0A6L8LFI5-F1-MODEL\_V4 \| 1.0 \| 1.83e-23 \| 573 \| 0.177 \| 776 \| 357 \| 33 \| 1 \| 750 \| 212 \| 731 \| PriCT\_2 domain-containing protein \| PriCT\_2 domain-containing protein; afdb-uniprot50 \| AF-A0A085TW06-F1-MODEL\_V4 \| 1.0 \| 6.438e-22 \| 573 \| 0.17 \| 778 \| 360 \| 28 \| 1 \| 750 \| 215 \| 734 \| PriCT\_2 domain-containing protein \| PriCT\_2 domain-containing protein; afdb-uniprot50 \| AF-A0A3N7RH32-F1-MODEL\_V4 \| 1.0 \| 7.915e-30 \| 573 \| 0.192 \| 872 \| 478 \| 46 \| 8 \| 737 \| 2 \| 788 \| DUF927 domain-containing protein \| DUF927 domain-containing protein; afdb-uniprot50 \| AF-A0A0P9QTD8-F1-MODEL\_V4 \| 1.0 \| 6.114e-14 \| 573 \| 0.327 \| 214 \| 122 \| 8 \| 136 \| 346 \| 3 \| 197 \| LPD7 domain-containing protein \| LPD7 domain-containing protein; afdb-uniprot50 \| AF-A0A1H7HXF3-F1-MODEL\_V4 \| 1.0 \| 1.854e-17 \| 572 \| 0.211 \| 387 \| 265 \| 17 \| 391 \| 758 \| 2 \| 367 \| Virulence-associated protein E \| Virulence-associated protein E; afdb-uniprot50 \| AF-A0A6H9P0V4-F1-MODEL\_V4 \| 1.0 \| 2.25e-15 \| 571 \| 0.2 \| 300 \| 220 \| 13 \| 467 \| 757 \| 1 \| 289 \| Helicase \| Helicase; afdb-uniprot50 \| AF-A0A641UPF6-F1-MODEL\_V4 \| 1.0 \| 6.521e-16 \| 571 \| 0.208 \| 321 \| 231 \| 13 \| 381 \| 691 \| 61 \| 368 \| Helicase \| Helicase; afdb-uniprot50 \| AF-A0A5C7CZ33-F1-MODEL\_V4 \| 1.0 \| 4.316e-16 \| 571 \| 0.285 \| 284 \| 157 \| 11 \| 82 \| 358 \| 149 \| 393 \| Uncharacterized protein \| Uncharacterized protein; afdb-uniprot50 \| AF-A0A3M5UMZ6-F1-MODEL\_V4 \| 1.0 \| 8.173e-15 \| 571 \| 0.313 \| 255 \| 145 \| 11 \| 114 \| 358 \| 366 \| 600 \| DUF5710 domain-containing protein \| DUF5710 domain-containing protein; afdb-uniprot50 \| AF-A0A3D5KKU3-F1-MODEL\_V4 \| 1.0 \| 1.538e-16 \| 570 \| 0.185 \| 398 \| 274 \| 21 \| 384 \| 758 \| 20 \| 390 \| Uncharacterized protein \| Uncharacterized protein; afdb-uniprot50 \| AF-A0A2V4UFM1-F1-MODEL\_V4 \| 1.0 \| 1.807e-29 \| 570 \| 0.197 \| 789 \| 447 \| 40 \| 13 \| 744 \| 9 \| 667 \| Phage/plasmid primase-like uncharacterized protein \| Phage/plasmid primase-like uncharacterized protein; afdb-uniprot50 \| AF-A0A370NG87-F1-MODEL\_V4 \| 1.0 \| 1.284e-28 \| 570 \| 0.202 \| 771 \| 451 \| 43 \| 101 \| 768 \| 82 \| 791 \| SF3 helicase domain-containing protein \| SF3 helicase domain-containing protein; afdb-uniprot50 \| AF-Q1DD05-F1-MODEL\_V4 \| 1.0 \| 1.396e-21 \| 570 \| 0.21 \| 557 \| 348 \| 30 \| 239 \| 757 \| 339 \| 841 \| Virulence-associated protein E domain protein \| Virulence-associated protein E domain protein; afdb-uniprot50 \| AF-A0A1G4SVX6-F1-MODEL\_V4 \| 1.0 \| 2.193e-19 \| 569 \| 0.209 \| 526 \| 321 \| 22 \| 261 \| 762 \| 264 \| 718 \| Virulence-associated protein E \| Virulence-associated protein E; afdb-uniprot50 \| AF-A0A6I5Q3V8-F1-MODEL\_V4 \| 1.0 \| 1.058e-22 \| 569 \| 0.186 \| 634 \| 403 \| 34 \| 179 \| 751 \| 75 \| 656 \| Uncharacterized protein \| Uncharacterized protein; afdb-uniprot50 \| AF-A0A374TFD7-F1-MODEL\_V4 \| 1.0 \| 1.243e-19 \| 569 \| 0.164 \| 534 \| 339 \| 28 \| 261 \| 756 \| 269 \| 733 \| VirE\_N domain-containing protein \| VirE\_N domain-containing protein; afdb-uniprot50 \| AF-A0A0C1Y5L2-F1-MODEL\_V4 \| 1.0 \| 6.397e-17 \| 567 \| 0.205 \| 385 \| 261 \| 18 \| 393 \| 754 \| 45 \| 407 \| Uncharacterized protein \| Uncharacterized protein; afdb-uniprot50 \| AF-A0A841VQI7-F1-MODEL\_V4 \| 1.0 \| 1.251e-24 \| 567 \| 0.176 \| 713 \| 400 \| 42 \| 159 \| 769 \| 27 \| 653 \| Uncharacterized protein \| Uncharacterized protein; afdb-uniprot50 \| AF-A8SVC7-F1-MODEL\_V4 \| 1.0 \| 1.317e-16 \| 565 \| 0.224 \| 356 \| 233 \| 17 \| 441 \| 769 \| 2 \| 341 \| Virulence-associated protein E \| Virulence-associated protein E; afdb-uniprot50 \| AF-A0A2V2GG06-F1-MODEL\_V4 \| 1.0 \| 7.001e-15 \| 564 \| 0.203 \| 310 \| 222 \| 11 \| 453 \| 751 \| 17 \| 312 \| Helicase \| Helicase; afdb-uniprot50 \| AF-J9DI45-F1-MODEL\_V4 \| 1.0 \| 4.344e-21 \| 564 \| 0.172 \| 625 \| 378 \| 32 \| 183 \| 751 \| 119 \| 660 \| Prim-Pol domain-containing protein \| Prim-Pol domain-containing protein; afdb-uniprot50 \| AF-A0A829YB94-F1-MODEL\_V4 \| 1.0 \| 3.65e-22 \| 564 \| 0.16 \| 790 \| 366 \| 25 \| 1 \| 757 \| 230 \| 754 \| Uncharacterized protein \| Uncharacterized protein; afdb-uniprot50 \| AF-A0A2W6ZVY5-F1-MODEL\_V4 \| 1.0 \| 1.651e-23 \| 564 \| 0.21 \| 657 \| 395 \| 34 \| 179 \| 755 \| 164 \| 776 \| Prim\_Zn\_Ribbon domain-containing protein \| Prim\_Zn\_Ribbon domain-containing protein; afdb-uniprot50 \| AF-A0A1H9F653-F1-MODEL\_V4 \| 1.0 \| 1.219e-28 \| 564 \| 0.206 \| 794 \| 449 \| 39 \| 102 \| 769 \| 126 \| 864 \| Uncharacterized domain associated with phage/plasmid primase \| Uncharacterized domain associated with phage/plasmid primase; afdb-uniprot50 \| AF-A0A359D3R9-F1-MODEL\_V4 \| 1.0 \| 9.239e-22 \| 563 \| 0.144 \| 763 \| 389 \| 29 \| 12 \| 757 \| 189 \| 704 \| Uncharacterized protein \| Uncharacterized protein; afdb-uniprot50 \| AF-A0A3E4N9U9-F1-MODEL\_V4 \| 1.0 \| 4.847e-18 \| 562 \| 0.17 \| 441 \| 336 \| 12 \| 330 \| 756 \| 21 \| 445 \| Helicase \| Helicase; afdb-uniprot50 \| AF-A0A5C7Q0U9-F1-MODEL\_V4 \| 1.0 \| 8.017e-24 \| 562 \| 0.178 \| 796 \| 426 \| 40 \| 2 \| 757 \| 216 \| 823 \| PriCT\_2 domain-containing protein \| PriCT\_2 domain-containing protein; afdb-uniprot50 \| AF-A0A1F1G801-F1-MODEL\_V4 \| 1.0 \| 3.208e-18 \| 561 \| 0.204 \| 411 \| 266 \| 19 \| 407 \| 769 \| 158 \| 555 \| Uncharacterized protein \| Uncharacterized protein; afdb-uniprot50 \| AF-A0A5C7M7G3-F1-MODEL\_V4 \| 1.0 \| 1.683e-22 \| 561 \| 0.172 \| 685 \| 416 \| 37 \| 143 \| 760 \| 86 \| 686 \| Prim-Pol domain-containing protein \| Prim-Pol domain-containing protein; afdb-uniprot50 \| AF-A0A844DJS6-F1-MODEL\_V4 \| 1.0 \| 1.018e-16 \| 560 \| 0.218 \| 376 \| 246 \| 18 \| 423 \| 769 \| 3 \| 359 \| Conjugal transfer protein TraC \| Conjugal transfer protein TraC; afdb-uniprot50 \| AF-A0A560FLU5-F1-MODEL\_V4 \| 1.0 \| 4.457e-17 \| 560 \| 0.2 \| 429 \| 291 \| 21 \| 379 \| 769 \| 4 \| 418 \| Virulence-associated protein E \| Virulence-associated protein E; afdb-uniprot50 \| AF-R4WX46-F1-MODEL\_V4 \| 1.0 \| 2.463e-29 \| 560 \| 0.196 \| 870 \| 472 \| 50 \| 6 \| 737 \| 1 \| 781 \| Uncharacterized protein \| Uncharacterized protein; afdb-uniprot50 \| AF-A0A433W3D6-F1-MODEL\_V4 \| 1.0 \| 1.181e-19 \| 559 \| 0.197 \| 526 \| 312 \| 23 \| 261 \| 757 \| 221 \| 665 \| RepB\_primase domain-containing protein \| RepB\_primase domain-containing protein; afdb-uniprot50 \| AF-A0A1Q4BTB8-F1-MODEL\_V4 \| 1.0 \| 1.423e-20 \| 559 \| 0.186 \| 568 \| 343 \| 32 \| 246 \| 769 \| 266 \| 758 \| PriCT\_2 domain-containing protein \| PriCT\_2 domain-containing protein; afdb-uniprot50 \| AF-A0A7X3VTX1-F1-MODEL\_V4 \| 1.0 \| 2.165e-17 \| 558 \| 0.223 \| 421 \| 272 \| 22 \| 370 \| 753 \| 32 \| 434 \| Uncharacterized protein \| Uncharacterized protein; afdb-uniprot50 \| AF-A0A6L8GF99-F1-MODEL\_V4 \| 1.0 \| 1.672e-17 \| 558 \| 0.179 \| 423 \| 283 \| 23 \| 379 \| 753 \| 71 \| 477 \| Uncharacterized protein \| Uncharacterized protein; afdb-uniprot50 \| AF-A0A435FA41-F1-MODEL\_V4 \| 1.0 \| 2.931e-20 \| 558 \| 0.193 \| 543 \| 331 \| 33 \| 260 \| 751 \| 141 \| 627 \| Uncharacterized protein \| Uncharacterized protein; afdb-uniprot50 \| AF-A0A3D5P9B6-F1-MODEL\_V4 \| 1.0 \| 1.528e-19 \| 558 \| 0.194 \| 567 \| 346 \| 34 \| 257 \| 757 \| 290 \| 811 \| Virulence-associated protein E \| Virulence-associated protein E; afdb-uniprot50 \| AF-A0A6A7WE66-F1-MODEL\_V4 \| 1.0 \| 1.086e-18 \| 557 \| 0.145 \| 530 \| 351 \| 29 \| 261 \| 755 \| 275 \| 737 \| VirE\_N domain-containing protein \| VirE\_N domain-containing protein; afdb-uniprot50 \| AF-A0A094YPP6-F1-MODEL\_V4 \| 1.0 \| 7.914e-22 \| 557 \| 0.165 \| 778 \| 356 \| 30 \| 12 \| 752 \| 243 \| 763 \| Virulence-associated E \| Virulence-associated E; afdb-uniprot50 \| AF-A0A4P5SJT8-F1-MODEL\_V4 \| 1.0 \| 1.965e-14 \| 556 \| 0.327 \| 235 \| 122 \| 11 \| 129 \| 346 \| 229 \| 444 \| Toprim domain-containing protein \| Toprim domain-containing protein; afdb-uniprot50 \| AF-A0A357LIR7-F1-MODEL\_V4 \| 1.0 \| 5.844e-19 \| 556 \| 0.176 \| 567 \| 337 \| 32 \| 239 \| 767 \| 32 \| 506 \| Uncharacterized protein \| Uncharacterized protein; afdb-uniprot50 \| AF-E0U6Z0-F1-MODEL\_V4 \| 1.0 \| 2.495e-23 \| 556 \| 0.176 \| 680 \| 403 \| 36 \| 177 \| 757 \| 64 \| 685 \| Virulence-associated E family protein \| Virulence-associated E family protein; afdb-uniprot50 \| AF-D6E493-F1-MODEL\_V4 \| 1.0 \| 8.279e-17 \| 555 \| 0.214 \| 401 \| 270 \| 18 \| 394 \| 769 \| 2 \| 382 \| Predicted P-loop ATPase and inactivated derivatives \| Predicted P-loop ATPase and inactivated derivatives; afdb-uniprot50 \| AF-A0A7G7KQ79-F1-MODEL\_V4 \| 1.0 \| 9.48e-18 \| 554 \| 0.197 \| 476 \| 327 \| 27 \| 318 \| 769 \| 1 \| 445 \| Uncharacterized protein \| Uncharacterized protein; afdb-uniprot50 \| AF-A0A2E4ID67-F1-MODEL\_V4 \| 1.0 \| 2.527e-17 \| 553 \| 0.184 \| 422 \| 302 \| 17 \| 370 \| 768 \| 29 \| 431 \| Uncharacterized protein \| Uncharacterized protein; afdb-uniprot50 \| AF-A0A1I5CW14-F1-MODEL\_V4 \| 1.0 \| 5.444e-20 \| 553 \| 0.185 \| 538 \| 355 \| 27 \| 246 \| 757 \| 116 \| 596 \| Predicted P-loop ATPase and inactivated derivatives \| Predicted P-loop ATPase and inactivated derivatives; afdb-uniprot50 \| AF-A0A7J5ET25-F1-MODEL\_V4 \| 1.0 \| 1.489e-23 \| 553 \| 0.168 \| 830 \| 421 \| 34 \| 1 \| 752 \| 222 \| 859 \| Prim-Pol domain-containing protein \| Prim-Pol domain-containing protein; afdb-uniprot50 \| AF-A0A7Z2G310-F1-MODEL\_V4 \| 1.0 \| 4.515e-19 \| 552 \| 0.192 \| 519 \| 355 \| 22 \| 264 \| 756 \| 266 \| 746 \| Uncharacterized protein \| Uncharacterized protein; afdb-uniprot50 \| AF-G5QKM5-F1-MODEL\_V4 \| 1.0 \| 8.173e-15 \| 552 \| 0.273 \| 274 \| 162 \| 10 \| 99 \| 358 \| 138 \| 388 \| DNA primase traC \| DNA primase traC; afdb-uniprot50 \| AF-A0A255SPK4-F1-MODEL\_V4 \| 1.0 \| 1.334e-18 \| 550 \| 0.208 \| 418 \| 265 \| 16 \| 263 \| 668 \| 241 \| 604 \| VirE\_N domain-containing protein \| VirE\_N domain-containing protein; afdb-uniprot50 \| AF-A0A843FDY4-F1-MODEL\_V4 \| 1.0 \| 2.096e-16 \| 549 \| 0.178 \| 410 \| 273 \| 18 \| 396 \| 769 \| 2 \| 383 \| Uncharacterized protein \| Uncharacterized protein; afdb-uniprot50 \| AF-D6YV80-F1-MODEL\_V4 \| 1.0 \| 1.738e-15 \| 549 \| 0.307 \| 267 \| 151 \| 13 \| 92 \| 347 \| 82 \| 325 \| Uncharacterized protein \| Uncharacterized protein; afdb-uniprot50 \| AF-A0A2V2SG54-F1-MODEL\_V4 \| 1.0 \| 1.079e-21 \| 549 \| 0.165 \| 791 \| 378 \| 28 \| 1 \| 757 \| 223 \| 765 \| Prim-Pol domain-containing protein \| Prim-Pol domain-containing protein; afdb-uniprot50 \| AF-A0A542BG12-F1-MODEL\_V4 \| 1.0 \| 1.196e-13 \| 549 \| 0.316 \| 237 \| 138 \| 10 \| 114 \| 346 \| 160 \| 376 \| Putative DNA primase/helicase \| Putative DNA primase/helicase; afdb-uniprot50 \| AF-A0A7J0AZE4-F1-MODEL\_V4 \| 1.0 \| 1.204e-18 \| 548 \| 0.207 \| 482 \| 277 \| 29 \| 370 \| 769 \| 134 \| 592 \| Uncharacterized protein \| Uncharacterized protein; afdb-uniprot50 \| AF-A0A081RVY4-F1-MODEL\_V4 \| 1.0 \| 1.284e-20 \| 548 \| 0.199 \| 547 \| 333 \| 31 \| 261 \| 753 \| 247 \| 742 \| Putative P-loop ATPase \| Putative P-loop ATPase; afdb-uniprot50 \| AF-A0A828NJX6-F1-MODEL\_V4 \| 1.0 \| 1.173e-14 \| 548 \| 0.272 \| 268 \| 163 \| 11 \| 90 \| 346 \| 126 \| 372 \| DNA primase \| DNA primase; afdb-uniprot50 \| AF-A0A1Y6D1S4-F1-MODEL\_V4 \| 1.0 \| 2.221e-29 \| 548 \| 0.195 \| 883 \| 482 \| 51 \| 10 \| 739 \| 52 \| 858 \| Putative DNA primase/helicase \| Putative DNA primase/helicase; afdb-uniprot50 \| AF-A0A4Q7EAM6-F1-MODEL\_V4 \| 1.0 \| 3.008e-16 \| 547 \| 0.206 \| 383 \| 259 \| 18 \| 395 \| 754 \| 133 \| 493 \| Uncharacterized protein \| Uncharacterized protein; afdb-uniprot50 \| AF-A0A5C4LBI4-F1-MODEL\_V4 \| 1.0 \| 1.196e-21 \| 547 \| 0.162 \| 700 \| 402 \| 31 \| 150 \| 752 \| 74 \| 685 \| Prim-Pol domain-containing protein \| Prim-Pol domain-containing protein; afdb-uniprot50 \| AF-A0A2N3BD34-F1-MODEL\_V4 \| 1.0 \| 6.397e-25 \| 547 \| 0.181 \| 834 \| 432 \| 45 \| 1 \| 671 \| 7 \| 752 \| P-loop ATPase \| P-loop ATPase; afdb-uniprot50 \| AF-A0A1H6V3P5-F1-MODEL\_V4 \| 1.0 \| 1.599e-22 \| 547 \| 0.165 \| 792 \| 367 \| 31 \| 1 \| 755 \| 235 \| 769 \| Primase C terminal 2 (PriCT-2) \| Primase C terminal 2 (PriCT-2); afdb-uniprot50 \| AF-A0A1Q4GUC4-F1-MODEL\_V4 \| 1.0 \| 4.126e-29 \| 547 \| 0.193 \| 888 \| 418 \| 48 \| 11 \| 731 \| 1 \| 756 \| PriCT\_2 domain-containing protein \| PriCT\_2 domain-containing protein; afdb-uniprot50 \| AF-A0A6P0ZFJ0-F1-MODEL\_V4 \| 1.0 \| 1.705e-24 \| 546 \| 0.172 \| 703 \| 432 \| 40 \| 143 \| 762 \| 18 \| 653 \| DUF3854 domain-containing protein \| DUF3854 domain-containing protein; afdb-uniprot50 \| AF-A0A7L9BQY4-F1-MODEL\_V4 \| 1.0 \| 1.578e-20 \| 546 \| 0.202 \| 568 \| 345 \| 31 \| 260 \| 762 \| 230 \| 754 \| Uncharacterized protein \| Uncharacterized protein; afdb-uniprot50 \| AF-A0A1H6K2S3-F1-MODEL\_V4 \| 1.0 \| 4.099e-16 \| 545 \| 0.299 \| 297 \| 149 \| 13 \| 68 \| 347 \| 488 \| 742 \| Uncharacterized domain associated with phage/plasmid primase \| Uncharacterized domain associated with phage/plasmid primase; afdb-uniprot50 \| AF-A0A371XDT2-F1-MODEL\_V4 \| 1.0 \| 3.489e-19 \| 545 \| 0.197 \| 532 \| 330 \| 22 \| 261 \| 756 \| 295 \| 765 \| Uncharacterized protein \| Uncharacterized protein; afdb-uniprot50 \| AF-A0A1H0EEJ0-F1-MODEL\_V4 \| 1.0 \| 7.467e-17 \| 544 \| 0.188 \| 409 \| 275 \| 23 \| 382 \| 755 \| 24 \| 410 \| Virulence-associated protein E \| Virulence-associated protein E; afdb-uniprot50 \| AF-A0A3Y9C6Z3-F1-MODEL\_V4 \| 1.0 \| 1.3e-14 \| 544 \| 0.279 \| 265 \| 153 \| 10 \| 98 \| 346 \| 134 \| 376 \| DUF5710 domain-containing protein \| DUF5710 domain-containing protein; afdb-uniprot50 \| AF-A0A367RLN6-F1-MODEL\_V4 \| 1.0 \| 1.761e-17 \| 544 \| 0.202 \| 449 \| 268 \| 19 \| 381 \| 758 \| 100 \| 529 \| Uncharacterized protein \| Uncharacterized protein; afdb-uniprot50 \| AF-A0A1I4C3P1-F1-MODEL\_V4 \| 1.0 \| 2.731e-29 \| 544 \| 0.19 \| 765 \| 421 \| 39 \| 91 \| 751 \| 72 \| 741 \| Putative DNA primase/helicase \| Putative DNA primase/helicase; afdb-uniprot50 \| AF-W2CQM4-F1-MODEL\_V4 \| 1.0 \| 1.489e-15 \| 543 \| 0.236 \| 288 \| 196 \| 12 \| 379 \| 652 \| 4 \| 281 \| Virulence-associated protein E \| Virulence-associated protein E; afdb-uniprot50 \| AF-A0A098C1H9-F1-MODEL\_V4 \| 1.0 \| 2.369e-15 \| 543 \| 0.233 \| 304 \| 206 \| 13 \| 384 \| 674 \| 14 \| 303 \| Virulence-associated E family protein \| Virulence-associated E family protein; afdb-uniprot50 \| AF-A0A4V1YW25-F1-MODEL\_V4 \| 1.0 \| 2.627e-15 \| 543 \| 0.281 \| 288 \| 187 \| 9 \| 370 \| 647 \| 31 \| 308 \| Virulence-associated E family protein \| Virulence-associated E family protein; afdb-uniprot50 \| AF-A0A239R0C2-F1-MODEL\_V4 \| 1.0 \| 7.092e-17 \| 543 \| 0.189 \| 412 \| 276 \| 16 \| 384 \| 755 \| 21 \| 414 \| DUF3874 domain-containing protein \| DUF3874 domain-containing protein; afdb-uniprot50 \| AF-A0A160UEW2-F1-MODEL\_V4 \| 1.0 \| 3.292e-14 \| 542 \| 0.248 \| 274 \| 188 \| 8 \| 484 \| 752 \| 11 \| 271 \| Virulenceassociated E \| Virulenceassociated E; afdb-uniprot50 \| AF-A0A4Q3J348-F1-MODEL\_V4 \| 1.0 \| 1.414e-15 \| 542 \| 0.25 \| 279 \| 187 \| 10 \| 381 \| 649 \| 108 \| 374 \| Uncharacterized protein \| Uncharacterized protein; afdb-uniprot50 \| AF-A0A2W7HY36-F1-MODEL\_V4 \| 1.0 \| 6.397e-17 \| 542 \| 0.291 \| 319 \| 150 \| 14 \| 38 \| 349 \| 70 \| 319 \| Putative DNA primase/helicase \| Putative DNA primase/helicase; afdb-uniprot50 \| AF-G5SSC0-F1-MODEL\_V4 \| 1.0 \| 3.208e-18 \| 542 \| 0.185 \| 464 \| 317 \| 22 \| 316 \| 755 \| 16 \| 442 \| DUF3874 domain-containing protein \| DUF3874 domain-containing protein; afdb-uniprot50 \| AF-A0A6G1TK16-F1-MODEL\_V4 \| 1.0 \| 3.943e-18 \| 542 \| 0.177 \| 518 \| 315 \| 23 \| 243 \| 730 \| 54 \| 490 \| Uncharacterized protein \| Uncharacterized protein; afdb-uniprot50 \| AF-X5DJW4-F1-MODEL\_V4 \| 1.0 \| 3.626e-17 \| 542 \| 0.172 \| 446 \| 310 \| 27 \| 326 \| 752 \| 277 \| 682 \| Bifunctional DNA primase/polymerase, N-terminal \| Bifunctional DNA primase/polymerase, N-terminal; afdb-uniprot50 \| AF-A0A557QLQ1-F1-MODEL\_V4 \| 1.0 \| 1.1e-28 \| 542 \| 0.212 \| 791 \| 427 \| 45 \| 92 \| 753 \| 64 \| 787 \| Toprim domain-containing protein \| Toprim domain-containing protein; afdb-uniprot50 \| AF-A0A7Y0DXY3-F1-MODEL\_V4 \| 1.0 \| 1.396e-13 \| 541 \| 0.361 \| 202 \| 108 \| 7 \| 156 \| 355 \| 2 \| 184 \| Toprim domain-containing protein \| Toprim domain-containing protein; afdb-uniprot50 \| AF-A0A3A6CGK6-F1-MODEL\_V4 \| 1.0 \| 4.544e-16 \| 541 \| 0.2 \| 359 \| 243 \| 15 \| 439 \| 769 \| 6 \| 348 \| Virulence-associated protein E \| Virulence-associated protein E; afdb-uniprot50 \| AF-A0A533RGC8-F1-MODEL\_V4 \| 1.0 \| 1.284e-28 \| 541 \| 0.191 \| 761 \| 437 \| 49 \| 92 \| 752 \| 78 \| 759 \| DNA primase \| DNA primase; afdb-uniprot50 \| AF-A0A844GG09-F1-MODEL\_V4 \| 1.0 \| 6.565e-29 \| 540 \| 0.21 \| 835 \| 433 \| 45 \| 12 \| 684 \| 5 \| 774 \| DUF927 domain-containing protein \| DUF927 domain-containing protein; afdb-uniprot50 \| AF-A0A5N3P4R6-F1-MODEL\_V4 \| 1.0 \| 9.541e-15 \| 539 \| 0.241 \| 307 \| 207 \| 15 \| 460 \| 755 \| 2 \| 293 \| Uncharacterized protein \| Uncharacterized protein; afdb-uniprot50 \| AF-A0A2T1EHM2-F1-MODEL\_V4 \| 1.0 \| 1.173e-22 \| 539 \| 0.197 \| 648 \| 402 \| 31 \| 166 \| 755 \| 107 \| 694 \| Uncharacterized protein \| Uncharacterized protein; afdb-uniprot50 \| AF-A0A562C7R2-F1-MODEL\_V4 \| 1.0 \| 7.565e-27 \| 539 \| 0.187 \| 762 \| 443 \| 45 \| 93 \| 767 \| 82 \| 753 \| Putative DNA primase/helicase \| Putative DNA primase/helicase; afdb-uniprot50 \| AF-A0A7C1H7I6-F1-MODEL\_V4 \| 1.0 \| 4.02e-17 \| 538 \| 0.272 \| 348 \| 186 \| 16 \| 93 \| 431 \| 113 \| 402 \| Toprim domain-containing protein \| Toprim domain-containing protein; afdb-uniprot50 \| AF-A0A1M3E3M9-F1-MODEL\_V4 \| 1.0 \| 2.463e-21 \| 537 \| 0.162 \| 787 \| 377 \| 34 \| 1 \| 757 \| 180 \| 713 \| Uncharacterized protein \| Uncharacterized protein; afdb-uniprot50 \| AF-W2CS72-F1-MODEL\_V4 \| 1.0 \| 8.173e-15 \| 536 \| 0.252 \| 261 \| 180 \| 10 \| 505 \| 756 \| 1 \| 255 \| Virulence-associated protein E \| Virulence-associated protein E; afdb-uniprot50 \| AF-A0A658JSB4-F1-MODEL\_V4 \| 1.0 \| 2.766e-15 \| 536 \| 0.33 \| 254 \| 151 \| 8 \| 380 \| 624 \| 381 \| 624 \| Virulence-associated E family protein \| Virulence-associated E family protein; afdb-uniprot50 \| AF-A0A158BCW7-F1-MODEL\_V4 \| 1.0 \| 9.421e-29 \| 536 \| 0.195 \| 985 \| 503 \| 56 \| 1 \| 756 \| 1 \| 924 \| Inner membrane protein \| Inner membrane protein; afdb-uniprot50 \| AF-A0A3D0G2H0-F1-MODEL\_V4 \| 1.0 \| 1.89e-16 \| 536 \| 0.199 \| 462 \| 284 \| 23 \| 370 \| 753 \| 464 \| 917 \| Uncharacterized protein \| Uncharacterized protein; afdb-uniprot50 \| AF-E0NRI5-F1-MODEL\_V4 \| 1.0 \| 1.598e-14 \| 535 \| 0.23 \| 265 \| 197 \| 5 \| 499 \| 757 \| 2 \| 265 \| Uncharacterized protein \| Uncharacterized protein; afdb-uniprot50 \| AF-A0A849TJR6-F1-MODEL\_V4 \| 1.0 \| 7.278e-29 \| 535 \| 0.185 \| 878 \| 441 \| 49 \| 1 \| 731 \| 19 \| 768 \| DUF3987 domain-containing protein \| DUF3987 domain-containing protein; afdb-uniprot50 \| AF-A0A0T7BQB0-F1-MODEL\_V4 \| 1.0 \| 3.843e-22 \| 535 \| 0.187 \| 645 \| 393 \| 35 \| 178 \| 757 \| 73 \| 651 \| DUF3854 domain-containing protein \| DUF3854 domain-containing protein; afdb-uniprot50 \| AF-A0A5T2E3M0-F1-MODEL\_V4 \| 1.0 \| 1.3e-14 \| 534 \| 0.273 \| 278 \| 162 \| 10 \| 99 \| 358 \| 138 \| 393 \| Uncharacterized protein \| Uncharacterized protein; afdb-uniprot50 \| AF-A0A154VF00-F1-MODEL\_V4 \| 1.0 \| 2.462e-13 \| 533 \| 0.343 \| 195 \| 106 \| 7 \| 155 \| 347 \| 2 \| 176 \| Toprim domain-containing protein \| Toprim domain-containing protein; afdb-uniprot50 \| AF-A0A0S2TFP5-F1-MODEL\_V4 \| 1.0 \| 2.193e-27 \| 533 \| 0.206 \| 770 \| 428 \| 43 \| 98 \| 752 \| 106 \| 807 \| Uncharacterized protein \| Uncharacterized protein; afdb-uniprot50 \| AF-I3Y9A5-F1-MODEL\_V4 \| 1.0 \| 1.807e-29 \| 533 \| 0.204 \| 884 \| 473 \| 47 \| 12 \| 738 \| 131 \| 940 \| DNA/RNA helicase, superfamily II \| DNA/RNA helicase, superfamily II; afdb-uniprot50 \| AF-I2PZX8-F1-MODEL\_V4 \| 1.0 \| 2.838e-27 \| 532 \| 0.175 \| 826 \| 423 \| 48 \| 102 \| 753 \| 77 \| 818 \| Toprim domain-containing protein \| Toprim domain-containing protein; afdb-uniprot50 \| AF-A0A4P2QP84-F1-MODEL\_V4 \| 1.0 \| 3.843e-22 \| 532 \| 0.177 \| 727 \| 410 \| 34 \| 179 \| 751 \| 79 \| 771 \| DUF3854 domain-containing protein \| DUF3854 domain-containing protein; afdb-uniprot50 \| AF-A0A8B2TSA0-F1-MODEL\_V4 \| 1.0 \| 1.94e-20 \| 531 \| 0.139 \| 773 \| 365 \| 27 \| 2 \| 751 \| 229 \| 723 \| Uncharacterized protein \| Uncharacterized protein; afdb-uniprot50 \| AF-A0A6P0YCZ3-F1-MODEL\_V4 \| 1.0 \| 7.002e-23 \| 531 \| 0.177 \| 716 \| 439 \| 42 \| 132 \| 762 \| 5 \| 655 \| Uncharacterized protein \| Uncharacterized protein; afdb-uniprot50 \| AF-A0A1L5KUQ4-F1-MODEL\_V4 \| 1.0 \| 4.344e-13 \| 530 \| 0.233 \| 240 \| 169 \| 10 \| 453 \| 687 \| 11 \| 240 \| Virulence-associated protein E \| Virulence-associated protein E; afdb-uniprot50 \| AF-A0A1H0XV22-F1-MODEL\_V4 \| 1.0 \| 8.121e-18 \| 529 \| 0.154 \| 555 \| 380 \| 28 \| 259 \| 769 \| 8 \| 516 \| Predicted P-loop ATPase and inactivated derivatives \| Predicted P-loop ATPase and inactivated derivatives; afdb-uniprot50 \| AF-K9SHA4-F1-MODEL\_V4 \| 1.0 \| 4.429e-20 \| 529 \| 0.188 \| 577 \| 348 \| 30 \| 224 \| 757 \| 243 \| 741 \| Virulence-associated E family protein \| Virulence-associated E family protein; afdb-uniprot50 \| AF-A0A0Q6VCR0-F1-MODEL\_V4 \| 1.0 \| 8.069e-21 \| 529 \| 0.159 \| 784 \| 373 \| 28 \| 1 \| 757 \| 220 \| 744 \| Uncharacterized protein \| Uncharacterized protein; afdb-uniprot50 \| AF-A0A7C1TYP0-F1-MODEL\_V4 \| 1.0 \| 2.931e-28 \| 529 \| 0.197 \| 785 \| 426 \| 49 \| 92 \| 753 \| 64 \| 767 \| Toprim domain-containing protein \| Toprim domain-containing protein; afdb-uniprot50 \| AF-A0A250DKS7-F1-MODEL\_V4 \| 1.0 \| 2.369e-23 \| 529 \| 0.184 \| 732 \| 433 \| 45 \| 167 \| 767 \| 167 \| 865 \| Virulence protein E \| Virulence protein E; afdb-uniprot50 \| AF-A0A0G1YTK1-F1-MODEL\_V4 \| 1.0 \| 6.315e-23 \| 529 \| 0.181 \| 837 \| 496 \| 41 \| 6 \| 767 \| 232 \| 953 \| Virulence-associated protein E domain protein \| Virulence-associated protein E domain protein; afdb-uniprot50 \| AF-A0A1H4HT56-F1-MODEL\_V4 \| 1.0 \| 1.284e-20 \| 528 \| 0.142 \| 788 \| 374 \| 29 \| 1 \| 753 \| 215 \| 735 \| Primase C terminal 2 (PriCT-2) \| Primase C terminal 2 (PriCT-2); afdb-uniprot50 \| AF-A0A0A2CPT5-F1-MODEL\_V4 \| 1.0 \| 4.515e-19 \| 528 \| 0.186 \| 551 \| 351 \| 28 \| 237 \| 757 \| 260 \| 742 \| PriCT\_1 domain-containing protein \| PriCT\_1 domain-containing protein; afdb-uniprot50 \| AF-A0A1Z4JI45-F1-MODEL\_V4 \| 1.0 \| 2.179e-22 \| 528 \| 0.164 \| 837 \| 436 \| 34 \| 1 \| 769 \| 305 \| 945 \| Uncharacterized protein \| Uncharacterized protein; afdb-uniprot50 \| AF-A0A639X8R3-F1-MODEL\_V4 \| 1.0 \| 6.314e-15 \| 528 \| 0.278 \| 287 \| 162 \| 12 \| 97 \| 360 \| 134 \| 398 \| Uncharacterized protein \| Uncharacterized protein; afdb-uniprot50 \| AF-A0A252D898-F1-MODEL\_V4 \| 1.0 \| 2.695e-19 \| 527 \| 0.192 \| 539 \| 324 \| 20 \| 239 \| 759 \| 288 \| 732 \| Prim-Pol domain-containing protein \| Prim-Pol domain-containing protein; afdb-uniprot50 \| AF-A0A5Y8ZF30-F1-MODEL\_V4 \| 1.0 \| 5.237e-14 \| 526 \| 0.285 \| 263 \| 147 \| 14 \| 105 \| 347 \| 461 \| 702 \| DUF1738 domain-containing protein \| DUF1738 domain-containing protein; afdb-uniprot50 \| AF-A0A369XMD4-F1-MODEL\_V4 \| 1.0 \| 3.868e-27 \| 526 \| 0.208 \| 798 \| 393 \| 51 \| 93 \| 740 \| 116 \| 824 \| DUF927 domain-containing protein \| DUF927 domain-containing protein; afdb-uniprot50 \| AF-A0A414T7G4-F1-MODEL\_V4 \| 1.0 \| 4.4e-15 \| 525 \| 0.189 \| 316 \| 216 \| 12 \| 485 \| 769 \| 6 \| 312 \| Uncharacterized protein \| Uncharacterized protein; afdb-uniprot50 \| AF-A0A2I8A4T1-F1-MODEL\_V4 \| 1.0 \| 1.181e-19 \| 525 \| 0.175 \| 558 \| 341 \| 30 \| 226 \| 753 \| 214 \| 682 \| Virulence-associated E \| Virulence-associated E; afdb-uniprot50 \| AF-A0A2E9L0S9-F1-MODEL\_V4 \| 1.0 \| 1.378e-19 \| 525 \| 0.195 \| 548 \| 353 \| 21 \| 243 \| 755 \| 234 \| 728 \| VirE\_N domain-containing protein \| VirE\_N domain-containing protein; afdb-uniprot50 \| AF-A0A094K1B3-F1-MODEL\_V4 \| 1.0 \| 1.196e-13 \| 524 \| 0.29 \| 251 \| 146 \| 9 \| 92 \| 335 \| 64 \| 289 \| Toprim domain-containing protein \| Toprim domain-containing protein; afdb-uniprot50 \| AF-A0A352X4W4-F1-MODEL\_V4 \| 1.0 \| 3.511e-24 \| 524 \| 0.165 \| 839 \| 480 \| 50 \| 46 \| 757 \| 84 \| 828 \| Toprim domain-containing protein \| Toprim domain-containing protein; afdb-uniprot50 \| AF-A0A6N6T6R6-F1-MODEL\_V4 \| 1.0 \| 4.516e-27 \| 522 \| 0.228 \| 761 \| 374 \| 43 \| 12 \| 615 \| 5 \| 709 \| DUF3987 domain-containing protein \| DUF3987 domain-containing protein; afdb-uniprot50 \| AF-A0A7T1MLF3-F1-MODEL\_V4 \| 1.0 \| 3.187e-21 \| 522 \| 0.177 \| 683 \| 419 \| 38 \| 167 \| 762 \| 40 \| 666 \| Uncharacterized protein \| Uncharacterized protein; afdb-uniprot50 \| AF-A0A6I1RJD4-F1-MODEL\_V4 \| 1.0 \| 1.211e-15 \| 521 \| 0.186 \| 344 \| 242 \| 17 \| 331 \| 658 \| 29 \| 350 \| Uncharacterized protein \| Uncharacterized protein; afdb-uniprot50 \| AF-Q3M8I3-F1-MODEL\_V4 \| 1.0 \| 2.931e-20 \| 521 \| 0.201 \| 605 \| 348 \| 31 \| 202 \| 769 \| 146 \| 652 \| Virulence-associated E \| Virulence-associated E; afdb-uniprot50 \| AF-A0A2E2RUS7-F1-MODEL\_V4 \| 1.0 \| 1.63e-21 \| 521 \| 0.185 \| 705 \| 395 \| 39 \| 151 \| 766 \| 94 \| 707 \| Prim-Pol domain-containing protein \| Prim-Pol domain-containing protein; afdb-uniprot50 \| AF-A0A2N8MB77-F1-MODEL\_V4 \| 1.0 \| 2.309e-27 \| 521 \| 0.193 \| 754 \| 399 \| 40 \| 99 \| 750 \| 98 \| 744 \| DNA primase \| DNA primase; afdb-uniprot50 \| AF-A0A0S3PXG1-F1-MODEL\_V4 \| 1.0 \| 3.77e-23 \| 521 \| 0.151 \| 816 \| 462 \| 46 \| 1 \| 768 \| 189 \| 821 \| Virulence-associated protein E \| Virulence-associated protein E; afdb-uniprot50 \| AF-I2UTG7-F1-MODEL\_V4 \| 1.0 \| 1.1e-12 \| 521 \| 0.317 \| 217 \| 126 \| 9 \| 133 \| 346 \| 4 \| 201 \| LPD7 domain-containing protein \| LPD7 domain-containing protein; afdb-uniprot50 \| AF-A0A2C6Z4N4-F1-MODEL\_V4 \| 1.0 \| 1.423e-12 \| 520 \| 0.282 \| 202 \| 125 \| 6 \| 139 \| 337 \| 1 \| 185 \| Uncharacterized protein \| Uncharacterized protein; afdb-uniprot50 \| AF-A0A6I4YYG7-F1-MODEL\_V4 \| 1.0 \| 3.027e-21 \| 520 \| 0.139 \| 801 \| 393 \| 31 \| 1 \| 757 \| 253 \| 800 \| Uncharacterized protein \| Uncharacterized protein; afdb-uniprot50 \| AF-A0A522SU55-F1-MODEL\_V4 \| 1.0 \| 2.73e-21 \| 519 \| 0.159 \| 771 \| 383 \| 30 \| 1 \| 751 \| 183 \| 708 \| Uncharacterized protein \| Uncharacterized protein; afdb-uniprot50 \| AF-W0DZN8-F1-MODEL\_V4 \| 1.0 \| 1.451e-27 \| 519 \| 0.197 \| 789 \| 426 \| 47 \| 93 \| 762 \| 77 \| 776 \| Toprim domain-containing protein \| Toprim domain-containing protein; afdb-uniprot50 \| AF-A0A1Q3NZT7-F1-MODEL\_V4 \| 1.0 \| 5.272e-27 \| 519 \| 0.175 \| 802 \| 464 \| 50 \| 102 \| 757 \| 84 \| 833 \| SF3 helicase domain-containing protein \| SF3 helicase domain-containing protein; afdb-uniprot50 \| AF-R4WFQ0-F1-MODEL\_V4 \| 1.0 \| 1.558e-26 \| 518 \| 0.18 \| 854 \| 454 \| 45 \| 107 \| 769 \| 85 \| 883 \| Toprim domain-containing protein \| Toprim domain-containing protein; afdb-uniprot50 \| AF-A0A228H1M8-F1-MODEL\_V4 \| 1.0 \| 4.126e-21 \| 516 \| 0.143 \| 793 \| 377 \| 34 \| 1 \| 755 \| 247 \| 774 \| PriCT\_2 domain-containing protein \| PriCT\_2 domain-containing protein; afdb-uniprot50 \| AF-A0A0P1I0V3-F1-MODEL\_V4 \| 1.0 \| 7.278e-21 \| 514 \| 0.151 \| 803 \| 393 \| 40 \| 2 \| 756 \| 241 \| 802 \| Putative P-loop ATPase \| Putative P-loop ATPase; afdb-uniprot50 \| AF-A0A1F1XH87-F1-MODEL\_V4 \| 1.0 \| 2.802e-17 \| 513 \| 0.186 \| 472 \| 295 \| 23 \| 370 \| 768 \| 17 \| 472 \| Uncharacterized protein \| Uncharacterized protein; afdb-uniprot50 \| AF-A0A4D9CEY9-F1-MODEL\_V4 \| 1.0 \| 3.047e-18 \| 513 \| 0.2 \| 545 \| 316 \| 26 \| 261 \| 769 \| 274 \| 734 \| RepB\_primase domain-containing protein \| RepB\_primase domain-containing protein; afdb-uniprot50 \| AF-A0A5P3VML8-F1-MODEL\_V4 \| 1.0 \| 8.551e-18 \| 513 \| 0.2 \| 518 \| 333 \| 28 \| 285 \| 762 \| 328 \| 804 \| Prim-Pol domain-containing protein \| Prim-Pol domain-containing protein; afdb-uniprot50 \| AF-K9ULE3-F1-MODEL\_V4 \| 1.0 \| 2.354e-18 \| 513 \| 0.187 \| 566 \| 351 \| 30 \| 237 \| 751 \| 234 \| 741 \| Putative P-loop ATPase \| Putative P-loop ATPase; afdb-uniprot50 \| AF-A0A1M4TMH4-F1-MODEL\_V4 \| 1.0 \| 3.843e-14 \| 512 \| 0.236 \| 262 \| 189 \| 8 \| 501 \| 757 \| 2 \| 257 \| Virulence-associated protein E \| Virulence-associated protein E; afdb-uniprot50 \| AF-A0A1Z4TD68-F1-MODEL\_V4 \| 1.0 \| 9.604e-20 \| 512 \| 0.183 \| 588 \| 356 \| 26 \| 233 \| 762 \| 71 \| 592 \| Uncharacterized protein \| Uncharacterized protein; afdb-uniprot50 \| AF-A0A3A0FPI8-F1-MODEL\_V4 \| 1.0 \| 4.574e-21 \| 512 \| 0.157 \| 768 \| 344 \| 30 \| 4 \| 735 \| 243 \| 743 \| Prim-Pol domain-containing protein \| Prim-Pol domain-containing protein; afdb-uniprot50 \| AF-A0A5C7L6B0-F1-MODEL\_V4 \| 1.0 \| 2.678e-14 \| 511 \| 0.29 \| 275 \| 155 \| 9 \| 59 \| 333 \| 74 \| 308 \| Toprim domain-containing protein \| Toprim domain-containing protein; afdb-uniprot50 \| AF-N0B2Q9-F1-MODEL\_V4 \| 1.0 \| 1.284e-20 \| 511 \| 0.185 \| 648 \| 365 \| 36 \| 182 \| 753 \| 123 \| 683 \| PriCT\_1 domain-containing protein \| PriCT\_1 domain-containing protein; afdb-uniprot50 \| AF-A0A1J5R0E3-F1-MODEL\_V4 \| 1.0 \| 1.335e-26 \| 511 \| 0.184 \| 752 \| 418 \| 40 \| 100 \| 750 \| 75 \| 731 \| DNA primase TraC \| DNA primase TraC; afdb-uniprot50 \| AF-A0A5C7PWQ8-F1-MODEL\_V4 \| 1.0 \| 2.415e-14 \| 510 \| 0.239 \| 317 \| 203 \| 16 \| 455 \| 757 \| 16 \| 308 \| Uncharacterized protein \| Uncharacterized protein; afdb-uniprot50 \| AF-A0A3B9BTJ8-F1-MODEL\_V4 \| 1.0 \| 6.606e-18 \| 509 \| 0.2 \| 493 \| 287 \| 24 \| 239 \| 691 \| 186 \| 611 \| Prim-Pol domain-containing protein \| Prim-Pol domain-containing protein; afdb-uniprot50 \| AF-A0A5C7JLL6-F1-MODEL\_V4 \| 1.0 \| 3.086e-20 \| 509 \| 0.142 \| 785 \| 409 \| 34 \| 4 \| 769 \| 188 \| 726 \| Uncharacterized protein \| Uncharacterized protein; afdb-uniprot50 \| AF-A0A838W082-F1-MODEL\_V4 \| 1.0 \| 1.044e-12 \| 508 \| 0.235 \| 221 \| 147 \| 9 \| 131 \| 347 \| 8 \| 210 \| Toprim domain-containing protein \| Toprim domain-containing protein; afdb-uniprot50 \| AF-A0A2L2NLQ2-F1-MODEL\_V4 \| 1.0 \| 3.511e-16 \| 508 \| 0.189 \| 455 \| 256 \| 24 \| 381 \| 751 \| 88 \| 513 \| Virulence-associated E family protein \| Virulence-associated E family protein; afdb-uniprot50 \| AF-A0A7X7WZM3-F1-MODEL\_V4 \| 1.0 \| 5.104e-18 \| 508 \| 0.145 \| 516 \| 350 \| 26 \| 264 \| 756 \| 273 \| 720 \| VirE\_N domain-containing protein \| VirE\_N domain-containing protein; afdb-uniprot50 \| AF-G1UZP2-F1-MODEL\_V4 \| 1.0 \| 1.227e-17 \| 507 \| 0.251 \| 414 \| 204 \| 20 \| 91 \| 454 \| 70 \| 427 \| Uncharacterized protein \| Uncharacterized protein; afdb-uniprot50 \| AF-A0A6M2A343-F1-MODEL\_V4 \| 1.0 \| 6.957e-26 \| 506 \| 0.163 \| 801 \| 447 \| 49 \| 98 \| 755 \| 117 \| 837 \| DNA primase TraC \| DNA primase TraC; afdb-uniprot50 \| AF-A0A3C0NTJ6-F1-MODEL\_V4 \| 1.0 \| 1.058e-14 \| 505 \| 0.252 \| 356 \| 222 \| 17 \| 436 \| 769 \| 2 \| 335 \| Conjugal transfer protein TraC \| Conjugal transfer protein TraC; afdb-uniprot50 \| AF-A0A7U8BI53-F1-MODEL\_V4 \| 1.0 \| 2.339e-13 \| 504 \| 0.248 \| 274 \| 168 \| 10 \| 92 \| 346 \| 69 \| 323 \| Toprim domain-containing protein \| Toprim domain-containing protein; afdb-uniprot50 \| AF-A0A1I1CGG0-F1-MODEL\_V4 \| 1.0 \| 4.878e-15 \| 504 \| 0.221 \| 380 \| 239 \| 19 \| 436 \| 769 \| 1 \| 369 \| Virulence-associated protein E \| Virulence-associated protein E; afdb-uniprot50 \| AF-A0A525WLH8-F1-MODEL\_V4 \| 1.0 \| 9.666e-25 \| 504 \| 0.184 \| 750 \| 374 \| 36 \| 100 \| 769 \| 77 \| 668 \| DUF3631 domain-containing protein \| DUF3631 domain-containing protein; afdb-uniprot50 \| AF-A0A827WY38-F1-MODEL\_V4 \| 1.0 \| 1.284e-12 \| 503 \| 0.313 \| 217 \| 128 \| 8 \| 115 \| 328 \| 160 \| 358 \| Uncharacterized protein \| Uncharacterized protein; afdb-uniprot50 \| AF-A0A2V6F860-F1-MODEL\_V4 \| 1.0 \| 1.136e-21 \| 502 \| 0.233 \| 531 \| 283 \| 26 \| 113 \| 607 \| 84 \| 526 \| Toprim domain-containing protein \| Toprim domain-containing protein; afdb-uniprot50 \| AF-A0A0E4FRT6-F1-MODEL\_V4 \| 1.0 \| 3.918e-21 \| 502 \| 0.142 \| 710 \| 430 \| 38 \| 143 \| 750 \| 73 \| 705 \| Prim-Pol domain-containing protein \| Prim-Pol domain-containing protein; afdb-uniprot50 \| AF-A0A6M3JPC8-F1-MODEL\_V4 \| 1.0 \| 1.165e-17 \| 502 \| 0.12 \| 555 \| 356 \| 31 \| 299 \| 756 \| 266 \| 785 \| Putative DNA primase \| Putative DNA primase; afdb-uniprot50 \| AF-A0A5T0QKQ6-F1-MODEL\_V4 \| 1.0 \| 7.662e-13 \| 501 \| 0.258 \| 259 \| 159 \| 9 \| 107 \| 347 \| 13 \| 256 \| Toprim domain-containing protein \| Toprim domain-containing protein; afdb-uniprot50 \| AF-A0A561PL40-F1-MODEL\_V4 \| 1.0 \| 6.692e-20 \| 501 \| 0.155 \| 785 \| 393 \| 34 \| 1 \| 760 \| 176 \| 715 \| Primase-like protein \| Primase-like protein; afdb-uniprot50 \| AF-A0A4P7L1H0-F1-MODEL\_V4 \| 1.0 \| 2.838e-19 \| 500 \| 0.181 \| 573 \| 370 \| 38 \| 240 \| 764 \| 79 \| 600 \| Virulence-associated protein E \| Virulence-associated protein E; afdb-uniprot50 \| AF-A0A2H6G363-F1-MODEL\_V4 \| 1.0 \| 1.031e-26 \| 500 \| 0.178 \| 730 \| 409 \| 41 \| 115 \| 751 \| 2 \| 633 \| DNA primase TraC \| DNA primase TraC; afdb-uniprot50 \| AF-A0A0Q7SIY5-F1-MODEL\_V4 \| 1.0 \| 2.748e-26 \| 500 \| 0.208 \| 783 \| 425 \| 51 \| 90 \| 753 \| 78 \| 784 \| Uncharacterized protein \| Uncharacterized protein; afdb-uniprot50 \| AF-A0A6P0U2Y5-F1-MODEL\_V4 \| 1.0 \| 3.769e-15 \| 499 \| 0.223 \| 372 \| 243 \| 16 \| 412 \| 751 \| 6 \| 363 \| Uncharacterized protein \| Uncharacterized protein; afdb-uniprot50 \| AF-A0A845X7V7-F1-MODEL\_V4 \| 1.0 \| 7.047e-20 \| 499 \| 0.201 \| 591 \| 345 \| 35 \| 239 \| 762 \| 236 \| 766 \| Uncharacterized protein \| Uncharacterized protein; afdb-uniprot50 \| AF-A0A558BT14-F1-MODEL\_V4 \| 1.0 \| 2.384e-20 \| 498 \| 0.149 \| 791 \| 422 \| 32 \| 1 \| 757 \| 17 \| 590 \| PriCT\_2 domain-containing protein \| PriCT\_2 domain-containing protein; afdb-uniprot50 \| AF-A0A0F9U4J5-F1-MODEL\_V4 \| 1.0 \| 1.031e-18 \| 498 \| 0.157 \| 787 \| 363 \| 32 \| 1 \| 757 \| 163 \| 679 \| Prim-Pol domain-containing protein \| Prim-Pol domain-containing protein; afdb-uniprot50 \| AF-A0A5J4QFG3-F1-MODEL\_V4 \| 1.0 \| 4.26e-14 \| 497 \| 0.233 \| 266 \| 186 \| 10 \| 382 \| 638 \| 20 \| 276 \| Uncharacterized protein \| Uncharacterized protein; afdb-uniprot50 \| AF-A0A2N0V8M1-F1-MODEL\_V4 \| 1.0 \| 6.155e-27 \| 497 \| 0.202 \| 749 \| 425 \| 44 \| 100 \| 757 \| 72 \| 738 \| SF3 helicase domain-containing protein \| SF3 helicase domain-containing protein; afdb-uniprot50 \| AF-A0A2M7MUG5-F1-MODEL\_V4 \| 1.0 \| 3.745e-26 \| 497 \| 0.198 \| 906 \| 410 \| 48 \| 12 \| 736 \| 5 \| 774 \| Uncharacterized protein \| Uncharacterized protein; afdb-uniprot50 \| AF-A0A2P7U2R3-F1-MODEL\_V4 \| 1.0 \| 2.766e-23 \| 496 \| 0.196 \| 849 \| 406 \| 42 \| 1 \| 744 \| 2 \| 678 \| Toprim domain-containing protein \| Toprim domain-containing protein; afdb-uniprot50 \| AF-A0A1F4HLP3-F1-MODEL\_V4 \| 1.0 \| 3.745e-26 \| 495 \| 0.218 \| 801 \| 410 \| 46 \| 36 \| 679 \| 1 \| 742 \| Uncharacterized protein \| Uncharacterized protein; afdb-uniprot50 \| AF-A0A126NQG0-F1-MODEL\_V4 \| 1.0 \| 1.878e-19 \| 495 \| 0.186 \| 568 \| 340 \| 32 \| 261 \| 757 \| 242 \| 758 \| Prim-Pol domain-containing protein \| Prim-Pol domain-containing protein; afdb-uniprot50 \| AF-A0A5U3F134-F1-MODEL\_V4 \| 1.0 \| 6.649e-15 \| 494 \| 0.267 \| 299 \| 158 \| 15 \| 68 \| 355 \| 140 \| 388 \| DNA primase \| DNA primase; afdb-uniprot50 \| AF-A0A2V5PJC8-F1-MODEL\_V4 \| 1.0 \| 1.107e-25 \| 493 \| 0.189 \| 803 \| 407 \| 47 \| 92 \| 752 \| 64 \| 764 \| Uncharacterized protein \| Uncharacterized protein; afdb-uniprot50 \| AF-A0A7Z0B884-F1-MODEL\_V4 \| 1.0 \| 1.267e-26 \| 493 \| 0.185 \| 762 \| 425 \| 40 \| 102 \| 750 \| 80 \| 758 \| Putative DNA primase/helicase \| Putative DNA primase/helicase; afdb-uniprot50 \| AF-A0A1I0JA19-F1-MODEL\_V4 \| 1.0 \| 3.534e-21 \| 492 \| 0.138 \| 777 \| 438 \| 34 \| 2 \| 755 \| 200 \| 767 \| Bifunctional DNA primase/polymerase, N-terminal \| Bifunctional DNA primase/polymerase, N-terminal; afdb-uniprot50 \| AF-A0A1G5SD75-F1-MODEL\_V4 \| 1.0 \| 2.003e-13 \| 491 \| 0.294 \| 258 \| 136 \| 9 \| 91 \| 348 \| 103 \| 314 \| Toprim protein \| Toprim protein; afdb-uniprot50 \| AF-E0UET4-F1-MODEL\_V4 \| 1.0 \| 3.626e-17 \| 491 \| 0.189 \| 496 \| 294 \| 24 \| 290 \| 753 \| 57 \| 476 \| Virulence-associated E family protein \| Virulence-associated E family protein; afdb-uniprot50 \| AF-B9KGK4-F1-MODEL\_V4 \| 1.0 \| 3.027e-13 \| 490 \| 0.259 \| 274 \| 160 \| 12 \| 91 \| 346 \| 87 \| 335 \| DNA primase TraC \| DNA primase TraC; afdb-uniprot50 \| AF-R5P0F0-F1-MODEL\_V4 \| 1.0 \| 2.137e-15 \| 490 \| 0.265 \| 358 \| 196 \| 18 \| 457 \| 760 \| 2 \| 346 \| Uncharacterized protein \| Uncharacterized protein; afdb-uniprot50 \| AF-A0A5N1JF43-F1-MODEL\_V4 \| 1.0 \| 4.429e-20 \| 490 \| 0.151 \| 804 \| 396 \| 35 \| 1 \| 769 \| 237 \| 788 \| Uncharacterized protein \| Uncharacterized protein; afdb-uniprot50 \| AF-A0A5C7ZG73-F1-MODEL\_V4 \| 1.0 \| 2.802e-25 \| 488 \| 0.183 \| 861 \| 422 \| 45 \| 13 \| 753 \| 6 \| 705 \| Topoisomerase \| Topoisomerase; afdb-uniprot50 \| AF-G1UVN8-F1-MODEL\_V4 \| 1.0 \| 6.036e-20 \| 485 \| 0.223 \| 536 \| 203 \| 24 \| 102 \| 614 \| 102 \| 447 \| Toprim domain-containing protein \| Toprim domain-containing protein; afdb-uniprot50 \| AF-A0A5J4SK06-F1-MODEL\_V4 \| 1.0 \| 1.1e-12 \| 484 \| 0.262 \| 213 \| 143 \| 6 \| 490 \| 699 \| 10 \| 211 \| Uncharacterized protein \| Uncharacterized protein; afdb-uniprot50 \| AF-A0A0H4B4X1-F1-MODEL\_V4 \| 1.0 \| 1.114e-14 \| 484 \| 0.204 \| 391 \| 250 \| 17 \| 412 \| 766 \| 1 \| 366 \| Uncharacterized protein \| Uncharacterized protein; afdb-uniprot50 \| AF-A0A2V4UF45-F1-MODEL\_V4 \| 1.0 \| 1.378e-19 \| 484 \| 0.231 \| 513 \| 266 \| 21 \| 102 \| 508 \| 73 \| 563 \| Phage/plasmid primase-like uncharacterized protein \| Phage/plasmid primase-like uncharacterized protein; afdb-uniprot50 \| AF-A0A4R5P3B1-F1-MODEL\_V4 \| 1.0 \| 5.732e-20 \| 484 \| 0.143 \| 675 \| 450 \| 39 \| 151 \| 757 \| 142 \| 756 \| Uncharacterized protein \| Uncharacterized protein; afdb-uniprot50 \| AF-A0A7U4DN83-F1-MODEL\_V4 \| 1.0 \| 8.122e-26 \| 484 \| 0.169 \| 784 \| 445 \| 42 \| 64 \| 750 \| 69 \| 742 \| Uncharacterized protein \| Uncharacterized protein; afdb-uniprot50 \| AF-A0A158DR21-F1-MODEL\_V4 \| 1.0 \| 7.713e-26 \| 483 \| 0.184 \| 755 \| 423 \| 43 \| 102 \| 750 \| 79 \| 746 \| TOPRIM domain-containing protein \| TOPRIM domain-containing protein; afdb-uniprot50 \| AF-A0A2N8MBZ7-F1-MODEL\_V4 \| 1.0 \| 5.48e-25 \| 483 \| 0.211 \| 705 \| 372 \| 43 \| 99 \| 689 \| 69 \| 703 \| Uncharacterized protein \| Uncharacterized protein; afdb-uniprot50 \| AF-A0A0S4I2H5-F1-MODEL\_V4 \| 1.0 \| 1.661e-12 \| 481 \| 0.338 \| 204 \| 117 \| 6 \| 92 \| 287 \| 61 \| 254 \| DNA primase TraC \| DNA primase TraC; afdb-uniprot50 \| AF-A1SXJ4-F1-MODEL\_V4 \| 1.0 \| 6.035e-12 \| 481 \| 0.241 \| 228 \| 149 \| 8 \| 115 \| 335 \| 86 \| 296 \| Toprim domain-containing protein \| Toprim domain-containing protein; afdb-uniprot50 \| AF-A0A7G8DD21-F1-MODEL\_V4 \| 1.0 \| 1.317e-16 \| 481 \| 0.153 \| 489 \| 329 \| 25 \| 316 \| 769 \| 1 \| 439 \| Virulence-associated E family protein \| Virulence-associated E family protein; afdb-uniprot50 \| AF-A0A1Z9AXB4-F1-MODEL\_V4 \| 1.0 \| 8.387e-19 \| 481 \| 0.184 \| 573 \| 335 \| 30 \| 264 \| 750 \| 194 \| 719 \| Uncharacterized protein \| Uncharacterized protein; afdb-uniprot50 \| AF-A0A845Z468-F1-MODEL\_V4 \| 1.0 \| 3.422e-20 \| 480 \| 0.185 \| 599 \| 364 \| 32 \| 170 \| 690 \| 78 \| 630 \| Uncharacterized protein \| Uncharacterized protein; afdb-uniprot50 \| AF-A0A158LQY7-F1-MODEL\_V4 \| 1.0 \| 1.952e-25 \| 479 \| 0.17 \| 947 \| 495 \| 52 \| 3 \| 751 \| 1 \| 854 \| Prim\_Zn\_Ribbon domain-containing protein \| Prim\_Zn\_Ribbon domain-containing protein; afdb-uniprot50 \| AF-A0A3G3H284-F1-MODEL\_V4 \| 1.0 \| 8.552e-26 \| 478 \| 0.178 \| 764 \| 408 \| 46 \| 93 \| 737 \| 74 \| 736 \| Uncharacterized protein \| Uncharacterized protein; afdb-uniprot50 \| AF-A0A6L3IIZ1-F1-MODEL\_V4 \| 1.0 \| 6.233e-13 \| 477 \| 0.242 \| 252 \| 180 \| 8 \| 512 \| 758 \| 7 \| 252 \| Uncharacterized protein \| Uncharacterized protein; afdb-uniprot50 \| AF-A0A7K0GNH6-F1-MODEL\_V4 \| 1.0 \| 2.593e-21 \| 477 \| 0.164 \| 737 \| 394 \| 36 \| 1 \| 684 \| 1 \| 568 \| Uncharacterized protein \| Uncharacterized protein; afdb-uniprot50 \| AF-A0A433VJU3-F1-MODEL\_V4 \| 1.0 \| 4.816e-21 \| 477 \| 0.171 \| 684 \| 426 \| 33 \| 161 \| 756 \| 82 \| 712 \| Uncharacterized protein \| Uncharacterized protein; afdb-uniprot50 \| AF-A0A850R2Y2-F1-MODEL\_V4 \| 1.0 \| 5.306e-24 \| 477 \| 0.175 \| 822 \| 459 \| 53 \| 93 \| 762 \| 74 \| 828 \| Toprim domain-containing protein \| Toprim domain-containing protein; afdb-uniprot50 \| AF-A0A011PBN5-F1-MODEL\_V4 \| 1.0 \| 2.875e-13 \| 476 \| 0.226 \| 296 \| 203 \| 15 \| 468 \| 751 \| 1 \| 282 \| Putative P-loop ATPase \| Putative P-loop ATPase; afdb-uniprot50 \| AF-K9TV34-F1-MODEL\_V4 \| 1.0 \| 5.959e-18 \| 476 \| 0.186 \| 569 \| 349 \| 33 \| 227 \| 769 \| 225 \| 705 \| Virulence-associated E family protein \| Virulence-associated E family protein; afdb-uniprot50 \| AF-A0A1I5Z275-F1-MODEL\_V4 \| 1.0 \| 1.267e-18 \| 475 \| 0.122 \| 810 \| 446 \| 45 \| 1 \| 757 \| 26 \| 623 \| Primase C terminal 2 (PriCT-2) \| Primase C terminal 2 (PriCT-2); afdb-uniprot50 \| AF-A0A080LT75-F1-MODEL\_V4 \| 1.0 \| 9.18e-25 \| 475 \| 0.187 \| 768 \| 464 \| 42 \| 102 \| 769 \| 81 \| 788 \| DNA primase TraC \| DNA primase TraC; afdb-uniprot50 \| AF-A0A328V111-F1-MODEL\_V4 \| 1.0 \| 2.123e-18 \| 474 \| 0.183 \| 562 \| 333 \| 29 \| 237 \| 756 \| 220 \| 697 \| Prim-Pol domain-containing protein \| Prim-Pol domain-containing protein; afdb-uniprot50 \| AF-D2QEV8-F1-MODEL\_V4 \| 1.0 \| 4.663e-20 \| 474 \| 0.145 \| 796 \| 390 \| 38 \| 2 \| 757 \| 204 \| 748 \| p-loop ATPase and inactivated derivatives-like protein \| p-loop ATPase and inactivated derivatives-like protein; afdb-uniprot50 \| AF-A0A7H0HGN8-F1-MODEL\_V4 \| 1.0 \| 2.627e-23 \| 472 \| 0.226 \| 626 \| 321 \| 37 \| 89 \| 645 \| 61 \| 591 \| DUF3987 domain-containing protein \| DUF3987 domain-containing protein; afdb-uniprot50 \| AF-A0A3N7INQ9-F1-MODEL\_V4 \| 1.0 \| 6.233e-13 \| 471 \| 0.264 \| 253 \| 170 \| 8 \| 526 \| 768 \| 1 \| 247 \| Virulence-associated E family protein \| Virulence-associated E family protein; afdb-uniprot50 \| AF-A0A350H4D5-F1-MODEL\_V4 \| 1.0 \| 1.568e-15 \| 471 \| 0.206 \| 392 \| 249 \| 18 \| 381 \| 743 \| 91 \| 449 \| Uncharacterized protein \| Uncharacterized protein; afdb-uniprot50 \| AF-C4K524-F1-MODEL\_V4 \| 1.0 \| 3.106e-17 \| 471 \| 0.219 \| 438 \| 230 \| 19 \| 99 \| 447 \| 137 \| 551 \| DNA primase TraC \| DNA primase TraC; afdb-uniprot50 \| AF-A0A5F0YSJ3-F1-MODEL\_V4 \| 1.0 \| 3.4e-15 \| 470 \| 0.229 \| 366 \| 207 \| 18 \| 100 \| 446 \| 128 \| 437 \| Toprim domain-containing protein \| Toprim domain-containing protein; afdb-uniprot50 \| AF-A0A5C7LJD7-F1-MODEL\_V4 \| 1.0 \| 6.606e-18 \| 470 \| 0.165 \| 579 \| 353 \| 32 \| 238 \| 757 \| 216 \| 723 \| Prim-Pol domain-containing protein \| Prim-Pol domain-containing protein; afdb-uniprot50 \| AF-A0A133PTK4-F1-MODEL\_V4 \| 1.0 \| 6.114e-14 \| 469 \| 0.228 \| 284 \| 192 \| 13 \| 369 \| 638 \| 43 \| 313 \| Uncharacterized protein \| Uncharacterized protein; afdb-uniprot50 \| AF-A0A414BQ09-F1-MODEL\_V4 \| 1.0 \| 1.369e-14 \| 469 \| 0.179 \| 351 \| 253 \| 13 \| 330 \| 659 \| 256 \| 592 \| Virulence protein E \| Virulence protein E; afdb-uniprot50 \| AF-A0A1L3SNY3-F1-MODEL\_V4 \| 1.0 \| 6.564e-21 \| 469 \| 0.162 \| 880 \| 439 \| 47 \| 1 \| 756 \| 149 \| 854 \| Uncharacterized protein \| Uncharacterized protein; afdb-uniprot50 \| AF-A0A844LJA6-F1-MODEL\_V4 \| 1.0 \| 2.627e-23 \| 468 \| 0.158 \| 896 \| 469 \| 44 \| 1 \| 745 \| 1 \| 762 \| Uncharacterized protein \| Uncharacterized protein; afdb-uniprot50 \| AF-A0A8A8P149-F1-MODEL\_V4 \| 1.0 \| 2.783e-12 \| 467 \| 0.296 \| 243 \| 143 \| 11 \| 99 \| 336 \| 113 \| 332 \| Uncharacterized protein \| Uncharacterized protein; afdb-uniprot50 \| AF-A0A367RNI6-F1-MODEL\_V4 \| 1.0 \| 9.061e-15 \| 467 \| 0.16 \| 448 \| 280 \| 20 \| 381 \| 751 \| 70 \| 498 \| Uncharacterized protein \| Uncharacterized protein; afdb-uniprot50 \| AF-D6Z5I8-F1-MODEL\_V4 \| 1.0 \| 1.292e-25 \| 465 \| 0.188 \| 879 \| 425 \| 54 \| 100 \| 751 \| 73 \| 889 \| Uncharacterized protein \| Uncharacterized protein; afdb-uniprot50 \| AF-A0A7U4BPL7-F1-MODEL\_V4 \| 1.0 \| 6.114e-14 \| 464 \| 0.272 \| 279 \| 160 \| 11 \| 58 \| 322 \| 67 \| 316 \| Prim\_Zn\_Ribbon domain-containing protein \| Prim\_Zn\_Ribbon domain-containing protein; afdb-uniprot50 \| AF-A0A416MI66-F1-MODEL\_V4 \| 1.0 \| 1.396e-13 \| 464 \| 0.207 \| 318 \| 211 \| 15 \| 477 \| 769 \| 2 \| 303 \| Virulence-associated protein E \| Virulence-associated protein E; afdb-uniprot50 \| AF-A0A7K1M057-F1-MODEL\_V4 \| 1.0 \| 4.4e-15 \| 464 \| 0.19 \| 394 \| 274 \| 20 \| 286 \| 659 \| 86 \| 454 \| Uncharacterized protein \| Uncharacterized protein; afdb-uniprot50 \| AF-A0A1V8M6R2-F1-MODEL\_V4 \| 1.0 \| 2.056e-25 \| 464 \| 0.174 \| 785 \| 446 \| 41 \| 91 \| 740 \| 77 \| 794 \| DUF927 domain-containing protein \| DUF927 domain-containing protein; afdb-uniprot50 \| AF-A0A2V2BHY3-F1-MODEL\_V4 \| 1.0 \| 6.867e-16 \| 463 \| 0.244 \| 356 \| 199 \| 14 \| 118 \| 449 \| 166 \| 475 \| Putative DNA primase/helicase \| Putative DNA primase/helicase; afdb-uniprot50 \| AF-A0A1I4TDG0-F1-MODEL\_V4 \| 1.0 \| 4.785e-16 \| 461 \| 0.18 \| 411 \| 253 \| 20 \| 264 \| 661 \| 288 \| 627 \| Virulence-associated protein E \| Virulence-associated protein E; afdb-uniprot50 \| AF-A0A7U7G979-F1-MODEL\_V4 \| 1.0 \| 5.732e-12 \| 460 \| 0.32 \| 231 \| 129 \| 10 \| 103 \| 319 \| 79 \| 295 \| Uncharacterized protein \| Uncharacterized protein; afdb-uniprot50 \| AF-A0A832X2K7-F1-MODEL\_V4 \| 1.0 \| 3.378e-18 \| 459 \| 0.224 \| 489 \| 240 \| 25 \| 1 \| 431 \| 16 \| 423 \| AAA family ATPase \| AAA family ATPase; afdb-uniprot50 \| AF-A0A328ZM50-F1-MODEL\_V4 \| 1.0 \| 3.603e-20 \| 459 \| 0.222 \| 562 \| 298 \| 33 \| 99 \| 615 \| 76 \| 543 \| Phage/plasmid primase-like uncharacterized protein \| Phage/plasmid primase-like uncharacterized protein; afdb-uniprot50 \| AF-A0A1J1LKJ5-F1-MODEL\_V4 \| 1.0 \| 8.387e-19 \| 459 \| 0.184 \| 592 \| 356 \| 28 \| 231 \| 769 \| 230 \| 747 \| Prim-Pol domain-containing protein \| Prim-Pol domain-containing protein; afdb-uniprot50 \| AF-A0A1S9D3E4-F1-MODEL\_V4 \| 1.0 \| 4.401e-23 \| 457 \| 0.187 \| 874 \| 417 \| 44 \| 12 \| 762 \| 6 \| 708 \| DNA primase TraC \| DNA primase TraC; afdb-uniprot50 \| AF-B6WV28-F1-MODEL\_V4 \| 1.0 \| 9.666e-25 \| 456 \| 0.204 \| 745 \| 400 \| 44 \| 102 \| 736 \| 76 \| 737 \| Toprim domain protein \| Toprim domain protein; afdb-uniprot50 \| AF-R6M6C9-F1-MODEL\_V4 \| 1.0 \| 5.622e-13 \| 455 \| 0.197 \| 248 \| 189 \| 7 \| 512 \| 755 \| 2 \| 243 \| VirE N-terminal domain protein \| VirE N-terminal domain protein; afdb-uniprot50 \| AF-A0A163VSB7-F1-MODEL\_V4 \| 1.0 \| 7.712e-18 \| 454 \| 0.187 \| 528 \| 333 \| 30 \| 288 \| 759 \| 285 \| 772 \| Virulence-associated protein E \| Virulence-associated protein E; afdb-uniprot50 \| AF-A0A127Q8R1-F1-MODEL\_V4 \| 1.0 \| 9.666e-25 \| 454 \| 0.188 \| 765 \| 411 \| 43 \| 102 \| 737 \| 107 \| 790 \| Toprim domain protein \| Toprim domain protein; afdb-uniprot50 \| AF-A0A6L8Z9G2-F1-MODEL\_V4 \| 1.0 \| 7.863e-17 \| 453 \| 0.178 \| 447 \| 266 \| 23 \| 240 \| 661 \| 242 \| 612 \| Virulence protein E \| Virulence protein E; afdb-uniprot50 \| AF-A5KIU4-F1-MODEL\_V4 \| 1.0 \| 4.125e-13 \| 451 \| 0.231 \| 298 \| 198 \| 14 \| 492 \| 769 \| 1 \| 287 \| Virulence-associated protein E \| Virulence-associated protein E; afdb-uniprot50 \| AF-A0A1Z4S646-F1-MODEL\_V4 \| 1.0 \| 2.294e-14 \| 450 \| 0.177 \| 435 \| 290 \| 20 \| 372 \| 758 \| 93 \| 507 \| Virulence-associated E family protein \| Virulence-associated E family protein; afdb-uniprot50 \| AF-A0A090TZ23-F1-MODEL\_V4 \| 1.0 \| 1.518e-22 \| 450 \| 0.15 \| 806 \| 447 \| 49 \| 119 \| 760 \| 3 \| 734 \| DNA primase phage-associated \| DNA primase phage-associated; afdb-uniprot50 \| AF-A0A2D3TEK2-F1-MODEL\_V4 \| 1.0 \| 1.072e-16 \| 448 \| 0.244 \| 413 \| 210 \| 17 \| 107 \| 447 \| 155 \| 537 \| Uncharacterized protein \| Uncharacterized protein; afdb-uniprot50 \| AF-A0A2X1NEA3-F1-MODEL\_V4 \| 1.0 \| 3.292e-14 \| 446 \| 0.221 \| 347 \| 188 \| 12 \| 73 \| 416 \| 79 \| 346 \| DNA primase TraC \| DNA primase TraC; afdb-uniprot50 \| AF-A0A742HTC7-F1-MODEL\_V4 \| 1.0 \| 1.114e-14 \| 446 \| 0.238 \| 361 \| 200 \| 15 \| 98 \| 442 \| 135 \| 436 \| DUF5710 domain-containing protein \| DUF5710 domain-containing protein; afdb-uniprot50 \| AF-A0A7G7VVW2-F1-MODEL\_V4 \| 1.0 \| 1.528e-19 \| 446 \| 0.165 \| 664 \| 380 \| 35 \| 178 \| 756 \| 123 \| 697 \| Bifunctional DNA primase/polymerase \| Bifunctional DNA primase/polymerase; afdb-uniprot50 \| AF-G4FLD9-F1-MODEL\_V4 \| 1.0 \| 6.521e-16 \| 445 \| 0.151 \| 568 \| 355 \| 30 \| 261 \| 764 \| 26 \| 530 \| Virulence-associated E family protein \| Virulence-associated E family protein; afdb-uniprot50 \| AF-A0A2X0RDQ2-F1-MODEL\_V4 \| 1.0 \| 1.915e-18 \| 445 \| 0.219 \| 519 \| 267 \| 26 \| 98 \| 539 \| 87 \| 544 \| Toprim domain-containing protein \| Toprim domain-containing protein; afdb-uniprot50 \| AF-A4CPW0-F1-MODEL\_V4 \| 1.0 \| 4.604e-18 \| 444 \| 0.155 \| 800 \| 407 \| 33 \| 1 \| 769 \| 183 \| 744 \| Uncharacterized protein \| Uncharacterized protein; afdb-uniprot50 \| AF-A0A7W2RV62-F1-MODEL\_V4 \| 1.0 \| 2.643e-12 \| 443 \| 0.247 \| 263 \| 153 \| 13 \| 91 \| 346 \| 69 \| 293 \| Toprim domain-containing protein \| Toprim domain-containing protein; afdb-uniprot50 \| AF-A0A847UTT6-F1-MODEL\_V4 \| 1.0 \| 1.599e-22 \| 443 \| 0.181 \| 876 \| 432 \| 58 \| 1 \| 753 \| 22 \| 735 \| AAA family ATPase \| AAA family ATPase; afdb-uniprot50 \| AF-K9P6A2-F1-MODEL\_V4 \| 1.0 \| 3.794e-20 \| 443 \| 0.138 \| 800 \| 415 \| 33 \| 1 \| 759 \| 230 \| 795 \| Putative P-loop ATPase \| Putative P-loop ATPase; afdb-uniprot50 \| AF-B5INY9-F1-MODEL\_V4 \| 1.0 \| 1.143e-18 \| 443 \| 0.146 \| 783 \| 351 \| 28 \| 1 \| 743 \| 196 \| 700 \| Primase C terminal 2 family \| Primase C terminal 2 family; afdb-uniprot50 \| AF-A0A128ESL8-F1-MODEL\_V4 \| 1.0 \| 2.109e-21 \| 439 \| 0.158 \| 787 \| 451 \| 41 \| 122 \| 756 \| 92 \| 818 \| DNA primase TraC \| DNA primase TraC; afdb-uniprot50 \| AF-X0WPY2-F1-MODEL\_V4 \| 1.0 \| 7.812e-12 \| 437 \| 0.209 \| 262 \| 189 \| 11 \| 410 \| 663 \| 6 \| 257 \| Uncharacterized protein \| Uncharacterized protein; afdb-uniprot50 \| AF-A0A7L8A2Q6-F1-MODEL\_V4 \| 1.0 \| 2.494e-15 \| 436 \| 0.21 \| 404 \| 227 \| 20 \| 102 \| 485 \| 69 \| 400 \| Toprim domain-containing protein \| Toprim domain-containing protein; afdb-uniprot50 \| AF-A0A1I4I4F4-F1-MODEL\_V4 \| 1.0 \| 7.001e-15 \| 436 \| 0.158 \| 453 \| 289 \| 23 \| 368 \| 744 \| 65 \| 501 \| Virulence-associated protein E \| Virulence-associated protein E; afdb-uniprot50 \| AF-A0A2S9KFP8-F1-MODEL\_V4 \| 1.0 \| 4.545e-24 \| 434 \| 0.207 \| 772 \| 405 \| 47 \| 93 \| 739 \| 81 \| 770 \| Uncharacterized protein \| Uncharacterized protein; afdb-uniprot50 \| AF-A0A415EYP9-F1-MODEL\_V4 \| 1.0 \| 1.011e-11 \| 433 \| 0.2 \| 265 \| 192 \| 9 \| 407 \| 659 \| 10 \| 266 \| Uncharacterized protein \| Uncharacterized protein; afdb-uniprot50 \| AF-A0A380Z8N6-F1-MODEL\_V4 \| 1.0 \| 1.275e-15 \| 433 \| 0.194 \| 436 \| 260 \| 25 \| 261 \| 659 \| 91 \| 472 \| Predicted P-loop ATPase and inactivated derivatives \| Predicted P-loop ATPase and inactivated derivatives; afdb-uniprot50 \| AF-A0A0P0M208-F1-MODEL\_V4 \| 1.0 \| 3.086e-12 \| 432 \| 0.172 \| 266 \| 193 \| 11 \| 505 \| 757 \| 3 \| 254 \| Putative helicase \| Putative helicase; afdb-uniprot50 \| AF-A0A6A7Z0Y2-F1-MODEL\_V4 \| 1.0 \| 4.206e-12 \| 432 \| 0.263 \| 254 \| 155 \| 10 \| 488 \| 722 \| 29 \| 269 \| Uncharacterized protein \| Uncharacterized protein; afdb-uniprot50 \| AF-A0A1F0MEI0-F1-MODEL\_V4 \| 1.0 \| 1.196e-13 \| 430 \| 0.213 \| 314 \| 201 \| 14 \| 370 \| 654 \| 60 \| 356 \| Uncharacterized protein \| Uncharacterized protein; afdb-uniprot50 \| AF-A0A6L7TTJ7-F1-MODEL\_V4 \| 1.0 \| 4.724e-14 \| 430 \| 0.193 \| 429 \| 288 \| 22 \| 387 \| 763 \| 362 \| 784 \| VirE\_N domain-containing protein \| VirE\_N domain-containing protein; afdb-uniprot50 \| AF-A0A412XP58-F1-MODEL\_V4 \| 1.0 \| 4.663e-12 \| 429 \| 0.245 \| 232 \| 169 \| 4 \| 528 \| 756 \| 2 \| 230 \| Virulence protein E \| Virulence protein E; afdb-uniprot50 \| AF-A0A5K7ZQE0-F1-MODEL\_V4 \| 1.0 \| 5.374e-18 \| 427 \| 0.211 \| 576 \| 247 \| 25 \| 93 \| 624 \| 69 \| 481 \| Toprim domain-containing protein \| Toprim domain-containing protein; afdb-uniprot50 \| AF-B2IUW1-F1-MODEL\_V4 \| 1.0 \| 2.165e-17 \| 426 \| 0.185 \| 555 \| 323 \| 32 \| 261 \| 769 \| 287 \| 758 \| Virulence-associated E family protein \| Virulence-associated E family protein; afdb-uniprot50 \| AF-A0A521VJ17-F1-MODEL\_V4 \| 1.0 \| 1.3e-22 \| 424 \| 0.189 \| 873 \| 373 \| 47 \| 92 \| 762 \| 90 \| 829 \| DUF3987 domain-containing protein \| DUF3987 domain-containing protein; afdb-uniprot50 \| AF-A0A1T4M124-F1-MODEL\_V4 \| 1.0 \| 2.783e-12 \| 423 \| 0.167 \| 352 \| 245 \| 15 \| 441 \| 756 \| 2 \| 341 \| Virulence-associated protein E \| Virulence-associated protein E; afdb-uniprot50 \| AF-A0A3A6TG99-F1-MODEL\_V4 \| 1.0 \| 1.89e-16 \| 423 \| 0.209 \| 444 \| 248 \| 24 \| 117 \| 495 \| 168 \| 573 \| Uncharacterized protein \| Uncharacterized protein; afdb-uniprot50 \| AF-A0A7G7KLA7-F1-MODEL\_V4 \| 1.0 \| 2.875e-13 \| 421 \| 0.269 \| 286 \| 174 \| 14 \| 488 \| 757 \| 4 \| 270 \| Uncharacterized protein \| Uncharacterized protein; afdb-uniprot50 \| AF-A0A6G1V8R3-F1-MODEL\_V4 \| 1.0 \| 2.431e-11 \| 420 \| 0.199 \| 241 \| 183 \| 6 \| 517 \| 751 \| 2 \| 238 \| Helicase \| Helicase; afdb-uniprot50 \| AF-I9HUA8-F1-MODEL\_V4 \| 1.0 \| 1.528e-11 \| 418 \| 0.181 \| 265 \| 205 \| 8 \| 498 \| 756 \| 2 \| 260 \| DUF3874 domain-containing protein \| DUF3874 domain-containing protein; afdb-uniprot50 \| AF-A0A1Z4FZF4-F1-MODEL\_V4 \| 1.0 \| 3.603e-12 \| 418 \| 0.244 \| 262 \| 181 \| 7 \| 513 \| 769 \| 2 \| 251 \| Virulence-associated E family protein \| Virulence-associated E family protein; afdb-uniprot50 \| AF-K5DAG2-F1-MODEL\_V4 \| 1.0 \| 9.79e-11 \| 417 \| 0.295 \| 176 \| 120 \| 3 \| 514 \| 688 \| 23 \| 195 \| Uncharacterized protein \| Uncharacterized protein; afdb-uniprot50 \| AF-A0A1Q8QRC0-F1-MODEL\_V4 \| 1.0 \| 4.288e-19 \| 414 \| 0.196 \| 601 \| 258 \| 30 \| 102 \| 643 \| 82 \| 516 \| DNA primase \| DNA primase; afdb-uniprot50 \| AF-A0A7W0X7N0-F1-MODEL\_V4 \| 1.0 \| 1.065e-19 \| 414 \| 0.193 \| 676 \| 320 \| 35 \| 102 \| 673 \| 76 \| 629 \| DUF3987 domain-containing protein \| DUF3987 domain-containing protein; afdb-uniprot50 \| AF-A0A1H6F9A3-F1-MODEL\_V4 \| 1.0 \| 1.093e-23 \| 414 \| 0.176 \| 860 \| 438 \| 53 \| 12 \| 731 \| 8 \| 736 \| DNA primase TraC \| DNA primase TraC; afdb-uniprot50 \| AF-A0A240TVL1-F1-MODEL\_V4 \| 1.0 \| 2.463e-21 \| 413 \| 0.186 \| 733 \| 348 \| 37 \| 102 \| 728 \| 78 \| 667 \| Toprim domain-containing protein \| Toprim domain-containing protein; afdb-uniprot50 \| AF-A0A7U8B3W4-F1-MODEL\_V4 \| 1.0 \| 2.559e-11 \| 412 \| 0.22 \| 268 \| 167 \| 10 \| 98 \| 343 \| 126 \| 373 \| DNA primase \| DNA primase; afdb-uniprot50 \| AF-A0A3M5N5A8-F1-MODEL\_V4 \| 1.0 \| 1.651e-15 \| 412 \| 0.231 \| 428 \| 222 \| 18 \| 114 \| 469 \| 354 \| 746 \| DNA primase TraC \| DNA primase TraC; afdb-uniprot50 \| AF-U2LAS6-F1-MODEL\_V4 \| 1.0 \| 1.284e-12 \| 411 \| 0.202 \| 301 \| 197 \| 11 \| 370 \| 638 \| 51 \| 340 \| Virulence-associated protein E \| Virulence-associated protein E; afdb-uniprot50 \| AF-A0A524BHU0-F1-MODEL\_V4 \| 1.0 \| 6.035e-12 \| 408 \| 0.248 \| 249 \| 173 \| 7 \| 519 \| 762 \| 2 \| 241 \| Uncharacterized protein \| Uncharacterized protein; afdb-uniprot50 \| AF-A0A2L2NAA1-F1-MODEL\_V4 \| 1.0 \| 6.779e-14 \| 407 \| 0.189 \| 460 \| 236 \| 20 \| 260 \| 671 \| 25 \| 395 \| Virulence-associated E family protein \| Virulence-associated E family protein; afdb-uniprot50 \| AF-A0A3A9HBK2-F1-MODEL\_V4 \| 1.0 \| 4.515e-11 \| 406 \| 0.217 \| 262 \| 168 \| 12 \| 488 \| 728 \| 10 \| 255 \| Virulence-associated protein E \| Virulence-associated protein E; afdb-uniprot50 \| AF-G2H1Q2-F1-MODEL\_V4 \| 1.0 \| 3.843e-22 \| 406 \| 0.151 \| 917 \| 493 \| 52 \| 1 \| 736 \| 29 \| 840 \| Primase C \| Primase C; afdb-uniprot50 \| AF-A0A558HMQ4-F1-MODEL\_V4 \| 1.0 \| 5.271e-11 \| 405 \| 0.18 \| 277 \| 195 \| 13 \| 80 \| 344 \| 120 \| 376 \| Uncharacterized protein \| Uncharacterized protein; afdb-uniprot50 \| AF-A0A7K1M065-F1-MODEL\_V4 \| 1.0 \| 2.354e-10 \| 404 \| 0.222 \| 198 \| 141 \| 7 \| 466 \| 653 \| 3 \| 197 \| Uncharacterized protein \| Uncharacterized protein; afdb-uniprot50 \| AF-A0A5K1IZ44-F1-MODEL\_V4 \| 1.0 \| 6.735e-17 \| 404 \| 0.149 \| 616 \| 368 \| 35 \| 226 \| 759 \| 199 \| 740 \| Virulence-associated protein E \| Virulence-associated protein E; afdb-uniprot50 \| AF-A0A5J4PAM3-F1-MODEL\_V4 \| 1.0 \| 1.727e-10 \| 403 \| 0.282 \| 191 \| 129 \| 6 \| 514 \| 699 \| 2 \| 189 \| Uncharacterized protein \| Uncharacterized protein; afdb-uniprot50 \| AF-A0A6M9PYL6-F1-MODEL\_V4 \| 1.0 \| 1.079e-21 \| 403 \| 0.199 \| 723 \| 382 \| 45 \| 89 \| 732 \| 61 \| 665 \| Toprim domain-containing protein \| Toprim domain-containing protein; afdb-uniprot50 \| AF-A0A6S6S6H1-F1-MODEL\_V4 \| 1.0 \| 1.085e-10 \| 402 \| 0.287 \| 205 \| 126 \| 6 \| 91 \| 295 \| 432 \| 616 \| DNA primase (EC) \| DNA primase (EC); afdb-uniprot50 \| AF-A0A3P1ZSQ5-F1-MODEL\_V4 \| 1.0 \| 6.779e-22 \| 402 \| 0.196 \| 789 \| 412 \| 43 \| 92 \| 731 \| 71 \| 786 \| DUF927 domain-containing protein \| DUF927 domain-containing protein; afdb-uniprot50 \| AF-A0A257SN45-F1-MODEL\_V4 \| 1.0 \| 5.514e-14 \| 401 \| 0.206 \| 398 \| 222 \| 20 \| 102 \| 454 \| 86 \| 434 \| Toprim domain-containing protein \| Toprim domain-containing protein; afdb-uniprot50 \| AF-A0A1Z5HDV9-F1-MODEL\_V4 \| 1.0 \| 5.844e-11 \| 400 \| 0.269 \| 219 \| 134 \| 9 \| 466 \| 666 \| 12 \| 222 \| Uncharacterized protein \| Uncharacterized protein; afdb-uniprot50 \| AF-A0A496KDJ9-F1-MODEL\_V4 \| 1.0 \| 1.807e-13 \| 399 \| 0.203 \| 374 \| 199 \| 20 \| 98 \| 447 \| 75 \| 373 \| Toprim domain-containing protein \| Toprim domain-containing protein; afdb-uniprot50 \| AF-F1W2P6-F1-MODEL\_V4 \| 1.0 \| 1.044e-20 \| 398 \| 0.157 \| 808 \| 385 \| 46 \| 100 \| 755 \| 78 \| 741 \| Toprim domain-containing protein \| Toprim domain-containing protein; afdb-uniprot50 \| AF-A0A535HSC6-F1-MODEL\_V4 \| 1.0 \| 1.079e-21 \| 396 \| 0.176 \| 825 \| 407 \| 42 \| 106 \| 738 \| 1 \| 744 \| DUF927 domain-containing protein \| DUF927 domain-containing protein; afdb-uniprot50 \| AF-A0A0F7KH82-F1-MODEL\_V4 \| 1.0 \| 1.369e-22 \| 395 \| 0.202 \| 761 \| 386 \| 47 \| 92 \| 737 \| 71 \| 725 \| Putative DNA primase/helicase \| Putative DNA primase/helicase; afdb-uniprot50 \| AF-A0A5E4TKB0-F1-MODEL\_V4 \| 1.0 \| 1.058e-22 \| 395 \| 0.186 \| 757 \| 388 \| 45 \| 119 \| 739 \| 2 \| 666 \| DNA primase \| DNA primase; afdb-uniprot50 \| AF-W2CNN4-F1-MODEL\_V4 \| 1.0 \| 3.556e-10 \| 394 \| 0.258 \| 186 \| 128 \| 4 \| 471 \| 655 \| 4 \| 180 \| Virulence-associated protein E \| Virulence-associated protein E; afdb-uniprot50 \| AF-A0A1T5CZF5-F1-MODEL\_V4 \| 1.0 \| 1.818e-10 \| 394 \| 0.248 \| 225 \| 161 \| 6 \| 534 \| 755 \| 1 \| 220 \| Virulence-associated protein E \| Virulence-associated protein E; afdb-uniprot50 \| AF-A0A533QNK2-F1-MODEL\_V4 \| 1.0 \| 3.868e-19 \| 392 \| 0.183 \| 763 \| 340 \| 43 \| 98 \| 750 \| 73 \| 662 \| DNA primase TraC \| DNA primase TraC; afdb-uniprot50 \| AF-A0A858ZQH3-F1-MODEL\_V4 \| 1.0 \| 1.121e-19 \| 388 \| 0.178 \| 794 \| 372 \| 40 \| 93 \| 769 \| 75 \| 704 \| AAA family ATPase \| AAA family ATPase; afdb-uniprot50 \| AF-A0A2E5Y8U8-F1-MODEL\_V4 \| 1.0 \| 1.352e-20 \| 388 \| 0.152 \| 854 \| 385 \| 49 \| 102 \| 753 \| 75 \| 791 \| DNA primase \| DNA primase; afdb-uniprot50 \| AF-X1J433-F1-MODEL\_V4 \| 1.0 \| 3.943e-10 \| 387 \| 0.2 \| 274 \| 196 \| 14 \| 407 \| 669 \| 2 \| 263 \| Uncharacterized protein \| Uncharacterized protein; afdb-uniprot50 \| AF-A0A6N9A9R1-F1-MODEL\_V4 \| 1.0 \| 7.002e-23 \| 383 \| 0.165 \| 816 \| 453 \| 55 \| 54 \| 745 \| 59 \| 770 \| Uncharacterized protein \| Uncharacterized protein; afdb-uniprot50 \| AF-A0A349EI79-F1-MODEL\_V4 \| 1.0 \| 3.378e-18 \| 378 \| 0.181 \| 793 \| 316 \| 39 \| 92 \| 753 \| 72 \| 662 \| Toprim domain-containing protein \| Toprim domain-containing protein; afdb-uniprot50 \| AF-A0A4R6Y3D7-F1-MODEL\_V4 \| 1.0 \| 7.965e-19 \| 378 \| 0.197 \| 769 \| 356 \| 42 \| 91 \| 758 \| 70 \| 677 \| Putative DNA primase/helicase \| Putative DNA primase/helicase; afdb-uniprot50 \| AF-A0A3R6E138-F1-MODEL\_V4 \| 1.0 \| 3.443e-09 \| 374 \| 0.225 \| 173 \| 129 \| 4 \| 482 \| 652 \| 3 \| 172 \| Virulence protein E \| Virulence protein E; afdb-uniprot50 \| AF-K1RXF7-F1-MODEL\_V4 \| 1.0 \| 7.711e-10 \| 372 \| 0.229 \| 187 \| 129 \| 5 \| 511 \| 684 \| 8 \| 192 \| Uncharacterized protein \| Uncharacterized protein; afdb-uniprot50 \| AF-A0A090BV70-F1-MODEL\_V4 \| 1.0 \| 6.564e-21 \| 372 \| 0.165 \| 779 \| 410 \| 43 \| 92 \| 747 \| 67 \| 728 \| Toprim domain-containing protein \| Toprim domain-containing protein; afdb-uniprot50 \| AF-A0A0H5Q210-F1-MODEL\_V4 \| 1.0 \| 8.121e-18 \| 370 \| 0.168 \| 740 \| 331 \| 35 \| 102 \| 744 \| 85 \| 636 \| Toprim domain-containing protein \| Toprim domain-containing protein; afdb-uniprot50 \| AF-A0A660N8U3-F1-MODEL\_V4 \| 1.0 \| 2.415e-14 \| 370 \| 0.216 \| 458 \| 223 \| 23 \| 87 \| 477 \| 2 \| 390 \| Toprim domain-containing protein \| Toprim domain-containing protein; afdb-uniprot50 \| AF-A0A142JNG9-F1-MODEL\_V4 \| 1.0 \| 4.344e-13 \| 368 \| 0.17 \| 482 \| 285 \| 31 \| 368 \| 759 \| 100 \| 556 \| Uncharacterized protein \| Uncharacterized protein; afdb-uniprot50 \| AF-A0A5C7LNP3-F1-MODEL\_V4 \| 1.0 \| 1.235e-14 \| 368 \| 0.151 \| 541 \| 357 \| 28 \| 290 \| 769 \| 181 \| 680 \| ZnF\_CHCC domain-containing protein \| ZnF\_CHCC domain-containing protein; afdb-uniprot50 \| AF-A0A1F9LYL6-F1-MODEL\_V4 \| 1.0 \| 1.609e-19 \| 368 \| 0.173 \| 776 \| 348 \| 38 \| 102 \| 731 \| 107 \| 734 \| Toprim domain-containing protein \| Toprim domain-containing protein; afdb-uniprot50 \| AF-A0A5E5AI01-F1-MODEL\_V4 \| 1.0 \| 4.428e-12 \| 367 \| 0.177 \| 434 \| 280 \| 25 \| 377 \| 750 \| 109 \| 525 \| Virulence protein E \| Virulence protein E; afdb-uniprot50 \| AF-A0A6J4G0U8-F1-MODEL\_V4 \| 1.0 \| 1.651e-07 \| 365 \| 0.431 \| 116 \| 62 \| 2 \| 234 \| 347 \| 22 \| 135 \| Toprim domain-containing protein \| Toprim domain-containing protein; afdb-uniprot50 \| AF-A0A6I7WYX8-F1-MODEL\_V4 \| 1.0 \| 2.695e-11 \| 365 \| 0.244 \| 229 \| 156 \| 7 \| 506 \| 724 \| 9 \| 230 \| Uncharacterized protein \| Uncharacterized protein; afdb-uniprot50 \| AF-A0A5E4TGU1-F1-MODEL\_V4 \| 1.0 \| 7.662e-13 \| 364 \| 0.165 \| 448 \| 270 \| 25 \| 377 \| 751 \| 204 \| 620 \| Virulence protein E \| Virulence protein E; afdb-uniprot50 \| AF-A0A133Y5T5-F1-MODEL\_V4 \| 1.0 \| 3.046e-10 \| 363 \| 0.219 \| 251 \| 180 \| 10 \| 380 \| 622 \| 8 \| 250 \| Uncharacterized protein \| Uncharacterized protein; afdb-uniprot50 \| AF-A0A651FY11-F1-MODEL\_V4 \| 1.0 \| 7.712e-18 \| 363 \| 0.173 \| 743 \| 317 \| 43 \| 102 \| 743 \| 78 \| 624 \| Toprim domain-containing protein \| Toprim domain-containing protein; afdb-uniprot50 \| AF-A0A412FI27-F1-MODEL\_V4 \| 1.0 \| 2.235e-10 \| 361 \| 0.227 \| 255 \| 181 \| 9 \| 379 \| 624 \| 16 \| 263 \| Virulence protein \| Virulence protein; afdb-uniprot50 \| AF-I5D485-F1-MODEL\_V4 \| 1.0 \| 1.024e-13 \| 360 \| 0.166 \| 451 \| 276 \| 31 \| 377 \| 754 \| 94 \| 517 \| Virulence-associated E family protein \| Virulence-associated E family protein; afdb-uniprot50 \| AF-A0A1G7B4P0-F1-MODEL\_V4 \| 1.0 \| 3.378e-18 \| 358 \| 0.166 \| 792 \| 329 \| 42 \| 101 \| 757 \| 93 \| 687 \| Uncharacterized domain associated with phage/plasmid primase \| Uncharacterized domain associated with phage/plasmid primase; afdb-uniprot50 \| AF-A0A4Z0F4C3-F1-MODEL\_V4 \| 1.0 \| 2.15e-12 \| 355 \| 0.173 \| 374 \| 247 \| 20 \| 318 \| 650 \| 125 \| 477 \| Uncharacterized protein \| Uncharacterized protein; afdb-uniprot50 \| AF-A0A134BA49-F1-MODEL\_V4 \| 1.0 \| 1.705e-08 \| 353 \| 0.293 \| 150 \| 103 \| 2 \| 490 \| 638 \| 9 \| 156 \| Uncharacterized protein \| Uncharacterized protein; afdb-uniprot50 \| AF-A0A845UU64-F1-MODEL\_V4 \| 1.0 \| 1.107e-17 \| 352 \| 0.171 \| 762 \| 337 \| 43 \| 99 \| 757 \| 88 \| 657 \| AAA family ATPase \| AAA family ATPase; afdb-uniprot50 \| AF-A0A1V0QD50-F1-MODEL\_V4 \| 1.0 \| 3.106e-09 \| 351 \| 0.186 \| 230 \| 161 \| 9 \| 463 \| 687 \| 3 \| 211 \| Uncharacterized protein \| Uncharacterized protein; afdb-uniprot50 \| AF-A0A4R1VW43-F1-MODEL\_V4 \| 1.0 \| 5.305e-16 \| 350 \| 0.202 \| 588 \| 291 \| 35 \| 77 \| 609 \| 81 \| 545 \| Phage/plasmid primase-like uncharacterized protein \| Phage/plasmid primase-like uncharacterized protein; afdb-uniprot50 \| AF-A0A5S9QFP7-F1-MODEL\_V4 \| 1.0 \| 8.226e-20 \| 350 \| 0.154 \| 861 \| 458 \| 51 \| 93 \| 765 \| 88 \| 866 \| DNA primase TraC \| DNA primase TraC; afdb-uniprot50 \| AF-A0A3C0I2Y8-F1-MODEL\_V4 \| 1.0 \| 5.038e-16 \| 349 \| 0.162 \| 566 \| 278 \| 27 \| 99 \| 618 \| 94 \| 509 \| Uncharacterized protein \| Uncharacterized protein; afdb-uniprot50 \| AF-A0A2M8TYS1-F1-MODEL\_V4 \| 1.0 \| 5.204e-17 \| 348 \| 0.19 \| 661 \| 341 \| 38 \| 100 \| 636 \| 68 \| 658 \| Toprim domain-containing protein \| Toprim domain-containing protein; afdb-uniprot50 \| AF-A0A1E5Q402-F1-MODEL\_V4 \| 1.0 \| 1.727e-18 \| 346 \| 0.167 \| 728 \| 386 \| 49 \| 100 \| 759 \| 115 \| 690 \| Toprim domain-containing protein \| Toprim domain-containing protein; afdb-uniprot50 \| AF-A0A0H3A6N2-F1-MODEL\_V4 \| 1.0 \| 9.121e-20 \| 344 \| 0.186 \| 805 \| 376 \| 49 \| 112 \| 752 \| 85 \| 774 \| Toprim sub domain protein \| Toprim sub domain protein; afdb-uniprot50 \| AF-A0A2J6MQS6-F1-MODEL\_V4 \| 1.0 \| 6.691e-12 \| 343 \| 0.16 \| 449 \| 271 \| 26 \| 377 \| 751 \| 151 \| 567 \| Uncharacterized protein \| Uncharacterized protein; afdb-uniprot50 \| AF-A0A4R7DVC6-F1-MODEL\_V4 \| 1.0 \| 2.354e-18 \| 343 \| 0.177 \| 767 \| 377 \| 52 \| 88 \| 756 \| 60 \| 670 \| Putative DNA primase/helicase \| Putative DNA primase/helicase; afdb-uniprot50 \| AF-A0A4R5QFG5-F1-MODEL\_V4 \| 1.0 \| 2.893e-18 \| 343 \| 0.168 \| 835 \| 411 \| 43 \| 1 \| 743 \| 1 \| 643 \| Toprim domain-containing protein \| Toprim domain-containing protein; afdb-uniprot50 \| AF-A0A7V2GZR5-F1-MODEL\_V4 \| 1.0 \| 2.056e-17 \| 336 \| 0.186 \| 723 \| 317 \| 46 \| 132 \| 757 \| 3 \| 550 \| Toprim domain-containing protein \| Toprim domain-containing protein; afdb-uniprot50 \| AF-A0A847UVH2-F1-MODEL\_V4 \| 1.0 \| 1.405e-10 \| 335 \| 0.31 \| 216 \| 126 \| 9 \| 13 \| 212 \| 2 \| 210 \| Uncharacterized protein \| Uncharacterized protein; afdb-uniprot50 \| AF-A0A0H3ZTP3-F1-MODEL\_V4 \| 1.0 \| 1.334e-18 \| 335 \| 0.128 \| 944 \| 513 \| 53 \| 6 \| 745 \| 2 \| 839 \| DNA primase \| DNA primase; afdb-uniprot50 \| AF-A0A257DJ96-F1-MODEL\_V4 \| 1.0 \| 1.15e-15 \| 333 \| 0.2 \| 588 \| 299 \| 33 \| 76 \| 609 \| 80 \| 550 \| ZnF\_CHCC domain-containing protein \| ZnF\_CHCC domain-containing protein; afdb-uniprot50 \| AF-A0A1F7FGC5-F1-MODEL\_V4 \| 1.0 \| 3.334e-16 \| 332 \| 0.12 \| 724 \| 431 \| 51 \| 162 \| 755 \| 89 \| 736 \| SF3 helicase domain-containing protein \| SF3 helicase domain-containing protein; afdb-uniprot50 \| AF-A0A258SRK8-F1-MODEL\_V4 \| 1.0 \| 7.612e-08 \| 329 \| 0.315 \| 130 \| 87 \| 1 \| 362 \| 489 \| 57 \| 186 \| Uncharacterized protein \| Uncharacterized protein; afdb-uniprot50 \| AF-A0A5C7K204-F1-MODEL\_V4 \| 1.0 \| 3.066e-15 \| 321 \| 0.188 \| 574 \| 297 \| 36 \| 115 \| 614 \| 114 \| 592 \| Herpes\_ori\_bp domain-containing protein \| Herpes\_ori\_bp domain-containing protein; afdb-uniprot50 \| AF-A0A6J4FPK4-F1-MODEL\_V4 \| 1.0 \| 3.943e-18 \| 315 \| 0.198 \| 740 \| 311 \| 38 \| 125 \| 754 \| 102 \| 669 \| Toprim domain-containing protein \| Toprim domain-containing protein; afdb-uniprot50 \| AF-A0A1M6LF95-F1-MODEL\_V4 \| 1.0 \| 1.46e-08 \| 312 \| 0.182 \| 236 \| 180 \| 8 \| 534 \| 762 \| 1 \| 230 \| Virulence-associated protein E \| Virulence-associated protein E; afdb-uniprot50 \| AF-A0A4Z1QPV4-F1-MODEL\_V4 \| 1.0 \| 1.878e-11 \| 311 \| 0.134 \| 492 \| 330 \| 28 \| 328 \| 769 \| 53 \| 498 \| Uncharacterized protein \| Uncharacterized protein; afdb-uniprot50 \| AF-A0A4E0QLS9-F1-MODEL\_V4 \| 1.0 \| 1.114e-14 \| 311 \| 0.129 \| 632 \| 376 \| 34 \| 81 \| 628 \| 98 \| 639 \| Uncharacterized protein \| Uncharacterized protein; afdb-uniprot50 \| AF-A0A7X1KDD5-F1-MODEL\_V4 \| 1.0 \| 1.508e-17 \| 305 \| 0.188 \| 749 \| 320 \| 36 \| 125 \| 754 \| 312 \| 891 \| Bifunctional DNA primase/polymerase \| Bifunctional DNA primase/polymerase; afdb-uniprot50 \| AF-A0A7J0A256-F1-MODEL\_V4 \| 1.0 \| 3.007e-08 \| 301 \| 0.243 \| 197 \| 144 \| 3 \| 563 \| 756 \| 5 \| 199 \| Uncharacterized protein \| Uncharacterized protein; afdb-uniprot50 \| AF-A0A6I2NXZ8-F1-MODEL\_V4 \| 1.0 \| 2.4e-09 \| 296 \| 0.189 \| 274 \| 193 \| 11 \| 503 \| 762 \| 20 \| 278 \| DUF3874 domain-containing protein \| DUF3874 domain-containing protein; afdb-uniprot50 \| AF-A0A6N1AJ04-F1-MODEL\_V4 \| 1.0 \| 1.334e-10 \| 294 \| 0.148 \| 466 \| 291 \| 28 \| 376 \| 757 \| 83 \| 526 \| Uncharacterized protein \| Uncharacterized protein; afdb-uniprot50 \| AF-A0A7G8I5M2-F1-MODEL\_V4 \| 1.0 \| 4.02e-09 \| 293 \| 0.147 \| 278 \| 203 \| 9 \| 497 \| 751 \| 3 \| 269 \| Virulence-associated E family protein \| Virulence-associated E family protein; afdb-uniprot50 \| AF-A0A160JDL1-F1-MODEL\_V4 \| 1.0 \| 5.55e-11 \| 292 \| 0.126 \| 567 \| 365 \| 32 \| 239 \| 769 \| 107 \| 578 \| Uncharacterized protein \| Uncharacterized protein; afdb-uniprot50 \| AF-A0A1S7U2B0-F1-MODEL\_V4 \| 1.0 \| 6.606e-10 \| 287 \| 0.129 \| 409 \| 279 \| 21 \| 381 \| 744 \| 98 \| 474 \| Uncharacterized protein \| Uncharacterized protein; afdb-uniprot50 \| AF-K1SV85-F1-MODEL\_V4 \| 1.0 \| 1.058e-06 \| 277 \| 0.305 \| 134 \| 91 \| 2 \| 509 \| 641 \| 4 \| 136 \| Virulence-associated protein E \| Virulence-associated protein E; afdb-uniprot50 \| AF-A0A6N2RAJ0-F1-MODEL\_V4 \| 1.0 \| 1.761e-09 \| 262 \| 0.153 \| 405 \| 242 \| 13 \| 376 \| 767 \| 44 \| 360 \| DUF based on B. Theta Gene description \| DUF based on B. Theta Gene description; afdb-uniprot50 \| AF-A0A4U1BRV7-F1-MODEL\_V4 \| 1.0 \| 1.1e-12 \| 258 \| 0.146 \| 520 \| 251 \| 25 \| 151 \| 616 \| 4 \| 384 \| Toprim domain-containing protein \| Toprim domain-containing protein; afdb-uniprot50 \| AF-A0A358XVY2-F1-MODEL\_V4 \| 1.0 \| 0.0003421 \| 257 \| 0.325 \| 83 \| 56 \| 0 \| 12 \| 94 \| 194 \| 276 \| PriCT\_2 domain-containing protein \| PriCT\_2 domain-containing protein; afdb-uniprot50 \| AF-A0A5C5UXX3-F1-MODEL\_V4 \| 1.0 \| 3.603e-12 \| 251 \| 0.13 \| 758 \| 447 \| 56 \| 150 \| 769 \| 133 \| 816 \| SF3 helicase domain-containing protein \| SF3 helicase domain-containing protein; afdb-uniprot50 \| AF-A0A7Y8UR69-F1-MODEL\_V4 \| 1.0 \| 4.152e-10 \| 248 \| 0.145 \| 569 \| 350 \| 32 \| 287 \| 769 \| 21 \| 539 \| Uncharacterized protein \| Uncharacterized protein; afdb-uniprot50 \| AF-A0A1L9QN36-F1-MODEL\_V4 \| 1.0 \| 4.663e-12 \| 228 \| 0.126 \| 721 \| 428 \| 40 \| 180 \| 751 \| 58 \| 725 \| SF3 helicase domain-containing protein \| SF3 helicase domain-containing protein; afdb-uniprot50 \| AF-A0A1V5NKL3-F1-MODEL\_V4 \| 1.0 \| 2.912e-15 \| 226 \| 0.151 \| 711 \| 339 \| 34 \| 215 \| 769 \| 8 \| 609 \| DNA primase TraC \| DNA primase TraC; afdb-uniprot50 \| AF-A0A174TGT0-F1-MODEL\_V4 \| 1.0 \| 1.058e-06 \| 224 \| 0.217 \| 207 \| 152 \| 7 \| 561 \| 762 \| 18 \| 219 \| Predicted P-loop ATPase and inactivated derivatives \| Predicted P-loop ATPase and inactivated derivatives; afdb-uniprot50 \| AF-R5NEB3-F1-MODEL\_V4 \| 1.0 \| 2.221e-05 \| 223 \| 0.191 \| 209 \| 162 \| 5 \| 549 \| 753 \| 2 \| 207 \| SWIRM domain-containing protein \| SWIRM domain-containing protein; afdb-uniprot50 \| AF-A0A7X2MAP5-F1-MODEL\_V4 \| 1.0 \| 4.457e-09 \| 223 \| 0.125 \| 447 \| 273 \| 28 \| 381 \| 753 \| 229 \| 631 \| DNA primase \| DNA primase; afdb-uniprot50 \| AF-A0A0J6VN82-F1-MODEL\_V4 \| 1.0 \| 1.165e-09 \| 223 \| 0.12 \| 557 \| 348 \| 32 \| 240 \| 721 \| 174 \| 663 \| Uncharacterized protein \| Uncharacterized protein; afdb-uniprot50 \| AF-A0A2E4HGQ7-F1-MODEL\_V4 \| 1.0 \| 3.867e-11 \| 221 \| 0.12 \| 844 \| 403 \| 46 \| 9 \| 751 \| 224 \| 828 \| PriCT\_2 domain-containing protein \| PriCT\_2 domain-containing protein; afdb-uniprot50 \| AF-A0A2P7THR1-F1-MODEL\_V4 \| 1.0 \| 4.344e-13 \| 218 \| 0.099 \| 905 \| 508 \| 55 \| 60 \| 750 \| 86 \| 897 \| Toprim domain-containing protein \| Toprim domain-containing protein; afdb-uniprot50 \| AF-A0A480BUQ6-F1-MODEL\_V4 \| 1.0 \| 1.915e-10 \| 207 \| 0.161 \| 595 \| 336 \| 38 \| 267 \| 755 \| 3 \| 540 \| Uncharacterized protein \| Uncharacterized protein; afdb-uniprot50 \| AF-A0A0D6AFU3-F1-MODEL\_V4 \| 1.0 \| 2.95e-09 \| 192 \| 0.107 \| 612 \| 358 \| 35 \| 261 \| 768 \| 373 \| 899 \| SF3 helicase domain-containing protein \| SF3 helicase domain-containing protein; afdb-uniprot50 \| AF-A0A5C8BGE8-F1-MODEL\_V4 \| 1.0 \| 2.136e-07 \| 188 \| 0.112 \| 505 \| 309 \| 30 \| 327 \| 751 \| 18 \| 463 \| SF3 helicase domain-containing protein \| SF3 helicase domain-containing protein; afdb-uniprot50 \| AF-A0A200I1W5-F1-MODEL\_V4 \| 1.0 \| 1.211e-07 \| 187 \| 0.142 \| 449 \| 271 \| 32 \| 376 \| 745 \| 58 \| 471 \| SF3 helicase domain-containing protein \| SF3 helicase domain-containing protein; afdb-uniprot50 \| AF-A0A401LNS6-F1-MODEL\_V4 \| 1.0 \| 0.0002783 \| 185 \| 0.223 \| 134 \| 103 \| 1 \| 556 \| 689 \| 57 \| 189 \| Uncharacterized protein \| Uncharacterized protein; afdb-uniprot50 \| AF-A0A0B0H9C4-F1-MODEL\_V4 \| 1.0 \| 6.314e-07 \| 184 \| 0.13 \| 491 \| 292 \| 34 \| 374 \| 769 \| 158 \| 608 \| Phage/plasmid primase \| Phage/plasmid primase; afdb-uniprot50 \| AF-A0A3A5AP02-F1-MODEL\_V4 \| 1.0 \| 6.521e-08 \| 184 \| 0.124 \| 564 \| 340 \| 40 \| 289 \| 757 \| 259 \| 763 \| SF3 helicase domain-containing protein \| SF3 helicase domain-containing protein; afdb-uniprot50 \| AF-A0A1L9QT25-F1-MODEL\_V4 \| 1.0 \| 1.114e-06 \| 174 \| 0.104 \| 439 \| 293 \| 26 \| 394 \| 757 \| 8 \| 421 \| SF3 helicase domain-containing protein \| SF3 helicase domain-containing protein; afdb-uniprot50 \| AF-A0A8A3PZQ0-F1-MODEL\_V4 \| 1.0 \| 6.193e-08 \| 174 \| 0.134 \| 461 \| 273 \| 32 \| 383 \| 753 \| 350 \| 774 \| Uncharacterized protein \| Uncharacterized protein; afdb-uniprot50 \| AF-A0A2S9SYH7-F1-MODEL\_V4 \| 1.0 \| 2.294e-06 \| 173 \| 0.095 \| 460 \| 298 \| 22 \| 384 \| 751 \| 222 \| 655 \| Uncharacterized protein \| Uncharacterized protein; afdb-uniprot50 \| AF-A0A1W2G805-F1-MODEL\_V4 \| 1.0 \| 5.958e-10 \| 171 \| 0.096 \| 799 \| 432 \| 54 \| 145 \| 744 \| 5 \| 712 \| Uncharacterized protein \| Uncharacterized protein; afdb-uniprot50 \| AF-A0A1M6M5E6-F1-MODEL\_V4 \| 1.0 \| 3.399e-07 \| 171 \| 0.119 \| 484 \| 277 \| 33 \| 383 \| 753 \| 679 \| 1126 \| Uncharacterized protein \| Uncharacterized protein; afdb-uniprot50 \| AF-A0A661W849-F1-MODEL\_V4 \| 1.0 \| 2.003e-05 \| 169 \| 0.157 \| 209 \| 148 \| 12 \| 473 \| 675 \| 2 \| 188 \| Uncharacterized protein \| Uncharacterized protein; afdb-uniprot50 \| AF-L0GXI9-F1-MODEL\_V4 \| 1.0 \| 4.91e-12 \| 169 \| 0.13 \| 943 \| 379 \| 53 \| 1 \| 757 \| 20 \| 707 \| Primase C terminal 2 (PriCT-2) \| Primase C terminal 2 (PriCT-2); afdb-uniprot50 \| AF-A0A7X3HIV0-F1-MODEL\_V4 \| 1.0 \| 8.887e-08 \| 168 \| 0.128 \| 451 \| 271 \| 29 \| 384 \| 762 \| 30 \| 430 \| Uncharacterized protein \| Uncharacterized protein; afdb-uniprot50 \| AF-A0A2A8H4K1-F1-MODEL\_V4 \| 1.0 \| 2.178e-06 \| 160 \| 0.113 \| 457 \| 288 \| 28 \| 384 \| 751 \| 373 \| 801 \| DNA primase \| DNA primase; afdb-uniprot50 \| AF-A0A3D3BMH9-F1-MODEL\_V4 \| 1.0 \| 5.237e-06 \| 155 \| 0.141 \| 346 \| 212 \| 25 \| 463 \| 752 \| 44 \| 360 \| Uncharacterized protein \| Uncharacterized protein; afdb-uniprot50 \| AF-A0A2D7PG15-F1-MODEL\_V4 \| 1.0 \| 8.172e-07 \| 154 \| 0.111 \| 423 \| 251 \| 25 \| 440 \| 753 \| 519 \| 925 \| ZnF\_CHCC domain-containing protein \| ZnF\_CHCC domain-containing protein; afdb-uniprot50 \| AF-A0A843HQ44-F1-MODEL\_V4 \| 1.0 \| 5.136e-07 \| 153 \| 0.109 \| 702 \| 380 \| 48 \| 153 \| 753 \| 62 \| 619 \| Uncharacterized protein \| Uncharacterized protein; afdb-uniprot50 \| AF-A0A7U3YKR5-F1-MODEL\_V4 \| 1.0 \| 1.716e-05 \| 152 \| 0.113 \| 474 \| 290 \| 30 \| 384 \| 767 \| 115 \| 548 \| Phage/plasmid primase, P4 family \| Phage/plasmid primase, P4 family; afdb-uniprot50 \| AF-A0A149RZ98-F1-MODEL\_V4 \| 1.0 \| 1.058e-06 \| 150 \| 0.12 \| 489 \| 297 \| 35 \| 384 \| 769 \| 37 \| 495 \| Uncharacterized protein \| Uncharacterized protein; afdb-uniprot50 \| AF-A0A0R1LZ24-F1-MODEL\_V4 \| 1.0 \| 4.046e-06 \| 146 \| 0.111 \| 485 \| 296 \| 34 \| 370 \| 755 \| 62 \| 510 \| SF3 helicase domain-containing protein \| SF3 helicase domain-containing protein; afdb-uniprot50 \| AF-A0A495V3N2-F1-MODEL\_V4 \| 1.0 \| 1.83e-07 \| 146 \| 0.102 \| 831 \| 440 \| 52 \| 60 \| 757 \| 240 \| 897 \| P4 family phage/plasmid primase-like protein \| P4 family phage/plasmid primase-like protein; afdb-uniprot50 \| AF-A0A0F9LV59-F1-MODEL\_V4 \| 1.0 \| 2.462e-05 \| 145 \| 0.125 \| 366 \| 226 \| 21 \| 451 \| 753 \| 416 \| 750 \| Uncharacterized protein \| Uncharacterized protein; afdb-uniprot50 \| AF-A0A7X2P799-F1-MODEL\_V4 \| 1.0 \| 1.259e-05 \| 142 \| 0.113 \| 467 \| 265 \| 30 \| 384 \| 757 \| 256 \| 666 \| DNA primase \| DNA primase; afdb-uniprot50 \| AF-A0A5M1BU63-F1-MODEL\_V4 \| 1.0 \| 1.965e-06 \| 142 \| 0.1 \| 587 \| 347 \| 43 \| 243 \| 734 \| 231 \| 731 \| DNA primase \| DNA primase; afdb-uniprot50 \| AF-A0A0Q0ER42-F1-MODEL\_V4 \| 1.0 \| 0.0003086 \| 141 \| 0.119 \| 317 \| 230 \| 20 \| 463 \| 751 \| 20 \| 315 \| Bifunctional DNA primase/polymerase \| Bifunctional DNA primase/polymerase; afdb-uniprot50 \| AF-A0A6H1NSJ1-F1-MODEL\_V4 \| 1.0 \| 0.0001283 \| 141 \| 0.118 \| 303 \| 208 \| 20 \| 471 \| 739 \| 214 \| 491 \| DNA primase \| DNA primase; afdb-uniprot50 \| AF-R7LTZ8-F1-MODEL\_V4 \| 1.0 \| 8.015e-08 \| 139 \| 0.103 \| 722 \| 379 \| 43 \| 77 \| 737 \| 78 \| 591 \| DNA primase \| DNA primase; afdb-uniprot50 \| AF-A0A517YE19-F1-MODEL\_V4 \| 1.0 \| 7.761e-07 \| 139 \| 0.112 \| 639 \| 350 \| 38 \| 259 \| 757 \| 80 \| 641 \| SF3 helicase domain-containing protein \| SF3 helicase domain-containing protein; afdb-uniprot50 \| AF-A0A2T4U2M5-F1-MODEL\_V4 \| 1.0 \| 2.178e-06 \| 136 \| 0.114 \| 570 \| 340 \| 40 \| 261 \| 737 \| 216 \| 713 \| DNA primase \| DNA primase; afdb-uniprot50 \| AF-A0A2G6B4P4-F1-MODEL\_V4 \| 1.0 \| 1.683e-06 \| 135 \| 0.098 \| 591 \| 346 \| 39 \| 243 \| 731 \| 141 \| 646 \| DNA primase \| DNA primase; afdb-uniprot50 \| AF-A0A844G9M1-F1-MODEL\_V4 \| 1.0 \| 7.515e-06 \| 134 \| 0.112 \| 559 \| 313 \| 38 \| 337 \| 755 \| 344 \| 859 \| Uncharacterized protein \| Uncharacterized protein; afdb-uniprot50 \| AF-A0A496Z8V0-F1-MODEL\_V4 \| 1.0 \| 2.712e-08 \| 127 \| 0.134 \| 719 \| 312 \| 38 \| 9 \| 683 \| 230 \| 681 \| Uncharacterized protein \| Uncharacterized protein; afdb-uniprot50 \| AF-A0A317LFU4-F1-MODEL\_V4 \| 1.0 \| 1.518e-06 \| 121 \| 0.114 \| 912 \| 340 \| 46 \| 1 \| 769 \| 198 \| 784 \| Prim-Pol domain-containing protein \| Prim-Pol domain-containing protein; afdb-uniprot50 \| AF-A0A5B8XK45-F1-MODEL\_V4 \| 1.0 \| 4.26e-06 \| 115 \| 0.107 \| 605 \| 328 \| 44 \| 260 \| 761 \| 44 \| 539 \| Uncharacterized protein \| Uncharacterized protein; afdb-uniprot50 \| AF-A0A0Q6S7Q0-F1-MODEL\_V4 \| 1.0 \| 3.291e-06 \| 114 \| 0.16 \| 437 \| 180 \| 23 \| 1 \| 320 \| 135 \| 501 \| Uncharacterized protein \| Uncharacterized protein |
| Top keywords  (threshold 1.00e-02 (evalue)) | **DNA, domain\_containing, E, Virulence\_associated, primase, replication, factor, licensing, helicase, Toprim** |
| Output files | ../../similar\_structures/54\_FANPEZAQ\_CDS\_0054\_afdb-proteome\_foldseek.tsv ../../similar\_structures/54\_FANPEZAQ\_CDS\_0054\_afdb-uniprot50\_foldseek.tsv ../../similar\_structures/54\_FANPEZAQ\_CDS\_0054\_merged.svg ../../similar\_structures/54\_FANPEZAQ\_CDS\_0054\_pdb\_foldseek.tsv |

  
  
  

Return to summary | Go to previous | Go to next

  


---

**Sequence/structure alignments coloring**  
Each object in the alignment figures is colored according to its E-value following this color coding:

1e-100
10
