## Supplementary material for "Completing the BASEL phage collection to unlock hidden diversity for systematic exploration of phage-host interactions": Data S2: 57.html

FANPEZAQ\_CDS\_0057


Return to summary | Go to previous | Go to next

|  |  |
| --- | --- |
| FANPEZAQ\_CDS\_0057  Page creation date: 02 Sep 2024, 12:00  Project folder: n/a  Input sequences file: Escherichia\_virus\_HeidiAbel.gb | domain\_containing duf551 hypothetical phage eaa eaa1 putative duf550 duf3850 ead lar restriction alleviation ea22\_like fragment p181809 vi\_03505 p158937 vi\_06901 conserved 3'\_5' exoribonuclease molecular chaperone dnaj prophage asch pua dtdp\_6\_deoxy\_l\_hexose 3\_o\_methyltransferase bacteriophage replication restart dna helicase pria |

### Sequence information

|  |  |
| --- | --- |
| Name | FANPEZAQ\_CDS\_0057  57\_FANPEZAQ\_CDS\_0057 (pipeline id) |
| Imported annotations | Escherichia\_virus\_HeidiAbel Bas97 |
| Protein sequence | MKQVTIDTVRERIAELEMVNSHGELSLRGEFELACLRMLADSMGGVDDDVRNVVVLLENN EWAEHCTNTVLGSRLESEITRLVGHVQPAPVASDAEPSFDAMMRALDAFYADDDVPELAM LAAFRILLADVRNQPAPVVIDDVMEEINRWATDRSLVGRNNYTQQQNALIDVYIGLTEPQ VPVAVPDEVTSEQAYQIGDYYGDPIDVFARGANWMRSKIIGNMLLADMDRAATRKVQELT TCNTAQQFEALTTNGKSEDLLNAERIAREAKALSDRLNDTSAVHPRVSGWSKRQQKVISY SLWRFMHSAYEMAKDAAQKGTKEDFDRFMQDAKCAEEIREEFTSAGSGKP |
| Number of residues | 350 |
| Molecular weight (Da) | 39171.61 |
| Output files | ../../query\_sequences/57\_FANPEZAQ\_CDS\_0057.fasta |

### Putative domain architecture and protein family

#### Search results (HHblits)^1^

|  |  |
| --- | --- |
| Domain family databases searched | Pfam, Ncbi-cd, Cath, Phrogs |
| Results, scheme(s)  (Top layers only; threshold 1.00e-03 (evalue)) | xml version="1.0" encoding="utf-8" standalone="no"?       2024-09-02T21:08:24.990238 image/svg+xml   Matplotlib v3.7.2, https://matplotlib.org/ |
| Results, table  (E-value ≤ 1.00e-03 (evalue)) | db \| id \| prob \| evalue \| pvalue \| score \| cols \| query \| query\_len \| template \| template\_len \| name \| description; phrogs \| 20174 \| 99.8 \| 4.5e-24 \| 5e-28 \| 183.9 \| 65 \| (25, 94) \| 350 \| (4, 68) \| 221 \| NA \| NA; Category: unknown function; p181809 VI\_03505; phrogs \| 19524 \| 99.7 \| 2.9e-22 \| 3.2e-26 \| 162.4 \| 60 \| (25, 89) \| 350 \| (4, 63) \| 127 \| NA \| NA; Category: unknown function; p158937 VI\_06901 |
| Top keywords  (threshold 1.00e-03 (evalue)) | **p181809, VI\_03505, p158937, VI\_06901** |
| Output files | ../../domain\_architecture/57\_FANPEZAQ\_CDS\_0057\_cath.hhr ../../domain\_architecture/57\_FANPEZAQ\_CDS\_0057\_merged.svg ../../domain\_architecture/57\_FANPEZAQ\_CDS\_0057\_ncbi-cd.hhr ../../domain\_architecture/57\_FANPEZAQ\_CDS\_0057\_pfam.hhr ../../domain\_architecture/57\_FANPEZAQ\_CDS\_0057\_phrogs.hhr |

### Identical protein sequences/structures

#### Search results

|  |  |
| --- | --- |
| Protein sequence databases searched | Pdb, Swissprot, Refseq |
| Identical proteins found | -- |
| Top keywords | -- |
| Output files | -- |

### Similar protein sequences/structures

#### Sequence similarity search results (HHblits)^1^

|  |  |
| --- | --- |
| Sequence databases searched | Uniclust, Pdb70 |
| Results, scheme(s)  (Top layers only, threshold 1.00e-03 (evalue)) | xml version="1.0" encoding="utf-8" standalone="no"?       2024-09-02T21:08:53.048347 image/svg+xml   Matplotlib v3.7.2, https://matplotlib.org/ |
| Results, table(s)  (threshold 1.00e-03 (evalue)) | db \| id \| prob \| evalue \| pvalue \| score \| cols \| query \| query\_len \| template \| template\_len \| name \| description; uniclust \| UniRef100\_A0A088FQU9 \| 99.9 \| 4.4e-25 \| 8.2e-31 \| 204.7 \| 142 \| (1, 142) \| 350 \| (1, 142) \| 275 \| DUF551 domain-containing protein \| DUF551 domain-containing protein; uniclust \| UniRef100\_UPI000AB3930E \| 99.8 \| 3.9e-22 \| 8e-28 \| 190.0 \| 158 \| (36, 231) \| 350 \| (92, 266) \| 269 \| DUF551 domain-containing protein \| DUF551 domain-containing protein; uniclust \| UniRef100\_A0A4U9I001 \| 99.8 \| 9.1e-22 \| 2.3e-27 \| 199.1 \| 153 \| (77, 232) \| 350 \| (118, 310) \| 362 \| DUF551 domain-containing protein \| DUF551 domain-containing protein; uniclust \| UniRef100\_A0A6S5JYC3 \| 99.7 \| 9.9e-21 \| 2.4e-26 \| 186.2 \| 166 \| (62, 237) \| 350 \| (73, 266) \| 289 \| DUF551 domain-containing protein \| DUF551 domain-containing protein; uniclust \| UniRef100\_A0A482MEI6 \| 99.7 \| 3.6e-20 \| 7.7e-26 \| 182.8 \| 155 \| (76, 232) \| 350 \| (74, 269) \| 353 \| DUF551 domain-containing protein \| DUF551 domain-containing protein; uniclust \| UniRef100\_UPI001CA50654 \| 99.6 \| 6.5e-19 \| 1.4e-24 \| 164.9 \| 125 \| (80, 231) \| 350 \| (68, 202) \| 206 \| hypothetical protein \| hypothetical protein; uniclust \| UniRef100\_A0A3R0VRQ8 \| 99.6 \| 1.1e-18 \| 2.5e-24 \| 170.2 \| 167 \| (36, 231) \| 350 \| (85, 286) \| 290 \| DUF551 domain-containing protein \| DUF551 domain-containing protein; uniclust \| UniRef100\_UPI0020CDA783 \| 99.6 \| 1.9e-18 \| 3.4e-24 \| 155.7 \| 139 \| (76, 231) \| 350 \| (48, 192) \| 195 \| DUF550 domain-containing protein \| DUF550 domain-containing protein; uniclust \| UniRef100\_A0A0J1LUR9 \| 99.5 \| 5.9e-17 \| 1.4e-22 \| 157.0 \| 78 \| (148, 234) \| 350 \| (148, 230) \| 251 \| Phage protein \| Phage protein; uniclust \| UniRef100\_UPI0007CC47B1 \| 99.5 \| 8.5e-17 \| 1.9e-22 \| 157.9 \| 105 \| (132, 236) \| 350 \| (151, 282) \| 293 \| hypothetical protein \| hypothetical protein; uniclust \| UniRef100\_A0A0H3CRI4 \| 99.5 \| 1.3e-16 \| 3.7e-22 \| 160.9 \| 63 \| (168, 234) \| 350 \| (198, 261) \| 307 \| DUF551 domain-containing protein \| DUF551 domain-containing protein; uniclust \| UniRef100\_A0A3R0PNL9 \| 99.4 \| 2.2e-16 \| 5.4e-22 \| 156.2 \| 73 \| (24, 96) \| 350 \| (63, 139) \| 284 \| DUF551 domain-containing protein \| DUF551 domain-containing protein; uniclust \| UniRef100\_A0A2H4FNA7 \| 99.4 \| 6.4e-16 \| 1.5e-21 \| 146.9 \| 76 \| (21, 96) \| 350 \| (69, 149) \| 213 \| Eaa protein \| Eaa protein; uniclust \| UniRef100\_UPI0007CD0D32 \| 99.3 \| 3.6e-15 \| 6.6e-21 \| 138.2 \| 141 \| (78, 231) \| 350 \| (43, 227) \| 230 \| hypothetical protein \| hypothetical protein; uniclust \| UniRef100\_A0A447R0Q5 \| 99.3 \| 3.5e-15 \| 7e-21 \| 136.6 \| 62 \| (36, 97) \| 350 \| (84, 150) \| 186 \| Phage protein \| Phage protein; uniclust \| UniRef100\_A0A3J4L5J7 \| 99.3 \| 5.8e-15 \| 1.2e-20 \| 142.3 \| 85 \| (41, 126) \| 350 \| (103, 189) \| 262 \| Eaa protein \| Eaa protein; uniclust \| UniRef100\_A0A6L3Y1N6 \| 99.3 \| 7.2e-15 \| 1.6e-20 \| 148.1 \| 73 \| (156, 237) \| 350 \| (261, 334) \| 359 \| DUF551 domain-containing protein \| DUF551 domain-containing protein; uniclust \| UniRef100\_A0A411E019 \| 99.3 \| 8.6e-15 \| 1.6e-20 \| 120.3 \| 86 \| (1, 90) \| 350 \| (1, 88) \| 92 \| Uncharacterized protein \| Uncharacterized protein; uniclust \| UniRef100\_D5LH12 \| 99.3 \| 1.9e-14 \| 3.5e-20 \| 136.4 \| 86 \| (1, 86) \| 350 \| (1, 160) \| 274 \| Conserved phage protein \| Conserved phage protein; uniclust \| UniRef100\_UPI00064338B8 \| 99.3 \| 2e-14 \| 3.9e-20 \| 138.1 \| 59 \| (168, 230) \| 350 \| (208, 267) \| 270 \| hypothetical protein \| hypothetical protein; uniclust \| UniRef100\_UPI00094A7D01 \| 99.2 \| 5.6e-14 \| 1e-19 \| 124.7 \| 102 \| (127, 232) \| 350 \| (42, 156) \| 159 \| hypothetical protein \| hypothetical protein; uniclust \| UniRef100\_UPI0007B3E07F \| 99.2 \| 6e-14 \| 1.2e-19 \| 123.8 \| 70 \| (153, 231) \| 350 \| (65, 135) \| 138 \| hypothetical protein \| hypothetical protein; uniclust \| UniRef100\_UPI0020766232 \| 99.2 \| 1.6e-13 \| 2.9e-19 \| 124.6 \| 60 \| (168, 231) \| 350 \| (125, 185) \| 187 \| hypothetical protein \| hypothetical protein; uniclust \| UniRef100\_UPI0007F92665 \| 99.1 \| 4.3e-13 \| 8.5e-19 \| 128.9 \| 60 \| (168, 231) \| 350 \| (189, 249) \| 252 \| hypothetical protein \| hypothetical protein; uniclust \| UniRef100\_A0A2J0PF48 \| 99.1 \| 5.8e-13 \| 1.1e-18 \| 126.2 \| 68 \| (159, 230) \| 350 \| (121, 189) \| 261 \| Uncharacterized protein \| Uncharacterized protein; uniclust \| UniRef100\_UPI001FFE00E5 \| 99.1 \| 8.1e-13 \| 1.5e-18 \| 126.0 \| 65 \| (168, 236) \| 350 \| (174, 239) \| 267 \| 3'-5' exoribonuclease \| 3'-5' exoribonuclease; uniclust \| UniRef100\_UPI001F2AB062 \| 99.0 \| 1.2e-12 \| 2.3e-18 \| 126.5 \| 60 \| (168, 231) \| 350 \| (249, 309) \| 312 \| hypothetical protein \| hypothetical protein; uniclust \| UniRef100\_A0A447R477 \| 99.0 \| 1.2e-12 \| 2.3e-18 \| 119.6 \| 81 \| (16, 96) \| 350 \| (50, 140) \| 186 \| Phage protein \| Phage protein; uniclust \| UniRef100\_UPI001E5CA50A \| 99.0 \| 4.1e-12 \| 7.5e-18 \| 107.8 \| 60 \| (168, 231) \| 350 \| (49, 109) \| 111 \| hypothetical protein \| hypothetical protein; uniclust \| UniRef100\_UPI001F0C3520 \| 98.9 \| 1.1e-11 \| 2.2e-17 \| 120.2 \| 60 \| (170, 233) \| 350 \| (203, 263) \| 285 \| hypothetical protein \| hypothetical protein; uniclust \| UniRef100\_A0A5B8RN22 \| 98.9 \| 1.6e-11 \| 3.2e-17 \| 121.2 \| 169 \| (60, 228) \| 350 \| (110, 307) \| 311 \| Molecular chaperone DnaJ \| Molecular chaperone DnaJ; uniclust \| UniRef100\_A0A4Q4ADR6 \| 98.8 \| 2.6e-11 \| 5.4e-17 \| 116.0 \| 58 \| (170, 231) \| 350 \| (164, 222) \| 226 \| Uncharacterized protein \| Uncharacterized protein; uniclust \| UniRef100\_UPI0009D42707 \| 98.8 \| 3.5e-11 \| 6.9e-17 \| 122.9 \| 157 \| (61, 228) \| 350 \| (235, 420) \| 424 \| hypothetical protein \| hypothetical protein; uniclust \| UniRef100\_UPI000E2B2B70 \| 98.8 \| 3.3e-11 \| 8e-17 \| 127.3 \| 60 \| (168, 233) \| 350 \| (315, 376) \| 436 \| hypothetical protein \| hypothetical protein; uniclust \| UniRef100\_UPI0021B4286A \| 98.8 \| 5.1e-11 \| 9.4e-17 \| 110.3 \| 61 \| (36, 96) \| 350 \| (55, 119) \| 202 \| hypothetical protein \| hypothetical protein; uniclust \| UniRef100\_A0A636KA63 \| 98.8 \| 5.9e-11 \| 1.1e-16 \| 111.4 \| 57 \| (41, 97) \| 350 \| (131, 187) \| 225 \| Eaa protein \| Eaa protein; uniclust \| UniRef100\_UPI000F2AB43C \| 98.8 \| 6.1e-11 \| 1.2e-16 \| 112.4 \| 52 \| (180, 231) \| 350 \| (171, 223) \| 226 \| hypothetical protein \| hypothetical protein; uniclust \| UniRef100\_UPI002032BBFA \| 98.8 \| 6.7e-11 \| 1.2e-16 \| 111.7 \| 65 \| (158, 231) \| 350 \| (169, 234) \| 236 \| hypothetical protein \| hypothetical protein; uniclust \| UniRef100\_UPI001908214B \| 98.7 \| 8e-11 \| 1.5e-16 \| 106.9 \| 57 \| (176, 232) \| 350 \| (86, 143) \| 172 \| hypothetical protein \| hypothetical protein; uniclust \| UniRef100\_UPI002005553F \| 98.7 \| 9.3e-11 \| 1.7e-16 \| 104.1 \| 60 \| (168, 231) \| 350 \| (85, 145) \| 146 \| hypothetical protein \| hypothetical protein; uniclust \| UniRef100\_UPI001F4431F5 \| 98.7 \| 1.7e-10 \| 3.1e-16 \| 102.9 \| 59 \| (168, 230) \| 350 \| (88, 147) \| 149 \| hypothetical protein \| hypothetical protein; uniclust \| UniRef100\_A0A8S7XWP4 \| 98.7 \| 2.2e-10 \| 4.2e-16 \| 106.8 \| 55 \| (41, 95) \| 350 \| (15, 69) \| 189 \| Uncharacterized protein \| Uncharacterized protein; uniclust \| UniRef100\_UPI0009AEFA08 \| 98.6 \| 3.4e-10 \| 6.2e-16 \| 108.1 \| 61 \| (36, 96) \| 350 \| (113, 177) \| 253 \| hypothetical protein \| hypothetical protein; uniclust \| UniRef100\_UPI000A5CD1B2 \| 98.6 \| 4.4e-10 \| 8e-16 \| 105.8 \| 51 \| (181, 231) \| 350 \| (171, 222) \| 223 \| Lar family restriction alleviation protein \| Lar family restriction alleviation protein; uniclust \| UniRef100\_UPI00044908D2 \| 98.6 \| 3.3e-10 \| 8.3e-16 \| 115.0 \| 59 \| (80, 142) \| 350 \| (78, 140) \| 292 \| DUF3850 domain-containing protein \| DUF3850 domain-containing protein; uniclust \| UniRef100\_UPI0007356114 \| 98.6 \| 4.4e-10 \| 9.3e-16 \| 113.6 \| 63 \| (168, 235) \| 350 \| (232, 296) \| 318 \| hypothetical protein \| hypothetical protein; uniclust \| UniRef100\_UPI000DCAFC4D \| 98.6 \| 5.3e-10 \| 9.9e-16 \| 105.0 \| 56 \| (176, 231) \| 350 \| (155, 211) \| 213 \| hypothetical protein \| hypothetical protein; uniclust \| UniRef100\_UPI0003ED00DA \| 98.6 \| 5.2e-10 \| 1.3e-15 \| 114.1 \| 60 \| (168, 231) \| 350 \| (243, 304) \| 307 \| hypothetical protein \| hypothetical protein; uniclust \| UniRef100\_A0A5V5DTZ8 \| 98.6 \| 6.9e-10 \| 1.4e-15 \| 111.1 \| 60 \| (168, 231) \| 350 \| (246, 316) \| 318 \| DUF551 domain-containing protein \| DUF551 domain-containing protein; uniclust \| UniRef100\_UPI00083CDA3F \| 98.5 \| 8.8e-10 \| 1.6e-15 \| 104.0 \| 85 \| (1, 86) \| 350 \| (1, 85) \| 224 \| hypothetical protein \| hypothetical protein; uniclust \| UniRef100\_UPI001F14D20D \| 98.5 \| 9.8e-10 \| 1.8e-15 \| 101.7 \| 76 \| (15, 90) \| 350 \| (58, 143) \| 192 \| hypothetical protein \| hypothetical protein; uniclust \| UniRef100\_UPI0009ACDD1E \| 98.5 \| 9.9e-10 \| 1.9e-15 \| 105.3 \| 126 \| (17, 144) \| 350 \| (51, 202) \| 228 \| hypothetical protein \| hypothetical protein; uniclust \| UniRef100\_G9Z9C5 \| 98.5 \| 1.3e-09 \| 2.4e-15 \| 103.7 \| 59 \| (173, 231) \| 350 \| (177, 236) \| 239 \| Uncharacterized protein \| Uncharacterized protein; uniclust \| UniRef100\_UPI001F118484 \| 98.5 \| 1.3e-09 \| 2.4e-15 \| 109.8 \| 56 \| (176, 231) \| 350 \| (354, 410) \| 413 \| hypothetical protein \| hypothetical protein; uniclust \| UniRef100\_A0A376JIE3 \| 98.4 \| 2e-09 \| 4.8e-15 \| 110.9 \| 74 \| (168, 245) \| 350 \| (223, 306) \| 330 \| Putative prophage protein \| Putative prophage protein; uniclust \| UniRef100\_UPI0001BDC4F6 \| 98.4 \| 3.6e-09 \| 7e-15 \| 99.7 \| 56 \| (170, 229) \| 350 \| (129, 190) \| 200 \| DUF551 domain-containing protein \| DUF551 domain-containing protein; uniclust \| UniRef100\_UPI0020177EC0 \| 98.4 \| 4.4e-09 \| 8.4e-15 \| 103.7 \| 57 \| (176, 232) \| 350 \| (229, 286) \| 289 \| hypothetical protein \| hypothetical protein; uniclust \| UniRef100\_UPI001CF306BC \| 98.4 \| 4.8e-09 \| 1e-14 \| 106.8 \| 138 \| (82, 232) \| 350 \| (73, 244) \| 334 \| DUF551 domain-containing protein \| DUF551 domain-containing protein; uniclust \| UniRef100\_A0A8B3JTV7 \| 98.3 \| 6.4e-09 \| 1.2e-14 \| 102.2 \| 52 \| (180, 231) \| 350 \| (254, 306) \| 309 \| Uncharacterized protein \| Uncharacterized protein; uniclust \| UniRef100\_UPI002073060C \| 98.3 \| 6.8e-09 \| 1.2e-14 \| 100.1 \| 52 \| (180, 231) \| 350 \| (204, 256) \| 258 \| hypothetical protein \| hypothetical protein; uniclust \| UniRef100\_A0A024HVZ1 \| 98.3 \| 5.1e-09 \| 1.3e-14 \| 109.4 \| 25 \| (73, 97) \| 350 \| (112, 136) \| 345 \| DUF551 domain-containing protein \| DUF551 domain-containing protein; uniclust \| UniRef100\_UPI0021CE8872 \| 98.3 \| 8.5e-09 \| 1.6e-14 \| 102.4 \| 52 \| (180, 231) \| 350 \| (283, 335) \| 337 \| hypothetical protein \| hypothetical protein; uniclust \| UniRef100\_A0A7I7AAG6 \| 98.3 \| 9.2e-09 \| 2.3e-14 \| 103.2 \| 76 \| (153, 232) \| 350 \| (143, 225) \| 247 \| DUF551 domain-containing protein \| DUF551 domain-containing protein; uniclust \| UniRef100\_UPI001D009ECB \| 98.2 \| 1.5e-08 \| 2.7e-14 \| 94.4 \| 52 \| (180, 231) \| 350 \| (137, 189) \| 190 \| hypothetical protein \| hypothetical protein; uniclust \| UniRef100\_A0A2S4QDE0 \| 98.2 \| 1e-08 \| 2.7e-14 \| 105.8 \| 66 \| (158, 234) \| 350 \| (221, 290) \| 308 \| DUF551 domain-containing protein \| DUF551 domain-containing protein; uniclust \| UniRef100\_UPI00036C2CC1 \| 98.2 \| 1.3e-08 \| 2.8e-14 \| 100.9 \| 150 \| (74, 231) \| 350 \| (71, 245) \| 250 \| DUF551 domain-containing protein \| DUF551 domain-containing protein; uniclust \| UniRef100\_UPI001D0D4EBB \| 98.2 \| 1.6e-08 \| 3.1e-14 \| 97.4 \| 147 \| (80, 232) \| 350 \| (26, 230) \| 233 \| DUF551 domain-containing protein \| DUF551 domain-containing protein; uniclust \| UniRef100\_UPI00073553D8 \| 98.2 \| 1.7e-08 \| 3.2e-14 \| 98.4 \| 55 \| (176, 230) \| 350 \| (194, 249) \| 251 \| hypothetical protein \| hypothetical protein; uniclust \| UniRef100\_A0A0A0FB64 \| 98.2 \| 1.2e-08 \| 3.3e-14 \| 104.9 \| 71 \| (155, 234) \| 350 \| (193, 267) \| 297 \| Uncharacterized protein \| Uncharacterized protein; uniclust \| UniRef100\_A0A0F1A3D6 \| 98.2 \| 1.8e-08 \| 4.1e-14 \| 98.5 \| 176 \| (1, 232) \| 350 \| (1, 183) \| 211 \| Eaa protein \| Eaa protein; uniclust \| UniRef100\_A0A482J7X3 \| 98.2 \| 1.6e-08 \| 4.6e-14 \| 102.0 \| 39 \| (153, 195) \| 350 \| (138, 177) \| 247 \| Eaa1 \| Eaa1; uniclust \| UniRef100\_A0A837KEC1 \| 98.2 \| 2.1e-08 \| 4.6e-14 \| 101.0 \| 152 \| (75, 235) \| 350 \| (97, 271) \| 283 \| DUF551 domain-containing protein \| DUF551 domain-containing protein; uniclust \| UniRef100\_UPI0004D9BA72 \| 98.2 \| 2.4e-08 \| 5.7e-14 \| 104.7 \| 77 \| (153, 233) \| 350 \| (261, 348) \| 362 \| DUF551 domain-containing protein \| DUF551 domain-containing protein; uniclust \| UniRef100\_UPI001E39654B \| 98.2 \| 2.9e-08 \| 5.8e-14 \| 99.1 \| 63 \| (155, 227) \| 350 \| (178, 242) \| 286 \| hypothetical protein \| hypothetical protein; uniclust \| UniRef100\_A0A4C9H8G9 \| 98.2 \| 2.5e-08 \| 6.1e-14 \| 102.5 \| 36 \| (155, 194) \| 350 \| (137, 173) \| 295 \| Putative phage protein \| Putative phage protein; uniclust \| UniRef100\_UPI0021F4EEC0 \| 98.1 \| 4.8e-08 \| 8.9e-14 \| 98.7 \| 48 \| (182, 229) \| 350 \| (334, 382) \| 384 \| ASCH/PUA domain-containing protein \| ASCH/PUA domain-containing protein; uniclust \| UniRef100\_UPI0022F09B78 \| 98.1 \| 5.3e-08 \| 9.8e-14 \| 90.9 \| 60 \| (153, 221) \| 350 \| (125, 186) \| 189 \| hypothetical protein \| hypothetical protein; uniclust \| UniRef100\_UPI00069C98EA \| 98.1 \| 4.8e-08 \| 1e-13 \| 96.4 \| 75 \| (153, 231) \| 350 \| (152, 236) \| 240 \| DUF551 domain-containing protein \| DUF551 domain-containing protein; uniclust \| UniRef100\_A0A0T9RK48 \| 98.1 \| 3.9e-08 \| 1.1e-13 \| 102.3 \| 42 \| (190, 231) \| 350 \| (263, 305) \| 307 \| Uncharacterized protein \| Uncharacterized protein; uniclust \| UniRef100\_A0A0D8VD89 \| 98.0 \| 6.2e-08 \| 1.5e-13 \| 102.0 \| 129 \| (82, 233) \| 350 \| (130, 270) \| 354 \| DUF551 domain-containing protein \| DUF551 domain-containing protein; uniclust \| UniRef100\_A0A0L1BMB8 \| 98.0 \| 7.4e-08 \| 1.9e-13 \| 97.1 \| 85 \| (78, 193) \| 350 \| (103, 188) \| 241 \| DUF551 domain-containing protein \| DUF551 domain-containing protein; uniclust \| UniRef100\_A0A1E7Z2Q0 \| 98.0 \| 7.6e-08 \| 1.9e-13 \| 97.3 \| 76 \| (153, 233) \| 350 \| (123, 206) \| 249 \| DUF551 domain-containing protein \| DUF551 domain-containing protein; uniclust \| UniRef100\_A0A377C8S0 \| 98.0 \| 8.9e-08 \| 2.3e-13 \| 99.5 \| 67 \| (156, 233) \| 350 \| (237, 309) \| 311 \| Eaa protein \| Eaa protein; uniclust \| UniRef100\_UPI00058238E0 \| 98.0 \| 1e-07 \| 2.4e-13 \| 100.1 \| 70 \| (160, 238) \| 350 \| (260, 337) \| 360 \| DUF551 domain-containing protein \| DUF551 domain-containing protein; uniclust \| UniRef100\_UPI0022BF4606 \| 98.0 \| 1.4e-07 \| 2.5e-13 \| 84.5 \| 56 \| (176, 231) \| 350 \| (75, 131) \| 136 \| hypothetical protein \| hypothetical protein; uniclust \| UniRef100\_A0A7H5A556 \| 98.0 \| 1e-07 \| 2.7e-13 \| 94.8 \| 23 \| (75, 97) \| 350 \| (86, 108) \| 218 \| Eaa protein \| Eaa protein; uniclust \| UniRef100\_A0A1Z3N2X6 \| 97.9 \| 1.7e-07 \| 3.4e-13 \| 93.2 \| 57 \| (34, 90) \| 350 \| (95, 152) \| 263 \| Ead/Ea22-like family protein \| Ead/Ea22-like family protein; uniclust \| UniRef100\_A0A348D9R6 \| 97.9 \| 1.9e-07 \| 3.5e-13 \| 91.1 \| 54 \| (176, 229) \| 350 \| (209, 263) \| 265 \| Phage protein \| Phage protein; uniclust \| UniRef100\_A0A0F6TJ66 \| 97.9 \| 1.7e-07 \| 3.9e-13 \| 96.5 \| 78 \| (158, 244) \| 350 \| (159, 245) \| 295 \| DUF551 domain-containing protein \| DUF551 domain-containing protein; uniclust \| UniRef100\_A0A0J1LU56 \| 97.9 \| 1.8e-07 \| 4e-13 \| 97.5 \| 73 \| (153, 235) \| 350 \| (257, 333) \| 357 \| Phage protein \| Phage protein; uniclust \| UniRef100\_UPI000A3D7A99 \| 97.9 \| 2.1e-07 \| 4.5e-13 \| 95.5 \| 62 \| (168, 233) \| 350 \| (232, 299) \| 320 \| DUF550 domain-containing protein \| DUF550 domain-containing protein; uniclust \| UniRef100\_UPI00068B16CB \| 97.9 \| 2.1e-07 \| 5.2e-13 \| 93.1 \| 94 \| (132, 234) \| 350 \| (95, 211) \| 232 \| DUF551 domain-containing protein \| DUF551 domain-containing protein; uniclust \| UniRef100\_A0A096XER3 \| 97.9 \| 2.3e-07 \| 6e-13 \| 97.9 \| 37 \| (155, 200) \| 350 \| (254, 290) \| 345 \| dTDP-6-deoxy-L-hexose 3-O-methyltransferase \| dTDP-6-deoxy-L-hexose 3-O-methyltransferase; uniclust \| UniRef100\_A0A7Z1J7P7 \| 97.9 \| 2.8e-07 \| 6.2e-13 \| 90.1 \| 63 \| (168, 234) \| 350 \| (112, 181) \| 207 \| DUF551 domain-containing protein \| DUF551 domain-containing protein; uniclust \| UniRef100\_A0A0J1LWP4 \| 97.8 \| 2.7e-07 \| 6.9e-13 \| 92.5 \| 67 \| (153, 232) \| 350 \| (137, 206) \| 231 \| Lar family restriction alleviation protein \| Lar family restriction alleviation protein; uniclust \| UniRef100\_A0A2X3CAE9 \| 97.8 \| 2.6e-07 \| 7.2e-13 \| 95.9 \| 61 \| (168, 233) \| 350 \| (190, 260) \| 295 \| DUF551 domain-containing protein \| DUF551 domain-containing protein; uniclust \| UniRef100\_A0A0E1CIM4 \| 97.8 \| 3.7e-07 \| 8.8e-13 \| 91.8 \| 23 \| (168, 194) \| 350 \| (173, 195) \| 241 \| DUF551 domain-containing protein \| DUF551 domain-containing protein; uniclust \| UniRef100\_UPI001F23EE67 \| 97.8 \| 5.1e-07 \| 9.4e-13 \| 93.1 \| 56 \| (176, 231) \| 350 \| (375, 431) \| 436 \| hypothetical protein \| hypothetical protein; uniclust \| UniRef100\_UPI000CDDAC1C \| 97.8 \| 6e-07 \| 1.1e-12 \| 85.6 \| 51 \| (181, 231) \| 350 \| (143, 194) \| 196 \| hypothetical protein \| hypothetical protein; uniclust \| UniRef100\_A0A377YIT5 \| 97.7 \| 5.1e-07 \| 1.4e-12 \| 92.1 \| 15 \| (81, 95) \| 350 \| (92, 106) \| 255 \| Eaa1 \| Eaa1; uniclust \| UniRef100\_UPI001F2E8690 \| 97.7 \| 8.4e-07 \| 1.6e-12 \| 71.1 \| 59 \| (25, 88) \| 350 \| (4, 62) \| 66 \| hypothetical protein \| hypothetical protein; uniclust \| UniRef100\_A0A2X4T8E3 \| 97.7 \| 7.6e-07 \| 1.6e-12 \| 87.8 \| 70 \| (153, 231) \| 350 \| (137, 217) \| 221 \| Phage protein \| Phage protein; uniclust \| UniRef100\_UPI000501F9A2 \| 97.7 \| 7.3e-07 \| 1.6e-12 \| 91.6 \| 68 \| (156, 232) \| 350 \| (194, 270) \| 293 \| hypothetical protein \| hypothetical protein; uniclust \| UniRef100\_A0A485A5N0 \| 97.7 \| 7e-07 \| 1.8e-12 \| 89.5 \| 33 \| (60, 95) \| 350 \| (67, 99) \| 223 \| DUF551 domain-containing protein \| DUF551 domain-containing protein; uniclust \| UniRef100\_A0A378CDL6 \| 97.7 \| 8.2e-07 \| 1.9e-12 \| 91.8 \| 137 \| (73, 233) \| 350 \| (136, 292) \| 306 \| Protein of uncharacterized function (DUF551) \| Protein of uncharacterized function (DUF551); uniclust \| UniRef100\_UPI000297C7C1 \| 97.7 \| 8.2e-07 \| 1.9e-12 \| 87.4 \| 87 \| (79, 177) \| 350 \| (70, 165) \| 205 \| hypothetical protein \| hypothetical protein; uniclust \| UniRef100\_A0A2S1Y6Q1 \| 97.7 \| 1e-06 \| 2.3e-12 \| 86.2 \| 75 \| (155, 234) \| 350 \| (92, 183) \| 196 \| DUF551 domain-containing protein \| DUF551 domain-containing protein; uniclust \| UniRef100\_A0A5T7YCH7 \| 97.7 \| 1.2e-06 \| 2.5e-12 \| 80.1 \| 66 \| (158, 232) \| 350 \| (56, 132) \| 133 \| DUF551 domain-containing protein \| DUF551 domain-containing protein; uniclust \| UniRef100\_A0A1S0ZHQ4 \| 97.7 \| 1.1e-06 \| 2.5e-12 \| 88.2 \| 78 \| (79, 178) \| 350 \| (91, 171) \| 229 \| DUF551 domain-containing protein \| DUF551 domain-containing protein; uniclust \| UniRef100\_UPI000E2D2FA1 \| 97.6 \| 1.3e-06 \| 2.6e-12 \| 92.0 \| 78 \| (85, 164) \| 350 \| (128, 234) \| 395 \| DUF551 domain-containing protein \| DUF551 domain-containing protein; uniclust \| UniRef100\_UPI0008A4C7D4 \| 97.6 \| 1.4e-06 \| 2.9e-12 \| 89.9 \| 102 \| (77, 178) \| 350 \| (125, 279) \| 347 \| DUF3850 domain-containing protein \| DUF3850 domain-containing protein; uniclust \| UniRef100\_UPI0002413D61 \| 97.6 \| 1.8e-06 \| 3.6e-12 \| 84.9 \| 66 \| (157, 231) \| 350 \| (139, 216) \| 221 \| DUF551 domain-containing protein \| DUF551 domain-containing protein; uniclust \| UniRef100\_A0A2T8SVC1 \| 97.6 \| 1.6e-06 \| 3.6e-12 \| 79.7 \| 75 \| (149, 232) \| 350 \| (32, 112) \| 126 \| DUF551 domain-containing protein (Fragment) \| DUF551 domain-containing protein (Fragment); uniclust \| UniRef100\_A0A7T8LMQ9 \| 97.5 \| 2.6e-06 \| 4.9e-12 \| 82.9 \| 54 \| (42, 97) \| 350 \| (30, 88) \| 219 \| DUF551 domain-containing protein \| DUF551 domain-containing protein; uniclust \| UniRef100\_A0A0K4M1A2 \| 97.5 \| 2.8e-06 \| 5.2e-12 \| 76.2 \| 66 \| (25, 96) \| 350 \| (4, 69) \| 127 \| Phage protein \| Phage protein; uniclust \| UniRef100\_A0A0F7JC87 \| 97.5 \| 2.2e-06 \| 5.2e-12 \| 89.1 \| 86 \| (1, 96) \| 350 \| (42, 136) \| 303 \| Eaa protein \| Eaa protein; uniclust \| UniRef100\_UPI001CC98A5F \| 97.5 \| 2.6e-06 \| 5.2e-12 \| 86.6 \| 59 \| (73, 131) \| 350 \| (76, 138) \| 293 \| DUF551 domain-containing protein \| DUF551 domain-containing protein; uniclust \| UniRef100\_UPI00028301C4 \| 97.5 \| 2.4e-06 \| 5.5e-12 \| 91.4 \| 76 \| (153, 232) \| 350 \| (290, 377) \| 392 \| DUF551 domain-containing protein \| DUF551 domain-containing protein; uniclust \| UniRef100\_A0A702BB40 \| 97.5 \| 2.4e-06 \| 5.7e-12 \| 85.1 \| 86 \| (77, 187) \| 350 \| (77, 163) \| 214 \| Eaa protein \| Eaa protein; uniclust \| UniRef100\_UPI001D125E14 \| 97.5 \| 3.5e-06 \| 6.8e-12 \| 84.0 \| 137 \| (89, 231) \| 350 \| (80, 257) \| 260 \| DUF551 domain-containing protein \| DUF551 domain-containing protein; uniclust \| UniRef100\_UPI001F14CB3B \| 97.5 \| 3.7e-06 \| 6.8e-12 \| 77.4 \| 64 \| (168, 235) \| 350 \| (68, 134) \| 154 \| DUF551 domain-containing protein \| DUF551 domain-containing protein; uniclust \| UniRef100\_A0A088F853 \| 97.5 \| 3.9e-06 \| 7.9e-12 \| 73.8 \| 40 \| (1, 40) \| 350 \| (12, 53) \| 100 \| Uncharacterized protein \| Uncharacterized protein; uniclust \| UniRef100\_B5Y0H2 \| 97.5 \| 3.2e-06 \| 8e-12 \| 90.8 \| 58 \| (170, 232) \| 350 \| (325, 384) \| 387 \| DUF551 domain-containing protein \| DUF551 domain-containing protein; uniclust \| UniRef100\_A0A6C9E065 \| 97.5 \| 4.3e-06 \| 8.3e-12 \| 76.0 \| 49 \| (183, 231) \| 350 \| (78, 127) \| 129 \| Uncharacterized protein \| Uncharacterized protein; uniclust \| UniRef100\_A0A3Q9UB38 \| 97.5 \| 4e-06 \| 8.5e-12 \| 85.3 \| 97 \| (72, 178) \| 350 \| (81, 184) \| 271 \| Uncharacterized protein \| Uncharacterized protein; uniclust \| UniRef100\_A0A1X3LMB4 \| 97.4 \| 4.2e-06 \| 1e-11 \| 84.6 \| 28 \| (157, 193) \| 350 \| (134, 161) \| 230 \| Eaa protein \| Eaa protein; uniclust \| UniRef100\_A0A7U9Q9Z2 \| 97.4 \| 5.3e-06 \| 1.2e-11 \| 88.1 \| 23 \| (168, 194) \| 350 \| (297, 319) \| 369 \| Putative phage protein \| Putative phage protein; uniclust \| UniRef100\_A0A5U9SIS9 \| 97.4 \| 6.9e-06 \| 1.4e-11 \| 80.5 \| 55 \| (42, 96) \| 350 \| (26, 90) \| 207 \| Eaa protein \| Eaa protein; uniclust \| UniRef100\_A0A9E7BH61 \| 97.4 \| 6.4e-06 \| 1.4e-11 \| 83.7 \| 53 \| (183, 236) \| 350 \| (174, 236) \| 255 \| Uncharacterized protein \| Uncharacterized protein; uniclust \| UniRef100\_A0A7I7AAS7 \| 97.4 \| 6.2e-06 \| 1.5e-11 \| 82.5 \| 89 \| (1, 97) \| 350 \| (3, 93) \| 210 \| Uncharacterized protein \| Uncharacterized protein; uniclust \| UniRef100\_A0A2U3EYJ7 \| 97.3 \| 8.1e-06 \| 1.6e-11 \| 76.7 \| 67 \| (156, 231) \| 350 \| (81, 152) \| 155 \| DUF551 domain-containing protein \| DUF551 domain-containing protein; uniclust \| UniRef100\_A0A2X3IQU6 \| 97.3 \| 5.8e-06 \| 1.7e-11 \| 89.8 \| 65 \| (168, 236) \| 350 \| (274, 344) \| 395 \| Eaa1 \| Eaa1; uniclust \| UniRef100\_A0A403N2V7 \| 97.3 \| 8.2e-06 \| 1.7e-11 \| 83.1 \| 21 \| (153, 178) \| 350 \| (187, 207) \| 274 \| Uncharacterized protein \| Uncharacterized protein; uniclust \| UniRef100\_A0A2W5GG89 \| 97.3 \| 8.2e-06 \| 1.8e-11 \| 74.8 \| 50 \| (155, 213) \| 350 \| (35, 84) \| 119 \| Uncharacterized protein \| Uncharacterized protein; uniclust \| UniRef100\_UPI00069F21E5 \| 97.3 \| 8.5e-06 \| 1.9e-11 \| 74.4 \| 63 \| (159, 231) \| 350 \| (48, 114) \| 116 \| DUF551 domain-containing protein \| DUF551 domain-containing protein; uniclust \| UniRef100\_UPI001D0F47EE \| 97.3 \| 9.8e-06 \| 1.9e-11 \| 77.1 \| 18 \| (79, 96) \| 350 \| (15, 32) \| 176 \| DUF551 domain-containing protein \| DUF551 domain-containing protein; uniclust \| UniRef100\_A0A0M2KI41 \| 97.3 \| 8.5e-06 \| 2e-11 \| 79.2 \| 69 \| (159, 236) \| 350 \| (99, 171) \| 172 \| DUF551 domain-containing protein \| DUF551 domain-containing protein; uniclust \| UniRef100\_UPI001BDF9013 \| 97.3 \| 1.1e-05 \| 2.1e-11 \| 72.0 \| 68 \| (154, 231) \| 350 \| (50, 121) \| 122 \| DUF551 domain-containing protein \| DUF551 domain-containing protein; uniclust \| UniRef100\_A0A5Q9LHJ5 \| 97.3 \| 7.8e-06 \| 2.2e-11 \| 86.2 \| 75 \| (153, 232) \| 350 \| (222, 303) \| 306 \| Uncharacterized protein \| Uncharacterized protein; uniclust \| UniRef100\_A0A3F3IH09 \| 97.3 \| 1.2e-05 \| 2.6e-11 \| 76.5 \| 73 \| (152, 231) \| 350 \| (71, 155) \| 158 \| Protein Eaa \| Protein Eaa; uniclust \| UniRef100\_UPI000B422C02 \| 97.3 \| 1.4e-05 \| 2.7e-11 \| 70.8 \| 44 \| (182, 225) \| 350 \| (71, 115) \| 115 \| hypothetical protein \| hypothetical protein; uniclust \| UniRef100\_A0A085ASF4 \| 97.3 \| 1.1e-05 \| 2.7e-11 \| 87.5 \| 32 \| (56, 96) \| 350 \| (187, 218) \| 403 \| DUF551 domain-containing protein \| DUF551 domain-containing protein; uniclust \| UniRef100\_UPI000CFB54C0 \| 97.2 \| 1.5e-05 \| 3e-11 \| 78.6 \| 105 \| (55, 194) \| 350 \| (64, 171) \| 221 \| DUF551 domain-containing protein \| DUF551 domain-containing protein; uniclust \| UniRef100\_UPI000AED1C5F \| 97.2 \| 1.7e-05 \| 3.1e-11 \| 75.1 \| 129 \| (78, 230) \| 350 \| (34, 180) \| 182 \| DUF550 domain-containing protein \| DUF550 domain-containing protein; uniclust \| UniRef100\_A0A377WN62 \| 97.2 \| 1.2e-05 \| 3.3e-11 \| 88.2 \| 75 \| (153, 231) \| 350 \| (300, 383) \| 422 \| Eaa1 \| Eaa1; uniclust \| UniRef100\_A0A1X3LI39 \| 97.2 \| 1.5e-05 \| 3.4e-11 \| 84.1 \| 61 \| (168, 232) \| 350 \| (203, 276) \| 343 \| Putative bacteriophage protein \| Putative bacteriophage protein; uniclust \| UniRef100\_UPI0020D2028B \| 97.2 \| 2.2e-05 \| 4e-11 \| 79.8 \| 59 \| (169, 231) \| 350 \| (260, 329) \| 330 \| DUF551 domain-containing protein \| DUF551 domain-containing protein; uniclust \| UniRef100\_A0A7I6XQ80 \| 97.1 \| 2.5e-05 \| 5.6e-11 \| 81.7 \| 61 \| (168, 237) \| 350 \| (215, 281) \| 307 \| DUF551 domain-containing protein \| DUF551 domain-containing protein; uniclust \| UniRef100\_A0A7H9JWU5 \| 97.1 \| 2.7e-05 \| 6.2e-11 \| 81.3 \| 26 \| (153, 178) \| 350 \| (195, 236) \| 299 \| DUF551 domain-containing protein \| DUF551 domain-containing protein; uniclust \| UniRef100\_A0A641J2S1 \| 97.1 \| 2.8e-05 \| 6.2e-11 \| 79.5 \| 30 \| (168, 201) \| 350 \| (185, 214) \| 256 \| DUF551 domain-containing protein \| DUF551 domain-containing protein; uniclust \| UniRef100\_UPI001ABB5B0F \| 97.1 \| 3.5e-05 \| 6.5e-11 \| 79.2 \| 52 \| (170, 225) \| 350 \| (301, 353) \| 360 \| hypothetical protein \| hypothetical protein; uniclust \| UniRef100\_UPI001B3928F5 \| 97.1 \| 3.5e-05 \| 6.5e-11 \| 78.8 \| 60 \| (37, 96) \| 350 \| (137, 201) \| 343 \| hypothetical protein \| hypothetical protein; uniclust \| UniRef100\_A0A7G3ES10 \| 97.1 \| 2.7e-05 \| 6.7e-11 \| 78.4 \| 85 \| (1, 96) \| 350 \| (1, 88) \| 213 \| DUF551 domain-containing protein \| DUF551 domain-containing protein; uniclust \| UniRef100\_UPI000E4CA42D \| 97.1 \| 4e-05 \| 7.4e-11 \| 78.7 \| 50 \| (181, 230) \| 350 \| (203, 253) \| 353 \| hypothetical protein \| hypothetical protein; uniclust \| UniRef100\_UPI001D0D554A \| 97.0 \| 4.5e-05 \| 8.3e-11 \| 73.4 \| 64 \| (156, 230) \| 350 \| (133, 197) \| 199 \| DUF551 domain-containing protein \| DUF551 domain-containing protein; uniclust \| UniRef100\_UPI0007EBD3DF \| 97.0 \| 3.8e-05 \| 8.4e-11 \| 80.7 \| 81 \| (80, 178) \| 350 \| (157, 240) \| 315 \| hypothetical protein \| hypothetical protein; uniclust \| UniRef100\_UPI001CF3EDA8 \| 97.0 \| 4.6e-05 \| 8.5e-11 \| 70.1 \| 67 \| (155, 230) \| 350 \| (63, 140) \| 143 \| DUF551 domain-containing protein \| DUF551 domain-containing protein; uniclust \| UniRef100\_UPI001587EB33 \| 97.0 \| 4.9e-05 \| 9e-11 \| 68.3 \| 51 \| (176, 226) \| 350 \| (62, 113) \| 121 \| hypothetical protein \| hypothetical protein; uniclust \| UniRef100\_UPI000808C4DE \| 97.0 \| 4.4e-05 \| 9.1e-11 \| 80.4 \| 144 \| (76, 236) \| 350 \| (175, 331) \| 342 \| DUF551 domain-containing protein \| DUF551 domain-containing protein; uniclust \| UniRef100\_UPI00079C252A \| 96.9 \| 6.6e-05 \| 1.3e-10 \| 71.1 \| 88 \| (1, 96) \| 350 \| (1, 90) \| 150 \| hypothetical protein \| hypothetical protein; uniclust \| UniRef100\_UPI0006597CA0 \| 96.9 \| 6.8e-05 \| 1.4e-10 \| 77.6 \| 84 \| (79, 164) \| 350 \| (142, 240) \| 287 \| DUF551 domain-containing protein \| DUF551 domain-containing protein; uniclust \| UniRef100\_A0A443WQQ1 \| 96.9 \| 5.8e-05 \| 1.5e-10 \| 78.8 \| 85 \| (78, 178) \| 350 \| (109, 212) \| 270 \| Uncharacterized protein \| Uncharacterized protein; uniclust \| UniRef100\_A0A2T3SVJ2 \| 96.9 \| 7.4e-05 \| 1.6e-10 \| 77.0 \| 60 \| (168, 231) \| 350 \| (200, 271) \| 275 \| DUF551 domain-containing protein \| DUF551 domain-containing protein; uniclust \| UniRef100\_UPI000AC50A34 \| 96.9 \| 7.6e-05 \| 1.7e-10 \| 77.2 \| 58 \| (169, 232) \| 350 \| (203, 265) \| 268 \| DUF551 domain-containing protein \| DUF551 domain-containing protein; uniclust \| UniRef100\_UPI001E47FED3 \| 96.9 \| 0.00011 \| 2e-10 \| 68.8 \| 70 \| (153, 231) \| 350 \| (74, 154) \| 156 \| DUF551 domain-containing protein \| DUF551 domain-containing protein; uniclust \| UniRef100\_A0A3T2UUV1 \| 96.8 \| 9.8e-05 \| 2.2e-10 \| 75.5 \| 65 \| (158, 232) \| 350 \| (167, 239) \| 242 \| DUF551 domain-containing protein \| DUF551 domain-containing protein; uniclust \| UniRef100\_UPI00080AA30A \| 96.8 \| 0.00012 \| 2.2e-10 \| 76.2 \| 52 \| (176, 227) \| 350 \| (275, 327) \| 384 \| hypothetical protein \| hypothetical protein; uniclust \| UniRef100\_UPI000DCD3321 \| 96.8 \| 0.00011 \| 2.3e-10 \| 73.9 \| 80 \| (148, 232) \| 350 \| (128, 222) \| 226 \| hypothetical protein \| hypothetical protein; uniclust \| UniRef100\_A0A5C7BMG0 \| 96.8 \| 0.00011 \| 2.3e-10 \| 74.1 \| 24 \| (168, 195) \| 350 \| (134, 157) \| 214 \| DUF551 domain-containing protein \| DUF551 domain-containing protein; uniclust \| UniRef100\_UPI001D114CB1 \| 96.8 \| 0.00012 \| 2.4e-10 \| 78.1 \| 141 \| (77, 231) \| 350 \| (136, 314) \| 383 \| DUF3850 domain-containing protein \| DUF3850 domain-containing protein; uniclust \| UniRef100\_S5MU58 \| 96.8 \| 0.00013 \| 2.4e-10 \| 74.1 \| 55 \| (42, 98) \| 350 \| (115, 174) \| 303 \| Phage EaD protein \| Phage EaD protein; uniclust \| UniRef100\_A0A2T4Y165 \| 96.8 \| 8.9e-05 \| 2.4e-10 \| 79.8 \| 22 \| (169, 194) \| 350 \| (248, 269) \| 337 \| DUF551 domain-containing protein \| DUF551 domain-containing protein; uniclust \| UniRef100\_A0A3V3HL53 \| 96.8 \| 0.00015 \| 2.8e-10 \| 71.7 \| 33 \| (153, 195) \| 350 \| (156, 188) \| 240 \| DUF551 domain-containing protein \| DUF551 domain-containing protein; uniclust \| UniRef100\_UPI00168151F6 \| 96.8 \| 0.00015 \| 2.8e-10 \| 69.9 \| 44 \| (176, 219) \| 350 \| (140, 183) \| 195 \| hypothetical protein \| hypothetical protein; uniclust \| UniRef100\_UPI000D745A46 \| 96.8 \| 0.00015 \| 3e-10 \| 71.9 \| 113 \| (80, 205) \| 350 \| (37, 161) \| 197 \| hypothetical protein \| hypothetical protein; uniclust \| UniRef100\_UPI000685C591 \| 96.7 \| 0.00015 \| 3.2e-10 \| 71.8 \| 56 \| (41, 96) \| 350 \| (98, 154) \| 193 \| hypothetical protein \| hypothetical protein; uniclust \| UniRef100\_UPI0015E53E8C \| 96.7 \| 0.00019 \| 3.4e-10 \| 62.1 \| 39 \| (1, 39) \| 350 \| (1, 39) \| 91 \| hypothetical protein \| hypothetical protein; uniclust \| UniRef100\_UPI001920B9A5 \| 96.7 \| 0.00019 \| 3.5e-10 \| 71.5 \| 54 \| (42, 95) \| 350 \| (84, 137) \| 254 \| hypothetical protein \| hypothetical protein; uniclust \| UniRef100\_A0A377X5V4 \| 96.7 \| 0.00014 \| 3.6e-10 \| 81.4 \| 76 \| (153, 232) \| 350 \| (371, 453) \| 475 \| Protein of uncharacterized function (DUF551) \| Protein of uncharacterized function (DUF551); uniclust \| UniRef100\_A0A636IGV6 \| 96.7 \| 0.00019 \| 3.7e-10 \| 74.1 \| 52 \| (43, 96) \| 350 \| (90, 146) \| 312 \| DUF551 domain-containing protein \| DUF551 domain-containing protein; uniclust \| UniRef100\_A0A3Y7SQI3 \| 96.7 \| 0.00019 \| 3.8e-10 \| 64.3 \| 29 \| (155, 192) \| 350 \| (21, 49) \| 100 \| DUF551 domain-containing protein \| DUF551 domain-containing protein; uniclust \| UniRef100\_A0A2H4FNA9 \| 96.7 \| 0.00019 \| 3.9e-10 \| 73.2 \| 78 \| (153, 234) \| 350 \| (155, 240) \| 250 \| DUF551 domain-containing protein \| DUF551 domain-containing protein; uniclust \| UniRef100\_A0A286P6T1 \| 96.7 \| 0.0002 \| 3.9e-10 \| 70.1 \| 52 \| (168, 223) \| 350 \| (106, 158) \| 197 \| Uncharacterized protein \| Uncharacterized protein; uniclust \| UniRef100\_A0A4R7FZ57 \| 96.7 \| 0.0002 \| 4.1e-10 \| 73.9 \| 48 \| (176, 223) \| 350 \| (225, 273) \| 287 \| Replication restart DNA helicase PriA \| Replication restart DNA helicase PriA; uniclust \| UniRef100\_A0A4P8YP68 \| 96.7 \| 0.00021 \| 4.5e-10 \| 64.0 \| 41 \| (147, 192) \| 350 \| (7, 48) \| 99 \| DUF551 domain-containing protein \| DUF551 domain-containing protein; uniclust \| UniRef100\_A0A0D6ZJB3 \| 96.6 \| 0.00024 \| 5.7e-10 \| 71.1 \| 40 \| (153, 196) \| 350 \| (97, 141) \| 193 \| DUF551 domain-containing protein \| DUF551 domain-containing protein; uniclust \| UniRef100\_UPI0003BBF7AD \| 96.6 \| 0.00032 \| 5.8e-10 \| 74.0 \| 59 \| (168, 230) \| 350 \| (346, 415) \| 417 \| DUF551 domain-containing protein \| DUF551 domain-containing protein; uniclust \| UniRef100\_UPI001B2924A7 \| 96.6 \| 0.00032 \| 5.9e-10 \| 69.1 \| 40 \| (187, 226) \| 350 \| (182, 222) \| 223 \| hypothetical protein \| hypothetical protein; uniclust \| UniRef100\_UPI000AAEB9B5 \| 96.6 \| 0.00034 \| 6.3e-10 \| 69.3 \| 48 \| (135, 191) \| 350 \| (122, 175) \| 235 \| DUF551 domain-containing protein \| DUF551 domain-containing protein; uniclust \| UniRef100\_UPI0005B32617 \| 96.6 \| 0.00035 \| 6.6e-10 \| 71.9 \| 48 \| (176, 223) \| 350 \| (244, 292) \| 305 \| hypothetical protein \| hypothetical protein; uniclust \| UniRef100\_UPI00038F7A69 \| 96.6 \| 0.00038 \| 6.9e-10 \| 67.5 \| 55 \| (42, 97) \| 350 \| (84, 138) \| 195 \| hypothetical protein \| hypothetical protein; uniclust \| UniRef100\_A0A5I4QGP7 \| 96.5 \| 0.00033 \| 7.4e-10 \| 72.2 \| 72 \| (155, 235) \| 350 \| (149, 229) \| 246 \| DUF551 domain-containing protein \| DUF551 domain-containing protein; uniclust \| UniRef100\_A0A2D2W4X9 \| 96.5 \| 0.00041 \| 7.5e-10 \| 60.6 \| 40 \| (1, 40) \| 350 \| (1, 42) \| 95 \| Uncharacterized protein \| Uncharacterized protein; uniclust \| UniRef100\_A0A543Q1M0 \| 96.5 \| 0.00038 \| 7.5e-10 \| 59.2 \| 36 \| (189, 224) \| 350 \| (27, 63) \| 73 \| Uncharacterized protein \| Uncharacterized protein; uniclust \| UniRef100\_UPI000A38D79B \| 96.5 \| 0.00037 \| 7.6e-10 \| 73.1 \| 80 \| (153, 236) \| 350 \| (190, 294) \| 313 \| DUF551 domain-containing protein \| DUF551 domain-containing protein; uniclust \| UniRef100\_A0A7U6G891 \| 96.5 \| 0.00043 \| 8.2e-10 \| 70.4 \| 59 \| (168, 230) \| 350 \| (199, 264) \| 267 \| DUF551 domain-containing protein \| DUF551 domain-containing protein; uniclust \| UniRef100\_A0A5N3CXX3 \| 96.5 \| 0.00045 \| 8.4e-10 \| 72.0 \| 54 \| (42, 95) \| 350 \| (87, 140) \| 355 \| Phage protein \| Phage protein; uniclust \| UniRef100\_A0A0E1M694 \| 96.5 \| 0.00037 \| 8.7e-10 \| 73.7 \| 36 \| (158, 198) \| 350 \| (204, 239) \| 290 \| DUF551 domain-containing protein \| DUF551 domain-containing protein; uniclust \| UniRef100\_UPI001E2EB100 \| 96.5 \| 0.00048 \| 9.1e-10 \| 69.5 \| 17 \| (80, 96) \| 350 \| (93, 109) \| 254 \| DUF551 domain-containing protein \| DUF551 domain-containing protein; uniclust \| UniRef100\_UPI001D0E6CF5 \| 96.5 \| 0.00048 \| 9.2e-10 \| 71.1 \| 117 \| (77, 204) \| 350 \| (120, 258) \| 302 \| DUF551 domain-containing protein \| DUF551 domain-containing protein; uniclust \| UniRef100\_A0A481W6W9 \| 96.5 \| 0.00052 \| 9.6e-10 \| 55.7 \| 40 \| (1, 40) \| 350 \| (1, 42) \| 61 \| Uncharacterized protein \| Uncharacterized protein; uniclust \| UniRef100\_UPI001C8BD278 \| 96.5 \| 0.00053 \| 9.7e-10 \| 64.7 \| 40 \| (182, 221) \| 350 \| (115, 154) \| 158 \| hypothetical protein \| hypothetical protein; uniclust \| UniRef100\_D8A596 \| 96.5 \| 0.00052 \| 1e-09 \| 64.1 \| 37 \| (158, 203) \| 350 \| (65, 101) \| 136 \| DUF551 domain-containing protein (Fragment) \| DUF551 domain-containing protein (Fragment); uniclust \| UniRef100\_A0A822WF58 \| 96.4 \| 0.00058 \| 1.2e-09 \| 68.0 \| 22 \| (153, 179) \| 350 \| (149, 170) \| 202 \| DUF551 domain-containing protein \| DUF551 domain-containing protein; uniclust \| UniRef100\_UPI000D697C96 \| 96.4 \| 0.00064 \| 1.2e-09 \| 62.1 \| 38 \| (187, 224) \| 350 \| (80, 118) \| 119 \| hypothetical protein \| hypothetical protein; uniclust \| UniRef100\_UPI0019290DBF \| 96.4 \| 0.00066 \| 1.2e-09 \| 64.5 \| 54 \| (176, 229) \| 350 \| (105, 159) \| 164 \| hypothetical protein \| hypothetical protein; uniclust \| UniRef100\_A0A2G2PQ67 \| 96.4 \| 0.00057 \| 1.2e-09 \| 64.1 \| 38 \| (168, 209) \| 350 \| (51, 88) \| 125 \| DUF551 domain-containing protein \| DUF551 domain-containing protein; uniclust \| UniRef100\_A0A076LNT5 \| 96.4 \| 0.00068 \| 1.3e-09 \| 62.3 \| 46 \| (42, 87) \| 350 \| (77, 122) \| 130 \| Ead/Ea22-like family protein \| Ead/Ea22-like family protein; uniclust \| UniRef100\_A0A855F7K2 \| 96.4 \| 0.00059 \| 1.4e-09 \| 72.2 \| 24 \| (72, 95) \| 350 \| (76, 99) \| 286 \| DUF551 domain-containing protein \| DUF551 domain-containing protein; uniclust \| UniRef100\_A0A2I8SEA0 \| 96.4 \| 0.00063 \| 1.4e-09 \| 63.8 \| 31 \| (155, 195) \| 350 \| (39, 69) \| 123 \| DUF551 domain-containing protein \| DUF551 domain-containing protein; uniclust \| UniRef100\_A0A1C3H9H7 \| 96.4 \| 0.00057 \| 1.4e-09 \| 71.7 \| 24 \| (168, 195) \| 350 \| (159, 182) \| 263 \| Eaa1 \| Eaa1; uniclust \| UniRef100\_A0A0P0ZCX1 \| 96.3 \| 0.00074 \| 1.5e-09 \| 66.3 \| 72 \| (155, 235) \| 350 \| (93, 176) \| 187 \| DUF551 domain-containing protein \| DUF551 domain-containing protein; uniclust \| UniRef100\_UPI000799D453 \| 96.3 \| 0.00074 \| 1.5e-09 \| 64.9 \| 62 \| (168, 233) \| 350 \| (67, 141) \| 155 \| DUF551 domain-containing protein \| DUF551 domain-containing protein; uniclust \| UniRef100\_A0A4Q2QYQ8 \| 96.3 \| 0.00087 \| 1.6e-09 \| 66.3 \| 69 \| (153, 230) \| 350 \| (51, 134) \| 221 \| DUF551 domain-containing protein \| DUF551 domain-containing protein; uniclust \| UniRef100\_UPI00069C164F \| 96.3 \| 0.00074 \| 1.7e-09 \| 69.8 \| 21 \| (153, 178) \| 350 \| (145, 165) \| 238 \| DUF551 domain-containing protein \| DUF551 domain-containing protein; uniclust \| UniRef100\_UPI00040D00C3 \| 96.3 \| 0.00076 \| 1.7e-09 \| 71.3 \| 24 \| (73, 96) \| 350 \| (136, 159) \| 293 \| hypothetical protein \| hypothetical protein; uniclust \| UniRef100\_A0A4D6DY86 \| 96.3 \| 0.00094 \| 1.7e-09 \| 59.2 \| 40 \| (1, 40) \| 350 \| (4, 45) \| 101 \| Uncharacterized protein \| Uncharacterized protein; uniclust \| UniRef100\_UPI00202223FF \| 96.3 \| 0.001 \| 1.9e-09 \| 64.1 \| 26 \| (153, 187) \| 350 \| (74, 99) \| 179 \| DUF551 domain-containing protein \| DUF551 domain-containing protein; uniclust \| UniRef100\_A0A0H2Z1D9 \| 96.2 \| 0.00076 \| 2.1e-09 \| 70.2 \| 64 \| (158, 231) \| 350 \| (167, 236) \| 241 \| DUF551 domain-containing protein \| DUF551 domain-containing protein; uniclust \| UniRef100\_UPI0007CA5FDA \| 96.2 \| 0.00092 \| 2.2e-09 \| 69.1 \| 21 \| (75, 96) \| 350 \| (91, 111) \| 232 \| hypothetical protein \| hypothetical protein; uniclust \| UniRef100\_A0A447RMB8 \| 96.2 \| 0.00086 \| 2.3e-09 \| 72.3 \| 77 \| (153, 236) \| 350 \| (222, 302) \| 313 \| Eaa1 \| Eaa1; uniclust \| UniRef100\_A0A486MKY1 \| 96.2 \| 0.001 \| 2.4e-09 \| 68.5 \| 82 \| (80, 175) \| 350 \| (83, 170) \| 228 \| Uncharacterized protein \| Uncharacterized protein |
| Top keywords  (threshold 1.00e-03 (evalue)) | **domain\_containing, DUF551, hypothetical, Phage, Eaa, Eaa1, Putative, DUF550, DUF3850, Ead** |
| Output files | ../../similar\_sequences/57\_FANPEZAQ\_CDS\_0057\_merged.svg ../../similar\_sequences/57\_FANPEZAQ\_CDS\_0057\_pdb70.a3m ../../similar\_sequences/57\_FANPEZAQ\_CDS\_0057\_pdb70.hhr ../../similar\_sequences/57\_FANPEZAQ\_CDS\_0057\_uniclust.a3m ../../similar\_sequences/57\_FANPEZAQ\_CDS\_0057\_uniclust.hhr |

#### Structure prediction (AlphaFold)^2^

|  |  |
| --- | --- |
| Stats | xml version="1.0" encoding="utf-8" standalone="no"?       2024-09-02T21:09:53.750450 image/svg+xml   Matplotlib v3.7.2, https://matplotlib.org/ |
| Predicted structure | **NGL Viewer Controls:**  - Center: *Left-Click* - Rotate: *Left-Click + Drag* - Translate: *Right-Click + Drag* - Zoom: *Shift + Left-Click + Drag* |
| Output files | ../../predicted\_structures/57\_FANPEZAQ\_CDS\_0057/features.pkl ../../predicted\_structures/57\_FANPEZAQ\_CDS\_0057/ranked\_0.pdb ../../predicted\_structures/57\_FANPEZAQ\_CDS\_0057/ranked\_0\_plots.svg ../../predicted\_structures/57\_FANPEZAQ\_CDS\_0057/result\_model\_1\_ptm\_pred\_0.pkl |

#### Structure similarity search results (Foldseek)^3^

|  |  |
| --- | --- |
| Structure databases searched | Pdb, Afdb-proteome, Afdb-uniprot50 |
| Results, scheme(s)  (Top layers only, threshold 1.00e-02 (evalue)) | xml version="1.0" encoding="utf-8" standalone="no"?       2024-09-02T21:11:28.848085 image/svg+xml   Matplotlib v3.7.2, https://matplotlib.org/ |
| Results, table  (threshold 1.00e-02 (evalue)) | -- |
| Top keywords  (threshold 1.00e-02 (evalue)) | -- |
| Output files | ../../similar\_structures/57\_FANPEZAQ\_CDS\_0057\_afdb-proteome\_foldseek.tsv ../../similar\_structures/57\_FANPEZAQ\_CDS\_0057\_afdb-uniprot50\_foldseek.tsv ../../similar\_structures/57\_FANPEZAQ\_CDS\_0057\_merged.svg ../../similar\_structures/57\_FANPEZAQ\_CDS\_0057\_pdb\_foldseek.tsv |

  
  
  

Return to summary | Go to previous | Go to next

  


---

**Sequence/structure alignments coloring**  
Each object in the alignment figures is colored according to its E-value following this color coding:

1e-100
10
