## Supplementary material for "Completing the BASEL phage collection to unlock hidden diversity for systematic exploration of phage-host interactions": Data S2: 61.html

FANPEZAQ\_CDS\_0061


Return to summary | Go to previous | Go to next

|  |  |
| --- | --- |
| FANPEZAQ\_CDS\_0061  Page creation date: 02 Sep 2024, 12:00  Project folder: n/a  Input sequences file: Escherichia\_virus\_HeidiAbel.gb | mor domain\_containing transcription activator regulator transcriptional helix\_turn\_helix response dna hypothetical binding and hth regulatory dna\_binding operon homeodomain\_like rna regulation middle nucleotide metabolism system two\_component putative fragment conserved resolvase gol activator\_like factor quorum luxr\_type narl winged terminase head packaging repressor mse complex sensing fixj |

### Sequence information

|  |  |
| --- | --- |
| Name | FANPEZAQ\_CDS\_0061  61\_FANPEZAQ\_CDS\_0061 (pipeline id) |
| Imported annotations | Escherichia\_virus\_HeidiAbel Bas97 |
| Protein sequence | MELPRTVQEIADVIGRERALFLIGQLPKLWVPSQQYHKVILYVPKRIRPNDPLVQILGWQ DASKMVQHFGGEMLHPANCEYIYRHFIHRSIKRMHSEGMDAKAIAELLDVSERTVKRHCT DKPHKDTSPANDNTPQRLLTAMGQ |
| Number of residues | 144 |
| Molecular weight (Da) | 16644.11 |
| Output files | ../../query\_sequences/61\_FANPEZAQ\_CDS\_0061.fasta |

### Putative domain architecture and protein family

#### Search results (HHblits)^1^

|  |  |
| --- | --- |
| Domain family databases searched | Pfam, Ncbi-cd, Cath, Phrogs |
| Results, scheme(s)  (Top layers only; threshold 1.00e-03 (evalue)) | xml version="1.0" encoding="utf-8" standalone="no"?       2024-09-02T21:08:25.687149 image/svg+xml   Matplotlib v3.7.2, https://matplotlib.org/ |
| Results, table  (E-value ≤ 1.00e-03 (evalue)) | db \| id \| prob \| evalue \| pvalue \| score \| cols \| query \| query\_len \| template \| template\_len \| name \| description; pfam \| PF08765 \| 99.4 \| 1.5e-17 \| 3e-21 \| 108.3 \| 97 \| (2, 124) \| 144 \| (2, 98) \| 108 \| Mor \| Mor transcription activator family; pfam \| PF10654 \| 97.0 \| 5.3e-07 \| 1e-10 \| 58.2 \| 54 \| (84, 137) \| 144 \| (66, 119) \| 126 \| DUF2481 \| Protein of unknown function (DUF2481); pfam \| PF11242 \| 96.7 \| 1.8e-06 \| 3.3e-10 \| 49.9 \| 37 \| (88, 124) \| 144 \| (4, 40) \| 64 \| DUF2774 \| Protein of unknown function (DUF2774); pfam \| PF05344 \| 95.9 \| 2.7e-05 \| 5.3e-09 \| 42.6 \| 39 \| (89, 127) \| 144 \| (3, 41) \| 62 \| DUF746 \| Domain of Unknown Function (DUF746); pfam \| PF18010 \| 93.7 \| 0.00097 \| 2e-07 \| 29.4 \| 22 \| (97, 118) \| 144 \| (7, 28) \| 29 \| HTH\_49 \| Cry35Ab1 HTH C-terminal domain; phrogs \| 836 \| 99.9 \| 9.6e-33 \| 1.3e-36 \| 203.8 \| 115 \| (1, 125) \| 144 \| (21, 137) \| 149 \| late transcriptional activator \| late transcriptional activator; Category: transcription regulation; p206917 VI\_10124; phrogs \| 532 \| 99.4 \| 7.8e-18 \| 1e-21 \| 120.7 \| 76 \| (51, 127) \| 144 \| (34, 109) \| 132 \| late transcriptional activator \| late transcriptional activator; Category: transcription regulation; p318767 VI\_01623; phrogs \| 4151 \| 97.8 \| 6.3e-09 \| 7.8e-13 \| 75.8 \| 52 \| (86, 137) \| 144 \| (8, 59) \| 168 \| DNA binding protein \| DNA binding protein; Category: DNA, RNA and nucleotide metabolism; p284557 VI\_05952; phrogs \| 517 \| 97.4 \| 8.2e-08 \| 1e-11 \| 64.9 \| 44 \| (80, 123) \| 144 \| (62, 105) \| 107 \| HTH DNA binding protein \| HTH DNA binding protein; Category: DNA, RNA and nucleotide metabolism; MF668278\_p45; phrogs \| 664 \| 97.4 \| 7.6e-08 \| 1e-11 \| 65.7 \| 55 \| (81, 135) \| 144 \| (55, 109) \| 117 \| DNA binding protein \| DNA binding protein; Category: DNA, RNA and nucleotide metabolism; MF072690\_p140; phrogs \| 6156 \| 97.4 \| 1.1e-07 \| 1.4e-11 \| 60.6 \| 58 \| (76, 133) \| 144 \| (11, 69) \| 78 \| endonuclease \| endonuclease; Category: DNA, RNA and nucleotide metabolism; KU160668\_p36; phrogs \| 20467 \| 97.1 \| 6.1e-07 \| 7e-11 \| 60.3 \| 50 \| (76, 126) \| 144 \| (66, 115) \| 116 \| HNH endonuclease \| HNH endonuclease; Category: DNA, RNA and nucleotide metabolism; NC\_026590\_p30; phrogs \| 7067 \| 96.8 \| 2e-06 \| 2.3e-10 \| 50.8 \| 49 \| (68, 121) \| 144 \| (5, 53) \| 55 \| terminase small subunit \| terminase small subunit; Category: head and packaging; NC\_034627\_p52; phrogs \| 6831 \| 96.8 \| 2.1e-06 \| 2.5e-10 \| 53.0 \| 37 \| (88, 124) \| 144 \| (5, 41) \| 66 \| transcriptional regulator \| transcriptional regulator; Category: transcription regulation; p431926 VI\_12195; phrogs \| 4494 \| 96.8 \| 2.2e-06 \| 2.5e-10 \| 55.5 \| 52 \| (87, 138) \| 144 \| (6, 57) \| 88 \| NA \| NA; Category: unknown function; p409661 VI\_06907; phrogs \| 3855 \| 96.7 \| 2.9e-06 \| 3.6e-10 \| 57.7 \| 72 \| (55, 126) \| 144 \| (35, 107) \| 114 \| HTH DNA binding protein \| HTH DNA binding protein; Category: DNA, RNA and nucleotide metabolism; KY290952\_p65; phrogs \| 5536 \| 96.6 \| 3.9e-06 \| 4.8e-10 \| 60.1 \| 39 \| (83, 121) \| 144 \| (5, 43) \| 165 \| NA \| NA; Category: unknown function; p166286 VI\_04146; phrogs \| 6223 \| 96.3 \| 1.4e-05 \| 1.7e-09 \| 50.5 \| 35 \| (95, 129) \| 144 \| (14, 48) \| 77 \| transcriptional repressor \| transcriptional repressor; Category: transcription regulation; p192116 VI\_09192; phrogs \| 1157 \| 96.1 \| 2.9e-05 \| 3.6e-09 \| 60.0 \| 37 \| (86, 122) \| 144 \| (5, 41) \| 291 \| transposase \| transposase; Category: integration and excision; p237322 VI\_08188; phrogs \| 1373 \| 96.0 \| 3.2e-05 \| 4.2e-09 \| 50.4 \| 34 \| (95, 128) \| 144 \| (43, 76) \| 95 \| transcriptional regulator \| transcriptional regulator; Category: transcription regulation; NC\_005857\_p48; phrogs \| 2308 \| 96.0 \| 3.9e-05 \| 4.7e-09 \| 54.0 \| 41 \| (83, 123) \| 144 \| (1, 41) \| 151 \| NA \| NA; Category: unknown function; p149266 VI\_05209; phrogs \| 11376 \| 95.8 \| 6.5e-05 \| 7.6e-09 \| 51.4 \| 55 \| (78, 132) \| 144 \| (38, 92) \| 134 \| NA \| NA; Category: unknown function; p46013 VI\_01297; phrogs \| 4779 \| 95.7 \| 8.1e-05 \| 1e-08 \| 46.9 \| 30 \| (95, 124) \| 144 \| (11, 40) \| 78 \| transcriptional regulator \| transcriptional regulator; Category: transcription regulation; p225361 VI\_06249; phrogs \| 7061 \| 95.5 \| 0.00012 \| 1.5e-08 \| 45.4 \| 35 \| (89, 123) \| 144 \| (38, 72) \| 73 \| HTH DNA binding protein \| HTH DNA binding protein; Category: DNA, RNA and nucleotide metabolism; KU160668\_p38; phrogs \| 15481 \| 95.2 \| 0.00025 \| 2.7e-08 \| 38.7 \| 39 \| (82, 120) \| 144 \| (3, 41) \| 43 \| NA \| NA; Category: unknown function; KY683735\_p262; phrogs \| 2538 \| 95.1 \| 0.00024 \| 3.1e-08 \| 47.5 \| 35 \| (90, 124) \| 144 \| (8, 42) \| 113 \| terminase small subunit \| terminase small subunit; Category: head and packaging; p133796 VI\_05051; phrogs \| 151 \| 95.1 \| 0.00025 \| 3.4e-08 \| 44.9 \| 37 \| (91, 127) \| 144 \| (15, 51) \| 91 \| transposase \| transposase; Category: integration and excision; p268658 VI\_04551; phrogs \| 7847 \| 95.0 \| 0.00031 \| 3.6e-08 \| 51.6 \| 44 \| (88, 131) \| 144 \| (7, 50) \| 247 \| NA \| NA; Category: unknown function; p386826 VI\_07743; phrogs \| 6119 \| 95.0 \| 0.00035 \| 3.9e-08 \| 43.8 \| 64 \| (61, 124) \| 144 \| (19, 87) \| 94 \| NA \| NA; Category: unknown function; p232261 VI\_06268; phrogs \| 1381 \| 95.0 \| 0.00032 \| 4e-08 \| 45.9 \| 34 \| (98, 131) \| 144 \| (19, 52) \| 99 \| plasmid antitoxin with HTH domain \| plasmid antitoxin with HTH domain; Category: moron, auxiliary metabolic gene and host takeover; p79485 VI\_11488; phrogs \| 12823 \| 94.9 \| 0.00039 \| 4.4e-08 \| 46.1 \| 57 \| (87, 143) \| 144 \| (17, 75) \| 117 \| NA \| NA; Category: unknown function; MF919537\_p145; phrogs \| 1337 \| 94.9 \| 0.00037 \| 4.7e-08 \| 41.7 \| 33 \| (94, 126) \| 144 \| (11, 43) \| 61 \| DNA binding protein \| DNA binding protein; Category: DNA, RNA and nucleotide metabolism; p177899 VI\_00920; phrogs \| 25961 \| 94.9 \| 0.00042 \| 4.7e-08 \| 40.9 \| 34 \| (49, 82) \| 144 \| (32, 65) \| 66 \| NA \| NA; Category: unknown function; p32746 VI\_10942; phrogs \| 5751 \| 94.9 \| 0.0004 \| 4.7e-08 \| 48.9 \| 45 \| (88, 132) \| 144 \| (14, 58) \| 164 \| terminase large subunit \| terminase large subunit; Category: head and packaging; p273217 VI\_06917; phrogs \| 1468 \| 94.8 \| 0.00039 \| 4.8e-08 \| 42.7 \| 37 \| (94, 130) \| 144 \| (19, 55) \| 69 \| transcriptional repressor \| transcriptional repressor; Category: transcription regulation; p307668 VI\_06058; phrogs \| 27475 \| 94.8 \| 0.00047 \| 5.3e-08 \| 54.9 \| 49 \| (84, 132) \| 144 \| (3, 51) \| 570 \| NA \| NA; Category: unknown function; p89643 VI\_08714; phrogs \| 4291 \| 94.6 \| 0.00055 \| 6.5e-08 \| 58.2 \| 70 \| (50, 119) \| 144 \| (833, 905) \| 906 \| RNA-dependent RNA polymerase \| RNA-dependent RNA polymerase; Category: DNA, RNA and nucleotide metabolism; p46481 VI\_01201; phrogs \| 11138 \| 94.6 \| 0.00053 \| 6.5e-08 \| 46.7 \| 58 \| (57, 125) \| 144 \| (67, 124) \| 129 \| NA \| NA; Category: unknown function; p272681 VI\_04059; phrogs \| 323 \| 94.5 \| 0.00055 \| 7.5e-08 \| 41.9 \| 47 \| (82, 129) \| 144 \| (6, 52) \| 78 \| transcriptional regulator, D5-like \| transcriptional regulator, D5-like; Category: transcription regulation; p118890 VI\_11329; phrogs \| 7676 \| 94.4 \| 0.00077 \| 9e-08 \| 41.5 \| 31 \| (95, 125) \| 144 \| (13, 43) \| 75 \| NA \| NA; Category: unknown function; NC\_023688\_p227; phrogs \| 59 \| 94.2 \| 0.00079 \| 1.1e-07 \| 46.4 \| 48 \| (78, 125) \| 144 \| (111, 159) \| 161 \| HNH endonuclease \| HNH endonuclease; Category: DNA, RNA and nucleotide metabolism; MG812495\_p46; phrogs \| 609 \| 94.2 \| 0.00091 \| 1.1e-07 \| 53.3 \| 43 \| (83, 125) \| 144 \| (6, 48) \| 405 \| RIIB lysis inhibitor \| RIIB lysis inhibitor; Category: lysis; MG696114\_p269; phrogs \| 2104 \| 94.2 \| 0.00092 \| 1.1e-07 \| 43.1 \| 49 \| (81, 131) \| 144 \| (28, 76) \| 92 \| NA \| NA; Category: unknown function; p72070 VI\_03178; phrogs \| 1206 \| 94.1 \| 0.001 \| 1.3e-07 \| 48.8 \| 46 \| (85, 130) \| 144 \| (16, 62) \| 212 \| terminase small subunit \| terminase small subunit; Category: head and packaging; p287057 VI\_06574; phrogs \| 309 \| 94.1 \| 0.00094 \| 1.3e-07 \| 46.1 \| 39 \| (84, 122) \| 144 \| (9, 47) \| 153 \| terminase small subunit \| terminase small subunit; Category: head and packaging; p301521 VI\_00938 |
| Top keywords  (threshold 1.00e-03 (evalue)) | **and, DNA, RNA, nucleotide, metabolism, transcription, transcriptional, regulation, binding, HTH** |
| Output files | ../../domain\_architecture/61\_FANPEZAQ\_CDS\_0061\_cath.hhr ../../domain\_architecture/61\_FANPEZAQ\_CDS\_0061\_merged.svg ../../domain\_architecture/61\_FANPEZAQ\_CDS\_0061\_ncbi-cd.hhr ../../domain\_architecture/61\_FANPEZAQ\_CDS\_0061\_pfam.hhr ../../domain\_architecture/61\_FANPEZAQ\_CDS\_0061\_phrogs.hhr |

### Identical protein sequences/structures

#### Search results

|  |  |
| --- | --- |
| Protein sequence databases searched | Pdb, Swissprot, Refseq |
| Identical proteins found | -- |
| Top keywords | -- |
| Output files | -- |

### Similar protein sequences/structures

#### Sequence similarity search results (HHblits)^1^

|  |  |
| --- | --- |
| Sequence databases searched | Uniclust, Pdb70 |
| Results, scheme(s)  (Top layers only, threshold 1.00e-03 (evalue)) | xml version="1.0" encoding="utf-8" standalone="no"?       2024-09-02T21:08:54.119180 image/svg+xml   Matplotlib v3.7.2, https://matplotlib.org/ |
| Results, table(s)  (threshold 1.00e-03 (evalue)) | db \| id \| prob \| evalue \| pvalue \| score \| cols \| query \| query\_len \| template \| template\_len \| name \| description; uniclust \| UniRef100\_A0A0F7KCJ2 \| 99.9 \| 1.5e-31 \| 3.3e-37 \| 197.2 \| 110 \| (2, 121) \| 144 \| (60, 169) \| 196 \| Mor family transcriptional regulator \| Mor family transcriptional regulator; uniclust \| UniRef100\_A0A0J6SFM2 \| 99.9 \| 8e-31 \| 1.8e-36 \| 190.2 \| 114 \| (2, 125) \| 144 \| (26, 141) \| 164 \| Mor transcription activator domain-containing protein \| Mor transcription activator domain-containing protein; uniclust \| UniRef100\_A0A143DDS5 \| 99.9 \| 9.2e-31 \| 1.9e-36 \| 190.1 \| 135 \| (1, 135) \| 144 \| (25, 161) \| 197 \| Mor transcription activator domain-containing protein \| Mor transcription activator domain-containing protein; uniclust \| UniRef100\_A0A088FQJ8 \| 99.9 \| 2.1e-30 \| 4.3e-36 \| 185.2 \| 144 \| (1, 144) \| 144 \| (21, 167) \| 169 \| Uncharacterized protein \| Uncharacterized protein; uniclust \| UniRef100\_A0A1E7HXZ1 \| 99.9 \| 1.8e-29 \| 3.9e-35 \| 179.1 \| 108 \| (3, 122) \| 144 \| (17, 124) \| 146 \| Mor transcription activator domain-containing protein \| Mor transcription activator domain-containing protein; uniclust \| UniRef100\_A0A075CDW0 \| 99.9 \| 1.1e-28 \| 2.4e-34 \| 182.4 \| 112 \| (2, 123) \| 144 \| (33, 157) \| 191 \| Mor transcription activator domain-containing protein \| Mor transcription activator domain-containing protein; uniclust \| UniRef100\_A0A011TVI4 \| 99.9 \| 1.6e-28 \| 3.5e-34 \| 174.3 \| 111 \| (2, 122) \| 144 \| (15, 128) \| 145 \| HTH luxR-type domain-containing protein \| HTH luxR-type domain-containing protein; uniclust \| UniRef100\_A0A212KJX9 \| 99.9 \| 3e-28 \| 6.4e-34 \| 174.7 \| 116 \| (2, 127) \| 144 \| (25, 143) \| 157 \| Mor transcription activator domain-containing protein \| Mor transcription activator domain-containing protein; uniclust \| UniRef100\_A0A0D8KFG8 \| 99.9 \| 9.3e-27 \| 1.9e-32 \| 164.2 \| 110 \| (2, 123) \| 144 \| (23, 132) \| 148 \| Mor transcription activator domain-containing protein \| Mor transcription activator domain-containing protein; uniclust \| UniRef100\_A0A0X8JK61 \| 99.9 \| 1.3e-26 \| 2.7e-32 \| 166.7 \| 115 \| (2, 126) \| 144 \| (36, 163) \| 174 \| Mor transcription activator domain-containing protein \| Mor transcription activator domain-containing protein; uniclust \| UniRef100\_A0A081B6E8 \| 99.9 \| 1.4e-26 \| 3e-32 \| 165.4 \| 110 \| (2, 123) \| 144 \| (11, 120) \| 147 \| Mor, transcription regulation \| Mor, transcription regulation; uniclust \| UniRef100\_A0A0W1G863 \| 99.9 \| 1.8e-26 \| 3.9e-32 \| 162.7 \| 106 \| (5, 125) \| 144 \| (18, 124) \| 140 \| Mor transcription activator domain-containing protein \| Mor transcription activator domain-containing protein; uniclust \| UniRef100\_A0A063BCU2 \| 99.9 \| 4.6e-26 \| 9.9e-32 \| 165.8 \| 112 \| (2, 123) \| 144 \| (26, 149) \| 176 \| Mor transcription activator domain protein \| Mor transcription activator domain protein; uniclust \| UniRef100\_A0A0J7J645 \| 99.9 \| 1.5e-25 \| 2.9e-31 \| 153.7 \| 115 \| (2, 126) \| 144 \| (9, 123) \| 135 \| Mor transcription activator family \| Mor transcription activator family; uniclust \| UniRef100\_A0A070A9J5 \| 99.8 \| 6.1e-25 \| 1.3e-30 \| 157.0 \| 104 \| (2, 123) \| 144 \| (18, 121) \| 148 \| Mor transcription activator domain-containing protein \| Mor transcription activator domain-containing protein; uniclust \| UniRef100\_A0A0D5LXL7 \| 99.8 \| 6.4e-25 \| 1.4e-30 \| 156.7 \| 106 \| (3, 125) \| 144 \| (17, 122) \| 147 \| Mor transcription activator domain-containing protein \| Mor transcription activator domain-containing protein; uniclust \| UniRef100\_A0A077FBH5 \| 99.8 \| 8.3e-25 \| 1.7e-30 \| 160.7 \| 114 \| (2, 126) \| 144 \| (47, 160) \| 195 \| Mor transcription activator domain-containing protein \| Mor transcription activator domain-containing protein; uniclust \| UniRef100\_A0A0Q3EQP9 \| 99.8 \| 8.9e-25 \| 1.9e-30 \| 155.0 \| 112 \| (2, 123) \| 144 \| (8, 119) \| 151 \| Mor transcription activator domain-containing protein \| Mor transcription activator domain-containing protein; uniclust \| UniRef100\_A0A0Q7I488 \| 99.8 \| 1.7e-24 \| 3.5e-30 \| 153.0 \| 109 \| (2, 120) \| 144 \| (13, 128) \| 147 \| Mor transcription activator domain-containing protein \| Mor transcription activator domain-containing protein; uniclust \| UniRef100\_A0A1H8FWD2 \| 99.8 \| 2e-24 \| 4.1e-30 \| 150.6 \| 112 \| (3, 125) \| 144 \| (19, 130) \| 138 \| Mor transcription activator family protein \| Mor transcription activator family protein; uniclust \| UniRef100\_A0A6P1ZJ74 \| 99.8 \| 2.4e-24 \| 4.7e-30 \| 151.1 \| 125 \| (2, 136) \| 144 \| (32, 156) \| 161 \| Mor transcription activator domain-containing protein \| Mor transcription activator domain-containing protein; uniclust \| UniRef100\_A0A106BHZ3 \| 99.8 \| 2.8e-24 \| 5.9e-30 \| 154.3 \| 110 \| (3, 122) \| 144 \| (7, 116) \| 156 \| Mor transcription activator domain-containing protein \| Mor transcription activator domain-containing protein; uniclust \| UniRef100\_A0A0N7I1N9 \| 99.8 \| 4e-24 \| 8.3e-30 \| 153.4 \| 114 \| (1, 124) \| 144 \| (21, 138) \| 164 \| Two-component system, NarL family, nitrate/nitrite response regulator NarP \| Two-component system, NarL family, nitrate/nitrite response regulator NarP; uniclust \| UniRef100\_A0A0E4G398 \| 99.8 \| 4.4e-24 \| 8.9e-30 \| 148.4 \| 109 \| (3, 123) \| 144 \| (11, 123) \| 141 \| Mor transcription activator domain-containing protein \| Mor transcription activator domain-containing protein; uniclust \| UniRef100\_A0A5E4RYD4 \| 99.8 \| 2.7e-23 \| 5.3e-29 \| 141.6 \| 111 \| (2, 122) \| 144 \| (11, 121) \| 132 \| Mor transcription activator domain-containing protein \| Mor transcription activator domain-containing protein; uniclust \| UniRef100\_A0A077P2T4 \| 99.8 \| 2.7e-23 \| 5.8e-29 \| 154.9 \| 113 \| (2, 124) \| 144 \| (42, 167) \| 200 \| Mor transcription activator domain-containing protein \| Mor transcription activator domain-containing protein; uniclust \| UniRef100\_A0A1G0CQ82 \| 99.8 \| 2.9e-23 \| 5.9e-29 \| 148.5 \| 113 \| (1, 123) \| 144 \| (28, 150) \| 166 \| Mor transcription activator domain-containing protein \| Mor transcription activator domain-containing protein; uniclust \| UniRef100\_A0A0A8F3J8 \| 99.8 \| 6.6e-23 \| 1.4e-28 \| 147.5 \| 115 \| (2, 126) \| 144 \| (24, 149) \| 167 \| Mor transcription activator domain-containing protein \| Mor transcription activator domain-containing protein; uniclust \| UniRef100\_A0A011P6X2 \| 99.8 \| 8.8e-23 \| 1.8e-28 \| 149.5 \| 116 \| (2, 127) \| 144 \| (37, 166) \| 183 \| Mor transcription activator family protein \| Mor transcription activator family protein; uniclust \| UniRef100\_A0A1Y6D0G8 \| 99.8 \| 2.6e-22 \| 5.1e-28 \| 140.4 \| 112 \| (2, 123) \| 144 \| (14, 127) \| 145 \| Mor transcription activator family protein \| Mor transcription activator family protein; uniclust \| UniRef100\_A0A2G2I3T3 \| 99.8 \| 2.5e-22 \| 5.3e-28 \| 145.6 \| 112 \| (2, 124) \| 144 \| (19, 130) \| 166 \| Mor transcription activator domain-containing protein \| Mor transcription activator domain-containing protein; uniclust \| UniRef100\_A0A2N1ANB5 \| 99.8 \| 6.3e-22 \| 1.2e-27 \| 140.7 \| 117 \| (1, 121) \| 144 \| (1, 119) \| 174 \| Mor transcription activator domain-containing protein \| Mor transcription activator domain-containing protein; uniclust \| UniRef100\_A0A0H2MCL8 \| 99.8 \| 1e-21 \| 1.9e-27 \| 138.0 \| 112 \| (2, 123) \| 144 \| (14, 125) \| 163 \| Mor transcription activator family protein \| Mor transcription activator family protein; uniclust \| UniRef100\_A0A4D7B5F0 \| 99.8 \| 9.6e-22 \| 2e-27 \| 142.3 \| 103 \| (2, 116) \| 144 \| (23, 125) \| 173 \| Helix-turn-helix domain-containing protein \| Helix-turn-helix domain-containing protein; uniclust \| UniRef100\_A0A4D7DAD3 \| 99.8 \| 1e-21 \| 2.1e-27 \| 136.8 \| 104 \| (6, 124) \| 144 \| (14, 117) \| 135 \| Mor transcription activator domain-containing protein \| Mor transcription activator domain-containing protein; uniclust \| UniRef100\_A0A0B4BIH4 \| 99.7 \| 1.3e-21 \| 2.6e-27 \| 145.0 \| 118 \| (2, 122) \| 144 \| (75, 192) \| 225 \| Uncharacterized protein \| Uncharacterized protein; uniclust \| UniRef100\_A0A0M7AQF8 \| 99.7 \| 1.9e-21 \| 3.9e-27 \| 140.4 \| 110 \| (2, 121) \| 144 \| (31, 150) \| 176 \| Uncharacterized protein \| Uncharacterized protein; uniclust \| UniRef100\_A0A009ZMY9 \| 99.7 \| 3.7e-21 \| 7.3e-27 \| 134.1 \| 116 \| (2, 127) \| 144 \| (9, 130) \| 146 \| Mor transcription activator family protein \| Mor transcription activator family protein; uniclust \| UniRef100\_A0A2T5J3R6 \| 99.7 \| 6.6e-21 \| 1.3e-26 \| 132.7 \| 110 \| (5, 124) \| 144 \| (23, 139) \| 149 \| Mor transcription activator family protein \| Mor transcription activator family protein; uniclust \| UniRef100\_A0A7D7IUH2 \| 99.7 \| 7.3e-21 \| 1.4e-26 \| 131.8 \| 134 \| (7, 140) \| 144 \| (1, 138) \| 151 \| Mor transcription activator domain-containing protein \| Mor transcription activator domain-containing protein; uniclust \| UniRef100\_A0A4Q5L708 \| 99.7 \| 7.5e-21 \| 1.4e-26 \| 131.8 \| 115 \| (2, 126) \| 144 \| (33, 147) \| 154 \| Mor transcription activator domain-containing protein \| Mor transcription activator domain-containing protein; uniclust \| UniRef100\_A0A1H9CJ13 \| 99.7 \| 7.3e-21 \| 1.4e-26 \| 130.9 \| 105 \| (5, 121) \| 144 \| (8, 112) \| 136 \| Mor transcription activator family protein \| Mor transcription activator family protein; uniclust \| UniRef100\_A0A350X8X0 \| 99.7 \| 1.2e-20 \| 2.3e-26 \| 131.1 \| 113 \| (4, 127) \| 144 \| (24, 136) \| 144 \| Mor transcription activator domain-containing protein \| Mor transcription activator domain-containing protein; uniclust \| UniRef100\_UPI000400307B \| 99.7 \| 1.9e-20 \| 3.5e-26 \| 129.2 \| 114 \| (2, 125) \| 144 \| (24, 138) \| 163 \| Mor transcription activator family protein \| Mor transcription activator family protein; uniclust \| UniRef100\_A0A241VCE8 \| 99.7 \| 2.1e-20 \| 4.2e-26 \| 129.8 \| 122 \| (2, 123) \| 144 \| (6, 131) \| 136 \| Mor transcription activator domain-containing protein \| Mor transcription activator domain-containing protein; uniclust \| UniRef100\_A0A0D8CPA5 \| 99.7 \| 2.1e-20 \| 4.3e-26 \| 130.4 \| 105 \| (8, 125) \| 144 \| (20, 124) \| 134 \| Mor transcription activator domain-containing protein \| Mor transcription activator domain-containing protein; uniclust \| UniRef100\_A0A845TMQ7 \| 99.7 \| 4.6e-20 \| 8.7e-26 \| 125.3 \| 118 \| (1, 128) \| 144 \| (1, 125) \| 131 \| Mor transcription activator domain-containing protein \| Mor transcription activator domain-containing protein; uniclust \| UniRef100\_A0A1D9CY07 \| 99.7 \| 5.5e-20 \| 1.1e-25 \| 127.7 \| 111 \| (4, 124) \| 144 \| (7, 119) \| 140 \| Putative bacteriophage transcriptional regulator \| Putative bacteriophage transcriptional regulator; uniclust \| UniRef100\_A0A1S0V4H7 \| 99.7 \| 7e-20 \| 1.3e-25 \| 131.4 \| 124 \| (1, 124) \| 144 \| (10, 141) \| 186 \| Mor transcription activator domain-containing protein (Fragment) \| Mor transcription activator domain-containing protein (Fragment); uniclust \| UniRef100\_A0A2S5CLI8 \| 99.7 \| 7.7e-20 \| 1.5e-25 \| 125.9 \| 116 \| (2, 127) \| 144 \| (11, 126) \| 141 \| Mor transcription activator domain-containing protein \| Mor transcription activator domain-containing protein; uniclust \| UniRef100\_A0A258P502 \| 99.7 \| 9.4e-20 \| 1.8e-25 \| 129.8 \| 110 \| (2, 121) \| 144 \| (57, 170) \| 173 \| Mor transcription activator domain-containing protein \| Mor transcription activator domain-containing protein; uniclust \| UniRef100\_A0A0H2X8T4 \| 99.7 \| 9.8e-20 \| 1.9e-25 \| 130.2 \| 123 \| (2, 125) \| 144 \| (17, 147) \| 166 \| Mor transcription activator domain-containing protein \| Mor transcription activator domain-containing protein; uniclust \| UniRef100\_A0A062GXL3 \| 99.7 \| 1.2e-19 \| 2.2e-25 \| 127.5 \| 113 \| (2, 124) \| 144 \| (22, 136) \| 154 \| CENP-B N-terminal DNA-binding domain protein \| CENP-B N-terminal DNA-binding domain protein; uniclust \| UniRef100\_A0A0F2N9K9 \| 99.7 \| 1.8e-19 \| 3.7e-25 \| 128.5 \| 107 \| (8, 125) \| 144 \| (15, 121) \| 149 \| Mor transcription activator domain-containing protein \| Mor transcription activator domain-containing protein; uniclust \| UniRef100\_A0A401JFQ9 \| 99.7 \| 2.2e-19 \| 4e-25 \| 120.5 \| 108 \| (5, 122) \| 144 \| (24, 131) \| 133 \| Mor transcription activator domain-containing protein \| Mor transcription activator domain-containing protein; uniclust \| UniRef100\_A0A2E3ES45 \| 99.6 \| 2.4e-19 \| 4.6e-25 \| 122.4 \| 112 \| (2, 125) \| 144 \| (5, 116) \| 126 \| Mor transcription activator domain-containing protein \| Mor transcription activator domain-containing protein; uniclust \| UniRef100\_A0A090DT46 \| 99.6 \| 2.7e-19 \| 5.4e-25 \| 128.3 \| 102 \| (6, 125) \| 144 \| (23, 124) \| 159 \| Mor transcription activator domain-containing protein \| Mor transcription activator domain-containing protein; uniclust \| UniRef100\_B2FRB5 \| 99.6 \| 3.3e-19 \| 6e-25 \| 120.9 \| 130 \| (2, 131) \| 144 \| (3, 132) \| 142 \| Mor transcription activator domain-containing protein \| Mor transcription activator domain-containing protein; uniclust \| UniRef100\_A0A1G4ZYZ4 \| 99.6 \| 3.8e-19 \| 7.1e-25 \| 125.8 \| 111 \| (2, 122) \| 144 \| (58, 172) \| 175 \| Mor transcription activator family protein \| Mor transcription activator family protein; uniclust \| UniRef100\_UPI000F8E7E3B \| 99.6 \| 4.9e-19 \| 9e-25 \| 121.2 \| 130 \| (2, 134) \| 144 \| (5, 134) \| 151 \| hypothetical protein \| hypothetical protein; uniclust \| UniRef100\_A0A6I2UT81 \| 99.6 \| 4.9e-19 \| 9.3e-25 \| 127.7 \| 117 \| (2, 121) \| 144 \| (81, 197) \| 202 \| Helix-turn-helix domain-containing protein \| Helix-turn-helix domain-containing protein; uniclust \| UniRef100\_A0A060H6I6 \| 99.6 \| 5.2e-19 \| 9.5e-25 \| 127.8 \| 121 \| (2, 122) \| 144 \| (50, 170) \| 219 \| Uncharacterized protein \| Uncharacterized protein; uniclust \| UniRef100\_A0A5C7QTA1 \| 99.6 \| 6.8e-19 \| 1.3e-24 \| 122.6 \| 119 \| (2, 131) \| 144 \| (18, 138) \| 143 \| Mor transcription activator domain-containing protein \| Mor transcription activator domain-containing protein; uniclust \| UniRef100\_A0A522VJQ9 \| 99.6 \| 7.7e-19 \| 1.4e-24 \| 121.4 \| 112 \| (2, 123) \| 144 \| (44, 156) \| 161 \| Mor transcription activator domain-containing protein \| Mor transcription activator domain-containing protein; uniclust \| UniRef100\_A0A109W8W4 \| 99.6 \| 7e-19 \| 1.4e-24 \| 132.1 \| 113 \| (2, 124) \| 144 \| (61, 184) \| 215 \| Mor transcription activator domain-containing protein \| Mor transcription activator domain-containing protein; uniclust \| UniRef100\_A0A8T3KWY8 \| 99.6 \| 8.8e-19 \| 1.6e-24 \| 121.6 \| 120 \| (2, 131) \| 144 \| (28, 152) \| 166 \| Helix-turn-helix domain-containing protein \| Helix-turn-helix domain-containing protein; uniclust \| UniRef100\_A0A5C7Q033 \| 99.6 \| 3.1e-18 \| 6.1e-24 \| 120.7 \| 113 \| (5, 126) \| 144 \| (6, 118) \| 156 \| Winged helix-turn-helix transcriptional regulator \| Winged helix-turn-helix transcriptional regulator; uniclust \| UniRef100\_A0A068Z4A5 \| 99.6 \| 3.4e-18 \| 6.7e-24 \| 122.8 \| 113 \| (2, 124) \| 144 \| (18, 144) \| 168 \| Mor transcription activator domain-containing protein \| Mor transcription activator domain-containing protein; uniclust \| UniRef100\_I3YBF0 \| 99.6 \| 4e-18 \| 7.5e-24 \| 121.2 \| 113 \| (2, 125) \| 144 \| (30, 142) \| 184 \| Mor transcription activator-like protein \| Mor transcription activator-like protein; uniclust \| UniRef100\_A0A6N4DE39 \| 99.6 \| 4e-18 \| 7.8e-24 \| 116.2 \| 108 \| (2, 123) \| 144 \| (6, 113) \| 123 \| Mor transcription activator domain-containing protein \| Mor transcription activator domain-containing protein; uniclust \| UniRef100\_A0A1Z4C0F4 \| 99.6 \| 4.2e-18 \| 7.9e-24 \| 120.2 \| 114 \| (2, 125) \| 144 \| (35, 148) \| 169 \| Mor transcription activator domain-containing protein \| Mor transcription activator domain-containing protein; uniclust \| UniRef100\_A0A961ZJA7 \| 99.6 \| 4.5e-18 \| 8.3e-24 \| 118.4 \| 111 \| (2, 122) \| 144 \| (25, 138) \| 168 \| Uncharacterized protein \| Uncharacterized protein; uniclust \| UniRef100\_A0A2K9NFC4 \| 99.6 \| 6.3e-18 \| 1.2e-23 \| 116.7 \| 106 \| (3, 123) \| 144 \| (22, 127) \| 137 \| Mor transcription activator domain-containing protein \| Mor transcription activator domain-containing protein; uniclust \| UniRef100\_A0A0C1G7F9 \| 99.6 \| 6.6e-18 \| 1.3e-23 \| 120.0 \| 112 \| (2, 126) \| 144 \| (9, 120) \| 148 \| Mor transcription activator domain-containing protein \| Mor transcription activator domain-containing protein; uniclust \| UniRef100\_UPI001D0F25A5 \| 99.6 \| 8.4e-18 \| 1.5e-23 \| 115.8 \| 111 \| (2, 122) \| 144 \| (42, 152) \| 155 \| Mor transcription activator family protein \| Mor transcription activator family protein; uniclust \| UniRef100\_A0A081NY94 \| 99.6 \| 8.3e-18 \| 1.8e-23 \| 117.8 \| 77 \| (47, 124) \| 144 \| (22, 98) \| 125 \| Mor transcription activator domain-containing protein \| Mor transcription activator domain-containing protein; uniclust \| UniRef100\_A0A139DN11 \| 99.6 \| 9.2e-18 \| 1.8e-23 \| 122.0 \| 122 \| (6, 138) \| 144 \| (34, 155) \| 181 \| Mor transcription activator domain-containing protein \| Mor transcription activator domain-containing protein; uniclust \| UniRef100\_A0A0B3S273 \| 99.5 \| 1.6e-17 \| 3.3e-23 \| 119.4 \| 100 \| (3, 119) \| 144 \| (27, 126) \| 152 \| Mor transcription activator domain-containing protein \| Mor transcription activator domain-containing protein; uniclust \| UniRef100\_A0A0P1FJJ4 \| 99.5 \| 1.7e-17 \| 3.7e-23 \| 119.9 \| 104 \| (3, 123) \| 144 \| (23, 126) \| 150 \| Helix-turn-helix domain-containing protein \| Helix-turn-helix domain-containing protein; uniclust \| UniRef100\_A0A3A6NWI4 \| 99.5 \| 2.1e-17 \| 3.9e-23 \| 113.8 \| 113 \| (2, 124) \| 144 \| (16, 133) \| 154 \| Mor transcription activator domain-containing protein \| Mor transcription activator domain-containing protein; uniclust \| UniRef100\_A0A0A2TST3 \| 99.5 \| 1.9e-17 \| 4.1e-23 \| 117.7 \| 80 \| (47, 127) \| 144 \| (28, 107) \| 133 \| Mor transcription activator domain-containing protein \| Mor transcription activator domain-containing protein; uniclust \| UniRef100\_A0A929AAA7 \| 99.5 \| 2.4e-17 \| 4.4e-23 \| 110.3 \| 105 \| (5, 121) \| 144 \| (21, 125) \| 127 \| Mor transcription activator domain-containing protein \| Mor transcription activator domain-containing protein; uniclust \| UniRef100\_A0A017RUI4 \| 99.5 \| 2.3e-17 \| 4.9e-23 \| 116.9 \| 75 \| (48, 123) \| 144 \| (26, 100) \| 130 \| Mor transcription activator domain-containing protein \| Mor transcription activator domain-containing protein; uniclust \| UniRef100\_A0A1V4WIM0 \| 99.5 \| 3e-17 \| 6.2e-23 \| 113.3 \| 80 \| (47, 127) \| 144 \| (25, 104) \| 118 \| Mor transcription activator family protein \| Mor transcription activator family protein; uniclust \| UniRef100\_A0A1E3H4F5 \| 99.5 \| 4e-17 \| 7.9e-23 \| 118.9 \| 111 \| (2, 125) \| 144 \| (23, 135) \| 183 \| Mor transcription activator family protein \| Mor transcription activator family protein; uniclust \| UniRef100\_A0A1Z3U8U4 \| 99.5 \| 4.1e-17 \| 8e-23 \| 113.7 \| 105 \| (2, 116) \| 144 \| (11, 117) \| 140 \| Uncharacterized protein \| Uncharacterized protein; uniclust \| UniRef100\_A0A0U3JVX5 \| 99.5 \| 4.3e-17 \| 8.7e-23 \| 124.2 \| 121 \| (2, 132) \| 144 \| (106, 239) \| 244 \| Uncharacterized protein \| Uncharacterized protein; uniclust \| UniRef100\_A0A022PC28 \| 99.5 \| 4e-17 \| 9.2e-23 \| 120.3 \| 74 \| (51, 125) \| 144 \| (43, 116) \| 159 \| Mor transcription activator domain-containing protein \| Mor transcription activator domain-containing protein; uniclust \| UniRef100\_A0A2N9YH57 \| 99.5 \| 6.6e-17 \| 1.2e-22 \| 111.3 \| 112 \| (2, 123) \| 144 \| (10, 127) \| 139 \| Mor transcription activator domain-containing protein \| Mor transcription activator domain-containing protein; uniclust \| UniRef100\_UPI0009301444 \| 99.5 \| 7.1e-17 \| 1.3e-22 \| 115.7 \| 104 \| (5, 123) \| 144 \| (97, 201) \| 203 \| helix-turn-helix domain-containing protein \| helix-turn-helix domain-containing protein; uniclust \| UniRef100\_A0A0C9PS36 \| 99.5 \| 7.6e-17 \| 1.6e-22 \| 112.4 \| 79 \| (49, 128) \| 144 \| (21, 99) \| 123 \| Uncharacterized conserved protein \| Uncharacterized conserved protein; uniclust \| UniRef100\_A0A109W455 \| 99.5 \| 7.9e-17 \| 1.7e-22 \| 113.5 \| 78 \| (52, 130) \| 144 \| (34, 111) \| 128 \| Mor transcription activator domain-containing protein \| Mor transcription activator domain-containing protein; uniclust \| UniRef100\_A0A7V8CQF7 \| 99.5 \| 1.2e-16 \| 2.2e-22 \| 110.5 \| 115 \| (2, 126) \| 144 \| (7, 131) \| 141 \| Mor transcription activator domain-containing protein \| Mor transcription activator domain-containing protein; uniclust \| UniRef100\_A0A0N0M130 \| 99.5 \| 1.2e-16 \| 2.4e-22 \| 112.3 \| 111 \| (2, 122) \| 144 \| (22, 133) \| 147 \| Mor transcription activator domain-containing protein \| Mor transcription activator domain-containing protein; uniclust \| UniRef100\_A0A0F6AFK3 \| 99.5 \| 1.3e-16 \| 2.5e-22 \| 112.1 \| 110 \| (2, 122) \| 144 \| (23, 132) \| 151 \| Mor transcription activator domain-containing protein \| Mor transcription activator domain-containing protein; uniclust \| UniRef100\_A0A011MI82 \| 99.5 \| 1.9e-16 \| 3.9e-22 \| 111.4 \| 77 \| (51, 128) \| 144 \| (39, 115) \| 135 \| Transcriptional regulator \| Transcriptional regulator; uniclust \| UniRef100\_A0A228J1B3 \| 99.4 \| 3.1e-16 \| 6e-22 \| 111.4 \| 109 \| (7, 125) \| 144 \| (52, 160) \| 163 \| Mor transcription activator domain-containing protein \| Mor transcription activator domain-containing protein; uniclust \| UniRef100\_UPI00188FAA08 \| 99.4 \| 3.4e-16 \| 6.3e-22 \| 108.5 \| 113 \| (3, 126) \| 144 \| (20, 136) \| 158 \| Mor transcription activator family protein \| Mor transcription activator family protein; uniclust \| UniRef100\_A0A0U5MJB7 \| 99.4 \| 3.3e-16 \| 6.4e-22 \| 110.2 \| 116 \| (2, 127) \| 144 \| (22, 140) \| 149 \| Helix-turn-helix domain-containing protein \| Helix-turn-helix domain-containing protein; uniclust \| UniRef100\_A0A2X3CYD8 \| 99.4 \| 3.3e-16 \| 6.4e-22 \| 105.9 \| 85 \| (2, 96) \| 144 \| (10, 99) \| 114 \| Mor transcription activator family \| Mor transcription activator family; uniclust \| UniRef100\_A0A0F1AW83 \| 99.4 \| 4.4e-16 \| 8.6e-22 \| 108.4 \| 113 \| (2, 124) \| 144 \| (3, 128) \| 138 \| Mor transcription activator domain-containing protein \| Mor transcription activator domain-containing protein; uniclust \| UniRef100\_A0A2W5KVX1 \| 99.4 \| 5.2e-16 \| 1e-21 \| 102.8 \| 89 \| (41, 129) \| 144 \| (2, 92) \| 100 \| Resolvase HTH domain-containing protein (Fragment) \| Resolvase HTH domain-containing protein (Fragment); uniclust \| UniRef100\_UPI00035FAD56 \| 99.4 \| 5.3e-16 \| 1e-21 \| 105.4 \| 110 \| (4, 127) \| 144 \| (4, 113) \| 119 \| hypothetical protein \| hypothetical protein; uniclust \| UniRef100\_A0A4D7B5Y7 \| 99.4 \| 6e-16 \| 1.1e-21 \| 109.1 \| 113 \| (2, 124) \| 144 \| (47, 159) \| 177 \| Helix-turn-helix domain-containing protein \| Helix-turn-helix domain-containing protein; uniclust \| UniRef100\_UPI001FAB3CB7 \| 99.4 \| 8.4e-16 \| 1.6e-21 \| 108.1 \| 115 \| (3, 127) \| 144 \| (54, 168) \| 174 \| Mor transcription activator family protein \| Mor transcription activator family protein; uniclust \| UniRef100\_A0A2M6UU58 \| 99.4 \| 8.4e-16 \| 1.6e-21 \| 106.2 \| 111 \| (2, 125) \| 144 \| (9, 123) \| 141 \| Helix-turn-helix domain-containing protein \| Helix-turn-helix domain-containing protein; uniclust \| UniRef100\_A0A286GYP9 \| 99.4 \| 8.9e-16 \| 1.7e-21 \| 109.5 \| 114 \| (2, 125) \| 144 \| (19, 137) \| 155 \| Homeodomain-like domain-containing protein \| Homeodomain-like domain-containing protein; uniclust \| UniRef100\_K9RSU5 \| 99.4 \| 9.7e-16 \| 1.8e-21 \| 101.5 \| 100 \| (7, 119) \| 144 \| (13, 113) \| 116 \| Mor transcription activator-like protein \| Mor transcription activator-like protein; uniclust \| UniRef100\_UPI000695E811 \| 99.4 \| 9.7e-16 \| 1.8e-21 \| 116.9 \| 127 \| (1, 127) \| 144 \| (5, 136) \| 320 \| Mor transcription activator family protein \| Mor transcription activator family protein; uniclust \| UniRef100\_A0A956STM1 \| 99.4 \| 9.9e-16 \| 1.8e-21 \| 103.1 \| 112 \| (2, 123) \| 144 \| (12, 124) \| 129 \| Uncharacterized protein \| Uncharacterized protein; uniclust \| UniRef100\_A0A6L8HVT0 \| 99.4 \| 1.2e-15 \| 2.2e-21 \| 106.6 \| 102 \| (8, 121) \| 144 \| (63, 164) \| 165 \| Mor transcription activator domain-containing protein \| Mor transcription activator domain-containing protein; uniclust \| UniRef100\_A0A1J4X7E1 \| 99.4 \| 1.9e-15 \| 3.5e-21 \| 103.1 \| 111 \| (2, 124) \| 144 \| (18, 130) \| 140 \| Mor transcription activator domain-containing protein \| Mor transcription activator domain-containing protein; uniclust \| UniRef100\_A0A839IQE8 \| 99.4 \| 1.9e-15 \| 3.5e-21 \| 104.1 \| 114 \| (2, 126) \| 144 \| (19, 132) \| 149 \| Transcriptional regulator \| Transcriptional regulator; uniclust \| UniRef100\_A0A0A8VVE1 \| 99.4 \| 2.1e-15 \| 4.3e-21 \| 105.9 \| 77 \| (48, 125) \| 144 \| (33, 109) \| 131 \| Transcriptional regulator \| Transcriptional regulator; uniclust \| UniRef100\_A0A9D1UAM0 \| 99.4 \| 2.3e-15 \| 4.3e-21 \| 106.5 \| 110 \| (6, 125) \| 144 \| (28, 150) \| 170 \| Uncharacterized protein \| Uncharacterized protein; uniclust \| UniRef100\_A0A4R3L9E1 \| 99.4 \| 2.6e-15 \| 4.9e-21 \| 107.7 \| 107 \| (6, 122) \| 144 \| (11, 123) \| 172 \| Mor transcription activator family protein \| Mor transcription activator family protein; uniclust \| UniRef100\_A0A4Q8MMR8 \| 99.4 \| 2.8e-15 \| 5.4e-21 \| 108.5 \| 108 \| (6, 123) \| 144 \| (20, 131) \| 186 \| Mor transcription activator domain-containing protein \| Mor transcription activator domain-containing protein; uniclust \| UniRef100\_A0A0P8Y9C8 \| 99.3 \| 3.3e-15 \| 6.1e-21 \| 107.2 \| 107 \| (7, 123) \| 144 \| (27, 133) \| 198 \| Mor transcription activator domain-containing protein \| Mor transcription activator domain-containing protein; uniclust \| UniRef100\_A0A1V0PQE7 \| 99.3 \| 3.5e-15 \| 6.9e-21 \| 103.8 \| 109 \| (6, 126) \| 144 \| (9, 118) \| 131 \| Mor transcription activator domain-containing protein \| Mor transcription activator domain-containing protein; uniclust \| UniRef100\_UPI001FAB7209 \| 99.3 \| 3.8e-15 \| 7e-21 \| 105.7 \| 113 \| (2, 124) \| 144 \| (6, 131) \| 182 \| Mor transcription activator family protein \| Mor transcription activator family protein; uniclust \| UniRef100\_A0A0C9NCV1 \| 99.3 \| 3.5e-15 \| 7.2e-21 \| 113.0 \| 114 \| (3, 127) \| 144 \| (66, 180) \| 217 \| DNA, contig: SP630 \| DNA, contig: SP630; uniclust \| UniRef100\_A0A1Z8PYJ8 \| 99.3 \| 4.4e-15 \| 8.3e-21 \| 101.0 \| 109 \| (4, 127) \| 144 \| (14, 122) \| 126 \| Mor transcription activator domain-containing protein \| Mor transcription activator domain-containing protein; uniclust \| UniRef100\_A0A1H5GMI8 \| 99.3 \| 5.2e-15 \| 9.6e-21 \| 103.3 \| 110 \| (2, 122) \| 144 \| (48, 157) \| 162 \| Transcriptional regulator, Middle operon regulator (Mor) family \| Transcriptional regulator, Middle operon regulator (Mor) family; uniclust \| UniRef100\_A0A081CS68 \| 99.3 \| 4.8e-15 \| 1e-20 \| 113.3 \| 109 \| (2, 123) \| 144 \| (74, 187) \| 216 \| Uncharacterized protein \| Uncharacterized protein; uniclust \| UniRef100\_A0A0D8J127 \| 99.3 \| 5.5e-15 \| 1.1e-20 \| 103.7 \| 77 \| (48, 125) \| 144 \| (18, 95) \| 128 \| Mor transcription activator domain-containing protein \| Mor transcription activator domain-containing protein; uniclust \| UniRef100\_A0A2S5JEN6 \| 99.3 \| 6.5e-15 \| 1.2e-20 \| 100.2 \| 116 \| (1, 126) \| 144 \| (1, 122) \| 136 \| Mor transcription activator family protein \| Mor transcription activator family protein; uniclust \| UniRef100\_A0A1H6HI74 \| 99.3 \| 6.7e-15 \| 1.3e-20 \| 101.3 \| 111 \| (6, 131) \| 144 \| (7, 120) \| 134 \| Helix-turn-helix domain of resolvase \| Helix-turn-helix domain of resolvase; uniclust \| UniRef100\_A0A8I0VL83 \| 99.3 \| 8.8e-15 \| 1.6e-20 \| 98.9 \| 113 \| (2, 123) \| 144 \| (8, 120) \| 130 \| Mor transcription activator domain-containing protein \| Mor transcription activator domain-containing protein; uniclust \| UniRef100\_A0A3T0N1F4 \| 99.3 \| 9.3e-15 \| 1.8e-20 \| 101.6 \| 115 \| (1, 125) \| 144 \| (21, 136) \| 139 \| Mor transcription activator domain-containing protein \| Mor transcription activator domain-containing protein; uniclust \| UniRef100\_A0A3B9L6A9 \| 99.3 \| 9.9e-15 \| 1.8e-20 \| 102.4 \| 107 \| (2, 121) \| 144 \| (48, 154) \| 167 \| Mor transcription activator domain-containing protein \| Mor transcription activator domain-containing protein; uniclust \| UniRef100\_A0A3F3GTU9 \| 99.3 \| 1.1e-14 \| 2e-20 \| 104.2 \| 116 \| (2, 127) \| 144 \| (42, 162) \| 177 \| Putative DNA transposition protein \| Putative DNA transposition protein; uniclust \| UniRef100\_A0A0U2B4P9 \| 99.3 \| 9.9e-15 \| 2.1e-20 \| 103.0 \| 75 \| (51, 126) \| 144 \| (34, 108) \| 127 \| Mor transcription activator domain-containing protein \| Mor transcription activator domain-containing protein; uniclust \| UniRef100\_UPI0019340A10 \| 99.3 \| 1.2e-14 \| 2.3e-20 \| 101.2 \| 106 \| (7, 129) \| 144 \| (17, 123) \| 139 \| MarR family transcriptional regulator \| MarR family transcriptional regulator; uniclust \| UniRef100\_A0A962X3A9 \| 99.3 \| 1.3e-14 \| 2.4e-20 \| 99.1 \| 112 \| (2, 123) \| 144 \| (8, 119) \| 139 \| Uncharacterized protein \| Uncharacterized protein; uniclust \| UniRef100\_A0A1E3UZN7 \| 99.3 \| 1.6e-14 \| 2.9e-20 \| 98.1 \| 107 \| (2, 123) \| 144 \| (15, 122) \| 133 \| Mor transcription activator domain-containing protein \| Mor transcription activator domain-containing protein; uniclust \| UniRef100\_A0A7X3YCW7 \| 99.3 \| 1.8e-14 \| 3.3e-20 \| 94.2 \| 90 \| (19, 120) \| 144 \| (15, 104) \| 105 \| Mor transcription activator domain-containing protein \| Mor transcription activator domain-containing protein; uniclust \| UniRef100\_UPI001EEB33AC \| 99.3 \| 1.8e-14 \| 3.4e-20 \| 98.5 \| 120 \| (2, 124) \| 144 \| (13, 133) \| 140 \| hypothetical protein \| hypothetical protein; uniclust \| UniRef100\_UPI0009EC8F65 \| 99.3 \| 1.9e-14 \| 3.6e-20 \| 104.4 \| 110 \| (2, 121) \| 144 \| (9, 129) \| 213 \| Mor transcription activator family protein \| Mor transcription activator family protein; uniclust \| UniRef100\_UPI0012BB55E9 \| 99.3 \| 2.5e-14 \| 4.5e-20 \| 99.2 \| 100 \| (9, 120) \| 144 \| (6, 105) \| 148 \| Mor transcription activator family protein \| Mor transcription activator family protein; uniclust \| UniRef100\_A0A2I1RGQ4 \| 99.3 \| 2.5e-14 \| 4.6e-20 \| 100.6 \| 114 \| (2, 125) \| 144 \| (23, 149) \| 168 \| Mor transcription activator domain-containing protein \| Mor transcription activator domain-containing protein; uniclust \| UniRef100\_A0A9E4GYE6 \| 99.3 \| 2.8e-14 \| 5.3e-20 \| 97.5 \| 107 \| (2, 124) \| 144 \| (13, 121) \| 125 \| Helix-turn-helix domain-containing protein \| Helix-turn-helix domain-containing protein; uniclust \| UniRef100\_A0A562HZL9 \| 99.2 \| 2.8e-14 \| 5.4e-20 \| 102.3 \| 113 \| (2, 124) \| 144 \| (23, 148) \| 162 \| Mor transcription activator family protein \| Mor transcription activator family protein; uniclust \| UniRef100\_A0A3L6M032 \| 99.2 \| 3e-14 \| 6e-20 \| 104.4 \| 113 \| (2, 124) \| 144 \| (32, 160) \| 180 \| Mor transcription activator domain-containing protein \| Mor transcription activator domain-containing protein; uniclust \| UniRef100\_A0A345CP23 \| 99.2 \| 3.1e-14 \| 6e-20 \| 97.3 \| 84 \| (2, 95) \| 144 \| (14, 102) \| 117 \| Mor transcription activator domain-containing protein \| Mor transcription activator domain-containing protein; uniclust \| UniRef100\_A0A1I5W2T6 \| 99.2 \| 3.1e-14 \| 6.1e-20 \| 98.3 \| 110 \| (3, 123) \| 144 \| (4, 113) \| 127 \| Homeodomain-like domain-containing protein \| Homeodomain-like domain-containing protein; uniclust \| UniRef100\_A0A1B8Q513 \| 99.2 \| 3.2e-14 \| 6.7e-20 \| 100.9 \| 78 \| (50, 128) \| 144 \| (41, 118) \| 131 \| Mor transcription activator domain-containing protein \| Mor transcription activator domain-containing protein; uniclust \| UniRef100\_UPI000A949961 \| 99.2 \| 3.6e-14 \| 7.2e-20 \| 102.5 \| 115 \| (2, 126) \| 144 \| (25, 156) \| 165 \| hypothetical protein \| hypothetical protein; uniclust \| UniRef100\_UPI001FAB374C \| 99.2 \| 4.2e-14 \| 7.7e-20 \| 101.2 \| 115 \| (2, 126) \| 144 \| (14, 128) \| 190 \| Mor transcription activator family protein \| Mor transcription activator family protein; uniclust \| UniRef100\_A0A140L4X4 \| 99.2 \| 4.1e-14 \| 8.4e-20 \| 94.4 \| 75 \| (48, 122) \| 144 \| (15, 90) \| 92 \| Mor transcription activator domain-containing protein \| Mor transcription activator domain-containing protein; uniclust \| UniRef100\_A0A066Q1X8 \| 99.2 \| 3.8e-14 \| 8.5e-20 \| 104.9 \| 74 \| (52, 126) \| 144 \| (70, 143) \| 159 \| Mor transcription activator family protein \| Mor transcription activator family protein; uniclust \| UniRef100\_A0A099GE87 \| 99.2 \| 4.6e-14 \| 9.4e-20 \| 101.9 \| 111 \| (5, 126) \| 144 \| (17, 133) \| 155 \| Mor transcription activator family protein \| Mor transcription activator family protein; uniclust \| UniRef100\_A0A5Q0EN91 \| 99.2 \| 5.4e-14 \| 1e-19 \| 94.1 \| 105 \| (8, 122) \| 144 \| (6, 116) \| 120 \| Mor transcription activator domain-containing protein \| Mor transcription activator domain-containing protein; uniclust \| UniRef100\_A0A952RM33 \| 99.2 \| 5.6e-14 \| 1e-19 \| 97.1 \| 111 \| (2, 124) \| 144 \| (20, 132) \| 148 \| Uncharacterized protein \| Uncharacterized protein; uniclust \| UniRef100\_A0A192IDQ6 \| 99.2 \| 5.9e-14 \| 1.1e-19 \| 96.2 \| 105 \| (2, 125) \| 144 \| (7, 112) \| 122 \| Helix-turn-helix domain-containing protein \| Helix-turn-helix domain-containing protein; uniclust \| UniRef100\_UPI001412D10D \| 99.2 \| 6.4e-14 \| 1.2e-19 \| 96.1 \| 104 \| (6, 124) \| 144 \| (8, 111) \| 129 \| helix-turn-helix domain-containing protein \| helix-turn-helix domain-containing protein; uniclust \| UniRef100\_A0A5C7NL60 \| 99.2 \| 7.2e-14 \| 1.3e-19 \| 98.5 \| 109 \| (3, 125) \| 144 \| (52, 160) \| 170 \| Mor transcription activator domain-containing protein \| Mor transcription activator domain-containing protein; uniclust \| UniRef100\_A0A1L9C9I9 \| 99.2 \| 7.4e-14 \| 1.5e-19 \| 104.5 \| 105 \| (6, 127) \| 144 \| (27, 138) \| 200 \| Helix-turn-helix domain-containing protein \| Helix-turn-helix domain-containing protein; uniclust \| UniRef100\_A0A963FRH9 \| 99.2 \| 8.9e-14 \| 1.6e-19 \| 99.8 \| 108 \| (5, 122) \| 144 \| (6, 113) \| 193 \| Uncharacterized protein \| Uncharacterized protein; uniclust \| UniRef100\_A0A4R2KJZ0 \| 99.2 \| 9.9e-14 \| 1.9e-19 \| 100.2 \| 106 \| (7, 125) \| 144 \| (30, 137) \| 180 \| Mor transcription activator family protein \| Mor transcription activator family protein; uniclust \| UniRef100\_A0A3A9HIU9 \| 99.2 \| 1e-13 \| 2e-19 \| 92.3 \| 74 \| (48, 123) \| 144 \| (13, 86) \| 98 \| Mor transcription activator domain-containing protein \| Mor transcription activator domain-containing protein; uniclust \| UniRef100\_A0A0A0DCH4 \| 99.2 \| 9.9e-14 \| 2e-19 \| 98.9 \| 109 \| (2, 125) \| 144 \| (11, 119) \| 144 \| Mor transcription activator domain-containing protein \| Mor transcription activator domain-containing protein; uniclust \| UniRef100\_B0RUR6 \| 99.2 \| 1.1e-13 \| 2e-19 \| 106.3 \| 125 \| (2, 127) \| 144 \| (126, 258) \| 296 \| Mor transcription activator domain-containing protein \| Mor transcription activator domain-containing protein; uniclust \| UniRef100\_UPI0012DBD419 \| 99.2 \| 1.2e-13 \| 2.2e-19 \| 96.0 \| 110 \| (2, 121) \| 144 \| (32, 151) \| 153 \| Mor transcription activator family protein \| Mor transcription activator family protein; uniclust \| UniRef100\_A0A2E0QVY5 \| 99.2 \| 1.1e-13 \| 2.3e-19 \| 99.3 \| 105 \| (3, 124) \| 144 \| (22, 126) \| 151 \| Resolvase HTH domain-containing protein \| Resolvase HTH domain-containing protein; uniclust \| UniRef100\_A0A951T787 \| 99.2 \| 1.3e-13 \| 2.4e-19 \| 93.5 \| 108 \| (10, 127) \| 144 \| (11, 118) \| 130 \| Helix-turn-helix domain-containing protein \| Helix-turn-helix domain-containing protein; uniclust \| UniRef100\_A0A1M5RUN6 \| 99.2 \| 1.2e-13 \| 2.4e-19 \| 93.1 \| 78 \| (48, 125) \| 144 \| (17, 94) \| 111 \| Mor transcription activator family protein \| Mor transcription activator family protein; uniclust \| UniRef100\_UPI0016882A40 \| 99.2 \| 1.4e-13 \| 2.5e-19 \| 99.9 \| 110 \| (7, 126) \| 144 \| (52, 162) \| 208 \| Mor transcription activator family protein \| Mor transcription activator family protein; uniclust \| UniRef100\_A0A0F5EYF4 \| 99.2 \| 1.3e-13 \| 2.8e-19 \| 95.7 \| 65 \| (56, 121) \| 144 \| (37, 101) \| 113 \| Uncharacterized conserved protein \| Uncharacterized conserved protein; uniclust \| UniRef100\_A0A0J9BG83 \| 99.2 \| 1.5e-13 \| 3e-19 \| 94.8 \| 80 \| (48, 128) \| 144 \| (21, 100) \| 116 \| Mor transcription activator domain-containing protein \| Mor transcription activator domain-containing protein; uniclust \| UniRef100\_K5Z2H2 \| 99.1 \| 2.1e-13 \| 3.9e-19 \| 100.0 \| 121 \| (2, 122) \| 144 \| (95, 220) \| 225 \| Uncharacterized protein \| Uncharacterized protein; uniclust \| UniRef100\_A0A191ZDV3 \| 99.1 \| 2e-13 \| 4e-19 \| 96.7 \| 100 \| (4, 129) \| 144 \| (22, 121) \| 137 \| Mor transcription activator domain-containing protein \| Mor transcription activator domain-containing protein; uniclust \| UniRef100\_A0A127K415 \| 99.1 \| 1.9e-13 \| 4.1e-19 \| 99.5 \| 76 \| (50, 126) \| 144 \| (49, 124) \| 145 \| Mor transcription activator domain-containing protein \| Mor transcription activator domain-containing protein; uniclust \| UniRef100\_UPI001F2B9EB0 \| 99.1 \| 2.2e-13 \| 4.2e-19 \| 89.7 \| 91 \| (22, 124) \| 144 \| (1, 91) \| 100 \| helix-turn-helix domain-containing protein \| helix-turn-helix domain-containing protein; uniclust \| UniRef100\_UPI00233FAF5B \| 99.1 \| 2.3e-13 \| 4.3e-19 \| 90.7 \| 101 \| (14, 124) \| 144 \| (1, 103) \| 116 \| Mor transcription activator family protein \| Mor transcription activator family protein; uniclust \| UniRef100\_A0A560II84 \| 99.1 \| 2.4e-13 \| 4.4e-19 \| 94.4 \| 113 \| (2, 127) \| 144 \| (8, 120) \| 151 \| Mor transcription activator family protein \| Mor transcription activator family protein; uniclust \| UniRef100\_A0A858Q8R5 \| 99.1 \| 2.5e-13 \| 4.6e-19 \| 87.1 \| 76 \| (48, 123) \| 144 \| (7, 82) \| 92 \| Mor transcription activator domain-containing protein \| Mor transcription activator domain-containing protein; uniclust \| UniRef100\_A0A2T6B1K6 \| 99.1 \| 2.5e-13 \| 5e-19 \| 95.8 \| 107 \| (4, 127) \| 144 \| (12, 118) \| 130 \| Homeodomain-like domain-containing protein \| Homeodomain-like domain-containing protein; uniclust \| UniRef100\_A0A076PVY6 \| 99.1 \| 2.9e-13 \| 5.4e-19 \| 96.2 \| 113 \| (2, 124) \| 144 \| (11, 126) \| 177 \| Mor transcription activator domain-containing protein \| Mor transcription activator domain-containing protein; uniclust \| UniRef100\_A0A1X7P996 \| 99.1 \| 2.9e-13 \| 5.7e-19 \| 96.6 \| 113 \| (5, 127) \| 144 \| (21, 135) \| 157 \| Homeodomain-like domain-containing protein \| Homeodomain-like domain-containing protein; uniclust \| UniRef100\_A0A257EV10 \| 99.1 \| 2.7e-13 \| 5.7e-19 \| 99.2 \| 117 \| (1, 132) \| 144 \| (23, 141) \| 152 \| Uncharacterized protein \| Uncharacterized protein; uniclust \| UniRef100\_A0A0H5SSP8 \| 99.1 \| 2.7e-13 \| 5.7e-19 \| 97.3 \| 72 \| (57, 129) \| 144 \| (57, 128) \| 138 \| Middle operon regulator-related protein \| Middle operon regulator-related protein; uniclust \| UniRef100\_A0A315EKC3 \| 99.1 \| 3.4e-13 \| 6.2e-19 \| 87.2 \| 81 \| (40, 120) \| 144 \| (15, 95) \| 96 \| Mor transcription activator domain-containing protein \| Mor transcription activator domain-containing protein; uniclust \| UniRef100\_UPI001F22401B \| 99.1 \| 3.4e-13 \| 6.2e-19 \| 91.9 \| 107 \| (2, 124) \| 144 \| (7, 113) \| 133 \| Mor transcription activator family protein \| Mor transcription activator family protein; uniclust \| UniRef100\_A0A4R8HMW6 \| 99.1 \| 3.7e-13 \| 6.7e-19 \| 93.9 \| 113 \| (3, 126) \| 144 \| (24, 140) \| 155 \| Mor transcription activator family protein \| Mor transcription activator family protein; uniclust \| UniRef100\_A0A062XBM9 \| 99.1 \| 3.2e-13 \| 7e-19 \| 96.9 \| 78 \| (49, 128) \| 144 \| (34, 111) \| 129 \| Phage-associated protein, BcepMu gp16 family \| Phage-associated protein, BcepMu gp16 family; uniclust \| UniRef100\_A0A2G6DFJ6 \| 99.1 \| 4e-13 \| 7.3e-19 \| 95.6 \| 110 \| (5, 124) \| 144 \| (52, 163) \| 178 \| Mor transcription activator domain-containing protein \| Mor transcription activator domain-containing protein; uniclust \| UniRef100\_A0A241XT80 \| 99.1 \| 3.8e-13 \| 7.3e-19 \| 95.8 \| 111 \| (2, 122) \| 144 \| (30, 153) \| 161 \| Putative integral membrane protein \| Putative integral membrane protein; uniclust \| UniRef100\_UPI0018A8FC44 \| 99.1 \| 3.7e-13 \| 7.3e-19 \| 92.0 \| 86 \| (41, 129) \| 144 \| (24, 109) \| 114 \| Mor transcription activator family protein \| Mor transcription activator family protein; uniclust \| UniRef100\_A0A149SVR6 \| 99.1 \| 3.9e-13 \| 7.3e-19 \| 91.3 \| 110 \| (5, 123) \| 144 \| (4, 113) \| 120 \| Resolvase HTH domain-containing protein \| Resolvase HTH domain-containing protein; uniclust \| UniRef100\_A0A142BHK6 \| 99.1 \| 4.6e-13 \| 8.5e-19 \| 94.4 \| 103 \| (8, 122) \| 144 \| (5, 107) \| 158 \| Mor transcription activator domain-containing protein \| Mor transcription activator domain-containing protein; uniclust \| UniRef100\_A0A9E7AG12 \| 99.1 \| 4.7e-13 \| 8.6e-19 \| 93.5 \| 107 \| (3, 126) \| 144 \| (41, 147) \| 156 \| Uncharacterized protein \| Uncharacterized protein; uniclust \| UniRef100\_A0A099KN35 \| 99.1 \| 4.6e-13 \| 8.6e-19 \| 94.3 \| 116 \| (2, 128) \| 144 \| (19, 135) \| 150 \| Mor transcription activator domain protein \| Mor transcription activator domain protein; uniclust \| UniRef100\_A0A7T0FZL0 \| 99.1 \| 4.4e-13 \| 8.8e-19 \| 94.6 \| 83 \| (46, 129) \| 144 \| (35, 117) \| 131 \| Mor transcription activator domain-containing protein \| Mor transcription activator domain-containing protein; uniclust \| UniRef100\_A0A1G0EZ32 \| 99.1 \| 4.9e-13 \| 8.9e-19 \| 88.1 \| 104 \| (7, 120) \| 144 \| (2, 105) \| 107 \| Mor transcription activator domain-containing protein \| Mor transcription activator domain-containing protein; uniclust \| UniRef100\_A0A1Y6CRI6 \| 99.1 \| 5.1e-13 \| 9.5e-19 \| 93.3 \| 111 \| (2, 126) \| 144 \| (20, 131) \| 145 \| Homeodomain-like domain-containing protein \| Homeodomain-like domain-containing protein; uniclust \| UniRef100\_A0A973XNX3 \| 99.1 \| 5.4e-13 \| 9.9e-19 \| 102.9 \| 110 \| (2, 122) \| 144 \| (5, 123) \| 337 \| Helix-turn-helix domain-containing protein \| Helix-turn-helix domain-containing protein; uniclust \| UniRef100\_A0A1Y4LTI1 \| 99.1 \| 5.7e-13 \| 1e-18 \| 86.4 \| 76 \| (51, 126) \| 144 \| (16, 91) \| 97 \| Mor transcription activator domain-containing protein \| Mor transcription activator domain-containing protein; uniclust \| UniRef100\_A0A071LTN5 \| 99.1 \| 5e-13 \| 1.1e-18 \| 101.9 \| 71 \| (55, 126) \| 144 \| (99, 169) \| 198 \| Middle operon regulator \| Middle operon regulator; uniclust \| UniRef100\_A0A022G338 \| 99.1 \| 5.2e-13 \| 1.1e-18 \| 95.0 \| 99 \| (13, 128) \| 144 \| (18, 116) \| 129 \| DNA-binding protein \| DNA-binding protein; uniclust \| UniRef100\_UPI001BD169B3 \| 99.1 \| 7.1e-13 \| 1.3e-18 \| 86.1 \| 84 \| (21, 122) \| 144 \| (2, 85) \| 98 \| helix-turn-helix domain-containing protein \| helix-turn-helix domain-containing protein; uniclust \| UniRef100\_A0A1E7PY91 \| 99.1 \| 6.1e-13 \| 1.3e-18 \| 94.0 \| 69 \| (52, 124) \| 144 \| (39, 107) \| 117 \| Mor transcription activator domain-containing protein \| Mor transcription activator domain-containing protein; uniclust \| UniRef100\_E2CN54 \| 99.1 \| 8.8e-13 \| 1.6e-18 \| 90.8 \| 110 \| (3, 122) \| 144 \| (17, 128) \| 141 \| Resolvase HTH domain-containing protein \| Resolvase HTH domain-containing protein; uniclust \| UniRef100\_A0A367WL54 \| 99.1 \| 8.8e-13 \| 1.6e-18 \| 98.2 \| 111 \| (5, 126) \| 144 \| (22, 133) \| 249 \| Mor transcription activator domain-containing protein \| Mor transcription activator domain-containing protein; uniclust \| UniRef100\_A0A395D1X7 \| 99.1 \| 1.1e-12 \| 2e-18 \| 91.5 \| 117 \| (1, 127) \| 144 \| (1, 123) \| 153 \| Helix-turn-helix domain-containing protein \| Helix-turn-helix domain-containing protein; uniclust \| UniRef100\_A0A1G8B086 \| 99.1 \| 1.1e-12 \| 2e-18 \| 85.0 \| 72 \| (48, 120) \| 144 \| (4, 76) \| 89 \| Mor transcription activator family protein \| Mor transcription activator family protein; uniclust \| UniRef100\_UPI000A927094 \| 99.1 \| 1.1e-12 \| 2e-18 \| 94.5 \| 106 \| (8, 124) \| 144 \| (20, 127) \| 174 \| helix-turn-helix domain-containing protein \| helix-turn-helix domain-containing protein; uniclust \| UniRef100\_A0A521YLK1 \| 99.0 \| 1.2e-12 \| 2.2e-18 \| 97.7 \| 107 \| (5, 122) \| 144 \| (121, 229) \| 251 \| Mor transcription activator domain-containing protein \| Mor transcription activator domain-containing protein; uniclust \| UniRef100\_A4JDC4 \| 99.0 \| 1.6e-12 \| 3e-18 \| 103.0 \| 110 \| (3, 122) \| 144 \| (165, 286) \| 428 \| Mor transcription activator domain-containing protein \| Mor transcription activator domain-containing protein; uniclust \| UniRef100\_A0A1F4QW68 \| 99.0 \| 1.5e-12 \| 3.1e-18 \| 90.6 \| 88 \| (41, 129) \| 144 \| (12, 99) \| 112 \| Mor transcription activator domain-containing protein \| Mor transcription activator domain-containing protein; uniclust \| UniRef100\_A0A4Q0YJD9 \| 99.0 \| 1.7e-12 \| 3.1e-18 \| 91.5 \| 98 \| (6, 115) \| 144 \| (3, 106) \| 164 \| Uncharacterized protein \| Uncharacterized protein; uniclust \| UniRef100\_A0A291IG12 \| 99.0 \| 1.7e-12 \| 3.2e-18 \| 83.8 \| 80 \| (2, 91) \| 144 \| (14, 93) \| 94 \| DnaA N-terminal domain-containing protein \| DnaA N-terminal domain-containing protein; uniclust \| UniRef100\_A0A158E8H2 \| 99.0 \| 1.9e-12 \| 3.4e-18 \| 90.5 \| 81 \| (1, 81) \| 144 \| (28, 111) \| 154 \| Uncharacterized protein \| Uncharacterized protein; uniclust \| UniRef100\_A0A439VN72 \| 99.0 \| 2.3e-12 \| 4.2e-18 \| 95.0 \| 113 \| (3, 125) \| 144 \| (107, 219) \| 227 \| Helix-turn-helix domain-containing protein \| Helix-turn-helix domain-containing protein; uniclust \| UniRef100\_A0A4P6I322 \| 99.0 \| 2.3e-12 \| 4.2e-18 \| 92.4 \| 111 \| (4, 124) \| 144 \| (20, 139) \| 174 \| Mor transcription activator domain-containing protein \| Mor transcription activator domain-containing protein; uniclust \| UniRef100\_A0A1D8IVF8 \| 99.0 \| 2.3e-12 \| 4.2e-18 \| 93.7 \| 112 \| (5, 125) \| 144 \| (6, 117) \| 204 \| Mor transcription activator domain-containing protein \| Mor transcription activator domain-containing protein; uniclust \| UniRef100\_UPI0013A6E23F \| 99.0 \| 2.4e-12 \| 4.3e-18 \| 91.4 \| 115 \| (2, 119) \| 144 \| (52, 166) \| 172 \| hypothetical protein \| hypothetical protein; uniclust \| UniRef100\_A0A3M1PQC8 \| 99.0 \| 2.4e-12 \| 4.5e-18 \| 88.5 \| 105 \| (8, 125) \| 144 \| (7, 111) \| 138 \| Helix-turn-helix domain-containing protein \| Helix-turn-helix domain-containing protein; uniclust \| UniRef100\_A0A3T0N8M3 \| 99.0 \| 2.4e-12 \| 4.5e-18 \| 83.9 \| 82 \| (40, 121) \| 144 \| (7, 88) \| 99 \| Helix-turn-helix domain-containing protein \| Helix-turn-helix domain-containing protein; uniclust \| UniRef100\_A0A0F3KGH2 \| 99.0 \| 2.3e-12 \| 4.9e-18 \| 93.5 \| 76 \| (57, 132) \| 144 \| (58, 133) \| 143 \| Mor transcription activator domain-containing protein \| Mor transcription activator domain-containing protein; uniclust \| UniRef100\_A0A511VBD2 \| 99.0 \| 2.7e-12 \| 5.2e-18 \| 89.1 \| 82 \| (48, 129) \| 144 \| (45, 126) \| 135 \| Mor transcription activator domain-containing protein \| Mor transcription activator domain-containing protein; uniclust \| UniRef100\_N1ZFG9 \| 99.0 \| 2.9e-12 \| 5.3e-18 \| 80.7 \| 75 \| (50, 125) \| 144 \| (5, 79) \| 81 \| Mor transcription activator domain-containing protein \| Mor transcription activator domain-containing protein; uniclust \| UniRef100\_A0A942LWW7 \| 99.0 \| 2.9e-12 \| 5.4e-18 \| 91.2 \| 113 \| (3, 126) \| 144 \| (1, 114) \| 175 \| Uncharacterized protein \| Uncharacterized protein; uniclust \| UniRef100\_A0A2E9SLK3 \| 99.0 \| 3.6e-12 \| 6.5e-18 \| 86.6 \| 107 \| (6, 123) \| 144 \| (3, 109) \| 127 \| Mor transcription activator domain-containing protein \| Mor transcription activator domain-containing protein; uniclust \| UniRef100\_A0A1M5SM93 \| 99.0 \| 3.8e-12 \| 7.1e-18 \| 89.6 \| 110 \| (5, 125) \| 144 \| (23, 132) \| 147 \| Mor transcription activator family protein \| Mor transcription activator family protein; uniclust \| UniRef100\_A0A522VTV5 \| 99.0 \| 3.9e-12 \| 7.1e-18 \| 87.5 \| 112 \| (3, 124) \| 144 \| (14, 126) \| 138 \| Mor transcription activator domain-containing protein \| Mor transcription activator domain-containing protein; uniclust \| UniRef100\_UPI00113DD0F0 \| 99.0 \| 4e-12 \| 7.7e-18 \| 88.5 \| 104 \| (7, 127) \| 144 \| (19, 127) \| 134 \| hypothetical protein \| hypothetical protein; uniclust \| UniRef100\_UPI001BFF8277 \| 99.0 \| 4.5e-12 \| 8.3e-18 \| 87.2 \| 113 \| (2, 124) \| 144 \| (19, 131) \| 138 \| Mor transcription activator family protein \| Mor transcription activator family protein; uniclust \| UniRef100\_A0A2G1LMB9 \| 99.0 \| 4.5e-12 \| 8.5e-18 \| 91.2 \| 104 \| (9, 129) \| 144 \| (8, 113) \| 166 \| Transcriptional regulator \| Transcriptional regulator; uniclust \| UniRef100\_UPI001FFD560B \| 99.0 \| 4.7e-12 \| 8.6e-18 \| 89.2 \| 114 \| (4, 127) \| 144 \| (4, 124) \| 161 \| helix-turn-helix domain-containing protein \| helix-turn-helix domain-containing protein; uniclust \| UniRef100\_UPI001F4838F6 \| 98.9 \| 6.1e-12 \| 1.1e-17 \| 82.4 \| 88 \| (12, 110) \| 144 \| (2, 93) \| 101 \| hypothetical protein \| hypothetical protein; uniclust \| UniRef100\_A0A1X7QDV6 \| 98.9 \| 6e-12 \| 1.1e-17 \| 91.2 \| 105 \| (2, 119) \| 144 \| (13, 119) \| 178 \| Uncharacterized protein \| Uncharacterized protein; uniclust \| UniRef100\_T2JY53 \| 98.9 \| 6.5e-12 \| 1.2e-17 \| 84.5 \| 108 \| (2, 119) \| 144 \| (11, 118) \| 119 \| Mor transcription activator domain-containing protein \| Mor transcription activator domain-containing protein; uniclust \| UniRef100\_R6HUG7 \| 98.9 \| 6.6e-12 \| 1.2e-17 \| 86.7 \| 117 \| (6, 132) \| 144 \| (10, 127) \| 141 \| Mor transcription activator domain-containing protein \| Mor transcription activator domain-containing protein; uniclust \| UniRef100\_A0A2N2RB11 \| 98.9 \| 6.5e-12 \| 1.2e-17 \| 88.4 \| 118 \| (7, 136) \| 144 \| (14, 133) \| 145 \| Uncharacterized protein \| Uncharacterized protein; uniclust \| UniRef100\_A0A1H8U5F3 \| 98.9 \| 6.5e-12 \| 1.3e-17 \| 88.0 \| 75 \| (50, 125) \| 144 \| (38, 112) \| 127 \| Mor transcription activator family protein \| Mor transcription activator family protein; uniclust \| UniRef100\_A0A7H4K6A5 \| 98.9 \| 7.2e-12 \| 1.3e-17 \| 87.6 \| 102 \| (8, 127) \| 144 \| (43, 144) \| 153 \| Mor transcription activator domain-containing protein \| Mor transcription activator domain-containing protein; uniclust \| UniRef100\_A0A564WHC9 \| 98.9 \| 7.6e-12 \| 1.4e-17 \| 88.4 \| 111 \| (2, 122) \| 144 \| (51, 162) \| 164 \| Mor transcription activator domain-containing protein \| Mor transcription activator domain-containing protein; uniclust \| UniRef100\_A0A345DE69 \| 98.9 \| 8e-12 \| 1.5e-17 \| 87.6 \| 109 \| (6, 124) \| 144 \| (3, 111) \| 155 \| Mor transcription activator domain-containing protein \| Mor transcription activator domain-containing protein; uniclust \| UniRef100\_UPI000755AEB5 \| 98.9 \| 9.1e-12 \| 1.7e-17 \| 84.6 \| 80 \| (2, 81) \| 144 \| (3, 84) \| 126 \| hypothetical protein \| hypothetical protein; uniclust \| UniRef100\_UPI0002631E43 \| 98.9 \| 9.1e-12 \| 1.7e-17 \| 89.9 \| 104 \| (6, 126) \| 144 \| (13, 116) \| 190 \| hypothetical protein \| hypothetical protein; uniclust \| UniRef100\_A0A1H7X2Y5 \| 98.9 \| 8.6e-12 \| 1.8e-17 \| 90.5 \| 72 \| (55, 127) \| 144 \| (59, 130) \| 146 \| Transcriptional regulator, Middle operon regulator (Mor) family \| Transcriptional regulator, Middle operon regulator (Mor) family; uniclust \| UniRef100\_A0A2G1LMC0 \| 98.9 \| 1e-11 \| 1.8e-17 \| 86.3 \| 105 \| (6, 127) \| 144 \| (2, 106) \| 146 \| Mor transcription activator domain-containing protein \| Mor transcription activator domain-containing protein; uniclust \| UniRef100\_UPI001F5430B7 \| 98.9 \| 1.2e-11 \| 2.2e-17 \| 86.1 \| 116 \| (5, 120) \| 144 \| (18, 143) \| 147 \| hypothetical protein \| hypothetical protein; uniclust \| UniRef100\_UPI001C0BED20 \| 98.9 \| 1.2e-11 \| 2.2e-17 \| 85.1 \| 98 \| (10, 124) \| 144 \| (34, 131) \| 137 \| hypothetical protein \| hypothetical protein; uniclust \| UniRef100\_UPI0009DAAD1D \| 98.9 \| 1.2e-11 \| 2.4e-17 \| 84.0 \| 109 \| (5, 121) \| 144 \| (4, 112) \| 115 \| helix-turn-helix domain-containing protein \| helix-turn-helix domain-containing protein; uniclust \| UniRef100\_A0A5U2RGL3 \| 98.9 \| 1.3e-11 \| 2.4e-17 \| 89.9 \| 121 \| (2, 126) \| 144 \| (57, 185) \| 190 \| Mor transcription activator family protein \| Mor transcription activator family protein; uniclust \| UniRef100\_A0A135L136 \| 98.9 \| 1.2e-11 \| 2.4e-17 \| 80.4 \| 71 \| (51, 121) \| 144 \| (8, 81) \| 83 \| Mor transcription activator domain-containing protein \| Mor transcription activator domain-containing protein; uniclust \| UniRef100\_A0A081RZJ1 \| 98.9 \| 1.2e-11 \| 2.4e-17 \| 81.4 \| 65 \| (64, 129) \| 144 \| (4, 68) \| 86 \| Mor transcription activator-like protein \| Mor transcription activator-like protein; uniclust \| UniRef100\_A0A6N9P271 \| 98.9 \| 1.4e-11 \| 2.6e-17 \| 86.1 \| 81 \| (50, 130) \| 144 \| (58, 138) \| 152 \| Mor transcription activator domain-containing protein \| Mor transcription activator domain-containing protein; uniclust \| UniRef100\_A0A4V2V2Q2 \| 98.9 \| 1.5e-11 \| 2.7e-17 \| 83.8 \| 99 \| (14, 122) \| 144 \| (13, 116) \| 121 \| Homeodomain-like domain-containing protein \| Homeodomain-like domain-containing protein; uniclust \| UniRef100\_A0A0F2KP01 \| 98.9 \| 1.5e-11 \| 2.8e-17 \| 90.2 \| 110 \| (2, 124) \| 144 \| (10, 121) \| 194 \| Uncharacterized protein \| Uncharacterized protein; uniclust \| UniRef100\_A0A1I3FMF4 \| 98.9 \| 1.6e-11 \| 2.8e-17 \| 86.5 \| 106 \| (6, 123) \| 144 \| (19, 127) \| 159 \| Regulatory protein, luxR family \| Regulatory protein, luxR family; uniclust \| UniRef100\_UPI001FCE4F32 \| 98.9 \| 1.6e-11 \| 2.9e-17 \| 84.7 \| 106 \| (5, 128) \| 144 \| (13, 118) \| 138 \| hypothetical protein \| hypothetical protein; uniclust \| UniRef100\_A0A0M2DU47 \| 98.9 \| 1.4e-11 \| 2.9e-17 \| 89.9 \| 74 \| (52, 126) \| 144 \| (59, 132) \| 154 \| Mor transcription activator domain-containing protein \| Mor transcription activator domain-containing protein; uniclust \| UniRef100\_UPI000AA29095 \| 98.9 \| 1.6e-11 \| 2.9e-17 \| 79.5 \| 87 \| (8, 106) \| 144 \| (5, 91) \| 93 \| hypothetical protein \| hypothetical protein; uniclust \| UniRef100\_UPI0022EB6F38 \| 98.9 \| 1.7e-11 \| 3.1e-17 \| 94.8 \| 116 \| (2, 120) \| 144 \| (201, 316) \| 328 \| DUF1064 domain-containing protein \| DUF1064 domain-containing protein; uniclust \| UniRef100\_A0A4Q7BE38 \| 98.9 \| 1.7e-11 \| 3.2e-17 \| 87.8 \| 120 \| (2, 121) \| 144 \| (48, 170) \| 180 \| Mor transcription activator domain-containing protein \| Mor transcription activator domain-containing protein; uniclust \| UniRef100\_A0A072T170 \| 98.9 \| 1.6e-11 \| 3.4e-17 \| 91.5 \| 105 \| (3, 128) \| 144 \| (41, 147) \| 172 \| Mor transcription activator domain-containing protein \| Mor transcription activator domain-containing protein; uniclust \| UniRef100\_A0A1Y6CHR8 \| 98.9 \| 2e-11 \| 3.7e-17 \| 85.4 \| 108 \| (5, 123) \| 144 \| (39, 146) \| 152 \| Mor transcription activator family protein \| Mor transcription activator family protein; uniclust \| UniRef100\_A0A1G7LRH3 \| 98.9 \| 2e-11 \| 3.7e-17 \| 87.2 \| 103 \| (9, 123) \| 144 \| (22, 127) \| 160 \| Uncharacterized protein \| Uncharacterized protein; uniclust \| UniRef100\_A0A062XDH3 \| 98.9 \| 1.8e-11 \| 3.8e-17 \| 86.7 \| 79 \| (48, 128) \| 144 \| (33, 111) \| 120 \| RNA polymerase sigma factor, sigma-70 family \| RNA polymerase sigma factor, sigma-70 family; uniclust \| UniRef100\_UPI001905C364 \| 98.9 \| 2e-11 \| 3.9e-17 \| 86.0 \| 110 \| (2, 121) \| 144 \| (20, 142) \| 146 \| hypothetical protein \| hypothetical protein; uniclust \| UniRef100\_A0A0K1NDC6 \| 98.8 \| 2.2e-11 \| 4.6e-17 \| 85.5 \| 64 \| (61, 125) \| 144 \| (45, 108) \| 118 \| Mor transcription activator domain-containing protein \| Mor transcription activator domain-containing protein; uniclust \| UniRef100\_A0A450WDE4 \| 98.8 \| 2.5e-11 \| 4.6e-17 \| 85.3 \| 98 \| (9, 128) \| 144 \| (26, 123) \| 156 \| Transcriptional regulator, Middle operon regulator (Mor) family \| Transcriptional regulator, Middle operon regulator (Mor) family; uniclust \| UniRef100\_UPI0015754E79 \| 98.8 \| 2.5e-11 \| 4.6e-17 \| 89.1 \| 101 \| (8, 125) \| 144 \| (109, 209) \| 213 \| hypothetical protein \| hypothetical protein; uniclust \| UniRef100\_A0A3B9PNW9 \| 98.8 \| 2.8e-11 \| 5.2e-17 \| 94.4 \| 114 \| (2, 125) \| 144 \| (201, 325) \| 350 \| Mor transcription activator domain-containing protein \| Mor transcription activator domain-containing protein; uniclust \| UniRef100\_A0A3C1SU58 \| 98.8 \| 2.7e-11 \| 5.3e-17 \| 80.5 \| 64 \| (57, 123) \| 144 \| (25, 88) \| 91 \| Mor transcription activator domain-containing protein (Fragment) \| Mor transcription activator domain-containing protein (Fragment); uniclust \| UniRef100\_A0A084Z6E6 \| 98.8 \| 2.8e-11 \| 5.4e-17 \| 83.5 \| 67 \| (58, 125) \| 144 \| (23, 89) \| 116 \| Positive regulator (Fragment) \| Positive regulator (Fragment); uniclust \| UniRef100\_UPI001BAA3F89 \| 98.8 \| 3.1e-11 \| 5.6e-17 \| 84.9 \| 111 \| (5, 125) \| 144 \| (35, 149) \| 156 \| helix-turn-helix domain-containing protein \| helix-turn-helix domain-containing protein; uniclust \| UniRef100\_A0A109KKA2 \| 98.8 \| 2.8e-11 \| 5.8e-17 \| 88.2 \| 105 \| (13, 131) \| 144 \| (20, 125) \| 149 \| Mor transcription activator family protein \| Mor transcription activator family protein; uniclust \| UniRef100\_A0A3S4VXS7 \| 98.8 \| 3.5e-11 \| 6.7e-17 \| 75.5 \| 66 \| (61, 127) \| 144 \| (3, 68) \| 70 \| Uncharacterized conserved protein \| Uncharacterized conserved protein; uniclust \| UniRef100\_UPI000F4B675E \| 98.8 \| 3.7e-11 \| 6.8e-17 \| 80.1 \| 76 \| (50, 126) \| 144 \| (11, 86) \| 110 \| Mor transcription activator family protein \| Mor transcription activator family protein; uniclust \| UniRef100\_UPI0015D4FDA7 \| 98.8 \| 4.1e-11 \| 7.6e-17 \| 87.9 \| 111 \| (2, 125) \| 144 \| (8, 118) \| 210 \| hypothetical protein \| hypothetical protein; uniclust \| UniRef100\_A0A0J7J773 \| 98.8 \| 3.8e-11 \| 7.7e-17 \| 88.1 \| 68 \| (57, 125) \| 144 \| (76, 143) \| 159 \| Transcriptional regulator, Middle operon regulator (Mor) family \| Transcriptional regulator, Middle operon regulator (Mor) family; uniclust \| UniRef100\_A0A0N1BSW6 \| 98.8 \| 4.6e-11 \| 8.6e-17 \| 86.1 \| 113 \| (2, 127) \| 144 \| (46, 158) \| 173 \| Transposase IS30-like HTH domain-containing protein \| Transposase IS30-like HTH domain-containing protein; uniclust \| UniRef100\_UPI00145E8AB1 \| 98.8 \| 4.7e-11 \| 8.7e-17 \| 85.0 \| 108 \| (6, 123) \| 144 \| (50, 157) \| 169 \| hypothetical protein \| hypothetical protein; uniclust \| UniRef100\_A0A7Z0N040 \| 98.8 \| 5.1e-11 \| 9.7e-17 \| 86.3 \| 105 \| (5, 119) \| 144 \| (13, 117) \| 172 \| Mor transcription activator domain-containing protein \| Mor transcription activator domain-containing protein; uniclust \| UniRef100\_A0A927JUI3 \| 98.8 \| 5.7e-11 \| 1e-16 \| 85.6 \| 106 \| (10, 125) \| 144 \| (13, 128) \| 183 \| Helix-turn-helix domain-containing protein \| Helix-turn-helix domain-containing protein; uniclust \| UniRef100\_A0A2M7Q369 \| 98.8 \| 6.3e-11 \| 1.1e-16 \| 82.4 \| 111 \| (2, 125) \| 144 \| (19, 136) \| 143 \| Mor transcription activator domain-containing protein \| Mor transcription activator domain-containing protein; uniclust \| UniRef100\_R6CNA9 \| 98.8 \| 6.5e-11 \| 1.2e-16 \| 79.1 \| 73 \| (51, 123) \| 144 \| (26, 98) \| 109 \| Mor transcription activator domain-containing protein \| Mor transcription activator domain-containing protein; uniclust \| UniRef100\_A0A9D2KRI3 \| 98.8 \| 6.9e-11 \| 1.3e-16 \| 87.1 \| 113 \| (3, 125) \| 144 \| (81, 204) \| 217 \| Uncharacterized protein \| Uncharacterized protein; uniclust \| UniRef100\_A0A2W4TD25 \| 98.8 \| 7.2e-11 \| 1.3e-16 \| 85.4 \| 112 \| (3, 126) \| 144 \| (72, 183) \| 189 \| Helix-turn-helix domain-containing protein \| Helix-turn-helix domain-containing protein; uniclust \| UniRef100\_A0A2M7G6P5 \| 98.8 \| 7.1e-11 \| 1.4e-16 \| 83.3 \| 110 \| (4, 123) \| 144 \| (20, 140) \| 141 \| Mor transcription activator domain-containing protein \| Mor transcription activator domain-containing protein; uniclust \| UniRef100\_A0A7Z7WCS0 \| 98.8 \| 7.8e-11 \| 1.4e-16 \| 90.0 \| 108 \| (5, 122) \| 144 \| (164, 283) \| 287 \| Uncharacterized conserved protein \| Uncharacterized conserved protein; uniclust \| UniRef100\_A0A344J3T3 \| 98.8 \| 7e-11 \| 1.4e-16 \| 87.8 \| 72 \| (53, 125) \| 144 \| (75, 146) \| 169 \| Transcriptional regulator \| Transcriptional regulator; uniclust \| UniRef100\_UPI00147BFF21 \| 98.8 \| 7.8e-11 \| 1.4e-16 \| 81.9 \| 104 \| (8, 122) \| 144 \| (2, 107) \| 143 \| hypothetical protein \| hypothetical protein; uniclust \| UniRef100\_E9I3C0 \| 98.7 \| 7.9e-11 \| 1.5e-16 \| 82.9 \| 114 \| (7, 130) \| 144 \| (6, 119) \| 156 \| Uncharacterized protein \| Uncharacterized protein; uniclust \| UniRef100\_Q31HV4 \| 98.7 \| 8e-11 \| 1.5e-16 \| 81.9 \| 108 \| (6, 123) \| 144 \| (13, 120) \| 144 \| Mor transcription activator domain-containing protein \| Mor transcription activator domain-containing protein; uniclust \| UniRef100\_A0A317DTH4 \| 98.7 \| 8.1e-11 \| 1.5e-16 \| 87.3 \| 111 \| (2, 125) \| 144 \| (61, 177) \| 226 \| Helix-turn-helix domain-containing protein \| Helix-turn-helix domain-containing protein; uniclust \| UniRef100\_UPI00036465FF \| 98.7 \| 8.5e-11 \| 1.6e-16 \| 83.6 \| 111 \| (2, 122) \| 144 \| (22, 145) \| 148 \| Mor transcription activator family protein \| Mor transcription activator family protein; uniclust \| UniRef100\_UPI000381B40D \| 98.7 \| 9.1e-11 \| 1.7e-16 \| 73.6 \| 63 \| (6, 78) \| 144 \| (3, 65) \| 71 \| hypothetical protein \| hypothetical protein; uniclust \| UniRef100\_A0A1I1UI57 \| 98.7 \| 9.6e-11 \| 1.8e-16 \| 80.2 \| 106 \| (8, 123) \| 144 \| (2, 111) \| 129 \| Homeodomain-like domain-containing protein \| Homeodomain-like domain-containing protein; uniclust \| UniRef100\_UPI0011E59512 \| 98.7 \| 1.1e-10 \| 2.1e-16 \| 85.0 \| 114 \| (6, 131) \| 144 \| (5, 120) \| 197 \| hypothetical protein \| hypothetical protein; uniclust \| UniRef100\_A0A440JH48 \| 98.7 \| 1.1e-10 \| 2.1e-16 \| 88.3 \| 100 \| (8, 125) \| 144 \| (150, 249) \| 264 \| Winged helix-turn-helix domain-containing protein \| Winged helix-turn-helix domain-containing protein; uniclust \| UniRef100\_A0A1I5P0R3 \| 98.7 \| 1.2e-10 \| 2.2e-16 \| 83.6 \| 120 \| (2, 121) \| 144 \| (32, 157) \| 177 \| Mor transcription activator family protein \| Mor transcription activator family protein; uniclust \| UniRef100\_UPI002351D03E \| 98.7 \| 1.2e-10 \| 2.2e-16 \| 77.0 \| 89 \| (22, 120) \| 144 \| (3, 104) \| 104 \| Mor transcription activator family protein \| Mor transcription activator family protein; uniclust \| UniRef100\_A0A031WGA1 \| 98.7 \| 1.2e-10 \| 2.2e-16 \| 77.0 \| 74 \| (41, 122) \| 144 \| (18, 91) \| 93 \| Putative transcriptional regulator, activator Mor \| Putative transcriptional regulator, activator Mor; uniclust \| UniRef100\_A0A961S296 \| 98.7 \| 1.3e-10 \| 2.4e-16 \| 81.2 \| 117 \| (2, 129) \| 144 \| (14, 133) \| 147 \| Helix-turn-helix domain-containing protein \| Helix-turn-helix domain-containing protein; uniclust \| UniRef100\_A0A7Z7WCS0 \| 98.7 \| 1.4e-10 \| 2.5e-16 \| 88.7 \| 113 \| (2, 124) \| 144 \| (5, 129) \| 287 \| Uncharacterized conserved protein \| Uncharacterized conserved protein; uniclust \| UniRef100\_K2JSJ1 \| 98.7 \| 1.4e-10 \| 2.6e-16 \| 80.3 \| 114 \| (2, 126) \| 144 \| (10, 124) \| 138 \| Mor transcription activator domain-containing protein \| Mor transcription activator domain-containing protein; uniclust \| UniRef100\_A0A7Y6R9Q8 \| 98.7 \| 1.4e-10 \| 2.7e-16 \| 83.4 \| 118 \| (2, 122) \| 144 \| (10, 129) \| 160 \| HDOD domain-containing protein \| HDOD domain-containing protein; uniclust \| UniRef100\_A0A0Q2YXZ0 \| 98.7 \| 1.5e-10 \| 2.8e-16 \| 79.8 \| 109 \| (5, 126) \| 144 \| (9, 118) \| 134 \| Mor transcription activator domain-containing protein \| Mor transcription activator domain-containing protein; uniclust \| UniRef100\_A0A650EP32 \| 98.7 \| 1.4e-10 \| 2.8e-16 \| 85.6 \| 98 \| (18, 126) \| 144 \| (35, 146) \| 164 \| Mor transcription activator domain-containing protein \| Mor transcription activator domain-containing protein; uniclust \| UniRef100\_A0A7C1YCC8 \| 98.7 \| 1.5e-10 \| 2.8e-16 \| 77.3 \| 71 \| (51, 122) \| 144 \| (27, 97) \| 99 \| Mor transcription activator domain-containing protein \| Mor transcription activator domain-containing protein; uniclust \| UniRef100\_A0A748ASM0 \| 98.7 \| 1.5e-10 \| 2.8e-16 \| 82.9 \| 111 \| (2, 122) \| 144 \| (33, 157) \| 174 \| Mor transcription activator domain-containing protein \| Mor transcription activator domain-containing protein; uniclust \| UniRef100\_A0A1Y3SP53 \| 98.7 \| 1.5e-10 \| 2.9e-16 \| 75.8 \| 73 \| (50, 122) \| 144 \| (10, 83) \| 85 \| Mor transcription activator domain-containing protein \| Mor transcription activator domain-containing protein; uniclust \| UniRef100\_A0A1U9VJQ1 \| 98.7 \| 1.5e-10 \| 3.1e-16 \| 85.0 \| 108 \| (2, 124) \| 144 \| (37, 149) \| 158 \| DNA-binding protein \| DNA-binding protein; uniclust \| UniRef100\_A0A1G9KD85 \| 98.7 \| 1.6e-10 \| 3.1e-16 \| 78.6 \| 72 \| (49, 120) \| 144 \| (22, 93) \| 105 \| Homeodomain-like domain-containing protein \| Homeodomain-like domain-containing protein; uniclust \| UniRef100\_A0A847URR4 \| 98.7 \| 1.7e-10 \| 3.2e-16 \| 83.2 \| 104 \| (7, 122) \| 144 \| (8, 111) \| 164 \| Helix-turn-helix domain-containing protein \| Helix-turn-helix domain-containing protein; uniclust \| UniRef100\_A0A2X4N2M8 \| 98.7 \| 1.6e-10 \| 3.2e-16 \| 82.4 \| 81 \| (49, 131) \| 144 \| (34, 114) \| 136 \| Uncharacterized conserved protein \| Uncharacterized conserved protein; uniclust \| UniRef100\_A0A5C7NTW8 \| 98.7 \| 2.1e-10 \| 3.9e-16 \| 79.5 \| 109 \| (3, 121) \| 144 \| (12, 136) \| 138 \| Mor transcription activator domain-containing protein \| Mor transcription activator domain-containing protein; uniclust \| UniRef100\_A0A450WGZ3 \| 98.7 \| 2.3e-10 \| 4.3e-16 \| 78.1 \| 81 \| (48, 129) \| 144 \| (35, 115) \| 126 \| Transcriptional regulator, Middle operon regulator (Mor) family \| Transcriptional regulator, Middle operon regulator (Mor) family; uniclust \| UniRef100\_A0A1X7N693 \| 98.7 \| 2.4e-10 \| 4.4e-16 \| 80.8 \| 112 \| (2, 126) \| 144 \| (17, 130) \| 158 \| Mor transcription activator domain-containing protein \| Mor transcription activator domain-containing protein; uniclust \| UniRef100\_UPI001358B363 \| 98.7 \| 2.4e-10 \| 4.5e-16 \| 75.9 \| 92 \| (4, 105) \| 144 \| (4, 104) \| 106 \| hypothetical protein \| hypothetical protein; uniclust \| UniRef100\_A0A1V6M161 \| 98.7 \| 2.5e-10 \| 4.5e-16 \| 79.5 \| 117 \| (2, 129) \| 144 \| (13, 135) \| 142 \| Mor transcription activator domain-containing protein \| Mor transcription activator domain-containing protein; uniclust \| UniRef100\_A0A1Q9A418 \| 98.7 \| 2.3e-10 \| 4.7e-16 \| 85.6 \| 102 \| (5, 123) \| 144 \| (33, 136) \| 180 \| Uncharacterized protein \| Uncharacterized protein; uniclust \| UniRef100\_UPI001884F6D1 \| 98.6 \| 2.6e-10 \| 4.8e-16 \| 81.2 \| 101 \| (3, 121) \| 144 \| (42, 142) \| 165 \| helix-turn-helix domain-containing protein \| helix-turn-helix domain-containing protein; uniclust \| UniRef100\_A0A0S9S3A9 \| 98.6 \| 2.7e-10 \| 5e-16 \| 75.9 \| 95 \| (10, 121) \| 144 \| (4, 98) \| 105 \| Resolvase HTH domain-containing protein \| Resolvase HTH domain-containing protein; uniclust \| UniRef100\_A0A935K439 \| 98.6 \| 2.9e-10 \| 5.3e-16 \| 75.8 \| 98 \| (11, 121) \| 144 \| (3, 105) \| 108 \| Uncharacterized protein \| Uncharacterized protein; uniclust \| UniRef100\_A0A1V5UYJ7 \| 98.6 \| 2.6e-10 \| 5.3e-16 \| 77.3 \| 75 \| (50, 126) \| 144 \| (14, 88) \| 97 \| Mor transcription activator family protein \| Mor transcription activator family protein; uniclust \| UniRef100\_A0A9C9NCZ5 \| 98.6 \| 3e-10 \| 5.6e-16 \| 82.8 \| 108 \| (4, 125) \| 144 \| (31, 138) \| 195 \| Uncharacterized protein \| Uncharacterized protein; uniclust \| UniRef100\_A0A2C8FCX0 \| 98.6 \| 3.2e-10 \| 5.9e-16 \| 70.5 \| 69 \| (50, 121) \| 144 \| (3, 71) \| 72 \| Mor transcription activator domain-containing protein \| Mor transcription activator domain-containing protein; uniclust \| UniRef100\_A0A0M2KGX3 \| 98.6 \| 3.2e-10 \| 6.1e-16 \| 83.4 \| 77 \| (54, 130) \| 144 \| (92, 168) \| 177 \| Regulator of pectin lyase production \| Regulator of pectin lyase production; uniclust \| UniRef100\_UPI0022351324 \| 98.6 \| 3.4e-10 \| 6.3e-16 \| 72.5 \| 80 \| (43, 122) \| 144 \| (3, 82) \| 85 \| Mor transcription activator family protein \| Mor transcription activator family protein; uniclust \| UniRef100\_A0A378WIV1 \| 98.6 \| 3.5e-10 \| 6.4e-16 \| 78.9 \| 108 \| (3, 120) \| 144 \| (20, 142) \| 143 \| Phage protein \| Phage protein; uniclust \| UniRef100\_A0A0A0HG49 \| 98.6 \| 3.5e-10 \| 6.5e-16 \| 78.5 \| 110 \| (4, 125) \| 144 \| (5, 114) \| 129 \| Mor transcription activator family \| Mor transcription activator family; uniclust \| UniRef100\_UPI001EDCF87F \| 98.6 \| 3.7e-10 \| 6.8e-16 \| 84.4 \| 124 \| (2, 125) \| 144 \| (13, 143) \| 236 \| hypothetical protein \| hypothetical protein; uniclust \| UniRef100\_A0A0E3C1B1 \| 98.6 \| 3.4e-10 \| 6.9e-16 \| 80.6 \| 64 \| (63, 127) \| 144 \| (60, 123) \| 130 \| HTH iclR-type domain-containing protein \| HTH iclR-type domain-containing protein; uniclust \| UniRef100\_A0A8T3N508 \| 98.6 \| 4.3e-10 \| 7.9e-16 \| 73.9 \| 86 \| (11, 122) \| 144 \| (2, 87) \| 98 \| Mor transcription activator domain-containing protein \| Mor transcription activator domain-containing protein; uniclust \| UniRef100\_A0A024P8A0 \| 98.6 \| 3.7e-10 \| 8.1e-16 \| 82.2 \| 81 \| (52, 132) \| 144 \| (28, 120) \| 135 \| Mor transcription activator family protein \| Mor transcription activator family protein; uniclust \| UniRef100\_A0A547Q840 \| 98.6 \| 4.8e-10 \| 9.1e-16 \| 78.5 \| 110 \| (4, 123) \| 144 \| (8, 119) \| 132 \| Mor transcription activator domain-containing protein \| Mor transcription activator domain-containing protein; uniclust \| UniRef100\_UPI0015754E79 \| 98.6 \| 5e-10 \| 9.2e-16 \| 82.7 \| 102 \| (8, 126) \| 144 \| (5, 106) \| 213 \| hypothetical protein \| hypothetical protein; uniclust \| UniRef100\_A0A8S5NR56 \| 98.6 \| 5.1e-10 \| 9.4e-16 \| 71.3 \| 70 \| (51, 120) \| 144 \| (9, 79) \| 82 \| Mor transcription activator family \| Mor transcription activator family; uniclust \| UniRef100\_UPI000D19E10B \| 98.6 \| 5.2e-10 \| 9.6e-16 \| 77.5 \| 109 \| (2, 121) \| 144 \| (20, 131) \| 137 \| helix-turn-helix domain-containing protein \| helix-turn-helix domain-containing protein; uniclust \| UniRef100\_A0A257PH65 \| 98.6 \| 5.2e-10 \| 9.6e-16 \| 75.3 \| 97 \| (9, 122) \| 144 \| (6, 102) \| 114 \| Mor transcription activator domain-containing protein \| Mor transcription activator domain-containing protein; uniclust \| UniRef100\_UPI00111B3C11 \| 98.6 \| 5.4e-10 \| 1e-15 \| 81.3 \| 96 \| (2, 107) \| 144 \| (11, 115) \| 191 \| DUF1834 family protein \| DUF1834 family protein; uniclust \| UniRef100\_A0A0L0UL23 \| 98.6 \| 5.3e-10 \| 1e-15 \| 74.7 \| 76 \| (51, 127) \| 144 \| (3, 78) \| 92 \| Mor transcription activator domain-containing protein \| Mor transcription activator domain-containing protein; uniclust \| UniRef100\_A0A1Y2K7L6 \| 98.6 \| 6.3e-10 \| 1.2e-15 \| 76.0 \| 101 \| (5, 123) \| 144 \| (14, 114) \| 124 \| Helix-turn-helix domain-containing protein \| Helix-turn-helix domain-containing protein; uniclust \| UniRef100\_A0A0B3W048 \| 98.6 \| 6.2e-10 \| 1.2e-15 \| 73.1 \| 72 \| (49, 121) \| 144 \| (11, 82) \| 84 \| Transcriptional regulator \| Transcriptional regulator; uniclust \| UniRef100\_A0A357ANT8 \| 98.6 \| 6.6e-10 \| 1.2e-15 \| 74.1 \| 80 \| (48, 127) \| 144 \| (24, 103) \| 107 \| Mor transcription activator domain-containing protein \| Mor transcription activator domain-containing protein; uniclust \| UniRef100\_UPI0020B8F391 \| 98.6 \| 7.1e-10 \| 1.3e-15 \| 81.0 \| 112 \| (7, 128) \| 144 \| (16, 130) \| 196 \| hypothetical protein \| hypothetical protein; uniclust \| UniRef100\_A0A1V5V371 \| 98.6 \| 6.9e-10 \| 1.3e-15 \| 77.1 \| 75 \| (52, 126) \| 144 \| (24, 103) \| 117 \| Mor transcription activator family protein \| Mor transcription activator family protein; uniclust \| UniRef100\_N6VLK0 \| 98.6 \| 7.3e-10 \| 1.3e-15 \| 70.6 \| 68 \| (2, 79) \| 144 \| (8, 79) \| 82 \| Uncharacterized protein \| Uncharacterized protein; uniclust \| UniRef100\_A0A090SPL0 \| 98.6 \| 7.4e-10 \| 1.4e-15 \| 77.3 \| 99 \| (2, 114) \| 144 \| (9, 108) \| 143 \| Transcriptional regulator \| Transcriptional regulator; uniclust \| UniRef100\_A0A1J5INX3 \| 98.6 \| 7.3e-10 \| 1.4e-15 \| 71.2 \| 69 \| (54, 124) \| 144 \| (2, 70) \| 80 \| Mor transcription activator domain-containing protein \| Mor transcription activator domain-containing protein; uniclust \| UniRef100\_A0A316MS27 \| 98.6 \| 7.4e-10 \| 1.4e-15 \| 75.7 \| 74 \| (52, 126) \| 144 \| (31, 104) \| 119 \| DNA-binding protein \| DNA-binding protein; uniclust \| UniRef100\_A0A1B8TFA3 \| 98.6 \| 7e-10 \| 1.4e-15 \| 78.6 \| 76 \| (52, 128) \| 144 \| (34, 109) \| 129 \| Mor transcription activator-like protein \| Mor transcription activator-like protein; uniclust \| UniRef100\_UPI00045E7B36 \| 98.5 \| 8.1e-10 \| 1.5e-15 \| 71.9 \| 77 \| (44, 120) \| 144 \| (2, 78) \| 92 \| hypothetical protein \| hypothetical protein; uniclust \| UniRef100\_A0A1A9RK94 \| 98.5 \| 7.1e-10 \| 1.5e-15 \| 85.8 \| 109 \| (1, 142) \| 144 \| (80, 194) \| 205 \| Uncharacterized protein \| Uncharacterized protein; uniclust \| UniRef100\_A0A345DDH0 \| 98.5 \| 8.2e-10 \| 1.5e-15 \| 76.4 \| 111 \| (3, 123) \| 144 \| (9, 120) \| 134 \| Mor transcription activator domain-containing protein \| Mor transcription activator domain-containing protein; uniclust \| UniRef100\_UPI0016AEF5FB \| 98.5 \| 8.3e-10 \| 1.5e-15 \| 71.5 \| 75 \| (2, 79) \| 144 \| (6, 82) \| 90 \| hypothetical protein \| hypothetical protein; uniclust \| UniRef100\_A0A950CSA0 \| 98.5 \| 8.8e-10 \| 1.6e-15 \| 71.8 \| 63 \| (6, 78) \| 144 \| (31, 93) \| 93 \| Uncharacterized protein (Fragment) \| Uncharacterized protein (Fragment); uniclust \| UniRef100\_A0A1S2CM03 \| 98.5 \| 8.3e-10 \| 1.6e-15 \| 78.6 \| 79 \| (50, 129) \| 144 \| (44, 122) \| 134 \| Mor transcription activator domain-containing protein \| Mor transcription activator domain-containing protein; uniclust \| UniRef100\_A0A2A2JZL6 \| 98.5 \| 8.9e-10 \| 1.6e-15 \| 87.6 \| 93 \| (2, 105) \| 144 \| (69, 161) \| 392 \| Uncharacterized protein \| Uncharacterized protein; uniclust \| UniRef100\_A0A1M5PL34 \| 98.5 \| 8.5e-10 \| 1.7e-15 \| 75.7 \| 74 \| (49, 126) \| 144 \| (17, 90) \| 107 \| Mor transcription activator family protein \| Mor transcription activator family protein; uniclust \| UniRef100\_A0A072TCN7 \| 98.5 \| 9.6e-10 \| 1.8e-15 \| 88.7 \| 76 \| (53, 129) \| 144 \| (362, 437) \| 450 \| Mor family transcriptional regulator \| Mor family transcriptional regulator; uniclust \| UniRef100\_A0A0B1TT43 \| 98.5 \| 1.1e-09 \| 2e-15 \| 73.7 \| 74 \| (51, 124) \| 144 \| (8, 85) \| 101 \| Mor transcription activator domain-containing protein \| Mor transcription activator domain-containing protein; uniclust \| UniRef100\_A0A846UIR1 \| 98.5 \| 1.1e-09 \| 2e-15 \| 69.7 \| 67 \| (52, 122) \| 144 \| (14, 80) \| 81 \| Mor family transcriptional regulator \| Mor family transcriptional regulator; uniclust \| UniRef100\_A0A1Q6QCR6 \| 98.5 \| 1.2e-09 \| 2.1e-15 \| 68.0 \| 68 \| (54, 122) \| 144 \| (2, 69) \| 71 \| Mor transcription activator domain-containing protein \| Mor transcription activator domain-containing protein; uniclust \| UniRef100\_A0A147H032 \| 98.5 \| 1.2e-09 \| 2.1e-15 \| 69.0 \| 60 \| (61, 121) \| 144 \| (17, 76) \| 77 \| Mor transcription activator domain-containing protein \| Mor transcription activator domain-containing protein; uniclust \| UniRef100\_A0A1H0G9V9 \| 98.5 \| 1.2e-09 \| 2.3e-15 \| 70.8 \| 59 \| (62, 125) \| 144 \| (18, 76) \| 81 \| Mor transcription activator family protein \| Mor transcription activator family protein; uniclust \| UniRef100\_A0A3N9RYD7 \| 98.5 \| 1.4e-09 \| 2.5e-15 \| 81.5 \| 112 \| (2, 124) \| 144 \| (54, 166) \| 234 \| Uncharacterized protein \| Uncharacterized protein; uniclust \| UniRef100\_UPI0018E781FA \| 98.5 \| 1.4e-09 \| 2.6e-15 \| 77.5 \| 103 \| (6, 122) \| 144 \| (31, 136) \| 162 \| LuxR C-terminal-related transcriptional regulator \| LuxR C-terminal-related transcriptional regulator; uniclust \| UniRef100\_A0A522LCM6 \| 98.5 \| 1.4e-09 \| 2.6e-15 \| 81.1 \| 120 \| (3, 132) \| 144 \| (30, 164) \| 226 \| Mor transcription activator domain-containing protein \| Mor transcription activator domain-containing protein; uniclust \| UniRef100\_A0A2M8EEU5 \| 98.5 \| 1.4e-09 \| 2.6e-15 \| 80.0 \| 113 \| (2, 124) \| 144 \| (30, 164) \| 181 \| Mor transcription activator domain-containing protein \| Mor transcription activator domain-containing protein; uniclust \| UniRef100\_A0A265Q2W8 \| 98.5 \| 1.4e-09 \| 2.7e-15 \| 72.5 \| 75 \| (49, 125) \| 144 \| (15, 89) \| 93 \| Mor transcription activator domain-containing protein \| Mor transcription activator domain-containing protein; uniclust \| UniRef100\_A0A8K2AGT6 \| 98.5 \| 1.5e-09 \| 2.7e-15 \| 78.1 \| 105 \| (4, 120) \| 144 \| (4, 108) \| 173 \| Mor transcription activator domain-containing protein \| Mor transcription activator domain-containing protein; uniclust \| UniRef100\_A0A0C5E2Q8 \| 98.5 \| 1.5e-09 \| 2.9e-15 \| 76.5 \| 76 \| (50, 126) \| 144 \| (45, 120) \| 138 \| Mor transcription activator domain-containing protein \| Mor transcription activator domain-containing protein; uniclust \| UniRef100\_UPI000695E811 \| 98.5 \| 1.6e-09 \| 3e-15 \| 84.6 \| 120 \| (3, 122) \| 144 \| (146, 270) \| 320 \| Mor transcription activator family protein \| Mor transcription activator family protein; uniclust \| UniRef100\_UPI0021D601D8 \| 98.5 \| 1.8e-09 \| 3.3e-15 \| 70.7 \| 71 \| (54, 124) \| 144 \| (2, 81) \| 95 \| Mor transcription activator family protein \| Mor transcription activator family protein; uniclust \| UniRef100\_A0A2U2AMQ7 \| 98.5 \| 1.8e-09 \| 3.4e-15 \| 81.5 \| 115 \| (2, 126) \| 144 \| (115, 240) \| 249 \| Mor transcription activator domain-containing protein \| Mor transcription activator domain-containing protein; uniclust \| UniRef100\_A0A3D5S739 \| 98.5 \| 2e-09 \| 3.8e-15 \| 73.4 \| 101 \| (10, 120) \| 144 \| (1, 117) \| 121 \| Mor transcription activator domain-containing protein \| Mor transcription activator domain-containing protein; uniclust \| UniRef100\_A0A840C4F7 \| 98.4 \| 2.4e-09 \| 4.4e-15 \| 76.6 \| 116 \| (1, 125) \| 144 \| (37, 160) \| 165 \| Mor transcription activator domain-containing protein \| Mor transcription activator domain-containing protein; uniclust \| UniRef100\_A0A1S1HV25 \| 98.4 \| 2.4e-09 \| 4.4e-15 \| 72.8 \| 76 \| (50, 125) \| 144 \| (30, 105) \| 119 \| Mor transcription activator domain-containing protein \| Mor transcription activator domain-containing protein; uniclust \| UniRef100\_B6IMZ5 \| 98.4 \| 2.5e-09 \| 4.6e-15 \| 77.3 \| 114 \| (3, 126) \| 144 \| (53, 172) \| 178 \| Uncharacterized protein \| Uncharacterized protein; uniclust \| UniRef100\_A0A7Y6TZ55 \| 98.4 \| 2.5e-09 \| 4.6e-15 \| 72.8 \| 70 \| (60, 129) \| 144 \| (41, 110) \| 119 \| Mor transcription activator domain-containing protein \| Mor transcription activator domain-containing protein; uniclust \| UniRef100\_UPI00215AD7B1 \| 98.4 \| 2.5e-09 \| 4.6e-15 \| 73.6 \| 78 \| (46, 123) \| 144 \| (15, 92) \| 128 \| Mor transcription activator family protein \| Mor transcription activator family protein; uniclust \| UniRef100\_A0A0J6KKT4 \| 98.4 \| 2.3e-09 \| 4.7e-15 \| 78.9 \| 79 \| (52, 130) \| 144 \| (50, 128) \| 153 \| Mor transcription activator domain-containing protein \| Mor transcription activator domain-containing protein; uniclust \| UniRef100\_A0A1M5S655 \| 98.4 \| 2.4e-09 \| 4.9e-15 \| 76.8 \| 66 \| (57, 122) \| 144 \| (46, 112) \| 130 \| Mor transcription activator family protein \| Mor transcription activator family protein; uniclust \| UniRef100\_A0A1G3LYB2 \| 98.4 \| 2.7e-09 \| 4.9e-15 \| 71.0 \| 75 \| (49, 125) \| 144 \| (18, 93) \| 103 \| Mor transcription activator domain-containing protein \| Mor transcription activator domain-containing protein; uniclust \| UniRef100\_A0A258LJQ1 \| 98.4 \| 2.9e-09 \| 5.3e-15 \| 70.3 \| 64 \| (7, 80) \| 144 \| (3, 66) \| 99 \| Transcriptional regulator \| Transcriptional regulator; uniclust \| UniRef100\_UPI0012EC8521 \| 98.4 \| 3.1e-09 \| 5.7e-15 \| 68.9 \| 81 \| (43, 128) \| 144 \| (2, 83) \| 89 \| Mor transcription activator family protein \| Mor transcription activator family protein; uniclust \| UniRef100\_A0A9E3YQJ6 \| 98.4 \| 3.2e-09 \| 5.9e-15 \| 72.3 \| 109 \| (3, 121) \| 144 \| (2, 116) \| 119 \| Uncharacterized protein \| Uncharacterized protein; uniclust \| UniRef100\_A0A6N6T8X9 \| 98.4 \| 3.2e-09 \| 6e-15 \| 69.4 \| 65 \| (61, 126) \| 144 \| (17, 81) \| 85 \| Mor transcription activator domain-containing protein \| Mor transcription activator domain-containing protein; uniclust \| UniRef100\_A0A6F9X1I1 \| 98.4 \| 3.3e-09 \| 6e-15 \| 75.1 \| 114 \| (3, 127) \| 144 \| (19, 141) \| 153 \| Mor transcription activator domain-containing protein \| Mor transcription activator domain-containing protein; uniclust \| UniRef100\_UPI000428175B \| 98.4 \| 3.2e-09 \| 6.2e-15 \| 72.7 \| 70 \| (57, 127) \| 144 \| (28, 97) \| 108 \| Mor transcription activator family protein \| Mor transcription activator family protein; uniclust \| UniRef100\_A0A317E9Z1 \| 98.4 \| 3.5e-09 \| 6.5e-15 \| 76.2 \| 110 \| (2, 125) \| 144 \| (10, 123) \| 172 \| Helix-turn-helix domain-containing protein \| Helix-turn-helix domain-containing protein; uniclust \| UniRef100\_A0A292RV70 \| 98.4 \| 3.7e-09 \| 6.7e-15 \| 70.0 \| 72 \| (51, 122) \| 144 \| (27, 98) \| 100 \| Mor transcription activator domain-containing protein \| Mor transcription activator domain-containing protein; uniclust \| UniRef100\_A0A1H3CN96 \| 98.4 \| 3.9e-09 \| 7.5e-15 \| 73.3 \| 73 \| (55, 128) \| 144 \| (36, 110) \| 115 \| Transcriptional regulator, Middle operon regulator (Mor) family \| Transcriptional regulator, Middle operon regulator (Mor) family; uniclust \| UniRef100\_UPI001C5AB0C2 \| 98.4 \| 4.2e-09 \| 7.7e-15 \| 72.0 \| 102 \| (14, 125) \| 144 \| (4, 116) \| 122 \| Mor transcription activator family protein \| Mor transcription activator family protein; uniclust \| UniRef100\_G1UW96 \| 98.4 \| 4.3e-09 \| 7.9e-15 \| 81.3 \| 93 \| (20, 123) \| 144 \| (50, 151) \| 293 \| Mor transcription activator domain-containing protein \| Mor transcription activator domain-containing protein; uniclust \| UniRef100\_A0A9D5VAY5 \| 98.4 \| 4.5e-09 \| 8.2e-15 \| 79.3 \| 110 \| (3, 123) \| 144 \| (9, 119) \| 242 \| Uncharacterized protein \| Uncharacterized protein; uniclust \| UniRef100\_A0A1E4LD59 \| 98.4 \| 4.5e-09 \| 8.5e-15 \| 72.5 \| 77 \| (49, 125) \| 144 \| (28, 104) \| 116 \| Mor transcription activator domain-containing protein \| Mor transcription activator domain-containing protein; uniclust \| UniRef100\_A0A0R1UPE2 \| 98.4 \| 4.4e-09 \| 8.5e-15 \| 68.4 \| 74 \| (57, 132) \| 144 \| (2, 75) \| 78 \| Mor transcription activator domain-containing protein \| Mor transcription activator domain-containing protein; uniclust \| UniRef100\_UPI0022B816E3 \| 98.4 \| 4.9e-09 \| 9e-15 \| 73.0 \| 101 \| (10, 127) \| 144 \| (28, 128) \| 137 \| hypothetical protein \| hypothetical protein; uniclust \| UniRef100\_A0A7D5LKB6 \| 98.4 \| 4.8e-09 \| 9.3e-15 \| 73.8 \| 69 \| (57, 125) \| 144 \| (47, 115) \| 127 \| Mor transcription activator domain-containing protein \| Mor transcription activator domain-containing protein; uniclust \| UniRef100\_A0A7W7YWM5 \| 98.4 \| 5.1e-09 \| 9.4e-15 \| 74.6 \| 95 \| (9, 121) \| 144 \| (57, 151) \| 159 \| Mor transcription activator domain-containing protein \| Mor transcription activator domain-containing protein; uniclust \| UniRef100\_A0A0H3A7L4 \| 98.4 \| 4.6e-09 \| 9.5e-15 \| 74.2 \| 77 \| (49, 125) \| 144 \| (21, 107) \| 115 \| Mor transcription activator domain-containing protein \| Mor transcription activator domain-containing protein; uniclust \| UniRef100\_A0A069DBT8 \| 98.4 \| 4.7e-09 \| 9.6e-15 \| 76.0 \| 73 \| (51, 123) \| 144 \| (22, 106) \| 133 \| Mor transcription activator domain-containing protein \| Mor transcription activator domain-containing protein; uniclust \| UniRef100\_A0A1X7QCW3 \| 98.4 \| 5.5e-09 \| 1e-14 \| 79.0 \| 112 \| (2, 126) \| 144 \| (52, 165) \| 245 \| Helix-turn-helix domain-containing protein \| Helix-turn-helix domain-containing protein; uniclust \| UniRef100\_UPI0022745754 \| 98.4 \| 5.5e-09 \| 1e-14 \| 71.3 \| 105 \| (3, 124) \| 144 \| (12, 116) \| 120 \| hypothetical protein \| hypothetical protein; uniclust \| UniRef100\_A0A3R8L878 \| 98.3 \| 5.8e-09 \| 1.1e-14 \| 79.7 \| 109 \| (2, 122) \| 144 \| (84, 195) \| 249 \| DNA-binding protein \| DNA-binding protein; uniclust \| UniRef100\_A0A0C5E5W8 \| 98.3 \| 5.5e-09 \| 1.1e-14 \| 76.8 \| 72 \| (57, 128) \| 144 \| (67, 139) \| 156 \| Mor transcription activator domain-containing protein \| Mor transcription activator domain-containing protein; uniclust \| UniRef100\_A0A2S7CGT1 \| 98.3 \| 5.4e-09 \| 1.1e-14 \| 75.2 \| 74 \| (51, 124) \| 144 \| (40, 118) \| 131 \| Mor transcription activator domain-containing protein \| Mor transcription activator domain-containing protein; uniclust \| UniRef100\_A0A1V8NVX7 \| 98.3 \| 5.7e-09 \| 1.1e-14 \| 80.9 \| 75 \| (51, 126) \| 144 \| (52, 126) \| 225 \| Mor transcription activator domain-containing protein \| Mor transcription activator domain-containing protein; uniclust \| UniRef100\_A0A2S6H5A2 \| 98.3 \| 6.1e-09 \| 1.1e-14 \| 67.5 \| 58 \| (60, 120) \| 144 \| (24, 81) \| 82 \| Mor transcription activator family protein \| Mor transcription activator family protein; uniclust \| UniRef100\_UPI000B1C3237 \| 98.3 \| 6.4e-09 \| 1.2e-14 \| 86.3 \| 85 \| (2, 96) \| 144 \| (9, 98) \| 576 \| DNA circularization N-terminal domain-containing protein \| DNA circularization N-terminal domain-containing protein; uniclust \| UniRef100\_A0A0P7M6A8 \| 98.3 \| 6.5e-09 \| 1.2e-14 \| 70.8 \| 81 \| (52, 132) \| 144 \| (33, 113) \| 118 \| Mor transcription activator domain-containing protein \| Mor transcription activator domain-containing protein; uniclust \| UniRef100\_A0A3D1VPS2 \| 98.3 \| 6.7e-09 \| 1.3e-14 \| 67.4 \| 69 \| (52, 121) \| 144 \| (9, 77) \| 83 \| Mor transcription activator domain-containing protein \| Mor transcription activator domain-containing protein; uniclust \| UniRef100\_UPI00215ADC39 \| 98.3 \| 7e-09 \| 1.3e-14 \| 73.5 \| 83 \| (41, 124) \| 144 \| (60, 151) \| 153 \| Mor transcription activator family protein \| Mor transcription activator family protein; uniclust \| UniRef100\_A0A4Y6UA46 \| 98.3 \| 7e-09 \| 1.3e-14 \| 74.8 \| 109 \| (5, 118) \| 144 \| (59, 168) \| 172 \| Uncharacterized protein \| Uncharacterized protein; uniclust \| UniRef100\_A0A942QUK7 \| 98.3 \| 6.8e-09 \| 1.3e-14 \| 71.1 \| 68 \| (51, 122) \| 144 \| (40, 107) \| 110 \| Mor transcription activator domain-containing protein \| Mor transcription activator domain-containing protein; uniclust \| UniRef100\_A0A2G4RG90 \| 98.3 \| 7.6e-09 \| 1.4e-14 \| 70.6 \| 102 \| (6, 124) \| 144 \| (8, 115) \| 119 \| Mor transcription activator domain-containing protein \| Mor transcription activator domain-containing protein; uniclust \| UniRef100\_UPI0009F57BC5 \| 98.3 \| 7.5e-09 \| 1.4e-14 \| 68.9 \| 69 \| (56, 125) \| 144 \| (15, 83) \| 93 \| Mor transcription activator family protein \| Mor transcription activator family protein; uniclust \| UniRef100\_A0A3S5A9Q7 \| 98.3 \| 7.3e-09 \| 1.4e-14 \| 72.3 \| 69 \| (58, 126) \| 144 \| (40, 109) \| 114 \| Putative bacteriophage transcriptional regulator \| Putative bacteriophage transcriptional regulator; uniclust \| UniRef100\_A0A953CGI5 \| 98.3 \| 8.3e-09 \| 1.5e-14 \| 77.0 \| 109 \| (7, 125) \| 144 \| (19, 130) \| 220 \| Uncharacterized protein \| Uncharacterized protein; uniclust \| UniRef100\_UPI0015D49757 \| 98.3 \| 8.3e-09 \| 1.5e-14 \| 74.6 \| 107 \| (2, 122) \| 144 \| (15, 134) \| 175 \| hypothetical protein \| hypothetical protein; uniclust \| UniRef100\_A0A2H6APG1 \| 98.3 \| 7.8e-09 \| 1.6e-14 \| 71.7 \| 74 \| (50, 126) \| 144 \| (25, 98) \| 106 \| Mor transcription activator domain-containing protein \| Mor transcription activator domain-containing protein; uniclust \| UniRef100\_A0A2D8HLU1 \| 98.3 \| 8.6e-09 \| 1.6e-14 \| 73.7 \| 111 \| (3, 123) \| 144 \| (35, 149) \| 161 \| Mor transcription activator domain-containing protein \| Mor transcription activator domain-containing protein; uniclust \| UniRef100\_A0A3N0ZBC0 \| 98.3 \| 8.8e-09 \| 1.7e-14 \| 68.2 \| 70 \| (49, 119) \| 144 \| (19, 88) \| 91 \| Helix-turn-helix domain-containing protein \| Helix-turn-helix domain-containing protein; uniclust \| UniRef100\_A4JDC4 \| 98.3 \| 9.5e-09 \| 1.7e-14 \| 82.9 \| 109 \| (6, 124) \| 144 \| (16, 137) \| 428 \| Mor transcription activator domain-containing protein \| Mor transcription activator domain-containing protein; uniclust \| UniRef100\_B6IMZ4 \| 98.3 \| 9.8e-09 \| 1.8e-14 \| 72.7 \| 110 \| (6, 127) \| 144 \| (25, 135) \| 151 \| Mor transcription activator domain-containing protein \| Mor transcription activator domain-containing protein; uniclust \| UniRef100\_A0A0F4QDL7 \| 98.3 \| 9.4e-09 \| 1.8e-14 \| 73.5 \| 75 \| (52, 127) \| 144 \| (38, 112) \| 137 \| Transcriptional regulator \| Transcriptional regulator; uniclust \| UniRef100\_A0A3S1BGX4 \| 98.3 \| 1e-08 \| 1.9e-14 \| 74.4 \| 69 \| (57, 126) \| 144 \| (96, 164) \| 178 \| Mor transcription activator domain-containing protein \| Mor transcription activator domain-containing protein; uniclust \| UniRef100\_UPI0010BF961F \| 98.3 \| 1e-08 \| 1.9e-14 \| 71.9 \| 105 \| (10, 124) \| 144 \| (15, 122) \| 142 \| helix-turn-helix domain-containing protein \| helix-turn-helix domain-containing protein; uniclust \| UniRef100\_A0A0Q5ZD56 \| 98.3 \| 1e-08 \| 2.1e-14 \| 71.5 \| 62 \| (63, 124) \| 144 \| (41, 102) \| 105 \| Resolvase HTH domain-containing protein \| Resolvase HTH domain-containing protein; uniclust \| UniRef100\_A0A923AS64 \| 98.3 \| 1.2e-08 \| 2.3e-14 \| 76.7 \| 111 \| (2, 122) \| 144 \| (91, 218) \| 232 \| Mor transcription activator domain-containing protein \| Mor transcription activator domain-containing protein; uniclust \| UniRef100\_UPI00102DFDF1 \| 98.3 \| 1.2e-08 \| 2.3e-14 \| 67.9 \| 70 \| (54, 124) \| 144 \| (29, 98) \| 102 \| Mor transcription activator family protein \| Mor transcription activator family protein; uniclust \| UniRef100\_T0IRN1 \| 98.3 \| 1.3e-08 \| 2.3e-14 \| 72.0 \| 108 \| (11, 127) \| 144 \| (27, 135) \| 149 \| Mor transcription activator domain-containing protein \| Mor transcription activator domain-containing protein; uniclust \| UniRef100\_UPI00117FD392 \| 98.3 \| 1.2e-08 \| 2.3e-14 \| 75.0 \| 113 \| (2, 119) \| 144 \| (73, 189) \| 190 \| hypothetical protein \| hypothetical protein; uniclust \| UniRef100\_A0A1G6EHB1 \| 98.3 \| 1.2e-08 \| 2.3e-14 \| 73.6 \| 78 \| (49, 128) \| 144 \| (57, 134) \| 138 \| Mor transcription activator family protein \| Mor transcription activator family protein; uniclust \| UniRef100\_A0A511V899 \| 98.3 \| 1.3e-08 \| 2.3e-14 \| 68.1 \| 75 \| (48, 122) \| 144 \| (20, 94) \| 104 \| Mor transcription activator domain-containing protein \| Mor transcription activator domain-containing protein; uniclust \| UniRef100\_A0A014NJ40 \| 98.3 \| 1.2e-08 \| 2.3e-14 \| 73.3 \| 69 \| (57, 126) \| 144 \| (57, 125) \| 141 \| Mor transcription activator domain-containing protein \| Mor transcription activator domain-containing protein; uniclust \| UniRef100\_UPI0021C4554C \| 98.3 \| 1.3e-08 \| 2.4e-14 \| 64.7 \| 55 \| (67, 122) \| 144 \| (3, 57) \| 78 \| Mor transcription activator family protein \| Mor transcription activator family protein; uniclust \| UniRef100\_A0A2C5TWB0 \| 98.3 \| 1.4e-08 \| 2.5e-14 \| 71.4 \| 90 \| (20, 119) \| 144 \| (1, 103) \| 143 \| Mor transcription activator domain-containing protein \| Mor transcription activator domain-containing protein; uniclust \| UniRef100\_A0A0J1GQ97 \| 98.2 \| 1.5e-08 \| 2.9e-14 \| 72.3 \| 76 \| (52, 127) \| 144 \| (20, 107) \| 123 \| Mor transcription activator domain-containing protein \| Mor transcription activator domain-containing protein; uniclust \| UniRef100\_A0A3R1ADR8 \| 98.2 \| 1.5e-08 \| 2.9e-14 \| 71.9 \| 76 \| (52, 127) \| 144 \| (43, 118) \| 130 \| Mor transcription activator domain-containing protein \| Mor transcription activator domain-containing protein; uniclust \| UniRef100\_A0A1E3ZIA4 \| 98.2 \| 1.6e-08 \| 3e-14 \| 69.3 \| 80 \| (44, 128) \| 144 \| (26, 105) \| 121 \| Mor transcription activator domain-containing protein \| Mor transcription activator domain-containing protein; uniclust \| UniRef100\_A0A1A9WW73 \| 98.2 \| 1.7e-08 \| 3.1e-14 \| 72.9 \| 78 \| (50, 128) \| 144 \| (79, 156) \| 171 \| Mor transcription activator domain-containing protein \| Mor transcription activator domain-containing protein; uniclust \| UniRef100\_A0A0A8H9R4 \| 98.2 \| 1.5e-08 \| 3.1e-14 \| 71.5 \| 62 \| (60, 124) \| 144 \| (34, 103) \| 112 \| Transcriptional regulator, Mor family \| Transcriptional regulator, Mor family; uniclust \| UniRef100\_A0A2V2EJZ8 \| 98.2 \| 1.7e-08 \| 3.1e-14 \| 69.1 \| 73 \| (49, 122) \| 144 \| (46, 118) \| 120 \| Mor transcription activator domain-containing protein \| Mor transcription activator domain-containing protein; uniclust \| UniRef100\_UPI000B853F07 \| 98.2 \| 1.7e-08 \| 3.1e-14 \| 71.7 \| 100 \| (16, 125) \| 144 \| (45, 144) \| 153 \| Mor transcription activator family protein \| Mor transcription activator family protein; uniclust \| UniRef100\_A0A847LF34 \| 98.2 \| 1.9e-08 \| 3.6e-14 \| 75.2 \| 113 \| (1, 125) \| 144 \| (72, 184) \| 221 \| Mor transcription activator domain-containing protein \| Mor transcription activator domain-containing protein; uniclust \| UniRef100\_UPI0013C2DCD9 \| 98.2 \| 2.1e-08 \| 3.8e-14 \| 74.4 \| 122 \| (2, 123) \| 144 \| (10, 138) \| 207 \| AlpA family phage regulatory protein \| AlpA family phage regulatory protein; uniclust \| UniRef100\_UPI000424C323 \| 98.2 \| 2.2e-08 \| 4e-14 \| 67.6 \| 78 \| (47, 124) \| 144 \| (21, 98) \| 110 \| Mor transcription activator family protein \| Mor transcription activator family protein; uniclust \| UniRef100\_A0A4T5K8K5 \| 98.2 \| 2.1e-08 \| 4e-14 \| 72.5 \| 72 \| (58, 129) \| 144 \| (64, 135) \| 142 \| Transcriptional regulator \| Transcriptional regulator; uniclust \| UniRef100\_A0A2D8KBN5 \| 98.2 \| 2.2e-08 \| 4.1e-14 \| 71.1 \| 111 \| (6, 126) \| 144 \| (14, 132) \| 152 \| Helix-turn-helix domain-containing protein \| Helix-turn-helix domain-containing protein; uniclust \| UniRef100\_UPI000AAE4DE3 \| 98.2 \| 2.2e-08 \| 4.1e-14 \| 71.3 \| 69 \| (53, 122) \| 144 \| (85, 153) \| 155 \| Mor transcription activator family protein \| Mor transcription activator family protein; uniclust \| UniRef100\_A0A127QUE9 \| 98.2 \| 2e-08 \| 4.1e-14 \| 73.5 \| 94 \| (1, 127) \| 144 \| (39, 139) \| 140 \| Uncharacterized protein \| Uncharacterized protein; uniclust \| UniRef100\_R5R2B3 \| 98.2 \| 2.6e-08 \| 4.8e-14 \| 65.9 \| 81 \| (41, 121) \| 144 \| (13, 94) \| 97 \| Mor transcription activator domain-containing protein \| Mor transcription activator domain-containing protein; uniclust \| UniRef100\_UPI001FD7F15E \| 98.2 \| 2.7e-08 \| 4.9e-14 \| 75.1 \| 77 \| (50, 126) \| 144 \| (45, 129) \| 235 \| Mor transcription activator family protein \| Mor transcription activator family protein; uniclust \| UniRef100\_A0A091ATJ0 \| 98.2 \| 2.5e-08 \| 5.1e-14 \| 71.2 \| 68 \| (57, 127) \| 144 \| (40, 107) \| 124 \| Mor transcription activator domain-containing protein \| Mor transcription activator domain-containing protein; uniclust \| UniRef100\_A0A270BEU5 \| 98.2 \| 2.9e-08 \| 5.3e-14 \| 69.4 \| 108 \| (8, 128) \| 144 \| (14, 121) \| 136 \| MarR family transcriptional regulator \| MarR family transcriptional regulator; uniclust \| UniRef100\_A0A522DPM1 \| 98.2 \| 3.1e-08 \| 5.7e-14 \| 65.6 \| 75 \| (52, 126) \| 144 \| (7, 81) \| 97 \| Mor transcription activator domain-containing protein \| Mor transcription activator domain-containing protein; uniclust \| UniRef100\_UPI001F1206BA \| 98.2 \| 3.1e-08 \| 5.8e-14 \| 65.2 \| 74 \| (53, 127) \| 144 \| (10, 85) \| 94 \| Mor transcription activator family protein \| Mor transcription activator family protein; uniclust \| UniRef100\_UPI0012F83529 \| 98.2 \| 3.2e-08 \| 5.8e-14 \| 69.4 \| 98 \| (15, 122) \| 144 \| (29, 126) \| 139 \| hypothetical protein \| hypothetical protein; uniclust \| UniRef100\_UPI002112227B \| 98.2 \| 3.2e-08 \| 5.9e-14 \| 63.4 \| 72 \| (50, 121) \| 144 \| (7, 78) \| 80 \| Mor transcription activator family protein \| Mor transcription activator family protein; uniclust \| UniRef100\_A0A540V321 \| 98.2 \| 3.2e-08 \| 5.9e-14 \| 59.5 \| 51 \| (67, 121) \| 144 \| (4, 54) \| 57 \| DNA-binding protein (Fragment) \| DNA-binding protein (Fragment); uniclust \| UniRef100\_UPI000DA609CB \| 98.2 \| 3.2e-08 \| 6e-14 \| 58.6 \| 51 \| (67, 121) \| 144 \| (2, 52) \| 53 \| Mor transcription activator family protein \| Mor transcription activator family protein; uniclust \| UniRef100\_A0A921TC53 \| 98.2 \| 3.4e-08 \| 6.2e-14 \| 75.3 \| 114 \| (7, 131) \| 144 \| (36, 149) \| 254 \| N-acetylmuramoyl-L-alanine amidase \| N-acetylmuramoyl-L-alanine amidase; uniclust \| UniRef100\_A0A3S0MTD3 \| 98.2 \| 3.2e-08 \| 6.3e-14 \| 70.6 \| 64 \| (64, 128) \| 144 \| (57, 120) \| 129 \| Mor transcription activator domain-containing protein \| Mor transcription activator domain-containing protein; uniclust \| UniRef100\_A0A4Q8XRF6 \| 98.1 \| 3.4e-08 \| 6.4e-14 \| 72.3 \| 100 \| (3, 120) \| 144 \| (18, 122) \| 165 \| Uncharacterized protein \| Uncharacterized protein; uniclust \| UniRef100\_A0A1T4L4A5 \| 98.1 \| 3.5e-08 \| 6.5e-14 \| 65.5 \| 71 \| (50, 121) \| 144 \| (18, 88) \| 99 \| Mor transcription activator family protein \| Mor transcription activator family protein; uniclust \| UniRef100\_A0A4Q3PAH2 \| 98.1 \| 3.5e-08 \| 6.9e-14 \| 70.0 \| 72 \| (48, 123) \| 144 \| (39, 110) \| 126 \| Mor transcription activator domain-containing protein \| Mor transcription activator domain-containing protein; uniclust \| UniRef100\_UPI00041F2CD4 \| 98.1 \| 3.7e-08 \| 6.9e-14 \| 66.3 \| 71 \| (52, 122) \| 144 \| (4, 82) \| 98 \| Mor transcription activator family protein \| Mor transcription activator family protein; uniclust \| UniRef100\_A0A194AFZ1 \| 98.1 \| 3.8e-08 \| 7e-14 \| 64.1 \| 76 \| (50, 125) \| 144 \| (6, 82) \| 88 \| Mor transcription activator domain-containing protein \| Mor transcription activator domain-containing protein; uniclust \| UniRef100\_A0A6L3AN94 \| 98.1 \| 3.9e-08 \| 7.1e-14 \| 64.2 \| 69 \| (51, 120) \| 144 \| (15, 87) \| 89 \| Mor transcription activator domain-containing protein \| Mor transcription activator domain-containing protein; uniclust \| UniRef100\_A0A2D8KBQ5 \| 98.1 \| 3.9e-08 \| 7.1e-14 \| 72.2 \| 104 \| (3, 124) \| 144 \| (57, 161) \| 190 \| HTH luxR-type domain-containing protein \| HTH luxR-type domain-containing protein; uniclust \| UniRef100\_A0A1X7C3R7 \| 98.1 \| 4e-08 \| 7.4e-14 \| 63.9 \| 70 \| (50, 121) \| 144 \| (15, 85) \| 87 \| Mor transcription activator family protein \| Mor transcription activator family protein; uniclust \| UniRef100\_UPI0012EBF5E3 \| 98.1 \| 4.1e-08 \| 7.5e-14 \| 65.0 \| 72 \| (50, 121) \| 144 \| (19, 90) \| 97 \| helix-turn-helix domain-containing protein \| helix-turn-helix domain-containing protein; uniclust \| UniRef100\_A3YAG8 \| 98.1 \| 4e-08 \| 7.6e-14 \| 68.8 \| 72 \| (56, 127) \| 144 \| (48, 119) \| 123 \| Mor transcription activator domain-containing protein \| Mor transcription activator domain-containing protein; uniclust \| UniRef100\_A0A3C0QNU7 \| 98.1 \| 4.2e-08 \| 7.7e-14 \| 68.7 \| 116 \| (3, 120) \| 144 \| (9, 136) \| 137 \| Mor transcription activator domain-containing protein \| Mor transcription activator domain-containing protein; uniclust \| UniRef100\_A4EV43 \| 98.1 \| 4.3e-08 \| 7.8e-14 \| 69.2 \| 98 \| (7, 121) \| 144 \| (45, 142) \| 144 \| Helix-turn-helix domain-containing protein \| Helix-turn-helix domain-containing protein; uniclust \| UniRef100\_A0A0F5P8U2 \| 98.1 \| 4.5e-08 \| 8.2e-14 \| 71.2 \| 104 \| (7, 120) \| 144 \| (1, 106) \| 176 \| Mor transcription activator domain-containing protein \| Mor transcription activator domain-containing protein; uniclust \| UniRef100\_A0A660E659 \| 98.1 \| 4.2e-08 \| 8.4e-14 \| 67.2 \| 78 \| (49, 127) \| 144 \| (14, 91) \| 96 \| Mor transcription activator domain-containing protein \| Mor transcription activator domain-containing protein; uniclust \| UniRef100\_UPI001145015F \| 98.1 \| 4.9e-08 \| 9e-14 \| 67.2 \| 107 \| (3, 121) \| 144 \| (11, 121) \| 122 \| helix-turn-helix domain-containing protein \| helix-turn-helix domain-containing protein; uniclust \| UniRef100\_A0A8J6NST4 \| 98.1 \| 4.9e-08 \| 9e-14 \| 59.5 \| 57 \| (64, 121) \| 144 \| (4, 60) \| 61 \| Mor transcription activator domain-containing protein \| Mor transcription activator domain-containing protein; uniclust \| UniRef100\_UPI000AAA50FC \| 98.1 \| 5e-08 \| 9.1e-14 \| 63.3 \| 66 \| (58, 123) \| 144 \| (2, 75) \| 86 \| Mor transcription activator family protein \| Mor transcription activator family protein; uniclust \| UniRef100\_A0A427CJC7 \| 98.1 \| 5e-08 \| 9.1e-14 \| 69.2 \| 74 \| (51, 124) \| 144 \| (60, 133) \| 148 \| Mor transcription activator-like protein \| Mor transcription activator-like protein; uniclust \| UniRef100\_A0A095V3V3 \| 98.1 \| 5e-08 \| 9.1e-14 \| 71.7 \| 108 \| (2, 120) \| 144 \| (21, 135) \| 190 \| Transposase \| Transposase; uniclust \| UniRef100\_UPI001157E51F \| 98.1 \| 5.2e-08 \| 9.5e-14 \| 71.8 \| 108 \| (2, 122) \| 144 \| (27, 142) \| 193 \| helix-turn-helix domain-containing protein \| helix-turn-helix domain-containing protein; uniclust \| UniRef100\_A0A653XI87 \| 98.1 \| 5.3e-08 \| 9.7e-14 \| 64.3 \| 73 \| (54, 126) \| 144 \| (19, 91) \| 95 \| Mor transcription activator domain-containing protein \| Mor transcription activator domain-containing protein; uniclust \| UniRef100\_A0A1C6AG62 \| 98.1 \| 5.3e-08 \| 1e-13 \| 65.6 \| 77 \| (50, 126) \| 144 \| (17, 94) \| 99 \| Uncharacterized conserved protein \| Uncharacterized conserved protein; uniclust \| UniRef100\_A0A443K8S6 \| 98.1 \| 5.6e-08 \| 1e-13 \| 70.2 \| 101 \| (13, 125) \| 144 \| (35, 138) \| 167 \| Helix-turn-helix domain-containing protein \| Helix-turn-helix domain-containing protein; uniclust \| UniRef100\_A0A921TDK8 \| 98.1 \| 5.8e-08 \| 1.1e-13 \| 75.1 \| 112 \| (6, 128) \| 144 \| (167, 278) \| 282 \| Uncharacterized protein (Fragment) \| Uncharacterized protein (Fragment); uniclust \| UniRef100\_A0A0Q2XPP4 \| 98.1 \| 5.8e-08 \| 1.1e-13 \| 73.6 \| 106 \| (6, 125) \| 144 \| (9, 118) \| 238 \| Mor transcription activator domain-containing protein \| Mor transcription activator domain-containing protein; uniclust \| UniRef100\_UPI001C0C1CEE \| 98.1 \| 6.4e-08 \| 1.2e-13 \| 69.3 \| 118 \| (2, 132) \| 144 \| (23, 146) \| 157 \| hypothetical protein \| hypothetical protein; uniclust \| UniRef100\_UPI001EE639DA \| 98.1 \| 6.5e-08 \| 1.2e-13 \| 65.5 \| 77 \| (49, 125) \| 144 \| (19, 103) \| 110 \| Mor transcription activator family protein \| Mor transcription activator family protein; uniclust \| UniRef100\_A0A099VBY3 \| 98.1 \| 6.2e-08 \| 1.2e-13 \| 64.1 \| 62 \| (61, 127) \| 144 \| (18, 79) \| 84 \| Mor transcription activator domain-containing protein \| Mor transcription activator domain-containing protein; uniclust \| UniRef100\_UPI0016685DF9 \| 98.1 \| 6.6e-08 \| 1.2e-13 \| 69.6 \| 119 \| (4, 122) \| 144 \| (23, 147) \| 164 \| helix-turn-helix domain-containing protein \| helix-turn-helix domain-containing protein; uniclust \| UniRef100\_A0A351VJ38 \| 98.1 \| 6.5e-08 \| 1.3e-13 \| 64.9 \| 65 \| (57, 122) \| 144 \| (24, 88) \| 91 \| Mor transcription activator domain-containing protein \| Mor transcription activator domain-containing protein; uniclust \| UniRef100\_UPI001FE71F71 \| 98.1 \| 6.9e-08 \| 1.3e-13 \| 60.6 \| 58 \| (69, 127) \| 144 \| (2, 59) \| 71 \| Mor transcription activator family protein \| Mor transcription activator family protein; uniclust \| UniRef100\_UPI00223A6E0C \| 98.1 \| 6.8e-08 \| 1.3e-13 \| 63.5 \| 66 \| (60, 125) \| 144 \| (13, 78) \| 90 \| Mor transcription activator family protein \| Mor transcription activator family protein; uniclust \| UniRef100\_A0A3E2N6R8 \| 98.1 \| 6e-08 \| 1.3e-13 \| 72.2 \| 76 \| (49, 127) \| 144 \| (55, 130) \| 148 \| Mor transcription activator family protein \| Mor transcription activator family protein; uniclust \| UniRef100\_A0A1C3FDH3 \| 98.1 \| 6e-08 \| 1.3e-13 \| 68.6 \| 74 \| (49, 125) \| 144 \| (25, 98) \| 108 \| Mor transcription activator domain-containing protein \| Mor transcription activator domain-containing protein; uniclust \| UniRef100\_UPI00226DF99C \| 98.1 \| 7.1e-08 \| 1.3e-13 \| 65.7 \| 65 \| (59, 124) \| 144 \| (36, 100) \| 114 \| Mor transcription activator family protein \| Mor transcription activator family protein; uniclust \| UniRef100\_A0A5C7LYQ9 \| 98.1 \| 7.3e-08 \| 1.3e-13 \| 65.5 \| 78 \| (50, 127) \| 144 \| (27, 104) \| 112 \| Mor transcription activator domain-containing protein \| Mor transcription activator domain-containing protein; uniclust \| UniRef100\_A0A962WFG0 \| 98.1 \| 7.4e-08 \| 1.4e-13 \| 67.1 \| 115 \| (3, 126) \| 144 \| (4, 120) \| 131 \| Uncharacterized protein \| Uncharacterized protein; uniclust \| UniRef100\_UPI0012EB89D4 \| 98.0 \| 8.4e-08 \| 1.5e-13 \| 67.0 \| 102 \| (6, 124) \| 144 \| (26, 127) \| 133 \| hypothetical protein \| hypothetical protein; uniclust \| UniRef100\_UPI00055724BA \| 98.0 \| 8.8e-08 \| 1.6e-13 \| 63.4 \| 76 \| (52, 127) \| 144 \| (10, 87) \| 91 \| Mor transcription activator family protein \| Mor transcription activator family protein; pdb70 \| 1RR7\_A \| 99.2 \| 2.2e-15 \| 1.8e-19 \| 100.9 \| 93 \| (2, 125) \| 144 \| (27, 119) \| 129 \| Middle operon regulator \| 1RR7\_A Middle operon regulator Mor, TRANSCRIPTION; pdb70 \| 6EO2\_A \| 96.5 \| 8e-06 \| 7.5e-10 \| 53.1 \| 61 \| (79, 142) \| 144 \| (146, 206) \| 210 \| Transcriptional regulatory protein RcsB \| 6EO2\_A Transcriptional regulatory protein RcsB Transcriptional factor, DNA BINDING PROTEIN; pdb70 \| 5O8Z\_A \| 96.5 \| 8.3e-06 \| 7.8e-10 \| 53.4 \| 61 \| (79, 142) \| 144 \| (146, 206) \| 216 \| Transcriptional regulatory protein RcsB \| 5O8Z\_A Transcriptional regulatory protein RcsB Response regulator Transcriptional factor, TRANSCRIPTION; pdb70 \| 5U8K\_A \| 96.4 \| 1.4e-05 \| 1.3e-09 \| 52.9 \| 61 \| (80, 143) \| 144 \| (155, 223) \| 231 \| Response regulator \| 5U8K\_A Response regulator Repressor of iron transporter, Aspartate-less HET: MSE; pdb70 \| 5XT2\_C \| 96.4 \| 1.5e-05 \| 1.4e-09 \| 52.2 \| 60 \| (81, 143) \| 144 \| (150, 209) \| 215 \| Response regulator FixJ \| 5XT2\_C Response regulator FixJ Response Regulater, FixJ/NarL Family, DNA HET: GOL, FMT; pdb70 \| 5XSO\_A \| 96.3 \| 2.1e-05 \| 2e-09 \| 51.4 \| 60 \| (81, 143) \| 144 \| (150, 209) \| 215 \| Response regulator FixJ \| 5XSO\_A Response regulator FixJ Response Regulater, FixJ/NarL Family, DNA HET: FMT, GOL; pdb70 \| 5U8K\_B \| 96.3 \| 2.1e-05 \| 2e-09 \| 51.9 \| 61 \| (80, 143) \| 144 \| (155, 223) \| 231 \| Response regulator \| 5U8K\_B Response regulator Repressor of iron transporter, Aspartate-less HET: MSE; pdb70 \| 1A04\_B \| 96.2 \| 2.2e-05 \| 2.1e-09 \| 51.1 \| 59 \| (82, 143) \| 144 \| (153, 211) \| 215 \| NITRATE/NITRITE RESPONSE REGULATOR PROTEIN NARL \| 1A04\_B NITRATE/NITRITE RESPONSE REGULATOR PROTEIN NARL SIGNAL TRANSDUCTION PROTEIN, RESPONSE REGULATORS; pdb70 \| 6MWZ\_A \| 96.2 \| 2.5e-05 \| 2.2e-09 \| 54.5 \| 61 \| (78, 141) \| 144 \| (171, 231) \| 239 \| Transcriptional regulator LasR, ALA-HIS-HIS-HIS-HIS-ALA \| 6MWZ\_A Transcriptional regulator LasR, ALA-HIS-HIS-HIS-HIS-ALA Transcriptional activator, TRANSCRIPTION HET: K5M; pdb70 \| 1RNL\_A \| 96.2 \| 2.4e-05 \| 2.2e-09 \| 51.0 \| 59 \| (82, 143) \| 144 \| (153, 211) \| 215 \| NITRATE/NITRITE RESPONSE REGULATOR PROTEIN NARL \| 1RNL\_A NITRATE/NITRITE RESPONSE REGULATOR PROTEIN NARL RESPONSE REGULATORS, TWO-COMPONENT SYSTEMS, SIGNAL HET: GOL; pdb70 \| 5VFA\_A \| 96.2 \| 2.5e-05 \| 2.3e-09 \| 51.6 \| 61 \| (80, 143) \| 144 \| (154, 222) \| 230 \| Response regulator \| 5VFA\_A Response regulator Repressor of iron transporter, Aspartate-less; pdb70 \| 4ZMR\_A \| 96.2 \| 2.6e-05 \| 2.4e-09 \| 50.4 \| 61 \| (80, 143) \| 144 \| (143, 203) \| 207 \| Response regulator \| 4ZMR\_A Response regulator Response regulator, DNA BINDING PROTEIN; pdb70 \| 4ZMS\_A \| 96.2 \| 2.6e-05 \| 2.4e-09 \| 50.4 \| 61 \| (80, 143) \| 144 \| (143, 203) \| 207 \| Response regulator \| 4ZMS\_A Response regulator Response regulator, DNA BINDING PROTEIN HET: 4QT; pdb70 \| 4YN8\_A \| 96.2 \| 2.7e-05 \| 2.5e-09 \| 50.2 \| 60 \| (81, 143) \| 144 \| (140, 199) \| 203 \| ChrA \| 4YN8\_A ChrA DNA-binding, helix-turn-helix, DNA BINDING PROTEIN HET: SO4; pdb70 \| 2HQR\_A \| 96.2 \| 2.8e-05 \| 2.6e-09 \| 51.4 \| 60 \| (81, 143) \| 144 \| (141, 209) \| 223 \| Putative TRANSCRIPTIONAL REGULATOR \| 2HQR\_A Putative TRANSCRIPTIONAL REGULATOR Phosporylation-Independent Response Regulator, H. pylori; pdb70 \| 1L3L\_A \| 96.2 \| 3e-05 \| 2.6e-09 \| 54.0 \| 61 \| (79, 142) \| 144 \| (169, 229) \| 234 \| Transcriptional activator protein traR/DNA Complex \| 1L3L\_A Transcriptional activator protein traR/DNA Complex helix-turn-helix DNA binding motif, alpha/beta/alpha HET: LAE, MSE; pdb70 \| 4KFC\_A \| 96.1 \| 3e-05 \| 2.8e-09 \| 51.3 \| 63 \| (79, 143) \| 144 \| (149, 220) \| 227 \| KDP operon transcriptional regulatory protein \| 4KFC\_A KDP operon transcriptional regulatory protein receiver domain, DNA-binding domain, TRANSCRIPTION; pdb70 \| 4LFU\_A \| 96.1 \| 3.2e-05 \| 2.8e-09 \| 54.5 \| 61 \| (79, 142) \| 144 \| (175, 235) \| 248 \| Regulatory protein SdiA \| 4LFU\_A Regulatory protein SdiA LuxR-type quorum sensing receptor, transcription HET: PG4; pdb70 \| 5F64\_C \| 96.1 \| 3.1e-05 \| 2.9e-09 \| 49.6 \| 60 \| (81, 143) \| 144 \| (144, 203) \| 207 \| Positive transcription regulator EvgA \| 5F64\_C Positive transcription regulator EvgA transcription regulator, sensor, EvgS, structural; pdb70 \| 1YS7\_B \| 96.1 \| 3.3e-05 \| 3.1e-09 \| 51.2 \| 62 \| (79, 143) \| 144 \| (155, 225) \| 233 \| Transcriptional regulatory protein prrA \| 1YS7\_B Transcriptional regulatory protein prrA Response regulator, DNA binding domain HET: GOL; pdb70 \| 2Q0O\_A \| 96.1 \| 3.7e-05 \| 3.3e-09 \| 53.5 \| 62 \| (78, 142) \| 144 \| (170, 231) \| 236 \| quorum sensing transcription factor TraR \| 2Q0O\_A quorum sensing transcription factor TraR helix-turn-helix, two-helix coiled coil, TRANSCRIPTION HET: LAE; pdb70 \| 3Q9S\_A \| 96.1 \| 3.8e-05 \| 3.5e-09 \| 52.4 \| 62 \| (79, 143) \| 144 \| (178, 248) \| 249 \| DNA-binding response regulator \| 3Q9S\_A DNA-binding response regulator response regulator, DNA BINDING PROTEIN; pdb70 \| 1YS6\_B \| 96.1 \| 3.9e-05 \| 3.7e-09 \| 50.6 \| 61 \| (80, 143) \| 144 \| (156, 225) \| 233 \| Transcriptional regulatory protein prrA \| 1YS6\_B Transcriptional regulatory protein prrA Response regulator protein, Dna binding; pdb70 \| 6MWL\_A \| 96.0 \| 4.3e-05 \| 3.8e-09 \| 53.3 \| 60 \| (79, 141) \| 144 \| (172, 231) \| 239 \| Transcriptional regulator LasR \| 6MWL\_A Transcriptional regulator LasR Transcriptional activator, TRANSCRIPTION HET: K5G; pdb70 \| 4Y13\_A \| 96.0 \| 4.4e-05 \| 3.8e-09 \| 53.8 \| 59 \| (80, 141) \| 144 \| (176, 234) \| 246 \| SdiA in complex with octanoyl-rac-glycerol \| 4Y13\_A SdiA in complex with octanoyl-rac-glycerol Quorum sensor, DNA, DNA BINDING HET: SO4, MSE, GOL, 480; pdb70 \| 3SZT\_A \| 96.0 \| 4.4e-05 \| 3.9e-09 \| 53.2 \| 61 \| (78, 141) \| 144 \| (170, 230) \| 237 \| PhzR \| 3SZT\_A PhzR quorum sensing acyl-homoserine lactone, Helix-turn-Helix HET: OHN; pdb70 \| 6CBQ\_A \| 96.0 \| 4.4e-05 \| 3.9e-09 \| 53.2 \| 61 \| (78, 141) \| 144 \| (170, 230) \| 237 \| Chromatin assembly factor 1 subunit \| 6CBQ\_A Chromatin assembly factor 1 subunit LuxR-type AHL receptor, Pseudomonas aeruginosa HET: EVY; pdb70 \| 6CC0\_B \| 96.0 \| 4.4e-05 \| 3.9e-09 \| 53.2 \| 61 \| (78, 141) \| 144 \| (170, 230) \| 237 \| Chromatin assembly factor 1 subunit \| 6CC0\_B Chromatin assembly factor 1 subunit LuxR-type AHL receptor, Pseudomonas aeruginosa HET: EWM; pdb70 \| 4GVP\_C \| 96.0 \| 4.2e-05 \| 4e-09 \| 49.3 \| 58 \| (83, 143) \| 144 \| (146, 203) \| 208 \| Response regulator protein vraR \| 4GVP\_C Response regulator protein vraR Response regulator, Two-component system, Bacterial; pdb70 \| 4IF4\_A \| 96.0 \| 4.2e-05 \| 4e-09 \| 49.3 \| 58 \| (83, 143) \| 144 \| (146, 203) \| 208 \| Response regulator protein VraR \| 4IF4\_A Response regulator protein VraR Response regulator, Two-component system, Bacterial HET: BEF, SO4; pdb70 \| 5HEV\_C \| 96.0 \| 4.2e-05 \| 4e-09 \| 49.3 \| 59 \| (82, 143) \| 144 \| (147, 205) \| 210 \| Response regulator protein VraR \| 5HEV\_C Response regulator protein VraR Enterococcus faecium, LiaR, response regulator HET: BEF; pdb70 \| 4HYE\_B \| 96.0 \| 4.3e-05 \| 4e-09 \| 50.2 \| 59 \| (82, 143) \| 144 \| (158, 216) \| 220 \| Response regulator \| 4HYE\_B Response regulator Cytosol, TRANSCRIPTION ACTIVATOR; pdb70 \| 3QP6\_A \| 96.0 \| 4.8e-05 \| 4.2e-09 \| 54.2 \| 59 \| (80, 141) \| 144 \| (194, 252) \| 265 \| CviR transcriptional regulator \| 3QP6\_A CviR transcriptional regulator quorum sensing, agonist, antagonist, LuxR HET: HL6; pdb70 \| 5HEV\_B \| 96.0 \| 4.5e-05 \| 4.3e-09 \| 49.1 \| 59 \| (82, 143) \| 144 \| (147, 205) \| 210 \| Response regulator protein VraR \| 5HEV\_B Response regulator protein VraR Enterococcus faecium, LiaR, response regulator HET: BEF; pdb70 \| 1KGS\_A \| 95.9 \| 6.1e-05 \| 5.8e-09 \| 49.3 \| 60 \| (81, 143) \| 144 \| (149, 217) \| 225 \| DNA BINDING RESPONSE REGULATOR D \| 1KGS\_A DNA BINDING RESPONSE REGULATOR D DNA-binding protein, alph-beta sandwich, winged-helix HET: SCN; pdb70 \| 1YIO\_A \| 95.9 \| 6.3e-05 \| 5.9e-09 \| 48.8 \| 56 \| (84, 142) \| 144 \| (143, 198) \| 208 \| response regulator StyR \| 1YIO\_A response regulator StyR Transcription regulation, DNA BINDING PROTEIN; pdb70 \| 2OQR\_A \| 95.9 \| 6.6e-05 \| 6.1e-09 \| 49.6 \| 50 \| (81, 130) \| 144 \| (154, 212) \| 230 \| Sensory transduction protein regX3 \| 2OQR\_A Sensory transduction protein regX3 response regulator, winged-helix-turn-helix, dna-binding, 3D; pdb70 \| 4KNY\_B \| 95.8 \| 7.2e-05 \| 6.7e-09 \| 49.3 \| 61 \| (81, 143) \| 144 \| (151, 220) \| 227 \| KDP operon transcriptional regulatory protein \| 4KNY\_B KDP operon transcriptional regulatory protein receiver domain, DNA-binding domain, TRANSCRIPTION; pdb70 \| 2GWR\_A \| 95.8 \| 8.2e-05 \| 7.6e-09 \| 49.9 \| 52 \| (79, 130) \| 144 \| (149, 209) \| 238 \| DNA-binding response regulator mtrA \| 2GWR\_A DNA-binding response regulator mtrA Two-component regulatory system, Transcription regulation HET: GOL; pdb70 \| 3R0J\_A \| 95.7 \| 9.7e-05 \| 9.1e-09 \| 49.6 \| 59 \| (81, 143) \| 144 \| (174, 241) \| 250 \| POSSIBLE TWO COMPONENT SYSTEM RESPONSE \| 3R0J\_A POSSIBLE TWO COMPONENT SYSTEM RESPONSE beta-alpha fold, winged helix-turn-helix, transcription HET: SO4, MSE, MSO; pdb70 \| 5ED4\_B \| 95.7 \| 0.00011 \| 9.8e-09 \| 49.5 \| 59 \| (80, 142) \| 144 \| (173, 240) \| 250 \| Response regulator/DNA Complex \| 5ED4\_B Response regulator/DNA Complex protein-DNA complex, winged helix-turn-helix, direct HET: EDO; pdb70 \| 3KLO\_A \| 95.5 \| 0.00014 \| 1.4e-08 \| 47.7 \| 59 \| (80, 141) \| 144 \| (156, 214) \| 225 \| Transcriptional regulator VpsT \| 3KLO\_A Transcriptional regulator VpsT REC domain, HTH domain, DNA-binding HET: TAR, C2E; pdb70 \| 1P2F\_A \| 95.5 \| 0.00016 \| 1.5e-08 \| 47.5 \| 61 \| (81, 143) \| 144 \| (143, 210) \| 220 \| Response Regulator \| 1P2F\_A Response Regulator response regulator, DrrB, OmpR/PhoB, Transcription; pdb70 \| 6KJU\_B \| 95.3 \| 0.00024 \| 2.1e-08 \| 49.6 \| 57 \| (82, 141) \| 144 \| (154, 210) \| 254 \| Transcriptional regulator LuxR family/DNA Complex \| 6KJU\_B Transcriptional regulator LuxR family/DNA Complex Vibrio cholera, Quorum sensing, transcription HET: A1U; pdb70 \| 4LDZ\_A \| 95.3 \| 0.00025 \| 2.4e-08 \| 45.4 \| 59 \| (82, 143) \| 144 \| (140, 198) \| 203 \| Transcriptional regulatory protein DesR \| 4LDZ\_A Transcriptional regulatory protein DesR Response regulator, two-component system, transcription HET: GOL; pdb70 \| 4B09\_E \| 94.8 \| 0.00053 \| 5e-08 \| 45.2 \| 50 \| (81, 130) \| 144 \| (158, 216) \| 240 \| TRANSCRIPTIONAL REGULATORY PROTEIN BAER \| 4B09\_E TRANSCRIPTIONAL REGULATORY PROTEIN BAER TRANSCRIPTION, RESPONSE REGULATOR, DNA BINDING HET: TBR, CAS; pdb70 \| 4B09\_H \| 94.8 \| 0.00053 \| 5e-08 \| 45.2 \| 50 \| (81, 130) \| 144 \| (158, 216) \| 240 \| TRANSCRIPTIONAL REGULATORY PROTEIN BAER \| 4B09\_H TRANSCRIPTIONAL REGULATORY PROTEIN BAER TRANSCRIPTION, RESPONSE REGULATOR, DNA BINDING HET: TBR, CAS; pdb70 \| 3C3W\_B \| 94.5 \| 0.00089 \| 8.2e-08 \| 43.9 \| 48 \| (83, 130) \| 144 \| (149, 196) \| 225 \| TWO COMPONENT TRANSCRIPTIONAL REGULATORY PROTEIN \| 3C3W\_B TWO COMPONENT TRANSCRIPTIONAL REGULATORY PROTEIN RESPONSE REGULATOR, TWO-COMPONENT REGULATORY SYSTEM; pdb70 \| 3C3W\_A \| 94.4 \| 0.00094 \| 8.7e-08 \| 43.8 \| 48 \| (84, 131) \| 144 \| (150, 197) \| 225 \| TWO COMPONENT TRANSCRIPTIONAL REGULATORY PROTEIN \| 3C3W\_A TWO COMPONENT TRANSCRIPTIONAL REGULATORY PROTEIN RESPONSE REGULATOR, TWO-COMPONENT REGULATORY SYSTEM; pdb70 \| 4S05\_A \| 94.3 \| 0.001 \| 9.7e-08 \| 43.8 \| 50 \| (80, 129) \| 144 \| (146, 204) \| 232 \| DNA-binding transcriptional regulator BasR \| 4S05\_A DNA-binding transcriptional regulator BasR Two-Component System, response regulator, PmrA |
| Top keywords  (threshold 1.00e-03 (evalue)) | **transcription, Mor, activator, domain\_containing, regulator, Helix\_turn\_helix, Transcriptional, Response, hypothetical, DNA** |
| Output files | ../../similar\_sequences/61\_FANPEZAQ\_CDS\_0061\_merged.svg ../../similar\_sequences/61\_FANPEZAQ\_CDS\_0061\_pdb70.a3m ../../similar\_sequences/61\_FANPEZAQ\_CDS\_0061\_pdb70.hhr ../../similar\_sequences/61\_FANPEZAQ\_CDS\_0061\_uniclust.a3m ../../similar\_sequences/61\_FANPEZAQ\_CDS\_0061\_uniclust.hhr |

#### Structure prediction (AlphaFold)^2^

|  |  |
| --- | --- |
| Stats | xml version="1.0" encoding="utf-8" standalone="no"?       2024-09-02T21:09:56.983254 image/svg+xml   Matplotlib v3.7.2, https://matplotlib.org/ |
| Predicted structure | **NGL Viewer Controls:**  - Center: *Left-Click* - Rotate: *Left-Click + Drag* - Translate: *Right-Click + Drag* - Zoom: *Shift + Left-Click + Drag* |
| Output files | ../../predicted\_structures/61\_FANPEZAQ\_CDS\_0061/features.pkl ../../predicted\_structures/61\_FANPEZAQ\_CDS\_0061/ranked\_0.pdb ../../predicted\_structures/61\_FANPEZAQ\_CDS\_0061/ranked\_0\_plots.svg ../../predicted\_structures/61\_FANPEZAQ\_CDS\_0061/result\_model\_1\_ptm\_pred\_0.pkl |

#### Structure similarity search results (Foldseek)^3^

|  |  |
| --- | --- |
| Structure databases searched | Pdb, Afdb-proteome, Afdb-uniprot50 |
| Results, scheme(s)  (Top layers only, threshold 1.00e-02 (evalue)) | xml version="1.0" encoding="utf-8" standalone="no"?       2024-09-02T21:11:29.367064 image/svg+xml   Matplotlib v3.7.2, https://matplotlib.org/ |
| Results, table  (threshold 1.00e-02 (evalue)) | db \| id \| prob \| evalue \| bits \| fident \| alnlen \| mismatch \| gapopen \| qstart \| qend \| tstart \| tend \| name \| description; afdb-uniprot50 \| AF-A0A8A6KH34-F1-MODEL\_V4 \| 1.0 \| 9.589e-13 \| 518 \| 0.438 \| 121 \| 66 \| 1 \| 1 \| 119 \| 4 \| 124 \| Uncharacterized protein \| Uncharacterized protein; afdb-uniprot50 \| AF-A0A6M4YR27-F1-MODEL\_V4 \| 1.0 \| 8.436e-14 \| 509 \| 0.471 \| 140 \| 68 \| 2 \| 2 \| 135 \| 7 \| 146 \| Uncharacterized protein \| Uncharacterized protein; afdb-uniprot50 \| AF-A0A1S0V4H7-F1-MODEL\_V4 \| 1.0 \| 3.799e-13 \| 505 \| 0.419 \| 143 \| 83 \| 0 \| 1 \| 143 \| 3 \| 145 \| Uncharacterized protein \| Uncharacterized protein; afdb-uniprot50 \| AF-A0A653KZR3-F1-MODEL\_V4 \| 1.0 \| 7.962e-14 \| 498 \| 0.456 \| 149 \| 73 \| 3 \| 2 \| 144 \| 7 \| 153 \| Uncharacterized protein \| Uncharacterized protein; afdb-uniprot50 \| AF-A0A060HA06-F1-MODEL\_V4 \| 1.0 \| 3.193e-13 \| 495 \| 0.412 \| 148 \| 80 \| 2 \| 1 \| 144 \| 44 \| 188 \| Uncharacterized protein \| Uncharacterized protein; afdb-uniprot50 \| AF-A0A6L6JDU6-F1-MODEL\_V4 \| 1.0 \| 1.944e-11 \| 494 \| 0.443 \| 124 \| 66 \| 1 \| 1 \| 124 \| 80 \| 200 \| Uncharacterized protein \| Uncharacterized protein; afdb-uniprot50 \| AF-A0A315D133-F1-MODEL\_V4 \| 1.0 \| 4.519e-13 \| 489 \| 0.458 \| 133 \| 65 \| 2 \| 2 \| 127 \| 4 \| 136 \| Uncharacterized protein \| Uncharacterized protein; afdb-uniprot50 \| AF-K5Z2H2-F1-MODEL\_V4 \| 1.0 \| 1.029e-11 \| 479 \| 0.447 \| 123 \| 63 \| 2 \| 1 \| 118 \| 94 \| 216 \| Uncharacterized protein \| Uncharacterized protein; afdb-uniprot50 \| AF-A0A1H2R5R9-F1-MODEL\_V4 \| 1.0 \| 1.944e-11 \| 478 \| 0.461 \| 117 \| 60 \| 2 \| 2 \| 118 \| 8 \| 121 \| Uncharacterized protein \| Uncharacterized protein; afdb-uniprot50 \| AF-A0A291LZN5-F1-MODEL\_V4 \| 1.0 \| 8.261e-11 \| 455 \| 0.418 \| 122 \| 68 \| 1 \| 1 \| 122 \| 13 \| 131 \| Uncharacterized protein \| Uncharacterized protein; afdb-uniprot50 \| AF-A0A660NN87-F1-MODEL\_V4 \| 1.0 \| 1.835e-11 \| 455 \| 0.428 \| 119 \| 66 \| 1 \| 2 \| 118 \| 17 \| 135 \| Uncharacterized protein \| Uncharacterized protein; afdb-uniprot50 \| AF-A0A2S0MNB5-F1-MODEL\_V4 \| 1.0 \| 6.185e-11 \| 444 \| 0.373 \| 126 \| 75 \| 2 \| 1 \| 125 \| 7 \| 129 \| Uncharacterized protein \| Uncharacterized protein; afdb-uniprot50 \| AF-A0A7L9Q1W2-F1-MODEL\_V4 \| 1.0 \| 2.284e-12 \| 419 \| 0.473 \| 133 \| 58 \| 4 \| 1 \| 121 \| 1 \| 133 \| Uncharacterized protein \| Uncharacterized protein; afdb-uniprot50 \| AF-A0A241VCE8-F1-MODEL\_V4 \| 1.0 \| 4.176e-10 \| 416 \| 0.352 \| 125 \| 78 \| 2 \| 1 \| 122 \| 4 \| 128 \| Uncharacterized protein \| Uncharacterized protein; afdb-uniprot50 \| AF-A0A143DGK1-F1-MODEL\_V4 \| 1.0 \| 1.254e-09 \| 413 \| 0.406 \| 118 \| 67 \| 2 \| 1 \| 118 \| 1 \| 115 \| Uncharacterized protein \| Uncharacterized protein; afdb-uniprot50 \| AF-A0A4Q3Z6S0-F1-MODEL\_V4 \| 1.0 \| 4.176e-10 \| 403 \| 0.386 \| 119 \| 69 \| 2 \| 1 \| 118 \| 6 \| 121 \| Uncharacterized protein \| Uncharacterized protein; afdb-uniprot50 \| AF-A0A410UF43-F1-MODEL\_V4 \| 1.0 \| 1.654e-10 \| 396 \| 0.338 \| 118 \| 78 \| 0 \| 1 \| 118 \| 17 \| 134 \| Uncharacterized protein \| Uncharacterized protein; afdb-uniprot50 \| AF-A0A150HM30-F1-MODEL\_V4 \| 1.0 \| 2.628e-10 \| 391 \| 0.314 \| 124 \| 82 \| 2 \| 1 \| 121 \| 6 \| 129 \| Uncharacterized protein \| Uncharacterized protein; afdb-uniprot50 \| AF-A0A7T7R8M9-F1-MODEL\_V4 \| 1.0 \| 1.968e-10 \| 380 \| 0.354 \| 124 \| 77 \| 2 \| 3 \| 123 \| 8 \| 131 \| Uncharacterized protein \| Uncharacterized protein; afdb-uniprot50 \| AF-T1XCZ5-F1-MODEL\_V4 \| 1.0 \| 1.944e-11 \| 380 \| 0.428 \| 133 \| 68 \| 4 \| 1 \| 125 \| 1 \| 133 \| Uncharacterized protein \| Uncharacterized protein; afdb-uniprot50 \| AF-A0A4R7UHV5-F1-MODEL\_V4 \| 1.0 \| 7.03e-10 \| 373 \| 0.324 \| 145 \| 88 \| 2 \| 3 \| 144 \| 4 \| 141 \| Helix-turn-helix protein \| Helix-turn-helix protein; afdb-uniprot50 \| AF-A0A1I1F6V4-F1-MODEL\_V4 \| 1.0 \| 1.753e-10 \| 370 \| 0.419 \| 124 \| 67 \| 3 \| 1 \| 122 \| 3 \| 123 \| Uncharacterized protein \| Uncharacterized protein; afdb-uniprot50 \| AF-A0A3G6WB74-F1-MODEL\_V4 \| 1.0 \| 1.198e-08 \| 337 \| 0.273 \| 128 \| 90 \| 1 \| 1 \| 128 \| 62 \| 186 \| Uncharacterized protein \| Uncharacterized protein; afdb-uniprot50 \| AF-A0A3A1P024-F1-MODEL\_V4 \| 1.0 \| 7.893e-10 \| 334 \| 0.343 \| 128 \| 81 \| 2 \| 1 \| 125 \| 1 \| 128 \| Mor domain-containing protein \| Mor domain-containing protein; afdb-uniprot50 \| AF-A0A7V8FKC4-F1-MODEL\_V4 \| 1.0 \| 1.6e-08 \| 328 \| 0.343 \| 131 \| 75 \| 2 \| 2 \| 128 \| 6 \| 129 \| Mor domain-containing protein \| Mor domain-containing protein; afdb-uniprot50 \| AF-A0A373B2W6-F1-MODEL\_V4 \| 1.0 \| 4.228e-09 \| 324 \| 0.347 \| 115 \| 72 \| 2 \| 7 \| 118 \| 11 \| 125 \| Mor domain-containing protein \| Mor domain-containing protein; afdb-uniprot50 \| AF-A0A4Q7BE38-F1-MODEL\_V4 \| 1.0 \| 1.007e-08 \| 324 \| 0.314 \| 121 \| 78 \| 3 \| 1 \| 118 \| 49 \| 167 \| Mor domain-containing protein \| Mor domain-containing protein; afdb-uniprot50 \| AF-A0A291FP91-F1-MODEL\_V4 \| 1.0 \| 2.292e-07 \| 310 \| 0.282 \| 117 \| 81 \| 1 \| 2 \| 118 \| 63 \| 176 \| Uncharacterized protein \| Uncharacterized protein; afdb-uniprot50 \| AF-A0A7V7G2S6-F1-MODEL\_V4 \| 1.0 \| 3.354e-09 \| 305 \| 0.285 \| 175 \| 81 \| 5 \| 1 \| 144 \| 1 \| 162 \| Uncharacterized protein \| Uncharacterized protein; afdb-uniprot50 \| AF-A0A1I5P0R3-F1-MODEL\_V4 \| 1.0 \| 1.88e-09 \| 303 \| 0.337 \| 148 \| 90 \| 3 \| 2 \| 143 \| 32 \| 177 \| Uncharacterized protein \| Uncharacterized protein; afdb-uniprot50 \| AF-A0A158S129-F1-MODEL\_V4 \| 1.0 \| 2.854e-08 \| 290 \| 0.411 \| 107 \| 57 \| 2 \| 24 \| 124 \| 2 \| 108 \| Uncharacterized protein \| Uncharacterized protein; afdb-uniprot50 \| AF-A0A7J0BLE3-F1-MODEL\_V4 \| 1.0 \| 7.729e-07 \| 290 \| 0.35 \| 117 \| 66 \| 2 \| 2 \| 118 \| 5 \| 111 \| Uncharacterized protein \| Uncharacterized protein; afdb-uniprot50 \| AF-A0A158E8H2-F1-MODEL\_V4 \| 1.0 \| 1.548e-06 \| 278 \| 0.433 \| 83 \| 44 \| 1 \| 1 \| 80 \| 28 \| 110 \| Uncharacterized protein \| Uncharacterized protein; afdb-uniprot50 \| AF-A0A4T2A620-F1-MODEL\_V4 \| 1.0 \| 7.729e-07 \| 271 \| 0.294 \| 119 \| 70 \| 2 \| 2 \| 118 \| 4 \| 110 \| HTH luxR-type domain-containing protein \| HTH luxR-type domain-containing protein; afdb-uniprot50 \| AF-A0A7W9SAC2-F1-MODEL\_V4 \| 1.0 \| 1.951e-06 \| 265 \| 0.233 \| 124 \| 85 \| 1 \| 3 \| 126 \| 8 \| 121 \| Mor family transcriptional regulator \| Mor family transcriptional regulator; afdb-uniprot50 \| AF-A0A4Y6UA46-F1-MODEL\_V4 \| 1.0 \| 2.727e-07 \| 264 \| 0.218 \| 119 \| 87 \| 3 \| 2 \| 119 \| 56 \| 169 \| Uncharacterized protein \| Uncharacterized protein; afdb-uniprot50 \| AF-A0A7M3MKQ0-F1-MODEL\_V4 \| 1.0 \| 6.497e-07 \| 263 \| 0.294 \| 119 \| 74 \| 2 \| 3 \| 121 \| 40 \| 148 \| Mor domain-containing protein \| Mor domain-containing protein; afdb-uniprot50 \| AF-A0A1F6G643-F1-MODEL\_V4 \| 1.0 \| 2.761e-06 \| 254 \| 0.225 \| 124 \| 85 \| 2 \| 7 \| 129 \| 2 \| 115 \| Mor domain-containing protein \| Mor domain-containing protein; afdb-uniprot50 \| AF-A0A1J5T9J9-F1-MODEL\_V4 \| 1.0 \| 4.386e-06 \| 253 \| 0.314 \| 121 \| 70 \| 3 \| 4 \| 122 \| 13 \| 122 \| Mor transcription activator family protein \| Mor transcription activator family protein; afdb-uniprot50 \| AF-A0A7C8HXL4-F1-MODEL\_V4 \| 1.0 \| 9.308e-06 \| 246 \| 0.308 \| 120 \| 74 \| 2 \| 2 \| 119 \| 6 \| 118 \| Sigma70\_r4\_2 domain-containing protein \| Sigma70\_r4\_2 domain-containing protein; afdb-uniprot50 \| AF-A0A1F6GLC2-F1-MODEL\_V4 \| 1.0 \| 5.529e-06 \| 244 \| 0.26 \| 119 \| 78 \| 1 \| 7 \| 125 \| 2 \| 110 \| Mor domain-containing protein \| Mor domain-containing protein; afdb-uniprot50 \| AF-A0A7C8HVY2-F1-MODEL\_V4 \| 1.0 \| 6.577e-06 \| 244 \| 0.288 \| 125 \| 77 \| 3 \| 1 \| 120 \| 3 \| 120 \| Uncharacterized protein \| Uncharacterized protein; afdb-uniprot50 \| AF-A0A557QXB9-F1-MODEL\_V4 \| 1.0 \| 8.784e-06 \| 244 \| 0.25 \| 116 \| 77 \| 1 \| 3 \| 118 \| 24 \| 129 \| Mor domain-containing protein \| Mor domain-containing protein; afdb-uniprot50 \| AF-A0A7Y8NKK1-F1-MODEL\_V4 \| 1.0 \| 1.738e-06 \| 243 \| 0.242 \| 140 \| 91 \| 5 \| 3 \| 131 \| 15 \| 150 \| Mor domain-containing protein \| Mor domain-containing protein; afdb-uniprot50 \| AF-I3YGT9-F1-MODEL\_V4 \| 1.0 \| 1.567e-05 \| 241 \| 0.302 \| 119 \| 71 \| 2 \| 3 \| 119 \| 24 \| 132 \| Mor transcription activator-like protein \| Mor transcription activator-like protein; afdb-uniprot50 \| AF-A0A423PRX3-F1-MODEL\_V4 \| 1.0 \| 1.173e-05 \| 240 \| 0.29 \| 117 \| 73 \| 1 \| 3 \| 119 \| 10 \| 116 \| Mor domain-containing protein \| Mor domain-containing protein; afdb-uniprot50 \| AF-A0A1T2L9D4-F1-MODEL\_V4 \| 1.0 \| 6.577e-06 \| 239 \| 0.198 \| 116 \| 81 \| 2 \| 4 \| 118 \| 9 \| 113 \| Mor domain-containing protein \| Mor domain-containing protein; afdb-uniprot50 \| AF-A0A7W6S2N3-F1-MODEL\_V4 \| 1.0 \| 3.907e-06 \| 238 \| 0.246 \| 130 \| 79 \| 4 \| 1 \| 125 \| 1 \| 116 \| DNA-binding transcriptional ArsR family regulator \| DNA-binding transcriptional ArsR family regulator; afdb-uniprot50 \| AF-F9ZD98-F1-MODEL\_V4 \| 1.0 \| 3.48e-06 \| 234 \| 0.241 \| 124 \| 83 \| 4 \| 3 \| 125 \| 10 \| 123 \| Mor transcription activator domain protein \| Mor transcription activator domain protein; afdb-uniprot50 \| AF-A0A1N6I1K7-F1-MODEL\_V4 \| 1.0 \| 1.243e-05 \| 232 \| 0.276 \| 123 \| 74 \| 2 \| 3 \| 120 \| 8 \| 120 \| Mor transcription activator family protein \| Mor transcription activator family protein; afdb-uniprot50 \| AF-A0A1Y0N3Z3-F1-MODEL\_V4 \| 1.0 \| 6.207e-06 \| 232 \| 0.218 \| 142 \| 93 \| 2 \| 3 \| 144 \| 15 \| 138 \| Uncharacterized protein \| Uncharacterized protein; afdb-uniprot50 \| AF-A0A3B9L6A9-F1-MODEL\_V4 \| 1.0 \| 3.325e-05 \| 229 \| 0.205 \| 117 \| 80 \| 2 \| 3 \| 119 \| 49 \| 152 \| Mor domain-containing protein \| Mor domain-containing protein; afdb-uniprot50 \| AF-A0A2Z3IDQ1-F1-MODEL\_V4 \| 1.0 \| 1.951e-06 \| 228 \| 0.437 \| 96 \| 52 \| 1 \| 41 \| 134 \| 2 \| 97 \| Uncharacterized protein \| Uncharacterized protein; afdb-uniprot50 \| AF-A0A832XIX8-F1-MODEL\_V4 \| 1.0 \| 2.067e-06 \| 227 \| 0.246 \| 126 \| 82 \| 2 \| 1 \| 123 \| 10 \| 125 \| Uncharacterized protein \| Uncharacterized protein; afdb-uniprot50 \| AF-A0A4Z0WD49-F1-MODEL\_V4 \| 1.0 \| 1.094e-06 \| 226 \| 0.281 \| 142 \| 82 \| 3 \| 3 \| 144 \| 12 \| 133 \| Mor domain-containing protein \| Mor domain-containing protein; afdb-uniprot50 \| AF-B0RUR6-F1-MODEL\_V4 \| 1.0 \| 5.218e-06 \| 226 \| 0.263 \| 133 \| 85 \| 7 \| 3 \| 125 \| 119 \| 248 \| Uncharacterized protein \| Uncharacterized protein; afdb-uniprot50 \| AF-A0A4D7B5Y7-F1-MODEL\_V4 \| 1.0 \| 9.742e-07 \| 223 \| 0.293 \| 133 \| 73 \| 5 \| 3 \| 125 \| 48 \| 169 \| Uncharacterized protein \| Uncharacterized protein; afdb-uniprot50 \| AF-A0A106BHZ3-F1-MODEL\_V4 \| 1.0 \| 1.317e-05 \| 222 \| 0.212 \| 113 \| 79 \| 1 \| 6 \| 118 \| 7 \| 109 \| Uncharacterized protein \| Uncharacterized protein; afdb-uniprot50 \| AF-A0A512HA32-F1-MODEL\_V4 \| 1.0 \| 9.862e-06 \| 222 \| 0.238 \| 126 \| 83 \| 3 \| 3 \| 128 \| 11 \| 123 \| Mor domain-containing protein \| Mor domain-containing protein; afdb-uniprot50 \| AF-A0A5S9PH66-F1-MODEL\_V4 \| 1.0 \| 2.321e-06 \| 220 \| 0.263 \| 129 \| 80 \| 2 \| 2 \| 125 \| 7 \| 125 \| Mor domain-containing protein \| Mor domain-containing protein; afdb-uniprot50 \| AF-A0A522X0T2-F1-MODEL\_V4 \| 1.0 \| 8.393e-05 \| 219 \| 0.198 \| 116 \| 83 \| 1 \| 3 \| 118 \| 36 \| 141 \| Mor domain-containing protein \| Mor domain-containing protein; afdb-uniprot50 \| AF-A0A2E2T381-F1-MODEL\_V4 \| 1.0 \| 5.529e-06 \| 218 \| 0.273 \| 117 \| 75 \| 1 \| 3 \| 119 \| 8 \| 114 \| Mor domain-containing protein \| Mor domain-containing protein; afdb-uniprot50 \| AF-A0A4Q5L708-F1-MODEL\_V4 \| 1.0 \| 6.207e-06 \| 218 \| 0.169 \| 118 \| 88 \| 1 \| 4 \| 121 \| 35 \| 142 \| Mor domain-containing protein \| Mor domain-containing protein; afdb-uniprot50 \| AF-A0A349MJQ0-F1-MODEL\_V4 \| 1.0 \| 2.925e-06 \| 217 \| 0.228 \| 127 \| 88 \| 2 \| 2 \| 128 \| 17 \| 133 \| Mor domain-containing protein \| Mor domain-containing protein; afdb-uniprot50 \| AF-A0A7Y4W2P4-F1-MODEL\_V4 \| 1.0 \| 8.893e-05 \| 215 \| 0.172 \| 116 \| 86 \| 1 \| 3 \| 118 \| 6 \| 111 \| Mor domain-containing protein \| Mor domain-containing protein; afdb-uniprot50 \| AF-A0A7Y7LP38-F1-MODEL\_V4 \| 1.0 \| 1.975e-05 \| 215 \| 0.231 \| 138 \| 87 \| 4 \| 3 \| 139 \| 15 \| 134 \| Mor family transcriptional regulator \| Mor family transcriptional regulator; afdb-uniprot50 \| AF-A0A1Y5STF4-F1-MODEL\_V4 \| 1.0 \| 2.093e-05 \| 215 \| 0.254 \| 118 \| 78 \| 1 \| 3 \| 120 \| 12 \| 119 \| Mor transcription activator family protein \| Mor transcription activator family protein; afdb-uniprot50 \| AF-A0A1I7CKY0-F1-MODEL\_V4 \| 1.0 \| 4.986e-05 \| 214 \| 0.194 \| 118 \| 84 \| 2 \| 2 \| 119 \| 27 \| 133 \| Mor transcription activator family protein \| Mor transcription activator family protein; afdb-uniprot50 \| AF-A0A2G6DFJ6-F1-MODEL\_V4 \| 1.0 \| 0.0001333 \| 214 \| 0.177 \| 118 \| 85 \| 2 \| 5 \| 120 \| 52 \| 159 \| Mor domain-containing protein \| Mor domain-containing protein; afdb-uniprot50 \| AF-A0A850RKP1-F1-MODEL\_V4 \| 1.0 \| 2.795e-05 \| 213 \| 0.255 \| 129 \| 84 \| 2 \| 3 \| 129 \| 20 \| 138 \| Transcriptional regulator \| Transcriptional regulator; afdb-uniprot50 \| AF-A0A212KJX9-F1-MODEL\_V4 \| 1.0 \| 2.925e-06 \| 211 \| 0.223 \| 130 \| 88 \| 3 \| 1 \| 127 \| 2 \| 121 \| Mor domain-containing protein \| Mor domain-containing protein; afdb-uniprot50 \| AF-M4NQN5-F1-MODEL\_V4 \| 1.0 \| 4.191e-05 \| 211 \| 0.173 \| 144 \| 98 \| 3 \| 3 \| 144 \| 13 \| 137 \| Uncharacterized protein \| Uncharacterized protein; afdb-uniprot50 \| AF-X5MEB9-F1-MODEL\_V4 \| 1.0 \| 2.925e-06 \| 210 \| 0.256 \| 117 \| 75 \| 4 \| 3 \| 118 \| 8 \| 113 \| Uncharacterized protein \| Uncharacterized protein; afdb-uniprot50 \| AF-A0A7D5NCC3-F1-MODEL\_V4 \| 1.0 \| 2.638e-05 \| 210 \| 0.217 \| 129 \| 85 \| 3 \| 3 \| 127 \| 5 \| 121 \| Mor domain-containing protein \| Mor domain-containing protein; afdb-uniprot50 \| AF-A0A522VNP1-F1-MODEL\_V4 \| 1.0 \| 5.931e-05 \| 209 \| 0.279 \| 118 \| 72 \| 4 \| 3 \| 118 \| 12 \| 118 \| Mor domain-containing protein \| Mor domain-containing protein; afdb-uniprot50 \| AF-A0A178HNW5-F1-MODEL\_V4 \| 1.0 \| 1.759e-05 \| 209 \| 0.246 \| 126 \| 83 \| 3 \| 2 \| 126 \| 12 \| 126 \| Uncharacterized protein \| Uncharacterized protein; afdb-uniprot50 \| AF-T2G7I7-F1-MODEL\_V4 \| 1.0 \| 4.705e-05 \| 209 \| 0.184 \| 119 \| 87 \| 1 \| 3 \| 121 \| 25 \| 133 \| Uncharacterized protein \| Uncharacterized protein; afdb-uniprot50 \| AF-A0A4R7ABL8-F1-MODEL\_V4 \| 1.0 \| 2.925e-06 \| 209 \| 0.219 \| 146 \| 90 \| 4 \| 3 \| 144 \| 6 \| 131 \| Homeodomain-like domain-containing protein \| Homeodomain-like domain-containing protein; afdb-uniprot50 \| AF-A0A149SVR6-F1-MODEL\_V4 \| 1.0 \| 9.308e-06 \| 208 \| 0.243 \| 111 \| 75 \| 4 \| 8 \| 118 \| 2 \| 103 \| HTH\_7 domain-containing protein \| HTH\_7 domain-containing protein; afdb-uniprot50 \| AF-A0A1Y6D0G8-F1-MODEL\_V4 \| 1.0 \| 3.907e-06 \| 208 \| 0.232 \| 129 \| 86 \| 3 \| 2 \| 127 \| 13 \| 131 \| Mor transcription activator family protein \| Mor transcription activator family protein; afdb-uniprot50 \| AF-A0A840G8N0-F1-MODEL\_V4 \| 1.0 \| 1.975e-05 \| 208 \| 0.232 \| 116 \| 79 \| 1 \| 3 \| 118 \| 20 \| 125 \| Mor family transcriptional regulator \| Mor family transcriptional regulator; afdb-uniprot50 \| AF-A0A349MJR3-F1-MODEL\_V4 \| 1.0 \| 6.659e-05 \| 207 \| 0.233 \| 124 \| 81 \| 3 \| 1 \| 122 \| 1 \| 112 \| Mor domain-containing protein \| Mor domain-containing protein; afdb-uniprot50 \| AF-A0A2D3T993-F1-MODEL\_V4 \| 1.0 \| 5.529e-06 \| 207 \| 0.269 \| 130 \| 78 \| 4 \| 1 \| 125 \| 6 \| 123 \| Mor domain-containing protein \| Mor domain-containing protein; afdb-uniprot50 \| AF-A0A1G0CQ82-F1-MODEL\_V4 \| 1.0 \| 5.597e-05 \| 207 \| 0.226 \| 119 \| 79 \| 3 \| 4 \| 119 \| 13 \| 121 \| Mor domain-containing protein \| Mor domain-containing protein; afdb-uniprot50 \| AF-A0A0M0FPC9-F1-MODEL\_V4 \| 1.0 \| 4.191e-05 \| 207 \| 0.252 \| 119 \| 77 \| 3 \| 1 \| 118 \| 7 \| 114 \| Mor domain-containing protein \| Mor domain-containing protein; afdb-uniprot50 \| AF-D5CUC0-F1-MODEL\_V4 \| 1.0 \| 5.597e-05 \| 206 \| 0.213 \| 117 \| 82 \| 2 \| 3 \| 119 \| 11 \| 117 \| Mor domain-containing protein \| Mor domain-containing protein; afdb-uniprot50 \| AF-A0A1W9JSH1-F1-MODEL\_V4 \| 1.0 \| 2.217e-05 \| 206 \| 0.219 \| 132 \| 86 \| 4 \| 1 \| 125 \| 3 \| 124 \| Mor domain-containing protein \| Mor domain-containing protein; afdb-uniprot50 \| AF-A9ID21-F1-MODEL\_V4 \| 1.0 \| 1.317e-05 \| 206 \| 0.204 \| 142 \| 93 \| 2 \| 3 \| 144 \| 12 \| 133 \| Mor domain-containing protein \| Mor domain-containing protein; afdb-uniprot50 \| AF-A0A3M1WU79-F1-MODEL\_V4 \| 1.0 \| 4.986e-05 \| 206 \| 0.215 \| 139 \| 87 \| 2 \| 3 \| 129 \| 12 \| 140 \| DNA transposition protein \| DNA transposition protein; afdb-uniprot50 \| AF-N6Y808-F1-MODEL\_V4 \| 1.0 \| 5.283e-05 \| 205 \| 0.251 \| 135 \| 80 \| 4 \| 4 \| 129 \| 26 \| 148 \| Uncharacterized protein \| Uncharacterized protein; afdb-uniprot50 \| AF-I3YBF0-F1-MODEL\_V4 \| 1.0 \| 2.093e-05 \| 205 \| 0.214 \| 135 \| 86 \| 3 \| 3 \| 127 \| 18 \| 142 \| Mor transcription activator-like protein \| Mor transcription activator-like protein; afdb-uniprot50 \| AF-A8I7L4-F1-MODEL\_V4 \| 1.0 \| 1.045e-05 \| 202 \| 0.201 \| 119 \| 85 \| 1 \| 1 \| 119 \| 5 \| 113 \| Uncharacterized protein \| Uncharacterized protein; afdb-uniprot50 \| AF-A0A1H6U3N5-F1-MODEL\_V4 \| 1.0 \| 7.294e-07 \| 202 \| 0.429 \| 121 \| 59 \| 2 \| 14 \| 125 \| 2 \| 121 \| Winged helix-turn helix \| Winged helix-turn helix; afdb-uniprot50 \| AF-A0A350X8X0-F1-MODEL\_V4 \| 1.0 \| 8.784e-06 \| 202 \| 0.243 \| 115 \| 76 \| 2 \| 7 \| 121 \| 14 \| 117 \| Mor domain-containing protein \| Mor domain-containing protein; afdb-uniprot50 \| AF-A0A1V3HDH7-F1-MODEL\_V4 \| 1.0 \| 8.893e-05 \| 201 \| 0.232 \| 116 \| 78 \| 2 \| 3 \| 118 \| 20 \| 124 \| HTH psq-type domain-containing protein \| HTH psq-type domain-containing protein; afdb-uniprot50 \| AF-A0A2S5CLI8-F1-MODEL\_V4 \| 1.0 \| 4.191e-05 \| 201 \| 0.194 \| 118 \| 81 \| 3 \| 3 \| 118 \| 12 \| 117 \| Mor domain-containing protein \| Mor domain-containing protein; afdb-uniprot50 \| AF-A9C2C1-F1-MODEL\_V4 \| 1.0 \| 1.567e-05 \| 201 \| 0.2 \| 150 \| 100 \| 3 \| 1 \| 139 \| 2 \| 142 \| Uncharacterized protein \| Uncharacterized protein; afdb-uniprot50 \| AF-A0A4R3J9E2-F1-MODEL\_V4 \| 1.0 \| 2.638e-05 \| 200 \| 0.246 \| 134 \| 83 \| 4 \| 1 \| 128 \| 1 \| 122 \| Homeodomain-like domain-containing protein \| Homeodomain-like domain-containing protein; afdb-uniprot50 \| AF-A0A315EKC3-F1-MODEL\_V4 \| 1.0 \| 0.0002119 \| 199 \| 0.273 \| 84 \| 61 \| 0 \| 38 \| 121 \| 13 \| 96 \| Mor domain-containing protein \| Mor domain-containing protein; afdb-uniprot50 \| AF-A0A1I2FJZ9-F1-MODEL\_V4 \| 1.0 \| 1.567e-05 \| 199 \| 0.211 \| 123 \| 86 \| 2 \| 3 \| 125 \| 10 \| 121 \| Mor transcription activator family protein \| Mor transcription activator family protein; afdb-uniprot50 \| AF-A0A143Z940-F1-MODEL\_V4 \| 1.0 \| 1.759e-05 \| 199 \| 0.248 \| 125 \| 84 \| 1 \| 3 \| 127 \| 13 \| 127 \| Homeodomain-like domain-containing protein \| Homeodomain-like domain-containing protein; afdb-uniprot50 \| AF-A0A375GAU5-F1-MODEL\_V4 \| 1.0 \| 7.921e-05 \| 199 \| 0.21 \| 114 \| 80 \| 1 \| 7 \| 120 \| 11 \| 114 \| Uncharacterized protein \| Uncharacterized protein; afdb-uniprot50 \| AF-A0A368TMP4-F1-MODEL\_V4 \| 1.0 \| 6.207e-06 \| 198 \| 0.235 \| 153 \| 92 \| 6 \| 1 \| 144 \| 1 \| 137 \| Uncharacterized protein \| Uncharacterized protein; afdb-uniprot50 \| AF-A0A1I5IF11-F1-MODEL\_V4 \| 1.0 \| 0.000267 \| 198 \| 0.157 \| 121 \| 88 \| 2 \| 3 \| 119 \| 59 \| 169 \| Mor transcription activator family protein \| Mor transcription activator family protein; afdb-uniprot50 \| AF-A0A1I7AGL9-F1-MODEL\_V4 \| 1.0 \| 3.523e-05 \| 197 \| 0.216 \| 120 \| 84 \| 1 \| 1 \| 120 \| 8 \| 117 \| Transcriptional regulator, Middle operon regulator (Mor) family \| Transcriptional regulator, Middle operon regulator (Mor) family; afdb-uniprot50 \| AF-A0A0J7J645-F1-MODEL\_V4 \| 1.0 \| 6.969e-06 \| 197 \| 0.22 \| 127 \| 88 \| 2 \| 3 \| 128 \| 9 \| 125 \| Mor transcription activator family \| Mor transcription activator family; afdb-uniprot50 \| AF-A0A5C7Q033-F1-MODEL\_V4 \| 1.0 \| 2.638e-05 \| 197 \| 0.219 \| 141 \| 94 \| 3 \| 1 \| 134 \| 2 \| 133 \| Winged helix-turn-helix transcriptional regulator \| Winged helix-turn-helix transcriptional regulator; afdb-uniprot50 \| AF-A0A6H3FDI8-F1-MODEL\_V4 \| 1.0 \| 1.864e-05 \| 196 \| 0.231 \| 147 \| 85 \| 7 \| 3 \| 139 \| 41 \| 169 \| Mor domain-containing protein \| Mor domain-containing protein; afdb-uniprot50 \| AF-A0A3A0EQS8-F1-MODEL\_V4 \| 1.0 \| 4.191e-05 \| 194 \| 0.226 \| 119 \| 82 \| 1 \| 3 \| 121 \| 14 \| 122 \| Mor domain-containing protein \| Mor domain-containing protein; afdb-uniprot50 \| AF-A0A849SXZ0-F1-MODEL\_V4 \| 1.0 \| 0.0001333 \| 194 \| 0.198 \| 131 \| 88 \| 3 \| 3 \| 128 \| 32 \| 150 \| Uncharacterized protein \| Uncharacterized protein; afdb-uniprot50 \| AF-A0A258P502-F1-MODEL\_V4 \| 1.0 \| 7.921e-05 \| 194 \| 0.22 \| 127 \| 81 \| 3 \| 3 \| 123 \| 58 \| 172 \| Mor domain-containing protein \| Mor domain-containing protein; afdb-uniprot50 \| AF-A0A1Q6UEQ9-F1-MODEL\_V4 \| 1.0 \| 5.283e-05 \| 192 \| 0.254 \| 118 \| 74 \| 4 \| 7 \| 122 \| 8 \| 113 \| Uncharacterized protein \| Uncharacterized protein; afdb-uniprot50 \| AF-A0A3R9ZQF3-F1-MODEL\_V4 \| 1.0 \| 7.921e-05 \| 192 \| 0.221 \| 122 \| 81 \| 2 \| 2 \| 119 \| 28 \| 139 \| Mor domain-containing protein \| Mor domain-containing protein; afdb-uniprot50 \| AF-A0A142BHK6-F1-MODEL\_V4 \| 1.0 \| 1.567e-05 \| 192 \| 0.179 \| 139 \| 100 \| 3 \| 8 \| 144 \| 5 \| 131 \| Mor transcription activator domain-containing protein \| Mor transcription activator domain-containing protein; afdb-uniprot50 \| AF-A0A5C7KLE1-F1-MODEL\_V4 \| 1.0 \| 9.862e-06 \| 191 \| 0.262 \| 141 \| 82 \| 6 \| 4 \| 144 \| 11 \| 129 \| Mor domain-containing protein \| Mor domain-containing protein; afdb-uniprot50 \| AF-A0A345DE69-F1-MODEL\_V4 \| 1.0 \| 4.191e-05 \| 191 \| 0.197 \| 137 \| 85 \| 3 \| 7 \| 128 \| 4 \| 130 \| Uncharacterized protein \| Uncharacterized protein; afdb-uniprot50 \| AF-A0A3R8L878-F1-MODEL\_V4 \| 1.0 \| 5.597e-05 \| 191 \| 0.196 \| 132 \| 90 \| 4 \| 3 \| 128 \| 85 \| 206 \| Uncharacterized protein \| Uncharacterized protein; afdb-uniprot50 \| AF-A0A2P5M266-F1-MODEL\_V4 \| 1.0 \| 7.921e-05 \| 190 \| 0.206 \| 126 \| 89 \| 3 \| 3 \| 127 \| 11 \| 126 \| Mor domain-containing protein \| Mor domain-containing protein; afdb-uniprot50 \| AF-A0A7U5K534-F1-MODEL\_V4 \| 1.0 \| 2.962e-05 \| 190 \| 0.241 \| 124 \| 83 \| 2 \| 3 \| 125 \| 12 \| 125 \| Uncharacterized protein \| Uncharacterized protein; afdb-uniprot50 \| AF-A0A1P8JXX9-F1-MODEL\_V4 \| 1.0 \| 0.0003779 \| 190 \| 0.186 \| 118 \| 84 \| 3 \| 3 \| 119 \| 13 \| 119 \| Mor domain-containing protein \| Mor domain-containing protein; afdb-uniprot50 \| AF-A0A522VTV5-F1-MODEL\_V4 \| 1.0 \| 0.0001681 \| 190 \| 0.193 \| 129 \| 91 \| 4 \| 3 \| 128 \| 14 \| 132 \| Mor domain-containing protein \| Mor domain-containing protein; afdb-uniprot50 \| AF-A0A6G9Q100-F1-MODEL\_V4 \| 1.0 \| 0.0001121 \| 190 \| 0.203 \| 128 \| 89 \| 3 \| 3 \| 129 \| 19 \| 134 \| Helix-turn-helix domain-containing protein \| Helix-turn-helix domain-containing protein; afdb-uniprot50 \| AF-A0A439VN72-F1-MODEL\_V4 \| 1.0 \| 3.733e-05 \| 190 \| 0.247 \| 117 \| 72 \| 4 \| 5 \| 118 \| 109 \| 212 \| Uncharacterized protein \| Uncharacterized protein; afdb-uniprot50 \| AF-A0A286AC00-F1-MODEL\_V4 \| 1.0 \| 0.0003177 \| 189 \| 0.222 \| 117 \| 79 \| 3 \| 3 \| 118 \| 10 \| 115 \| Mor transcription activator family protein \| Mor transcription activator family protein; afdb-uniprot50 \| AF-A0A1C3EL69-F1-MODEL\_V4 \| 1.0 \| 4.986e-05 \| 188 \| 0.26 \| 123 \| 71 \| 5 \| 6 \| 121 \| 3 \| 112 \| Mor domain-containing protein \| Mor domain-containing protein; afdb-uniprot50 \| AF-H1SDM0-F1-MODEL\_V4 \| 1.0 \| 0.0001681 \| 188 \| 0.17 \| 129 \| 88 \| 3 \| 7 \| 128 \| 20 \| 136 \| Mor domain-containing protein \| Mor domain-containing protein; afdb-uniprot50 \| AF-A0A272EMK5-F1-MODEL\_V4 \| 1.0 \| 3.955e-05 \| 188 \| 0.214 \| 121 \| 82 \| 2 \| 1 \| 118 \| 2 \| 112 \| Uncharacterized protein \| Uncharacterized protein; afdb-uniprot50 \| AF-A0A7W9QXS3-F1-MODEL\_V4 \| 1.0 \| 9.308e-06 \| 187 \| 0.22 \| 136 \| 79 \| 5 \| 1 \| 124 \| 6 \| 126 \| Uncharacterized protein \| Uncharacterized protein; afdb-uniprot50 \| AF-A0A2E6KN34-F1-MODEL\_V4 \| 1.0 \| 4.705e-05 \| 186 \| 0.196 \| 117 \| 79 \| 2 \| 2 \| 118 \| 10 \| 111 \| Uncharacterized protein \| Uncharacterized protein; afdb-uniprot50 \| AF-A0A6G9Q124-F1-MODEL\_V4 \| 1.0 \| 0.0002245 \| 186 \| 0.195 \| 123 \| 86 \| 3 \| 3 \| 124 \| 19 \| 129 \| Helix-turn-helix domain-containing protein \| Helix-turn-helix domain-containing protein; afdb-uniprot50 \| AF-M5DM24-F1-MODEL\_V4 \| 1.0 \| 2.49e-05 \| 186 \| 0.223 \| 121 \| 73 \| 3 \| 3 \| 119 \| 10 \| 113 \| Mor domain-containing protein \| Mor domain-containing protein; afdb-uniprot50 \| AF-A0A1N6X0E0-F1-MODEL\_V4 \| 1.0 \| 9.984e-05 \| 186 \| 0.208 \| 134 \| 84 \| 4 \| 3 \| 128 \| 13 \| 132 \| Mor transcription activator family protein \| Mor transcription activator family protein; afdb-uniprot50 \| AF-A0A521YLK1-F1-MODEL\_V4 \| 1.0 \| 8.393e-05 \| 186 \| 0.214 \| 140 \| 87 \| 4 \| 1 \| 128 \| 117 \| 245 \| Mor domain-containing protein \| Mor domain-containing protein; afdb-uniprot50 \| AF-F5RBY1-F1-MODEL\_V4 \| 1.0 \| 6.659e-05 \| 185 \| 0.223 \| 143 \| 86 \| 5 \| 1 \| 128 \| 1 \| 133 \| Bacteriophage DNA transposition protein B \| Bacteriophage DNA transposition protein B; afdb-uniprot50 \| AF-N9R432-F1-MODEL\_V4 \| 1.0 \| 9.423e-05 \| 185 \| 0.198 \| 141 \| 96 \| 5 \| 3 \| 141 \| 23 \| 148 \| Mor domain-containing protein \| Mor domain-containing protein; afdb-uniprot50 \| AF-A0A5C7L0Z4-F1-MODEL\_V4 \| 1.0 \| 7.921e-05 \| 184 \| 0.169 \| 112 \| 83 \| 1 \| 7 \| 118 \| 6 \| 107 \| Mor domain-containing protein \| Mor domain-containing protein; afdb-uniprot50 \| AF-A0A4Q0YJD9-F1-MODEL\_V4 \| 1.0 \| 5.931e-05 \| 184 \| 0.258 \| 120 \| 71 \| 4 \| 6 \| 119 \| 3 \| 110 \| Uncharacterized protein \| Uncharacterized protein; afdb-uniprot50 \| AF-A0A2N9YH57-F1-MODEL\_V4 \| 1.0 \| 0.0003779 \| 183 \| 0.191 \| 120 \| 82 \| 4 \| 3 \| 121 \| 10 \| 115 \| Mor domain-containing protein \| Mor domain-containing protein; afdb-uniprot50 \| AF-A0A6N4DE39-F1-MODEL\_V4 \| 1.0 \| 0.0004004 \| 183 \| 0.25 \| 116 \| 73 \| 2 \| 3 \| 118 \| 7 \| 108 \| Uncharacterized protein \| Uncharacterized protein; afdb-uniprot50 \| AF-A0A3D9Q0H6-F1-MODEL\_V4 \| 1.0 \| 1.759e-05 \| 183 \| 0.268 \| 134 \| 80 \| 3 \| 3 \| 128 \| 10 \| 133 \| Uncharacterized protein \| Uncharacterized protein; afdb-uniprot50 \| AF-A0A3P1ZRA8-F1-MODEL\_V4 \| 1.0 \| 2.638e-05 \| 183 \| 0.166 \| 156 \| 106 \| 6 \| 3 \| 144 \| 55 \| 200 \| Mor domain-containing protein \| Mor domain-containing protein; afdb-uniprot50 \| AF-Q1QMW9-F1-MODEL\_V4 \| 1.0 \| 8.393e-05 \| 182 \| 0.277 \| 119 \| 74 \| 3 \| 1 \| 118 \| 3 \| 110 \| Uncharacterized protein \| Uncharacterized protein; afdb-uniprot50 \| AF-A0A5P9EX33-F1-MODEL\_V4 \| 1.0 \| 0.0001681 \| 182 \| 0.216 \| 125 \| 86 \| 3 \| 2 \| 125 \| 18 \| 131 \| Mor transcription activator family protein \| Mor transcription activator family protein; afdb-uniprot50 \| AF-A0A2A2B555-F1-MODEL\_V4 \| 1.0 \| 0.0003779 \| 182 \| 0.206 \| 116 \| 80 \| 2 \| 5 \| 118 \| 8 \| 113 \| Mor domain-containing protein \| Mor domain-containing protein; afdb-uniprot50 \| AF-A0A833MQ40-F1-MODEL\_V4 \| 1.0 \| 1.479e-05 \| 182 \| 0.274 \| 131 \| 79 \| 2 \| 1 \| 125 \| 12 \| 132 \| Uncharacterized protein \| Uncharacterized protein; afdb-uniprot50 \| AF-A0A2W4TD25-F1-MODEL\_V4 \| 1.0 \| 6.659e-05 \| 182 \| 0.256 \| 121 \| 76 \| 3 \| 7 \| 125 \| 76 \| 184 \| Uncharacterized protein \| Uncharacterized protein; afdb-uniprot50 \| AF-G2HDC1-F1-MODEL\_V4 \| 1.0 \| 8.893e-05 \| 181 \| 0.202 \| 138 \| 83 \| 4 \| 7 \| 144 \| 9 \| 119 \| Uncharacterized protein \| Uncharacterized protein; afdb-uniprot50 \| AF-A0A2T5J3R6-F1-MODEL\_V4 \| 1.0 \| 0.0006004 \| 181 \| 0.185 \| 124 \| 88 \| 3 \| 7 \| 127 \| 5 \| 118 \| Mor transcription activator family protein \| Mor transcription activator family protein; afdb-uniprot50 \| AF-A0A6N7JS93-F1-MODEL\_V4 \| 1.0 \| 3.325e-05 \| 181 \| 0.246 \| 130 \| 73 \| 6 \| 4 \| 125 \| 12 \| 124 \| Uncharacterized protein \| Uncharacterized protein; afdb-uniprot50 \| AF-A0A2U2N1B0-F1-MODEL\_V4 \| 1.0 \| 0.0004763 \| 181 \| 0.294 \| 119 \| 70 \| 5 \| 3 \| 118 \| 9 \| 116 \| Uncharacterized protein \| Uncharacterized protein; afdb-uniprot50 \| AF-A0A0U5FC51-F1-MODEL\_V4 \| 1.0 \| 1.567e-05 \| 181 \| 0.195 \| 133 \| 92 \| 5 \| 3 \| 124 \| 13 \| 141 \| Uncharacterized protein \| Uncharacterized protein; afdb-uniprot50 \| AF-A0A1J4X7E1-F1-MODEL\_V4 \| 1.0 \| 0.0002998 \| 180 \| 0.21 \| 133 \| 86 \| 6 \| 2 \| 129 \| 18 \| 136 \| Mor domain-containing protein \| Mor domain-containing protein; afdb-uniprot50 \| AF-A0A1G8EZ96-F1-MODEL\_V4 \| 1.0 \| 1.864e-05 \| 180 \| 0.214 \| 135 \| 85 \| 3 \| 1 \| 127 \| 10 \| 131 \| Homeodomain-like domain-containing protein \| Homeodomain-like domain-containing protein; afdb-uniprot50 \| AF-A0A1D2X552-F1-MODEL\_V4 \| 1.0 \| 0.0002119 \| 180 \| 0.195 \| 123 \| 85 \| 3 \| 8 \| 128 \| 5 \| 115 \| Mor domain-containing protein \| Mor domain-containing protein; afdb-uniprot50 \| AF-A0A1I5W2T6-F1-MODEL\_V4 \| 1.0 \| 0.0001497 \| 179 \| 0.184 \| 119 \| 84 \| 3 \| 1 \| 119 \| 1 \| 106 \| Uncharacterized protein \| Uncharacterized protein; afdb-uniprot50 \| AF-A0A357V9S2-F1-MODEL\_V4 \| 1.0 \| 0.0001497 \| 179 \| 0.216 \| 134 \| 85 \| 2 \| 1 \| 124 \| 1 \| 124 \| Uncharacterized protein \| Uncharacterized protein; afdb-uniprot50 \| AF-K0D2L3-F1-MODEL\_V4 \| 1.0 \| 0.0004004 \| 179 \| 0.184 \| 119 \| 82 \| 5 \| 8 \| 125 \| 14 \| 118 \| Mor domain-containing protein \| Mor domain-containing protein; afdb-uniprot50 \| AF-A0A3D3KCN7-F1-MODEL\_V4 \| 1.0 \| 0.0005667 \| 179 \| 0.173 \| 121 \| 88 \| 2 \| 1 \| 119 \| 1 \| 111 \| Mor domain-containing protein \| Mor domain-containing protein; afdb-uniprot50 \| AF-A0A401JFQ9-F1-MODEL\_V4 \| 1.0 \| 2.093e-05 \| 179 \| 0.219 \| 132 \| 85 \| 2 \| 1 \| 124 \| 12 \| 133 \| Mor domain-containing protein \| Mor domain-containing protein; afdb-uniprot50 \| AF-A0A0D2GJW6-F1-MODEL\_V4 \| 1.0 \| 7.055e-05 \| 177 \| 0.201 \| 114 \| 78 \| 2 \| 5 \| 118 \| 4 \| 104 \| Uncharacterized protein \| Uncharacterized protein; afdb-uniprot50 \| AF-A0A5C7VFA7-F1-MODEL\_V4 \| 1.0 \| 0.0001058 \| 177 \| 0.162 \| 123 \| 90 \| 3 \| 1 \| 123 \| 12 \| 121 \| Mor domain-containing protein \| Mor domain-containing protein; afdb-uniprot50 \| AF-A0A0W1G863-F1-MODEL\_V4 \| 1.0 \| 1.66e-05 \| 177 \| 0.258 \| 143 \| 86 \| 6 \| 1 \| 133 \| 1 \| 133 \| Uncharacterized protein \| Uncharacterized protein; afdb-uniprot50 \| AF-A0A5K6V8A9-F1-MODEL\_V4 \| 1.0 \| 7.055e-05 \| 176 \| 0.254 \| 114 \| 68 \| 5 \| 11 \| 119 \| 2 \| 103 \| Mor domain-containing protein \| Mor domain-containing protein; afdb-uniprot50 \| AF-A0A1N7JLA1-F1-MODEL\_V4 \| 1.0 \| 0.0001887 \| 176 \| 0.26 \| 123 \| 71 \| 4 \| 2 \| 118 \| 5 \| 113 \| Uncharacterized protein \| Uncharacterized protein; afdb-uniprot50 \| AF-A0A2W5N6E1-F1-MODEL\_V4 \| 1.0 \| 0.0001121 \| 176 \| 0.247 \| 109 \| 67 \| 2 \| 10 \| 118 \| 19 \| 112 \| Uncharacterized protein \| Uncharacterized protein; afdb-uniprot50 \| AF-A0A1I1UI57-F1-MODEL\_V4 \| 1.0 \| 3.138e-05 \| 176 \| 0.26 \| 119 \| 66 \| 4 \| 8 \| 118 \| 2 \| 106 \| Uncharacterized protein \| Uncharacterized protein; afdb-uniprot50 \| AF-A0A3M1PQC8-F1-MODEL\_V4 \| 1.0 \| 5.931e-05 \| 176 \| 0.187 \| 139 \| 97 \| 4 \| 7 \| 144 \| 6 \| 129 \| Uncharacterized protein \| Uncharacterized protein; afdb-uniprot50 \| AF-A0A3T0W468-F1-MODEL\_V4 \| 1.0 \| 0.0001681 \| 176 \| 0.162 \| 117 \| 88 \| 1 \| 3 \| 119 \| 14 \| 120 \| Mor domain-containing protein \| Mor domain-containing protein; afdb-uniprot50 \| AF-A0A7W8C0C3-F1-MODEL\_V4 \| 1.0 \| 4.986e-05 \| 176 \| 0.192 \| 161 \| 94 \| 7 \| 3 \| 144 \| 63 \| 206 \| Mor family transcriptional regulator \| Mor family transcriptional regulator; afdb-uniprot50 \| AF-A0A4P9VQV6-F1-MODEL\_V4 \| 1.0 \| 0.0001999 \| 175 \| 0.201 \| 119 \| 80 \| 5 \| 3 \| 119 \| 14 \| 119 \| Transcriptional regulator \| Transcriptional regulator; afdb-uniprot50 \| AF-A0A4D7DET1-F1-MODEL\_V4 \| 1.0 \| 2.49e-05 \| 174 \| 0.23 \| 126 \| 76 \| 5 \| 6 \| 125 \| 11 \| 121 \| Uncharacterized protein \| Uncharacterized protein; afdb-uniprot50 \| AF-A0A1W9J765-F1-MODEL\_V4 \| 1.0 \| 0.0002379 \| 174 \| 0.224 \| 116 \| 80 \| 1 \| 3 \| 118 \| 12 \| 117 \| Mor domain-containing protein \| Mor domain-containing protein; afdb-uniprot50 \| AF-A0A6L5XLQ5-F1-MODEL\_V4 \| 1.0 \| 5.597e-05 \| 174 \| 0.18 \| 150 \| 100 \| 6 \| 3 \| 139 \| 48 \| 187 \| Mor domain-containing protein \| Mor domain-containing protein; afdb-uniprot50 \| AF-A0A286GYP9-F1-MODEL\_V4 \| 1.0 \| 0.0002245 \| 173 \| 0.188 \| 122 \| 83 \| 2 \| 3 \| 118 \| 12 \| 123 \| Uncharacterized protein \| Uncharacterized protein; afdb-uniprot50 \| AF-B2FRB5-F1-MODEL\_V4 \| 1.0 \| 1.396e-05 \| 172 \| 0.301 \| 126 \| 72 \| 4 \| 27 \| 143 \| 1 \| 119 \| Uncharacterized protein \| Uncharacterized protein; afdb-uniprot50 \| AF-A0A089WQT6-F1-MODEL\_V4 \| 1.0 \| 0.0005047 \| 170 \| 0.194 \| 118 \| 82 \| 3 \| 3 \| 119 \| 14 \| 119 \| Transcriptional regulator \| Transcriptional regulator; afdb-uniprot50 \| AF-A0A7K3NNL5-F1-MODEL\_V4 \| 1.0 \| 6.284e-05 \| 170 \| 0.255 \| 133 \| 81 \| 4 \| 1 \| 125 \| 7 \| 129 \| Uncharacterized protein \| Uncharacterized protein; afdb-uniprot50 \| AF-A0A2W4T512-F1-MODEL\_V4 \| 1.0 \| 3.523e-05 \| 169 \| 0.23 \| 139 \| 80 \| 5 \| 3 \| 128 \| 11 \| 135 \| Uncharacterized protein \| Uncharacterized protein; afdb-uniprot50 \| AF-A0A1X7NE24-F1-MODEL\_V4 \| 1.0 \| 0.0009003 \| 168 \| 0.204 \| 122 \| 82 \| 5 \| 2 \| 121 \| 4 \| 112 \| Uncharacterized protein \| Uncharacterized protein; afdb-uniprot50 \| AF-A0A0J6SFM2-F1-MODEL\_V4 \| 1.0 \| 3.138e-05 \| 168 \| 0.196 \| 132 \| 92 \| 3 \| 1 \| 128 \| 8 \| 129 \| Uncharacterized protein \| Uncharacterized protein; afdb-uniprot50 \| AF-A0A6H3FAD0-F1-MODEL\_V4 \| 1.0 \| 0.0001586 \| 167 \| 0.206 \| 121 \| 81 \| 4 \| 7 \| 125 \| 5 \| 112 \| Uncharacterized protein \| Uncharacterized protein; afdb-uniprot50 \| AF-A0A5C7NL60-F1-MODEL\_V4 \| 1.0 \| 3.523e-05 \| 167 \| 0.257 \| 136 \| 82 \| 5 \| 4 \| 139 \| 53 \| 169 \| Mor domain-containing protein \| Mor domain-containing protein; afdb-uniprot50 \| AF-A0A1X7QCW3-F1-MODEL\_V4 \| 1.0 \| 7.055e-05 \| 167 \| 0.207 \| 169 \| 96 \| 7 \| 1 \| 144 \| 51 \| 206 \| Uncharacterized protein \| Uncharacterized protein; afdb-uniprot50 \| AF-A0A554WZV0-F1-MODEL\_V4 \| 1.0 \| 0.0006362 \| 166 \| 0.203 \| 123 \| 85 \| 3 \| 7 \| 128 \| 3 \| 113 \| Mor transcription activator family protein \| Mor transcription activator family protein; afdb-uniprot50 \| AF-A0A7T3E5I3-F1-MODEL\_V4 \| 1.0 \| 9.984e-05 \| 166 \| 0.194 \| 139 \| 86 \| 4 \| 6 \| 143 \| 10 \| 123 \| Uncharacterized protein \| Uncharacterized protein; afdb-uniprot50 \| AF-A0A1Z4C0F4-F1-MODEL\_V4 \| 1.0 \| 0.002408 \| 166 \| 0.196 \| 117 \| 82 \| 3 \| 3 \| 118 \| 13 \| 118 \| Mor domain-containing protein \| Mor domain-containing protein; afdb-uniprot50 \| AF-E2CFN9-F1-MODEL\_V4 \| 1.0 \| 0.0001058 \| 166 \| 0.197 \| 137 \| 83 \| 6 \| 3 \| 128 \| 7 \| 127 \| Uncharacterized protein \| Uncharacterized protein; afdb-uniprot50 \| AF-A0A839IQE8-F1-MODEL\_V4 \| 1.0 \| 0.0007143 \| 166 \| 0.198 \| 116 \| 82 \| 2 \| 3 \| 118 \| 20 \| 124 \| Transcriptional regulator \| Transcriptional regulator; afdb-uniprot50 \| AF-A0A3B9NYV6-F1-MODEL\_V4 \| 1.0 \| 0.0001781 \| 165 \| 0.201 \| 129 \| 89 \| 3 \| 3 \| 128 \| 15 \| 132 \| Uncharacterized protein \| Uncharacterized protein; afdb-uniprot50 \| AF-A0A2S5JEN6-F1-MODEL\_V4 \| 1.0 \| 0.0005047 \| 165 \| 0.242 \| 128 \| 77 \| 4 \| 1 \| 120 \| 1 \| 116 \| Uncharacterized protein \| Uncharacterized protein; afdb-uniprot50 \| AF-A0A139DN11-F1-MODEL\_V4 \| 1.0 \| 0.0009539 \| 165 \| 0.188 \| 122 \| 81 \| 4 \| 3 \| 119 \| 31 \| 139 \| Mor domain-containing protein \| Mor domain-containing protein; afdb-uniprot50 \| AF-A0A2D3TET5-F1-MODEL\_V4 \| 1.0 \| 4.986e-05 \| 164 \| 0.242 \| 140 \| 88 \| 5 \| 11 \| 144 \| 2 \| 129 \| Mor domain-containing protein \| Mor domain-containing protein; afdb-uniprot50 \| AF-A0A2W5H275-F1-MODEL\_V4 \| 1.0 \| 0.001071 \| 164 \| 0.21 \| 119 \| 80 \| 4 \| 1 \| 118 \| 1 \| 106 \| Mor domain-containing protein \| Mor domain-containing protein; afdb-uniprot50 \| AF-E9I3C0-F1-MODEL\_V4 \| 1.0 \| 2.35e-05 \| 164 \| 0.177 \| 152 \| 101 \| 3 \| 1 \| 139 \| 1 \| 141 \| Uncharacterized protein \| Uncharacterized protein; afdb-uniprot50 \| AF-A0A3B9PP60-F1-MODEL\_V4 \| 1.0 \| 0.0001497 \| 164 \| 0.253 \| 150 \| 85 \| 5 \| 3 \| 144 \| 19 \| 149 \| Mor domain-containing protein \| Mor domain-containing protein; afdb-uniprot50 \| AF-A0A1H6GGU4-F1-MODEL\_V4 \| 1.0 \| 8.893e-05 \| 163 \| 0.201 \| 129 \| 85 \| 3 \| 3 \| 127 \| 16 \| 130 \| Uncharacterized protein \| Uncharacterized protein; afdb-uniprot50 \| AF-A0A2M8DST2-F1-MODEL\_V4 \| 1.0 \| 0.0003366 \| 163 \| 0.2 \| 120 \| 82 \| 3 \| 4 \| 119 \| 27 \| 136 \| Mor domain-containing protein \| Mor domain-containing protein; afdb-uniprot50 \| AF-A0A418W4I4-F1-MODEL\_V4 \| 1.0 \| 3.733e-05 \| 163 \| 0.24 \| 150 \| 88 \| 6 \| 3 \| 141 \| 14 \| 148 \| Uncharacterized protein \| Uncharacterized protein; afdb-uniprot50 \| AF-A0A853HSH7-F1-MODEL\_V4 \| 1.0 \| 7.921e-05 \| 163 \| 0.18 \| 155 \| 98 \| 5 \| 1 \| 139 \| 1 \| 142 \| Uncharacterized protein \| Uncharacterized protein; afdb-uniprot50 \| AF-A0A2T6M6E5-F1-MODEL\_V4 \| 1.0 \| 0.000267 \| 162 \| 0.196 \| 122 \| 84 \| 3 \| 2 \| 119 \| 3 \| 114 \| Uncharacterized protein \| Uncharacterized protein; afdb-uniprot50 \| AF-A0A2K1DE74-F1-MODEL\_V4 \| 1.0 \| 0.0005348 \| 162 \| 0.184 \| 130 \| 87 \| 4 \| 2 \| 124 \| 18 \| 135 \| Transcriptional regulator \| Transcriptional regulator; afdb-uniprot50 \| AF-A0A1H5GMI8-F1-MODEL\_V4 \| 1.0 \| 0.00143 \| 162 \| 0.205 \| 117 \| 82 \| 3 \| 3 \| 119 \| 49 \| 154 \| Transcriptional regulator, Middle operon regulator (Mor) family \| Transcriptional regulator, Middle operon regulator (Mor) family; afdb-uniprot50 \| AF-A0A6L8HVT0-F1-MODEL\_V4 \| 1.0 \| 0.0003177 \| 162 \| 0.206 \| 116 \| 72 \| 5 \| 7 \| 118 \| 62 \| 161 \| Mor domain-containing protein \| Mor domain-containing protein; afdb-uniprot50 \| AF-A0A1G0EZ32-F1-MODEL\_V4 \| 1.0 \| 0.00135 \| 161 \| 0.157 \| 121 \| 82 \| 3 \| 7 \| 122 \| 2 \| 107 \| Uncharacterized protein \| Uncharacterized protein; afdb-uniprot50 \| AF-A0A2N2RB11-F1-MODEL\_V4 \| 1.0 \| 0.0002245 \| 161 \| 0.234 \| 115 \| 72 \| 4 \| 7 \| 118 \| 13 \| 114 \| Uncharacterized protein \| Uncharacterized protein; afdb-uniprot50 \| AF-A0A451AQL9-F1-MODEL\_V4 \| 1.0 \| 0.0006741 \| 161 \| 0.164 \| 134 \| 94 \| 2 \| 3 \| 128 \| 12 \| 135 \| Mor transcription activator family protein \| Mor transcription activator family protein; afdb-uniprot50 \| AF-T2JY53-F1-MODEL\_V4 \| 1.0 \| 0.001071 \| 160 \| 0.211 \| 118 \| 83 \| 1 \| 3 \| 120 \| 12 \| 119 \| Uncharacterized protein \| Uncharacterized protein; afdb-uniprot50 \| AF-A0A2S6N2X3-F1-MODEL\_V4 \| 1.0 \| 0.0007568 \| 160 \| 0.186 \| 118 \| 84 \| 2 \| 3 \| 118 \| 5 \| 112 \| Uncharacterized protein \| Uncharacterized protein; afdb-uniprot50 \| AF-A0A1H1G473-F1-MODEL\_V4 \| 1.0 \| 0.0004496 \| 160 \| 0.21 \| 128 \| 81 \| 3 \| 1 \| 118 \| 2 \| 119 \| Helix-turn-helix domain-containing protein \| Helix-turn-helix domain-containing protein; afdb-uniprot50 \| AF-A0A4Y8W9I1-F1-MODEL\_V4 \| 1.0 \| 0.0005667 \| 160 \| 0.218 \| 119 \| 76 \| 4 \| 3 \| 118 \| 22 \| 126 \| Uncharacterized protein \| Uncharacterized protein; afdb-uniprot50 \| AF-A0A4V2V2Q2-F1-MODEL\_V4 \| 1.0 \| 0.0005667 \| 159 \| 0.225 \| 124 \| 78 \| 5 \| 1 \| 118 \| 1 \| 112 \| Homeodomain-like domain-containing protein \| Homeodomain-like domain-containing protein; afdb-uniprot50 \| AF-A0A5K1I944-F1-MODEL\_V4 \| 1.0 \| 0.0003177 \| 159 \| 0.232 \| 112 \| 76 \| 2 \| 7 \| 118 \| 8 \| 109 \| Mor transcription activator family protein \| Mor transcription activator family protein; afdb-uniprot50 \| AF-V4NRA8-F1-MODEL\_V4 \| 1.0 \| 0.0002998 \| 159 \| 0.213 \| 122 \| 76 \| 4 \| 1 \| 118 \| 11 \| 116 \| Uncharacterized protein \| Uncharacterized protein; afdb-uniprot50 \| AF-A0A4S4AP65-F1-MODEL\_V4 \| 1.0 \| 0.0003779 \| 159 \| 0.19 \| 152 \| 92 \| 4 \| 3 \| 144 \| 43 \| 173 \| Mor domain-containing protein \| Mor domain-containing protein; afdb-uniprot50 \| AF-A0A2Z6DXP8-F1-MODEL\_V4 \| 1.0 \| 0.0009003 \| 158 \| 0.226 \| 128 \| 83 \| 4 \| 1 \| 124 \| 1 \| 116 \| Mor domain-containing protein \| Mor domain-containing protein; afdb-uniprot50 \| AF-A0A3F3GTU9-F1-MODEL\_V4 \| 1.0 \| 0.0007568 \| 158 \| 0.197 \| 152 \| 97 \| 4 \| 3 \| 144 \| 13 \| 149 \| Putative DNA transposition protein \| Putative DNA transposition protein; afdb-uniprot50 \| AF-A0A845TMQ7-F1-MODEL\_V4 \| 1.0 \| 0.0003779 \| 156 \| 0.17 \| 129 \| 84 \| 4 \| 1 \| 119 \| 1 \| 116 \| Uncharacterized protein \| Uncharacterized protein; afdb-uniprot50 \| AF-A0A522VKF2-F1-MODEL\_V4 \| 1.0 \| 9.984e-05 \| 156 \| 0.186 \| 145 \| 95 \| 5 \| 3 \| 144 \| 14 \| 138 \| Mor domain-containing protein \| Mor domain-containing protein; afdb-uniprot50 \| AF-A0A2D9F5H7-F1-MODEL\_V4 \| 1.0 \| 0.0003567 \| 155 \| 0.203 \| 123 \| 76 \| 4 \| 2 \| 118 \| 9 \| 115 \| Mor domain-containing protein \| Mor domain-containing protein; afdb-uniprot50 \| AF-A0A1H3FGD2-F1-MODEL\_V4 \| 1.0 \| 0.0006362 \| 155 \| 0.23 \| 117 \| 78 \| 4 \| 3 \| 118 \| 9 \| 114 \| Mor transcription activator family protein \| Mor transcription activator family protein; afdb-uniprot50 \| AF-A0A1B8Q882-F1-MODEL\_V4 \| 1.0 \| 0.0002245 \| 155 \| 0.188 \| 127 \| 83 \| 4 \| 3 \| 119 \| 19 \| 135 \| Uncharacterized protein \| Uncharacterized protein; afdb-uniprot50 \| AF-A0A7Z0N040-F1-MODEL\_V4 \| 1.0 \| 0.003408 \| 155 \| 0.151 \| 112 \| 85 \| 2 \| 7 \| 118 \| 14 \| 115 \| Uncharacterized protein \| Uncharacterized protein; afdb-uniprot50 \| AF-F4QGB6-F1-MODEL\_V4 \| 1.0 \| 0.0004004 \| 154 \| 0.204 \| 122 \| 78 \| 3 \| 8 \| 125 \| 2 \| 108 \| Uncharacterized protein \| Uncharacterized protein; afdb-uniprot50 \| AF-A0A7X3YCW7-F1-MODEL\_V4 \| 1.0 \| 0.0006004 \| 153 \| 0.181 \| 116 \| 82 \| 2 \| 6 \| 121 \| 3 \| 105 \| Mor domain-containing protein \| Mor domain-containing protein; afdb-uniprot50 \| AF-A0A1G8GH27-F1-MODEL\_V4 \| 1.0 \| 0.000252 \| 153 \| 0.221 \| 122 \| 75 \| 5 \| 3 \| 118 \| 10 \| 117 \| Uncharacterized protein \| Uncharacterized protein; afdb-uniprot50 \| AF-A0A6A7Y6F2-F1-MODEL\_V4 \| 1.0 \| 0.0001121 \| 153 \| 0.239 \| 138 \| 87 \| 7 \| 3 \| 134 \| 12 \| 137 \| Uncharacterized protein \| Uncharacterized protein; afdb-uniprot50 \| AF-A0A843YXE8-F1-MODEL\_V4 \| 1.0 \| 0.0001887 \| 153 \| 0.217 \| 138 \| 81 \| 5 \| 7 \| 129 \| 11 \| 136 \| Uncharacterized protein \| Uncharacterized protein; afdb-uniprot50 \| AF-A0A2W5WW80-F1-MODEL\_V4 \| 1.0 \| 0.0006741 \| 152 \| 0.198 \| 141 \| 92 \| 4 \| 1 \| 136 \| 2 \| 126 \| Helix-turn-helix domain-containing protein \| Helix-turn-helix domain-containing protein; afdb-uniprot50 \| AF-T0IRN1-F1-MODEL\_V4 \| 1.0 \| 0.000267 \| 152 \| 0.184 \| 130 \| 88 \| 3 \| 6 \| 127 \| 22 \| 141 \| Uncharacterized protein \| Uncharacterized protein; afdb-uniprot50 \| AF-A0A848C494-F1-MODEL\_V4 \| 1.0 \| 0.0006362 \| 152 \| 0.165 \| 151 \| 97 \| 6 \| 3 \| 143 \| 31 \| 162 \| Uncharacterized protein \| Uncharacterized protein; afdb-uniprot50 \| AF-A0A7W5ZUY6-F1-MODEL\_V4 \| 1.0 \| 0.0007143 \| 151 \| 0.156 \| 134 \| 90 \| 3 \| 1 \| 128 \| 1 \| 117 \| Uncharacterized protein \| Uncharacterized protein; afdb-uniprot50 \| AF-A0A510UGA1-F1-MODEL\_V4 \| 1.0 \| 0.00143 \| 151 \| 0.176 \| 119 \| 85 \| 3 \| 3 \| 120 \| 21 \| 127 \| Transcriptional regulator \| Transcriptional regulator; afdb-uniprot50 \| AF-A0A291IMH9-F1-MODEL\_V4 \| 1.0 \| 0.0004496 \| 151 \| 0.208 \| 134 \| 88 \| 5 \| 7 \| 137 \| 10 \| 128 \| Mor domain-containing protein \| Mor domain-containing protein; afdb-uniprot50 \| AF-A0A6N8BMD9-F1-MODEL\_V4 \| 1.0 \| 0.0008019 \| 151 \| 0.146 \| 157 \| 108 \| 7 \| 3 \| 144 \| 62 \| 207 \| Mor domain-containing protein \| Mor domain-containing protein; afdb-uniprot50 \| AF-A0A1D7Z023-F1-MODEL\_V4 \| 1.0 \| 0.002551 \| 150 \| 0.213 \| 117 \| 79 \| 3 \| 2 \| 118 \| 20 \| 123 \| Mor domain-containing protein \| Mor domain-containing protein; afdb-uniprot50 \| AF-A0A4D7DA93-F1-MODEL\_V4 \| 1.0 \| 0.0004004 \| 150 \| 0.195 \| 148 \| 103 \| 5 \| 2 \| 144 \| 23 \| 159 \| Uncharacterized protein \| Uncharacterized protein; afdb-uniprot50 \| AF-A0A564WHC9-F1-MODEL\_V4 \| 1.0 \| 0.0006362 \| 150 \| 0.211 \| 118 \| 82 \| 3 \| 2 \| 118 \| 51 \| 158 \| Uncharacterized protein \| Uncharacterized protein; afdb-uniprot50 \| AF-T2GC33-F1-MODEL\_V4 \| 1.0 \| 0.001702 \| 149 \| 0.205 \| 112 \| 79 \| 1 \| 7 \| 118 \| 14 \| 115 \| Uncharacterized protein \| Uncharacterized protein; afdb-uniprot50 \| AF-A0A2E9SLK3-F1-MODEL\_V4 \| 1.0 \| 0.000252 \| 149 \| 0.226 \| 119 \| 81 \| 3 \| 7 \| 125 \| 4 \| 111 \| Uncharacterized protein \| Uncharacterized protein; afdb-uniprot50 \| AF-A0A3N9RYX3-F1-MODEL\_V4 \| 1.0 \| 0.002703 \| 149 \| 0.2 \| 120 \| 82 \| 4 \| 5 \| 122 \| 24 \| 131 \| Uncharacterized protein \| Uncharacterized protein; afdb-uniprot50 \| AF-A0A1G3LVZ0-F1-MODEL\_V4 \| 1.0 \| 0.0002379 \| 148 \| 0.175 \| 137 \| 87 \| 5 \| 7 \| 129 \| 3 \| 127 \| Uncharacterized protein \| Uncharacterized protein; afdb-uniprot50 \| AF-A0A853IG23-F1-MODEL\_V4 \| 1.0 \| 0.000267 \| 147 \| 0.172 \| 133 \| 86 \| 5 \| 7 \| 127 \| 4 \| 124 \| Uncharacterized protein \| Uncharacterized protein; afdb-uniprot50 \| AF-A0A420ZY38-F1-MODEL\_V4 \| 1.0 \| 0.0007568 \| 147 \| 0.236 \| 114 \| 75 \| 3 \| 7 \| 119 \| 83 \| 185 \| Mor transcription activator family protein \| Mor transcription activator family protein; afdb-uniprot50 \| AF-A0A3N9RYD7-F1-MODEL\_V4 \| 1.0 \| 0.0006004 \| 147 \| 0.201 \| 139 \| 96 \| 5 \| 4 \| 139 \| 56 \| 182 \| Uncharacterized protein \| Uncharacterized protein; afdb-uniprot50 \| AF-A0A1H9CJ13-F1-MODEL\_V4 \| 1.0 \| 0.002551 \| 146 \| 0.181 \| 116 \| 77 \| 3 \| 7 \| 119 \| 2 \| 102 \| Mor transcription activator family protein \| Mor transcription activator family protein; afdb-uniprot50 \| AF-A0A7W9RCT4-F1-MODEL\_V4 \| 1.0 \| 0.0001999 \| 145 \| 0.224 \| 125 \| 75 \| 5 \| 3 \| 119 \| 8 \| 118 \| Uncharacterized protein \| Uncharacterized protein; afdb-uniprot50 \| AF-E2CN54-F1-MODEL\_V4 \| 1.0 \| 0.0008497 \| 145 \| 0.227 \| 132 \| 83 \| 3 \| 3 \| 125 \| 17 \| 138 \| HTH\_7 domain-containing protein \| HTH\_7 domain-containing protein; afdb-uniprot50 \| AF-R6HUG7-F1-MODEL\_V4 \| 1.0 \| 0.002551 \| 144 \| 0.185 \| 113 \| 81 \| 2 \| 7 \| 118 \| 11 \| 113 \| Uncharacterized protein \| Uncharacterized protein; afdb-uniprot50 \| AF-A0A522VJQ9-F1-MODEL\_V4 \| 1.0 \| 0.0005348 \| 143 \| 0.22 \| 127 \| 85 \| 4 \| 3 \| 125 \| 45 \| 161 \| Mor domain-containing protein \| Mor domain-containing protein; afdb-uniprot50 \| AF-A0A554X2Y4-F1-MODEL\_V4 \| 1.0 \| 0.0008497 \| 143 \| 0.197 \| 162 \| 100 \| 5 \| 1 \| 144 \| 2 \| 151 \| Mor transcription activator family protein \| Mor transcription activator family protein; afdb-uniprot50 \| AF-A0A0Q2YXZ0-F1-MODEL\_V4 \| 1.0 \| 0.0009003 \| 142 \| 0.202 \| 148 \| 94 \| 6 \| 2 \| 144 \| 6 \| 134 \| Uncharacterized protein \| Uncharacterized protein; afdb-uniprot50 \| AF-A0A0U5MJB7-F1-MODEL\_V4 \| 1.0 \| 0.0003779 \| 142 \| 0.223 \| 130 \| 83 \| 5 \| 6 \| 128 \| 16 \| 134 \| Uncharacterized protein \| Uncharacterized protein; afdb-uniprot50 \| AF-A0A5E4XGN4-F1-MODEL\_V4 \| 1.0 \| 0.0004496 \| 141 \| 0.205 \| 136 \| 84 \| 4 \| 1 \| 124 \| 1 \| 124 \| Mor domain-containing protein \| Mor domain-containing protein; afdb-uniprot50 \| AF-A0A7C1YRL4-F1-MODEL\_V4 \| 1.0 \| 0.002408 \| 137 \| 0.169 \| 118 \| 88 \| 1 \| 2 \| 119 \| 10 \| 117 \| Uncharacterized protein \| Uncharacterized protein; afdb-uniprot50 \| AF-B8GL06-F1-MODEL\_V4 \| 1.0 \| 0.009114 \| 136 \| 0.156 \| 128 \| 92 \| 5 \| 3 \| 127 \| 16 \| 130 \| Mor domain-containing protein \| Mor domain-containing protein; afdb-uniprot50 \| AF-A0A1H8FWD2-F1-MODEL\_V4 \| 1.0 \| 0.00191 \| 135 \| 0.173 \| 121 \| 90 \| 2 \| 4 \| 124 \| 6 \| 116 \| Mor transcription activator family protein \| Mor transcription activator family protein; afdb-uniprot50 \| AF-A0A2D3TE22-F1-MODEL\_V4 \| 1.0 \| 0.0004004 \| 135 \| 0.234 \| 132 \| 70 \| 5 \| 1 \| 131 \| 23 \| 124 \| Uncharacterized protein \| Uncharacterized protein; afdb-uniprot50 \| AF-A0A6A4RCI3-F1-MODEL\_V4 \| 1.0 \| 0.001516 \| 134 \| 0.248 \| 133 \| 80 \| 5 \| 3 \| 127 \| 9 \| 129 \| Uncharacterized protein \| Uncharacterized protein; afdb-uniprot50 \| AF-A0A4Q8UBK0-F1-MODEL\_V4 \| 1.0 \| 0.0007568 \| 134 \| 0.116 \| 146 \| 109 \| 6 \| 1 \| 140 \| 1 \| 132 \| Mor domain-containing protein \| Mor domain-containing protein; afdb-uniprot50 \| AF-N9PZH4-F1-MODEL\_V4 \| 1.0 \| 0.001702 \| 134 \| 0.185 \| 140 \| 90 \| 4 \| 2 \| 127 \| 9 \| 138 \| Mor domain-containing protein \| Mor domain-containing protein; afdb-uniprot50 \| AF-A0A6L6WQR3-F1-MODEL\_V4 \| 1.0 \| 0.003035 \| 132 \| 0.176 \| 119 \| 87 \| 2 \| 1 \| 118 \| 6 \| 114 \| Uncharacterized protein \| Uncharacterized protein; afdb-uniprot50 \| AF-A0A5E4ZWP6-F1-MODEL\_V4 \| 1.0 \| 0.004551 \| 131 \| 0.178 \| 112 \| 82 \| 1 \| 8 \| 119 \| 22 \| 123 \| Mor domain-containing protein \| Mor domain-containing protein; afdb-uniprot50 \| AF-Q31HV4-F1-MODEL\_V4 \| 1.0 \| 0.004295 \| 131 \| 0.166 \| 114 \| 85 \| 2 \| 5 \| 118 \| 12 \| 115 \| Uncharacterized protein \| Uncharacterized protein; afdb-uniprot50 \| AF-A0A858Q8R5-F1-MODEL\_V4 \| 1.0 \| 0.004822 \| 130 \| 0.247 \| 85 \| 62 \| 1 \| 46 \| 128 \| 5 \| 89 \| Uncharacterized protein \| Uncharacterized protein; afdb-uniprot50 \| AF-A0A847G9G6-F1-MODEL\_V4 \| 1.0 \| 0.001202 \| 130 \| 0.183 \| 142 \| 100 \| 5 \| 1 \| 139 \| 1 \| 129 \| Uncharacterized protein \| Uncharacterized protein; afdb-uniprot50 \| AF-A0A850KBD1-F1-MODEL\_V4 \| 1.0 \| 0.000252 \| 130 \| 0.207 \| 140 \| 96 \| 4 \| 7 \| 143 \| 8 \| 135 \| Helix-turn-helix domain-containing protein \| Helix-turn-helix domain-containing protein; afdb-uniprot50 \| AF-A0A3D9Q2J0-F1-MODEL\_V4 \| 1.0 \| 0.0004763 \| 130 \| 0.219 \| 146 \| 85 \| 6 \| 2 \| 128 \| 6 \| 141 \| Mor transcription activator family protein \| Mor transcription activator family protein; afdb-uniprot50 \| AF-A0A6L7X7N5-F1-MODEL\_V4 \| 1.0 \| 0.001011 \| 129 \| 0.186 \| 134 \| 86 \| 6 \| 1 \| 124 \| 3 \| 123 \| Mor domain-containing protein \| Mor domain-containing protein; afdb-uniprot50 \| AF-A0A3T0N3Z2-F1-MODEL\_V4 \| 1.0 \| 0.009114 \| 129 \| 0.168 \| 119 \| 88 \| 2 \| 1 \| 118 \| 21 \| 129 \| Uncharacterized protein \| Uncharacterized protein; afdb-uniprot50 \| AF-A0A2G1LMC0-F1-MODEL\_V4 \| 1.0 \| 0.0007143 \| 128 \| 0.154 \| 155 \| 97 \| 5 \| 7 \| 144 \| 3 \| 140 \| Uncharacterized protein \| Uncharacterized protein; afdb-uniprot50 \| AF-A0A7W4G402-F1-MODEL\_V4 \| 1.0 \| 0.0009003 \| 127 \| 0.1 \| 119 \| 96 \| 2 \| 3 \| 120 \| 11 \| 119 \| Mor domain-containing protein \| Mor domain-containing protein; afdb-uniprot50 \| AF-Q3IGJ2-F1-MODEL\_V4 \| 1.0 \| 0.0009003 \| 127 \| 0.1 \| 119 \| 96 \| 2 \| 3 \| 120 \| 11 \| 119 \| Putative orphan protein \| Putative orphan protein; afdb-uniprot50 \| AF-A0A5Q0EN91-F1-MODEL\_V4 \| 1.0 \| 0.002145 \| 126 \| 0.261 \| 134 \| 67 \| 10 \| 1 \| 121 \| 1 \| 115 \| Mor domain-containing protein \| Mor domain-containing protein; afdb-uniprot50 \| AF-A0A7W6S3Y5-F1-MODEL\_V4 \| 1.0 \| 0.004822 \| 126 \| 0.209 \| 124 \| 80 \| 4 \| 2 \| 118 \| 16 \| 128 \| DNA-binding NarL/FixJ family response regulator \| DNA-binding NarL/FixJ family response regulator; afdb-uniprot50 \| AF-A0A270BEU5-F1-MODEL\_V4 \| 1.0 \| 0.002865 \| 125 \| 0.193 \| 119 \| 84 \| 3 \| 1 \| 118 \| 4 \| 111 \| Uncharacterized protein \| Uncharacterized protein; afdb-uniprot50 \| AF-A0A0P8Y9C8-F1-MODEL\_V4 \| 1.0 \| 0.002145 \| 125 \| 0.186 \| 145 \| 100 \| 3 \| 1 \| 142 \| 21 \| 150 \| Mor domain-containing protein \| Mor domain-containing protein; afdb-uniprot50 \| AF-A0A0F2RXJ4-F1-MODEL\_V4 \| 1.0 \| 0.00191 \| 124 \| 0.161 \| 118 \| 87 \| 2 \| 1 \| 118 \| 2 \| 107 \| Uncharacterized protein \| Uncharacterized protein; afdb-uniprot50 \| AF-A0A656HD19-F1-MODEL\_V4 \| 1.0 \| 0.006441 \| 123 \| 0.168 \| 119 \| 83 \| 5 \| 3 \| 118 \| 7 \| 112 \| Mor transcription activator domain protein \| Mor transcription activator domain protein; afdb-uniprot50 \| AF-B8CUZ8-F1-MODEL\_V4 \| 1.0 \| 0.002551 \| 122 \| 0.172 \| 139 \| 95 \| 7 \| 3 \| 139 \| 22 \| 142 \| Mor domain-containing protein \| Mor domain-containing protein; afdb-uniprot50 \| AF-A0A7C5KXW5-F1-MODEL\_V4 \| 1.0 \| 0.006441 \| 122 \| 0.176 \| 130 \| 88 \| 4 \| 3 \| 125 \| 15 \| 132 \| Mor domain-containing protein \| Mor domain-containing protein; afdb-uniprot50 \| AF-A0A5P8XH25-F1-MODEL\_V4 \| 1.0 \| 0.008118 \| 116 \| 0.214 \| 107 \| 71 \| 4 \| 39 \| 139 \| 10 \| 109 \| Mor domain-containing protein \| Mor domain-containing protein; afdb-uniprot50 \| AF-A0A3N1PMJ1-F1-MODEL\_V4 \| 1.0 \| 0.00511 \| 110 \| 0.175 \| 131 \| 87 \| 3 \| 3 \| 124 \| 15 \| 133 \| Mor transcription activator family protein \| Mor transcription activator family protein; afdb-uniprot50 \| AF-A0A2M7G6P5-F1-MODEL\_V4 \| 1.0 \| 0.009657 \| 103 \| 0.183 \| 120 \| 74 \| 7 \| 11 \| 118 \| 28 \| 135 \| Mor domain-containing protein \| Mor domain-containing protein |
| Top keywords  (threshold 1.00e-02 (evalue)) | **Mor, domain\_containing, transcription, activator, regulator, Transcriptional, Helix\_turn\_helix, Homeodomain\_like, DNA, transposition** |
| Output files | ../../similar\_structures/61\_FANPEZAQ\_CDS\_0061\_afdb-proteome\_foldseek.tsv ../../similar\_structures/61\_FANPEZAQ\_CDS\_0061\_afdb-uniprot50\_foldseek.tsv ../../similar\_structures/61\_FANPEZAQ\_CDS\_0061\_merged.svg ../../similar\_structures/61\_FANPEZAQ\_CDS\_0061\_pdb\_foldseek.tsv |

  
  
  

Return to summary | Go to previous | Go to next

  


---

**Sequence/structure alignments coloring**  
Each object in the alignment figures is colored according to its E-value following this color coding:

1e-100
10
