## Supplementary material for "Completing the BASEL phage collection to unlock hidden diversity for systematic exploration of phage-host interactions": Data S2: 62.html

FANPEZAQ\_CDS\_0062


Return to summary | Go to previous | Go to next

|  |  |
| --- | --- |
| FANPEZAQ\_CDS\_0062  Page creation date: 02 Sep 2024, 12:00  Project folder: n/a  Input sequences file: Escherichia\_virus\_HeidiAbel.gb | holin hypothetical putative phage membrane fragment transporter mfs lysis peptidase m48 ste24p nc\_023006\_p63 p387102 vi\_09157 p363326 vi\_04187 p127786 vi\_03659 antiholin lambda holin\_like ii conserved n\_acetylmuramidase duf1097 domain\_containing prophage hol44 superfamily v contig\_80 whole genome shotgun sequence ethanolamine permease |

### Sequence information

|  |  |
| --- | --- |
| Name | FANPEZAQ\_CDS\_0062  62\_FANPEZAQ\_CDS\_0062 (pipeline id) |
| Imported annotations | Escherichia\_virus\_HeidiAbel Bas97 |
| Protein sequence | MEFFQKFLDNIGWGIAGIMGSLVSLPFHDDIKSVKARIWFVCTGAICAYFLTDLVSLYFK IDRGLAGSVGFLLGAFGGSLLAAVIRAIKAADIWQLLKNRFGGGNA |
| Number of residues | 106 |
| Molecular weight (Da) | 11463.38 |
| Output files | ../../query\_sequences/62\_FANPEZAQ\_CDS\_0062.fasta |

### Putative domain architecture and protein family

#### Search results (HHblits)^1^

|  |  |
| --- | --- |
| Domain family databases searched | Pfam, Ncbi-cd, Cath, Phrogs |
| Results, scheme(s)  (Top layers only; threshold 1.00e-03 (evalue)) | xml version="1.0" encoding="utf-8" standalone="no"?       2024-09-02T21:08:25.903983 image/svg+xml   Matplotlib v3.7.2, https://matplotlib.org/ |
| Results, table  (E-value ≤ 1.00e-03 (evalue)) | db \| id \| prob \| evalue \| pvalue \| score \| cols \| query \| query\_len \| template \| template\_len \| name \| description; phrogs \| 2359 \| 99.8 \| 6.3e-25 \| 7.9e-29 \| 143.6 \| 105 \| (1, 105) \| 106 \| (1, 105) \| 107 \| holin \| holin; Category: lysis; NC\_023006\_p63; phrogs \| 198 \| 99.7 \| 1.4e-21 \| 1.8e-25 \| 129.1 \| 86 \| (9, 97) \| 106 \| (23, 108) \| 116 \| holin \| holin; Category: lysis; p387102 VI\_09157; phrogs \| 13918 \| 97.9 \| 4.1e-09 \| 4.8e-13 \| 68.5 \| 80 \| (9, 89) \| 106 \| (21, 101) \| 127 \| NA \| NA; Category: unknown function; p363326 VI\_04187; phrogs \| 21855 \| 95.0 \| 0.00031 \| 3.6e-08 \| 38.3 \| 29 \| (9, 38) \| 106 \| (27, 55) \| 56 \| NA \| NA; Category: unknown function; p127786 VI\_03659 |
| Top keywords  (threshold 1.00e-03 (evalue)) | **holin, lysis, NC\_023006\_p63, p387102, VI\_09157, p363326, VI\_04187, p127786, VI\_03659** |
| Output files | ../../domain\_architecture/62\_FANPEZAQ\_CDS\_0062\_cath.hhr ../../domain\_architecture/62\_FANPEZAQ\_CDS\_0062\_merged.svg ../../domain\_architecture/62\_FANPEZAQ\_CDS\_0062\_ncbi-cd.hhr ../../domain\_architecture/62\_FANPEZAQ\_CDS\_0062\_pfam.hhr ../../domain\_architecture/62\_FANPEZAQ\_CDS\_0062\_phrogs.hhr |

### Identical protein sequences/structures

#### Search results

|  |  |
| --- | --- |
| Protein sequence databases searched | Pdb, Swissprot, Refseq |
| Identical proteins found | -- |
| Top keywords | -- |
| Output files | -- |

### Similar protein sequences/structures

#### Sequence similarity search results (HHblits)^1^

|  |  |
| --- | --- |
| Sequence databases searched | Uniclust, Pdb70 |
| Results, scheme(s)  (Top layers only, threshold 1.00e-03 (evalue)) | xml version="1.0" encoding="utf-8" standalone="no"?       2024-09-02T21:08:54.385043 image/svg+xml   Matplotlib v3.7.2, https://matplotlib.org/ |
| Results, table(s)  (threshold 1.00e-03 (evalue)) | db \| id \| prob \| evalue \| pvalue \| score \| cols \| query \| query\_len \| template \| template\_len \| name \| description; uniclust \| UniRef100\_A0A059KTJ6 \| 99.9 \| 1.9e-26 \| 4e-32 \| 154.2 \| 105 \| (1, 105) \| 106 \| (6, 112) \| 115 \| MFS transporter \| MFS transporter; uniclust \| UniRef100\_A0A077LKE0 \| 99.8 \| 2.3e-23 \| 4.3e-29 \| 145.4 \| 105 \| (1, 105) \| 106 \| (13, 122) \| 173 \| MFS transporter \| MFS transporter; uniclust \| UniRef100\_A0A083UGU3 \| 99.6 \| 3e-18 \| 6.8e-24 \| 121.1 \| 95 \| (8, 105) \| 106 \| (37, 131) \| 146 \| Phage holin \| Phage holin; uniclust \| UniRef100\_A0A127QUD9 \| 99.6 \| 3.7e-18 \| 7.5e-24 \| 119.5 \| 95 \| (10, 105) \| 106 \| (45, 141) \| 149 \| Holin \| Holin; uniclust \| UniRef100\_A0A345L6P1 \| 99.5 \| 1.2e-17 \| 2.4e-23 \| 111.7 \| 104 \| (1, 104) \| 106 \| (1, 105) \| 108 \| Holin/antiholin \| Holin/antiholin; uniclust \| UniRef100\_A0A0M7GTX4 \| 99.5 \| 1.2e-17 \| 2.6e-23 \| 114.4 \| 92 \| (8, 104) \| 106 \| (17, 111) \| 123 \| Phage holin, lambda family \| Phage holin, lambda family; uniclust \| UniRef100\_A0A1G9MGV6 \| 99.4 \| 4.3e-16 \| 8.6e-22 \| 106.7 \| 90 \| (1, 91) \| 106 \| (9, 98) \| 128 \| Holin \| Holin; uniclust \| UniRef100\_A0A087N8V0 \| 99.4 \| 4.6e-16 \| 1e-21 \| 107.6 \| 100 \| (5, 105) \| 106 \| (14, 115) \| 124 \| Holin \| Holin; uniclust \| UniRef100\_A0A0Q5DKQ6 \| 99.4 \| 8.3e-16 \| 1.8e-21 \| 108.1 \| 84 \| (7, 91) \| 106 \| (19, 110) \| 140 \| Peptidase M48, Ste24p \| Peptidase M48, Ste24p; uniclust \| UniRef100\_A0A009PIY2 \| 99.4 \| 1e-15 \| 2e-21 \| 105.0 \| 94 \| (9, 105) \| 106 \| (30, 123) \| 125 \| Putative membrane protein \| Putative membrane protein; uniclust \| UniRef100\_A0A011V762 \| 99.4 \| 9.8e-16 \| 2.2e-21 \| 109.8 \| 88 \| (3, 91) \| 106 \| (27, 116) \| 158 \| Phage holin family protein \| Phage holin family protein; uniclust \| UniRef100\_A0A059V0X5 \| 99.4 \| 1.2e-15 \| 2.9e-21 \| 107.8 \| 87 \| (9, 98) \| 106 \| (38, 124) \| 138 \| Membrane protein \| Membrane protein; uniclust \| UniRef100\_A0A010SYK4 \| 99.3 \| 5.4e-15 \| 1.1e-20 \| 99.5 \| 84 \| (6, 90) \| 106 \| (11, 98) \| 107 \| Holin \| Holin; uniclust \| UniRef100\_A0A246JHP1 \| 99.3 \| 8.4e-15 \| 1.8e-20 \| 98.7 \| 86 \| (12, 98) \| 106 \| (15, 101) \| 107 \| Holin \| Holin; uniclust \| UniRef100\_A0A021X861 \| 99.3 \| 2e-14 \| 4.1e-20 \| 100.8 \| 88 \| (2, 90) \| 106 \| (18, 113) \| 141 \| Holin \| Holin; uniclust \| UniRef100\_A0A0Q4GP77 \| 99.2 \| 3e-14 \| 6.5e-20 \| 97.7 \| 102 \| (2, 104) \| 106 \| (7, 110) \| 116 \| Uncharacterized protein \| Uncharacterized protein; uniclust \| UniRef100\_A0A088U760 \| 99.2 \| 3.6e-14 \| 7.6e-20 \| 100.7 \| 97 \| (7, 104) \| 106 \| (21, 121) \| 145 \| Putative membrane protein \| Putative membrane protein; uniclust \| UniRef100\_A0A076YME0 \| 99.2 \| 4.9e-14 \| 9.9e-20 \| 96.0 \| 94 \| (1, 94) \| 106 \| (1, 96) \| 116 \| Holin-like class II protein \| Holin-like class II protein; uniclust \| UniRef100\_A0A0U3F4J3 \| 99.2 \| 5.3e-14 \| 1.2e-19 \| 96.6 \| 88 \| (1, 89) \| 106 \| (4, 92) \| 112 \| Holin \| Holin; uniclust \| UniRef100\_A0A0Q8XZH4 \| 99.2 \| 9.1e-14 \| 1.9e-19 \| 97.7 \| 92 \| (10, 104) \| 106 \| (38, 129) \| 139 \| Holin \| Holin; uniclust \| UniRef100\_A0A0A8TGY1 \| 99.1 \| 3.2e-13 \| 6.4e-19 \| 92.8 \| 99 \| (6, 104) \| 106 \| (12, 120) \| 123 \| Holin \| Holin; uniclust \| UniRef100\_A0A166F525 \| 99.1 \| 3.9e-13 \| 7.8e-19 \| 91.5 \| 90 \| (2, 91) \| 106 \| (3, 98) \| 115 \| Phage holin family 2 \| Phage holin family 2; uniclust \| UniRef100\_A0A109J7U2 \| 99.1 \| 4.6e-13 \| 8.9e-19 \| 94.5 \| 90 \| (3, 93) \| 106 \| (15, 107) \| 155 \| Holin \| Holin; uniclust \| UniRef100\_A0A0E0URI3 \| 99.1 \| 4.7e-13 \| 9.7e-19 \| 93.2 \| 92 \| (11, 103) \| 106 \| (23, 124) \| 129 \| Holin \| Holin; uniclust \| UniRef100\_E7EKS8 \| 99.1 \| 6.2e-13 \| 1.1e-18 \| 89.9 \| 95 \| (1, 95) \| 106 \| (15, 110) \| 127 \| Putative holin (Fragment) \| Putative holin (Fragment); uniclust \| UniRef100\_A0A6G6Y576 \| 99.1 \| 6e-13 \| 1.2e-18 \| 89.4 \| 94 \| (11, 105) \| 106 \| (14, 107) \| 108 \| Uncharacterized protein \| Uncharacterized protein; uniclust \| UniRef100\_A0A2N7QRA9 \| 99.1 \| 9.5e-13 \| 1.9e-18 \| 89.4 \| 86 \| (4, 90) \| 106 \| (10, 95) \| 116 \| Holin \| Holin; uniclust \| UniRef100\_A0A149SVI2 \| 99.1 \| 1e-12 \| 2e-18 \| 87.9 \| 92 \| (10, 105) \| 106 \| (12, 103) \| 104 \| Holin \| Holin; uniclust \| UniRef100\_A0A085FNY7 \| 99.0 \| 2e-12 \| 4.1e-18 \| 89.9 \| 85 \| (6, 91) \| 106 \| (14, 106) \| 128 \| Uncharacterized protein \| Uncharacterized protein; uniclust \| UniRef100\_A0A345BNC1 \| 99.0 \| 2.7e-12 \| 5e-18 \| 88.9 \| 98 \| (8, 105) \| 106 \| (49, 146) \| 148 \| Putative holin \| Putative holin; uniclust \| UniRef100\_UPI002013492F \| 99.0 \| 3.1e-12 \| 5.7e-18 \| 93.0 \| 95 \| (10, 105) \| 106 \| (15, 111) \| 203 \| hypothetical protein \| hypothetical protein; uniclust \| UniRef100\_A0A0A8IL65 \| 99.0 \| 2.6e-12 \| 5.9e-18 \| 90.2 \| 89 \| (1, 91) \| 106 \| (11, 107) \| 127 \| Holin \| Holin; uniclust \| UniRef100\_A0A127Q5M9 \| 98.9 \| 9.8e-12 \| 1.9e-17 \| 80.7 \| 76 \| (29, 105) \| 106 \| (8, 84) \| 89 \| Holin \| Holin; uniclust \| UniRef100\_A0A853FDE8 \| 98.9 \| 1.6e-11 \| 3e-17 \| 82.7 \| 94 \| (3, 97) \| 106 \| (3, 97) \| 110 \| Holin \| Holin; uniclust \| UniRef100\_UPI0018D78283 \| 98.8 \| 3.7e-11 \| 6.9e-17 \| 76.9 \| 79 \| (25, 103) \| 106 \| (3, 81) \| 86 \| hypothetical protein \| hypothetical protein; uniclust \| UniRef100\_A0A0D0KIG3 \| 98.8 \| 4.1e-11 \| 8.9e-17 \| 83.0 \| 87 \| (1, 89) \| 106 \| (1, 96) \| 115 \| Holin \| Holin; uniclust \| UniRef100\_UPI000BAFFC2F \| 98.7 \| 1.9e-10 \| 3.6e-16 \| 76.5 \| 96 \| (7, 102) \| 106 \| (5, 101) \| 107 \| hypothetical protein \| hypothetical protein; uniclust \| UniRef100\_A0A0E3BDL1 \| 98.6 \| 2.5e-10 \| 5.1e-16 \| 78.4 \| 91 \| (3, 102) \| 106 \| (10, 104) \| 115 \| Holin \| Holin; uniclust \| UniRef100\_A0A090EBA1 \| 98.6 \| 3e-10 \| 5.7e-16 \| 80.7 \| 88 \| (2, 90) \| 106 \| (37, 133) \| 153 \| Holin \| Holin; uniclust \| UniRef100\_A0A2N1F928 \| 98.6 \| 7e-10 \| 1.4e-15 \| 75.1 \| 93 \| (1, 94) \| 106 \| (5, 97) \| 105 \| Holin \| Holin; uniclust \| UniRef100\_A0A2Z6EUY1 \| 98.5 \| 7.7e-10 \| 1.5e-15 \| 75.2 \| 96 \| (8, 104) \| 106 \| (17, 114) \| 115 \| Holin protein \| Holin protein; uniclust \| UniRef100\_A0A6G5Y843 \| 98.5 \| 1.1e-09 \| 2e-15 \| 71.7 \| 86 \| (10, 96) \| 106 \| (4, 90) \| 91 \| Holin \| Holin; uniclust \| UniRef100\_A0A0Q5M7J0 \| 98.5 \| 1.1e-09 \| 2.2e-15 \| 76.6 \| 88 \| (2, 90) \| 106 \| (9, 96) \| 130 \| Holin \| Holin; uniclust \| UniRef100\_A0A1R0FA65 \| 98.5 \| 1.2e-09 \| 2.3e-15 \| 78.1 \| 88 \| (2, 90) \| 106 \| (34, 135) \| 145 \| Uncharacterized protein \| Uncharacterized protein; uniclust \| UniRef100\_A0A0K2FIU1 \| 98.5 \| 1.3e-09 \| 2.6e-15 \| 73.8 \| 85 \| (10, 94) \| 106 \| (20, 105) \| 108 \| Holin \| Holin; uniclust \| UniRef100\_A0A0F2S303 \| 98.5 \| 1.3e-09 \| 2.9e-15 \| 78.3 \| 78 \| (12, 90) \| 106 \| (41, 123) \| 140 \| Uncharacterized protein \| Uncharacterized protein; uniclust \| UniRef100\_A0A6J5MI17 \| 98.5 \| 1.8e-09 \| 3.6e-15 \| 72.1 \| 86 \| (4, 90) \| 106 \| (3, 89) \| 96 \| Holin \| Holin; uniclust \| UniRef100\_A0A0Q6UK20 \| 98.4 \| 2.3e-09 \| 4.7e-15 \| 75.7 \| 95 \| (8, 103) \| 106 \| (14, 117) \| 128 \| Holin \| Holin; uniclust \| UniRef100\_W0LM32 \| 98.4 \| 4.1e-09 \| 7.5e-15 \| 69.3 \| 77 \| (19, 95) \| 106 \| (1, 78) \| 95 \| Putative holin \| Putative holin; uniclust \| UniRef100\_A0A239F4Z5 \| 98.3 \| 6.1e-09 \| 1.1e-14 \| 67.7 \| 87 \| (14, 103) \| 106 \| (2, 88) \| 89 \| Phage holin \| Phage holin; uniclust \| UniRef100\_A0A5E7QE83 \| 98.3 \| 7.4e-09 \| 1.4e-14 \| 69.0 \| 89 \| (11, 99) \| 106 \| (11, 99) \| 103 \| Holin \| Holin; uniclust \| UniRef100\_A0A959GDS2 \| 98.3 \| 1.1e-08 \| 1.9e-14 \| 71.8 \| 76 \| (7, 83) \| 106 \| (33, 108) \| 143 \| Uncharacterized protein \| Uncharacterized protein; uniclust \| UniRef100\_UPI001EE52D85 \| 98.3 \| 1.1e-08 \| 2.1e-14 \| 68.2 \| 88 \| (12, 100) \| 106 \| (14, 102) \| 103 \| hypothetical protein \| hypothetical protein; uniclust \| UniRef100\_A0A2S5TKI1 \| 98.2 \| 1.6e-08 \| 2.9e-14 \| 68.0 \| 94 \| (6, 100) \| 106 \| (3, 101) \| 107 \| Holin \| Holin; uniclust \| UniRef100\_A0A1D3JUZ2 \| 98.2 \| 2.2e-08 \| 4.2e-14 \| 69.2 \| 94 \| (9, 105) \| 106 \| (25, 121) \| 122 \| Conserved hypothetical membrane protein \| Conserved hypothetical membrane protein; uniclust \| UniRef100\_A0A0F0E234 \| 98.2 \| 2.6e-08 \| 5e-14 \| 68.6 \| 82 \| (7, 90) \| 106 \| (10, 92) \| 114 \| Holin \| Holin; uniclust \| UniRef100\_UPI0018A7A48E \| 98.2 \| 2.9e-08 \| 5.3e-14 \| 65.3 \| 91 \| (12, 104) \| 106 \| (3, 93) \| 93 \| hypothetical protein \| hypothetical protein; uniclust \| UniRef100\_A0A8T8CVW9 \| 98.1 \| 7e-08 \| 1.3e-13 \| 70.7 \| 89 \| (2, 91) \| 106 \| (81, 177) \| 192 \| N-acetylmuramidase family protein \| N-acetylmuramidase family protein; uniclust \| UniRef100\_UPI001900EF55 \| 98.0 \| 9.4e-08 \| 1.8e-13 \| 63.5 \| 76 \| (29, 104) \| 106 \| (7, 91) \| 93 \| hypothetical protein \| hypothetical protein; uniclust \| UniRef100\_A0A366F9C6 \| 98.0 \| 1.1e-07 \| 1.9e-13 \| 59.4 \| 61 \| (42, 105) \| 106 \| (3, 66) \| 67 \| Holin \| Holin; uniclust \| UniRef100\_A0A1I1D0C4 \| 97.9 \| 2.9e-07 \| 6.4e-13 \| 66.9 \| 93 \| (10, 105) \| 106 \| (32, 130) \| 138 \| Putative phage holin \| Putative phage holin; uniclust \| UniRef100\_A0A1U7D4W0 \| 97.8 \| 4.4e-07 \| 9e-13 \| 63.0 \| 91 \| (13, 105) \| 106 \| (14, 106) \| 106 \| Holin \| Holin; uniclust \| UniRef100\_A0A3M4HEX7 \| 97.8 \| 4.9e-07 \| 9.3e-13 \| 55.2 \| 49 \| (1, 49) \| 106 \| (1, 53) \| 55 \| Uncharacterized protein \| Uncharacterized protein; uniclust \| UniRef100\_A0A519X5A4 \| 97.8 \| 5.1e-07 \| 9.6e-13 \| 61.7 \| 83 \| (2, 85) \| 106 \| (11, 93) \| 108 \| Holin \| Holin; uniclust \| UniRef100\_A0A366DXW5 \| 97.8 \| 5.4e-07 \| 1e-12 \| 59.2 \| 59 \| (2, 61) \| 106 \| (5, 63) \| 83 \| Uncharacterized protein \| Uncharacterized protein; uniclust \| UniRef100\_UPI001678847F \| 97.8 \| 5.8e-07 \| 1.1e-12 \| 62.5 \| 90 \| (13, 105) \| 106 \| (30, 119) \| 121 \| putative holin \| putative holin; uniclust \| UniRef100\_A0A806CKK2 \| 97.7 \| 7.5e-07 \| 1.4e-12 \| 60.9 \| 103 \| (2, 105) \| 106 \| (7, 111) \| 112 \| Holin \| Holin; uniclust \| UniRef100\_H8L657 \| 97.7 \| 8.9e-07 \| 1.6e-12 \| 59.4 \| 88 \| (2, 90) \| 106 \| (4, 91) \| 99 \| Phage holin \| Phage holin; uniclust \| UniRef100\_A0A0D0J2P5 \| 97.7 \| 1e-06 \| 2.2e-12 \| 62.9 \| 76 \| (10, 86) \| 106 \| (29, 107) \| 122 \| Phage holin \| Phage holin; uniclust \| UniRef100\_A0A7Y5IYH3 \| 97.6 \| 1.5e-06 \| 2.8e-12 \| 60.1 \| 77 \| (10, 87) \| 106 \| (21, 97) \| 120 \| Holin \| Holin; uniclust \| UniRef100\_A0A5C8PAN5 \| 97.6 \| 1.5e-06 \| 2.9e-12 \| 60.6 \| 88 \| (3, 90) \| 106 \| (6, 97) \| 122 \| Phage holin family protein \| Phage holin family protein; uniclust \| UniRef100\_UPI0010C024CC \| 97.6 \| 2.5e-06 \| 4.8e-12 \| 64.4 \| 87 \| (10, 99) \| 106 \| (28, 114) \| 211 \| putative holin \| putative holin; uniclust \| UniRef100\_A0A089YNB7 \| 97.5 \| 3e-06 \| 5.6e-12 \| 60.4 \| 90 \| (10, 102) \| 106 \| (42, 131) \| 132 \| Holin \| Holin; uniclust \| UniRef100\_A0A6J5QP20 \| 97.5 \| 3.1e-06 \| 5.8e-12 \| 58.0 \| 80 \| (8, 87) \| 106 \| (8, 88) \| 111 \| Holin \| Holin; uniclust \| UniRef100\_UPI000319E467 \| 97.5 \| 3.5e-06 \| 6.5e-12 \| 53.9 \| 60 \| (1, 60) \| 106 \| (1, 64) \| 73 \| hypothetical protein \| hypothetical protein; uniclust \| UniRef100\_A0A241XK97 \| 97.5 \| 4e-06 \| 7.4e-12 \| 59.7 \| 86 \| (10, 98) \| 106 \| (38, 123) \| 140 \| Holin \| Holin; uniclust \| UniRef100\_UPI00207B5E30 \| 97.5 \| 4.8e-06 \| 8.8e-12 \| 62.2 \| 87 \| (10, 99) \| 106 \| (99, 185) \| 200 \| putative holin \| putative holin; uniclust \| UniRef100\_A0A102EQF3 \| 97.4 \| 4.9e-06 \| 9e-12 \| 51.9 \| 59 \| (46, 104) \| 106 \| (2, 61) \| 63 \| Uncharacterized protein \| Uncharacterized protein; uniclust \| UniRef100\_UPI00217D4006 \| 97.4 \| 6e-06 \| 1.1e-11 \| 53.8 \| 77 \| (1, 77) \| 106 \| (1, 79) \| 80 \| hypothetical protein \| hypothetical protein; uniclust \| UniRef100\_A0A0K1JXH2 \| 97.4 \| 6.5e-06 \| 1.4e-11 \| 62.3 \| 82 \| (9, 91) \| 106 \| (64, 153) \| 173 \| Uncharacterized protein \| Uncharacterized protein; uniclust \| UniRef100\_UPI001E5DDA53 \| 97.4 \| 7.8e-06 \| 1.4e-11 \| 56.8 \| 85 \| (4, 90) \| 106 \| (11, 102) \| 119 \| hypothetical protein \| hypothetical protein; uniclust \| UniRef100\_UPI00217CC2F6 \| 97.4 \| 8e-06 \| 1.5e-11 \| 56.5 \| 85 \| (10, 94) \| 106 \| (7, 92) \| 115 \| hypothetical protein \| hypothetical protein; uniclust \| UniRef100\_A0A2M9P829 \| 97.4 \| 8e-06 \| 1.5e-11 \| 54.9 \| 85 \| (16, 102) \| 106 \| (10, 95) \| 96 \| Phage holin family protein \| Phage holin family protein; uniclust \| UniRef100\_A0A1Y6CU26 \| 97.4 \| 8.6e-06 \| 1.6e-11 \| 56.3 \| 87 \| (7, 94) \| 106 \| (11, 105) \| 114 \| Uncharacterized protein \| Uncharacterized protein; uniclust \| UniRef100\_A0A1I5MXA9 \| 97.3 \| 9.9e-06 \| 1.8e-11 \| 55.8 \| 88 \| (2, 91) \| 106 \| (4, 91) \| 111 \| Holin \| Holin; uniclust \| UniRef100\_UPI0019D322D2 \| 97.3 \| 1.3e-05 \| 2.4e-11 \| 51.0 \| 55 \| (7, 62) \| 106 \| (10, 64) \| 69 \| hypothetical protein \| hypothetical protein; uniclust \| UniRef100\_A0A2A2CFL2 \| 97.3 \| 1.4e-05 \| 2.6e-11 \| 56.0 \| 88 \| (10, 100) \| 106 \| (30, 117) \| 117 \| Membrane transporter protein \| Membrane transporter protein; uniclust \| UniRef100\_UPI001F180A34 \| 97.2 \| 2.1e-05 \| 4e-11 \| 56.7 \| 93 \| (9, 102) \| 106 \| (21, 121) \| 144 \| hypothetical protein \| hypothetical protein; uniclust \| UniRef100\_A0A959HN31 \| 97.2 \| 2.6e-05 \| 4.7e-11 \| 54.1 \| 73 \| (11, 86) \| 106 \| (12, 87) \| 114 \| Uncharacterized protein \| Uncharacterized protein; uniclust \| UniRef100\_A0A3S9U7J0 \| 97.1 \| 2.8e-05 \| 5.2e-11 \| 53.2 \| 86 \| (9, 94) \| 106 \| (15, 101) \| 105 \| Holin \| Holin; uniclust \| UniRef100\_F3C3E1 \| 97.1 \| 3.8e-05 \| 6.9e-11 \| 48.4 \| 59 \| (46, 104) \| 106 \| (2, 61) \| 64 \| MFS transporter (Fragment) \| MFS transporter (Fragment); uniclust \| UniRef100\_J8RRD5 \| 97.0 \| 5e-05 \| 9.2e-11 \| 47.9 \| 60 \| (42, 103) \| 106 \| (4, 63) \| 64 \| Uncharacterized protein \| Uncharacterized protein; uniclust \| UniRef100\_A0A3S1MB93 \| 97.0 \| 4.9e-05 \| 9.3e-11 \| 51.1 \| 57 \| (33, 89) \| 106 \| (11, 76) \| 85 \| Uncharacterized protein (Fragment) \| Uncharacterized protein (Fragment); uniclust \| UniRef100\_UPI001157799A \| 97.0 \| 5.4e-05 \| 1e-10 \| 54.6 \| 87 \| (3, 90) \| 106 \| (20, 111) \| 146 \| hypothetical protein \| hypothetical protein; uniclust \| UniRef100\_A0A5B0DZY6 \| 97.0 \| 5.4e-05 \| 1e-10 \| 52.6 \| 85 \| (2, 88) \| 106 \| (2, 95) \| 113 \| Holin \| Holin; uniclust \| UniRef100\_A0A077K9X1 \| 97.0 \| 4.7e-05 \| 1.1e-10 \| 56.2 \| 80 \| (10, 90) \| 106 \| (29, 113) \| 134 \| Holin \| Holin; uniclust \| UniRef100\_A0A2V4JFC9 \| 97.0 \| 5.6e-05 \| 1.1e-10 \| 51.8 \| 62 \| (10, 74) \| 106 \| (28, 89) \| 90 \| DUF1097 domain-containing protein (Fragment) \| DUF1097 domain-containing protein (Fragment); uniclust \| UniRef100\_A0A1B1INP0 \| 97.0 \| 5.5e-05 \| 1.2e-10 \| 53.9 \| 83 \| (7, 90) \| 106 \| (9, 95) \| 110 \| Holin \| Holin; uniclust \| UniRef100\_UPI00216732F2 \| 97.0 \| 6.9e-05 \| 1.3e-10 \| 48.0 \| 64 \| (40, 103) \| 106 \| (2, 67) \| 69 \| hypothetical protein \| hypothetical protein; uniclust \| UniRef100\_A0A066T1C6 \| 96.9 \| 5.8e-05 \| 1.3e-10 \| 57.2 \| 81 \| (11, 92) \| 106 \| (54, 141) \| 159 \| Uncharacterized protein \| Uncharacterized protein; uniclust \| UniRef100\_A0A0U2C0W7 \| 96.9 \| 6.6e-05 \| 1.4e-10 \| 54.0 \| 88 \| (17, 105) \| 106 \| (13, 103) \| 115 \| Holin \| Holin; uniclust \| UniRef100\_A0A024HES2 \| 96.9 \| 8.2e-05 \| 1.5e-10 \| 50.2 \| 78 \| (11, 89) \| 106 \| (7, 90) \| 93 \| Uncharacterized protein \| Uncharacterized protein; uniclust \| UniRef100\_A0A085G117 \| 96.9 \| 7.9e-05 \| 1.6e-10 \| 55.7 \| 94 \| (10, 104) \| 106 \| (41, 139) \| 151 \| Putative prophage membrane protein \| Putative prophage membrane protein; uniclust \| UniRef100\_UPI00234D9A13 \| 96.9 \| 0.00011 \| 1.9e-10 \| 50.6 \| 76 \| (10, 88) \| 106 \| (9, 89) \| 104 \| hypothetical protein \| hypothetical protein; uniclust \| UniRef100\_A0A8E4ZL07 \| 96.8 \| 0.00012 \| 2.1e-10 \| 50.7 \| 89 \| (10, 98) \| 106 \| (6, 100) \| 108 \| Holin \| Holin; uniclust \| UniRef100\_A0A081J3M5 \| 96.8 \| 0.00014 \| 2.6e-10 \| 48.4 \| 67 \| (33, 102) \| 106 \| (6, 76) \| 85 \| Uncharacterized protein \| Uncharacterized protein; uniclust \| UniRef100\_A0A017HBY4 \| 96.8 \| 0.00014 \| 3e-10 \| 52.8 \| 73 \| (13, 86) \| 106 \| (32, 108) \| 119 \| Uncharacterized protein \| Uncharacterized protein; uniclust \| UniRef100\_UPI0015B67E84 \| 96.7 \| 0.00019 \| 3.6e-10 \| 47.2 \| 61 \| (1, 61) \| 106 \| (1, 63) \| 74 \| hypothetical protein \| hypothetical protein; uniclust \| UniRef100\_A0A944HBG9 \| 96.7 \| 0.00019 \| 3.7e-10 \| 50.8 \| 99 \| (4, 105) \| 106 \| (13, 113) \| 114 \| Uncharacterized protein \| Uncharacterized protein; uniclust \| UniRef100\_UPI001848C271 \| 96.7 \| 0.0002 \| 3.8e-10 \| 47.3 \| 48 \| (11, 59) \| 106 \| (27, 74) \| 75 \| putative holin \| putative holin; uniclust \| UniRef100\_A0A2P7RER1 \| 96.7 \| 0.00022 \| 4.1e-10 \| 47.5 \| 66 \| (24, 90) \| 106 \| (3, 76) \| 84 \| Uncharacterized protein (Fragment) \| Uncharacterized protein (Fragment); uniclust \| UniRef100\_UPI00094B3F0A \| 96.7 \| 0.00022 \| 4.1e-10 \| 47.4 \| 59 \| (1, 61) \| 106 \| (1, 60) \| 83 \| hypothetical protein \| hypothetical protein; uniclust \| UniRef100\_A0A2W6XWQ2 \| 96.7 \| 0.00023 \| 4.2e-10 \| 45.1 \| 52 \| (2, 54) \| 106 \| (10, 62) \| 63 \| Uncharacterized protein (Fragment) \| Uncharacterized protein (Fragment); uniclust \| UniRef100\_UPI001571687F \| 96.7 \| 0.00025 \| 4.6e-10 \| 50.6 \| 89 \| (1, 90) \| 106 \| (1, 91) \| 129 \| hypothetical protein \| hypothetical protein; uniclust \| UniRef100\_A0A3G3GJ83 \| 96.6 \| 0.00025 \| 4.8e-10 \| 52.3 \| 100 \| (5, 104) \| 106 \| (29, 135) \| 152 \| Holin \| Holin; uniclust \| UniRef100\_A0A2S0PEK2 \| 96.6 \| 0.00024 \| 5.2e-10 \| 54.7 \| 93 \| (10, 103) \| 106 \| (62, 155) \| 178 \| Uncharacterized protein \| Uncharacterized protein; uniclust \| UniRef100\_A0A0A1W9E3 \| 96.6 \| 0.0003 \| 6.2e-10 \| 50.3 \| 85 \| (5, 90) \| 106 \| (7, 97) \| 108 \| Uncharacterized protein \| Uncharacterized protein; uniclust \| UniRef100\_A0A0P6RA73 \| 96.6 \| 0.00028 \| 6.2e-10 \| 51.5 \| 89 \| (14, 105) \| 106 \| (29, 118) \| 122 \| Holin \| Holin; uniclust \| UniRef100\_UPI00037B93FB \| 96.6 \| 0.00035 \| 6.4e-10 \| 50.2 \| 91 \| (12, 103) \| 106 \| (13, 109) \| 134 \| hypothetical protein \| hypothetical protein; uniclust \| UniRef100\_A0A971Y2T2 \| 96.5 \| 0.00051 \| 9.4e-10 \| 47.6 \| 78 \| (19, 101) \| 106 \| (20, 97) \| 104 \| Uncharacterized protein \| Uncharacterized protein; uniclust \| UniRef100\_A0A1G7B018 \| 96.5 \| 0.00052 \| 9.9e-10 \| 48.1 \| 85 \| (1, 88) \| 106 \| (1, 86) \| 104 \| Phage holin family Hol44, holin superfamily V \| Phage holin family Hol44, holin superfamily V; uniclust \| UniRef100\_A0A848MF35 \| 96.4 \| 0.00057 \| 1e-09 \| 46.6 \| 87 \| (11, 100) \| 106 \| (5, 91) \| 94 \| Holin \| Holin; uniclust \| UniRef100\_A0A6N6ZDS7 \| 96.4 \| 0.00058 \| 1.1e-09 \| 48.8 \| 78 \| (7, 85) \| 106 \| (9, 87) \| 123 \| Peptidase M48, Ste24p \| Peptidase M48, Ste24p; uniclust \| UniRef100\_A0A071M1T4 \| 96.4 \| 0.00053 \| 1.1e-09 \| 52.4 \| 85 \| (10, 98) \| 106 \| (49, 137) \| 169 \| Membrane protein \| Membrane protein; uniclust \| UniRef100\_A0A0Q7XKN5 \| 96.4 \| 0.00076 \| 1.4e-09 \| 49.0 \| 78 \| (10, 89) \| 106 \| (44, 129) \| 140 \| Holin \| Holin; uniclust \| UniRef100\_A0A2P5K7X0 \| 96.4 \| 0.0007 \| 1.4e-09 \| 49.9 \| 76 \| (12, 91) \| 106 \| (25, 100) \| 131 \| Putative phage holin \| Putative phage holin; uniclust \| UniRef100\_A0A068CCL5 \| 96.3 \| 0.00078 \| 1.5e-09 \| 48.8 \| 88 \| (1, 92) \| 106 \| (4, 96) \| 121 \| Holin \| Holin; uniclust \| UniRef100\_UPI00193FE6FD \| 96.3 \| 0.00084 \| 1.6e-09 \| 47.3 \| 90 \| (10, 100) \| 106 \| (15, 105) \| 114 \| hypothetical protein \| hypothetical protein; uniclust \| UniRef100\_A0A212DP65 \| 96.3 \| 0.00092 \| 1.7e-09 \| 45.9 \| 53 \| (38, 90) \| 106 \| (2, 63) \| 97 \| Uncharacterized protein \| Uncharacterized protein; uniclust \| UniRef100\_W0V419 \| 96.3 \| 0.001 \| 1.9e-09 \| 45.5 \| 79 \| (11, 90) \| 106 \| (3, 90) \| 95 \| Putative membrane protein \| Putative membrane protein |
| Top keywords  (threshold 1.00e-03 (evalue)) | **Holin, hypothetical, Putative, Phage, membrane, Fragment, transporter, MFS, Peptidase, M48** |
| Output files | ../../similar\_sequences/62\_FANPEZAQ\_CDS\_0062\_merged.svg ../../similar\_sequences/62\_FANPEZAQ\_CDS\_0062\_pdb70.a3m ../../similar\_sequences/62\_FANPEZAQ\_CDS\_0062\_pdb70.hhr ../../similar\_sequences/62\_FANPEZAQ\_CDS\_0062\_uniclust.a3m ../../similar\_sequences/62\_FANPEZAQ\_CDS\_0062\_uniclust.hhr |

#### Structure prediction (AlphaFold)^2^

|  |  |
| --- | --- |
| Stats | xml version="1.0" encoding="utf-8" standalone="no"?       2024-09-02T21:09:57.713479 image/svg+xml   Matplotlib v3.7.2, https://matplotlib.org/ |
| Predicted structure | **NGL Viewer Controls:**  - Center: *Left-Click* - Rotate: *Left-Click + Drag* - Translate: *Right-Click + Drag* - Zoom: *Shift + Left-Click + Drag* |
| Output files | ../../predicted\_structures/62\_FANPEZAQ\_CDS\_0062/features.pkl ../../predicted\_structures/62\_FANPEZAQ\_CDS\_0062/ranked\_0.pdb ../../predicted\_structures/62\_FANPEZAQ\_CDS\_0062/ranked\_0\_plots.svg ../../predicted\_structures/62\_FANPEZAQ\_CDS\_0062/result\_model\_1\_ptm\_pred\_0.pkl |

#### Structure similarity search results (Foldseek)^3^

|  |  |
| --- | --- |
| Structure databases searched | Pdb, Afdb-proteome, Afdb-uniprot50 |
| Results, scheme(s)  (Top layers only, threshold 1.00e-02 (evalue)) | xml version="1.0" encoding="utf-8" standalone="no"?       2024-09-02T21:11:29.532784 image/svg+xml   Matplotlib v3.7.2, https://matplotlib.org/ |
| Results, table  (threshold 1.00e-02 (evalue)) | db \| id \| prob \| evalue \| bits \| fident \| alnlen \| mismatch \| gapopen \| qstart \| qend \| tstart \| tend \| name \| description; afdb-uniprot50 \| AF-A0A3S1A8T4-F1-MODEL\_V4 \| 1.0 \| 1.828e-08 \| 432 \| 0.49 \| 106 \| 54 \| 0 \| 1 \| 106 \| 1 \| 106 \| MFS transporter \| MFS transporter; afdb-uniprot50 \| AF-A0A653KX91-F1-MODEL\_V4 \| 1.0 \| 3.559e-07 \| 349 \| 0.472 \| 108 \| 54 \| 1 \| 1 \| 105 \| 1 \| 108 \| Contig\_80, whole genome shotgun sequence \| Contig\_80, whole genome shotgun sequence; afdb-uniprot50 \| AF-A0A5E7NBM4-F1-MODEL\_V4 \| 1.0 \| 1.276e-07 \| 340 \| 0.472 \| 108 \| 55 \| 2 \| 1 \| 106 \| 1 \| 108 \| Uncharacterized protein \| Uncharacterized protein; afdb-uniprot50 \| AF-A0A158E8Z8-F1-MODEL\_V4 \| 1.0 \| 2.868e-07 \| 330 \| 0.49 \| 104 \| 53 \| 0 \| 2 \| 105 \| 3 \| 106 \| Uncharacterized protein \| Uncharacterized protein; afdb-uniprot50 \| AF-A0A3M6EJ76-F1-MODEL\_V4 \| 1.0 \| 0.0001866 \| 240 \| 0.376 \| 85 \| 51 \| 2 \| 1 \| 83 \| 1 \| 85 \| Uncharacterized protein \| Uncharacterized protein; afdb-uniprot50 \| AF-A0A0A8TGY1-F1-MODEL\_V4 \| 1.0 \| 0.0004672 \| 233 \| 0.354 \| 93 \| 59 \| 1 \| 2 \| 93 \| 8 \| 100 \| Uncharacterized protein \| Uncharacterized protein; afdb-uniprot50 \| AF-A0A1B5DBB0-F1-MODEL\_V4 \| 1.0 \| 3.317e-05 \| 227 \| 0.423 \| 111 \| 56 \| 4 \| 1 \| 106 \| 1 \| 108 \| Ethanolamine permease \| Ethanolamine permease; afdb-uniprot50 \| AF-A0A150HN48-F1-MODEL\_V4 \| 1.0 \| 0.000646 \| 205 \| 0.27 \| 111 \| 71 \| 2 \| 2 \| 102 \| 8 \| 118 \| Uncharacterized protein \| Uncharacterized protein; afdb-uniprot50 \| AF-A0A6G6Y576-F1-MODEL\_V4 \| 1.0 \| 0.0002874 \| 203 \| 0.302 \| 109 \| 65 \| 2 \| 5 \| 103 \| 8 \| 115 \| Uncharacterized protein \| Uncharacterized protein; afdb-uniprot50 \| AF-A0A149SVI2-F1-MODEL\_V4 \| 1.0 \| 0.0009426 \| 194 \| 0.257 \| 101 \| 72 \| 1 \| 5 \| 105 \| 4 \| 101 \| Uncharacterized protein \| Uncharacterized protein; afdb-uniprot50 \| AF-A0A1I7FZH2-F1-MODEL\_V4 \| 1.0 \| 9.764e-05 \| 191 \| 0.438 \| 105 \| 54 \| 3 \| 1 \| 100 \| 1 \| 105 \| Uncharacterized protein \| Uncharacterized protein; afdb-uniprot50 \| AF-A0A7U9JL14-F1-MODEL\_V4 \| 1.0 \| 0.002118 \| 181 \| 0.329 \| 94 \| 62 \| 1 \| 13 \| 105 \| 18 \| 111 \| Holin \| Holin; afdb-uniprot50 \| AF-A0A5E7QE83-F1-MODEL\_V4 \| 1.0 \| 0.00105 \| 177 \| 0.303 \| 89 \| 60 \| 2 \| 12 \| 99 \| 12 \| 99 \| Uncharacterized protein \| Uncharacterized protein; afdb-uniprot50 \| AF-A0A853FDE8-F1-MODEL\_V4 \| 1.0 \| 0.009099 \| 174 \| 0.241 \| 91 \| 68 \| 1 \| 9 \| 99 \| 10 \| 99 \| Uncharacterized protein \| Uncharacterized protein; afdb-uniprot50 \| AF-A0A845A3H7-F1-MODEL\_V4 \| 1.0 \| 0.007739 \| 172 \| 0.202 \| 99 \| 78 \| 1 \| 5 \| 103 \| 8 \| 105 \| Uncharacterized protein \| Uncharacterized protein; afdb-uniprot50 \| AF-A0A2S5TKI1-F1-MODEL\_V4 \| 1.0 \| 0.003091 \| 163 \| 0.233 \| 103 \| 76 \| 2 \| 6 \| 106 \| 3 \| 104 \| Uncharacterized protein \| Uncharacterized protein; afdb-uniprot50 \| AF-A0A7V8Q7L9-F1-MODEL\_V4 \| 1.0 \| 0.009604 \| 150 \| 0.152 \| 131 \| 84 \| 3 \| 3 \| 106 \| 9 \| 139 \| Uncharacterized protein \| Uncharacterized protein |
| Top keywords  (threshold 1.00e-02 (evalue)) | **MFS, transporter, Contig\_80, whole, genome, shotgun, sequence, Ethanolamine, permease, Holin** |
| Output files | ../../similar\_structures/62\_FANPEZAQ\_CDS\_0062\_afdb-proteome\_foldseek.tsv ../../similar\_structures/62\_FANPEZAQ\_CDS\_0062\_afdb-uniprot50\_foldseek.tsv ../../similar\_structures/62\_FANPEZAQ\_CDS\_0062\_merged.svg ../../similar\_structures/62\_FANPEZAQ\_CDS\_0062\_pdb\_foldseek.tsv |

  
  
  

Return to summary | Go to previous | Go to next

  


---

**Sequence/structure alignments coloring**  
Each object in the alignment figures is colored according to its E-value following this color coding:

1e-100
10
