## Supplementary material for "Completing the BASEL phage collection to unlock hidden diversity for systematic exploration of phage-host interactions": Data S2: 63.html

FANPEZAQ\_CDS\_0063


Return to summary | Go to previous | Go to next

|  |  |
| --- | --- |
| FANPEZAQ\_CDS\_0063  Page creation date: 02 Sep 2024, 12:00  Project folder: n/a  Input sequences file: Escherichia\_virus\_HeidiAbel.gb | holin hypothetical putative holin\_like i membrane lysis nc\_019419\_p7 rhs repeat\_associated core domain\_containing feob\_associated cys\_rich |

### Sequence information

|  |  |
| --- | --- |
| Name | FANPEZAQ\_CDS\_0063  63\_FANPEZAQ\_CDS\_0063 (pipeline id) |
| Imported annotations | Escherichia\_virus\_HeidiAbel Bas97 |
| Protein sequence | MLLTVINFISVLVIFCWSTWCVLHHGVRDGIVGKILFSCVAISALAVILHEVQGEYLTRS TVTLHATIAAVAVRHFFVNTVWKPFFSKYFHE |
| Number of residues | 92 |
| Molecular weight (Da) | 10420.25 |
| Output files | ../../query\_sequences/63\_FANPEZAQ\_CDS\_0063.fasta |

### Putative domain architecture and protein family

#### Search results (HHblits)^1^

|  |  |
| --- | --- |
| Domain family databases searched | Pfam, Ncbi-cd, Cath, Phrogs |
| Results, scheme(s)  (Top layers only; threshold 1.00e-03 (evalue)) | xml version="1.0" encoding="utf-8" standalone="no"?       2024-09-02T21:08:26.100168 image/svg+xml   Matplotlib v3.7.2, https://matplotlib.org/ |
| Results, table  (E-value ≤ 1.00e-03 (evalue)) | db \| id \| prob \| evalue \| pvalue \| score \| cols \| query \| query\_len \| template \| template\_len \| name \| description; phrogs \| 2508 \| 100.0 \| 1.2e-35 \| 1.5e-39 \| 194.9 \| 80 \| (5, 86) \| 92 \| (2, 81) \| 81 \| holin \| holin; Category: lysis; NC\_019419\_p7 |
| Top keywords  (threshold 1.00e-03 (evalue)) | **holin, lysis, NC\_019419\_p7** |
| Output files | ../../domain\_architecture/63\_FANPEZAQ\_CDS\_0063\_cath.hhr ../../domain\_architecture/63\_FANPEZAQ\_CDS\_0063\_merged.svg ../../domain\_architecture/63\_FANPEZAQ\_CDS\_0063\_ncbi-cd.hhr ../../domain\_architecture/63\_FANPEZAQ\_CDS\_0063\_pfam.hhr ../../domain\_architecture/63\_FANPEZAQ\_CDS\_0063\_phrogs.hhr |

### Identical protein sequences/structures

#### Search results

|  |  |
| --- | --- |
| Protein sequence databases searched | Pdb, Swissprot, Refseq |
| Identical proteins found | -- |
| Top keywords | -- |
| Output files | -- |

### Similar protein sequences/structures

#### Sequence similarity search results (HHblits)^1^

|  |  |
| --- | --- |
| Sequence databases searched | Uniclust, Pdb70 |
| Results, scheme(s)  (Top layers only, threshold 1.00e-03 (evalue)) | xml version="1.0" encoding="utf-8" standalone="no"?       2024-09-02T21:08:54.594164 image/svg+xml   Matplotlib v3.7.2, https://matplotlib.org/ |
| Results, table(s)  (threshold 1.00e-03 (evalue)) | db \| id \| prob \| evalue \| pvalue \| score \| cols \| query \| query\_len \| template \| template\_len \| name \| description; uniclust \| UniRef100\_A0A059KTZ2 \| 100.0 \| 8.3e-41 \| 1.8e-46 \| 240.9 \| 87 \| (1, 87) \| 92 \| (2, 89) \| 126 \| Holin \| Holin; uniclust \| UniRef100\_A0A0D0PMH9 \| 99.9 \| 1.3e-28 \| 3e-34 \| 173.4 \| 84 \| (1, 89) \| 92 \| (2, 85) \| 101 \| Uncharacterized protein \| Uncharacterized protein; uniclust \| UniRef100\_A0A193GYR7 \| 99.9 \| 2.5e-25 \| 5.1e-31 \| 154.7 \| 79 \| (1, 79) \| 92 \| (1, 79) \| 93 \| Holin \| Holin; uniclust \| UniRef100\_UPI001032C2BF \| 99.8 \| 6.8e-25 \| 1.3e-30 \| 159.0 \| 86 \| (1, 86) \| 92 \| (1, 87) \| 131 \| hypothetical protein \| hypothetical protein; uniclust \| UniRef100\_A0A0K2FI78 \| 99.7 \| 5.2e-20 \| 1.2e-25 \| 131.8 \| 80 \| (3, 82) \| 92 \| (7, 86) \| 104 \| Putative holin-like class I \| Putative holin-like class I; uniclust \| UniRef100\_A0A158E9V0 \| 99.6 \| 1.7e-18 \| 3.5e-24 \| 121.5 \| 82 \| (1, 86) \| 92 \| (1, 82) \| 92 \| Uncharacterized protein \| Uncharacterized protein; uniclust \| UniRef100\_A0A127R0E2 \| 99.4 \| 4.3e-16 \| 9.3e-22 \| 109.9 \| 73 \| (1, 73) \| 92 \| (1, 74) \| 90 \| Putative membrane protein \| Putative membrane protein; uniclust \| UniRef100\_A0A093S375 \| 99.2 \| 1.2e-13 \| 2.1e-19 \| 95.5 \| 72 \| (6, 80) \| 92 \| (2, 74) \| 80 \| Holin \| Holin; uniclust \| UniRef100\_UPI00209AFE34 \| 99.0 \| 3.1e-12 \| 5.7e-18 \| 98.5 \| 53 \| (1, 53) \| 92 \| (1, 53) \| 170 \| RHS repeat-associated core domain-containing protein \| RHS repeat-associated core domain-containing protein; uniclust \| UniRef100\_A0A853FDB7 \| 98.9 \| 5.6e-12 \| 1.1e-17 \| 90.5 \| 82 \| (3, 87) \| 92 \| (2, 87) \| 96 \| Holin \| Holin; uniclust \| UniRef100\_G0AAG9 \| 98.7 \| 1.1e-10 \| 2e-16 \| 86.6 \| 72 \| (2, 73) \| 92 \| (35, 107) \| 122 \| Uncharacterized protein \| Uncharacterized protein; uniclust \| UniRef100\_UPI001B8D6944 \| 98.3 \| 1.1e-08 \| 2e-14 \| 67.8 \| 53 \| (2, 54) \| 92 \| (1, 53) \| 55 \| hypothetical protein \| hypothetical protein; uniclust \| UniRef100\_A0A2N5F659 \| 98.2 \| 1.9e-08 \| 3.5e-14 \| 67.7 \| 31 \| (1, 31) \| 92 \| (1, 31) \| 59 \| FeoB-associated Cys-rich membrane protein \| FeoB-associated Cys-rich membrane protein; uniclust \| UniRef100\_A0A076YQQ2 \| 98.1 \| 5.8e-08 \| 1.2e-13 \| 71.1 \| 87 \| (4, 90) \| 92 \| (9, 95) \| 96 \| Putative holin-like class I protein \| Putative holin-like class I protein; uniclust \| UniRef100\_A0A7S9XD84 \| 98.0 \| 8.5e-08 \| 1.6e-13 \| 70.4 \| 78 \| (6, 83) \| 92 \| (12, 89) \| 102 \| Putative holin-like class I \| Putative holin-like class I; uniclust \| UniRef100\_UPI0014302FF6 \| 97.9 \| 2.6e-07 \| 4.8e-13 \| 64.6 \| 59 \| (23, 81) \| 92 \| (1, 60) \| 72 \| hypothetical protein \| hypothetical protein; uniclust \| UniRef100\_A0A022GLN2 \| 97.8 \| 4.9e-07 \| 1.1e-12 \| 66.3 \| 86 \| (1, 88) \| 92 \| (1, 86) \| 92 \| Membrane protein \| Membrane protein; uniclust \| UniRef100\_UPI001375FE1C \| 97.7 \| 1.3e-06 \| 2.4e-12 \| 58.5 \| 27 \| (1, 27) \| 92 \| (1, 27) \| 54 \| hypothetical protein \| hypothetical protein; uniclust \| UniRef100\_A0A4D6DYF3 \| 97.5 \| 3.8e-06 \| 7e-12 \| 64.3 \| 80 \| (12, 91) \| 92 \| (47, 126) \| 126 \| Class I holin-like protein \| Class I holin-like protein; uniclust \| UniRef100\_A0A102EQN0 \| 97.3 \| 1.1e-05 \| 2.5e-11 \| 60.1 \| 85 \| (1, 85) \| 92 \| (3, 88) \| 96 \| Holin \| Holin; uniclust \| UniRef100\_A0A0F4TAU4 \| 97.0 \| 5.9e-05 \| 1.1e-10 \| 53.3 \| 54 \| (23, 76) \| 92 \| (4, 60) \| 69 \| Uncharacterized protein \| Uncharacterized protein |
| Top keywords  (threshold 1.00e-03 (evalue)) | **Holin, hypothetical, Putative, holin\_like, I, membrane, RHS, repeat\_associated, core, domain\_containing** |
| Output files | ../../similar\_sequences/63\_FANPEZAQ\_CDS\_0063\_merged.svg ../../similar\_sequences/63\_FANPEZAQ\_CDS\_0063\_pdb70.a3m ../../similar\_sequences/63\_FANPEZAQ\_CDS\_0063\_pdb70.hhr ../../similar\_sequences/63\_FANPEZAQ\_CDS\_0063\_uniclust.a3m ../../similar\_sequences/63\_FANPEZAQ\_CDS\_0063\_uniclust.hhr |

#### Structure prediction (AlphaFold)^2^

|  |  |
| --- | --- |
| Stats | xml version="1.0" encoding="utf-8" standalone="no"?       2024-09-02T21:09:58.372452 image/svg+xml   Matplotlib v3.7.2, https://matplotlib.org/ |
| Predicted structure | **NGL Viewer Controls:**  - Center: *Left-Click* - Rotate: *Left-Click + Drag* - Translate: *Right-Click + Drag* - Zoom: *Shift + Left-Click + Drag* |
| Output files | ../../predicted\_structures/63\_FANPEZAQ\_CDS\_0063/features.pkl ../../predicted\_structures/63\_FANPEZAQ\_CDS\_0063/ranked\_0.pdb ../../predicted\_structures/63\_FANPEZAQ\_CDS\_0063/ranked\_0\_plots.svg ../../predicted\_structures/63\_FANPEZAQ\_CDS\_0063/result\_model\_1\_ptm\_pred\_0.pkl |

#### Structure similarity search results (Foldseek)^3^

|  |  |
| --- | --- |
| Structure databases searched | Pdb, Afdb-proteome, Afdb-uniprot50 |
| Results, scheme(s)  (Top layers only, threshold 1.00e-02 (evalue)) | xml version="1.0" encoding="utf-8" standalone="no"?       2024-09-02T21:11:29.705335 image/svg+xml   Matplotlib v3.7.2, https://matplotlib.org/ |
| Results, table  (threshold 1.00e-02 (evalue)) | db \| id \| prob \| evalue \| bits \| fident \| alnlen \| mismatch \| gapopen \| qstart \| qend \| tstart \| tend \| name \| description; afdb-uniprot50 \| AF-A0A5E7Q5Q9-F1-MODEL\_V4 \| 1.0 \| 3.868e-05 \| 272 \| 0.362 \| 91 \| 57 \| 1 \| 1 \| 90 \| 1 \| 91 \| Uncharacterized protein \| Uncharacterized protein; afdb-uniprot50 \| AF-A0A0C5RPM8-F1-MODEL\_V4 \| 1.0 \| 0.000604 \| 223 \| 0.364 \| 85 \| 53 \| 1 \| 1 \| 84 \| 1 \| 85 \| Uncharacterized protein \| Uncharacterized protein; afdb-uniprot50 \| AF-A0A3G7X3J1-F1-MODEL\_V4 \| 1.0 \| 0.001854 \| 204 \| 0.359 \| 89 \| 54 \| 2 \| 1 \| 87 \| 1 \| 88 \| Uncharacterized protein \| Uncharacterized protein; afdb-uniprot50 \| AF-A0A436ZDM6-F1-MODEL\_V4 \| 1.0 \| 0.001854 \| 194 \| 0.281 \| 96 \| 65 \| 2 \| 1 \| 92 \| 1 \| 96 \| Uncharacterized protein \| Uncharacterized protein; afdb-uniprot50 \| AF-A0A0Q5F466-F1-MODEL\_V4 \| 1.0 \| 0.002455 \| 185 \| 0.318 \| 91 \| 55 \| 3 \| 1 \| 87 \| 1 \| 88 \| Uncharacterized protein \| Uncharacterized protein; afdb-uniprot50 \| AF-A0A158E9V0-F1-MODEL\_V4 \| 1.0 \| 0.006368 \| 155 \| 0.239 \| 92 \| 62 \| 2 \| 1 \| 92 \| 1 \| 84 \| Uncharacterized protein \| Uncharacterized protein |
| Top keywords  (threshold 1.00e-02 (evalue)) | -- |
| Output files | ../../similar\_structures/63\_FANPEZAQ\_CDS\_0063\_afdb-proteome\_foldseek.tsv ../../similar\_structures/63\_FANPEZAQ\_CDS\_0063\_afdb-uniprot50\_foldseek.tsv ../../similar\_structures/63\_FANPEZAQ\_CDS\_0063\_merged.svg ../../similar\_structures/63\_FANPEZAQ\_CDS\_0063\_pdb\_foldseek.tsv |

  
  
  

Return to summary | Go to previous | Go to next

  


---

**Sequence/structure alignments coloring**  
Each object in the alignment figures is colored according to its E-value following this color coding:

1e-100
10
